# The landscape of antibody binding in SARS-CoV-2 infection

**DOI:** 10.1101/2020.10.10.334292

**Authors:** Anna S. Heffron, Sean J. McIlwain, Maya F. Amjadi, David A. Baker, Saniya Khullar, Ajay K. Sethi, Ann C. Palmenberg, Miriam A. Shelef, David H. O’Connor, Irene M. Ong

**Affiliations:** Department of Pathology and Laboratory Medicine, University of Wisconsin-Madison, Madison, WI, United States of America; Department of Biostatistics and Medical Informatics, University of Wisconsin-Madison, Madison, WI, United States of America; University of Wisconsin Carbone Comprehensive Cancer Center, University of Wisconsin-Madison, Madison, WI, United States of America; Department of Medicine, University of Wisconsin-Madison, Madison, WI, United States of America; Department of Population Health Sciences, University of Wisconsin-Madison, Madison, WI, United States of America; Department of Biochemistry, Institute for Molecular Virology, University of Wisconsin–Madison, Madison, WI, United States of America; William S. Middleton Memorial Veterans Hospital, Madison, WI, United States of America; Wisconsin National Primate Research Center, University of Wisconsin-Madison, Madison, Wisconsin, United States of America; Department of Obstetrics and Gynecology, University of Wisconsin-Madison, Madison, WI, United States of America

## Abstract

The search for potential antibody-based diagnostics, vaccines, and therapeutics for pandemic severe acute respiratory syndrome coronavirus 2 (SARS-CoV-2) has focused almost exclusively on the spike (S) and nucleocapsid (N) proteins. Coronavirus membrane (M), ORF3a, and ORF8 proteins are humoral immunogens in other coronaviruses (CoVs) but remain largely uninvestigated for SARS-CoV-2. Here we use ultradense peptide microarray mapping to show that SARS-CoV-2 infection induces robust antibody responses to epitopes throughout the SARS-CoV-2 proteome, particularly in M, in which one epitope achieved excellent diagnostic accuracy. We map 79 B cell epitopes throughout the SARS-CoV-2 proteome and demonstrate that antibodies that develop in response to SARS-CoV-2 infection bind homologous peptide sequences in the six other known human CoVs. We also confirm reactivity against four of our top-ranking epitopes by enzyme-linked immunosorbent assay (ELISA). Illness severity correlated with increased reactivity to nine SARS-CoV-2 epitopes in S, M, N, and ORF3a in our population. Our results demonstrate previously unknown, highly reactive B cell epitopes throughout the full proteome of SARS-CoV-2 and other CoV proteins.

## Introduction

Antibodies correlate with protection from coronaviruses (CoVs) including severe acute respiratory syndrome coronavirus 2 (SARS-CoV-2) [1–8], severe acute respiratory syndrome coronavirus (SARS-CoV) [8–12] and Middle Eastern respiratory syndrome coronavirus (MERS-CoV) [8, 13–16]. All CoVs encode four main structural proteins, spike (S), envelope (E), membrane (M), and nucleocapsid (N), as well as multiple non-structural proteins and accessory proteins [17]. In SARS-CoV-2, anti-S and anti-N antibodies have received the most attention to date [1–8], including in serology-based diagnostic tests [1–5] and vaccine candidates [6–8]. The immunogenicity of S-based vaccines is variable [18, 19], so better representation of the breadth of antibody reactivity in vaccines, therapeutics, and diagnostics will be important as the pandemic continues especially as new variants emerge. Prior reports observed that not all individuals infected with SARS-CoV-2 produce detectable antibodies against S or N [1– 5], indicating a need for expanded antibody-based options.

Much less is known about antibody responses to other SARS-CoV-2 proteins, though data from other CoVs suggest they may be important. Antibodies against SARS-CoV M can be more potent than antibodies against SARS-CoV S [20–22], and some experimental SARS-CoV and MERS-CoV vaccines elicit responses to M, E, and ORF8 [8]. Additionally, previous work has demonstrated humoral cross-reactivity between CoVs [7, 11, 23–26] and suggested it could be protective [26, 27], although full-proteome cross-reactivity has not been investigated.

We designed a peptide microarray tiling the proteomes of SARS-CoV-2 and eight other human and animal CoVs in order to assess antibody epitope specificity and potential cross-reactivity with other CoVs. We examined IgG antibody responses in 40 COVID-19 convalescent patients and 20 SARS-CoV-2-naïve controls. Independent ELISAs confirm four of the highest-performing epitopes. We detected antibody responses to epitopes throughout the SARS-CoV-2 proteome, with several antibodies exhibiting apparent cross-reactive binding to homologous epitopes in multiple other CoVs.

## Results

### SARS-CoV-2-naïve controls show consistent binding in “common cold” CoVs and limited binding in SARS-CoV-2, SARS-CoV, and MERS-CoV

Greater than 90% of adult humans are seropositive for the human “common cold” CoVs (CCCoVs: HCoV-HKU1, HCoV-OC43, HCoV-NL63, and HCoV-229E) [28, 29], but the effect of these pre-existing antibodies upon immune responses to SARS-CoV-2 or other CoVs remains uncertain. We measured IgG reactivity in sera from 20 SARS-CoV-2-naïve control subjects to CoV linear peptides, considering reactivity that was >3.00 standard deviations above the mean for the log_2_-quantile normalized array data to be indicative of antibody binding [30]. All sera (SARS-CoV-2-naïve and COVID-19-convalescent) exhibited binding in known epitopes of at least one of the control non-CoV strains (poliovirus vaccine and rhinovirus; Fig. 1, Extended data 1) and all were collected in Wisconsin, USA, where exposure to SARS-CoV or MERS-CoV was extremely unlikely. We found that at least one epitope in structural or accessory proteins showed binding in 100% of controls for HCoV-HKU1, 85% of controls for HCoV-OC43, 65% for HCoV-NL63, and 55% for HCoV-229E (Fig. 2, Extended data 1). The apparent cross-reactive binding was observed in 45% of controls for MERS-CoV, 50% for SARS-CoV, and 50% for SARS-CoV-2.

**Figure 1.**
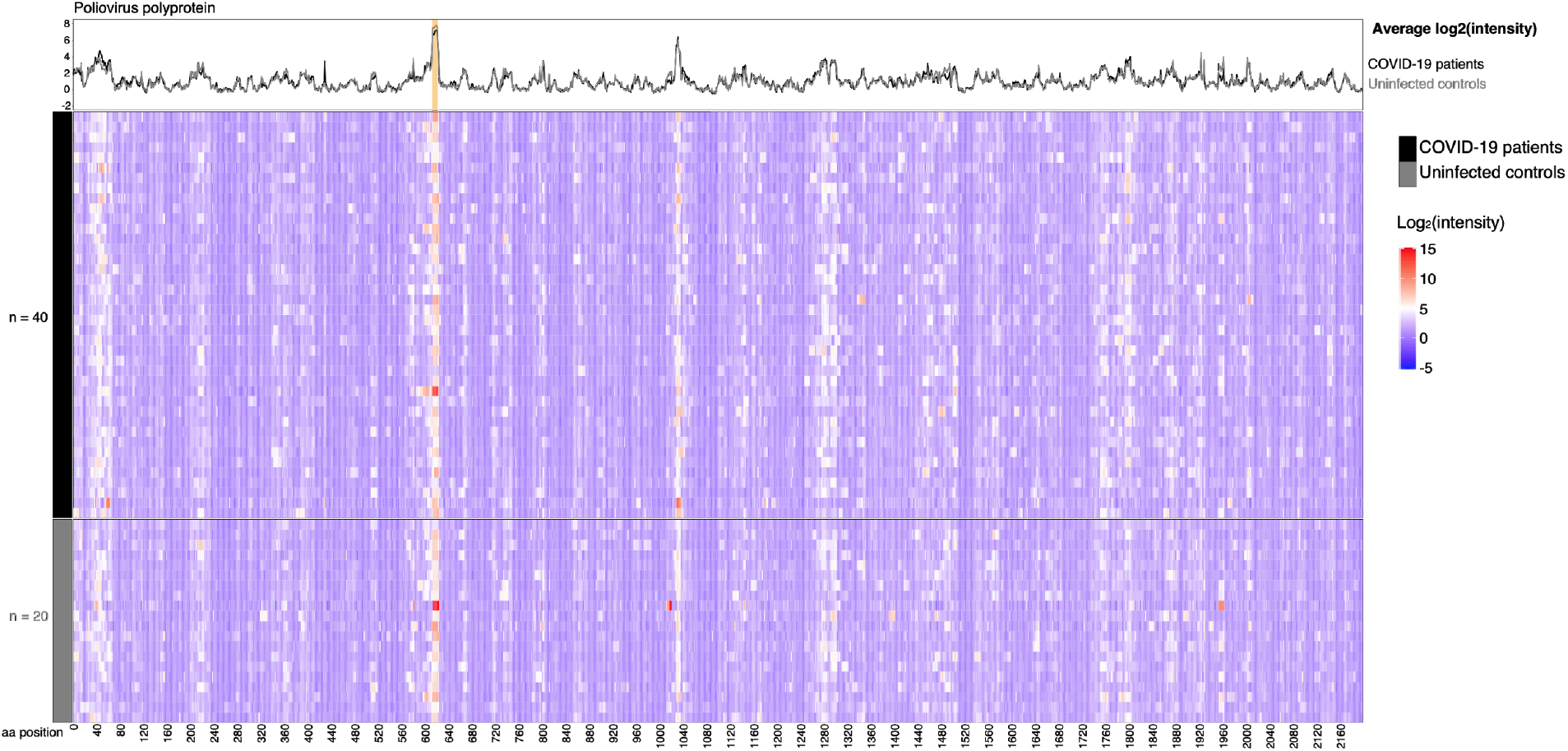
Patients and control subjects show reactivity to a poliovirus control. Sera from 20 control subjects collected before 2019 were assayed for IgG binding to the full proteome of human poliovirus 1 on a peptide microarray. Binding was measured as reactivity that was >3.00 standard deviations above the mean for the log_2_-quantile normalized array data. Patients and controls alike showed reactivity to a well-documented linear poliovirus epitope (start position 613 [IEDB.org]; orange shading in line plot).

**Figure 2.**
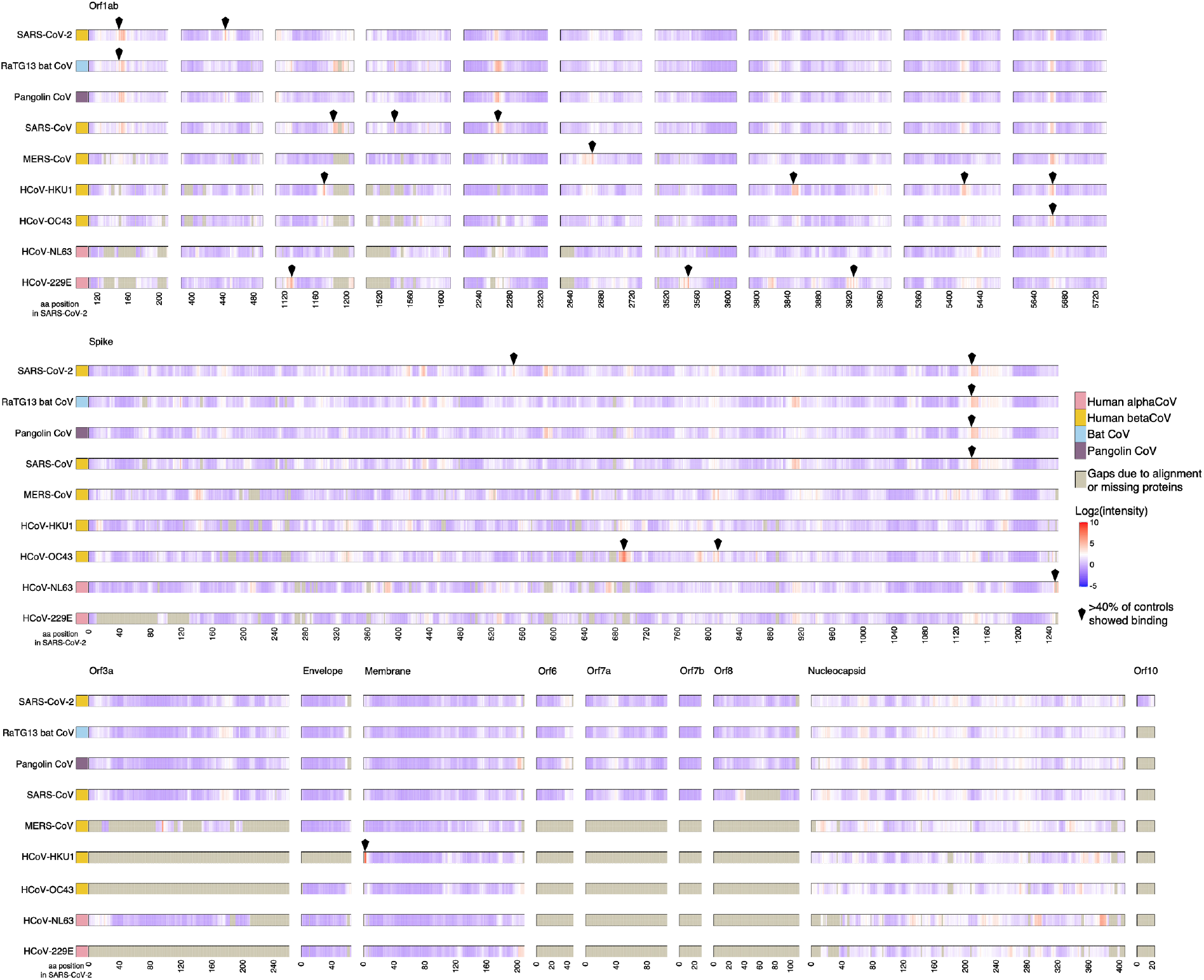
Control sera show reactivity to CCCoVs and to SARS-CoV, MERS-CoV, and SARS-CoV-2. Sera from 20 control subjects collected before 2019 were assayed for IgG binding to the full proteomes of nine CoVs on a peptide microarray. Viral proteins are shown aligned to the SARS-CoV-2 proteome with each virus having an individual panel; SARS-CoV-2 amino acid (aa) position is represented on the x-axis. Binding was measured as reactivity that was >3.00 standard deviations above the mean for the log_2_-quantile normalized array data. Peptides for which >40% of the controls showed binding are indicated by a diamond.

### SARS-CoV-2 infection induces antibodies binding throughout the proteome

We aimed to map the full breadth of IgG binding induced by SARS-CoV-2 infection and to rank the identified epitopes in terms of likelihood of immunodominance. We defined epitope recognition as antibody binding to contiguous peptides in which the average log_2_-normalized intensity for patients was at least 2-fold greater than for controls with *t-*test statistics yielding adjusted *p*-values <0.1. We chose these criteria, rather than the 3.00 standard deviation cut-off (Extended data 2), in order to ensure that binding detected would be greater than background binding seen in controls (2-fold greater) and to remove regions of binding that were not at least weakly significantly different from controls (adjusted *p*<0.1).

These criteria identified 79 B cell epitopes (Fig. 3, Table 1) in S, M, N, ORF1ab, ORF3a, ORF6, and ORF8. We ranked these epitopes by minimum adjusted *p*-value for any 16-mer in the epitope in order to determine the greatest likelihood of difference from controls as a proxy for likelihood of immunodominance. The highest-ranking epitope occurred in the N-terminus of M (1-M-24). Patient sera showed high-magnitude reactivity (up to an average of 6.7 fluorescence intensity units) in other epitopes in S, M, N, and ORF3a, with lower-magnitude reactivity (average of <3.3 fluorescence intensity units) epitopes in other proteins. The epitopes with the greatest reactivity in S were located in the S2 subunit of the protein (residues 686–1273) rather than the S1 subunit (residues 14-685) [6] (Fig. 3). The greatest reactivity in S occurred in the fusion peptide (residues 788-806) and at the base of the extracellular portion of the protein (between the heptad repeat 1 and heptad repeat 2, roughly residues 984-1163) (Fig. 3, Fig. 4). The highest magnitude antibody binding (red sites in Fig. 4) on S are below the flexible head region that must be in the “up” position for ACE2 binding to occur. Notably less reactivity occurred in the receptor-binding domain (residues 319-541) [6]. Four detected epitopes (553-S-26, 624-S-23, 807-S-26, and 1140-S-25) have previously been shown to be potently neutralizing [31–33], and all four of these were ranked within the top 10 of our 79 epitopes. Forty-two of our detected epitopes (including 1-M-24, 553-S-26, 624-S-23, 807-S-26, and 1140-S-25; Table 1) confirm bioinformatic predictions of antigenicity based on SARS-CoV and MERS-CoV [7, 8, 34–36], including each of the 12 topranking epitopes.

**Figure 3.**
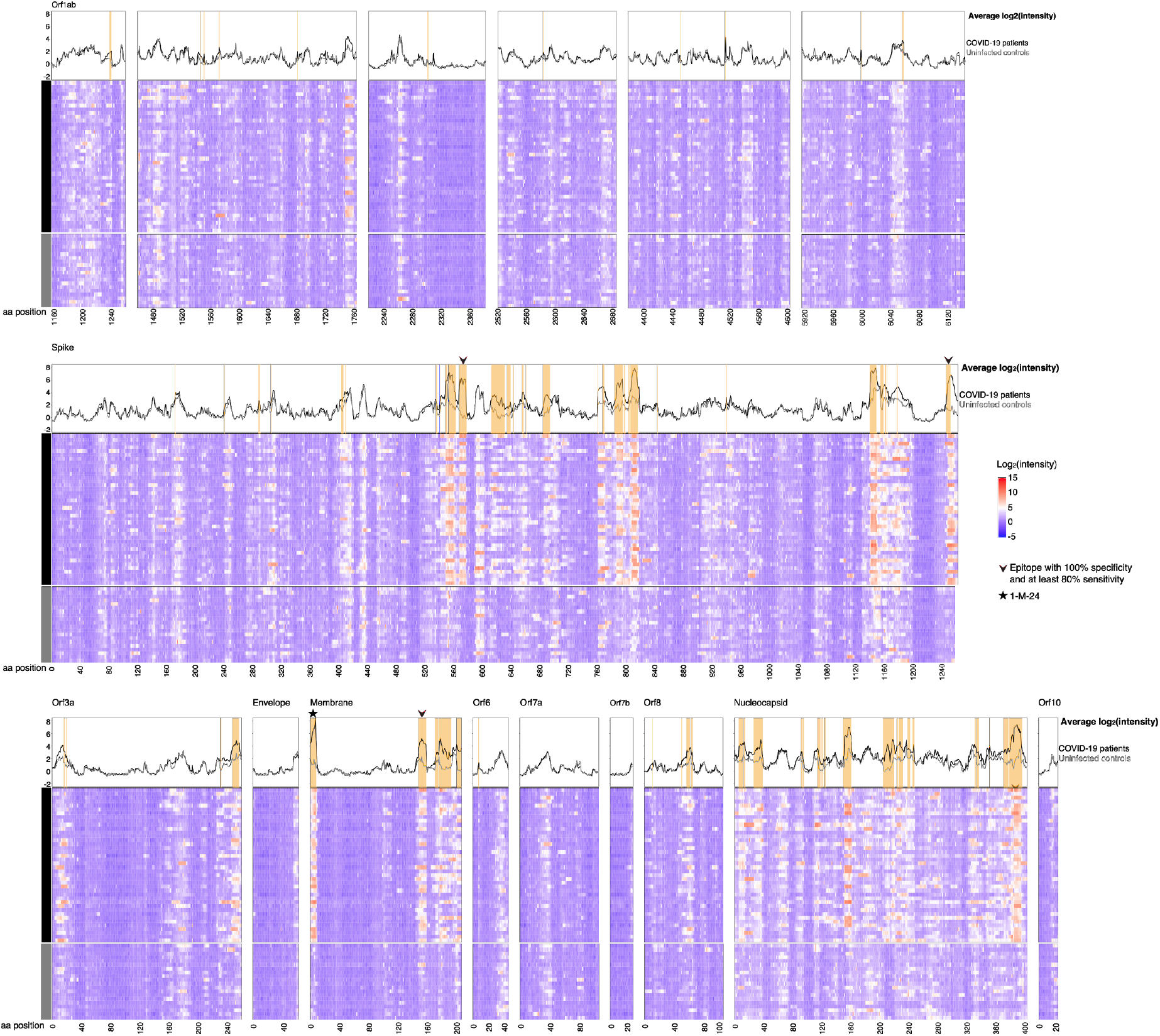
Anti-SARS-CoV-2 antibodies bind throughout the viral proteome. Sera from 40 COVID-19 convalescent subjects were assayed for IgG binding to the full SARS-CoV-2 proteome on a peptide microarray. B cell epitopes were defined as peptides in which patients’ average log_2_-normalized intensity (black lines in line plots) is 2-fold greater than controls’ (gray lines in line plots) and *ŕ*-test statistics yield adjusted *p-*values < 0.1; epitopes are identified by orange shading in the line plots.

**Table 1.**
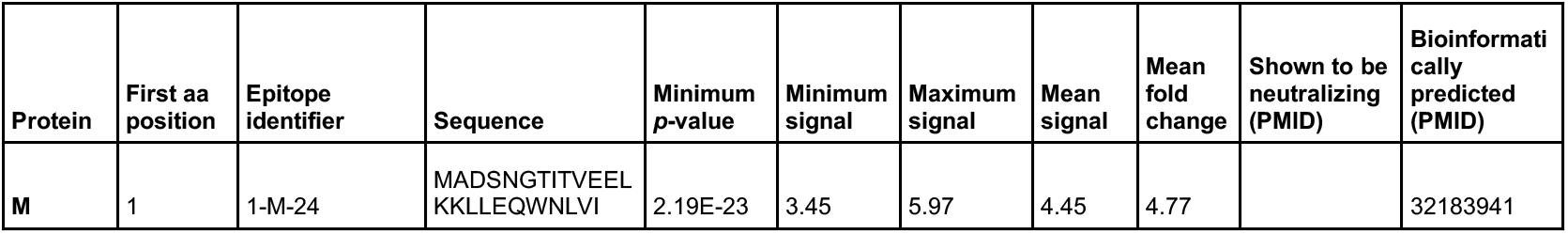
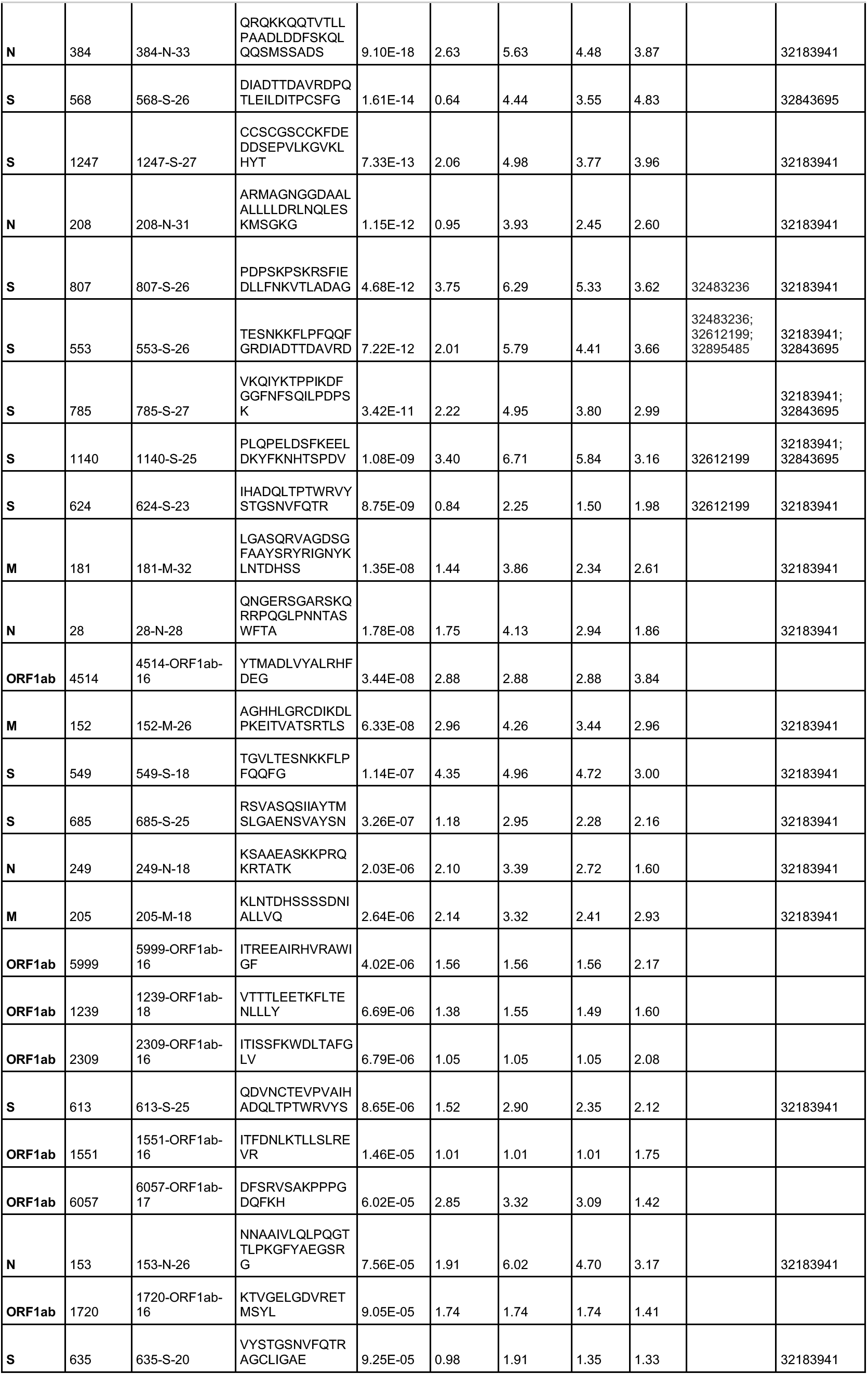
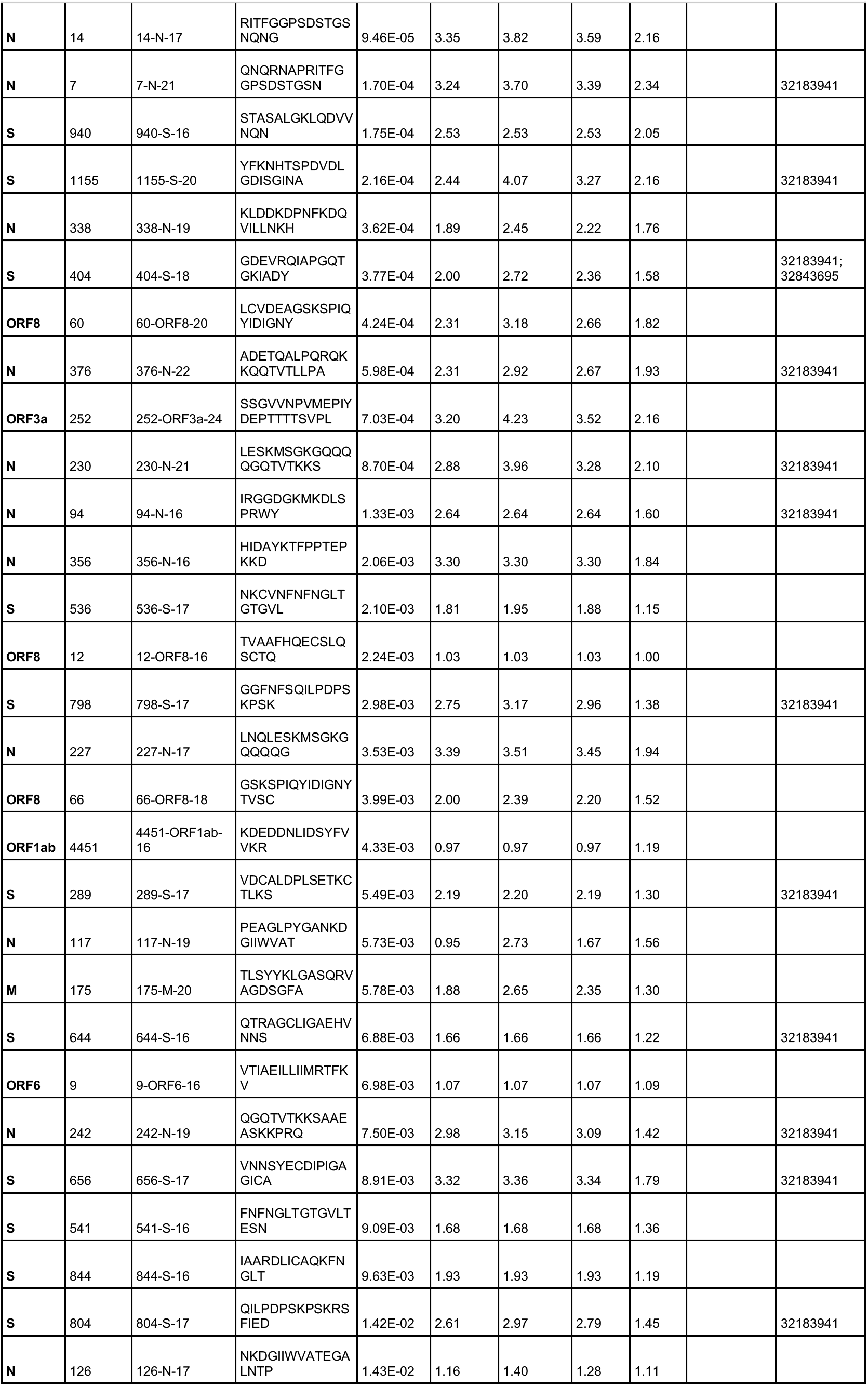
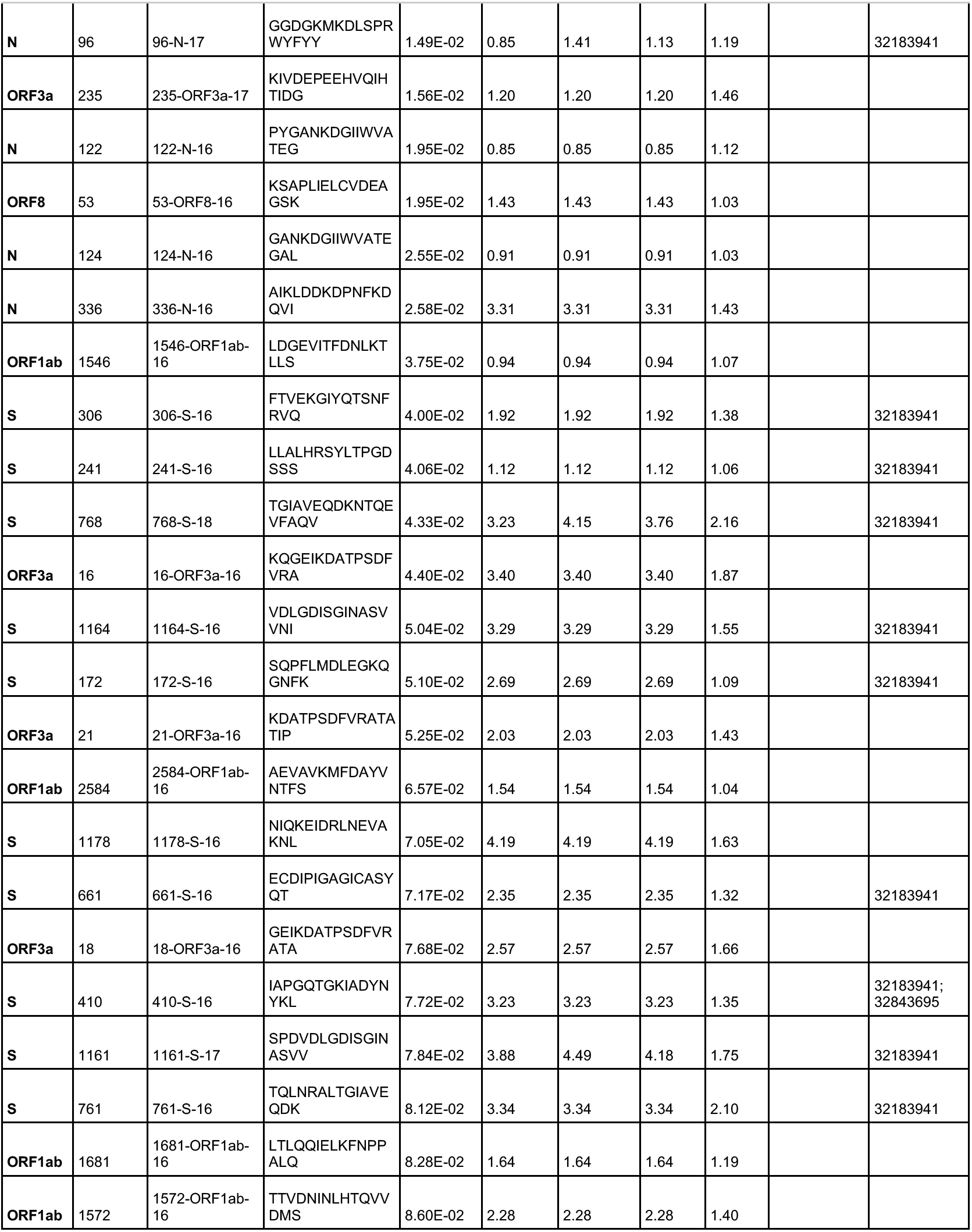
Profiling antibody binding in 40 COVID-19 convalescent patients compared to 20 naïve controls identifies B cell epitopes in SARS-CoV-2 (all data is log_2_-normalized).

**Figure 4.**
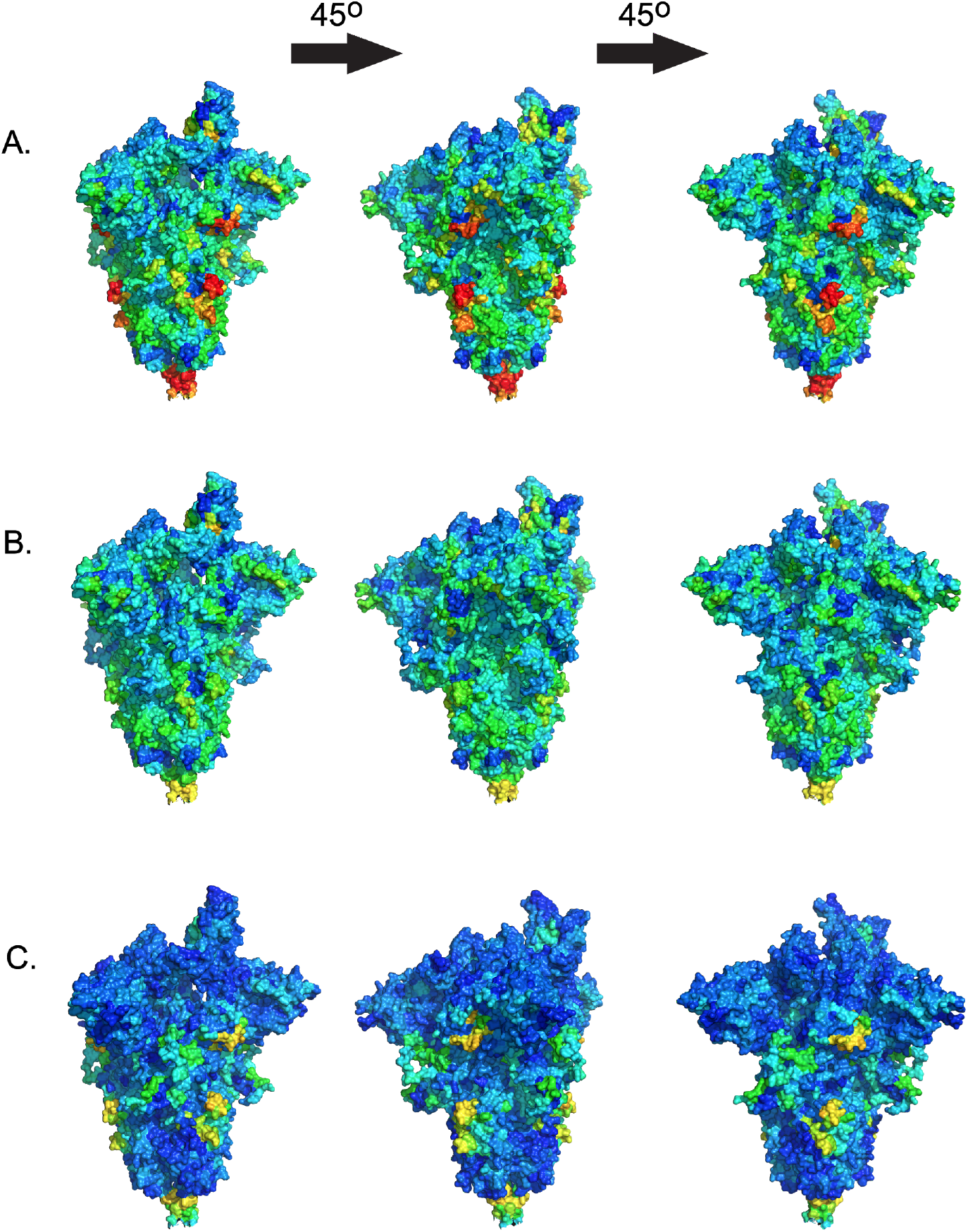
Anti-SARS-CoV-2 antibodies to S protein show the highest binding in the fusion cleavage site. Binding reactivities for COVID-19 convalescent patients (A), naïve controls (B), and the difference between patients and controls (C) were localized on a coordinate file for a trimer of the SARS-CoV-2 S protein using a dark blue (low, 0.00 fluorescence intensity) to red (high, 9.00 fluorescence intensity) color scale. The highest reactivity occurred in the fusion peptide (aa 788-806) and at the base of the extracellular portion of the molecule (aa 984-1163), with lower reactivity in the receptor-binding domain (aa 319-541).

The highest specificity (100%) and sensitivity (98%), determined by linear discriminant analysis leave-one-out cross-validation, for any individual peptide was observed for a 16-mer within the 1-M-24 epitope: ITVEELKKLLEQWNLV (Extended data 3). Fifteen additional individual peptides in M, S, and N had 100% measured specificity and at least 80% sensitivity (Table 2). Combinations of 1-M-24 with one of five other epitopes (384-N-33, 807-S-26, 6057-ORF1ab-17, 227-N-17, 4451-ORF1ab-16) yielded an area under the curve receiver operating characteristic of 1.00 (Extended data 4) based on linear discriminant analysis leave-one-out-cross-validation.

**Table 2.**
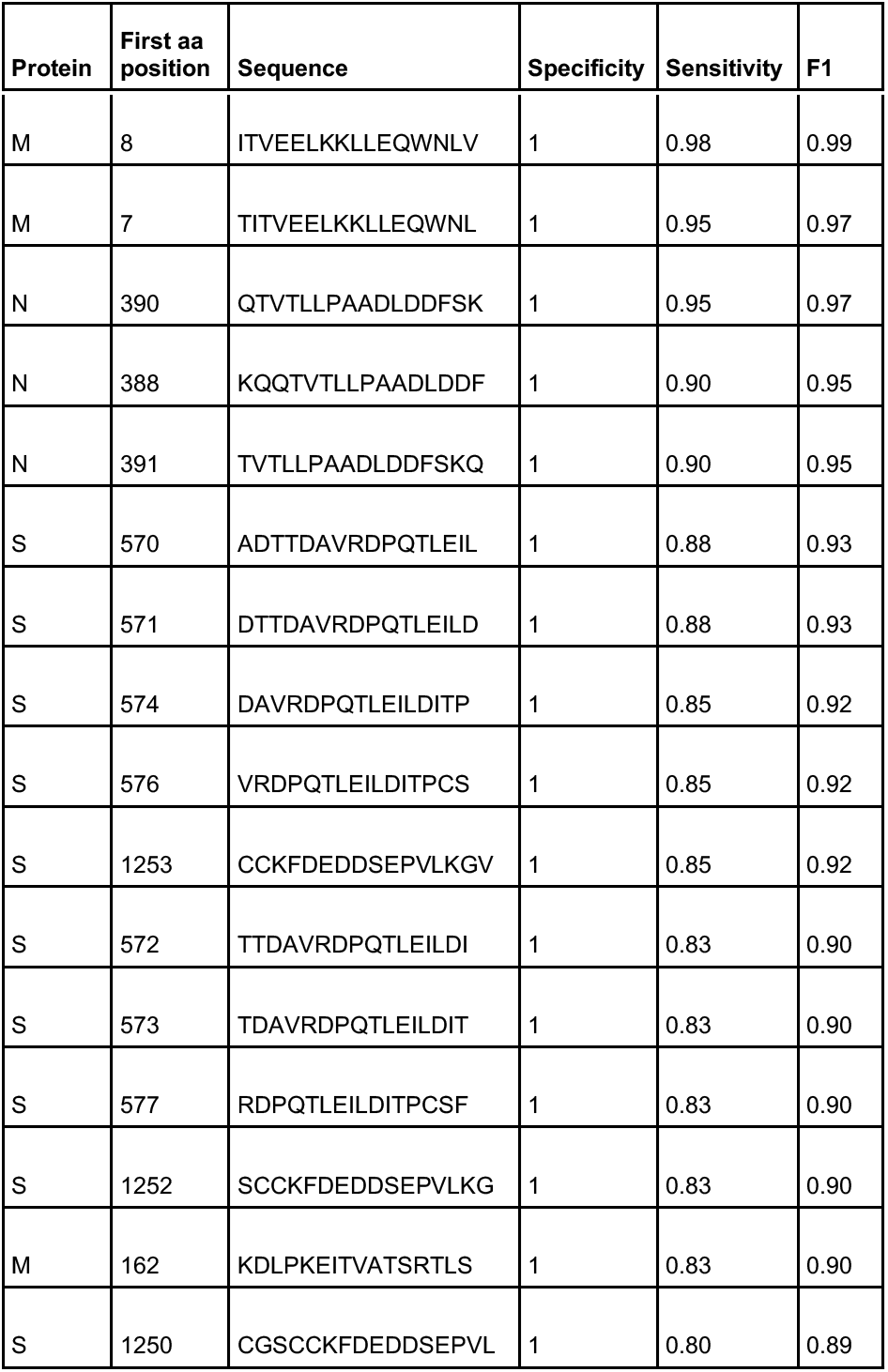
Sixteen peptides in the SARS-CoV-2 proteome had 100% specificity and at least 80% sensitivity for SARS-CoV-2 infection in 40 COVID-19 convalescent patients compared to 20 naïve controls

### Anti-SARS-CoV-2 antibodies may cross-reactively bind peptides in other CoVs

We determined epitopes bound by anti-SARS-CoV-2 antibodies in non-SARS-CoV-2 CoVs by the same criteria we used to determine epitopes in SARS-CoV-2. Epitopes in these viruses were defined as binding by antibodies in COVID-19 convalescent sera to peptides at an average log_2_-normalized intensity at least 2-fold greater than in controls with *t*-test statistics yielding adjusted *p*-values <0.1. Some of these epitopes were identical sequences with SARS-CoV-2, particularly in the RaTG13 bat betacoronavirus (β-CoV), the closest known relative of SARS-CoV-2 (96% nucleotide identity) [37, 38], the pangolin CoV (85% nucleotide identity with SARS-CoV-2) [39], and SARS-CoV(78% identity) [37]. Cross-reactivity of an antibody is typically determined by evaluating a pure preparation of specific antibodies or by competition assays. However, since our Wisconsin subjects are almost certainly naïve to MERS-CoV, SARS-CoV, and bat and pangolin CoVs, we can make predictions about cross-reactivity (as opposed to binding due to sequence identity).

Antibodies in COVID-19-convalescent sera appeared to be cross-reactive with identical or homologous epitopes in S, M, N, ORF1ab, ORF3, ORF6, and ORF8 in other CoVs (Fig. 5, Extended data 5, Extended data 6, Extended data 7). Overall, the greatest number of epitopes in any non-SARS-CoV-2 CoV occurred in the RaTG13 bat betacoronavirus (β-CoV) at 75 epitopes (60 identical to SARS-CoV-2, 10 homologous non-identical, five non-homologous non-identical). The second greatest number, 60 epitopes, occurred in the pangolin CoV (23 identical to SARS-CoV-2, 28 homologous non-identical, nine non-homologous non-identical), and third SARS-CoV with 45 epitopes, (10 identical to SARS-CoV-2, 30 homologous non-identical, five non-homologous non-identical) (Extended data 6, Extended data 7). One region, corresponding to SARS-CoV-2 epitope 807-S-26, showed binding or potential crossreactivity across all CoVs, and one, corresponding to SARS-CoV-2 epitope 1140-S-25, showed binding or potential cross-reactivity across all β-CoVs (Fig. 5). Epitope 807-S-26 includes the CoV S fusion peptide, and 1140-S-25 is immediately adjacent to the heptad repeat region 2, both of which are involved in membrane fusion [40].

**Figure 5.**
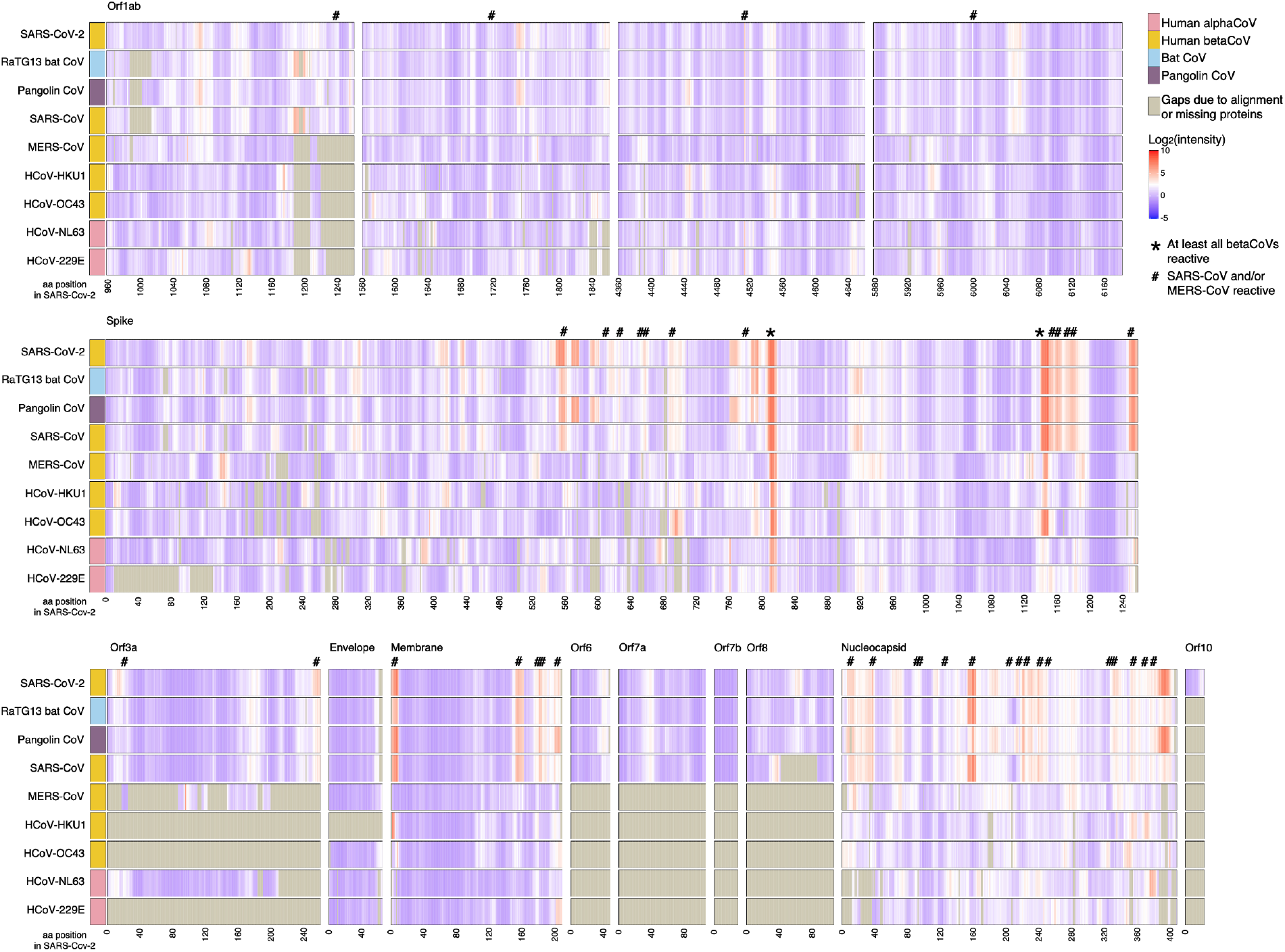
Anti-SARS-CoV-2 antibodies may cross-react with other CoVs. Sera from 40 COVID-19 convalescent patients were assayed for IgG binding to 9 CoVs on a peptide microarray; averages for all 40 are shown. Viral proteins are aligned to the SARS-CoV-2 proteome; SARS-CoV-2 amino acid (aa) position is represented on the x-axis. Regions that may be cross-reactive across all β-CoVs (*) or cross-reactive for SARS-CoV or MERS-CoV (#) are indicated. Gray shading indicates gaps due to alignment or lacking homologous proteins. Cross-reactive binding is defined as peptides in which patients’ average log_2_-normalized intensity is 2-fold greater than controls’ and *t-*test statistics yield adjusted *p*-values < 0.1.

### Enzyme-linked immunosorbent assays (ELISAs) confirm peptide microarray findings

Having determined reactivity and apparent cross-reactivity by peptide array, we aimed to independently confirm and validate these findings by ELISA. We selected four peptides for ELISA evaluation (1253-S-16, 814-S-16, 8-M-16, and 390-N-16) from those in our top 10 ranked epitopes, considering diversity among the proteins represented, neutralizing capacity and potential cross-reactivity across multiple CoVs, and using the 16-mer in each epitope that most correctly discriminated between patients and controls. All four SARS-CoV-2 peptides had higher IgG binding in COVID-19 convalescent sera than in controls (Fig. 6). Peptide 8-M-16 showed the greatest discrimination between COVID-19 convalescent and control sera with only three COVID-19 convalescent samples having values similar to controls. Both peptides 1253-S-16 and 814-S-16 showed greater binding in controls than either 8-M-16 or 390-N-16, confirming our findings of greater potential cross-reactivity among epitopes found in S.

**Figure 6.**
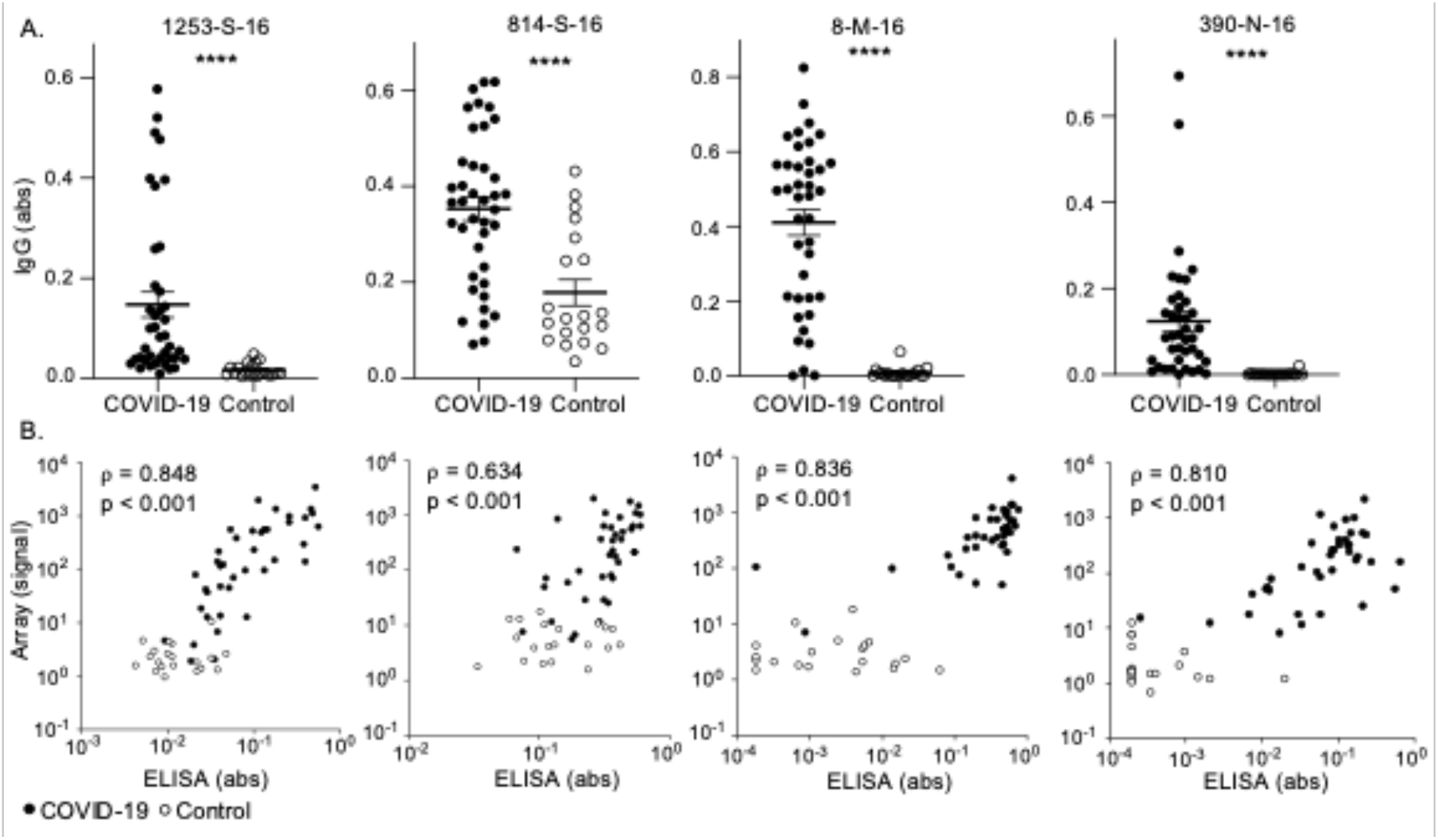
Higher IgG binding to SARS-CoV-2 peptides in COVID-19 convalescent subjects compared to controls by ELISA. (A) IgG binding to SARS-CoV-2 peptides in COVID-19 convalescent (n=40) and naïve control (n=20) sera was measured by ELISA. Bars indicate mean absorbance (abs) +/- SEM and \*\*\*\**p*<0.0001 by *t*-test. (B) Anti-SARS-CoV-2 peptide IgG detected by ELISA was compared to array findings by Spearman’s rank-order correlation (Spearman correlation coefficient, ρ) for COVID-19 convalescent (n=40, closed circles) and control (n=20, open circles) sera.

### Reactivity in some epitopes correlates with disease severity

Increased antibody titer and duration have been associated with increased severity of illness due to infection with SARS-CoV-2 [41–45] and other CoVs [46], though data on epitope-level differences by severity is lacking [47]. We compared reactivity in patients within our cohort whose COVID-19 course required intubation and mechanical ventilation (n=8) with reactivity in COVID-19 convalescent patients who never required hospitalization (n=25) using multilinear regression accounting for age, sex, immunocompromising conditions, and Charlson comorbidity index score [48] to determine epitope-level resolution of differences in reactivity. Nine epitopes in S, M, N, and ORF3a showed statistically significant (*p*<0.05) increases in reactivity for intubated patients relative to never-hospitalized patients (Fig. 7a, Extended data 8).

**Figure 7.**
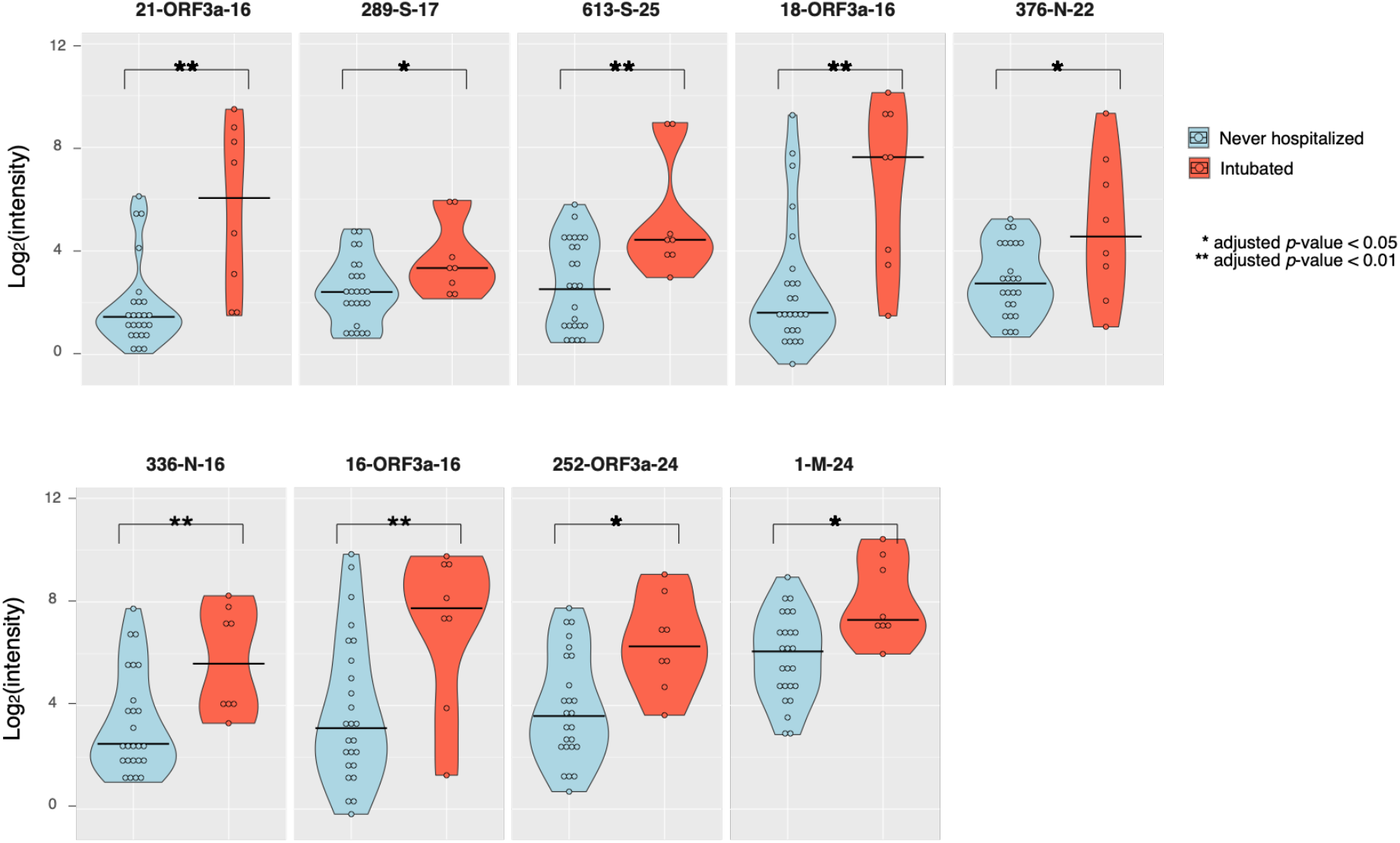
Disease severity correlates with increased antibody binding in specific SARS-CoV-2 epitopes. IgG reactivity against SARS-CoV-2 epitopes identified by peptide microarray in COVID-19 convalescent patients who were never hospitalized versus intubated patients showed statistically significant increases in reactivity in intubated patients for 11 epitopes.

## Discussion

In our analysis of antibody binding to the full proteome of SARS-CoV-2, the highest magnitude binding of anti-SARS-CoV-2 antibodies from human sera occurred for an epitope in the N-terminus of M protein, with high specificity and sensitivity. Antibodies produced after infection with SARS-CoV-2 reacted with epitopes throughout the proteomes of other human and non-human CoVs, recognizing homologous regions across all CoVs. Taken together, these results confirm that humans mount strong, broad antibody responses to SARS-CoV-2 proteins in addition to S and N, and implicate M epitopes as highly relevant to diagnostic and potentially to vaccine design.

M proteins are the most abundant proteins in CoV virions [17]. The N-terminus of M is known in other CoVs to be a small, glycosylated ectodomain that protrudes outside the virion and interacts with S, N, and E [17], while the rest of M resides within the viral particle. Full-length SARS-CoV M has been shown to induce protective antibodies [20, 49], and patterns of antibodies binding to SARS-CoV M are similar to those we found in SARS-CoV-2 [34]. SARS-CoV anti-M antibodies can synergize with anti-S and anti-N antibodies for improved neutralization [20, 49], and M has been used in protective SARS-CoV and MERS-CoV vaccines [8]. However, the mechanism of protection of anti-M antibodies remains unknown, and this protein remains largely understudied and underutilized as an antigen. Other groups have not previously identified the high magnitude binding we observed for M, though that may be due to using earlier sample timepoints or different techniques, populations, or computational algorithms [50, 51]. Notably, some of the highest binding we observed in the S protein occurred at the base of the extracellular portion of the protein, which would be the site of the putative interaction between SARS-CoV-2 S and M. The ACE2 binding site and the S helix in extended fusion are not as immunodominant as expected, suggesting that other, less-investigated epitopes may be playing a larger role in immunity to SARS-CoV-2 than is currently appreciated. Our results, in concert with prior knowledge of anti-SARS-CoV antibodies, strongly suggest that epitopes in M, particularly the 1-M-24 epitope as well as other novel epitopes we identified, should be investigated further as potential targets in SARS-CoV-2 diagnostics, vaccines, and therapeutics. Interestingly, we found antibodies bind three of the non-S mutations, in ORF8 and N, in the B.1.1.7 variant of SARS-CoV-2 that has recently emerged in the United Kingdom and which is considered to potentially be more transmissible than previous known variants [52].

We also found that antibodies produced in response to SARS-CoV-2 infection appear to bind peptides representing homologous epitopes throughout the proteomes of other human and non-human CoVs. Hundreds of CoVs have been discovered in bats and other species [27, 37–39, 53, 54], making future spillovers inevitable. The potential broad cross-reactivity we observed in some homologous peptide sequences may help guide the development of pan-CoV vaccines [15], especially given that antibodies binding to 807-S-26 and 1140-S-25, which showed potential cross-reactivity across all CoVs and all β-CoVs, respectively, are known to be potently neutralizing [31, 32]. A caveat is that our methods cannot discern whether the increased IgG binding to CCCoVs in COVID-19 convalescent sera is due to newly developed cross-reactive antibodies or due to the stimulation of a memory response against the original CCCoV antigens. However, cross-reactivity of anti-SARS-CoV-2 antibodies with SARS-CoV or MERS-CoV is likely real, since our population was very unlikely to have been exposed to those viruses. A more stringent assessment of cross-reactivity as well as functional investigations into these cross-reactive antibodies will be vital in determining their capacity for cross-protection. Further, our methods efficiently detect antibody binding to linear epitopes [55], but their sensitivity for detecting parts of conformational epitopes is unknown, and additional analyses will be required to determine whether epitopes identified induce neutralizing or otherwise protective antibodies.

Finally, we demonstrated that more severely ill patients have significantly greater reactivity to certain epitopes in S, M, N, and ORF3a. The nine epitopes with significantly higher magnitude reactivity in intubated patients may play a role in the overaggressive immune response known to characterize severe COVID-19 [7, 56], suggesting that they may be targets for treatment in or prevention of severe disease. Alternatively, the antibody response in general may be higher in very sick patients, expanding the repertoire of antibody reactivity. Future studies should investigate whether these differences can be detected early in the disease course to determine their potential utility as predictive markers of disease severity. The correlation of reactivity to CCCoVs with reactivity to SARS-CoV-2 in all subjects including uninfected controls suggests preexisting antibodies to CCCoVs may be relevant to an individual’s capacity to effectively produce anti-SARS-CoV-2 antibodies, especially given that pre-existing anti-CoV antibodies are more common in children and adolescents [57].

Many questions remain regarding the biology and immunology related to SARS-CoV-2. Our extensive profiling of epitope-level resolution antibody reactivity in COVID-19 convalescent subjects, confirmed by independent assays, provides new epitopes that could serve as important targets in the development of improved diagnostics, vaccines, and therapeutics against SARS-CoV-2 and dangerous human CoVs that may emerge in the future.

## Extended data

**Extended data 1.**Percentages of the 40 COVID-19 convalescent patients and 20 naïve controls reacted to known epitopes in at least one control virus (rhinovirus and poliovirus strains).

**Extended data 2.**Percentages and individual data for the 40 COVID-19 convalescent patients and 20 naïve controls showing log_2_-normalized fluorescence intensity at least 3.00 standard deviations above the mean for the array for nine species of CoVs.

**Extended data 3.**Specificity and sensitivity for past SARS-CoV-2 infection in 40 COVID-19 convalescent patients compared to 20 naïve controls of individual 16-mer peptides comprising epitopes throughout the full SARS-CoV-2 proteome.

**Extended data 4.**Epitopes paired with the 1-M-24 epitope obtained an area under the receiver operating characteristic curve (AUC-ROC) of 1.0 for SARS-CoV-2 infection in 40 COVID-19 convalescent patients and 20 naïve controls using leave-one-out cross validation with linear discriminant analysis.

**Extended data 5.**Alignment of epitopes in human and animal CoVs for which antibodies in sera from 40 COVID-19 convalescent patients showed apparent crossreactive binding. Alignments were performed in Geneious Prime 2020.1.2 (Auckland, New Zealand).

**Extended data 6.**Cross-reactive binding of antibodies against other CoVs in 40 COVID-19 convalescent patients compared to 20 naïve controls.

**Extended data 7.**Cross-reactive binding of antibodies in 40 COVID-19 convalescent patients compared to 20 naïve controls in protein motifs in other CoVs aligned to SARS-CoV-2.

**Extended data 8.**Comparison of antibody binding in SARS-CoV-2 B cell epitopes in 8 intubated COVID-19 convalescent patients compared to 25 symptomatic but never hospitalized COVID-19 convalescent patients compared by multilinear regression accounting for age, sex, immunocompromising conditions, and Charlson comorbidity index score.

## Acknowledgments

The authors are grateful to Dr. Christina Newman, Dr. Nathan Sherer, Dr. Thomas Friedrich, Dr. Amelia Haj, Dr. James Gern, Dr. Christine Seroogy, and Gage Moreno for their thoughtful comments and helpful discussions in preparing this manuscript. The authors are grateful to Dr. Robert Kirchdoerfer for generously providing the chimeric pdb file used for structural representations of SARS-CoV-2 spike protein in this work.

## Author contributions

A.S.H., S.J.M., D.A.B., M.F.A., M.A.S., D.H.O. and I.M.O. conceptualized this study.

A.S.H., D.A.B., and I.M.O. created the array design. M.F.A. selected patient and control samples and performed ELISA assays under the guidance of M.A.S.. A.S.H. and M.F.A. collected clinical and demographic information by medical record chart review under the guidance of M.A.S.. A.S.H., S.J.M., M.F.A., D.A.B., S.K., A.C.P., and I.M.O. performed data normalizations, analyses, and validations, and created graphical data visualizations. A.S.H., S.J.M., M.F.A., D.A.B., A.K.S., and I.M.O. performed formal statistical analyses. S.J.M., D.A.B., S.K., and I.M.O. wrote the custom software scripts used. A.S.H. and M.F.A. wrote the original draft of the manuscript with input from M.A.S, D.H.O., and I.M.O.. A.S.H., S.J.M., D.A.B., M.F.A., and I.M.O. wrote sections of the methods. All authors contributed to reviewing and editing.

The authors declare the following competing interests: A.S.H., S.J.M., D.A.B., M.F.A., S.K., M.A.S., D.H.O., and I.M.O are listed as the inventors on a patent filed that is related to findings in this study. Application: 63/080568, 63/083671. Title: IDENTIFICATION OF SARS-COV-2 EPITOPES DISCRIMINATING COVID-19 INFECTION FROM CONTROL AND METHODS OF USE. Application type: Provisional. Status: Filed. Country: United States. Filing date: September 18, 2020, September 25, 2020.

## Methods

### Peptide microarray design and synthesis

Viral protein sequences were selected and submitted to Nimble Therapeutics (Madison, WI, USA) for development into a peptide microarray [55]. Sequences represented include proteomes of all seven coronaviruses known to infect humans, proteomes of closely related coronaviruses found in bats and pangolins, and spike proteins from other coronaviruses (accession numbers and replicates per peptide shown in **Supplementary Table 1**). A number of proteins were included as controls, including poliovirus, seven strains of human rhinovirus, and human cytomegalovirus 65kDa phosphoprotein. We chose these controls given that we expect most human adults will have antibody reactivity to at least one of these proteins and proteomes. Accession numbers used to represent each viral protein are listed in the supplemental material (accession numbers and replicates per peptide shown in **Supplementary Table 1**). All proteins were tiled as 16 amino acid peptides overlapping by 15 amino acids. All unique peptides were tiled in a lawn of thousands of copies, with each unique peptide represented in at least 3 and up to 5 replicates (**Supplementary Table 1**). The peptide sequences were synthesized in situ with a Nimble Therapeutics Maskless Array Synthesizer (MAS) by light-directed solid-phase peptide synthesis using an amino-functionalized support (Geiner Bio-One) coupled with a 6-aminohexanoic acid linker and amino acid derivatives carrying a photosensitive 2-(2-nitrophenyl) propyloxycarbonyl (NPPOC) protection group (Orgentis Chemicals). Unique peptides were synthesized in random positions on the array to minimize impact of positional bias. Each array consists of twelve subarrays, where each subarray can process one sample and each subarray contains up to 389,000 unique peptide sequences.

**Supplementary Table 1.** Proteins represented on the peptide microarray

|  | Protein(s) | GenBank accession number(s) | Number of replicates of each unique peptide |
| --- | --- | --- | --- |
| **Coronavirus proteins** | Severe acute respiratory syndrome coronavirus 2 proteome | NC_045512.2 | 4-5 |
|  | Severe acute respiratory syndrome coronavirus proteome | NC_004718.3 | 3 |
|  | Middle Eastern respiratory syndrome coronavirus proteome | NC_019843.3 | 3 |
|  | Human coronavirus HKU1 proteome | NC_006577.2 | 3 |
|  | Human coronavirus OC43 proteome | NC_006213.1 | 3 |
|  | Human coronavirus 229E proteome | NC_002645.1 | 3 |
|  | Human coronavirus NL63 proteome | NC_005831.2 | 3 |
|  | Bat coronavirus (RaTG13 isolate) proteome | MN996532.1 | 3 |
|  | Pangolin coronavirus proteome | MT072864.1 | 3 |
| **Control proteins** | Human rhinovirus A1 polyprotein | NC_038311.1 | 3 |
|  | Human rhinovirus A7 polyprotein | DQ473503.1 | 3 |
|  | Human rhinovirus A16 polyprotein | L24917.1 | 3 |
|  | Human rhinovirus A36 polyprotein | JX074050.1 | 3 |
|  | Human rhinovirus C2 polyprotein | EF077280.1 | 3 |
|  | Human rhinovirus C15 polyprotein | GU219984.1 | 3 |
|  | Human rhinovirus C41 polyprotein | KY189321.1 | 3 |
|  | Human poliovirus 1 polyprotein | ANA67904.1 | 3 |

### Human subjects

The study was conducted in accordance with the Declaration of Helsinki and approved by the Institutional Review Board of the University of Wisconsin-Madison. Clinical data and sera from subjects infected with SARS-CoV-2 were obtained from the University of Wisconsin (UW) COVID-19 Convalescent Biobank and from control subjects (sera collected prior to 2019) from the UW Rheumatology Biobank [58]. All subjects were 18 years of age or older at the time of recruitment and provided informed consent. COVID-19 convalescent subjects had a positive SARS-COV-2 PCR test at UW Health with sera collected 5-6 weeks after self-reported COVID-19 symptom resolution except blood was collected for one subject after 9 weeks. Age, sex, medications, and medical problems were abstracted from UW Health’s electronic medical record (EMR). Race and ethnicity were self-reported. Hospitalization and intubation for COVID-19 and smoking status at the time of blood collection (controls) or COVID-19 were obtained by EMR abstraction and self-report and were in complete agreement. Two thirds of COVID-19 convalescent subjects and all controls had a primary care appointment at UW Health within 2 years of the blood draw as an indicator of the completeness of the medical information. Subjects were considered to have an immunocompromising condition if they met any of the following criteria: immunosuppressing medications, systemic inflammatory or autoimmune disease, cancer not in remission, uncontrolled diabetes (secondary manifestations or hemoglobin A1c >7.0%), or congenital or acquired immunodeficiency. Control and COVID-19 subjects were similar in regard to demographics and health (**Supplementary Table 2**), and subjects who were not hospitalized, were hospitalized, or were hospitalized and intubated also were compared (**Supplementary Table 3**). No subjects were current smokers.

**Supplementary Table 2.**
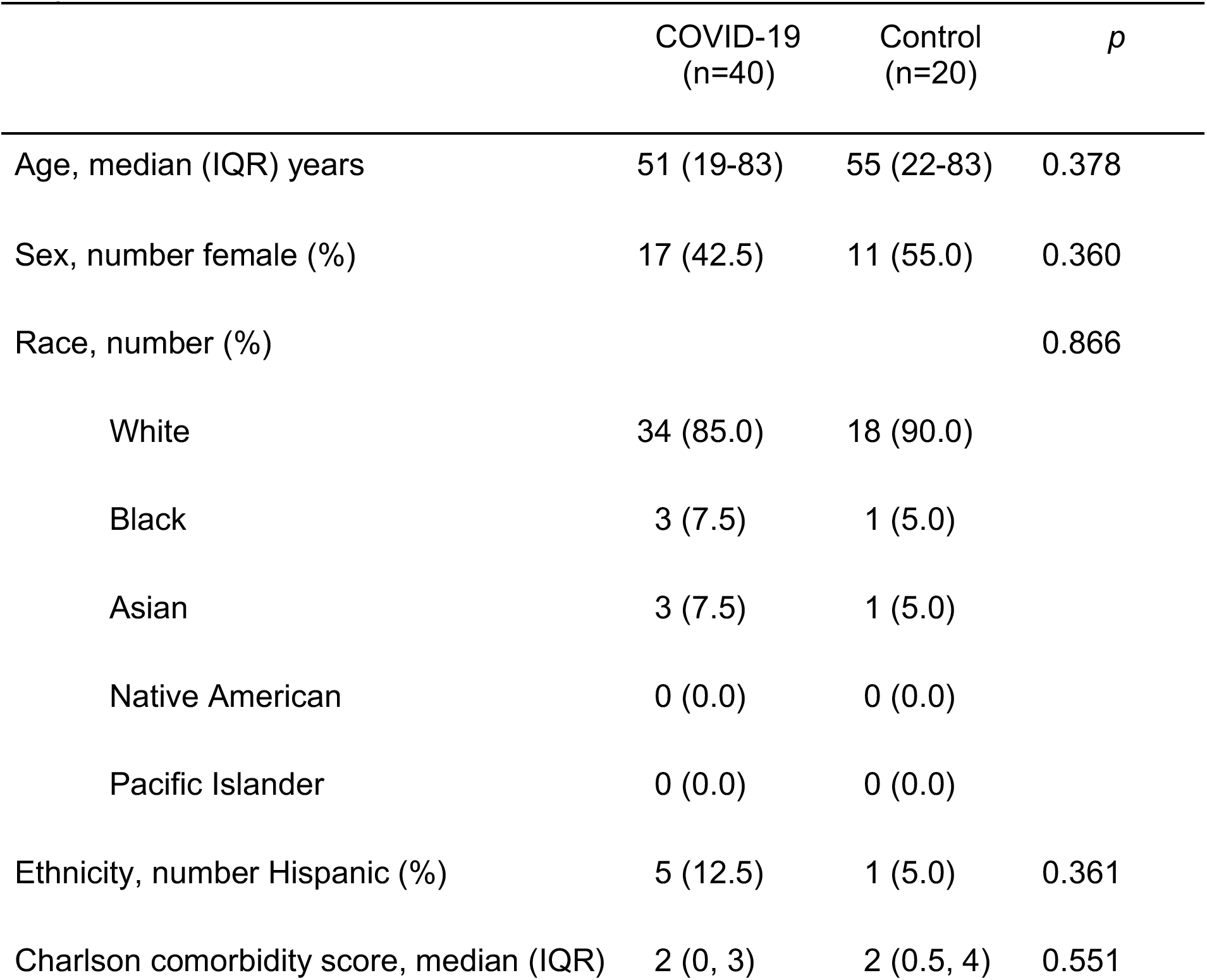
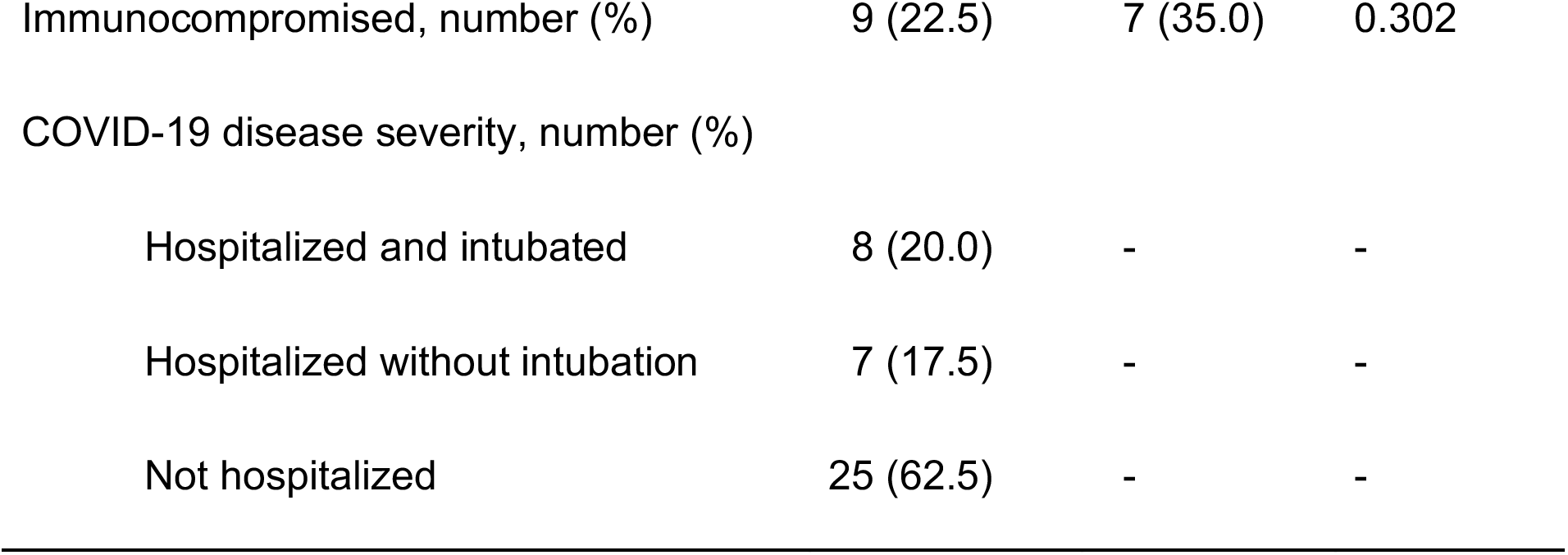
Characteristics of COVID-19 Convalescent and Control Subjects

|  | COVID-19<br>(n=40) | Control<br>(n=20) | *p* |
| --- | --- | --- | --- |
| Age, median (IQR) years | 51 (19-83) | 55 (22-83) | 0.378 |
| Sex, number female (%) | 17 (42.5) | 11 (55.0) | 0.360 |
| Race, number (%) |  |  | 0.866 |
| White | 34 (85.0) | 18 (90.0) |  |
| Black | 3 (7.5) | 1 (5.0) |  |
| Asian | 3 (7.5) | 1 (5.0) |  |
| Native American | 0 (0.0) | 0 (0.0) |  |
| Pacific Islander | 0 (0.0) | 0 (0.0) |  |
| Ethnicity, number Hispanic (%) | 5 (12.5) | 1 (5.0) | 0.361 |
| Charlson comorbidity score, median (IQR) | 2 (0, 3) | 2 (0.5, 4) | 0.551 |
| Immunocompromised, number (%) | 9 (22.5) | 7 (35.0) | 0.302 |
| COVID-19 disease severity, number (%) |  |  |  |
| Hospitalized and intubated | 8 (20.0) | - | - |
| Hospitalized without intubation | 7 (17.5) | - | - |
| Not hospitalized | 25 (62.5) | - | - |

**Supplementary Table 3.** Characteristics of COVID-19 Convalescent Subjects According to Hospitalization Status

| Characteristic | Not<br>hospitalize<br>d (n=25) | Hospitalized<br>without<br>intubation<br>(n=7) | Hospitalized<br>and<br>intubated<br>(n=8) | *p* |
| --- | --- | --- | --- | --- |
| Age, median (IQR) years | 49 (30, 56) | 66 (48, 83) | 63 (58, 68) | 0.013 |
| Sex, number female (%) | 12 (48.0) | 3 (42.9) | 2 (25.0) | 0.519 |
| Race, number (%) |  |  |  | 0.537 |
| White | 20 (80.0) | 6 (85.7) | 8 (100.0) |  |
| Black | 3 (12.0) | 0 (0.0) | 0 (0.0) |  |
| Asian | 2 (8.0) | 1 (14.3) | 0 (0.0) |  |
| Native American | 0 (0.0) | 0 (0.0) | 0 (0.0) |  |
| Pacific Islander | 0 (0.0) | 0 (0.0) | 0 (0.0) |  |
| Ethnicity, number Hispanic (%) | 5 (20.0) | 0 (0.0) | 0 (0.0) | 0.180 |
| Charlson comorbidity score, median (IQR) | 1 (0, 2) | 2 (0, 6) | 2.5 (2, 4) | 0.008 |
| Immunocompromised, number (%) | 5 (20.0) | 2 (28.6) | 2 (25.0) | 0.875 |

### Peptide array sample binding

Samples were diluted 1:100 in binding buffer (0.01M Tris-Cl, pH 7.4, 1% alkali-soluble casein, 0.05% Tween-20) and bound to arrays overnight at 4°C. After sample binding, the arrays were washed 3× in wash buffer (1× TBS, 0.05% Tween-20), 10 min per wash. Primary sample binding was detected via Alexa Fluor^®^ 647-conjugated goat antihuman IgG secondary antibody (Jackson ImmunoResearch). The secondary antibody was diluted 1:10,000 (final concentration 0.1 ng/μl) in secondary binding buffer (1× TBS, 1% alkali-soluble casein, 0.05% Tween-20). Arrays were incubated with secondary antibody for 3 h at room temperature, then washed 3× in wash buffer (10 min per wash), washed for 30 sec in reagent-grade water, and then dried by spinning in a microcentrifuge equipped with an array holder. The fluorescent signal of the secondary antibody was detected by scanning at 635 nm at 2 μm resolution using an Innopsys 910AL microarray scanner. Scanned array images were analyzed with proprietary Nimble Therapeutics software to extract fluorescence intensity values for each peptide.

### Peptide microarray findings validation

We included sequences on the array of viruses which we expected all adult humans to be likely to have been exposed to as positive controls: one poliovirus strain (measuring vaccine exposure), and seven rhinovirus strains. Any subject whose sera did not react to at least one positive control would be considered a failed run and removed from the analysis. All subjects in this analysis reacted to epitopes in at least one control strain (Fig. 1, Extended data 1).

### Peptide microarray data analysis and data availability

The raw fluorescence signal intensity values were log_2_ transformed. Clusters of fluorescence intensity of statistically unlikely magnitude, indicating array defects, were identified and removed. Local and large area spatial corrections were applied, and the median transformed intensity of the peptide replicates was determined. The resulting median data was cross-normalized using quantile normalization. All peptide microarray datasets and code used in these analyses can be downloaded from https://github.com/Ong-Research/Ong_UW_Adult_Covid-19.git.

### Protein structures

The SARS-CoV-2 S-chimera.pdb used to make S protein structures was built by Robert Kirchdoerfer using 6VYB.pdb, 5X4S.pdb and 6LZG coordinates and filling in internal unresolved residues from known (presumably) analogous sites determined for SARS-CoV S from 6CRV.pdb. Additional unmodeled regions were generated using Modeller [59]. C-proximal HR2 regions were modeled as single helices (Phe1148-Leu1211) in Coot [60].

The data2bfactor Python script written by Robert L. Campbell, Thomas Holder, and Suguru Asai (downloaded from http://pldserver1.biochem.queensu.ca/~rlc/work/pymol/) was used to substitute peptide array data onto this structure in place of the B factor in PyMol (The PyMOL Molecular Graphics System, Version 2.0 Schrödinger, LLC) using a dark blue (low) to red (high) color scale. Data used for these visualizations was the average reactivity in the 40 COVID-19 convalescent patients, the average reactivity in the 20 naïve controls, and the difference between averages for the patients and for the controls.

### Enzyme-linked immunosorbent assays (ELISAs)

Costar 96-well high binding plates (Corning, Corning, USA) were incubated at 4°C overnight with 5μg/ml streptavidin (Thermo Fisher Scientific, Waltham, USA) in PBS (Corning). Plates were washed twice with PBS and incubated at room temperature for 1 hour with 0.5mM of the following peptides (Biomatik, Kitchener, Canada) in PBS: 814-S-16 (KRSFIEDLLFNKVTLA-K-biotin), 1253-S-16 (CCKFDEDDSEPVLKGV-K-biotin), 390-N-16 (QTVTLLPAADLDDFSK-K-biotin), 8-M-16 (ITVEELKKLLEQWNLV-K-biotin). Plates were washed thrice with wash buffer (0.2% Tween-20 in PBS), then incubated for 1 hour in blocking solution (5% nonfat dry milk in wash buffer) at room temperature, incubated overnight at 4°C with sera at 1:200 in blocking solution, washed four times with wash buffer, incubated for 1 hour at room temperature with mouse anti-human IgG conjugated to horse radish peroxidase (Southern Biotech, Birmingham, USA) diluted 1:5000 in blocking solution, washed four times with wash buffer, and incubated with tetramethyl benzidine substrate solution (Thermo Fisher Scientific) for 5 minutes followed by 0.18M sulfuric acid. Absorbance was read on a FilterMax F3 Multi-mode Microplate reader (Molecular Devices, San Jose, USA) at 450 and 562nm. Background signal from 562nm absorbance and wells with no peptide and no serum were subtracted. Plates were normalized using a pooled serum sample on every plate. Absorbance values of zero were plotted as 0.0002 to allow a log scale for graphs. Samples were run in duplicate.

### Statistical analysis

Statistical analyses were performed in R (v 4.0.2) using in-house scripts. For each peptide, a *p*-value from a two-sided *t*-test with unequal variance between sets of patient and control responses were calculated and adjusted using the Benjamini-Hochberg (BH) algorithm. To determine whether the peptide was in an epitope (in SARS-CoV-2 proteins) or cross-reactive for anti-SARS-CoV-2 antibodies (in non-SARS-CoV-2 proteins), we used an adjusted *p*-value cutoff of <0.1 (based on multiple hypothesis testing correction for all 119,487 unique sequences on the array) and a fold-change of greater than or equal to 2 and grouped consecutive peptides as a represented epitope. Linear discriminant analysis leave-one-out cross validation was used to determine specificity and sensitivity on each peptide and from each epitope using the average signal of the component peptides.

To identify cross-reactive epitopes, we used each SARS-CoV-2 epitope sequence as a query, searched the database of proteins from the sequences in the peptide array using blastp (-word-size 2, num-targets 4000) to find homologous sequences in the bat, pangolin, and other human CoV strains, then determined whether the average log_2_-normalized intensity for these sequences in patients was at least 2-fold greater than in controls with *t*-test statistics yielding adjusted *p*-values <0.1. Each blast hit was then mapped back to the corresponding probe ranges.

The clinical and demographic characteristics of convalescent subjects were compared to those of the controls using *χ^2^* tests for categorical variables and Wilcoxon rank-sum tests for non-normally distributed continuous measures.

Heatmaps were created using the gridtext [61] and complexheatmap [62] packages in R. Alignments for heatmaps were created using MUSCLE [63].

## References

1. Deeks JJ, Dinnes J, Takwoingi Y et al. Antibody tests for identification of current and past infection with SARS-CoV-2. Cochrane Database Syst Rev. 2020;6:CD013652.

2. Liu W, Liu L, Kou G et al. Evaluation of Nucleocapsid and Spike Protein-Based Enzyme-Linked Immunosorbent Assays for Detecting Antibodies against SARS-CoV-2. J Clin Microbiol. 2020;58

3. Tré-Hardy M, Wilmet A, Beukinga I et al. Analytical and clinical validation of an ELISA for specific SARS-CoV-2 IgG, IgA, and IgM antibodies. J Med Virol. 2020

4. Lisboa Bastos M, Tavaziva G, Abidi SK et al. Diagnostic accuracy of serological tests for covid-19: systematic review and meta-analysis. BMJ.2020;370:m2516.

5. Ayouba A, Thaurignac G, Morquin D et al. Multiplex detection and dynamics of IgG antibodies to SARS-CoV2 and the highly pathogenic human coronaviruses SARS-CoV and MERS-CoV. J Clin Virol. 2020;129:104521.

6. Huang Y, Yang C, Xu XF, Xu W, Liu SW. Structural and functional properties of SARS-CoV-2 spike protein: potential antivirus drug development for COVID-19. Acta Pharmacol Sin. 2020;41:1141–1149.

7. Chen W. Promise and challenges in the development of COVID-19 vaccines. Hum Vaccin Immunother. 20201–5.

8. Ong E, Wong MU, Huffman A, He Y. COVID-19 Coronavirus Vaccine Design Using Reverse Vaccinology and Machine Learning. Front Immunol. 2020;11:1581.

9. Sui J, Li W, Murakami A et al. Potent neutralization of severe acute respiratory syndrome (SARS) coronavirus by a human mAb to S1 protein that blocks receptor association. Proc Natl Acad Sci U S A. 2004;101:2536–2541.

10. ter Meulen J, Bakker AB, van den Brink EN et al. Human monoclonal antibody as prophylaxis for SARS coronavirus infection in ferrets. Lancet. 2004;363:2139–2141.

11. Rockx B, Corti D, Donaldson E et al. Structural basis for potent cross-neutralizing human monoclonal antibody protection against lethal human and zoonotic severe acute respiratory syndrome coronavirus challenge. J Virol. 2008;82:3220–3235.

12. Martin JE, Louder MK, Holman LA et al. A SARS DNA vaccine induces neutralizing antibody and cellular immune responses in healthy adults in a Phase I clinical trial. Vaccine. 2008;26:6338–6343.

13. Jiang L, Wang N, Zuo T et al. Potent neutralization of MERS-CoV by human neutralizing monoclonal antibodies to the viral spike glycoprotein. Sci Transl Med. 2014;6:234ra59.

14. Chen Z, Bao L, Chen C et al. Human Neutralizing Monoclonal Antibody Inhibition of Middle East Respiratory Syndrome Coronavirus Replication in the Common Marmoset. J Infect Dis. 2017;215:1807–1815.

15. Burton DR, Walker LM. Rational Vaccine Design in the Time of COVID-19. Cell Host Microbe. 2020;27:695–698.

16. Zhou Y, Jiang S, Du L. Prospects for a MERS-CoV spike vaccine. Expert Rev Vaccines. 2018;17:677–686.

17. Maier HJ, Bickerton E, Britton P. Coronaviruses: methods and protocols. New York: Humana Press; Springer; 2015:xi, 285 pages.

18. Ravichandran S, Coyle EM, Klenow L et al. Antibody signature induced by SARS-CoV-2 spike protein immunogens in rabbits. Sci Transl Med. 2020;12

19. Knoll MD Wonodi C. Oxford-AstraZeneca COVID-19 vaccine efficacy. Lancet. 2020

20. Pang H, Liu Y, Han X et al. Protective humoral responses to severe acute respiratory syndrome-associated coronavirus: implications for the design of an effective protein-based vaccine. J Gen Virol. 2004;85:3109–3113.

21. Chow SC, Ho CY, Tam TT et al. Specific epitopes of the structural and hypothetical proteins elicit variable humoral responses in SARS patients. J Clin Pathol. 2006;59:468–476.

22. He Y, Zhou Y, Siddiqui P, Niu J, Jiang S. Identification of immunodominant epitopes on the membrane protein of the severe acute respiratory syndrome-associated coronavirus. J Clin Microbiol. 2005;43:3718–3726.

23. Tian X, Li C, Huang A et al. Potent binding of 2019 novel coronavirus spike protein by a SARS coronavirus-specific human monoclonal antibody.[letter]. Emerg Microbes Infect 2020;9(1):382–385.

24. Lv H, Wu NC, Tsang OT et al. Cross-reactive Antibody Response between SARS-CoV-2 and SARS-CoV Infections. Cell Rep. 2020;31:107725.

25. Pinto D, Park YJ, Beltramello M et al. Cross-neutralization of SARS-CoV-2 by a human monoclonal SARS-CoV antibody. Nature. 2020;583:290–295.

26. Nickbakhsh S, Ho A, Marques DFP, McMenamin J, Gunson RN, Murcia PR. Epidemiology of Seasonal Coronaviruses: Establishing the Context for the Emergence of Coronavirus Disease 2019. J Infect Dis. 2020;222:17–25.

27. Morens DM, Fauci AS. Emerging Pandemic Diseases: How We Got to COVID-19. Cell. 2020;182:1077–1092.

28. Gorse GJ, Patel GB, Vitale JN, O’Connor TZ. Prevalence of antibodies to four human coronaviruses is lower in nasal secretions than in serum. Clin Vaccine Immunol. 2010;17:1875–1880.

29. Premkumar L, Segovia-Chumbez B, Jadi R et al. The receptor binding domain of the viral spike protein is an immunodominant and highly specific target of antibodies in SARS-CoV-2 patients. Sci Immunol. 2020;5

30. Valentini D, Rao M, Rane L et al. Peptide microarray-based characterization of antibody responses to host proteins after bacille Calmette-Guérin vaccination. Int J Infect Dis. 2017;56:140–154.

31. Poh CM, Carissimo G, Wang B et al. Two linear epitopes on the SARS-CoV-2 spike protein that elicit neutralising antibodies in COVID-19 patients. Nat Commun. 2020;11:2806.

32. Zhang BZ, Hu YF, Chen LL et al. Mining of epitopes on spike protein of SARS-CoV-2 from COVID-19 patients.[letter]. Cell Res 2020;30(8):702–704.

33. Li Y, Lai DY, Zhang HN et al. Linear epitopes of SARS-CoV-2 spike protein elicit neutralizing antibodies in COVID-19 patients. Cell Mol Immunol. 2020;17:1095–1097.

34. Grifoni A, Sidney J, Zhang Y, Scheuermann RH, Peters B, Sette A. A Sequence Homology and Bioinformatic Approach Can Predict Candidate Targets for Immune Responses to SARS-CoV-2. Cell Host Microbe. 2020;27:671–680.e2.

35. Ahmed SF, Quadeer AA, McKay MR. Preliminary Identification of Potential Vaccine Targets for the COVID-19 Coronavirus (SARS-CoV-2) Based on SARS-CoV Immunological Studies. Viruses. 2020;12

36. Crooke SN, Ovsyannikova IG, Kennedy RB, Poland GA. Immunoinformatic identification of B cell and T cell epitopes in the SARS-CoV-2 proteome. Sci Rep. 2020;10:14179.

37. Zhou P, Yang XL, Wang XG et al. A pneumonia outbreak associated with a new coronavirus of probable bat origin. Nature. 2020;579:270–273.

38. Ge XY, Wang N, Zhang W et al. Coexistence of multiple coronaviruses in several bat colonies in an abandoned mineshaft. Virol Sin. 2016;31:31–40.

39. Xiao K, Zhai J, Feng Y et al. Isolation of SARS-CoV-2-related coronavirus from Malayan pangolins. Nature. 2020;583:286–289.

40. Tang T, Bidon M, Jaimes JA, Whittaker GR, Daniel S. Coronavirus membrane fusion mechanism offers a potential target for antiviral development. Antiviral Res. 2020;178:104792.

41. Behrens GMN, Cossmann A, Stankov MV et al. Perceived versus proven SARS-CoV-2-specific immune responses in health-care professionals. Infection. 2020;48:631–634.

42. Caturegli G, Materi J, Howard BM, Caturegli P. Clinical Validity of Serum Antibodies to SARS-CoV-2: A Case-Control Study. Ann Intern Med. 2020

43. Long QX, Tang XJ, Shi QL et al. Clinical and immunological assessment of asymptomatic SARS-CoV-2 infections. Nat Med. 2020;26:1200–1204.

44. Ko JH, Joo EJ, Park SJ et al. Neutralizing Antibody Production in Asymptomatic and Mild COVID-19 Patients, in Comparison with Pneumonic COVID-19 Patients. J Clin Med. 2020;9

45. Kirkcaldy RD, King BA, Brooks JT. COVID-19 and Postinfection Immunity: Limited Evidence, Many Remaining Questions. JAMA. 2020;323:2245–2246.

46. Huang AT, Garcia-Carreras B, Hitchings MDT et al. A systematic review of antibody mediated immunity to coronaviruses: kinetics, correlates of protection, and association with severity. Nat Commun. 2020;11:4704.

47. Juno JA, Tan HX, Lee WS et al. Humoral and circulating follicular helper T cell responses in recovered patients with COVID-19. Nat Med. 2020;26:1428–1434.

48. Charlson ME, Pompei P, Ales KL, MacKenzie CR. A new method of classifying prognostic comorbidity in longitudinal studies: development and validation. J Chronic Dis. 1987;40:373–383.

49. Shi SQ, Peng JP, Li YC et al. The expression of membrane protein augments the specific responses induced by SARS-CoV nucleocapsid DNA immunization. Mol Immunol. 2006;43:1791–1798.

50. Shrock E, Fujimura E, Kula T et al. Viral epitope profiling of COVID-19 patients reveals cross-reactivity and correlates of severity. Science. 2020

51. Mishra N, Huang X, Joshi S et al. Immunoreactive peptide maps of SARS-CoV-2 and other human coronaviruses. bioRxiv. 2020. https://www.biorxiv.org/content/10.1101/2020.08.13.249953v1.full.pdf.

52. ARTIC Network. Preliminary genomic characterisation of an emergent SARS-CoV-2 lineage in the UK defined by a novel set of spike mutations. 2020. https://virological.org/t/preliminary-genomic-characterisation-of-an-emergent-sars-cov-2-lineage-in-the-uk-defined-by-a-novel-set-of-spike-mutations/563

53. Woo PC, Lau SK, Huang Y, Yuen KY. Coronavirus diversity, phylogeny and interspecies jumping. Exp Biol Med (Maywood). 2009;234:1117–1127.

54. Anthony SJ, Johnson CK, Greig DJ et al. Global patterns in coronavirus diversity. Virus Evol. 2017;3:vex012.

55. Heffron AS, Mohr EL, Baker D et al. Antibody responses to Zika virus proteins in pregnant and non-pregnant macaques. PLoS Negl Trop Dis. 2018;12:e0006903.

56. Moore JB, June CH. Cytokine release syndrome in severe COVID-19. Science. 2020;368:473–474.

57. Ng KW, Faulkner N, Cornish GH et al. Preexisting and de novo humoral immunity to SARS-CoV-2 in humans. Science. 2020

58. Holmes CL, Peyton CG, Bier AM et al. Reduced IgG titers against pertussis in rheumatoid arthritis: Evidence for a citrulline-biased immune response and medication effects. PLoS One. 2019;14:e0217221.

59. Webb B, Sali A. Comparative Protein Structure Modeling Using MODELLER. Curr Protoc Bioinformatics. 2016;54:5.6.1-5.6.37.

60. Emsley P, Lohkamp B, Scott WG, Cowtan K. Features and development of Coot. Acta Crystallogr D Biol Crystallogr. 2010;66:486–501.

61. Wilke CO. gridtext: Improved Text Rendering Support for ‘Grid’ Graphics. R package version 0.1.1.2020

62. Gu Z, Eils R, Schlesner M. Complex heatmaps reveal patterns and correlations in multidimensional genomic data. Bioinformatics. 2016;32:2847–2849.

63. Edgar RC. MUSCLE: multiple sequence alignment with high accuracy and high throughput. Nucleic Acids Res. 2004;32:1792–1797.

