## Extended data 1 for "The landscape of antibody binding in SARS-CoV-2 infection"

| Extended data 1 | | | | | | |
|---|---|---|---|---|---|---|
| **Protein** | **Start position (peptide)** | **Stop position (peptide)** | **Number of peptides** | **Minimum fraction (per peptide) of control subjects with reactivity >3 sd above average** | **Maximum fraction (per peptide) of control subjects with reactivity >3 sd above average** | **Mean fraction (over all peptides in this contiguous stretch) of controlsubjects with reactivity >3 sd above average** |
| ANA67904.1_human-poliovirus-1 | 1 | 17 | 17 | 0.05 | 0.2 | 0.111764706 |
| ANA67904.1_human-poliovirus-1 | 24 | 24 | 1 | 0.05 | 0.05 | 0.05 |
| ANA67904.1_human-poliovirus-1 | 26 | 56 | 31 | 0.05 | 0.45 | 0.209677419 |
| ANA67904.1_human-poliovirus-1 | 58 | 65 | 8 | 0.05 | 0.35 | 0.175 |
| ANA67904.1_human-poliovirus-1 | 83 | 83 | 1 | 0.05 | 0.05 | 0.05 |
| ANA67904.1_human-poliovirus-1 | 97 | 97 | 1 | 0.05 | 0.05 | 0.05 |
| ANA67904.1_human-poliovirus-1 | 117 | 117 | 1 | 0.1 | 0.1 | 0.1 |
| ANA67904.1_human-poliovirus-1 | 149 | 149 | 1 | 0.05 | 0.05 | 0.05 |
| ANA67904.1_human-poliovirus-1 | 151 | 151 | 1 | 0.15 | 0.15 | 0.15 |
| ANA67904.1_human-poliovirus-1 | 153 | 156 | 4 | 0.05 | 0.1 | 0.0875 |
| ANA67904.1_human-poliovirus-1 | 182 | 182 | 1 | 0.1 | 0.1 | 0.1 |
| ANA67904.1_human-poliovirus-1 | 201 | 201 | 1 | 0.1 | 0.1 | 0.1 |
| ANA67904.1_human-poliovirus-1 | 203 | 205 | 3 | 0.05 | 0.05 | 0.05 |
| ANA67904.1_human-poliovirus-1 | 208 | 210 | 3 | 0.05 | 0.1 | 0.083333333 |
| ANA67904.1_human-poliovirus-1 | 212 | 222 | 11 | 0.05 | 0.3 | 0.163636364 |
| ANA67904.1_human-poliovirus-1 | 238 | 239 | 2 | 0.05 | 0.05 | 0.05 |
| ANA67904.1_human-poliovirus-1 | 280 | 280 | 1 | 0.05 | 0.05 | 0.05 |
| ANA67904.1_human-poliovirus-1 | 304 | 304 | 1 | 0.05 | 0.05 | 0.05 |
| ANA67904.1_human-poliovirus-1 | 316 | 317 | 2 | 0.05 | 0.05 | 0.05 |
| ANA67904.1_human-poliovirus-1 | 319 | 330 | 12 | 0.05 | 0.05 | 0.05 |
| ANA67904.1_human-poliovirus-1 | 344 | 344 | 1 | 0.05 | 0.05 | 0.05 |
| ANA67904.1_human-poliovirus-1 | 348 | 348 | 1 | 0.05 | 0.05 | 0.05 |
| ANA67904.1_human-poliovirus-1 | 351 | 351 | 1 | 0.05 | 0.05 | 0.05 |
| ANA67904.1_human-poliovirus-1 | 358 | 360 | 3 | 0.1 | 0.1 | 0.1 |
| ANA67904.1_human-poliovirus-1 | 362 | 366 | 5 | 0.05 | 0.15 | 0.11 |
| ANA67904.1_human-poliovirus-1 | 385 | 389 | 5 | 0.05 | 0.05 | 0.05 |
| ANA67904.1_human-poliovirus-1 | 391 | 393 | 3 | 0.05 | 0.05 | 0.05 |
| ANA67904.1_human-poliovirus-1 | 397 | 397 | 1 | 0.05 | 0.05 | 0.05 |
| ANA67904.1_human-poliovirus-1 | 399 | 399 | 1 | 0.05 | 0.05 | 0.05 |
| ANA67904.1_human-poliovirus-1 | 407 | 407 | 1 | 0.05 | 0.05 | 0.05 |
| ANA67904.1_human-poliovirus-1 | 466 | 466 | 1 | 0.05 | 0.05 | 0.05 |
| ANA67904.1_human-poliovirus-1 | 480 | 481 | 2 | 0.05 | 0.1 | 0.075 |
| ANA67904.1_human-poliovirus-1 | 496 | 496 | 1 | 0.05 | 0.05 | 0.05 |
| ANA67904.1_human-poliovirus-1 | 508 | 518 | 11 | 0.05 | 0.2 | 0.122727273 |
| ANA67904.1_human-poliovirus-1 | 523 | 523 | 1 | 0.05 | 0.05 | 0.05 |
| ANA67904.1_human-poliovirus-1 | 570 | 586 | 17 | 0.05 | 0.35 | 0.129411765 |
| ANA67904.1_human-poliovirus-1 | 588 | 624 | 37 | 0.05 | 1 | 0.417567568 |

| Extended data 1 | | | | | | |
|---|---|---|---|---|---|---|
| Protein | Start position (peptide) | Stop position (peptide) | Number of peptides | Minimum fraction (per peptide) of control subjects with reactivity >3 sd above average | Maximum fraction (per peptide) of control subjects with reactivity >3 sd above average | Mean fraction (over all peptides in this contiguous stretch) of controlsubjects with reactivity >3 sd above average |
| ANA67904.1_human-poliovirus-1 | 630 | 630 | 1 | 0.05 | 0.05 | 0.05 |
| ANA67904.1_human-poliovirus-1 | 657 | 657 | 1 | 0.05 | 0.05 | 0.05 |
| ANA67904.1_human-poliovirus-1 | 661 | 670 | 10 | 0.05 | 0.15 | 0.115 |
| ANA67904.1_human-poliovirus-1 | 715 | 715 | 1 | 0.05 | 0.05 | 0.05 |
| ANA67904.1_human-poliovirus-1 | 718 | 718 | 1 | 0.15 | 0.15 | 0.15 |
| ANA67904.1_human-poliovirus-1 | 722 | 722 | 1 | 0.05 | 0.05 | 0.05 |
| ANA67904.1_human-poliovirus-1 | 728 | 743 | 16 | 0.05 | 0.25 | 0.115625 |
| ANA67904.1_human-poliovirus-1 | 746 | 746 | 1 | 0.05 | 0.05 | 0.05 |
| ANA67904.1_human-poliovirus-1 | 754 | 754 | 1 | 0.05 | 0.05 | 0.05 |
| ANA67904.1_human-poliovirus-1 | 780 | 780 | 1 | 0.05 | 0.05 | 0.05 |
| ANA67904.1_human-poliovirus-1 | 787 | 798 | 12 | 0.05 | 0.35 | 0.1 |
| ANA67904.1_human-poliovirus-1 | 800 | 803 | 4 | 0.2 | 0.35 | 0.275 |
| ANA67904.1_human-poliovirus-1 | 812 | 812 | 1 | 0.25 | 0.25 | 0.25 |
| ANA67904.1_human-poliovirus-1 | 849 | 850 | 2 | 0.05 | 0.05 | 0.05 |
| ANA67904.1_human-poliovirus-1 | 852 | 859 | 8 | 0.05 | 0.1 | 0.0625 |
| ANA67904.1_human-poliovirus-1 | 862 | 862 | 1 | 0.1 | 0.1 | 0.1 |
| ANA67904.1_human-poliovirus-1 | 865 | 865 | 1 | 0.05 | 0.05 | 0.05 |
| ANA67904.1_human-poliovirus-1 | 898 | 899 | 2 | 0.05 | 0.05 | 0.05 |
| ANA67904.1_human-poliovirus-1 | 901 | 901 | 1 | 0.05 | 0.05 | 0.05 |
| ANA67904.1_human-poliovirus-1 | 905 | 905 | 1 | 0.05 | 0.05 | 0.05 |
| ANA67904.1_human-poliovirus-1 | 924 | 924 | 1 | 0.1 | 0.1 | 0.1 |
| ANA67904.1_human-poliovirus-1 | 928 | 928 | 1 | 0.05 | 0.05 | 0.05 |
| ANA67904.1_human-poliovirus-1 | 945 | 946 | 2 | 0.05 | 0.1 | 0.075 |
| ANA67904.1_human-poliovirus-1 | 958 | 958 | 1 | 0.05 | 0.05 | 0.05 |
| ANA67904.1_human-poliovirus-1 | 977 | 977 | 1 | 0.05 | 0.05 | 0.05 |
| ANA67904.1_human-poliovirus-1 | 1012 | 1020 | 9 | 0.05 | 0.1 | 0.061111111 |
| ANA67904.1_human-poliovirus-1 | 1023 | 1023 | 1 | 0.05 | 0.05 | 0.05 |
| ANA67904.1_human-poliovirus-1 | 1026 | 1035 | 10 | 0.1 | 0.85 | 0.65 |
| ANA67904.1_human-poliovirus-1 | 1041 | 1042 | 2 | 0.05 | 0.05 | 0.05 |
| ANA67904.1_human-poliovirus-1 | 1089 | 1089 | 1 | 0.05 | 0.05 | 0.05 |
| ANA67904.1_human-poliovirus-1 | 1107 | 1107 | 1 | 0.05 | 0.05 | 0.05 |
| ANA67904.1_human-poliovirus-1 | 1109 | 1110 | 2 | 0.05 | 0.05 | 0.05 |
| ANA67904.1_human-poliovirus-1 | 1125 | 1134 | 10 | 0.05 | 0.3 | 0.1 |
| ANA67904.1_human-poliovirus-1 | 1138 | 1138 | 1 | 0.05 | 0.05 | 0.05 |
| ANA67904.1_human-poliovirus-1 | 1142 | 1146 | 5 | 0.15 | 0.2 | 0.19 |
| ANA67904.1_human-poliovirus-1 | 1148 | 1148 | 1 | 0.05 | 0.05 | 0.05 |
| ANA67904.1_human-poliovirus-1 | 1150 | 1150 | 1 | 0.05 | 0.05 | 0.05 |

| Extended data 1 | | | | | | |
|---|---|---|---|---|---|---|
| Protein | Start position (peptide) | Stop position (peptide) | Number of peptides | Minimum fraction (per peptide) of control subjects with reactivity >3 sd above average | Maximum fraction (per peptide) of control subjects with reactivity >3 sd above average | Mean fraction (over all peptides in this contiguous stretch) of controlsubjects with reactivity >3 sd above average |
| ANA67904.1_human-poliovirus-1 | 1155 | 1164 | 10 | 0.05 | 0.05 | 0.05 |
| ANA67904.1_human-poliovirus-1 | 1166 | 1174 | 9 | 0.05 | 0.15 | 0.061111111 |
| ANA67904.1_human-poliovirus-1 | 1178 | 1178 | 1 | 0.05 | 0.05 | 0.05 |
| ANA67904.1_human-poliovirus-1 | 1183 | 1183 | 1 | 0.05 | 0.05 | 0.05 |
| ANA67904.1_human-poliovirus-1 | 1214 | 1214 | 1 | 0.05 | 0.05 | 0.05 |
| ANA67904.1_human-poliovirus-1 | 1222 | 1222 | 1 | 0.05 | 0.05 | 0.05 |
| ANA67904.1_human-poliovirus-1 | 1224 | 1224 | 1 | 0.1 | 0.1 | 0.1 |
| ANA67904.1_human-poliovirus-1 | 1250 | 1250 | 1 | 0.05 | 0.05 | 0.05 |
| ANA67904.1_human-poliovirus-1 | 1265 | 1265 | 1 | 0.05 | 0.05 | 0.05 |
| ANA67904.1_human-poliovirus-1 | 1271 | 1271 | 1 | 0.05 | 0.05 | 0.05 |
| ANA67904.1_human-poliovirus-1 | 1274 | 1285 | 12 | 0.05 | 0.4 | 0.2625 |
| ANA67904.1_human-poliovirus-1 | 1288 | 1288 | 1 | 0.05 | 0.05 | 0.05 |
| ANA67904.1_human-poliovirus-1 | 1291 | 1306 | 16 | 0.05 | 0.3 | 0.203125 |
| ANA67904.1_human-poliovirus-1 | 1311 | 1319 | 9 | 0.05 | 0.05 | 0.05 |
| ANA67904.1_human-poliovirus-1 | 1321 | 1321 | 1 | 0.1 | 0.1 | 0.1 |
| ANA67904.1_human-poliovirus-1 | 1325 | 1325 | 1 | 0.05 | 0.05 | 0.05 |
| ANA67904.1_human-poliovirus-1 | 1337 | 1349 | 13 | 0.05 | 0.15 | 0.084615385 |
| ANA67904.1_human-poliovirus-1 | 1375 | 1379 | 5 | 0.05 | 0.05 | 0.05 |
| ANA67904.1_human-poliovirus-1 | 1391 | 1400 | 10 | 0.05 | 0.05 | 0.05 |
| ANA67904.1_human-poliovirus-1 | 1408 | 1409 | 2 | 0.1 | 0.1 | 0.1 |
| ANA67904.1_human-poliovirus-1 | 1417 | 1419 | 3 | 0.05 | 0.15 | 0.083333333 |
| ANA67904.1_human-poliovirus-1 | 1421 | 1421 | 1 | 0.05 | 0.05 | 0.05 |
| ANA67904.1_human-poliovirus-1 | 1429 | 1429 | 1 | 0.05 | 0.05 | 0.05 |
| ANA67904.1_human-poliovirus-1 | 1436 | 1441 | 6 | 0.05 | 0.1 | 0.058333333 |
| ANA67904.1_human-poliovirus-1 | 1443 | 1443 | 1 | 0.05 | 0.05 | 0.05 |
| ANA67904.1_human-poliovirus-1 | 1450 | 1453 | 4 | 0.05 | 0.1 | 0.0625 |
| ANA67904.1_human-poliovirus-1 | 1456 | 1457 | 2 | 0.05 | 0.2 | 0.125 |
| ANA67904.1_human-poliovirus-1 | 1463 | 1467 | 5 | 0.05 | 0.15 | 0.09 |
| ANA67904.1_human-poliovirus-1 | 1469 | 1475 | 7 | 0.05 | 0.25 | 0.171428571 |
| ANA67904.1_human-poliovirus-1 | 1477 | 1477 | 1 | 0.05 | 0.05 | 0.05 |
| ANA67904.1_human-poliovirus-1 | 1480 | 1495 | 16 | 0.05 | 0.25 | 0.115625 |
| ANA67904.1_human-poliovirus-1 | 1499 | 1506 | 8 | 0.05 | 0.15 | 0.10625 |
| ANA67904.1_human-poliovirus-1 | 1537 | 1546 | 10 | 0.05 | 0.1 | 0.055 |
| ANA67904.1_human-poliovirus-1 | 1557 | 1565 | 9 | 0.05 | 0.2 | 0.077777778 |
| ANA67904.1_human-poliovirus-1 | 1567 | 1575 | 9 | 0.05 | 0.2 | 0.116666667 |
| ANA67904.1_human-poliovirus-1 | 1577 | 1577 | 1 | 0.1 | 0.1 | 0.1 |
| ANA67904.1_human-poliovirus-1 | 1597 | 1597 | 1 | 0.05 | 0.05 | 0.05 |

| Extended data 1 | | | | | | |
|---|---|---|---|---|---|---|
| Protein | Start position (peptide) | Stop position (peptide) | Number of peptides | Minimum fraction (per peptide) of control subjects with reactivity >3 sd above average | Maximum fraction (per peptide) of control subjects with reactivity >3 sd above average | Mean fraction (over all peptides in this contiguous stretch) of controlsubjects with reactivity >3 sd above average |
| ANA67904.1_human-poliovirus-1 | 1607 | 1607 | 1 | 0.05 | 0.05 | 0.05 |
| ANA67904.1_human-poliovirus-1 | 1618 | 1618 | 1 | 0.05 | 0.05 | 0.05 |
| ANA67904.1_human-poliovirus-1 | 1622 | 1625 | 4 | 0.05 | 0.05 | 0.05 |
| ANA67904.1_human-poliovirus-1 | 1627 | 1627 | 1 | 0.05 | 0.05 | 0.05 |
| ANA67904.1_human-poliovirus-1 | 1636 | 1636 | 1 | 0.05 | 0.05 | 0.05 |
| ANA67904.1_human-poliovirus-1 | 1638 | 1647 | 10 | 0.05 | 0.25 | 0.105 |
| ANA67904.1_human-poliovirus-1 | 1658 | 1663 | 6 | 0.05 | 0.05 | 0.05 |
| ANA67904.1_human-poliovirus-1 | 1670 | 1670 | 1 | 0.05 | 0.05 | 0.05 |
| ANA67904.1_human-poliovirus-1 | 1678 | 1679 | 2 | 0.05 | 0.05 | 0.05 |
| ANA67904.1_human-poliovirus-1 | 1683 | 1683 | 1 | 0.1 | 0.1 | 0.1 |
| ANA67904.1_human-poliovirus-1 | 1722 | 1723 | 2 | 0.05 | 0.05 | 0.05 |
| ANA67904.1_human-poliovirus-1 | 1743 | 1743 | 1 | 0.05 | 0.05 | 0.05 |
| ANA67904.1_human-poliovirus-1 | 1750 | 1764 | 15 | 0.05 | 0.3 | 0.183333333 |
| ANA67904.1_human-poliovirus-1 | 1769 | 1771 | 3 | 0.05 | 0.05 | 0.05 |
| ANA67904.1_human-poliovirus-1 | 1782 | 1804 | 23 | 0.05 | 0.35 | 0.130434783 |
| ANA67904.1_human-poliovirus-1 | 1806 | 1806 | 1 | 0.05 | 0.05 | 0.05 |
| ANA67904.1_human-poliovirus-1 | 1808 | 1808 | 1 | 0.05 | 0.05 | 0.05 |
| ANA67904.1_human-poliovirus-1 | 1810 | 1810 | 1 | 0.05 | 0.05 | 0.05 |
| ANA67904.1_human-poliovirus-1 | 1816 | 1817 | 2 | 0.05 | 0.05 | 0.05 |
| ANA67904.1_human-poliovirus-1 | 1821 | 1821 | 1 | 0.05 | 0.05 | 0.05 |
| ANA67904.1_human-poliovirus-1 | 1828 | 1828 | 1 | 0.05 | 0.05 | 0.05 |
| ANA67904.1_human-poliovirus-1 | 1834 | 1836 | 3 | 0.05 | 0.1 | 0.066666667 |
| ANA67904.1_human-poliovirus-1 | 1838 | 1839 | 2 | 0.05 | 0.05 | 0.05 |
| ANA67904.1_human-poliovirus-1 | 1844 | 1845 | 2 | 0.05 | 0.05 | 0.05 |
| ANA67904.1_human-poliovirus-1 | 1854 | 1854 | 1 | 0.15 | 0.15 | 0.15 |
| ANA67904.1_human-poliovirus-1 | 1858 | 1875 | 18 | 0.05 | 0.3 | 0.1 |
| ANA67904.1_human-poliovirus-1 | 1877 | 1880 | 4 | 0.05 | 0.15 | 0.0875 |
| ANA67904.1_human-poliovirus-1 | 1883 | 1883 | 1 | 0.05 | 0.05 | 0.05 |
| ANA67904.1_human-poliovirus-1 | 1885 | 1885 | 1 | 0.05 | 0.05 | 0.05 |
| ANA67904.1_human-poliovirus-1 | 1892 | 1901 | 10 | 0.05 | 0.1 | 0.055 |
| ANA67904.1_human-poliovirus-1 | 1914 | 1916 | 3 | 0.05 | 0.05 | 0.05 |
| ANA67904.1_human-poliovirus-1 | 1923 | 1923 | 1 | 0.6 | 0.6 | 0.6 |
| ANA67904.1_human-poliovirus-1 | 1927 | 1928 | 2 | 0.1 | 0.45 | 0.275 |
| ANA67904.1_human-poliovirus-1 | 1930 | 1930 | 1 | 0.05 | 0.05 | 0.05 |
| ANA67904.1_human-poliovirus-1 | 1940 | 1943 | 4 | 0.05 | 0.05 | 0.05 |
| ANA67904.1_human-poliovirus-1 | 1953 | 1962 | 10 | 0.1 | 0.4 | 0.275 |
| ANA67904.1_human-poliovirus-1 | 1969 | 1969 | 1 | 0.05 | 0.05 | 0.05 |

| Extended data 1 | | | | | | |
|---|---|---|---|---|---|---|
| Protein | Start position (peptide) | Stop position (peptide) | Number of peptides | Minimum fraction (per peptide) of control subjects with reactivity >3 sd above average | Maximum fraction (per peptide) of control subjects with reactivity >3 sd above average | Mean fraction (over all peptides in this contiguous stretch) of controlsubjects with reactivity >3 sd above average |
| ANA67904.1_human-poliovirus-1 | 1975 | 1976 | 2 | 0.1 | 0.15 | 0.125 |
| ANA67904.1_human-poliovirus-1 | 1997 | 1997 | 1 | 0.05 | 0.05 | 0.05 |
| ANA67904.1_human-poliovirus-1 | 2002 | 2008 | 7 | 0.05 | 0.4 | 0.192857143 |
| ANA67904.1_human-poliovirus-1 | 2023 | 2025 | 3 | 0.05 | 0.05 | 0.05 |
| ANA67904.1_human-poliovirus-1 | 2058 | 2058 | 1 | 0.1 | 0.1 | 0.1 |
| ANA67904.1_human-poliovirus-1 | 2063 | 2063 | 1 | 0.1 | 0.1 | 0.1 |
| ANA67904.1_human-poliovirus-1 | 2089 | 2092 | 4 | 0.05 | 0.1 | 0.075 |
| ANA67904.1_human-poliovirus-1 | 2095 | 2096 | 2 | 0.05 | 0.1 | 0.075 |
| ANA67904.1_human-poliovirus-1 | 2123 | 2125 | 3 | 0.05 | 0.1 | 0.066666667 |
| ANA67904.1_human-poliovirus-1 | 2147 | 2147 | 1 | 0.2 | 0.2 | 0.2 |
| ANA67904.1_human-poliovirus-1 | 2173 | 2173 | 1 | 0.05 | 0.05 | 0.05 |
| DQ473503.1;ABF51197.1;Human_rhinovirus_A7_com | 6 | 16 | 11 | 0.05 | 0.2 | 0.118181818 |
| DQ473503.1;ABF51197.1;Human_rhinovirus_A7_com | 20 | 64 | 45 | 0.05 | 0.6 | 0.301111111 |
| DQ473503.1;ABF51197.1;Human_rhinovirus_A7_com | 77 | 88 | 12 | 0.05 | 0.15 | 0.058333333 |
| DQ473503.1;ABF51197.1;Human_rhinovirus_A7_com | 95 | 95 | 1 | 0.05 | 0.05 | 0.05 |
| DQ473503.1;ABF51197.1;Human_rhinovirus_A7_com | 103 | 125 | 23 | 0.05 | 0.45 | 0.219565217 |
| DQ473503.1;ABF51197.1;Human_rhinovirus_A7_com | 128 | 129 | 2 | 0.05 | 0.05 | 0.05 |
| DQ473503.1;ABF51197.1;Human_rhinovirus_A7_com | 149 | 149 | 1 | 0.05 | 0.05 | 0.05 |
| DQ473503.1;ABF51197.1;Human_rhinovirus_A7_com | 151 | 151 | 1 | 0.05 | 0.05 | 0.05 |
| DQ473503.1;ABF51197.1;Human_rhinovirus_A7_com | 191 | 201 | 11 | 0.05 | 0.05 | 0.05 |
| DQ473503.1;ABF51197.1;Human_rhinovirus_A7_com | 203 | 217 | 15 | 0.05 | 0.7 | 0.486666667 |
| DQ473503.1;ABF51197.1;Human_rhinovirus_A7_com | 221 | 223 | 3 | 0.05 | 0.05 | 0.05 |
| DQ473503.1;ABF51197.1;Human_rhinovirus_A7_com | 229 | 229 | 1 | 0.05 | 0.05 | 0.05 |
| DQ473503.1;ABF51197.1;Human_rhinovirus_A7_com | 239 | 240 | 2 | 0.05 | 0.05 | 0.05 |
| DQ473503.1;ABF51197.1;Human_rhinovirus_A7_com | 297 | 297 | 1 | 0.05 | 0.05 | 0.05 |
| DQ473503.1;ABF51197.1;Human_rhinovirus_A7_com | 301 | 301 | 1 | 0.05 | 0.05 | 0.05 |
| DQ473503.1;ABF51197.1;Human_rhinovirus_A7_com | 310 | 310 | 1 | 0.1 | 0.1 | 0.1 |
| DQ473503.1;ABF51197.1;Human_rhinovirus_A7_com | 316 | 317 | 2 | 0.05 | 0.05 | 0.05 |
| DQ473503.1;ABF51197.1;Human_rhinovirus_A7_com | 332 | 332 | 1 | 0.05 | 0.05 | 0.05 |
| DQ473503.1;ABF51197.1;Human_rhinovirus_A7_com | 334 | 334 | 1 | 0.05 | 0.05 | 0.05 |
| DQ473503.1;ABF51197.1;Human_rhinovirus_A7_com | 340 | 340 | 1 | 0.1 | 0.1 | 0.1 |
| DQ473503.1;ABF51197.1;Human_rhinovirus_A7_com | 351 | 353 | 3 | 0.05 | 0.15 | 0.083333333 |
| DQ473503.1;ABF51197.1;Human_rhinovirus_A7_com | 369 | 369 | 1 | 0.05 | 0.05 | 0.05 |
| DQ473503.1;ABF51197.1;Human_rhinovirus_A7_com | 378 | 391 | 14 | 0.05 | 0.45 | 0.264285714 |
| DQ473503.1;ABF51197.1;Human_rhinovirus_A7_com | 393 | 393 | 1 | 0.05 | 0.05 | 0.05 |
| DQ473503.1;ABF51197.1;Human_rhinovirus_A7_com | 397 | 408 | 12 | 0.05 | 0.1 | 0.054166667 |
| DQ473503.1;ABF51197.1;Human_rhinovirus_A7_com | 422 | 422 | 1 | 0.05 | 0.05 | 0.05 |

| Extended data 1 | | | | | | |
|---|---|---|---|---|---|---|
| Protein | Start position (peptide) | Stop position (peptide) | Number of peptides | Minimum fraction (per peptide) of control subjects with reactivity >3 sd above average | Maximum fraction (per peptide) of control subjects with reactivity >3 sd above average | Mean fraction (over all peptides in this contiguous stretch) of controlsubjects with reactivity >3 sd above average |
| DQ473503.1;ABF51197.1;Human_rhinovirus_A7_com | 489 | 489 | 1 | 0.05 | 0.05 | 0.05 |
| DQ473503.1;ABF51197.1;Human_rhinovirus_A7_com | 502 | 502 | 1 | 0.05 | 0.05 | 0.05 |
| DQ473503.1;ABF51197.1;Human_rhinovirus_A7_com | 506 | 507 | 2 | 0.05 | 0.05 | 0.05 |
| DQ473503.1;ABF51197.1;Human_rhinovirus_A7_com | 529 | 530 | 2 | 0.05 | 0.05 | 0.05 |
| DQ473503.1;ABF51197.1;Human_rhinovirus_A7_com | 532 | 532 | 1 | 0.05 | 0.05 | 0.05 |
| DQ473503.1;ABF51197.1;Human_rhinovirus_A7_com | 560 | 560 | 1 | 0.05 | 0.05 | 0.05 |
| DQ473503.1;ABF51197.1;Human_rhinovirus_A7_com | 565 | 588 | 24 | 0.15 | 1 | 0.69375 |
| DQ473503.1;ABF51197.1;Human_rhinovirus_A7_com | 590 | 590 | 1 | 0.1 | 0.1 | 0.1 |
| DQ473503.1;ABF51197.1;Human_rhinovirus_A7_com | 594 | 611 | 18 | 0.05 | 0.95 | 0.616666667 |
| DQ473503.1;ABF51197.1;Human_rhinovirus_A7_com | 613 | 615 | 3 | 0.05 | 0.05 | 0.05 |
| DQ473503.1;ABF51197.1;Human_rhinovirus_A7_com | 619 | 619 | 1 | 0.05 | 0.05 | 0.05 |
| DQ473503.1;ABF51197.1;Human_rhinovirus_A7_com | 621 | 625 | 5 | 0.05 | 0.05 | 0.05 |
| DQ473503.1;ABF51197.1;Human_rhinovirus_A7_com | 645 | 645 | 1 | 0.05 | 0.05 | 0.05 |
| DQ473503.1;ABF51197.1;Human_rhinovirus_A7_com | 679 | 679 | 1 | 0.05 | 0.05 | 0.05 |
| DQ473503.1;ABF51197.1;Human_rhinovirus_A7_com | 691 | 692 | 2 | 0.05 | 0.05 | 0.05 |
| DQ473503.1;ABF51197.1;Human_rhinovirus_A7_com | 711 | 726 | 16 | 0.05 | 0.1 | 0.065625 |
| DQ473503.1;ABF51197.1;Human_rhinovirus_A7_com | 728 | 731 | 4 | 0.05 | 0.25 | 0.1625 |
| DQ473503.1;ABF51197.1;Human_rhinovirus_A7_com | 765 | 777 | 13 | 0.05 | 0.2 | 0.15 |
| DQ473503.1;ABF51197.1;Human_rhinovirus_A7_com | 826 | 841 | 16 | 0.05 | 0.25 | 0.11875 |
| DQ473503.1;ABF51197.1;Human_rhinovirus_A7_com | 844 | 853 | 10 | 0.05 | 0.3 | 0.16 |
| DQ473503.1;ABF51197.1;Human_rhinovirus_A7_com | 860 | 861 | 2 | 0.05 | 0.05 | 0.05 |
| DQ473503.1;ABF51197.1;Human_rhinovirus_A7_com | 895 | 903 | 9 | 0.05 | 0.1 | 0.055555556 |
| DQ473503.1;ABF51197.1;Human_rhinovirus_A7_com | 905 | 905 | 1 | 0.05 | 0.05 | 0.05 |
| DQ473503.1;ABF51197.1;Human_rhinovirus_A7_com | 940 | 940 | 1 | 0.05 | 0.05 | 0.05 |
| DQ473503.1;ABF51197.1;Human_rhinovirus_A7_com | 942 | 943 | 2 | 0.05 | 0.05 | 0.05 |
| DQ473503.1;ABF51197.1;Human_rhinovirus_A7_com | 955 | 955 | 1 | 0.05 | 0.05 | 0.05 |
| DQ473503.1;ABF51197.1;Human_rhinovirus_A7_com | 975 | 975 | 1 | 0.05 | 0.05 | 0.05 |
| DQ473503.1;ABF51197.1;Human_rhinovirus_A7_com | 995 | 995 | 1 | 0.05 | 0.05 | 0.05 |
| DQ473503.1;ABF51197.1;Human_rhinovirus_A7_com | 1004 | 1008 | 5 | 0.15 | 0.9 | 0.56 |
| DQ473503.1;ABF51197.1;Human_rhinovirus_A7_com | 1013 | 1020 | 8 | 0.1 | 0.2 | 0.15625 |
| DQ473503.1;ABF51197.1;Human_rhinovirus_A7_com | 1022 | 1022 | 1 | 0.05 | 0.05 | 0.05 |
| DQ473503.1;ABF51197.1;Human_rhinovirus_A7_com | 1025 | 1025 | 1 | 0.1 | 0.1 | 0.1 |
| DQ473503.1;ABF51197.1;Human_rhinovirus_A7_com | 1029 | 1029 | 1 | 0.05 | 0.05 | 0.05 |
| DQ473503.1;ABF51197.1;Human_rhinovirus_A7_com | 1087 | 1097 | 11 | 0.05 | 0.1 | 0.063636364 |
| DQ473503.1;ABF51197.1;Human_rhinovirus_A7_com | 1099 | 1108 | 10 | 0.05 | 0.2 | 0.085 |
| DQ473503.1;ABF51197.1;Human_rhinovirus_A7_com | 1116 | 1118 | 3 | 0.05 | 0.1 | 0.083333333 |
| DQ473503.1;ABF51197.1;Human_rhinovirus_A7_com | 1127 | 1142 | 16 | 0.05 | 0.25 | 0.1125 |

| Extended data 1 | | | | | | |
|---|---|---|---|---|---|---|
| Protein | Start position (peptide) | Stop position (peptide) | Number of peptides | Minimum fraction (per peptide) of control subjects with reactivity >3 sd above average | Maximum fraction (per peptide) of control subjects with reactivity >3 sd above average | Mean fraction (over all peptides in this contiguous stretch) of controlsubjects with reactivity >3 sd above average |
| DQ473503.1;ABF51197.1;Human_rhinovirus_A7_com | 1145 | 1146 | 2 | 0.05 | 0.05 | 0.05 |
| DQ473503.1;ABF51197.1;Human_rhinovirus_A7_com | 1151 | 1151 | 1 | 0.05 | 0.05 | 0.05 |
| DQ473503.1;ABF51197.1;Human_rhinovirus_A7_com | 1157 | 1159 | 3 | 0.05 | 0.05 | 0.05 |
| DQ473503.1;ABF51197.1;Human_rhinovirus_A7_com | 1164 | 1166 | 3 | 0.05 | 0.05 | 0.05 |
| DQ473503.1;ABF51197.1;Human_rhinovirus_A7_com | 1168 | 1170 | 3 | 0.05 | 0.05 | 0.05 |
| DQ473503.1;ABF51197.1;Human_rhinovirus_A7_com | 1172 | 1172 | 1 | 0.05 | 0.05 | 0.05 |
| DQ473503.1;ABF51197.1;Human_rhinovirus_A7_com | 1182 | 1182 | 1 | 0.05 | 0.05 | 0.05 |
| DQ473503.1;ABF51197.1;Human_rhinovirus_A7_com | 1185 | 1187 | 3 | 0.05 | 0.1 | 0.066666667 |
| DQ473503.1;ABF51197.1;Human_rhinovirus_A7_com | 1189 | 1191 | 3 | 0.05 | 0.1 | 0.083333333 |
| DQ473503.1;ABF51197.1;Human_rhinovirus_A7_com | 1193 | 1194 | 2 | 0.05 | 0.05 | 0.05 |
| DQ473503.1;ABF51197.1;Human_rhinovirus_A7_com | 1196 | 1196 | 1 | 0.05 | 0.05 | 0.05 |
| DQ473503.1;ABF51197.1;Human_rhinovirus_A7_com | 1202 | 1202 | 1 | 0.05 | 0.05 | 0.05 |
| DQ473503.1;ABF51197.1;Human_rhinovirus_A7_com | 1205 | 1205 | 1 | 0.05 | 0.05 | 0.05 |
| DQ473503.1;ABF51197.1;Human_rhinovirus_A7_com | 1240 | 1255 | 16 | 0.05 | 0.7 | 0.44375 |
| DQ473503.1;ABF51197.1;Human_rhinovirus_A7_com | 1259 | 1259 | 1 | 0.05 | 0.05 | 0.05 |
| DQ473503.1;ABF51197.1;Human_rhinovirus_A7_com | 1268 | 1268 | 1 | 0.05 | 0.05 | 0.05 |
| DQ473503.1;ABF51197.1;Human_rhinovirus_A7_com | 1272 | 1273 | 2 | 0.05 | 0.1 | 0.075 |
| DQ473503.1;ABF51197.1;Human_rhinovirus_A7_com | 1275 | 1275 | 1 | 0.05 | 0.05 | 0.05 |
| DQ473503.1;ABF51197.1;Human_rhinovirus_A7_com | 1285 | 1297 | 13 | 0.15 | 1 | 0.696153846 |
| DQ473503.1;ABF51197.1;Human_rhinovirus_A7_com | 1313 | 1321 | 9 | 0.05 | 0.1 | 0.094444444 |
| DQ473503.1;ABF51197.1;Human_rhinovirus_A7_com | 1323 | 1323 | 1 | 0.05 | 0.05 | 0.05 |
| DQ473503.1;ABF51197.1;Human_rhinovirus_A7_com | 1339 | 1350 | 12 | 0.1 | 0.25 | 0.166666667 |
| DQ473503.1;ABF51197.1;Human_rhinovirus_A7_com | 1353 | 1353 | 1 | 0.1 | 0.1 | 0.1 |
| DQ473503.1;ABF51197.1;Human_rhinovirus_A7_com | 1360 | 1364 | 5 | 0.05 | 0.1 | 0.06 |
| DQ473503.1;ABF51197.1;Human_rhinovirus_A7_com | 1371 | 1381 | 11 | 0.05 | 0.1 | 0.077272727 |
| DQ473503.1;ABF51197.1;Human_rhinovirus_A7_com | 1399 | 1400 | 2 | 0.05 | 0.1 | 0.075 |
| DQ473503.1;ABF51197.1;Human_rhinovirus_A7_com | 1402 | 1404 | 3 | 0.05 | 0.05 | 0.05 |
| DQ473503.1;ABF51197.1;Human_rhinovirus_A7_com | 1406 | 1410 | 5 | 0.05 | 0.05 | 0.05 |
| DQ473503.1;ABF51197.1;Human_rhinovirus_A7_com | 1412 | 1412 | 1 | 0.05 | 0.05 | 0.05 |
| DQ473503.1;ABF51197.1;Human_rhinovirus_A7_com | 1416 | 1416 | 1 | 0.05 | 0.05 | 0.05 |
| DQ473503.1;ABF51197.1;Human_rhinovirus_A7_com | 1418 | 1418 | 1 | 0.05 | 0.05 | 0.05 |
| DQ473503.1;ABF51197.1;Human_rhinovirus_A7_com | 1422 | 1454 | 33 | 0.05 | 0.8 | 0.303030303 |
| DQ473503.1;ABF51197.1;Human_rhinovirus_A7_com | 1456 | 1457 | 2 | 0.05 | 0.15 | 0.1 |
| DQ473503.1;ABF51197.1;Human_rhinovirus_A7_com | 1464 | 1464 | 1 | 0.15 | 0.15 | 0.15 |
| DQ473503.1;ABF51197.1;Human_rhinovirus_A7_com | 1508 | 1509 | 2 | 0.05 | 0.1 | 0.075 |
| DQ473503.1;ABF51197.1;Human_rhinovirus_A7_com | 1512 | 1513 | 2 | 0.05 | 0.05 | 0.05 |
| DQ473503.1;ABF51197.1;Human_rhinovirus_A7_com | 1515 | 1515 | 1 | 0.05 | 0.05 | 0.05 |

| Extended data 1 | | | | | | |
|---|---|---|---|---|---|---|
| Protein | Start position (peptide) | Stop position (peptide) | Number of peptides | Minimum fraction (per peptide) of control subjects with reactivity >3 sd above average | Maximum fraction (per peptide) of control subjects with reactivity >3 sd above average | Mean fraction (over all peptides in this contiguous stretch) of controlsubjects with reactivity >3 sd above average |
| DQ473503.1;ABF51197.1;Human_rhinovirus_A7_com | 1520 | 1520 | 1 | 0.1 | 0.1 | 0.1 |
| DQ473503.1;ABF51197.1;Human_rhinovirus_A7_com | 1530 | 1535 | 6 | 0.1 | 0.15 | 0.133333333 |
| DQ473503.1;ABF51197.1;Human_rhinovirus_A7_com | 1538 | 1538 | 1 | 0.05 | 0.05 | 0.05 |
| DQ473503.1;ABF51197.1;Human_rhinovirus_A7_com | 1548 | 1548 | 1 | 0.05 | 0.05 | 0.05 |
| DQ473503.1;ABF51197.1;Human_rhinovirus_A7_com | 1553 | 1553 | 1 | 0.05 | 0.05 | 0.05 |
| DQ473503.1;ABF51197.1;Human_rhinovirus_A7_com | 1556 | 1557 | 2 | 0.05 | 0.05 | 0.05 |
| DQ473503.1;ABF51197.1;Human_rhinovirus_A7_com | 1589 | 1601 | 13 | 0.05 | 0.25 | 0.142307692 |
| DQ473503.1;ABF51197.1;Human_rhinovirus_A7_com | 1606 | 1606 | 1 | 0.05 | 0.05 | 0.05 |
| DQ473503.1;ABF51197.1;Human_rhinovirus_A7_com | 1621 | 1621 | 1 | 0.1 | 0.1 | 0.1 |
| DQ473503.1;ABF51197.1;Human_rhinovirus_A7_com | 1625 | 1625 | 1 | 0.05 | 0.05 | 0.05 |
| DQ473503.1;ABF51197.1;Human_rhinovirus_A7_com | 1627 | 1628 | 2 | 0.05 | 0.05 | 0.05 |
| DQ473503.1;ABF51197.1;Human_rhinovirus_A7_com | 1647 | 1647 | 1 | 0.05 | 0.05 | 0.05 |
| DQ473503.1;ABF51197.1;Human_rhinovirus_A7_com | 1654 | 1654 | 1 | 0.05 | 0.05 | 0.05 |
| DQ473503.1;ABF51197.1;Human_rhinovirus_A7_com | 1688 | 1690 | 3 | 0.05 | 0.05 | 0.05 |
| DQ473503.1;ABF51197.1;Human_rhinovirus_A7_com | 1695 | 1695 | 1 | 0.05 | 0.05 | 0.05 |
| DQ473503.1;ABF51197.1;Human_rhinovirus_A7_com | 1697 | 1711 | 15 | 0.05 | 0.3 | 0.076666667 |
| DQ473503.1;ABF51197.1;Human_rhinovirus_A7_com | 1717 | 1717 | 1 | 0.05 | 0.05 | 0.05 |
| DQ473503.1;ABF51197.1;Human_rhinovirus_A7_com | 1719 | 1746 | 28 | 0.05 | 0.4 | 0.2125 |
| DQ473503.1;ABF51197.1;Human_rhinovirus_A7_com | 1752 | 1753 | 2 | 0.05 | 0.1 | 0.075 |
| DQ473503.1;ABF51197.1;Human_rhinovirus_A7_com | 1755 | 1755 | 1 | 0.05 | 0.05 | 0.05 |
| DQ473503.1;ABF51197.1;Human_rhinovirus_A7_com | 1757 | 1757 | 1 | 0.1 | 0.1 | 0.1 |
| DQ473503.1;ABF51197.1;Human_rhinovirus_A7_com | 1766 | 1766 | 1 | 0.05 | 0.05 | 0.05 |
| DQ473503.1;ABF51197.1;Human_rhinovirus_A7_com | 1770 | 1770 | 1 | 0.05 | 0.05 | 0.05 |
| DQ473503.1;ABF51197.1;Human_rhinovirus_A7_com | 1782 | 1782 | 1 | 0.05 | 0.05 | 0.05 |
| DQ473503.1;ABF51197.1;Human_rhinovirus_A7_com | 1790 | 1805 | 16 | 0.05 | 0.15 | 0.0875 |
| DQ473503.1;ABF51197.1;Human_rhinovirus_A7_com | 1807 | 1809 | 3 | 0.05 | 0.15 | 0.1 |
| DQ473503.1;ABF51197.1;Human_rhinovirus_A7_com | 1812 | 1824 | 13 | 0.05 | 0.1 | 0.080769231 |
| DQ473503.1;ABF51197.1;Human_rhinovirus_A7_com | 1826 | 1826 | 1 | 0.05 | 0.05 | 0.05 |
| DQ473503.1;ABF51197.1;Human_rhinovirus_A7_com | 1828 | 1832 | 5 | 0.1 | 0.3 | 0.14 |
| DQ473503.1;ABF51197.1;Human_rhinovirus_A7_com | 1838 | 1854 | 17 | 0.05 | 0.2 | 0.091176471 |
| DQ473503.1;ABF51197.1;Human_rhinovirus_A7_com | 1870 | 1871 | 2 | 0.05 | 0.05 | 0.05 |
| DQ473503.1;ABF51197.1;Human_rhinovirus_A7_com | 1873 | 1874 | 2 | 0.05 | 0.05 | 0.05 |
| DQ473503.1;ABF51197.1;Human_rhinovirus_A7_com | 1882 | 1882 | 1 | 0.05 | 0.05 | 0.05 |
| DQ473503.1;ABF51197.1;Human_rhinovirus_A7_com | 1893 | 1896 | 4 | 0.05 | 0.05 | 0.05 |
| DQ473503.1;ABF51197.1;Human_rhinovirus_A7_com | 1924 | 1934 | 11 | 0.05 | 0.1 | 0.054545455 |
| DQ473503.1;ABF51197.1;Human_rhinovirus_A7_com | 1937 | 1937 | 1 | 0.05 | 0.05 | 0.05 |
| DQ473503.1;ABF51197.1;Human_rhinovirus_A7_com | 1945 | 1945 | 1 | 0.05 | 0.05 | 0.05 |

| Extended data 1 | | | | | | |
|---|---|---|---|---|---|---|
| **Protein** | **Start position (peptide)** | **Stop position (peptide)** | **Number of peptides** | **Minimum fraction (per peptide) of control subjects with reactivity >3 sd above average** | **Maximum fraction (per peptide) of control subjects with reactivity >3 sd above average** | **Mean fraction (over all peptides in this contiguous stretch) of controlsubjects with reactivity >3 sd above average** |
| DQ473503.1;ABF51197.1;Human_rhinovirus_A7_com | 1949 | 1950 | 2 | 0.05 | 0.05 | 0.05 |
| DQ473503.1;ABF51197.1;Human_rhinovirus_A7_com | 1965 | 1965 | 1 | 0.1 | 0.1 | 0.1 |
| DQ473503.1;ABF51197.1;Human_rhinovirus_A7_com | 1971 | 1971 | 1 | 0.05 | 0.05 | 0.05 |
| DQ473503.1;ABF51197.1;Human_rhinovirus_A7_com | 1975 | 1975 | 1 | 0.05 | 0.05 | 0.05 |
| DQ473503.1;ABF51197.1;Human_rhinovirus_A7_com | 1981 | 1981 | 1 | 0.05 | 0.05 | 0.05 |
| DQ473503.1;ABF51197.1;Human_rhinovirus_A7_com | 1984 | 1984 | 1 | 0.05 | 0.05 | 0.05 |
| DQ473503.1;ABF51197.1;Human_rhinovirus_A7_com | 1987 | 1987 | 1 | 0.05 | 0.05 | 0.05 |
| DQ473503.1;ABF51197.1;Human_rhinovirus_A7_com | 2011 | 2011 | 1 | 0.05 | 0.05 | 0.05 |
| DQ473503.1;ABF51197.1;Human_rhinovirus_A7_com | 2015 | 2015 | 1 | 0.05 | 0.05 | 0.05 |
| DQ473503.1;ABF51197.1;Human_rhinovirus_A7_com | 2033 | 2033 | 1 | 0.05 | 0.05 | 0.05 |
| DQ473503.1;ABF51197.1;Human_rhinovirus_A7_com | 2046 | 2046 | 1 | 0.05 | 0.05 | 0.05 |
| DQ473503.1;ABF51197.1;Human_rhinovirus_A7_com | 2050 | 2050 | 1 | 0.05 | 0.05 | 0.05 |
| DQ473503.1;ABF51197.1;Human_rhinovirus_A7_com | 2073 | 2073 | 1 | 0.05 | 0.05 | 0.05 |
| DQ473503.1;ABF51197.1;Human_rhinovirus_A7_com | 2093 | 2097 | 5 | 0.05 | 0.05 | 0.05 |
| DQ473503.1;ABF51197.1;Human_rhinovirus_A7_com | 2100 | 2100 | 1 | 0.1 | 0.1 | 0.1 |
| DQ473503.1;ABF51197.1;Human_rhinovirus_A7_com | 2117 | 2119 | 3 | 0.05 | 0.05 | 0.05 |
| DQ473503.1;ABF51197.1;Human_rhinovirus_A7_com | 2121 | 2121 | 1 | 0.05 | 0.05 | 0.05 |
| DQ473503.1;ABF51197.1;Human_rhinovirus_A7_com | 2148 | 2149 | 2 | 0.05 | 0.1 | 0.075 |
| EF077280.1;ABQ51392.1;Human_rhinovirus_C2_NAT | 1 | 3 | 3 | 0.05 | 0.05 | 0.05 |
| EF077280.1;ABQ51392.1;Human_rhinovirus_C2_NAT | 7 | 7 | 1 | 0.05 | 0.05 | 0.05 |
| EF077280.1;ABQ51392.1;Human_rhinovirus_C2_NAT | 21 | 21 | 1 | 0.05 | 0.05 | 0.05 |
| EF077280.1;ABQ51392.1;Human_rhinovirus_C2_NAT | 23 | 28 | 6 | 0.05 | 0.1 | 0.058333333 |
| EF077280.1;ABQ51392.1;Human_rhinovirus_C2_NAT | 31 | 33 | 3 | 0.05 | 0.05 | 0.05 |
| EF077280.1;ABQ51392.1;Human_rhinovirus_C2_NAT | 35 | 52 | 18 | 0.05 | 0.35 | 0.155555556 |
| EF077280.1;ABQ51392.1;Human_rhinovirus_C2_NAT | 55 | 55 | 1 | 0.1 | 0.1 | 0.1 |
| EF077280.1;ABQ51392.1;Human_rhinovirus_C2_NAT | 73 | 74 | 2 | 0.05 | 0.1 | 0.075 |
| EF077280.1;ABQ51392.1;Human_rhinovirus_C2_NAT | 80 | 81 | 2 | 0.05 | 0.15 | 0.1 |
| EF077280.1;ABQ51392.1;Human_rhinovirus_C2_NAT | 83 | 95 | 13 | 0.05 | 0.05 | 0.05 |
| EF077280.1;ABQ51392.1;Human_rhinovirus_C2_NAT | 100 | 100 | 1 | 0.05 | 0.05 | 0.05 |
| EF077280.1;ABQ51392.1;Human_rhinovirus_C2_NAT | 104 | 114 | 11 | 0.1 | 0.3 | 0.204545455 |
| EF077280.1;ABQ51392.1;Human_rhinovirus_C2_NAT | 141 | 141 | 1 | 0.05 | 0.05 | 0.05 |
| EF077280.1;ABQ51392.1;Human_rhinovirus_C2_NAT | 153 | 153 | 1 | 0.05 | 0.05 | 0.05 |
| EF077280.1;ABQ51392.1;Human_rhinovirus_C2_NAT | 191 | 191 | 1 | 0.05 | 0.05 | 0.05 |
| EF077280.1;ABQ51392.1;Human_rhinovirus_C2_NAT | 193 | 195 | 3 | 0.05 | 0.05 | 0.05 |
| EF077280.1;ABQ51392.1;Human_rhinovirus_C2_NAT | 197 | 198 | 2 | 0.05 | 0.05 | 0.05 |
| EF077280.1;ABQ51392.1;Human_rhinovirus_C2_NAT | 201 | 214 | 14 | 0.1 | 0.45 | 0.335714286 |
| EF077280.1;ABQ51392.1;Human_rhinovirus_C2_NAT | 218 | 218 | 1 | 0.05 | 0.05 | 0.05 |

| Extended data 1 | | | | | | |
|---|---|---|---|---|---|---|
| Protein | Start position (peptide) | Stop position (peptide) | Number of peptides | Minimum fraction (per peptide) of control subjects with reactivity >3 sd above average | Maximum fraction (per peptide) of control subjects with reactivity >3 sd above average | Mean fraction (over all peptides in this contiguous stretch) of controlsubjects with reactivity >3 sd above average |
| EF077280.1;ABQ51392.1;Human_rhinovirus_C2_NAT | 220 | 220 | 1 | 0.05 | 0.05 | 0.05 |
| EF077280.1;ABQ51392.1;Human_rhinovirus_C2_NAT | 223 | 225 | 3 | 0.05 | 0.1 | 0.066666667 |
| EF077280.1;ABQ51392.1;Human_rhinovirus_C2_NAT | 230 | 230 | 1 | 0.05 | 0.05 | 0.05 |
| EF077280.1;ABQ51392.1;Human_rhinovirus_C2_NAT | 253 | 253 | 1 | 0.05 | 0.05 | 0.05 |
| EF077280.1;ABQ51392.1;Human_rhinovirus_C2_NAT | 256 | 256 | 1 | 0.05 | 0.05 | 0.05 |
| EF077280.1;ABQ51392.1;Human_rhinovirus_C2_NAT | 271 | 271 | 1 | 0.05 | 0.05 | 0.05 |
| EF077280.1;ABQ51392.1;Human_rhinovirus_C2_NAT | 288 | 288 | 1 | 0.05 | 0.05 | 0.05 |
| EF077280.1;ABQ51392.1;Human_rhinovirus_C2_NAT | 295 | 295 | 1 | 0.05 | 0.05 | 0.05 |
| EF077280.1;ABQ51392.1;Human_rhinovirus_C2_NAT | 299 | 300 | 2 | 0.05 | 0.1 | 0.075 |
| EF077280.1;ABQ51392.1;Human_rhinovirus_C2_NAT | 346 | 346 | 1 | 0.05 | 0.05 | 0.05 |
| EF077280.1;ABQ51392.1;Human_rhinovirus_C2_NAT | 358 | 358 | 1 | 0.05 | 0.05 | 0.05 |
| EF077280.1;ABQ51392.1;Human_rhinovirus_C2_NAT | 377 | 377 | 1 | 0.05 | 0.05 | 0.05 |
| EF077280.1;ABQ51392.1;Human_rhinovirus_C2_NAT | 387 | 387 | 1 | 0.05 | 0.05 | 0.05 |
| EF077280.1;ABQ51392.1;Human_rhinovirus_C2_NAT | 396 | 396 | 1 | 0.05 | 0.05 | 0.05 |
| EF077280.1;ABQ51392.1;Human_rhinovirus_C2_NAT | 403 | 404 | 2 | 0.05 | 0.05 | 0.05 |
| EF077280.1;ABQ51392.1;Human_rhinovirus_C2_NAT | 450 | 451 | 2 | 0.1 | 0.15 | 0.125 |
| EF077280.1;ABQ51392.1;Human_rhinovirus_C2_NAT | 453 | 453 | 1 | 0.05 | 0.05 | 0.05 |
| EF077280.1;ABQ51392.1;Human_rhinovirus_C2_NAT | 457 | 457 | 1 | 0.05 | 0.05 | 0.05 |
| EF077280.1;ABQ51392.1;Human_rhinovirus_C2_NAT | 463 | 465 | 3 | 0.05 | 0.05 | 0.05 |
| EF077280.1;ABQ51392.1;Human_rhinovirus_C2_NAT | 483 | 483 | 1 | 0.05 | 0.05 | 0.05 |
| EF077280.1;ABQ51392.1;Human_rhinovirus_C2_NAT | 501 | 501 | 1 | 0.05 | 0.05 | 0.05 |
| EF077280.1;ABQ51392.1;Human_rhinovirus_C2_NAT | 518 | 518 | 1 | 0.05 | 0.05 | 0.05 |
| EF077280.1;ABQ51392.1;Human_rhinovirus_C2_NAT | 522 | 524 | 3 | 0.05 | 0.05 | 0.05 |
| EF077280.1;ABQ51392.1;Human_rhinovirus_C2_NAT | 549 | 549 | 1 | 0.1 | 0.1 | 0.1 |
| EF077280.1;ABQ51392.1;Human_rhinovirus_C2_NAT | 553 | 553 | 1 | 0.1 | 0.1 | 0.1 |
| EF077280.1;ABQ51392.1;Human_rhinovirus_C2_NAT | 556 | 556 | 1 | 0.05 | 0.05 | 0.05 |
| EF077280.1;ABQ51392.1;Human_rhinovirus_C2_NAT | 558 | 592 | 35 | 0.05 | 0.8 | 0.45 |
| EF077280.1;ABQ51392.1;Human_rhinovirus_C2_NAT | 604 | 604 | 1 | 0.05 | 0.05 | 0.05 |
| EF077280.1;ABQ51392.1;Human_rhinovirus_C2_NAT | 616 | 624 | 9 | 0.05 | 0.15 | 0.088888889 |
| EF077280.1;ABQ51392.1;Human_rhinovirus_C2_NAT | 641 | 643 | 3 | 0.05 | 0.05 | 0.05 |
| EF077280.1;ABQ51392.1;Human_rhinovirus_C2_NAT | 645 | 647 | 3 | 0.05 | 0.1 | 0.083333333 |
| EF077280.1;ABQ51392.1;Human_rhinovirus_C2_NAT | 650 | 652 | 3 | 0.05 | 0.05 | 0.05 |
| EF077280.1;ABQ51392.1;Human_rhinovirus_C2_NAT | 694 | 694 | 1 | 0.05 | 0.05 | 0.05 |
| EF077280.1;ABQ51392.1;Human_rhinovirus_C2_NAT | 703 | 705 | 3 | 0.05 | 0.4 | 0.183333333 |
| EF077280.1;ABQ51392.1;Human_rhinovirus_C2_NAT | 712 | 713 | 2 | 0.05 | 0.05 | 0.05 |
| EF077280.1;ABQ51392.1;Human_rhinovirus_C2_NAT | 715 | 724 | 10 | 0.05 | 0.15 | 0.105 |
| EF077280.1;ABQ51392.1;Human_rhinovirus_C2_NAT | 742 | 742 | 1 | 0.05 | 0.05 | 0.05 |

| Extended data 1 | | | | | | |
|---|---|---|---|---|---|---|
| Protein | Start position (peptide) | Stop position (peptide) | Number of peptides | Minimum fraction (per peptide) of control subjects with reactivity >3 sd above average | Maximum fraction (per peptide) of control subjects with reactivity >3 sd above average | Mean fraction (over all peptides in this contiguous stretch) of controlsubjects with reactivity >3 sd above average |
| EF077280.1;ABQ51392.1;Human_rhinovirus_C2_NAT | 749 | 754 | 6 | 0.05 | 0.05 | 0.05 |
| EF077280.1;ABQ51392.1;Human_rhinovirus_C2_NAT | 769 | 771 | 3 | 0.05 | 0.1 | 0.066666667 |
| EF077280.1;ABQ51392.1;Human_rhinovirus_C2_NAT | 773 | 773 | 1 | 0.1 | 0.1 | 0.1 |
| EF077280.1;ABQ51392.1;Human_rhinovirus_C2_NAT | 815 | 831 | 17 | 0.05 | 0.35 | 0.185294118 |
| EF077280.1;ABQ51392.1;Human_rhinovirus_C2_NAT | 834 | 834 | 1 | 0.05 | 0.05 | 0.05 |
| EF077280.1;ABQ51392.1;Human_rhinovirus_C2_NAT | 838 | 838 | 1 | 0.15 | 0.15 | 0.15 |
| EF077280.1;ABQ51392.1;Human_rhinovirus_C2_NAT | 843 | 843 | 1 | 0.05 | 0.05 | 0.05 |
| EF077280.1;ABQ51392.1;Human_rhinovirus_C2_NAT | 845 | 852 | 8 | 0.05 | 0.1 | 0.075 |
| EF077280.1;ABQ51392.1;Human_rhinovirus_C2_NAT | 872 | 872 | 1 | 0.05 | 0.05 | 0.05 |
| EF077280.1;ABQ51392.1;Human_rhinovirus_C2_NAT | 899 | 899 | 1 | 0.05 | 0.05 | 0.05 |
| EF077280.1;ABQ51392.1;Human_rhinovirus_C2_NAT | 904 | 904 | 1 | 0.05 | 0.05 | 0.05 |
| EF077280.1;ABQ51392.1;Human_rhinovirus_C2_NAT | 906 | 909 | 4 | 0.05 | 0.05 | 0.05 |
| EF077280.1;ABQ51392.1;Human_rhinovirus_C2_NAT | 912 | 912 | 1 | 0.05 | 0.05 | 0.05 |
| EF077280.1;ABQ51392.1;Human_rhinovirus_C2_NAT | 914 | 914 | 1 | 0.05 | 0.05 | 0.05 |
| EF077280.1;ABQ51392.1;Human_rhinovirus_C2_NAT | 968 | 968 | 1 | 0.05 | 0.05 | 0.05 |
| EF077280.1;ABQ51392.1;Human_rhinovirus_C2_NAT | 973 | 996 | 24 | 0.05 | 0.8 | 0.325 |
| EF077280.1;ABQ51392.1;Human_rhinovirus_C2_NAT | 1002 | 1002 | 1 | 0.05 | 0.05 | 0.05 |
| EF077280.1;ABQ51392.1;Human_rhinovirus_C2_NAT | 1012 | 1012 | 1 | 0.05 | 0.05 | 0.05 |
| EF077280.1;ABQ51392.1;Human_rhinovirus_C2_NAT | 1059 | 1059 | 1 | 0.05 | 0.05 | 0.05 |
| EF077280.1;ABQ51392.1;Human_rhinovirus_C2_NAT | 1078 | 1079 | 2 | 0.05 | 0.2 | 0.125 |
| EF077280.1;ABQ51392.1;Human_rhinovirus_C2_NAT | 1087 | 1089 | 3 | 0.05 | 0.05 | 0.05 |
| EF077280.1;ABQ51392.1;Human_rhinovirus_C2_NAT | 1093 | 1093 | 1 | 0.05 | 0.05 | 0.05 |
| EF077280.1;ABQ51392.1;Human_rhinovirus_C2_NAT | 1097 | 1097 | 1 | 0.05 | 0.05 | 0.05 |
| EF077280.1;ABQ51392.1;Human_rhinovirus_C2_NAT | 1101 | 1102 | 2 | 0.05 | 0.05 | 0.05 |
| EF077280.1;ABQ51392.1;Human_rhinovirus_C2_NAT | 1130 | 1130 | 1 | 0.05 | 0.05 | 0.05 |
| EF077280.1;ABQ51392.1;Human_rhinovirus_C2_NAT | 1164 | 1173 | 10 | 0.05 | 0.1 | 0.065 |
| EF077280.1;ABQ51392.1;Human_rhinovirus_C2_NAT | 1183 | 1185 | 3 | 0.05 | 0.05 | 0.05 |
| EF077280.1;ABQ51392.1;Human_rhinovirus_C2_NAT | 1221 | 1235 | 15 | 0.05 | 0.45 | 0.296666667 |
| EF077280.1;ABQ51392.1;Human_rhinovirus_C2_NAT | 1244 | 1255 | 12 | 0.05 | 0.05 | 0.05 |
| EF077280.1;ABQ51392.1;Human_rhinovirus_C2_NAT | 1257 | 1257 | 1 | 0.05 | 0.05 | 0.05 |
| EF077280.1;ABQ51392.1;Human_rhinovirus_C2_NAT | 1264 | 1264 | 1 | 0.05 | 0.05 | 0.05 |
| EF077280.1;ABQ51392.1;Human_rhinovirus_C2_NAT | 1273 | 1273 | 1 | 0.05 | 0.05 | 0.05 |
| EF077280.1;ABQ51392.1;Human_rhinovirus_C2_NAT | 1275 | 1279 | 5 | 0.05 | 0.05 | 0.05 |
| EF077280.1;ABQ51392.1;Human_rhinovirus_C2_NAT | 1285 | 1285 | 1 | 0.05 | 0.05 | 0.05 |
| EF077280.1;ABQ51392.1;Human_rhinovirus_C2_NAT | 1290 | 1290 | 1 | 0.05 | 0.05 | 0.05 |
| EF077280.1;ABQ51392.1;Human_rhinovirus_C2_NAT | 1292 | 1293 | 2 | 0.05 | 0.05 | 0.05 |
| EF077280.1;ABQ51392.1;Human_rhinovirus_C2_NAT | 1295 | 1298 | 4 | 0.05 | 0.05 | 0.05 |

| Extended data 1 | | | | | | |
|---|---|---|---|---|---|---|
| Protein | Start position (peptide) | Stop position (peptide) | Number of peptides | Minimum fraction (per peptide) of control subjects with reactivity >3 sd above average | Maximum fraction (per peptide) of control subjects with reactivity >3 sd above average | Mean fraction (over all peptides in this contiguous stretch) of controlsubjects with reactivity >3 sd above average |
| EF077280.1;ABQ51392.1;Human_rhinovirus_C2_NAT | 1300 | 1300 | 1 | 0.1 | 0.1 | 0.1 |
| EF077280.1;ABQ51392.1;Human_rhinovirus_C2_NAT | 1308 | 1308 | 1 | 0.05 | 0.05 | 0.05 |
| EF077280.1;ABQ51392.1;Human_rhinovirus_C2_NAT | 1313 | 1313 | 1 | 0.05 | 0.05 | 0.05 |
| EF077280.1;ABQ51392.1;Human_rhinovirus_C2_NAT | 1336 | 1336 | 1 | 0.05 | 0.05 | 0.05 |
| EF077280.1;ABQ51392.1;Human_rhinovirus_C2_NAT | 1368 | 1368 | 1 | 0.05 | 0.05 | 0.05 |
| EF077280.1;ABQ51392.1;Human_rhinovirus_C2_NAT | 1375 | 1379 | 5 | 0.05 | 0.05 | 0.05 |
| EF077280.1;ABQ51392.1;Human_rhinovirus_C2_NAT | 1386 | 1386 | 1 | 0.05 | 0.05 | 0.05 |
| EF077280.1;ABQ51392.1;Human_rhinovirus_C2_NAT | 1403 | 1415 | 13 | 0.05 | 0.7 | 0.338461538 |
| EF077280.1;ABQ51392.1;Human_rhinovirus_C2_NAT | 1417 | 1424 | 8 | 0.05 | 0.15 | 0.0875 |
| EF077280.1;ABQ51392.1;Human_rhinovirus_C2_NAT | 1517 | 1517 | 1 | 0.05 | 0.05 | 0.05 |
| EF077280.1;ABQ51392.1;Human_rhinovirus_C2_NAT | 1544 | 1545 | 2 | 0.05 | 0.05 | 0.05 |
| EF077280.1;ABQ51392.1;Human_rhinovirus_C2_NAT | 1547 | 1548 | 2 | 0.1 | 0.1 | 0.1 |
| EF077280.1;ABQ51392.1;Human_rhinovirus_C2_NAT | 1574 | 1583 | 10 | 0.05 | 0.3 | 0.125 |
| EF077280.1;ABQ51392.1;Human_rhinovirus_C2_NAT | 1605 | 1607 | 3 | 0.05 | 0.1 | 0.083333333 |
| EF077280.1;ABQ51392.1;Human_rhinovirus_C2_NAT | 1609 | 1609 | 1 | 0.05 | 0.05 | 0.05 |
| EF077280.1;ABQ51392.1;Human_rhinovirus_C2_NAT | 1612 | 1612 | 1 | 0.1 | 0.1 | 0.1 |
| EF077280.1;ABQ51392.1;Human_rhinovirus_C2_NAT | 1614 | 1614 | 1 | 0.05 | 0.05 | 0.05 |
| EF077280.1;ABQ51392.1;Human_rhinovirus_C2_NAT | 1620 | 1620 | 1 | 0.05 | 0.05 | 0.05 |
| EF077280.1;ABQ51392.1;Human_rhinovirus_C2_NAT | 1635 | 1635 | 1 | 0.05 | 0.05 | 0.05 |
| EF077280.1;ABQ51392.1;Human_rhinovirus_C2_NAT | 1639 | 1639 | 1 | 0.05 | 0.05 | 0.05 |
| EF077280.1;ABQ51392.1;Human_rhinovirus_C2_NAT | 1650 | 1650 | 1 | 0.05 | 0.05 | 0.05 |
| EF077280.1;ABQ51392.1;Human_rhinovirus_C2_NAT | 1659 | 1659 | 1 | 0.1 | 0.1 | 0.1 |
| EF077280.1;ABQ51392.1;Human_rhinovirus_C2_NAT | 1690 | 1716 | 27 | 0.05 | 0.35 | 0.116666667 |
| EF077280.1;ABQ51392.1;Human_rhinovirus_C2_NAT | 1718 | 1722 | 5 | 0.05 | 0.1 | 0.07 |
| EF077280.1;ABQ51392.1;Human_rhinovirus_C2_NAT | 1724 | 1728 | 5 | 0.05 | 0.35 | 0.16 |
| EF077280.1;ABQ51392.1;Human_rhinovirus_C2_NAT | 1730 | 1754 | 25 | 0.05 | 0.35 | 0.212 |
| EF077280.1;ABQ51392.1;Human_rhinovirus_C2_NAT | 1770 | 1770 | 1 | 0.05 | 0.05 | 0.05 |
| EF077280.1;ABQ51392.1;Human_rhinovirus_C2_NAT | 1772 | 1780 | 9 | 0.05 | 0.15 | 0.105555556 |
| EF077280.1;ABQ51392.1;Human_rhinovirus_C2_NAT | 1795 | 1795 | 1 | 0.05 | 0.05 | 0.05 |
| EF077280.1;ABQ51392.1;Human_rhinovirus_C2_NAT | 1802 | 1803 | 2 | 0.05 | 0.05 | 0.05 |
| EF077280.1;ABQ51392.1;Human_rhinovirus_C2_NAT | 1810 | 1811 | 2 | 0.05 | 0.1 | 0.075 |
| EF077280.1;ABQ51392.1;Human_rhinovirus_C2_NAT | 1814 | 1818 | 5 | 0.05 | 0.1 | 0.06 |
| EF077280.1;ABQ51392.1;Human_rhinovirus_C2_NAT | 1822 | 1829 | 8 | 0.05 | 0.1 | 0.075 |
| EF077280.1;ABQ51392.1;Human_rhinovirus_C2_NAT | 1832 | 1836 | 5 | 0.05 | 0.05 | 0.05 |
| EF077280.1;ABQ51392.1;Human_rhinovirus_C2_NAT | 1844 | 1852 | 9 | 0.05 | 0.05 | 0.05 |
| EF077280.1;ABQ51392.1;Human_rhinovirus_C2_NAT | 1854 | 1854 | 1 | 0.05 | 0.05 | 0.05 |
| EF077280.1;ABQ51392.1;Human_rhinovirus_C2_NAT | 1859 | 1859 | 1 | 0.25 | 0.25 | 0.25 |

| Extended data 1 | | | | | | |
|---|---|---|---|---|---|---|
| Protein | Start position (peptide) | Stop position (peptide) | Number of peptides | Minimum fraction (per peptide) of control subjects with reactivity >3 sd above average | Maximum fraction (per peptide) of control subjects with reactivity >3 sd above average | Mean fraction (over all peptides in this contiguous stretch) of controlsubjects with reactivity >3 sd above average |
| EF077280.1;ABQ51392.1;Human_rhinovirus_C2_NAT | 1864 | 1864 | 1 | 0.3 | 0.3 | 0.3 |
| EF077280.1;ABQ51392.1;Human_rhinovirus_C2_NAT | 1896 | 1896 | 1 | 0.05 | 0.05 | 0.05 |
| EF077280.1;ABQ51392.1;Human_rhinovirus_C2_NAT | 1907 | 1915 | 9 | 0.05 | 0.05 | 0.05 |
| EF077280.1;ABQ51392.1;Human_rhinovirus_C2_NAT | 1950 | 1952 | 3 | 0.05 | 0.05 | 0.05 |
| EF077280.1;ABQ51392.1;Human_rhinovirus_C2_NAT | 1956 | 1958 | 3 | 0.05 | 0.1 | 0.066666667 |
| EF077280.1;ABQ51392.1;Human_rhinovirus_C2_NAT | 1961 | 1961 | 1 | 0.05 | 0.05 | 0.05 |
| EF077280.1;ABQ51392.1;Human_rhinovirus_C2_NAT | 1965 | 1965 | 1 | 0.05 | 0.05 | 0.05 |
| EF077280.1;ABQ51392.1;Human_rhinovirus_C2_NAT | 1970 | 1970 | 1 | 0.05 | 0.05 | 0.05 |
| EF077280.1;ABQ51392.1;Human_rhinovirus_C2_NAT | 1992 | 1992 | 1 | 0.05 | 0.05 | 0.05 |
| EF077280.1;ABQ51392.1;Human_rhinovirus_C2_NAT | 2007 | 2023 | 17 | 0.05 | 0.15 | 0.094117647 |
| EF077280.1;ABQ51392.1;Human_rhinovirus_C2_NAT | 2029 | 2029 | 1 | 0.15 | 0.15 | 0.15 |
| EF077280.1;ABQ51392.1;Human_rhinovirus_C2_NAT | 2031 | 2039 | 9 | 0.05 | 0.1 | 0.072222222 |
| EF077280.1;ABQ51392.1;Human_rhinovirus_C2_NAT | 2049 | 2061 | 13 | 0.05 | 0.1 | 0.065384615 |
| EF077280.1;ABQ51392.1;Human_rhinovirus_C2_NAT | 2078 | 2081 | 4 | 0.05 | 0.2 | 0.0875 |
| EF077280.1;ABQ51392.1;Human_rhinovirus_C2_NAT | 2100 | 2103 | 4 | 0.05 | 0.2 | 0.1 |
| EF077280.1;ABQ51392.1;Human_rhinovirus_C2_NAT | 2108 | 2108 | 1 | 0.05 | 0.05 | 0.05 |
| EF077280.1;ABQ51392.1;Human_rhinovirus_C2_NAT | 2113 | 2114 | 2 | 0.05 | 0.05 | 0.05 |
| GU219984.1;ACZ67658.1;Human_rhinovirus_C15_st | 3 | 3 | 1 | 0.05 | 0.05 | 0.05 |
| GU219984.1;ACZ67658.1;Human_rhinovirus_C15_st | 5 | 16 | 12 | 0.05 | 0.2 | 0.1125 |
| GU219984.1;ACZ67658.1;Human_rhinovirus_C15_st | 20 | 21 | 2 | 0.05 | 0.05 | 0.05 |
| GU219984.1;ACZ67658.1;Human_rhinovirus_C15_st | 23 | 28 | 6 | 0.05 | 0.15 | 0.083333333 |
| GU219984.1;ACZ67658.1;Human_rhinovirus_C15_st | 35 | 47 | 13 | 0.05 | 0.25 | 0.180769231 |
| GU219984.1;ACZ67658.1;Human_rhinovirus_C15_st | 49 | 49 | 1 | 0.05 | 0.05 | 0.05 |
| GU219984.1;ACZ67658.1;Human_rhinovirus_C15_st | 51 | 51 | 1 | 0.05 | 0.05 | 0.05 |
| GU219984.1;ACZ67658.1;Human_rhinovirus_C15_st | 72 | 72 | 1 | 0.1 | 0.1 | 0.1 |
| GU219984.1;ACZ67658.1;Human_rhinovirus_C15_st | 74 | 74 | 1 | 0.1 | 0.1 | 0.1 |
| GU219984.1;ACZ67658.1;Human_rhinovirus_C15_st | 80 | 81 | 2 | 0.05 | 0.15 | 0.1 |
| GU219984.1;ACZ67658.1;Human_rhinovirus_C15_st | 104 | 114 | 11 | 0.1 | 0.2 | 0.15 |
| GU219984.1;ACZ67658.1;Human_rhinovirus_C15_st | 117 | 117 | 1 | 0.1 | 0.1 | 0.1 |
| GU219984.1;ACZ67658.1;Human_rhinovirus_C15_st | 120 | 121 | 2 | 0.1 | 0.25 | 0.175 |
| GU219984.1;ACZ67658.1;Human_rhinovirus_C15_st | 146 | 149 | 4 | 0.05 | 0.1 | 0.075 |
| GU219984.1;ACZ67658.1;Human_rhinovirus_C15_st | 151 | 151 | 1 | 0.05 | 0.05 | 0.05 |
| GU219984.1;ACZ67658.1;Human_rhinovirus_C15_st | 153 | 154 | 2 | 0.05 | 0.1 | 0.075 |
| GU219984.1;ACZ67658.1;Human_rhinovirus_C15_st | 201 | 213 | 13 | 0.05 | 0.45 | 0.311538462 |
| GU219984.1;ACZ67658.1;Human_rhinovirus_C15_st | 217 | 217 | 1 | 0.05 | 0.05 | 0.05 |
| GU219984.1;ACZ67658.1;Human_rhinovirus_C15_st | 220 | 220 | 1 | 0.05 | 0.05 | 0.05 |
| GU219984.1;ACZ67658.1;Human_rhinovirus_C15_st | 224 | 224 | 1 | 0.05 | 0.05 | 0.05 |

| Extended data 1 | | | | | | |
|---|---|---|---|---|---|---|
| Protein | Start position (peptide) | Stop position (peptide) | Number of peptides | Minimum fraction (per peptide) of control subjects with reactivity >3 sd above average | Maximum fraction (per peptide) of control subjects with reactivity >3 sd above average | Mean fraction (over all peptides in this contiguous stretch) of controlsubjects with reactivity >3 sd above average |
| GU219984.1;ACZ67658.1;Human_rhinovirus_C15_st | 226 | 236 | 11 | 0.05 | 0.1 | 0.054545455 |
| GU219984.1;ACZ67658.1;Human_rhinovirus_C15_st | 260 | 260 | 1 | 0.05 | 0.05 | 0.05 |
| GU219984.1;ACZ67658.1;Human_rhinovirus_C15_st | 274 | 274 | 1 | 0.05 | 0.05 | 0.05 |
| GU219984.1;ACZ67658.1;Human_rhinovirus_C15_st | 277 | 278 | 2 | 0.05 | 0.05 | 0.05 |
| GU219984.1;ACZ67658.1;Human_rhinovirus_C15_st | 296 | 297 | 2 | 0.05 | 0.05 | 0.05 |
| GU219984.1;ACZ67658.1;Human_rhinovirus_C15_st | 310 | 310 | 1 | 0.05 | 0.05 | 0.05 |
| GU219984.1;ACZ67658.1;Human_rhinovirus_C15_st | 314 | 316 | 3 | 0.05 | 0.05 | 0.05 |
| GU219984.1;ACZ67658.1;Human_rhinovirus_C15_st | 319 | 319 | 1 | 0.05 | 0.05 | 0.05 |
| GU219984.1;ACZ67658.1;Human_rhinovirus_C15_st | 321 | 321 | 1 | 0.05 | 0.05 | 0.05 |
| GU219984.1;ACZ67658.1;Human_rhinovirus_C15_st | 326 | 327 | 2 | 0.05 | 0.15 | 0.1 |
| GU219984.1;ACZ67658.1;Human_rhinovirus_C15_st | 349 | 349 | 1 | 0.05 | 0.05 | 0.05 |
| GU219984.1;ACZ67658.1;Human_rhinovirus_C15_st | 373 | 373 | 1 | 0.1 | 0.1 | 0.1 |
| GU219984.1;ACZ67658.1;Human_rhinovirus_C15_st | 382 | 382 | 1 | 0.15 | 0.15 | 0.15 |
| GU219984.1;ACZ67658.1;Human_rhinovirus_C15_st | 411 | 411 | 1 | 0.05 | 0.05 | 0.05 |
| GU219984.1;ACZ67658.1;Human_rhinovirus_C15_st | 469 | 469 | 1 | 0.05 | 0.05 | 0.05 |
| GU219984.1;ACZ67658.1;Human_rhinovirus_C15_st | 473 | 473 | 1 | 0.05 | 0.05 | 0.05 |
| GU219984.1;ACZ67658.1;Human_rhinovirus_C15_st | 485 | 485 | 1 | 0.05 | 0.05 | 0.05 |
| GU219984.1;ACZ67658.1;Human_rhinovirus_C15_st | 500 | 500 | 1 | 0.05 | 0.05 | 0.05 |
| GU219984.1;ACZ67658.1;Human_rhinovirus_C15_st | 503 | 503 | 1 | 0.05 | 0.05 | 0.05 |
| GU219984.1;ACZ67658.1;Human_rhinovirus_C15_st | 512 | 512 | 1 | 0.05 | 0.05 | 0.05 |
| GU219984.1;ACZ67658.1;Human_rhinovirus_C15_st | 520 | 520 | 1 | 0.05 | 0.05 | 0.05 |
| GU219984.1;ACZ67658.1;Human_rhinovirus_C15_st | 559 | 559 | 1 | 0.05 | 0.05 | 0.05 |
| GU219984.1;ACZ67658.1;Human_rhinovirus_C15_st | 561 | 563 | 3 | 0.05 | 0.05 | 0.05 |
| GU219984.1;ACZ67658.1;Human_rhinovirus_C15_st | 566 | 604 | 39 | 0.05 | 0.75 | 0.330769231 |
| GU219984.1;ACZ67658.1;Human_rhinovirus_C15_st | 607 | 611 | 5 | 0.05 | 0.15 | 0.07 |
| GU219984.1;ACZ67658.1;Human_rhinovirus_C15_st | 613 | 613 | 1 | 0.05 | 0.05 | 0.05 |
| GU219984.1;ACZ67658.1;Human_rhinovirus_C15_st | 616 | 629 | 14 | 0.05 | 0.55 | 0.182142857 |
| GU219984.1;ACZ67658.1;Human_rhinovirus_C15_st | 681 | 683 | 3 | 0.05 | 0.05 | 0.05 |
| GU219984.1;ACZ67658.1;Human_rhinovirus_C15_st | 692 | 694 | 3 | 0.05 | 0.1 | 0.066666667 |
| GU219984.1;ACZ67658.1;Human_rhinovirus_C15_st | 696 | 705 | 10 | 0.05 | 0.15 | 0.09 |
| GU219984.1;ACZ67658.1;Human_rhinovirus_C15_st | 712 | 712 | 1 | 0.05 | 0.05 | 0.05 |
| GU219984.1;ACZ67658.1;Human_rhinovirus_C15_st | 717 | 717 | 1 | 0.05 | 0.05 | 0.05 |
| GU219984.1;ACZ67658.1;Human_rhinovirus_C15_st | 719 | 720 | 2 | 0.05 | 0.15 | 0.1 |
| GU219984.1;ACZ67658.1;Human_rhinovirus_C15_st | 722 | 722 | 1 | 0.1 | 0.1 | 0.1 |
| GU219984.1;ACZ67658.1;Human_rhinovirus_C15_st | 724 | 728 | 5 | 0.05 | 0.1 | 0.09 |
| GU219984.1;ACZ67658.1;Human_rhinovirus_C15_st | 747 | 748 | 2 | 0.05 | 0.1 | 0.075 |
| GU219984.1;ACZ67658.1;Human_rhinovirus_C15_st | 750 | 750 | 1 | 0.05 | 0.05 | 0.05 |

| Extended data 1 | | | | | | |
|---|---|---|---|---|---|---|
| Protein | Start position (peptide) | Stop position (peptide) | Number of peptides | Minimum fraction (per peptide) of control subjects with reactivity >3 sd above average | Maximum fraction (per peptide) of control subjects with reactivity >3 sd above average | Mean fraction (over all peptides in this contiguous stretch) of controlsubjects with reactivity >3 sd above average |
| GU219984.1;ACZ67658.1;Human_rhinovirus_C15_st | 754 | 754 | 1 | 0.05 | 0.05 | 0.05 |
| GU219984.1;ACZ67658.1;Human_rhinovirus_C15_st | 773 | 775 | 3 | 0.05 | 0.15 | 0.1 |
| GU219984.1;ACZ67658.1;Human_rhinovirus_C15_st | 780 | 780 | 1 | 0.05 | 0.05 | 0.05 |
| GU219984.1;ACZ67658.1;Human_rhinovirus_C15_st | 790 | 790 | 1 | 0.05 | 0.05 | 0.05 |
| GU219984.1;ACZ67658.1;Human_rhinovirus_C15_st | 805 | 812 | 8 | 0.05 | 0.15 | 0.075 |
| GU219984.1;ACZ67658.1;Human_rhinovirus_C15_st | 817 | 819 | 3 | 0.05 | 0.1 | 0.066666667 |
| GU219984.1;ACZ67658.1;Human_rhinovirus_C15_st | 822 | 840 | 19 | 0.05 | 0.55 | 0.202631579 |
| GU219984.1;ACZ67658.1;Human_rhinovirus_C15_st | 844 | 844 | 1 | 0.05 | 0.05 | 0.05 |
| GU219984.1;ACZ67658.1;Human_rhinovirus_C15_st | 847 | 847 | 1 | 0.05 | 0.05 | 0.05 |
| GU219984.1;ACZ67658.1;Human_rhinovirus_C15_st | 856 | 859 | 4 | 0.05 | 0.05 | 0.05 |
| GU219984.1;ACZ67658.1;Human_rhinovirus_C15_st | 881 | 883 | 3 | 0.05 | 0.1 | 0.083333333 |
| GU219984.1;ACZ67658.1;Human_rhinovirus_C15_st | 886 | 886 | 1 | 0.05 | 0.05 | 0.05 |
| GU219984.1;ACZ67658.1;Human_rhinovirus_C15_st | 901 | 901 | 1 | 0.05 | 0.05 | 0.05 |
| GU219984.1;ACZ67658.1;Human_rhinovirus_C15_st | 954 | 954 | 1 | 0.05 | 0.05 | 0.05 |
| GU219984.1;ACZ67658.1;Human_rhinovirus_C15_st | 960 | 960 | 1 | 0.05 | 0.05 | 0.05 |
| GU219984.1;ACZ67658.1;Human_rhinovirus_C15_st | 967 | 967 | 1 | 0.05 | 0.05 | 0.05 |
| GU219984.1;ACZ67658.1;Human_rhinovirus_C15_st | 975 | 979 | 5 | 0.05 | 0.05 | 0.05 |
| GU219984.1;ACZ67658.1;Human_rhinovirus_C15_st | 981 | 982 | 2 | 0.05 | 0.05 | 0.05 |
| GU219984.1;ACZ67658.1;Human_rhinovirus_C15_st | 984 | 1002 | 19 | 0.05 | 0.9 | 0.352631579 |
| GU219984.1;ACZ67658.1;Human_rhinovirus_C15_st | 1011 | 1012 | 2 | 0.05 | 0.1 | 0.075 |
| GU219984.1;ACZ67658.1;Human_rhinovirus_C15_st | 1042 | 1042 | 1 | 0.05 | 0.05 | 0.05 |
| GU219984.1;ACZ67658.1;Human_rhinovirus_C15_st | 1071 | 1080 | 10 | 0.05 | 0.05 | 0.05 |
| GU219984.1;ACZ67658.1;Human_rhinovirus_C15_st | 1087 | 1088 | 2 | 0.05 | 0.05 | 0.05 |
| GU219984.1;ACZ67658.1;Human_rhinovirus_C15_st | 1114 | 1114 | 1 | 0.05 | 0.05 | 0.05 |
| GU219984.1;ACZ67658.1;Human_rhinovirus_C15_st | 1121 | 1121 | 1 | 0.05 | 0.05 | 0.05 |
| GU219984.1;ACZ67658.1;Human_rhinovirus_C15_st | 1130 | 1130 | 1 | 0.2 | 0.2 | 0.2 |
| GU219984.1;ACZ67658.1;Human_rhinovirus_C15_st | 1134 | 1139 | 6 | 0.05 | 0.05 | 0.05 |
| GU219984.1;ACZ67658.1;Human_rhinovirus_C15_st | 1149 | 1149 | 1 | 0.05 | 0.05 | 0.05 |
| GU219984.1;ACZ67658.1;Human_rhinovirus_C15_st | 1170 | 1177 | 8 | 0.05 | 0.35 | 0.09375 |
| GU219984.1;ACZ67658.1;Human_rhinovirus_C15_st | 1194 | 1194 | 1 | 0.05 | 0.05 | 0.05 |
| GU219984.1;ACZ67658.1;Human_rhinovirus_C15_st | 1198 | 1198 | 1 | 0.05 | 0.05 | 0.05 |
| GU219984.1;ACZ67658.1;Human_rhinovirus_C15_st | 1230 | 1244 | 15 | 0.1 | 0.65 | 0.446666667 |
| GU219984.1;ACZ67658.1;Human_rhinovirus_C15_st | 1248 | 1248 | 1 | 0.05 | 0.05 | 0.05 |
| GU219984.1;ACZ67658.1;Human_rhinovirus_C15_st | 1251 | 1263 | 13 | 0.05 | 0.2 | 0.103846154 |
| GU219984.1;ACZ67658.1;Human_rhinovirus_C15_st | 1280 | 1282 | 3 | 0.05 | 0.05 | 0.05 |
| GU219984.1;ACZ67658.1;Human_rhinovirus_C15_st | 1286 | 1286 | 1 | 0.15 | 0.15 | 0.15 |
| GU219984.1;ACZ67658.1;Human_rhinovirus_C15_st | 1301 | 1301 | 1 | 0.05 | 0.05 | 0.05 |

| Extended data 1 | | | | | | |
|---|---|---|---|---|---|---|
| Protein | Start position (peptide) | Stop position (peptide) | Number of peptides | Minimum fraction (per peptide) of control subjects with reactivity >3 sd above average | Maximum fraction (per peptide) of control subjects with reactivity >3 sd above average | Mean fraction (over all peptides in this contiguous stretch) of controlsubjects with reactivity >3 sd above average |
| GU219984.1;ACZ67658.1;Human_rhinovirus_C15_st | 1305 | 1305 | 1 | 0.1 | 0.1 | 0.1 |
| GU219984.1;ACZ67658.1;Human_rhinovirus_C15_st | 1307 | 1307 | 1 | 0.05 | 0.05 | 0.05 |
| GU219984.1;ACZ67658.1;Human_rhinovirus_C15_st | 1314 | 1319 | 6 | 0.05 | 0.05 | 0.05 |
| GU219984.1;ACZ67658.1;Human_rhinovirus_C15_st | 1321 | 1321 | 1 | 0.1 | 0.1 | 0.1 |
| GU219984.1;ACZ67658.1;Human_rhinovirus_C15_st | 1325 | 1328 | 4 | 0.05 | 0.1 | 0.0875 |
| GU219984.1;ACZ67658.1;Human_rhinovirus_C15_st | 1330 | 1330 | 1 | 0.05 | 0.05 | 0.05 |
| GU219984.1;ACZ67658.1;Human_rhinovirus_C15_st | 1345 | 1345 | 1 | 0.05 | 0.05 | 0.05 |
| GU219984.1;ACZ67658.1;Human_rhinovirus_C15_st | 1359 | 1359 | 1 | 0.05 | 0.05 | 0.05 |
| GU219984.1;ACZ67658.1;Human_rhinovirus_C15_st | 1375 | 1383 | 9 | 0.05 | 0.15 | 0.1 |
| GU219984.1;ACZ67658.1;Human_rhinovirus_C15_st | 1386 | 1387 | 2 | 0.05 | 0.05 | 0.05 |
| GU219984.1;ACZ67658.1;Human_rhinovirus_C15_st | 1397 | 1398 | 2 | 0.05 | 0.05 | 0.05 |
| GU219984.1;ACZ67658.1;Human_rhinovirus_C15_st | 1403 | 1409 | 7 | 0.05 | 0.1 | 0.078571429 |
| GU219984.1;ACZ67658.1;Human_rhinovirus_C15_st | 1412 | 1424 | 13 | 0.05 | 0.8 | 0.423076923 |
| GU219984.1;ACZ67658.1;Human_rhinovirus_C15_st | 1426 | 1429 | 4 | 0.05 | 0.1 | 0.0625 |
| GU219984.1;ACZ67658.1;Human_rhinovirus_C15_st | 1431 | 1431 | 1 | 0.05 | 0.05 | 0.05 |
| GU219984.1;ACZ67658.1;Human_rhinovirus_C15_st | 1433 | 1437 | 5 | 0.05 | 0.1 | 0.06 |
| GU219984.1;ACZ67658.1;Human_rhinovirus_C15_st | 1440 | 1440 | 1 | 0.1 | 0.1 | 0.1 |
| GU219984.1;ACZ67658.1;Human_rhinovirus_C15_st | 1442 | 1449 | 8 | 0.05 | 0.1 | 0.05625 |
| GU219984.1;ACZ67658.1;Human_rhinovirus_C15_st | 1454 | 1454 | 1 | 0.05 | 0.05 | 0.05 |
| GU219984.1;ACZ67658.1;Human_rhinovirus_C15_st | 1484 | 1484 | 1 | 0.05 | 0.05 | 0.05 |
| GU219984.1;ACZ67658.1;Human_rhinovirus_C15_st | 1486 | 1491 | 6 | 0.05 | 0.35 | 0.116666667 |
| GU219984.1;ACZ67658.1;Human_rhinovirus_C15_st | 1528 | 1530 | 3 | 0.05 | 0.1 | 0.066666667 |
| GU219984.1;ACZ67658.1;Human_rhinovirus_C15_st | 1582 | 1592 | 11 | 0.05 | 0.25 | 0.122727273 |
| GU219984.1;ACZ67658.1;Human_rhinovirus_C15_st | 1606 | 1606 | 1 | 0.1 | 0.1 | 0.1 |
| GU219984.1;ACZ67658.1;Human_rhinovirus_C15_st | 1610 | 1614 | 5 | 0.05 | 0.2 | 0.09 |
| GU219984.1;ACZ67658.1;Human_rhinovirus_C15_st | 1620 | 1620 | 1 | 0.05 | 0.05 | 0.05 |
| GU219984.1;ACZ67658.1;Human_rhinovirus_C15_st | 1624 | 1627 | 4 | 0.05 | 0.05 | 0.05 |
| GU219984.1;ACZ67658.1;Human_rhinovirus_C15_st | 1651 | 1651 | 1 | 0.05 | 0.05 | 0.05 |
| GU219984.1;ACZ67658.1;Human_rhinovirus_C15_st | 1653 | 1654 | 2 | 0.1 | 0.1 | 0.1 |
| GU219984.1;ACZ67658.1;Human_rhinovirus_C15_st | 1661 | 1662 | 2 | 0.1 | 0.1 | 0.1 |
| GU219984.1;ACZ67658.1;Human_rhinovirus_C15_st | 1668 | 1668 | 1 | 0.1 | 0.1 | 0.1 |
| GU219984.1;ACZ67658.1;Human_rhinovirus_C15_st | 1684 | 1684 | 1 | 0.05 | 0.05 | 0.05 |
| GU219984.1;ACZ67658.1;Human_rhinovirus_C15_st | 1700 | 1740 | 41 | 0.05 | 0.6 | 0.185365854 |
| GU219984.1;ACZ67658.1;Human_rhinovirus_C15_st | 1745 | 1755 | 11 | 0.05 | 0.2 | 0.113636364 |
| GU219984.1;ACZ67658.1;Human_rhinovirus_C15_st | 1760 | 1760 | 1 | 0.05 | 0.05 | 0.05 |
| GU219984.1;ACZ67658.1;Human_rhinovirus_C15_st | 1763 | 1763 | 1 | 0.05 | 0.05 | 0.05 |
| GU219984.1;ACZ67658.1;Human_rhinovirus_C15_st | 1775 | 1775 | 1 | 0.05 | 0.05 | 0.05 |

| Extended data 1 | | | | | | |
|---|---|---|---|---|---|---|
| **Protein** | **Start position (peptide)** | **Stop position (peptide)** | **Number of peptides** | **Minimum fraction (per peptide) of control subjects with reactivity >3 sd above average** | **Maximum fraction (per peptide) of control subjects with reactivity >3 sd above average** | **Mean fraction (over all peptides in this contiguous stretch) of controlsubjects with reactivity >3 sd above average** |
| GU219984.1;ACZ67658.1;Human_rhinovirus_C15_st | 1782 | 1789 | 8 | 0.05 | 0.1 | 0.05625 |
| GU219984.1;ACZ67658.1;Human_rhinovirus_C15_st | 1796 | 1796 | 1 | 0.25 | 0.25 | 0.25 |
| GU219984.1;ACZ67658.1;Human_rhinovirus_C15_st | 1820 | 1820 | 1 | 0.05 | 0.05 | 0.05 |
| GU219984.1;ACZ67658.1;Human_rhinovirus_C15_st | 1829 | 1845 | 17 | 0.05 | 0.15 | 0.088235294 |
| GU219984.1;ACZ67658.1;Human_rhinovirus_C15_st | 1860 | 1861 | 2 | 0.05 | 0.1 | 0.075 |
| GU219984.1;ACZ67658.1;Human_rhinovirus_C15_st | 1863 | 1863 | 1 | 0.05 | 0.05 | 0.05 |
| GU219984.1;ACZ67658.1;Human_rhinovirus_C15_st | 1865 | 1865 | 1 | 0.05 | 0.05 | 0.05 |
| GU219984.1;ACZ67658.1;Human_rhinovirus_C15_st | 1868 | 1868 | 1 | 0.1 | 0.1 | 0.1 |
| GU219984.1;ACZ67658.1;Human_rhinovirus_C15_st | 1870 | 1870 | 1 | 0.05 | 0.05 | 0.05 |
| GU219984.1;ACZ67658.1;Human_rhinovirus_C15_st | 1873 | 1873 | 1 | 0.2 | 0.2 | 0.2 |
| GU219984.1;ACZ67658.1;Human_rhinovirus_C15_st | 1889 | 1889 | 1 | 0.05 | 0.05 | 0.05 |
| GU219984.1;ACZ67658.1;Human_rhinovirus_C15_st | 1896 | 1896 | 1 | 0.05 | 0.05 | 0.05 |
| GU219984.1;ACZ67658.1;Human_rhinovirus_C15_st | 1914 | 1924 | 11 | 0.05 | 0.1 | 0.054545455 |
| GU219984.1;ACZ67658.1;Human_rhinovirus_C15_st | 1926 | 1926 | 1 | 0.05 | 0.05 | 0.05 |
| GU219984.1;ACZ67658.1;Human_rhinovirus_C15_st | 1938 | 1940 | 3 | 0.05 | 0.1 | 0.066666667 |
| GU219984.1;ACZ67658.1;Human_rhinovirus_C15_st | 1942 | 1944 | 3 | 0.05 | 0.05 | 0.05 |
| GU219984.1;ACZ67658.1;Human_rhinovirus_C15_st | 1961 | 1970 | 10 | 0.05 | 0.15 | 0.135 |
| GU219984.1;ACZ67658.1;Human_rhinovirus_C15_st | 1974 | 1974 | 1 | 0.05 | 0.05 | 0.05 |
| GU219984.1;ACZ67658.1;Human_rhinovirus_C15_st | 1999 | 1999 | 1 | 0.05 | 0.05 | 0.05 |
| GU219984.1;ACZ67658.1;Human_rhinovirus_C15_st | 2005 | 2005 | 1 | 0.05 | 0.05 | 0.05 |
| GU219984.1;ACZ67658.1;Human_rhinovirus_C15_st | 2044 | 2049 | 6 | 0.05 | 0.15 | 0.091666667 |
| GU219984.1;ACZ67658.1;Human_rhinovirus_C15_st | 2059 | 2063 | 5 | 0.05 | 0.1 | 0.06 |
| GU219984.1;ACZ67658.1;Human_rhinovirus_C15_st | 2065 | 2068 | 4 | 0.05 | 0.05 | 0.05 |
| GU219984.1;ACZ67658.1;Human_rhinovirus_C15_st | 2070 | 2070 | 1 | 0.05 | 0.05 | 0.05 |
| GU219984.1;ACZ67658.1;Human_rhinovirus_C15_st | 2085 | 2090 | 6 | 0.05 | 0.2 | 0.075 |
| GU219984.1;ACZ67658.1;Human_rhinovirus_C15_st | 2094 | 2094 | 1 | 0.05 | 0.05 | 0.05 |
| GU219984.1;ACZ67658.1;Human_rhinovirus_C15_st | 2108 | 2109 | 2 | 0.05 | 0.05 | 0.05 |
| GU219984.1;ACZ67658.1;Human_rhinovirus_C15_st | 2111 | 2113 | 3 | 0.05 | 0.05 | 0.05 |
| JX074050.1;AFK65737.1;Human_rhinovirus_A36_st | 5 | 5 | 1 | 0.1 | 0.1 | 0.1 |
| JX074050.1;AFK65737.1;Human_rhinovirus_A36_st | 7 | 9 | 3 | 0.05 | 0.05 | 0.05 |
| JX074050.1;AFK65737.1;Human_rhinovirus_A36_st | 11 | 11 | 1 | 0.05 | 0.05 | 0.05 |
| JX074050.1;AFK65737.1;Human_rhinovirus_A36_st | 14 | 15 | 2 | 0.05 | 0.15 | 0.1 |
| JX074050.1;AFK65737.1;Human_rhinovirus_A36_st | 20 | 64 | 45 | 0.05 | 0.6 | 0.301111111 |
| JX074050.1;AFK65737.1;Human_rhinovirus_A36_st | 77 | 77 | 1 | 0.05 | 0.05 | 0.05 |
| JX074050.1;AFK65737.1;Human_rhinovirus_A36_st | 79 | 88 | 10 | 0.05 | 0.05 | 0.05 |
| JX074050.1;AFK65737.1;Human_rhinovirus_A36_st | 95 | 95 | 1 | 0.05 | 0.05 | 0.05 |
| JX074050.1;AFK65737.1;Human_rhinovirus_A36_st | 103 | 191 | 89 | 0.05 | 0.5 | 0.097752809 |

| Extended data 1 | | | | | | |
|---|---|---|---|---|---|---|
| Protein | Start position (peptide) | Stop position (peptide) | Number of peptides | Minimum fraction (per peptide) of control subjects with reactivity >3 sd above average | Maximum fraction (per peptide) of control subjects with reactivity >3 sd above average | Mean fraction (over all peptides in this contiguous stretch) of controlsubjects with reactivity >3 sd above average |
| JX074050.1;AFK65737.1;Human_rhinovirus_A36_st | 203 | 217 | 15 | 0.05 | 0.7 | 0.493333333 |
| JX074050.1;AFK65737.1;Human_rhinovirus_A36_st | 223 | 224 | 2 | 0.05 | 0.05 | 0.05 |
| JX074050.1;AFK65737.1;Human_rhinovirus_A36_st | 228 | 228 | 1 | 0.05 | 0.05 | 0.05 |
| JX074050.1;AFK65737.1;Human_rhinovirus_A36_st | 230 | 231 | 2 | 0.05 | 0.05 | 0.05 |
| JX074050.1;AFK65737.1;Human_rhinovirus_A36_st | 235 | 235 | 1 | 0.05 | 0.05 | 0.05 |
| JX074050.1;AFK65737.1;Human_rhinovirus_A36_st | 239 | 240 | 2 | 0.05 | 0.05 | 0.05 |
| JX074050.1;AFK65737.1;Human_rhinovirus_A36_st | 291 | 291 | 1 | 0.05 | 0.05 | 0.05 |
| JX074050.1;AFK65737.1;Human_rhinovirus_A36_st | 294 | 294 | 1 | 0.05 | 0.05 | 0.05 |
| JX074050.1;AFK65737.1;Human_rhinovirus_A36_st | 317 | 317 | 1 | 0.05 | 0.05 | 0.05 |
| JX074050.1;AFK65737.1;Human_rhinovirus_A36_st | 319 | 319 | 1 | 0.05 | 0.05 | 0.05 |
| JX074050.1;AFK65737.1;Human_rhinovirus_A36_st | 321 | 321 | 1 | 0.05 | 0.05 | 0.05 |
| JX074050.1;AFK65737.1;Human_rhinovirus_A36_st | 324 | 334 | 11 | 0.05 | 0.1 | 0.059090909 |
| JX074050.1;AFK65737.1;Human_rhinovirus_A36_st | 342 | 342 | 1 | 0.1 | 0.1 | 0.1 |
| JX074050.1;AFK65737.1;Human_rhinovirus_A36_st | 353 | 355 | 3 | 0.05 | 0.15 | 0.083333333 |
| JX074050.1;AFK65737.1;Human_rhinovirus_A36_st | 371 | 371 | 1 | 0.05 | 0.05 | 0.05 |
| JX074050.1;AFK65737.1;Human_rhinovirus_A36_st | 377 | 573 | 197 | 0.05 | 0.1 | 0.050253807 |
| JX074050.1;AFK65737.1;Human_rhinovirus_A36_st | 578 | 589 | 12 | 0.15 | 0.5 | 0.383333333 |
| JX074050.1;AFK65737.1;Human_rhinovirus_A36_st | 591 | 592 | 2 | 0.05 | 0.05 | 0.05 |
| JX074050.1;AFK65737.1;Human_rhinovirus_A36_st | 594 | 612 | 19 | 0.1 | 0.9 | 0.560526316 |
| JX074050.1;AFK65737.1;Human_rhinovirus_A36_st | 614 | 616 | 3 | 0.05 | 0.05 | 0.05 |
| JX074050.1;AFK65737.1;Human_rhinovirus_A36_st | 620 | 620 | 1 | 0.05 | 0.05 | 0.05 |
| JX074050.1;AFK65737.1;Human_rhinovirus_A36_st | 622 | 626 | 5 | 0.05 | 0.05 | 0.05 |
| JX074050.1;AFK65737.1;Human_rhinovirus_A36_st | 653 | 653 | 1 | 0.05 | 0.05 | 0.05 |
| JX074050.1;AFK65737.1;Human_rhinovirus_A36_st | 693 | 693 | 1 | 0.05 | 0.05 | 0.05 |
| JX074050.1;AFK65737.1;Human_rhinovirus_A36_st | 713 | 728 | 16 | 0.05 | 0.2 | 0.0875 |
| JX074050.1;AFK65737.1;Human_rhinovirus_A36_st | 730 | 733 | 4 | 0.05 | 0.2 | 0.15 |
| JX074050.1;AFK65737.1;Human_rhinovirus_A36_st | 742 | 742 | 1 | 0.05 | 0.05 | 0.05 |
| JX074050.1;AFK65737.1;Human_rhinovirus_A36_st | 767 | 767 | 1 | 0.05 | 0.05 | 0.05 |
| JX074050.1;AFK65737.1;Human_rhinovirus_A36_st | 769 | 779 | 11 | 0.05 | 0.2 | 0.109090909 |
| JX074050.1;AFK65737.1;Human_rhinovirus_A36_st | 799 | 800 | 2 | 0.1 | 0.1 | 0.1 |
| JX074050.1;AFK65737.1;Human_rhinovirus_A36_st | 802 | 802 | 1 | 0.05 | 0.05 | 0.05 |
| JX074050.1;AFK65737.1;Human_rhinovirus_A36_st | 828 | 842 | 15 | 0.05 | 0.3 | 0.136666667 |
| JX074050.1;AFK65737.1;Human_rhinovirus_A36_st | 846 | 855 | 10 | 0.05 | 0.55 | 0.315 |
| JX074050.1;AFK65737.1;Human_rhinovirus_A36_st | 895 | 902 | 8 | 0.05 | 0.1 | 0.05625 |
| JX074050.1;AFK65737.1;Human_rhinovirus_A36_st | 939 | 939 | 1 | 0.05 | 0.05 | 0.05 |
| JX074050.1;AFK65737.1;Human_rhinovirus_A36_st | 941 | 941 | 1 | 0.05 | 0.05 | 0.05 |
| JX074050.1;AFK65737.1;Human_rhinovirus_A36_st | 972 | 972 | 1 | 0.05 | 0.05 | 0.05 |

| Extended data 1 | | | | | | |
|---|---|---|---|---|---|---|
| **Protein** | **Start position (peptide)** | **Stop position (peptide)** | **Number of peptides** | **Minimum fraction (per peptide) of control subjects with reactivity >3 sd above average** | **Maximum fraction (per peptide) of control subjects with reactivity >3 sd above average** | **Mean fraction (over all peptides in this contiguous stretch) of controlsubjects with reactivity >3 sd above average** |
| JX074050.1;AFK65737.1;Human_rhinovirus_A36_st | 977 | 978 | 2 | 0.05 | 0.05 | 0.05 |
| JX074050.1;AFK65737.1;Human_rhinovirus_A36_st | 997 | 998 | 2 | 0.05 | 0.05 | 0.05 |
| JX074050.1;AFK65737.1;Human_rhinovirus_A36_st | 1003 | 1007 | 5 | 0.05 | 0.5 | 0.26 |
| JX074050.1;AFK65737.1;Human_rhinovirus_A36_st | 1009 | 1009 | 1 | 0.05 | 0.05 | 0.05 |
| JX074050.1;AFK65737.1;Human_rhinovirus_A36_st | 1011 | 1019 | 9 | 0.05 | 0.2 | 0.127777778 |
| JX074050.1;AFK65737.1;Human_rhinovirus_A36_st | 1021 | 1021 | 1 | 0.05 | 0.05 | 0.05 |
| JX074050.1;AFK65737.1;Human_rhinovirus_A36_st | 1024 | 1024 | 1 | 0.1 | 0.1 | 0.1 |
| JX074050.1;AFK65737.1;Human_rhinovirus_A36_st | 1028 | 1028 | 1 | 0.05 | 0.05 | 0.05 |
| JX074050.1;AFK65737.1;Human_rhinovirus_A36_st | 1098 | 1107 | 10 | 0.05 | 0.2 | 0.085 |
| JX074050.1;AFK65737.1;Human_rhinovirus_A36_st | 1115 | 1117 | 3 | 0.05 | 0.1 | 0.083333333 |
| JX074050.1;AFK65737.1;Human_rhinovirus_A36_st | 1126 | 1133 | 8 | 0.05 | 0.3 | 0.15625 |
| JX074050.1;AFK65737.1;Human_rhinovirus_A36_st | 1142 | 1142 | 1 | 0.05 | 0.05 | 0.05 |
| JX074050.1;AFK65737.1;Human_rhinovirus_A36_st | 1145 | 1145 | 1 | 0.05 | 0.05 | 0.05 |
| JX074050.1;AFK65737.1;Human_rhinovirus_A36_st | 1160 | 1160 | 1 | 0.1 | 0.1 | 0.1 |
| JX074050.1;AFK65737.1;Human_rhinovirus_A36_st | 1164 | 1164 | 1 | 0.25 | 0.25 | 0.25 |
| JX074050.1;AFK65737.1;Human_rhinovirus_A36_st | 1181 | 1181 | 1 | 0.05 | 0.05 | 0.05 |
| JX074050.1;AFK65737.1;Human_rhinovirus_A36_st | 1184 | 1186 | 3 | 0.05 | 0.05 | 0.05 |
| JX074050.1;AFK65737.1;Human_rhinovirus_A36_st | 1204 | 1204 | 1 | 0.05 | 0.05 | 0.05 |
| JX074050.1;AFK65737.1;Human_rhinovirus_A36_st | 1239 | 1254 | 16 | 0.05 | 0.7 | 0.44375 |
| JX074050.1;AFK65737.1;Human_rhinovirus_A36_st | 1258 | 1258 | 1 | 0.05 | 0.05 | 0.05 |
| JX074050.1;AFK65737.1;Human_rhinovirus_A36_st | 1266 | 1273 | 8 | 0.05 | 0.1 | 0.05625 |
| JX074050.1;AFK65737.1;Human_rhinovirus_A36_st | 1276 | 1276 | 1 | 0.05 | 0.05 | 0.05 |
| JX074050.1;AFK65737.1;Human_rhinovirus_A36_st | 1284 | 1296 | 13 | 0.15 | 1 | 0.696153846 |
| JX074050.1;AFK65737.1;Human_rhinovirus_A36_st | 1312 | 1320 | 9 | 0.1 | 0.15 | 0.105555556 |
| JX074050.1;AFK65737.1;Human_rhinovirus_A36_st | 1322 | 1322 | 1 | 0.05 | 0.05 | 0.05 |
| JX074050.1;AFK65737.1;Human_rhinovirus_A36_st | 1339 | 1340 | 2 | 0.05 | 0.1 | 0.075 |
| JX074050.1;AFK65737.1;Human_rhinovirus_A36_st | 1342 | 1342 | 1 | 0.05 | 0.05 | 0.05 |
| JX074050.1;AFK65737.1;Human_rhinovirus_A36_st | 1344 | 1348 | 5 | 0.1 | 0.25 | 0.16 |
| JX074050.1;AFK65737.1;Human_rhinovirus_A36_st | 1351 | 1352 | 2 | 0.05 | 0.05 | 0.05 |
| JX074050.1;AFK65737.1;Human_rhinovirus_A36_st | 1370 | 1381 | 12 | 0.05 | 0.1 | 0.079166667 |
| JX074050.1;AFK65737.1;Human_rhinovirus_A36_st | 1397 | 1398 | 2 | 0.05 | 0.1 | 0.075 |
| JX074050.1;AFK65737.1;Human_rhinovirus_A36_st | 1407 | 1409 | 3 | 0.05 | 0.05 | 0.05 |
| JX074050.1;AFK65737.1;Human_rhinovirus_A36_st | 1415 | 1415 | 1 | 0.05 | 0.05 | 0.05 |
| JX074050.1;AFK65737.1;Human_rhinovirus_A36_st | 1417 | 1417 | 1 | 0.05 | 0.05 | 0.05 |
| JX074050.1;AFK65737.1;Human_rhinovirus_A36_st | 1419 | 1419 | 1 | 0.05 | 0.05 | 0.05 |
| JX074050.1;AFK65737.1;Human_rhinovirus_A36_st | 1421 | 1432 | 12 | 0.15 | 0.8 | 0.6375 |
| JX074050.1;AFK65737.1;Human_rhinovirus_A36_st | 1434 | 1454 | 21 | 0.05 | 0.2 | 0.097619048 |

| Extended data 1 | | | | | | |
|---|---|---|---|---|---|---|
| Protein | Start position (peptide) | Stop position (peptide) | Number of peptides | Minimum fraction (per peptide) of control subjects with reactivity >3 sd above average | Maximum fraction (per peptide) of control subjects with reactivity >3 sd above average | Mean fraction (over all peptides in this contiguous stretch) of controlsubjects with reactivity >3 sd above average |
| JX074050.1;AFK65737.1;Human_rhinovirus_A36_st | 1456 | 1456 | 1 | 0.15 | 0.15 | 0.15 |
| JX074050.1;AFK65737.1;Human_rhinovirus_A36_st | 1463 | 1463 | 1 | 0.15 | 0.15 | 0.15 |
| JX074050.1;AFK65737.1;Human_rhinovirus_A36_st | 1503 | 1503 | 1 | 0.05 | 0.05 | 0.05 |
| JX074050.1;AFK65737.1;Human_rhinovirus_A36_st | 1506 | 1506 | 1 | 0.05 | 0.05 | 0.05 |
| JX074050.1;AFK65737.1;Human_rhinovirus_A36_st | 1511 | 1512 | 2 | 0.05 | 0.05 | 0.05 |
| JX074050.1;AFK65737.1;Human_rhinovirus_A36_st | 1514 | 1514 | 1 | 0.05 | 0.05 | 0.05 |
| JX074050.1;AFK65737.1;Human_rhinovirus_A36_st | 1517 | 1517 | 1 | 0.05 | 0.05 | 0.05 |
| JX074050.1;AFK65737.1;Human_rhinovirus_A36_st | 1529 | 1538 | 10 | 0.05 | 0.15 | 0.125 |
| JX074050.1;AFK65737.1;Human_rhinovirus_A36_st | 1545 | 1545 | 1 | 0.05 | 0.05 | 0.05 |
| JX074050.1;AFK65737.1;Human_rhinovirus_A36_st | 1547 | 1547 | 1 | 0.15 | 0.15 | 0.15 |
| JX074050.1;AFK65737.1;Human_rhinovirus_A36_st | 1553 | 1553 | 1 | 0.05 | 0.05 | 0.05 |
| JX074050.1;AFK65737.1;Human_rhinovirus_A36_st | 1555 | 1556 | 2 | 0.05 | 0.05 | 0.05 |
| JX074050.1;AFK65737.1;Human_rhinovirus_A36_st | 1571 | 1571 | 1 | 0.05 | 0.05 | 0.05 |
| JX074050.1;AFK65737.1;Human_rhinovirus_A36_st | 1573 | 1573 | 1 | 0.05 | 0.05 | 0.05 |
| JX074050.1;AFK65737.1;Human_rhinovirus_A36_st | 1605 | 1606 | 2 | 0.05 | 0.05 | 0.05 |
| JX074050.1;AFK65737.1;Human_rhinovirus_A36_st | 1612 | 1612 | 1 | 0.1 | 0.1 | 0.1 |
| JX074050.1;AFK65737.1;Human_rhinovirus_A36_st | 1615 | 1615 | 1 | 0.05 | 0.05 | 0.05 |
| JX074050.1;AFK65737.1;Human_rhinovirus_A36_st | 1628 | 1628 | 1 | 0.05 | 0.05 | 0.05 |
| JX074050.1;AFK65737.1;Human_rhinovirus_A36_st | 1636 | 1636 | 1 | 0.05 | 0.05 | 0.05 |
| JX074050.1;AFK65737.1;Human_rhinovirus_A36_st | 1638 | 1638 | 1 | 0.1 | 0.1 | 0.1 |
| JX074050.1;AFK65737.1;Human_rhinovirus_A36_st | 1646 | 1646 | 1 | 0.05 | 0.05 | 0.05 |
| JX074050.1;AFK65737.1;Human_rhinovirus_A36_st | 1653 | 1653 | 1 | 0.05 | 0.05 | 0.05 |
| JX074050.1;AFK65737.1;Human_rhinovirus_A36_st | 1696 | 1697 | 2 | 0.05 | 0.15 | 0.1 |
| JX074050.1;AFK65737.1;Human_rhinovirus_A36_st | 1707 | 1709 | 3 | 0.05 | 0.1 | 0.066666667 |
| JX074050.1;AFK65737.1;Human_rhinovirus_A36_st | 1716 | 1716 | 1 | 0.05 | 0.05 | 0.05 |
| JX074050.1;AFK65737.1;Human_rhinovirus_A36_st | 1718 | 1745 | 28 | 0.05 | 0.5 | 0.251785714 |
| JX074050.1;AFK65737.1;Human_rhinovirus_A36_st | 1748 | 1748 | 1 | 0.05 | 0.05 | 0.05 |
| JX074050.1;AFK65737.1;Human_rhinovirus_A36_st | 1751 | 1753 | 3 | 0.05 | 0.1 | 0.066666667 |
| JX074050.1;AFK65737.1;Human_rhinovirus_A36_st | 1765 | 1765 | 1 | 0.05 | 0.05 | 0.05 |
| JX074050.1;AFK65737.1;Human_rhinovirus_A36_st | 1769 | 1769 | 1 | 0.05 | 0.05 | 0.05 |
| JX074050.1;AFK65737.1;Human_rhinovirus_A36_st | 1781 | 1781 | 1 | 0.05 | 0.05 | 0.05 |
| JX074050.1;AFK65737.1;Human_rhinovirus_A36_st | 1785 | 1857 | 73 | 0.05 | 0.1 | 0.053424658 |
| JX074050.1;AFK65737.1;Human_rhinovirus_A36_st | 1867 | 1867 | 1 | 0.05 | 0.05 | 0.05 |
| JX074050.1;AFK65737.1;Human_rhinovirus_A36_st | 1870 | 1870 | 1 | 0.05 | 0.05 | 0.05 |
| JX074050.1;AFK65737.1;Human_rhinovirus_A36_st | 1872 | 1872 | 1 | 0.05 | 0.05 | 0.05 |
| JX074050.1;AFK65737.1;Human_rhinovirus_A36_st | 1923 | 1933 | 11 | 0.05 | 0.05 | 0.05 |
| JX074050.1;AFK65737.1;Human_rhinovirus_A36_st | 1936 | 1937 | 2 | 0.05 | 0.05 | 0.05 |

| Extended data 1 | | | | | | |
|---|---|---|---|---|---|---|
| Protein | Start position (peptide) | Stop position (peptide) | Number of peptides | Minimum fraction (per peptide) of control subjects with reactivity >3 sd above average | Maximum fraction (per peptide) of control subjects with reactivity >3 sd above average | Mean fraction (over all peptides in this contiguous stretch) of controlsubjects with reactivity >3 sd above average |
| JX074050.1;AFK65737.1;Human_rhinovirus_A36_st | 1948 | 1948 | 1 | 0.05 | 0.05 | 0.05 |
| JX074050.1;AFK65737.1;Human_rhinovirus_A36_st | 1955 | 1955 | 1 | 0.05 | 0.05 | 0.05 |
| JX074050.1;AFK65737.1;Human_rhinovirus_A36_st | 1964 | 1964 | 1 | 0.05 | 0.05 | 0.05 |
| JX074050.1;AFK65737.1;Human_rhinovirus_A36_st | 1970 | 1970 | 1 | 0.05 | 0.05 | 0.05 |
| JX074050.1;AFK65737.1;Human_rhinovirus_A36_st | 1974 | 1974 | 1 | 0.05 | 0.05 | 0.05 |
| JX074050.1;AFK65737.1;Human_rhinovirus_A36_st | 1980 | 1980 | 1 | 0.05 | 0.05 | 0.05 |
| JX074050.1;AFK65737.1;Human_rhinovirus_A36_st | 1983 | 1983 | 1 | 0.05 | 0.05 | 0.05 |
| JX074050.1;AFK65737.1;Human_rhinovirus_A36_st | 1986 | 1986 | 1 | 0.05 | 0.05 | 0.05 |
| JX074050.1;AFK65737.1;Human_rhinovirus_A36_st | 1988 | 1988 | 1 | 0.05 | 0.05 | 0.05 |
| JX074050.1;AFK65737.1;Human_rhinovirus_A36_st | 2005 | 2006 | 2 | 0.05 | 0.1 | 0.075 |
| JX074050.1;AFK65737.1;Human_rhinovirus_A36_st | 2010 | 2010 | 1 | 0.05 | 0.05 | 0.05 |
| JX074050.1;AFK65737.1;Human_rhinovirus_A36_st | 2014 | 2014 | 1 | 0.05 | 0.05 | 0.05 |
| JX074050.1;AFK65737.1;Human_rhinovirus_A36_st | 2039 | 2039 | 1 | 0.05 | 0.05 | 0.05 |
| JX074050.1;AFK65737.1;Human_rhinovirus_A36_st | 2048 | 2048 | 1 | 0.05 | 0.05 | 0.05 |
| JX074050.1;AFK65737.1;Human_rhinovirus_A36_st | 2058 | 2058 | 1 | 0.05 | 0.05 | 0.05 |
| JX074050.1;AFK65737.1;Human_rhinovirus_A36_st | 2079 | 2080 | 2 | 0.05 | 0.05 | 0.05 |
| JX074050.1;AFK65737.1;Human_rhinovirus_A36_st | 2099 | 2099 | 1 | 0.1 | 0.1 | 0.1 |
| KY189321.1;ARE30269.1;Rhinovirus_C41_strain_S | 1 | 3 | 3 | 0.05 | 0.05 | 0.05 |
| KY189321.1;ARE30269.1;Rhinovirus_C41_strain_S | 11 | 12 | 2 | 0.05 | 0.05 | 0.05 |
| KY189321.1;ARE30269.1;Rhinovirus_C41_strain_S | 23 | 28 | 6 | 0.05 | 0.15 | 0.091666667 |
| KY189321.1;ARE30269.1;Rhinovirus_C41_strain_S | 35 | 47 | 13 | 0.05 | 0.25 | 0.180769231 |
| KY189321.1;ARE30269.1;Rhinovirus_C41_strain_S | 49 | 49 | 1 | 0.05 | 0.05 | 0.05 |
| KY189321.1;ARE30269.1;Rhinovirus_C41_strain_S | 51 | 51 | 1 | 0.05 | 0.05 | 0.05 |
| KY189321.1;ARE30269.1;Rhinovirus_C41_strain_S | 58 | 58 | 1 | 0.05 | 0.05 | 0.05 |
| KY189321.1;ARE30269.1;Rhinovirus_C41_strain_S | 73 | 74 | 2 | 0.05 | 0.1 | 0.075 |
| KY189321.1;ARE30269.1;Rhinovirus_C41_strain_S | 80 | 81 | 2 | 0.05 | 0.15 | 0.1 |
| KY189321.1;ARE30269.1;Rhinovirus_C41_strain_S | 83 | 95 | 13 | 0.05 | 0.1 | 0.053846154 |
| KY189321.1;ARE30269.1;Rhinovirus_C41_strain_S | 104 | 114 | 11 | 0.1 | 0.2 | 0.15 |
| KY189321.1;ARE30269.1;Rhinovirus_C41_strain_S | 117 | 117 | 1 | 0.1 | 0.1 | 0.1 |
| KY189321.1;ARE30269.1;Rhinovirus_C41_strain_S | 159 | 159 | 1 | 0.05 | 0.05 | 0.05 |
| KY189321.1;ARE30269.1;Rhinovirus_C41_strain_S | 201 | 214 | 14 | 0.05 | 0.55 | 0.353571429 |
| KY189321.1;ARE30269.1;Rhinovirus_C41_strain_S | 220 | 220 | 1 | 0.05 | 0.05 | 0.05 |
| KY189321.1;ARE30269.1;Rhinovirus_C41_strain_S | 223 | 224 | 2 | 0.05 | 0.05 | 0.05 |
| KY189321.1;ARE30269.1;Rhinovirus_C41_strain_S | 274 | 274 | 1 | 0.05 | 0.05 | 0.05 |
| KY189321.1;ARE30269.1;Rhinovirus_C41_strain_S | 276 | 278 | 3 | 0.05 | 0.05 | 0.05 |
| KY189321.1;ARE30269.1;Rhinovirus_C41_strain_S | 299 | 299 | 1 | 0.05 | 0.05 | 0.05 |
| KY189321.1;ARE30269.1;Rhinovirus_C41_strain_S | 307 | 307 | 1 | 0.05 | 0.05 | 0.05 |

| Extended data 1 | | | | | | |
|---|---|---|---|---|---|---|
| Protein | Start position (peptide) | Stop position (peptide) | Number of peptides | Minimum fraction (per peptide) of control subjects with reactivity >3 sd above average | Maximum fraction (per peptide) of control subjects with reactivity >3 sd above average | Mean fraction (over all peptides in this contiguous stretch) of controlsubjects with reactivity >3 sd above average |
| KY189321.1;ARE30269.1;Rhinovirus_C41_strain_S | 310 | 311 | 2 | 0.05 | 0.15 | 0.1 |
| KY189321.1;ARE30269.1;Rhinovirus_C41_strain_S | 314 | 316 | 3 | 0.05 | 0.1 | 0.066666667 |
| KY189321.1;ARE30269.1;Rhinovirus_C41_strain_S | 319 | 319 | 1 | 0.05 | 0.05 | 0.05 |
| KY189321.1;ARE30269.1;Rhinovirus_C41_strain_S | 321 | 321 | 1 | 0.05 | 0.05 | 0.05 |
| KY189321.1;ARE30269.1;Rhinovirus_C41_strain_S | 326 | 327 | 2 | 0.05 | 0.05 | 0.05 |
| KY189321.1;ARE30269.1;Rhinovirus_C41_strain_S | 349 | 349 | 1 | 0.1 | 0.1 | 0.1 |
| KY189321.1;ARE30269.1;Rhinovirus_C41_strain_S | 372 | 373 | 2 | 0.05 | 0.1 | 0.075 |
| KY189321.1;ARE30269.1;Rhinovirus_C41_strain_S | 377 | 379 | 3 | 0.05 | 0.05 | 0.05 |
| KY189321.1;ARE30269.1;Rhinovirus_C41_strain_S | 382 | 382 | 1 | 0.2 | 0.2 | 0.2 |
| KY189321.1;ARE30269.1;Rhinovirus_C41_strain_S | 396 | 407 | 12 | 0.05 | 0.1 | 0.058333333 |
| KY189321.1;ARE30269.1;Rhinovirus_C41_strain_S | 463 | 471 | 9 | 0.05 | 0.05 | 0.05 |
| KY189321.1;ARE30269.1;Rhinovirus_C41_strain_S | 485 | 485 | 1 | 0.05 | 0.05 | 0.05 |
| KY189321.1;ARE30269.1;Rhinovirus_C41_strain_S | 501 | 502 | 2 | 0.05 | 0.05 | 0.05 |
| KY189321.1;ARE30269.1;Rhinovirus_C41_strain_S | 505 | 505 | 1 | 0.05 | 0.05 | 0.05 |
| KY189321.1;ARE30269.1;Rhinovirus_C41_strain_S | 520 | 520 | 1 | 0.05 | 0.05 | 0.05 |
| KY189321.1;ARE30269.1;Rhinovirus_C41_strain_S | 525 | 525 | 1 | 0.05 | 0.05 | 0.05 |
| KY189321.1;ARE30269.1;Rhinovirus_C41_strain_S | 554 | 554 | 1 | 0.05 | 0.05 | 0.05 |
| KY189321.1;ARE30269.1;Rhinovirus_C41_strain_S | 557 | 557 | 1 | 0.05 | 0.05 | 0.05 |
| KY189321.1;ARE30269.1;Rhinovirus_C41_strain_S | 560 | 560 | 1 | 0.05 | 0.05 | 0.05 |
| KY189321.1;ARE30269.1;Rhinovirus_C41_strain_S | 564 | 602 | 39 | 0.05 | 0.65 | 0.319230769 |
| KY189321.1;ARE30269.1;Rhinovirus_C41_strain_S | 607 | 607 | 1 | 0.05 | 0.05 | 0.05 |
| KY189321.1;ARE30269.1;Rhinovirus_C41_strain_S | 616 | 627 | 12 | 0.05 | 0.4 | 0.166666667 |
| KY189321.1;ARE30269.1;Rhinovirus_C41_strain_S | 646 | 646 | 1 | 0.1 | 0.1 | 0.1 |
| KY189321.1;ARE30269.1;Rhinovirus_C41_strain_S | 650 | 655 | 6 | 0.05 | 0.2 | 0.141666667 |
| KY189321.1;ARE30269.1;Rhinovirus_C41_strain_S | 680 | 682 | 3 | 0.05 | 0.05 | 0.05 |
| KY189321.1;ARE30269.1;Rhinovirus_C41_strain_S | 694 | 695 | 2 | 0.05 | 0.1 | 0.075 |
| KY189321.1;ARE30269.1;Rhinovirus_C41_strain_S | 697 | 700 | 4 | 0.05 | 0.05 | 0.05 |
| KY189321.1;ARE30269.1;Rhinovirus_C41_strain_S | 702 | 708 | 7 | 0.05 | 0.05 | 0.05 |
| KY189321.1;ARE30269.1;Rhinovirus_C41_strain_S | 715 | 716 | 2 | 0.05 | 0.1 | 0.075 |
| KY189321.1;ARE30269.1;Rhinovirus_C41_strain_S | 718 | 727 | 10 | 0.05 | 0.15 | 0.105 |
| KY189321.1;ARE30269.1;Rhinovirus_C41_strain_S | 743 | 750 | 8 | 0.05 | 0.25 | 0.18125 |
| KY189321.1;ARE30269.1;Rhinovirus_C41_strain_S | 753 | 756 | 4 | 0.05 | 0.1 | 0.0625 |
| KY189321.1;ARE30269.1;Rhinovirus_C41_strain_S | 769 | 775 | 7 | 0.05 | 0.15 | 0.085714286 |
| KY189321.1;ARE30269.1;Rhinovirus_C41_strain_S | 805 | 819 | 15 | 0.05 | 0.2 | 0.08 |
| KY189321.1;ARE30269.1;Rhinovirus_C41_strain_S | 824 | 833 | 10 | 0.05 | 0.15 | 0.075 |
| KY189321.1;ARE30269.1;Rhinovirus_C41_strain_S | 842 | 842 | 1 | 0.05 | 0.05 | 0.05 |
| KY189321.1;ARE30269.1;Rhinovirus_C41_strain_S | 869 | 869 | 1 | 0.05 | 0.05 | 0.05 |

| Extended data 1 | | | | | | |
|---|---|---|---|---|---|---|
| **Protein** | **Start position (peptide)** | **Stop position (peptide)** | **Number of peptides** | **Minimum fraction (per peptide) of control subjects with reactivity >3 sd above average** | **Maximum fraction (per peptide) of control subjects with reactivity >3 sd above average** | **Mean fraction (over all peptides in this contiguous stretch) of controlsubjects with reactivity >3 sd above average** |
| KY189321.1;ARE30269.1;Rhinovirus_C41_strain_S | 871 | 871 | 1 | 0.05 | 0.05 | 0.05 |
| KY189321.1;ARE30269.1;Rhinovirus_C41_strain_S | 878 | 878 | 1 | 0.05 | 0.05 | 0.05 |
| KY189321.1;ARE30269.1;Rhinovirus_C41_strain_S | 880 | 881 | 2 | 0.05 | 0.05 | 0.05 |
| KY189321.1;ARE30269.1;Rhinovirus_C41_strain_S | 916 | 916 | 1 | 0.05 | 0.05 | 0.05 |
| KY189321.1;ARE30269.1;Rhinovirus_C41_strain_S | 918 | 918 | 1 | 0.05 | 0.05 | 0.05 |
| KY189321.1;ARE30269.1;Rhinovirus_C41_strain_S | 930 | 935 | 6 | 0.05 | 0.1 | 0.058333333 |
| KY189321.1;ARE30269.1;Rhinovirus_C41_strain_S | 937 | 937 | 1 | 0.05 | 0.05 | 0.05 |
| KY189321.1;ARE30269.1;Rhinovirus_C41_strain_S | 949 | 949 | 1 | 0.05 | 0.05 | 0.05 |
| KY189321.1;ARE30269.1;Rhinovirus_C41_strain_S | 955 | 955 | 1 | 0.05 | 0.05 | 0.05 |
| KY189321.1;ARE30269.1;Rhinovirus_C41_strain_S | 970 | 974 | 5 | 0.05 | 0.05 | 0.05 |
| KY189321.1;ARE30269.1;Rhinovirus_C41_strain_S | 979 | 997 | 19 | 0.05 | 0.9 | 0.352631579 |
| KY189321.1;ARE30269.1;Rhinovirus_C41_strain_S | 1037 | 1037 | 1 | 0.05 | 0.05 | 0.05 |
| KY189321.1;ARE30269.1;Rhinovirus_C41_strain_S | 1075 | 1075 | 1 | 0.05 | 0.05 | 0.05 |
| KY189321.1;ARE30269.1;Rhinovirus_C41_strain_S | 1082 | 1083 | 2 | 0.05 | 0.1 | 0.075 |
| KY189321.1;ARE30269.1;Rhinovirus_C41_strain_S | 1091 | 1091 | 1 | 0.05 | 0.05 | 0.05 |
| KY189321.1;ARE30269.1;Rhinovirus_C41_strain_S | 1099 | 1099 | 1 | 0.05 | 0.05 | 0.05 |
| KY189321.1;ARE30269.1;Rhinovirus_C41_strain_S | 1101 | 1101 | 1 | 0.05 | 0.05 | 0.05 |
| KY189321.1;ARE30269.1;Rhinovirus_C41_strain_S | 1113 | 1113 | 1 | 0.05 | 0.05 | 0.05 |
| KY189321.1;ARE30269.1;Rhinovirus_C41_strain_S | 1125 | 1125 | 1 | 0.2 | 0.2 | 0.2 |
| KY189321.1;ARE30269.1;Rhinovirus_C41_strain_S | 1129 | 1134 | 6 | 0.05 | 0.05 | 0.05 |
| KY189321.1;ARE30269.1;Rhinovirus_C41_strain_S | 1166 | 1168 | 3 | 0.05 | 0.4 | 0.166666667 |
| KY189321.1;ARE30269.1;Rhinovirus_C41_strain_S | 1173 | 1179 | 7 | 0.05 | 0.1 | 0.071428571 |
| KY189321.1;ARE30269.1;Rhinovirus_C41_strain_S | 1182 | 1182 | 1 | 0.05 | 0.05 | 0.05 |
| KY189321.1;ARE30269.1;Rhinovirus_C41_strain_S | 1208 | 1208 | 1 | 0.05 | 0.05 | 0.05 |
| KY189321.1;ARE30269.1;Rhinovirus_C41_strain_S | 1224 | 1239 | 16 | 0.05 | 0.6 | 0.3625 |
| KY189321.1;ARE30269.1;Rhinovirus_C41_strain_S | 1243 | 1243 | 1 | 0.05 | 0.05 | 0.05 |
| KY189321.1;ARE30269.1;Rhinovirus_C41_strain_S | 1245 | 1245 | 1 | 0.05 | 0.05 | 0.05 |
| KY189321.1;ARE30269.1;Rhinovirus_C41_strain_S | 1247 | 1247 | 1 | 0.05 | 0.05 | 0.05 |
| KY189321.1;ARE30269.1;Rhinovirus_C41_strain_S | 1249 | 1253 | 5 | 0.05 | 0.1 | 0.07 |
| KY189321.1;ARE30269.1;Rhinovirus_C41_strain_S | 1255 | 1256 | 2 | 0.05 | 0.05 | 0.05 |
| KY189321.1;ARE30269.1;Rhinovirus_C41_strain_S | 1258 | 1258 | 1 | 0.05 | 0.05 | 0.05 |
| KY189321.1;ARE30269.1;Rhinovirus_C41_strain_S | 1268 | 1268 | 1 | 0.05 | 0.05 | 0.05 |
| KY189321.1;ARE30269.1;Rhinovirus_C41_strain_S | 1273 | 1277 | 5 | 0.05 | 0.05 | 0.05 |
| KY189321.1;ARE30269.1;Rhinovirus_C41_strain_S | 1281 | 1281 | 1 | 0.05 | 0.05 | 0.05 |
| KY189321.1;ARE30269.1;Rhinovirus_C41_strain_S | 1296 | 1296 | 1 | 0.05 | 0.05 | 0.05 |
| KY189321.1;ARE30269.1;Rhinovirus_C41_strain_S | 1298 | 1298 | 1 | 0.05 | 0.05 | 0.05 |
| KY189321.1;ARE30269.1;Rhinovirus_C41_strain_S | 1300 | 1300 | 1 | 0.05 | 0.05 | 0.05 |

| Extended data 1 | | | | | | |
|---|---|---|---|---|---|---|
| Protein | Start position (peptide) | Stop position (peptide) | Number of peptides | Minimum fraction (per peptide) of control subjects with reactivity >3 sd above average | Maximum fraction (per peptide) of control subjects with reactivity >3 sd above average | Mean fraction (over all peptides in this contiguous stretch) of controlsubjects with reactivity >3 sd above average |
| KY189321.1;ARE30269.1;Rhinovirus_C41_strain_S | 1302 | 1302 | 1 | 0.05 | 0.05 | 0.05 |
| KY189321.1;ARE30269.1;Rhinovirus_C41_strain_S | 1309 | 1312 | 4 | 0.05 | 0.05 | 0.05 |
| KY189321.1;ARE30269.1;Rhinovirus_C41_strain_S | 1328 | 1338 | 11 | 0.05 | 0.2 | 0.113636364 |
| KY189321.1;ARE30269.1;Rhinovirus_C41_strain_S | 1368 | 1378 | 11 | 0.05 | 0.15 | 0.095454545 |
| KY189321.1;ARE30269.1;Rhinovirus_C41_strain_S | 1381 | 1382 | 2 | 0.05 | 0.05 | 0.05 |
| KY189321.1;ARE30269.1;Rhinovirus_C41_strain_S | 1389 | 1389 | 1 | 0.05 | 0.05 | 0.05 |
| KY189321.1;ARE30269.1;Rhinovirus_C41_strain_S | 1406 | 1419 | 14 | 0.05 | 0.8 | 0.378571429 |
| KY189321.1;ARE30269.1;Rhinovirus_C41_strain_S | 1421 | 1424 | 4 | 0.05 | 0.1 | 0.0625 |
| KY189321.1;ARE30269.1;Rhinovirus_C41_strain_S | 1426 | 1426 | 1 | 0.05 | 0.05 | 0.05 |
| KY189321.1;ARE30269.1;Rhinovirus_C41_strain_S | 1428 | 1432 | 5 | 0.05 | 0.05 | 0.05 |
| KY189321.1;ARE30269.1;Rhinovirus_C41_strain_S | 1449 | 1449 | 1 | 0.05 | 0.05 | 0.05 |
| KY189321.1;ARE30269.1;Rhinovirus_C41_strain_S | 1482 | 1482 | 1 | 0.1 | 0.1 | 0.1 |
| KY189321.1;ARE30269.1;Rhinovirus_C41_strain_S | 1484 | 1484 | 1 | 0.15 | 0.15 | 0.15 |
| KY189321.1;ARE30269.1;Rhinovirus_C41_strain_S | 1516 | 1517 | 2 | 0.05 | 0.05 | 0.05 |
| KY189321.1;ARE30269.1;Rhinovirus_C41_strain_S | 1543 | 1547 | 5 | 0.05 | 0.05 | 0.05 |
| KY189321.1;ARE30269.1;Rhinovirus_C41_strain_S | 1549 | 1550 | 2 | 0.05 | 0.1 | 0.075 |
| KY189321.1;ARE30269.1;Rhinovirus_C41_strain_S | 1559 | 1563 | 5 | 0.05 | 0.05 | 0.05 |
| KY189321.1;ARE30269.1;Rhinovirus_C41_strain_S | 1577 | 1587 | 11 | 0.05 | 0.25 | 0.127272727 |
| KY189321.1;ARE30269.1;Rhinovirus_C41_strain_S | 1599 | 1605 | 7 | 0.05 | 0.1 | 0.057142857 |
| KY189321.1;ARE30269.1;Rhinovirus_C41_strain_S | 1607 | 1609 | 3 | 0.05 | 0.1 | 0.066666667 |
| KY189321.1;ARE30269.1;Rhinovirus_C41_strain_S | 1620 | 1620 | 1 | 0.05 | 0.05 | 0.05 |
| KY189321.1;ARE30269.1;Rhinovirus_C41_strain_S | 1623 | 1623 | 1 | 0.05 | 0.05 | 0.05 |
| KY189321.1;ARE30269.1;Rhinovirus_C41_strain_S | 1639 | 1639 | 1 | 0.05 | 0.05 | 0.05 |
| KY189321.1;ARE30269.1;Rhinovirus_C41_strain_S | 1646 | 1646 | 1 | 0.05 | 0.05 | 0.05 |
| KY189321.1;ARE30269.1;Rhinovirus_C41_strain_S | 1648 | 1649 | 2 | 0.1 | 0.1 | 0.1 |
| KY189321.1;ARE30269.1;Rhinovirus_C41_strain_S | 1656 | 1657 | 2 | 0.1 | 0.1 | 0.1 |
| KY189321.1;ARE30269.1;Rhinovirus_C41_strain_S | 1692 | 1692 | 1 | 0.05 | 0.05 | 0.05 |
| KY189321.1;ARE30269.1;Rhinovirus_C41_strain_S | 1694 | 1694 | 1 | 0.05 | 0.05 | 0.05 |
| KY189321.1;ARE30269.1;Rhinovirus_C41_strain_S | 1698 | 1735 | 38 | 0.05 | 0.55 | 0.209210526 |
| KY189321.1;ARE30269.1;Rhinovirus_C41_strain_S | 1740 | 1742 | 3 | 0.05 | 0.1 | 0.083333333 |
| KY189321.1;ARE30269.1;Rhinovirus_C41_strain_S | 1745 | 1752 | 8 | 0.05 | 0.1 | 0.0625 |
| KY189321.1;ARE30269.1;Rhinovirus_C41_strain_S | 1756 | 1757 | 2 | 0.05 | 0.2 | 0.125 |
| KY189321.1;ARE30269.1;Rhinovirus_C41_strain_S | 1760 | 1760 | 1 | 0.05 | 0.05 | 0.05 |
| KY189321.1;ARE30269.1;Rhinovirus_C41_strain_S | 1764 | 1764 | 1 | 0.05 | 0.05 | 0.05 |
| KY189321.1;ARE30269.1;Rhinovirus_C41_strain_S | 1781 | 1782 | 2 | 0.05 | 0.05 | 0.05 |
| KY189321.1;ARE30269.1;Rhinovirus_C41_strain_S | 1799 | 1800 | 2 | 0.05 | 0.05 | 0.05 |
| KY189321.1;ARE30269.1;Rhinovirus_C41_strain_S | 1802 | 1806 | 5 | 0.05 | 0.05 | 0.05 |

| Extended data 1 | | | | | | |
|---|---|---|---|---|---|---|
| Protein | Start position (peptide) | Stop position (peptide) | Number of peptides | Minimum fraction (per peptide) of control subjects with reactivity >3 sd above average | Maximum fraction (per peptide) of control subjects with reactivity >3 sd above average | Mean fraction (over all peptides in this contiguous stretch) of controlsubjects with reactivity >3 sd above average |
| KY189321.1;ARE30269.1;Rhinovirus_C41_strain_S | 1811 | 1812 | 2 | 0.05 | 0.05 | 0.05 |
| KY189321.1;ARE30269.1;Rhinovirus_C41_strain_S | 1814 | 1820 | 7 | 0.05 | 0.15 | 0.1 |
| KY189321.1;ARE30269.1;Rhinovirus_C41_strain_S | 1824 | 1840 | 17 | 0.05 | 0.15 | 0.088235294 |
| KY189321.1;ARE30269.1;Rhinovirus_C41_strain_S | 1855 | 1856 | 2 | 0.05 | 0.1 | 0.075 |
| KY189321.1;ARE30269.1;Rhinovirus_C41_strain_S | 1859 | 1860 | 2 | 0.05 | 0.05 | 0.05 |
| KY189321.1;ARE30269.1;Rhinovirus_C41_strain_S | 1862 | 1863 | 2 | 0.05 | 0.1 | 0.075 |
| KY189321.1;ARE30269.1;Rhinovirus_C41_strain_S | 1865 | 1865 | 1 | 0.05 | 0.05 | 0.05 |
| KY189321.1;ARE30269.1;Rhinovirus_C41_strain_S | 1868 | 1868 | 1 | 0.35 | 0.35 | 0.35 |
| KY189321.1;ARE30269.1;Rhinovirus_C41_strain_S | 1870 | 1870 | 1 | 0.05 | 0.05 | 0.05 |
| KY189321.1;ARE30269.1;Rhinovirus_C41_strain_S | 1901 | 1901 | 1 | 0.05 | 0.05 | 0.05 |
| KY189321.1;ARE30269.1;Rhinovirus_C41_strain_S | 1909 | 1919 | 11 | 0.05 | 0.1 | 0.054545455 |
| KY189321.1;ARE30269.1;Rhinovirus_C41_strain_S | 1936 | 1936 | 1 | 0.05 | 0.05 | 0.05 |
| KY189321.1;ARE30269.1;Rhinovirus_C41_strain_S | 1952 | 1965 | 14 | 0.05 | 0.25 | 0.139285714 |
| KY189321.1;ARE30269.1;Rhinovirus_C41_strain_S | 1969 | 1969 | 1 | 0.05 | 0.05 | 0.05 |
| KY189321.1;ARE30269.1;Rhinovirus_C41_strain_S | 1991 | 1992 | 2 | 0.05 | 0.05 | 0.05 |
| KY189321.1;ARE30269.1;Rhinovirus_C41_strain_S | 1999 | 2002 | 4 | 0.05 | 0.05 | 0.05 |
| KY189321.1;ARE30269.1;Rhinovirus_C41_strain_S | 2010 | 2028 | 19 | 0.05 | 0.15 | 0.086842105 |
| KY189321.1;ARE30269.1;Rhinovirus_C41_strain_S | 2030 | 2030 | 1 | 0.05 | 0.05 | 0.05 |
| KY189321.1;ARE30269.1;Rhinovirus_C41_strain_S | 2061 | 2061 | 1 | 0.05 | 0.05 | 0.05 |
| KY189321.1;ARE30269.1;Rhinovirus_C41_strain_S | 2079 | 2079 | 1 | 0.05 | 0.05 | 0.05 |
| KY189321.1;ARE30269.1;Rhinovirus_C41_strain_S | 2082 | 2085 | 4 | 0.05 | 0.2 | 0.1 |
| KY189321.1;ARE30269.1;Rhinovirus_C41_strain_S | 2105 | 2110 | 6 | 0.05 | 0.05 | 0.05 |
| L24917.1;AAA69862.1;Human_rhinovirus_type_A16 | 5 | 5 | 1 | 0.05 | 0.05 | 0.05 |
| L24917.1;AAA69862.1;Human_rhinovirus_type_A16 | 14 | 14 | 1 | 0.05 | 0.05 | 0.05 |
| L24917.1;AAA69862.1;Human_rhinovirus_type_A16 | 20 | 64 | 45 | 0.05 | 0.6 | 0.3 |
| L24917.1;AAA69862.1;Human_rhinovirus_type_A16 | 80 | 88 | 9 | 0.05 | 0.1 | 0.055555556 |
| L24917.1;AAA69862.1;Human_rhinovirus_type_A16 | 102 | 125 | 24 | 0.05 | 0.45 | 0.20625 |
| L24917.1;AAA69862.1;Human_rhinovirus_type_A16 | 128 | 141 | 14 | 0.05 | 0.15 | 0.071428571 |
| L24917.1;AAA69862.1;Human_rhinovirus_type_A16 | 148 | 149 | 2 | 0.05 | 0.05 | 0.05 |
| L24917.1;AAA69862.1;Human_rhinovirus_type_A16 | 151 | 151 | 1 | 0.05 | 0.05 | 0.05 |
| L24917.1;AAA69862.1;Human_rhinovirus_type_A16 | 155 | 156 | 2 | 0.05 | 0.05 | 0.05 |
| L24917.1;AAA69862.1;Human_rhinovirus_type_A16 | 182 | 182 | 1 | 0.05 | 0.05 | 0.05 |
| L24917.1;AAA69862.1;Human_rhinovirus_type_A16 | 198 | 198 | 1 | 0.15 | 0.15 | 0.15 |
| L24917.1;AAA69862.1;Human_rhinovirus_type_A16 | 200 | 216 | 17 | 0.05 | 0.75 | 0.491176471 |
| L24917.1;AAA69862.1;Human_rhinovirus_type_A16 | 220 | 220 | 1 | 0.1 | 0.1 | 0.1 |
| L24917.1;AAA69862.1;Human_rhinovirus_type_A16 | 222 | 233 | 12 | 0.1 | 0.3 | 0.225 |
| L24917.1;AAA69862.1;Human_rhinovirus_type_A16 | 270 | 272 | 3 | 0.05 | 0.15 | 0.083333333 |

| Extended data 1 | | | | | | |
|---|---|---|---|---|---|---|
| **Protein** | **Start position (peptide)** | **Stop position (peptide)** | **Number of peptides** | **Minimum fraction (per peptide) of control subjects with reactivity >3 sd above average** | **Maximum fraction (per peptide) of control subjects with reactivity >3 sd above average** | **Mean fraction (over all peptides in this contiguous stretch) of controlsubjects with reactivity >3 sd above average** |
| L24917.1;AAA69862.1;Human_rhinovirus_type_A16 | 291 | 297 | 7 | 0.05 | 0.3 | 0.142857143 |
| L24917.1;AAA69862.1;Human_rhinovirus_type_A16 | 299 | 299 | 1 | 0.05 | 0.05 | 0.05 |
| L24917.1;AAA69862.1;Human_rhinovirus_type_A16 | 307 | 307 | 1 | 0.05 | 0.05 | 0.05 |
| L24917.1;AAA69862.1;Human_rhinovirus_type_A16 | 309 | 314 | 6 | 0.05 | 0.1 | 0.066666667 |
| L24917.1;AAA69862.1;Human_rhinovirus_type_A16 | 347 | 350 | 4 | 0.05 | 0.1 | 0.0625 |
| L24917.1;AAA69862.1;Human_rhinovirus_type_A16 | 353 | 357 | 5 | 0.05 | 0.05 | 0.05 |
| L24917.1;AAA69862.1;Human_rhinovirus_type_A16 | 374 | 375 | 2 | 0.05 | 0.05 | 0.05 |
| L24917.1;AAA69862.1;Human_rhinovirus_type_A16 | 379 | 379 | 1 | 0.05 | 0.05 | 0.05 |
| L24917.1;AAA69862.1;Human_rhinovirus_type_A16 | 381 | 385 | 5 | 0.05 | 0.15 | 0.08 |
| L24917.1;AAA69862.1;Human_rhinovirus_type_A16 | 443 | 448 | 6 | 0.05 | 0.1 | 0.075 |
| L24917.1;AAA69862.1;Human_rhinovirus_type_A16 | 450 | 453 | 4 | 0.05 | 0.05 | 0.05 |
| L24917.1;AAA69862.1;Human_rhinovirus_type_A16 | 484 | 484 | 1 | 0.05 | 0.05 | 0.05 |
| L24917.1;AAA69862.1;Human_rhinovirus_type_A16 | 499 | 500 | 2 | 0.1 | 0.1 | 0.1 |
| L24917.1;AAA69862.1;Human_rhinovirus_type_A16 | 502 | 502 | 1 | 0.15 | 0.15 | 0.15 |
| L24917.1;AAA69862.1;Human_rhinovirus_type_A16 | 555 | 555 | 1 | 0.15 | 0.15 | 0.15 |
| L24917.1;AAA69862.1;Human_rhinovirus_type_A16 | 560 | 616 | 57 | 0.05 | 1 | 0.50877193 |
| L24917.1;AAA69862.1;Human_rhinovirus_type_A16 | 642 | 653 | 12 | 0.05 | 0.3 | 0.1875 |
| L24917.1;AAA69862.1;Human_rhinovirus_type_A16 | 656 | 661 | 6 | 0.05 | 0.1 | 0.058333333 |
| L24917.1;AAA69862.1;Human_rhinovirus_type_A16 | 670 | 670 | 1 | 0.05 | 0.05 | 0.05 |
| L24917.1;AAA69862.1;Human_rhinovirus_type_A16 | 679 | 679 | 1 | 0.05 | 0.05 | 0.05 |
| L24917.1;AAA69862.1;Human_rhinovirus_type_A16 | 681 | 684 | 4 | 0.05 | 0.1 | 0.075 |
| L24917.1;AAA69862.1;Human_rhinovirus_type_A16 | 688 | 688 | 1 | 0.05 | 0.05 | 0.05 |
| L24917.1;AAA69862.1;Human_rhinovirus_type_A16 | 702 | 717 | 16 | 0.05 | 0.3 | 0.1 |
| L24917.1;AAA69862.1;Human_rhinovirus_type_A16 | 719 | 722 | 4 | 0.05 | 0.2 | 0.125 |
| L24917.1;AAA69862.1;Human_rhinovirus_type_A16 | 725 | 725 | 1 | 0.05 | 0.05 | 0.05 |
| L24917.1;AAA69862.1;Human_rhinovirus_type_A16 | 727 | 731 | 5 | 0.05 | 0.1 | 0.06 |
| L24917.1;AAA69862.1;Human_rhinovirus_type_A16 | 756 | 762 | 7 | 0.05 | 0.05 | 0.05 |
| L24917.1;AAA69862.1;Human_rhinovirus_type_A16 | 767 | 767 | 1 | 0.05 | 0.05 | 0.05 |
| L24917.1;AAA69862.1;Human_rhinovirus_type_A16 | 799 | 799 | 1 | 0.05 | 0.05 | 0.05 |
| L24917.1;AAA69862.1;Human_rhinovirus_type_A16 | 817 | 828 | 12 | 0.1 | 0.25 | 0.1875 |
| L24917.1;AAA69862.1;Human_rhinovirus_type_A16 | 830 | 834 | 5 | 0.05 | 0.05 | 0.05 |
| L24917.1;AAA69862.1;Human_rhinovirus_type_A16 | 836 | 845 | 10 | 0.05 | 0.25 | 0.105 |
| L24917.1;AAA69862.1;Human_rhinovirus_type_A16 | 847 | 847 | 1 | 0.05 | 0.05 | 0.05 |
| L24917.1;AAA69862.1;Human_rhinovirus_type_A16 | 903 | 905 | 3 | 0.05 | 0.05 | 0.05 |
| L24917.1;AAA69862.1;Human_rhinovirus_type_A16 | 907 | 907 | 1 | 0.05 | 0.05 | 0.05 |
| L24917.1;AAA69862.1;Human_rhinovirus_type_A16 | 909 | 909 | 1 | 0.05 | 0.05 | 0.05 |
| L24917.1;AAA69862.1;Human_rhinovirus_type_A16 | 914 | 914 | 1 | 0.05 | 0.05 | 0.05 |

| Extended data 1 | | | | | | |
|---|---|---|---|---|---|---|
| Protein | Start position (peptide) | Stop position (peptide) | Number of peptides | Minimum fraction (per peptide) of control subjects with reactivity >3 sd above average | Maximum fraction (per peptide) of control subjects with reactivity >3 sd above average | Mean fraction (over all peptides in this contiguous stretch) of controlsubjects with reactivity >3 sd above average |
| L24917.1;AAA69862.1;Human_rhinovirus_type_A16 | 929 | 929 | 1 | 0.05 | 0.05 | 0.05 |
| L24917.1;AAA69862.1;Human_rhinovirus_type_A16 | 931 | 931 | 1 | 0.05 | 0.05 | 0.05 |
| L24917.1;AAA69862.1;Human_rhinovirus_type_A16 | 988 | 989 | 2 | 0.05 | 0.05 | 0.05 |
| L24917.1;AAA69862.1;Human_rhinovirus_type_A16 | 991 | 1012 | 22 | 0.1 | 0.95 | 0.447727273 |
| L24917.1;AAA69862.1;Human_rhinovirus_type_A16 | 1014 | 1018 | 5 | 0.05 | 0.1 | 0.07 |
| L24917.1;AAA69862.1;Human_rhinovirus_type_A16 | 1024 | 1024 | 1 | 0.05 | 0.05 | 0.05 |
| L24917.1;AAA69862.1;Human_rhinovirus_type_A16 | 1074 | 1086 | 13 | 0.05 | 0.3 | 0.115384615 |
| L24917.1;AAA69862.1;Human_rhinovirus_type_A16 | 1089 | 1092 | 4 | 0.05 | 0.2 | 0.1 |
| L24917.1;AAA69862.1;Human_rhinovirus_type_A16 | 1100 | 1103 | 4 | 0.05 | 0.05 | 0.05 |
| L24917.1;AAA69862.1;Human_rhinovirus_type_A16 | 1105 | 1108 | 4 | 0.2 | 0.3 | 0.2375 |
| L24917.1;AAA69862.1;Human_rhinovirus_type_A16 | 1113 | 1113 | 1 | 0.1 | 0.1 | 0.1 |
| L24917.1;AAA69862.1;Human_rhinovirus_type_A16 | 1115 | 1123 | 9 | 0.05 | 0.2 | 0.127777778 |
| L24917.1;AAA69862.1;Human_rhinovirus_type_A16 | 1135 | 1138 | 4 | 0.05 | 0.05 | 0.05 |
| L24917.1;AAA69862.1;Human_rhinovirus_type_A16 | 1141 | 1142 | 2 | 0.05 | 0.05 | 0.05 |
| L24917.1;AAA69862.1;Human_rhinovirus_type_A16 | 1146 | 1147 | 2 | 0.05 | 0.05 | 0.05 |
| L24917.1;AAA69862.1;Human_rhinovirus_type_A16 | 1150 | 1151 | 2 | 0.05 | 0.1 | 0.075 |
| L24917.1;AAA69862.1;Human_rhinovirus_type_A16 | 1153 | 1156 | 4 | 0.05 | 0.1 | 0.075 |
| L24917.1;AAA69862.1;Human_rhinovirus_type_A16 | 1171 | 1176 | 6 | 0.05 | 0.05 | 0.05 |
| L24917.1;AAA69862.1;Human_rhinovirus_type_A16 | 1221 | 1221 | 1 | 0.05 | 0.05 | 0.05 |
| L24917.1;AAA69862.1;Human_rhinovirus_type_A16 | 1226 | 1245 | 20 | 0.05 | 0.7 | 0.3975 |
| L24917.1;AAA69862.1;Human_rhinovirus_type_A16 | 1248 | 1252 | 5 | 0.05 | 0.1 | 0.06 |
| L24917.1;AAA69862.1;Human_rhinovirus_type_A16 | 1266 | 1266 | 1 | 0.05 | 0.05 | 0.05 |
| L24917.1;AAA69862.1;Human_rhinovirus_type_A16 | 1274 | 1286 | 13 | 0.1 | 1 | 0.7 |
| L24917.1;AAA69862.1;Human_rhinovirus_type_A16 | 1303 | 1310 | 8 | 0.05 | 0.15 | 0.125 |
| L24917.1;AAA69862.1;Human_rhinovirus_type_A16 | 1328 | 1335 | 8 | 0.05 | 0.15 | 0.09375 |
| L24917.1;AAA69862.1;Human_rhinovirus_type_A16 | 1337 | 1337 | 1 | 0.05 | 0.05 | 0.05 |
| L24917.1;AAA69862.1;Human_rhinovirus_type_A16 | 1354 | 1355 | 2 | 0.1 | 0.15 | 0.125 |
| L24917.1;AAA69862.1;Human_rhinovirus_type_A16 | 1362 | 1362 | 1 | 0.05 | 0.05 | 0.05 |
| L24917.1;AAA69862.1;Human_rhinovirus_type_A16 | 1372 | 1377 | 6 | 0.05 | 0.05 | 0.05 |
| L24917.1;AAA69862.1;Human_rhinovirus_type_A16 | 1380 | 1381 | 2 | 0.05 | 0.05 | 0.05 |
| L24917.1;AAA69862.1;Human_rhinovirus_type_A16 | 1390 | 1390 | 1 | 0.05 | 0.05 | 0.05 |
| L24917.1;AAA69862.1;Human_rhinovirus_type_A16 | 1392 | 1392 | 1 | 0.05 | 0.05 | 0.05 |
| L24917.1;AAA69862.1;Human_rhinovirus_type_A16 | 1395 | 1405 | 11 | 0.05 | 0.2 | 0.127272727 |
| L24917.1;AAA69862.1;Human_rhinovirus_type_A16 | 1409 | 1445 | 37 | 0.05 | 0.8 | 0.406756757 |
| L24917.1;AAA69862.1;Human_rhinovirus_type_A16 | 1455 | 1455 | 1 | 0.05 | 0.05 | 0.05 |
| L24917.1;AAA69862.1;Human_rhinovirus_type_A16 | 1503 | 1504 | 2 | 0.05 | 0.05 | 0.05 |
| L24917.1;AAA69862.1;Human_rhinovirus_type_A16 | 1511 | 1511 | 1 | 0.05 | 0.05 | 0.05 |

| Extended data 1 | | | | | | |
|---|---|---|---|---|---|---|
| Protein | Start position (peptide) | Stop position (peptide) | Number of peptides | Minimum fraction (per peptide) of control subjects with reactivity >3 sd above average | Maximum fraction (per peptide) of control subjects with reactivity >3 sd above average | Mean fraction (over all peptides in this contiguous stretch) of controlsubjects with reactivity >3 sd above average |
| L24917.1;AAA69862.1;Human_rhinovirus_type_A16 | 1539 | 1539 | 1 | 0.05 | 0.05 | 0.05 |
| L24917.1;AAA69862.1;Human_rhinovirus_type_A16 | 1544 | 1544 | 1 | 0.05 | 0.05 | 0.05 |
| L24917.1;AAA69862.1;Human_rhinovirus_type_A16 | 1546 | 1548 | 3 | 0.05 | 0.1 | 0.066666667 |
| L24917.1;AAA69862.1;Human_rhinovirus_type_A16 | 1550 | 1553 | 4 | 0.05 | 0.1 | 0.0625 |
| L24917.1;AAA69862.1;Human_rhinovirus_type_A16 | 1556 | 1556 | 1 | 0.05 | 0.05 | 0.05 |
| L24917.1;AAA69862.1;Human_rhinovirus_type_A16 | 1559 | 1567 | 9 | 0.05 | 0.2 | 0.1 |
| L24917.1;AAA69862.1;Human_rhinovirus_type_A16 | 1580 | 1592 | 13 | 0.05 | 0.3 | 0.146153846 |
| L24917.1;AAA69862.1;Human_rhinovirus_type_A16 | 1604 | 1604 | 1 | 0.15 | 0.15 | 0.15 |
| L24917.1;AAA69862.1;Human_rhinovirus_type_A16 | 1687 | 1687 | 1 | 0.1 | 0.1 | 0.1 |
| L24917.1;AAA69862.1;Human_rhinovirus_type_A16 | 1695 | 1695 | 1 | 0.05 | 0.05 | 0.05 |
| L24917.1;AAA69862.1;Human_rhinovirus_type_A16 | 1701 | 1724 | 24 | 0.05 | 0.5 | 0.2125 |
| L24917.1;AAA69862.1;Human_rhinovirus_type_A16 | 1727 | 1740 | 14 | 0.05 | 0.2 | 0.089285714 |
| L24917.1;AAA69862.1;Human_rhinovirus_type_A16 | 1743 | 1743 | 1 | 0.05 | 0.05 | 0.05 |
| L24917.1;AAA69862.1;Human_rhinovirus_type_A16 | 1745 | 1750 | 6 | 0.05 | 0.15 | 0.083333333 |
| L24917.1;AAA69862.1;Human_rhinovirus_type_A16 | 1752 | 1753 | 2 | 0.05 | 0.05 | 0.05 |
| L24917.1;AAA69862.1;Human_rhinovirus_type_A16 | 1755 | 1755 | 1 | 0.1 | 0.1 | 0.1 |
| L24917.1;AAA69862.1;Human_rhinovirus_type_A16 | 1762 | 1762 | 1 | 0.05 | 0.05 | 0.05 |
| L24917.1;AAA69862.1;Human_rhinovirus_type_A16 | 1773 | 1774 | 2 | 0.05 | 0.05 | 0.05 |
| L24917.1;AAA69862.1;Human_rhinovirus_type_A16 | 1778 | 1778 | 1 | 0.05 | 0.05 | 0.05 |
| L24917.1;AAA69862.1;Human_rhinovirus_type_A16 | 1780 | 1786 | 7 | 0.05 | 0.05 | 0.05 |
| L24917.1;AAA69862.1;Human_rhinovirus_type_A16 | 1792 | 1792 | 1 | 0.05 | 0.05 | 0.05 |
| L24917.1;AAA69862.1;Human_rhinovirus_type_A16 | 1795 | 1796 | 2 | 0.05 | 0.05 | 0.05 |
| L24917.1;AAA69862.1;Human_rhinovirus_type_A16 | 1798 | 1800 | 3 | 0.05 | 0.1 | 0.066666667 |
| L24917.1;AAA69862.1;Human_rhinovirus_type_A16 | 1803 | 1804 | 2 | 0.05 | 0.05 | 0.05 |
| L24917.1;AAA69862.1;Human_rhinovirus_type_A16 | 1806 | 1812 | 7 | 0.05 | 0.1 | 0.078571429 |
| L24917.1;AAA69862.1;Human_rhinovirus_type_A16 | 1817 | 1817 | 1 | 0.05 | 0.05 | 0.05 |
| L24917.1;AAA69862.1;Human_rhinovirus_type_A16 | 1820 | 1820 | 1 | 0.05 | 0.05 | 0.05 |
| L24917.1;AAA69862.1;Human_rhinovirus_type_A16 | 1829 | 1830 | 2 | 0.1 | 0.1 | 0.1 |
| L24917.1;AAA69862.1;Human_rhinovirus_type_A16 | 1832 | 1832 | 1 | 0.05 | 0.05 | 0.05 |
| L24917.1;AAA69862.1;Human_rhinovirus_type_A16 | 1837 | 1845 | 9 | 0.05 | 0.2 | 0.116666667 |
| L24917.1;AAA69862.1;Human_rhinovirus_type_A16 | 1853 | 1855 | 3 | 0.05 | 0.05 | 0.05 |
| L24917.1;AAA69862.1;Human_rhinovirus_type_A16 | 1857 | 1863 | 7 | 0.05 | 0.15 | 0.071428571 |
| L24917.1;AAA69862.1;Human_rhinovirus_type_A16 | 1873 | 1873 | 1 | 0.05 | 0.05 | 0.05 |
| L24917.1;AAA69862.1;Human_rhinovirus_type_A16 | 1898 | 1898 | 1 | 0.1 | 0.1 | 0.1 |
| L24917.1;AAA69862.1;Human_rhinovirus_type_A16 | 1900 | 1900 | 1 | 0.1 | 0.1 | 0.1 |
| L24917.1;AAA69862.1;Human_rhinovirus_type_A16 | 1902 | 1902 | 1 | 0.05 | 0.05 | 0.05 |
| L24917.1;AAA69862.1;Human_rhinovirus_type_A16 | 1904 | 1907 | 4 | 0.05 | 0.4 | 0.1875 |

| Extended data 1 | | | | | | |
|---|---|---|---|---|---|---|
| **Protein** | **Start position (peptide)** | **Stop position (peptide)** | **Number of peptides** | **Minimum fraction (per peptide) of control subjects with reactivity >3 sd above average** | **Maximum fraction (per peptide) of control subjects with reactivity >3 sd above average** | **Mean fraction (over all peptides in this contiguous stretch) of controlsubjects with reactivity >3 sd above average** |
| L24917.1;AAA69862.1;Human_rhinovirus_type_A16 | 1915 | 1925 | 11 | 0.05 | 0.1 | 0.054545455 |
| L24917.1;AAA69862.1;Human_rhinovirus_type_A16 | 1927 | 1927 | 1 | 0.05 | 0.05 | 0.05 |
| L24917.1;AAA69862.1;Human_rhinovirus_type_A16 | 1938 | 1938 | 1 | 0.05 | 0.05 | 0.05 |
| L24917.1;AAA69862.1;Human_rhinovirus_type_A16 | 1940 | 1941 | 2 | 0.05 | 0.1 | 0.075 |
| L24917.1;AAA69862.1;Human_rhinovirus_type_A16 | 1962 | 1962 | 1 | 0.05 | 0.05 | 0.05 |
| L24917.1;AAA69862.1;Human_rhinovirus_type_A16 | 1968 | 1970 | 3 | 0.05 | 0.05 | 0.05 |
| L24917.1;AAA69862.1;Human_rhinovirus_type_A16 | 1972 | 1973 | 2 | 0.05 | 0.05 | 0.05 |
| L24917.1;AAA69862.1;Human_rhinovirus_type_A16 | 1975 | 1975 | 1 | 0.05 | 0.05 | 0.05 |
| L24917.1;AAA69862.1;Human_rhinovirus_type_A16 | 2018 | 2018 | 1 | 0.05 | 0.05 | 0.05 |
| L24917.1;AAA69862.1;Human_rhinovirus_type_A16 | 2024 | 2024 | 1 | 0.05 | 0.05 | 0.05 |
| L24917.1;AAA69862.1;Human_rhinovirus_type_A16 | 2028 | 2035 | 8 | 0.05 | 0.05 | 0.05 |
| L24917.1;AAA69862.1;Human_rhinovirus_type_A16 | 2038 | 2039 | 2 | 0.05 | 0.1 | 0.075 |
| L24917.1;AAA69862.1;Human_rhinovirus_type_A16 | 2041 | 2043 | 3 | 0.05 | 0.05 | 0.05 |
| L24917.1;AAA69862.1;Human_rhinovirus_type_A16 | 2046 | 2048 | 3 | 0.05 | 0.05 | 0.05 |
| L24917.1;AAA69862.1;Human_rhinovirus_type_A16 | 2068 | 2072 | 5 | 0.05 | 0.15 | 0.08 |
| L24917.1;AAA69862.1;Human_rhinovirus_type_A16 | 2084 | 2084 | 1 | 0.05 | 0.05 | 0.05 |
| L24917.1;AAA69862.1;Human_rhinovirus_type_A16 | 2086 | 2088 | 3 | 0.05 | 0.1 | 0.066666667 |
| L24917.1;AAA69862.1;Human_rhinovirus_type_A16 | 2090 | 2092 | 3 | 0.05 | 0.2 | 0.1 |
| L24917.1;AAA69862.1;Human_rhinovirus_type_A16 | 2095 | 2095 | 1 | 0.05 | 0.05 | 0.05 |
| L24917.1;AAA69862.1;Human_rhinovirus_type_A16 | 2111 | 2119 | 9 | 0.05 | 0.2 | 0.094444444 |
| NC_002645.1;NP_073549.1;HCoV-229E_1ab | 1 | 1 | 1 | 0.05 | 0.05 | 0.05 |
| NC_002645.1;NP_073549.1;HCoV-229E_1ab | 8 | 8 | 1 | 0.05 | 0.05 | 0.05 |
| NC_002645.1;NP_073549.1;HCoV-229E_1ab | 20 | 21 | 2 | 0.05 | 0.05 | 0.05 |
| NC_002645.1;NP_073549.1;HCoV-229E_1ab | 46 | 46 | 1 | 0.05 | 0.05 | 0.05 |
| NC_002645.1;NP_073549.1;HCoV-229E_1ab | 55 | 55 | 1 | 0.2 | 0.2 | 0.2 |
| NC_002645.1;NP_073549.1;HCoV-229E_1ab | 58 | 58 | 1 | 0.05 | 0.05 | 0.05 |
| NC_002645.1;NP_073549.1;HCoV-229E_1ab | 89 | 90 | 2 | 0.05 | 0.2 | 0.125 |
| NC_002645.1;NP_073549.1;HCoV-229E_1ab | 95 | 97 | 3 | 0.05 | 0.1 | 0.066666667 |
| NC_002645.1;NP_073549.1;HCoV-229E_1ab | 99 | 99 | 1 | 0.05 | 0.05 | 0.05 |
| NC_002645.1;NP_073549.1;HCoV-229E_1ab | 103 | 103 | 1 | 0.05 | 0.05 | 0.05 |
| NC_002645.1;NP_073549.1;HCoV-229E_1ab | 107 | 107 | 1 | 0.05 | 0.05 | 0.05 |
| NC_002645.1;NP_073549.1;HCoV-229E_1ab | 113 | 121 | 9 | 0.05 | 0.05 | 0.05 |
| NC_002645.1;NP_073549.1;HCoV-229E_1ab | 156 | 156 | 1 | 0.05 | 0.05 | 0.05 |
| NC_002645.1;NP_073549.1;HCoV-229E_1ab | 163 | 164 | 2 | 0.05 | 0.05 | 0.05 |
| NC_002645.1;NP_073549.1;HCoV-229E_1ab | 166 | 168 | 3 | 0.05 | 0.05 | 0.05 |
| NC_002645.1;NP_073549.1;HCoV-229E_1ab | 170 | 171 | 2 | 0.05 | 0.05 | 0.05 |
| NC_002645.1;NP_073549.1;HCoV-229E_1ab | 189 | 189 | 1 | 0.15 | 0.15 | 0.15 |

| Extended data 1 | | | | | | |
|---|---|---|---|---|---|---|
| Protein | Start position (peptide) | Stop position (peptide) | Number of peptides | Minimum fraction (per peptide) of control subjects with reactivity >3 sd above average | Maximum fraction (per peptide) of control subjects with reactivity >3 sd above average | Mean fraction (over all peptides in this contiguous stretch) of controlsubjects with reactivity >3 sd above average |
| NC_002645.1;NP_073549.1;HCoV-229E_1ab | 195 | 195 | 1 | 0.05 | 0.05 | 0.05 |
| NC_002645.1;NP_073549.1;HCoV-229E_1ab | 203 | 203 | 1 | 0.05 | 0.05 | 0.05 |
| NC_002645.1;NP_073549.1;HCoV-229E_1ab | 207 | 207 | 1 | 0.05 | 0.05 | 0.05 |
| NC_002645.1;NP_073549.1;HCoV-229E_1ab | 210 | 210 | 1 | 0.05 | 0.05 | 0.05 |
| NC_002645.1;NP_073549.1;HCoV-229E_1ab | 222 | 222 | 1 | 0.05 | 0.05 | 0.05 |
| NC_002645.1;NP_073549.1;HCoV-229E_1ab | 243 | 245 | 3 | 0.05 | 0.05 | 0.05 |
| NC_002645.1;NP_073549.1;HCoV-229E_1ab | 247 | 251 | 5 | 0.05 | 0.05 | 0.05 |
| NC_002645.1;NP_073549.1;HCoV-229E_1ab | 269 | 276 | 8 | 0.05 | 0.15 | 0.09375 |
| NC_002645.1;NP_073549.1;HCoV-229E_1ab | 278 | 278 | 1 | 0.05 | 0.05 | 0.05 |
| NC_002645.1;NP_073549.1;HCoV-229E_1ab | 361 | 361 | 1 | 0.05 | 0.05 | 0.05 |
| NC_002645.1;NP_073549.1;HCoV-229E_1ab | 364 | 365 | 2 | 0.05 | 0.05 | 0.05 |
| NC_002645.1;NP_073549.1;HCoV-229E_1ab | 384 | 384 | 1 | 0.05 | 0.05 | 0.05 |
| NC_002645.1;NP_073549.1;HCoV-229E_1ab | 387 | 395 | 9 | 0.05 | 0.1 | 0.061111111 |
| NC_002645.1;NP_073549.1;HCoV-229E_1ab | 405 | 406 | 2 | 0.05 | 0.05 | 0.05 |
| NC_002645.1;NP_073549.1;HCoV-229E_1ab | 413 | 416 | 4 | 0.1 | 0.1 | 0.1 |
| NC_002645.1;NP_073549.1;HCoV-229E_1ab | 443 | 443 | 1 | 0.05 | 0.05 | 0.05 |
| NC_002645.1;NP_073549.1;HCoV-229E_1ab | 448 | 452 | 5 | 0.05 | 0.05 | 0.05 |
| NC_002645.1;NP_073549.1;HCoV-229E_1ab | 474 | 475 | 2 | 0.05 | 0.05 | 0.05 |
| NC_002645.1;NP_073549.1;HCoV-229E_1ab | 477 | 478 | 2 | 0.15 | 0.2 | 0.175 |
| NC_002645.1;NP_073549.1;HCoV-229E_1ab | 544 | 544 | 1 | 0.1 | 0.1 | 0.1 |
| NC_002645.1;NP_073549.1;HCoV-229E_1ab | 592 | 592 | 1 | 0.1 | 0.1 | 0.1 |
| NC_002645.1;NP_073549.1;HCoV-229E_1ab | 594 | 594 | 1 | 0.05 | 0.05 | 0.05 |
| NC_002645.1;NP_073549.1;HCoV-229E_1ab | 610 | 618 | 9 | 0.05 | 0.05 | 0.05 |
| NC_002645.1;NP_073549.1;HCoV-229E_1ab | 627 | 629 | 3 | 0.05 | 0.05 | 0.05 |
| NC_002645.1;NP_073549.1;HCoV-229E_1ab | 631 | 643 | 13 | 0.05 | 0.5 | 0.288461538 |
| NC_002645.1;NP_073549.1;HCoV-229E_1ab | 653 | 653 | 1 | 0.05 | 0.05 | 0.05 |
| NC_002645.1;NP_073549.1;HCoV-229E_1ab | 656 | 656 | 1 | 0.05 | 0.05 | 0.05 |
| NC_002645.1;NP_073549.1;HCoV-229E_1ab | 658 | 659 | 2 | 0.05 | 0.05 | 0.05 |
| NC_002645.1;NP_073549.1;HCoV-229E_1ab | 662 | 662 | 1 | 0.05 | 0.05 | 0.05 |
| NC_002645.1;NP_073549.1;HCoV-229E_1ab | 665 | 672 | 8 | 0.05 | 0.1 | 0.0625 |
| NC_002645.1;NP_073549.1;HCoV-229E_1ab | 681 | 681 | 1 | 0.05 | 0.05 | 0.05 |
| NC_002645.1;NP_073549.1;HCoV-229E_1ab | 683 | 684 | 2 | 0.05 | 0.1 | 0.075 |
| NC_002645.1;NP_073549.1;HCoV-229E_1ab | 686 | 695 | 10 | 0.05 | 0.1 | 0.065 |
| NC_002645.1;NP_073549.1;HCoV-229E_1ab | 698 | 700 | 3 | 0.05 | 0.15 | 0.116666667 |
| NC_002645.1;NP_073549.1;HCoV-229E_1ab | 704 | 704 | 1 | 0.05 | 0.05 | 0.05 |
| NC_002645.1;NP_073549.1;HCoV-229E_1ab | 723 | 723 | 1 | 0.05 | 0.05 | 0.05 |
| NC_002645.1;NP_073549.1;HCoV-229E_1ab | 738 | 738 | 1 | 0.05 | 0.05 | 0.05 |

| Extended data 1 | | | | | | |
|---|---|---|---|---|---|---|
| Protein | Start position (peptide) | Stop position (peptide) | Number of peptides | Minimum fraction (per peptide) of control subjects with reactivity >3 sd above average | Maximum fraction (per peptide) of control subjects with reactivity >3 sd above average | Mean fraction (over all peptides in this contiguous stretch) of controlsubjects with reactivity >3 sd above average |
| NC_002645.1;NP_073549.1;HCoV-229E_1ab | 797 | 797 | 1 | 0.05 | 0.05 | 0.05 |
| NC_002645.1;NP_073549.1;HCoV-229E_1ab | 824 | 827 | 4 | 0.05 | 0.1 | 0.075 |
| NC_002645.1;NP_073549.1;HCoV-229E_1ab | 831 | 831 | 1 | 0.1 | 0.1 | 0.1 |
| NC_002645.1;NP_073549.1;HCoV-229E_1ab | 852 | 852 | 1 | 0.05 | 0.05 | 0.05 |
| NC_002645.1;NP_073549.1;HCoV-229E_1ab | 865 | 876 | 12 | 0.05 | 0.2 | 0.075 |
| NC_002645.1;NP_073549.1;HCoV-229E_1ab | 879 | 881 | 3 | 0.05 | 0.05 | 0.05 |
| NC_002645.1;NP_073549.1;HCoV-229E_1ab | 883 | 883 | 1 | 0.05 | 0.05 | 0.05 |
| NC_002645.1;NP_073549.1;HCoV-229E_1ab | 900 | 901 | 2 | 0.05 | 0.1 | 0.075 |
| NC_002645.1;NP_073549.1;HCoV-229E_1ab | 903 | 903 | 1 | 0.1 | 0.1 | 0.1 |
| NC_002645.1;NP_073549.1;HCoV-229E_1ab | 906 | 906 | 1 | 0.05 | 0.05 | 0.05 |
| NC_002645.1;NP_073549.1;HCoV-229E_1ab | 924 | 924 | 1 | 0.05 | 0.05 | 0.05 |
| NC_002645.1;NP_073549.1;HCoV-229E_1ab | 972 | 972 | 1 | 0.05 | 0.05 | 0.05 |
| NC_002645.1;NP_073549.1;HCoV-229E_1ab | 974 | 975 | 2 | 0.05 | 0.1 | 0.075 |
| NC_002645.1;NP_073549.1;HCoV-229E_1ab | 983 | 983 | 1 | 0.05 | 0.05 | 0.05 |
| NC_002645.1;NP_073549.1;HCoV-229E_1ab | 989 | 989 | 1 | 0.05 | 0.05 | 0.05 |
| NC_002645.1;NP_073549.1;HCoV-229E_1ab | 997 | 1002 | 6 | 0.05 | 0.05 | 0.05 |
| NC_002645.1;NP_073549.1;HCoV-229E_1ab | 1005 | 1009 | 5 | 0.05 | 0.2 | 0.12 |
| NC_002645.1;NP_073549.1;HCoV-229E_1ab | 1011 | 1019 | 9 | 0.05 | 0.15 | 0.105555556 |
| NC_002645.1;NP_073549.1;HCoV-229E_1ab | 1022 | 1022 | 1 | 0.05 | 0.05 | 0.05 |
| NC_002645.1;NP_073549.1;HCoV-229E_1ab | 1024 | 1031 | 8 | 0.05 | 0.1 | 0.05625 |
| NC_002645.1;NP_073549.1;HCoV-229E_1ab | 1038 | 1038 | 1 | 0.05 | 0.05 | 0.05 |
| NC_002645.1;NP_073549.1;HCoV-229E_1ab | 1045 | 1045 | 1 | 0.05 | 0.05 | 0.05 |
| NC_002645.1;NP_073549.1;HCoV-229E_1ab | 1064 | 1067 | 4 | 0.05 | 0.1 | 0.0625 |
| NC_002645.1;NP_073549.1;HCoV-229E_1ab | 1088 | 1088 | 1 | 0.05 | 0.05 | 0.05 |
| NC_002645.1;NP_073549.1;HCoV-229E_1ab | 1110 | 1110 | 1 | 0.05 | 0.05 | 0.05 |
| NC_002645.1;NP_073549.1;HCoV-229E_1ab | 1112 | 1112 | 1 | 0.05 | 0.05 | 0.05 |
| NC_002645.1;NP_073549.1;HCoV-229E_1ab | 1149 | 1149 | 1 | 0.05 | 0.05 | 0.05 |
| NC_002645.1;NP_073549.1;HCoV-229E_1ab | 1173 | 1174 | 2 | 0.05 | 0.1 | 0.075 |
| NC_002645.1;NP_073549.1;HCoV-229E_1ab | 1183 | 1183 | 1 | 0.05 | 0.05 | 0.05 |
| NC_002645.1;NP_073549.1;HCoV-229E_1ab | 1208 | 1208 | 1 | 0.05 | 0.05 | 0.05 |
| NC_002645.1;NP_073549.1;HCoV-229E_1ab | 1215 | 1227 | 13 | 0.05 | 0.15 | 0.065384615 |
| NC_002645.1;NP_073549.1;HCoV-229E_1ab | 1231 | 1242 | 12 | 0.05 | 0.1 | 0.070833333 |
| NC_002645.1;NP_073549.1;HCoV-229E_1ab | 1250 | 1250 | 1 | 0.05 | 0.05 | 0.05 |
| NC_002645.1;NP_073549.1;HCoV-229E_1ab | 1257 | 1266 | 10 | 0.05 | 0.2 | 0.09 |
| NC_002645.1;NP_073549.1;HCoV-229E_1ab | 1272 | 1288 | 17 | 0.05 | 0.15 | 0.061764706 |
| NC_002645.1;NP_073549.1;HCoV-229E_1ab | 1292 | 1292 | 1 | 0.05 | 0.05 | 0.05 |
| NC_002645.1;NP_073549.1;HCoV-229E_1ab | 1294 | 1302 | 9 | 0.05 | 0.3 | 0.083333333 |

| Extended data 1 | | | | | | |
|---|---|---|---|---|---|---|
| Protein | Start position (peptide) | Stop position (peptide) | Number of peptides | Minimum fraction (per peptide) of control subjects with reactivity >3 sd above average | Maximum fraction (per peptide) of control subjects with reactivity >3 sd above average | Mean fraction (over all peptides in this contiguous stretch) of controlsubjects with reactivity >3 sd above average |
| NC_002645.1;NP_073549.1;HCoV-229E_1ab | 1325 | 1326 | 2 | 0.05 | 0.05 | 0.05 |
| NC_002645.1;NP_073549.1;HCoV-229E_1ab | 1365 | 1375 | 11 | 0.1 | 0.55 | 0.313636364 |
| NC_002645.1;NP_073549.1;HCoV-229E_1ab | 1429 | 1429 | 1 | 0.05 | 0.05 | 0.05 |
| NC_002645.1;NP_073549.1;HCoV-229E_1ab | 1432 | 1445 | 14 | 0.05 | 0.2 | 0.128571429 |
| NC_002645.1;NP_073549.1;HCoV-229E_1ab | 1467 | 1469 | 3 | 0.05 | 0.05 | 0.05 |
| NC_002645.1;NP_073549.1;HCoV-229E_1ab | 1490 | 1490 | 1 | 0.05 | 0.05 | 0.05 |
| NC_002645.1;NP_073549.1;HCoV-229E_1ab | 1492 | 1497 | 6 | 0.05 | 0.05 | 0.05 |
| NC_002645.1;NP_073549.1;HCoV-229E_1ab | 1506 | 1506 | 1 | 0.1 | 0.1 | 0.1 |
| NC_002645.1;NP_073549.1;HCoV-229E_1ab | 1510 | 1511 | 2 | 0.05 | 0.05 | 0.05 |
| NC_002645.1;NP_073549.1;HCoV-229E_1ab | 1514 | 1514 | 1 | 0.1 | 0.1 | 0.1 |
| NC_002645.1;NP_073549.1;HCoV-229E_1ab | 1516 | 1516 | 1 | 0.05 | 0.05 | 0.05 |
| NC_002645.1;NP_073549.1;HCoV-229E_1ab | 1519 | 1519 | 1 | 0.05 | 0.05 | 0.05 |
| NC_002645.1;NP_073549.1;HCoV-229E_1ab | 1521 | 1521 | 1 | 0.1 | 0.1 | 0.1 |
| NC_002645.1;NP_073549.1;HCoV-229E_1ab | 1583 | 1583 | 1 | 0.05 | 0.05 | 0.05 |
| NC_002645.1;NP_073549.1;HCoV-229E_1ab | 1586 | 1589 | 4 | 0.05 | 0.05 | 0.05 |
| NC_002645.1;NP_073549.1;HCoV-229E_1ab | 1607 | 1607 | 1 | 0.05 | 0.05 | 0.05 |
| NC_002645.1;NP_073549.1;HCoV-229E_1ab | 1611 | 1611 | 1 | 0.05 | 0.05 | 0.05 |
| NC_002645.1;NP_073549.1;HCoV-229E_1ab | 1627 | 1627 | 1 | 0.05 | 0.05 | 0.05 |
| NC_002645.1;NP_073549.1;HCoV-229E_1ab | 1631 | 1635 | 5 | 0.05 | 0.1 | 0.06 |
| NC_002645.1;NP_073549.1;HCoV-229E_1ab | 1684 | 1684 | 1 | 0.05 | 0.05 | 0.05 |
| NC_002645.1;NP_073549.1;HCoV-229E_1ab | 1690 | 1690 | 1 | 0.05 | 0.05 | 0.05 |
| NC_002645.1;NP_073549.1;HCoV-229E_1ab | 1710 | 1710 | 1 | 0.05 | 0.05 | 0.05 |
| NC_002645.1;NP_073549.1;HCoV-229E_1ab | 1718 | 1721 | 4 | 0.05 | 0.05 | 0.05 |
| NC_002645.1;NP_073549.1;HCoV-229E_1ab | 1741 | 1741 | 1 | 0.1 | 0.1 | 0.1 |
| NC_002645.1;NP_073549.1;HCoV-229E_1ab | 1753 | 1760 | 8 | 0.05 | 0.1 | 0.075 |
| NC_002645.1;NP_073549.1;HCoV-229E_1ab | 1762 | 1762 | 1 | 0.05 | 0.05 | 0.05 |
| NC_002645.1;NP_073549.1;HCoV-229E_1ab | 1764 | 1764 | 1 | 0.05 | 0.05 | 0.05 |
| NC_002645.1;NP_073549.1;HCoV-229E_1ab | 1777 | 1778 | 2 | 0.05 | 0.05 | 0.05 |
| NC_002645.1;NP_073549.1;HCoV-229E_1ab | 1783 | 1786 | 4 | 0.1 | 0.15 | 0.1375 |
| NC_002645.1;NP_073549.1;HCoV-229E_1ab | 1797 | 1798 | 2 | 0.05 | 0.05 | 0.05 |
| NC_002645.1;NP_073549.1;HCoV-229E_1ab | 1802 | 1802 | 1 | 0.05 | 0.05 | 0.05 |
| NC_002645.1;NP_073549.1;HCoV-229E_1ab | 1804 | 1804 | 1 | 0.2 | 0.2 | 0.2 |
| NC_002645.1;NP_073549.1;HCoV-229E_1ab | 1822 | 1826 | 5 | 0.05 | 0.1 | 0.07 |
| NC_002645.1;NP_073549.1;HCoV-229E_1ab | 1832 | 1832 | 1 | 0.05 | 0.05 | 0.05 |
| NC_002645.1;NP_073549.1;HCoV-229E_1ab | 1836 | 1836 | 1 | 0.05 | 0.05 | 0.05 |
| NC_002645.1;NP_073549.1;HCoV-229E_1ab | 1849 | 1851 | 3 | 0.05 | 0.05 | 0.05 |
| NC_002645.1;NP_073549.1;HCoV-229E_1ab | 1855 | 1855 | 1 | 0.05 | 0.05 | 0.05 |

| Extended data 1 | | | | | | |
|---|---|---|---|---|---|---|
| Protein | Start position (peptide) | Stop position (peptide) | Number of peptides | Minimum fraction (per peptide) of control subjects with reactivity >3 sd above average | Maximum fraction (per peptide) of control subjects with reactivity >3 sd above average | Mean fraction (over all peptides in this contiguous stretch) of controlsubjects with reactivity >3 sd above average |
| NC_002645.1;NP_073549.1;HCoV-229E_1ab | 1866 | 1866 | 1 | 0.05 | 0.05 | 0.05 |
| NC_002645.1;NP_073549.1;HCoV-229E_1ab | 1868 | 1868 | 1 | 0.15 | 0.15 | 0.15 |
| NC_002645.1;NP_073549.1;HCoV-229E_1ab | 1870 | 1870 | 1 | 0.05 | 0.05 | 0.05 |
| NC_002645.1;NP_073549.1;HCoV-229E_1ab | 1872 | 1872 | 1 | 0.05 | 0.05 | 0.05 |
| NC_002645.1;NP_073549.1;HCoV-229E_1ab | 1899 | 1899 | 1 | 0.05 | 0.05 | 0.05 |
| NC_002645.1;NP_073549.1;HCoV-229E_1ab | 1901 | 1901 | 1 | 0.05 | 0.05 | 0.05 |
| NC_002645.1;NP_073549.1;HCoV-229E_1ab | 1903 | 1903 | 1 | 0.05 | 0.05 | 0.05 |
| NC_002645.1;NP_073549.1;HCoV-229E_1ab | 1907 | 1907 | 1 | 0.05 | 0.05 | 0.05 |
| NC_002645.1;NP_073549.1;HCoV-229E_1ab | 1909 | 1917 | 9 | 0.05 | 0.2 | 0.116666667 |
| NC_002645.1;NP_073549.1;HCoV-229E_1ab | 1919 | 1919 | 1 | 0.05 | 0.05 | 0.05 |
| NC_002645.1;NP_073549.1;HCoV-229E_1ab | 1942 | 1943 | 2 | 0.05 | 0.05 | 0.05 |
| NC_002645.1;NP_073549.1;HCoV-229E_1ab | 1950 | 1950 | 1 | 0.05 | 0.05 | 0.05 |
| NC_002645.1;NP_073549.1;HCoV-229E_1ab | 1968 | 1968 | 1 | 0.05 | 0.05 | 0.05 |
| NC_002645.1;NP_073549.1;HCoV-229E_1ab | 2018 | 2019 | 2 | 0.05 | 0.05 | 0.05 |
| NC_002645.1;NP_073549.1;HCoV-229E_1ab | 2022 | 2025 | 4 | 0.05 | 0.1 | 0.0625 |
| NC_002645.1;NP_073549.1;HCoV-229E_1ab | 2036 | 2036 | 1 | 0.25 | 0.25 | 0.25 |
| NC_002645.1;NP_073549.1;HCoV-229E_1ab | 2039 | 2040 | 2 | 0.1 | 0.1 | 0.1 |
| NC_002645.1;NP_073549.1;HCoV-229E_1ab | 2103 | 2104 | 2 | 0.05 | 0.05 | 0.05 |
| NC_002645.1;NP_073549.1;HCoV-229E_1ab | 2151 | 2151 | 1 | 0.05 | 0.05 | 0.05 |
| NC_002645.1;NP_073549.1;HCoV-229E_1ab | 2173 | 2173 | 1 | 0.05 | 0.05 | 0.05 |
| NC_002645.1;NP_073549.1;HCoV-229E_1ab | 2175 | 2175 | 1 | 0.05 | 0.05 | 0.05 |
| NC_002645.1;NP_073549.1;HCoV-229E_1ab | 2208 | 2210 | 3 | 0.05 | 0.05 | 0.05 |
| NC_002645.1;NP_073549.1;HCoV-229E_1ab | 2216 | 2224 | 9 | 0.05 | 0.15 | 0.061111111 |
| NC_002645.1;NP_073549.1;HCoV-229E_1ab | 2228 | 2228 | 1 | 0.1 | 0.1 | 0.1 |
| NC_002645.1;NP_073549.1;HCoV-229E_1ab | 2232 | 2234 | 3 | 0.05 | 0.3 | 0.15 |
| NC_002645.1;NP_073549.1;HCoV-229E_1ab | 2255 | 2255 | 1 | 0.05 | 0.05 | 0.05 |
| NC_002645.1;NP_073549.1;HCoV-229E_1ab | 2259 | 2261 | 3 | 0.05 | 0.1 | 0.066666667 |
| NC_002645.1;NP_073549.1;HCoV-229E_1ab | 2266 | 2267 | 2 | 0.05 | 0.05 | 0.05 |
| NC_002645.1;NP_073549.1;HCoV-229E_1ab | 2281 | 2293 | 13 | 0.05 | 0.25 | 0.134615385 |
| NC_002645.1;NP_073549.1;HCoV-229E_1ab | 2314 | 2314 | 1 | 0.05 | 0.05 | 0.05 |
| NC_002645.1;NP_073549.1;HCoV-229E_1ab | 2336 | 2347 | 12 | 0.05 | 0.1 | 0.066666667 |
| NC_002645.1;NP_073549.1;HCoV-229E_1ab | 2355 | 2358 | 4 | 0.05 | 0.2 | 0.0875 |
| NC_002645.1;NP_073549.1;HCoV-229E_1ab | 2363 | 2363 | 1 | 0.05 | 0.05 | 0.05 |
| NC_002645.1;NP_073549.1;HCoV-229E_1ab | 2374 | 2374 | 1 | 0.05 | 0.05 | 0.05 |
| NC_002645.1;NP_073549.1;HCoV-229E_1ab | 2383 | 2383 | 1 | 0.05 | 0.05 | 0.05 |
| NC_002645.1;NP_073549.1;HCoV-229E_1ab | 2386 | 2386 | 1 | 0.1 | 0.1 | 0.1 |
| NC_002645.1;NP_073549.1;HCoV-229E_1ab | 2388 | 2390 | 3 | 0.05 | 0.1 | 0.066666667 |

| Extended data 1 | | | | | | |
|---|---|---|---|---|---|---|
| Protein | Start position (peptide) | Stop position (peptide) | Number of peptides | Minimum fraction (per peptide) of control subjects with reactivity >3 sd above average | Maximum fraction (per peptide) of control subjects with reactivity >3 sd above average | Mean fraction (over all peptides in this contiguous stretch) of controlsubjects with reactivity >3 sd above average |
| NC_002645.1;NP_073549.1;HCoV-229E_1ab | 2392 | 2392 | 1 | 0.05 | 0.05 | 0.05 |
| NC_002645.1;NP_073549.1;HCoV-229E_1ab | 2415 | 2416 | 2 | 0.05 | 0.05 | 0.05 |
| NC_002645.1;NP_073549.1;HCoV-229E_1ab | 2418 | 2418 | 1 | 0.05 | 0.05 | 0.05 |
| NC_002645.1;NP_073549.1;HCoV-229E_1ab | 2427 | 2430 | 4 | 0.05 | 0.1 | 0.0625 |
| NC_002645.1;NP_073549.1;HCoV-229E_1ab | 2432 | 2436 | 5 | 0.05 | 0.15 | 0.09 |
| NC_002645.1;NP_073549.1;HCoV-229E_1ab | 2522 | 2537 | 16 | 0.05 | 0.2 | 0.090625 |
| NC_002645.1;NP_073549.1;HCoV-229E_1ab | 2539 | 2540 | 2 | 0.05 | 0.1 | 0.075 |
| NC_002645.1;NP_073549.1;HCoV-229E_1ab | 2545 | 2547 | 3 | 0.05 | 0.1 | 0.066666667 |
| NC_002645.1;NP_073549.1;HCoV-229E_1ab | 2549 | 2549 | 1 | 0.1 | 0.1 | 0.1 |
| NC_002645.1;NP_073549.1;HCoV-229E_1ab | 2576 | 2577 | 2 | 0.1 | 0.15 | 0.125 |
| NC_002645.1;NP_073549.1;HCoV-229E_1ab | 2581 | 2581 | 1 | 0.05 | 0.05 | 0.05 |
| NC_002645.1;NP_073549.1;HCoV-229E_1ab | 2623 | 2623 | 1 | 0.05 | 0.05 | 0.05 |
| NC_002645.1;NP_073549.1;HCoV-229E_1ab | 2655 | 2658 | 4 | 0.1 | 0.2 | 0.15 |
| NC_002645.1;NP_073549.1;HCoV-229E_1ab | 2661 | 2667 | 7 | 0.05 | 0.15 | 0.078571429 |
| NC_002645.1;NP_073549.1;HCoV-229E_1ab | 2692 | 2696 | 5 | 0.05 | 0.05 | 0.05 |
| NC_002645.1;NP_073549.1;HCoV-229E_1ab | 2702 | 2702 | 1 | 0.05 | 0.05 | 0.05 |
| NC_002645.1;NP_073549.1;HCoV-229E_1ab | 2707 | 2707 | 1 | 0.05 | 0.05 | 0.05 |
| NC_002645.1;NP_073549.1;HCoV-229E_1ab | 2709 | 2709 | 1 | 0.05 | 0.05 | 0.05 |
| NC_002645.1;NP_073549.1;HCoV-229E_1ab | 2712 | 2712 | 1 | 0.05 | 0.05 | 0.05 |
| NC_002645.1;NP_073549.1;HCoV-229E_1ab | 2715 | 2715 | 1 | 0.05 | 0.05 | 0.05 |
| NC_002645.1;NP_073549.1;HCoV-229E_1ab | 2735 | 2735 | 1 | 0.05 | 0.05 | 0.05 |
| NC_002645.1;NP_073549.1;HCoV-229E_1ab | 2859 | 2859 | 1 | 0.05 | 0.05 | 0.05 |
| NC_002645.1;NP_073549.1;HCoV-229E_1ab | 2861 | 2867 | 7 | 0.05 | 0.15 | 0.085714286 |
| NC_002645.1;NP_073549.1;HCoV-229E_1ab | 2869 | 2870 | 2 | 0.05 | 0.05 | 0.05 |
| NC_002645.1;NP_073549.1;HCoV-229E_1ab | 2889 | 2891 | 3 | 0.05 | 0.05 | 0.05 |
| NC_002645.1;NP_073549.1;HCoV-229E_1ab | 2895 | 2897 | 3 | 0.05 | 0.05 | 0.05 |
| NC_002645.1;NP_073549.1;HCoV-229E_1ab | 2903 | 2903 | 1 | 0.05 | 0.05 | 0.05 |
| NC_002645.1;NP_073549.1;HCoV-229E_1ab | 2913 | 2914 | 2 | 0.05 | 0.25 | 0.15 |
| NC_002645.1;NP_073549.1;HCoV-229E_1ab | 2917 | 2918 | 2 | 0.05 | 0.05 | 0.05 |
| NC_002645.1;NP_073549.1;HCoV-229E_1ab | 2920 | 2920 | 1 | 0.05 | 0.05 | 0.05 |
| NC_002645.1;NP_073549.1;HCoV-229E_1ab | 2925 | 2925 | 1 | 0.05 | 0.05 | 0.05 |
| NC_002645.1;NP_073549.1;HCoV-229E_1ab | 2935 | 2936 | 2 | 0.05 | 0.05 | 0.05 |
| NC_002645.1;NP_073549.1;HCoV-229E_1ab | 2940 | 2940 | 1 | 0.05 | 0.05 | 0.05 |
| NC_002645.1;NP_073549.1;HCoV-229E_1ab | 3062 | 3062 | 1 | 0.05 | 0.05 | 0.05 |
| NC_002645.1;NP_073549.1;HCoV-229E_1ab | 3102 | 3111 | 10 | 0.05 | 0.1 | 0.055 |
| NC_002645.1;NP_073549.1;HCoV-229E_1ab | 3125 | 3125 | 1 | 0.05 | 0.05 | 0.05 |
| NC_002645.1;NP_073549.1;HCoV-229E_1ab | 3133 | 3133 | 1 | 0.1 | 0.1 | 0.1 |

| Extended data 1 | | | | | | |
|---|---|---|---|---|---|---|
| Protein | Start position (peptide) | Stop position (peptide) | Number of peptides | Minimum fraction (per peptide) of control subjects with reactivity >3 sd above average | Maximum fraction (per peptide) of control subjects with reactivity >3 sd above average | Mean fraction (over all peptides in this contiguous stretch) of controlsubjects with reactivity >3 sd above average |
| NC_002645.1;NP_073549.1;HCoV-229E_1ab | 3136 | 3148 | 13 | 0.05 | 0.05 | 0.05 |
| NC_002645.1;NP_073549.1;HCoV-229E_1ab | 3155 | 3156 | 2 | 0.05 | 0.05 | 0.05 |
| NC_002645.1;NP_073549.1;HCoV-229E_1ab | 3173 | 3173 | 1 | 0.05 | 0.05 | 0.05 |
| NC_002645.1;NP_073549.1;HCoV-229E_1ab | 3193 | 3193 | 1 | 0.05 | 0.05 | 0.05 |
| NC_002645.1;NP_073549.1;HCoV-229E_1ab | 3202 | 3202 | 1 | 0.05 | 0.05 | 0.05 |
| NC_002645.1;NP_073549.1;HCoV-229E_1ab | 3224 | 3224 | 1 | 0.1 | 0.1 | 0.1 |
| NC_002645.1;NP_073549.1;HCoV-229E_1ab | 3229 | 3229 | 1 | 0.05 | 0.05 | 0.05 |
| NC_002645.1;NP_073549.1;HCoV-229E_1ab | 3236 | 3238 | 3 | 0.05 | 0.2 | 0.133333333 |
| NC_002645.1;NP_073549.1;HCoV-229E_1ab | 3240 | 3243 | 4 | 0.05 | 0.4 | 0.225 |
| NC_002645.1;NP_073549.1;HCoV-229E_1ab | 3245 | 3248 | 4 | 0.05 | 0.55 | 0.225 |
| NC_002645.1;NP_073549.1;HCoV-229E_1ab | 3252 | 3252 | 1 | 0.05 | 0.05 | 0.05 |
| NC_002645.1;NP_073549.1;HCoV-229E_1ab | 3254 | 3254 | 1 | 0.05 | 0.05 | 0.05 |
| NC_002645.1;NP_073549.1;HCoV-229E_1ab | 3263 | 3263 | 1 | 0.05 | 0.05 | 0.05 |
| NC_002645.1;NP_073549.1;HCoV-229E_1ab | 3350 | 3362 | 13 | 0.05 | 0.15 | 0.084615385 |
| NC_002645.1;NP_073549.1;HCoV-229E_1ab | 3455 | 3455 | 1 | 0.05 | 0.05 | 0.05 |
| NC_002645.1;NP_073549.1;HCoV-229E_1ab | 3496 | 3497 | 2 | 0.05 | 0.05 | 0.05 |
| NC_002645.1;NP_073549.1;HCoV-229E_1ab | 3502 | 3511 | 10 | 0.05 | 0.35 | 0.135 |
| NC_002645.1;NP_073549.1;HCoV-229E_1ab | 3533 | 3541 | 9 | 0.05 | 0.1 | 0.055555556 |
| NC_002645.1;NP_073549.1;HCoV-229E_1ab | 3548 | 3549 | 2 | 0.1 | 0.15 | 0.125 |
| NC_002645.1;NP_073549.1;HCoV-229E_1ab | 3555 | 3555 | 1 | 0.05 | 0.05 | 0.05 |
| NC_002645.1;NP_073549.1;HCoV-229E_1ab | 3559 | 3564 | 6 | 0.05 | 0.1 | 0.075 |
| NC_002645.1;NP_073549.1;HCoV-229E_1ab | 3567 | 3570 | 4 | 0.05 | 0.1 | 0.0875 |
| NC_002645.1;NP_073549.1;HCoV-229E_1ab | 3579 | 3579 | 1 | 0.05 | 0.05 | 0.05 |
| NC_002645.1;NP_073549.1;HCoV-229E_1ab | 3602 | 3617 | 16 | 0.05 | 0.55 | 0.1875 |
| NC_002645.1;NP_073549.1;HCoV-229E_1ab | 3631 | 3641 | 11 | 0.05 | 0.1 | 0.077272727 |
| NC_002645.1;NP_073549.1;HCoV-229E_1ab | 3645 | 3645 | 1 | 0.05 | 0.05 | 0.05 |
| NC_002645.1;NP_073549.1;HCoV-229E_1ab | 3669 | 3670 | 2 | 0.05 | 0.05 | 0.05 |
| NC_002645.1;NP_073549.1;HCoV-229E_1ab | 3688 | 3693 | 6 | 0.05 | 0.15 | 0.075 |
| NC_002645.1;NP_073549.1;HCoV-229E_1ab | 3731 | 3733 | 3 | 0.05 | 0.15 | 0.083333333 |
| NC_002645.1;NP_073549.1;HCoV-229E_1ab | 3740 | 3740 | 1 | 0.05 | 0.05 | 0.05 |
| NC_002645.1;NP_073549.1;HCoV-229E_1ab | 3758 | 3758 | 1 | 0.05 | 0.05 | 0.05 |
| NC_002645.1;NP_073549.1;HCoV-229E_1ab | 3765 | 3765 | 1 | 0.1 | 0.1 | 0.1 |
| NC_002645.1;NP_073549.1;HCoV-229E_1ab | 3788 | 3789 | 2 | 0.05 | 0.05 | 0.05 |
| NC_002645.1;NP_073549.1;HCoV-229E_1ab | 3798 | 3800 | 3 | 0.05 | 0.05 | 0.05 |
| NC_002645.1;NP_073549.1;HCoV-229E_1ab | 3820 | 3820 | 1 | 0.05 | 0.05 | 0.05 |
| NC_002645.1;NP_073549.1;HCoV-229E_1ab | 3841 | 3845 | 5 | 0.05 | 0.15 | 0.08 |
| NC_002645.1;NP_073549.1;HCoV-229E_1ab | 3851 | 3851 | 1 | 0.05 | 0.05 | 0.05 |

| Extended data 1 | | | | | | |
|---|---|---|---|---|---|---|
| Protein | Start position (peptide) | Stop position (peptide) | Number of peptides | Minimum fraction (per peptide) of control subjects with reactivity >3 sd above average | Maximum fraction (per peptide) of control subjects with reactivity >3 sd above average | Mean fraction (over all peptides in this contiguous stretch) of controlsubjects with reactivity >3 sd above average |
| NC_002645.1;NP_073549.1;HCoV-229E_1ab | 3857 | 3857 | 1 | 0.05 | 0.05 | 0.05 |
| NC_002645.1;NP_073549.1;HCoV-229E_1ab | 3866 | 3869 | 4 | 0.05 | 0.05 | 0.05 |
| NC_002645.1;NP_073549.1;HCoV-229E_1ab | 3887 | 3887 | 1 | 0.05 | 0.05 | 0.05 |
| NC_002645.1;NP_073549.1;HCoV-229E_1ab | 3895 | 3895 | 1 | 0.05 | 0.05 | 0.05 |
| NC_002645.1;NP_073549.1;HCoV-229E_1ab | 3923 | 3928 | 6 | 0.05 | 0.05 | 0.05 |
| NC_002645.1;NP_073549.1;HCoV-229E_1ab | 3943 | 3944 | 2 | 0.05 | 0.05 | 0.05 |
| NC_002645.1;NP_073549.1;HCoV-229E_1ab | 3947 | 3958 | 12 | 0.05 | 0.2 | 0.1 |
| NC_002645.1;NP_073549.1;HCoV-229E_1ab | 3962 | 3963 | 2 | 0.05 | 0.05 | 0.05 |
| NC_002645.1;NP_073549.1;HCoV-229E_1ab | 3971 | 3971 | 1 | 0.05 | 0.05 | 0.05 |
| NC_002645.1;NP_073549.1;HCoV-229E_1ab | 3986 | 3986 | 1 | 0.05 | 0.05 | 0.05 |
| NC_002645.1;NP_073549.1;HCoV-229E_1ab | 3988 | 3988 | 1 | 0.1 | 0.1 | 0.1 |
| NC_002645.1;NP_073549.1;HCoV-229E_1ab | 3990 | 3990 | 1 | 0.05 | 0.05 | 0.05 |
| NC_002645.1;NP_073549.1;HCoV-229E_1ab | 3993 | 3993 | 1 | 0.05 | 0.05 | 0.05 |
| NC_002645.1;NP_073549.1;HCoV-229E_1ab | 4023 | 4024 | 2 | 0.05 | 0.05 | 0.05 |
| NC_002645.1;NP_073549.1;HCoV-229E_1ab | 4031 | 4032 | 2 | 0.05 | 0.05 | 0.05 |
| NC_002645.1;NP_073549.1;HCoV-229E_1ab | 4034 | 4034 | 1 | 0.05 | 0.05 | 0.05 |
| NC_002645.1;NP_073549.1;HCoV-229E_1ab | 4061 | 4063 | 3 | 0.05 | 0.05 | 0.05 |
| NC_002645.1;NP_073549.1;HCoV-229E_1ab | 4068 | 4069 | 2 | 0.05 | 0.05 | 0.05 |
| NC_002645.1;NP_073549.1;HCoV-229E_1ab | 4073 | 4074 | 2 | 0.05 | 0.05 | 0.05 |
| NC_002645.1;NP_073549.1;HCoV-229E_1ab | 4092 | 4092 | 1 | 0.05 | 0.05 | 0.05 |
| NC_002645.1;NP_073549.1;HCoV-229E_1ab | 4098 | 4107 | 10 | 0.05 | 0.15 | 0.075 |
| NC_002645.1;NP_073549.1;HCoV-229E_1ab | 4114 | 4117 | 4 | 0.05 | 0.1 | 0.0625 |
| NC_002645.1;NP_073549.1;HCoV-229E_1ab | 4120 | 4121 | 2 | 0.05 | 0.05 | 0.05 |
| NC_002645.1;NP_073549.1;HCoV-229E_1ab | 4132 | 4132 | 1 | 0.05 | 0.05 | 0.05 |
| NC_002645.1;NP_073549.1;HCoV-229E_1ab | 4135 | 4135 | 1 | 0.1 | 0.1 | 0.1 |
| NC_002645.1;NP_073549.1;HCoV-229E_1ab | 4138 | 4138 | 1 | 0.05 | 0.05 | 0.05 |
| NC_002645.1;NP_073549.1;HCoV-229E_1ab | 4140 | 4140 | 1 | 0.05 | 0.05 | 0.05 |
| NC_002645.1;NP_073549.1;HCoV-229E_1ab | 4185 | 4185 | 1 | 0.1 | 0.1 | 0.1 |
| NC_002645.1;NP_073549.1;HCoV-229E_1ab | 4190 | 4193 | 4 | 0.05 | 0.1 | 0.0625 |
| NC_002645.1;NP_073549.1;HCoV-229E_1ab | 4197 | 4197 | 1 | 0.05 | 0.05 | 0.05 |
| NC_002645.1;NP_073549.1;HCoV-229E_1ab | 4207 | 4216 | 10 | 0.15 | 0.2 | 0.17 |
| NC_002645.1;NP_073549.1;HCoV-229E_1ab | 4218 | 4218 | 1 | 0.05 | 0.05 | 0.05 |
| NC_002645.1;NP_073549.1;HCoV-229E_1ab | 4224 | 4224 | 1 | 0.05 | 0.05 | 0.05 |
| NC_002645.1;NP_073549.1;HCoV-229E_1ab | 4233 | 4233 | 1 | 0.05 | 0.05 | 0.05 |
| NC_002645.1;NP_073549.1;HCoV-229E_1ab | 4235 | 4235 | 1 | 0.1 | 0.1 | 0.1 |
| NC_002645.1;NP_073549.1;HCoV-229E_1ab | 4249 | 4251 | 3 | 0.05 | 0.05 | 0.05 |
| NC_002645.1;NP_073549.1;HCoV-229E_1ab | 4261 | 4261 | 1 | 0.05 | 0.05 | 0.05 |

| Extended data 1 | | | | | | |
|---|---|---|---|---|---|---|
| Protein | Start position (peptide) | Stop position (peptide) | Number of peptides | Minimum fraction (per peptide) of control subjects with reactivity >3 sd above average | Maximum fraction (per peptide) of control subjects with reactivity >3 sd above average | Mean fraction (over all peptides in this contiguous stretch) of controlsubjects with reactivity >3 sd above average |
| NC_002645.1;NP_073549.1;HCoV-229E_1ab | 4265 | 4265 | 1 | 0.1 | 0.1 | 0.1 |
| NC_002645.1;NP_073549.1;HCoV-229E_1ab | 4313 | 4320 | 8 | 0.1 | 0.2 | 0.11875 |
| NC_002645.1;NP_073549.1;HCoV-229E_1ab | 4325 | 4325 | 1 | 0.05 | 0.05 | 0.05 |
| NC_002645.1;NP_073549.1;HCoV-229E_1ab | 4333 | 4335 | 3 | 0.05 | 0.1 | 0.066666667 |
| NC_002645.1;NP_073549.1;HCoV-229E_1ab | 4412 | 4412 | 1 | 0.05 | 0.05 | 0.05 |
| NC_002645.1;NP_073549.1;HCoV-229E_1ab | 4448 | 4448 | 1 | 0.05 | 0.05 | 0.05 |
| NC_002645.1;NP_073549.1;HCoV-229E_1ab | 4463 | 4463 | 1 | 0.1 | 0.1 | 0.1 |
| NC_002645.1;NP_073549.1;HCoV-229E_1ab | 4467 | 4476 | 10 | 0.15 | 0.25 | 0.18 |
| NC_002645.1;NP_073549.1;HCoV-229E_1ab | 4478 | 4478 | 1 | 0.05 | 0.05 | 0.05 |
| NC_002645.1;NP_073549.1;HCoV-229E_1ab | 4486 | 4486 | 1 | 0.05 | 0.05 | 0.05 |
| NC_002645.1;NP_073549.1;HCoV-229E_1ab | 4488 | 4490 | 3 | 0.05 | 0.1 | 0.066666667 |
| NC_002645.1;NP_073549.1;HCoV-229E_1ab | 4497 | 4497 | 1 | 0.05 | 0.05 | 0.05 |
| NC_002645.1;NP_073549.1;HCoV-229E_1ab | 4507 | 4507 | 1 | 0.05 | 0.05 | 0.05 |
| NC_002645.1;NP_073549.1;HCoV-229E_1ab | 4510 | 4511 | 2 | 0.05 | 0.15 | 0.1 |
| NC_002645.1;NP_073549.1;HCoV-229E_1ab | 4513 | 4515 | 3 | 0.05 | 0.1 | 0.083333333 |
| NC_002645.1;NP_073549.1;HCoV-229E_1ab | 4520 | 4520 | 1 | 0.05 | 0.05 | 0.05 |
| NC_002645.1;NP_073549.1;HCoV-229E_1ab | 4525 | 4525 | 1 | 0.05 | 0.05 | 0.05 |
| NC_002645.1;NP_073549.1;HCoV-229E_1ab | 4547 | 4563 | 17 | 0.05 | 0.25 | 0.117647059 |
| NC_002645.1;NP_073549.1;HCoV-229E_1ab | 4565 | 4565 | 1 | 0.05 | 0.05 | 0.05 |
| NC_002645.1;NP_073549.1;HCoV-229E_1ab | 4580 | 4580 | 1 | 0.05 | 0.05 | 0.05 |
| NC_002645.1;NP_073549.1;HCoV-229E_1ab | 4582 | 4584 | 3 | 0.05 | 0.05 | 0.05 |
| NC_002645.1;NP_073549.1;HCoV-229E_1ab | 4586 | 4586 | 1 | 0.05 | 0.05 | 0.05 |
| NC_002645.1;NP_073549.1;HCoV-229E_1ab | 4594 | 4595 | 2 | 0.05 | 0.15 | 0.1 |
| NC_002645.1;NP_073549.1;HCoV-229E_1ab | 4600 | 4602 | 3 | 0.05 | 0.05 | 0.05 |
| NC_002645.1;NP_073549.1;HCoV-229E_1ab | 4611 | 4611 | 1 | 0.1 | 0.1 | 0.1 |
| NC_002645.1;NP_073549.1;HCoV-229E_1ab | 4652 | 4652 | 1 | 0.05 | 0.05 | 0.05 |
| NC_002645.1;NP_073549.1;HCoV-229E_1ab | 4656 | 4658 | 3 | 0.05 | 0.15 | 0.083333333 |
| NC_002645.1;NP_073549.1;HCoV-229E_1ab | 4663 | 4674 | 12 | 0.05 | 0.15 | 0.066666667 |
| NC_002645.1;NP_073549.1;HCoV-229E_1ab | 4701 | 4701 | 1 | 0.1 | 0.1 | 0.1 |
| NC_002645.1;NP_073549.1;HCoV-229E_1ab | 4711 | 4711 | 1 | 0.05 | 0.05 | 0.05 |
| NC_002645.1;NP_073549.1;HCoV-229E_1ab | 4720 | 4720 | 1 | 0.05 | 0.05 | 0.05 |
| NC_002645.1;NP_073549.1;HCoV-229E_1ab | 4731 | 4732 | 2 | 0.05 | 0.05 | 0.05 |
| NC_002645.1;NP_073549.1;HCoV-229E_1ab | 4737 | 4743 | 7 | 0.05 | 0.05 | 0.05 |
| NC_002645.1;NP_073549.1;HCoV-229E_1ab | 4762 | 4762 | 1 | 0.15 | 0.15 | 0.15 |
| NC_002645.1;NP_073549.1;HCoV-229E_1ab | 4767 | 4774 | 8 | 0.05 | 0.2 | 0.1125 |
| NC_002645.1;NP_073549.1;HCoV-229E_1ab | 4778 | 4779 | 2 | 0.05 | 0.05 | 0.05 |
| NC_002645.1;NP_073549.1;HCoV-229E_1ab | 4795 | 4796 | 2 | 0.05 | 0.05 | 0.05 |

| Extended data 1 | | | | | | |
|---|---|---|---|---|---|---|
| Protein | Start position (peptide) | Stop position (peptide) | Number of peptides | Minimum fraction (per peptide) of control subjects with reactivity >3 sd above average | Maximum fraction (per peptide) of control subjects with reactivity >3 sd above average | Mean fraction (over all peptides in this contiguous stretch) of controlsubjects with reactivity >3 sd above average |
| NC_002645.1;NP_073549.1;HCoV-229E_1ab | 4798 | 4803 | 6 | 0.05 | 0.1 | 0.066666667 |
| NC_002645.1;NP_073549.1;HCoV-229E_1ab | 4818 | 4823 | 6 | 0.05 | 0.05 | 0.05 |
| NC_002645.1;NP_073549.1;HCoV-229E_1ab | 4829 | 4829 | 1 | 0.05 | 0.05 | 0.05 |
| NC_002645.1;NP_073549.1;HCoV-229E_1ab | 4832 | 4832 | 1 | 0.05 | 0.05 | 0.05 |
| NC_002645.1;NP_073549.1;HCoV-229E_1ab | 4866 | 4866 | 1 | 0.05 | 0.05 | 0.05 |
| NC_002645.1;NP_073549.1;HCoV-229E_1ab | 4876 | 4876 | 1 | 0.05 | 0.05 | 0.05 |
| NC_002645.1;NP_073549.1;HCoV-229E_1ab | 4878 | 4878 | 1 | 0.05 | 0.05 | 0.05 |
| NC_002645.1;NP_073549.1;HCoV-229E_1ab | 4881 | 4881 | 1 | 0.05 | 0.05 | 0.05 |
| NC_002645.1;NP_073549.1;HCoV-229E_1ab | 4883 | 4883 | 1 | 0.05 | 0.05 | 0.05 |
| NC_002645.1;NP_073549.1;HCoV-229E_1ab | 4897 | 4903 | 7 | 0.05 | 0.15 | 0.085714286 |
| NC_002645.1;NP_073549.1;HCoV-229E_1ab | 4908 | 4908 | 1 | 0.05 | 0.05 | 0.05 |
| NC_002645.1;NP_073549.1;HCoV-229E_1ab | 4944 | 4954 | 11 | 0.1 | 0.25 | 0.154545455 |
| NC_002645.1;NP_073549.1;HCoV-229E_1ab | 4967 | 4967 | 1 | 0.05 | 0.05 | 0.05 |
| NC_002645.1;NP_073549.1;HCoV-229E_1ab | 4969 | 4969 | 1 | 0.05 | 0.05 | 0.05 |
| NC_002645.1;NP_073549.1;HCoV-229E_1ab | 4975 | 4976 | 2 | 0.05 | 0.05 | 0.05 |
| NC_002645.1;NP_073549.1;HCoV-229E_1ab | 4980 | 4980 | 1 | 0.05 | 0.05 | 0.05 |
| NC_002645.1;NP_073549.1;HCoV-229E_1ab | 4982 | 4982 | 1 | 0.05 | 0.05 | 0.05 |
| NC_002645.1;NP_073549.1;HCoV-229E_1ab | 5042 | 5047 | 6 | 0.05 | 0.1 | 0.058333333 |
| NC_002645.1;NP_073549.1;HCoV-229E_1ab | 5082 | 5097 | 16 | 0.05 | 0.15 | 0.075 |
| NC_002645.1;NP_073549.1;HCoV-229E_1ab | 5106 | 5115 | 10 | 0.05 | 0.2 | 0.125 |
| NC_002645.1;NP_073549.1;HCoV-229E_1ab | 5122 | 5123 | 2 | 0.05 | 0.05 | 0.05 |
| NC_002645.1;NP_073549.1;HCoV-229E_1ab | 5130 | 5130 | 1 | 0.1 | 0.1 | 0.1 |
| NC_002645.1;NP_073549.1;HCoV-229E_1ab | 5132 | 5134 | 3 | 0.05 | 0.05 | 0.05 |
| NC_002645.1;NP_073549.1;HCoV-229E_1ab | 5136 | 5137 | 2 | 0.05 | 0.1 | 0.075 |
| NC_002645.1;NP_073549.1;HCoV-229E_1ab | 5141 | 5141 | 1 | 0.05 | 0.05 | 0.05 |
| NC_002645.1;NP_073549.1;HCoV-229E_1ab | 5145 | 5145 | 1 | 0.05 | 0.05 | 0.05 |
| NC_002645.1;NP_073549.1;HCoV-229E_1ab | 5163 | 5163 | 1 | 0.05 | 0.05 | 0.05 |
| NC_002645.1;NP_073549.1;HCoV-229E_1ab | 5190 | 5193 | 4 | 0.05 | 0.05 | 0.05 |
| NC_002645.1;NP_073549.1;HCoV-229E_1ab | 5197 | 5198 | 2 | 0.05 | 0.05 | 0.05 |
| NC_002645.1;NP_073549.1;HCoV-229E_1ab | 5205 | 5206 | 2 | 0.05 | 0.05 | 0.05 |
| NC_002645.1;NP_073549.1;HCoV-229E_1ab | 5229 | 5230 | 2 | 0.05 | 0.1 | 0.075 |
| NC_002645.1;NP_073549.1;HCoV-229E_1ab | 5267 | 5267 | 1 | 0.05 | 0.05 | 0.05 |
| NC_002645.1;NP_073549.1;HCoV-229E_1ab | 5271 | 5271 | 1 | 0.05 | 0.05 | 0.05 |
| NC_002645.1;NP_073549.1;HCoV-229E_1ab | 5294 | 5294 | 1 | 0.05 | 0.05 | 0.05 |
| NC_002645.1;NP_073549.1;HCoV-229E_1ab | 5331 | 5331 | 1 | 0.05 | 0.05 | 0.05 |
| NC_002645.1;NP_073549.1;HCoV-229E_1ab | 5336 | 5345 | 10 | 0.05 | 0.1 | 0.065 |
| NC_002645.1;NP_073549.1;HCoV-229E_1ab | 5348 | 5349 | 2 | 0.05 | 0.1 | 0.075 |

| Extended data 1 | | | | | | |
|---|---|---|---|---|---|---|
| Protein | Start position (peptide) | Stop position (peptide) | Number of peptides | Minimum fraction (per peptide) of control subjects with reactivity >3 sd above average | Maximum fraction (per peptide) of control subjects with reactivity >3 sd above average | Mean fraction (over all peptides in this contiguous stretch) of controlsubjects with reactivity >3 sd above average |
| NC_002645.1;NP_073549.1;HCoV-229E_1ab | 5352 | 5352 | 1 | 0.05 | 0.05 | 0.05 |
| NC_002645.1;NP_073549.1;HCoV-229E_1ab | 5355 | 5363 | 9 | 0.05 | 0.1 | 0.072222222 |
| NC_002645.1;NP_073549.1;HCoV-229E_1ab | 5366 | 5369 | 4 | 0.05 | 0.05 | 0.05 |
| NC_002645.1;NP_073549.1;HCoV-229E_1ab | 5408 | 5408 | 1 | 0.05 | 0.05 | 0.05 |
| NC_002645.1;NP_073549.1;HCoV-229E_1ab | 5413 | 5413 | 1 | 0.05 | 0.05 | 0.05 |
| NC_002645.1;NP_073549.1;HCoV-229E_1ab | 5442 | 5442 | 1 | 0.1 | 0.1 | 0.1 |
| NC_002645.1;NP_073549.1;HCoV-229E_1ab | 5444 | 5445 | 2 | 0.05 | 0.05 | 0.05 |
| NC_002645.1;NP_073549.1;HCoV-229E_1ab | 5448 | 5448 | 1 | 0.05 | 0.05 | 0.05 |
| NC_002645.1;NP_073549.1;HCoV-229E_1ab | 5476 | 5476 | 1 | 0.1 | 0.1 | 0.1 |
| NC_002645.1;NP_073549.1;HCoV-229E_1ab | 5479 | 5479 | 1 | 0.05 | 0.05 | 0.05 |
| NC_002645.1;NP_073549.1;HCoV-229E_1ab | 5500 | 5500 | 1 | 0.05 | 0.05 | 0.05 |
| NC_002645.1;NP_073549.1;HCoV-229E_1ab | 5503 | 5506 | 4 | 0.05 | 0.1 | 0.075 |
| NC_002645.1;NP_073549.1;HCoV-229E_1ab | 5523 | 5525 | 3 | 0.05 | 0.05 | 0.05 |
| NC_002645.1;NP_073549.1;HCoV-229E_1ab | 5543 | 5543 | 1 | 0.05 | 0.05 | 0.05 |
| NC_002645.1;NP_073549.1;HCoV-229E_1ab | 5545 | 5545 | 1 | 0.05 | 0.05 | 0.05 |
| NC_002645.1;NP_073549.1;HCoV-229E_1ab | 5554 | 5555 | 2 | 0.05 | 0.05 | 0.05 |
| NC_002645.1;NP_073549.1;HCoV-229E_1ab | 5589 | 5589 | 1 | 0.05 | 0.05 | 0.05 |
| NC_002645.1;NP_073549.1;HCoV-229E_1ab | 5596 | 5596 | 1 | 0.05 | 0.05 | 0.05 |
| NC_002645.1;NP_073549.1;HCoV-229E_1ab | 5598 | 5598 | 1 | 0.05 | 0.05 | 0.05 |
| NC_002645.1;NP_073549.1;HCoV-229E_1ab | 5603 | 5603 | 1 | 0.05 | 0.05 | 0.05 |
| NC_002645.1;NP_073549.1;HCoV-229E_1ab | 5606 | 5606 | 1 | 0.05 | 0.05 | 0.05 |
| NC_002645.1;NP_073549.1;HCoV-229E_1ab | 5618 | 5630 | 13 | 0.05 | 0.2 | 0.115384615 |
| NC_002645.1;NP_073549.1;HCoV-229E_1ab | 5683 | 5691 | 9 | 0.05 | 0.1 | 0.061111111 |
| NC_002645.1;NP_073549.1;HCoV-229E_1ab | 5695 | 5695 | 1 | 0.05 | 0.05 | 0.05 |
| NC_002645.1;NP_073549.1;HCoV-229E_1ab | 5699 | 5702 | 4 | 0.05 | 0.1 | 0.075 |
| NC_002645.1;NP_073549.1;HCoV-229E_1ab | 5706 | 5706 | 1 | 0.15 | 0.15 | 0.15 |
| NC_002645.1;NP_073549.1;HCoV-229E_1ab | 5720 | 5720 | 1 | 0.05 | 0.05 | 0.05 |
| NC_002645.1;NP_073549.1;HCoV-229E_1ab | 5727 | 5727 | 1 | 0.05 | 0.05 | 0.05 |
| NC_002645.1;NP_073549.1;HCoV-229E_1ab | 5756 | 5756 | 1 | 0.05 | 0.05 | 0.05 |
| NC_002645.1;NP_073549.1;HCoV-229E_1ab | 5807 | 5808 | 2 | 0.05 | 0.05 | 0.05 |
| NC_002645.1;NP_073549.1;HCoV-229E_1ab | 5812 | 5812 | 1 | 0.05 | 0.05 | 0.05 |
| NC_002645.1;NP_073549.1;HCoV-229E_1ab | 5846 | 5846 | 1 | 0.05 | 0.05 | 0.05 |
| NC_002645.1;NP_073549.1;HCoV-229E_1ab | 5877 | 5891 | 15 | 0.05 | 0.15 | 0.083333333 |
| NC_002645.1;NP_073549.1;HCoV-229E_1ab | 5907 | 5907 | 1 | 0.05 | 0.05 | 0.05 |
| NC_002645.1;NP_073549.1;HCoV-229E_1ab | 5919 | 5920 | 2 | 0.05 | 0.05 | 0.05 |
| NC_002645.1;NP_073549.1;HCoV-229E_1ab | 5923 | 5923 | 1 | 0.05 | 0.05 | 0.05 |
| NC_002645.1;NP_073549.1;HCoV-229E_1ab | 5933 | 5933 | 1 | 0.05 | 0.05 | 0.05 |

| Extended data 1 | | | | | | |
|---|---|---|---|---|---|---|
| Protein | Start position (peptide) | Stop position (peptide) | Number of peptides | Minimum fraction (per peptide) of control subjects with reactivity >3 sd above average | Maximum fraction (per peptide) of control subjects with reactivity >3 sd above average | Mean fraction (over all peptides in this contiguous stretch) of controlsubjects with reactivity >3 sd above average |
| NC_002645.1;NP_073549.1;HCoV-229E_1ab | 5935 | 5935 | 1 | 0.05 | 0.05 | 0.05 |
| NC_002645.1;NP_073549.1;HCoV-229E_1ab | 5938 | 5953 | 16 | 0.05 | 0.15 | 0.0625 |
| NC_002645.1;NP_073549.1;HCoV-229E_1ab | 5987 | 5987 | 1 | 0.1 | 0.1 | 0.1 |
| NC_002645.1;NP_073549.1;HCoV-229E_1ab | 5992 | 5993 | 2 | 0.05 | 0.1 | 0.075 |
| NC_002645.1;NP_073549.1;HCoV-229E_1ab | 6005 | 6010 | 6 | 0.05 | 0.05 | 0.05 |
| NC_002645.1;NP_073549.1;HCoV-229E_1ab | 6012 | 6013 | 2 | 0.05 | 0.05 | 0.05 |
| NC_002645.1;NP_073549.1;HCoV-229E_1ab | 6015 | 6015 | 1 | 0.05 | 0.05 | 0.05 |
| NC_002645.1;NP_073549.1;HCoV-229E_1ab | 6029 | 6030 | 2 | 0.05 | 0.05 | 0.05 |
| NC_002645.1;NP_073549.1;HCoV-229E_1ab | 6032 | 6032 | 1 | 0.05 | 0.05 | 0.05 |
| NC_002645.1;NP_073549.1;HCoV-229E_1ab | 6036 | 6036 | 1 | 0.1 | 0.1 | 0.1 |
| NC_002645.1;NP_073549.1;HCoV-229E_1ab | 6057 | 6057 | 1 | 0.05 | 0.05 | 0.05 |
| NC_002645.1;NP_073549.1;HCoV-229E_1ab | 6103 | 6103 | 1 | 0.05 | 0.05 | 0.05 |
| NC_002645.1;NP_073549.1;HCoV-229E_1ab | 6111 | 6111 | 1 | 0.2 | 0.2 | 0.2 |
| NC_002645.1;NP_073549.1;HCoV-229E_1ab | 6116 | 6124 | 9 | 0.05 | 0.1 | 0.055555556 |
| NC_002645.1;NP_073549.1;HCoV-229E_1ab | 6127 | 6127 | 1 | 0.05 | 0.05 | 0.05 |
| NC_002645.1;NP_073549.1;HCoV-229E_1ab | 6129 | 6129 | 1 | 0.05 | 0.05 | 0.05 |
| NC_002645.1;NP_073549.1;HCoV-229E_1ab | 6133 | 6133 | 1 | 0.05 | 0.05 | 0.05 |
| NC_002645.1;NP_073549.1;HCoV-229E_1ab | 6135 | 6135 | 1 | 0.05 | 0.05 | 0.05 |
| NC_002645.1;NP_073549.1;HCoV-229E_1ab | 6141 | 6149 | 9 | 0.05 | 0.05 | 0.05 |
| NC_002645.1;NP_073549.1;HCoV-229E_1ab | 6151 | 6151 | 1 | 0.1 | 0.1 | 0.1 |
| NC_002645.1;NP_073549.1;HCoV-229E_1ab | 6153 | 6153 | 1 | 0.05 | 0.05 | 0.05 |
| NC_002645.1;NP_073549.1;HCoV-229E_1ab | 6170 | 6170 | 1 | 0.15 | 0.15 | 0.15 |
| NC_002645.1;NP_073549.1;HCoV-229E_1ab | 6193 | 6193 | 1 | 0.05 | 0.05 | 0.05 |
| NC_002645.1;NP_073549.1;HCoV-229E_1ab | 6213 | 6213 | 1 | 0.1 | 0.1 | 0.1 |
| NC_002645.1;NP_073549.1;HCoV-229E_1ab | 6224 | 6224 | 1 | 0.05 | 0.05 | 0.05 |
| NC_002645.1;NP_073549.1;HCoV-229E_1ab | 6270 | 6274 | 5 | 0.05 | 0.25 | 0.11 |
| NC_002645.1;NP_073549.1;HCoV-229E_1ab | 6288 | 6290 | 3 | 0.05 | 0.05 | 0.05 |
| NC_002645.1;NP_073549.1;HCoV-229E_1ab | 6297 | 6297 | 1 | 0.05 | 0.05 | 0.05 |
| NC_002645.1;NP_073549.1;HCoV-229E_1ab | 6305 | 6307 | 3 | 0.05 | 0.05 | 0.05 |
| NC_002645.1;NP_073549.1;HCoV-229E_1ab | 6311 | 6311 | 1 | 0.05 | 0.05 | 0.05 |
| NC_002645.1;NP_073549.1;HCoV-229E_1ab | 6333 | 6333 | 1 | 0.05 | 0.05 | 0.05 |
| NC_002645.1;NP_073549.1;HCoV-229E_1ab | 6337 | 6338 | 2 | 0.1 | 0.1 | 0.1 |
| NC_002645.1;NP_073549.1;HCoV-229E_1ab | 6343 | 6343 | 1 | 0.05 | 0.05 | 0.05 |
| NC_002645.1;NP_073549.1;HCoV-229E_1ab | 6360 | 6368 | 9 | 0.05 | 0.1 | 0.061111111 |
| NC_002645.1;NP_073549.1;HCoV-229E_1ab | 6370 | 6370 | 1 | 0.1 | 0.1 | 0.1 |
| NC_002645.1;NP_073549.1;HCoV-229E_1ab | 6378 | 6378 | 1 | 0.1 | 0.1 | 0.1 |
| NC_002645.1;NP_073549.1;HCoV-229E_1ab | 6407 | 6407 | 1 | 0.05 | 0.05 | 0.05 |

| Extended data 1 | | | | | | |
|---|---|---|---|---|---|---|
| Protein | Start position (peptide) | Stop position (peptide) | Number of peptides | Minimum fraction (per peptide) of control subjects with reactivity >3 sd above average | Maximum fraction (per peptide) of control subjects with reactivity >3 sd above average | Mean fraction (over all peptides in this contiguous stretch) of controlsubjects with reactivity >3 sd above average |
| NC_002645.1;NP_073549.1;HCoV-229E_1ab | 6422 | 6422 | 1 | 0.05 | 0.05 | 0.05 |
| NC_002645.1;NP_073549.1;HCoV-229E_1ab | 6434 | 6434 | 1 | 0.05 | 0.05 | 0.05 |
| NC_002645.1;NP_073549.1;HCoV-229E_1ab | 6444 | 6444 | 1 | 0.05 | 0.05 | 0.05 |
| NC_002645.1;NP_073549.1;HCoV-229E_1ab | 6446 | 6446 | 1 | 0.05 | 0.05 | 0.05 |
| NC_002645.1;NP_073549.1;HCoV-229E_1ab | 6453 | 6454 | 2 | 0.05 | 0.05 | 0.05 |
| NC_002645.1;NP_073549.1;HCoV-229E_1ab | 6506 | 6509 | 4 | 0.05 | 0.05 | 0.05 |
| NC_002645.1;NP_073549.1;HCoV-229E_1ab | 6521 | 6528 | 8 | 0.05 | 0.1 | 0.0625 |
| NC_002645.1;NP_073549.1;HCoV-229E_1ab | 6548 | 6549 | 2 | 0.05 | 0.05 | 0.05 |
| NC_002645.1;NP_073549.1;HCoV-229E_1ab | 6576 | 6580 | 5 | 0.05 | 0.35 | 0.2 |
| NC_002645.1;NP_073549.1;HCoV-229E_1ab | 6584 | 6584 | 1 | 0.05 | 0.05 | 0.05 |
| NC_002645.1;NP_073549.1;HCoV-229E_1ab | 6592 | 6594 | 3 | 0.05 | 0.05 | 0.05 |
| NC_002645.1;NP_073549.1;HCoV-229E_1ab | 6596 | 6597 | 2 | 0.05 | 0.05 | 0.05 |
| NC_002645.1;NP_073549.1;HCoV-229E_1ab | 6626 | 6627 | 2 | 0.05 | 0.05 | 0.05 |
| NC_002645.1;NP_073549.1;HCoV-229E_1ab | 6665 | 6665 | 1 | 0.05 | 0.05 | 0.05 |
| NC_002645.1;NP_073549.1;HCoV-229E_1ab | 6671 | 6674 | 4 | 0.05 | 0.1 | 0.0625 |
| NC_002645.1;NP_073550.1;HCoV-229E_1a | 1 | 1 | 1 | 0.05 | 0.05 | 0.05 |
| NC_002645.1;NP_073550.1;HCoV-229E_1a | 8 | 8 | 1 | 0.05 | 0.05 | 0.05 |
| NC_002645.1;NP_073550.1;HCoV-229E_1a | 20 | 21 | 2 | 0.05 | 0.05 | 0.05 |
| NC_002645.1;NP_073550.1;HCoV-229E_1a | 46 | 46 | 1 | 0.05 | 0.05 | 0.05 |
| NC_002645.1;NP_073550.1;HCoV-229E_1a | 55 | 55 | 1 | 0.2 | 0.2 | 0.2 |
| NC_002645.1;NP_073550.1;HCoV-229E_1a | 58 | 58 | 1 | 0.05 | 0.05 | 0.05 |
| NC_002645.1;NP_073550.1;HCoV-229E_1a | 89 | 90 | 2 | 0.05 | 0.2 | 0.125 |
| NC_002645.1;NP_073550.1;HCoV-229E_1a | 95 | 97 | 3 | 0.05 | 0.1 | 0.066666667 |
| NC_002645.1;NP_073550.1;HCoV-229E_1a | 99 | 99 | 1 | 0.05 | 0.05 | 0.05 |
| NC_002645.1;NP_073550.1;HCoV-229E_1a | 103 | 103 | 1 | 0.05 | 0.05 | 0.05 |
| NC_002645.1;NP_073550.1;HCoV-229E_1a | 107 | 107 | 1 | 0.05 | 0.05 | 0.05 |
| NC_002645.1;NP_073550.1;HCoV-229E_1a | 113 | 121 | 9 | 0.05 | 0.05 | 0.05 |
| NC_002645.1;NP_073550.1;HCoV-229E_1a | 156 | 156 | 1 | 0.05 | 0.05 | 0.05 |
| NC_002645.1;NP_073550.1;HCoV-229E_1a | 163 | 164 | 2 | 0.05 | 0.05 | 0.05 |
| NC_002645.1;NP_073550.1;HCoV-229E_1a | 166 | 168 | 3 | 0.05 | 0.05 | 0.05 |
| NC_002645.1;NP_073550.1;HCoV-229E_1a | 170 | 171 | 2 | 0.05 | 0.05 | 0.05 |
| NC_002645.1;NP_073550.1;HCoV-229E_1a | 189 | 189 | 1 | 0.15 | 0.15 | 0.15 |
| NC_002645.1;NP_073550.1;HCoV-229E_1a | 195 | 195 | 1 | 0.05 | 0.05 | 0.05 |
| NC_002645.1;NP_073550.1;HCoV-229E_1a | 203 | 203 | 1 | 0.05 | 0.05 | 0.05 |
| NC_002645.1;NP_073550.1;HCoV-229E_1a | 207 | 207 | 1 | 0.05 | 0.05 | 0.05 |
| NC_002645.1;NP_073550.1;HCoV-229E_1a | 210 | 210 | 1 | 0.05 | 0.05 | 0.05 |
| NC_002645.1;NP_073550.1;HCoV-229E_1a | 222 | 222 | 1 | 0.05 | 0.05 | 0.05 |

| Extended data 1 | | | | | | |
|---|---|---|---|---|---|---|
| **Protein** | **Start position (peptide)** | **Stop position (peptide)** | **Number of peptides** | **Minimum fraction (per peptide) of control subjects with reactivity >3 sd above average** | **Maximum fraction (per peptide) of control subjects with reactivity >3 sd above average** | **Mean fraction (over all peptides in this contiguous stretch) of controlsubjects with reactivity >3 sd above average** |
| NC_002645.1;NP_073550.1;HCoV-229E_1a | 243 | 245 | 3 | 0.05 | 0.05 | 0.05 |
| NC_002645.1;NP_073550.1;HCoV-229E_1a | 247 | 251 | 5 | 0.05 | 0.05 | 0.05 |
| NC_002645.1;NP_073550.1;HCoV-229E_1a | 269 | 276 | 8 | 0.05 | 0.15 | 0.09375 |
| NC_002645.1;NP_073550.1;HCoV-229E_1a | 278 | 278 | 1 | 0.05 | 0.05 | 0.05 |
| NC_002645.1;NP_073550.1;HCoV-229E_1a | 361 | 361 | 1 | 0.05 | 0.05 | 0.05 |
| NC_002645.1;NP_073550.1;HCoV-229E_1a | 364 | 365 | 2 | 0.05 | 0.05 | 0.05 |
| NC_002645.1;NP_073550.1;HCoV-229E_1a | 384 | 384 | 1 | 0.05 | 0.05 | 0.05 |
| NC_002645.1;NP_073550.1;HCoV-229E_1a | 387 | 395 | 9 | 0.05 | 0.1 | 0.061111111 |
| NC_002645.1;NP_073550.1;HCoV-229E_1a | 405 | 406 | 2 | 0.05 | 0.05 | 0.05 |
| NC_002645.1;NP_073550.1;HCoV-229E_1a | 413 | 416 | 4 | 0.1 | 0.1 | 0.1 |
| NC_002645.1;NP_073550.1;HCoV-229E_1a | 443 | 443 | 1 | 0.05 | 0.05 | 0.05 |
| NC_002645.1;NP_073550.1;HCoV-229E_1a | 448 | 452 | 5 | 0.05 | 0.05 | 0.05 |
| NC_002645.1;NP_073550.1;HCoV-229E_1a | 474 | 475 | 2 | 0.05 | 0.05 | 0.05 |
| NC_002645.1;NP_073550.1;HCoV-229E_1a | 477 | 478 | 2 | 0.15 | 0.2 | 0.175 |
| NC_002645.1;NP_073550.1;HCoV-229E_1a | 544 | 544 | 1 | 0.1 | 0.1 | 0.1 |
| NC_002645.1;NP_073550.1;HCoV-229E_1a | 592 | 592 | 1 | 0.1 | 0.1 | 0.1 |
| NC_002645.1;NP_073550.1;HCoV-229E_1a | 594 | 594 | 1 | 0.05 | 0.05 | 0.05 |
| NC_002645.1;NP_073550.1;HCoV-229E_1a | 610 | 618 | 9 | 0.05 | 0.05 | 0.05 |
| NC_002645.1;NP_073550.1;HCoV-229E_1a | 627 | 629 | 3 | 0.05 | 0.05 | 0.05 |
| NC_002645.1;NP_073550.1;HCoV-229E_1a | 631 | 643 | 13 | 0.05 | 0.5 | 0.288461538 |
| NC_002645.1;NP_073550.1;HCoV-229E_1a | 653 | 653 | 1 | 0.05 | 0.05 | 0.05 |
| NC_002645.1;NP_073550.1;HCoV-229E_1a | 656 | 656 | 1 | 0.05 | 0.05 | 0.05 |
| NC_002645.1;NP_073550.1;HCoV-229E_1a | 658 | 659 | 2 | 0.05 | 0.05 | 0.05 |
| NC_002645.1;NP_073550.1;HCoV-229E_1a | 662 | 662 | 1 | 0.05 | 0.05 | 0.05 |
| NC_002645.1;NP_073550.1;HCoV-229E_1a | 665 | 672 | 8 | 0.05 | 0.1 | 0.0625 |
| NC_002645.1;NP_073550.1;HCoV-229E_1a | 681 | 681 | 1 | 0.05 | 0.05 | 0.05 |
| NC_002645.1;NP_073550.1;HCoV-229E_1a | 683 | 684 | 2 | 0.05 | 0.1 | 0.075 |
| NC_002645.1;NP_073550.1;HCoV-229E_1a | 686 | 695 | 10 | 0.05 | 0.1 | 0.065 |
| NC_002645.1;NP_073550.1;HCoV-229E_1a | 698 | 700 | 3 | 0.05 | 0.15 | 0.116666667 |
| NC_002645.1;NP_073550.1;HCoV-229E_1a | 704 | 704 | 1 | 0.05 | 0.05 | 0.05 |
| NC_002645.1;NP_073550.1;HCoV-229E_1a | 723 | 723 | 1 | 0.05 | 0.05 | 0.05 |
| NC_002645.1;NP_073550.1;HCoV-229E_1a | 738 | 738 | 1 | 0.05 | 0.05 | 0.05 |
| NC_002645.1;NP_073550.1;HCoV-229E_1a | 797 | 797 | 1 | 0.05 | 0.05 | 0.05 |
| NC_002645.1;NP_073550.1;HCoV-229E_1a | 824 | 827 | 4 | 0.05 | 0.1 | 0.075 |
| NC_002645.1;NP_073550.1;HCoV-229E_1a | 831 | 831 | 1 | 0.1 | 0.1 | 0.1 |
| NC_002645.1;NP_073550.1;HCoV-229E_1a | 852 | 852 | 1 | 0.05 | 0.05 | 0.05 |
| NC_002645.1;NP_073550.1;HCoV-229E_1a | 865 | 876 | 12 | 0.05 | 0.2 | 0.075 |

| Extended data 1 | | | | | | |
|---|---|---|---|---|---|---|
| Protein | Start position (peptide) | Stop position (peptide) | Number of peptides | Minimum fraction (per peptide) of control subjects with reactivity >3 sd above average | Maximum fraction (per peptide) of control subjects with reactivity >3 sd above average | Mean fraction (over all peptides in this contiguous stretch) of controlsubjects with reactivity >3 sd above average |
| NC_002645.1;NP_073550.1;HCoV-229E_1a | 879 | 881 | 3 | 0.05 | 0.05 | 0.05 |
| NC_002645.1;NP_073550.1;HCoV-229E_1a | 883 | 883 | 1 | 0.05 | 0.05 | 0.05 |
| NC_002645.1;NP_073550.1;HCoV-229E_1a | 900 | 901 | 2 | 0.05 | 0.1 | 0.075 |
| NC_002645.1;NP_073550.1;HCoV-229E_1a | 903 | 903 | 1 | 0.1 | 0.1 | 0.1 |
| NC_002645.1;NP_073550.1;HCoV-229E_1a | 906 | 906 | 1 | 0.05 | 0.05 | 0.05 |
| NC_002645.1;NP_073550.1;HCoV-229E_1a | 924 | 924 | 1 | 0.05 | 0.05 | 0.05 |
| NC_002645.1;NP_073550.1;HCoV-229E_1a | 972 | 972 | 1 | 0.05 | 0.05 | 0.05 |
| NC_002645.1;NP_073550.1;HCoV-229E_1a | 974 | 975 | 2 | 0.05 | 0.1 | 0.075 |
| NC_002645.1;NP_073550.1;HCoV-229E_1a | 983 | 983 | 1 | 0.05 | 0.05 | 0.05 |
| NC_002645.1;NP_073550.1;HCoV-229E_1a | 989 | 989 | 1 | 0.05 | 0.05 | 0.05 |
| NC_002645.1;NP_073550.1;HCoV-229E_1a | 997 | 1002 | 6 | 0.05 | 0.05 | 0.05 |
| NC_002645.1;NP_073550.1;HCoV-229E_1a | 1005 | 1009 | 5 | 0.05 | 0.2 | 0.12 |
| NC_002645.1;NP_073550.1;HCoV-229E_1a | 1011 | 1019 | 9 | 0.05 | 0.15 | 0.105555556 |
| NC_002645.1;NP_073550.1;HCoV-229E_1a | 1022 | 1022 | 1 | 0.05 | 0.05 | 0.05 |
| NC_002645.1;NP_073550.1;HCoV-229E_1a | 1024 | 1031 | 8 | 0.05 | 0.1 | 0.05625 |
| NC_002645.1;NP_073550.1;HCoV-229E_1a | 1038 | 1038 | 1 | 0.05 | 0.05 | 0.05 |
| NC_002645.1;NP_073550.1;HCoV-229E_1a | 1045 | 1045 | 1 | 0.05 | 0.05 | 0.05 |
| NC_002645.1;NP_073550.1;HCoV-229E_1a | 1064 | 1067 | 4 | 0.05 | 0.1 | 0.0625 |
| NC_002645.1;NP_073550.1;HCoV-229E_1a | 1088 | 1088 | 1 | 0.05 | 0.05 | 0.05 |
| NC_002645.1;NP_073550.1;HCoV-229E_1a | 1110 | 1110 | 1 | 0.05 | 0.05 | 0.05 |
| NC_002645.1;NP_073550.1;HCoV-229E_1a | 1112 | 1112 | 1 | 0.05 | 0.05 | 0.05 |
| NC_002645.1;NP_073550.1;HCoV-229E_1a | 1149 | 1149 | 1 | 0.05 | 0.05 | 0.05 |
| NC_002645.1;NP_073550.1;HCoV-229E_1a | 1173 | 1174 | 2 | 0.05 | 0.1 | 0.075 |
| NC_002645.1;NP_073550.1;HCoV-229E_1a | 1183 | 1183 | 1 | 0.05 | 0.05 | 0.05 |
| NC_002645.1;NP_073550.1;HCoV-229E_1a | 1208 | 1208 | 1 | 0.05 | 0.05 | 0.05 |
| NC_002645.1;NP_073550.1;HCoV-229E_1a | 1215 | 1227 | 13 | 0.05 | 0.15 | 0.065384615 |
| NC_002645.1;NP_073550.1;HCoV-229E_1a | 1231 | 1242 | 12 | 0.05 | 0.1 | 0.070833333 |
| NC_002645.1;NP_073550.1;HCoV-229E_1a | 1250 | 1250 | 1 | 0.05 | 0.05 | 0.05 |
| NC_002645.1;NP_073550.1;HCoV-229E_1a | 1257 | 1266 | 10 | 0.05 | 0.2 | 0.09 |
| NC_002645.1;NP_073550.1;HCoV-229E_1a | 1272 | 1288 | 17 | 0.05 | 0.15 | 0.061764706 |
| NC_002645.1;NP_073550.1;HCoV-229E_1a | 1292 | 1292 | 1 | 0.05 | 0.05 | 0.05 |
| NC_002645.1;NP_073550.1;HCoV-229E_1a | 1294 | 1302 | 9 | 0.05 | 0.3 | 0.083333333 |
| NC_002645.1;NP_073550.1;HCoV-229E_1a | 1325 | 1326 | 2 | 0.05 | 0.05 | 0.05 |
| NC_002645.1;NP_073550.1;HCoV-229E_1a | 1365 | 1375 | 11 | 0.1 | 0.55 | 0.313636364 |
| NC_002645.1;NP_073550.1;HCoV-229E_1a | 1429 | 1429 | 1 | 0.05 | 0.05 | 0.05 |
| NC_002645.1;NP_073550.1;HCoV-229E_1a | 1432 | 1445 | 14 | 0.05 | 0.2 | 0.128571429 |
| NC_002645.1;NP_073550.1;HCoV-229E_1a | 1467 | 1469 | 3 | 0.05 | 0.05 | 0.05 |

| Extended data 1 | | | | | | |
|---|---|---|---|---|---|---|
| Protein | Start position (peptide) | Stop position (peptide) | Number of peptides | Minimum fraction (per peptide) of control subjects with reactivity >3 sd above average | Maximum fraction (per peptide) of control subjects with reactivity >3 sd above average | Mean fraction (over all peptides in this contiguous stretch) of controlsubjects with reactivity >3 sd above average |
| NC_002645.1;NP_073550.1;HCoV-229E_1a | 1490 | 1490 | 1 | 0.05 | 0.05 | 0.05 |
| NC_002645.1;NP_073550.1;HCoV-229E_1a | 1492 | 1497 | 6 | 0.05 | 0.05 | 0.05 |
| NC_002645.1;NP_073550.1;HCoV-229E_1a | 1506 | 1506 | 1 | 0.1 | 0.1 | 0.1 |
| NC_002645.1;NP_073550.1;HCoV-229E_1a | 1510 | 1511 | 2 | 0.05 | 0.05 | 0.05 |
| NC_002645.1;NP_073550.1;HCoV-229E_1a | 1514 | 1514 | 1 | 0.1 | 0.1 | 0.1 |
| NC_002645.1;NP_073550.1;HCoV-229E_1a | 1516 | 1516 | 1 | 0.05 | 0.05 | 0.05 |
| NC_002645.1;NP_073550.1;HCoV-229E_1a | 1519 | 1519 | 1 | 0.05 | 0.05 | 0.05 |
| NC_002645.1;NP_073550.1;HCoV-229E_1a | 1521 | 1521 | 1 | 0.1 | 0.1 | 0.1 |
| NC_002645.1;NP_073550.1;HCoV-229E_1a | 1583 | 1583 | 1 | 0.05 | 0.05 | 0.05 |
| NC_002645.1;NP_073550.1;HCoV-229E_1a | 1586 | 1589 | 4 | 0.05 | 0.05 | 0.05 |
| NC_002645.1;NP_073550.1;HCoV-229E_1a | 1607 | 1607 | 1 | 0.05 | 0.05 | 0.05 |
| NC_002645.1;NP_073550.1;HCoV-229E_1a | 1611 | 1611 | 1 | 0.05 | 0.05 | 0.05 |
| NC_002645.1;NP_073550.1;HCoV-229E_1a | 1627 | 1627 | 1 | 0.05 | 0.05 | 0.05 |
| NC_002645.1;NP_073550.1;HCoV-229E_1a | 1631 | 1635 | 5 | 0.05 | 0.1 | 0.06 |
| NC_002645.1;NP_073550.1;HCoV-229E_1a | 1684 | 1684 | 1 | 0.05 | 0.05 | 0.05 |
| NC_002645.1;NP_073550.1;HCoV-229E_1a | 1690 | 1690 | 1 | 0.05 | 0.05 | 0.05 |
| NC_002645.1;NP_073550.1;HCoV-229E_1a | 1710 | 1710 | 1 | 0.05 | 0.05 | 0.05 |
| NC_002645.1;NP_073550.1;HCoV-229E_1a | 1718 | 1721 | 4 | 0.05 | 0.05 | 0.05 |
| NC_002645.1;NP_073550.1;HCoV-229E_1a | 1741 | 1741 | 1 | 0.1 | 0.1 | 0.1 |
| NC_002645.1;NP_073550.1;HCoV-229E_1a | 1753 | 1760 | 8 | 0.05 | 0.1 | 0.075 |
| NC_002645.1;NP_073550.1;HCoV-229E_1a | 1762 | 1762 | 1 | 0.05 | 0.05 | 0.05 |
| NC_002645.1;NP_073550.1;HCoV-229E_1a | 1764 | 1764 | 1 | 0.05 | 0.05 | 0.05 |
| NC_002645.1;NP_073550.1;HCoV-229E_1a | 1777 | 1778 | 2 | 0.05 | 0.05 | 0.05 |
| NC_002645.1;NP_073550.1;HCoV-229E_1a | 1783 | 1786 | 4 | 0.1 | 0.15 | 0.1375 |
| NC_002645.1;NP_073550.1;HCoV-229E_1a | 1797 | 1798 | 2 | 0.05 | 0.05 | 0.05 |
| NC_002645.1;NP_073550.1;HCoV-229E_1a | 1802 | 1802 | 1 | 0.05 | 0.05 | 0.05 |
| NC_002645.1;NP_073550.1;HCoV-229E_1a | 1804 | 1804 | 1 | 0.2 | 0.2 | 0.2 |
| NC_002645.1;NP_073550.1;HCoV-229E_1a | 1822 | 1826 | 5 | 0.05 | 0.1 | 0.07 |
| NC_002645.1;NP_073550.1;HCoV-229E_1a | 1832 | 1832 | 1 | 0.05 | 0.05 | 0.05 |
| NC_002645.1;NP_073550.1;HCoV-229E_1a | 1836 | 1836 | 1 | 0.05 | 0.05 | 0.05 |
| NC_002645.1;NP_073550.1;HCoV-229E_1a | 1849 | 1851 | 3 | 0.05 | 0.05 | 0.05 |
| NC_002645.1;NP_073550.1;HCoV-229E_1a | 1855 | 1855 | 1 | 0.05 | 0.05 | 0.05 |
| NC_002645.1;NP_073550.1;HCoV-229E_1a | 1866 | 1866 | 1 | 0.05 | 0.05 | 0.05 |
| NC_002645.1;NP_073550.1;HCoV-229E_1a | 1868 | 1868 | 1 | 0.15 | 0.15 | 0.15 |
| NC_002645.1;NP_073550.1;HCoV-229E_1a | 1870 | 1870 | 1 | 0.05 | 0.05 | 0.05 |
| NC_002645.1;NP_073550.1;HCoV-229E_1a | 1872 | 1872 | 1 | 0.05 | 0.05 | 0.05 |
| NC_002645.1;NP_073550.1;HCoV-229E_1a | 1899 | 1899 | 1 | 0.05 | 0.05 | 0.05 |

| Extended data 1 | | | | | | |
|---|---|---|---|---|---|---|
| **Protein** | **Start position (peptide)** | **Stop position (peptide)** | **Number of peptides** | **Minimum fraction (per peptide) of control subjects with reactivity >3 sd above average** | **Maximum fraction (per peptide) of control subjects with reactivity >3 sd above average** | **Mean fraction (over all peptides in this contiguous stretch) of controlsubjects with reactivity >3 sd above average** |
| NC_002645.1;NP_073550.1;HCoV-229E_1a | 1901 | 1901 | 1 | 0.05 | 0.05 | 0.05 |
| NC_002645.1;NP_073550.1;HCoV-229E_1a | 1903 | 1903 | 1 | 0.05 | 0.05 | 0.05 |
| NC_002645.1;NP_073550.1;HCoV-229E_1a | 1907 | 1907 | 1 | 0.05 | 0.05 | 0.05 |
| NC_002645.1;NP_073550.1;HCoV-229E_1a | 1909 | 1917 | 9 | 0.05 | 0.2 | 0.116666667 |
| NC_002645.1;NP_073550.1;HCoV-229E_1a | 1919 | 1919 | 1 | 0.05 | 0.05 | 0.05 |
| NC_002645.1;NP_073550.1;HCoV-229E_1a | 1942 | 1943 | 2 | 0.05 | 0.05 | 0.05 |
| NC_002645.1;NP_073550.1;HCoV-229E_1a | 1950 | 1950 | 1 | 0.05 | 0.05 | 0.05 |
| NC_002645.1;NP_073550.1;HCoV-229E_1a | 1968 | 1968 | 1 | 0.05 | 0.05 | 0.05 |
| NC_002645.1;NP_073550.1;HCoV-229E_1a | 2018 | 2019 | 2 | 0.05 | 0.05 | 0.05 |
| NC_002645.1;NP_073550.1;HCoV-229E_1a | 2022 | 2025 | 4 | 0.05 | 0.1 | 0.0625 |
| NC_002645.1;NP_073550.1;HCoV-229E_1a | 2036 | 2036 | 1 | 0.25 | 0.25 | 0.25 |
| NC_002645.1;NP_073550.1;HCoV-229E_1a | 2039 | 2040 | 2 | 0.1 | 0.1 | 0.1 |
| NC_002645.1;NP_073550.1;HCoV-229E_1a | 2103 | 2104 | 2 | 0.05 | 0.05 | 0.05 |
| NC_002645.1;NP_073550.1;HCoV-229E_1a | 2151 | 2151 | 1 | 0.05 | 0.05 | 0.05 |
| NC_002645.1;NP_073550.1;HCoV-229E_1a | 2173 | 2173 | 1 | 0.05 | 0.05 | 0.05 |
| NC_002645.1;NP_073550.1;HCoV-229E_1a | 2175 | 2175 | 1 | 0.05 | 0.05 | 0.05 |
| NC_002645.1;NP_073550.1;HCoV-229E_1a | 2208 | 2210 | 3 | 0.05 | 0.05 | 0.05 |
| NC_002645.1;NP_073550.1;HCoV-229E_1a | 2216 | 2224 | 9 | 0.05 | 0.15 | 0.061111111 |
| NC_002645.1;NP_073550.1;HCoV-229E_1a | 2228 | 2228 | 1 | 0.1 | 0.1 | 0.1 |
| NC_002645.1;NP_073550.1;HCoV-229E_1a | 2232 | 2234 | 3 | 0.05 | 0.3 | 0.15 |
| NC_002645.1;NP_073550.1;HCoV-229E_1a | 2255 | 2255 | 1 | 0.05 | 0.05 | 0.05 |
| NC_002645.1;NP_073550.1;HCoV-229E_1a | 2259 | 2261 | 3 | 0.05 | 0.1 | 0.066666667 |
| NC_002645.1;NP_073550.1;HCoV-229E_1a | 2266 | 2267 | 2 | 0.05 | 0.05 | 0.05 |
| NC_002645.1;NP_073550.1;HCoV-229E_1a | 2281 | 2293 | 13 | 0.05 | 0.25 | 0.134615385 |
| NC_002645.1;NP_073550.1;HCoV-229E_1a | 2314 | 2314 | 1 | 0.05 | 0.05 | 0.05 |
| NC_002645.1;NP_073550.1;HCoV-229E_1a | 2336 | 2347 | 12 | 0.05 | 0.1 | 0.066666667 |
| NC_002645.1;NP_073550.1;HCoV-229E_1a | 2355 | 2358 | 4 | 0.05 | 0.2 | 0.0875 |
| NC_002645.1;NP_073550.1;HCoV-229E_1a | 2363 | 2363 | 1 | 0.05 | 0.05 | 0.05 |
| NC_002645.1;NP_073550.1;HCoV-229E_1a | 2374 | 2374 | 1 | 0.05 | 0.05 | 0.05 |
| NC_002645.1;NP_073550.1;HCoV-229E_1a | 2383 | 2383 | 1 | 0.05 | 0.05 | 0.05 |
| NC_002645.1;NP_073550.1;HCoV-229E_1a | 2386 | 2386 | 1 | 0.1 | 0.1 | 0.1 |
| NC_002645.1;NP_073550.1;HCoV-229E_1a | 2388 | 2390 | 3 | 0.05 | 0.1 | 0.066666667 |
| NC_002645.1;NP_073550.1;HCoV-229E_1a | 2392 | 2392 | 1 | 0.05 | 0.05 | 0.05 |
| NC_002645.1;NP_073550.1;HCoV-229E_1a | 2415 | 2416 | 2 | 0.05 | 0.05 | 0.05 |
| NC_002645.1;NP_073550.1;HCoV-229E_1a | 2418 | 2418 | 1 | 0.05 | 0.05 | 0.05 |
| NC_002645.1;NP_073550.1;HCoV-229E_1a | 2427 | 2430 | 4 | 0.05 | 0.1 | 0.0625 |
| NC_002645.1;NP_073550.1;HCoV-229E_1a | 2432 | 2436 | 5 | 0.05 | 0.15 | 0.09 |

| Extended data 1 | | | | | | |
|---|---|---|---|---|---|---|
| Protein | Start position (peptide) | Stop position (peptide) | Number of peptides | Minimum fraction (per peptide) of control subjects with reactivity >3 sd above average | Maximum fraction (per peptide) of control subjects with reactivity >3 sd above average | Mean fraction (over all peptides in this contiguous stretch) of controlsubjects with reactivity >3 sd above average |
| NC_002645.1;NP_073550.1;HCoV-229E_1a | 2522 | 2537 | 16 | 0.05 | 0.2 | 0.090625 |
| NC_002645.1;NP_073550.1;HCoV-229E_1a | 2539 | 2540 | 2 | 0.05 | 0.1 | 0.075 |
| NC_002645.1;NP_073550.1;HCoV-229E_1a | 2545 | 2547 | 3 | 0.05 | 0.1 | 0.066666667 |
| NC_002645.1;NP_073550.1;HCoV-229E_1a | 2549 | 2549 | 1 | 0.1 | 0.1 | 0.1 |
| NC_002645.1;NP_073550.1;HCoV-229E_1a | 2576 | 2577 | 2 | 0.1 | 0.15 | 0.125 |
| NC_002645.1;NP_073550.1;HCoV-229E_1a | 2581 | 2581 | 1 | 0.05 | 0.05 | 0.05 |
| NC_002645.1;NP_073550.1;HCoV-229E_1a | 2623 | 2623 | 1 | 0.05 | 0.05 | 0.05 |
| NC_002645.1;NP_073550.1;HCoV-229E_1a | 2655 | 2658 | 4 | 0.1 | 0.2 | 0.15 |
| NC_002645.1;NP_073550.1;HCoV-229E_1a | 2661 | 2667 | 7 | 0.05 | 0.15 | 0.078571429 |
| NC_002645.1;NP_073550.1;HCoV-229E_1a | 2692 | 2696 | 5 | 0.05 | 0.05 | 0.05 |
| NC_002645.1;NP_073550.1;HCoV-229E_1a | 2702 | 2702 | 1 | 0.05 | 0.05 | 0.05 |
| NC_002645.1;NP_073550.1;HCoV-229E_1a | 2707 | 2707 | 1 | 0.05 | 0.05 | 0.05 |
| NC_002645.1;NP_073550.1;HCoV-229E_1a | 2709 | 2709 | 1 | 0.05 | 0.05 | 0.05 |
| NC_002645.1;NP_073550.1;HCoV-229E_1a | 2712 | 2712 | 1 | 0.05 | 0.05 | 0.05 |
| NC_002645.1;NP_073550.1;HCoV-229E_1a | 2715 | 2715 | 1 | 0.05 | 0.05 | 0.05 |
| NC_002645.1;NP_073550.1;HCoV-229E_1a | 2735 | 2735 | 1 | 0.05 | 0.05 | 0.05 |
| NC_002645.1;NP_073550.1;HCoV-229E_1a | 2859 | 2859 | 1 | 0.05 | 0.05 | 0.05 |
| NC_002645.1;NP_073550.1;HCoV-229E_1a | 2861 | 2867 | 7 | 0.05 | 0.15 | 0.085714286 |
| NC_002645.1;NP_073550.1;HCoV-229E_1a | 2869 | 2870 | 2 | 0.05 | 0.05 | 0.05 |
| NC_002645.1;NP_073550.1;HCoV-229E_1a | 2889 | 2891 | 3 | 0.05 | 0.05 | 0.05 |
| NC_002645.1;NP_073550.1;HCoV-229E_1a | 2895 | 2897 | 3 | 0.05 | 0.05 | 0.05 |
| NC_002645.1;NP_073550.1;HCoV-229E_1a | 2903 | 2903 | 1 | 0.05 | 0.05 | 0.05 |
| NC_002645.1;NP_073550.1;HCoV-229E_1a | 2913 | 2914 | 2 | 0.05 | 0.25 | 0.15 |
| NC_002645.1;NP_073550.1;HCoV-229E_1a | 2917 | 2918 | 2 | 0.05 | 0.05 | 0.05 |
| NC_002645.1;NP_073550.1;HCoV-229E_1a | 2920 | 2920 | 1 | 0.05 | 0.05 | 0.05 |
| NC_002645.1;NP_073550.1;HCoV-229E_1a | 2925 | 2925 | 1 | 0.05 | 0.05 | 0.05 |
| NC_002645.1;NP_073550.1;HCoV-229E_1a | 2935 | 2936 | 2 | 0.05 | 0.05 | 0.05 |
| NC_002645.1;NP_073550.1;HCoV-229E_1a | 2940 | 2940 | 1 | 0.05 | 0.05 | 0.05 |
| NC_002645.1;NP_073550.1;HCoV-229E_1a | 3062 | 3062 | 1 | 0.05 | 0.05 | 0.05 |
| NC_002645.1;NP_073550.1;HCoV-229E_1a | 3102 | 3111 | 10 | 0.05 | 0.1 | 0.055 |
| NC_002645.1;NP_073550.1;HCoV-229E_1a | 3125 | 3125 | 1 | 0.05 | 0.05 | 0.05 |
| NC_002645.1;NP_073550.1;HCoV-229E_1a | 3133 | 3133 | 1 | 0.1 | 0.1 | 0.1 |
| NC_002645.1;NP_073550.1;HCoV-229E_1a | 3136 | 3148 | 13 | 0.05 | 0.05 | 0.05 |
| NC_002645.1;NP_073550.1;HCoV-229E_1a | 3155 | 3156 | 2 | 0.05 | 0.05 | 0.05 |
| NC_002645.1;NP_073550.1;HCoV-229E_1a | 3173 | 3173 | 1 | 0.05 | 0.05 | 0.05 |
| NC_002645.1;NP_073550.1;HCoV-229E_1a | 3193 | 3193 | 1 | 0.05 | 0.05 | 0.05 |
| NC_002645.1;NP_073550.1;HCoV-229E_1a | 3202 | 3202 | 1 | 0.05 | 0.05 | 0.05 |

| Extended data 1 | | | | | | |
|---|---|---|---|---|---|---|
| **Protein** | **Start position (peptide)** | **Stop position (peptide)** | **Number of peptides** | **Minimum fraction (per peptide) of control subjects with reactivity >3 sd above average** | **Maximum fraction (per peptide) of control subjects with reactivity >3 sd above average** | **Mean fraction (over all peptides in this contiguous stretch) of controlsubjects with reactivity >3 sd above average** |
| NC_002645.1;NP_073550.1;HCoV-229E_1a | 3224 | 3224 | 1 | 0.1 | 0.1 | 0.1 |
| NC_002645.1;NP_073550.1;HCoV-229E_1a | 3229 | 3229 | 1 | 0.05 | 0.05 | 0.05 |
| NC_002645.1;NP_073550.1;HCoV-229E_1a | 3236 | 3238 | 3 | 0.05 | 0.2 | 0.133333333 |
| NC_002645.1;NP_073550.1;HCoV-229E_1a | 3240 | 3243 | 4 | 0.05 | 0.4 | 0.225 |
| NC_002645.1;NP_073550.1;HCoV-229E_1a | 3245 | 3248 | 4 | 0.05 | 0.55 | 0.225 |
| NC_002645.1;NP_073550.1;HCoV-229E_1a | 3252 | 3252 | 1 | 0.05 | 0.05 | 0.05 |
| NC_002645.1;NP_073550.1;HCoV-229E_1a | 3254 | 3254 | 1 | 0.05 | 0.05 | 0.05 |
| NC_002645.1;NP_073550.1;HCoV-229E_1a | 3263 | 3263 | 1 | 0.05 | 0.05 | 0.05 |
| NC_002645.1;NP_073550.1;HCoV-229E_1a | 3350 | 3362 | 13 | 0.05 | 0.15 | 0.084615385 |
| NC_002645.1;NP_073550.1;HCoV-229E_1a | 3455 | 3455 | 1 | 0.05 | 0.05 | 0.05 |
| NC_002645.1;NP_073550.1;HCoV-229E_1a | 3496 | 3497 | 2 | 0.05 | 0.05 | 0.05 |
| NC_002645.1;NP_073550.1;HCoV-229E_1a | 3502 | 3511 | 10 | 0.05 | 0.35 | 0.135 |
| NC_002645.1;NP_073550.1;HCoV-229E_1a | 3533 | 3541 | 9 | 0.05 | 0.1 | 0.055555556 |
| NC_002645.1;NP_073550.1;HCoV-229E_1a | 3548 | 3549 | 2 | 0.1 | 0.15 | 0.125 |
| NC_002645.1;NP_073550.1;HCoV-229E_1a | 3555 | 3555 | 1 | 0.05 | 0.05 | 0.05 |
| NC_002645.1;NP_073550.1;HCoV-229E_1a | 3559 | 3564 | 6 | 0.05 | 0.1 | 0.075 |
| NC_002645.1;NP_073550.1;HCoV-229E_1a | 3567 | 3570 | 4 | 0.05 | 0.1 | 0.0875 |
| NC_002645.1;NP_073550.1;HCoV-229E_1a | 3579 | 3579 | 1 | 0.05 | 0.05 | 0.05 |
| NC_002645.1;NP_073550.1;HCoV-229E_1a | 3602 | 3617 | 16 | 0.05 | 0.55 | 0.1875 |
| NC_002645.1;NP_073550.1;HCoV-229E_1a | 3631 | 3641 | 11 | 0.05 | 0.1 | 0.077272727 |
| NC_002645.1;NP_073550.1;HCoV-229E_1a | 3645 | 3645 | 1 | 0.05 | 0.05 | 0.05 |
| NC_002645.1;NP_073550.1;HCoV-229E_1a | 3669 | 3670 | 2 | 0.05 | 0.05 | 0.05 |
| NC_002645.1;NP_073550.1;HCoV-229E_1a | 3688 | 3693 | 6 | 0.05 | 0.15 | 0.075 |
| NC_002645.1;NP_073550.1;HCoV-229E_1a | 3731 | 3733 | 3 | 0.05 | 0.15 | 0.083333333 |
| NC_002645.1;NP_073550.1;HCoV-229E_1a | 3740 | 3740 | 1 | 0.05 | 0.05 | 0.05 |
| NC_002645.1;NP_073550.1;HCoV-229E_1a | 3758 | 3758 | 1 | 0.05 | 0.05 | 0.05 |
| NC_002645.1;NP_073550.1;HCoV-229E_1a | 3765 | 3765 | 1 | 0.1 | 0.1 | 0.1 |
| NC_002645.1;NP_073550.1;HCoV-229E_1a | 3788 | 3789 | 2 | 0.05 | 0.05 | 0.05 |
| NC_002645.1;NP_073550.1;HCoV-229E_1a | 3798 | 3800 | 3 | 0.05 | 0.05 | 0.05 |
| NC_002645.1;NP_073550.1;HCoV-229E_1a | 3820 | 3820 | 1 | 0.05 | 0.05 | 0.05 |
| NC_002645.1;NP_073550.1;HCoV-229E_1a | 3841 | 3845 | 5 | 0.05 | 0.15 | 0.08 |
| NC_002645.1;NP_073550.1;HCoV-229E_1a | 3851 | 3851 | 1 | 0.05 | 0.05 | 0.05 |
| NC_002645.1;NP_073550.1;HCoV-229E_1a | 3857 | 3857 | 1 | 0.05 | 0.05 | 0.05 |
| NC_002645.1;NP_073550.1;HCoV-229E_1a | 3866 | 3869 | 4 | 0.05 | 0.05 | 0.05 |
| NC_002645.1;NP_073550.1;HCoV-229E_1a | 3887 | 3887 | 1 | 0.05 | 0.05 | 0.05 |
| NC_002645.1;NP_073550.1;HCoV-229E_1a | 3895 | 3895 | 1 | 0.05 | 0.05 | 0.05 |
| NC_002645.1;NP_073550.1;HCoV-229E_1a | 3923 | 3928 | 6 | 0.05 | 0.05 | 0.05 |

| Extended data 1 | | | | | | |
|---|---|---|---|---|---|---|
| **Protein** | **Start position (peptide)** | **Stop position (peptide)** | **Number of peptides** | **Minimum fraction (per peptide) of control subjects with reactivity >3 sd above average** | **Maximum fraction (per peptide) of control subjects with reactivity >3 sd above average** | **Mean fraction (over all peptides in this contiguous stretch) of controlsubjects with reactivity >3 sd above average** |
| NC_002645.1;NP_073550.1;HCoV-229E_1a | 3943 | 3944 | 2 | 0.05 | 0.05 | 0.05 |
| NC_002645.1;NP_073550.1;HCoV-229E_1a | 3947 | 3958 | 12 | 0.05 | 0.2 | 0.1 |
| NC_002645.1;NP_073550.1;HCoV-229E_1a | 3962 | 3963 | 2 | 0.05 | 0.05 | 0.05 |
| NC_002645.1;NP_073550.1;HCoV-229E_1a | 3971 | 3971 | 1 | 0.05 | 0.05 | 0.05 |
| NC_002645.1;NP_073550.1;HCoV-229E_1a | 3986 | 3986 | 1 | 0.05 | 0.05 | 0.05 |
| NC_002645.1;NP_073550.1;HCoV-229E_1a | 3988 | 3988 | 1 | 0.1 | 0.1 | 0.1 |
| NC_002645.1;NP_073550.1;HCoV-229E_1a | 3990 | 3990 | 1 | 0.05 | 0.05 | 0.05 |
| NC_002645.1;NP_073550.1;HCoV-229E_1a | 3993 | 3993 | 1 | 0.05 | 0.05 | 0.05 |
| NC_002645.1;NP_073550.1;HCoV-229E_1a | 4023 | 4024 | 2 | 0.05 | 0.05 | 0.05 |
| NC_002645.1;NP_073550.1;HCoV-229E_1a | 4031 | 4032 | 2 | 0.05 | 0.05 | 0.05 |
| NC_002645.1;NP_073550.1;HCoV-229E_1a | 4034 | 4034 | 1 | 0.05 | 0.05 | 0.05 |
| NC_002645.1;NP_073550.1;HCoV-229E_1a | 4061 | 4061 | 1 | 0.05 | 0.05 | 0.05 |
| NC_002645.1;NP_073550.1;HCoV-229E_1a | 4064 | 4065 | 2 | 0.05 | 0.05 | 0.05 |
| NC_002645.1;NP_073551.1;HCoV-229E_surface | 9 | 9 | 1 | 0.05 | 0.05 | 0.05 |
| NC_002645.1;NP_073551.1;HCoV-229E_surface | 14 | 17 | 4 | 0.05 | 0.05 | 0.05 |
| NC_002645.1;NP_073551.1;HCoV-229E_surface | 31 | 31 | 1 | 0.05 | 0.05 | 0.05 |
| NC_002645.1;NP_073551.1;HCoV-229E_surface | 37 | 38 | 2 | 0.05 | 0.05 | 0.05 |
| NC_002645.1;NP_073551.1;HCoV-229E_surface | 40 | 43 | 4 | 0.05 | 0.05 | 0.05 |
| NC_002645.1;NP_073551.1;HCoV-229E_surface | 54 | 54 | 1 | 0.05 | 0.05 | 0.05 |
| NC_002645.1;NP_073551.1;HCoV-229E_surface | 58 | 61 | 4 | 0.05 | 0.1 | 0.075 |
| NC_002645.1;NP_073551.1;HCoV-229E_surface | 112 | 112 | 1 | 0.05 | 0.05 | 0.05 |
| NC_002645.1;NP_073551.1;HCoV-229E_surface | 143 | 152 | 10 | 0.05 | 0.15 | 0.105 |
| NC_002645.1;NP_073551.1;HCoV-229E_surface | 169 | 169 | 1 | 0.05 | 0.05 | 0.05 |
| NC_002645.1;NP_073551.1;HCoV-229E_surface | 174 | 175 | 2 | 0.05 | 0.05 | 0.05 |
| NC_002645.1;NP_073551.1;HCoV-229E_surface | 207 | 207 | 1 | 0.05 | 0.05 | 0.05 |
| NC_002645.1;NP_073551.1;HCoV-229E_surface | 211 | 211 | 1 | 0.05 | 0.05 | 0.05 |
| NC_002645.1;NP_073551.1;HCoV-229E_surface | 258 | 264 | 7 | 0.05 | 0.05 | 0.05 |
| NC_002645.1;NP_073551.1;HCoV-229E_surface | 300 | 300 | 1 | 0.05 | 0.05 | 0.05 |
| NC_002645.1;NP_073551.1;HCoV-229E_surface | 304 | 305 | 2 | 0.05 | 0.05 | 0.05 |
| NC_002645.1;NP_073551.1;HCoV-229E_surface | 313 | 313 | 1 | 0.1 | 0.1 | 0.1 |
| NC_002645.1;NP_073551.1;HCoV-229E_surface | 321 | 333 | 13 | 0.05 | 0.15 | 0.076923077 |
| NC_002645.1;NP_073551.1;HCoV-229E_surface | 358 | 358 | 1 | 0.05 | 0.05 | 0.05 |
| NC_002645.1;NP_073551.1;HCoV-229E_surface | 362 | 364 | 3 | 0.05 | 0.1 | 0.066666667 |
| NC_002645.1;NP_073551.1;HCoV-229E_surface | 390 | 390 | 1 | 0.05 | 0.05 | 0.05 |
| NC_002645.1;NP_073551.1;HCoV-229E_surface | 408 | 408 | 1 | 0.05 | 0.05 | 0.05 |
| NC_002645.1;NP_073551.1;HCoV-229E_surface | 410 | 418 | 9 | 0.05 | 0.05 | 0.05 |
| NC_002645.1;NP_073551.1;HCoV-229E_surface | 430 | 430 | 1 | 0.05 | 0.05 | 0.05 |

| Extended data 1 | | | | | | |
|---|---|---|---|---|---|---|
| **Protein** | **Start position (peptide)** | **Stop position (peptide)** | **Number of peptides** | **Minimum fraction (per peptide) of control subjects with reactivity >3 sd above average** | **Maximum fraction (per peptide) of control subjects with reactivity >3 sd above average** | **Mean fraction (over all peptides in this contiguous stretch) of controlsubjects with reactivity >3 sd above average** |
| NC_002645.1;NP_073551.1;HCoV-229E_surface | 437 | 448 | 12 | 0.05 | 0.35 | 0.204166667 |
| NC_002645.1;NP_073551.1;HCoV-229E_surface | 470 | 473 | 4 | 0.05 | 0.15 | 0.1 |
| NC_002645.1;NP_073551.1;HCoV-229E_surface | 500 | 500 | 1 | 0.05 | 0.05 | 0.05 |
| NC_002645.1;NP_073551.1;HCoV-229E_surface | 509 | 519 | 11 | 0.05 | 0.15 | 0.072727273 |
| NC_002645.1;NP_073551.1;HCoV-229E_surface | 522 | 529 | 8 | 0.05 | 0.15 | 0.0875 |
| NC_002645.1;NP_073551.1;HCoV-229E_surface | 543 | 545 | 3 | 0.05 | 0.05 | 0.05 |
| NC_002645.1;NP_073551.1;HCoV-229E_surface | 562 | 562 | 1 | 0.1 | 0.1 | 0.1 |
| NC_002645.1;NP_073551.1;HCoV-229E_surface | 565 | 565 | 1 | 0.05 | 0.05 | 0.05 |
| NC_002645.1;NP_073551.1;HCoV-229E_surface | 576 | 578 | 3 | 0.05 | 0.05 | 0.05 |
| NC_002645.1;NP_073551.1;HCoV-229E_surface | 610 | 610 | 1 | 0.05 | 0.05 | 0.05 |
| NC_002645.1;NP_073551.1;HCoV-229E_surface | 612 | 612 | 1 | 0.05 | 0.05 | 0.05 |
| NC_002645.1;NP_073551.1;HCoV-229E_surface | 627 | 627 | 1 | 0.05 | 0.05 | 0.05 |
| NC_002645.1;NP_073551.1;HCoV-229E_surface | 629 | 632 | 4 | 0.05 | 0.05 | 0.05 |
| NC_002645.1;NP_073551.1;HCoV-229E_surface | 641 | 651 | 11 | 0.05 | 0.2 | 0.077272727 |
| NC_002645.1;NP_073551.1;HCoV-229E_surface | 656 | 656 | 1 | 0.05 | 0.05 | 0.05 |
| NC_002645.1;NP_073551.1;HCoV-229E_surface | 663 | 663 | 1 | 0.05 | 0.05 | 0.05 |
| NC_002645.1;NP_073551.1;HCoV-229E_surface | 665 | 667 | 3 | 0.05 | 0.05 | 0.05 |
| NC_002645.1;NP_073551.1;HCoV-229E_surface | 669 | 669 | 1 | 0.15 | 0.15 | 0.15 |
| NC_002645.1;NP_073551.1;HCoV-229E_surface | 684 | 684 | 1 | 0.05 | 0.05 | 0.05 |
| NC_002645.1;NP_073551.1;HCoV-229E_surface | 691 | 691 | 1 | 0.15 | 0.15 | 0.15 |
| NC_002645.1;NP_073551.1;HCoV-229E_surface | 694 | 696 | 3 | 0.05 | 0.05 | 0.05 |
| NC_002645.1;NP_073551.1;HCoV-229E_surface | 698 | 698 | 1 | 0.05 | 0.05 | 0.05 |
| NC_002645.1;NP_073551.1;HCoV-229E_surface | 700 | 701 | 2 | 0.1 | 0.1 | 0.1 |
| NC_002645.1;NP_073551.1;HCoV-229E_surface | 704 | 704 | 1 | 0.05 | 0.05 | 0.05 |
| NC_002645.1;NP_073551.1;HCoV-229E_surface | 708 | 717 | 10 | 0.05 | 0.15 | 0.12 |
| NC_002645.1;NP_073551.1;HCoV-229E_surface | 719 | 719 | 1 | 0.05 | 0.05 | 0.05 |
| NC_002645.1;NP_073551.1;HCoV-229E_surface | 725 | 725 | 1 | 0.05 | 0.05 | 0.05 |
| NC_002645.1;NP_073551.1;HCoV-229E_surface | 727 | 727 | 1 | 0.05 | 0.05 | 0.05 |
| NC_002645.1;NP_073551.1;HCoV-229E_surface | 778 | 778 | 1 | 0.05 | 0.05 | 0.05 |
| NC_002645.1;NP_073551.1;HCoV-229E_surface | 781 | 781 | 1 | 0.05 | 0.05 | 0.05 |
| NC_002645.1;NP_073551.1;HCoV-229E_surface | 784 | 796 | 13 | 0.05 | 0.3 | 0.165384615 |
| NC_002645.1;NP_073551.1;HCoV-229E_surface | 798 | 801 | 4 | 0.05 | 0.05 | 0.05 |
| NC_002645.1;NP_073551.1;HCoV-229E_surface | 803 | 804 | 2 | 0.05 | 0.1 | 0.075 |
| NC_002645.1;NP_073551.1;HCoV-229E_surface | 807 | 807 | 1 | 0.05 | 0.05 | 0.05 |
| NC_002645.1;NP_073551.1;HCoV-229E_surface | 811 | 811 | 1 | 0.05 | 0.05 | 0.05 |
| NC_002645.1;NP_073551.1;HCoV-229E_surface | 817 | 827 | 11 | 0.05 | 0.15 | 0.109090909 |
| NC_002645.1;NP_073551.1;HCoV-229E_surface | 829 | 839 | 11 | 0.05 | 0.1 | 0.059090909 |

| Extended data 1 | | | | | | |
|---|---|---|---|---|---|---|
| Protein | Start position (peptide) | Stop position (peptide) | Number of peptides | Minimum fraction (per peptide) of control subjects with reactivity >3 sd above average | Maximum fraction (per peptide) of control subjects with reactivity >3 sd above average | Mean fraction (over all peptides in this contiguous stretch) of controlsubjects with reactivity >3 sd above average |
| NC_002645.1;NP_073551.1;HCoV-229E_surface | 857 | 857 | 1 | 0.05 | 0.05 | 0.05 |
| NC_002645.1;NP_073551.1;HCoV-229E_surface | 860 | 862 | 3 | 0.05 | 0.05 | 0.05 |
| NC_002645.1;NP_073551.1;HCoV-229E_surface | 864 | 865 | 2 | 0.05 | 0.15 | 0.1 |
| NC_002645.1;NP_073551.1;HCoV-229E_surface | 873 | 874 | 2 | 0.05 | 0.05 | 0.05 |
| NC_002645.1;NP_073551.1;HCoV-229E_surface | 887 | 888 | 2 | 0.05 | 0.05 | 0.05 |
| NC_002645.1;NP_073551.1;HCoV-229E_surface | 898 | 899 | 2 | 0.1 | 0.1 | 0.1 |
| NC_002645.1;NP_073551.1;HCoV-229E_surface | 907 | 907 | 1 | 0.1 | 0.1 | 0.1 |
| NC_002645.1;NP_073551.1;HCoV-229E_surface | 975 | 975 | 1 | 0.05 | 0.05 | 0.05 |
| NC_002645.1;NP_073551.1;HCoV-229E_surface | 991 | 991 | 1 | 0.05 | 0.05 | 0.05 |
| NC_002645.1;NP_073551.1;HCoV-229E_surface | 1010 | 1013 | 4 | 0.05 | 0.05 | 0.05 |
| NC_002645.1;NP_073551.1;HCoV-229E_surface | 1017 | 1019 | 3 | 0.05 | 0.05 | 0.05 |
| NC_002645.1;NP_073551.1;HCoV-229E_surface | 1021 | 1026 | 6 | 0.05 | 0.1 | 0.058333333 |
| NC_002645.1;NP_073551.1;HCoV-229E_surface | 1032 | 1032 | 1 | 0.05 | 0.05 | 0.05 |
| NC_002645.1;NP_073551.1;HCoV-229E_surface | 1036 | 1036 | 1 | 0.05 | 0.05 | 0.05 |
| NC_002645.1;NP_073551.1;HCoV-229E_surface | 1039 | 1053 | 15 | 0.05 | 0.35 | 0.166666667 |
| NC_002645.1;NP_073551.1;HCoV-229E_surface | 1056 | 1057 | 2 | 0.05 | 0.05 | 0.05 |
| NC_002645.1;NP_073551.1;HCoV-229E_surface | 1072 | 1075 | 4 | 0.05 | 0.05 | 0.05 |
| NC_002645.1;NP_073551.1;HCoV-229E_surface | 1083 | 1087 | 5 | 0.05 | 0.05 | 0.05 |
| NC_002645.1;NP_073551.1;HCoV-229E_surface | 1091 | 1095 | 5 | 0.05 | 0.15 | 0.07 |
| NC_002645.1;NP_073551.1;HCoV-229E_surface | 1098 | 1099 | 2 | 0.05 | 0.1 | 0.075 |
| NC_002645.1;NP_073551.1;HCoV-229E_surface | 1106 | 1106 | 1 | 0.05 | 0.05 | 0.05 |
| NC_002645.1;NP_073551.1;HCoV-229E_surface | 1154 | 1161 | 8 | 0.05 | 0.1 | 0.05625 |
| NC_002645.1;NP_073552.1;HCoV-229E_4a | 4 | 6 | 3 | 0.05 | 0.1 | 0.083333333 |
| NC_002645.1;NP_073552.1;HCoV-229E_4a | 8 | 8 | 1 | 0.05 | 0.05 | 0.05 |
| NC_002645.1;NP_073552.1;HCoV-229E_4a | 10 | 13 | 4 | 0.05 | 0.1 | 0.0625 |
| NC_002645.1;NP_073552.1;HCoV-229E_4a | 18 | 32 | 15 | 0.05 | 0.2 | 0.136666667 |
| NC_002645.1;NP_073553.1;HCoV-229E_4b | 37 | 37 | 1 | 0.05 | 0.05 | 0.05 |
| NC_002645.1;NP_073553.1;HCoV-229E_4b | 40 | 40 | 1 | 0.05 | 0.05 | 0.05 |
| NC_002645.1;NP_073553.1;HCoV-229E_4b | 69 | 77 | 9 | 0.05 | 0.05 | 0.05 |
| NC_002645.1;NP_073555.1;HCoV-229E_membrane | 146 | 146 | 1 | 0.05 | 0.05 | 0.05 |
| NC_002645.1;NP_073555.1;HCoV-229E_membrane | 156 | 156 | 1 | 0.05 | 0.05 | 0.05 |
| NC_002645.1;NP_073555.1;HCoV-229E_membrane | 180 | 180 | 1 | 0.05 | 0.05 | 0.05 |
| NC_002645.1;NP_073555.1;HCoV-229E_membrane | 203 | 214 | 12 | 0.1 | 0.35 | 0.229166667 |
| NC_002645.1;NP_073556.1;HCoV-229E_nucleocapsi | 1 | 1 | 1 | 0.05 | 0.05 | 0.05 |
| NC_002645.1;NP_073556.1;HCoV-229E_nucleocapsi | 4 | 14 | 11 | 0.05 | 0.1 | 0.059090909 |
| NC_002645.1;NP_073556.1;HCoV-229E_nucleocapsi | 23 | 23 | 1 | 0.05 | 0.05 | 0.05 |
| NC_002645.1;NP_073556.1;HCoV-229E_nucleocapsi | 28 | 28 | 1 | 0.05 | 0.05 | 0.05 |

| Extended data 1 | | | | | | |
|---|---|---|---|---|---|---|
| **Protein** | **Start position (peptide)** | **Stop position (peptide)** | **Number of peptides** | **Minimum fraction (per peptide) of control subjects with reactivity >3 sd above average** | **Maximum fraction (per peptide) of control subjects with reactivity >3 sd above average** | **Mean fraction (over all peptides in this contiguous stretch) of controlsubjects with reactivity >3 sd above average** |
| NC_002645.1;NP_073556.1;HCoV-229E_nucleocapsi | 31 | 32 | 2 | 0.05 | 0.05 | 0.05 |
| NC_002645.1;NP_073556.1;HCoV-229E_nucleocapsi | 55 | 60 | 6 | 0.05 | 0.05 | 0.05 |
| NC_002645.1;NP_073556.1;HCoV-229E_nucleocapsi | 63 | 63 | 1 | 0.05 | 0.05 | 0.05 |
| NC_002645.1;NP_073556.1;HCoV-229E_nucleocapsi | 95 | 95 | 1 | 0.05 | 0.05 | 0.05 |
| NC_002645.1;NP_073556.1;HCoV-229E_nucleocapsi | 103 | 103 | 1 | 0.05 | 0.05 | 0.05 |
| NC_002645.1;NP_073556.1;HCoV-229E_nucleocapsi | 117 | 117 | 1 | 0.05 | 0.05 | 0.05 |
| NC_002645.1;NP_073556.1;HCoV-229E_nucleocapsi | 121 | 121 | 1 | 0.05 | 0.05 | 0.05 |
| NC_002645.1;NP_073556.1;HCoV-229E_nucleocapsi | 132 | 141 | 10 | 0.05 | 0.05 | 0.05 |
| NC_002645.1;NP_073556.1;HCoV-229E_nucleocapsi | 144 | 144 | 1 | 0.05 | 0.05 | 0.05 |
| NC_002645.1;NP_073556.1;HCoV-229E_nucleocapsi | 148 | 148 | 1 | 0.05 | 0.05 | 0.05 |
| NC_002645.1;NP_073556.1;HCoV-229E_nucleocapsi | 162 | 162 | 1 | 0.1 | 0.1 | 0.1 |
| NC_002645.1;NP_073556.1;HCoV-229E_nucleocapsi | 173 | 178 | 6 | 0.05 | 0.05 | 0.05 |
| NC_002645.1;NP_073556.1;HCoV-229E_nucleocapsi | 181 | 207 | 27 | 0.05 | 0.1 | 0.053703704 |
| NC_002645.1;NP_073556.1;HCoV-229E_nucleocapsi | 213 | 213 | 1 | 0.05 | 0.05 | 0.05 |
| NC_002645.1;NP_073556.1;HCoV-229E_nucleocapsi | 219 | 219 | 1 | 0.05 | 0.05 | 0.05 |
| NC_002645.1;NP_073556.1;HCoV-229E_nucleocapsi | 232 | 234 | 3 | 0.05 | 0.1 | 0.083333333 |
| NC_002645.1;NP_073556.1;HCoV-229E_nucleocapsi | 241 | 241 | 1 | 0.05 | 0.05 | 0.05 |
| NC_002645.1;NP_073556.1;HCoV-229E_nucleocapsi | 243 | 256 | 14 | 0.05 | 0.2 | 0.1 |
| NC_002645.1;NP_073556.1;HCoV-229E_nucleocapsi | 258 | 260 | 3 | 0.05 | 0.1 | 0.066666667 |
| NC_002645.1;NP_073556.1;HCoV-229E_nucleocapsi | 266 | 266 | 1 | 0.05 | 0.05 | 0.05 |
| NC_002645.1;NP_073556.1;HCoV-229E_nucleocapsi | 268 | 268 | 1 | 0.05 | 0.05 | 0.05 |
| NC_002645.1;NP_073556.1;HCoV-229E_nucleocapsi | 279 | 289 | 11 | 0.05 | 0.2 | 0.090909091 |
| NC_002645.1;NP_073556.1;HCoV-229E_nucleocapsi | 300 | 308 | 9 | 0.05 | 0.15 | 0.077777778 |
| NC_002645.1;NP_073556.1;HCoV-229E_nucleocapsi | 326 | 327 | 2 | 0.05 | 0.05 | 0.05 |
| NC_002645.1;NP_073556.1;HCoV-229E_nucleocapsi | 331 | 378 | 48 | 0.05 | 0.15 | 0.060416667 |
| NC_004718.3;NP_828849.2;SARS-CoV_orf1ab | 1 | 1 | 1 | 0.05 | 0.05 | 0.05 |
| NC_004718.3;NP_828849.2;SARS-CoV_orf1ab | 4 | 6 | 3 | 0.05 | 0.1 | 0.066666667 |
| NC_004718.3;NP_828849.2;SARS-CoV_orf1ab | 29 | 30 | 2 | 0.1 | 0.25 | 0.175 |
| NC_004718.3;NP_828849.2;SARS-CoV_orf1ab | 32 | 34 | 3 | 0.05 | 0.05 | 0.05 |
| NC_004718.3;NP_828849.2;SARS-CoV_orf1ab | 44 | 44 | 1 | 0.05 | 0.05 | 0.05 |
| NC_004718.3;NP_828849.2;SARS-CoV_orf1ab | 52 | 52 | 1 | 0.05 | 0.05 | 0.05 |
| NC_004718.3;NP_828849.2;SARS-CoV_orf1ab | 65 | 65 | 1 | 0.05 | 0.05 | 0.05 |
| NC_004718.3;NP_828849.2;SARS-CoV_orf1ab | 71 | 72 | 2 | 0.05 | 0.05 | 0.05 |
| NC_004718.3;NP_828849.2;SARS-CoV_orf1ab | 90 | 91 | 2 | 0.05 | 0.2 | 0.125 |
| NC_004718.3;NP_828849.2;SARS-CoV_orf1ab | 116 | 121 | 6 | 0.05 | 0.1 | 0.058333333 |
| NC_004718.3;NP_828849.2;SARS-CoV_orf1ab | 124 | 125 | 2 | 0.05 | 0.15 | 0.1 |
| NC_004718.3;NP_828849.2;SARS-CoV_orf1ab | 131 | 133 | 3 | 0.05 | 0.1 | 0.066666667 |

| Extended data 1 | | | | | | |
|---|---|---|---|---|---|---|
| Protein | Start position (peptide) | Stop position (peptide) | Number of peptides | Minimum fraction (per peptide) of control subjects with reactivity >3 sd above average | Maximum fraction (per peptide) of control subjects with reactivity >3 sd above average | Mean fraction (over all peptides in this contiguous stretch) of controlsubjects with reactivity >3 sd above average |
| NC_004718.3;NP_828849.2;SARS-CoV_orf1ab | 139 | 140 | 2 | 0.05 | 0.05 | 0.05 |
| NC_004718.3;NP_828849.2;SARS-CoV_orf1ab | 145 | 145 | 1 | 0.2 | 0.2 | 0.2 |
| NC_004718.3;NP_828849.2;SARS-CoV_orf1ab | 149 | 157 | 9 | 0.05 | 0.25 | 0.133333333 |
| NC_004718.3;NP_828849.2;SARS-CoV_orf1ab | 165 | 165 | 1 | 0.05 | 0.05 | 0.05 |
| NC_004718.3;NP_828849.2;SARS-CoV_orf1ab | 168 | 168 | 1 | 0.2 | 0.2 | 0.2 |
| NC_004718.3;NP_828849.2;SARS-CoV_orf1ab | 173 | 173 | 1 | 0.05 | 0.05 | 0.05 |
| NC_004718.3;NP_828849.2;SARS-CoV_orf1ab | 179 | 180 | 2 | 0.05 | 0.15 | 0.1 |
| NC_004718.3;NP_828849.2;SARS-CoV_orf1ab | 189 | 189 | 1 | 0.05 | 0.05 | 0.05 |
| NC_004718.3;NP_828849.2;SARS-CoV_orf1ab | 204 | 204 | 1 | 0.05 | 0.05 | 0.05 |
| NC_004718.3;NP_828849.2;SARS-CoV_orf1ab | 214 | 218 | 5 | 0.05 | 0.15 | 0.1 |
| NC_004718.3;NP_828849.2;SARS-CoV_orf1ab | 220 | 223 | 4 | 0.05 | 0.15 | 0.1 |
| NC_004718.3;NP_828849.2;SARS-CoV_orf1ab | 225 | 225 | 1 | 0.05 | 0.05 | 0.05 |
| NC_004718.3;NP_828849.2;SARS-CoV_orf1ab | 240 | 241 | 2 | 0.05 | 0.05 | 0.05 |
| NC_004718.3;NP_828849.2;SARS-CoV_orf1ab | 254 | 256 | 3 | 0.05 | 0.05 | 0.05 |
| NC_004718.3;NP_828849.2;SARS-CoV_orf1ab | 262 | 263 | 2 | 0.05 | 0.1 | 0.075 |
| NC_004718.3;NP_828849.2;SARS-CoV_orf1ab | 271 | 271 | 1 | 0.05 | 0.05 | 0.05 |
| NC_004718.3;NP_828849.2;SARS-CoV_orf1ab | 276 | 285 | 10 | 0.05 | 0.15 | 0.06 |
| NC_004718.3;NP_828849.2;SARS-CoV_orf1ab | 296 | 301 | 6 | 0.05 | 0.05 | 0.05 |
| NC_004718.3;NP_828849.2;SARS-CoV_orf1ab | 315 | 315 | 1 | 0.1 | 0.1 | 0.1 |
| NC_004718.3;NP_828849.2;SARS-CoV_orf1ab | 324 | 324 | 1 | 0.05 | 0.05 | 0.05 |
| NC_004718.3;NP_828849.2;SARS-CoV_orf1ab | 356 | 358 | 3 | 0.05 | 0.05 | 0.05 |
| NC_004718.3;NP_828849.2;SARS-CoV_orf1ab | 362 | 362 | 1 | 0.05 | 0.05 | 0.05 |
| NC_004718.3;NP_828849.2;SARS-CoV_orf1ab | 366 | 366 | 1 | 0.05 | 0.05 | 0.05 |
| NC_004718.3;NP_828849.2;SARS-CoV_orf1ab | 368 | 375 | 8 | 0.05 | 0.1 | 0.05625 |
| NC_004718.3;NP_828849.2;SARS-CoV_orf1ab | 377 | 377 | 1 | 0.1 | 0.1 | 0.1 |
| NC_004718.3;NP_828849.2;SARS-CoV_orf1ab | 383 | 383 | 1 | 0.05 | 0.05 | 0.05 |
| NC_004718.3;NP_828849.2;SARS-CoV_orf1ab | 425 | 425 | 1 | 0.05 | 0.05 | 0.05 |
| NC_004718.3;NP_828849.2;SARS-CoV_orf1ab | 428 | 428 | 1 | 0.05 | 0.05 | 0.05 |
| NC_004718.3;NP_828849.2;SARS-CoV_orf1ab | 430 | 430 | 1 | 0.05 | 0.05 | 0.05 |
| NC_004718.3;NP_828849.2;SARS-CoV_orf1ab | 438 | 438 | 1 | 0.05 | 0.05 | 0.05 |
| NC_004718.3;NP_828849.2;SARS-CoV_orf1ab | 444 | 448 | 5 | 0.05 | 0.1 | 0.06 |
| NC_004718.3;NP_828849.2;SARS-CoV_orf1ab | 453 | 453 | 1 | 0.05 | 0.05 | 0.05 |
| NC_004718.3;NP_828849.2;SARS-CoV_orf1ab | 457 | 458 | 2 | 0.05 | 0.05 | 0.05 |
| NC_004718.3;NP_828849.2;SARS-CoV_orf1ab | 477 | 484 | 8 | 0.05 | 0.1 | 0.0625 |
| NC_004718.3;NP_828849.2;SARS-CoV_orf1ab | 497 | 497 | 1 | 0.05 | 0.05 | 0.05 |
| NC_004718.3;NP_828849.2;SARS-CoV_orf1ab | 526 | 526 | 1 | 0.05 | 0.05 | 0.05 |
| NC_004718.3;NP_828849.2;SARS-CoV_orf1ab | 531 | 532 | 2 | 0.05 | 0.05 | 0.05 |

| Extended data 1 | | | | | | |
|---|---|---|---|---|---|---|
| Protein | Start position (peptide) | Stop position (peptide) | Number of peptides | Minimum fraction (per peptide) of control subjects with reactivity >3 sd above average | Maximum fraction (per peptide) of control subjects with reactivity >3 sd above average | Mean fraction (over all peptides in this contiguous stretch) of controlsubjects with reactivity >3 sd above average |
| NC_004718.3;NP_828849.2;SARS-CoV_orf1ab | 545 | 551 | 7 | 0.05 | 0.1 | 0.057142857 |
| NC_004718.3;NP_828849.2;SARS-CoV_orf1ab | 553 | 554 | 2 | 0.05 | 0.05 | 0.05 |
| NC_004718.3;NP_828849.2;SARS-CoV_orf1ab | 572 | 572 | 1 | 0.05 | 0.05 | 0.05 |
| NC_004718.3;NP_828849.2;SARS-CoV_orf1ab | 589 | 589 | 1 | 0.05 | 0.05 | 0.05 |
| NC_004718.3;NP_828849.2;SARS-CoV_orf1ab | 605 | 605 | 1 | 0.1 | 0.1 | 0.1 |
| NC_004718.3;NP_828849.2;SARS-CoV_orf1ab | 619 | 622 | 4 | 0.05 | 0.05 | 0.05 |
| NC_004718.3;NP_828849.2;SARS-CoV_orf1ab | 630 | 630 | 1 | 0.05 | 0.05 | 0.05 |
| NC_004718.3;NP_828849.2;SARS-CoV_orf1ab | 633 | 635 | 3 | 0.05 | 0.05 | 0.05 |
| NC_004718.3;NP_828849.2;SARS-CoV_orf1ab | 655 | 655 | 1 | 0.05 | 0.05 | 0.05 |
| NC_004718.3;NP_828849.2;SARS-CoV_orf1ab | 660 | 660 | 1 | 0.05 | 0.05 | 0.05 |
| NC_004718.3;NP_828849.2;SARS-CoV_orf1ab | 666 | 666 | 1 | 0.05 | 0.05 | 0.05 |
| NC_004718.3;NP_828849.2;SARS-CoV_orf1ab | 682 | 684 | 3 | 0.05 | 0.05 | 0.05 |
| NC_004718.3;NP_828849.2;SARS-CoV_orf1ab | 694 | 694 | 1 | 0.05 | 0.05 | 0.05 |
| NC_004718.3;NP_828849.2;SARS-CoV_orf1ab | 700 | 701 | 2 | 0.05 | 0.05 | 0.05 |
| NC_004718.3;NP_828849.2;SARS-CoV_orf1ab | 741 | 741 | 1 | 0.05 | 0.05 | 0.05 |
| NC_004718.3;NP_828849.2;SARS-CoV_orf1ab | 744 | 746 | 3 | 0.05 | 0.2 | 0.1 |
| NC_004718.3;NP_828849.2;SARS-CoV_orf1ab | 748 | 755 | 8 | 0.05 | 0.1 | 0.05625 |
| NC_004718.3;NP_828849.2;SARS-CoV_orf1ab | 770 | 771 | 2 | 0.05 | 0.1 | 0.075 |
| NC_004718.3;NP_828849.2;SARS-CoV_orf1ab | 782 | 782 | 1 | 0.05 | 0.05 | 0.05 |
| NC_004718.3;NP_828849.2;SARS-CoV_orf1ab | 789 | 789 | 1 | 0.05 | 0.05 | 0.05 |
| NC_004718.3;NP_828849.2;SARS-CoV_orf1ab | 801 | 801 | 1 | 0.05 | 0.05 | 0.05 |
| NC_004718.3;NP_828849.2;SARS-CoV_orf1ab | 816 | 816 | 1 | 0.05 | 0.05 | 0.05 |
| NC_004718.3;NP_828849.2;SARS-CoV_orf1ab | 826 | 826 | 1 | 0.05 | 0.05 | 0.05 |
| NC_004718.3;NP_828849.2;SARS-CoV_orf1ab | 828 | 828 | 1 | 0.1 | 0.1 | 0.1 |
| NC_004718.3;NP_828849.2;SARS-CoV_orf1ab | 832 | 832 | 1 | 0.05 | 0.05 | 0.05 |
| NC_004718.3;NP_828849.2;SARS-CoV_orf1ab | 837 | 837 | 1 | 0.05 | 0.05 | 0.05 |
| NC_004718.3;NP_828849.2;SARS-CoV_orf1ab | 842 | 844 | 3 | 0.05 | 0.05 | 0.05 |
| NC_004718.3;NP_828849.2;SARS-CoV_orf1ab | 850 | 855 | 6 | 0.05 | 0.15 | 0.116666667 |
| NC_004718.3;NP_828849.2;SARS-CoV_orf1ab | 864 | 864 | 1 | 0.05 | 0.05 | 0.05 |
| NC_004718.3;NP_828849.2;SARS-CoV_orf1ab | 878 | 878 | 1 | 0.05 | 0.05 | 0.05 |
| NC_004718.3;NP_828849.2;SARS-CoV_orf1ab | 885 | 885 | 1 | 0.15 | 0.15 | 0.15 |
| NC_004718.3;NP_828849.2;SARS-CoV_orf1ab | 889 | 889 | 1 | 0.05 | 0.05 | 0.05 |
| NC_004718.3;NP_828849.2;SARS-CoV_orf1ab | 933 | 933 | 1 | 0.25 | 0.25 | 0.25 |
| NC_004718.3;NP_828849.2;SARS-CoV_orf1ab | 950 | 955 | 6 | 0.05 | 0.15 | 0.066666667 |
| NC_004718.3;NP_828849.2;SARS-CoV_orf1ab | 979 | 979 | 1 | 0.05 | 0.05 | 0.05 |
| NC_004718.3;NP_828849.2;SARS-CoV_orf1ab | 992 | 994 | 3 | 0.05 | 0.05 | 0.05 |
| NC_004718.3;NP_828849.2;SARS-CoV_orf1ab | 996 | 996 | 1 | 0.05 | 0.05 | 0.05 |

| Extended data 1 | | | | | | |
|---|---|---|---|---|---|---|
| Protein | Start position (peptide) | Stop position (peptide) | Number of peptides | Minimum fraction (per peptide) of control subjects with reactivity >3 sd above average | Maximum fraction (per peptide) of control subjects with reactivity >3 sd above average | Mean fraction (over all peptides in this contiguous stretch) of controlsubjects with reactivity >3 sd above average |
| NC_004718.3;NP_828849.2;SARS-CoV_orf1ab | 1001 | 1008 | 8 | 0.05 | 0.05 | 0.05 |
| NC_004718.3;NP_828849.2;SARS-CoV_orf1ab | 1010 | 1011 | 2 | 0.05 | 0.05 | 0.05 |
| NC_004718.3;NP_828849.2;SARS-CoV_orf1ab | 1024 | 1025 | 2 | 0.05 | 0.05 | 0.05 |
| NC_004718.3;NP_828849.2;SARS-CoV_orf1ab | 1033 | 1033 | 1 | 0.05 | 0.05 | 0.05 |
| NC_004718.3;NP_828849.2;SARS-CoV_orf1ab | 1035 | 1035 | 1 | 0.1 | 0.1 | 0.1 |
| NC_004718.3;NP_828849.2;SARS-CoV_orf1ab | 1045 | 1045 | 1 | 0.05 | 0.05 | 0.05 |
| NC_004718.3;NP_828849.2;SARS-CoV_orf1ab | 1047 | 1047 | 1 | 0.05 | 0.05 | 0.05 |
| NC_004718.3;NP_828849.2;SARS-CoV_orf1ab | 1049 | 1065 | 17 | 0.05 | 0.2 | 0.147058824 |
| NC_004718.3;NP_828849.2;SARS-CoV_orf1ab | 1090 | 1091 | 2 | 0.05 | 0.05 | 0.05 |
| NC_004718.3;NP_828849.2;SARS-CoV_orf1ab | 1093 | 1093 | 1 | 0.05 | 0.05 | 0.05 |
| NC_004718.3;NP_828849.2;SARS-CoV_orf1ab | 1095 | 1097 | 3 | 0.05 | 0.05 | 0.05 |
| NC_004718.3;NP_828849.2;SARS-CoV_orf1ab | 1102 | 1104 | 3 | 0.05 | 0.1 | 0.066666667 |
| NC_004718.3;NP_828849.2;SARS-CoV_orf1ab | 1110 | 1110 | 1 | 0.1 | 0.1 | 0.1 |
| NC_004718.3;NP_828849.2;SARS-CoV_orf1ab | 1158 | 1158 | 1 | 0.05 | 0.05 | 0.05 |
| NC_004718.3;NP_828849.2;SARS-CoV_orf1ab | 1164 | 1173 | 10 | 0.05 | 0.45 | 0.37 |
| NC_004718.3;NP_828849.2;SARS-CoV_orf1ab | 1190 | 1192 | 3 | 0.05 | 0.1 | 0.066666667 |
| NC_004718.3;NP_828849.2;SARS-CoV_orf1ab | 1196 | 1201 | 6 | 0.05 | 0.15 | 0.091666667 |
| NC_004718.3;NP_828849.2;SARS-CoV_orf1ab | 1213 | 1213 | 1 | 0.05 | 0.05 | 0.05 |
| NC_004718.3;NP_828849.2;SARS-CoV_orf1ab | 1231 | 1234 | 4 | 0.05 | 0.15 | 0.0875 |
| NC_004718.3;NP_828849.2;SARS-CoV_orf1ab | 1240 | 1240 | 1 | 0.05 | 0.05 | 0.05 |
| NC_004718.3;NP_828849.2;SARS-CoV_orf1ab | 1250 | 1250 | 1 | 0.05 | 0.05 | 0.05 |
| NC_004718.3;NP_828849.2;SARS-CoV_orf1ab | 1272 | 1278 | 7 | 0.05 | 0.05 | 0.05 |
| NC_004718.3;NP_828849.2;SARS-CoV_orf1ab | 1287 | 1287 | 1 | 0.05 | 0.05 | 0.05 |
| NC_004718.3;NP_828849.2;SARS-CoV_orf1ab | 1295 | 1297 | 3 | 0.05 | 0.1 | 0.066666667 |
| NC_004718.3;NP_828849.2;SARS-CoV_orf1ab | 1300 | 1300 | 1 | 0.05 | 0.05 | 0.05 |
| NC_004718.3;NP_828849.2;SARS-CoV_orf1ab | 1335 | 1335 | 1 | 0.05 | 0.05 | 0.05 |
| NC_004718.3;NP_828849.2;SARS-CoV_orf1ab | 1341 | 1341 | 1 | 0.05 | 0.05 | 0.05 |
| NC_004718.3;NP_828849.2;SARS-CoV_orf1ab | 1358 | 1359 | 2 | 0.05 | 0.05 | 0.05 |
| NC_004718.3;NP_828849.2;SARS-CoV_orf1ab | 1372 | 1382 | 11 | 0.05 | 0.15 | 0.090909091 |
| NC_004718.3;NP_828849.2;SARS-CoV_orf1ab | 1390 | 1390 | 1 | 0.05 | 0.05 | 0.05 |
| NC_004718.3;NP_828849.2;SARS-CoV_orf1ab | 1406 | 1406 | 1 | 0.05 | 0.05 | 0.05 |
| NC_004718.3;NP_828849.2;SARS-CoV_orf1ab | 1409 | 1409 | 1 | 0.05 | 0.05 | 0.05 |
| NC_004718.3;NP_828849.2;SARS-CoV_orf1ab | 1411 | 1413 | 3 | 0.05 | 0.15 | 0.1 |
| NC_004718.3;NP_828849.2;SARS-CoV_orf1ab | 1416 | 1416 | 1 | 0.1 | 0.1 | 0.1 |
| NC_004718.3;NP_828849.2;SARS-CoV_orf1ab | 1424 | 1424 | 1 | 0.05 | 0.05 | 0.05 |
| NC_004718.3;NP_828849.2;SARS-CoV_orf1ab | 1434 | 1434 | 1 | 0.05 | 0.05 | 0.05 |
| NC_004718.3;NP_828849.2;SARS-CoV_orf1ab | 1448 | 1448 | 1 | 0.05 | 0.05 | 0.05 |

| Extended data 1 | | | | | | |
|---|---|---|---|---|---|---|
| Protein | Start position (peptide) | Stop position (peptide) | Number of peptides | Minimum fraction (per peptide) of control subjects with reactivity >3 sd above average | Maximum fraction (per peptide) of control subjects with reactivity >3 sd above average | Mean fraction (over all peptides in this contiguous stretch) of controlsubjects with reactivity >3 sd above average |
| NC_004718.3;NP_828849.2;SARS-CoV_orf1ab | 1456 | 1466 | 11 | 0.05 | 0.1 | 0.059090909 |
| NC_004718.3;NP_828849.2;SARS-CoV_orf1ab | 1468 | 1471 | 4 | 0.05 | 0.05 | 0.05 |
| NC_004718.3;NP_828849.2;SARS-CoV_orf1ab | 1485 | 1485 | 1 | 0.05 | 0.05 | 0.05 |
| NC_004718.3;NP_828849.2;SARS-CoV_orf1ab | 1503 | 1506 | 4 | 0.05 | 0.05 | 0.05 |
| NC_004718.3;NP_828849.2;SARS-CoV_orf1ab | 1508 | 1509 | 2 | 0.05 | 0.05 | 0.05 |
| NC_004718.3;NP_828849.2;SARS-CoV_orf1ab | 1516 | 1518 | 3 | 0.05 | 0.45 | 0.216666667 |
| NC_004718.3;NP_828849.2;SARS-CoV_orf1ab | 1542 | 1542 | 1 | 0.05 | 0.05 | 0.05 |
| NC_004718.3;NP_828849.2;SARS-CoV_orf1ab | 1545 | 1546 | 2 | 0.05 | 0.05 | 0.05 |
| NC_004718.3;NP_828849.2;SARS-CoV_orf1ab | 1560 | 1560 | 1 | 0.2 | 0.2 | 0.2 |
| NC_004718.3;NP_828849.2;SARS-CoV_orf1ab | 1572 | 1572 | 1 | 0.15 | 0.15 | 0.15 |
| NC_004718.3;NP_828849.2;SARS-CoV_orf1ab | 1574 | 1574 | 1 | 0.05 | 0.05 | 0.05 |
| NC_004718.3;NP_828849.2;SARS-CoV_orf1ab | 1579 | 1579 | 1 | 0.05 | 0.05 | 0.05 |
| NC_004718.3;NP_828849.2;SARS-CoV_orf1ab | 1608 | 1616 | 9 | 0.05 | 0.05 | 0.05 |
| NC_004718.3;NP_828849.2;SARS-CoV_orf1ab | 1625 | 1625 | 1 | 0.05 | 0.05 | 0.05 |
| NC_004718.3;NP_828849.2;SARS-CoV_orf1ab | 1634 | 1641 | 8 | 0.05 | 0.05 | 0.05 |
| NC_004718.3;NP_828849.2;SARS-CoV_orf1ab | 1661 | 1662 | 2 | 0.05 | 0.1 | 0.075 |
| NC_004718.3;NP_828849.2;SARS-CoV_orf1ab | 1664 | 1664 | 1 | 0.1 | 0.1 | 0.1 |
| NC_004718.3;NP_828849.2;SARS-CoV_orf1ab | 1668 | 1672 | 5 | 0.05 | 0.1 | 0.06 |
| NC_004718.3;NP_828849.2;SARS-CoV_orf1ab | 1690 | 1692 | 3 | 0.05 | 0.15 | 0.1 |
| NC_004718.3;NP_828849.2;SARS-CoV_orf1ab | 1694 | 1694 | 1 | 0.05 | 0.05 | 0.05 |
| NC_004718.3;NP_828849.2;SARS-CoV_orf1ab | 1700 | 1700 | 1 | 0.05 | 0.05 | 0.05 |
| NC_004718.3;NP_828849.2;SARS-CoV_orf1ab | 1703 | 1703 | 1 | 0.05 | 0.05 | 0.05 |
| NC_004718.3;NP_828849.2;SARS-CoV_orf1ab | 1719 | 1719 | 1 | 0.1 | 0.1 | 0.1 |
| NC_004718.3;NP_828849.2;SARS-CoV_orf1ab | 1722 | 1729 | 8 | 0.05 | 0.1 | 0.06875 |
| NC_004718.3;NP_828849.2;SARS-CoV_orf1ab | 1733 | 1734 | 2 | 0.05 | 0.05 | 0.05 |
| NC_004718.3;NP_828849.2;SARS-CoV_orf1ab | 1738 | 1751 | 14 | 0.05 | 0.2 | 0.121428571 |
| NC_004718.3;NP_828849.2;SARS-CoV_orf1ab | 1764 | 1764 | 1 | 0.05 | 0.05 | 0.05 |
| NC_004718.3;NP_828849.2;SARS-CoV_orf1ab | 1772 | 1772 | 1 | 0.05 | 0.05 | 0.05 |
| NC_004718.3;NP_828849.2;SARS-CoV_orf1ab | 1774 | 1774 | 1 | 0.1 | 0.1 | 0.1 |
| NC_004718.3;NP_828849.2;SARS-CoV_orf1ab | 1781 | 1781 | 1 | 0.05 | 0.05 | 0.05 |
| NC_004718.3;NP_828849.2;SARS-CoV_orf1ab | 1806 | 1806 | 1 | 0.1 | 0.1 | 0.1 |
| NC_004718.3;NP_828849.2;SARS-CoV_orf1ab | 1808 | 1808 | 1 | 0.05 | 0.05 | 0.05 |
| NC_004718.3;NP_828849.2;SARS-CoV_orf1ab | 1811 | 1816 | 6 | 0.05 | 0.05 | 0.05 |
| NC_004718.3;NP_828849.2;SARS-CoV_orf1ab | 1818 | 1818 | 1 | 0.05 | 0.05 | 0.05 |
| NC_004718.3;NP_828849.2;SARS-CoV_orf1ab | 1838 | 1838 | 1 | 0.05 | 0.05 | 0.05 |
| NC_004718.3;NP_828849.2;SARS-CoV_orf1ab | 1843 | 1844 | 2 | 0.05 | 0.05 | 0.05 |
| NC_004718.3;NP_828849.2;SARS-CoV_orf1ab | 1846 | 1846 | 1 | 0.05 | 0.05 | 0.05 |

| Extended data 1 | | | | | | |
|---|---|---|---|---|---|---|
| Protein | Start position (peptide) | Stop position (peptide) | Number of peptides | Minimum fraction (per peptide) of control subjects with reactivity >3 sd above average | Maximum fraction (per peptide) of control subjects with reactivity >3 sd above average | Mean fraction (over all peptides in this contiguous stretch) of controlsubjects with reactivity >3 sd above average |
| NC_004718.3;NP_828849.2;SARS-CoV_orf1ab | 1850 | 1850 | 1 | 0.05 | 0.05 | 0.05 |
| NC_004718.3;NP_828849.2;SARS-CoV_orf1ab | 1857 | 1860 | 4 | 0.05 | 0.05 | 0.05 |
| NC_004718.3;NP_828849.2;SARS-CoV_orf1ab | 1863 | 1863 | 1 | 0.2 | 0.2 | 0.2 |
| NC_004718.3;NP_828849.2;SARS-CoV_orf1ab | 1868 | 1868 | 1 | 0.05 | 0.05 | 0.05 |
| NC_004718.3;NP_828849.2;SARS-CoV_orf1ab | 1870 | 1898 | 29 | 0.05 | 0.15 | 0.072413793 |
| NC_004718.3;NP_828849.2;SARS-CoV_orf1ab | 1904 | 1915 | 12 | 0.05 | 0.4 | 0.195833333 |
| NC_004718.3;NP_828849.2;SARS-CoV_orf1ab | 1942 | 1942 | 1 | 0.05 | 0.05 | 0.05 |
| NC_004718.3;NP_828849.2;SARS-CoV_orf1ab | 1951 | 1951 | 1 | 0.05 | 0.05 | 0.05 |
| NC_004718.3;NP_828849.2;SARS-CoV_orf1ab | 1967 | 1975 | 9 | 0.05 | 0.1 | 0.066666667 |
| NC_004718.3;NP_828849.2;SARS-CoV_orf1ab | 1986 | 1991 | 6 | 0.05 | 0.15 | 0.091666667 |
| NC_004718.3;NP_828849.2;SARS-CoV_orf1ab | 1994 | 1996 | 3 | 0.05 | 0.15 | 0.083333333 |
| NC_004718.3;NP_828849.2;SARS-CoV_orf1ab | 2000 | 2000 | 1 | 0.05 | 0.05 | 0.05 |
| NC_004718.3;NP_828849.2;SARS-CoV_orf1ab | 2005 | 2006 | 2 | 0.05 | 0.1 | 0.075 |
| NC_004718.3;NP_828849.2;SARS-CoV_orf1ab | 2008 | 2008 | 1 | 0.15 | 0.15 | 0.15 |
| NC_004718.3;NP_828849.2;SARS-CoV_orf1ab | 2031 | 2031 | 1 | 0.05 | 0.05 | 0.05 |
| NC_004718.3;NP_828849.2;SARS-CoV_orf1ab | 2060 | 2060 | 1 | 0.05 | 0.05 | 0.05 |
| NC_004718.3;NP_828849.2;SARS-CoV_orf1ab | 2068 | 2068 | 1 | 0.25 | 0.25 | 0.25 |
| NC_004718.3;NP_828849.2;SARS-CoV_orf1ab | 2074 | 2078 | 5 | 0.05 | 0.15 | 0.09 |
| NC_004718.3;NP_828849.2;SARS-CoV_orf1ab | 2091 | 2091 | 1 | 0.05 | 0.05 | 0.05 |
| NC_004718.3;NP_828849.2;SARS-CoV_orf1ab | 2100 | 2107 | 8 | 0.05 | 0.05 | 0.05 |
| NC_004718.3;NP_828849.2;SARS-CoV_orf1ab | 2109 | 2110 | 2 | 0.05 | 0.05 | 0.05 |
| NC_004718.3;NP_828849.2;SARS-CoV_orf1ab | 2117 | 2117 | 1 | 0.05 | 0.05 | 0.05 |
| NC_004718.3;NP_828849.2;SARS-CoV_orf1ab | 2119 | 2120 | 2 | 0.05 | 0.1 | 0.075 |
| NC_004718.3;NP_828849.2;SARS-CoV_orf1ab | 2127 | 2127 | 1 | 0.1 | 0.1 | 0.1 |
| NC_004718.3;NP_828849.2;SARS-CoV_orf1ab | 2158 | 2158 | 1 | 0.05 | 0.05 | 0.05 |
| NC_004718.3;NP_828849.2;SARS-CoV_orf1ab | 2165 | 2165 | 1 | 0.05 | 0.05 | 0.05 |
| NC_004718.3;NP_828849.2;SARS-CoV_orf1ab | 2178 | 2185 | 8 | 0.05 | 0.05 | 0.05 |
| NC_004718.3;NP_828849.2;SARS-CoV_orf1ab | 2187 | 2189 | 3 | 0.05 | 0.1 | 0.066666667 |
| NC_004718.3;NP_828849.2;SARS-CoV_orf1ab | 2231 | 2231 | 1 | 0.05 | 0.05 | 0.05 |
| NC_004718.3;NP_828849.2;SARS-CoV_orf1ab | 2238 | 2248 | 11 | 0.05 | 0.45 | 0.245454545 |
| NC_004718.3;NP_828849.2;SARS-CoV_orf1ab | 2251 | 2251 | 1 | 0.05 | 0.05 | 0.05 |
| NC_004718.3;NP_828849.2;SARS-CoV_orf1ab | 2268 | 2268 | 1 | 0.05 | 0.05 | 0.05 |
| NC_004718.3;NP_828849.2;SARS-CoV_orf1ab | 2270 | 2273 | 4 | 0.05 | 0.05 | 0.05 |
| NC_004718.3;NP_828849.2;SARS-CoV_orf1ab | 2287 | 2287 | 1 | 0.05 | 0.05 | 0.05 |
| NC_004718.3;NP_828849.2;SARS-CoV_orf1ab | 2446 | 2448 | 3 | 0.05 | 0.05 | 0.05 |
| NC_004718.3;NP_828849.2;SARS-CoV_orf1ab | 2461 | 2461 | 1 | 0.05 | 0.05 | 0.05 |
| NC_004718.3;NP_828849.2;SARS-CoV_orf1ab | 2499 | 2502 | 4 | 0.05 | 0.05 | 0.05 |

| Extended data 1 | | | | | | |
|---|---|---|---|---|---|---|
| Protein | Start position (peptide) | Stop position (peptide) | Number of peptides | Minimum fraction (per peptide) of control subjects with reactivity >3 sd above average | Maximum fraction (per peptide) of control subjects with reactivity >3 sd above average | Mean fraction (over all peptides in this contiguous stretch) of controlsubjects with reactivity >3 sd above average |
| NC_004718.3;NP_828849.2;SARS-CoV_orf1ab | 2505 | 2505 | 1 | 0.05 | 0.05 | 0.05 |
| NC_004718.3;NP_828849.2;SARS-CoV_orf1ab | 2513 | 2516 | 4 | 0.05 | 0.05 | 0.05 |
| NC_004718.3;NP_828849.2;SARS-CoV_orf1ab | 2518 | 2520 | 3 | 0.05 | 0.15 | 0.083333333 |
| NC_004718.3;NP_828849.2;SARS-CoV_orf1ab | 2525 | 2525 | 1 | 0.05 | 0.05 | 0.05 |
| NC_004718.3;NP_828849.2;SARS-CoV_orf1ab | 2527 | 2528 | 2 | 0.05 | 0.05 | 0.05 |
| NC_004718.3;NP_828849.2;SARS-CoV_orf1ab | 2548 | 2548 | 1 | 0.1 | 0.1 | 0.1 |
| NC_004718.3;NP_828849.2;SARS-CoV_orf1ab | 2563 | 2564 | 2 | 0.05 | 0.05 | 0.05 |
| NC_004718.3;NP_828849.2;SARS-CoV_orf1ab | 2572 | 2573 | 2 | 0.05 | 0.1 | 0.075 |
| NC_004718.3;NP_828849.2;SARS-CoV_orf1ab | 2586 | 2596 | 11 | 0.1 | 0.25 | 0.177272727 |
| NC_004718.3;NP_828849.2;SARS-CoV_orf1ab | 2599 | 2599 | 1 | 0.05 | 0.05 | 0.05 |
| NC_004718.3;NP_828849.2;SARS-CoV_orf1ab | 2605 | 2605 | 1 | 0.05 | 0.05 | 0.05 |
| NC_004718.3;NP_828849.2;SARS-CoV_orf1ab | 2615 | 2616 | 2 | 0.05 | 0.05 | 0.05 |
| NC_004718.3;NP_828849.2;SARS-CoV_orf1ab | 2638 | 2638 | 1 | 0.05 | 0.05 | 0.05 |
| NC_004718.3;NP_828849.2;SARS-CoV_orf1ab | 2640 | 2641 | 2 | 0.05 | 0.05 | 0.05 |
| NC_004718.3;NP_828849.2;SARS-CoV_orf1ab | 2652 | 2654 | 3 | 0.05 | 0.05 | 0.05 |
| NC_004718.3;NP_828849.2;SARS-CoV_orf1ab | 2661 | 2662 | 2 | 0.05 | 0.05 | 0.05 |
| NC_004718.3;NP_828849.2;SARS-CoV_orf1ab | 2666 | 2666 | 1 | 0.05 | 0.05 | 0.05 |
| NC_004718.3;NP_828849.2;SARS-CoV_orf1ab | 2668 | 2668 | 1 | 0.05 | 0.05 | 0.05 |
| NC_004718.3;NP_828849.2;SARS-CoV_orf1ab | 2670 | 2670 | 1 | 0.05 | 0.05 | 0.05 |
| NC_004718.3;NP_828849.2;SARS-CoV_orf1ab | 2679 | 2679 | 1 | 0.05 | 0.05 | 0.05 |
| NC_004718.3;NP_828849.2;SARS-CoV_orf1ab | 2692 | 2692 | 1 | 0.05 | 0.05 | 0.05 |
| NC_004718.3;NP_828849.2;SARS-CoV_orf1ab | 2695 | 2695 | 1 | 0.05 | 0.05 | 0.05 |
| NC_004718.3;NP_828849.2;SARS-CoV_orf1ab | 2705 | 2706 | 2 | 0.05 | 0.05 | 0.05 |
| NC_004718.3;NP_828849.2;SARS-CoV_orf1ab | 2767 | 2768 | 2 | 0.05 | 0.05 | 0.05 |
| NC_004718.3;NP_828849.2;SARS-CoV_orf1ab | 2770 | 2770 | 1 | 0.05 | 0.05 | 0.05 |
| NC_004718.3;NP_828849.2;SARS-CoV_orf1ab | 2773 | 2774 | 2 | 0.05 | 0.1 | 0.075 |
| NC_004718.3;NP_828849.2;SARS-CoV_orf1ab | 2776 | 2784 | 9 | 0.05 | 0.15 | 0.094444444 |
| NC_004718.3;NP_828849.2;SARS-CoV_orf1ab | 2788 | 2788 | 1 | 0.1 | 0.1 | 0.1 |
| NC_004718.3;NP_828849.2;SARS-CoV_orf1ab | 2792 | 2792 | 1 | 0.15 | 0.15 | 0.15 |
| NC_004718.3;NP_828849.2;SARS-CoV_orf1ab | 2798 | 2799 | 2 | 0.05 | 0.05 | 0.05 |
| NC_004718.3;NP_828849.2;SARS-CoV_orf1ab | 2805 | 2805 | 1 | 0.05 | 0.05 | 0.05 |
| NC_004718.3;NP_828849.2;SARS-CoV_orf1ab | 2811 | 2812 | 2 | 0.05 | 0.05 | 0.05 |
| NC_004718.3;NP_828849.2;SARS-CoV_orf1ab | 2814 | 2815 | 2 | 0.05 | 0.05 | 0.05 |
| NC_004718.3;NP_828849.2;SARS-CoV_orf1ab | 2818 | 2821 | 4 | 0.05 | 0.1 | 0.075 |
| NC_004718.3;NP_828849.2;SARS-CoV_orf1ab | 2842 | 2843 | 2 | 0.05 | 0.05 | 0.05 |
| NC_004718.3;NP_828849.2;SARS-CoV_orf1ab | 2850 | 2850 | 1 | 0.05 | 0.05 | 0.05 |
| NC_004718.3;NP_828849.2;SARS-CoV_orf1ab | 2872 | 2876 | 5 | 0.05 | 0.05 | 0.05 |

| Extended data 1 | | | | | | |
|---|---|---|---|---|---|---|
| Protein | Start position (peptide) | Stop position (peptide) | Number of peptides | Minimum fraction (per peptide) of control subjects with reactivity >3 sd above average | Maximum fraction (per peptide) of control subjects with reactivity >3 sd above average | Mean fraction (over all peptides in this contiguous stretch) of controlsubjects with reactivity >3 sd above average |
| NC_004718.3;NP_828849.2;SARS-CoV_orf1ab | 2878 | 2881 | 4 | 0.05 | 0.05 | 0.05 |
| NC_004718.3;NP_828849.2;SARS-CoV_orf1ab | 2916 | 2926 | 11 | 0.05 | 0.25 | 0.109090909 |
| NC_004718.3;NP_828849.2;SARS-CoV_orf1ab | 2948 | 2948 | 1 | 0.05 | 0.05 | 0.05 |
| NC_004718.3;NP_828849.2;SARS-CoV_orf1ab | 2992 | 2992 | 1 | 0.05 | 0.05 | 0.05 |
| NC_004718.3;NP_828849.2;SARS-CoV_orf1ab | 2997 | 2998 | 2 | 0.05 | 0.05 | 0.05 |
| NC_004718.3;NP_828849.2;SARS-CoV_orf1ab | 3013 | 3013 | 1 | 0.05 | 0.05 | 0.05 |
| NC_004718.3;NP_828849.2;SARS-CoV_orf1ab | 3042 | 3042 | 1 | 0.05 | 0.05 | 0.05 |
| NC_004718.3;NP_828849.2;SARS-CoV_orf1ab | 3094 | 3094 | 1 | 0.05 | 0.05 | 0.05 |
| NC_004718.3;NP_828849.2;SARS-CoV_orf1ab | 3137 | 3137 | 1 | 0.05 | 0.05 | 0.05 |
| NC_004718.3;NP_828849.2;SARS-CoV_orf1ab | 3141 | 3141 | 1 | 0.05 | 0.05 | 0.05 |
| NC_004718.3;NP_828849.2;SARS-CoV_orf1ab | 3192 | 3192 | 1 | 0.05 | 0.05 | 0.05 |
| NC_004718.3;NP_828849.2;SARS-CoV_orf1ab | 3207 | 3207 | 1 | 0.05 | 0.05 | 0.05 |
| NC_004718.3;NP_828849.2;SARS-CoV_orf1ab | 3214 | 3216 | 3 | 0.05 | 0.3 | 0.166666667 |
| NC_004718.3;NP_828849.2;SARS-CoV_orf1ab | 3218 | 3218 | 1 | 0.1 | 0.1 | 0.1 |
| NC_004718.3;NP_828849.2;SARS-CoV_orf1ab | 3227 | 3227 | 1 | 0.05 | 0.05 | 0.05 |
| NC_004718.3;NP_828849.2;SARS-CoV_orf1ab | 3247 | 3247 | 1 | 0.05 | 0.05 | 0.05 |
| NC_004718.3;NP_828849.2;SARS-CoV_orf1ab | 3279 | 3279 | 1 | 0.05 | 0.05 | 0.05 |
| NC_004718.3;NP_828849.2;SARS-CoV_orf1ab | 3330 | 3330 | 1 | 0.15 | 0.15 | 0.15 |
| NC_004718.3;NP_828849.2;SARS-CoV_orf1ab | 3371 | 3371 | 1 | 0.05 | 0.05 | 0.05 |
| NC_004718.3;NP_828849.2;SARS-CoV_orf1ab | 3403 | 3414 | 12 | 0.05 | 0.25 | 0.175 |
| NC_004718.3;NP_828849.2;SARS-CoV_orf1ab | 3417 | 3421 | 5 | 0.05 | 0.1 | 0.07 |
| NC_004718.3;NP_828849.2;SARS-CoV_orf1ab | 3423 | 3423 | 1 | 0.05 | 0.05 | 0.05 |
| NC_004718.3;NP_828849.2;SARS-CoV_orf1ab | 3425 | 3425 | 1 | 0.05 | 0.05 | 0.05 |
| NC_004718.3;NP_828849.2;SARS-CoV_orf1ab | 3456 | 3467 | 12 | 0.05 | 0.25 | 0.075 |
| NC_004718.3;NP_828849.2;SARS-CoV_orf1ab | 3469 | 3469 | 1 | 0.05 | 0.05 | 0.05 |
| NC_004718.3;NP_828849.2;SARS-CoV_orf1ab | 3473 | 3473 | 1 | 0.05 | 0.05 | 0.05 |
| NC_004718.3;NP_828849.2;SARS-CoV_orf1ab | 3480 | 3480 | 1 | 0.05 | 0.05 | 0.05 |
| NC_004718.3;NP_828849.2;SARS-CoV_orf1ab | 3482 | 3492 | 11 | 0.05 | 0.15 | 0.068181818 |
| NC_004718.3;NP_828849.2;SARS-CoV_orf1ab | 3498 | 3498 | 1 | 0.05 | 0.05 | 0.05 |
| NC_004718.3;NP_828849.2;SARS-CoV_orf1ab | 3528 | 3528 | 1 | 0.1 | 0.1 | 0.1 |
| NC_004718.3;NP_828849.2;SARS-CoV_orf1ab | 3536 | 3543 | 8 | 0.05 | 0.05 | 0.05 |
| NC_004718.3;NP_828849.2;SARS-CoV_orf1ab | 3545 | 3547 | 3 | 0.05 | 0.05 | 0.05 |
| NC_004718.3;NP_828849.2;SARS-CoV_orf1ab | 3695 | 3699 | 5 | 0.05 | 0.05 | 0.05 |
| NC_004718.3;NP_828849.2;SARS-CoV_orf1ab | 3786 | 3786 | 1 | 0.25 | 0.25 | 0.25 |
| NC_004718.3;NP_828849.2;SARS-CoV_orf1ab | 3795 | 3796 | 2 | 0.05 | 0.05 | 0.05 |
| NC_004718.3;NP_828849.2;SARS-CoV_orf1ab | 3801 | 3806 | 6 | 0.05 | 0.2 | 0.125 |
| NC_004718.3;NP_828849.2;SARS-CoV_orf1ab | 3823 | 3829 | 7 | 0.05 | 0.1 | 0.057142857 |

| Extended data 1 | | | | | | |
|---|---|---|---|---|---|---|
| Protein | Start position (peptide) | Stop position (peptide) | Number of peptides | Minimum fraction (per peptide) of control subjects with reactivity >3 sd above average | Maximum fraction (per peptide) of control subjects with reactivity >3 sd above average | Mean fraction (over all peptides in this contiguous stretch) of controlsubjects with reactivity >3 sd above average |
| NC_004718.3;NP_828849.2;SARS-CoV_orf1ab | 3831 | 3834 | 4 | 0.05 | 0.05 | 0.05 |
| NC_004718.3;NP_828849.2;SARS-CoV_orf1ab | 3865 | 3867 | 3 | 0.05 | 0.05 | 0.05 |
| NC_004718.3;NP_828849.2;SARS-CoV_orf1ab | 3894 | 3900 | 7 | 0.05 | 0.1 | 0.078571429 |
| NC_004718.3;NP_828849.2;SARS-CoV_orf1ab | 3906 | 3908 | 3 | 0.05 | 0.05 | 0.05 |
| NC_004718.3;NP_828849.2;SARS-CoV_orf1ab | 3910 | 3910 | 1 | 0.05 | 0.05 | 0.05 |
| NC_004718.3;NP_828849.2;SARS-CoV_orf1ab | 3918 | 3918 | 1 | 0.05 | 0.05 | 0.05 |
| NC_004718.3;NP_828849.2;SARS-CoV_orf1ab | 3922 | 3922 | 1 | 0.05 | 0.05 | 0.05 |
| NC_004718.3;NP_828849.2;SARS-CoV_orf1ab | 3928 | 3928 | 1 | 0.05 | 0.05 | 0.05 |
| NC_004718.3;NP_828849.2;SARS-CoV_orf1ab | 3930 | 3934 | 5 | 0.05 | 0.1 | 0.06 |
| NC_004718.3;NP_828849.2;SARS-CoV_orf1ab | 3937 | 3937 | 1 | 0.05 | 0.05 | 0.05 |
| NC_004718.3;NP_828849.2;SARS-CoV_orf1ab | 3957 | 3958 | 2 | 0.05 | 0.05 | 0.05 |
| NC_004718.3;NP_828849.2;SARS-CoV_orf1ab | 3960 | 3960 | 1 | 0.05 | 0.05 | 0.05 |
| NC_004718.3;NP_828849.2;SARS-CoV_orf1ab | 4008 | 4018 | 11 | 0.05 | 0.4 | 0.227272727 |
| NC_004718.3;NP_828849.2;SARS-CoV_orf1ab | 4020 | 4021 | 2 | 0.05 | 0.1 | 0.075 |
| NC_004718.3;NP_828849.2;SARS-CoV_orf1ab | 4023 | 4024 | 2 | 0.05 | 0.05 | 0.05 |
| NC_004718.3;NP_828849.2;SARS-CoV_orf1ab | 4030 | 4031 | 2 | 0.05 | 0.05 | 0.05 |
| NC_004718.3;NP_828849.2;SARS-CoV_orf1ab | 4040 | 4050 | 11 | 0.05 | 0.1 | 0.054545455 |
| NC_004718.3;NP_828849.2;SARS-CoV_orf1ab | 4052 | 4052 | 1 | 0.05 | 0.05 | 0.05 |
| NC_004718.3;NP_828849.2;SARS-CoV_orf1ab | 4069 | 4070 | 2 | 0.05 | 0.05 | 0.05 |
| NC_004718.3;NP_828849.2;SARS-CoV_orf1ab | 4080 | 4080 | 1 | 0.15 | 0.15 | 0.15 |
| NC_004718.3;NP_828849.2;SARS-CoV_orf1ab | 4109 | 4111 | 3 | 0.05 | 0.1 | 0.066666667 |
| NC_004718.3;NP_828849.2;SARS-CoV_orf1ab | 4114 | 4114 | 1 | 0.05 | 0.05 | 0.05 |
| NC_004718.3;NP_828849.2;SARS-CoV_orf1ab | 4118 | 4118 | 1 | 0.05 | 0.05 | 0.05 |
| NC_004718.3;NP_828849.2;SARS-CoV_orf1ab | 4138 | 4141 | 4 | 0.05 | 0.25 | 0.15 |
| NC_004718.3;NP_828849.2;SARS-CoV_orf1ab | 4180 | 4180 | 1 | 0.2 | 0.2 | 0.2 |
| NC_004718.3;NP_828849.2;SARS-CoV_orf1ab | 4182 | 4185 | 4 | 0.05 | 0.05 | 0.05 |
| NC_004718.3;NP_828849.2;SARS-CoV_orf1ab | 4192 | 4192 | 1 | 0.05 | 0.05 | 0.05 |
| NC_004718.3;NP_828849.2;SARS-CoV_orf1ab | 4206 | 4212 | 7 | 0.05 | 0.25 | 0.107142857 |
| NC_004718.3;NP_828849.2;SARS-CoV_orf1ab | 4224 | 4224 | 1 | 0.05 | 0.05 | 0.05 |
| NC_004718.3;NP_828849.2;SARS-CoV_orf1ab | 4244 | 4251 | 8 | 0.05 | 0.15 | 0.08125 |
| NC_004718.3;NP_828849.2;SARS-CoV_orf1ab | 4256 | 4257 | 2 | 0.05 | 0.15 | 0.1 |
| NC_004718.3;NP_828849.2;SARS-CoV_orf1ab | 4285 | 4285 | 1 | 0.05 | 0.05 | 0.05 |
| NC_004718.3;NP_828849.2;SARS-CoV_orf1ab | 4287 | 4287 | 1 | 0.05 | 0.05 | 0.05 |
| NC_004718.3;NP_828849.2;SARS-CoV_orf1ab | 4321 | 4325 | 5 | 0.05 | 0.05 | 0.05 |
| NC_004718.3;NP_828849.2;SARS-CoV_orf1ab | 4363 | 4363 | 1 | 0.05 | 0.05 | 0.05 |
| NC_004718.3;NP_828849.2;SARS-CoV_orf1ab | 4365 | 4365 | 1 | 0.05 | 0.05 | 0.05 |
| NC_004718.3;NP_828849.2;SARS-CoV_orf1ab | 4371 | 4371 | 1 | 0.05 | 0.05 | 0.05 |

| Extended data 1 | | | | | | |
|---|---|---|---|---|---|---|
| Protein | Start position (peptide) | Stop position (peptide) | Number of peptides | Minimum fraction (per peptide) of control subjects with reactivity >3 sd above average | Maximum fraction (per peptide) of control subjects with reactivity >3 sd above average | Mean fraction (over all peptides in this contiguous stretch) of controlsubjects with reactivity >3 sd above average |
| NC_004718.3;NP_828849.2;SARS-CoV_orf1ab | 4395 | 4397 | 3 | 0.05 | 0.1 | 0.083333333 |
| NC_004718.3;NP_828849.2;SARS-CoV_orf1ab | 4418 | 4425 | 8 | 0.05 | 0.1 | 0.075 |
| NC_004718.3;NP_828849.2;SARS-CoV_orf1ab | 4427 | 4427 | 1 | 0.05 | 0.05 | 0.05 |
| NC_004718.3;NP_828849.2;SARS-CoV_orf1ab | 4447 | 4447 | 1 | 0.05 | 0.05 | 0.05 |
| NC_004718.3;NP_828849.2;SARS-CoV_orf1ab | 4454 | 4456 | 3 | 0.05 | 0.05 | 0.05 |
| NC_004718.3;NP_828849.2;SARS-CoV_orf1ab | 4497 | 4497 | 1 | 0.05 | 0.05 | 0.05 |
| NC_004718.3;NP_828849.2;SARS-CoV_orf1ab | 4522 | 4523 | 2 | 0.05 | 0.2 | 0.125 |
| NC_004718.3;NP_828849.2;SARS-CoV_orf1ab | 4526 | 4528 | 3 | 0.05 | 0.1 | 0.066666667 |
| NC_004718.3;NP_828849.2;SARS-CoV_orf1ab | 4534 | 4536 | 3 | 0.05 | 0.05 | 0.05 |
| NC_004718.3;NP_828849.2;SARS-CoV_orf1ab | 4542 | 4542 | 1 | 0.05 | 0.05 | 0.05 |
| NC_004718.3;NP_828849.2;SARS-CoV_orf1ab | 4549 | 4549 | 1 | 0.05 | 0.05 | 0.05 |
| NC_004718.3;NP_828849.2;SARS-CoV_orf1ab | 4565 | 4566 | 2 | 0.1 | 0.1 | 0.1 |
| NC_004718.3;NP_828849.2;SARS-CoV_orf1ab | 4568 | 4568 | 1 | 0.05 | 0.05 | 0.05 |
| NC_004718.3;NP_828849.2;SARS-CoV_orf1ab | 4572 | 4572 | 1 | 0.05 | 0.05 | 0.05 |
| NC_004718.3;NP_828849.2;SARS-CoV_orf1ab | 4582 | 4582 | 1 | 0.05 | 0.05 | 0.05 |
| NC_004718.3;NP_828849.2;SARS-CoV_orf1ab | 4586 | 4587 | 2 | 0.05 | 0.1 | 0.075 |
| NC_004718.3;NP_828849.2;SARS-CoV_orf1ab | 4612 | 4612 | 1 | 0.05 | 0.05 | 0.05 |
| NC_004718.3;NP_828849.2;SARS-CoV_orf1ab | 4618 | 4618 | 1 | 0.05 | 0.05 | 0.05 |
| NC_004718.3;NP_828849.2;SARS-CoV_orf1ab | 4621 | 4629 | 9 | 0.05 | 0.15 | 0.066666667 |
| NC_004718.3;NP_828849.2;SARS-CoV_orf1ab | 4631 | 4636 | 6 | 0.05 | 0.1 | 0.058333333 |
| NC_004718.3;NP_828849.2;SARS-CoV_orf1ab | 4638 | 4641 | 4 | 0.05 | 0.05 | 0.05 |
| NC_004718.3;NP_828849.2;SARS-CoV_orf1ab | 4647 | 4647 | 1 | 0.05 | 0.05 | 0.05 |
| NC_004718.3;NP_828849.2;SARS-CoV_orf1ab | 4726 | 4726 | 1 | 0.05 | 0.05 | 0.05 |
| NC_004718.3;NP_828849.2;SARS-CoV_orf1ab | 4734 | 4737 | 4 | 0.05 | 0.1 | 0.075 |
| NC_004718.3;NP_828849.2;SARS-CoV_orf1ab | 4754 | 4754 | 1 | 0.05 | 0.05 | 0.05 |
| NC_004718.3;NP_828849.2;SARS-CoV_orf1ab | 4768 | 4769 | 2 | 0.05 | 0.05 | 0.05 |
| NC_004718.3;NP_828849.2;SARS-CoV_orf1ab | 4776 | 4782 | 7 | 0.05 | 0.2 | 0.1 |
| NC_004718.3;NP_828849.2;SARS-CoV_orf1ab | 4785 | 4787 | 3 | 0.05 | 0.05 | 0.05 |
| NC_004718.3;NP_828849.2;SARS-CoV_orf1ab | 4794 | 4794 | 1 | 0.05 | 0.05 | 0.05 |
| NC_004718.3;NP_828849.2;SARS-CoV_orf1ab | 4803 | 4804 | 2 | 0.05 | 0.05 | 0.05 |
| NC_004718.3;NP_828849.2;SARS-CoV_orf1ab | 4807 | 4807 | 1 | 0.05 | 0.05 | 0.05 |
| NC_004718.3;NP_828849.2;SARS-CoV_orf1ab | 4813 | 4821 | 9 | 0.05 | 0.25 | 0.15 |
| NC_004718.3;NP_828849.2;SARS-CoV_orf1ab | 4832 | 4832 | 1 | 0.05 | 0.05 | 0.05 |
| NC_004718.3;NP_828849.2;SARS-CoV_orf1ab | 4845 | 4848 | 4 | 0.05 | 0.05 | 0.05 |
| NC_004718.3;NP_828849.2;SARS-CoV_orf1ab | 4850 | 4851 | 2 | 0.05 | 0.05 | 0.05 |
| NC_004718.3;NP_828849.2;SARS-CoV_orf1ab | 4853 | 4853 | 1 | 0.05 | 0.05 | 0.05 |
| NC_004718.3;NP_828849.2;SARS-CoV_orf1ab | 4855 | 4857 | 3 | 0.05 | 0.1 | 0.066666667 |

| Extended data 1 | | | | | | |
|---|---|---|---|---|---|---|
| Protein | Start position (peptide) | Stop position (peptide) | Number of peptides | Minimum fraction (per peptide) of control subjects with reactivity >3 sd above average | Maximum fraction (per peptide) of control subjects with reactivity >3 sd above average | Mean fraction (over all peptides in this contiguous stretch) of controlsubjects with reactivity >3 sd above average |
| NC_004718.3;NP_828849.2;SARS-CoV_orf1ab | 4859 | 4859 | 1 | 0.15 | 0.15 | 0.15 |
| NC_004718.3;NP_828849.2;SARS-CoV_orf1ab | 4861 | 4862 | 2 | 0.05 | 0.15 | 0.1 |
| NC_004718.3;NP_828849.2;SARS-CoV_orf1ab | 4874 | 4874 | 1 | 0.05 | 0.05 | 0.05 |
| NC_004718.3;NP_828849.2;SARS-CoV_orf1ab | 4876 | 4876 | 1 | 0.05 | 0.05 | 0.05 |
| NC_004718.3;NP_828849.2;SARS-CoV_orf1ab | 4886 | 4894 | 9 | 0.05 | 0.2 | 0.077777778 |
| NC_004718.3;NP_828849.2;SARS-CoV_orf1ab | 4900 | 4901 | 2 | 0.05 | 0.1 | 0.075 |
| NC_004718.3;NP_828849.2;SARS-CoV_orf1ab | 4906 | 4906 | 1 | 0.1 | 0.1 | 0.1 |
| NC_004718.3;NP_828849.2;SARS-CoV_orf1ab | 4940 | 4949 | 10 | 0.15 | 0.4 | 0.26 |
| NC_004718.3;NP_828849.2;SARS-CoV_orf1ab | 5034 | 5034 | 1 | 0.05 | 0.05 | 0.05 |
| NC_004718.3;NP_828849.2;SARS-CoV_orf1ab | 5046 | 5047 | 2 | 0.05 | 0.1 | 0.075 |
| NC_004718.3;NP_828849.2;SARS-CoV_orf1ab | 5049 | 5049 | 1 | 0.05 | 0.05 | 0.05 |
| NC_004718.3;NP_828849.2;SARS-CoV_orf1ab | 5071 | 5071 | 1 | 0.05 | 0.05 | 0.05 |
| NC_004718.3;NP_828849.2;SARS-CoV_orf1ab | 5074 | 5074 | 1 | 0.05 | 0.05 | 0.05 |
| NC_004718.3;NP_828849.2;SARS-CoV_orf1ab | 5078 | 5078 | 1 | 0.05 | 0.05 | 0.05 |
| NC_004718.3;NP_828849.2;SARS-CoV_orf1ab | 5090 | 5090 | 1 | 0.05 | 0.05 | 0.05 |
| NC_004718.3;NP_828849.2;SARS-CoV_orf1ab | 5101 | 5101 | 1 | 0.05 | 0.05 | 0.05 |
| NC_004718.3;NP_828849.2;SARS-CoV_orf1ab | 5108 | 5108 | 1 | 0.05 | 0.05 | 0.05 |
| NC_004718.3;NP_828849.2;SARS-CoV_orf1ab | 5116 | 5116 | 1 | 0.05 | 0.05 | 0.05 |
| NC_004718.3;NP_828849.2;SARS-CoV_orf1ab | 5126 | 5127 | 2 | 0.05 | 0.05 | 0.05 |
| NC_004718.3;NP_828849.2;SARS-CoV_orf1ab | 5131 | 5132 | 2 | 0.05 | 0.1 | 0.075 |
| NC_004718.3;NP_828849.2;SARS-CoV_orf1ab | 5152 | 5152 | 1 | 0.05 | 0.05 | 0.05 |
| NC_004718.3;NP_828849.2;SARS-CoV_orf1ab | 5161 | 5161 | 1 | 0.05 | 0.05 | 0.05 |
| NC_004718.3;NP_828849.2;SARS-CoV_orf1ab | 5163 | 5163 | 1 | 0.05 | 0.05 | 0.05 |
| NC_004718.3;NP_828849.2;SARS-CoV_orf1ab | 5166 | 5169 | 4 | 0.05 | 0.05 | 0.05 |
| NC_004718.3;NP_828849.2;SARS-CoV_orf1ab | 5183 | 5184 | 2 | 0.05 | 0.05 | 0.05 |
| NC_004718.3;NP_828849.2;SARS-CoV_orf1ab | 5186 | 5186 | 1 | 0.05 | 0.05 | 0.05 |
| NC_004718.3;NP_828849.2;SARS-CoV_orf1ab | 5188 | 5188 | 1 | 0.1 | 0.1 | 0.1 |
| NC_004718.3;NP_828849.2;SARS-CoV_orf1ab | 5206 | 5206 | 1 | 0.05 | 0.05 | 0.05 |
| NC_004718.3;NP_828849.2;SARS-CoV_orf1ab | 5279 | 5279 | 1 | 0.05 | 0.05 | 0.05 |
| NC_004718.3;NP_828849.2;SARS-CoV_orf1ab | 5322 | 5322 | 1 | 0.05 | 0.05 | 0.05 |
| NC_004718.3;NP_828849.2;SARS-CoV_orf1ab | 5347 | 5350 | 4 | 0.05 | 0.05 | 0.05 |
| NC_004718.3;NP_828849.2;SARS-CoV_orf1ab | 5355 | 5355 | 1 | 0.05 | 0.05 | 0.05 |
| NC_004718.3;NP_828849.2;SARS-CoV_orf1ab | 5392 | 5402 | 11 | 0.05 | 0.2 | 0.086363636 |
| NC_004718.3;NP_828849.2;SARS-CoV_orf1ab | 5418 | 5418 | 1 | 0.05 | 0.05 | 0.05 |
| NC_004718.3;NP_828849.2;SARS-CoV_orf1ab | 5429 | 5431 | 3 | 0.05 | 0.05 | 0.05 |
| NC_004718.3;NP_828849.2;SARS-CoV_orf1ab | 5436 | 5436 | 1 | 0.05 | 0.05 | 0.05 |
| NC_004718.3;NP_828849.2;SARS-CoV_orf1ab | 5443 | 5443 | 1 | 0.05 | 0.05 | 0.05 |

| Extended data 1 | | | | | | |
|---|---|---|---|---|---|---|
| Protein | Start position (peptide) | Stop position (peptide) | Number of peptides | Minimum fraction (per peptide) of control subjects with reactivity >3 sd above average | Maximum fraction (per peptide) of control subjects with reactivity >3 sd above average | Mean fraction (over all peptides in this contiguous stretch) of controlsubjects with reactivity >3 sd above average |
| NC_004718.3;NP_828849.2;SARS-CoV_orf1ab | 5447 | 5452 | 6 | 0.05 | 0.05 | 0.05 |
| NC_004718.3;NP_828849.2;SARS-CoV_orf1ab | 5460 | 5460 | 1 | 0.05 | 0.05 | 0.05 |
| NC_004718.3;NP_828849.2;SARS-CoV_orf1ab | 5483 | 5496 | 14 | 0.05 | 0.1 | 0.071428571 |
| NC_004718.3;NP_828849.2;SARS-CoV_orf1ab | 5499 | 5499 | 1 | 0.1 | 0.1 | 0.1 |
| NC_004718.3;NP_828849.2;SARS-CoV_orf1ab | 5504 | 5504 | 1 | 0.05 | 0.05 | 0.05 |
| NC_004718.3;NP_828849.2;SARS-CoV_orf1ab | 5524 | 5524 | 1 | 0.05 | 0.05 | 0.05 |
| NC_004718.3;NP_828849.2;SARS-CoV_orf1ab | 5548 | 5557 | 10 | 0.05 | 0.1 | 0.055 |
| NC_004718.3;NP_828849.2;SARS-CoV_orf1ab | 5559 | 5559 | 1 | 0.05 | 0.05 | 0.05 |
| NC_004718.3;NP_828849.2;SARS-CoV_orf1ab | 5561 | 5562 | 2 | 0.1 | 0.1 | 0.1 |
| NC_004718.3;NP_828849.2;SARS-CoV_orf1ab | 5564 | 5564 | 1 | 0.05 | 0.05 | 0.05 |
| NC_004718.3;NP_828849.2;SARS-CoV_orf1ab | 5575 | 5575 | 1 | 0.05 | 0.05 | 0.05 |
| NC_004718.3;NP_828849.2;SARS-CoV_orf1ab | 5615 | 5615 | 1 | 0.05 | 0.05 | 0.05 |
| NC_004718.3;NP_828849.2;SARS-CoV_orf1ab | 5636 | 5636 | 1 | 0.05 | 0.05 | 0.05 |
| NC_004718.3;NP_828849.2;SARS-CoV_orf1ab | 5638 | 5644 | 7 | 0.05 | 0.4 | 0.214285714 |
| NC_004718.3;NP_828849.2;SARS-CoV_orf1ab | 5707 | 5715 | 9 | 0.05 | 0.1 | 0.088888889 |
| NC_004718.3;NP_828849.2;SARS-CoV_orf1ab | 5739 | 5750 | 12 | 0.1 | 0.15 | 0.116666667 |
| NC_004718.3;NP_828849.2;SARS-CoV_orf1ab | 5754 | 5754 | 1 | 0.05 | 0.05 | 0.05 |
| NC_004718.3;NP_828849.2;SARS-CoV_orf1ab | 5756 | 5756 | 1 | 0.05 | 0.05 | 0.05 |
| NC_004718.3;NP_828849.2;SARS-CoV_orf1ab | 5762 | 5762 | 1 | 0.05 | 0.05 | 0.05 |
| NC_004718.3;NP_828849.2;SARS-CoV_orf1ab | 5779 | 5781 | 3 | 0.05 | 0.05 | 0.05 |
| NC_004718.3;NP_828849.2;SARS-CoV_orf1ab | 5784 | 5784 | 1 | 0.05 | 0.05 | 0.05 |
| NC_004718.3;NP_828849.2;SARS-CoV_orf1ab | 5805 | 5805 | 1 | 0.05 | 0.05 | 0.05 |
| NC_004718.3;NP_828849.2;SARS-CoV_orf1ab | 5809 | 5810 | 2 | 0.05 | 0.15 | 0.1 |
| NC_004718.3;NP_828849.2;SARS-CoV_orf1ab | 5814 | 5816 | 3 | 0.05 | 0.1 | 0.083333333 |
| NC_004718.3;NP_828849.2;SARS-CoV_orf1ab | 5823 | 5823 | 1 | 0.05 | 0.05 | 0.05 |
| NC_004718.3;NP_828849.2;SARS-CoV_orf1ab | 5840 | 5845 | 6 | 0.05 | 0.05 | 0.05 |
| NC_004718.3;NP_828849.2;SARS-CoV_orf1ab | 5848 | 5850 | 3 | 0.05 | 0.05 | 0.05 |
| NC_004718.3;NP_828849.2;SARS-CoV_orf1ab | 5874 | 5876 | 3 | 0.05 | 0.1 | 0.066666667 |
| NC_004718.3;NP_828849.2;SARS-CoV_orf1ab | 5889 | 5895 | 7 | 0.05 | 0.15 | 0.078571429 |
| NC_004718.3;NP_828849.2;SARS-CoV_orf1ab | 5898 | 5898 | 1 | 0.05 | 0.05 | 0.05 |
| NC_004718.3;NP_828849.2;SARS-CoV_orf1ab | 5911 | 5911 | 1 | 0.05 | 0.05 | 0.05 |
| NC_004718.3;NP_828849.2;SARS-CoV_orf1ab | 5929 | 5930 | 2 | 0.05 | 0.05 | 0.05 |
| NC_004718.3;NP_828849.2;SARS-CoV_orf1ab | 5959 | 5961 | 3 | 0.05 | 0.1 | 0.066666667 |
| NC_004718.3;NP_828849.2;SARS-CoV_orf1ab | 5966 | 5966 | 1 | 0.05 | 0.05 | 0.05 |
| NC_004718.3;NP_828849.2;SARS-CoV_orf1ab | 6024 | 6025 | 2 | 0.05 | 0.1 | 0.075 |
| NC_004718.3;NP_828849.2;SARS-CoV_orf1ab | 6027 | 6027 | 1 | 0.2 | 0.2 | 0.2 |
| NC_004718.3;NP_828849.2;SARS-CoV_orf1ab | 6029 | 6029 | 1 | 0.1 | 0.1 | 0.1 |

| Extended data 1 | | | | | | |
|---|---|---|---|---|---|---|
| Protein | Start position (peptide) | Stop position (peptide) | Number of peptides | Minimum fraction (per peptide) of control subjects with reactivity >3 sd above average | Maximum fraction (per peptide) of control subjects with reactivity >3 sd above average | Mean fraction (over all peptides in this contiguous stretch) of controlsubjects with reactivity >3 sd above average |
| NC_004718.3;NP_828849.2;SARS-CoV_orf1ab | 6033 | 6034 | 2 | 0.05 | 0.1 | 0.075 |
| NC_004718.3;NP_828849.2;SARS-CoV_orf1ab | 6036 | 6037 | 2 | 0.05 | 0.1 | 0.075 |
| NC_004718.3;NP_828849.2;SARS-CoV_orf1ab | 6093 | 6093 | 1 | 0.05 | 0.05 | 0.05 |
| NC_004718.3;NP_828849.2;SARS-CoV_orf1ab | 6103 | 6104 | 2 | 0.05 | 0.05 | 0.05 |
| NC_004718.3;NP_828849.2;SARS-CoV_orf1ab | 6110 | 6110 | 1 | 0.05 | 0.05 | 0.05 |
| NC_004718.3;NP_828849.2;SARS-CoV_orf1ab | 6112 | 6112 | 1 | 0.05 | 0.05 | 0.05 |
| NC_004718.3;NP_828849.2;SARS-CoV_orf1ab | 6156 | 6157 | 2 | 0.05 | 0.05 | 0.05 |
| NC_004718.3;NP_828849.2;SARS-CoV_orf1ab | 6190 | 6190 | 1 | 0.05 | 0.05 | 0.05 |
| NC_004718.3;NP_828849.2;SARS-CoV_orf1ab | 6201 | 6201 | 1 | 0.1 | 0.1 | 0.1 |
| NC_004718.3;NP_828849.2;SARS-CoV_orf1ab | 6206 | 6206 | 1 | 0.05 | 0.05 | 0.05 |
| NC_004718.3;NP_828849.2;SARS-CoV_orf1ab | 6232 | 6233 | 2 | 0.05 | 0.05 | 0.05 |
| NC_004718.3;NP_828849.2;SARS-CoV_orf1ab | 6236 | 6237 | 2 | 0.05 | 0.05 | 0.05 |
| NC_004718.3;NP_828849.2;SARS-CoV_orf1ab | 6247 | 6249 | 3 | 0.05 | 0.1 | 0.066666667 |
| NC_004718.3;NP_828849.2;SARS-CoV_orf1ab | 6251 | 6251 | 1 | 0.05 | 0.05 | 0.05 |
| NC_004718.3;NP_828849.2;SARS-CoV_orf1ab | 6265 | 6270 | 6 | 0.05 | 0.25 | 0.125 |
| NC_004718.3;NP_828849.2;SARS-CoV_orf1ab | 6283 | 6283 | 1 | 0.05 | 0.05 | 0.05 |
| NC_004718.3;NP_828849.2;SARS-CoV_orf1ab | 6304 | 6308 | 5 | 0.05 | 0.1 | 0.06 |
| NC_004718.3;NP_828849.2;SARS-CoV_orf1ab | 6327 | 6335 | 9 | 0.05 | 0.1 | 0.055555556 |
| NC_004718.3;NP_828849.2;SARS-CoV_orf1ab | 6359 | 6359 | 1 | 0.05 | 0.05 | 0.05 |
| NC_004718.3;NP_828849.2;SARS-CoV_orf1ab | 6388 | 6388 | 1 | 0.05 | 0.05 | 0.05 |
| NC_004718.3;NP_828849.2;SARS-CoV_orf1ab | 6425 | 6425 | 1 | 0.05 | 0.05 | 0.05 |
| NC_004718.3;NP_828849.2;SARS-CoV_orf1ab | 6427 | 6427 | 1 | 0.05 | 0.05 | 0.05 |
| NC_004718.3;NP_828849.2;SARS-CoV_orf1ab | 6430 | 6431 | 2 | 0.05 | 0.05 | 0.05 |
| NC_004718.3;NP_828849.2;SARS-CoV_orf1ab | 6436 | 6436 | 1 | 0.05 | 0.05 | 0.05 |
| NC_004718.3;NP_828849.2;SARS-CoV_orf1ab | 6438 | 6438 | 1 | 0.05 | 0.05 | 0.05 |
| NC_004718.3;NP_828849.2;SARS-CoV_orf1ab | 6446 | 6446 | 1 | 0.05 | 0.05 | 0.05 |
| NC_004718.3;NP_828849.2;SARS-CoV_orf1ab | 6452 | 6452 | 1 | 0.05 | 0.05 | 0.05 |
| NC_004718.3;NP_828849.2;SARS-CoV_orf1ab | 6455 | 6455 | 1 | 0.05 | 0.05 | 0.05 |
| NC_004718.3;NP_828849.2;SARS-CoV_orf1ab | 6466 | 6466 | 1 | 0.1 | 0.1 | 0.1 |
| NC_004718.3;NP_828849.2;SARS-CoV_orf1ab | 6468 | 6468 | 1 | 0.1 | 0.1 | 0.1 |
| NC_004718.3;NP_828849.2;SARS-CoV_orf1ab | 6473 | 6473 | 1 | 0.05 | 0.05 | 0.05 |
| NC_004718.3;NP_828849.2;SARS-CoV_orf1ab | 6482 | 6482 | 1 | 0.05 | 0.05 | 0.05 |
| NC_004718.3;NP_828849.2;SARS-CoV_orf1ab | 6485 | 6485 | 1 | 0.1 | 0.1 | 0.1 |
| NC_004718.3;NP_828849.2;SARS-CoV_orf1ab | 6488 | 6489 | 2 | 0.05 | 0.1 | 0.075 |
| NC_004718.3;NP_828849.2;SARS-CoV_orf1ab | 6491 | 6493 | 3 | 0.15 | 0.3 | 0.233333333 |
| NC_004718.3;NP_828849.2;SARS-CoV_orf1ab | 6495 | 6495 | 1 | 0.05 | 0.05 | 0.05 |
| NC_004718.3;NP_828849.2;SARS-CoV_orf1ab | 6505 | 6505 | 1 | 0.05 | 0.05 | 0.05 |

| Extended data 1 | | | | | | |
|---|---|---|---|---|---|---|
| Protein | Start position (peptide) | Stop position (peptide) | Number of peptides | Minimum fraction (per peptide) of control subjects with reactivity >3 sd above average | Maximum fraction (per peptide) of control subjects with reactivity >3 sd above average | Mean fraction (over all peptides in this contiguous stretch) of controlsubjects with reactivity >3 sd above average |
| NC_004718.3;NP_828849.2;SARS-CoV_orf1ab | 6513 | 6514 | 2 | 0.05 | 0.05 | 0.05 |
| NC_004718.3;NP_828849.2;SARS-CoV_orf1ab | 6524 | 6534 | 11 | 0.05 | 0.1 | 0.063636364 |
| NC_004718.3;NP_828849.2;SARS-CoV_orf1ab | 6553 | 6553 | 1 | 0.05 | 0.05 | 0.05 |
| NC_004718.3;NP_828849.2;SARS-CoV_orf1ab | 6575 | 6575 | 1 | 0.05 | 0.05 | 0.05 |
| NC_004718.3;NP_828849.2;SARS-CoV_orf1ab | 6581 | 6581 | 1 | 0.05 | 0.05 | 0.05 |
| NC_004718.3;NP_828849.2;SARS-CoV_orf1ab | 6598 | 6599 | 2 | 0.1 | 0.1 | 0.1 |
| NC_004718.3;NP_828849.2;SARS-CoV_orf1ab | 6606 | 6608 | 3 | 0.05 | 0.1 | 0.083333333 |
| NC_004718.3;NP_828849.2;SARS-CoV_orf1ab | 6610 | 6610 | 1 | 0.05 | 0.05 | 0.05 |
| NC_004718.3;NP_828849.2;SARS-CoV_orf1ab | 6620 | 6629 | 10 | 0.05 | 0.2 | 0.08 |
| NC_004718.3;NP_828849.2;SARS-CoV_orf1ab | 6685 | 6685 | 1 | 0.05 | 0.05 | 0.05 |
| NC_004718.3;NP_828849.2;SARS-CoV_orf1ab | 6687 | 6689 | 3 | 0.05 | 0.05 | 0.05 |
| NC_004718.3;NP_828849.2;SARS-CoV_orf1ab | 6692 | 6694 | 3 | 0.05 | 0.15 | 0.083333333 |
| NC_004718.3;NP_828849.2;SARS-CoV_orf1ab | 6696 | 6698 | 3 | 0.05 | 0.1 | 0.083333333 |
| NC_004718.3;NP_828849.2;SARS-CoV_orf1ab | 6700 | 6705 | 6 | 0.05 | 0.1 | 0.075 |
| NC_004718.3;NP_828849.2;SARS-CoV_orf1ab | 6722 | 6727 | 6 | 0.05 | 0.1 | 0.058333333 |
| NC_004718.3;NP_828849.2;SARS-CoV_orf1ab | 6732 | 6732 | 1 | 0.05 | 0.05 | 0.05 |
| NC_004718.3;NP_828849.2;SARS-CoV_orf1ab | 6770 | 6770 | 1 | 0.1 | 0.1 | 0.1 |
| NC_004718.3;NP_828849.2;SARS-CoV_orf1ab | 6772 | 6772 | 1 | 0.05 | 0.05 | 0.05 |
| NC_004718.3;NP_828849.2;SARS-CoV_orf1ab | 6774 | 6774 | 1 | 0.05 | 0.05 | 0.05 |
| NC_004718.3;NP_828849.2;SARS-CoV_orf1ab | 6794 | 6794 | 1 | 0.05 | 0.05 | 0.05 |
| NC_004718.3;NP_828849.2;SARS-CoV_orf1ab | 6797 | 6799 | 3 | 0.05 | 0.05 | 0.05 |
| NC_004718.3;NP_828849.2;SARS-CoV_orf1ab | 6801 | 6805 | 5 | 0.05 | 0.05 | 0.05 |
| NC_004718.3;NP_828849.2;SARS-CoV_orf1ab | 6839 | 6839 | 1 | 0.05 | 0.05 | 0.05 |
| NC_004718.3;NP_828849.2;SARS-CoV_orf1ab | 6847 | 6848 | 2 | 0.05 | 0.05 | 0.05 |
| NC_004718.3;NP_828849.2;SARS-CoV_orf1ab | 6868 | 6870 | 3 | 0.05 | 0.05 | 0.05 |
| NC_004718.3;NP_828849.2;SARS-CoV_orf1ab | 6873 | 6873 | 1 | 0.1 | 0.1 | 0.1 |
| NC_004718.3;NP_828849.2;SARS-CoV_orf1ab | 6876 | 6882 | 7 | 0.05 | 0.05 | 0.05 |
| NC_004718.3;NP_828849.2;SARS-CoV_orf1ab | 6885 | 6885 | 1 | 0.05 | 0.05 | 0.05 |
| NC_004718.3;NP_828849.2;SARS-CoV_orf1ab | 6896 | 6896 | 1 | 0.1 | 0.1 | 0.1 |
| NC_004718.3;NP_828849.2;SARS-CoV_orf1ab | 6899 | 6900 | 2 | 0.05 | 0.05 | 0.05 |
| NC_004718.3;NP_828849.2;SARS-CoV_orf1ab | 6903 | 6905 | 3 | 0.05 | 0.05 | 0.05 |
| NC_004718.3;NP_828849.2;SARS-CoV_orf1ab | 6907 | 6913 | 7 | 0.05 | 0.05 | 0.05 |
| NC_004718.3;NP_828849.2;SARS-CoV_orf1ab | 6967 | 6967 | 1 | 0.05 | 0.05 | 0.05 |
| NC_004718.3;NP_828849.2;SARS-CoV_orf1ab | 6970 | 6970 | 1 | 0.05 | 0.05 | 0.05 |
| NC_004718.3;NP_828849.2;SARS-CoV_orf1ab | 6977 | 6978 | 2 | 0.05 | 0.05 | 0.05 |
| NC_004718.3;NP_828849.2;SARS-CoV_orf1ab | 6980 | 6986 | 7 | 0.05 | 0.25 | 0.092857143 |
| NC_004718.3;NP_828849.2;SARS-CoV_orf1ab | 6992 | 6992 | 1 | 0.05 | 0.05 | 0.05 |

| Extended data 1 | | | | | | |
|---|---|---|---|---|---|---|
| Protein | Start position (peptide) | Stop position (peptide) | Number of peptides | Minimum fraction (per peptide) of control subjects with reactivity >3 sd above average | Maximum fraction (per peptide) of control subjects with reactivity >3 sd above average | Mean fraction (over all peptides in this contiguous stretch) of controlsubjects with reactivity >3 sd above average |
| NC_004718.3;NP_828849.2;SARS-CoV_orf1ab | 7009 | 7015 | 7 | 0.05 | 0.05 | 0.05 |
| NC_004718.3;NP_828849.2;SARS-CoV_orf1ab | 7026 | 7026 | 1 | 0.05 | 0.05 | 0.05 |
| NC_004718.3;NP_828850.1;SARS-CoV_orf1a | 1 | 1 | 1 | 0.05 | 0.05 | 0.05 |
| NC_004718.3;NP_828850.1;SARS-CoV_orf1a | 4 | 6 | 3 | 0.05 | 0.1 | 0.066666667 |
| NC_004718.3;NP_828850.1;SARS-CoV_orf1a | 29 | 30 | 2 | 0.1 | 0.25 | 0.175 |
| NC_004718.3;NP_828850.1;SARS-CoV_orf1a | 32 | 34 | 3 | 0.05 | 0.05 | 0.05 |
| NC_004718.3;NP_828850.1;SARS-CoV_orf1a | 44 | 44 | 1 | 0.05 | 0.05 | 0.05 |
| NC_004718.3;NP_828850.1;SARS-CoV_orf1a | 52 | 52 | 1 | 0.05 | 0.05 | 0.05 |
| NC_004718.3;NP_828850.1;SARS-CoV_orf1a | 65 | 65 | 1 | 0.05 | 0.05 | 0.05 |
| NC_004718.3;NP_828850.1;SARS-CoV_orf1a | 71 | 72 | 2 | 0.05 | 0.05 | 0.05 |
| NC_004718.3;NP_828850.1;SARS-CoV_orf1a | 90 | 91 | 2 | 0.05 | 0.2 | 0.125 |
| NC_004718.3;NP_828850.1;SARS-CoV_orf1a | 116 | 121 | 6 | 0.05 | 0.1 | 0.058333333 |
| NC_004718.3;NP_828850.1;SARS-CoV_orf1a | 124 | 125 | 2 | 0.05 | 0.15 | 0.1 |
| NC_004718.3;NP_828850.1;SARS-CoV_orf1a | 131 | 133 | 3 | 0.05 | 0.1 | 0.066666667 |
| NC_004718.3;NP_828850.1;SARS-CoV_orf1a | 139 | 140 | 2 | 0.05 | 0.05 | 0.05 |
| NC_004718.3;NP_828850.1;SARS-CoV_orf1a | 145 | 145 | 1 | 0.2 | 0.2 | 0.2 |
| NC_004718.3;NP_828850.1;SARS-CoV_orf1a | 149 | 157 | 9 | 0.05 | 0.25 | 0.133333333 |
| NC_004718.3;NP_828850.1;SARS-CoV_orf1a | 165 | 165 | 1 | 0.05 | 0.05 | 0.05 |
| NC_004718.3;NP_828850.1;SARS-CoV_orf1a | 168 | 168 | 1 | 0.2 | 0.2 | 0.2 |
| NC_004718.3;NP_828850.1;SARS-CoV_orf1a | 173 | 173 | 1 | 0.05 | 0.05 | 0.05 |
| NC_004718.3;NP_828850.1;SARS-CoV_orf1a | 179 | 180 | 2 | 0.05 | 0.15 | 0.1 |
| NC_004718.3;NP_828850.1;SARS-CoV_orf1a | 189 | 189 | 1 | 0.05 | 0.05 | 0.05 |
| NC_004718.3;NP_828850.1;SARS-CoV_orf1a | 204 | 204 | 1 | 0.05 | 0.05 | 0.05 |
| NC_004718.3;NP_828850.1;SARS-CoV_orf1a | 214 | 218 | 5 | 0.05 | 0.15 | 0.1 |
| NC_004718.3;NP_828851.1;SARS-CoV_E2-glycopr | 8 | 11 | 4 | 0.05 | 0.05 | 0.05 |
| NC_004718.3;NP_828851.1;SARS-CoV_E2-glycopr | 17 | 23 | 7 | 0.05 | 0.05 | 0.05 |
| NC_004718.3;NP_828851.1;SARS-CoV_E2-glycopr | 30 | 30 | 1 | 0.05 | 0.05 | 0.05 |
| NC_004718.3;NP_828851.1;SARS-CoV_E2-glycopr | 76 | 76 | 1 | 0.05 | 0.05 | 0.05 |
| NC_004718.3;NP_828851.1;SARS-CoV_E2-glycopr | 83 | 83 | 1 | 0.05 | 0.05 | 0.05 |
| NC_004718.3;NP_828851.1;SARS-CoV_E2-glycopr | 87 | 90 | 4 | 0.05 | 0.05 | 0.05 |
| NC_004718.3;NP_828851.1;SARS-CoV_E2-glycopr | 93 | 93 | 1 | 0.05 | 0.05 | 0.05 |
| NC_004718.3;NP_828851.1;SARS-CoV_E2-glycopr | 107 | 118 | 12 | 0.05 | 0.25 | 0.091666667 |
| NC_004718.3;NP_828851.1;SARS-CoV_E2-glycopr | 121 | 121 | 1 | 0.05 | 0.05 | 0.05 |
| NC_004718.3;NP_828851.1;SARS-CoV_E2-glycopr | 134 | 135 | 2 | 0.05 | 0.05 | 0.05 |
| NC_004718.3;NP_828851.1;SARS-CoV_E2-glycopr | 168 | 170 | 3 | 0.05 | 0.15 | 0.1 |
| NC_004718.3;NP_828851.1;SARS-CoV_E2-glycopr | 176 | 176 | 1 | 0.05 | 0.05 | 0.05 |
| NC_004718.3;NP_828851.1;SARS-CoV_E2-glycopr | 179 | 181 | 3 | 0.05 | 0.05 | 0.05 |

| Extended data 1 | | | | | | |
|---|---|---|---|---|---|---|
| Protein | Start position (peptide) | Stop position (peptide) | Number of peptides | Minimum fraction (per peptide) of control subjects with reactivity >3 sd above average | Maximum fraction (per peptide) of control subjects with reactivity >3 sd above average | Mean fraction (over all peptides in this contiguous stretch) of controlsubjects with reactivity >3 sd above average |
| NC_004718.3;NP_828851.1;SARS-CoV_E2-glycopr | 193 | 201 | 9 | 0.05 | 0.2 | 0.111111111 |
| NC_004718.3;NP_828851.1;SARS-CoV_E2-glycopr | 204 | 204 | 1 | 0.05 | 0.05 | 0.05 |
| NC_004718.3;NP_828851.1;SARS-CoV_E2-glycopr | 235 | 236 | 2 | 0.05 | 0.05 | 0.05 |
| NC_004718.3;NP_828851.1;SARS-CoV_E2-glycopr | 255 | 255 | 1 | 0.05 | 0.05 | 0.05 |
| NC_004718.3;NP_828851.1;SARS-CoV_E2-glycopr | 258 | 261 | 4 | 0.05 | 0.05 | 0.05 |
| NC_004718.3;NP_828851.1;SARS-CoV_E2-glycopr | 284 | 284 | 1 | 0.05 | 0.05 | 0.05 |
| NC_004718.3;NP_828851.1;SARS-CoV_E2-glycopr | 286 | 291 | 6 | 0.05 | 0.15 | 0.091666667 |
| NC_004718.3;NP_828851.1;SARS-CoV_E2-glycopr | 294 | 294 | 1 | 0.05 | 0.05 | 0.05 |
| NC_004718.3;NP_828851.1;SARS-CoV_E2-glycopr | 297 | 299 | 3 | 0.05 | 0.1 | 0.066666667 |
| NC_004718.3;NP_828851.1;SARS-CoV_E2-glycopr | 302 | 302 | 1 | 0.05 | 0.05 | 0.05 |
| NC_004718.3;NP_828851.1;SARS-CoV_E2-glycopr | 306 | 306 | 1 | 0.1 | 0.1 | 0.1 |
| NC_004718.3;NP_828851.1;SARS-CoV_E2-glycopr | 313 | 313 | 1 | 0.05 | 0.05 | 0.05 |
| NC_004718.3;NP_828851.1;SARS-CoV_E2-glycopr | 316 | 324 | 9 | 0.05 | 0.05 | 0.05 |
| NC_004718.3;NP_828851.1;SARS-CoV_E2-glycopr | 330 | 330 | 1 | 0.1 | 0.1 | 0.1 |
| NC_004718.3;NP_828851.1;SARS-CoV_E2-glycopr | 340 | 341 | 2 | 0.05 | 0.05 | 0.05 |
| NC_004718.3;NP_828851.1;SARS-CoV_E2-glycopr | 378 | 380 | 3 | 0.05 | 0.05 | 0.05 |
| NC_004718.3;NP_828851.1;SARS-CoV_E2-glycopr | 384 | 385 | 2 | 0.1 | 0.1 | 0.1 |
| NC_004718.3;NP_828851.1;SARS-CoV_E2-glycopr | 396 | 397 | 2 | 0.05 | 0.05 | 0.05 |
| NC_004718.3;NP_828851.1;SARS-CoV_E2-glycopr | 400 | 403 | 4 | 0.05 | 0.1 | 0.0625 |
| NC_004718.3;NP_828851.1;SARS-CoV_E2-glycopr | 406 | 407 | 2 | 0.05 | 0.05 | 0.05 |
| NC_004718.3;NP_828851.1;SARS-CoV_E2-glycopr | 419 | 428 | 10 | 0.05 | 0.25 | 0.125 |
| NC_004718.3;NP_828851.1;SARS-CoV_E2-glycopr | 444 | 444 | 1 | 0.05 | 0.05 | 0.05 |
| NC_004718.3;NP_828851.1;SARS-CoV_E2-glycopr | 446 | 446 | 1 | 0.05 | 0.05 | 0.05 |
| NC_004718.3;NP_828851.1;SARS-CoV_E2-glycopr | 448 | 449 | 2 | 0.05 | 0.1 | 0.075 |
| NC_004718.3;NP_828851.1;SARS-CoV_E2-glycopr | 452 | 452 | 1 | 0.05 | 0.05 | 0.05 |
| NC_004718.3;NP_828851.1;SARS-CoV_E2-glycopr | 455 | 456 | 2 | 0.05 | 0.1 | 0.075 |
| NC_004718.3;NP_828851.1;SARS-CoV_E2-glycopr | 506 | 511 | 6 | 0.05 | 0.15 | 0.066666667 |
| NC_004718.3;NP_828851.1;SARS-CoV_E2-glycopr | 516 | 516 | 1 | 0.1 | 0.1 | 0.1 |
| NC_004718.3;NP_828851.1;SARS-CoV_E2-glycopr | 518 | 518 | 1 | 0.05 | 0.05 | 0.05 |
| NC_004718.3;NP_828851.1;SARS-CoV_E2-glycopr | 520 | 521 | 2 | 0.05 | 0.05 | 0.05 |
| NC_004718.3;NP_828851.1;SARS-CoV_E2-glycopr | 537 | 537 | 1 | 0.05 | 0.05 | 0.05 |
| NC_004718.3;NP_828851.1;SARS-CoV_E2-glycopr | 542 | 542 | 1 | 0.05 | 0.05 | 0.05 |
| NC_004718.3;NP_828851.1;SARS-CoV_E2-glycopr | 549 | 557 | 9 | 0.05 | 0.15 | 0.077777778 |
| NC_004718.3;NP_828851.1;SARS-CoV_E2-glycopr | 561 | 562 | 2 | 0.05 | 0.05 | 0.05 |
| NC_004718.3;NP_828851.1;SARS-CoV_E2-glycopr | 576 | 578 | 3 | 0.05 | 0.1 | 0.066666667 |
| NC_004718.3;NP_828851.1;SARS-CoV_E2-glycopr | 609 | 609 | 1 | 0.05 | 0.05 | 0.05 |
| NC_004718.3;NP_828851.1;SARS-CoV_E2-glycopr | 613 | 622 | 10 | 0.05 | 0.05 | 0.05 |

| Extended data 1 | | | | | | |
|---|---|---|---|---|---|---|
| Protein | Start position (peptide) | Stop position (peptide) | Number of peptides | Minimum fraction (per peptide) of control subjects with reactivity >3 sd above average | Maximum fraction (per peptide) of control subjects with reactivity >3 sd above average | Mean fraction (over all peptides in this contiguous stretch) of controlsubjects with reactivity >3 sd above average |
| NC_004718.3;NP_828851.1;SARS-CoV_E2-glycopr | 629 | 629 | 1 | 0.05 | 0.05 | 0.05 |
| NC_004718.3;NP_828851.1;SARS-CoV_E2-glycopr | 642 | 645 | 4 | 0.05 | 0.1 | 0.0625 |
| NC_004718.3;NP_828851.1;SARS-CoV_E2-glycopr | 676 | 689 | 14 | 0.05 | 0.2 | 0.085714286 |
| NC_004718.3;NP_828851.1;SARS-CoV_E2-glycopr | 717 | 725 | 9 | 0.05 | 0.1 | 0.072222222 |
| NC_004718.3;NP_828851.1;SARS-CoV_E2-glycopr | 739 | 739 | 1 | 0.05 | 0.05 | 0.05 |
| NC_004718.3;NP_828851.1;SARS-CoV_E2-glycopr | 743 | 751 | 9 | 0.05 | 0.05 | 0.05 |
| NC_004718.3;NP_828851.1;SARS-CoV_E2-glycopr | 753 | 761 | 9 | 0.05 | 0.1 | 0.055555556 |
| NC_004718.3;NP_828851.1;SARS-CoV_E2-glycopr | 779 | 780 | 2 | 0.05 | 0.05 | 0.05 |
| NC_004718.3;NP_828851.1;SARS-CoV_E2-glycopr | 783 | 795 | 13 | 0.05 | 0.1 | 0.061538462 |
| NC_004718.3;NP_828851.1;SARS-CoV_E2-glycopr | 799 | 800 | 2 | 0.05 | 0.1 | 0.075 |
| NC_004718.3;NP_828851.1;SARS-CoV_E2-glycopr | 810 | 810 | 1 | 0.1 | 0.1 | 0.1 |
| NC_004718.3;NP_828851.1;SARS-CoV_E2-glycopr | 814 | 814 | 1 | 0.05 | 0.05 | 0.05 |
| NC_004718.3;NP_828851.1;SARS-CoV_E2-glycopr | 830 | 830 | 1 | 0.05 | 0.05 | 0.05 |
| NC_004718.3;NP_828851.1;SARS-CoV_E2-glycopr | 843 | 843 | 1 | 0.05 | 0.05 | 0.05 |
| NC_004718.3;NP_828851.1;SARS-CoV_E2-glycopr | 860 | 860 | 1 | 0.1 | 0.1 | 0.1 |
| NC_004718.3;NP_828851.1;SARS-CoV_E2-glycopr | 874 | 883 | 10 | 0.05 | 0.15 | 0.07 |
| NC_004718.3;NP_828851.1;SARS-CoV_E2-glycopr | 887 | 887 | 1 | 0.05 | 0.05 | 0.05 |
| NC_004718.3;NP_828851.1;SARS-CoV_E2-glycopr | 890 | 892 | 3 | 0.05 | 0.05 | 0.05 |
| NC_004718.3;NP_828851.1;SARS-CoV_E2-glycopr | 894 | 906 | 13 | 0.05 | 0.35 | 0.203846154 |
| NC_004718.3;NP_828851.1;SARS-CoV_E2-glycopr | 912 | 912 | 1 | 0.05 | 0.05 | 0.05 |
| NC_004718.3;NP_828851.1;SARS-CoV_E2-glycopr | 928 | 928 | 1 | 0.1 | 0.1 | 0.1 |
| NC_004718.3;NP_828851.1;SARS-CoV_E2-glycopr | 933 | 933 | 1 | 0.05 | 0.05 | 0.05 |
| NC_004718.3;NP_828851.1;SARS-CoV_E2-glycopr | 954 | 954 | 1 | 0.05 | 0.05 | 0.05 |
| NC_004718.3;NP_828851.1;SARS-CoV_E2-glycopr | 956 | 958 | 3 | 0.05 | 0.05 | 0.05 |
| NC_004718.3;NP_828851.1;SARS-CoV_E2-glycopr | 961 | 962 | 2 | 0.05 | 0.05 | 0.05 |
| NC_004718.3;NP_828851.1;SARS-CoV_E2-glycopr | 970 | 970 | 1 | 0.05 | 0.05 | 0.05 |
| NC_004718.3;NP_828851.1;SARS-CoV_E2-glycopr | 993 | 993 | 1 | 0.05 | 0.05 | 0.05 |
| NC_004718.3;NP_828851.1;SARS-CoV_E2-glycopr | 998 | 1000 | 3 | 0.05 | 0.05 | 0.05 |
| NC_004718.3;NP_828851.1;SARS-CoV_E2-glycopr | 1023 | 1023 | 1 | 0.05 | 0.05 | 0.05 |
| NC_004718.3;NP_828851.1;SARS-CoV_E2-glycopr | 1048 | 1049 | 2 | 0.1 | 0.2 | 0.15 |
| NC_004718.3;NP_828851.1;SARS-CoV_E2-glycopr | 1052 | 1053 | 2 | 0.05 | 0.2 | 0.125 |
| NC_004718.3;NP_828851.1;SARS-CoV_E2-glycopr | 1059 | 1062 | 4 | 0.05 | 0.05 | 0.05 |
| NC_004718.3;NP_828851.1;SARS-CoV_E2-glycopr | 1090 | 1098 | 9 | 0.05 | 0.05 | 0.05 |
| NC_004718.3;NP_828851.1;SARS-CoV_E2-glycopr | 1123 | 1136 | 14 | 0.05 | 0.5 | 0.314285714 |
| NC_004718.3;NP_828851.1;SARS-CoV_E2-glycopr | 1140 | 1143 | 4 | 0.05 | 0.2 | 0.1125 |
| NC_004718.3;NP_828851.1;SARS-CoV_E2-glycopr | 1149 | 1176 | 28 | 0.05 | 0.2 | 0.078571429 |
| NC_004718.3;NP_828851.1;SARS-CoV_E2-glycopr | 1236 | 1243 | 8 | 0.05 | 0.1 | 0.0625 |

| Extended data 1 | | | | | | |
|---|---|---|---|---|---|---|
| Protein | Start position (peptide) | Stop position (peptide) | Number of peptides | Minimum fraction (per peptide) of control subjects with reactivity >3 sd above average | Maximum fraction (per peptide) of control subjects with reactivity >3 sd above average | Mean fraction (over all peptides in this contiguous stretch) of controlsubjects with reactivity >3 sd above average |
| NC_004718.3;NP_828852.2;SARS-CoV_3a | 8 | 8 | 1 | 0.05 | 0.05 | 0.05 |
| NC_004718.3;NP_828852.2;SARS-CoV_3a | 13 | 19 | 7 | 0.05 | 0.1 | 0.071428571 |
| NC_004718.3;NP_828852.2;SARS-CoV_3a | 36 | 36 | 1 | 0.05 | 0.05 | 0.05 |
| NC_004718.3;NP_828852.2;SARS-CoV_3a | 129 | 129 | 1 | 0.05 | 0.05 | 0.05 |
| NC_004718.3;NP_828852.2;SARS-CoV_3a | 171 | 189 | 19 | 0.05 | 0.25 | 0.139473684 |
| NC_004718.3;NP_828852.2;SARS-CoV_3a | 196 | 196 | 1 | 0.1 | 0.1 | 0.1 |
| NC_004718.3;NP_828852.2;SARS-CoV_3a | 209 | 210 | 2 | 0.05 | 0.05 | 0.05 |
| NC_004718.3;NP_828852.2;SARS-CoV_3a | 212 | 213 | 2 | 0.05 | 0.05 | 0.05 |
| NC_004718.3;NP_828852.2;SARS-CoV_3a | 235 | 235 | 1 | 0.05 | 0.05 | 0.05 |
| NC_004718.3;NP_828852.2;SARS-CoV_3a | 239 | 249 | 11 | 0.05 | 0.1 | 0.054545455 |
| NC_004718.3;NP_828852.2;SARS-CoV_3a | 251 | 251 | 1 | 0.05 | 0.05 | 0.05 |
| NC_004718.3;NP_828852.2;SARS-CoV_3a | 257 | 260 | 4 | 0.05 | 0.05 | 0.05 |
| NC_004718.3;NP_828853.1;SARS-CoV_3b | 130 | 130 | 1 | 0.05 | 0.05 | 0.05 |
| NC_004718.3;NP_828853.1;SARS-CoV_3b | 132 | 132 | 1 | 0.05 | 0.05 | 0.05 |
| NC_004718.3;NP_828854.1;SARS-CoV_E | 57 | 57 | 1 | 0.05 | 0.05 | 0.05 |
| NC_004718.3;NP_828854.1;SARS-CoV_E | 60 | 61 | 2 | 0.05 | 0.1 | 0.075 |
| NC_004718.3;NP_828855.1;SARS-CoV_matrix | 3 | 4 | 2 | 0.05 | 0.05 | 0.05 |
| NC_004718.3;NP_828855.1;SARS-CoV_matrix | 13 | 13 | 1 | 0.05 | 0.05 | 0.05 |
| NC_004718.3;NP_828855.1;SARS-CoV_matrix | 38 | 39 | 2 | 0.05 | 0.05 | 0.05 |
| NC_004718.3;NP_828855.1;SARS-CoV_matrix | 100 | 100 | 1 | 0.05 | 0.05 | 0.05 |
| NC_004718.3;NP_828855.1;SARS-CoV_matrix | 116 | 116 | 1 | 0.05 | 0.05 | 0.05 |
| NC_004718.3;NP_828855.1;SARS-CoV_matrix | 147 | 157 | 11 | 0.05 | 0.1 | 0.054545455 |
| NC_004718.3;NP_828855.1;SARS-CoV_matrix | 179 | 179 | 1 | 0.05 | 0.05 | 0.05 |
| NC_004718.3;NP_828855.1;SARS-CoV_matrix | 199 | 200 | 2 | 0.05 | 0.1 | 0.075 |
| NC_004718.3;NP_828856.1;SARS-CoV_6 | 39 | 40 | 2 | 0.05 | 0.05 | 0.05 |
| NC_004718.3;NP_828856.1;SARS-CoV_6 | 48 | 50 | 3 | 0.05 | 0.05 | 0.05 |
| NC_004718.3;NP_828857.1;SARS-CoV_7 | 23 | 25 | 3 | 0.05 | 0.05 | 0.05 |
| NC_004718.3;NP_828857.1;SARS-CoV_7 | 79 | 85 | 7 | 0.05 | 0.1 | 0.078571429 |
| NC_004718.3;NP_828857.1;SARS-CoV_7 | 88 | 88 | 1 | 0.05 | 0.05 | 0.05 |
| NC_004718.3;NP_828858.1;SARS-CoV_nucleocapsid | 3 | 3 | 1 | 0.05 | 0.05 | 0.05 |
| NC_004718.3;NP_828858.1;SARS-CoV_nucleocapsid | 6 | 15 | 10 | 0.05 | 0.1 | 0.06 |
| NC_004718.3;NP_828858.1;SARS-CoV_nucleocapsid | 17 | 28 | 12 | 0.05 | 0.2 | 0.104166667 |
| NC_004718.3;NP_828858.1;SARS-CoV_nucleocapsid | 30 | 30 | 1 | 0.05 | 0.05 | 0.05 |
| NC_004718.3;NP_828858.1;SARS-CoV_nucleocapsid | 37 | 38 | 2 | 0.05 | 0.05 | 0.05 |
| NC_004718.3;NP_828858.1;SARS-CoV_nucleocapsid | 63 | 75 | 13 | 0.05 | 0.2 | 0.1 |
| NC_004718.3;NP_828858.1;SARS-CoV_nucleocapsid | 79 | 81 | 3 | 0.05 | 0.05 | 0.05 |
| NC_004718.3;NP_828858.1;SARS-CoV_nucleocapsid | 84 | 86 | 3 | 0.05 | 0.05 | 0.05 |

| Extended data 1 | | | | | | |
|---|---|---|---|---|---|---|
| Protein | Start position (peptide) | Stop position (peptide) | Number of peptides | Minimum fraction (per peptide) of control subjects with reactivity >3 sd above average | Maximum fraction (per peptide) of control subjects with reactivity >3 sd above average | Mean fraction (over all peptides in this contiguous stretch) of controlsubjects with reactivity >3 sd above average |
| NC_004718.3;NP_828858.1;SARS-CoV_nucleocapsid | 113 | 114 | 2 | 0.05 | 0.1 | 0.075 |
| NC_004718.3;NP_828858.1;SARS-CoV_nucleocapsid | 139 | 144 | 6 | 0.05 | 0.15 | 0.091666667 |
| NC_004718.3;NP_828858.1;SARS-CoV_nucleocapsid | 149 | 151 | 3 | 0.1 | 0.2 | 0.166666667 |
| NC_004718.3;NP_828858.1;SARS-CoV_nucleocapsid | 155 | 165 | 11 | 0.05 | 0.2 | 0.127272727 |
| NC_004718.3;NP_828858.1;SARS-CoV_nucleocapsid | 169 | 169 | 1 | 0.05 | 0.05 | 0.05 |
| NC_004718.3;NP_828858.1;SARS-CoV_nucleocapsid | 197 | 203 | 7 | 0.05 | 0.15 | 0.085714286 |
| NC_004718.3;NP_828858.1;SARS-CoV_nucleocapsid | 219 | 226 | 8 | 0.05 | 0.1 | 0.05625 |
| NC_004718.3;NP_828858.1;SARS-CoV_nucleocapsid | 230 | 230 | 1 | 0.05 | 0.05 | 0.05 |
| NC_004718.3;NP_828858.1;SARS-CoV_nucleocapsid | 232 | 232 | 1 | 0.05 | 0.05 | 0.05 |
| NC_004718.3;NP_828858.1;SARS-CoV_nucleocapsid | 239 | 242 | 4 | 0.05 | 0.25 | 0.1 |
| NC_004718.3;NP_828858.1;SARS-CoV_nucleocapsid | 248 | 248 | 1 | 0.05 | 0.05 | 0.05 |
| NC_004718.3;NP_828858.1;SARS-CoV_nucleocapsid | 253 | 253 | 1 | 0.05 | 0.05 | 0.05 |
| NC_004718.3;NP_828858.1;SARS-CoV_nucleocapsid | 259 | 289 | 31 | 0.05 | 0.15 | 0.077419355 |
| NC_004718.3;NP_828858.1;SARS-CoV_nucleocapsid | 295 | 295 | 1 | 0.05 | 0.05 | 0.05 |
| NC_004718.3;NP_828858.1;SARS-CoV_nucleocapsid | 298 | 298 | 1 | 0.15 | 0.15 | 0.15 |
| NC_004718.3;NP_828858.1;SARS-CoV_nucleocapsid | 311 | 311 | 1 | 0.05 | 0.05 | 0.05 |
| NC_004718.3;NP_828858.1;SARS-CoV_nucleocapsid | 314 | 315 | 2 | 0.05 | 0.05 | 0.05 |
| NC_004718.3;NP_828858.1;SARS-CoV_nucleocapsid | 333 | 333 | 1 | 0.05 | 0.05 | 0.05 |
| NC_004718.3;NP_828858.1;SARS-CoV_nucleocapsid | 336 | 336 | 1 | 0.05 | 0.05 | 0.05 |
| NC_004718.3;NP_828858.1;SARS-CoV_nucleocapsid | 338 | 338 | 1 | 0.05 | 0.05 | 0.05 |
| NC_004718.3;NP_828858.1;SARS-CoV_nucleocapsid | 342 | 342 | 1 | 0.05 | 0.05 | 0.05 |
| NC_004718.3;NP_828858.1;SARS-CoV_nucleocapsid | 346 | 346 | 1 | 0.05 | 0.05 | 0.05 |
| NC_004718.3;NP_828858.1;SARS-CoV_nucleocapsid | 354 | 354 | 1 | 0.05 | 0.05 | 0.05 |
| NC_004718.3;NP_828858.1;SARS-CoV_nucleocapsid | 358 | 359 | 2 | 0.05 | 0.05 | 0.05 |
| NC_004718.3;NP_828858.1;SARS-CoV_nucleocapsid | 363 | 365 | 3 | 0.05 | 0.05 | 0.05 |
| NC_004718.3;NP_828858.1;SARS-CoV_nucleocapsid | 370 | 370 | 1 | 0.05 | 0.05 | 0.05 |
| NC_004718.3;NP_828858.1;SARS-CoV_nucleocapsid | 372 | 374 | 3 | 0.05 | 0.1 | 0.066666667 |
| NC_004718.3;NP_828858.1;SARS-CoV_nucleocapsid | 376 | 379 | 4 | 0.05 | 0.05 | 0.05 |
| NC_004718.3;NP_828858.1;SARS-CoV_nucleocapsid | 384 | 384 | 1 | 0.05 | 0.05 | 0.05 |
| NC_004718.3;NP_828858.1;SARS-CoV_nucleocapsid | 402 | 402 | 1 | 0.05 | 0.05 | 0.05 |
| NC_004718.3;NP_828858.1;SARS-CoV_nucleocapsid | 410 | 411 | 2 | 0.05 | 0.05 | 0.05 |
| NC_004718.3;NP_828859.1;SARS-CoV_9b | 1 | 6 | 6 | 0.05 | 0.2 | 0.116666667 |
| NC_004718.3;NP_828859.1;SARS-CoV_9b | 50 | 50 | 1 | 0.05 | 0.05 | 0.05 |
| NC_004718.3;NP_828859.1;SARS-CoV_9b | 52 | 61 | 10 | 0.05 | 0.1 | 0.085 |
| NC_004718.3;NP_828859.1;SARS-CoV_9b | 78 | 78 | 1 | 0.05 | 0.05 | 0.05 |
| NC_004718.3;NP_828859.1;SARS-CoV_9b | 83 | 83 | 1 | 0.05 | 0.05 | 0.05 |
| NC_004718.3;NP_828859.1;SARS-CoV_9b | 85 | 87 | 3 | 0.05 | 0.05 | 0.05 |

| Extended data 1 | | | | | | |
|---|---|---|---|---|---|---|
| Protein | Start position (peptide) | Stop position (peptide) | Number of peptides | Minimum fraction (per peptide) of control subjects with reactivity >3 sd above average | Maximum fraction (per peptide) of control subjects with reactivity >3 sd above average | Mean fraction (over all peptides in this contiguous stretch) of controlsubjects with reactivity >3 sd above average |
| NC_004718.3;NP_849176.1;SARS-CoV_8a | 17 | 17 | 1 | 0.05 | 0.05 | 0.05 |
| NC_004718.3;NP_849176.1;SARS-CoV_8a | 19 | 25 | 7 | 0.05 | 0.1 | 0.085714286 |
| NC_004718.3;NP_849177.1;SARS-CoV_8b | 32 | 43 | 12 | 0.05 | 0.2 | 0.1375 |
| NC_005831.2;YP_003766.2;HCoV-NL63_1ab | 1 | 1 | 1 | 0.05 | 0.05 | 0.05 |
| NC_005831.2;YP_003766.2;HCoV-NL63_1ab | 7 | 8 | 2 | 0.05 | 0.05 | 0.05 |
| NC_005831.2;YP_003766.2;HCoV-NL63_1ab | 12 | 12 | 1 | 0.05 | 0.05 | 0.05 |
| NC_005831.2;YP_003766.2;HCoV-NL63_1ab | 20 | 21 | 2 | 0.05 | 0.05 | 0.05 |
| NC_005831.2;YP_003766.2;HCoV-NL63_1ab | 55 | 55 | 1 | 0.1 | 0.1 | 0.1 |
| NC_005831.2;YP_003766.2;HCoV-NL63_1ab | 89 | 89 | 1 | 0.05 | 0.05 | 0.05 |
| NC_005831.2;YP_003766.2;HCoV-NL63_1ab | 113 | 113 | 1 | 0.05 | 0.05 | 0.05 |
| NC_005831.2;YP_003766.2;HCoV-NL63_1ab | 122 | 122 | 1 | 0.05 | 0.05 | 0.05 |
| NC_005831.2;YP_003766.2;HCoV-NL63_1ab | 125 | 128 | 4 | 0.05 | 0.05 | 0.05 |
| NC_005831.2;YP_003766.2;HCoV-NL63_1ab | 130 | 130 | 1 | 0.1 | 0.1 | 0.1 |
| NC_005831.2;YP_003766.2;HCoV-NL63_1ab | 150 | 150 | 1 | 0.05 | 0.05 | 0.05 |
| NC_005831.2;YP_003766.2;HCoV-NL63_1ab | 154 | 155 | 2 | 0.05 | 0.05 | 0.05 |
| NC_005831.2;YP_003766.2;HCoV-NL63_1ab | 158 | 158 | 1 | 0.05 | 0.05 | 0.05 |
| NC_005831.2;YP_003766.2;HCoV-NL63_1ab | 238 | 241 | 4 | 0.05 | 0.05 | 0.05 |
| NC_005831.2;YP_003766.2;HCoV-NL63_1ab | 243 | 244 | 2 | 0.05 | 0.05 | 0.05 |
| NC_005831.2;YP_003766.2;HCoV-NL63_1ab | 255 | 255 | 1 | 0.05 | 0.05 | 0.05 |
| NC_005831.2;YP_003766.2;HCoV-NL63_1ab | 267 | 268 | 2 | 0.05 | 0.05 | 0.05 |
| NC_005831.2;YP_003766.2;HCoV-NL63_1ab | 270 | 274 | 5 | 0.05 | 0.1 | 0.06 |
| NC_005831.2;YP_003766.2;HCoV-NL63_1ab | 281 | 281 | 1 | 0.05 | 0.05 | 0.05 |
| NC_005831.2;YP_003766.2;HCoV-NL63_1ab | 294 | 294 | 1 | 0.05 | 0.05 | 0.05 |
| NC_005831.2;YP_003766.2;HCoV-NL63_1ab | 315 | 315 | 1 | 0.05 | 0.05 | 0.05 |
| NC_005831.2;YP_003766.2;HCoV-NL63_1ab | 328 | 328 | 1 | 0.05 | 0.05 | 0.05 |
| NC_005831.2;YP_003766.2;HCoV-NL63_1ab | 333 | 333 | 1 | 0.05 | 0.05 | 0.05 |
| NC_005831.2;YP_003766.2;HCoV-NL63_1ab | 338 | 338 | 1 | 0.05 | 0.05 | 0.05 |
| NC_005831.2;YP_003766.2;HCoV-NL63_1ab | 341 | 341 | 1 | 0.05 | 0.05 | 0.05 |
| NC_005831.2;YP_003766.2;HCoV-NL63_1ab | 351 | 351 | 1 | 0.1 | 0.1 | 0.1 |
| NC_005831.2;YP_003766.2;HCoV-NL63_1ab | 355 | 368 | 14 | 0.05 | 0.4 | 0.225 |
| NC_005831.2;YP_003766.2;HCoV-NL63_1ab | 385 | 386 | 2 | 0.05 | 0.05 | 0.05 |
| NC_005831.2;YP_003766.2;HCoV-NL63_1ab | 391 | 391 | 1 | 0.05 | 0.05 | 0.05 |
| NC_005831.2;YP_003766.2;HCoV-NL63_1ab | 393 | 393 | 1 | 0.05 | 0.05 | 0.05 |
| NC_005831.2;YP_003766.2;HCoV-NL63_1ab | 398 | 405 | 8 | 0.05 | 0.1 | 0.05625 |
| NC_005831.2;YP_003766.2;HCoV-NL63_1ab | 412 | 412 | 1 | 0.05 | 0.05 | 0.05 |
| NC_005831.2;YP_003766.2;HCoV-NL63_1ab | 447 | 447 | 1 | 0.05 | 0.05 | 0.05 |
| NC_005831.2;YP_003766.2;HCoV-NL63_1ab | 471 | 485 | 15 | 0.05 | 0.2 | 0.083333333 |

| Extended data 1 | | | | | | |
|---|---|---|---|---|---|---|
| **Protein** | **Start position (peptide)** | **Stop position (peptide)** | **Number of peptides** | **Minimum fraction (per peptide) of control subjects with reactivity >3 sd above average** | **Maximum fraction (per peptide) of control subjects with reactivity >3 sd above average** | **Mean fraction (over all peptides in this contiguous stretch) of controlsubjects with reactivity >3 sd above average** |
| NC_005831.2;YP_003766.2;HCoV-NL63_1ab | 516 | 520 | 5 | 0.05 | 0.05 | 0.05 |
| NC_005831.2;YP_003766.2;HCoV-NL63_1ab | 522 | 522 | 1 | 0.05 | 0.05 | 0.05 |
| NC_005831.2;YP_003766.2;HCoV-NL63_1ab | 526 | 526 | 1 | 0.05 | 0.05 | 0.05 |
| NC_005831.2;YP_003766.2;HCoV-NL63_1ab | 578 | 578 | 1 | 0.05 | 0.05 | 0.05 |
| NC_005831.2;YP_003766.2;HCoV-NL63_1ab | 641 | 644 | 4 | 0.05 | 0.05 | 0.05 |
| NC_005831.2;YP_003766.2;HCoV-NL63_1ab | 680 | 681 | 2 | 0.05 | 0.05 | 0.05 |
| NC_005831.2;YP_003766.2;HCoV-NL63_1ab | 686 | 687 | 2 | 0.05 | 0.1 | 0.075 |
| NC_005831.2;YP_003766.2;HCoV-NL63_1ab | 689 | 689 | 1 | 0.05 | 0.05 | 0.05 |
| NC_005831.2;YP_003766.2;HCoV-NL63_1ab | 695 | 695 | 1 | 0.05 | 0.05 | 0.05 |
| NC_005831.2;YP_003766.2;HCoV-NL63_1ab | 702 | 702 | 1 | 0.1 | 0.1 | 0.1 |
| NC_005831.2;YP_003766.2;HCoV-NL63_1ab | 708 | 708 | 1 | 0.05 | 0.05 | 0.05 |
| NC_005831.2;YP_003766.2;HCoV-NL63_1ab | 720 | 720 | 1 | 0.05 | 0.05 | 0.05 |
| NC_005831.2;YP_003766.2;HCoV-NL63_1ab | 829 | 829 | 1 | 0.05 | 0.05 | 0.05 |
| NC_005831.2;YP_003766.2;HCoV-NL63_1ab | 835 | 835 | 1 | 0.05 | 0.05 | 0.05 |
| NC_005831.2;YP_003766.2;HCoV-NL63_1ab | 875 | 875 | 1 | 0.05 | 0.05 | 0.05 |
| NC_005831.2;YP_003766.2;HCoV-NL63_1ab | 877 | 877 | 1 | 0.1 | 0.1 | 0.1 |
| NC_005831.2;YP_003766.2;HCoV-NL63_1ab | 884 | 884 | 1 | 0.05 | 0.05 | 0.05 |
| NC_005831.2;YP_003766.2;HCoV-NL63_1ab | 901 | 901 | 1 | 0.05 | 0.05 | 0.05 |
| NC_005831.2;YP_003766.2;HCoV-NL63_1ab | 933 | 934 | 2 | 0.05 | 0.1 | 0.075 |
| NC_005831.2;YP_003766.2;HCoV-NL63_1ab | 949 | 949 | 1 | 0.05 | 0.05 | 0.05 |
| NC_005831.2;YP_003766.2;HCoV-NL63_1ab | 968 | 970 | 3 | 0.05 | 0.05 | 0.05 |
| NC_005831.2;YP_003766.2;HCoV-NL63_1ab | 976 | 976 | 1 | 0.05 | 0.05 | 0.05 |
| NC_005831.2;YP_003766.2;HCoV-NL63_1ab | 982 | 989 | 8 | 0.05 | 0.15 | 0.075 |
| NC_005831.2;YP_003766.2;HCoV-NL63_1ab | 1015 | 1025 | 11 | 0.05 | 0.05 | 0.05 |
| NC_005831.2;YP_003766.2;HCoV-NL63_1ab | 1042 | 1047 | 6 | 0.05 | 0.2 | 0.091666667 |
| NC_005831.2;YP_003766.2;HCoV-NL63_1ab | 1051 | 1051 | 1 | 0.05 | 0.05 | 0.05 |
| NC_005831.2;YP_003766.2;HCoV-NL63_1ab | 1054 | 1054 | 1 | 0.05 | 0.05 | 0.05 |
| NC_005831.2;YP_003766.2;HCoV-NL63_1ab | 1078 | 1078 | 1 | 0.05 | 0.05 | 0.05 |
| NC_005831.2;YP_003766.2;HCoV-NL63_1ab | 1080 | 1080 | 1 | 0.05 | 0.05 | 0.05 |
| NC_005831.2;YP_003766.2;HCoV-NL63_1ab | 1109 | 1109 | 1 | 0.05 | 0.05 | 0.05 |
| NC_005831.2;YP_003766.2;HCoV-NL63_1ab | 1128 | 1128 | 1 | 0.05 | 0.05 | 0.05 |
| NC_005831.2;YP_003766.2;HCoV-NL63_1ab | 1130 | 1131 | 2 | 0.05 | 0.05 | 0.05 |
| NC_005831.2;YP_003766.2;HCoV-NL63_1ab | 1135 | 1135 | 1 | 0.05 | 0.05 | 0.05 |
| NC_005831.2;YP_003766.2;HCoV-NL63_1ab | 1137 | 1137 | 1 | 0.05 | 0.05 | 0.05 |
| NC_005831.2;YP_003766.2;HCoV-NL63_1ab | 1146 | 1146 | 1 | 0.05 | 0.05 | 0.05 |
| NC_005831.2;YP_003766.2;HCoV-NL63_1ab | 1148 | 1151 | 4 | 0.05 | 0.05 | 0.05 |
| NC_005831.2;YP_003766.2;HCoV-NL63_1ab | 1185 | 1187 | 3 | 0.05 | 0.05 | 0.05 |

| Extended data 1 | | | | | | |
|---|---|---|---|---|---|---|
| Protein | Start position (peptide) | Stop position (peptide) | Number of peptides | Minimum fraction (per peptide) of control subjects with reactivity >3 sd above average | Maximum fraction (per peptide) of control subjects with reactivity >3 sd above average | Mean fraction (over all peptides in this contiguous stretch) of controlsubjects with reactivity >3 sd above average |
| NC_005831.2;YP_003766.2;HCoV-NL63_1ab | 1206 | 1206 | 1 | 0.05 | 0.05 | 0.05 |
| NC_005831.2;YP_003766.2;HCoV-NL63_1ab | 1210 | 1216 | 7 | 0.05 | 0.1 | 0.078571429 |
| NC_005831.2;YP_003766.2;HCoV-NL63_1ab | 1218 | 1220 | 3 | 0.05 | 0.05 | 0.05 |
| NC_005831.2;YP_003766.2;HCoV-NL63_1ab | 1222 | 1232 | 11 | 0.05 | 0.45 | 0.181818182 |
| NC_005831.2;YP_003766.2;HCoV-NL63_1ab | 1244 | 1251 | 8 | 0.05 | 0.05 | 0.05 |
| NC_005831.2;YP_003766.2;HCoV-NL63_1ab | 1283 | 1283 | 1 | 0.05 | 0.05 | 0.05 |
| NC_005831.2;YP_003766.2;HCoV-NL63_1ab | 1285 | 1285 | 1 | 0.05 | 0.05 | 0.05 |
| NC_005831.2;YP_003766.2;HCoV-NL63_1ab | 1296 | 1296 | 1 | 0.05 | 0.05 | 0.05 |
| NC_005831.2;YP_003766.2;HCoV-NL63_1ab | 1301 | 1301 | 1 | 0.05 | 0.05 | 0.05 |
| NC_005831.2;YP_003766.2;HCoV-NL63_1ab | 1305 | 1316 | 12 | 0.05 | 0.2 | 0.1125 |
| NC_005831.2;YP_003766.2;HCoV-NL63_1ab | 1318 | 1318 | 1 | 0.05 | 0.05 | 0.05 |
| NC_005831.2;YP_003766.2;HCoV-NL63_1ab | 1320 | 1321 | 2 | 0.05 | 0.05 | 0.05 |
| NC_005831.2;YP_003766.2;HCoV-NL63_1ab | 1325 | 1325 | 1 | 0.05 | 0.05 | 0.05 |
| NC_005831.2;YP_003766.2;HCoV-NL63_1ab | 1328 | 1328 | 1 | 0.05 | 0.05 | 0.05 |
| NC_005831.2;YP_003766.2;HCoV-NL63_1ab | 1330 | 1334 | 5 | 0.05 | 0.05 | 0.05 |
| NC_005831.2;YP_003766.2;HCoV-NL63_1ab | 1336 | 1337 | 2 | 0.05 | 0.1 | 0.075 |
| NC_005831.2;YP_003766.2;HCoV-NL63_1ab | 1339 | 1339 | 1 | 0.05 | 0.05 | 0.05 |
| NC_005831.2;YP_003766.2;HCoV-NL63_1ab | 1342 | 1343 | 2 | 0.05 | 0.05 | 0.05 |
| NC_005831.2;YP_003766.2;HCoV-NL63_1ab | 1357 | 1359 | 3 | 0.1 | 0.1 | 0.1 |
| NC_005831.2;YP_003766.2;HCoV-NL63_1ab | 1403 | 1404 | 2 | 0.05 | 0.05 | 0.05 |
| NC_005831.2;YP_003766.2;HCoV-NL63_1ab | 1416 | 1416 | 1 | 0.05 | 0.05 | 0.05 |
| NC_005831.2;YP_003766.2;HCoV-NL63_1ab | 1439 | 1442 | 4 | 0.05 | 0.1 | 0.0625 |
| NC_005831.2;YP_003766.2;HCoV-NL63_1ab | 1449 | 1449 | 1 | 0.05 | 0.05 | 0.05 |
| NC_005831.2;YP_003766.2;HCoV-NL63_1ab | 1457 | 1474 | 18 | 0.05 | 0.25 | 0.141666667 |
| NC_005831.2;YP_003766.2;HCoV-NL63_1ab | 1478 | 1478 | 1 | 0.05 | 0.05 | 0.05 |
| NC_005831.2;YP_003766.2;HCoV-NL63_1ab | 1510 | 1517 | 8 | 0.05 | 0.35 | 0.1125 |
| NC_005831.2;YP_003766.2;HCoV-NL63_1ab | 1538 | 1539 | 2 | 0.05 | 0.05 | 0.05 |
| NC_005831.2;YP_003766.2;HCoV-NL63_1ab | 1563 | 1568 | 6 | 0.05 | 0.1 | 0.058333333 |
| NC_005831.2;YP_003766.2;HCoV-NL63_1ab | 1570 | 1570 | 1 | 0.05 | 0.05 | 0.05 |
| NC_005831.2;YP_003766.2;HCoV-NL63_1ab | 1577 | 1577 | 1 | 0.05 | 0.05 | 0.05 |
| NC_005831.2;YP_003766.2;HCoV-NL63_1ab | 1579 | 1579 | 1 | 0.05 | 0.05 | 0.05 |
| NC_005831.2;YP_003766.2;HCoV-NL63_1ab | 1592 | 1602 | 11 | 0.05 | 0.1 | 0.059090909 |
| NC_005831.2;YP_003766.2;HCoV-NL63_1ab | 1613 | 1616 | 4 | 0.05 | 0.1 | 0.075 |
| NC_005831.2;YP_003766.2;HCoV-NL63_1ab | 1625 | 1625 | 1 | 0.05 | 0.05 | 0.05 |
| NC_005831.2;YP_003766.2;HCoV-NL63_1ab | 1633 | 1633 | 1 | 0.05 | 0.05 | 0.05 |
| NC_005831.2;YP_003766.2;HCoV-NL63_1ab | 1651 | 1652 | 2 | 0.1 | 0.3 | 0.2 |
| NC_005831.2;YP_003766.2;HCoV-NL63_1ab | 1695 | 1695 | 1 | 0.05 | 0.05 | 0.05 |

| Extended data 1 | | | | | | |
|---|---|---|---|---|---|---|
| Protein | Start position (peptide) | Stop position (peptide) | Number of peptides | Minimum fraction (per peptide) of control subjects with reactivity >3 sd above average | Maximum fraction (per peptide) of control subjects with reactivity >3 sd above average | Mean fraction (over all peptides in this contiguous stretch) of controlsubjects with reactivity >3 sd above average |
| NC_005831.2;YP_003766.2;HCoV-NL63_1ab | 1727 | 1727 | 1 | 0.05 | 0.05 | 0.05 |
| NC_005831.2;YP_003766.2;HCoV-NL63_1ab | 1739 | 1739 | 1 | 0.05 | 0.05 | 0.05 |
| NC_005831.2;YP_003766.2;HCoV-NL63_1ab | 1765 | 1765 | 1 | 0.05 | 0.05 | 0.05 |
| NC_005831.2;YP_003766.2;HCoV-NL63_1ab | 1806 | 1806 | 1 | 0.05 | 0.05 | 0.05 |
| NC_005831.2;YP_003766.2;HCoV-NL63_1ab | 1810 | 1810 | 1 | 0.05 | 0.05 | 0.05 |
| NC_005831.2;YP_003766.2;HCoV-NL63_1ab | 1837 | 1839 | 3 | 0.05 | 0.1 | 0.066666667 |
| NC_005831.2;YP_003766.2;HCoV-NL63_1ab | 1842 | 1842 | 1 | 0.05 | 0.05 | 0.05 |
| NC_005831.2;YP_003766.2;HCoV-NL63_1ab | 1845 | 1845 | 1 | 0.05 | 0.05 | 0.05 |
| NC_005831.2;YP_003766.2;HCoV-NL63_1ab | 1890 | 1890 | 1 | 0.1 | 0.1 | 0.1 |
| NC_005831.2;YP_003766.2;HCoV-NL63_1ab | 1895 | 1895 | 1 | 0.05 | 0.05 | 0.05 |
| NC_005831.2;YP_003766.2;HCoV-NL63_1ab | 1992 | 1993 | 2 | 0.05 | 0.05 | 0.05 |
| NC_005831.2;YP_003766.2;HCoV-NL63_1ab | 2003 | 2007 | 5 | 0.05 | 0.1 | 0.06 |
| NC_005831.2;YP_003766.2;HCoV-NL63_1ab | 2029 | 2029 | 1 | 0.05 | 0.05 | 0.05 |
| NC_005831.2;YP_003766.2;HCoV-NL63_1ab | 2129 | 2129 | 1 | 0.05 | 0.05 | 0.05 |
| NC_005831.2;YP_003766.2;HCoV-NL63_1ab | 2134 | 2134 | 1 | 0.05 | 0.05 | 0.05 |
| NC_005831.2;YP_003766.2;HCoV-NL63_1ab | 2153 | 2154 | 2 | 0.05 | 0.05 | 0.05 |
| NC_005831.2;YP_003766.2;HCoV-NL63_1ab | 2159 | 2159 | 1 | 0.05 | 0.05 | 0.05 |
| NC_005831.2;YP_003766.2;HCoV-NL63_1ab | 2179 | 2185 | 7 | 0.05 | 0.15 | 0.064285714 |
| NC_005831.2;YP_003766.2;HCoV-NL63_1ab | 2198 | 2209 | 12 | 0.05 | 0.1 | 0.091666667 |
| NC_005831.2;YP_003766.2;HCoV-NL63_1ab | 2232 | 2235 | 4 | 0.05 | 0.05 | 0.05 |
| NC_005831.2;YP_003766.2;HCoV-NL63_1ab | 2237 | 2239 | 3 | 0.05 | 0.1 | 0.083333333 |
| NC_005831.2;YP_003766.2;HCoV-NL63_1ab | 2244 | 2245 | 2 | 0.05 | 0.05 | 0.05 |
| NC_005831.2;YP_003766.2;HCoV-NL63_1ab | 2257 | 2266 | 10 | 0.05 | 0.1 | 0.055 |
| NC_005831.2;YP_003766.2;HCoV-NL63_1ab | 2292 | 2292 | 1 | 0.05 | 0.05 | 0.05 |
| NC_005831.2;YP_003766.2;HCoV-NL63_1ab | 2320 | 2320 | 1 | 0.05 | 0.05 | 0.05 |
| NC_005831.2;YP_003766.2;HCoV-NL63_1ab | 2364 | 2364 | 1 | 0.05 | 0.05 | 0.05 |
| NC_005831.2;YP_003766.2;HCoV-NL63_1ab | 2378 | 2378 | 1 | 0.05 | 0.05 | 0.05 |
| NC_005831.2;YP_003766.2;HCoV-NL63_1ab | 2381 | 2381 | 1 | 0.05 | 0.05 | 0.05 |
| NC_005831.2;YP_003766.2;HCoV-NL63_1ab | 2407 | 2408 | 2 | 0.1 | 0.15 | 0.125 |
| NC_005831.2;YP_003766.2;HCoV-NL63_1ab | 2412 | 2412 | 1 | 0.15 | 0.15 | 0.15 |
| NC_005831.2;YP_003766.2;HCoV-NL63_1ab | 2443 | 2443 | 1 | 0.05 | 0.05 | 0.05 |
| NC_005831.2;YP_003766.2;HCoV-NL63_1ab | 2451 | 2459 | 9 | 0.05 | 0.05 | 0.05 |
| NC_005831.2;YP_003766.2;HCoV-NL63_1ab | 2482 | 2484 | 3 | 0.05 | 0.05 | 0.05 |
| NC_005831.2;YP_003766.2;HCoV-NL63_1ab | 2495 | 2497 | 3 | 0.05 | 0.05 | 0.05 |
| NC_005831.2;YP_003766.2;HCoV-NL63_1ab | 2501 | 2502 | 2 | 0.05 | 0.05 | 0.05 |
| NC_005831.2;YP_003766.2;HCoV-NL63_1ab | 2513 | 2514 | 2 | 0.05 | 0.05 | 0.05 |
| NC_005831.2;YP_003766.2;HCoV-NL63_1ab | 2532 | 2533 | 2 | 0.05 | 0.05 | 0.05 |

| Extended data 1 | | | | | | |
|---|---|---|---|---|---|---|
| Protein | Start position (peptide) | Stop position (peptide) | Number of peptides | Minimum fraction (per peptide) of control subjects with reactivity >3 sd above average | Maximum fraction (per peptide) of control subjects with reactivity >3 sd above average | Mean fraction (over all peptides in this contiguous stretch) of controlsubjects with reactivity >3 sd above average |
| NC_005831.2;YP_003766.2;HCoV-NL63_1ab | 2550 | 2551 | 2 | 0.05 | 0.05 | 0.05 |
| NC_005831.2;YP_003766.2;HCoV-NL63_1ab | 2553 | 2553 | 1 | 0.05 | 0.05 | 0.05 |
| NC_005831.2;YP_003766.2;HCoV-NL63_1ab | 2580 | 2580 | 1 | 0.05 | 0.05 | 0.05 |
| NC_005831.2;YP_003766.2;HCoV-NL63_1ab | 2582 | 2589 | 8 | 0.05 | 0.05 | 0.05 |
| NC_005831.2;YP_003766.2;HCoV-NL63_1ab | 2597 | 2597 | 1 | 0.1 | 0.1 | 0.1 |
| NC_005831.2;YP_003766.2;HCoV-NL63_1ab | 2606 | 2607 | 2 | 0.05 | 0.05 | 0.05 |
| NC_005831.2;YP_003766.2;HCoV-NL63_1ab | 2610 | 2610 | 1 | 0.05 | 0.05 | 0.05 |
| NC_005831.2;YP_003766.2;HCoV-NL63_1ab | 2622 | 2624 | 3 | 0.1 | 0.25 | 0.166666667 |
| NC_005831.2;YP_003766.2;HCoV-NL63_1ab | 2626 | 2631 | 6 | 0.05 | 0.15 | 0.083333333 |
| NC_005831.2;YP_003766.2;HCoV-NL63_1ab | 2638 | 2639 | 2 | 0.05 | 0.1 | 0.075 |
| NC_005831.2;YP_003766.2;HCoV-NL63_1ab | 2683 | 2683 | 1 | 0.05 | 0.05 | 0.05 |
| NC_005831.2;YP_003766.2;HCoV-NL63_1ab | 2699 | 2699 | 1 | 0.05 | 0.05 | 0.05 |
| NC_005831.2;YP_003766.2;HCoV-NL63_1ab | 2824 | 2824 | 1 | 0.05 | 0.05 | 0.05 |
| NC_005831.2;YP_003766.2;HCoV-NL63_1ab | 2836 | 2841 | 6 | 0.05 | 0.15 | 0.108333333 |
| NC_005831.2;YP_003766.2;HCoV-NL63_1ab | 2843 | 2844 | 2 | 0.05 | 0.05 | 0.05 |
| NC_005831.2;YP_003766.2;HCoV-NL63_1ab | 2864 | 2866 | 3 | 0.05 | 0.05 | 0.05 |
| NC_005831.2;YP_003766.2;HCoV-NL63_1ab | 2868 | 2868 | 1 | 0.05 | 0.05 | 0.05 |
| NC_005831.2;YP_003766.2;HCoV-NL63_1ab | 2870 | 2871 | 2 | 0.05 | 0.05 | 0.05 |
| NC_005831.2;YP_003766.2;HCoV-NL63_1ab | 2881 | 2881 | 1 | 0.05 | 0.05 | 0.05 |
| NC_005831.2;YP_003766.2;HCoV-NL63_1ab | 2887 | 2887 | 1 | 0.05 | 0.05 | 0.05 |
| NC_005831.2;YP_003766.2;HCoV-NL63_1ab | 2891 | 2892 | 2 | 0.05 | 0.05 | 0.05 |
| NC_005831.2;YP_003766.2;HCoV-NL63_1ab | 2898 | 2898 | 1 | 0.05 | 0.05 | 0.05 |
| NC_005831.2;YP_003766.2;HCoV-NL63_1ab | 2920 | 2922 | 3 | 0.05 | 0.1 | 0.083333333 |
| NC_005831.2;YP_003766.2;HCoV-NL63_1ab | 2926 | 2926 | 1 | 0.05 | 0.05 | 0.05 |
| NC_005831.2;YP_003766.2;HCoV-NL63_1ab | 2930 | 2930 | 1 | 0.05 | 0.05 | 0.05 |
| NC_005831.2;YP_003766.2;HCoV-NL63_1ab | 2953 | 2953 | 1 | 0.05 | 0.05 | 0.05 |
| NC_005831.2;YP_003766.2;HCoV-NL63_1ab | 3028 | 3029 | 2 | 0.05 | 0.15 | 0.1 |
| NC_005831.2;YP_003766.2;HCoV-NL63_1ab | 3048 | 3048 | 1 | 0.1 | 0.1 | 0.1 |
| NC_005831.2;YP_003766.2;HCoV-NL63_1ab | 3083 | 3083 | 1 | 0.05 | 0.05 | 0.05 |
| NC_005831.2;YP_003766.2;HCoV-NL63_1ab | 3097 | 3102 | 6 | 0.05 | 0.05 | 0.05 |
| NC_005831.2;YP_003766.2;HCoV-NL63_1ab | 3112 | 3112 | 1 | 0.05 | 0.05 | 0.05 |
| NC_005831.2;YP_003766.2;HCoV-NL63_1ab | 3114 | 3124 | 11 | 0.1 | 0.15 | 0.104545455 |
| NC_005831.2;YP_003766.2;HCoV-NL63_1ab | 3126 | 3126 | 1 | 0.05 | 0.05 | 0.05 |
| NC_005831.2;YP_003766.2;HCoV-NL63_1ab | 3130 | 3131 | 2 | 0.05 | 0.2 | 0.125 |
| NC_005831.2;YP_003766.2;HCoV-NL63_1ab | 3160 | 3168 | 9 | 0.05 | 0.15 | 0.088888889 |
| NC_005831.2;YP_003766.2;HCoV-NL63_1ab | 3171 | 3174 | 4 | 0.05 | 0.05 | 0.05 |
| NC_005831.2;YP_003766.2;HCoV-NL63_1ab | 3199 | 3199 | 1 | 0.05 | 0.05 | 0.05 |

| Extended data 1 | | | | | | |
|---|---|---|---|---|---|---|
| Protein | Start position (peptide) | Stop position (peptide) | Number of peptides | Minimum fraction (per peptide) of control subjects with reactivity >3 sd above average | Maximum fraction (per peptide) of control subjects with reactivity >3 sd above average | Mean fraction (over all peptides in this contiguous stretch) of controlsubjects with reactivity >3 sd above average |
| NC_005831.2;YP_003766.2;HCoV-NL63_1ab | 3204 | 3207 | 4 | 0.05 | 0.05 | 0.05 |
| NC_005831.2;YP_003766.2;HCoV-NL63_1ab | 3210 | 3216 | 7 | 0.05 | 0.05 | 0.05 |
| NC_005831.2;YP_003766.2;HCoV-NL63_1ab | 3220 | 3222 | 3 | 0.05 | 0.1 | 0.083333333 |
| NC_005831.2;YP_003766.2;HCoV-NL63_1ab | 3224 | 3224 | 1 | 0.05 | 0.05 | 0.05 |
| NC_005831.2;YP_003766.2;HCoV-NL63_1ab | 3229 | 3230 | 2 | 0.05 | 0.05 | 0.05 |
| NC_005831.2;YP_003766.2;HCoV-NL63_1ab | 3330 | 3330 | 1 | 0.05 | 0.05 | 0.05 |
| NC_005831.2;YP_003766.2;HCoV-NL63_1ab | 3426 | 3427 | 2 | 0.05 | 0.1 | 0.075 |
| NC_005831.2;YP_003766.2;HCoV-NL63_1ab | 3465 | 3465 | 1 | 0.05 | 0.05 | 0.05 |
| NC_005831.2;YP_003766.2;HCoV-NL63_1ab | 3474 | 3476 | 3 | 0.05 | 0.1 | 0.083333333 |
| NC_005831.2;YP_003766.2;HCoV-NL63_1ab | 3478 | 3481 | 4 | 0.05 | 0.15 | 0.075 |
| NC_005831.2;YP_003766.2;HCoV-NL63_1ab | 3483 | 3483 | 1 | 0.05 | 0.05 | 0.05 |
| NC_005831.2;YP_003766.2;HCoV-NL63_1ab | 3507 | 3517 | 11 | 0.05 | 0.1 | 0.054545455 |
| NC_005831.2;YP_003766.2;HCoV-NL63_1ab | 3530 | 3530 | 1 | 0.1 | 0.1 | 0.1 |
| NC_005831.2;YP_003766.2;HCoV-NL63_1ab | 3544 | 3548 | 5 | 0.05 | 0.15 | 0.09 |
| NC_005831.2;YP_003766.2;HCoV-NL63_1ab | 3553 | 3554 | 2 | 0.05 | 0.1 | 0.075 |
| NC_005831.2;YP_003766.2;HCoV-NL63_1ab | 3578 | 3584 | 7 | 0.05 | 0.1 | 0.057142857 |
| NC_005831.2;YP_003766.2;HCoV-NL63_1ab | 3588 | 3589 | 2 | 0.05 | 0.05 | 0.05 |
| NC_005831.2;YP_003766.2;HCoV-NL63_1ab | 3592 | 3592 | 1 | 0.1 | 0.1 | 0.1 |
| NC_005831.2;YP_003766.2;HCoV-NL63_1ab | 3594 | 3594 | 1 | 0.05 | 0.05 | 0.05 |
| NC_005831.2;YP_003766.2;HCoV-NL63_1ab | 3612 | 3618 | 7 | 0.05 | 0.15 | 0.078571429 |
| NC_005831.2;YP_003766.2;HCoV-NL63_1ab | 3633 | 3634 | 2 | 0.05 | 0.05 | 0.05 |
| NC_005831.2;YP_003766.2;HCoV-NL63_1ab | 3645 | 3645 | 1 | 0.05 | 0.05 | 0.05 |
| NC_005831.2;YP_003766.2;HCoV-NL63_1ab | 3655 | 3655 | 1 | 0.05 | 0.05 | 0.05 |
| NC_005831.2;YP_003766.2;HCoV-NL63_1ab | 3679 | 3679 | 1 | 0.05 | 0.05 | 0.05 |
| NC_005831.2;YP_003766.2;HCoV-NL63_1ab | 3704 | 3704 | 1 | 0.1 | 0.1 | 0.1 |
| NC_005831.2;YP_003766.2;HCoV-NL63_1ab | 3715 | 3718 | 4 | 0.05 | 0.2 | 0.1125 |
| NC_005831.2;YP_003766.2;HCoV-NL63_1ab | 3756 | 3756 | 1 | 0.05 | 0.05 | 0.05 |
| NC_005831.2;YP_003766.2;HCoV-NL63_1ab | 3758 | 3760 | 3 | 0.05 | 0.15 | 0.083333333 |
| NC_005831.2;YP_003766.2;HCoV-NL63_1ab | 3768 | 3768 | 1 | 0.05 | 0.05 | 0.05 |
| NC_005831.2;YP_003766.2;HCoV-NL63_1ab | 3770 | 3770 | 1 | 0.05 | 0.05 | 0.05 |
| NC_005831.2;YP_003766.2;HCoV-NL63_1ab | 3793 | 3793 | 1 | 0.05 | 0.05 | 0.05 |
| NC_005831.2;YP_003766.2;HCoV-NL63_1ab | 3802 | 3802 | 1 | 0.05 | 0.05 | 0.05 |
| NC_005831.2;YP_003766.2;HCoV-NL63_1ab | 3819 | 3826 | 8 | 0.05 | 0.2 | 0.0875 |
| NC_005831.2;YP_003766.2;HCoV-NL63_1ab | 3832 | 3832 | 1 | 0.05 | 0.05 | 0.05 |
| NC_005831.2;YP_003766.2;HCoV-NL63_1ab | 3840 | 3843 | 4 | 0.05 | 0.15 | 0.1 |
| NC_005831.2;YP_003766.2;HCoV-NL63_1ab | 3845 | 3845 | 1 | 0.05 | 0.05 | 0.05 |
| NC_005831.2;YP_003766.2;HCoV-NL63_1ab | 3850 | 3850 | 1 | 0.1 | 0.1 | 0.1 |

| Extended data 1 | | | | | | |
|---|---|---|---|---|---|---|
| **Protein** | **Start position (peptide)** | **Stop position (peptide)** | **Number of peptides** | **Minimum fraction (per peptide) of control subjects with reactivity >3 sd above average** | **Maximum fraction (per peptide) of control subjects with reactivity >3 sd above average** | **Mean fraction (over all peptides in this contiguous stretch) of controlsubjects with reactivity >3 sd above average** |
| NC_005831.2;YP_003766.2;HCoV-NL63_1ab | 3852 | 3852 | 1 | 0.05 | 0.05 | 0.05 |
| NC_005831.2;YP_003766.2;HCoV-NL63_1ab | 3865 | 3865 | 1 | 0.05 | 0.05 | 0.05 |
| NC_005831.2;YP_003766.2;HCoV-NL63_1ab | 3867 | 3867 | 1 | 0.05 | 0.05 | 0.05 |
| NC_005831.2;YP_003766.2;HCoV-NL63_1ab | 3899 | 3903 | 5 | 0.05 | 0.05 | 0.05 |
| NC_005831.2;YP_003766.2;HCoV-NL63_1ab | 3920 | 3920 | 1 | 0.05 | 0.05 | 0.05 |
| NC_005831.2;YP_003766.2;HCoV-NL63_1ab | 3945 | 3947 | 3 | 0.05 | 0.05 | 0.05 |
| NC_005831.2;YP_003766.2;HCoV-NL63_1ab | 3950 | 3950 | 1 | 0.05 | 0.05 | 0.05 |
| NC_005831.2;YP_003766.2;HCoV-NL63_1ab | 3962 | 3963 | 2 | 0.05 | 0.1 | 0.075 |
| NC_005831.2;YP_003766.2;HCoV-NL63_1ab | 4031 | 4031 | 1 | 0.05 | 0.05 | 0.05 |
| NC_005831.2;YP_003766.2;HCoV-NL63_1ab | 4033 | 4034 | 2 | 0.05 | 0.05 | 0.05 |
| NC_005831.2;YP_003766.2;HCoV-NL63_1ab | 4044 | 4044 | 1 | 0.05 | 0.05 | 0.05 |
| NC_005831.2;YP_003766.2;HCoV-NL63_1ab | 4048 | 4049 | 2 | 0.05 | 0.05 | 0.05 |
| NC_005831.2;YP_003766.2;HCoV-NL63_1ab | 4053 | 4053 | 1 | 0.05 | 0.05 | 0.05 |
| NC_005831.2;YP_003766.2;HCoV-NL63_1ab | 4077 | 4081 | 5 | 0.05 | 0.05 | 0.05 |
| NC_005831.2;YP_003766.2;HCoV-NL63_1ab | 4084 | 4085 | 2 | 0.05 | 0.05 | 0.05 |
| NC_005831.2;YP_003766.2;HCoV-NL63_1ab | 4147 | 4150 | 4 | 0.05 | 0.05 | 0.05 |
| NC_005831.2;YP_003766.2;HCoV-NL63_1ab | 4158 | 4165 | 8 | 0.05 | 0.1 | 0.05625 |
| NC_005831.2;YP_003766.2;HCoV-NL63_1ab | 4168 | 4172 | 5 | 0.05 | 0.15 | 0.07 |
| NC_005831.2;YP_003766.2;HCoV-NL63_1ab | 4195 | 4196 | 2 | 0.05 | 0.05 | 0.05 |
| NC_005831.2;YP_003766.2;HCoV-NL63_1ab | 4201 | 4201 | 1 | 0.05 | 0.05 | 0.05 |
| NC_005831.2;YP_003766.2;HCoV-NL63_1ab | 4203 | 4203 | 1 | 0.05 | 0.05 | 0.05 |
| NC_005831.2;YP_003766.2;HCoV-NL63_1ab | 4208 | 4208 | 1 | 0.05 | 0.05 | 0.05 |
| NC_005831.2;YP_003766.2;HCoV-NL63_1ab | 4236 | 4236 | 1 | 0.05 | 0.05 | 0.05 |
| NC_005831.2;YP_003766.2;HCoV-NL63_1ab | 4240 | 4240 | 1 | 0.1 | 0.1 | 0.1 |
| NC_005831.2;YP_003766.2;HCoV-NL63_1ab | 4254 | 4254 | 1 | 0.05 | 0.05 | 0.05 |
| NC_005831.2;YP_003766.2;HCoV-NL63_1ab | 4287 | 4295 | 9 | 0.05 | 0.15 | 0.105555556 |
| NC_005831.2;YP_003766.2;HCoV-NL63_1ab | 4307 | 4310 | 4 | 0.05 | 0.15 | 0.1 |
| NC_005831.2;YP_003766.2;HCoV-NL63_1ab | 4333 | 4339 | 7 | 0.05 | 0.05 | 0.05 |
| NC_005831.2;YP_003766.2;HCoV-NL63_1ab | 4423 | 4423 | 1 | 0.05 | 0.05 | 0.05 |
| NC_005831.2;YP_003766.2;HCoV-NL63_1ab | 4426 | 4426 | 1 | 0.1 | 0.1 | 0.1 |
| NC_005831.2;YP_003766.2;HCoV-NL63_1ab | 4438 | 4438 | 1 | 0.1 | 0.1 | 0.1 |
| NC_005831.2;YP_003766.2;HCoV-NL63_1ab | 4442 | 4450 | 9 | 0.05 | 0.2 | 0.127777778 |
| NC_005831.2;YP_003766.2;HCoV-NL63_1ab | 4453 | 4453 | 1 | 0.05 | 0.05 | 0.05 |
| NC_005831.2;YP_003766.2;HCoV-NL63_1ab | 4461 | 4461 | 1 | 0.05 | 0.05 | 0.05 |
| NC_005831.2;YP_003766.2;HCoV-NL63_1ab | 4465 | 4465 | 1 | 0.1 | 0.1 | 0.1 |
| NC_005831.2;YP_003766.2;HCoV-NL63_1ab | 4482 | 4490 | 9 | 0.05 | 0.2 | 0.144444444 |
| NC_005831.2;YP_003766.2;HCoV-NL63_1ab | 4521 | 4538 | 18 | 0.05 | 0.25 | 0.105555556 |

| Extended data 1 | | | | | | |
|---|---|---|---|---|---|---|
| Protein | Start position (peptide) | Stop position (peptide) | Number of peptides | Minimum fraction (per peptide) of control subjects with reactivity >3 sd above average | Maximum fraction (per peptide) of control subjects with reactivity >3 sd above average | Mean fraction (over all peptides in this contiguous stretch) of controlsubjects with reactivity >3 sd above average |
| NC_005831.2;YP_003766.2;HCoV-NL63_1ab | 4569 | 4570 | 2 | 0.15 | 0.2 | 0.175 |
| NC_005831.2;YP_003766.2;HCoV-NL63_1ab | 4575 | 4577 | 3 | 0.05 | 0.05 | 0.05 |
| NC_005831.2;YP_003766.2;HCoV-NL63_1ab | 4586 | 4586 | 1 | 0.1 | 0.1 | 0.1 |
| NC_005831.2;YP_003766.2;HCoV-NL63_1ab | 4609 | 4617 | 9 | 0.05 | 0.05 | 0.05 |
| NC_005831.2;YP_003766.2;HCoV-NL63_1ab | 4632 | 4633 | 2 | 0.05 | 0.05 | 0.05 |
| NC_005831.2;YP_003766.2;HCoV-NL63_1ab | 4635 | 4635 | 1 | 0.05 | 0.05 | 0.05 |
| NC_005831.2;YP_003766.2;HCoV-NL63_1ab | 4641 | 4641 | 1 | 0.05 | 0.05 | 0.05 |
| NC_005831.2;YP_003766.2;HCoV-NL63_1ab | 4645 | 4645 | 1 | 0.05 | 0.05 | 0.05 |
| NC_005831.2;YP_003766.2;HCoV-NL63_1ab | 4695 | 4695 | 1 | 0.05 | 0.05 | 0.05 |
| NC_005831.2;YP_003766.2;HCoV-NL63_1ab | 4706 | 4707 | 2 | 0.05 | 0.05 | 0.05 |
| NC_005831.2;YP_003766.2;HCoV-NL63_1ab | 4712 | 4713 | 2 | 0.05 | 0.05 | 0.05 |
| NC_005831.2;YP_003766.2;HCoV-NL63_1ab | 4718 | 4718 | 1 | 0.05 | 0.05 | 0.05 |
| NC_005831.2;YP_003766.2;HCoV-NL63_1ab | 4734 | 4734 | 1 | 0.05 | 0.05 | 0.05 |
| NC_005831.2;YP_003766.2;HCoV-NL63_1ab | 4736 | 4738 | 3 | 0.05 | 0.2 | 0.1 |
| NC_005831.2;YP_003766.2;HCoV-NL63_1ab | 4743 | 4743 | 1 | 0.05 | 0.05 | 0.05 |
| NC_005831.2;YP_003766.2;HCoV-NL63_1ab | 4751 | 4751 | 1 | 0.05 | 0.05 | 0.05 |
| NC_005831.2;YP_003766.2;HCoV-NL63_1ab | 4772 | 4775 | 4 | 0.05 | 0.1 | 0.0625 |
| NC_005831.2;YP_003766.2;HCoV-NL63_1ab | 4777 | 4778 | 2 | 0.05 | 0.1 | 0.075 |
| NC_005831.2;YP_003766.2;HCoV-NL63_1ab | 4796 | 4796 | 1 | 0.05 | 0.05 | 0.05 |
| NC_005831.2;YP_003766.2;HCoV-NL63_1ab | 4806 | 4807 | 2 | 0.1 | 0.1 | 0.1 |
| NC_005831.2;YP_003766.2;HCoV-NL63_1ab | 4832 | 4832 | 1 | 0.05 | 0.05 | 0.05 |
| NC_005831.2;YP_003766.2;HCoV-NL63_1ab | 4838 | 4838 | 1 | 0.05 | 0.05 | 0.05 |
| NC_005831.2;YP_003766.2;HCoV-NL63_1ab | 4841 | 4841 | 1 | 0.05 | 0.05 | 0.05 |
| NC_005831.2;YP_003766.2;HCoV-NL63_1ab | 4846 | 4847 | 2 | 0.05 | 0.2 | 0.125 |
| NC_005831.2;YP_003766.2;HCoV-NL63_1ab | 4857 | 4857 | 1 | 0.05 | 0.05 | 0.05 |
| NC_005831.2;YP_003766.2;HCoV-NL63_1ab | 4902 | 4903 | 2 | 0.05 | 0.05 | 0.05 |
| NC_005831.2;YP_003766.2;HCoV-NL63_1ab | 4919 | 4926 | 8 | 0.05 | 0.2 | 0.125 |
| NC_005831.2;YP_003766.2;HCoV-NL63_1ab | 4940 | 4940 | 1 | 0.05 | 0.05 | 0.05 |
| NC_005831.2;YP_003766.2;HCoV-NL63_1ab | 4957 | 4957 | 1 | 0.05 | 0.05 | 0.05 |
| NC_005831.2;YP_003766.2;HCoV-NL63_1ab | 4959 | 4959 | 1 | 0.05 | 0.05 | 0.05 |
| NC_005831.2;YP_003766.2;HCoV-NL63_1ab | 5019 | 5019 | 1 | 0.05 | 0.05 | 0.05 |
| NC_005831.2;YP_003766.2;HCoV-NL63_1ab | 5026 | 5026 | 1 | 0.1 | 0.1 | 0.1 |
| NC_005831.2;YP_003766.2;HCoV-NL63_1ab | 5053 | 5067 | 15 | 0.05 | 0.1 | 0.063333333 |
| NC_005831.2;YP_003766.2;HCoV-NL63_1ab | 5081 | 5090 | 10 | 0.05 | 0.25 | 0.155 |
| NC_005831.2;YP_003766.2;HCoV-NL63_1ab | 5097 | 5097 | 1 | 0.1 | 0.1 | 0.1 |
| NC_005831.2;YP_003766.2;HCoV-NL63_1ab | 5099 | 5099 | 1 | 0.05 | 0.05 | 0.05 |
| NC_005831.2;YP_003766.2;HCoV-NL63_1ab | 5108 | 5108 | 1 | 0.05 | 0.05 | 0.05 |

| Extended data 1 | | | | | | |
|---|---|---|---|---|---|---|
| **Protein** | **Start position (peptide)** | **Stop position (peptide)** | **Number of peptides** | **Minimum fraction (per peptide) of control subjects with reactivity >3 sd above average** | **Maximum fraction (per peptide) of control subjects with reactivity >3 sd above average** | **Mean fraction (over all peptides in this contiguous stretch) of controlsubjects with reactivity >3 sd above average** |
| NC_005831.2;YP_003766.2;HCoV-NL63_1ab | 5111 | 5112 | 2 | 0.05 | 0.05 | 0.05 |
| NC_005831.2;YP_003766.2;HCoV-NL63_1ab | 5116 | 5117 | 2 | 0.05 | 0.05 | 0.05 |
| NC_005831.2;YP_003766.2;HCoV-NL63_1ab | 5119 | 5120 | 2 | 0.1 | 0.1 | 0.1 |
| NC_005831.2;YP_003766.2;HCoV-NL63_1ab | 5124 | 5124 | 1 | 0.05 | 0.05 | 0.05 |
| NC_005831.2;YP_003766.2;HCoV-NL63_1ab | 5133 | 5133 | 1 | 0.05 | 0.05 | 0.05 |
| NC_005831.2;YP_003766.2;HCoV-NL63_1ab | 5138 | 5138 | 1 | 0.05 | 0.05 | 0.05 |
| NC_005831.2;YP_003766.2;HCoV-NL63_1ab | 5157 | 5157 | 1 | 0.05 | 0.05 | 0.05 |
| NC_005831.2;YP_003766.2;HCoV-NL63_1ab | 5164 | 5164 | 1 | 0.05 | 0.05 | 0.05 |
| NC_005831.2;YP_003766.2;HCoV-NL63_1ab | 5181 | 5181 | 1 | 0.05 | 0.05 | 0.05 |
| NC_005831.2;YP_003766.2;HCoV-NL63_1ab | 5203 | 5205 | 3 | 0.05 | 0.05 | 0.05 |
| NC_005831.2;YP_003766.2;HCoV-NL63_1ab | 5209 | 5209 | 1 | 0.05 | 0.05 | 0.05 |
| NC_005831.2;YP_003766.2;HCoV-NL63_1ab | 5306 | 5306 | 1 | 0.05 | 0.05 | 0.05 |
| NC_005831.2;YP_003766.2;HCoV-NL63_1ab | 5309 | 5315 | 7 | 0.05 | 0.25 | 0.128571429 |
| NC_005831.2;YP_003766.2;HCoV-NL63_1ab | 5317 | 5317 | 1 | 0.1 | 0.1 | 0.1 |
| NC_005831.2;YP_003766.2;HCoV-NL63_1ab | 5321 | 5324 | 4 | 0.05 | 0.05 | 0.05 |
| NC_005831.2;YP_003766.2;HCoV-NL63_1ab | 5328 | 5328 | 1 | 0.05 | 0.05 | 0.05 |
| NC_005831.2;YP_003766.2;HCoV-NL63_1ab | 5330 | 5338 | 9 | 0.05 | 0.1 | 0.072222222 |
| NC_005831.2;YP_003766.2;HCoV-NL63_1ab | 5341 | 5344 | 4 | 0.05 | 0.05 | 0.05 |
| NC_005831.2;YP_003766.2;HCoV-NL63_1ab | 5383 | 5383 | 1 | 0.05 | 0.05 | 0.05 |
| NC_005831.2;YP_003766.2;HCoV-NL63_1ab | 5388 | 5388 | 1 | 0.05 | 0.05 | 0.05 |
| NC_005831.2;YP_003766.2;HCoV-NL63_1ab | 5417 | 5417 | 1 | 0.1 | 0.1 | 0.1 |
| NC_005831.2;YP_003766.2;HCoV-NL63_1ab | 5419 | 5420 | 2 | 0.05 | 0.05 | 0.05 |
| NC_005831.2;YP_003766.2;HCoV-NL63_1ab | 5424 | 5424 | 1 | 0.05 | 0.05 | 0.05 |
| NC_005831.2;YP_003766.2;HCoV-NL63_1ab | 5429 | 5429 | 1 | 0.05 | 0.05 | 0.05 |
| NC_005831.2;YP_003766.2;HCoV-NL63_1ab | 5439 | 5447 | 9 | 0.05 | 0.1 | 0.072222222 |
| NC_005831.2;YP_003766.2;HCoV-NL63_1ab | 5450 | 5452 | 3 | 0.05 | 0.1 | 0.066666667 |
| NC_005831.2;YP_003766.2;HCoV-NL63_1ab | 5475 | 5475 | 1 | 0.15 | 0.15 | 0.15 |
| NC_005831.2;YP_003766.2;HCoV-NL63_1ab | 5478 | 5479 | 2 | 0.05 | 0.05 | 0.05 |
| NC_005831.2;YP_003766.2;HCoV-NL63_1ab | 5491 | 5501 | 11 | 0.05 | 0.15 | 0.104545455 |
| NC_005831.2;YP_003766.2;HCoV-NL63_1ab | 5518 | 5520 | 3 | 0.05 | 0.05 | 0.05 |
| NC_005831.2;YP_003766.2;HCoV-NL63_1ab | 5529 | 5530 | 2 | 0.05 | 0.05 | 0.05 |
| NC_005831.2;YP_003766.2;HCoV-NL63_1ab | 5554 | 5554 | 1 | 0.05 | 0.05 | 0.05 |
| NC_005831.2;YP_003766.2;HCoV-NL63_1ab | 5592 | 5601 | 10 | 0.05 | 0.15 | 0.075 |
| NC_005831.2;YP_003766.2;HCoV-NL63_1ab | 5609 | 5609 | 1 | 0.05 | 0.05 | 0.05 |
| NC_005831.2;YP_003766.2;HCoV-NL63_1ab | 5658 | 5668 | 11 | 0.05 | 0.05 | 0.05 |
| NC_005831.2;YP_003766.2;HCoV-NL63_1ab | 5670 | 5670 | 1 | 0.05 | 0.05 | 0.05 |
| NC_005831.2;YP_003766.2;HCoV-NL63_1ab | 5766 | 5766 | 1 | 0.05 | 0.05 | 0.05 |

| Extended data 1 | | | | | | |
|---|---|---|---|---|---|---|
| Protein | Start position (peptide) | Stop position (peptide) | Number of peptides | Minimum fraction (per peptide) of control subjects with reactivity >3 sd above average | Maximum fraction (per peptide) of control subjects with reactivity >3 sd above average | Mean fraction (over all peptides in this contiguous stretch) of controlsubjects with reactivity >3 sd above average |
| NC_005831.2;YP_003766.2;HCoV-NL63_1ab | 5773 | 5776 | 4 | 0.05 | 0.05 | 0.05 |
| NC_005831.2;YP_003766.2;HCoV-NL63_1ab | 5863 | 5867 | 5 | 0.05 | 0.1 | 0.06 |
| NC_005831.2;YP_003766.2;HCoV-NL63_1ab | 5871 | 5871 | 1 | 0.05 | 0.05 | 0.05 |
| NC_005831.2;YP_003766.2;HCoV-NL63_1ab | 5908 | 5908 | 1 | 0.05 | 0.05 | 0.05 |
| NC_005831.2;YP_003766.2;HCoV-NL63_1ab | 5914 | 5918 | 5 | 0.05 | 0.05 | 0.05 |
| NC_005831.2;YP_003766.2;HCoV-NL63_1ab | 5920 | 5921 | 2 | 0.05 | 0.05 | 0.05 |
| NC_005831.2;YP_003766.2;HCoV-NL63_1ab | 5923 | 5928 | 6 | 0.05 | 0.1 | 0.066666667 |
| NC_005831.2;YP_003766.2;HCoV-NL63_1ab | 5960 | 5960 | 1 | 0.05 | 0.05 | 0.05 |
| NC_005831.2;YP_003766.2;HCoV-NL63_1ab | 5962 | 5963 | 2 | 0.05 | 0.05 | 0.05 |
| NC_005831.2;YP_003766.2;HCoV-NL63_1ab | 5966 | 5966 | 1 | 0.1 | 0.1 | 0.1 |
| NC_005831.2;YP_003766.2;HCoV-NL63_1ab | 5988 | 5988 | 1 | 0.05 | 0.05 | 0.05 |
| NC_005831.2;YP_003766.2;HCoV-NL63_1ab | 6003 | 6017 | 15 | 0.05 | 0.2 | 0.066666667 |
| NC_005831.2;YP_003766.2;HCoV-NL63_1ab | 6022 | 6023 | 2 | 0.15 | 0.15 | 0.15 |
| NC_005831.2;YP_003766.2;HCoV-NL63_1ab | 6031 | 6032 | 2 | 0.05 | 0.05 | 0.05 |
| NC_005831.2;YP_003766.2;HCoV-NL63_1ab | 6045 | 6045 | 1 | 0.05 | 0.05 | 0.05 |
| NC_005831.2;YP_003766.2;HCoV-NL63_1ab | 6048 | 6048 | 1 | 0.05 | 0.05 | 0.05 |
| NC_005831.2;YP_003766.2;HCoV-NL63_1ab | 6051 | 6052 | 2 | 0.05 | 0.05 | 0.05 |
| NC_005831.2;YP_003766.2;HCoV-NL63_1ab | 6072 | 6072 | 1 | 0.05 | 0.05 | 0.05 |
| NC_005831.2;YP_003766.2;HCoV-NL63_1ab | 6078 | 6079 | 2 | 0.05 | 0.1 | 0.075 |
| NC_005831.2;YP_003766.2;HCoV-NL63_1ab | 6081 | 6081 | 1 | 0.05 | 0.05 | 0.05 |
| NC_005831.2;YP_003766.2;HCoV-NL63_1ab | 6085 | 6092 | 8 | 0.05 | 0.05 | 0.05 |
| NC_005831.2;YP_003766.2;HCoV-NL63_1ab | 6094 | 6095 | 2 | 0.05 | 0.05 | 0.05 |
| NC_005831.2;YP_003766.2;HCoV-NL63_1ab | 6100 | 6107 | 8 | 0.05 | 0.1 | 0.09375 |
| NC_005831.2;YP_003766.2;HCoV-NL63_1ab | 6109 | 6109 | 1 | 0.05 | 0.05 | 0.05 |
| NC_005831.2;YP_003766.2;HCoV-NL63_1ab | 6116 | 6128 | 13 | 0.05 | 0.1 | 0.057692308 |
| NC_005831.2;YP_003766.2;HCoV-NL63_1ab | 6136 | 6136 | 1 | 0.05 | 0.05 | 0.05 |
| NC_005831.2;YP_003766.2;HCoV-NL63_1ab | 6168 | 6172 | 5 | 0.05 | 0.1 | 0.06 |
| NC_005831.2;YP_003766.2;HCoV-NL63_1ab | 6226 | 6226 | 1 | 0.05 | 0.05 | 0.05 |
| NC_005831.2;YP_003766.2;HCoV-NL63_1ab | 6228 | 6228 | 1 | 0.05 | 0.05 | 0.05 |
| NC_005831.2;YP_003766.2;HCoV-NL63_1ab | 6230 | 6230 | 1 | 0.05 | 0.05 | 0.05 |
| NC_005831.2;YP_003766.2;HCoV-NL63_1ab | 6258 | 6264 | 7 | 0.05 | 0.1 | 0.071428571 |
| NC_005831.2;YP_003766.2;HCoV-NL63_1ab | 6272 | 6273 | 2 | 0.05 | 0.05 | 0.05 |
| NC_005831.2;YP_003766.2;HCoV-NL63_1ab | 6275 | 6280 | 6 | 0.05 | 0.05 | 0.05 |
| NC_005831.2;YP_003766.2;HCoV-NL63_1ab | 6282 | 6283 | 2 | 0.05 | 0.05 | 0.05 |
| NC_005831.2;YP_003766.2;HCoV-NL63_1ab | 6285 | 6285 | 1 | 0.05 | 0.05 | 0.05 |
| NC_005831.2;YP_003766.2;HCoV-NL63_1ab | 6288 | 6288 | 1 | 0.05 | 0.05 | 0.05 |
| NC_005831.2;YP_003766.2;HCoV-NL63_1ab | 6300 | 6302 | 3 | 0.05 | 0.05 | 0.05 |

| Extended data 1 | | | | | | |
|---|---|---|---|---|---|---|
| Protein | Start position (peptide) | Stop position (peptide) | Number of peptides | Minimum fraction (per peptide) of control subjects with reactivity >3 sd above average | Maximum fraction (per peptide) of control subjects with reactivity >3 sd above average | Mean fraction (over all peptides in this contiguous stretch) of controlsubjects with reactivity >3 sd above average |
| NC_005831.2;YP_003766.2;HCoV-NL63_1ab | 6308 | 6309 | 2 | 0.1 | 0.1 | 0.1 |
| NC_005831.2;YP_003766.2;HCoV-NL63_1ab | 6314 | 6314 | 1 | 0.05 | 0.05 | 0.05 |
| NC_005831.2;YP_003766.2;HCoV-NL63_1ab | 6331 | 6341 | 11 | 0.05 | 0.1 | 0.077272727 |
| NC_005831.2;YP_003766.2;HCoV-NL63_1ab | 6343 | 6345 | 3 | 0.05 | 0.05 | 0.05 |
| NC_005831.2;YP_003766.2;HCoV-NL63_1ab | 6393 | 6393 | 1 | 0.1 | 0.1 | 0.1 |
| NC_005831.2;YP_003766.2;HCoV-NL63_1ab | 6405 | 6405 | 1 | 0.05 | 0.05 | 0.05 |
| NC_005831.2;YP_003766.2;HCoV-NL63_1ab | 6417 | 6417 | 1 | 0.05 | 0.05 | 0.05 |
| NC_005831.2;YP_003766.2;HCoV-NL63_1ab | 6426 | 6426 | 1 | 0.05 | 0.05 | 0.05 |
| NC_005831.2;YP_003766.2;HCoV-NL63_1ab | 6474 | 6481 | 8 | 0.05 | 0.05 | 0.05 |
| NC_005831.2;YP_003766.2;HCoV-NL63_1ab | 6487 | 6487 | 1 | 0.05 | 0.05 | 0.05 |
| NC_005831.2;YP_003766.2;HCoV-NL63_1ab | 6492 | 6493 | 2 | 0.05 | 0.05 | 0.05 |
| NC_005831.2;YP_003766.2;HCoV-NL63_1ab | 6497 | 6497 | 1 | 0.05 | 0.05 | 0.05 |
| NC_005831.2;YP_003766.2;HCoV-NL63_1ab | 6501 | 6501 | 1 | 0.05 | 0.05 | 0.05 |
| NC_005831.2;YP_003766.2;HCoV-NL63_1ab | 6514 | 6514 | 1 | 0.05 | 0.05 | 0.05 |
| NC_005831.2;YP_003766.2;HCoV-NL63_1ab | 6516 | 6516 | 1 | 0.05 | 0.05 | 0.05 |
| NC_005831.2;YP_003766.2;HCoV-NL63_1ab | 6547 | 6550 | 4 | 0.1 | 0.35 | 0.225 |
| NC_005831.2;YP_003766.2;HCoV-NL63_1ab | 6564 | 6564 | 1 | 0.05 | 0.05 | 0.05 |
| NC_005831.2;YP_003766.2;HCoV-NL63_1ab | 6568 | 6570 | 3 | 0.05 | 0.05 | 0.05 |
| NC_005831.2;YP_003766.2;HCoV-NL63_1ab | 6581 | 6581 | 1 | 0.05 | 0.05 | 0.05 |
| NC_005831.2;YP_003767.1;HCoV-NL63_spike | 18 | 18 | 1 | 0.1 | 0.1 | 0.1 |
| NC_005831.2;YP_003767.1;HCoV-NL63_spike | 87 | 87 | 1 | 0.1 | 0.1 | 0.1 |
| NC_005831.2;YP_003767.1;HCoV-NL63_spike | 117 | 117 | 1 | 0.05 | 0.05 | 0.05 |
| NC_005831.2;YP_003767.1;HCoV-NL63_spike | 150 | 150 | 1 | 0.05 | 0.05 | 0.05 |
| NC_005831.2;YP_003767.1;HCoV-NL63_spike | 184 | 184 | 1 | 0.05 | 0.05 | 0.05 |
| NC_005831.2;YP_003767.1;HCoV-NL63_spike | 186 | 186 | 1 | 0.05 | 0.05 | 0.05 |
| NC_005831.2;YP_003767.1;HCoV-NL63_spike | 212 | 212 | 1 | 0.3 | 0.3 | 0.3 |
| NC_005831.2;YP_003767.1;HCoV-NL63_spike | 217 | 220 | 4 | 0.05 | 0.1 | 0.0625 |
| NC_005831.2;YP_003767.1;HCoV-NL63_spike | 244 | 244 | 1 | 0.1 | 0.1 | 0.1 |
| NC_005831.2;YP_003767.1;HCoV-NL63_spike | 267 | 277 | 11 | 0.05 | 0.1 | 0.054545455 |
| NC_005831.2;YP_003767.1;HCoV-NL63_spike | 291 | 291 | 1 | 0.05 | 0.05 | 0.05 |
| NC_005831.2;YP_003767.1;HCoV-NL63_spike | 293 | 296 | 4 | 0.05 | 0.2 | 0.125 |
| NC_005831.2;YP_003767.1;HCoV-NL63_spike | 298 | 300 | 3 | 0.05 | 0.1 | 0.083333333 |
| NC_005831.2;YP_003767.1;HCoV-NL63_spike | 329 | 334 | 6 | 0.05 | 0.1 | 0.075 |
| NC_005831.2;YP_003767.1;HCoV-NL63_spike | 343 | 349 | 7 | 0.05 | 0.05 | 0.05 |
| NC_005831.2;YP_003767.1;HCoV-NL63_spike | 352 | 353 | 2 | 0.05 | 0.05 | 0.05 |
| NC_005831.2;YP_003767.1;HCoV-NL63_spike | 395 | 399 | 5 | 0.05 | 0.15 | 0.09 |
| NC_005831.2;YP_003767.1;HCoV-NL63_spike | 416 | 416 | 1 | 0.05 | 0.05 | 0.05 |

| Extended data 1 | | | | | | |
|---|---|---|---|---|---|---|
| **Protein** | **Start position (peptide)** | **Stop position (peptide)** | **Number of peptides** | **Minimum fraction (per peptide) of control subjects with reactivity >3 sd above average** | **Maximum fraction (per peptide) of control subjects with reactivity >3 sd above average** | **Mean fraction (over all peptides in this contiguous stretch) of controlsubjects with reactivity >3 sd above average** |
| NC_005831.2;YP_003767.1;HCoV-NL63_spike | 432 | 433 | 2 | 0.05 | 0.05 | 0.05 |
| NC_005831.2;YP_003767.1;HCoV-NL63_spike | 435 | 440 | 6 | 0.05 | 0.1 | 0.066666667 |
| NC_005831.2;YP_003767.1;HCoV-NL63_spike | 450 | 460 | 11 | 0.05 | 0.4 | 0.268181818 |
| NC_005831.2;YP_003767.1;HCoV-NL63_spike | 480 | 480 | 1 | 0.05 | 0.05 | 0.05 |
| NC_005831.2;YP_003767.1;HCoV-NL63_spike | 495 | 505 | 11 | 0.05 | 0.1 | 0.059090909 |
| NC_005831.2;YP_003767.1;HCoV-NL63_spike | 508 | 508 | 1 | 0.05 | 0.05 | 0.05 |
| NC_005831.2;YP_003767.1;HCoV-NL63_spike | 526 | 528 | 3 | 0.1 | 0.15 | 0.116666667 |
| NC_005831.2;YP_003767.1;HCoV-NL63_spike | 538 | 538 | 1 | 0.05 | 0.05 | 0.05 |
| NC_005831.2;YP_003767.1;HCoV-NL63_spike | 544 | 546 | 3 | 0.05 | 0.05 | 0.05 |
| NC_005831.2;YP_003767.1;HCoV-NL63_spike | 599 | 600 | 2 | 0.1 | 0.2 | 0.15 |
| NC_005831.2;YP_003767.1;HCoV-NL63_spike | 613 | 616 | 4 | 0.05 | 0.2 | 0.0875 |
| NC_005831.2;YP_003767.1;HCoV-NL63_spike | 618 | 642 | 25 | 0.05 | 0.2 | 0.096 |
| NC_005831.2;YP_003767.1;HCoV-NL63_spike | 644 | 644 | 1 | 0.05 | 0.05 | 0.05 |
| NC_005831.2;YP_003767.1;HCoV-NL63_spike | 651 | 654 | 4 | 0.05 | 0.2 | 0.0875 |
| NC_005831.2;YP_003767.1;HCoV-NL63_spike | 656 | 656 | 1 | 0.15 | 0.15 | 0.15 |
| NC_005831.2;YP_003767.1;HCoV-NL63_spike | 678 | 689 | 12 | 0.05 | 0.15 | 0.079166667 |
| NC_005831.2;YP_003767.1;HCoV-NL63_spike | 693 | 693 | 1 | 0.05 | 0.05 | 0.05 |
| NC_005831.2;YP_003767.1;HCoV-NL63_spike | 698 | 698 | 1 | 0.05 | 0.05 | 0.05 |
| NC_005831.2;YP_003767.1;HCoV-NL63_spike | 700 | 700 | 1 | 0.15 | 0.15 | 0.15 |
| NC_005831.2;YP_003767.1;HCoV-NL63_spike | 706 | 706 | 1 | 0.05 | 0.05 | 0.05 |
| NC_005831.2;YP_003767.1;HCoV-NL63_spike | 738 | 747 | 10 | 0.05 | 0.4 | 0.245 |
| NC_005831.2;YP_003767.1;HCoV-NL63_spike | 757 | 757 | 1 | 0.05 | 0.05 | 0.05 |
| NC_005831.2;YP_003767.1;HCoV-NL63_spike | 759 | 759 | 1 | 0.05 | 0.05 | 0.05 |
| NC_005831.2;YP_003767.1;HCoV-NL63_spike | 800 | 801 | 2 | 0.05 | 0.05 | 0.05 |
| NC_005831.2;YP_003767.1;HCoV-NL63_spike | 836 | 837 | 2 | 0.05 | 0.05 | 0.05 |
| NC_005831.2;YP_003767.1;HCoV-NL63_spike | 844 | 845 | 2 | 0.05 | 0.05 | 0.05 |
| NC_005831.2;YP_003767.1;HCoV-NL63_spike | 851 | 851 | 1 | 0.05 | 0.05 | 0.05 |
| NC_005831.2;YP_003767.1;HCoV-NL63_spike | 854 | 854 | 1 | 0.05 | 0.05 | 0.05 |
| NC_005831.2;YP_003767.1;HCoV-NL63_spike | 856 | 856 | 1 | 0.05 | 0.05 | 0.05 |
| NC_005831.2;YP_003767.1;HCoV-NL63_spike | 859 | 859 | 1 | 0.05 | 0.05 | 0.05 |
| NC_005831.2;YP_003767.1;HCoV-NL63_spike | 863 | 865 | 3 | 0.05 | 0.1 | 0.066666667 |
| NC_005831.2;YP_003767.1;HCoV-NL63_spike | 872 | 872 | 1 | 0.1 | 0.1 | 0.1 |
| NC_005831.2;YP_003767.1;HCoV-NL63_spike | 875 | 877 | 3 | 0.05 | 0.05 | 0.05 |
| NC_005831.2;YP_003767.1;HCoV-NL63_spike | 879 | 879 | 1 | 0.05 | 0.05 | 0.05 |
| NC_005831.2;YP_003767.1;HCoV-NL63_spike | 890 | 898 | 9 | 0.1 | 0.15 | 0.116666667 |
| NC_005831.2;YP_003767.1;HCoV-NL63_spike | 900 | 900 | 1 | 0.05 | 0.05 | 0.05 |
| NC_005831.2;YP_003767.1;HCoV-NL63_spike | 906 | 906 | 1 | 0.05 | 0.05 | 0.05 |

| Extended data 1 | | | | | | |
|---|---|---|---|---|---|---|
| Protein | Start position (peptide) | Stop position (peptide) | Number of peptides | Minimum fraction (per peptide) of control subjects with reactivity >3 sd above average | Maximum fraction (per peptide) of control subjects with reactivity >3 sd above average | Mean fraction (over all peptides in this contiguous stretch) of controlsubjects with reactivity >3 sd above average |
| NC_005831.2;YP_003767.1;HCoV-NL63_spike | 908 | 908 | 1 | 0.05 | 0.05 | 0.05 |
| NC_005831.2;YP_003767.1;HCoV-NL63_spike | 926 | 926 | 1 | 0.1 | 0.1 | 0.1 |
| NC_005831.2;YP_003767.1;HCoV-NL63_spike | 931 | 932 | 2 | 0.05 | 0.05 | 0.05 |
| NC_005831.2;YP_003767.1;HCoV-NL63_spike | 959 | 959 | 1 | 0.05 | 0.05 | 0.05 |
| NC_005831.2;YP_003767.1;HCoV-NL63_spike | 962 | 962 | 1 | 0.05 | 0.05 | 0.05 |
| NC_005831.2;YP_003767.1;HCoV-NL63_spike | 965 | 981 | 17 | 0.05 | 0.3 | 0.170588235 |
| NC_005831.2;YP_003767.1;HCoV-NL63_spike | 987 | 993 | 7 | 0.05 | 0.15 | 0.064285714 |
| NC_005831.2;YP_003767.1;HCoV-NL63_spike | 1002 | 1004 | 3 | 0.05 | 0.05 | 0.05 |
| NC_005831.2;YP_003767.1;HCoV-NL63_spike | 1011 | 1011 | 1 | 0.05 | 0.05 | 0.05 |
| NC_005831.2;YP_003767.1;HCoV-NL63_spike | 1038 | 1046 | 9 | 0.05 | 0.2 | 0.077777778 |
| NC_005831.2;YP_003767.1;HCoV-NL63_spike | 1054 | 1055 | 2 | 0.05 | 0.05 | 0.05 |
| NC_005831.2;YP_003767.1;HCoV-NL63_spike | 1083 | 1083 | 1 | 0.05 | 0.05 | 0.05 |
| NC_005831.2;YP_003767.1;HCoV-NL63_spike | 1086 | 1086 | 1 | 0.05 | 0.05 | 0.05 |
| NC_005831.2;YP_003767.1;HCoV-NL63_spike | 1097 | 1097 | 1 | 0.05 | 0.05 | 0.05 |
| NC_005831.2;YP_003767.1;HCoV-NL63_spike | 1100 | 1103 | 4 | 0.05 | 0.05 | 0.05 |
| NC_005831.2;YP_003767.1;HCoV-NL63_spike | 1135 | 1136 | 2 | 0.05 | 0.05 | 0.05 |
| NC_005831.2;YP_003767.1;HCoV-NL63_spike | 1172 | 1172 | 1 | 0.05 | 0.05 | 0.05 |
| NC_005831.2;YP_003767.1;HCoV-NL63_spike | 1199 | 1199 | 1 | 0.05 | 0.05 | 0.05 |
| NC_005831.2;YP_003767.1;HCoV-NL63_spike | 1209 | 1209 | 1 | 0.05 | 0.05 | 0.05 |
| NC_005831.2;YP_003767.1;HCoV-NL63_spike | 1211 | 1217 | 7 | 0.05 | 0.25 | 0.157142857 |
| NC_005831.2;YP_003767.1;HCoV-NL63_spike | 1220 | 1229 | 10 | 0.05 | 0.1 | 0.06 |
| NC_005831.2;YP_003767.1;HCoV-NL63_spike | 1232 | 1232 | 1 | 0.05 | 0.05 | 0.05 |
| NC_005831.2;YP_003767.1;HCoV-NL63_spike | 1246 | 1253 | 8 | 0.05 | 0.15 | 0.0875 |
| NC_005831.2;YP_003767.1;HCoV-NL63_spike | 1271 | 1273 | 3 | 0.05 | 0.3 | 0.183333333 |
| NC_005831.2;YP_003767.1;HCoV-NL63_spike | 1277 | 1277 | 1 | 0.05 | 0.05 | 0.05 |
| NC_005831.2;YP_003767.1;HCoV-NL63_spike | 1281 | 1281 | 1 | 0.05 | 0.05 | 0.05 |
| NC_005831.2;YP_003767.1;HCoV-NL63_spike | 1336 | 1345 | 10 | 0.05 | 0.45 | 0.345 |
| NC_005831.2;YP_003768.1;HCoV-NL63_3 | 2 | 2 | 1 | 0.05 | 0.05 | 0.05 |
| NC_005831.2;YP_003768.1;HCoV-NL63_3 | 5 | 5 | 1 | 0.05 | 0.05 | 0.05 |
| NC_005831.2;YP_003768.1;HCoV-NL63_3 | 10 | 31 | 22 | 0.05 | 0.3 | 0.106818182 |
| NC_005831.2;YP_003768.1;HCoV-NL63_3 | 136 | 136 | 1 | 0.05 | 0.05 | 0.05 |
| NC_005831.2;YP_003768.1;HCoV-NL63_3 | 172 | 173 | 2 | 0.05 | 0.05 | 0.05 |
| NC_005831.2;YP_003768.1;HCoV-NL63_3 | 182 | 182 | 1 | 0.05 | 0.05 | 0.05 |
| NC_005831.2;YP_003768.1;HCoV-NL63_3 | 184 | 185 | 2 | 0.05 | 0.05 | 0.05 |
| NC_005831.2;YP_003768.1;HCoV-NL63_3 | 213 | 214 | 2 | 0.05 | 0.05 | 0.05 |
| NC_005831.2;YP_003769.1;HCoV-NL63_envelope | 58 | 58 | 1 | 0.05 | 0.05 | 0.05 |
| NC_005831.2;YP_003769.1;HCoV-NL63_envelope | 60 | 61 | 2 | 0.05 | 0.15 | 0.1 |

| Extended data 1 | | | | | | |
|---|---|---|---|---|---|---|
| **Protein** | **Start position (peptide)** | **Stop position (peptide)** | **Number of peptides** | **Minimum fraction (per peptide) of control subjects with reactivity >3 sd above average** | **Maximum fraction (per peptide) of control subjects with reactivity >3 sd above average** | **Mean fraction (over all peptides in this contiguous stretch) of controlsubjects with reactivity >3 sd above average** |
| NC_005831.2;YP_003769.1;HCoV-NL63_envelope | 65 | 66 | 2 | 0.05 | 0.05 | 0.05 |
| NC_005831.2;YP_003770.1;HCoV-NL63_membrane | 1 | 2 | 2 | 0.1 | 0.15 | 0.125 |
| NC_005831.2;YP_003770.1;HCoV-NL63_membrane | 121 | 121 | 1 | 0.05 | 0.05 | 0.05 |
| NC_005831.2;YP_003770.1;HCoV-NL63_membrane | 177 | 183 | 7 | 0.05 | 0.05 | 0.05 |
| NC_005831.2;YP_003770.1;HCoV-NL63_membrane | 195 | 195 | 1 | 0.05 | 0.05 | 0.05 |
| NC_005831.2;YP_003770.1;HCoV-NL63_membrane | 204 | 210 | 7 | 0.05 | 0.05 | 0.05 |
| NC_005831.2;YP_003770.1;HCoV-NL63_membrane | 214 | 215 | 2 | 0.05 | 0.05 | 0.05 |
| NC_005831.2;YP_003771.1;HCoV-NL63_nucleocap | 1 | 6 | 6 | 0.05 | 0.1 | 0.058333333 |
| NC_005831.2;YP_003771.1;HCoV-NL63_nucleocap | 16 | 26 | 11 | 0.05 | 0.1 | 0.059090909 |
| NC_005831.2;YP_003771.1;HCoV-NL63_nucleocap | 35 | 36 | 2 | 0.05 | 0.1 | 0.075 |
| NC_005831.2;YP_003771.1;HCoV-NL63_nucleocap | 38 | 38 | 1 | 0.05 | 0.05 | 0.05 |
| NC_005831.2;YP_003771.1;HCoV-NL63_nucleocap | 40 | 42 | 3 | 0.05 | 0.05 | 0.05 |
| NC_005831.2;YP_003771.1;HCoV-NL63_nucleocap | 46 | 46 | 1 | 0.05 | 0.05 | 0.05 |
| NC_005831.2;YP_003771.1;HCoV-NL63_nucleocap | 49 | 49 | 1 | 0.05 | 0.05 | 0.05 |
| NC_005831.2;YP_003771.1;HCoV-NL63_nucleocap | 58 | 62 | 5 | 0.05 | 0.15 | 0.09 |
| NC_005831.2;YP_003771.1;HCoV-NL63_nucleocap | 93 | 110 | 18 | 0.05 | 0.3 | 0.119444444 |
| NC_005831.2;YP_003771.1;HCoV-NL63_nucleocap | 116 | 117 | 2 | 0.05 | 0.05 | 0.05 |
| NC_005831.2;YP_003771.1;HCoV-NL63_nucleocap | 119 | 120 | 2 | 0.1 | 0.1 | 0.1 |
| NC_005831.2;YP_003771.1;HCoV-NL63_nucleocap | 134 | 141 | 8 | 0.05 | 0.1 | 0.08125 |
| NC_005831.2;YP_003771.1;HCoV-NL63_nucleocap | 152 | 153 | 2 | 0.05 | 0.05 | 0.05 |
| NC_005831.2;YP_003771.1;HCoV-NL63_nucleocap | 155 | 162 | 8 | 0.05 | 0.05 | 0.05 |
| NC_005831.2;YP_003771.1;HCoV-NL63_nucleocap | 171 | 173 | 3 | 0.05 | 0.05 | 0.05 |
| NC_005831.2;YP_003771.1;HCoV-NL63_nucleocap | 177 | 177 | 1 | 0.05 | 0.05 | 0.05 |
| NC_005831.2;YP_003771.1;HCoV-NL63_nucleocap | 186 | 215 | 30 | 0.05 | 0.25 | 0.088333333 |
| NC_005831.2;YP_003771.1;HCoV-NL63_nucleocap | 229 | 235 | 7 | 0.05 | 0.1 | 0.078571429 |
| NC_005831.2;YP_003771.1;HCoV-NL63_nucleocap | 239 | 239 | 1 | 0.05 | 0.05 | 0.05 |
| NC_005831.2;YP_003771.1;HCoV-NL63_nucleocap | 261 | 272 | 12 | 0.05 | 0.4 | 0.266666667 |
| NC_005831.2;YP_003771.1;HCoV-NL63_nucleocap | 278 | 288 | 11 | 0.05 | 0.1 | 0.059090909 |
| NC_005831.2;YP_003771.1;HCoV-NL63_nucleocap | 294 | 294 | 1 | 0.05 | 0.05 | 0.05 |
| NC_005831.2;YP_003771.1;HCoV-NL63_nucleocap | 296 | 296 | 1 | 0.05 | 0.05 | 0.05 |
| NC_005831.2;YP_003771.1;HCoV-NL63_nucleocap | 324 | 324 | 1 | 0.15 | 0.15 | 0.15 |
| NC_005831.2;YP_003771.1;HCoV-NL63_nucleocap | 327 | 337 | 11 | 0.05 | 0.05 | 0.05 |
| NC_005831.2;YP_003771.1;HCoV-NL63_nucleocap | 339 | 339 | 1 | 0.05 | 0.05 | 0.05 |
| NC_005831.2;YP_003771.1;HCoV-NL63_nucleocap | 341 | 341 | 1 | 0.1 | 0.1 | 0.1 |
| NC_005831.2;YP_003771.1;HCoV-NL63_nucleocap | 343 | 360 | 18 | 0.05 | 0.65 | 0.344444444 |
| NC_006213.1;YP_009555238.1;HCoV-OC43_orf1ab | 6 | 6 | 1 | 0.05 | 0.05 | 0.05 |
| NC_006213.1;YP_009555238.1;HCoV-OC43_orf1ab | 14 | 15 | 2 | 0.05 | 0.05 | 0.05 |

| Extended data 1 | | | | | | |
|---|---|---|---|---|---|---|
| Protein | Start position (peptide) | Stop position (peptide) | Number of peptides | Minimum fraction (per peptide) of control subjects with reactivity >3 sd above average | Maximum fraction (per peptide) of control subjects with reactivity >3 sd above average | Mean fraction (over all peptides in this contiguous stretch) of controlsubjects with reactivity >3 sd above average |
| NC_006213.1;YP_009555238.1;HCoV-OC43_orf1ab | 19 | 23 | 5 | 0.05 | 0.35 | 0.11 |
| NC_006213.1;YP_009555238.1;HCoV-OC43_orf1ab | 25 | 26 | 2 | 0.05 | 0.05 | 0.05 |
| NC_006213.1;YP_009555238.1;HCoV-OC43_orf1ab | 31 | 31 | 1 | 0.05 | 0.05 | 0.05 |
| NC_006213.1;YP_009555238.1;HCoV-OC43_orf1ab | 43 | 44 | 2 | 0.05 | 0.05 | 0.05 |
| NC_006213.1;YP_009555238.1;HCoV-OC43_orf1ab | 46 | 48 | 3 | 0.05 | 0.2 | 0.116666667 |
| NC_006213.1;YP_009555238.1;HCoV-OC43_orf1ab | 73 | 88 | 16 | 0.05 | 0.1 | 0.059375 |
| NC_006213.1;YP_009555238.1;HCoV-OC43_orf1ab | 92 | 92 | 1 | 0.05 | 0.05 | 0.05 |
| NC_006213.1;YP_009555238.1;HCoV-OC43_orf1ab | 189 | 189 | 1 | 0.05 | 0.05 | 0.05 |
| NC_006213.1;YP_009555238.1;HCoV-OC43_orf1ab | 191 | 191 | 1 | 0.05 | 0.05 | 0.05 |
| NC_006213.1;YP_009555238.1;HCoV-OC43_orf1ab | 194 | 194 | 1 | 0.05 | 0.05 | 0.05 |
| NC_006213.1;YP_009555238.1;HCoV-OC43_orf1ab | 202 | 202 | 1 | 0.05 | 0.05 | 0.05 |
| NC_006213.1;YP_009555238.1;HCoV-OC43_orf1ab | 204 | 204 | 1 | 0.05 | 0.05 | 0.05 |
| NC_006213.1;YP_009555238.1;HCoV-OC43_orf1ab | 206 | 207 | 2 | 0.05 | 0.05 | 0.05 |
| NC_006213.1;YP_009555238.1;HCoV-OC43_orf1ab | 210 | 210 | 1 | 0.1 | 0.1 | 0.1 |
| NC_006213.1;YP_009555238.1;HCoV-OC43_orf1ab | 218 | 218 | 1 | 0.05 | 0.05 | 0.05 |
| NC_006213.1;YP_009555238.1;HCoV-OC43_orf1ab | 259 | 270 | 12 | 0.05 | 0.1 | 0.0625 |
| NC_006213.1;YP_009555238.1;HCoV-OC43_orf1ab | 291 | 302 | 12 | 0.1 | 0.15 | 0.133333333 |
| NC_006213.1;YP_009555238.1;HCoV-OC43_orf1ab | 317 | 317 | 1 | 0.05 | 0.05 | 0.05 |
| NC_006213.1;YP_009555238.1;HCoV-OC43_orf1ab | 324 | 324 | 1 | 0.05 | 0.05 | 0.05 |
| NC_006213.1;YP_009555238.1;HCoV-OC43_orf1ab | 355 | 355 | 1 | 0.05 | 0.05 | 0.05 |
| NC_006213.1;YP_009555238.1;HCoV-OC43_orf1ab | 378 | 378 | 1 | 0.05 | 0.05 | 0.05 |
| NC_006213.1;YP_009555238.1;HCoV-OC43_orf1ab | 398 | 400 | 3 | 0.05 | 0.15 | 0.1 |
| NC_006213.1;YP_009555238.1;HCoV-OC43_orf1ab | 409 | 419 | 11 | 0.05 | 0.05 | 0.05 |
| NC_006213.1;YP_009555238.1;HCoV-OC43_orf1ab | 421 | 421 | 1 | 0.05 | 0.05 | 0.05 |
| NC_006213.1;YP_009555238.1;HCoV-OC43_orf1ab | 427 | 437 | 11 | 0.05 | 0.15 | 0.068181818 |
| NC_006213.1;YP_009555238.1;HCoV-OC43_orf1ab | 497 | 497 | 1 | 0.05 | 0.05 | 0.05 |
| NC_006213.1;YP_009555238.1;HCoV-OC43_orf1ab | 518 | 523 | 6 | 0.05 | 0.05 | 0.05 |
| NC_006213.1;YP_009555238.1;HCoV-OC43_orf1ab | 534 | 534 | 1 | 0.05 | 0.05 | 0.05 |
| NC_006213.1;YP_009555238.1;HCoV-OC43_orf1ab | 546 | 548 | 3 | 0.05 | 0.1 | 0.083333333 |
| NC_006213.1;YP_009555238.1;HCoV-OC43_orf1ab | 553 | 555 | 3 | 0.05 | 0.05 | 0.05 |
| NC_006213.1;YP_009555238.1;HCoV-OC43_orf1ab | 592 | 594 | 3 | 0.05 | 0.1 | 0.066666667 |
| NC_006213.1;YP_009555238.1;HCoV-OC43_orf1ab | 596 | 596 | 1 | 0.05 | 0.05 | 0.05 |
| NC_006213.1;YP_009555238.1;HCoV-OC43_orf1ab | 600 | 604 | 5 | 0.05 | 0.05 | 0.05 |
| NC_006213.1;YP_009555238.1;HCoV-OC43_orf1ab | 607 | 608 | 2 | 0.05 | 0.05 | 0.05 |
| NC_006213.1;YP_009555238.1;HCoV-OC43_orf1ab | 615 | 615 | 1 | 0.1 | 0.1 | 0.1 |
| NC_006213.1;YP_009555238.1;HCoV-OC43_orf1ab | 620 | 620 | 1 | 0.05 | 0.05 | 0.05 |
| NC_006213.1;YP_009555238.1;HCoV-OC43_orf1ab | 628 | 629 | 2 | 0.05 | 0.05 | 0.05 |

| Extended data 1 | | | | | | |
|---|---|---|---|---|---|---|
| **Protein** | **Start position (peptide)** | **Stop position (peptide)** | **Number of peptides** | **Minimum fraction (per peptide) of control subjects with reactivity >3 sd above average** | **Maximum fraction (per peptide) of control subjects with reactivity >3 sd above average** | **Mean fraction (over all peptides in this contiguous stretch) of controlsubjects with reactivity >3 sd above average** |
| NC_006213.1;YP_009555238.1;HCoV-OC43_orf1ab | 689 | 689 | 1 | 0.05 | 0.05 | 0.05 |
| NC_006213.1;YP_009555238.1;HCoV-OC43_orf1ab | 693 | 694 | 2 | 0.05 | 0.05 | 0.05 |
| NC_006213.1;YP_009555238.1;HCoV-OC43_orf1ab | 696 | 696 | 1 | 0.05 | 0.05 | 0.05 |
| NC_006213.1;YP_009555238.1;HCoV-OC43_orf1ab | 700 | 700 | 1 | 0.05 | 0.05 | 0.05 |
| NC_006213.1;YP_009555238.1;HCoV-OC43_orf1ab | 743 | 747 | 5 | 0.05 | 0.1 | 0.08 |
| NC_006213.1;YP_009555238.1;HCoV-OC43_orf1ab | 749 | 755 | 7 | 0.05 | 0.1 | 0.057142857 |
| NC_006213.1;YP_009555238.1;HCoV-OC43_orf1ab | 772 | 780 | 9 | 0.05 | 0.1 | 0.055555556 |
| NC_006213.1;YP_009555238.1;HCoV-OC43_orf1ab | 797 | 801 | 5 | 0.05 | 0.05 | 0.05 |
| NC_006213.1;YP_009555238.1;HCoV-OC43_orf1ab | 815 | 816 | 2 | 0.05 | 0.05 | 0.05 |
| NC_006213.1;YP_009555238.1;HCoV-OC43_orf1ab | 818 | 818 | 1 | 0.15 | 0.15 | 0.15 |
| NC_006213.1;YP_009555238.1;HCoV-OC43_orf1ab | 821 | 822 | 2 | 0.05 | 0.05 | 0.05 |
| NC_006213.1;YP_009555238.1;HCoV-OC43_orf1ab | 841 | 841 | 1 | 0.05 | 0.05 | 0.05 |
| NC_006213.1;YP_009555238.1;HCoV-OC43_orf1ab | 848 | 848 | 1 | 0.1 | 0.1 | 0.1 |
| NC_006213.1;YP_009555238.1;HCoV-OC43_orf1ab | 877 | 878 | 2 | 0.05 | 0.05 | 0.05 |
| NC_006213.1;YP_009555238.1;HCoV-OC43_orf1ab | 911 | 911 | 1 | 0.05 | 0.05 | 0.05 |
| NC_006213.1;YP_009555238.1;HCoV-OC43_orf1ab | 926 | 929 | 4 | 0.05 | 0.05 | 0.05 |
| NC_006213.1;YP_009555238.1;HCoV-OC43_orf1ab | 935 | 937 | 3 | 0.05 | 0.05 | 0.05 |
| NC_006213.1;YP_009555238.1;HCoV-OC43_orf1ab | 939 | 942 | 4 | 0.05 | 0.1 | 0.075 |
| NC_006213.1;YP_009555238.1;HCoV-OC43_orf1ab | 944 | 944 | 1 | 0.05 | 0.05 | 0.05 |
| NC_006213.1;YP_009555238.1;HCoV-OC43_orf1ab | 965 | 965 | 1 | 0.1 | 0.1 | 0.1 |
| NC_006213.1;YP_009555238.1;HCoV-OC43_orf1ab | 977 | 978 | 2 | 0.05 | 0.15 | 0.1 |
| NC_006213.1;YP_009555238.1;HCoV-OC43_orf1ab | 1002 | 1009 | 8 | 0.05 | 0.1 | 0.05625 |
| NC_006213.1;YP_009555238.1;HCoV-OC43_orf1ab | 1027 | 1036 | 10 | 0.05 | 0.1 | 0.085 |
| NC_006213.1;YP_009555238.1;HCoV-OC43_orf1ab | 1084 | 1085 | 2 | 0.05 | 0.05 | 0.05 |
| NC_006213.1;YP_009555238.1;HCoV-OC43_orf1ab | 1093 | 1093 | 1 | 0.05 | 0.05 | 0.05 |
| NC_006213.1;YP_009555238.1;HCoV-OC43_orf1ab | 1108 | 1108 | 1 | 0.05 | 0.05 | 0.05 |
| NC_006213.1;YP_009555238.1;HCoV-OC43_orf1ab | 1111 | 1111 | 1 | 0.1 | 0.1 | 0.1 |
| NC_006213.1;YP_009555238.1;HCoV-OC43_orf1ab | 1152 | 1153 | 2 | 0.05 | 0.05 | 0.05 |
| NC_006213.1;YP_009555238.1;HCoV-OC43_orf1ab | 1187 | 1187 | 1 | 0.05 | 0.05 | 0.05 |
| NC_006213.1;YP_009555238.1;HCoV-OC43_orf1ab | 1214 | 1214 | 1 | 0.05 | 0.05 | 0.05 |
| NC_006213.1;YP_009555238.1;HCoV-OC43_orf1ab | 1225 | 1226 | 2 | 0.05 | 0.05 | 0.05 |
| NC_006213.1;YP_009555238.1;HCoV-OC43_orf1ab | 1232 | 1236 | 5 | 0.05 | 0.05 | 0.05 |
| NC_006213.1;YP_009555238.1;HCoV-OC43_orf1ab | 1240 | 1246 | 7 | 0.05 | 0.15 | 0.078571429 |
| NC_006213.1;YP_009555238.1;HCoV-OC43_orf1ab | 1283 | 1283 | 1 | 0.05 | 0.05 | 0.05 |
| NC_006213.1;YP_009555238.1;HCoV-OC43_orf1ab | 1290 | 1290 | 1 | 0.05 | 0.05 | 0.05 |
| NC_006213.1;YP_009555238.1;HCoV-OC43_orf1ab | 1293 | 1294 | 2 | 0.15 | 0.15 | 0.15 |
| NC_006213.1;YP_009555238.1;HCoV-OC43_orf1ab | 1296 | 1304 | 9 | 0.05 | 0.1 | 0.055555556 |

| Extended data 1 | | | | | | |
|---|---|---|---|---|---|---|
| **Protein** | **Start position (peptide)** | **Stop position (peptide)** | **Number of peptides** | **Minimum fraction (per peptide) of control subjects with reactivity >3 sd above average** | **Maximum fraction (per peptide) of control subjects with reactivity >3 sd above average** | **Mean fraction (over all peptides in this contiguous stretch) of controlsubjects with reactivity >3 sd above average** |
| NC_006213.1;YP_009555238.1;HCoV-OC43_orf1ab | 1332 | 1332 | 1 | 0.05 | 0.05 | 0.05 |
| NC_006213.1;YP_009555238.1;HCoV-OC43_orf1ab | 1361 | 1361 | 1 | 0.05 | 0.05 | 0.05 |
| NC_006213.1;YP_009555238.1;HCoV-OC43_orf1ab | 1372 | 1374 | 3 | 0.05 | 0.1 | 0.066666667 |
| NC_006213.1;YP_009555238.1;HCoV-OC43_orf1ab | 1397 | 1397 | 1 | 0.05 | 0.05 | 0.05 |
| NC_006213.1;YP_009555238.1;HCoV-OC43_orf1ab | 1409 | 1411 | 3 | 0.05 | 0.05 | 0.05 |
| NC_006213.1;YP_009555238.1;HCoV-OC43_orf1ab | 1415 | 1417 | 3 | 0.05 | 0.15 | 0.1 |
| NC_006213.1;YP_009555238.1;HCoV-OC43_orf1ab | 1422 | 1425 | 4 | 0.05 | 0.15 | 0.1125 |
| NC_006213.1;YP_009555238.1;HCoV-OC43_orf1ab | 1428 | 1428 | 1 | 0.05 | 0.05 | 0.05 |
| NC_006213.1;YP_009555238.1;HCoV-OC43_orf1ab | 1446 | 1446 | 1 | 0.05 | 0.05 | 0.05 |
| NC_006213.1;YP_009555238.1;HCoV-OC43_orf1ab | 1449 | 1449 | 1 | 0.05 | 0.05 | 0.05 |
| NC_006213.1;YP_009555238.1;HCoV-OC43_orf1ab | 1454 | 1454 | 1 | 0.05 | 0.05 | 0.05 |
| NC_006213.1;YP_009555238.1;HCoV-OC43_orf1ab | 1476 | 1476 | 1 | 0.15 | 0.15 | 0.15 |
| NC_006213.1;YP_009555238.1;HCoV-OC43_orf1ab | 1482 | 1482 | 1 | 0.05 | 0.05 | 0.05 |
| NC_006213.1;YP_009555238.1;HCoV-OC43_orf1ab | 1496 | 1496 | 1 | 0.05 | 0.05 | 0.05 |
| NC_006213.1;YP_009555238.1;HCoV-OC43_orf1ab | 1498 | 1499 | 2 | 0.05 | 0.1 | 0.075 |
| NC_006213.1;YP_009555238.1;HCoV-OC43_orf1ab | 1504 | 1505 | 2 | 0.05 | 0.05 | 0.05 |
| NC_006213.1;YP_009555238.1;HCoV-OC43_orf1ab | 1531 | 1531 | 1 | 0.05 | 0.05 | 0.05 |
| NC_006213.1;YP_009555238.1;HCoV-OC43_orf1ab | 1536 | 1545 | 10 | 0.05 | 0.15 | 0.115 |
| NC_006213.1;YP_009555238.1;HCoV-OC43_orf1ab | 1551 | 1553 | 3 | 0.05 | 0.05 | 0.05 |
| NC_006213.1;YP_009555238.1;HCoV-OC43_orf1ab | 1555 | 1555 | 1 | 0.05 | 0.05 | 0.05 |
| NC_006213.1;YP_009555238.1;HCoV-OC43_orf1ab | 1558 | 1558 | 1 | 0.05 | 0.05 | 0.05 |
| NC_006213.1;YP_009555238.1;HCoV-OC43_orf1ab | 1563 | 1563 | 1 | 0.05 | 0.05 | 0.05 |
| NC_006213.1;YP_009555238.1;HCoV-OC43_orf1ab | 1568 | 1574 | 7 | 0.05 | 0.2 | 0.114285714 |
| NC_006213.1;YP_009555238.1;HCoV-OC43_orf1ab | 1600 | 1602 | 3 | 0.05 | 0.05 | 0.05 |
| NC_006213.1;YP_009555238.1;HCoV-OC43_orf1ab | 1611 | 1612 | 2 | 0.05 | 0.05 | 0.05 |
| NC_006213.1;YP_009555238.1;HCoV-OC43_orf1ab | 1616 | 1616 | 1 | 0.05 | 0.05 | 0.05 |
| NC_006213.1;YP_009555238.1;HCoV-OC43_orf1ab | 1622 | 1624 | 3 | 0.05 | 0.1 | 0.066666667 |
| NC_006213.1;YP_009555238.1;HCoV-OC43_orf1ab | 1628 | 1628 | 1 | 0.1 | 0.1 | 0.1 |
| NC_006213.1;YP_009555238.1;HCoV-OC43_orf1ab | 1630 | 1632 | 3 | 0.05 | 0.1 | 0.066666667 |
| NC_006213.1;YP_009555238.1;HCoV-OC43_orf1ab | 1634 | 1634 | 1 | 0.05 | 0.05 | 0.05 |
| NC_006213.1;YP_009555238.1;HCoV-OC43_orf1ab | 1637 | 1637 | 1 | 0.05 | 0.05 | 0.05 |
| NC_006213.1;YP_009555238.1;HCoV-OC43_orf1ab | 1645 | 1645 | 1 | 0.05 | 0.05 | 0.05 |
| NC_006213.1;YP_009555238.1;HCoV-OC43_orf1ab | 1655 | 1666 | 12 | 0.05 | 0.1 | 0.0625 |
| NC_006213.1;YP_009555238.1;HCoV-OC43_orf1ab | 1706 | 1716 | 11 | 0.05 | 0.25 | 0.122727273 |
| NC_006213.1;YP_009555238.1;HCoV-OC43_orf1ab | 1720 | 1720 | 1 | 0.05 | 0.05 | 0.05 |
| NC_006213.1;YP_009555238.1;HCoV-OC43_orf1ab | 1722 | 1722 | 1 | 0.05 | 0.05 | 0.05 |
| NC_006213.1;YP_009555238.1;HCoV-OC43_orf1ab | 1756 | 1756 | 1 | 0.05 | 0.05 | 0.05 |

| Extended data 1 | | | | | | |
|---|---|---|---|---|---|---|
| Protein | Start position (peptide) | Stop position (peptide) | Number of peptides | Minimum fraction (per peptide) of control subjects with reactivity >3 sd above average | Maximum fraction (per peptide) of control subjects with reactivity >3 sd above average | Mean fraction (over all peptides in this contiguous stretch) of controlsubjects with reactivity >3 sd above average |
| NC_006213.1;YP_009555238.1;HCoV-OC43_orf1ab | 1760 | 1760 | 1 | 0.05 | 0.05 | 0.05 |
| NC_006213.1;YP_009555238.1;HCoV-OC43_orf1ab | 1775 | 1776 | 2 | 0.05 | 0.05 | 0.05 |
| NC_006213.1;YP_009555238.1;HCoV-OC43_orf1ab | 1803 | 1803 | 1 | 0.05 | 0.05 | 0.05 |
| NC_006213.1;YP_009555238.1;HCoV-OC43_orf1ab | 1811 | 1812 | 2 | 0.05 | 0.05 | 0.05 |
| NC_006213.1;YP_009555238.1;HCoV-OC43_orf1ab | 1832 | 1841 | 10 | 0.05 | 0.3 | 0.09 |
| NC_006213.1;YP_009555238.1;HCoV-OC43_orf1ab | 1850 | 1850 | 1 | 0.05 | 0.05 | 0.05 |
| NC_006213.1;YP_009555238.1;HCoV-OC43_orf1ab | 1860 | 1860 | 1 | 0.05 | 0.05 | 0.05 |
| NC_006213.1;YP_009555238.1;HCoV-OC43_orf1ab | 1870 | 1871 | 2 | 0.05 | 0.1 | 0.075 |
| NC_006213.1;YP_009555238.1;HCoV-OC43_orf1ab | 1873 | 1877 | 5 | 0.05 | 0.1 | 0.07 |
| NC_006213.1;YP_009555238.1;HCoV-OC43_orf1ab | 1879 | 1879 | 1 | 0.1 | 0.1 | 0.1 |
| NC_006213.1;YP_009555238.1;HCoV-OC43_orf1ab | 1887 | 1887 | 1 | 0.1 | 0.1 | 0.1 |
| NC_006213.1;YP_009555238.1;HCoV-OC43_orf1ab | 1904 | 1904 | 1 | 0.05 | 0.05 | 0.05 |
| NC_006213.1;YP_009555238.1;HCoV-OC43_orf1ab | 1909 | 1910 | 2 | 0.05 | 0.05 | 0.05 |
| NC_006213.1;YP_009555238.1;HCoV-OC43_orf1ab | 1912 | 1913 | 2 | 0.05 | 0.05 | 0.05 |
| NC_006213.1;YP_009555238.1;HCoV-OC43_orf1ab | 1921 | 1921 | 1 | 0.1 | 0.1 | 0.1 |
| NC_006213.1;YP_009555238.1;HCoV-OC43_orf1ab | 1926 | 1933 | 8 | 0.05 | 0.1 | 0.06875 |
| NC_006213.1;YP_009555238.1;HCoV-OC43_orf1ab | 1935 | 1937 | 3 | 0.05 | 0.05 | 0.05 |
| NC_006213.1;YP_009555238.1;HCoV-OC43_orf1ab | 1941 | 1941 | 1 | 0.05 | 0.05 | 0.05 |
| NC_006213.1;YP_009555238.1;HCoV-OC43_orf1ab | 1943 | 1945 | 3 | 0.05 | 0.1 | 0.066666667 |
| NC_006213.1;YP_009555238.1;HCoV-OC43_orf1ab | 1957 | 1957 | 1 | 0.05 | 0.05 | 0.05 |
| NC_006213.1;YP_009555238.1;HCoV-OC43_orf1ab | 1993 | 1993 | 1 | 0.05 | 0.05 | 0.05 |
| NC_006213.1;YP_009555238.1;HCoV-OC43_orf1ab | 1995 | 2004 | 10 | 0.05 | 0.1 | 0.07 |
| NC_006213.1;YP_009555238.1;HCoV-OC43_orf1ab | 2008 | 2008 | 1 | 0.05 | 0.05 | 0.05 |
| NC_006213.1;YP_009555238.1;HCoV-OC43_orf1ab | 2017 | 2017 | 1 | 0.05 | 0.05 | 0.05 |
| NC_006213.1;YP_009555238.1;HCoV-OC43_orf1ab | 2040 | 2040 | 1 | 0.05 | 0.05 | 0.05 |
| NC_006213.1;YP_009555238.1;HCoV-OC43_orf1ab | 2042 | 2042 | 1 | 0.05 | 0.05 | 0.05 |
| NC_006213.1;YP_009555238.1;HCoV-OC43_orf1ab | 2045 | 2045 | 1 | 0.05 | 0.05 | 0.05 |
| NC_006213.1;YP_009555238.1;HCoV-OC43_orf1ab | 2055 | 2055 | 1 | 0.05 | 0.05 | 0.05 |
| NC_006213.1;YP_009555238.1;HCoV-OC43_orf1ab | 2111 | 2114 | 4 | 0.05 | 0.15 | 0.075 |
| NC_006213.1;YP_009555238.1;HCoV-OC43_orf1ab | 2157 | 2157 | 1 | 0.05 | 0.05 | 0.05 |
| NC_006213.1;YP_009555238.1;HCoV-OC43_orf1ab | 2214 | 2223 | 10 | 0.05 | 0.05 | 0.05 |
| NC_006213.1;YP_009555238.1;HCoV-OC43_orf1ab | 2253 | 2258 | 6 | 0.05 | 0.15 | 0.083333333 |
| NC_006213.1;YP_009555238.1;HCoV-OC43_orf1ab | 2271 | 2272 | 2 | 0.05 | 0.05 | 0.05 |
| NC_006213.1;YP_009555238.1;HCoV-OC43_orf1ab | 2274 | 2277 | 4 | 0.05 | 0.05 | 0.05 |
| NC_006213.1;YP_009555238.1;HCoV-OC43_orf1ab | 2287 | 2290 | 4 | 0.05 | 0.15 | 0.075 |
| NC_006213.1;YP_009555238.1;HCoV-OC43_orf1ab | 2356 | 2356 | 1 | 0.2 | 0.2 | 0.2 |
| NC_006213.1;YP_009555238.1;HCoV-OC43_orf1ab | 2414 | 2414 | 1 | 0.05 | 0.05 | 0.05 |

| Extended data 1 | | | | | | |
|---|---|---|---|---|---|---|
| Protein | Start position (peptide) | Stop position (peptide) | Number of peptides | Minimum fraction (per peptide) of control subjects with reactivity >3 sd above average | Maximum fraction (per peptide) of control subjects with reactivity >3 sd above average | Mean fraction (over all peptides in this contiguous stretch) of controlsubjects with reactivity >3 sd above average |
| NC_006213.1;YP_009555238.1;HCoV-OC43_orf1ab | 2430 | 2430 | 1 | 0.05 | 0.05 | 0.05 |
| NC_006213.1;YP_009555238.1;HCoV-OC43_orf1ab | 2437 | 2437 | 1 | 0.05 | 0.05 | 0.05 |
| NC_006213.1;YP_009555238.1;HCoV-OC43_orf1ab | 2451 | 2455 | 5 | 0.05 | 0.05 | 0.05 |
| NC_006213.1;YP_009555238.1;HCoV-OC43_orf1ab | 2472 | 2474 | 3 | 0.05 | 0.1 | 0.066666667 |
| NC_006213.1;YP_009555238.1;HCoV-OC43_orf1ab | 2479 | 2479 | 1 | 0.05 | 0.05 | 0.05 |
| NC_006213.1;YP_009555238.1;HCoV-OC43_orf1ab | 2490 | 2495 | 6 | 0.05 | 0.15 | 0.1 |
| NC_006213.1;YP_009555238.1;HCoV-OC43_orf1ab | 2509 | 2510 | 2 | 0.15 | 0.15 | 0.15 |
| NC_006213.1;YP_009555238.1;HCoV-OC43_orf1ab | 2512 | 2512 | 1 | 0.05 | 0.05 | 0.05 |
| NC_006213.1;YP_009555238.1;HCoV-OC43_orf1ab | 2517 | 2517 | 1 | 0.05 | 0.05 | 0.05 |
| NC_006213.1;YP_009555238.1;HCoV-OC43_orf1ab | 2522 | 2522 | 1 | 0.05 | 0.05 | 0.05 |
| NC_006213.1;YP_009555238.1;HCoV-OC43_orf1ab | 2524 | 2531 | 8 | 0.05 | 0.1 | 0.06875 |
| NC_006213.1;YP_009555238.1;HCoV-OC43_orf1ab | 2534 | 2534 | 1 | 0.05 | 0.05 | 0.05 |
| NC_006213.1;YP_009555238.1;HCoV-OC43_orf1ab | 2555 | 2559 | 5 | 0.05 | 0.05 | 0.05 |
| NC_006213.1;YP_009555238.1;HCoV-OC43_orf1ab | 2581 | 2588 | 8 | 0.1 | 0.15 | 0.1125 |
| NC_006213.1;YP_009555238.1;HCoV-OC43_orf1ab | 2602 | 2603 | 2 | 0.05 | 0.05 | 0.05 |
| NC_006213.1;YP_009555238.1;HCoV-OC43_orf1ab | 2641 | 2641 | 1 | 0.05 | 0.05 | 0.05 |
| NC_006213.1;YP_009555238.1;HCoV-OC43_orf1ab | 2652 | 2655 | 4 | 0.05 | 0.1 | 0.0625 |
| NC_006213.1;YP_009555238.1;HCoV-OC43_orf1ab | 2657 | 2658 | 2 | 0.05 | 0.05 | 0.05 |
| NC_006213.1;YP_009555238.1;HCoV-OC43_orf1ab | 2662 | 2662 | 1 | 0.05 | 0.05 | 0.05 |
| NC_006213.1;YP_009555238.1;HCoV-OC43_orf1ab | 2673 | 2673 | 1 | 0.05 | 0.05 | 0.05 |
| NC_006213.1;YP_009555238.1;HCoV-OC43_orf1ab | 2678 | 2678 | 1 | 0.05 | 0.05 | 0.05 |
| NC_006213.1;YP_009555238.1;HCoV-OC43_orf1ab | 2695 | 2695 | 1 | 0.05 | 0.05 | 0.05 |
| NC_006213.1;YP_009555238.1;HCoV-OC43_orf1ab | 2699 | 2704 | 6 | 0.05 | 0.2 | 0.091666667 |
| NC_006213.1;YP_009555238.1;HCoV-OC43_orf1ab | 2721 | 2722 | 2 | 0.05 | 0.05 | 0.05 |
| NC_006213.1;YP_009555238.1;HCoV-OC43_orf1ab | 2772 | 2782 | 11 | 0.05 | 0.1 | 0.077272727 |
| NC_006213.1;YP_009555238.1;HCoV-OC43_orf1ab | 2788 | 2788 | 1 | 0.05 | 0.05 | 0.05 |
| NC_006213.1;YP_009555238.1;HCoV-OC43_orf1ab | 2799 | 2799 | 1 | 0.05 | 0.05 | 0.05 |
| NC_006213.1;YP_009555238.1;HCoV-OC43_orf1ab | 2804 | 2805 | 2 | 0.05 | 0.1 | 0.075 |
| NC_006213.1;YP_009555238.1;HCoV-OC43_orf1ab | 2812 | 2812 | 1 | 0.05 | 0.05 | 0.05 |
| NC_006213.1;YP_009555238.1;HCoV-OC43_orf1ab | 2843 | 2843 | 1 | 0.05 | 0.05 | 0.05 |
| NC_006213.1;YP_009555238.1;HCoV-OC43_orf1ab | 2845 | 2845 | 1 | 0.05 | 0.05 | 0.05 |
| NC_006213.1;YP_009555238.1;HCoV-OC43_orf1ab | 2900 | 2900 | 1 | 0.05 | 0.05 | 0.05 |
| NC_006213.1;YP_009555238.1;HCoV-OC43_orf1ab | 2920 | 2920 | 1 | 0.05 | 0.05 | 0.05 |
| NC_006213.1;YP_009555238.1;HCoV-OC43_orf1ab | 2973 | 2973 | 1 | 0.05 | 0.05 | 0.05 |
| NC_006213.1;YP_009555238.1;HCoV-OC43_orf1ab | 3145 | 3145 | 1 | 0.05 | 0.05 | 0.05 |
| NC_006213.1;YP_009555238.1;HCoV-OC43_orf1ab | 3172 | 3172 | 1 | 0.05 | 0.05 | 0.05 |
| NC_006213.1;YP_009555238.1;HCoV-OC43_orf1ab | 3192 | 3192 | 1 | 0.05 | 0.05 | 0.05 |

| Extended data 1 | | | | | | |
|---|---|---|---|---|---|---|
| Protein | Start position (peptide) | Stop position (peptide) | Number of peptides | Minimum fraction (per peptide) of control subjects with reactivity >3 sd above average | Maximum fraction (per peptide) of control subjects with reactivity >3 sd above average | Mean fraction (over all peptides in this contiguous stretch) of controlsubjects with reactivity >3 sd above average |
| NC_006213.1;YP_009555238.1;HCoV-OC43_orf1ab | 3196 | 3197 | 2 | 0.05 | 0.1 | 0.075 |
| NC_006213.1;YP_009555238.1;HCoV-OC43_orf1ab | 3202 | 3202 | 1 | 0.05 | 0.05 | 0.05 |
| NC_006213.1;YP_009555238.1;HCoV-OC43_orf1ab | 3209 | 3209 | 1 | 0.05 | 0.05 | 0.05 |
| NC_006213.1;YP_009555238.1;HCoV-OC43_orf1ab | 3211 | 3213 | 3 | 0.05 | 0.1 | 0.066666667 |
| NC_006213.1;YP_009555238.1;HCoV-OC43_orf1ab | 3220 | 3222 | 3 | 0.05 | 0.25 | 0.133333333 |
| NC_006213.1;YP_009555238.1;HCoV-OC43_orf1ab | 3224 | 3224 | 1 | 0.05 | 0.05 | 0.05 |
| NC_006213.1;YP_009555238.1;HCoV-OC43_orf1ab | 3226 | 3226 | 1 | 0.05 | 0.05 | 0.05 |
| NC_006213.1;YP_009555238.1;HCoV-OC43_orf1ab | 3228 | 3229 | 2 | 0.05 | 0.05 | 0.05 |
| NC_006213.1;YP_009555238.1;HCoV-OC43_orf1ab | 3233 | 3233 | 1 | 0.1 | 0.1 | 0.1 |
| NC_006213.1;YP_009555238.1;HCoV-OC43_orf1ab | 3248 | 3248 | 1 | 0.05 | 0.05 | 0.05 |
| NC_006213.1;YP_009555238.1;HCoV-OC43_orf1ab | 3259 | 3270 | 12 | 0.05 | 0.1 | 0.066666667 |
| NC_006213.1;YP_009555238.1;HCoV-OC43_orf1ab | 3288 | 3289 | 2 | 0.05 | 0.05 | 0.05 |
| NC_006213.1;YP_009555238.1;HCoV-OC43_orf1ab | 3292 | 3292 | 1 | 0.05 | 0.05 | 0.05 |
| NC_006213.1;YP_009555238.1;HCoV-OC43_orf1ab | 3297 | 3299 | 3 | 0.05 | 0.05 | 0.05 |
| NC_006213.1;YP_009555238.1;HCoV-OC43_orf1ab | 3348 | 3348 | 1 | 0.05 | 0.05 | 0.05 |
| NC_006213.1;YP_009555238.1;HCoV-OC43_orf1ab | 3362 | 3363 | 2 | 0.05 | 0.05 | 0.05 |
| NC_006213.1;YP_009555238.1;HCoV-OC43_orf1ab | 3391 | 3393 | 3 | 0.05 | 0.1 | 0.066666667 |
| NC_006213.1;YP_009555238.1;HCoV-OC43_orf1ab | 3395 | 3395 | 1 | 0.1 | 0.1 | 0.1 |
| NC_006213.1;YP_009555238.1;HCoV-OC43_orf1ab | 3397 | 3400 | 4 | 0.05 | 0.1 | 0.075 |
| NC_006213.1;YP_009555238.1;HCoV-OC43_orf1ab | 3417 | 3417 | 1 | 0.05 | 0.05 | 0.05 |
| NC_006213.1;YP_009555238.1;HCoV-OC43_orf1ab | 3419 | 3422 | 4 | 0.05 | 0.1 | 0.0625 |
| NC_006213.1;YP_009555238.1;HCoV-OC43_orf1ab | 3456 | 3456 | 1 | 0.05 | 0.05 | 0.05 |
| NC_006213.1;YP_009555238.1;HCoV-OC43_orf1ab | 3467 | 3467 | 1 | 0.05 | 0.05 | 0.05 |
| NC_006213.1;YP_009555238.1;HCoV-OC43_orf1ab | 3472 | 3472 | 1 | 0.1 | 0.1 | 0.1 |
| NC_006213.1;YP_009555238.1;HCoV-OC43_orf1ab | 3480 | 3480 | 1 | 0.05 | 0.05 | 0.05 |
| NC_006213.1;YP_009555238.1;HCoV-OC43_orf1ab | 3484 | 3485 | 2 | 0.05 | 0.05 | 0.05 |
| NC_006213.1;YP_009555238.1;HCoV-OC43_orf1ab | 3490 | 3490 | 1 | 0.05 | 0.05 | 0.05 |
| NC_006213.1;YP_009555238.1;HCoV-OC43_orf1ab | 3496 | 3496 | 1 | 0.05 | 0.05 | 0.05 |
| NC_006213.1;YP_009555238.1;HCoV-OC43_orf1ab | 3512 | 3512 | 1 | 0.05 | 0.05 | 0.05 |
| NC_006213.1;YP_009555238.1;HCoV-OC43_orf1ab | 3518 | 3518 | 1 | 0.05 | 0.05 | 0.05 |
| NC_006213.1;YP_009555238.1;HCoV-OC43_orf1ab | 3526 | 3527 | 2 | 0.05 | 0.05 | 0.05 |
| NC_006213.1;YP_009555238.1;HCoV-OC43_orf1ab | 3529 | 3529 | 1 | 0.05 | 0.05 | 0.05 |
| NC_006213.1;YP_009555238.1;HCoV-OC43_orf1ab | 3531 | 3531 | 1 | 0.1 | 0.1 | 0.1 |
| NC_006213.1;YP_009555238.1;HCoV-OC43_orf1ab | 3647 | 3647 | 1 | 0.05 | 0.05 | 0.05 |
| NC_006213.1;YP_009555238.1;HCoV-OC43_orf1ab | 3702 | 3702 | 1 | 0.05 | 0.05 | 0.05 |
| NC_006213.1;YP_009555238.1;HCoV-OC43_orf1ab | 3704 | 3704 | 1 | 0.05 | 0.05 | 0.05 |
| NC_006213.1;YP_009555238.1;HCoV-OC43_orf1ab | 3792 | 3792 | 1 | 0.05 | 0.05 | 0.05 |

| Extended data 1 | | | | | | |
|---|---|---|---|---|---|---|
| Protein | Start position (peptide) | Stop position (peptide) | Number of peptides | Minimum fraction (per peptide) of control subjects with reactivity >3 sd above average | Maximum fraction (per peptide) of control subjects with reactivity >3 sd above average | Mean fraction (over all peptides in this contiguous stretch) of controlsubjects with reactivity >3 sd above average |
| NC_006213.1;YP_009555238.1;HCoV-OC43_orf1ab | 3796 | 3796 | 1 | 0.1 | 0.1 | 0.1 |
| NC_006213.1;YP_009555238.1;HCoV-OC43_orf1ab | 3802 | 3802 | 1 | 0.05 | 0.05 | 0.05 |
| NC_006213.1;YP_009555238.1;HCoV-OC43_orf1ab | 3826 | 3831 | 6 | 0.05 | 0.1 | 0.066666667 |
| NC_006213.1;YP_009555238.1;HCoV-OC43_orf1ab | 3838 | 3839 | 2 | 0.05 | 0.15 | 0.1 |
| NC_006213.1;YP_009555238.1;HCoV-OC43_orf1ab | 3850 | 3855 | 6 | 0.05 | 0.05 | 0.05 |
| NC_006213.1;YP_009555238.1;HCoV-OC43_orf1ab | 3858 | 3859 | 2 | 0.05 | 0.15 | 0.1 |
| NC_006213.1;YP_009555238.1;HCoV-OC43_orf1ab | 3869 | 3869 | 1 | 0.05 | 0.05 | 0.05 |
| NC_006213.1;YP_009555238.1;HCoV-OC43_orf1ab | 3909 | 3910 | 2 | 0.05 | 0.05 | 0.05 |
| NC_006213.1;YP_009555238.1;HCoV-OC43_orf1ab | 3930 | 3931 | 2 | 0.05 | 0.05 | 0.05 |
| NC_006213.1;YP_009555238.1;HCoV-OC43_orf1ab | 3933 | 3941 | 9 | 0.05 | 0.1 | 0.061111111 |
| NC_006213.1;YP_009555238.1;HCoV-OC43_orf1ab | 3946 | 3946 | 1 | 0.05 | 0.05 | 0.05 |
| NC_006213.1;YP_009555238.1;HCoV-OC43_orf1ab | 3958 | 3958 | 1 | 0.05 | 0.05 | 0.05 |
| NC_006213.1;YP_009555238.1;HCoV-OC43_orf1ab | 3965 | 3965 | 1 | 0.05 | 0.05 | 0.05 |
| NC_006213.1;YP_009555238.1;HCoV-OC43_orf1ab | 3967 | 3968 | 2 | 0.05 | 0.15 | 0.1 |
| NC_006213.1;YP_009555238.1;HCoV-OC43_orf1ab | 3977 | 3977 | 1 | 0.05 | 0.05 | 0.05 |
| NC_006213.1;YP_009555238.1;HCoV-OC43_orf1ab | 3986 | 3986 | 1 | 0.05 | 0.05 | 0.05 |
| NC_006213.1;YP_009555238.1;HCoV-OC43_orf1ab | 3990 | 3990 | 1 | 0.05 | 0.05 | 0.05 |
| NC_006213.1;YP_009555238.1;HCoV-OC43_orf1ab | 3999 | 3999 | 1 | 0.05 | 0.05 | 0.05 |
| NC_006213.1;YP_009555238.1;HCoV-OC43_orf1ab | 4015 | 4031 | 17 | 0.05 | 0.35 | 0.161764706 |
| NC_006213.1;YP_009555238.1;HCoV-OC43_orf1ab | 4034 | 4034 | 1 | 0.15 | 0.15 | 0.15 |
| NC_006213.1;YP_009555238.1;HCoV-OC43_orf1ab | 4036 | 4036 | 1 | 0.05 | 0.05 | 0.05 |
| NC_006213.1;YP_009555238.1;HCoV-OC43_orf1ab | 4038 | 4038 | 1 | 0.1 | 0.1 | 0.1 |
| NC_006213.1;YP_009555238.1;HCoV-OC43_orf1ab | 4040 | 4042 | 3 | 0.05 | 0.1 | 0.066666667 |
| NC_006213.1;YP_009555238.1;HCoV-OC43_orf1ab | 4044 | 4059 | 16 | 0.05 | 0.25 | 0.096875 |
| NC_006213.1;YP_009555238.1;HCoV-OC43_orf1ab | 4074 | 4080 | 7 | 0.05 | 0.15 | 0.121428571 |
| NC_006213.1;YP_009555238.1;HCoV-OC43_orf1ab | 4083 | 4083 | 1 | 0.05 | 0.05 | 0.05 |
| NC_006213.1;YP_009555238.1;HCoV-OC43_orf1ab | 4085 | 4093 | 9 | 0.05 | 0.1 | 0.072222222 |
| NC_006213.1;YP_009555238.1;HCoV-OC43_orf1ab | 4109 | 4110 | 2 | 0.1 | 0.1 | 0.1 |
| NC_006213.1;YP_009555238.1;HCoV-OC43_orf1ab | 4118 | 4120 | 3 | 0.05 | 0.1 | 0.083333333 |
| NC_006213.1;YP_009555238.1;HCoV-OC43_orf1ab | 4133 | 4133 | 1 | 0.05 | 0.05 | 0.05 |
| NC_006213.1;YP_009555238.1;HCoV-OC43_orf1ab | 4141 | 4152 | 12 | 0.05 | 0.4 | 0.145833333 |
| NC_006213.1;YP_009555238.1;HCoV-OC43_orf1ab | 4159 | 4159 | 1 | 0.05 | 0.05 | 0.05 |
| NC_006213.1;YP_009555238.1;HCoV-OC43_orf1ab | 4167 | 4178 | 12 | 0.05 | 0.3 | 0.179166667 |
| NC_006213.1;YP_009555238.1;HCoV-OC43_orf1ab | 4185 | 4190 | 6 | 0.05 | 0.05 | 0.05 |
| NC_006213.1;YP_009555238.1;HCoV-OC43_orf1ab | 4193 | 4193 | 1 | 0.05 | 0.05 | 0.05 |
| NC_006213.1;YP_009555238.1;HCoV-OC43_orf1ab | 4223 | 4229 | 7 | 0.05 | 0.1 | 0.064285714 |
| NC_006213.1;YP_009555238.1;HCoV-OC43_orf1ab | 4231 | 4231 | 1 | 0.1 | 0.1 | 0.1 |

| Extended data 1 | | | | | | |
|---|---|---|---|---|---|---|
| Protein | Start position (peptide) | Stop position (peptide) | Number of peptides | Minimum fraction (per peptide) of control subjects with reactivity >3 sd above average | Maximum fraction (per peptide) of control subjects with reactivity >3 sd above average | Mean fraction (over all peptides in this contiguous stretch) of controlsubjects with reactivity >3 sd above average |
| NC_006213.1;YP_009555238.1;HCoV-OC43_orf1ab | 4240 | 4240 | 1 | 0.05 | 0.05 | 0.05 |
| NC_006213.1;YP_009555238.1;HCoV-OC43_orf1ab | 4250 | 4253 | 4 | 0.05 | 0.1 | 0.075 |
| NC_006213.1;YP_009555238.1;HCoV-OC43_orf1ab | 4261 | 4261 | 1 | 0.05 | 0.05 | 0.05 |
| NC_006213.1;YP_009555238.1;HCoV-OC43_orf1ab | 4285 | 4285 | 1 | 0.05 | 0.05 | 0.05 |
| NC_006213.1;YP_009555238.1;HCoV-OC43_orf1ab | 4287 | 4287 | 1 | 0.05 | 0.05 | 0.05 |
| NC_006213.1;YP_009555238.1;HCoV-OC43_orf1ab | 4326 | 4327 | 2 | 0.05 | 0.05 | 0.05 |
| NC_006213.1;YP_009555238.1;HCoV-OC43_orf1ab | 4356 | 4370 | 15 | 0.05 | 0.2 | 0.093333333 |
| NC_006213.1;YP_009555238.1;HCoV-OC43_orf1ab | 4388 | 4388 | 1 | 0.2 | 0.2 | 0.2 |
| NC_006213.1;YP_009555238.1;HCoV-OC43_orf1ab | 4396 | 4397 | 2 | 0.05 | 0.05 | 0.05 |
| NC_006213.1;YP_009555238.1;HCoV-OC43_orf1ab | 4420 | 4425 | 6 | 0.05 | 0.15 | 0.066666667 |
| NC_006213.1;YP_009555238.1;HCoV-OC43_orf1ab | 4442 | 4442 | 1 | 0.1 | 0.1 | 0.1 |
| NC_006213.1;YP_009555238.1;HCoV-OC43_orf1ab | 4457 | 4457 | 1 | 0.05 | 0.05 | 0.05 |
| NC_006213.1;YP_009555238.1;HCoV-OC43_orf1ab | 4471 | 4471 | 1 | 0.05 | 0.05 | 0.05 |
| NC_006213.1;YP_009555238.1;HCoV-OC43_orf1ab | 4522 | 4523 | 2 | 0.05 | 0.2 | 0.125 |
| NC_006213.1;YP_009555238.1;HCoV-OC43_orf1ab | 4526 | 4528 | 3 | 0.05 | 0.1 | 0.066666667 |
| NC_006213.1;YP_009555238.1;HCoV-OC43_orf1ab | 4530 | 4531 | 2 | 0.05 | 0.05 | 0.05 |
| NC_006213.1;YP_009555238.1;HCoV-OC43_orf1ab | 4533 | 4533 | 1 | 0.05 | 0.05 | 0.05 |
| NC_006213.1;YP_009555238.1;HCoV-OC43_orf1ab | 4568 | 4568 | 1 | 0.05 | 0.05 | 0.05 |
| NC_006213.1;YP_009555238.1;HCoV-OC43_orf1ab | 4572 | 4572 | 1 | 0.15 | 0.15 | 0.15 |
| NC_006213.1;YP_009555238.1;HCoV-OC43_orf1ab | 4574 | 4574 | 1 | 0.05 | 0.05 | 0.05 |
| NC_006213.1;YP_009555238.1;HCoV-OC43_orf1ab | 4601 | 4601 | 1 | 0.05 | 0.05 | 0.05 |
| NC_006213.1;YP_009555238.1;HCoV-OC43_orf1ab | 4629 | 4632 | 4 | 0.05 | 0.05 | 0.05 |
| NC_006213.1;YP_009555238.1;HCoV-OC43_orf1ab | 4634 | 4637 | 4 | 0.05 | 0.1 | 0.075 |
| NC_006213.1;YP_009555238.1;HCoV-OC43_orf1ab | 4650 | 4651 | 2 | 0.05 | 0.1 | 0.075 |
| NC_006213.1;YP_009555238.1;HCoV-OC43_orf1ab | 4712 | 4712 | 1 | 0.05 | 0.05 | 0.05 |
| NC_006213.1;YP_009555238.1;HCoV-OC43_orf1ab | 4750 | 4750 | 1 | 0.05 | 0.05 | 0.05 |
| NC_006213.1;YP_009555238.1;HCoV-OC43_orf1ab | 4764 | 4766 | 3 | 0.05 | 0.05 | 0.05 |
| NC_006213.1;YP_009555238.1;HCoV-OC43_orf1ab | 4772 | 4773 | 2 | 0.05 | 0.15 | 0.1 |
| NC_006213.1;YP_009555238.1;HCoV-OC43_orf1ab | 4776 | 4788 | 13 | 0.05 | 0.2 | 0.084615385 |
| NC_006213.1;YP_009555238.1;HCoV-OC43_orf1ab | 4792 | 4792 | 1 | 0.05 | 0.05 | 0.05 |
| NC_006213.1;YP_009555238.1;HCoV-OC43_orf1ab | 4799 | 4799 | 1 | 0.05 | 0.05 | 0.05 |
| NC_006213.1;YP_009555238.1;HCoV-OC43_orf1ab | 4801 | 4801 | 1 | 0.05 | 0.05 | 0.05 |
| NC_006213.1;YP_009555238.1;HCoV-OC43_orf1ab | 4809 | 4818 | 10 | 0.05 | 0.25 | 0.195 |
| NC_006213.1;YP_009555238.1;HCoV-OC43_orf1ab | 4852 | 4853 | 2 | 0.05 | 0.05 | 0.05 |
| NC_006213.1;YP_009555238.1;HCoV-OC43_orf1ab | 4855 | 4865 | 11 | 0.05 | 0.25 | 0.127272727 |
| NC_006213.1;YP_009555238.1;HCoV-OC43_orf1ab | 4896 | 4897 | 2 | 0.05 | 0.2 | 0.125 |
| NC_006213.1;YP_009555238.1;HCoV-OC43_orf1ab | 4960 | 4966 | 7 | 0.05 | 0.2 | 0.078571429 |

| Extended data 1 | | | | | | |
|---|---|---|---|---|---|---|
| Protein | Start position (peptide) | Stop position (peptide) | Number of peptides | Minimum fraction (per peptide) of control subjects with reactivity >3 sd above average | Maximum fraction (per peptide) of control subjects with reactivity >3 sd above average | Mean fraction (over all peptides in this contiguous stretch) of controlsubjects with reactivity >3 sd above average |
| NC_006213.1;YP_009555238.1;HCoV-OC43_orf1ab | 4968 | 4972 | 5 | 0.1 | 0.2 | 0.12 |
| NC_006213.1;YP_009555238.1;HCoV-OC43_orf1ab | 5007 | 5013 | 7 | 0.05 | 0.1 | 0.057142857 |
| NC_006213.1;YP_009555238.1;HCoV-OC43_orf1ab | 5039 | 5045 | 7 | 0.05 | 0.05 | 0.05 |
| NC_006213.1;YP_009555238.1;HCoV-OC43_orf1ab | 5064 | 5064 | 1 | 0.05 | 0.05 | 0.05 |
| NC_006213.1;YP_009555238.1;HCoV-OC43_orf1ab | 5071 | 5071 | 1 | 0.05 | 0.05 | 0.05 |
| NC_006213.1;YP_009555238.1;HCoV-OC43_orf1ab | 5073 | 5073 | 1 | 0.05 | 0.05 | 0.05 |
| NC_006213.1;YP_009555238.1;HCoV-OC43_orf1ab | 5104 | 5104 | 1 | 0.05 | 0.05 | 0.05 |
| NC_006213.1;YP_009555238.1;HCoV-OC43_orf1ab | 5106 | 5106 | 1 | 0.05 | 0.05 | 0.05 |
| NC_006213.1;YP_009555238.1;HCoV-OC43_orf1ab | 5112 | 5112 | 1 | 0.05 | 0.05 | 0.05 |
| NC_006213.1;YP_009555238.1;HCoV-OC43_orf1ab | 5128 | 5128 | 1 | 0.05 | 0.05 | 0.05 |
| NC_006213.1;YP_009555238.1;HCoV-OC43_orf1ab | 5130 | 5130 | 1 | 0.05 | 0.05 | 0.05 |
| NC_006213.1;YP_009555238.1;HCoV-OC43_orf1ab | 5148 | 5148 | 1 | 0.05 | 0.05 | 0.05 |
| NC_006213.1;YP_009555238.1;HCoV-OC43_orf1ab | 5158 | 5159 | 2 | 0.05 | 0.1 | 0.075 |
| NC_006213.1;YP_009555238.1;HCoV-OC43_orf1ab | 5178 | 5178 | 1 | 0.05 | 0.05 | 0.05 |
| NC_006213.1;YP_009555238.1;HCoV-OC43_orf1ab | 5182 | 5186 | 5 | 0.1 | 0.15 | 0.13 |
| NC_006213.1;YP_009555238.1;HCoV-OC43_orf1ab | 5202 | 5203 | 2 | 0.05 | 0.05 | 0.05 |
| NC_006213.1;YP_009555238.1;HCoV-OC43_orf1ab | 5231 | 5231 | 1 | 0.05 | 0.05 | 0.05 |
| NC_006213.1;YP_009555238.1;HCoV-OC43_orf1ab | 5249 | 5255 | 7 | 0.05 | 0.15 | 0.1 |
| NC_006213.1;YP_009555238.1;HCoV-OC43_orf1ab | 5267 | 5267 | 1 | 0.05 | 0.05 | 0.05 |
| NC_006213.1;YP_009555238.1;HCoV-OC43_orf1ab | 5270 | 5276 | 7 | 0.05 | 0.05 | 0.05 |
| NC_006213.1;YP_009555238.1;HCoV-OC43_orf1ab | 5341 | 5341 | 1 | 0.05 | 0.05 | 0.05 |
| NC_006213.1;YP_009555238.1;HCoV-OC43_orf1ab | 5344 | 5344 | 1 | 0.05 | 0.05 | 0.05 |
| NC_006213.1;YP_009555238.1;HCoV-OC43_orf1ab | 5346 | 5346 | 1 | 0.05 | 0.05 | 0.05 |
| NC_006213.1;YP_009555238.1;HCoV-OC43_orf1ab | 5350 | 5351 | 2 | 0.05 | 0.05 | 0.05 |
| NC_006213.1;YP_009555238.1;HCoV-OC43_orf1ab | 5364 | 5365 | 2 | 0.05 | 0.05 | 0.05 |
| NC_006213.1;YP_009555238.1;HCoV-OC43_orf1ab | 5389 | 5389 | 1 | 0.05 | 0.05 | 0.05 |
| NC_006213.1;YP_009555238.1;HCoV-OC43_orf1ab | 5392 | 5393 | 2 | 0.05 | 0.1 | 0.075 |
| NC_006213.1;YP_009555238.1;HCoV-OC43_orf1ab | 5395 | 5395 | 1 | 0.05 | 0.05 | 0.05 |
| NC_006213.1;YP_009555238.1;HCoV-OC43_orf1ab | 5414 | 5415 | 2 | 0.05 | 0.05 | 0.05 |
| NC_006213.1;YP_009555238.1;HCoV-OC43_orf1ab | 5417 | 5417 | 1 | 0.05 | 0.05 | 0.05 |
| NC_006213.1;YP_009555238.1;HCoV-OC43_orf1ab | 5423 | 5423 | 1 | 0.05 | 0.05 | 0.05 |
| NC_006213.1;YP_009555238.1;HCoV-OC43_orf1ab | 5425 | 5425 | 1 | 0.05 | 0.05 | 0.05 |
| NC_006213.1;YP_009555238.1;HCoV-OC43_orf1ab | 5433 | 5434 | 2 | 0.05 | 0.05 | 0.05 |
| NC_006213.1;YP_009555238.1;HCoV-OC43_orf1ab | 5456 | 5457 | 2 | 0.05 | 0.05 | 0.05 |
| NC_006213.1;YP_009555238.1;HCoV-OC43_orf1ab | 5461 | 5463 | 3 | 0.05 | 0.05 | 0.05 |
| NC_006213.1;YP_009555238.1;HCoV-OC43_orf1ab | 5465 | 5467 | 3 | 0.05 | 0.1 | 0.066666667 |
| NC_006213.1;YP_009555238.1;HCoV-OC43_orf1ab | 5494 | 5494 | 1 | 0.05 | 0.05 | 0.05 |

| Extended data 1 | | | | | | |
|---|---|---|---|---|---|---|
| Protein | Start position (peptide) | Stop position (peptide) | Number of peptides | Minimum fraction (per peptide) of control subjects with reactivity >3 sd above average | Maximum fraction (per peptide) of control subjects with reactivity >3 sd above average | Mean fraction (over all peptides in this contiguous stretch) of controlsubjects with reactivity >3 sd above average |
| NC_006213.1;YP_009555238.1;HCoV-OC43_orf1ab | 5500 | 5500 | 1 | 0.05 | 0.05 | 0.05 |
| NC_006213.1;YP_009555238.1;HCoV-OC43_orf1ab | 5520 | 5520 | 1 | 0.05 | 0.05 | 0.05 |
| NC_006213.1;YP_009555238.1;HCoV-OC43_orf1ab | 5529 | 5531 | 3 | 0.05 | 0.05 | 0.05 |
| NC_006213.1;YP_009555238.1;HCoV-OC43_orf1ab | 5535 | 5538 | 4 | 0.05 | 0.05 | 0.05 |
| NC_006213.1;YP_009555238.1;HCoV-OC43_orf1ab | 5594 | 5594 | 1 | 0.05 | 0.05 | 0.05 |
| NC_006213.1;YP_009555238.1;HCoV-OC43_orf1ab | 5609 | 5610 | 2 | 0.05 | 0.05 | 0.05 |
| NC_006213.1;YP_009555238.1;HCoV-OC43_orf1ab | 5612 | 5612 | 1 | 0.05 | 0.05 | 0.05 |
| NC_006213.1;YP_009555238.1;HCoV-OC43_orf1ab | 5614 | 5614 | 1 | 0.05 | 0.05 | 0.05 |
| NC_006213.1;YP_009555238.1;HCoV-OC43_orf1ab | 5616 | 5617 | 2 | 0.05 | 0.05 | 0.05 |
| NC_006213.1;YP_009555238.1;HCoV-OC43_orf1ab | 5631 | 5639 | 9 | 0.05 | 0.5 | 0.211111111 |
| NC_006213.1;YP_009555238.1;HCoV-OC43_orf1ab | 5648 | 5649 | 2 | 0.05 | 0.05 | 0.05 |
| NC_006213.1;YP_009555238.1;HCoV-OC43_orf1ab | 5658 | 5658 | 1 | 0.05 | 0.05 | 0.05 |
| NC_006213.1;YP_009555238.1;HCoV-OC43_orf1ab | 5660 | 5668 | 9 | 0.05 | 0.1 | 0.055555556 |
| NC_006213.1;YP_009555238.1;HCoV-OC43_orf1ab | 5671 | 5671 | 1 | 0.05 | 0.05 | 0.05 |
| NC_006213.1;YP_009555238.1;HCoV-OC43_orf1ab | 5699 | 5700 | 2 | 0.05 | 0.05 | 0.05 |
| NC_006213.1;YP_009555238.1;HCoV-OC43_orf1ab | 5707 | 5707 | 1 | 0.05 | 0.05 | 0.05 |
| NC_006213.1;YP_009555238.1;HCoV-OC43_orf1ab | 5716 | 5716 | 1 | 0.05 | 0.05 | 0.05 |
| NC_006213.1;YP_009555238.1;HCoV-OC43_orf1ab | 5734 | 5745 | 12 | 0.05 | 0.15 | 0.104166667 |
| NC_006213.1;YP_009555238.1;HCoV-OC43_orf1ab | 5779 | 5780 | 2 | 0.05 | 0.05 | 0.05 |
| NC_006213.1;YP_009555238.1;HCoV-OC43_orf1ab | 5789 | 5790 | 2 | 0.05 | 0.05 | 0.05 |
| NC_006213.1;YP_009555238.1;HCoV-OC43_orf1ab | 5794 | 5794 | 1 | 0.05 | 0.05 | 0.05 |
| NC_006213.1;YP_009555238.1;HCoV-OC43_orf1ab | 5803 | 5803 | 1 | 0.1 | 0.1 | 0.1 |
| NC_006213.1;YP_009555238.1;HCoV-OC43_orf1ab | 5806 | 5806 | 1 | 0.05 | 0.05 | 0.05 |
| NC_006213.1;YP_009555238.1;HCoV-OC43_orf1ab | 5808 | 5809 | 2 | 0.05 | 0.05 | 0.05 |
| NC_006213.1;YP_009555238.1;HCoV-OC43_orf1ab | 5818 | 5819 | 2 | 0.05 | 0.15 | 0.1 |
| NC_006213.1;YP_009555238.1;HCoV-OC43_orf1ab | 5823 | 5825 | 3 | 0.05 | 0.05 | 0.05 |
| NC_006213.1;YP_009555238.1;HCoV-OC43_orf1ab | 5836 | 5836 | 1 | 0.05 | 0.05 | 0.05 |
| NC_006213.1;YP_009555238.1;HCoV-OC43_orf1ab | 5843 | 5844 | 2 | 0.05 | 0.1 | 0.075 |
| NC_006213.1;YP_009555238.1;HCoV-OC43_orf1ab | 5854 | 5857 | 4 | 0.05 | 0.05 | 0.05 |
| NC_006213.1;YP_009555238.1;HCoV-OC43_orf1ab | 5883 | 5891 | 9 | 0.05 | 0.1 | 0.066666667 |
| NC_006213.1;YP_009555238.1;HCoV-OC43_orf1ab | 5904 | 5904 | 1 | 0.05 | 0.05 | 0.05 |
| NC_006213.1;YP_009555238.1;HCoV-OC43_orf1ab | 5909 | 5909 | 1 | 0.05 | 0.05 | 0.05 |
| NC_006213.1;YP_009555238.1;HCoV-OC43_orf1ab | 5918 | 5925 | 8 | 0.05 | 0.2 | 0.13125 |
| NC_006213.1;YP_009555238.1;HCoV-OC43_orf1ab | 5933 | 5933 | 1 | 0.05 | 0.05 | 0.05 |
| NC_006213.1;YP_009555238.1;HCoV-OC43_orf1ab | 5942 | 5942 | 1 | 0.05 | 0.05 | 0.05 |
| NC_006213.1;YP_009555238.1;HCoV-OC43_orf1ab | 5990 | 6000 | 11 | 0.05 | 0.05 | 0.05 |
| NC_006213.1;YP_009555238.1;HCoV-OC43_orf1ab | 6154 | 6154 | 1 | 0.05 | 0.05 | 0.05 |

| Extended data 1 | | | | | | |
|---|---|---|---|---|---|---|
| Protein | Start position (peptide) | Stop position (peptide) | Number of peptides | Minimum fraction (per peptide) of control subjects with reactivity >3 sd above average | Maximum fraction (per peptide) of control subjects with reactivity >3 sd above average | Mean fraction (over all peptides in this contiguous stretch) of controlsubjects with reactivity >3 sd above average |
| NC_006213.1;YP_009555238.1;HCoV-OC43_orf1ab | 6186 | 6188 | 3 | 0.05 | 0.05 | 0.05 |
| NC_006213.1;YP_009555238.1;HCoV-OC43_orf1ab | 6192 | 6192 | 1 | 0.05 | 0.05 | 0.05 |
| NC_006213.1;YP_009555238.1;HCoV-OC43_orf1ab | 6196 | 6196 | 1 | 0.05 | 0.05 | 0.05 |
| NC_006213.1;YP_009555238.1;HCoV-OC43_orf1ab | 6238 | 6238 | 1 | 0.05 | 0.05 | 0.05 |
| NC_006213.1;YP_009555238.1;HCoV-OC43_orf1ab | 6240 | 6244 | 5 | 0.05 | 0.3 | 0.12 |
| NC_006213.1;YP_009555238.1;HCoV-OC43_orf1ab | 6246 | 6246 | 1 | 0.05 | 0.05 | 0.05 |
| NC_006213.1;YP_009555238.1;HCoV-OC43_orf1ab | 6259 | 6268 | 10 | 0.05 | 0.1 | 0.065 |
| NC_006213.1;YP_009555238.1;HCoV-OC43_orf1ab | 6270 | 6272 | 3 | 0.05 | 0.05 | 0.05 |
| NC_006213.1;YP_009555238.1;HCoV-OC43_orf1ab | 6275 | 6276 | 2 | 0.05 | 0.05 | 0.05 |
| NC_006213.1;YP_009555238.1;HCoV-OC43_orf1ab | 6291 | 6301 | 11 | 0.05 | 0.2 | 0.095454545 |
| NC_006213.1;YP_009555238.1;HCoV-OC43_orf1ab | 6382 | 6386 | 5 | 0.05 | 0.15 | 0.11 |
| NC_006213.1;YP_009555238.1;HCoV-OC43_orf1ab | 6427 | 6427 | 1 | 0.05 | 0.05 | 0.05 |
| NC_006213.1;YP_009555238.1;HCoV-OC43_orf1ab | 6430 | 6432 | 3 | 0.05 | 0.1 | 0.066666667 |
| NC_006213.1;YP_009555238.1;HCoV-OC43_orf1ab | 6436 | 6444 | 9 | 0.05 | 0.15 | 0.072222222 |
| NC_006213.1;YP_009555238.1;HCoV-OC43_orf1ab | 6446 | 6447 | 2 | 0.05 | 0.1 | 0.075 |
| NC_006213.1;YP_009555238.1;HCoV-OC43_orf1ab | 6462 | 6464 | 3 | 0.05 | 0.1 | 0.083333333 |
| NC_006213.1;YP_009555238.1;HCoV-OC43_orf1ab | 6472 | 6472 | 1 | 0.05 | 0.05 | 0.05 |
| NC_006213.1;YP_009555238.1;HCoV-OC43_orf1ab | 6516 | 6521 | 6 | 0.05 | 0.05 | 0.05 |
| NC_006213.1;YP_009555238.1;HCoV-OC43_orf1ab | 6540 | 6540 | 1 | 0.05 | 0.05 | 0.05 |
| NC_006213.1;YP_009555238.1;HCoV-OC43_orf1ab | 6552 | 6553 | 2 | 0.05 | 0.05 | 0.05 |
| NC_006213.1;YP_009555238.1;HCoV-OC43_orf1ab | 6569 | 6569 | 1 | 0.05 | 0.05 | 0.05 |
| NC_006213.1;YP_009555238.1;HCoV-OC43_orf1ab | 6571 | 6571 | 1 | 0.05 | 0.05 | 0.05 |
| NC_006213.1;YP_009555238.1;HCoV-OC43_orf1ab | 6599 | 6609 | 11 | 0.05 | 0.2 | 0.122727273 |
| NC_006213.1;YP_009555238.1;HCoV-OC43_orf1ab | 6611 | 6613 | 3 | 0.05 | 0.15 | 0.083333333 |
| NC_006213.1;YP_009555238.1;HCoV-OC43_orf1ab | 6615 | 6617 | 3 | 0.05 | 0.05 | 0.05 |
| NC_006213.1;YP_009555238.1;HCoV-OC43_orf1ab | 6620 | 6627 | 8 | 0.05 | 0.1 | 0.09375 |
| NC_006213.1;YP_009555238.1;HCoV-OC43_orf1ab | 6652 | 6661 | 10 | 0.05 | 0.1 | 0.08 |
| NC_006213.1;YP_009555238.1;HCoV-OC43_orf1ab | 6664 | 6669 | 6 | 0.05 | 0.15 | 0.083333333 |
| NC_006213.1;YP_009555238.1;HCoV-OC43_orf1ab | 6671 | 6671 | 1 | 0.05 | 0.05 | 0.05 |
| NC_006213.1;YP_009555238.1;HCoV-OC43_orf1ab | 6674 | 6674 | 1 | 0.05 | 0.05 | 0.05 |
| NC_006213.1;YP_009555238.1;HCoV-OC43_orf1ab | 6676 | 6676 | 1 | 0.1 | 0.1 | 0.1 |
| NC_006213.1;YP_009555238.1;HCoV-OC43_orf1ab | 6722 | 6722 | 1 | 0.05 | 0.05 | 0.05 |
| NC_006213.1;YP_009555238.1;HCoV-OC43_orf1ab | 6766 | 6767 | 2 | 0.05 | 0.05 | 0.05 |
| NC_006213.1;YP_009555238.1;HCoV-OC43_orf1ab | 6772 | 6774 | 3 | 0.05 | 0.1 | 0.066666667 |
| NC_006213.1;YP_009555238.1;HCoV-OC43_orf1ab | 6783 | 6784 | 2 | 0.05 | 0.05 | 0.05 |
| NC_006213.1;YP_009555238.1;HCoV-OC43_orf1ab | 6792 | 6793 | 2 | 0.05 | 0.05 | 0.05 |
| NC_006213.1;YP_009555238.1;HCoV-OC43_orf1ab | 6810 | 6811 | 2 | 0.05 | 0.05 | 0.05 |

| Extended data 1 | | | | | | |
|---|---|---|---|---|---|---|
| Protein | Start position (peptide) | Stop position (peptide) | Number of peptides | Minimum fraction (per peptide) of control subjects with reactivity >3 sd above average | Maximum fraction (per peptide) of control subjects with reactivity >3 sd above average | Mean fraction (over all peptides in this contiguous stretch) of controlsubjects with reactivity >3 sd above average |
| NC_006213.1;YP_009555238.1;HCoV-OC43_orf1ab | 6819 | 6819 | 1 | 0.05 | 0.05 | 0.05 |
| NC_006213.1;YP_009555238.1;HCoV-OC43_orf1ab | 6826 | 6826 | 1 | 0.05 | 0.05 | 0.05 |
| NC_006213.1;YP_009555238.1;HCoV-OC43_orf1ab | 6839 | 6839 | 1 | 0.05 | 0.05 | 0.05 |
| NC_006213.1;YP_009555238.1;HCoV-OC43_orf1ab | 6843 | 6846 | 4 | 0.05 | 0.05 | 0.05 |
| NC_006213.1;YP_009555238.1;HCoV-OC43_orf1ab | 6858 | 6858 | 1 | 0.05 | 0.05 | 0.05 |
| NC_006213.1;YP_009555238.1;HCoV-OC43_orf1ab | 6860 | 6860 | 1 | 0.05 | 0.05 | 0.05 |
| NC_006213.1;YP_009555238.1;HCoV-OC43_orf1ab | 6868 | 6868 | 1 | 0.05 | 0.05 | 0.05 |
| NC_006213.1;YP_009555238.1;HCoV-OC43_orf1ab | 6924 | 6935 | 12 | 0.05 | 0.2 | 0.1 |
| NC_006213.1;YP_009555238.1;HCoV-OC43_orf1ab | 6949 | 6949 | 1 | 0.05 | 0.05 | 0.05 |
| NC_006213.1;YP_009555238.1;HCoV-OC43_orf1ab | 6991 | 6991 | 1 | 0.05 | 0.05 | 0.05 |
| NC_006213.1;YP_009555238.1;HCoV-OC43_orf1ab | 7007 | 7011 | 5 | 0.05 | 0.05 | 0.05 |
| NC_006213.1;YP_009555238.1;HCoV-OC43_orf1ab | 7013 | 7014 | 2 | 0.25 | 0.25 | 0.25 |
| NC_006213.1;YP_009555238.1;HCoV-OC43_orf1ab | 7030 | 7037 | 8 | 0.05 | 0.05 | 0.05 |
| NC_006213.1;YP_009555238.1;HCoV-OC43_orf1ab | 7054 | 7055 | 2 | 0.05 | 0.05 | 0.05 |
| NC_006213.1;YP_009555238.1;HCoV-OC43_orf1ab | 7061 | 7062 | 2 | 0.05 | 0.05 | 0.05 |
| NC_006213.1;YP_009555238.1;HCoV-OC43_orf1ab | 7065 | 7067 | 3 | 0.05 | 0.05 | 0.05 |
| NC_006213.1;YP_009555238.1;HCoV-OC43_orf1ab | 7069 | 7076 | 8 | 0.05 | 0.25 | 0.1875 |
| NC_006213.1;YP_009555239.1;HCoV-OC43_ns2 | 3 | 4 | 2 | 0.05 | 0.05 | 0.05 |
| NC_006213.1;YP_009555239.1;HCoV-OC43_ns2 | 23 | 23 | 1 | 0.05 | 0.05 | 0.05 |
| NC_006213.1;YP_009555239.1;HCoV-OC43_ns2 | 54 | 54 | 1 | 0.05 | 0.05 | 0.05 |
| NC_006213.1;YP_009555239.1;HCoV-OC43_ns2 | 58 | 64 | 7 | 0.05 | 0.15 | 0.078571429 |
| NC_006213.1;YP_009555239.1;HCoV-OC43_ns2 | 66 | 70 | 5 | 0.05 | 0.1 | 0.06 |
| NC_006213.1;YP_009555239.1;HCoV-OC43_ns2 | 101 | 101 | 1 | 0.45 | 0.45 | 0.45 |
| NC_006213.1;YP_009555239.1;HCoV-OC43_ns2 | 105 | 105 | 1 | 0.05 | 0.05 | 0.05 |
| NC_006213.1;YP_009555239.1;HCoV-OC43_ns2 | 107 | 108 | 2 | 0.1 | 0.25 | 0.175 |
| NC_006213.1;YP_009555239.1;HCoV-OC43_ns2 | 112 | 114 | 3 | 0.05 | 0.05 | 0.05 |
| NC_006213.1;YP_009555239.1;HCoV-OC43_ns2 | 119 | 120 | 2 | 0.05 | 0.05 | 0.05 |
| NC_006213.1;YP_009555239.1;HCoV-OC43_ns2 | 140 | 140 | 1 | 0.05 | 0.05 | 0.05 |
| NC_006213.1;YP_009555239.1;HCoV-OC43_ns2 | 142 | 144 | 3 | 0.1 | 0.15 | 0.116666667 |
| NC_006213.1;YP_009555239.1;HCoV-OC43_ns2 | 151 | 151 | 1 | 0.05 | 0.05 | 0.05 |
| NC_006213.1;YP_009555239.1;HCoV-OC43_ns2 | 154 | 155 | 2 | 0.05 | 0.05 | 0.05 |
| NC_006213.1;YP_009555239.1;HCoV-OC43_ns2 | 157 | 165 | 9 | 0.05 | 0.1 | 0.061111111 |
| NC_006213.1;YP_009555239.1;HCoV-OC43_ns2 | 168 | 173 | 6 | 0.05 | 0.1 | 0.066666667 |
| NC_006213.1;YP_009555239.1;HCoV-OC43_ns2 | 177 | 177 | 1 | 0.05 | 0.05 | 0.05 |
| NC_006213.1;YP_009555239.1;HCoV-OC43_ns2 | 187 | 187 | 1 | 0.05 | 0.05 | 0.05 |
| NC_006213.1;YP_009555239.1;HCoV-OC43_ns2 | 189 | 194 | 6 | 0.05 | 0.15 | 0.083333333 |
| NC_006213.1;YP_009555239.1;HCoV-OC43_ns2 | 206 | 211 | 6 | 0.05 | 0.05 | 0.05 |

| Extended data 1 | | | | | | |
|---|---|---|---|---|---|---|
| Protein | Start position (peptide) | Stop position (peptide) | Number of peptides | Minimum fraction (per peptide) of control subjects with reactivity >3 sd above average | Maximum fraction (per peptide) of control subjects with reactivity >3 sd above average | Mean fraction (over all peptides in this contiguous stretch) of controlsubjects with reactivity >3 sd above average |
| NC_006213.1;YP_009555240.1;HCoV-OC43_HA-E | 13 | 13 | 1 | 0.1 | 0.1 | 0.1 |
| NC_006213.1;YP_009555240.1;HCoV-OC43_HA-E | 16 | 20 | 5 | 0.05 | 0.1 | 0.06 |
| NC_006213.1;YP_009555240.1;HCoV-OC43_HA-E | 24 | 24 | 1 | 0.05 | 0.05 | 0.05 |
| NC_006213.1;YP_009555240.1;HCoV-OC43_HA-E | 36 | 36 | 1 | 0.05 | 0.05 | 0.05 |
| NC_006213.1;YP_009555240.1;HCoV-OC43_HA-E | 39 | 39 | 1 | 0.05 | 0.05 | 0.05 |
| NC_006213.1;YP_009555240.1;HCoV-OC43_HA-E | 46 | 48 | 3 | 0.05 | 0.1 | 0.066666667 |
| NC_006213.1;YP_009555240.1;HCoV-OC43_HA-E | 51 | 51 | 1 | 0.05 | 0.05 | 0.05 |
| NC_006213.1;YP_009555240.1;HCoV-OC43_HA-E | 54 | 55 | 2 | 0.05 | 0.05 | 0.05 |
| NC_006213.1;YP_009555240.1;HCoV-OC43_HA-E | 63 | 64 | 2 | 0.05 | 0.05 | 0.05 |
| NC_006213.1;YP_009555240.1;HCoV-OC43_HA-E | 80 | 81 | 2 | 0.05 | 0.05 | 0.05 |
| NC_006213.1;YP_009555240.1;HCoV-OC43_HA-E | 104 | 118 | 15 | 0.05 | 0.45 | 0.133333333 |
| NC_006213.1;YP_009555240.1;HCoV-OC43_HA-E | 149 | 149 | 1 | 0.05 | 0.05 | 0.05 |
| NC_006213.1;YP_009555240.1;HCoV-OC43_HA-E | 156 | 158 | 3 | 0.05 | 0.1 | 0.083333333 |
| NC_006213.1;YP_009555240.1;HCoV-OC43_HA-E | 162 | 164 | 3 | 0.05 | 0.25 | 0.15 |
| NC_006213.1;YP_009555240.1;HCoV-OC43_HA-E | 167 | 170 | 4 | 0.05 | 0.1 | 0.0625 |
| NC_006213.1;YP_009555240.1;HCoV-OC43_HA-E | 175 | 175 | 1 | 0.1 | 0.1 | 0.1 |
| NC_006213.1;YP_009555240.1;HCoV-OC43_HA-E | 178 | 178 | 1 | 0.05 | 0.05 | 0.05 |
| NC_006213.1;YP_009555240.1;HCoV-OC43_HA-E | 187 | 187 | 1 | 0.05 | 0.05 | 0.05 |
| NC_006213.1;YP_009555240.1;HCoV-OC43_HA-E | 204 | 210 | 7 | 0.05 | 0.35 | 0.228571429 |
| NC_006213.1;YP_009555240.1;HCoV-OC43_HA-E | 212 | 217 | 6 | 0.05 | 0.15 | 0.116666667 |
| NC_006213.1;YP_009555240.1;HCoV-OC43_HA-E | 219 | 219 | 1 | 0.1 | 0.1 | 0.1 |
| NC_006213.1;YP_009555240.1;HCoV-OC43_HA-E | 227 | 230 | 4 | 0.05 | 0.05 | 0.05 |
| NC_006213.1;YP_009555240.1;HCoV-OC43_HA-E | 233 | 233 | 1 | 0.05 | 0.05 | 0.05 |
| NC_006213.1;YP_009555240.1;HCoV-OC43_HA-E | 243 | 243 | 1 | 0.05 | 0.05 | 0.05 |
| NC_006213.1;YP_009555240.1;HCoV-OC43_HA-E | 269 | 271 | 3 | 0.05 | 0.05 | 0.05 |
| NC_006213.1;YP_009555240.1;HCoV-OC43_HA-E | 285 | 287 | 3 | 0.05 | 0.05 | 0.05 |
| NC_006213.1;YP_009555240.1;HCoV-OC43_HA-E | 290 | 292 | 3 | 0.05 | 0.15 | 0.083333333 |
| NC_006213.1;YP_009555240.1;HCoV-OC43_HA-E | 297 | 297 | 1 | 0.05 | 0.05 | 0.05 |
| NC_006213.1;YP_009555240.1;HCoV-OC43_HA-E | 305 | 309 | 5 | 0.1 | 0.1 | 0.1 |
| NC_006213.1;YP_009555240.1;HCoV-OC43_HA-E | 311 | 316 | 6 | 0.05 | 0.2 | 0.125 |
| NC_006213.1;YP_009555240.1;HCoV-OC43_HA-E | 323 | 323 | 1 | 0.05 | 0.05 | 0.05 |
| NC_006213.1;YP_009555240.1;HCoV-OC43_HA-E | 360 | 360 | 1 | 0.05 | 0.05 | 0.05 |
| NC_006213.1;YP_009555240.1;HCoV-OC43_HA-E | 362 | 371 | 10 | 0.1 | 0.45 | 0.165 |
| NC_006213.1;YP_009555241.1;HCoV-OC43_spike | 10 | 20 | 11 | 0.05 | 0.05 | 0.05 |
| NC_006213.1;YP_009555241.1;HCoV-OC43_spike | 23 | 23 | 1 | 0.05 | 0.05 | 0.05 |
| NC_006213.1;YP_009555241.1;HCoV-OC43_spike | 30 | 34 | 5 | 0.05 | 0.1 | 0.06 |
| NC_006213.1;YP_009555241.1;HCoV-OC43_spike | 36 | 37 | 2 | 0.05 | 0.1 | 0.075 |

| Extended data 1 | | | | | | |
|---|---|---|---|---|---|---|
| Protein | Start position (peptide) | Stop position (peptide) | Number of peptides | Minimum fraction (per peptide) of control subjects with reactivity >3 sd above average | Maximum fraction (per peptide) of control subjects with reactivity >3 sd above average | Mean fraction (over all peptides in this contiguous stretch) of controlsubjects with reactivity >3 sd above average |
| NC_006213.1;YP_009555241.1;HCoV-OC43_spike | 58 | 58 | 1 | 0.05 | 0.05 | 0.05 |
| NC_006213.1;YP_009555241.1;HCoV-OC43_spike | 64 | 64 | 1 | 0.05 | 0.05 | 0.05 |
| NC_006213.1;YP_009555241.1;HCoV-OC43_spike | 97 | 98 | 2 | 0.05 | 0.05 | 0.05 |
| NC_006213.1;YP_009555241.1;HCoV-OC43_spike | 107 | 113 | 7 | 0.05 | 0.05 | 0.05 |
| NC_006213.1;YP_009555241.1;HCoV-OC43_spike | 132 | 133 | 2 | 0.05 | 0.05 | 0.05 |
| NC_006213.1;YP_009555241.1;HCoV-OC43_spike | 137 | 143 | 7 | 0.05 | 0.1 | 0.064285714 |
| NC_006213.1;YP_009555241.1;HCoV-OC43_spike | 157 | 157 | 1 | 0.05 | 0.05 | 0.05 |
| NC_006213.1;YP_009555241.1;HCoV-OC43_spike | 170 | 171 | 2 | 0.05 | 0.05 | 0.05 |
| NC_006213.1;YP_009555241.1;HCoV-OC43_spike | 173 | 174 | 2 | 0.05 | 0.05 | 0.05 |
| NC_006213.1;YP_009555241.1;HCoV-OC43_spike | 176 | 178 | 3 | 0.05 | 0.05 | 0.05 |
| NC_006213.1;YP_009555241.1;HCoV-OC43_spike | 200 | 200 | 1 | 0.05 | 0.05 | 0.05 |
| NC_006213.1;YP_009555241.1;HCoV-OC43_spike | 205 | 207 | 3 | 0.05 | 0.05 | 0.05 |
| NC_006213.1;YP_009555241.1;HCoV-OC43_spike | 273 | 277 | 5 | 0.05 | 0.05 | 0.05 |
| NC_006213.1;YP_009555241.1;HCoV-OC43_spike | 286 | 286 | 1 | 0.05 | 0.05 | 0.05 |
| NC_006213.1;YP_009555241.1;HCoV-OC43_spike | 292 | 293 | 2 | 0.05 | 0.05 | 0.05 |
| NC_006213.1;YP_009555241.1;HCoV-OC43_spike | 301 | 301 | 1 | 0.05 | 0.05 | 0.05 |
| NC_006213.1;YP_009555241.1;HCoV-OC43_spike | 306 | 311 | 6 | 0.05 | 0.1 | 0.058333333 |
| NC_006213.1;YP_009555241.1;HCoV-OC43_spike | 320 | 320 | 1 | 0.05 | 0.05 | 0.05 |
| NC_006213.1;YP_009555241.1;HCoV-OC43_spike | 329 | 330 | 2 | 0.1 | 0.15 | 0.125 |
| NC_006213.1;YP_009555241.1;HCoV-OC43_spike | 332 | 338 | 7 | 0.05 | 0.25 | 0.135714286 |
| NC_006213.1;YP_009555241.1;HCoV-OC43_spike | 340 | 343 | 4 | 0.05 | 0.1 | 0.0625 |
| NC_006213.1;YP_009555241.1;HCoV-OC43_spike | 345 | 346 | 2 | 0.05 | 0.1 | 0.075 |
| NC_006213.1;YP_009555241.1;HCoV-OC43_spike | 348 | 349 | 2 | 0.05 | 0.1 | 0.075 |
| NC_006213.1;YP_009555241.1;HCoV-OC43_spike | 353 | 353 | 1 | 0.05 | 0.05 | 0.05 |
| NC_006213.1;YP_009555241.1;HCoV-OC43_spike | 360 | 360 | 1 | 0.05 | 0.05 | 0.05 |
| NC_006213.1;YP_009555241.1;HCoV-OC43_spike | 369 | 369 | 1 | 0.05 | 0.05 | 0.05 |
| NC_006213.1;YP_009555241.1;HCoV-OC43_spike | 373 | 377 | 5 | 0.05 | 0.15 | 0.07 |
| NC_006213.1;YP_009555241.1;HCoV-OC43_spike | 394 | 394 | 1 | 0.05 | 0.05 | 0.05 |
| NC_006213.1;YP_009555241.1;HCoV-OC43_spike | 400 | 400 | 1 | 0.05 | 0.05 | 0.05 |
| NC_006213.1;YP_009555241.1;HCoV-OC43_spike | 425 | 426 | 2 | 0.05 | 0.1 | 0.075 |
| NC_006213.1;YP_009555241.1;HCoV-OC43_spike | 447 | 447 | 1 | 0.05 | 0.05 | 0.05 |
| NC_006213.1;YP_009555241.1;HCoV-OC43_spike | 449 | 459 | 11 | 0.05 | 0.2 | 0.122727273 |
| NC_006213.1;YP_009555241.1;HCoV-OC43_spike | 465 | 465 | 1 | 0.05 | 0.05 | 0.05 |
| NC_006213.1;YP_009555241.1;HCoV-OC43_spike | 470 | 479 | 10 | 0.05 | 0.05 | 0.05 |
| NC_006213.1;YP_009555241.1;HCoV-OC43_spike | 491 | 491 | 1 | 0.05 | 0.05 | 0.05 |
| NC_006213.1;YP_009555241.1;HCoV-OC43_spike | 493 | 493 | 1 | 0.05 | 0.05 | 0.05 |
| NC_006213.1;YP_009555241.1;HCoV-OC43_spike | 495 | 506 | 12 | 0.05 | 0.1 | 0.054166667 |

| Extended data 1 | | | | | | |
|---|---|---|---|---|---|---|
| Protein | Start position (peptide) | Stop position (peptide) | Number of peptides | Minimum fraction (per peptide) of control subjects with reactivity >3 sd above average | Maximum fraction (per peptide) of control subjects with reactivity >3 sd above average | Mean fraction (over all peptides in this contiguous stretch) of controlsubjects with reactivity >3 sd above average |
| NC_006213.1;YP_009555241.1;HCoV-OC43_spike | 535 | 535 | 1 | 0.05 | 0.05 | 0.05 |
| NC_006213.1;YP_009555241.1;HCoV-OC43_spike | 555 | 555 | 1 | 0.05 | 0.05 | 0.05 |
| NC_006213.1;YP_009555241.1;HCoV-OC43_spike | 566 | 566 | 1 | 0.05 | 0.05 | 0.05 |
| NC_006213.1;YP_009555241.1;HCoV-OC43_spike | 587 | 587 | 1 | 0.05 | 0.05 | 0.05 |
| NC_006213.1;YP_009555241.1;HCoV-OC43_spike | 591 | 592 | 2 | 0.05 | 0.05 | 0.05 |
| NC_006213.1;YP_009555241.1;HCoV-OC43_spike | 595 | 602 | 8 | 0.05 | 0.1 | 0.05625 |
| NC_006213.1;YP_009555241.1;HCoV-OC43_spike | 642 | 642 | 1 | 0.1 | 0.1 | 0.1 |
| NC_006213.1;YP_009555241.1;HCoV-OC43_spike | 650 | 650 | 1 | 0.05 | 0.05 | 0.05 |
| NC_006213.1;YP_009555241.1;HCoV-OC43_spike | 652 | 652 | 1 | 0.05 | 0.05 | 0.05 |
| NC_006213.1;YP_009555241.1;HCoV-OC43_spike | 667 | 667 | 1 | 0.05 | 0.05 | 0.05 |
| NC_006213.1;YP_009555241.1;HCoV-OC43_spike | 703 | 704 | 2 | 0.05 | 0.05 | 0.05 |
| NC_006213.1;YP_009555241.1;HCoV-OC43_spike | 714 | 714 | 1 | 0.05 | 0.05 | 0.05 |
| NC_006213.1;YP_009555241.1;HCoV-OC43_spike | 717 | 718 | 2 | 0.05 | 0.05 | 0.05 |
| NC_006213.1;YP_009555241.1;HCoV-OC43_spike | 747 | 749 | 3 | 0.05 | 0.1 | 0.083333333 |
| NC_006213.1;YP_009555241.1;HCoV-OC43_spike | 755 | 774 | 20 | 0.05 | 0.85 | 0.4375 |
| NC_006213.1;YP_009555241.1;HCoV-OC43_spike | 776 | 776 | 1 | 0.25 | 0.25 | 0.25 |
| NC_006213.1;YP_009555241.1;HCoV-OC43_spike | 797 | 799 | 3 | 0.05 | 0.05 | 0.05 |
| NC_006213.1;YP_009555241.1;HCoV-OC43_spike | 803 | 803 | 1 | 0.05 | 0.05 | 0.05 |
| NC_006213.1;YP_009555241.1;HCoV-OC43_spike | 811 | 811 | 1 | 0.05 | 0.05 | 0.05 |
| NC_006213.1;YP_009555241.1;HCoV-OC43_spike | 837 | 838 | 2 | 0.05 | 0.05 | 0.05 |
| NC_006213.1;YP_009555241.1;HCoV-OC43_spike | 846 | 847 | 2 | 0.05 | 0.05 | 0.05 |
| NC_006213.1;YP_009555241.1;HCoV-OC43_spike | 852 | 852 | 1 | 0.05 | 0.05 | 0.05 |
| NC_006213.1;YP_009555241.1;HCoV-OC43_spike | 865 | 865 | 1 | 0.05 | 0.05 | 0.05 |
| NC_006213.1;YP_009555241.1;HCoV-OC43_spike | 867 | 877 | 11 | 0.05 | 0.3 | 0.163636364 |
| NC_006213.1;YP_009555241.1;HCoV-OC43_spike | 879 | 881 | 3 | 0.05 | 0.1 | 0.083333333 |
| NC_006213.1;YP_009555241.1;HCoV-OC43_spike | 886 | 886 | 1 | 0.05 | 0.05 | 0.05 |
| NC_006213.1;YP_009555241.1;HCoV-OC43_spike | 896 | 901 | 6 | 0.05 | 0.2 | 0.116666667 |
| NC_006213.1;YP_009555241.1;HCoV-OC43_spike | 903 | 905 | 3 | 0.05 | 0.45 | 0.2 |
| NC_006213.1;YP_009555241.1;HCoV-OC43_spike | 913 | 915 | 3 | 0.05 | 0.05 | 0.05 |
| NC_006213.1;YP_009555241.1;HCoV-OC43_spike | 917 | 917 | 1 | 0.05 | 0.05 | 0.05 |
| NC_006213.1;YP_009555241.1;HCoV-OC43_spike | 930 | 930 | 1 | 0.05 | 0.05 | 0.05 |
| NC_006213.1;YP_009555241.1;HCoV-OC43_spike | 933 | 937 | 5 | 0.05 | 0.05 | 0.05 |
| NC_006213.1;YP_009555241.1;HCoV-OC43_spike | 939 | 943 | 5 | 0.05 | 0.1 | 0.06 |
| NC_006213.1;YP_009555241.1;HCoV-OC43_spike | 945 | 948 | 4 | 0.05 | 0.05 | 0.05 |
| NC_006213.1;YP_009555241.1;HCoV-OC43_spike | 950 | 950 | 1 | 0.05 | 0.05 | 0.05 |
| NC_006213.1;YP_009555241.1;HCoV-OC43_spike | 954 | 955 | 2 | 0.2 | 0.2 | 0.2 |
| NC_006213.1;YP_009555241.1;HCoV-OC43_spike | 958 | 958 | 1 | 0.05 | 0.05 | 0.05 |

| Extended data 1 | | | | | | |
|---|---|---|---|---|---|---|
| **Protein** | **Start position (peptide)** | **Stop position (peptide)** | **Number of peptides** | **Minimum fraction (per peptide) of control subjects with reactivity >3 sd above average** | **Maximum fraction (per peptide) of control subjects with reactivity >3 sd above average** | **Mean fraction (over all peptides in this contiguous stretch) of controlsubjects with reactivity >3 sd above average** |
| NC_006213.1;YP_009555241.1;HCoV-OC43_spike | 962 | 963 | 2 | 0.05 | 0.1 | 0.075 |
| NC_006213.1;YP_009555241.1;HCoV-OC43_spike | 965 | 967 | 3 | 0.05 | 0.1 | 0.066666667 |
| NC_006213.1;YP_009555241.1;HCoV-OC43_spike | 997 | 999 | 3 | 0.05 | 0.05 | 0.05 |
| NC_006213.1;YP_009555241.1;HCoV-OC43_spike | 1005 | 1005 | 1 | 0.05 | 0.05 | 0.05 |
| NC_006213.1;YP_009555241.1;HCoV-OC43_spike | 1011 | 1019 | 9 | 0.05 | 0.1 | 0.066666667 |
| NC_006213.1;YP_009555241.1;HCoV-OC43_spike | 1024 | 1024 | 1 | 0.05 | 0.05 | 0.05 |
| NC_006213.1;YP_009555241.1;HCoV-OC43_spike | 1030 | 1035 | 6 | 0.05 | 0.1 | 0.066666667 |
| NC_006213.1;YP_009555241.1;HCoV-OC43_spike | 1040 | 1040 | 1 | 0.05 | 0.05 | 0.05 |
| NC_006213.1;YP_009555241.1;HCoV-OC43_spike | 1046 | 1047 | 2 | 0.05 | 0.1 | 0.075 |
| NC_006213.1;YP_009555241.1;HCoV-OC43_spike | 1050 | 1051 | 2 | 0.05 | 0.05 | 0.05 |
| NC_006213.1;YP_009555241.1;HCoV-OC43_spike | 1057 | 1057 | 1 | 0.05 | 0.05 | 0.05 |
| NC_006213.1;YP_009555241.1;HCoV-OC43_spike | 1064 | 1064 | 1 | 0.05 | 0.05 | 0.05 |
| NC_006213.1;YP_009555241.1;HCoV-OC43_spike | 1066 | 1066 | 1 | 0.1 | 0.1 | 0.1 |
| NC_006213.1;YP_009555241.1;HCoV-OC43_spike | 1070 | 1070 | 1 | 0.05 | 0.05 | 0.05 |
| NC_006213.1;YP_009555241.1;HCoV-OC43_spike | 1072 | 1074 | 3 | 0.05 | 0.1 | 0.066666667 |
| NC_006213.1;YP_009555241.1;HCoV-OC43_spike | 1086 | 1092 | 7 | 0.05 | 0.05 | 0.05 |
| NC_006213.1;YP_009555241.1;HCoV-OC43_spike | 1110 | 1111 | 2 | 0.05 | 0.05 | 0.05 |
| NC_006213.1;YP_009555241.1;HCoV-OC43_spike | 1160 | 1160 | 1 | 0.05 | 0.05 | 0.05 |
| NC_006213.1;YP_009555241.1;HCoV-OC43_spike | 1166 | 1166 | 1 | 0.05 | 0.05 | 0.05 |
| NC_006213.1;YP_009555241.1;HCoV-OC43_spike | 1173 | 1173 | 1 | 0.05 | 0.05 | 0.05 |
| NC_006213.1;YP_009555241.1;HCoV-OC43_spike | 1179 | 1180 | 2 | 0.05 | 0.05 | 0.05 |
| NC_006213.1;YP_009555241.1;HCoV-OC43_spike | 1190 | 1191 | 2 | 0.05 | 0.05 | 0.05 |
| NC_006213.1;YP_009555241.1;HCoV-OC43_spike | 1193 | 1193 | 1 | 0.05 | 0.05 | 0.05 |
| NC_006213.1;YP_009555241.1;HCoV-OC43_spike | 1198 | 1198 | 1 | 0.05 | 0.05 | 0.05 |
| NC_006213.1;YP_009555241.1;HCoV-OC43_spike | 1217 | 1218 | 2 | 0.05 | 0.05 | 0.05 |
| NC_006213.1;YP_009555241.1;HCoV-OC43_spike | 1221 | 1221 | 1 | 0.05 | 0.05 | 0.05 |
| NC_006213.1;YP_009555241.1;HCoV-OC43_spike | 1228 | 1230 | 3 | 0.05 | 0.05 | 0.05 |
| NC_006213.1;YP_009555241.1;HCoV-OC43_spike | 1232 | 1235 | 4 | 0.05 | 0.15 | 0.0875 |
| NC_006213.1;YP_009555241.1;HCoV-OC43_spike | 1238 | 1239 | 2 | 0.05 | 0.1 | 0.075 |
| NC_006213.1;YP_009555241.1;HCoV-OC43_spike | 1272 | 1276 | 5 | 0.05 | 0.05 | 0.05 |
| NC_006213.1;YP_009555241.1;HCoV-OC43_spike | 1284 | 1284 | 1 | 0.05 | 0.05 | 0.05 |
| NC_006213.1;YP_009555241.1;HCoV-OC43_spike | 1329 | 1342 | 14 | 0.05 | 0.3 | 0.125 |
| NC_006213.1;YP_009555242.1;HCoV-OC43_ns12.9 | 6 | 21 | 16 | 0.05 | 0.15 | 0.1 |
| NC_006213.1;YP_009555242.1;HCoV-OC43_ns12.9 | 23 | 24 | 2 | 0.05 | 0.05 | 0.05 |
| NC_006213.1;YP_009555242.1;HCoV-OC43_ns12.9 | 29 | 29 | 1 | 0.05 | 0.05 | 0.05 |
| NC_006213.1;YP_009555242.1;HCoV-OC43_ns12.9 | 74 | 83 | 10 | 0.05 | 0.1 | 0.06 |
| NC_006213.1;YP_009555243.1;HCoV-OC43_envelope | 62 | 63 | 2 | 0.05 | 0.05 | 0.05 |

| Extended data 1 | | | | | | |
|---|---|---|---|---|---|---|
| **Protein** | **Start position (peptide)** | **Stop position (peptide)** | **Number of peptides** | **Minimum fraction (per peptide) of control subjects with reactivity >3 sd above average** | **Maximum fraction (per peptide) of control subjects with reactivity >3 sd above average** | **Mean fraction (over all peptides in this contiguous stretch) of controlsubjects with reactivity >3 sd above average** |
| NC_006213.1;YP_009555243.1;HCoV-OC43_envelope | 65 | 70 | 6 | 0.05 | 0.1 | 0.083333333 |
| NC_006213.1;YP_009555244.1;HCoV-OC43_membrane | 1 | 7 | 7 | 0.05 | 0.05 | 0.05 |
| NC_006213.1;YP_009555244.1;HCoV-OC43_membrane | 9 | 13 | 5 | 0.1 | 0.15 | 0.13 |
| NC_006213.1;YP_009555244.1;HCoV-OC43_membrane | 22 | 22 | 1 | 0.05 | 0.05 | 0.05 |
| NC_006213.1;YP_009555244.1;HCoV-OC43_membrane | 115 | 120 | 6 | 0.05 | 0.1 | 0.058333333 |
| NC_006213.1;YP_009555244.1;HCoV-OC43_membrane | 188 | 188 | 1 | 0.05 | 0.05 | 0.05 |
| NC_006213.1;YP_009555244.1;HCoV-OC43_membrane | 202 | 202 | 1 | 0.05 | 0.05 | 0.05 |
| NC_006213.1;YP_009555244.1;HCoV-OC43_membrane | 207 | 207 | 1 | 0.05 | 0.05 | 0.05 |
| NC_006213.1;YP_009555244.1;HCoV-OC43_membrane | 210 | 217 | 8 | 0.1 | 0.2 | 0.15 |
| NC_006213.1;YP_009555245.1;HCoV-OC43_nucleoca | 1 | 7 | 7 | 0.05 | 0.05 | 0.05 |
| NC_006213.1;YP_009555245.1;HCoV-OC43_nucleoca | 13 | 13 | 1 | 0.05 | 0.05 | 0.05 |
| NC_006213.1;YP_009555245.1;HCoV-OC43_nucleoca | 19 | 25 | 7 | 0.05 | 0.35 | 0.192857143 |
| NC_006213.1;YP_009555245.1;HCoV-OC43_nucleoca | 35 | 46 | 12 | 0.05 | 0.15 | 0.079166667 |
| NC_006213.1;YP_009555245.1;HCoV-OC43_nucleoca | 68 | 68 | 1 | 0.1 | 0.1 | 0.1 |
| NC_006213.1;YP_009555245.1;HCoV-OC43_nucleoca | 72 | 72 | 1 | 0.1 | 0.1 | 0.1 |
| NC_006213.1;YP_009555245.1;HCoV-OC43_nucleoca | 75 | 87 | 13 | 0.05 | 0.15 | 0.092307692 |
| NC_006213.1;YP_009555245.1;HCoV-OC43_nucleoca | 125 | 128 | 4 | 0.05 | 0.1 | 0.0625 |
| NC_006213.1;YP_009555245.1;HCoV-OC43_nucleoca | 139 | 139 | 1 | 0.05 | 0.05 | 0.05 |
| NC_006213.1;YP_009555245.1;HCoV-OC43_nucleoca | 141 | 141 | 1 | 0.05 | 0.05 | 0.05 |
| NC_006213.1;YP_009555245.1;HCoV-OC43_nucleoca | 149 | 149 | 1 | 0.05 | 0.05 | 0.05 |
| NC_006213.1;YP_009555245.1;HCoV-OC43_nucleoca | 151 | 151 | 1 | 0.05 | 0.05 | 0.05 |
| NC_006213.1;YP_009555245.1;HCoV-OC43_nucleoca | 157 | 157 | 1 | 0.05 | 0.05 | 0.05 |
| NC_006213.1;YP_009555245.1;HCoV-OC43_nucleoca | 160 | 160 | 1 | 0.05 | 0.05 | 0.05 |
| NC_006213.1;YP_009555245.1;HCoV-OC43_nucleoca | 165 | 165 | 1 | 0.05 | 0.05 | 0.05 |
| NC_006213.1;YP_009555245.1;HCoV-OC43_nucleoca | 170 | 170 | 1 | 0.05 | 0.05 | 0.05 |
| NC_006213.1;YP_009555245.1;HCoV-OC43_nucleoca | 181 | 181 | 1 | 0.05 | 0.05 | 0.05 |
| NC_006213.1;YP_009555245.1;HCoV-OC43_nucleoca | 187 | 187 | 1 | 0.05 | 0.05 | 0.05 |
| NC_006213.1;YP_009555245.1;HCoV-OC43_nucleoca | 195 | 195 | 1 | 0.05 | 0.05 | 0.05 |
| NC_006213.1;YP_009555245.1;HCoV-OC43_nucleoca | 200 | 200 | 1 | 0.05 | 0.05 | 0.05 |
| NC_006213.1;YP_009555245.1;HCoV-OC43_nucleoca | 203 | 206 | 4 | 0.05 | 0.05 | 0.05 |
| NC_006213.1;YP_009555245.1;HCoV-OC43_nucleoca | 208 | 210 | 3 | 0.05 | 0.1 | 0.066666667 |
| NC_006213.1;YP_009555245.1;HCoV-OC43_nucleoca | 212 | 213 | 2 | 0.05 | 0.1 | 0.075 |
| NC_006213.1;YP_009555245.1;HCoV-OC43_nucleoca | 216 | 218 | 3 | 0.05 | 0.1 | 0.083333333 |
| NC_006213.1;YP_009555245.1;HCoV-OC43_nucleoca | 237 | 241 | 5 | 0.05 | 0.1 | 0.08 |
| NC_006213.1;YP_009555245.1;HCoV-OC43_nucleoca | 243 | 244 | 2 | 0.05 | 0.05 | 0.05 |
| NC_006213.1;YP_009555245.1;HCoV-OC43_nucleoca | 246 | 248 | 3 | 0.05 | 0.35 | 0.15 |
| NC_006213.1;YP_009555245.1;HCoV-OC43_nucleoca | 253 | 264 | 12 | 0.05 | 0.05 | 0.05 |

| Extended data 1 | | | | | | |
|---|---|---|---|---|---|---|
| **Protein** | **Start position (peptide)** | **Stop position (peptide)** | **Number of peptides** | **Minimum fraction (per peptide) of control subjects with reactivity >3 sd above average** | **Maximum fraction (per peptide) of control subjects with reactivity >3 sd above average** | **Mean fraction (over all peptides in this contiguous stretch) of controlsubjects with reactivity >3 sd above average** |
| NC_006213.1;YP_009555245.1;HCoV-OC43_nucleoca | 270 | 270 | 1 | 0.05 | 0.05 | 0.05 |
| NC_006213.1;YP_009555245.1;HCoV-OC43_nucleoca | 275 | 275 | 1 | 0.05 | 0.05 | 0.05 |
| NC_006213.1;YP_009555245.1;HCoV-OC43_nucleoca | 282 | 286 | 5 | 0.05 | 0.35 | 0.17 |
| NC_006213.1;YP_009555245.1;HCoV-OC43_nucleoca | 289 | 289 | 1 | 0.05 | 0.05 | 0.05 |
| NC_006213.1;YP_009555245.1;HCoV-OC43_nucleoca | 303 | 308 | 6 | 0.05 | 0.1 | 0.058333333 |
| NC_006213.1;YP_009555245.1;HCoV-OC43_nucleoca | 310 | 311 | 2 | 0.05 | 0.05 | 0.05 |
| NC_006213.1;YP_009555245.1;HCoV-OC43_nucleoca | 328 | 341 | 14 | 0.05 | 0.2 | 0.128571429 |
| NC_006213.1;YP_009555245.1;HCoV-OC43_nucleoca | 350 | 350 | 1 | 0.05 | 0.05 | 0.05 |
| NC_006213.1;YP_009555245.1;HCoV-OC43_nucleoca | 352 | 353 | 2 | 0.05 | 0.05 | 0.05 |
| NC_006213.1;YP_009555245.1;HCoV-OC43_nucleoca | 355 | 358 | 4 | 0.05 | 0.05 | 0.05 |
| NC_006213.1;YP_009555245.1;HCoV-OC43_nucleoca | 363 | 363 | 1 | 0.05 | 0.05 | 0.05 |
| NC_006213.1;YP_009555245.1;HCoV-OC43_nucleoca | 365 | 374 | 10 | 0.05 | 0.25 | 0.105 |
| NC_006213.1;YP_009555245.1;HCoV-OC43_nucleoca | 393 | 393 | 1 | 0.05 | 0.05 | 0.05 |
| NC_006213.1;YP_009555245.1;HCoV-OC43_nucleoca | 396 | 397 | 2 | 0.05 | 0.1 | 0.075 |
| NC_006213.1;YP_009555245.1;HCoV-OC43_nucleoca | 399 | 405 | 7 | 0.05 | 0.1 | 0.085714286 |
| NC_006213.1;YP_009555245.1;HCoV-OC43_nucleoca | 410 | 416 | 7 | 0.05 | 0.1 | 0.057142857 |
| NC_006213.1;YP_009555245.1;HCoV-OC43_nucleoca | 420 | 427 | 8 | 0.05 | 0.15 | 0.075 |
| NC_006213.1;YP_009555245.1;HCoV-OC43_nucleoca | 429 | 429 | 1 | 0.05 | 0.05 | 0.05 |
| NC_006577.2;YP_173236.1;HCoV-HKU1_orf1ab | 3 | 3 | 1 | 0.05 | 0.05 | 0.05 |
| NC_006577.2;YP_173236.1;HCoV-HKU1_orf1ab | 18 | 19 | 2 | 0.05 | 0.05 | 0.05 |
| NC_006577.2;YP_173236.1;HCoV-HKU1_orf1ab | 26 | 36 | 11 | 0.05 | 0.1 | 0.054545455 |
| NC_006577.2;YP_173236.1;HCoV-HKU1_orf1ab | 47 | 47 | 1 | 0.05 | 0.05 | 0.05 |
| NC_006577.2;YP_173236.1;HCoV-HKU1_orf1ab | 71 | 71 | 1 | 0.05 | 0.05 | 0.05 |
| NC_006577.2;YP_173236.1;HCoV-HKU1_orf1ab | 83 | 87 | 5 | 0.05 | 0.1 | 0.09 |
| NC_006577.2;YP_173236.1;HCoV-HKU1_orf1ab | 89 | 91 | 3 | 0.05 | 0.05 | 0.05 |
| NC_006577.2;YP_173236.1;HCoV-HKU1_orf1ab | 93 | 94 | 2 | 0.1 | 0.1 | 0.1 |
| NC_006577.2;YP_173236.1;HCoV-HKU1_orf1ab | 96 | 98 | 3 | 0.1 | 0.1 | 0.1 |
| NC_006577.2;YP_173236.1;HCoV-HKU1_orf1ab | 182 | 183 | 2 | 0.05 | 0.1 | 0.075 |
| NC_006577.2;YP_173236.1;HCoV-HKU1_orf1ab | 191 | 191 | 1 | 0.05 | 0.05 | 0.05 |
| NC_006577.2;YP_173236.1;HCoV-HKU1_orf1ab | 193 | 194 | 2 | 0.05 | 0.05 | 0.05 |
| NC_006577.2;YP_173236.1;HCoV-HKU1_orf1ab | 227 | 227 | 1 | 0.05 | 0.05 | 0.05 |
| NC_006577.2;YP_173236.1;HCoV-HKU1_orf1ab | 233 | 233 | 1 | 0.05 | 0.05 | 0.05 |
| NC_006577.2;YP_173236.1;HCoV-HKU1_orf1ab | 235 | 237 | 3 | 0.05 | 0.05 | 0.05 |
| NC_006577.2;YP_173236.1;HCoV-HKU1_orf1ab | 252 | 256 | 5 | 0.05 | 0.05 | 0.05 |
| NC_006577.2;YP_173236.1;HCoV-HKU1_orf1ab | 294 | 294 | 1 | 0.1 | 0.1 | 0.1 |
| NC_006577.2;YP_173236.1;HCoV-HKU1_orf1ab | 300 | 300 | 1 | 0.05 | 0.05 | 0.05 |
| NC_006577.2;YP_173236.1;HCoV-HKU1_orf1ab | 303 | 303 | 1 | 0.05 | 0.05 | 0.05 |

| Extended data 1 | | | | | | |
|---|---|---|---|---|---|---|
| Protein | Start position (peptide) | Stop position (peptide) | Number of peptides | Minimum fraction (per peptide) of control subjects with reactivity >3 sd above average | Maximum fraction (per peptide) of control subjects with reactivity >3 sd above average | Mean fraction (over all peptides in this contiguous stretch) of controlsubjects with reactivity >3 sd above average |
| NC_006577.2;YP_173236.1;HCoV-HKU1_orf1ab | 327 | 327 | 1 | 0.05 | 0.05 | 0.05 |
| NC_006577.2;YP_173236.1;HCoV-HKU1_orf1ab | 356 | 356 | 1 | 0.05 | 0.05 | 0.05 |
| NC_006577.2;YP_173236.1;HCoV-HKU1_orf1ab | 373 | 374 | 2 | 0.05 | 0.05 | 0.05 |
| NC_006577.2;YP_173236.1;HCoV-HKU1_orf1ab | 393 | 400 | 8 | 0.05 | 0.05 | 0.05 |
| NC_006577.2;YP_173236.1;HCoV-HKU1_orf1ab | 402 | 402 | 1 | 0.05 | 0.05 | 0.05 |
| NC_006577.2;YP_173236.1;HCoV-HKU1_orf1ab | 404 | 404 | 1 | 0.05 | 0.05 | 0.05 |
| NC_006577.2;YP_173236.1;HCoV-HKU1_orf1ab | 409 | 409 | 1 | 0.05 | 0.05 | 0.05 |
| NC_006577.2;YP_173236.1;HCoV-HKU1_orf1ab | 411 | 416 | 6 | 0.05 | 0.05 | 0.05 |
| NC_006577.2;YP_173236.1;HCoV-HKU1_orf1ab | 423 | 423 | 1 | 0.1 | 0.1 | 0.1 |
| NC_006577.2;YP_173236.1;HCoV-HKU1_orf1ab | 488 | 488 | 1 | 0.05 | 0.05 | 0.05 |
| NC_006577.2;YP_173236.1;HCoV-HKU1_orf1ab | 491 | 492 | 2 | 0.05 | 0.1 | 0.075 |
| NC_006577.2;YP_173236.1;HCoV-HKU1_orf1ab | 512 | 512 | 1 | 0.05 | 0.05 | 0.05 |
| NC_006577.2;YP_173236.1;HCoV-HKU1_orf1ab | 516 | 524 | 9 | 0.05 | 0.05 | 0.05 |
| NC_006577.2;YP_173236.1;HCoV-HKU1_orf1ab | 527 | 527 | 1 | 0.05 | 0.05 | 0.05 |
| NC_006577.2;YP_173236.1;HCoV-HKU1_orf1ab | 534 | 534 | 1 | 0.05 | 0.05 | 0.05 |
| NC_006577.2;YP_173236.1;HCoV-HKU1_orf1ab | 551 | 551 | 1 | 0.05 | 0.05 | 0.05 |
| NC_006577.2;YP_173236.1;HCoV-HKU1_orf1ab | 570 | 570 | 1 | 0.05 | 0.05 | 0.05 |
| NC_006577.2;YP_173236.1;HCoV-HKU1_orf1ab | 619 | 619 | 1 | 0.05 | 0.05 | 0.05 |
| NC_006577.2;YP_173236.1;HCoV-HKU1_orf1ab | 621 | 625 | 5 | 0.1 | 0.15 | 0.13 |
| NC_006577.2;YP_173236.1;HCoV-HKU1_orf1ab | 652 | 653 | 2 | 0.05 | 0.1 | 0.075 |
| NC_006577.2;YP_173236.1;HCoV-HKU1_orf1ab | 743 | 743 | 1 | 0.05 | 0.05 | 0.05 |
| NC_006577.2;YP_173236.1;HCoV-HKU1_orf1ab | 746 | 746 | 1 | 0.1 | 0.1 | 0.1 |
| NC_006577.2;YP_173236.1;HCoV-HKU1_orf1ab | 749 | 750 | 2 | 0.05 | 0.05 | 0.05 |
| NC_006577.2;YP_173236.1;HCoV-HKU1_orf1ab | 754 | 754 | 1 | 0.1 | 0.1 | 0.1 |
| NC_006577.2;YP_173236.1;HCoV-HKU1_orf1ab | 757 | 757 | 1 | 0.1 | 0.1 | 0.1 |
| NC_006577.2;YP_173236.1;HCoV-HKU1_orf1ab | 764 | 764 | 1 | 0.05 | 0.05 | 0.05 |
| NC_006577.2;YP_173236.1;HCoV-HKU1_orf1ab | 768 | 768 | 1 | 0.05 | 0.05 | 0.05 |
| NC_006577.2;YP_173236.1;HCoV-HKU1_orf1ab | 773 | 775 | 3 | 0.05 | 0.05 | 0.05 |
| NC_006577.2;YP_173236.1;HCoV-HKU1_orf1ab | 778 | 778 | 1 | 0.05 | 0.05 | 0.05 |
| NC_006577.2;YP_173236.1;HCoV-HKU1_orf1ab | 782 | 784 | 3 | 0.05 | 0.05 | 0.05 |
| NC_006577.2;YP_173236.1;HCoV-HKU1_orf1ab | 789 | 789 | 1 | 0.05 | 0.05 | 0.05 |
| NC_006577.2;YP_173236.1;HCoV-HKU1_orf1ab | 799 | 799 | 1 | 0.05 | 0.05 | 0.05 |
| NC_006577.2;YP_173236.1;HCoV-HKU1_orf1ab | 808 | 808 | 1 | 0.1 | 0.1 | 0.1 |
| NC_006577.2;YP_173236.1;HCoV-HKU1_orf1ab | 812 | 812 | 1 | 0.1 | 0.1 | 0.1 |
| NC_006577.2;YP_173236.1;HCoV-HKU1_orf1ab | 814 | 814 | 1 | 0.05 | 0.05 | 0.05 |
| NC_006577.2;YP_173236.1;HCoV-HKU1_orf1ab | 830 | 830 | 1 | 0.05 | 0.05 | 0.05 |
| NC_006577.2;YP_173236.1;HCoV-HKU1_orf1ab | 837 | 838 | 2 | 0.05 | 0.05 | 0.05 |

| Extended data 1 | | | | | | |
|---|---|---|---|---|---|---|
| Protein | Start position (peptide) | Stop position (peptide) | Number of peptides | Minimum fraction (per peptide) of control subjects with reactivity >3 sd above average | Maximum fraction (per peptide) of control subjects with reactivity >3 sd above average | Mean fraction (over all peptides in this contiguous stretch) of controlsubjects with reactivity >3 sd above average |
| NC_006577.2;YP_173236.1;HCoV-HKU1_orf1ab | 842 | 842 | 1 | 0.05 | 0.05 | 0.05 |
| NC_006577.2;YP_173236.1;HCoV-HKU1_orf1ab | 864 | 866 | 3 | 0.05 | 0.05 | 0.05 |
| NC_006577.2;YP_173236.1;HCoV-HKU1_orf1ab | 868 | 869 | 2 | 0.05 | 0.1 | 0.075 |
| NC_006577.2;YP_173236.1;HCoV-HKU1_orf1ab | 872 | 874 | 3 | 0.05 | 0.05 | 0.05 |
| NC_006577.2;YP_173236.1;HCoV-HKU1_orf1ab | 884 | 884 | 1 | 0.05 | 0.05 | 0.05 |
| NC_006577.2;YP_173236.1;HCoV-HKU1_orf1ab | 886 | 886 | 1 | 0.05 | 0.05 | 0.05 |
| NC_006577.2;YP_173236.1;HCoV-HKU1_orf1ab | 889 | 889 | 1 | 0.05 | 0.05 | 0.05 |
| NC_006577.2;YP_173236.1;HCoV-HKU1_orf1ab | 893 | 894 | 2 | 0.05 | 0.1 | 0.075 |
| NC_006577.2;YP_173236.1;HCoV-HKU1_orf1ab | 896 | 900 | 5 | 0.05 | 0.05 | 0.05 |
| NC_006577.2;YP_173236.1;HCoV-HKU1_orf1ab | 919 | 919 | 1 | 0.1 | 0.1 | 0.1 |
| NC_006577.2;YP_173236.1;HCoV-HKU1_orf1ab | 923 | 924 | 2 | 0.05 | 0.05 | 0.05 |
| NC_006577.2;YP_173236.1;HCoV-HKU1_orf1ab | 930 | 935 | 6 | 0.05 | 0.2 | 0.108333333 |
| NC_006577.2;YP_173236.1;HCoV-HKU1_orf1ab | 937 | 1091 | 155 | 0.05 | 0.25 | 0.087419355 |
| NC_006577.2;YP_173236.1;HCoV-HKU1_orf1ab | 1095 | 1095 | 1 | 0.05 | 0.05 | 0.05 |
| NC_006577.2;YP_173236.1;HCoV-HKU1_orf1ab | 1101 | 1112 | 12 | 0.05 | 0.25 | 0.095833333 |
| NC_006577.2;YP_173236.1;HCoV-HKU1_orf1ab | 1132 | 1132 | 1 | 0.05 | 0.05 | 0.05 |
| NC_006577.2;YP_173236.1;HCoV-HKU1_orf1ab | 1134 | 1135 | 2 | 0.05 | 0.05 | 0.05 |
| NC_006577.2;YP_173236.1;HCoV-HKU1_orf1ab | 1137 | 1137 | 1 | 0.05 | 0.05 | 0.05 |
| NC_006577.2;YP_173236.1;HCoV-HKU1_orf1ab | 1140 | 1140 | 1 | 0.05 | 0.05 | 0.05 |
| NC_006577.2;YP_173236.1;HCoV-HKU1_orf1ab | 1170 | 1170 | 1 | 0.1 | 0.1 | 0.1 |
| NC_006577.2;YP_173236.1;HCoV-HKU1_orf1ab | 1182 | 1182 | 1 | 0.05 | 0.05 | 0.05 |
| NC_006577.2;YP_173236.1;HCoV-HKU1_orf1ab | 1234 | 1234 | 1 | 0.05 | 0.05 | 0.05 |
| NC_006577.2;YP_173236.1;HCoV-HKU1_orf1ab | 1260 | 1260 | 1 | 0.05 | 0.05 | 0.05 |
| NC_006577.2;YP_173236.1;HCoV-HKU1_orf1ab | 1295 | 1295 | 1 | 0.05 | 0.05 | 0.05 |
| NC_006577.2;YP_173236.1;HCoV-HKU1_orf1ab | 1300 | 1300 | 1 | 0.05 | 0.05 | 0.05 |
| NC_006577.2;YP_173236.1;HCoV-HKU1_orf1ab | 1309 | 1311 | 3 | 0.05 | 0.05 | 0.05 |
| NC_006577.2;YP_173236.1;HCoV-HKU1_orf1ab | 1319 | 1329 | 11 | 0.05 | 0.15 | 0.095454545 |
| NC_006577.2;YP_173236.1;HCoV-HKU1_orf1ab | 1333 | 1333 | 1 | 0.05 | 0.05 | 0.05 |
| NC_006577.2;YP_173236.1;HCoV-HKU1_orf1ab | 1346 | 1346 | 1 | 0.05 | 0.05 | 0.05 |
| NC_006577.2;YP_173236.1;HCoV-HKU1_orf1ab | 1379 | 1379 | 1 | 0.05 | 0.05 | 0.05 |
| NC_006577.2;YP_173236.1;HCoV-HKU1_orf1ab | 1381 | 1381 | 1 | 0.05 | 0.05 | 0.05 |
| NC_006577.2;YP_173236.1;HCoV-HKU1_orf1ab | 1383 | 1385 | 3 | 0.05 | 0.05 | 0.05 |
| NC_006577.2;YP_173236.1;HCoV-HKU1_orf1ab | 1387 | 1388 | 2 | 0.05 | 0.05 | 0.05 |
| NC_006577.2;YP_173236.1;HCoV-HKU1_orf1ab | 1390 | 1391 | 2 | 0.05 | 0.05 | 0.05 |
| NC_006577.2;YP_173236.1;HCoV-HKU1_orf1ab | 1419 | 1419 | 1 | 0.05 | 0.05 | 0.05 |
| NC_006577.2;YP_173236.1;HCoV-HKU1_orf1ab | 1440 | 1443 | 4 | 0.05 | 0.05 | 0.05 |
| NC_006577.2;YP_173236.1;HCoV-HKU1_orf1ab | 1458 | 1458 | 1 | 0.1 | 0.1 | 0.1 |

| Extended data 1 | | | | | | |
|---|---|---|---|---|---|---|
| Protein | Start position (peptide) | Stop position (peptide) | Number of peptides | Minimum fraction (per peptide) of control subjects with reactivity >3 sd above average | Maximum fraction (per peptide) of control subjects with reactivity >3 sd above average | Mean fraction (over all peptides in this contiguous stretch) of controlsubjects with reactivity >3 sd above average |
| NC_006577.2;YP_173236.1;HCoV-HKU1_orf1ab | 1484 | 1484 | 1 | 0.05 | 0.05 | 0.05 |
| NC_006577.2;YP_173236.1;HCoV-HKU1_orf1ab | 1494 | 1497 | 4 | 0.05 | 0.05 | 0.05 |
| NC_006577.2;YP_173236.1;HCoV-HKU1_orf1ab | 1500 | 1504 | 5 | 0.05 | 0.6 | 0.22 |
| NC_006577.2;YP_173236.1;HCoV-HKU1_orf1ab | 1510 | 1510 | 1 | 0.05 | 0.05 | 0.05 |
| NC_006577.2;YP_173236.1;HCoV-HKU1_orf1ab | 1533 | 1534 | 2 | 0.05 | 0.05 | 0.05 |
| NC_006577.2;YP_173236.1;HCoV-HKU1_orf1ab | 1550 | 1553 | 4 | 0.1 | 0.2 | 0.1375 |
| NC_006577.2;YP_173236.1;HCoV-HKU1_orf1ab | 1559 | 1560 | 2 | 0.05 | 0.05 | 0.05 |
| NC_006577.2;YP_173236.1;HCoV-HKU1_orf1ab | 1566 | 1566 | 1 | 0.05 | 0.05 | 0.05 |
| NC_006577.2;YP_173236.1;HCoV-HKU1_orf1ab | 1569 | 1569 | 1 | 0.05 | 0.05 | 0.05 |
| NC_006577.2;YP_173236.1;HCoV-HKU1_orf1ab | 1575 | 1581 | 7 | 0.05 | 0.15 | 0.064285714 |
| NC_006577.2;YP_173236.1;HCoV-HKU1_orf1ab | 1583 | 1583 | 1 | 0.05 | 0.05 | 0.05 |
| NC_006577.2;YP_173236.1;HCoV-HKU1_orf1ab | 1588 | 1592 | 5 | 0.05 | 0.1 | 0.09 |
| NC_006577.2;YP_173236.1;HCoV-HKU1_orf1ab | 1594 | 1594 | 1 | 0.05 | 0.05 | 0.05 |
| NC_006577.2;YP_173236.1;HCoV-HKU1_orf1ab | 1597 | 1597 | 1 | 0.05 | 0.05 | 0.05 |
| NC_006577.2;YP_173236.1;HCoV-HKU1_orf1ab | 1624 | 1624 | 1 | 0.05 | 0.05 | 0.05 |
| NC_006577.2;YP_173236.1;HCoV-HKU1_orf1ab | 1638 | 1638 | 1 | 0.05 | 0.05 | 0.05 |
| NC_006577.2;YP_173236.1;HCoV-HKU1_orf1ab | 1648 | 1648 | 1 | 0.05 | 0.05 | 0.05 |
| NC_006577.2;YP_173236.1;HCoV-HKU1_orf1ab | 1650 | 1652 | 3 | 0.05 | 0.05 | 0.05 |
| NC_006577.2;YP_173236.1;HCoV-HKU1_orf1ab | 1677 | 1679 | 3 | 0.05 | 0.05 | 0.05 |
| NC_006577.2;YP_173236.1;HCoV-HKU1_orf1ab | 1685 | 1686 | 2 | 0.05 | 0.05 | 0.05 |
| NC_006577.2;YP_173236.1;HCoV-HKU1_orf1ab | 1697 | 1698 | 2 | 0.05 | 0.1 | 0.075 |
| NC_006577.2;YP_173236.1;HCoV-HKU1_orf1ab | 1700 | 1700 | 1 | 0.05 | 0.05 | 0.05 |
| NC_006577.2;YP_173236.1;HCoV-HKU1_orf1ab | 1702 | 1703 | 2 | 0.05 | 0.05 | 0.05 |
| NC_006577.2;YP_173236.1;HCoV-HKU1_orf1ab | 1709 | 1709 | 1 | 0.05 | 0.05 | 0.05 |
| NC_006577.2;YP_173236.1;HCoV-HKU1_orf1ab | 1715 | 1715 | 1 | 0.1 | 0.1 | 0.1 |
| NC_006577.2;YP_173236.1;HCoV-HKU1_orf1ab | 1742 | 1743 | 2 | 0.05 | 0.05 | 0.05 |
| NC_006577.2;YP_173236.1;HCoV-HKU1_orf1ab | 1747 | 1748 | 2 | 0.05 | 0.15 | 0.1 |
| NC_006577.2;YP_173236.1;HCoV-HKU1_orf1ab | 1751 | 1751 | 1 | 0.05 | 0.05 | 0.05 |
| NC_006577.2;YP_173236.1;HCoV-HKU1_orf1ab | 1772 | 1772 | 1 | 0.05 | 0.05 | 0.05 |
| NC_006577.2;YP_173236.1;HCoV-HKU1_orf1ab | 1795 | 1800 | 6 | 0.05 | 0.3 | 0.166666667 |
| NC_006577.2;YP_173236.1;HCoV-HKU1_orf1ab | 1830 | 1840 | 11 | 0.05 | 0.1 | 0.054545455 |
| NC_006577.2;YP_173236.1;HCoV-HKU1_orf1ab | 1842 | 1842 | 1 | 0.05 | 0.05 | 0.05 |
| NC_006577.2;YP_173236.1;HCoV-HKU1_orf1ab | 1855 | 1855 | 1 | 0.05 | 0.05 | 0.05 |
| NC_006577.2;YP_173236.1;HCoV-HKU1_orf1ab | 1884 | 1895 | 12 | 0.05 | 0.2 | 0.104166667 |
| NC_006577.2;YP_173236.1;HCoV-HKU1_orf1ab | 1962 | 1962 | 1 | 0.05 | 0.05 | 0.05 |
| NC_006577.2;YP_173236.1;HCoV-HKU1_orf1ab | 1968 | 1968 | 1 | 0.05 | 0.05 | 0.05 |
| NC_006577.2;YP_173236.1;HCoV-HKU1_orf1ab | 1970 | 1972 | 3 | 0.05 | 0.1 | 0.066666667 |

| Extended data 1 | | | | | | |
|---|---|---|---|---|---|---|
| Protein | Start position (peptide) | Stop position (peptide) | Number of peptides | Minimum fraction (per peptide) of control subjects with reactivity >3 sd above average | Maximum fraction (per peptide) of control subjects with reactivity >3 sd above average | Mean fraction (over all peptides in this contiguous stretch) of controlsubjects with reactivity >3 sd above average |
| NC_006577.2;YP_173236.1;HCoV-HKU1_orf1ab | 1975 | 1975 | 1 | 0.1 | 0.1 | 0.1 |
| NC_006577.2;YP_173236.1;HCoV-HKU1_orf1ab | 1982 | 1982 | 1 | 0.15 | 0.15 | 0.15 |
| NC_006577.2;YP_173236.1;HCoV-HKU1_orf1ab | 1988 | 1988 | 1 | 0.05 | 0.05 | 0.05 |
| NC_006577.2;YP_173236.1;HCoV-HKU1_orf1ab | 1990 | 1990 | 1 | 0.05 | 0.05 | 0.05 |
| NC_006577.2;YP_173236.1;HCoV-HKU1_orf1ab | 2000 | 2007 | 8 | 0.05 | 0.05 | 0.05 |
| NC_006577.2;YP_173236.1;HCoV-HKU1_orf1ab | 2018 | 2018 | 1 | 0.05 | 0.05 | 0.05 |
| NC_006577.2;YP_173236.1;HCoV-HKU1_orf1ab | 2038 | 2038 | 1 | 0.05 | 0.05 | 0.05 |
| NC_006577.2;YP_173236.1;HCoV-HKU1_orf1ab | 2043 | 2043 | 1 | 0.05 | 0.05 | 0.05 |
| NC_006577.2;YP_173236.1;HCoV-HKU1_orf1ab | 2050 | 2050 | 1 | 0.05 | 0.05 | 0.05 |
| NC_006577.2;YP_173236.1;HCoV-HKU1_orf1ab | 2084 | 2084 | 1 | 0.05 | 0.05 | 0.05 |
| NC_006577.2;YP_173236.1;HCoV-HKU1_orf1ab | 2086 | 2086 | 1 | 0.05 | 0.05 | 0.05 |
| NC_006577.2;YP_173236.1;HCoV-HKU1_orf1ab | 2088 | 2102 | 15 | 0.05 | 0.25 | 0.126666667 |
| NC_006577.2;YP_173236.1;HCoV-HKU1_orf1ab | 2105 | 2106 | 2 | 0.05 | 0.05 | 0.05 |
| NC_006577.2;YP_173236.1;HCoV-HKU1_orf1ab | 2113 | 2114 | 2 | 0.05 | 0.05 | 0.05 |
| NC_006577.2;YP_173236.1;HCoV-HKU1_orf1ab | 2136 | 2140 | 5 | 0.05 | 0.1 | 0.07 |
| NC_006577.2;YP_173236.1;HCoV-HKU1_orf1ab | 2211 | 2211 | 1 | 0.05 | 0.05 | 0.05 |
| NC_006577.2;YP_173236.1;HCoV-HKU1_orf1ab | 2360 | 2360 | 1 | 0.05 | 0.05 | 0.05 |
| NC_006577.2;YP_173236.1;HCoV-HKU1_orf1ab | 2502 | 2502 | 1 | 0.05 | 0.05 | 0.05 |
| NC_006577.2;YP_173236.1;HCoV-HKU1_orf1ab | 2505 | 2505 | 1 | 0.1 | 0.1 | 0.1 |
| NC_006577.2;YP_173236.1;HCoV-HKU1_orf1ab | 2518 | 2518 | 1 | 0.05 | 0.05 | 0.05 |
| NC_006577.2;YP_173236.1;HCoV-HKU1_orf1ab | 2536 | 2537 | 2 | 0.05 | 0.05 | 0.05 |
| NC_006577.2;YP_173236.1;HCoV-HKU1_orf1ab | 2539 | 2542 | 4 | 0.05 | 0.05 | 0.05 |
| NC_006577.2;YP_173236.1;HCoV-HKU1_orf1ab | 2549 | 2549 | 1 | 0.05 | 0.05 | 0.05 |
| NC_006577.2;YP_173236.1;HCoV-HKU1_orf1ab | 2551 | 2551 | 1 | 0.05 | 0.05 | 0.05 |
| NC_006577.2;YP_173236.1;HCoV-HKU1_orf1ab | 2553 | 2553 | 1 | 0.05 | 0.05 | 0.05 |
| NC_006577.2;YP_173236.1;HCoV-HKU1_orf1ab | 2567 | 2567 | 1 | 0.05 | 0.05 | 0.05 |
| NC_006577.2;YP_173236.1;HCoV-HKU1_orf1ab | 2575 | 2579 | 5 | 0.05 | 0.05 | 0.05 |
| NC_006577.2;YP_173236.1;HCoV-HKU1_orf1ab | 2583 | 2583 | 1 | 0.05 | 0.05 | 0.05 |
| NC_006577.2;YP_173236.1;HCoV-HKU1_orf1ab | 2599 | 2599 | 1 | 0.1 | 0.1 | 0.1 |
| NC_006577.2;YP_173236.1;HCoV-HKU1_orf1ab | 2601 | 2601 | 1 | 0.05 | 0.05 | 0.05 |
| NC_006577.2;YP_173236.1;HCoV-HKU1_orf1ab | 2605 | 2605 | 1 | 0.05 | 0.05 | 0.05 |
| NC_006577.2;YP_173236.1;HCoV-HKU1_orf1ab | 2617 | 2617 | 1 | 0.05 | 0.05 | 0.05 |
| NC_006577.2;YP_173236.1;HCoV-HKU1_orf1ab | 2644 | 2644 | 1 | 0.05 | 0.05 | 0.05 |
| NC_006577.2;YP_173236.1;HCoV-HKU1_orf1ab | 2670 | 2670 | 1 | 0.05 | 0.05 | 0.05 |
| NC_006577.2;YP_173236.1;HCoV-HKU1_orf1ab | 2672 | 2673 | 2 | 0.05 | 0.05 | 0.05 |
| NC_006577.2;YP_173236.1;HCoV-HKU1_orf1ab | 2721 | 2721 | 1 | 0.05 | 0.05 | 0.05 |
| NC_006577.2;YP_173236.1;HCoV-HKU1_orf1ab | 2733 | 2733 | 1 | 0.05 | 0.05 | 0.05 |

| Extended data 1 | | | | | | |
|---|---|---|---|---|---|---|
| Protein | Start position (peptide) | Stop position (peptide) | Number of peptides | Minimum fraction (per peptide) of control subjects with reactivity >3 sd above average | Maximum fraction (per peptide) of control subjects with reactivity >3 sd above average | Mean fraction (over all peptides in this contiguous stretch) of controlsubjects with reactivity >3 sd above average |
| NC_006577.2;YP_173236.1;HCoV-HKU1_orf1ab | 2736 | 2736 | 1 | 0.05 | 0.05 | 0.05 |
| NC_006577.2;YP_173236.1;HCoV-HKU1_orf1ab | 2738 | 2739 | 2 | 0.05 | 0.05 | 0.05 |
| NC_006577.2;YP_173236.1;HCoV-HKU1_orf1ab | 2742 | 2743 | 2 | 0.1 | 0.1 | 0.1 |
| NC_006577.2;YP_173236.1;HCoV-HKU1_orf1ab | 2745 | 2746 | 2 | 0.05 | 0.1 | 0.075 |
| NC_006577.2;YP_173236.1;HCoV-HKU1_orf1ab | 2750 | 2750 | 1 | 0.05 | 0.05 | 0.05 |
| NC_006577.2;YP_173236.1;HCoV-HKU1_orf1ab | 2761 | 2761 | 1 | 0.05 | 0.05 | 0.05 |
| NC_006577.2;YP_173236.1;HCoV-HKU1_orf1ab | 2813 | 2821 | 9 | 0.05 | 0.05 | 0.05 |
| NC_006577.2;YP_173236.1;HCoV-HKU1_orf1ab | 2879 | 2879 | 1 | 0.05 | 0.05 | 0.05 |
| NC_006577.2;YP_173236.1;HCoV-HKU1_orf1ab | 2887 | 2887 | 1 | 0.3 | 0.3 | 0.3 |
| NC_006577.2;YP_173236.1;HCoV-HKU1_orf1ab | 2892 | 2893 | 2 | 0.05 | 0.2 | 0.125 |
| NC_006577.2;YP_173236.1;HCoV-HKU1_orf1ab | 2932 | 2932 | 1 | 0.05 | 0.05 | 0.05 |
| NC_006577.2;YP_173236.1;HCoV-HKU1_orf1ab | 2989 | 2989 | 1 | 0.05 | 0.05 | 0.05 |
| NC_006577.2;YP_173236.1;HCoV-HKU1_orf1ab | 2996 | 2996 | 1 | 0.05 | 0.05 | 0.05 |
| NC_006577.2;YP_173236.1;HCoV-HKU1_orf1ab | 3001 | 3006 | 6 | 0.05 | 0.15 | 0.1 |
| NC_006577.2;YP_173236.1;HCoV-HKU1_orf1ab | 3008 | 3010 | 3 | 0.05 | 0.1 | 0.066666667 |
| NC_006577.2;YP_173236.1;HCoV-HKU1_orf1ab | 3029 | 3030 | 2 | 0.05 | 0.05 | 0.05 |
| NC_006577.2;YP_173236.1;HCoV-HKU1_orf1ab | 3231 | 3232 | 2 | 0.05 | 0.05 | 0.05 |
| NC_006577.2;YP_173236.1;HCoV-HKU1_orf1ab | 3256 | 3266 | 11 | 0.05 | 0.15 | 0.1 |
| NC_006577.2;YP_173236.1;HCoV-HKU1_orf1ab | 3268 | 3268 | 1 | 0.05 | 0.05 | 0.05 |
| NC_006577.2;YP_173236.1;HCoV-HKU1_orf1ab | 3280 | 3280 | 1 | 0.05 | 0.05 | 0.05 |
| NC_006577.2;YP_173236.1;HCoV-HKU1_orf1ab | 3284 | 3285 | 2 | 0.05 | 0.1 | 0.075 |
| NC_006577.2;YP_173236.1;HCoV-HKU1_orf1ab | 3290 | 3290 | 1 | 0.05 | 0.05 | 0.05 |
| NC_006577.2;YP_173236.1;HCoV-HKU1_orf1ab | 3296 | 3297 | 2 | 0.05 | 0.05 | 0.05 |
| NC_006577.2;YP_173236.1;HCoV-HKU1_orf1ab | 3299 | 3299 | 1 | 0.05 | 0.05 | 0.05 |
| NC_006577.2;YP_173236.1;HCoV-HKU1_orf1ab | 3301 | 3302 | 2 | 0.05 | 0.1 | 0.075 |
| NC_006577.2;YP_173236.1;HCoV-HKU1_orf1ab | 3309 | 3312 | 4 | 0.05 | 0.1 | 0.0625 |
| NC_006577.2;YP_173236.1;HCoV-HKU1_orf1ab | 3314 | 3314 | 1 | 0.05 | 0.05 | 0.05 |
| NC_006577.2;YP_173236.1;HCoV-HKU1_orf1ab | 3316 | 3317 | 2 | 0.05 | 0.05 | 0.05 |
| NC_006577.2;YP_173236.1;HCoV-HKU1_orf1ab | 3321 | 3321 | 1 | 0.1 | 0.1 | 0.1 |
| NC_006577.2;YP_173236.1;HCoV-HKU1_orf1ab | 3332 | 3339 | 8 | 0.05 | 0.1 | 0.0875 |
| NC_006577.2;YP_173236.1;HCoV-HKU1_orf1ab | 3342 | 3342 | 1 | 0.05 | 0.05 | 0.05 |
| NC_006577.2;YP_173236.1;HCoV-HKU1_orf1ab | 3375 | 3375 | 1 | 0.05 | 0.05 | 0.05 |
| NC_006577.2;YP_173236.1;HCoV-HKU1_orf1ab | 3377 | 3377 | 1 | 0.1 | 0.1 | 0.1 |
| NC_006577.2;YP_173236.1;HCoV-HKU1_orf1ab | 3429 | 3433 | 5 | 0.05 | 0.15 | 0.1 |
| NC_006577.2;YP_173236.1;HCoV-HKU1_orf1ab | 3436 | 3436 | 1 | 0.05 | 0.05 | 0.05 |
| NC_006577.2;YP_173236.1;HCoV-HKU1_orf1ab | 3478 | 3487 | 10 | 0.1 | 0.1 | 0.1 |
| NC_006577.2;YP_173236.1;HCoV-HKU1_orf1ab | 3500 | 3508 | 9 | 0.05 | 0.05 | 0.05 |

| Extended data 1 | | | | | | |
|---|---|---|---|---|---|---|
| Protein | Start position (peptide) | Stop position (peptide) | Number of peptides | Minimum fraction (per peptide) of control subjects with reactivity >3 sd above average | Maximum fraction (per peptide) of control subjects with reactivity >3 sd above average | Mean fraction (over all peptides in this contiguous stretch) of controlsubjects with reactivity >3 sd above average |
| NC_006577.2;YP_173236.1;HCoV-HKU1_orf1ab | 3520 | 3520 | 1 | 0.05 | 0.05 | 0.05 |
| NC_006577.2;YP_173236.1;HCoV-HKU1_orf1ab | 3560 | 3560 | 1 | 0.05 | 0.05 | 0.05 |
| NC_006577.2;YP_173236.1;HCoV-HKU1_orf1ab | 3563 | 3563 | 1 | 0.1 | 0.1 | 0.1 |
| NC_006577.2;YP_173236.1;HCoV-HKU1_orf1ab | 3571 | 3573 | 3 | 0.05 | 0.05 | 0.05 |
| NC_006577.2;YP_173236.1;HCoV-HKU1_orf1ab | 3584 | 3584 | 1 | 0.05 | 0.05 | 0.05 |
| NC_006577.2;YP_173236.1;HCoV-HKU1_orf1ab | 3613 | 3615 | 3 | 0.05 | 0.05 | 0.05 |
| NC_006577.2;YP_173236.1;HCoV-HKU1_orf1ab | 3617 | 3617 | 1 | 0.1 | 0.1 | 0.1 |
| NC_006577.2;YP_173236.1;HCoV-HKU1_orf1ab | 3619 | 3619 | 1 | 0.1 | 0.1 | 0.1 |
| NC_006577.2;YP_173236.1;HCoV-HKU1_orf1ab | 3633 | 3634 | 2 | 0.05 | 0.05 | 0.05 |
| NC_006577.2;YP_173236.1;HCoV-HKU1_orf1ab | 3880 | 3880 | 1 | 0.05 | 0.05 | 0.05 |
| NC_006577.2;YP_173236.1;HCoV-HKU1_orf1ab | 3884 | 3884 | 1 | 0.1 | 0.1 | 0.1 |
| NC_006577.2;YP_173236.1;HCoV-HKU1_orf1ab | 3890 | 3890 | 1 | 0.05 | 0.05 | 0.05 |
| NC_006577.2;YP_173236.1;HCoV-HKU1_orf1ab | 3910 | 3920 | 11 | 0.05 | 0.5 | 0.345454545 |
| NC_006577.2;YP_173236.1;HCoV-HKU1_orf1ab | 3922 | 3922 | 1 | 0.05 | 0.05 | 0.05 |
| NC_006577.2;YP_173236.1;HCoV-HKU1_orf1ab | 3926 | 3927 | 2 | 0.05 | 0.15 | 0.1 |
| NC_006577.2;YP_173236.1;HCoV-HKU1_orf1ab | 3938 | 3947 | 10 | 0.05 | 0.1 | 0.055 |
| NC_006577.2;YP_173236.1;HCoV-HKU1_orf1ab | 3957 | 3957 | 1 | 0.05 | 0.05 | 0.05 |
| NC_006577.2;YP_173236.1;HCoV-HKU1_orf1ab | 3987 | 3988 | 2 | 0.05 | 0.15 | 0.1 |
| NC_006577.2;YP_173236.1;HCoV-HKU1_orf1ab | 3997 | 3997 | 1 | 0.05 | 0.05 | 0.05 |
| NC_006577.2;YP_173236.1;HCoV-HKU1_orf1ab | 4002 | 4004 | 3 | 0.05 | 0.1 | 0.066666667 |
| NC_006577.2;YP_173236.1;HCoV-HKU1_orf1ab | 4008 | 4008 | 1 | 0.05 | 0.05 | 0.05 |
| NC_006577.2;YP_173236.1;HCoV-HKU1_orf1ab | 4018 | 4019 | 2 | 0.05 | 0.05 | 0.05 |
| NC_006577.2;YP_173236.1;HCoV-HKU1_orf1ab | 4021 | 4032 | 12 | 0.05 | 0.15 | 0.075 |
| NC_006577.2;YP_173236.1;HCoV-HKU1_orf1ab | 4034 | 4038 | 5 | 0.05 | 0.15 | 0.08 |
| NC_006577.2;YP_173236.1;HCoV-HKU1_orf1ab | 4054 | 4055 | 2 | 0.05 | 0.05 | 0.05 |
| NC_006577.2;YP_173236.1;HCoV-HKU1_orf1ab | 4065 | 4066 | 2 | 0.05 | 0.05 | 0.05 |
| NC_006577.2;YP_173236.1;HCoV-HKU1_orf1ab | 4074 | 4074 | 1 | 0.05 | 0.05 | 0.05 |
| NC_006577.2;YP_173236.1;HCoV-HKU1_orf1ab | 4078 | 4078 | 1 | 0.05 | 0.05 | 0.05 |
| NC_006577.2;YP_173236.1;HCoV-HKU1_orf1ab | 4087 | 4087 | 1 | 0.05 | 0.05 | 0.05 |
| NC_006577.2;YP_173236.1;HCoV-HKU1_orf1ab | 4103 | 4118 | 16 | 0.05 | 0.35 | 0.16875 |
| NC_006577.2;YP_173236.1;HCoV-HKU1_orf1ab | 4122 | 4122 | 1 | 0.15 | 0.15 | 0.15 |
| NC_006577.2;YP_173236.1;HCoV-HKU1_orf1ab | 4124 | 4124 | 1 | 0.05 | 0.05 | 0.05 |
| NC_006577.2;YP_173236.1;HCoV-HKU1_orf1ab | 4126 | 4126 | 1 | 0.05 | 0.05 | 0.05 |
| NC_006577.2;YP_173236.1;HCoV-HKU1_orf1ab | 4138 | 4138 | 1 | 0.05 | 0.05 | 0.05 |
| NC_006577.2;YP_173236.1;HCoV-HKU1_orf1ab | 4141 | 4148 | 8 | 0.05 | 0.1 | 0.06875 |
| NC_006577.2;YP_173236.1;HCoV-HKU1_orf1ab | 4160 | 4175 | 16 | 0.05 | 0.2 | 0.115625 |
| NC_006577.2;YP_173236.1;HCoV-HKU1_orf1ab | 4179 | 4179 | 1 | 0.05 | 0.05 | 0.05 |

| Extended data 1 | | | | | | |
|---|---|---|---|---|---|---|
| Protein | Start position (peptide) | Stop position (peptide) | Number of peptides | Minimum fraction (per peptide) of control subjects with reactivity >3 sd above average | Maximum fraction (per peptide) of control subjects with reactivity >3 sd above average | Mean fraction (over all peptides in this contiguous stretch) of controlsubjects with reactivity >3 sd above average |
| NC_006577.2;YP_173236.1;HCoV-HKU1_orf1ab | 4189 | 4189 | 1 | 0.05 | 0.05 | 0.05 |
| NC_006577.2;YP_173236.1;HCoV-HKU1_orf1ab | 4194 | 4194 | 1 | 0.3 | 0.3 | 0.3 |
| NC_006577.2;YP_173236.1;HCoV-HKU1_orf1ab | 4197 | 4198 | 2 | 0.1 | 0.15 | 0.125 |
| NC_006577.2;YP_173236.1;HCoV-HKU1_orf1ab | 4208 | 4208 | 1 | 0.05 | 0.05 | 0.05 |
| NC_006577.2;YP_173236.1;HCoV-HKU1_orf1ab | 4214 | 4214 | 1 | 0.05 | 0.05 | 0.05 |
| NC_006577.2;YP_173236.1;HCoV-HKU1_orf1ab | 4221 | 4221 | 1 | 0.05 | 0.05 | 0.05 |
| NC_006577.2;YP_173236.1;HCoV-HKU1_orf1ab | 4226 | 4226 | 1 | 0.05 | 0.05 | 0.05 |
| NC_006577.2;YP_173236.1;HCoV-HKU1_orf1ab | 4233 | 4234 | 2 | 0.05 | 0.05 | 0.05 |
| NC_006577.2;YP_173236.1;HCoV-HKU1_orf1ab | 4236 | 4236 | 1 | 0.05 | 0.05 | 0.05 |
| NC_006577.2;YP_173236.1;HCoV-HKU1_orf1ab | 4247 | 4247 | 1 | 0.05 | 0.05 | 0.05 |
| NC_006577.2;YP_173236.1;HCoV-HKU1_orf1ab | 4250 | 4250 | 1 | 0.05 | 0.05 | 0.05 |
| NC_006577.2;YP_173236.1;HCoV-HKU1_orf1ab | 4255 | 4266 | 12 | 0.05 | 0.2 | 0.125 |
| NC_006577.2;YP_173236.1;HCoV-HKU1_orf1ab | 4284 | 4284 | 1 | 0.05 | 0.05 | 0.05 |
| NC_006577.2;YP_173236.1;HCoV-HKU1_orf1ab | 4314 | 4314 | 1 | 0.05 | 0.05 | 0.05 |
| NC_006577.2;YP_173236.1;HCoV-HKU1_orf1ab | 4328 | 4328 | 1 | 0.05 | 0.05 | 0.05 |
| NC_006577.2;YP_173236.1;HCoV-HKU1_orf1ab | 4337 | 4342 | 6 | 0.05 | 0.1 | 0.066666667 |
| NC_006577.2;YP_173236.1;HCoV-HKU1_orf1ab | 4375 | 4375 | 1 | 0.05 | 0.05 | 0.05 |
| NC_006577.2;YP_173236.1;HCoV-HKU1_orf1ab | 4380 | 4380 | 1 | 0.05 | 0.05 | 0.05 |
| NC_006577.2;YP_173236.1;HCoV-HKU1_orf1ab | 4414 | 4414 | 1 | 0.05 | 0.05 | 0.05 |
| NC_006577.2;YP_173236.1;HCoV-HKU1_orf1ab | 4440 | 4440 | 1 | 0.05 | 0.05 | 0.05 |
| NC_006577.2;YP_173236.1;HCoV-HKU1_orf1ab | 4442 | 4442 | 1 | 0.05 | 0.05 | 0.05 |
| NC_006577.2;YP_173236.1;HCoV-HKU1_orf1ab | 4448 | 4448 | 1 | 0.05 | 0.05 | 0.05 |
| NC_006577.2;YP_173236.1;HCoV-HKU1_orf1ab | 4476 | 4476 | 1 | 0.2 | 0.2 | 0.2 |
| NC_006577.2;YP_173236.1;HCoV-HKU1_orf1ab | 4508 | 4515 | 8 | 0.05 | 0.35 | 0.15625 |
| NC_006577.2;YP_173236.1;HCoV-HKU1_orf1ab | 4518 | 4518 | 1 | 0.15 | 0.15 | 0.15 |
| NC_006577.2;YP_173236.1;HCoV-HKU1_orf1ab | 4520 | 4531 | 12 | 0.05 | 0.15 | 0.070833333 |
| NC_006577.2;YP_173236.1;HCoV-HKU1_orf1ab | 4533 | 4533 | 1 | 0.05 | 0.05 | 0.05 |
| NC_006577.2;YP_173236.1;HCoV-HKU1_orf1ab | 4570 | 4571 | 2 | 0.05 | 0.05 | 0.05 |
| NC_006577.2;YP_173236.1;HCoV-HKU1_orf1ab | 4610 | 4611 | 2 | 0.05 | 0.25 | 0.15 |
| NC_006577.2;YP_173236.1;HCoV-HKU1_orf1ab | 4614 | 4616 | 3 | 0.05 | 0.1 | 0.083333333 |
| NC_006577.2;YP_173236.1;HCoV-HKU1_orf1ab | 4674 | 4674 | 1 | 0.1 | 0.1 | 0.1 |
| NC_006577.2;YP_173236.1;HCoV-HKU1_orf1ab | 4681 | 4681 | 1 | 0.05 | 0.05 | 0.05 |
| NC_006577.2;YP_173236.1;HCoV-HKU1_orf1ab | 4688 | 4690 | 3 | 0.05 | 0.05 | 0.05 |
| NC_006577.2;YP_173236.1;HCoV-HKU1_orf1ab | 4710 | 4710 | 1 | 0.05 | 0.05 | 0.05 |
| NC_006577.2;YP_173236.1;HCoV-HKU1_orf1ab | 4713 | 4713 | 1 | 0.05 | 0.05 | 0.05 |
| NC_006577.2;YP_173236.1;HCoV-HKU1_orf1ab | 4717 | 4720 | 4 | 0.05 | 0.05 | 0.05 |
| NC_006577.2;YP_173236.1;HCoV-HKU1_orf1ab | 4722 | 4725 | 4 | 0.05 | 0.1 | 0.075 |

| Extended data 1 | | | | | | |
|---|---|---|---|---|---|---|
| Protein | Start position (peptide) | Stop position (peptide) | Number of peptides | Minimum fraction (per peptide) of control subjects with reactivity >3 sd above average | Maximum fraction (per peptide) of control subjects with reactivity >3 sd above average | Mean fraction (over all peptides in this contiguous stretch) of controlsubjects with reactivity >3 sd above average |
| NC_006577.2;YP_173236.1;HCoV-HKU1_orf1ab | 4740 | 4741 | 2 | 0.05 | 0.1 | 0.075 |
| NC_006577.2;YP_173236.1;HCoV-HKU1_orf1ab | 4744 | 4745 | 2 | 0.05 | 0.05 | 0.05 |
| NC_006577.2;YP_173236.1;HCoV-HKU1_orf1ab | 4800 | 4802 | 3 | 0.05 | 0.05 | 0.05 |
| NC_006577.2;YP_173236.1;HCoV-HKU1_orf1ab | 4825 | 4825 | 1 | 0.05 | 0.05 | 0.05 |
| NC_006577.2;YP_173236.1;HCoV-HKU1_orf1ab | 4852 | 4854 | 3 | 0.05 | 0.05 | 0.05 |
| NC_006577.2;YP_173236.1;HCoV-HKU1_orf1ab | 4860 | 4866 | 7 | 0.05 | 0.15 | 0.078571429 |
| NC_006577.2;YP_173236.1;HCoV-HKU1_orf1ab | 4870 | 4870 | 1 | 0.05 | 0.05 | 0.05 |
| NC_006577.2;YP_173236.1;HCoV-HKU1_orf1ab | 4874 | 4875 | 2 | 0.05 | 0.05 | 0.05 |
| NC_006577.2;YP_173236.1;HCoV-HKU1_orf1ab | 4887 | 4887 | 1 | 0.05 | 0.05 | 0.05 |
| NC_006577.2;YP_173236.1;HCoV-HKU1_orf1ab | 4889 | 4889 | 1 | 0.05 | 0.05 | 0.05 |
| NC_006577.2;YP_173236.1;HCoV-HKU1_orf1ab | 4897 | 4906 | 10 | 0.05 | 0.25 | 0.195 |
| NC_006577.2;YP_173236.1;HCoV-HKU1_orf1ab | 4940 | 4941 | 2 | 0.05 | 0.05 | 0.05 |
| NC_006577.2;YP_173236.1;HCoV-HKU1_orf1ab | 4943 | 4953 | 11 | 0.05 | 0.25 | 0.127272727 |
| NC_006577.2;YP_173236.1;HCoV-HKU1_orf1ab | 4970 | 4970 | 1 | 0.05 | 0.05 | 0.05 |
| NC_006577.2;YP_173236.1;HCoV-HKU1_orf1ab | 4984 | 4985 | 2 | 0.05 | 0.2 | 0.125 |
| NC_006577.2;YP_173236.1;HCoV-HKU1_orf1ab | 5046 | 5048 | 3 | 0.05 | 0.05 | 0.05 |
| NC_006577.2;YP_173236.1;HCoV-HKU1_orf1ab | 5050 | 5050 | 1 | 0.15 | 0.15 | 0.15 |
| NC_006577.2;YP_173236.1;HCoV-HKU1_orf1ab | 5054 | 5054 | 1 | 0.1 | 0.1 | 0.1 |
| NC_006577.2;YP_173236.1;HCoV-HKU1_orf1ab | 5056 | 5060 | 5 | 0.05 | 0.1 | 0.08 |
| NC_006577.2;YP_173236.1;HCoV-HKU1_orf1ab | 5096 | 5101 | 6 | 0.05 | 0.05 | 0.05 |
| NC_006577.2;YP_173236.1;HCoV-HKU1_orf1ab | 5127 | 5133 | 7 | 0.05 | 0.05 | 0.05 |
| NC_006577.2;YP_173236.1;HCoV-HKU1_orf1ab | 5174 | 5174 | 1 | 0.05 | 0.05 | 0.05 |
| NC_006577.2;YP_173236.1;HCoV-HKU1_orf1ab | 5181 | 5182 | 2 | 0.05 | 0.05 | 0.05 |
| NC_006577.2;YP_173236.1;HCoV-HKU1_orf1ab | 5194 | 5194 | 1 | 0.05 | 0.05 | 0.05 |
| NC_006577.2;YP_173236.1;HCoV-HKU1_orf1ab | 5216 | 5216 | 1 | 0.05 | 0.05 | 0.05 |
| NC_006577.2;YP_173236.1;HCoV-HKU1_orf1ab | 5218 | 5218 | 1 | 0.05 | 0.05 | 0.05 |
| NC_006577.2;YP_173236.1;HCoV-HKU1_orf1ab | 5236 | 5236 | 1 | 0.05 | 0.05 | 0.05 |
| NC_006577.2;YP_173236.1;HCoV-HKU1_orf1ab | 5243 | 5253 | 11 | 0.05 | 0.1 | 0.054545455 |
| NC_006577.2;YP_173236.1;HCoV-HKU1_orf1ab | 5256 | 5256 | 1 | 0.05 | 0.05 | 0.05 |
| NC_006577.2;YP_173236.1;HCoV-HKU1_orf1ab | 5290 | 5291 | 2 | 0.05 | 0.05 | 0.05 |
| NC_006577.2;YP_173236.1;HCoV-HKU1_orf1ab | 5323 | 5323 | 1 | 0.05 | 0.05 | 0.05 |
| NC_006577.2;YP_173236.1;HCoV-HKU1_orf1ab | 5339 | 5342 | 4 | 0.05 | 0.1 | 0.0625 |
| NC_006577.2;YP_173236.1;HCoV-HKU1_orf1ab | 5356 | 5356 | 1 | 0.05 | 0.05 | 0.05 |
| NC_006577.2;YP_173236.1;HCoV-HKU1_orf1ab | 5360 | 5372 | 13 | 0.05 | 0.2 | 0.107692308 |
| NC_006577.2;YP_173236.1;HCoV-HKU1_orf1ab | 5374 | 5374 | 1 | 0.05 | 0.05 | 0.05 |
| NC_006577.2;YP_173236.1;HCoV-HKU1_orf1ab | 5410 | 5413 | 4 | 0.05 | 0.05 | 0.05 |
| NC_006577.2;YP_173236.1;HCoV-HKU1_orf1ab | 5432 | 5432 | 1 | 0.05 | 0.05 | 0.05 |

| Extended data 1 | | | | | | |
|---|---|---|---|---|---|---|
| Protein | Start position (peptide) | Stop position (peptide) | Number of peptides | Minimum fraction (per peptide) of control subjects with reactivity >3 sd above average | Maximum fraction (per peptide) of control subjects with reactivity >3 sd above average | Mean fraction (over all peptides in this contiguous stretch) of controlsubjects with reactivity >3 sd above average |
| NC_006577.2;YP_173236.1;HCoV-HKU1_orf1ab | 5434 | 5434 | 1 | 0.05 | 0.05 | 0.05 |
| NC_006577.2;YP_173236.1;HCoV-HKU1_orf1ab | 5450 | 5451 | 2 | 0.05 | 0.05 | 0.05 |
| NC_006577.2;YP_173236.1;HCoV-HKU1_orf1ab | 5477 | 5489 | 13 | 0.05 | 0.5 | 0.157692308 |
| NC_006577.2;YP_173236.1;HCoV-HKU1_orf1ab | 5491 | 5491 | 1 | 0.05 | 0.05 | 0.05 |
| NC_006577.2;YP_173236.1;HCoV-HKU1_orf1ab | 5505 | 5505 | 1 | 0.05 | 0.05 | 0.05 |
| NC_006577.2;YP_173236.1;HCoV-HKU1_orf1ab | 5512 | 5513 | 2 | 0.05 | 0.05 | 0.05 |
| NC_006577.2;YP_173236.1;HCoV-HKU1_orf1ab | 5521 | 5522 | 2 | 0.05 | 0.05 | 0.05 |
| NC_006577.2;YP_173236.1;HCoV-HKU1_orf1ab | 5548 | 5552 | 5 | 0.05 | 0.15 | 0.08 |
| NC_006577.2;YP_173236.1;HCoV-HKU1_orf1ab | 5554 | 5555 | 2 | 0.05 | 0.05 | 0.05 |
| NC_006577.2;YP_173236.1;HCoV-HKU1_orf1ab | 5582 | 5582 | 1 | 0.05 | 0.05 | 0.05 |
| NC_006577.2;YP_173236.1;HCoV-HKU1_orf1ab | 5588 | 5588 | 1 | 0.05 | 0.05 | 0.05 |
| NC_006577.2;YP_173236.1;HCoV-HKU1_orf1ab | 5593 | 5593 | 1 | 0.05 | 0.05 | 0.05 |
| NC_006577.2;YP_173236.1;HCoV-HKU1_orf1ab | 5608 | 5608 | 1 | 0.05 | 0.05 | 0.05 |
| NC_006577.2;YP_173236.1;HCoV-HKU1_orf1ab | 5618 | 5619 | 2 | 0.05 | 0.1 | 0.075 |
| NC_006577.2;YP_173236.1;HCoV-HKU1_orf1ab | 5628 | 5628 | 1 | 0.05 | 0.05 | 0.05 |
| NC_006577.2;YP_173236.1;HCoV-HKU1_orf1ab | 5637 | 5637 | 1 | 0.05 | 0.05 | 0.05 |
| NC_006577.2;YP_173236.1;HCoV-HKU1_orf1ab | 5641 | 5641 | 1 | 0.05 | 0.05 | 0.05 |
| NC_006577.2;YP_173236.1;HCoV-HKU1_orf1ab | 5682 | 5682 | 1 | 0.05 | 0.05 | 0.05 |
| NC_006577.2;YP_173236.1;HCoV-HKU1_orf1ab | 5697 | 5698 | 2 | 0.05 | 0.05 | 0.05 |
| NC_006577.2;YP_173236.1;HCoV-HKU1_orf1ab | 5700 | 5700 | 1 | 0.05 | 0.05 | 0.05 |
| NC_006577.2;YP_173236.1;HCoV-HKU1_orf1ab | 5702 | 5704 | 3 | 0.05 | 0.05 | 0.05 |
| NC_006577.2;YP_173236.1;HCoV-HKU1_orf1ab | 5719 | 5728 | 10 | 0.05 | 0.5 | 0.21 |
| NC_006577.2;YP_173236.1;HCoV-HKU1_orf1ab | 5748 | 5756 | 9 | 0.05 | 0.05 | 0.05 |
| NC_006577.2;YP_173236.1;HCoV-HKU1_orf1ab | 5759 | 5759 | 1 | 0.05 | 0.05 | 0.05 |
| NC_006577.2;YP_173236.1;HCoV-HKU1_orf1ab | 5770 | 5771 | 2 | 0.05 | 0.05 | 0.05 |
| NC_006577.2;YP_173236.1;HCoV-HKU1_orf1ab | 5773 | 5773 | 1 | 0.05 | 0.05 | 0.05 |
| NC_006577.2;YP_173236.1;HCoV-HKU1_orf1ab | 5787 | 5787 | 1 | 0.05 | 0.05 | 0.05 |
| NC_006577.2;YP_173236.1;HCoV-HKU1_orf1ab | 5796 | 5796 | 1 | 0.05 | 0.05 | 0.05 |
| NC_006577.2;YP_173236.1;HCoV-HKU1_orf1ab | 5822 | 5825 | 4 | 0.05 | 0.15 | 0.1125 |
| NC_006577.2;YP_173236.1;HCoV-HKU1_orf1ab | 5828 | 5832 | 5 | 0.05 | 0.15 | 0.07 |
| NC_006577.2;YP_173236.1;HCoV-HKU1_orf1ab | 5837 | 5837 | 1 | 0.1 | 0.1 | 0.1 |
| NC_006577.2;YP_173236.1;HCoV-HKU1_orf1ab | 5854 | 5857 | 4 | 0.05 | 0.3 | 0.1375 |
| NC_006577.2;YP_173236.1;HCoV-HKU1_orf1ab | 5882 | 5882 | 1 | 0.05 | 0.05 | 0.05 |
| NC_006577.2;YP_173236.1;HCoV-HKU1_orf1ab | 5888 | 5888 | 1 | 0.05 | 0.05 | 0.05 |
| NC_006577.2;YP_173236.1;HCoV-HKU1_orf1ab | 5891 | 5891 | 1 | 0.05 | 0.05 | 0.05 |
| NC_006577.2;YP_173236.1;HCoV-HKU1_orf1ab | 5896 | 5896 | 1 | 0.05 | 0.05 | 0.05 |
| NC_006577.2;YP_173236.1;HCoV-HKU1_orf1ab | 5904 | 5904 | 1 | 0.1 | 0.1 | 0.1 |

| Extended data 1 | | | | | | |
|---|---|---|---|---|---|---|
| Protein | Start position (peptide) | Stop position (peptide) | Number of peptides | Minimum fraction (per peptide) of control subjects with reactivity >3 sd above average | Maximum fraction (per peptide) of control subjects with reactivity >3 sd above average | Mean fraction (over all peptides in this contiguous stretch) of controlsubjects with reactivity >3 sd above average |
| NC_006577.2;YP_173236.1;HCoV-HKU1_orf1ab | 5907 | 5907 | 1 | 0.05 | 0.05 | 0.05 |
| NC_006577.2;YP_173236.1;HCoV-HKU1_orf1ab | 5912 | 5913 | 2 | 0.05 | 0.05 | 0.05 |
| NC_006577.2;YP_173236.1;HCoV-HKU1_orf1ab | 5924 | 5924 | 1 | 0.05 | 0.05 | 0.05 |
| NC_006577.2;YP_173236.1;HCoV-HKU1_orf1ab | 5931 | 5932 | 2 | 0.05 | 0.1 | 0.075 |
| NC_006577.2;YP_173236.1;HCoV-HKU1_orf1ab | 5942 | 5943 | 2 | 0.05 | 0.05 | 0.05 |
| NC_006577.2;YP_173236.1;HCoV-HKU1_orf1ab | 5977 | 5978 | 2 | 0.05 | 0.05 | 0.05 |
| NC_006577.2;YP_173236.1;HCoV-HKU1_orf1ab | 5981 | 5981 | 1 | 0.05 | 0.05 | 0.05 |
| NC_006577.2;YP_173236.1;HCoV-HKU1_orf1ab | 5983 | 5983 | 1 | 0.05 | 0.05 | 0.05 |
| NC_006577.2;YP_173236.1;HCoV-HKU1_orf1ab | 6006 | 6015 | 10 | 0.05 | 0.2 | 0.16 |
| NC_006577.2;YP_173236.1;HCoV-HKU1_orf1ab | 6064 | 6065 | 2 | 0.05 | 0.05 | 0.05 |
| NC_006577.2;YP_173236.1;HCoV-HKU1_orf1ab | 6069 | 6069 | 1 | 0.05 | 0.05 | 0.05 |
| NC_006577.2;YP_173236.1;HCoV-HKU1_orf1ab | 6078 | 6078 | 1 | 0.05 | 0.05 | 0.05 |
| NC_006577.2;YP_173236.1;HCoV-HKU1_orf1ab | 6086 | 6086 | 1 | 0.05 | 0.05 | 0.05 |
| NC_006577.2;YP_173236.1;HCoV-HKU1_orf1ab | 6088 | 6088 | 1 | 0.05 | 0.05 | 0.05 |
| NC_006577.2;YP_173236.1;HCoV-HKU1_orf1ab | 6123 | 6124 | 2 | 0.05 | 0.1 | 0.075 |
| NC_006577.2;YP_173236.1;HCoV-HKU1_orf1ab | 6127 | 6129 | 3 | 0.05 | 0.05 | 0.05 |
| NC_006577.2;YP_173236.1;HCoV-HKU1_orf1ab | 6197 | 6198 | 2 | 0.05 | 0.1 | 0.075 |
| NC_006577.2;YP_173236.1;HCoV-HKU1_orf1ab | 6239 | 6239 | 1 | 0.05 | 0.05 | 0.05 |
| NC_006577.2;YP_173236.1;HCoV-HKU1_orf1ab | 6242 | 6242 | 1 | 0.05 | 0.05 | 0.05 |
| NC_006577.2;YP_173236.1;HCoV-HKU1_orf1ab | 6247 | 6247 | 1 | 0.05 | 0.05 | 0.05 |
| NC_006577.2;YP_173236.1;HCoV-HKU1_orf1ab | 6274 | 6276 | 3 | 0.05 | 0.15 | 0.1 |
| NC_006577.2;YP_173236.1;HCoV-HKU1_orf1ab | 6281 | 6281 | 1 | 0.05 | 0.05 | 0.05 |
| NC_006577.2;YP_173236.1;HCoV-HKU1_orf1ab | 6284 | 6284 | 1 | 0.05 | 0.05 | 0.05 |
| NC_006577.2;YP_173236.1;HCoV-HKU1_orf1ab | 6322 | 6322 | 1 | 0.05 | 0.05 | 0.05 |
| NC_006577.2;YP_173236.1;HCoV-HKU1_orf1ab | 6327 | 6327 | 1 | 0.05 | 0.05 | 0.05 |
| NC_006577.2;YP_173236.1;HCoV-HKU1_orf1ab | 6336 | 6336 | 1 | 0.05 | 0.05 | 0.05 |
| NC_006577.2;YP_173236.1;HCoV-HKU1_orf1ab | 6346 | 6355 | 10 | 0.05 | 0.15 | 0.09 |
| NC_006577.2;YP_173236.1;HCoV-HKU1_orf1ab | 6359 | 6360 | 2 | 0.05 | 0.05 | 0.05 |
| NC_006577.2;YP_173236.1;HCoV-HKU1_orf1ab | 6363 | 6363 | 1 | 0.1 | 0.1 | 0.1 |
| NC_006577.2;YP_173236.1;HCoV-HKU1_orf1ab | 6410 | 6410 | 1 | 0.05 | 0.05 | 0.05 |
| NC_006577.2;YP_173236.1;HCoV-HKU1_orf1ab | 6412 | 6412 | 1 | 0.05 | 0.05 | 0.05 |
| NC_006577.2;YP_173236.1;HCoV-HKU1_orf1ab | 6428 | 6430 | 3 | 0.05 | 0.05 | 0.05 |
| NC_006577.2;YP_173236.1;HCoV-HKU1_orf1ab | 6435 | 6441 | 7 | 0.05 | 0.2 | 0.078571429 |
| NC_006577.2;YP_173236.1;HCoV-HKU1_orf1ab | 6501 | 6503 | 3 | 0.05 | 0.05 | 0.05 |
| NC_006577.2;YP_173236.1;HCoV-HKU1_orf1ab | 6517 | 6517 | 1 | 0.05 | 0.05 | 0.05 |
| NC_006577.2;YP_173236.1;HCoV-HKU1_orf1ab | 6541 | 6542 | 2 | 0.05 | 0.05 | 0.05 |
| NC_006577.2;YP_173236.1;HCoV-HKU1_orf1ab | 6544 | 6548 | 5 | 0.05 | 0.2 | 0.09 |

| Extended data 1 | | | | | | |
|---|---|---|---|---|---|---|
| Protein | Start position (peptide) | Stop position (peptide) | Number of peptides | Minimum fraction (per peptide) of control subjects with reactivity >3 sd above average | Maximum fraction (per peptide) of control subjects with reactivity >3 sd above average | Mean fraction (over all peptides in this contiguous stretch) of controlsubjects with reactivity >3 sd above average |
| NC_006577.2;YP_173236.1;HCoV-HKU1_orf1ab | 6550 | 6551 | 2 | 0.05 | 0.05 | 0.05 |
| NC_006577.2;YP_173236.1;HCoV-HKU1_orf1ab | 6570 | 6570 | 1 | 0.05 | 0.05 | 0.05 |
| NC_006577.2;YP_173236.1;HCoV-HKU1_orf1ab | 6572 | 6572 | 1 | 0.05 | 0.05 | 0.05 |
| NC_006577.2;YP_173236.1;HCoV-HKU1_orf1ab | 6579 | 6579 | 1 | 0.05 | 0.05 | 0.05 |
| NC_006577.2;YP_173236.1;HCoV-HKU1_orf1ab | 6605 | 6608 | 4 | 0.05 | 0.15 | 0.075 |
| NC_006577.2;YP_173236.1;HCoV-HKU1_orf1ab | 6625 | 6625 | 1 | 0.05 | 0.05 | 0.05 |
| NC_006577.2;YP_173236.1;HCoV-HKU1_orf1ab | 6659 | 6660 | 2 | 0.05 | 0.05 | 0.05 |
| NC_006577.2;YP_173236.1;HCoV-HKU1_orf1ab | 6662 | 6662 | 1 | 0.05 | 0.05 | 0.05 |
| NC_006577.2;YP_173236.1;HCoV-HKU1_orf1ab | 6664 | 6664 | 1 | 0.05 | 0.05 | 0.05 |
| NC_006577.2;YP_173236.1;HCoV-HKU1_orf1ab | 6694 | 6694 | 1 | 0.05 | 0.05 | 0.05 |
| NC_006577.2;YP_173236.1;HCoV-HKU1_orf1ab | 6697 | 6699 | 3 | 0.05 | 0.1 | 0.066666667 |
| NC_006577.2;YP_173236.1;HCoV-HKU1_orf1ab | 6701 | 6705 | 5 | 0.05 | 0.15 | 0.1 |
| NC_006577.2;YP_173236.1;HCoV-HKU1_orf1ab | 6707 | 6715 | 9 | 0.05 | 0.15 | 0.083333333 |
| NC_006577.2;YP_173236.1;HCoV-HKU1_orf1ab | 6735 | 6735 | 1 | 0.05 | 0.05 | 0.05 |
| NC_006577.2;YP_173236.1;HCoV-HKU1_orf1ab | 6745 | 6745 | 1 | 0.05 | 0.05 | 0.05 |
| NC_006577.2;YP_173236.1;HCoV-HKU1_orf1ab | 6747 | 6747 | 1 | 0.05 | 0.05 | 0.05 |
| NC_006577.2;YP_173236.1;HCoV-HKU1_orf1ab | 6752 | 6752 | 1 | 0.05 | 0.05 | 0.05 |
| NC_006577.2;YP_173236.1;HCoV-HKU1_orf1ab | 6763 | 6763 | 1 | 0.1 | 0.1 | 0.1 |
| NC_006577.2;YP_173236.1;HCoV-HKU1_orf1ab | 6796 | 6798 | 3 | 0.05 | 0.05 | 0.05 |
| NC_006577.2;YP_173236.1;HCoV-HKU1_orf1ab | 6813 | 6815 | 3 | 0.05 | 0.05 | 0.05 |
| NC_006577.2;YP_173236.1;HCoV-HKU1_orf1ab | 6850 | 6852 | 3 | 0.05 | 0.05 | 0.05 |
| NC_006577.2;YP_173236.1;HCoV-HKU1_orf1ab | 6854 | 6854 | 1 | 0.05 | 0.05 | 0.05 |
| NC_006577.2;YP_173236.1;HCoV-HKU1_orf1ab | 6859 | 6862 | 4 | 0.05 | 0.1 | 0.0875 |
| NC_006577.2;YP_173236.1;HCoV-HKU1_orf1ab | 6880 | 6881 | 2 | 0.05 | 0.05 | 0.05 |
| NC_006577.2;YP_173236.1;HCoV-HKU1_orf1ab | 6906 | 6906 | 1 | 0.05 | 0.05 | 0.05 |
| NC_006577.2;YP_173236.1;HCoV-HKU1_orf1ab | 6913 | 6913 | 1 | 0.05 | 0.05 | 0.05 |
| NC_006577.2;YP_173236.1;HCoV-HKU1_orf1ab | 6926 | 6926 | 1 | 0.05 | 0.05 | 0.05 |
| NC_006577.2;YP_173236.1;HCoV-HKU1_orf1ab | 6930 | 6933 | 4 | 0.05 | 0.05 | 0.05 |
| NC_006577.2;YP_173236.1;HCoV-HKU1_orf1ab | 6956 | 6956 | 1 | 0.05 | 0.05 | 0.05 |
| NC_006577.2;YP_173236.1;HCoV-HKU1_orf1ab | 7010 | 7021 | 12 | 0.05 | 0.2 | 0.095833333 |
| NC_006577.2;YP_173236.1;HCoV-HKU1_orf1ab | 7036 | 7036 | 1 | 0.05 | 0.05 | 0.05 |
| NC_006577.2;YP_173236.1;HCoV-HKU1_orf1ab | 7056 | 7056 | 1 | 0.05 | 0.05 | 0.05 |
| NC_006577.2;YP_173236.1;HCoV-HKU1_orf1ab | 7078 | 7078 | 1 | 0.1 | 0.1 | 0.1 |
| NC_006577.2;YP_173236.1;HCoV-HKU1_orf1ab | 7092 | 7092 | 1 | 0.05 | 0.05 | 0.05 |
| NC_006577.2;YP_173236.1;HCoV-HKU1_orf1ab | 7094 | 7098 | 5 | 0.05 | 0.2 | 0.1 |
| NC_006577.2;YP_173236.1;HCoV-HKU1_orf1ab | 7100 | 7101 | 2 | 0.15 | 0.25 | 0.2 |
| NC_006577.2;YP_173236.1;HCoV-HKU1_orf1ab | 7117 | 7119 | 3 | 0.05 | 0.05 | 0.05 |

| Extended data 1 | | | | | | |
|---|---|---|---|---|---|---|
| Protein | Start position (peptide) | Stop position (peptide) | Number of peptides | Minimum fraction (per peptide) of control subjects with reactivity >3 sd above average | Maximum fraction (per peptide) of control subjects with reactivity >3 sd above average | Mean fraction (over all peptides in this contiguous stretch) of controlsubjects with reactivity >3 sd above average |
| NC_006577.2;YP_173236.1;HCoV-HKU1_orf1ab | 7156 | 7160 | 5 | 0.05 | 0.15 | 0.1 |
| NC_006577.2;YP_173237.1;HCoV-HKU1_HA-E-glycop | 13 | 13 | 1 | 0.05 | 0.05 | 0.05 |
| NC_006577.2;YP_173237.1;HCoV-HKU1_HA-E-glycop | 29 | 31 | 3 | 0.05 | 0.1 | 0.083333333 |
| NC_006577.2;YP_173237.1;HCoV-HKU1_HA-E-glycop | 34 | 34 | 1 | 0.05 | 0.05 | 0.05 |
| NC_006577.2;YP_173237.1;HCoV-HKU1_HA-E-glycop | 36 | 37 | 2 | 0.05 | 0.1 | 0.075 |
| NC_006577.2;YP_173237.1;HCoV-HKU1_HA-E-glycop | 41 | 41 | 1 | 0.05 | 0.05 | 0.05 |
| NC_006577.2;YP_173237.1;HCoV-HKU1_HA-E-glycop | 46 | 47 | 2 | 0.05 | 0.05 | 0.05 |
| NC_006577.2;YP_173237.1;HCoV-HKU1_HA-E-glycop | 105 | 108 | 4 | 0.05 | 0.05 | 0.05 |
| NC_006577.2;YP_173237.1;HCoV-HKU1_HA-E-glycop | 142 | 142 | 1 | 0.05 | 0.05 | 0.05 |
| NC_006577.2;YP_173237.1;HCoV-HKU1_HA-E-glycop | 151 | 151 | 1 | 0.05 | 0.05 | 0.05 |
| NC_006577.2;YP_173237.1;HCoV-HKU1_HA-E-glycop | 153 | 153 | 1 | 0.05 | 0.05 | 0.05 |
| NC_006577.2;YP_173237.1;HCoV-HKU1_HA-E-glycop | 160 | 160 | 1 | 0.05 | 0.05 | 0.05 |
| NC_006577.2;YP_173237.1;HCoV-HKU1_HA-E-glycop | 191 | 191 | 1 | 0.05 | 0.05 | 0.05 |
| NC_006577.2;YP_173237.1;HCoV-HKU1_HA-E-glycop | 203 | 204 | 2 | 0.05 | 0.05 | 0.05 |
| NC_006577.2;YP_173237.1;HCoV-HKU1_HA-E-glycop | 221 | 221 | 1 | 0.05 | 0.05 | 0.05 |
| NC_006577.2;YP_173237.1;HCoV-HKU1_HA-E-glycop | 227 | 227 | 1 | 0.05 | 0.05 | 0.05 |
| NC_006577.2;YP_173237.1;HCoV-HKU1_HA-E-glycop | 250 | 250 | 1 | 0.05 | 0.05 | 0.05 |
| NC_006577.2;YP_173237.1;HCoV-HKU1_HA-E-glycop | 254 | 255 | 2 | 0.05 | 0.05 | 0.05 |
| NC_006577.2;YP_173237.1;HCoV-HKU1_HA-E-glycop | 257 | 257 | 1 | 0.1 | 0.1 | 0.1 |
| NC_006577.2;YP_173237.1;HCoV-HKU1_HA-E-glycop | 261 | 261 | 1 | 0.05 | 0.05 | 0.05 |
| NC_006577.2;YP_173237.1;HCoV-HKU1_HA-E-glycop | 281 | 287 | 7 | 0.05 | 0.2 | 0.1 |
| NC_006577.2;YP_173237.1;HCoV-HKU1_HA-E-glycop | 291 | 291 | 1 | 0.05 | 0.05 | 0.05 |
| NC_006577.2;YP_173238.1;HCoV-HKU1_spike | 14 | 17 | 4 | 0.05 | 0.3 | 0.1125 |
| NC_006577.2;YP_173238.1;HCoV-HKU1_spike | 19 | 24 | 6 | 0.05 | 0.1 | 0.075 |
| NC_006577.2;YP_173238.1;HCoV-HKU1_spike | 64 | 64 | 1 | 0.05 | 0.05 | 0.05 |
| NC_006577.2;YP_173238.1;HCoV-HKU1_spike | 86 | 86 | 1 | 0.05 | 0.05 | 0.05 |
| NC_006577.2;YP_173238.1;HCoV-HKU1_spike | 105 | 105 | 1 | 0.05 | 0.05 | 0.05 |
| NC_006577.2;YP_173238.1;HCoV-HKU1_spike | 165 | 169 | 5 | 0.05 | 0.15 | 0.09 |
| NC_006577.2;YP_173238.1;HCoV-HKU1_spike | 171 | 171 | 1 | 0.05 | 0.05 | 0.05 |
| NC_006577.2;YP_173238.1;HCoV-HKU1_spike | 177 | 183 | 7 | 0.05 | 0.05 | 0.05 |
| NC_006577.2;YP_173238.1;HCoV-HKU1_spike | 186 | 186 | 1 | 0.05 | 0.05 | 0.05 |
| NC_006577.2;YP_173238.1;HCoV-HKU1_spike | 236 | 237 | 2 | 0.1 | 0.1 | 0.1 |
| NC_006577.2;YP_173238.1;HCoV-HKU1_spike | 241 | 241 | 1 | 0.05 | 0.05 | 0.05 |
| NC_006577.2;YP_173238.1;HCoV-HKU1_spike | 244 | 245 | 2 | 0.05 | 0.15 | 0.1 |
| NC_006577.2;YP_173238.1;HCoV-HKU1_spike | 266 | 269 | 4 | 0.05 | 0.1 | 0.0875 |
| NC_006577.2;YP_173238.1;HCoV-HKU1_spike | 276 | 276 | 1 | 0.05 | 0.05 | 0.05 |
| NC_006577.2;YP_173238.1;HCoV-HKU1_spike | 278 | 279 | 2 | 0.05 | 0.05 | 0.05 |

| Extended data 1 | | | | | | |
|---|---|---|---|---|---|---|
| Protein | Start position (peptide) | Stop position (peptide) | Number of peptides | Minimum fraction (per peptide) of control subjects with reactivity >3 sd above average | Maximum fraction (per peptide) of control subjects with reactivity >3 sd above average | Mean fraction (over all peptides in this contiguous stretch) of controlsubjects with reactivity >3 sd above average |
| NC_006577.2;YP_173238.1;HCoV-HKU1_spike | 287 | 287 | 1 | 0.1 | 0.1 | 0.1 |
| NC_006577.2;YP_173238.1;HCoV-HKU1_spike | 289 | 292 | 4 | 0.05 | 0.05 | 0.05 |
| NC_006577.2;YP_173238.1;HCoV-HKU1_spike | 294 | 294 | 1 | 0.05 | 0.05 | 0.05 |
| NC_006577.2;YP_173238.1;HCoV-HKU1_spike | 300 | 300 | 1 | 0.1 | 0.1 | 0.1 |
| NC_006577.2;YP_173238.1;HCoV-HKU1_spike | 322 | 334 | 13 | 0.05 | 0.2 | 0.1 |
| NC_006577.2;YP_173238.1;HCoV-HKU1_spike | 338 | 338 | 1 | 0.05 | 0.05 | 0.05 |
| NC_006577.2;YP_173238.1;HCoV-HKU1_spike | 345 | 345 | 1 | 0.05 | 0.05 | 0.05 |
| NC_006577.2;YP_173238.1;HCoV-HKU1_spike | 365 | 370 | 6 | 0.05 | 0.2 | 0.108333333 |
| NC_006577.2;YP_173238.1;HCoV-HKU1_spike | 386 | 387 | 2 | 0.05 | 0.05 | 0.05 |
| NC_006577.2;YP_173238.1;HCoV-HKU1_spike | 404 | 411 | 8 | 0.05 | 0.2 | 0.1 |
| NC_006577.2;YP_173238.1;HCoV-HKU1_spike | 415 | 416 | 2 | 0.05 | 0.05 | 0.05 |
| NC_006577.2;YP_173238.1;HCoV-HKU1_spike | 422 | 433 | 12 | 0.05 | 0.1 | 0.0875 |
| NC_006577.2;YP_173238.1;HCoV-HKU1_spike | 468 | 468 | 1 | 0.05 | 0.05 | 0.05 |
| NC_006577.2;YP_173238.1;HCoV-HKU1_spike | 471 | 473 | 3 | 0.05 | 0.05 | 0.05 |
| NC_006577.2;YP_173238.1;HCoV-HKU1_spike | 479 | 481 | 3 | 0.05 | 0.2 | 0.1 |
| NC_006577.2;YP_173238.1;HCoV-HKU1_spike | 491 | 491 | 1 | 0.05 | 0.05 | 0.05 |
| NC_006577.2;YP_173238.1;HCoV-HKU1_spike | 497 | 497 | 1 | 0.05 | 0.05 | 0.05 |
| NC_006577.2;YP_173238.1;HCoV-HKU1_spike | 506 | 506 | 1 | 0.05 | 0.05 | 0.05 |
| NC_006577.2;YP_173238.1;HCoV-HKU1_spike | 515 | 515 | 1 | 0.05 | 0.05 | 0.05 |
| NC_006577.2;YP_173238.1;HCoV-HKU1_spike | 529 | 529 | 1 | 0.05 | 0.05 | 0.05 |
| NC_006577.2;YP_173238.1;HCoV-HKU1_spike | 548 | 549 | 2 | 0.05 | 0.15 | 0.1 |
| NC_006577.2;YP_173238.1;HCoV-HKU1_spike | 583 | 583 | 1 | 0.05 | 0.05 | 0.05 |
| NC_006577.2;YP_173238.1;HCoV-HKU1_spike | 587 | 587 | 1 | 0.05 | 0.05 | 0.05 |
| NC_006577.2;YP_173238.1;HCoV-HKU1_spike | 593 | 593 | 1 | 0.05 | 0.05 | 0.05 |
| NC_006577.2;YP_173238.1;HCoV-HKU1_spike | 595 | 595 | 1 | 0.1 | 0.1 | 0.1 |
| NC_006577.2;YP_173238.1;HCoV-HKU1_spike | 597 | 597 | 1 | 0.05 | 0.05 | 0.05 |
| NC_006577.2;YP_173238.1;HCoV-HKU1_spike | 600 | 602 | 3 | 0.05 | 0.05 | 0.05 |
| NC_006577.2;YP_173238.1;HCoV-HKU1_spike | 604 | 604 | 1 | 0.05 | 0.05 | 0.05 |
| NC_006577.2;YP_173238.1;HCoV-HKU1_spike | 606 | 606 | 1 | 0.05 | 0.05 | 0.05 |
| NC_006577.2;YP_173238.1;HCoV-HKU1_spike | 642 | 644 | 3 | 0.1 | 0.15 | 0.133333333 |
| NC_006577.2;YP_173238.1;HCoV-HKU1_spike | 647 | 647 | 1 | 0.15 | 0.15 | 0.15 |
| NC_006577.2;YP_173238.1;HCoV-HKU1_spike | 649 | 650 | 2 | 0.05 | 0.1 | 0.075 |
| NC_006577.2;YP_173238.1;HCoV-HKU1_spike | 655 | 659 | 5 | 0.05 | 0.15 | 0.07 |
| NC_006577.2;YP_173238.1;HCoV-HKU1_spike | 661 | 662 | 2 | 0.05 | 0.05 | 0.05 |
| NC_006577.2;YP_173238.1;HCoV-HKU1_spike | 664 | 665 | 2 | 0.05 | 0.05 | 0.05 |
| NC_006577.2;YP_173238.1;HCoV-HKU1_spike | 667 | 667 | 1 | 0.05 | 0.05 | 0.05 |
| NC_006577.2;YP_173238.1;HCoV-HKU1_spike | 713 | 723 | 11 | 0.05 | 0.05 | 0.05 |

| Extended data 1 | | | | | | |
|---|---|---|---|---|---|---|
| Protein | Start position (peptide) | Stop position (peptide) | Number of peptides | Minimum fraction (per peptide) of control subjects with reactivity >3 sd above average | Maximum fraction (per peptide) of control subjects with reactivity >3 sd above average | Mean fraction (over all peptides in this contiguous stretch) of controlsubjects with reactivity >3 sd above average |
| NC_006577.2;YP_173238.1;HCoV-HKU1_spike | 740 | 740 | 1 | 0.05 | 0.05 | 0.05 |
| NC_006577.2;YP_173238.1;HCoV-HKU1_spike | 750 | 750 | 1 | 0.1 | 0.1 | 0.1 |
| NC_006577.2;YP_173238.1;HCoV-HKU1_spike | 758 | 773 | 16 | 0.05 | 0.25 | 0.11875 |
| NC_006577.2;YP_173238.1;HCoV-HKU1_spike | 775 | 775 | 1 | 0.05 | 0.05 | 0.05 |
| NC_006577.2;YP_173238.1;HCoV-HKU1_spike | 778 | 778 | 1 | 0.05 | 0.05 | 0.05 |
| NC_006577.2;YP_173238.1;HCoV-HKU1_spike | 782 | 782 | 1 | 0.05 | 0.05 | 0.05 |
| NC_006577.2;YP_173238.1;HCoV-HKU1_spike | 798 | 805 | 8 | 0.05 | 0.15 | 0.075 |
| NC_006577.2;YP_173238.1;HCoV-HKU1_spike | 813 | 813 | 1 | 0.05 | 0.05 | 0.05 |
| NC_006577.2;YP_173238.1;HCoV-HKU1_spike | 820 | 820 | 1 | 0.1 | 0.1 | 0.1 |
| NC_006577.2;YP_173238.1;HCoV-HKU1_spike | 831 | 831 | 1 | 0.05 | 0.05 | 0.05 |
| NC_006577.2;YP_173238.1;HCoV-HKU1_spike | 833 | 841 | 9 | 0.05 | 0.15 | 0.094444444 |
| NC_006577.2;YP_173238.1;HCoV-HKU1_spike | 843 | 844 | 2 | 0.05 | 0.1 | 0.075 |
| NC_006577.2;YP_173238.1;HCoV-HKU1_spike | 846 | 856 | 11 | 0.05 | 0.15 | 0.072727273 |
| NC_006577.2;YP_173238.1;HCoV-HKU1_spike | 859 | 868 | 10 | 0.05 | 0.1 | 0.055 |
| NC_006577.2;YP_173238.1;HCoV-HKU1_spike | 871 | 877 | 7 | 0.05 | 0.15 | 0.107142857 |
| NC_006577.2;YP_173238.1;HCoV-HKU1_spike | 882 | 882 | 1 | 0.05 | 0.05 | 0.05 |
| NC_006577.2;YP_173238.1;HCoV-HKU1_spike | 898 | 902 | 5 | 0.05 | 0.15 | 0.08 |
| NC_006577.2;YP_173238.1;HCoV-HKU1_spike | 914 | 916 | 3 | 0.05 | 0.05 | 0.05 |
| NC_006577.2;YP_173238.1;HCoV-HKU1_spike | 918 | 918 | 1 | 0.05 | 0.05 | 0.05 |
| NC_006577.2;YP_173238.1;HCoV-HKU1_spike | 931 | 931 | 1 | 0.05 | 0.05 | 0.05 |
| NC_006577.2;YP_173238.1;HCoV-HKU1_spike | 951 | 951 | 1 | 0.05 | 0.05 | 0.05 |
| NC_006577.2;YP_173238.1;HCoV-HKU1_spike | 954 | 955 | 2 | 0.05 | 0.05 | 0.05 |
| NC_006577.2;YP_173238.1;HCoV-HKU1_spike | 963 | 966 | 4 | 0.05 | 0.05 | 0.05 |
| NC_006577.2;YP_173238.1;HCoV-HKU1_spike | 984 | 984 | 1 | 0.05 | 0.05 | 0.05 |
| NC_006577.2;YP_173238.1;HCoV-HKU1_spike | 999 | 999 | 1 | 0.1 | 0.1 | 0.1 |
| NC_006577.2;YP_173238.1;HCoV-HKU1_spike | 1010 | 1010 | 1 | 0.05 | 0.05 | 0.05 |
| NC_006577.2;YP_173238.1;HCoV-HKU1_spike | 1014 | 1014 | 1 | 0.05 | 0.05 | 0.05 |
| NC_006577.2;YP_173238.1;HCoV-HKU1_spike | 1025 | 1025 | 1 | 0.05 | 0.05 | 0.05 |
| NC_006577.2;YP_173238.1;HCoV-HKU1_spike | 1031 | 1031 | 1 | 0.05 | 0.05 | 0.05 |
| NC_006577.2;YP_173238.1;HCoV-HKU1_spike | 1033 | 1036 | 4 | 0.05 | 0.15 | 0.0875 |
| NC_006577.2;YP_173238.1;HCoV-HKU1_spike | 1057 | 1057 | 1 | 0.05 | 0.05 | 0.05 |
| NC_006577.2;YP_173238.1;HCoV-HKU1_spike | 1065 | 1066 | 2 | 0.05 | 0.1 | 0.075 |
| NC_006577.2;YP_173238.1;HCoV-HKU1_spike | 1071 | 1071 | 1 | 0.05 | 0.05 | 0.05 |
| NC_006577.2;YP_173238.1;HCoV-HKU1_spike | 1073 | 1073 | 1 | 0.05 | 0.05 | 0.05 |
| NC_006577.2;YP_173238.1;HCoV-HKU1_spike | 1093 | 1093 | 1 | 0.05 | 0.05 | 0.05 |
| NC_006577.2;YP_173238.1;HCoV-HKU1_spike | 1098 | 1098 | 1 | 0.05 | 0.05 | 0.05 |
| NC_006577.2;YP_173238.1;HCoV-HKU1_spike | 1101 | 1104 | 4 | 0.05 | 0.05 | 0.05 |

| Extended data 1 | | | | | | |
|---|---|---|---|---|---|---|
| **Protein** | **Start position (peptide)** | **Stop position (peptide)** | **Number of peptides** | **Minimum fraction (per peptide) of control subjects with reactivity >3 sd above average** | **Maximum fraction (per peptide) of control subjects with reactivity >3 sd above average** | **Mean fraction (over all peptides in this contiguous stretch) of controlsubjects with reactivity >3 sd above average** |
| NC_006577.2;YP_173238.1;HCoV-HKU1_spike | 1167 | 1167 | 1 | 0.05 | 0.05 | 0.05 |
| NC_006577.2;YP_173238.1;HCoV-HKU1_spike | 1171 | 1171 | 1 | 0.05 | 0.05 | 0.05 |
| NC_006577.2;YP_173238.1;HCoV-HKU1_spike | 1195 | 1201 | 7 | 0.05 | 0.1 | 0.057142857 |
| NC_006577.2;YP_173238.1;HCoV-HKU1_spike | 1221 | 1221 | 1 | 0.05 | 0.05 | 0.05 |
| NC_006577.2;YP_173238.1;HCoV-HKU1_spike | 1228 | 1236 | 9 | 0.05 | 0.1 | 0.061111111 |
| NC_006577.2;YP_173238.1;HCoV-HKU1_spike | 1240 | 1250 | 11 | 0.05 | 0.1 | 0.068181818 |
| NC_006577.2;YP_173238.1;HCoV-HKU1_spike | 1269 | 1269 | 1 | 0.05 | 0.05 | 0.05 |
| NC_006577.2;YP_173238.1;HCoV-HKU1_spike | 1271 | 1280 | 10 | 0.05 | 0.3 | 0.14 |
| NC_006577.2;YP_173238.1;HCoV-HKU1_spike | 1287 | 1287 | 1 | 0.05 | 0.05 | 0.05 |
| NC_006577.2;YP_173238.1;HCoV-HKU1_spike | 1341 | 1345 | 5 | 0.15 | 0.15 | 0.15 |
| NC_006577.2;YP_173239.1;HCoV-HKU1_non-structu | 12 | 20 | 9 | 0.05 | 0.15 | 0.077777778 |
| NC_006577.2;YP_173239.1;HCoV-HKU1_non-structu | 60 | 60 | 1 | 0.05 | 0.05 | 0.05 |
| NC_006577.2;YP_173239.1;HCoV-HKU1_non-structu | 74 | 74 | 1 | 0.05 | 0.05 | 0.05 |
| NC_006577.2;YP_173239.1;HCoV-HKU1_non-structu | 77 | 77 | 1 | 0.05 | 0.05 | 0.05 |
| NC_006577.2;YP_173240.1;HCoV-HKU1_small-membr | 57 | 57 | 1 | 0.05 | 0.05 | 0.05 |
| NC_006577.2;YP_173240.1;HCoV-HKU1_small-membr | 60 | 60 | 1 | 0.05 | 0.05 | 0.05 |
| NC_006577.2;YP_173240.1;HCoV-HKU1_small-membr | 66 | 71 | 6 | 0.05 | 0.05 | 0.05 |
| NC_006577.2;YP_173241.1;HCoV-HKU1_membrane | 1 | 9 | 9 | 0.05 | 1 | 0.355555556 |
| NC_006577.2;YP_173241.1;HCoV-HKU1_membrane | 111 | 115 | 5 | 0.05 | 0.1 | 0.06 |
| NC_006577.2;YP_173241.1;HCoV-HKU1_membrane | 123 | 123 | 1 | 0.05 | 0.05 | 0.05 |
| NC_006577.2;YP_173241.1;HCoV-HKU1_membrane | 153 | 153 | 1 | 0.05 | 0.05 | 0.05 |
| NC_006577.2;YP_173241.1;HCoV-HKU1_membrane | 179 | 179 | 1 | 0.05 | 0.05 | 0.05 |
| NC_006577.2;YP_173241.1;HCoV-HKU1_membrane | 182 | 182 | 1 | 0.05 | 0.05 | 0.05 |
| NC_006577.2;YP_173242.1;HCoV-HKU1_nucleocapsi | 2 | 2 | 1 | 0.05 | 0.05 | 0.05 |
| NC_006577.2;YP_173242.1;HCoV-HKU1_nucleocapsi | 4 | 4 | 1 | 0.05 | 0.05 | 0.05 |
| NC_006577.2;YP_173242.1;HCoV-HKU1_nucleocapsi | 16 | 23 | 8 | 0.05 | 0.05 | 0.05 |
| NC_006577.2;YP_173242.1;HCoV-HKU1_nucleocapsi | 25 | 26 | 2 | 0.05 | 0.15 | 0.1 |
| NC_006577.2;YP_173242.1;HCoV-HKU1_nucleocapsi | 67 | 67 | 1 | 0.05 | 0.05 | 0.05 |
| NC_006577.2;YP_173242.1;HCoV-HKU1_nucleocapsi | 71 | 71 | 1 | 0.1 | 0.1 | 0.1 |
| NC_006577.2;YP_173242.1;HCoV-HKU1_nucleocapsi | 77 | 77 | 1 | 0.05 | 0.05 | 0.05 |
| NC_006577.2;YP_173242.1;HCoV-HKU1_nucleocapsi | 111 | 111 | 1 | 0.05 | 0.05 | 0.05 |
| NC_006577.2;YP_173242.1;HCoV-HKU1_nucleocapsi | 128 | 129 | 2 | 0.05 | 0.1 | 0.075 |
| NC_006577.2;YP_173242.1;HCoV-HKU1_nucleocapsi | 139 | 151 | 13 | 0.05 | 0.05 | 0.05 |
| NC_006577.2;YP_173242.1;HCoV-HKU1_nucleocapsi | 155 | 156 | 2 | 0.05 | 0.05 | 0.05 |
| NC_006577.2;YP_173242.1;HCoV-HKU1_nucleocapsi | 158 | 164 | 7 | 0.05 | 0.1 | 0.057142857 |
| NC_006577.2;YP_173242.1;HCoV-HKU1_nucleocapsi | 169 | 169 | 1 | 0.05 | 0.05 | 0.05 |
| NC_006577.2;YP_173242.1;HCoV-HKU1_nucleocapsi | 180 | 180 | 1 | 0.1 | 0.1 | 0.1 |

| Extended data 1 | | | | | | |
|---|---|---|---|---|---|---|
| **Protein** | **Start position (peptide)** | **Stop position (peptide)** | **Number of peptides** | **Minimum fraction (per peptide) of control subjects with reactivity >3 sd above average** | **Maximum fraction (per peptide) of control subjects with reactivity >3 sd above average** | **Mean fraction (over all peptides in this contiguous stretch) of controlsubjects with reactivity >3 sd above average** |
| NC_006577.2;YP_173242.1;HCoV-HKU1_nucleocapsi | 186 | 187 | 2 | 0.05 | 0.05 | 0.05 |
| NC_006577.2;YP_173242.1;HCoV-HKU1_nucleocapsi | 194 | 194 | 1 | 0.1 | 0.1 | 0.1 |
| NC_006577.2;YP_173242.1;HCoV-HKU1_nucleocapsi | 205 | 205 | 1 | 0.05 | 0.05 | 0.05 |
| NC_006577.2;YP_173242.1;HCoV-HKU1_nucleocapsi | 212 | 213 | 2 | 0.05 | 0.05 | 0.05 |
| NC_006577.2;YP_173242.1;HCoV-HKU1_nucleocapsi | 216 | 216 | 1 | 0.05 | 0.05 | 0.05 |
| NC_006577.2;YP_173242.1;HCoV-HKU1_nucleocapsi | 232 | 248 | 17 | 0.05 | 0.1 | 0.052941176 |
| NC_006577.2;YP_173242.1;HCoV-HKU1_nucleocapsi | 250 | 255 | 6 | 0.05 | 0.05 | 0.05 |
| NC_006577.2;YP_173242.1;HCoV-HKU1_nucleocapsi | 268 | 268 | 1 | 0.05 | 0.05 | 0.05 |
| NC_006577.2;YP_173242.1;HCoV-HKU1_nucleocapsi | 280 | 286 | 7 | 0.05 | 0.05 | 0.05 |
| NC_006577.2;YP_173242.1;HCoV-HKU1_nucleocapsi | 301 | 305 | 5 | 0.05 | 0.1 | 0.06 |
| NC_006577.2;YP_173242.1;HCoV-HKU1_nucleocapsi | 309 | 309 | 1 | 0.05 | 0.05 | 0.05 |
| NC_006577.2;YP_173242.1;HCoV-HKU1_nucleocapsi | 326 | 326 | 1 | 0.05 | 0.05 | 0.05 |
| NC_006577.2;YP_173242.1;HCoV-HKU1_nucleocapsi | 331 | 331 | 1 | 0.1 | 0.1 | 0.1 |
| NC_006577.2;YP_173242.1;HCoV-HKU1_nucleocapsi | 336 | 337 | 2 | 0.05 | 0.05 | 0.05 |
| NC_006577.2;YP_173242.1;HCoV-HKU1_nucleocapsi | 350 | 352 | 3 | 0.05 | 0.1 | 0.083333333 |
| NC_006577.2;YP_173242.1;HCoV-HKU1_nucleocapsi | 360 | 360 | 1 | 0.05 | 0.05 | 0.05 |
| NC_006577.2;YP_173242.1;HCoV-HKU1_nucleocapsi | 366 | 366 | 1 | 0.05 | 0.05 | 0.05 |
| NC_006577.2;YP_173242.1;HCoV-HKU1_nucleocapsi | 368 | 372 | 5 | 0.05 | 0.2 | 0.1 |
| NC_006577.2;YP_173242.1;HCoV-HKU1_nucleocapsi | 374 | 376 | 3 | 0.05 | 0.1 | 0.066666667 |
| NC_006577.2;YP_173242.1;HCoV-HKU1_nucleocapsi | 391 | 391 | 1 | 0.05 | 0.05 | 0.05 |
| NC_006577.2;YP_173242.1;HCoV-HKU1_nucleocapsi | 394 | 404 | 11 | 0.1 | 0.35 | 0.254545455 |
| NC_006577.2;YP_173242.1;HCoV-HKU1_nucleocapsi | 406 | 406 | 1 | 0.1 | 0.1 | 0.1 |
| NC_006577.2;YP_173242.1;HCoV-HKU1_nucleocapsi | 409 | 409 | 1 | 0.05 | 0.05 | 0.05 |
| NC_006577.2;YP_173242.1;HCoV-HKU1_nucleocapsi | 417 | 417 | 1 | 0.1 | 0.1 | 0.1 |
| NC_006577.2;YP_173242.1;HCoV-HKU1_nucleocapsi | 426 | 430 | 5 | 0.05 | 0.1 | 0.06 |
| NC_006577.2;YP_173243.1;HCoV-HKU1_nucleocap2 | 10 | 11 | 2 | 0.05 | 0.2 | 0.125 |
| NC_006577.2;YP_173243.1;HCoV-HKU1_nucleocap2 | 13 | 13 | 1 | 0.1 | 0.1 | 0.1 |
| NC_006577.2;YP_173243.1;HCoV-HKU1_nucleocap2 | 21 | 29 | 9 | 0.05 | 0.15 | 0.083333333 |
| NC_006577.2;YP_173243.1;HCoV-HKU1_nucleocap2 | 46 | 46 | 1 | 0.05 | 0.05 | 0.05 |
| NC_006577.2;YP_173243.1;HCoV-HKU1_nucleocap2 | 49 | 49 | 1 | 0.05 | 0.05 | 0.05 |
| NC_006577.2;YP_173243.1;HCoV-HKU1_nucleocap2 | 57 | 58 | 2 | 0.05 | 0.05 | 0.05 |
| NC_006577.2;YP_173243.1;HCoV-HKU1_nucleocap2 | 62 | 64 | 3 | 0.05 | 0.15 | 0.083333333 |
| NC_006577.2;YP_173243.1;HCoV-HKU1_nucleocap2 | 67 | 67 | 1 | 0.05 | 0.05 | 0.05 |
| NC_006577.2;YP_173243.1;HCoV-HKU1_nucleocap2 | 73 | 73 | 1 | 0.05 | 0.05 | 0.05 |
| NC_006577.2;YP_173243.1;HCoV-HKU1_nucleocap2 | 76 | 79 | 4 | 0.05 | 0.05 | 0.05 |
| NC_006577.2;YP_173243.1;HCoV-HKU1_nucleocap2 | 89 | 89 | 1 | 0.05 | 0.05 | 0.05 |
| NC_006577.2;YP_173243.1;HCoV-HKU1_nucleocap2 | 103 | 103 | 1 | 0.05 | 0.05 | 0.05 |

| Extended data 1 | | | | | | |
|---|---|---|---|---|---|---|
| **Protein** | **Start position (peptide)** | **Stop position (peptide)** | **Number of peptides** | **Minimum fraction (per peptide) of control subjects with reactivity >3 sd above average** | **Maximum fraction (per peptide) of control subjects with reactivity >3 sd above average** | **Mean fraction (over all peptides in this contiguous stretch) of controlsubjects with reactivity >3 sd above average** |
| NC_006577.2;YP_173243.1;HCoV-HKU1_nucleocap2 | 107 | 107 | 1 | 0.05 | 0.05 | 0.05 |
| NC_006577.2;YP_173243.1;HCoV-HKU1_nucleocap2 | 112 | 112 | 1 | 0.1 | 0.1 | 0.1 |
| NC_006577.2;YP_173243.1;HCoV-HKU1_nucleocap2 | 188 | 189 | 2 | 0.05 | 0.15 | 0.1 |
| NC_019843.3;YP_009047202.1;MERS_1ab | 4 | 4 | 1 | 0.05 | 0.05 | 0.05 |
| NC_019843.3;YP_009047202.1;MERS_1ab | 8 | 8 | 1 | 0.05 | 0.05 | 0.05 |
| NC_019843.3;YP_009047202.1;MERS_1ab | 14 | 14 | 1 | 0.05 | 0.05 | 0.05 |
| NC_019843.3;YP_009047202.1;MERS_1ab | 16 | 16 | 1 | 0.05 | 0.05 | 0.05 |
| NC_019843.3;YP_009047202.1;MERS_1ab | 19 | 19 | 1 | 0.1 | 0.1 | 0.1 |
| NC_019843.3;YP_009047202.1;MERS_1ab | 29 | 39 | 11 | 0.05 | 0.15 | 0.059090909 |
| NC_019843.3;YP_009047202.1;MERS_1ab | 46 | 46 | 1 | 0.05 | 0.05 | 0.05 |
| NC_019843.3;YP_009047202.1;MERS_1ab | 92 | 92 | 1 | 0.1 | 0.1 | 0.1 |
| NC_019843.3;YP_009047202.1;MERS_1ab | 98 | 98 | 1 | 0.05 | 0.05 | 0.05 |
| NC_019843.3;YP_009047202.1;MERS_1ab | 101 | 104 | 4 | 0.05 | 0.05 | 0.05 |
| NC_019843.3;YP_009047202.1;MERS_1ab | 107 | 107 | 1 | 0.05 | 0.05 | 0.05 |
| NC_019843.3;YP_009047202.1;MERS_1ab | 120 | 120 | 1 | 0.05 | 0.05 | 0.05 |
| NC_019843.3;YP_009047202.1;MERS_1ab | 126 | 128 | 3 | 0.05 | 0.1 | 0.066666667 |
| NC_019843.3;YP_009047202.1;MERS_1ab | 163 | 170 | 8 | 0.05 | 0.1 | 0.09375 |
| NC_019843.3;YP_009047202.1;MERS_1ab | 176 | 176 | 1 | 0.05 | 0.05 | 0.05 |
| NC_019843.3;YP_009047202.1;MERS_1ab | 184 | 184 | 1 | 0.05 | 0.05 | 0.05 |
| NC_019843.3;YP_009047202.1;MERS_1ab | 186 | 188 | 3 | 0.05 | 0.1 | 0.083333333 |
| NC_019843.3;YP_009047202.1;MERS_1ab | 196 | 197 | 2 | 0.05 | 0.1 | 0.075 |
| NC_019843.3;YP_009047202.1;MERS_1ab | 199 | 199 | 1 | 0.05 | 0.05 | 0.05 |
| NC_019843.3;YP_009047202.1;MERS_1ab | 219 | 219 | 1 | 0.05 | 0.05 | 0.05 |
| NC_019843.3;YP_009047202.1;MERS_1ab | 232 | 232 | 1 | 0.05 | 0.05 | 0.05 |
| NC_019843.3;YP_009047202.1;MERS_1ab | 259 | 260 | 2 | 0.05 | 0.05 | 0.05 |
| NC_019843.3;YP_009047202.1;MERS_1ab | 267 | 267 | 1 | 0.05 | 0.05 | 0.05 |
| NC_019843.3;YP_009047202.1;MERS_1ab | 304 | 304 | 1 | 0.05 | 0.05 | 0.05 |
| NC_019843.3;YP_009047202.1;MERS_1ab | 334 | 334 | 1 | 0.05 | 0.05 | 0.05 |
| NC_019843.3;YP_009047202.1;MERS_1ab | 340 | 340 | 1 | 0.05 | 0.05 | 0.05 |
| NC_019843.3;YP_009047202.1;MERS_1ab | 347 | 347 | 1 | 0.05 | 0.05 | 0.05 |
| NC_019843.3;YP_009047202.1;MERS_1ab | 356 | 356 | 1 | 0.05 | 0.05 | 0.05 |
| NC_019843.3;YP_009047202.1;MERS_1ab | 388 | 388 | 1 | 0.05 | 0.05 | 0.05 |
| NC_019843.3;YP_009047202.1;MERS_1ab | 392 | 392 | 1 | 0.05 | 0.05 | 0.05 |
| NC_019843.3;YP_009047202.1;MERS_1ab | 405 | 408 | 4 | 0.05 | 0.1 | 0.075 |
| NC_019843.3;YP_009047202.1;MERS_1ab | 411 | 411 | 1 | 0.05 | 0.05 | 0.05 |
| NC_019843.3;YP_009047202.1;MERS_1ab | 467 | 467 | 1 | 0.05 | 0.05 | 0.05 |
| NC_019843.3;YP_009047202.1;MERS_1ab | 489 | 489 | 1 | 0.05 | 0.05 | 0.05 |

| Extended data 1 | | | | | | |
|---|---|---|---|---|---|---|
| Protein | Start position (peptide) | Stop position (peptide) | Number of peptides | Minimum fraction (per peptide) of control subjects with reactivity >3 sd above average | Maximum fraction (per peptide) of control subjects with reactivity >3 sd above average | Mean fraction (over all peptides in this contiguous stretch) of controlsubjects with reactivity >3 sd above average |
| NC_019843.3;YP_009047202.1;MERS_1ab | 497 | 502 | 6 | 0.05 | 0.05 | 0.05 |
| NC_019843.3;YP_009047202.1;MERS_1ab | 505 | 505 | 1 | 0.05 | 0.05 | 0.05 |
| NC_019843.3;YP_009047202.1;MERS_1ab | 512 | 519 | 8 | 0.05 | 0.1 | 0.06875 |
| NC_019843.3;YP_009047202.1;MERS_1ab | 521 | 521 | 1 | 0.05 | 0.05 | 0.05 |
| NC_019843.3;YP_009047202.1;MERS_1ab | 526 | 526 | 1 | 0.05 | 0.05 | 0.05 |
| NC_019843.3;YP_009047202.1;MERS_1ab | 529 | 530 | 2 | 0.05 | 0.05 | 0.05 |
| NC_019843.3;YP_009047202.1;MERS_1ab | 535 | 536 | 2 | 0.05 | 0.05 | 0.05 |
| NC_019843.3;YP_009047202.1;MERS_1ab | 539 | 539 | 1 | 0.1 | 0.1 | 0.1 |
| NC_019843.3;YP_009047202.1;MERS_1ab | 564 | 564 | 1 | 0.05 | 0.05 | 0.05 |
| NC_019843.3;YP_009047202.1;MERS_1ab | 567 | 567 | 1 | 0.05 | 0.05 | 0.05 |
| NC_019843.3;YP_009047202.1;MERS_1ab | 574 | 574 | 1 | 0.05 | 0.05 | 0.05 |
| NC_019843.3;YP_009047202.1;MERS_1ab | 594 | 598 | 5 | 0.05 | 0.15 | 0.07 |
| NC_019843.3;YP_009047202.1;MERS_1ab | 692 | 692 | 1 | 0.05 | 0.05 | 0.05 |
| NC_019843.3;YP_009047202.1;MERS_1ab | 701 | 703 | 3 | 0.1 | 0.15 | 0.116666667 |
| NC_019843.3;YP_009047202.1;MERS_1ab | 729 | 729 | 1 | 0.05 | 0.05 | 0.05 |
| NC_019843.3;YP_009047202.1;MERS_1ab | 745 | 746 | 2 | 0.05 | 0.05 | 0.05 |
| NC_019843.3;YP_009047202.1;MERS_1ab | 749 | 749 | 1 | 0.05 | 0.05 | 0.05 |
| NC_019843.3;YP_009047202.1;MERS_1ab | 751 | 751 | 1 | 0.1 | 0.1 | 0.1 |
| NC_019843.3;YP_009047202.1;MERS_1ab | 757 | 757 | 1 | 0.05 | 0.05 | 0.05 |
| NC_019843.3;YP_009047202.1;MERS_1ab | 766 | 766 | 1 | 0.05 | 0.05 | 0.05 |
| NC_019843.3;YP_009047202.1;MERS_1ab | 771 | 772 | 2 | 0.05 | 0.05 | 0.05 |
| NC_019843.3;YP_009047202.1;MERS_1ab | 778 | 781 | 4 | 0.05 | 0.1 | 0.075 |
| NC_019843.3;YP_009047202.1;MERS_1ab | 783 | 787 | 5 | 0.05 | 0.05 | 0.05 |
| NC_019843.3;YP_009047202.1;MERS_1ab | 790 | 790 | 1 | 0.05 | 0.05 | 0.05 |
| NC_019843.3;YP_009047202.1;MERS_1ab | 817 | 818 | 2 | 0.05 | 0.05 | 0.05 |
| NC_019843.3;YP_009047202.1;MERS_1ab | 820 | 822 | 3 | 0.05 | 0.1 | 0.066666667 |
| NC_019843.3;YP_009047202.1;MERS_1ab | 824 | 828 | 5 | 0.05 | 0.1 | 0.06 |
| NC_019843.3;YP_009047202.1;MERS_1ab | 852 | 852 | 1 | 0.05 | 0.05 | 0.05 |
| NC_019843.3;YP_009047202.1;MERS_1ab | 856 | 856 | 1 | 0.05 | 0.05 | 0.05 |
| NC_019843.3;YP_009047202.1;MERS_1ab | 881 | 881 | 1 | 0.05 | 0.05 | 0.05 |
| NC_019843.3;YP_009047202.1;MERS_1ab | 889 | 890 | 2 | 0.05 | 0.05 | 0.05 |
| NC_019843.3;YP_009047202.1;MERS_1ab | 895 | 895 | 1 | 0.05 | 0.05 | 0.05 |
| NC_019843.3;YP_009047202.1;MERS_1ab | 915 | 915 | 1 | 0.05 | 0.05 | 0.05 |
| NC_019843.3;YP_009047202.1;MERS_1ab | 920 | 928 | 9 | 0.05 | 0.15 | 0.083333333 |
| NC_019843.3;YP_009047202.1;MERS_1ab | 930 | 931 | 2 | 0.05 | 0.05 | 0.05 |
| NC_019843.3;YP_009047202.1;MERS_1ab | 959 | 959 | 1 | 0.05 | 0.05 | 0.05 |
| NC_019843.3;YP_009047202.1;MERS_1ab | 982 | 982 | 1 | 0.05 | 0.05 | 0.05 |

| Extended data 1 | | | | | | |
|---|---|---|---|---|---|---|
| Protein | Start position (peptide) | Stop position (peptide) | Number of peptides | Minimum fraction (per peptide) of control subjects with reactivity >3 sd above average | Maximum fraction (per peptide) of control subjects with reactivity >3 sd above average | Mean fraction (over all peptides in this contiguous stretch) of controlsubjects with reactivity >3 sd above average |
| NC_019843.3;YP_009047202.1;MERS_1ab | 992 | 993 | 2 | 0.05 | 0.05 | 0.05 |
| NC_019843.3;YP_009047202.1;MERS_1ab | 995 | 1010 | 16 | 0.05 | 0.15 | 0.08125 |
| NC_019843.3;YP_009047202.1;MERS_1ab | 1019 | 1019 | 1 | 0.1 | 0.1 | 0.1 |
| NC_019843.3;YP_009047202.1;MERS_1ab | 1034 | 1035 | 2 | 0.05 | 0.05 | 0.05 |
| NC_019843.3;YP_009047202.1;MERS_1ab | 1040 | 1044 | 5 | 0.05 | 0.05 | 0.05 |
| NC_019843.3;YP_009047202.1;MERS_1ab | 1051 | 1051 | 1 | 0.05 | 0.05 | 0.05 |
| NC_019843.3;YP_009047202.1;MERS_1ab | 1056 | 1060 | 5 | 0.05 | 0.1 | 0.06 |
| NC_019843.3;YP_009047202.1;MERS_1ab | 1062 | 1063 | 2 | 0.05 | 0.1 | 0.075 |
| NC_019843.3;YP_009047202.1;MERS_1ab | 1066 | 1066 | 1 | 0.05 | 0.05 | 0.05 |
| NC_019843.3;YP_009047202.1;MERS_1ab | 1082 | 1094 | 13 | 0.05 | 0.2 | 0.103846154 |
| NC_019843.3;YP_009047202.1;MERS_1ab | 1099 | 1099 | 1 | 0.05 | 0.05 | 0.05 |
| NC_019843.3;YP_009047202.1;MERS_1ab | 1105 | 1106 | 2 | 0.05 | 0.15 | 0.1 |
| NC_019843.3;YP_009047202.1;MERS_1ab | 1126 | 1126 | 1 | 0.05 | 0.05 | 0.05 |
| NC_019843.3;YP_009047202.1;MERS_1ab | 1131 | 1131 | 1 | 0.05 | 0.05 | 0.05 |
| NC_019843.3;YP_009047202.1;MERS_1ab | 1139 | 1140 | 2 | 0.05 | 0.05 | 0.05 |
| NC_019843.3;YP_009047202.1;MERS_1ab | 1142 | 1142 | 1 | 0.4 | 0.4 | 0.4 |
| NC_019843.3;YP_009047202.1;MERS_1ab | 1148 | 1148 | 1 | 0.05 | 0.05 | 0.05 |
| NC_019843.3;YP_009047202.1;MERS_1ab | 1153 | 1162 | 10 | 0.05 | 0.1 | 0.055 |
| NC_019843.3;YP_009047202.1;MERS_1ab | 1167 | 1168 | 2 | 0.05 | 0.05 | 0.05 |
| NC_019843.3;YP_009047202.1;MERS_1ab | 1170 | 1170 | 1 | 0.05 | 0.05 | 0.05 |
| NC_019843.3;YP_009047202.1;MERS_1ab | 1172 | 1172 | 1 | 0.1 | 0.1 | 0.1 |
| NC_019843.3;YP_009047202.1;MERS_1ab | 1203 | 1210 | 8 | 0.05 | 0.1 | 0.05625 |
| NC_019843.3;YP_009047202.1;MERS_1ab | 1289 | 1289 | 1 | 0.05 | 0.05 | 0.05 |
| NC_019843.3;YP_009047202.1;MERS_1ab | 1306 | 1306 | 1 | 0.1 | 0.1 | 0.1 |
| NC_019843.3;YP_009047202.1;MERS_1ab | 1318 | 1319 | 2 | 0.05 | 0.1 | 0.075 |
| NC_019843.3;YP_009047202.1;MERS_1ab | 1336 | 1340 | 5 | 0.1 | 0.15 | 0.13 |
| NC_019843.3;YP_009047202.1;MERS_1ab | 1343 | 1346 | 4 | 0.05 | 0.1 | 0.075 |
| NC_019843.3;YP_009047202.1;MERS_1ab | 1378 | 1381 | 4 | 0.05 | 0.05 | 0.05 |
| NC_019843.3;YP_009047202.1;MERS_1ab | 1394 | 1395 | 2 | 0.05 | 0.05 | 0.05 |
| NC_019843.3;YP_009047202.1;MERS_1ab | 1398 | 1398 | 1 | 0.05 | 0.05 | 0.05 |
| NC_019843.3;YP_009047202.1;MERS_1ab | 1401 | 1401 | 1 | 0.05 | 0.05 | 0.05 |
| NC_019843.3;YP_009047202.1;MERS_1ab | 1403 | 1404 | 2 | 0.05 | 0.4 | 0.225 |
| NC_019843.3;YP_009047202.1;MERS_1ab | 1424 | 1424 | 1 | 0.05 | 0.05 | 0.05 |
| NC_019843.3;YP_009047202.1;MERS_1ab | 1437 | 1437 | 1 | 0.05 | 0.05 | 0.05 |
| NC_019843.3;YP_009047202.1;MERS_1ab | 1473 | 1473 | 1 | 0.05 | 0.05 | 0.05 |
| NC_019843.3;YP_009047202.1;MERS_1ab | 1490 | 1490 | 1 | 0.1 | 0.1 | 0.1 |
| NC_019843.3;YP_009047202.1;MERS_1ab | 1514 | 1516 | 3 | 0.05 | 0.05 | 0.05 |

| Extended data 1 | | | | | | |
|---|---|---|---|---|---|---|
| Protein | Start position (peptide) | Stop position (peptide) | Number of peptides | Minimum fraction (per peptide) of control subjects with reactivity >3 sd above average | Maximum fraction (per peptide) of control subjects with reactivity >3 sd above average | Mean fraction (over all peptides in this contiguous stretch) of controlsubjects with reactivity >3 sd above average |
| NC_019843.3;YP_009047202.1;MERS_1ab | 1518 | 1518 | 1 | 0.05 | 0.05 | 0.05 |
| NC_019843.3;YP_009047202.1;MERS_1ab | 1520 | 1521 | 2 | 0.05 | 0.05 | 0.05 |
| NC_019843.3;YP_009047202.1;MERS_1ab | 1523 | 1523 | 1 | 0.05 | 0.05 | 0.05 |
| NC_019843.3;YP_009047202.1;MERS_1ab | 1530 | 1545 | 16 | 0.05 | 0.15 | 0.084375 |
| NC_019843.3;YP_009047202.1;MERS_1ab | 1603 | 1604 | 2 | 0.05 | 0.15 | 0.1 |
| NC_019843.3;YP_009047202.1;MERS_1ab | 1613 | 1617 | 5 | 0.05 | 0.1 | 0.07 |
| NC_019843.3;YP_009047202.1;MERS_1ab | 1633 | 1641 | 9 | 0.05 | 0.15 | 0.088888889 |
| NC_019843.3;YP_009047202.1;MERS_1ab | 1645 | 1645 | 1 | 0.15 | 0.15 | 0.15 |
| NC_019843.3;YP_009047202.1;MERS_1ab | 1676 | 1676 | 1 | 0.15 | 0.15 | 0.15 |
| NC_019843.3;YP_009047202.1;MERS_1ab | 1752 | 1752 | 1 | 0.1 | 0.1 | 0.1 |
| NC_019843.3;YP_009047202.1;MERS_1ab | 1773 | 1774 | 2 | 0.05 | 0.05 | 0.05 |
| NC_019843.3;YP_009047202.1;MERS_1ab | 1787 | 1787 | 1 | 0.05 | 0.05 | 0.05 |
| NC_019843.3;YP_009047202.1;MERS_1ab | 1789 | 1789 | 1 | 0.05 | 0.05 | 0.05 |
| NC_019843.3;YP_009047202.1;MERS_1ab | 1803 | 1803 | 1 | 0.2 | 0.2 | 0.2 |
| NC_019843.3;YP_009047202.1;MERS_1ab | 1805 | 1810 | 6 | 0.05 | 0.1 | 0.066666667 |
| NC_019843.3;YP_009047202.1;MERS_1ab | 1815 | 1825 | 11 | 0.05 | 0.3 | 0.181818182 |
| NC_019843.3;YP_009047202.1;MERS_1ab | 1828 | 1828 | 1 | 0.05 | 0.05 | 0.05 |
| NC_019843.3;YP_009047202.1;MERS_1ab | 1850 | 1850 | 1 | 0.1 | 0.1 | 0.1 |
| NC_019843.3;YP_009047202.1;MERS_1ab | 1858 | 1858 | 1 | 0.05 | 0.05 | 0.05 |
| NC_019843.3;YP_009047202.1;MERS_1ab | 1863 | 1863 | 1 | 0.05 | 0.05 | 0.05 |
| NC_019843.3;YP_009047202.1;MERS_1ab | 1868 | 1868 | 1 | 0.05 | 0.05 | 0.05 |
| NC_019843.3;YP_009047202.1;MERS_1ab | 1871 | 1871 | 1 | 0.05 | 0.05 | 0.05 |
| NC_019843.3;YP_009047202.1;MERS_1ab | 1873 | 1874 | 2 | 0.05 | 0.05 | 0.05 |
| NC_019843.3;YP_009047202.1;MERS_1ab | 1888 | 1889 | 2 | 0.05 | 0.05 | 0.05 |
| NC_019843.3;YP_009047202.1;MERS_1ab | 1895 | 1895 | 1 | 0.05 | 0.05 | 0.05 |
| NC_019843.3;YP_009047202.1;MERS_1ab | 1901 | 1901 | 1 | 0.05 | 0.05 | 0.05 |
| NC_019843.3;YP_009047202.1;MERS_1ab | 1905 | 1905 | 1 | 0.05 | 0.05 | 0.05 |
| NC_019843.3;YP_009047202.1;MERS_1ab | 1920 | 1922 | 3 | 0.05 | 0.1 | 0.083333333 |
| NC_019843.3;YP_009047202.1;MERS_1ab | 1924 | 1930 | 7 | 0.05 | 0.15 | 0.107142857 |
| NC_019843.3;YP_009047202.1;MERS_1ab | 1932 | 1932 | 1 | 0.05 | 0.05 | 0.05 |
| NC_019843.3;YP_009047202.1;MERS_1ab | 1938 | 1938 | 1 | 0.05 | 0.05 | 0.05 |
| NC_019843.3;YP_009047202.1;MERS_1ab | 1941 | 1949 | 9 | 0.05 | 0.3 | 0.133333333 |
| NC_019843.3;YP_009047202.1;MERS_1ab | 1955 | 1955 | 1 | 0.05 | 0.05 | 0.05 |
| NC_019843.3;YP_009047202.1;MERS_1ab | 1964 | 1973 | 10 | 0.05 | 0.05 | 0.05 |
| NC_019843.3;YP_009047202.1;MERS_1ab | 1979 | 1980 | 2 | 0.05 | 0.05 | 0.05 |
| NC_019843.3;YP_009047202.1;MERS_1ab | 1982 | 1988 | 7 | 0.05 | 0.1 | 0.078571429 |
| NC_019843.3;YP_009047202.1;MERS_1ab | 1992 | 1992 | 1 | 0.05 | 0.05 | 0.05 |

| Extended data 1 | | | | | | |
|---|---|---|---|---|---|---|
| Protein | Start position (peptide) | Stop position (peptide) | Number of peptides | Minimum fraction (per peptide) of control subjects with reactivity >3 sd above average | Maximum fraction (per peptide) of control subjects with reactivity >3 sd above average | Mean fraction (over all peptides in this contiguous stretch) of controlsubjects with reactivity >3 sd above average |
| NC_019843.3;YP_009047202.1;MERS_1ab | 2001 | 2008 | 8 | 0.05 | 0.05 | 0.05 |
| NC_019843.3;YP_009047202.1;MERS_1ab | 2013 | 2013 | 1 | 0.05 | 0.05 | 0.05 |
| NC_019843.3;YP_009047202.1;MERS_1ab | 2015 | 2015 | 1 | 0.05 | 0.05 | 0.05 |
| NC_019843.3;YP_009047202.1;MERS_1ab | 2042 | 2042 | 1 | 0.05 | 0.05 | 0.05 |
| NC_019843.3;YP_009047202.1;MERS_1ab | 2119 | 2121 | 3 | 0.05 | 0.05 | 0.05 |
| NC_019843.3;YP_009047202.1;MERS_1ab | 2143 | 2143 | 1 | 0.05 | 0.05 | 0.05 |
| NC_019843.3;YP_009047202.1;MERS_1ab | 2208 | 2208 | 1 | 0.05 | 0.05 | 0.05 |
| NC_019843.3;YP_009047202.1;MERS_1ab | 2220 | 2220 | 1 | 0.05 | 0.05 | 0.05 |
| NC_019843.3;YP_009047202.1;MERS_1ab | 2222 | 2222 | 1 | 0.05 | 0.05 | 0.05 |
| NC_019843.3;YP_009047202.1;MERS_1ab | 2228 | 2235 | 8 | 0.05 | 0.15 | 0.11875 |
| NC_019843.3;YP_009047202.1;MERS_1ab | 2239 | 2239 | 1 | 0.05 | 0.05 | 0.05 |
| NC_019843.3;YP_009047202.1;MERS_1ab | 2259 | 2259 | 1 | 0.15 | 0.15 | 0.15 |
| NC_019843.3;YP_009047202.1;MERS_1ab | 2333 | 2333 | 1 | 0.05 | 0.05 | 0.05 |
| NC_019843.3;YP_009047202.1;MERS_1ab | 2437 | 2437 | 1 | 0.05 | 0.05 | 0.05 |
| NC_019843.3;YP_009047202.1;MERS_1ab | 2439 | 2447 | 9 | 0.05 | 0.1 | 0.055555556 |
| NC_019843.3;YP_009047202.1;MERS_1ab | 2483 | 2483 | 1 | 0.05 | 0.05 | 0.05 |
| NC_019843.3;YP_009047202.1;MERS_1ab | 2498 | 2499 | 2 | 0.05 | 0.05 | 0.05 |
| NC_019843.3;YP_009047202.1;MERS_1ab | 2544 | 2544 | 1 | 0.25 | 0.25 | 0.25 |
| NC_019843.3;YP_009047202.1;MERS_1ab | 2565 | 2565 | 1 | 0.05 | 0.05 | 0.05 |
| NC_019843.3;YP_009047202.1;MERS_1ab | 2577 | 2581 | 5 | 0.05 | 0.05 | 0.05 |
| NC_019843.3;YP_009047202.1;MERS_1ab | 2588 | 2589 | 2 | 0.05 | 0.05 | 0.05 |
| NC_019843.3;YP_009047202.1;MERS_1ab | 2608 | 2608 | 1 | 0.05 | 0.05 | 0.05 |
| NC_019843.3;YP_009047202.1;MERS_1ab | 2610 | 2616 | 7 | 0.05 | 0.1 | 0.057142857 |
| NC_019843.3;YP_009047202.1;MERS_1ab | 2626 | 2634 | 9 | 0.05 | 0.2 | 0.138888889 |
| NC_019843.3;YP_009047202.1;MERS_1ab | 2637 | 2639 | 3 | 0.1 | 0.1 | 0.1 |
| NC_019843.3;YP_009047202.1;MERS_1ab | 2641 | 2644 | 4 | 0.05 | 0.45 | 0.1875 |
| NC_019843.3;YP_009047202.1;MERS_1ab | 2648 | 2648 | 1 | 0.05 | 0.05 | 0.05 |
| NC_019843.3;YP_009047202.1;MERS_1ab | 2657 | 2657 | 1 | 0.05 | 0.05 | 0.05 |
| NC_019843.3;YP_009047202.1;MERS_1ab | 2676 | 2676 | 1 | 0.05 | 0.05 | 0.05 |
| NC_019843.3;YP_009047202.1;MERS_1ab | 2682 | 2684 | 3 | 0.05 | 0.05 | 0.05 |
| NC_019843.3;YP_009047202.1;MERS_1ab | 2686 | 2686 | 1 | 0.05 | 0.05 | 0.05 |
| NC_019843.3;YP_009047202.1;MERS_1ab | 2715 | 2715 | 1 | 0.05 | 0.05 | 0.05 |
| NC_019843.3;YP_009047202.1;MERS_1ab | 2731 | 2731 | 1 | 0.05 | 0.05 | 0.05 |
| NC_019843.3;YP_009047202.1;MERS_1ab | 2780 | 2781 | 2 | 0.05 | 0.2 | 0.125 |
| NC_019843.3;YP_009047202.1;MERS_1ab | 2788 | 2789 | 2 | 0.05 | 0.05 | 0.05 |
| NC_019843.3;YP_009047202.1;MERS_1ab | 2794 | 2794 | 1 | 0.05 | 0.05 | 0.05 |
| NC_019843.3;YP_009047202.1;MERS_1ab | 2815 | 2819 | 5 | 0.05 | 0.1 | 0.06 |

| Extended data 1 | | | | | | |
|---|---|---|---|---|---|---|
| Protein | Start position (peptide) | Stop position (peptide) | Number of peptides | Minimum fraction (per peptide) of control subjects with reactivity >3 sd above average | Maximum fraction (per peptide) of control subjects with reactivity >3 sd above average | Mean fraction (over all peptides in this contiguous stretch) of controlsubjects with reactivity >3 sd above average |
| NC_019843.3;YP_009047202.1;MERS_1ab | 2821 | 2823 | 3 | 0.05 | 0.05 | 0.05 |
| NC_019843.3;YP_009047202.1;MERS_1ab | 2840 | 2842 | 3 | 0.05 | 0.15 | 0.083333333 |
| NC_019843.3;YP_009047202.1;MERS_1ab | 2844 | 2844 | 1 | 0.05 | 0.05 | 0.05 |
| NC_019843.3;YP_009047202.1;MERS_1ab | 2847 | 2849 | 3 | 0.05 | 0.1 | 0.066666667 |
| NC_019843.3;YP_009047202.1;MERS_1ab | 2870 | 2870 | 1 | 0.05 | 0.05 | 0.05 |
| NC_019843.3;YP_009047202.1;MERS_1ab | 2879 | 2880 | 2 | 0.05 | 0.05 | 0.05 |
| NC_019843.3;YP_009047202.1;MERS_1ab | 2884 | 2884 | 1 | 0.05 | 0.05 | 0.05 |
| NC_019843.3;YP_009047202.1;MERS_1ab | 2897 | 2906 | 10 | 0.05 | 0.05 | 0.05 |
| NC_019843.3;YP_009047202.1;MERS_1ab | 2910 | 2910 | 1 | 0.05 | 0.05 | 0.05 |
| NC_019843.3;YP_009047202.1;MERS_1ab | 2912 | 2912 | 1 | 0.05 | 0.05 | 0.05 |
| NC_019843.3;YP_009047202.1;MERS_1ab | 2918 | 2918 | 1 | 0.1 | 0.1 | 0.1 |
| NC_019843.3;YP_009047202.1;MERS_1ab | 2924 | 2924 | 1 | 0.05 | 0.05 | 0.05 |
| NC_019843.3;YP_009047202.1;MERS_1ab | 2930 | 2930 | 1 | 0.05 | 0.05 | 0.05 |
| NC_019843.3;YP_009047202.1;MERS_1ab | 2935 | 2938 | 4 | 0.05 | 0.05 | 0.05 |
| NC_019843.3;YP_009047202.1;MERS_1ab | 3046 | 3047 | 2 | 0.05 | 0.05 | 0.05 |
| NC_019843.3;YP_009047202.1;MERS_1ab | 3142 | 3143 | 2 | 0.05 | 0.05 | 0.05 |
| NC_019843.3;YP_009047202.1;MERS_1ab | 3150 | 3150 | 1 | 0.05 | 0.05 | 0.05 |
| NC_019843.3;YP_009047202.1;MERS_1ab | 3152 | 3155 | 4 | 0.05 | 0.05 | 0.05 |
| NC_019843.3;YP_009047202.1;MERS_1ab | 3163 | 3163 | 1 | 0.05 | 0.05 | 0.05 |
| NC_019843.3;YP_009047202.1;MERS_1ab | 3171 | 3171 | 1 | 0.05 | 0.05 | 0.05 |
| NC_019843.3;YP_009047202.1;MERS_1ab | 3173 | 3173 | 1 | 0.05 | 0.05 | 0.05 |
| NC_019843.3;YP_009047202.1;MERS_1ab | 3181 | 3181 | 1 | 0.05 | 0.05 | 0.05 |
| NC_019843.3;YP_009047202.1;MERS_1ab | 3194 | 3194 | 1 | 0.05 | 0.05 | 0.05 |
| NC_019843.3;YP_009047202.1;MERS_1ab | 3199 | 3200 | 2 | 0.05 | 0.05 | 0.05 |
| NC_019843.3;YP_009047202.1;MERS_1ab | 3214 | 3224 | 11 | 0.05 | 0.2 | 0.072727273 |
| NC_019843.3;YP_009047202.1;MERS_1ab | 3229 | 3229 | 1 | 0.05 | 0.05 | 0.05 |
| NC_019843.3;YP_009047202.1;MERS_1ab | 3234 | 3234 | 1 | 0.05 | 0.05 | 0.05 |
| NC_019843.3;YP_009047202.1;MERS_1ab | 3244 | 3247 | 4 | 0.05 | 0.1 | 0.0625 |
| NC_019843.3;YP_009047202.1;MERS_1ab | 3249 | 3249 | 1 | 0.05 | 0.05 | 0.05 |
| NC_019843.3;YP_009047202.1;MERS_1ab | 3283 | 3283 | 1 | 0.05 | 0.05 | 0.05 |
| NC_019843.3;YP_009047202.1;MERS_1ab | 3306 | 3318 | 13 | 0.05 | 0.15 | 0.107692308 |
| NC_019843.3;YP_009047202.1;MERS_1ab | 3341 | 3341 | 1 | 0.05 | 0.05 | 0.05 |
| NC_019843.3;YP_009047202.1;MERS_1ab | 3345 | 3346 | 2 | 0.05 | 0.1 | 0.075 |
| NC_019843.3;YP_009047202.1;MERS_1ab | 3389 | 3393 | 5 | 0.05 | 0.1 | 0.08 |
| NC_019843.3;YP_009047202.1;MERS_1ab | 3400 | 3400 | 1 | 0.05 | 0.05 | 0.05 |
| NC_019843.3;YP_009047202.1;MERS_1ab | 3413 | 3421 | 9 | 0.05 | 0.05 | 0.05 |
| NC_019843.3;YP_009047202.1;MERS_1ab | 3423 | 3423 | 1 | 0.05 | 0.05 | 0.05 |

| Extended data 1 | | | | | | |
|---|---|---|---|---|---|---|
| Protein | Start position (peptide) | Stop position (peptide) | Number of peptides | Minimum fraction (per peptide) of control subjects with reactivity >3 sd above average | Maximum fraction (per peptide) of control subjects with reactivity >3 sd above average | Mean fraction (over all peptides in this contiguous stretch) of controlsubjects with reactivity >3 sd above average |
| NC_019843.3;YP_009047202.1;MERS_1ab | 3491 | 3491 | 1 | 0.05 | 0.05 | 0.05 |
| NC_019843.3;YP_009047202.1;MERS_1ab | 3535 | 3535 | 1 | 0.1 | 0.1 | 0.1 |
| NC_019843.3;YP_009047202.1;MERS_1ab | 3537 | 3537 | 1 | 0.05 | 0.05 | 0.05 |
| NC_019843.3;YP_009047202.1;MERS_1ab | 3550 | 3550 | 1 | 0.05 | 0.05 | 0.05 |
| NC_019843.3;YP_009047202.1;MERS_1ab | 3646 | 3657 | 12 | 0.05 | 0.35 | 0.191666667 |
| NC_019843.3;YP_009047202.1;MERS_1ab | 3699 | 3705 | 7 | 0.05 | 0.1 | 0.057142857 |
| NC_019843.3;YP_009047202.1;MERS_1ab | 3805 | 3805 | 1 | 0.05 | 0.05 | 0.05 |
| NC_019843.3;YP_009047202.1;MERS_1ab | 3809 | 3809 | 1 | 0.05 | 0.05 | 0.05 |
| NC_019843.3;YP_009047202.1;MERS_1ab | 3811 | 3811 | 1 | 0.15 | 0.15 | 0.15 |
| NC_019843.3;YP_009047202.1;MERS_1ab | 3814 | 3815 | 2 | 0.05 | 0.05 | 0.05 |
| NC_019843.3;YP_009047202.1;MERS_1ab | 3817 | 3817 | 1 | 0.05 | 0.05 | 0.05 |
| NC_019843.3;YP_009047202.1;MERS_1ab | 3832 | 3840 | 9 | 0.05 | 0.1 | 0.055555556 |
| NC_019843.3;YP_009047202.1;MERS_1ab | 3872 | 3872 | 1 | 0.05 | 0.05 | 0.05 |
| NC_019843.3;YP_009047202.1;MERS_1ab | 3917 | 3917 | 1 | 0.05 | 0.05 | 0.05 |
| NC_019843.3;YP_009047202.1;MERS_1ab | 3936 | 3936 | 1 | 0.05 | 0.05 | 0.05 |
| NC_019843.3;YP_009047202.1;MERS_1ab | 3938 | 3942 | 5 | 0.05 | 0.05 | 0.05 |
| NC_019843.3;YP_009047202.1;MERS_1ab | 3966 | 3971 | 6 | 0.05 | 0.1 | 0.066666667 |
| NC_019843.3;YP_009047202.1;MERS_1ab | 3981 | 3981 | 1 | 0.05 | 0.05 | 0.05 |
| NC_019843.3;YP_009047202.1;MERS_1ab | 3992 | 3996 | 5 | 0.05 | 0.1 | 0.06 |
| NC_019843.3;YP_009047202.1;MERS_1ab | 4018 | 4034 | 17 | 0.05 | 0.3 | 0.144117647 |
| NC_019843.3;YP_009047202.1;MERS_1ab | 4036 | 4036 | 1 | 0.05 | 0.05 | 0.05 |
| NC_019843.3;YP_009047202.1;MERS_1ab | 4049 | 4050 | 2 | 0.05 | 0.05 | 0.05 |
| NC_019843.3;YP_009047202.1;MERS_1ab | 4053 | 4053 | 1 | 0.05 | 0.05 | 0.05 |
| NC_019843.3;YP_009047202.1;MERS_1ab | 4075 | 4075 | 1 | 0.05 | 0.05 | 0.05 |
| NC_019843.3;YP_009047202.1;MERS_1ab | 4077 | 4079 | 3 | 0.05 | 0.05 | 0.05 |
| NC_019843.3;YP_009047202.1;MERS_1ab | 4094 | 4094 | 1 | 0.05 | 0.05 | 0.05 |
| NC_019843.3;YP_009047202.1;MERS_1ab | 4096 | 4098 | 3 | 0.05 | 0.1 | 0.066666667 |
| NC_019843.3;YP_009047202.1;MERS_1ab | 4114 | 4116 | 3 | 0.05 | 0.05 | 0.05 |
| NC_019843.3;YP_009047202.1;MERS_1ab | 4126 | 4126 | 1 | 0.05 | 0.05 | 0.05 |
| NC_019843.3;YP_009047202.1;MERS_1ab | 4159 | 4160 | 2 | 0.05 | 0.05 | 0.05 |
| NC_019843.3;YP_009047202.1;MERS_1ab | 4163 | 4163 | 1 | 0.05 | 0.05 | 0.05 |
| NC_019843.3;YP_009047202.1;MERS_1ab | 4172 | 4177 | 6 | 0.05 | 0.15 | 0.075 |
| NC_019843.3;YP_009047202.1;MERS_1ab | 4180 | 4183 | 4 | 0.05 | 0.05 | 0.05 |
| NC_019843.3;YP_009047202.1;MERS_1ab | 4217 | 4217 | 1 | 0.1 | 0.1 | 0.1 |
| NC_019843.3;YP_009047202.1;MERS_1ab | 4231 | 4231 | 1 | 0.1 | 0.1 | 0.1 |
| NC_019843.3;YP_009047202.1;MERS_1ab | 4236 | 4236 | 1 | 0.05 | 0.05 | 0.05 |
| NC_019843.3;YP_009047202.1;MERS_1ab | 4245 | 4245 | 1 | 0.1 | 0.1 | 0.1 |

| Extended data 1 | | | | | | |
|---|---|---|---|---|---|---|
| **Protein** | **Start position (peptide)** | **Stop position (peptide)** | **Number of peptides** | **Minimum fraction (per peptide) of control subjects with reactivity >3 sd above average** | **Maximum fraction (per peptide) of control subjects with reactivity >3 sd above average** | **Mean fraction (over all peptides in this contiguous stretch) of controlsubjects with reactivity >3 sd above average** |
| NC_019843.3;YP_009047202.1;MERS_1ab | 4254 | 4263 | 10 | 0.05 | 0.15 | 0.1 |
| NC_019843.3;YP_009047202.1;MERS_1ab | 4273 | 4273 | 1 | 0.05 | 0.05 | 0.05 |
| NC_019843.3;YP_009047202.1;MERS_1ab | 4276 | 4276 | 1 | 0.05 | 0.05 | 0.05 |
| NC_019843.3;YP_009047202.1;MERS_1ab | 4278 | 4278 | 1 | 0.05 | 0.05 | 0.05 |
| NC_019843.3;YP_009047202.1;MERS_1ab | 4283 | 4285 | 3 | 0.05 | 0.05 | 0.05 |
| NC_019843.3;YP_009047202.1;MERS_1ab | 4329 | 4329 | 1 | 0.05 | 0.05 | 0.05 |
| NC_019843.3;YP_009047202.1;MERS_1ab | 4331 | 4332 | 2 | 0.05 | 0.05 | 0.05 |
| NC_019843.3;YP_009047202.1;MERS_1ab | 4343 | 4343 | 1 | 0.05 | 0.05 | 0.05 |
| NC_019843.3;YP_009047202.1;MERS_1ab | 4359 | 4359 | 1 | 0.05 | 0.05 | 0.05 |
| NC_019843.3;YP_009047202.1;MERS_1ab | 4368 | 4368 | 1 | 0.1 | 0.1 | 0.1 |
| NC_019843.3;YP_009047202.1;MERS_1ab | 4370 | 4380 | 11 | 0.05 | 0.15 | 0.095454545 |
| NC_019843.3;YP_009047202.1;MERS_1ab | 4400 | 4400 | 1 | 0.05 | 0.05 | 0.05 |
| NC_019843.3;YP_009047202.1;MERS_1ab | 4427 | 4433 | 7 | 0.05 | 0.15 | 0.071428571 |
| NC_019843.3;YP_009047202.1;MERS_1ab | 4446 | 4446 | 1 | 0.05 | 0.05 | 0.05 |
| NC_019843.3;YP_009047202.1;MERS_1ab | 4460 | 4460 | 1 | 0.05 | 0.05 | 0.05 |
| NC_019843.3;YP_009047202.1;MERS_1ab | 4523 | 4525 | 3 | 0.05 | 0.2 | 0.1 |
| NC_019843.3;YP_009047202.1;MERS_1ab | 4531 | 4531 | 1 | 0.05 | 0.05 | 0.05 |
| NC_019843.3;YP_009047202.1;MERS_1ab | 4535 | 4535 | 1 | 0.05 | 0.05 | 0.05 |
| NC_019843.3;YP_009047202.1;MERS_1ab | 4558 | 4558 | 1 | 0.05 | 0.05 | 0.05 |
| NC_019843.3;YP_009047202.1;MERS_1ab | 4577 | 4577 | 1 | 0.05 | 0.05 | 0.05 |
| NC_019843.3;YP_009047202.1;MERS_1ab | 4581 | 4581 | 1 | 0.15 | 0.15 | 0.15 |
| NC_019843.3;YP_009047202.1;MERS_1ab | 4583 | 4583 | 1 | 0.05 | 0.05 | 0.05 |
| NC_019843.3;YP_009047202.1;MERS_1ab | 4595 | 4595 | 1 | 0.1 | 0.1 | 0.1 |
| NC_019843.3;YP_009047202.1;MERS_1ab | 4616 | 4616 | 1 | 0.05 | 0.05 | 0.05 |
| NC_019843.3;YP_009047202.1;MERS_1ab | 4638 | 4639 | 2 | 0.05 | 0.05 | 0.05 |
| NC_019843.3;YP_009047202.1;MERS_1ab | 4642 | 4650 | 9 | 0.05 | 0.15 | 0.077777778 |
| NC_019843.3;YP_009047202.1;MERS_1ab | 4666 | 4666 | 1 | 0.05 | 0.05 | 0.05 |
| NC_019843.3;YP_009047202.1;MERS_1ab | 4725 | 4725 | 1 | 0.05 | 0.05 | 0.05 |
| NC_019843.3;YP_009047202.1;MERS_1ab | 4730 | 4731 | 2 | 0.05 | 0.05 | 0.05 |
| NC_019843.3;YP_009047202.1;MERS_1ab | 4733 | 4744 | 12 | 0.05 | 0.15 | 0.108333333 |
| NC_019843.3;YP_009047202.1;MERS_1ab | 4749 | 4760 | 12 | 0.05 | 0.1 | 0.0625 |
| NC_019843.3;YP_009047202.1;MERS_1ab | 4763 | 4763 | 1 | 0.05 | 0.05 | 0.05 |
| NC_019843.3;YP_009047202.1;MERS_1ab | 4778 | 4778 | 1 | 0.05 | 0.05 | 0.05 |
| NC_019843.3;YP_009047202.1;MERS_1ab | 4781 | 4782 | 2 | 0.05 | 0.05 | 0.05 |
| NC_019843.3;YP_009047202.1;MERS_1ab | 4784 | 4790 | 7 | 0.05 | 0.2 | 0.107142857 |
| NC_019843.3;YP_009047202.1;MERS_1ab | 4795 | 4795 | 1 | 0.05 | 0.05 | 0.05 |
| NC_019843.3;YP_009047202.1;MERS_1ab | 4812 | 4812 | 1 | 0.05 | 0.05 | 0.05 |

| Extended data 1 | | | | | | |
|---|---|---|---|---|---|---|
| Protein | Start position (peptide) | Stop position (peptide) | Number of peptides | Minimum fraction (per peptide) of control subjects with reactivity >3 sd above average | Maximum fraction (per peptide) of control subjects with reactivity >3 sd above average | Mean fraction (over all peptides in this contiguous stretch) of controlsubjects with reactivity >3 sd above average |
| NC_019843.3;YP_009047202.1;MERS_1ab | 4819 | 4819 | 1 | 0.05 | 0.05 | 0.05 |
| NC_019843.3;YP_009047202.1;MERS_1ab | 4822 | 4822 | 1 | 0.05 | 0.05 | 0.05 |
| NC_019843.3;YP_009047202.1;MERS_1ab | 4824 | 4824 | 1 | 0.05 | 0.05 | 0.05 |
| NC_019843.3;YP_009047202.1;MERS_1ab | 4826 | 4830 | 5 | 0.05 | 0.1 | 0.08 |
| NC_019843.3;YP_009047202.1;MERS_1ab | 4842 | 4842 | 1 | 0.05 | 0.05 | 0.05 |
| NC_019843.3;YP_009047202.1;MERS_1ab | 4865 | 4865 | 1 | 0.05 | 0.05 | 0.05 |
| NC_019843.3;YP_009047202.1;MERS_1ab | 4868 | 4868 | 1 | 0.05 | 0.05 | 0.05 |
| NC_019843.3;YP_009047202.1;MERS_1ab | 4871 | 4871 | 1 | 0.05 | 0.05 | 0.05 |
| NC_019843.3;YP_009047202.1;MERS_1ab | 4895 | 4895 | 1 | 0.05 | 0.05 | 0.05 |
| NC_019843.3;YP_009047202.1;MERS_1ab | 4902 | 4904 | 3 | 0.05 | 0.05 | 0.05 |
| NC_019843.3;YP_009047202.1;MERS_1ab | 4909 | 4910 | 2 | 0.05 | 0.1 | 0.075 |
| NC_019843.3;YP_009047202.1;MERS_1ab | 4944 | 4944 | 1 | 0.05 | 0.05 | 0.05 |
| NC_019843.3;YP_009047202.1;MERS_1ab | 4949 | 4958 | 10 | 0.05 | 0.4 | 0.225 |
| NC_019843.3;YP_009047202.1;MERS_1ab | 4975 | 4975 | 1 | 0.05 | 0.05 | 0.05 |
| NC_019843.3;YP_009047202.1;MERS_1ab | 4985 | 4985 | 1 | 0.05 | 0.05 | 0.05 |
| NC_019843.3;YP_009047202.1;MERS_1ab | 5024 | 5026 | 3 | 0.05 | 0.1 | 0.066666667 |
| NC_019843.3;YP_009047202.1;MERS_1ab | 5035 | 5035 | 1 | 0.05 | 0.05 | 0.05 |
| NC_019843.3;YP_009047202.1;MERS_1ab | 5042 | 5042 | 1 | 0.05 | 0.05 | 0.05 |
| NC_019843.3;YP_009047202.1;MERS_1ab | 5050 | 5050 | 1 | 0.05 | 0.05 | 0.05 |
| NC_019843.3;YP_009047202.1;MERS_1ab | 5055 | 5056 | 2 | 0.05 | 0.1 | 0.075 |
| NC_019843.3;YP_009047202.1;MERS_1ab | 5058 | 5058 | 1 | 0.05 | 0.05 | 0.05 |
| NC_019843.3;YP_009047202.1;MERS_1ab | 5072 | 5073 | 2 | 0.05 | 0.05 | 0.05 |
| NC_019843.3;YP_009047202.1;MERS_1ab | 5077 | 5088 | 12 | 0.05 | 0.1 | 0.0625 |
| NC_019843.3;YP_009047202.1;MERS_1ab | 5106 | 5106 | 1 | 0.1 | 0.1 | 0.1 |
| NC_019843.3;YP_009047202.1;MERS_1ab | 5108 | 5110 | 3 | 0.05 | 0.1 | 0.066666667 |
| NC_019843.3;YP_009047202.1;MERS_1ab | 5114 | 5114 | 1 | 0.05 | 0.05 | 0.05 |
| NC_019843.3;YP_009047202.1;MERS_1ab | 5125 | 5125 | 1 | 0.05 | 0.05 | 0.05 |
| NC_019843.3;YP_009047202.1;MERS_1ab | 5135 | 5137 | 3 | 0.05 | 0.05 | 0.05 |
| NC_019843.3;YP_009047202.1;MERS_1ab | 5148 | 5162 | 15 | 0.05 | 0.15 | 0.06 |
| NC_019843.3;YP_009047202.1;MERS_1ab | 5172 | 5172 | 1 | 0.05 | 0.05 | 0.05 |
| NC_019843.3;YP_009047202.1;MERS_1ab | 5179 | 5179 | 1 | 0.05 | 0.05 | 0.05 |
| NC_019843.3;YP_009047202.1;MERS_1ab | 5199 | 5199 | 1 | 0.05 | 0.05 | 0.05 |
| NC_019843.3;YP_009047202.1;MERS_1ab | 5215 | 5215 | 1 | 0.05 | 0.05 | 0.05 |
| NC_019843.3;YP_009047202.1;MERS_1ab | 5260 | 5260 | 1 | 0.05 | 0.05 | 0.05 |
| NC_019843.3;YP_009047202.1;MERS_1ab | 5263 | 5263 | 1 | 0.05 | 0.05 | 0.05 |
| NC_019843.3;YP_009047202.1;MERS_1ab | 5265 | 5265 | 1 | 0.05 | 0.05 | 0.05 |
| NC_019843.3;YP_009047202.1;MERS_1ab | 5277 | 5277 | 1 | 0.05 | 0.05 | 0.05 |

| Extended data 1 | | | | | | |
|---|---|---|---|---|---|---|
| Protein | Start position (peptide) | Stop position (peptide) | Number of peptides | Minimum fraction (per peptide) of control subjects with reactivity >3 sd above average | Maximum fraction (per peptide) of control subjects with reactivity >3 sd above average | Mean fraction (over all peptides in this contiguous stretch) of controlsubjects with reactivity >3 sd above average |
| NC_019843.3;YP_009047202.1;MERS_1ab | 5285 | 5285 | 1 | 0.05 | 0.05 | 0.05 |
| NC_019843.3;YP_009047202.1;MERS_1ab | 5289 | 5289 | 1 | 0.05 | 0.05 | 0.05 |
| NC_019843.3;YP_009047202.1;MERS_1ab | 5357 | 5357 | 1 | 0.05 | 0.05 | 0.05 |
| NC_019843.3;YP_009047202.1;MERS_1ab | 5366 | 5366 | 1 | 0.05 | 0.05 | 0.05 |
| NC_019843.3;YP_009047202.1;MERS_1ab | 5427 | 5428 | 2 | 0.05 | 0.05 | 0.05 |
| NC_019843.3;YP_009047202.1;MERS_1ab | 5430 | 5430 | 1 | 0.05 | 0.05 | 0.05 |
| NC_019843.3;YP_009047202.1;MERS_1ab | 5465 | 5465 | 1 | 0.05 | 0.05 | 0.05 |
| NC_019843.3;YP_009047202.1;MERS_1ab | 5470 | 5470 | 1 | 0.05 | 0.05 | 0.05 |
| NC_019843.3;YP_009047202.1;MERS_1ab | 5475 | 5475 | 1 | 0.05 | 0.05 | 0.05 |
| NC_019843.3;YP_009047202.1;MERS_1ab | 5492 | 5494 | 3 | 0.05 | 0.1 | 0.083333333 |
| NC_019843.3;YP_009047202.1;MERS_1ab | 5497 | 5497 | 1 | 0.05 | 0.05 | 0.05 |
| NC_019843.3;YP_009047202.1;MERS_1ab | 5507 | 5509 | 3 | 0.05 | 0.1 | 0.066666667 |
| NC_019843.3;YP_009047202.1;MERS_1ab | 5512 | 5513 | 2 | 0.05 | 0.05 | 0.05 |
| NC_019843.3;YP_009047202.1;MERS_1ab | 5519 | 5519 | 1 | 0.05 | 0.05 | 0.05 |
| NC_019843.3;YP_009047202.1;MERS_1ab | 5546 | 5546 | 1 | 0.05 | 0.05 | 0.05 |
| NC_019843.3;YP_009047202.1;MERS_1ab | 5579 | 5581 | 3 | 0.05 | 0.05 | 0.05 |
| NC_019843.3;YP_009047202.1;MERS_1ab | 5608 | 5608 | 1 | 0.05 | 0.05 | 0.05 |
| NC_019843.3;YP_009047202.1;MERS_1ab | 5645 | 5645 | 1 | 0.05 | 0.05 | 0.05 |
| NC_019843.3;YP_009047202.1;MERS_1ab | 5647 | 5662 | 16 | 0.05 | 0.4 | 0.178125 |
| NC_019843.3;YP_009047202.1;MERS_1ab | 5669 | 5677 | 9 | 0.05 | 0.1 | 0.072222222 |
| NC_019843.3;YP_009047202.1;MERS_1ab | 5682 | 5682 | 1 | 0.05 | 0.05 | 0.05 |
| NC_019843.3;YP_009047202.1;MERS_1ab | 5696 | 5696 | 1 | 0.05 | 0.05 | 0.05 |
| NC_019843.3;YP_009047202.1;MERS_1ab | 5699 | 5699 | 1 | 0.05 | 0.05 | 0.05 |
| NC_019843.3;YP_009047202.1;MERS_1ab | 5718 | 5718 | 1 | 0.05 | 0.05 | 0.05 |
| NC_019843.3;YP_009047202.1;MERS_1ab | 5720 | 5722 | 3 | 0.05 | 0.05 | 0.05 |
| NC_019843.3;YP_009047202.1;MERS_1ab | 5748 | 5750 | 3 | 0.05 | 0.05 | 0.05 |
| NC_019843.3;YP_009047202.1;MERS_1ab | 5754 | 5754 | 1 | 0.05 | 0.05 | 0.05 |
| NC_019843.3;YP_009047202.1;MERS_1ab | 5767 | 5767 | 1 | 0.05 | 0.05 | 0.05 |
| NC_019843.3;YP_009047202.1;MERS_1ab | 5769 | 5769 | 1 | 0.05 | 0.05 | 0.05 |
| NC_019843.3;YP_009047202.1;MERS_1ab | 5786 | 5787 | 2 | 0.05 | 0.05 | 0.05 |
| NC_019843.3;YP_009047202.1;MERS_1ab | 5792 | 5793 | 2 | 0.05 | 0.2 | 0.125 |
| NC_019843.3;YP_009047202.1;MERS_1ab | 5814 | 5814 | 1 | 0.15 | 0.15 | 0.15 |
| NC_019843.3;YP_009047202.1;MERS_1ab | 5817 | 5817 | 1 | 0.1 | 0.1 | 0.1 |
| NC_019843.3;YP_009047202.1;MERS_1ab | 5819 | 5819 | 1 | 0.05 | 0.05 | 0.05 |
| NC_019843.3;YP_009047202.1;MERS_1ab | 5822 | 5824 | 3 | 0.05 | 0.15 | 0.083333333 |
| NC_019843.3;YP_009047202.1;MERS_1ab | 5852 | 5852 | 1 | 0.05 | 0.05 | 0.05 |
| NC_019843.3;YP_009047202.1;MERS_1ab | 5854 | 5866 | 13 | 0.05 | 0.2 | 0.111538462 |

| Extended data 1 | | | | | | |
|---|---|---|---|---|---|---|
| Protein | Start position (peptide) | Stop position (peptide) | Number of peptides | Minimum fraction (per peptide) of control subjects with reactivity >3 sd above average | Maximum fraction (per peptide) of control subjects with reactivity >3 sd above average | Mean fraction (over all peptides in this contiguous stretch) of controlsubjects with reactivity >3 sd above average |
| NC_019843.3;YP_009047202.1;MERS_1ab | 5881 | 5882 | 2 | 0.05 | 0.05 | 0.05 |
| NC_019843.3;YP_009047202.1;MERS_1ab | 5917 | 5917 | 1 | 0.05 | 0.05 | 0.05 |
| NC_019843.3;YP_009047202.1;MERS_1ab | 5919 | 5919 | 1 | 0.05 | 0.05 | 0.05 |
| NC_019843.3;YP_009047202.1;MERS_1ab | 5924 | 5932 | 9 | 0.05 | 0.15 | 0.083333333 |
| NC_019843.3;YP_009047202.1;MERS_1ab | 5934 | 5936 | 3 | 0.05 | 0.05 | 0.05 |
| NC_019843.3;YP_009047202.1;MERS_1ab | 5941 | 5942 | 2 | 0.05 | 0.05 | 0.05 |
| NC_019843.3;YP_009047202.1;MERS_1ab | 5961 | 5961 | 1 | 0.05 | 0.05 | 0.05 |
| NC_019843.3;YP_009047202.1;MERS_1ab | 5964 | 5964 | 1 | 0.05 | 0.05 | 0.05 |
| NC_019843.3;YP_009047202.1;MERS_1ab | 6031 | 6031 | 1 | 0.1 | 0.1 | 0.1 |
| NC_019843.3;YP_009047202.1;MERS_1ab | 6051 | 6051 | 1 | 0.05 | 0.05 | 0.05 |
| NC_019843.3;YP_009047202.1;MERS_1ab | 6069 | 6070 | 2 | 0.05 | 0.05 | 0.05 |
| NC_019843.3;YP_009047202.1;MERS_1ab | 6099 | 6099 | 1 | 0.05 | 0.05 | 0.05 |
| NC_019843.3;YP_009047202.1;MERS_1ab | 6104 | 6104 | 1 | 0.05 | 0.05 | 0.05 |
| NC_019843.3;YP_009047202.1;MERS_1ab | 6109 | 6109 | 1 | 0.05 | 0.05 | 0.05 |
| NC_019843.3;YP_009047202.1;MERS_1ab | 6163 | 6163 | 1 | 0.05 | 0.05 | 0.05 |
| NC_019843.3;YP_009047202.1;MERS_1ab | 6168 | 6170 | 3 | 0.05 | 0.05 | 0.05 |
| NC_019843.3;YP_009047202.1;MERS_1ab | 6196 | 6203 | 8 | 0.05 | 0.15 | 0.0875 |
| NC_019843.3;YP_009047202.1;MERS_1ab | 6213 | 6213 | 1 | 0.05 | 0.05 | 0.05 |
| NC_019843.3;YP_009047202.1;MERS_1ab | 6230 | 6242 | 13 | 0.05 | 0.15 | 0.073076923 |
| NC_019843.3;YP_009047202.1;MERS_1ab | 6251 | 6251 | 1 | 0.05 | 0.05 | 0.05 |
| NC_019843.3;YP_009047202.1;MERS_1ab | 6261 | 6261 | 1 | 0.05 | 0.05 | 0.05 |
| NC_019843.3;YP_009047202.1;MERS_1ab | 6263 | 6263 | 1 | 0.05 | 0.05 | 0.05 |
| NC_019843.3;YP_009047202.1;MERS_1ab | 6275 | 6275 | 1 | 0.05 | 0.05 | 0.05 |
| NC_019843.3;YP_009047202.1;MERS_1ab | 6281 | 6281 | 1 | 0.05 | 0.05 | 0.05 |
| NC_019843.3;YP_009047202.1;MERS_1ab | 6330 | 6330 | 1 | 0.1 | 0.1 | 0.1 |
| NC_019843.3;YP_009047202.1;MERS_1ab | 6332 | 6332 | 1 | 0.05 | 0.05 | 0.05 |
| NC_019843.3;YP_009047202.1;MERS_1ab | 6357 | 6357 | 1 | 0.05 | 0.05 | 0.05 |
| NC_019843.3;YP_009047202.1;MERS_1ab | 6361 | 6362 | 2 | 0.05 | 0.05 | 0.05 |
| NC_019843.3;YP_009047202.1;MERS_1ab | 6382 | 6388 | 7 | 0.05 | 0.1 | 0.071428571 |
| NC_019843.3;YP_009047202.1;MERS_1ab | 6393 | 6393 | 1 | 0.05 | 0.05 | 0.05 |
| NC_019843.3;YP_009047202.1;MERS_1ab | 6433 | 6433 | 1 | 0.1 | 0.1 | 0.1 |
| NC_019843.3;YP_009047202.1;MERS_1ab | 6440 | 6442 | 3 | 0.05 | 0.05 | 0.05 |
| NC_019843.3;YP_009047202.1;MERS_1ab | 6447 | 6454 | 8 | 0.05 | 0.15 | 0.1 |
| NC_019843.3;YP_009047202.1;MERS_1ab | 6468 | 6469 | 2 | 0.05 | 0.35 | 0.2 |
| NC_019843.3;YP_009047202.1;MERS_1ab | 6471 | 6474 | 4 | 0.05 | 0.1 | 0.075 |
| NC_019843.3;YP_009047202.1;MERS_1ab | 6476 | 6476 | 1 | 0.1 | 0.1 | 0.1 |
| NC_019843.3;YP_009047202.1;MERS_1ab | 6489 | 6490 | 2 | 0.05 | 0.05 | 0.05 |

| Extended data 1 | | | | | | |
|---|---|---|---|---|---|---|
| Protein | Start position (peptide) | Stop position (peptide) | Number of peptides | Minimum fraction (per peptide) of control subjects with reactivity >3 sd above average | Maximum fraction (per peptide) of control subjects with reactivity >3 sd above average | Mean fraction (over all peptides in this contiguous stretch) of controlsubjects with reactivity >3 sd above average |
| NC_019843.3;YP_009047202.1;MERS_1ab | 6497 | 6497 | 1 | 0.05 | 0.05 | 0.05 |
| NC_019843.3;YP_009047202.1;MERS_1ab | 6499 | 6499 | 1 | 0.05 | 0.05 | 0.05 |
| NC_019843.3;YP_009047202.1;MERS_1ab | 6528 | 6528 | 1 | 0.05 | 0.05 | 0.05 |
| NC_019843.3;YP_009047202.1;MERS_1ab | 6550 | 6551 | 2 | 0.05 | 0.05 | 0.05 |
| NC_019843.3;YP_009047202.1;MERS_1ab | 6560 | 6567 | 8 | 0.05 | 0.1 | 0.05625 |
| NC_019843.3;YP_009047202.1;MERS_1ab | 6590 | 6590 | 1 | 0.05 | 0.05 | 0.05 |
| NC_019843.3;YP_009047202.1;MERS_1ab | 6602 | 6602 | 1 | 0.05 | 0.05 | 0.05 |
| NC_019843.3;YP_009047202.1;MERS_1ab | 6605 | 6611 | 7 | 0.05 | 0.35 | 0.192857143 |
| NC_019843.3;YP_009047202.1;MERS_1ab | 6626 | 6627 | 2 | 0.05 | 0.05 | 0.05 |
| NC_019843.3;YP_009047202.1;MERS_1ab | 6638 | 6638 | 1 | 0.05 | 0.05 | 0.05 |
| NC_019843.3;YP_009047202.1;MERS_1ab | 6643 | 6643 | 1 | 0.05 | 0.05 | 0.05 |
| NC_019843.3;YP_009047202.1;MERS_1ab | 6655 | 6655 | 1 | 0.05 | 0.05 | 0.05 |
| NC_019843.3;YP_009047202.1;MERS_1ab | 6682 | 6682 | 1 | 0.05 | 0.05 | 0.05 |
| NC_019843.3;YP_009047202.1;MERS_1ab | 6684 | 6684 | 1 | 0.05 | 0.05 | 0.05 |
| NC_019843.3;YP_009047202.1;MERS_1ab | 6691 | 6691 | 1 | 0.05 | 0.05 | 0.05 |
| NC_019843.3;YP_009047202.1;MERS_1ab | 6701 | 6701 | 1 | 0.05 | 0.05 | 0.05 |
| NC_019843.3;YP_009047202.1;MERS_1ab | 6704 | 6704 | 1 | 0.05 | 0.05 | 0.05 |
| NC_019843.3;YP_009047202.1;MERS_1ab | 6722 | 6723 | 2 | 0.05 | 0.05 | 0.05 |
| NC_019843.3;YP_009047202.1;MERS_1ab | 6727 | 6727 | 1 | 0.05 | 0.05 | 0.05 |
| NC_019843.3;YP_009047202.1;MERS_1ab | 6754 | 6754 | 1 | 0.05 | 0.05 | 0.05 |
| NC_019843.3;YP_009047202.1;MERS_1ab | 6762 | 6763 | 2 | 0.05 | 0.05 | 0.05 |
| NC_019843.3;YP_009047202.1;MERS_1ab | 6772 | 6773 | 2 | 0.05 | 0.05 | 0.05 |
| NC_019843.3;YP_009047202.1;MERS_1ab | 6790 | 6790 | 1 | 0.05 | 0.05 | 0.05 |
| NC_019843.3;YP_009047202.1;MERS_1ab | 6792 | 6792 | 1 | 0.05 | 0.05 | 0.05 |
| NC_019843.3;YP_009047202.1;MERS_1ab | 6798 | 6798 | 1 | 0.05 | 0.05 | 0.05 |
| NC_019843.3;YP_009047202.1;MERS_1ab | 6810 | 6810 | 1 | 0.05 | 0.05 | 0.05 |
| NC_019843.3;YP_009047202.1;MERS_1ab | 6848 | 6848 | 1 | 0.05 | 0.05 | 0.05 |
| NC_019843.3;YP_009047202.1;MERS_1ab | 6866 | 6869 | 4 | 0.05 | 0.15 | 0.075 |
| NC_019843.3;YP_009047202.1;MERS_1ab | 6871 | 6871 | 1 | 0.05 | 0.05 | 0.05 |
| NC_019843.3;YP_009047202.1;MERS_1ab | 6877 | 6877 | 1 | 0.05 | 0.05 | 0.05 |
| NC_019843.3;YP_009047202.1;MERS_1ab | 6885 | 6885 | 1 | 0.05 | 0.05 | 0.05 |
| NC_019843.3;YP_009047202.1;MERS_1ab | 6899 | 6912 | 14 | 0.05 | 0.3 | 0.171428571 |
| NC_019843.3;YP_009047202.1;MERS_1ab | 6928 | 6928 | 1 | 0.05 | 0.05 | 0.05 |
| NC_019843.3;YP_009047202.1;MERS_1ab | 6970 | 6970 | 1 | 0.05 | 0.05 | 0.05 |
| NC_019843.3;YP_009047202.1;MERS_1ab | 6983 | 6983 | 1 | 0.05 | 0.05 | 0.05 |
| NC_019843.3;YP_009047202.1;MERS_1ab | 7030 | 7031 | 2 | 0.05 | 0.05 | 0.05 |
| NC_019843.3;YP_009047202.1;MERS_1ab | 7035 | 7035 | 1 | 0.05 | 0.05 | 0.05 |

| Extended data 1 | | | | | | |
|---|---|---|---|---|---|---|
| Protein | Start position (peptide) | Stop position (peptide) | Number of peptides | Minimum fraction (per peptide) of control subjects with reactivity >3 sd above average | Maximum fraction (per peptide) of control subjects with reactivity >3 sd above average | Mean fraction (over all peptides in this contiguous stretch) of controlsubjects with reactivity >3 sd above average |
| NC_019843.3;YP_009047202.1;MERS_1ab | 7047 | 7054 | 8 | 0.05 | 0.15 | 0.08125 |
| NC_019843.3;YP_009047203.1;MERS_1a | 4 | 4 | 1 | 0.05 | 0.05 | 0.05 |
| NC_019843.3;YP_009047203.1;MERS_1a | 8 | 8 | 1 | 0.05 | 0.05 | 0.05 |
| NC_019843.3;YP_009047203.1;MERS_1a | 14 | 14 | 1 | 0.05 | 0.05 | 0.05 |
| NC_019843.3;YP_009047203.1;MERS_1a | 16 | 16 | 1 | 0.05 | 0.05 | 0.05 |
| NC_019843.3;YP_009047203.1;MERS_1a | 19 | 19 | 1 | 0.1 | 0.1 | 0.1 |
| NC_019843.3;YP_009047203.1;MERS_1a | 29 | 39 | 11 | 0.05 | 0.15 | 0.059090909 |
| NC_019843.3;YP_009047203.1;MERS_1a | 46 | 46 | 1 | 0.05 | 0.05 | 0.05 |
| NC_019843.3;YP_009047203.1;MERS_1a | 92 | 92 | 1 | 0.1 | 0.1 | 0.1 |
| NC_019843.3;YP_009047203.1;MERS_1a | 98 | 98 | 1 | 0.05 | 0.05 | 0.05 |
| NC_019843.3;YP_009047203.1;MERS_1a | 101 | 104 | 4 | 0.05 | 0.05 | 0.05 |
| NC_019843.3;YP_009047203.1;MERS_1a | 107 | 107 | 1 | 0.05 | 0.05 | 0.05 |
| NC_019843.3;YP_009047203.1;MERS_1a | 120 | 120 | 1 | 0.05 | 0.05 | 0.05 |
| NC_019843.3;YP_009047203.1;MERS_1a | 126 | 128 | 3 | 0.05 | 0.1 | 0.066666667 |
| NC_019843.3;YP_009047203.1;MERS_1a | 163 | 170 | 8 | 0.05 | 0.1 | 0.09375 |
| NC_019843.3;YP_009047203.1;MERS_1a | 176 | 176 | 1 | 0.05 | 0.05 | 0.05 |
| NC_019843.3;YP_009047203.1;MERS_1a | 184 | 184 | 1 | 0.05 | 0.05 | 0.05 |
| NC_019843.3;YP_009047203.1;MERS_1a | 186 | 188 | 3 | 0.05 | 0.1 | 0.083333333 |
| NC_019843.3;YP_009047203.1;MERS_1a | 196 | 197 | 2 | 0.05 | 0.1 | 0.075 |
| NC_019843.3;YP_009047203.1;MERS_1a | 199 | 199 | 1 | 0.05 | 0.05 | 0.05 |
| NC_019843.3;YP_009047203.1;MERS_1a | 219 | 219 | 1 | 0.05 | 0.05 | 0.05 |
| NC_019843.3;YP_009047203.1;MERS_1a | 232 | 232 | 1 | 0.05 | 0.05 | 0.05 |
| NC_019843.3;YP_009047203.1;MERS_1a | 259 | 260 | 2 | 0.05 | 0.05 | 0.05 |
| NC_019843.3;YP_009047203.1;MERS_1a | 267 | 267 | 1 | 0.05 | 0.05 | 0.05 |
| NC_019843.3;YP_009047203.1;MERS_1a | 304 | 304 | 1 | 0.05 | 0.05 | 0.05 |
| NC_019843.3;YP_009047203.1;MERS_1a | 334 | 334 | 1 | 0.05 | 0.05 | 0.05 |
| NC_019843.3;YP_009047203.1;MERS_1a | 340 | 340 | 1 | 0.05 | 0.05 | 0.05 |
| NC_019843.3;YP_009047203.1;MERS_1a | 347 | 347 | 1 | 0.05 | 0.05 | 0.05 |
| NC_019843.3;YP_009047203.1;MERS_1a | 356 | 356 | 1 | 0.05 | 0.05 | 0.05 |
| NC_019843.3;YP_009047203.1;MERS_1a | 388 | 388 | 1 | 0.05 | 0.05 | 0.05 |
| NC_019843.3;YP_009047203.1;MERS_1a | 392 | 392 | 1 | 0.05 | 0.05 | 0.05 |
| NC_019843.3;YP_009047203.1;MERS_1a | 405 | 408 | 4 | 0.05 | 0.1 | 0.075 |
| NC_019843.3;YP_009047203.1;MERS_1a | 411 | 411 | 1 | 0.05 | 0.05 | 0.05 |
| NC_019843.3;YP_009047203.1;MERS_1a | 467 | 467 | 1 | 0.05 | 0.05 | 0.05 |
| NC_019843.3;YP_009047203.1;MERS_1a | 489 | 489 | 1 | 0.05 | 0.05 | 0.05 |
| NC_019843.3;YP_009047203.1;MERS_1a | 497 | 502 | 6 | 0.05 | 0.05 | 0.05 |
| NC_019843.3;YP_009047203.1;MERS_1a | 505 | 505 | 1 | 0.05 | 0.05 | 0.05 |

| Extended data 1 | | | | | | |
|---|---|---|---|---|---|---|
| Protein | Start position (peptide) | Stop position (peptide) | Number of peptides | Minimum fraction (per peptide) of control subjects with reactivity >3 sd above average | Maximum fraction (per peptide) of control subjects with reactivity >3 sd above average | Mean fraction (over all peptides in this contiguous stretch) of controlsubjects with reactivity >3 sd above average |
| NC_019843.3;YP_009047203.1;MERS_1a | 512 | 519 | 8 | 0.05 | 0.1 | 0.06875 |
| NC_019843.3;YP_009047203.1;MERS_1a | 521 | 521 | 1 | 0.05 | 0.05 | 0.05 |
| NC_019843.3;YP_009047203.1;MERS_1a | 526 | 526 | 1 | 0.05 | 0.05 | 0.05 |
| NC_019843.3;YP_009047203.1;MERS_1a | 529 | 530 | 2 | 0.05 | 0.05 | 0.05 |
| NC_019843.3;YP_009047203.1;MERS_1a | 535 | 536 | 2 | 0.05 | 0.05 | 0.05 |
| NC_019843.3;YP_009047203.1;MERS_1a | 539 | 539 | 1 | 0.1 | 0.1 | 0.1 |
| NC_019843.3;YP_009047203.1;MERS_1a | 564 | 564 | 1 | 0.05 | 0.05 | 0.05 |
| NC_019843.3;YP_009047203.1;MERS_1a | 567 | 567 | 1 | 0.05 | 0.05 | 0.05 |
| NC_019843.3;YP_009047203.1;MERS_1a | 574 | 574 | 1 | 0.05 | 0.05 | 0.05 |
| NC_019843.3;YP_009047203.1;MERS_1a | 594 | 598 | 5 | 0.05 | 0.15 | 0.07 |
| NC_019843.3;YP_009047203.1;MERS_1a | 692 | 692 | 1 | 0.05 | 0.05 | 0.05 |
| NC_019843.3;YP_009047203.1;MERS_1a | 701 | 703 | 3 | 0.1 | 0.15 | 0.116666667 |
| NC_019843.3;YP_009047203.1;MERS_1a | 729 | 729 | 1 | 0.05 | 0.05 | 0.05 |
| NC_019843.3;YP_009047203.1;MERS_1a | 745 | 746 | 2 | 0.05 | 0.05 | 0.05 |
| NC_019843.3;YP_009047203.1;MERS_1a | 749 | 749 | 1 | 0.05 | 0.05 | 0.05 |
| NC_019843.3;YP_009047203.1;MERS_1a | 751 | 751 | 1 | 0.1 | 0.1 | 0.1 |
| NC_019843.3;YP_009047203.1;MERS_1a | 757 | 757 | 1 | 0.05 | 0.05 | 0.05 |
| NC_019843.3;YP_009047203.1;MERS_1a | 766 | 766 | 1 | 0.05 | 0.05 | 0.05 |
| NC_019843.3;YP_009047203.1;MERS_1a | 771 | 772 | 2 | 0.05 | 0.05 | 0.05 |
| NC_019843.3;YP_009047203.1;MERS_1a | 778 | 781 | 4 | 0.05 | 0.1 | 0.075 |
| NC_019843.3;YP_009047203.1;MERS_1a | 783 | 787 | 5 | 0.05 | 0.05 | 0.05 |
| NC_019843.3;YP_009047203.1;MERS_1a | 790 | 790 | 1 | 0.05 | 0.05 | 0.05 |
| NC_019843.3;YP_009047203.1;MERS_1a | 817 | 818 | 2 | 0.05 | 0.05 | 0.05 |
| NC_019843.3;YP_009047203.1;MERS_1a | 820 | 822 | 3 | 0.05 | 0.1 | 0.066666667 |
| NC_019843.3;YP_009047203.1;MERS_1a | 824 | 828 | 5 | 0.05 | 0.1 | 0.06 |
| NC_019843.3;YP_009047203.1;MERS_1a | 852 | 852 | 1 | 0.05 | 0.05 | 0.05 |
| NC_019843.3;YP_009047203.1;MERS_1a | 856 | 856 | 1 | 0.05 | 0.05 | 0.05 |
| NC_019843.3;YP_009047203.1;MERS_1a | 881 | 881 | 1 | 0.05 | 0.05 | 0.05 |
| NC_019843.3;YP_009047203.1;MERS_1a | 889 | 890 | 2 | 0.05 | 0.05 | 0.05 |
| NC_019843.3;YP_009047203.1;MERS_1a | 895 | 895 | 1 | 0.05 | 0.05 | 0.05 |
| NC_019843.3;YP_009047203.1;MERS_1a | 915 | 915 | 1 | 0.05 | 0.05 | 0.05 |
| NC_019843.3;YP_009047203.1;MERS_1a | 920 | 928 | 9 | 0.05 | 0.15 | 0.083333333 |
| NC_019843.3;YP_009047203.1;MERS_1a | 930 | 931 | 2 | 0.05 | 0.05 | 0.05 |
| NC_019843.3;YP_009047203.1;MERS_1a | 959 | 959 | 1 | 0.05 | 0.05 | 0.05 |
| NC_019843.3;YP_009047203.1;MERS_1a | 982 | 982 | 1 | 0.05 | 0.05 | 0.05 |
| NC_019843.3;YP_009047203.1;MERS_1a | 992 | 993 | 2 | 0.05 | 0.05 | 0.05 |
| NC_019843.3;YP_009047203.1;MERS_1a | 995 | 1010 | 16 | 0.05 | 0.15 | 0.08125 |

| Extended data 1 | | | | | | |
|---|---|---|---|---|---|---|
| Protein | Start position (peptide) | Stop position (peptide) | Number of peptides | Minimum fraction (per peptide) of control subjects with reactivity >3 sd above average | Maximum fraction (per peptide) of control subjects with reactivity >3 sd above average | Mean fraction (over all peptides in this contiguous stretch) of controlsubjects with reactivity >3 sd above average |
| NC_019843.3;YP_009047203.1;MERS_1a | 1019 | 1019 | 1 | 0.1 | 0.1 | 0.1 |
| NC_019843.3;YP_009047203.1;MERS_1a | 1034 | 1035 | 2 | 0.05 | 0.05 | 0.05 |
| NC_019843.3;YP_009047203.1;MERS_1a | 1040 | 1044 | 5 | 0.05 | 0.05 | 0.05 |
| NC_019843.3;YP_009047203.1;MERS_1a | 1051 | 1051 | 1 | 0.05 | 0.05 | 0.05 |
| NC_019843.3;YP_009047203.1;MERS_1a | 1056 | 1060 | 5 | 0.05 | 0.1 | 0.06 |
| NC_019843.3;YP_009047203.1;MERS_1a | 1062 | 1063 | 2 | 0.05 | 0.1 | 0.075 |
| NC_019843.3;YP_009047203.1;MERS_1a | 1066 | 1066 | 1 | 0.05 | 0.05 | 0.05 |
| NC_019843.3;YP_009047203.1;MERS_1a | 1082 | 1094 | 13 | 0.05 | 0.2 | 0.103846154 |
| NC_019843.3;YP_009047203.1;MERS_1a | 1099 | 1099 | 1 | 0.05 | 0.05 | 0.05 |
| NC_019843.3;YP_009047203.1;MERS_1a | 1105 | 1106 | 2 | 0.05 | 0.15 | 0.1 |
| NC_019843.3;YP_009047203.1;MERS_1a | 1126 | 1126 | 1 | 0.05 | 0.05 | 0.05 |
| NC_019843.3;YP_009047203.1;MERS_1a | 1131 | 1131 | 1 | 0.05 | 0.05 | 0.05 |
| NC_019843.3;YP_009047203.1;MERS_1a | 1139 | 1140 | 2 | 0.05 | 0.05 | 0.05 |
| NC_019843.3;YP_009047203.1;MERS_1a | 1142 | 1142 | 1 | 0.4 | 0.4 | 0.4 |
| NC_019843.3;YP_009047203.1;MERS_1a | 1148 | 1148 | 1 | 0.05 | 0.05 | 0.05 |
| NC_019843.3;YP_009047203.1;MERS_1a | 1153 | 1162 | 10 | 0.05 | 0.1 | 0.055 |
| NC_019843.3;YP_009047203.1;MERS_1a | 1167 | 1168 | 2 | 0.05 | 0.05 | 0.05 |
| NC_019843.3;YP_009047203.1;MERS_1a | 1170 | 1170 | 1 | 0.05 | 0.05 | 0.05 |
| NC_019843.3;YP_009047203.1;MERS_1a | 1172 | 1172 | 1 | 0.1 | 0.1 | 0.1 |
| NC_019843.3;YP_009047203.1;MERS_1a | 1203 | 1210 | 8 | 0.05 | 0.1 | 0.05625 |
| NC_019843.3;YP_009047203.1;MERS_1a | 1289 | 1289 | 1 | 0.05 | 0.05 | 0.05 |
| NC_019843.3;YP_009047203.1;MERS_1a | 1306 | 1306 | 1 | 0.1 | 0.1 | 0.1 |
| NC_019843.3;YP_009047203.1;MERS_1a | 1318 | 1319 | 2 | 0.05 | 0.1 | 0.075 |
| NC_019843.3;YP_009047203.1;MERS_1a | 1336 | 1340 | 5 | 0.1 | 0.15 | 0.13 |
| NC_019843.3;YP_009047203.1;MERS_1a | 1343 | 1346 | 4 | 0.05 | 0.1 | 0.075 |
| NC_019843.3;YP_009047203.1;MERS_1a | 1378 | 1381 | 4 | 0.05 | 0.05 | 0.05 |
| NC_019843.3;YP_009047203.1;MERS_1a | 1394 | 1395 | 2 | 0.05 | 0.05 | 0.05 |
| NC_019843.3;YP_009047203.1;MERS_1a | 1398 | 1398 | 1 | 0.05 | 0.05 | 0.05 |
| NC_019843.3;YP_009047203.1;MERS_1a | 1401 | 1401 | 1 | 0.05 | 0.05 | 0.05 |
| NC_019843.3;YP_009047203.1;MERS_1a | 1403 | 1404 | 2 | 0.05 | 0.4 | 0.225 |
| NC_019843.3;YP_009047203.1;MERS_1a | 1424 | 1424 | 1 | 0.05 | 0.05 | 0.05 |
| NC_019843.3;YP_009047203.1;MERS_1a | 1437 | 1437 | 1 | 0.05 | 0.05 | 0.05 |
| NC_019843.3;YP_009047203.1;MERS_1a | 1473 | 1473 | 1 | 0.05 | 0.05 | 0.05 |
| NC_019843.3;YP_009047203.1;MERS_1a | 1490 | 1490 | 1 | 0.1 | 0.1 | 0.1 |
| NC_019843.3;YP_009047203.1;MERS_1a | 1514 | 1516 | 3 | 0.05 | 0.05 | 0.05 |
| NC_019843.3;YP_009047203.1;MERS_1a | 1518 | 1518 | 1 | 0.05 | 0.05 | 0.05 |
| NC_019843.3;YP_009047203.1;MERS_1a | 1520 | 1521 | 2 | 0.05 | 0.05 | 0.05 |

| Extended data 1 | | | | | | |
|---|---|---|---|---|---|---|
| Protein | Start position (peptide) | Stop position (peptide) | Number of peptides | Minimum fraction (per peptide) of control subjects with reactivity >3 sd above average | Maximum fraction (per peptide) of control subjects with reactivity >3 sd above average | Mean fraction (over all peptides in this contiguous stretch) of controlsubjects with reactivity >3 sd above average |
| NC_019843.3;YP_009047203.1;MERS_1a | 1523 | 1523 | 1 | 0.05 | 0.05 | 0.05 |
| NC_019843.3;YP_009047203.1;MERS_1a | 1530 | 1545 | 16 | 0.05 | 0.15 | 0.084375 |
| NC_019843.3;YP_009047203.1;MERS_1a | 1603 | 1604 | 2 | 0.05 | 0.15 | 0.1 |
| NC_019843.3;YP_009047203.1;MERS_1a | 1613 | 1617 | 5 | 0.05 | 0.1 | 0.07 |
| NC_019843.3;YP_009047203.1;MERS_1a | 1633 | 1641 | 9 | 0.05 | 0.15 | 0.088888889 |
| NC_019843.3;YP_009047203.1;MERS_1a | 1645 | 1645 | 1 | 0.15 | 0.15 | 0.15 |
| NC_019843.3;YP_009047203.1;MERS_1a | 1676 | 1676 | 1 | 0.15 | 0.15 | 0.15 |
| NC_019843.3;YP_009047203.1;MERS_1a | 1752 | 1752 | 1 | 0.1 | 0.1 | 0.1 |
| NC_019843.3;YP_009047203.1;MERS_1a | 1773 | 1774 | 2 | 0.05 | 0.05 | 0.05 |
| NC_019843.3;YP_009047203.1;MERS_1a | 1787 | 1787 | 1 | 0.05 | 0.05 | 0.05 |
| NC_019843.3;YP_009047203.1;MERS_1a | 1789 | 1789 | 1 | 0.05 | 0.05 | 0.05 |
| NC_019843.3;YP_009047203.1;MERS_1a | 1803 | 1803 | 1 | 0.2 | 0.2 | 0.2 |
| NC_019843.3;YP_009047203.1;MERS_1a | 1805 | 1810 | 6 | 0.05 | 0.1 | 0.066666667 |
| NC_019843.3;YP_009047203.1;MERS_1a | 1815 | 1825 | 11 | 0.05 | 0.3 | 0.181818182 |
| NC_019843.3;YP_009047203.1;MERS_1a | 1828 | 1828 | 1 | 0.05 | 0.05 | 0.05 |
| NC_019843.3;YP_009047203.1;MERS_1a | 1850 | 1850 | 1 | 0.1 | 0.1 | 0.1 |
| NC_019843.3;YP_009047203.1;MERS_1a | 1858 | 1858 | 1 | 0.05 | 0.05 | 0.05 |
| NC_019843.3;YP_009047203.1;MERS_1a | 1863 | 1863 | 1 | 0.05 | 0.05 | 0.05 |
| NC_019843.3;YP_009047203.1;MERS_1a | 1868 | 1868 | 1 | 0.05 | 0.05 | 0.05 |
| NC_019843.3;YP_009047203.1;MERS_1a | 1871 | 1871 | 1 | 0.05 | 0.05 | 0.05 |
| NC_019843.3;YP_009047203.1;MERS_1a | 1873 | 1874 | 2 | 0.05 | 0.05 | 0.05 |
| NC_019843.3;YP_009047203.1;MERS_1a | 1888 | 1889 | 2 | 0.05 | 0.05 | 0.05 |
| NC_019843.3;YP_009047203.1;MERS_1a | 1895 | 1895 | 1 | 0.05 | 0.05 | 0.05 |
| NC_019843.3;YP_009047203.1;MERS_1a | 1901 | 1901 | 1 | 0.05 | 0.05 | 0.05 |
| NC_019843.3;YP_009047203.1;MERS_1a | 1905 | 1905 | 1 | 0.05 | 0.05 | 0.05 |
| NC_019843.3;YP_009047203.1;MERS_1a | 1920 | 1922 | 3 | 0.05 | 0.1 | 0.083333333 |
| NC_019843.3;YP_009047203.1;MERS_1a | 1924 | 1930 | 7 | 0.05 | 0.15 | 0.107142857 |
| NC_019843.3;YP_009047203.1;MERS_1a | 1932 | 1932 | 1 | 0.05 | 0.05 | 0.05 |
| NC_019843.3;YP_009047203.1;MERS_1a | 1938 | 1938 | 1 | 0.05 | 0.05 | 0.05 |
| NC_019843.3;YP_009047203.1;MERS_1a | 1941 | 1949 | 9 | 0.05 | 0.3 | 0.133333333 |
| NC_019843.3;YP_009047203.1;MERS_1a | 1955 | 1955 | 1 | 0.05 | 0.05 | 0.05 |
| NC_019843.3;YP_009047203.1;MERS_1a | 1964 | 1973 | 10 | 0.05 | 0.05 | 0.05 |
| NC_019843.3;YP_009047203.1;MERS_1a | 1979 | 1980 | 2 | 0.05 | 0.05 | 0.05 |
| NC_019843.3;YP_009047203.1;MERS_1a | 1982 | 1988 | 7 | 0.05 | 0.1 | 0.078571429 |
| NC_019843.3;YP_009047203.1;MERS_1a | 1992 | 1992 | 1 | 0.05 | 0.05 | 0.05 |
| NC_019843.3;YP_009047203.1;MERS_1a | 2001 | 2008 | 8 | 0.05 | 0.05 | 0.05 |
| NC_019843.3;YP_009047203.1;MERS_1a | 2013 | 2013 | 1 | 0.05 | 0.05 | 0.05 |

| Extended data 1 | | | | | | |
|---|---|---|---|---|---|---|
| **Protein** | **Start position (peptide)** | **Stop position (peptide)** | **Number of peptides** | **Minimum fraction (per peptide) of control subjects with reactivity >3 sd above average** | **Maximum fraction (per peptide) of control subjects with reactivity >3 sd above average** | **Mean fraction (over all peptides in this contiguous stretch) of controlsubjects with reactivity >3 sd above average** |
| NC_019843.3;YP_009047203.1;MERS_1a | 2015 | 2015 | 1 | 0.05 | 0.05 | 0.05 |
| NC_019843.3;YP_009047203.1;MERS_1a | 2042 | 2042 | 1 | 0.05 | 0.05 | 0.05 |
| NC_019843.3;YP_009047203.1;MERS_1a | 2119 | 2121 | 3 | 0.05 | 0.05 | 0.05 |
| NC_019843.3;YP_009047203.1;MERS_1a | 2143 | 2143 | 1 | 0.05 | 0.05 | 0.05 |
| NC_019843.3;YP_009047203.1;MERS_1a | 2208 | 2208 | 1 | 0.05 | 0.05 | 0.05 |
| NC_019843.3;YP_009047203.1;MERS_1a | 2220 | 2220 | 1 | 0.05 | 0.05 | 0.05 |
| NC_019843.3;YP_009047203.1;MERS_1a | 2222 | 2222 | 1 | 0.05 | 0.05 | 0.05 |
| NC_019843.3;YP_009047203.1;MERS_1a | 2228 | 2235 | 8 | 0.05 | 0.15 | 0.11875 |
| NC_019843.3;YP_009047203.1;MERS_1a | 2239 | 2239 | 1 | 0.05 | 0.05 | 0.05 |
| NC_019843.3;YP_009047203.1;MERS_1a | 2259 | 2259 | 1 | 0.15 | 0.15 | 0.15 |
| NC_019843.3;YP_009047203.1;MERS_1a | 2333 | 2333 | 1 | 0.05 | 0.05 | 0.05 |
| NC_019843.3;YP_009047203.1;MERS_1a | 2437 | 2437 | 1 | 0.05 | 0.05 | 0.05 |
| NC_019843.3;YP_009047203.1;MERS_1a | 2439 | 2447 | 9 | 0.05 | 0.1 | 0.055555556 |
| NC_019843.3;YP_009047203.1;MERS_1a | 2483 | 2483 | 1 | 0.05 | 0.05 | 0.05 |
| NC_019843.3;YP_009047203.1;MERS_1a | 2498 | 2499 | 2 | 0.05 | 0.05 | 0.05 |
| NC_019843.3;YP_009047203.1;MERS_1a | 2544 | 2544 | 1 | 0.25 | 0.25 | 0.25 |
| NC_019843.3;YP_009047203.1;MERS_1a | 2565 | 2565 | 1 | 0.05 | 0.05 | 0.05 |
| NC_019843.3;YP_009047203.1;MERS_1a | 2577 | 2581 | 5 | 0.05 | 0.05 | 0.05 |
| NC_019843.3;YP_009047203.1;MERS_1a | 2588 | 2589 | 2 | 0.05 | 0.05 | 0.05 |
| NC_019843.3;YP_009047203.1;MERS_1a | 2608 | 2608 | 1 | 0.05 | 0.05 | 0.05 |
| NC_019843.3;YP_009047203.1;MERS_1a | 2610 | 2616 | 7 | 0.05 | 0.1 | 0.057142857 |
| NC_019843.3;YP_009047203.1;MERS_1a | 2626 | 2634 | 9 | 0.05 | 0.2 | 0.138888889 |
| NC_019843.3;YP_009047203.1;MERS_1a | 2637 | 2639 | 3 | 0.1 | 0.1 | 0.1 |
| NC_019843.3;YP_009047203.1;MERS_1a | 2641 | 2644 | 4 | 0.05 | 0.45 | 0.1875 |
| NC_019843.3;YP_009047203.1;MERS_1a | 2648 | 2648 | 1 | 0.05 | 0.05 | 0.05 |
| NC_019843.3;YP_009047203.1;MERS_1a | 2657 | 2657 | 1 | 0.05 | 0.05 | 0.05 |
| NC_019843.3;YP_009047203.1;MERS_1a | 2676 | 2676 | 1 | 0.05 | 0.05 | 0.05 |
| NC_019843.3;YP_009047203.1;MERS_1a | 2682 | 2684 | 3 | 0.05 | 0.05 | 0.05 |
| NC_019843.3;YP_009047203.1;MERS_1a | 2686 | 2686 | 1 | 0.05 | 0.05 | 0.05 |
| NC_019843.3;YP_009047203.1;MERS_1a | 2715 | 2715 | 1 | 0.05 | 0.05 | 0.05 |
| NC_019843.3;YP_009047203.1;MERS_1a | 2731 | 2731 | 1 | 0.05 | 0.05 | 0.05 |
| NC_019843.3;YP_009047203.1;MERS_1a | 2780 | 2781 | 2 | 0.05 | 0.2 | 0.125 |
| NC_019843.3;YP_009047203.1;MERS_1a | 2788 | 2789 | 2 | 0.05 | 0.05 | 0.05 |
| NC_019843.3;YP_009047203.1;MERS_1a | 2794 | 2794 | 1 | 0.05 | 0.05 | 0.05 |
| NC_019843.3;YP_009047203.1;MERS_1a | 2815 | 2819 | 5 | 0.05 | 0.1 | 0.06 |
| NC_019843.3;YP_009047203.1;MERS_1a | 2821 | 2823 | 3 | 0.05 | 0.05 | 0.05 |
| NC_019843.3;YP_009047203.1;MERS_1a | 2840 | 2842 | 3 | 0.05 | 0.15 | 0.083333333 |

| Extended data 1 | | | | | | |
|---|---|---|---|---|---|---|
| Protein | Start position (peptide) | Stop position (peptide) | Number of peptides | Minimum fraction (per peptide) of control subjects with reactivity >3 sd above average | Maximum fraction (per peptide) of control subjects with reactivity >3 sd above average | Mean fraction (over all peptides in this contiguous stretch) of controlsubjects with reactivity >3 sd above average |
| NC_019843.3;YP_009047203.1;MERS_1a | 2844 | 2844 | 1 | 0.05 | 0.05 | 0.05 |
| NC_019843.3;YP_009047203.1;MERS_1a | 2847 | 2849 | 3 | 0.05 | 0.1 | 0.066666667 |
| NC_019843.3;YP_009047203.1;MERS_1a | 2870 | 2870 | 1 | 0.05 | 0.05 | 0.05 |
| NC_019843.3;YP_009047203.1;MERS_1a | 2879 | 2880 | 2 | 0.05 | 0.05 | 0.05 |
| NC_019843.3;YP_009047203.1;MERS_1a | 2884 | 2884 | 1 | 0.05 | 0.05 | 0.05 |
| NC_019843.3;YP_009047203.1;MERS_1a | 2897 | 2906 | 10 | 0.05 | 0.05 | 0.05 |
| NC_019843.3;YP_009047203.1;MERS_1a | 2910 | 2910 | 1 | 0.05 | 0.05 | 0.05 |
| NC_019843.3;YP_009047203.1;MERS_1a | 2912 | 2912 | 1 | 0.05 | 0.05 | 0.05 |
| NC_019843.3;YP_009047203.1;MERS_1a | 2918 | 2918 | 1 | 0.1 | 0.1 | 0.1 |
| NC_019843.3;YP_009047203.1;MERS_1a | 2924 | 2924 | 1 | 0.05 | 0.05 | 0.05 |
| NC_019843.3;YP_009047203.1;MERS_1a | 2930 | 2930 | 1 | 0.05 | 0.05 | 0.05 |
| NC_019843.3;YP_009047203.1;MERS_1a | 2935 | 2938 | 4 | 0.05 | 0.05 | 0.05 |
| NC_019843.3;YP_009047203.1;MERS_1a | 3046 | 3047 | 2 | 0.05 | 0.05 | 0.05 |
| NC_019843.3;YP_009047203.1;MERS_1a | 3142 | 3143 | 2 | 0.05 | 0.05 | 0.05 |
| NC_019843.3;YP_009047203.1;MERS_1a | 3150 | 3150 | 1 | 0.05 | 0.05 | 0.05 |
| NC_019843.3;YP_009047203.1;MERS_1a | 3152 | 3155 | 4 | 0.05 | 0.05 | 0.05 |
| NC_019843.3;YP_009047203.1;MERS_1a | 3163 | 3163 | 1 | 0.05 | 0.05 | 0.05 |
| NC_019843.3;YP_009047203.1;MERS_1a | 3171 | 3171 | 1 | 0.05 | 0.05 | 0.05 |
| NC_019843.3;YP_009047203.1;MERS_1a | 3173 | 3173 | 1 | 0.05 | 0.05 | 0.05 |
| NC_019843.3;YP_009047203.1;MERS_1a | 3181 | 3181 | 1 | 0.05 | 0.05 | 0.05 |
| NC_019843.3;YP_009047203.1;MERS_1a | 3194 | 3194 | 1 | 0.05 | 0.05 | 0.05 |
| NC_019843.3;YP_009047203.1;MERS_1a | 3199 | 3200 | 2 | 0.05 | 0.05 | 0.05 |
| NC_019843.3;YP_009047203.1;MERS_1a | 3214 | 3224 | 11 | 0.05 | 0.2 | 0.072727273 |
| NC_019843.3;YP_009047203.1;MERS_1a | 3229 | 3229 | 1 | 0.05 | 0.05 | 0.05 |
| NC_019843.3;YP_009047203.1;MERS_1a | 3234 | 3234 | 1 | 0.05 | 0.05 | 0.05 |
| NC_019843.3;YP_009047203.1;MERS_1a | 3244 | 3247 | 4 | 0.05 | 0.1 | 0.0625 |
| NC_019843.3;YP_009047203.1;MERS_1a | 3249 | 3249 | 1 | 0.05 | 0.05 | 0.05 |
| NC_019843.3;YP_009047203.1;MERS_1a | 3283 | 3283 | 1 | 0.05 | 0.05 | 0.05 |
| NC_019843.3;YP_009047203.1;MERS_1a | 3306 | 3318 | 13 | 0.05 | 0.15 | 0.107692308 |
| NC_019843.3;YP_009047203.1;MERS_1a | 3341 | 3341 | 1 | 0.05 | 0.05 | 0.05 |
| NC_019843.3;YP_009047203.1;MERS_1a | 3345 | 3346 | 2 | 0.05 | 0.1 | 0.075 |
| NC_019843.3;YP_009047203.1;MERS_1a | 3389 | 3393 | 5 | 0.05 | 0.1 | 0.08 |
| NC_019843.3;YP_009047203.1;MERS_1a | 3400 | 3400 | 1 | 0.05 | 0.05 | 0.05 |
| NC_019843.3;YP_009047203.1;MERS_1a | 3413 | 3421 | 9 | 0.05 | 0.05 | 0.05 |
| NC_019843.3;YP_009047203.1;MERS_1a | 3423 | 3423 | 1 | 0.05 | 0.05 | 0.05 |
| NC_019843.3;YP_009047203.1;MERS_1a | 3491 | 3491 | 1 | 0.05 | 0.05 | 0.05 |
| NC_019843.3;YP_009047203.1;MERS_1a | 3535 | 3535 | 1 | 0.1 | 0.1 | 0.1 |

| Extended data 1 | | | | | | |
|---|---|---|---|---|---|---|
| Protein | Start position (peptide) | Stop position (peptide) | Number of peptides | Minimum fraction (per peptide) of control subjects with reactivity >3 sd above average | Maximum fraction (per peptide) of control subjects with reactivity >3 sd above average | Mean fraction (over all peptides in this contiguous stretch) of controlsubjects with reactivity >3 sd above average |
| NC_019843.3;YP_009047203.1;MERS_1a | 3537 | 3537 | 1 | 0.05 | 0.05 | 0.05 |
| NC_019843.3;YP_009047203.1;MERS_1a | 3550 | 3550 | 1 | 0.05 | 0.05 | 0.05 |
| NC_019843.3;YP_009047203.1;MERS_1a | 3646 | 3657 | 12 | 0.05 | 0.35 | 0.191666667 |
| NC_019843.3;YP_009047203.1;MERS_1a | 3699 | 3705 | 7 | 0.05 | 0.1 | 0.057142857 |
| NC_019843.3;YP_009047203.1;MERS_1a | 3805 | 3805 | 1 | 0.05 | 0.05 | 0.05 |
| NC_019843.3;YP_009047203.1;MERS_1a | 3809 | 3809 | 1 | 0.05 | 0.05 | 0.05 |
| NC_019843.3;YP_009047203.1;MERS_1a | 3811 | 3811 | 1 | 0.15 | 0.15 | 0.15 |
| NC_019843.3;YP_009047203.1;MERS_1a | 3814 | 3815 | 2 | 0.05 | 0.05 | 0.05 |
| NC_019843.3;YP_009047203.1;MERS_1a | 3817 | 3817 | 1 | 0.05 | 0.05 | 0.05 |
| NC_019843.3;YP_009047203.1;MERS_1a | 3832 | 3840 | 9 | 0.05 | 0.1 | 0.055555556 |
| NC_019843.3;YP_009047203.1;MERS_1a | 3872 | 3872 | 1 | 0.05 | 0.05 | 0.05 |
| NC_019843.3;YP_009047203.1;MERS_1a | 3917 | 3917 | 1 | 0.05 | 0.05 | 0.05 |
| NC_019843.3;YP_009047203.1;MERS_1a | 3936 | 3936 | 1 | 0.05 | 0.05 | 0.05 |
| NC_019843.3;YP_009047203.1;MERS_1a | 3938 | 3942 | 5 | 0.05 | 0.05 | 0.05 |
| NC_019843.3;YP_009047203.1;MERS_1a | 3966 | 3971 | 6 | 0.05 | 0.1 | 0.066666667 |
| NC_019843.3;YP_009047203.1;MERS_1a | 3981 | 3981 | 1 | 0.05 | 0.05 | 0.05 |
| NC_019843.3;YP_009047203.1;MERS_1a | 3992 | 3996 | 5 | 0.05 | 0.1 | 0.06 |
| NC_019843.3;YP_009047203.1;MERS_1a | 4018 | 4034 | 17 | 0.05 | 0.3 | 0.144117647 |
| NC_019843.3;YP_009047203.1;MERS_1a | 4036 | 4036 | 1 | 0.05 | 0.05 | 0.05 |
| NC_019843.3;YP_009047203.1;MERS_1a | 4049 | 4050 | 2 | 0.05 | 0.05 | 0.05 |
| NC_019843.3;YP_009047203.1;MERS_1a | 4053 | 4053 | 1 | 0.05 | 0.05 | 0.05 |
| NC_019843.3;YP_009047203.1;MERS_1a | 4075 | 4075 | 1 | 0.05 | 0.05 | 0.05 |
| NC_019843.3;YP_009047203.1;MERS_1a | 4077 | 4079 | 3 | 0.05 | 0.05 | 0.05 |
| NC_019843.3;YP_009047203.1;MERS_1a | 4094 | 4094 | 1 | 0.05 | 0.05 | 0.05 |
| NC_019843.3;YP_009047203.1;MERS_1a | 4096 | 4098 | 3 | 0.05 | 0.1 | 0.066666667 |
| NC_019843.3;YP_009047203.1;MERS_1a | 4114 | 4116 | 3 | 0.05 | 0.05 | 0.05 |
| NC_019843.3;YP_009047203.1;MERS_1a | 4126 | 4126 | 1 | 0.05 | 0.05 | 0.05 |
| NC_019843.3;YP_009047203.1;MERS_1a | 4159 | 4160 | 2 | 0.05 | 0.05 | 0.05 |
| NC_019843.3;YP_009047203.1;MERS_1a | 4163 | 4163 | 1 | 0.05 | 0.05 | 0.05 |
| NC_019843.3;YP_009047203.1;MERS_1a | 4172 | 4177 | 6 | 0.05 | 0.15 | 0.075 |
| NC_019843.3;YP_009047203.1;MERS_1a | 4180 | 4183 | 4 | 0.05 | 0.05 | 0.05 |
| NC_019843.3;YP_009047203.1;MERS_1a | 4217 | 4217 | 1 | 0.1 | 0.1 | 0.1 |
| NC_019843.3;YP_009047203.1;MERS_1a | 4231 | 4231 | 1 | 0.1 | 0.1 | 0.1 |
| NC_019843.3;YP_009047203.1;MERS_1a | 4236 | 4236 | 1 | 0.05 | 0.05 | 0.05 |
| NC_019843.3;YP_009047203.1;MERS_1a | 4245 | 4245 | 1 | 0.1 | 0.1 | 0.1 |
| NC_019843.3;YP_009047203.1;MERS_1a | 4254 | 4263 | 10 | 0.05 | 0.15 | 0.1 |
| NC_019843.3;YP_009047203.1;MERS_1a | 4273 | 4273 | 1 | 0.05 | 0.05 | 0.05 |

| Extended data 1 | | | | | | |
|---|---|---|---|---|---|---|
| **Protein** | **Start position (peptide)** | **Stop position (peptide)** | **Number of peptides** | **Minimum fraction (per peptide) of control subjects with reactivity >3 sd above average** | **Maximum fraction (per peptide) of control subjects with reactivity >3 sd above average** | **Mean fraction (over all peptides in this contiguous stretch) of controlsubjects with reactivity >3 sd above average** |
| NC_019843.3;YP_009047203.1;MERS_1a | 4276 | 4276 | 1 | 0.05 | 0.05 | 0.05 |
| NC_019843.3;YP_009047203.1;MERS_1a | 4278 | 4278 | 1 | 0.05 | 0.05 | 0.05 |
| NC_019843.3;YP_009047203.1;MERS_1a | 4283 | 4285 | 3 | 0.05 | 0.05 | 0.05 |
| NC_019843.3;YP_009047203.1;MERS_1a | 4329 | 4329 | 1 | 0.05 | 0.05 | 0.05 |
| NC_019843.3;YP_009047203.1;MERS_1a | 4331 | 4332 | 2 | 0.05 | 0.05 | 0.05 |
| NC_019843.3;YP_009047203.1;MERS_1a | 4343 | 4343 | 1 | 0.05 | 0.05 | 0.05 |
| NC_019843.3;YP_009047203.1;MERS_1a | 4359 | 4359 | 1 | 0.05 | 0.05 | 0.05 |
| NC_019843.3;YP_009047203.1;MERS_1a | 4368 | 4368 | 1 | 0.1 | 0.1 | 0.1 |
| NC_019843.3;YP_009047203.1;MERS_1a | 4370 | 4380 | 11 | 0.05 | 0.25 | 0.145454545 |
| NC_019843.3;YP_009047204.1;MERS_spike | 12 | 13 | 2 | 0.05 | 0.1 | 0.075 |
| NC_019843.3;YP_009047204.1;MERS_spike | 20 | 20 | 1 | 0.05 | 0.05 | 0.05 |
| NC_019843.3;YP_009047204.1;MERS_spike | 37 | 42 | 6 | 0.05 | 0.1 | 0.066666667 |
| NC_019843.3;YP_009047204.1;MERS_spike | 45 | 47 | 3 | 0.05 | 0.1 | 0.066666667 |
| NC_019843.3;YP_009047204.1;MERS_spike | 51 | 52 | 2 | 0.05 | 0.05 | 0.05 |
| NC_019843.3;YP_009047204.1;MERS_spike | 70 | 70 | 1 | 0.05 | 0.05 | 0.05 |
| NC_019843.3;YP_009047204.1;MERS_spike | 84 | 88 | 5 | 0.05 | 0.05 | 0.05 |
| NC_019843.3;YP_009047204.1;MERS_spike | 99 | 99 | 1 | 0.05 | 0.05 | 0.05 |
| NC_019843.3;YP_009047204.1;MERS_spike | 101 | 102 | 2 | 0.1 | 0.1 | 0.1 |
| NC_019843.3;YP_009047204.1;MERS_spike | 108 | 109 | 2 | 0.05 | 0.05 | 0.05 |
| NC_019843.3;YP_009047204.1;MERS_spike | 111 | 118 | 8 | 0.05 | 0.1 | 0.08125 |
| NC_019843.3;YP_009047204.1;MERS_spike | 123 | 123 | 1 | 0.05 | 0.05 | 0.05 |
| NC_019843.3;YP_009047204.1;MERS_spike | 189 | 198 | 10 | 0.05 | 0.1 | 0.06 |
| NC_019843.3;YP_009047204.1;MERS_spike | 205 | 205 | 1 | 0.05 | 0.05 | 0.05 |
| NC_019843.3;YP_009047204.1;MERS_spike | 208 | 222 | 15 | 0.05 | 0.45 | 0.166666667 |
| NC_019843.3;YP_009047204.1;MERS_spike | 226 | 227 | 2 | 0.05 | 0.1 | 0.075 |
| NC_019843.3;YP_009047204.1;MERS_spike | 232 | 232 | 1 | 0.05 | 0.05 | 0.05 |
| NC_019843.3;YP_009047204.1;MERS_spike | 246 | 250 | 5 | 0.05 | 0.15 | 0.07 |
| NC_019843.3;YP_009047204.1;MERS_spike | 265 | 272 | 8 | 0.05 | 0.1 | 0.075 |
| NC_019843.3;YP_009047204.1;MERS_spike | 336 | 336 | 1 | 0.05 | 0.05 | 0.05 |
| NC_019843.3;YP_009047204.1;MERS_spike | 340 | 340 | 1 | 0.05 | 0.05 | 0.05 |
| NC_019843.3;YP_009047204.1;MERS_spike | 357 | 359 | 3 | 0.05 | 0.1 | 0.066666667 |
| NC_019843.3;YP_009047204.1;MERS_spike | 363 | 363 | 1 | 0.05 | 0.05 | 0.05 |
| NC_019843.3;YP_009047204.1;MERS_spike | 380 | 380 | 1 | 0.1 | 0.1 | 0.1 |
| NC_019843.3;YP_009047204.1;MERS_spike | 417 | 417 | 1 | 0.1 | 0.1 | 0.1 |
| NC_019843.3;YP_009047204.1;MERS_spike | 419 | 419 | 1 | 0.15 | 0.15 | 0.15 |
| NC_019843.3;YP_009047204.1;MERS_spike | 445 | 446 | 2 | 0.05 | 0.05 | 0.05 |
| NC_019843.3;YP_009047204.1;MERS_spike | 453 | 453 | 1 | 0.1 | 0.1 | 0.1 |

| Extended data 1 | | | | | | |
|---|---|---|---|---|---|---|
| **Protein** | **Start position (peptide)** | **Stop position (peptide)** | **Number of peptides** | **Minimum fraction (per peptide) of control subjects with reactivity >3 sd above average** | **Maximum fraction (per peptide) of control subjects with reactivity >3 sd above average** | **Mean fraction (over all peptides in this contiguous stretch) of controlsubjects with reactivity >3 sd above average** |
| NC_019843.3;YP_009047204.1;MERS_spike | 462 | 462 | 1 | 0.35 | 0.35 | 0.35 |
| NC_019843.3;YP_009047204.1;MERS_spike | 479 | 483 | 5 | 0.05 | 0.05 | 0.05 |
| NC_019843.3;YP_009047204.1;MERS_spike | 496 | 496 | 1 | 0.05 | 0.05 | 0.05 |
| NC_019843.3;YP_009047204.1;MERS_spike | 498 | 500 | 3 | 0.05 | 0.05 | 0.05 |
| NC_019843.3;YP_009047204.1;MERS_spike | 505 | 505 | 1 | 0.05 | 0.05 | 0.05 |
| NC_019843.3;YP_009047204.1;MERS_spike | 517 | 517 | 1 | 0.25 | 0.25 | 0.25 |
| NC_019843.3;YP_009047204.1;MERS_spike | 531 | 531 | 1 | 0.05 | 0.05 | 0.05 |
| NC_019843.3;YP_009047204.1;MERS_spike | 535 | 539 | 5 | 0.05 | 0.35 | 0.15 |
| NC_019843.3;YP_009047204.1;MERS_spike | 543 | 543 | 1 | 0.05 | 0.05 | 0.05 |
| NC_019843.3;YP_009047204.1;MERS_spike | 549 | 549 | 1 | 0.05 | 0.05 | 0.05 |
| NC_019843.3;YP_009047204.1;MERS_spike | 571 | 590 | 20 | 0.05 | 0.3 | 0.1 |
| NC_019843.3;YP_009047204.1;MERS_spike | 592 | 592 | 1 | 0.05 | 0.05 | 0.05 |
| NC_019843.3;YP_009047204.1;MERS_spike | 613 | 613 | 1 | 0.05 | 0.05 | 0.05 |
| NC_019843.3;YP_009047204.1;MERS_spike | 688 | 689 | 2 | 0.05 | 0.05 | 0.05 |
| NC_019843.3;YP_009047204.1;MERS_spike | 691 | 691 | 1 | 0.05 | 0.05 | 0.05 |
| NC_019843.3;YP_009047204.1;MERS_spike | 693 | 699 | 7 | 0.05 | 0.3 | 0.171428571 |
| NC_019843.3;YP_009047204.1;MERS_spike | 717 | 717 | 1 | 0.2 | 0.2 | 0.2 |
| NC_019843.3;YP_009047204.1;MERS_spike | 720 | 721 | 2 | 0.05 | 0.05 | 0.05 |
| NC_019843.3;YP_009047204.1;MERS_spike | 724 | 724 | 1 | 0.05 | 0.05 | 0.05 |
| NC_019843.3;YP_009047204.1;MERS_spike | 727 | 729 | 3 | 0.05 | 0.15 | 0.083333333 |
| NC_019843.3;YP_009047204.1;MERS_spike | 735 | 737 | 3 | 0.05 | 0.05 | 0.05 |
| NC_019843.3;YP_009047204.1;MERS_spike | 739 | 745 | 7 | 0.05 | 0.1 | 0.057142857 |
| NC_019843.3;YP_009047204.1;MERS_spike | 759 | 763 | 5 | 0.05 | 0.1 | 0.08 |
| NC_019843.3;YP_009047204.1;MERS_spike | 770 | 770 | 1 | 0.05 | 0.05 | 0.05 |
| NC_019843.3;YP_009047204.1;MERS_spike | 787 | 787 | 1 | 0.05 | 0.05 | 0.05 |
| NC_019843.3;YP_009047204.1;MERS_spike | 831 | 831 | 1 | 0.05 | 0.05 | 0.05 |
| NC_019843.3;YP_009047204.1;MERS_spike | 838 | 838 | 1 | 0.05 | 0.05 | 0.05 |
| NC_019843.3;YP_009047204.1;MERS_spike | 852 | 852 | 1 | 0.05 | 0.05 | 0.05 |
| NC_019843.3;YP_009047204.1;MERS_spike | 863 | 863 | 1 | 0.05 | 0.05 | 0.05 |
| NC_019843.3;YP_009047204.1;MERS_spike | 875 | 875 | 1 | 0.05 | 0.05 | 0.05 |
| NC_019843.3;YP_009047204.1;MERS_spike | 878 | 884 | 7 | 0.05 | 0.35 | 0.171428571 |
| NC_019843.3;YP_009047204.1;MERS_spike | 888 | 890 | 3 | 0.1 | 0.35 | 0.183333333 |
| NC_019843.3;YP_009047204.1;MERS_spike | 902 | 902 | 1 | 0.05 | 0.05 | 0.05 |
| NC_019843.3;YP_009047204.1;MERS_spike | 906 | 917 | 12 | 0.05 | 0.1 | 0.066666667 |
| NC_019843.3;YP_009047204.1;MERS_spike | 931 | 931 | 1 | 0.05 | 0.05 | 0.05 |
| NC_019843.3;YP_009047204.1;MERS_spike | 954 | 954 | 1 | 0.05 | 0.05 | 0.05 |
| NC_019843.3;YP_009047204.1;MERS_spike | 984 | 984 | 1 | 0.1 | 0.1 | 0.1 |

| Extended data 1 | | | | | | |
|---|---|---|---|---|---|---|
| Protein | Start position (peptide) | Stop position (peptide) | Number of peptides | Minimum fraction (per peptide) of control subjects with reactivity >3 sd above average | Maximum fraction (per peptide) of control subjects with reactivity >3 sd above average | Mean fraction (over all peptides in this contiguous stretch) of controlsubjects with reactivity >3 sd above average |
| NC_019843.3;YP_009047204.1;MERS_spike | 994 | 994 | 1 | 0.05 | 0.05 | 0.05 |
| NC_019843.3;YP_009047204.1;MERS_spike | 996 | 1012 | 17 | 0.05 | 0.2 | 0.097058824 |
| NC_019843.3;YP_009047204.1;MERS_spike | 1014 | 1031 | 18 | 0.05 | 0.2 | 0.086111111 |
| NC_019843.3;YP_009047204.1;MERS_spike | 1035 | 1035 | 1 | 0.05 | 0.05 | 0.05 |
| NC_019843.3;YP_009047204.1;MERS_spike | 1045 | 1045 | 1 | 0.05 | 0.05 | 0.05 |
| NC_019843.3;YP_009047204.1;MERS_spike | 1050 | 1050 | 1 | 0.1 | 0.1 | 0.1 |
| NC_019843.3;YP_009047204.1;MERS_spike | 1059 | 1059 | 1 | 0.05 | 0.05 | 0.05 |
| NC_019843.3;YP_009047204.1;MERS_spike | 1063 | 1064 | 2 | 0.05 | 0.05 | 0.05 |
| NC_019843.3;YP_009047204.1;MERS_spike | 1078 | 1078 | 1 | 0.05 | 0.05 | 0.05 |
| NC_019843.3;YP_009047204.1;MERS_spike | 1095 | 1096 | 2 | 0.05 | 0.1 | 0.075 |
| NC_019843.3;YP_009047204.1;MERS_spike | 1155 | 1155 | 1 | 0.1 | 0.1 | 0.1 |
| NC_019843.3;YP_009047204.1;MERS_spike | 1187 | 1195 | 9 | 0.05 | 0.15 | 0.094444444 |
| NC_019843.3;YP_009047204.1;MERS_spike | 1198 | 1199 | 2 | 0.05 | 0.05 | 0.05 |
| NC_019843.3;YP_009047204.1;MERS_spike | 1202 | 1213 | 12 | 0.05 | 0.35 | 0.166666667 |
| NC_019843.3;YP_009047204.1;MERS_spike | 1228 | 1231 | 4 | 0.05 | 0.05 | 0.05 |
| NC_019843.3;YP_009047204.1;MERS_spike | 1233 | 1233 | 1 | 0.05 | 0.05 | 0.05 |
| NC_019843.3;YP_009047204.1;MERS_spike | 1236 | 1236 | 1 | 0.1 | 0.1 | 0.1 |
| NC_019843.3;YP_009047204.1;MERS_spike | 1241 | 1241 | 1 | 0.05 | 0.05 | 0.05 |
| NC_019843.3;YP_009047204.1;MERS_spike | 1267 | 1276 | 10 | 0.05 | 0.2 | 0.12 |
| NC_019843.3;YP_009047204.1;MERS_spike | 1278 | 1278 | 1 | 0.05 | 0.05 | 0.05 |
| NC_019843.3;YP_009047205.1;MERS_ns3 | 17 | 22 | 6 | 0.05 | 0.8 | 0.191666667 |
| NC_019843.3;YP_009047205.1;MERS_ns3 | 36 | 48 | 13 | 0.05 | 0.05 | 0.05 |
| NC_019843.3;YP_009047205.1;MERS_ns3 | 52 | 52 | 1 | 0.05 | 0.05 | 0.05 |
| NC_019843.3;YP_009047205.1;MERS_ns3 | 54 | 54 | 1 | 0.05 | 0.05 | 0.05 |
| NC_019843.3;YP_009047205.1;MERS_ns3 | 79 | 80 | 2 | 0.05 | 0.05 | 0.05 |
| NC_019843.3;YP_009047205.1;MERS_ns3 | 84 | 90 | 7 | 0.05 | 0.05 | 0.05 |
| NC_019843.3;YP_009047206.1;MERS_ns4a | 24 | 24 | 1 | 0.05 | 0.05 | 0.05 |
| NC_019843.3;YP_009047206.1;MERS_ns4a | 28 | 28 | 1 | 0.05 | 0.05 | 0.05 |
| NC_019843.3;YP_009047206.1;MERS_ns4a | 48 | 51 | 4 | 0.05 | 0.15 | 0.0875 |
| NC_019843.3;YP_009047206.1;MERS_ns4a | 53 | 56 | 4 | 0.1 | 0.15 | 0.125 |
| NC_019843.3;YP_009047206.1;MERS_ns4a | 67 | 67 | 1 | 0.05 | 0.05 | 0.05 |
| NC_019843.3;YP_009047207.1;MERS_ns4b | 3 | 4 | 2 | 0.05 | 0.05 | 0.05 |
| NC_019843.3;YP_009047207.1;MERS_ns4b | 20 | 20 | 1 | 0.05 | 0.05 | 0.05 |
| NC_019843.3;YP_009047207.1;MERS_ns4b | 57 | 57 | 1 | 0.05 | 0.05 | 0.05 |
| NC_019843.3;YP_009047207.1;MERS_ns4b | 102 | 102 | 1 | 0.05 | 0.05 | 0.05 |
| NC_019843.3;YP_009047207.1;MERS_ns4b | 136 | 136 | 1 | 0.05 | 0.05 | 0.05 |
| NC_019843.3;YP_009047207.1;MERS_ns4b | 152 | 152 | 1 | 0.05 | 0.05 | 0.05 |

| Extended data 1 | | | | | | |
|---|---|---|---|---|---|---|
| Protein | Start position (peptide) | Stop position (peptide) | Number of peptides | Minimum fraction (per peptide) of control subjects with reactivity >3 sd above average | Maximum fraction (per peptide) of control subjects with reactivity >3 sd above average | Mean fraction (over all peptides in this contiguous stretch) of controlsubjects with reactivity >3 sd above average |
| NC_019843.3;YP_009047207.1;MERS_ns4b | 155 | 155 | 1 | 0.05 | 0.05 | 0.05 |
| NC_019843.3;YP_009047207.1;MERS_ns4b | 177 | 184 | 8 | 0.05 | 0.05 | 0.05 |
| NC_019843.3;YP_009047207.1;MERS_ns4b | 192 | 192 | 1 | 0.05 | 0.05 | 0.05 |
| NC_019843.3;YP_009047207.1;MERS_ns4b | 208 | 208 | 1 | 0.05 | 0.05 | 0.05 |
| NC_019843.3;YP_009047207.1;MERS_ns4b | 215 | 216 | 2 | 0.05 | 0.05 | 0.05 |
| NC_019843.3;YP_009047207.1;MERS_ns4b | 218 | 218 | 1 | 0.1 | 0.1 | 0.1 |
| NC_019843.3;YP_009047207.1;MERS_ns4b | 222 | 226 | 5 | 0.05 | 0.2 | 0.09 |
| NC_019843.3;YP_009047207.1;MERS_ns4b | 233 | 233 | 1 | 0.05 | 0.05 | 0.05 |
| NC_019843.3;YP_009047208.1;MERS_ns5 | 3 | 6 | 4 | 0.05 | 0.05 | 0.05 |
| NC_019843.3;YP_009047208.1;MERS_ns5 | 9 | 9 | 1 | 0.05 | 0.05 | 0.05 |
| NC_019843.3;YP_009047208.1;MERS_ns5 | 11 | 13 | 3 | 0.05 | 0.05 | 0.05 |
| NC_019843.3;YP_009047208.1;MERS_ns5 | 25 | 25 | 1 | 0.05 | 0.05 | 0.05 |
| NC_019843.3;YP_009047208.1;MERS_ns5 | 27 | 27 | 1 | 0.05 | 0.05 | 0.05 |
| NC_019843.3;YP_009047208.1;MERS_ns5 | 135 | 144 | 10 | 0.05 | 0.05 | 0.05 |
| NC_019843.3;YP_009047208.1;MERS_ns5 | 189 | 189 | 1 | 0.05 | 0.05 | 0.05 |
| NC_019843.3;YP_009047208.1;MERS_ns5 | 195 | 195 | 1 | 0.05 | 0.05 | 0.05 |
| NC_019843.3;YP_009047208.1;MERS_ns5 | 198 | 198 | 1 | 0.05 | 0.05 | 0.05 |
| NC_019843.3;YP_009047209.1;MERS_envelope | 40 | 40 | 1 | 0.05 | 0.05 | 0.05 |
| NC_019843.3;YP_009047209.1;MERS_envelope | 63 | 63 | 1 | 0.05 | 0.05 | 0.05 |
| NC_019843.3;YP_009047209.1;MERS_envelope | 67 | 71 | 5 | 0.05 | 0.1 | 0.09 |
| NC_019843.3;YP_009047210.1;MERS_membrane | 70 | 70 | 1 | 0.05 | 0.05 | 0.05 |
| NC_019843.3;YP_009047210.1;MERS_membrane | 110 | 110 | 1 | 0.05 | 0.05 | 0.05 |
| NC_019843.3;YP_009047210.1;MERS_membrane | 132 | 132 | 1 | 0.05 | 0.05 | 0.05 |
| NC_019843.3;YP_009047210.1;MERS_membrane | 134 | 134 | 1 | 0.05 | 0.05 | 0.05 |
| NC_019843.3;YP_009047210.1;MERS_membrane | 146 | 146 | 1 | 0.05 | 0.05 | 0.05 |
| NC_019843.3;YP_009047210.1;MERS_membrane | 155 | 162 | 8 | 0.05 | 0.1 | 0.075 |
| NC_019843.3;YP_009047210.1;MERS_membrane | 178 | 178 | 1 | 0.05 | 0.05 | 0.05 |
| NC_019843.3;YP_009047210.1;MERS_membrane | 198 | 200 | 3 | 0.05 | 0.05 | 0.05 |
| NC_019843.3;YP_009047211.1;MERS_nucleoprote | 1 | 21 | 21 | 0.05 | 0.2 | 0.088095238 |
| NC_019843.3;YP_009047211.1;MERS_nucleoprote | 23 | 23 | 1 | 0.05 | 0.05 | 0.05 |
| NC_019843.3;YP_009047211.1;MERS_nucleoprote | 36 | 36 | 1 | 0.05 | 0.05 | 0.05 |
| NC_019843.3;YP_009047211.1;MERS_nucleoprote | 49 | 49 | 1 | 0.05 | 0.05 | 0.05 |
| NC_019843.3;YP_009047211.1;MERS_nucleoprote | 55 | 55 | 1 | 0.15 | 0.15 | 0.15 |
| NC_019843.3;YP_009047211.1;MERS_nucleoprote | 60 | 64 | 5 | 0.05 | 0.1 | 0.08 |
| NC_019843.3;YP_009047211.1;MERS_nucleoprote | 67 | 67 | 1 | 0.05 | 0.05 | 0.05 |
| NC_019843.3;YP_009047211.1;MERS_nucleoprote | 69 | 83 | 15 | 0.05 | 0.1 | 0.06 |
| NC_019843.3;YP_009047211.1;MERS_nucleoprote | 87 | 87 | 1 | 0.05 | 0.05 | 0.05 |

| Extended data 1 | | | | | | |
|---|---|---|---|---|---|---|
| Protein | Start position (peptide) | Stop position (peptide) | Number of peptides | Minimum fraction (per peptide) of control subjects with reactivity >3 sd above average | Maximum fraction (per peptide) of control subjects with reactivity >3 sd above average | Mean fraction (over all peptides in this contiguous stretch) of controlsubjects with reactivity >3 sd above average |
| NC_019843.3;YP_009047211.1;MERS_nucleoprote | 95 | 95 | 1 | 0.05 | 0.05 | 0.05 |
| NC_019843.3;YP_009047211.1;MERS_nucleoprote | 128 | 137 | 10 | 0.05 | 0.2 | 0.07 |
| NC_019843.3;YP_009047211.1;MERS_nucleoprote | 139 | 139 | 1 | 0.05 | 0.05 | 0.05 |
| NC_019843.3;YP_009047211.1;MERS_nucleoprote | 149 | 150 | 2 | 0.05 | 0.05 | 0.05 |
| NC_019843.3;YP_009047211.1;MERS_nucleoprote | 152 | 152 | 1 | 0.05 | 0.05 | 0.05 |
| NC_019843.3;YP_009047211.1;MERS_nucleoprote | 156 | 157 | 2 | 0.05 | 0.05 | 0.05 |
| NC_019843.3;YP_009047211.1;MERS_nucleoprote | 163 | 173 | 11 | 0.05 | 0.2 | 0.063636364 |
| NC_019843.3;YP_009047211.1;MERS_nucleoprote | 193 | 193 | 1 | 0.1 | 0.1 | 0.1 |
| NC_019843.3;YP_009047211.1;MERS_nucleoprote | 196 | 196 | 1 | 0.1 | 0.1 | 0.1 |
| NC_019843.3;YP_009047211.1;MERS_nucleoprote | 208 | 209 | 2 | 0.05 | 0.05 | 0.05 |
| NC_019843.3;YP_009047211.1;MERS_nucleoprote | 211 | 211 | 1 | 0.05 | 0.05 | 0.05 |
| NC_019843.3;YP_009047211.1;MERS_nucleoprote | 214 | 214 | 1 | 0.1 | 0.1 | 0.1 |
| NC_019843.3;YP_009047211.1;MERS_nucleoprote | 225 | 232 | 8 | 0.05 | 0.05 | 0.05 |
| NC_019843.3;YP_009047211.1;MERS_nucleoprote | 240 | 240 | 1 | 0.05 | 0.05 | 0.05 |
| NC_019843.3;YP_009047211.1;MERS_nucleoprote | 261 | 276 | 16 | 0.05 | 0.15 | 0.0875 |
| NC_019843.3;YP_009047211.1;MERS_nucleoprote | 278 | 278 | 1 | 0.05 | 0.05 | 0.05 |
| NC_019843.3;YP_009047211.1;MERS_nucleoprote | 282 | 284 | 3 | 0.05 | 0.05 | 0.05 |
| NC_019843.3;YP_009047211.1;MERS_nucleoprote | 289 | 289 | 1 | 0.05 | 0.05 | 0.05 |
| NC_019843.3;YP_009047211.1;MERS_nucleoprote | 291 | 291 | 1 | 0.05 | 0.05 | 0.05 |
| NC_019843.3;YP_009047211.1;MERS_nucleoprote | 332 | 340 | 9 | 0.05 | 0.25 | 0.111111111 |
| NC_019843.3;YP_009047211.1;MERS_nucleoprote | 344 | 344 | 1 | 0.05 | 0.05 | 0.05 |
| NC_019843.3;YP_009047211.1;MERS_nucleoprote | 348 | 353 | 6 | 0.05 | 0.1 | 0.066666667 |
| NC_019843.3;YP_009047211.1;MERS_nucleoprote | 355 | 358 | 4 | 0.05 | 0.05 | 0.05 |
| NC_019843.3;YP_009047211.1;MERS_nucleoprote | 365 | 365 | 1 | 0.05 | 0.05 | 0.05 |
| NC_019843.3;YP_009047211.1;MERS_nucleoprote | 370 | 370 | 1 | 0.05 | 0.05 | 0.05 |
| NC_019843.3;YP_009047211.1;MERS_nucleoprote | 375 | 375 | 1 | 0.05 | 0.05 | 0.05 |
| NC_019843.3;YP_009047211.1;MERS_nucleoprote | 380 | 381 | 2 | 0.05 | 0.1 | 0.075 |
| NC_019843.3;YP_009047212.1;MERS_orf8b | 25 | 25 | 1 | 0.05 | 0.05 | 0.05 |
| NC_019843.3;YP_009047212.1;MERS_orf8b | 49 | 49 | 1 | 0.05 | 0.05 | 0.05 |
| NC_038311.1;YP_009505608.1;Rhinovirus-A1 | 5 | 5 | 1 | 0.1 | 0.1 | 0.1 |
| NC_038311.1;YP_009505608.1;Rhinovirus-A1 | 7 | 9 | 3 | 0.05 | 0.05 | 0.05 |
| NC_038311.1;YP_009505608.1;Rhinovirus-A1 | 11 | 11 | 1 | 0.05 | 0.05 | 0.05 |
| NC_038311.1;YP_009505608.1;Rhinovirus-A1 | 14 | 15 | 2 | 0.05 | 0.15 | 0.1 |
| NC_038311.1;YP_009505608.1;Rhinovirus-A1 | 20 | 64 | 45 | 0.05 | 0.6 | 0.3 |
| NC_038311.1;YP_009505608.1;Rhinovirus-A1 | 77 | 88 | 12 | 0.05 | 0.1 | 0.054166667 |
| NC_038311.1;YP_009505608.1;Rhinovirus-A1 | 102 | 125 | 24 | 0.05 | 0.45 | 0.208333333 |
| NC_038311.1;YP_009505608.1;Rhinovirus-A1 | 128 | 129 | 2 | 0.05 | 0.05 | 0.05 |

| Extended data 1 | | | | | | |
|---|---|---|---|---|---|---|
| Protein | Start position (peptide) | Stop position (peptide) | Number of peptides | Minimum fraction (per peptide) of control subjects with reactivity >3 sd above average | Maximum fraction (per peptide) of control subjects with reactivity >3 sd above average | Mean fraction (over all peptides in this contiguous stretch) of controlsubjects with reactivity >3 sd above average |
| NC_038311.1;YP_009505608.1;Rhinovirus-A1 | 143 | 143 | 1 | 0.05 | 0.05 | 0.05 |
| NC_038311.1;YP_009505608.1;Rhinovirus-A1 | 148 | 149 | 2 | 0.05 | 0.05 | 0.05 |
| NC_038311.1;YP_009505608.1;Rhinovirus-A1 | 151 | 151 | 1 | 0.05 | 0.05 | 0.05 |
| NC_038311.1;YP_009505608.1;Rhinovirus-A1 | 155 | 156 | 2 | 0.05 | 0.05 | 0.05 |
| NC_038311.1;YP_009505608.1;Rhinovirus-A1 | 203 | 216 | 14 | 0.1 | 0.65 | 0.507142857 |
| NC_038311.1;YP_009505608.1;Rhinovirus-A1 | 224 | 229 | 6 | 0.05 | 0.15 | 0.083333333 |
| NC_038311.1;YP_009505608.1;Rhinovirus-A1 | 238 | 238 | 1 | 0.05 | 0.05 | 0.05 |
| NC_038311.1;YP_009505608.1;Rhinovirus-A1 | 271 | 272 | 2 | 0.05 | 0.05 | 0.05 |
| NC_038311.1;YP_009505608.1;Rhinovirus-A1 | 292 | 302 | 11 | 0.05 | 0.15 | 0.059090909 |
| NC_038311.1;YP_009505608.1;Rhinovirus-A1 | 309 | 309 | 1 | 0.05 | 0.05 | 0.05 |
| NC_038311.1;YP_009505608.1;Rhinovirus-A1 | 311 | 320 | 10 | 0.05 | 0.1 | 0.06 |
| NC_038311.1;YP_009505608.1;Rhinovirus-A1 | 324 | 324 | 1 | 0.05 | 0.05 | 0.05 |
| NC_038311.1;YP_009505608.1;Rhinovirus-A1 | 326 | 326 | 1 | 0.05 | 0.05 | 0.05 |
| NC_038311.1;YP_009505608.1;Rhinovirus-A1 | 349 | 359 | 11 | 0.05 | 0.1 | 0.054545455 |
| NC_038311.1;YP_009505608.1;Rhinovirus-A1 | 377 | 390 | 14 | 0.05 | 0.35 | 0.157142857 |
| NC_038311.1;YP_009505608.1;Rhinovirus-A1 | 399 | 399 | 1 | 0.05 | 0.05 | 0.05 |
| NC_038311.1;YP_009505608.1;Rhinovirus-A1 | 413 | 413 | 1 | 0.05 | 0.05 | 0.05 |
| NC_038311.1;YP_009505608.1;Rhinovirus-A1 | 445 | 450 | 6 | 0.05 | 0.1 | 0.066666667 |
| NC_038311.1;YP_009505608.1;Rhinovirus-A1 | 452 | 455 | 4 | 0.05 | 0.05 | 0.05 |
| NC_038311.1;YP_009505608.1;Rhinovirus-A1 | 461 | 461 | 1 | 0.05 | 0.05 | 0.05 |
| NC_038311.1;YP_009505608.1;Rhinovirus-A1 | 464 | 465 | 2 | 0.05 | 0.1 | 0.075 |
| NC_038311.1;YP_009505608.1;Rhinovirus-A1 | 469 | 469 | 1 | 0.05 | 0.05 | 0.05 |
| NC_038311.1;YP_009505608.1;Rhinovirus-A1 | 486 | 486 | 1 | 0.05 | 0.05 | 0.05 |
| NC_038311.1;YP_009505608.1;Rhinovirus-A1 | 499 | 499 | 1 | 0.05 | 0.05 | 0.05 |
| NC_038311.1;YP_009505608.1;Rhinovirus-A1 | 527 | 527 | 1 | 0.05 | 0.05 | 0.05 |
| NC_038311.1;YP_009505608.1;Rhinovirus-A1 | 562 | 608 | 47 | 0.1 | 1 | 0.590425532 |
| NC_038311.1;YP_009505608.1;Rhinovirus-A1 | 610 | 612 | 3 | 0.05 | 0.1 | 0.066666667 |
| NC_038311.1;YP_009505608.1;Rhinovirus-A1 | 615 | 616 | 2 | 0.05 | 0.05 | 0.05 |
| NC_038311.1;YP_009505608.1;Rhinovirus-A1 | 645 | 645 | 1 | 0.05 | 0.05 | 0.05 |
| NC_038311.1;YP_009505608.1;Rhinovirus-A1 | 649 | 657 | 9 | 0.05 | 0.3 | 0.122222222 |
| NC_038311.1;YP_009505608.1;Rhinovirus-A1 | 660 | 662 | 3 | 0.05 | 0.1 | 0.083333333 |
| NC_038311.1;YP_009505608.1;Rhinovirus-A1 | 707 | 721 | 15 | 0.05 | 0.25 | 0.083333333 |
| NC_038311.1;YP_009505608.1;Rhinovirus-A1 | 724 | 727 | 4 | 0.05 | 0.15 | 0.1 |
| NC_038311.1;YP_009505608.1;Rhinovirus-A1 | 731 | 736 | 6 | 0.05 | 0.1 | 0.091666667 |
| NC_038311.1;YP_009505608.1;Rhinovirus-A1 | 763 | 767 | 5 | 0.05 | 0.1 | 0.07 |
| NC_038311.1;YP_009505608.1;Rhinovirus-A1 | 770 | 770 | 1 | 0.05 | 0.05 | 0.05 |
| NC_038311.1;YP_009505608.1;Rhinovirus-A1 | 773 | 773 | 1 | 0.05 | 0.05 | 0.05 |

| Extended data 1 | | | | | | |
|---|---|---|---|---|---|---|
| Protein | Start position (peptide) | Stop position (peptide) | Number of peptides | Minimum fraction (per peptide) of control subjects with reactivity >3 sd above average | Maximum fraction (per peptide) of control subjects with reactivity >3 sd above average | Mean fraction (over all peptides in this contiguous stretch) of controlsubjects with reactivity >3 sd above average |
| NC_038311.1;YP_009505608.1;Rhinovirus-A1 | 822 | 838 | 17 | 0.05 | 0.3 | 0.158823529 |
| NC_038311.1;YP_009505608.1;Rhinovirus-A1 | 840 | 851 | 12 | 0.05 | 0.2 | 0.116666667 |
| NC_038311.1;YP_009505608.1;Rhinovirus-A1 | 853 | 855 | 3 | 0.05 | 0.05 | 0.05 |
| NC_038311.1;YP_009505608.1;Rhinovirus-A1 | 858 | 858 | 1 | 0.05 | 0.05 | 0.05 |
| NC_038311.1;YP_009505608.1;Rhinovirus-A1 | 889 | 896 | 8 | 0.05 | 0.25 | 0.10625 |
| NC_038311.1;YP_009505608.1;Rhinovirus-A1 | 933 | 933 | 1 | 0.05 | 0.05 | 0.05 |
| NC_038311.1;YP_009505608.1;Rhinovirus-A1 | 935 | 935 | 1 | 0.05 | 0.05 | 0.05 |
| NC_038311.1;YP_009505608.1;Rhinovirus-A1 | 993 | 993 | 1 | 0.05 | 0.05 | 0.05 |
| NC_038311.1;YP_009505608.1;Rhinovirus-A1 | 995 | 1024 | 30 | 0.05 | 0.95 | 0.335 |
| NC_038311.1;YP_009505608.1;Rhinovirus-A1 | 1078 | 1090 | 13 | 0.05 | 0.3 | 0.115384615 |
| NC_038311.1;YP_009505608.1;Rhinovirus-A1 | 1093 | 1096 | 4 | 0.05 | 0.2 | 0.1 |
| NC_038311.1;YP_009505608.1;Rhinovirus-A1 | 1104 | 1107 | 4 | 0.05 | 0.05 | 0.05 |
| NC_038311.1;YP_009505608.1;Rhinovirus-A1 | 1109 | 1112 | 4 | 0.2 | 0.3 | 0.2375 |
| NC_038311.1;YP_009505608.1;Rhinovirus-A1 | 1117 | 1117 | 1 | 0.1 | 0.1 | 0.1 |
| NC_038311.1;YP_009505608.1;Rhinovirus-A1 | 1119 | 1127 | 9 | 0.05 | 0.2 | 0.138888889 |
| NC_038311.1;YP_009505608.1;Rhinovirus-A1 | 1139 | 1140 | 2 | 0.05 | 0.05 | 0.05 |
| NC_038311.1;YP_009505608.1;Rhinovirus-A1 | 1145 | 1145 | 1 | 0.1 | 0.1 | 0.1 |
| NC_038311.1;YP_009505608.1;Rhinovirus-A1 | 1147 | 1151 | 5 | 0.05 | 0.05 | 0.05 |
| NC_038311.1;YP_009505608.1;Rhinovirus-A1 | 1153 | 1161 | 9 | 0.05 | 0.05 | 0.05 |
| NC_038311.1;YP_009505608.1;Rhinovirus-A1 | 1199 | 1199 | 1 | 0.05 | 0.05 | 0.05 |
| NC_038311.1;YP_009505608.1;Rhinovirus-A1 | 1225 | 1226 | 2 | 0.05 | 0.05 | 0.05 |
| NC_038311.1;YP_009505608.1;Rhinovirus-A1 | 1230 | 1249 | 20 | 0.05 | 0.7 | 0.3975 |
| NC_038311.1;YP_009505608.1;Rhinovirus-A1 | 1252 | 1256 | 5 | 0.05 | 0.1 | 0.06 |
| NC_038311.1;YP_009505608.1;Rhinovirus-A1 | 1258 | 1268 | 11 | 0.05 | 0.05 | 0.05 |
| NC_038311.1;YP_009505608.1;Rhinovirus-A1 | 1278 | 1290 | 13 | 0.1 | 1 | 0.696153846 |
| NC_038311.1;YP_009505608.1;Rhinovirus-A1 | 1307 | 1314 | 8 | 0.05 | 0.15 | 0.125 |
| NC_038311.1;YP_009505608.1;Rhinovirus-A1 | 1317 | 1317 | 1 | 0.05 | 0.05 | 0.05 |
| NC_038311.1;YP_009505608.1;Rhinovirus-A1 | 1328 | 1330 | 3 | 0.05 | 0.05 | 0.05 |
| NC_038311.1;YP_009505608.1;Rhinovirus-A1 | 1335 | 1337 | 3 | 0.05 | 0.1 | 0.066666667 |
| NC_038311.1;YP_009505608.1;Rhinovirus-A1 | 1344 | 1347 | 4 | 0.05 | 0.05 | 0.05 |
| NC_038311.1;YP_009505608.1;Rhinovirus-A1 | 1349 | 1349 | 1 | 0.05 | 0.05 | 0.05 |
| NC_038311.1;YP_009505608.1;Rhinovirus-A1 | 1355 | 1355 | 1 | 0.05 | 0.05 | 0.05 |
| NC_038311.1;YP_009505608.1;Rhinovirus-A1 | 1391 | 1391 | 1 | 0.05 | 0.05 | 0.05 |
| NC_038311.1;YP_009505608.1;Rhinovirus-A1 | 1395 | 1396 | 2 | 0.05 | 0.05 | 0.05 |
| NC_038311.1;YP_009505608.1;Rhinovirus-A1 | 1399 | 1409 | 11 | 0.05 | 0.2 | 0.113636364 |
| NC_038311.1;YP_009505608.1;Rhinovirus-A1 | 1413 | 1414 | 2 | 0.05 | 0.05 | 0.05 |
| NC_038311.1;YP_009505608.1;Rhinovirus-A1 | 1416 | 1450 | 35 | 0.05 | 0.85 | 0.47 |

| Extended data 1 | | | | | | |
|---|---|---|---|---|---|---|
| Protein | Start position (peptide) | Stop position (peptide) | Number of peptides | Minimum fraction (per peptide) of control subjects with reactivity >3 sd above average | Maximum fraction (per peptide) of control subjects with reactivity >3 sd above average | Mean fraction (over all peptides in this contiguous stretch) of controlsubjects with reactivity >3 sd above average |
| NC_038311.1;YP_009505608.1;Rhinovirus-A1 | 1459 | 1459 | 1 | 0.05 | 0.05 | 0.05 |
| NC_038311.1;YP_009505608.1;Rhinovirus-A1 | 1507 | 1508 | 2 | 0.05 | 0.05 | 0.05 |
| NC_038311.1;YP_009505608.1;Rhinovirus-A1 | 1510 | 1510 | 1 | 0.05 | 0.05 | 0.05 |
| NC_038311.1;YP_009505608.1;Rhinovirus-A1 | 1515 | 1515 | 1 | 0.1 | 0.1 | 0.1 |
| NC_038311.1;YP_009505608.1;Rhinovirus-A1 | 1531 | 1531 | 1 | 0.1 | 0.1 | 0.1 |
| NC_038311.1;YP_009505608.1;Rhinovirus-A1 | 1564 | 1564 | 1 | 0.05 | 0.05 | 0.05 |
| NC_038311.1;YP_009505608.1;Rhinovirus-A1 | 1568 | 1571 | 4 | 0.05 | 0.2 | 0.1 |
| NC_038311.1;YP_009505608.1;Rhinovirus-A1 | 1585 | 1596 | 12 | 0.1 | 0.3 | 0.166666667 |
| NC_038311.1;YP_009505608.1;Rhinovirus-A1 | 1604 | 1604 | 1 | 0.05 | 0.05 | 0.05 |
| NC_038311.1;YP_009505608.1;Rhinovirus-A1 | 1608 | 1609 | 2 | 0.05 | 0.2 | 0.125 |
| NC_038311.1;YP_009505608.1;Rhinovirus-A1 | 1634 | 1634 | 1 | 0.05 | 0.05 | 0.05 |
| NC_038311.1;YP_009505608.1;Rhinovirus-A1 | 1638 | 1638 | 1 | 0.05 | 0.05 | 0.05 |
| NC_038311.1;YP_009505608.1;Rhinovirus-A1 | 1685 | 1694 | 10 | 0.05 | 0.1 | 0.07 |
| NC_038311.1;YP_009505608.1;Rhinovirus-A1 | 1704 | 1729 | 26 | 0.05 | 0.55 | 0.223076923 |
| NC_038311.1;YP_009505608.1;Rhinovirus-A1 | 1731 | 1737 | 7 | 0.05 | 0.2 | 0.128571429 |
| NC_038311.1;YP_009505608.1;Rhinovirus-A1 | 1740 | 1741 | 2 | 0.05 | 0.05 | 0.05 |
| NC_038311.1;YP_009505608.1;Rhinovirus-A1 | 1743 | 1743 | 1 | 0.05 | 0.05 | 0.05 |
| NC_038311.1;YP_009505608.1;Rhinovirus-A1 | 1749 | 1760 | 12 | 0.05 | 0.15 | 0.091666667 |
| NC_038311.1;YP_009505608.1;Rhinovirus-A1 | 1777 | 1777 | 1 | 0.05 | 0.05 | 0.05 |
| NC_038311.1;YP_009505608.1;Rhinovirus-A1 | 1788 | 1788 | 1 | 0.05 | 0.05 | 0.05 |
| NC_038311.1;YP_009505608.1;Rhinovirus-A1 | 1790 | 1791 | 2 | 0.05 | 0.05 | 0.05 |
| NC_038311.1;YP_009505608.1;Rhinovirus-A1 | 1795 | 1796 | 2 | 0.05 | 0.05 | 0.05 |
| NC_038311.1;YP_009505608.1;Rhinovirus-A1 | 1799 | 1800 | 2 | 0.05 | 0.25 | 0.15 |
| NC_038311.1;YP_009505608.1;Rhinovirus-A1 | 1802 | 1803 | 2 | 0.05 | 0.15 | 0.1 |
| NC_038311.1;YP_009505608.1;Rhinovirus-A1 | 1805 | 1805 | 1 | 0.05 | 0.05 | 0.05 |
| NC_038311.1;YP_009505608.1;Rhinovirus-A1 | 1807 | 1807 | 1 | 0.05 | 0.05 | 0.05 |
| NC_038311.1;YP_009505608.1;Rhinovirus-A1 | 1813 | 1824 | 12 | 0.1 | 0.25 | 0.1375 |
| NC_038311.1;YP_009505608.1;Rhinovirus-A1 | 1832 | 1850 | 19 | 0.05 | 0.25 | 0.131578947 |
| NC_038311.1;YP_009505608.1;Rhinovirus-A1 | 1854 | 1862 | 9 | 0.05 | 0.05 | 0.05 |
| NC_038311.1;YP_009505608.1;Rhinovirus-A1 | 1864 | 1867 | 4 | 0.05 | 0.15 | 0.075 |
| NC_038311.1;YP_009505608.1;Rhinovirus-A1 | 1877 | 1877 | 1 | 0.1 | 0.1 | 0.1 |
| NC_038311.1;YP_009505608.1;Rhinovirus-A1 | 1893 | 1893 | 1 | 0.05 | 0.05 | 0.05 |
| NC_038311.1;YP_009505608.1;Rhinovirus-A1 | 1902 | 1902 | 1 | 0.1 | 0.1 | 0.1 |
| NC_038311.1;YP_009505608.1;Rhinovirus-A1 | 1904 | 1911 | 8 | 0.05 | 0.4 | 0.1625 |
| NC_038311.1;YP_009505608.1;Rhinovirus-A1 | 1919 | 1929 | 11 | 0.05 | 0.05 | 0.05 |
| NC_038311.1;YP_009505608.1;Rhinovirus-A1 | 1947 | 1949 | 3 | 0.05 | 0.05 | 0.05 |
| NC_038311.1;YP_009505608.1;Rhinovirus-A1 | 1951 | 1952 | 2 | 0.05 | 0.05 | 0.05 |

| Extended data 1 | | | | | | |
|---|---|---|---|---|---|---|
| Protein | Start position (peptide) | Stop position (peptide) | Number of peptides | Minimum fraction (per peptide) of control subjects with reactivity >3 sd above average | Maximum fraction (per peptide) of control subjects with reactivity >3 sd above average | Mean fraction (over all peptides in this contiguous stretch) of controlsubjects with reactivity >3 sd above average |
| NC_038311.1;YP_009505608.1;Rhinovirus-A1 | 1966 | 1966 | 1 | 0.05 | 0.05 | 0.05 |
| NC_038311.1;YP_009505608.1;Rhinovirus-A1 | 1982 | 1982 | 1 | 0.05 | 0.05 | 0.05 |
| NC_038311.1;YP_009505608.1;Rhinovirus-A1 | 2001 | 2002 | 2 | 0.05 | 0.1 | 0.075 |
| NC_038311.1;YP_009505608.1;Rhinovirus-A1 | 2006 | 2006 | 1 | 0.05 | 0.05 | 0.05 |
| NC_038311.1;YP_009505608.1;Rhinovirus-A1 | 2010 | 2010 | 1 | 0.05 | 0.05 | 0.05 |
| NC_038311.1;YP_009505608.1;Rhinovirus-A1 | 2030 | 2034 | 5 | 0.05 | 0.15 | 0.07 |
| NC_038311.1;YP_009505608.1;Rhinovirus-A1 | 2036 | 2036 | 1 | 0.05 | 0.05 | 0.05 |
| NC_038311.1;YP_009505608.1;Rhinovirus-A1 | 2045 | 2045 | 1 | 0.05 | 0.05 | 0.05 |
| NC_038311.1;YP_009505608.1;Rhinovirus-A1 | 2048 | 2054 | 7 | 0.05 | 0.05 | 0.05 |
| NC_038311.1;YP_009505608.1;Rhinovirus-A1 | 2061 | 2061 | 1 | 0.05 | 0.05 | 0.05 |
| NC_038311.1;YP_009505608.1;Rhinovirus-A1 | 2063 | 2063 | 1 | 0.05 | 0.05 | 0.05 |
| NC_038311.1;YP_009505608.1;Rhinovirus-A1 | 2068 | 2068 | 1 | 0.05 | 0.05 | 0.05 |
| NC_038311.1;YP_009505608.1;Rhinovirus-A1 | 2088 | 2088 | 1 | 0.05 | 0.05 | 0.05 |
| NC_038311.1;YP_009505608.1;Rhinovirus-A1 | 2094 | 2096 | 3 | 0.05 | 0.2 | 0.1 |
| NC_045512.2;YP_009724389.1;Wu1-SARS2_orf1ab | 29 | 30 | 2 | 0.1 | 0.25 | 0.175 |
| NC_045512.2;YP_009724389.1;Wu1-SARS2_orf1ab | 33 | 34 | 2 | 0.05 | 0.1 | 0.075 |
| NC_045512.2;YP_009724389.1;Wu1-SARS2_orf1ab | 44 | 44 | 1 | 0.05 | 0.05 | 0.05 |
| NC_045512.2;YP_009724389.1;Wu1-SARS2_orf1ab | 52 | 52 | 1 | 0.05 | 0.05 | 0.05 |
| NC_045512.2;YP_009724389.1;Wu1-SARS2_orf1ab | 54 | 54 | 1 | 0.05 | 0.05 | 0.05 |
| NC_045512.2;YP_009724389.1;Wu1-SARS2_orf1ab | 71 | 71 | 1 | 0.05 | 0.05 | 0.05 |
| NC_045512.2;YP_009724389.1;Wu1-SARS2_orf1ab | 118 | 118 | 1 | 0.05 | 0.05 | 0.05 |
| NC_045512.2;YP_009724389.1;Wu1-SARS2_orf1ab | 121 | 121 | 1 | 0.05 | 0.05 | 0.05 |
| NC_045512.2;YP_009724389.1;Wu1-SARS2_orf1ab | 125 | 125 | 1 | 0.1 | 0.1 | 0.1 |
| NC_045512.2;YP_009724389.1;Wu1-SARS2_orf1ab | 131 | 133 | 3 | 0.05 | 0.1 | 0.066666667 |
| NC_045512.2;YP_009724389.1;Wu1-SARS2_orf1ab | 135 | 135 | 1 | 0.05 | 0.05 | 0.05 |
| NC_045512.2;YP_009724389.1;Wu1-SARS2_orf1ab | 139 | 140 | 2 | 0.05 | 0.05 | 0.05 |
| NC_045512.2;YP_009724389.1;Wu1-SARS2_orf1ab | 143 | 143 | 1 | 0.05 | 0.05 | 0.05 |
| NC_045512.2;YP_009724389.1;Wu1-SARS2_orf1ab | 145 | 146 | 2 | 0.05 | 0.2 | 0.125 |
| NC_045512.2;YP_009724389.1;Wu1-SARS2_orf1ab | 148 | 158 | 11 | 0.05 | 0.5 | 0.290909091 |
| NC_045512.2;YP_009724389.1;Wu1-SARS2_orf1ab | 165 | 166 | 2 | 0.05 | 0.1 | 0.075 |
| NC_045512.2;YP_009724389.1;Wu1-SARS2_orf1ab | 168 | 168 | 1 | 0.1 | 0.1 | 0.1 |
| NC_045512.2;YP_009724389.1;Wu1-SARS2_orf1ab | 174 | 174 | 1 | 0.05 | 0.05 | 0.05 |
| NC_045512.2;YP_009724389.1;Wu1-SARS2_orf1ab | 197 | 197 | 1 | 0.05 | 0.05 | 0.05 |
| NC_045512.2;YP_009724389.1;Wu1-SARS2_orf1ab | 204 | 204 | 1 | 0.05 | 0.05 | 0.05 |
| NC_045512.2;YP_009724389.1;Wu1-SARS2_orf1ab | 211 | 211 | 1 | 0.05 | 0.05 | 0.05 |
| NC_045512.2;YP_009724389.1;Wu1-SARS2_orf1ab | 214 | 215 | 2 | 0.1 | 0.1 | 0.1 |
| NC_045512.2;YP_009724389.1;Wu1-SARS2_orf1ab | 220 | 220 | 1 | 0.05 | 0.05 | 0.05 |

| Extended data 1 | | | | | | |
|---|---|---|---|---|---|---|
| Protein | Start position (peptide) | Stop position (peptide) | Number of peptides | Minimum fraction (per peptide) of control subjects with reactivity >3 sd above average | Maximum fraction (per peptide) of control subjects with reactivity >3 sd above average | Mean fraction (over all peptides in this contiguous stretch) of controlsubjects with reactivity >3 sd above average |
| NC_045512.2;YP_009724389.1;Wu1-SARS2_orf1ab | 225 | 225 | 1 | 0.05 | 0.05 | 0.05 |
| NC_045512.2;YP_009724389.1;Wu1-SARS2_orf1ab | 252 | 253 | 2 | 0.05 | 0.05 | 0.05 |
| NC_045512.2;YP_009724389.1;Wu1-SARS2_orf1ab | 256 | 256 | 1 | 0.1 | 0.1 | 0.1 |
| NC_045512.2;YP_009724389.1;Wu1-SARS2_orf1ab | 258 | 261 | 4 | 0.05 | 0.05 | 0.05 |
| NC_045512.2;YP_009724389.1;Wu1-SARS2_orf1ab | 276 | 286 | 11 | 0.05 | 0.1 | 0.063636364 |
| NC_045512.2;YP_009724389.1;Wu1-SARS2_orf1ab | 291 | 291 | 1 | 0.05 | 0.05 | 0.05 |
| NC_045512.2;YP_009724389.1;Wu1-SARS2_orf1ab | 296 | 305 | 10 | 0.05 | 0.1 | 0.085 |
| NC_045512.2;YP_009724389.1;Wu1-SARS2_orf1ab | 324 | 324 | 1 | 0.05 | 0.05 | 0.05 |
| NC_045512.2;YP_009724389.1;Wu1-SARS2_orf1ab | 347 | 347 | 1 | 0.05 | 0.05 | 0.05 |
| NC_045512.2;YP_009724389.1;Wu1-SARS2_orf1ab | 349 | 349 | 1 | 0.05 | 0.05 | 0.05 |
| NC_045512.2;YP_009724389.1;Wu1-SARS2_orf1ab | 379 | 379 | 1 | 0.15 | 0.15 | 0.15 |
| NC_045512.2;YP_009724389.1;Wu1-SARS2_orf1ab | 385 | 386 | 2 | 0.05 | 0.15 | 0.1 |
| NC_045512.2;YP_009724389.1;Wu1-SARS2_orf1ab | 424 | 425 | 2 | 0.05 | 0.05 | 0.05 |
| NC_045512.2;YP_009724389.1;Wu1-SARS2_orf1ab | 443 | 448 | 6 | 0.05 | 0.5 | 0.15 |
| NC_045512.2;YP_009724389.1;Wu1-SARS2_orf1ab | 452 | 452 | 1 | 0.05 | 0.05 | 0.05 |
| NC_045512.2;YP_009724389.1;Wu1-SARS2_orf1ab | 456 | 458 | 3 | 0.05 | 0.05 | 0.05 |
| NC_045512.2;YP_009724389.1;Wu1-SARS2_orf1ab | 489 | 489 | 1 | 0.05 | 0.05 | 0.05 |
| NC_045512.2;YP_009724389.1;Wu1-SARS2_orf1ab | 493 | 494 | 2 | 0.05 | 0.05 | 0.05 |
| NC_045512.2;YP_009724389.1;Wu1-SARS2_orf1ab | 518 | 518 | 1 | 0.1 | 0.1 | 0.1 |
| NC_045512.2;YP_009724389.1;Wu1-SARS2_orf1ab | 525 | 526 | 2 | 0.05 | 0.05 | 0.05 |
| NC_045512.2;YP_009724389.1;Wu1-SARS2_orf1ab | 551 | 551 | 1 | 0.05 | 0.05 | 0.05 |
| NC_045512.2;YP_009724389.1;Wu1-SARS2_orf1ab | 553 | 553 | 1 | 0.05 | 0.05 | 0.05 |
| NC_045512.2;YP_009724389.1;Wu1-SARS2_orf1ab | 555 | 555 | 1 | 0.05 | 0.05 | 0.05 |
| NC_045512.2;YP_009724389.1;Wu1-SARS2_orf1ab | 572 | 572 | 1 | 0.1 | 0.1 | 0.1 |
| NC_045512.2;YP_009724389.1;Wu1-SARS2_orf1ab | 589 | 589 | 1 | 0.05 | 0.05 | 0.05 |
| NC_045512.2;YP_009724389.1;Wu1-SARS2_orf1ab | 617 | 621 | 5 | 0.05 | 0.1 | 0.06 |
| NC_045512.2;YP_009724389.1;Wu1-SARS2_orf1ab | 638 | 638 | 1 | 0.05 | 0.05 | 0.05 |
| NC_045512.2;YP_009724389.1;Wu1-SARS2_orf1ab | 712 | 712 | 1 | 0.05 | 0.05 | 0.05 |
| NC_045512.2;YP_009724389.1;Wu1-SARS2_orf1ab | 726 | 726 | 1 | 0.05 | 0.05 | 0.05 |
| NC_045512.2;YP_009724389.1;Wu1-SARS2_orf1ab | 744 | 759 | 16 | 0.05 | 0.15 | 0.0875 |
| NC_045512.2;YP_009724389.1;Wu1-SARS2_orf1ab | 764 | 767 | 4 | 0.05 | 0.05 | 0.05 |
| NC_045512.2;YP_009724389.1;Wu1-SARS2_orf1ab | 775 | 775 | 1 | 0.05 | 0.05 | 0.05 |
| NC_045512.2;YP_009724389.1;Wu1-SARS2_orf1ab | 782 | 782 | 1 | 0.05 | 0.05 | 0.05 |
| NC_045512.2;YP_009724389.1;Wu1-SARS2_orf1ab | 789 | 789 | 1 | 0.05 | 0.05 | 0.05 |
| NC_045512.2;YP_009724389.1;Wu1-SARS2_orf1ab | 797 | 798 | 2 | 0.05 | 0.05 | 0.05 |
| NC_045512.2;YP_009724389.1;Wu1-SARS2_orf1ab | 801 | 801 | 1 | 0.05 | 0.05 | 0.05 |
| NC_045512.2;YP_009724389.1;Wu1-SARS2_orf1ab | 803 | 806 | 4 | 0.05 | 0.1 | 0.0625 |

| Extended data 1 | | | | | | |
|---|---|---|---|---|---|---|
| Protein | Start position (peptide) | Stop position (peptide) | Number of peptides | Minimum fraction (per peptide) of control subjects with reactivity >3 sd above average | Maximum fraction (per peptide) of control subjects with reactivity >3 sd above average | Mean fraction (over all peptides in this contiguous stretch) of controlsubjects with reactivity >3 sd above average |
| NC_045512.2;YP_009724389.1;Wu1-SARS2_orf1ab | 813 | 813 | 1 | 0.05 | 0.05 | 0.05 |
| NC_045512.2;YP_009724389.1;Wu1-SARS2_orf1ab | 819 | 820 | 2 | 0.05 | 0.05 | 0.05 |
| NC_045512.2;YP_009724389.1;Wu1-SARS2_orf1ab | 827 | 827 | 1 | 0.1 | 0.1 | 0.1 |
| NC_045512.2;YP_009724389.1;Wu1-SARS2_orf1ab | 831 | 831 | 1 | 0.05 | 0.05 | 0.05 |
| NC_045512.2;YP_009724389.1;Wu1-SARS2_orf1ab | 836 | 837 | 2 | 0.05 | 0.05 | 0.05 |
| NC_045512.2;YP_009724389.1;Wu1-SARS2_orf1ab | 840 | 844 | 5 | 0.05 | 0.25 | 0.16 |
| NC_045512.2;YP_009724389.1;Wu1-SARS2_orf1ab | 876 | 876 | 1 | 0.05 | 0.05 | 0.05 |
| NC_045512.2;YP_009724389.1;Wu1-SARS2_orf1ab | 881 | 881 | 1 | 0.05 | 0.05 | 0.05 |
| NC_045512.2;YP_009724389.1;Wu1-SARS2_orf1ab | 884 | 884 | 1 | 0.05 | 0.05 | 0.05 |
| NC_045512.2;YP_009724389.1;Wu1-SARS2_orf1ab | 900 | 900 | 1 | 0.1 | 0.1 | 0.1 |
| NC_045512.2;YP_009724389.1;Wu1-SARS2_orf1ab | 906 | 906 | 1 | 0.05 | 0.05 | 0.05 |
| NC_045512.2;YP_009724389.1;Wu1-SARS2_orf1ab | 927 | 929 | 3 | 0.05 | 0.35 | 0.15 |
| NC_045512.2;YP_009724389.1;Wu1-SARS2_orf1ab | 933 | 933 | 1 | 0.05 | 0.05 | 0.05 |
| NC_045512.2;YP_009724389.1;Wu1-SARS2_orf1ab | 940 | 940 | 1 | 0.05 | 0.05 | 0.05 |
| NC_045512.2;YP_009724389.1;Wu1-SARS2_orf1ab | 952 | 953 | 2 | 0.1 | 0.15 | 0.125 |
| NC_045512.2;YP_009724389.1;Wu1-SARS2_orf1ab | 977 | 977 | 1 | 0.1 | 0.1 | 0.1 |
| NC_045512.2;YP_009724389.1;Wu1-SARS2_orf1ab | 1015 | 1015 | 1 | 0.1 | 0.1 | 0.1 |
| NC_045512.2;YP_009724389.1;Wu1-SARS2_orf1ab | 1023 | 1034 | 12 | 0.05 | 0.2 | 0.070833333 |
| NC_045512.2;YP_009724389.1;Wu1-SARS2_orf1ab | 1048 | 1048 | 1 | 0.05 | 0.05 | 0.05 |
| NC_045512.2;YP_009724389.1;Wu1-SARS2_orf1ab | 1055 | 1055 | 1 | 0.05 | 0.05 | 0.05 |
| NC_045512.2;YP_009724389.1;Wu1-SARS2_orf1ab | 1057 | 1057 | 1 | 0.05 | 0.05 | 0.05 |
| NC_045512.2;YP_009724389.1;Wu1-SARS2_orf1ab | 1063 | 1063 | 1 | 0.05 | 0.05 | 0.05 |
| NC_045512.2;YP_009724389.1;Wu1-SARS2_orf1ab | 1067 | 1075 | 9 | 0.05 | 0.2 | 0.111111111 |
| NC_045512.2;YP_009724389.1;Wu1-SARS2_orf1ab | 1078 | 1079 | 2 | 0.05 | 0.05 | 0.05 |
| NC_045512.2;YP_009724389.1;Wu1-SARS2_orf1ab | 1087 | 1088 | 2 | 0.05 | 0.15 | 0.1 |
| NC_045512.2;YP_009724389.1;Wu1-SARS2_orf1ab | 1111 | 1115 | 5 | 0.05 | 0.15 | 0.07 |
| NC_045512.2;YP_009724389.1;Wu1-SARS2_orf1ab | 1117 | 1122 | 6 | 0.05 | 0.2 | 0.1 |
| NC_045512.2;YP_009724389.1;Wu1-SARS2_orf1ab | 1125 | 1126 | 2 | 0.05 | 0.05 | 0.05 |
| NC_045512.2;YP_009724389.1;Wu1-SARS2_orf1ab | 1144 | 1144 | 1 | 0.05 | 0.05 | 0.05 |
| NC_045512.2;YP_009724389.1;Wu1-SARS2_orf1ab | 1173 | 1175 | 3 | 0.05 | 0.05 | 0.05 |
| NC_045512.2;YP_009724389.1;Wu1-SARS2_orf1ab | 1177 | 1179 | 3 | 0.05 | 0.1 | 0.066666667 |
| NC_045512.2;YP_009724389.1;Wu1-SARS2_orf1ab | 1185 | 1190 | 6 | 0.05 | 0.1 | 0.058333333 |
| NC_045512.2;YP_009724389.1;Wu1-SARS2_orf1ab | 1192 | 1192 | 1 | 0.05 | 0.05 | 0.05 |
| NC_045512.2;YP_009724389.1;Wu1-SARS2_orf1ab | 1194 | 1201 | 8 | 0.05 | 0.1 | 0.06875 |
| NC_045512.2;YP_009724389.1;Wu1-SARS2_orf1ab | 1204 | 1218 | 15 | 0.05 | 0.1 | 0.066666667 |
| NC_045512.2;YP_009724389.1;Wu1-SARS2_orf1ab | 1220 | 1226 | 7 | 0.05 | 0.05 | 0.05 |
| NC_045512.2;YP_009724389.1;Wu1-SARS2_orf1ab | 1255 | 1256 | 2 | 0.05 | 0.1 | 0.075 |

| Extended data 1 | | | | | | |
|---|---|---|---|---|---|---|
| Protein | Start position (peptide) | Stop position (peptide) | Number of peptides | Minimum fraction (per peptide) of control subjects with reactivity >3 sd above average | Maximum fraction (per peptide) of control subjects with reactivity >3 sd above average | Mean fraction (over all peptides in this contiguous stretch) of controlsubjects with reactivity >3 sd above average |
| NC_045512.2;YP_009724389.1;Wu1-SARS2_orf1ab | 1272 | 1272 | 1 | 0.05 | 0.05 | 0.05 |
| NC_045512.2;YP_009724389.1;Wu1-SARS2_orf1ab | 1296 | 1302 | 7 | 0.05 | 0.05 | 0.05 |
| NC_045512.2;YP_009724389.1;Wu1-SARS2_orf1ab | 1309 | 1309 | 1 | 0.05 | 0.05 | 0.05 |
| NC_045512.2;YP_009724389.1;Wu1-SARS2_orf1ab | 1316 | 1318 | 3 | 0.05 | 0.05 | 0.05 |
| NC_045512.2;YP_009724389.1;Wu1-SARS2_orf1ab | 1323 | 1323 | 1 | 0.05 | 0.05 | 0.05 |
| NC_045512.2;YP_009724389.1;Wu1-SARS2_orf1ab | 1328 | 1329 | 2 | 0.05 | 0.05 | 0.05 |
| NC_045512.2;YP_009724389.1;Wu1-SARS2_orf1ab | 1354 | 1354 | 1 | 0.05 | 0.05 | 0.05 |
| NC_045512.2;YP_009724389.1;Wu1-SARS2_orf1ab | 1357 | 1357 | 1 | 0.05 | 0.05 | 0.05 |
| NC_045512.2;YP_009724389.1;Wu1-SARS2_orf1ab | 1359 | 1359 | 1 | 0.05 | 0.05 | 0.05 |
| NC_045512.2;YP_009724389.1;Wu1-SARS2_orf1ab | 1365 | 1365 | 1 | 0.05 | 0.05 | 0.05 |
| NC_045512.2;YP_009724389.1;Wu1-SARS2_orf1ab | 1382 | 1383 | 2 | 0.05 | 0.05 | 0.05 |
| NC_045512.2;YP_009724389.1;Wu1-SARS2_orf1ab | 1396 | 1406 | 11 | 0.05 | 0.1 | 0.090909091 |
| NC_045512.2;YP_009724389.1;Wu1-SARS2_orf1ab | 1414 | 1414 | 1 | 0.05 | 0.05 | 0.05 |
| NC_045512.2;YP_009724389.1;Wu1-SARS2_orf1ab | 1427 | 1442 | 16 | 0.05 | 0.35 | 0.1875 |
| NC_045512.2;YP_009724389.1;Wu1-SARS2_orf1ab | 1446 | 1446 | 1 | 0.05 | 0.05 | 0.05 |
| NC_045512.2;YP_009724389.1;Wu1-SARS2_orf1ab | 1448 | 1451 | 4 | 0.05 | 0.1 | 0.0625 |
| NC_045512.2;YP_009724389.1;Wu1-SARS2_orf1ab | 1460 | 1462 | 3 | 0.05 | 0.15 | 0.1 |
| NC_045512.2;YP_009724389.1;Wu1-SARS2_orf1ab | 1473 | 1473 | 1 | 0.05 | 0.05 | 0.05 |
| NC_045512.2;YP_009724389.1;Wu1-SARS2_orf1ab | 1480 | 1493 | 14 | 0.05 | 0.25 | 0.117857143 |
| NC_045512.2;YP_009724389.1;Wu1-SARS2_orf1ab | 1506 | 1506 | 1 | 0.05 | 0.05 | 0.05 |
| NC_045512.2;YP_009724389.1;Wu1-SARS2_orf1ab | 1509 | 1509 | 1 | 0.05 | 0.05 | 0.05 |
| NC_045512.2;YP_009724389.1;Wu1-SARS2_orf1ab | 1524 | 1524 | 1 | 0.05 | 0.05 | 0.05 |
| NC_045512.2;YP_009724389.1;Wu1-SARS2_orf1ab | 1526 | 1526 | 1 | 0.05 | 0.05 | 0.05 |
| NC_045512.2;YP_009724389.1;Wu1-SARS2_orf1ab | 1573 | 1573 | 1 | 0.05 | 0.05 | 0.05 |
| NC_045512.2;YP_009724389.1;Wu1-SARS2_orf1ab | 1595 | 1595 | 1 | 0.15 | 0.15 | 0.15 |
| NC_045512.2;YP_009724389.1;Wu1-SARS2_orf1ab | 1602 | 1603 | 2 | 0.05 | 0.05 | 0.05 |
| NC_045512.2;YP_009724389.1;Wu1-SARS2_orf1ab | 1611 | 1616 | 6 | 0.05 | 0.05 | 0.05 |
| NC_045512.2;YP_009724389.1;Wu1-SARS2_orf1ab | 1631 | 1634 | 4 | 0.05 | 0.05 | 0.05 |
| NC_045512.2;YP_009724389.1;Wu1-SARS2_orf1ab | 1639 | 1639 | 1 | 0.05 | 0.05 | 0.05 |
| NC_045512.2;YP_009724389.1;Wu1-SARS2_orf1ab | 1657 | 1664 | 8 | 0.05 | 0.05 | 0.05 |
| NC_045512.2;YP_009724389.1;Wu1-SARS2_orf1ab | 1679 | 1679 | 1 | 0.05 | 0.05 | 0.05 |
| NC_045512.2;YP_009724389.1;Wu1-SARS2_orf1ab | 1692 | 1697 | 6 | 0.05 | 0.15 | 0.075 |
| NC_045512.2;YP_009724389.1;Wu1-SARS2_orf1ab | 1723 | 1724 | 2 | 0.05 | 0.05 | 0.05 |
| NC_045512.2;YP_009724389.1;Wu1-SARS2_orf1ab | 1726 | 1726 | 1 | 0.05 | 0.05 | 0.05 |
| NC_045512.2;YP_009724389.1;Wu1-SARS2_orf1ab | 1748 | 1757 | 10 | 0.05 | 0.2 | 0.1 |
| NC_045512.2;YP_009724389.1;Wu1-SARS2_orf1ab | 1761 | 1775 | 15 | 0.05 | 0.15 | 0.116666667 |
| NC_045512.2;YP_009724389.1;Wu1-SARS2_orf1ab | 1786 | 1786 | 1 | 0.05 | 0.05 | 0.05 |

| Extended data 1 | | | | | | |
|---|---|---|---|---|---|---|
| **Protein** | **Start position (peptide)** | **Stop position (peptide)** | **Number of peptides** | **Minimum fraction (per peptide) of control subjects with reactivity >3 sd above average** | **Maximum fraction (per peptide) of control subjects with reactivity >3 sd above average** | **Mean fraction (over all peptides in this contiguous stretch) of controlsubjects with reactivity >3 sd above average** |
| NC_045512.2;YP_009724389.1;Wu1-SARS2_orf1ab | 1790 | 1790 | 1 | 0.05 | 0.05 | 0.05 |
| NC_045512.2;YP_009724389.1;Wu1-SARS2_orf1ab | 1793 | 1793 | 1 | 0.05 | 0.05 | 0.05 |
| NC_045512.2;YP_009724389.1;Wu1-SARS2_orf1ab | 1795 | 1795 | 1 | 0.05 | 0.05 | 0.05 |
| NC_045512.2;YP_009724389.1;Wu1-SARS2_orf1ab | 1797 | 1797 | 1 | 0.1 | 0.1 | 0.1 |
| NC_045512.2;YP_009724389.1;Wu1-SARS2_orf1ab | 1810 | 1810 | 1 | 0.05 | 0.05 | 0.05 |
| NC_045512.2;YP_009724389.1;Wu1-SARS2_orf1ab | 1817 | 1817 | 1 | 0.1 | 0.1 | 0.1 |
| NC_045512.2;YP_009724389.1;Wu1-SARS2_orf1ab | 1831 | 1832 | 2 | 0.05 | 0.05 | 0.05 |
| NC_045512.2;YP_009724389.1;Wu1-SARS2_orf1ab | 1854 | 1861 | 8 | 0.05 | 0.25 | 0.09375 |
| NC_045512.2;YP_009724389.1;Wu1-SARS2_orf1ab | 1865 | 1865 | 1 | 0.1 | 0.1 | 0.1 |
| NC_045512.2;YP_009724389.1;Wu1-SARS2_orf1ab | 1868 | 1868 | 1 | 0.05 | 0.05 | 0.05 |
| NC_045512.2;YP_009724389.1;Wu1-SARS2_orf1ab | 1886 | 1886 | 1 | 0.05 | 0.05 | 0.05 |
| NC_045512.2;YP_009724389.1;Wu1-SARS2_orf1ab | 1890 | 1901 | 12 | 0.05 | 0.35 | 0.191666667 |
| NC_045512.2;YP_009724389.1;Wu1-SARS2_orf1ab | 1903 | 1908 | 6 | 0.05 | 0.15 | 0.075 |
| NC_045512.2;YP_009724389.1;Wu1-SARS2_orf1ab | 1910 | 1910 | 1 | 0.2 | 0.2 | 0.2 |
| NC_045512.2;YP_009724389.1;Wu1-SARS2_orf1ab | 1914 | 1921 | 8 | 0.05 | 0.05 | 0.05 |
| NC_045512.2;YP_009724389.1;Wu1-SARS2_orf1ab | 1924 | 1924 | 1 | 0.05 | 0.05 | 0.05 |
| NC_045512.2;YP_009724389.1;Wu1-SARS2_orf1ab | 1927 | 1940 | 14 | 0.05 | 0.3 | 0.146428571 |
| NC_045512.2;YP_009724389.1;Wu1-SARS2_orf1ab | 1959 | 1965 | 7 | 0.05 | 0.05 | 0.05 |
| NC_045512.2;YP_009724389.1;Wu1-SARS2_orf1ab | 1984 | 1985 | 2 | 0.05 | 0.05 | 0.05 |
| NC_045512.2;YP_009724389.1;Wu1-SARS2_orf1ab | 1987 | 1997 | 11 | 0.05 | 0.25 | 0.104545455 |
| NC_045512.2;YP_009724389.1;Wu1-SARS2_orf1ab | 2009 | 2020 | 12 | 0.05 | 0.35 | 0.15 |
| NC_045512.2;YP_009724389.1;Wu1-SARS2_orf1ab | 2023 | 2024 | 2 | 0.05 | 0.05 | 0.05 |
| NC_045512.2;YP_009724389.1;Wu1-SARS2_orf1ab | 2068 | 2069 | 2 | 0.05 | 0.05 | 0.05 |
| NC_045512.2;YP_009724389.1;Wu1-SARS2_orf1ab | 2083 | 2083 | 1 | 0.05 | 0.05 | 0.05 |
| NC_045512.2;YP_009724389.1;Wu1-SARS2_orf1ab | 2091 | 2091 | 1 | 0.05 | 0.05 | 0.05 |
| NC_045512.2;YP_009724389.1;Wu1-SARS2_orf1ab | 2097 | 2099 | 3 | 0.05 | 0.25 | 0.116666667 |
| NC_045512.2;YP_009724389.1;Wu1-SARS2_orf1ab | 2114 | 2115 | 2 | 0.05 | 0.1 | 0.075 |
| NC_045512.2;YP_009724389.1;Wu1-SARS2_orf1ab | 2118 | 2119 | 2 | 0.05 | 0.05 | 0.05 |
| NC_045512.2;YP_009724389.1;Wu1-SARS2_orf1ab | 2124 | 2144 | 21 | 0.05 | 0.2 | 0.080952381 |
| NC_045512.2;YP_009724389.1;Wu1-SARS2_orf1ab | 2180 | 2181 | 2 | 0.05 | 0.1 | 0.075 |
| NC_045512.2;YP_009724389.1;Wu1-SARS2_orf1ab | 2185 | 2186 | 2 | 0.05 | 0.05 | 0.05 |
| NC_045512.2;YP_009724389.1;Wu1-SARS2_orf1ab | 2212 | 2212 | 1 | 0.05 | 0.05 | 0.05 |
| NC_045512.2;YP_009724389.1;Wu1-SARS2_orf1ab | 2260 | 2270 | 11 | 0.05 | 0.3 | 0.140909091 |
| NC_045512.2;YP_009724389.1;Wu1-SARS2_orf1ab | 2274 | 2274 | 1 | 0.05 | 0.05 | 0.05 |
| NC_045512.2;YP_009724389.1;Wu1-SARS2_orf1ab | 2291 | 2291 | 1 | 0.05 | 0.05 | 0.05 |
| NC_045512.2;YP_009724389.1;Wu1-SARS2_orf1ab | 2294 | 2295 | 2 | 0.05 | 0.05 | 0.05 |
| NC_045512.2;YP_009724389.1;Wu1-SARS2_orf1ab | 2469 | 2471 | 3 | 0.05 | 0.05 | 0.05 |

| Extended data 1 | | | | | | |
|---|---|---|---|---|---|---|
| Protein | Start position (peptide) | Stop position (peptide) | Number of peptides | Minimum fraction (per peptide) of control subjects with reactivity >3 sd above average | Maximum fraction (per peptide) of control subjects with reactivity >3 sd above average | Mean fraction (over all peptides in this contiguous stretch) of controlsubjects with reactivity >3 sd above average |
| NC_045512.2;YP_009724389.1;Wu1-SARS2_orf1ab | 2482 | 2482 | 1 | 0.05 | 0.05 | 0.05 |
| NC_045512.2;YP_009724389.1;Wu1-SARS2_orf1ab | 2504 | 2504 | 1 | 0.05 | 0.05 | 0.05 |
| NC_045512.2;YP_009724389.1;Wu1-SARS2_orf1ab | 2522 | 2525 | 4 | 0.05 | 0.05 | 0.05 |
| NC_045512.2;YP_009724389.1;Wu1-SARS2_orf1ab | 2528 | 2528 | 1 | 0.05 | 0.05 | 0.05 |
| NC_045512.2;YP_009724389.1;Wu1-SARS2_orf1ab | 2536 | 2538 | 3 | 0.05 | 0.05 | 0.05 |
| NC_045512.2;YP_009724389.1;Wu1-SARS2_orf1ab | 2543 | 2543 | 1 | 0.1 | 0.1 | 0.1 |
| NC_045512.2;YP_009724389.1;Wu1-SARS2_orf1ab | 2548 | 2548 | 1 | 0.05 | 0.05 | 0.05 |
| NC_045512.2;YP_009724389.1;Wu1-SARS2_orf1ab | 2551 | 2551 | 1 | 0.05 | 0.05 | 0.05 |
| NC_045512.2;YP_009724389.1;Wu1-SARS2_orf1ab | 2571 | 2571 | 1 | 0.05 | 0.05 | 0.05 |
| NC_045512.2;YP_009724389.1;Wu1-SARS2_orf1ab | 2577 | 2577 | 1 | 0.05 | 0.05 | 0.05 |
| NC_045512.2;YP_009724389.1;Wu1-SARS2_orf1ab | 2579 | 2579 | 1 | 0.05 | 0.05 | 0.05 |
| NC_045512.2;YP_009724389.1;Wu1-SARS2_orf1ab | 2593 | 2596 | 4 | 0.05 | 0.15 | 0.075 |
| NC_045512.2;YP_009724389.1;Wu1-SARS2_orf1ab | 2616 | 2619 | 4 | 0.05 | 0.1 | 0.0625 |
| NC_045512.2;YP_009724389.1;Wu1-SARS2_orf1ab | 2622 | 2622 | 1 | 0.05 | 0.05 | 0.05 |
| NC_045512.2;YP_009724389.1;Wu1-SARS2_orf1ab | 2661 | 2661 | 1 | 0.05 | 0.05 | 0.05 |
| NC_045512.2;YP_009724389.1;Wu1-SARS2_orf1ab | 2663 | 2664 | 2 | 0.05 | 0.05 | 0.05 |
| NC_045512.2;YP_009724389.1;Wu1-SARS2_orf1ab | 2668 | 2669 | 2 | 0.05 | 0.1 | 0.075 |
| NC_045512.2;YP_009724389.1;Wu1-SARS2_orf1ab | 2672 | 2672 | 1 | 0.05 | 0.05 | 0.05 |
| NC_045512.2;YP_009724389.1;Wu1-SARS2_orf1ab | 2675 | 2677 | 3 | 0.05 | 0.05 | 0.05 |
| NC_045512.2;YP_009724389.1;Wu1-SARS2_orf1ab | 2684 | 2685 | 2 | 0.05 | 0.05 | 0.05 |
| NC_045512.2;YP_009724389.1;Wu1-SARS2_orf1ab | 2689 | 2689 | 1 | 0.05 | 0.05 | 0.05 |
| NC_045512.2;YP_009724389.1;Wu1-SARS2_orf1ab | 2691 | 2691 | 1 | 0.05 | 0.05 | 0.05 |
| NC_045512.2;YP_009724389.1;Wu1-SARS2_orf1ab | 2718 | 2718 | 1 | 0.05 | 0.05 | 0.05 |
| NC_045512.2;YP_009724389.1;Wu1-SARS2_orf1ab | 2729 | 2729 | 1 | 0.05 | 0.05 | 0.05 |
| NC_045512.2;YP_009724389.1;Wu1-SARS2_orf1ab | 2796 | 2796 | 1 | 0.05 | 0.05 | 0.05 |
| NC_045512.2;YP_009724389.1;Wu1-SARS2_orf1ab | 2798 | 2806 | 9 | 0.05 | 0.1 | 0.055555556 |
| NC_045512.2;YP_009724389.1;Wu1-SARS2_orf1ab | 2818 | 2818 | 1 | 0.05 | 0.05 | 0.05 |
| NC_045512.2;YP_009724389.1;Wu1-SARS2_orf1ab | 2821 | 2822 | 2 | 0.05 | 0.1 | 0.075 |
| NC_045512.2;YP_009724389.1;Wu1-SARS2_orf1ab | 2828 | 2828 | 1 | 0.1 | 0.1 | 0.1 |
| NC_045512.2;YP_009724389.1;Wu1-SARS2_orf1ab | 2835 | 2835 | 1 | 0.05 | 0.05 | 0.05 |
| NC_045512.2;YP_009724389.1;Wu1-SARS2_orf1ab | 2839 | 2839 | 1 | 0.05 | 0.05 | 0.05 |
| NC_045512.2;YP_009724389.1;Wu1-SARS2_orf1ab | 2841 | 2841 | 1 | 0.05 | 0.05 | 0.05 |
| NC_045512.2;YP_009724389.1;Wu1-SARS2_orf1ab | 2891 | 2892 | 2 | 0.05 | 0.05 | 0.05 |
| NC_045512.2;YP_009724389.1;Wu1-SARS2_orf1ab | 2895 | 2898 | 4 | 0.05 | 0.05 | 0.05 |
| NC_045512.2;YP_009724389.1;Wu1-SARS2_orf1ab | 2903 | 2903 | 1 | 0.05 | 0.05 | 0.05 |
| NC_045512.2;YP_009724389.1;Wu1-SARS2_orf1ab | 2913 | 2918 | 6 | 0.05 | 0.15 | 0.083333333 |
| NC_045512.2;YP_009724389.1;Wu1-SARS2_orf1ab | 2920 | 2921 | 2 | 0.05 | 0.1 | 0.075 |

| Extended data 1 | | | | | | |
|---|---|---|---|---|---|---|
| Protein | Start position (peptide) | Stop position (peptide) | Number of peptides | Minimum fraction (per peptide) of control subjects with reactivity >3 sd above average | Maximum fraction (per peptide) of control subjects with reactivity >3 sd above average | Mean fraction (over all peptides in this contiguous stretch) of controlsubjects with reactivity >3 sd above average |
| NC_045512.2;YP_009724389.1;Wu1-SARS2_orf1ab | 2936 | 2936 | 1 | 0.1 | 0.1 | 0.1 |
| NC_045512.2;YP_009724389.1;Wu1-SARS2_orf1ab | 2939 | 2949 | 11 | 0.05 | 0.15 | 0.1 |
| NC_045512.2;YP_009724389.1;Wu1-SARS2_orf1ab | 2971 | 2971 | 1 | 0.05 | 0.05 | 0.05 |
| NC_045512.2;YP_009724389.1;Wu1-SARS2_orf1ab | 3010 | 3021 | 12 | 0.1 | 0.15 | 0.104166667 |
| NC_045512.2;YP_009724389.1;Wu1-SARS2_orf1ab | 3025 | 3025 | 1 | 0.05 | 0.05 | 0.05 |
| NC_045512.2;YP_009724389.1;Wu1-SARS2_orf1ab | 3027 | 3032 | 6 | 0.05 | 0.05 | 0.05 |
| NC_045512.2;YP_009724389.1;Wu1-SARS2_orf1ab | 3036 | 3036 | 1 | 0.05 | 0.05 | 0.05 |
| NC_045512.2;YP_009724389.1;Wu1-SARS2_orf1ab | 3164 | 3164 | 1 | 0.05 | 0.05 | 0.05 |
| NC_045512.2;YP_009724389.1;Wu1-SARS2_orf1ab | 3182 | 3192 | 11 | 0.05 | 0.15 | 0.095454545 |
| NC_045512.2;YP_009724389.1;Wu1-SARS2_orf1ab | 3215 | 3215 | 1 | 0.05 | 0.05 | 0.05 |
| NC_045512.2;YP_009724389.1;Wu1-SARS2_orf1ab | 3230 | 3230 | 1 | 0.05 | 0.05 | 0.05 |
| NC_045512.2;YP_009724389.1;Wu1-SARS2_orf1ab | 3234 | 3234 | 1 | 0.05 | 0.05 | 0.05 |
| NC_045512.2;YP_009724389.1;Wu1-SARS2_orf1ab | 3237 | 3241 | 5 | 0.05 | 0.3 | 0.13 |
| NC_045512.2;YP_009724389.1;Wu1-SARS2_orf1ab | 3250 | 3250 | 1 | 0.05 | 0.05 | 0.05 |
| NC_045512.2;YP_009724389.1;Wu1-SARS2_orf1ab | 3270 | 3270 | 1 | 0.05 | 0.05 | 0.05 |
| NC_045512.2;YP_009724389.1;Wu1-SARS2_orf1ab | 3313 | 3313 | 1 | 0.05 | 0.05 | 0.05 |
| NC_045512.2;YP_009724389.1;Wu1-SARS2_orf1ab | 3343 | 3343 | 1 | 0.05 | 0.05 | 0.05 |
| NC_045512.2;YP_009724389.1;Wu1-SARS2_orf1ab | 3392 | 3392 | 1 | 0.05 | 0.05 | 0.05 |
| NC_045512.2;YP_009724389.1;Wu1-SARS2_orf1ab | 3394 | 3394 | 1 | 0.05 | 0.05 | 0.05 |
| NC_045512.2;YP_009724389.1;Wu1-SARS2_orf1ab | 3426 | 3444 | 19 | 0.05 | 0.3 | 0.163157895 |
| NC_045512.2;YP_009724389.1;Wu1-SARS2_orf1ab | 3446 | 3446 | 1 | 0.05 | 0.05 | 0.05 |
| NC_045512.2;YP_009724389.1;Wu1-SARS2_orf1ab | 3448 | 3448 | 1 | 0.05 | 0.05 | 0.05 |
| NC_045512.2;YP_009724389.1;Wu1-SARS2_orf1ab | 3479 | 3490 | 12 | 0.05 | 0.25 | 0.075 |
| NC_045512.2;YP_009724389.1;Wu1-SARS2_orf1ab | 3492 | 3492 | 1 | 0.05 | 0.05 | 0.05 |
| NC_045512.2;YP_009724389.1;Wu1-SARS2_orf1ab | 3496 | 3496 | 1 | 0.05 | 0.05 | 0.05 |
| NC_045512.2;YP_009724389.1;Wu1-SARS2_orf1ab | 3503 | 3503 | 1 | 0.05 | 0.05 | 0.05 |
| NC_045512.2;YP_009724389.1;Wu1-SARS2_orf1ab | 3505 | 3515 | 11 | 0.05 | 0.15 | 0.063636364 |
| NC_045512.2;YP_009724389.1;Wu1-SARS2_orf1ab | 3521 | 3521 | 1 | 0.05 | 0.05 | 0.05 |
| NC_045512.2;YP_009724389.1;Wu1-SARS2_orf1ab | 3527 | 3529 | 3 | 0.05 | 0.05 | 0.05 |
| NC_045512.2;YP_009724389.1;Wu1-SARS2_orf1ab | 3547 | 3548 | 2 | 0.05 | 0.05 | 0.05 |
| NC_045512.2;YP_009724389.1;Wu1-SARS2_orf1ab | 3551 | 3551 | 1 | 0.1 | 0.1 | 0.1 |
| NC_045512.2;YP_009724389.1;Wu1-SARS2_orf1ab | 3718 | 3722 | 5 | 0.05 | 0.05 | 0.05 |
| NC_045512.2;YP_009724389.1;Wu1-SARS2_orf1ab | 3809 | 3809 | 1 | 0.25 | 0.25 | 0.25 |
| NC_045512.2;YP_009724389.1;Wu1-SARS2_orf1ab | 3818 | 3819 | 2 | 0.05 | 0.15 | 0.1 |
| NC_045512.2;YP_009724389.1;Wu1-SARS2_orf1ab | 3822 | 3831 | 10 | 0.05 | 0.3 | 0.15 |
| NC_045512.2;YP_009724389.1;Wu1-SARS2_orf1ab | 3846 | 3852 | 7 | 0.05 | 0.1 | 0.057142857 |
| NC_045512.2;YP_009724389.1;Wu1-SARS2_orf1ab | 3854 | 3857 | 4 | 0.05 | 0.05 | 0.05 |

| Extended data 1 | | | | | | |
|---|---|---|---|---|---|---|
| **Protein** | **Start position (peptide)** | **Stop position (peptide)** | **Number of peptides** | **Minimum fraction (per peptide) of control subjects with reactivity >3 sd above average** | **Maximum fraction (per peptide) of control subjects with reactivity >3 sd above average** | **Mean fraction (over all peptides in this contiguous stretch) of controlsubjects with reactivity >3 sd above average** |
| NC_045512.2;YP_009724389.1;Wu1-SARS2_orf1ab | 3888 | 3890 | 3 | 0.05 | 0.05 | 0.05 |
| NC_045512.2;YP_009724389.1;Wu1-SARS2_orf1ab | 3919 | 3923 | 5 | 0.05 | 0.1 | 0.06 |
| NC_045512.2;YP_009724389.1;Wu1-SARS2_orf1ab | 3929 | 3931 | 3 | 0.05 | 0.05 | 0.05 |
| NC_045512.2;YP_009724389.1;Wu1-SARS2_orf1ab | 3933 | 3933 | 1 | 0.05 | 0.05 | 0.05 |
| NC_045512.2;YP_009724389.1;Wu1-SARS2_orf1ab | 3941 | 3941 | 1 | 0.05 | 0.05 | 0.05 |
| NC_045512.2;YP_009724389.1;Wu1-SARS2_orf1ab | 3947 | 3951 | 5 | 0.05 | 0.1 | 0.06 |
| NC_045512.2;YP_009724389.1;Wu1-SARS2_orf1ab | 3953 | 3957 | 5 | 0.05 | 0.1 | 0.06 |
| NC_045512.2;YP_009724389.1;Wu1-SARS2_orf1ab | 3960 | 3960 | 1 | 0.05 | 0.05 | 0.05 |
| NC_045512.2;YP_009724389.1;Wu1-SARS2_orf1ab | 3980 | 3981 | 2 | 0.05 | 0.05 | 0.05 |
| NC_045512.2;YP_009724389.1;Wu1-SARS2_orf1ab | 3983 | 3983 | 1 | 0.05 | 0.05 | 0.05 |
| NC_045512.2;YP_009724389.1;Wu1-SARS2_orf1ab | 4031 | 4041 | 11 | 0.05 | 0.4 | 0.227272727 |
| NC_045512.2;YP_009724389.1;Wu1-SARS2_orf1ab | 4043 | 4044 | 2 | 0.05 | 0.1 | 0.075 |
| NC_045512.2;YP_009724389.1;Wu1-SARS2_orf1ab | 4046 | 4047 | 2 | 0.05 | 0.05 | 0.05 |
| NC_045512.2;YP_009724389.1;Wu1-SARS2_orf1ab | 4053 | 4054 | 2 | 0.05 | 0.05 | 0.05 |
| NC_045512.2;YP_009724389.1;Wu1-SARS2_orf1ab | 4063 | 4063 | 1 | 0.05 | 0.05 | 0.05 |
| NC_045512.2;YP_009724389.1;Wu1-SARS2_orf1ab | 4070 | 4076 | 7 | 0.05 | 0.2 | 0.092857143 |
| NC_045512.2;YP_009724389.1;Wu1-SARS2_orf1ab | 4092 | 4093 | 2 | 0.05 | 0.05 | 0.05 |
| NC_045512.2;YP_009724389.1;Wu1-SARS2_orf1ab | 4103 | 4103 | 1 | 0.05 | 0.05 | 0.05 |
| NC_045512.2;YP_009724389.1;Wu1-SARS2_orf1ab | 4132 | 4134 | 3 | 0.05 | 0.1 | 0.066666667 |
| NC_045512.2;YP_009724389.1;Wu1-SARS2_orf1ab | 4137 | 4137 | 1 | 0.05 | 0.05 | 0.05 |
| NC_045512.2;YP_009724389.1;Wu1-SARS2_orf1ab | 4141 | 4141 | 1 | 0.05 | 0.05 | 0.05 |
| NC_045512.2;YP_009724389.1;Wu1-SARS2_orf1ab | 4161 | 4163 | 3 | 0.05 | 0.2 | 0.116666667 |
| NC_045512.2;YP_009724389.1;Wu1-SARS2_orf1ab | 4182 | 4182 | 1 | 0.05 | 0.05 | 0.05 |
| NC_045512.2;YP_009724389.1;Wu1-SARS2_orf1ab | 4184 | 4188 | 5 | 0.05 | 0.3 | 0.14 |
| NC_045512.2;YP_009724389.1;Wu1-SARS2_orf1ab | 4203 | 4203 | 1 | 0.2 | 0.2 | 0.2 |
| NC_045512.2;YP_009724389.1;Wu1-SARS2_orf1ab | 4205 | 4208 | 4 | 0.05 | 0.05 | 0.05 |
| NC_045512.2;YP_009724389.1;Wu1-SARS2_orf1ab | 4215 | 4215 | 1 | 0.05 | 0.05 | 0.05 |
| NC_045512.2;YP_009724389.1;Wu1-SARS2_orf1ab | 4229 | 4235 | 7 | 0.05 | 0.25 | 0.107142857 |
| NC_045512.2;YP_009724389.1;Wu1-SARS2_orf1ab | 4247 | 4247 | 1 | 0.05 | 0.05 | 0.05 |
| NC_045512.2;YP_009724389.1;Wu1-SARS2_orf1ab | 4268 | 4268 | 1 | 0.05 | 0.05 | 0.05 |
| NC_045512.2;YP_009724389.1;Wu1-SARS2_orf1ab | 4271 | 4275 | 5 | 0.05 | 0.2 | 0.11 |
| NC_045512.2;YP_009724389.1;Wu1-SARS2_orf1ab | 4279 | 4280 | 2 | 0.05 | 0.15 | 0.1 |
| NC_045512.2;YP_009724389.1;Wu1-SARS2_orf1ab | 4308 | 4308 | 1 | 0.05 | 0.05 | 0.05 |
| NC_045512.2;YP_009724389.1;Wu1-SARS2_orf1ab | 4310 | 4310 | 1 | 0.05 | 0.05 | 0.05 |
| NC_045512.2;YP_009724389.1;Wu1-SARS2_orf1ab | 4344 | 4348 | 5 | 0.05 | 0.05 | 0.05 |
| NC_045512.2;YP_009724389.1;Wu1-SARS2_orf1ab | 4353 | 4353 | 1 | 0.05 | 0.05 | 0.05 |
| NC_045512.2;YP_009724389.1;Wu1-SARS2_orf1ab | 4386 | 4388 | 3 | 0.05 | 0.15 | 0.083333333 |

| Extended data 1 | | | | | | |
|---|---|---|---|---|---|---|
| Protein | Start position (peptide) | Stop position (peptide) | Number of peptides | Minimum fraction (per peptide) of control subjects with reactivity >3 sd above average | Maximum fraction (per peptide) of control subjects with reactivity >3 sd above average | Mean fraction (over all peptides in this contiguous stretch) of controlsubjects with reactivity >3 sd above average |
| NC_045512.2;YP_009724389.1;Wu1-SARS2_orf1ab | 4390 | 4390 | 1 | 0.05 | 0.05 | 0.05 |
| NC_045512.2;YP_009724389.1;Wu1-SARS2_orf1ab | 4394 | 4394 | 1 | 0.05 | 0.05 | 0.05 |
| NC_045512.2;YP_009724389.1;Wu1-SARS2_orf1ab | 4419 | 4421 | 3 | 0.05 | 0.1 | 0.066666667 |
| NC_045512.2;YP_009724389.1;Wu1-SARS2_orf1ab | 4423 | 4423 | 1 | 0.05 | 0.05 | 0.05 |
| NC_045512.2;YP_009724389.1;Wu1-SARS2_orf1ab | 4446 | 4446 | 1 | 0.05 | 0.05 | 0.05 |
| NC_045512.2;YP_009724389.1;Wu1-SARS2_orf1ab | 4469 | 4469 | 1 | 0.05 | 0.05 | 0.05 |
| NC_045512.2;YP_009724389.1;Wu1-SARS2_orf1ab | 4520 | 4520 | 1 | 0.05 | 0.05 | 0.05 |
| NC_045512.2;YP_009724389.1;Wu1-SARS2_orf1ab | 4545 | 4546 | 2 | 0.05 | 0.2 | 0.125 |
| NC_045512.2;YP_009724389.1;Wu1-SARS2_orf1ab | 4549 | 4551 | 3 | 0.05 | 0.1 | 0.066666667 |
| NC_045512.2;YP_009724389.1;Wu1-SARS2_orf1ab | 4557 | 4559 | 3 | 0.05 | 0.05 | 0.05 |
| NC_045512.2;YP_009724389.1;Wu1-SARS2_orf1ab | 4565 | 4565 | 1 | 0.05 | 0.05 | 0.05 |
| NC_045512.2;YP_009724389.1;Wu1-SARS2_orf1ab | 4572 | 4572 | 1 | 0.05 | 0.05 | 0.05 |
| NC_045512.2;YP_009724389.1;Wu1-SARS2_orf1ab | 4591 | 4591 | 1 | 0.05 | 0.05 | 0.05 |
| NC_045512.2;YP_009724389.1;Wu1-SARS2_orf1ab | 4595 | 4595 | 1 | 0.05 | 0.05 | 0.05 |
| NC_045512.2;YP_009724389.1;Wu1-SARS2_orf1ab | 4609 | 4609 | 1 | 0.1 | 0.1 | 0.1 |
| NC_045512.2;YP_009724389.1;Wu1-SARS2_orf1ab | 4635 | 4635 | 1 | 0.05 | 0.05 | 0.05 |
| NC_045512.2;YP_009724389.1;Wu1-SARS2_orf1ab | 4641 | 4644 | 4 | 0.05 | 0.05 | 0.05 |
| NC_045512.2;YP_009724389.1;Wu1-SARS2_orf1ab | 4646 | 4646 | 1 | 0.05 | 0.05 | 0.05 |
| NC_045512.2;YP_009724389.1;Wu1-SARS2_orf1ab | 4649 | 4650 | 2 | 0.05 | 0.1 | 0.075 |
| NC_045512.2;YP_009724389.1;Wu1-SARS2_orf1ab | 4656 | 4665 | 10 | 0.05 | 0.1 | 0.06 |
| NC_045512.2;YP_009724389.1;Wu1-SARS2_orf1ab | 4749 | 4749 | 1 | 0.05 | 0.05 | 0.05 |
| NC_045512.2;YP_009724389.1;Wu1-SARS2_orf1ab | 4757 | 4760 | 4 | 0.05 | 0.1 | 0.075 |
| NC_045512.2;YP_009724389.1;Wu1-SARS2_orf1ab | 4777 | 4777 | 1 | 0.05 | 0.05 | 0.05 |
| NC_045512.2;YP_009724389.1;Wu1-SARS2_orf1ab | 4791 | 4792 | 2 | 0.05 | 0.05 | 0.05 |
| NC_045512.2;YP_009724389.1;Wu1-SARS2_orf1ab | 4799 | 4805 | 7 | 0.05 | 0.2 | 0.1 |
| NC_045512.2;YP_009724389.1;Wu1-SARS2_orf1ab | 4808 | 4810 | 3 | 0.05 | 0.05 | 0.05 |
| NC_045512.2;YP_009724389.1;Wu1-SARS2_orf1ab | 4817 | 4817 | 1 | 0.05 | 0.05 | 0.05 |
| NC_045512.2;YP_009724389.1;Wu1-SARS2_orf1ab | 4826 | 4827 | 2 | 0.05 | 0.05 | 0.05 |
| NC_045512.2;YP_009724389.1;Wu1-SARS2_orf1ab | 4830 | 4830 | 1 | 0.05 | 0.05 | 0.05 |
| NC_045512.2;YP_009724389.1;Wu1-SARS2_orf1ab | 4836 | 4844 | 9 | 0.05 | 0.25 | 0.15 |
| NC_045512.2;YP_009724389.1;Wu1-SARS2_orf1ab | 4855 | 4855 | 1 | 0.05 | 0.05 | 0.05 |
| NC_045512.2;YP_009724389.1;Wu1-SARS2_orf1ab | 4868 | 4871 | 4 | 0.05 | 0.05 | 0.05 |
| NC_045512.2;YP_009724389.1;Wu1-SARS2_orf1ab | 4873 | 4874 | 2 | 0.05 | 0.05 | 0.05 |
| NC_045512.2;YP_009724389.1;Wu1-SARS2_orf1ab | 4876 | 4876 | 1 | 0.05 | 0.05 | 0.05 |
| NC_045512.2;YP_009724389.1;Wu1-SARS2_orf1ab | 4878 | 4880 | 3 | 0.05 | 0.1 | 0.066666667 |
| NC_045512.2;YP_009724389.1;Wu1-SARS2_orf1ab | 4882 | 4882 | 1 | 0.15 | 0.15 | 0.15 |
| NC_045512.2;YP_009724389.1;Wu1-SARS2_orf1ab | 4884 | 4885 | 2 | 0.05 | 0.15 | 0.1 |

| Extended data 1 | | | | | | |
|---|---|---|---|---|---|---|
| Protein | Start position (peptide) | Stop position (peptide) | Number of peptides | Minimum fraction (per peptide) of control subjects with reactivity >3 sd above average | Maximum fraction (per peptide) of control subjects with reactivity >3 sd above average | Mean fraction (over all peptides in this contiguous stretch) of controlsubjects with reactivity >3 sd above average |
| NC_045512.2;YP_009724389.1;Wu1-SARS2_orf1ab | 4897 | 4897 | 1 | 0.05 | 0.05 | 0.05 |
| NC_045512.2;YP_009724389.1;Wu1-SARS2_orf1ab | 4899 | 4899 | 1 | 0.05 | 0.05 | 0.05 |
| NC_045512.2;YP_009724389.1;Wu1-SARS2_orf1ab | 4909 | 4917 | 9 | 0.05 | 0.2 | 0.077777778 |
| NC_045512.2;YP_009724389.1;Wu1-SARS2_orf1ab | 4923 | 4924 | 2 | 0.05 | 0.1 | 0.075 |
| NC_045512.2;YP_009724389.1;Wu1-SARS2_orf1ab | 4929 | 4929 | 1 | 0.1 | 0.1 | 0.1 |
| NC_045512.2;YP_009724389.1;Wu1-SARS2_orf1ab | 4963 | 4972 | 10 | 0.15 | 0.4 | 0.26 |
| NC_045512.2;YP_009724389.1;Wu1-SARS2_orf1ab | 5057 | 5057 | 1 | 0.05 | 0.05 | 0.05 |
| NC_045512.2;YP_009724389.1;Wu1-SARS2_orf1ab | 5069 | 5070 | 2 | 0.05 | 0.1 | 0.075 |
| NC_045512.2;YP_009724389.1;Wu1-SARS2_orf1ab | 5072 | 5072 | 1 | 0.05 | 0.05 | 0.05 |
| NC_045512.2;YP_009724389.1;Wu1-SARS2_orf1ab | 5094 | 5094 | 1 | 0.05 | 0.05 | 0.05 |
| NC_045512.2;YP_009724389.1;Wu1-SARS2_orf1ab | 5097 | 5097 | 1 | 0.05 | 0.05 | 0.05 |
| NC_045512.2;YP_009724389.1;Wu1-SARS2_orf1ab | 5101 | 5101 | 1 | 0.05 | 0.05 | 0.05 |
| NC_045512.2;YP_009724389.1;Wu1-SARS2_orf1ab | 5113 | 5113 | 1 | 0.05 | 0.05 | 0.05 |
| NC_045512.2;YP_009724389.1;Wu1-SARS2_orf1ab | 5124 | 5124 | 1 | 0.05 | 0.05 | 0.05 |
| NC_045512.2;YP_009724389.1;Wu1-SARS2_orf1ab | 5131 | 5131 | 1 | 0.05 | 0.05 | 0.05 |
| NC_045512.2;YP_009724389.1;Wu1-SARS2_orf1ab | 5133 | 5133 | 1 | 0.05 | 0.05 | 0.05 |
| NC_045512.2;YP_009724389.1;Wu1-SARS2_orf1ab | 5139 | 5139 | 1 | 0.05 | 0.05 | 0.05 |
| NC_045512.2;YP_009724389.1;Wu1-SARS2_orf1ab | 5154 | 5154 | 1 | 0.05 | 0.05 | 0.05 |
| NC_045512.2;YP_009724389.1;Wu1-SARS2_orf1ab | 5175 | 5175 | 1 | 0.05 | 0.05 | 0.05 |
| NC_045512.2;YP_009724389.1;Wu1-SARS2_orf1ab | 5184 | 5184 | 1 | 0.05 | 0.05 | 0.05 |
| NC_045512.2;YP_009724389.1;Wu1-SARS2_orf1ab | 5186 | 5186 | 1 | 0.05 | 0.05 | 0.05 |
| NC_045512.2;YP_009724389.1;Wu1-SARS2_orf1ab | 5189 | 5192 | 4 | 0.05 | 0.05 | 0.05 |
| NC_045512.2;YP_009724389.1;Wu1-SARS2_orf1ab | 5206 | 5207 | 2 | 0.05 | 0.05 | 0.05 |
| NC_045512.2;YP_009724389.1;Wu1-SARS2_orf1ab | 5209 | 5209 | 1 | 0.05 | 0.05 | 0.05 |
| NC_045512.2;YP_009724389.1;Wu1-SARS2_orf1ab | 5211 | 5211 | 1 | 0.1 | 0.1 | 0.1 |
| NC_045512.2;YP_009724389.1;Wu1-SARS2_orf1ab | 5229 | 5229 | 1 | 0.05 | 0.05 | 0.05 |
| NC_045512.2;YP_009724389.1;Wu1-SARS2_orf1ab | 5302 | 5302 | 1 | 0.05 | 0.05 | 0.05 |
| NC_045512.2;YP_009724389.1;Wu1-SARS2_orf1ab | 5345 | 5345 | 1 | 0.05 | 0.05 | 0.05 |
| NC_045512.2;YP_009724389.1;Wu1-SARS2_orf1ab | 5370 | 5373 | 4 | 0.05 | 0.05 | 0.05 |
| NC_045512.2;YP_009724389.1;Wu1-SARS2_orf1ab | 5378 | 5378 | 1 | 0.05 | 0.05 | 0.05 |
| NC_045512.2;YP_009724389.1;Wu1-SARS2_orf1ab | 5415 | 5425 | 11 | 0.05 | 0.2 | 0.086363636 |
| NC_045512.2;YP_009724389.1;Wu1-SARS2_orf1ab | 5441 | 5441 | 1 | 0.05 | 0.05 | 0.05 |
| NC_045512.2;YP_009724389.1;Wu1-SARS2_orf1ab | 5452 | 5454 | 3 | 0.05 | 0.05 | 0.05 |
| NC_045512.2;YP_009724389.1;Wu1-SARS2_orf1ab | 5459 | 5459 | 1 | 0.05 | 0.05 | 0.05 |
| NC_045512.2;YP_009724389.1;Wu1-SARS2_orf1ab | 5466 | 5466 | 1 | 0.05 | 0.05 | 0.05 |
| NC_045512.2;YP_009724389.1;Wu1-SARS2_orf1ab | 5470 | 5475 | 6 | 0.05 | 0.05 | 0.05 |
| NC_045512.2;YP_009724389.1;Wu1-SARS2_orf1ab | 5483 | 5483 | 1 | 0.05 | 0.05 | 0.05 |

| Extended data 1 | | | | | | |
|---|---|---|---|---|---|---|
| **Protein** | **Start position (peptide)** | **Stop position (peptide)** | **Number of peptides** | **Minimum fraction (per peptide) of control subjects with reactivity >3 sd above average** | **Maximum fraction (per peptide) of control subjects with reactivity >3 sd above average** | **Mean fraction (over all peptides in this contiguous stretch) of controlsubjects with reactivity >3 sd above average** |
| NC_045512.2;YP_009724389.1;Wu1-SARS2_orf1ab | 5506 | 5519 | 14 | 0.05 | 0.1 | 0.071428571 |
| NC_045512.2;YP_009724389.1;Wu1-SARS2_orf1ab | 5522 | 5522 | 1 | 0.1 | 0.1 | 0.1 |
| NC_045512.2;YP_009724389.1;Wu1-SARS2_orf1ab | 5527 | 5527 | 1 | 0.05 | 0.05 | 0.05 |
| NC_045512.2;YP_009724389.1;Wu1-SARS2_orf1ab | 5547 | 5547 | 1 | 0.05 | 0.05 | 0.05 |
| NC_045512.2;YP_009724389.1;Wu1-SARS2_orf1ab | 5571 | 5580 | 10 | 0.05 | 0.1 | 0.055 |
| NC_045512.2;YP_009724389.1;Wu1-SARS2_orf1ab | 5582 | 5582 | 1 | 0.05 | 0.05 | 0.05 |
| NC_045512.2;YP_009724389.1;Wu1-SARS2_orf1ab | 5584 | 5585 | 2 | 0.1 | 0.1 | 0.1 |
| NC_045512.2;YP_009724389.1;Wu1-SARS2_orf1ab | 5587 | 5587 | 1 | 0.05 | 0.05 | 0.05 |
| NC_045512.2;YP_009724389.1;Wu1-SARS2_orf1ab | 5598 | 5598 | 1 | 0.05 | 0.05 | 0.05 |
| NC_045512.2;YP_009724389.1;Wu1-SARS2_orf1ab | 5638 | 5638 | 1 | 0.05 | 0.05 | 0.05 |
| NC_045512.2;YP_009724389.1;Wu1-SARS2_orf1ab | 5659 | 5659 | 1 | 0.05 | 0.05 | 0.05 |
| NC_045512.2;YP_009724389.1;Wu1-SARS2_orf1ab | 5661 | 5667 | 7 | 0.05 | 0.4 | 0.214285714 |
| NC_045512.2;YP_009724389.1;Wu1-SARS2_orf1ab | 5730 | 5738 | 9 | 0.05 | 0.1 | 0.088888889 |
| NC_045512.2;YP_009724389.1;Wu1-SARS2_orf1ab | 5762 | 5773 | 12 | 0.1 | 0.15 | 0.116666667 |
| NC_045512.2;YP_009724389.1;Wu1-SARS2_orf1ab | 5777 | 5777 | 1 | 0.05 | 0.05 | 0.05 |
| NC_045512.2;YP_009724389.1;Wu1-SARS2_orf1ab | 5779 | 5779 | 1 | 0.05 | 0.05 | 0.05 |
| NC_045512.2;YP_009724389.1;Wu1-SARS2_orf1ab | 5785 | 5785 | 1 | 0.05 | 0.05 | 0.05 |
| NC_045512.2;YP_009724389.1;Wu1-SARS2_orf1ab | 5802 | 5804 | 3 | 0.05 | 0.05 | 0.05 |
| NC_045512.2;YP_009724389.1;Wu1-SARS2_orf1ab | 5807 | 5807 | 1 | 0.05 | 0.05 | 0.05 |
| NC_045512.2;YP_009724389.1;Wu1-SARS2_orf1ab | 5828 | 5828 | 1 | 0.05 | 0.05 | 0.05 |
| NC_045512.2;YP_009724389.1;Wu1-SARS2_orf1ab | 5832 | 5833 | 2 | 0.05 | 0.15 | 0.1 |
| NC_045512.2;YP_009724389.1;Wu1-SARS2_orf1ab | 5837 | 5839 | 3 | 0.05 | 0.1 | 0.083333333 |
| NC_045512.2;YP_009724389.1;Wu1-SARS2_orf1ab | 5846 | 5846 | 1 | 0.05 | 0.05 | 0.05 |
| NC_045512.2;YP_009724389.1;Wu1-SARS2_orf1ab | 5863 | 5868 | 6 | 0.05 | 0.05 | 0.05 |
| NC_045512.2;YP_009724389.1;Wu1-SARS2_orf1ab | 5871 | 5873 | 3 | 0.05 | 0.05 | 0.05 |
| NC_045512.2;YP_009724389.1;Wu1-SARS2_orf1ab | 5897 | 5899 | 3 | 0.05 | 0.1 | 0.066666667 |
| NC_045512.2;YP_009724389.1;Wu1-SARS2_orf1ab | 5912 | 5918 | 7 | 0.05 | 0.15 | 0.078571429 |
| NC_045512.2;YP_009724389.1;Wu1-SARS2_orf1ab | 5921 | 5921 | 1 | 0.05 | 0.05 | 0.05 |
| NC_045512.2;YP_009724389.1;Wu1-SARS2_orf1ab | 5934 | 5934 | 1 | 0.05 | 0.05 | 0.05 |
| NC_045512.2;YP_009724389.1;Wu1-SARS2_orf1ab | 5982 | 5984 | 3 | 0.05 | 0.1 | 0.066666667 |
| NC_045512.2;YP_009724389.1;Wu1-SARS2_orf1ab | 5989 | 5989 | 1 | 0.05 | 0.05 | 0.05 |
| NC_045512.2;YP_009724389.1;Wu1-SARS2_orf1ab | 6043 | 6045 | 3 | 0.05 | 0.1 | 0.066666667 |
| NC_045512.2;YP_009724389.1;Wu1-SARS2_orf1ab | 6047 | 6048 | 2 | 0.05 | 0.15 | 0.1 |
| NC_045512.2;YP_009724389.1;Wu1-SARS2_orf1ab | 6050 | 6050 | 1 | 0.05 | 0.05 | 0.05 |
| NC_045512.2;YP_009724389.1;Wu1-SARS2_orf1ab | 6052 | 6052 | 1 | 0.05 | 0.05 | 0.05 |
| NC_045512.2;YP_009724389.1;Wu1-SARS2_orf1ab | 6054 | 6054 | 1 | 0.05 | 0.05 | 0.05 |
| NC_045512.2;YP_009724389.1;Wu1-SARS2_orf1ab | 6116 | 6116 | 1 | 0.05 | 0.05 | 0.05 |

| Extended data 1 | | | | | | |
|---|---|---|---|---|---|---|
| **Protein** | **Start position (peptide)** | **Stop position (peptide)** | **Number of peptides** | **Minimum fraction (per peptide) of control subjects with reactivity >3 sd above average** | **Maximum fraction (per peptide) of control subjects with reactivity >3 sd above average** | **Mean fraction (over all peptides in this contiguous stretch) of controlsubjects with reactivity >3 sd above average** |
| NC_045512.2;YP_009724389.1;Wu1-SARS2_orf1ab | 6132 | 6136 | 5 | 0.05 | 0.15 | 0.07 |
| NC_045512.2;YP_009724389.1;Wu1-SARS2_orf1ab | 6180 | 6180 | 1 | 0.05 | 0.05 | 0.05 |
| NC_045512.2;YP_009724389.1;Wu1-SARS2_orf1ab | 6214 | 6224 | 11 | 0.05 | 0.1 | 0.086363636 |
| NC_045512.2;YP_009724389.1;Wu1-SARS2_orf1ab | 6242 | 6243 | 2 | 0.05 | 0.05 | 0.05 |
| NC_045512.2;YP_009724389.1;Wu1-SARS2_orf1ab | 6255 | 6255 | 1 | 0.05 | 0.05 | 0.05 |
| NC_045512.2;YP_009724389.1;Wu1-SARS2_orf1ab | 6262 | 6266 | 5 | 0.05 | 0.05 | 0.05 |
| NC_045512.2;YP_009724389.1;Wu1-SARS2_orf1ab | 6269 | 6272 | 4 | 0.05 | 0.1 | 0.0875 |
| NC_045512.2;YP_009724389.1;Wu1-SARS2_orf1ab | 6274 | 6274 | 1 | 0.05 | 0.05 | 0.05 |
| NC_045512.2;YP_009724389.1;Wu1-SARS2_orf1ab | 6289 | 6293 | 5 | 0.1 | 0.2 | 0.13 |
| NC_045512.2;YP_009724389.1;Wu1-SARS2_orf1ab | 6327 | 6331 | 5 | 0.05 | 0.1 | 0.06 |
| NC_045512.2;YP_009724389.1;Wu1-SARS2_orf1ab | 6351 | 6358 | 8 | 0.05 | 0.05 | 0.05 |
| NC_045512.2;YP_009724389.1;Wu1-SARS2_orf1ab | 6382 | 6382 | 1 | 0.05 | 0.05 | 0.05 |
| NC_045512.2;YP_009724389.1;Wu1-SARS2_orf1ab | 6445 | 6445 | 1 | 0.05 | 0.05 | 0.05 |
| NC_045512.2;YP_009724389.1;Wu1-SARS2_orf1ab | 6454 | 6455 | 2 | 0.05 | 0.05 | 0.05 |
| NC_045512.2;YP_009724389.1;Wu1-SARS2_orf1ab | 6457 | 6459 | 3 | 0.05 | 0.1 | 0.066666667 |
| NC_045512.2;YP_009724389.1;Wu1-SARS2_orf1ab | 6461 | 6461 | 1 | 0.05 | 0.05 | 0.05 |
| NC_045512.2;YP_009724389.1;Wu1-SARS2_orf1ab | 6475 | 6475 | 1 | 0.05 | 0.05 | 0.05 |
| NC_045512.2;YP_009724389.1;Wu1-SARS2_orf1ab | 6496 | 6496 | 1 | 0.05 | 0.05 | 0.05 |
| NC_045512.2;YP_009724389.1;Wu1-SARS2_orf1ab | 6505 | 6505 | 1 | 0.05 | 0.05 | 0.05 |
| NC_045512.2;YP_009724389.1;Wu1-SARS2_orf1ab | 6507 | 6516 | 10 | 0.05 | 0.4 | 0.175 |
| NC_045512.2;YP_009724389.1;Wu1-SARS2_orf1ab | 6518 | 6518 | 1 | 0.05 | 0.05 | 0.05 |
| NC_045512.2;YP_009724389.1;Wu1-SARS2_orf1ab | 6528 | 6528 | 1 | 0.05 | 0.05 | 0.05 |
| NC_045512.2;YP_009724389.1;Wu1-SARS2_orf1ab | 6537 | 6537 | 1 | 0.05 | 0.05 | 0.05 |
| NC_045512.2;YP_009724389.1;Wu1-SARS2_orf1ab | 6541 | 6541 | 1 | 0.15 | 0.15 | 0.15 |
| NC_045512.2;YP_009724389.1;Wu1-SARS2_orf1ab | 6546 | 6557 | 12 | 0.05 | 0.15 | 0.0875 |
| NC_045512.2;YP_009724389.1;Wu1-SARS2_orf1ab | 6573 | 6573 | 1 | 0.05 | 0.05 | 0.05 |
| NC_045512.2;YP_009724389.1;Wu1-SARS2_orf1ab | 6579 | 6579 | 1 | 0.05 | 0.05 | 0.05 |
| NC_045512.2;YP_009724389.1;Wu1-SARS2_orf1ab | 6592 | 6594 | 3 | 0.05 | 0.05 | 0.05 |
| NC_045512.2;YP_009724389.1;Wu1-SARS2_orf1ab | 6596 | 6596 | 1 | 0.05 | 0.05 | 0.05 |
| NC_045512.2;YP_009724389.1;Wu1-SARS2_orf1ab | 6598 | 6598 | 1 | 0.05 | 0.05 | 0.05 |
| NC_045512.2;YP_009724389.1;Wu1-SARS2_orf1ab | 6600 | 6600 | 1 | 0.05 | 0.05 | 0.05 |
| NC_045512.2;YP_009724389.1;Wu1-SARS2_orf1ab | 6619 | 6619 | 1 | 0.05 | 0.05 | 0.05 |
| NC_045512.2;YP_009724389.1;Wu1-SARS2_orf1ab | 6621 | 6621 | 1 | 0.05 | 0.05 | 0.05 |
| NC_045512.2;YP_009724389.1;Wu1-SARS2_orf1ab | 6625 | 6627 | 3 | 0.05 | 0.1 | 0.066666667 |
| NC_045512.2;YP_009724389.1;Wu1-SARS2_orf1ab | 6629 | 6633 | 5 | 0.05 | 0.1 | 0.07 |
| NC_045512.2;YP_009724389.1;Wu1-SARS2_orf1ab | 6642 | 6642 | 1 | 0.05 | 0.05 | 0.05 |
| NC_045512.2;YP_009724389.1;Wu1-SARS2_orf1ab | 6663 | 6665 | 3 | 0.05 | 0.05 | 0.05 |

| Extended data 1 | | | | | | |
|---|---|---|---|---|---|---|
| Protein | Start position (peptide) | Stop position (peptide) | Number of peptides | Minimum fraction (per peptide) of control subjects with reactivity >3 sd above average | Maximum fraction (per peptide) of control subjects with reactivity >3 sd above average | Mean fraction (over all peptides in this contiguous stretch) of controlsubjects with reactivity >3 sd above average |
| NC_045512.2;YP_009724389.1;Wu1-SARS2_orf1ab | 6668 | 6668 | 1 | 0.05 | 0.05 | 0.05 |
| NC_045512.2;YP_009724389.1;Wu1-SARS2_orf1ab | 6701 | 6708 | 8 | 0.05 | 0.15 | 0.08125 |
| NC_045512.2;YP_009724389.1;Wu1-SARS2_orf1ab | 6711 | 6712 | 2 | 0.05 | 0.15 | 0.1 |
| NC_045512.2;YP_009724389.1;Wu1-SARS2_orf1ab | 6715 | 6717 | 3 | 0.05 | 0.15 | 0.083333333 |
| NC_045512.2;YP_009724389.1;Wu1-SARS2_orf1ab | 6719 | 6721 | 3 | 0.05 | 0.1 | 0.083333333 |
| NC_045512.2;YP_009724389.1;Wu1-SARS2_orf1ab | 6723 | 6728 | 6 | 0.05 | 0.1 | 0.075 |
| NC_045512.2;YP_009724389.1;Wu1-SARS2_orf1ab | 6745 | 6750 | 6 | 0.05 | 0.1 | 0.058333333 |
| NC_045512.2;YP_009724389.1;Wu1-SARS2_orf1ab | 6793 | 6793 | 1 | 0.1 | 0.1 | 0.1 |
| NC_045512.2;YP_009724389.1;Wu1-SARS2_orf1ab | 6821 | 6828 | 8 | 0.05 | 0.1 | 0.0875 |
| NC_045512.2;YP_009724389.1;Wu1-SARS2_orf1ab | 6830 | 6830 | 1 | 0.25 | 0.25 | 0.25 |
| NC_045512.2;YP_009724389.1;Wu1-SARS2_orf1ab | 6862 | 6862 | 1 | 0.05 | 0.05 | 0.05 |
| NC_045512.2;YP_009724389.1;Wu1-SARS2_orf1ab | 6870 | 6871 | 2 | 0.05 | 0.05 | 0.05 |
| NC_045512.2;YP_009724389.1;Wu1-SARS2_orf1ab | 6891 | 6893 | 3 | 0.05 | 0.05 | 0.05 |
| NC_045512.2;YP_009724389.1;Wu1-SARS2_orf1ab | 6896 | 6896 | 1 | 0.1 | 0.1 | 0.1 |
| NC_045512.2;YP_009724389.1;Wu1-SARS2_orf1ab | 6899 | 6905 | 7 | 0.05 | 0.05 | 0.05 |
| NC_045512.2;YP_009724389.1;Wu1-SARS2_orf1ab | 6908 | 6908 | 1 | 0.05 | 0.05 | 0.05 |
| NC_045512.2;YP_009724389.1;Wu1-SARS2_orf1ab | 6919 | 6919 | 1 | 0.15 | 0.15 | 0.15 |
| NC_045512.2;YP_009724389.1;Wu1-SARS2_orf1ab | 6923 | 6936 | 14 | 0.05 | 0.25 | 0.121428571 |
| NC_045512.2;YP_009724389.1;Wu1-SARS2_orf1ab | 6990 | 6990 | 1 | 0.05 | 0.05 | 0.05 |
| NC_045512.2;YP_009724389.1;Wu1-SARS2_orf1ab | 7003 | 7004 | 2 | 0.05 | 0.05 | 0.05 |
| NC_045512.2;YP_009724389.1;Wu1-SARS2_orf1ab | 7006 | 7010 | 5 | 0.05 | 0.1 | 0.09 |
| NC_045512.2;YP_009724389.1;Wu1-SARS2_orf1ab | 7015 | 7015 | 1 | 0.05 | 0.05 | 0.05 |
| NC_045512.2;YP_009724389.1;Wu1-SARS2_orf1ab | 7032 | 7038 | 7 | 0.05 | 0.05 | 0.05 |
| NC_045512.2;YP_009724389.1;Wu1-SARS2_orf1ab | 7053 | 7057 | 5 | 0.05 | 0.1 | 0.06 |
| NC_045512.2;YP_009724390.1;Wu1-SARS2_surface | 8 | 8 | 1 | 0.05 | 0.05 | 0.05 |
| NC_045512.2;YP_009724390.1;Wu1-SARS2_surface | 27 | 27 | 1 | 0.05 | 0.05 | 0.05 |
| NC_045512.2;YP_009724390.1;Wu1-SARS2_surface | 39 | 39 | 1 | 0.05 | 0.05 | 0.05 |
| NC_045512.2;YP_009724390.1;Wu1-SARS2_surface | 74 | 74 | 1 | 0.05 | 0.05 | 0.05 |
| NC_045512.2;YP_009724390.1;Wu1-SARS2_surface | 82 | 92 | 11 | 0.05 | 0.05 | 0.05 |
| NC_045512.2;YP_009724390.1;Wu1-SARS2_surface | 111 | 114 | 4 | 0.05 | 0.05 | 0.05 |
| NC_045512.2;YP_009724390.1;Wu1-SARS2_surface | 119 | 120 | 2 | 0.05 | 0.05 | 0.05 |
| NC_045512.2;YP_009724390.1;Wu1-SARS2_surface | 124 | 124 | 1 | 0.05 | 0.05 | 0.05 |
| NC_045512.2;YP_009724390.1;Wu1-SARS2_surface | 139 | 144 | 6 | 0.05 | 0.1 | 0.058333333 |
| NC_045512.2;YP_009724390.1;Wu1-SARS2_surface | 146 | 149 | 4 | 0.05 | 0.05 | 0.05 |
| NC_045512.2;YP_009724390.1;Wu1-SARS2_surface | 158 | 158 | 1 | 0.05 | 0.05 | 0.05 |
| NC_045512.2;YP_009724390.1;Wu1-SARS2_surface | 169 | 169 | 1 | 0.05 | 0.05 | 0.05 |
| NC_045512.2;YP_009724390.1;Wu1-SARS2_surface | 171 | 171 | 1 | 0.05 | 0.05 | 0.05 |

| Extended data 1 | | | | | | |
| --- | --- | --- | --- | --- | --- | --- |
| **Protein** | **Start position (peptide)** | **Stop position (peptide)** | **Number of peptides** | **Minimum fraction (per peptide) of control subjects with reactivity >3 sd above average** | **Maximum fraction (per peptide) of control subjects with reactivity >3 sd above average** | **Mean fraction (over all peptides in this contiguous stretch) of controlsubjects with reactivity >3 sd above average** |
| NC_045512.2;YP_009724390.1;Wu1-SARS2_surface | 173 | 180 | 8 | 0.05 | 0.15 | 0.08125 |
| NC_045512.2;YP_009724390.1;Wu1-SARS2_surface | 183 | 185 | 3 | 0.05 | 0.1 | 0.083333333 |
| NC_045512.2;YP_009724390.1;Wu1-SARS2_surface | 224 | 224 | 1 | 0.05 | 0.05 | 0.05 |
| NC_045512.2;YP_009724390.1;Wu1-SARS2_surface | 248 | 249 | 2 | 0.1 | 0.15 | 0.125 |
| NC_045512.2;YP_009724390.1;Wu1-SARS2_surface | 272 | 274 | 3 | 0.05 | 0.05 | 0.05 |
| NC_045512.2;YP_009724390.1;Wu1-SARS2_surface | 277 | 278 | 2 | 0.05 | 0.05 | 0.05 |
| NC_045512.2;YP_009724390.1;Wu1-SARS2_surface | 295 | 296 | 2 | 0.05 | 0.1 | 0.075 |
| NC_045512.2;YP_009724390.1;Wu1-SARS2_surface | 302 | 302 | 1 | 0.05 | 0.05 | 0.05 |
| NC_045512.2;YP_009724390.1;Wu1-SARS2_surface | 308 | 312 | 5 | 0.05 | 0.1 | 0.07 |
| NC_045512.2;YP_009724390.1;Wu1-SARS2_surface | 320 | 320 | 1 | 0.05 | 0.05 | 0.05 |
| NC_045512.2;YP_009724390.1;Wu1-SARS2_surface | 324 | 324 | 1 | 0.05 | 0.05 | 0.05 |
| NC_045512.2;YP_009724390.1;Wu1-SARS2_surface | 329 | 330 | 2 | 0.05 | 0.05 | 0.05 |
| NC_045512.2;YP_009724390.1;Wu1-SARS2_surface | 334 | 335 | 2 | 0.05 | 0.05 | 0.05 |
| NC_045512.2;YP_009724390.1;Wu1-SARS2_surface | 349 | 349 | 1 | 0.05 | 0.05 | 0.05 |
| NC_045512.2;YP_009724390.1;Wu1-SARS2_surface | 363 | 363 | 1 | 0.05 | 0.05 | 0.05 |
| NC_045512.2;YP_009724390.1;Wu1-SARS2_surface | 369 | 369 | 1 | 0.05 | 0.05 | 0.05 |
| NC_045512.2;YP_009724390.1;Wu1-SARS2_surface | 374 | 375 | 2 | 0.05 | 0.05 | 0.05 |
| NC_045512.2;YP_009724390.1;Wu1-SARS2_surface | 393 | 393 | 1 | 0.05 | 0.05 | 0.05 |
| NC_045512.2;YP_009724390.1;Wu1-SARS2_surface | 397 | 398 | 2 | 0.05 | 0.15 | 0.1 |
| NC_045512.2;YP_009724390.1;Wu1-SARS2_surface | 408 | 409 | 2 | 0.05 | 0.15 | 0.1 |
| NC_045512.2;YP_009724390.1;Wu1-SARS2_surface | 412 | 417 | 6 | 0.05 | 0.25 | 0.141666667 |
| NC_045512.2;YP_009724390.1;Wu1-SARS2_surface | 419 | 419 | 1 | 0.05 | 0.05 | 0.05 |
| NC_045512.2;YP_009724390.1;Wu1-SARS2_surface | 430 | 438 | 9 | 0.05 | 0.3 | 0.161111111 |
| NC_045512.2;YP_009724390.1;Wu1-SARS2_surface | 452 | 458 | 7 | 0.05 | 0.1 | 0.071428571 |
| NC_045512.2;YP_009724390.1;Wu1-SARS2_surface | 462 | 462 | 1 | 0.05 | 0.05 | 0.05 |
| NC_045512.2;YP_009724390.1;Wu1-SARS2_surface | 465 | 465 | 1 | 0.05 | 0.05 | 0.05 |
| NC_045512.2;YP_009724390.1;Wu1-SARS2_surface | 481 | 481 | 1 | 0.05 | 0.05 | 0.05 |
| NC_045512.2;YP_009724390.1;Wu1-SARS2_surface | 489 | 489 | 1 | 0.05 | 0.05 | 0.05 |
| NC_045512.2;YP_009724390.1;Wu1-SARS2_surface | 518 | 518 | 1 | 0.05 | 0.05 | 0.05 |
| NC_045512.2;YP_009724390.1;Wu1-SARS2_surface | 542 | 542 | 1 | 0.05 | 0.05 | 0.05 |
| NC_045512.2;YP_009724390.1;Wu1-SARS2_surface | 546 | 546 | 1 | 0.05 | 0.05 | 0.05 |
| NC_045512.2;YP_009724390.1;Wu1-SARS2_surface | 548 | 548 | 1 | 0.05 | 0.05 | 0.05 |
| NC_045512.2;YP_009724390.1;Wu1-SARS2_surface | 550 | 555 | 6 | 0.05 | 0.5 | 0.183333333 |
| NC_045512.2;YP_009724390.1;Wu1-SARS2_surface | 564 | 565 | 2 | 0.05 | 0.05 | 0.05 |
| NC_045512.2;YP_009724390.1;Wu1-SARS2_surface | 569 | 569 | 1 | 0.05 | 0.05 | 0.05 |
| NC_045512.2;YP_009724390.1;Wu1-SARS2_surface | 590 | 602 | 13 | 0.1 | 0.4 | 0.246153846 |
| NC_045512.2;YP_009724390.1;Wu1-SARS2_surface | 623 | 623 | 1 | 0.05 | 0.05 | 0.05 |

| Extended data 1 | | | | | | |
|---|---|---|---|---|---|---|
| Protein | Start position (peptide) | Stop position (peptide) | Number of peptides | Minimum fraction (per peptide) of control subjects with reactivity >3 sd above average | Maximum fraction (per peptide) of control subjects with reactivity >3 sd above average | Mean fraction (over all peptides in this contiguous stretch) of controlsubjects with reactivity >3 sd above average |
| NC_045512.2;YP_009724390.1;Wu1-SARS2_surface | 634 | 634 | 1 | 0.05 | 0.05 | 0.05 |
| NC_045512.2;YP_009724390.1;Wu1-SARS2_surface | 655 | 660 | 6 | 0.05 | 0.1 | 0.058333333 |
| NC_045512.2;YP_009724390.1;Wu1-SARS2_surface | 675 | 679 | 5 | 0.05 | 0.1 | 0.07 |
| NC_045512.2;YP_009724390.1;Wu1-SARS2_surface | 681 | 681 | 1 | 0.05 | 0.05 | 0.05 |
| NC_045512.2;YP_009724390.1;Wu1-SARS2_surface | 691 | 691 | 1 | 0.1 | 0.1 | 0.1 |
| NC_045512.2;YP_009724390.1;Wu1-SARS2_surface | 696 | 696 | 1 | 0.05 | 0.05 | 0.05 |
| NC_045512.2;YP_009724390.1;Wu1-SARS2_surface | 698 | 708 | 11 | 0.05 | 0.2 | 0.113636364 |
| NC_045512.2;YP_009724390.1;Wu1-SARS2_surface | 736 | 738 | 3 | 0.05 | 0.05 | 0.05 |
| NC_045512.2;YP_009724390.1;Wu1-SARS2_surface | 740 | 743 | 4 | 0.05 | 0.05 | 0.05 |
| NC_045512.2;YP_009724390.1;Wu1-SARS2_surface | 761 | 769 | 9 | 0.1 | 0.2 | 0.144444444 |
| NC_045512.2;YP_009724390.1;Wu1-SARS2_surface | 771 | 774 | 4 | 0.1 | 0.1 | 0.1 |
| NC_045512.2;YP_009724390.1;Wu1-SARS2_surface | 777 | 779 | 3 | 0.05 | 0.05 | 0.05 |
| NC_045512.2;YP_009724390.1;Wu1-SARS2_surface | 794 | 794 | 1 | 0.05 | 0.05 | 0.05 |
| NC_045512.2;YP_009724390.1;Wu1-SARS2_surface | 797 | 797 | 1 | 0.05 | 0.05 | 0.05 |
| NC_045512.2;YP_009724390.1;Wu1-SARS2_surface | 806 | 813 | 8 | 0.05 | 0.1 | 0.0625 |
| NC_045512.2;YP_009724390.1;Wu1-SARS2_surface | 817 | 818 | 2 | 0.05 | 0.1 | 0.075 |
| NC_045512.2;YP_009724390.1;Wu1-SARS2_surface | 827 | 828 | 2 | 0.05 | 0.1 | 0.075 |
| NC_045512.2;YP_009724390.1;Wu1-SARS2_surface | 831 | 831 | 1 | 0.05 | 0.05 | 0.05 |
| NC_045512.2;YP_009724390.1;Wu1-SARS2_surface | 835 | 835 | 1 | 0.05 | 0.05 | 0.05 |
| NC_045512.2;YP_009724390.1;Wu1-SARS2_surface | 840 | 840 | 1 | 0.05 | 0.05 | 0.05 |
| NC_045512.2;YP_009724390.1;Wu1-SARS2_surface | 842 | 843 | 2 | 0.05 | 0.1 | 0.075 |
| NC_045512.2;YP_009724390.1;Wu1-SARS2_surface | 848 | 848 | 1 | 0.05 | 0.05 | 0.05 |
| NC_045512.2;YP_009724390.1;Wu1-SARS2_surface | 878 | 878 | 1 | 0.1 | 0.1 | 0.1 |
| NC_045512.2;YP_009724390.1;Wu1-SARS2_surface | 892 | 901 | 10 | 0.05 | 0.15 | 0.07 |
| NC_045512.2;YP_009724390.1;Wu1-SARS2_surface | 905 | 905 | 1 | 0.05 | 0.05 | 0.05 |
| NC_045512.2;YP_009724390.1;Wu1-SARS2_surface | 907 | 907 | 1 | 0.05 | 0.05 | 0.05 |
| NC_045512.2;YP_009724390.1;Wu1-SARS2_surface | 909 | 910 | 2 | 0.05 | 0.05 | 0.05 |
| NC_045512.2;YP_009724390.1;Wu1-SARS2_surface | 913 | 913 | 1 | 0.05 | 0.05 | 0.05 |
| NC_045512.2;YP_009724390.1;Wu1-SARS2_surface | 916 | 916 | 1 | 0.05 | 0.05 | 0.05 |
| NC_045512.2;YP_009724390.1;Wu1-SARS2_surface | 918 | 918 | 1 | 0.05 | 0.05 | 0.05 |
| NC_045512.2;YP_009724390.1;Wu1-SARS2_surface | 920 | 928 | 9 | 0.05 | 0.1 | 0.061111111 |
| NC_045512.2;YP_009724390.1;Wu1-SARS2_surface | 932 | 932 | 1 | 0.05 | 0.05 | 0.05 |
| NC_045512.2;YP_009724390.1;Wu1-SARS2_surface | 946 | 946 | 1 | 0.1 | 0.1 | 0.1 |
| NC_045512.2;YP_009724390.1;Wu1-SARS2_surface | 951 | 951 | 1 | 0.05 | 0.05 | 0.05 |
| NC_045512.2;YP_009724390.1;Wu1-SARS2_surface | 972 | 972 | 1 | 0.05 | 0.05 | 0.05 |
| NC_045512.2;YP_009724390.1;Wu1-SARS2_surface | 974 | 976 | 3 | 0.05 | 0.05 | 0.05 |
| NC_045512.2;YP_009724390.1;Wu1-SARS2_surface | 979 | 980 | 2 | 0.05 | 0.05 | 0.05 |

| Extended data 1 | | | | | | |
|---|---|---|---|---|---|---|
| Protein | Start position (peptide) | Stop position (peptide) | Number of peptides | Minimum fraction (per peptide) of control subjects with reactivity >3 sd above average | Maximum fraction (per peptide) of control subjects with reactivity >3 sd above average | Mean fraction (over all peptides in this contiguous stretch) of controlsubjects with reactivity >3 sd above average |
| NC_045512.2;YP_009724390.1;Wu1-SARS2_surface | 988 | 988 | 1 | 0.05 | 0.05 | 0.05 |
| NC_045512.2;YP_009724390.1;Wu1-SARS2_surface | 1011 | 1011 | 1 | 0.05 | 0.05 | 0.05 |
| NC_045512.2;YP_009724390.1;Wu1-SARS2_surface | 1016 | 1018 | 3 | 0.05 | 0.05 | 0.05 |
| NC_045512.2;YP_009724390.1;Wu1-SARS2_surface | 1042 | 1042 | 1 | 0.05 | 0.05 | 0.05 |
| NC_045512.2;YP_009724390.1;Wu1-SARS2_surface | 1064 | 1067 | 4 | 0.05 | 0.1 | 0.0625 |
| NC_045512.2;YP_009724390.1;Wu1-SARS2_surface | 1071 | 1071 | 1 | 0.1 | 0.1 | 0.1 |
| NC_045512.2;YP_009724390.1;Wu1-SARS2_surface | 1078 | 1079 | 2 | 0.05 | 0.05 | 0.05 |
| NC_045512.2;YP_009724390.1;Wu1-SARS2_surface | 1081 | 1081 | 1 | 0.05 | 0.05 | 0.05 |
| NC_045512.2;YP_009724390.1;Wu1-SARS2_surface | 1104 | 1106 | 3 | 0.05 | 0.1 | 0.066666667 |
| NC_045512.2;YP_009724390.1;Wu1-SARS2_surface | 1108 | 1116 | 9 | 0.05 | 0.1 | 0.055555556 |
| NC_045512.2;YP_009724390.1;Wu1-SARS2_surface | 1141 | 1154 | 14 | 0.05 | 0.5 | 0.314285714 |
| NC_045512.2;YP_009724390.1;Wu1-SARS2_surface | 1158 | 1161 | 4 | 0.05 | 0.2 | 0.1125 |
| NC_045512.2;YP_009724390.1;Wu1-SARS2_surface | 1167 | 1194 | 28 | 0.05 | 0.2 | 0.078571429 |
| NC_045512.2;YP_009724390.1;Wu1-SARS2_surface | 1254 | 1261 | 8 | 0.05 | 0.1 | 0.0625 |
| NC_045512.2;YP_009724391.1;Wu1-SARS2_orf3a | 8 | 8 | 1 | 0.05 | 0.05 | 0.05 |
| NC_045512.2;YP_009724391.1;Wu1-SARS2_orf3a | 10 | 10 | 1 | 0.05 | 0.05 | 0.05 |
| NC_045512.2;YP_009724391.1;Wu1-SARS2_orf3a | 12 | 15 | 4 | 0.05 | 0.05 | 0.05 |
| NC_045512.2;YP_009724391.1;Wu1-SARS2_orf3a | 36 | 36 | 1 | 0.05 | 0.05 | 0.05 |
| NC_045512.2;YP_009724391.1;Wu1-SARS2_orf3a | 153 | 153 | 1 | 0.05 | 0.05 | 0.05 |
| NC_045512.2;YP_009724391.1;Wu1-SARS2_orf3a | 176 | 185 | 10 | 0.05 | 0.1 | 0.06 |
| NC_045512.2;YP_009724391.1;Wu1-SARS2_orf3a | 209 | 209 | 1 | 0.05 | 0.05 | 0.05 |
| NC_045512.2;YP_009724391.1;Wu1-SARS2_orf3a | 220 | 220 | 1 | 0.05 | 0.05 | 0.05 |
| NC_045512.2;YP_009724391.1;Wu1-SARS2_orf3a | 231 | 231 | 1 | 0.05 | 0.05 | 0.05 |
| NC_045512.2;YP_009724391.1;Wu1-SARS2_orf3a | 241 | 252 | 12 | 0.05 | 0.05 | 0.05 |
| NC_045512.2;YP_009724391.1;Wu1-SARS2_orf3a | 257 | 261 | 5 | 0.05 | 0.05 | 0.05 |
| NC_045512.2;YP_009724392.1;Wu1-SARS2_envelope | 60 | 64 | 5 | 0.05 | 0.1 | 0.08 |
| NC_045512.2;YP_009724393.1;Wu1-SARS2_membrane | 11 | 11 | 1 | 0.05 | 0.05 | 0.05 |
| NC_045512.2;YP_009724393.1;Wu1-SARS2_membrane | 101 | 101 | 1 | 0.05 | 0.05 | 0.05 |
| NC_045512.2;YP_009724393.1;Wu1-SARS2_membrane | 148 | 158 | 11 | 0.05 | 0.1 | 0.054545455 |
| NC_045512.2;YP_009724393.1;Wu1-SARS2_membrane | 180 | 180 | 1 | 0.05 | 0.05 | 0.05 |
| NC_045512.2;YP_009724393.1;Wu1-SARS2_membrane | 200 | 205 | 6 | 0.05 | 0.1 | 0.058333333 |
| NC_045512.2;YP_009724394.1;Wu1-SARS2_orf6 | 31 | 46 | 16 | 0.05 | 0.1 | 0.065625 |
| NC_045512.2;YP_009724395.1;Wu1-SARS2_orf7a | 23 | 25 | 3 | 0.05 | 0.05 | 0.05 |
| NC_045512.2;YP_009724395.1;Wu1-SARS2_orf7a | 36 | 36 | 1 | 0.05 | 0.05 | 0.05 |
| NC_045512.2;YP_009724395.1;Wu1-SARS2_orf7a | 79 | 81 | 3 | 0.05 | 0.15 | 0.083333333 |
| NC_045512.2;YP_009724395.1;Wu1-SARS2_orf7a | 84 | 85 | 2 | 0.05 | 0.05 | 0.05 |
| NC_045512.2;YP_009724396.1;Wu1-SARS2_orf8 | 62 | 62 | 1 | 0.05 | 0.05 | 0.05 |

| Extended data 1 | | | | | | |
|---|---|---|---|---|---|---|
| **Protein** | **Start position (peptide)** | **Stop position (peptide)** | **Number of peptides** | **Minimum fraction (per peptide) of control subjects with reactivity >3 sd above average** | **Maximum fraction (per peptide) of control subjects with reactivity >3 sd above average** | **Mean fraction (over all peptides in this contiguous stretch) of controlsubjects with reactivity >3 sd above average** |
| NC_045512.2;YP_009724396.1;Wu1-SARS2_orf8 | 83 | 85 | 3 | 0.05 | 0.05 | 0.05 |
| NC_045512.2;YP_009724397.2;Wu1-SARS2_nucleoca | 1 | 13 | 13 | 0.05 | 0.15 | 0.065384615 |
| NC_045512.2;YP_009724397.2;Wu1-SARS2_nucleoca | 19 | 19 | 1 | 0.05 | 0.05 | 0.05 |
| NC_045512.2;YP_009724397.2;Wu1-SARS2_nucleoca | 21 | 22 | 2 | 0.05 | 0.05 | 0.05 |
| NC_045512.2;YP_009724397.2;Wu1-SARS2_nucleoca | 62 | 62 | 1 | 0.05 | 0.05 | 0.05 |
| NC_045512.2;YP_009724397.2;Wu1-SARS2_nucleoca | 64 | 64 | 1 | 0.1 | 0.1 | 0.1 |
| NC_045512.2;YP_009724397.2;Wu1-SARS2_nucleoca | 67 | 67 | 1 | 0.1 | 0.1 | 0.1 |
| NC_045512.2;YP_009724397.2;Wu1-SARS2_nucleoca | 73 | 73 | 1 | 0.15 | 0.15 | 0.15 |
| NC_045512.2;YP_009724397.2;Wu1-SARS2_nucleoca | 78 | 79 | 2 | 0.05 | 0.05 | 0.05 |
| NC_045512.2;YP_009724397.2;Wu1-SARS2_nucleoca | 85 | 85 | 1 | 0.05 | 0.05 | 0.05 |
| NC_045512.2;YP_009724397.2;Wu1-SARS2_nucleoca | 111 | 112 | 2 | 0.05 | 0.1 | 0.075 |
| NC_045512.2;YP_009724397.2;Wu1-SARS2_nucleoca | 138 | 138 | 1 | 0.05 | 0.05 | 0.05 |
| NC_045512.2;YP_009724397.2;Wu1-SARS2_nucleoca | 141 | 141 | 1 | 0.05 | 0.05 | 0.05 |
| NC_045512.2;YP_009724397.2;Wu1-SARS2_nucleoca | 154 | 164 | 11 | 0.05 | 0.2 | 0.122727273 |
| NC_045512.2;YP_009724397.2;Wu1-SARS2_nucleoca | 168 | 168 | 1 | 0.05 | 0.05 | 0.05 |
| NC_045512.2;YP_009724397.2;Wu1-SARS2_nucleoca | 181 | 181 | 1 | 0.05 | 0.05 | 0.05 |
| NC_045512.2;YP_009724397.2;Wu1-SARS2_nucleoca | 183 | 191 | 9 | 0.05 | 0.2 | 0.133333333 |
| NC_045512.2;YP_009724397.2;Wu1-SARS2_nucleoca | 197 | 197 | 1 | 0.05 | 0.05 | 0.05 |
| NC_045512.2;YP_009724397.2;Wu1-SARS2_nucleoca | 200 | 203 | 4 | 0.05 | 0.05 | 0.05 |
| NC_045512.2;YP_009724397.2;Wu1-SARS2_nucleoca | 218 | 226 | 9 | 0.05 | 0.1 | 0.055555556 |
| NC_045512.2;YP_009724397.2;Wu1-SARS2_nucleoca | 229 | 229 | 1 | 0.05 | 0.05 | 0.05 |
| NC_045512.2;YP_009724397.2;Wu1-SARS2_nucleoca | 231 | 231 | 1 | 0.05 | 0.05 | 0.05 |
| NC_045512.2;YP_009724397.2;Wu1-SARS2_nucleoca | 238 | 241 | 4 | 0.05 | 0.25 | 0.1 |
| NC_045512.2;YP_009724397.2;Wu1-SARS2_nucleoca | 247 | 247 | 1 | 0.05 | 0.05 | 0.05 |
| NC_045512.2;YP_009724397.2;Wu1-SARS2_nucleoca | 258 | 258 | 1 | 0.05 | 0.05 | 0.05 |
| NC_045512.2;YP_009724397.2;Wu1-SARS2_nucleoca | 263 | 280 | 18 | 0.05 | 0.15 | 0.077777778 |
| NC_045512.2;YP_009724397.2;Wu1-SARS2_nucleoca | 294 | 294 | 1 | 0.05 | 0.05 | 0.05 |
| NC_045512.2;YP_009724397.2;Wu1-SARS2_nucleoca | 297 | 297 | 1 | 0.15 | 0.15 | 0.15 |
| NC_045512.2;YP_009724397.2;Wu1-SARS2_nucleoca | 310 | 310 | 1 | 0.05 | 0.05 | 0.05 |
| NC_045512.2;YP_009724397.2;Wu1-SARS2_nucleoca | 313 | 314 | 2 | 0.05 | 0.05 | 0.05 |
| NC_045512.2;YP_009724397.2;Wu1-SARS2_nucleoca | 332 | 333 | 2 | 0.05 | 0.1 | 0.075 |
| NC_045512.2;YP_009724397.2;Wu1-SARS2_nucleoca | 335 | 335 | 1 | 0.1 | 0.1 | 0.1 |
| NC_045512.2;YP_009724397.2;Wu1-SARS2_nucleoca | 337 | 337 | 1 | 0.05 | 0.05 | 0.05 |
| NC_045512.2;YP_009724397.2;Wu1-SARS2_nucleoca | 343 | 345 | 3 | 0.05 | 0.05 | 0.05 |
| NC_045512.2;YP_009724397.2;Wu1-SARS2_nucleoca | 353 | 353 | 1 | 0.05 | 0.05 | 0.05 |
| NC_045512.2;YP_009724397.2;Wu1-SARS2_nucleoca | 357 | 358 | 2 | 0.05 | 0.05 | 0.05 |
| NC_045512.2;YP_009724397.2;Wu1-SARS2_nucleoca | 362 | 364 | 3 | 0.05 | 0.05 | 0.05 |

| Extended data 1 | | | | | | |
|---|---|---|---|---|---|---|
| Protein | Start position (peptide) | Stop position (peptide) | Number of peptides | Minimum fraction (per peptide) of control subjects with reactivity >3 sd above average | Maximum fraction (per peptide) of control subjects with reactivity >3 sd above average | Mean fraction (over all peptides in this contiguous stretch) of controlsubjects with reactivity >3 sd above average |
| NC_045512.2;YP_009724397.2;Wu1-SARS2_nucleoca | 377 | 377 | 1 | 0.05 | 0.05 | 0.05 |
| NC_045512.2;YP_009724397.2;Wu1-SARS2_nucleoca | 383 | 383 | 1 | 0.05 | 0.05 | 0.05 |
| NC_045512.2;YP_009724397.2;Wu1-SARS2_nucleoca | 395 | 395 | 1 | 0.05 | 0.05 | 0.05 |
| NC_045512.2;YP_009724397.2;Wu1-SARS2_nucleoca | 397 | 398 | 2 | 0.1 | 0.25 | 0.175 |
| NC_045512.2;YP_009725255.1;Wu1-SARS2_orf10 | 16 | 19 | 4 | 0.05 | 0.15 | 0.075 |
| NC_045512.2;YP_009725255.1;Wu1-SARS2_orf10 | 21 | 21 | 1 | 0.1 | 0.1 | 0.1 |
| NC_045512.2;YP_009725255.1;Wu1-SARS2_orf10 | 26 | 27 | 2 | 0.05 | 0.05 | 0.05 |
