## Extended data 2 for "The landscape of antibody binding in SARS-CoV-2 infection"

| Peptide | Fraction of control subjects with reactivity >3 sd above average | Fraction of COVID-19 convalescent subjects with reactivity >3 sd above average |
|---|---|---|
| NC_045512.2;YP_009724389.1;Wu1-SARS2_orf1ab;1 | 0 | 0 |
| NC_045512.2;YP_009724389.1;Wu1-SARS2_orf1ab;2 | 0 | 0.025 |
| NC_045512.2;YP_009724389.1;Wu1-SARS2_orf1ab;3 | 0 | 0 |
| NC_045512.2;YP_009724389.1;Wu1-SARS2_orf1ab;4 | 0 | 0.025 |
| NC_045512.2;YP_009724389.1;Wu1-SARS2_orf1ab;5 | 0 | 0 |
| NC_045512.2;YP_009724389.1;Wu1-SARS2_orf1ab;6 | 0 | 0 |
| NC_045512.2;YP_009724389.1;Wu1-SARS2_orf1ab;7 | 0 | 0 |
| NC_045512.2;YP_009724389.1;Wu1-SARS2_orf1ab;8 | 0 | 0 |
| NC_045512.2;YP_009724389.1;Wu1-SARS2_orf1ab;9 | 0 | 0 |
| NC_045512.2;YP_009724389.1;Wu1-SARS2_orf1ab;10 | 0 | 0 |
| NC_045512.2;YP_009724389.1;Wu1-SARS2_orf1ab;11 | 0 | 0 |
| NC_045512.2;YP_009724389.1;Wu1-SARS2_orf1ab;12 | 0 | 0 |
| NC_045512.2;YP_009724389.1;Wu1-SARS2_orf1ab;13 | 0 | 0 |
| NC_045512.2;YP_009724389.1;Wu1-SARS2_orf1ab;14 | 0 | 0 |
| NC_045512.2;YP_009724389.1;Wu1-SARS2_orf1ab;15 | 0 | 0 |
| NC_045512.2;YP_009724389.1;Wu1-SARS2_orf1ab;16 | 0 | 0 |
| NC_045512.2;YP_009724389.1;Wu1-SARS2_orf1ab;17 | 0 | 0 |
| NC_045512.2;YP_009724389.1;Wu1-SARS2_orf1ab;18 | 0 | 0 |
| NC_045512.2;YP_009724389.1;Wu1-SARS2_orf1ab;19 | 0 | 0 |
| NC_045512.2;YP_009724389.1;Wu1-SARS2_orf1ab;20 | 0 | 0 |
| NC_045512.2;YP_009724389.1;Wu1-SARS2_orf1ab;21 | 0 | 0 |
| NC_045512.2;YP_009724389.1;Wu1-SARS2_orf1ab;22 | 0 | 0 |
| NC_045512.2;YP_009724389.1;Wu1-SARS2_orf1ab;23 | 0 | 0 |
| NC_045512.2;YP_009724389.1;Wu1-SARS2_orf1ab;24 | 0 | 0 |
| NC_045512.2;YP_009724389.1;Wu1-SARS2_orf1ab;25 | 0 | 0.025 |
| NC_045512.2;YP_009724389.1;Wu1-SARS2_orf1ab;26 | 0 | 0.025 |
| NC_045512.2;YP_009724389.1;Wu1-SARS2_orf1ab;27 | 0 | 0.025 |
| NC_045512.2;YP_009724389.1;Wu1-SARS2_orf1ab;28 | 0 | 0.025 |
| NC_045512.2;YP_009724389.1;Wu1-SARS2_orf1ab;29 | 0.1 | 0.025 |
| NC_045512.2;YP_009724389.1;Wu1-SARS2_orf1ab;30 | 0.25 | 0.15 |
| NC_045512.2;YP_009724389.1;Wu1-SARS2_orf1ab;31 | 0 | 0.025 |
| NC_045512.2;YP_009724389.1;Wu1-SARS2_orf1ab;32 | 0 | 0.05 |
| NC_045512.2;YP_009724389.1;Wu1-SARS2_orf1ab;33 | 0.05 | 0 |

| | | |
|---|---|---|
| NC_045512.2;YP_009724389.1;Wu1-SARS2_orf1ab;34 | 0.1 | 0 |
| NC_045512.2;YP_009724389.1;Wu1-SARS2_orf1ab;35 | 0 | 0.025 |
| NC_045512.2;YP_009724389.1;Wu1-SARS2_orf1ab;36 | 0 | 0.025 |
| NC_045512.2;YP_009724389.1;Wu1-SARS2_orf1ab;37 | 0 | 0 |
| NC_045512.2;YP_009724389.1;Wu1-SARS2_orf1ab;38 | 0 | 0 |
| NC_045512.2;YP_009724389.1;Wu1-SARS2_orf1ab;39 | 0 | 0 |
| NC_045512.2;YP_009724389.1;Wu1-SARS2_orf1ab;40 | 0 | 0 |
| NC_045512.2;YP_009724389.1;Wu1-SARS2_orf1ab;41 | 0 | 0 |
| NC_045512.2;YP_009724389.1;Wu1-SARS2_orf1ab;42 | 0 | 0 |
| NC_045512.2;YP_009724389.1;Wu1-SARS2_orf1ab;43 | 0 | 0 |
| NC_045512.2;YP_009724389.1;Wu1-SARS2_orf1ab;44 | 0.05 | 0 |
| NC_045512.2;YP_009724389.1;Wu1-SARS2_orf1ab;45 | 0 | 0 |
| NC_045512.2;YP_009724389.1;Wu1-SARS2_orf1ab;46 | 0 | 0 |
| NC_045512.2;YP_009724389.1;Wu1-SARS2_orf1ab;47 | 0 | 0 |
| NC_045512.2;YP_009724389.1;Wu1-SARS2_orf1ab;48 | 0 | 0 |
| NC_045512.2;YP_009724389.1;Wu1-SARS2_orf1ab;49 | 0 | 0 |
| NC_045512.2;YP_009724389.1;Wu1-SARS2_orf1ab;50 | 0 | 0 |
| NC_045512.2;YP_009724389.1;Wu1-SARS2_orf1ab;51 | 0 | 0 |
| NC_045512.2;YP_009724389.1;Wu1-SARS2_orf1ab;52 | 0.05 | 0.025 |
| NC_045512.2;YP_009724389.1;Wu1-SARS2_orf1ab;53 | 0 | 0.025 |
| NC_045512.2;YP_009724389.1;Wu1-SARS2_orf1ab;54 | 0.05 | 0.025 |
| NC_045512.2;YP_009724389.1;Wu1-SARS2_orf1ab;55 | 0 | 0.025 |
| NC_045512.2;YP_009724389.1;Wu1-SARS2_orf1ab;56 | 0 | 0.025 |
| NC_045512.2;YP_009724389.1;Wu1-SARS2_orf1ab;57 | 0 | 0.025 |
| NC_045512.2;YP_009724389.1;Wu1-SARS2_orf1ab;58 | 0 | 0.025 |
| NC_045512.2;YP_009724389.1;Wu1-SARS2_orf1ab;59 | 0 | 0 |
| NC_045512.2;YP_009724389.1;Wu1-SARS2_orf1ab;60 | 0 | 0 |
| NC_045512.2;YP_009724389.1;Wu1-SARS2_orf1ab;61 | 0 | 0 |
| NC_045512.2;YP_009724389.1;Wu1-SARS2_orf1ab;62 | 0 | 0 |
| NC_045512.2;YP_009724389.1;Wu1-SARS2_orf1ab;63 | 0 | 0 |
| NC_045512.2;YP_009724389.1;Wu1-SARS2_orf1ab;64 | 0 | 0 |
| NC_045512.2;YP_009724389.1;Wu1-SARS2_orf1ab;65 | 0 | 0 |
| NC_045512.2;YP_009724389.1;Wu1-SARS2_orf1ab;66 | 0 | 0 |
| NC_045512.2;YP_009724389.1;Wu1-SARS2_orf1ab;67 | 0 | 0 |
| NC_045512.2;YP_009724389.1;Wu1-SARS2_orf1ab;68 | 0 | 0 |
| NC_045512.2;YP_009724389.1;Wu1-SARS2_orf1ab;69 | 0 | 0 |
| NC_045512.2;YP_009724389.1;Wu1-SARS2_orf1ab;70 | 0 | 0 |
| NC_045512.2;YP_009724389.1;Wu1-SARS2_orf1ab;71 | 0.05 | 0 |
| NC_045512.2;YP_009724389.1;Wu1-SARS2_orf1ab;72 | 0 | 0 |
| NC_045512.2;YP_009724389.1;Wu1-SARS2_orf1ab;73 | 0 | 0 |
| NC_045512.2;YP_009724389.1;Wu1-SARS2_orf1ab;74 | 0 | 0 |
| NC_045512.2;YP_009724389.1;Wu1-SARS2_orf1ab;75 | 0 | 0.025 |
| NC_045512.2;YP_009724389.1;Wu1-SARS2_orf1ab;76 | 0 | 0 |

| | | |
|---|---|---|
| NC_045512.2;YP_009724389.1;Wu1-SARS2_orf1ab;77 | 0 | 0 |
| NC_045512.2;YP_009724389.1;Wu1-SARS2_orf1ab;78 | 0 | 0 |
| NC_045512.2;YP_009724389.1;Wu1-SARS2_orf1ab;79 | 0 | 0 |
| NC_045512.2;YP_009724389.1;Wu1-SARS2_orf1ab;80 | 0 | 0 |
| NC_045512.2;YP_009724389.1;Wu1-SARS2_orf1ab;81 | 0 | 0 |
| NC_045512.2;YP_009724389.1;Wu1-SARS2_orf1ab;82 | 0 | 0 |
| NC_045512.2;YP_009724389.1;Wu1-SARS2_orf1ab;83 | 0 | 0 |
| NC_045512.2;YP_009724389.1;Wu1-SARS2_orf1ab;84 | 0 | 0 |
| NC_045512.2;YP_009724389.1;Wu1-SARS2_orf1ab;85 | 0 | 0 |
| NC_045512.2;YP_009724389.1;Wu1-SARS2_orf1ab;86 | 0 | 0.025 |
| NC_045512.2;YP_009724389.1;Wu1-SARS2_orf1ab;87 | 0 | 0 |
| NC_045512.2;YP_009724389.1;Wu1-SARS2_orf1ab;88 | 0 | 0 |
| NC_045512.2;YP_009724389.1;Wu1-SARS2_orf1ab;89 | 0 | 0 |
| NC_045512.2;YP_009724389.1;Wu1-SARS2_orf1ab;90 | 0 | 0 |
| NC_045512.2;YP_009724389.1;Wu1-SARS2_orf1ab;91 | 0 | 0 |
| NC_045512.2;YP_009724389.1;Wu1-SARS2_orf1ab;92 | 0 | 0 |
| NC_045512.2;YP_009724389.1;Wu1-SARS2_orf1ab;93 | 0 | 0 |
| NC_045512.2;YP_009724389.1;Wu1-SARS2_orf1ab;94 | 0 | 0.025 |
| NC_045512.2;YP_009724389.1;Wu1-SARS2_orf1ab;95 | 0 | 0 |
| NC_045512.2;YP_009724389.1;Wu1-SARS2_orf1ab;96 | 0 | 0 |
| NC_045512.2;YP_009724389.1;Wu1-SARS2_orf1ab;97 | 0 | 0 |
| NC_045512.2;YP_009724389.1;Wu1-SARS2_orf1ab;98 | 0 | 0 |
| NC_045512.2;YP_009724389.1;Wu1-SARS2_orf1ab;99 | 0 | 0 |
| NC_045512.2;YP_009724389.1;Wu1-SARS2_orf1ab;100 | 0 | 0 |
| NC_045512.2;YP_009724389.1;Wu1-SARS2_orf1ab;101 | 0 | 0 |
| NC_045512.2;YP_009724389.1;Wu1-SARS2_orf1ab;102 | 0 | 0 |
| NC_045512.2;YP_009724389.1;Wu1-SARS2_orf1ab;103 | 0 | 0 |
| NC_045512.2;YP_009724389.1;Wu1-SARS2_orf1ab;104 | 0 | 0 |
| NC_045512.2;YP_009724389.1;Wu1-SARS2_orf1ab;105 | 0 | 0 |
| NC_045512.2;YP_009724389.1;Wu1-SARS2_orf1ab;106 | 0 | 0 |
| NC_045512.2;YP_009724389.1;Wu1-SARS2_orf1ab;107 | 0 | 0 |
| NC_045512.2;YP_009724389.1;Wu1-SARS2_orf1ab;108 | 0 | 0 |
| NC_045512.2;YP_009724389.1;Wu1-SARS2_orf1ab;109 | 0 | 0 |
| NC_045512.2;YP_009724389.1;Wu1-SARS2_orf1ab;110 | 0 | 0 |
| NC_045512.2;YP_009724389.1;Wu1-SARS2_orf1ab;111 | 0 | 0 |
| NC_045512.2;YP_009724389.1;Wu1-SARS2_orf1ab;112 | 0 | 0 |
| NC_045512.2;YP_009724389.1;Wu1-SARS2_orf1ab;113 | 0 | 0 |
| NC_045512.2;YP_009724389.1;Wu1-SARS2_orf1ab;114 | 0 | 0.025 |
| NC_045512.2;YP_009724389.1;Wu1-SARS2_orf1ab;115 | 0 | 0.025 |
| NC_045512.2;YP_009724389.1;Wu1-SARS2_orf1ab;116 | 0 | 0.025 |
| NC_045512.2;YP_009724389.1;Wu1-SARS2_orf1ab;117 | 0 | 0.025 |
| NC_045512.2;YP_009724389.1;Wu1-SARS2_orf1ab;118 | 0.05 | 0.025 |
| NC_045512.2;YP_009724389.1;Wu1-SARS2_orf1ab;119 | 0 | 0.025 |

| | | |
|---|---|---|
| NC_045512.2;YP_009724389.1;Wu1-SARS2_orf1ab;120 | 0 | 0.025 |
| NC_045512.2;YP_009724389.1;Wu1-SARS2_orf1ab;121 | 0.05 | 0.025 |
| NC_045512.2;YP_009724389.1;Wu1-SARS2_orf1ab;122 | 0 | 0.025 |
| NC_045512.2;YP_009724389.1;Wu1-SARS2_orf1ab;123 | 0 | 0.025 |
| NC_045512.2;YP_009724389.1;Wu1-SARS2_orf1ab;124 | 0 | 0.05 |
| NC_045512.2;YP_009724389.1;Wu1-SARS2_orf1ab;125 | 0.1 | 0.025 |
| NC_045512.2;YP_009724389.1;Wu1-SARS2_orf1ab;126 | 0 | 0 |
| NC_045512.2;YP_009724389.1;Wu1-SARS2_orf1ab;127 | 0 | 0 |
| NC_045512.2;YP_009724389.1;Wu1-SARS2_orf1ab;128 | 0 | 0.025 |
| NC_045512.2;YP_009724389.1;Wu1-SARS2_orf1ab;129 | 0 | 0.05 |
| NC_045512.2;YP_009724389.1;Wu1-SARS2_orf1ab;130 | 0 | 0.025 |
| NC_045512.2;YP_009724389.1;Wu1-SARS2_orf1ab;131 | 0.1 | 0.025 |
| NC_045512.2;YP_009724389.1;Wu1-SARS2_orf1ab;132 | 0.05 | 0.025 |
| NC_045512.2;YP_009724389.1;Wu1-SARS2_orf1ab;133 | 0.05 | 0 |
| NC_045512.2;YP_009724389.1;Wu1-SARS2_orf1ab;134 | 0 | 0 |
| NC_045512.2;YP_009724389.1;Wu1-SARS2_orf1ab;135 | 0.05 | 0 |
| NC_045512.2;YP_009724389.1;Wu1-SARS2_orf1ab;136 | 0 | 0.025 |
| NC_045512.2;YP_009724389.1;Wu1-SARS2_orf1ab;137 | 0 | 0 |
| NC_045512.2;YP_009724389.1;Wu1-SARS2_orf1ab;138 | 0 | 0 |
| NC_045512.2;YP_009724389.1;Wu1-SARS2_orf1ab;139 | 0.05 | 0 |
| NC_045512.2;YP_009724389.1;Wu1-SARS2_orf1ab;140 | 0.05 | 0 |
| NC_045512.2;YP_009724389.1;Wu1-SARS2_orf1ab;141 | 0 | 0 |
| NC_045512.2;YP_009724389.1;Wu1-SARS2_orf1ab;142 | 0 | 0 |
| NC_045512.2;YP_009724389.1;Wu1-SARS2_orf1ab;143 | 0.05 | 0 |
| NC_045512.2;YP_009724389.1;Wu1-SARS2_orf1ab;144 | 0 | 0.025 |
| NC_045512.2;YP_009724389.1;Wu1-SARS2_orf1ab;145 | 0.2 | 0.1 |
| NC_045512.2;YP_009724389.1;Wu1-SARS2_orf1ab;146 | 0.05 | 0 |
| NC_045512.2;YP_009724389.1;Wu1-SARS2_orf1ab;147 | 0 | 0 |
| NC_045512.2;YP_009724389.1;Wu1-SARS2_orf1ab;148 | 0.05 | 0 |
| NC_045512.2;YP_009724389.1;Wu1-SARS2_orf1ab;149 | 0.3 | 0.2 |
| NC_045512.2;YP_009724389.1;Wu1-SARS2_orf1ab;150 | 0.25 | 0.075 |
| NC_045512.2;YP_009724389.1;Wu1-SARS2_orf1ab;151 | 0.3 | 0.15 |
| NC_045512.2;YP_009724389.1;Wu1-SARS2_orf1ab;152 | 0.45 | 0.225 |
| NC_045512.2;YP_009724389.1;Wu1-SARS2_orf1ab;153 | 0.35 | 0.2 |
| NC_045512.2;YP_009724389.1;Wu1-SARS2_orf1ab;154 | 0.45 | 0.35 |
| NC_045512.2;YP_009724389.1;Wu1-SARS2_orf1ab;155 | 0.5 | 0.325 |
| NC_045512.2;YP_009724389.1;Wu1-SARS2_orf1ab;156 | 0.3 | 0.1 |
| NC_045512.2;YP_009724389.1;Wu1-SARS2_orf1ab;157 | 0.2 | 0.05 |
| NC_045512.2;YP_009724389.1;Wu1-SARS2_orf1ab;158 | 0.05 | 0 |
| NC_045512.2;YP_009724389.1;Wu1-SARS2_orf1ab;159 | 0 | 0 |
| NC_045512.2;YP_009724389.1;Wu1-SARS2_orf1ab;160 | 0 | 0 |
| NC_045512.2;YP_009724389.1;Wu1-SARS2_orf1ab;161 | 0 | 0 |
| NC_045512.2;YP_009724389.1;Wu1-SARS2_orf1ab;162 | 0 | 0 |

| | | |
|---|---|---|
| NC_045512.2;YP_009724389.1;Wu1-SARS2_orf1ab;163 | 0 | 0.025 |
| NC_045512.2;YP_009724389.1;Wu1-SARS2_orf1ab;164 | 0 | 0 |
| NC_045512.2;YP_009724389.1;Wu1-SARS2_orf1ab;165 | 0.1 | 0 |
| NC_045512.2;YP_009724389.1;Wu1-SARS2_orf1ab;166 | 0.05 | 0.025 |
| NC_045512.2;YP_009724389.1;Wu1-SARS2_orf1ab;167 | 0 | 0.025 |
| NC_045512.2;YP_009724389.1;Wu1-SARS2_orf1ab;168 | 0.1 | 0.025 |
| NC_045512.2;YP_009724389.1;Wu1-SARS2_orf1ab;169 | 0 | 0 |
| NC_045512.2;YP_009724389.1;Wu1-SARS2_orf1ab;170 | 0 | 0.025 |
| NC_045512.2;YP_009724389.1;Wu1-SARS2_orf1ab;171 | 0 | 0 |
| NC_045512.2;YP_009724389.1;Wu1-SARS2_orf1ab;172 | 0 | 0 |
| NC_045512.2;YP_009724389.1;Wu1-SARS2_orf1ab;173 | 0 | 0 |
| NC_045512.2;YP_009724389.1;Wu1-SARS2_orf1ab;174 | 0.05 | 0.025 |
| NC_045512.2;YP_009724389.1;Wu1-SARS2_orf1ab;175 | 0 | 0 |
| NC_045512.2;YP_009724389.1;Wu1-SARS2_orf1ab;176 | 0 | 0 |
| NC_045512.2;YP_009724389.1;Wu1-SARS2_orf1ab;177 | 0 | 0 |
| NC_045512.2;YP_009724389.1;Wu1-SARS2_orf1ab;178 | 0 | 0 |
| NC_045512.2;YP_009724389.1;Wu1-SARS2_orf1ab;179 | 0 | 0 |
| NC_045512.2;YP_009724389.1;Wu1-SARS2_orf1ab;180 | 0 | 0 |
| NC_045512.2;YP_009724389.1;Wu1-SARS2_orf1ab;181 | 0 | 0.025 |
| NC_045512.2;YP_009724389.1;Wu1-SARS2_orf1ab;182 | 0 | 0 |
| NC_045512.2;YP_009724389.1;Wu1-SARS2_orf1ab;183 | 0 | 0 |
| NC_045512.2;YP_009724389.1;Wu1-SARS2_orf1ab;184 | 0 | 0.025 |
| NC_045512.2;YP_009724389.1;Wu1-SARS2_orf1ab;185 | 0 | 0.025 |
| NC_045512.2;YP_009724389.1;Wu1-SARS2_orf1ab;186 | 0 | 0.025 |
| NC_045512.2;YP_009724389.1;Wu1-SARS2_orf1ab;187 | 0 | 0 |
| NC_045512.2;YP_009724389.1;Wu1-SARS2_orf1ab;188 | 0 | 0 |
| NC_045512.2;YP_009724389.1;Wu1-SARS2_orf1ab;189 | 0 | 0 |
| NC_045512.2;YP_009724389.1;Wu1-SARS2_orf1ab;190 | 0 | 0 |
| NC_045512.2;YP_009724389.1;Wu1-SARS2_orf1ab;191 | 0 | 0 |
| NC_045512.2;YP_009724389.1;Wu1-SARS2_orf1ab;192 | 0 | 0 |
| NC_045512.2;YP_009724389.1;Wu1-SARS2_orf1ab;193 | 0 | 0 |
| NC_045512.2;YP_009724389.1;Wu1-SARS2_orf1ab;194 | 0 | 0 |
| NC_045512.2;YP_009724389.1;Wu1-SARS2_orf1ab;195 | 0 | 0 |
| NC_045512.2;YP_009724389.1;Wu1-SARS2_orf1ab;196 | 0 | 0 |
| NC_045512.2;YP_009724389.1;Wu1-SARS2_orf1ab;197 | 0.05 | 0 |
| NC_045512.2;YP_009724389.1;Wu1-SARS2_orf1ab;198 | 0 | 0 |
| NC_045512.2;YP_009724389.1;Wu1-SARS2_orf1ab;199 | 0 | 0 |
| NC_045512.2;YP_009724389.1;Wu1-SARS2_orf1ab;200 | 0 | 0 |
| NC_045512.2;YP_009724389.1;Wu1-SARS2_orf1ab;201 | 0 | 0 |
| NC_045512.2;YP_009724389.1;Wu1-SARS2_orf1ab;202 | 0 | 0 |
| NC_045512.2;YP_009724389.1;Wu1-SARS2_orf1ab;203 | 0 | 0 |
| NC_045512.2;YP_009724389.1;Wu1-SARS2_orf1ab;204 | 0.05 | 0 |
| NC_045512.2;YP_009724389.1;Wu1-SARS2_orf1ab;205 | 0 | 0 |

| | | |
|---|---|---|
| NC_045512.2;YP_009724389.1;Wu1-SARS2_orf1ab;206 | 0 | 0 |
| NC_045512.2;YP_009724389.1;Wu1-SARS2_orf1ab;207 | 0 | 0 |
| NC_045512.2;YP_009724389.1;Wu1-SARS2_orf1ab;208 | 0 | 0 |
| NC_045512.2;YP_009724389.1;Wu1-SARS2_orf1ab;209 | 0 | 0 |
| NC_045512.2;YP_009724389.1;Wu1-SARS2_orf1ab;210 | 0 | 0 |
| NC_045512.2;YP_009724389.1;Wu1-SARS2_orf1ab;211 | 0.05 | 0.075 |
| NC_045512.2;YP_009724389.1;Wu1-SARS2_orf1ab;212 | 0 | 0.1 |
| NC_045512.2;YP_009724389.1;Wu1-SARS2_orf1ab;213 | 0 | 0.1 |
| NC_045512.2;YP_009724389.1;Wu1-SARS2_orf1ab;214 | 0.1 | 0.075 |
| NC_045512.2;YP_009724389.1;Wu1-SARS2_orf1ab;215 | 0.1 | 0.075 |
| NC_045512.2;YP_009724389.1;Wu1-SARS2_orf1ab;216 | 0 | 0.075 |
| NC_045512.2;YP_009724389.1;Wu1-SARS2_orf1ab;217 | 0 | 0.025 |
| NC_045512.2;YP_009724389.1;Wu1-SARS2_orf1ab;218 | 0 | 0.025 |
| NC_045512.2;YP_009724389.1;Wu1-SARS2_orf1ab;219 | 0 | 0.025 |
| NC_045512.2;YP_009724389.1;Wu1-SARS2_orf1ab;220 | 0.05 | 0.075 |
| NC_045512.2;YP_009724389.1;Wu1-SARS2_orf1ab;221 | 0 | 0.025 |
| NC_045512.2;YP_009724389.1;Wu1-SARS2_orf1ab;222 | 0 | 0.025 |
| NC_045512.2;YP_009724389.1;Wu1-SARS2_orf1ab;223 | 0 | 0.025 |
| NC_045512.2;YP_009724389.1;Wu1-SARS2_orf1ab;224 | 0 | 0 |
| NC_045512.2;YP_009724389.1;Wu1-SARS2_orf1ab;225 | 0.05 | 0 |
| NC_045512.2;YP_009724389.1;Wu1-SARS2_orf1ab;226 | 0 | 0 |
| NC_045512.2;YP_009724389.1;Wu1-SARS2_orf1ab;227 | 0 | 0 |
| NC_045512.2;YP_009724389.1;Wu1-SARS2_orf1ab;228 | 0 | 0 |
| NC_045512.2;YP_009724389.1;Wu1-SARS2_orf1ab;229 | 0 | 0 |
| NC_045512.2;YP_009724389.1;Wu1-SARS2_orf1ab;230 | 0 | 0 |
| NC_045512.2;YP_009724389.1;Wu1-SARS2_orf1ab;231 | 0 | 0 |
| NC_045512.2;YP_009724389.1;Wu1-SARS2_orf1ab;232 | 0 | 0 |
| NC_045512.2;YP_009724389.1;Wu1-SARS2_orf1ab;233 | 0 | 0 |
| NC_045512.2;YP_009724389.1;Wu1-SARS2_orf1ab;234 | 0 | 0 |
| NC_045512.2;YP_009724389.1;Wu1-SARS2_orf1ab;235 | 0 | 0 |
| NC_045512.2;YP_009724389.1;Wu1-SARS2_orf1ab;236 | 0 | 0 |
| NC_045512.2;YP_009724389.1;Wu1-SARS2_orf1ab;237 | 0 | 0 |
| NC_045512.2;YP_009724389.1;Wu1-SARS2_orf1ab;238 | 0 | 0 |
| NC_045512.2;YP_009724389.1;Wu1-SARS2_orf1ab;239 | 0 | 0 |
| NC_045512.2;YP_009724389.1;Wu1-SARS2_orf1ab;240 | 0 | 0.025 |
| NC_045512.2;YP_009724389.1;Wu1-SARS2_orf1ab;241 | 0 | 0.025 |
| NC_045512.2;YP_009724389.1;Wu1-SARS2_orf1ab;242 | 0 | 0.05 |
| NC_045512.2;YP_009724389.1;Wu1-SARS2_orf1ab;243 | 0 | 0.05 |
| NC_045512.2;YP_009724389.1;Wu1-SARS2_orf1ab;244 | 0 | 0.05 |
| NC_045512.2;YP_009724389.1;Wu1-SARS2_orf1ab;245 | 0 | 0.05 |
| NC_045512.2;YP_009724389.1;Wu1-SARS2_orf1ab;246 | 0 | 0.075 |
| NC_045512.2;YP_009724389.1;Wu1-SARS2_orf1ab;247 | 0 | 0.05 |
| NC_045512.2;YP_009724389.1;Wu1-SARS2_orf1ab;248 | 0 | 0.025 |

| | | |
|---|---|---|
| NC_045512.2;YP_009724389.1;Wu1-SARS2_orf1ab;249 | 0 | 0.05 |
| NC_045512.2;YP_009724389.1;Wu1-SARS2_orf1ab;250 | 0 | 0.025 |
| NC_045512.2;YP_009724389.1;Wu1-SARS2_orf1ab;251 | 0 | 0.025 |
| NC_045512.2;YP_009724389.1;Wu1-SARS2_orf1ab;252 | 0.05 | 0 |
| NC_045512.2;YP_009724389.1;Wu1-SARS2_orf1ab;253 | 0.05 | 0 |
| NC_045512.2;YP_009724389.1;Wu1-SARS2_orf1ab;254 | 0 | 0 |
| NC_045512.2;YP_009724389.1;Wu1-SARS2_orf1ab;255 | 0 | 0.05 |
| NC_045512.2;YP_009724389.1;Wu1-SARS2_orf1ab;256 | 0.1 | 0.1 |
| NC_045512.2;YP_009724389.1;Wu1-SARS2_orf1ab;257 | 0 | 0 |
| NC_045512.2;YP_009724389.1;Wu1-SARS2_orf1ab;258 | 0.05 | 0 |
| NC_045512.2;YP_009724389.1;Wu1-SARS2_orf1ab;259 | 0.05 | 0.05 |
| NC_045512.2;YP_009724389.1;Wu1-SARS2_orf1ab;260 | 0.05 | 0.025 |
| NC_045512.2;YP_009724389.1;Wu1-SARS2_orf1ab;261 | 0.05 | 0.025 |
| NC_045512.2;YP_009724389.1;Wu1-SARS2_orf1ab;262 | 0 | 0 |
| NC_045512.2;YP_009724389.1;Wu1-SARS2_orf1ab;263 | 0 | 0.025 |
| NC_045512.2;YP_009724389.1;Wu1-SARS2_orf1ab;264 | 0 | 0.025 |
| NC_045512.2;YP_009724389.1;Wu1-SARS2_orf1ab;265 | 0 | 0 |
| NC_045512.2;YP_009724389.1;Wu1-SARS2_orf1ab;266 | 0 | 0 |
| NC_045512.2;YP_009724389.1;Wu1-SARS2_orf1ab;267 | 0 | 0 |
| NC_045512.2;YP_009724389.1;Wu1-SARS2_orf1ab;268 | 0 | 0 |
| NC_045512.2;YP_009724389.1;Wu1-SARS2_orf1ab;269 | 0 | 0 |
| NC_045512.2;YP_009724389.1;Wu1-SARS2_orf1ab;270 | 0 | 0 |
| NC_045512.2;YP_009724389.1;Wu1-SARS2_orf1ab;271 | 0 | 0 |
| NC_045512.2;YP_009724389.1;Wu1-SARS2_orf1ab;272 | 0 | 0 |
| NC_045512.2;YP_009724389.1;Wu1-SARS2_orf1ab;273 | 0 | 0 |
| NC_045512.2;YP_009724389.1;Wu1-SARS2_orf1ab;274 | 0 | 0 |
| NC_045512.2;YP_009724389.1;Wu1-SARS2_orf1ab;275 | 0 | 0.025 |
| NC_045512.2;YP_009724389.1;Wu1-SARS2_orf1ab;276 | 0.1 | 0.025 |
| NC_045512.2;YP_009724389.1;Wu1-SARS2_orf1ab;277 | 0.05 | 0 |
| NC_045512.2;YP_009724389.1;Wu1-SARS2_orf1ab;278 | 0.05 | 0 |
| NC_045512.2;YP_009724389.1;Wu1-SARS2_orf1ab;279 | 0.05 | 0 |
| NC_045512.2;YP_009724389.1;Wu1-SARS2_orf1ab;280 | 0.05 | 0 |
| NC_045512.2;YP_009724389.1;Wu1-SARS2_orf1ab;281 | 0.1 | 0 |
| NC_045512.2;YP_009724389.1;Wu1-SARS2_orf1ab;282 | 0.05 | 0 |
| NC_045512.2;YP_009724389.1;Wu1-SARS2_orf1ab;283 | 0.1 | 0.025 |
| NC_045512.2;YP_009724389.1;Wu1-SARS2_orf1ab;284 | 0.05 | 0.025 |
| NC_045512.2;YP_009724389.1;Wu1-SARS2_orf1ab;285 | 0.05 | 0.025 |
| NC_045512.2;YP_009724389.1;Wu1-SARS2_orf1ab;286 | 0.05 | 0.025 |
| NC_045512.2;YP_009724389.1;Wu1-SARS2_orf1ab;287 | 0 | 0 |
| NC_045512.2;YP_009724389.1;Wu1-SARS2_orf1ab;288 | 0 | 0 |
| NC_045512.2;YP_009724389.1;Wu1-SARS2_orf1ab;289 | 0 | 0 |
| NC_045512.2;YP_009724389.1;Wu1-SARS2_orf1ab;290 | 0 | 0 |
| NC_045512.2;YP_009724389.1;Wu1-SARS2_orf1ab;291 | 0.05 | 0 |

| | | |
|---|---|---|
| NC_045512.2;YP_009724389.1;Wu1-SARS2_orf1ab;292 | 0 | 0 |
| NC_045512.2;YP_009724389.1;Wu1-SARS2_orf1ab;293 | 0 | 0 |
| NC_045512.2;YP_009724389.1;Wu1-SARS2_orf1ab;294 | 0 | 0 |
| NC_045512.2;YP_009724389.1;Wu1-SARS2_orf1ab;295 | 0 | 0 |
| NC_045512.2;YP_009724389.1;Wu1-SARS2_orf1ab;296 | 0.1 | 0 |
| NC_045512.2;YP_009724389.1;Wu1-SARS2_orf1ab;297 | 0.1 | 0 |
| NC_045512.2;YP_009724389.1;Wu1-SARS2_orf1ab;298 | 0.1 | 0 |
| NC_045512.2;YP_009724389.1;Wu1-SARS2_orf1ab;299 | 0.1 | 0 |
| NC_045512.2;YP_009724389.1;Wu1-SARS2_orf1ab;300 | 0.1 | 0 |
| NC_045512.2;YP_009724389.1;Wu1-SARS2_orf1ab;301 | 0.1 | 0.1 |
| NC_045512.2;YP_009724389.1;Wu1-SARS2_orf1ab;302 | 0.05 | 0 |
| NC_045512.2;YP_009724389.1;Wu1-SARS2_orf1ab;303 | 0.05 | 0 |
| NC_045512.2;YP_009724389.1;Wu1-SARS2_orf1ab;304 | 0.1 | 0 |
| NC_045512.2;YP_009724389.1;Wu1-SARS2_orf1ab;305 | 0.05 | 0 |
| NC_045512.2;YP_009724389.1;Wu1-SARS2_orf1ab;306 | 0 | 0 |
| NC_045512.2;YP_009724389.1;Wu1-SARS2_orf1ab;307 | 0 | 0 |
| NC_045512.2;YP_009724389.1;Wu1-SARS2_orf1ab;308 | 0 | 0 |
| NC_045512.2;YP_009724389.1;Wu1-SARS2_orf1ab;309 | 0 | 0 |
| NC_045512.2;YP_009724389.1;Wu1-SARS2_orf1ab;310 | 0 | 0 |
| NC_045512.2;YP_009724389.1;Wu1-SARS2_orf1ab;311 | 0 | 0 |
| NC_045512.2;YP_009724389.1;Wu1-SARS2_orf1ab;312 | 0 | 0 |
| NC_045512.2;YP_009724389.1;Wu1-SARS2_orf1ab;313 | 0 | 0 |
| NC_045512.2;YP_009724389.1;Wu1-SARS2_orf1ab;314 | 0 | 0 |
| NC_045512.2;YP_009724389.1;Wu1-SARS2_orf1ab;315 | 0 | 0 |
| NC_045512.2;YP_009724389.1;Wu1-SARS2_orf1ab;316 | 0 | 0 |
| NC_045512.2;YP_009724389.1;Wu1-SARS2_orf1ab;317 | 0 | 0 |
| NC_045512.2;YP_009724389.1;Wu1-SARS2_orf1ab;318 | 0 | 0 |
| NC_045512.2;YP_009724389.1;Wu1-SARS2_orf1ab;319 | 0 | 0 |
| NC_045512.2;YP_009724389.1;Wu1-SARS2_orf1ab;320 | 0 | 0 |
| NC_045512.2;YP_009724389.1;Wu1-SARS2_orf1ab;321 | 0 | 0 |
| NC_045512.2;YP_009724389.1;Wu1-SARS2_orf1ab;322 | 0 | 0 |
| NC_045512.2;YP_009724389.1;Wu1-SARS2_orf1ab;323 | 0 | 0 |
| NC_045512.2;YP_009724389.1;Wu1-SARS2_orf1ab;324 | 0.05 | 0 |
| NC_045512.2;YP_009724389.1;Wu1-SARS2_orf1ab;325 | 0 | 0.05 |
| NC_045512.2;YP_009724389.1;Wu1-SARS2_orf1ab;326 | 0 | 0 |
| NC_045512.2;YP_009724389.1;Wu1-SARS2_orf1ab;327 | 0 | 0 |
| NC_045512.2;YP_009724389.1;Wu1-SARS2_orf1ab;328 | 0 | 0 |
| NC_045512.2;YP_009724389.1;Wu1-SARS2_orf1ab;329 | 0 | 0 |
| NC_045512.2;YP_009724389.1;Wu1-SARS2_orf1ab;330 | 0 | 0 |
| NC_045512.2;YP_009724389.1;Wu1-SARS2_orf1ab;331 | 0 | 0 |
| NC_045512.2;YP_009724389.1;Wu1-SARS2_orf1ab;332 | 0 | 0 |
| NC_045512.2;YP_009724389.1;Wu1-SARS2_orf1ab;333 | 0 | 0 |
| NC_045512.2;YP_009724389.1;Wu1-SARS2_orf1ab;334 | 0 | 0 |

| | | |
|---|---|---|
| NC_045512.2;YP_009724389.1;Wu1-SARS2_orf1ab;335 | 0 | 0 |
| NC_045512.2;YP_009724389.1;Wu1-SARS2_orf1ab;336 | 0 | 0 |
| NC_045512.2;YP_009724389.1;Wu1-SARS2_orf1ab;337 | 0 | 0 |
| NC_045512.2;YP_009724389.1;Wu1-SARS2_orf1ab;338 | 0 | 0 |
| NC_045512.2;YP_009724389.1;Wu1-SARS2_orf1ab;339 | 0 | 0 |
| NC_045512.2;YP_009724389.1;Wu1-SARS2_orf1ab;340 | 0 | 0 |
| NC_045512.2;YP_009724389.1;Wu1-SARS2_orf1ab;341 | 0 | 0 |
| NC_045512.2;YP_009724389.1;Wu1-SARS2_orf1ab;342 | 0 | 0 |
| NC_045512.2;YP_009724389.1;Wu1-SARS2_orf1ab;343 | 0 | 0 |
| NC_045512.2;YP_009724389.1;Wu1-SARS2_orf1ab;344 | 0 | 0 |
| NC_045512.2;YP_009724389.1;Wu1-SARS2_orf1ab;345 | 0 | 0 |
| NC_045512.2;YP_009724389.1;Wu1-SARS2_orf1ab;346 | 0 | 0 |
| NC_045512.2;YP_009724389.1;Wu1-SARS2_orf1ab;347 | 0.05 | 0 |
| NC_045512.2;YP_009724389.1;Wu1-SARS2_orf1ab;348 | 0 | 0 |
| NC_045512.2;YP_009724389.1;Wu1-SARS2_orf1ab;349 | 0.05 | 0 |
| NC_045512.2;YP_009724389.1;Wu1-SARS2_orf1ab;350 | 0 | 0 |
| NC_045512.2;YP_009724389.1;Wu1-SARS2_orf1ab;351 | 0 | 0 |
| NC_045512.2;YP_009724389.1;Wu1-SARS2_orf1ab;352 | 0 | 0.025 |
| NC_045512.2;YP_009724389.1;Wu1-SARS2_orf1ab;353 | 0 | 0 |
| NC_045512.2;YP_009724389.1;Wu1-SARS2_orf1ab;354 | 0 | 0 |
| NC_045512.2;YP_009724389.1;Wu1-SARS2_orf1ab;355 | 0 | 0 |
| NC_045512.2;YP_009724389.1;Wu1-SARS2_orf1ab;356 | 0 | 0 |
| NC_045512.2;YP_009724389.1;Wu1-SARS2_orf1ab;357 | 0 | 0 |
| NC_045512.2;YP_009724389.1;Wu1-SARS2_orf1ab;358 | 0 | 0 |
| NC_045512.2;YP_009724389.1;Wu1-SARS2_orf1ab;359 | 0 | 0 |
| NC_045512.2;YP_009724389.1;Wu1-SARS2_orf1ab;360 | 0 | 0 |
| NC_045512.2;YP_009724389.1;Wu1-SARS2_orf1ab;361 | 0 | 0 |
| NC_045512.2;YP_009724389.1;Wu1-SARS2_orf1ab;362 | 0 | 0 |
| NC_045512.2;YP_009724389.1;Wu1-SARS2_orf1ab;363 | 0 | 0 |
| NC_045512.2;YP_009724389.1;Wu1-SARS2_orf1ab;364 | 0 | 0 |
| NC_045512.2;YP_009724389.1;Wu1-SARS2_orf1ab;365 | 0 | 0 |
| NC_045512.2;YP_009724389.1;Wu1-SARS2_orf1ab;366 | 0 | 0 |
| NC_045512.2;YP_009724389.1;Wu1-SARS2_orf1ab;367 | 0 | 0 |
| NC_045512.2;YP_009724389.1;Wu1-SARS2_orf1ab;368 | 0 | 0 |
| NC_045512.2;YP_009724389.1;Wu1-SARS2_orf1ab;369 | 0 | 0 |
| NC_045512.2;YP_009724389.1;Wu1-SARS2_orf1ab;370 | 0 | 0 |
| NC_045512.2;YP_009724389.1;Wu1-SARS2_orf1ab;371 | 0 | 0 |
| NC_045512.2;YP_009724389.1;Wu1-SARS2_orf1ab;372 | 0 | 0 |
| NC_045512.2;YP_009724389.1;Wu1-SARS2_orf1ab;373 | 0 | 0 |
| NC_045512.2;YP_009724389.1;Wu1-SARS2_orf1ab;374 | 0 | 0.025 |
| NC_045512.2;YP_009724389.1;Wu1-SARS2_orf1ab;375 | 0 | 0 |
| NC_045512.2;YP_009724389.1;Wu1-SARS2_orf1ab;376 | 0 | 0 |
| NC_045512.2;YP_009724389.1;Wu1-SARS2_orf1ab;377 | 0 | 0 |

| | | |
|---|---|---|
| NC_045512.2;YP_009724389.1;Wu1-SARS2_orf1ab;378 | 0 | 0 |
| NC_045512.2;YP_009724389.1;Wu1-SARS2_orf1ab;379 | 0.15 | 0.05 |
| NC_045512.2;YP_009724389.1;Wu1-SARS2_orf1ab;380 | 0 | 0.025 |
| NC_045512.2;YP_009724389.1;Wu1-SARS2_orf1ab;381 | 0 | 0.025 |
| NC_045512.2;YP_009724389.1;Wu1-SARS2_orf1ab;382 | 0 | 0.025 |
| NC_045512.2;YP_009724389.1;Wu1-SARS2_orf1ab;383 | 0 | 0.025 |
| NC_045512.2;YP_009724389.1;Wu1-SARS2_orf1ab;384 | 0 | 0.025 |
| NC_045512.2;YP_009724389.1;Wu1-SARS2_orf1ab;385 | 0.15 | 0.075 |
| NC_045512.2;YP_009724389.1;Wu1-SARS2_orf1ab;386 | 0.05 | 0.075 |
| NC_045512.2;YP_009724389.1;Wu1-SARS2_orf1ab;387 | 0 | 0 |
| NC_045512.2;YP_009724389.1;Wu1-SARS2_orf1ab;388 | 0 | 0 |
| NC_045512.2;YP_009724389.1;Wu1-SARS2_orf1ab;389 | 0 | 0 |
| NC_045512.2;YP_009724389.1;Wu1-SARS2_orf1ab;390 | 0 | 0 |
| NC_045512.2;YP_009724389.1;Wu1-SARS2_orf1ab;391 | 0 | 0 |
| NC_045512.2;YP_009724389.1;Wu1-SARS2_orf1ab;392 | 0 | 0 |
| NC_045512.2;YP_009724389.1;Wu1-SARS2_orf1ab;393 | 0 | 0 |
| NC_045512.2;YP_009724389.1;Wu1-SARS2_orf1ab;394 | 0 | 0 |
| NC_045512.2;YP_009724389.1;Wu1-SARS2_orf1ab;395 | 0 | 0 |
| NC_045512.2;YP_009724389.1;Wu1-SARS2_orf1ab;396 | 0 | 0 |
| NC_045512.2;YP_009724389.1;Wu1-SARS2_orf1ab;397 | 0 | 0.025 |
| NC_045512.2;YP_009724389.1;Wu1-SARS2_orf1ab;398 | 0 | 0 |
| NC_045512.2;YP_009724389.1;Wu1-SARS2_orf1ab;399 | 0 | 0 |
| NC_045512.2;YP_009724389.1;Wu1-SARS2_orf1ab;400 | 0 | 0 |
| NC_045512.2;YP_009724389.1;Wu1-SARS2_orf1ab;401 | 0 | 0 |
| NC_045512.2;YP_009724389.1;Wu1-SARS2_orf1ab;402 | 0 | 0 |
| NC_045512.2;YP_009724389.1;Wu1-SARS2_orf1ab;403 | 0 | 0 |
| NC_045512.2;YP_009724389.1;Wu1-SARS2_orf1ab;404 | 0 | 0 |
| NC_045512.2;YP_009724389.1;Wu1-SARS2_orf1ab;405 | 0 | 0 |
| NC_045512.2;YP_009724389.1;Wu1-SARS2_orf1ab;406 | 0 | 0 |
| NC_045512.2;YP_009724389.1;Wu1-SARS2_orf1ab;407 | 0 | 0 |
| NC_045512.2;YP_009724389.1;Wu1-SARS2_orf1ab;408 | 0 | 0 |
| NC_045512.2;YP_009724389.1;Wu1-SARS2_orf1ab;409 | 0 | 0 |
| NC_045512.2;YP_009724389.1;Wu1-SARS2_orf1ab;410 | 0 | 0 |
| NC_045512.2;YP_009724389.1;Wu1-SARS2_orf1ab;411 | 0 | 0 |
| NC_045512.2;YP_009724389.1;Wu1-SARS2_orf1ab;412 | 0 | 0 |
| NC_045512.2;YP_009724389.1;Wu1-SARS2_orf1ab;413 | 0 | 0 |
| NC_045512.2;YP_009724389.1;Wu1-SARS2_orf1ab;414 | 0 | 0 |
| NC_045512.2;YP_009724389.1;Wu1-SARS2_orf1ab;415 | 0 | 0 |
| NC_045512.2;YP_009724389.1;Wu1-SARS2_orf1ab;416 | 0 | 0 |
| NC_045512.2;YP_009724389.1;Wu1-SARS2_orf1ab;417 | 0 | 0 |
| NC_045512.2;YP_009724389.1;Wu1-SARS2_orf1ab;418 | 0 | 0 |
| NC_045512.2;YP_009724389.1;Wu1-SARS2_orf1ab;419 | 0 | 0 |
| NC_045512.2;YP_009724389.1;Wu1-SARS2_orf1ab;420 | 0 | 0 |

| | | |
|---|---|---|
| NC_045512.2;YP_009724389.1;Wu1-SARS2_orf1ab;421 | 0 | 0 |
| NC_045512.2;YP_009724389.1;Wu1-SARS2_orf1ab;422 | 0 | 0 |
| NC_045512.2;YP_009724389.1;Wu1-SARS2_orf1ab;423 | 0 | 0 |
| NC_045512.2;YP_009724389.1;Wu1-SARS2_orf1ab;424 | 0.05 | 0.025 |
| NC_045512.2;YP_009724389.1;Wu1-SARS2_orf1ab;425 | 0.05 | 0.025 |
| NC_045512.2;YP_009724389.1;Wu1-SARS2_orf1ab;426 | 0 | 0 |
| NC_045512.2;YP_009724389.1;Wu1-SARS2_orf1ab;427 | 0 | 0.025 |
| NC_045512.2;YP_009724389.1;Wu1-SARS2_orf1ab;428 | 0 | 0 |
| NC_045512.2;YP_009724389.1;Wu1-SARS2_orf1ab;429 | 0 | 0 |
| NC_045512.2;YP_009724389.1;Wu1-SARS2_orf1ab;430 | 0 | 0 |
| NC_045512.2;YP_009724389.1;Wu1-SARS2_orf1ab;431 | 0 | 0 |
| NC_045512.2;YP_009724389.1;Wu1-SARS2_orf1ab;432 | 0 | 0 |
| NC_045512.2;YP_009724389.1;Wu1-SARS2_orf1ab;433 | 0 | 0 |
| NC_045512.2;YP_009724389.1;Wu1-SARS2_orf1ab;434 | 0 | 0 |
| NC_045512.2;YP_009724389.1;Wu1-SARS2_orf1ab;435 | 0 | 0 |
| NC_045512.2;YP_009724389.1;Wu1-SARS2_orf1ab;436 | 0 | 0 |
| NC_045512.2;YP_009724389.1;Wu1-SARS2_orf1ab;437 | 0 | 0 |
| NC_045512.2;YP_009724389.1;Wu1-SARS2_orf1ab;438 | 0 | 0 |
| NC_045512.2;YP_009724389.1;Wu1-SARS2_orf1ab;439 | 0 | 0 |
| NC_045512.2;YP_009724389.1;Wu1-SARS2_orf1ab;440 | 0 | 0 |
| NC_045512.2;YP_009724389.1;Wu1-SARS2_orf1ab;441 | 0 | 0 |
| NC_045512.2;YP_009724389.1;Wu1-SARS2_orf1ab;442 | 0 | 0 |
| NC_045512.2;YP_009724389.1;Wu1-SARS2_orf1ab;443 | 0.05 | 0 |
| NC_045512.2;YP_009724389.1;Wu1-SARS2_orf1ab;444 | 0.2 | 0.025 |
| NC_045512.2;YP_009724389.1;Wu1-SARS2_orf1ab;445 | 0.5 | 0.475 |
| NC_045512.2;YP_009724389.1;Wu1-SARS2_orf1ab;446 | 0.05 | 0.05 |
| NC_045512.2;YP_009724389.1;Wu1-SARS2_orf1ab;447 | 0.05 | 0.05 |
| NC_045512.2;YP_009724389.1;Wu1-SARS2_orf1ab;448 | 0.05 | 0.05 |
| NC_045512.2;YP_009724389.1;Wu1-SARS2_orf1ab;449 | 0 | 0.025 |
| NC_045512.2;YP_009724389.1;Wu1-SARS2_orf1ab;450 | 0 | 0 |
| NC_045512.2;YP_009724389.1;Wu1-SARS2_orf1ab;451 | 0 | 0 |
| NC_045512.2;YP_009724389.1;Wu1-SARS2_orf1ab;452 | 0.05 | 0 |
| NC_045512.2;YP_009724389.1;Wu1-SARS2_orf1ab;453 | 0 | 0 |
| NC_045512.2;YP_009724389.1;Wu1-SARS2_orf1ab;454 | 0 | 0 |
| NC_045512.2;YP_009724389.1;Wu1-SARS2_orf1ab;455 | 0 | 0 |
| NC_045512.2;YP_009724389.1;Wu1-SARS2_orf1ab;456 | 0.05 | 0 |
| NC_045512.2;YP_009724389.1;Wu1-SARS2_orf1ab;457 | 0.05 | 0 |
| NC_045512.2;YP_009724389.1;Wu1-SARS2_orf1ab;458 | 0.05 | 0 |
| NC_045512.2;YP_009724389.1;Wu1-SARS2_orf1ab;459 | 0 | 0 |
| NC_045512.2;YP_009724389.1;Wu1-SARS2_orf1ab;460 | 0 | 0 |
| NC_045512.2;YP_009724389.1;Wu1-SARS2_orf1ab;461 | 0 | 0 |
| NC_045512.2;YP_009724389.1;Wu1-SARS2_orf1ab;462 | 0 | 0 |
| NC_045512.2;YP_009724389.1;Wu1-SARS2_orf1ab;463 | 0 | 0 |

| | | |
|---|---|---|
| NC_045512.2;YP_009724389.1;Wu1-SARS2_orf1ab;464 | 0 | 0 |
| NC_045512.2;YP_009724389.1;Wu1-SARS2_orf1ab;465 | 0 | 0 |
| NC_045512.2;YP_009724389.1;Wu1-SARS2_orf1ab;466 | 0 | 0 |
| NC_045512.2;YP_009724389.1;Wu1-SARS2_orf1ab;467 | 0 | 0 |
| NC_045512.2;YP_009724389.1;Wu1-SARS2_orf1ab;468 | 0 | 0 |
| NC_045512.2;YP_009724389.1;Wu1-SARS2_orf1ab;469 | 0 | 0 |
| NC_045512.2;YP_009724389.1;Wu1-SARS2_orf1ab;470 | 0 | 0 |
| NC_045512.2;YP_009724389.1;Wu1-SARS2_orf1ab;471 | 0 | 0 |
| NC_045512.2;YP_009724389.1;Wu1-SARS2_orf1ab;472 | 0 | 0 |
| NC_045512.2;YP_009724389.1;Wu1-SARS2_orf1ab;473 | 0 | 0 |
| NC_045512.2;YP_009724389.1;Wu1-SARS2_orf1ab;474 | 0 | 0 |
| NC_045512.2;YP_009724389.1;Wu1-SARS2_orf1ab;475 | 0 | 0 |
| NC_045512.2;YP_009724389.1;Wu1-SARS2_orf1ab;476 | 0 | 0 |
| NC_045512.2;YP_009724389.1;Wu1-SARS2_orf1ab;477 | 0 | 0 |
| NC_045512.2;YP_009724389.1;Wu1-SARS2_orf1ab;478 | 0 | 0 |
| NC_045512.2;YP_009724389.1;Wu1-SARS2_orf1ab;479 | 0 | 0 |
| NC_045512.2;YP_009724389.1;Wu1-SARS2_orf1ab;480 | 0 | 0.05 |
| NC_045512.2;YP_009724389.1;Wu1-SARS2_orf1ab;481 | 0 | 0.05 |
| NC_045512.2;YP_009724389.1;Wu1-SARS2_orf1ab;482 | 0 | 0.025 |
| NC_045512.2;YP_009724389.1;Wu1-SARS2_orf1ab;483 | 0 | 0.025 |
| NC_045512.2;YP_009724389.1;Wu1-SARS2_orf1ab;484 | 0 | 0 |
| NC_045512.2;YP_009724389.1;Wu1-SARS2_orf1ab;485 | 0 | 0 |
| NC_045512.2;YP_009724389.1;Wu1-SARS2_orf1ab;486 | 0 | 0.025 |
| NC_045512.2;YP_009724389.1;Wu1-SARS2_orf1ab;487 | 0 | 0 |
| NC_045512.2;YP_009724389.1;Wu1-SARS2_orf1ab;488 | 0 | 0 |
| NC_045512.2;YP_009724389.1;Wu1-SARS2_orf1ab;489 | 0.05 | 0.025 |
| NC_045512.2;YP_009724389.1;Wu1-SARS2_orf1ab;490 | 0 | 0 |
| NC_045512.2;YP_009724389.1;Wu1-SARS2_orf1ab;491 | 0 | 0 |
| NC_045512.2;YP_009724389.1;Wu1-SARS2_orf1ab;492 | 0 | 0 |
| NC_045512.2;YP_009724389.1;Wu1-SARS2_orf1ab;493 | 0.05 | 0.05 |
| NC_045512.2;YP_009724389.1;Wu1-SARS2_orf1ab;494 | 0.05 | 0.025 |
| NC_045512.2;YP_009724389.1;Wu1-SARS2_orf1ab;495 | 0 | 0 |
| NC_045512.2;YP_009724389.1;Wu1-SARS2_orf1ab;496 | 0 | 0 |
| NC_045512.2;YP_009724389.1;Wu1-SARS2_orf1ab;497 | 0 | 0 |
| NC_045512.2;YP_009724389.1;Wu1-SARS2_orf1ab;498 | 0 | 0 |
| NC_045512.2;YP_009724389.1;Wu1-SARS2_orf1ab;499 | 0 | 0 |
| NC_045512.2;YP_009724389.1;Wu1-SARS2_orf1ab;500 | 0 | 0 |
| NC_045512.2;YP_009724389.1;Wu1-SARS2_orf1ab;501 | 0 | 0 |
| NC_045512.2;YP_009724389.1;Wu1-SARS2_orf1ab;502 | 0 | 0 |
| NC_045512.2;YP_009724389.1;Wu1-SARS2_orf1ab;503 | 0 | 0 |
| NC_045512.2;YP_009724389.1;Wu1-SARS2_orf1ab;504 | 0 | 0 |
| NC_045512.2;YP_009724389.1;Wu1-SARS2_orf1ab;505 | 0 | 0 |
| NC_045512.2;YP_009724389.1;Wu1-SARS2_orf1ab;506 | 0 | 0 |

| | | |
|---|---|---|
| NC_045512.2;YP_009724389.1;Wu1-SARS2_orf1ab;507 | 0 | 0 |
| NC_045512.2;YP_009724389.1;Wu1-SARS2_orf1ab;508 | 0 | 0 |
| NC_045512.2;YP_009724389.1;Wu1-SARS2_orf1ab;509 | 0 | 0 |
| NC_045512.2;YP_009724389.1;Wu1-SARS2_orf1ab;510 | 0 | 0 |
| NC_045512.2;YP_009724389.1;Wu1-SARS2_orf1ab;511 | 0 | 0 |
| NC_045512.2;YP_009724389.1;Wu1-SARS2_orf1ab;512 | 0 | 0 |
| NC_045512.2;YP_009724389.1;Wu1-SARS2_orf1ab;513 | 0 | 0.025 |
| NC_045512.2;YP_009724389.1;Wu1-SARS2_orf1ab;514 | 0 | 0.025 |
| NC_045512.2;YP_009724389.1;Wu1-SARS2_orf1ab;515 | 0 | 0.025 |
| NC_045512.2;YP_009724389.1;Wu1-SARS2_orf1ab;516 | 0 | 0.025 |
| NC_045512.2;YP_009724389.1;Wu1-SARS2_orf1ab;517 | 0 | 0 |
| NC_045512.2;YP_009724389.1;Wu1-SARS2_orf1ab;518 | 0.1 | 0.025 |
| NC_045512.2;YP_009724389.1;Wu1-SARS2_orf1ab;519 | 0 | 0 |
| NC_045512.2;YP_009724389.1;Wu1-SARS2_orf1ab;520 | 0 | 0 |
| NC_045512.2;YP_009724389.1;Wu1-SARS2_orf1ab;521 | 0 | 0 |
| NC_045512.2;YP_009724389.1;Wu1-SARS2_orf1ab;522 | 0 | 0 |
| NC_045512.2;YP_009724389.1;Wu1-SARS2_orf1ab;523 | 0 | 0 |
| NC_045512.2;YP_009724389.1;Wu1-SARS2_orf1ab;524 | 0 | 0 |
| NC_045512.2;YP_009724389.1;Wu1-SARS2_orf1ab;525 | 0.05 | 0 |
| NC_045512.2;YP_009724389.1;Wu1-SARS2_orf1ab;526 | 0.05 | 0.025 |
| NC_045512.2;YP_009724389.1;Wu1-SARS2_orf1ab;527 | 0 | 0 |
| NC_045512.2;YP_009724389.1;Wu1-SARS2_orf1ab;528 | 0 | 0 |
| NC_045512.2;YP_009724389.1;Wu1-SARS2_orf1ab;529 | 0 | 0 |
| NC_045512.2;YP_009724389.1;Wu1-SARS2_orf1ab;530 | 0 | 0 |
| NC_045512.2;YP_009724389.1;Wu1-SARS2_orf1ab;531 | 0 | 0 |
| NC_045512.2;YP_009724389.1;Wu1-SARS2_orf1ab;532 | 0 | 0 |
| NC_045512.2;YP_009724389.1;Wu1-SARS2_orf1ab;533 | 0 | 0 |
| NC_045512.2;YP_009724389.1;Wu1-SARS2_orf1ab;534 | 0 | 0 |
| NC_045512.2;YP_009724389.1;Wu1-SARS2_orf1ab;535 | 0 | 0 |
| NC_045512.2;YP_009724389.1;Wu1-SARS2_orf1ab;536 | 0 | 0 |
| NC_045512.2;YP_009724389.1;Wu1-SARS2_orf1ab;537 | 0 | 0 |
| NC_045512.2;YP_009724389.1;Wu1-SARS2_orf1ab;538 | 0 | 0 |
| NC_045512.2;YP_009724389.1;Wu1-SARS2_orf1ab;539 | 0 | 0 |
| NC_045512.2;YP_009724389.1;Wu1-SARS2_orf1ab;540 | 0 | 0 |
| NC_045512.2;YP_009724389.1;Wu1-SARS2_orf1ab;541 | 0 | 0 |
| NC_045512.2;YP_009724389.1;Wu1-SARS2_orf1ab;542 | 0 | 0 |
| NC_045512.2;YP_009724389.1;Wu1-SARS2_orf1ab;543 | 0 | 0 |
| NC_045512.2;YP_009724389.1;Wu1-SARS2_orf1ab;544 | 0 | 0 |
| NC_045512.2;YP_009724389.1;Wu1-SARS2_orf1ab;545 | 0 | 0 |
| NC_045512.2;YP_009724389.1;Wu1-SARS2_orf1ab;546 | 0 | 0.025 |
| NC_045512.2;YP_009724389.1;Wu1-SARS2_orf1ab;547 | 0 | 0 |
| NC_045512.2;YP_009724389.1;Wu1-SARS2_orf1ab;548 | 0 | 0 |
| NC_045512.2;YP_009724389.1;Wu1-SARS2_orf1ab;549 | 0 | 0 |

| | | |
|---|---|---|
| NC_045512.2;YP_009724389.1;Wu1-SARS2_orf1ab;550 | 0 | 0 |
| NC_045512.2;YP_009724389.1;Wu1-SARS2_orf1ab;551 | 0.05 | 0 |
| NC_045512.2;YP_009724389.1;Wu1-SARS2_orf1ab;552 | 0 | 0 |
| NC_045512.2;YP_009724389.1;Wu1-SARS2_orf1ab;553 | 0.05 | 0 |
| NC_045512.2;YP_009724389.1;Wu1-SARS2_orf1ab;554 | 0 | 0 |
| NC_045512.2;YP_009724389.1;Wu1-SARS2_orf1ab;555 | 0.05 | 0 |
| NC_045512.2;YP_009724389.1;Wu1-SARS2_orf1ab;556 | 0 | 0 |
| NC_045512.2;YP_009724389.1;Wu1-SARS2_orf1ab;557 | 0 | 0 |
| NC_045512.2;YP_009724389.1;Wu1-SARS2_orf1ab;558 | 0 | 0 |
| NC_045512.2;YP_009724389.1;Wu1-SARS2_orf1ab;559 | 0 | 0 |
| NC_045512.2;YP_009724389.1;Wu1-SARS2_orf1ab;560 | 0 | 0 |
| NC_045512.2;YP_009724389.1;Wu1-SARS2_orf1ab;561 | 0 | 0 |
| NC_045512.2;YP_009724389.1;Wu1-SARS2_orf1ab;562 | 0 | 0 |
| NC_045512.2;YP_009724389.1;Wu1-SARS2_orf1ab;563 | 0 | 0 |
| NC_045512.2;YP_009724389.1;Wu1-SARS2_orf1ab;564 | 0 | 0 |
| NC_045512.2;YP_009724389.1;Wu1-SARS2_orf1ab;565 | 0 | 0.025 |
| NC_045512.2;YP_009724389.1;Wu1-SARS2_orf1ab;566 | 0 | 0 |
| NC_045512.2;YP_009724389.1;Wu1-SARS2_orf1ab;567 | 0 | 0 |
| NC_045512.2;YP_009724389.1;Wu1-SARS2_orf1ab;568 | 0 | 0 |
| NC_045512.2;YP_009724389.1;Wu1-SARS2_orf1ab;569 | 0 | 0.025 |
| NC_045512.2;YP_009724389.1;Wu1-SARS2_orf1ab;570 | 0 | 0.025 |
| NC_045512.2;YP_009724389.1;Wu1-SARS2_orf1ab;571 | 0 | 0.025 |
| NC_045512.2;YP_009724389.1;Wu1-SARS2_orf1ab;572 | 0.1 | 0.05 |
| NC_045512.2;YP_009724389.1;Wu1-SARS2_orf1ab;573 | 0 | 0.025 |
| NC_045512.2;YP_009724389.1;Wu1-SARS2_orf1ab;574 | 0 | 0 |
| NC_045512.2;YP_009724389.1;Wu1-SARS2_orf1ab;575 | 0 | 0 |
| NC_045512.2;YP_009724389.1;Wu1-SARS2_orf1ab;576 | 0 | 0 |
| NC_045512.2;YP_009724389.1;Wu1-SARS2_orf1ab;577 | 0 | 0 |
| NC_045512.2;YP_009724389.1;Wu1-SARS2_orf1ab;578 | 0 | 0.025 |
| NC_045512.2;YP_009724389.1;Wu1-SARS2_orf1ab;579 | 0 | 0 |
| NC_045512.2;YP_009724389.1;Wu1-SARS2_orf1ab;580 | 0 | 0 |
| NC_045512.2;YP_009724389.1;Wu1-SARS2_orf1ab;581 | 0 | 0 |
| NC_045512.2;YP_009724389.1;Wu1-SARS2_orf1ab;582 | 0 | 0 |
| NC_045512.2;YP_009724389.1;Wu1-SARS2_orf1ab;583 | 0 | 0 |
| NC_045512.2;YP_009724389.1;Wu1-SARS2_orf1ab;584 | 0 | 0 |
| NC_045512.2;YP_009724389.1;Wu1-SARS2_orf1ab;585 | 0 | 0 |
| NC_045512.2;YP_009724389.1;Wu1-SARS2_orf1ab;586 | 0 | 0 |
| NC_045512.2;YP_009724389.1;Wu1-SARS2_orf1ab;587 | 0 | 0 |
| NC_045512.2;YP_009724389.1;Wu1-SARS2_orf1ab;588 | 0 | 0 |
| NC_045512.2;YP_009724389.1;Wu1-SARS2_orf1ab;589 | 0.05 | 0 |
| NC_045512.2;YP_009724389.1;Wu1-SARS2_orf1ab;590 | 0 | 0 |
| NC_045512.2;YP_009724389.1;Wu1-SARS2_orf1ab;591 | 0 | 0 |
| NC_045512.2;YP_009724389.1;Wu1-SARS2_orf1ab;592 | 0 | 0 |

| | | |
|---|---|---|
| NC_045512.2;YP_009724389.1;Wu1-SARS2_orf1ab;593 | 0 | 0 |
| NC_045512.2;YP_009724389.1;Wu1-SARS2_orf1ab;594 | 0 | 0 |
| NC_045512.2;YP_009724389.1;Wu1-SARS2_orf1ab;595 | 0 | 0 |
| NC_045512.2;YP_009724389.1;Wu1-SARS2_orf1ab;596 | 0 | 0 |
| NC_045512.2;YP_009724389.1;Wu1-SARS2_orf1ab;597 | 0 | 0 |
| NC_045512.2;YP_009724389.1;Wu1-SARS2_orf1ab;598 | 0 | 0 |
| NC_045512.2;YP_009724389.1;Wu1-SARS2_orf1ab;599 | 0 | 0 |
| NC_045512.2;YP_009724389.1;Wu1-SARS2_orf1ab;600 | 0 | 0 |
| NC_045512.2;YP_009724389.1;Wu1-SARS2_orf1ab;601 | 0 | 0 |
| NC_045512.2;YP_009724389.1;Wu1-SARS2_orf1ab;602 | 0 | 0 |
| NC_045512.2;YP_009724389.1;Wu1-SARS2_orf1ab;603 | 0 | 0 |
| NC_045512.2;YP_009724389.1;Wu1-SARS2_orf1ab;604 | 0 | 0 |
| NC_045512.2;YP_009724389.1;Wu1-SARS2_orf1ab;605 | 0 | 0 |
| NC_045512.2;YP_009724389.1;Wu1-SARS2_orf1ab;606 | 0 | 0 |
| NC_045512.2;YP_009724389.1;Wu1-SARS2_orf1ab;607 | 0 | 0 |
| NC_045512.2;YP_009724389.1;Wu1-SARS2_orf1ab;608 | 0 | 0 |
| NC_045512.2;YP_009724389.1;Wu1-SARS2_orf1ab;609 | 0 | 0 |
| NC_045512.2;YP_009724389.1;Wu1-SARS2_orf1ab;610 | 0 | 0 |
| NC_045512.2;YP_009724389.1;Wu1-SARS2_orf1ab;611 | 0 | 0 |
| NC_045512.2;YP_009724389.1;Wu1-SARS2_orf1ab;612 | 0 | 0 |
| NC_045512.2;YP_009724389.1;Wu1-SARS2_orf1ab;613 | 0 | 0 |
| NC_045512.2;YP_009724389.1;Wu1-SARS2_orf1ab;614 | 0 | 0.05 |
| NC_045512.2;YP_009724389.1;Wu1-SARS2_orf1ab;615 | 0 | 0 |
| NC_045512.2;YP_009724389.1;Wu1-SARS2_orf1ab;616 | 0 | 0 |
| NC_045512.2;YP_009724389.1;Wu1-SARS2_orf1ab;617 | 0.05 | 0.025 |
| NC_045512.2;YP_009724389.1;Wu1-SARS2_orf1ab;618 | 0.05 | 0.05 |
| NC_045512.2;YP_009724389.1;Wu1-SARS2_orf1ab;619 | 0.05 | 0.025 |
| NC_045512.2;YP_009724389.1;Wu1-SARS2_orf1ab;620 | 0.05 | 0.025 |
| NC_045512.2;YP_009724389.1;Wu1-SARS2_orf1ab;621 | 0.1 | 0.025 |
| NC_045512.2;YP_009724389.1;Wu1-SARS2_orf1ab;622 | 0 | 0.025 |
| NC_045512.2;YP_009724389.1;Wu1-SARS2_orf1ab;623 | 0 | 0.025 |
| NC_045512.2;YP_009724389.1;Wu1-SARS2_orf1ab;624 | 0 | 0.05 |
| NC_045512.2;YP_009724389.1;Wu1-SARS2_orf1ab;625 | 0 | 0 |
| NC_045512.2;YP_009724389.1;Wu1-SARS2_orf1ab;626 | 0 | 0 |
| NC_045512.2;YP_009724389.1;Wu1-SARS2_orf1ab;627 | 0 | 0 |
| NC_045512.2;YP_009724389.1;Wu1-SARS2_orf1ab;628 | 0 | 0.025 |
| NC_045512.2;YP_009724389.1;Wu1-SARS2_orf1ab;629 | 0 | 0.025 |
| NC_045512.2;YP_009724389.1;Wu1-SARS2_orf1ab;630 | 0 | 0.025 |
| NC_045512.2;YP_009724389.1;Wu1-SARS2_orf1ab;631 | 0 | 0.05 |
| NC_045512.2;YP_009724389.1;Wu1-SARS2_orf1ab;632 | 0 | 0.05 |
| NC_045512.2;YP_009724389.1;Wu1-SARS2_orf1ab;633 | 0 | 0.025 |
| NC_045512.2;YP_009724389.1;Wu1-SARS2_orf1ab;634 | 0 | 0 |
| NC_045512.2;YP_009724389.1;Wu1-SARS2_orf1ab;635 | 0 | 0.025 |

| | | |
|---|---|---|
| NC_045512.2;YP_009724389.1;Wu1-SARS2_orf1ab;636 | 0 | 0.05 |
| NC_045512.2;YP_009724389.1;Wu1-SARS2_orf1ab;637 | 0 | 0 |
| NC_045512.2;YP_009724389.1;Wu1-SARS2_orf1ab;638 | 0.05 | 0.025 |
| NC_045512.2;YP_009724389.1;Wu1-SARS2_orf1ab;639 | 0 | 0 |
| NC_045512.2;YP_009724389.1;Wu1-SARS2_orf1ab;640 | 0 | 0 |
| NC_045512.2;YP_009724389.1;Wu1-SARS2_orf1ab;641 | 0 | 0 |
| NC_045512.2;YP_009724389.1;Wu1-SARS2_orf1ab;642 | 0 | 0 |
| NC_045512.2;YP_009724389.1;Wu1-SARS2_orf1ab;643 | 0 | 0 |
| NC_045512.2;YP_009724389.1;Wu1-SARS2_orf1ab;644 | 0 | 0 |
| NC_045512.2;YP_009724389.1;Wu1-SARS2_orf1ab;645 | 0 | 0 |
| NC_045512.2;YP_009724389.1;Wu1-SARS2_orf1ab;646 | 0 | 0 |
| NC_045512.2;YP_009724389.1;Wu1-SARS2_orf1ab;647 | 0 | 0 |
| NC_045512.2;YP_009724389.1;Wu1-SARS2_orf1ab;648 | 0 | 0 |
| NC_045512.2;YP_009724389.1;Wu1-SARS2_orf1ab;649 | 0 | 0 |
| NC_045512.2;YP_009724389.1;Wu1-SARS2_orf1ab;650 | 0 | 0 |
| NC_045512.2;YP_009724389.1;Wu1-SARS2_orf1ab;651 | 0 | 0 |
| NC_045512.2;YP_009724389.1;Wu1-SARS2_orf1ab;652 | 0 | 0 |
| NC_045512.2;YP_009724389.1;Wu1-SARS2_orf1ab;653 | 0 | 0 |
| NC_045512.2;YP_009724389.1;Wu1-SARS2_orf1ab;654 | 0 | 0 |
| NC_045512.2;YP_009724389.1;Wu1-SARS2_orf1ab;655 | 0 | 0 |
| NC_045512.2;YP_009724389.1;Wu1-SARS2_orf1ab;656 | 0 | 0 |
| NC_045512.2;YP_009724389.1;Wu1-SARS2_orf1ab;657 | 0 | 0.025 |
| NC_045512.2;YP_009724389.1;Wu1-SARS2_orf1ab;658 | 0 | 0.05 |
| NC_045512.2;YP_009724389.1;Wu1-SARS2_orf1ab;659 | 0 | 0.025 |
| NC_045512.2;YP_009724389.1;Wu1-SARS2_orf1ab;660 | 0 | 0.025 |
| NC_045512.2;YP_009724389.1;Wu1-SARS2_orf1ab;661 | 0 | 0.025 |
| NC_045512.2;YP_009724389.1;Wu1-SARS2_orf1ab;662 | 0 | 0.025 |
| NC_045512.2;YP_009724389.1;Wu1-SARS2_orf1ab;663 | 0 | 0.025 |
| NC_045512.2;YP_009724389.1;Wu1-SARS2_orf1ab;664 | 0 | 0.025 |
| NC_045512.2;YP_009724389.1;Wu1-SARS2_orf1ab;665 | 0 | 0.025 |
| NC_045512.2;YP_009724389.1;Wu1-SARS2_orf1ab;666 | 0 | 0.025 |
| NC_045512.2;YP_009724389.1;Wu1-SARS2_orf1ab;667 | 0 | 0.025 |
| NC_045512.2;YP_009724389.1;Wu1-SARS2_orf1ab;668 | 0 | 0 |
| NC_045512.2;YP_009724389.1;Wu1-SARS2_orf1ab;669 | 0 | 0 |
| NC_045512.2;YP_009724389.1;Wu1-SARS2_orf1ab;670 | 0 | 0 |
| NC_045512.2;YP_009724389.1;Wu1-SARS2_orf1ab;671 | 0 | 0 |
| NC_045512.2;YP_009724389.1;Wu1-SARS2_orf1ab;672 | 0 | 0 |
| NC_045512.2;YP_009724389.1;Wu1-SARS2_orf1ab;673 | 0 | 0 |
| NC_045512.2;YP_009724389.1;Wu1-SARS2_orf1ab;674 | 0 | 0 |
| NC_045512.2;YP_009724389.1;Wu1-SARS2_orf1ab;675 | 0 | 0 |
| NC_045512.2;YP_009724389.1;Wu1-SARS2_orf1ab;676 | 0 | 0 |
| NC_045512.2;YP_009724389.1;Wu1-SARS2_orf1ab;677 | 0 | 0 |
| NC_045512.2;YP_009724389.1;Wu1-SARS2_orf1ab;678 | 0 | 0 |

| | | |
|---|---|---|
| NC_045512.2;YP_009724389.1;Wu1-SARS2_orf1ab;679 | 0 | 0 |
| NC_045512.2;YP_009724389.1;Wu1-SARS2_orf1ab;680 | 0 | 0 |
| NC_045512.2;YP_009724389.1;Wu1-SARS2_orf1ab;681 | 0 | 0 |
| NC_045512.2;YP_009724389.1;Wu1-SARS2_orf1ab;682 | 0 | 0 |
| NC_045512.2;YP_009724389.1;Wu1-SARS2_orf1ab;683 | 0 | 0 |
| NC_045512.2;YP_009724389.1;Wu1-SARS2_orf1ab;684 | 0 | 0 |
| NC_045512.2;YP_009724389.1;Wu1-SARS2_orf1ab;685 | 0 | 0 |
| NC_045512.2;YP_009724389.1;Wu1-SARS2_orf1ab;686 | 0 | 0 |
| NC_045512.2;YP_009724389.1;Wu1-SARS2_orf1ab;687 | 0 | 0 |
| NC_045512.2;YP_009724389.1;Wu1-SARS2_orf1ab;688 | 0 | 0 |
| NC_045512.2;YP_009724389.1;Wu1-SARS2_orf1ab;689 | 0 | 0 |
| NC_045512.2;YP_009724389.1;Wu1-SARS2_orf1ab;690 | 0 | 0 |
| NC_045512.2;YP_009724389.1;Wu1-SARS2_orf1ab;691 | 0 | 0 |
| NC_045512.2;YP_009724389.1;Wu1-SARS2_orf1ab;692 | 0 | 0 |
| NC_045512.2;YP_009724389.1;Wu1-SARS2_orf1ab;693 | 0 | 0.025 |
| NC_045512.2;YP_009724389.1;Wu1-SARS2_orf1ab;694 | 0 | 0 |
| NC_045512.2;YP_009724389.1;Wu1-SARS2_orf1ab;695 | 0 | 0 |
| NC_045512.2;YP_009724389.1;Wu1-SARS2_orf1ab;696 | 0 | 0.025 |
| NC_045512.2;YP_009724389.1;Wu1-SARS2_orf1ab;697 | 0 | 0 |
| NC_045512.2;YP_009724389.1;Wu1-SARS2_orf1ab;698 | 0 | 0 |
| NC_045512.2;YP_009724389.1;Wu1-SARS2_orf1ab;699 | 0 | 0 |
| NC_045512.2;YP_009724389.1;Wu1-SARS2_orf1ab;700 | 0 | 0 |
| NC_045512.2;YP_009724389.1;Wu1-SARS2_orf1ab;701 | 0 | 0 |
| NC_045512.2;YP_009724389.1;Wu1-SARS2_orf1ab;702 | 0 | 0 |
| NC_045512.2;YP_009724389.1;Wu1-SARS2_orf1ab;703 | 0 | 0 |
| NC_045512.2;YP_009724389.1;Wu1-SARS2_orf1ab;704 | 0 | 0 |
| NC_045512.2;YP_009724389.1;Wu1-SARS2_orf1ab;705 | 0 | 0 |
| NC_045512.2;YP_009724389.1;Wu1-SARS2_orf1ab;706 | 0 | 0 |
| NC_045512.2;YP_009724389.1;Wu1-SARS2_orf1ab;707 | 0 | 0 |
| NC_045512.2;YP_009724389.1;Wu1-SARS2_orf1ab;708 | 0 | 0 |
| NC_045512.2;YP_009724389.1;Wu1-SARS2_orf1ab;709 | 0 | 0 |
| NC_045512.2;YP_009724389.1;Wu1-SARS2_orf1ab;710 | 0 | 0 |
| NC_045512.2;YP_009724389.1;Wu1-SARS2_orf1ab;711 | 0 | 0 |
| NC_045512.2;YP_009724389.1;Wu1-SARS2_orf1ab;712 | 0.05 | 0 |
| NC_045512.2;YP_009724389.1;Wu1-SARS2_orf1ab;713 | 0 | 0 |
| NC_045512.2;YP_009724389.1;Wu1-SARS2_orf1ab;714 | 0 | 0.025 |
| NC_045512.2;YP_009724389.1;Wu1-SARS2_orf1ab;715 | 0 | 0 |
| NC_045512.2;YP_009724389.1;Wu1-SARS2_orf1ab;716 | 0 | 0 |
| NC_045512.2;YP_009724389.1;Wu1-SARS2_orf1ab;717 | 0 | 0 |
| NC_045512.2;YP_009724389.1;Wu1-SARS2_orf1ab;718 | 0 | 0.025 |
| NC_045512.2;YP_009724389.1;Wu1-SARS2_orf1ab;719 | 0 | 0 |
| NC_045512.2;YP_009724389.1;Wu1-SARS2_orf1ab;720 | 0 | 0 |
| NC_045512.2;YP_009724389.1;Wu1-SARS2_orf1ab;721 | 0 | 0.05 |

| | | |
|---|---|---|
| NC_045512.2;YP_009724389.1;Wu1-SARS2_orf1ab;722 | 0 | 0.025 |
| NC_045512.2;YP_009724389.1;Wu1-SARS2_orf1ab;723 | 0 | 0 |
| NC_045512.2;YP_009724389.1;Wu1-SARS2_orf1ab;724 | 0 | 0 |
| NC_045512.2;YP_009724389.1;Wu1-SARS2_orf1ab;725 | 0 | 0 |
| NC_045512.2;YP_009724389.1;Wu1-SARS2_orf1ab;726 | 0.05 | 0 |
| NC_045512.2;YP_009724389.1;Wu1-SARS2_orf1ab;727 | 0 | 0 |
| NC_045512.2;YP_009724389.1;Wu1-SARS2_orf1ab;728 | 0 | 0 |
| NC_045512.2;YP_009724389.1;Wu1-SARS2_orf1ab;729 | 0 | 0 |
| NC_045512.2;YP_009724389.1;Wu1-SARS2_orf1ab;730 | 0 | 0 |
| NC_045512.2;YP_009724389.1;Wu1-SARS2_orf1ab;731 | 0 | 0 |
| NC_045512.2;YP_009724389.1;Wu1-SARS2_orf1ab;732 | 0 | 0 |
| NC_045512.2;YP_009724389.1;Wu1-SARS2_orf1ab;733 | 0 | 0 |
| NC_045512.2;YP_009724389.1;Wu1-SARS2_orf1ab;734 | 0 | 0 |
| NC_045512.2;YP_009724389.1;Wu1-SARS2_orf1ab;735 | 0 | 0 |
| NC_045512.2;YP_009724389.1;Wu1-SARS2_orf1ab;736 | 0 | 0 |
| NC_045512.2;YP_009724389.1;Wu1-SARS2_orf1ab;737 | 0 | 0 |
| NC_045512.2;YP_009724389.1;Wu1-SARS2_orf1ab;738 | 0 | 0 |
| NC_045512.2;YP_009724389.1;Wu1-SARS2_orf1ab;739 | 0 | 0 |
| NC_045512.2;YP_009724389.1;Wu1-SARS2_orf1ab;740 | 0 | 0 |
| NC_045512.2;YP_009724389.1;Wu1-SARS2_orf1ab;741 | 0 | 0 |
| NC_045512.2;YP_009724389.1;Wu1-SARS2_orf1ab;742 | 0 | 0 |
| NC_045512.2;YP_009724389.1;Wu1-SARS2_orf1ab;743 | 0 | 0 |
| NC_045512.2;YP_009724389.1;Wu1-SARS2_orf1ab;744 | 0.15 | 0.05 |
| NC_045512.2;YP_009724389.1;Wu1-SARS2_orf1ab;745 | 0.1 | 0.05 |
| NC_045512.2;YP_009724389.1;Wu1-SARS2_orf1ab;746 | 0.15 | 0.025 |
| NC_045512.2;YP_009724389.1;Wu1-SARS2_orf1ab;747 | 0.1 | 0.025 |
| NC_045512.2;YP_009724389.1;Wu1-SARS2_orf1ab;748 | 0.1 | 0.05 |
| NC_045512.2;YP_009724389.1;Wu1-SARS2_orf1ab;749 | 0.15 | 0.075 |
| NC_045512.2;YP_009724389.1;Wu1-SARS2_orf1ab;750 | 0.15 | 0.05 |
| NC_045512.2;YP_009724389.1;Wu1-SARS2_orf1ab;751 | 0.05 | 0.05 |
| NC_045512.2;YP_009724389.1;Wu1-SARS2_orf1ab;752 | 0.05 | 0.05 |
| NC_045512.2;YP_009724389.1;Wu1-SARS2_orf1ab;753 | 0.05 | 0.075 |
| NC_045512.2;YP_009724389.1;Wu1-SARS2_orf1ab;754 | 0.1 | 0.075 |
| NC_045512.2;YP_009724389.1;Wu1-SARS2_orf1ab;755 | 0.05 | 0.05 |
| NC_045512.2;YP_009724389.1;Wu1-SARS2_orf1ab;756 | 0.05 | 0.05 |
| NC_045512.2;YP_009724389.1;Wu1-SARS2_orf1ab;757 | 0.05 | 0.05 |
| NC_045512.2;YP_009724389.1;Wu1-SARS2_orf1ab;758 | 0.05 | 0.05 |
| NC_045512.2;YP_009724389.1;Wu1-SARS2_orf1ab;759 | 0.05 | 0.075 |
| NC_045512.2;YP_009724389.1;Wu1-SARS2_orf1ab;760 | 0 | 0.025 |
| NC_045512.2;YP_009724389.1;Wu1-SARS2_orf1ab;761 | 0 | 0.025 |
| NC_045512.2;YP_009724389.1;Wu1-SARS2_orf1ab;762 | 0 | 0.025 |
| NC_045512.2;YP_009724389.1;Wu1-SARS2_orf1ab;763 | 0 | 0.025 |
| NC_045512.2;YP_009724389.1;Wu1-SARS2_orf1ab;764 | 0.05 | 0.025 |

| | | |
|---|---|---|
| NC_045512.2;YP_009724389.1;Wu1-SARS2_orf1ab;765 | 0.05 | 0.025 |
| NC_045512.2;YP_009724389.1;Wu1-SARS2_orf1ab;766 | 0.05 | 0.075 |
| NC_045512.2;YP_009724389.1;Wu1-SARS2_orf1ab;767 | 0.05 | 0.05 |
| NC_045512.2;YP_009724389.1;Wu1-SARS2_orf1ab;768 | 0 | 0.075 |
| NC_045512.2;YP_009724389.1;Wu1-SARS2_orf1ab;769 | 0 | 0 |
| NC_045512.2;YP_009724389.1;Wu1-SARS2_orf1ab;770 | 0 | 0 |
| NC_045512.2;YP_009724389.1;Wu1-SARS2_orf1ab;771 | 0 | 0 |
| NC_045512.2;YP_009724389.1;Wu1-SARS2_orf1ab;772 | 0 | 0 |
| NC_045512.2;YP_009724389.1;Wu1-SARS2_orf1ab;773 | 0 | 0 |
| NC_045512.2;YP_009724389.1;Wu1-SARS2_orf1ab;774 | 0 | 0 |
| NC_045512.2;YP_009724389.1;Wu1-SARS2_orf1ab;775 | 0.05 | 0 |
| NC_045512.2;YP_009724389.1;Wu1-SARS2_orf1ab;776 | 0 | 0 |
| NC_045512.2;YP_009724389.1;Wu1-SARS2_orf1ab;777 | 0 | 0 |
| NC_045512.2;YP_009724389.1;Wu1-SARS2_orf1ab;778 | 0 | 0 |
| NC_045512.2;YP_009724389.1;Wu1-SARS2_orf1ab;779 | 0 | 0 |
| NC_045512.2;YP_009724389.1;Wu1-SARS2_orf1ab;780 | 0 | 0.025 |
| NC_045512.2;YP_009724389.1;Wu1-SARS2_orf1ab;781 | 0 | 0 |
| NC_045512.2;YP_009724389.1;Wu1-SARS2_orf1ab;782 | 0.05 | 0 |
| NC_045512.2;YP_009724389.1;Wu1-SARS2_orf1ab;783 | 0 | 0 |
| NC_045512.2;YP_009724389.1;Wu1-SARS2_orf1ab;784 | 0 | 0 |
| NC_045512.2;YP_009724389.1;Wu1-SARS2_orf1ab;785 | 0 | 0 |
| NC_045512.2;YP_009724389.1;Wu1-SARS2_orf1ab;786 | 0 | 0 |
| NC_045512.2;YP_009724389.1;Wu1-SARS2_orf1ab;787 | 0 | 0.025 |
| NC_045512.2;YP_009724389.1;Wu1-SARS2_orf1ab;788 | 0 | 0.025 |
| NC_045512.2;YP_009724389.1;Wu1-SARS2_orf1ab;789 | 0.05 | 0 |
| NC_045512.2;YP_009724389.1;Wu1-SARS2_orf1ab;790 | 0 | 0.025 |
| NC_045512.2;YP_009724389.1;Wu1-SARS2_orf1ab;791 | 0 | 0.05 |
| NC_045512.2;YP_009724389.1;Wu1-SARS2_orf1ab;792 | 0 | 0.05 |
| NC_045512.2;YP_009724389.1;Wu1-SARS2_orf1ab;793 | 0 | 0.025 |
| NC_045512.2;YP_009724389.1;Wu1-SARS2_orf1ab;794 | 0 | 0 |
| NC_045512.2;YP_009724389.1;Wu1-SARS2_orf1ab;795 | 0 | 0 |
| NC_045512.2;YP_009724389.1;Wu1-SARS2_orf1ab;796 | 0 | 0 |
| NC_045512.2;YP_009724389.1;Wu1-SARS2_orf1ab;797 | 0.05 | 0.05 |
| NC_045512.2;YP_009724389.1;Wu1-SARS2_orf1ab;798 | 0.05 | 0 |
| NC_045512.2;YP_009724389.1;Wu1-SARS2_orf1ab;799 | 0 | 0 |
| NC_045512.2;YP_009724389.1;Wu1-SARS2_orf1ab;800 | 0 | 0 |
| NC_045512.2;YP_009724389.1;Wu1-SARS2_orf1ab;801 | 0.05 | 0 |
| NC_045512.2;YP_009724389.1;Wu1-SARS2_orf1ab;802 | 0 | 0 |
| NC_045512.2;YP_009724389.1;Wu1-SARS2_orf1ab;803 | 0.1 | 0 |
| NC_045512.2;YP_009724389.1;Wu1-SARS2_orf1ab;804 | 0.05 | 0 |
| NC_045512.2;YP_009724389.1;Wu1-SARS2_orf1ab;805 | 0.05 | 0 |
| NC_045512.2;YP_009724389.1;Wu1-SARS2_orf1ab;806 | 0.05 | 0 |
| NC_045512.2;YP_009724389.1;Wu1-SARS2_orf1ab;807 | 0 | 0.025 |

| | | |
|---|---|---|
| NC_045512.2;YP_009724389.1;Wu1-SARS2_orf1ab;808 | 0 | 0 |
| NC_045512.2;YP_009724389.1;Wu1-SARS2_orf1ab;809 | 0 | 0 |
| NC_045512.2;YP_009724389.1;Wu1-SARS2_orf1ab;810 | 0 | 0 |
| NC_045512.2;YP_009724389.1;Wu1-SARS2_orf1ab;811 | 0 | 0 |
| NC_045512.2;YP_009724389.1;Wu1-SARS2_orf1ab;812 | 0 | 0 |
| NC_045512.2;YP_009724389.1;Wu1-SARS2_orf1ab;813 | 0.05 | 0 |
| NC_045512.2;YP_009724389.1;Wu1-SARS2_orf1ab;814 | 0 | 0.025 |
| NC_045512.2;YP_009724389.1;Wu1-SARS2_orf1ab;815 | 0 | 0 |
| NC_045512.2;YP_009724389.1;Wu1-SARS2_orf1ab;816 | 0 | 0 |
| NC_045512.2;YP_009724389.1;Wu1-SARS2_orf1ab;817 | 0 | 0 |
| NC_045512.2;YP_009724389.1;Wu1-SARS2_orf1ab;818 | 0 | 0 |
| NC_045512.2;YP_009724389.1;Wu1-SARS2_orf1ab;819 | 0.05 | 0 |
| NC_045512.2;YP_009724389.1;Wu1-SARS2_orf1ab;820 | 0.05 | 0 |
| NC_045512.2;YP_009724389.1;Wu1-SARS2_orf1ab;821 | 0 | 0 |
| NC_045512.2;YP_009724389.1;Wu1-SARS2_orf1ab;822 | 0 | 0.025 |
| NC_045512.2;YP_009724389.1;Wu1-SARS2_orf1ab;823 | 0 | 0.025 |
| NC_045512.2;YP_009724389.1;Wu1-SARS2_orf1ab;824 | 0 | 0.025 |
| NC_045512.2;YP_009724389.1;Wu1-SARS2_orf1ab;825 | 0 | 0.025 |
| NC_045512.2;YP_009724389.1;Wu1-SARS2_orf1ab;826 | 0 | 0 |
| NC_045512.2;YP_009724389.1;Wu1-SARS2_orf1ab;827 | 0.1 | 0 |
| NC_045512.2;YP_009724389.1;Wu1-SARS2_orf1ab;828 | 0 | 0 |
| NC_045512.2;YP_009724389.1;Wu1-SARS2_orf1ab;829 | 0 | 0 |
| NC_045512.2;YP_009724389.1;Wu1-SARS2_orf1ab;830 | 0 | 0 |
| NC_045512.2;YP_009724389.1;Wu1-SARS2_orf1ab;831 | 0.05 | 0.05 |
| NC_045512.2;YP_009724389.1;Wu1-SARS2_orf1ab;832 | 0 | 0 |
| NC_045512.2;YP_009724389.1;Wu1-SARS2_orf1ab;833 | 0 | 0 |
| NC_045512.2;YP_009724389.1;Wu1-SARS2_orf1ab;834 | 0 | 0 |
| NC_045512.2;YP_009724389.1;Wu1-SARS2_orf1ab;835 | 0 | 0 |
| NC_045512.2;YP_009724389.1;Wu1-SARS2_orf1ab;836 | 0.05 | 0 |
| NC_045512.2;YP_009724389.1;Wu1-SARS2_orf1ab;837 | 0.05 | 0 |
| NC_045512.2;YP_009724389.1;Wu1-SARS2_orf1ab;838 | 0 | 0 |
| NC_045512.2;YP_009724389.1;Wu1-SARS2_orf1ab;839 | 0 | 0.025 |
| NC_045512.2;YP_009724389.1;Wu1-SARS2_orf1ab;840 | 0.2 | 0.1 |
| NC_045512.2;YP_009724389.1;Wu1-SARS2_orf1ab;841 | 0.1 | 0.15 |
| NC_045512.2;YP_009724389.1;Wu1-SARS2_orf1ab;842 | 0.2 | 0.1 |
| NC_045512.2;YP_009724389.1;Wu1-SARS2_orf1ab;843 | 0.25 | 0.1 |
| NC_045512.2;YP_009724389.1;Wu1-SARS2_orf1ab;844 | 0.05 | 0.05 |
| NC_045512.2;YP_009724389.1;Wu1-SARS2_orf1ab;845 | 0 | 0.05 |
| NC_045512.2;YP_009724389.1;Wu1-SARS2_orf1ab;846 | 0 | 0.05 |
| NC_045512.2;YP_009724389.1;Wu1-SARS2_orf1ab;847 | 0 | 0.025 |
| NC_045512.2;YP_009724389.1;Wu1-SARS2_orf1ab;848 | 0 | 0 |
| NC_045512.2;YP_009724389.1;Wu1-SARS2_orf1ab;849 | 0 | 0 |
| NC_045512.2;YP_009724389.1;Wu1-SARS2_orf1ab;850 | 0 | 0 |

| | | |
|---|---|---|
| NC_045512.2;YP_009724389.1;Wu1-SARS2_orf1ab;851 | 0 | 0 |
| NC_045512.2;YP_009724389.1;Wu1-SARS2_orf1ab;852 | 0 | 0 |
| NC_045512.2;YP_009724389.1;Wu1-SARS2_orf1ab;853 | 0 | 0 |
| NC_045512.2;YP_009724389.1;Wu1-SARS2_orf1ab;854 | 0 | 0 |
| NC_045512.2;YP_009724389.1;Wu1-SARS2_orf1ab;855 | 0 | 0 |
| NC_045512.2;YP_009724389.1;Wu1-SARS2_orf1ab;856 | 0 | 0 |
| NC_045512.2;YP_009724389.1;Wu1-SARS2_orf1ab;857 | 0 | 0 |
| NC_045512.2;YP_009724389.1;Wu1-SARS2_orf1ab;858 | 0 | 0 |
| NC_045512.2;YP_009724389.1;Wu1-SARS2_orf1ab;859 | 0 | 0.025 |
| NC_045512.2;YP_009724389.1;Wu1-SARS2_orf1ab;860 | 0 | 0 |
| NC_045512.2;YP_009724389.1;Wu1-SARS2_orf1ab;861 | 0 | 0 |
| NC_045512.2;YP_009724389.1;Wu1-SARS2_orf1ab;862 | 0 | 0 |
| NC_045512.2;YP_009724389.1;Wu1-SARS2_orf1ab;863 | 0 | 0 |
| NC_045512.2;YP_009724389.1;Wu1-SARS2_orf1ab;864 | 0 | 0 |
| NC_045512.2;YP_009724389.1;Wu1-SARS2_orf1ab;865 | 0 | 0 |
| NC_045512.2;YP_009724389.1;Wu1-SARS2_orf1ab;866 | 0 | 0 |
| NC_045512.2;YP_009724389.1;Wu1-SARS2_orf1ab;867 | 0 | 0 |
| NC_045512.2;YP_009724389.1;Wu1-SARS2_orf1ab;868 | 0 | 0 |
| NC_045512.2;YP_009724389.1;Wu1-SARS2_orf1ab;869 | 0 | 0 |
| NC_045512.2;YP_009724389.1;Wu1-SARS2_orf1ab;870 | 0 | 0 |
| NC_045512.2;YP_009724389.1;Wu1-SARS2_orf1ab;871 | 0 | 0 |
| NC_045512.2;YP_009724389.1;Wu1-SARS2_orf1ab;872 | 0 | 0 |
| NC_045512.2;YP_009724389.1;Wu1-SARS2_orf1ab;873 | 0 | 0 |
| NC_045512.2;YP_009724389.1;Wu1-SARS2_orf1ab;874 | 0 | 0 |
| NC_045512.2;YP_009724389.1;Wu1-SARS2_orf1ab;875 | 0 | 0.025 |
| NC_045512.2;YP_009724389.1;Wu1-SARS2_orf1ab;876 | 0.05 | 0 |
| NC_045512.2;YP_009724389.1;Wu1-SARS2_orf1ab;877 | 0 | 0.025 |
| NC_045512.2;YP_009724389.1;Wu1-SARS2_orf1ab;878 | 0 | 0.05 |
| NC_045512.2;YP_009724389.1;Wu1-SARS2_orf1ab;879 | 0 | 0.025 |
| NC_045512.2;YP_009724389.1;Wu1-SARS2_orf1ab;880 | 0 | 0.025 |
| NC_045512.2;YP_009724389.1;Wu1-SARS2_orf1ab;881 | 0.05 | 0.025 |
| NC_045512.2;YP_009724389.1;Wu1-SARS2_orf1ab;882 | 0 | 0.025 |
| NC_045512.2;YP_009724389.1;Wu1-SARS2_orf1ab;883 | 0 | 0 |
| NC_045512.2;YP_009724389.1;Wu1-SARS2_orf1ab;884 | 0.05 | 0 |
| NC_045512.2;YP_009724389.1;Wu1-SARS2_orf1ab;885 | 0 | 0 |
| NC_045512.2;YP_009724389.1;Wu1-SARS2_orf1ab;886 | 0 | 0 |
| NC_045512.2;YP_009724389.1;Wu1-SARS2_orf1ab;887 | 0 | 0 |
| NC_045512.2;YP_009724389.1;Wu1-SARS2_orf1ab;888 | 0 | 0 |
| NC_045512.2;YP_009724389.1;Wu1-SARS2_orf1ab;889 | 0 | 0 |
| NC_045512.2;YP_009724389.1;Wu1-SARS2_orf1ab;890 | 0 | 0 |
| NC_045512.2;YP_009724389.1;Wu1-SARS2_orf1ab;891 | 0 | 0 |
| NC_045512.2;YP_009724389.1;Wu1-SARS2_orf1ab;892 | 0 | 0 |
| NC_045512.2;YP_009724389.1;Wu1-SARS2_orf1ab;893 | 0 | 0 |

| | | |
|---|---|---|
| NC_045512.2;YP_009724389.1;Wu1-SARS2_orf1ab;894 | 0 | 0 |
| NC_045512.2;YP_009724389.1;Wu1-SARS2_orf1ab;895 | 0 | 0 |
| NC_045512.2;YP_009724389.1;Wu1-SARS2_orf1ab;896 | 0 | 0 |
| NC_045512.2;YP_009724389.1;Wu1-SARS2_orf1ab;897 | 0 | 0 |
| NC_045512.2;YP_009724389.1;Wu1-SARS2_orf1ab;898 | 0 | 0 |
| NC_045512.2;YP_009724389.1;Wu1-SARS2_orf1ab;899 | 0 | 0 |
| NC_045512.2;YP_009724389.1;Wu1-SARS2_orf1ab;900 | 0.1 | 0 |
| NC_045512.2;YP_009724389.1;Wu1-SARS2_orf1ab;901 | 0 | 0 |
| NC_045512.2;YP_009724389.1;Wu1-SARS2_orf1ab;902 | 0 | 0 |
| NC_045512.2;YP_009724389.1;Wu1-SARS2_orf1ab;903 | 0 | 0 |
| NC_045512.2;YP_009724389.1;Wu1-SARS2_orf1ab;904 | 0 | 0 |
| NC_045512.2;YP_009724389.1;Wu1-SARS2_orf1ab;905 | 0 | 0 |
| NC_045512.2;YP_009724389.1;Wu1-SARS2_orf1ab;906 | 0.05 | 0 |
| NC_045512.2;YP_009724389.1;Wu1-SARS2_orf1ab;907 | 0 | 0 |
| NC_045512.2;YP_009724389.1;Wu1-SARS2_orf1ab;908 | 0 | 0 |
| NC_045512.2;YP_009724389.1;Wu1-SARS2_orf1ab;909 | 0 | 0 |
| NC_045512.2;YP_009724389.1;Wu1-SARS2_orf1ab;910 | 0 | 0 |
| NC_045512.2;YP_009724389.1;Wu1-SARS2_orf1ab;911 | 0 | 0 |
| NC_045512.2;YP_009724389.1;Wu1-SARS2_orf1ab;912 | 0 | 0 |
| NC_045512.2;YP_009724389.1;Wu1-SARS2_orf1ab;913 | 0 | 0 |
| NC_045512.2;YP_009724389.1;Wu1-SARS2_orf1ab;914 | 0 | 0 |
| NC_045512.2;YP_009724389.1;Wu1-SARS2_orf1ab;915 | 0 | 0 |
| NC_045512.2;YP_009724389.1;Wu1-SARS2_orf1ab;916 | 0 | 0.025 |
| NC_045512.2;YP_009724389.1;Wu1-SARS2_orf1ab;917 | 0 | 0 |
| NC_045512.2;YP_009724389.1;Wu1-SARS2_orf1ab;918 | 0 | 0.025 |
| NC_045512.2;YP_009724389.1;Wu1-SARS2_orf1ab;919 | 0 | 0.05 |
| NC_045512.2;YP_009724389.1;Wu1-SARS2_orf1ab;920 | 0 | 0.075 |
| NC_045512.2;YP_009724389.1;Wu1-SARS2_orf1ab;921 | 0 | 0.1 |
| NC_045512.2;YP_009724389.1;Wu1-SARS2_orf1ab;922 | 0 | 0.1 |
| NC_045512.2;YP_009724389.1;Wu1-SARS2_orf1ab;923 | 0 | 0.1 |
| NC_045512.2;YP_009724389.1;Wu1-SARS2_orf1ab;924 | 0 | 0.125 |
| NC_045512.2;YP_009724389.1;Wu1-SARS2_orf1ab;925 | 0 | 0.1 |
| NC_045512.2;YP_009724389.1;Wu1-SARS2_orf1ab;926 | 0 | 0.025 |
| NC_045512.2;YP_009724389.1;Wu1-SARS2_orf1ab;927 | 0.05 | 0 |
| NC_045512.2;YP_009724389.1;Wu1-SARS2_orf1ab;928 | 0.05 | 0.025 |
| NC_045512.2;YP_009724389.1;Wu1-SARS2_orf1ab;929 | 0.35 | 0.25 |
| NC_045512.2;YP_009724389.1;Wu1-SARS2_orf1ab;930 | 0 | 0 |
| NC_045512.2;YP_009724389.1;Wu1-SARS2_orf1ab;931 | 0 | 0 |
| NC_045512.2;YP_009724389.1;Wu1-SARS2_orf1ab;932 | 0 | 0 |
| NC_045512.2;YP_009724389.1;Wu1-SARS2_orf1ab;933 | 0.05 | 0 |
| NC_045512.2;YP_009724389.1;Wu1-SARS2_orf1ab;934 | 0 | 0 |
| NC_045512.2;YP_009724389.1;Wu1-SARS2_orf1ab;935 | 0 | 0 |
| NC_045512.2;YP_009724389.1;Wu1-SARS2_orf1ab;936 | 0 | 0 |

| | | |
|---|---|---|
| NC_045512.2;YP_009724389.1;Wu1-SARS2_orf1ab;937 | 0 | 0 |
| NC_045512.2;YP_009724389.1;Wu1-SARS2_orf1ab;938 | 0 | 0 |
| NC_045512.2;YP_009724389.1;Wu1-SARS2_orf1ab;939 | 0 | 0 |
| NC_045512.2;YP_009724389.1;Wu1-SARS2_orf1ab;940 | 0.05 | 0.05 |
| NC_045512.2;YP_009724389.1;Wu1-SARS2_orf1ab;941 | 0 | 0.025 |
| NC_045512.2;YP_009724389.1;Wu1-SARS2_orf1ab;942 | 0 | 0 |
| NC_045512.2;YP_009724389.1;Wu1-SARS2_orf1ab;943 | 0 | 0 |
| NC_045512.2;YP_009724389.1;Wu1-SARS2_orf1ab;944 | 0 | 0 |
| NC_045512.2;YP_009724389.1;Wu1-SARS2_orf1ab;945 | 0 | 0 |
| NC_045512.2;YP_009724389.1;Wu1-SARS2_orf1ab;946 | 0 | 0.025 |
| NC_045512.2;YP_009724389.1;Wu1-SARS2_orf1ab;947 | 0 | 0.025 |
| NC_045512.2;YP_009724389.1;Wu1-SARS2_orf1ab;948 | 0 | 0 |
| NC_045512.2;YP_009724389.1;Wu1-SARS2_orf1ab;949 | 0 | 0.05 |
| NC_045512.2;YP_009724389.1;Wu1-SARS2_orf1ab;950 | 0 | 0.075 |
| NC_045512.2;YP_009724389.1;Wu1-SARS2_orf1ab;951 | 0 | 0 |
| NC_045512.2;YP_009724389.1;Wu1-SARS2_orf1ab;952 | 0.1 | 0 |
| NC_045512.2;YP_009724389.1;Wu1-SARS2_orf1ab;953 | 0.15 | 0.025 |
| NC_045512.2;YP_009724389.1;Wu1-SARS2_orf1ab;954 | 0 | 0 |
| NC_045512.2;YP_009724389.1;Wu1-SARS2_orf1ab;955 | 0 | 0 |
| NC_045512.2;YP_009724389.1;Wu1-SARS2_orf1ab;956 | 0 | 0 |
| NC_045512.2;YP_009724389.1;Wu1-SARS2_orf1ab;957 | 0 | 0 |
| NC_045512.2;YP_009724389.1;Wu1-SARS2_orf1ab;958 | 0 | 0 |
| NC_045512.2;YP_009724389.1;Wu1-SARS2_orf1ab;959 | 0 | 0 |
| NC_045512.2;YP_009724389.1;Wu1-SARS2_orf1ab;960 | 0 | 0 |
| NC_045512.2;YP_009724389.1;Wu1-SARS2_orf1ab;961 | 0 | 0 |
| NC_045512.2;YP_009724389.1;Wu1-SARS2_orf1ab;962 | 0 | 0 |
| NC_045512.2;YP_009724389.1;Wu1-SARS2_orf1ab;963 | 0 | 0 |
| NC_045512.2;YP_009724389.1;Wu1-SARS2_orf1ab;964 | 0 | 0.025 |
| NC_045512.2;YP_009724389.1;Wu1-SARS2_orf1ab;965 | 0 | 0.025 |
| NC_045512.2;YP_009724389.1;Wu1-SARS2_orf1ab;966 | 0 | 0.025 |
| NC_045512.2;YP_009724389.1;Wu1-SARS2_orf1ab;967 | 0 | 0.025 |
| NC_045512.2;YP_009724389.1;Wu1-SARS2_orf1ab;968 | 0 | 0.025 |
| NC_045512.2;YP_009724389.1;Wu1-SARS2_orf1ab;969 | 0 | 0.025 |
| NC_045512.2;YP_009724389.1;Wu1-SARS2_orf1ab;970 | 0 | 0.025 |
| NC_045512.2;YP_009724389.1;Wu1-SARS2_orf1ab;971 | 0 | 0.025 |
| NC_045512.2;YP_009724389.1;Wu1-SARS2_orf1ab;972 | 0 | 0.05 |
| NC_045512.2;YP_009724389.1;Wu1-SARS2_orf1ab;973 | 0 | 0.025 |
| NC_045512.2;YP_009724389.1;Wu1-SARS2_orf1ab;974 | 0 | 0.025 |
| NC_045512.2;YP_009724389.1;Wu1-SARS2_orf1ab;975 | 0 | 0.025 |
| NC_045512.2;YP_009724389.1;Wu1-SARS2_orf1ab;976 | 0 | 0.075 |
| NC_045512.2;YP_009724389.1;Wu1-SARS2_orf1ab;977 | 0.1 | 0.025 |
| NC_045512.2;YP_009724389.1;Wu1-SARS2_orf1ab;978 | 0 | 0.025 |
| NC_045512.2;YP_009724389.1;Wu1-SARS2_orf1ab;979 | 0 | 0.025 |

| | | |
|---|---|---|
| NC_045512.2;YP_009724389.1;Wu1-SARS2_orf1ab;980 | 0 | 0 |
| NC_045512.2;YP_009724389.1;Wu1-SARS2_orf1ab;981 | 0 | 0 |
| NC_045512.2;YP_009724389.1;Wu1-SARS2_orf1ab;982 | 0 | 0 |
| NC_045512.2;YP_009724389.1;Wu1-SARS2_orf1ab;983 | 0 | 0 |
| NC_045512.2;YP_009724389.1;Wu1-SARS2_orf1ab;984 | 0 | 0.025 |
| NC_045512.2;YP_009724389.1;Wu1-SARS2_orf1ab;985 | 0 | 0 |
| NC_045512.2;YP_009724389.1;Wu1-SARS2_orf1ab;986 | 0 | 0 |
| NC_045512.2;YP_009724389.1;Wu1-SARS2_orf1ab;987 | 0 | 0 |
| NC_045512.2;YP_009724389.1;Wu1-SARS2_orf1ab;988 | 0 | 0 |
| NC_045512.2;YP_009724389.1;Wu1-SARS2_orf1ab;989 | 0 | 0 |
| NC_045512.2;YP_009724389.1;Wu1-SARS2_orf1ab;990 | 0 | 0 |
| NC_045512.2;YP_009724389.1;Wu1-SARS2_orf1ab;991 | 0 | 0 |
| NC_045512.2;YP_009724389.1;Wu1-SARS2_orf1ab;992 | 0 | 0 |
| NC_045512.2;YP_009724389.1;Wu1-SARS2_orf1ab;993 | 0 | 0 |
| NC_045512.2;YP_009724389.1;Wu1-SARS2_orf1ab;994 | 0 | 0 |
| NC_045512.2;YP_009724389.1;Wu1-SARS2_orf1ab;995 | 0 | 0 |
| NC_045512.2;YP_009724389.1;Wu1-SARS2_orf1ab;996 | 0 | 0.025 |
| NC_045512.2;YP_009724389.1;Wu1-SARS2_orf1ab;997 | 0 | 0 |
| NC_045512.2;YP_009724389.1;Wu1-SARS2_orf1ab;998 | 0 | 0 |
| NC_045512.2;YP_009724389.1;Wu1-SARS2_orf1ab;999 | 0 | 0 |
| NC_045512.2;YP_009724389.1;Wu1-SARS2_orf1ab;1000 | 0 | 0 |
| NC_045512.2;YP_009724389.1;Wu1-SARS2_orf1ab;1001 | 0 | 0 |
| NC_045512.2;YP_009724389.1;Wu1-SARS2_orf1ab;1002 | 0 | 0 |
| NC_045512.2;YP_009724389.1;Wu1-SARS2_orf1ab;1003 | 0 | 0 |
| NC_045512.2;YP_009724389.1;Wu1-SARS2_orf1ab;1004 | 0 | 0 |
| NC_045512.2;YP_009724389.1;Wu1-SARS2_orf1ab;1005 | 0 | 0 |
| NC_045512.2;YP_009724389.1;Wu1-SARS2_orf1ab;1006 | 0 | 0 |
| NC_045512.2;YP_009724389.1;Wu1-SARS2_orf1ab;1007 | 0 | 0 |
| NC_045512.2;YP_009724389.1;Wu1-SARS2_orf1ab;1008 | 0 | 0 |
| NC_045512.2;YP_009724389.1;Wu1-SARS2_orf1ab;1009 | 0 | 0 |
| NC_045512.2;YP_009724389.1;Wu1-SARS2_orf1ab;1010 | 0 | 0 |
| NC_045512.2;YP_009724389.1;Wu1-SARS2_orf1ab;1011 | 0 | 0 |
| NC_045512.2;YP_009724389.1;Wu1-SARS2_orf1ab;1012 | 0 | 0 |
| NC_045512.2;YP_009724389.1;Wu1-SARS2_orf1ab;1013 | 0 | 0 |
| NC_045512.2;YP_009724389.1;Wu1-SARS2_orf1ab;1014 | 0 | 0.05 |
| NC_045512.2;YP_009724389.1;Wu1-SARS2_orf1ab;1015 | 0.1 | 0.05 |
| NC_045512.2;YP_009724389.1;Wu1-SARS2_orf1ab;1016 | 0 | 0.05 |
| NC_045512.2;YP_009724389.1;Wu1-SARS2_orf1ab;1017 | 0 | 0.025 |
| NC_045512.2;YP_009724389.1;Wu1-SARS2_orf1ab;1018 | 0 | 0 |
| NC_045512.2;YP_009724389.1;Wu1-SARS2_orf1ab;1019 | 0 | 0 |
| NC_045512.2;YP_009724389.1;Wu1-SARS2_orf1ab;1020 | 0 | 0 |
| NC_045512.2;YP_009724389.1;Wu1-SARS2_orf1ab;1021 | 0 | 0 |
| NC_045512.2;YP_009724389.1;Wu1-SARS2_orf1ab;1022 | 0 | 0 |

| | | |
|---|---|---|
| NC_045512.2;YP_009724389.1;Wu1-SARS2_orf1ab;1023 | 0.05 | 0 |
| NC_045512.2;YP_009724389.1;Wu1-SARS2_orf1ab;1024 | 0.05 | 0 |
| NC_045512.2;YP_009724389.1;Wu1-SARS2_orf1ab;1025 | 0.05 | 0 |
| NC_045512.2;YP_009724389.1;Wu1-SARS2_orf1ab;1026 | 0.05 | 0 |
| NC_045512.2;YP_009724389.1;Wu1-SARS2_orf1ab;1027 | 0.05 | 0 |
| NC_045512.2;YP_009724389.1;Wu1-SARS2_orf1ab;1028 | 0.05 | 0 |
| NC_045512.2;YP_009724389.1;Wu1-SARS2_orf1ab;1029 | 0.05 | 0 |
| NC_045512.2;YP_009724389.1;Wu1-SARS2_orf1ab;1030 | 0.05 | 0.025 |
| NC_045512.2;YP_009724389.1;Wu1-SARS2_orf1ab;1031 | 0.05 | 0 |
| NC_045512.2;YP_009724389.1;Wu1-SARS2_orf1ab;1032 | 0.15 | 0.025 |
| NC_045512.2;YP_009724389.1;Wu1-SARS2_orf1ab;1033 | 0.2 | 0.025 |
| NC_045512.2;YP_009724389.1;Wu1-SARS2_orf1ab;1034 | 0.05 | 0.025 |
| NC_045512.2;YP_009724389.1;Wu1-SARS2_orf1ab;1035 | 0 | 0.05 |
| NC_045512.2;YP_009724389.1;Wu1-SARS2_orf1ab;1036 | 0 | 0.025 |
| NC_045512.2;YP_009724389.1;Wu1-SARS2_orf1ab;1037 | 0 | 0 |
| NC_045512.2;YP_009724389.1;Wu1-SARS2_orf1ab;1038 | 0 | 0 |
| NC_045512.2;YP_009724389.1;Wu1-SARS2_orf1ab;1039 | 0 | 0 |
| NC_045512.2;YP_009724389.1;Wu1-SARS2_orf1ab;1040 | 0 | 0 |
| NC_045512.2;YP_009724389.1;Wu1-SARS2_orf1ab;1041 | 0 | 0 |
| NC_045512.2;YP_009724389.1;Wu1-SARS2_orf1ab;1042 | 0 | 0 |
| NC_045512.2;YP_009724389.1;Wu1-SARS2_orf1ab;1043 | 0 | 0.025 |
| NC_045512.2;YP_009724389.1;Wu1-SARS2_orf1ab;1044 | 0 | 0 |
| NC_045512.2;YP_009724389.1;Wu1-SARS2_orf1ab;1045 | 0 | 0 |
| NC_045512.2;YP_009724389.1;Wu1-SARS2_orf1ab;1046 | 0 | 0.025 |
| NC_045512.2;YP_009724389.1;Wu1-SARS2_orf1ab;1047 | 0 | 0.025 |
| NC_045512.2;YP_009724389.1;Wu1-SARS2_orf1ab;1048 | 0.05 | 0 |
| NC_045512.2;YP_009724389.1;Wu1-SARS2_orf1ab;1049 | 0 | 0 |
| NC_045512.2;YP_009724389.1;Wu1-SARS2_orf1ab;1050 | 0 | 0 |
| NC_045512.2;YP_009724389.1;Wu1-SARS2_orf1ab;1051 | 0 | 0 |
| NC_045512.2;YP_009724389.1;Wu1-SARS2_orf1ab;1052 | 0 | 0 |
| NC_045512.2;YP_009724389.1;Wu1-SARS2_orf1ab;1053 | 0 | 0 |
| NC_045512.2;YP_009724389.1;Wu1-SARS2_orf1ab;1054 | 0 | 0 |
| NC_045512.2;YP_009724389.1;Wu1-SARS2_orf1ab;1055 | 0.05 | 0 |
| NC_045512.2;YP_009724389.1;Wu1-SARS2_orf1ab;1056 | 0 | 0 |
| NC_045512.2;YP_009724389.1;Wu1-SARS2_orf1ab;1057 | 0.05 | 0.05 |
| NC_045512.2;YP_009724389.1;Wu1-SARS2_orf1ab;1058 | 0 | 0 |
| NC_045512.2;YP_009724389.1;Wu1-SARS2_orf1ab;1059 | 0 | 0 |
| NC_045512.2;YP_009724389.1;Wu1-SARS2_orf1ab;1060 | 0 | 0 |
| NC_045512.2;YP_009724389.1;Wu1-SARS2_orf1ab;1061 | 0 | 0 |
| NC_045512.2;YP_009724389.1;Wu1-SARS2_orf1ab;1062 | 0 | 0 |
| NC_045512.2;YP_009724389.1;Wu1-SARS2_orf1ab;1063 | 0.05 | 0 |
| NC_045512.2;YP_009724389.1;Wu1-SARS2_orf1ab;1064 | 0 | 0 |
| NC_045512.2;YP_009724389.1;Wu1-SARS2_orf1ab;1065 | 0 | 0 |

| | | |
|---|---|---|
| NC_045512.2;YP_009724389.1;Wu1-SARS2_orf1ab;1066 | 0 | 0.025 |
| NC_045512.2;YP_009724389.1;Wu1-SARS2_orf1ab;1067 | 0.1 | 0.025 |
| NC_045512.2;YP_009724389.1;Wu1-SARS2_orf1ab;1068 | 0.15 | 0.025 |
| NC_045512.2;YP_009724389.1;Wu1-SARS2_orf1ab;1069 | 0.05 | 0.025 |
| NC_045512.2;YP_009724389.1;Wu1-SARS2_orf1ab;1070 | 0.05 | 0.025 |
| NC_045512.2;YP_009724389.1;Wu1-SARS2_orf1ab;1071 | 0.15 | 0.2 |
| NC_045512.2;YP_009724389.1;Wu1-SARS2_orf1ab;1072 | 0.2 | 0.2 |
| NC_045512.2;YP_009724389.1;Wu1-SARS2_orf1ab;1073 | 0.2 | 0.2 |
| NC_045512.2;YP_009724389.1;Wu1-SARS2_orf1ab;1074 | 0.05 | 0.15 |
| NC_045512.2;YP_009724389.1;Wu1-SARS2_orf1ab;1075 | 0.05 | 0.15 |
| NC_045512.2;YP_009724389.1;Wu1-SARS2_orf1ab;1076 | 0 | 0.025 |
| NC_045512.2;YP_009724389.1;Wu1-SARS2_orf1ab;1077 | 0 | 0.025 |
| NC_045512.2;YP_009724389.1;Wu1-SARS2_orf1ab;1078 | 0.05 | 0.025 |
| NC_045512.2;YP_009724389.1;Wu1-SARS2_orf1ab;1079 | 0.05 | 0 |
| NC_045512.2;YP_009724389.1;Wu1-SARS2_orf1ab;1080 | 0 | 0 |
| NC_045512.2;YP_009724389.1;Wu1-SARS2_orf1ab;1081 | 0 | 0 |
| NC_045512.2;YP_009724389.1;Wu1-SARS2_orf1ab;1082 | 0 | 0.025 |
| NC_045512.2;YP_009724389.1;Wu1-SARS2_orf1ab;1083 | 0 | 0 |
| NC_045512.2;YP_009724389.1;Wu1-SARS2_orf1ab;1084 | 0 | 0 |
| NC_045512.2;YP_009724389.1;Wu1-SARS2_orf1ab;1085 | 0 | 0.025 |
| NC_045512.2;YP_009724389.1;Wu1-SARS2_orf1ab;1086 | 0 | 0.05 |
| NC_045512.2;YP_009724389.1;Wu1-SARS2_orf1ab;1087 | 0.15 | 0.05 |
| NC_045512.2;YP_009724389.1;Wu1-SARS2_orf1ab;1088 | 0.05 | 0.05 |
| NC_045512.2;YP_009724389.1;Wu1-SARS2_orf1ab;1089 | 0 | 0.025 |
| NC_045512.2;YP_009724389.1;Wu1-SARS2_orf1ab;1090 | 0 | 0 |
| NC_045512.2;YP_009724389.1;Wu1-SARS2_orf1ab;1091 | 0 | 0 |
| NC_045512.2;YP_009724389.1;Wu1-SARS2_orf1ab;1092 | 0 | 0 |
| NC_045512.2;YP_009724389.1;Wu1-SARS2_orf1ab;1093 | 0 | 0 |
| NC_045512.2;YP_009724389.1;Wu1-SARS2_orf1ab;1094 | 0 | 0 |
| NC_045512.2;YP_009724389.1;Wu1-SARS2_orf1ab;1095 | 0 | 0 |
| NC_045512.2;YP_009724389.1;Wu1-SARS2_orf1ab;1096 | 0 | 0 |
| NC_045512.2;YP_009724389.1;Wu1-SARS2_orf1ab;1097 | 0 | 0 |
| NC_045512.2;YP_009724389.1;Wu1-SARS2_orf1ab;1098 | 0 | 0 |
| NC_045512.2;YP_009724389.1;Wu1-SARS2_orf1ab;1099 | 0 | 0 |
| NC_045512.2;YP_009724389.1;Wu1-SARS2_orf1ab;1100 | 0 | 0 |
| NC_045512.2;YP_009724389.1;Wu1-SARS2_orf1ab;1101 | 0 | 0 |
| NC_045512.2;YP_009724389.1;Wu1-SARS2_orf1ab;1102 | 0 | 0 |
| NC_045512.2;YP_009724389.1;Wu1-SARS2_orf1ab;1103 | 0 | 0 |
| NC_045512.2;YP_009724389.1;Wu1-SARS2_orf1ab;1104 | 0 | 0 |
| NC_045512.2;YP_009724389.1;Wu1-SARS2_orf1ab;1105 | 0 | 0 |
| NC_045512.2;YP_009724389.1;Wu1-SARS2_orf1ab;1106 | 0 | 0 |
| NC_045512.2;YP_009724389.1;Wu1-SARS2_orf1ab;1107 | 0 | 0 |
| NC_045512.2;YP_009724389.1;Wu1-SARS2_orf1ab;1108 | 0 | 0 |

| | | |
|---|---|---|
| NC_045512.2;YP_009724389.1;Wu1-SARS2_orf1ab;1109 | 0 | 0 |
| NC_045512.2;YP_009724389.1;Wu1-SARS2_orf1ab;1110 | 0 | 0 |
| NC_045512.2;YP_009724389.1;Wu1-SARS2_orf1ab;1111 | 0.05 | 0 |
| NC_045512.2;YP_009724389.1;Wu1-SARS2_orf1ab;1112 | 0.05 | 0 |
| NC_045512.2;YP_009724389.1;Wu1-SARS2_orf1ab;1113 | 0.05 | 0.075 |
| NC_045512.2;YP_009724389.1;Wu1-SARS2_orf1ab;1114 | 0.05 | 0.125 |
| NC_045512.2;YP_009724389.1;Wu1-SARS2_orf1ab;1115 | 0.15 | 0.075 |
| NC_045512.2;YP_009724389.1;Wu1-SARS2_orf1ab;1116 | 0 | 0.075 |
| NC_045512.2;YP_009724389.1;Wu1-SARS2_orf1ab;1117 | 0.2 | 0.1 |
| NC_045512.2;YP_009724389.1;Wu1-SARS2_orf1ab;1118 | 0.2 | 0.125 |
| NC_045512.2;YP_009724389.1;Wu1-SARS2_orf1ab;1119 | 0.05 | 0.075 |
| NC_045512.2;YP_009724389.1;Wu1-SARS2_orf1ab;1120 | 0.05 | 0.05 |
| NC_045512.2;YP_009724389.1;Wu1-SARS2_orf1ab;1121 | 0.05 | 0.025 |
| NC_045512.2;YP_009724389.1;Wu1-SARS2_orf1ab;1122 | 0.05 | 0.05 |
| NC_045512.2;YP_009724389.1;Wu1-SARS2_orf1ab;1123 | 0 | 0.025 |
| NC_045512.2;YP_009724389.1;Wu1-SARS2_orf1ab;1124 | 0 | 0.025 |
| NC_045512.2;YP_009724389.1;Wu1-SARS2_orf1ab;1125 | 0.05 | 0.025 |
| NC_045512.2;YP_009724389.1;Wu1-SARS2_orf1ab;1126 | 0.05 | 0.025 |
| NC_045512.2;YP_009724389.1;Wu1-SARS2_orf1ab;1127 | 0 | 0 |
| NC_045512.2;YP_009724389.1;Wu1-SARS2_orf1ab;1128 | 0 | 0 |
| NC_045512.2;YP_009724389.1;Wu1-SARS2_orf1ab;1129 | 0 | 0 |
| NC_045512.2;YP_009724389.1;Wu1-SARS2_orf1ab;1130 | 0 | 0 |
| NC_045512.2;YP_009724389.1;Wu1-SARS2_orf1ab;1131 | 0 | 0.025 |
| NC_045512.2;YP_009724389.1;Wu1-SARS2_orf1ab;1132 | 0 | 0.05 |
| NC_045512.2;YP_009724389.1;Wu1-SARS2_orf1ab;1133 | 0 | 0 |
| NC_045512.2;YP_009724389.1;Wu1-SARS2_orf1ab;1134 | 0 | 0 |
| NC_045512.2;YP_009724389.1;Wu1-SARS2_orf1ab;1135 | 0 | 0 |
| NC_045512.2;YP_009724389.1;Wu1-SARS2_orf1ab;1136 | 0 | 0 |
| NC_045512.2;YP_009724389.1;Wu1-SARS2_orf1ab;1137 | 0 | 0 |
| NC_045512.2;YP_009724389.1;Wu1-SARS2_orf1ab;1138 | 0 | 0 |
| NC_045512.2;YP_009724389.1;Wu1-SARS2_orf1ab;1139 | 0 | 0 |
| NC_045512.2;YP_009724389.1;Wu1-SARS2_orf1ab;1140 | 0 | 0 |
| NC_045512.2;YP_009724389.1;Wu1-SARS2_orf1ab;1141 | 0 | 0 |
| NC_045512.2;YP_009724389.1;Wu1-SARS2_orf1ab;1142 | 0 | 0 |
| NC_045512.2;YP_009724389.1;Wu1-SARS2_orf1ab;1143 | 0 | 0 |
| NC_045512.2;YP_009724389.1;Wu1-SARS2_orf1ab;1144 | 0.05 | 0.025 |
| NC_045512.2;YP_009724389.1;Wu1-SARS2_orf1ab;1145 | 0 | 0 |
| NC_045512.2;YP_009724389.1;Wu1-SARS2_orf1ab;1146 | 0 | 0 |
| NC_045512.2;YP_009724389.1;Wu1-SARS2_orf1ab;1147 | 0 | 0.025 |
| NC_045512.2;YP_009724389.1;Wu1-SARS2_orf1ab;1148 | 0 | 0 |
| NC_045512.2;YP_009724389.1;Wu1-SARS2_orf1ab;1149 | 0 | 0.025 |
| NC_045512.2;YP_009724389.1;Wu1-SARS2_orf1ab;1150 | 0 | 0 |
| NC_045512.2;YP_009724389.1;Wu1-SARS2_orf1ab;1151 | 0 | 0 |

| | | |
|---|---|---|
| NC_045512.2;YP_009724389.1;Wu1-SARS2_orf1ab;1152 | 0 | 0 |
| NC_045512.2;YP_009724389.1;Wu1-SARS2_orf1ab;1153 | 0 | 0 |
| NC_045512.2;YP_009724389.1;Wu1-SARS2_orf1ab;1154 | 0 | 0 |
| NC_045512.2;YP_009724389.1;Wu1-SARS2_orf1ab;1155 | 0 | 0 |
| NC_045512.2;YP_009724389.1;Wu1-SARS2_orf1ab;1156 | 0 | 0 |
| NC_045512.2;YP_009724389.1;Wu1-SARS2_orf1ab;1157 | 0 | 0 |
| NC_045512.2;YP_009724389.1;Wu1-SARS2_orf1ab;1158 | 0 | 0 |
| NC_045512.2;YP_009724389.1;Wu1-SARS2_orf1ab;1159 | 0 | 0 |
| NC_045512.2;YP_009724389.1;Wu1-SARS2_orf1ab;1160 | 0 | 0 |
| NC_045512.2;YP_009724389.1;Wu1-SARS2_orf1ab;1161 | 0 | 0 |
| NC_045512.2;YP_009724389.1;Wu1-SARS2_orf1ab;1162 | 0 | 0 |
| NC_045512.2;YP_009724389.1;Wu1-SARS2_orf1ab;1163 | 0 | 0 |
| NC_045512.2;YP_009724389.1;Wu1-SARS2_orf1ab;1164 | 0 | 0 |
| NC_045512.2;YP_009724389.1;Wu1-SARS2_orf1ab;1165 | 0 | 0 |
| NC_045512.2;YP_009724389.1;Wu1-SARS2_orf1ab;1166 | 0 | 0.05 |
| NC_045512.2;YP_009724389.1;Wu1-SARS2_orf1ab;1167 | 0 | 0 |
| NC_045512.2;YP_009724389.1;Wu1-SARS2_orf1ab;1168 | 0 | 0 |
| NC_045512.2;YP_009724389.1;Wu1-SARS2_orf1ab;1169 | 0 | 0 |
| NC_045512.2;YP_009724389.1;Wu1-SARS2_orf1ab;1170 | 0 | 0 |
| NC_045512.2;YP_009724389.1;Wu1-SARS2_orf1ab;1171 | 0 | 0 |
| NC_045512.2;YP_009724389.1;Wu1-SARS2_orf1ab;1172 | 0 | 0 |
| NC_045512.2;YP_009724389.1;Wu1-SARS2_orf1ab;1173 | 0.05 | 0 |
| NC_045512.2;YP_009724389.1;Wu1-SARS2_orf1ab;1174 | 0.05 | 0 |
| NC_045512.2;YP_009724389.1;Wu1-SARS2_orf1ab;1175 | 0.05 | 0 |
| NC_045512.2;YP_009724389.1;Wu1-SARS2_orf1ab;1176 | 0 | 0 |
| NC_045512.2;YP_009724389.1;Wu1-SARS2_orf1ab;1177 | 0.05 | 0 |
| NC_045512.2;YP_009724389.1;Wu1-SARS2_orf1ab;1178 | 0.05 | 0 |
| NC_045512.2;YP_009724389.1;Wu1-SARS2_orf1ab;1179 | 0.1 | 0 |
| NC_045512.2;YP_009724389.1;Wu1-SARS2_orf1ab;1180 | 0 | 0 |
| NC_045512.2;YP_009724389.1;Wu1-SARS2_orf1ab;1181 | 0 | 0 |
| NC_045512.2;YP_009724389.1;Wu1-SARS2_orf1ab;1182 | 0 | 0 |
| NC_045512.2;YP_009724389.1;Wu1-SARS2_orf1ab;1183 | 0 | 0 |
| NC_045512.2;YP_009724389.1;Wu1-SARS2_orf1ab;1184 | 0 | 0 |
| NC_045512.2;YP_009724389.1;Wu1-SARS2_orf1ab;1185 | 0.05 | 0 |
| NC_045512.2;YP_009724389.1;Wu1-SARS2_orf1ab;1186 | 0.05 | 0 |
| NC_045512.2;YP_009724389.1;Wu1-SARS2_orf1ab;1187 | 0.1 | 0.025 |
| NC_045512.2;YP_009724389.1;Wu1-SARS2_orf1ab;1188 | 0.05 | 0.025 |
| NC_045512.2;YP_009724389.1;Wu1-SARS2_orf1ab;1189 | 0.05 | 0.025 |
| NC_045512.2;YP_009724389.1;Wu1-SARS2_orf1ab;1190 | 0.05 | 0.075 |
| NC_045512.2;YP_009724389.1;Wu1-SARS2_orf1ab;1191 | 0 | 0.05 |
| NC_045512.2;YP_009724389.1;Wu1-SARS2_orf1ab;1192 | 0.05 | 0.025 |
| NC_045512.2;YP_009724389.1;Wu1-SARS2_orf1ab;1193 | 0 | 0 |
| NC_045512.2;YP_009724389.1;Wu1-SARS2_orf1ab;1194 | 0.05 | 0 |

| | | |
|---|---|---|
| NC_045512.2;YP_009724389.1;Wu1-SARS2_orf1ab;1195 | 0.05 | 0.05 |
| NC_045512.2;YP_009724389.1;Wu1-SARS2_orf1ab;1196 | 0.05 | 0.05 |
| NC_045512.2;YP_009724389.1;Wu1-SARS2_orf1ab;1197 | 0.1 | 0.025 |
| NC_045512.2;YP_009724389.1;Wu1-SARS2_orf1ab;1198 | 0.05 | 0.025 |
| NC_045512.2;YP_009724389.1;Wu1-SARS2_orf1ab;1199 | 0.05 | 0 |
| NC_045512.2;YP_009724389.1;Wu1-SARS2_orf1ab;1200 | 0.1 | 0.025 |
| NC_045512.2;YP_009724389.1;Wu1-SARS2_orf1ab;1201 | 0.1 | 0.025 |
| NC_045512.2;YP_009724389.1;Wu1-SARS2_orf1ab;1202 | 0 | 0.05 |
| NC_045512.2;YP_009724389.1;Wu1-SARS2_orf1ab;1203 | 0 | 0.05 |
| NC_045512.2;YP_009724389.1;Wu1-SARS2_orf1ab;1204 | 0.05 | 0.05 |
| NC_045512.2;YP_009724389.1;Wu1-SARS2_orf1ab;1205 | 0.05 | 0.05 |
| NC_045512.2;YP_009724389.1;Wu1-SARS2_orf1ab;1206 | 0.05 | 0.025 |
| NC_045512.2;YP_009724389.1;Wu1-SARS2_orf1ab;1207 | 0.1 | 0.025 |
| NC_045512.2;YP_009724389.1;Wu1-SARS2_orf1ab;1208 | 0.05 | 0 |
| NC_045512.2;YP_009724389.1;Wu1-SARS2_orf1ab;1209 | 0.05 | 0 |
| NC_045512.2;YP_009724389.1;Wu1-SARS2_orf1ab;1210 | 0.1 | 0.05 |
| NC_045512.2;YP_009724389.1;Wu1-SARS2_orf1ab;1211 | 0.1 | 0.025 |
| NC_045512.2;YP_009724389.1;Wu1-SARS2_orf1ab;1212 | 0.05 | 0.025 |
| NC_045512.2;YP_009724389.1;Wu1-SARS2_orf1ab;1213 | 0.1 | 0.075 |
| NC_045512.2;YP_009724389.1;Wu1-SARS2_orf1ab;1214 | 0.05 | 0.05 |
| NC_045512.2;YP_009724389.1;Wu1-SARS2_orf1ab;1215 | 0.1 | 0.025 |
| NC_045512.2;YP_009724389.1;Wu1-SARS2_orf1ab;1216 | 0.05 | 0.025 |
| NC_045512.2;YP_009724389.1;Wu1-SARS2_orf1ab;1217 | 0.05 | 0.125 |
| NC_045512.2;YP_009724389.1;Wu1-SARS2_orf1ab;1218 | 0.05 | 0 |
| NC_045512.2;YP_009724389.1;Wu1-SARS2_orf1ab;1219 | 0 | 0 |
| NC_045512.2;YP_009724389.1;Wu1-SARS2_orf1ab;1220 | 0.05 | 0 |
| NC_045512.2;YP_009724389.1;Wu1-SARS2_orf1ab;1221 | 0.05 | 0 |
| NC_045512.2;YP_009724389.1;Wu1-SARS2_orf1ab;1222 | 0.05 | 0 |
| NC_045512.2;YP_009724389.1;Wu1-SARS2_orf1ab;1223 | 0.05 | 0 |
| NC_045512.2;YP_009724389.1;Wu1-SARS2_orf1ab;1224 | 0.05 | 0 |
| NC_045512.2;YP_009724389.1;Wu1-SARS2_orf1ab;1225 | 0.05 | 0.025 |
| NC_045512.2;YP_009724389.1;Wu1-SARS2_orf1ab;1226 | 0.05 | 0.025 |
| NC_045512.2;YP_009724389.1;Wu1-SARS2_orf1ab;1227 | 0 | 0.025 |
| NC_045512.2;YP_009724389.1;Wu1-SARS2_orf1ab;1228 | 0 | 0.025 |
| NC_045512.2;YP_009724389.1;Wu1-SARS2_orf1ab;1229 | 0 | 0.025 |
| NC_045512.2;YP_009724389.1;Wu1-SARS2_orf1ab;1230 | 0 | 0.025 |
| NC_045512.2;YP_009724389.1;Wu1-SARS2_orf1ab;1231 | 0 | 0.025 |
| NC_045512.2;YP_009724389.1;Wu1-SARS2_orf1ab;1232 | 0 | 0.025 |
| NC_045512.2;YP_009724389.1;Wu1-SARS2_orf1ab;1233 | 0 | 0.025 |
| NC_045512.2;YP_009724389.1;Wu1-SARS2_orf1ab;1234 | 0 | 0.025 |
| NC_045512.2;YP_009724389.1;Wu1-SARS2_orf1ab;1235 | 0 | 0.025 |
| NC_045512.2;YP_009724389.1;Wu1-SARS2_orf1ab;1236 | 0 | 0 |
| NC_045512.2;YP_009724389.1;Wu1-SARS2_orf1ab;1237 | 0 | 0.075 |

| | | |
|---|---|---|
| NC_045512.2;YP_009724389.1;Wu1-SARS2_orf1ab;1238 | 0 | 0.05 |
| NC_045512.2;YP_009724389.1;Wu1-SARS2_orf1ab;1239 | 0 | 0.025 |
| NC_045512.2;YP_009724389.1;Wu1-SARS2_orf1ab;1240 | 0 | 0.025 |
| NC_045512.2;YP_009724389.1;Wu1-SARS2_orf1ab;1241 | 0 | 0.05 |
| NC_045512.2;YP_009724389.1;Wu1-SARS2_orf1ab;1242 | 0 | 0 |
| NC_045512.2;YP_009724389.1;Wu1-SARS2_orf1ab;1243 | 0 | 0 |
| NC_045512.2;YP_009724389.1;Wu1-SARS2_orf1ab;1244 | 0 | 0 |
| NC_045512.2;YP_009724389.1;Wu1-SARS2_orf1ab;1245 | 0 | 0 |
| NC_045512.2;YP_009724389.1;Wu1-SARS2_orf1ab;1246 | 0 | 0 |
| NC_045512.2;YP_009724389.1;Wu1-SARS2_orf1ab;1247 | 0 | 0 |
| NC_045512.2;YP_009724389.1;Wu1-SARS2_orf1ab;1248 | 0 | 0 |
| NC_045512.2;YP_009724389.1;Wu1-SARS2_orf1ab;1249 | 0 | 0 |
| NC_045512.2;YP_009724389.1;Wu1-SARS2_orf1ab;1250 | 0 | 0 |
| NC_045512.2;YP_009724389.1;Wu1-SARS2_orf1ab;1251 | 0 | 0 |
| NC_045512.2;YP_009724389.1;Wu1-SARS2_orf1ab;1252 | 0 | 0 |
| NC_045512.2;YP_009724389.1;Wu1-SARS2_orf1ab;1253 | 0 | 0 |
| NC_045512.2;YP_009724389.1;Wu1-SARS2_orf1ab;1254 | 0 | 0.025 |
| NC_045512.2;YP_009724389.1;Wu1-SARS2_orf1ab;1255 | 0.05 | 0.075 |
| NC_045512.2;YP_009724389.1;Wu1-SARS2_orf1ab;1256 | 0.1 | 0.075 |
| NC_045512.2;YP_009724389.1;Wu1-SARS2_orf1ab;1257 | 0 | 0 |
| NC_045512.2;YP_009724389.1;Wu1-SARS2_orf1ab;1258 | 0 | 0 |
| NC_045512.2;YP_009724389.1;Wu1-SARS2_orf1ab;1259 | 0 | 0 |
| NC_045512.2;YP_009724389.1;Wu1-SARS2_orf1ab;1260 | 0 | 0 |
| NC_045512.2;YP_009724389.1;Wu1-SARS2_orf1ab;1261 | 0 | 0 |
| NC_045512.2;YP_009724389.1;Wu1-SARS2_orf1ab;1262 | 0 | 0 |
| NC_045512.2;YP_009724389.1;Wu1-SARS2_orf1ab;1263 | 0 | 0 |
| NC_045512.2;YP_009724389.1;Wu1-SARS2_orf1ab;1264 | 0 | 0 |
| NC_045512.2;YP_009724389.1;Wu1-SARS2_orf1ab;1265 | 0 | 0 |
| NC_045512.2;YP_009724389.1;Wu1-SARS2_orf1ab;1266 | 0 | 0.025 |
| NC_045512.2;YP_009724389.1;Wu1-SARS2_orf1ab;1267 | 0 | 0.025 |
| NC_045512.2;YP_009724389.1;Wu1-SARS2_orf1ab;1268 | 0 | 0.025 |
| NC_045512.2;YP_009724389.1;Wu1-SARS2_orf1ab;1269 | 0 | 0.05 |
| NC_045512.2;YP_009724389.1;Wu1-SARS2_orf1ab;1270 | 0 | 0.075 |
| NC_045512.2;YP_009724389.1;Wu1-SARS2_orf1ab;1271 | 0 | 0.075 |
| NC_045512.2;YP_009724389.1;Wu1-SARS2_orf1ab;1272 | 0.05 | 0.075 |
| NC_045512.2;YP_009724389.1;Wu1-SARS2_orf1ab;1273 | 0 | 0.025 |
| NC_045512.2;YP_009724389.1;Wu1-SARS2_orf1ab;1274 | 0 | 0.025 |
| NC_045512.2;YP_009724389.1;Wu1-SARS2_orf1ab;1275 | 0 | 0.025 |
| NC_045512.2;YP_009724389.1;Wu1-SARS2_orf1ab;1276 | 0 | 0 |
| NC_045512.2;YP_009724389.1;Wu1-SARS2_orf1ab;1277 | 0 | 0 |
| NC_045512.2;YP_009724389.1;Wu1-SARS2_orf1ab;1278 | 0 | 0 |
| NC_045512.2;YP_009724389.1;Wu1-SARS2_orf1ab;1279 | 0 | 0 |
| NC_045512.2;YP_009724389.1;Wu1-SARS2_orf1ab;1280 | 0 | 0 |

| | | |
|---|---|---|
| NC_045512.2;YP_009724389.1;Wu1-SARS2_orf1ab;1281 | 0 | 0 |
| NC_045512.2;YP_009724389.1;Wu1-SARS2_orf1ab;1282 | 0 | 0 |
| NC_045512.2;YP_009724389.1;Wu1-SARS2_orf1ab;1283 | 0 | 0 |
| NC_045512.2;YP_009724389.1;Wu1-SARS2_orf1ab;1284 | 0 | 0 |
| NC_045512.2;YP_009724389.1;Wu1-SARS2_orf1ab;1285 | 0 | 0 |
| NC_045512.2;YP_009724389.1;Wu1-SARS2_orf1ab;1286 | 0 | 0 |
| NC_045512.2;YP_009724389.1;Wu1-SARS2_orf1ab;1287 | 0 | 0 |
| NC_045512.2;YP_009724389.1;Wu1-SARS2_orf1ab;1288 | 0 | 0.025 |
| NC_045512.2;YP_009724389.1;Wu1-SARS2_orf1ab;1289 | 0 | 0 |
| NC_045512.2;YP_009724389.1;Wu1-SARS2_orf1ab;1290 | 0 | 0 |
| NC_045512.2;YP_009724389.1;Wu1-SARS2_orf1ab;1291 | 0 | 0 |
| NC_045512.2;YP_009724389.1;Wu1-SARS2_orf1ab;1292 | 0 | 0 |
| NC_045512.2;YP_009724389.1;Wu1-SARS2_orf1ab;1293 | 0 | 0 |
| NC_045512.2;YP_009724389.1;Wu1-SARS2_orf1ab;1294 | 0 | 0 |
| NC_045512.2;YP_009724389.1;Wu1-SARS2_orf1ab;1295 | 0 | 0 |
| NC_045512.2;YP_009724389.1;Wu1-SARS2_orf1ab;1296 | 0.05 | 0 |
| NC_045512.2;YP_009724389.1;Wu1-SARS2_orf1ab;1297 | 0.05 | 0.025 |
| NC_045512.2;YP_009724389.1;Wu1-SARS2_orf1ab;1298 | 0.05 | 0.025 |
| NC_045512.2;YP_009724389.1;Wu1-SARS2_orf1ab;1299 | 0.05 | 0.025 |
| NC_045512.2;YP_009724389.1;Wu1-SARS2_orf1ab;1300 | 0.05 | 0 |
| NC_045512.2;YP_009724389.1;Wu1-SARS2_orf1ab;1301 | 0.05 | 0 |
| NC_045512.2;YP_009724389.1;Wu1-SARS2_orf1ab;1302 | 0.05 | 0 |
| NC_045512.2;YP_009724389.1;Wu1-SARS2_orf1ab;1303 | 0 | 0 |
| NC_045512.2;YP_009724389.1;Wu1-SARS2_orf1ab;1304 | 0 | 0 |
| NC_045512.2;YP_009724389.1;Wu1-SARS2_orf1ab;1305 | 0 | 0 |
| NC_045512.2;YP_009724389.1;Wu1-SARS2_orf1ab;1306 | 0 | 0 |
| NC_045512.2;YP_009724389.1;Wu1-SARS2_orf1ab;1307 | 0 | 0 |
| NC_045512.2;YP_009724389.1;Wu1-SARS2_orf1ab;1308 | 0 | 0 |
| NC_045512.2;YP_009724389.1;Wu1-SARS2_orf1ab;1309 | 0.05 | 0 |
| NC_045512.2;YP_009724389.1;Wu1-SARS2_orf1ab;1310 | 0 | 0 |
| NC_045512.2;YP_009724389.1;Wu1-SARS2_orf1ab;1311 | 0 | 0 |
| NC_045512.2;YP_009724389.1;Wu1-SARS2_orf1ab;1312 | 0 | 0 |
| NC_045512.2;YP_009724389.1;Wu1-SARS2_orf1ab;1313 | 0 | 0 |
| NC_045512.2;YP_009724389.1;Wu1-SARS2_orf1ab;1314 | 0 | 0 |
| NC_045512.2;YP_009724389.1;Wu1-SARS2_orf1ab;1315 | 0 | 0 |
| NC_045512.2;YP_009724389.1;Wu1-SARS2_orf1ab;1316 | 0.05 | 0 |
| NC_045512.2;YP_009724389.1;Wu1-SARS2_orf1ab;1317 | 0.05 | 0 |
| NC_045512.2;YP_009724389.1;Wu1-SARS2_orf1ab;1318 | 0.05 | 0.025 |
| NC_045512.2;YP_009724389.1;Wu1-SARS2_orf1ab;1319 | 0 | 0 |
| NC_045512.2;YP_009724389.1;Wu1-SARS2_orf1ab;1320 | 0 | 0.05 |
| NC_045512.2;YP_009724389.1;Wu1-SARS2_orf1ab;1321 | 0 | 0 |
| NC_045512.2;YP_009724389.1;Wu1-SARS2_orf1ab;1322 | 0 | 0 |
| NC_045512.2;YP_009724389.1;Wu1-SARS2_orf1ab;1323 | 0.05 | 0 |

| | | |
|---|---|---|
| NC_045512.2;YP_009724389.1;Wu1-SARS2_orf1ab;1324 | 0 | 0 |
| NC_045512.2;YP_009724389.1;Wu1-SARS2_orf1ab;1325 | 0 | 0 |
| NC_045512.2;YP_009724389.1;Wu1-SARS2_orf1ab;1326 | 0 | 0 |
| NC_045512.2;YP_009724389.1;Wu1-SARS2_orf1ab;1327 | 0 | 0 |
| NC_045512.2;YP_009724389.1;Wu1-SARS2_orf1ab;1328 | 0.05 | 0 |
| NC_045512.2;YP_009724389.1;Wu1-SARS2_orf1ab;1329 | 0.05 | 0 |
| NC_045512.2;YP_009724389.1;Wu1-SARS2_orf1ab;1330 | 0 | 0 |
| NC_045512.2;YP_009724389.1;Wu1-SARS2_orf1ab;1331 | 0 | 0.025 |
| NC_045512.2;YP_009724389.1;Wu1-SARS2_orf1ab;1332 | 0 | 0.05 |
| NC_045512.2;YP_009724389.1;Wu1-SARS2_orf1ab;1333 | 0 | 0 |
| NC_045512.2;YP_009724389.1;Wu1-SARS2_orf1ab;1334 | 0 | 0 |
| NC_045512.2;YP_009724389.1;Wu1-SARS2_orf1ab;1335 | 0 | 0 |
| NC_045512.2;YP_009724389.1;Wu1-SARS2_orf1ab;1336 | 0 | 0.025 |
| NC_045512.2;YP_009724389.1;Wu1-SARS2_orf1ab;1337 | 0 | 0.05 |
| NC_045512.2;YP_009724389.1;Wu1-SARS2_orf1ab;1338 | 0 | 0 |
| NC_045512.2;YP_009724389.1;Wu1-SARS2_orf1ab;1339 | 0 | 0 |
| NC_045512.2;YP_009724389.1;Wu1-SARS2_orf1ab;1340 | 0 | 0 |
| NC_045512.2;YP_009724389.1;Wu1-SARS2_orf1ab;1341 | 0 | 0 |
| NC_045512.2;YP_009724389.1;Wu1-SARS2_orf1ab;1342 | 0 | 0 |
| NC_045512.2;YP_009724389.1;Wu1-SARS2_orf1ab;1343 | 0 | 0 |
| NC_045512.2;YP_009724389.1;Wu1-SARS2_orf1ab;1344 | 0 | 0 |
| NC_045512.2;YP_009724389.1;Wu1-SARS2_orf1ab;1345 | 0 | 0 |
| NC_045512.2;YP_009724389.1;Wu1-SARS2_orf1ab;1346 | 0 | 0 |
| NC_045512.2;YP_009724389.1;Wu1-SARS2_orf1ab;1347 | 0 | 0 |
| NC_045512.2;YP_009724389.1;Wu1-SARS2_orf1ab;1348 | 0 | 0 |
| NC_045512.2;YP_009724389.1;Wu1-SARS2_orf1ab;1349 | 0 | 0 |
| NC_045512.2;YP_009724389.1;Wu1-SARS2_orf1ab;1350 | 0 | 0.025 |
| NC_045512.2;YP_009724389.1;Wu1-SARS2_orf1ab;1351 | 0 | 0 |
| NC_045512.2;YP_009724389.1;Wu1-SARS2_orf1ab;1352 | 0 | 0 |
| NC_045512.2;YP_009724389.1;Wu1-SARS2_orf1ab;1353 | 0 | 0 |
| NC_045512.2;YP_009724389.1;Wu1-SARS2_orf1ab;1354 | 0.05 | 0 |
| NC_045512.2;YP_009724389.1;Wu1-SARS2_orf1ab;1355 | 0 | 0 |
| NC_045512.2;YP_009724389.1;Wu1-SARS2_orf1ab;1356 | 0 | 0 |
| NC_045512.2;YP_009724389.1;Wu1-SARS2_orf1ab;1357 | 0.05 | 0.025 |
| NC_045512.2;YP_009724389.1;Wu1-SARS2_orf1ab;1358 | 0 | 0 |
| NC_045512.2;YP_009724389.1;Wu1-SARS2_orf1ab;1359 | 0.05 | 0 |
| NC_045512.2;YP_009724389.1;Wu1-SARS2_orf1ab;1360 | 0 | 0 |
| NC_045512.2;YP_009724389.1;Wu1-SARS2_orf1ab;1361 | 0 | 0 |
| NC_045512.2;YP_009724389.1;Wu1-SARS2_orf1ab;1362 | 0 | 0 |
| NC_045512.2;YP_009724389.1;Wu1-SARS2_orf1ab;1363 | 0 | 0 |
| NC_045512.2;YP_009724389.1;Wu1-SARS2_orf1ab;1364 | 0 | 0 |
| NC_045512.2;YP_009724389.1;Wu1-SARS2_orf1ab;1365 | 0.05 | 0 |
| NC_045512.2;YP_009724389.1;Wu1-SARS2_orf1ab;1366 | 0 | 0 |

| | | |
|---|---|---|
| NC_045512.2;YP_009724389.1;Wu1-SARS2_orf1ab;1367 | 0 | 0 |
| NC_045512.2;YP_009724389.1;Wu1-SARS2_orf1ab;1368 | 0 | 0 |
| NC_045512.2;YP_009724389.1;Wu1-SARS2_orf1ab;1369 | 0 | 0.025 |
| NC_045512.2;YP_009724389.1;Wu1-SARS2_orf1ab;1370 | 0 | 0 |
| NC_045512.2;YP_009724389.1;Wu1-SARS2_orf1ab;1371 | 0 | 0 |
| NC_045512.2;YP_009724389.1;Wu1-SARS2_orf1ab;1372 | 0 | 0.05 |
| NC_045512.2;YP_009724389.1;Wu1-SARS2_orf1ab;1373 | 0 | 0.025 |
| NC_045512.2;YP_009724389.1;Wu1-SARS2_orf1ab;1374 | 0 | 0 |
| NC_045512.2;YP_009724389.1;Wu1-SARS2_orf1ab;1375 | 0 | 0.025 |
| NC_045512.2;YP_009724389.1;Wu1-SARS2_orf1ab;1376 | 0 | 0 |
| NC_045512.2;YP_009724389.1;Wu1-SARS2_orf1ab;1377 | 0 | 0 |
| NC_045512.2;YP_009724389.1;Wu1-SARS2_orf1ab;1378 | 0 | 0 |
| NC_045512.2;YP_009724389.1;Wu1-SARS2_orf1ab;1379 | 0 | 0 |
| NC_045512.2;YP_009724389.1;Wu1-SARS2_orf1ab;1380 | 0 | 0 |
| NC_045512.2;YP_009724389.1;Wu1-SARS2_orf1ab;1381 | 0 | 0 |
| NC_045512.2;YP_009724389.1;Wu1-SARS2_orf1ab;1382 | 0.05 | 0.025 |
| NC_045512.2;YP_009724389.1;Wu1-SARS2_orf1ab;1383 | 0.05 | 0.025 |
| NC_045512.2;YP_009724389.1;Wu1-SARS2_orf1ab;1384 | 0 | 0 |
| NC_045512.2;YP_009724389.1;Wu1-SARS2_orf1ab;1385 | 0 | 0 |
| NC_045512.2;YP_009724389.1;Wu1-SARS2_orf1ab;1386 | 0 | 0 |
| NC_045512.2;YP_009724389.1;Wu1-SARS2_orf1ab;1387 | 0 | 0 |
| NC_045512.2;YP_009724389.1;Wu1-SARS2_orf1ab;1388 | 0 | 0 |
| NC_045512.2;YP_009724389.1;Wu1-SARS2_orf1ab;1389 | 0 | 0 |
| NC_045512.2;YP_009724389.1;Wu1-SARS2_orf1ab;1390 | 0 | 0 |
| NC_045512.2;YP_009724389.1;Wu1-SARS2_orf1ab;1391 | 0 | 0 |
| NC_045512.2;YP_009724389.1;Wu1-SARS2_orf1ab;1392 | 0 | 0 |
| NC_045512.2;YP_009724389.1;Wu1-SARS2_orf1ab;1393 | 0 | 0 |
| NC_045512.2;YP_009724389.1;Wu1-SARS2_orf1ab;1394 | 0 | 0 |
| NC_045512.2;YP_009724389.1;Wu1-SARS2_orf1ab;1395 | 0 | 0 |
| NC_045512.2;YP_009724389.1;Wu1-SARS2_orf1ab;1396 | 0.05 | 0 |
| NC_045512.2;YP_009724389.1;Wu1-SARS2_orf1ab;1397 | 0.05 | 0 |
| NC_045512.2;YP_009724389.1;Wu1-SARS2_orf1ab;1398 | 0.1 | 0 |
| NC_045512.2;YP_009724389.1;Wu1-SARS2_orf1ab;1399 | 0.1 | 0 |
| NC_045512.2;YP_009724389.1;Wu1-SARS2_orf1ab;1400 | 0.1 | 0 |
| NC_045512.2;YP_009724389.1;Wu1-SARS2_orf1ab;1401 | 0.1 | 0 |
| NC_045512.2;YP_009724389.1;Wu1-SARS2_orf1ab;1402 | 0.1 | 0 |
| NC_045512.2;YP_009724389.1;Wu1-SARS2_orf1ab;1403 | 0.1 | 0.025 |
| NC_045512.2;YP_009724389.1;Wu1-SARS2_orf1ab;1404 | 0.1 | 0.025 |
| NC_045512.2;YP_009724389.1;Wu1-SARS2_orf1ab;1405 | 0.1 | 0.025 |
| NC_045512.2;YP_009724389.1;Wu1-SARS2_orf1ab;1406 | 0.1 | 0.025 |
| NC_045512.2;YP_009724389.1;Wu1-SARS2_orf1ab;1407 | 0 | 0.025 |
| NC_045512.2;YP_009724389.1;Wu1-SARS2_orf1ab;1408 | 0 | 0 |
| NC_045512.2;YP_009724389.1;Wu1-SARS2_orf1ab;1409 | 0 | 0 |

| | | |
|---|---|---|
| NC_045512.2;YP_009724389.1;Wu1-SARS2_orf1ab;1410 | 0 | 0 |
| NC_045512.2;YP_009724389.1;Wu1-SARS2_orf1ab;1411 | 0 | 0 |
| NC_045512.2;YP_009724389.1;Wu1-SARS2_orf1ab;1412 | 0 | 0 |
| NC_045512.2;YP_009724389.1;Wu1-SARS2_orf1ab;1413 | 0 | 0 |
| NC_045512.2;YP_009724389.1;Wu1-SARS2_orf1ab;1414 | 0.05 | 0.1 |
| NC_045512.2;YP_009724389.1;Wu1-SARS2_orf1ab;1415 | 0 | 0 |
| NC_045512.2;YP_009724389.1;Wu1-SARS2_orf1ab;1416 | 0 | 0 |
| NC_045512.2;YP_009724389.1;Wu1-SARS2_orf1ab;1417 | 0 | 0 |
| NC_045512.2;YP_009724389.1;Wu1-SARS2_orf1ab;1418 | 0 | 0 |
| NC_045512.2;YP_009724389.1;Wu1-SARS2_orf1ab;1419 | 0 | 0 |
| NC_045512.2;YP_009724389.1;Wu1-SARS2_orf1ab;1420 | 0 | 0 |
| NC_045512.2;YP_009724389.1;Wu1-SARS2_orf1ab;1421 | 0 | 0 |
| NC_045512.2;YP_009724389.1;Wu1-SARS2_orf1ab;1422 | 0 | 0 |
| NC_045512.2;YP_009724389.1;Wu1-SARS2_orf1ab;1423 | 0 | 0 |
| NC_045512.2;YP_009724389.1;Wu1-SARS2_orf1ab;1424 | 0 | 0.025 |
| NC_045512.2;YP_009724389.1;Wu1-SARS2_orf1ab;1425 | 0 | 0.05 |
| NC_045512.2;YP_009724389.1;Wu1-SARS2_orf1ab;1426 | 0 | 0.025 |
| NC_045512.2;YP_009724389.1;Wu1-SARS2_orf1ab;1427 | 0.2 | 0.1 |
| NC_045512.2;YP_009724389.1;Wu1-SARS2_orf1ab;1428 | 0.25 | 0.125 |
| NC_045512.2;YP_009724389.1;Wu1-SARS2_orf1ab;1429 | 0.15 | 0.25 |
| NC_045512.2;YP_009724389.1;Wu1-SARS2_orf1ab;1430 | 0.2 | 0.25 |
| NC_045512.2;YP_009724389.1;Wu1-SARS2_orf1ab;1431 | 0.3 | 0.35 |
| NC_045512.2;YP_009724389.1;Wu1-SARS2_orf1ab;1432 | 0.2 | 0.225 |
| NC_045512.2;YP_009724389.1;Wu1-SARS2_orf1ab;1433 | 0.2 | 0.175 |
| NC_045512.2;YP_009724389.1;Wu1-SARS2_orf1ab;1434 | 0.2 | 0.225 |
| NC_045512.2;YP_009724389.1;Wu1-SARS2_orf1ab;1435 | 0.25 | 0.2 |
| NC_045512.2;YP_009724389.1;Wu1-SARS2_orf1ab;1436 | 0.2 | 0.2 |
| NC_045512.2;YP_009724389.1;Wu1-SARS2_orf1ab;1437 | 0.35 | 0.3 |
| NC_045512.2;YP_009724389.1;Wu1-SARS2_orf1ab;1438 | 0.15 | 0.15 |
| NC_045512.2;YP_009724389.1;Wu1-SARS2_orf1ab;1439 | 0.1 | 0.1 |
| NC_045512.2;YP_009724389.1;Wu1-SARS2_orf1ab;1440 | 0.1 | 0.1 |
| NC_045512.2;YP_009724389.1;Wu1-SARS2_orf1ab;1441 | 0.1 | 0.1 |
| NC_045512.2;YP_009724389.1;Wu1-SARS2_orf1ab;1442 | 0.05 | 0.075 |
| NC_045512.2;YP_009724389.1;Wu1-SARS2_orf1ab;1443 | 0 | 0 |
| NC_045512.2;YP_009724389.1;Wu1-SARS2_orf1ab;1444 | 0 | 0 |
| NC_045512.2;YP_009724389.1;Wu1-SARS2_orf1ab;1445 | 0 | 0 |
| NC_045512.2;YP_009724389.1;Wu1-SARS2_orf1ab;1446 | 0.05 | 0 |
| NC_045512.2;YP_009724389.1;Wu1-SARS2_orf1ab;1447 | 0 | 0 |
| NC_045512.2;YP_009724389.1;Wu1-SARS2_orf1ab;1448 | 0.1 | 0.05 |
| NC_045512.2;YP_009724389.1;Wu1-SARS2_orf1ab;1449 | 0.05 | 0.025 |
| NC_045512.2;YP_009724389.1;Wu1-SARS2_orf1ab;1450 | 0.05 | 0.025 |
| NC_045512.2;YP_009724389.1;Wu1-SARS2_orf1ab;1451 | 0.05 | 0.025 |
| NC_045512.2;YP_009724389.1;Wu1-SARS2_orf1ab;1452 | 0 | 0 |

| | | |
|---|---|---|
| NC_045512.2;YP_009724389.1;Wu1-SARS2_orf1ab;1453 | 0 | 0 |
| NC_045512.2;YP_009724389.1;Wu1-SARS2_orf1ab;1454 | 0 | 0 |
| NC_045512.2;YP_009724389.1;Wu1-SARS2_orf1ab;1455 | 0 | 0 |
| NC_045512.2;YP_009724389.1;Wu1-SARS2_orf1ab;1456 | 0 | 0.025 |
| NC_045512.2;YP_009724389.1;Wu1-SARS2_orf1ab;1457 | 0 | 0 |
| NC_045512.2;YP_009724389.1;Wu1-SARS2_orf1ab;1458 | 0 | 0 |
| NC_045512.2;YP_009724389.1;Wu1-SARS2_orf1ab;1459 | 0 | 0.075 |
| NC_045512.2;YP_009724389.1;Wu1-SARS2_orf1ab;1460 | 0.1 | 0.075 |
| NC_045512.2;YP_009724389.1;Wu1-SARS2_orf1ab;1461 | 0.15 | 0.125 |
| NC_045512.2;YP_009724389.1;Wu1-SARS2_orf1ab;1462 | 0.05 | 0 |
| NC_045512.2;YP_009724389.1;Wu1-SARS2_orf1ab;1463 | 0 | 0.025 |
| NC_045512.2;YP_009724389.1;Wu1-SARS2_orf1ab;1464 | 0 | 0 |
| NC_045512.2;YP_009724389.1;Wu1-SARS2_orf1ab;1465 | 0 | 0 |
| NC_045512.2;YP_009724389.1;Wu1-SARS2_orf1ab;1466 | 0 | 0 |
| NC_045512.2;YP_009724389.1;Wu1-SARS2_orf1ab;1467 | 0 | 0 |
| NC_045512.2;YP_009724389.1;Wu1-SARS2_orf1ab;1468 | 0 | 0 |
| NC_045512.2;YP_009724389.1;Wu1-SARS2_orf1ab;1469 | 0 | 0 |
| NC_045512.2;YP_009724389.1;Wu1-SARS2_orf1ab;1470 | 0 | 0 |
| NC_045512.2;YP_009724389.1;Wu1-SARS2_orf1ab;1471 | 0 | 0 |
| NC_045512.2;YP_009724389.1;Wu1-SARS2_orf1ab;1472 | 0 | 0.05 |
| NC_045512.2;YP_009724389.1;Wu1-SARS2_orf1ab;1473 | 0.05 | 0.025 |
| NC_045512.2;YP_009724389.1;Wu1-SARS2_orf1ab;1474 | 0 | 0.1 |
| NC_045512.2;YP_009724389.1;Wu1-SARS2_orf1ab;1475 | 0 | 0.025 |
| NC_045512.2;YP_009724389.1;Wu1-SARS2_orf1ab;1476 | 0 | 0 |
| NC_045512.2;YP_009724389.1;Wu1-SARS2_orf1ab;1477 | 0 | 0 |
| NC_045512.2;YP_009724389.1;Wu1-SARS2_orf1ab;1478 | 0 | 0 |
| NC_045512.2;YP_009724389.1;Wu1-SARS2_orf1ab;1479 | 0 | 0 |
| NC_045512.2;YP_009724389.1;Wu1-SARS2_orf1ab;1480 | 0.05 | 0.05 |
| NC_045512.2;YP_009724389.1;Wu1-SARS2_orf1ab;1481 | 0.05 | 0.075 |
| NC_045512.2;YP_009724389.1;Wu1-SARS2_orf1ab;1482 | 0.1 | 0.05 |
| NC_045512.2;YP_009724389.1;Wu1-SARS2_orf1ab;1483 | 0.15 | 0.1 |
| NC_045512.2;YP_009724389.1;Wu1-SARS2_orf1ab;1484 | 0.1 | 0.15 |
| NC_045512.2;YP_009724389.1;Wu1-SARS2_orf1ab;1485 | 0.1 | 0.125 |
| NC_045512.2;YP_009724389.1;Wu1-SARS2_orf1ab;1486 | 0.15 | 0.125 |
| NC_045512.2;YP_009724389.1;Wu1-SARS2_orf1ab;1487 | 0.2 | 0.125 |
| NC_045512.2;YP_009724389.1;Wu1-SARS2_orf1ab;1488 | 0.25 | 0.1 |
| NC_045512.2;YP_009724389.1;Wu1-SARS2_orf1ab;1489 | 0.2 | 0.1 |
| NC_045512.2;YP_009724389.1;Wu1-SARS2_orf1ab;1490 | 0.15 | 0.025 |
| NC_045512.2;YP_009724389.1;Wu1-SARS2_orf1ab;1491 | 0.05 | 0 |
| NC_045512.2;YP_009724389.1;Wu1-SARS2_orf1ab;1492 | 0.05 | 0 |
| NC_045512.2;YP_009724389.1;Wu1-SARS2_orf1ab;1493 | 0.05 | 0 |
| NC_045512.2;YP_009724389.1;Wu1-SARS2_orf1ab;1494 | 0 | 0 |
| NC_045512.2;YP_009724389.1;Wu1-SARS2_orf1ab;1495 | 0 | 0 |

| | | |
|---|---|---|
| NC_045512.2;YP_009724389.1;Wu1-SARS2_orf1ab;1496 | 0 | 0 |
| NC_045512.2;YP_009724389.1;Wu1-SARS2_orf1ab;1497 | 0 | 0 |
| NC_045512.2;YP_009724389.1;Wu1-SARS2_orf1ab;1498 | 0 | 0 |
| NC_045512.2;YP_009724389.1;Wu1-SARS2_orf1ab;1499 | 0 | 0 |
| NC_045512.2;YP_009724389.1;Wu1-SARS2_orf1ab;1500 | 0 | 0 |
| NC_045512.2;YP_009724389.1;Wu1-SARS2_orf1ab;1501 | 0 | 0 |
| NC_045512.2;YP_009724389.1;Wu1-SARS2_orf1ab;1502 | 0 | 0 |
| NC_045512.2;YP_009724389.1;Wu1-SARS2_orf1ab;1503 | 0 | 0 |
| NC_045512.2;YP_009724389.1;Wu1-SARS2_orf1ab;1504 | 0 | 0 |
| NC_045512.2;YP_009724389.1;Wu1-SARS2_orf1ab;1505 | 0 | 0 |
| NC_045512.2;YP_009724389.1;Wu1-SARS2_orf1ab;1506 | 0.05 | 0.05 |
| NC_045512.2;YP_009724389.1;Wu1-SARS2_orf1ab;1507 | 0 | 0.05 |
| NC_045512.2;YP_009724389.1;Wu1-SARS2_orf1ab;1508 | 0 | 0.025 |
| NC_045512.2;YP_009724389.1;Wu1-SARS2_orf1ab;1509 | 0.05 | 0 |
| NC_045512.2;YP_009724389.1;Wu1-SARS2_orf1ab;1510 | 0 | 0 |
| NC_045512.2;YP_009724389.1;Wu1-SARS2_orf1ab;1511 | 0 | 0.025 |
| NC_045512.2;YP_009724389.1;Wu1-SARS2_orf1ab;1512 | 0 | 0 |
| NC_045512.2;YP_009724389.1;Wu1-SARS2_orf1ab;1513 | 0 | 0 |
| NC_045512.2;YP_009724389.1;Wu1-SARS2_orf1ab;1514 | 0 | 0 |
| NC_045512.2;YP_009724389.1;Wu1-SARS2_orf1ab;1515 | 0 | 0 |
| NC_045512.2;YP_009724389.1;Wu1-SARS2_orf1ab;1516 | 0 | 0 |
| NC_045512.2;YP_009724389.1;Wu1-SARS2_orf1ab;1517 | 0 | 0 |
| NC_045512.2;YP_009724389.1;Wu1-SARS2_orf1ab;1518 | 0 | 0.075 |
| NC_045512.2;YP_009724389.1;Wu1-SARS2_orf1ab;1519 | 0 | 0.075 |
| NC_045512.2;YP_009724389.1;Wu1-SARS2_orf1ab;1520 | 0 | 0.075 |
| NC_045512.2;YP_009724389.1;Wu1-SARS2_orf1ab;1521 | 0 | 0.075 |
| NC_045512.2;YP_009724389.1;Wu1-SARS2_orf1ab;1522 | 0 | 0.05 |
| NC_045512.2;YP_009724389.1;Wu1-SARS2_orf1ab;1523 | 0 | 0.05 |
| NC_045512.2;YP_009724389.1;Wu1-SARS2_orf1ab;1524 | 0.05 | 0.075 |
| NC_045512.2;YP_009724389.1;Wu1-SARS2_orf1ab;1525 | 0 | 0.05 |
| NC_045512.2;YP_009724389.1;Wu1-SARS2_orf1ab;1526 | 0.05 | 0.075 |
| NC_045512.2;YP_009724389.1;Wu1-SARS2_orf1ab;1527 | 0 | 0.025 |
| NC_045512.2;YP_009724389.1;Wu1-SARS2_orf1ab;1528 | 0 | 0.025 |
| NC_045512.2;YP_009724389.1;Wu1-SARS2_orf1ab;1529 | 0 | 0.025 |
| NC_045512.2;YP_009724389.1;Wu1-SARS2_orf1ab;1530 | 0 | 0 |
| NC_045512.2;YP_009724389.1;Wu1-SARS2_orf1ab;1531 | 0 | 0 |
| NC_045512.2;YP_009724389.1;Wu1-SARS2_orf1ab;1532 | 0 | 0 |
| NC_045512.2;YP_009724389.1;Wu1-SARS2_orf1ab;1533 | 0 | 0 |
| NC_045512.2;YP_009724389.1;Wu1-SARS2_orf1ab;1534 | 0 | 0 |
| NC_045512.2;YP_009724389.1;Wu1-SARS2_orf1ab;1535 | 0 | 0 |
| NC_045512.2;YP_009724389.1;Wu1-SARS2_orf1ab;1536 | 0 | 0 |
| NC_045512.2;YP_009724389.1;Wu1-SARS2_orf1ab;1537 | 0 | 0 |
| NC_045512.2;YP_009724389.1;Wu1-SARS2_orf1ab;1538 | 0 | 0 |

| | | |
|---|---|---|
| NC_045512.2;YP_009724389.1;Wu1-SARS2_orf1ab;1539 | 0 | 0 |
| NC_045512.2;YP_009724389.1;Wu1-SARS2_orf1ab;1540 | 0 | 0 |
| NC_045512.2;YP_009724389.1;Wu1-SARS2_orf1ab;1541 | 0 | 0 |
| NC_045512.2;YP_009724389.1;Wu1-SARS2_orf1ab;1542 | 0 | 0.025 |
| NC_045512.2;YP_009724389.1;Wu1-SARS2_orf1ab;1543 | 0 | 0.025 |
| NC_045512.2;YP_009724389.1;Wu1-SARS2_orf1ab;1544 | 0 | 0.025 |
| NC_045512.2;YP_009724389.1;Wu1-SARS2_orf1ab;1545 | 0 | 0.05 |
| NC_045512.2;YP_009724389.1;Wu1-SARS2_orf1ab;1546 | 0 | 0.05 |
| NC_045512.2;YP_009724389.1;Wu1-SARS2_orf1ab;1547 | 0 | 0.025 |
| NC_045512.2;YP_009724389.1;Wu1-SARS2_orf1ab;1548 | 0 | 0 |
| NC_045512.2;YP_009724389.1;Wu1-SARS2_orf1ab;1549 | 0 | 0.025 |
| NC_045512.2;YP_009724389.1;Wu1-SARS2_orf1ab;1550 | 0 | 0 |
| NC_045512.2;YP_009724389.1;Wu1-SARS2_orf1ab;1551 | 0 | 0.05 |
| NC_045512.2;YP_009724389.1;Wu1-SARS2_orf1ab;1552 | 0 | 0 |
| NC_045512.2;YP_009724389.1;Wu1-SARS2_orf1ab;1553 | 0 | 0 |
| NC_045512.2;YP_009724389.1;Wu1-SARS2_orf1ab;1554 | 0 | 0 |
| NC_045512.2;YP_009724389.1;Wu1-SARS2_orf1ab;1555 | 0 | 0 |
| NC_045512.2;YP_009724389.1;Wu1-SARS2_orf1ab;1556 | 0 | 0 |
| NC_045512.2;YP_009724389.1;Wu1-SARS2_orf1ab;1557 | 0 | 0 |
| NC_045512.2;YP_009724389.1;Wu1-SARS2_orf1ab;1558 | 0 | 0 |
| NC_045512.2;YP_009724389.1;Wu1-SARS2_orf1ab;1559 | 0 | 0 |
| NC_045512.2;YP_009724389.1;Wu1-SARS2_orf1ab;1560 | 0 | 0 |
| NC_045512.2;YP_009724389.1;Wu1-SARS2_orf1ab;1561 | 0 | 0 |
| NC_045512.2;YP_009724389.1;Wu1-SARS2_orf1ab;1562 | 0 | 0 |
| NC_045512.2;YP_009724389.1;Wu1-SARS2_orf1ab;1563 | 0 | 0.025 |
| NC_045512.2;YP_009724389.1;Wu1-SARS2_orf1ab;1564 | 0 | 0 |
| NC_045512.2;YP_009724389.1;Wu1-SARS2_orf1ab;1565 | 0 | 0 |
| NC_045512.2;YP_009724389.1;Wu1-SARS2_orf1ab;1566 | 0 | 0 |
| NC_045512.2;YP_009724389.1;Wu1-SARS2_orf1ab;1567 | 0 | 0.025 |
| NC_045512.2;YP_009724389.1;Wu1-SARS2_orf1ab;1568 | 0 | 0.025 |
| NC_045512.2;YP_009724389.1;Wu1-SARS2_orf1ab;1569 | 0 | 0.025 |
| NC_045512.2;YP_009724389.1;Wu1-SARS2_orf1ab;1570 | 0 | 0.025 |
| NC_045512.2;YP_009724389.1;Wu1-SARS2_orf1ab;1571 | 0 | 0.025 |
| NC_045512.2;YP_009724389.1;Wu1-SARS2_orf1ab;1572 | 0 | 0.025 |
| NC_045512.2;YP_009724389.1;Wu1-SARS2_orf1ab;1573 | 0.05 | 0.125 |
| NC_045512.2;YP_009724389.1;Wu1-SARS2_orf1ab;1574 | 0 | 0.075 |
| NC_045512.2;YP_009724389.1;Wu1-SARS2_orf1ab;1575 | 0 | 0.025 |
| NC_045512.2;YP_009724389.1;Wu1-SARS2_orf1ab;1576 | 0 | 0.025 |
| NC_045512.2;YP_009724389.1;Wu1-SARS2_orf1ab;1577 | 0 | 0.025 |
| NC_045512.2;YP_009724389.1;Wu1-SARS2_orf1ab;1578 | 0 | 0.025 |
| NC_045512.2;YP_009724389.1;Wu1-SARS2_orf1ab;1579 | 0 | 0.025 |
| NC_045512.2;YP_009724389.1;Wu1-SARS2_orf1ab;1580 | 0 | 0 |
| NC_045512.2;YP_009724389.1;Wu1-SARS2_orf1ab;1581 | 0 | 0 |

| | | |
|---|---|---|
| NC_045512.2;YP_009724389.1;Wu1-SARS2_orf1ab;1582 | 0 | 0 |
| NC_045512.2;YP_009724389.1;Wu1-SARS2_orf1ab;1583 | 0 | 0 |
| NC_045512.2;YP_009724389.1;Wu1-SARS2_orf1ab;1584 | 0 | 0 |
| NC_045512.2;YP_009724389.1;Wu1-SARS2_orf1ab;1585 | 0 | 0 |
| NC_045512.2;YP_009724389.1;Wu1-SARS2_orf1ab;1586 | 0 | 0 |
| NC_045512.2;YP_009724389.1;Wu1-SARS2_orf1ab;1587 | 0 | 0 |
| NC_045512.2;YP_009724389.1;Wu1-SARS2_orf1ab;1588 | 0 | 0 |
| NC_045512.2;YP_009724389.1;Wu1-SARS2_orf1ab;1589 | 0 | 0 |
| NC_045512.2;YP_009724389.1;Wu1-SARS2_orf1ab;1590 | 0 | 0 |
| NC_045512.2;YP_009724389.1;Wu1-SARS2_orf1ab;1591 | 0 | 0.025 |
| NC_045512.2;YP_009724389.1;Wu1-SARS2_orf1ab;1592 | 0 | 0.025 |
| NC_045512.2;YP_009724389.1;Wu1-SARS2_orf1ab;1593 | 0 | 0.025 |
| NC_045512.2;YP_009724389.1;Wu1-SARS2_orf1ab;1594 | 0 | 0.025 |
| NC_045512.2;YP_009724389.1;Wu1-SARS2_orf1ab;1595 | 0.15 | 0.125 |
| NC_045512.2;YP_009724389.1;Wu1-SARS2_orf1ab;1596 | 0 | 0.025 |
| NC_045512.2;YP_009724389.1;Wu1-SARS2_orf1ab;1597 | 0 | 0.025 |
| NC_045512.2;YP_009724389.1;Wu1-SARS2_orf1ab;1598 | 0 | 0 |
| NC_045512.2;YP_009724389.1;Wu1-SARS2_orf1ab;1599 | 0 | 0 |
| NC_045512.2;YP_009724389.1;Wu1-SARS2_orf1ab;1600 | 0 | 0 |
| NC_045512.2;YP_009724389.1;Wu1-SARS2_orf1ab;1601 | 0 | 0.025 |
| NC_045512.2;YP_009724389.1;Wu1-SARS2_orf1ab;1602 | 0.05 | 0 |
| NC_045512.2;YP_009724389.1;Wu1-SARS2_orf1ab;1603 | 0.05 | 0 |
| NC_045512.2;YP_009724389.1;Wu1-SARS2_orf1ab;1604 | 0 | 0 |
| NC_045512.2;YP_009724389.1;Wu1-SARS2_orf1ab;1605 | 0 | 0 |
| NC_045512.2;YP_009724389.1;Wu1-SARS2_orf1ab;1606 | 0 | 0 |
| NC_045512.2;YP_009724389.1;Wu1-SARS2_orf1ab;1607 | 0 | 0 |
| NC_045512.2;YP_009724389.1;Wu1-SARS2_orf1ab;1608 | 0 | 0 |
| NC_045512.2;YP_009724389.1;Wu1-SARS2_orf1ab;1609 | 0 | 0 |
| NC_045512.2;YP_009724389.1;Wu1-SARS2_orf1ab;1610 | 0 | 0 |
| NC_045512.2;YP_009724389.1;Wu1-SARS2_orf1ab;1611 | 0.05 | 0 |
| NC_045512.2;YP_009724389.1;Wu1-SARS2_orf1ab;1612 | 0.05 | 0 |
| NC_045512.2;YP_009724389.1;Wu1-SARS2_orf1ab;1613 | 0.05 | 0 |
| NC_045512.2;YP_009724389.1;Wu1-SARS2_orf1ab;1614 | 0.05 | 0 |
| NC_045512.2;YP_009724389.1;Wu1-SARS2_orf1ab;1615 | 0.05 | 0 |
| NC_045512.2;YP_009724389.1;Wu1-SARS2_orf1ab;1616 | 0.05 | 0 |
| NC_045512.2;YP_009724389.1;Wu1-SARS2_orf1ab;1617 | 0 | 0 |
| NC_045512.2;YP_009724389.1;Wu1-SARS2_orf1ab;1618 | 0 | 0 |
| NC_045512.2;YP_009724389.1;Wu1-SARS2_orf1ab;1619 | 0 | 0 |
| NC_045512.2;YP_009724389.1;Wu1-SARS2_orf1ab;1620 | 0 | 0 |
| NC_045512.2;YP_009724389.1;Wu1-SARS2_orf1ab;1621 | 0 | 0 |
| NC_045512.2;YP_009724389.1;Wu1-SARS2_orf1ab;1622 | 0 | 0 |
| NC_045512.2;YP_009724389.1;Wu1-SARS2_orf1ab;1623 | 0 | 0 |
| NC_045512.2;YP_009724389.1;Wu1-SARS2_orf1ab;1624 | 0 | 0 |

| | | |
|---|---|---|
| NC_045512.2;YP_009724389.1;Wu1-SARS2_orf1ab;1625 | 0 | 0 |
| NC_045512.2;YP_009724389.1;Wu1-SARS2_orf1ab;1626 | 0 | 0 |
| NC_045512.2;YP_009724389.1;Wu1-SARS2_orf1ab;1627 | 0 | 0 |
| NC_045512.2;YP_009724389.1;Wu1-SARS2_orf1ab;1628 | 0 | 0 |
| NC_045512.2;YP_009724389.1;Wu1-SARS2_orf1ab;1629 | 0 | 0 |
| NC_045512.2;YP_009724389.1;Wu1-SARS2_orf1ab;1630 | 0 | 0 |
| NC_045512.2;YP_009724389.1;Wu1-SARS2_orf1ab;1631 | 0.05 | 0 |
| NC_045512.2;YP_009724389.1;Wu1-SARS2_orf1ab;1632 | 0.05 | 0 |
| NC_045512.2;YP_009724389.1;Wu1-SARS2_orf1ab;1633 | 0.05 | 0 |
| NC_045512.2;YP_009724389.1;Wu1-SARS2_orf1ab;1634 | 0.05 | 0 |
| NC_045512.2;YP_009724389.1;Wu1-SARS2_orf1ab;1635 | 0 | 0 |
| NC_045512.2;YP_009724389.1;Wu1-SARS2_orf1ab;1636 | 0 | 0 |
| NC_045512.2;YP_009724389.1;Wu1-SARS2_orf1ab;1637 | 0 | 0 |
| NC_045512.2;YP_009724389.1;Wu1-SARS2_orf1ab;1638 | 0 | 0 |
| NC_045512.2;YP_009724389.1;Wu1-SARS2_orf1ab;1639 | 0.05 | 0 |
| NC_045512.2;YP_009724389.1;Wu1-SARS2_orf1ab;1640 | 0 | 0 |
| NC_045512.2;YP_009724389.1;Wu1-SARS2_orf1ab;1641 | 0 | 0 |
| NC_045512.2;YP_009724389.1;Wu1-SARS2_orf1ab;1642 | 0 | 0 |
| NC_045512.2;YP_009724389.1;Wu1-SARS2_orf1ab;1643 | 0 | 0 |
| NC_045512.2;YP_009724389.1;Wu1-SARS2_orf1ab;1644 | 0 | 0 |
| NC_045512.2;YP_009724389.1;Wu1-SARS2_orf1ab;1645 | 0 | 0 |
| NC_045512.2;YP_009724389.1;Wu1-SARS2_orf1ab;1646 | 0 | 0 |
| NC_045512.2;YP_009724389.1;Wu1-SARS2_orf1ab;1647 | 0 | 0 |
| NC_045512.2;YP_009724389.1;Wu1-SARS2_orf1ab;1648 | 0 | 0 |
| NC_045512.2;YP_009724389.1;Wu1-SARS2_orf1ab;1649 | 0 | 0 |
| NC_045512.2;YP_009724389.1;Wu1-SARS2_orf1ab;1650 | 0 | 0 |
| NC_045512.2;YP_009724389.1;Wu1-SARS2_orf1ab;1651 | 0 | 0 |
| NC_045512.2;YP_009724389.1;Wu1-SARS2_orf1ab;1652 | 0 | 0 |
| NC_045512.2;YP_009724389.1;Wu1-SARS2_orf1ab;1653 | 0 | 0 |
| NC_045512.2;YP_009724389.1;Wu1-SARS2_orf1ab;1654 | 0 | 0 |
| NC_045512.2;YP_009724389.1;Wu1-SARS2_orf1ab;1655 | 0 | 0 |
| NC_045512.2;YP_009724389.1;Wu1-SARS2_orf1ab;1656 | 0 | 0 |
| NC_045512.2;YP_009724389.1;Wu1-SARS2_orf1ab;1657 | 0.05 | 0.025 |
| NC_045512.2;YP_009724389.1;Wu1-SARS2_orf1ab;1658 | 0.05 | 0.05 |
| NC_045512.2;YP_009724389.1;Wu1-SARS2_orf1ab;1659 | 0.05 | 0.025 |
| NC_045512.2;YP_009724389.1;Wu1-SARS2_orf1ab;1660 | 0.05 | 0 |
| NC_045512.2;YP_009724389.1;Wu1-SARS2_orf1ab;1661 | 0.05 | 0 |
| NC_045512.2;YP_009724389.1;Wu1-SARS2_orf1ab;1662 | 0.05 | 0 |
| NC_045512.2;YP_009724389.1;Wu1-SARS2_orf1ab;1663 | 0.05 | 0 |
| NC_045512.2;YP_009724389.1;Wu1-SARS2_orf1ab;1664 | 0.05 | 0 |
| NC_045512.2;YP_009724389.1;Wu1-SARS2_orf1ab;1665 | 0 | 0 |
| NC_045512.2;YP_009724389.1;Wu1-SARS2_orf1ab;1666 | 0 | 0 |
| NC_045512.2;YP_009724389.1;Wu1-SARS2_orf1ab;1667 | 0 | 0 |

| | | |
|---|---|---|
| NC_045512.2;YP_009724389.1;Wu1-SARS2_orf1ab;1668 | 0 | 0 |
| NC_045512.2;YP_009724389.1;Wu1-SARS2_orf1ab;1669 | 0 | 0 |
| NC_045512.2;YP_009724389.1;Wu1-SARS2_orf1ab;1670 | 0 | 0 |
| NC_045512.2;YP_009724389.1;Wu1-SARS2_orf1ab;1671 | 0 | 0 |
| NC_045512.2;YP_009724389.1;Wu1-SARS2_orf1ab;1672 | 0 | 0 |
| NC_045512.2;YP_009724389.1;Wu1-SARS2_orf1ab;1673 | 0 | 0 |
| NC_045512.2;YP_009724389.1;Wu1-SARS2_orf1ab;1674 | 0 | 0 |
| NC_045512.2;YP_009724389.1;Wu1-SARS2_orf1ab;1675 | 0 | 0 |
| NC_045512.2;YP_009724389.1;Wu1-SARS2_orf1ab;1676 | 0 | 0 |
| NC_045512.2;YP_009724389.1;Wu1-SARS2_orf1ab;1677 | 0 | 0 |
| NC_045512.2;YP_009724389.1;Wu1-SARS2_orf1ab;1678 | 0 | 0 |
| NC_045512.2;YP_009724389.1;Wu1-SARS2_orf1ab;1679 | 0.05 | 0 |
| NC_045512.2;YP_009724389.1;Wu1-SARS2_orf1ab;1680 | 0 | 0 |
| NC_045512.2;YP_009724389.1;Wu1-SARS2_orf1ab;1681 | 0 | 0 |
| NC_045512.2;YP_009724389.1;Wu1-SARS2_orf1ab;1682 | 0 | 0.025 |
| NC_045512.2;YP_009724389.1;Wu1-SARS2_orf1ab;1683 | 0 | 0 |
| NC_045512.2;YP_009724389.1;Wu1-SARS2_orf1ab;1684 | 0 | 0 |
| NC_045512.2;YP_009724389.1;Wu1-SARS2_orf1ab;1685 | 0 | 0 |
| NC_045512.2;YP_009724389.1;Wu1-SARS2_orf1ab;1686 | 0 | 0.05 |
| NC_045512.2;YP_009724389.1;Wu1-SARS2_orf1ab;1687 | 0 | 0.025 |
| NC_045512.2;YP_009724389.1;Wu1-SARS2_orf1ab;1688 | 0 | 0 |
| NC_045512.2;YP_009724389.1;Wu1-SARS2_orf1ab;1689 | 0 | 0 |
| NC_045512.2;YP_009724389.1;Wu1-SARS2_orf1ab;1690 | 0 | 0 |
| NC_045512.2;YP_009724389.1;Wu1-SARS2_orf1ab;1691 | 0 | 0 |
| NC_045512.2;YP_009724389.1;Wu1-SARS2_orf1ab;1692 | 0.05 | 0.025 |
| NC_045512.2;YP_009724389.1;Wu1-SARS2_orf1ab;1693 | 0.1 | 0.075 |
| NC_045512.2;YP_009724389.1;Wu1-SARS2_orf1ab;1694 | 0.05 | 0.025 |
| NC_045512.2;YP_009724389.1;Wu1-SARS2_orf1ab;1695 | 0.15 | 0.025 |
| NC_045512.2;YP_009724389.1;Wu1-SARS2_orf1ab;1696 | 0.05 | 0.025 |
| NC_045512.2;YP_009724389.1;Wu1-SARS2_orf1ab;1697 | 0.05 | 0 |
| NC_045512.2;YP_009724389.1;Wu1-SARS2_orf1ab;1698 | 0 | 0 |
| NC_045512.2;YP_009724389.1;Wu1-SARS2_orf1ab;1699 | 0 | 0 |
| NC_045512.2;YP_009724389.1;Wu1-SARS2_orf1ab;1700 | 0 | 0 |
| NC_045512.2;YP_009724389.1;Wu1-SARS2_orf1ab;1701 | 0 | 0 |
| NC_045512.2;YP_009724389.1;Wu1-SARS2_orf1ab;1702 | 0 | 0 |
| NC_045512.2;YP_009724389.1;Wu1-SARS2_orf1ab;1703 | 0 | 0 |
| NC_045512.2;YP_009724389.1;Wu1-SARS2_orf1ab;1704 | 0 | 0 |
| NC_045512.2;YP_009724389.1;Wu1-SARS2_orf1ab;1705 | 0 | 0 |
| NC_045512.2;YP_009724389.1;Wu1-SARS2_orf1ab;1706 | 0 | 0 |
| NC_045512.2;YP_009724389.1;Wu1-SARS2_orf1ab;1707 | 0 | 0 |
| NC_045512.2;YP_009724389.1;Wu1-SARS2_orf1ab;1708 | 0 | 0 |
| NC_045512.2;YP_009724389.1;Wu1-SARS2_orf1ab;1709 | 0 | 0 |
| NC_045512.2;YP_009724389.1;Wu1-SARS2_orf1ab;1710 | 0 | 0 |

| | | |
|---|---|---|
| NC_045512.2;YP_009724389.1;Wu1-SARS2_orf1ab;1711 | 0 | 0.025 |
| NC_045512.2;YP_009724389.1;Wu1-SARS2_orf1ab;1712 | 0 | 0.025 |
| NC_045512.2;YP_009724389.1;Wu1-SARS2_orf1ab;1713 | 0 | 0.05 |
| NC_045512.2;YP_009724389.1;Wu1-SARS2_orf1ab;1714 | 0 | 0.025 |
| NC_045512.2;YP_009724389.1;Wu1-SARS2_orf1ab;1715 | 0 | 0 |
| NC_045512.2;YP_009724389.1;Wu1-SARS2_orf1ab;1716 | 0 | 0 |
| NC_045512.2;YP_009724389.1;Wu1-SARS2_orf1ab;1717 | 0 | 0 |
| NC_045512.2;YP_009724389.1;Wu1-SARS2_orf1ab;1718 | 0 | 0 |
| NC_045512.2;YP_009724389.1;Wu1-SARS2_orf1ab;1719 | 0 | 0.025 |
| NC_045512.2;YP_009724389.1;Wu1-SARS2_orf1ab;1720 | 0 | 0.025 |
| NC_045512.2;YP_009724389.1;Wu1-SARS2_orf1ab;1721 | 0 | 0 |
| NC_045512.2;YP_009724389.1;Wu1-SARS2_orf1ab;1722 | 0 | 0 |
| NC_045512.2;YP_009724389.1;Wu1-SARS2_orf1ab;1723 | 0.05 | 0.025 |
| NC_045512.2;YP_009724389.1;Wu1-SARS2_orf1ab;1724 | 0.05 | 0 |
| NC_045512.2;YP_009724389.1;Wu1-SARS2_orf1ab;1725 | 0 | 0 |
| NC_045512.2;YP_009724389.1;Wu1-SARS2_orf1ab;1726 | 0.05 | 0 |
| NC_045512.2;YP_009724389.1;Wu1-SARS2_orf1ab;1727 | 0 | 0 |
| NC_045512.2;YP_009724389.1;Wu1-SARS2_orf1ab;1728 | 0 | 0 |
| NC_045512.2;YP_009724389.1;Wu1-SARS2_orf1ab;1729 | 0 | 0 |
| NC_045512.2;YP_009724389.1;Wu1-SARS2_orf1ab;1730 | 0 | 0 |
| NC_045512.2;YP_009724389.1;Wu1-SARS2_orf1ab;1731 | 0 | 0 |
| NC_045512.2;YP_009724389.1;Wu1-SARS2_orf1ab;1732 | 0 | 0 |
| NC_045512.2;YP_009724389.1;Wu1-SARS2_orf1ab;1733 | 0 | 0 |
| NC_045512.2;YP_009724389.1;Wu1-SARS2_orf1ab;1734 | 0 | 0 |
| NC_045512.2;YP_009724389.1;Wu1-SARS2_orf1ab;1735 | 0 | 0 |
| NC_045512.2;YP_009724389.1;Wu1-SARS2_orf1ab;1736 | 0 | 0 |
| NC_045512.2;YP_009724389.1;Wu1-SARS2_orf1ab;1737 | 0 | 0 |
| NC_045512.2;YP_009724389.1;Wu1-SARS2_orf1ab;1738 | 0 | 0 |
| NC_045512.2;YP_009724389.1;Wu1-SARS2_orf1ab;1739 | 0 | 0 |
| NC_045512.2;YP_009724389.1;Wu1-SARS2_orf1ab;1740 | 0 | 0 |
| NC_045512.2;YP_009724389.1;Wu1-SARS2_orf1ab;1741 | 0 | 0 |
| NC_045512.2;YP_009724389.1;Wu1-SARS2_orf1ab;1742 | 0 | 0 |
| NC_045512.2;YP_009724389.1;Wu1-SARS2_orf1ab;1743 | 0 | 0 |
| NC_045512.2;YP_009724389.1;Wu1-SARS2_orf1ab;1744 | 0 | 0 |
| NC_045512.2;YP_009724389.1;Wu1-SARS2_orf1ab;1745 | 0 | 0.075 |
| NC_045512.2;YP_009724389.1;Wu1-SARS2_orf1ab;1746 | 0 | 0.025 |
| NC_045512.2;YP_009724389.1;Wu1-SARS2_orf1ab;1747 | 0 | 0.025 |
| NC_045512.2;YP_009724389.1;Wu1-SARS2_orf1ab;1748 | 0.1 | 0.325 |
| NC_045512.2;YP_009724389.1;Wu1-SARS2_orf1ab;1749 | 0.1 | 0.3 |
| NC_045512.2;YP_009724389.1;Wu1-SARS2_orf1ab;1750 | 0.1 | 0.3 |
| NC_045512.2;YP_009724389.1;Wu1-SARS2_orf1ab;1751 | 0.15 | 0.325 |
| NC_045512.2;YP_009724389.1;Wu1-SARS2_orf1ab;1752 | 0.2 | 0.325 |
| NC_045512.2;YP_009724389.1;Wu1-SARS2_orf1ab;1753 | 0.1 | 0.35 |

| | | |
|---|---|---|
| NC_045512.2;YP_009724389.1;Wu1-SARS2_orf1ab;1754 | 0.05 | 0.275 |
| NC_045512.2;YP_009724389.1;Wu1-SARS2_orf1ab;1755 | 0.05 | 0.2 |
| NC_045512.2;YP_009724389.1;Wu1-SARS2_orf1ab;1756 | 0.1 | 0.25 |
| NC_045512.2;YP_009724389.1;Wu1-SARS2_orf1ab;1757 | 0.05 | 0.2 |
| NC_045512.2;YP_009724389.1;Wu1-SARS2_orf1ab;1758 | 0 | 0.125 |
| NC_045512.2;YP_009724389.1;Wu1-SARS2_orf1ab;1759 | 0 | 0 |
| NC_045512.2;YP_009724389.1;Wu1-SARS2_orf1ab;1760 | 0 | 0 |
| NC_045512.2;YP_009724389.1;Wu1-SARS2_orf1ab;1761 | 0.1 | 0 |
| NC_045512.2;YP_009724389.1;Wu1-SARS2_orf1ab;1762 | 0.15 | 0 |
| NC_045512.2;YP_009724389.1;Wu1-SARS2_orf1ab;1763 | 0.15 | 0 |
| NC_045512.2;YP_009724389.1;Wu1-SARS2_orf1ab;1764 | 0.1 | 0 |
| NC_045512.2;YP_009724389.1;Wu1-SARS2_orf1ab;1765 | 0.1 | 0.05 |
| NC_045512.2;YP_009724389.1;Wu1-SARS2_orf1ab;1766 | 0.1 | 0.05 |
| NC_045512.2;YP_009724389.1;Wu1-SARS2_orf1ab;1767 | 0.05 | 0.025 |
| NC_045512.2;YP_009724389.1;Wu1-SARS2_orf1ab;1768 | 0.05 | 0.025 |
| NC_045512.2;YP_009724389.1;Wu1-SARS2_orf1ab;1769 | 0.15 | 0.05 |
| NC_045512.2;YP_009724389.1;Wu1-SARS2_orf1ab;1770 | 0.05 | 0.05 |
| NC_045512.2;YP_009724389.1;Wu1-SARS2_orf1ab;1771 | 0.15 | 0.05 |
| NC_045512.2;YP_009724389.1;Wu1-SARS2_orf1ab;1772 | 0.15 | 0.075 |
| NC_045512.2;YP_009724389.1;Wu1-SARS2_orf1ab;1773 | 0.15 | 0.05 |
| NC_045512.2;YP_009724389.1;Wu1-SARS2_orf1ab;1774 | 0.15 | 0.075 |
| NC_045512.2;YP_009724389.1;Wu1-SARS2_orf1ab;1775 | 0.15 | 0.1 |
| NC_045512.2;YP_009724389.1;Wu1-SARS2_orf1ab;1776 | 0 | 0 |
| NC_045512.2;YP_009724389.1;Wu1-SARS2_orf1ab;1777 | 0 | 0 |
| NC_045512.2;YP_009724389.1;Wu1-SARS2_orf1ab;1778 | 0 | 0 |
| NC_045512.2;YP_009724389.1;Wu1-SARS2_orf1ab;1779 | 0 | 0 |
| NC_045512.2;YP_009724389.1;Wu1-SARS2_orf1ab;1780 | 0 | 0.025 |
| NC_045512.2;YP_009724389.1;Wu1-SARS2_orf1ab;1781 | 0 | 0.025 |
| NC_045512.2;YP_009724389.1;Wu1-SARS2_orf1ab;1782 | 0 | 0.025 |
| NC_045512.2;YP_009724389.1;Wu1-SARS2_orf1ab;1783 | 0 | 0.025 |
| NC_045512.2;YP_009724389.1;Wu1-SARS2_orf1ab;1784 | 0 | 0.025 |
| NC_045512.2;YP_009724389.1;Wu1-SARS2_orf1ab;1785 | 0 | 0.025 |
| NC_045512.2;YP_009724389.1;Wu1-SARS2_orf1ab;1786 | 0.05 | 0.075 |
| NC_045512.2;YP_009724389.1;Wu1-SARS2_orf1ab;1787 | 0 | 0.075 |
| NC_045512.2;YP_009724389.1;Wu1-SARS2_orf1ab;1788 | 0 | 0.05 |
| NC_045512.2;YP_009724389.1;Wu1-SARS2_orf1ab;1789 | 0 | 0.05 |
| NC_045512.2;YP_009724389.1;Wu1-SARS2_orf1ab;1790 | 0.05 | 0.05 |
| NC_045512.2;YP_009724389.1;Wu1-SARS2_orf1ab;1791 | 0 | 0.025 |
| NC_045512.2;YP_009724389.1;Wu1-SARS2_orf1ab;1792 | 0 | 0.025 |
| NC_045512.2;YP_009724389.1;Wu1-SARS2_orf1ab;1793 | 0.05 | 0.05 |
| NC_045512.2;YP_009724389.1;Wu1-SARS2_orf1ab;1794 | 0 | 0.05 |
| NC_045512.2;YP_009724389.1;Wu1-SARS2_orf1ab;1795 | 0.05 | 0.025 |
| NC_045512.2;YP_009724389.1;Wu1-SARS2_orf1ab;1796 | 0 | 0 |

| | | |
|---|---|---|
| NC_045512.2;YP_009724389.1;Wu1-SARS2_orf1ab;1797 | 0.1 | 0 |
| NC_045512.2;YP_009724389.1;Wu1-SARS2_orf1ab;1798 | 0 | 0.025 |
| NC_045512.2;YP_009724389.1;Wu1-SARS2_orf1ab;1799 | 0 | 0 |
| NC_045512.2;YP_009724389.1;Wu1-SARS2_orf1ab;1800 | 0 | 0 |
| NC_045512.2;YP_009724389.1;Wu1-SARS2_orf1ab;1801 | 0 | 0.025 |
| NC_045512.2;YP_009724389.1;Wu1-SARS2_orf1ab;1802 | 0 | 0 |
| NC_045512.2;YP_009724389.1;Wu1-SARS2_orf1ab;1803 | 0 | 0 |
| NC_045512.2;YP_009724389.1;Wu1-SARS2_orf1ab;1804 | 0 | 0 |
| NC_045512.2;YP_009724389.1;Wu1-SARS2_orf1ab;1805 | 0 | 0 |
| NC_045512.2;YP_009724389.1;Wu1-SARS2_orf1ab;1806 | 0 | 0 |
| NC_045512.2;YP_009724389.1;Wu1-SARS2_orf1ab;1807 | 0 | 0 |
| NC_045512.2;YP_009724389.1;Wu1-SARS2_orf1ab;1808 | 0 | 0 |
| NC_045512.2;YP_009724389.1;Wu1-SARS2_orf1ab;1809 | 0 | 0 |
| NC_045512.2;YP_009724389.1;Wu1-SARS2_orf1ab;1810 | 0.05 | 0.025 |
| NC_045512.2;YP_009724389.1;Wu1-SARS2_orf1ab;1811 | 0 | 0 |
| NC_045512.2;YP_009724389.1;Wu1-SARS2_orf1ab;1812 | 0 | 0 |
| NC_045512.2;YP_009724389.1;Wu1-SARS2_orf1ab;1813 | 0 | 0 |
| NC_045512.2;YP_009724389.1;Wu1-SARS2_orf1ab;1814 | 0 | 0 |
| NC_045512.2;YP_009724389.1;Wu1-SARS2_orf1ab;1815 | 0 | 0 |
| NC_045512.2;YP_009724389.1;Wu1-SARS2_orf1ab;1816 | 0 | 0.025 |
| NC_045512.2;YP_009724389.1;Wu1-SARS2_orf1ab;1817 | 0.1 | 0.075 |
| NC_045512.2;YP_009724389.1;Wu1-SARS2_orf1ab;1818 | 0 | 0 |
| NC_045512.2;YP_009724389.1;Wu1-SARS2_orf1ab;1819 | 0 | 0 |
| NC_045512.2;YP_009724389.1;Wu1-SARS2_orf1ab;1820 | 0 | 0 |
| NC_045512.2;YP_009724389.1;Wu1-SARS2_orf1ab;1821 | 0 | 0 |
| NC_045512.2;YP_009724389.1;Wu1-SARS2_orf1ab;1822 | 0 | 0 |
| NC_045512.2;YP_009724389.1;Wu1-SARS2_orf1ab;1823 | 0 | 0 |
| NC_045512.2;YP_009724389.1;Wu1-SARS2_orf1ab;1824 | 0 | 0 |
| NC_045512.2;YP_009724389.1;Wu1-SARS2_orf1ab;1825 | 0 | 0 |
| NC_045512.2;YP_009724389.1;Wu1-SARS2_orf1ab;1826 | 0 | 0.15 |
| NC_045512.2;YP_009724389.1;Wu1-SARS2_orf1ab;1827 | 0 | 0 |
| NC_045512.2;YP_009724389.1;Wu1-SARS2_orf1ab;1828 | 0 | 0 |
| NC_045512.2;YP_009724389.1;Wu1-SARS2_orf1ab;1829 | 0 | 0 |
| NC_045512.2;YP_009724389.1;Wu1-SARS2_orf1ab;1830 | 0 | 0 |
| NC_045512.2;YP_009724389.1;Wu1-SARS2_orf1ab;1831 | 0.05 | 0 |
| NC_045512.2;YP_009724389.1;Wu1-SARS2_orf1ab;1832 | 0.05 | 0 |
| NC_045512.2;YP_009724389.1;Wu1-SARS2_orf1ab;1833 | 0 | 0 |
| NC_045512.2;YP_009724389.1;Wu1-SARS2_orf1ab;1834 | 0 | 0 |
| NC_045512.2;YP_009724389.1;Wu1-SARS2_orf1ab;1835 | 0 | 0 |
| NC_045512.2;YP_009724389.1;Wu1-SARS2_orf1ab;1836 | 0 | 0 |
| NC_045512.2;YP_009724389.1;Wu1-SARS2_orf1ab;1837 | 0 | 0 |
| NC_045512.2;YP_009724389.1;Wu1-SARS2_orf1ab;1838 | 0 | 0 |
| NC_045512.2;YP_009724389.1;Wu1-SARS2_orf1ab;1839 | 0 | 0 |

| | | |
|---|---|---|
| NC_045512.2;YP_009724389.1;Wu1-SARS2_orf1ab;1840 | 0 | 0 |
| NC_045512.2;YP_009724389.1;Wu1-SARS2_orf1ab;1841 | 0 | 0 |
| NC_045512.2;YP_009724389.1;Wu1-SARS2_orf1ab;1842 | 0 | 0 |
| NC_045512.2;YP_009724389.1;Wu1-SARS2_orf1ab;1843 | 0 | 0 |
| NC_045512.2;YP_009724389.1;Wu1-SARS2_orf1ab;1844 | 0 | 0 |
| NC_045512.2;YP_009724389.1;Wu1-SARS2_orf1ab;1845 | 0 | 0 |
| NC_045512.2;YP_009724389.1;Wu1-SARS2_orf1ab;1846 | 0 | 0 |
| NC_045512.2;YP_009724389.1;Wu1-SARS2_orf1ab;1847 | 0 | 0.025 |
| NC_045512.2;YP_009724389.1;Wu1-SARS2_orf1ab;1848 | 0 | 0 |
| NC_045512.2;YP_009724389.1;Wu1-SARS2_orf1ab;1849 | 0 | 0.025 |
| NC_045512.2;YP_009724389.1;Wu1-SARS2_orf1ab;1850 | 0 | 0.075 |
| NC_045512.2;YP_009724389.1;Wu1-SARS2_orf1ab;1851 | 0 | 0.05 |
| NC_045512.2;YP_009724389.1;Wu1-SARS2_orf1ab;1852 | 0 | 0.025 |
| NC_045512.2;YP_009724389.1;Wu1-SARS2_orf1ab;1853 | 0 | 0.025 |
| NC_045512.2;YP_009724389.1;Wu1-SARS2_orf1ab;1854 | 0.05 | 0.075 |
| NC_045512.2;YP_009724389.1;Wu1-SARS2_orf1ab;1855 | 0.05 | 0.1 |
| NC_045512.2;YP_009724389.1;Wu1-SARS2_orf1ab;1856 | 0.1 | 0.125 |
| NC_045512.2;YP_009724389.1;Wu1-SARS2_orf1ab;1857 | 0.1 | 0.1 |
| NC_045512.2;YP_009724389.1;Wu1-SARS2_orf1ab;1858 | 0.1 | 0.05 |
| NC_045512.2;YP_009724389.1;Wu1-SARS2_orf1ab;1859 | 0.05 | 0.075 |
| NC_045512.2;YP_009724389.1;Wu1-SARS2_orf1ab;1860 | 0.05 | 0.1 |
| NC_045512.2;YP_009724389.1;Wu1-SARS2_orf1ab;1861 | 0.25 | 0.15 |
| NC_045512.2;YP_009724389.1;Wu1-SARS2_orf1ab;1862 | 0 | 0 |
| NC_045512.2;YP_009724389.1;Wu1-SARS2_orf1ab;1863 | 0 | 0.025 |
| NC_045512.2;YP_009724389.1;Wu1-SARS2_orf1ab;1864 | 0 | 0.025 |
| NC_045512.2;YP_009724389.1;Wu1-SARS2_orf1ab;1865 | 0.1 | 0.025 |
| NC_045512.2;YP_009724389.1;Wu1-SARS2_orf1ab;1866 | 0 | 0.025 |
| NC_045512.2;YP_009724389.1;Wu1-SARS2_orf1ab;1867 | 0 | 0.025 |
| NC_045512.2;YP_009724389.1;Wu1-SARS2_orf1ab;1868 | 0.05 | 0.025 |
| NC_045512.2;YP_009724389.1;Wu1-SARS2_orf1ab;1869 | 0 | 0.025 |
| NC_045512.2;YP_009724389.1;Wu1-SARS2_orf1ab;1870 | 0 | 0.025 |
| NC_045512.2;YP_009724389.1;Wu1-SARS2_orf1ab;1871 | 0 | 0.025 |
| NC_045512.2;YP_009724389.1;Wu1-SARS2_orf1ab;1872 | 0 | 0.025 |
| NC_045512.2;YP_009724389.1;Wu1-SARS2_orf1ab;1873 | 0 | 0 |
| NC_045512.2;YP_009724389.1;Wu1-SARS2_orf1ab;1874 | 0 | 0 |
| NC_045512.2;YP_009724389.1;Wu1-SARS2_orf1ab;1875 | 0 | 0 |
| NC_045512.2;YP_009724389.1;Wu1-SARS2_orf1ab;1876 | 0 | 0 |
| NC_045512.2;YP_009724389.1;Wu1-SARS2_orf1ab;1877 | 0 | 0 |
| NC_045512.2;YP_009724389.1;Wu1-SARS2_orf1ab;1878 | 0 | 0 |
| NC_045512.2;YP_009724389.1;Wu1-SARS2_orf1ab;1879 | 0 | 0 |
| NC_045512.2;YP_009724389.1;Wu1-SARS2_orf1ab;1880 | 0 | 0 |
| NC_045512.2;YP_009724389.1;Wu1-SARS2_orf1ab;1881 | 0 | 0.025 |
| NC_045512.2;YP_009724389.1;Wu1-SARS2_orf1ab;1882 | 0 | 0 |

| | | |
|---|---|---|
| NC_045512.2;YP_009724389.1;Wu1-SARS2_orf1ab;1883 | 0 | 0 |
| NC_045512.2;YP_009724389.1;Wu1-SARS2_orf1ab;1884 | 0 | 0 |
| NC_045512.2;YP_009724389.1;Wu1-SARS2_orf1ab;1885 | 0 | 0 |
| NC_045512.2;YP_009724389.1;Wu1-SARS2_orf1ab;1886 | 0.05 | 0.125 |
| NC_045512.2;YP_009724389.1;Wu1-SARS2_orf1ab;1887 | 0 | 0.1 |
| NC_045512.2;YP_009724389.1;Wu1-SARS2_orf1ab;1888 | 0 | 0.15 |
| NC_045512.2;YP_009724389.1;Wu1-SARS2_orf1ab;1889 | 0 | 0.175 |
| NC_045512.2;YP_009724389.1;Wu1-SARS2_orf1ab;1890 | 0.2 | 0.25 |
| NC_045512.2;YP_009724389.1;Wu1-SARS2_orf1ab;1891 | 0.15 | 0.35 |
| NC_045512.2;YP_009724389.1;Wu1-SARS2_orf1ab;1892 | 0.1 | 0.225 |
| NC_045512.2;YP_009724389.1;Wu1-SARS2_orf1ab;1893 | 0.05 | 0.175 |
| NC_045512.2;YP_009724389.1;Wu1-SARS2_orf1ab;1894 | 0.3 | 0.175 |
| NC_045512.2;YP_009724389.1;Wu1-SARS2_orf1ab;1895 | 0.35 | 0.175 |
| NC_045512.2;YP_009724389.1;Wu1-SARS2_orf1ab;1896 | 0.2 | 0.175 |
| NC_045512.2;YP_009724389.1;Wu1-SARS2_orf1ab;1897 | 0.3 | 0.1 |
| NC_045512.2;YP_009724389.1;Wu1-SARS2_orf1ab;1898 | 0.15 | 0.075 |
| NC_045512.2;YP_009724389.1;Wu1-SARS2_orf1ab;1899 | 0.2 | 0.05 |
| NC_045512.2;YP_009724389.1;Wu1-SARS2_orf1ab;1900 | 0.15 | 0.05 |
| NC_045512.2;YP_009724389.1;Wu1-SARS2_orf1ab;1901 | 0.15 | 0.15 |
| NC_045512.2;YP_009724389.1;Wu1-SARS2_orf1ab;1902 | 0 | 0.025 |
| NC_045512.2;YP_009724389.1;Wu1-SARS2_orf1ab;1903 | 0.05 | 0 |
| NC_045512.2;YP_009724389.1;Wu1-SARS2_orf1ab;1904 | 0.05 | 0 |
| NC_045512.2;YP_009724389.1;Wu1-SARS2_orf1ab;1905 | 0.15 | 0.2 |
| NC_045512.2;YP_009724389.1;Wu1-SARS2_orf1ab;1906 | 0.05 | 0 |
| NC_045512.2;YP_009724389.1;Wu1-SARS2_orf1ab;1907 | 0.05 | 0 |
| NC_045512.2;YP_009724389.1;Wu1-SARS2_orf1ab;1908 | 0.1 | 0 |
| NC_045512.2;YP_009724389.1;Wu1-SARS2_orf1ab;1909 | 0 | 0.025 |
| NC_045512.2;YP_009724389.1;Wu1-SARS2_orf1ab;1910 | 0.2 | 0.1 |
| NC_045512.2;YP_009724389.1;Wu1-SARS2_orf1ab;1911 | 0 | 0.075 |
| NC_045512.2;YP_009724389.1;Wu1-SARS2_orf1ab;1912 | 0 | 0.05 |
| NC_045512.2;YP_009724389.1;Wu1-SARS2_orf1ab;1913 | 0 | 0.05 |
| NC_045512.2;YP_009724389.1;Wu1-SARS2_orf1ab;1914 | 0.05 | 0.075 |
| NC_045512.2;YP_009724389.1;Wu1-SARS2_orf1ab;1915 | 0.05 | 0.05 |
| NC_045512.2;YP_009724389.1;Wu1-SARS2_orf1ab;1916 | 0.05 | 0.025 |
| NC_045512.2;YP_009724389.1;Wu1-SARS2_orf1ab;1917 | 0.05 | 0.025 |
| NC_045512.2;YP_009724389.1;Wu1-SARS2_orf1ab;1918 | 0.05 | 0.025 |
| NC_045512.2;YP_009724389.1;Wu1-SARS2_orf1ab;1919 | 0.05 | 0.025 |
| NC_045512.2;YP_009724389.1;Wu1-SARS2_orf1ab;1920 | 0.05 | 0 |
| NC_045512.2;YP_009724389.1;Wu1-SARS2_orf1ab;1921 | 0.05 | 0 |
| NC_045512.2;YP_009724389.1;Wu1-SARS2_orf1ab;1922 | 0 | 0 |
| NC_045512.2;YP_009724389.1;Wu1-SARS2_orf1ab;1923 | 0 | 0.025 |
| NC_045512.2;YP_009724389.1;Wu1-SARS2_orf1ab;1924 | 0.05 | 0 |
| NC_045512.2;YP_009724389.1;Wu1-SARS2_orf1ab;1925 | 0 | 0 |

| | | |
|---|---|---|
| NC_045512.2;YP_009724389.1;Wu1-SARS2_orf1ab;1926 | 0 | 0 |
| NC_045512.2;YP_009724389.1;Wu1-SARS2_orf1ab;1927 | 0.2 | 0.025 |
| NC_045512.2;YP_009724389.1;Wu1-SARS2_orf1ab;1928 | 0.2 | 0 |
| NC_045512.2;YP_009724389.1;Wu1-SARS2_orf1ab;1929 | 0.2 | 0.05 |
| NC_045512.2;YP_009724389.1;Wu1-SARS2_orf1ab;1930 | 0.3 | 0.025 |
| NC_045512.2;YP_009724389.1;Wu1-SARS2_orf1ab;1931 | 0.2 | 0 |
| NC_045512.2;YP_009724389.1;Wu1-SARS2_orf1ab;1932 | 0.25 | 0 |
| NC_045512.2;YP_009724389.1;Wu1-SARS2_orf1ab;1933 | 0.15 | 0.075 |
| NC_045512.2;YP_009724389.1;Wu1-SARS2_orf1ab;1934 | 0.1 | 0.025 |
| NC_045512.2;YP_009724389.1;Wu1-SARS2_orf1ab;1935 | 0.1 | 0.05 |
| NC_045512.2;YP_009724389.1;Wu1-SARS2_orf1ab;1936 | 0.1 | 0 |
| NC_045512.2;YP_009724389.1;Wu1-SARS2_orf1ab;1937 | 0.05 | 0 |
| NC_045512.2;YP_009724389.1;Wu1-SARS2_orf1ab;1938 | 0.1 | 0.025 |
| NC_045512.2;YP_009724389.1;Wu1-SARS2_orf1ab;1939 | 0.05 | 0.05 |
| NC_045512.2;YP_009724389.1;Wu1-SARS2_orf1ab;1940 | 0.05 | 0 |
| NC_045512.2;YP_009724389.1;Wu1-SARS2_orf1ab;1941 | 0 | 0 |
| NC_045512.2;YP_009724389.1;Wu1-SARS2_orf1ab;1942 | 0 | 0 |
| NC_045512.2;YP_009724389.1;Wu1-SARS2_orf1ab;1943 | 0 | 0 |
| NC_045512.2;YP_009724389.1;Wu1-SARS2_orf1ab;1944 | 0 | 0 |
| NC_045512.2;YP_009724389.1;Wu1-SARS2_orf1ab;1945 | 0 | 0 |
| NC_045512.2;YP_009724389.1;Wu1-SARS2_orf1ab;1946 | 0 | 0 |
| NC_045512.2;YP_009724389.1;Wu1-SARS2_orf1ab;1947 | 0 | 0 |
| NC_045512.2;YP_009724389.1;Wu1-SARS2_orf1ab;1948 | 0 | 0 |
| NC_045512.2;YP_009724389.1;Wu1-SARS2_orf1ab;1949 | 0 | 0 |
| NC_045512.2;YP_009724389.1;Wu1-SARS2_orf1ab;1950 | 0 | 0 |
| NC_045512.2;YP_009724389.1;Wu1-SARS2_orf1ab;1951 | 0 | 0 |
| NC_045512.2;YP_009724389.1;Wu1-SARS2_orf1ab;1952 | 0 | 0.025 |
| NC_045512.2;YP_009724389.1;Wu1-SARS2_orf1ab;1953 | 0 | 0.025 |
| NC_045512.2;YP_009724389.1;Wu1-SARS2_orf1ab;1954 | 0 | 0.05 |
| NC_045512.2;YP_009724389.1;Wu1-SARS2_orf1ab;1955 | 0 | 0 |
| NC_045512.2;YP_009724389.1;Wu1-SARS2_orf1ab;1956 | 0 | 0.025 |
| NC_045512.2;YP_009724389.1;Wu1-SARS2_orf1ab;1957 | 0 | 0 |
| NC_045512.2;YP_009724389.1;Wu1-SARS2_orf1ab;1958 | 0 | 0.025 |
| NC_045512.2;YP_009724389.1;Wu1-SARS2_orf1ab;1959 | 0.05 | 0.025 |
| NC_045512.2;YP_009724389.1;Wu1-SARS2_orf1ab;1960 | 0.05 | 0.05 |
| NC_045512.2;YP_009724389.1;Wu1-SARS2_orf1ab;1961 | 0.05 | 0.075 |
| NC_045512.2;YP_009724389.1;Wu1-SARS2_orf1ab;1962 | 0.05 | 0.075 |
| NC_045512.2;YP_009724389.1;Wu1-SARS2_orf1ab;1963 | 0.05 | 0.025 |
| NC_045512.2;YP_009724389.1;Wu1-SARS2_orf1ab;1964 | 0.05 | 0.025 |
| NC_045512.2;YP_009724389.1;Wu1-SARS2_orf1ab;1965 | 0.05 | 0.075 |
| NC_045512.2;YP_009724389.1;Wu1-SARS2_orf1ab;1966 | 0 | 0.025 |
| NC_045512.2;YP_009724389.1;Wu1-SARS2_orf1ab;1967 | 0 | 0.025 |
| NC_045512.2;YP_009724389.1;Wu1-SARS2_orf1ab;1968 | 0 | 0.05 |

| | | |
|---|---|---|
| NC_045512.2;YP_009724389.1;Wu1-SARS2_orf1ab;1969 | 0 | 0.075 |
| NC_045512.2;YP_009724389.1;Wu1-SARS2_orf1ab;1970 | 0 | 0 |
| NC_045512.2;YP_009724389.1;Wu1-SARS2_orf1ab;1971 | 0 | 0 |
| NC_045512.2;YP_009724389.1;Wu1-SARS2_orf1ab;1972 | 0 | 0 |
| NC_045512.2;YP_009724389.1;Wu1-SARS2_orf1ab;1973 | 0 | 0 |
| NC_045512.2;YP_009724389.1;Wu1-SARS2_orf1ab;1974 | 0 | 0 |
| NC_045512.2;YP_009724389.1;Wu1-SARS2_orf1ab;1975 | 0 | 0.025 |
| NC_045512.2;YP_009724389.1;Wu1-SARS2_orf1ab;1976 | 0 | 0 |
| NC_045512.2;YP_009724389.1;Wu1-SARS2_orf1ab;1977 | 0 | 0 |
| NC_045512.2;YP_009724389.1;Wu1-SARS2_orf1ab;1978 | 0 | 0 |
| NC_045512.2;YP_009724389.1;Wu1-SARS2_orf1ab;1979 | 0 | 0 |
| NC_045512.2;YP_009724389.1;Wu1-SARS2_orf1ab;1980 | 0 | 0 |
| NC_045512.2;YP_009724389.1;Wu1-SARS2_orf1ab;1981 | 0 | 0 |
| NC_045512.2;YP_009724389.1;Wu1-SARS2_orf1ab;1982 | 0 | 0 |
| NC_045512.2;YP_009724389.1;Wu1-SARS2_orf1ab;1983 | 0 | 0 |
| NC_045512.2;YP_009724389.1;Wu1-SARS2_orf1ab;1984 | 0.05 | 0 |
| NC_045512.2;YP_009724389.1;Wu1-SARS2_orf1ab;1985 | 0.05 | 0 |
| NC_045512.2;YP_009724389.1;Wu1-SARS2_orf1ab;1986 | 0 | 0 |
| NC_045512.2;YP_009724389.1;Wu1-SARS2_orf1ab;1987 | 0.05 | 0 |
| NC_045512.2;YP_009724389.1;Wu1-SARS2_orf1ab;1988 | 0.05 | 0.05 |
| NC_045512.2;YP_009724389.1;Wu1-SARS2_orf1ab;1989 | 0.1 | 0.025 |
| NC_045512.2;YP_009724389.1;Wu1-SARS2_orf1ab;1990 | 0.05 | 0.075 |
| NC_045512.2;YP_009724389.1;Wu1-SARS2_orf1ab;1991 | 0.1 | 0.1 |
| NC_045512.2;YP_009724389.1;Wu1-SARS2_orf1ab;1992 | 0.15 | 0.15 |
| NC_045512.2;YP_009724389.1;Wu1-SARS2_orf1ab;1993 | 0.15 | 0.15 |
| NC_045512.2;YP_009724389.1;Wu1-SARS2_orf1ab;1994 | 0.25 | 0.1 |
| NC_045512.2;YP_009724389.1;Wu1-SARS2_orf1ab;1995 | 0.15 | 0.075 |
| NC_045512.2;YP_009724389.1;Wu1-SARS2_orf1ab;1996 | 0.05 | 0.025 |
| NC_045512.2;YP_009724389.1;Wu1-SARS2_orf1ab;1997 | 0.05 | 0.025 |
| NC_045512.2;YP_009724389.1;Wu1-SARS2_orf1ab;1998 | 0 | 0 |
| NC_045512.2;YP_009724389.1;Wu1-SARS2_orf1ab;1999 | 0 | 0 |
| NC_045512.2;YP_009724389.1;Wu1-SARS2_orf1ab;2000 | 0 | 0 |
| NC_045512.2;YP_009724389.1;Wu1-SARS2_orf1ab;2001 | 0 | 0 |
| NC_045512.2;YP_009724389.1;Wu1-SARS2_orf1ab;2002 | 0 | 0 |
| NC_045512.2;YP_009724389.1;Wu1-SARS2_orf1ab;2003 | 0 | 0 |
| NC_045512.2;YP_009724389.1;Wu1-SARS2_orf1ab;2004 | 0 | 0 |
| NC_045512.2;YP_009724389.1;Wu1-SARS2_orf1ab;2005 | 0 | 0 |
| NC_045512.2;YP_009724389.1;Wu1-SARS2_orf1ab;2006 | 0 | 0 |
| NC_045512.2;YP_009724389.1;Wu1-SARS2_orf1ab;2007 | 0 | 0 |
| NC_045512.2;YP_009724389.1;Wu1-SARS2_orf1ab;2008 | 0 | 0 |
| NC_045512.2;YP_009724389.1;Wu1-SARS2_orf1ab;2009 | 0.05 | 0 |
| NC_045512.2;YP_009724389.1;Wu1-SARS2_orf1ab;2010 | 0.1 | 0.075 |
| NC_045512.2;YP_009724389.1;Wu1-SARS2_orf1ab;2011 | 0.2 | 0.1 |

| | | |
|---|---|---|
| NC_045512.2;YP_009724389.1;Wu1-SARS2_orf1ab;2012 | 0.1 | 0.05 |
| NC_045512.2;YP_009724389.1;Wu1-SARS2_orf1ab;2013 | 0.15 | 0.075 |
| NC_045512.2;YP_009724389.1;Wu1-SARS2_orf1ab;2014 | 0.35 | 0.1 |
| NC_045512.2;YP_009724389.1;Wu1-SARS2_orf1ab;2015 | 0.25 | 0.1 |
| NC_045512.2;YP_009724389.1;Wu1-SARS2_orf1ab;2016 | 0.1 | 0.05 |
| NC_045512.2;YP_009724389.1;Wu1-SARS2_orf1ab;2017 | 0.1 | 0.05 |
| NC_045512.2;YP_009724389.1;Wu1-SARS2_orf1ab;2018 | 0.15 | 0.125 |
| NC_045512.2;YP_009724389.1;Wu1-SARS2_orf1ab;2019 | 0.2 | 0.05 |
| NC_045512.2;YP_009724389.1;Wu1-SARS2_orf1ab;2020 | 0.05 | 0 |
| NC_045512.2;YP_009724389.1;Wu1-SARS2_orf1ab;2021 | 0 | 0.025 |
| NC_045512.2;YP_009724389.1;Wu1-SARS2_orf1ab;2022 | 0 | 0.025 |
| NC_045512.2;YP_009724389.1;Wu1-SARS2_orf1ab;2023 | 0.05 | 0.025 |
| NC_045512.2;YP_009724389.1;Wu1-SARS2_orf1ab;2024 | 0.05 | 0.025 |
| NC_045512.2;YP_009724389.1;Wu1-SARS2_orf1ab;2025 | 0 | 0.025 |
| NC_045512.2;YP_009724389.1;Wu1-SARS2_orf1ab;2026 | 0 | 0.025 |
| NC_045512.2;YP_009724389.1;Wu1-SARS2_orf1ab;2027 | 0 | 0.025 |
| NC_045512.2;YP_009724389.1;Wu1-SARS2_orf1ab;2028 | 0 | 0.025 |
| NC_045512.2;YP_009724389.1;Wu1-SARS2_orf1ab;2029 | 0 | 0.025 |
| NC_045512.2;YP_009724389.1;Wu1-SARS2_orf1ab;2030 | 0 | 0 |
| NC_045512.2;YP_009724389.1;Wu1-SARS2_orf1ab;2031 | 0 | 0 |
| NC_045512.2;YP_009724389.1;Wu1-SARS2_orf1ab;2032 | 0 | 0 |
| NC_045512.2;YP_009724389.1;Wu1-SARS2_orf1ab;2033 | 0 | 0 |
| NC_045512.2;YP_009724389.1;Wu1-SARS2_orf1ab;2034 | 0 | 0 |
| NC_045512.2;YP_009724389.1;Wu1-SARS2_orf1ab;2035 | 0 | 0 |
| NC_045512.2;YP_009724389.1;Wu1-SARS2_orf1ab;2036 | 0 | 0 |
| NC_045512.2;YP_009724389.1;Wu1-SARS2_orf1ab;2037 | 0 | 0 |
| NC_045512.2;YP_009724389.1;Wu1-SARS2_orf1ab;2038 | 0 | 0 |
| NC_045512.2;YP_009724389.1;Wu1-SARS2_orf1ab;2039 | 0 | 0 |
| NC_045512.2;YP_009724389.1;Wu1-SARS2_orf1ab;2040 | 0 | 0 |
| NC_045512.2;YP_009724389.1;Wu1-SARS2_orf1ab;2041 | 0 | 0 |
| NC_045512.2;YP_009724389.1;Wu1-SARS2_orf1ab;2042 | 0 | 0.025 |
| NC_045512.2;YP_009724389.1;Wu1-SARS2_orf1ab;2043 | 0 | 0 |
| NC_045512.2;YP_009724389.1;Wu1-SARS2_orf1ab;2044 | 0 | 0.05 |
| NC_045512.2;YP_009724389.1;Wu1-SARS2_orf1ab;2045 | 0 | 0.05 |
| NC_045512.2;YP_009724389.1;Wu1-SARS2_orf1ab;2046 | 0 | 0.05 |
| NC_045512.2;YP_009724389.1;Wu1-SARS2_orf1ab;2047 | 0 | 0.05 |
| NC_045512.2;YP_009724389.1;Wu1-SARS2_orf1ab;2048 | 0 | 0.05 |
| NC_045512.2;YP_009724389.1;Wu1-SARS2_orf1ab;2049 | 0 | 0.05 |
| NC_045512.2;YP_009724389.1;Wu1-SARS2_orf1ab;2050 | 0 | 0.05 |
| NC_045512.2;YP_009724389.1;Wu1-SARS2_orf1ab;2051 | 0 | 0.05 |
| NC_045512.2;YP_009724389.1;Wu1-SARS2_orf1ab;2052 | 0 | 0.05 |
| NC_045512.2;YP_009724389.1;Wu1-SARS2_orf1ab;2053 | 0 | 0.025 |
| NC_045512.2;YP_009724389.1;Wu1-SARS2_orf1ab;2054 | 0 | 0 |

| | | |
|---|---|---|
| NC_045512.2;YP_009724389.1;Wu1-SARS2_orf1ab;2055 | 0 | 0 |
| NC_045512.2;YP_009724389.1;Wu1-SARS2_orf1ab;2056 | 0 | 0 |
| NC_045512.2;YP_009724389.1;Wu1-SARS2_orf1ab;2057 | 0 | 0 |
| NC_045512.2;YP_009724389.1;Wu1-SARS2_orf1ab;2058 | 0 | 0 |
| NC_045512.2;YP_009724389.1;Wu1-SARS2_orf1ab;2059 | 0 | 0 |
| NC_045512.2;YP_009724389.1;Wu1-SARS2_orf1ab;2060 | 0 | 0 |
| NC_045512.2;YP_009724389.1;Wu1-SARS2_orf1ab;2061 | 0 | 0 |
| NC_045512.2;YP_009724389.1;Wu1-SARS2_orf1ab;2062 | 0 | 0 |
| NC_045512.2;YP_009724389.1;Wu1-SARS2_orf1ab;2063 | 0 | 0 |
| NC_045512.2;YP_009724389.1;Wu1-SARS2_orf1ab;2064 | 0 | 0 |
| NC_045512.2;YP_009724389.1;Wu1-SARS2_orf1ab;2065 | 0 | 0 |
| NC_045512.2;YP_009724389.1;Wu1-SARS2_orf1ab;2066 | 0 | 0 |
| NC_045512.2;YP_009724389.1;Wu1-SARS2_orf1ab;2067 | 0 | 0 |
| NC_045512.2;YP_009724389.1;Wu1-SARS2_orf1ab;2068 | 0.05 | 0.025 |
| NC_045512.2;YP_009724389.1;Wu1-SARS2_orf1ab;2069 | 0.05 | 0.05 |
| NC_045512.2;YP_009724389.1;Wu1-SARS2_orf1ab;2070 | 0 | 0 |
| NC_045512.2;YP_009724389.1;Wu1-SARS2_orf1ab;2071 | 0 | 0 |
| NC_045512.2;YP_009724389.1;Wu1-SARS2_orf1ab;2072 | 0 | 0 |
| NC_045512.2;YP_009724389.1;Wu1-SARS2_orf1ab;2073 | 0 | 0.025 |
| NC_045512.2;YP_009724389.1;Wu1-SARS2_orf1ab;2074 | 0 | 0 |
| NC_045512.2;YP_009724389.1;Wu1-SARS2_orf1ab;2075 | 0 | 0 |
| NC_045512.2;YP_009724389.1;Wu1-SARS2_orf1ab;2076 | 0 | 0 |
| NC_045512.2;YP_009724389.1;Wu1-SARS2_orf1ab;2077 | 0 | 0 |
| NC_045512.2;YP_009724389.1;Wu1-SARS2_orf1ab;2078 | 0 | 0 |
| NC_045512.2;YP_009724389.1;Wu1-SARS2_orf1ab;2079 | 0 | 0 |
| NC_045512.2;YP_009724389.1;Wu1-SARS2_orf1ab;2080 | 0 | 0 |
| NC_045512.2;YP_009724389.1;Wu1-SARS2_orf1ab;2081 | 0 | 0 |
| NC_045512.2;YP_009724389.1;Wu1-SARS2_orf1ab;2082 | 0 | 0 |
| NC_045512.2;YP_009724389.1;Wu1-SARS2_orf1ab;2083 | 0.05 | 0.025 |
| NC_045512.2;YP_009724389.1;Wu1-SARS2_orf1ab;2084 | 0 | 0 |
| NC_045512.2;YP_009724389.1;Wu1-SARS2_orf1ab;2085 | 0 | 0 |
| NC_045512.2;YP_009724389.1;Wu1-SARS2_orf1ab;2086 | 0 | 0 |
| NC_045512.2;YP_009724389.1;Wu1-SARS2_orf1ab;2087 | 0 | 0 |
| NC_045512.2;YP_009724389.1;Wu1-SARS2_orf1ab;2088 | 0 | 0 |
| NC_045512.2;YP_009724389.1;Wu1-SARS2_orf1ab;2089 | 0 | 0 |
| NC_045512.2;YP_009724389.1;Wu1-SARS2_orf1ab;2090 | 0 | 0 |
| NC_045512.2;YP_009724389.1;Wu1-SARS2_orf1ab;2091 | 0.05 | 0.05 |
| NC_045512.2;YP_009724389.1;Wu1-SARS2_orf1ab;2092 | 0 | 0.05 |
| NC_045512.2;YP_009724389.1;Wu1-SARS2_orf1ab;2093 | 0 | 0.025 |
| NC_045512.2;YP_009724389.1;Wu1-SARS2_orf1ab;2094 | 0 | 0 |
| NC_045512.2;YP_009724389.1;Wu1-SARS2_orf1ab;2095 | 0 | 0.025 |
| NC_045512.2;YP_009724389.1;Wu1-SARS2_orf1ab;2096 | 0 | 0.025 |
| NC_045512.2;YP_009724389.1;Wu1-SARS2_orf1ab;2097 | 0.05 | 0.025 |

| | | |
|---|---|---|
| NC_045512.2;YP_009724389.1;Wu1-SARS2_orf1ab;2098 | 0.05 | 0.05 |
| NC_045512.2;YP_009724389.1;Wu1-SARS2_orf1ab;2099 | 0.25 | 0.125 |
| NC_045512.2;YP_009724389.1;Wu1-SARS2_orf1ab;2100 | 0 | 0.05 |
| NC_045512.2;YP_009724389.1;Wu1-SARS2_orf1ab;2101 | 0 | 0.05 |
| NC_045512.2;YP_009724389.1;Wu1-SARS2_orf1ab;2102 | 0 | 0 |
| NC_045512.2;YP_009724389.1;Wu1-SARS2_orf1ab;2103 | 0 | 0 |
| NC_045512.2;YP_009724389.1;Wu1-SARS2_orf1ab;2104 | 0 | 0 |
| NC_045512.2;YP_009724389.1;Wu1-SARS2_orf1ab;2105 | 0 | 0 |
| NC_045512.2;YP_009724389.1;Wu1-SARS2_orf1ab;2106 | 0 | 0 |
| NC_045512.2;YP_009724389.1;Wu1-SARS2_orf1ab;2107 | 0 | 0 |
| NC_045512.2;YP_009724389.1;Wu1-SARS2_orf1ab;2108 | 0 | 0 |
| NC_045512.2;YP_009724389.1;Wu1-SARS2_orf1ab;2109 | 0 | 0 |
| NC_045512.2;YP_009724389.1;Wu1-SARS2_orf1ab;2110 | 0 | 0 |
| NC_045512.2;YP_009724389.1;Wu1-SARS2_orf1ab;2111 | 0 | 0 |
| NC_045512.2;YP_009724389.1;Wu1-SARS2_orf1ab;2112 | 0 | 0 |
| NC_045512.2;YP_009724389.1;Wu1-SARS2_orf1ab;2113 | 0 | 0 |
| NC_045512.2;YP_009724389.1;Wu1-SARS2_orf1ab;2114 | 0.1 | 0.025 |
| NC_045512.2;YP_009724389.1;Wu1-SARS2_orf1ab;2115 | 0.05 | 0 |
| NC_045512.2;YP_009724389.1;Wu1-SARS2_orf1ab;2116 | 0 | 0 |
| NC_045512.2;YP_009724389.1;Wu1-SARS2_orf1ab;2117 | 0 | 0 |
| NC_045512.2;YP_009724389.1;Wu1-SARS2_orf1ab;2118 | 0.05 | 0 |
| NC_045512.2;YP_009724389.1;Wu1-SARS2_orf1ab;2119 | 0.05 | 0 |
| NC_045512.2;YP_009724389.1;Wu1-SARS2_orf1ab;2120 | 0 | 0 |
| NC_045512.2;YP_009724389.1;Wu1-SARS2_orf1ab;2121 | 0 | 0 |
| NC_045512.2;YP_009724389.1;Wu1-SARS2_orf1ab;2122 | 0 | 0 |
| NC_045512.2;YP_009724389.1;Wu1-SARS2_orf1ab;2123 | 0 | 0 |
| NC_045512.2;YP_009724389.1;Wu1-SARS2_orf1ab;2124 | 0.05 | 0 |
| NC_045512.2;YP_009724389.1;Wu1-SARS2_orf1ab;2125 | 0.05 | 0 |
| NC_045512.2;YP_009724389.1;Wu1-SARS2_orf1ab;2126 | 0.05 | 0.025 |
| NC_045512.2;YP_009724389.1;Wu1-SARS2_orf1ab;2127 | 0.05 | 0 |
| NC_045512.2;YP_009724389.1;Wu1-SARS2_orf1ab;2128 | 0.1 | 0 |
| NC_045512.2;YP_009724389.1;Wu1-SARS2_orf1ab;2129 | 0.15 | 0.025 |
| NC_045512.2;YP_009724389.1;Wu1-SARS2_orf1ab;2130 | 0.1 | 0 |
| NC_045512.2;YP_009724389.1;Wu1-SARS2_orf1ab;2131 | 0.05 | 0 |
| NC_045512.2;YP_009724389.1;Wu1-SARS2_orf1ab;2132 | 0.1 | 0 |
| NC_045512.2;YP_009724389.1;Wu1-SARS2_orf1ab;2133 | 0.1 | 0.05 |
| NC_045512.2;YP_009724389.1;Wu1-SARS2_orf1ab;2134 | 0.1 | 0.025 |
| NC_045512.2;YP_009724389.1;Wu1-SARS2_orf1ab;2135 | 0.1 | 0.025 |
| NC_045512.2;YP_009724389.1;Wu1-SARS2_orf1ab;2136 | 0.2 | 0.075 |
| NC_045512.2;YP_009724389.1;Wu1-SARS2_orf1ab;2137 | 0.05 | 0.025 |
| NC_045512.2;YP_009724389.1;Wu1-SARS2_orf1ab;2138 | 0.05 | 0.05 |
| NC_045512.2;YP_009724389.1;Wu1-SARS2_orf1ab;2139 | 0.1 | 0.05 |
| NC_045512.2;YP_009724389.1;Wu1-SARS2_orf1ab;2140 | 0.05 | 0.05 |

| | | |
|---|---|---|
| NC_045512.2;YP_009724389.1;Wu1-SARS2_orf1ab;2141 | 0.05 | 0.05 |
| NC_045512.2;YP_009724389.1;Wu1-SARS2_orf1ab;2142 | 0.05 | 0.05 |
| NC_045512.2;YP_009724389.1;Wu1-SARS2_orf1ab;2143 | 0.1 | 0.025 |
| NC_045512.2;YP_009724389.1;Wu1-SARS2_orf1ab;2144 | 0.05 | 0 |
| NC_045512.2;YP_009724389.1;Wu1-SARS2_orf1ab;2145 | 0 | 0 |
| NC_045512.2;YP_009724389.1;Wu1-SARS2_orf1ab;2146 | 0 | 0 |
| NC_045512.2;YP_009724389.1;Wu1-SARS2_orf1ab;2147 | 0 | 0 |
| NC_045512.2;YP_009724389.1;Wu1-SARS2_orf1ab;2148 | 0 | 0 |
| NC_045512.2;YP_009724389.1;Wu1-SARS2_orf1ab;2149 | 0 | 0 |
| NC_045512.2;YP_009724389.1;Wu1-SARS2_orf1ab;2150 | 0 | 0 |
| NC_045512.2;YP_009724389.1;Wu1-SARS2_orf1ab;2151 | 0 | 0 |
| NC_045512.2;YP_009724389.1;Wu1-SARS2_orf1ab;2152 | 0 | 0 |
| NC_045512.2;YP_009724389.1;Wu1-SARS2_orf1ab;2153 | 0 | 0 |
| NC_045512.2;YP_009724389.1;Wu1-SARS2_orf1ab;2154 | 0 | 0 |
| NC_045512.2;YP_009724389.1;Wu1-SARS2_orf1ab;2155 | 0 | 0 |
| NC_045512.2;YP_009724389.1;Wu1-SARS2_orf1ab;2156 | 0 | 0 |
| NC_045512.2;YP_009724389.1;Wu1-SARS2_orf1ab;2157 | 0 | 0 |
| NC_045512.2;YP_009724389.1;Wu1-SARS2_orf1ab;2158 | 0 | 0 |
| NC_045512.2;YP_009724389.1;Wu1-SARS2_orf1ab;2159 | 0 | 0 |
| NC_045512.2;YP_009724389.1;Wu1-SARS2_orf1ab;2160 | 0 | 0 |
| NC_045512.2;YP_009724389.1;Wu1-SARS2_orf1ab;2161 | 0 | 0 |
| NC_045512.2;YP_009724389.1;Wu1-SARS2_orf1ab;2162 | 0 | 0 |
| NC_045512.2;YP_009724389.1;Wu1-SARS2_orf1ab;2163 | 0 | 0 |
| NC_045512.2;YP_009724389.1;Wu1-SARS2_orf1ab;2164 | 0 | 0 |
| NC_045512.2;YP_009724389.1;Wu1-SARS2_orf1ab;2165 | 0 | 0 |
| NC_045512.2;YP_009724389.1;Wu1-SARS2_orf1ab;2166 | 0 | 0 |
| NC_045512.2;YP_009724389.1;Wu1-SARS2_orf1ab;2167 | 0 | 0 |
| NC_045512.2;YP_009724389.1;Wu1-SARS2_orf1ab;2168 | 0 | 0 |
| NC_045512.2;YP_009724389.1;Wu1-SARS2_orf1ab;2169 | 0 | 0 |
| NC_045512.2;YP_009724389.1;Wu1-SARS2_orf1ab;2170 | 0 | 0 |
| NC_045512.2;YP_009724389.1;Wu1-SARS2_orf1ab;2171 | 0 | 0 |
| NC_045512.2;YP_009724389.1;Wu1-SARS2_orf1ab;2172 | 0 | 0 |
| NC_045512.2;YP_009724389.1;Wu1-SARS2_orf1ab;2173 | 0 | 0 |
| NC_045512.2;YP_009724389.1;Wu1-SARS2_orf1ab;2174 | 0 | 0 |
| NC_045512.2;YP_009724389.1;Wu1-SARS2_orf1ab;2175 | 0 | 0 |
| NC_045512.2;YP_009724389.1;Wu1-SARS2_orf1ab;2176 | 0 | 0 |
| NC_045512.2;YP_009724389.1;Wu1-SARS2_orf1ab;2177 | 0 | 0 |
| NC_045512.2;YP_009724389.1;Wu1-SARS2_orf1ab;2178 | 0 | 0 |
| NC_045512.2;YP_009724389.1;Wu1-SARS2_orf1ab;2179 | 0 | 0 |
| NC_045512.2;YP_009724389.1;Wu1-SARS2_orf1ab;2180 | 0.05 | 0 |
| NC_045512.2;YP_009724389.1;Wu1-SARS2_orf1ab;2181 | 0.1 | 0 |
| NC_045512.2;YP_009724389.1;Wu1-SARS2_orf1ab;2182 | 0 | 0 |
| NC_045512.2;YP_009724389.1;Wu1-SARS2_orf1ab;2183 | 0 | 0.025 |

| | | |
|---|---|---|
| NC_045512.2;YP_009724389.1;Wu1-SARS2_orf1ab;2184 | 0 | 0 |
| NC_045512.2;YP_009724389.1;Wu1-SARS2_orf1ab;2185 | 0.05 | 0 |
| NC_045512.2;YP_009724389.1;Wu1-SARS2_orf1ab;2186 | 0.05 | 0 |
| NC_045512.2;YP_009724389.1;Wu1-SARS2_orf1ab;2187 | 0 | 0 |
| NC_045512.2;YP_009724389.1;Wu1-SARS2_orf1ab;2188 | 0 | 0 |
| NC_045512.2;YP_009724389.1;Wu1-SARS2_orf1ab;2189 | 0 | 0 |
| NC_045512.2;YP_009724389.1;Wu1-SARS2_orf1ab;2190 | 0 | 0 |
| NC_045512.2;YP_009724389.1;Wu1-SARS2_orf1ab;2191 | 0 | 0 |
| NC_045512.2;YP_009724389.1;Wu1-SARS2_orf1ab;2192 | 0 | 0 |
| NC_045512.2;YP_009724389.1;Wu1-SARS2_orf1ab;2193 | 0 | 0 |
| NC_045512.2;YP_009724389.1;Wu1-SARS2_orf1ab;2194 | 0 | 0 |
| NC_045512.2;YP_009724389.1;Wu1-SARS2_orf1ab;2195 | 0 | 0 |
| NC_045512.2;YP_009724389.1;Wu1-SARS2_orf1ab;2196 | 0 | 0 |
| NC_045512.2;YP_009724389.1;Wu1-SARS2_orf1ab;2197 | 0 | 0 |
| NC_045512.2;YP_009724389.1;Wu1-SARS2_orf1ab;2198 | 0 | 0 |
| NC_045512.2;YP_009724389.1;Wu1-SARS2_orf1ab;2199 | 0 | 0 |
| NC_045512.2;YP_009724389.1;Wu1-SARS2_orf1ab;2200 | 0 | 0 |
| NC_045512.2;YP_009724389.1;Wu1-SARS2_orf1ab;2201 | 0 | 0 |
| NC_045512.2;YP_009724389.1;Wu1-SARS2_orf1ab;2202 | 0 | 0 |
| NC_045512.2;YP_009724389.1;Wu1-SARS2_orf1ab;2203 | 0 | 0 |
| NC_045512.2;YP_009724389.1;Wu1-SARS2_orf1ab;2204 | 0 | 0 |
| NC_045512.2;YP_009724389.1;Wu1-SARS2_orf1ab;2205 | 0 | 0 |
| NC_045512.2;YP_009724389.1;Wu1-SARS2_orf1ab;2206 | 0 | 0 |
| NC_045512.2;YP_009724389.1;Wu1-SARS2_orf1ab;2207 | 0 | 0 |
| NC_045512.2;YP_009724389.1;Wu1-SARS2_orf1ab;2208 | 0 | 0 |
| NC_045512.2;YP_009724389.1;Wu1-SARS2_orf1ab;2209 | 0 | 0 |
| NC_045512.2;YP_009724389.1;Wu1-SARS2_orf1ab;2210 | 0 | 0 |
| NC_045512.2;YP_009724389.1;Wu1-SARS2_orf1ab;2211 | 0 | 0 |
| NC_045512.2;YP_009724389.1;Wu1-SARS2_orf1ab;2212 | 0.05 | 0 |
| NC_045512.2;YP_009724389.1;Wu1-SARS2_orf1ab;2213 | 0 | 0 |
| NC_045512.2;YP_009724389.1;Wu1-SARS2_orf1ab;2214 | 0 | 0 |
| NC_045512.2;YP_009724389.1;Wu1-SARS2_orf1ab;2215 | 0 | 0 |
| NC_045512.2;YP_009724389.1;Wu1-SARS2_orf1ab;2216 | 0 | 0 |
| NC_045512.2;YP_009724389.1;Wu1-SARS2_orf1ab;2217 | 0 | 0 |
| NC_045512.2;YP_009724389.1;Wu1-SARS2_orf1ab;2218 | 0 | 0 |
| NC_045512.2;YP_009724389.1;Wu1-SARS2_orf1ab;2219 | 0 | 0 |
| NC_045512.2;YP_009724389.1;Wu1-SARS2_orf1ab;2220 | 0 | 0 |
| NC_045512.2;YP_009724389.1;Wu1-SARS2_orf1ab;2221 | 0 | 0 |
| NC_045512.2;YP_009724389.1;Wu1-SARS2_orf1ab;2222 | 0 | 0 |
| NC_045512.2;YP_009724389.1;Wu1-SARS2_orf1ab;2223 | 0 | 0 |
| NC_045512.2;YP_009724389.1;Wu1-SARS2_orf1ab;2224 | 0 | 0 |
| NC_045512.2;YP_009724389.1;Wu1-SARS2_orf1ab;2225 | 0 | 0 |
| NC_045512.2;YP_009724389.1;Wu1-SARS2_orf1ab;2226 | 0 | 0 |

| | | |
|---|---|---|
| NC_045512.2;YP_009724389.1;Wu1-SARS2_orf1ab;2227 | 0 | 0 |
| NC_045512.2;YP_009724389.1;Wu1-SARS2_orf1ab;2228 | 0 | 0 |
| NC_045512.2;YP_009724389.1;Wu1-SARS2_orf1ab;2229 | 0 | 0 |
| NC_045512.2;YP_009724389.1;Wu1-SARS2_orf1ab;2230 | 0 | 0 |
| NC_045512.2;YP_009724389.1;Wu1-SARS2_orf1ab;2231 | 0 | 0 |
| NC_045512.2;YP_009724389.1;Wu1-SARS2_orf1ab;2232 | 0 | 0 |
| NC_045512.2;YP_009724389.1;Wu1-SARS2_orf1ab;2233 | 0 | 0 |
| NC_045512.2;YP_009724389.1;Wu1-SARS2_orf1ab;2234 | 0 | 0 |
| NC_045512.2;YP_009724389.1;Wu1-SARS2_orf1ab;2235 | 0 | 0 |
| NC_045512.2;YP_009724389.1;Wu1-SARS2_orf1ab;2236 | 0 | 0 |
| NC_045512.2;YP_009724389.1;Wu1-SARS2_orf1ab;2237 | 0 | 0 |
| NC_045512.2;YP_009724389.1;Wu1-SARS2_orf1ab;2238 | 0 | 0 |
| NC_045512.2;YP_009724389.1;Wu1-SARS2_orf1ab;2239 | 0 | 0 |
| NC_045512.2;YP_009724389.1;Wu1-SARS2_orf1ab;2240 | 0 | 0 |
| NC_045512.2;YP_009724389.1;Wu1-SARS2_orf1ab;2241 | 0 | 0 |
| NC_045512.2;YP_009724389.1;Wu1-SARS2_orf1ab;2242 | 0 | 0 |
| NC_045512.2;YP_009724389.1;Wu1-SARS2_orf1ab;2243 | 0 | 0 |
| NC_045512.2;YP_009724389.1;Wu1-SARS2_orf1ab;2244 | 0 | 0 |
| NC_045512.2;YP_009724389.1;Wu1-SARS2_orf1ab;2245 | 0 | 0 |
| NC_045512.2;YP_009724389.1;Wu1-SARS2_orf1ab;2246 | 0 | 0 |
| NC_045512.2;YP_009724389.1;Wu1-SARS2_orf1ab;2247 | 0 | 0 |
| NC_045512.2;YP_009724389.1;Wu1-SARS2_orf1ab;2248 | 0 | 0.025 |
| NC_045512.2;YP_009724389.1;Wu1-SARS2_orf1ab;2249 | 0 | 0.025 |
| NC_045512.2;YP_009724389.1;Wu1-SARS2_orf1ab;2250 | 0 | 0 |
| NC_045512.2;YP_009724389.1;Wu1-SARS2_orf1ab;2251 | 0 | 0 |
| NC_045512.2;YP_009724389.1;Wu1-SARS2_orf1ab;2252 | 0 | 0 |
| NC_045512.2;YP_009724389.1;Wu1-SARS2_orf1ab;2253 | 0 | 0 |
| NC_045512.2;YP_009724389.1;Wu1-SARS2_orf1ab;2254 | 0 | 0 |
| NC_045512.2;YP_009724389.1;Wu1-SARS2_orf1ab;2255 | 0 | 0 |
| NC_045512.2;YP_009724389.1;Wu1-SARS2_orf1ab;2256 | 0 | 0 |
| NC_045512.2;YP_009724389.1;Wu1-SARS2_orf1ab;2257 | 0 | 0 |
| NC_045512.2;YP_009724389.1;Wu1-SARS2_orf1ab;2258 | 0 | 0.05 |
| NC_045512.2;YP_009724389.1;Wu1-SARS2_orf1ab;2259 | 0 | 0 |
| NC_045512.2;YP_009724389.1;Wu1-SARS2_orf1ab;2260 | 0.2 | 0.125 |
| NC_045512.2;YP_009724389.1;Wu1-SARS2_orf1ab;2261 | 0.15 | 0.1 |
| NC_045512.2;YP_009724389.1;Wu1-SARS2_orf1ab;2262 | 0.3 | 0.275 |
| NC_045512.2;YP_009724389.1;Wu1-SARS2_orf1ab;2263 | 0.2 | 0.15 |
| NC_045512.2;YP_009724389.1;Wu1-SARS2_orf1ab;2264 | 0.25 | 0.175 |
| NC_045512.2;YP_009724389.1;Wu1-SARS2_orf1ab;2265 | 0.2 | 0.1 |
| NC_045512.2;YP_009724389.1;Wu1-SARS2_orf1ab;2266 | 0.05 | 0 |
| NC_045512.2;YP_009724389.1;Wu1-SARS2_orf1ab;2267 | 0.05 | 0.025 |
| NC_045512.2;YP_009724389.1;Wu1-SARS2_orf1ab;2268 | 0.05 | 0.025 |
| NC_045512.2;YP_009724389.1;Wu1-SARS2_orf1ab;2269 | 0.05 | 0 |

| | | |
|---|---|---|
| NC_045512.2;YP_009724389.1;Wu1-SARS2_orf1ab;2270 | 0.05 | 0 |
| NC_045512.2;YP_009724389.1;Wu1-SARS2_orf1ab;2271 | 0 | 0 |
| NC_045512.2;YP_009724389.1;Wu1-SARS2_orf1ab;2272 | 0 | 0 |
| NC_045512.2;YP_009724389.1;Wu1-SARS2_orf1ab;2273 | 0 | 0 |
| NC_045512.2;YP_009724389.1;Wu1-SARS2_orf1ab;2274 | 0.05 | 0 |
| NC_045512.2;YP_009724389.1;Wu1-SARS2_orf1ab;2275 | 0 | 0 |
| NC_045512.2;YP_009724389.1;Wu1-SARS2_orf1ab;2276 | 0 | 0 |
| NC_045512.2;YP_009724389.1;Wu1-SARS2_orf1ab;2277 | 0 | 0 |
| NC_045512.2;YP_009724389.1;Wu1-SARS2_orf1ab;2278 | 0 | 0 |
| NC_045512.2;YP_009724389.1;Wu1-SARS2_orf1ab;2279 | 0 | 0 |
| NC_045512.2;YP_009724389.1;Wu1-SARS2_orf1ab;2280 | 0 | 0 |
| NC_045512.2;YP_009724389.1;Wu1-SARS2_orf1ab;2281 | 0 | 0 |
| NC_045512.2;YP_009724389.1;Wu1-SARS2_orf1ab;2282 | 0 | 0 |
| NC_045512.2;YP_009724389.1;Wu1-SARS2_orf1ab;2283 | 0 | 0 |
| NC_045512.2;YP_009724389.1;Wu1-SARS2_orf1ab;2284 | 0 | 0 |
| NC_045512.2;YP_009724389.1;Wu1-SARS2_orf1ab;2285 | 0 | 0 |
| NC_045512.2;YP_009724389.1;Wu1-SARS2_orf1ab;2286 | 0 | 0 |
| NC_045512.2;YP_009724389.1;Wu1-SARS2_orf1ab;2287 | 0 | 0 |
| NC_045512.2;YP_009724389.1;Wu1-SARS2_orf1ab;2288 | 0 | 0.025 |
| NC_045512.2;YP_009724389.1;Wu1-SARS2_orf1ab;2289 | 0 | 0 |
| NC_045512.2;YP_009724389.1;Wu1-SARS2_orf1ab;2290 | 0 | 0 |
| NC_045512.2;YP_009724389.1;Wu1-SARS2_orf1ab;2291 | 0.05 | 0.025 |
| NC_045512.2;YP_009724389.1;Wu1-SARS2_orf1ab;2292 | 0 | 0 |
| NC_045512.2;YP_009724389.1;Wu1-SARS2_orf1ab;2293 | 0 | 0 |
| NC_045512.2;YP_009724389.1;Wu1-SARS2_orf1ab;2294 | 0.05 | 0.025 |
| NC_045512.2;YP_009724389.1;Wu1-SARS2_orf1ab;2295 | 0.05 | 0 |
| NC_045512.2;YP_009724389.1;Wu1-SARS2_orf1ab;2296 | 0 | 0 |
| NC_045512.2;YP_009724389.1;Wu1-SARS2_orf1ab;2297 | 0 | 0 |
| NC_045512.2;YP_009724389.1;Wu1-SARS2_orf1ab;2298 | 0 | 0 |
| NC_045512.2;YP_009724389.1;Wu1-SARS2_orf1ab;2299 | 0 | 0 |
| NC_045512.2;YP_009724389.1;Wu1-SARS2_orf1ab;2300 | 0 | 0 |
| NC_045512.2;YP_009724389.1;Wu1-SARS2_orf1ab;2301 | 0 | 0 |
| NC_045512.2;YP_009724389.1;Wu1-SARS2_orf1ab;2302 | 0 | 0 |
| NC_045512.2;YP_009724389.1;Wu1-SARS2_orf1ab;2303 | 0 | 0 |
| NC_045512.2;YP_009724389.1;Wu1-SARS2_orf1ab;2304 | 0 | 0 |
| NC_045512.2;YP_009724389.1;Wu1-SARS2_orf1ab;2305 | 0 | 0 |
| NC_045512.2;YP_009724389.1;Wu1-SARS2_orf1ab;2306 | 0 | 0 |
| NC_045512.2;YP_009724389.1;Wu1-SARS2_orf1ab;2307 | 0 | 0 |
| NC_045512.2;YP_009724389.1;Wu1-SARS2_orf1ab;2308 | 0 | 0 |
| NC_045512.2;YP_009724389.1;Wu1-SARS2_orf1ab;2309 | 0 | 0.075 |
| NC_045512.2;YP_009724389.1;Wu1-SARS2_orf1ab;2310 | 0 | 0 |
| NC_045512.2;YP_009724389.1;Wu1-SARS2_orf1ab;2311 | 0 | 0 |
| NC_045512.2;YP_009724389.1;Wu1-SARS2_orf1ab;2312 | 0 | 0 |

| | | |
|---|---|---|
| NC_045512.2;YP_009724389.1;Wu1-SARS2_orf1ab;2313 | 0 | 0 |
| NC_045512.2;YP_009724389.1;Wu1-SARS2_orf1ab;2314 | 0 | 0 |
| NC_045512.2;YP_009724389.1;Wu1-SARS2_orf1ab;2315 | 0 | 0 |
| NC_045512.2;YP_009724389.1;Wu1-SARS2_orf1ab;2316 | 0 | 0 |
| NC_045512.2;YP_009724389.1;Wu1-SARS2_orf1ab;2317 | 0 | 0 |
| NC_045512.2;YP_009724389.1;Wu1-SARS2_orf1ab;2318 | 0 | 0 |
| NC_045512.2;YP_009724389.1;Wu1-SARS2_orf1ab;2319 | 0 | 0 |
| NC_045512.2;YP_009724389.1;Wu1-SARS2_orf1ab;2320 | 0 | 0 |
| NC_045512.2;YP_009724389.1;Wu1-SARS2_orf1ab;2321 | 0 | 0 |
| NC_045512.2;YP_009724389.1;Wu1-SARS2_orf1ab;2322 | 0 | 0 |
| NC_045512.2;YP_009724389.1;Wu1-SARS2_orf1ab;2323 | 0 | 0 |
| NC_045512.2;YP_009724389.1;Wu1-SARS2_orf1ab;2324 | 0 | 0 |
| NC_045512.2;YP_009724389.1;Wu1-SARS2_orf1ab;2325 | 0 | 0 |
| NC_045512.2;YP_009724389.1;Wu1-SARS2_orf1ab;2326 | 0 | 0 |
| NC_045512.2;YP_009724389.1;Wu1-SARS2_orf1ab;2327 | 0 | 0 |
| NC_045512.2;YP_009724389.1;Wu1-SARS2_orf1ab;2328 | 0 | 0 |
| NC_045512.2;YP_009724389.1;Wu1-SARS2_orf1ab;2329 | 0 | 0 |
| NC_045512.2;YP_009724389.1;Wu1-SARS2_orf1ab;2330 | 0 | 0 |
| NC_045512.2;YP_009724389.1;Wu1-SARS2_orf1ab;2331 | 0 | 0 |
| NC_045512.2;YP_009724389.1;Wu1-SARS2_orf1ab;2332 | 0 | 0 |
| NC_045512.2;YP_009724389.1;Wu1-SARS2_orf1ab;2333 | 0 | 0 |
| NC_045512.2;YP_009724389.1;Wu1-SARS2_orf1ab;2334 | 0 | 0 |
| NC_045512.2;YP_009724389.1;Wu1-SARS2_orf1ab;2335 | 0 | 0 |
| NC_045512.2;YP_009724389.1;Wu1-SARS2_orf1ab;2336 | 0 | 0 |
| NC_045512.2;YP_009724389.1;Wu1-SARS2_orf1ab;2337 | 0 | 0 |
| NC_045512.2;YP_009724389.1;Wu1-SARS2_orf1ab;2338 | 0 | 0 |
| NC_045512.2;YP_009724389.1;Wu1-SARS2_orf1ab;2339 | 0 | 0 |
| NC_045512.2;YP_009724389.1;Wu1-SARS2_orf1ab;2340 | 0 | 0 |
| NC_045512.2;YP_009724389.1;Wu1-SARS2_orf1ab;2341 | 0 | 0 |
| NC_045512.2;YP_009724389.1;Wu1-SARS2_orf1ab;2342 | 0 | 0 |
| NC_045512.2;YP_009724389.1;Wu1-SARS2_orf1ab;2343 | 0 | 0 |
| NC_045512.2;YP_009724389.1;Wu1-SARS2_orf1ab;2344 | 0 | 0 |
| NC_045512.2;YP_009724389.1;Wu1-SARS2_orf1ab;2345 | 0 | 0 |
| NC_045512.2;YP_009724389.1;Wu1-SARS2_orf1ab;2346 | 0 | 0 |
| NC_045512.2;YP_009724389.1;Wu1-SARS2_orf1ab;2347 | 0 | 0 |
| NC_045512.2;YP_009724389.1;Wu1-SARS2_orf1ab;2348 | 0 | 0 |
| NC_045512.2;YP_009724389.1;Wu1-SARS2_orf1ab;2349 | 0 | 0 |
| NC_045512.2;YP_009724389.1;Wu1-SARS2_orf1ab;2350 | 0 | 0 |
| NC_045512.2;YP_009724389.1;Wu1-SARS2_orf1ab;2351 | 0 | 0 |
| NC_045512.2;YP_009724389.1;Wu1-SARS2_orf1ab;2352 | 0 | 0 |
| NC_045512.2;YP_009724389.1;Wu1-SARS2_orf1ab;2353 | 0 | 0 |
| NC_045512.2;YP_009724389.1;Wu1-SARS2_orf1ab;2354 | 0 | 0 |
| NC_045512.2;YP_009724389.1;Wu1-SARS2_orf1ab;2355 | 0 | 0 |

| | | |
|---|---|---|
| NC_045512.2;YP_009724389.1;Wu1-SARS2_orf1ab;2356 | 0 | 0 |
| NC_045512.2;YP_009724389.1;Wu1-SARS2_orf1ab;2357 | 0 | 0 |
| NC_045512.2;YP_009724389.1;Wu1-SARS2_orf1ab;2358 | 0 | 0 |
| NC_045512.2;YP_009724389.1;Wu1-SARS2_orf1ab;2359 | 0 | 0 |
| NC_045512.2;YP_009724389.1;Wu1-SARS2_orf1ab;2360 | 0 | 0 |
| NC_045512.2;YP_009724389.1;Wu1-SARS2_orf1ab;2361 | 0 | 0 |
| NC_045512.2;YP_009724389.1;Wu1-SARS2_orf1ab;2362 | 0 | 0 |
| NC_045512.2;YP_009724389.1;Wu1-SARS2_orf1ab;2363 | 0 | 0 |
| NC_045512.2;YP_009724389.1;Wu1-SARS2_orf1ab;2364 | 0 | 0 |
| NC_045512.2;YP_009724389.1;Wu1-SARS2_orf1ab;2365 | 0 | 0 |
| NC_045512.2;YP_009724389.1;Wu1-SARS2_orf1ab;2366 | 0 | 0 |
| NC_045512.2;YP_009724389.1;Wu1-SARS2_orf1ab;2367 | 0 | 0 |
| NC_045512.2;YP_009724389.1;Wu1-SARS2_orf1ab;2368 | 0 | 0 |
| NC_045512.2;YP_009724389.1;Wu1-SARS2_orf1ab;2369 | 0 | 0 |
| NC_045512.2;YP_009724389.1;Wu1-SARS2_orf1ab;2370 | 0 | 0 |
| NC_045512.2;YP_009724389.1;Wu1-SARS2_orf1ab;2371 | 0 | 0 |
| NC_045512.2;YP_009724389.1;Wu1-SARS2_orf1ab;2372 | 0 | 0 |
| NC_045512.2;YP_009724389.1;Wu1-SARS2_orf1ab;2373 | 0 | 0 |
| NC_045512.2;YP_009724389.1;Wu1-SARS2_orf1ab;2374 | 0 | 0 |
| NC_045512.2;YP_009724389.1;Wu1-SARS2_orf1ab;2375 | 0 | 0 |
| NC_045512.2;YP_009724389.1;Wu1-SARS2_orf1ab;2376 | 0 | 0 |
| NC_045512.2;YP_009724389.1;Wu1-SARS2_orf1ab;2377 | 0 | 0 |
| NC_045512.2;YP_009724389.1;Wu1-SARS2_orf1ab;2378 | 0 | 0 |
| NC_045512.2;YP_009724389.1;Wu1-SARS2_orf1ab;2379 | 0 | 0 |
| NC_045512.2;YP_009724389.1;Wu1-SARS2_orf1ab;2380 | 0 | 0 |
| NC_045512.2;YP_009724389.1;Wu1-SARS2_orf1ab;2381 | 0 | 0 |
| NC_045512.2;YP_009724389.1;Wu1-SARS2_orf1ab;2382 | 0 | 0 |
| NC_045512.2;YP_009724389.1;Wu1-SARS2_orf1ab;2383 | 0 | 0 |
| NC_045512.2;YP_009724389.1;Wu1-SARS2_orf1ab;2384 | 0 | 0 |
| NC_045512.2;YP_009724389.1;Wu1-SARS2_orf1ab;2385 | 0 | 0 |
| NC_045512.2;YP_009724389.1;Wu1-SARS2_orf1ab;2386 | 0 | 0 |
| NC_045512.2;YP_009724389.1;Wu1-SARS2_orf1ab;2387 | 0 | 0 |
| NC_045512.2;YP_009724389.1;Wu1-SARS2_orf1ab;2388 | 0 | 0 |
| NC_045512.2;YP_009724389.1;Wu1-SARS2_orf1ab;2389 | 0 | 0 |
| NC_045512.2;YP_009724389.1;Wu1-SARS2_orf1ab;2390 | 0 | 0 |
| NC_045512.2;YP_009724389.1;Wu1-SARS2_orf1ab;2391 | 0 | 0 |
| NC_045512.2;YP_009724389.1;Wu1-SARS2_orf1ab;2392 | 0 | 0 |
| NC_045512.2;YP_009724389.1;Wu1-SARS2_orf1ab;2393 | 0 | 0 |
| NC_045512.2;YP_009724389.1;Wu1-SARS2_orf1ab;2394 | 0 | 0 |
| NC_045512.2;YP_009724389.1;Wu1-SARS2_orf1ab;2395 | 0 | 0 |
| NC_045512.2;YP_009724389.1;Wu1-SARS2_orf1ab;2396 | 0 | 0 |
| NC_045512.2;YP_009724389.1;Wu1-SARS2_orf1ab;2397 | 0 | 0 |
| NC_045512.2;YP_009724389.1;Wu1-SARS2_orf1ab;2398 | 0 | 0 |

| | | |
|---|---|---|
| NC_045512.2;YP_009724389.1;Wu1-SARS2_orf1ab;2399 | 0 | 0 |
| NC_045512.2;YP_009724389.1;Wu1-SARS2_orf1ab;2400 | 0 | 0 |
| NC_045512.2;YP_009724389.1;Wu1-SARS2_orf1ab;2401 | 0 | 0 |
| NC_045512.2;YP_009724389.1;Wu1-SARS2_orf1ab;2402 | 0 | 0 |
| NC_045512.2;YP_009724389.1;Wu1-SARS2_orf1ab;2403 | 0 | 0 |
| NC_045512.2;YP_009724389.1;Wu1-SARS2_orf1ab;2404 | 0 | 0 |
| NC_045512.2;YP_009724389.1;Wu1-SARS2_orf1ab;2405 | 0 | 0 |
| NC_045512.2;YP_009724389.1;Wu1-SARS2_orf1ab;2406 | 0 | 0 |
| NC_045512.2;YP_009724389.1;Wu1-SARS2_orf1ab;2407 | 0 | 0 |
| NC_045512.2;YP_009724389.1;Wu1-SARS2_orf1ab;2408 | 0 | 0 |
| NC_045512.2;YP_009724389.1;Wu1-SARS2_orf1ab;2409 | 0 | 0 |
| NC_045512.2;YP_009724389.1;Wu1-SARS2_orf1ab;2410 | 0 | 0 |
| NC_045512.2;YP_009724389.1;Wu1-SARS2_orf1ab;2411 | 0 | 0 |
| NC_045512.2;YP_009724389.1;Wu1-SARS2_orf1ab;2412 | 0 | 0 |
| NC_045512.2;YP_009724389.1;Wu1-SARS2_orf1ab;2413 | 0 | 0 |
| NC_045512.2;YP_009724389.1;Wu1-SARS2_orf1ab;2414 | 0 | 0 |
| NC_045512.2;YP_009724389.1;Wu1-SARS2_orf1ab;2415 | 0 | 0 |
| NC_045512.2;YP_009724389.1;Wu1-SARS2_orf1ab;2416 | 0 | 0.025 |
| NC_045512.2;YP_009724389.1;Wu1-SARS2_orf1ab;2417 | 0 | 0 |
| NC_045512.2;YP_009724389.1;Wu1-SARS2_orf1ab;2418 | 0 | 0 |
| NC_045512.2;YP_009724389.1;Wu1-SARS2_orf1ab;2419 | 0 | 0 |
| NC_045512.2;YP_009724389.1;Wu1-SARS2_orf1ab;2420 | 0 | 0 |
| NC_045512.2;YP_009724389.1;Wu1-SARS2_orf1ab;2421 | 0 | 0 |
| NC_045512.2;YP_009724389.1;Wu1-SARS2_orf1ab;2422 | 0 | 0 |
| NC_045512.2;YP_009724389.1;Wu1-SARS2_orf1ab;2423 | 0 | 0 |
| NC_045512.2;YP_009724389.1;Wu1-SARS2_orf1ab;2424 | 0 | 0 |
| NC_045512.2;YP_009724389.1;Wu1-SARS2_orf1ab;2425 | 0 | 0 |
| NC_045512.2;YP_009724389.1;Wu1-SARS2_orf1ab;2426 | 0 | 0 |
| NC_045512.2;YP_009724389.1;Wu1-SARS2_orf1ab;2427 | 0 | 0 |
| NC_045512.2;YP_009724389.1;Wu1-SARS2_orf1ab;2428 | 0 | 0 |
| NC_045512.2;YP_009724389.1;Wu1-SARS2_orf1ab;2429 | 0 | 0 |
| NC_045512.2;YP_009724389.1;Wu1-SARS2_orf1ab;2430 | 0 | 0 |
| NC_045512.2;YP_009724389.1;Wu1-SARS2_orf1ab;2431 | 0 | 0 |
| NC_045512.2;YP_009724389.1;Wu1-SARS2_orf1ab;2432 | 0 | 0 |
| NC_045512.2;YP_009724389.1;Wu1-SARS2_orf1ab;2433 | 0 | 0 |
| NC_045512.2;YP_009724389.1;Wu1-SARS2_orf1ab;2434 | 0 | 0 |
| NC_045512.2;YP_009724389.1;Wu1-SARS2_orf1ab;2435 | 0 | 0 |
| NC_045512.2;YP_009724389.1;Wu1-SARS2_orf1ab;2436 | 0 | 0 |
| NC_045512.2;YP_009724389.1;Wu1-SARS2_orf1ab;2437 | 0 | 0 |
| NC_045512.2;YP_009724389.1;Wu1-SARS2_orf1ab;2438 | 0 | 0 |
| NC_045512.2;YP_009724389.1;Wu1-SARS2_orf1ab;2439 | 0 | 0 |
| NC_045512.2;YP_009724389.1;Wu1-SARS2_orf1ab;2440 | 0 | 0 |
| NC_045512.2;YP_009724389.1;Wu1-SARS2_orf1ab;2441 | 0 | 0.025 |

| | | |
|---|---|---|
| NC_045512.2;YP_009724389.1;Wu1-SARS2_orf1ab;2442 | 0 | 0.025 |
| NC_045512.2;YP_009724389.1;Wu1-SARS2_orf1ab;2443 | 0 | 0 |
| NC_045512.2;YP_009724389.1;Wu1-SARS2_orf1ab;2444 | 0 | 0 |
| NC_045512.2;YP_009724389.1;Wu1-SARS2_orf1ab;2445 | 0 | 0 |
| NC_045512.2;YP_009724389.1;Wu1-SARS2_orf1ab;2446 | 0 | 0 |
| NC_045512.2;YP_009724389.1;Wu1-SARS2_orf1ab;2447 | 0 | 0 |
| NC_045512.2;YP_009724389.1;Wu1-SARS2_orf1ab;2448 | 0 | 0 |
| NC_045512.2;YP_009724389.1;Wu1-SARS2_orf1ab;2449 | 0 | 0 |
| NC_045512.2;YP_009724389.1;Wu1-SARS2_orf1ab;2450 | 0 | 0 |
| NC_045512.2;YP_009724389.1;Wu1-SARS2_orf1ab;2451 | 0 | 0 |
| NC_045512.2;YP_009724389.1;Wu1-SARS2_orf1ab;2452 | 0 | 0 |
| NC_045512.2;YP_009724389.1;Wu1-SARS2_orf1ab;2453 | 0 | 0 |
| NC_045512.2;YP_009724389.1;Wu1-SARS2_orf1ab;2454 | 0 | 0 |
| NC_045512.2;YP_009724389.1;Wu1-SARS2_orf1ab;2455 | 0 | 0 |
| NC_045512.2;YP_009724389.1;Wu1-SARS2_orf1ab;2456 | 0 | 0 |
| NC_045512.2;YP_009724389.1;Wu1-SARS2_orf1ab;2457 | 0 | 0 |
| NC_045512.2;YP_009724389.1;Wu1-SARS2_orf1ab;2458 | 0 | 0 |
| NC_045512.2;YP_009724389.1;Wu1-SARS2_orf1ab;2459 | 0 | 0 |
| NC_045512.2;YP_009724389.1;Wu1-SARS2_orf1ab;2460 | 0 | 0 |
| NC_045512.2;YP_009724389.1;Wu1-SARS2_orf1ab;2461 | 0 | 0 |
| NC_045512.2;YP_009724389.1;Wu1-SARS2_orf1ab;2462 | 0 | 0 |
| NC_045512.2;YP_009724389.1;Wu1-SARS2_orf1ab;2463 | 0 | 0 |
| NC_045512.2;YP_009724389.1;Wu1-SARS2_orf1ab;2464 | 0 | 0 |
| NC_045512.2;YP_009724389.1;Wu1-SARS2_orf1ab;2465 | 0 | 0 |
| NC_045512.2;YP_009724389.1;Wu1-SARS2_orf1ab;2466 | 0 | 0 |
| NC_045512.2;YP_009724389.1;Wu1-SARS2_orf1ab;2467 | 0 | 0 |
| NC_045512.2;YP_009724389.1;Wu1-SARS2_orf1ab;2468 | 0 | 0 |
| NC_045512.2;YP_009724389.1;Wu1-SARS2_orf1ab;2469 | 0.05 | 0 |
| NC_045512.2;YP_009724389.1;Wu1-SARS2_orf1ab;2470 | 0.05 | 0.025 |
| NC_045512.2;YP_009724389.1;Wu1-SARS2_orf1ab;2471 | 0.05 | 0.025 |
| NC_045512.2;YP_009724389.1;Wu1-SARS2_orf1ab;2472 | 0 | 0 |
| NC_045512.2;YP_009724389.1;Wu1-SARS2_orf1ab;2473 | 0 | 0 |
| NC_045512.2;YP_009724389.1;Wu1-SARS2_orf1ab;2474 | 0 | 0 |
| NC_045512.2;YP_009724389.1;Wu1-SARS2_orf1ab;2475 | 0 | 0.025 |
| NC_045512.2;YP_009724389.1;Wu1-SARS2_orf1ab;2476 | 0 | 0 |
| NC_045512.2;YP_009724389.1;Wu1-SARS2_orf1ab;2477 | 0 | 0 |
| NC_045512.2;YP_009724389.1;Wu1-SARS2_orf1ab;2478 | 0 | 0.05 |
| NC_045512.2;YP_009724389.1;Wu1-SARS2_orf1ab;2479 | 0 | 0 |
| NC_045512.2;YP_009724389.1;Wu1-SARS2_orf1ab;2480 | 0 | 0 |
| NC_045512.2;YP_009724389.1;Wu1-SARS2_orf1ab;2481 | 0 | 0 |
| NC_045512.2;YP_009724389.1;Wu1-SARS2_orf1ab;2482 | 0.05 | 0 |
| NC_045512.2;YP_009724389.1;Wu1-SARS2_orf1ab;2483 | 0 | 0 |
| NC_045512.2;YP_009724389.1;Wu1-SARS2_orf1ab;2484 | 0 | 0 |

| | | |
|---|---|---|
| NC_045512.2;YP_009724389.1;Wu1-SARS2_orf1ab;2485 | 0 | 0 |
| NC_045512.2;YP_009724389.1;Wu1-SARS2_orf1ab;2486 | 0 | 0 |
| NC_045512.2;YP_009724389.1;Wu1-SARS2_orf1ab;2487 | 0 | 0 |
| NC_045512.2;YP_009724389.1;Wu1-SARS2_orf1ab;2488 | 0 | 0 |
| NC_045512.2;YP_009724389.1;Wu1-SARS2_orf1ab;2489 | 0 | 0 |
| NC_045512.2;YP_009724389.1;Wu1-SARS2_orf1ab;2490 | 0 | 0 |
| NC_045512.2;YP_009724389.1;Wu1-SARS2_orf1ab;2491 | 0 | 0 |
| NC_045512.2;YP_009724389.1;Wu1-SARS2_orf1ab;2492 | 0 | 0 |
| NC_045512.2;YP_009724389.1;Wu1-SARS2_orf1ab;2493 | 0 | 0 |
| NC_045512.2;YP_009724389.1;Wu1-SARS2_orf1ab;2494 | 0 | 0 |
| NC_045512.2;YP_009724389.1;Wu1-SARS2_orf1ab;2495 | 0 | 0 |
| NC_045512.2;YP_009724389.1;Wu1-SARS2_orf1ab;2496 | 0 | 0 |
| NC_045512.2;YP_009724389.1;Wu1-SARS2_orf1ab;2497 | 0 | 0 |
| NC_045512.2;YP_009724389.1;Wu1-SARS2_orf1ab;2498 | 0 | 0 |
| NC_045512.2;YP_009724389.1;Wu1-SARS2_orf1ab;2499 | 0 | 0 |
| NC_045512.2;YP_009724389.1;Wu1-SARS2_orf1ab;2500 | 0 | 0 |
| NC_045512.2;YP_009724389.1;Wu1-SARS2_orf1ab;2501 | 0 | 0 |
| NC_045512.2;YP_009724389.1;Wu1-SARS2_orf1ab;2502 | 0 | 0 |
| NC_045512.2;YP_009724389.1;Wu1-SARS2_orf1ab;2503 | 0 | 0 |
| NC_045512.2;YP_009724389.1;Wu1-SARS2_orf1ab;2504 | 0.05 | 0.05 |
| NC_045512.2;YP_009724389.1;Wu1-SARS2_orf1ab;2505 | 0 | 0 |
| NC_045512.2;YP_009724389.1;Wu1-SARS2_orf1ab;2506 | 0 | 0 |
| NC_045512.2;YP_009724389.1;Wu1-SARS2_orf1ab;2507 | 0 | 0 |
| NC_045512.2;YP_009724389.1;Wu1-SARS2_orf1ab;2508 | 0 | 0 |
| NC_045512.2;YP_009724389.1;Wu1-SARS2_orf1ab;2509 | 0 | 0 |
| NC_045512.2;YP_009724389.1;Wu1-SARS2_orf1ab;2510 | 0 | 0 |
| NC_045512.2;YP_009724389.1;Wu1-SARS2_orf1ab;2511 | 0 | 0 |
| NC_045512.2;YP_009724389.1;Wu1-SARS2_orf1ab;2512 | 0 | 0 |
| NC_045512.2;YP_009724389.1;Wu1-SARS2_orf1ab;2513 | 0 | 0 |
| NC_045512.2;YP_009724389.1;Wu1-SARS2_orf1ab;2514 | 0 | 0 |
| NC_045512.2;YP_009724389.1;Wu1-SARS2_orf1ab;2515 | 0 | 0 |
| NC_045512.2;YP_009724389.1;Wu1-SARS2_orf1ab;2516 | 0 | 0 |
| NC_045512.2;YP_009724389.1;Wu1-SARS2_orf1ab;2517 | 0 | 0 |
| NC_045512.2;YP_009724389.1;Wu1-SARS2_orf1ab;2518 | 0 | 0 |
| NC_045512.2;YP_009724389.1;Wu1-SARS2_orf1ab;2519 | 0 | 0 |
| NC_045512.2;YP_009724389.1;Wu1-SARS2_orf1ab;2520 | 0 | 0 |
| NC_045512.2;YP_009724389.1;Wu1-SARS2_orf1ab;2521 | 0 | 0.025 |
| NC_045512.2;YP_009724389.1;Wu1-SARS2_orf1ab;2522 | 0.05 | 0.025 |
| NC_045512.2;YP_009724389.1;Wu1-SARS2_orf1ab;2523 | 0.05 | 0.025 |
| NC_045512.2;YP_009724389.1;Wu1-SARS2_orf1ab;2524 | 0.05 | 0.025 |
| NC_045512.2;YP_009724389.1;Wu1-SARS2_orf1ab;2525 | 0.05 | 0.025 |
| NC_045512.2;YP_009724389.1;Wu1-SARS2_orf1ab;2526 | 0 | 0 |
| NC_045512.2;YP_009724389.1;Wu1-SARS2_orf1ab;2527 | 0 | 0 |

| | | |
|---|---|---|
| NC_045512.2;YP_009724389.1;Wu1-SARS2_orf1ab;2528 | 0.05 | 0 |
| NC_045512.2;YP_009724389.1;Wu1-SARS2_orf1ab;2529 | 0 | 0 |
| NC_045512.2;YP_009724389.1;Wu1-SARS2_orf1ab;2530 | 0 | 0 |
| NC_045512.2;YP_009724389.1;Wu1-SARS2_orf1ab;2531 | 0 | 0 |
| NC_045512.2;YP_009724389.1;Wu1-SARS2_orf1ab;2532 | 0 | 0 |
| NC_045512.2;YP_009724389.1;Wu1-SARS2_orf1ab;2533 | 0 | 0 |
| NC_045512.2;YP_009724389.1;Wu1-SARS2_orf1ab;2534 | 0 | 0.025 |
| NC_045512.2;YP_009724389.1;Wu1-SARS2_orf1ab;2535 | 0 | 0.025 |
| NC_045512.2;YP_009724389.1;Wu1-SARS2_orf1ab;2536 | 0.05 | 0.025 |
| NC_045512.2;YP_009724389.1;Wu1-SARS2_orf1ab;2537 | 0.05 | 0.025 |
| NC_045512.2;YP_009724389.1;Wu1-SARS2_orf1ab;2538 | 0.05 | 0.025 |
| NC_045512.2;YP_009724389.1;Wu1-SARS2_orf1ab;2539 | 0 | 0.025 |
| NC_045512.2;YP_009724389.1;Wu1-SARS2_orf1ab;2540 | 0 | 0.025 |
| NC_045512.2;YP_009724389.1;Wu1-SARS2_orf1ab;2541 | 0 | 0.025 |
| NC_045512.2;YP_009724389.1;Wu1-SARS2_orf1ab;2542 | 0 | 0.025 |
| NC_045512.2;YP_009724389.1;Wu1-SARS2_orf1ab;2543 | 0.1 | 0.05 |
| NC_045512.2;YP_009724389.1;Wu1-SARS2_orf1ab;2544 | 0 | 0 |
| NC_045512.2;YP_009724389.1;Wu1-SARS2_orf1ab;2545 | 0 | 0 |
| NC_045512.2;YP_009724389.1;Wu1-SARS2_orf1ab;2546 | 0 | 0 |
| NC_045512.2;YP_009724389.1;Wu1-SARS2_orf1ab;2547 | 0 | 0 |
| NC_045512.2;YP_009724389.1;Wu1-SARS2_orf1ab;2548 | 0.05 | 0 |
| NC_045512.2;YP_009724389.1;Wu1-SARS2_orf1ab;2549 | 0 | 0 |
| NC_045512.2;YP_009724389.1;Wu1-SARS2_orf1ab;2550 | 0 | 0 |
| NC_045512.2;YP_009724389.1;Wu1-SARS2_orf1ab;2551 | 0.05 | 0 |
| NC_045512.2;YP_009724389.1;Wu1-SARS2_orf1ab;2552 | 0 | 0 |
| NC_045512.2;YP_009724389.1;Wu1-SARS2_orf1ab;2553 | 0 | 0 |
| NC_045512.2;YP_009724389.1;Wu1-SARS2_orf1ab;2554 | 0 | 0 |
| NC_045512.2;YP_009724389.1;Wu1-SARS2_orf1ab;2555 | 0 | 0 |
| NC_045512.2;YP_009724389.1;Wu1-SARS2_orf1ab;2556 | 0 | 0 |
| NC_045512.2;YP_009724389.1;Wu1-SARS2_orf1ab;2557 | 0 | 0.025 |
| NC_045512.2;YP_009724389.1;Wu1-SARS2_orf1ab;2558 | 0 | 0 |
| NC_045512.2;YP_009724389.1;Wu1-SARS2_orf1ab;2559 | 0 | 0 |
| NC_045512.2;YP_009724389.1;Wu1-SARS2_orf1ab;2560 | 0 | 0 |
| NC_045512.2;YP_009724389.1;Wu1-SARS2_orf1ab;2561 | 0 | 0 |
| NC_045512.2;YP_009724389.1;Wu1-SARS2_orf1ab;2562 | 0 | 0 |
| NC_045512.2;YP_009724389.1;Wu1-SARS2_orf1ab;2563 | 0 | 0 |
| NC_045512.2;YP_009724389.1;Wu1-SARS2_orf1ab;2564 | 0 | 0 |
| NC_045512.2;YP_009724389.1;Wu1-SARS2_orf1ab;2565 | 0 | 0 |
| NC_045512.2;YP_009724389.1;Wu1-SARS2_orf1ab;2566 | 0 | 0 |
| NC_045512.2;YP_009724389.1;Wu1-SARS2_orf1ab;2567 | 0 | 0 |
| NC_045512.2;YP_009724389.1;Wu1-SARS2_orf1ab;2568 | 0 | 0 |
| NC_045512.2;YP_009724389.1;Wu1-SARS2_orf1ab;2569 | 0 | 0.025 |
| NC_045512.2;YP_009724389.1;Wu1-SARS2_orf1ab;2570 | 0 | 0.025 |

| | | |
|---|---|---|
| NC_045512.2;YP_009724389.1;Wu1-SARS2_orf1ab;2571 | 0.05 | 0.1 |
| NC_045512.2;YP_009724389.1;Wu1-SARS2_orf1ab;2572 | 0 | 0.05 |
| NC_045512.2;YP_009724389.1;Wu1-SARS2_orf1ab;2573 | 0 | 0 |
| NC_045512.2;YP_009724389.1;Wu1-SARS2_orf1ab;2574 | 0 | 0 |
| NC_045512.2;YP_009724389.1;Wu1-SARS2_orf1ab;2575 | 0 | 0 |
| NC_045512.2;YP_009724389.1;Wu1-SARS2_orf1ab;2576 | 0 | 0 |
| NC_045512.2;YP_009724389.1;Wu1-SARS2_orf1ab;2577 | 0.05 | 0 |
| NC_045512.2;YP_009724389.1;Wu1-SARS2_orf1ab;2578 | 0 | 0 |
| NC_045512.2;YP_009724389.1;Wu1-SARS2_orf1ab;2579 | 0.05 | 0 |
| NC_045512.2;YP_009724389.1;Wu1-SARS2_orf1ab;2580 | 0 | 0 |
| NC_045512.2;YP_009724389.1;Wu1-SARS2_orf1ab;2581 | 0 | 0 |
| NC_045512.2;YP_009724389.1;Wu1-SARS2_orf1ab;2582 | 0 | 0 |
| NC_045512.2;YP_009724389.1;Wu1-SARS2_orf1ab;2583 | 0 | 0.025 |
| NC_045512.2;YP_009724389.1;Wu1-SARS2_orf1ab;2584 | 0 | 0.025 |
| NC_045512.2;YP_009724389.1;Wu1-SARS2_orf1ab;2585 | 0 | 0 |
| NC_045512.2;YP_009724389.1;Wu1-SARS2_orf1ab;2586 | 0 | 0 |
| NC_045512.2;YP_009724389.1;Wu1-SARS2_orf1ab;2587 | 0 | 0 |
| NC_045512.2;YP_009724389.1;Wu1-SARS2_orf1ab;2588 | 0 | 0 |
| NC_045512.2;YP_009724389.1;Wu1-SARS2_orf1ab;2589 | 0 | 0 |
| NC_045512.2;YP_009724389.1;Wu1-SARS2_orf1ab;2590 | 0 | 0.025 |
| NC_045512.2;YP_009724389.1;Wu1-SARS2_orf1ab;2591 | 0 | 0 |
| NC_045512.2;YP_009724389.1;Wu1-SARS2_orf1ab;2592 | 0 | 0 |
| NC_045512.2;YP_009724389.1;Wu1-SARS2_orf1ab;2593 | 0.05 | 0.05 |
| NC_045512.2;YP_009724389.1;Wu1-SARS2_orf1ab;2594 | 0.05 | 0.025 |
| NC_045512.2;YP_009724389.1;Wu1-SARS2_orf1ab;2595 | 0.15 | 0.025 |
| NC_045512.2;YP_009724389.1;Wu1-SARS2_orf1ab;2596 | 0.05 | 0 |
| NC_045512.2;YP_009724389.1;Wu1-SARS2_orf1ab;2597 | 0 | 0 |
| NC_045512.2;YP_009724389.1;Wu1-SARS2_orf1ab;2598 | 0 | 0 |
| NC_045512.2;YP_009724389.1;Wu1-SARS2_orf1ab;2599 | 0 | 0.025 |
| NC_045512.2;YP_009724389.1;Wu1-SARS2_orf1ab;2600 | 0 | 0 |
| NC_045512.2;YP_009724389.1;Wu1-SARS2_orf1ab;2601 | 0 | 0 |
| NC_045512.2;YP_009724389.1;Wu1-SARS2_orf1ab;2602 | 0 | 0 |
| NC_045512.2;YP_009724389.1;Wu1-SARS2_orf1ab;2603 | 0 | 0 |
| NC_045512.2;YP_009724389.1;Wu1-SARS2_orf1ab;2604 | 0 | 0 |
| NC_045512.2;YP_009724389.1;Wu1-SARS2_orf1ab;2605 | 0 | 0 |
| NC_045512.2;YP_009724389.1;Wu1-SARS2_orf1ab;2606 | 0 | 0 |
| NC_045512.2;YP_009724389.1;Wu1-SARS2_orf1ab;2607 | 0 | 0.05 |
| NC_045512.2;YP_009724389.1;Wu1-SARS2_orf1ab;2608 | 0 | 0 |
| NC_045512.2;YP_009724389.1;Wu1-SARS2_orf1ab;2609 | 0 | 0 |
| NC_045512.2;YP_009724389.1;Wu1-SARS2_orf1ab;2610 | 0 | 0 |
| NC_045512.2;YP_009724389.1;Wu1-SARS2_orf1ab;2611 | 0 | 0 |
| NC_045512.2;YP_009724389.1;Wu1-SARS2_orf1ab;2612 | 0 | 0 |
| NC_045512.2;YP_009724389.1;Wu1-SARS2_orf1ab;2613 | 0 | 0 |

| | | |
|---|---|---|
| NC_045512.2;YP_009724389.1;Wu1-SARS2_orf1ab;2614 | 0 | 0.025 |
| NC_045512.2;YP_009724389.1;Wu1-SARS2_orf1ab;2615 | 0 | 0.05 |
| NC_045512.2;YP_009724389.1;Wu1-SARS2_orf1ab;2616 | 0.05 | 0.025 |
| NC_045512.2;YP_009724389.1;Wu1-SARS2_orf1ab;2617 | 0.1 | 0.05 |
| NC_045512.2;YP_009724389.1;Wu1-SARS2_orf1ab;2618 | 0.05 | 0.025 |
| NC_045512.2;YP_009724389.1;Wu1-SARS2_orf1ab;2619 | 0.05 | 0 |
| NC_045512.2;YP_009724389.1;Wu1-SARS2_orf1ab;2620 | 0 | 0 |
| NC_045512.2;YP_009724389.1;Wu1-SARS2_orf1ab;2621 | 0 | 0 |
| NC_045512.2;YP_009724389.1;Wu1-SARS2_orf1ab;2622 | 0.05 | 0 |
| NC_045512.2;YP_009724389.1;Wu1-SARS2_orf1ab;2623 | 0 | 0 |
| NC_045512.2;YP_009724389.1;Wu1-SARS2_orf1ab;2624 | 0 | 0 |
| NC_045512.2;YP_009724389.1;Wu1-SARS2_orf1ab;2625 | 0 | 0 |
| NC_045512.2;YP_009724389.1;Wu1-SARS2_orf1ab;2626 | 0 | 0 |
| NC_045512.2;YP_009724389.1;Wu1-SARS2_orf1ab;2627 | 0 | 0 |
| NC_045512.2;YP_009724389.1;Wu1-SARS2_orf1ab;2628 | 0 | 0 |
| NC_045512.2;YP_009724389.1;Wu1-SARS2_orf1ab;2629 | 0 | 0 |
| NC_045512.2;YP_009724389.1;Wu1-SARS2_orf1ab;2630 | 0 | 0 |
| NC_045512.2;YP_009724389.1;Wu1-SARS2_orf1ab;2631 | 0 | 0 |
| NC_045512.2;YP_009724389.1;Wu1-SARS2_orf1ab;2632 | 0 | 0 |
| NC_045512.2;YP_009724389.1;Wu1-SARS2_orf1ab;2633 | 0 | 0 |
| NC_045512.2;YP_009724389.1;Wu1-SARS2_orf1ab;2634 | 0 | 0 |
| NC_045512.2;YP_009724389.1;Wu1-SARS2_orf1ab;2635 | 0 | 0 |
| NC_045512.2;YP_009724389.1;Wu1-SARS2_orf1ab;2636 | 0 | 0 |
| NC_045512.2;YP_009724389.1;Wu1-SARS2_orf1ab;2637 | 0 | 0 |
| NC_045512.2;YP_009724389.1;Wu1-SARS2_orf1ab;2638 | 0 | 0 |
| NC_045512.2;YP_009724389.1;Wu1-SARS2_orf1ab;2639 | 0 | 0 |
| NC_045512.2;YP_009724389.1;Wu1-SARS2_orf1ab;2640 | 0 | 0 |
| NC_045512.2;YP_009724389.1;Wu1-SARS2_orf1ab;2641 | 0 | 0 |
| NC_045512.2;YP_009724389.1;Wu1-SARS2_orf1ab;2642 | 0 | 0 |
| NC_045512.2;YP_009724389.1;Wu1-SARS2_orf1ab;2643 | 0 | 0 |
| NC_045512.2;YP_009724389.1;Wu1-SARS2_orf1ab;2644 | 0 | 0 |
| NC_045512.2;YP_009724389.1;Wu1-SARS2_orf1ab;2645 | 0 | 0 |
| NC_045512.2;YP_009724389.1;Wu1-SARS2_orf1ab;2646 | 0 | 0 |
| NC_045512.2;YP_009724389.1;Wu1-SARS2_orf1ab;2647 | 0 | 0.025 |
| NC_045512.2;YP_009724389.1;Wu1-SARS2_orf1ab;2648 | 0 | 0 |
| NC_045512.2;YP_009724389.1;Wu1-SARS2_orf1ab;2649 | 0 | 0 |
| NC_045512.2;YP_009724389.1;Wu1-SARS2_orf1ab;2650 | 0 | 0 |
| NC_045512.2;YP_009724389.1;Wu1-SARS2_orf1ab;2651 | 0 | 0 |
| NC_045512.2;YP_009724389.1;Wu1-SARS2_orf1ab;2652 | 0 | 0 |
| NC_045512.2;YP_009724389.1;Wu1-SARS2_orf1ab;2653 | 0 | 0 |
| NC_045512.2;YP_009724389.1;Wu1-SARS2_orf1ab;2654 | 0 | 0 |
| NC_045512.2;YP_009724389.1;Wu1-SARS2_orf1ab;2655 | 0 | 0 |
| NC_045512.2;YP_009724389.1;Wu1-SARS2_orf1ab;2656 | 0 | 0 |

| | | |
|---|---|---|
| NC_045512.2;YP_009724389.1;Wu1-SARS2_orf1ab;2657 | 0 | 0 |
| NC_045512.2;YP_009724389.1;Wu1-SARS2_orf1ab;2658 | 0 | 0 |
| NC_045512.2;YP_009724389.1;Wu1-SARS2_orf1ab;2659 | 0 | 0 |
| NC_045512.2;YP_009724389.1;Wu1-SARS2_orf1ab;2660 | 0 | 0 |
| NC_045512.2;YP_009724389.1;Wu1-SARS2_orf1ab;2661 | 0.05 | 0 |
| NC_045512.2;YP_009724389.1;Wu1-SARS2_orf1ab;2662 | 0 | 0 |
| NC_045512.2;YP_009724389.1;Wu1-SARS2_orf1ab;2663 | 0.05 | 0.05 |
| NC_045512.2;YP_009724389.1;Wu1-SARS2_orf1ab;2664 | 0.05 | 0.05 |
| NC_045512.2;YP_009724389.1;Wu1-SARS2_orf1ab;2665 | 0 | 0 |
| NC_045512.2;YP_009724389.1;Wu1-SARS2_orf1ab;2666 | 0 | 0 |
| NC_045512.2;YP_009724389.1;Wu1-SARS2_orf1ab;2667 | 0 | 0.025 |
| NC_045512.2;YP_009724389.1;Wu1-SARS2_orf1ab;2668 | 0.1 | 0.05 |
| NC_045512.2;YP_009724389.1;Wu1-SARS2_orf1ab;2669 | 0.05 | 0 |
| NC_045512.2;YP_009724389.1;Wu1-SARS2_orf1ab;2670 | 0 | 0 |
| NC_045512.2;YP_009724389.1;Wu1-SARS2_orf1ab;2671 | 0 | 0 |
| NC_045512.2;YP_009724389.1;Wu1-SARS2_orf1ab;2672 | 0.05 | 0 |
| NC_045512.2;YP_009724389.1;Wu1-SARS2_orf1ab;2673 | 0 | 0 |
| NC_045512.2;YP_009724389.1;Wu1-SARS2_orf1ab;2674 | 0 | 0.025 |
| NC_045512.2;YP_009724389.1;Wu1-SARS2_orf1ab;2675 | 0.05 | 0.075 |
| NC_045512.2;YP_009724389.1;Wu1-SARS2_orf1ab;2676 | 0.05 | 0.025 |
| NC_045512.2;YP_009724389.1;Wu1-SARS2_orf1ab;2677 | 0.05 | 0.05 |
| NC_045512.2;YP_009724389.1;Wu1-SARS2_orf1ab;2678 | 0 | 0 |
| NC_045512.2;YP_009724389.1;Wu1-SARS2_orf1ab;2679 | 0 | 0 |
| NC_045512.2;YP_009724389.1;Wu1-SARS2_orf1ab;2680 | 0 | 0 |
| NC_045512.2;YP_009724389.1;Wu1-SARS2_orf1ab;2681 | 0 | 0 |
| NC_045512.2;YP_009724389.1;Wu1-SARS2_orf1ab;2682 | 0 | 0 |
| NC_045512.2;YP_009724389.1;Wu1-SARS2_orf1ab;2683 | 0 | 0 |
| NC_045512.2;YP_009724389.1;Wu1-SARS2_orf1ab;2684 | 0.05 | 0 |
| NC_045512.2;YP_009724389.1;Wu1-SARS2_orf1ab;2685 | 0.05 | 0 |
| NC_045512.2;YP_009724389.1;Wu1-SARS2_orf1ab;2686 | 0 | 0 |
| NC_045512.2;YP_009724389.1;Wu1-SARS2_orf1ab;2687 | 0 | 0 |
| NC_045512.2;YP_009724389.1;Wu1-SARS2_orf1ab;2688 | 0 | 0 |
| NC_045512.2;YP_009724389.1;Wu1-SARS2_orf1ab;2689 | 0.05 | 0.05 |
| NC_045512.2;YP_009724389.1;Wu1-SARS2_orf1ab;2690 | 0 | 0 |
| NC_045512.2;YP_009724389.1;Wu1-SARS2_orf1ab;2691 | 0.05 | 0 |
| NC_045512.2;YP_009724389.1;Wu1-SARS2_orf1ab;2692 | 0 | 0 |
| NC_045512.2;YP_009724389.1;Wu1-SARS2_orf1ab;2693 | 0 | 0 |
| NC_045512.2;YP_009724389.1;Wu1-SARS2_orf1ab;2694 | 0 | 0 |
| NC_045512.2;YP_009724389.1;Wu1-SARS2_orf1ab;2695 | 0 | 0 |
| NC_045512.2;YP_009724389.1;Wu1-SARS2_orf1ab;2696 | 0 | 0 |
| NC_045512.2;YP_009724389.1;Wu1-SARS2_orf1ab;2697 | 0 | 0 |
| NC_045512.2;YP_009724389.1;Wu1-SARS2_orf1ab;2698 | 0 | 0 |
| NC_045512.2;YP_009724389.1;Wu1-SARS2_orf1ab;2699 | 0 | 0 |

| | | |
|---|---|---|
| NC_045512.2;YP_009724389.1;Wu1-SARS2_orf1ab;2700 | 0 | 0 |
| NC_045512.2;YP_009724389.1;Wu1-SARS2_orf1ab;2701 | 0 | 0 |
| NC_045512.2;YP_009724389.1;Wu1-SARS2_orf1ab;2702 | 0 | 0 |
| NC_045512.2;YP_009724389.1;Wu1-SARS2_orf1ab;2703 | 0 | 0 |
| NC_045512.2;YP_009724389.1;Wu1-SARS2_orf1ab;2704 | 0 | 0 |
| NC_045512.2;YP_009724389.1;Wu1-SARS2_orf1ab;2705 | 0 | 0 |
| NC_045512.2;YP_009724389.1;Wu1-SARS2_orf1ab;2706 | 0 | 0 |
| NC_045512.2;YP_009724389.1;Wu1-SARS2_orf1ab;2707 | 0 | 0 |
| NC_045512.2;YP_009724389.1;Wu1-SARS2_orf1ab;2708 | 0 | 0 |
| NC_045512.2;YP_009724389.1;Wu1-SARS2_orf1ab;2709 | 0 | 0 |
| NC_045512.2;YP_009724389.1;Wu1-SARS2_orf1ab;2710 | 0 | 0 |
| NC_045512.2;YP_009724389.1;Wu1-SARS2_orf1ab;2711 | 0 | 0 |
| NC_045512.2;YP_009724389.1;Wu1-SARS2_orf1ab;2712 | 0 | 0 |
| NC_045512.2;YP_009724389.1;Wu1-SARS2_orf1ab;2713 | 0 | 0 |
| NC_045512.2;YP_009724389.1;Wu1-SARS2_orf1ab;2714 | 0 | 0 |
| NC_045512.2;YP_009724389.1;Wu1-SARS2_orf1ab;2715 | 0 | 0 |
| NC_045512.2;YP_009724389.1;Wu1-SARS2_orf1ab;2716 | 0 | 0 |
| NC_045512.2;YP_009724389.1;Wu1-SARS2_orf1ab;2717 | 0 | 0 |
| NC_045512.2;YP_009724389.1;Wu1-SARS2_orf1ab;2718 | 0.05 | 0 |
| NC_045512.2;YP_009724389.1;Wu1-SARS2_orf1ab;2719 | 0 | 0 |
| NC_045512.2;YP_009724389.1;Wu1-SARS2_orf1ab;2720 | 0 | 0 |
| NC_045512.2;YP_009724389.1;Wu1-SARS2_orf1ab;2721 | 0 | 0 |
| NC_045512.2;YP_009724389.1;Wu1-SARS2_orf1ab;2722 | 0 | 0.025 |
| NC_045512.2;YP_009724389.1;Wu1-SARS2_orf1ab;2723 | 0 | 0 |
| NC_045512.2;YP_009724389.1;Wu1-SARS2_orf1ab;2724 | 0 | 0 |
| NC_045512.2;YP_009724389.1;Wu1-SARS2_orf1ab;2725 | 0 | 0 |
| NC_045512.2;YP_009724389.1;Wu1-SARS2_orf1ab;2726 | 0 | 0.025 |
| NC_045512.2;YP_009724389.1;Wu1-SARS2_orf1ab;2727 | 0 | 0.025 |
| NC_045512.2;YP_009724389.1;Wu1-SARS2_orf1ab;2728 | 0 | 0.025 |
| NC_045512.2;YP_009724389.1;Wu1-SARS2_orf1ab;2729 | 0.05 | 0.025 |
| NC_045512.2;YP_009724389.1;Wu1-SARS2_orf1ab;2730 | 0 | 0.025 |
| NC_045512.2;YP_009724389.1;Wu1-SARS2_orf1ab;2731 | 0 | 0.025 |
| NC_045512.2;YP_009724389.1;Wu1-SARS2_orf1ab;2732 | 0 | 0.025 |
| NC_045512.2;YP_009724389.1;Wu1-SARS2_orf1ab;2733 | 0 | 0.025 |
| NC_045512.2;YP_009724389.1;Wu1-SARS2_orf1ab;2734 | 0 | 0 |
| NC_045512.2;YP_009724389.1;Wu1-SARS2_orf1ab;2735 | 0 | 0 |
| NC_045512.2;YP_009724389.1;Wu1-SARS2_orf1ab;2736 | 0 | 0 |
| NC_045512.2;YP_009724389.1;Wu1-SARS2_orf1ab;2737 | 0 | 0 |
| NC_045512.2;YP_009724389.1;Wu1-SARS2_orf1ab;2738 | 0 | 0.025 |
| NC_045512.2;YP_009724389.1;Wu1-SARS2_orf1ab;2739 | 0 | 0 |
| NC_045512.2;YP_009724389.1;Wu1-SARS2_orf1ab;2740 | 0 | 0 |
| NC_045512.2;YP_009724389.1;Wu1-SARS2_orf1ab;2741 | 0 | 0.025 |
| NC_045512.2;YP_009724389.1;Wu1-SARS2_orf1ab;2742 | 0 | 0 |

| | | |
|---|---|---|
| NC_045512.2;YP_009724389.1;Wu1-SARS2_orf1ab;2743 | 0 | 0.025 |
| NC_045512.2;YP_009724389.1;Wu1-SARS2_orf1ab;2744 | 0 | 0.025 |
| NC_045512.2;YP_009724389.1;Wu1-SARS2_orf1ab;2745 | 0 | 0 |
| NC_045512.2;YP_009724389.1;Wu1-SARS2_orf1ab;2746 | 0 | 0 |
| NC_045512.2;YP_009724389.1;Wu1-SARS2_orf1ab;2747 | 0 | 0 |
| NC_045512.2;YP_009724389.1;Wu1-SARS2_orf1ab;2748 | 0 | 0 |
| NC_045512.2;YP_009724389.1;Wu1-SARS2_orf1ab;2749 | 0 | 0 |
| NC_045512.2;YP_009724389.1;Wu1-SARS2_orf1ab;2750 | 0 | 0 |
| NC_045512.2;YP_009724389.1;Wu1-SARS2_orf1ab;2751 | 0 | 0 |
| NC_045512.2;YP_009724389.1;Wu1-SARS2_orf1ab;2752 | 0 | 0 |
| NC_045512.2;YP_009724389.1;Wu1-SARS2_orf1ab;2753 | 0 | 0 |
| NC_045512.2;YP_009724389.1;Wu1-SARS2_orf1ab;2754 | 0 | 0 |
| NC_045512.2;YP_009724389.1;Wu1-SARS2_orf1ab;2755 | 0 | 0 |
| NC_045512.2;YP_009724389.1;Wu1-SARS2_orf1ab;2756 | 0 | 0 |
| NC_045512.2;YP_009724389.1;Wu1-SARS2_orf1ab;2757 | 0 | 0 |
| NC_045512.2;YP_009724389.1;Wu1-SARS2_orf1ab;2758 | 0 | 0 |
| NC_045512.2;YP_009724389.1;Wu1-SARS2_orf1ab;2759 | 0 | 0 |
| NC_045512.2;YP_009724389.1;Wu1-SARS2_orf1ab;2760 | 0 | 0 |
| NC_045512.2;YP_009724389.1;Wu1-SARS2_orf1ab;2761 | 0 | 0 |
| NC_045512.2;YP_009724389.1;Wu1-SARS2_orf1ab;2762 | 0 | 0 |
| NC_045512.2;YP_009724389.1;Wu1-SARS2_orf1ab;2763 | 0 | 0 |
| NC_045512.2;YP_009724389.1;Wu1-SARS2_orf1ab;2764 | 0 | 0 |
| NC_045512.2;YP_009724389.1;Wu1-SARS2_orf1ab;2765 | 0 | 0 |
| NC_045512.2;YP_009724389.1;Wu1-SARS2_orf1ab;2766 | 0 | 0 |
| NC_045512.2;YP_009724389.1;Wu1-SARS2_orf1ab;2767 | 0 | 0 |
| NC_045512.2;YP_009724389.1;Wu1-SARS2_orf1ab;2768 | 0 | 0 |
| NC_045512.2;YP_009724389.1;Wu1-SARS2_orf1ab;2769 | 0 | 0 |
| NC_045512.2;YP_009724389.1;Wu1-SARS2_orf1ab;2770 | 0 | 0 |
| NC_045512.2;YP_009724389.1;Wu1-SARS2_orf1ab;2771 | 0 | 0 |
| NC_045512.2;YP_009724389.1;Wu1-SARS2_orf1ab;2772 | 0 | 0 |
| NC_045512.2;YP_009724389.1;Wu1-SARS2_orf1ab;2773 | 0 | 0 |
| NC_045512.2;YP_009724389.1;Wu1-SARS2_orf1ab;2774 | 0 | 0 |
| NC_045512.2;YP_009724389.1;Wu1-SARS2_orf1ab;2775 | 0 | 0 |
| NC_045512.2;YP_009724389.1;Wu1-SARS2_orf1ab;2776 | 0 | 0 |
| NC_045512.2;YP_009724389.1;Wu1-SARS2_orf1ab;2777 | 0 | 0 |
| NC_045512.2;YP_009724389.1;Wu1-SARS2_orf1ab;2778 | 0 | 0 |
| NC_045512.2;YP_009724389.1;Wu1-SARS2_orf1ab;2779 | 0 | 0 |
| NC_045512.2;YP_009724389.1;Wu1-SARS2_orf1ab;2780 | 0 | 0 |
| NC_045512.2;YP_009724389.1;Wu1-SARS2_orf1ab;2781 | 0 | 0 |
| NC_045512.2;YP_009724389.1;Wu1-SARS2_orf1ab;2782 | 0 | 0 |
| NC_045512.2;YP_009724389.1;Wu1-SARS2_orf1ab;2783 | 0 | 0 |
| NC_045512.2;YP_009724389.1;Wu1-SARS2_orf1ab;2784 | 0 | 0 |
| NC_045512.2;YP_009724389.1;Wu1-SARS2_orf1ab;2785 | 0 | 0 |

| | | |
|---|---|---|
| NC_045512.2;YP_009724389.1;Wu1-SARS2_orf1ab;2786 | 0 | 0 |
| NC_045512.2;YP_009724389.1;Wu1-SARS2_orf1ab;2787 | 0 | 0 |
| NC_045512.2;YP_009724389.1;Wu1-SARS2_orf1ab;2788 | 0 | 0 |
| NC_045512.2;YP_009724389.1;Wu1-SARS2_orf1ab;2789 | 0 | 0 |
| NC_045512.2;YP_009724389.1;Wu1-SARS2_orf1ab;2790 | 0 | 0 |
| NC_045512.2;YP_009724389.1;Wu1-SARS2_orf1ab;2791 | 0 | 0 |
| NC_045512.2;YP_009724389.1;Wu1-SARS2_orf1ab;2792 | 0 | 0 |
| NC_045512.2;YP_009724389.1;Wu1-SARS2_orf1ab;2793 | 0 | 0 |
| NC_045512.2;YP_009724389.1;Wu1-SARS2_orf1ab;2794 | 0 | 0 |
| NC_045512.2;YP_009724389.1;Wu1-SARS2_orf1ab;2795 | 0 | 0 |
| NC_045512.2;YP_009724389.1;Wu1-SARS2_orf1ab;2796 | 0.05 | 0 |
| NC_045512.2;YP_009724389.1;Wu1-SARS2_orf1ab;2797 | 0 | 0.025 |
| NC_045512.2;YP_009724389.1;Wu1-SARS2_orf1ab;2798 | 0.05 | 0.025 |
| NC_045512.2;YP_009724389.1;Wu1-SARS2_orf1ab;2799 | 0.05 | 0.025 |
| NC_045512.2;YP_009724389.1;Wu1-SARS2_orf1ab;2800 | 0.05 | 0.025 |
| NC_045512.2;YP_009724389.1;Wu1-SARS2_orf1ab;2801 | 0.05 | 0.025 |
| NC_045512.2;YP_009724389.1;Wu1-SARS2_orf1ab;2802 | 0.1 | 0.025 |
| NC_045512.2;YP_009724389.1;Wu1-SARS2_orf1ab;2803 | 0.05 | 0.025 |
| NC_045512.2;YP_009724389.1;Wu1-SARS2_orf1ab;2804 | 0.05 | 0.075 |
| NC_045512.2;YP_009724389.1;Wu1-SARS2_orf1ab;2805 | 0.05 | 0.05 |
| NC_045512.2;YP_009724389.1;Wu1-SARS2_orf1ab;2806 | 0.05 | 0 |
| NC_045512.2;YP_009724389.1;Wu1-SARS2_orf1ab;2807 | 0 | 0 |
| NC_045512.2;YP_009724389.1;Wu1-SARS2_orf1ab;2808 | 0 | 0 |
| NC_045512.2;YP_009724389.1;Wu1-SARS2_orf1ab;2809 | 0 | 0 |
| NC_045512.2;YP_009724389.1;Wu1-SARS2_orf1ab;2810 | 0 | 0 |
| NC_045512.2;YP_009724389.1;Wu1-SARS2_orf1ab;2811 | 0 | 0 |
| NC_045512.2;YP_009724389.1;Wu1-SARS2_orf1ab;2812 | 0 | 0 |
| NC_045512.2;YP_009724389.1;Wu1-SARS2_orf1ab;2813 | 0 | 0 |
| NC_045512.2;YP_009724389.1;Wu1-SARS2_orf1ab;2814 | 0 | 0 |
| NC_045512.2;YP_009724389.1;Wu1-SARS2_orf1ab;2815 | 0 | 0.05 |
| NC_045512.2;YP_009724389.1;Wu1-SARS2_orf1ab;2816 | 0 | 0 |
| NC_045512.2;YP_009724389.1;Wu1-SARS2_orf1ab;2817 | 0 | 0.025 |
| NC_045512.2;YP_009724389.1;Wu1-SARS2_orf1ab;2818 | 0.05 | 0 |
| NC_045512.2;YP_009724389.1;Wu1-SARS2_orf1ab;2819 | 0 | 0 |
| NC_045512.2;YP_009724389.1;Wu1-SARS2_orf1ab;2820 | 0 | 0 |
| NC_045512.2;YP_009724389.1;Wu1-SARS2_orf1ab;2821 | 0.05 | 0 |
| NC_045512.2;YP_009724389.1;Wu1-SARS2_orf1ab;2822 | 0.1 | 0 |
| NC_045512.2;YP_009724389.1;Wu1-SARS2_orf1ab;2823 | 0 | 0 |
| NC_045512.2;YP_009724389.1;Wu1-SARS2_orf1ab;2824 | 0 | 0 |
| NC_045512.2;YP_009724389.1;Wu1-SARS2_orf1ab;2825 | 0 | 0 |
| NC_045512.2;YP_009724389.1;Wu1-SARS2_orf1ab;2826 | 0 | 0 |
| NC_045512.2;YP_009724389.1;Wu1-SARS2_orf1ab;2827 | 0 | 0 |
| NC_045512.2;YP_009724389.1;Wu1-SARS2_orf1ab;2828 | 0.1 | 0.025 |

| | | |
|---|---|---|
| NC_045512.2;YP_009724389.1;Wu1-SARS2_orf1ab;2829 | 0 | 0 |
| NC_045512.2;YP_009724389.1;Wu1-SARS2_orf1ab;2830 | 0 | 0 |
| NC_045512.2;YP_009724389.1;Wu1-SARS2_orf1ab;2831 | 0 | 0 |
| NC_045512.2;YP_009724389.1;Wu1-SARS2_orf1ab;2832 | 0 | 0 |
| NC_045512.2;YP_009724389.1;Wu1-SARS2_orf1ab;2833 | 0 | 0 |
| NC_045512.2;YP_009724389.1;Wu1-SARS2_orf1ab;2834 | 0 | 0 |
| NC_045512.2;YP_009724389.1;Wu1-SARS2_orf1ab;2835 | 0.05 | 0 |
| NC_045512.2;YP_009724389.1;Wu1-SARS2_orf1ab;2836 | 0 | 0.05 |
| NC_045512.2;YP_009724389.1;Wu1-SARS2_orf1ab;2837 | 0 | 0 |
| NC_045512.2;YP_009724389.1;Wu1-SARS2_orf1ab;2838 | 0 | 0 |
| NC_045512.2;YP_009724389.1;Wu1-SARS2_orf1ab;2839 | 0.05 | 0 |
| NC_045512.2;YP_009724389.1;Wu1-SARS2_orf1ab;2840 | 0 | 0.025 |
| NC_045512.2;YP_009724389.1;Wu1-SARS2_orf1ab;2841 | 0.05 | 0.025 |
| NC_045512.2;YP_009724389.1;Wu1-SARS2_orf1ab;2842 | 0 | 0 |
| NC_045512.2;YP_009724389.1;Wu1-SARS2_orf1ab;2843 | 0 | 0 |
| NC_045512.2;YP_009724389.1;Wu1-SARS2_orf1ab;2844 | 0 | 0.05 |
| NC_045512.2;YP_009724389.1;Wu1-SARS2_orf1ab;2845 | 0 | 0 |
| NC_045512.2;YP_009724389.1;Wu1-SARS2_orf1ab;2846 | 0 | 0 |
| NC_045512.2;YP_009724389.1;Wu1-SARS2_orf1ab;2847 | 0 | 0 |
| NC_045512.2;YP_009724389.1;Wu1-SARS2_orf1ab;2848 | 0 | 0 |
| NC_045512.2;YP_009724389.1;Wu1-SARS2_orf1ab;2849 | 0 | 0 |
| NC_045512.2;YP_009724389.1;Wu1-SARS2_orf1ab;2850 | 0 | 0 |
| NC_045512.2;YP_009724389.1;Wu1-SARS2_orf1ab;2851 | 0 | 0 |
| NC_045512.2;YP_009724389.1;Wu1-SARS2_orf1ab;2852 | 0 | 0 |
| NC_045512.2;YP_009724389.1;Wu1-SARS2_orf1ab;2853 | 0 | 0 |
| NC_045512.2;YP_009724389.1;Wu1-SARS2_orf1ab;2854 | 0 | 0 |
| NC_045512.2;YP_009724389.1;Wu1-SARS2_orf1ab;2855 | 0 | 0 |
| NC_045512.2;YP_009724389.1;Wu1-SARS2_orf1ab;2856 | 0 | 0 |
| NC_045512.2;YP_009724389.1;Wu1-SARS2_orf1ab;2857 | 0 | 0.025 |
| NC_045512.2;YP_009724389.1;Wu1-SARS2_orf1ab;2858 | 0 | 0.025 |
| NC_045512.2;YP_009724389.1;Wu1-SARS2_orf1ab;2859 | 0 | 0.025 |
| NC_045512.2;YP_009724389.1;Wu1-SARS2_orf1ab;2860 | 0 | 0 |
| NC_045512.2;YP_009724389.1;Wu1-SARS2_orf1ab;2861 | 0 | 0 |
| NC_045512.2;YP_009724389.1;Wu1-SARS2_orf1ab;2862 | 0 | 0 |
| NC_045512.2;YP_009724389.1;Wu1-SARS2_orf1ab;2863 | 0 | 0 |
| NC_045512.2;YP_009724389.1;Wu1-SARS2_orf1ab;2864 | 0 | 0 |
| NC_045512.2;YP_009724389.1;Wu1-SARS2_orf1ab;2865 | 0 | 0 |
| NC_045512.2;YP_009724389.1;Wu1-SARS2_orf1ab;2866 | 0 | 0 |
| NC_045512.2;YP_009724389.1;Wu1-SARS2_orf1ab;2867 | 0 | 0 |
| NC_045512.2;YP_009724389.1;Wu1-SARS2_orf1ab;2868 | 0 | 0 |
| NC_045512.2;YP_009724389.1;Wu1-SARS2_orf1ab;2869 | 0 | 0 |
| NC_045512.2;YP_009724389.1;Wu1-SARS2_orf1ab;2870 | 0 | 0 |
| NC_045512.2;YP_009724389.1;Wu1-SARS2_orf1ab;2871 | 0 | 0 |

| | | |
|---|---|---|
| NC_045512.2;YP_009724389.1;Wu1-SARS2_orf1ab;2872 | 0 | 0 |
| NC_045512.2;YP_009724389.1;Wu1-SARS2_orf1ab;2873 | 0 | 0 |
| NC_045512.2;YP_009724389.1;Wu1-SARS2_orf1ab;2874 | 0 | 0 |
| NC_045512.2;YP_009724389.1;Wu1-SARS2_orf1ab;2875 | 0 | 0 |
| NC_045512.2;YP_009724389.1;Wu1-SARS2_orf1ab;2876 | 0 | 0 |
| NC_045512.2;YP_009724389.1;Wu1-SARS2_orf1ab;2877 | 0 | 0 |
| NC_045512.2;YP_009724389.1;Wu1-SARS2_orf1ab;2878 | 0 | 0 |
| NC_045512.2;YP_009724389.1;Wu1-SARS2_orf1ab;2879 | 0 | 0 |
| NC_045512.2;YP_009724389.1;Wu1-SARS2_orf1ab;2880 | 0 | 0 |
| NC_045512.2;YP_009724389.1;Wu1-SARS2_orf1ab;2881 | 0 | 0 |
| NC_045512.2;YP_009724389.1;Wu1-SARS2_orf1ab;2882 | 0 | 0 |
| NC_045512.2;YP_009724389.1;Wu1-SARS2_orf1ab;2883 | 0 | 0 |
| NC_045512.2;YP_009724389.1;Wu1-SARS2_orf1ab;2884 | 0 | 0 |
| NC_045512.2;YP_009724389.1;Wu1-SARS2_orf1ab;2885 | 0 | 0 |
| NC_045512.2;YP_009724389.1;Wu1-SARS2_orf1ab;2886 | 0 | 0 |
| NC_045512.2;YP_009724389.1;Wu1-SARS2_orf1ab;2887 | 0 | 0 |
| NC_045512.2;YP_009724389.1;Wu1-SARS2_orf1ab;2888 | 0 | 0 |
| NC_045512.2;YP_009724389.1;Wu1-SARS2_orf1ab;2889 | 0 | 0 |
| NC_045512.2;YP_009724389.1;Wu1-SARS2_orf1ab;2890 | 0 | 0 |
| NC_045512.2;YP_009724389.1;Wu1-SARS2_orf1ab;2891 | 0.05 | 0 |
| NC_045512.2;YP_009724389.1;Wu1-SARS2_orf1ab;2892 | 0.05 | 0.025 |
| NC_045512.2;YP_009724389.1;Wu1-SARS2_orf1ab;2893 | 0 | 0.025 |
| NC_045512.2;YP_009724389.1;Wu1-SARS2_orf1ab;2894 | 0 | 0.025 |
| NC_045512.2;YP_009724389.1;Wu1-SARS2_orf1ab;2895 | 0.05 | 0 |
| NC_045512.2;YP_009724389.1;Wu1-SARS2_orf1ab;2896 | 0.05 | 0.05 |
| NC_045512.2;YP_009724389.1;Wu1-SARS2_orf1ab;2897 | 0.05 | 0.025 |
| NC_045512.2;YP_009724389.1;Wu1-SARS2_orf1ab;2898 | 0.05 | 0 |
| NC_045512.2;YP_009724389.1;Wu1-SARS2_orf1ab;2899 | 0 | 0 |
| NC_045512.2;YP_009724389.1;Wu1-SARS2_orf1ab;2900 | 0 | 0 |
| NC_045512.2;YP_009724389.1;Wu1-SARS2_orf1ab;2901 | 0 | 0 |
| NC_045512.2;YP_009724389.1;Wu1-SARS2_orf1ab;2902 | 0 | 0 |
| NC_045512.2;YP_009724389.1;Wu1-SARS2_orf1ab;2903 | 0.05 | 0 |
| NC_045512.2;YP_009724389.1;Wu1-SARS2_orf1ab;2904 | 0 | 0 |
| NC_045512.2;YP_009724389.1;Wu1-SARS2_orf1ab;2905 | 0 | 0 |
| NC_045512.2;YP_009724389.1;Wu1-SARS2_orf1ab;2906 | 0 | 0 |
| NC_045512.2;YP_009724389.1;Wu1-SARS2_orf1ab;2907 | 0 | 0 |
| NC_045512.2;YP_009724389.1;Wu1-SARS2_orf1ab;2908 | 0 | 0 |
| NC_045512.2;YP_009724389.1;Wu1-SARS2_orf1ab;2909 | 0 | 0 |
| NC_045512.2;YP_009724389.1;Wu1-SARS2_orf1ab;2910 | 0 | 0 |
| NC_045512.2;YP_009724389.1;Wu1-SARS2_orf1ab;2911 | 0 | 0 |
| NC_045512.2;YP_009724389.1;Wu1-SARS2_orf1ab;2912 | 0 | 0 |
| NC_045512.2;YP_009724389.1;Wu1-SARS2_orf1ab;2913 | 0.05 | 0.025 |
| NC_045512.2;YP_009724389.1;Wu1-SARS2_orf1ab;2914 | 0.05 | 0.025 |

| | | |
|---|---|---|
| NC_045512.2;YP_009724389.1;Wu1-SARS2_orf1ab;2915 | 0.05 | 0.125 |
| NC_045512.2;YP_009724389.1;Wu1-SARS2_orf1ab;2916 | 0.15 | 0.15 |
| NC_045512.2;YP_009724389.1;Wu1-SARS2_orf1ab;2917 | 0.1 | 0.05 |
| NC_045512.2;YP_009724389.1;Wu1-SARS2_orf1ab;2918 | 0.1 | 0.05 |
| NC_045512.2;YP_009724389.1;Wu1-SARS2_orf1ab;2919 | 0 | 0.025 |
| NC_045512.2;YP_009724389.1;Wu1-SARS2_orf1ab;2920 | 0.1 | 0.025 |
| NC_045512.2;YP_009724389.1;Wu1-SARS2_orf1ab;2921 | 0.05 | 0.025 |
| NC_045512.2;YP_009724389.1;Wu1-SARS2_orf1ab;2922 | 0 | 0.025 |
| NC_045512.2;YP_009724389.1;Wu1-SARS2_orf1ab;2923 | 0 | 0.025 |
| NC_045512.2;YP_009724389.1;Wu1-SARS2_orf1ab;2924 | 0 | 0 |
| NC_045512.2;YP_009724389.1;Wu1-SARS2_orf1ab;2925 | 0 | 0 |
| NC_045512.2;YP_009724389.1;Wu1-SARS2_orf1ab;2926 | 0 | 0 |
| NC_045512.2;YP_009724389.1;Wu1-SARS2_orf1ab;2927 | 0 | 0 |
| NC_045512.2;YP_009724389.1;Wu1-SARS2_orf1ab;2928 | 0 | 0 |
| NC_045512.2;YP_009724389.1;Wu1-SARS2_orf1ab;2929 | 0 | 0.025 |
| NC_045512.2;YP_009724389.1;Wu1-SARS2_orf1ab;2930 | 0 | 0 |
| NC_045512.2;YP_009724389.1;Wu1-SARS2_orf1ab;2931 | 0 | 0 |
| NC_045512.2;YP_009724389.1;Wu1-SARS2_orf1ab;2932 | 0 | 0 |
| NC_045512.2;YP_009724389.1;Wu1-SARS2_orf1ab;2933 | 0 | 0 |
| NC_045512.2;YP_009724389.1;Wu1-SARS2_orf1ab;2934 | 0 | 0 |
| NC_045512.2;YP_009724389.1;Wu1-SARS2_orf1ab;2935 | 0 | 0 |
| NC_045512.2;YP_009724389.1;Wu1-SARS2_orf1ab;2936 | 0.1 | 0 |
| NC_045512.2;YP_009724389.1;Wu1-SARS2_orf1ab;2937 | 0 | 0.025 |
| NC_045512.2;YP_009724389.1;Wu1-SARS2_orf1ab;2938 | 0 | 0.025 |
| NC_045512.2;YP_009724389.1;Wu1-SARS2_orf1ab;2939 | 0.15 | 0.15 |
| NC_045512.2;YP_009724389.1;Wu1-SARS2_orf1ab;2940 | 0.15 | 0.15 |
| NC_045512.2;YP_009724389.1;Wu1-SARS2_orf1ab;2941 | 0.1 | 0.025 |
| NC_045512.2;YP_009724389.1;Wu1-SARS2_orf1ab;2942 | 0.1 | 0.05 |
| NC_045512.2;YP_009724389.1;Wu1-SARS2_orf1ab;2943 | 0.05 | 0.025 |
| NC_045512.2;YP_009724389.1;Wu1-SARS2_orf1ab;2944 | 0.15 | 0.075 |
| NC_045512.2;YP_009724389.1;Wu1-SARS2_orf1ab;2945 | 0.1 | 0.1 |
| NC_045512.2;YP_009724389.1;Wu1-SARS2_orf1ab;2946 | 0.15 | 0.025 |
| NC_045512.2;YP_009724389.1;Wu1-SARS2_orf1ab;2947 | 0.05 | 0.025 |
| NC_045512.2;YP_009724389.1;Wu1-SARS2_orf1ab;2948 | 0.05 | 0.05 |
| NC_045512.2;YP_009724389.1;Wu1-SARS2_orf1ab;2949 | 0.05 | 0.025 |
| NC_045512.2;YP_009724389.1;Wu1-SARS2_orf1ab;2950 | 0 | 0 |
| NC_045512.2;YP_009724389.1;Wu1-SARS2_orf1ab;2951 | 0 | 0 |
| NC_045512.2;YP_009724389.1;Wu1-SARS2_orf1ab;2952 | 0 | 0 |
| NC_045512.2;YP_009724389.1;Wu1-SARS2_orf1ab;2953 | 0 | 0 |
| NC_045512.2;YP_009724389.1;Wu1-SARS2_orf1ab;2954 | 0 | 0 |
| NC_045512.2;YP_009724389.1;Wu1-SARS2_orf1ab;2955 | 0 | 0 |
| NC_045512.2;YP_009724389.1;Wu1-SARS2_orf1ab;2956 | 0 | 0 |
| NC_045512.2;YP_009724389.1;Wu1-SARS2_orf1ab;2957 | 0 | 0 |

| | | |
|---|---|---|
| NC_045512.2;YP_009724389.1;Wu1-SARS2_orf1ab;2958 | 0 | 0 |
| NC_045512.2;YP_009724389.1;Wu1-SARS2_orf1ab;2959 | 0 | 0 |
| NC_045512.2;YP_009724389.1;Wu1-SARS2_orf1ab;2960 | 0 | 0.025 |
| NC_045512.2;YP_009724389.1;Wu1-SARS2_orf1ab;2961 | 0 | 0 |
| NC_045512.2;YP_009724389.1;Wu1-SARS2_orf1ab;2962 | 0 | 0 |
| NC_045512.2;YP_009724389.1;Wu1-SARS2_orf1ab;2963 | 0 | 0 |
| NC_045512.2;YP_009724389.1;Wu1-SARS2_orf1ab;2964 | 0 | 0 |
| NC_045512.2;YP_009724389.1;Wu1-SARS2_orf1ab;2965 | 0 | 0 |
| NC_045512.2;YP_009724389.1;Wu1-SARS2_orf1ab;2966 | 0 | 0 |
| NC_045512.2;YP_009724389.1;Wu1-SARS2_orf1ab;2967 | 0 | 0 |
| NC_045512.2;YP_009724389.1;Wu1-SARS2_orf1ab;2968 | 0 | 0 |
| NC_045512.2;YP_009724389.1;Wu1-SARS2_orf1ab;2969 | 0 | 0 |
| NC_045512.2;YP_009724389.1;Wu1-SARS2_orf1ab;2970 | 0 | 0 |
| NC_045512.2;YP_009724389.1;Wu1-SARS2_orf1ab;2971 | 0.05 | 0.1 |
| NC_045512.2;YP_009724389.1;Wu1-SARS2_orf1ab;2972 | 0 | 0.05 |
| NC_045512.2;YP_009724389.1;Wu1-SARS2_orf1ab;2973 | 0 | 0.075 |
| NC_045512.2;YP_009724389.1;Wu1-SARS2_orf1ab;2974 | 0 | 0.075 |
| NC_045512.2;YP_009724389.1;Wu1-SARS2_orf1ab;2975 | 0 | 0.05 |
| NC_045512.2;YP_009724389.1;Wu1-SARS2_orf1ab;2976 | 0 | 0.05 |
| NC_045512.2;YP_009724389.1;Wu1-SARS2_orf1ab;2977 | 0 | 0.05 |
| NC_045512.2;YP_009724389.1;Wu1-SARS2_orf1ab;2978 | 0 | 0.05 |
| NC_045512.2;YP_009724389.1;Wu1-SARS2_orf1ab;2979 | 0 | 0.05 |
| NC_045512.2;YP_009724389.1;Wu1-SARS2_orf1ab;2980 | 0 | 0 |
| NC_045512.2;YP_009724389.1;Wu1-SARS2_orf1ab;2981 | 0 | 0 |
| NC_045512.2;YP_009724389.1;Wu1-SARS2_orf1ab;2982 | 0 | 0 |
| NC_045512.2;YP_009724389.1;Wu1-SARS2_orf1ab;2983 | 0 | 0 |
| NC_045512.2;YP_009724389.1;Wu1-SARS2_orf1ab;2984 | 0 | 0 |
| NC_045512.2;YP_009724389.1;Wu1-SARS2_orf1ab;2985 | 0 | 0 |
| NC_045512.2;YP_009724389.1;Wu1-SARS2_orf1ab;2986 | 0 | 0 |
| NC_045512.2;YP_009724389.1;Wu1-SARS2_orf1ab;2987 | 0 | 0 |
| NC_045512.2;YP_009724389.1;Wu1-SARS2_orf1ab;2988 | 0 | 0 |
| NC_045512.2;YP_009724389.1;Wu1-SARS2_orf1ab;2989 | 0 | 0 |
| NC_045512.2;YP_009724389.1;Wu1-SARS2_orf1ab;2990 | 0 | 0 |
| NC_045512.2;YP_009724389.1;Wu1-SARS2_orf1ab;2991 | 0 | 0 |
| NC_045512.2;YP_009724389.1;Wu1-SARS2_orf1ab;2992 | 0 | 0 |
| NC_045512.2;YP_009724389.1;Wu1-SARS2_orf1ab;2993 | 0 | 0 |
| NC_045512.2;YP_009724389.1;Wu1-SARS2_orf1ab;2994 | 0 | 0 |
| NC_045512.2;YP_009724389.1;Wu1-SARS2_orf1ab;2995 | 0 | 0 |
| NC_045512.2;YP_009724389.1;Wu1-SARS2_orf1ab;2996 | 0 | 0 |
| NC_045512.2;YP_009724389.1;Wu1-SARS2_orf1ab;2997 | 0 | 0 |
| NC_045512.2;YP_009724389.1;Wu1-SARS2_orf1ab;2998 | 0 | 0 |
| NC_045512.2;YP_009724389.1;Wu1-SARS2_orf1ab;2999 | 0 | 0 |
| NC_045512.2;YP_009724389.1;Wu1-SARS2_orf1ab;3000 | 0 | 0 |

| | | |
|---|---|---|
| NC_045512.2;YP_009724389.1;Wu1-SARS2_orf1ab;3001 | 0 | 0 |
| NC_045512.2;YP_009724389.1;Wu1-SARS2_orf1ab;3002 | 0 | 0 |
| NC_045512.2;YP_009724389.1;Wu1-SARS2_orf1ab;3003 | 0 | 0 |
| NC_045512.2;YP_009724389.1;Wu1-SARS2_orf1ab;3004 | 0 | 0 |
| NC_045512.2;YP_009724389.1;Wu1-SARS2_orf1ab;3005 | 0 | 0 |
| NC_045512.2;YP_009724389.1;Wu1-SARS2_orf1ab;3006 | 0 | 0 |
| NC_045512.2;YP_009724389.1;Wu1-SARS2_orf1ab;3007 | 0 | 0 |
| NC_045512.2;YP_009724389.1;Wu1-SARS2_orf1ab;3008 | 0 | 0 |
| NC_045512.2;YP_009724389.1;Wu1-SARS2_orf1ab;3009 | 0 | 0 |
| NC_045512.2;YP_009724389.1;Wu1-SARS2_orf1ab;3010 | 0.1 | 0.15 |
| NC_045512.2;YP_009724389.1;Wu1-SARS2_orf1ab;3011 | 0.1 | 0.15 |
| NC_045512.2;YP_009724389.1;Wu1-SARS2_orf1ab;3012 | 0.1 | 0.125 |
| NC_045512.2;YP_009724389.1;Wu1-SARS2_orf1ab;3013 | 0.1 | 0.15 |
| NC_045512.2;YP_009724389.1;Wu1-SARS2_orf1ab;3014 | 0.1 | 0.15 |
| NC_045512.2;YP_009724389.1;Wu1-SARS2_orf1ab;3015 | 0.1 | 0.15 |
| NC_045512.2;YP_009724389.1;Wu1-SARS2_orf1ab;3016 | 0.1 | 0.15 |
| NC_045512.2;YP_009724389.1;Wu1-SARS2_orf1ab;3017 | 0.1 | 0.15 |
| NC_045512.2;YP_009724389.1;Wu1-SARS2_orf1ab;3018 | 0.1 | 0.15 |
| NC_045512.2;YP_009724389.1;Wu1-SARS2_orf1ab;3019 | 0.1 | 0.15 |
| NC_045512.2;YP_009724389.1;Wu1-SARS2_orf1ab;3020 | 0.15 | 0.225 |
| NC_045512.2;YP_009724389.1;Wu1-SARS2_orf1ab;3021 | 0.1 | 0.15 |
| NC_045512.2;YP_009724389.1;Wu1-SARS2_orf1ab;3022 | 0 | 0 |
| NC_045512.2;YP_009724389.1;Wu1-SARS2_orf1ab;3023 | 0 | 0 |
| NC_045512.2;YP_009724389.1;Wu1-SARS2_orf1ab;3024 | 0 | 0 |
| NC_045512.2;YP_009724389.1;Wu1-SARS2_orf1ab;3025 | 0.05 | 0 |
| NC_045512.2;YP_009724389.1;Wu1-SARS2_orf1ab;3026 | 0 | 0 |
| NC_045512.2;YP_009724389.1;Wu1-SARS2_orf1ab;3027 | 0.05 | 0 |
| NC_045512.2;YP_009724389.1;Wu1-SARS2_orf1ab;3028 | 0.05 | 0 |
| NC_045512.2;YP_009724389.1;Wu1-SARS2_orf1ab;3029 | 0.05 | 0 |
| NC_045512.2;YP_009724389.1;Wu1-SARS2_orf1ab;3030 | 0.05 | 0 |
| NC_045512.2;YP_009724389.1;Wu1-SARS2_orf1ab;3031 | 0.05 | 0 |
| NC_045512.2;YP_009724389.1;Wu1-SARS2_orf1ab;3032 | 0.05 | 0 |
| NC_045512.2;YP_009724389.1;Wu1-SARS2_orf1ab;3033 | 0 | 0 |
| NC_045512.2;YP_009724389.1;Wu1-SARS2_orf1ab;3034 | 0 | 0 |
| NC_045512.2;YP_009724389.1;Wu1-SARS2_orf1ab;3035 | 0 | 0 |
| NC_045512.2;YP_009724389.1;Wu1-SARS2_orf1ab;3036 | 0.05 | 0 |
| NC_045512.2;YP_009724389.1;Wu1-SARS2_orf1ab;3037 | 0 | 0 |
| NC_045512.2;YP_009724389.1;Wu1-SARS2_orf1ab;3038 | 0 | 0 |
| NC_045512.2;YP_009724389.1;Wu1-SARS2_orf1ab;3039 | 0 | 0.025 |
| NC_045512.2;YP_009724389.1;Wu1-SARS2_orf1ab;3040 | 0 | 0 |
| NC_045512.2;YP_009724389.1;Wu1-SARS2_orf1ab;3041 | 0 | 0 |
| NC_045512.2;YP_009724389.1;Wu1-SARS2_orf1ab;3042 | 0 | 0 |
| NC_045512.2;YP_009724389.1;Wu1-SARS2_orf1ab;3043 | 0 | 0 |

| | | |
|---|---|---|
| NC_045512.2;YP_009724389.1;Wu1-SARS2_orf1ab;3044 | 0 | 0 |
| NC_045512.2;YP_009724389.1;Wu1-SARS2_orf1ab;3045 | 0 | 0 |
| NC_045512.2;YP_009724389.1;Wu1-SARS2_orf1ab;3046 | 0 | 0 |
| NC_045512.2;YP_009724389.1;Wu1-SARS2_orf1ab;3047 | 0 | 0 |
| NC_045512.2;YP_009724389.1;Wu1-SARS2_orf1ab;3048 | 0 | 0 |
| NC_045512.2;YP_009724389.1;Wu1-SARS2_orf1ab;3049 | 0 | 0 |
| NC_045512.2;YP_009724389.1;Wu1-SARS2_orf1ab;3050 | 0 | 0 |
| NC_045512.2;YP_009724389.1;Wu1-SARS2_orf1ab;3051 | 0 | 0 |
| NC_045512.2;YP_009724389.1;Wu1-SARS2_orf1ab;3052 | 0 | 0 |
| NC_045512.2;YP_009724389.1;Wu1-SARS2_orf1ab;3053 | 0 | 0 |
| NC_045512.2;YP_009724389.1;Wu1-SARS2_orf1ab;3054 | 0 | 0 |
| NC_045512.2;YP_009724389.1;Wu1-SARS2_orf1ab;3055 | 0 | 0 |
| NC_045512.2;YP_009724389.1;Wu1-SARS2_orf1ab;3056 | 0 | 0 |
| NC_045512.2;YP_009724389.1;Wu1-SARS2_orf1ab;3057 | 0 | 0 |
| NC_045512.2;YP_009724389.1;Wu1-SARS2_orf1ab;3058 | 0 | 0 |
| NC_045512.2;YP_009724389.1;Wu1-SARS2_orf1ab;3059 | 0 | 0 |
| NC_045512.2;YP_009724389.1;Wu1-SARS2_orf1ab;3060 | 0 | 0 |
| NC_045512.2;YP_009724389.1;Wu1-SARS2_orf1ab;3061 | 0 | 0 |
| NC_045512.2;YP_009724389.1;Wu1-SARS2_orf1ab;3062 | 0 | 0 |
| NC_045512.2;YP_009724389.1;Wu1-SARS2_orf1ab;3063 | 0 | 0 |
| NC_045512.2;YP_009724389.1;Wu1-SARS2_orf1ab;3064 | 0 | 0 |
| NC_045512.2;YP_009724389.1;Wu1-SARS2_orf1ab;3065 | 0 | 0 |
| NC_045512.2;YP_009724389.1;Wu1-SARS2_orf1ab;3066 | 0 | 0 |
| NC_045512.2;YP_009724389.1;Wu1-SARS2_orf1ab;3067 | 0 | 0 |
| NC_045512.2;YP_009724389.1;Wu1-SARS2_orf1ab;3068 | 0 | 0 |
| NC_045512.2;YP_009724389.1;Wu1-SARS2_orf1ab;3069 | 0 | 0 |
| NC_045512.2;YP_009724389.1;Wu1-SARS2_orf1ab;3070 | 0 | 0 |
| NC_045512.2;YP_009724389.1;Wu1-SARS2_orf1ab;3071 | 0 | 0 |
| NC_045512.2;YP_009724389.1;Wu1-SARS2_orf1ab;3072 | 0 | 0 |
| NC_045512.2;YP_009724389.1;Wu1-SARS2_orf1ab;3073 | 0 | 0 |
| NC_045512.2;YP_009724389.1;Wu1-SARS2_orf1ab;3074 | 0 | 0 |
| NC_045512.2;YP_009724389.1;Wu1-SARS2_orf1ab;3075 | 0 | 0 |
| NC_045512.2;YP_009724389.1;Wu1-SARS2_orf1ab;3076 | 0 | 0 |
| NC_045512.2;YP_009724389.1;Wu1-SARS2_orf1ab;3077 | 0 | 0 |
| NC_045512.2;YP_009724389.1;Wu1-SARS2_orf1ab;3078 | 0 | 0 |
| NC_045512.2;YP_009724389.1;Wu1-SARS2_orf1ab;3079 | 0 | 0 |
| NC_045512.2;YP_009724389.1;Wu1-SARS2_orf1ab;3080 | 0 | 0 |
| NC_045512.2;YP_009724389.1;Wu1-SARS2_orf1ab;3081 | 0 | 0 |
| NC_045512.2;YP_009724389.1;Wu1-SARS2_orf1ab;3082 | 0 | 0 |
| NC_045512.2;YP_009724389.1;Wu1-SARS2_orf1ab;3083 | 0 | 0 |
| NC_045512.2;YP_009724389.1;Wu1-SARS2_orf1ab;3084 | 0 | 0 |
| NC_045512.2;YP_009724389.1;Wu1-SARS2_orf1ab;3085 | 0 | 0 |
| NC_045512.2;YP_009724389.1;Wu1-SARS2_orf1ab;3086 | 0 | 0 |

| | | |
|---|---|---|
| NC_045512.2;YP_009724389.1;Wu1-SARS2_orf1ab;3087 | 0 | 0 |
| NC_045512.2;YP_009724389.1;Wu1-SARS2_orf1ab;3088 | 0 | 0 |
| NC_045512.2;YP_009724389.1;Wu1-SARS2_orf1ab;3089 | 0 | 0 |
| NC_045512.2;YP_009724389.1;Wu1-SARS2_orf1ab;3090 | 0 | 0 |
| NC_045512.2;YP_009724389.1;Wu1-SARS2_orf1ab;3091 | 0 | 0 |
| NC_045512.2;YP_009724389.1;Wu1-SARS2_orf1ab;3092 | 0 | 0 |
| NC_045512.2;YP_009724389.1;Wu1-SARS2_orf1ab;3093 | 0 | 0 |
| NC_045512.2;YP_009724389.1;Wu1-SARS2_orf1ab;3094 | 0 | 0 |
| NC_045512.2;YP_009724389.1;Wu1-SARS2_orf1ab;3095 | 0 | 0 |
| NC_045512.2;YP_009724389.1;Wu1-SARS2_orf1ab;3096 | 0 | 0 |
| NC_045512.2;YP_009724389.1;Wu1-SARS2_orf1ab;3097 | 0 | 0 |
| NC_045512.2;YP_009724389.1;Wu1-SARS2_orf1ab;3098 | 0 | 0 |
| NC_045512.2;YP_009724389.1;Wu1-SARS2_orf1ab;3099 | 0 | 0 |
| NC_045512.2;YP_009724389.1;Wu1-SARS2_orf1ab;3100 | 0 | 0 |
| NC_045512.2;YP_009724389.1;Wu1-SARS2_orf1ab;3101 | 0 | 0 |
| NC_045512.2;YP_009724389.1;Wu1-SARS2_orf1ab;3102 | 0 | 0 |
| NC_045512.2;YP_009724389.1;Wu1-SARS2_orf1ab;3103 | 0 | 0 |
| NC_045512.2;YP_009724389.1;Wu1-SARS2_orf1ab;3104 | 0 | 0 |
| NC_045512.2;YP_009724389.1;Wu1-SARS2_orf1ab;3105 | 0 | 0 |
| NC_045512.2;YP_009724389.1;Wu1-SARS2_orf1ab;3106 | 0 | 0 |
| NC_045512.2;YP_009724389.1;Wu1-SARS2_orf1ab;3107 | 0 | 0 |
| NC_045512.2;YP_009724389.1;Wu1-SARS2_orf1ab;3108 | 0 | 0 |
| NC_045512.2;YP_009724389.1;Wu1-SARS2_orf1ab;3109 | 0 | 0 |
| NC_045512.2;YP_009724389.1;Wu1-SARS2_orf1ab;3110 | 0 | 0 |
| NC_045512.2;YP_009724389.1;Wu1-SARS2_orf1ab;3111 | 0 | 0 |
| NC_045512.2;YP_009724389.1;Wu1-SARS2_orf1ab;3112 | 0 | 0 |
| NC_045512.2;YP_009724389.1;Wu1-SARS2_orf1ab;3113 | 0 | 0 |
| NC_045512.2;YP_009724389.1;Wu1-SARS2_orf1ab;3114 | 0 | 0 |
| NC_045512.2;YP_009724389.1;Wu1-SARS2_orf1ab;3115 | 0 | 0 |
| NC_045512.2;YP_009724389.1;Wu1-SARS2_orf1ab;3116 | 0 | 0 |
| NC_045512.2;YP_009724389.1;Wu1-SARS2_orf1ab;3117 | 0 | 0 |
| NC_045512.2;YP_009724389.1;Wu1-SARS2_orf1ab;3118 | 0 | 0 |
| NC_045512.2;YP_009724389.1;Wu1-SARS2_orf1ab;3119 | 0 | 0 |
| NC_045512.2;YP_009724389.1;Wu1-SARS2_orf1ab;3120 | 0 | 0 |
| NC_045512.2;YP_009724389.1;Wu1-SARS2_orf1ab;3121 | 0 | 0 |
| NC_045512.2;YP_009724389.1;Wu1-SARS2_orf1ab;3122 | 0 | 0 |
| NC_045512.2;YP_009724389.1;Wu1-SARS2_orf1ab;3123 | 0 | 0 |
| NC_045512.2;YP_009724389.1;Wu1-SARS2_orf1ab;3124 | 0 | 0 |
| NC_045512.2;YP_009724389.1;Wu1-SARS2_orf1ab;3125 | 0 | 0 |
| NC_045512.2;YP_009724389.1;Wu1-SARS2_orf1ab;3126 | 0 | 0 |
| NC_045512.2;YP_009724389.1;Wu1-SARS2_orf1ab;3127 | 0 | 0 |
| NC_045512.2;YP_009724389.1;Wu1-SARS2_orf1ab;3128 | 0 | 0 |
| NC_045512.2;YP_009724389.1;Wu1-SARS2_orf1ab;3129 | 0 | 0 |

| | | |
|---|---|---|
| NC_045512.2;YP_009724389.1;Wu1-SARS2_orf1ab;3130 | 0 | 0 |
| NC_045512.2;YP_009724389.1;Wu1-SARS2_orf1ab;3131 | 0 | 0 |
| NC_045512.2;YP_009724389.1;Wu1-SARS2_orf1ab;3132 | 0 | 0 |
| NC_045512.2;YP_009724389.1;Wu1-SARS2_orf1ab;3133 | 0 | 0 |
| NC_045512.2;YP_009724389.1;Wu1-SARS2_orf1ab;3134 | 0 | 0 |
| NC_045512.2;YP_009724389.1;Wu1-SARS2_orf1ab;3135 | 0 | 0 |
| NC_045512.2;YP_009724389.1;Wu1-SARS2_orf1ab;3136 | 0 | 0 |
| NC_045512.2;YP_009724389.1;Wu1-SARS2_orf1ab;3137 | 0 | 0 |
| NC_045512.2;YP_009724389.1;Wu1-SARS2_orf1ab;3138 | 0 | 0 |
| NC_045512.2;YP_009724389.1;Wu1-SARS2_orf1ab;3139 | 0 | 0 |
| NC_045512.2;YP_009724389.1;Wu1-SARS2_orf1ab;3140 | 0 | 0 |
| NC_045512.2;YP_009724389.1;Wu1-SARS2_orf1ab;3141 | 0 | 0 |
| NC_045512.2;YP_009724389.1;Wu1-SARS2_orf1ab;3142 | 0 | 0 |
| NC_045512.2;YP_009724389.1;Wu1-SARS2_orf1ab;3143 | 0 | 0 |
| NC_045512.2;YP_009724389.1;Wu1-SARS2_orf1ab;3144 | 0 | 0 |
| NC_045512.2;YP_009724389.1;Wu1-SARS2_orf1ab;3145 | 0 | 0 |
| NC_045512.2;YP_009724389.1;Wu1-SARS2_orf1ab;3146 | 0 | 0 |
| NC_045512.2;YP_009724389.1;Wu1-SARS2_orf1ab;3147 | 0 | 0 |
| NC_045512.2;YP_009724389.1;Wu1-SARS2_orf1ab;3148 | 0 | 0 |
| NC_045512.2;YP_009724389.1;Wu1-SARS2_orf1ab;3149 | 0 | 0 |
| NC_045512.2;YP_009724389.1;Wu1-SARS2_orf1ab;3150 | 0 | 0 |
| NC_045512.2;YP_009724389.1;Wu1-SARS2_orf1ab;3151 | 0 | 0 |
| NC_045512.2;YP_009724389.1;Wu1-SARS2_orf1ab;3152 | 0 | 0 |
| NC_045512.2;YP_009724389.1;Wu1-SARS2_orf1ab;3153 | 0 | 0 |
| NC_045512.2;YP_009724389.1;Wu1-SARS2_orf1ab;3154 | 0 | 0 |
| NC_045512.2;YP_009724389.1;Wu1-SARS2_orf1ab;3155 | 0 | 0 |
| NC_045512.2;YP_009724389.1;Wu1-SARS2_orf1ab;3156 | 0 | 0 |
| NC_045512.2;YP_009724389.1;Wu1-SARS2_orf1ab;3157 | 0 | 0 |
| NC_045512.2;YP_009724389.1;Wu1-SARS2_orf1ab;3158 | 0 | 0 |
| NC_045512.2;YP_009724389.1;Wu1-SARS2_orf1ab;3159 | 0 | 0 |
| NC_045512.2;YP_009724389.1;Wu1-SARS2_orf1ab;3160 | 0 | 0 |
| NC_045512.2;YP_009724389.1;Wu1-SARS2_orf1ab;3161 | 0 | 0 |
| NC_045512.2;YP_009724389.1;Wu1-SARS2_orf1ab;3162 | 0 | 0.075 |
| NC_045512.2;YP_009724389.1;Wu1-SARS2_orf1ab;3163 | 0 | 0 |
| NC_045512.2;YP_009724389.1;Wu1-SARS2_orf1ab;3164 | 0.05 | 0.025 |
| NC_045512.2;YP_009724389.1;Wu1-SARS2_orf1ab;3165 | 0 | 0 |
| NC_045512.2;YP_009724389.1;Wu1-SARS2_orf1ab;3166 | 0 | 0 |
| NC_045512.2;YP_009724389.1;Wu1-SARS2_orf1ab;3167 | 0 | 0 |
| NC_045512.2;YP_009724389.1;Wu1-SARS2_orf1ab;3168 | 0 | 0 |
| NC_045512.2;YP_009724389.1;Wu1-SARS2_orf1ab;3169 | 0 | 0 |
| NC_045512.2;YP_009724389.1;Wu1-SARS2_orf1ab;3170 | 0 | 0 |
| NC_045512.2;YP_009724389.1;Wu1-SARS2_orf1ab;3171 | 0 | 0 |
| NC_045512.2;YP_009724389.1;Wu1-SARS2_orf1ab;3172 | 0 | 0 |

| | | |
|---|---|---|
| NC_045512.2;YP_009724389.1;Wu1-SARS2_orf1ab;3173 | 0 | 0 |
| NC_045512.2;YP_009724389.1;Wu1-SARS2_orf1ab;3174 | 0 | 0 |
| NC_045512.2;YP_009724389.1;Wu1-SARS2_orf1ab;3175 | 0 | 0 |
| NC_045512.2;YP_009724389.1;Wu1-SARS2_orf1ab;3176 | 0 | 0 |
| NC_045512.2;YP_009724389.1;Wu1-SARS2_orf1ab;3177 | 0 | 0 |
| NC_045512.2;YP_009724389.1;Wu1-SARS2_orf1ab;3178 | 0 | 0 |
| NC_045512.2;YP_009724389.1;Wu1-SARS2_orf1ab;3179 | 0 | 0 |
| NC_045512.2;YP_009724389.1;Wu1-SARS2_orf1ab;3180 | 0 | 0 |
| NC_045512.2;YP_009724389.1;Wu1-SARS2_orf1ab;3181 | 0 | 0 |
| NC_045512.2;YP_009724389.1;Wu1-SARS2_orf1ab;3182 | 0.05 | 0 |
| NC_045512.2;YP_009724389.1;Wu1-SARS2_orf1ab;3183 | 0.1 | 0 |
| NC_045512.2;YP_009724389.1;Wu1-SARS2_orf1ab;3184 | 0.1 | 0 |
| NC_045512.2;YP_009724389.1;Wu1-SARS2_orf1ab;3185 | 0.1 | 0 |
| NC_045512.2;YP_009724389.1;Wu1-SARS2_orf1ab;3186 | 0.1 | 0 |
| NC_045512.2;YP_009724389.1;Wu1-SARS2_orf1ab;3187 | 0.1 | 0 |
| NC_045512.2;YP_009724389.1;Wu1-SARS2_orf1ab;3188 | 0.15 | 0 |
| NC_045512.2;YP_009724389.1;Wu1-SARS2_orf1ab;3189 | 0.1 | 0 |
| NC_045512.2;YP_009724389.1;Wu1-SARS2_orf1ab;3190 | 0.1 | 0 |
| NC_045512.2;YP_009724389.1;Wu1-SARS2_orf1ab;3191 | 0.1 | 0.05 |
| NC_045512.2;YP_009724389.1;Wu1-SARS2_orf1ab;3192 | 0.05 | 0.025 |
| NC_045512.2;YP_009724389.1;Wu1-SARS2_orf1ab;3193 | 0 | 0.025 |
| NC_045512.2;YP_009724389.1;Wu1-SARS2_orf1ab;3194 | 0 | 0.025 |
| NC_045512.2;YP_009724389.1;Wu1-SARS2_orf1ab;3195 | 0 | 0 |
| NC_045512.2;YP_009724389.1;Wu1-SARS2_orf1ab;3196 | 0 | 0 |
| NC_045512.2;YP_009724389.1;Wu1-SARS2_orf1ab;3197 | 0 | 0 |
| NC_045512.2;YP_009724389.1;Wu1-SARS2_orf1ab;3198 | 0 | 0 |
| NC_045512.2;YP_009724389.1;Wu1-SARS2_orf1ab;3199 | 0 | 0 |
| NC_045512.2;YP_009724389.1;Wu1-SARS2_orf1ab;3200 | 0 | 0 |
| NC_045512.2;YP_009724389.1;Wu1-SARS2_orf1ab;3201 | 0 | 0 |
| NC_045512.2;YP_009724389.1;Wu1-SARS2_orf1ab;3202 | 0 | 0 |
| NC_045512.2;YP_009724389.1;Wu1-SARS2_orf1ab;3203 | 0 | 0 |
| NC_045512.2;YP_009724389.1;Wu1-SARS2_orf1ab;3204 | 0 | 0 |
| NC_045512.2;YP_009724389.1;Wu1-SARS2_orf1ab;3205 | 0 | 0 |
| NC_045512.2;YP_009724389.1;Wu1-SARS2_orf1ab;3206 | 0 | 0 |
| NC_045512.2;YP_009724389.1;Wu1-SARS2_orf1ab;3207 | 0 | 0 |
| NC_045512.2;YP_009724389.1;Wu1-SARS2_orf1ab;3208 | 0 | 0 |
| NC_045512.2;YP_009724389.1;Wu1-SARS2_orf1ab;3209 | 0 | 0.025 |
| NC_045512.2;YP_009724389.1;Wu1-SARS2_orf1ab;3210 | 0 | 0.025 |
| NC_045512.2;YP_009724389.1;Wu1-SARS2_orf1ab;3211 | 0 | 0.025 |
| NC_045512.2;YP_009724389.1;Wu1-SARS2_orf1ab;3212 | 0 | 0 |
| NC_045512.2;YP_009724389.1;Wu1-SARS2_orf1ab;3213 | 0 | 0 |
| NC_045512.2;YP_009724389.1;Wu1-SARS2_orf1ab;3214 | 0 | 0 |
| NC_045512.2;YP_009724389.1;Wu1-SARS2_orf1ab;3215 | 0.05 | 0.075 |

| | | |
|---|---|---|
| NC_045512.2;YP_009724389.1;Wu1-SARS2_orf1ab;3216 | 0 | 0 |
| NC_045512.2;YP_009724389.1;Wu1-SARS2_orf1ab;3217 | 0 | 0 |
| NC_045512.2;YP_009724389.1;Wu1-SARS2_orf1ab;3218 | 0 | 0 |
| NC_045512.2;YP_009724389.1;Wu1-SARS2_orf1ab;3219 | 0 | 0.025 |
| NC_045512.2;YP_009724389.1;Wu1-SARS2_orf1ab;3220 | 0 | 0 |
| NC_045512.2;YP_009724389.1;Wu1-SARS2_orf1ab;3221 | 0 | 0 |
| NC_045512.2;YP_009724389.1;Wu1-SARS2_orf1ab;3222 | 0 | 0 |
| NC_045512.2;YP_009724389.1;Wu1-SARS2_orf1ab;3223 | 0 | 0 |
| NC_045512.2;YP_009724389.1;Wu1-SARS2_orf1ab;3224 | 0 | 0 |
| NC_045512.2;YP_009724389.1;Wu1-SARS2_orf1ab;3225 | 0 | 0.075 |
| NC_045512.2;YP_009724389.1;Wu1-SARS2_orf1ab;3226 | 0 | 0.025 |
| NC_045512.2;YP_009724389.1;Wu1-SARS2_orf1ab;3227 | 0 | 0.025 |
| NC_045512.2;YP_009724389.1;Wu1-SARS2_orf1ab;3228 | 0 | 0.025 |
| NC_045512.2;YP_009724389.1;Wu1-SARS2_orf1ab;3229 | 0 | 0.025 |
| NC_045512.2;YP_009724389.1;Wu1-SARS2_orf1ab;3230 | 0.05 | 0.025 |
| NC_045512.2;YP_009724389.1;Wu1-SARS2_orf1ab;3231 | 0 | 0.025 |
| NC_045512.2;YP_009724389.1;Wu1-SARS2_orf1ab;3232 | 0 | 0.025 |
| NC_045512.2;YP_009724389.1;Wu1-SARS2_orf1ab;3233 | 0 | 0.025 |
| NC_045512.2;YP_009724389.1;Wu1-SARS2_orf1ab;3234 | 0.05 | 0 |
| NC_045512.2;YP_009724389.1;Wu1-SARS2_orf1ab;3235 | 0 | 0 |
| NC_045512.2;YP_009724389.1;Wu1-SARS2_orf1ab;3236 | 0 | 0 |
| NC_045512.2;YP_009724389.1;Wu1-SARS2_orf1ab;3237 | 0.15 | 0.05 |
| NC_045512.2;YP_009724389.1;Wu1-SARS2_orf1ab;3238 | 0.3 | 0.05 |
| NC_045512.2;YP_009724389.1;Wu1-SARS2_orf1ab;3239 | 0.05 | 0 |
| NC_045512.2;YP_009724389.1;Wu1-SARS2_orf1ab;3240 | 0.05 | 0.05 |
| NC_045512.2;YP_009724389.1;Wu1-SARS2_orf1ab;3241 | 0.1 | 0.1 |
| NC_045512.2;YP_009724389.1;Wu1-SARS2_orf1ab;3242 | 0 | 0.05 |
| NC_045512.2;YP_009724389.1;Wu1-SARS2_orf1ab;3243 | 0 | 0.05 |
| NC_045512.2;YP_009724389.1;Wu1-SARS2_orf1ab;3244 | 0 | 0 |
| NC_045512.2;YP_009724389.1;Wu1-SARS2_orf1ab;3245 | 0 | 0.025 |
| NC_045512.2;YP_009724389.1;Wu1-SARS2_orf1ab;3246 | 0 | 0 |
| NC_045512.2;YP_009724389.1;Wu1-SARS2_orf1ab;3247 | 0 | 0 |
| NC_045512.2;YP_009724389.1;Wu1-SARS2_orf1ab;3248 | 0 | 0 |
| NC_045512.2;YP_009724389.1;Wu1-SARS2_orf1ab;3249 | 0 | 0 |
| NC_045512.2;YP_009724389.1;Wu1-SARS2_orf1ab;3250 | 0.05 | 0 |
| NC_045512.2;YP_009724389.1;Wu1-SARS2_orf1ab;3251 | 0 | 0 |
| NC_045512.2;YP_009724389.1;Wu1-SARS2_orf1ab;3252 | 0 | 0 |
| NC_045512.2;YP_009724389.1;Wu1-SARS2_orf1ab;3253 | 0 | 0 |
| NC_045512.2;YP_009724389.1;Wu1-SARS2_orf1ab;3254 | 0 | 0 |
| NC_045512.2;YP_009724389.1;Wu1-SARS2_orf1ab;3255 | 0 | 0 |
| NC_045512.2;YP_009724389.1;Wu1-SARS2_orf1ab;3256 | 0 | 0 |
| NC_045512.2;YP_009724389.1;Wu1-SARS2_orf1ab;3257 | 0 | 0 |
| NC_045512.2;YP_009724389.1;Wu1-SARS2_orf1ab;3258 | 0 | 0 |

| | | |
|---|---|---|
| NC_045512.2;YP_009724389.1;Wu1-SARS2_orf1ab;3259 | 0 | 0 |
| NC_045512.2;YP_009724389.1;Wu1-SARS2_orf1ab;3260 | 0 | 0 |
| NC_045512.2;YP_009724389.1;Wu1-SARS2_orf1ab;3261 | 0 | 0 |
| NC_045512.2;YP_009724389.1;Wu1-SARS2_orf1ab;3262 | 0 | 0 |
| NC_045512.2;YP_009724389.1;Wu1-SARS2_orf1ab;3263 | 0 | 0 |
| NC_045512.2;YP_009724389.1;Wu1-SARS2_orf1ab;3264 | 0 | 0 |
| NC_045512.2;YP_009724389.1;Wu1-SARS2_orf1ab;3265 | 0 | 0 |
| NC_045512.2;YP_009724389.1;Wu1-SARS2_orf1ab;3266 | 0 | 0 |
| NC_045512.2;YP_009724389.1;Wu1-SARS2_orf1ab;3267 | 0 | 0 |
| NC_045512.2;YP_009724389.1;Wu1-SARS2_orf1ab;3268 | 0 | 0.025 |
| NC_045512.2;YP_009724389.1;Wu1-SARS2_orf1ab;3269 | 0 | 0 |
| NC_045512.2;YP_009724389.1;Wu1-SARS2_orf1ab;3270 | 0.05 | 0 |
| NC_045512.2;YP_009724389.1;Wu1-SARS2_orf1ab;3271 | 0 | 0 |
| NC_045512.2;YP_009724389.1;Wu1-SARS2_orf1ab;3272 | 0 | 0 |
| NC_045512.2;YP_009724389.1;Wu1-SARS2_orf1ab;3273 | 0 | 0 |
| NC_045512.2;YP_009724389.1;Wu1-SARS2_orf1ab;3274 | 0 | 0 |
| NC_045512.2;YP_009724389.1;Wu1-SARS2_orf1ab;3275 | 0 | 0 |
| NC_045512.2;YP_009724389.1;Wu1-SARS2_orf1ab;3276 | 0 | 0 |
| NC_045512.2;YP_009724389.1;Wu1-SARS2_orf1ab;3277 | 0 | 0 |
| NC_045512.2;YP_009724389.1;Wu1-SARS2_orf1ab;3278 | 0 | 0 |
| NC_045512.2;YP_009724389.1;Wu1-SARS2_orf1ab;3279 | 0 | 0 |
| NC_045512.2;YP_009724389.1;Wu1-SARS2_orf1ab;3280 | 0 | 0 |
| NC_045512.2;YP_009724389.1;Wu1-SARS2_orf1ab;3281 | 0 | 0 |
| NC_045512.2;YP_009724389.1;Wu1-SARS2_orf1ab;3282 | 0 | 0 |
| NC_045512.2;YP_009724389.1;Wu1-SARS2_orf1ab;3283 | 0 | 0 |
| NC_045512.2;YP_009724389.1;Wu1-SARS2_orf1ab;3284 | 0 | 0 |
| NC_045512.2;YP_009724389.1;Wu1-SARS2_orf1ab;3285 | 0 | 0 |
| NC_045512.2;YP_009724389.1;Wu1-SARS2_orf1ab;3286 | 0 | 0 |
| NC_045512.2;YP_009724389.1;Wu1-SARS2_orf1ab;3287 | 0 | 0 |
| NC_045512.2;YP_009724389.1;Wu1-SARS2_orf1ab;3288 | 0 | 0 |
| NC_045512.2;YP_009724389.1;Wu1-SARS2_orf1ab;3289 | 0 | 0 |
| NC_045512.2;YP_009724389.1;Wu1-SARS2_orf1ab;3290 | 0 | 0 |
| NC_045512.2;YP_009724389.1;Wu1-SARS2_orf1ab;3291 | 0 | 0 |
| NC_045512.2;YP_009724389.1;Wu1-SARS2_orf1ab;3292 | 0 | 0 |
| NC_045512.2;YP_009724389.1;Wu1-SARS2_orf1ab;3293 | 0 | 0 |
| NC_045512.2;YP_009724389.1;Wu1-SARS2_orf1ab;3294 | 0 | 0 |
| NC_045512.2;YP_009724389.1;Wu1-SARS2_orf1ab;3295 | 0 | 0 |
| NC_045512.2;YP_009724389.1;Wu1-SARS2_orf1ab;3296 | 0 | 0 |
| NC_045512.2;YP_009724389.1;Wu1-SARS2_orf1ab;3297 | 0 | 0 |
| NC_045512.2;YP_009724389.1;Wu1-SARS2_orf1ab;3298 | 0 | 0 |
| NC_045512.2;YP_009724389.1;Wu1-SARS2_orf1ab;3299 | 0 | 0.025 |
| NC_045512.2;YP_009724389.1;Wu1-SARS2_orf1ab;3300 | 0 | 0 |
| NC_045512.2;YP_009724389.1;Wu1-SARS2_orf1ab;3301 | 0 | 0 |

| | | |
|---|---|---|
| NC_045512.2;YP_009724389.1;Wu1-SARS2_orf1ab;3302 | 0 | 0 |
| NC_045512.2;YP_009724389.1;Wu1-SARS2_orf1ab;3303 | 0 | 0 |
| NC_045512.2;YP_009724389.1;Wu1-SARS2_orf1ab;3304 | 0 | 0 |
| NC_045512.2;YP_009724389.1;Wu1-SARS2_orf1ab;3305 | 0 | 0 |
| NC_045512.2;YP_009724389.1;Wu1-SARS2_orf1ab;3306 | 0 | 0.025 |
| NC_045512.2;YP_009724389.1;Wu1-SARS2_orf1ab;3307 | 0 | 0 |
| NC_045512.2;YP_009724389.1;Wu1-SARS2_orf1ab;3308 | 0 | 0 |
| NC_045512.2;YP_009724389.1;Wu1-SARS2_orf1ab;3309 | 0 | 0 |
| NC_045512.2;YP_009724389.1;Wu1-SARS2_orf1ab;3310 | 0 | 0.025 |
| NC_045512.2;YP_009724389.1;Wu1-SARS2_orf1ab;3311 | 0 | 0.025 |
| NC_045512.2;YP_009724389.1;Wu1-SARS2_orf1ab;3312 | 0 | 0 |
| NC_045512.2;YP_009724389.1;Wu1-SARS2_orf1ab;3313 | 0.05 | 0 |
| NC_045512.2;YP_009724389.1;Wu1-SARS2_orf1ab;3314 | 0 | 0.05 |
| NC_045512.2;YP_009724389.1;Wu1-SARS2_orf1ab;3315 | 0 | 0 |
| NC_045512.2;YP_009724389.1;Wu1-SARS2_orf1ab;3316 | 0 | 0 |
| NC_045512.2;YP_009724389.1;Wu1-SARS2_orf1ab;3317 | 0 | 0 |
| NC_045512.2;YP_009724389.1;Wu1-SARS2_orf1ab;3318 | 0 | 0 |
| NC_045512.2;YP_009724389.1;Wu1-SARS2_orf1ab;3319 | 0 | 0 |
| NC_045512.2;YP_009724389.1;Wu1-SARS2_orf1ab;3320 | 0 | 0 |
| NC_045512.2;YP_009724389.1;Wu1-SARS2_orf1ab;3321 | 0 | 0 |
| NC_045512.2;YP_009724389.1;Wu1-SARS2_orf1ab;3322 | 0 | 0 |
| NC_045512.2;YP_009724389.1;Wu1-SARS2_orf1ab;3323 | 0 | 0 |
| NC_045512.2;YP_009724389.1;Wu1-SARS2_orf1ab;3324 | 0 | 0 |
| NC_045512.2;YP_009724389.1;Wu1-SARS2_orf1ab;3325 | 0 | 0 |
| NC_045512.2;YP_009724389.1;Wu1-SARS2_orf1ab;3326 | 0 | 0 |
| NC_045512.2;YP_009724389.1;Wu1-SARS2_orf1ab;3327 | 0 | 0 |
| NC_045512.2;YP_009724389.1;Wu1-SARS2_orf1ab;3328 | 0 | 0 |
| NC_045512.2;YP_009724389.1;Wu1-SARS2_orf1ab;3329 | 0 | 0 |
| NC_045512.2;YP_009724389.1;Wu1-SARS2_orf1ab;3330 | 0 | 0 |
| NC_045512.2;YP_009724389.1;Wu1-SARS2_orf1ab;3331 | 0 | 0 |
| NC_045512.2;YP_009724389.1;Wu1-SARS2_orf1ab;3332 | 0 | 0 |
| NC_045512.2;YP_009724389.1;Wu1-SARS2_orf1ab;3333 | 0 | 0 |
| NC_045512.2;YP_009724389.1;Wu1-SARS2_orf1ab;3334 | 0 | 0 |
| NC_045512.2;YP_009724389.1;Wu1-SARS2_orf1ab;3335 | 0 | 0 |
| NC_045512.2;YP_009724389.1;Wu1-SARS2_orf1ab;3336 | 0 | 0 |
| NC_045512.2;YP_009724389.1;Wu1-SARS2_orf1ab;3337 | 0 | 0 |
| NC_045512.2;YP_009724389.1;Wu1-SARS2_orf1ab;3338 | 0 | 0 |
| NC_045512.2;YP_009724389.1;Wu1-SARS2_orf1ab;3339 | 0 | 0 |
| NC_045512.2;YP_009724389.1;Wu1-SARS2_orf1ab;3340 | 0 | 0 |
| NC_045512.2;YP_009724389.1;Wu1-SARS2_orf1ab;3341 | 0 | 0 |
| NC_045512.2;YP_009724389.1;Wu1-SARS2_orf1ab;3342 | 0 | 0.025 |
| NC_045512.2;YP_009724389.1;Wu1-SARS2_orf1ab;3343 | 0.05 | 0 |
| NC_045512.2;YP_009724389.1;Wu1-SARS2_orf1ab;3344 | 0 | 0 |

| | | |
|---|---|---|
| NC_045512.2;YP_009724389.1;Wu1-SARS2_orf1ab;3345 | 0 | 0 |
| NC_045512.2;YP_009724389.1;Wu1-SARS2_orf1ab;3346 | 0 | 0 |
| NC_045512.2;YP_009724389.1;Wu1-SARS2_orf1ab;3347 | 0 | 0 |
| NC_045512.2;YP_009724389.1;Wu1-SARS2_orf1ab;3348 | 0 | 0 |
| NC_045512.2;YP_009724389.1;Wu1-SARS2_orf1ab;3349 | 0 | 0 |
| NC_045512.2;YP_009724389.1;Wu1-SARS2_orf1ab;3350 | 0 | 0 |
| NC_045512.2;YP_009724389.1;Wu1-SARS2_orf1ab;3351 | 0 | 0 |
| NC_045512.2;YP_009724389.1;Wu1-SARS2_orf1ab;3352 | 0 | 0.025 |
| NC_045512.2;YP_009724389.1;Wu1-SARS2_orf1ab;3353 | 0 | 0 |
| NC_045512.2;YP_009724389.1;Wu1-SARS2_orf1ab;3354 | 0 | 0 |
| NC_045512.2;YP_009724389.1;Wu1-SARS2_orf1ab;3355 | 0 | 0 |
| NC_045512.2;YP_009724389.1;Wu1-SARS2_orf1ab;3356 | 0 | 0 |
| NC_045512.2;YP_009724389.1;Wu1-SARS2_orf1ab;3357 | 0 | 0 |
| NC_045512.2;YP_009724389.1;Wu1-SARS2_orf1ab;3358 | 0 | 0 |
| NC_045512.2;YP_009724389.1;Wu1-SARS2_orf1ab;3359 | 0 | 0 |
| NC_045512.2;YP_009724389.1;Wu1-SARS2_orf1ab;3360 | 0 | 0 |
| NC_045512.2;YP_009724389.1;Wu1-SARS2_orf1ab;3361 | 0 | 0 |
| NC_045512.2;YP_009724389.1;Wu1-SARS2_orf1ab;3362 | 0 | 0 |
| NC_045512.2;YP_009724389.1;Wu1-SARS2_orf1ab;3363 | 0 | 0 |
| NC_045512.2;YP_009724389.1;Wu1-SARS2_orf1ab;3364 | 0 | 0 |
| NC_045512.2;YP_009724389.1;Wu1-SARS2_orf1ab;3365 | 0 | 0 |
| NC_045512.2;YP_009724389.1;Wu1-SARS2_orf1ab;3366 | 0 | 0 |
| NC_045512.2;YP_009724389.1;Wu1-SARS2_orf1ab;3367 | 0 | 0 |
| NC_045512.2;YP_009724389.1;Wu1-SARS2_orf1ab;3368 | 0 | 0 |
| NC_045512.2;YP_009724389.1;Wu1-SARS2_orf1ab;3369 | 0 | 0 |
| NC_045512.2;YP_009724389.1;Wu1-SARS2_orf1ab;3370 | 0 | 0.025 |
| NC_045512.2;YP_009724389.1;Wu1-SARS2_orf1ab;3371 | 0 | 0 |
| NC_045512.2;YP_009724389.1;Wu1-SARS2_orf1ab;3372 | 0 | 0 |
| NC_045512.2;YP_009724389.1;Wu1-SARS2_orf1ab;3373 | 0 | 0 |
| NC_045512.2;YP_009724389.1;Wu1-SARS2_orf1ab;3374 | 0 | 0 |
| NC_045512.2;YP_009724389.1;Wu1-SARS2_orf1ab;3375 | 0 | 0 |
| NC_045512.2;YP_009724389.1;Wu1-SARS2_orf1ab;3376 | 0 | 0 |
| NC_045512.2;YP_009724389.1;Wu1-SARS2_orf1ab;3377 | 0 | 0 |
| NC_045512.2;YP_009724389.1;Wu1-SARS2_orf1ab;3378 | 0 | 0 |
| NC_045512.2;YP_009724389.1;Wu1-SARS2_orf1ab;3379 | 0 | 0 |
| NC_045512.2;YP_009724389.1;Wu1-SARS2_orf1ab;3380 | 0 | 0 |
| NC_045512.2;YP_009724389.1;Wu1-SARS2_orf1ab;3381 | 0 | 0.025 |
| NC_045512.2;YP_009724389.1;Wu1-SARS2_orf1ab;3382 | 0 | 0 |
| NC_045512.2;YP_009724389.1;Wu1-SARS2_orf1ab;3383 | 0 | 0 |
| NC_045512.2;YP_009724389.1;Wu1-SARS2_orf1ab;3384 | 0 | 0 |
| NC_045512.2;YP_009724389.1;Wu1-SARS2_orf1ab;3385 | 0 | 0 |
| NC_045512.2;YP_009724389.1;Wu1-SARS2_orf1ab;3386 | 0 | 0 |
| NC_045512.2;YP_009724389.1;Wu1-SARS2_orf1ab;3387 | 0 | 0.025 |

| | | |
|---|---|---|
| NC_045512.2;YP_009724389.1;Wu1-SARS2_orf1ab;3388 | 0 | 0 |
| NC_045512.2;YP_009724389.1;Wu1-SARS2_orf1ab;3389 | 0 | 0 |
| NC_045512.2;YP_009724389.1;Wu1-SARS2_orf1ab;3390 | 0 | 0 |
| NC_045512.2;YP_009724389.1;Wu1-SARS2_orf1ab;3391 | 0 | 0 |
| NC_045512.2;YP_009724389.1;Wu1-SARS2_orf1ab;3392 | 0.05 | 0 |
| NC_045512.2;YP_009724389.1;Wu1-SARS2_orf1ab;3393 | 0 | 0 |
| NC_045512.2;YP_009724389.1;Wu1-SARS2_orf1ab;3394 | 0.05 | 0 |
| NC_045512.2;YP_009724389.1;Wu1-SARS2_orf1ab;3395 | 0 | 0 |
| NC_045512.2;YP_009724389.1;Wu1-SARS2_orf1ab;3396 | 0 | 0 |
| NC_045512.2;YP_009724389.1;Wu1-SARS2_orf1ab;3397 | 0 | 0 |
| NC_045512.2;YP_009724389.1;Wu1-SARS2_orf1ab;3398 | 0 | 0 |
| NC_045512.2;YP_009724389.1;Wu1-SARS2_orf1ab;3399 | 0 | 0 |
| NC_045512.2;YP_009724389.1;Wu1-SARS2_orf1ab;3400 | 0 | 0.025 |
| NC_045512.2;YP_009724389.1;Wu1-SARS2_orf1ab;3401 | 0 | 0.025 |
| NC_045512.2;YP_009724389.1;Wu1-SARS2_orf1ab;3402 | 0 | 0.025 |
| NC_045512.2;YP_009724389.1;Wu1-SARS2_orf1ab;3403 | 0 | 0.05 |
| NC_045512.2;YP_009724389.1;Wu1-SARS2_orf1ab;3404 | 0 | 0.05 |
| NC_045512.2;YP_009724389.1;Wu1-SARS2_orf1ab;3405 | 0 | 0.025 |
| NC_045512.2;YP_009724389.1;Wu1-SARS2_orf1ab;3406 | 0 | 0.025 |
| NC_045512.2;YP_009724389.1;Wu1-SARS2_orf1ab;3407 | 0 | 0 |
| NC_045512.2;YP_009724389.1;Wu1-SARS2_orf1ab;3408 | 0 | 0 |
| NC_045512.2;YP_009724389.1;Wu1-SARS2_orf1ab;3409 | 0 | 0 |
| NC_045512.2;YP_009724389.1;Wu1-SARS2_orf1ab;3410 | 0 | 0 |
| NC_045512.2;YP_009724389.1;Wu1-SARS2_orf1ab;3411 | 0 | 0 |
| NC_045512.2;YP_009724389.1;Wu1-SARS2_orf1ab;3412 | 0 | 0 |
| NC_045512.2;YP_009724389.1;Wu1-SARS2_orf1ab;3413 | 0 | 0 |
| NC_045512.2;YP_009724389.1;Wu1-SARS2_orf1ab;3414 | 0 | 0 |
| NC_045512.2;YP_009724389.1;Wu1-SARS2_orf1ab;3415 | 0 | 0 |
| NC_045512.2;YP_009724389.1;Wu1-SARS2_orf1ab;3416 | 0 | 0 |
| NC_045512.2;YP_009724389.1;Wu1-SARS2_orf1ab;3417 | 0 | 0 |
| NC_045512.2;YP_009724389.1;Wu1-SARS2_orf1ab;3418 | 0 | 0 |
| NC_045512.2;YP_009724389.1;Wu1-SARS2_orf1ab;3419 | 0 | 0 |
| NC_045512.2;YP_009724389.1;Wu1-SARS2_orf1ab;3420 | 0 | 0 |
| NC_045512.2;YP_009724389.1;Wu1-SARS2_orf1ab;3421 | 0 | 0 |
| NC_045512.2;YP_009724389.1;Wu1-SARS2_orf1ab;3422 | 0 | 0 |
| NC_045512.2;YP_009724389.1;Wu1-SARS2_orf1ab;3423 | 0 | 0 |
| NC_045512.2;YP_009724389.1;Wu1-SARS2_orf1ab;3424 | 0 | 0 |
| NC_045512.2;YP_009724389.1;Wu1-SARS2_orf1ab;3425 | 0 | 0 |
| NC_045512.2;YP_009724389.1;Wu1-SARS2_orf1ab;3426 | 0.15 | 0.025 |
| NC_045512.2;YP_009724389.1;Wu1-SARS2_orf1ab;3427 | 0.1 | 0.05 |
| NC_045512.2;YP_009724389.1;Wu1-SARS2_orf1ab;3428 | 0.05 | 0.05 |
| NC_045512.2;YP_009724389.1;Wu1-SARS2_orf1ab;3429 | 0.2 | 0.075 |
| NC_045512.2;YP_009724389.1;Wu1-SARS2_orf1ab;3430 | 0.25 | 0.125 |

| | | |
|---|---|---|
| NC_045512.2;YP_009724389.1;Wu1-SARS2_orf1ab;3431 | 0.25 | 0.125 |
| NC_045512.2;YP_009724389.1;Wu1-SARS2_orf1ab;3432 | 0.3 | 0.125 |
| NC_045512.2;YP_009724389.1;Wu1-SARS2_orf1ab;3433 | 0.3 | 0.15 |
| NC_045512.2;YP_009724389.1;Wu1-SARS2_orf1ab;3434 | 0.2 | 0.075 |
| NC_045512.2;YP_009724389.1;Wu1-SARS2_orf1ab;3435 | 0.25 | 0.1 |
| NC_045512.2;YP_009724389.1;Wu1-SARS2_orf1ab;3436 | 0.25 | 0.075 |
| NC_045512.2;YP_009724389.1;Wu1-SARS2_orf1ab;3437 | 0.25 | 0.2 |
| NC_045512.2;YP_009724389.1;Wu1-SARS2_orf1ab;3438 | 0.1 | 0.025 |
| NC_045512.2;YP_009724389.1;Wu1-SARS2_orf1ab;3439 | 0.1 | 0.025 |
| NC_045512.2;YP_009724389.1;Wu1-SARS2_orf1ab;3440 | 0.05 | 0.025 |
| NC_045512.2;YP_009724389.1;Wu1-SARS2_orf1ab;3441 | 0.05 | 0 |
| NC_045512.2;YP_009724389.1;Wu1-SARS2_orf1ab;3442 | 0.1 | 0.025 |
| NC_045512.2;YP_009724389.1;Wu1-SARS2_orf1ab;3443 | 0.05 | 0 |
| NC_045512.2;YP_009724389.1;Wu1-SARS2_orf1ab;3444 | 0.1 | 0 |
| NC_045512.2;YP_009724389.1;Wu1-SARS2_orf1ab;3445 | 0 | 0 |
| NC_045512.2;YP_009724389.1;Wu1-SARS2_orf1ab;3446 | 0.05 | 0 |
| NC_045512.2;YP_009724389.1;Wu1-SARS2_orf1ab;3447 | 0 | 0 |
| NC_045512.2;YP_009724389.1;Wu1-SARS2_orf1ab;3448 | 0.05 | 0.025 |
| NC_045512.2;YP_009724389.1;Wu1-SARS2_orf1ab;3449 | 0 | 0 |
| NC_045512.2;YP_009724389.1;Wu1-SARS2_orf1ab;3450 | 0 | 0 |
| NC_045512.2;YP_009724389.1;Wu1-SARS2_orf1ab;3451 | 0 | 0 |
| NC_045512.2;YP_009724389.1;Wu1-SARS2_orf1ab;3452 | 0 | 0 |
| NC_045512.2;YP_009724389.1;Wu1-SARS2_orf1ab;3453 | 0 | 0 |
| NC_045512.2;YP_009724389.1;Wu1-SARS2_orf1ab;3454 | 0 | 0 |
| NC_045512.2;YP_009724389.1;Wu1-SARS2_orf1ab;3455 | 0 | 0 |
| NC_045512.2;YP_009724389.1;Wu1-SARS2_orf1ab;3456 | 0 | 0 |
| NC_045512.2;YP_009724389.1;Wu1-SARS2_orf1ab;3457 | 0 | 0 |
| NC_045512.2;YP_009724389.1;Wu1-SARS2_orf1ab;3458 | 0 | 0 |
| NC_045512.2;YP_009724389.1;Wu1-SARS2_orf1ab;3459 | 0 | 0 |
| NC_045512.2;YP_009724389.1;Wu1-SARS2_orf1ab;3460 | 0 | 0 |
| NC_045512.2;YP_009724389.1;Wu1-SARS2_orf1ab;3461 | 0 | 0 |
| NC_045512.2;YP_009724389.1;Wu1-SARS2_orf1ab;3462 | 0 | 0 |
| NC_045512.2;YP_009724389.1;Wu1-SARS2_orf1ab;3463 | 0 | 0 |
| NC_045512.2;YP_009724389.1;Wu1-SARS2_orf1ab;3464 | 0 | 0 |
| NC_045512.2;YP_009724389.1;Wu1-SARS2_orf1ab;3465 | 0 | 0 |
| NC_045512.2;YP_009724389.1;Wu1-SARS2_orf1ab;3466 | 0 | 0 |
| NC_045512.2;YP_009724389.1;Wu1-SARS2_orf1ab;3467 | 0 | 0 |
| NC_045512.2;YP_009724389.1;Wu1-SARS2_orf1ab;3468 | 0 | 0 |
| NC_045512.2;YP_009724389.1;Wu1-SARS2_orf1ab;3469 | 0 | 0 |
| NC_045512.2;YP_009724389.1;Wu1-SARS2_orf1ab;3470 | 0 | 0 |
| NC_045512.2;YP_009724389.1;Wu1-SARS2_orf1ab;3471 | 0 | 0 |
| NC_045512.2;YP_009724389.1;Wu1-SARS2_orf1ab;3472 | 0 | 0 |
| NC_045512.2;YP_009724389.1;Wu1-SARS2_orf1ab;3473 | 0 | 0 |

| | | |
|---|---|---|
| NC_045512.2;YP_009724389.1;Wu1-SARS2_orf1ab;3474 | 0 | 0 |
| NC_045512.2;YP_009724389.1;Wu1-SARS2_orf1ab;3475 | 0 | 0 |
| NC_045512.2;YP_009724389.1;Wu1-SARS2_orf1ab;3476 | 0 | 0 |
| NC_045512.2;YP_009724389.1;Wu1-SARS2_orf1ab;3477 | 0 | 0 |
| NC_045512.2;YP_009724389.1;Wu1-SARS2_orf1ab;3478 | 0 | 0 |
| NC_045512.2;YP_009724389.1;Wu1-SARS2_orf1ab;3479 | 0.05 | 0 |
| NC_045512.2;YP_009724389.1;Wu1-SARS2_orf1ab;3480 | 0.05 | 0 |
| NC_045512.2;YP_009724389.1;Wu1-SARS2_orf1ab;3481 | 0.05 | 0 |
| NC_045512.2;YP_009724389.1;Wu1-SARS2_orf1ab;3482 | 0.05 | 0.025 |
| NC_045512.2;YP_009724389.1;Wu1-SARS2_orf1ab;3483 | 0.1 | 0 |
| NC_045512.2;YP_009724389.1;Wu1-SARS2_orf1ab;3484 | 0.05 | 0 |
| NC_045512.2;YP_009724389.1;Wu1-SARS2_orf1ab;3485 | 0.1 | 0 |
| NC_045512.2;YP_009724389.1;Wu1-SARS2_orf1ab;3486 | 0.05 | 0 |
| NC_045512.2;YP_009724389.1;Wu1-SARS2_orf1ab;3487 | 0.05 | 0.025 |
| NC_045512.2;YP_009724389.1;Wu1-SARS2_orf1ab;3488 | 0.05 | 0.15 |
| NC_045512.2;YP_009724389.1;Wu1-SARS2_orf1ab;3489 | 0.25 | 0.1 |
| NC_045512.2;YP_009724389.1;Wu1-SARS2_orf1ab;3490 | 0.05 | 0 |
| NC_045512.2;YP_009724389.1;Wu1-SARS2_orf1ab;3491 | 0 | 0 |
| NC_045512.2;YP_009724389.1;Wu1-SARS2_orf1ab;3492 | 0.05 | 0 |
| NC_045512.2;YP_009724389.1;Wu1-SARS2_orf1ab;3493 | 0 | 0 |
| NC_045512.2;YP_009724389.1;Wu1-SARS2_orf1ab;3494 | 0 | 0.025 |
| NC_045512.2;YP_009724389.1;Wu1-SARS2_orf1ab;3495 | 0 | 0.025 |
| NC_045512.2;YP_009724389.1;Wu1-SARS2_orf1ab;3496 | 0.05 | 0 |
| NC_045512.2;YP_009724389.1;Wu1-SARS2_orf1ab;3497 | 0 | 0 |
| NC_045512.2;YP_009724389.1;Wu1-SARS2_orf1ab;3498 | 0 | 0 |
| NC_045512.2;YP_009724389.1;Wu1-SARS2_orf1ab;3499 | 0 | 0 |
| NC_045512.2;YP_009724389.1;Wu1-SARS2_orf1ab;3500 | 0 | 0 |
| NC_045512.2;YP_009724389.1;Wu1-SARS2_orf1ab;3501 | 0 | 0 |
| NC_045512.2;YP_009724389.1;Wu1-SARS2_orf1ab;3502 | 0 | 0 |
| NC_045512.2;YP_009724389.1;Wu1-SARS2_orf1ab;3503 | 0.05 | 0 |
| NC_045512.2;YP_009724389.1;Wu1-SARS2_orf1ab;3504 | 0 | 0 |
| NC_045512.2;YP_009724389.1;Wu1-SARS2_orf1ab;3505 | 0.05 | 0.025 |
| NC_045512.2;YP_009724389.1;Wu1-SARS2_orf1ab;3506 | 0.15 | 0.1 |
| NC_045512.2;YP_009724389.1;Wu1-SARS2_orf1ab;3507 | 0.05 | 0.05 |
| NC_045512.2;YP_009724389.1;Wu1-SARS2_orf1ab;3508 | 0.1 | 0.15 |
| NC_045512.2;YP_009724389.1;Wu1-SARS2_orf1ab;3509 | 0.05 | 0.025 |
| NC_045512.2;YP_009724389.1;Wu1-SARS2_orf1ab;3510 | 0.05 | 0 |
| NC_045512.2;YP_009724389.1;Wu1-SARS2_orf1ab;3511 | 0.05 | 0 |
| NC_045512.2;YP_009724389.1;Wu1-SARS2_orf1ab;3512 | 0.05 | 0 |
| NC_045512.2;YP_009724389.1;Wu1-SARS2_orf1ab;3513 | 0.05 | 0 |
| NC_045512.2;YP_009724389.1;Wu1-SARS2_orf1ab;3514 | 0.05 | 0 |
| NC_045512.2;YP_009724389.1;Wu1-SARS2_orf1ab;3515 | 0.05 | 0 |
| NC_045512.2;YP_009724389.1;Wu1-SARS2_orf1ab;3516 | 0 | 0 |

| | | |
|---|---|---|
| NC_045512.2;YP_009724389.1;Wu1-SARS2_orf1ab;3517 | 0 | 0 |
| NC_045512.2;YP_009724389.1;Wu1-SARS2_orf1ab;3518 | 0 | 0 |
| NC_045512.2;YP_009724389.1;Wu1-SARS2_orf1ab;3519 | 0 | 0 |
| NC_045512.2;YP_009724389.1;Wu1-SARS2_orf1ab;3520 | 0 | 0 |
| NC_045512.2;YP_009724389.1;Wu1-SARS2_orf1ab;3521 | 0.05 | 0 |
| NC_045512.2;YP_009724389.1;Wu1-SARS2_orf1ab;3522 | 0 | 0 |
| NC_045512.2;YP_009724389.1;Wu1-SARS2_orf1ab;3523 | 0 | 0 |
| NC_045512.2;YP_009724389.1;Wu1-SARS2_orf1ab;3524 | 0 | 0 |
| NC_045512.2;YP_009724389.1;Wu1-SARS2_orf1ab;3525 | 0 | 0 |
| NC_045512.2;YP_009724389.1;Wu1-SARS2_orf1ab;3526 | 0 | 0.025 |
| NC_045512.2;YP_009724389.1;Wu1-SARS2_orf1ab;3527 | 0.05 | 0 |
| NC_045512.2;YP_009724389.1;Wu1-SARS2_orf1ab;3528 | 0.05 | 0 |
| NC_045512.2;YP_009724389.1;Wu1-SARS2_orf1ab;3529 | 0.05 | 0.025 |
| NC_045512.2;YP_009724389.1;Wu1-SARS2_orf1ab;3530 | 0 | 0.025 |
| NC_045512.2;YP_009724389.1;Wu1-SARS2_orf1ab;3531 | 0 | 0.025 |
| NC_045512.2;YP_009724389.1;Wu1-SARS2_orf1ab;3532 | 0 | 0.025 |
| NC_045512.2;YP_009724389.1;Wu1-SARS2_orf1ab;3533 | 0 | 0 |
| NC_045512.2;YP_009724389.1;Wu1-SARS2_orf1ab;3534 | 0 | 0 |
| NC_045512.2;YP_009724389.1;Wu1-SARS2_orf1ab;3535 | 0 | 0 |
| NC_045512.2;YP_009724389.1;Wu1-SARS2_orf1ab;3536 | 0 | 0 |
| NC_045512.2;YP_009724389.1;Wu1-SARS2_orf1ab;3537 | 0 | 0 |
| NC_045512.2;YP_009724389.1;Wu1-SARS2_orf1ab;3538 | 0 | 0 |
| NC_045512.2;YP_009724389.1;Wu1-SARS2_orf1ab;3539 | 0 | 0.025 |
| NC_045512.2;YP_009724389.1;Wu1-SARS2_orf1ab;3540 | 0 | 0 |
| NC_045512.2;YP_009724389.1;Wu1-SARS2_orf1ab;3541 | 0 | 0.025 |
| NC_045512.2;YP_009724389.1;Wu1-SARS2_orf1ab;3542 | 0 | 0.025 |
| NC_045512.2;YP_009724389.1;Wu1-SARS2_orf1ab;3543 | 0 | 0.05 |
| NC_045512.2;YP_009724389.1;Wu1-SARS2_orf1ab;3544 | 0 | 0.05 |
| NC_045512.2;YP_009724389.1;Wu1-SARS2_orf1ab;3545 | 0 | 0.025 |
| NC_045512.2;YP_009724389.1;Wu1-SARS2_orf1ab;3546 | 0 | 0.025 |
| NC_045512.2;YP_009724389.1;Wu1-SARS2_orf1ab;3547 | 0.05 | 0.05 |
| NC_045512.2;YP_009724389.1;Wu1-SARS2_orf1ab;3548 | 0.05 | 0.05 |
| NC_045512.2;YP_009724389.1;Wu1-SARS2_orf1ab;3549 | 0 | 0.025 |
| NC_045512.2;YP_009724389.1;Wu1-SARS2_orf1ab;3550 | 0 | 0.05 |
| NC_045512.2;YP_009724389.1;Wu1-SARS2_orf1ab;3551 | 0.1 | 0.025 |
| NC_045512.2;YP_009724389.1;Wu1-SARS2_orf1ab;3552 | 0 | 0 |
| NC_045512.2;YP_009724389.1;Wu1-SARS2_orf1ab;3553 | 0 | 0 |
| NC_045512.2;YP_009724389.1;Wu1-SARS2_orf1ab;3554 | 0 | 0 |
| NC_045512.2;YP_009724389.1;Wu1-SARS2_orf1ab;3555 | 0 | 0 |
| NC_045512.2;YP_009724389.1;Wu1-SARS2_orf1ab;3556 | 0 | 0 |
| NC_045512.2;YP_009724389.1;Wu1-SARS2_orf1ab;3557 | 0 | 0 |
| NC_045512.2;YP_009724389.1;Wu1-SARS2_orf1ab;3558 | 0 | 0 |
| NC_045512.2;YP_009724389.1;Wu1-SARS2_orf1ab;3559 | 0 | 0 |

| | | |
|---|---|---|
| NC_045512.2;YP_009724389.1;Wu1-SARS2_orf1ab;3560 | 0 | 0 |
| NC_045512.2;YP_009724389.1;Wu1-SARS2_orf1ab;3561 | 0 | 0 |
| NC_045512.2;YP_009724389.1;Wu1-SARS2_orf1ab;3562 | 0 | 0.025 |
| NC_045512.2;YP_009724389.1;Wu1-SARS2_orf1ab;3563 | 0 | 0.025 |
| NC_045512.2;YP_009724389.1;Wu1-SARS2_orf1ab;3564 | 0 | 0 |
| NC_045512.2;YP_009724389.1;Wu1-SARS2_orf1ab;3565 | 0 | 0 |
| NC_045512.2;YP_009724389.1;Wu1-SARS2_orf1ab;3566 | 0 | 0 |
| NC_045512.2;YP_009724389.1;Wu1-SARS2_orf1ab;3567 | 0 | 0 |
| NC_045512.2;YP_009724389.1;Wu1-SARS2_orf1ab;3568 | 0 | 0 |
| NC_045512.2;YP_009724389.1;Wu1-SARS2_orf1ab;3569 | 0 | 0 |
| NC_045512.2;YP_009724389.1;Wu1-SARS2_orf1ab;3570 | 0 | 0 |
| NC_045512.2;YP_009724389.1;Wu1-SARS2_orf1ab;3571 | 0 | 0 |
| NC_045512.2;YP_009724389.1;Wu1-SARS2_orf1ab;3572 | 0 | 0 |
| NC_045512.2;YP_009724389.1;Wu1-SARS2_orf1ab;3573 | 0 | 0 |
| NC_045512.2;YP_009724389.1;Wu1-SARS2_orf1ab;3574 | 0 | 0 |
| NC_045512.2;YP_009724389.1;Wu1-SARS2_orf1ab;3575 | 0 | 0 |
| NC_045512.2;YP_009724389.1;Wu1-SARS2_orf1ab;3576 | 0 | 0 |
| NC_045512.2;YP_009724389.1;Wu1-SARS2_orf1ab;3577 | 0 | 0 |
| NC_045512.2;YP_009724389.1;Wu1-SARS2_orf1ab;3578 | 0 | 0 |
| NC_045512.2;YP_009724389.1;Wu1-SARS2_orf1ab;3579 | 0 | 0 |
| NC_045512.2;YP_009724389.1;Wu1-SARS2_orf1ab;3580 | 0 | 0 |
| NC_045512.2;YP_009724389.1;Wu1-SARS2_orf1ab;3581 | 0 | 0 |
| NC_045512.2;YP_009724389.1;Wu1-SARS2_orf1ab;3582 | 0 | 0 |
| NC_045512.2;YP_009724389.1;Wu1-SARS2_orf1ab;3583 | 0 | 0 |
| NC_045512.2;YP_009724389.1;Wu1-SARS2_orf1ab;3584 | 0 | 0 |
| NC_045512.2;YP_009724389.1;Wu1-SARS2_orf1ab;3585 | 0 | 0 |
| NC_045512.2;YP_009724389.1;Wu1-SARS2_orf1ab;3586 | 0 | 0 |
| NC_045512.2;YP_009724389.1;Wu1-SARS2_orf1ab;3587 | 0 | 0 |
| NC_045512.2;YP_009724389.1;Wu1-SARS2_orf1ab;3588 | 0 | 0 |
| NC_045512.2;YP_009724389.1;Wu1-SARS2_orf1ab;3589 | 0 | 0 |
| NC_045512.2;YP_009724389.1;Wu1-SARS2_orf1ab;3590 | 0 | 0 |
| NC_045512.2;YP_009724389.1;Wu1-SARS2_orf1ab;3591 | 0 | 0 |
| NC_045512.2;YP_009724389.1;Wu1-SARS2_orf1ab;3592 | 0 | 0 |
| NC_045512.2;YP_009724389.1;Wu1-SARS2_orf1ab;3593 | 0 | 0 |
| NC_045512.2;YP_009724389.1;Wu1-SARS2_orf1ab;3594 | 0 | 0 |
| NC_045512.2;YP_009724389.1;Wu1-SARS2_orf1ab;3595 | 0 | 0 |
| NC_045512.2;YP_009724389.1;Wu1-SARS2_orf1ab;3596 | 0 | 0 |
| NC_045512.2;YP_009724389.1;Wu1-SARS2_orf1ab;3597 | 0 | 0 |
| NC_045512.2;YP_009724389.1;Wu1-SARS2_orf1ab;3598 | 0 | 0 |
| NC_045512.2;YP_009724389.1;Wu1-SARS2_orf1ab;3599 | 0 | 0 |
| NC_045512.2;YP_009724389.1;Wu1-SARS2_orf1ab;3600 | 0 | 0 |
| NC_045512.2;YP_009724389.1;Wu1-SARS2_orf1ab;3601 | 0 | 0 |
| NC_045512.2;YP_009724389.1;Wu1-SARS2_orf1ab;3602 | 0 | 0 |

| | | |
|---|---|---|
| NC_045512.2;YP_009724389.1;Wu1-SARS2_orf1ab;3603 | 0 | 0 |
| NC_045512.2;YP_009724389.1;Wu1-SARS2_orf1ab;3604 | 0 | 0 |
| NC_045512.2;YP_009724389.1;Wu1-SARS2_orf1ab;3605 | 0 | 0 |
| NC_045512.2;YP_009724389.1;Wu1-SARS2_orf1ab;3606 | 0 | 0 |
| NC_045512.2;YP_009724389.1;Wu1-SARS2_orf1ab;3607 | 0 | 0 |
| NC_045512.2;YP_009724389.1;Wu1-SARS2_orf1ab;3608 | 0 | 0 |
| NC_045512.2;YP_009724389.1;Wu1-SARS2_orf1ab;3609 | 0 | 0 |
| NC_045512.2;YP_009724389.1;Wu1-SARS2_orf1ab;3610 | 0 | 0 |
| NC_045512.2;YP_009724389.1;Wu1-SARS2_orf1ab;3611 | 0 | 0 |
| NC_045512.2;YP_009724389.1;Wu1-SARS2_orf1ab;3612 | 0 | 0 |
| NC_045512.2;YP_009724389.1;Wu1-SARS2_orf1ab;3613 | 0 | 0 |
| NC_045512.2;YP_009724389.1;Wu1-SARS2_orf1ab;3614 | 0 | 0 |
| NC_045512.2;YP_009724389.1;Wu1-SARS2_orf1ab;3615 | 0 | 0 |
| NC_045512.2;YP_009724389.1;Wu1-SARS2_orf1ab;3616 | 0 | 0 |
| NC_045512.2;YP_009724389.1;Wu1-SARS2_orf1ab;3617 | 0 | 0 |
| NC_045512.2;YP_009724389.1;Wu1-SARS2_orf1ab;3618 | 0 | 0 |
| NC_045512.2;YP_009724389.1;Wu1-SARS2_orf1ab;3619 | 0 | 0 |
| NC_045512.2;YP_009724389.1;Wu1-SARS2_orf1ab;3620 | 0 | 0 |
| NC_045512.2;YP_009724389.1;Wu1-SARS2_orf1ab;3621 | 0 | 0 |
| NC_045512.2;YP_009724389.1;Wu1-SARS2_orf1ab;3622 | 0 | 0 |
| NC_045512.2;YP_009724389.1;Wu1-SARS2_orf1ab;3623 | 0 | 0 |
| NC_045512.2;YP_009724389.1;Wu1-SARS2_orf1ab;3624 | 0 | 0 |
| NC_045512.2;YP_009724389.1;Wu1-SARS2_orf1ab;3625 | 0 | 0 |
| NC_045512.2;YP_009724389.1;Wu1-SARS2_orf1ab;3626 | 0 | 0 |
| NC_045512.2;YP_009724389.1;Wu1-SARS2_orf1ab;3627 | 0 | 0 |
| NC_045512.2;YP_009724389.1;Wu1-SARS2_orf1ab;3628 | 0 | 0 |
| NC_045512.2;YP_009724389.1;Wu1-SARS2_orf1ab;3629 | 0 | 0 |
| NC_045512.2;YP_009724389.1;Wu1-SARS2_orf1ab;3630 | 0 | 0 |
| NC_045512.2;YP_009724389.1;Wu1-SARS2_orf1ab;3631 | 0 | 0 |
| NC_045512.2;YP_009724389.1;Wu1-SARS2_orf1ab;3632 | 0 | 0 |
| NC_045512.2;YP_009724389.1;Wu1-SARS2_orf1ab;3633 | 0 | 0 |
| NC_045512.2;YP_009724389.1;Wu1-SARS2_orf1ab;3634 | 0 | 0 |
| NC_045512.2;YP_009724389.1;Wu1-SARS2_orf1ab;3635 | 0 | 0 |
| NC_045512.2;YP_009724389.1;Wu1-SARS2_orf1ab;3636 | 0 | 0 |
| NC_045512.2;YP_009724389.1;Wu1-SARS2_orf1ab;3637 | 0 | 0 |
| NC_045512.2;YP_009724389.1;Wu1-SARS2_orf1ab;3638 | 0 | 0 |
| NC_045512.2;YP_009724389.1;Wu1-SARS2_orf1ab;3639 | 0 | 0 |
| NC_045512.2;YP_009724389.1;Wu1-SARS2_orf1ab;3640 | 0 | 0 |
| NC_045512.2;YP_009724389.1;Wu1-SARS2_orf1ab;3641 | 0 | 0 |
| NC_045512.2;YP_009724389.1;Wu1-SARS2_orf1ab;3642 | 0 | 0 |
| NC_045512.2;YP_009724389.1;Wu1-SARS2_orf1ab;3643 | 0 | 0 |
| NC_045512.2;YP_009724389.1;Wu1-SARS2_orf1ab;3644 | 0 | 0 |
| NC_045512.2;YP_009724389.1;Wu1-SARS2_orf1ab;3645 | 0 | 0 |

| | | |
|---|---|---|
| NC_045512.2;YP_009724389.1;Wu1-SARS2_orf1ab;3646 | 0 | 0 |
| NC_045512.2;YP_009724389.1;Wu1-SARS2_orf1ab;3647 | 0 | 0 |
| NC_045512.2;YP_009724389.1;Wu1-SARS2_orf1ab;3648 | 0 | 0 |
| NC_045512.2;YP_009724389.1;Wu1-SARS2_orf1ab;3649 | 0 | 0 |
| NC_045512.2;YP_009724389.1;Wu1-SARS2_orf1ab;3650 | 0 | 0 |
| NC_045512.2;YP_009724389.1;Wu1-SARS2_orf1ab;3651 | 0 | 0 |
| NC_045512.2;YP_009724389.1;Wu1-SARS2_orf1ab;3652 | 0 | 0 |
| NC_045512.2;YP_009724389.1;Wu1-SARS2_orf1ab;3653 | 0 | 0 |
| NC_045512.2;YP_009724389.1;Wu1-SARS2_orf1ab;3654 | 0 | 0 |
| NC_045512.2;YP_009724389.1;Wu1-SARS2_orf1ab;3655 | 0 | 0 |
| NC_045512.2;YP_009724389.1;Wu1-SARS2_orf1ab;3656 | 0 | 0 |
| NC_045512.2;YP_009724389.1;Wu1-SARS2_orf1ab;3657 | 0 | 0 |
| NC_045512.2;YP_009724389.1;Wu1-SARS2_orf1ab;3658 | 0 | 0 |
| NC_045512.2;YP_009724389.1;Wu1-SARS2_orf1ab;3659 | 0 | 0 |
| NC_045512.2;YP_009724389.1;Wu1-SARS2_orf1ab;3660 | 0 | 0 |
| NC_045512.2;YP_009724389.1;Wu1-SARS2_orf1ab;3661 | 0 | 0 |
| NC_045512.2;YP_009724389.1;Wu1-SARS2_orf1ab;3662 | 0 | 0 |
| NC_045512.2;YP_009724389.1;Wu1-SARS2_orf1ab;3663 | 0 | 0 |
| NC_045512.2;YP_009724389.1;Wu1-SARS2_orf1ab;3664 | 0 | 0.025 |
| NC_045512.2;YP_009724389.1;Wu1-SARS2_orf1ab;3665 | 0 | 0.025 |
| NC_045512.2;YP_009724389.1;Wu1-SARS2_orf1ab;3666 | 0 | 0.05 |
| NC_045512.2;YP_009724389.1;Wu1-SARS2_orf1ab;3667 | 0 | 0.025 |
| NC_045512.2;YP_009724389.1;Wu1-SARS2_orf1ab;3668 | 0 | 0.025 |
| NC_045512.2;YP_009724389.1;Wu1-SARS2_orf1ab;3669 | 0 | 0.025 |
| NC_045512.2;YP_009724389.1;Wu1-SARS2_orf1ab;3670 | 0 | 0.025 |
| NC_045512.2;YP_009724389.1;Wu1-SARS2_orf1ab;3671 | 0 | 0.025 |
| NC_045512.2;YP_009724389.1;Wu1-SARS2_orf1ab;3672 | 0 | 0.025 |
| NC_045512.2;YP_009724389.1;Wu1-SARS2_orf1ab;3673 | 0 | 0.025 |
| NC_045512.2;YP_009724389.1;Wu1-SARS2_orf1ab;3674 | 0 | 0 |
| NC_045512.2;YP_009724389.1;Wu1-SARS2_orf1ab;3675 | 0 | 0 |
| NC_045512.2;YP_009724389.1;Wu1-SARS2_orf1ab;3676 | 0 | 0 |
| NC_045512.2;YP_009724389.1;Wu1-SARS2_orf1ab;3677 | 0 | 0 |
| NC_045512.2;YP_009724389.1;Wu1-SARS2_orf1ab;3678 | 0 | 0 |
| NC_045512.2;YP_009724389.1;Wu1-SARS2_orf1ab;3679 | 0 | 0 |
| NC_045512.2;YP_009724389.1;Wu1-SARS2_orf1ab;3680 | 0 | 0 |
| NC_045512.2;YP_009724389.1;Wu1-SARS2_orf1ab;3681 | 0 | 0 |
| NC_045512.2;YP_009724389.1;Wu1-SARS2_orf1ab;3682 | 0 | 0 |
| NC_045512.2;YP_009724389.1;Wu1-SARS2_orf1ab;3683 | 0 | 0 |
| NC_045512.2;YP_009724389.1;Wu1-SARS2_orf1ab;3684 | 0 | 0 |
| NC_045512.2;YP_009724389.1;Wu1-SARS2_orf1ab;3685 | 0 | 0 |
| NC_045512.2;YP_009724389.1;Wu1-SARS2_orf1ab;3686 | 0 | 0 |
| NC_045512.2;YP_009724389.1;Wu1-SARS2_orf1ab;3687 | 0 | 0 |
| NC_045512.2;YP_009724389.1;Wu1-SARS2_orf1ab;3688 | 0 | 0 |

| | | |
|---|---|---|
| NC_045512.2;YP_009724389.1;Wu1-SARS2_orf1ab;3689 | 0 | 0 |
| NC_045512.2;YP_009724389.1;Wu1-SARS2_orf1ab;3690 | 0 | 0 |
| NC_045512.2;YP_009724389.1;Wu1-SARS2_orf1ab;3691 | 0 | 0 |
| NC_045512.2;YP_009724389.1;Wu1-SARS2_orf1ab;3692 | 0 | 0 |
| NC_045512.2;YP_009724389.1;Wu1-SARS2_orf1ab;3693 | 0 | 0 |
| NC_045512.2;YP_009724389.1;Wu1-SARS2_orf1ab;3694 | 0 | 0 |
| NC_045512.2;YP_009724389.1;Wu1-SARS2_orf1ab;3695 | 0 | 0 |
| NC_045512.2;YP_009724389.1;Wu1-SARS2_orf1ab;3696 | 0 | 0 |
| NC_045512.2;YP_009724389.1;Wu1-SARS2_orf1ab;3697 | 0 | 0 |
| NC_045512.2;YP_009724389.1;Wu1-SARS2_orf1ab;3698 | 0 | 0.025 |
| NC_045512.2;YP_009724389.1;Wu1-SARS2_orf1ab;3699 | 0 | 0 |
| NC_045512.2;YP_009724389.1;Wu1-SARS2_orf1ab;3700 | 0 | 0 |
| NC_045512.2;YP_009724389.1;Wu1-SARS2_orf1ab;3701 | 0 | 0 |
| NC_045512.2;YP_009724389.1;Wu1-SARS2_orf1ab;3702 | 0 | 0 |
| NC_045512.2;YP_009724389.1;Wu1-SARS2_orf1ab;3703 | 0 | 0 |
| NC_045512.2;YP_009724389.1;Wu1-SARS2_orf1ab;3704 | 0 | 0 |
| NC_045512.2;YP_009724389.1;Wu1-SARS2_orf1ab;3705 | 0 | 0 |
| NC_045512.2;YP_009724389.1;Wu1-SARS2_orf1ab;3706 | 0 | 0 |
| NC_045512.2;YP_009724389.1;Wu1-SARS2_orf1ab;3707 | 0 | 0 |
| NC_045512.2;YP_009724389.1;Wu1-SARS2_orf1ab;3708 | 0 | 0 |
| NC_045512.2;YP_009724389.1;Wu1-SARS2_orf1ab;3709 | 0 | 0 |
| NC_045512.2;YP_009724389.1;Wu1-SARS2_orf1ab;3710 | 0 | 0 |
| NC_045512.2;YP_009724389.1;Wu1-SARS2_orf1ab;3711 | 0 | 0 |
| NC_045512.2;YP_009724389.1;Wu1-SARS2_orf1ab;3712 | 0 | 0 |
| NC_045512.2;YP_009724389.1;Wu1-SARS2_orf1ab;3713 | 0 | 0 |
| NC_045512.2;YP_009724389.1;Wu1-SARS2_orf1ab;3714 | 0 | 0 |
| NC_045512.2;YP_009724389.1;Wu1-SARS2_orf1ab;3715 | 0 | 0 |
| NC_045512.2;YP_009724389.1;Wu1-SARS2_orf1ab;3716 | 0 | 0 |
| NC_045512.2;YP_009724389.1;Wu1-SARS2_orf1ab;3717 | 0 | 0 |
| NC_045512.2;YP_009724389.1;Wu1-SARS2_orf1ab;3718 | 0.05 | 0 |
| NC_045512.2;YP_009724389.1;Wu1-SARS2_orf1ab;3719 | 0.05 | 0 |
| NC_045512.2;YP_009724389.1;Wu1-SARS2_orf1ab;3720 | 0.05 | 0 |
| NC_045512.2;YP_009724389.1;Wu1-SARS2_orf1ab;3721 | 0.05 | 0.025 |
| NC_045512.2;YP_009724389.1;Wu1-SARS2_orf1ab;3722 | 0.05 | 0 |
| NC_045512.2;YP_009724389.1;Wu1-SARS2_orf1ab;3723 | 0 | 0 |
| NC_045512.2;YP_009724389.1;Wu1-SARS2_orf1ab;3724 | 0 | 0 |
| NC_045512.2;YP_009724389.1;Wu1-SARS2_orf1ab;3725 | 0 | 0 |
| NC_045512.2;YP_009724389.1;Wu1-SARS2_orf1ab;3726 | 0 | 0 |
| NC_045512.2;YP_009724389.1;Wu1-SARS2_orf1ab;3727 | 0 | 0 |
| NC_045512.2;YP_009724389.1;Wu1-SARS2_orf1ab;3728 | 0 | 0 |
| NC_045512.2;YP_009724389.1;Wu1-SARS2_orf1ab;3729 | 0 | 0 |
| NC_045512.2;YP_009724389.1;Wu1-SARS2_orf1ab;3730 | 0 | 0 |
| NC_045512.2;YP_009724389.1;Wu1-SARS2_orf1ab;3731 | 0 | 0 |

| | | |
|---|---|---|
| NC_045512.2;YP_009724389.1;Wu1-SARS2_orf1ab;3732 | 0 | 0 |
| NC_045512.2;YP_009724389.1;Wu1-SARS2_orf1ab;3733 | 0 | 0 |
| NC_045512.2;YP_009724389.1;Wu1-SARS2_orf1ab;3734 | 0 | 0 |
| NC_045512.2;YP_009724389.1;Wu1-SARS2_orf1ab;3735 | 0 | 0 |
| NC_045512.2;YP_009724389.1;Wu1-SARS2_orf1ab;3736 | 0 | 0 |
| NC_045512.2;YP_009724389.1;Wu1-SARS2_orf1ab;3737 | 0 | 0 |
| NC_045512.2;YP_009724389.1;Wu1-SARS2_orf1ab;3738 | 0 | 0 |
| NC_045512.2;YP_009724389.1;Wu1-SARS2_orf1ab;3739 | 0 | 0 |
| NC_045512.2;YP_009724389.1;Wu1-SARS2_orf1ab;3740 | 0 | 0 |
| NC_045512.2;YP_009724389.1;Wu1-SARS2_orf1ab;3741 | 0 | 0 |
| NC_045512.2;YP_009724389.1;Wu1-SARS2_orf1ab;3742 | 0 | 0 |
| NC_045512.2;YP_009724389.1;Wu1-SARS2_orf1ab;3743 | 0 | 0 |
| NC_045512.2;YP_009724389.1;Wu1-SARS2_orf1ab;3744 | 0 | 0 |
| NC_045512.2;YP_009724389.1;Wu1-SARS2_orf1ab;3745 | 0 | 0 |
| NC_045512.2;YP_009724389.1;Wu1-SARS2_orf1ab;3746 | 0 | 0 |
| NC_045512.2;YP_009724389.1;Wu1-SARS2_orf1ab;3747 | 0 | 0 |
| NC_045512.2;YP_009724389.1;Wu1-SARS2_orf1ab;3748 | 0 | 0 |
| NC_045512.2;YP_009724389.1;Wu1-SARS2_orf1ab;3749 | 0 | 0 |
| NC_045512.2;YP_009724389.1;Wu1-SARS2_orf1ab;3750 | 0 | 0 |
| NC_045512.2;YP_009724389.1;Wu1-SARS2_orf1ab;3751 | 0 | 0 |
| NC_045512.2;YP_009724389.1;Wu1-SARS2_orf1ab;3752 | 0 | 0 |
| NC_045512.2;YP_009724389.1;Wu1-SARS2_orf1ab;3753 | 0 | 0 |
| NC_045512.2;YP_009724389.1;Wu1-SARS2_orf1ab;3754 | 0 | 0 |
| NC_045512.2;YP_009724389.1;Wu1-SARS2_orf1ab;3755 | 0 | 0 |
| NC_045512.2;YP_009724389.1;Wu1-SARS2_orf1ab;3756 | 0 | 0 |
| NC_045512.2;YP_009724389.1;Wu1-SARS2_orf1ab;3757 | 0 | 0 |
| NC_045512.2;YP_009724389.1;Wu1-SARS2_orf1ab;3758 | 0 | 0 |
| NC_045512.2;YP_009724389.1;Wu1-SARS2_orf1ab;3759 | 0 | 0 |
| NC_045512.2;YP_009724389.1;Wu1-SARS2_orf1ab;3760 | 0 | 0 |
| NC_045512.2;YP_009724389.1;Wu1-SARS2_orf1ab;3761 | 0 | 0 |
| NC_045512.2;YP_009724389.1;Wu1-SARS2_orf1ab;3762 | 0 | 0 |
| NC_045512.2;YP_009724389.1;Wu1-SARS2_orf1ab;3763 | 0 | 0 |
| NC_045512.2;YP_009724389.1;Wu1-SARS2_orf1ab;3764 | 0 | 0 |
| NC_045512.2;YP_009724389.1;Wu1-SARS2_orf1ab;3765 | 0 | 0 |
| NC_045512.2;YP_009724389.1;Wu1-SARS2_orf1ab;3766 | 0 | 0 |
| NC_045512.2;YP_009724389.1;Wu1-SARS2_orf1ab;3767 | 0 | 0 |
| NC_045512.2;YP_009724389.1;Wu1-SARS2_orf1ab;3768 | 0 | 0 |
| NC_045512.2;YP_009724389.1;Wu1-SARS2_orf1ab;3769 | 0 | 0 |
| NC_045512.2;YP_009724389.1;Wu1-SARS2_orf1ab;3770 | 0 | 0 |
| NC_045512.2;YP_009724389.1;Wu1-SARS2_orf1ab;3771 | 0 | 0 |
| NC_045512.2;YP_009724389.1;Wu1-SARS2_orf1ab;3772 | 0 | 0 |
| NC_045512.2;YP_009724389.1;Wu1-SARS2_orf1ab;3773 | 0 | 0 |
| NC_045512.2;YP_009724389.1;Wu1-SARS2_orf1ab;3774 | 0 | 0 |

| | | |
|---|---|---|
| NC_045512.2;YP_009724389.1;Wu1-SARS2_orf1ab;3775 | 0 | 0 |
| NC_045512.2;YP_009724389.1;Wu1-SARS2_orf1ab;3776 | 0 | 0 |
| NC_045512.2;YP_009724389.1;Wu1-SARS2_orf1ab;3777 | 0 | 0 |
| NC_045512.2;YP_009724389.1;Wu1-SARS2_orf1ab;3778 | 0 | 0 |
| NC_045512.2;YP_009724389.1;Wu1-SARS2_orf1ab;3779 | 0 | 0 |
| NC_045512.2;YP_009724389.1;Wu1-SARS2_orf1ab;3780 | 0 | 0 |
| NC_045512.2;YP_009724389.1;Wu1-SARS2_orf1ab;3781 | 0 | 0 |
| NC_045512.2;YP_009724389.1;Wu1-SARS2_orf1ab;3782 | 0 | 0 |
| NC_045512.2;YP_009724389.1;Wu1-SARS2_orf1ab;3783 | 0 | 0 |
| NC_045512.2;YP_009724389.1;Wu1-SARS2_orf1ab;3784 | 0 | 0 |
| NC_045512.2;YP_009724389.1;Wu1-SARS2_orf1ab;3785 | 0 | 0 |
| NC_045512.2;YP_009724389.1;Wu1-SARS2_orf1ab;3786 | 0 | 0 |
| NC_045512.2;YP_009724389.1;Wu1-SARS2_orf1ab;3787 | 0 | 0 |
| NC_045512.2;YP_009724389.1;Wu1-SARS2_orf1ab;3788 | 0 | 0 |
| NC_045512.2;YP_009724389.1;Wu1-SARS2_orf1ab;3789 | 0 | 0 |
| NC_045512.2;YP_009724389.1;Wu1-SARS2_orf1ab;3790 | 0 | 0 |
| NC_045512.2;YP_009724389.1;Wu1-SARS2_orf1ab;3791 | 0 | 0 |
| NC_045512.2;YP_009724389.1;Wu1-SARS2_orf1ab;3792 | 0 | 0 |
| NC_045512.2;YP_009724389.1;Wu1-SARS2_orf1ab;3793 | 0 | 0 |
| NC_045512.2;YP_009724389.1;Wu1-SARS2_orf1ab;3794 | 0 | 0 |
| NC_045512.2;YP_009724389.1;Wu1-SARS2_orf1ab;3795 | 0 | 0 |
| NC_045512.2;YP_009724389.1;Wu1-SARS2_orf1ab;3796 | 0 | 0 |
| NC_045512.2;YP_009724389.1;Wu1-SARS2_orf1ab;3797 | 0 | 0 |
| NC_045512.2;YP_009724389.1;Wu1-SARS2_orf1ab;3798 | 0 | 0 |
| NC_045512.2;YP_009724389.1;Wu1-SARS2_orf1ab;3799 | 0 | 0 |
| NC_045512.2;YP_009724389.1;Wu1-SARS2_orf1ab;3800 | 0 | 0 |
| NC_045512.2;YP_009724389.1;Wu1-SARS2_orf1ab;3801 | 0 | 0 |
| NC_045512.2;YP_009724389.1;Wu1-SARS2_orf1ab;3802 | 0 | 0 |
| NC_045512.2;YP_009724389.1;Wu1-SARS2_orf1ab;3803 | 0 | 0 |
| NC_045512.2;YP_009724389.1;Wu1-SARS2_orf1ab;3804 | 0 | 0 |
| NC_045512.2;YP_009724389.1;Wu1-SARS2_orf1ab;3805 | 0 | 0 |
| NC_045512.2;YP_009724389.1;Wu1-SARS2_orf1ab;3806 | 0 | 0 |
| NC_045512.2;YP_009724389.1;Wu1-SARS2_orf1ab;3807 | 0 | 0 |
| NC_045512.2;YP_009724389.1;Wu1-SARS2_orf1ab;3808 | 0 | 0 |
| NC_045512.2;YP_009724389.1;Wu1-SARS2_orf1ab;3809 | 0.25 | 0.175 |
| NC_045512.2;YP_009724389.1;Wu1-SARS2_orf1ab;3810 | 0 | 0 |
| NC_045512.2;YP_009724389.1;Wu1-SARS2_orf1ab;3811 | 0 | 0 |
| NC_045512.2;YP_009724389.1;Wu1-SARS2_orf1ab;3812 | 0 | 0 |
| NC_045512.2;YP_009724389.1;Wu1-SARS2_orf1ab;3813 | 0 | 0 |
| NC_045512.2;YP_009724389.1;Wu1-SARS2_orf1ab;3814 | 0 | 0 |
| NC_045512.2;YP_009724389.1;Wu1-SARS2_orf1ab;3815 | 0 | 0 |
| NC_045512.2;YP_009724389.1;Wu1-SARS2_orf1ab;3816 | 0 | 0 |
| NC_045512.2;YP_009724389.1;Wu1-SARS2_orf1ab;3817 | 0 | 0.05 |

| | | |
|---|---|---|
| NC_045512.2;YP_009724389.1;Wu1-SARS2_orf1ab;3818 | 0.05 | 0.025 |
| NC_045512.2;YP_009724389.1;Wu1-SARS2_orf1ab;3819 | 0.15 | 0.075 |
| NC_045512.2;YP_009724389.1;Wu1-SARS2_orf1ab;3820 | 0 | 0.025 |
| NC_045512.2;YP_009724389.1;Wu1-SARS2_orf1ab;3821 | 0 | 0.025 |
| NC_045512.2;YP_009724389.1;Wu1-SARS2_orf1ab;3822 | 0.05 | 0.125 |
| NC_045512.2;YP_009724389.1;Wu1-SARS2_orf1ab;3823 | 0.1 | 0.05 |
| NC_045512.2;YP_009724389.1;Wu1-SARS2_orf1ab;3824 | 0.3 | 0.1 |
| NC_045512.2;YP_009724389.1;Wu1-SARS2_orf1ab;3825 | 0.25 | 0.1 |
| NC_045512.2;YP_009724389.1;Wu1-SARS2_orf1ab;3826 | 0.25 | 0.1 |
| NC_045512.2;YP_009724389.1;Wu1-SARS2_orf1ab;3827 | 0.2 | 0.1 |
| NC_045512.2;YP_009724389.1;Wu1-SARS2_orf1ab;3828 | 0.15 | 0.075 |
| NC_045512.2;YP_009724389.1;Wu1-SARS2_orf1ab;3829 | 0.1 | 0.075 |
| NC_045512.2;YP_009724389.1;Wu1-SARS2_orf1ab;3830 | 0.05 | 0.075 |
| NC_045512.2;YP_009724389.1;Wu1-SARS2_orf1ab;3831 | 0.05 | 0.05 |
| NC_045512.2;YP_009724389.1;Wu1-SARS2_orf1ab;3832 | 0 | 0.05 |
| NC_045512.2;YP_009724389.1;Wu1-SARS2_orf1ab;3833 | 0 | 0 |
| NC_045512.2;YP_009724389.1;Wu1-SARS2_orf1ab;3834 | 0 | 0 |
| NC_045512.2;YP_009724389.1;Wu1-SARS2_orf1ab;3835 | 0 | 0 |
| NC_045512.2;YP_009724389.1;Wu1-SARS2_orf1ab;3836 | 0 | 0 |
| NC_045512.2;YP_009724389.1;Wu1-SARS2_orf1ab;3837 | 0 | 0 |
| NC_045512.2;YP_009724389.1;Wu1-SARS2_orf1ab;3838 | 0 | 0 |
| NC_045512.2;YP_009724389.1;Wu1-SARS2_orf1ab;3839 | 0 | 0 |
| NC_045512.2;YP_009724389.1;Wu1-SARS2_orf1ab;3840 | 0 | 0 |
| NC_045512.2;YP_009724389.1;Wu1-SARS2_orf1ab;3841 | 0 | 0 |
| NC_045512.2;YP_009724389.1;Wu1-SARS2_orf1ab;3842 | 0 | 0.025 |
| NC_045512.2;YP_009724389.1;Wu1-SARS2_orf1ab;3843 | 0 | 0 |
| NC_045512.2;YP_009724389.1;Wu1-SARS2_orf1ab;3844 | 0 | 0.025 |
| NC_045512.2;YP_009724389.1;Wu1-SARS2_orf1ab;3845 | 0 | 0.025 |
| NC_045512.2;YP_009724389.1;Wu1-SARS2_orf1ab;3846 | 0.05 | 0.05 |
| NC_045512.2;YP_009724389.1;Wu1-SARS2_orf1ab;3847 | 0.05 | 0.05 |
| NC_045512.2;YP_009724389.1;Wu1-SARS2_orf1ab;3848 | 0.05 | 0.05 |
| NC_045512.2;YP_009724389.1;Wu1-SARS2_orf1ab;3849 | 0.1 | 0.1 |
| NC_045512.2;YP_009724389.1;Wu1-SARS2_orf1ab;3850 | 0.05 | 0.075 |
| NC_045512.2;YP_009724389.1;Wu1-SARS2_orf1ab;3851 | 0.05 | 0.05 |
| NC_045512.2;YP_009724389.1;Wu1-SARS2_orf1ab;3852 | 0.05 | 0.025 |
| NC_045512.2;YP_009724389.1;Wu1-SARS2_orf1ab;3853 | 0 | 0.025 |
| NC_045512.2;YP_009724389.1;Wu1-SARS2_orf1ab;3854 | 0.05 | 0.025 |
| NC_045512.2;YP_009724389.1;Wu1-SARS2_orf1ab;3855 | 0.05 | 0.025 |
| NC_045512.2;YP_009724389.1;Wu1-SARS2_orf1ab;3856 | 0.05 | 0.025 |
| NC_045512.2;YP_009724389.1;Wu1-SARS2_orf1ab;3857 | 0.05 | 0 |
| NC_045512.2;YP_009724389.1;Wu1-SARS2_orf1ab;3858 | 0 | 0 |
| NC_045512.2;YP_009724389.1;Wu1-SARS2_orf1ab;3859 | 0 | 0 |
| NC_045512.2;YP_009724389.1;Wu1-SARS2_orf1ab;3860 | 0 | 0 |

| | | |
|---|---|---|
| NC_045512.2;YP_009724389.1;Wu1-SARS2_orf1ab;3861 | 0 | 0 |
| NC_045512.2;YP_009724389.1;Wu1-SARS2_orf1ab;3862 | 0 | 0 |
| NC_045512.2;YP_009724389.1;Wu1-SARS2_orf1ab;3863 | 0 | 0 |
| NC_045512.2;YP_009724389.1;Wu1-SARS2_orf1ab;3864 | 0 | 0 |
| NC_045512.2;YP_009724389.1;Wu1-SARS2_orf1ab;3865 | 0 | 0 |
| NC_045512.2;YP_009724389.1;Wu1-SARS2_orf1ab;3866 | 0 | 0 |
| NC_045512.2;YP_009724389.1;Wu1-SARS2_orf1ab;3867 | 0 | 0 |
| NC_045512.2;YP_009724389.1;Wu1-SARS2_orf1ab;3868 | 0 | 0 |
| NC_045512.2;YP_009724389.1;Wu1-SARS2_orf1ab;3869 | 0 | 0 |
| NC_045512.2;YP_009724389.1;Wu1-SARS2_orf1ab;3870 | 0 | 0 |
| NC_045512.2;YP_009724389.1;Wu1-SARS2_orf1ab;3871 | 0 | 0 |
| NC_045512.2;YP_009724389.1;Wu1-SARS2_orf1ab;3872 | 0 | 0 |
| NC_045512.2;YP_009724389.1;Wu1-SARS2_orf1ab;3873 | 0 | 0 |
| NC_045512.2;YP_009724389.1;Wu1-SARS2_orf1ab;3874 | 0 | 0 |
| NC_045512.2;YP_009724389.1;Wu1-SARS2_orf1ab;3875 | 0 | 0 |
| NC_045512.2;YP_009724389.1;Wu1-SARS2_orf1ab;3876 | 0 | 0 |
| NC_045512.2;YP_009724389.1;Wu1-SARS2_orf1ab;3877 | 0 | 0 |
| NC_045512.2;YP_009724389.1;Wu1-SARS2_orf1ab;3878 | 0 | 0 |
| NC_045512.2;YP_009724389.1;Wu1-SARS2_orf1ab;3879 | 0 | 0 |
| NC_045512.2;YP_009724389.1;Wu1-SARS2_orf1ab;3880 | 0 | 0 |
| NC_045512.2;YP_009724389.1;Wu1-SARS2_orf1ab;3881 | 0 | 0 |
| NC_045512.2;YP_009724389.1;Wu1-SARS2_orf1ab;3882 | 0 | 0.025 |
| NC_045512.2;YP_009724389.1;Wu1-SARS2_orf1ab;3883 | 0 | 0 |
| NC_045512.2;YP_009724389.1;Wu1-SARS2_orf1ab;3884 | 0 | 0.025 |
| NC_045512.2;YP_009724389.1;Wu1-SARS2_orf1ab;3885 | 0 | 0 |
| NC_045512.2;YP_009724389.1;Wu1-SARS2_orf1ab;3886 | 0 | 0 |
| NC_045512.2;YP_009724389.1;Wu1-SARS2_orf1ab;3887 | 0 | 0 |
| NC_045512.2;YP_009724389.1;Wu1-SARS2_orf1ab;3888 | 0.05 | 0 |
| NC_045512.2;YP_009724389.1;Wu1-SARS2_orf1ab;3889 | 0.05 | 0 |
| NC_045512.2;YP_009724389.1;Wu1-SARS2_orf1ab;3890 | 0.05 | 0 |
| NC_045512.2;YP_009724389.1;Wu1-SARS2_orf1ab;3891 | 0 | 0 |
| NC_045512.2;YP_009724389.1;Wu1-SARS2_orf1ab;3892 | 0 | 0 |
| NC_045512.2;YP_009724389.1;Wu1-SARS2_orf1ab;3893 | 0 | 0 |
| NC_045512.2;YP_009724389.1;Wu1-SARS2_orf1ab;3894 | 0 | 0 |
| NC_045512.2;YP_009724389.1;Wu1-SARS2_orf1ab;3895 | 0 | 0 |
| NC_045512.2;YP_009724389.1;Wu1-SARS2_orf1ab;3896 | 0 | 0 |
| NC_045512.2;YP_009724389.1;Wu1-SARS2_orf1ab;3897 | 0 | 0 |
| NC_045512.2;YP_009724389.1;Wu1-SARS2_orf1ab;3898 | 0 | 0 |
| NC_045512.2;YP_009724389.1;Wu1-SARS2_orf1ab;3899 | 0 | 0 |
| NC_045512.2;YP_009724389.1;Wu1-SARS2_orf1ab;3900 | 0 | 0 |
| NC_045512.2;YP_009724389.1;Wu1-SARS2_orf1ab;3901 | 0 | 0 |
| NC_045512.2;YP_009724389.1;Wu1-SARS2_orf1ab;3902 | 0 | 0 |
| NC_045512.2;YP_009724389.1;Wu1-SARS2_orf1ab;3903 | 0 | 0 |

| | | |
|---|---|---|
| NC_045512.2;YP_009724389.1;Wu1-SARS2_orf1ab;3904 | 0 | 0 |
| NC_045512.2;YP_009724389.1;Wu1-SARS2_orf1ab;3905 | 0 | 0 |
| NC_045512.2;YP_009724389.1;Wu1-SARS2_orf1ab;3906 | 0 | 0 |
| NC_045512.2;YP_009724389.1;Wu1-SARS2_orf1ab;3907 | 0 | 0 |
| NC_045512.2;YP_009724389.1;Wu1-SARS2_orf1ab;3908 | 0 | 0 |
| NC_045512.2;YP_009724389.1;Wu1-SARS2_orf1ab;3909 | 0 | 0 |
| NC_045512.2;YP_009724389.1;Wu1-SARS2_orf1ab;3910 | 0 | 0 |
| NC_045512.2;YP_009724389.1;Wu1-SARS2_orf1ab;3911 | 0 | 0 |
| NC_045512.2;YP_009724389.1;Wu1-SARS2_orf1ab;3912 | 0 | 0 |
| NC_045512.2;YP_009724389.1;Wu1-SARS2_orf1ab;3913 | 0 | 0 |
| NC_045512.2;YP_009724389.1;Wu1-SARS2_orf1ab;3914 | 0 | 0.05 |
| NC_045512.2;YP_009724389.1;Wu1-SARS2_orf1ab;3915 | 0 | 0.025 |
| NC_045512.2;YP_009724389.1;Wu1-SARS2_orf1ab;3916 | 0 | 0.025 |
| NC_045512.2;YP_009724389.1;Wu1-SARS2_orf1ab;3917 | 0 | 0.025 |
| NC_045512.2;YP_009724389.1;Wu1-SARS2_orf1ab;3918 | 0 | 0.025 |
| NC_045512.2;YP_009724389.1;Wu1-SARS2_orf1ab;3919 | 0.05 | 0.05 |
| NC_045512.2;YP_009724389.1;Wu1-SARS2_orf1ab;3920 | 0.05 | 0.05 |
| NC_045512.2;YP_009724389.1;Wu1-SARS2_orf1ab;3921 | 0.05 | 0.075 |
| NC_045512.2;YP_009724389.1;Wu1-SARS2_orf1ab;3922 | 0.05 | 0.05 |
| NC_045512.2;YP_009724389.1;Wu1-SARS2_orf1ab;3923 | 0.1 | 0.025 |
| NC_045512.2;YP_009724389.1;Wu1-SARS2_orf1ab;3924 | 0 | 0.025 |
| NC_045512.2;YP_009724389.1;Wu1-SARS2_orf1ab;3925 | 0 | 0.05 |
| NC_045512.2;YP_009724389.1;Wu1-SARS2_orf1ab;3926 | 0 | 0.025 |
| NC_045512.2;YP_009724389.1;Wu1-SARS2_orf1ab;3927 | 0 | 0 |
| NC_045512.2;YP_009724389.1;Wu1-SARS2_orf1ab;3928 | 0 | 0 |
| NC_045512.2;YP_009724389.1;Wu1-SARS2_orf1ab;3929 | 0.05 | 0 |
| NC_045512.2;YP_009724389.1;Wu1-SARS2_orf1ab;3930 | 0.05 | 0 |
| NC_045512.2;YP_009724389.1;Wu1-SARS2_orf1ab;3931 | 0.05 | 0 |
| NC_045512.2;YP_009724389.1;Wu1-SARS2_orf1ab;3932 | 0 | 0 |
| NC_045512.2;YP_009724389.1;Wu1-SARS2_orf1ab;3933 | 0.05 | 0 |
| NC_045512.2;YP_009724389.1;Wu1-SARS2_orf1ab;3934 | 0 | 0 |
| NC_045512.2;YP_009724389.1;Wu1-SARS2_orf1ab;3935 | 0 | 0 |
| NC_045512.2;YP_009724389.1;Wu1-SARS2_orf1ab;3936 | 0 | 0 |
| NC_045512.2;YP_009724389.1;Wu1-SARS2_orf1ab;3937 | 0 | 0 |
| NC_045512.2;YP_009724389.1;Wu1-SARS2_orf1ab;3938 | 0 | 0 |
| NC_045512.2;YP_009724389.1;Wu1-SARS2_orf1ab;3939 | 0 | 0 |
| NC_045512.2;YP_009724389.1;Wu1-SARS2_orf1ab;3940 | 0 | 0 |
| NC_045512.2;YP_009724389.1;Wu1-SARS2_orf1ab;3941 | 0.05 | 0.05 |
| NC_045512.2;YP_009724389.1;Wu1-SARS2_orf1ab;3942 | 0 | 0.025 |
| NC_045512.2;YP_009724389.1;Wu1-SARS2_orf1ab;3943 | 0 | 0 |
| NC_045512.2;YP_009724389.1;Wu1-SARS2_orf1ab;3944 | 0 | 0 |
| NC_045512.2;YP_009724389.1;Wu1-SARS2_orf1ab;3945 | 0 | 0 |
| NC_045512.2;YP_009724389.1;Wu1-SARS2_orf1ab;3946 | 0 | 0.05 |

| | | |
|---|---|---|
| NC_045512.2;YP_009724389.1;Wu1-SARS2_orf1ab;3947 | 0.05 | 0 |
| NC_045512.2;YP_009724389.1;Wu1-SARS2_orf1ab;3948 | 0.05 | 0 |
| NC_045512.2;YP_009724389.1;Wu1-SARS2_orf1ab;3949 | 0.05 | 0 |
| NC_045512.2;YP_009724389.1;Wu1-SARS2_orf1ab;3950 | 0.05 | 0.05 |
| NC_045512.2;YP_009724389.1;Wu1-SARS2_orf1ab;3951 | 0.1 | 0.05 |
| NC_045512.2;YP_009724389.1;Wu1-SARS2_orf1ab;3952 | 0 | 0.025 |
| NC_045512.2;YP_009724389.1;Wu1-SARS2_orf1ab;3953 | 0.1 | 0.05 |
| NC_045512.2;YP_009724389.1;Wu1-SARS2_orf1ab;3954 | 0.05 | 0.125 |
| NC_045512.2;YP_009724389.1;Wu1-SARS2_orf1ab;3955 | 0.05 | 0.075 |
| NC_045512.2;YP_009724389.1;Wu1-SARS2_orf1ab;3956 | 0.05 | 0.15 |
| NC_045512.2;YP_009724389.1;Wu1-SARS2_orf1ab;3957 | 0.05 | 0.05 |
| NC_045512.2;YP_009724389.1;Wu1-SARS2_orf1ab;3958 | 0 | 0.025 |
| NC_045512.2;YP_009724389.1;Wu1-SARS2_orf1ab;3959 | 0 | 0 |
| NC_045512.2;YP_009724389.1;Wu1-SARS2_orf1ab;3960 | 0.05 | 0 |
| NC_045512.2;YP_009724389.1;Wu1-SARS2_orf1ab;3961 | 0 | 0 |
| NC_045512.2;YP_009724389.1;Wu1-SARS2_orf1ab;3962 | 0 | 0 |
| NC_045512.2;YP_009724389.1;Wu1-SARS2_orf1ab;3963 | 0 | 0 |
| NC_045512.2;YP_009724389.1;Wu1-SARS2_orf1ab;3964 | 0 | 0 |
| NC_045512.2;YP_009724389.1;Wu1-SARS2_orf1ab;3965 | 0 | 0 |
| NC_045512.2;YP_009724389.1;Wu1-SARS2_orf1ab;3966 | 0 | 0 |
| NC_045512.2;YP_009724389.1;Wu1-SARS2_orf1ab;3967 | 0 | 0 |
| NC_045512.2;YP_009724389.1;Wu1-SARS2_orf1ab;3968 | 0 | 0 |
| NC_045512.2;YP_009724389.1;Wu1-SARS2_orf1ab;3969 | 0 | 0 |
| NC_045512.2;YP_009724389.1;Wu1-SARS2_orf1ab;3970 | 0 | 0 |
| NC_045512.2;YP_009724389.1;Wu1-SARS2_orf1ab;3971 | 0 | 0 |
| NC_045512.2;YP_009724389.1;Wu1-SARS2_orf1ab;3972 | 0 | 0 |
| NC_045512.2;YP_009724389.1;Wu1-SARS2_orf1ab;3973 | 0 | 0 |
| NC_045512.2;YP_009724389.1;Wu1-SARS2_orf1ab;3974 | 0 | 0 |
| NC_045512.2;YP_009724389.1;Wu1-SARS2_orf1ab;3975 | 0 | 0 |
| NC_045512.2;YP_009724389.1;Wu1-SARS2_orf1ab;3976 | 0 | 0 |
| NC_045512.2;YP_009724389.1;Wu1-SARS2_orf1ab;3977 | 0 | 0.025 |
| NC_045512.2;YP_009724389.1;Wu1-SARS2_orf1ab;3978 | 0 | 0.05 |
| NC_045512.2;YP_009724389.1;Wu1-SARS2_orf1ab;3979 | 0 | 0.025 |
| NC_045512.2;YP_009724389.1;Wu1-SARS2_orf1ab;3980 | 0.05 | 0.025 |
| NC_045512.2;YP_009724389.1;Wu1-SARS2_orf1ab;3981 | 0.05 | 0.025 |
| NC_045512.2;YP_009724389.1;Wu1-SARS2_orf1ab;3982 | 0 | 0.05 |
| NC_045512.2;YP_009724389.1;Wu1-SARS2_orf1ab;3983 | 0.05 | 0.025 |
| NC_045512.2;YP_009724389.1;Wu1-SARS2_orf1ab;3984 | 0 | 0.025 |
| NC_045512.2;YP_009724389.1;Wu1-SARS2_orf1ab;3985 | 0 | 0 |
| NC_045512.2;YP_009724389.1;Wu1-SARS2_orf1ab;3986 | 0 | 0 |
| NC_045512.2;YP_009724389.1;Wu1-SARS2_orf1ab;3987 | 0 | 0 |
| NC_045512.2;YP_009724389.1;Wu1-SARS2_orf1ab;3988 | 0 | 0 |
| NC_045512.2;YP_009724389.1;Wu1-SARS2_orf1ab;3989 | 0 | 0 |

| | | |
|---|---|---|
| NC_045512.2;YP_009724389.1;Wu1-SARS2_orf1ab;3990 | 0 | 0 |
| NC_045512.2;YP_009724389.1;Wu1-SARS2_orf1ab;3991 | 0 | 0 |
| NC_045512.2;YP_009724389.1;Wu1-SARS2_orf1ab;3992 | 0 | 0 |
| NC_045512.2;YP_009724389.1;Wu1-SARS2_orf1ab;3993 | 0 | 0 |
| NC_045512.2;YP_009724389.1;Wu1-SARS2_orf1ab;3994 | 0 | 0.025 |
| NC_045512.2;YP_009724389.1;Wu1-SARS2_orf1ab;3995 | 0 | 0 |
| NC_045512.2;YP_009724389.1;Wu1-SARS2_orf1ab;3996 | 0 | 0 |
| NC_045512.2;YP_009724389.1;Wu1-SARS2_orf1ab;3997 | 0 | 0 |
| NC_045512.2;YP_009724389.1;Wu1-SARS2_orf1ab;3998 | 0 | 0.05 |
| NC_045512.2;YP_009724389.1;Wu1-SARS2_orf1ab;3999 | 0 | 0.025 |
| NC_045512.2;YP_009724389.1;Wu1-SARS2_orf1ab;4000 | 0 | 0 |
| NC_045512.2;YP_009724389.1;Wu1-SARS2_orf1ab;4001 | 0 | 0 |
| NC_045512.2;YP_009724389.1;Wu1-SARS2_orf1ab;4002 | 0 | 0 |
| NC_045512.2;YP_009724389.1;Wu1-SARS2_orf1ab;4003 | 0 | 0 |
| NC_045512.2;YP_009724389.1;Wu1-SARS2_orf1ab;4004 | 0 | 0 |
| NC_045512.2;YP_009724389.1;Wu1-SARS2_orf1ab;4005 | 0 | 0 |
| NC_045512.2;YP_009724389.1;Wu1-SARS2_orf1ab;4006 | 0 | 0.1 |
| NC_045512.2;YP_009724389.1;Wu1-SARS2_orf1ab;4007 | 0 | 0.075 |
| NC_045512.2;YP_009724389.1;Wu1-SARS2_orf1ab;4008 | 0 | 0.05 |
| NC_045512.2;YP_009724389.1;Wu1-SARS2_orf1ab;4009 | 0 | 0.05 |
| NC_045512.2;YP_009724389.1;Wu1-SARS2_orf1ab;4010 | 0 | 0.05 |
| NC_045512.2;YP_009724389.1;Wu1-SARS2_orf1ab;4011 | 0 | 0 |
| NC_045512.2;YP_009724389.1;Wu1-SARS2_orf1ab;4012 | 0 | 0 |
| NC_045512.2;YP_009724389.1;Wu1-SARS2_orf1ab;4013 | 0 | 0 |
| NC_045512.2;YP_009724389.1;Wu1-SARS2_orf1ab;4014 | 0 | 0 |
| NC_045512.2;YP_009724389.1;Wu1-SARS2_orf1ab;4015 | 0 | 0 |
| NC_045512.2;YP_009724389.1;Wu1-SARS2_orf1ab;4016 | 0 | 0 |
| NC_045512.2;YP_009724389.1;Wu1-SARS2_orf1ab;4017 | 0 | 0 |
| NC_045512.2;YP_009724389.1;Wu1-SARS2_orf1ab;4018 | 0 | 0 |
| NC_045512.2;YP_009724389.1;Wu1-SARS2_orf1ab;4019 | 0 | 0 |
| NC_045512.2;YP_009724389.1;Wu1-SARS2_orf1ab;4020 | 0 | 0 |
| NC_045512.2;YP_009724389.1;Wu1-SARS2_orf1ab;4021 | 0 | 0 |
| NC_045512.2;YP_009724389.1;Wu1-SARS2_orf1ab;4022 | 0 | 0 |
| NC_045512.2;YP_009724389.1;Wu1-SARS2_orf1ab;4023 | 0 | 0 |
| NC_045512.2;YP_009724389.1;Wu1-SARS2_orf1ab;4024 | 0 | 0 |
| NC_045512.2;YP_009724389.1;Wu1-SARS2_orf1ab;4025 | 0 | 0 |
| NC_045512.2;YP_009724389.1;Wu1-SARS2_orf1ab;4026 | 0 | 0 |
| NC_045512.2;YP_009724389.1;Wu1-SARS2_orf1ab;4027 | 0 | 0 |
| NC_045512.2;YP_009724389.1;Wu1-SARS2_orf1ab;4028 | 0 | 0 |
| NC_045512.2;YP_009724389.1;Wu1-SARS2_orf1ab;4029 | 0 | 0 |
| NC_045512.2;YP_009724389.1;Wu1-SARS2_orf1ab;4030 | 0 | 0 |
| NC_045512.2;YP_009724389.1;Wu1-SARS2_orf1ab;4031 | 0.15 | 0.1 |
| NC_045512.2;YP_009724389.1;Wu1-SARS2_orf1ab;4032 | 0.2 | 0.1 |

| | | |
|---|---|---|
| NC_045512.2;YP_009724389.1;Wu1-SARS2_orf1ab;4033 | 0.2 | 0.075 |
| NC_045512.2;YP_009724389.1;Wu1-SARS2_orf1ab;4034 | 0.25 | 0.1 |
| NC_045512.2;YP_009724389.1;Wu1-SARS2_orf1ab;4035 | 0.25 | 0.275 |
| NC_045512.2;YP_009724389.1;Wu1-SARS2_orf1ab;4036 | 0.3 | 0.3 |
| NC_045512.2;YP_009724389.1;Wu1-SARS2_orf1ab;4037 | 0.35 | 0.4 |
| NC_045512.2;YP_009724389.1;Wu1-SARS2_orf1ab;4038 | 0.25 | 0.25 |
| NC_045512.2;YP_009724389.1;Wu1-SARS2_orf1ab;4039 | 0.4 | 0.5 |
| NC_045512.2;YP_009724389.1;Wu1-SARS2_orf1ab;4040 | 0.1 | 0.025 |
| NC_045512.2;YP_009724389.1;Wu1-SARS2_orf1ab;4041 | 0.05 | 0.025 |
| NC_045512.2;YP_009724389.1;Wu1-SARS2_orf1ab;4042 | 0 | 0 |
| NC_045512.2;YP_009724389.1;Wu1-SARS2_orf1ab;4043 | 0.05 | 0.1 |
| NC_045512.2;YP_009724389.1;Wu1-SARS2_orf1ab;4044 | 0.1 | 0.075 |
| NC_045512.2;YP_009724389.1;Wu1-SARS2_orf1ab;4045 | 0 | 0.075 |
| NC_045512.2;YP_009724389.1;Wu1-SARS2_orf1ab;4046 | 0.05 | 0.075 |
| NC_045512.2;YP_009724389.1;Wu1-SARS2_orf1ab;4047 | 0.05 | 0 |
| NC_045512.2;YP_009724389.1;Wu1-SARS2_orf1ab;4048 | 0 | 0.05 |
| NC_045512.2;YP_009724389.1;Wu1-SARS2_orf1ab;4049 | 0 | 0 |
| NC_045512.2;YP_009724389.1;Wu1-SARS2_orf1ab;4050 | 0 | 0 |
| NC_045512.2;YP_009724389.1;Wu1-SARS2_orf1ab;4051 | 0 | 0 |
| NC_045512.2;YP_009724389.1;Wu1-SARS2_orf1ab;4052 | 0 | 0 |
| NC_045512.2;YP_009724389.1;Wu1-SARS2_orf1ab;4053 | 0.05 | 0.025 |
| NC_045512.2;YP_009724389.1;Wu1-SARS2_orf1ab;4054 | 0.05 | 0.025 |
| NC_045512.2;YP_009724389.1;Wu1-SARS2_orf1ab;4055 | 0 | 0 |
| NC_045512.2;YP_009724389.1;Wu1-SARS2_orf1ab;4056 | 0 | 0 |
| NC_045512.2;YP_009724389.1;Wu1-SARS2_orf1ab;4057 | 0 | 0 |
| NC_045512.2;YP_009724389.1;Wu1-SARS2_orf1ab;4058 | 0 | 0 |
| NC_045512.2;YP_009724389.1;Wu1-SARS2_orf1ab;4059 | 0 | 0 |
| NC_045512.2;YP_009724389.1;Wu1-SARS2_orf1ab;4060 | 0 | 0 |
| NC_045512.2;YP_009724389.1;Wu1-SARS2_orf1ab;4061 | 0 | 0 |
| NC_045512.2;YP_009724389.1;Wu1-SARS2_orf1ab;4062 | 0 | 0.025 |
| NC_045512.2;YP_009724389.1;Wu1-SARS2_orf1ab;4063 | 0.05 | 0 |
| NC_045512.2;YP_009724389.1;Wu1-SARS2_orf1ab;4064 | 0 | 0.025 |
| NC_045512.2;YP_009724389.1;Wu1-SARS2_orf1ab;4065 | 0 | 0 |
| NC_045512.2;YP_009724389.1;Wu1-SARS2_orf1ab;4066 | 0 | 0 |
| NC_045512.2;YP_009724389.1;Wu1-SARS2_orf1ab;4067 | 0 | 0 |
| NC_045512.2;YP_009724389.1;Wu1-SARS2_orf1ab;4068 | 0 | 0.025 |
| NC_045512.2;YP_009724389.1;Wu1-SARS2_orf1ab;4069 | 0 | 0 |
| NC_045512.2;YP_009724389.1;Wu1-SARS2_orf1ab;4070 | 0.1 | 0 |
| NC_045512.2;YP_009724389.1;Wu1-SARS2_orf1ab;4071 | 0.1 | 0.025 |
| NC_045512.2;YP_009724389.1;Wu1-SARS2_orf1ab;4072 | 0.2 | 0.025 |
| NC_045512.2;YP_009724389.1;Wu1-SARS2_orf1ab;4073 | 0.05 | 0.025 |
| NC_045512.2;YP_009724389.1;Wu1-SARS2_orf1ab;4074 | 0.1 | 0.05 |
| NC_045512.2;YP_009724389.1;Wu1-SARS2_orf1ab;4075 | 0.05 | 0.025 |

| | | |
|---|---|---|
| NC_045512.2;YP_009724389.1;Wu1-SARS2_orf1ab;4076 | 0.05 | 0 |
| NC_045512.2;YP_009724389.1;Wu1-SARS2_orf1ab;4077 | 0 | 0 |
| NC_045512.2;YP_009724389.1;Wu1-SARS2_orf1ab;4078 | 0 | 0 |
| NC_045512.2;YP_009724389.1;Wu1-SARS2_orf1ab;4079 | 0 | 0 |
| NC_045512.2;YP_009724389.1;Wu1-SARS2_orf1ab;4080 | 0 | 0 |
| NC_045512.2;YP_009724389.1;Wu1-SARS2_orf1ab;4081 | 0 | 0 |
| NC_045512.2;YP_009724389.1;Wu1-SARS2_orf1ab;4082 | 0 | 0 |
| NC_045512.2;YP_009724389.1;Wu1-SARS2_orf1ab;4083 | 0 | 0 |
| NC_045512.2;YP_009724389.1;Wu1-SARS2_orf1ab;4084 | 0 | 0 |
| NC_045512.2;YP_009724389.1;Wu1-SARS2_orf1ab;4085 | 0 | 0 |
| NC_045512.2;YP_009724389.1;Wu1-SARS2_orf1ab;4086 | 0 | 0 |
| NC_045512.2;YP_009724389.1;Wu1-SARS2_orf1ab;4087 | 0 | 0 |
| NC_045512.2;YP_009724389.1;Wu1-SARS2_orf1ab;4088 | 0 | 0 |
| NC_045512.2;YP_009724389.1;Wu1-SARS2_orf1ab;4089 | 0 | 0 |
| NC_045512.2;YP_009724389.1;Wu1-SARS2_orf1ab;4090 | 0 | 0 |
| NC_045512.2;YP_009724389.1;Wu1-SARS2_orf1ab;4091 | 0 | 0 |
| NC_045512.2;YP_009724389.1;Wu1-SARS2_orf1ab;4092 | 0.05 | 0.025 |
| NC_045512.2;YP_009724389.1;Wu1-SARS2_orf1ab;4093 | 0.05 | 0.05 |
| NC_045512.2;YP_009724389.1;Wu1-SARS2_orf1ab;4094 | 0 | 0 |
| NC_045512.2;YP_009724389.1;Wu1-SARS2_orf1ab;4095 | 0 | 0.025 |
| NC_045512.2;YP_009724389.1;Wu1-SARS2_orf1ab;4096 | 0 | 0 |
| NC_045512.2;YP_009724389.1;Wu1-SARS2_orf1ab;4097 | 0 | 0 |
| NC_045512.2;YP_009724389.1;Wu1-SARS2_orf1ab;4098 | 0 | 0 |
| NC_045512.2;YP_009724389.1;Wu1-SARS2_orf1ab;4099 | 0 | 0.025 |
| NC_045512.2;YP_009724389.1;Wu1-SARS2_orf1ab;4100 | 0 | 0 |
| NC_045512.2;YP_009724389.1;Wu1-SARS2_orf1ab;4101 | 0 | 0 |
| NC_045512.2;YP_009724389.1;Wu1-SARS2_orf1ab;4102 | 0 | 0 |
| NC_045512.2;YP_009724389.1;Wu1-SARS2_orf1ab;4103 | 0.05 | 0 |
| NC_045512.2;YP_009724389.1;Wu1-SARS2_orf1ab;4104 | 0 | 0 |
| NC_045512.2;YP_009724389.1;Wu1-SARS2_orf1ab;4105 | 0 | 0 |
| NC_045512.2;YP_009724389.1;Wu1-SARS2_orf1ab;4106 | 0 | 0 |
| NC_045512.2;YP_009724389.1;Wu1-SARS2_orf1ab;4107 | 0 | 0 |
| NC_045512.2;YP_009724389.1;Wu1-SARS2_orf1ab;4108 | 0 | 0.025 |
| NC_045512.2;YP_009724389.1;Wu1-SARS2_orf1ab;4109 | 0 | 0 |
| NC_045512.2;YP_009724389.1;Wu1-SARS2_orf1ab;4110 | 0 | 0 |
| NC_045512.2;YP_009724389.1;Wu1-SARS2_orf1ab;4111 | 0 | 0 |
| NC_045512.2;YP_009724389.1;Wu1-SARS2_orf1ab;4112 | 0 | 0 |
| NC_045512.2;YP_009724389.1;Wu1-SARS2_orf1ab;4113 | 0 | 0 |
| NC_045512.2;YP_009724389.1;Wu1-SARS2_orf1ab;4114 | 0 | 0 |
| NC_045512.2;YP_009724389.1;Wu1-SARS2_orf1ab;4115 | 0 | 0 |
| NC_045512.2;YP_009724389.1;Wu1-SARS2_orf1ab;4116 | 0 | 0 |
| NC_045512.2;YP_009724389.1;Wu1-SARS2_orf1ab;4117 | 0 | 0 |
| NC_045512.2;YP_009724389.1;Wu1-SARS2_orf1ab;4118 | 0 | 0 |

| | | |
|---|---|---|
| NC_045512.2;YP_009724389.1;Wu1-SARS2_orf1ab;4119 | 0 | 0 |
| NC_045512.2;YP_009724389.1;Wu1-SARS2_orf1ab;4120 | 0 | 0 |
| NC_045512.2;YP_009724389.1;Wu1-SARS2_orf1ab;4121 | 0 | 0 |
| NC_045512.2;YP_009724389.1;Wu1-SARS2_orf1ab;4122 | 0 | 0 |
| NC_045512.2;YP_009724389.1;Wu1-SARS2_orf1ab;4123 | 0 | 0 |
| NC_045512.2;YP_009724389.1;Wu1-SARS2_orf1ab;4124 | 0 | 0 |
| NC_045512.2;YP_009724389.1;Wu1-SARS2_orf1ab;4125 | 0 | 0 |
| NC_045512.2;YP_009724389.1;Wu1-SARS2_orf1ab;4126 | 0 | 0 |
| NC_045512.2;YP_009724389.1;Wu1-SARS2_orf1ab;4127 | 0 | 0.025 |
| NC_045512.2;YP_009724389.1;Wu1-SARS2_orf1ab;4128 | 0 | 0.075 |
| NC_045512.2;YP_009724389.1;Wu1-SARS2_orf1ab;4129 | 0 | 0.05 |
| NC_045512.2;YP_009724389.1;Wu1-SARS2_orf1ab;4130 | 0 | 0.025 |
| NC_045512.2;YP_009724389.1;Wu1-SARS2_orf1ab;4131 | 0 | 0.025 |
| NC_045512.2;YP_009724389.1;Wu1-SARS2_orf1ab;4132 | 0.05 | 0.025 |
| NC_045512.2;YP_009724389.1;Wu1-SARS2_orf1ab;4133 | 0.1 | 0.025 |
| NC_045512.2;YP_009724389.1;Wu1-SARS2_orf1ab;4134 | 0.05 | 0.025 |
| NC_045512.2;YP_009724389.1;Wu1-SARS2_orf1ab;4135 | 0 | 0.025 |
| NC_045512.2;YP_009724389.1;Wu1-SARS2_orf1ab;4136 | 0 | 0 |
| NC_045512.2;YP_009724389.1;Wu1-SARS2_orf1ab;4137 | 0.05 | 0.025 |
| NC_045512.2;YP_009724389.1;Wu1-SARS2_orf1ab;4138 | 0 | 0 |
| NC_045512.2;YP_009724389.1;Wu1-SARS2_orf1ab;4139 | 0 | 0 |
| NC_045512.2;YP_009724389.1;Wu1-SARS2_orf1ab;4140 | 0 | 0.025 |
| NC_045512.2;YP_009724389.1;Wu1-SARS2_orf1ab;4141 | 0.05 | 0.05 |
| NC_045512.2;YP_009724389.1;Wu1-SARS2_orf1ab;4142 | 0 | 0 |
| NC_045512.2;YP_009724389.1;Wu1-SARS2_orf1ab;4143 | 0 | 0 |
| NC_045512.2;YP_009724389.1;Wu1-SARS2_orf1ab;4144 | 0 | 0 |
| NC_045512.2;YP_009724389.1;Wu1-SARS2_orf1ab;4145 | 0 | 0 |
| NC_045512.2;YP_009724389.1;Wu1-SARS2_orf1ab;4146 | 0 | 0 |
| NC_045512.2;YP_009724389.1;Wu1-SARS2_orf1ab;4147 | 0 | 0 |
| NC_045512.2;YP_009724389.1;Wu1-SARS2_orf1ab;4148 | 0 | 0 |
| NC_045512.2;YP_009724389.1;Wu1-SARS2_orf1ab;4149 | 0 | 0 |
| NC_045512.2;YP_009724389.1;Wu1-SARS2_orf1ab;4150 | 0 | 0 |
| NC_045512.2;YP_009724389.1;Wu1-SARS2_orf1ab;4151 | 0 | 0 |
| NC_045512.2;YP_009724389.1;Wu1-SARS2_orf1ab;4152 | 0 | 0 |
| NC_045512.2;YP_009724389.1;Wu1-SARS2_orf1ab;4153 | 0 | 0 |
| NC_045512.2;YP_009724389.1;Wu1-SARS2_orf1ab;4154 | 0 | 0 |
| NC_045512.2;YP_009724389.1;Wu1-SARS2_orf1ab;4155 | 0 | 0 |
| NC_045512.2;YP_009724389.1;Wu1-SARS2_orf1ab;4156 | 0 | 0 |
| NC_045512.2;YP_009724389.1;Wu1-SARS2_orf1ab;4157 | 0 | 0 |
| NC_045512.2;YP_009724389.1;Wu1-SARS2_orf1ab;4158 | 0 | 0 |
| NC_045512.2;YP_009724389.1;Wu1-SARS2_orf1ab;4159 | 0 | 0 |
| NC_045512.2;YP_009724389.1;Wu1-SARS2_orf1ab;4160 | 0 | 0 |
| NC_045512.2;YP_009724389.1;Wu1-SARS2_orf1ab;4161 | 0.05 | 0.05 |

| | | |
|---|---|---|
| NC_045512.2;YP_009724389.1;Wu1-SARS2_orf1ab;4162 | 0.1 | 0.05 |
| NC_045512.2;YP_009724389.1;Wu1-SARS2_orf1ab;4163 | 0.2 | 0.05 |
| NC_045512.2;YP_009724389.1;Wu1-SARS2_orf1ab;4164 | 0 | 0 |
| NC_045512.2;YP_009724389.1;Wu1-SARS2_orf1ab;4165 | 0 | 0 |
| NC_045512.2;YP_009724389.1;Wu1-SARS2_orf1ab;4166 | 0 | 0 |
| NC_045512.2;YP_009724389.1;Wu1-SARS2_orf1ab;4167 | 0 | 0 |
| NC_045512.2;YP_009724389.1;Wu1-SARS2_orf1ab;4168 | 0 | 0 |
| NC_045512.2;YP_009724389.1;Wu1-SARS2_orf1ab;4169 | 0 | 0 |
| NC_045512.2;YP_009724389.1;Wu1-SARS2_orf1ab;4170 | 0 | 0 |
| NC_045512.2;YP_009724389.1;Wu1-SARS2_orf1ab;4171 | 0 | 0 |
| NC_045512.2;YP_009724389.1;Wu1-SARS2_orf1ab;4172 | 0 | 0 |
| NC_045512.2;YP_009724389.1;Wu1-SARS2_orf1ab;4173 | 0 | 0 |
| NC_045512.2;YP_009724389.1;Wu1-SARS2_orf1ab;4174 | 0 | 0 |
| NC_045512.2;YP_009724389.1;Wu1-SARS2_orf1ab;4175 | 0 | 0 |
| NC_045512.2;YP_009724389.1;Wu1-SARS2_orf1ab;4176 | 0 | 0 |
| NC_045512.2;YP_009724389.1;Wu1-SARS2_orf1ab;4177 | 0 | 0 |
| NC_045512.2;YP_009724389.1;Wu1-SARS2_orf1ab;4178 | 0 | 0 |
| NC_045512.2;YP_009724389.1;Wu1-SARS2_orf1ab;4179 | 0 | 0 |
| NC_045512.2;YP_009724389.1;Wu1-SARS2_orf1ab;4180 | 0 | 0 |
| NC_045512.2;YP_009724389.1;Wu1-SARS2_orf1ab;4181 | 0 | 0 |
| NC_045512.2;YP_009724389.1;Wu1-SARS2_orf1ab;4182 | 0.05 | 0 |
| NC_045512.2;YP_009724389.1;Wu1-SARS2_orf1ab;4183 | 0 | 0 |
| NC_045512.2;YP_009724389.1;Wu1-SARS2_orf1ab;4184 | 0.05 | 0 |
| NC_045512.2;YP_009724389.1;Wu1-SARS2_orf1ab;4185 | 0.05 | 0.05 |
| NC_045512.2;YP_009724389.1;Wu1-SARS2_orf1ab;4186 | 0.15 | 0.1 |
| NC_045512.2;YP_009724389.1;Wu1-SARS2_orf1ab;4187 | 0.3 | 0.25 |
| NC_045512.2;YP_009724389.1;Wu1-SARS2_orf1ab;4188 | 0.15 | 0.05 |
| NC_045512.2;YP_009724389.1;Wu1-SARS2_orf1ab;4189 | 0 | 0 |
| NC_045512.2;YP_009724389.1;Wu1-SARS2_orf1ab;4190 | 0 | 0 |
| NC_045512.2;YP_009724389.1;Wu1-SARS2_orf1ab;4191 | 0 | 0 |
| NC_045512.2;YP_009724389.1;Wu1-SARS2_orf1ab;4192 | 0 | 0 |
| NC_045512.2;YP_009724389.1;Wu1-SARS2_orf1ab;4193 | 0 | 0 |
| NC_045512.2;YP_009724389.1;Wu1-SARS2_orf1ab;4194 | 0 | 0 |
| NC_045512.2;YP_009724389.1;Wu1-SARS2_orf1ab;4195 | 0 | 0 |
| NC_045512.2;YP_009724389.1;Wu1-SARS2_orf1ab;4196 | 0 | 0 |
| NC_045512.2;YP_009724389.1;Wu1-SARS2_orf1ab;4197 | 0 | 0 |
| NC_045512.2;YP_009724389.1;Wu1-SARS2_orf1ab;4198 | 0 | 0 |
| NC_045512.2;YP_009724389.1;Wu1-SARS2_orf1ab;4199 | 0 | 0 |
| NC_045512.2;YP_009724389.1;Wu1-SARS2_orf1ab;4200 | 0 | 0 |
| NC_045512.2;YP_009724389.1;Wu1-SARS2_orf1ab;4201 | 0 | 0.025 |
| NC_045512.2;YP_009724389.1;Wu1-SARS2_orf1ab;4202 | 0 | 0 |
| NC_045512.2;YP_009724389.1;Wu1-SARS2_orf1ab;4203 | 0.2 | 0.15 |
| NC_045512.2;YP_009724389.1;Wu1-SARS2_orf1ab;4204 | 0 | 0 |

| | | |
|---|---|---|
| NC_045512.2;YP_009724389.1;Wu1-SARS2_orf1ab;4205 | 0.05 | 0.15 |
| NC_045512.2;YP_009724389.1;Wu1-SARS2_orf1ab;4206 | 0.05 | 0.075 |
| NC_045512.2;YP_009724389.1;Wu1-SARS2_orf1ab;4207 | 0.05 | 0.05 |
| NC_045512.2;YP_009724389.1;Wu1-SARS2_orf1ab;4208 | 0.05 | 0.025 |
| NC_045512.2;YP_009724389.1;Wu1-SARS2_orf1ab;4209 | 0 | 0 |
| NC_045512.2;YP_009724389.1;Wu1-SARS2_orf1ab;4210 | 0 | 0.025 |
| NC_045512.2;YP_009724389.1;Wu1-SARS2_orf1ab;4211 | 0 | 0 |
| NC_045512.2;YP_009724389.1;Wu1-SARS2_orf1ab;4212 | 0 | 0 |
| NC_045512.2;YP_009724389.1;Wu1-SARS2_orf1ab;4213 | 0 | 0 |
| NC_045512.2;YP_009724389.1;Wu1-SARS2_orf1ab;4214 | 0 | 0 |
| NC_045512.2;YP_009724389.1;Wu1-SARS2_orf1ab;4215 | 0.05 | 0 |
| NC_045512.2;YP_009724389.1;Wu1-SARS2_orf1ab;4216 | 0 | 0 |
| NC_045512.2;YP_009724389.1;Wu1-SARS2_orf1ab;4217 | 0 | 0 |
| NC_045512.2;YP_009724389.1;Wu1-SARS2_orf1ab;4218 | 0 | 0 |
| NC_045512.2;YP_009724389.1;Wu1-SARS2_orf1ab;4219 | 0 | 0 |
| NC_045512.2;YP_009724389.1;Wu1-SARS2_orf1ab;4220 | 0 | 0 |
| NC_045512.2;YP_009724389.1;Wu1-SARS2_orf1ab;4221 | 0 | 0 |
| NC_045512.2;YP_009724389.1;Wu1-SARS2_orf1ab;4222 | 0 | 0 |
| NC_045512.2;YP_009724389.1;Wu1-SARS2_orf1ab;4223 | 0 | 0 |
| NC_045512.2;YP_009724389.1;Wu1-SARS2_orf1ab;4224 | 0 | 0 |
| NC_045512.2;YP_009724389.1;Wu1-SARS2_orf1ab;4225 | 0 | 0 |
| NC_045512.2;YP_009724389.1;Wu1-SARS2_orf1ab;4226 | 0 | 0 |
| NC_045512.2;YP_009724389.1;Wu1-SARS2_orf1ab;4227 | 0 | 0 |
| NC_045512.2;YP_009724389.1;Wu1-SARS2_orf1ab;4228 | 0 | 0 |
| NC_045512.2;YP_009724389.1;Wu1-SARS2_orf1ab;4229 | 0.05 | 0.025 |
| NC_045512.2;YP_009724389.1;Wu1-SARS2_orf1ab;4230 | 0.1 | 0.075 |
| NC_045512.2;YP_009724389.1;Wu1-SARS2_orf1ab;4231 | 0.15 | 0.1 |
| NC_045512.2;YP_009724389.1;Wu1-SARS2_orf1ab;4232 | 0.25 | 0.1 |
| NC_045512.2;YP_009724389.1;Wu1-SARS2_orf1ab;4233 | 0.1 | 0.1 |
| NC_045512.2;YP_009724389.1;Wu1-SARS2_orf1ab;4234 | 0.05 | 0.05 |
| NC_045512.2;YP_009724389.1;Wu1-SARS2_orf1ab;4235 | 0.05 | 0.025 |
| NC_045512.2;YP_009724389.1;Wu1-SARS2_orf1ab;4236 | 0 | 0 |
| NC_045512.2;YP_009724389.1;Wu1-SARS2_orf1ab;4237 | 0 | 0 |
| NC_045512.2;YP_009724389.1;Wu1-SARS2_orf1ab;4238 | 0 | 0 |
| NC_045512.2;YP_009724389.1;Wu1-SARS2_orf1ab;4239 | 0 | 0 |
| NC_045512.2;YP_009724389.1;Wu1-SARS2_orf1ab;4240 | 0 | 0 |
| NC_045512.2;YP_009724389.1;Wu1-SARS2_orf1ab;4241 | 0 | 0 |
| NC_045512.2;YP_009724389.1;Wu1-SARS2_orf1ab;4242 | 0 | 0 |
| NC_045512.2;YP_009724389.1;Wu1-SARS2_orf1ab;4243 | 0 | 0 |
| NC_045512.2;YP_009724389.1;Wu1-SARS2_orf1ab;4244 | 0 | 0.025 |
| NC_045512.2;YP_009724389.1;Wu1-SARS2_orf1ab;4245 | 0 | 0 |
| NC_045512.2;YP_009724389.1;Wu1-SARS2_orf1ab;4246 | 0 | 0.025 |
| NC_045512.2;YP_009724389.1;Wu1-SARS2_orf1ab;4247 | 0.05 | 0.025 |

| | | |
|---|---|---|
| NC_045512.2;YP_009724389.1;Wu1-SARS2_orf1ab;4248 | 0 | 0.025 |
| NC_045512.2;YP_009724389.1;Wu1-SARS2_orf1ab;4249 | 0 | 0 |
| NC_045512.2;YP_009724389.1;Wu1-SARS2_orf1ab;4250 | 0 | 0 |
| NC_045512.2;YP_009724389.1;Wu1-SARS2_orf1ab;4251 | 0 | 0.025 |
| NC_045512.2;YP_009724389.1;Wu1-SARS2_orf1ab;4252 | 0 | 0.025 |
| NC_045512.2;YP_009724389.1;Wu1-SARS2_orf1ab;4253 | 0 | 0 |
| NC_045512.2;YP_009724389.1;Wu1-SARS2_orf1ab;4254 | 0 | 0 |
| NC_045512.2;YP_009724389.1;Wu1-SARS2_orf1ab;4255 | 0 | 0 |
| NC_045512.2;YP_009724389.1;Wu1-SARS2_orf1ab;4256 | 0 | 0 |
| NC_045512.2;YP_009724389.1;Wu1-SARS2_orf1ab;4257 | 0 | 0 |
| NC_045512.2;YP_009724389.1;Wu1-SARS2_orf1ab;4258 | 0 | 0 |
| NC_045512.2;YP_009724389.1;Wu1-SARS2_orf1ab;4259 | 0 | 0 |
| NC_045512.2;YP_009724389.1;Wu1-SARS2_orf1ab;4260 | 0 | 0 |
| NC_045512.2;YP_009724389.1;Wu1-SARS2_orf1ab;4261 | 0 | 0 |
| NC_045512.2;YP_009724389.1;Wu1-SARS2_orf1ab;4262 | 0 | 0 |
| NC_045512.2;YP_009724389.1;Wu1-SARS2_orf1ab;4263 | 0 | 0 |
| NC_045512.2;YP_009724389.1;Wu1-SARS2_orf1ab;4264 | 0 | 0 |
| NC_045512.2;YP_009724389.1;Wu1-SARS2_orf1ab;4265 | 0 | 0 |
| NC_045512.2;YP_009724389.1;Wu1-SARS2_orf1ab;4266 | 0 | 0 |
| NC_045512.2;YP_009724389.1;Wu1-SARS2_orf1ab;4267 | 0 | 0 |
| NC_045512.2;YP_009724389.1;Wu1-SARS2_orf1ab;4268 | 0.05 | 0 |
| NC_045512.2;YP_009724389.1;Wu1-SARS2_orf1ab;4269 | 0 | 0 |
| NC_045512.2;YP_009724389.1;Wu1-SARS2_orf1ab;4270 | 0 | 0 |
| NC_045512.2;YP_009724389.1;Wu1-SARS2_orf1ab;4271 | 0.05 | 0 |
| NC_045512.2;YP_009724389.1;Wu1-SARS2_orf1ab;4272 | 0.05 | 0 |
| NC_045512.2;YP_009724389.1;Wu1-SARS2_orf1ab;4273 | 0.2 | 0.05 |
| NC_045512.2;YP_009724389.1;Wu1-SARS2_orf1ab;4274 | 0.15 | 0.025 |
| NC_045512.2;YP_009724389.1;Wu1-SARS2_orf1ab;4275 | 0.1 | 0.025 |
| NC_045512.2;YP_009724389.1;Wu1-SARS2_orf1ab;4276 | 0 | 0.025 |
| NC_045512.2;YP_009724389.1;Wu1-SARS2_orf1ab;4277 | 0 | 0.05 |
| NC_045512.2;YP_009724389.1;Wu1-SARS2_orf1ab;4278 | 0 | 0.05 |
| NC_045512.2;YP_009724389.1;Wu1-SARS2_orf1ab;4279 | 0.15 | 0.05 |
| NC_045512.2;YP_009724389.1;Wu1-SARS2_orf1ab;4280 | 0.05 | 0.05 |
| NC_045512.2;YP_009724389.1;Wu1-SARS2_orf1ab;4281 | 0 | 0 |
| NC_045512.2;YP_009724389.1;Wu1-SARS2_orf1ab;4282 | 0 | 0 |
| NC_045512.2;YP_009724389.1;Wu1-SARS2_orf1ab;4283 | 0 | 0 |
| NC_045512.2;YP_009724389.1;Wu1-SARS2_orf1ab;4284 | 0 | 0 |
| NC_045512.2;YP_009724389.1;Wu1-SARS2_orf1ab;4285 | 0 | 0 |
| NC_045512.2;YP_009724389.1;Wu1-SARS2_orf1ab;4286 | 0 | 0 |
| NC_045512.2;YP_009724389.1;Wu1-SARS2_orf1ab;4287 | 0 | 0 |
| NC_045512.2;YP_009724389.1;Wu1-SARS2_orf1ab;4288 | 0 | 0 |
| NC_045512.2;YP_009724389.1;Wu1-SARS2_orf1ab;4289 | 0 | 0.025 |
| NC_045512.2;YP_009724389.1;Wu1-SARS2_orf1ab;4290 | 0 | 0 |

| | | |
|---|---|---|
| NC_045512.2;YP_009724389.1;Wu1-SARS2_orf1ab;4291 | 0 | 0.025 |
| NC_045512.2;YP_009724389.1;Wu1-SARS2_orf1ab;4292 | 0 | 0.025 |
| NC_045512.2;YP_009724389.1;Wu1-SARS2_orf1ab;4293 | 0 | 0 |
| NC_045512.2;YP_009724389.1;Wu1-SARS2_orf1ab;4294 | 0 | 0 |
| NC_045512.2;YP_009724389.1;Wu1-SARS2_orf1ab;4295 | 0 | 0 |
| NC_045512.2;YP_009724389.1;Wu1-SARS2_orf1ab;4296 | 0 | 0 |
| NC_045512.2;YP_009724389.1;Wu1-SARS2_orf1ab;4297 | 0 | 0 |
| NC_045512.2;YP_009724389.1;Wu1-SARS2_orf1ab;4298 | 0 | 0 |
| NC_045512.2;YP_009724389.1;Wu1-SARS2_orf1ab;4299 | 0 | 0 |
| NC_045512.2;YP_009724389.1;Wu1-SARS2_orf1ab;4300 | 0 | 0 |
| NC_045512.2;YP_009724389.1;Wu1-SARS2_orf1ab;4301 | 0 | 0 |
| NC_045512.2;YP_009724389.1;Wu1-SARS2_orf1ab;4302 | 0 | 0.025 |
| NC_045512.2;YP_009724389.1;Wu1-SARS2_orf1ab;4303 | 0 | 0.025 |
| NC_045512.2;YP_009724389.1;Wu1-SARS2_orf1ab;4304 | 0 | 0.025 |
| NC_045512.2;YP_009724389.1;Wu1-SARS2_orf1ab;4305 | 0 | 0.025 |
| NC_045512.2;YP_009724389.1;Wu1-SARS2_orf1ab;4306 | 0 | 0.025 |
| NC_045512.2;YP_009724389.1;Wu1-SARS2_orf1ab;4307 | 0 | 0.025 |
| NC_045512.2;YP_009724389.1;Wu1-SARS2_orf1ab;4308 | 0.05 | 0.025 |
| NC_045512.2;YP_009724389.1;Wu1-SARS2_orf1ab;4309 | 0 | 0.025 |
| NC_045512.2;YP_009724389.1;Wu1-SARS2_orf1ab;4310 | 0.05 | 0.025 |
| NC_045512.2;YP_009724389.1;Wu1-SARS2_orf1ab;4311 | 0 | 0 |
| NC_045512.2;YP_009724389.1;Wu1-SARS2_orf1ab;4312 | 0 | 0 |
| NC_045512.2;YP_009724389.1;Wu1-SARS2_orf1ab;4313 | 0 | 0 |
| NC_045512.2;YP_009724389.1;Wu1-SARS2_orf1ab;4314 | 0 | 0 |
| NC_045512.2;YP_009724389.1;Wu1-SARS2_orf1ab;4315 | 0 | 0 |
| NC_045512.2;YP_009724389.1;Wu1-SARS2_orf1ab;4316 | 0 | 0 |
| NC_045512.2;YP_009724389.1;Wu1-SARS2_orf1ab;4317 | 0 | 0 |
| NC_045512.2;YP_009724389.1;Wu1-SARS2_orf1ab;4318 | 0 | 0 |
| NC_045512.2;YP_009724389.1;Wu1-SARS2_orf1ab;4319 | 0 | 0 |
| NC_045512.2;YP_009724389.1;Wu1-SARS2_orf1ab;4320 | 0 | 0 |
| NC_045512.2;YP_009724389.1;Wu1-SARS2_orf1ab;4321 | 0 | 0 |
| NC_045512.2;YP_009724389.1;Wu1-SARS2_orf1ab;4322 | 0 | 0 |
| NC_045512.2;YP_009724389.1;Wu1-SARS2_orf1ab;4323 | 0 | 0 |
| NC_045512.2;YP_009724389.1;Wu1-SARS2_orf1ab;4324 | 0 | 0 |
| NC_045512.2;YP_009724389.1;Wu1-SARS2_orf1ab;4325 | 0 | 0 |
| NC_045512.2;YP_009724389.1;Wu1-SARS2_orf1ab;4326 | 0 | 0 |
| NC_045512.2;YP_009724389.1;Wu1-SARS2_orf1ab;4327 | 0 | 0 |
| NC_045512.2;YP_009724389.1;Wu1-SARS2_orf1ab;4328 | 0 | 0 |
| NC_045512.2;YP_009724389.1;Wu1-SARS2_orf1ab;4329 | 0 | 0.025 |
| NC_045512.2;YP_009724389.1;Wu1-SARS2_orf1ab;4330 | 0 | 0.025 |
| NC_045512.2;YP_009724389.1;Wu1-SARS2_orf1ab;4331 | 0 | 0.025 |
| NC_045512.2;YP_009724389.1;Wu1-SARS2_orf1ab;4332 | 0 | 0.025 |
| NC_045512.2;YP_009724389.1;Wu1-SARS2_orf1ab;4333 | 0 | 0.025 |

| | | |
|---|---|---|
| NC_045512.2;YP_009724389.1;Wu1-SARS2_orf1ab;4334 | 0 | 0.025 |
| NC_045512.2;YP_009724389.1;Wu1-SARS2_orf1ab;4335 | 0 | 0 |
| NC_045512.2;YP_009724389.1;Wu1-SARS2_orf1ab;4336 | 0 | 0 |
| NC_045512.2;YP_009724389.1;Wu1-SARS2_orf1ab;4337 | 0 | 0 |
| NC_045512.2;YP_009724389.1;Wu1-SARS2_orf1ab;4338 | 0 | 0 |
| NC_045512.2;YP_009724389.1;Wu1-SARS2_orf1ab;4339 | 0 | 0 |
| NC_045512.2;YP_009724389.1;Wu1-SARS2_orf1ab;4340 | 0 | 0.05 |
| NC_045512.2;YP_009724389.1;Wu1-SARS2_orf1ab;4341 | 0 | 0.05 |
| NC_045512.2;YP_009724389.1;Wu1-SARS2_orf1ab;4342 | 0 | 0 |
| NC_045512.2;YP_009724389.1;Wu1-SARS2_orf1ab;4343 | 0 | 0 |
| NC_045512.2;YP_009724389.1;Wu1-SARS2_orf1ab;4344 | 0.05 | 0 |
| NC_045512.2;YP_009724389.1;Wu1-SARS2_orf1ab;4345 | 0.05 | 0.025 |
| NC_045512.2;YP_009724389.1;Wu1-SARS2_orf1ab;4346 | 0.05 | 0.05 |
| NC_045512.2;YP_009724389.1;Wu1-SARS2_orf1ab;4347 | 0.05 | 0 |
| NC_045512.2;YP_009724389.1;Wu1-SARS2_orf1ab;4348 | 0.05 | 0 |
| NC_045512.2;YP_009724389.1;Wu1-SARS2_orf1ab;4349 | 0 | 0 |
| NC_045512.2;YP_009724389.1;Wu1-SARS2_orf1ab;4350 | 0 | 0.025 |
| NC_045512.2;YP_009724389.1;Wu1-SARS2_orf1ab;4351 | 0 | 0.025 |
| NC_045512.2;YP_009724389.1;Wu1-SARS2_orf1ab;4352 | 0 | 0.025 |
| NC_045512.2;YP_009724389.1;Wu1-SARS2_orf1ab;4353 | 0.05 | 0.05 |
| NC_045512.2;YP_009724389.1;Wu1-SARS2_orf1ab;4354 | 0 | 0.025 |
| NC_045512.2;YP_009724389.1;Wu1-SARS2_orf1ab;4355 | 0 | 0.025 |
| NC_045512.2;YP_009724389.1;Wu1-SARS2_orf1ab;4356 | 0 | 0.025 |
| NC_045512.2;YP_009724389.1;Wu1-SARS2_orf1ab;4357 | 0 | 0.025 |
| NC_045512.2;YP_009724389.1;Wu1-SARS2_orf1ab;4358 | 0 | 0 |
| NC_045512.2;YP_009724389.1;Wu1-SARS2_orf1ab;4359 | 0 | 0 |
| NC_045512.2;YP_009724389.1;Wu1-SARS2_orf1ab;4360 | 0 | 0 |
| NC_045512.2;YP_009724389.1;Wu1-SARS2_orf1ab;4361 | 0 | 0 |
| NC_045512.2;YP_009724389.1;Wu1-SARS2_orf1ab;4362 | 0 | 0 |
| NC_045512.2;YP_009724389.1;Wu1-SARS2_orf1ab;4363 | 0 | 0 |
| NC_045512.2;YP_009724389.1;Wu1-SARS2_orf1ab;4364 | 0 | 0 |
| NC_045512.2;YP_009724389.1;Wu1-SARS2_orf1ab;4365 | 0 | 0 |
| NC_045512.2;YP_009724389.1;Wu1-SARS2_orf1ab;4366 | 0 | 0 |
| NC_045512.2;YP_009724389.1;Wu1-SARS2_orf1ab;4367 | 0 | 0 |
| NC_045512.2;YP_009724389.1;Wu1-SARS2_orf1ab;4368 | 0 | 0 |
| NC_045512.2;YP_009724389.1;Wu1-SARS2_orf1ab;4369 | 0 | 0 |
| NC_045512.2;YP_009724389.1;Wu1-SARS2_orf1ab;4370 | 0 | 0 |
| NC_045512.2;YP_009724389.1;Wu1-SARS2_orf1ab;4371 | 0 | 0 |
| NC_045512.2;YP_009724389.1;Wu1-SARS2_orf1ab;4372 | 0 | 0 |
| NC_045512.2;YP_009724389.1;Wu1-SARS2_orf1ab;4373 | 0 | 0 |
| NC_045512.2;YP_009724389.1;Wu1-SARS2_orf1ab;4374 | 0 | 0 |
| NC_045512.2;YP_009724389.1;Wu1-SARS2_orf1ab;4375 | 0 | 0.075 |
| NC_045512.2;YP_009724389.1;Wu1-SARS2_orf1ab;4376 | 0 | 0 |

| | | |
|---|---|---|
| NC_045512.2;YP_009724389.1;Wu1-SARS2_orf1ab;4377 | 0 | 0.025 |
| NC_045512.2;YP_009724389.1;Wu1-SARS2_orf1ab;4378 | 0 | 0 |
| NC_045512.2;YP_009724389.1;Wu1-SARS2_orf1ab;4379 | 0 | 0 |
| NC_045512.2;YP_009724389.1;Wu1-SARS2_orf1ab;4380 | 0 | 0 |
| NC_045512.2;YP_009724389.1;Wu1-SARS2_orf1ab;4381 | 0 | 0 |
| NC_045512.2;YP_009724389.1;Wu1-SARS2_orf1ab;4382 | 0 | 0 |
| NC_045512.2;YP_009724389.1;Wu1-SARS2_orf1ab;4383 | 0 | 0 |
| NC_045512.2;YP_009724389.1;Wu1-SARS2_orf1ab;4384 | 0 | 0 |
| NC_045512.2;YP_009724389.1;Wu1-SARS2_orf1ab;4385 | 0 | 0 |
| NC_045512.2;YP_009724389.1;Wu1-SARS2_orf1ab;4386 | 0.05 | 0 |
| NC_045512.2;YP_009724389.1;Wu1-SARS2_orf1ab;4387 | 0.05 | 0.025 |
| NC_045512.2;YP_009724389.1;Wu1-SARS2_orf1ab;4388 | 0.15 | 0.025 |
| NC_045512.2;YP_009724389.1;Wu1-SARS2_orf1ab;4389 | 0 | 0 |
| NC_045512.2;YP_009724389.1;Wu1-SARS2_orf1ab;4390 | 0.05 | 0 |
| NC_045512.2;YP_009724389.1;Wu1-SARS2_orf1ab;4391 | 0 | 0 |
| NC_045512.2;YP_009724389.1;Wu1-SARS2_orf1ab;4392 | 0 | 0 |
| NC_045512.2;YP_009724389.1;Wu1-SARS2_orf1ab;4393 | 0 | 0 |
| NC_045512.2;YP_009724389.1;Wu1-SARS2_orf1ab;4394 | 0.05 | 0 |
| NC_045512.2;YP_009724389.1;Wu1-SARS2_orf1ab;4395 | 0 | 0 |
| NC_045512.2;YP_009724389.1;Wu1-SARS2_orf1ab;4396 | 0 | 0 |
| NC_045512.2;YP_009724389.1;Wu1-SARS2_orf1ab;4397 | 0 | 0 |
| NC_045512.2;YP_009724389.1;Wu1-SARS2_orf1ab;4398 | 0 | 0 |
| NC_045512.2;YP_009724389.1;Wu1-SARS2_orf1ab;4399 | 0 | 0 |
| NC_045512.2;YP_009724389.1;Wu1-SARS2_orf1ab;4400 | 0 | 0 |
| NC_045512.2;YP_009724389.1;Wu1-SARS2_orf1ab;4401 | 0 | 0 |
| NC_045512.2;YP_009724389.1;Wu1-SARS2_orf1ab;4402 | 0 | 0 |
| NC_045512.2;YP_009724389.1;Wu1-SARS2_orf1ab;4403 | 0 | 0 |
| NC_045512.2;YP_009724389.1;Wu1-SARS2_orf1ab;4404 | 0 | 0 |
| NC_045512.2;YP_009724389.1;Wu1-SARS2_orf1ab;4405 | 0 | 0 |
| NC_045512.2;YP_009724389.1;Wu1-SARS2_orf1ab;4406 | 0 | 0 |
| NC_045512.2;YP_009724389.1;Wu1-SARS2_orf1ab;4407 | 0 | 0.025 |
| NC_045512.2;YP_009724389.1;Wu1-SARS2_orf1ab;4408 | 0 | 0 |
| NC_045512.2;YP_009724389.1;Wu1-SARS2_orf1ab;4409 | 0 | 0 |
| NC_045512.2;YP_009724389.1;Wu1-SARS2_orf1ab;4410 | 0 | 0 |
| NC_045512.2;YP_009724389.1;Wu1-SARS2_orf1ab;4411 | 0 | 0 |
| NC_045512.2;YP_009724389.1;Wu1-SARS2_orf1ab;4412 | 0 | 0 |
| NC_045512.2;YP_009724389.1;Wu1-SARS2_orf1ab;4413 | 0 | 0 |
| NC_045512.2;YP_009724389.1;Wu1-SARS2_orf1ab;4414 | 0 | 0 |
| NC_045512.2;YP_009724389.1;Wu1-SARS2_orf1ab;4415 | 0 | 0 |
| NC_045512.2;YP_009724389.1;Wu1-SARS2_orf1ab;4416 | 0 | 0 |
| NC_045512.2;YP_009724389.1;Wu1-SARS2_orf1ab;4417 | 0 | 0 |
| NC_045512.2;YP_009724389.1;Wu1-SARS2_orf1ab;4418 | 0 | 0.025 |
| NC_045512.2;YP_009724389.1;Wu1-SARS2_orf1ab;4419 | 0.1 | 0 |

| | | |
|---|---|---|
| NC_045512.2;YP_009724389.1;Wu1-SARS2_orf1ab;4420 | 0.05 | 0.05 |
| NC_045512.2;YP_009724389.1;Wu1-SARS2_orf1ab;4421 | 0.05 | 0 |
| NC_045512.2;YP_009724389.1;Wu1-SARS2_orf1ab;4422 | 0 | 0 |
| NC_045512.2;YP_009724389.1;Wu1-SARS2_orf1ab;4423 | 0.05 | 0 |
| NC_045512.2;YP_009724389.1;Wu1-SARS2_orf1ab;4424 | 0 | 0.025 |
| NC_045512.2;YP_009724389.1;Wu1-SARS2_orf1ab;4425 | 0 | 0 |
| NC_045512.2;YP_009724389.1;Wu1-SARS2_orf1ab;4426 | 0 | 0 |
| NC_045512.2;YP_009724389.1;Wu1-SARS2_orf1ab;4427 | 0 | 0 |
| NC_045512.2;YP_009724389.1;Wu1-SARS2_orf1ab;4428 | 0 | 0 |
| NC_045512.2;YP_009724389.1;Wu1-SARS2_orf1ab;4429 | 0 | 0 |
| NC_045512.2;YP_009724389.1;Wu1-SARS2_orf1ab;4430 | 0 | 0 |
| NC_045512.2;YP_009724389.1;Wu1-SARS2_orf1ab;4431 | 0 | 0 |
| NC_045512.2;YP_009724389.1;Wu1-SARS2_orf1ab;4432 | 0 | 0 |
| NC_045512.2;YP_009724389.1;Wu1-SARS2_orf1ab;4433 | 0 | 0 |
| NC_045512.2;YP_009724389.1;Wu1-SARS2_orf1ab;4434 | 0 | 0 |
| NC_045512.2;YP_009724389.1;Wu1-SARS2_orf1ab;4435 | 0 | 0 |
| NC_045512.2;YP_009724389.1;Wu1-SARS2_orf1ab;4436 | 0 | 0 |
| NC_045512.2;YP_009724389.1;Wu1-SARS2_orf1ab;4437 | 0 | 0 |
| NC_045512.2;YP_009724389.1;Wu1-SARS2_orf1ab;4438 | 0 | 0.025 |
| NC_045512.2;YP_009724389.1;Wu1-SARS2_orf1ab;4439 | 0 | 0 |
| NC_045512.2;YP_009724389.1;Wu1-SARS2_orf1ab;4440 | 0 | 0 |
| NC_045512.2;YP_009724389.1;Wu1-SARS2_orf1ab;4441 | 0 | 0.025 |
| NC_045512.2;YP_009724389.1;Wu1-SARS2_orf1ab;4442 | 0 | 0 |
| NC_045512.2;YP_009724389.1;Wu1-SARS2_orf1ab;4443 | 0 | 0 |
| NC_045512.2;YP_009724389.1;Wu1-SARS2_orf1ab;4444 | 0 | 0.05 |
| NC_045512.2;YP_009724389.1;Wu1-SARS2_orf1ab;4445 | 0 | 0.05 |
| NC_045512.2;YP_009724389.1;Wu1-SARS2_orf1ab;4446 | 0.05 | 0.025 |
| NC_045512.2;YP_009724389.1;Wu1-SARS2_orf1ab;4447 | 0 | 0.05 |
| NC_045512.2;YP_009724389.1;Wu1-SARS2_orf1ab;4448 | 0 | 0.025 |
| NC_045512.2;YP_009724389.1;Wu1-SARS2_orf1ab;4449 | 0 | 0 |
| NC_045512.2;YP_009724389.1;Wu1-SARS2_orf1ab;4450 | 0 | 0.025 |
| NC_045512.2;YP_009724389.1;Wu1-SARS2_orf1ab;4451 | 0 | 0.025 |
| NC_045512.2;YP_009724389.1;Wu1-SARS2_orf1ab;4452 | 0 | 0.025 |
| NC_045512.2;YP_009724389.1;Wu1-SARS2_orf1ab;4453 | 0 | 0 |
| NC_045512.2;YP_009724389.1;Wu1-SARS2_orf1ab;4454 | 0 | 0 |
| NC_045512.2;YP_009724389.1;Wu1-SARS2_orf1ab;4455 | 0 | 0 |
| NC_045512.2;YP_009724389.1;Wu1-SARS2_orf1ab;4456 | 0 | 0 |
| NC_045512.2;YP_009724389.1;Wu1-SARS2_orf1ab;4457 | 0 | 0 |
| NC_045512.2;YP_009724389.1;Wu1-SARS2_orf1ab;4458 | 0 | 0 |
| NC_045512.2;YP_009724389.1;Wu1-SARS2_orf1ab;4459 | 0 | 0 |
| NC_045512.2;YP_009724389.1;Wu1-SARS2_orf1ab;4460 | 0 | 0 |
| NC_045512.2;YP_009724389.1;Wu1-SARS2_orf1ab;4461 | 0 | 0 |
| NC_045512.2;YP_009724389.1;Wu1-SARS2_orf1ab;4462 | 0 | 0 |

| | | |
|---|---|---|
| NC_045512.2;YP_009724389.1;Wu1-SARS2_orf1ab;4463 | 0 | 0 |
| NC_045512.2;YP_009724389.1;Wu1-SARS2_orf1ab;4464 | 0 | 0 |
| NC_045512.2;YP_009724389.1;Wu1-SARS2_orf1ab;4465 | 0 | 0 |
| NC_045512.2;YP_009724389.1;Wu1-SARS2_orf1ab;4466 | 0 | 0 |
| NC_045512.2;YP_009724389.1;Wu1-SARS2_orf1ab;4467 | 0 | 0 |
| NC_045512.2;YP_009724389.1;Wu1-SARS2_orf1ab;4468 | 0 | 0 |
| NC_045512.2;YP_009724389.1;Wu1-SARS2_orf1ab;4469 | 0.05 | 0.05 |
| NC_045512.2;YP_009724389.1;Wu1-SARS2_orf1ab;4470 | 0 | 0 |
| NC_045512.2;YP_009724389.1;Wu1-SARS2_orf1ab;4471 | 0 | 0 |
| NC_045512.2;YP_009724389.1;Wu1-SARS2_orf1ab;4472 | 0 | 0 |
| NC_045512.2;YP_009724389.1;Wu1-SARS2_orf1ab;4473 | 0 | 0 |
| NC_045512.2;YP_009724389.1;Wu1-SARS2_orf1ab;4474 | 0 | 0 |
| NC_045512.2;YP_009724389.1;Wu1-SARS2_orf1ab;4475 | 0 | 0 |
| NC_045512.2;YP_009724389.1;Wu1-SARS2_orf1ab;4476 | 0 | 0 |
| NC_045512.2;YP_009724389.1;Wu1-SARS2_orf1ab;4477 | 0 | 0.025 |
| NC_045512.2;YP_009724389.1;Wu1-SARS2_orf1ab;4478 | 0 | 0 |
| NC_045512.2;YP_009724389.1;Wu1-SARS2_orf1ab;4479 | 0 | 0.025 |
| NC_045512.2;YP_009724389.1;Wu1-SARS2_orf1ab;4480 | 0 | 0 |
| NC_045512.2;YP_009724389.1;Wu1-SARS2_orf1ab;4481 | 0 | 0.025 |
| NC_045512.2;YP_009724389.1;Wu1-SARS2_orf1ab;4482 | 0 | 0 |
| NC_045512.2;YP_009724389.1;Wu1-SARS2_orf1ab;4483 | 0 | 0 |
| NC_045512.2;YP_009724389.1;Wu1-SARS2_orf1ab;4484 | 0 | 0.025 |
| NC_045512.2;YP_009724389.1;Wu1-SARS2_orf1ab;4485 | 0 | 0.025 |
| NC_045512.2;YP_009724389.1;Wu1-SARS2_orf1ab;4486 | 0 | 0.05 |
| NC_045512.2;YP_009724389.1;Wu1-SARS2_orf1ab;4487 | 0 | 0.05 |
| NC_045512.2;YP_009724389.1;Wu1-SARS2_orf1ab;4488 | 0 | 0.05 |
| NC_045512.2;YP_009724389.1;Wu1-SARS2_orf1ab;4489 | 0 | 0.05 |
| NC_045512.2;YP_009724389.1;Wu1-SARS2_orf1ab;4490 | 0 | 0.025 |
| NC_045512.2;YP_009724389.1;Wu1-SARS2_orf1ab;4491 | 0 | 0.025 |
| NC_045512.2;YP_009724389.1;Wu1-SARS2_orf1ab;4492 | 0 | 0.025 |
| NC_045512.2;YP_009724389.1;Wu1-SARS2_orf1ab;4493 | 0 | 0 |
| NC_045512.2;YP_009724389.1;Wu1-SARS2_orf1ab;4494 | 0 | 0 |
| NC_045512.2;YP_009724389.1;Wu1-SARS2_orf1ab;4495 | 0 | 0 |
| NC_045512.2;YP_009724389.1;Wu1-SARS2_orf1ab;4496 | 0 | 0 |
| NC_045512.2;YP_009724389.1;Wu1-SARS2_orf1ab;4497 | 0 | 0 |
| NC_045512.2;YP_009724389.1;Wu1-SARS2_orf1ab;4498 | 0 | 0 |
| NC_045512.2;YP_009724389.1;Wu1-SARS2_orf1ab;4499 | 0 | 0 |
| NC_045512.2;YP_009724389.1;Wu1-SARS2_orf1ab;4500 | 0 | 0 |
| NC_045512.2;YP_009724389.1;Wu1-SARS2_orf1ab;4501 | 0 | 0 |
| NC_045512.2;YP_009724389.1;Wu1-SARS2_orf1ab;4502 | 0 | 0 |
| NC_045512.2;YP_009724389.1;Wu1-SARS2_orf1ab;4503 | 0 | 0 |
| NC_045512.2;YP_009724389.1;Wu1-SARS2_orf1ab;4504 | 0 | 0 |
| NC_045512.2;YP_009724389.1;Wu1-SARS2_orf1ab;4505 | 0 | 0 |

| | | |
|---|---|---|
| NC_045512.2;YP_009724389.1;Wu1-SARS2_orf1ab;4506 | 0 | 0 |
| NC_045512.2;YP_009724389.1;Wu1-SARS2_orf1ab;4507 | 0 | 0 |
| NC_045512.2;YP_009724389.1;Wu1-SARS2_orf1ab;4508 | 0 | 0 |
| NC_045512.2;YP_009724389.1;Wu1-SARS2_orf1ab;4509 | 0 | 0 |
| NC_045512.2;YP_009724389.1;Wu1-SARS2_orf1ab;4510 | 0 | 0 |
| NC_045512.2;YP_009724389.1;Wu1-SARS2_orf1ab;4511 | 0 | 0 |
| NC_045512.2;YP_009724389.1;Wu1-SARS2_orf1ab;4512 | 0 | 0 |
| NC_045512.2;YP_009724389.1;Wu1-SARS2_orf1ab;4513 | 0 | 0 |
| NC_045512.2;YP_009724389.1;Wu1-SARS2_orf1ab;4514 | 0 | 0.375 |
| NC_045512.2;YP_009724389.1;Wu1-SARS2_orf1ab;4515 | 0 | 0 |
| NC_045512.2;YP_009724389.1;Wu1-SARS2_orf1ab;4516 | 0 | 0 |
| NC_045512.2;YP_009724389.1;Wu1-SARS2_orf1ab;4517 | 0 | 0 |
| NC_045512.2;YP_009724389.1;Wu1-SARS2_orf1ab;4518 | 0 | 0 |
| NC_045512.2;YP_009724389.1;Wu1-SARS2_orf1ab;4519 | 0 | 0 |
| NC_045512.2;YP_009724389.1;Wu1-SARS2_orf1ab;4520 | 0.05 | 0.1 |
| NC_045512.2;YP_009724389.1;Wu1-SARS2_orf1ab;4521 | 0 | 0.025 |
| NC_045512.2;YP_009724389.1;Wu1-SARS2_orf1ab;4522 | 0 | 0 |
| NC_045512.2;YP_009724389.1;Wu1-SARS2_orf1ab;4523 | 0 | 0 |
| NC_045512.2;YP_009724389.1;Wu1-SARS2_orf1ab;4524 | 0 | 0 |
| NC_045512.2;YP_009724389.1;Wu1-SARS2_orf1ab;4525 | 0 | 0 |
| NC_045512.2;YP_009724389.1;Wu1-SARS2_orf1ab;4526 | 0 | 0.025 |
| NC_045512.2;YP_009724389.1;Wu1-SARS2_orf1ab;4527 | 0 | 0 |
| NC_045512.2;YP_009724389.1;Wu1-SARS2_orf1ab;4528 | 0 | 0 |
| NC_045512.2;YP_009724389.1;Wu1-SARS2_orf1ab;4529 | 0 | 0 |
| NC_045512.2;YP_009724389.1;Wu1-SARS2_orf1ab;4530 | 0 | 0 |
| NC_045512.2;YP_009724389.1;Wu1-SARS2_orf1ab;4531 | 0 | 0 |
| NC_045512.2;YP_009724389.1;Wu1-SARS2_orf1ab;4532 | 0 | 0 |
| NC_045512.2;YP_009724389.1;Wu1-SARS2_orf1ab;4533 | 0 | 0.025 |
| NC_045512.2;YP_009724389.1;Wu1-SARS2_orf1ab;4534 | 0 | 0 |
| NC_045512.2;YP_009724389.1;Wu1-SARS2_orf1ab;4535 | 0 | 0 |
| NC_045512.2;YP_009724389.1;Wu1-SARS2_orf1ab;4536 | 0 | 0 |
| NC_045512.2;YP_009724389.1;Wu1-SARS2_orf1ab;4537 | 0 | 0 |
| NC_045512.2;YP_009724389.1;Wu1-SARS2_orf1ab;4538 | 0 | 0.025 |
| NC_045512.2;YP_009724389.1;Wu1-SARS2_orf1ab;4539 | 0 | 0 |
| NC_045512.2;YP_009724389.1;Wu1-SARS2_orf1ab;4540 | 0 | 0 |
| NC_045512.2;YP_009724389.1;Wu1-SARS2_orf1ab;4541 | 0 | 0.025 |
| NC_045512.2;YP_009724389.1;Wu1-SARS2_orf1ab;4542 | 0 | 0.025 |
| NC_045512.2;YP_009724389.1;Wu1-SARS2_orf1ab;4543 | 0 | 0 |
| NC_045512.2;YP_009724389.1;Wu1-SARS2_orf1ab;4544 | 0 | 0 |
| NC_045512.2;YP_009724389.1;Wu1-SARS2_orf1ab;4545 | 0.2 | 0.025 |
| NC_045512.2;YP_009724389.1;Wu1-SARS2_orf1ab;4546 | 0.05 | 0 |
| NC_045512.2;YP_009724389.1;Wu1-SARS2_orf1ab;4547 | 0 | 0.025 |
| NC_045512.2;YP_009724389.1;Wu1-SARS2_orf1ab;4548 | 0 | 0.075 |

| | | |
|---|---|---|
| NC_045512.2;YP_009724389.1;Wu1-SARS2_orf1ab;4549 | 0.05 | 0.075 |
| NC_045512.2;YP_009724389.1;Wu1-SARS2_orf1ab;4550 | 0.05 | 0.075 |
| NC_045512.2;YP_009724389.1;Wu1-SARS2_orf1ab;4551 | 0.1 | 0.05 |
| NC_045512.2;YP_009724389.1;Wu1-SARS2_orf1ab;4552 | 0 | 0.025 |
| NC_045512.2;YP_009724389.1;Wu1-SARS2_orf1ab;4553 | 0 | 0 |
| NC_045512.2;YP_009724389.1;Wu1-SARS2_orf1ab;4554 | 0 | 0 |
| NC_045512.2;YP_009724389.1;Wu1-SARS2_orf1ab;4555 | 0 | 0 |
| NC_045512.2;YP_009724389.1;Wu1-SARS2_orf1ab;4556 | 0 | 0 |
| NC_045512.2;YP_009724389.1;Wu1-SARS2_orf1ab;4557 | 0.05 | 0 |
| NC_045512.2;YP_009724389.1;Wu1-SARS2_orf1ab;4558 | 0.05 | 0.025 |
| NC_045512.2;YP_009724389.1;Wu1-SARS2_orf1ab;4559 | 0.05 | 0 |
| NC_045512.2;YP_009724389.1;Wu1-SARS2_orf1ab;4560 | 0 | 0 |
| NC_045512.2;YP_009724389.1;Wu1-SARS2_orf1ab;4561 | 0 | 0 |
| NC_045512.2;YP_009724389.1;Wu1-SARS2_orf1ab;4562 | 0 | 0 |
| NC_045512.2;YP_009724389.1;Wu1-SARS2_orf1ab;4563 | 0 | 0 |
| NC_045512.2;YP_009724389.1;Wu1-SARS2_orf1ab;4564 | 0 | 0 |
| NC_045512.2;YP_009724389.1;Wu1-SARS2_orf1ab;4565 | 0.05 | 0 |
| NC_045512.2;YP_009724389.1;Wu1-SARS2_orf1ab;4566 | 0 | 0 |
| NC_045512.2;YP_009724389.1;Wu1-SARS2_orf1ab;4567 | 0 | 0 |
| NC_045512.2;YP_009724389.1;Wu1-SARS2_orf1ab;4568 | 0 | 0 |
| NC_045512.2;YP_009724389.1;Wu1-SARS2_orf1ab;4569 | 0 | 0 |
| NC_045512.2;YP_009724389.1;Wu1-SARS2_orf1ab;4570 | 0 | 0 |
| NC_045512.2;YP_009724389.1;Wu1-SARS2_orf1ab;4571 | 0 | 0 |
| NC_045512.2;YP_009724389.1;Wu1-SARS2_orf1ab;4572 | 0.05 | 0 |
| NC_045512.2;YP_009724389.1;Wu1-SARS2_orf1ab;4573 | 0 | 0 |
| NC_045512.2;YP_009724389.1;Wu1-SARS2_orf1ab;4574 | 0 | 0 |
| NC_045512.2;YP_009724389.1;Wu1-SARS2_orf1ab;4575 | 0 | 0 |
| NC_045512.2;YP_009724389.1;Wu1-SARS2_orf1ab;4576 | 0 | 0 |
| NC_045512.2;YP_009724389.1;Wu1-SARS2_orf1ab;4577 | 0 | 0 |
| NC_045512.2;YP_009724389.1;Wu1-SARS2_orf1ab;4578 | 0 | 0 |
| NC_045512.2;YP_009724389.1;Wu1-SARS2_orf1ab;4579 | 0 | 0 |
| NC_045512.2;YP_009724389.1;Wu1-SARS2_orf1ab;4580 | 0 | 0 |
| NC_045512.2;YP_009724389.1;Wu1-SARS2_orf1ab;4581 | 0 | 0 |
| NC_045512.2;YP_009724389.1;Wu1-SARS2_orf1ab;4582 | 0 | 0 |
| NC_045512.2;YP_009724389.1;Wu1-SARS2_orf1ab;4583 | 0 | 0.025 |
| NC_045512.2;YP_009724389.1;Wu1-SARS2_orf1ab;4584 | 0 | 0 |
| NC_045512.2;YP_009724389.1;Wu1-SARS2_orf1ab;4585 | 0 | 0 |
| NC_045512.2;YP_009724389.1;Wu1-SARS2_orf1ab;4586 | 0 | 0 |
| NC_045512.2;YP_009724389.1;Wu1-SARS2_orf1ab;4587 | 0 | 0 |
| NC_045512.2;YP_009724389.1;Wu1-SARS2_orf1ab;4588 | 0 | 0 |
| NC_045512.2;YP_009724389.1;Wu1-SARS2_orf1ab;4589 | 0 | 0 |
| NC_045512.2;YP_009724389.1;Wu1-SARS2_orf1ab;4590 | 0 | 0 |
| NC_045512.2;YP_009724389.1;Wu1-SARS2_orf1ab;4591 | 0.05 | 0 |

| | | |
|---|---|---|
| NC_045512.2;YP_009724389.1;Wu1-SARS2_orf1ab;4592 | 0 | 0 |
| NC_045512.2;YP_009724389.1;Wu1-SARS2_orf1ab;4593 | 0 | 0 |
| NC_045512.2;YP_009724389.1;Wu1-SARS2_orf1ab;4594 | 0 | 0 |
| NC_045512.2;YP_009724389.1;Wu1-SARS2_orf1ab;4595 | 0.05 | 0 |
| NC_045512.2;YP_009724389.1;Wu1-SARS2_orf1ab;4596 | 0 | 0 |
| NC_045512.2;YP_009724389.1;Wu1-SARS2_orf1ab;4597 | 0 | 0 |
| NC_045512.2;YP_009724389.1;Wu1-SARS2_orf1ab;4598 | 0 | 0 |
| NC_045512.2;YP_009724389.1;Wu1-SARS2_orf1ab;4599 | 0 | 0 |
| NC_045512.2;YP_009724389.1;Wu1-SARS2_orf1ab;4600 | 0 | 0 |
| NC_045512.2;YP_009724389.1;Wu1-SARS2_orf1ab;4601 | 0 | 0 |
| NC_045512.2;YP_009724389.1;Wu1-SARS2_orf1ab;4602 | 0 | 0.025 |
| NC_045512.2;YP_009724389.1;Wu1-SARS2_orf1ab;4603 | 0 | 0 |
| NC_045512.2;YP_009724389.1;Wu1-SARS2_orf1ab;4604 | 0 | 0 |
| NC_045512.2;YP_009724389.1;Wu1-SARS2_orf1ab;4605 | 0 | 0.025 |
| NC_045512.2;YP_009724389.1;Wu1-SARS2_orf1ab;4606 | 0 | 0 |
| NC_045512.2;YP_009724389.1;Wu1-SARS2_orf1ab;4607 | 0 | 0 |
| NC_045512.2;YP_009724389.1;Wu1-SARS2_orf1ab;4608 | 0 | 0 |
| NC_045512.2;YP_009724389.1;Wu1-SARS2_orf1ab;4609 | 0.1 | 0.05 |
| NC_045512.2;YP_009724389.1;Wu1-SARS2_orf1ab;4610 | 0 | 0 |
| NC_045512.2;YP_009724389.1;Wu1-SARS2_orf1ab;4611 | 0 | 0 |
| NC_045512.2;YP_009724389.1;Wu1-SARS2_orf1ab;4612 | 0 | 0 |
| NC_045512.2;YP_009724389.1;Wu1-SARS2_orf1ab;4613 | 0 | 0 |
| NC_045512.2;YP_009724389.1;Wu1-SARS2_orf1ab;4614 | 0 | 0 |
| NC_045512.2;YP_009724389.1;Wu1-SARS2_orf1ab;4615 | 0 | 0 |
| NC_045512.2;YP_009724389.1;Wu1-SARS2_orf1ab;4616 | 0 | 0.025 |
| NC_045512.2;YP_009724389.1;Wu1-SARS2_orf1ab;4617 | 0 | 0 |
| NC_045512.2;YP_009724389.1;Wu1-SARS2_orf1ab;4618 | 0 | 0 |
| NC_045512.2;YP_009724389.1;Wu1-SARS2_orf1ab;4619 | 0 | 0 |
| NC_045512.2;YP_009724389.1;Wu1-SARS2_orf1ab;4620 | 0 | 0 |
| NC_045512.2;YP_009724389.1;Wu1-SARS2_orf1ab;4621 | 0 | 0 |
| NC_045512.2;YP_009724389.1;Wu1-SARS2_orf1ab;4622 | 0 | 0 |
| NC_045512.2;YP_009724389.1;Wu1-SARS2_orf1ab;4623 | 0 | 0 |
| NC_045512.2;YP_009724389.1;Wu1-SARS2_orf1ab;4624 | 0 | 0 |
| NC_045512.2;YP_009724389.1;Wu1-SARS2_orf1ab;4625 | 0 | 0 |
| NC_045512.2;YP_009724389.1;Wu1-SARS2_orf1ab;4626 | 0 | 0 |
| NC_045512.2;YP_009724389.1;Wu1-SARS2_orf1ab;4627 | 0 | 0 |
| NC_045512.2;YP_009724389.1;Wu1-SARS2_orf1ab;4628 | 0 | 0 |
| NC_045512.2;YP_009724389.1;Wu1-SARS2_orf1ab;4629 | 0 | 0 |
| NC_045512.2;YP_009724389.1;Wu1-SARS2_orf1ab;4630 | 0 | 0 |
| NC_045512.2;YP_009724389.1;Wu1-SARS2_orf1ab;4631 | 0 | 0 |
| NC_045512.2;YP_009724389.1;Wu1-SARS2_orf1ab;4632 | 0 | 0 |
| NC_045512.2;YP_009724389.1;Wu1-SARS2_orf1ab;4633 | 0 | 0 |
| NC_045512.2;YP_009724389.1;Wu1-SARS2_orf1ab;4634 | 0 | 0 |

| | | |
|---|---|---|
| NC_045512.2;YP_009724389.1;Wu1-SARS2_orf1ab;4635 | 0.05 | 0 |
| NC_045512.2;YP_009724389.1;Wu1-SARS2_orf1ab;4636 | 0 | 0 |
| NC_045512.2;YP_009724389.1;Wu1-SARS2_orf1ab;4637 | 0 | 0 |
| NC_045512.2;YP_009724389.1;Wu1-SARS2_orf1ab;4638 | 0 | 0 |
| NC_045512.2;YP_009724389.1;Wu1-SARS2_orf1ab;4639 | 0 | 0 |
| NC_045512.2;YP_009724389.1;Wu1-SARS2_orf1ab;4640 | 0 | 0 |
| NC_045512.2;YP_009724389.1;Wu1-SARS2_orf1ab;4641 | 0.05 | 0 |
| NC_045512.2;YP_009724389.1;Wu1-SARS2_orf1ab;4642 | 0.05 | 0 |
| NC_045512.2;YP_009724389.1;Wu1-SARS2_orf1ab;4643 | 0.05 | 0 |
| NC_045512.2;YP_009724389.1;Wu1-SARS2_orf1ab;4644 | 0.05 | 0 |
| NC_045512.2;YP_009724389.1;Wu1-SARS2_orf1ab;4645 | 0 | 0.025 |
| NC_045512.2;YP_009724389.1;Wu1-SARS2_orf1ab;4646 | 0.05 | 0.075 |
| NC_045512.2;YP_009724389.1;Wu1-SARS2_orf1ab;4647 | 0 | 0.025 |
| NC_045512.2;YP_009724389.1;Wu1-SARS2_orf1ab;4648 | 0 | 0 |
| NC_045512.2;YP_009724389.1;Wu1-SARS2_orf1ab;4649 | 0.05 | 0.05 |
| NC_045512.2;YP_009724389.1;Wu1-SARS2_orf1ab;4650 | 0.1 | 0 |
| NC_045512.2;YP_009724389.1;Wu1-SARS2_orf1ab;4651 | 0 | 0 |
| NC_045512.2;YP_009724389.1;Wu1-SARS2_orf1ab;4652 | 0 | 0 |
| NC_045512.2;YP_009724389.1;Wu1-SARS2_orf1ab;4653 | 0 | 0 |
| NC_045512.2;YP_009724389.1;Wu1-SARS2_orf1ab;4654 | 0 | 0 |
| NC_045512.2;YP_009724389.1;Wu1-SARS2_orf1ab;4655 | 0 | 0 |
| NC_045512.2;YP_009724389.1;Wu1-SARS2_orf1ab;4656 | 0.05 | 0 |
| NC_045512.2;YP_009724389.1;Wu1-SARS2_orf1ab;4657 | 0.05 | 0 |
| NC_045512.2;YP_009724389.1;Wu1-SARS2_orf1ab;4658 | 0.1 | 0 |
| NC_045512.2;YP_009724389.1;Wu1-SARS2_orf1ab;4659 | 0.05 | 0.05 |
| NC_045512.2;YP_009724389.1;Wu1-SARS2_orf1ab;4660 | 0.1 | 0.025 |
| NC_045512.2;YP_009724389.1;Wu1-SARS2_orf1ab;4661 | 0.05 | 0.025 |
| NC_045512.2;YP_009724389.1;Wu1-SARS2_orf1ab;4662 | 0.05 | 0.05 |
| NC_045512.2;YP_009724389.1;Wu1-SARS2_orf1ab;4663 | 0.05 | 0.075 |
| NC_045512.2;YP_009724389.1;Wu1-SARS2_orf1ab;4664 | 0.05 | 0.175 |
| NC_045512.2;YP_009724389.1;Wu1-SARS2_orf1ab;4665 | 0.05 | 0.125 |
| NC_045512.2;YP_009724389.1;Wu1-SARS2_orf1ab;4666 | 0 | 0 |
| NC_045512.2;YP_009724389.1;Wu1-SARS2_orf1ab;4667 | 0 | 0 |
| NC_045512.2;YP_009724389.1;Wu1-SARS2_orf1ab;4668 | 0 | 0 |
| NC_045512.2;YP_009724389.1;Wu1-SARS2_orf1ab;4669 | 0 | 0 |
| NC_045512.2;YP_009724389.1;Wu1-SARS2_orf1ab;4670 | 0 | 0.025 |
| NC_045512.2;YP_009724389.1;Wu1-SARS2_orf1ab;4671 | 0 | 0 |
| NC_045512.2;YP_009724389.1;Wu1-SARS2_orf1ab;4672 | 0 | 0 |
| NC_045512.2;YP_009724389.1;Wu1-SARS2_orf1ab;4673 | 0 | 0 |
| NC_045512.2;YP_009724389.1;Wu1-SARS2_orf1ab;4674 | 0 | 0.025 |
| NC_045512.2;YP_009724389.1;Wu1-SARS2_orf1ab;4675 | 0 | 0 |
| NC_045512.2;YP_009724389.1;Wu1-SARS2_orf1ab;4676 | 0 | 0.025 |
| NC_045512.2;YP_009724389.1;Wu1-SARS2_orf1ab;4677 | 0 | 0.025 |

| | | |
|---|---|---|
| NC_045512.2;YP_009724389.1;Wu1-SARS2_orf1ab;4678 | 0 | 0.05 |
| NC_045512.2;YP_009724389.1;Wu1-SARS2_orf1ab;4679 | 0 | 0.075 |
| NC_045512.2;YP_009724389.1;Wu1-SARS2_orf1ab;4680 | 0 | 0.1 |
| NC_045512.2;YP_009724389.1;Wu1-SARS2_orf1ab;4681 | 0 | 0.05 |
| NC_045512.2;YP_009724389.1;Wu1-SARS2_orf1ab;4682 | 0 | 0.025 |
| NC_045512.2;YP_009724389.1;Wu1-SARS2_orf1ab;4683 | 0 | 0.05 |
| NC_045512.2;YP_009724389.1;Wu1-SARS2_orf1ab;4684 | 0 | 0.05 |
| NC_045512.2;YP_009724389.1;Wu1-SARS2_orf1ab;4685 | 0 | 0.025 |
| NC_045512.2;YP_009724389.1;Wu1-SARS2_orf1ab;4686 | 0 | 0.025 |
| NC_045512.2;YP_009724389.1;Wu1-SARS2_orf1ab;4687 | 0 | 0.025 |
| NC_045512.2;YP_009724389.1;Wu1-SARS2_orf1ab;4688 | 0 | 0 |
| NC_045512.2;YP_009724389.1;Wu1-SARS2_orf1ab;4689 | 0 | 0 |
| NC_045512.2;YP_009724389.1;Wu1-SARS2_orf1ab;4690 | 0 | 0 |
| NC_045512.2;YP_009724389.1;Wu1-SARS2_orf1ab;4691 | 0 | 0 |
| NC_045512.2;YP_009724389.1;Wu1-SARS2_orf1ab;4692 | 0 | 0 |
| NC_045512.2;YP_009724389.1;Wu1-SARS2_orf1ab;4693 | 0 | 0 |
| NC_045512.2;YP_009724389.1;Wu1-SARS2_orf1ab;4694 | 0 | 0 |
| NC_045512.2;YP_009724389.1;Wu1-SARS2_orf1ab;4695 | 0 | 0 |
| NC_045512.2;YP_009724389.1;Wu1-SARS2_orf1ab;4696 | 0 | 0 |
| NC_045512.2;YP_009724389.1;Wu1-SARS2_orf1ab;4697 | 0 | 0 |
| NC_045512.2;YP_009724389.1;Wu1-SARS2_orf1ab;4698 | 0 | 0 |
| NC_045512.2;YP_009724389.1;Wu1-SARS2_orf1ab;4699 | 0 | 0 |
| NC_045512.2;YP_009724389.1;Wu1-SARS2_orf1ab;4700 | 0 | 0 |
| NC_045512.2;YP_009724389.1;Wu1-SARS2_orf1ab;4701 | 0 | 0 |
| NC_045512.2;YP_009724389.1;Wu1-SARS2_orf1ab;4702 | 0 | 0 |
| NC_045512.2;YP_009724389.1;Wu1-SARS2_orf1ab;4703 | 0 | 0 |
| NC_045512.2;YP_009724389.1;Wu1-SARS2_orf1ab;4704 | 0 | 0 |
| NC_045512.2;YP_009724389.1;Wu1-SARS2_orf1ab;4705 | 0 | 0 |
| NC_045512.2;YP_009724389.1;Wu1-SARS2_orf1ab;4706 | 0 | 0 |
| NC_045512.2;YP_009724389.1;Wu1-SARS2_orf1ab;4707 | 0 | 0 |
| NC_045512.2;YP_009724389.1;Wu1-SARS2_orf1ab;4708 | 0 | 0 |
| NC_045512.2;YP_009724389.1;Wu1-SARS2_orf1ab;4709 | 0 | 0 |
| NC_045512.2;YP_009724389.1;Wu1-SARS2_orf1ab;4710 | 0 | 0 |
| NC_045512.2;YP_009724389.1;Wu1-SARS2_orf1ab;4711 | 0 | 0 |
| NC_045512.2;YP_009724389.1;Wu1-SARS2_orf1ab;4712 | 0 | 0 |
| NC_045512.2;YP_009724389.1;Wu1-SARS2_orf1ab;4713 | 0 | 0 |
| NC_045512.2;YP_009724389.1;Wu1-SARS2_orf1ab;4714 | 0 | 0 |
| NC_045512.2;YP_009724389.1;Wu1-SARS2_orf1ab;4715 | 0 | 0 |
| NC_045512.2;YP_009724389.1;Wu1-SARS2_orf1ab;4716 | 0 | 0 |
| NC_045512.2;YP_009724389.1;Wu1-SARS2_orf1ab;4717 | 0 | 0 |
| NC_045512.2;YP_009724389.1;Wu1-SARS2_orf1ab;4718 | 0 | 0 |
| NC_045512.2;YP_009724389.1;Wu1-SARS2_orf1ab;4719 | 0 | 0 |
| NC_045512.2;YP_009724389.1;Wu1-SARS2_orf1ab;4720 | 0 | 0 |

| | | |
|---|---|---|
| NC_045512.2;YP_009724389.1;Wu1-SARS2_orf1ab;4721 | 0 | 0 |
| NC_045512.2;YP_009724389.1;Wu1-SARS2_orf1ab;4722 | 0 | 0 |
| NC_045512.2;YP_009724389.1;Wu1-SARS2_orf1ab;4723 | 0 | 0 |
| NC_045512.2;YP_009724389.1;Wu1-SARS2_orf1ab;4724 | 0 | 0 |
| NC_045512.2;YP_009724389.1;Wu1-SARS2_orf1ab;4725 | 0 | 0 |
| NC_045512.2;YP_009724389.1;Wu1-SARS2_orf1ab;4726 | 0 | 0 |
| NC_045512.2;YP_009724389.1;Wu1-SARS2_orf1ab;4727 | 0 | 0 |
| NC_045512.2;YP_009724389.1;Wu1-SARS2_orf1ab;4728 | 0 | 0 |
| NC_045512.2;YP_009724389.1;Wu1-SARS2_orf1ab;4729 | 0 | 0 |
| NC_045512.2;YP_009724389.1;Wu1-SARS2_orf1ab;4730 | 0 | 0 |
| NC_045512.2;YP_009724389.1;Wu1-SARS2_orf1ab;4731 | 0 | 0 |
| NC_045512.2;YP_009724389.1;Wu1-SARS2_orf1ab;4732 | 0 | 0 |
| NC_045512.2;YP_009724389.1;Wu1-SARS2_orf1ab;4733 | 0 | 0 |
| NC_045512.2;YP_009724389.1;Wu1-SARS2_orf1ab;4734 | 0 | 0 |
| NC_045512.2;YP_009724389.1;Wu1-SARS2_orf1ab;4735 | 0 | 0 |
| NC_045512.2;YP_009724389.1;Wu1-SARS2_orf1ab;4736 | 0 | 0 |
| NC_045512.2;YP_009724389.1;Wu1-SARS2_orf1ab;4737 | 0 | 0 |
| NC_045512.2;YP_009724389.1;Wu1-SARS2_orf1ab;4738 | 0 | 0 |
| NC_045512.2;YP_009724389.1;Wu1-SARS2_orf1ab;4739 | 0 | 0 |
| NC_045512.2;YP_009724389.1;Wu1-SARS2_orf1ab;4740 | 0 | 0 |
| NC_045512.2;YP_009724389.1;Wu1-SARS2_orf1ab;4741 | 0 | 0 |
| NC_045512.2;YP_009724389.1;Wu1-SARS2_orf1ab;4742 | 0 | 0 |
| NC_045512.2;YP_009724389.1;Wu1-SARS2_orf1ab;4743 | 0 | 0 |
| NC_045512.2;YP_009724389.1;Wu1-SARS2_orf1ab;4744 | 0 | 0 |
| NC_045512.2;YP_009724389.1;Wu1-SARS2_orf1ab;4745 | 0 | 0 |
| NC_045512.2;YP_009724389.1;Wu1-SARS2_orf1ab;4746 | 0 | 0 |
| NC_045512.2;YP_009724389.1;Wu1-SARS2_orf1ab;4747 | 0 | 0.025 |
| NC_045512.2;YP_009724389.1;Wu1-SARS2_orf1ab;4748 | 0 | 0.075 |
| NC_045512.2;YP_009724389.1;Wu1-SARS2_orf1ab;4749 | 0.05 | 0.025 |
| NC_045512.2;YP_009724389.1;Wu1-SARS2_orf1ab;4750 | 0 | 0 |
| NC_045512.2;YP_009724389.1;Wu1-SARS2_orf1ab;4751 | 0 | 0 |
| NC_045512.2;YP_009724389.1;Wu1-SARS2_orf1ab;4752 | 0 | 0 |
| NC_045512.2;YP_009724389.1;Wu1-SARS2_orf1ab;4753 | 0 | 0 |
| NC_045512.2;YP_009724389.1;Wu1-SARS2_orf1ab;4754 | 0 | 0 |
| NC_045512.2;YP_009724389.1;Wu1-SARS2_orf1ab;4755 | 0 | 0 |
| NC_045512.2;YP_009724389.1;Wu1-SARS2_orf1ab;4756 | 0 | 0 |
| NC_045512.2;YP_009724389.1;Wu1-SARS2_orf1ab;4757 | 0.05 | 0 |
| NC_045512.2;YP_009724389.1;Wu1-SARS2_orf1ab;4758 | 0.1 | 0.025 |
| NC_045512.2;YP_009724389.1;Wu1-SARS2_orf1ab;4759 | 0.1 | 0.025 |
| NC_045512.2;YP_009724389.1;Wu1-SARS2_orf1ab;4760 | 0.05 | 0.025 |
| NC_045512.2;YP_009724389.1;Wu1-SARS2_orf1ab;4761 | 0 | 0 |
| NC_045512.2;YP_009724389.1;Wu1-SARS2_orf1ab;4762 | 0 | 0 |
| NC_045512.2;YP_009724389.1;Wu1-SARS2_orf1ab;4763 | 0 | 0 |

| | | |
|---|---|---|
| NC_045512.2;YP_009724389.1;Wu1-SARS2_orf1ab;4764 | 0 | 0.025 |
| NC_045512.2;YP_009724389.1;Wu1-SARS2_orf1ab;4765 | 0 | 0.025 |
| NC_045512.2;YP_009724389.1;Wu1-SARS2_orf1ab;4766 | 0 | 0 |
| NC_045512.2;YP_009724389.1;Wu1-SARS2_orf1ab;4767 | 0 | 0 |
| NC_045512.2;YP_009724389.1;Wu1-SARS2_orf1ab;4768 | 0 | 0 |
| NC_045512.2;YP_009724389.1;Wu1-SARS2_orf1ab;4769 | 0 | 0 |
| NC_045512.2;YP_009724389.1;Wu1-SARS2_orf1ab;4770 | 0 | 0 |
| NC_045512.2;YP_009724389.1;Wu1-SARS2_orf1ab;4771 | 0 | 0 |
| NC_045512.2;YP_009724389.1;Wu1-SARS2_orf1ab;4772 | 0 | 0 |
| NC_045512.2;YP_009724389.1;Wu1-SARS2_orf1ab;4773 | 0 | 0 |
| NC_045512.2;YP_009724389.1;Wu1-SARS2_orf1ab;4774 | 0 | 0 |
| NC_045512.2;YP_009724389.1;Wu1-SARS2_orf1ab;4775 | 0 | 0 |
| NC_045512.2;YP_009724389.1;Wu1-SARS2_orf1ab;4776 | 0 | 0 |
| NC_045512.2;YP_009724389.1;Wu1-SARS2_orf1ab;4777 | 0.05 | 0 |
| NC_045512.2;YP_009724389.1;Wu1-SARS2_orf1ab;4778 | 0 | 0 |
| NC_045512.2;YP_009724389.1;Wu1-SARS2_orf1ab;4779 | 0 | 0 |
| NC_045512.2;YP_009724389.1;Wu1-SARS2_orf1ab;4780 | 0 | 0 |
| NC_045512.2;YP_009724389.1;Wu1-SARS2_orf1ab;4781 | 0 | 0 |
| NC_045512.2;YP_009724389.1;Wu1-SARS2_orf1ab;4782 | 0 | 0 |
| NC_045512.2;YP_009724389.1;Wu1-SARS2_orf1ab;4783 | 0 | 0 |
| NC_045512.2;YP_009724389.1;Wu1-SARS2_orf1ab;4784 | 0 | 0 |
| NC_045512.2;YP_009724389.1;Wu1-SARS2_orf1ab;4785 | 0 | 0 |
| NC_045512.2;YP_009724389.1;Wu1-SARS2_orf1ab;4786 | 0 | 0 |
| NC_045512.2;YP_009724389.1;Wu1-SARS2_orf1ab;4787 | 0 | 0 |
| NC_045512.2;YP_009724389.1;Wu1-SARS2_orf1ab;4788 | 0 | 0 |
| NC_045512.2;YP_009724389.1;Wu1-SARS2_orf1ab;4789 | 0 | 0 |
| NC_045512.2;YP_009724389.1;Wu1-SARS2_orf1ab;4790 | 0 | 0 |
| NC_045512.2;YP_009724389.1;Wu1-SARS2_orf1ab;4791 | 0.05 | 0 |
| NC_045512.2;YP_009724389.1;Wu1-SARS2_orf1ab;4792 | 0.05 | 0.05 |
| NC_045512.2;YP_009724389.1;Wu1-SARS2_orf1ab;4793 | 0 | 0 |
| NC_045512.2;YP_009724389.1;Wu1-SARS2_orf1ab;4794 | 0 | 0 |
| NC_045512.2;YP_009724389.1;Wu1-SARS2_orf1ab;4795 | 0 | 0.025 |
| NC_045512.2;YP_009724389.1;Wu1-SARS2_orf1ab;4796 | 0 | 0.025 |
| NC_045512.2;YP_009724389.1;Wu1-SARS2_orf1ab;4797 | 0 | 0 |
| NC_045512.2;YP_009724389.1;Wu1-SARS2_orf1ab;4798 | 0 | 0 |
| NC_045512.2;YP_009724389.1;Wu1-SARS2_orf1ab;4799 | 0.05 | 0 |
| NC_045512.2;YP_009724389.1;Wu1-SARS2_orf1ab;4800 | 0.05 | 0.025 |
| NC_045512.2;YP_009724389.1;Wu1-SARS2_orf1ab;4801 | 0.1 | 0.025 |
| NC_045512.2;YP_009724389.1;Wu1-SARS2_orf1ab;4802 | 0.1 | 0.025 |
| NC_045512.2;YP_009724389.1;Wu1-SARS2_orf1ab;4803 | 0.1 | 0.025 |
| NC_045512.2;YP_009724389.1;Wu1-SARS2_orf1ab;4804 | 0.2 | 0.025 |
| NC_045512.2;YP_009724389.1;Wu1-SARS2_orf1ab;4805 | 0.1 | 0 |
| NC_045512.2;YP_009724389.1;Wu1-SARS2_orf1ab;4806 | 0 | 0 |

| | | |
|---|---|---|
| NC_045512.2;YP_009724389.1;Wu1-SARS2_orf1ab;4807 | 0 | 0 |
| NC_045512.2;YP_009724389.1;Wu1-SARS2_orf1ab;4808 | 0.05 | 0 |
| NC_045512.2;YP_009724389.1;Wu1-SARS2_orf1ab;4809 | 0.05 | 0 |
| NC_045512.2;YP_009724389.1;Wu1-SARS2_orf1ab;4810 | 0.05 | 0 |
| NC_045512.2;YP_009724389.1;Wu1-SARS2_orf1ab;4811 | 0 | 0 |
| NC_045512.2;YP_009724389.1;Wu1-SARS2_orf1ab;4812 | 0 | 0 |
| NC_045512.2;YP_009724389.1;Wu1-SARS2_orf1ab;4813 | 0 | 0 |
| NC_045512.2;YP_009724389.1;Wu1-SARS2_orf1ab;4814 | 0 | 0 |
| NC_045512.2;YP_009724389.1;Wu1-SARS2_orf1ab;4815 | 0 | 0 |
| NC_045512.2;YP_009724389.1;Wu1-SARS2_orf1ab;4816 | 0 | 0 |
| NC_045512.2;YP_009724389.1;Wu1-SARS2_orf1ab;4817 | 0.05 | 0 |
| NC_045512.2;YP_009724389.1;Wu1-SARS2_orf1ab;4818 | 0 | 0.025 |
| NC_045512.2;YP_009724389.1;Wu1-SARS2_orf1ab;4819 | 0 | 0 |
| NC_045512.2;YP_009724389.1;Wu1-SARS2_orf1ab;4820 | 0 | 0 |
| NC_045512.2;YP_009724389.1;Wu1-SARS2_orf1ab;4821 | 0 | 0 |
| NC_045512.2;YP_009724389.1;Wu1-SARS2_orf1ab;4822 | 0 | 0 |
| NC_045512.2;YP_009724389.1;Wu1-SARS2_orf1ab;4823 | 0 | 0 |
| NC_045512.2;YP_009724389.1;Wu1-SARS2_orf1ab;4824 | 0 | 0 |
| NC_045512.2;YP_009724389.1;Wu1-SARS2_orf1ab;4825 | 0 | 0 |
| NC_045512.2;YP_009724389.1;Wu1-SARS2_orf1ab;4826 | 0.05 | 0 |
| NC_045512.2;YP_009724389.1;Wu1-SARS2_orf1ab;4827 | 0.05 | 0 |
| NC_045512.2;YP_009724389.1;Wu1-SARS2_orf1ab;4828 | 0 | 0 |
| NC_045512.2;YP_009724389.1;Wu1-SARS2_orf1ab;4829 | 0 | 0 |
| NC_045512.2;YP_009724389.1;Wu1-SARS2_orf1ab;4830 | 0.05 | 0 |
| NC_045512.2;YP_009724389.1;Wu1-SARS2_orf1ab;4831 | 0 | 0 |
| NC_045512.2;YP_009724389.1;Wu1-SARS2_orf1ab;4832 | 0 | 0 |
| NC_045512.2;YP_009724389.1;Wu1-SARS2_orf1ab;4833 | 0 | 0 |
| NC_045512.2;YP_009724389.1;Wu1-SARS2_orf1ab;4834 | 0 | 0.025 |
| NC_045512.2;YP_009724389.1;Wu1-SARS2_orf1ab;4835 | 0 | 0 |
| NC_045512.2;YP_009724389.1;Wu1-SARS2_orf1ab;4836 | 0.1 | 0.025 |
| NC_045512.2;YP_009724389.1;Wu1-SARS2_orf1ab;4837 | 0.05 | 0.025 |
| NC_045512.2;YP_009724389.1;Wu1-SARS2_orf1ab;4838 | 0.15 | 0 |
| NC_045512.2;YP_009724389.1;Wu1-SARS2_orf1ab;4839 | 0.2 | 0 |
| NC_045512.2;YP_009724389.1;Wu1-SARS2_orf1ab;4840 | 0.15 | 0.075 |
| NC_045512.2;YP_009724389.1;Wu1-SARS2_orf1ab;4841 | 0.25 | 0.025 |
| NC_045512.2;YP_009724389.1;Wu1-SARS2_orf1ab;4842 | 0.15 | 0 |
| NC_045512.2;YP_009724389.1;Wu1-SARS2_orf1ab;4843 | 0.15 | 0 |
| NC_045512.2;YP_009724389.1;Wu1-SARS2_orf1ab;4844 | 0.15 | 0 |
| NC_045512.2;YP_009724389.1;Wu1-SARS2_orf1ab;4845 | 0 | 0 |
| NC_045512.2;YP_009724389.1;Wu1-SARS2_orf1ab;4846 | 0 | 0 |
| NC_045512.2;YP_009724389.1;Wu1-SARS2_orf1ab;4847 | 0 | 0 |
| NC_045512.2;YP_009724389.1;Wu1-SARS2_orf1ab;4848 | 0 | 0 |
| NC_045512.2;YP_009724389.1;Wu1-SARS2_orf1ab;4849 | 0 | 0 |

| | | |
|---|---|---|
| NC_045512.2;YP_009724389.1;Wu1-SARS2_orf1ab;4850 | 0 | 0 |
| NC_045512.2;YP_009724389.1;Wu1-SARS2_orf1ab;4851 | 0 | 0 |
| NC_045512.2;YP_009724389.1;Wu1-SARS2_orf1ab;4852 | 0 | 0 |
| NC_045512.2;YP_009724389.1;Wu1-SARS2_orf1ab;4853 | 0 | 0 |
| NC_045512.2;YP_009724389.1;Wu1-SARS2_orf1ab;4854 | 0 | 0 |
| NC_045512.2;YP_009724389.1;Wu1-SARS2_orf1ab;4855 | 0.05 | 0.025 |
| NC_045512.2;YP_009724389.1;Wu1-SARS2_orf1ab;4856 | 0 | 0 |
| NC_045512.2;YP_009724389.1;Wu1-SARS2_orf1ab;4857 | 0 | 0 |
| NC_045512.2;YP_009724389.1;Wu1-SARS2_orf1ab;4858 | 0 | 0.025 |
| NC_045512.2;YP_009724389.1;Wu1-SARS2_orf1ab;4859 | 0 | 0 |
| NC_045512.2;YP_009724389.1;Wu1-SARS2_orf1ab;4860 | 0 | 0 |
| NC_045512.2;YP_009724389.1;Wu1-SARS2_orf1ab;4861 | 0 | 0 |
| NC_045512.2;YP_009724389.1;Wu1-SARS2_orf1ab;4862 | 0 | 0 |
| NC_045512.2;YP_009724389.1;Wu1-SARS2_orf1ab;4863 | 0 | 0 |
| NC_045512.2;YP_009724389.1;Wu1-SARS2_orf1ab;4864 | 0 | 0.025 |
| NC_045512.2;YP_009724389.1;Wu1-SARS2_orf1ab;4865 | 0 | 0.025 |
| NC_045512.2;YP_009724389.1;Wu1-SARS2_orf1ab;4866 | 0 | 0 |
| NC_045512.2;YP_009724389.1;Wu1-SARS2_orf1ab;4867 | 0 | 0 |
| NC_045512.2;YP_009724389.1;Wu1-SARS2_orf1ab;4868 | 0.05 | 0 |
| NC_045512.2;YP_009724389.1;Wu1-SARS2_orf1ab;4869 | 0.05 | 0.075 |
| NC_045512.2;YP_009724389.1;Wu1-SARS2_orf1ab;4870 | 0.05 | 0.025 |
| NC_045512.2;YP_009724389.1;Wu1-SARS2_orf1ab;4871 | 0.05 | 0 |
| NC_045512.2;YP_009724389.1;Wu1-SARS2_orf1ab;4872 | 0 | 0.025 |
| NC_045512.2;YP_009724389.1;Wu1-SARS2_orf1ab;4873 | 0.05 | 0 |
| NC_045512.2;YP_009724389.1;Wu1-SARS2_orf1ab;4874 | 0.05 | 0 |
| NC_045512.2;YP_009724389.1;Wu1-SARS2_orf1ab;4875 | 0 | 0 |
| NC_045512.2;YP_009724389.1;Wu1-SARS2_orf1ab;4876 | 0.05 | 0.025 |
| NC_045512.2;YP_009724389.1;Wu1-SARS2_orf1ab;4877 | 0 | 0.025 |
| NC_045512.2;YP_009724389.1;Wu1-SARS2_orf1ab;4878 | 0.05 | 0.025 |
| NC_045512.2;YP_009724389.1;Wu1-SARS2_orf1ab;4879 | 0.05 | 0.025 |
| NC_045512.2;YP_009724389.1;Wu1-SARS2_orf1ab;4880 | 0.1 | 0.025 |
| NC_045512.2;YP_009724389.1;Wu1-SARS2_orf1ab;4881 | 0 | 0.025 |
| NC_045512.2;YP_009724389.1;Wu1-SARS2_orf1ab;4882 | 0.15 | 0.1 |
| NC_045512.2;YP_009724389.1;Wu1-SARS2_orf1ab;4883 | 0 | 0.05 |
| NC_045512.2;YP_009724389.1;Wu1-SARS2_orf1ab;4884 | 0.15 | 0.075 |
| NC_045512.2;YP_009724389.1;Wu1-SARS2_orf1ab;4885 | 0.05 | 0.05 |
| NC_045512.2;YP_009724389.1;Wu1-SARS2_orf1ab;4886 | 0 | 0.025 |
| NC_045512.2;YP_009724389.1;Wu1-SARS2_orf1ab;4887 | 0 | 0 |
| NC_045512.2;YP_009724389.1;Wu1-SARS2_orf1ab;4888 | 0 | 0 |
| NC_045512.2;YP_009724389.1;Wu1-SARS2_orf1ab;4889 | 0 | 0 |
| NC_045512.2;YP_009724389.1;Wu1-SARS2_orf1ab;4890 | 0 | 0 |
| NC_045512.2;YP_009724389.1;Wu1-SARS2_orf1ab;4891 | 0 | 0 |
| NC_045512.2;YP_009724389.1;Wu1-SARS2_orf1ab;4892 | 0 | 0 |

| | | |
|---|---|---|
| NC_045512.2;YP_009724389.1;Wu1-SARS2_orf1ab;4893 | 0 | 0 |
| NC_045512.2;YP_009724389.1;Wu1-SARS2_orf1ab;4894 | 0 | 0 |
| NC_045512.2;YP_009724389.1;Wu1-SARS2_orf1ab;4895 | 0 | 0 |
| NC_045512.2;YP_009724389.1;Wu1-SARS2_orf1ab;4896 | 0 | 0 |
| NC_045512.2;YP_009724389.1;Wu1-SARS2_orf1ab;4897 | 0.05 | 0 |
| NC_045512.2;YP_009724389.1;Wu1-SARS2_orf1ab;4898 | 0 | 0 |
| NC_045512.2;YP_009724389.1;Wu1-SARS2_orf1ab;4899 | 0.05 | 0 |
| NC_045512.2;YP_009724389.1;Wu1-SARS2_orf1ab;4900 | 0 | 0 |
| NC_045512.2;YP_009724389.1;Wu1-SARS2_orf1ab;4901 | 0 | 0 |
| NC_045512.2;YP_009724389.1;Wu1-SARS2_orf1ab;4902 | 0 | 0 |
| NC_045512.2;YP_009724389.1;Wu1-SARS2_orf1ab;4903 | 0 | 0 |
| NC_045512.2;YP_009724389.1;Wu1-SARS2_orf1ab;4904 | 0 | 0 |
| NC_045512.2;YP_009724389.1;Wu1-SARS2_orf1ab;4905 | 0 | 0 |
| NC_045512.2;YP_009724389.1;Wu1-SARS2_orf1ab;4906 | 0 | 0 |
| NC_045512.2;YP_009724389.1;Wu1-SARS2_orf1ab;4907 | 0 | 0 |
| NC_045512.2;YP_009724389.1;Wu1-SARS2_orf1ab;4908 | 0 | 0 |
| NC_045512.2;YP_009724389.1;Wu1-SARS2_orf1ab;4909 | 0.1 | 0.025 |
| NC_045512.2;YP_009724389.1;Wu1-SARS2_orf1ab;4910 | 0.05 | 0.025 |
| NC_045512.2;YP_009724389.1;Wu1-SARS2_orf1ab;4911 | 0.05 | 0 |
| NC_045512.2;YP_009724389.1;Wu1-SARS2_orf1ab;4912 | 0.1 | 0.05 |
| NC_045512.2;YP_009724389.1;Wu1-SARS2_orf1ab;4913 | 0.05 | 0 |
| NC_045512.2;YP_009724389.1;Wu1-SARS2_orf1ab;4914 | 0.2 | 0.025 |
| NC_045512.2;YP_009724389.1;Wu1-SARS2_orf1ab;4915 | 0.05 | 0.025 |
| NC_045512.2;YP_009724389.1;Wu1-SARS2_orf1ab;4916 | 0.05 | 0 |
| NC_045512.2;YP_009724389.1;Wu1-SARS2_orf1ab;4917 | 0.05 | 0 |
| NC_045512.2;YP_009724389.1;Wu1-SARS2_orf1ab;4918 | 0 | 0 |
| NC_045512.2;YP_009724389.1;Wu1-SARS2_orf1ab;4919 | 0 | 0 |
| NC_045512.2;YP_009724389.1;Wu1-SARS2_orf1ab;4920 | 0 | 0 |
| NC_045512.2;YP_009724389.1;Wu1-SARS2_orf1ab;4921 | 0 | 0 |
| NC_045512.2;YP_009724389.1;Wu1-SARS2_orf1ab;4922 | 0 | 0 |
| NC_045512.2;YP_009724389.1;Wu1-SARS2_orf1ab;4923 | 0.05 | 0 |
| NC_045512.2;YP_009724389.1;Wu1-SARS2_orf1ab;4924 | 0.1 | 0.15 |
| NC_045512.2;YP_009724389.1;Wu1-SARS2_orf1ab;4925 | 0 | 0 |
| NC_045512.2;YP_009724389.1;Wu1-SARS2_orf1ab;4926 | 0 | 0 |
| NC_045512.2;YP_009724389.1;Wu1-SARS2_orf1ab;4927 | 0 | 0 |
| NC_045512.2;YP_009724389.1;Wu1-SARS2_orf1ab;4928 | 0 | 0.025 |
| NC_045512.2;YP_009724389.1;Wu1-SARS2_orf1ab;4929 | 0.1 | 0.05 |
| NC_045512.2;YP_009724389.1;Wu1-SARS2_orf1ab;4930 | 0 | 0.025 |
| NC_045512.2;YP_009724389.1;Wu1-SARS2_orf1ab;4931 | 0 | 0.025 |
| NC_045512.2;YP_009724389.1;Wu1-SARS2_orf1ab;4932 | 0 | 0.025 |
| NC_045512.2;YP_009724389.1;Wu1-SARS2_orf1ab;4933 | 0 | 0.025 |
| NC_045512.2;YP_009724389.1;Wu1-SARS2_orf1ab;4934 | 0 | 0 |
| NC_045512.2;YP_009724389.1;Wu1-SARS2_orf1ab;4935 | 0 | 0 |

| | | |
|---|---|---|
| NC_045512.2;YP_009724389.1;Wu1-SARS2_orf1ab;4936 | 0 | 0 |
| NC_045512.2;YP_009724389.1;Wu1-SARS2_orf1ab;4937 | 0 | 0 |
| NC_045512.2;YP_009724389.1;Wu1-SARS2_orf1ab;4938 | 0 | 0 |
| NC_045512.2;YP_009724389.1;Wu1-SARS2_orf1ab;4939 | 0 | 0 |
| NC_045512.2;YP_009724389.1;Wu1-SARS2_orf1ab;4940 | 0 | 0 |
| NC_045512.2;YP_009724389.1;Wu1-SARS2_orf1ab;4941 | 0 | 0 |
| NC_045512.2;YP_009724389.1;Wu1-SARS2_orf1ab;4942 | 0 | 0 |
| NC_045512.2;YP_009724389.1;Wu1-SARS2_orf1ab;4943 | 0 | 0 |
| NC_045512.2;YP_009724389.1;Wu1-SARS2_orf1ab;4944 | 0 | 0 |
| NC_045512.2;YP_009724389.1;Wu1-SARS2_orf1ab;4945 | 0 | 0 |
| NC_045512.2;YP_009724389.1;Wu1-SARS2_orf1ab;4946 | 0 | 0 |
| NC_045512.2;YP_009724389.1;Wu1-SARS2_orf1ab;4947 | 0 | 0 |
| NC_045512.2;YP_009724389.1;Wu1-SARS2_orf1ab;4948 | 0 | 0 |
| NC_045512.2;YP_009724389.1;Wu1-SARS2_orf1ab;4949 | 0 | 0 |
| NC_045512.2;YP_009724389.1;Wu1-SARS2_orf1ab;4950 | 0 | 0 |
| NC_045512.2;YP_009724389.1;Wu1-SARS2_orf1ab;4951 | 0 | 0 |
| NC_045512.2;YP_009724389.1;Wu1-SARS2_orf1ab;4952 | 0 | 0 |
| NC_045512.2;YP_009724389.1;Wu1-SARS2_orf1ab;4953 | 0 | 0 |
| NC_045512.2;YP_009724389.1;Wu1-SARS2_orf1ab;4954 | 0 | 0 |
| NC_045512.2;YP_009724389.1;Wu1-SARS2_orf1ab;4955 | 0 | 0 |
| NC_045512.2;YP_009724389.1;Wu1-SARS2_orf1ab;4956 | 0 | 0 |
| NC_045512.2;YP_009724389.1;Wu1-SARS2_orf1ab;4957 | 0 | 0 |
| NC_045512.2;YP_009724389.1;Wu1-SARS2_orf1ab;4958 | 0 | 0 |
| NC_045512.2;YP_009724389.1;Wu1-SARS2_orf1ab;4959 | 0 | 0 |
| NC_045512.2;YP_009724389.1;Wu1-SARS2_orf1ab;4960 | 0 | 0 |
| NC_045512.2;YP_009724389.1;Wu1-SARS2_orf1ab;4961 | 0 | 0 |
| NC_045512.2;YP_009724389.1;Wu1-SARS2_orf1ab;4962 | 0 | 0 |
| NC_045512.2;YP_009724389.1;Wu1-SARS2_orf1ab;4963 | 0.3 | 0.175 |
| NC_045512.2;YP_009724389.1;Wu1-SARS2_orf1ab;4964 | 0.4 | 0.275 |
| NC_045512.2;YP_009724389.1;Wu1-SARS2_orf1ab;4965 | 0.35 | 0.275 |
| NC_045512.2;YP_009724389.1;Wu1-SARS2_orf1ab;4966 | 0.25 | 0.025 |
| NC_045512.2;YP_009724389.1;Wu1-SARS2_orf1ab;4967 | 0.25 | 0.075 |
| NC_045512.2;YP_009724389.1;Wu1-SARS2_orf1ab;4968 | 0.25 | 0.025 |
| NC_045512.2;YP_009724389.1;Wu1-SARS2_orf1ab;4969 | 0.25 | 0.05 |
| NC_045512.2;YP_009724389.1;Wu1-SARS2_orf1ab;4970 | 0.25 | 0.025 |
| NC_045512.2;YP_009724389.1;Wu1-SARS2_orf1ab;4971 | 0.15 | 0.025 |
| NC_045512.2;YP_009724389.1;Wu1-SARS2_orf1ab;4972 | 0.15 | 0.025 |
| NC_045512.2;YP_009724389.1;Wu1-SARS2_orf1ab;4973 | 0 | 0 |
| NC_045512.2;YP_009724389.1;Wu1-SARS2_orf1ab;4974 | 0 | 0 |
| NC_045512.2;YP_009724389.1;Wu1-SARS2_orf1ab;4975 | 0 | 0 |
| NC_045512.2;YP_009724389.1;Wu1-SARS2_orf1ab;4976 | 0 | 0 |
| NC_045512.2;YP_009724389.1;Wu1-SARS2_orf1ab;4977 | 0 | 0 |
| NC_045512.2;YP_009724389.1;Wu1-SARS2_orf1ab;4978 | 0 | 0 |

| | | |
|---|---|---|
| NC_045512.2;YP_009724389.1;Wu1-SARS2_orf1ab;4979 | 0 | 0 |
| NC_045512.2;YP_009724389.1;Wu1-SARS2_orf1ab;4980 | 0 | 0 |
| NC_045512.2;YP_009724389.1;Wu1-SARS2_orf1ab;4981 | 0 | 0 |
| NC_045512.2;YP_009724389.1;Wu1-SARS2_orf1ab;4982 | 0 | 0 |
| NC_045512.2;YP_009724389.1;Wu1-SARS2_orf1ab;4983 | 0 | 0 |
| NC_045512.2;YP_009724389.1;Wu1-SARS2_orf1ab;4984 | 0 | 0 |
| NC_045512.2;YP_009724389.1;Wu1-SARS2_orf1ab;4985 | 0 | 0.05 |
| NC_045512.2;YP_009724389.1;Wu1-SARS2_orf1ab;4986 | 0 | 0 |
| NC_045512.2;YP_009724389.1;Wu1-SARS2_orf1ab;4987 | 0 | 0 |
| NC_045512.2;YP_009724389.1;Wu1-SARS2_orf1ab;4988 | 0 | 0 |
| NC_045512.2;YP_009724389.1;Wu1-SARS2_orf1ab;4989 | 0 | 0 |
| NC_045512.2;YP_009724389.1;Wu1-SARS2_orf1ab;4990 | 0 | 0 |
| NC_045512.2;YP_009724389.1;Wu1-SARS2_orf1ab;4991 | 0 | 0 |
| NC_045512.2;YP_009724389.1;Wu1-SARS2_orf1ab;4992 | 0 | 0 |
| NC_045512.2;YP_009724389.1;Wu1-SARS2_orf1ab;4993 | 0 | 0 |
| NC_045512.2;YP_009724389.1;Wu1-SARS2_orf1ab;4994 | 0 | 0 |
| NC_045512.2;YP_009724389.1;Wu1-SARS2_orf1ab;4995 | 0 | 0 |
| NC_045512.2;YP_009724389.1;Wu1-SARS2_orf1ab;4996 | 0 | 0 |
| NC_045512.2;YP_009724389.1;Wu1-SARS2_orf1ab;4997 | 0 | 0.075 |
| NC_045512.2;YP_009724389.1;Wu1-SARS2_orf1ab;4998 | 0 | 0.025 |
| NC_045512.2;YP_009724389.1;Wu1-SARS2_orf1ab;4999 | 0 | 0.025 |
| NC_045512.2;YP_009724389.1;Wu1-SARS2_orf1ab;5000 | 0 | 0.025 |
| NC_045512.2;YP_009724389.1;Wu1-SARS2_orf1ab;5001 | 0 | 0.025 |
| NC_045512.2;YP_009724389.1;Wu1-SARS2_orf1ab;5002 | 0 | 0.025 |
| NC_045512.2;YP_009724389.1;Wu1-SARS2_orf1ab;5003 | 0 | 0.025 |
| NC_045512.2;YP_009724389.1;Wu1-SARS2_orf1ab;5004 | 0 | 0.025 |
| NC_045512.2;YP_009724389.1;Wu1-SARS2_orf1ab;5005 | 0 | 0.025 |
| NC_045512.2;YP_009724389.1;Wu1-SARS2_orf1ab;5006 | 0 | 0.025 |
| NC_045512.2;YP_009724389.1;Wu1-SARS2_orf1ab;5007 | 0 | 0.025 |
| NC_045512.2;YP_009724389.1;Wu1-SARS2_orf1ab;5008 | 0 | 0.025 |
| NC_045512.2;YP_009724389.1;Wu1-SARS2_orf1ab;5009 | 0 | 0 |
| NC_045512.2;YP_009724389.1;Wu1-SARS2_orf1ab;5010 | 0 | 0 |
| NC_045512.2;YP_009724389.1;Wu1-SARS2_orf1ab;5011 | 0 | 0 |
| NC_045512.2;YP_009724389.1;Wu1-SARS2_orf1ab;5012 | 0 | 0 |
| NC_045512.2;YP_009724389.1;Wu1-SARS2_orf1ab;5013 | 0 | 0 |
| NC_045512.2;YP_009724389.1;Wu1-SARS2_orf1ab;5014 | 0 | 0 |
| NC_045512.2;YP_009724389.1;Wu1-SARS2_orf1ab;5015 | 0 | 0 |
| NC_045512.2;YP_009724389.1;Wu1-SARS2_orf1ab;5016 | 0 | 0 |
| NC_045512.2;YP_009724389.1;Wu1-SARS2_orf1ab;5017 | 0 | 0 |
| NC_045512.2;YP_009724389.1;Wu1-SARS2_orf1ab;5018 | 0 | 0 |
| NC_045512.2;YP_009724389.1;Wu1-SARS2_orf1ab;5019 | 0 | 0 |
| NC_045512.2;YP_009724389.1;Wu1-SARS2_orf1ab;5020 | 0 | 0 |
| NC_045512.2;YP_009724389.1;Wu1-SARS2_orf1ab;5021 | 0 | 0 |

| | | |
|---|---|---|
| NC_045512.2;YP_009724389.1;Wu1-SARS2_orf1ab;5022 | 0 | 0 |
| NC_045512.2;YP_009724389.1;Wu1-SARS2_orf1ab;5023 | 0 | 0 |
| NC_045512.2;YP_009724389.1;Wu1-SARS2_orf1ab;5024 | 0 | 0 |
| NC_045512.2;YP_009724389.1;Wu1-SARS2_orf1ab;5025 | 0 | 0 |
| NC_045512.2;YP_009724389.1;Wu1-SARS2_orf1ab;5026 | 0 | 0 |
| NC_045512.2;YP_009724389.1;Wu1-SARS2_orf1ab;5027 | 0 | 0 |
| NC_045512.2;YP_009724389.1;Wu1-SARS2_orf1ab;5028 | 0 | 0 |
| NC_045512.2;YP_009724389.1;Wu1-SARS2_orf1ab;5029 | 0 | 0 |
| NC_045512.2;YP_009724389.1;Wu1-SARS2_orf1ab;5030 | 0 | 0 |
| NC_045512.2;YP_009724389.1;Wu1-SARS2_orf1ab;5031 | 0 | 0 |
| NC_045512.2;YP_009724389.1;Wu1-SARS2_orf1ab;5032 | 0 | 0 |
| NC_045512.2;YP_009724389.1;Wu1-SARS2_orf1ab;5033 | 0 | 0 |
| NC_045512.2;YP_009724389.1;Wu1-SARS2_orf1ab;5034 | 0 | 0 |
| NC_045512.2;YP_009724389.1;Wu1-SARS2_orf1ab;5035 | 0 | 0 |
| NC_045512.2;YP_009724389.1;Wu1-SARS2_orf1ab;5036 | 0 | 0 |
| NC_045512.2;YP_009724389.1;Wu1-SARS2_orf1ab;5037 | 0 | 0 |
| NC_045512.2;YP_009724389.1;Wu1-SARS2_orf1ab;5038 | 0 | 0 |
| NC_045512.2;YP_009724389.1;Wu1-SARS2_orf1ab;5039 | 0 | 0 |
| NC_045512.2;YP_009724389.1;Wu1-SARS2_orf1ab;5040 | 0 | 0 |
| NC_045512.2;YP_009724389.1;Wu1-SARS2_orf1ab;5041 | 0 | 0 |
| NC_045512.2;YP_009724389.1;Wu1-SARS2_orf1ab;5042 | 0 | 0 |
| NC_045512.2;YP_009724389.1;Wu1-SARS2_orf1ab;5043 | 0 | 0 |
| NC_045512.2;YP_009724389.1;Wu1-SARS2_orf1ab;5044 | 0 | 0 |
| NC_045512.2;YP_009724389.1;Wu1-SARS2_orf1ab;5045 | 0 | 0 |
| NC_045512.2;YP_009724389.1;Wu1-SARS2_orf1ab;5046 | 0 | 0 |
| NC_045512.2;YP_009724389.1;Wu1-SARS2_orf1ab;5047 | 0 | 0 |
| NC_045512.2;YP_009724389.1;Wu1-SARS2_orf1ab;5048 | 0 | 0 |
| NC_045512.2;YP_009724389.1;Wu1-SARS2_orf1ab;5049 | 0 | 0 |
| NC_045512.2;YP_009724389.1;Wu1-SARS2_orf1ab;5050 | 0 | 0 |
| NC_045512.2;YP_009724389.1;Wu1-SARS2_orf1ab;5051 | 0 | 0 |
| NC_045512.2;YP_009724389.1;Wu1-SARS2_orf1ab;5052 | 0 | 0 |
| NC_045512.2;YP_009724389.1;Wu1-SARS2_orf1ab;5053 | 0 | 0 |
| NC_045512.2;YP_009724389.1;Wu1-SARS2_orf1ab;5054 | 0 | 0 |
| NC_045512.2;YP_009724389.1;Wu1-SARS2_orf1ab;5055 | 0 | 0 |
| NC_045512.2;YP_009724389.1;Wu1-SARS2_orf1ab;5056 | 0 | 0 |
| NC_045512.2;YP_009724389.1;Wu1-SARS2_orf1ab;5057 | 0.05 | 0 |
| NC_045512.2;YP_009724389.1;Wu1-SARS2_orf1ab;5058 | 0 | 0 |
| NC_045512.2;YP_009724389.1;Wu1-SARS2_orf1ab;5059 | 0 | 0 |
| NC_045512.2;YP_009724389.1;Wu1-SARS2_orf1ab;5060 | 0 | 0 |
| NC_045512.2;YP_009724389.1;Wu1-SARS2_orf1ab;5061 | 0 | 0.025 |
| NC_045512.2;YP_009724389.1;Wu1-SARS2_orf1ab;5062 | 0 | 0.025 |
| NC_045512.2;YP_009724389.1;Wu1-SARS2_orf1ab;5063 | 0 | 0.025 |
| NC_045512.2;YP_009724389.1;Wu1-SARS2_orf1ab;5064 | 0 | 0.025 |

| | | |
|---|---|---|
| NC_045512.2;YP_009724389.1;Wu1-SARS2_orf1ab;5065 | 0 | 0.05 |
| NC_045512.2;YP_009724389.1;Wu1-SARS2_orf1ab;5066 | 0 | 0.025 |
| NC_045512.2;YP_009724389.1;Wu1-SARS2_orf1ab;5067 | 0 | 0.025 |
| NC_045512.2;YP_009724389.1;Wu1-SARS2_orf1ab;5068 | 0 | 0.05 |
| NC_045512.2;YP_009724389.1;Wu1-SARS2_orf1ab;5069 | 0.05 | 0.025 |
| NC_045512.2;YP_009724389.1;Wu1-SARS2_orf1ab;5070 | 0.1 | 0.05 |
| NC_045512.2;YP_009724389.1;Wu1-SARS2_orf1ab;5071 | 0 | 0 |
| NC_045512.2;YP_009724389.1;Wu1-SARS2_orf1ab;5072 | 0.05 | 0 |
| NC_045512.2;YP_009724389.1;Wu1-SARS2_orf1ab;5073 | 0 | 0 |
| NC_045512.2;YP_009724389.1;Wu1-SARS2_orf1ab;5074 | 0 | 0 |
| NC_045512.2;YP_009724389.1;Wu1-SARS2_orf1ab;5075 | 0 | 0 |
| NC_045512.2;YP_009724389.1;Wu1-SARS2_orf1ab;5076 | 0 | 0 |
| NC_045512.2;YP_009724389.1;Wu1-SARS2_orf1ab;5077 | 0 | 0 |
| NC_045512.2;YP_009724389.1;Wu1-SARS2_orf1ab;5078 | 0 | 0 |
| NC_045512.2;YP_009724389.1;Wu1-SARS2_orf1ab;5079 | 0 | 0 |
| NC_045512.2;YP_009724389.1;Wu1-SARS2_orf1ab;5080 | 0 | 0.025 |
| NC_045512.2;YP_009724389.1;Wu1-SARS2_orf1ab;5081 | 0 | 0 |
| NC_045512.2;YP_009724389.1;Wu1-SARS2_orf1ab;5082 | 0 | 0.025 |
| NC_045512.2;YP_009724389.1;Wu1-SARS2_orf1ab;5083 | 0 | 0 |
| NC_045512.2;YP_009724389.1;Wu1-SARS2_orf1ab;5084 | 0 | 0 |
| NC_045512.2;YP_009724389.1;Wu1-SARS2_orf1ab;5085 | 0 | 0 |
| NC_045512.2;YP_009724389.1;Wu1-SARS2_orf1ab;5086 | 0 | 0 |
| NC_045512.2;YP_009724389.1;Wu1-SARS2_orf1ab;5087 | 0 | 0 |
| NC_045512.2;YP_009724389.1;Wu1-SARS2_orf1ab;5088 | 0 | 0 |
| NC_045512.2;YP_009724389.1;Wu1-SARS2_orf1ab;5089 | 0 | 0.025 |
| NC_045512.2;YP_009724389.1;Wu1-SARS2_orf1ab;5090 | 0 | 0 |
| NC_045512.2;YP_009724389.1;Wu1-SARS2_orf1ab;5091 | 0 | 0.025 |
| NC_045512.2;YP_009724389.1;Wu1-SARS2_orf1ab;5092 | 0 | 0.025 |
| NC_045512.2;YP_009724389.1;Wu1-SARS2_orf1ab;5093 | 0 | 0.025 |
| NC_045512.2;YP_009724389.1;Wu1-SARS2_orf1ab;5094 | 0.05 | 0.025 |
| NC_045512.2;YP_009724389.1;Wu1-SARS2_orf1ab;5095 | 0 | 0 |
| NC_045512.2;YP_009724389.1;Wu1-SARS2_orf1ab;5096 | 0 | 0.075 |
| NC_045512.2;YP_009724389.1;Wu1-SARS2_orf1ab;5097 | 0.05 | 0.05 |
| NC_045512.2;YP_009724389.1;Wu1-SARS2_orf1ab;5098 | 0 | 0.025 |
| NC_045512.2;YP_009724389.1;Wu1-SARS2_orf1ab;5099 | 0 | 0.05 |
| NC_045512.2;YP_009724389.1;Wu1-SARS2_orf1ab;5100 | 0 | 0.05 |
| NC_045512.2;YP_009724389.1;Wu1-SARS2_orf1ab;5101 | 0.05 | 0.025 |
| NC_045512.2;YP_009724389.1;Wu1-SARS2_orf1ab;5102 | 0 | 0 |
| NC_045512.2;YP_009724389.1;Wu1-SARS2_orf1ab;5103 | 0 | 0 |
| NC_045512.2;YP_009724389.1;Wu1-SARS2_orf1ab;5104 | 0 | 0 |
| NC_045512.2;YP_009724389.1;Wu1-SARS2_orf1ab;5105 | 0 | 0 |
| NC_045512.2;YP_009724389.1;Wu1-SARS2_orf1ab;5106 | 0 | 0 |
| NC_045512.2;YP_009724389.1;Wu1-SARS2_orf1ab;5107 | 0 | 0 |

| | | |
|---|---|---|
| NC_045512.2;YP_009724389.1;Wu1-SARS2_orf1ab;5108 | 0 | 0 |
| NC_045512.2;YP_009724389.1;Wu1-SARS2_orf1ab;5109 | 0 | 0 |
| NC_045512.2;YP_009724389.1;Wu1-SARS2_orf1ab;5110 | 0 | 0 |
| NC_045512.2;YP_009724389.1;Wu1-SARS2_orf1ab;5111 | 0 | 0 |
| NC_045512.2;YP_009724389.1;Wu1-SARS2_orf1ab;5112 | 0 | 0 |
| NC_045512.2;YP_009724389.1;Wu1-SARS2_orf1ab;5113 | 0.05 | 0 |
| NC_045512.2;YP_009724389.1;Wu1-SARS2_orf1ab;5114 | 0 | 0 |
| NC_045512.2;YP_009724389.1;Wu1-SARS2_orf1ab;5115 | 0 | 0 |
| NC_045512.2;YP_009724389.1;Wu1-SARS2_orf1ab;5116 | 0 | 0 |
| NC_045512.2;YP_009724389.1;Wu1-SARS2_orf1ab;5117 | 0 | 0 |
| NC_045512.2;YP_009724389.1;Wu1-SARS2_orf1ab;5118 | 0 | 0 |
| NC_045512.2;YP_009724389.1;Wu1-SARS2_orf1ab;5119 | 0 | 0 |
| NC_045512.2;YP_009724389.1;Wu1-SARS2_orf1ab;5120 | 0 | 0 |
| NC_045512.2;YP_009724389.1;Wu1-SARS2_orf1ab;5121 | 0 | 0 |
| NC_045512.2;YP_009724389.1;Wu1-SARS2_orf1ab;5122 | 0 | 0 |
| NC_045512.2;YP_009724389.1;Wu1-SARS2_orf1ab;5123 | 0 | 0.05 |
| NC_045512.2;YP_009724389.1;Wu1-SARS2_orf1ab;5124 | 0.05 | 0.05 |
| NC_045512.2;YP_009724389.1;Wu1-SARS2_orf1ab;5125 | 0 | 0.05 |
| NC_045512.2;YP_009724389.1;Wu1-SARS2_orf1ab;5126 | 0 | 0 |
| NC_045512.2;YP_009724389.1;Wu1-SARS2_orf1ab;5127 | 0 | 0.025 |
| NC_045512.2;YP_009724389.1;Wu1-SARS2_orf1ab;5128 | 0 | 0 |
| NC_045512.2;YP_009724389.1;Wu1-SARS2_orf1ab;5129 | 0 | 0.025 |
| NC_045512.2;YP_009724389.1;Wu1-SARS2_orf1ab;5130 | 0 | 0 |
| NC_045512.2;YP_009724389.1;Wu1-SARS2_orf1ab;5131 | 0.05 | 0.025 |
| NC_045512.2;YP_009724389.1;Wu1-SARS2_orf1ab;5132 | 0 | 0 |
| NC_045512.2;YP_009724389.1;Wu1-SARS2_orf1ab;5133 | 0.05 | 0.025 |
| NC_045512.2;YP_009724389.1;Wu1-SARS2_orf1ab;5134 | 0 | 0 |
| NC_045512.2;YP_009724389.1;Wu1-SARS2_orf1ab;5135 | 0 | 0 |
| NC_045512.2;YP_009724389.1;Wu1-SARS2_orf1ab;5136 | 0 | 0 |
| NC_045512.2;YP_009724389.1;Wu1-SARS2_orf1ab;5137 | 0 | 0 |
| NC_045512.2;YP_009724389.1;Wu1-SARS2_orf1ab;5138 | 0 | 0 |
| NC_045512.2;YP_009724389.1;Wu1-SARS2_orf1ab;5139 | 0.05 | 0 |
| NC_045512.2;YP_009724389.1;Wu1-SARS2_orf1ab;5140 | 0 | 0 |
| NC_045512.2;YP_009724389.1;Wu1-SARS2_orf1ab;5141 | 0 | 0 |
| NC_045512.2;YP_009724389.1;Wu1-SARS2_orf1ab;5142 | 0 | 0 |
| NC_045512.2;YP_009724389.1;Wu1-SARS2_orf1ab;5143 | 0 | 0 |
| NC_045512.2;YP_009724389.1;Wu1-SARS2_orf1ab;5144 | 0 | 0 |
| NC_045512.2;YP_009724389.1;Wu1-SARS2_orf1ab;5145 | 0 | 0 |
| NC_045512.2;YP_009724389.1;Wu1-SARS2_orf1ab;5146 | 0 | 0 |
| NC_045512.2;YP_009724389.1;Wu1-SARS2_orf1ab;5147 | 0 | 0 |
| NC_045512.2;YP_009724389.1;Wu1-SARS2_orf1ab;5148 | 0 | 0 |
| NC_045512.2;YP_009724389.1;Wu1-SARS2_orf1ab;5149 | 0 | 0 |
| NC_045512.2;YP_009724389.1;Wu1-SARS2_orf1ab;5150 | 0 | 0 |

| | | |
|---|---|---|
| NC_045512.2;YP_009724389.1;Wu1-SARS2_orf1ab;5151 | 0 | 0 |
| NC_045512.2;YP_009724389.1;Wu1-SARS2_orf1ab;5152 | 0 | 0 |
| NC_045512.2;YP_009724389.1;Wu1-SARS2_orf1ab;5153 | 0 | 0 |
| NC_045512.2;YP_009724389.1;Wu1-SARS2_orf1ab;5154 | 0.05 | 0.05 |
| NC_045512.2;YP_009724389.1;Wu1-SARS2_orf1ab;5155 | 0 | 0 |
| NC_045512.2;YP_009724389.1;Wu1-SARS2_orf1ab;5156 | 0 | 0 |
| NC_045512.2;YP_009724389.1;Wu1-SARS2_orf1ab;5157 | 0 | 0 |
| NC_045512.2;YP_009724389.1;Wu1-SARS2_orf1ab;5158 | 0 | 0 |
| NC_045512.2;YP_009724389.1;Wu1-SARS2_orf1ab;5159 | 0 | 0 |
| NC_045512.2;YP_009724389.1;Wu1-SARS2_orf1ab;5160 | 0 | 0 |
| NC_045512.2;YP_009724389.1;Wu1-SARS2_orf1ab;5161 | 0 | 0 |
| NC_045512.2;YP_009724389.1;Wu1-SARS2_orf1ab;5162 | 0 | 0 |
| NC_045512.2;YP_009724389.1;Wu1-SARS2_orf1ab;5163 | 0 | 0 |
| NC_045512.2;YP_009724389.1;Wu1-SARS2_orf1ab;5164 | 0 | 0 |
| NC_045512.2;YP_009724389.1;Wu1-SARS2_orf1ab;5165 | 0 | 0 |
| NC_045512.2;YP_009724389.1;Wu1-SARS2_orf1ab;5166 | 0 | 0 |
| NC_045512.2;YP_009724389.1;Wu1-SARS2_orf1ab;5167 | 0 | 0 |
| NC_045512.2;YP_009724389.1;Wu1-SARS2_orf1ab;5168 | 0 | 0 |
| NC_045512.2;YP_009724389.1;Wu1-SARS2_orf1ab;5169 | 0 | 0 |
| NC_045512.2;YP_009724389.1;Wu1-SARS2_orf1ab;5170 | 0 | 0 |
| NC_045512.2;YP_009724389.1;Wu1-SARS2_orf1ab;5171 | 0 | 0 |
| NC_045512.2;YP_009724389.1;Wu1-SARS2_orf1ab;5172 | 0 | 0 |
| NC_045512.2;YP_009724389.1;Wu1-SARS2_orf1ab;5173 | 0 | 0 |
| NC_045512.2;YP_009724389.1;Wu1-SARS2_orf1ab;5174 | 0 | 0 |
| NC_045512.2;YP_009724389.1;Wu1-SARS2_orf1ab;5175 | 0.05 | 0 |
| NC_045512.2;YP_009724389.1;Wu1-SARS2_orf1ab;5176 | 0 | 0 |
| NC_045512.2;YP_009724389.1;Wu1-SARS2_orf1ab;5177 | 0 | 0 |
| NC_045512.2;YP_009724389.1;Wu1-SARS2_orf1ab;5178 | 0 | 0 |
| NC_045512.2;YP_009724389.1;Wu1-SARS2_orf1ab;5179 | 0 | 0.025 |
| NC_045512.2;YP_009724389.1;Wu1-SARS2_orf1ab;5180 | 0 | 0.025 |
| NC_045512.2;YP_009724389.1;Wu1-SARS2_orf1ab;5181 | 0 | 0.025 |
| NC_045512.2;YP_009724389.1;Wu1-SARS2_orf1ab;5182 | 0 | 0.025 |
| NC_045512.2;YP_009724389.1;Wu1-SARS2_orf1ab;5183 | 0 | 0.025 |
| NC_045512.2;YP_009724389.1;Wu1-SARS2_orf1ab;5184 | 0.05 | 0.025 |
| NC_045512.2;YP_009724389.1;Wu1-SARS2_orf1ab;5185 | 0 | 0.025 |
| NC_045512.2;YP_009724389.1;Wu1-SARS2_orf1ab;5186 | 0.05 | 0.025 |
| NC_045512.2;YP_009724389.1;Wu1-SARS2_orf1ab;5187 | 0 | 0.025 |
| NC_045512.2;YP_009724389.1;Wu1-SARS2_orf1ab;5188 | 0 | 0.025 |
| NC_045512.2;YP_009724389.1;Wu1-SARS2_orf1ab;5189 | 0.05 | 0.025 |
| NC_045512.2;YP_009724389.1;Wu1-SARS2_orf1ab;5190 | 0.05 | 0.025 |
| NC_045512.2;YP_009724389.1;Wu1-SARS2_orf1ab;5191 | 0.05 | 0.025 |
| NC_045512.2;YP_009724389.1;Wu1-SARS2_orf1ab;5192 | 0.05 | 0.025 |
| NC_045512.2;YP_009724389.1;Wu1-SARS2_orf1ab;5193 | 0 | 0 |

| | | |
|---|---|---|
| NC_045512.2;YP_009724389.1;Wu1-SARS2_orf1ab;5194 | 0 | 0 |
| NC_045512.2;YP_009724389.1;Wu1-SARS2_orf1ab;5195 | 0 | 0 |
| NC_045512.2;YP_009724389.1;Wu1-SARS2_orf1ab;5196 | 0 | 0 |
| NC_045512.2;YP_009724389.1;Wu1-SARS2_orf1ab;5197 | 0 | 0 |
| NC_045512.2;YP_009724389.1;Wu1-SARS2_orf1ab;5198 | 0 | 0 |
| NC_045512.2;YP_009724389.1;Wu1-SARS2_orf1ab;5199 | 0 | 0 |
| NC_045512.2;YP_009724389.1;Wu1-SARS2_orf1ab;5200 | 0 | 0 |
| NC_045512.2;YP_009724389.1;Wu1-SARS2_orf1ab;5201 | 0 | 0 |
| NC_045512.2;YP_009724389.1;Wu1-SARS2_orf1ab;5202 | 0 | 0 |
| NC_045512.2;YP_009724389.1;Wu1-SARS2_orf1ab;5203 | 0 | 0.025 |
| NC_045512.2;YP_009724389.1;Wu1-SARS2_orf1ab;5204 | 0 | 0.025 |
| NC_045512.2;YP_009724389.1;Wu1-SARS2_orf1ab;5205 | 0 | 0.025 |
| NC_045512.2;YP_009724389.1;Wu1-SARS2_orf1ab;5206 | 0.05 | 0 |
| NC_045512.2;YP_009724389.1;Wu1-SARS2_orf1ab;5207 | 0.05 | 0 |
| NC_045512.2;YP_009724389.1;Wu1-SARS2_orf1ab;5208 | 0 | 0 |
| NC_045512.2;YP_009724389.1;Wu1-SARS2_orf1ab;5209 | 0.05 | 0 |
| NC_045512.2;YP_009724389.1;Wu1-SARS2_orf1ab;5210 | 0 | 0 |
| NC_045512.2;YP_009724389.1;Wu1-SARS2_orf1ab;5211 | 0.1 | 0.1 |
| NC_045512.2;YP_009724389.1;Wu1-SARS2_orf1ab;5212 | 0 | 0.025 |
| NC_045512.2;YP_009724389.1;Wu1-SARS2_orf1ab;5213 | 0 | 0.025 |
| NC_045512.2;YP_009724389.1;Wu1-SARS2_orf1ab;5214 | 0 | 0 |
| NC_045512.2;YP_009724389.1;Wu1-SARS2_orf1ab;5215 | 0 | 0 |
| NC_045512.2;YP_009724389.1;Wu1-SARS2_orf1ab;5216 | 0 | 0 |
| NC_045512.2;YP_009724389.1;Wu1-SARS2_orf1ab;5217 | 0 | 0 |
| NC_045512.2;YP_009724389.1;Wu1-SARS2_orf1ab;5218 | 0 | 0 |
| NC_045512.2;YP_009724389.1;Wu1-SARS2_orf1ab;5219 | 0 | 0 |
| NC_045512.2;YP_009724389.1;Wu1-SARS2_orf1ab;5220 | 0 | 0 |
| NC_045512.2;YP_009724389.1;Wu1-SARS2_orf1ab;5221 | 0 | 0 |
| NC_045512.2;YP_009724389.1;Wu1-SARS2_orf1ab;5222 | 0 | 0 |
| NC_045512.2;YP_009724389.1;Wu1-SARS2_orf1ab;5223 | 0 | 0 |
| NC_045512.2;YP_009724389.1;Wu1-SARS2_orf1ab;5224 | 0 | 0 |
| NC_045512.2;YP_009724389.1;Wu1-SARS2_orf1ab;5225 | 0 | 0 |
| NC_045512.2;YP_009724389.1;Wu1-SARS2_orf1ab;5226 | 0 | 0 |
| NC_045512.2;YP_009724389.1;Wu1-SARS2_orf1ab;5227 | 0 | 0 |
| NC_045512.2;YP_009724389.1;Wu1-SARS2_orf1ab;5228 | 0 | 0 |
| NC_045512.2;YP_009724389.1;Wu1-SARS2_orf1ab;5229 | 0.05 | 0 |
| NC_045512.2;YP_009724389.1;Wu1-SARS2_orf1ab;5230 | 0 | 0 |
| NC_045512.2;YP_009724389.1;Wu1-SARS2_orf1ab;5231 | 0 | 0 |
| NC_045512.2;YP_009724389.1;Wu1-SARS2_orf1ab;5232 | 0 | 0 |
| NC_045512.2;YP_009724389.1;Wu1-SARS2_orf1ab;5233 | 0 | 0 |
| NC_045512.2;YP_009724389.1;Wu1-SARS2_orf1ab;5234 | 0 | 0 |
| NC_045512.2;YP_009724389.1;Wu1-SARS2_orf1ab;5235 | 0 | 0 |
| NC_045512.2;YP_009724389.1;Wu1-SARS2_orf1ab;5236 | 0 | 0 |

| | | |
|---|---|---|
| NC_045512.2;YP_009724389.1;Wu1-SARS2_orf1ab;5237 | 0 | 0 |
| NC_045512.2;YP_009724389.1;Wu1-SARS2_orf1ab;5238 | 0 | 0 |
| NC_045512.2;YP_009724389.1;Wu1-SARS2_orf1ab;5239 | 0 | 0 |
| NC_045512.2;YP_009724389.1;Wu1-SARS2_orf1ab;5240 | 0 | 0 |
| NC_045512.2;YP_009724389.1;Wu1-SARS2_orf1ab;5241 | 0 | 0 |
| NC_045512.2;YP_009724389.1;Wu1-SARS2_orf1ab;5242 | 0 | 0 |
| NC_045512.2;YP_009724389.1;Wu1-SARS2_orf1ab;5243 | 0 | 0 |
| NC_045512.2;YP_009724389.1;Wu1-SARS2_orf1ab;5244 | 0 | 0 |
| NC_045512.2;YP_009724389.1;Wu1-SARS2_orf1ab;5245 | 0 | 0 |
| NC_045512.2;YP_009724389.1;Wu1-SARS2_orf1ab;5246 | 0 | 0 |
| NC_045512.2;YP_009724389.1;Wu1-SARS2_orf1ab;5247 | 0 | 0 |
| NC_045512.2;YP_009724389.1;Wu1-SARS2_orf1ab;5248 | 0 | 0 |
| NC_045512.2;YP_009724389.1;Wu1-SARS2_orf1ab;5249 | 0 | 0 |
| NC_045512.2;YP_009724389.1;Wu1-SARS2_orf1ab;5250 | 0 | 0 |
| NC_045512.2;YP_009724389.1;Wu1-SARS2_orf1ab;5251 | 0 | 0 |
| NC_045512.2;YP_009724389.1;Wu1-SARS2_orf1ab;5252 | 0 | 0 |
| NC_045512.2;YP_009724389.1;Wu1-SARS2_orf1ab;5253 | 0 | 0 |
| NC_045512.2;YP_009724389.1;Wu1-SARS2_orf1ab;5254 | 0 | 0 |
| NC_045512.2;YP_009724389.1;Wu1-SARS2_orf1ab;5255 | 0 | 0 |
| NC_045512.2;YP_009724389.1;Wu1-SARS2_orf1ab;5256 | 0 | 0 |
| NC_045512.2;YP_009724389.1;Wu1-SARS2_orf1ab;5257 | 0 | 0 |
| NC_045512.2;YP_009724389.1;Wu1-SARS2_orf1ab;5258 | 0 | 0 |
| NC_045512.2;YP_009724389.1;Wu1-SARS2_orf1ab;5259 | 0 | 0 |
| NC_045512.2;YP_009724389.1;Wu1-SARS2_orf1ab;5260 | 0 | 0 |
| NC_045512.2;YP_009724389.1;Wu1-SARS2_orf1ab;5261 | 0 | 0 |
| NC_045512.2;YP_009724389.1;Wu1-SARS2_orf1ab;5262 | 0 | 0 |
| NC_045512.2;YP_009724389.1;Wu1-SARS2_orf1ab;5263 | 0 | 0 |
| NC_045512.2;YP_009724389.1;Wu1-SARS2_orf1ab;5264 | 0 | 0 |
| NC_045512.2;YP_009724389.1;Wu1-SARS2_orf1ab;5265 | 0 | 0 |
| NC_045512.2;YP_009724389.1;Wu1-SARS2_orf1ab;5266 | 0 | 0 |
| NC_045512.2;YP_009724389.1;Wu1-SARS2_orf1ab;5267 | 0 | 0 |
| NC_045512.2;YP_009724389.1;Wu1-SARS2_orf1ab;5268 | 0 | 0 |
| NC_045512.2;YP_009724389.1;Wu1-SARS2_orf1ab;5269 | 0 | 0 |
| NC_045512.2;YP_009724389.1;Wu1-SARS2_orf1ab;5270 | 0 | 0 |
| NC_045512.2;YP_009724389.1;Wu1-SARS2_orf1ab;5271 | 0 | 0 |
| NC_045512.2;YP_009724389.1;Wu1-SARS2_orf1ab;5272 | 0 | 0 |
| NC_045512.2;YP_009724389.1;Wu1-SARS2_orf1ab;5273 | 0 | 0 |
| NC_045512.2;YP_009724389.1;Wu1-SARS2_orf1ab;5274 | 0 | 0 |
| NC_045512.2;YP_009724389.1;Wu1-SARS2_orf1ab;5275 | 0 | 0 |
| NC_045512.2;YP_009724389.1;Wu1-SARS2_orf1ab;5276 | 0 | 0 |
| NC_045512.2;YP_009724389.1;Wu1-SARS2_orf1ab;5277 | 0 | 0 |
| NC_045512.2;YP_009724389.1;Wu1-SARS2_orf1ab;5278 | 0 | 0 |
| NC_045512.2;YP_009724389.1;Wu1-SARS2_orf1ab;5279 | 0 | 0 |

| | | |
|---|---|---|
| NC_045512.2;YP_009724389.1;Wu1-SARS2_orf1ab;5280 | 0 | 0 |
| NC_045512.2;YP_009724389.1;Wu1-SARS2_orf1ab;5281 | 0 | 0.075 |
| NC_045512.2;YP_009724389.1;Wu1-SARS2_orf1ab;5282 | 0 | 0.025 |
| NC_045512.2;YP_009724389.1;Wu1-SARS2_orf1ab;5283 | 0 | 0 |
| NC_045512.2;YP_009724389.1;Wu1-SARS2_orf1ab;5284 | 0 | 0 |
| NC_045512.2;YP_009724389.1;Wu1-SARS2_orf1ab;5285 | 0 | 0 |
| NC_045512.2;YP_009724389.1;Wu1-SARS2_orf1ab;5286 | 0 | 0 |
| NC_045512.2;YP_009724389.1;Wu1-SARS2_orf1ab;5287 | 0 | 0 |
| NC_045512.2;YP_009724389.1;Wu1-SARS2_orf1ab;5288 | 0 | 0 |
| NC_045512.2;YP_009724389.1;Wu1-SARS2_orf1ab;5289 | 0 | 0 |
| NC_045512.2;YP_009724389.1;Wu1-SARS2_orf1ab;5290 | 0 | 0 |
| NC_045512.2;YP_009724389.1;Wu1-SARS2_orf1ab;5291 | 0 | 0 |
| NC_045512.2;YP_009724389.1;Wu1-SARS2_orf1ab;5292 | 0 | 0 |
| NC_045512.2;YP_009724389.1;Wu1-SARS2_orf1ab;5293 | 0 | 0 |
| NC_045512.2;YP_009724389.1;Wu1-SARS2_orf1ab;5294 | 0 | 0 |
| NC_045512.2;YP_009724389.1;Wu1-SARS2_orf1ab;5295 | 0 | 0 |
| NC_045512.2;YP_009724389.1;Wu1-SARS2_orf1ab;5296 | 0 | 0.025 |
| NC_045512.2;YP_009724389.1;Wu1-SARS2_orf1ab;5297 | 0 | 0 |
| NC_045512.2;YP_009724389.1;Wu1-SARS2_orf1ab;5298 | 0 | 0 |
| NC_045512.2;YP_009724389.1;Wu1-SARS2_orf1ab;5299 | 0 | 0 |
| NC_045512.2;YP_009724389.1;Wu1-SARS2_orf1ab;5300 | 0 | 0 |
| NC_045512.2;YP_009724389.1;Wu1-SARS2_orf1ab;5301 | 0 | 0 |
| NC_045512.2;YP_009724389.1;Wu1-SARS2_orf1ab;5302 | 0.05 | 0 |
| NC_045512.2;YP_009724389.1;Wu1-SARS2_orf1ab;5303 | 0 | 0 |
| NC_045512.2;YP_009724389.1;Wu1-SARS2_orf1ab;5304 | 0 | 0 |
| NC_045512.2;YP_009724389.1;Wu1-SARS2_orf1ab;5305 | 0 | 0 |
| NC_045512.2;YP_009724389.1;Wu1-SARS2_orf1ab;5306 | 0 | 0 |
| NC_045512.2;YP_009724389.1;Wu1-SARS2_orf1ab;5307 | 0 | 0 |
| NC_045512.2;YP_009724389.1;Wu1-SARS2_orf1ab;5308 | 0 | 0 |
| NC_045512.2;YP_009724389.1;Wu1-SARS2_orf1ab;5309 | 0 | 0 |
| NC_045512.2;YP_009724389.1;Wu1-SARS2_orf1ab;5310 | 0 | 0 |
| NC_045512.2;YP_009724389.1;Wu1-SARS2_orf1ab;5311 | 0 | 0 |
| NC_045512.2;YP_009724389.1;Wu1-SARS2_orf1ab;5312 | 0 | 0 |
| NC_045512.2;YP_009724389.1;Wu1-SARS2_orf1ab;5313 | 0 | 0 |
| NC_045512.2;YP_009724389.1;Wu1-SARS2_orf1ab;5314 | 0 | 0 |
| NC_045512.2;YP_009724389.1;Wu1-SARS2_orf1ab;5315 | 0 | 0 |
| NC_045512.2;YP_009724389.1;Wu1-SARS2_orf1ab;5316 | 0 | 0 |
| NC_045512.2;YP_009724389.1;Wu1-SARS2_orf1ab;5317 | 0 | 0 |
| NC_045512.2;YP_009724389.1;Wu1-SARS2_orf1ab;5318 | 0 | 0 |
| NC_045512.2;YP_009724389.1;Wu1-SARS2_orf1ab;5319 | 0 | 0 |
| NC_045512.2;YP_009724389.1;Wu1-SARS2_orf1ab;5320 | 0 | 0 |
| NC_045512.2;YP_009724389.1;Wu1-SARS2_orf1ab;5321 | 0 | 0 |
| NC_045512.2;YP_009724389.1;Wu1-SARS2_orf1ab;5322 | 0 | 0 |

| | | |
|---|---|---|
| NC_045512.2;YP_009724389.1;Wu1-SARS2_orf1ab;5323 | 0 | 0 |
| NC_045512.2;YP_009724389.1;Wu1-SARS2_orf1ab;5324 | 0 | 0 |
| NC_045512.2;YP_009724389.1;Wu1-SARS2_orf1ab;5325 | 0 | 0 |
| NC_045512.2;YP_009724389.1;Wu1-SARS2_orf1ab;5326 | 0 | 0 |
| NC_045512.2;YP_009724389.1;Wu1-SARS2_orf1ab;5327 | 0 | 0 |
| NC_045512.2;YP_009724389.1;Wu1-SARS2_orf1ab;5328 | 0 | 0 |
| NC_045512.2;YP_009724389.1;Wu1-SARS2_orf1ab;5329 | 0 | 0 |
| NC_045512.2;YP_009724389.1;Wu1-SARS2_orf1ab;5330 | 0 | 0 |
| NC_045512.2;YP_009724389.1;Wu1-SARS2_orf1ab;5331 | 0 | 0 |
| NC_045512.2;YP_009724389.1;Wu1-SARS2_orf1ab;5332 | 0 | 0 |
| NC_045512.2;YP_009724389.1;Wu1-SARS2_orf1ab;5333 | 0 | 0 |
| NC_045512.2;YP_009724389.1;Wu1-SARS2_orf1ab;5334 | 0 | 0 |
| NC_045512.2;YP_009724389.1;Wu1-SARS2_orf1ab;5335 | 0 | 0 |
| NC_045512.2;YP_009724389.1;Wu1-SARS2_orf1ab;5336 | 0 | 0 |
| NC_045512.2;YP_009724389.1;Wu1-SARS2_orf1ab;5337 | 0 | 0 |
| NC_045512.2;YP_009724389.1;Wu1-SARS2_orf1ab;5338 | 0 | 0 |
| NC_045512.2;YP_009724389.1;Wu1-SARS2_orf1ab;5339 | 0 | 0 |
| NC_045512.2;YP_009724389.1;Wu1-SARS2_orf1ab;5340 | 0 | 0 |
| NC_045512.2;YP_009724389.1;Wu1-SARS2_orf1ab;5341 | 0 | 0 |
| NC_045512.2;YP_009724389.1;Wu1-SARS2_orf1ab;5342 | 0 | 0 |
| NC_045512.2;YP_009724389.1;Wu1-SARS2_orf1ab;5343 | 0 | 0 |
| NC_045512.2;YP_009724389.1;Wu1-SARS2_orf1ab;5344 | 0 | 0 |
| NC_045512.2;YP_009724389.1;Wu1-SARS2_orf1ab;5345 | 0.05 | 0 |
| NC_045512.2;YP_009724389.1;Wu1-SARS2_orf1ab;5346 | 0 | 0 |
| NC_045512.2;YP_009724389.1;Wu1-SARS2_orf1ab;5347 | 0 | 0 |
| NC_045512.2;YP_009724389.1;Wu1-SARS2_orf1ab;5348 | 0 | 0 |
| NC_045512.2;YP_009724389.1;Wu1-SARS2_orf1ab;5349 | 0 | 0 |
| NC_045512.2;YP_009724389.1;Wu1-SARS2_orf1ab;5350 | 0 | 0 |
| NC_045512.2;YP_009724389.1;Wu1-SARS2_orf1ab;5351 | 0 | 0 |
| NC_045512.2;YP_009724389.1;Wu1-SARS2_orf1ab;5352 | 0 | 0 |
| NC_045512.2;YP_009724389.1;Wu1-SARS2_orf1ab;5353 | 0 | 0 |
| NC_045512.2;YP_009724389.1;Wu1-SARS2_orf1ab;5354 | 0 | 0 |
| NC_045512.2;YP_009724389.1;Wu1-SARS2_orf1ab;5355 | 0 | 0.025 |
| NC_045512.2;YP_009724389.1;Wu1-SARS2_orf1ab;5356 | 0 | 0 |
| NC_045512.2;YP_009724389.1;Wu1-SARS2_orf1ab;5357 | 0 | 0 |
| NC_045512.2;YP_009724389.1;Wu1-SARS2_orf1ab;5358 | 0 | 0 |
| NC_045512.2;YP_009724389.1;Wu1-SARS2_orf1ab;5359 | 0 | 0 |
| NC_045512.2;YP_009724389.1;Wu1-SARS2_orf1ab;5360 | 0 | 0 |
| NC_045512.2;YP_009724389.1;Wu1-SARS2_orf1ab;5361 | 0 | 0 |
| NC_045512.2;YP_009724389.1;Wu1-SARS2_orf1ab;5362 | 0 | 0 |
| NC_045512.2;YP_009724389.1;Wu1-SARS2_orf1ab;5363 | 0 | 0 |
| NC_045512.2;YP_009724389.1;Wu1-SARS2_orf1ab;5364 | 0 | 0 |
| NC_045512.2;YP_009724389.1;Wu1-SARS2_orf1ab;5365 | 0 | 0 |

| | | |
|---|---|---|
| NC_045512.2;YP_009724389.1;Wu1-SARS2_orf1ab;5366 | 0 | 0.075 |
| NC_045512.2;YP_009724389.1;Wu1-SARS2_orf1ab;5367 | 0 | 0 |
| NC_045512.2;YP_009724389.1;Wu1-SARS2_orf1ab;5368 | 0 | 0 |
| NC_045512.2;YP_009724389.1;Wu1-SARS2_orf1ab;5369 | 0 | 0 |
| NC_045512.2;YP_009724389.1;Wu1-SARS2_orf1ab;5370 | 0.05 | 0 |
| NC_045512.2;YP_009724389.1;Wu1-SARS2_orf1ab;5371 | 0.05 | 0 |
| NC_045512.2;YP_009724389.1;Wu1-SARS2_orf1ab;5372 | 0.05 | 0 |
| NC_045512.2;YP_009724389.1;Wu1-SARS2_orf1ab;5373 | 0.05 | 0 |
| NC_045512.2;YP_009724389.1;Wu1-SARS2_orf1ab;5374 | 0 | 0 |
| NC_045512.2;YP_009724389.1;Wu1-SARS2_orf1ab;5375 | 0 | 0 |
| NC_045512.2;YP_009724389.1;Wu1-SARS2_orf1ab;5376 | 0 | 0 |
| NC_045512.2;YP_009724389.1;Wu1-SARS2_orf1ab;5377 | 0 | 0 |
| NC_045512.2;YP_009724389.1;Wu1-SARS2_orf1ab;5378 | 0.05 | 0 |
| NC_045512.2;YP_009724389.1;Wu1-SARS2_orf1ab;5379 | 0 | 0 |
| NC_045512.2;YP_009724389.1;Wu1-SARS2_orf1ab;5380 | 0 | 0 |
| NC_045512.2;YP_009724389.1;Wu1-SARS2_orf1ab;5381 | 0 | 0 |
| NC_045512.2;YP_009724389.1;Wu1-SARS2_orf1ab;5382 | 0 | 0 |
| NC_045512.2;YP_009724389.1;Wu1-SARS2_orf1ab;5383 | 0 | 0 |
| NC_045512.2;YP_009724389.1;Wu1-SARS2_orf1ab;5384 | 0 | 0 |
| NC_045512.2;YP_009724389.1;Wu1-SARS2_orf1ab;5385 | 0 | 0 |
| NC_045512.2;YP_009724389.1;Wu1-SARS2_orf1ab;5386 | 0 | 0 |
| NC_045512.2;YP_009724389.1;Wu1-SARS2_orf1ab;5387 | 0 | 0 |
| NC_045512.2;YP_009724389.1;Wu1-SARS2_orf1ab;5388 | 0 | 0 |
| NC_045512.2;YP_009724389.1;Wu1-SARS2_orf1ab;5389 | 0 | 0 |
| NC_045512.2;YP_009724389.1;Wu1-SARS2_orf1ab;5390 | 0 | 0 |
| NC_045512.2;YP_009724389.1;Wu1-SARS2_orf1ab;5391 | 0 | 0 |
| NC_045512.2;YP_009724389.1;Wu1-SARS2_orf1ab;5392 | 0 | 0 |
| NC_045512.2;YP_009724389.1;Wu1-SARS2_orf1ab;5393 | 0 | 0 |
| NC_045512.2;YP_009724389.1;Wu1-SARS2_orf1ab;5394 | 0 | 0 |
| NC_045512.2;YP_009724389.1;Wu1-SARS2_orf1ab;5395 | 0 | 0 |
| NC_045512.2;YP_009724389.1;Wu1-SARS2_orf1ab;5396 | 0 | 0 |
| NC_045512.2;YP_009724389.1;Wu1-SARS2_orf1ab;5397 | 0 | 0 |
| NC_045512.2;YP_009724389.1;Wu1-SARS2_orf1ab;5398 | 0 | 0 |
| NC_045512.2;YP_009724389.1;Wu1-SARS2_orf1ab;5399 | 0 | 0 |
| NC_045512.2;YP_009724389.1;Wu1-SARS2_orf1ab;5400 | 0 | 0 |
| NC_045512.2;YP_009724389.1;Wu1-SARS2_orf1ab;5401 | 0 | 0.025 |
| NC_045512.2;YP_009724389.1;Wu1-SARS2_orf1ab;5402 | 0 | 0 |
| NC_045512.2;YP_009724389.1;Wu1-SARS2_orf1ab;5403 | 0 | 0 |
| NC_045512.2;YP_009724389.1;Wu1-SARS2_orf1ab;5404 | 0 | 0 |
| NC_045512.2;YP_009724389.1;Wu1-SARS2_orf1ab;5405 | 0 | 0 |
| NC_045512.2;YP_009724389.1;Wu1-SARS2_orf1ab;5406 | 0 | 0 |
| NC_045512.2;YP_009724389.1;Wu1-SARS2_orf1ab;5407 | 0 | 0 |
| NC_045512.2;YP_009724389.1;Wu1-SARS2_orf1ab;5408 | 0 | 0 |

| | | |
|---|---|---|
| NC_045512.2;YP_009724389.1;Wu1-SARS2_orf1ab;5409 | 0 | 0 |
| NC_045512.2;YP_009724389.1;Wu1-SARS2_orf1ab;5410 | 0 | 0 |
| NC_045512.2;YP_009724389.1;Wu1-SARS2_orf1ab;5411 | 0 | 0 |
| NC_045512.2;YP_009724389.1;Wu1-SARS2_orf1ab;5412 | 0 | 0 |
| NC_045512.2;YP_009724389.1;Wu1-SARS2_orf1ab;5413 | 0 | 0 |
| NC_045512.2;YP_009724389.1;Wu1-SARS2_orf1ab;5414 | 0 | 0 |
| NC_045512.2;YP_009724389.1;Wu1-SARS2_orf1ab;5415 | 0.05 | 0 |
| NC_045512.2;YP_009724389.1;Wu1-SARS2_orf1ab;5416 | 0.15 | 0.125 |
| NC_045512.2;YP_009724389.1;Wu1-SARS2_orf1ab;5417 | 0.15 | 0.125 |
| NC_045512.2;YP_009724389.1;Wu1-SARS2_orf1ab;5418 | 0.2 | 0.15 |
| NC_045512.2;YP_009724389.1;Wu1-SARS2_orf1ab;5419 | 0.1 | 0.175 |
| NC_045512.2;YP_009724389.1;Wu1-SARS2_orf1ab;5420 | 0.05 | 0.1 |
| NC_045512.2;YP_009724389.1;Wu1-SARS2_orf1ab;5421 | 0.05 | 0.075 |
| NC_045512.2;YP_009724389.1;Wu1-SARS2_orf1ab;5422 | 0.05 | 0.2 |
| NC_045512.2;YP_009724389.1;Wu1-SARS2_orf1ab;5423 | 0.05 | 0.075 |
| NC_045512.2;YP_009724389.1;Wu1-SARS2_orf1ab;5424 | 0.05 | 0.025 |
| NC_045512.2;YP_009724389.1;Wu1-SARS2_orf1ab;5425 | 0.05 | 0.05 |
| NC_045512.2;YP_009724389.1;Wu1-SARS2_orf1ab;5426 | 0 | 0.025 |
| NC_045512.2;YP_009724389.1;Wu1-SARS2_orf1ab;5427 | 0 | 0 |
| NC_045512.2;YP_009724389.1;Wu1-SARS2_orf1ab;5428 | 0 | 0 |
| NC_045512.2;YP_009724389.1;Wu1-SARS2_orf1ab;5429 | 0 | 0 |
| NC_045512.2;YP_009724389.1;Wu1-SARS2_orf1ab;5430 | 0 | 0 |
| NC_045512.2;YP_009724389.1;Wu1-SARS2_orf1ab;5431 | 0 | 0 |
| NC_045512.2;YP_009724389.1;Wu1-SARS2_orf1ab;5432 | 0 | 0 |
| NC_045512.2;YP_009724389.1;Wu1-SARS2_orf1ab;5433 | 0 | 0 |
| NC_045512.2;YP_009724389.1;Wu1-SARS2_orf1ab;5434 | 0 | 0 |
| NC_045512.2;YP_009724389.1;Wu1-SARS2_orf1ab;5435 | 0 | 0 |
| NC_045512.2;YP_009724389.1;Wu1-SARS2_orf1ab;5436 | 0 | 0 |
| NC_045512.2;YP_009724389.1;Wu1-SARS2_orf1ab;5437 | 0 | 0 |
| NC_045512.2;YP_009724389.1;Wu1-SARS2_orf1ab;5438 | 0 | 0 |
| NC_045512.2;YP_009724389.1;Wu1-SARS2_orf1ab;5439 | 0 | 0 |
| NC_045512.2;YP_009724389.1;Wu1-SARS2_orf1ab;5440 | 0 | 0 |
| NC_045512.2;YP_009724389.1;Wu1-SARS2_orf1ab;5441 | 0.05 | 0 |
| NC_045512.2;YP_009724389.1;Wu1-SARS2_orf1ab;5442 | 0 | 0 |
| NC_045512.2;YP_009724389.1;Wu1-SARS2_orf1ab;5443 | 0 | 0 |
| NC_045512.2;YP_009724389.1;Wu1-SARS2_orf1ab;5444 | 0 | 0 |
| NC_045512.2;YP_009724389.1;Wu1-SARS2_orf1ab;5445 | 0 | 0 |
| NC_045512.2;YP_009724389.1;Wu1-SARS2_orf1ab;5446 | 0 | 0 |
| NC_045512.2;YP_009724389.1;Wu1-SARS2_orf1ab;5447 | 0 | 0 |
| NC_045512.2;YP_009724389.1;Wu1-SARS2_orf1ab;5448 | 0 | 0 |
| NC_045512.2;YP_009724389.1;Wu1-SARS2_orf1ab;5449 | 0 | 0.025 |
| NC_045512.2;YP_009724389.1;Wu1-SARS2_orf1ab;5450 | 0 | 0 |
| NC_045512.2;YP_009724389.1;Wu1-SARS2_orf1ab;5451 | 0 | 0 |

| | | |
|---|---|---|
| NC_045512.2;YP_009724389.1;Wu1-SARS2_orf1ab;5452 | 0.05 | 0 |
| NC_045512.2;YP_009724389.1;Wu1-SARS2_orf1ab;5453 | 0.05 | 0 |
| NC_045512.2;YP_009724389.1;Wu1-SARS2_orf1ab;5454 | 0.05 | 0 |
| NC_045512.2;YP_009724389.1;Wu1-SARS2_orf1ab;5455 | 0 | 0 |
| NC_045512.2;YP_009724389.1;Wu1-SARS2_orf1ab;5456 | 0 | 0 |
| NC_045512.2;YP_009724389.1;Wu1-SARS2_orf1ab;5457 | 0 | 0 |
| NC_045512.2;YP_009724389.1;Wu1-SARS2_orf1ab;5458 | 0 | 0 |
| NC_045512.2;YP_009724389.1;Wu1-SARS2_orf1ab;5459 | 0.05 | 0.025 |
| NC_045512.2;YP_009724389.1;Wu1-SARS2_orf1ab;5460 | 0 | 0 |
| NC_045512.2;YP_009724389.1;Wu1-SARS2_orf1ab;5461 | 0 | 0.025 |
| NC_045512.2;YP_009724389.1;Wu1-SARS2_orf1ab;5462 | 0 | 0 |
| NC_045512.2;YP_009724389.1;Wu1-SARS2_orf1ab;5463 | 0 | 0 |
| NC_045512.2;YP_009724389.1;Wu1-SARS2_orf1ab;5464 | 0 | 0 |
| NC_045512.2;YP_009724389.1;Wu1-SARS2_orf1ab;5465 | 0 | 0 |
| NC_045512.2;YP_009724389.1;Wu1-SARS2_orf1ab;5466 | 0.05 | 0 |
| NC_045512.2;YP_009724389.1;Wu1-SARS2_orf1ab;5467 | 0 | 0 |
| NC_045512.2;YP_009724389.1;Wu1-SARS2_orf1ab;5468 | 0 | 0 |
| NC_045512.2;YP_009724389.1;Wu1-SARS2_orf1ab;5469 | 0 | 0 |
| NC_045512.2;YP_009724389.1;Wu1-SARS2_orf1ab;5470 | 0.05 | 0 |
| NC_045512.2;YP_009724389.1;Wu1-SARS2_orf1ab;5471 | 0.05 | 0 |
| NC_045512.2;YP_009724389.1;Wu1-SARS2_orf1ab;5472 | 0.05 | 0 |
| NC_045512.2;YP_009724389.1;Wu1-SARS2_orf1ab;5473 | 0.05 | 0.05 |
| NC_045512.2;YP_009724389.1;Wu1-SARS2_orf1ab;5474 | 0.05 | 0.05 |
| NC_045512.2;YP_009724389.1;Wu1-SARS2_orf1ab;5475 | 0.05 | 0 |
| NC_045512.2;YP_009724389.1;Wu1-SARS2_orf1ab;5476 | 0 | 0.05 |
| NC_045512.2;YP_009724389.1;Wu1-SARS2_orf1ab;5477 | 0 | 0 |
| NC_045512.2;YP_009724389.1;Wu1-SARS2_orf1ab;5478 | 0 | 0 |
| NC_045512.2;YP_009724389.1;Wu1-SARS2_orf1ab;5479 | 0 | 0 |
| NC_045512.2;YP_009724389.1;Wu1-SARS2_orf1ab;5480 | 0 | 0.05 |
| NC_045512.2;YP_009724389.1;Wu1-SARS2_orf1ab;5481 | 0 | 0.05 |
| NC_045512.2;YP_009724389.1;Wu1-SARS2_orf1ab;5482 | 0 | 0.05 |
| NC_045512.2;YP_009724389.1;Wu1-SARS2_orf1ab;5483 | 0.05 | 0.075 |
| NC_045512.2;YP_009724389.1;Wu1-SARS2_orf1ab;5484 | 0 | 0 |
| NC_045512.2;YP_009724389.1;Wu1-SARS2_orf1ab;5485 | 0 | 0 |
| NC_045512.2;YP_009724389.1;Wu1-SARS2_orf1ab;5486 | 0 | 0 |
| NC_045512.2;YP_009724389.1;Wu1-SARS2_orf1ab;5487 | 0 | 0 |
| NC_045512.2;YP_009724389.1;Wu1-SARS2_orf1ab;5488 | 0 | 0 |
| NC_045512.2;YP_009724389.1;Wu1-SARS2_orf1ab;5489 | 0 | 0 |
| NC_045512.2;YP_009724389.1;Wu1-SARS2_orf1ab;5490 | 0 | 0 |
| NC_045512.2;YP_009724389.1;Wu1-SARS2_orf1ab;5491 | 0 | 0 |
| NC_045512.2;YP_009724389.1;Wu1-SARS2_orf1ab;5492 | 0 | 0 |
| NC_045512.2;YP_009724389.1;Wu1-SARS2_orf1ab;5493 | 0 | 0 |
| NC_045512.2;YP_009724389.1;Wu1-SARS2_orf1ab;5494 | 0 | 0 |

| | | |
|---|---|---|
| NC_045512.2;YP_009724389.1;Wu1-SARS2_orf1ab;5495 | 0 | 0 |
| NC_045512.2;YP_009724389.1;Wu1-SARS2_orf1ab;5496 | 0 | 0 |
| NC_045512.2;YP_009724389.1;Wu1-SARS2_orf1ab;5497 | 0 | 0 |
| NC_045512.2;YP_009724389.1;Wu1-SARS2_orf1ab;5498 | 0 | 0 |
| NC_045512.2;YP_009724389.1;Wu1-SARS2_orf1ab;5499 | 0 | 0 |
| NC_045512.2;YP_009724389.1;Wu1-SARS2_orf1ab;5500 | 0 | 0 |
| NC_045512.2;YP_009724389.1;Wu1-SARS2_orf1ab;5501 | 0 | 0 |
| NC_045512.2;YP_009724389.1;Wu1-SARS2_orf1ab;5502 | 0 | 0 |
| NC_045512.2;YP_009724389.1;Wu1-SARS2_orf1ab;5503 | 0 | 0 |
| NC_045512.2;YP_009724389.1;Wu1-SARS2_orf1ab;5504 | 0 | 0 |
| NC_045512.2;YP_009724389.1;Wu1-SARS2_orf1ab;5505 | 0 | 0 |
| NC_045512.2;YP_009724389.1;Wu1-SARS2_orf1ab;5506 | 0.1 | 0.025 |
| NC_045512.2;YP_009724389.1;Wu1-SARS2_orf1ab;5507 | 0.1 | 0.025 |
| NC_045512.2;YP_009724389.1;Wu1-SARS2_orf1ab;5508 | 0.1 | 0.025 |
| NC_045512.2;YP_009724389.1;Wu1-SARS2_orf1ab;5509 | 0.1 | 0 |
| NC_045512.2;YP_009724389.1;Wu1-SARS2_orf1ab;5510 | 0.05 | 0.05 |
| NC_045512.2;YP_009724389.1;Wu1-SARS2_orf1ab;5511 | 0.05 | 0 |
| NC_045512.2;YP_009724389.1;Wu1-SARS2_orf1ab;5512 | 0.1 | 0 |
| NC_045512.2;YP_009724389.1;Wu1-SARS2_orf1ab;5513 | 0.05 | 0 |
| NC_045512.2;YP_009724389.1;Wu1-SARS2_orf1ab;5514 | 0.05 | 0.025 |
| NC_045512.2;YP_009724389.1;Wu1-SARS2_orf1ab;5515 | 0.05 | 0.025 |
| NC_045512.2;YP_009724389.1;Wu1-SARS2_orf1ab;5516 | 0.05 | 0.025 |
| NC_045512.2;YP_009724389.1;Wu1-SARS2_orf1ab;5517 | 0.05 | 0.025 |
| NC_045512.2;YP_009724389.1;Wu1-SARS2_orf1ab;5518 | 0.1 | 0.025 |
| NC_045512.2;YP_009724389.1;Wu1-SARS2_orf1ab;5519 | 0.05 | 0.025 |
| NC_045512.2;YP_009724389.1;Wu1-SARS2_orf1ab;5520 | 0 | 0 |
| NC_045512.2;YP_009724389.1;Wu1-SARS2_orf1ab;5521 | 0 | 0.025 |
| NC_045512.2;YP_009724389.1;Wu1-SARS2_orf1ab;5522 | 0.1 | 0 |
| NC_045512.2;YP_009724389.1;Wu1-SARS2_orf1ab;5523 | 0 | 0.025 |
| NC_045512.2;YP_009724389.1;Wu1-SARS2_orf1ab;5524 | 0 | 0.025 |
| NC_045512.2;YP_009724389.1;Wu1-SARS2_orf1ab;5525 | 0 | 0.025 |
| NC_045512.2;YP_009724389.1;Wu1-SARS2_orf1ab;5526 | 0 | 0.025 |
| NC_045512.2;YP_009724389.1;Wu1-SARS2_orf1ab;5527 | 0.05 | 0 |
| NC_045512.2;YP_009724389.1;Wu1-SARS2_orf1ab;5528 | 0 | 0 |
| NC_045512.2;YP_009724389.1;Wu1-SARS2_orf1ab;5529 | 0 | 0 |
| NC_045512.2;YP_009724389.1;Wu1-SARS2_orf1ab;5530 | 0 | 0 |
| NC_045512.2;YP_009724389.1;Wu1-SARS2_orf1ab;5531 | 0 | 0 |
| NC_045512.2;YP_009724389.1;Wu1-SARS2_orf1ab;5532 | 0 | 0 |
| NC_045512.2;YP_009724389.1;Wu1-SARS2_orf1ab;5533 | 0 | 0 |
| NC_045512.2;YP_009724389.1;Wu1-SARS2_orf1ab;5534 | 0 | 0 |
| NC_045512.2;YP_009724389.1;Wu1-SARS2_orf1ab;5535 | 0 | 0 |
| NC_045512.2;YP_009724389.1;Wu1-SARS2_orf1ab;5536 | 0 | 0 |
| NC_045512.2;YP_009724389.1;Wu1-SARS2_orf1ab;5537 | 0 | 0 |

| | | |
|---|---|---|
| NC_045512.2;YP_009724389.1;Wu1-SARS2_orf1ab;5538 | 0 | 0 |
| NC_045512.2;YP_009724389.1;Wu1-SARS2_orf1ab;5539 | 0 | 0 |
| NC_045512.2;YP_009724389.1;Wu1-SARS2_orf1ab;5540 | 0 | 0 |
| NC_045512.2;YP_009724389.1;Wu1-SARS2_orf1ab;5541 | 0 | 0 |
| NC_045512.2;YP_009724389.1;Wu1-SARS2_orf1ab;5542 | 0 | 0 |
| NC_045512.2;YP_009724389.1;Wu1-SARS2_orf1ab;5543 | 0 | 0 |
| NC_045512.2;YP_009724389.1;Wu1-SARS2_orf1ab;5544 | 0 | 0 |
| NC_045512.2;YP_009724389.1;Wu1-SARS2_orf1ab;5545 | 0 | 0 |
| NC_045512.2;YP_009724389.1;Wu1-SARS2_orf1ab;5546 | 0 | 0 |
| NC_045512.2;YP_009724389.1;Wu1-SARS2_orf1ab;5547 | 0.05 | 0 |
| NC_045512.2;YP_009724389.1;Wu1-SARS2_orf1ab;5548 | 0 | 0 |
| NC_045512.2;YP_009724389.1;Wu1-SARS2_orf1ab;5549 | 0 | 0 |
| NC_045512.2;YP_009724389.1;Wu1-SARS2_orf1ab;5550 | 0 | 0 |
| NC_045512.2;YP_009724389.1;Wu1-SARS2_orf1ab;5551 | 0 | 0.025 |
| NC_045512.2;YP_009724389.1;Wu1-SARS2_orf1ab;5552 | 0 | 0 |
| NC_045512.2;YP_009724389.1;Wu1-SARS2_orf1ab;5553 | 0 | 0 |
| NC_045512.2;YP_009724389.1;Wu1-SARS2_orf1ab;5554 | 0 | 0 |
| NC_045512.2;YP_009724389.1;Wu1-SARS2_orf1ab;5555 | 0 | 0.025 |
| NC_045512.2;YP_009724389.1;Wu1-SARS2_orf1ab;5556 | 0 | 0.025 |
| NC_045512.2;YP_009724389.1;Wu1-SARS2_orf1ab;5557 | 0 | 0.025 |
| NC_045512.2;YP_009724389.1;Wu1-SARS2_orf1ab;5558 | 0 | 0.025 |
| NC_045512.2;YP_009724389.1;Wu1-SARS2_orf1ab;5559 | 0 | 0.025 |
| NC_045512.2;YP_009724389.1;Wu1-SARS2_orf1ab;5560 | 0 | 0.025 |
| NC_045512.2;YP_009724389.1;Wu1-SARS2_orf1ab;5561 | 0 | 0.025 |
| NC_045512.2;YP_009724389.1;Wu1-SARS2_orf1ab;5562 | 0 | 0 |
| NC_045512.2;YP_009724389.1;Wu1-SARS2_orf1ab;5563 | 0 | 0 |
| NC_045512.2;YP_009724389.1;Wu1-SARS2_orf1ab;5564 | 0 | 0.025 |
| NC_045512.2;YP_009724389.1;Wu1-SARS2_orf1ab;5565 | 0 | 0.025 |
| NC_045512.2;YP_009724389.1;Wu1-SARS2_orf1ab;5566 | 0 | 0 |
| NC_045512.2;YP_009724389.1;Wu1-SARS2_orf1ab;5567 | 0 | 0.025 |
| NC_045512.2;YP_009724389.1;Wu1-SARS2_orf1ab;5568 | 0 | 0 |
| NC_045512.2;YP_009724389.1;Wu1-SARS2_orf1ab;5569 | 0 | 0 |
| NC_045512.2;YP_009724389.1;Wu1-SARS2_orf1ab;5570 | 0 | 0 |
| NC_045512.2;YP_009724389.1;Wu1-SARS2_orf1ab;5571 | 0.05 | 0 |
| NC_045512.2;YP_009724389.1;Wu1-SARS2_orf1ab;5572 | 0.05 | 0.025 |
| NC_045512.2;YP_009724389.1;Wu1-SARS2_orf1ab;5573 | 0.05 | 0.025 |
| NC_045512.2;YP_009724389.1;Wu1-SARS2_orf1ab;5574 | 0.05 | 0.025 |
| NC_045512.2;YP_009724389.1;Wu1-SARS2_orf1ab;5575 | 0.05 | 0.025 |
| NC_045512.2;YP_009724389.1;Wu1-SARS2_orf1ab;5576 | 0.05 | 0.075 |
| NC_045512.2;YP_009724389.1;Wu1-SARS2_orf1ab;5577 | 0.05 | 0.175 |
| NC_045512.2;YP_009724389.1;Wu1-SARS2_orf1ab;5578 | 0.05 | 0.075 |
| NC_045512.2;YP_009724389.1;Wu1-SARS2_orf1ab;5579 | 0.1 | 0.125 |
| NC_045512.2;YP_009724389.1;Wu1-SARS2_orf1ab;5580 | 0.05 | 0.075 |

| | | |
|---|---|---|
| NC_045512.2;YP_009724389.1;Wu1-SARS2_orf1ab;5581 | 0 | 0.05 |
| NC_045512.2;YP_009724389.1;Wu1-SARS2_orf1ab;5582 | 0.05 | 0.025 |
| NC_045512.2;YP_009724389.1;Wu1-SARS2_orf1ab;5583 | 0 | 0 |
| NC_045512.2;YP_009724389.1;Wu1-SARS2_orf1ab;5584 | 0.1 | 0.1 |
| NC_045512.2;YP_009724389.1;Wu1-SARS2_orf1ab;5585 | 0.1 | 0.025 |
| NC_045512.2;YP_009724389.1;Wu1-SARS2_orf1ab;5586 | 0 | 0 |
| NC_045512.2;YP_009724389.1;Wu1-SARS2_orf1ab;5587 | 0.05 | 0.05 |
| NC_045512.2;YP_009724389.1;Wu1-SARS2_orf1ab;5588 | 0 | 0.025 |
| NC_045512.2;YP_009724389.1;Wu1-SARS2_orf1ab;5589 | 0 | 0.025 |
| NC_045512.2;YP_009724389.1;Wu1-SARS2_orf1ab;5590 | 0 | 0.025 |
| NC_045512.2;YP_009724389.1;Wu1-SARS2_orf1ab;5591 | 0 | 0.025 |
| NC_045512.2;YP_009724389.1;Wu1-SARS2_orf1ab;5592 | 0 | 0.025 |
| NC_045512.2;YP_009724389.1;Wu1-SARS2_orf1ab;5593 | 0 | 0.075 |
| NC_045512.2;YP_009724389.1;Wu1-SARS2_orf1ab;5594 | 0 | 0.025 |
| NC_045512.2;YP_009724389.1;Wu1-SARS2_orf1ab;5595 | 0 | 0.025 |
| NC_045512.2;YP_009724389.1;Wu1-SARS2_orf1ab;5596 | 0 | 0 |
| NC_045512.2;YP_009724389.1;Wu1-SARS2_orf1ab;5597 | 0 | 0 |
| NC_045512.2;YP_009724389.1;Wu1-SARS2_orf1ab;5598 | 0.05 | 0.025 |
| NC_045512.2;YP_009724389.1;Wu1-SARS2_orf1ab;5599 | 0 | 0 |
| NC_045512.2;YP_009724389.1;Wu1-SARS2_orf1ab;5600 | 0 | 0 |
| NC_045512.2;YP_009724389.1;Wu1-SARS2_orf1ab;5601 | 0 | 0.025 |
| NC_045512.2;YP_009724389.1;Wu1-SARS2_orf1ab;5602 | 0 | 0 |
| NC_045512.2;YP_009724389.1;Wu1-SARS2_orf1ab;5603 | 0 | 0 |
| NC_045512.2;YP_009724389.1;Wu1-SARS2_orf1ab;5604 | 0 | 0 |
| NC_045512.2;YP_009724389.1;Wu1-SARS2_orf1ab;5605 | 0 | 0 |
| NC_045512.2;YP_009724389.1;Wu1-SARS2_orf1ab;5606 | 0 | 0 |
| NC_045512.2;YP_009724389.1;Wu1-SARS2_orf1ab;5607 | 0 | 0 |
| NC_045512.2;YP_009724389.1;Wu1-SARS2_orf1ab;5608 | 0 | 0 |
| NC_045512.2;YP_009724389.1;Wu1-SARS2_orf1ab;5609 | 0 | 0 |
| NC_045512.2;YP_009724389.1;Wu1-SARS2_orf1ab;5610 | 0 | 0 |
| NC_045512.2;YP_009724389.1;Wu1-SARS2_orf1ab;5611 | 0 | 0 |
| NC_045512.2;YP_009724389.1;Wu1-SARS2_orf1ab;5612 | 0 | 0 |
| NC_045512.2;YP_009724389.1;Wu1-SARS2_orf1ab;5613 | 0 | 0 |
| NC_045512.2;YP_009724389.1;Wu1-SARS2_orf1ab;5614 | 0 | 0 |
| NC_045512.2;YP_009724389.1;Wu1-SARS2_orf1ab;5615 | 0 | 0 |
| NC_045512.2;YP_009724389.1;Wu1-SARS2_orf1ab;5616 | 0 | 0 |
| NC_045512.2;YP_009724389.1;Wu1-SARS2_orf1ab;5617 | 0 | 0 |
| NC_045512.2;YP_009724389.1;Wu1-SARS2_orf1ab;5618 | 0 | 0 |
| NC_045512.2;YP_009724389.1;Wu1-SARS2_orf1ab;5619 | 0 | 0 |
| NC_045512.2;YP_009724389.1;Wu1-SARS2_orf1ab;5620 | 0 | 0 |
| NC_045512.2;YP_009724389.1;Wu1-SARS2_orf1ab;5621 | 0 | 0 |
| NC_045512.2;YP_009724389.1;Wu1-SARS2_orf1ab;5622 | 0 | 0 |
| NC_045512.2;YP_009724389.1;Wu1-SARS2_orf1ab;5623 | 0 | 0 |

| | | |
|---|---|---|
| NC_045512.2;YP_009724389.1;Wu1-SARS2_orf1ab;5624 | 0 | 0.025 |
| NC_045512.2;YP_009724389.1;Wu1-SARS2_orf1ab;5625 | 0 | 0.025 |
| NC_045512.2;YP_009724389.1;Wu1-SARS2_orf1ab;5626 | 0 | 0.05 |
| NC_045512.2;YP_009724389.1;Wu1-SARS2_orf1ab;5627 | 0 | 0 |
| NC_045512.2;YP_009724389.1;Wu1-SARS2_orf1ab;5628 | 0 | 0 |
| NC_045512.2;YP_009724389.1;Wu1-SARS2_orf1ab;5629 | 0 | 0 |
| NC_045512.2;YP_009724389.1;Wu1-SARS2_orf1ab;5630 | 0 | 0 |
| NC_045512.2;YP_009724389.1;Wu1-SARS2_orf1ab;5631 | 0 | 0 |
| NC_045512.2;YP_009724389.1;Wu1-SARS2_orf1ab;5632 | 0 | 0 |
| NC_045512.2;YP_009724389.1;Wu1-SARS2_orf1ab;5633 | 0 | 0 |
| NC_045512.2;YP_009724389.1;Wu1-SARS2_orf1ab;5634 | 0 | 0 |
| NC_045512.2;YP_009724389.1;Wu1-SARS2_orf1ab;5635 | 0 | 0 |
| NC_045512.2;YP_009724389.1;Wu1-SARS2_orf1ab;5636 | 0 | 0 |
| NC_045512.2;YP_009724389.1;Wu1-SARS2_orf1ab;5637 | 0 | 0 |
| NC_045512.2;YP_009724389.1;Wu1-SARS2_orf1ab;5638 | 0.05 | 0 |
| NC_045512.2;YP_009724389.1;Wu1-SARS2_orf1ab;5639 | 0 | 0 |
| NC_045512.2;YP_009724389.1;Wu1-SARS2_orf1ab;5640 | 0 | 0 |
| NC_045512.2;YP_009724389.1;Wu1-SARS2_orf1ab;5641 | 0 | 0 |
| NC_045512.2;YP_009724389.1;Wu1-SARS2_orf1ab;5642 | 0 | 0 |
| NC_045512.2;YP_009724389.1;Wu1-SARS2_orf1ab;5643 | 0 | 0 |
| NC_045512.2;YP_009724389.1;Wu1-SARS2_orf1ab;5644 | 0 | 0 |
| NC_045512.2;YP_009724389.1;Wu1-SARS2_orf1ab;5645 | 0 | 0 |
| NC_045512.2;YP_009724389.1;Wu1-SARS2_orf1ab;5646 | 0 | 0 |
| NC_045512.2;YP_009724389.1;Wu1-SARS2_orf1ab;5647 | 0 | 0 |
| NC_045512.2;YP_009724389.1;Wu1-SARS2_orf1ab;5648 | 0 | 0 |
| NC_045512.2;YP_009724389.1;Wu1-SARS2_orf1ab;5649 | 0 | 0 |
| NC_045512.2;YP_009724389.1;Wu1-SARS2_orf1ab;5650 | 0 | 0 |
| NC_045512.2;YP_009724389.1;Wu1-SARS2_orf1ab;5651 | 0 | 0 |
| NC_045512.2;YP_009724389.1;Wu1-SARS2_orf1ab;5652 | 0 | 0 |
| NC_045512.2;YP_009724389.1;Wu1-SARS2_orf1ab;5653 | 0 | 0 |
| NC_045512.2;YP_009724389.1;Wu1-SARS2_orf1ab;5654 | 0 | 0 |
| NC_045512.2;YP_009724389.1;Wu1-SARS2_orf1ab;5655 | 0 | 0 |
| NC_045512.2;YP_009724389.1;Wu1-SARS2_orf1ab;5656 | 0 | 0 |
| NC_045512.2;YP_009724389.1;Wu1-SARS2_orf1ab;5657 | 0 | 0.025 |
| NC_045512.2;YP_009724389.1;Wu1-SARS2_orf1ab;5658 | 0 | 0.025 |
| NC_045512.2;YP_009724389.1;Wu1-SARS2_orf1ab;5659 | 0.05 | 0.025 |
| NC_045512.2;YP_009724389.1;Wu1-SARS2_orf1ab;5660 | 0 | 0.025 |
| NC_045512.2;YP_009724389.1;Wu1-SARS2_orf1ab;5661 | 0.05 | 0.075 |
| NC_045512.2;YP_009724389.1;Wu1-SARS2_orf1ab;5662 | 0.25 | 0.075 |
| NC_045512.2;YP_009724389.1;Wu1-SARS2_orf1ab;5663 | 0.4 | 0.125 |
| NC_045512.2;YP_009724389.1;Wu1-SARS2_orf1ab;5664 | 0.35 | 0.075 |
| NC_045512.2;YP_009724389.1;Wu1-SARS2_orf1ab;5665 | 0.3 | 0.075 |
| NC_045512.2;YP_009724389.1;Wu1-SARS2_orf1ab;5666 | 0.1 | 0.025 |

| | | |
|---|---|---|
| NC_045512.2;YP_009724389.1;Wu1-SARS2_orf1ab;5667 | 0.05 | 0.025 |
| NC_045512.2;YP_009724389.1;Wu1-SARS2_orf1ab;5668 | 0 | 0.025 |
| NC_045512.2;YP_009724389.1;Wu1-SARS2_orf1ab;5669 | 0 | 0 |
| NC_045512.2;YP_009724389.1;Wu1-SARS2_orf1ab;5670 | 0 | 0 |
| NC_045512.2;YP_009724389.1;Wu1-SARS2_orf1ab;5671 | 0 | 0 |
| NC_045512.2;YP_009724389.1;Wu1-SARS2_orf1ab;5672 | 0 | 0 |
| NC_045512.2;YP_009724389.1;Wu1-SARS2_orf1ab;5673 | 0 | 0 |
| NC_045512.2;YP_009724389.1;Wu1-SARS2_orf1ab;5674 | 0 | 0 |
| NC_045512.2;YP_009724389.1;Wu1-SARS2_orf1ab;5675 | 0 | 0 |
| NC_045512.2;YP_009724389.1;Wu1-SARS2_orf1ab;5676 | 0 | 0.025 |
| NC_045512.2;YP_009724389.1;Wu1-SARS2_orf1ab;5677 | 0 | 0 |
| NC_045512.2;YP_009724389.1;Wu1-SARS2_orf1ab;5678 | 0 | 0 |
| NC_045512.2;YP_009724389.1;Wu1-SARS2_orf1ab;5679 | 0 | 0 |
| NC_045512.2;YP_009724389.1;Wu1-SARS2_orf1ab;5680 | 0 | 0 |
| NC_045512.2;YP_009724389.1;Wu1-SARS2_orf1ab;5681 | 0 | 0 |
| NC_045512.2;YP_009724389.1;Wu1-SARS2_orf1ab;5682 | 0 | 0 |
| NC_045512.2;YP_009724389.1;Wu1-SARS2_orf1ab;5683 | 0 | 0 |
| NC_045512.2;YP_009724389.1;Wu1-SARS2_orf1ab;5684 | 0 | 0 |
| NC_045512.2;YP_009724389.1;Wu1-SARS2_orf1ab;5685 | 0 | 0 |
| NC_045512.2;YP_009724389.1;Wu1-SARS2_orf1ab;5686 | 0 | 0 |
| NC_045512.2;YP_009724389.1;Wu1-SARS2_orf1ab;5687 | 0 | 0 |
| NC_045512.2;YP_009724389.1;Wu1-SARS2_orf1ab;5688 | 0 | 0.025 |
| NC_045512.2;YP_009724389.1;Wu1-SARS2_orf1ab;5689 | 0 | 0.025 |
| NC_045512.2;YP_009724389.1;Wu1-SARS2_orf1ab;5690 | 0 | 0 |
| NC_045512.2;YP_009724389.1;Wu1-SARS2_orf1ab;5691 | 0 | 0 |
| NC_045512.2;YP_009724389.1;Wu1-SARS2_orf1ab;5692 | 0 | 0.075 |
| NC_045512.2;YP_009724389.1;Wu1-SARS2_orf1ab;5693 | 0 | 0 |
| NC_045512.2;YP_009724389.1;Wu1-SARS2_orf1ab;5694 | 0 | 0.025 |
| NC_045512.2;YP_009724389.1;Wu1-SARS2_orf1ab;5695 | 0 | 0.025 |
| NC_045512.2;YP_009724389.1;Wu1-SARS2_orf1ab;5696 | 0 | 0.05 |
| NC_045512.2;YP_009724389.1;Wu1-SARS2_orf1ab;5697 | 0 | 0.05 |
| NC_045512.2;YP_009724389.1;Wu1-SARS2_orf1ab;5698 | 0 | 0.025 |
| NC_045512.2;YP_009724389.1;Wu1-SARS2_orf1ab;5699 | 0 | 0.025 |
| NC_045512.2;YP_009724389.1;Wu1-SARS2_orf1ab;5700 | 0 | 0.025 |
| NC_045512.2;YP_009724389.1;Wu1-SARS2_orf1ab;5701 | 0 | 0.025 |
| NC_045512.2;YP_009724389.1;Wu1-SARS2_orf1ab;5702 | 0 | 0 |
| NC_045512.2;YP_009724389.1;Wu1-SARS2_orf1ab;5703 | 0 | 0 |
| NC_045512.2;YP_009724389.1;Wu1-SARS2_orf1ab;5704 | 0 | 0 |
| NC_045512.2;YP_009724389.1;Wu1-SARS2_orf1ab;5705 | 0 | 0.025 |
| NC_045512.2;YP_009724389.1;Wu1-SARS2_orf1ab;5706 | 0 | 0.025 |
| NC_045512.2;YP_009724389.1;Wu1-SARS2_orf1ab;5707 | 0 | 0.025 |
| NC_045512.2;YP_009724389.1;Wu1-SARS2_orf1ab;5708 | 0 | 0 |
| NC_045512.2;YP_009724389.1;Wu1-SARS2_orf1ab;5709 | 0 | 0 |

| | | |
|---|---|---|
| NC_045512.2;YP_009724389.1;Wu1-SARS2_orf1ab;5710 | 0 | 0.025 |
| NC_045512.2;YP_009724389.1;Wu1-SARS2_orf1ab;5711 | 0 | 0.025 |
| NC_045512.2;YP_009724389.1;Wu1-SARS2_orf1ab;5712 | 0 | 0.025 |
| NC_045512.2;YP_009724389.1;Wu1-SARS2_orf1ab;5713 | 0 | 0.025 |
| NC_045512.2;YP_009724389.1;Wu1-SARS2_orf1ab;5714 | 0 | 0.025 |
| NC_045512.2;YP_009724389.1;Wu1-SARS2_orf1ab;5715 | 0 | 0.025 |
| NC_045512.2;YP_009724389.1;Wu1-SARS2_orf1ab;5716 | 0 | 0.025 |
| NC_045512.2;YP_009724389.1;Wu1-SARS2_orf1ab;5717 | 0 | 0 |
| NC_045512.2;YP_009724389.1;Wu1-SARS2_orf1ab;5718 | 0 | 0 |
| NC_045512.2;YP_009724389.1;Wu1-SARS2_orf1ab;5719 | 0 | 0 |
| NC_045512.2;YP_009724389.1;Wu1-SARS2_orf1ab;5720 | 0 | 0 |
| NC_045512.2;YP_009724389.1;Wu1-SARS2_orf1ab;5721 | 0 | 0.025 |
| NC_045512.2;YP_009724389.1;Wu1-SARS2_orf1ab;5722 | 0 | 0 |
| NC_045512.2;YP_009724389.1;Wu1-SARS2_orf1ab;5723 | 0 | 0.025 |
| NC_045512.2;YP_009724389.1;Wu1-SARS2_orf1ab;5724 | 0 | 0.025 |
| NC_045512.2;YP_009724389.1;Wu1-SARS2_orf1ab;5725 | 0 | 0.025 |
| NC_045512.2;YP_009724389.1;Wu1-SARS2_orf1ab;5726 | 0 | 0 |
| NC_045512.2;YP_009724389.1;Wu1-SARS2_orf1ab;5727 | 0 | 0.025 |
| NC_045512.2;YP_009724389.1;Wu1-SARS2_orf1ab;5728 | 0 | 0.075 |
| NC_045512.2;YP_009724389.1;Wu1-SARS2_orf1ab;5729 | 0 | 0.075 |
| NC_045512.2;YP_009724389.1;Wu1-SARS2_orf1ab;5730 | 0.05 | 0.1 |
| NC_045512.2;YP_009724389.1;Wu1-SARS2_orf1ab;5731 | 0.1 | 0.075 |
| NC_045512.2;YP_009724389.1;Wu1-SARS2_orf1ab;5732 | 0.1 | 0.075 |
| NC_045512.2;YP_009724389.1;Wu1-SARS2_orf1ab;5733 | 0.1 | 0.05 |
| NC_045512.2;YP_009724389.1;Wu1-SARS2_orf1ab;5734 | 0.1 | 0.05 |
| NC_045512.2;YP_009724389.1;Wu1-SARS2_orf1ab;5735 | 0.1 | 0.05 |
| NC_045512.2;YP_009724389.1;Wu1-SARS2_orf1ab;5736 | 0.1 | 0.05 |
| NC_045512.2;YP_009724389.1;Wu1-SARS2_orf1ab;5737 | 0.1 | 0 |
| NC_045512.2;YP_009724389.1;Wu1-SARS2_orf1ab;5738 | 0.05 | 0.025 |
| NC_045512.2;YP_009724389.1;Wu1-SARS2_orf1ab;5739 | 0 | 0 |
| NC_045512.2;YP_009724389.1;Wu1-SARS2_orf1ab;5740 | 0 | 0 |
| NC_045512.2;YP_009724389.1;Wu1-SARS2_orf1ab;5741 | 0 | 0 |
| NC_045512.2;YP_009724389.1;Wu1-SARS2_orf1ab;5742 | 0 | 0 |
| NC_045512.2;YP_009724389.1;Wu1-SARS2_orf1ab;5743 | 0 | 0 |
| NC_045512.2;YP_009724389.1;Wu1-SARS2_orf1ab;5744 | 0 | 0 |
| NC_045512.2;YP_009724389.1;Wu1-SARS2_orf1ab;5745 | 0 | 0 |
| NC_045512.2;YP_009724389.1;Wu1-SARS2_orf1ab;5746 | 0 | 0 |
| NC_045512.2;YP_009724389.1;Wu1-SARS2_orf1ab;5747 | 0 | 0 |
| NC_045512.2;YP_009724389.1;Wu1-SARS2_orf1ab;5748 | 0 | 0 |
| NC_045512.2;YP_009724389.1;Wu1-SARS2_orf1ab;5749 | 0 | 0.025 |
| NC_045512.2;YP_009724389.1;Wu1-SARS2_orf1ab;5750 | 0 | 0 |
| NC_045512.2;YP_009724389.1;Wu1-SARS2_orf1ab;5751 | 0 | 0 |
| NC_045512.2;YP_009724389.1;Wu1-SARS2_orf1ab;5752 | 0 | 0 |

| | | |
|---|---|---|
| NC_045512.2;YP_009724389.1;Wu1-SARS2_orf1ab;5753 | 0 | 0 |
| NC_045512.2;YP_009724389.1;Wu1-SARS2_orf1ab;5754 | 0 | 0 |
| NC_045512.2;YP_009724389.1;Wu1-SARS2_orf1ab;5755 | 0 | 0 |
| NC_045512.2;YP_009724389.1;Wu1-SARS2_orf1ab;5756 | 0 | 0 |
| NC_045512.2;YP_009724389.1;Wu1-SARS2_orf1ab;5757 | 0 | 0 |
| NC_045512.2;YP_009724389.1;Wu1-SARS2_orf1ab;5758 | 0 | 0 |
| NC_045512.2;YP_009724389.1;Wu1-SARS2_orf1ab;5759 | 0 | 0 |
| NC_045512.2;YP_009724389.1;Wu1-SARS2_orf1ab;5760 | 0 | 0 |
| NC_045512.2;YP_009724389.1;Wu1-SARS2_orf1ab;5761 | 0 | 0 |
| NC_045512.2;YP_009724389.1;Wu1-SARS2_orf1ab;5762 | 0.1 | 0.05 |
| NC_045512.2;YP_009724389.1;Wu1-SARS2_orf1ab;5763 | 0.1 | 0.05 |
| NC_045512.2;YP_009724389.1;Wu1-SARS2_orf1ab;5764 | 0.15 | 0.125 |
| NC_045512.2;YP_009724389.1;Wu1-SARS2_orf1ab;5765 | 0.15 | 0.15 |
| NC_045512.2;YP_009724389.1;Wu1-SARS2_orf1ab;5766 | 0.1 | 0.05 |
| NC_045512.2;YP_009724389.1;Wu1-SARS2_orf1ab;5767 | 0.1 | 0.025 |
| NC_045512.2;YP_009724389.1;Wu1-SARS2_orf1ab;5768 | 0.1 | 0.075 |
| NC_045512.2;YP_009724389.1;Wu1-SARS2_orf1ab;5769 | 0.1 | 0.15 |
| NC_045512.2;YP_009724389.1;Wu1-SARS2_orf1ab;5770 | 0.15 | 0.225 |
| NC_045512.2;YP_009724389.1;Wu1-SARS2_orf1ab;5771 | 0.1 | 0.05 |
| NC_045512.2;YP_009724389.1;Wu1-SARS2_orf1ab;5772 | 0.1 | 0.075 |
| NC_045512.2;YP_009724389.1;Wu1-SARS2_orf1ab;5773 | 0.15 | 0.15 |
| NC_045512.2;YP_009724389.1;Wu1-SARS2_orf1ab;5774 | 0 | 0 |
| NC_045512.2;YP_009724389.1;Wu1-SARS2_orf1ab;5775 | 0 | 0 |
| NC_045512.2;YP_009724389.1;Wu1-SARS2_orf1ab;5776 | 0 | 0 |
| NC_045512.2;YP_009724389.1;Wu1-SARS2_orf1ab;5777 | 0.05 | 0 |
| NC_045512.2;YP_009724389.1;Wu1-SARS2_orf1ab;5778 | 0 | 0 |
| NC_045512.2;YP_009724389.1;Wu1-SARS2_orf1ab;5779 | 0.05 | 0.025 |
| NC_045512.2;YP_009724389.1;Wu1-SARS2_orf1ab;5780 | 0 | 0.025 |
| NC_045512.2;YP_009724389.1;Wu1-SARS2_orf1ab;5781 | 0 | 0.025 |
| NC_045512.2;YP_009724389.1;Wu1-SARS2_orf1ab;5782 | 0 | 0.025 |
| NC_045512.2;YP_009724389.1;Wu1-SARS2_orf1ab;5783 | 0 | 0.025 |
| NC_045512.2;YP_009724389.1;Wu1-SARS2_orf1ab;5784 | 0 | 0.025 |
| NC_045512.2;YP_009724389.1;Wu1-SARS2_orf1ab;5785 | 0.05 | 0 |
| NC_045512.2;YP_009724389.1;Wu1-SARS2_orf1ab;5786 | 0 | 0 |
| NC_045512.2;YP_009724389.1;Wu1-SARS2_orf1ab;5787 | 0 | 0 |
| NC_045512.2;YP_009724389.1;Wu1-SARS2_orf1ab;5788 | 0 | 0 |
| NC_045512.2;YP_009724389.1;Wu1-SARS2_orf1ab;5789 | 0 | 0 |
| NC_045512.2;YP_009724389.1;Wu1-SARS2_orf1ab;5790 | 0 | 0 |
| NC_045512.2;YP_009724389.1;Wu1-SARS2_orf1ab;5791 | 0 | 0 |
| NC_045512.2;YP_009724389.1;Wu1-SARS2_orf1ab;5792 | 0 | 0 |
| NC_045512.2;YP_009724389.1;Wu1-SARS2_orf1ab;5793 | 0 | 0 |
| NC_045512.2;YP_009724389.1;Wu1-SARS2_orf1ab;5794 | 0 | 0 |
| NC_045512.2;YP_009724389.1;Wu1-SARS2_orf1ab;5795 | 0 | 0 |

| | | |
|---|---|---|
| NC_045512.2;YP_009724389.1;Wu1-SARS2_orf1ab;5796 | 0 | 0 |
| NC_045512.2;YP_009724389.1;Wu1-SARS2_orf1ab;5797 | 0 | 0 |
| NC_045512.2;YP_009724389.1;Wu1-SARS2_orf1ab;5798 | 0 | 0 |
| NC_045512.2;YP_009724389.1;Wu1-SARS2_orf1ab;5799 | 0 | 0 |
| NC_045512.2;YP_009724389.1;Wu1-SARS2_orf1ab;5800 | 0 | 0 |
| NC_045512.2;YP_009724389.1;Wu1-SARS2_orf1ab;5801 | 0 | 0.025 |
| NC_045512.2;YP_009724389.1;Wu1-SARS2_orf1ab;5802 | 0.05 | 0 |
| NC_045512.2;YP_009724389.1;Wu1-SARS2_orf1ab;5803 | 0.05 | 0 |
| NC_045512.2;YP_009724389.1;Wu1-SARS2_orf1ab;5804 | 0.05 | 0 |
| NC_045512.2;YP_009724389.1;Wu1-SARS2_orf1ab;5805 | 0 | 0 |
| NC_045512.2;YP_009724389.1;Wu1-SARS2_orf1ab;5806 | 0 | 0 |
| NC_045512.2;YP_009724389.1;Wu1-SARS2_orf1ab;5807 | 0.05 | 0 |
| NC_045512.2;YP_009724389.1;Wu1-SARS2_orf1ab;5808 | 0 | 0 |
| NC_045512.2;YP_009724389.1;Wu1-SARS2_orf1ab;5809 | 0 | 0 |
| NC_045512.2;YP_009724389.1;Wu1-SARS2_orf1ab;5810 | 0 | 0 |
| NC_045512.2;YP_009724389.1;Wu1-SARS2_orf1ab;5811 | 0 | 0 |
| NC_045512.2;YP_009724389.1;Wu1-SARS2_orf1ab;5812 | 0 | 0 |
| NC_045512.2;YP_009724389.1;Wu1-SARS2_orf1ab;5813 | 0 | 0 |
| NC_045512.2;YP_009724389.1;Wu1-SARS2_orf1ab;5814 | 0 | 0 |
| NC_045512.2;YP_009724389.1;Wu1-SARS2_orf1ab;5815 | 0 | 0 |
| NC_045512.2;YP_009724389.1;Wu1-SARS2_orf1ab;5816 | 0 | 0 |
| NC_045512.2;YP_009724389.1;Wu1-SARS2_orf1ab;5817 | 0 | 0 |
| NC_045512.2;YP_009724389.1;Wu1-SARS2_orf1ab;5818 | 0 | 0.025 |
| NC_045512.2;YP_009724389.1;Wu1-SARS2_orf1ab;5819 | 0 | 0 |
| NC_045512.2;YP_009724389.1;Wu1-SARS2_orf1ab;5820 | 0 | 0 |
| NC_045512.2;YP_009724389.1;Wu1-SARS2_orf1ab;5821 | 0 | 0 |
| NC_045512.2;YP_009724389.1;Wu1-SARS2_orf1ab;5822 | 0 | 0 |
| NC_045512.2;YP_009724389.1;Wu1-SARS2_orf1ab;5823 | 0 | 0 |
| NC_045512.2;YP_009724389.1;Wu1-SARS2_orf1ab;5824 | 0 | 0 |
| NC_045512.2;YP_009724389.1;Wu1-SARS2_orf1ab;5825 | 0 | 0 |
| NC_045512.2;YP_009724389.1;Wu1-SARS2_orf1ab;5826 | 0 | 0.05 |
| NC_045512.2;YP_009724389.1;Wu1-SARS2_orf1ab;5827 | 0 | 0 |
| NC_045512.2;YP_009724389.1;Wu1-SARS2_orf1ab;5828 | 0.05 | 0 |
| NC_045512.2;YP_009724389.1;Wu1-SARS2_orf1ab;5829 | 0 | 0 |
| NC_045512.2;YP_009724389.1;Wu1-SARS2_orf1ab;5830 | 0 | 0 |
| NC_045512.2;YP_009724389.1;Wu1-SARS2_orf1ab;5831 | 0 | 0 |
| NC_045512.2;YP_009724389.1;Wu1-SARS2_orf1ab;5832 | 0.15 | 0.025 |
| NC_045512.2;YP_009724389.1;Wu1-SARS2_orf1ab;5833 | 0.05 | 0 |
| NC_045512.2;YP_009724389.1;Wu1-SARS2_orf1ab;5834 | 0 | 0 |
| NC_045512.2;YP_009724389.1;Wu1-SARS2_orf1ab;5835 | 0 | 0.025 |
| NC_045512.2;YP_009724389.1;Wu1-SARS2_orf1ab;5836 | 0 | 0.025 |
| NC_045512.2;YP_009724389.1;Wu1-SARS2_orf1ab;5837 | 0.1 | 0 |
| NC_045512.2;YP_009724389.1;Wu1-SARS2_orf1ab;5838 | 0.1 | 0.05 |

| | | |
|---|---|---|
| NC_045512.2;YP_009724389.1;Wu1-SARS2_orf1ab;5839 | 0.05 | 0 |
| NC_045512.2;YP_009724389.1;Wu1-SARS2_orf1ab;5840 | 0 | 0 |
| NC_045512.2;YP_009724389.1;Wu1-SARS2_orf1ab;5841 | 0 | 0 |
| NC_045512.2;YP_009724389.1;Wu1-SARS2_orf1ab;5842 | 0 | 0.05 |
| NC_045512.2;YP_009724389.1;Wu1-SARS2_orf1ab;5843 | 0 | 0 |
| NC_045512.2;YP_009724389.1;Wu1-SARS2_orf1ab;5844 | 0 | 0.025 |
| NC_045512.2;YP_009724389.1;Wu1-SARS2_orf1ab;5845 | 0 | 0 |
| NC_045512.2;YP_009724389.1;Wu1-SARS2_orf1ab;5846 | 0.05 | 0 |
| NC_045512.2;YP_009724389.1;Wu1-SARS2_orf1ab;5847 | 0 | 0 |
| NC_045512.2;YP_009724389.1;Wu1-SARS2_orf1ab;5848 | 0 | 0 |
| NC_045512.2;YP_009724389.1;Wu1-SARS2_orf1ab;5849 | 0 | 0.05 |
| NC_045512.2;YP_009724389.1;Wu1-SARS2_orf1ab;5850 | 0 | 0.025 |
| NC_045512.2;YP_009724389.1;Wu1-SARS2_orf1ab;5851 | 0 | 0.025 |
| NC_045512.2;YP_009724389.1;Wu1-SARS2_orf1ab;5852 | 0 | 0.025 |
| NC_045512.2;YP_009724389.1;Wu1-SARS2_orf1ab;5853 | 0 | 0.025 |
| NC_045512.2;YP_009724389.1;Wu1-SARS2_orf1ab;5854 | 0 | 0 |
| NC_045512.2;YP_009724389.1;Wu1-SARS2_orf1ab;5855 | 0 | 0 |
| NC_045512.2;YP_009724389.1;Wu1-SARS2_orf1ab;5856 | 0 | 0.025 |
| NC_045512.2;YP_009724389.1;Wu1-SARS2_orf1ab;5857 | 0 | 0.025 |
| NC_045512.2;YP_009724389.1;Wu1-SARS2_orf1ab;5858 | 0 | 0 |
| NC_045512.2;YP_009724389.1;Wu1-SARS2_orf1ab;5859 | 0 | 0 |
| NC_045512.2;YP_009724389.1;Wu1-SARS2_orf1ab;5860 | 0 | 0 |
| NC_045512.2;YP_009724389.1;Wu1-SARS2_orf1ab;5861 | 0 | 0 |
| NC_045512.2;YP_009724389.1;Wu1-SARS2_orf1ab;5862 | 0 | 0 |
| NC_045512.2;YP_009724389.1;Wu1-SARS2_orf1ab;5863 | 0.05 | 0 |
| NC_045512.2;YP_009724389.1;Wu1-SARS2_orf1ab;5864 | 0.05 | 0.025 |
| NC_045512.2;YP_009724389.1;Wu1-SARS2_orf1ab;5865 | 0.05 | 0 |
| NC_045512.2;YP_009724389.1;Wu1-SARS2_orf1ab;5866 | 0.05 | 0 |
| NC_045512.2;YP_009724389.1;Wu1-SARS2_orf1ab;5867 | 0.05 | 0 |
| NC_045512.2;YP_009724389.1;Wu1-SARS2_orf1ab;5868 | 0.05 | 0 |
| NC_045512.2;YP_009724389.1;Wu1-SARS2_orf1ab;5869 | 0 | 0 |
| NC_045512.2;YP_009724389.1;Wu1-SARS2_orf1ab;5870 | 0 | 0 |
| NC_045512.2;YP_009724389.1;Wu1-SARS2_orf1ab;5871 | 0.05 | 0 |
| NC_045512.2;YP_009724389.1;Wu1-SARS2_orf1ab;5872 | 0.05 | 0 |
| NC_045512.2;YP_009724389.1;Wu1-SARS2_orf1ab;5873 | 0.05 | 0.05 |
| NC_045512.2;YP_009724389.1;Wu1-SARS2_orf1ab;5874 | 0 | 0 |
| NC_045512.2;YP_009724389.1;Wu1-SARS2_orf1ab;5875 | 0 | 0 |
| NC_045512.2;YP_009724389.1;Wu1-SARS2_orf1ab;5876 | 0 | 0 |
| NC_045512.2;YP_009724389.1;Wu1-SARS2_orf1ab;5877 | 0 | 0 |
| NC_045512.2;YP_009724389.1;Wu1-SARS2_orf1ab;5878 | 0 | 0 |
| NC_045512.2;YP_009724389.1;Wu1-SARS2_orf1ab;5879 | 0 | 0 |
| NC_045512.2;YP_009724389.1;Wu1-SARS2_orf1ab;5880 | 0 | 0 |
| NC_045512.2;YP_009724389.1;Wu1-SARS2_orf1ab;5881 | 0 | 0 |

| | | |
|---|---|---|
| NC_045512.2;YP_009724389.1;Wu1-SARS2_orf1ab;5882 | 0 | 0 |
| NC_045512.2;YP_009724389.1;Wu1-SARS2_orf1ab;5883 | 0 | 0 |
| NC_045512.2;YP_009724389.1;Wu1-SARS2_orf1ab;5884 | 0 | 0 |
| NC_045512.2;YP_009724389.1;Wu1-SARS2_orf1ab;5885 | 0 | 0 |
| NC_045512.2;YP_009724389.1;Wu1-SARS2_orf1ab;5886 | 0 | 0 |
| NC_045512.2;YP_009724389.1;Wu1-SARS2_orf1ab;5887 | 0 | 0 |
| NC_045512.2;YP_009724389.1;Wu1-SARS2_orf1ab;5888 | 0 | 0 |
| NC_045512.2;YP_009724389.1;Wu1-SARS2_orf1ab;5889 | 0 | 0 |
| NC_045512.2;YP_009724389.1;Wu1-SARS2_orf1ab;5890 | 0 | 0 |
| NC_045512.2;YP_009724389.1;Wu1-SARS2_orf1ab;5891 | 0 | 0 |
| NC_045512.2;YP_009724389.1;Wu1-SARS2_orf1ab;5892 | 0 | 0 |
| NC_045512.2;YP_009724389.1;Wu1-SARS2_orf1ab;5893 | 0 | 0.025 |
| NC_045512.2;YP_009724389.1;Wu1-SARS2_orf1ab;5894 | 0 | 0 |
| NC_045512.2;YP_009724389.1;Wu1-SARS2_orf1ab;5895 | 0 | 0.05 |
| NC_045512.2;YP_009724389.1;Wu1-SARS2_orf1ab;5896 | 0 | 0.05 |
| NC_045512.2;YP_009724389.1;Wu1-SARS2_orf1ab;5897 | 0.05 | 0.05 |
| NC_045512.2;YP_009724389.1;Wu1-SARS2_orf1ab;5898 | 0.05 | 0.05 |
| NC_045512.2;YP_009724389.1;Wu1-SARS2_orf1ab;5899 | 0.1 | 0.075 |
| NC_045512.2;YP_009724389.1;Wu1-SARS2_orf1ab;5900 | 0 | 0.025 |
| NC_045512.2;YP_009724389.1;Wu1-SARS2_orf1ab;5901 | 0 | 0 |
| NC_045512.2;YP_009724389.1;Wu1-SARS2_orf1ab;5902 | 0 | 0 |
| NC_045512.2;YP_009724389.1;Wu1-SARS2_orf1ab;5903 | 0 | 0.025 |
| NC_045512.2;YP_009724389.1;Wu1-SARS2_orf1ab;5904 | 0 | 0.025 |
| NC_045512.2;YP_009724389.1;Wu1-SARS2_orf1ab;5905 | 0 | 0 |
| NC_045512.2;YP_009724389.1;Wu1-SARS2_orf1ab;5906 | 0 | 0 |
| NC_045512.2;YP_009724389.1;Wu1-SARS2_orf1ab;5907 | 0 | 0 |
| NC_045512.2;YP_009724389.1;Wu1-SARS2_orf1ab;5908 | 0 | 0.025 |
| NC_045512.2;YP_009724389.1;Wu1-SARS2_orf1ab;5909 | 0 | 0.025 |
| NC_045512.2;YP_009724389.1;Wu1-SARS2_orf1ab;5910 | 0 | 0.05 |
| NC_045512.2;YP_009724389.1;Wu1-SARS2_orf1ab;5911 | 0 | 0.05 |
| NC_045512.2;YP_009724389.1;Wu1-SARS2_orf1ab;5912 | 0.05 | 0.075 |
| NC_045512.2;YP_009724389.1;Wu1-SARS2_orf1ab;5913 | 0.1 | 0.05 |
| NC_045512.2;YP_009724389.1;Wu1-SARS2_orf1ab;5914 | 0.05 | 0.075 |
| NC_045512.2;YP_009724389.1;Wu1-SARS2_orf1ab;5915 | 0.05 | 0.05 |
| NC_045512.2;YP_009724389.1;Wu1-SARS2_orf1ab;5916 | 0.1 | 0.1 |
| NC_045512.2;YP_009724389.1;Wu1-SARS2_orf1ab;5917 | 0.15 | 0.025 |
| NC_045512.2;YP_009724389.1;Wu1-SARS2_orf1ab;5918 | 0.05 | 0 |
| NC_045512.2;YP_009724389.1;Wu1-SARS2_orf1ab;5919 | 0 | 0 |
| NC_045512.2;YP_009724389.1;Wu1-SARS2_orf1ab;5920 | 0 | 0 |
| NC_045512.2;YP_009724389.1;Wu1-SARS2_orf1ab;5921 | 0.05 | 0 |
| NC_045512.2;YP_009724389.1;Wu1-SARS2_orf1ab;5922 | 0 | 0 |
| NC_045512.2;YP_009724389.1;Wu1-SARS2_orf1ab;5923 | 0 | 0 |
| NC_045512.2;YP_009724389.1;Wu1-SARS2_orf1ab;5924 | 0 | 0 |

| | | |
|---|---|---|
| NC_045512.2;YP_009724389.1;Wu1-SARS2_orf1ab;5925 | 0 | 0 |
| NC_045512.2;YP_009724389.1;Wu1-SARS2_orf1ab;5926 | 0 | 0 |
| NC_045512.2;YP_009724389.1;Wu1-SARS2_orf1ab;5927 | 0 | 0 |
| NC_045512.2;YP_009724389.1;Wu1-SARS2_orf1ab;5928 | 0 | 0 |
| NC_045512.2;YP_009724389.1;Wu1-SARS2_orf1ab;5929 | 0 | 0 |
| NC_045512.2;YP_009724389.1;Wu1-SARS2_orf1ab;5930 | 0 | 0 |
| NC_045512.2;YP_009724389.1;Wu1-SARS2_orf1ab;5931 | 0 | 0 |
| NC_045512.2;YP_009724389.1;Wu1-SARS2_orf1ab;5932 | 0 | 0 |
| NC_045512.2;YP_009724389.1;Wu1-SARS2_orf1ab;5933 | 0 | 0 |
| NC_045512.2;YP_009724389.1;Wu1-SARS2_orf1ab;5934 | 0.05 | 0 |
| NC_045512.2;YP_009724389.1;Wu1-SARS2_orf1ab;5935 | 0 | 0 |
| NC_045512.2;YP_009724389.1;Wu1-SARS2_orf1ab;5936 | 0 | 0 |
| NC_045512.2;YP_009724389.1;Wu1-SARS2_orf1ab;5937 | 0 | 0 |
| NC_045512.2;YP_009724389.1;Wu1-SARS2_orf1ab;5938 | 0 | 0 |
| NC_045512.2;YP_009724389.1;Wu1-SARS2_orf1ab;5939 | 0 | 0 |
| NC_045512.2;YP_009724389.1;Wu1-SARS2_orf1ab;5940 | 0 | 0 |
| NC_045512.2;YP_009724389.1;Wu1-SARS2_orf1ab;5941 | 0 | 0 |
| NC_045512.2;YP_009724389.1;Wu1-SARS2_orf1ab;5942 | 0 | 0 |
| NC_045512.2;YP_009724389.1;Wu1-SARS2_orf1ab;5943 | 0 | 0 |
| NC_045512.2;YP_009724389.1;Wu1-SARS2_orf1ab;5944 | 0 | 0 |
| NC_045512.2;YP_009724389.1;Wu1-SARS2_orf1ab;5945 | 0 | 0 |
| NC_045512.2;YP_009724389.1;Wu1-SARS2_orf1ab;5946 | 0 | 0 |
| NC_045512.2;YP_009724389.1;Wu1-SARS2_orf1ab;5947 | 0 | 0 |
| NC_045512.2;YP_009724389.1;Wu1-SARS2_orf1ab;5948 | 0 | 0 |
| NC_045512.2;YP_009724389.1;Wu1-SARS2_orf1ab;5949 | 0 | 0 |
| NC_045512.2;YP_009724389.1;Wu1-SARS2_orf1ab;5950 | 0 | 0 |
| NC_045512.2;YP_009724389.1;Wu1-SARS2_orf1ab;5951 | 0 | 0 |
| NC_045512.2;YP_009724389.1;Wu1-SARS2_orf1ab;5952 | 0 | 0.025 |
| NC_045512.2;YP_009724389.1;Wu1-SARS2_orf1ab;5953 | 0 | 0 |
| NC_045512.2;YP_009724389.1;Wu1-SARS2_orf1ab;5954 | 0 | 0 |
| NC_045512.2;YP_009724389.1;Wu1-SARS2_orf1ab;5955 | 0 | 0 |
| NC_045512.2;YP_009724389.1;Wu1-SARS2_orf1ab;5956 | 0 | 0 |
| NC_045512.2;YP_009724389.1;Wu1-SARS2_orf1ab;5957 | 0 | 0 |
| NC_045512.2;YP_009724389.1;Wu1-SARS2_orf1ab;5958 | 0 | 0 |
| NC_045512.2;YP_009724389.1;Wu1-SARS2_orf1ab;5959 | 0 | 0 |
| NC_045512.2;YP_009724389.1;Wu1-SARS2_orf1ab;5960 | 0 | 0 |
| NC_045512.2;YP_009724389.1;Wu1-SARS2_orf1ab;5961 | 0 | 0 |
| NC_045512.2;YP_009724389.1;Wu1-SARS2_orf1ab;5962 | 0 | 0 |
| NC_045512.2;YP_009724389.1;Wu1-SARS2_orf1ab;5963 | 0 | 0 |
| NC_045512.2;YP_009724389.1;Wu1-SARS2_orf1ab;5964 | 0 | 0 |
| NC_045512.2;YP_009724389.1;Wu1-SARS2_orf1ab;5965 | 0 | 0 |
| NC_045512.2;YP_009724389.1;Wu1-SARS2_orf1ab;5966 | 0 | 0.025 |
| NC_045512.2;YP_009724389.1;Wu1-SARS2_orf1ab;5967 | 0 | 0 |

| | | |
|---|---|---|
| NC_045512.2;YP_009724389.1;Wu1-SARS2_orf1ab;5968 | 0 | 0 |
| NC_045512.2;YP_009724389.1;Wu1-SARS2_orf1ab;5969 | 0 | 0 |
| NC_045512.2;YP_009724389.1;Wu1-SARS2_orf1ab;5970 | 0 | 0 |
| NC_045512.2;YP_009724389.1;Wu1-SARS2_orf1ab;5971 | 0 | 0 |
| NC_045512.2;YP_009724389.1;Wu1-SARS2_orf1ab;5972 | 0 | 0 |
| NC_045512.2;YP_009724389.1;Wu1-SARS2_orf1ab;5973 | 0 | 0 |
| NC_045512.2;YP_009724389.1;Wu1-SARS2_orf1ab;5974 | 0 | 0 |
| NC_045512.2;YP_009724389.1;Wu1-SARS2_orf1ab;5975 | 0 | 0 |
| NC_045512.2;YP_009724389.1;Wu1-SARS2_orf1ab;5976 | 0 | 0 |
| NC_045512.2;YP_009724389.1;Wu1-SARS2_orf1ab;5977 | 0 | 0 |
| NC_045512.2;YP_009724389.1;Wu1-SARS2_orf1ab;5978 | 0 | 0 |
| NC_045512.2;YP_009724389.1;Wu1-SARS2_orf1ab;5979 | 0 | 0.025 |
| NC_045512.2;YP_009724389.1;Wu1-SARS2_orf1ab;5980 | 0 | 0.05 |
| NC_045512.2;YP_009724389.1;Wu1-SARS2_orf1ab;5981 | 0 | 0.05 |
| NC_045512.2;YP_009724389.1;Wu1-SARS2_orf1ab;5982 | 0.05 | 0.05 |
| NC_045512.2;YP_009724389.1;Wu1-SARS2_orf1ab;5983 | 0.1 | 0.05 |
| NC_045512.2;YP_009724389.1;Wu1-SARS2_orf1ab;5984 | 0.05 | 0.05 |
| NC_045512.2;YP_009724389.1;Wu1-SARS2_orf1ab;5985 | 0 | 0.05 |
| NC_045512.2;YP_009724389.1;Wu1-SARS2_orf1ab;5986 | 0 | 0.025 |
| NC_045512.2;YP_009724389.1;Wu1-SARS2_orf1ab;5987 | 0 | 0 |
| NC_045512.2;YP_009724389.1;Wu1-SARS2_orf1ab;5988 | 0 | 0 |
| NC_045512.2;YP_009724389.1;Wu1-SARS2_orf1ab;5989 | 0.05 | 0 |
| NC_045512.2;YP_009724389.1;Wu1-SARS2_orf1ab;5990 | 0 | 0 |
| NC_045512.2;YP_009724389.1;Wu1-SARS2_orf1ab;5991 | 0 | 0 |
| NC_045512.2;YP_009724389.1;Wu1-SARS2_orf1ab;5992 | 0 | 0 |
| NC_045512.2;YP_009724389.1;Wu1-SARS2_orf1ab;5993 | 0 | 0 |
| NC_045512.2;YP_009724389.1;Wu1-SARS2_orf1ab;5994 | 0 | 0 |
| NC_045512.2;YP_009724389.1;Wu1-SARS2_orf1ab;5995 | 0 | 0 |
| NC_045512.2;YP_009724389.1;Wu1-SARS2_orf1ab;5996 | 0 | 0 |
| NC_045512.2;YP_009724389.1;Wu1-SARS2_orf1ab;5997 | 0 | 0 |
| NC_045512.2;YP_009724389.1;Wu1-SARS2_orf1ab;5998 | 0 | 0 |
| NC_045512.2;YP_009724389.1;Wu1-SARS2_orf1ab;5999 | 0 | 0.025 |
| NC_045512.2;YP_009724389.1;Wu1-SARS2_orf1ab;6000 | 0 | 0 |
| NC_045512.2;YP_009724389.1;Wu1-SARS2_orf1ab;6001 | 0 | 0 |
| NC_045512.2;YP_009724389.1;Wu1-SARS2_orf1ab;6002 | 0 | 0 |
| NC_045512.2;YP_009724389.1;Wu1-SARS2_orf1ab;6003 | 0 | 0 |
| NC_045512.2;YP_009724389.1;Wu1-SARS2_orf1ab;6004 | 0 | 0 |
| NC_045512.2;YP_009724389.1;Wu1-SARS2_orf1ab;6005 | 0 | 0 |
| NC_045512.2;YP_009724389.1;Wu1-SARS2_orf1ab;6006 | 0 | 0 |
| NC_045512.2;YP_009724389.1;Wu1-SARS2_orf1ab;6007 | 0 | 0 |
| NC_045512.2;YP_009724389.1;Wu1-SARS2_orf1ab;6008 | 0 | 0 |
| NC_045512.2;YP_009724389.1;Wu1-SARS2_orf1ab;6009 | 0 | 0 |
| NC_045512.2;YP_009724389.1;Wu1-SARS2_orf1ab;6010 | 0 | 0 |

| | | |
|---|---|---|
| NC_045512.2;YP_009724389.1;Wu1-SARS2_orf1ab;6011 | 0 | 0 |
| NC_045512.2;YP_009724389.1;Wu1-SARS2_orf1ab;6012 | 0 | 0 |
| NC_045512.2;YP_009724389.1;Wu1-SARS2_orf1ab;6013 | 0 | 0 |
| NC_045512.2;YP_009724389.1;Wu1-SARS2_orf1ab;6014 | 0 | 0 |
| NC_045512.2;YP_009724389.1;Wu1-SARS2_orf1ab;6015 | 0 | 0 |
| NC_045512.2;YP_009724389.1;Wu1-SARS2_orf1ab;6016 | 0 | 0 |
| NC_045512.2;YP_009724389.1;Wu1-SARS2_orf1ab;6017 | 0 | 0.025 |
| NC_045512.2;YP_009724389.1;Wu1-SARS2_orf1ab;6018 | 0 | 0.025 |
| NC_045512.2;YP_009724389.1;Wu1-SARS2_orf1ab;6019 | 0 | 0 |
| NC_045512.2;YP_009724389.1;Wu1-SARS2_orf1ab;6020 | 0 | 0 |
| NC_045512.2;YP_009724389.1;Wu1-SARS2_orf1ab;6021 | 0 | 0 |
| NC_045512.2;YP_009724389.1;Wu1-SARS2_orf1ab;6022 | 0 | 0.025 |
| NC_045512.2;YP_009724389.1;Wu1-SARS2_orf1ab;6023 | 0 | 0 |
| NC_045512.2;YP_009724389.1;Wu1-SARS2_orf1ab;6024 | 0 | 0 |
| NC_045512.2;YP_009724389.1;Wu1-SARS2_orf1ab;6025 | 0 | 0.025 |
| NC_045512.2;YP_009724389.1;Wu1-SARS2_orf1ab;6026 | 0 | 0 |
| NC_045512.2;YP_009724389.1;Wu1-SARS2_orf1ab;6027 | 0 | 0 |
| NC_045512.2;YP_009724389.1;Wu1-SARS2_orf1ab;6028 | 0 | 0 |
| NC_045512.2;YP_009724389.1;Wu1-SARS2_orf1ab;6029 | 0 | 0.025 |
| NC_045512.2;YP_009724389.1;Wu1-SARS2_orf1ab;6030 | 0 | 0 |
| NC_045512.2;YP_009724389.1;Wu1-SARS2_orf1ab;6031 | 0 | 0 |
| NC_045512.2;YP_009724389.1;Wu1-SARS2_orf1ab;6032 | 0 | 0 |
| NC_045512.2;YP_009724389.1;Wu1-SARS2_orf1ab;6033 | 0 | 0 |
| NC_045512.2;YP_009724389.1;Wu1-SARS2_orf1ab;6034 | 0 | 0 |
| NC_045512.2;YP_009724389.1;Wu1-SARS2_orf1ab;6035 | 0 | 0 |
| NC_045512.2;YP_009724389.1;Wu1-SARS2_orf1ab;6036 | 0 | 0 |
| NC_045512.2;YP_009724389.1;Wu1-SARS2_orf1ab;6037 | 0 | 0 |
| NC_045512.2;YP_009724389.1;Wu1-SARS2_orf1ab;6038 | 0 | 0 |
| NC_045512.2;YP_009724389.1;Wu1-SARS2_orf1ab;6039 | 0 | 0 |
| NC_045512.2;YP_009724389.1;Wu1-SARS2_orf1ab;6040 | 0 | 0 |
| NC_045512.2;YP_009724389.1;Wu1-SARS2_orf1ab;6041 | 0 | 0.05 |
| NC_045512.2;YP_009724389.1;Wu1-SARS2_orf1ab;6042 | 0 | 0.075 |
| NC_045512.2;YP_009724389.1;Wu1-SARS2_orf1ab;6043 | 0.05 | 0.075 |
| NC_045512.2;YP_009724389.1;Wu1-SARS2_orf1ab;6044 | 0.05 | 0.025 |
| NC_045512.2;YP_009724389.1;Wu1-SARS2_orf1ab;6045 | 0.1 | 0.025 |
| NC_045512.2;YP_009724389.1;Wu1-SARS2_orf1ab;6046 | 0 | 0 |
| NC_045512.2;YP_009724389.1;Wu1-SARS2_orf1ab;6047 | 0.15 | 0.075 |
| NC_045512.2;YP_009724389.1;Wu1-SARS2_orf1ab;6048 | 0.05 | 0.025 |
| NC_045512.2;YP_009724389.1;Wu1-SARS2_orf1ab;6049 | 0 | 0.05 |
| NC_045512.2;YP_009724389.1;Wu1-SARS2_orf1ab;6050 | 0.05 | 0.075 |
| NC_045512.2;YP_009724389.1;Wu1-SARS2_orf1ab;6051 | 0 | 0.05 |
| NC_045512.2;YP_009724389.1;Wu1-SARS2_orf1ab;6052 | 0.05 | 0.025 |
| NC_045512.2;YP_009724389.1;Wu1-SARS2_orf1ab;6053 | 0 | 0.025 |

| | | |
|---|---|---|
| NC_045512.2;YP_009724389.1;Wu1-SARS2_orf1ab;6054 | 0.05 | 0.075 |
| NC_045512.2;YP_009724389.1;Wu1-SARS2_orf1ab;6055 | 0 | 0.025 |
| NC_045512.2;YP_009724389.1;Wu1-SARS2_orf1ab;6056 | 0 | 0.075 |
| NC_045512.2;YP_009724389.1;Wu1-SARS2_orf1ab;6057 | 0 | 0.1 |
| NC_045512.2;YP_009724389.1;Wu1-SARS2_orf1ab;6058 | 0 | 0.1 |
| NC_045512.2;YP_009724389.1;Wu1-SARS2_orf1ab;6059 | 0 | 0 |
| NC_045512.2;YP_009724389.1;Wu1-SARS2_orf1ab;6060 | 0 | 0 |
| NC_045512.2;YP_009724389.1;Wu1-SARS2_orf1ab;6061 | 0 | 0 |
| NC_045512.2;YP_009724389.1;Wu1-SARS2_orf1ab;6062 | 0 | 0 |
| NC_045512.2;YP_009724389.1;Wu1-SARS2_orf1ab;6063 | 0 | 0 |
| NC_045512.2;YP_009724389.1;Wu1-SARS2_orf1ab;6064 | 0 | 0 |
| NC_045512.2;YP_009724389.1;Wu1-SARS2_orf1ab;6065 | 0 | 0 |
| NC_045512.2;YP_009724389.1;Wu1-SARS2_orf1ab;6066 | 0 | 0 |
| NC_045512.2;YP_009724389.1;Wu1-SARS2_orf1ab;6067 | 0 | 0 |
| NC_045512.2;YP_009724389.1;Wu1-SARS2_orf1ab;6068 | 0 | 0 |
| NC_045512.2;YP_009724389.1;Wu1-SARS2_orf1ab;6069 | 0 | 0 |
| NC_045512.2;YP_009724389.1;Wu1-SARS2_orf1ab;6070 | 0 | 0 |
| NC_045512.2;YP_009724389.1;Wu1-SARS2_orf1ab;6071 | 0 | 0 |
| NC_045512.2;YP_009724389.1;Wu1-SARS2_orf1ab;6072 | 0 | 0 |
| NC_045512.2;YP_009724389.1;Wu1-SARS2_orf1ab;6073 | 0 | 0 |
| NC_045512.2;YP_009724389.1;Wu1-SARS2_orf1ab;6074 | 0 | 0 |
| NC_045512.2;YP_009724389.1;Wu1-SARS2_orf1ab;6075 | 0 | 0 |
| NC_045512.2;YP_009724389.1;Wu1-SARS2_orf1ab;6076 | 0 | 0 |
| NC_045512.2;YP_009724389.1;Wu1-SARS2_orf1ab;6077 | 0 | 0 |
| NC_045512.2;YP_009724389.1;Wu1-SARS2_orf1ab;6078 | 0 | 0 |
| NC_045512.2;YP_009724389.1;Wu1-SARS2_orf1ab;6079 | 0 | 0 |
| NC_045512.2;YP_009724389.1;Wu1-SARS2_orf1ab;6080 | 0 | 0 |
| NC_045512.2;YP_009724389.1;Wu1-SARS2_orf1ab;6081 | 0 | 0 |
| NC_045512.2;YP_009724389.1;Wu1-SARS2_orf1ab;6082 | 0 | 0 |
| NC_045512.2;YP_009724389.1;Wu1-SARS2_orf1ab;6083 | 0 | 0 |
| NC_045512.2;YP_009724389.1;Wu1-SARS2_orf1ab;6084 | 0 | 0 |
| NC_045512.2;YP_009724389.1;Wu1-SARS2_orf1ab;6085 | 0 | 0 |
| NC_045512.2;YP_009724389.1;Wu1-SARS2_orf1ab;6086 | 0 | 0 |
| NC_045512.2;YP_009724389.1;Wu1-SARS2_orf1ab;6087 | 0 | 0 |
| NC_045512.2;YP_009724389.1;Wu1-SARS2_orf1ab;6088 | 0 | 0 |
| NC_045512.2;YP_009724389.1;Wu1-SARS2_orf1ab;6089 | 0 | 0 |
| NC_045512.2;YP_009724389.1;Wu1-SARS2_orf1ab;6090 | 0 | 0 |
| NC_045512.2;YP_009724389.1;Wu1-SARS2_orf1ab;6091 | 0 | 0.025 |
| NC_045512.2;YP_009724389.1;Wu1-SARS2_orf1ab;6092 | 0 | 0 |
| NC_045512.2;YP_009724389.1;Wu1-SARS2_orf1ab;6093 | 0 | 0.025 |
| NC_045512.2;YP_009724389.1;Wu1-SARS2_orf1ab;6094 | 0 | 0.025 |
| NC_045512.2;YP_009724389.1;Wu1-SARS2_orf1ab;6095 | 0 | 0 |
| NC_045512.2;YP_009724389.1;Wu1-SARS2_orf1ab;6096 | 0 | 0 |

| | | |
|---|---|---|
| NC_045512.2;YP_009724389.1;Wu1-SARS2_orf1ab;6097 | 0 | 0 |
| NC_045512.2;YP_009724389.1;Wu1-SARS2_orf1ab;6098 | 0 | 0 |
| NC_045512.2;YP_009724389.1;Wu1-SARS2_orf1ab;6099 | 0 | 0 |
| NC_045512.2;YP_009724389.1;Wu1-SARS2_orf1ab;6100 | 0 | 0 |
| NC_045512.2;YP_009724389.1;Wu1-SARS2_orf1ab;6101 | 0 | 0 |
| NC_045512.2;YP_009724389.1;Wu1-SARS2_orf1ab;6102 | 0 | 0 |
| NC_045512.2;YP_009724389.1;Wu1-SARS2_orf1ab;6103 | 0 | 0 |
| NC_045512.2;YP_009724389.1;Wu1-SARS2_orf1ab;6104 | 0 | 0 |
| NC_045512.2;YP_009724389.1;Wu1-SARS2_orf1ab;6105 | 0 | 0 |
| NC_045512.2;YP_009724389.1;Wu1-SARS2_orf1ab;6106 | 0 | 0 |
| NC_045512.2;YP_009724389.1;Wu1-SARS2_orf1ab;6107 | 0 | 0 |
| NC_045512.2;YP_009724389.1;Wu1-SARS2_orf1ab;6108 | 0 | 0 |
| NC_045512.2;YP_009724389.1;Wu1-SARS2_orf1ab;6109 | 0 | 0 |
| NC_045512.2;YP_009724389.1;Wu1-SARS2_orf1ab;6110 | 0 | 0 |
| NC_045512.2;YP_009724389.1;Wu1-SARS2_orf1ab;6111 | 0 | 0 |
| NC_045512.2;YP_009724389.1;Wu1-SARS2_orf1ab;6112 | 0 | 0 |
| NC_045512.2;YP_009724389.1;Wu1-SARS2_orf1ab;6113 | 0 | 0 |
| NC_045512.2;YP_009724389.1;Wu1-SARS2_orf1ab;6114 | 0 | 0 |
| NC_045512.2;YP_009724389.1;Wu1-SARS2_orf1ab;6115 | 0 | 0 |
| NC_045512.2;YP_009724389.1;Wu1-SARS2_orf1ab;6116 | 0.05 | 0 |
| NC_045512.2;YP_009724389.1;Wu1-SARS2_orf1ab;6117 | 0 | 0 |
| NC_045512.2;YP_009724389.1;Wu1-SARS2_orf1ab;6118 | 0 | 0 |
| NC_045512.2;YP_009724389.1;Wu1-SARS2_orf1ab;6119 | 0 | 0 |
| NC_045512.2;YP_009724389.1;Wu1-SARS2_orf1ab;6120 | 0 | 0 |
| NC_045512.2;YP_009724389.1;Wu1-SARS2_orf1ab;6121 | 0 | 0 |
| NC_045512.2;YP_009724389.1;Wu1-SARS2_orf1ab;6122 | 0 | 0 |
| NC_045512.2;YP_009724389.1;Wu1-SARS2_orf1ab;6123 | 0 | 0 |
| NC_045512.2;YP_009724389.1;Wu1-SARS2_orf1ab;6124 | 0 | 0.025 |
| NC_045512.2;YP_009724389.1;Wu1-SARS2_orf1ab;6125 | 0 | 0 |
| NC_045512.2;YP_009724389.1;Wu1-SARS2_orf1ab;6126 | 0 | 0 |
| NC_045512.2;YP_009724389.1;Wu1-SARS2_orf1ab;6127 | 0 | 0 |
| NC_045512.2;YP_009724389.1;Wu1-SARS2_orf1ab;6128 | 0 | 0 |
| NC_045512.2;YP_009724389.1;Wu1-SARS2_orf1ab;6129 | 0 | 0.025 |
| NC_045512.2;YP_009724389.1;Wu1-SARS2_orf1ab;6130 | 0 | 0 |
| NC_045512.2;YP_009724389.1;Wu1-SARS2_orf1ab;6131 | 0 | 0 |
| NC_045512.2;YP_009724389.1;Wu1-SARS2_orf1ab;6132 | 0.05 | 0 |
| NC_045512.2;YP_009724389.1;Wu1-SARS2_orf1ab;6133 | 0.15 | 0 |
| NC_045512.2;YP_009724389.1;Wu1-SARS2_orf1ab;6134 | 0.05 | 0.025 |
| NC_045512.2;YP_009724389.1;Wu1-SARS2_orf1ab;6135 | 0.05 | 0.025 |
| NC_045512.2;YP_009724389.1;Wu1-SARS2_orf1ab;6136 | 0.05 | 0 |
| NC_045512.2;YP_009724389.1;Wu1-SARS2_orf1ab;6137 | 0 | 0 |
| NC_045512.2;YP_009724389.1;Wu1-SARS2_orf1ab;6138 | 0 | 0 |
| NC_045512.2;YP_009724389.1;Wu1-SARS2_orf1ab;6139 | 0 | 0 |

| | | |
|---|---|---|
| NC_045512.2;YP_009724389.1;Wu1-SARS2_orf1ab;6140 | 0 | 0 |
| NC_045512.2;YP_009724389.1;Wu1-SARS2_orf1ab;6141 | 0 | 0 |
| NC_045512.2;YP_009724389.1;Wu1-SARS2_orf1ab;6142 | 0 | 0 |
| NC_045512.2;YP_009724389.1;Wu1-SARS2_orf1ab;6143 | 0 | 0 |
| NC_045512.2;YP_009724389.1;Wu1-SARS2_orf1ab;6144 | 0 | 0 |
| NC_045512.2;YP_009724389.1;Wu1-SARS2_orf1ab;6145 | 0 | 0 |
| NC_045512.2;YP_009724389.1;Wu1-SARS2_orf1ab;6146 | 0 | 0 |
| NC_045512.2;YP_009724389.1;Wu1-SARS2_orf1ab;6147 | 0 | 0 |
| NC_045512.2;YP_009724389.1;Wu1-SARS2_orf1ab;6148 | 0 | 0 |
| NC_045512.2;YP_009724389.1;Wu1-SARS2_orf1ab;6149 | 0 | 0 |
| NC_045512.2;YP_009724389.1;Wu1-SARS2_orf1ab;6150 | 0 | 0 |
| NC_045512.2;YP_009724389.1;Wu1-SARS2_orf1ab;6151 | 0 | 0 |
| NC_045512.2;YP_009724389.1;Wu1-SARS2_orf1ab;6152 | 0 | 0 |
| NC_045512.2;YP_009724389.1;Wu1-SARS2_orf1ab;6153 | 0 | 0 |
| NC_045512.2;YP_009724389.1;Wu1-SARS2_orf1ab;6154 | 0 | 0 |
| NC_045512.2;YP_009724389.1;Wu1-SARS2_orf1ab;6155 | 0 | 0 |
| NC_045512.2;YP_009724389.1;Wu1-SARS2_orf1ab;6156 | 0 | 0 |
| NC_045512.2;YP_009724389.1;Wu1-SARS2_orf1ab;6157 | 0 | 0.05 |
| NC_045512.2;YP_009724389.1;Wu1-SARS2_orf1ab;6158 | 0 | 0 |
| NC_045512.2;YP_009724389.1;Wu1-SARS2_orf1ab;6159 | 0 | 0 |
| NC_045512.2;YP_009724389.1;Wu1-SARS2_orf1ab;6160 | 0 | 0 |
| NC_045512.2;YP_009724389.1;Wu1-SARS2_orf1ab;6161 | 0 | 0 |
| NC_045512.2;YP_009724389.1;Wu1-SARS2_orf1ab;6162 | 0 | 0 |
| NC_045512.2;YP_009724389.1;Wu1-SARS2_orf1ab;6163 | 0 | 0 |
| NC_045512.2;YP_009724389.1;Wu1-SARS2_orf1ab;6164 | 0 | 0 |
| NC_045512.2;YP_009724389.1;Wu1-SARS2_orf1ab;6165 | 0 | 0 |
| NC_045512.2;YP_009724389.1;Wu1-SARS2_orf1ab;6166 | 0 | 0 |
| NC_045512.2;YP_009724389.1;Wu1-SARS2_orf1ab;6167 | 0 | 0 |
| NC_045512.2;YP_009724389.1;Wu1-SARS2_orf1ab;6168 | 0 | 0 |
| NC_045512.2;YP_009724389.1;Wu1-SARS2_orf1ab;6169 | 0 | 0 |
| NC_045512.2;YP_009724389.1;Wu1-SARS2_orf1ab;6170 | 0 | 0 |
| NC_045512.2;YP_009724389.1;Wu1-SARS2_orf1ab;6171 | 0 | 0 |
| NC_045512.2;YP_009724389.1;Wu1-SARS2_orf1ab;6172 | 0 | 0 |
| NC_045512.2;YP_009724389.1;Wu1-SARS2_orf1ab;6173 | 0 | 0 |
| NC_045512.2;YP_009724389.1;Wu1-SARS2_orf1ab;6174 | 0 | 0 |
| NC_045512.2;YP_009724389.1;Wu1-SARS2_orf1ab;6175 | 0 | 0 |
| NC_045512.2;YP_009724389.1;Wu1-SARS2_orf1ab;6176 | 0 | 0 |
| NC_045512.2;YP_009724389.1;Wu1-SARS2_orf1ab;6177 | 0 | 0.025 |
| NC_045512.2;YP_009724389.1;Wu1-SARS2_orf1ab;6178 | 0 | 0 |
| NC_045512.2;YP_009724389.1;Wu1-SARS2_orf1ab;6179 | 0 | 0 |
| NC_045512.2;YP_009724389.1;Wu1-SARS2_orf1ab;6180 | 0.05 | 0.05 |
| NC_045512.2;YP_009724389.1;Wu1-SARS2_orf1ab;6181 | 0 | 0 |
| NC_045512.2;YP_009724389.1;Wu1-SARS2_orf1ab;6182 | 0 | 0 |

| | | |
|---|---|---|
| NC_045512.2;YP_009724389.1;Wu1-SARS2_orf1ab;6183 | 0 | 0 |
| NC_045512.2;YP_009724389.1;Wu1-SARS2_orf1ab;6184 | 0 | 0 |
| NC_045512.2;YP_009724389.1;Wu1-SARS2_orf1ab;6185 | 0 | 0 |
| NC_045512.2;YP_009724389.1;Wu1-SARS2_orf1ab;6186 | 0 | 0 |
| NC_045512.2;YP_009724389.1;Wu1-SARS2_orf1ab;6187 | 0 | 0 |
| NC_045512.2;YP_009724389.1;Wu1-SARS2_orf1ab;6188 | 0 | 0 |
| NC_045512.2;YP_009724389.1;Wu1-SARS2_orf1ab;6189 | 0 | 0 |
| NC_045512.2;YP_009724389.1;Wu1-SARS2_orf1ab;6190 | 0 | 0 |
| NC_045512.2;YP_009724389.1;Wu1-SARS2_orf1ab;6191 | 0 | 0 |
| NC_045512.2;YP_009724389.1;Wu1-SARS2_orf1ab;6192 | 0 | 0 |
| NC_045512.2;YP_009724389.1;Wu1-SARS2_orf1ab;6193 | 0 | 0 |
| NC_045512.2;YP_009724389.1;Wu1-SARS2_orf1ab;6194 | 0 | 0 |
| NC_045512.2;YP_009724389.1;Wu1-SARS2_orf1ab;6195 | 0 | 0 |
| NC_045512.2;YP_009724389.1;Wu1-SARS2_orf1ab;6196 | 0 | 0 |
| NC_045512.2;YP_009724389.1;Wu1-SARS2_orf1ab;6197 | 0 | 0 |
| NC_045512.2;YP_009724389.1;Wu1-SARS2_orf1ab;6198 | 0 | 0 |
| NC_045512.2;YP_009724389.1;Wu1-SARS2_orf1ab;6199 | 0 | 0 |
| NC_045512.2;YP_009724389.1;Wu1-SARS2_orf1ab;6200 | 0 | 0 |
| NC_045512.2;YP_009724389.1;Wu1-SARS2_orf1ab;6201 | 0 | 0 |
| NC_045512.2;YP_009724389.1;Wu1-SARS2_orf1ab;6202 | 0 | 0 |
| NC_045512.2;YP_009724389.1;Wu1-SARS2_orf1ab;6203 | 0 | 0 |
| NC_045512.2;YP_009724389.1;Wu1-SARS2_orf1ab;6204 | 0 | 0 |
| NC_045512.2;YP_009724389.1;Wu1-SARS2_orf1ab;6205 | 0 | 0 |
| NC_045512.2;YP_009724389.1;Wu1-SARS2_orf1ab;6206 | 0 | 0 |
| NC_045512.2;YP_009724389.1;Wu1-SARS2_orf1ab;6207 | 0 | 0 |
| NC_045512.2;YP_009724389.1;Wu1-SARS2_orf1ab;6208 | 0 | 0 |
| NC_045512.2;YP_009724389.1;Wu1-SARS2_orf1ab;6209 | 0 | 0 |
| NC_045512.2;YP_009724389.1;Wu1-SARS2_orf1ab;6210 | 0 | 0 |
| NC_045512.2;YP_009724389.1;Wu1-SARS2_orf1ab;6211 | 0 | 0 |
| NC_045512.2;YP_009724389.1;Wu1-SARS2_orf1ab;6212 | 0 | 0.025 |
| NC_045512.2;YP_009724389.1;Wu1-SARS2_orf1ab;6213 | 0 | 0.025 |
| NC_045512.2;YP_009724389.1;Wu1-SARS2_orf1ab;6214 | 0.1 | 0.075 |
| NC_045512.2;YP_009724389.1;Wu1-SARS2_orf1ab;6215 | 0.1 | 0.05 |
| NC_045512.2;YP_009724389.1;Wu1-SARS2_orf1ab;6216 | 0.1 | 0.05 |
| NC_045512.2;YP_009724389.1;Wu1-SARS2_orf1ab;6217 | 0.1 | 0.05 |
| NC_045512.2;YP_009724389.1;Wu1-SARS2_orf1ab;6218 | 0.1 | 0.025 |
| NC_045512.2;YP_009724389.1;Wu1-SARS2_orf1ab;6219 | 0.1 | 0.025 |
| NC_045512.2;YP_009724389.1;Wu1-SARS2_orf1ab;6220 | 0.1 | 0.05 |
| NC_045512.2;YP_009724389.1;Wu1-SARS2_orf1ab;6221 | 0.05 | 0 |
| NC_045512.2;YP_009724389.1;Wu1-SARS2_orf1ab;6222 | 0.05 | 0 |
| NC_045512.2;YP_009724389.1;Wu1-SARS2_orf1ab;6223 | 0.05 | 0 |
| NC_045512.2;YP_009724389.1;Wu1-SARS2_orf1ab;6224 | 0.1 | 0 |
| NC_045512.2;YP_009724389.1;Wu1-SARS2_orf1ab;6225 | 0 | 0 |

| | | |
|---|---|---|
| NC_045512.2;YP_009724389.1;Wu1-SARS2_orf1ab;6226 | 0 | 0 |
| NC_045512.2;YP_009724389.1;Wu1-SARS2_orf1ab;6227 | 0 | 0 |
| NC_045512.2;YP_009724389.1;Wu1-SARS2_orf1ab;6228 | 0 | 0 |
| NC_045512.2;YP_009724389.1;Wu1-SARS2_orf1ab;6229 | 0 | 0 |
| NC_045512.2;YP_009724389.1;Wu1-SARS2_orf1ab;6230 | 0 | 0 |
| NC_045512.2;YP_009724389.1;Wu1-SARS2_orf1ab;6231 | 0 | 0 |
| NC_045512.2;YP_009724389.1;Wu1-SARS2_orf1ab;6232 | 0 | 0 |
| NC_045512.2;YP_009724389.1;Wu1-SARS2_orf1ab;6233 | 0 | 0 |
| NC_045512.2;YP_009724389.1;Wu1-SARS2_orf1ab;6234 | 0 | 0 |
| NC_045512.2;YP_009724389.1;Wu1-SARS2_orf1ab;6235 | 0 | 0 |
| NC_045512.2;YP_009724389.1;Wu1-SARS2_orf1ab;6236 | 0 | 0 |
| NC_045512.2;YP_009724389.1;Wu1-SARS2_orf1ab;6237 | 0 | 0 |
| NC_045512.2;YP_009724389.1;Wu1-SARS2_orf1ab;6238 | 0 | 0 |
| NC_045512.2;YP_009724389.1;Wu1-SARS2_orf1ab;6239 | 0 | 0 |
| NC_045512.2;YP_009724389.1;Wu1-SARS2_orf1ab;6240 | 0 | 0 |
| NC_045512.2;YP_009724389.1;Wu1-SARS2_orf1ab;6241 | 0 | 0 |
| NC_045512.2;YP_009724389.1;Wu1-SARS2_orf1ab;6242 | 0.05 | 0 |
| NC_045512.2;YP_009724389.1;Wu1-SARS2_orf1ab;6243 | 0.05 | 0.025 |
| NC_045512.2;YP_009724389.1;Wu1-SARS2_orf1ab;6244 | 0 | 0 |
| NC_045512.2;YP_009724389.1;Wu1-SARS2_orf1ab;6245 | 0 | 0.025 |
| NC_045512.2;YP_009724389.1;Wu1-SARS2_orf1ab;6246 | 0 | 0 |
| NC_045512.2;YP_009724389.1;Wu1-SARS2_orf1ab;6247 | 0 | 0 |
| NC_045512.2;YP_009724389.1;Wu1-SARS2_orf1ab;6248 | 0 | 0 |
| NC_045512.2;YP_009724389.1;Wu1-SARS2_orf1ab;6249 | 0 | 0 |
| NC_045512.2;YP_009724389.1;Wu1-SARS2_orf1ab;6250 | 0 | 0 |
| NC_045512.2;YP_009724389.1;Wu1-SARS2_orf1ab;6251 | 0 | 0 |
| NC_045512.2;YP_009724389.1;Wu1-SARS2_orf1ab;6252 | 0 | 0 |
| NC_045512.2;YP_009724389.1;Wu1-SARS2_orf1ab;6253 | 0 | 0 |
| NC_045512.2;YP_009724389.1;Wu1-SARS2_orf1ab;6254 | 0 | 0 |
| NC_045512.2;YP_009724389.1;Wu1-SARS2_orf1ab;6255 | 0.05 | 0 |
| NC_045512.2;YP_009724389.1;Wu1-SARS2_orf1ab;6256 | 0 | 0 |
| NC_045512.2;YP_009724389.1;Wu1-SARS2_orf1ab;6257 | 0 | 0 |
| NC_045512.2;YP_009724389.1;Wu1-SARS2_orf1ab;6258 | 0 | 0 |
| NC_045512.2;YP_009724389.1;Wu1-SARS2_orf1ab;6259 | 0 | 0 |
| NC_045512.2;YP_009724389.1;Wu1-SARS2_orf1ab;6260 | 0 | 0 |
| NC_045512.2;YP_009724389.1;Wu1-SARS2_orf1ab;6261 | 0 | 0 |
| NC_045512.2;YP_009724389.1;Wu1-SARS2_orf1ab;6262 | 0.05 | 0.025 |
| NC_045512.2;YP_009724389.1;Wu1-SARS2_orf1ab;6263 | 0.05 | 0.025 |
| NC_045512.2;YP_009724389.1;Wu1-SARS2_orf1ab;6264 | 0.05 | 0.025 |
| NC_045512.2;YP_009724389.1;Wu1-SARS2_orf1ab;6265 | 0.05 | 0.025 |
| NC_045512.2;YP_009724389.1;Wu1-SARS2_orf1ab;6266 | 0.05 | 0.025 |
| NC_045512.2;YP_009724389.1;Wu1-SARS2_orf1ab;6267 | 0 | 0.025 |
| NC_045512.2;YP_009724389.1;Wu1-SARS2_orf1ab;6268 | 0 | 0.025 |

| | | |
|---|---|---|
| NC_045512.2;YP_009724389.1;Wu1-SARS2_orf1ab;6269 | 0.1 | 0 |
| NC_045512.2;YP_009724389.1;Wu1-SARS2_orf1ab;6270 | 0.1 | 0.025 |
| NC_045512.2;YP_009724389.1;Wu1-SARS2_orf1ab;6271 | 0.1 | 0.025 |
| NC_045512.2;YP_009724389.1;Wu1-SARS2_orf1ab;6272 | 0.05 | 0.05 |
| NC_045512.2;YP_009724389.1;Wu1-SARS2_orf1ab;6273 | 0 | 0 |
| NC_045512.2;YP_009724389.1;Wu1-SARS2_orf1ab;6274 | 0.05 | 0.025 |
| NC_045512.2;YP_009724389.1;Wu1-SARS2_orf1ab;6275 | 0 | 0 |
| NC_045512.2;YP_009724389.1;Wu1-SARS2_orf1ab;6276 | 0 | 0 |
| NC_045512.2;YP_009724389.1;Wu1-SARS2_orf1ab;6277 | 0 | 0 |
| NC_045512.2;YP_009724389.1;Wu1-SARS2_orf1ab;6278 | 0 | 0.025 |
| NC_045512.2;YP_009724389.1;Wu1-SARS2_orf1ab;6279 | 0 | 0.025 |
| NC_045512.2;YP_009724389.1;Wu1-SARS2_orf1ab;6280 | 0 | 0 |
| NC_045512.2;YP_009724389.1;Wu1-SARS2_orf1ab;6281 | 0 | 0 |
| NC_045512.2;YP_009724389.1;Wu1-SARS2_orf1ab;6282 | 0 | 0 |
| NC_045512.2;YP_009724389.1;Wu1-SARS2_orf1ab;6283 | 0 | 0 |
| NC_045512.2;YP_009724389.1;Wu1-SARS2_orf1ab;6284 | 0 | 0 |
| NC_045512.2;YP_009724389.1;Wu1-SARS2_orf1ab;6285 | 0 | 0 |
| NC_045512.2;YP_009724389.1;Wu1-SARS2_orf1ab;6286 | 0 | 0 |
| NC_045512.2;YP_009724389.1;Wu1-SARS2_orf1ab;6287 | 0 | 0 |
| NC_045512.2;YP_009724389.1;Wu1-SARS2_orf1ab;6288 | 0 | 0 |
| NC_045512.2;YP_009724389.1;Wu1-SARS2_orf1ab;6289 | 0.1 | 0.025 |
| NC_045512.2;YP_009724389.1;Wu1-SARS2_orf1ab;6290 | 0.15 | 0.025 |
| NC_045512.2;YP_009724389.1;Wu1-SARS2_orf1ab;6291 | 0.1 | 0.025 |
| NC_045512.2;YP_009724389.1;Wu1-SARS2_orf1ab;6292 | 0.2 | 0.025 |
| NC_045512.2;YP_009724389.1;Wu1-SARS2_orf1ab;6293 | 0.1 | 0.025 |
| NC_045512.2;YP_009724389.1;Wu1-SARS2_orf1ab;6294 | 0 | 0 |
| NC_045512.2;YP_009724389.1;Wu1-SARS2_orf1ab;6295 | 0 | 0 |
| NC_045512.2;YP_009724389.1;Wu1-SARS2_orf1ab;6296 | 0 | 0 |
| NC_045512.2;YP_009724389.1;Wu1-SARS2_orf1ab;6297 | 0 | 0 |
| NC_045512.2;YP_009724389.1;Wu1-SARS2_orf1ab;6298 | 0 | 0 |
| NC_045512.2;YP_009724389.1;Wu1-SARS2_orf1ab;6299 | 0 | 0 |
| NC_045512.2;YP_009724389.1;Wu1-SARS2_orf1ab;6300 | 0 | 0 |
| NC_045512.2;YP_009724389.1;Wu1-SARS2_orf1ab;6301 | 0 | 0 |
| NC_045512.2;YP_009724389.1;Wu1-SARS2_orf1ab;6302 | 0 | 0 |
| NC_045512.2;YP_009724389.1;Wu1-SARS2_orf1ab;6303 | 0 | 0 |
| NC_045512.2;YP_009724389.1;Wu1-SARS2_orf1ab;6304 | 0 | 0 |
| NC_045512.2;YP_009724389.1;Wu1-SARS2_orf1ab;6305 | 0 | 0 |
| NC_045512.2;YP_009724389.1;Wu1-SARS2_orf1ab;6306 | 0 | 0 |
| NC_045512.2;YP_009724389.1;Wu1-SARS2_orf1ab;6307 | 0 | 0 |
| NC_045512.2;YP_009724389.1;Wu1-SARS2_orf1ab;6308 | 0 | 0 |
| NC_045512.2;YP_009724389.1;Wu1-SARS2_orf1ab;6309 | 0 | 0 |
| NC_045512.2;YP_009724389.1;Wu1-SARS2_orf1ab;6310 | 0 | 0 |
| NC_045512.2;YP_009724389.1;Wu1-SARS2_orf1ab;6311 | 0 | 0 |

| | | |
|---|---|---|
| NC_045512.2;YP_009724389.1;Wu1-SARS2_orf1ab;6312 | 0 | 0 |
| NC_045512.2;YP_009724389.1;Wu1-SARS2_orf1ab;6313 | 0 | 0 |
| NC_045512.2;YP_009724389.1;Wu1-SARS2_orf1ab;6314 | 0 | 0 |
| NC_045512.2;YP_009724389.1;Wu1-SARS2_orf1ab;6315 | 0 | 0 |
| NC_045512.2;YP_009724389.1;Wu1-SARS2_orf1ab;6316 | 0 | 0 |
| NC_045512.2;YP_009724389.1;Wu1-SARS2_orf1ab;6317 | 0 | 0 |
| NC_045512.2;YP_009724389.1;Wu1-SARS2_orf1ab;6318 | 0 | 0 |
| NC_045512.2;YP_009724389.1;Wu1-SARS2_orf1ab;6319 | 0 | 0 |
| NC_045512.2;YP_009724389.1;Wu1-SARS2_orf1ab;6320 | 0 | 0 |
| NC_045512.2;YP_009724389.1;Wu1-SARS2_orf1ab;6321 | 0 | 0 |
| NC_045512.2;YP_009724389.1;Wu1-SARS2_orf1ab;6322 | 0 | 0 |
| NC_045512.2;YP_009724389.1;Wu1-SARS2_orf1ab;6323 | 0 | 0 |
| NC_045512.2;YP_009724389.1;Wu1-SARS2_orf1ab;6324 | 0 | 0 |
| NC_045512.2;YP_009724389.1;Wu1-SARS2_orf1ab;6325 | 0 | 0 |
| NC_045512.2;YP_009724389.1;Wu1-SARS2_orf1ab;6326 | 0 | 0 |
| NC_045512.2;YP_009724389.1;Wu1-SARS2_orf1ab;6327 | 0.1 | 0.025 |
| NC_045512.2;YP_009724389.1;Wu1-SARS2_orf1ab;6328 | 0.05 | 0.025 |
| NC_045512.2;YP_009724389.1;Wu1-SARS2_orf1ab;6329 | 0.05 | 0 |
| NC_045512.2;YP_009724389.1;Wu1-SARS2_orf1ab;6330 | 0.05 | 0 |
| NC_045512.2;YP_009724389.1;Wu1-SARS2_orf1ab;6331 | 0.05 | 0 |
| NC_045512.2;YP_009724389.1;Wu1-SARS2_orf1ab;6332 | 0 | 0.025 |
| NC_045512.2;YP_009724389.1;Wu1-SARS2_orf1ab;6333 | 0 | 0 |
| NC_045512.2;YP_009724389.1;Wu1-SARS2_orf1ab;6334 | 0 | 0 |
| NC_045512.2;YP_009724389.1;Wu1-SARS2_orf1ab;6335 | 0 | 0 |
| NC_045512.2;YP_009724389.1;Wu1-SARS2_orf1ab;6336 | 0 | 0 |
| NC_045512.2;YP_009724389.1;Wu1-SARS2_orf1ab;6337 | 0 | 0 |
| NC_045512.2;YP_009724389.1;Wu1-SARS2_orf1ab;6338 | 0 | 0 |
| NC_045512.2;YP_009724389.1;Wu1-SARS2_orf1ab;6339 | 0 | 0 |
| NC_045512.2;YP_009724389.1;Wu1-SARS2_orf1ab;6340 | 0 | 0 |
| NC_045512.2;YP_009724389.1;Wu1-SARS2_orf1ab;6341 | 0 | 0 |
| NC_045512.2;YP_009724389.1;Wu1-SARS2_orf1ab;6342 | 0 | 0 |
| NC_045512.2;YP_009724389.1;Wu1-SARS2_orf1ab;6343 | 0 | 0 |
| NC_045512.2;YP_009724389.1;Wu1-SARS2_orf1ab;6344 | 0 | 0 |
| NC_045512.2;YP_009724389.1;Wu1-SARS2_orf1ab;6345 | 0 | 0.025 |
| NC_045512.2;YP_009724389.1;Wu1-SARS2_orf1ab;6346 | 0 | 0.025 |
| NC_045512.2;YP_009724389.1;Wu1-SARS2_orf1ab;6347 | 0 | 0 |
| NC_045512.2;YP_009724389.1;Wu1-SARS2_orf1ab;6348 | 0 | 0.025 |
| NC_045512.2;YP_009724389.1;Wu1-SARS2_orf1ab;6349 | 0 | 0 |
| NC_045512.2;YP_009724389.1;Wu1-SARS2_orf1ab;6350 | 0 | 0.025 |
| NC_045512.2;YP_009724389.1;Wu1-SARS2_orf1ab;6351 | 0.05 | 0.025 |
| NC_045512.2;YP_009724389.1;Wu1-SARS2_orf1ab;6352 | 0.05 | 0.025 |
| NC_045512.2;YP_009724389.1;Wu1-SARS2_orf1ab;6353 | 0.05 | 0.025 |
| NC_045512.2;YP_009724389.1;Wu1-SARS2_orf1ab;6354 | 0.05 | 0.025 |

| | | |
|---|---|---|
| NC_045512.2;YP_009724389.1;Wu1-SARS2_orf1ab;6355 | 0.05 | 0.025 |
| NC_045512.2;YP_009724389.1;Wu1-SARS2_orf1ab;6356 | 0.05 | 0 |
| NC_045512.2;YP_009724389.1;Wu1-SARS2_orf1ab;6357 | 0.05 | 0.05 |
| NC_045512.2;YP_009724389.1;Wu1-SARS2_orf1ab;6358 | 0.05 | 0.025 |
| NC_045512.2;YP_009724389.1;Wu1-SARS2_orf1ab;6359 | 0 | 0 |
| NC_045512.2;YP_009724389.1;Wu1-SARS2_orf1ab;6360 | 0 | 0 |
| NC_045512.2;YP_009724389.1;Wu1-SARS2_orf1ab;6361 | 0 | 0.025 |
| NC_045512.2;YP_009724389.1;Wu1-SARS2_orf1ab;6362 | 0 | 0 |
| NC_045512.2;YP_009724389.1;Wu1-SARS2_orf1ab;6363 | 0 | 0 |
| NC_045512.2;YP_009724389.1;Wu1-SARS2_orf1ab;6364 | 0 | 0 |
| NC_045512.2;YP_009724389.1;Wu1-SARS2_orf1ab;6365 | 0 | 0 |
| NC_045512.2;YP_009724389.1;Wu1-SARS2_orf1ab;6366 | 0 | 0 |
| NC_045512.2;YP_009724389.1;Wu1-SARS2_orf1ab;6367 | 0 | 0 |
| NC_045512.2;YP_009724389.1;Wu1-SARS2_orf1ab;6368 | 0 | 0 |
| NC_045512.2;YP_009724389.1;Wu1-SARS2_orf1ab;6369 | 0 | 0 |
| NC_045512.2;YP_009724389.1;Wu1-SARS2_orf1ab;6370 | 0 | 0 |
| NC_045512.2;YP_009724389.1;Wu1-SARS2_orf1ab;6371 | 0 | 0.025 |
| NC_045512.2;YP_009724389.1;Wu1-SARS2_orf1ab;6372 | 0 | 0 |
| NC_045512.2;YP_009724389.1;Wu1-SARS2_orf1ab;6373 | 0 | 0 |
| NC_045512.2;YP_009724389.1;Wu1-SARS2_orf1ab;6374 | 0 | 0 |
| NC_045512.2;YP_009724389.1;Wu1-SARS2_orf1ab;6375 | 0 | 0 |
| NC_045512.2;YP_009724389.1;Wu1-SARS2_orf1ab;6376 | 0 | 0.05 |
| NC_045512.2;YP_009724389.1;Wu1-SARS2_orf1ab;6377 | 0 | 0.075 |
| NC_045512.2;YP_009724389.1;Wu1-SARS2_orf1ab;6378 | 0 | 0.05 |
| NC_045512.2;YP_009724389.1;Wu1-SARS2_orf1ab;6379 | 0 | 0.05 |
| NC_045512.2;YP_009724389.1;Wu1-SARS2_orf1ab;6380 | 0 | 0.075 |
| NC_045512.2;YP_009724389.1;Wu1-SARS2_orf1ab;6381 | 0 | 0.075 |
| NC_045512.2;YP_009724389.1;Wu1-SARS2_orf1ab;6382 | 0.05 | 0.05 |
| NC_045512.2;YP_009724389.1;Wu1-SARS2_orf1ab;6383 | 0 | 0.05 |
| NC_045512.2;YP_009724389.1;Wu1-SARS2_orf1ab;6384 | 0 | 0.025 |
| NC_045512.2;YP_009724389.1;Wu1-SARS2_orf1ab;6385 | 0 | 0.025 |
| NC_045512.2;YP_009724389.1;Wu1-SARS2_orf1ab;6386 | 0 | 0.025 |
| NC_045512.2;YP_009724389.1;Wu1-SARS2_orf1ab;6387 | 0 | 0 |
| NC_045512.2;YP_009724389.1;Wu1-SARS2_orf1ab;6388 | 0 | 0 |
| NC_045512.2;YP_009724389.1;Wu1-SARS2_orf1ab;6389 | 0 | 0 |
| NC_045512.2;YP_009724389.1;Wu1-SARS2_orf1ab;6390 | 0 | 0 |
| NC_045512.2;YP_009724389.1;Wu1-SARS2_orf1ab;6391 | 0 | 0 |
| NC_045512.2;YP_009724389.1;Wu1-SARS2_orf1ab;6392 | 0 | 0 |
| NC_045512.2;YP_009724389.1;Wu1-SARS2_orf1ab;6393 | 0 | 0 |
| NC_045512.2;YP_009724389.1;Wu1-SARS2_orf1ab;6394 | 0 | 0 |
| NC_045512.2;YP_009724389.1;Wu1-SARS2_orf1ab;6395 | 0 | 0 |
| NC_045512.2;YP_009724389.1;Wu1-SARS2_orf1ab;6396 | 0 | 0 |
| NC_045512.2;YP_009724389.1;Wu1-SARS2_orf1ab;6397 | 0 | 0 |

| | | |
|---|---|---|
| NC_045512.2;YP_009724389.1;Wu1-SARS2_orf1ab;6398 | 0 | 0 |
| NC_045512.2;YP_009724389.1;Wu1-SARS2_orf1ab;6399 | 0 | 0 |
| NC_045512.2;YP_009724389.1;Wu1-SARS2_orf1ab;6400 | 0 | 0 |
| NC_045512.2;YP_009724389.1;Wu1-SARS2_orf1ab;6401 | 0 | 0 |
| NC_045512.2;YP_009724389.1;Wu1-SARS2_orf1ab;6402 | 0 | 0 |
| NC_045512.2;YP_009724389.1;Wu1-SARS2_orf1ab;6403 | 0 | 0 |
| NC_045512.2;YP_009724389.1;Wu1-SARS2_orf1ab;6404 | 0 | 0 |
| NC_045512.2;YP_009724389.1;Wu1-SARS2_orf1ab;6405 | 0 | 0 |
| NC_045512.2;YP_009724389.1;Wu1-SARS2_orf1ab;6406 | 0 | 0 |
| NC_045512.2;YP_009724389.1;Wu1-SARS2_orf1ab;6407 | 0 | 0 |
| NC_045512.2;YP_009724389.1;Wu1-SARS2_orf1ab;6408 | 0 | 0 |
| NC_045512.2;YP_009724389.1;Wu1-SARS2_orf1ab;6409 | 0 | 0 |
| NC_045512.2;YP_009724389.1;Wu1-SARS2_orf1ab;6410 | 0 | 0.025 |
| NC_045512.2;YP_009724389.1;Wu1-SARS2_orf1ab;6411 | 0 | 0.05 |
| NC_045512.2;YP_009724389.1;Wu1-SARS2_orf1ab;6412 | 0 | 0.025 |
| NC_045512.2;YP_009724389.1;Wu1-SARS2_orf1ab;6413 | 0 | 0.025 |
| NC_045512.2;YP_009724389.1;Wu1-SARS2_orf1ab;6414 | 0 | 0.025 |
| NC_045512.2;YP_009724389.1;Wu1-SARS2_orf1ab;6415 | 0 | 0.025 |
| NC_045512.2;YP_009724389.1;Wu1-SARS2_orf1ab;6416 | 0 | 0.025 |
| NC_045512.2;YP_009724389.1;Wu1-SARS2_orf1ab;6417 | 0 | 0.025 |
| NC_045512.2;YP_009724389.1;Wu1-SARS2_orf1ab;6418 | 0 | 0.025 |
| NC_045512.2;YP_009724389.1;Wu1-SARS2_orf1ab;6419 | 0 | 0 |
| NC_045512.2;YP_009724389.1;Wu1-SARS2_orf1ab;6420 | 0 | 0 |
| NC_045512.2;YP_009724389.1;Wu1-SARS2_orf1ab;6421 | 0 | 0 |
| NC_045512.2;YP_009724389.1;Wu1-SARS2_orf1ab;6422 | 0 | 0 |
| NC_045512.2;YP_009724389.1;Wu1-SARS2_orf1ab;6423 | 0 | 0 |
| NC_045512.2;YP_009724389.1;Wu1-SARS2_orf1ab;6424 | 0 | 0 |
| NC_045512.2;YP_009724389.1;Wu1-SARS2_orf1ab;6425 | 0 | 0 |
| NC_045512.2;YP_009724389.1;Wu1-SARS2_orf1ab;6426 | 0 | 0 |
| NC_045512.2;YP_009724389.1;Wu1-SARS2_orf1ab;6427 | 0 | 0 |
| NC_045512.2;YP_009724389.1;Wu1-SARS2_orf1ab;6428 | 0 | 0 |
| NC_045512.2;YP_009724389.1;Wu1-SARS2_orf1ab;6429 | 0 | 0 |
| NC_045512.2;YP_009724389.1;Wu1-SARS2_orf1ab;6430 | 0 | 0 |
| NC_045512.2;YP_009724389.1;Wu1-SARS2_orf1ab;6431 | 0 | 0 |
| NC_045512.2;YP_009724389.1;Wu1-SARS2_orf1ab;6432 | 0 | 0 |
| NC_045512.2;YP_009724389.1;Wu1-SARS2_orf1ab;6433 | 0 | 0 |
| NC_045512.2;YP_009724389.1;Wu1-SARS2_orf1ab;6434 | 0 | 0 |
| NC_045512.2;YP_009724389.1;Wu1-SARS2_orf1ab;6435 | 0 | 0 |
| NC_045512.2;YP_009724389.1;Wu1-SARS2_orf1ab;6436 | 0 | 0 |
| NC_045512.2;YP_009724389.1;Wu1-SARS2_orf1ab;6437 | 0 | 0 |
| NC_045512.2;YP_009724389.1;Wu1-SARS2_orf1ab;6438 | 0 | 0 |
| NC_045512.2;YP_009724389.1;Wu1-SARS2_orf1ab;6439 | 0 | 0 |
| NC_045512.2;YP_009724389.1;Wu1-SARS2_orf1ab;6440 | 0 | 0 |

| | | |
|---|---|---|
| NC_045512.2;YP_009724389.1;Wu1-SARS2_orf1ab;6441 | 0 | 0 |
| NC_045512.2;YP_009724389.1;Wu1-SARS2_orf1ab;6442 | 0 | 0 |
| NC_045512.2;YP_009724389.1;Wu1-SARS2_orf1ab;6443 | 0 | 0 |
| NC_045512.2;YP_009724389.1;Wu1-SARS2_orf1ab;6444 | 0 | 0 |
| NC_045512.2;YP_009724389.1;Wu1-SARS2_orf1ab;6445 | 0.05 | 0 |
| NC_045512.2;YP_009724389.1;Wu1-SARS2_orf1ab;6446 | 0 | 0 |
| NC_045512.2;YP_009724389.1;Wu1-SARS2_orf1ab;6447 | 0 | 0 |
| NC_045512.2;YP_009724389.1;Wu1-SARS2_orf1ab;6448 | 0 | 0 |
| NC_045512.2;YP_009724389.1;Wu1-SARS2_orf1ab;6449 | 0 | 0 |
| NC_045512.2;YP_009724389.1;Wu1-SARS2_orf1ab;6450 | 0 | 0 |
| NC_045512.2;YP_009724389.1;Wu1-SARS2_orf1ab;6451 | 0 | 0 |
| NC_045512.2;YP_009724389.1;Wu1-SARS2_orf1ab;6452 | 0 | 0 |
| NC_045512.2;YP_009724389.1;Wu1-SARS2_orf1ab;6453 | 0 | 0 |
| NC_045512.2;YP_009724389.1;Wu1-SARS2_orf1ab;6454 | 0.05 | 0.025 |
| NC_045512.2;YP_009724389.1;Wu1-SARS2_orf1ab;6455 | 0.05 | 0.025 |
| NC_045512.2;YP_009724389.1;Wu1-SARS2_orf1ab;6456 | 0 | 0 |
| NC_045512.2;YP_009724389.1;Wu1-SARS2_orf1ab;6457 | 0.05 | 0 |
| NC_045512.2;YP_009724389.1;Wu1-SARS2_orf1ab;6458 | 0.1 | 0.025 |
| NC_045512.2;YP_009724389.1;Wu1-SARS2_orf1ab;6459 | 0.05 | 0.025 |
| NC_045512.2;YP_009724389.1;Wu1-SARS2_orf1ab;6460 | 0 | 0.025 |
| NC_045512.2;YP_009724389.1;Wu1-SARS2_orf1ab;6461 | 0.05 | 0.05 |
| NC_045512.2;YP_009724389.1;Wu1-SARS2_orf1ab;6462 | 0 | 0.025 |
| NC_045512.2;YP_009724389.1;Wu1-SARS2_orf1ab;6463 | 0 | 0.025 |
| NC_045512.2;YP_009724389.1;Wu1-SARS2_orf1ab;6464 | 0 | 0 |
| NC_045512.2;YP_009724389.1;Wu1-SARS2_orf1ab;6465 | 0 | 0 |
| NC_045512.2;YP_009724389.1;Wu1-SARS2_orf1ab;6466 | 0 | 0 |
| NC_045512.2;YP_009724389.1;Wu1-SARS2_orf1ab;6467 | 0 | 0 |
| NC_045512.2;YP_009724389.1;Wu1-SARS2_orf1ab;6468 | 0 | 0 |
| NC_045512.2;YP_009724389.1;Wu1-SARS2_orf1ab;6469 | 0 | 0 |
| NC_045512.2;YP_009724389.1;Wu1-SARS2_orf1ab;6470 | 0 | 0 |
| NC_045512.2;YP_009724389.1;Wu1-SARS2_orf1ab;6471 | 0 | 0.025 |
| NC_045512.2;YP_009724389.1;Wu1-SARS2_orf1ab;6472 | 0 | 0.025 |
| NC_045512.2;YP_009724389.1;Wu1-SARS2_orf1ab;6473 | 0 | 0.025 |
| NC_045512.2;YP_009724389.1;Wu1-SARS2_orf1ab;6474 | 0 | 0 |
| NC_045512.2;YP_009724389.1;Wu1-SARS2_orf1ab;6475 | 0.05 | 0 |
| NC_045512.2;YP_009724389.1;Wu1-SARS2_orf1ab;6476 | 0 | 0.05 |
| NC_045512.2;YP_009724389.1;Wu1-SARS2_orf1ab;6477 | 0 | 0.025 |
| NC_045512.2;YP_009724389.1;Wu1-SARS2_orf1ab;6478 | 0 | 0.025 |
| NC_045512.2;YP_009724389.1;Wu1-SARS2_orf1ab;6479 | 0 | 0.025 |
| NC_045512.2;YP_009724389.1;Wu1-SARS2_orf1ab;6480 | 0 | 0 |
| NC_045512.2;YP_009724389.1;Wu1-SARS2_orf1ab;6481 | 0 | 0 |
| NC_045512.2;YP_009724389.1;Wu1-SARS2_orf1ab;6482 | 0 | 0 |
| NC_045512.2;YP_009724389.1;Wu1-SARS2_orf1ab;6483 | 0 | 0 |

| | | |
|---|---|---|
| NC_045512.2;YP_009724389.1;Wu1-SARS2_orf1ab;6484 | 0 | 0.025 |
| NC_045512.2;YP_009724389.1;Wu1-SARS2_orf1ab;6485 | 0 | 0.05 |
| NC_045512.2;YP_009724389.1;Wu1-SARS2_orf1ab;6486 | 0 | 0.05 |
| NC_045512.2;YP_009724389.1;Wu1-SARS2_orf1ab;6487 | 0 | 0.025 |
| NC_045512.2;YP_009724389.1;Wu1-SARS2_orf1ab;6488 | 0 | 0.025 |
| NC_045512.2;YP_009724389.1;Wu1-SARS2_orf1ab;6489 | 0 | 0.05 |
| NC_045512.2;YP_009724389.1;Wu1-SARS2_orf1ab;6490 | 0 | 0.025 |
| NC_045512.2;YP_009724389.1;Wu1-SARS2_orf1ab;6491 | 0 | 0.075 |
| NC_045512.2;YP_009724389.1;Wu1-SARS2_orf1ab;6492 | 0 | 0.075 |
| NC_045512.2;YP_009724389.1;Wu1-SARS2_orf1ab;6493 | 0 | 0.05 |
| NC_045512.2;YP_009724389.1;Wu1-SARS2_orf1ab;6494 | 0 | 0.025 |
| NC_045512.2;YP_009724389.1;Wu1-SARS2_orf1ab;6495 | 0 | 0.025 |
| NC_045512.2;YP_009724389.1;Wu1-SARS2_orf1ab;6496 | 0.05 | 0.025 |
| NC_045512.2;YP_009724389.1;Wu1-SARS2_orf1ab;6497 | 0 | 0 |
| NC_045512.2;YP_009724389.1;Wu1-SARS2_orf1ab;6498 | 0 | 0 |
| NC_045512.2;YP_009724389.1;Wu1-SARS2_orf1ab;6499 | 0 | 0 |
| NC_045512.2;YP_009724389.1;Wu1-SARS2_orf1ab;6500 | 0 | 0 |
| NC_045512.2;YP_009724389.1;Wu1-SARS2_orf1ab;6501 | 0 | 0 |
| NC_045512.2;YP_009724389.1;Wu1-SARS2_orf1ab;6502 | 0 | 0 |
| NC_045512.2;YP_009724389.1;Wu1-SARS2_orf1ab;6503 | 0 | 0 |
| NC_045512.2;YP_009724389.1;Wu1-SARS2_orf1ab;6504 | 0 | 0 |
| NC_045512.2;YP_009724389.1;Wu1-SARS2_orf1ab;6505 | 0.05 | 0 |
| NC_045512.2;YP_009724389.1;Wu1-SARS2_orf1ab;6506 | 0 | 0 |
| NC_045512.2;YP_009724389.1;Wu1-SARS2_orf1ab;6507 | 0.05 | 0.1 |
| NC_045512.2;YP_009724389.1;Wu1-SARS2_orf1ab;6508 | 0.2 | 0.125 |
| NC_045512.2;YP_009724389.1;Wu1-SARS2_orf1ab;6509 | 0.05 | 0.1 |
| NC_045512.2;YP_009724389.1;Wu1-SARS2_orf1ab;6510 | 0.05 | 0.075 |
| NC_045512.2;YP_009724389.1;Wu1-SARS2_orf1ab;6511 | 0.1 | 0.075 |
| NC_045512.2;YP_009724389.1;Wu1-SARS2_orf1ab;6512 | 0.15 | 0.15 |
| NC_045512.2;YP_009724389.1;Wu1-SARS2_orf1ab;6513 | 0.15 | 0.125 |
| NC_045512.2;YP_009724389.1;Wu1-SARS2_orf1ab;6514 | 0.4 | 0.2 |
| NC_045512.2;YP_009724389.1;Wu1-SARS2_orf1ab;6515 | 0.4 | 0.2 |
| NC_045512.2;YP_009724389.1;Wu1-SARS2_orf1ab;6516 | 0.2 | 0.15 |
| NC_045512.2;YP_009724389.1;Wu1-SARS2_orf1ab;6517 | 0 | 0 |
| NC_045512.2;YP_009724389.1;Wu1-SARS2_orf1ab;6518 | 0.05 | 0 |
| NC_045512.2;YP_009724389.1;Wu1-SARS2_orf1ab;6519 | 0 | 0 |
| NC_045512.2;YP_009724389.1;Wu1-SARS2_orf1ab;6520 | 0 | 0 |
| NC_045512.2;YP_009724389.1;Wu1-SARS2_orf1ab;6521 | 0 | 0.025 |
| NC_045512.2;YP_009724389.1;Wu1-SARS2_orf1ab;6522 | 0 | 0 |
| NC_045512.2;YP_009724389.1;Wu1-SARS2_orf1ab;6523 | 0 | 0 |
| NC_045512.2;YP_009724389.1;Wu1-SARS2_orf1ab;6524 | 0 | 0 |
| NC_045512.2;YP_009724389.1;Wu1-SARS2_orf1ab;6525 | 0 | 0 |
| NC_045512.2;YP_009724389.1;Wu1-SARS2_orf1ab;6526 | 0 | 0 |

| | | |
|---|---|---|
| NC_045512.2;YP_009724389.1;Wu1-SARS2_orf1ab;6527 | 0 | 0 |
| NC_045512.2;YP_009724389.1;Wu1-SARS2_orf1ab;6528 | 0.05 | 0 |
| NC_045512.2;YP_009724389.1;Wu1-SARS2_orf1ab;6529 | 0 | 0.025 |
| NC_045512.2;YP_009724389.1;Wu1-SARS2_orf1ab;6530 | 0 | 0 |
| NC_045512.2;YP_009724389.1;Wu1-SARS2_orf1ab;6531 | 0 | 0.025 |
| NC_045512.2;YP_009724389.1;Wu1-SARS2_orf1ab;6532 | 0 | 0 |
| NC_045512.2;YP_009724389.1;Wu1-SARS2_orf1ab;6533 | 0 | 0 |
| NC_045512.2;YP_009724389.1;Wu1-SARS2_orf1ab;6534 | 0 | 0 |
| NC_045512.2;YP_009724389.1;Wu1-SARS2_orf1ab;6535 | 0 | 0 |
| NC_045512.2;YP_009724389.1;Wu1-SARS2_orf1ab;6536 | 0 | 0 |
| NC_045512.2;YP_009724389.1;Wu1-SARS2_orf1ab;6537 | 0.05 | 0 |
| NC_045512.2;YP_009724389.1;Wu1-SARS2_orf1ab;6538 | 0 | 0 |
| NC_045512.2;YP_009724389.1;Wu1-SARS2_orf1ab;6539 | 0 | 0 |
| NC_045512.2;YP_009724389.1;Wu1-SARS2_orf1ab;6540 | 0 | 0 |
| NC_045512.2;YP_009724389.1;Wu1-SARS2_orf1ab;6541 | 0.15 | 0.075 |
| NC_045512.2;YP_009724389.1;Wu1-SARS2_orf1ab;6542 | 0 | 0 |
| NC_045512.2;YP_009724389.1;Wu1-SARS2_orf1ab;6543 | 0 | 0 |
| NC_045512.2;YP_009724389.1;Wu1-SARS2_orf1ab;6544 | 0 | 0 |
| NC_045512.2;YP_009724389.1;Wu1-SARS2_orf1ab;6545 | 0 | 0 |
| NC_045512.2;YP_009724389.1;Wu1-SARS2_orf1ab;6546 | 0.05 | 0 |
| NC_045512.2;YP_009724389.1;Wu1-SARS2_orf1ab;6547 | 0.05 | 0 |
| NC_045512.2;YP_009724389.1;Wu1-SARS2_orf1ab;6548 | 0.1 | 0 |
| NC_045512.2;YP_009724389.1;Wu1-SARS2_orf1ab;6549 | 0.15 | 0.05 |
| NC_045512.2;YP_009724389.1;Wu1-SARS2_orf1ab;6550 | 0.15 | 0 |
| NC_045512.2;YP_009724389.1;Wu1-SARS2_orf1ab;6551 | 0.1 | 0 |
| NC_045512.2;YP_009724389.1;Wu1-SARS2_orf1ab;6552 | 0.15 | 0.025 |
| NC_045512.2;YP_009724389.1;Wu1-SARS2_orf1ab;6553 | 0.1 | 0.025 |
| NC_045512.2;YP_009724389.1;Wu1-SARS2_orf1ab;6554 | 0.05 | 0 |
| NC_045512.2;YP_009724389.1;Wu1-SARS2_orf1ab;6555 | 0.05 | 0.025 |
| NC_045512.2;YP_009724389.1;Wu1-SARS2_orf1ab;6556 | 0.05 | 0.05 |
| NC_045512.2;YP_009724389.1;Wu1-SARS2_orf1ab;6557 | 0.05 | 0.025 |
| NC_045512.2;YP_009724389.1;Wu1-SARS2_orf1ab;6558 | 0 | 0 |
| NC_045512.2;YP_009724389.1;Wu1-SARS2_orf1ab;6559 | 0 | 0 |
| NC_045512.2;YP_009724389.1;Wu1-SARS2_orf1ab;6560 | 0 | 0 |
| NC_045512.2;YP_009724389.1;Wu1-SARS2_orf1ab;6561 | 0 | 0 |
| NC_045512.2;YP_009724389.1;Wu1-SARS2_orf1ab;6562 | 0 | 0 |
| NC_045512.2;YP_009724389.1;Wu1-SARS2_orf1ab;6563 | 0 | 0 |
| NC_045512.2;YP_009724389.1;Wu1-SARS2_orf1ab;6564 | 0 | 0 |
| NC_045512.2;YP_009724389.1;Wu1-SARS2_orf1ab;6565 | 0 | 0 |
| NC_045512.2;YP_009724389.1;Wu1-SARS2_orf1ab;6566 | 0 | 0 |
| NC_045512.2;YP_009724389.1;Wu1-SARS2_orf1ab;6567 | 0 | 0 |
| NC_045512.2;YP_009724389.1;Wu1-SARS2_orf1ab;6568 | 0 | 0 |
| NC_045512.2;YP_009724389.1;Wu1-SARS2_orf1ab;6569 | 0 | 0 |

| | | |
|---|---|---|
| NC_045512.2;YP_009724389.1;Wu1-SARS2_orf1ab;6570 | 0 | 0 |
| NC_045512.2;YP_009724389.1;Wu1-SARS2_orf1ab;6571 | 0 | 0 |
| NC_045512.2;YP_009724389.1;Wu1-SARS2_orf1ab;6572 | 0 | 0 |
| NC_045512.2;YP_009724389.1;Wu1-SARS2_orf1ab;6573 | 0.05 | 0 |
| NC_045512.2;YP_009724389.1;Wu1-SARS2_orf1ab;6574 | 0 | 0 |
| NC_045512.2;YP_009724389.1;Wu1-SARS2_orf1ab;6575 | 0 | 0 |
| NC_045512.2;YP_009724389.1;Wu1-SARS2_orf1ab;6576 | 0 | 0 |
| NC_045512.2;YP_009724389.1;Wu1-SARS2_orf1ab;6577 | 0 | 0 |
| NC_045512.2;YP_009724389.1;Wu1-SARS2_orf1ab;6578 | 0 | 0 |
| NC_045512.2;YP_009724389.1;Wu1-SARS2_orf1ab;6579 | 0.05 | 0 |
| NC_045512.2;YP_009724389.1;Wu1-SARS2_orf1ab;6580 | 0 | 0 |
| NC_045512.2;YP_009724389.1;Wu1-SARS2_orf1ab;6581 | 0 | 0.025 |
| NC_045512.2;YP_009724389.1;Wu1-SARS2_orf1ab;6582 | 0 | 0 |
| NC_045512.2;YP_009724389.1;Wu1-SARS2_orf1ab;6583 | 0 | 0 |
| NC_045512.2;YP_009724389.1;Wu1-SARS2_orf1ab;6584 | 0 | 0 |
| NC_045512.2;YP_009724389.1;Wu1-SARS2_orf1ab;6585 | 0 | 0 |
| NC_045512.2;YP_009724389.1;Wu1-SARS2_orf1ab;6586 | 0 | 0 |
| NC_045512.2;YP_009724389.1;Wu1-SARS2_orf1ab;6587 | 0 | 0 |
| NC_045512.2;YP_009724389.1;Wu1-SARS2_orf1ab;6588 | 0 | 0 |
| NC_045512.2;YP_009724389.1;Wu1-SARS2_orf1ab;6589 | 0 | 0 |
| NC_045512.2;YP_009724389.1;Wu1-SARS2_orf1ab;6590 | 0 | 0 |
| NC_045512.2;YP_009724389.1;Wu1-SARS2_orf1ab;6591 | 0 | 0 |
| NC_045512.2;YP_009724389.1;Wu1-SARS2_orf1ab;6592 | 0.05 | 0.025 |
| NC_045512.2;YP_009724389.1;Wu1-SARS2_orf1ab;6593 | 0.05 | 0 |
| NC_045512.2;YP_009724389.1;Wu1-SARS2_orf1ab;6594 | 0.05 | 0 |
| NC_045512.2;YP_009724389.1;Wu1-SARS2_orf1ab;6595 | 0 | 0 |
| NC_045512.2;YP_009724389.1;Wu1-SARS2_orf1ab;6596 | 0.05 | 0 |
| NC_045512.2;YP_009724389.1;Wu1-SARS2_orf1ab;6597 | 0 | 0.025 |
| NC_045512.2;YP_009724389.1;Wu1-SARS2_orf1ab;6598 | 0.05 | 0.15 |
| NC_045512.2;YP_009724389.1;Wu1-SARS2_orf1ab;6599 | 0 | 0.025 |
| NC_045512.2;YP_009724389.1;Wu1-SARS2_orf1ab;6600 | 0.05 | 0 |
| NC_045512.2;YP_009724389.1;Wu1-SARS2_orf1ab;6601 | 0 | 0 |
| NC_045512.2;YP_009724389.1;Wu1-SARS2_orf1ab;6602 | 0 | 0.05 |
| NC_045512.2;YP_009724389.1;Wu1-SARS2_orf1ab;6603 | 0 | 0 |
| NC_045512.2;YP_009724389.1;Wu1-SARS2_orf1ab;6604 | 0 | 0 |
| NC_045512.2;YP_009724389.1;Wu1-SARS2_orf1ab;6605 | 0 | 0 |
| NC_045512.2;YP_009724389.1;Wu1-SARS2_orf1ab;6606 | 0 | 0 |
| NC_045512.2;YP_009724389.1;Wu1-SARS2_orf1ab;6607 | 0 | 0 |
| NC_045512.2;YP_009724389.1;Wu1-SARS2_orf1ab;6608 | 0 | 0 |
| NC_045512.2;YP_009724389.1;Wu1-SARS2_orf1ab;6609 | 0 | 0 |
| NC_045512.2;YP_009724389.1;Wu1-SARS2_orf1ab;6610 | 0 | 0 |
| NC_045512.2;YP_009724389.1;Wu1-SARS2_orf1ab;6611 | 0 | 0.025 |
| NC_045512.2;YP_009724389.1;Wu1-SARS2_orf1ab;6612 | 0 | 0.025 |

| | | |
|---|---|---|
| NC_045512.2;YP_009724389.1;Wu1-SARS2_orf1ab;6613 | 0 | 0.025 |
| NC_045512.2;YP_009724389.1;Wu1-SARS2_orf1ab;6614 | 0 | 0.025 |
| NC_045512.2;YP_009724389.1;Wu1-SARS2_orf1ab;6615 | 0 | 0.025 |
| NC_045512.2;YP_009724389.1;Wu1-SARS2_orf1ab;6616 | 0 | 0 |
| NC_045512.2;YP_009724389.1;Wu1-SARS2_orf1ab;6617 | 0 | 0 |
| NC_045512.2;YP_009724389.1;Wu1-SARS2_orf1ab;6618 | 0 | 0 |
| NC_045512.2;YP_009724389.1;Wu1-SARS2_orf1ab;6619 | 0.05 | 0 |
| NC_045512.2;YP_009724389.1;Wu1-SARS2_orf1ab;6620 | 0 | 0.05 |
| NC_045512.2;YP_009724389.1;Wu1-SARS2_orf1ab;6621 | 0.05 | 0.05 |
| NC_045512.2;YP_009724389.1;Wu1-SARS2_orf1ab;6622 | 0 | 0.075 |
| NC_045512.2;YP_009724389.1;Wu1-SARS2_orf1ab;6623 | 0 | 0.025 |
| NC_045512.2;YP_009724389.1;Wu1-SARS2_orf1ab;6624 | 0 | 0.025 |
| NC_045512.2;YP_009724389.1;Wu1-SARS2_orf1ab;6625 | 0.05 | 0.05 |
| NC_045512.2;YP_009724389.1;Wu1-SARS2_orf1ab;6626 | 0.05 | 0.05 |
| NC_045512.2;YP_009724389.1;Wu1-SARS2_orf1ab;6627 | 0.1 | 0.075 |
| NC_045512.2;YP_009724389.1;Wu1-SARS2_orf1ab;6628 | 0 | 0.075 |
| NC_045512.2;YP_009724389.1;Wu1-SARS2_orf1ab;6629 | 0.1 | 0.075 |
| NC_045512.2;YP_009724389.1;Wu1-SARS2_orf1ab;6630 | 0.05 | 0.1 |
| NC_045512.2;YP_009724389.1;Wu1-SARS2_orf1ab;6631 | 0.1 | 0.075 |
| NC_045512.2;YP_009724389.1;Wu1-SARS2_orf1ab;6632 | 0.05 | 0.025 |
| NC_045512.2;YP_009724389.1;Wu1-SARS2_orf1ab;6633 | 0.05 | 0.025 |
| NC_045512.2;YP_009724389.1;Wu1-SARS2_orf1ab;6634 | 0 | 0 |
| NC_045512.2;YP_009724389.1;Wu1-SARS2_orf1ab;6635 | 0 | 0 |
| NC_045512.2;YP_009724389.1;Wu1-SARS2_orf1ab;6636 | 0 | 0.025 |
| NC_045512.2;YP_009724389.1;Wu1-SARS2_orf1ab;6637 | 0 | 0 |
| NC_045512.2;YP_009724389.1;Wu1-SARS2_orf1ab;6638 | 0 | 0 |
| NC_045512.2;YP_009724389.1;Wu1-SARS2_orf1ab;6639 | 0 | 0.025 |
| NC_045512.2;YP_009724389.1;Wu1-SARS2_orf1ab;6640 | 0 | 0.05 |
| NC_045512.2;YP_009724389.1;Wu1-SARS2_orf1ab;6641 | 0 | 0.025 |
| NC_045512.2;YP_009724389.1;Wu1-SARS2_orf1ab;6642 | 0.05 | 0 |
| NC_045512.2;YP_009724389.1;Wu1-SARS2_orf1ab;6643 | 0 | 0 |
| NC_045512.2;YP_009724389.1;Wu1-SARS2_orf1ab;6644 | 0 | 0 |
| NC_045512.2;YP_009724389.1;Wu1-SARS2_orf1ab;6645 | 0 | 0 |
| NC_045512.2;YP_009724389.1;Wu1-SARS2_orf1ab;6646 | 0 | 0 |
| NC_045512.2;YP_009724389.1;Wu1-SARS2_orf1ab;6647 | 0 | 0 |
| NC_045512.2;YP_009724389.1;Wu1-SARS2_orf1ab;6648 | 0 | 0 |
| NC_045512.2;YP_009724389.1;Wu1-SARS2_orf1ab;6649 | 0 | 0 |
| NC_045512.2;YP_009724389.1;Wu1-SARS2_orf1ab;6650 | 0 | 0 |
| NC_045512.2;YP_009724389.1;Wu1-SARS2_orf1ab;6651 | 0 | 0 |
| NC_045512.2;YP_009724389.1;Wu1-SARS2_orf1ab;6652 | 0 | 0 |
| NC_045512.2;YP_009724389.1;Wu1-SARS2_orf1ab;6653 | 0 | 0 |
| NC_045512.2;YP_009724389.1;Wu1-SARS2_orf1ab;6654 | 0 | 0 |
| NC_045512.2;YP_009724389.1;Wu1-SARS2_orf1ab;6655 | 0 | 0 |

| | | |
|---|---|---|
| NC_045512.2;YP_009724389.1;Wu1-SARS2_orf1ab;6656 | 0 | 0 |
| NC_045512.2;YP_009724389.1;Wu1-SARS2_orf1ab;6657 | 0 | 0 |
| NC_045512.2;YP_009724389.1;Wu1-SARS2_orf1ab;6658 | 0 | 0.025 |
| NC_045512.2;YP_009724389.1;Wu1-SARS2_orf1ab;6659 | 0 | 0 |
| NC_045512.2;YP_009724389.1;Wu1-SARS2_orf1ab;6660 | 0 | 0 |
| NC_045512.2;YP_009724389.1;Wu1-SARS2_orf1ab;6661 | 0 | 0 |
| NC_045512.2;YP_009724389.1;Wu1-SARS2_orf1ab;6662 | 0 | 0 |
| NC_045512.2;YP_009724389.1;Wu1-SARS2_orf1ab;6663 | 0.05 | 0 |
| NC_045512.2;YP_009724389.1;Wu1-SARS2_orf1ab;6664 | 0.05 | 0 |
| NC_045512.2;YP_009724389.1;Wu1-SARS2_orf1ab;6665 | 0.05 | 0.025 |
| NC_045512.2;YP_009724389.1;Wu1-SARS2_orf1ab;6666 | 0 | 0.025 |
| NC_045512.2;YP_009724389.1;Wu1-SARS2_orf1ab;6667 | 0 | 0.025 |
| NC_045512.2;YP_009724389.1;Wu1-SARS2_orf1ab;6668 | 0.05 | 0.075 |
| NC_045512.2;YP_009724389.1;Wu1-SARS2_orf1ab;6669 | 0 | 0 |
| NC_045512.2;YP_009724389.1;Wu1-SARS2_orf1ab;6670 | 0 | 0 |
| NC_045512.2;YP_009724389.1;Wu1-SARS2_orf1ab;6671 | 0 | 0 |
| NC_045512.2;YP_009724389.1;Wu1-SARS2_orf1ab;6672 | 0 | 0 |
| NC_045512.2;YP_009724389.1;Wu1-SARS2_orf1ab;6673 | 0 | 0 |
| NC_045512.2;YP_009724389.1;Wu1-SARS2_orf1ab;6674 | 0 | 0 |
| NC_045512.2;YP_009724389.1;Wu1-SARS2_orf1ab;6675 | 0 | 0 |
| NC_045512.2;YP_009724389.1;Wu1-SARS2_orf1ab;6676 | 0 | 0 |
| NC_045512.2;YP_009724389.1;Wu1-SARS2_orf1ab;6677 | 0 | 0 |
| NC_045512.2;YP_009724389.1;Wu1-SARS2_orf1ab;6678 | 0 | 0 |
| NC_045512.2;YP_009724389.1;Wu1-SARS2_orf1ab;6679 | 0 | 0 |
| NC_045512.2;YP_009724389.1;Wu1-SARS2_orf1ab;6680 | 0 | 0 |
| NC_045512.2;YP_009724389.1;Wu1-SARS2_orf1ab;6681 | 0 | 0 |
| NC_045512.2;YP_009724389.1;Wu1-SARS2_orf1ab;6682 | 0 | 0 |
| NC_045512.2;YP_009724389.1;Wu1-SARS2_orf1ab;6683 | 0 | 0 |
| NC_045512.2;YP_009724389.1;Wu1-SARS2_orf1ab;6684 | 0 | 0 |
| NC_045512.2;YP_009724389.1;Wu1-SARS2_orf1ab;6685 | 0 | 0 |
| NC_045512.2;YP_009724389.1;Wu1-SARS2_orf1ab;6686 | 0 | 0 |
| NC_045512.2;YP_009724389.1;Wu1-SARS2_orf1ab;6687 | 0 | 0 |
| NC_045512.2;YP_009724389.1;Wu1-SARS2_orf1ab;6688 | 0 | 0 |
| NC_045512.2;YP_009724389.1;Wu1-SARS2_orf1ab;6689 | 0 | 0 |
| NC_045512.2;YP_009724389.1;Wu1-SARS2_orf1ab;6690 | 0 | 0 |
| NC_045512.2;YP_009724389.1;Wu1-SARS2_orf1ab;6691 | 0 | 0 |
| NC_045512.2;YP_009724389.1;Wu1-SARS2_orf1ab;6692 | 0 | 0 |
| NC_045512.2;YP_009724389.1;Wu1-SARS2_orf1ab;6693 | 0 | 0 |
| NC_045512.2;YP_009724389.1;Wu1-SARS2_orf1ab;6694 | 0 | 0 |
| NC_045512.2;YP_009724389.1;Wu1-SARS2_orf1ab;6695 | 0 | 0 |
| NC_045512.2;YP_009724389.1;Wu1-SARS2_orf1ab;6696 | 0 | 0 |
| NC_045512.2;YP_009724389.1;Wu1-SARS2_orf1ab;6697 | 0 | 0 |
| NC_045512.2;YP_009724389.1;Wu1-SARS2_orf1ab;6698 | 0 | 0 |

| | | |
|---|---|---|
| NC_045512.2;YP_009724389.1;Wu1-SARS2_orf1ab;6699 | 0 | 0 |
| NC_045512.2;YP_009724389.1;Wu1-SARS2_orf1ab;6700 | 0 | 0 |
| NC_045512.2;YP_009724389.1;Wu1-SARS2_orf1ab;6701 | 0.05 | 0 |
| NC_045512.2;YP_009724389.1;Wu1-SARS2_orf1ab;6702 | 0.05 | 0.025 |
| NC_045512.2;YP_009724389.1;Wu1-SARS2_orf1ab;6703 | 0.1 | 0.075 |
| NC_045512.2;YP_009724389.1;Wu1-SARS2_orf1ab;6704 | 0.05 | 0.075 |
| NC_045512.2;YP_009724389.1;Wu1-SARS2_orf1ab;6705 | 0.05 | 0.05 |
| NC_045512.2;YP_009724389.1;Wu1-SARS2_orf1ab;6706 | 0.05 | 0.075 |
| NC_045512.2;YP_009724389.1;Wu1-SARS2_orf1ab;6707 | 0.15 | 0.075 |
| NC_045512.2;YP_009724389.1;Wu1-SARS2_orf1ab;6708 | 0.15 | 0.075 |
| NC_045512.2;YP_009724389.1;Wu1-SARS2_orf1ab;6709 | 0 | 0 |
| NC_045512.2;YP_009724389.1;Wu1-SARS2_orf1ab;6710 | 0 | 0 |
| NC_045512.2;YP_009724389.1;Wu1-SARS2_orf1ab;6711 | 0.15 | 0 |
| NC_045512.2;YP_009724389.1;Wu1-SARS2_orf1ab;6712 | 0.05 | 0 |
| NC_045512.2;YP_009724389.1;Wu1-SARS2_orf1ab;6713 | 0 | 0 |
| NC_045512.2;YP_009724389.1;Wu1-SARS2_orf1ab;6714 | 0 | 0 |
| NC_045512.2;YP_009724389.1;Wu1-SARS2_orf1ab;6715 | 0.05 | 0 |
| NC_045512.2;YP_009724389.1;Wu1-SARS2_orf1ab;6716 | 0.05 | 0 |
| NC_045512.2;YP_009724389.1;Wu1-SARS2_orf1ab;6717 | 0.15 | 0 |
| NC_045512.2;YP_009724389.1;Wu1-SARS2_orf1ab;6718 | 0 | 0 |
| NC_045512.2;YP_009724389.1;Wu1-SARS2_orf1ab;6719 | 0.05 | 0 |
| NC_045512.2;YP_009724389.1;Wu1-SARS2_orf1ab;6720 | 0.1 | 0 |
| NC_045512.2;YP_009724389.1;Wu1-SARS2_orf1ab;6721 | 0.1 | 0.025 |
| NC_045512.2;YP_009724389.1;Wu1-SARS2_orf1ab;6722 | 0 | 0 |
| NC_045512.2;YP_009724389.1;Wu1-SARS2_orf1ab;6723 | 0.1 | 0 |
| NC_045512.2;YP_009724389.1;Wu1-SARS2_orf1ab;6724 | 0.05 | 0 |
| NC_045512.2;YP_009724389.1;Wu1-SARS2_orf1ab;6725 | 0.05 | 0 |
| NC_045512.2;YP_009724389.1;Wu1-SARS2_orf1ab;6726 | 0.1 | 0 |
| NC_045512.2;YP_009724389.1;Wu1-SARS2_orf1ab;6727 | 0.1 | 0 |
| NC_045512.2;YP_009724389.1;Wu1-SARS2_orf1ab;6728 | 0.05 | 0.025 |
| NC_045512.2;YP_009724389.1;Wu1-SARS2_orf1ab;6729 | 0 | 0 |
| NC_045512.2;YP_009724389.1;Wu1-SARS2_orf1ab;6730 | 0 | 0 |
| NC_045512.2;YP_009724389.1;Wu1-SARS2_orf1ab;6731 | 0 | 0 |
| NC_045512.2;YP_009724389.1;Wu1-SARS2_orf1ab;6732 | 0 | 0 |
| NC_045512.2;YP_009724389.1;Wu1-SARS2_orf1ab;6733 | 0 | 0 |
| NC_045512.2;YP_009724389.1;Wu1-SARS2_orf1ab;6734 | 0 | 0 |
| NC_045512.2;YP_009724389.1;Wu1-SARS2_orf1ab;6735 | 0 | 0 |
| NC_045512.2;YP_009724389.1;Wu1-SARS2_orf1ab;6736 | 0 | 0 |
| NC_045512.2;YP_009724389.1;Wu1-SARS2_orf1ab;6737 | 0 | 0 |
| NC_045512.2;YP_009724389.1;Wu1-SARS2_orf1ab;6738 | 0 | 0 |
| NC_045512.2;YP_009724389.1;Wu1-SARS2_orf1ab;6739 | 0 | 0 |
| NC_045512.2;YP_009724389.1;Wu1-SARS2_orf1ab;6740 | 0 | 0 |
| NC_045512.2;YP_009724389.1;Wu1-SARS2_orf1ab;6741 | 0 | 0 |

| | | |
|---|---|---|
| NC_045512.2;YP_009724389.1;Wu1-SARS2_orf1ab;6742 | 0 | 0 |
| NC_045512.2;YP_009724389.1;Wu1-SARS2_orf1ab;6743 | 0 | 0 |
| NC_045512.2;YP_009724389.1;Wu1-SARS2_orf1ab;6744 | 0 | 0 |
| NC_045512.2;YP_009724389.1;Wu1-SARS2_orf1ab;6745 | 0.05 | 0.025 |
| NC_045512.2;YP_009724389.1;Wu1-SARS2_orf1ab;6746 | 0.05 | 0 |
| NC_045512.2;YP_009724389.1;Wu1-SARS2_orf1ab;6747 | 0.05 | 0 |
| NC_045512.2;YP_009724389.1;Wu1-SARS2_orf1ab;6748 | 0.05 | 0 |
| NC_045512.2;YP_009724389.1;Wu1-SARS2_orf1ab;6749 | 0.1 | 0.125 |
| NC_045512.2;YP_009724389.1;Wu1-SARS2_orf1ab;6750 | 0.05 | 0 |
| NC_045512.2;YP_009724389.1;Wu1-SARS2_orf1ab;6751 | 0 | 0 |
| NC_045512.2;YP_009724389.1;Wu1-SARS2_orf1ab;6752 | 0 | 0 |
| NC_045512.2;YP_009724389.1;Wu1-SARS2_orf1ab;6753 | 0 | 0 |
| NC_045512.2;YP_009724389.1;Wu1-SARS2_orf1ab;6754 | 0 | 0 |
| NC_045512.2;YP_009724389.1;Wu1-SARS2_orf1ab;6755 | 0 | 0 |
| NC_045512.2;YP_009724389.1;Wu1-SARS2_orf1ab;6756 | 0 | 0 |
| NC_045512.2;YP_009724389.1;Wu1-SARS2_orf1ab;6757 | 0 | 0 |
| NC_045512.2;YP_009724389.1;Wu1-SARS2_orf1ab;6758 | 0 | 0 |
| NC_045512.2;YP_009724389.1;Wu1-SARS2_orf1ab;6759 | 0 | 0 |
| NC_045512.2;YP_009724389.1;Wu1-SARS2_orf1ab;6760 | 0 | 0 |
| NC_045512.2;YP_009724389.1;Wu1-SARS2_orf1ab;6761 | 0 | 0 |
| NC_045512.2;YP_009724389.1;Wu1-SARS2_orf1ab;6762 | 0 | 0 |
| NC_045512.2;YP_009724389.1;Wu1-SARS2_orf1ab;6763 | 0 | 0 |
| NC_045512.2;YP_009724389.1;Wu1-SARS2_orf1ab;6764 | 0 | 0 |
| NC_045512.2;YP_009724389.1;Wu1-SARS2_orf1ab;6765 | 0 | 0 |
| NC_045512.2;YP_009724389.1;Wu1-SARS2_orf1ab;6766 | 0 | 0 |
| NC_045512.2;YP_009724389.1;Wu1-SARS2_orf1ab;6767 | 0 | 0 |
| NC_045512.2;YP_009724389.1;Wu1-SARS2_orf1ab;6768 | 0 | 0 |
| NC_045512.2;YP_009724389.1;Wu1-SARS2_orf1ab;6769 | 0 | 0 |
| NC_045512.2;YP_009724389.1;Wu1-SARS2_orf1ab;6770 | 0 | 0 |
| NC_045512.2;YP_009724389.1;Wu1-SARS2_orf1ab;6771 | 0 | 0 |
| NC_045512.2;YP_009724389.1;Wu1-SARS2_orf1ab;6772 | 0 | 0.025 |
| NC_045512.2;YP_009724389.1;Wu1-SARS2_orf1ab;6773 | 0 | 0 |
| NC_045512.2;YP_009724389.1;Wu1-SARS2_orf1ab;6774 | 0 | 0 |
| NC_045512.2;YP_009724389.1;Wu1-SARS2_orf1ab;6775 | 0 | 0 |
| NC_045512.2;YP_009724389.1;Wu1-SARS2_orf1ab;6776 | 0 | 0 |
| NC_045512.2;YP_009724389.1;Wu1-SARS2_orf1ab;6777 | 0 | 0 |
| NC_045512.2;YP_009724389.1;Wu1-SARS2_orf1ab;6778 | 0 | 0 |
| NC_045512.2;YP_009724389.1;Wu1-SARS2_orf1ab;6779 | 0 | 0 |
| NC_045512.2;YP_009724389.1;Wu1-SARS2_orf1ab;6780 | 0 | 0 |
| NC_045512.2;YP_009724389.1;Wu1-SARS2_orf1ab;6781 | 0 | 0 |
| NC_045512.2;YP_009724389.1;Wu1-SARS2_orf1ab;6782 | 0 | 0 |
| NC_045512.2;YP_009724389.1;Wu1-SARS2_orf1ab;6783 | 0 | 0.025 |
| NC_045512.2;YP_009724389.1;Wu1-SARS2_orf1ab;6784 | 0 | 0.025 |

| | | |
|---|---|---|
| NC_045512.2;YP_009724389.1;Wu1-SARS2_orf1ab;6785 | 0 | 0.025 |
| NC_045512.2;YP_009724389.1;Wu1-SARS2_orf1ab;6786 | 0 | 0.025 |
| NC_045512.2;YP_009724389.1;Wu1-SARS2_orf1ab;6787 | 0 | 0.025 |
| NC_045512.2;YP_009724389.1;Wu1-SARS2_orf1ab;6788 | 0 | 0.025 |
| NC_045512.2;YP_009724389.1;Wu1-SARS2_orf1ab;6789 | 0 | 0.025 |
| NC_045512.2;YP_009724389.1;Wu1-SARS2_orf1ab;6790 | 0 | 0.025 |
| NC_045512.2;YP_009724389.1;Wu1-SARS2_orf1ab;6791 | 0 | 0.025 |
| NC_045512.2;YP_009724389.1;Wu1-SARS2_orf1ab;6792 | 0 | 0.025 |
| NC_045512.2;YP_009724389.1;Wu1-SARS2_orf1ab;6793 | 0.1 | 0.05 |
| NC_045512.2;YP_009724389.1;Wu1-SARS2_orf1ab;6794 | 0 | 0 |
| NC_045512.2;YP_009724389.1;Wu1-SARS2_orf1ab;6795 | 0 | 0 |
| NC_045512.2;YP_009724389.1;Wu1-SARS2_orf1ab;6796 | 0 | 0 |
| NC_045512.2;YP_009724389.1;Wu1-SARS2_orf1ab;6797 | 0 | 0 |
| NC_045512.2;YP_009724389.1;Wu1-SARS2_orf1ab;6798 | 0 | 0 |
| NC_045512.2;YP_009724389.1;Wu1-SARS2_orf1ab;6799 | 0 | 0 |
| NC_045512.2;YP_009724389.1;Wu1-SARS2_orf1ab;6800 | 0 | 0 |
| NC_045512.2;YP_009724389.1;Wu1-SARS2_orf1ab;6801 | 0 | 0 |
| NC_045512.2;YP_009724389.1;Wu1-SARS2_orf1ab;6802 | 0 | 0 |
| NC_045512.2;YP_009724389.1;Wu1-SARS2_orf1ab;6803 | 0 | 0 |
| NC_045512.2;YP_009724389.1;Wu1-SARS2_orf1ab;6804 | 0 | 0 |
| NC_045512.2;YP_009724389.1;Wu1-SARS2_orf1ab;6805 | 0 | 0 |
| NC_045512.2;YP_009724389.1;Wu1-SARS2_orf1ab;6806 | 0 | 0 |
| NC_045512.2;YP_009724389.1;Wu1-SARS2_orf1ab;6807 | 0 | 0 |
| NC_045512.2;YP_009724389.1;Wu1-SARS2_orf1ab;6808 | 0 | 0 |
| NC_045512.2;YP_009724389.1;Wu1-SARS2_orf1ab;6809 | 0 | 0.025 |
| NC_045512.2;YP_009724389.1;Wu1-SARS2_orf1ab;6810 | 0 | 0 |
| NC_045512.2;YP_009724389.1;Wu1-SARS2_orf1ab;6811 | 0 | 0 |
| NC_045512.2;YP_009724389.1;Wu1-SARS2_orf1ab;6812 | 0 | 0 |
| NC_045512.2;YP_009724389.1;Wu1-SARS2_orf1ab;6813 | 0 | 0 |
| NC_045512.2;YP_009724389.1;Wu1-SARS2_orf1ab;6814 | 0 | 0 |
| NC_045512.2;YP_009724389.1;Wu1-SARS2_orf1ab;6815 | 0 | 0 |
| NC_045512.2;YP_009724389.1;Wu1-SARS2_orf1ab;6816 | 0 | 0 |
| NC_045512.2;YP_009724389.1;Wu1-SARS2_orf1ab;6817 | 0 | 0 |
| NC_045512.2;YP_009724389.1;Wu1-SARS2_orf1ab;6818 | 0 | 0 |
| NC_045512.2;YP_009724389.1;Wu1-SARS2_orf1ab;6819 | 0 | 0 |
| NC_045512.2;YP_009724389.1;Wu1-SARS2_orf1ab;6820 | 0 | 0 |
| NC_045512.2;YP_009724389.1;Wu1-SARS2_orf1ab;6821 | 0.05 | 0.05 |
| NC_045512.2;YP_009724389.1;Wu1-SARS2_orf1ab;6822 | 0.1 | 0.075 |
| NC_045512.2;YP_009724389.1;Wu1-SARS2_orf1ab;6823 | 0.05 | 0 |
| NC_045512.2;YP_009724389.1;Wu1-SARS2_orf1ab;6824 | 0.1 | 0.025 |
| NC_045512.2;YP_009724389.1;Wu1-SARS2_orf1ab;6825 | 0.1 | 0.025 |
| NC_045512.2;YP_009724389.1;Wu1-SARS2_orf1ab;6826 | 0.1 | 0.025 |
| NC_045512.2;YP_009724389.1;Wu1-SARS2_orf1ab;6827 | 0.1 | 0.025 |

| | | |
|---|---|---|
| NC_045512.2;YP_009724389.1;Wu1-SARS2_orf1ab;6828 | 0.1 | 0.025 |
| NC_045512.2;YP_009724389.1;Wu1-SARS2_orf1ab;6829 | 0 | 0 |
| NC_045512.2;YP_009724389.1;Wu1-SARS2_orf1ab;6830 | 0.25 | 0.125 |
| NC_045512.2;YP_009724389.1;Wu1-SARS2_orf1ab;6831 | 0 | 0 |
| NC_045512.2;YP_009724389.1;Wu1-SARS2_orf1ab;6832 | 0 | 0 |
| NC_045512.2;YP_009724389.1;Wu1-SARS2_orf1ab;6833 | 0 | 0 |
| NC_045512.2;YP_009724389.1;Wu1-SARS2_orf1ab;6834 | 0 | 0 |
| NC_045512.2;YP_009724389.1;Wu1-SARS2_orf1ab;6835 | 0 | 0 |
| NC_045512.2;YP_009724389.1;Wu1-SARS2_orf1ab;6836 | 0 | 0.05 |
| NC_045512.2;YP_009724389.1;Wu1-SARS2_orf1ab;6837 | 0 | 0 |
| NC_045512.2;YP_009724389.1;Wu1-SARS2_orf1ab;6838 | 0 | 0 |
| NC_045512.2;YP_009724389.1;Wu1-SARS2_orf1ab;6839 | 0 | 0 |
| NC_045512.2;YP_009724389.1;Wu1-SARS2_orf1ab;6840 | 0 | 0 |
| NC_045512.2;YP_009724389.1;Wu1-SARS2_orf1ab;6841 | 0 | 0 |
| NC_045512.2;YP_009724389.1;Wu1-SARS2_orf1ab;6842 | 0 | 0 |
| NC_045512.2;YP_009724389.1;Wu1-SARS2_orf1ab;6843 | 0 | 0 |
| NC_045512.2;YP_009724389.1;Wu1-SARS2_orf1ab;6844 | 0 | 0 |
| NC_045512.2;YP_009724389.1;Wu1-SARS2_orf1ab;6845 | 0 | 0 |
| NC_045512.2;YP_009724389.1;Wu1-SARS2_orf1ab;6846 | 0 | 0 |
| NC_045512.2;YP_009724389.1;Wu1-SARS2_orf1ab;6847 | 0 | 0 |
| NC_045512.2;YP_009724389.1;Wu1-SARS2_orf1ab;6848 | 0 | 0 |
| NC_045512.2;YP_009724389.1;Wu1-SARS2_orf1ab;6849 | 0 | 0 |
| NC_045512.2;YP_009724389.1;Wu1-SARS2_orf1ab;6850 | 0 | 0 |
| NC_045512.2;YP_009724389.1;Wu1-SARS2_orf1ab;6851 | 0 | 0 |
| NC_045512.2;YP_009724389.1;Wu1-SARS2_orf1ab;6852 | 0 | 0 |
| NC_045512.2;YP_009724389.1;Wu1-SARS2_orf1ab;6853 | 0 | 0 |
| NC_045512.2;YP_009724389.1;Wu1-SARS2_orf1ab;6854 | 0 | 0 |
| NC_045512.2;YP_009724389.1;Wu1-SARS2_orf1ab;6855 | 0 | 0 |
| NC_045512.2;YP_009724389.1;Wu1-SARS2_orf1ab;6856 | 0 | 0 |
| NC_045512.2;YP_009724389.1;Wu1-SARS2_orf1ab;6857 | 0 | 0 |
| NC_045512.2;YP_009724389.1;Wu1-SARS2_orf1ab;6858 | 0 | 0 |
| NC_045512.2;YP_009724389.1;Wu1-SARS2_orf1ab;6859 | 0 | 0 |
| NC_045512.2;YP_009724389.1;Wu1-SARS2_orf1ab;6860 | 0 | 0 |
| NC_045512.2;YP_009724389.1;Wu1-SARS2_orf1ab;6861 | 0 | 0 |
| NC_045512.2;YP_009724389.1;Wu1-SARS2_orf1ab;6862 | 0.05 | 0.05 |
| NC_045512.2;YP_009724389.1;Wu1-SARS2_orf1ab;6863 | 0 | 0 |
| NC_045512.2;YP_009724389.1;Wu1-SARS2_orf1ab;6864 | 0 | 0 |
| NC_045512.2;YP_009724389.1;Wu1-SARS2_orf1ab;6865 | 0 | 0 |
| NC_045512.2;YP_009724389.1;Wu1-SARS2_orf1ab;6866 | 0 | 0 |
| NC_045512.2;YP_009724389.1;Wu1-SARS2_orf1ab;6867 | 0 | 0 |
| NC_045512.2;YP_009724389.1;Wu1-SARS2_orf1ab;6868 | 0 | 0 |
| NC_045512.2;YP_009724389.1;Wu1-SARS2_orf1ab;6869 | 0 | 0 |
| NC_045512.2;YP_009724389.1;Wu1-SARS2_orf1ab;6870 | 0.05 | 0 |

| | | |
|---|---|---|
| NC_045512.2;YP_009724389.1;Wu1-SARS2_orf1ab;6871 | 0.05 | 0 |
| NC_045512.2;YP_009724389.1;Wu1-SARS2_orf1ab;6872 | 0 | 0 |
| NC_045512.2;YP_009724389.1;Wu1-SARS2_orf1ab;6873 | 0 | 0 |
| NC_045512.2;YP_009724389.1;Wu1-SARS2_orf1ab;6874 | 0 | 0 |
| NC_045512.2;YP_009724389.1;Wu1-SARS2_orf1ab;6875 | 0 | 0 |
| NC_045512.2;YP_009724389.1;Wu1-SARS2_orf1ab;6876 | 0 | 0 |
| NC_045512.2;YP_009724389.1;Wu1-SARS2_orf1ab;6877 | 0 | 0 |
| NC_045512.2;YP_009724389.1;Wu1-SARS2_orf1ab;6878 | 0 | 0 |
| NC_045512.2;YP_009724389.1;Wu1-SARS2_orf1ab;6879 | 0 | 0 |
| NC_045512.2;YP_009724389.1;Wu1-SARS2_orf1ab;6880 | 0 | 0 |
| NC_045512.2;YP_009724389.1;Wu1-SARS2_orf1ab;6881 | 0 | 0 |
| NC_045512.2;YP_009724389.1;Wu1-SARS2_orf1ab;6882 | 0 | 0 |
| NC_045512.2;YP_009724389.1;Wu1-SARS2_orf1ab;6883 | 0 | 0 |
| NC_045512.2;YP_009724389.1;Wu1-SARS2_orf1ab;6884 | 0 | 0 |
| NC_045512.2;YP_009724389.1;Wu1-SARS2_orf1ab;6885 | 0 | 0 |
| NC_045512.2;YP_009724389.1;Wu1-SARS2_orf1ab;6886 | 0 | 0 |
| NC_045512.2;YP_009724389.1;Wu1-SARS2_orf1ab;6887 | 0 | 0.025 |
| NC_045512.2;YP_009724389.1;Wu1-SARS2_orf1ab;6888 | 0 | 0.025 |
| NC_045512.2;YP_009724389.1;Wu1-SARS2_orf1ab;6889 | 0 | 0.025 |
| NC_045512.2;YP_009724389.1;Wu1-SARS2_orf1ab;6890 | 0 | 0.025 |
| NC_045512.2;YP_009724389.1;Wu1-SARS2_orf1ab;6891 | 0.05 | 0.025 |
| NC_045512.2;YP_009724389.1;Wu1-SARS2_orf1ab;6892 | 0.05 | 0.05 |
| NC_045512.2;YP_009724389.1;Wu1-SARS2_orf1ab;6893 | 0.05 | 0.05 |
| NC_045512.2;YP_009724389.1;Wu1-SARS2_orf1ab;6894 | 0 | 0.025 |
| NC_045512.2;YP_009724389.1;Wu1-SARS2_orf1ab;6895 | 0 | 0.05 |
| NC_045512.2;YP_009724389.1;Wu1-SARS2_orf1ab;6896 | 0.1 | 0.05 |
| NC_045512.2;YP_009724389.1;Wu1-SARS2_orf1ab;6897 | 0 | 0.025 |
| NC_045512.2;YP_009724389.1;Wu1-SARS2_orf1ab;6898 | 0 | 0 |
| NC_045512.2;YP_009724389.1;Wu1-SARS2_orf1ab;6899 | 0.05 | 0 |
| NC_045512.2;YP_009724389.1;Wu1-SARS2_orf1ab;6900 | 0.05 | 0 |
| NC_045512.2;YP_009724389.1;Wu1-SARS2_orf1ab;6901 | 0.05 | 0 |
| NC_045512.2;YP_009724389.1;Wu1-SARS2_orf1ab;6902 | 0.05 | 0 |
| NC_045512.2;YP_009724389.1;Wu1-SARS2_orf1ab;6903 | 0.05 | 0 |
| NC_045512.2;YP_009724389.1;Wu1-SARS2_orf1ab;6904 | 0.05 | 0 |
| NC_045512.2;YP_009724389.1;Wu1-SARS2_orf1ab;6905 | 0.05 | 0 |
| NC_045512.2;YP_009724389.1;Wu1-SARS2_orf1ab;6906 | 0 | 0.025 |
| NC_045512.2;YP_009724389.1;Wu1-SARS2_orf1ab;6907 | 0 | 0 |
| NC_045512.2;YP_009724389.1;Wu1-SARS2_orf1ab;6908 | 0.05 | 0.175 |
| NC_045512.2;YP_009724389.1;Wu1-SARS2_orf1ab;6909 | 0 | 0.1 |
| NC_045512.2;YP_009724389.1;Wu1-SARS2_orf1ab;6910 | 0 | 0.025 |
| NC_045512.2;YP_009724389.1;Wu1-SARS2_orf1ab;6911 | 0 | 0 |
| NC_045512.2;YP_009724389.1;Wu1-SARS2_orf1ab;6912 | 0 | 0 |
| NC_045512.2;YP_009724389.1;Wu1-SARS2_orf1ab;6913 | 0 | 0 |

| | | |
|---|---|---|
| NC_045512.2;YP_009724389.1;Wu1-SARS2_orf1ab;6914 | 0 | 0.025 |
| NC_045512.2;YP_009724389.1;Wu1-SARS2_orf1ab;6915 | 0 | 0 |
| NC_045512.2;YP_009724389.1;Wu1-SARS2_orf1ab;6916 | 0 | 0 |
| NC_045512.2;YP_009724389.1;Wu1-SARS2_orf1ab;6917 | 0 | 0 |
| NC_045512.2;YP_009724389.1;Wu1-SARS2_orf1ab;6918 | 0 | 0 |
| NC_045512.2;YP_009724389.1;Wu1-SARS2_orf1ab;6919 | 0.15 | 0.05 |
| NC_045512.2;YP_009724389.1;Wu1-SARS2_orf1ab;6920 | 0 | 0 |
| NC_045512.2;YP_009724389.1;Wu1-SARS2_orf1ab;6921 | 0 | 0.025 |
| NC_045512.2;YP_009724389.1;Wu1-SARS2_orf1ab;6922 | 0 | 0 |
| NC_045512.2;YP_009724389.1;Wu1-SARS2_orf1ab;6923 | 0.05 | 0.025 |
| NC_045512.2;YP_009724389.1;Wu1-SARS2_orf1ab;6924 | 0.05 | 0 |
| NC_045512.2;YP_009724389.1;Wu1-SARS2_orf1ab;6925 | 0.05 | 0 |
| NC_045512.2;YP_009724389.1;Wu1-SARS2_orf1ab;6926 | 0.15 | 0.05 |
| NC_045512.2;YP_009724389.1;Wu1-SARS2_orf1ab;6927 | 0.2 | 0.075 |
| NC_045512.2;YP_009724389.1;Wu1-SARS2_orf1ab;6928 | 0.2 | 0.1 |
| NC_045512.2;YP_009724389.1;Wu1-SARS2_orf1ab;6929 | 0.1 | 0.05 |
| NC_045512.2;YP_009724389.1;Wu1-SARS2_orf1ab;6930 | 0.25 | 0.05 |
| NC_045512.2;YP_009724389.1;Wu1-SARS2_orf1ab;6931 | 0.15 | 0.05 |
| NC_045512.2;YP_009724389.1;Wu1-SARS2_orf1ab;6932 | 0.1 | 0.05 |
| NC_045512.2;YP_009724389.1;Wu1-SARS2_orf1ab;6933 | 0.05 | 0.075 |
| NC_045512.2;YP_009724389.1;Wu1-SARS2_orf1ab;6934 | 0.25 | 0.1 |
| NC_045512.2;YP_009724389.1;Wu1-SARS2_orf1ab;6935 | 0.05 | 0.025 |
| NC_045512.2;YP_009724389.1;Wu1-SARS2_orf1ab;6936 | 0.05 | 0.025 |
| NC_045512.2;YP_009724389.1;Wu1-SARS2_orf1ab;6937 | 0 | 0.025 |
| NC_045512.2;YP_009724389.1;Wu1-SARS2_orf1ab;6938 | 0 | 0.05 |
| NC_045512.2;YP_009724389.1;Wu1-SARS2_orf1ab;6939 | 0 | 0 |
| NC_045512.2;YP_009724389.1;Wu1-SARS2_orf1ab;6940 | 0 | 0 |
| NC_045512.2;YP_009724389.1;Wu1-SARS2_orf1ab;6941 | 0 | 0 |
| NC_045512.2;YP_009724389.1;Wu1-SARS2_orf1ab;6942 | 0 | 0 |
| NC_045512.2;YP_009724389.1;Wu1-SARS2_orf1ab;6943 | 0 | 0 |
| NC_045512.2;YP_009724389.1;Wu1-SARS2_orf1ab;6944 | 0 | 0 |
| NC_045512.2;YP_009724389.1;Wu1-SARS2_orf1ab;6945 | 0 | 0 |
| NC_045512.2;YP_009724389.1;Wu1-SARS2_orf1ab;6946 | 0 | 0 |
| NC_045512.2;YP_009724389.1;Wu1-SARS2_orf1ab;6947 | 0 | 0 |
| NC_045512.2;YP_009724389.1;Wu1-SARS2_orf1ab;6948 | 0 | 0 |
| NC_045512.2;YP_009724389.1;Wu1-SARS2_orf1ab;6949 | 0 | 0 |
| NC_045512.2;YP_009724389.1;Wu1-SARS2_orf1ab;6950 | 0 | 0 |
| NC_045512.2;YP_009724389.1;Wu1-SARS2_orf1ab;6951 | 0 | 0 |
| NC_045512.2;YP_009724389.1;Wu1-SARS2_orf1ab;6952 | 0 | 0 |
| NC_045512.2;YP_009724389.1;Wu1-SARS2_orf1ab;6953 | 0 | 0 |
| NC_045512.2;YP_009724389.1;Wu1-SARS2_orf1ab;6954 | 0 | 0 |
| NC_045512.2;YP_009724389.1;Wu1-SARS2_orf1ab;6955 | 0 | 0 |
| NC_045512.2;YP_009724389.1;Wu1-SARS2_orf1ab;6956 | 0 | 0 |

| | | |
|---|---|---|
| NC_045512.2;YP_009724389.1;Wu1-SARS2_orf1ab;6957 | 0 | 0 |
| NC_045512.2;YP_009724389.1;Wu1-SARS2_orf1ab;6958 | 0 | 0 |
| NC_045512.2;YP_009724389.1;Wu1-SARS2_orf1ab;6959 | 0 | 0 |
| NC_045512.2;YP_009724389.1;Wu1-SARS2_orf1ab;6960 | 0 | 0 |
| NC_045512.2;YP_009724389.1;Wu1-SARS2_orf1ab;6961 | 0 | 0 |
| NC_045512.2;YP_009724389.1;Wu1-SARS2_orf1ab;6962 | 0 | 0 |
| NC_045512.2;YP_009724389.1;Wu1-SARS2_orf1ab;6963 | 0 | 0 |
| NC_045512.2;YP_009724389.1;Wu1-SARS2_orf1ab;6964 | 0 | 0 |
| NC_045512.2;YP_009724389.1;Wu1-SARS2_orf1ab;6965 | 0 | 0.05 |
| NC_045512.2;YP_009724389.1;Wu1-SARS2_orf1ab;6966 | 0 | 0.025 |
| NC_045512.2;YP_009724389.1;Wu1-SARS2_orf1ab;6967 | 0 | 0.05 |
| NC_045512.2;YP_009724389.1;Wu1-SARS2_orf1ab;6968 | 0 | 0.025 |
| NC_045512.2;YP_009724389.1;Wu1-SARS2_orf1ab;6969 | 0 | 0.025 |
| NC_045512.2;YP_009724389.1;Wu1-SARS2_orf1ab;6970 | 0 | 0.025 |
| NC_045512.2;YP_009724389.1;Wu1-SARS2_orf1ab;6971 | 0 | 0.025 |
| NC_045512.2;YP_009724389.1;Wu1-SARS2_orf1ab;6972 | 0 | 0.025 |
| NC_045512.2;YP_009724389.1;Wu1-SARS2_orf1ab;6973 | 0 | 0.025 |
| NC_045512.2;YP_009724389.1;Wu1-SARS2_orf1ab;6974 | 0 | 0 |
| NC_045512.2;YP_009724389.1;Wu1-SARS2_orf1ab;6975 | 0 | 0 |
| NC_045512.2;YP_009724389.1;Wu1-SARS2_orf1ab;6976 | 0 | 0 |
| NC_045512.2;YP_009724389.1;Wu1-SARS2_orf1ab;6977 | 0 | 0 |
| NC_045512.2;YP_009724389.1;Wu1-SARS2_orf1ab;6978 | 0 | 0 |
| NC_045512.2;YP_009724389.1;Wu1-SARS2_orf1ab;6979 | 0 | 0 |
| NC_045512.2;YP_009724389.1;Wu1-SARS2_orf1ab;6980 | 0 | 0 |
| NC_045512.2;YP_009724389.1;Wu1-SARS2_orf1ab;6981 | 0 | 0 |
| NC_045512.2;YP_009724389.1;Wu1-SARS2_orf1ab;6982 | 0 | 0 |
| NC_045512.2;YP_009724389.1;Wu1-SARS2_orf1ab;6983 | 0 | 0 |
| NC_045512.2;YP_009724389.1;Wu1-SARS2_orf1ab;6984 | 0 | 0 |
| NC_045512.2;YP_009724389.1;Wu1-SARS2_orf1ab;6985 | 0 | 0 |
| NC_045512.2;YP_009724389.1;Wu1-SARS2_orf1ab;6986 | 0 | 0.025 |
| NC_045512.2;YP_009724389.1;Wu1-SARS2_orf1ab;6987 | 0 | 0.025 |
| NC_045512.2;YP_009724389.1;Wu1-SARS2_orf1ab;6988 | 0 | 0.05 |
| NC_045512.2;YP_009724389.1;Wu1-SARS2_orf1ab;6989 | 0 | 0 |
| NC_045512.2;YP_009724389.1;Wu1-SARS2_orf1ab;6990 | 0.05 | 0 |
| NC_045512.2;YP_009724389.1;Wu1-SARS2_orf1ab;6991 | 0 | 0 |
| NC_045512.2;YP_009724389.1;Wu1-SARS2_orf1ab;6992 | 0 | 0.05 |
| NC_045512.2;YP_009724389.1;Wu1-SARS2_orf1ab;6993 | 0 | 0 |
| NC_045512.2;YP_009724389.1;Wu1-SARS2_orf1ab;6994 | 0 | 0 |
| NC_045512.2;YP_009724389.1;Wu1-SARS2_orf1ab;6995 | 0 | 0 |
| NC_045512.2;YP_009724389.1;Wu1-SARS2_orf1ab;6996 | 0 | 0 |
| NC_045512.2;YP_009724389.1;Wu1-SARS2_orf1ab;6997 | 0 | 0 |
| NC_045512.2;YP_009724389.1;Wu1-SARS2_orf1ab;6998 | 0 | 0 |
| NC_045512.2;YP_009724389.1;Wu1-SARS2_orf1ab;6999 | 0 | 0 |

| | | |
|---|---|---|
| NC_045512.2;YP_009724389.1;Wu1-SARS2_orf1ab;7000 | 0 | 0 |
| NC_045512.2;YP_009724389.1;Wu1-SARS2_orf1ab;7001 | 0 | 0 |
| NC_045512.2;YP_009724389.1;Wu1-SARS2_orf1ab;7002 | 0 | 0 |
| NC_045512.2;YP_009724389.1;Wu1-SARS2_orf1ab;7003 | 0.05 | 0 |
| NC_045512.2;YP_009724389.1;Wu1-SARS2_orf1ab;7004 | 0.05 | 0 |
| NC_045512.2;YP_009724389.1;Wu1-SARS2_orf1ab;7005 | 0 | 0 |
| NC_045512.2;YP_009724389.1;Wu1-SARS2_orf1ab;7006 | 0.1 | 0 |
| NC_045512.2;YP_009724389.1;Wu1-SARS2_orf1ab;7007 | 0.1 | 0 |
| NC_045512.2;YP_009724389.1;Wu1-SARS2_orf1ab;7008 | 0.05 | 0 |
| NC_045512.2;YP_009724389.1;Wu1-SARS2_orf1ab;7009 | 0.1 | 0 |
| NC_045512.2;YP_009724389.1;Wu1-SARS2_orf1ab;7010 | 0.1 | 0.05 |
| NC_045512.2;YP_009724389.1;Wu1-SARS2_orf1ab;7011 | 0 | 0.05 |
| NC_045512.2;YP_009724389.1;Wu1-SARS2_orf1ab;7012 | 0 | 0.05 |
| NC_045512.2;YP_009724389.1;Wu1-SARS2_orf1ab;7013 | 0 | 0 |
| NC_045512.2;YP_009724389.1;Wu1-SARS2_orf1ab;7014 | 0 | 0 |
| NC_045512.2;YP_009724389.1;Wu1-SARS2_orf1ab;7015 | 0.05 | 0.05 |
| NC_045512.2;YP_009724389.1;Wu1-SARS2_orf1ab;7016 | 0 | 0 |
| NC_045512.2;YP_009724389.1;Wu1-SARS2_orf1ab;7017 | 0 | 0 |
| NC_045512.2;YP_009724389.1;Wu1-SARS2_orf1ab;7018 | 0 | 0 |
| NC_045512.2;YP_009724389.1;Wu1-SARS2_orf1ab;7019 | 0 | 0 |
| NC_045512.2;YP_009724389.1;Wu1-SARS2_orf1ab;7020 | 0 | 0 |
| NC_045512.2;YP_009724389.1;Wu1-SARS2_orf1ab;7021 | 0 | 0 |
| NC_045512.2;YP_009724389.1;Wu1-SARS2_orf1ab;7022 | 0 | 0 |
| NC_045512.2;YP_009724389.1;Wu1-SARS2_orf1ab;7023 | 0 | 0 |
| NC_045512.2;YP_009724389.1;Wu1-SARS2_orf1ab;7024 | 0 | 0 |
| NC_045512.2;YP_009724389.1;Wu1-SARS2_orf1ab;7025 | 0 | 0 |
| NC_045512.2;YP_009724389.1;Wu1-SARS2_orf1ab;7026 | 0 | 0 |
| NC_045512.2;YP_009724389.1;Wu1-SARS2_orf1ab;7027 | 0 | 0 |
| NC_045512.2;YP_009724389.1;Wu1-SARS2_orf1ab;7028 | 0 | 0 |
| NC_045512.2;YP_009724389.1;Wu1-SARS2_orf1ab;7029 | 0 | 0 |
| NC_045512.2;YP_009724389.1;Wu1-SARS2_orf1ab;7030 | 0 | 0 |
| NC_045512.2;YP_009724389.1;Wu1-SARS2_orf1ab;7031 | 0 | 0 |
| NC_045512.2;YP_009724389.1;Wu1-SARS2_orf1ab;7032 | 0.05 | 0.075 |
| NC_045512.2;YP_009724389.1;Wu1-SARS2_orf1ab;7033 | 0.05 | 0.075 |
| NC_045512.2;YP_009724389.1;Wu1-SARS2_orf1ab;7034 | 0.05 | 0.05 |
| NC_045512.2;YP_009724389.1;Wu1-SARS2_orf1ab;7035 | 0.05 | 0.05 |
| NC_045512.2;YP_009724389.1;Wu1-SARS2_orf1ab;7036 | 0.05 | 0.05 |
| NC_045512.2;YP_009724389.1;Wu1-SARS2_orf1ab;7037 | 0.05 | 0.075 |
| NC_045512.2;YP_009724389.1;Wu1-SARS2_orf1ab;7038 | 0.05 | 0.025 |
| NC_045512.2;YP_009724389.1;Wu1-SARS2_orf1ab;7039 | 0 | 0.025 |
| NC_045512.2;YP_009724389.1;Wu1-SARS2_orf1ab;7040 | 0 | 0 |
| NC_045512.2;YP_009724389.1;Wu1-SARS2_orf1ab;7041 | 0 | 0 |
| NC_045512.2;YP_009724389.1;Wu1-SARS2_orf1ab;7042 | 0 | 0 |

| | | |
|---|---|---|
| NC_045512.2;YP_009724389.1;Wu1-SARS2_orf1ab;7043 | 0 | 0 |
| NC_045512.2;YP_009724389.1;Wu1-SARS2_orf1ab;7044 | 0 | 0 |
| NC_045512.2;YP_009724389.1;Wu1-SARS2_orf1ab;7045 | 0 | 0 |
| NC_045512.2;YP_009724389.1;Wu1-SARS2_orf1ab;7046 | 0 | 0 |
| NC_045512.2;YP_009724389.1;Wu1-SARS2_orf1ab;7047 | 0 | 0 |
| NC_045512.2;YP_009724389.1;Wu1-SARS2_orf1ab;7048 | 0 | 0 |
| NC_045512.2;YP_009724389.1;Wu1-SARS2_orf1ab;7049 | 0 | 0 |
| NC_045512.2;YP_009724389.1;Wu1-SARS2_orf1ab;7050 | 0 | 0 |
| NC_045512.2;YP_009724389.1;Wu1-SARS2_orf1ab;7051 | 0 | 0 |
| NC_045512.2;YP_009724389.1;Wu1-SARS2_orf1ab;7052 | 0 | 0 |
| NC_045512.2;YP_009724389.1;Wu1-SARS2_orf1ab;7053 | 0.05 | 0 |
| NC_045512.2;YP_009724389.1;Wu1-SARS2_orf1ab;7054 | 0.05 | 0 |
| NC_045512.2;YP_009724389.1;Wu1-SARS2_orf1ab;7055 | 0.05 | 0 |
| NC_045512.2;YP_009724389.1;Wu1-SARS2_orf1ab;7056 | 0.1 | 0 |
| NC_045512.2;YP_009724389.1;Wu1-SARS2_orf1ab;7057 | 0.05 | 0 |
| NC_045512.2;YP_009724389.1;Wu1-SARS2_orf1ab;7058 | 0 | 0 |
| NC_045512.2;YP_009724389.1;Wu1-SARS2_orf1ab;7059 | 0 | 0 |
| NC_045512.2;YP_009724389.1;Wu1-SARS2_orf1ab;7060 | 0 | 0 |
| NC_045512.2;YP_009724389.1;Wu1-SARS2_orf1ab;7061 | 0 | 0 |
| NC_045512.2;YP_009724389.1;Wu1-SARS2_orf1ab;7062 | 0 | 0 |
| NC_045512.2;YP_009724389.1;Wu1-SARS2_orf1ab;7063 | 0 | 0 |
| NC_045512.2;YP_009724389.1;Wu1-SARS2_orf1ab;7064 | 0 | 0 |
| NC_045512.2;YP_009724389.1;Wu1-SARS2_orf1ab;7065 | 0 | 0 |
| NC_045512.2;YP_009724389.1;Wu1-SARS2_orf1ab;7066 | 0 | 0 |
| NC_045512.2;YP_009724389.1;Wu1-SARS2_orf1ab;7067 | 0 | 0 |
| NC_045512.2;YP_009724389.1;Wu1-SARS2_orf1ab;7068 | 0.05 | 0 |
| NC_045512.2;YP_009724389.1;Wu1-SARS2_orf1ab;7069 | 0 | 0 |
| NC_045512.2;YP_009724389.1;Wu1-SARS2_orf1ab;7070 | 0 | 0 |
| NC_045512.2;YP_009724389.1;Wu1-SARS2_orf1ab;7071 | 0 | 0.025 |
| NC_045512.2;YP_009724389.1;Wu1-SARS2_orf1ab;7072 | 0 | 0.025 |
| NC_045512.2;YP_009724389.1;Wu1-SARS2_orf1ab;7073 | 0 | 0.025 |
| NC_045512.2;YP_009724389.1;Wu1-SARS2_orf1ab;7074 | 0 | 0.025 |
| NC_045512.2;YP_009724389.1;Wu1-SARS2_orf1ab;7075 | 0 | 0.025 |
| NC_045512.2;YP_009724389.1;Wu1-SARS2_orf1ab;7076 | 0 | 0.025 |
| NC_045512.2;YP_009724389.1;Wu1-SARS2_orf1ab;7077 | 0 | 0.025 |
| NC_045512.2;YP_009724389.1;Wu1-SARS2_orf1ab;7078 | 0 | 0.025 |
| NC_045512.2;YP_009724389.1;Wu1-SARS2_orf1ab;7079 | 0 | 0.025 |
| NC_045512.2;YP_009724389.1;Wu1-SARS2_orf1ab;7080 | 0 | 0.025 |
| NC_045512.2;YP_009724389.1;Wu1-SARS2_orf1ab;7081 | 0 | 0.025 |
| NC_045512.2;YP_009724389.1;Wu1-SARS2_orf1ab;7082 | 0 | 0 |
| NC_045512.2;YP_009724389.1;Wu1-SARS2_orf1ab;7083 | 0 | 0 |
| NC_045512.2;YP_009724389.1;Wu1-SARS2_orf1ab;7084 | 0 | 0 |
| NC_045512.2;YP_009724389.1;Wu1-SARS2_orf1ab;7085 | 0 | 0 |

| | | |
|---|---|---|
| NC_045512.2;YP_009724390.1;Wu1-SARS2_surface;1 | 0 | 0 |
| NC_045512.2;YP_009724390.1;Wu1-SARS2_surface;2 | 0 | 0 |
| NC_045512.2;YP_009724390.1;Wu1-SARS2_surface;3 | 0 | 0 |
| NC_045512.2;YP_009724390.1;Wu1-SARS2_surface;4 | 0 | 0 |
| NC_045512.2;YP_009724390.1;Wu1-SARS2_surface;5 | 0 | 0 |
| NC_045512.2;YP_009724390.1;Wu1-SARS2_surface;6 | 0 | 0 |
| NC_045512.2;YP_009724390.1;Wu1-SARS2_surface;7 | 0 | 0 |
| NC_045512.2;YP_009724390.1;Wu1-SARS2_surface;8 | 0.05 | 0 |
| NC_045512.2;YP_009724390.1;Wu1-SARS2_surface;9 | 0 | 0 |
| NC_045512.2;YP_009724390.1;Wu1-SARS2_surface;10 | 0 | 0 |
| NC_045512.2;YP_009724390.1;Wu1-SARS2_surface;11 | 0 | 0 |
| NC_045512.2;YP_009724390.1;Wu1-SARS2_surface;12 | 0 | 0 |
| NC_045512.2;YP_009724390.1;Wu1-SARS2_surface;13 | 0 | 0 |
| NC_045512.2;YP_009724390.1;Wu1-SARS2_surface;14 | 0 | 0 |
| NC_045512.2;YP_009724390.1;Wu1-SARS2_surface;15 | 0 | 0 |
| NC_045512.2;YP_009724390.1;Wu1-SARS2_surface;16 | 0 | 0 |
| NC_045512.2;YP_009724390.1;Wu1-SARS2_surface;17 | 0 | 0 |
| NC_045512.2;YP_009724390.1;Wu1-SARS2_surface;18 | 0 | 0 |
| NC_045512.2;YP_009724390.1;Wu1-SARS2_surface;19 | 0 | 0 |
| NC_045512.2;YP_009724390.1;Wu1-SARS2_surface;20 | 0 | 0 |
| NC_045512.2;YP_009724390.1;Wu1-SARS2_surface;21 | 0 | 0 |
| NC_045512.2;YP_009724390.1;Wu1-SARS2_surface;22 | 0 | 0 |
| NC_045512.2;YP_009724390.1;Wu1-SARS2_surface;23 | 0 | 0 |
| NC_045512.2;YP_009724390.1;Wu1-SARS2_surface;24 | 0 | 0 |
| NC_045512.2;YP_009724390.1;Wu1-SARS2_surface;25 | 0 | 0.025 |
| NC_045512.2;YP_009724390.1;Wu1-SARS2_surface;26 | 0 | 0.025 |
| NC_045512.2;YP_009724390.1;Wu1-SARS2_surface;27 | 0.05 | 0 |
| NC_045512.2;YP_009724390.1;Wu1-SARS2_surface;28 | 0 | 0 |
| NC_045512.2;YP_009724390.1;Wu1-SARS2_surface;29 | 0 | 0 |
| NC_045512.2;YP_009724390.1;Wu1-SARS2_surface;30 | 0 | 0 |
| NC_045512.2;YP_009724390.1;Wu1-SARS2_surface;31 | 0 | 0 |
| NC_045512.2;YP_009724390.1;Wu1-SARS2_surface;32 | 0 | 0 |
| NC_045512.2;YP_009724390.1;Wu1-SARS2_surface;33 | 0 | 0 |
| NC_045512.2;YP_009724390.1;Wu1-SARS2_surface;34 | 0 | 0 |
| NC_045512.2;YP_009724390.1;Wu1-SARS2_surface;35 | 0 | 0 |
| NC_045512.2;YP_009724390.1;Wu1-SARS2_surface;36 | 0 | 0 |
| NC_045512.2;YP_009724390.1;Wu1-SARS2_surface;37 | 0 | 0 |
| NC_045512.2;YP_009724390.1;Wu1-SARS2_surface;38 | 0 | 0 |
| NC_045512.2;YP_009724390.1;Wu1-SARS2_surface;39 | 0.05 | 0 |
| NC_045512.2;YP_009724390.1;Wu1-SARS2_surface;40 | 0 | 0 |
| NC_045512.2;YP_009724390.1;Wu1-SARS2_surface;41 | 0 | 0 |
| NC_045512.2;YP_009724390.1;Wu1-SARS2_surface;42 | 0 | 0 |
| NC_045512.2;YP_009724390.1;Wu1-SARS2_surface;43 | 0 | 0 |

| | | |
|---|---|---|
| NC_045512.2;YP_009724390.1;Wu1-SARS2_surface;44 | 0 | 0 |
| NC_045512.2;YP_009724390.1;Wu1-SARS2_surface;45 | 0 | 0 |
| NC_045512.2;YP_009724390.1;Wu1-SARS2_surface;46 | 0 | 0 |
| NC_045512.2;YP_009724390.1;Wu1-SARS2_surface;47 | 0 | 0 |
| NC_045512.2;YP_009724390.1;Wu1-SARS2_surface;48 | 0 | 0 |
| NC_045512.2;YP_009724390.1;Wu1-SARS2_surface;49 | 0 | 0 |
| NC_045512.2;YP_009724390.1;Wu1-SARS2_surface;50 | 0 | 0 |
| NC_045512.2;YP_009724390.1;Wu1-SARS2_surface;51 | 0 | 0 |
| NC_045512.2;YP_009724390.1;Wu1-SARS2_surface;52 | 0 | 0 |
| NC_045512.2;YP_009724390.1;Wu1-SARS2_surface;53 | 0 | 0 |
| NC_045512.2;YP_009724390.1;Wu1-SARS2_surface;54 | 0 | 0 |
| NC_045512.2;YP_009724390.1;Wu1-SARS2_surface;55 | 0 | 0 |
| NC_045512.2;YP_009724390.1;Wu1-SARS2_surface;56 | 0 | 0 |
| NC_045512.2;YP_009724390.1;Wu1-SARS2_surface;57 | 0 | 0 |
| NC_045512.2;YP_009724390.1;Wu1-SARS2_surface;58 | 0 | 0 |
| NC_045512.2;YP_009724390.1;Wu1-SARS2_surface;59 | 0 | 0 |
| NC_045512.2;YP_009724390.1;Wu1-SARS2_surface;60 | 0 | 0 |
| NC_045512.2;YP_009724390.1;Wu1-SARS2_surface;61 | 0 | 0 |
| NC_045512.2;YP_009724390.1;Wu1-SARS2_surface;62 | 0 | 0 |
| NC_045512.2;YP_009724390.1;Wu1-SARS2_surface;63 | 0 | 0 |
| NC_045512.2;YP_009724390.1;Wu1-SARS2_surface;64 | 0 | 0 |
| NC_045512.2;YP_009724390.1;Wu1-SARS2_surface;65 | 0 | 0 |
| NC_045512.2;YP_009724390.1;Wu1-SARS2_surface;66 | 0 | 0 |
| NC_045512.2;YP_009724390.1;Wu1-SARS2_surface;67 | 0 | 0 |
| NC_045512.2;YP_009724390.1;Wu1-SARS2_surface;68 | 0 | 0 |
| NC_045512.2;YP_009724390.1;Wu1-SARS2_surface;69 | 0 | 0 |
| NC_045512.2;YP_009724390.1;Wu1-SARS2_surface;70 | 0 | 0 |
| NC_045512.2;YP_009724390.1;Wu1-SARS2_surface;71 | 0 | 0 |
| NC_045512.2;YP_009724390.1;Wu1-SARS2_surface;72 | 0 | 0 |
| NC_045512.2;YP_009724390.1;Wu1-SARS2_surface;73 | 0 | 0.025 |
| NC_045512.2;YP_009724390.1;Wu1-SARS2_surface;74 | 0.05 | 0.025 |
| NC_045512.2;YP_009724390.1;Wu1-SARS2_surface;75 | 0 | 0.025 |
| NC_045512.2;YP_009724390.1;Wu1-SARS2_surface;76 | 0 | 0.025 |
| NC_045512.2;YP_009724390.1;Wu1-SARS2_surface;77 | 0 | 0.025 |
| NC_045512.2;YP_009724390.1;Wu1-SARS2_surface;78 | 0 | 0 |
| NC_045512.2;YP_009724390.1;Wu1-SARS2_surface;79 | 0 | 0 |
| NC_045512.2;YP_009724390.1;Wu1-SARS2_surface;80 | 0 | 0 |
| NC_045512.2;YP_009724390.1;Wu1-SARS2_surface;81 | 0 | 0.025 |
| NC_045512.2;YP_009724390.1;Wu1-SARS2_surface;82 | 0.05 | 0.075 |
| NC_045512.2;YP_009724390.1;Wu1-SARS2_surface;83 | 0.05 | 0.1 |
| NC_045512.2;YP_009724390.1;Wu1-SARS2_surface;84 | 0.05 | 0.075 |
| NC_045512.2;YP_009724390.1;Wu1-SARS2_surface;85 | 0.05 | 0.05 |
| NC_045512.2;YP_009724390.1;Wu1-SARS2_surface;86 | 0.05 | 0.05 |

| | | |
|---|---|---|
| NC_045512.2;YP_009724390.1;Wu1-SARS2_surface;87 | 0.05 | 0 |
| NC_045512.2;YP_009724390.1;Wu1-SARS2_surface;88 | 0.05 | 0.025 |
| NC_045512.2;YP_009724390.1;Wu1-SARS2_surface;89 | 0.05 | 0 |
| NC_045512.2;YP_009724390.1;Wu1-SARS2_surface;90 | 0.05 | 0.025 |
| NC_045512.2;YP_009724390.1;Wu1-SARS2_surface;91 | 0.05 | 0 |
| NC_045512.2;YP_009724390.1;Wu1-SARS2_surface;92 | 0.05 | 0 |
| NC_045512.2;YP_009724390.1;Wu1-SARS2_surface;93 | 0 | 0 |
| NC_045512.2;YP_009724390.1;Wu1-SARS2_surface;94 | 0 | 0 |
| NC_045512.2;YP_009724390.1;Wu1-SARS2_surface;95 | 0 | 0 |
| NC_045512.2;YP_009724390.1;Wu1-SARS2_surface;96 | 0 | 0 |
| NC_045512.2;YP_009724390.1;Wu1-SARS2_surface;97 | 0 | 0 |
| NC_045512.2;YP_009724390.1;Wu1-SARS2_surface;98 | 0 | 0 |
| NC_045512.2;YP_009724390.1;Wu1-SARS2_surface;99 | 0 | 0 |
| NC_045512.2;YP_009724390.1;Wu1-SARS2_surface;100 | 0 | 0 |
| NC_045512.2;YP_009724390.1;Wu1-SARS2_surface;101 | 0 | 0 |
| NC_045512.2;YP_009724390.1;Wu1-SARS2_surface;102 | 0 | 0 |
| NC_045512.2;YP_009724390.1;Wu1-SARS2_surface;103 | 0 | 0 |
| NC_045512.2;YP_009724390.1;Wu1-SARS2_surface;104 | 0 | 0 |
| NC_045512.2;YP_009724390.1;Wu1-SARS2_surface;105 | 0 | 0 |
| NC_045512.2;YP_009724390.1;Wu1-SARS2_surface;106 | 0 | 0 |
| NC_045512.2;YP_009724390.1;Wu1-SARS2_surface;107 | 0 | 0 |
| NC_045512.2;YP_009724390.1;Wu1-SARS2_surface;108 | 0 | 0 |
| NC_045512.2;YP_009724390.1;Wu1-SARS2_surface;109 | 0 | 0.025 |
| NC_045512.2;YP_009724390.1;Wu1-SARS2_surface;110 | 0 | 0 |
| NC_045512.2;YP_009724390.1;Wu1-SARS2_surface;111 | 0.05 | 0.025 |
| NC_045512.2;YP_009724390.1;Wu1-SARS2_surface;112 | 0.05 | 0.025 |
| NC_045512.2;YP_009724390.1;Wu1-SARS2_surface;113 | 0.05 | 0 |
| NC_045512.2;YP_009724390.1;Wu1-SARS2_surface;114 | 0.05 | 0 |
| NC_045512.2;YP_009724390.1;Wu1-SARS2_surface;115 | 0 | 0 |
| NC_045512.2;YP_009724390.1;Wu1-SARS2_surface;116 | 0 | 0 |
| NC_045512.2;YP_009724390.1;Wu1-SARS2_surface;117 | 0 | 0 |
| NC_045512.2;YP_009724390.1;Wu1-SARS2_surface;118 | 0 | 0 |
| NC_045512.2;YP_009724390.1;Wu1-SARS2_surface;119 | 0.05 | 0 |
| NC_045512.2;YP_009724390.1;Wu1-SARS2_surface;120 | 0.05 | 0 |
| NC_045512.2;YP_009724390.1;Wu1-SARS2_surface;121 | 0 | 0 |
| NC_045512.2;YP_009724390.1;Wu1-SARS2_surface;122 | 0 | 0 |
| NC_045512.2;YP_009724390.1;Wu1-SARS2_surface;123 | 0 | 0 |
| NC_045512.2;YP_009724390.1;Wu1-SARS2_surface;124 | 0.05 | 0 |
| NC_045512.2;YP_009724390.1;Wu1-SARS2_surface;125 | 0 | 0 |
| NC_045512.2;YP_009724390.1;Wu1-SARS2_surface;126 | 0 | 0 |
| NC_045512.2;YP_009724390.1;Wu1-SARS2_surface;127 | 0 | 0 |
| NC_045512.2;YP_009724390.1;Wu1-SARS2_surface;128 | 0 | 0 |
| NC_045512.2;YP_009724390.1;Wu1-SARS2_surface;129 | 0 | 0 |

| | | |
|---|---|---|
| NC_045512.2;YP_009724390.1;Wu1-SARS2_surface;130 | 0 | 0 |
| NC_045512.2;YP_009724390.1;Wu1-SARS2_surface;131 | 0 | 0 |
| NC_045512.2;YP_009724390.1;Wu1-SARS2_surface;132 | 0 | 0 |
| NC_045512.2;YP_009724390.1;Wu1-SARS2_surface;133 | 0 | 0 |
| NC_045512.2;YP_009724390.1;Wu1-SARS2_surface;134 | 0 | 0 |
| NC_045512.2;YP_009724390.1;Wu1-SARS2_surface;135 | 0 | 0 |
| NC_045512.2;YP_009724390.1;Wu1-SARS2_surface;136 | 0 | 0 |
| NC_045512.2;YP_009724390.1;Wu1-SARS2_surface;137 | 0 | 0 |
| NC_045512.2;YP_009724390.1;Wu1-SARS2_surface;138 | 0 | 0 |
| NC_045512.2;YP_009724390.1;Wu1-SARS2_surface;139 | 0.05 | 0 |
| NC_045512.2;YP_009724390.1;Wu1-SARS2_surface;140 | 0.05 | 0.025 |
| NC_045512.2;YP_009724390.1;Wu1-SARS2_surface;141 | 0.05 | 0.075 |
| NC_045512.2;YP_009724390.1;Wu1-SARS2_surface;142 | 0.1 | 0.05 |
| NC_045512.2;YP_009724390.1;Wu1-SARS2_surface;143 | 0.05 | 0.025 |
| NC_045512.2;YP_009724390.1;Wu1-SARS2_surface;144 | 0.05 | 0 |
| NC_045512.2;YP_009724390.1;Wu1-SARS2_surface;145 | 0 | 0 |
| NC_045512.2;YP_009724390.1;Wu1-SARS2_surface;146 | 0.05 | 0 |
| NC_045512.2;YP_009724390.1;Wu1-SARS2_surface;147 | 0.05 | 0 |
| NC_045512.2;YP_009724390.1;Wu1-SARS2_surface;148 | 0.05 | 0 |
| NC_045512.2;YP_009724390.1;Wu1-SARS2_surface;149 | 0.05 | 0.025 |
| NC_045512.2;YP_009724390.1;Wu1-SARS2_surface;150 | 0 | 0.05 |
| NC_045512.2;YP_009724390.1;Wu1-SARS2_surface;151 | 0 | 0.025 |
| NC_045512.2;YP_009724390.1;Wu1-SARS2_surface;152 | 0 | 0.025 |
| NC_045512.2;YP_009724390.1;Wu1-SARS2_surface;153 | 0 | 0.025 |
| NC_045512.2;YP_009724390.1;Wu1-SARS2_surface;154 | 0 | 0.025 |
| NC_045512.2;YP_009724390.1;Wu1-SARS2_surface;155 | 0 | 0.025 |
| NC_045512.2;YP_009724390.1;Wu1-SARS2_surface;156 | 0 | 0 |
| NC_045512.2;YP_009724390.1;Wu1-SARS2_surface;157 | 0 | 0 |
| NC_045512.2;YP_009724390.1;Wu1-SARS2_surface;158 | 0.05 | 0.05 |
| NC_045512.2;YP_009724390.1;Wu1-SARS2_surface;159 | 0 | 0.1 |
| NC_045512.2;YP_009724390.1;Wu1-SARS2_surface;160 | 0 | 0 |
| NC_045512.2;YP_009724390.1;Wu1-SARS2_surface;161 | 0 | 0 |
| NC_045512.2;YP_009724390.1;Wu1-SARS2_surface;162 | 0 | 0.025 |
| NC_045512.2;YP_009724390.1;Wu1-SARS2_surface;163 | 0 | 0 |
| NC_045512.2;YP_009724390.1;Wu1-SARS2_surface;164 | 0 | 0 |
| NC_045512.2;YP_009724390.1;Wu1-SARS2_surface;165 | 0 | 0 |
| NC_045512.2;YP_009724390.1;Wu1-SARS2_surface;166 | 0 | 0 |
| NC_045512.2;YP_009724390.1;Wu1-SARS2_surface;167 | 0 | 0 |
| NC_045512.2;YP_009724390.1;Wu1-SARS2_surface;168 | 0 | 0 |
| NC_045512.2;YP_009724390.1;Wu1-SARS2_surface;169 | 0.05 | 0 |
| NC_045512.2;YP_009724390.1;Wu1-SARS2_surface;170 | 0 | 0 |
| NC_045512.2;YP_009724390.1;Wu1-SARS2_surface;171 | 0.05 | 0 |
| NC_045512.2;YP_009724390.1;Wu1-SARS2_surface;172 | 0 | 0.075 |

| | | |
|---|---|---|
| NC_045512.2;YP_009724390.1;Wu1-SARS2_surface;173 | 0.05 | 0.225 |
| NC_045512.2;YP_009724390.1;Wu1-SARS2_surface;174 | 0.1 | 0.125 |
| NC_045512.2;YP_009724390.1;Wu1-SARS2_surface;175 | 0.1 | 0.15 |
| NC_045512.2;YP_009724390.1;Wu1-SARS2_surface;176 | 0.1 | 0.175 |
| NC_045512.2;YP_009724390.1;Wu1-SARS2_surface;177 | 0.15 | 0.1 |
| NC_045512.2;YP_009724390.1;Wu1-SARS2_surface;178 | 0.05 | 0.2 |
| NC_045512.2;YP_009724390.1;Wu1-SARS2_surface;179 | 0.05 | 0.125 |
| NC_045512.2;YP_009724390.1;Wu1-SARS2_surface;180 | 0.05 | 0.05 |
| NC_045512.2;YP_009724390.1;Wu1-SARS2_surface;181 | 0 | 0.05 |
| NC_045512.2;YP_009724390.1;Wu1-SARS2_surface;182 | 0 | 0.025 |
| NC_045512.2;YP_009724390.1;Wu1-SARS2_surface;183 | 0.05 | 0.025 |
| NC_045512.2;YP_009724390.1;Wu1-SARS2_surface;184 | 0.1 | 0 |
| NC_045512.2;YP_009724390.1;Wu1-SARS2_surface;185 | 0.1 | 0 |
| NC_045512.2;YP_009724390.1;Wu1-SARS2_surface;186 | 0 | 0 |
| NC_045512.2;YP_009724390.1;Wu1-SARS2_surface;187 | 0 | 0 |
| NC_045512.2;YP_009724390.1;Wu1-SARS2_surface;188 | 0 | 0 |
| NC_045512.2;YP_009724390.1;Wu1-SARS2_surface;189 | 0 | 0 |
| NC_045512.2;YP_009724390.1;Wu1-SARS2_surface;190 | 0 | 0 |
| NC_045512.2;YP_009724390.1;Wu1-SARS2_surface;191 | 0 | 0 |
| NC_045512.2;YP_009724390.1;Wu1-SARS2_surface;192 | 0 | 0 |
| NC_045512.2;YP_009724390.1;Wu1-SARS2_surface;193 | 0 | 0 |
| NC_045512.2;YP_009724390.1;Wu1-SARS2_surface;194 | 0 | 0 |
| NC_045512.2;YP_009724390.1;Wu1-SARS2_surface;195 | 0 | 0 |
| NC_045512.2;YP_009724390.1;Wu1-SARS2_surface;196 | 0 | 0 |
| NC_045512.2;YP_009724390.1;Wu1-SARS2_surface;197 | 0 | 0 |
| NC_045512.2;YP_009724390.1;Wu1-SARS2_surface;198 | 0 | 0 |
| NC_045512.2;YP_009724390.1;Wu1-SARS2_surface;199 | 0 | 0 |
| NC_045512.2;YP_009724390.1;Wu1-SARS2_surface;200 | 0 | 0 |
| NC_045512.2;YP_009724390.1;Wu1-SARS2_surface;201 | 0 | 0 |
| NC_045512.2;YP_009724390.1;Wu1-SARS2_surface;202 | 0 | 0 |
| NC_045512.2;YP_009724390.1;Wu1-SARS2_surface;203 | 0 | 0 |
| NC_045512.2;YP_009724390.1;Wu1-SARS2_surface;204 | 0 | 0.025 |
| NC_045512.2;YP_009724390.1;Wu1-SARS2_surface;205 | 0 | 0 |
| NC_045512.2;YP_009724390.1;Wu1-SARS2_surface;206 | 0 | 0 |
| NC_045512.2;YP_009724390.1;Wu1-SARS2_surface;207 | 0 | 0 |
| NC_045512.2;YP_009724390.1;Wu1-SARS2_surface;208 | 0 | 0.05 |
| NC_045512.2;YP_009724390.1;Wu1-SARS2_surface;209 | 0 | 0 |
| NC_045512.2;YP_009724390.1;Wu1-SARS2_surface;210 | 0 | 0 |
| NC_045512.2;YP_009724390.1;Wu1-SARS2_surface;211 | 0 | 0 |
| NC_045512.2;YP_009724390.1;Wu1-SARS2_surface;212 | 0 | 0 |
| NC_045512.2;YP_009724390.1;Wu1-SARS2_surface;213 | 0 | 0 |
| NC_045512.2;YP_009724390.1;Wu1-SARS2_surface;214 | 0 | 0 |
| NC_045512.2;YP_009724390.1;Wu1-SARS2_surface;215 | 0 | 0 |

| | | |
|---|---|---|
| NC_045512.2;YP_009724390.1;Wu1-SARS2_surface;216 | 0 | 0 |
| NC_045512.2;YP_009724390.1;Wu1-SARS2_surface;217 | 0 | 0 |
| NC_045512.2;YP_009724390.1;Wu1-SARS2_surface;218 | 0 | 0 |
| NC_045512.2;YP_009724390.1;Wu1-SARS2_surface;219 | 0 | 0 |
| NC_045512.2;YP_009724390.1;Wu1-SARS2_surface;220 | 0 | 0 |
| NC_045512.2;YP_009724390.1;Wu1-SARS2_surface;221 | 0 | 0 |
| NC_045512.2;YP_009724390.1;Wu1-SARS2_surface;222 | 0 | 0 |
| NC_045512.2;YP_009724390.1;Wu1-SARS2_surface;223 | 0 | 0 |
| NC_045512.2;YP_009724390.1;Wu1-SARS2_surface;224 | 0.05 | 0 |
| NC_045512.2;YP_009724390.1;Wu1-SARS2_surface;225 | 0 | 0.025 |
| NC_045512.2;YP_009724390.1;Wu1-SARS2_surface;226 | 0 | 0 |
| NC_045512.2;YP_009724390.1;Wu1-SARS2_surface;227 | 0 | 0 |
| NC_045512.2;YP_009724390.1;Wu1-SARS2_surface;228 | 0 | 0 |
| NC_045512.2;YP_009724390.1;Wu1-SARS2_surface;229 | 0 | 0 |
| NC_045512.2;YP_009724390.1;Wu1-SARS2_surface;230 | 0 | 0 |
| NC_045512.2;YP_009724390.1;Wu1-SARS2_surface;231 | 0 | 0 |
| NC_045512.2;YP_009724390.1;Wu1-SARS2_surface;232 | 0 | 0 |
| NC_045512.2;YP_009724390.1;Wu1-SARS2_surface;233 | 0 | 0 |
| NC_045512.2;YP_009724390.1;Wu1-SARS2_surface;234 | 0 | 0 |
| NC_045512.2;YP_009724390.1;Wu1-SARS2_surface;235 | 0 | 0 |
| NC_045512.2;YP_009724390.1;Wu1-SARS2_surface;236 | 0 | 0 |
| NC_045512.2;YP_009724390.1;Wu1-SARS2_surface;237 | 0 | 0 |
| NC_045512.2;YP_009724390.1;Wu1-SARS2_surface;238 | 0 | 0 |
| NC_045512.2;YP_009724390.1;Wu1-SARS2_surface;239 | 0 | 0 |
| NC_045512.2;YP_009724390.1;Wu1-SARS2_surface;240 | 0 | 0.025 |
| NC_045512.2;YP_009724390.1;Wu1-SARS2_surface;241 | 0 | 0.025 |
| NC_045512.2;YP_009724390.1;Wu1-SARS2_surface;242 | 0 | 0.075 |
| NC_045512.2;YP_009724390.1;Wu1-SARS2_surface;243 | 0 | 0.025 |
| NC_045512.2;YP_009724390.1;Wu1-SARS2_surface;244 | 0 | 0.025 |
| NC_045512.2;YP_009724390.1;Wu1-SARS2_surface;245 | 0 | 0.05 |
| NC_045512.2;YP_009724390.1;Wu1-SARS2_surface;246 | 0 | 0.1 |
| NC_045512.2;YP_009724390.1;Wu1-SARS2_surface;247 | 0 | 0.1 |
| NC_045512.2;YP_009724390.1;Wu1-SARS2_surface;248 | 0.15 | 0.175 |
| NC_045512.2;YP_009724390.1;Wu1-SARS2_surface;249 | 0.1 | 0.15 |
| NC_045512.2;YP_009724390.1;Wu1-SARS2_surface;250 | 0 | 0 |
| NC_045512.2;YP_009724390.1;Wu1-SARS2_surface;251 | 0 | 0 |
| NC_045512.2;YP_009724390.1;Wu1-SARS2_surface;252 | 0 | 0.025 |
| NC_045512.2;YP_009724390.1;Wu1-SARS2_surface;253 | 0 | 0 |
| NC_045512.2;YP_009724390.1;Wu1-SARS2_surface;254 | 0 | 0 |
| NC_045512.2;YP_009724390.1;Wu1-SARS2_surface;255 | 0 | 0 |
| NC_045512.2;YP_009724390.1;Wu1-SARS2_surface;256 | 0 | 0 |
| NC_045512.2;YP_009724390.1;Wu1-SARS2_surface;257 | 0 | 0 |
| NC_045512.2;YP_009724390.1;Wu1-SARS2_surface;258 | 0 | 0 |

| | | |
|---|---|---|
| NC_045512.2;YP_009724390.1;Wu1-SARS2_surface;259 | 0 | 0 |
| NC_045512.2;YP_009724390.1;Wu1-SARS2_surface;260 | 0 | 0 |
| NC_045512.2;YP_009724390.1;Wu1-SARS2_surface;261 | 0 | 0 |
| NC_045512.2;YP_009724390.1;Wu1-SARS2_surface;262 | 0 | 0 |
| NC_045512.2;YP_009724390.1;Wu1-SARS2_surface;263 | 0 | 0 |
| NC_045512.2;YP_009724390.1;Wu1-SARS2_surface;264 | 0 | 0 |
| NC_045512.2;YP_009724390.1;Wu1-SARS2_surface;265 | 0 | 0 |
| NC_045512.2;YP_009724390.1;Wu1-SARS2_surface;266 | 0 | 0 |
| NC_045512.2;YP_009724390.1;Wu1-SARS2_surface;267 | 0 | 0 |
| NC_045512.2;YP_009724390.1;Wu1-SARS2_surface;268 | 0 | 0 |
| NC_045512.2;YP_009724390.1;Wu1-SARS2_surface;269 | 0 | 0 |
| NC_045512.2;YP_009724390.1;Wu1-SARS2_surface;270 | 0 | 0 |
| NC_045512.2;YP_009724390.1;Wu1-SARS2_surface;271 | 0 | 0 |
| NC_045512.2;YP_009724390.1;Wu1-SARS2_surface;272 | 0.05 | 0 |
| NC_045512.2;YP_009724390.1;Wu1-SARS2_surface;273 | 0.05 | 0 |
| NC_045512.2;YP_009724390.1;Wu1-SARS2_surface;274 | 0.05 | 0 |
| NC_045512.2;YP_009724390.1;Wu1-SARS2_surface;275 | 0 | 0 |
| NC_045512.2;YP_009724390.1;Wu1-SARS2_surface;276 | 0 | 0 |
| NC_045512.2;YP_009724390.1;Wu1-SARS2_surface;277 | 0.05 | 0 |
| NC_045512.2;YP_009724390.1;Wu1-SARS2_surface;278 | 0.05 | 0 |
| NC_045512.2;YP_009724390.1;Wu1-SARS2_surface;279 | 0 | 0 |
| NC_045512.2;YP_009724390.1;Wu1-SARS2_surface;280 | 0 | 0 |
| NC_045512.2;YP_009724390.1;Wu1-SARS2_surface;281 | 0 | 0 |
| NC_045512.2;YP_009724390.1;Wu1-SARS2_surface;282 | 0 | 0 |
| NC_045512.2;YP_009724390.1;Wu1-SARS2_surface;283 | 0 | 0 |
| NC_045512.2;YP_009724390.1;Wu1-SARS2_surface;284 | 0 | 0 |
| NC_045512.2;YP_009724390.1;Wu1-SARS2_surface;285 | 0 | 0 |
| NC_045512.2;YP_009724390.1;Wu1-SARS2_surface;286 | 0 | 0 |
| NC_045512.2;YP_009724390.1;Wu1-SARS2_surface;287 | 0 | 0 |
| NC_045512.2;YP_009724390.1;Wu1-SARS2_surface;288 | 0 | 0 |
| NC_045512.2;YP_009724390.1;Wu1-SARS2_surface;289 | 0 | 0 |
| NC_045512.2;YP_009724390.1;Wu1-SARS2_surface;290 | 0 | 0 |
| NC_045512.2;YP_009724390.1;Wu1-SARS2_surface;291 | 0 | 0 |
| NC_045512.2;YP_009724390.1;Wu1-SARS2_surface;292 | 0 | 0 |
| NC_045512.2;YP_009724390.1;Wu1-SARS2_surface;293 | 0 | 0 |
| NC_045512.2;YP_009724390.1;Wu1-SARS2_surface;294 | 0 | 0 |
| NC_045512.2;YP_009724390.1;Wu1-SARS2_surface;295 | 0.05 | 0 |
| NC_045512.2;YP_009724390.1;Wu1-SARS2_surface;296 | 0.1 | 0 |
| NC_045512.2;YP_009724390.1;Wu1-SARS2_surface;297 | 0 | 0 |
| NC_045512.2;YP_009724390.1;Wu1-SARS2_surface;298 | 0 | 0.025 |
| NC_045512.2;YP_009724390.1;Wu1-SARS2_surface;299 | 0 | 0 |
| NC_045512.2;YP_009724390.1;Wu1-SARS2_surface;300 | 0 | 0 |
| NC_045512.2;YP_009724390.1;Wu1-SARS2_surface;301 | 0 | 0 |

| | | |
|---|---|---|
| NC_045512.2;YP_009724390.1;Wu1-SARS2_surface;302 | 0.05 | 0.075 |
| NC_045512.2;YP_009724390.1;Wu1-SARS2_surface;303 | 0 | 0.1 |
| NC_045512.2;YP_009724390.1;Wu1-SARS2_surface;304 | 0 | 0.05 |
| NC_045512.2;YP_009724390.1;Wu1-SARS2_surface;305 | 0 | 0.05 |
| NC_045512.2;YP_009724390.1;Wu1-SARS2_surface;306 | 0 | 0.05 |
| NC_045512.2;YP_009724390.1;Wu1-SARS2_surface;307 | 0 | 0.1 |
| NC_045512.2;YP_009724390.1;Wu1-SARS2_surface;308 | 0.05 | 0.15 |
| NC_045512.2;YP_009724390.1;Wu1-SARS2_surface;309 | 0.1 | 0.25 |
| NC_045512.2;YP_009724390.1;Wu1-SARS2_surface;310 | 0.05 | 0.275 |
| NC_045512.2;YP_009724390.1;Wu1-SARS2_surface;311 | 0.1 | 0.025 |
| NC_045512.2;YP_009724390.1;Wu1-SARS2_surface;312 | 0.05 | 0.025 |
| NC_045512.2;YP_009724390.1;Wu1-SARS2_surface;313 | 0 | 0.025 |
| NC_045512.2;YP_009724390.1;Wu1-SARS2_surface;314 | 0 | 0.025 |
| NC_045512.2;YP_009724390.1;Wu1-SARS2_surface;315 | 0 | 0.025 |
| NC_045512.2;YP_009724390.1;Wu1-SARS2_surface;316 | 0 | 0.025 |
| NC_045512.2;YP_009724390.1;Wu1-SARS2_surface;317 | 0 | 0 |
| NC_045512.2;YP_009724390.1;Wu1-SARS2_surface;318 | 0 | 0 |
| NC_045512.2;YP_009724390.1;Wu1-SARS2_surface;319 | 0 | 0.025 |
| NC_045512.2;YP_009724390.1;Wu1-SARS2_surface;320 | 0.05 | 0 |
| NC_045512.2;YP_009724390.1;Wu1-SARS2_surface;321 | 0 | 0 |
| NC_045512.2;YP_009724390.1;Wu1-SARS2_surface;322 | 0 | 0 |
| NC_045512.2;YP_009724390.1;Wu1-SARS2_surface;323 | 0 | 0 |
| NC_045512.2;YP_009724390.1;Wu1-SARS2_surface;324 | 0.05 | 0 |
| NC_045512.2;YP_009724390.1;Wu1-SARS2_surface;325 | 0 | 0 |
| NC_045512.2;YP_009724390.1;Wu1-SARS2_surface;326 | 0 | 0 |
| NC_045512.2;YP_009724390.1;Wu1-SARS2_surface;327 | 0 | 0 |
| NC_045512.2;YP_009724390.1;Wu1-SARS2_surface;328 | 0 | 0 |
| NC_045512.2;YP_009724390.1;Wu1-SARS2_surface;329 | 0.05 | 0 |
| NC_045512.2;YP_009724390.1;Wu1-SARS2_surface;330 | 0.05 | 0.025 |
| NC_045512.2;YP_009724390.1;Wu1-SARS2_surface;331 | 0 | 0 |
| NC_045512.2;YP_009724390.1;Wu1-SARS2_surface;332 | 0 | 0 |
| NC_045512.2;YP_009724390.1;Wu1-SARS2_surface;333 | 0 | 0 |
| NC_045512.2;YP_009724390.1;Wu1-SARS2_surface;334 | 0.05 | 0 |
| NC_045512.2;YP_009724390.1;Wu1-SARS2_surface;335 | 0.05 | 0 |
| NC_045512.2;YP_009724390.1;Wu1-SARS2_surface;336 | 0 | 0 |
| NC_045512.2;YP_009724390.1;Wu1-SARS2_surface;337 | 0 | 0 |
| NC_045512.2;YP_009724390.1;Wu1-SARS2_surface;338 | 0 | 0 |
| NC_045512.2;YP_009724390.1;Wu1-SARS2_surface;339 | 0 | 0 |
| NC_045512.2;YP_009724390.1;Wu1-SARS2_surface;340 | 0 | 0 |
| NC_045512.2;YP_009724390.1;Wu1-SARS2_surface;341 | 0 | 0 |
| NC_045512.2;YP_009724390.1;Wu1-SARS2_surface;342 | 0 | 0 |
| NC_045512.2;YP_009724390.1;Wu1-SARS2_surface;343 | 0 | 0 |
| NC_045512.2;YP_009724390.1;Wu1-SARS2_surface;344 | 0 | 0 |

| | | |
|---|---|---|
| NC_045512.2;YP_009724390.1;Wu1-SARS2_surface;345 | 0 | 0 |
| NC_045512.2;YP_009724390.1;Wu1-SARS2_surface;346 | 0 | 0 |
| NC_045512.2;YP_009724390.1;Wu1-SARS2_surface;347 | 0 | 0.025 |
| NC_045512.2;YP_009724390.1;Wu1-SARS2_surface;348 | 0 | 0.025 |
| NC_045512.2;YP_009724390.1;Wu1-SARS2_surface;349 | 0.05 | 0.025 |
| NC_045512.2;YP_009724390.1;Wu1-SARS2_surface;350 | 0 | 0.025 |
| NC_045512.2;YP_009724390.1;Wu1-SARS2_surface;351 | 0 | 0.025 |
| NC_045512.2;YP_009724390.1;Wu1-SARS2_surface;352 | 0 | 0 |
| NC_045512.2;YP_009724390.1;Wu1-SARS2_surface;353 | 0 | 0 |
| NC_045512.2;YP_009724390.1;Wu1-SARS2_surface;354 | 0 | 0 |
| NC_045512.2;YP_009724390.1;Wu1-SARS2_surface;355 | 0 | 0 |
| NC_045512.2;YP_009724390.1;Wu1-SARS2_surface;356 | 0 | 0.025 |
| NC_045512.2;YP_009724390.1;Wu1-SARS2_surface;357 | 0 | 0 |
| NC_045512.2;YP_009724390.1;Wu1-SARS2_surface;358 | 0 | 0.05 |
| NC_045512.2;YP_009724390.1;Wu1-SARS2_surface;359 | 0 | 0.025 |
| NC_045512.2;YP_009724390.1;Wu1-SARS2_surface;360 | 0 | 0 |
| NC_045512.2;YP_009724390.1;Wu1-SARS2_surface;361 | 0 | 0 |
| NC_045512.2;YP_009724390.1;Wu1-SARS2_surface;362 | 0 | 0 |
| NC_045512.2;YP_009724390.1;Wu1-SARS2_surface;363 | 0.05 | 0 |
| NC_045512.2;YP_009724390.1;Wu1-SARS2_surface;364 | 0 | 0 |
| NC_045512.2;YP_009724390.1;Wu1-SARS2_surface;365 | 0 | 0 |
| NC_045512.2;YP_009724390.1;Wu1-SARS2_surface;366 | 0 | 0 |
| NC_045512.2;YP_009724390.1;Wu1-SARS2_surface;367 | 0 | 0 |
| NC_045512.2;YP_009724390.1;Wu1-SARS2_surface;368 | 0 | 0 |
| NC_045512.2;YP_009724390.1;Wu1-SARS2_surface;369 | 0.05 | 0 |
| NC_045512.2;YP_009724390.1;Wu1-SARS2_surface;370 | 0 | 0 |
| NC_045512.2;YP_009724390.1;Wu1-SARS2_surface;371 | 0 | 0 |
| NC_045512.2;YP_009724390.1;Wu1-SARS2_surface;372 | 0 | 0 |
| NC_045512.2;YP_009724390.1;Wu1-SARS2_surface;373 | 0 | 0 |
| NC_045512.2;YP_009724390.1;Wu1-SARS2_surface;374 | 0.05 | 0 |
| NC_045512.2;YP_009724390.1;Wu1-SARS2_surface;375 | 0.05 | 0 |
| NC_045512.2;YP_009724390.1;Wu1-SARS2_surface;376 | 0 | 0 |
| NC_045512.2;YP_009724390.1;Wu1-SARS2_surface;377 | 0 | 0 |
| NC_045512.2;YP_009724390.1;Wu1-SARS2_surface;378 | 0 | 0 |
| NC_045512.2;YP_009724390.1;Wu1-SARS2_surface;379 | 0 | 0 |
| NC_045512.2;YP_009724390.1;Wu1-SARS2_surface;380 | 0 | 0 |
| NC_045512.2;YP_009724390.1;Wu1-SARS2_surface;381 | 0 | 0 |
| NC_045512.2;YP_009724390.1;Wu1-SARS2_surface;382 | 0 | 0 |
| NC_045512.2;YP_009724390.1;Wu1-SARS2_surface;383 | 0 | 0.025 |
| NC_045512.2;YP_009724390.1;Wu1-SARS2_surface;384 | 0 | 0 |
| NC_045512.2;YP_009724390.1;Wu1-SARS2_surface;385 | 0 | 0.025 |
| NC_045512.2;YP_009724390.1;Wu1-SARS2_surface;386 | 0 | 0 |
| NC_045512.2;YP_009724390.1;Wu1-SARS2_surface;387 | 0 | 0 |

| | | |
|---|---|---|
| NC_045512.2;YP_009724390.1;Wu1-SARS2_surface;388 | 0 | 0 |
| NC_045512.2;YP_009724390.1;Wu1-SARS2_surface;389 | 0 | 0 |
| NC_045512.2;YP_009724390.1;Wu1-SARS2_surface;390 | 0 | 0 |
| NC_045512.2;YP_009724390.1;Wu1-SARS2_surface;391 | 0 | 0 |
| NC_045512.2;YP_009724390.1;Wu1-SARS2_surface;392 | 0 | 0.025 |
| NC_045512.2;YP_009724390.1;Wu1-SARS2_surface;393 | 0.05 | 0 |
| NC_045512.2;YP_009724390.1;Wu1-SARS2_surface;394 | 0 | 0 |
| NC_045512.2;YP_009724390.1;Wu1-SARS2_surface;395 | 0 | 0.025 |
| NC_045512.2;YP_009724390.1;Wu1-SARS2_surface;396 | 0 | 0 |
| NC_045512.2;YP_009724390.1;Wu1-SARS2_surface;397 | 0.05 | 0.025 |
| NC_045512.2;YP_009724390.1;Wu1-SARS2_surface;398 | 0.15 | 0.025 |
| NC_045512.2;YP_009724390.1;Wu1-SARS2_surface;399 | 0 | 0 |
| NC_045512.2;YP_009724390.1;Wu1-SARS2_surface;400 | 0 | 0 |
| NC_045512.2;YP_009724390.1;Wu1-SARS2_surface;401 | 0 | 0 |
| NC_045512.2;YP_009724390.1;Wu1-SARS2_surface;402 | 0 | 0.05 |
| NC_045512.2;YP_009724390.1;Wu1-SARS2_surface;403 | 0 | 0.05 |
| NC_045512.2;YP_009724390.1;Wu1-SARS2_surface;404 | 0 | 0.05 |
| NC_045512.2;YP_009724390.1;Wu1-SARS2_surface;405 | 0 | 0.05 |
| NC_045512.2;YP_009724390.1;Wu1-SARS2_surface;406 | 0 | 0.075 |
| NC_045512.2;YP_009724390.1;Wu1-SARS2_surface;407 | 0 | 0.075 |
| NC_045512.2;YP_009724390.1;Wu1-SARS2_surface;408 | 0.05 | 0.1 |
| NC_045512.2;YP_009724390.1;Wu1-SARS2_surface;409 | 0.15 | 0.125 |
| NC_045512.2;YP_009724390.1;Wu1-SARS2_surface;410 | 0 | 0.05 |
| NC_045512.2;YP_009724390.1;Wu1-SARS2_surface;411 | 0 | 0.05 |
| NC_045512.2;YP_009724390.1;Wu1-SARS2_surface;412 | 0.05 | 0.1 |
| NC_045512.2;YP_009724390.1;Wu1-SARS2_surface;413 | 0.15 | 0.075 |
| NC_045512.2;YP_009724390.1;Wu1-SARS2_surface;414 | 0.2 | 0.125 |
| NC_045512.2;YP_009724390.1;Wu1-SARS2_surface;415 | 0.1 | 0.125 |
| NC_045512.2;YP_009724390.1;Wu1-SARS2_surface;416 | 0.25 | 0.25 |
| NC_045512.2;YP_009724390.1;Wu1-SARS2_surface;417 | 0.1 | 0.05 |
| NC_045512.2;YP_009724390.1;Wu1-SARS2_surface;418 | 0 | 0.025 |
| NC_045512.2;YP_009724390.1;Wu1-SARS2_surface;419 | 0.05 | 0.025 |
| NC_045512.2;YP_009724390.1;Wu1-SARS2_surface;420 | 0 | 0.025 |
| NC_045512.2;YP_009724390.1;Wu1-SARS2_surface;421 | 0 | 0 |
| NC_045512.2;YP_009724390.1;Wu1-SARS2_surface;422 | 0 | 0 |
| NC_045512.2;YP_009724390.1;Wu1-SARS2_surface;423 | 0 | 0 |
| NC_045512.2;YP_009724390.1;Wu1-SARS2_surface;424 | 0 | 0 |
| NC_045512.2;YP_009724390.1;Wu1-SARS2_surface;425 | 0 | 0 |
| NC_045512.2;YP_009724390.1;Wu1-SARS2_surface;426 | 0 | 0 |
| NC_045512.2;YP_009724390.1;Wu1-SARS2_surface;427 | 0 | 0 |
| NC_045512.2;YP_009724390.1;Wu1-SARS2_surface;428 | 0 | 0 |
| NC_045512.2;YP_009724390.1;Wu1-SARS2_surface;429 | 0 | 0 |
| NC_045512.2;YP_009724390.1;Wu1-SARS2_surface;430 | 0.05 | 0 |

| | | |
|---|---|---|
| NC_045512.2;YP_009724390.1;Wu1-SARS2_surface;431 | 0.05 | 0 |
| NC_045512.2;YP_009724390.1;Wu1-SARS2_surface;432 | 0.15 | 0.025 |
| NC_045512.2;YP_009724390.1;Wu1-SARS2_surface;433 | 0.3 | 0.025 |
| NC_045512.2;YP_009724390.1;Wu1-SARS2_surface;434 | 0.2 | 0.125 |
| NC_045512.2;YP_009724390.1;Wu1-SARS2_surface;435 | 0.3 | 0.275 |
| NC_045512.2;YP_009724390.1;Wu1-SARS2_surface;436 | 0.3 | 0.225 |
| NC_045512.2;YP_009724390.1;Wu1-SARS2_surface;437 | 0.05 | 0.125 |
| NC_045512.2;YP_009724390.1;Wu1-SARS2_surface;438 | 0.05 | 0.025 |
| NC_045512.2;YP_009724390.1;Wu1-SARS2_surface;439 | 0 | 0.025 |
| NC_045512.2;YP_009724390.1;Wu1-SARS2_surface;440 | 0 | 0.025 |
| NC_045512.2;YP_009724390.1;Wu1-SARS2_surface;441 | 0 | 0.025 |
| NC_045512.2;YP_009724390.1;Wu1-SARS2_surface;442 | 0 | 0 |
| NC_045512.2;YP_009724390.1;Wu1-SARS2_surface;443 | 0 | 0 |
| NC_045512.2;YP_009724390.1;Wu1-SARS2_surface;444 | 0 | 0 |
| NC_045512.2;YP_009724390.1;Wu1-SARS2_surface;445 | 0 | 0 |
| NC_045512.2;YP_009724390.1;Wu1-SARS2_surface;446 | 0 | 0 |
| NC_045512.2;YP_009724390.1;Wu1-SARS2_surface;447 | 0 | 0 |
| NC_045512.2;YP_009724390.1;Wu1-SARS2_surface;448 | 0 | 0 |
| NC_045512.2;YP_009724390.1;Wu1-SARS2_surface;449 | 0 | 0 |
| NC_045512.2;YP_009724390.1;Wu1-SARS2_surface;450 | 0 | 0 |
| NC_045512.2;YP_009724390.1;Wu1-SARS2_surface;451 | 0 | 0 |
| NC_045512.2;YP_009724390.1;Wu1-SARS2_surface;452 | 0.05 | 0.025 |
| NC_045512.2;YP_009724390.1;Wu1-SARS2_surface;453 | 0.05 | 0.025 |
| NC_045512.2;YP_009724390.1;Wu1-SARS2_surface;454 | 0.05 | 0.025 |
| NC_045512.2;YP_009724390.1;Wu1-SARS2_surface;455 | 0.1 | 0 |
| NC_045512.2;YP_009724390.1;Wu1-SARS2_surface;456 | 0.1 | 0.025 |
| NC_045512.2;YP_009724390.1;Wu1-SARS2_surface;457 | 0.1 | 0.075 |
| NC_045512.2;YP_009724390.1;Wu1-SARS2_surface;458 | 0.05 | 0.05 |
| NC_045512.2;YP_009724390.1;Wu1-SARS2_surface;459 | 0 | 0 |
| NC_045512.2;YP_009724390.1;Wu1-SARS2_surface;460 | 0 | 0 |
| NC_045512.2;YP_009724390.1;Wu1-SARS2_surface;461 | 0 | 0 |
| NC_045512.2;YP_009724390.1;Wu1-SARS2_surface;462 | 0.05 | 0 |
| NC_045512.2;YP_009724390.1;Wu1-SARS2_surface;463 | 0 | 0 |
| NC_045512.2;YP_009724390.1;Wu1-SARS2_surface;464 | 0 | 0 |
| NC_045512.2;YP_009724390.1;Wu1-SARS2_surface;465 | 0.05 | 0 |
| NC_045512.2;YP_009724390.1;Wu1-SARS2_surface;466 | 0 | 0 |
| NC_045512.2;YP_009724390.1;Wu1-SARS2_surface;467 | 0 | 0 |
| NC_045512.2;YP_009724390.1;Wu1-SARS2_surface;468 | 0 | 0 |
| NC_045512.2;YP_009724390.1;Wu1-SARS2_surface;469 | 0 | 0 |
| NC_045512.2;YP_009724390.1;Wu1-SARS2_surface;470 | 0 | 0 |
| NC_045512.2;YP_009724390.1;Wu1-SARS2_surface;471 | 0 | 0 |
| NC_045512.2;YP_009724390.1;Wu1-SARS2_surface;472 | 0 | 0 |
| NC_045512.2;YP_009724390.1;Wu1-SARS2_surface;473 | 0 | 0 |

| | | |
|---|---|---|
| NC_045512.2;YP_009724390.1;Wu1-SARS2_surface;474 | 0 | 0 |
| NC_045512.2;YP_009724390.1;Wu1-SARS2_surface;475 | 0 | 0 |
| NC_045512.2;YP_009724390.1;Wu1-SARS2_surface;476 | 0 | 0 |
| NC_045512.2;YP_009724390.1;Wu1-SARS2_surface;477 | 0 | 0 |
| NC_045512.2;YP_009724390.1;Wu1-SARS2_surface;478 | 0 | 0 |
| NC_045512.2;YP_009724390.1;Wu1-SARS2_surface;479 | 0 | 0 |
| NC_045512.2;YP_009724390.1;Wu1-SARS2_surface;480 | 0 | 0 |
| NC_045512.2;YP_009724390.1;Wu1-SARS2_surface;481 | 0.05 | 0.075 |
| NC_045512.2;YP_009724390.1;Wu1-SARS2_surface;482 | 0 | 0.025 |
| NC_045512.2;YP_009724390.1;Wu1-SARS2_surface;483 | 0 | 0 |
| NC_045512.2;YP_009724390.1;Wu1-SARS2_surface;484 | 0 | 0 |
| NC_045512.2;YP_009724390.1;Wu1-SARS2_surface;485 | 0 | 0 |
| NC_045512.2;YP_009724390.1;Wu1-SARS2_surface;486 | 0 | 0 |
| NC_045512.2;YP_009724390.1;Wu1-SARS2_surface;487 | 0 | 0.025 |
| NC_045512.2;YP_009724390.1;Wu1-SARS2_surface;488 | 0 | 0.025 |
| NC_045512.2;YP_009724390.1;Wu1-SARS2_surface;489 | 0.05 | 0.05 |
| NC_045512.2;YP_009724390.1;Wu1-SARS2_surface;490 | 0 | 0.05 |
| NC_045512.2;YP_009724390.1;Wu1-SARS2_surface;491 | 0 | 0.075 |
| NC_045512.2;YP_009724390.1;Wu1-SARS2_surface;492 | 0 | 0.075 |
| NC_045512.2;YP_009724390.1;Wu1-SARS2_surface;493 | 0 | 0.05 |
| NC_045512.2;YP_009724390.1;Wu1-SARS2_surface;494 | 0 | 0.025 |
| NC_045512.2;YP_009724390.1;Wu1-SARS2_surface;495 | 0 | 0 |
| NC_045512.2;YP_009724390.1;Wu1-SARS2_surface;496 | 0 | 0 |
| NC_045512.2;YP_009724390.1;Wu1-SARS2_surface;497 | 0 | 0 |
| NC_045512.2;YP_009724390.1;Wu1-SARS2_surface;498 | 0 | 0 |
| NC_045512.2;YP_009724390.1;Wu1-SARS2_surface;499 | 0 | 0 |
| NC_045512.2;YP_009724390.1;Wu1-SARS2_surface;500 | 0 | 0 |
| NC_045512.2;YP_009724390.1;Wu1-SARS2_surface;501 | 0 | 0 |
| NC_045512.2;YP_009724390.1;Wu1-SARS2_surface;502 | 0 | 0 |
| NC_045512.2;YP_009724390.1;Wu1-SARS2_surface;503 | 0 | 0 |
| NC_045512.2;YP_009724390.1;Wu1-SARS2_surface;504 | 0 | 0 |
| NC_045512.2;YP_009724390.1;Wu1-SARS2_surface;505 | 0 | 0 |
| NC_045512.2;YP_009724390.1;Wu1-SARS2_surface;506 | 0 | 0 |
| NC_045512.2;YP_009724390.1;Wu1-SARS2_surface;507 | 0 | 0 |
| NC_045512.2;YP_009724390.1;Wu1-SARS2_surface;508 | 0 | 0 |
| NC_045512.2;YP_009724390.1;Wu1-SARS2_surface;509 | 0 | 0 |
| NC_045512.2;YP_009724390.1;Wu1-SARS2_surface;510 | 0 | 0 |
| NC_045512.2;YP_009724390.1;Wu1-SARS2_surface;511 | 0 | 0 |
| NC_045512.2;YP_009724390.1;Wu1-SARS2_surface;512 | 0 | 0 |
| NC_045512.2;YP_009724390.1;Wu1-SARS2_surface;513 | 0 | 0 |
| NC_045512.2;YP_009724390.1;Wu1-SARS2_surface;514 | 0 | 0 |
| NC_045512.2;YP_009724390.1;Wu1-SARS2_surface;515 | 0 | 0 |
| NC_045512.2;YP_009724390.1;Wu1-SARS2_surface;516 | 0 | 0 |

| | | |
|---|---|---|
| NC_045512.2;YP_009724390.1;Wu1-SARS2_surface;517 | 0 | 0 |
| NC_045512.2;YP_009724390.1;Wu1-SARS2_surface;518 | 0.05 | 0 |
| NC_045512.2;YP_009724390.1;Wu1-SARS2_surface;519 | 0 | 0 |
| NC_045512.2;YP_009724390.1;Wu1-SARS2_surface;520 | 0 | 0.025 |
| NC_045512.2;YP_009724390.1;Wu1-SARS2_surface;521 | 0 | 0.025 |
| NC_045512.2;YP_009724390.1;Wu1-SARS2_surface;522 | 0 | 0.025 |
| NC_045512.2;YP_009724390.1;Wu1-SARS2_surface;523 | 0 | 0.025 |
| NC_045512.2;YP_009724390.1;Wu1-SARS2_surface;524 | 0 | 0.025 |
| NC_045512.2;YP_009724390.1;Wu1-SARS2_surface;525 | 0 | 0.025 |
| NC_045512.2;YP_009724390.1;Wu1-SARS2_surface;526 | 0 | 0 |
| NC_045512.2;YP_009724390.1;Wu1-SARS2_surface;527 | 0 | 0.025 |
| NC_045512.2;YP_009724390.1;Wu1-SARS2_surface;528 | 0 | 0.075 |
| NC_045512.2;YP_009724390.1;Wu1-SARS2_surface;529 | 0 | 0.05 |
| NC_045512.2;YP_009724390.1;Wu1-SARS2_surface;530 | 0 | 0.075 |
| NC_045512.2;YP_009724390.1;Wu1-SARS2_surface;531 | 0 | 0 |
| NC_045512.2;YP_009724390.1;Wu1-SARS2_surface;532 | 0 | 0.025 |
| NC_045512.2;YP_009724390.1;Wu1-SARS2_surface;533 | 0 | 0.025 |
| NC_045512.2;YP_009724390.1;Wu1-SARS2_surface;534 | 0 | 0.05 |
| NC_045512.2;YP_009724390.1;Wu1-SARS2_surface;535 | 0 | 0.05 |
| NC_045512.2;YP_009724390.1;Wu1-SARS2_surface;536 | 0 | 0.025 |
| NC_045512.2;YP_009724390.1;Wu1-SARS2_surface;537 | 0 | 0.025 |
| NC_045512.2;YP_009724390.1;Wu1-SARS2_surface;538 | 0 | 0 |
| NC_045512.2;YP_009724390.1;Wu1-SARS2_surface;539 | 0 | 0 |
| NC_045512.2;YP_009724390.1;Wu1-SARS2_surface;540 | 0 | 0 |
| NC_045512.2;YP_009724390.1;Wu1-SARS2_surface;541 | 0 | 0.025 |
| NC_045512.2;YP_009724390.1;Wu1-SARS2_surface;542 | 0.05 | 0.125 |
| NC_045512.2;YP_009724390.1;Wu1-SARS2_surface;543 | 0 | 0.125 |
| NC_045512.2;YP_009724390.1;Wu1-SARS2_surface;544 | 0 | 0.125 |
| NC_045512.2;YP_009724390.1;Wu1-SARS2_surface;545 | 0 | 0.1 |
| NC_045512.2;YP_009724390.1;Wu1-SARS2_surface;546 | 0.05 | 0.125 |
| NC_045512.2;YP_009724390.1;Wu1-SARS2_surface;547 | 0 | 0.1 |
| NC_045512.2;YP_009724390.1;Wu1-SARS2_surface;548 | 0.05 | 0.125 |
| NC_045512.2;YP_009724390.1;Wu1-SARS2_surface;549 | 0 | 0.5 |
| NC_045512.2;YP_009724390.1;Wu1-SARS2_surface;550 | 0.25 | 0.475 |
| NC_045512.2;YP_009724390.1;Wu1-SARS2_surface;551 | 0.15 | 0.5 |
| NC_045512.2;YP_009724390.1;Wu1-SARS2_surface;552 | 0.5 | 0.75 |
| NC_045512.2;YP_009724390.1;Wu1-SARS2_surface;553 | 0.05 | 0.7 |
| NC_045512.2;YP_009724390.1;Wu1-SARS2_surface;554 | 0.1 | 0.775 |
| NC_045512.2;YP_009724390.1;Wu1-SARS2_surface;555 | 0.05 | 0.6 |
| NC_045512.2;YP_009724390.1;Wu1-SARS2_surface;556 | 0 | 0.7 |
| NC_045512.2;YP_009724390.1;Wu1-SARS2_surface;557 | 0 | 0.75 |
| NC_045512.2;YP_009724390.1;Wu1-SARS2_surface;558 | 0 | 0.725 |
| NC_045512.2;YP_009724390.1;Wu1-SARS2_surface;559 | 0 | 0.525 |

| | | |
|---|---|---|
| NC_045512.2;YP_009724390.1;Wu1-SARS2_surface;560 | 0 | 0.525 |
| NC_045512.2;YP_009724390.1;Wu1-SARS2_surface;561 | 0 | 0.275 |
| NC_045512.2;YP_009724390.1;Wu1-SARS2_surface;562 | 0 | 0.25 |
| NC_045512.2;YP_009724390.1;Wu1-SARS2_surface;563 | 0 | 0.025 |
| NC_045512.2;YP_009724390.1;Wu1-SARS2_surface;564 | 0.05 | 0.05 |
| NC_045512.2;YP_009724390.1;Wu1-SARS2_surface;565 | 0.05 | 0.075 |
| NC_045512.2;YP_009724390.1;Wu1-SARS2_surface;566 | 0 | 0.025 |
| NC_045512.2;YP_009724390.1;Wu1-SARS2_surface;567 | 0 | 0.05 |
| NC_045512.2;YP_009724390.1;Wu1-SARS2_surface;568 | 0 | 0.375 |
| NC_045512.2;YP_009724390.1;Wu1-SARS2_surface;569 | 0.05 | 0.6 |
| NC_045512.2;YP_009724390.1;Wu1-SARS2_surface;570 | 0 | 0.4 |
| NC_045512.2;YP_009724390.1;Wu1-SARS2_surface;571 | 0 | 0.675 |
| NC_045512.2;YP_009724390.1;Wu1-SARS2_surface;572 | 0 | 0.625 |
| NC_045512.2;YP_009724390.1;Wu1-SARS2_surface;573 | 0 | 0.475 |
| NC_045512.2;YP_009724390.1;Wu1-SARS2_surface;574 | 0 | 0.55 |
| NC_045512.2;YP_009724390.1;Wu1-SARS2_surface;575 | 0 | 0.625 |
| NC_045512.2;YP_009724390.1;Wu1-SARS2_surface;576 | 0 | 0.7 |
| NC_045512.2;YP_009724390.1;Wu1-SARS2_surface;577 | 0 | 0.325 |
| NC_045512.2;YP_009724390.1;Wu1-SARS2_surface;578 | 0 | 0 |
| NC_045512.2;YP_009724390.1;Wu1-SARS2_surface;579 | 0 | 0 |
| NC_045512.2;YP_009724390.1;Wu1-SARS2_surface;580 | 0 | 0 |
| NC_045512.2;YP_009724390.1;Wu1-SARS2_surface;581 | 0 | 0 |
| NC_045512.2;YP_009724390.1;Wu1-SARS2_surface;582 | 0 | 0.05 |
| NC_045512.2;YP_009724390.1;Wu1-SARS2_surface;583 | 0 | 0 |
| NC_045512.2;YP_009724390.1;Wu1-SARS2_surface;584 | 0 | 0 |
| NC_045512.2;YP_009724390.1;Wu1-SARS2_surface;585 | 0 | 0 |
| NC_045512.2;YP_009724390.1;Wu1-SARS2_surface;586 | 0 | 0 |
| NC_045512.2;YP_009724390.1;Wu1-SARS2_surface;587 | 0 | 0 |
| NC_045512.2;YP_009724390.1;Wu1-SARS2_surface;588 | 0 | 0 |
| NC_045512.2;YP_009724390.1;Wu1-SARS2_surface;589 | 0 | 0.025 |
| NC_045512.2;YP_009724390.1;Wu1-SARS2_surface;590 | 0.1 | 0.075 |
| NC_045512.2;YP_009724390.1;Wu1-SARS2_surface;591 | 0.25 | 0.125 |
| NC_045512.2;YP_009724390.1;Wu1-SARS2_surface;592 | 0.25 | 0.125 |
| NC_045512.2;YP_009724390.1;Wu1-SARS2_surface;593 | 0.4 | 0.25 |
| NC_045512.2;YP_009724390.1;Wu1-SARS2_surface;594 | 0.4 | 0.35 |
| NC_045512.2;YP_009724390.1;Wu1-SARS2_surface;595 | 0.4 | 0.375 |
| NC_045512.2;YP_009724390.1;Wu1-SARS2_surface;596 | 0.35 | 0.175 |
| NC_045512.2;YP_009724390.1;Wu1-SARS2_surface;597 | 0.15 | 0.075 |
| NC_045512.2;YP_009724390.1;Wu1-SARS2_surface;598 | 0.2 | 0.075 |
| NC_045512.2;YP_009724390.1;Wu1-SARS2_surface;599 | 0.2 | 0.15 |
| NC_045512.2;YP_009724390.1;Wu1-SARS2_surface;600 | 0.15 | 0.1 |
| NC_045512.2;YP_009724390.1;Wu1-SARS2_surface;601 | 0.25 | 0.175 |
| NC_045512.2;YP_009724390.1;Wu1-SARS2_surface;602 | 0.1 | 0.075 |

| | | |
|---|---|---|
| NC_045512.2;YP_009724390.1;Wu1-SARS2_surface;603 | 0 | 0.025 |
| NC_045512.2;YP_009724390.1;Wu1-SARS2_surface;604 | 0 | 0.025 |
| NC_045512.2;YP_009724390.1;Wu1-SARS2_surface;605 | 0 | 0.025 |
| NC_045512.2;YP_009724390.1;Wu1-SARS2_surface;606 | 0 | 0.025 |
| NC_045512.2;YP_009724390.1;Wu1-SARS2_surface;607 | 0 | 0.025 |
| NC_045512.2;YP_009724390.1;Wu1-SARS2_surface;608 | 0 | 0 |
| NC_045512.2;YP_009724390.1;Wu1-SARS2_surface;609 | 0 | 0 |
| NC_045512.2;YP_009724390.1;Wu1-SARS2_surface;610 | 0 | 0 |
| NC_045512.2;YP_009724390.1;Wu1-SARS2_surface;611 | 0 | 0 |
| NC_045512.2;YP_009724390.1;Wu1-SARS2_surface;612 | 0 | 0.075 |
| NC_045512.2;YP_009724390.1;Wu1-SARS2_surface;613 | 0 | 0.075 |
| NC_045512.2;YP_009724390.1;Wu1-SARS2_surface;614 | 0 | 0.1 |
| NC_045512.2;YP_009724390.1;Wu1-SARS2_surface;615 | 0 | 0.125 |
| NC_045512.2;YP_009724390.1;Wu1-SARS2_surface;616 | 0 | 0.175 |
| NC_045512.2;YP_009724390.1;Wu1-SARS2_surface;617 | 0 | 0.225 |
| NC_045512.2;YP_009724390.1;Wu1-SARS2_surface;618 | 0 | 0.125 |
| NC_045512.2;YP_009724390.1;Wu1-SARS2_surface;619 | 0 | 0.075 |
| NC_045512.2;YP_009724390.1;Wu1-SARS2_surface;620 | 0 | 0.05 |
| NC_045512.2;YP_009724390.1;Wu1-SARS2_surface;621 | 0 | 0.05 |
| NC_045512.2;YP_009724390.1;Wu1-SARS2_surface;622 | 0 | 0.075 |
| NC_045512.2;YP_009724390.1;Wu1-SARS2_surface;623 | 0.05 | 0.05 |
| NC_045512.2;YP_009724390.1;Wu1-SARS2_surface;624 | 0 | 0.025 |
| NC_045512.2;YP_009724390.1;Wu1-SARS2_surface;625 | 0 | 0.05 |
| NC_045512.2;YP_009724390.1;Wu1-SARS2_surface;626 | 0 | 0.075 |
| NC_045512.2;YP_009724390.1;Wu1-SARS2_surface;627 | 0 | 0.175 |
| NC_045512.2;YP_009724390.1;Wu1-SARS2_surface;628 | 0 | 0.05 |
| NC_045512.2;YP_009724390.1;Wu1-SARS2_surface;629 | 0 | 0.05 |
| NC_045512.2;YP_009724390.1;Wu1-SARS2_surface;630 | 0 | 0.05 |
| NC_045512.2;YP_009724390.1;Wu1-SARS2_surface;631 | 0 | 0 |
| NC_045512.2;YP_009724390.1;Wu1-SARS2_surface;632 | 0 | 0 |
| NC_045512.2;YP_009724390.1;Wu1-SARS2_surface;633 | 0 | 0.025 |
| NC_045512.2;YP_009724390.1;Wu1-SARS2_surface;634 | 0.05 | 0.025 |
| NC_045512.2;YP_009724390.1;Wu1-SARS2_surface;635 | 0 | 0.025 |
| NC_045512.2;YP_009724390.1;Wu1-SARS2_surface;636 | 0 | 0 |
| NC_045512.2;YP_009724390.1;Wu1-SARS2_surface;637 | 0 | 0 |
| NC_045512.2;YP_009724390.1;Wu1-SARS2_surface;638 | 0 | 0 |
| NC_045512.2;YP_009724390.1;Wu1-SARS2_surface;639 | 0 | 0 |
| NC_045512.2;YP_009724390.1;Wu1-SARS2_surface;640 | 0 | 0 |
| NC_045512.2;YP_009724390.1;Wu1-SARS2_surface;641 | 0 | 0 |
| NC_045512.2;YP_009724390.1;Wu1-SARS2_surface;642 | 0 | 0 |
| NC_045512.2;YP_009724390.1;Wu1-SARS2_surface;643 | 0 | 0.025 |
| NC_045512.2;YP_009724390.1;Wu1-SARS2_surface;644 | 0 | 0.025 |
| NC_045512.2;YP_009724390.1;Wu1-SARS2_surface;645 | 0 | 0.025 |

| | | |
|---|---|---|
| NC_045512.2;YP_009724390.1;Wu1-SARS2_surface;646 | 0 | 0.025 |
| NC_045512.2;YP_009724390.1;Wu1-SARS2_surface;647 | 0 | 0.025 |
| NC_045512.2;YP_009724390.1;Wu1-SARS2_surface;648 | 0 | 0.025 |
| NC_045512.2;YP_009724390.1;Wu1-SARS2_surface;649 | 0 | 0 |
| NC_045512.2;YP_009724390.1;Wu1-SARS2_surface;650 | 0 | 0.025 |
| NC_045512.2;YP_009724390.1;Wu1-SARS2_surface;651 | 0 | 0.025 |
| NC_045512.2;YP_009724390.1;Wu1-SARS2_surface;652 | 0 | 0.025 |
| NC_045512.2;YP_009724390.1;Wu1-SARS2_surface;653 | 0 | 0.075 |
| NC_045512.2;YP_009724390.1;Wu1-SARS2_surface;654 | 0 | 0.025 |
| NC_045512.2;YP_009724390.1;Wu1-SARS2_surface;655 | 0.05 | 0.05 |
| NC_045512.2;YP_009724390.1;Wu1-SARS2_surface;656 | 0.05 | 0.2 |
| NC_045512.2;YP_009724390.1;Wu1-SARS2_surface;657 | 0.05 | 0.2 |
| NC_045512.2;YP_009724390.1;Wu1-SARS2_surface;658 | 0.1 | 0.2 |
| NC_045512.2;YP_009724390.1;Wu1-SARS2_surface;659 | 0.05 | 0 |
| NC_045512.2;YP_009724390.1;Wu1-SARS2_surface;660 | 0.05 | 0 |
| NC_045512.2;YP_009724390.1;Wu1-SARS2_surface;661 | 0 | 0.025 |
| NC_045512.2;YP_009724390.1;Wu1-SARS2_surface;662 | 0 | 0.025 |
| NC_045512.2;YP_009724390.1;Wu1-SARS2_surface;663 | 0 | 0.025 |
| NC_045512.2;YP_009724390.1;Wu1-SARS2_surface;664 | 0 | 0.025 |
| NC_045512.2;YP_009724390.1;Wu1-SARS2_surface;665 | 0 | 0.025 |
| NC_045512.2;YP_009724390.1;Wu1-SARS2_surface;666 | 0 | 0.025 |
| NC_045512.2;YP_009724390.1;Wu1-SARS2_surface;667 | 0 | 0 |
| NC_045512.2;YP_009724390.1;Wu1-SARS2_surface;668 | 0 | 0 |
| NC_045512.2;YP_009724390.1;Wu1-SARS2_surface;669 | 0 | 0 |
| NC_045512.2;YP_009724390.1;Wu1-SARS2_surface;670 | 0 | 0 |
| NC_045512.2;YP_009724390.1;Wu1-SARS2_surface;671 | 0 | 0 |
| NC_045512.2;YP_009724390.1;Wu1-SARS2_surface;672 | 0 | 0 |
| NC_045512.2;YP_009724390.1;Wu1-SARS2_surface;673 | 0 | 0 |
| NC_045512.2;YP_009724390.1;Wu1-SARS2_surface;674 | 0 | 0 |
| NC_045512.2;YP_009724390.1;Wu1-SARS2_surface;675 | 0.1 | 0 |
| NC_045512.2;YP_009724390.1;Wu1-SARS2_surface;676 | 0.1 | 0.05 |
| NC_045512.2;YP_009724390.1;Wu1-SARS2_surface;677 | 0.05 | 0.025 |
| NC_045512.2;YP_009724390.1;Wu1-SARS2_surface;678 | 0.05 | 0 |
| NC_045512.2;YP_009724390.1;Wu1-SARS2_surface;679 | 0.05 | 0 |
| NC_045512.2;YP_009724390.1;Wu1-SARS2_surface;680 | 0 | 0 |
| NC_045512.2;YP_009724390.1;Wu1-SARS2_surface;681 | 0.05 | 0 |
| NC_045512.2;YP_009724390.1;Wu1-SARS2_surface;682 | 0 | 0 |
| NC_045512.2;YP_009724390.1;Wu1-SARS2_surface;683 | 0 | 0 |
| NC_045512.2;YP_009724390.1;Wu1-SARS2_surface;684 | 0 | 0.025 |
| NC_045512.2;YP_009724390.1;Wu1-SARS2_surface;685 | 0 | 0.025 |
| NC_045512.2;YP_009724390.1;Wu1-SARS2_surface;686 | 0 | 0.075 |
| NC_045512.2;YP_009724390.1;Wu1-SARS2_surface;687 | 0 | 0.125 |
| NC_045512.2;YP_009724390.1;Wu1-SARS2_surface;688 | 0 | 0.125 |

| | | |
|---|---|---|
| NC_045512.2;YP_009724390.1;Wu1-SARS2_surface;689 | 0 | 0.125 |
| NC_045512.2;YP_009724390.1;Wu1-SARS2_surface;690 | 0 | 0.125 |
| NC_045512.2;YP_009724390.1;Wu1-SARS2_surface;691 | 0.1 | 0.175 |
| NC_045512.2;YP_009724390.1;Wu1-SARS2_surface;692 | 0 | 0.125 |
| NC_045512.2;YP_009724390.1;Wu1-SARS2_surface;693 | 0 | 0.125 |
| NC_045512.2;YP_009724390.1;Wu1-SARS2_surface;694 | 0 | 0.1 |
| NC_045512.2;YP_009724390.1;Wu1-SARS2_surface;695 | 0 | 0.075 |
| NC_045512.2;YP_009724390.1;Wu1-SARS2_surface;696 | 0.05 | 0.025 |
| NC_045512.2;YP_009724390.1;Wu1-SARS2_surface;697 | 0 | 0.05 |
| NC_045512.2;YP_009724390.1;Wu1-SARS2_surface;698 | 0.05 | 0.1 |
| NC_045512.2;YP_009724390.1;Wu1-SARS2_surface;699 | 0.1 | 0.05 |
| NC_045512.2;YP_009724390.1;Wu1-SARS2_surface;700 | 0.1 | 0.1 |
| NC_045512.2;YP_009724390.1;Wu1-SARS2_surface;701 | 0.15 | 0.125 |
| NC_045512.2;YP_009724390.1;Wu1-SARS2_surface;702 | 0.15 | 0.05 |
| NC_045512.2;YP_009724390.1;Wu1-SARS2_surface;703 | 0.05 | 0.025 |
| NC_045512.2;YP_009724390.1;Wu1-SARS2_surface;704 | 0.2 | 0 |
| NC_045512.2;YP_009724390.1;Wu1-SARS2_surface;705 | 0.1 | 0.025 |
| NC_045512.2;YP_009724390.1;Wu1-SARS2_surface;706 | 0.2 | 0.125 |
| NC_045512.2;YP_009724390.1;Wu1-SARS2_surface;707 | 0.1 | 0 |
| NC_045512.2;YP_009724390.1;Wu1-SARS2_surface;708 | 0.05 | 0 |
| NC_045512.2;YP_009724390.1;Wu1-SARS2_surface;709 | 0 | 0 |
| NC_045512.2;YP_009724390.1;Wu1-SARS2_surface;710 | 0 | 0 |
| NC_045512.2;YP_009724390.1;Wu1-SARS2_surface;711 | 0 | 0.05 |
| NC_045512.2;YP_009724390.1;Wu1-SARS2_surface;712 | 0 | 0 |
| NC_045512.2;YP_009724390.1;Wu1-SARS2_surface;713 | 0 | 0.025 |
| NC_045512.2;YP_009724390.1;Wu1-SARS2_surface;714 | 0 | 0 |
| NC_045512.2;YP_009724390.1;Wu1-SARS2_surface;715 | 0 | 0 |
| NC_045512.2;YP_009724390.1;Wu1-SARS2_surface;716 | 0 | 0 |
| NC_045512.2;YP_009724390.1;Wu1-SARS2_surface;717 | 0 | 0 |
| NC_045512.2;YP_009724390.1;Wu1-SARS2_surface;718 | 0 | 0 |
| NC_045512.2;YP_009724390.1;Wu1-SARS2_surface;719 | 0 | 0 |
| NC_045512.2;YP_009724390.1;Wu1-SARS2_surface;720 | 0 | 0 |
| NC_045512.2;YP_009724390.1;Wu1-SARS2_surface;721 | 0 | 0 |
| NC_045512.2;YP_009724390.1;Wu1-SARS2_surface;722 | 0 | 0 |
| NC_045512.2;YP_009724390.1;Wu1-SARS2_surface;723 | 0 | 0 |
| NC_045512.2;YP_009724390.1;Wu1-SARS2_surface;724 | 0 | 0 |
| NC_045512.2;YP_009724390.1;Wu1-SARS2_surface;725 | 0 | 0 |
| NC_045512.2;YP_009724390.1;Wu1-SARS2_surface;726 | 0 | 0 |
| NC_045512.2;YP_009724390.1;Wu1-SARS2_surface;727 | 0 | 0 |
| NC_045512.2;YP_009724390.1;Wu1-SARS2_surface;728 | 0 | 0 |
| NC_045512.2;YP_009724390.1;Wu1-SARS2_surface;729 | 0 | 0 |
| NC_045512.2;YP_009724390.1;Wu1-SARS2_surface;730 | 0 | 0 |
| NC_045512.2;YP_009724390.1;Wu1-SARS2_surface;731 | 0 | 0 |

| | | |
|---|---|---|
| NC_045512.2;YP_009724390.1;Wu1-SARS2_surface;732 | 0 | 0 |
| NC_045512.2;YP_009724390.1;Wu1-SARS2_surface;733 | 0 | 0 |
| NC_045512.2;YP_009724390.1;Wu1-SARS2_surface;734 | 0 | 0 |
| NC_045512.2;YP_009724390.1;Wu1-SARS2_surface;735 | 0 | 0 |
| NC_045512.2;YP_009724390.1;Wu1-SARS2_surface;736 | 0.05 | 0 |
| NC_045512.2;YP_009724390.1;Wu1-SARS2_surface;737 | 0.05 | 0 |
| NC_045512.2;YP_009724390.1;Wu1-SARS2_surface;738 | 0.05 | 0 |
| NC_045512.2;YP_009724390.1;Wu1-SARS2_surface;739 | 0 | 0 |
| NC_045512.2;YP_009724390.1;Wu1-SARS2_surface;740 | 0.05 | 0 |
| NC_045512.2;YP_009724390.1;Wu1-SARS2_surface;741 | 0.05 | 0 |
| NC_045512.2;YP_009724390.1;Wu1-SARS2_surface;742 | 0.05 | 0 |
| NC_045512.2;YP_009724390.1;Wu1-SARS2_surface;743 | 0.05 | 0 |
| NC_045512.2;YP_009724390.1;Wu1-SARS2_surface;744 | 0 | 0 |
| NC_045512.2;YP_009724390.1;Wu1-SARS2_surface;745 | 0 | 0 |
| NC_045512.2;YP_009724390.1;Wu1-SARS2_surface;746 | 0 | 0 |
| NC_045512.2;YP_009724390.1;Wu1-SARS2_surface;747 | 0 | 0 |
| NC_045512.2;YP_009724390.1;Wu1-SARS2_surface;748 | 0 | 0 |
| NC_045512.2;YP_009724390.1;Wu1-SARS2_surface;749 | 0 | 0 |
| NC_045512.2;YP_009724390.1;Wu1-SARS2_surface;750 | 0 | 0 |
| NC_045512.2;YP_009724390.1;Wu1-SARS2_surface;751 | 0 | 0 |
| NC_045512.2;YP_009724390.1;Wu1-SARS2_surface;752 | 0 | 0 |
| NC_045512.2;YP_009724390.1;Wu1-SARS2_surface;753 | 0 | 0 |
| NC_045512.2;YP_009724390.1;Wu1-SARS2_surface;754 | 0 | 0 |
| NC_045512.2;YP_009724390.1;Wu1-SARS2_surface;755 | 0 | 0 |
| NC_045512.2;YP_009724390.1;Wu1-SARS2_surface;756 | 0 | 0 |
| NC_045512.2;YP_009724390.1;Wu1-SARS2_surface;757 | 0 | 0 |
| NC_045512.2;YP_009724390.1;Wu1-SARS2_surface;758 | 0 | 0 |
| NC_045512.2;YP_009724390.1;Wu1-SARS2_surface;759 | 0 | 0 |
| NC_045512.2;YP_009724390.1;Wu1-SARS2_surface;760 | 0 | 0 |
| NC_045512.2;YP_009724390.1;Wu1-SARS2_surface;761 | 0.1 | 0.275 |
| NC_045512.2;YP_009724390.1;Wu1-SARS2_surface;762 | 0.1 | 0.35 |
| NC_045512.2;YP_009724390.1;Wu1-SARS2_surface;763 | 0.15 | 0.35 |
| NC_045512.2;YP_009724390.1;Wu1-SARS2_surface;764 | 0.15 | 0.375 |
| NC_045512.2;YP_009724390.1;Wu1-SARS2_surface;765 | 0.15 | 0.4 |
| NC_045512.2;YP_009724390.1;Wu1-SARS2_surface;766 | 0.2 | 0.4 |
| NC_045512.2;YP_009724390.1;Wu1-SARS2_surface;767 | 0.15 | 0.425 |
| NC_045512.2;YP_009724390.1;Wu1-SARS2_surface;768 | 0.15 | 0.425 |
| NC_045512.2;YP_009724390.1;Wu1-SARS2_surface;769 | 0.15 | 0.4 |
| NC_045512.2;YP_009724390.1;Wu1-SARS2_surface;770 | 0 | 0.275 |
| NC_045512.2;YP_009724390.1;Wu1-SARS2_surface;771 | 0.1 | 0.225 |
| NC_045512.2;YP_009724390.1;Wu1-SARS2_surface;772 | 0.1 | 0.1 |
| NC_045512.2;YP_009724390.1;Wu1-SARS2_surface;773 | 0.1 | 0.15 |
| NC_045512.2;YP_009724390.1;Wu1-SARS2_surface;774 | 0.1 | 0.1 |

| | | |
|---|---|---|
| NC_045512.2;YP_009724390.1;Wu1-SARS2_surface;775 | 0 | 0.1 |
| NC_045512.2;YP_009724390.1;Wu1-SARS2_surface;776 | 0 | 0.05 |
| NC_045512.2;YP_009724390.1;Wu1-SARS2_surface;777 | 0.05 | 0.05 |
| NC_045512.2;YP_009724390.1;Wu1-SARS2_surface;778 | 0.05 | 0.05 |
| NC_045512.2;YP_009724390.1;Wu1-SARS2_surface;779 | 0.05 | 0.075 |
| NC_045512.2;YP_009724390.1;Wu1-SARS2_surface;780 | 0 | 0 |
| NC_045512.2;YP_009724390.1;Wu1-SARS2_surface;781 | 0 | 0 |
| NC_045512.2;YP_009724390.1;Wu1-SARS2_surface;782 | 0 | 0 |
| NC_045512.2;YP_009724390.1;Wu1-SARS2_surface;783 | 0 | 0 |
| NC_045512.2;YP_009724390.1;Wu1-SARS2_surface;784 | 0 | 0 |
| NC_045512.2;YP_009724390.1;Wu1-SARS2_surface;785 | 0 | 0.025 |
| NC_045512.2;YP_009724390.1;Wu1-SARS2_surface;786 | 0 | 0.025 |
| NC_045512.2;YP_009724390.1;Wu1-SARS2_surface;787 | 0 | 0.35 |
| NC_045512.2;YP_009724390.1;Wu1-SARS2_surface;788 | 0 | 0.475 |
| NC_045512.2;YP_009724390.1;Wu1-SARS2_surface;789 | 0 | 0.475 |
| NC_045512.2;YP_009724390.1;Wu1-SARS2_surface;790 | 0 | 0.425 |
| NC_045512.2;YP_009724390.1;Wu1-SARS2_surface;791 | 0 | 0.35 |
| NC_045512.2;YP_009724390.1;Wu1-SARS2_surface;792 | 0 | 0.45 |
| NC_045512.2;YP_009724390.1;Wu1-SARS2_surface;793 | 0 | 0.5 |
| NC_045512.2;YP_009724390.1;Wu1-SARS2_surface;794 | 0.05 | 0.55 |
| NC_045512.2;YP_009724390.1;Wu1-SARS2_surface;795 | 0 | 0.575 |
| NC_045512.2;YP_009724390.1;Wu1-SARS2_surface;796 | 0 | 0.2 |
| NC_045512.2;YP_009724390.1;Wu1-SARS2_surface;797 | 0.05 | 0.15 |
| NC_045512.2;YP_009724390.1;Wu1-SARS2_surface;798 | 0 | 0.175 |
| NC_045512.2;YP_009724390.1;Wu1-SARS2_surface;799 | 0 | 0.125 |
| NC_045512.2;YP_009724390.1;Wu1-SARS2_surface;800 | 0 | 0.05 |
| NC_045512.2;YP_009724390.1;Wu1-SARS2_surface;801 | 0 | 0.075 |
| NC_045512.2;YP_009724390.1;Wu1-SARS2_surface;802 | 0 | 0.075 |
| NC_045512.2;YP_009724390.1;Wu1-SARS2_surface;803 | 0 | 0.075 |
| NC_045512.2;YP_009724390.1;Wu1-SARS2_surface;804 | 0 | 0.15 |
| NC_045512.2;YP_009724390.1;Wu1-SARS2_surface;805 | 0 | 0.15 |
| NC_045512.2;YP_009724390.1;Wu1-SARS2_surface;806 | 0.1 | 0.25 |
| NC_045512.2;YP_009724390.1;Wu1-SARS2_surface;807 | 0.05 | 0.225 |
| NC_045512.2;YP_009724390.1;Wu1-SARS2_surface;808 | 0.05 | 0.625 |
| NC_045512.2;YP_009724390.1;Wu1-SARS2_surface;809 | 0.1 | 0.75 |
| NC_045512.2;YP_009724390.1;Wu1-SARS2_surface;810 | 0.05 | 0.85 |
| NC_045512.2;YP_009724390.1;Wu1-SARS2_surface;811 | 0.05 | 0.825 |
| NC_045512.2;YP_009724390.1;Wu1-SARS2_surface;812 | 0.05 | 0.875 |
| NC_045512.2;YP_009724390.1;Wu1-SARS2_surface;813 | 0.05 | 0.825 |
| NC_045512.2;YP_009724390.1;Wu1-SARS2_surface;814 | 0 | 0.7 |
| NC_045512.2;YP_009724390.1;Wu1-SARS2_surface;815 | 0 | 0.725 |
| NC_045512.2;YP_009724390.1;Wu1-SARS2_surface;816 | 0 | 0.35 |
| NC_045512.2;YP_009724390.1;Wu1-SARS2_surface;817 | 0.05 | 0.25 |

| | | |
|---|---|---|
| NC_045512.2;YP_009724390.1;Wu1-SARS2_surface;818 | 0.1 | 0.275 |
| NC_045512.2;YP_009724390.1;Wu1-SARS2_surface;819 | 0 | 0.125 |
| NC_045512.2;YP_009724390.1;Wu1-SARS2_surface;820 | 0 | 0 |
| NC_045512.2;YP_009724390.1;Wu1-SARS2_surface;821 | 0 | 0 |
| NC_045512.2;YP_009724390.1;Wu1-SARS2_surface;822 | 0 | 0 |
| NC_045512.2;YP_009724390.1;Wu1-SARS2_surface;823 | 0 | 0.025 |
| NC_045512.2;YP_009724390.1;Wu1-SARS2_surface;824 | 0 | 0.025 |
| NC_045512.2;YP_009724390.1;Wu1-SARS2_surface;825 | 0 | 0.025 |
| NC_045512.2;YP_009724390.1;Wu1-SARS2_surface;826 | 0 | 0.05 |
| NC_045512.2;YP_009724390.1;Wu1-SARS2_surface;827 | 0.05 | 0.025 |
| NC_045512.2;YP_009724390.1;Wu1-SARS2_surface;828 | 0.1 | 0.025 |
| NC_045512.2;YP_009724390.1;Wu1-SARS2_surface;829 | 0 | 0.025 |
| NC_045512.2;YP_009724390.1;Wu1-SARS2_surface;830 | 0 | 0.025 |
| NC_045512.2;YP_009724390.1;Wu1-SARS2_surface;831 | 0.05 | 0.025 |
| NC_045512.2;YP_009724390.1;Wu1-SARS2_surface;832 | 0 | 0.025 |
| NC_045512.2;YP_009724390.1;Wu1-SARS2_surface;833 | 0 | 0.025 |
| NC_045512.2;YP_009724390.1;Wu1-SARS2_surface;834 | 0 | 0.025 |
| NC_045512.2;YP_009724390.1;Wu1-SARS2_surface;835 | 0.05 | 0.025 |
| NC_045512.2;YP_009724390.1;Wu1-SARS2_surface;836 | 0 | 0 |
| NC_045512.2;YP_009724390.1;Wu1-SARS2_surface;837 | 0 | 0 |
| NC_045512.2;YP_009724390.1;Wu1-SARS2_surface;838 | 0 | 0 |
| NC_045512.2;YP_009724390.1;Wu1-SARS2_surface;839 | 0 | 0 |
| NC_045512.2;YP_009724390.1;Wu1-SARS2_surface;840 | 0.05 | 0.025 |
| NC_045512.2;YP_009724390.1;Wu1-SARS2_surface;841 | 0 | 0 |
| NC_045512.2;YP_009724390.1;Wu1-SARS2_surface;842 | 0.1 | 0.025 |
| NC_045512.2;YP_009724390.1;Wu1-SARS2_surface;843 | 0.05 | 0.025 |
| NC_045512.2;YP_009724390.1;Wu1-SARS2_surface;844 | 0 | 0.025 |
| NC_045512.2;YP_009724390.1;Wu1-SARS2_surface;845 | 0 | 0 |
| NC_045512.2;YP_009724390.1;Wu1-SARS2_surface;846 | 0 | 0 |
| NC_045512.2;YP_009724390.1;Wu1-SARS2_surface;847 | 0 | 0 |
| NC_045512.2;YP_009724390.1;Wu1-SARS2_surface;848 | 0.05 | 0 |
| NC_045512.2;YP_009724390.1;Wu1-SARS2_surface;849 | 0 | 0 |
| NC_045512.2;YP_009724390.1;Wu1-SARS2_surface;850 | 0 | 0 |
| NC_045512.2;YP_009724390.1;Wu1-SARS2_surface;851 | 0 | 0 |
| NC_045512.2;YP_009724390.1;Wu1-SARS2_surface;852 | 0 | 0 |
| NC_045512.2;YP_009724390.1;Wu1-SARS2_surface;853 | 0 | 0 |
| NC_045512.2;YP_009724390.1;Wu1-SARS2_surface;854 | 0 | 0 |
| NC_045512.2;YP_009724390.1;Wu1-SARS2_surface;855 | 0 | 0 |
| NC_045512.2;YP_009724390.1;Wu1-SARS2_surface;856 | 0 | 0 |
| NC_045512.2;YP_009724390.1;Wu1-SARS2_surface;857 | 0 | 0 |
| NC_045512.2;YP_009724390.1;Wu1-SARS2_surface;858 | 0 | 0 |
| NC_045512.2;YP_009724390.1;Wu1-SARS2_surface;859 | 0 | 0 |
| NC_045512.2;YP_009724390.1;Wu1-SARS2_surface;860 | 0 | 0 |

| | | |
|---|---|---|
| NC_045512.2;YP_009724390.1;Wu1-SARS2_surface;861 | 0 | 0 |
| NC_045512.2;YP_009724390.1;Wu1-SARS2_surface;862 | 0 | 0 |
| NC_045512.2;YP_009724390.1;Wu1-SARS2_surface;863 | 0 | 0 |
| NC_045512.2;YP_009724390.1;Wu1-SARS2_surface;864 | 0 | 0 |
| NC_045512.2;YP_009724390.1;Wu1-SARS2_surface;865 | 0 | 0 |
| NC_045512.2;YP_009724390.1;Wu1-SARS2_surface;866 | 0 | 0 |
| NC_045512.2;YP_009724390.1;Wu1-SARS2_surface;867 | 0 | 0 |
| NC_045512.2;YP_009724390.1;Wu1-SARS2_surface;868 | 0 | 0 |
| NC_045512.2;YP_009724390.1;Wu1-SARS2_surface;869 | 0 | 0.025 |
| NC_045512.2;YP_009724390.1;Wu1-SARS2_surface;870 | 0 | 0.025 |
| NC_045512.2;YP_009724390.1;Wu1-SARS2_surface;871 | 0 | 0 |
| NC_045512.2;YP_009724390.1;Wu1-SARS2_surface;872 | 0 | 0 |
| NC_045512.2;YP_009724390.1;Wu1-SARS2_surface;873 | 0 | 0 |
| NC_045512.2;YP_009724390.1;Wu1-SARS2_surface;874 | 0 | 0 |
| NC_045512.2;YP_009724390.1;Wu1-SARS2_surface;875 | 0 | 0 |
| NC_045512.2;YP_009724390.1;Wu1-SARS2_surface;876 | 0 | 0 |
| NC_045512.2;YP_009724390.1;Wu1-SARS2_surface;877 | 0 | 0.025 |
| NC_045512.2;YP_009724390.1;Wu1-SARS2_surface;878 | 0.1 | 0.05 |
| NC_045512.2;YP_009724390.1;Wu1-SARS2_surface;879 | 0 | 0 |
| NC_045512.2;YP_009724390.1;Wu1-SARS2_surface;880 | 0 | 0 |
| NC_045512.2;YP_009724390.1;Wu1-SARS2_surface;881 | 0 | 0 |
| NC_045512.2;YP_009724390.1;Wu1-SARS2_surface;882 | 0 | 0 |
| NC_045512.2;YP_009724390.1;Wu1-SARS2_surface;883 | 0 | 0 |
| NC_045512.2;YP_009724390.1;Wu1-SARS2_surface;884 | 0 | 0 |
| NC_045512.2;YP_009724390.1;Wu1-SARS2_surface;885 | 0 | 0 |
| NC_045512.2;YP_009724390.1;Wu1-SARS2_surface;886 | 0 | 0 |
| NC_045512.2;YP_009724390.1;Wu1-SARS2_surface;887 | 0 | 0 |
| NC_045512.2;YP_009724390.1;Wu1-SARS2_surface;888 | 0 | 0 |
| NC_045512.2;YP_009724390.1;Wu1-SARS2_surface;889 | 0 | 0.025 |
| NC_045512.2;YP_009724390.1;Wu1-SARS2_surface;890 | 0 | 0 |
| NC_045512.2;YP_009724390.1;Wu1-SARS2_surface;891 | 0 | 0 |
| NC_045512.2;YP_009724390.1;Wu1-SARS2_surface;892 | 0.05 | 0.025 |
| NC_045512.2;YP_009724390.1;Wu1-SARS2_surface;893 | 0.05 | 0.025 |
| NC_045512.2;YP_009724390.1;Wu1-SARS2_surface;894 | 0.05 | 0.025 |
| NC_045512.2;YP_009724390.1;Wu1-SARS2_surface;895 | 0.05 | 0.025 |
| NC_045512.2;YP_009724390.1;Wu1-SARS2_surface;896 | 0.05 | 0.025 |
| NC_045512.2;YP_009724390.1;Wu1-SARS2_surface;897 | 0.05 | 0.025 |
| NC_045512.2;YP_009724390.1;Wu1-SARS2_surface;898 | 0.05 | 0.025 |
| NC_045512.2;YP_009724390.1;Wu1-SARS2_surface;899 | 0.1 | 0.05 |
| NC_045512.2;YP_009724390.1;Wu1-SARS2_surface;900 | 0.15 | 0.05 |
| NC_045512.2;YP_009724390.1;Wu1-SARS2_surface;901 | 0.1 | 0.05 |
| NC_045512.2;YP_009724390.1;Wu1-SARS2_surface;902 | 0 | 0 |
| NC_045512.2;YP_009724390.1;Wu1-SARS2_surface;903 | 0 | 0 |

| | | |
|---|---|---|
| NC_045512.2;YP_009724390.1;Wu1-SARS2_surface;904 | 0 | 0 |
| NC_045512.2;YP_009724390.1;Wu1-SARS2_surface;905 | 0.05 | 0.025 |
| NC_045512.2;YP_009724390.1;Wu1-SARS2_surface;906 | 0 | 0.05 |
| NC_045512.2;YP_009724390.1;Wu1-SARS2_surface;907 | 0.05 | 0.05 |
| NC_045512.2;YP_009724390.1;Wu1-SARS2_surface;908 | 0 | 0.05 |
| NC_045512.2;YP_009724390.1;Wu1-SARS2_surface;909 | 0.05 | 0.075 |
| NC_045512.2;YP_009724390.1;Wu1-SARS2_surface;910 | 0.05 | 0.025 |
| NC_045512.2;YP_009724390.1;Wu1-SARS2_surface;911 | 0 | 0.05 |
| NC_045512.2;YP_009724390.1;Wu1-SARS2_surface;912 | 0 | 0.025 |
| NC_045512.2;YP_009724390.1;Wu1-SARS2_surface;913 | 0.05 | 0.05 |
| NC_045512.2;YP_009724390.1;Wu1-SARS2_surface;914 | 0 | 0 |
| NC_045512.2;YP_009724390.1;Wu1-SARS2_surface;915 | 0 | 0.05 |
| NC_045512.2;YP_009724390.1;Wu1-SARS2_surface;916 | 0.05 | 0.025 |
| NC_045512.2;YP_009724390.1;Wu1-SARS2_surface;917 | 0 | 0.05 |
| NC_045512.2;YP_009724390.1;Wu1-SARS2_surface;918 | 0.05 | 0.025 |
| NC_045512.2;YP_009724390.1;Wu1-SARS2_surface;919 | 0 | 0 |
| NC_045512.2;YP_009724390.1;Wu1-SARS2_surface;920 | 0.05 | 0 |
| NC_045512.2;YP_009724390.1;Wu1-SARS2_surface;921 | 0.05 | 0.025 |
| NC_045512.2;YP_009724390.1;Wu1-SARS2_surface;922 | 0.05 | 0.025 |
| NC_045512.2;YP_009724390.1;Wu1-SARS2_surface;923 | 0.05 | 0.05 |
| NC_045512.2;YP_009724390.1;Wu1-SARS2_surface;924 | 0.1 | 0.05 |
| NC_045512.2;YP_009724390.1;Wu1-SARS2_surface;925 | 0.05 | 0.025 |
| NC_045512.2;YP_009724390.1;Wu1-SARS2_surface;926 | 0.05 | 0.025 |
| NC_045512.2;YP_009724390.1;Wu1-SARS2_surface;927 | 0.05 | 0.05 |
| NC_045512.2;YP_009724390.1;Wu1-SARS2_surface;928 | 0.1 | 0.05 |
| NC_045512.2;YP_009724390.1;Wu1-SARS2_surface;929 | 0 | 0.025 |
| NC_045512.2;YP_009724390.1;Wu1-SARS2_surface;930 | 0 | 0.025 |
| NC_045512.2;YP_009724390.1;Wu1-SARS2_surface;931 | 0 | 0 |
| NC_045512.2;YP_009724390.1;Wu1-SARS2_surface;932 | 0.05 | 0.025 |
| NC_045512.2;YP_009724390.1;Wu1-SARS2_surface;933 | 0 | 0 |
| NC_045512.2;YP_009724390.1;Wu1-SARS2_surface;934 | 0 | 0 |
| NC_045512.2;YP_009724390.1;Wu1-SARS2_surface;935 | 0 | 0 |
| NC_045512.2;YP_009724390.1;Wu1-SARS2_surface;936 | 0 | 0 |
| NC_045512.2;YP_009724390.1;Wu1-SARS2_surface;937 | 0 | 0 |
| NC_045512.2;YP_009724390.1;Wu1-SARS2_surface;938 | 0 | 0 |
| NC_045512.2;YP_009724390.1;Wu1-SARS2_surface;939 | 0 | 0 |
| NC_045512.2;YP_009724390.1;Wu1-SARS2_surface;940 | 0 | 0.075 |
| NC_045512.2;YP_009724390.1;Wu1-SARS2_surface;941 | 0 | 0.025 |
| NC_045512.2;YP_009724390.1;Wu1-SARS2_surface;942 | 0 | 0.025 |
| NC_045512.2;YP_009724390.1;Wu1-SARS2_surface;943 | 0 | 0.025 |
| NC_045512.2;YP_009724390.1;Wu1-SARS2_surface;944 | 0 | 0.075 |
| NC_045512.2;YP_009724390.1;Wu1-SARS2_surface;945 | 0 | 0.125 |
| NC_045512.2;YP_009724390.1;Wu1-SARS2_surface;946 | 0.1 | 0.075 |

| | | |
|---|---|---|
| NC_045512.2;YP_009724390.1;Wu1-SARS2_surface;947 | 0 | 0.1 |
| NC_045512.2;YP_009724390.1;Wu1-SARS2_surface;948 | 0 | 0.075 |
| NC_045512.2;YP_009724390.1;Wu1-SARS2_surface;949 | 0 | 0.025 |
| NC_045512.2;YP_009724390.1;Wu1-SARS2_surface;950 | 0 | 0.05 |
| NC_045512.2;YP_009724390.1;Wu1-SARS2_surface;951 | 0.05 | 0 |
| NC_045512.2;YP_009724390.1;Wu1-SARS2_surface;952 | 0 | 0 |
| NC_045512.2;YP_009724390.1;Wu1-SARS2_surface;953 | 0 | 0.025 |
| NC_045512.2;YP_009724390.1;Wu1-SARS2_surface;954 | 0 | 0.025 |
| NC_045512.2;YP_009724390.1;Wu1-SARS2_surface;955 | 0 | 0 |
| NC_045512.2;YP_009724390.1;Wu1-SARS2_surface;956 | 0 | 0.025 |
| NC_045512.2;YP_009724390.1;Wu1-SARS2_surface;957 | 0 | 0.05 |
| NC_045512.2;YP_009724390.1;Wu1-SARS2_surface;958 | 0 | 0.025 |
| NC_045512.2;YP_009724390.1;Wu1-SARS2_surface;959 | 0 | 0 |
| NC_045512.2;YP_009724390.1;Wu1-SARS2_surface;960 | 0 | 0 |
| NC_045512.2;YP_009724390.1;Wu1-SARS2_surface;961 | 0 | 0 |
| NC_045512.2;YP_009724390.1;Wu1-SARS2_surface;962 | 0 | 0 |
| NC_045512.2;YP_009724390.1;Wu1-SARS2_surface;963 | 0 | 0 |
| NC_045512.2;YP_009724390.1;Wu1-SARS2_surface;964 | 0 | 0.025 |
| NC_045512.2;YP_009724390.1;Wu1-SARS2_surface;965 | 0 | 0 |
| NC_045512.2;YP_009724390.1;Wu1-SARS2_surface;966 | 0 | 0 |
| NC_045512.2;YP_009724390.1;Wu1-SARS2_surface;967 | 0 | 0 |
| NC_045512.2;YP_009724390.1;Wu1-SARS2_surface;968 | 0 | 0 |
| NC_045512.2;YP_009724390.1;Wu1-SARS2_surface;969 | 0 | 0 |
| NC_045512.2;YP_009724390.1;Wu1-SARS2_surface;970 | 0 | 0.025 |
| NC_045512.2;YP_009724390.1;Wu1-SARS2_surface;971 | 0 | 0.025 |
| NC_045512.2;YP_009724390.1;Wu1-SARS2_surface;972 | 0.05 | 0.05 |
| NC_045512.2;YP_009724390.1;Wu1-SARS2_surface;973 | 0 | 0.075 |
| NC_045512.2;YP_009724390.1;Wu1-SARS2_surface;974 | 0.05 | 0.075 |
| NC_045512.2;YP_009724390.1;Wu1-SARS2_surface;975 | 0.05 | 0.125 |
| NC_045512.2;YP_009724390.1;Wu1-SARS2_surface;976 | 0.05 | 0.1 |
| NC_045512.2;YP_009724390.1;Wu1-SARS2_surface;977 | 0 | 0.075 |
| NC_045512.2;YP_009724390.1;Wu1-SARS2_surface;978 | 0 | 0.05 |
| NC_045512.2;YP_009724390.1;Wu1-SARS2_surface;979 | 0.05 | 0.05 |
| NC_045512.2;YP_009724390.1;Wu1-SARS2_surface;980 | 0.05 | 0.025 |
| NC_045512.2;YP_009724390.1;Wu1-SARS2_surface;981 | 0 | 0 |
| NC_045512.2;YP_009724390.1;Wu1-SARS2_surface;982 | 0 | 0 |
| NC_045512.2;YP_009724390.1;Wu1-SARS2_surface;983 | 0 | 0 |
| NC_045512.2;YP_009724390.1;Wu1-SARS2_surface;984 | 0 | 0.025 |
| NC_045512.2;YP_009724390.1;Wu1-SARS2_surface;985 | 0 | 0 |
| NC_045512.2;YP_009724390.1;Wu1-SARS2_surface;986 | 0 | 0 |
| NC_045512.2;YP_009724390.1;Wu1-SARS2_surface;987 | 0 | 0 |
| NC_045512.2;YP_009724390.1;Wu1-SARS2_surface;988 | 0.05 | 0 |
| NC_045512.2;YP_009724390.1;Wu1-SARS2_surface;989 | 0 | 0 |

| | | |
|---|---|---|
| NC_045512.2;YP_009724390.1;Wu1-SARS2_surface;990 | 0 | 0 |
| NC_045512.2;YP_009724390.1;Wu1-SARS2_surface;991 | 0 | 0 |
| NC_045512.2;YP_009724390.1;Wu1-SARS2_surface;992 | 0 | 0 |
| NC_045512.2;YP_009724390.1;Wu1-SARS2_surface;993 | 0 | 0 |
| NC_045512.2;YP_009724390.1;Wu1-SARS2_surface;994 | 0 | 0 |
| NC_045512.2;YP_009724390.1;Wu1-SARS2_surface;995 | 0 | 0 |
| NC_045512.2;YP_009724390.1;Wu1-SARS2_surface;996 | 0 | 0 |
| NC_045512.2;YP_009724390.1;Wu1-SARS2_surface;997 | 0 | 0 |
| NC_045512.2;YP_009724390.1;Wu1-SARS2_surface;998 | 0 | 0 |
| NC_045512.2;YP_009724390.1;Wu1-SARS2_surface;999 | 0 | 0 |
| NC_045512.2;YP_009724390.1;Wu1-SARS2_surface;1000 | 0 | 0 |
| NC_045512.2;YP_009724390.1;Wu1-SARS2_surface;1001 | 0 | 0 |
| NC_045512.2;YP_009724390.1;Wu1-SARS2_surface;1002 | 0 | 0 |
| NC_045512.2;YP_009724390.1;Wu1-SARS2_surface;1003 | 0 | 0.025 |
| NC_045512.2;YP_009724390.1;Wu1-SARS2_surface;1004 | 0 | 0 |
| NC_045512.2;YP_009724390.1;Wu1-SARS2_surface;1005 | 0 | 0 |
| NC_045512.2;YP_009724390.1;Wu1-SARS2_surface;1006 | 0 | 0 |
| NC_045512.2;YP_009724390.1;Wu1-SARS2_surface;1007 | 0 | 0 |
| NC_045512.2;YP_009724390.1;Wu1-SARS2_surface;1008 | 0 | 0 |
| NC_045512.2;YP_009724390.1;Wu1-SARS2_surface;1009 | 0 | 0 |
| NC_045512.2;YP_009724390.1;Wu1-SARS2_surface;1010 | 0 | 0 |
| NC_045512.2;YP_009724390.1;Wu1-SARS2_surface;1011 | 0.05 | 0.025 |
| NC_045512.2;YP_009724390.1;Wu1-SARS2_surface;1012 | 0 | 0 |
| NC_045512.2;YP_009724390.1;Wu1-SARS2_surface;1013 | 0 | 0.025 |
| NC_045512.2;YP_009724390.1;Wu1-SARS2_surface;1014 | 0 | 0 |
| NC_045512.2;YP_009724390.1;Wu1-SARS2_surface;1015 | 0 | 0 |
| NC_045512.2;YP_009724390.1;Wu1-SARS2_surface;1016 | 0.05 | 0.025 |
| NC_045512.2;YP_009724390.1;Wu1-SARS2_surface;1017 | 0.05 | 0 |
| NC_045512.2;YP_009724390.1;Wu1-SARS2_surface;1018 | 0.05 | 0 |
| NC_045512.2;YP_009724390.1;Wu1-SARS2_surface;1019 | 0 | 0.025 |
| NC_045512.2;YP_009724390.1;Wu1-SARS2_surface;1020 | 0 | 0 |
| NC_045512.2;YP_009724390.1;Wu1-SARS2_surface;1021 | 0 | 0.025 |
| NC_045512.2;YP_009724390.1;Wu1-SARS2_surface;1022 | 0 | 0 |
| NC_045512.2;YP_009724390.1;Wu1-SARS2_surface;1023 | 0 | 0 |
| NC_045512.2;YP_009724390.1;Wu1-SARS2_surface;1024 | 0 | 0 |
| NC_045512.2;YP_009724390.1;Wu1-SARS2_surface;1025 | 0 | 0 |
| NC_045512.2;YP_009724390.1;Wu1-SARS2_surface;1026 | 0 | 0 |
| NC_045512.2;YP_009724390.1;Wu1-SARS2_surface;1027 | 0 | 0 |
| NC_045512.2;YP_009724390.1;Wu1-SARS2_surface;1028 | 0 | 0 |
| NC_045512.2;YP_009724390.1;Wu1-SARS2_surface;1029 | 0 | 0 |
| NC_045512.2;YP_009724390.1;Wu1-SARS2_surface;1030 | 0 | 0 |
| NC_045512.2;YP_009724390.1;Wu1-SARS2_surface;1031 | 0 | 0 |
| NC_045512.2;YP_009724390.1;Wu1-SARS2_surface;1032 | 0 | 0 |

| | | |
|---|---|---|
| NC_045512.2;YP_009724390.1;Wu1-SARS2_surface;1033 | 0 | 0 |
| NC_045512.2;YP_009724390.1;Wu1-SARS2_surface;1034 | 0 | 0 |
| NC_045512.2;YP_009724390.1;Wu1-SARS2_surface;1035 | 0 | 0 |
| NC_045512.2;YP_009724390.1;Wu1-SARS2_surface;1036 | 0 | 0 |
| NC_045512.2;YP_009724390.1;Wu1-SARS2_surface;1037 | 0 | 0 |
| NC_045512.2;YP_009724390.1;Wu1-SARS2_surface;1038 | 0 | 0 |
| NC_045512.2;YP_009724390.1;Wu1-SARS2_surface;1039 | 0 | 0 |
| NC_045512.2;YP_009724390.1;Wu1-SARS2_surface;1040 | 0 | 0 |
| NC_045512.2;YP_009724390.1;Wu1-SARS2_surface;1041 | 0 | 0.025 |
| NC_045512.2;YP_009724390.1;Wu1-SARS2_surface;1042 | 0.05 | 0 |
| NC_045512.2;YP_009724390.1;Wu1-SARS2_surface;1043 | 0 | 0 |
| NC_045512.2;YP_009724390.1;Wu1-SARS2_surface;1044 | 0 | 0.05 |
| NC_045512.2;YP_009724390.1;Wu1-SARS2_surface;1045 | 0 | 0.025 |
| NC_045512.2;YP_009724390.1;Wu1-SARS2_surface;1046 | 0 | 0 |
| NC_045512.2;YP_009724390.1;Wu1-SARS2_surface;1047 | 0 | 0 |
| NC_045512.2;YP_009724390.1;Wu1-SARS2_surface;1048 | 0 | 0 |
| NC_045512.2;YP_009724390.1;Wu1-SARS2_surface;1049 | 0 | 0 |
| NC_045512.2;YP_009724390.1;Wu1-SARS2_surface;1050 | 0 | 0 |
| NC_045512.2;YP_009724390.1;Wu1-SARS2_surface;1051 | 0 | 0 |
| NC_045512.2;YP_009724390.1;Wu1-SARS2_surface;1052 | 0 | 0 |
| NC_045512.2;YP_009724390.1;Wu1-SARS2_surface;1053 | 0 | 0 |
| NC_045512.2;YP_009724390.1;Wu1-SARS2_surface;1054 | 0 | 0 |
| NC_045512.2;YP_009724390.1;Wu1-SARS2_surface;1055 | 0 | 0 |
| NC_045512.2;YP_009724390.1;Wu1-SARS2_surface;1056 | 0 | 0 |
| NC_045512.2;YP_009724390.1;Wu1-SARS2_surface;1057 | 0 | 0 |
| NC_045512.2;YP_009724390.1;Wu1-SARS2_surface;1058 | 0 | 0 |
| NC_045512.2;YP_009724390.1;Wu1-SARS2_surface;1059 | 0 | 0 |
| NC_045512.2;YP_009724390.1;Wu1-SARS2_surface;1060 | 0 | 0 |
| NC_045512.2;YP_009724390.1;Wu1-SARS2_surface;1061 | 0 | 0 |
| NC_045512.2;YP_009724390.1;Wu1-SARS2_surface;1062 | 0 | 0 |
| NC_045512.2;YP_009724390.1;Wu1-SARS2_surface;1063 | 0 | 0 |
| NC_045512.2;YP_009724390.1;Wu1-SARS2_surface;1064 | 0.05 | 0 |
| NC_045512.2;YP_009724390.1;Wu1-SARS2_surface;1065 | 0.1 | 0 |
| NC_045512.2;YP_009724390.1;Wu1-SARS2_surface;1066 | 0.05 | 0 |
| NC_045512.2;YP_009724390.1;Wu1-SARS2_surface;1067 | 0.05 | 0 |
| NC_045512.2;YP_009724390.1;Wu1-SARS2_surface;1068 | 0 | 0 |
| NC_045512.2;YP_009724390.1;Wu1-SARS2_surface;1069 | 0 | 0 |
| NC_045512.2;YP_009724390.1;Wu1-SARS2_surface;1070 | 0 | 0 |
| NC_045512.2;YP_009724390.1;Wu1-SARS2_surface;1071 | 0.1 | 0 |
| NC_045512.2;YP_009724390.1;Wu1-SARS2_surface;1072 | 0 | 0 |
| NC_045512.2;YP_009724390.1;Wu1-SARS2_surface;1073 | 0 | 0.025 |
| NC_045512.2;YP_009724390.1;Wu1-SARS2_surface;1074 | 0 | 0 |
| NC_045512.2;YP_009724390.1;Wu1-SARS2_surface;1075 | 0 | 0 |

| | | |
|---|---|---|
| NC_045512.2;YP_009724390.1;Wu1-SARS2_surface;1076 | 0 | 0 |
| NC_045512.2;YP_009724390.1;Wu1-SARS2_surface;1077 | 0 | 0 |
| NC_045512.2;YP_009724390.1;Wu1-SARS2_surface;1078 | 0.05 | 0 |
| NC_045512.2;YP_009724390.1;Wu1-SARS2_surface;1079 | 0.05 | 0 |
| NC_045512.2;YP_009724390.1;Wu1-SARS2_surface;1080 | 0 | 0 |
| NC_045512.2;YP_009724390.1;Wu1-SARS2_surface;1081 | 0.05 | 0.025 |
| NC_045512.2;YP_009724390.1;Wu1-SARS2_surface;1082 | 0 | 0 |
| NC_045512.2;YP_009724390.1;Wu1-SARS2_surface;1083 | 0 | 0 |
| NC_045512.2;YP_009724390.1;Wu1-SARS2_surface;1084 | 0 | 0 |
| NC_045512.2;YP_009724390.1;Wu1-SARS2_surface;1085 | 0 | 0 |
| NC_045512.2;YP_009724390.1;Wu1-SARS2_surface;1086 | 0 | 0 |
| NC_045512.2;YP_009724390.1;Wu1-SARS2_surface;1087 | 0 | 0 |
| NC_045512.2;YP_009724390.1;Wu1-SARS2_surface;1088 | 0 | 0 |
| NC_045512.2;YP_009724390.1;Wu1-SARS2_surface;1089 | 0 | 0 |
| NC_045512.2;YP_009724390.1;Wu1-SARS2_surface;1090 | 0 | 0 |
| NC_045512.2;YP_009724390.1;Wu1-SARS2_surface;1091 | 0 | 0 |
| NC_045512.2;YP_009724390.1;Wu1-SARS2_surface;1092 | 0 | 0 |
| NC_045512.2;YP_009724390.1;Wu1-SARS2_surface;1093 | 0 | 0 |
| NC_045512.2;YP_009724390.1;Wu1-SARS2_surface;1094 | 0 | 0 |
| NC_045512.2;YP_009724390.1;Wu1-SARS2_surface;1095 | 0 | 0 |
| NC_045512.2;YP_009724390.1;Wu1-SARS2_surface;1096 | 0 | 0 |
| NC_045512.2;YP_009724390.1;Wu1-SARS2_surface;1097 | 0 | 0 |
| NC_045512.2;YP_009724390.1;Wu1-SARS2_surface;1098 | 0 | 0 |
| NC_045512.2;YP_009724390.1;Wu1-SARS2_surface;1099 | 0 | 0 |
| NC_045512.2;YP_009724390.1;Wu1-SARS2_surface;1100 | 0 | 0 |
| NC_045512.2;YP_009724390.1;Wu1-SARS2_surface;1101 | 0 | 0 |
| NC_045512.2;YP_009724390.1;Wu1-SARS2_surface;1102 | 0 | 0 |
| NC_045512.2;YP_009724390.1;Wu1-SARS2_surface;1103 | 0 | 0.025 |
| NC_045512.2;YP_009724390.1;Wu1-SARS2_surface;1104 | 0.1 | 0.025 |
| NC_045512.2;YP_009724390.1;Wu1-SARS2_surface;1105 | 0.05 | 0.025 |
| NC_045512.2;YP_009724390.1;Wu1-SARS2_surface;1106 | 0.05 | 0.025 |
| NC_045512.2;YP_009724390.1;Wu1-SARS2_surface;1107 | 0 | 0 |
| NC_045512.2;YP_009724390.1;Wu1-SARS2_surface;1108 | 0.05 | 0.025 |
| NC_045512.2;YP_009724390.1;Wu1-SARS2_surface;1109 | 0.05 | 0.025 |
| NC_045512.2;YP_009724390.1;Wu1-SARS2_surface;1110 | 0.05 | 0.1 |
| NC_045512.2;YP_009724390.1;Wu1-SARS2_surface;1111 | 0.1 | 0.025 |
| NC_045512.2;YP_009724390.1;Wu1-SARS2_surface;1112 | 0.05 | 0.05 |
| NC_045512.2;YP_009724390.1;Wu1-SARS2_surface;1113 | 0.05 | 0.075 |
| NC_045512.2;YP_009724390.1;Wu1-SARS2_surface;1114 | 0.05 | 0.05 |
| NC_045512.2;YP_009724390.1;Wu1-SARS2_surface;1115 | 0.05 | 0.025 |
| NC_045512.2;YP_009724390.1;Wu1-SARS2_surface;1116 | 0.05 | 0.05 |
| NC_045512.2;YP_009724390.1;Wu1-SARS2_surface;1117 | 0 | 0.025 |
| NC_045512.2;YP_009724390.1;Wu1-SARS2_surface;1118 | 0 | 0 |

| | | |
|---|---|---|
| NC_045512.2;YP_009724390.1;Wu1-SARS2_surface;1119 | 0 | 0 |
| NC_045512.2;YP_009724390.1;Wu1-SARS2_surface;1120 | 0 | 0 |
| NC_045512.2;YP_009724390.1;Wu1-SARS2_surface;1121 | 0 | 0 |
| NC_045512.2;YP_009724390.1;Wu1-SARS2_surface;1122 | 0 | 0 |
| NC_045512.2;YP_009724390.1;Wu1-SARS2_surface;1123 | 0 | 0 |
| NC_045512.2;YP_009724390.1;Wu1-SARS2_surface;1124 | 0 | 0 |
| NC_045512.2;YP_009724390.1;Wu1-SARS2_surface;1125 | 0 | 0 |
| NC_045512.2;YP_009724390.1;Wu1-SARS2_surface;1126 | 0 | 0 |
| NC_045512.2;YP_009724390.1;Wu1-SARS2_surface;1127 | 0 | 0 |
| NC_045512.2;YP_009724390.1;Wu1-SARS2_surface;1128 | 0 | 0 |
| NC_045512.2;YP_009724390.1;Wu1-SARS2_surface;1129 | 0 | 0 |
| NC_045512.2;YP_009724390.1;Wu1-SARS2_surface;1130 | 0 | 0.05 |
| NC_045512.2;YP_009724390.1;Wu1-SARS2_surface;1131 | 0 | 0 |
| NC_045512.2;YP_009724390.1;Wu1-SARS2_surface;1132 | 0 | 0 |
| NC_045512.2;YP_009724390.1;Wu1-SARS2_surface;1133 | 0 | 0.025 |
| NC_045512.2;YP_009724390.1;Wu1-SARS2_surface;1134 | 0 | 0 |
| NC_045512.2;YP_009724390.1;Wu1-SARS2_surface;1135 | 0 | 0.025 |
| NC_045512.2;YP_009724390.1;Wu1-SARS2_surface;1136 | 0 | 0 |
| NC_045512.2;YP_009724390.1;Wu1-SARS2_surface;1137 | 0 | 0 |
| NC_045512.2;YP_009724390.1;Wu1-SARS2_surface;1138 | 0 | 0 |
| NC_045512.2;YP_009724390.1;Wu1-SARS2_surface;1139 | 0 | 0.05 |
| NC_045512.2;YP_009724390.1;Wu1-SARS2_surface;1140 | 0 | 0.175 |
| NC_045512.2;YP_009724390.1;Wu1-SARS2_surface;1141 | 0.05 | 0.75 |
| NC_045512.2;YP_009724390.1;Wu1-SARS2_surface;1142 | 0.1 | 0.775 |
| NC_045512.2;YP_009724390.1;Wu1-SARS2_surface;1143 | 0.05 | 0.775 |
| NC_045512.2;YP_009724390.1;Wu1-SARS2_surface;1144 | 0.05 | 0.725 |
| NC_045512.2;YP_009724390.1;Wu1-SARS2_surface;1145 | 0.3 | 0.725 |
| NC_045512.2;YP_009724390.1;Wu1-SARS2_surface;1146 | 0.4 | 0.775 |
| NC_045512.2;YP_009724390.1;Wu1-SARS2_surface;1147 | 0.35 | 0.775 |
| NC_045512.2;YP_009724390.1;Wu1-SARS2_surface;1148 | 0.35 | 0.825 |
| NC_045512.2;YP_009724390.1;Wu1-SARS2_surface;1149 | 0.45 | 0.8 |
| NC_045512.2;YP_009724390.1;Wu1-SARS2_surface;1150 | 0.4 | 0.5 |
| NC_045512.2;YP_009724390.1;Wu1-SARS2_surface;1151 | 0.45 | 0.35 |
| NC_045512.2;YP_009724390.1;Wu1-SARS2_surface;1152 | 0.45 | 0.35 |
| NC_045512.2;YP_009724390.1;Wu1-SARS2_surface;1153 | 0.5 | 0.35 |
| NC_045512.2;YP_009724390.1;Wu1-SARS2_surface;1154 | 0.5 | 0.375 |
| NC_045512.2;YP_009724390.1;Wu1-SARS2_surface;1155 | 0 | 0.2 |
| NC_045512.2;YP_009724390.1;Wu1-SARS2_surface;1156 | 0 | 0.125 |
| NC_045512.2;YP_009724390.1;Wu1-SARS2_surface;1157 | 0 | 0.225 |
| NC_045512.2;YP_009724390.1;Wu1-SARS2_surface;1158 | 0.05 | 0.275 |
| NC_045512.2;YP_009724390.1;Wu1-SARS2_surface;1159 | 0.1 | 0.35 |
| NC_045512.2;YP_009724390.1;Wu1-SARS2_surface;1160 | 0.2 | 0.35 |
| NC_045512.2;YP_009724390.1;Wu1-SARS2_surface;1161 | 0.1 | 0.35 |

| | | |
|---|---|---|
| NC_045512.2;YP_009724390.1;Wu1-SARS2_surface;1162 | 0 | 0.3 |
| NC_045512.2;YP_009724390.1;Wu1-SARS2_surface;1163 | 0 | 0.325 |
| NC_045512.2;YP_009724390.1;Wu1-SARS2_surface;1164 | 0 | 0.15 |
| NC_045512.2;YP_009724390.1;Wu1-SARS2_surface;1165 | 0 | 0.2 |
| NC_045512.2;YP_009724390.1;Wu1-SARS2_surface;1166 | 0 | 0.05 |
| NC_045512.2;YP_009724390.1;Wu1-SARS2_surface;1167 | 0.05 | 0.1 |
| NC_045512.2;YP_009724390.1;Wu1-SARS2_surface;1168 | 0.15 | 0.05 |
| NC_045512.2;YP_009724390.1;Wu1-SARS2_surface;1169 | 0.05 | 0.125 |
| NC_045512.2;YP_009724390.1;Wu1-SARS2_surface;1170 | 0.05 | 0.05 |
| NC_045512.2;YP_009724390.1;Wu1-SARS2_surface;1171 | 0.05 | 0.1 |
| NC_045512.2;YP_009724390.1;Wu1-SARS2_surface;1172 | 0.05 | 0.175 |
| NC_045512.2;YP_009724390.1;Wu1-SARS2_surface;1173 | 0.05 | 0.225 |
| NC_045512.2;YP_009724390.1;Wu1-SARS2_surface;1174 | 0.1 | 0.225 |
| NC_045512.2;YP_009724390.1;Wu1-SARS2_surface;1175 | 0.1 | 0.275 |
| NC_045512.2;YP_009724390.1;Wu1-SARS2_surface;1176 | 0.1 | 0.275 |
| NC_045512.2;YP_009724390.1;Wu1-SARS2_surface;1177 | 0.1 | 0.3 |
| NC_045512.2;YP_009724390.1;Wu1-SARS2_surface;1178 | 0.05 | 0.3 |
| NC_045512.2;YP_009724390.1;Wu1-SARS2_surface;1179 | 0.2 | 0.325 |
| NC_045512.2;YP_009724390.1;Wu1-SARS2_surface;1180 | 0.15 | 0.3 |
| NC_045512.2;YP_009724390.1;Wu1-SARS2_surface;1181 | 0.1 | 0.175 |
| NC_045512.2;YP_009724390.1;Wu1-SARS2_surface;1182 | 0.1 | 0.15 |
| NC_045512.2;YP_009724390.1;Wu1-SARS2_surface;1183 | 0.15 | 0.125 |
| NC_045512.2;YP_009724390.1;Wu1-SARS2_surface;1184 | 0.05 | 0.125 |
| NC_045512.2;YP_009724390.1;Wu1-SARS2_surface;1185 | 0.05 | 0.125 |
| NC_045512.2;YP_009724390.1;Wu1-SARS2_surface;1186 | 0.05 | 0.1 |
| NC_045512.2;YP_009724390.1;Wu1-SARS2_surface;1187 | 0.05 | 0.125 |
| NC_045512.2;YP_009724390.1;Wu1-SARS2_surface;1188 | 0.05 | 0.1 |
| NC_045512.2;YP_009724390.1;Wu1-SARS2_surface;1189 | 0.05 | 0.125 |
| NC_045512.2;YP_009724390.1;Wu1-SARS2_surface;1190 | 0.05 | 0.125 |
| NC_045512.2;YP_009724390.1;Wu1-SARS2_surface;1191 | 0.05 | 0.075 |
| NC_045512.2;YP_009724390.1;Wu1-SARS2_surface;1192 | 0.05 | 0.125 |
| NC_045512.2;YP_009724390.1;Wu1-SARS2_surface;1193 | 0.1 | 0.15 |
| NC_045512.2;YP_009724390.1;Wu1-SARS2_surface;1194 | 0.05 | 0.125 |
| NC_045512.2;YP_009724390.1;Wu1-SARS2_surface;1195 | 0 | 0 |
| NC_045512.2;YP_009724390.1;Wu1-SARS2_surface;1196 | 0 | 0.05 |
| NC_045512.2;YP_009724390.1;Wu1-SARS2_surface;1197 | 0 | 0 |
| NC_045512.2;YP_009724390.1;Wu1-SARS2_surface;1198 | 0 | 0 |
| NC_045512.2;YP_009724390.1;Wu1-SARS2_surface;1199 | 0 | 0 |
| NC_045512.2;YP_009724390.1;Wu1-SARS2_surface;1200 | 0 | 0.025 |
| NC_045512.2;YP_009724390.1;Wu1-SARS2_surface;1201 | 0 | 0 |
| NC_045512.2;YP_009724390.1;Wu1-SARS2_surface;1202 | 0 | 0 |
| NC_045512.2;YP_009724390.1;Wu1-SARS2_surface;1203 | 0 | 0 |
| NC_045512.2;YP_009724390.1;Wu1-SARS2_surface;1204 | 0 | 0 |

| | | |
|---|---|---|
| NC_045512.2;YP_009724390.1;Wu1-SARS2_surface;1205 | 0 | 0 |
| NC_045512.2;YP_009724390.1;Wu1-SARS2_surface;1206 | 0 | 0.025 |
| NC_045512.2;YP_009724390.1;Wu1-SARS2_surface;1207 | 0 | 0 |
| NC_045512.2;YP_009724390.1;Wu1-SARS2_surface;1208 | 0 | 0 |
| NC_045512.2;YP_009724390.1;Wu1-SARS2_surface;1209 | 0 | 0 |
| NC_045512.2;YP_009724390.1;Wu1-SARS2_surface;1210 | 0 | 0 |
| NC_045512.2;YP_009724390.1;Wu1-SARS2_surface;1211 | 0 | 0 |
| NC_045512.2;YP_009724390.1;Wu1-SARS2_surface;1212 | 0 | 0 |
| NC_045512.2;YP_009724390.1;Wu1-SARS2_surface;1213 | 0 | 0 |
| NC_045512.2;YP_009724390.1;Wu1-SARS2_surface;1214 | 0 | 0 |
| NC_045512.2;YP_009724390.1;Wu1-SARS2_surface;1215 | 0 | 0 |
| NC_045512.2;YP_009724390.1;Wu1-SARS2_surface;1216 | 0 | 0 |
| NC_045512.2;YP_009724390.1;Wu1-SARS2_surface;1217 | 0 | 0 |
| NC_045512.2;YP_009724390.1;Wu1-SARS2_surface;1218 | 0 | 0 |
| NC_045512.2;YP_009724390.1;Wu1-SARS2_surface;1219 | 0 | 0 |
| NC_045512.2;YP_009724390.1;Wu1-SARS2_surface;1220 | 0 | 0 |
| NC_045512.2;YP_009724390.1;Wu1-SARS2_surface;1221 | 0 | 0 |
| NC_045512.2;YP_009724390.1;Wu1-SARS2_surface;1222 | 0 | 0 |
| NC_045512.2;YP_009724390.1;Wu1-SARS2_surface;1223 | 0 | 0 |
| NC_045512.2;YP_009724390.1;Wu1-SARS2_surface;1224 | 0 | 0 |
| NC_045512.2;YP_009724390.1;Wu1-SARS2_surface;1225 | 0 | 0 |
| NC_045512.2;YP_009724390.1;Wu1-SARS2_surface;1226 | 0 | 0 |
| NC_045512.2;YP_009724390.1;Wu1-SARS2_surface;1227 | 0 | 0 |
| NC_045512.2;YP_009724390.1;Wu1-SARS2_surface;1228 | 0 | 0 |
| NC_045512.2;YP_009724390.1;Wu1-SARS2_surface;1229 | 0 | 0 |
| NC_045512.2;YP_009724390.1;Wu1-SARS2_surface;1230 | 0 | 0 |
| NC_045512.2;YP_009724390.1;Wu1-SARS2_surface;1231 | 0 | 0 |
| NC_045512.2;YP_009724390.1;Wu1-SARS2_surface;1232 | 0 | 0 |
| NC_045512.2;YP_009724390.1;Wu1-SARS2_surface;1233 | 0 | 0 |
| NC_045512.2;YP_009724390.1;Wu1-SARS2_surface;1234 | 0 | 0 |
| NC_045512.2;YP_009724390.1;Wu1-SARS2_surface;1235 | 0 | 0 |
| NC_045512.2;YP_009724390.1;Wu1-SARS2_surface;1236 | 0 | 0 |
| NC_045512.2;YP_009724390.1;Wu1-SARS2_surface;1237 | 0 | 0 |
| NC_045512.2;YP_009724390.1;Wu1-SARS2_surface;1238 | 0 | 0 |
| NC_045512.2;YP_009724390.1;Wu1-SARS2_surface;1239 | 0 | 0 |
| NC_045512.2;YP_009724390.1;Wu1-SARS2_surface;1240 | 0 | 0 |
| NC_045512.2;YP_009724390.1;Wu1-SARS2_surface;1241 | 0 | 0 |
| NC_045512.2;YP_009724390.1;Wu1-SARS2_surface;1242 | 0 | 0 |
| NC_045512.2;YP_009724390.1;Wu1-SARS2_surface;1243 | 0 | 0 |
| NC_045512.2;YP_009724390.1;Wu1-SARS2_surface;1244 | 0 | 0 |
| NC_045512.2;YP_009724390.1;Wu1-SARS2_surface;1245 | 0 | 0.075 |
| NC_045512.2;YP_009724390.1;Wu1-SARS2_surface;1246 | 0 | 0.1 |
| NC_045512.2;YP_009724390.1;Wu1-SARS2_surface;1247 | 0 | 0.25 |

| | | |
|---|---|---|
| NC_045512.2;YP_009724390.1;Wu1-SARS2_surface;1248 | 0 | 0.5 |
| NC_045512.2;YP_009724390.1;Wu1-SARS2_surface;1249 | 0 | 0.55 |
| NC_045512.2;YP_009724390.1;Wu1-SARS2_surface;1250 | 0 | 0.575 |
| NC_045512.2;YP_009724390.1;Wu1-SARS2_surface;1251 | 0 | 0.65 |
| NC_045512.2;YP_009724390.1;Wu1-SARS2_surface;1252 | 0 | 0.65 |
| NC_045512.2;YP_009724390.1;Wu1-SARS2_surface;1253 | 0 | 0.675 |
| NC_045512.2;YP_009724390.1;Wu1-SARS2_surface;1254 | 0.1 | 0.65 |
| NC_045512.2;YP_009724390.1;Wu1-SARS2_surface;1255 | 0.05 | 0.6 |
| NC_045512.2;YP_009724390.1;Wu1-SARS2_surface;1256 | 0.05 | 0.45 |
| NC_045512.2;YP_009724390.1;Wu1-SARS2_surface;1257 | 0.05 | 0.475 |
| NC_045512.2;YP_009724390.1;Wu1-SARS2_surface;1258 | 0.1 | 0.3 |
| NC_045512.2;YP_009724390.1;Wu1-SARS2_surface;1259 | 0.05 | 0.225 |
| NC_045512.2;YP_009724390.1;Wu1-SARS2_surface;1260 | 0.05 | 0.225 |
| NC_045512.2;YP_009724390.1;Wu1-SARS2_surface;1261 | 0.05 | 0.175 |
| NC_045512.2;YP_009724390.1;Wu1-SARS2_surface;1262 | 0 | 0.175 |
| NC_045512.2;YP_009724391.1;Wu1-SARS2_orf3a;1 | 0 | 0 |
| NC_045512.2;YP_009724391.1;Wu1-SARS2_orf3a;2 | 0 | 0 |
| NC_045512.2;YP_009724391.1;Wu1-SARS2_orf3a;3 | 0 | 0 |
| NC_045512.2;YP_009724391.1;Wu1-SARS2_orf3a;4 | 0 | 0 |
| NC_045512.2;YP_009724391.1;Wu1-SARS2_orf3a;5 | 0 | 0 |
| NC_045512.2;YP_009724391.1;Wu1-SARS2_orf3a;6 | 0 | 0.025 |
| NC_045512.2;YP_009724391.1;Wu1-SARS2_orf3a;7 | 0 | 0.125 |
| NC_045512.2;YP_009724391.1;Wu1-SARS2_orf3a;8 | 0.05 | 0.1 |
| NC_045512.2;YP_009724391.1;Wu1-SARS2_orf3a;9 | 0 | 0.15 |
| NC_045512.2;YP_009724391.1;Wu1-SARS2_orf3a;10 | 0.05 | 0.175 |
| NC_045512.2;YP_009724391.1;Wu1-SARS2_orf3a;11 | 0 | 0.175 |
| NC_045512.2;YP_009724391.1;Wu1-SARS2_orf3a;12 | 0.05 | 0.175 |
| NC_045512.2;YP_009724391.1;Wu1-SARS2_orf3a;13 | 0.05 | 0.275 |
| NC_045512.2;YP_009724391.1;Wu1-SARS2_orf3a;14 | 0.05 | 0.275 |
| NC_045512.2;YP_009724391.1;Wu1-SARS2_orf3a;15 | 0.05 | 0.275 |
| NC_045512.2;YP_009724391.1;Wu1-SARS2_orf3a;16 | 0 | 0.3 |
| NC_045512.2;YP_009724391.1;Wu1-SARS2_orf3a;17 | 0 | 0.175 |
| NC_045512.2;YP_009724391.1;Wu1-SARS2_orf3a;18 | 0 | 0.2 |
| NC_045512.2;YP_009724391.1;Wu1-SARS2_orf3a;19 | 0 | 0.175 |
| NC_045512.2;YP_009724391.1;Wu1-SARS2_orf3a;20 | 0 | 0.15 |
| NC_045512.2;YP_009724391.1;Wu1-SARS2_orf3a;21 | 0 | 0.125 |
| NC_045512.2;YP_009724391.1;Wu1-SARS2_orf3a;22 | 0 | 0.075 |
| NC_045512.2;YP_009724391.1;Wu1-SARS2_orf3a;23 | 0 | 0.025 |
| NC_045512.2;YP_009724391.1;Wu1-SARS2_orf3a;24 | 0 | 0 |
| NC_045512.2;YP_009724391.1;Wu1-SARS2_orf3a;25 | 0 | 0 |
| NC_045512.2;YP_009724391.1;Wu1-SARS2_orf3a;26 | 0 | 0 |
| NC_045512.2;YP_009724391.1;Wu1-SARS2_orf3a;27 | 0 | 0 |
| NC_045512.2;YP_009724391.1;Wu1-SARS2_orf3a;28 | 0 | 0 |

| | | |
|---|---|---|
| NC_045512.2;YP_009724391.1;Wu1-SARS2_orf3a;29 | 0 | 0.025 |
| NC_045512.2;YP_009724391.1;Wu1-SARS2_orf3a;30 | 0 | 0.025 |
| NC_045512.2;YP_009724391.1;Wu1-SARS2_orf3a;31 | 0 | 0 |
| NC_045512.2;YP_009724391.1;Wu1-SARS2_orf3a;32 | 0 | 0 |
| NC_045512.2;YP_009724391.1;Wu1-SARS2_orf3a;33 | 0 | 0 |
| NC_045512.2;YP_009724391.1;Wu1-SARS2_orf3a;34 | 0 | 0 |
| NC_045512.2;YP_009724391.1;Wu1-SARS2_orf3a;35 | 0 | 0 |
| NC_045512.2;YP_009724391.1;Wu1-SARS2_orf3a;36 | 0.05 | 0 |
| NC_045512.2;YP_009724391.1;Wu1-SARS2_orf3a;37 | 0 | 0 |
| NC_045512.2;YP_009724391.1;Wu1-SARS2_orf3a;38 | 0 | 0 |
| NC_045512.2;YP_009724391.1;Wu1-SARS2_orf3a;39 | 0 | 0 |
| NC_045512.2;YP_009724391.1;Wu1-SARS2_orf3a;40 | 0 | 0 |
| NC_045512.2;YP_009724391.1;Wu1-SARS2_orf3a;41 | 0 | 0 |
| NC_045512.2;YP_009724391.1;Wu1-SARS2_orf3a;42 | 0 | 0 |
| NC_045512.2;YP_009724391.1;Wu1-SARS2_orf3a;43 | 0 | 0 |
| NC_045512.2;YP_009724391.1;Wu1-SARS2_orf3a;44 | 0 | 0 |
| NC_045512.2;YP_009724391.1;Wu1-SARS2_orf3a;45 | 0 | 0 |
| NC_045512.2;YP_009724391.1;Wu1-SARS2_orf3a;46 | 0 | 0 |
| NC_045512.2;YP_009724391.1;Wu1-SARS2_orf3a;47 | 0 | 0 |
| NC_045512.2;YP_009724391.1;Wu1-SARS2_orf3a;48 | 0 | 0 |
| NC_045512.2;YP_009724391.1;Wu1-SARS2_orf3a;49 | 0 | 0 |
| NC_045512.2;YP_009724391.1;Wu1-SARS2_orf3a;50 | 0 | 0 |
| NC_045512.2;YP_009724391.1;Wu1-SARS2_orf3a;51 | 0 | 0 |
| NC_045512.2;YP_009724391.1;Wu1-SARS2_orf3a;52 | 0 | 0 |
| NC_045512.2;YP_009724391.1;Wu1-SARS2_orf3a;53 | 0 | 0 |
| NC_045512.2;YP_009724391.1;Wu1-SARS2_orf3a;54 | 0 | 0 |
| NC_045512.2;YP_009724391.1;Wu1-SARS2_orf3a;55 | 0 | 0 |
| NC_045512.2;YP_009724391.1;Wu1-SARS2_orf3a;56 | 0 | 0 |
| NC_045512.2;YP_009724391.1;Wu1-SARS2_orf3a;57 | 0 | 0 |
| NC_045512.2;YP_009724391.1;Wu1-SARS2_orf3a;58 | 0 | 0 |
| NC_045512.2;YP_009724391.1;Wu1-SARS2_orf3a;59 | 0 | 0 |
| NC_045512.2;YP_009724391.1;Wu1-SARS2_orf3a;60 | 0 | 0 |
| NC_045512.2;YP_009724391.1;Wu1-SARS2_orf3a;61 | 0 | 0 |
| NC_045512.2;YP_009724391.1;Wu1-SARS2_orf3a;62 | 0 | 0 |
| NC_045512.2;YP_009724391.1;Wu1-SARS2_orf3a;63 | 0 | 0 |
| NC_045512.2;YP_009724391.1;Wu1-SARS2_orf3a;64 | 0 | 0 |
| NC_045512.2;YP_009724391.1;Wu1-SARS2_orf3a;65 | 0 | 0 |
| NC_045512.2;YP_009724391.1;Wu1-SARS2_orf3a;66 | 0 | 0 |
| NC_045512.2;YP_009724391.1;Wu1-SARS2_orf3a;67 | 0 | 0 |
| NC_045512.2;YP_009724391.1;Wu1-SARS2_orf3a;68 | 0 | 0 |
| NC_045512.2;YP_009724391.1;Wu1-SARS2_orf3a;69 | 0 | 0 |
| NC_045512.2;YP_009724391.1;Wu1-SARS2_orf3a;70 | 0 | 0 |
| NC_045512.2;YP_009724391.1;Wu1-SARS2_orf3a;71 | 0 | 0 |

| | | |
|---|---|---|
| NC_045512.2;YP_009724391.1;Wu1-SARS2_orf3a;72 | 0 | 0 |
| NC_045512.2;YP_009724391.1;Wu1-SARS2_orf3a;73 | 0 | 0 |
| NC_045512.2;YP_009724391.1;Wu1-SARS2_orf3a;74 | 0 | 0 |
| NC_045512.2;YP_009724391.1;Wu1-SARS2_orf3a;75 | 0 | 0 |
| NC_045512.2;YP_009724391.1;Wu1-SARS2_orf3a;76 | 0 | 0 |
| NC_045512.2;YP_009724391.1;Wu1-SARS2_orf3a;77 | 0 | 0 |
| NC_045512.2;YP_009724391.1;Wu1-SARS2_orf3a;78 | 0 | 0 |
| NC_045512.2;YP_009724391.1;Wu1-SARS2_orf3a;79 | 0 | 0 |
| NC_045512.2;YP_009724391.1;Wu1-SARS2_orf3a;80 | 0 | 0 |
| NC_045512.2;YP_009724391.1;Wu1-SARS2_orf3a;81 | 0 | 0 |
| NC_045512.2;YP_009724391.1;Wu1-SARS2_orf3a;82 | 0 | 0 |
| NC_045512.2;YP_009724391.1;Wu1-SARS2_orf3a;83 | 0 | 0 |
| NC_045512.2;YP_009724391.1;Wu1-SARS2_orf3a;84 | 0 | 0 |
| NC_045512.2;YP_009724391.1;Wu1-SARS2_orf3a;85 | 0 | 0 |
| NC_045512.2;YP_009724391.1;Wu1-SARS2_orf3a;86 | 0 | 0 |
| NC_045512.2;YP_009724391.1;Wu1-SARS2_orf3a;87 | 0 | 0 |
| NC_045512.2;YP_009724391.1;Wu1-SARS2_orf3a;88 | 0 | 0 |
| NC_045512.2;YP_009724391.1;Wu1-SARS2_orf3a;89 | 0 | 0 |
| NC_045512.2;YP_009724391.1;Wu1-SARS2_orf3a;90 | 0 | 0 |
| NC_045512.2;YP_009724391.1;Wu1-SARS2_orf3a;91 | 0 | 0 |
| NC_045512.2;YP_009724391.1;Wu1-SARS2_orf3a;92 | 0 | 0 |
| NC_045512.2;YP_009724391.1;Wu1-SARS2_orf3a;93 | 0 | 0 |
| NC_045512.2;YP_009724391.1;Wu1-SARS2_orf3a;94 | 0 | 0 |
| NC_045512.2;YP_009724391.1;Wu1-SARS2_orf3a;95 | 0 | 0 |
| NC_045512.2;YP_009724391.1;Wu1-SARS2_orf3a;96 | 0 | 0 |
| NC_045512.2;YP_009724391.1;Wu1-SARS2_orf3a;97 | 0 | 0 |
| NC_045512.2;YP_009724391.1;Wu1-SARS2_orf3a;98 | 0 | 0 |
| NC_045512.2;YP_009724391.1;Wu1-SARS2_orf3a;99 | 0 | 0 |
| NC_045512.2;YP_009724391.1;Wu1-SARS2_orf3a;100 | 0 | 0 |
| NC_045512.2;YP_009724391.1;Wu1-SARS2_orf3a;101 | 0 | 0 |
| NC_045512.2;YP_009724391.1;Wu1-SARS2_orf3a;102 | 0 | 0 |
| NC_045512.2;YP_009724391.1;Wu1-SARS2_orf3a;103 | 0 | 0 |
| NC_045512.2;YP_009724391.1;Wu1-SARS2_orf3a;104 | 0 | 0 |
| NC_045512.2;YP_009724391.1;Wu1-SARS2_orf3a;105 | 0 | 0 |
| NC_045512.2;YP_009724391.1;Wu1-SARS2_orf3a;106 | 0 | 0 |
| NC_045512.2;YP_009724391.1;Wu1-SARS2_orf3a;107 | 0 | 0 |
| NC_045512.2;YP_009724391.1;Wu1-SARS2_orf3a;108 | 0 | 0 |
| NC_045512.2;YP_009724391.1;Wu1-SARS2_orf3a;109 | 0 | 0 |
| NC_045512.2;YP_009724391.1;Wu1-SARS2_orf3a;110 | 0 | 0 |
| NC_045512.2;YP_009724391.1;Wu1-SARS2_orf3a;111 | 0 | 0 |
| NC_045512.2;YP_009724391.1;Wu1-SARS2_orf3a;112 | 0 | 0 |
| NC_045512.2;YP_009724391.1;Wu1-SARS2_orf3a;113 | 0 | 0 |
| NC_045512.2;YP_009724391.1;Wu1-SARS2_orf3a;114 | 0 | 0 |

| | | |
|---|---|---|
| NC_045512.2;YP_009724391.1;Wu1-SARS2_orf3a;115 | 0 | 0 |
| NC_045512.2;YP_009724391.1;Wu1-SARS2_orf3a;116 | 0 | 0 |
| NC_045512.2;YP_009724391.1;Wu1-SARS2_orf3a;117 | 0 | 0 |
| NC_045512.2;YP_009724391.1;Wu1-SARS2_orf3a;118 | 0 | 0 |
| NC_045512.2;YP_009724391.1;Wu1-SARS2_orf3a;119 | 0 | 0 |
| NC_045512.2;YP_009724391.1;Wu1-SARS2_orf3a;120 | 0 | 0 |
| NC_045512.2;YP_009724391.1;Wu1-SARS2_orf3a;121 | 0 | 0 |
| NC_045512.2;YP_009724391.1;Wu1-SARS2_orf3a;122 | 0 | 0 |
| NC_045512.2;YP_009724391.1;Wu1-SARS2_orf3a;123 | 0 | 0 |
| NC_045512.2;YP_009724391.1;Wu1-SARS2_orf3a;124 | 0 | 0 |
| NC_045512.2;YP_009724391.1;Wu1-SARS2_orf3a;125 | 0 | 0 |
| NC_045512.2;YP_009724391.1;Wu1-SARS2_orf3a;126 | 0 | 0 |
| NC_045512.2;YP_009724391.1;Wu1-SARS2_orf3a;127 | 0 | 0 |
| NC_045512.2;YP_009724391.1;Wu1-SARS2_orf3a;128 | 0 | 0 |
| NC_045512.2;YP_009724391.1;Wu1-SARS2_orf3a;129 | 0 | 0 |
| NC_045512.2;YP_009724391.1;Wu1-SARS2_orf3a;130 | 0 | 0 |
| NC_045512.2;YP_009724391.1;Wu1-SARS2_orf3a;131 | 0 | 0 |
| NC_045512.2;YP_009724391.1;Wu1-SARS2_orf3a;132 | 0 | 0 |
| NC_045512.2;YP_009724391.1;Wu1-SARS2_orf3a;133 | 0 | 0 |
| NC_045512.2;YP_009724391.1;Wu1-SARS2_orf3a;134 | 0 | 0 |
| NC_045512.2;YP_009724391.1;Wu1-SARS2_orf3a;135 | 0 | 0 |
| NC_045512.2;YP_009724391.1;Wu1-SARS2_orf3a;136 | 0 | 0 |
| NC_045512.2;YP_009724391.1;Wu1-SARS2_orf3a;137 | 0 | 0 |
| NC_045512.2;YP_009724391.1;Wu1-SARS2_orf3a;138 | 0 | 0 |
| NC_045512.2;YP_009724391.1;Wu1-SARS2_orf3a;139 | 0 | 0 |
| NC_045512.2;YP_009724391.1;Wu1-SARS2_orf3a;140 | 0 | 0 |
| NC_045512.2;YP_009724391.1;Wu1-SARS2_orf3a;141 | 0 | 0 |
| NC_045512.2;YP_009724391.1;Wu1-SARS2_orf3a;142 | 0 | 0 |
| NC_045512.2;YP_009724391.1;Wu1-SARS2_orf3a;143 | 0 | 0 |
| NC_045512.2;YP_009724391.1;Wu1-SARS2_orf3a;144 | 0 | 0 |
| NC_045512.2;YP_009724391.1;Wu1-SARS2_orf3a;145 | 0 | 0 |
| NC_045512.2;YP_009724391.1;Wu1-SARS2_orf3a;146 | 0 | 0 |
| NC_045512.2;YP_009724391.1;Wu1-SARS2_orf3a;147 | 0 | 0 |
| NC_045512.2;YP_009724391.1;Wu1-SARS2_orf3a;148 | 0 | 0 |
| NC_045512.2;YP_009724391.1;Wu1-SARS2_orf3a;149 | 0 | 0 |
| NC_045512.2;YP_009724391.1;Wu1-SARS2_orf3a;150 | 0 | 0 |
| NC_045512.2;YP_009724391.1;Wu1-SARS2_orf3a;151 | 0 | 0 |
| NC_045512.2;YP_009724391.1;Wu1-SARS2_orf3a;152 | 0 | 0 |
| NC_045512.2;YP_009724391.1;Wu1-SARS2_orf3a;153 | 0.05 | 0 |
| NC_045512.2;YP_009724391.1;Wu1-SARS2_orf3a;154 | 0 | 0 |
| NC_045512.2;YP_009724391.1;Wu1-SARS2_orf3a;155 | 0 | 0 |
| NC_045512.2;YP_009724391.1;Wu1-SARS2_orf3a;156 | 0 | 0 |
| NC_045512.2;YP_009724391.1;Wu1-SARS2_orf3a;157 | 0 | 0 |

| | | |
|---|---|---|
| NC_045512.2;YP_009724391.1;Wu1-SARS2_orf3a;158 | 0 | 0 |
| NC_045512.2;YP_009724391.1;Wu1-SARS2_orf3a;159 | 0 | 0 |
| NC_045512.2;YP_009724391.1;Wu1-SARS2_orf3a;160 | 0 | 0 |
| NC_045512.2;YP_009724391.1;Wu1-SARS2_orf3a;161 | 0 | 0 |
| NC_045512.2;YP_009724391.1;Wu1-SARS2_orf3a;162 | 0 | 0.025 |
| NC_045512.2;YP_009724391.1;Wu1-SARS2_orf3a;163 | 0 | 0 |
| NC_045512.2;YP_009724391.1;Wu1-SARS2_orf3a;164 | 0 | 0 |
| NC_045512.2;YP_009724391.1;Wu1-SARS2_orf3a;165 | 0 | 0 |
| NC_045512.2;YP_009724391.1;Wu1-SARS2_orf3a;166 | 0 | 0 |
| NC_045512.2;YP_009724391.1;Wu1-SARS2_orf3a;167 | 0 | 0.025 |
| NC_045512.2;YP_009724391.1;Wu1-SARS2_orf3a;168 | 0 | 0.025 |
| NC_045512.2;YP_009724391.1;Wu1-SARS2_orf3a;169 | 0 | 0.025 |
| NC_045512.2;YP_009724391.1;Wu1-SARS2_orf3a;170 | 0 | 0.025 |
| NC_045512.2;YP_009724391.1;Wu1-SARS2_orf3a;171 | 0 | 0.025 |
| NC_045512.2;YP_009724391.1;Wu1-SARS2_orf3a;172 | 0 | 0.025 |
| NC_045512.2;YP_009724391.1;Wu1-SARS2_orf3a;173 | 0 | 0.075 |
| NC_045512.2;YP_009724391.1;Wu1-SARS2_orf3a;174 | 0 | 0.025 |
| NC_045512.2;YP_009724391.1;Wu1-SARS2_orf3a;175 | 0 | 0.025 |
| NC_045512.2;YP_009724391.1;Wu1-SARS2_orf3a;176 | 0.05 | 0.025 |
| NC_045512.2;YP_009724391.1;Wu1-SARS2_orf3a;177 | 0.05 | 0.1 |
| NC_045512.2;YP_009724391.1;Wu1-SARS2_orf3a;178 | 0.05 | 0.075 |
| NC_045512.2;YP_009724391.1;Wu1-SARS2_orf3a;179 | 0.05 | 0.05 |
| NC_045512.2;YP_009724391.1;Wu1-SARS2_orf3a;180 | 0.05 | 0.05 |
| NC_045512.2;YP_009724391.1;Wu1-SARS2_orf3a;181 | 0.1 | 0.05 |
| NC_045512.2;YP_009724391.1;Wu1-SARS2_orf3a;182 | 0.05 | 0.05 |
| NC_045512.2;YP_009724391.1;Wu1-SARS2_orf3a;183 | 0.05 | 0.15 |
| NC_045512.2;YP_009724391.1;Wu1-SARS2_orf3a;184 | 0.1 | 0.075 |
| NC_045512.2;YP_009724391.1;Wu1-SARS2_orf3a;185 | 0.05 | 0.05 |
| NC_045512.2;YP_009724391.1;Wu1-SARS2_orf3a;186 | 0 | 0.05 |
| NC_045512.2;YP_009724391.1;Wu1-SARS2_orf3a;187 | 0 | 0 |
| NC_045512.2;YP_009724391.1;Wu1-SARS2_orf3a;188 | 0 | 0 |
| NC_045512.2;YP_009724391.1;Wu1-SARS2_orf3a;189 | 0 | 0 |
| NC_045512.2;YP_009724391.1;Wu1-SARS2_orf3a;190 | 0 | 0 |
| NC_045512.2;YP_009724391.1;Wu1-SARS2_orf3a;191 | 0 | 0 |
| NC_045512.2;YP_009724391.1;Wu1-SARS2_orf3a;192 | 0 | 0 |
| NC_045512.2;YP_009724391.1;Wu1-SARS2_orf3a;193 | 0 | 0 |
| NC_045512.2;YP_009724391.1;Wu1-SARS2_orf3a;194 | 0 | 0.025 |
| NC_045512.2;YP_009724391.1;Wu1-SARS2_orf3a;195 | 0 | 0 |
| NC_045512.2;YP_009724391.1;Wu1-SARS2_orf3a;196 | 0 | 0.05 |
| NC_045512.2;YP_009724391.1;Wu1-SARS2_orf3a;197 | 0 | 0 |
| NC_045512.2;YP_009724391.1;Wu1-SARS2_orf3a;198 | 0 | 0 |
| NC_045512.2;YP_009724391.1;Wu1-SARS2_orf3a;199 | 0 | 0 |
| NC_045512.2;YP_009724391.1;Wu1-SARS2_orf3a;200 | 0 | 0 |

| | | |
|---|---|---|
| NC_045512.2;YP_009724391.1;Wu1-SARS2_orf3a;201 | 0 | 0 |
| NC_045512.2;YP_009724391.1;Wu1-SARS2_orf3a;202 | 0 | 0 |
| NC_045512.2;YP_009724391.1;Wu1-SARS2_orf3a;203 | 0 | 0 |
| NC_045512.2;YP_009724391.1;Wu1-SARS2_orf3a;204 | 0 | 0 |
| NC_045512.2;YP_009724391.1;Wu1-SARS2_orf3a;205 | 0 | 0 |
| NC_045512.2;YP_009724391.1;Wu1-SARS2_orf3a;206 | 0 | 0 |
| NC_045512.2;YP_009724391.1;Wu1-SARS2_orf3a;207 | 0 | 0.025 |
| NC_045512.2;YP_009724391.1;Wu1-SARS2_orf3a;208 | 0 | 0.025 |
| NC_045512.2;YP_009724391.1;Wu1-SARS2_orf3a;209 | 0.05 | 0.025 |
| NC_045512.2;YP_009724391.1;Wu1-SARS2_orf3a;210 | 0 | 0.025 |
| NC_045512.2;YP_009724391.1;Wu1-SARS2_orf3a;211 | 0 | 0.025 |
| NC_045512.2;YP_009724391.1;Wu1-SARS2_orf3a;212 | 0 | 0.025 |
| NC_045512.2;YP_009724391.1;Wu1-SARS2_orf3a;213 | 0 | 0.025 |
| NC_045512.2;YP_009724391.1;Wu1-SARS2_orf3a;214 | 0 | 0.025 |
| NC_045512.2;YP_009724391.1;Wu1-SARS2_orf3a;215 | 0 | 0 |
| NC_045512.2;YP_009724391.1;Wu1-SARS2_orf3a;216 | 0 | 0 |
| NC_045512.2;YP_009724391.1;Wu1-SARS2_orf3a;217 | 0 | 0 |
| NC_045512.2;YP_009724391.1;Wu1-SARS2_orf3a;218 | 0 | 0 |
| NC_045512.2;YP_009724391.1;Wu1-SARS2_orf3a;219 | 0 | 0 |
| NC_045512.2;YP_009724391.1;Wu1-SARS2_orf3a;220 | 0.05 | 0 |
| NC_045512.2;YP_009724391.1;Wu1-SARS2_orf3a;221 | 0 | 0 |
| NC_045512.2;YP_009724391.1;Wu1-SARS2_orf3a;222 | 0 | 0 |
| NC_045512.2;YP_009724391.1;Wu1-SARS2_orf3a;223 | 0 | 0 |
| NC_045512.2;YP_009724391.1;Wu1-SARS2_orf3a;224 | 0 | 0 |
| NC_045512.2;YP_009724391.1;Wu1-SARS2_orf3a;225 | 0 | 0 |
| NC_045512.2;YP_009724391.1;Wu1-SARS2_orf3a;226 | 0 | 0 |
| NC_045512.2;YP_009724391.1;Wu1-SARS2_orf3a;227 | 0 | 0 |
| NC_045512.2;YP_009724391.1;Wu1-SARS2_orf3a;228 | 0 | 0 |
| NC_045512.2;YP_009724391.1;Wu1-SARS2_orf3a;229 | 0 | 0 |
| NC_045512.2;YP_009724391.1;Wu1-SARS2_orf3a;230 | 0 | 0.075 |
| NC_045512.2;YP_009724391.1;Wu1-SARS2_orf3a;231 | 0.05 | 0.025 |
| NC_045512.2;YP_009724391.1;Wu1-SARS2_orf3a;232 | 0 | 0.05 |
| NC_045512.2;YP_009724391.1;Wu1-SARS2_orf3a;233 | 0 | 0.075 |
| NC_045512.2;YP_009724391.1;Wu1-SARS2_orf3a;234 | 0 | 0 |
| NC_045512.2;YP_009724391.1;Wu1-SARS2_orf3a;235 | 0 | 0.05 |
| NC_045512.2;YP_009724391.1;Wu1-SARS2_orf3a;236 | 0 | 0.025 |
| NC_045512.2;YP_009724391.1;Wu1-SARS2_orf3a;237 | 0 | 0.05 |
| NC_045512.2;YP_009724391.1;Wu1-SARS2_orf3a;238 | 0 | 0.075 |
| NC_045512.2;YP_009724391.1;Wu1-SARS2_orf3a;239 | 0 | 0.05 |
| NC_045512.2;YP_009724391.1;Wu1-SARS2_orf3a;240 | 0 | 0.1 |
| NC_045512.2;YP_009724391.1;Wu1-SARS2_orf3a;241 | 0.05 | 0.05 |
| NC_045512.2;YP_009724391.1;Wu1-SARS2_orf3a;242 | 0.05 | 0.05 |
| NC_045512.2;YP_009724391.1;Wu1-SARS2_orf3a;243 | 0.05 | 0.1 |

| | | |
|---|---|---|
| NC_045512.2;YP_009724391.1;Wu1-SARS2_orf3a;244 | 0.05 | 0.075 |
| NC_045512.2;YP_009724391.1;Wu1-SARS2_orf3a;245 | 0.05 | 0.075 |
| NC_045512.2;YP_009724391.1;Wu1-SARS2_orf3a;246 | 0.05 | 0.05 |
| NC_045512.2;YP_009724391.1;Wu1-SARS2_orf3a;247 | 0.05 | 0.05 |
| NC_045512.2;YP_009724391.1;Wu1-SARS2_orf3a;248 | 0.05 | 0.075 |
| NC_045512.2;YP_009724391.1;Wu1-SARS2_orf3a;249 | 0.05 | 0.075 |
| NC_045512.2;YP_009724391.1;Wu1-SARS2_orf3a;250 | 0.05 | 0.15 |
| NC_045512.2;YP_009724391.1;Wu1-SARS2_orf3a;251 | 0.05 | 0.15 |
| NC_045512.2;YP_009724391.1;Wu1-SARS2_orf3a;252 | 0.05 | 0.275 |
| NC_045512.2;YP_009724391.1;Wu1-SARS2_orf3a;253 | 0 | 0.3 |
| NC_045512.2;YP_009724391.1;Wu1-SARS2_orf3a;254 | 0 | 0.35 |
| NC_045512.2;YP_009724391.1;Wu1-SARS2_orf3a;255 | 0 | 0.3 |
| NC_045512.2;YP_009724391.1;Wu1-SARS2_orf3a;256 | 0 | 0.35 |
| NC_045512.2;YP_009724391.1;Wu1-SARS2_orf3a;257 | 0.05 | 0.4 |
| NC_045512.2;YP_009724391.1;Wu1-SARS2_orf3a;258 | 0.05 | 0.275 |
| NC_045512.2;YP_009724391.1;Wu1-SARS2_orf3a;259 | 0.05 | 0.325 |
| NC_045512.2;YP_009724391.1;Wu1-SARS2_orf3a;260 | 0.05 | 0.35 |
| NC_045512.2;YP_009724391.1;Wu1-SARS2_orf3a;261 | 0.05 | 0.375 |
| NC_045512.2;YP_009724391.1;Wu1-SARS2_orf3a;262 | 0 | 0.075 |
| NC_045512.2;YP_009724391.1;Wu1-SARS2_orf3a;263 | 0.05 | 0.1 |
| NC_045512.2;YP_009724391.1;Wu1-SARS2_orf3a;264 | 0 | 0.05 |
| NC_045512.2;YP_009724392.1;Wu1-SARS2_envelope;1 | 0 | 0 |
| NC_045512.2;YP_009724392.1;Wu1-SARS2_envelope;2 | 0 | 0 |
| NC_045512.2;YP_009724392.1;Wu1-SARS2_envelope;3 | 0 | 0 |
| NC_045512.2;YP_009724392.1;Wu1-SARS2_envelope;4 | 0 | 0 |
| NC_045512.2;YP_009724392.1;Wu1-SARS2_envelope;5 | 0 | 0 |
| NC_045512.2;YP_009724392.1;Wu1-SARS2_envelope;6 | 0 | 0 |
| NC_045512.2;YP_009724392.1;Wu1-SARS2_envelope;7 | 0 | 0 |
| NC_045512.2;YP_009724392.1;Wu1-SARS2_envelope;8 | 0 | 0 |
| NC_045512.2;YP_009724392.1;Wu1-SARS2_envelope;9 | 0 | 0 |
| NC_045512.2;YP_009724392.1;Wu1-SARS2_envelope;10 | 0 | 0 |
| NC_045512.2;YP_009724392.1;Wu1-SARS2_envelope;11 | 0 | 0 |
| NC_045512.2;YP_009724392.1;Wu1-SARS2_envelope;12 | 0 | 0 |
| NC_045512.2;YP_009724392.1;Wu1-SARS2_envelope;13 | 0 | 0 |
| NC_045512.2;YP_009724392.1;Wu1-SARS2_envelope;14 | 0 | 0 |
| NC_045512.2;YP_009724392.1;Wu1-SARS2_envelope;15 | 0 | 0 |
| NC_045512.2;YP_009724392.1;Wu1-SARS2_envelope;16 | 0 | 0 |
| NC_045512.2;YP_009724392.1;Wu1-SARS2_envelope;17 | 0 | 0 |
| NC_045512.2;YP_009724392.1;Wu1-SARS2_envelope;18 | 0 | 0 |
| NC_045512.2;YP_009724392.1;Wu1-SARS2_envelope;19 | 0 | 0 |
| NC_045512.2;YP_009724392.1;Wu1-SARS2_envelope;20 | 0 | 0 |
| NC_045512.2;YP_009724392.1;Wu1-SARS2_envelope;21 | 0 | 0 |
| NC_045512.2;YP_009724392.1;Wu1-SARS2_envelope;22 | 0 | 0 |

| | | |
|---|---|---|
| NC_045512.2;YP_009724392.1;Wu1-SARS2_envelope;23 | 0 | 0 |
| NC_045512.2;YP_009724392.1;Wu1-SARS2_envelope;24 | 0 | 0 |
| NC_045512.2;YP_009724392.1;Wu1-SARS2_envelope;25 | 0 | 0 |
| NC_045512.2;YP_009724392.1;Wu1-SARS2_envelope;26 | 0 | 0 |
| NC_045512.2;YP_009724392.1;Wu1-SARS2_envelope;27 | 0 | 0 |
| NC_045512.2;YP_009724392.1;Wu1-SARS2_envelope;28 | 0 | 0 |
| NC_045512.2;YP_009724392.1;Wu1-SARS2_envelope;29 | 0 | 0 |
| NC_045512.2;YP_009724392.1;Wu1-SARS2_envelope;30 | 0 | 0 |
| NC_045512.2;YP_009724392.1;Wu1-SARS2_envelope;31 | 0 | 0 |
| NC_045512.2;YP_009724392.1;Wu1-SARS2_envelope;32 | 0 | 0 |
| NC_045512.2;YP_009724392.1;Wu1-SARS2_envelope;33 | 0 | 0 |
| NC_045512.2;YP_009724392.1;Wu1-SARS2_envelope;34 | 0 | 0 |
| NC_045512.2;YP_009724392.1;Wu1-SARS2_envelope;35 | 0 | 0 |
| NC_045512.2;YP_009724392.1;Wu1-SARS2_envelope;36 | 0 | 0 |
| NC_045512.2;YP_009724392.1;Wu1-SARS2_envelope;37 | 0 | 0 |
| NC_045512.2;YP_009724392.1;Wu1-SARS2_envelope;38 | 0 | 0 |
| NC_045512.2;YP_009724392.1;Wu1-SARS2_envelope;39 | 0 | 0 |
| NC_045512.2;YP_009724392.1;Wu1-SARS2_envelope;40 | 0 | 0.025 |
| NC_045512.2;YP_009724392.1;Wu1-SARS2_envelope;41 | 0 | 0 |
| NC_045512.2;YP_009724392.1;Wu1-SARS2_envelope;42 | 0 | 0 |
| NC_045512.2;YP_009724392.1;Wu1-SARS2_envelope;43 | 0 | 0 |
| NC_045512.2;YP_009724392.1;Wu1-SARS2_envelope;44 | 0 | 0 |
| NC_045512.2;YP_009724392.1;Wu1-SARS2_envelope;45 | 0 | 0 |
| NC_045512.2;YP_009724392.1;Wu1-SARS2_envelope;46 | 0 | 0 |
| NC_045512.2;YP_009724392.1;Wu1-SARS2_envelope;47 | 0 | 0 |
| NC_045512.2;YP_009724392.1;Wu1-SARS2_envelope;48 | 0 | 0 |
| NC_045512.2;YP_009724392.1;Wu1-SARS2_envelope;49 | 0 | 0 |
| NC_045512.2;YP_009724392.1;Wu1-SARS2_envelope;50 | 0 | 0 |
| NC_045512.2;YP_009724392.1;Wu1-SARS2_envelope;51 | 0 | 0 |
| NC_045512.2;YP_009724392.1;Wu1-SARS2_envelope;52 | 0 | 0 |
| NC_045512.2;YP_009724392.1;Wu1-SARS2_envelope;53 | 0 | 0 |
| NC_045512.2;YP_009724392.1;Wu1-SARS2_envelope;54 | 0 | 0 |
| NC_045512.2;YP_009724392.1;Wu1-SARS2_envelope;55 | 0 | 0 |
| NC_045512.2;YP_009724392.1;Wu1-SARS2_envelope;56 | 0 | 0 |
| NC_045512.2;YP_009724392.1;Wu1-SARS2_envelope;57 | 0 | 0 |
| NC_045512.2;YP_009724392.1;Wu1-SARS2_envelope;58 | 0 | 0.075 |
| NC_045512.2;YP_009724392.1;Wu1-SARS2_envelope;59 | 0 | 0 |
| NC_045512.2;YP_009724392.1;Wu1-SARS2_envelope;60 | 0.1 | 0 |
| NC_045512.2;YP_009724392.1;Wu1-SARS2_envelope;61 | 0.1 | 0 |
| NC_045512.2;YP_009724392.1;Wu1-SARS2_envelope;62 | 0.05 | 0.05 |
| NC_045512.2;YP_009724392.1;Wu1-SARS2_envelope;63 | 0.1 | 0 |
| NC_045512.2;YP_009724392.1;Wu1-SARS2_envelope;64 | 0.05 | 0.075 |
| NC_045512.2;YP_009724393.1;Wu1-SARS2_membrane;1 | 0 | 0.2 |

| | | |
|---|---|---|
| NC_045512.2;YP_009724393.1;Wu1-SARS2_membrane;2 | 0 | 0.25 |
| NC_045512.2;YP_009724393.1;Wu1-SARS2_membrane;3 | 0 | 0.475 |
| NC_045512.2;YP_009724393.1;Wu1-SARS2_membrane;4 | 0 | 0.6 |
| NC_045512.2;YP_009724393.1;Wu1-SARS2_membrane;5 | 0 | 0.725 |
| NC_045512.2;YP_009724393.1;Wu1-SARS2_membrane;6 | 0 | 0.65 |
| NC_045512.2;YP_009724393.1;Wu1-SARS2_membrane;7 | 0 | 0.775 |
| NC_045512.2;YP_009724393.1;Wu1-SARS2_membrane;8 | 0 | 0.9 |
| NC_045512.2;YP_009724393.1;Wu1-SARS2_membrane;9 | 0 | 0.4 |
| NC_045512.2;YP_009724393.1;Wu1-SARS2_membrane;10 | 0 | 0.025 |
| NC_045512.2;YP_009724393.1;Wu1-SARS2_membrane;11 | 0.05 | 0 |
| NC_045512.2;YP_009724393.1;Wu1-SARS2_membrane;12 | 0 | 0 |
| NC_045512.2;YP_009724393.1;Wu1-SARS2_membrane;13 | 0 | 0 |
| NC_045512.2;YP_009724393.1;Wu1-SARS2_membrane;14 | 0 | 0 |
| NC_045512.2;YP_009724393.1;Wu1-SARS2_membrane;15 | 0 | 0 |
| NC_045512.2;YP_009724393.1;Wu1-SARS2_membrane;16 | 0 | 0 |
| NC_045512.2;YP_009724393.1;Wu1-SARS2_membrane;17 | 0 | 0 |
| NC_045512.2;YP_009724393.1;Wu1-SARS2_membrane;18 | 0 | 0 |
| NC_045512.2;YP_009724393.1;Wu1-SARS2_membrane;19 | 0 | 0 |
| NC_045512.2;YP_009724393.1;Wu1-SARS2_membrane;20 | 0 | 0 |
| NC_045512.2;YP_009724393.1;Wu1-SARS2_membrane;21 | 0 | 0 |
| NC_045512.2;YP_009724393.1;Wu1-SARS2_membrane;22 | 0 | 0 |
| NC_045512.2;YP_009724393.1;Wu1-SARS2_membrane;23 | 0 | 0 |
| NC_045512.2;YP_009724393.1;Wu1-SARS2_membrane;24 | 0 | 0 |
| NC_045512.2;YP_009724393.1;Wu1-SARS2_membrane;25 | 0 | 0 |
| NC_045512.2;YP_009724393.1;Wu1-SARS2_membrane;26 | 0 | 0 |
| NC_045512.2;YP_009724393.1;Wu1-SARS2_membrane;27 | 0 | 0 |
| NC_045512.2;YP_009724393.1;Wu1-SARS2_membrane;28 | 0 | 0 |
| NC_045512.2;YP_009724393.1;Wu1-SARS2_membrane;29 | 0 | 0 |
| NC_045512.2;YP_009724393.1;Wu1-SARS2_membrane;30 | 0 | 0 |
| NC_045512.2;YP_009724393.1;Wu1-SARS2_membrane;31 | 0 | 0 |
| NC_045512.2;YP_009724393.1;Wu1-SARS2_membrane;32 | 0 | 0 |
| NC_045512.2;YP_009724393.1;Wu1-SARS2_membrane;33 | 0 | 0 |
| NC_045512.2;YP_009724393.1;Wu1-SARS2_membrane;34 | 0 | 0 |
| NC_045512.2;YP_009724393.1;Wu1-SARS2_membrane;35 | 0 | 0 |
| NC_045512.2;YP_009724393.1;Wu1-SARS2_membrane;36 | 0 | 0 |
| NC_045512.2;YP_009724393.1;Wu1-SARS2_membrane;37 | 0 | 0 |
| NC_045512.2;YP_009724393.1;Wu1-SARS2_membrane;38 | 0 | 0 |
| NC_045512.2;YP_009724393.1;Wu1-SARS2_membrane;39 | 0 | 0 |
| NC_045512.2;YP_009724393.1;Wu1-SARS2_membrane;40 | 0 | 0 |
| NC_045512.2;YP_009724393.1;Wu1-SARS2_membrane;41 | 0 | 0 |
| NC_045512.2;YP_009724393.1;Wu1-SARS2_membrane;42 | 0 | 0 |
| NC_045512.2;YP_009724393.1;Wu1-SARS2_membrane;43 | 0 | 0 |
| NC_045512.2;YP_009724393.1;Wu1-SARS2_membrane;44 | 0 | 0 |

| | | |
|---|---|---|
| NC_045512.2;YP_009724393.1;Wu1-SARS2_membrane;45 | 0 | 0 |
| NC_045512.2;YP_009724393.1;Wu1-SARS2_membrane;46 | 0 | 0 |
| NC_045512.2;YP_009724393.1;Wu1-SARS2_membrane;47 | 0 | 0 |
| NC_045512.2;YP_009724393.1;Wu1-SARS2_membrane;48 | 0 | 0 |
| NC_045512.2;YP_009724393.1;Wu1-SARS2_membrane;49 | 0 | 0 |
| NC_045512.2;YP_009724393.1;Wu1-SARS2_membrane;50 | 0 | 0 |
| NC_045512.2;YP_009724393.1;Wu1-SARS2_membrane;51 | 0 | 0 |
| NC_045512.2;YP_009724393.1;Wu1-SARS2_membrane;52 | 0 | 0 |
| NC_045512.2;YP_009724393.1;Wu1-SARS2_membrane;53 | 0 | 0 |
| NC_045512.2;YP_009724393.1;Wu1-SARS2_membrane;54 | 0 | 0 |
| NC_045512.2;YP_009724393.1;Wu1-SARS2_membrane;55 | 0 | 0 |
| NC_045512.2;YP_009724393.1;Wu1-SARS2_membrane;56 | 0 | 0 |
| NC_045512.2;YP_009724393.1;Wu1-SARS2_membrane;57 | 0 | 0 |
| NC_045512.2;YP_009724393.1;Wu1-SARS2_membrane;58 | 0 | 0 |
| NC_045512.2;YP_009724393.1;Wu1-SARS2_membrane;59 | 0 | 0 |
| NC_045512.2;YP_009724393.1;Wu1-SARS2_membrane;60 | 0 | 0 |
| NC_045512.2;YP_009724393.1;Wu1-SARS2_membrane;61 | 0 | 0 |
| NC_045512.2;YP_009724393.1;Wu1-SARS2_membrane;62 | 0 | 0 |
| NC_045512.2;YP_009724393.1;Wu1-SARS2_membrane;63 | 0 | 0 |
| NC_045512.2;YP_009724393.1;Wu1-SARS2_membrane;64 | 0 | 0 |
| NC_045512.2;YP_009724393.1;Wu1-SARS2_membrane;65 | 0 | 0 |
| NC_045512.2;YP_009724393.1;Wu1-SARS2_membrane;66 | 0 | 0 |
| NC_045512.2;YP_009724393.1;Wu1-SARS2_membrane;67 | 0 | 0 |
| NC_045512.2;YP_009724393.1;Wu1-SARS2_membrane;68 | 0 | 0 |
| NC_045512.2;YP_009724393.1;Wu1-SARS2_membrane;69 | 0 | 0 |
| NC_045512.2;YP_009724393.1;Wu1-SARS2_membrane;70 | 0 | 0 |
| NC_045512.2;YP_009724393.1;Wu1-SARS2_membrane;71 | 0 | 0 |
| NC_045512.2;YP_009724393.1;Wu1-SARS2_membrane;72 | 0 | 0 |
| NC_045512.2;YP_009724393.1;Wu1-SARS2_membrane;73 | 0 | 0 |
| NC_045512.2;YP_009724393.1;Wu1-SARS2_membrane;74 | 0 | 0 |
| NC_045512.2;YP_009724393.1;Wu1-SARS2_membrane;75 | 0 | 0 |
| NC_045512.2;YP_009724393.1;Wu1-SARS2_membrane;76 | 0 | 0 |
| NC_045512.2;YP_009724393.1;Wu1-SARS2_membrane;77 | 0 | 0 |
| NC_045512.2;YP_009724393.1;Wu1-SARS2_membrane;78 | 0 | 0 |
| NC_045512.2;YP_009724393.1;Wu1-SARS2_membrane;79 | 0 | 0 |
| NC_045512.2;YP_009724393.1;Wu1-SARS2_membrane;80 | 0 | 0 |
| NC_045512.2;YP_009724393.1;Wu1-SARS2_membrane;81 | 0 | 0 |
| NC_045512.2;YP_009724393.1;Wu1-SARS2_membrane;82 | 0 | 0 |
| NC_045512.2;YP_009724393.1;Wu1-SARS2_membrane;83 | 0 | 0 |
| NC_045512.2;YP_009724393.1;Wu1-SARS2_membrane;84 | 0 | 0 |
| NC_045512.2;YP_009724393.1;Wu1-SARS2_membrane;85 | 0 | 0 |
| NC_045512.2;YP_009724393.1;Wu1-SARS2_membrane;86 | 0 | 0 |
| NC_045512.2;YP_009724393.1;Wu1-SARS2_membrane;87 | 0 | 0 |

| | | |
|---|---|---|
| NC_045512.2;YP_009724393.1;Wu1-SARS2_membrane;88 | 0 | 0 |
| NC_045512.2;YP_009724393.1;Wu1-SARS2_membrane;89 | 0 | 0 |
| NC_045512.2;YP_009724393.1;Wu1-SARS2_membrane;90 | 0 | 0 |
| NC_045512.2;YP_009724393.1;Wu1-SARS2_membrane;91 | 0 | 0 |
| NC_045512.2;YP_009724393.1;Wu1-SARS2_membrane;92 | 0 | 0 |
| NC_045512.2;YP_009724393.1;Wu1-SARS2_membrane;93 | 0 | 0 |
| NC_045512.2;YP_009724393.1;Wu1-SARS2_membrane;94 | 0 | 0 |
| NC_045512.2;YP_009724393.1;Wu1-SARS2_membrane;95 | 0 | 0 |
| NC_045512.2;YP_009724393.1;Wu1-SARS2_membrane;96 | 0 | 0 |
| NC_045512.2;YP_009724393.1;Wu1-SARS2_membrane;97 | 0 | 0 |
| NC_045512.2;YP_009724393.1;Wu1-SARS2_membrane;98 | 0 | 0 |
| NC_045512.2;YP_009724393.1;Wu1-SARS2_membrane;99 | 0 | 0 |
| NC_045512.2;YP_009724393.1;Wu1-SARS2_membrane;100 | 0 | 0 |
| NC_045512.2;YP_009724393.1;Wu1-SARS2_membrane;101 | 0.05 | 0.025 |
| NC_045512.2;YP_009724393.1;Wu1-SARS2_membrane;102 | 0 | 0 |
| NC_045512.2;YP_009724393.1;Wu1-SARS2_membrane;103 | 0 | 0 |
| NC_045512.2;YP_009724393.1;Wu1-SARS2_membrane;104 | 0 | 0 |
| NC_045512.2;YP_009724393.1;Wu1-SARS2_membrane;105 | 0 | 0 |
| NC_045512.2;YP_009724393.1;Wu1-SARS2_membrane;106 | 0 | 0 |
| NC_045512.2;YP_009724393.1;Wu1-SARS2_membrane;107 | 0 | 0 |
| NC_045512.2;YP_009724393.1;Wu1-SARS2_membrane;108 | 0 | 0.025 |
| NC_045512.2;YP_009724393.1;Wu1-SARS2_membrane;109 | 0 | 0.025 |
| NC_045512.2;YP_009724393.1;Wu1-SARS2_membrane;110 | 0 | 0 |
| NC_045512.2;YP_009724393.1;Wu1-SARS2_membrane;111 | 0 | 0.025 |
| NC_045512.2;YP_009724393.1;Wu1-SARS2_membrane;112 | 0 | 0.025 |
| NC_045512.2;YP_009724393.1;Wu1-SARS2_membrane;113 | 0 | 0.025 |
| NC_045512.2;YP_009724393.1;Wu1-SARS2_membrane;114 | 0 | 0 |
| NC_045512.2;YP_009724393.1;Wu1-SARS2_membrane;115 | 0 | 0 |
| NC_045512.2;YP_009724393.1;Wu1-SARS2_membrane;116 | 0 | 0 |
| NC_045512.2;YP_009724393.1;Wu1-SARS2_membrane;117 | 0 | 0 |
| NC_045512.2;YP_009724393.1;Wu1-SARS2_membrane;118 | 0 | 0 |
| NC_045512.2;YP_009724393.1;Wu1-SARS2_membrane;119 | 0 | 0 |
| NC_045512.2;YP_009724393.1;Wu1-SARS2_membrane;120 | 0 | 0 |
| NC_045512.2;YP_009724393.1;Wu1-SARS2_membrane;121 | 0 | 0 |
| NC_045512.2;YP_009724393.1;Wu1-SARS2_membrane;122 | 0 | 0 |
| NC_045512.2;YP_009724393.1;Wu1-SARS2_membrane;123 | 0 | 0 |
| NC_045512.2;YP_009724393.1;Wu1-SARS2_membrane;124 | 0 | 0 |
| NC_045512.2;YP_009724393.1;Wu1-SARS2_membrane;125 | 0 | 0 |
| NC_045512.2;YP_009724393.1;Wu1-SARS2_membrane;126 | 0 | 0 |
| NC_045512.2;YP_009724393.1;Wu1-SARS2_membrane;127 | 0 | 0 |
| NC_045512.2;YP_009724393.1;Wu1-SARS2_membrane;128 | 0 | 0 |
| NC_045512.2;YP_009724393.1;Wu1-SARS2_membrane;129 | 0 | 0 |
| NC_045512.2;YP_009724393.1;Wu1-SARS2_membrane;130 | 0 | 0 |

| | | |
|---|---|---|
| NC_045512.2;YP_009724393.1;Wu1-SARS2_membrane;131 | 0 | 0 |
| NC_045512.2;YP_009724393.1;Wu1-SARS2_membrane;132 | 0 | 0 |
| NC_045512.2;YP_009724393.1;Wu1-SARS2_membrane;133 | 0 | 0 |
| NC_045512.2;YP_009724393.1;Wu1-SARS2_membrane;134 | 0 | 0 |
| NC_045512.2;YP_009724393.1;Wu1-SARS2_membrane;135 | 0 | 0 |
| NC_045512.2;YP_009724393.1;Wu1-SARS2_membrane;136 | 0 | 0 |
| NC_045512.2;YP_009724393.1;Wu1-SARS2_membrane;137 | 0 | 0 |
| NC_045512.2;YP_009724393.1;Wu1-SARS2_membrane;138 | 0 | 0 |
| NC_045512.2;YP_009724393.1;Wu1-SARS2_membrane;139 | 0 | 0 |
| NC_045512.2;YP_009724393.1;Wu1-SARS2_membrane;140 | 0 | 0 |
| NC_045512.2;YP_009724393.1;Wu1-SARS2_membrane;141 | 0 | 0 |
| NC_045512.2;YP_009724393.1;Wu1-SARS2_membrane;142 | 0 | 0 |
| NC_045512.2;YP_009724393.1;Wu1-SARS2_membrane;143 | 0 | 0 |
| NC_045512.2;YP_009724393.1;Wu1-SARS2_membrane;144 | 0 | 0 |
| NC_045512.2;YP_009724393.1;Wu1-SARS2_membrane;145 | 0 | 0 |
| NC_045512.2;YP_009724393.1;Wu1-SARS2_membrane;146 | 0 | 0 |
| NC_045512.2;YP_009724393.1;Wu1-SARS2_membrane;147 | 0 | 0 |
| NC_045512.2;YP_009724393.1;Wu1-SARS2_membrane;148 | 0.05 | 0.025 |
| NC_045512.2;YP_009724393.1;Wu1-SARS2_membrane;149 | 0.05 | 0.05 |
| NC_045512.2;YP_009724393.1;Wu1-SARS2_membrane;150 | 0.05 | 0.05 |
| NC_045512.2;YP_009724393.1;Wu1-SARS2_membrane;151 | 0.05 | 0.075 |
| NC_045512.2;YP_009724393.1;Wu1-SARS2_membrane;152 | 0.05 | 0.125 |
| NC_045512.2;YP_009724393.1;Wu1-SARS2_membrane;153 | 0.05 | 0.375 |
| NC_045512.2;YP_009724393.1;Wu1-SARS2_membrane;154 | 0.05 | 0.45 |
| NC_045512.2;YP_009724393.1;Wu1-SARS2_membrane;155 | 0.05 | 0.375 |
| NC_045512.2;YP_009724393.1;Wu1-SARS2_membrane;156 | 0.1 | 0.425 |
| NC_045512.2;YP_009724393.1;Wu1-SARS2_membrane;157 | 0.05 | 0.4 |
| NC_045512.2;YP_009724393.1;Wu1-SARS2_membrane;158 | 0.05 | 0.35 |
| NC_045512.2;YP_009724393.1;Wu1-SARS2_membrane;159 | 0 | 0.35 |
| NC_045512.2;YP_009724393.1;Wu1-SARS2_membrane;160 | 0 | 0.35 |
| NC_045512.2;YP_009724393.1;Wu1-SARS2_membrane;161 | 0 | 0.35 |
| NC_045512.2;YP_009724393.1;Wu1-SARS2_membrane;162 | 0 | 0.35 |
| NC_045512.2;YP_009724393.1;Wu1-SARS2_membrane;163 | 0 | 0.025 |
| NC_045512.2;YP_009724393.1;Wu1-SARS2_membrane;164 | 0 | 0 |
| NC_045512.2;YP_009724393.1;Wu1-SARS2_membrane;165 | 0 | 0 |
| NC_045512.2;YP_009724393.1;Wu1-SARS2_membrane;166 | 0 | 0 |
| NC_045512.2;YP_009724393.1;Wu1-SARS2_membrane;167 | 0 | 0 |
| NC_045512.2;YP_009724393.1;Wu1-SARS2_membrane;168 | 0 | 0 |
| NC_045512.2;YP_009724393.1;Wu1-SARS2_membrane;169 | 0 | 0 |
| NC_045512.2;YP_009724393.1;Wu1-SARS2_membrane;170 | 0 | 0 |
| NC_045512.2;YP_009724393.1;Wu1-SARS2_membrane;171 | 0 | 0 |
| NC_045512.2;YP_009724393.1;Wu1-SARS2_membrane;172 | 0 | 0 |
| NC_045512.2;YP_009724393.1;Wu1-SARS2_membrane;173 | 0 | 0 |

| | | |
|---|---|---|
| NC_045512.2;YP_009724393.1;Wu1-SARS2_membrane;174 | 0 | 0 |
| NC_045512.2;YP_009724393.1;Wu1-SARS2_membrane;175 | 0 | 0.025 |
| NC_045512.2;YP_009724393.1;Wu1-SARS2_membrane;176 | 0 | 0.075 |
| NC_045512.2;YP_009724393.1;Wu1-SARS2_membrane;177 | 0 | 0.1 |
| NC_045512.2;YP_009724393.1;Wu1-SARS2_membrane;178 | 0 | 0.05 |
| NC_045512.2;YP_009724393.1;Wu1-SARS2_membrane;179 | 0 | 0.075 |
| NC_045512.2;YP_009724393.1;Wu1-SARS2_membrane;180 | 0.05 | 0.125 |
| NC_045512.2;YP_009724393.1;Wu1-SARS2_membrane;181 | 0 | 0.35 |
| NC_045512.2;YP_009724393.1;Wu1-SARS2_membrane;182 | 0 | 0.5 |
| NC_045512.2;YP_009724393.1;Wu1-SARS2_membrane;183 | 0 | 0.4 |
| NC_045512.2;YP_009724393.1;Wu1-SARS2_membrane;184 | 0 | 0.325 |
| NC_045512.2;YP_009724393.1;Wu1-SARS2_membrane;185 | 0 | 0.275 |
| NC_045512.2;YP_009724393.1;Wu1-SARS2_membrane;186 | 0 | 0.125 |
| NC_045512.2;YP_009724393.1;Wu1-SARS2_membrane;187 | 0 | 0.175 |
| NC_045512.2;YP_009724393.1;Wu1-SARS2_membrane;188 | 0 | 0.25 |
| NC_045512.2;YP_009724393.1;Wu1-SARS2_membrane;189 | 0 | 0.225 |
| NC_045512.2;YP_009724393.1;Wu1-SARS2_membrane;190 | 0 | 0.2 |
| NC_045512.2;YP_009724393.1;Wu1-SARS2_membrane;191 | 0 | 0.1 |
| NC_045512.2;YP_009724393.1;Wu1-SARS2_membrane;192 | 0 | 0.1 |
| NC_045512.2;YP_009724393.1;Wu1-SARS2_membrane;193 | 0 | 0.075 |
| NC_045512.2;YP_009724393.1;Wu1-SARS2_membrane;194 | 0 | 0.15 |
| NC_045512.2;YP_009724393.1;Wu1-SARS2_membrane;195 | 0 | 0.075 |
| NC_045512.2;YP_009724393.1;Wu1-SARS2_membrane;196 | 0 | 0.1 |
| NC_045512.2;YP_009724393.1;Wu1-SARS2_membrane;197 | 0 | 0.125 |
| NC_045512.2;YP_009724393.1;Wu1-SARS2_membrane;198 | 0 | 0.125 |
| NC_045512.2;YP_009724393.1;Wu1-SARS2_membrane;199 | 0 | 0.125 |
| NC_045512.2;YP_009724393.1;Wu1-SARS2_membrane;200 | 0.05 | 0.2 |
| NC_045512.2;YP_009724393.1;Wu1-SARS2_membrane;201 | 0.1 | 0.125 |
| NC_045512.2;YP_009724393.1;Wu1-SARS2_membrane;202 | 0.05 | 0.125 |
| NC_045512.2;YP_009724393.1;Wu1-SARS2_membrane;203 | 0.05 | 0.125 |
| NC_045512.2;YP_009724393.1;Wu1-SARS2_membrane;204 | 0.05 | 0.175 |
| NC_045512.2;YP_009724393.1;Wu1-SARS2_membrane;205 | 0.05 | 0.225 |
| NC_045512.2;YP_009724393.1;Wu1-SARS2_membrane;206 | 0 | 0.2 |
| NC_045512.2;YP_009724393.1;Wu1-SARS2_membrane;207 | 0 | 0.2 |
| NC_045512.2;YP_009724393.1;Wu1-SARS2_membrane;208 | 0 | 0.225 |
| NC_045512.2;YP_009724393.1;Wu1-SARS2_membrane;209 | 0 | 0.175 |
| NC_045512.2;YP_009724393.1;Wu1-SARS2_membrane;210 | 0 | 0.175 |
| NC_045512.2;YP_009724393.1;Wu1-SARS2_membrane;211 | 0 | 0.175 |
| NC_045512.2;YP_009724394.1;Wu1-SARS2_orf6;1 | 0 | 0 |
| NC_045512.2;YP_009724394.1;Wu1-SARS2_orf6;2 | 0 | 0 |
| NC_045512.2;YP_009724394.1;Wu1-SARS2_orf6;3 | 0 | 0 |
| NC_045512.2;YP_009724394.1;Wu1-SARS2_orf6;4 | 0 | 0 |
| NC_045512.2;YP_009724394.1;Wu1-SARS2_orf6;5 | 0 | 0 |

| | | |
|---|---|---|
| NC_045512.2;YP_009724394.1;Wu1-SARS2_orf6;6 | 0 | 0 |
| NC_045512.2;YP_009724394.1;Wu1-SARS2_orf6;7 | 0 | 0 |
| NC_045512.2;YP_009724394.1;Wu1-SARS2_orf6;8 | 0 | 0 |
| NC_045512.2;YP_009724394.1;Wu1-SARS2_orf6;9 | 0 | 0.025 |
| NC_045512.2;YP_009724394.1;Wu1-SARS2_orf6;10 | 0 | 0 |
| NC_045512.2;YP_009724394.1;Wu1-SARS2_orf6;11 | 0 | 0 |
| NC_045512.2;YP_009724394.1;Wu1-SARS2_orf6;12 | 0 | 0 |
| NC_045512.2;YP_009724394.1;Wu1-SARS2_orf6;13 | 0 | 0 |
| NC_045512.2;YP_009724394.1;Wu1-SARS2_orf6;14 | 0 | 0 |
| NC_045512.2;YP_009724394.1;Wu1-SARS2_orf6;15 | 0 | 0 |
| NC_045512.2;YP_009724394.1;Wu1-SARS2_orf6;16 | 0 | 0 |
| NC_045512.2;YP_009724394.1;Wu1-SARS2_orf6;17 | 0 | 0 |
| NC_045512.2;YP_009724394.1;Wu1-SARS2_orf6;18 | 0 | 0 |
| NC_045512.2;YP_009724394.1;Wu1-SARS2_orf6;19 | 0 | 0 |
| NC_045512.2;YP_009724394.1;Wu1-SARS2_orf6;20 | 0 | 0 |
| NC_045512.2;YP_009724394.1;Wu1-SARS2_orf6;21 | 0 | 0 |
| NC_045512.2;YP_009724394.1;Wu1-SARS2_orf6;22 | 0 | 0 |
| NC_045512.2;YP_009724394.1;Wu1-SARS2_orf6;23 | 0 | 0 |
| NC_045512.2;YP_009724394.1;Wu1-SARS2_orf6;24 | 0 | 0 |
| NC_045512.2;YP_009724394.1;Wu1-SARS2_orf6;25 | 0 | 0 |
| NC_045512.2;YP_009724394.1;Wu1-SARS2_orf6;26 | 0 | 0 |
| NC_045512.2;YP_009724394.1;Wu1-SARS2_orf6;27 | 0 | 0 |
| NC_045512.2;YP_009724394.1;Wu1-SARS2_orf6;28 | 0 | 0 |
| NC_045512.2;YP_009724394.1;Wu1-SARS2_orf6;29 | 0 | 0 |
| NC_045512.2;YP_009724394.1;Wu1-SARS2_orf6;30 | 0 | 0 |
| NC_045512.2;YP_009724394.1;Wu1-SARS2_orf6;31 | 0.05 | 0 |
| NC_045512.2;YP_009724394.1;Wu1-SARS2_orf6;32 | 0.1 | 0 |
| NC_045512.2;YP_009724394.1;Wu1-SARS2_orf6;33 | 0.1 | 0 |
| NC_045512.2;YP_009724394.1;Wu1-SARS2_orf6;34 | 0.05 | 0 |
| NC_045512.2;YP_009724394.1;Wu1-SARS2_orf6;35 | 0.1 | 0 |
| NC_045512.2;YP_009724394.1;Wu1-SARS2_orf6;36 | 0.05 | 0.025 |
| NC_045512.2;YP_009724394.1;Wu1-SARS2_orf6;37 | 0.05 | 0 |
| NC_045512.2;YP_009724394.1;Wu1-SARS2_orf6;38 | 0.05 | 0.025 |
| NC_045512.2;YP_009724394.1;Wu1-SARS2_orf6;39 | 0.05 | 0.075 |
| NC_045512.2;YP_009724394.1;Wu1-SARS2_orf6;40 | 0.05 | 0.05 |
| NC_045512.2;YP_009724394.1;Wu1-SARS2_orf6;41 | 0.1 | 0.075 |
| NC_045512.2;YP_009724394.1;Wu1-SARS2_orf6;42 | 0.05 | 0.075 |
| NC_045512.2;YP_009724394.1;Wu1-SARS2_orf6;43 | 0.05 | 0.075 |
| NC_045512.2;YP_009724394.1;Wu1-SARS2_orf6;44 | 0.05 | 0.075 |
| NC_045512.2;YP_009724394.1;Wu1-SARS2_orf6;45 | 0.1 | 0.05 |
| NC_045512.2;YP_009724394.1;Wu1-SARS2_orf6;46 | 0.05 | 0.05 |
| NC_045512.2;YP_009724394.1;Wu1-SARS2_orf6;47 | 0 | 0 |
| NC_045512.2;YP_009724394.1;Wu1-SARS2_orf6;48 | 0 | 0 |

| | | |
|---|---|---|
| NC_045512.2;YP_009724394.1;Wu1-SARS2_orf6;49 | 0 | 0.025 |
| NC_045512.2;YP_009724394.1;Wu1-SARS2_orf6;50 | 0 | 0 |
| NC_045512.2;YP_009724395.1;Wu1-SARS2_orf7a;1 | 0 | 0 |
| NC_045512.2;YP_009724395.1;Wu1-SARS2_orf7a;2 | 0 | 0 |
| NC_045512.2;YP_009724395.1;Wu1-SARS2_orf7a;3 | 0 | 0 |
| NC_045512.2;YP_009724395.1;Wu1-SARS2_orf7a;4 | 0 | 0 |
| NC_045512.2;YP_009724395.1;Wu1-SARS2_orf7a;5 | 0 | 0 |
| NC_045512.2;YP_009724395.1;Wu1-SARS2_orf7a;6 | 0 | 0 |
| NC_045512.2;YP_009724395.1;Wu1-SARS2_orf7a;7 | 0 | 0 |
| NC_045512.2;YP_009724395.1;Wu1-SARS2_orf7a;8 | 0 | 0 |
| NC_045512.2;YP_009724395.1;Wu1-SARS2_orf7a;9 | 0 | 0 |
| NC_045512.2;YP_009724395.1;Wu1-SARS2_orf7a;10 | 0 | 0 |
| NC_045512.2;YP_009724395.1;Wu1-SARS2_orf7a;11 | 0 | 0 |
| NC_045512.2;YP_009724395.1;Wu1-SARS2_orf7a;12 | 0 | 0 |
| NC_045512.2;YP_009724395.1;Wu1-SARS2_orf7a;13 | 0 | 0 |
| NC_045512.2;YP_009724395.1;Wu1-SARS2_orf7a;14 | 0 | 0 |
| NC_045512.2;YP_009724395.1;Wu1-SARS2_orf7a;15 | 0 | 0 |
| NC_045512.2;YP_009724395.1;Wu1-SARS2_orf7a;16 | 0 | 0 |
| NC_045512.2;YP_009724395.1;Wu1-SARS2_orf7a;17 | 0 | 0 |
| NC_045512.2;YP_009724395.1;Wu1-SARS2_orf7a;18 | 0 | 0 |
| NC_045512.2;YP_009724395.1;Wu1-SARS2_orf7a;19 | 0 | 0 |
| NC_045512.2;YP_009724395.1;Wu1-SARS2_orf7a;20 | 0 | 0 |
| NC_045512.2;YP_009724395.1;Wu1-SARS2_orf7a;21 | 0 | 0 |
| NC_045512.2;YP_009724395.1;Wu1-SARS2_orf7a;22 | 0 | 0 |
| NC_045512.2;YP_009724395.1;Wu1-SARS2_orf7a;23 | 0.05 | 0 |
| NC_045512.2;YP_009724395.1;Wu1-SARS2_orf7a;24 | 0.05 | 0 |
| NC_045512.2;YP_009724395.1;Wu1-SARS2_orf7a;25 | 0.05 | 0 |
| NC_045512.2;YP_009724395.1;Wu1-SARS2_orf7a;26 | 0 | 0 |
| NC_045512.2;YP_009724395.1;Wu1-SARS2_orf7a;27 | 0 | 0 |
| NC_045512.2;YP_009724395.1;Wu1-SARS2_orf7a;28 | 0 | 0.025 |
| NC_045512.2;YP_009724395.1;Wu1-SARS2_orf7a;29 | 0 | 0.025 |
| NC_045512.2;YP_009724395.1;Wu1-SARS2_orf7a;30 | 0 | 0.025 |
| NC_045512.2;YP_009724395.1;Wu1-SARS2_orf7a;31 | 0 | 0 |
| NC_045512.2;YP_009724395.1;Wu1-SARS2_orf7a;32 | 0 | 0 |
| NC_045512.2;YP_009724395.1;Wu1-SARS2_orf7a;33 | 0 | 0 |
| NC_045512.2;YP_009724395.1;Wu1-SARS2_orf7a;34 | 0 | 0.025 |
| NC_045512.2;YP_009724395.1;Wu1-SARS2_orf7a;35 | 0 | 0.025 |
| NC_045512.2;YP_009724395.1;Wu1-SARS2_orf7a;36 | 0.05 | 0.025 |
| NC_045512.2;YP_009724395.1;Wu1-SARS2_orf7a;37 | 0 | 0.05 |
| NC_045512.2;YP_009724395.1;Wu1-SARS2_orf7a;38 | 0 | 0.05 |
| NC_045512.2;YP_009724395.1;Wu1-SARS2_orf7a;39 | 0 | 0.05 |
| NC_045512.2;YP_009724395.1;Wu1-SARS2_orf7a;40 | 0 | 0.05 |
| NC_045512.2;YP_009724395.1;Wu1-SARS2_orf7a;41 | 0 | 0.075 |

| | | |
|---|---|---|
| NC_045512.2;YP_009724395.1;Wu1-SARS2_orf7a;42 | 0 | 0.075 |
| NC_045512.2;YP_009724395.1;Wu1-SARS2_orf7a;43 | 0 | 0.05 |
| NC_045512.2;YP_009724395.1;Wu1-SARS2_orf7a;44 | 0 | 0.025 |
| NC_045512.2;YP_009724395.1;Wu1-SARS2_orf7a;45 | 0 | 0 |
| NC_045512.2;YP_009724395.1;Wu1-SARS2_orf7a;46 | 0 | 0 |
| NC_045512.2;YP_009724395.1;Wu1-SARS2_orf7a;47 | 0 | 0 |
| NC_045512.2;YP_009724395.1;Wu1-SARS2_orf7a;48 | 0 | 0 |
| NC_045512.2;YP_009724395.1;Wu1-SARS2_orf7a;49 | 0 | 0 |
| NC_045512.2;YP_009724395.1;Wu1-SARS2_orf7a;50 | 0 | 0 |
| NC_045512.2;YP_009724395.1;Wu1-SARS2_orf7a;51 | 0 | 0 |
| NC_045512.2;YP_009724395.1;Wu1-SARS2_orf7a;52 | 0 | 0 |
| NC_045512.2;YP_009724395.1;Wu1-SARS2_orf7a;53 | 0 | 0 |
| NC_045512.2;YP_009724395.1;Wu1-SARS2_orf7a;54 | 0 | 0 |
| NC_045512.2;YP_009724395.1;Wu1-SARS2_orf7a;55 | 0 | 0 |
| NC_045512.2;YP_009724395.1;Wu1-SARS2_orf7a;56 | 0 | 0 |
| NC_045512.2;YP_009724395.1;Wu1-SARS2_orf7a;57 | 0 | 0.025 |
| NC_045512.2;YP_009724395.1;Wu1-SARS2_orf7a;58 | 0 | 0 |
| NC_045512.2;YP_009724395.1;Wu1-SARS2_orf7a;59 | 0 | 0 |
| NC_045512.2;YP_009724395.1;Wu1-SARS2_orf7a;60 | 0 | 0.025 |
| NC_045512.2;YP_009724395.1;Wu1-SARS2_orf7a;61 | 0 | 0.025 |
| NC_045512.2;YP_009724395.1;Wu1-SARS2_orf7a;62 | 0 | 0 |
| NC_045512.2;YP_009724395.1;Wu1-SARS2_orf7a;63 | 0 | 0 |
| NC_045512.2;YP_009724395.1;Wu1-SARS2_orf7a;64 | 0 | 0 |
| NC_045512.2;YP_009724395.1;Wu1-SARS2_orf7a;65 | 0 | 0 |
| NC_045512.2;YP_009724395.1;Wu1-SARS2_orf7a;66 | 0 | 0 |
| NC_045512.2;YP_009724395.1;Wu1-SARS2_orf7a;67 | 0 | 0 |
| NC_045512.2;YP_009724395.1;Wu1-SARS2_orf7a;68 | 0 | 0 |
| NC_045512.2;YP_009724395.1;Wu1-SARS2_orf7a;69 | 0 | 0.025 |
| NC_045512.2;YP_009724395.1;Wu1-SARS2_orf7a;70 | 0 | 0 |
| NC_045512.2;YP_009724395.1;Wu1-SARS2_orf7a;71 | 0 | 0 |
| NC_045512.2;YP_009724395.1;Wu1-SARS2_orf7a;72 | 0 | 0 |
| NC_045512.2;YP_009724395.1;Wu1-SARS2_orf7a;73 | 0 | 0 |
| NC_045512.2;YP_009724395.1;Wu1-SARS2_orf7a;74 | 0 | 0 |
| NC_045512.2;YP_009724395.1;Wu1-SARS2_orf7a;75 | 0 | 0 |
| NC_045512.2;YP_009724395.1;Wu1-SARS2_orf7a;76 | 0 | 0 |
| NC_045512.2;YP_009724395.1;Wu1-SARS2_orf7a;77 | 0 | 0 |
| NC_045512.2;YP_009724395.1;Wu1-SARS2_orf7a;78 | 0 | 0 |
| NC_045512.2;YP_009724395.1;Wu1-SARS2_orf7a;79 | 0.05 | 0 |
| NC_045512.2;YP_009724395.1;Wu1-SARS2_orf7a;80 | 0.15 | 0 |
| NC_045512.2;YP_009724395.1;Wu1-SARS2_orf7a;81 | 0.05 | 0 |
| NC_045512.2;YP_009724395.1;Wu1-SARS2_orf7a;82 | 0 | 0 |
| NC_045512.2;YP_009724395.1;Wu1-SARS2_orf7a;83 | 0 | 0 |
| NC_045512.2;YP_009724395.1;Wu1-SARS2_orf7a;84 | 0.05 | 0 |

| | | |
|---|---|---|
| NC_045512.2;YP_009724395.1;Wu1-SARS2_orf7a;85 | 0.05 | 0 |
| NC_045512.2;YP_009724395.1;Wu1-SARS2_orf7a;86 | 0 | 0 |
| NC_045512.2;YP_009724395.1;Wu1-SARS2_orf7a;87 | 0 | 0 |
| NC_045512.2;YP_009724395.1;Wu1-SARS2_orf7a;88 | 0 | 0 |
| NC_045512.2;YP_009724395.1;Wu1-SARS2_orf7a;89 | 0 | 0 |
| NC_045512.2;YP_009724395.1;Wu1-SARS2_orf7a;90 | 0 | 0 |
| NC_045512.2;YP_009724395.1;Wu1-SARS2_orf7a;91 | 0 | 0 |
| NC_045512.2;YP_009724395.1;Wu1-SARS2_orf7a;92 | 0 | 0 |
| NC_045512.2;YP_009724395.1;Wu1-SARS2_orf7a;93 | 0 | 0 |
| NC_045512.2;YP_009724395.1;Wu1-SARS2_orf7a;94 | 0 | 0 |
| NC_045512.2;YP_009724395.1;Wu1-SARS2_orf7a;95 | 0 | 0 |
| NC_045512.2;YP_009724395.1;Wu1-SARS2_orf7a;96 | 0 | 0 |
| NC_045512.2;YP_009724395.1;Wu1-SARS2_orf7a;97 | 0 | 0 |
| NC_045512.2;YP_009724395.1;Wu1-SARS2_orf7a;98 | 0 | 0 |
| NC_045512.2;YP_009724395.1;Wu1-SARS2_orf7a;99 | 0 | 0 |
| NC_045512.2;YP_009724395.1;Wu1-SARS2_orf7a;100 | 0 | 0 |
| NC_045512.2;YP_009724395.1;Wu1-SARS2_orf7a;101 | 0 | 0 |
| NC_045512.2;YP_009724395.1;Wu1-SARS2_orf7a;102 | 0 | 0 |
| NC_045512.2;YP_009724395.1;Wu1-SARS2_orf7a;103 | 0 | 0 |
| NC_045512.2;YP_009724395.1;Wu1-SARS2_orf7a;104 | 0 | 0 |
| NC_045512.2;YP_009724395.1;Wu1-SARS2_orf7a;105 | 0 | 0 |
| NC_045512.2;YP_009724395.1;Wu1-SARS2_orf7a;106 | 0 | 0 |
| NC_045512.2;YP_009724395.1;Wu1-SARS2_orf7a;107 | 0 | 0 |
| NC_045512.2;YP_009724395.1;Wu1-SARS2_orf7a;108 | 0 | 0 |
| NC_045512.2;YP_009724395.1;Wu1-SARS2_orf7a;109 | 0 | 0 |
| NC_045512.2;YP_009724395.1;Wu1-SARS2_orf7a;110 | 0 | 0 |
| NC_045512.2;YP_009724396.1;Wu1-SARS2_orf8;1 | 0 | 0 |
| NC_045512.2;YP_009724396.1;Wu1-SARS2_orf8;2 | 0 | 0 |
| NC_045512.2;YP_009724396.1;Wu1-SARS2_orf8;3 | 0 | 0 |
| NC_045512.2;YP_009724396.1;Wu1-SARS2_orf8;4 | 0 | 0 |
| NC_045512.2;YP_009724396.1;Wu1-SARS2_orf8;5 | 0 | 0 |
| NC_045512.2;YP_009724396.1;Wu1-SARS2_orf8;6 | 0 | 0 |
| NC_045512.2;YP_009724396.1;Wu1-SARS2_orf8;7 | 0 | 0 |
| NC_045512.2;YP_009724396.1;Wu1-SARS2_orf8;8 | 0 | 0 |
| NC_045512.2;YP_009724396.1;Wu1-SARS2_orf8;9 | 0 | 0 |
| NC_045512.2;YP_009724396.1;Wu1-SARS2_orf8;10 | 0 | 0 |
| NC_045512.2;YP_009724396.1;Wu1-SARS2_orf8;11 | 0 | 0 |
| NC_045512.2;YP_009724396.1;Wu1-SARS2_orf8;12 | 0 | 0 |
| NC_045512.2;YP_009724396.1;Wu1-SARS2_orf8;13 | 0 | 0.025 |
| NC_045512.2;YP_009724396.1;Wu1-SARS2_orf8;14 | 0 | 0.025 |
| NC_045512.2;YP_009724396.1;Wu1-SARS2_orf8;15 | 0 | 0.025 |
| NC_045512.2;YP_009724396.1;Wu1-SARS2_orf8;16 | 0 | 0 |
| NC_045512.2;YP_009724396.1;Wu1-SARS2_orf8;17 | 0 | 0 |

| | | |
|---|---|---|
| NC_045512.2;YP_009724396.1;Wu1-SARS2_orf8;18 | 0 | 0 |
| NC_045512.2;YP_009724396.1;Wu1-SARS2_orf8;19 | 0 | 0 |
| NC_045512.2;YP_009724396.1;Wu1-SARS2_orf8;20 | 0 | 0 |
| NC_045512.2;YP_009724396.1;Wu1-SARS2_orf8;21 | 0 | 0 |
| NC_045512.2;YP_009724396.1;Wu1-SARS2_orf8;22 | 0 | 0 |
| NC_045512.2;YP_009724396.1;Wu1-SARS2_orf8;23 | 0 | 0 |
| NC_045512.2;YP_009724396.1;Wu1-SARS2_orf8;24 | 0 | 0 |
| NC_045512.2;YP_009724396.1;Wu1-SARS2_orf8;25 | 0 | 0 |
| NC_045512.2;YP_009724396.1;Wu1-SARS2_orf8;26 | 0 | 0 |
| NC_045512.2;YP_009724396.1;Wu1-SARS2_orf8;27 | 0 | 0 |
| NC_045512.2;YP_009724396.1;Wu1-SARS2_orf8;28 | 0 | 0 |
| NC_045512.2;YP_009724396.1;Wu1-SARS2_orf8;29 | 0 | 0 |
| NC_045512.2;YP_009724396.1;Wu1-SARS2_orf8;30 | 0 | 0 |
| NC_045512.2;YP_009724396.1;Wu1-SARS2_orf8;31 | 0 | 0 |
| NC_045512.2;YP_009724396.1;Wu1-SARS2_orf8;32 | 0 | 0 |
| NC_045512.2;YP_009724396.1;Wu1-SARS2_orf8;33 | 0 | 0 |
| NC_045512.2;YP_009724396.1;Wu1-SARS2_orf8;34 | 0 | 0 |
| NC_045512.2;YP_009724396.1;Wu1-SARS2_orf8;35 | 0 | 0 |
| NC_045512.2;YP_009724396.1;Wu1-SARS2_orf8;36 | 0 | 0 |
| NC_045512.2;YP_009724396.1;Wu1-SARS2_orf8;37 | 0 | 0 |
| NC_045512.2;YP_009724396.1;Wu1-SARS2_orf8;38 | 0 | 0 |
| NC_045512.2;YP_009724396.1;Wu1-SARS2_orf8;39 | 0 | 0 |
| NC_045512.2;YP_009724396.1;Wu1-SARS2_orf8;40 | 0 | 0 |
| NC_045512.2;YP_009724396.1;Wu1-SARS2_orf8;41 | 0 | 0 |
| NC_045512.2;YP_009724396.1;Wu1-SARS2_orf8;42 | 0 | 0 |
| NC_045512.2;YP_009724396.1;Wu1-SARS2_orf8;43 | 0 | 0 |
| NC_045512.2;YP_009724396.1;Wu1-SARS2_orf8;44 | 0 | 0 |
| NC_045512.2;YP_009724396.1;Wu1-SARS2_orf8;45 | 0 | 0 |
| NC_045512.2;YP_009724396.1;Wu1-SARS2_orf8;46 | 0 | 0 |
| NC_045512.2;YP_009724396.1;Wu1-SARS2_orf8;47 | 0 | 0.025 |
| NC_045512.2;YP_009724396.1;Wu1-SARS2_orf8;48 | 0 | 0 |
| NC_045512.2;YP_009724396.1;Wu1-SARS2_orf8;49 | 0 | 0 |
| NC_045512.2;YP_009724396.1;Wu1-SARS2_orf8;50 | 0 | 0.025 |
| NC_045512.2;YP_009724396.1;Wu1-SARS2_orf8;51 | 0 | 0.025 |
| NC_045512.2;YP_009724396.1;Wu1-SARS2_orf8;52 | 0 | 0.025 |
| NC_045512.2;YP_009724396.1;Wu1-SARS2_orf8;53 | 0 | 0.025 |
| NC_045512.2;YP_009724396.1;Wu1-SARS2_orf8;54 | 0 | 0.025 |
| NC_045512.2;YP_009724396.1;Wu1-SARS2_orf8;55 | 0 | 0.025 |
| NC_045512.2;YP_009724396.1;Wu1-SARS2_orf8;56 | 0 | 0.05 |
| NC_045512.2;YP_009724396.1;Wu1-SARS2_orf8;57 | 0 | 0.05 |
| NC_045512.2;YP_009724396.1;Wu1-SARS2_orf8;58 | 0 | 0.05 |
| NC_045512.2;YP_009724396.1;Wu1-SARS2_orf8;59 | 0 | 0.05 |
| NC_045512.2;YP_009724396.1;Wu1-SARS2_orf8;60 | 0 | 0.175 |

| | | |
|---|---|---|
| NC_045512.2;YP_009724396.1;Wu1-SARS2_orf8;61 | 0 | 0.15 |
| NC_045512.2;YP_009724396.1;Wu1-SARS2_orf8;62 | 0.05 | 0.1 |
| NC_045512.2;YP_009724396.1;Wu1-SARS2_orf8;63 | 0 | 0.125 |
| NC_045512.2;YP_009724396.1;Wu1-SARS2_orf8;64 | 0 | 0.2 |
| NC_045512.2;YP_009724396.1;Wu1-SARS2_orf8;65 | 0 | 0.2 |
| NC_045512.2;YP_009724396.1;Wu1-SARS2_orf8;66 | 0 | 0.025 |
| NC_045512.2;YP_009724396.1;Wu1-SARS2_orf8;67 | 0 | 0.075 |
| NC_045512.2;YP_009724396.1;Wu1-SARS2_orf8;68 | 0 | 0.025 |
| NC_045512.2;YP_009724396.1;Wu1-SARS2_orf8;69 | 0 | 0.05 |
| NC_045512.2;YP_009724396.1;Wu1-SARS2_orf8;70 | 0 | 0.025 |
| NC_045512.2;YP_009724396.1;Wu1-SARS2_orf8;71 | 0 | 0 |
| NC_045512.2;YP_009724396.1;Wu1-SARS2_orf8;72 | 0 | 0 |
| NC_045512.2;YP_009724396.1;Wu1-SARS2_orf8;73 | 0 | 0 |
| NC_045512.2;YP_009724396.1;Wu1-SARS2_orf8;74 | 0 | 0 |
| NC_045512.2;YP_009724396.1;Wu1-SARS2_orf8;75 | 0 | 0 |
| NC_045512.2;YP_009724396.1;Wu1-SARS2_orf8;76 | 0 | 0 |
| NC_045512.2;YP_009724396.1;Wu1-SARS2_orf8;77 | 0 | 0 |
| NC_045512.2;YP_009724396.1;Wu1-SARS2_orf8;78 | 0 | 0 |
| NC_045512.2;YP_009724396.1;Wu1-SARS2_orf8;79 | 0 | 0 |
| NC_045512.2;YP_009724396.1;Wu1-SARS2_orf8;80 | 0 | 0 |
| NC_045512.2;YP_009724396.1;Wu1-SARS2_orf8;81 | 0 | 0 |
| NC_045512.2;YP_009724396.1;Wu1-SARS2_orf8;82 | 0 | 0 |
| NC_045512.2;YP_009724396.1;Wu1-SARS2_orf8;83 | 0.05 | 0 |
| NC_045512.2;YP_009724396.1;Wu1-SARS2_orf8;84 | 0.05 | 0 |
| NC_045512.2;YP_009724396.1;Wu1-SARS2_orf8;85 | 0.05 | 0.05 |
| NC_045512.2;YP_009724396.1;Wu1-SARS2_orf8;86 | 0 | 0 |
| NC_045512.2;YP_009724396.1;Wu1-SARS2_orf8;87 | 0 | 0 |
| NC_045512.2;YP_009724396.1;Wu1-SARS2_orf8;88 | 0 | 0 |
| NC_045512.2;YP_009724396.1;Wu1-SARS2_orf8;89 | 0 | 0 |
| NC_045512.2;YP_009724396.1;Wu1-SARS2_orf8;90 | 0 | 0 |
| NC_045512.2;YP_009724396.1;Wu1-SARS2_orf8;91 | 0 | 0 |
| NC_045512.2;YP_009724396.1;Wu1-SARS2_orf8;92 | 0 | 0 |
| NC_045512.2;YP_009724396.1;Wu1-SARS2_orf8;93 | 0 | 0 |
| NC_045512.2;YP_009724396.1;Wu1-SARS2_orf8;94 | 0 | 0 |
| NC_045512.2;YP_009724396.1;Wu1-SARS2_orf8;95 | 0 | 0 |
| NC_045512.2;YP_009724396.1;Wu1-SARS2_orf8;96 | 0 | 0 |
| NC_045512.2;YP_009724396.1;Wu1-SARS2_orf8;97 | 0 | 0 |
| NC_045512.2;YP_009724396.1;Wu1-SARS2_orf8;98 | 0 | 0 |
| NC_045512.2;YP_009724396.1;Wu1-SARS2_orf8;99 | 0 | 0 |
| NC_045512.2;YP_009724396.1;Wu1-SARS2_orf8;100 | 0 | 0 |
| NC_045512.2;YP_009724396.1;Wu1-SARS2_orf8;101 | 0 | 0 |
| NC_045512.2;YP_009724396.1;Wu1-SARS2_orf8;102 | 0 | 0 |
| NC_045512.2;YP_009724396.1;Wu1-SARS2_orf8;103 | 0 | 0 |

| | | |
|---|---|---|
| NC_045512.2;YP_009724396.1;Wu1-SARS2_orf8;104 | 0 | 0 |
| NC_045512.2;YP_009724396.1;Wu1-SARS2_orf8;105 | 0 | 0 |
| NC_045512.2;YP_009724396.1;Wu1-SARS2_orf8;106 | 0 | 0 |
| NC_045512.2;YP_009724396.1;Wu1-SARS2_orf8;107 | 0 | 0 |
| NC_045512.2;YP_009724396.1;Wu1-SARS2_orf8;108 | 0 | 0 |
| NC_045512.2;YP_009724396.1;Wu1-SARS2_orf8;109 | 0 | 0 |
| NC_045512.2;YP_009724396.1;Wu1-SARS2_orf8;110 | 0 | 0 |
| NC_045512.2;YP_009724397.2;Wu1-SARS2_nucleoca;1 | 0.05 | 0.025 |
| NC_045512.2;YP_009724397.2;Wu1-SARS2_nucleoca;2 | 0.05 | 0.025 |
| NC_045512.2;YP_009724397.2;Wu1-SARS2_nucleoca;3 | 0.1 | 0.025 |
| NC_045512.2;YP_009724397.2;Wu1-SARS2_nucleoca;4 | 0.05 | 0.025 |
| NC_045512.2;YP_009724397.2;Wu1-SARS2_nucleoca;5 | 0.1 | 0.05 |
| NC_045512.2;YP_009724397.2;Wu1-SARS2_nucleoca;6 | 0.05 | 0.025 |
| NC_045512.2;YP_009724397.2;Wu1-SARS2_nucleoca;7 | 0.05 | 0.15 |
| NC_045512.2;YP_009724397.2;Wu1-SARS2_nucleoca;8 | 0.05 | 0.125 |
| NC_045512.2;YP_009724397.2;Wu1-SARS2_nucleoca;9 | 0.05 | 0.225 |
| NC_045512.2;YP_009724397.2;Wu1-SARS2_nucleoca;10 | 0.05 | 0.125 |
| NC_045512.2;YP_009724397.2;Wu1-SARS2_nucleoca;11 | 0.05 | 0.275 |
| NC_045512.2;YP_009724397.2;Wu1-SARS2_nucleoca;12 | 0.05 | 0.325 |
| NC_045512.2;YP_009724397.2;Wu1-SARS2_nucleoca;13 | 0.15 | 0.2 |
| NC_045512.2;YP_009724397.2;Wu1-SARS2_nucleoca;14 | 0 | 0.3 |
| NC_045512.2;YP_009724397.2;Wu1-SARS2_nucleoca;15 | 0 | 0.15 |
| NC_045512.2;YP_009724397.2;Wu1-SARS2_nucleoca;16 | 0 | 0.1 |
| NC_045512.2;YP_009724397.2;Wu1-SARS2_nucleoca;17 | 0 | 0.1 |
| NC_045512.2;YP_009724397.2;Wu1-SARS2_nucleoca;18 | 0 | 0.1 |
| NC_045512.2;YP_009724397.2;Wu1-SARS2_nucleoca;19 | 0.05 | 0.1 |
| NC_045512.2;YP_009724397.2;Wu1-SARS2_nucleoca;20 | 0 | 0.075 |
| NC_045512.2;YP_009724397.2;Wu1-SARS2_nucleoca;21 | 0.05 | 0.125 |
| NC_045512.2;YP_009724397.2;Wu1-SARS2_nucleoca;22 | 0.05 | 0.125 |
| NC_045512.2;YP_009724397.2;Wu1-SARS2_nucleoca;23 | 0 | 0.1 |
| NC_045512.2;YP_009724397.2;Wu1-SARS2_nucleoca;24 | 0 | 0.15 |
| NC_045512.2;YP_009724397.2;Wu1-SARS2_nucleoca;25 | 0 | 0.075 |
| NC_045512.2;YP_009724397.2;Wu1-SARS2_nucleoca;26 | 0 | 0.075 |
| NC_045512.2;YP_009724397.2;Wu1-SARS2_nucleoca;27 | 0 | 0.05 |
| NC_045512.2;YP_009724397.2;Wu1-SARS2_nucleoca;28 | 0 | 0.05 |
| NC_045512.2;YP_009724397.2;Wu1-SARS2_nucleoca;29 | 0 | 0.075 |
| NC_045512.2;YP_009724397.2;Wu1-SARS2_nucleoca;30 | 0 | 0.075 |
| NC_045512.2;YP_009724397.2;Wu1-SARS2_nucleoca;31 | 0 | 0.075 |
| NC_045512.2;YP_009724397.2;Wu1-SARS2_nucleoca;32 | 0 | 0.05 |
| NC_045512.2;YP_009724397.2;Wu1-SARS2_nucleoca;33 | 0 | 0.1 |
| NC_045512.2;YP_009724397.2;Wu1-SARS2_nucleoca;34 | 0 | 0.125 |
| NC_045512.2;YP_009724397.2;Wu1-SARS2_nucleoca;35 | 0 | 0.075 |
| NC_045512.2;YP_009724397.2;Wu1-SARS2_nucleoca;36 | 0 | 0.125 |

| | | |
|---|---|---|
| NC_045512.2;YP_009724397.2;Wu1-SARS2_nucleoca;37 | 0 | 0.425 |
| NC_045512.2;YP_009724397.2;Wu1-SARS2_nucleoca;38 | 0 | 0.225 |
| NC_045512.2;YP_009724397.2;Wu1-SARS2_nucleoca;39 | 0 | 0.15 |
| NC_045512.2;YP_009724397.2;Wu1-SARS2_nucleoca;40 | 0 | 0.125 |
| NC_045512.2;YP_009724397.2;Wu1-SARS2_nucleoca;41 | 0 | 0.025 |
| NC_045512.2;YP_009724397.2;Wu1-SARS2_nucleoca;42 | 0 | 0 |
| NC_045512.2;YP_009724397.2;Wu1-SARS2_nucleoca;43 | 0 | 0.025 |
| NC_045512.2;YP_009724397.2;Wu1-SARS2_nucleoca;44 | 0 | 0 |
| NC_045512.2;YP_009724397.2;Wu1-SARS2_nucleoca;45 | 0 | 0 |
| NC_045512.2;YP_009724397.2;Wu1-SARS2_nucleoca;46 | 0 | 0 |
| NC_045512.2;YP_009724397.2;Wu1-SARS2_nucleoca;47 | 0 | 0 |
| NC_045512.2;YP_009724397.2;Wu1-SARS2_nucleoca;48 | 0 | 0 |
| NC_045512.2;YP_009724397.2;Wu1-SARS2_nucleoca;49 | 0 | 0 |
| NC_045512.2;YP_009724397.2;Wu1-SARS2_nucleoca;50 | 0 | 0 |
| NC_045512.2;YP_009724397.2;Wu1-SARS2_nucleoca;51 | 0 | 0.025 |
| NC_045512.2;YP_009724397.2;Wu1-SARS2_nucleoca;52 | 0 | 0 |
| NC_045512.2;YP_009724397.2;Wu1-SARS2_nucleoca;53 | 0 | 0 |
| NC_045512.2;YP_009724397.2;Wu1-SARS2_nucleoca;54 | 0 | 0 |
| NC_045512.2;YP_009724397.2;Wu1-SARS2_nucleoca;55 | 0 | 0 |
| NC_045512.2;YP_009724397.2;Wu1-SARS2_nucleoca;56 | 0 | 0 |
| NC_045512.2;YP_009724397.2;Wu1-SARS2_nucleoca;57 | 0 | 0 |
| NC_045512.2;YP_009724397.2;Wu1-SARS2_nucleoca;58 | 0 | 0.025 |
| NC_045512.2;YP_009724397.2;Wu1-SARS2_nucleoca;59 | 0 | 0.025 |
| NC_045512.2;YP_009724397.2;Wu1-SARS2_nucleoca;60 | 0 | 0.075 |
| NC_045512.2;YP_009724397.2;Wu1-SARS2_nucleoca;61 | 0 | 0.05 |
| NC_045512.2;YP_009724397.2;Wu1-SARS2_nucleoca;62 | 0.05 | 0.125 |
| NC_045512.2;YP_009724397.2;Wu1-SARS2_nucleoca;63 | 0 | 0.1 |
| NC_045512.2;YP_009724397.2;Wu1-SARS2_nucleoca;64 | 0.1 | 0.1 |
| NC_045512.2;YP_009724397.2;Wu1-SARS2_nucleoca;65 | 0 | 0.1 |
| NC_045512.2;YP_009724397.2;Wu1-SARS2_nucleoca;66 | 0 | 0.175 |
| NC_045512.2;YP_009724397.2;Wu1-SARS2_nucleoca;67 | 0.1 | 0.1 |
| NC_045512.2;YP_009724397.2;Wu1-SARS2_nucleoca;68 | 0 | 0.1 |
| NC_045512.2;YP_009724397.2;Wu1-SARS2_nucleoca;69 | 0 | 0.075 |
| NC_045512.2;YP_009724397.2;Wu1-SARS2_nucleoca;70 | 0 | 0.075 |
| NC_045512.2;YP_009724397.2;Wu1-SARS2_nucleoca;71 | 0 | 0.05 |
| NC_045512.2;YP_009724397.2;Wu1-SARS2_nucleoca;72 | 0 | 0.025 |
| NC_045512.2;YP_009724397.2;Wu1-SARS2_nucleoca;73 | 0.15 | 0.1 |
| NC_045512.2;YP_009724397.2;Wu1-SARS2_nucleoca;74 | 0 | 0.025 |
| NC_045512.2;YP_009724397.2;Wu1-SARS2_nucleoca;75 | 0 | 0.025 |
| NC_045512.2;YP_009724397.2;Wu1-SARS2_nucleoca;76 | 0 | 0 |
| NC_045512.2;YP_009724397.2;Wu1-SARS2_nucleoca;77 | 0 | 0 |
| NC_045512.2;YP_009724397.2;Wu1-SARS2_nucleoca;78 | 0.05 | 0 |
| NC_045512.2;YP_009724397.2;Wu1-SARS2_nucleoca;79 | 0.05 | 0 |

| | | |
|---|---|---|
| NC_045512.2;YP_009724397.2;Wu1-SARS2_nucleoca;80 | 0 | 0 |
| NC_045512.2;YP_009724397.2;Wu1-SARS2_nucleoca;81 | 0 | 0 |
| NC_045512.2;YP_009724397.2;Wu1-SARS2_nucleoca;82 | 0 | 0 |
| NC_045512.2;YP_009724397.2;Wu1-SARS2_nucleoca;83 | 0 | 0 |
| NC_045512.2;YP_009724397.2;Wu1-SARS2_nucleoca;84 | 0 | 0 |
| NC_045512.2;YP_009724397.2;Wu1-SARS2_nucleoca;85 | 0.05 | 0 |
| NC_045512.2;YP_009724397.2;Wu1-SARS2_nucleoca;86 | 0 | 0 |
| NC_045512.2;YP_009724397.2;Wu1-SARS2_nucleoca;87 | 0 | 0.025 |
| NC_045512.2;YP_009724397.2;Wu1-SARS2_nucleoca;88 | 0 | 0 |
| NC_045512.2;YP_009724397.2;Wu1-SARS2_nucleoca;89 | 0 | 0.025 |
| NC_045512.2;YP_009724397.2;Wu1-SARS2_nucleoca;90 | 0 | 0.025 |
| NC_045512.2;YP_009724397.2;Wu1-SARS2_nucleoca;91 | 0 | 0.025 |
| NC_045512.2;YP_009724397.2;Wu1-SARS2_nucleoca;92 | 0 | 0.025 |
| NC_045512.2;YP_009724397.2;Wu1-SARS2_nucleoca;93 | 0 | 0.025 |
| NC_045512.2;YP_009724397.2;Wu1-SARS2_nucleoca;94 | 0 | 0.025 |
| NC_045512.2;YP_009724397.2;Wu1-SARS2_nucleoca;95 | 0 | 0.025 |
| NC_045512.2;YP_009724397.2;Wu1-SARS2_nucleoca;96 | 0 | 0.025 |
| NC_045512.2;YP_009724397.2;Wu1-SARS2_nucleoca;97 | 0 | 0.025 |
| NC_045512.2;YP_009724397.2;Wu1-SARS2_nucleoca;98 | 0 | 0.025 |
| NC_045512.2;YP_009724397.2;Wu1-SARS2_nucleoca;99 | 0 | 0 |
| NC_045512.2;YP_009724397.2;Wu1-SARS2_nucleoca;100 | 0 | 0 |
| NC_045512.2;YP_009724397.2;Wu1-SARS2_nucleoca;101 | 0 | 0 |
| NC_045512.2;YP_009724397.2;Wu1-SARS2_nucleoca;102 | 0 | 0 |
| NC_045512.2;YP_009724397.2;Wu1-SARS2_nucleoca;103 | 0 | 0 |
| NC_045512.2;YP_009724397.2;Wu1-SARS2_nucleoca;104 | 0 | 0 |
| NC_045512.2;YP_009724397.2;Wu1-SARS2_nucleoca;105 | 0 | 0 |
| NC_045512.2;YP_009724397.2;Wu1-SARS2_nucleoca;106 | 0 | 0 |
| NC_045512.2;YP_009724397.2;Wu1-SARS2_nucleoca;107 | 0 | 0 |
| NC_045512.2;YP_009724397.2;Wu1-SARS2_nucleoca;108 | 0 | 0 |
| NC_045512.2;YP_009724397.2;Wu1-SARS2_nucleoca;109 | 0 | 0 |
| NC_045512.2;YP_009724397.2;Wu1-SARS2_nucleoca;110 | 0 | 0 |
| NC_045512.2;YP_009724397.2;Wu1-SARS2_nucleoca;111 | 0.05 | 0 |
| NC_045512.2;YP_009724397.2;Wu1-SARS2_nucleoca;112 | 0.1 | 0.125 |
| NC_045512.2;YP_009724397.2;Wu1-SARS2_nucleoca;113 | 0 | 0.1 |
| NC_045512.2;YP_009724397.2;Wu1-SARS2_nucleoca;114 | 0 | 0.1 |
| NC_045512.2;YP_009724397.2;Wu1-SARS2_nucleoca;115 | 0 | 0.1 |
| NC_045512.2;YP_009724397.2;Wu1-SARS2_nucleoca;116 | 0 | 0.15 |
| NC_045512.2;YP_009724397.2;Wu1-SARS2_nucleoca;117 | 0 | 0.15 |
| NC_045512.2;YP_009724397.2;Wu1-SARS2_nucleoca;118 | 0 | 0.05 |
| NC_045512.2;YP_009724397.2;Wu1-SARS2_nucleoca;119 | 0 | 0.05 |
| NC_045512.2;YP_009724397.2;Wu1-SARS2_nucleoca;120 | 0 | 0.05 |
| NC_045512.2;YP_009724397.2;Wu1-SARS2_nucleoca;121 | 0 | 0.025 |
| NC_045512.2;YP_009724397.2;Wu1-SARS2_nucleoca;122 | 0 | 0.025 |

| | | |
|---|---|---|
| NC_045512.2;YP_009724397.2;Wu1-SARS2_nucleoca;123 | 0 | 0 |
| NC_045512.2;YP_009724397.2;Wu1-SARS2_nucleoca;124 | 0 | 0 |
| NC_045512.2;YP_009724397.2;Wu1-SARS2_nucleoca;125 | 0 | 0.025 |
| NC_045512.2;YP_009724397.2;Wu1-SARS2_nucleoca;126 | 0 | 0 |
| NC_045512.2;YP_009724397.2;Wu1-SARS2_nucleoca;127 | 0 | 0 |
| NC_045512.2;YP_009724397.2;Wu1-SARS2_nucleoca;128 | 0 | 0.05 |
| NC_045512.2;YP_009724397.2;Wu1-SARS2_nucleoca;129 | 0 | 0.075 |
| NC_045512.2;YP_009724397.2;Wu1-SARS2_nucleoca;130 | 0 | 0.05 |
| NC_045512.2;YP_009724397.2;Wu1-SARS2_nucleoca;131 | 0 | 0.075 |
| NC_045512.2;YP_009724397.2;Wu1-SARS2_nucleoca;132 | 0 | 0.075 |
| NC_045512.2;YP_009724397.2;Wu1-SARS2_nucleoca;133 | 0 | 0.05 |
| NC_045512.2;YP_009724397.2;Wu1-SARS2_nucleoca;134 | 0 | 0.025 |
| NC_045512.2;YP_009724397.2;Wu1-SARS2_nucleoca;135 | 0 | 0.025 |
| NC_045512.2;YP_009724397.2;Wu1-SARS2_nucleoca;136 | 0 | 0.025 |
| NC_045512.2;YP_009724397.2;Wu1-SARS2_nucleoca;137 | 0 | 0 |
| NC_045512.2;YP_009724397.2;Wu1-SARS2_nucleoca;138 | 0.05 | 0.05 |
| NC_045512.2;YP_009724397.2;Wu1-SARS2_nucleoca;139 | 0 | 0.05 |
| NC_045512.2;YP_009724397.2;Wu1-SARS2_nucleoca;140 | 0 | 0.075 |
| NC_045512.2;YP_009724397.2;Wu1-SARS2_nucleoca;141 | 0.05 | 0.125 |
| NC_045512.2;YP_009724397.2;Wu1-SARS2_nucleoca;142 | 0 | 0.05 |
| NC_045512.2;YP_009724397.2;Wu1-SARS2_nucleoca;143 | 0 | 0.05 |
| NC_045512.2;YP_009724397.2;Wu1-SARS2_nucleoca;144 | 0 | 0.025 |
| NC_045512.2;YP_009724397.2;Wu1-SARS2_nucleoca;145 | 0 | 0.05 |
| NC_045512.2;YP_009724397.2;Wu1-SARS2_nucleoca;146 | 0 | 0.025 |
| NC_045512.2;YP_009724397.2;Wu1-SARS2_nucleoca;147 | 0 | 0.025 |
| NC_045512.2;YP_009724397.2;Wu1-SARS2_nucleoca;148 | 0 | 0.075 |
| NC_045512.2;YP_009724397.2;Wu1-SARS2_nucleoca;149 | 0 | 0.05 |
| NC_045512.2;YP_009724397.2;Wu1-SARS2_nucleoca;150 | 0 | 0.05 |
| NC_045512.2;YP_009724397.2;Wu1-SARS2_nucleoca;151 | 0 | 0 |
| NC_045512.2;YP_009724397.2;Wu1-SARS2_nucleoca;152 | 0 | 0.05 |
| NC_045512.2;YP_009724397.2;Wu1-SARS2_nucleoca;153 | 0 | 0.075 |
| NC_045512.2;YP_009724397.2;Wu1-SARS2_nucleoca;154 | 0.15 | 0.45 |
| NC_045512.2;YP_009724397.2;Wu1-SARS2_nucleoca;155 | 0.15 | 0.5 |
| NC_045512.2;YP_009724397.2;Wu1-SARS2_nucleoca;156 | 0.1 | 0.45 |
| NC_045512.2;YP_009724397.2;Wu1-SARS2_nucleoca;157 | 0.1 | 0.45 |
| NC_045512.2;YP_009724397.2;Wu1-SARS2_nucleoca;158 | 0.1 | 0.5 |
| NC_045512.2;YP_009724397.2;Wu1-SARS2_nucleoca;159 | 0.2 | 0.625 |
| NC_045512.2;YP_009724397.2;Wu1-SARS2_nucleoca;160 | 0.2 | 0.625 |
| NC_045512.2;YP_009724397.2;Wu1-SARS2_nucleoca;161 | 0.1 | 0.6 |
| NC_045512.2;YP_009724397.2;Wu1-SARS2_nucleoca;162 | 0.1 | 0.475 |
| NC_045512.2;YP_009724397.2;Wu1-SARS2_nucleoca;163 | 0.1 | 0.45 |
| NC_045512.2;YP_009724397.2;Wu1-SARS2_nucleoca;164 | 0.05 | 0.1 |
| NC_045512.2;YP_009724397.2;Wu1-SARS2_nucleoca;165 | 0 | 0.05 |

| | | |
|---|---|---|
| NC_045512.2;YP_009724397.2;Wu1-SARS2_nucleoca;166 | 0 | 0.05 |
| NC_045512.2;YP_009724397.2;Wu1-SARS2_nucleoca;167 | 0 | 0 |
| NC_045512.2;YP_009724397.2;Wu1-SARS2_nucleoca;168 | 0.05 | 0.025 |
| NC_045512.2;YP_009724397.2;Wu1-SARS2_nucleoca;169 | 0 | 0.05 |
| NC_045512.2;YP_009724397.2;Wu1-SARS2_nucleoca;170 | 0 | 0 |
| NC_045512.2;YP_009724397.2;Wu1-SARS2_nucleoca;171 | 0 | 0 |
| NC_045512.2;YP_009724397.2;Wu1-SARS2_nucleoca;172 | 0 | 0 |
| NC_045512.2;YP_009724397.2;Wu1-SARS2_nucleoca;173 | 0 | 0.025 |
| NC_045512.2;YP_009724397.2;Wu1-SARS2_nucleoca;174 | 0 | 0.05 |
| NC_045512.2;YP_009724397.2;Wu1-SARS2_nucleoca;175 | 0 | 0.025 |
| NC_045512.2;YP_009724397.2;Wu1-SARS2_nucleoca;176 | 0 | 0.025 |
| NC_045512.2;YP_009724397.2;Wu1-SARS2_nucleoca;177 | 0 | 0.05 |
| NC_045512.2;YP_009724397.2;Wu1-SARS2_nucleoca;178 | 0 | 0.075 |
| NC_045512.2;YP_009724397.2;Wu1-SARS2_nucleoca;179 | 0 | 0.1 |
| NC_045512.2;YP_009724397.2;Wu1-SARS2_nucleoca;180 | 0 | 0.15 |
| NC_045512.2;YP_009724397.2;Wu1-SARS2_nucleoca;181 | 0.05 | 0.125 |
| NC_045512.2;YP_009724397.2;Wu1-SARS2_nucleoca;182 | 0 | 0.125 |
| NC_045512.2;YP_009724397.2;Wu1-SARS2_nucleoca;183 | 0.15 | 0.15 |
| NC_045512.2;YP_009724397.2;Wu1-SARS2_nucleoca;184 | 0.15 | 0.125 |
| NC_045512.2;YP_009724397.2;Wu1-SARS2_nucleoca;185 | 0.15 | 0.15 |
| NC_045512.2;YP_009724397.2;Wu1-SARS2_nucleoca;186 | 0.15 | 0.2 |
| NC_045512.2;YP_009724397.2;Wu1-SARS2_nucleoca;187 | 0.15 | 0.225 |
| NC_045512.2;YP_009724397.2;Wu1-SARS2_nucleoca;188 | 0.1 | 0.2 |
| NC_045512.2;YP_009724397.2;Wu1-SARS2_nucleoca;189 | 0.1 | 0.1 |
| NC_045512.2;YP_009724397.2;Wu1-SARS2_nucleoca;190 | 0.2 | 0.225 |
| NC_045512.2;YP_009724397.2;Wu1-SARS2_nucleoca;191 | 0.05 | 0.05 |
| NC_045512.2;YP_009724397.2;Wu1-SARS2_nucleoca;192 | 0 | 0.075 |
| NC_045512.2;YP_009724397.2;Wu1-SARS2_nucleoca;193 | 0 | 0.025 |
| NC_045512.2;YP_009724397.2;Wu1-SARS2_nucleoca;194 | 0 | 0.05 |
| NC_045512.2;YP_009724397.2;Wu1-SARS2_nucleoca;195 | 0 | 0.025 |
| NC_045512.2;YP_009724397.2;Wu1-SARS2_nucleoca;196 | 0 | 0.05 |
| NC_045512.2;YP_009724397.2;Wu1-SARS2_nucleoca;197 | 0.05 | 0.05 |
| NC_045512.2;YP_009724397.2;Wu1-SARS2_nucleoca;198 | 0 | 0.025 |
| NC_045512.2;YP_009724397.2;Wu1-SARS2_nucleoca;199 | 0 | 0.025 |
| NC_045512.2;YP_009724397.2;Wu1-SARS2_nucleoca;200 | 0.05 | 0.1 |
| NC_045512.2;YP_009724397.2;Wu1-SARS2_nucleoca;201 | 0.05 | 0.025 |
| NC_045512.2;YP_009724397.2;Wu1-SARS2_nucleoca;202 | 0.05 | 0.025 |
| NC_045512.2;YP_009724397.2;Wu1-SARS2_nucleoca;203 | 0.05 | 0.05 |
| NC_045512.2;YP_009724397.2;Wu1-SARS2_nucleoca;204 | 0 | 0.05 |
| NC_045512.2;YP_009724397.2;Wu1-SARS2_nucleoca;205 | 0 | 0.05 |
| NC_045512.2;YP_009724397.2;Wu1-SARS2_nucleoca;206 | 0 | 0.05 |
| NC_045512.2;YP_009724397.2;Wu1-SARS2_nucleoca;207 | 0 | 0 |
| NC_045512.2;YP_009724397.2;Wu1-SARS2_nucleoca;208 | 0 | 0.05 |

| | | |
|---|---|---|
| NC_045512.2;YP_009724397.2;Wu1-SARS2_nucleoca;209 | 0 | 0 |
| NC_045512.2;YP_009724397.2;Wu1-SARS2_nucleoca;210 | 0 | 0.05 |
| NC_045512.2;YP_009724397.2;Wu1-SARS2_nucleoca;211 | 0 | 0.1 |
| NC_045512.2;YP_009724397.2;Wu1-SARS2_nucleoca;212 | 0 | 0.125 |
| NC_045512.2;YP_009724397.2;Wu1-SARS2_nucleoca;213 | 0 | 0.125 |
| NC_045512.2;YP_009724397.2;Wu1-SARS2_nucleoca;214 | 0 | 0.2 |
| NC_045512.2;YP_009724397.2;Wu1-SARS2_nucleoca;215 | 0 | 0.125 |
| NC_045512.2;YP_009724397.2;Wu1-SARS2_nucleoca;216 | 0 | 0.05 |
| NC_045512.2;YP_009724397.2;Wu1-SARS2_nucleoca;217 | 0 | 0 |
| NC_045512.2;YP_009724397.2;Wu1-SARS2_nucleoca;218 | 0.05 | 0.05 |
| NC_045512.2;YP_009724397.2;Wu1-SARS2_nucleoca;219 | 0.05 | 0.1 |
| NC_045512.2;YP_009724397.2;Wu1-SARS2_nucleoca;220 | 0.05 | 0.225 |
| NC_045512.2;YP_009724397.2;Wu1-SARS2_nucleoca;221 | 0.05 | 0.4 |
| NC_045512.2;YP_009724397.2;Wu1-SARS2_nucleoca;222 | 0.1 | 0.325 |
| NC_045512.2;YP_009724397.2;Wu1-SARS2_nucleoca;223 | 0.05 | 0.175 |
| NC_045512.2;YP_009724397.2;Wu1-SARS2_nucleoca;224 | 0.05 | 0.075 |
| NC_045512.2;YP_009724397.2;Wu1-SARS2_nucleoca;225 | 0.05 | 0.1 |
| NC_045512.2;YP_009724397.2;Wu1-SARS2_nucleoca;226 | 0.05 | 0.175 |
| NC_045512.2;YP_009724397.2;Wu1-SARS2_nucleoca;227 | 0 | 0.25 |
| NC_045512.2;YP_009724397.2;Wu1-SARS2_nucleoca;228 | 0 | 0.275 |
| NC_045512.2;YP_009724397.2;Wu1-SARS2_nucleoca;229 | 0.05 | 0.275 |
| NC_045512.2;YP_009724397.2;Wu1-SARS2_nucleoca;230 | 0 | 0.275 |
| NC_045512.2;YP_009724397.2;Wu1-SARS2_nucleoca;231 | 0.05 | 0.375 |
| NC_045512.2;YP_009724397.2;Wu1-SARS2_nucleoca;232 | 0 | 0.275 |
| NC_045512.2;YP_009724397.2;Wu1-SARS2_nucleoca;233 | 0 | 0.25 |
| NC_045512.2;YP_009724397.2;Wu1-SARS2_nucleoca;234 | 0 | 0.1 |
| NC_045512.2;YP_009724397.2;Wu1-SARS2_nucleoca;235 | 0 | 0.1 |
| NC_045512.2;YP_009724397.2;Wu1-SARS2_nucleoca;236 | 0 | 0.05 |
| NC_045512.2;YP_009724397.2;Wu1-SARS2_nucleoca;237 | 0 | 0.025 |
| NC_045512.2;YP_009724397.2;Wu1-SARS2_nucleoca;238 | 0.05 | 0.25 |
| NC_045512.2;YP_009724397.2;Wu1-SARS2_nucleoca;239 | 0.05 | 0.175 |
| NC_045512.2;YP_009724397.2;Wu1-SARS2_nucleoca;240 | 0.25 | 0.225 |
| NC_045512.2;YP_009724397.2;Wu1-SARS2_nucleoca;241 | 0.05 | 0.075 |
| NC_045512.2;YP_009724397.2;Wu1-SARS2_nucleoca;242 | 0 | 0.025 |
| NC_045512.2;YP_009724397.2;Wu1-SARS2_nucleoca;243 | 0 | 0.075 |
| NC_045512.2;YP_009724397.2;Wu1-SARS2_nucleoca;244 | 0 | 0.05 |
| NC_045512.2;YP_009724397.2;Wu1-SARS2_nucleoca;245 | 0 | 0.075 |
| NC_045512.2;YP_009724397.2;Wu1-SARS2_nucleoca;246 | 0 | 0.025 |
| NC_045512.2;YP_009724397.2;Wu1-SARS2_nucleoca;247 | 0.05 | 0.025 |
| NC_045512.2;YP_009724397.2;Wu1-SARS2_nucleoca;248 | 0 | 0.075 |
| NC_045512.2;YP_009724397.2;Wu1-SARS2_nucleoca;249 | 0 | 0.075 |
| NC_045512.2;YP_009724397.2;Wu1-SARS2_nucleoca;250 | 0 | 0.3 |
| NC_045512.2;YP_009724397.2;Wu1-SARS2_nucleoca;251 | 0 | 0 |

| | | |
|---|---|---|
| NC_045512.2;YP_009724397.2;Wu1-SARS2_nucleoca;252 | 0 | 0 |
| NC_045512.2;YP_009724397.2;Wu1-SARS2_nucleoca;253 | 0 | 0 |
| NC_045512.2;YP_009724397.2;Wu1-SARS2_nucleoca;254 | 0 | 0 |
| NC_045512.2;YP_009724397.2;Wu1-SARS2_nucleoca;255 | 0 | 0 |
| NC_045512.2;YP_009724397.2;Wu1-SARS2_nucleoca;256 | 0 | 0.05 |
| NC_045512.2;YP_009724397.2;Wu1-SARS2_nucleoca;257 | 0 | 0.075 |
| NC_045512.2;YP_009724397.2;Wu1-SARS2_nucleoca;258 | 0.05 | 0.075 |
| NC_045512.2;YP_009724397.2;Wu1-SARS2_nucleoca;259 | 0 | 0.075 |
| NC_045512.2;YP_009724397.2;Wu1-SARS2_nucleoca;260 | 0 | 0.1 |
| NC_045512.2;YP_009724397.2;Wu1-SARS2_nucleoca;261 | 0 | 0.075 |
| NC_045512.2;YP_009724397.2;Wu1-SARS2_nucleoca;262 | 0 | 0.05 |
| NC_045512.2;YP_009724397.2;Wu1-SARS2_nucleoca;263 | 0.05 | 0.05 |
| NC_045512.2;YP_009724397.2;Wu1-SARS2_nucleoca;264 | 0.05 | 0.075 |
| NC_045512.2;YP_009724397.2;Wu1-SARS2_nucleoca;265 | 0.05 | 0.075 |
| NC_045512.2;YP_009724397.2;Wu1-SARS2_nucleoca;266 | 0.05 | 0.075 |
| NC_045512.2;YP_009724397.2;Wu1-SARS2_nucleoca;267 | 0.15 | 0 |
| NC_045512.2;YP_009724397.2;Wu1-SARS2_nucleoca;268 | 0.05 | 0 |
| NC_045512.2;YP_009724397.2;Wu1-SARS2_nucleoca;269 | 0.05 | 0 |
| NC_045512.2;YP_009724397.2;Wu1-SARS2_nucleoca;270 | 0.15 | 0.05 |
| NC_045512.2;YP_009724397.2;Wu1-SARS2_nucleoca;271 | 0.15 | 0.05 |
| NC_045512.2;YP_009724397.2;Wu1-SARS2_nucleoca;272 | 0.15 | 0.05 |
| NC_045512.2;YP_009724397.2;Wu1-SARS2_nucleoca;273 | 0.1 | 0.075 |
| NC_045512.2;YP_009724397.2;Wu1-SARS2_nucleoca;274 | 0.1 | 0.05 |
| NC_045512.2;YP_009724397.2;Wu1-SARS2_nucleoca;275 | 0.05 | 0.025 |
| NC_045512.2;YP_009724397.2;Wu1-SARS2_nucleoca;276 | 0.05 | 0.05 |
| NC_045512.2;YP_009724397.2;Wu1-SARS2_nucleoca;277 | 0.05 | 0.025 |
| NC_045512.2;YP_009724397.2;Wu1-SARS2_nucleoca;278 | 0.05 | 0 |
| NC_045512.2;YP_009724397.2;Wu1-SARS2_nucleoca;279 | 0.05 | 0.05 |
| NC_045512.2;YP_009724397.2;Wu1-SARS2_nucleoca;280 | 0.05 | 0.175 |
| NC_045512.2;YP_009724397.2;Wu1-SARS2_nucleoca;281 | 0 | 0.1 |
| NC_045512.2;YP_009724397.2;Wu1-SARS2_nucleoca;282 | 0 | 0.05 |
| NC_045512.2;YP_009724397.2;Wu1-SARS2_nucleoca;283 | 0 | 0.1 |
| NC_045512.2;YP_009724397.2;Wu1-SARS2_nucleoca;284 | 0 | 0.075 |
| NC_045512.2;YP_009724397.2;Wu1-SARS2_nucleoca;285 | 0 | 0.075 |
| NC_045512.2;YP_009724397.2;Wu1-SARS2_nucleoca;286 | 0 | 0.075 |
| NC_045512.2;YP_009724397.2;Wu1-SARS2_nucleoca;287 | 0 | 0.025 |
| NC_045512.2;YP_009724397.2;Wu1-SARS2_nucleoca;288 | 0 | 0 |
| NC_045512.2;YP_009724397.2;Wu1-SARS2_nucleoca;289 | 0 | 0 |
| NC_045512.2;YP_009724397.2;Wu1-SARS2_nucleoca;290 | 0 | 0.025 |
| NC_045512.2;YP_009724397.2;Wu1-SARS2_nucleoca;291 | 0 | 0 |
| NC_045512.2;YP_009724397.2;Wu1-SARS2_nucleoca;292 | 0 | 0 |
| NC_045512.2;YP_009724397.2;Wu1-SARS2_nucleoca;293 | 0 | 0 |
| NC_045512.2;YP_009724397.2;Wu1-SARS2_nucleoca;294 | 0.05 | 0 |

| | | |
|---|---|---|
| NC_045512.2;YP_009724397.2;Wu1-SARS2_nucleoca;295 | 0 | 0.05 |
| NC_045512.2;YP_009724397.2;Wu1-SARS2_nucleoca;296 | 0 | 0 |
| NC_045512.2;YP_009724397.2;Wu1-SARS2_nucleoca;297 | 0.15 | 0.025 |
| NC_045512.2;YP_009724397.2;Wu1-SARS2_nucleoca;298 | 0 | 0 |
| NC_045512.2;YP_009724397.2;Wu1-SARS2_nucleoca;299 | 0 | 0 |
| NC_045512.2;YP_009724397.2;Wu1-SARS2_nucleoca;300 | 0 | 0 |
| NC_045512.2;YP_009724397.2;Wu1-SARS2_nucleoca;301 | 0 | 0 |
| NC_045512.2;YP_009724397.2;Wu1-SARS2_nucleoca;302 | 0 | 0.025 |
| NC_045512.2;YP_009724397.2;Wu1-SARS2_nucleoca;303 | 0 | 0 |
| NC_045512.2;YP_009724397.2;Wu1-SARS2_nucleoca;304 | 0 | 0 |
| NC_045512.2;YP_009724397.2;Wu1-SARS2_nucleoca;305 | 0 | 0 |
| NC_045512.2;YP_009724397.2;Wu1-SARS2_nucleoca;306 | 0 | 0 |
| NC_045512.2;YP_009724397.2;Wu1-SARS2_nucleoca;307 | 0 | 0 |
| NC_045512.2;YP_009724397.2;Wu1-SARS2_nucleoca;308 | 0 | 0 |
| NC_045512.2;YP_009724397.2;Wu1-SARS2_nucleoca;309 | 0 | 0.025 |
| NC_045512.2;YP_009724397.2;Wu1-SARS2_nucleoca;310 | 0.05 | 0.025 |
| NC_045512.2;YP_009724397.2;Wu1-SARS2_nucleoca;311 | 0 | 0.05 |
| NC_045512.2;YP_009724397.2;Wu1-SARS2_nucleoca;312 | 0 | 0.075 |
| NC_045512.2;YP_009724397.2;Wu1-SARS2_nucleoca;313 | 0.05 | 0.05 |
| NC_045512.2;YP_009724397.2;Wu1-SARS2_nucleoca;314 | 0.05 | 0.05 |
| NC_045512.2;YP_009724397.2;Wu1-SARS2_nucleoca;315 | 0 | 0.05 |
| NC_045512.2;YP_009724397.2;Wu1-SARS2_nucleoca;316 | 0 | 0.025 |
| NC_045512.2;YP_009724397.2;Wu1-SARS2_nucleoca;317 | 0 | 0.025 |
| NC_045512.2;YP_009724397.2;Wu1-SARS2_nucleoca;318 | 0 | 0.025 |
| NC_045512.2;YP_009724397.2;Wu1-SARS2_nucleoca;319 | 0 | 0 |
| NC_045512.2;YP_009724397.2;Wu1-SARS2_nucleoca;320 | 0 | 0 |
| NC_045512.2;YP_009724397.2;Wu1-SARS2_nucleoca;321 | 0 | 0 |
| NC_045512.2;YP_009724397.2;Wu1-SARS2_nucleoca;322 | 0 | 0 |
| NC_045512.2;YP_009724397.2;Wu1-SARS2_nucleoca;323 | 0 | 0 |
| NC_045512.2;YP_009724397.2;Wu1-SARS2_nucleoca;324 | 0 | 0 |
| NC_045512.2;YP_009724397.2;Wu1-SARS2_nucleoca;325 | 0 | 0 |
| NC_045512.2;YP_009724397.2;Wu1-SARS2_nucleoca;326 | 0 | 0 |
| NC_045512.2;YP_009724397.2;Wu1-SARS2_nucleoca;327 | 0 | 0.075 |
| NC_045512.2;YP_009724397.2;Wu1-SARS2_nucleoca;328 | 0 | 0.05 |
| NC_045512.2;YP_009724397.2;Wu1-SARS2_nucleoca;329 | 0 | 0.075 |
| NC_045512.2;YP_009724397.2;Wu1-SARS2_nucleoca;330 | 0 | 0.1 |
| NC_045512.2;YP_009724397.2;Wu1-SARS2_nucleoca;331 | 0 | 0.175 |
| NC_045512.2;YP_009724397.2;Wu1-SARS2_nucleoca;332 | 0.1 | 0.175 |
| NC_045512.2;YP_009724397.2;Wu1-SARS2_nucleoca;333 | 0.05 | 0.25 |
| NC_045512.2;YP_009724397.2;Wu1-SARS2_nucleoca;334 | 0 | 0.175 |
| NC_045512.2;YP_009724397.2;Wu1-SARS2_nucleoca;335 | 0.1 | 0.225 |
| NC_045512.2;YP_009724397.2;Wu1-SARS2_nucleoca;336 | 0 | 0.2 |
| NC_045512.2;YP_009724397.2;Wu1-SARS2_nucleoca;337 | 0.05 | 0.175 |

| | | |
|---|---|---|
| NC_045512.2;YP_009724397.2;Wu1-SARS2_nucleoca;338 | 0 | 0.1 |
| NC_045512.2;YP_009724397.2;Wu1-SARS2_nucleoca;339 | 0 | 0.2 |
| NC_045512.2;YP_009724397.2;Wu1-SARS2_nucleoca;340 | 0 | 0.15 |
| NC_045512.2;YP_009724397.2;Wu1-SARS2_nucleoca;341 | 0 | 0.075 |
| NC_045512.2;YP_009724397.2;Wu1-SARS2_nucleoca;342 | 0 | 0.05 |
| NC_045512.2;YP_009724397.2;Wu1-SARS2_nucleoca;343 | 0.05 | 0.1 |
| NC_045512.2;YP_009724397.2;Wu1-SARS2_nucleoca;344 | 0.05 | 0.1 |
| NC_045512.2;YP_009724397.2;Wu1-SARS2_nucleoca;345 | 0.05 | 0.05 |
| NC_045512.2;YP_009724397.2;Wu1-SARS2_nucleoca;346 | 0 | 0.075 |
| NC_045512.2;YP_009724397.2;Wu1-SARS2_nucleoca;347 | 0 | 0.05 |
| NC_045512.2;YP_009724397.2;Wu1-SARS2_nucleoca;348 | 0 | 0 |
| NC_045512.2;YP_009724397.2;Wu1-SARS2_nucleoca;349 | 0 | 0 |
| NC_045512.2;YP_009724397.2;Wu1-SARS2_nucleoca;350 | 0 | 0 |
| NC_045512.2;YP_009724397.2;Wu1-SARS2_nucleoca;351 | 0 | 0.025 |
| NC_045512.2;YP_009724397.2;Wu1-SARS2_nucleoca;352 | 0 | 0.15 |
| NC_045512.2;YP_009724397.2;Wu1-SARS2_nucleoca;353 | 0.05 | 0.125 |
| NC_045512.2;YP_009724397.2;Wu1-SARS2_nucleoca;354 | 0 | 0.125 |
| NC_045512.2;YP_009724397.2;Wu1-SARS2_nucleoca;355 | 0 | 0.05 |
| NC_045512.2;YP_009724397.2;Wu1-SARS2_nucleoca;356 | 0 | 0.25 |
| NC_045512.2;YP_009724397.2;Wu1-SARS2_nucleoca;357 | 0.05 | 0.125 |
| NC_045512.2;YP_009724397.2;Wu1-SARS2_nucleoca;358 | 0.05 | 0.1 |
| NC_045512.2;YP_009724397.2;Wu1-SARS2_nucleoca;359 | 0 | 0.075 |
| NC_045512.2;YP_009724397.2;Wu1-SARS2_nucleoca;360 | 0 | 0.075 |
| NC_045512.2;YP_009724397.2;Wu1-SARS2_nucleoca;361 | 0 | 0.05 |
| NC_045512.2;YP_009724397.2;Wu1-SARS2_nucleoca;362 | 0.05 | 0.1 |
| NC_045512.2;YP_009724397.2;Wu1-SARS2_nucleoca;363 | 0.05 | 0.1 |
| NC_045512.2;YP_009724397.2;Wu1-SARS2_nucleoca;364 | 0.05 | 0.025 |
| NC_045512.2;YP_009724397.2;Wu1-SARS2_nucleoca;365 | 0 | 0 |
| NC_045512.2;YP_009724397.2;Wu1-SARS2_nucleoca;366 | 0 | 0 |
| NC_045512.2;YP_009724397.2;Wu1-SARS2_nucleoca;367 | 0 | 0.025 |
| NC_045512.2;YP_009724397.2;Wu1-SARS2_nucleoca;368 | 0 | 0.05 |
| NC_045512.2;YP_009724397.2;Wu1-SARS2_nucleoca;369 | 0 | 0.075 |
| NC_045512.2;YP_009724397.2;Wu1-SARS2_nucleoca;370 | 0 | 0.075 |
| NC_045512.2;YP_009724397.2;Wu1-SARS2_nucleoca;371 | 0 | 0.05 |
| NC_045512.2;YP_009724397.2;Wu1-SARS2_nucleoca;372 | 0 | 0.1 |
| NC_045512.2;YP_009724397.2;Wu1-SARS2_nucleoca;373 | 0 | 0.1 |
| NC_045512.2;YP_009724397.2;Wu1-SARS2_nucleoca;374 | 0 | 0.125 |
| NC_045512.2;YP_009724397.2;Wu1-SARS2_nucleoca;375 | 0 | 0.1 |
| NC_045512.2;YP_009724397.2;Wu1-SARS2_nucleoca;376 | 0 | 0.125 |
| NC_045512.2;YP_009724397.2;Wu1-SARS2_nucleoca;377 | 0.05 | 0.125 |
| NC_045512.2;YP_009724397.2;Wu1-SARS2_nucleoca;378 | 0 | 0.175 |
| NC_045512.2;YP_009724397.2;Wu1-SARS2_nucleoca;379 | 0 | 0.125 |
| NC_045512.2;YP_009724397.2;Wu1-SARS2_nucleoca;380 | 0 | 0.15 |

| | | |
|---|---|---|
| NC_045512.2;YP_009724397.2;Wu1-SARS2_nucleoca;381 | 0 | 0.225 |
| NC_045512.2;YP_009724397.2;Wu1-SARS2_nucleoca;382 | 0 | 0.225 |
| NC_045512.2;YP_009724397.2;Wu1-SARS2_nucleoca;383 | 0.05 | 0.125 |
| NC_045512.2;YP_009724397.2;Wu1-SARS2_nucleoca;384 | 0 | 0.125 |
| NC_045512.2;YP_009724397.2;Wu1-SARS2_nucleoca;385 | 0 | 0.15 |
| NC_045512.2;YP_009724397.2;Wu1-SARS2_nucleoca;386 | 0 | 0.4 |
| NC_045512.2;YP_009724397.2;Wu1-SARS2_nucleoca;387 | 0 | 0.5 |
| NC_045512.2;YP_009724397.2;Wu1-SARS2_nucleoca;388 | 0 | 0.575 |
| NC_045512.2;YP_009724397.2;Wu1-SARS2_nucleoca;389 | 0 | 0.3 |
| NC_045512.2;YP_009724397.2;Wu1-SARS2_nucleoca;390 | 0 | 0.7 |
| NC_045512.2;YP_009724397.2;Wu1-SARS2_nucleoca;391 | 0 | 0.75 |
| NC_045512.2;YP_009724397.2;Wu1-SARS2_nucleoca;392 | 0 | 0.775 |
| NC_045512.2;YP_009724397.2;Wu1-SARS2_nucleoca;393 | 0 | 0.8 |
| NC_045512.2;YP_009724397.2;Wu1-SARS2_nucleoca;394 | 0 | 0.825 |
| NC_045512.2;YP_009724397.2;Wu1-SARS2_nucleoca;395 | 0.05 | 0.775 |
| NC_045512.2;YP_009724397.2;Wu1-SARS2_nucleoca;396 | 0 | 0.725 |
| NC_045512.2;YP_009724397.2;Wu1-SARS2_nucleoca;397 | 0.1 | 0.7 |
| NC_045512.2;YP_009724397.2;Wu1-SARS2_nucleoca;398 | 0.25 | 0.55 |
| NC_045512.2;YP_009724397.2;Wu1-SARS2_nucleoca;399 | 0 | 0.675 |
| NC_045512.2;YP_009724397.2;Wu1-SARS2_nucleoca;400 | 0 | 0.2 |
| NC_045512.2;YP_009724397.2;Wu1-SARS2_nucleoca;401 | 0 | 0.075 |
| NC_045512.2;YP_009724397.2;Wu1-SARS2_nucleoca;402 | 0 | 0.075 |
| NC_045512.2;YP_009724397.2;Wu1-SARS2_nucleoca;403 | 0.05 | 0.1 |
| NC_045512.2;YP_009724397.2;Wu1-SARS2_nucleoca;404 | 0 | 0.05 |
| NC_045512.2;YP_009724397.2;Wu1-SARS2_nucleoca;405 | 0 | 0.025 |
| NC_045512.2;YP_009724397.2;Wu1-SARS2_nucleoca;406 | 0 | 0.05 |
| NC_045512.2;YP_009724397.2;Wu1-SARS2_nucleoca;407 | 0 | 0.075 |
| NC_045512.2;YP_009724397.2;Wu1-SARS2_nucleoca;408 | 0 | 0.025 |
| NC_045512.2;YP_009725255.1;Wu1-SARS2_orf10;1 | 0 | 0 |
| NC_045512.2;YP_009725255.1;Wu1-SARS2_orf10;2 | 0 | 0 |
| NC_045512.2;YP_009725255.1;Wu1-SARS2_orf10;3 | 0 | 0 |
| NC_045512.2;YP_009725255.1;Wu1-SARS2_orf10;4 | 0 | 0 |
| NC_045512.2;YP_009725255.1;Wu1-SARS2_orf10;5 | 0 | 0 |
| NC_045512.2;YP_009725255.1;Wu1-SARS2_orf10;6 | 0 | 0 |
| NC_045512.2;YP_009725255.1;Wu1-SARS2_orf10;7 | 0 | 0 |
| NC_045512.2;YP_009725255.1;Wu1-SARS2_orf10;8 | 0 | 0 |
| NC_045512.2;YP_009725255.1;Wu1-SARS2_orf10;9 | 0 | 0 |
| NC_045512.2;YP_009725255.1;Wu1-SARS2_orf10;10 | 0 | 0 |
| NC_045512.2;YP_009725255.1;Wu1-SARS2_orf10;11 | 0 | 0 |
| NC_045512.2;YP_009725255.1;Wu1-SARS2_orf10;12 | 0 | 0 |
| NC_045512.2;YP_009725255.1;Wu1-SARS2_orf10;13 | 0 | 0 |
| NC_045512.2;YP_009725255.1;Wu1-SARS2_orf10;14 | 0 | 0 |
| NC_045512.2;YP_009725255.1;Wu1-SARS2_orf10;15 | 0 | 0 |

| | | |
|---|---|---|
| NC_045512.2;YP_009725255.1;Wu1-SARS2_orf10;16 | 0.05 | 0.025 |
| NC_045512.2;YP_009725255.1;Wu1-SARS2_orf10;17 | 0.05 | 0.025 |
| NC_045512.2;YP_009725255.1;Wu1-SARS2_orf10;18 | 0.05 | 0.025 |
| NC_045512.2;YP_009725255.1;Wu1-SARS2_orf10;19 | 0.15 | 0.05 |
| NC_045512.2;YP_009725255.1;Wu1-SARS2_orf10;20 | 0 | 0.05 |
| NC_045512.2;YP_009725255.1;Wu1-SARS2_orf10;21 | 0.1 | 0.05 |
| NC_045512.2;YP_009725255.1;Wu1-SARS2_orf10;22 | 0 | 0 |
| NC_045512.2;YP_009725255.1;Wu1-SARS2_orf10;23 | 0 | 0 |
| NC_045512.2;YP_009725255.1;Wu1-SARS2_orf10;24 | 0 | 0 |
| NC_045512.2;YP_009725255.1;Wu1-SARS2_orf10;25 | 0 | 0 |
| NC_045512.2;YP_009725255.1;Wu1-SARS2_orf10;26 | 0.05 | 0 |
| NC_045512.2;YP_009725255.1;Wu1-SARS2_orf10;27 | 0.05 | 0.05 |
| NC_045512.2;YP_009725295.1;Wu1-SARS2_orf1a;1 | 0 | 0 |
| NC_045512.2;YP_009725295.1;Wu1-SARS2_orf1a;2 | 0 | 0.025 |
| NC_045512.2;YP_009725295.1;Wu1-SARS2_orf1a;3 | 0 | 0 |
| NC_045512.2;YP_009725295.1;Wu1-SARS2_orf1a;4 | 0 | 0.025 |
| NC_045512.2;YP_009725295.1;Wu1-SARS2_orf1a;5 | 0 | 0 |
| NC_045512.2;YP_009725295.1;Wu1-SARS2_orf1a;6 | 0 | 0 |
| NC_045512.2;YP_009725295.1;Wu1-SARS2_orf1a;7 | 0 | 0 |
| NC_045512.2;YP_009725295.1;Wu1-SARS2_orf1a;8 | 0 | 0 |
| NC_045512.2;YP_009725295.1;Wu1-SARS2_orf1a;9 | 0 | 0 |
| NC_045512.2;YP_009725295.1;Wu1-SARS2_orf1a;10 | 0 | 0 |
| NC_045512.2;YP_009725295.1;Wu1-SARS2_orf1a;11 | 0 | 0 |
| NC_045512.2;YP_009725295.1;Wu1-SARS2_orf1a;12 | 0 | 0 |
| NC_045512.2;YP_009725295.1;Wu1-SARS2_orf1a;13 | 0 | 0 |
| NC_045512.2;YP_009725295.1;Wu1-SARS2_orf1a;14 | 0 | 0 |
| NC_045512.2;YP_009725295.1;Wu1-SARS2_orf1a;15 | 0 | 0 |
| NC_045512.2;YP_009725295.1;Wu1-SARS2_orf1a;16 | 0 | 0 |
| NC_045512.2;YP_009725295.1;Wu1-SARS2_orf1a;17 | 0 | 0 |
| NC_045512.2;YP_009725295.1;Wu1-SARS2_orf1a;18 | 0 | 0 |
| NC_045512.2;YP_009725295.1;Wu1-SARS2_orf1a;19 | 0 | 0 |
| NC_045512.2;YP_009725295.1;Wu1-SARS2_orf1a;20 | 0 | 0 |
| NC_045512.2;YP_009725295.1;Wu1-SARS2_orf1a;21 | 0 | 0 |
| NC_045512.2;YP_009725295.1;Wu1-SARS2_orf1a;22 | 0 | 0 |
| NC_045512.2;YP_009725295.1;Wu1-SARS2_orf1a;23 | 0 | 0 |
| NC_045512.2;YP_009725295.1;Wu1-SARS2_orf1a;24 | 0 | 0 |
| NC_045512.2;YP_009725295.1;Wu1-SARS2_orf1a;25 | 0 | 0.025 |
| NC_045512.2;YP_009725295.1;Wu1-SARS2_orf1a;26 | 0 | 0.025 |
| NC_045512.2;YP_009725295.1;Wu1-SARS2_orf1a;27 | 0 | 0.025 |
| NC_045512.2;YP_009725295.1;Wu1-SARS2_orf1a;28 | 0 | 0.025 |
| NC_045512.2;YP_009725295.1;Wu1-SARS2_orf1a;29 | 0.1 | 0.025 |
| NC_045512.2;YP_009725295.1;Wu1-SARS2_orf1a;30 | 0.25 | 0.15 |
| NC_045512.2;YP_009725295.1;Wu1-SARS2_orf1a;31 | 0 | 0.025 |

| | | |
|---|---|---|
| NC_045512.2;YP_009725295.1;Wu1-SARS2_orf1a;32 | 0 | 0.05 |
| NC_045512.2;YP_009725295.1;Wu1-SARS2_orf1a;33 | 0.05 | 0 |
| NC_045512.2;YP_009725295.1;Wu1-SARS2_orf1a;34 | 0.1 | 0 |
| NC_045512.2;YP_009725295.1;Wu1-SARS2_orf1a;35 | 0 | 0.025 |
| NC_045512.2;YP_009725295.1;Wu1-SARS2_orf1a;36 | 0 | 0.025 |
| NC_045512.2;YP_009725295.1;Wu1-SARS2_orf1a;37 | 0 | 0 |
| NC_045512.2;YP_009725295.1;Wu1-SARS2_orf1a;38 | 0 | 0 |
| NC_045512.2;YP_009725295.1;Wu1-SARS2_orf1a;39 | 0 | 0 |
| NC_045512.2;YP_009725295.1;Wu1-SARS2_orf1a;40 | 0 | 0 |
| NC_045512.2;YP_009725295.1;Wu1-SARS2_orf1a;41 | 0 | 0 |
| NC_045512.2;YP_009725295.1;Wu1-SARS2_orf1a;42 | 0 | 0 |
| NC_045512.2;YP_009725295.1;Wu1-SARS2_orf1a;43 | 0 | 0 |
| NC_045512.2;YP_009725295.1;Wu1-SARS2_orf1a;44 | 0.05 | 0 |
| NC_045512.2;YP_009725295.1;Wu1-SARS2_orf1a;45 | 0 | 0 |
| NC_045512.2;YP_009725295.1;Wu1-SARS2_orf1a;46 | 0 | 0 |
| NC_045512.2;YP_009725295.1;Wu1-SARS2_orf1a;47 | 0 | 0 |
| NC_045512.2;YP_009725295.1;Wu1-SARS2_orf1a;48 | 0 | 0 |
| NC_045512.2;YP_009725295.1;Wu1-SARS2_orf1a;49 | 0 | 0 |
| NC_045512.2;YP_009725295.1;Wu1-SARS2_orf1a;50 | 0 | 0 |
| NC_045512.2;YP_009725295.1;Wu1-SARS2_orf1a;51 | 0 | 0 |
| NC_045512.2;YP_009725295.1;Wu1-SARS2_orf1a;52 | 0.05 | 0.025 |
| NC_045512.2;YP_009725295.1;Wu1-SARS2_orf1a;53 | 0 | 0.025 |
| NC_045512.2;YP_009725295.1;Wu1-SARS2_orf1a;54 | 0.05 | 0.025 |
| NC_045512.2;YP_009725295.1;Wu1-SARS2_orf1a;55 | 0 | 0.025 |
| NC_045512.2;YP_009725295.1;Wu1-SARS2_orf1a;56 | 0 | 0.025 |
| NC_045512.2;YP_009725295.1;Wu1-SARS2_orf1a;57 | 0 | 0.025 |
| NC_045512.2;YP_009725295.1;Wu1-SARS2_orf1a;58 | 0 | 0.025 |
| NC_045512.2;YP_009725295.1;Wu1-SARS2_orf1a;59 | 0 | 0 |
| NC_045512.2;YP_009725295.1;Wu1-SARS2_orf1a;60 | 0 | 0 |
| NC_045512.2;YP_009725295.1;Wu1-SARS2_orf1a;61 | 0 | 0 |
| NC_045512.2;YP_009725295.1;Wu1-SARS2_orf1a;62 | 0 | 0 |
| NC_045512.2;YP_009725295.1;Wu1-SARS2_orf1a;63 | 0 | 0 |
| NC_045512.2;YP_009725295.1;Wu1-SARS2_orf1a;64 | 0 | 0 |
| NC_045512.2;YP_009725295.1;Wu1-SARS2_orf1a;65 | 0 | 0 |
| NC_045512.2;YP_009725295.1;Wu1-SARS2_orf1a;66 | 0 | 0 |
| NC_045512.2;YP_009725295.1;Wu1-SARS2_orf1a;67 | 0 | 0 |
| NC_045512.2;YP_009725295.1;Wu1-SARS2_orf1a;68 | 0 | 0 |
| NC_045512.2;YP_009725295.1;Wu1-SARS2_orf1a;69 | 0 | 0 |
| NC_045512.2;YP_009725295.1;Wu1-SARS2_orf1a;70 | 0 | 0 |
| NC_045512.2;YP_009725295.1;Wu1-SARS2_orf1a;71 | 0.05 | 0 |
| NC_045512.2;YP_009725295.1;Wu1-SARS2_orf1a;72 | 0 | 0 |
| NC_045512.2;YP_009725295.1;Wu1-SARS2_orf1a;73 | 0 | 0 |
| NC_045512.2;YP_009725295.1;Wu1-SARS2_orf1a;74 | 0 | 0 |

| | | |
|---|---|---|
| NC_045512.2;YP_009725295.1;Wu1-SARS2_orf1a;75 | 0 | 0.025 |
| NC_045512.2;YP_009725295.1;Wu1-SARS2_orf1a;76 | 0 | 0 |
| NC_045512.2;YP_009725295.1;Wu1-SARS2_orf1a;77 | 0 | 0 |
| NC_045512.2;YP_009725295.1;Wu1-SARS2_orf1a;78 | 0 | 0 |
| NC_045512.2;YP_009725295.1;Wu1-SARS2_orf1a;79 | 0 | 0 |
| NC_045512.2;YP_009725295.1;Wu1-SARS2_orf1a;80 | 0 | 0 |
| NC_045512.2;YP_009725295.1;Wu1-SARS2_orf1a;81 | 0 | 0 |
| NC_045512.2;YP_009725295.1;Wu1-SARS2_orf1a;82 | 0 | 0 |
| NC_045512.2;YP_009725295.1;Wu1-SARS2_orf1a;83 | 0 | 0 |
| NC_045512.2;YP_009725295.1;Wu1-SARS2_orf1a;84 | 0 | 0 |
| NC_045512.2;YP_009725295.1;Wu1-SARS2_orf1a;85 | 0 | 0 |
| NC_045512.2;YP_009725295.1;Wu1-SARS2_orf1a;86 | 0 | 0.025 |
| NC_045512.2;YP_009725295.1;Wu1-SARS2_orf1a;87 | 0 | 0 |
| NC_045512.2;YP_009725295.1;Wu1-SARS2_orf1a;88 | 0 | 0 |
| NC_045512.2;YP_009725295.1;Wu1-SARS2_orf1a;89 | 0 | 0 |
| NC_045512.2;YP_009725295.1;Wu1-SARS2_orf1a;90 | 0 | 0 |
| NC_045512.2;YP_009725295.1;Wu1-SARS2_orf1a;91 | 0 | 0 |
| NC_045512.2;YP_009725295.1;Wu1-SARS2_orf1a;92 | 0 | 0 |
| NC_045512.2;YP_009725295.1;Wu1-SARS2_orf1a;93 | 0 | 0 |
| NC_045512.2;YP_009725295.1;Wu1-SARS2_orf1a;94 | 0 | 0.025 |
| NC_045512.2;YP_009725295.1;Wu1-SARS2_orf1a;95 | 0 | 0 |
| NC_045512.2;YP_009725295.1;Wu1-SARS2_orf1a;96 | 0 | 0 |
| NC_045512.2;YP_009725295.1;Wu1-SARS2_orf1a;97 | 0 | 0 |
| NC_045512.2;YP_009725295.1;Wu1-SARS2_orf1a;98 | 0 | 0 |
| NC_045512.2;YP_009725295.1;Wu1-SARS2_orf1a;99 | 0 | 0 |
| NC_045512.2;YP_009725295.1;Wu1-SARS2_orf1a;100 | 0 | 0 |
| NC_045512.2;YP_009725295.1;Wu1-SARS2_orf1a;101 | 0 | 0 |
| NC_045512.2;YP_009725295.1;Wu1-SARS2_orf1a;102 | 0 | 0 |
| NC_045512.2;YP_009725295.1;Wu1-SARS2_orf1a;103 | 0 | 0 |
| NC_045512.2;YP_009725295.1;Wu1-SARS2_orf1a;104 | 0 | 0 |
| NC_045512.2;YP_009725295.1;Wu1-SARS2_orf1a;105 | 0 | 0 |
| NC_045512.2;YP_009725295.1;Wu1-SARS2_orf1a;106 | 0 | 0 |
| NC_045512.2;YP_009725295.1;Wu1-SARS2_orf1a;107 | 0 | 0 |
| NC_045512.2;YP_009725295.1;Wu1-SARS2_orf1a;108 | 0 | 0 |
| NC_045512.2;YP_009725295.1;Wu1-SARS2_orf1a;109 | 0 | 0 |
| NC_045512.2;YP_009725295.1;Wu1-SARS2_orf1a;110 | 0 | 0 |
| NC_045512.2;YP_009725295.1;Wu1-SARS2_orf1a;111 | 0 | 0 |
| NC_045512.2;YP_009725295.1;Wu1-SARS2_orf1a;112 | 0 | 0 |
| NC_045512.2;YP_009725295.1;Wu1-SARS2_orf1a;113 | 0 | 0 |
| NC_045512.2;YP_009725295.1;Wu1-SARS2_orf1a;114 | 0 | 0.025 |
| NC_045512.2;YP_009725295.1;Wu1-SARS2_orf1a;115 | 0 | 0.025 |
| NC_045512.2;YP_009725295.1;Wu1-SARS2_orf1a;116 | 0 | 0.025 |
| NC_045512.2;YP_009725295.1;Wu1-SARS2_orf1a;117 | 0 | 0.025 |

| | | |
|---|---|---|
| NC_045512.2;YP_009725295.1;Wu1-SARS2_orf1a;118 | 0.05 | 0.025 |
| NC_045512.2;YP_009725295.1;Wu1-SARS2_orf1a;119 | 0 | 0.025 |
| NC_045512.2;YP_009725295.1;Wu1-SARS2_orf1a;120 | 0 | 0.025 |
| NC_045512.2;YP_009725295.1;Wu1-SARS2_orf1a;121 | 0.05 | 0.025 |
| NC_045512.2;YP_009725295.1;Wu1-SARS2_orf1a;122 | 0 | 0.025 |
| NC_045512.2;YP_009725295.1;Wu1-SARS2_orf1a;123 | 0 | 0.025 |
| NC_045512.2;YP_009725295.1;Wu1-SARS2_orf1a;124 | 0 | 0.05 |
| NC_045512.2;YP_009725295.1;Wu1-SARS2_orf1a;125 | 0.1 | 0.025 |
| NC_045512.2;YP_009725295.1;Wu1-SARS2_orf1a;126 | 0 | 0 |
| NC_045512.2;YP_009725295.1;Wu1-SARS2_orf1a;127 | 0 | 0 |
| NC_045512.2;YP_009725295.1;Wu1-SARS2_orf1a;128 | 0 | 0.025 |
| NC_045512.2;YP_009725295.1;Wu1-SARS2_orf1a;129 | 0 | 0.05 |
| NC_045512.2;YP_009725295.1;Wu1-SARS2_orf1a;130 | 0 | 0.025 |
| NC_045512.2;YP_009725295.1;Wu1-SARS2_orf1a;131 | 0.1 | 0.025 |
| NC_045512.2;YP_009725295.1;Wu1-SARS2_orf1a;132 | 0.05 | 0.025 |
| NC_045512.2;YP_009725295.1;Wu1-SARS2_orf1a;133 | 0.05 | 0 |
| NC_045512.2;YP_009725295.1;Wu1-SARS2_orf1a;134 | 0 | 0 |
| NC_045512.2;YP_009725295.1;Wu1-SARS2_orf1a;135 | 0.05 | 0 |
| NC_045512.2;YP_009725295.1;Wu1-SARS2_orf1a;136 | 0 | 0.025 |
| NC_045512.2;YP_009725295.1;Wu1-SARS2_orf1a;137 | 0 | 0 |
| NC_045512.2;YP_009725295.1;Wu1-SARS2_orf1a;138 | 0 | 0 |
| NC_045512.2;YP_009725295.1;Wu1-SARS2_orf1a;139 | 0.05 | 0 |
| NC_045512.2;YP_009725295.1;Wu1-SARS2_orf1a;140 | 0.05 | 0 |
| NC_045512.2;YP_009725295.1;Wu1-SARS2_orf1a;141 | 0 | 0 |
| NC_045512.2;YP_009725295.1;Wu1-SARS2_orf1a;142 | 0 | 0 |
| NC_045512.2;YP_009725295.1;Wu1-SARS2_orf1a;143 | 0.05 | 0 |
| NC_045512.2;YP_009725295.1;Wu1-SARS2_orf1a;144 | 0 | 0.025 |
| NC_045512.2;YP_009725295.1;Wu1-SARS2_orf1a;145 | 0.2 | 0.1 |
| NC_045512.2;YP_009725295.1;Wu1-SARS2_orf1a;146 | 0.05 | 0 |
| NC_045512.2;YP_009725295.1;Wu1-SARS2_orf1a;147 | 0 | 0 |
| NC_045512.2;YP_009725295.1;Wu1-SARS2_orf1a;148 | 0.05 | 0 |
| NC_045512.2;YP_009725295.1;Wu1-SARS2_orf1a;149 | 0.3 | 0.2 |
| NC_045512.2;YP_009725295.1;Wu1-SARS2_orf1a;150 | 0.25 | 0.075 |
| NC_045512.2;YP_009725295.1;Wu1-SARS2_orf1a;151 | 0.3 | 0.15 |
| NC_045512.2;YP_009725295.1;Wu1-SARS2_orf1a;152 | 0.45 | 0.225 |
| NC_045512.2;YP_009725295.1;Wu1-SARS2_orf1a;153 | 0.35 | 0.2 |
| NC_045512.2;YP_009725295.1;Wu1-SARS2_orf1a;154 | 0.45 | 0.35 |
| NC_045512.2;YP_009725295.1;Wu1-SARS2_orf1a;155 | 0.5 | 0.325 |
| NC_045512.2;YP_009725295.1;Wu1-SARS2_orf1a;156 | 0.3 | 0.1 |
| NC_045512.2;YP_009725295.1;Wu1-SARS2_orf1a;157 | 0.2 | 0.05 |
| NC_045512.2;YP_009725295.1;Wu1-SARS2_orf1a;158 | 0.05 | 0 |
| NC_045512.2;YP_009725295.1;Wu1-SARS2_orf1a;159 | 0 | 0 |
| NC_045512.2;YP_009725295.1;Wu1-SARS2_orf1a;160 | 0 | 0 |

| | | |
|---|---|---|
| NC_045512.2;YP_009725295.1;Wu1-SARS2_orf1a;161 | 0 | 0 |
| NC_045512.2;YP_009725295.1;Wu1-SARS2_orf1a;162 | 0 | 0 |
| NC_045512.2;YP_009725295.1;Wu1-SARS2_orf1a;163 | 0 | 0.025 |
| NC_045512.2;YP_009725295.1;Wu1-SARS2_orf1a;164 | 0 | 0 |
| NC_045512.2;YP_009725295.1;Wu1-SARS2_orf1a;165 | 0.1 | 0 |
| NC_045512.2;YP_009725295.1;Wu1-SARS2_orf1a;166 | 0.05 | 0.025 |
| NC_045512.2;YP_009725295.1;Wu1-SARS2_orf1a;167 | 0 | 0.025 |
| NC_045512.2;YP_009725295.1;Wu1-SARS2_orf1a;168 | 0.1 | 0.025 |
| NC_045512.2;YP_009725295.1;Wu1-SARS2_orf1a;169 | 0 | 0 |
| NC_045512.2;YP_009725295.1;Wu1-SARS2_orf1a;170 | 0 | 0.025 |
| NC_045512.2;YP_009725295.1;Wu1-SARS2_orf1a;171 | 0 | 0 |
| NC_045512.2;YP_009725295.1;Wu1-SARS2_orf1a;172 | 0 | 0 |
| NC_045512.2;YP_009725295.1;Wu1-SARS2_orf1a;173 | 0 | 0 |
| NC_045512.2;YP_009725295.1;Wu1-SARS2_orf1a;174 | 0.05 | 0.025 |
| NC_045512.2;YP_009725295.1;Wu1-SARS2_orf1a;175 | 0 | 0 |
| NC_045512.2;YP_009725295.1;Wu1-SARS2_orf1a;176 | 0 | 0 |
| NC_045512.2;YP_009725295.1;Wu1-SARS2_orf1a;177 | 0 | 0 |
| NC_045512.2;YP_009725295.1;Wu1-SARS2_orf1a;178 | 0 | 0 |
| NC_045512.2;YP_009725295.1;Wu1-SARS2_orf1a;179 | 0 | 0 |
| NC_045512.2;YP_009725295.1;Wu1-SARS2_orf1a;180 | 0 | 0 |
| NC_045512.2;YP_009725295.1;Wu1-SARS2_orf1a;181 | 0 | 0.025 |
| NC_045512.2;YP_009725295.1;Wu1-SARS2_orf1a;182 | 0 | 0 |
| NC_045512.2;YP_009725295.1;Wu1-SARS2_orf1a;183 | 0 | 0 |
| NC_045512.2;YP_009725295.1;Wu1-SARS2_orf1a;184 | 0 | 0.025 |
| NC_045512.2;YP_009725295.1;Wu1-SARS2_orf1a;185 | 0 | 0.025 |
| NC_045512.2;YP_009725295.1;Wu1-SARS2_orf1a;186 | 0 | 0.025 |
| NC_045512.2;YP_009725295.1;Wu1-SARS2_orf1a;187 | 0 | 0 |
| NC_045512.2;YP_009725295.1;Wu1-SARS2_orf1a;188 | 0 | 0 |
| NC_045512.2;YP_009725295.1;Wu1-SARS2_orf1a;189 | 0 | 0 |
| NC_045512.2;YP_009725295.1;Wu1-SARS2_orf1a;190 | 0 | 0 |
| NC_045512.2;YP_009725295.1;Wu1-SARS2_orf1a;191 | 0 | 0 |
| NC_045512.2;YP_009725295.1;Wu1-SARS2_orf1a;192 | 0 | 0 |
| NC_045512.2;YP_009725295.1;Wu1-SARS2_orf1a;193 | 0 | 0 |
| NC_045512.2;YP_009725295.1;Wu1-SARS2_orf1a;194 | 0 | 0 |
| NC_045512.2;YP_009725295.1;Wu1-SARS2_orf1a;195 | 0 | 0 |
| NC_045512.2;YP_009725295.1;Wu1-SARS2_orf1a;196 | 0 | 0 |
| NC_045512.2;YP_009725295.1;Wu1-SARS2_orf1a;197 | 0.05 | 0 |
| NC_045512.2;YP_009725295.1;Wu1-SARS2_orf1a;198 | 0 | 0 |
| NC_045512.2;YP_009725295.1;Wu1-SARS2_orf1a;199 | 0 | 0 |
| NC_045512.2;YP_009725295.1;Wu1-SARS2_orf1a;200 | 0 | 0 |
| NC_045512.2;YP_009725295.1;Wu1-SARS2_orf1a;201 | 0 | 0 |
| NC_045512.2;YP_009725295.1;Wu1-SARS2_orf1a;202 | 0 | 0 |
| NC_045512.2;YP_009725295.1;Wu1-SARS2_orf1a;203 | 0 | 0 |

| | | |
|---|---|---|
| NC_045512.2;YP_009725295.1;Wu1-SARS2_orf1a;204 | 0.05 | 0 |
| NC_045512.2;YP_009725295.1;Wu1-SARS2_orf1a;205 | 0 | 0 |
| NC_045512.2;YP_009725295.1;Wu1-SARS2_orf1a;206 | 0 | 0 |
| NC_045512.2;YP_009725295.1;Wu1-SARS2_orf1a;207 | 0 | 0 |
| NC_045512.2;YP_009725295.1;Wu1-SARS2_orf1a;208 | 0 | 0 |
| NC_045512.2;YP_009725295.1;Wu1-SARS2_orf1a;209 | 0 | 0 |
| NC_045512.2;YP_009725295.1;Wu1-SARS2_orf1a;210 | 0 | 0 |
| NC_045512.2;YP_009725295.1;Wu1-SARS2_orf1a;211 | 0.05 | 0.075 |
| NC_045512.2;YP_009725295.1;Wu1-SARS2_orf1a;212 | 0 | 0.1 |
| NC_045512.2;YP_009725295.1;Wu1-SARS2_orf1a;213 | 0 | 0.1 |
| NC_045512.2;YP_009725295.1;Wu1-SARS2_orf1a;214 | 0.1 | 0.075 |
| NC_045512.2;YP_009725295.1;Wu1-SARS2_orf1a;215 | 0.1 | 0.075 |
| NC_045512.2;YP_009725295.1;Wu1-SARS2_orf1a;216 | 0 | 0.075 |
| NC_045512.2;YP_009725295.1;Wu1-SARS2_orf1a;217 | 0 | 0.025 |
| NC_045512.2;YP_009725295.1;Wu1-SARS2_orf1a;218 | 0 | 0.025 |
| NC_045512.2;YP_009725295.1;Wu1-SARS2_orf1a;219 | 0 | 0.025 |
| NC_045512.2;YP_009725295.1;Wu1-SARS2_orf1a;220 | 0.05 | 0.075 |
| NC_045512.2;YP_009725295.1;Wu1-SARS2_orf1a;221 | 0 | 0.025 |
| NC_045512.2;YP_009725295.1;Wu1-SARS2_orf1a;222 | 0 | 0.025 |
| NC_045512.2;YP_009725295.1;Wu1-SARS2_orf1a;223 | 0 | 0.025 |
| NC_045512.2;YP_009725295.1;Wu1-SARS2_orf1a;224 | 0 | 0 |
| NC_045512.2;YP_009725295.1;Wu1-SARS2_orf1a;225 | 0.05 | 0 |
| NC_045512.2;YP_009725295.1;Wu1-SARS2_orf1a;226 | 0 | 0 |
| NC_045512.2;YP_009725295.1;Wu1-SARS2_orf1a;227 | 0 | 0 |
| NC_045512.2;YP_009725295.1;Wu1-SARS2_orf1a;228 | 0 | 0 |
| NC_045512.2;YP_009725295.1;Wu1-SARS2_orf1a;229 | 0 | 0 |
| NC_045512.2;YP_009725295.1;Wu1-SARS2_orf1a;230 | 0 | 0 |
| NC_045512.2;YP_009725295.1;Wu1-SARS2_orf1a;231 | 0 | 0 |
| NC_045512.2;YP_009725295.1;Wu1-SARS2_orf1a;232 | 0 | 0 |
| NC_045512.2;YP_009725295.1;Wu1-SARS2_orf1a;233 | 0 | 0 |
| NC_045512.2;YP_009725295.1;Wu1-SARS2_orf1a;234 | 0 | 0 |
| NC_045512.2;YP_009725295.1;Wu1-SARS2_orf1a;235 | 0 | 0 |
| NC_045512.2;YP_009725295.1;Wu1-SARS2_orf1a;236 | 0 | 0 |
| NC_045512.2;YP_009725295.1;Wu1-SARS2_orf1a;237 | 0 | 0 |
| NC_045512.2;YP_009725295.1;Wu1-SARS2_orf1a;238 | 0 | 0 |
| NC_045512.2;YP_009725295.1;Wu1-SARS2_orf1a;239 | 0 | 0 |
| NC_045512.2;YP_009725295.1;Wu1-SARS2_orf1a;240 | 0 | 0.025 |
| NC_045512.2;YP_009725295.1;Wu1-SARS2_orf1a;241 | 0 | 0.025 |
| NC_045512.2;YP_009725295.1;Wu1-SARS2_orf1a;242 | 0 | 0.05 |
| NC_045512.2;YP_009725295.1;Wu1-SARS2_orf1a;243 | 0 | 0.05 |
| NC_045512.2;YP_009725295.1;Wu1-SARS2_orf1a;244 | 0 | 0.05 |
| NC_045512.2;YP_009725295.1;Wu1-SARS2_orf1a;245 | 0 | 0.05 |
| NC_045512.2;YP_009725295.1;Wu1-SARS2_orf1a;246 | 0 | 0.075 |

| | | |
|---|---|---|
| NC_045512.2;YP_009725295.1;Wu1-SARS2_orf1a;247 | 0 | 0.05 |
| NC_045512.2;YP_009725295.1;Wu1-SARS2_orf1a;248 | 0 | 0.025 |
| NC_045512.2;YP_009725295.1;Wu1-SARS2_orf1a;249 | 0 | 0.05 |
| NC_045512.2;YP_009725295.1;Wu1-SARS2_orf1a;250 | 0 | 0.025 |
| NC_045512.2;YP_009725295.1;Wu1-SARS2_orf1a;251 | 0 | 0.025 |
| NC_045512.2;YP_009725295.1;Wu1-SARS2_orf1a;252 | 0.05 | 0 |
| NC_045512.2;YP_009725295.1;Wu1-SARS2_orf1a;253 | 0.05 | 0 |
| NC_045512.2;YP_009725295.1;Wu1-SARS2_orf1a;254 | 0 | 0 |
| NC_045512.2;YP_009725295.1;Wu1-SARS2_orf1a;255 | 0 | 0.05 |
| NC_045512.2;YP_009725295.1;Wu1-SARS2_orf1a;256 | 0.1 | 0.1 |
| NC_045512.2;YP_009725295.1;Wu1-SARS2_orf1a;257 | 0 | 0 |
| NC_045512.2;YP_009725295.1;Wu1-SARS2_orf1a;258 | 0.05 | 0 |
| NC_045512.2;YP_009725295.1;Wu1-SARS2_orf1a;259 | 0.05 | 0.05 |
| NC_045512.2;YP_009725295.1;Wu1-SARS2_orf1a;260 | 0.05 | 0.025 |
| NC_045512.2;YP_009725295.1;Wu1-SARS2_orf1a;261 | 0.05 | 0.025 |
| NC_045512.2;YP_009725295.1;Wu1-SARS2_orf1a;262 | 0 | 0 |
| NC_045512.2;YP_009725295.1;Wu1-SARS2_orf1a;263 | 0 | 0.025 |
| NC_045512.2;YP_009725295.1;Wu1-SARS2_orf1a;264 | 0 | 0.025 |
| NC_045512.2;YP_009725295.1;Wu1-SARS2_orf1a;265 | 0 | 0 |
| NC_045512.2;YP_009725295.1;Wu1-SARS2_orf1a;266 | 0 | 0 |
| NC_045512.2;YP_009725295.1;Wu1-SARS2_orf1a;267 | 0 | 0 |
| NC_045512.2;YP_009725295.1;Wu1-SARS2_orf1a;268 | 0 | 0 |
| NC_045512.2;YP_009725295.1;Wu1-SARS2_orf1a;269 | 0 | 0 |
| NC_045512.2;YP_009725295.1;Wu1-SARS2_orf1a;270 | 0 | 0 |
| NC_045512.2;YP_009725295.1;Wu1-SARS2_orf1a;271 | 0 | 0 |
| NC_045512.2;YP_009725295.1;Wu1-SARS2_orf1a;272 | 0 | 0 |
| NC_045512.2;YP_009725295.1;Wu1-SARS2_orf1a;273 | 0 | 0 |
| NC_045512.2;YP_009725295.1;Wu1-SARS2_orf1a;274 | 0 | 0 |
| NC_045512.2;YP_009725295.1;Wu1-SARS2_orf1a;275 | 0 | 0.025 |
| NC_045512.2;YP_009725295.1;Wu1-SARS2_orf1a;276 | 0.1 | 0.025 |
| NC_045512.2;YP_009725295.1;Wu1-SARS2_orf1a;277 | 0.05 | 0 |
| NC_045512.2;YP_009725295.1;Wu1-SARS2_orf1a;278 | 0.05 | 0 |
| NC_045512.2;YP_009725295.1;Wu1-SARS2_orf1a;279 | 0.05 | 0 |
| NC_045512.2;YP_009725295.1;Wu1-SARS2_orf1a;280 | 0.05 | 0 |
| NC_045512.2;YP_009725295.1;Wu1-SARS2_orf1a;281 | 0.1 | 0 |
| NC_045512.2;YP_009725295.1;Wu1-SARS2_orf1a;282 | 0.05 | 0 |
| NC_045512.2;YP_009725295.1;Wu1-SARS2_orf1a;283 | 0.1 | 0.025 |
| NC_045512.2;YP_009725295.1;Wu1-SARS2_orf1a;284 | 0.05 | 0.025 |
| NC_045512.2;YP_009725295.1;Wu1-SARS2_orf1a;285 | 0.05 | 0.025 |
| NC_045512.2;YP_009725295.1;Wu1-SARS2_orf1a;286 | 0.05 | 0.025 |
| NC_045512.2;YP_009725295.1;Wu1-SARS2_orf1a;287 | 0 | 0 |
| NC_045512.2;YP_009725295.1;Wu1-SARS2_orf1a;288 | 0 | 0 |
| NC_045512.2;YP_009725295.1;Wu1-SARS2_orf1a;289 | 0 | 0 |

| | | |
|---|---|---|
| NC_045512.2;YP_009725295.1;Wu1-SARS2_orf1a;290 | 0 | 0 |
| NC_045512.2;YP_009725295.1;Wu1-SARS2_orf1a;291 | 0.05 | 0 |
| NC_045512.2;YP_009725295.1;Wu1-SARS2_orf1a;292 | 0 | 0 |
| NC_045512.2;YP_009725295.1;Wu1-SARS2_orf1a;293 | 0 | 0 |
| NC_045512.2;YP_009725295.1;Wu1-SARS2_orf1a;294 | 0 | 0 |
| NC_045512.2;YP_009725295.1;Wu1-SARS2_orf1a;295 | 0 | 0 |
| NC_045512.2;YP_009725295.1;Wu1-SARS2_orf1a;296 | 0.1 | 0 |
| NC_045512.2;YP_009725295.1;Wu1-SARS2_orf1a;297 | 0.1 | 0 |
| NC_045512.2;YP_009725295.1;Wu1-SARS2_orf1a;298 | 0.1 | 0 |
| NC_045512.2;YP_009725295.1;Wu1-SARS2_orf1a;299 | 0.1 | 0 |
| NC_045512.2;YP_009725295.1;Wu1-SARS2_orf1a;300 | 0.1 | 0 |
| NC_045512.2;YP_009725295.1;Wu1-SARS2_orf1a;301 | 0.1 | 0.1 |
| NC_045512.2;YP_009725295.1;Wu1-SARS2_orf1a;302 | 0.05 | 0 |
| NC_045512.2;YP_009725295.1;Wu1-SARS2_orf1a;303 | 0.05 | 0 |
| NC_045512.2;YP_009725295.1;Wu1-SARS2_orf1a;304 | 0.1 | 0 |
| NC_045512.2;YP_009725295.1;Wu1-SARS2_orf1a;305 | 0.05 | 0 |
| NC_045512.2;YP_009725295.1;Wu1-SARS2_orf1a;306 | 0 | 0 |
| NC_045512.2;YP_009725295.1;Wu1-SARS2_orf1a;307 | 0 | 0 |
| NC_045512.2;YP_009725295.1;Wu1-SARS2_orf1a;308 | 0 | 0 |
| NC_045512.2;YP_009725295.1;Wu1-SARS2_orf1a;309 | 0 | 0 |
| NC_045512.2;YP_009725295.1;Wu1-SARS2_orf1a;310 | 0 | 0 |
| NC_045512.2;YP_009725295.1;Wu1-SARS2_orf1a;311 | 0 | 0 |
| NC_045512.2;YP_009725295.1;Wu1-SARS2_orf1a;312 | 0 | 0 |
| NC_045512.2;YP_009725295.1;Wu1-SARS2_orf1a;313 | 0 | 0 |
| NC_045512.2;YP_009725295.1;Wu1-SARS2_orf1a;314 | 0 | 0 |
| NC_045512.2;YP_009725295.1;Wu1-SARS2_orf1a;315 | 0 | 0 |
| NC_045512.2;YP_009725295.1;Wu1-SARS2_orf1a;316 | 0 | 0 |
| NC_045512.2;YP_009725295.1;Wu1-SARS2_orf1a;317 | 0 | 0 |
| NC_045512.2;YP_009725295.1;Wu1-SARS2_orf1a;318 | 0 | 0 |
| NC_045512.2;YP_009725295.1;Wu1-SARS2_orf1a;319 | 0 | 0 |
| NC_045512.2;YP_009725295.1;Wu1-SARS2_orf1a;320 | 0 | 0 |
| NC_045512.2;YP_009725295.1;Wu1-SARS2_orf1a;321 | 0 | 0 |
| NC_045512.2;YP_009725295.1;Wu1-SARS2_orf1a;322 | 0 | 0 |
| NC_045512.2;YP_009725295.1;Wu1-SARS2_orf1a;323 | 0 | 0 |
| NC_045512.2;YP_009725295.1;Wu1-SARS2_orf1a;324 | 0.05 | 0 |
| NC_045512.2;YP_009725295.1;Wu1-SARS2_orf1a;325 | 0 | 0.05 |
| NC_045512.2;YP_009725295.1;Wu1-SARS2_orf1a;326 | 0 | 0 |
| NC_045512.2;YP_009725295.1;Wu1-SARS2_orf1a;327 | 0 | 0 |
| NC_045512.2;YP_009725295.1;Wu1-SARS2_orf1a;328 | 0 | 0 |
| NC_045512.2;YP_009725295.1;Wu1-SARS2_orf1a;329 | 0 | 0 |
| NC_045512.2;YP_009725295.1;Wu1-SARS2_orf1a;330 | 0 | 0 |
| NC_045512.2;YP_009725295.1;Wu1-SARS2_orf1a;331 | 0 | 0 |
| NC_045512.2;YP_009725295.1;Wu1-SARS2_orf1a;332 | 0 | 0 |

| | | |
|---|---|---|
| NC_045512.2;YP_009725295.1;Wu1-SARS2_orf1a;333 | 0 | 0 |
| NC_045512.2;YP_009725295.1;Wu1-SARS2_orf1a;334 | 0 | 0 |
| NC_045512.2;YP_009725295.1;Wu1-SARS2_orf1a;335 | 0 | 0 |
| NC_045512.2;YP_009725295.1;Wu1-SARS2_orf1a;336 | 0 | 0 |
| NC_045512.2;YP_009725295.1;Wu1-SARS2_orf1a;337 | 0 | 0 |
| NC_045512.2;YP_009725295.1;Wu1-SARS2_orf1a;338 | 0 | 0 |
| NC_045512.2;YP_009725295.1;Wu1-SARS2_orf1a;339 | 0 | 0 |
| NC_045512.2;YP_009725295.1;Wu1-SARS2_orf1a;340 | 0 | 0 |
| NC_045512.2;YP_009725295.1;Wu1-SARS2_orf1a;341 | 0 | 0 |
| NC_045512.2;YP_009725295.1;Wu1-SARS2_orf1a;342 | 0 | 0 |
| NC_045512.2;YP_009725295.1;Wu1-SARS2_orf1a;343 | 0 | 0 |
| NC_045512.2;YP_009725295.1;Wu1-SARS2_orf1a;344 | 0 | 0 |
| NC_045512.2;YP_009725295.1;Wu1-SARS2_orf1a;345 | 0 | 0 |
| NC_045512.2;YP_009725295.1;Wu1-SARS2_orf1a;346 | 0 | 0 |
| NC_045512.2;YP_009725295.1;Wu1-SARS2_orf1a;347 | 0.05 | 0 |
| NC_045512.2;YP_009725295.1;Wu1-SARS2_orf1a;348 | 0 | 0 |
| NC_045512.2;YP_009725295.1;Wu1-SARS2_orf1a;349 | 0.05 | 0 |
| NC_045512.2;YP_009725295.1;Wu1-SARS2_orf1a;350 | 0 | 0 |
| NC_045512.2;YP_009725295.1;Wu1-SARS2_orf1a;351 | 0 | 0 |
| NC_045512.2;YP_009725295.1;Wu1-SARS2_orf1a;352 | 0 | 0.025 |
| NC_045512.2;YP_009725295.1;Wu1-SARS2_orf1a;353 | 0 | 0 |
| NC_045512.2;YP_009725295.1;Wu1-SARS2_orf1a;354 | 0 | 0 |
| NC_045512.2;YP_009725295.1;Wu1-SARS2_orf1a;355 | 0 | 0 |
| NC_045512.2;YP_009725295.1;Wu1-SARS2_orf1a;356 | 0 | 0 |
| NC_045512.2;YP_009725295.1;Wu1-SARS2_orf1a;357 | 0 | 0 |
| NC_045512.2;YP_009725295.1;Wu1-SARS2_orf1a;358 | 0 | 0 |
| NC_045512.2;YP_009725295.1;Wu1-SARS2_orf1a;359 | 0 | 0 |
| NC_045512.2;YP_009725295.1;Wu1-SARS2_orf1a;360 | 0 | 0 |
| NC_045512.2;YP_009725295.1;Wu1-SARS2_orf1a;361 | 0 | 0 |
| NC_045512.2;YP_009725295.1;Wu1-SARS2_orf1a;362 | 0 | 0 |
| NC_045512.2;YP_009725295.1;Wu1-SARS2_orf1a;363 | 0 | 0 |
| NC_045512.2;YP_009725295.1;Wu1-SARS2_orf1a;364 | 0 | 0 |
| NC_045512.2;YP_009725295.1;Wu1-SARS2_orf1a;365 | 0 | 0 |
| NC_045512.2;YP_009725295.1;Wu1-SARS2_orf1a;366 | 0 | 0 |
| NC_045512.2;YP_009725295.1;Wu1-SARS2_orf1a;367 | 0 | 0 |
| NC_045512.2;YP_009725295.1;Wu1-SARS2_orf1a;368 | 0 | 0 |
| NC_045512.2;YP_009725295.1;Wu1-SARS2_orf1a;369 | 0 | 0 |
| NC_045512.2;YP_009725295.1;Wu1-SARS2_orf1a;370 | 0 | 0 |
| NC_045512.2;YP_009725295.1;Wu1-SARS2_orf1a;371 | 0 | 0 |
| NC_045512.2;YP_009725295.1;Wu1-SARS2_orf1a;372 | 0 | 0 |
| NC_045512.2;YP_009725295.1;Wu1-SARS2_orf1a;373 | 0 | 0 |
| NC_045512.2;YP_009725295.1;Wu1-SARS2_orf1a;374 | 0 | 0.025 |
| NC_045512.2;YP_009725295.1;Wu1-SARS2_orf1a;375 | 0 | 0 |

| | | |
|---|---|---|
| NC_045512.2;YP_009725295.1;Wu1-SARS2_orf1a;376 | 0 | 0 |
| NC_045512.2;YP_009725295.1;Wu1-SARS2_orf1a;377 | 0 | 0 |
| NC_045512.2;YP_009725295.1;Wu1-SARS2_orf1a;378 | 0 | 0 |
| NC_045512.2;YP_009725295.1;Wu1-SARS2_orf1a;379 | 0.15 | 0.05 |
| NC_045512.2;YP_009725295.1;Wu1-SARS2_orf1a;380 | 0 | 0.025 |
| NC_045512.2;YP_009725295.1;Wu1-SARS2_orf1a;381 | 0 | 0.025 |
| NC_045512.2;YP_009725295.1;Wu1-SARS2_orf1a;382 | 0 | 0.025 |
| NC_045512.2;YP_009725295.1;Wu1-SARS2_orf1a;383 | 0 | 0.025 |
| NC_045512.2;YP_009725295.1;Wu1-SARS2_orf1a;384 | 0 | 0.025 |
| NC_045512.2;YP_009725295.1;Wu1-SARS2_orf1a;385 | 0.15 | 0.075 |
| NC_045512.2;YP_009725295.1;Wu1-SARS2_orf1a;386 | 0.05 | 0.075 |
| NC_045512.2;YP_009725295.1;Wu1-SARS2_orf1a;387 | 0 | 0 |
| NC_045512.2;YP_009725295.1;Wu1-SARS2_orf1a;388 | 0 | 0 |
| NC_045512.2;YP_009725295.1;Wu1-SARS2_orf1a;389 | 0 | 0 |
| NC_045512.2;YP_009725295.1;Wu1-SARS2_orf1a;390 | 0 | 0 |
| NC_045512.2;YP_009725295.1;Wu1-SARS2_orf1a;391 | 0 | 0 |
| NC_045512.2;YP_009725295.1;Wu1-SARS2_orf1a;392 | 0 | 0 |
| NC_045512.2;YP_009725295.1;Wu1-SARS2_orf1a;393 | 0 | 0 |
| NC_045512.2;YP_009725295.1;Wu1-SARS2_orf1a;394 | 0 | 0 |
| NC_045512.2;YP_009725295.1;Wu1-SARS2_orf1a;395 | 0 | 0 |
| NC_045512.2;YP_009725295.1;Wu1-SARS2_orf1a;396 | 0 | 0 |
| NC_045512.2;YP_009725295.1;Wu1-SARS2_orf1a;397 | 0 | 0.025 |
| NC_045512.2;YP_009725295.1;Wu1-SARS2_orf1a;398 | 0 | 0 |
| NC_045512.2;YP_009725295.1;Wu1-SARS2_orf1a;399 | 0 | 0 |
| NC_045512.2;YP_009725295.1;Wu1-SARS2_orf1a;400 | 0 | 0 |
| NC_045512.2;YP_009725295.1;Wu1-SARS2_orf1a;401 | 0 | 0 |
| NC_045512.2;YP_009725295.1;Wu1-SARS2_orf1a;402 | 0 | 0 |
| NC_045512.2;YP_009725295.1;Wu1-SARS2_orf1a;403 | 0 | 0 |
| NC_045512.2;YP_009725295.1;Wu1-SARS2_orf1a;404 | 0 | 0 |
| NC_045512.2;YP_009725295.1;Wu1-SARS2_orf1a;405 | 0 | 0 |
| NC_045512.2;YP_009725295.1;Wu1-SARS2_orf1a;406 | 0 | 0 |
| NC_045512.2;YP_009725295.1;Wu1-SARS2_orf1a;407 | 0 | 0 |
| NC_045512.2;YP_009725295.1;Wu1-SARS2_orf1a;408 | 0 | 0 |
| NC_045512.2;YP_009725295.1;Wu1-SARS2_orf1a;409 | 0 | 0 |
| NC_045512.2;YP_009725295.1;Wu1-SARS2_orf1a;410 | 0 | 0 |
| NC_045512.2;YP_009725295.1;Wu1-SARS2_orf1a;411 | 0 | 0 |
| NC_045512.2;YP_009725295.1;Wu1-SARS2_orf1a;412 | 0 | 0 |
| NC_045512.2;YP_009725295.1;Wu1-SARS2_orf1a;413 | 0 | 0 |
| NC_045512.2;YP_009725295.1;Wu1-SARS2_orf1a;414 | 0 | 0 |
| NC_045512.2;YP_009725295.1;Wu1-SARS2_orf1a;415 | 0 | 0 |
| NC_045512.2;YP_009725295.1;Wu1-SARS2_orf1a;416 | 0 | 0 |
| NC_045512.2;YP_009725295.1;Wu1-SARS2_orf1a;417 | 0 | 0 |
| NC_045512.2;YP_009725295.1;Wu1-SARS2_orf1a;418 | 0 | 0 |

| | | |
|---|---|---|
| NC_045512.2;YP_009725295.1;Wu1-SARS2_orf1a;419 | 0 | 0 |
| NC_045512.2;YP_009725295.1;Wu1-SARS2_orf1a;420 | 0 | 0 |
| NC_045512.2;YP_009725295.1;Wu1-SARS2_orf1a;421 | 0 | 0 |
| NC_045512.2;YP_009725295.1;Wu1-SARS2_orf1a;422 | 0 | 0 |
| NC_045512.2;YP_009725295.1;Wu1-SARS2_orf1a;423 | 0 | 0 |
| NC_045512.2;YP_009725295.1;Wu1-SARS2_orf1a;424 | 0.05 | 0.025 |
| NC_045512.2;YP_009725295.1;Wu1-SARS2_orf1a;425 | 0.05 | 0.025 |
| NC_045512.2;YP_009725295.1;Wu1-SARS2_orf1a;426 | 0 | 0 |
| NC_045512.2;YP_009725295.1;Wu1-SARS2_orf1a;427 | 0 | 0.025 |
| NC_045512.2;YP_009725295.1;Wu1-SARS2_orf1a;428 | 0 | 0 |
| NC_045512.2;YP_009725295.1;Wu1-SARS2_orf1a;429 | 0 | 0 |
| NC_045512.2;YP_009725295.1;Wu1-SARS2_orf1a;430 | 0 | 0 |
| NC_045512.2;YP_009725295.1;Wu1-SARS2_orf1a;431 | 0 | 0 |
| NC_045512.2;YP_009725295.1;Wu1-SARS2_orf1a;432 | 0 | 0 |
| NC_045512.2;YP_009725295.1;Wu1-SARS2_orf1a;433 | 0 | 0 |
| NC_045512.2;YP_009725295.1;Wu1-SARS2_orf1a;434 | 0 | 0 |
| NC_045512.2;YP_009725295.1;Wu1-SARS2_orf1a;435 | 0 | 0 |
| NC_045512.2;YP_009725295.1;Wu1-SARS2_orf1a;436 | 0 | 0 |
| NC_045512.2;YP_009725295.1;Wu1-SARS2_orf1a;437 | 0 | 0 |
| NC_045512.2;YP_009725295.1;Wu1-SARS2_orf1a;438 | 0 | 0 |
| NC_045512.2;YP_009725295.1;Wu1-SARS2_orf1a;439 | 0 | 0 |
| NC_045512.2;YP_009725295.1;Wu1-SARS2_orf1a;440 | 0 | 0 |
| NC_045512.2;YP_009725295.1;Wu1-SARS2_orf1a;441 | 0 | 0 |
| NC_045512.2;YP_009725295.1;Wu1-SARS2_orf1a;442 | 0 | 0 |
| NC_045512.2;YP_009725295.1;Wu1-SARS2_orf1a;443 | 0.05 | 0 |
| NC_045512.2;YP_009725295.1;Wu1-SARS2_orf1a;444 | 0.2 | 0.025 |
| NC_045512.2;YP_009725295.1;Wu1-SARS2_orf1a;445 | 0.5 | 0.475 |
| NC_045512.2;YP_009725295.1;Wu1-SARS2_orf1a;446 | 0.05 | 0.05 |
| NC_045512.2;YP_009725295.1;Wu1-SARS2_orf1a;447 | 0.05 | 0.05 |
| NC_045512.2;YP_009725295.1;Wu1-SARS2_orf1a;448 | 0.05 | 0.05 |
| NC_045512.2;YP_009725295.1;Wu1-SARS2_orf1a;449 | 0 | 0.025 |
| NC_045512.2;YP_009725295.1;Wu1-SARS2_orf1a;450 | 0 | 0 |
| NC_045512.2;YP_009725295.1;Wu1-SARS2_orf1a;451 | 0 | 0 |
| NC_045512.2;YP_009725295.1;Wu1-SARS2_orf1a;452 | 0.05 | 0 |
| NC_045512.2;YP_009725295.1;Wu1-SARS2_orf1a;453 | 0 | 0 |
| NC_045512.2;YP_009725295.1;Wu1-SARS2_orf1a;454 | 0 | 0 |
| NC_045512.2;YP_009725295.1;Wu1-SARS2_orf1a;455 | 0 | 0 |
| NC_045512.2;YP_009725295.1;Wu1-SARS2_orf1a;456 | 0.05 | 0 |
| NC_045512.2;YP_009725295.1;Wu1-SARS2_orf1a;457 | 0.05 | 0 |
| NC_045512.2;YP_009725295.1;Wu1-SARS2_orf1a;458 | 0.05 | 0 |
| NC_045512.2;YP_009725295.1;Wu1-SARS2_orf1a;459 | 0 | 0 |
| NC_045512.2;YP_009725295.1;Wu1-SARS2_orf1a;460 | 0 | 0 |
| NC_045512.2;YP_009725295.1;Wu1-SARS2_orf1a;461 | 0 | 0 |

| | | |
|---|---|---|
| NC_045512.2;YP_009725295.1;Wu1-SARS2_orf1a;462 | 0 | 0 |
| NC_045512.2;YP_009725295.1;Wu1-SARS2_orf1a;463 | 0 | 0 |
| NC_045512.2;YP_009725295.1;Wu1-SARS2_orf1a;464 | 0 | 0 |
| NC_045512.2;YP_009725295.1;Wu1-SARS2_orf1a;465 | 0 | 0 |
| NC_045512.2;YP_009725295.1;Wu1-SARS2_orf1a;466 | 0 | 0 |
| NC_045512.2;YP_009725295.1;Wu1-SARS2_orf1a;467 | 0 | 0 |
| NC_045512.2;YP_009725295.1;Wu1-SARS2_orf1a;468 | 0 | 0 |
| NC_045512.2;YP_009725295.1;Wu1-SARS2_orf1a;469 | 0 | 0 |
| NC_045512.2;YP_009725295.1;Wu1-SARS2_orf1a;470 | 0 | 0 |
| NC_045512.2;YP_009725295.1;Wu1-SARS2_orf1a;471 | 0 | 0 |
| NC_045512.2;YP_009725295.1;Wu1-SARS2_orf1a;472 | 0 | 0 |
| NC_045512.2;YP_009725295.1;Wu1-SARS2_orf1a;473 | 0 | 0 |
| NC_045512.2;YP_009725295.1;Wu1-SARS2_orf1a;474 | 0 | 0 |
| NC_045512.2;YP_009725295.1;Wu1-SARS2_orf1a;475 | 0 | 0 |
| NC_045512.2;YP_009725295.1;Wu1-SARS2_orf1a;476 | 0 | 0 |
| NC_045512.2;YP_009725295.1;Wu1-SARS2_orf1a;477 | 0 | 0 |
| NC_045512.2;YP_009725295.1;Wu1-SARS2_orf1a;478 | 0 | 0 |
| NC_045512.2;YP_009725295.1;Wu1-SARS2_orf1a;479 | 0 | 0 |
| NC_045512.2;YP_009725295.1;Wu1-SARS2_orf1a;480 | 0 | 0.05 |
| NC_045512.2;YP_009725295.1;Wu1-SARS2_orf1a;481 | 0 | 0.05 |
| NC_045512.2;YP_009725295.1;Wu1-SARS2_orf1a;482 | 0 | 0.025 |
| NC_045512.2;YP_009725295.1;Wu1-SARS2_orf1a;483 | 0 | 0.025 |
| NC_045512.2;YP_009725295.1;Wu1-SARS2_orf1a;484 | 0 | 0 |
| NC_045512.2;YP_009725295.1;Wu1-SARS2_orf1a;485 | 0 | 0 |
| NC_045512.2;YP_009725295.1;Wu1-SARS2_orf1a;486 | 0 | 0.025 |
| NC_045512.2;YP_009725295.1;Wu1-SARS2_orf1a;487 | 0 | 0 |
| NC_045512.2;YP_009725295.1;Wu1-SARS2_orf1a;488 | 0 | 0 |
| NC_045512.2;YP_009725295.1;Wu1-SARS2_orf1a;489 | 0.05 | 0.025 |
| NC_045512.2;YP_009725295.1;Wu1-SARS2_orf1a;490 | 0 | 0 |
| NC_045512.2;YP_009725295.1;Wu1-SARS2_orf1a;491 | 0 | 0 |
| NC_045512.2;YP_009725295.1;Wu1-SARS2_orf1a;492 | 0 | 0 |
| NC_045512.2;YP_009725295.1;Wu1-SARS2_orf1a;493 | 0.05 | 0.05 |
| NC_045512.2;YP_009725295.1;Wu1-SARS2_orf1a;494 | 0.05 | 0.025 |
| NC_045512.2;YP_009725295.1;Wu1-SARS2_orf1a;495 | 0 | 0 |
| NC_045512.2;YP_009725295.1;Wu1-SARS2_orf1a;496 | 0 | 0 |
| NC_045512.2;YP_009725295.1;Wu1-SARS2_orf1a;497 | 0 | 0 |
| NC_045512.2;YP_009725295.1;Wu1-SARS2_orf1a;498 | 0 | 0 |
| NC_045512.2;YP_009725295.1;Wu1-SARS2_orf1a;499 | 0 | 0 |
| NC_045512.2;YP_009725295.1;Wu1-SARS2_orf1a;500 | 0 | 0 |
| NC_045512.2;YP_009725295.1;Wu1-SARS2_orf1a;501 | 0 | 0 |
| NC_045512.2;YP_009725295.1;Wu1-SARS2_orf1a;502 | 0 | 0 |
| NC_045512.2;YP_009725295.1;Wu1-SARS2_orf1a;503 | 0 | 0 |
| NC_045512.2;YP_009725295.1;Wu1-SARS2_orf1a;504 | 0 | 0 |

| | | |
|---|---|---|
| NC_045512.2;YP_009725295.1;Wu1-SARS2_orf1a;505 | 0 | 0 |
| NC_045512.2;YP_009725295.1;Wu1-SARS2_orf1a;506 | 0 | 0 |
| NC_045512.2;YP_009725295.1;Wu1-SARS2_orf1a;507 | 0 | 0 |
| NC_045512.2;YP_009725295.1;Wu1-SARS2_orf1a;508 | 0 | 0 |
| NC_045512.2;YP_009725295.1;Wu1-SARS2_orf1a;509 | 0 | 0 |
| NC_045512.2;YP_009725295.1;Wu1-SARS2_orf1a;510 | 0 | 0 |
| NC_045512.2;YP_009725295.1;Wu1-SARS2_orf1a;511 | 0 | 0 |
| NC_045512.2;YP_009725295.1;Wu1-SARS2_orf1a;512 | 0 | 0 |
| NC_045512.2;YP_009725295.1;Wu1-SARS2_orf1a;513 | 0 | 0.025 |
| NC_045512.2;YP_009725295.1;Wu1-SARS2_orf1a;514 | 0 | 0.025 |
| NC_045512.2;YP_009725295.1;Wu1-SARS2_orf1a;515 | 0 | 0.025 |
| NC_045512.2;YP_009725295.1;Wu1-SARS2_orf1a;516 | 0 | 0.025 |
| NC_045512.2;YP_009725295.1;Wu1-SARS2_orf1a;517 | 0 | 0 |
| NC_045512.2;YP_009725295.1;Wu1-SARS2_orf1a;518 | 0.1 | 0.025 |
| NC_045512.2;YP_009725295.1;Wu1-SARS2_orf1a;519 | 0 | 0 |
| NC_045512.2;YP_009725295.1;Wu1-SARS2_orf1a;520 | 0 | 0 |
| NC_045512.2;YP_009725295.1;Wu1-SARS2_orf1a;521 | 0 | 0 |
| NC_045512.2;YP_009725295.1;Wu1-SARS2_orf1a;522 | 0 | 0 |
| NC_045512.2;YP_009725295.1;Wu1-SARS2_orf1a;523 | 0 | 0 |
| NC_045512.2;YP_009725295.1;Wu1-SARS2_orf1a;524 | 0 | 0 |
| NC_045512.2;YP_009725295.1;Wu1-SARS2_orf1a;525 | 0.05 | 0 |
| NC_045512.2;YP_009725295.1;Wu1-SARS2_orf1a;526 | 0.05 | 0.025 |
| NC_045512.2;YP_009725295.1;Wu1-SARS2_orf1a;527 | 0 | 0 |
| NC_045512.2;YP_009725295.1;Wu1-SARS2_orf1a;528 | 0 | 0 |
| NC_045512.2;YP_009725295.1;Wu1-SARS2_orf1a;529 | 0 | 0 |
| NC_045512.2;YP_009725295.1;Wu1-SARS2_orf1a;530 | 0 | 0 |
| NC_045512.2;YP_009725295.1;Wu1-SARS2_orf1a;531 | 0 | 0 |
| NC_045512.2;YP_009725295.1;Wu1-SARS2_orf1a;532 | 0 | 0 |
| NC_045512.2;YP_009725295.1;Wu1-SARS2_orf1a;533 | 0 | 0 |
| NC_045512.2;YP_009725295.1;Wu1-SARS2_orf1a;534 | 0 | 0 |
| NC_045512.2;YP_009725295.1;Wu1-SARS2_orf1a;535 | 0 | 0 |
| NC_045512.2;YP_009725295.1;Wu1-SARS2_orf1a;536 | 0 | 0 |
| NC_045512.2;YP_009725295.1;Wu1-SARS2_orf1a;537 | 0 | 0 |
| NC_045512.2;YP_009725295.1;Wu1-SARS2_orf1a;538 | 0 | 0 |
| NC_045512.2;YP_009725295.1;Wu1-SARS2_orf1a;539 | 0 | 0 |
| NC_045512.2;YP_009725295.1;Wu1-SARS2_orf1a;540 | 0 | 0 |
| NC_045512.2;YP_009725295.1;Wu1-SARS2_orf1a;541 | 0 | 0 |
| NC_045512.2;YP_009725295.1;Wu1-SARS2_orf1a;542 | 0 | 0 |
| NC_045512.2;YP_009725295.1;Wu1-SARS2_orf1a;543 | 0 | 0 |
| NC_045512.2;YP_009725295.1;Wu1-SARS2_orf1a;544 | 0 | 0 |
| NC_045512.2;YP_009725295.1;Wu1-SARS2_orf1a;545 | 0 | 0 |
| NC_045512.2;YP_009725295.1;Wu1-SARS2_orf1a;546 | 0 | 0.025 |
| NC_045512.2;YP_009725295.1;Wu1-SARS2_orf1a;547 | 0 | 0 |

| | | |
|---|---|---|
| NC_045512.2;YP_009725295.1;Wu1-SARS2_orf1a;548 | 0 | 0 |
| NC_045512.2;YP_009725295.1;Wu1-SARS2_orf1a;549 | 0 | 0 |
| NC_045512.2;YP_009725295.1;Wu1-SARS2_orf1a;550 | 0 | 0 |
| NC_045512.2;YP_009725295.1;Wu1-SARS2_orf1a;551 | 0.05 | 0 |
| NC_045512.2;YP_009725295.1;Wu1-SARS2_orf1a;552 | 0 | 0 |
| NC_045512.2;YP_009725295.1;Wu1-SARS2_orf1a;553 | 0.05 | 0 |
| NC_045512.2;YP_009725295.1;Wu1-SARS2_orf1a;554 | 0 | 0 |
| NC_045512.2;YP_009725295.1;Wu1-SARS2_orf1a;555 | 0.05 | 0 |
| NC_045512.2;YP_009725295.1;Wu1-SARS2_orf1a;556 | 0 | 0 |
| NC_045512.2;YP_009725295.1;Wu1-SARS2_orf1a;557 | 0 | 0 |
| NC_045512.2;YP_009725295.1;Wu1-SARS2_orf1a;558 | 0 | 0 |
| NC_045512.2;YP_009725295.1;Wu1-SARS2_orf1a;559 | 0 | 0 |
| NC_045512.2;YP_009725295.1;Wu1-SARS2_orf1a;560 | 0 | 0 |
| NC_045512.2;YP_009725295.1;Wu1-SARS2_orf1a;561 | 0 | 0 |
| NC_045512.2;YP_009725295.1;Wu1-SARS2_orf1a;562 | 0 | 0 |
| NC_045512.2;YP_009725295.1;Wu1-SARS2_orf1a;563 | 0 | 0 |
| NC_045512.2;YP_009725295.1;Wu1-SARS2_orf1a;564 | 0 | 0 |
| NC_045512.2;YP_009725295.1;Wu1-SARS2_orf1a;565 | 0 | 0.025 |
| NC_045512.2;YP_009725295.1;Wu1-SARS2_orf1a;566 | 0 | 0 |
| NC_045512.2;YP_009725295.1;Wu1-SARS2_orf1a;567 | 0 | 0 |
| NC_045512.2;YP_009725295.1;Wu1-SARS2_orf1a;568 | 0 | 0 |
| NC_045512.2;YP_009725295.1;Wu1-SARS2_orf1a;569 | 0 | 0.025 |
| NC_045512.2;YP_009725295.1;Wu1-SARS2_orf1a;570 | 0 | 0.025 |
| NC_045512.2;YP_009725295.1;Wu1-SARS2_orf1a;571 | 0 | 0.025 |
| NC_045512.2;YP_009725295.1;Wu1-SARS2_orf1a;572 | 0.1 | 0.05 |
| NC_045512.2;YP_009725295.1;Wu1-SARS2_orf1a;573 | 0 | 0.025 |
| NC_045512.2;YP_009725295.1;Wu1-SARS2_orf1a;574 | 0 | 0 |
| NC_045512.2;YP_009725295.1;Wu1-SARS2_orf1a;575 | 0 | 0 |
| NC_045512.2;YP_009725295.1;Wu1-SARS2_orf1a;576 | 0 | 0 |
| NC_045512.2;YP_009725295.1;Wu1-SARS2_orf1a;577 | 0 | 0 |
| NC_045512.2;YP_009725295.1;Wu1-SARS2_orf1a;578 | 0 | 0.025 |
| NC_045512.2;YP_009725295.1;Wu1-SARS2_orf1a;579 | 0 | 0 |
| NC_045512.2;YP_009725295.1;Wu1-SARS2_orf1a;580 | 0 | 0 |
| NC_045512.2;YP_009725295.1;Wu1-SARS2_orf1a;581 | 0 | 0 |
| NC_045512.2;YP_009725295.1;Wu1-SARS2_orf1a;582 | 0 | 0 |
| NC_045512.2;YP_009725295.1;Wu1-SARS2_orf1a;583 | 0 | 0 |
| NC_045512.2;YP_009725295.1;Wu1-SARS2_orf1a;584 | 0 | 0 |
| NC_045512.2;YP_009725295.1;Wu1-SARS2_orf1a;585 | 0 | 0 |
| NC_045512.2;YP_009725295.1;Wu1-SARS2_orf1a;586 | 0 | 0 |
| NC_045512.2;YP_009725295.1;Wu1-SARS2_orf1a;587 | 0 | 0 |
| NC_045512.2;YP_009725295.1;Wu1-SARS2_orf1a;588 | 0 | 0 |
| NC_045512.2;YP_009725295.1;Wu1-SARS2_orf1a;589 | 0.05 | 0 |
| NC_045512.2;YP_009725295.1;Wu1-SARS2_orf1a;590 | 0 | 0 |

| | | |
|---|---|---|
| NC_045512.2;YP_009725295.1;Wu1-SARS2_orf1a;591 | 0 | 0 |
| NC_045512.2;YP_009725295.1;Wu1-SARS2_orf1a;592 | 0 | 0 |
| NC_045512.2;YP_009725295.1;Wu1-SARS2_orf1a;593 | 0 | 0 |
| NC_045512.2;YP_009725295.1;Wu1-SARS2_orf1a;594 | 0 | 0 |
| NC_045512.2;YP_009725295.1;Wu1-SARS2_orf1a;595 | 0 | 0 |
| NC_045512.2;YP_009725295.1;Wu1-SARS2_orf1a;596 | 0 | 0 |
| NC_045512.2;YP_009725295.1;Wu1-SARS2_orf1a;597 | 0 | 0 |
| NC_045512.2;YP_009725295.1;Wu1-SARS2_orf1a;598 | 0 | 0 |
| NC_045512.2;YP_009725295.1;Wu1-SARS2_orf1a;599 | 0 | 0 |
| NC_045512.2;YP_009725295.1;Wu1-SARS2_orf1a;600 | 0 | 0 |
| NC_045512.2;YP_009725295.1;Wu1-SARS2_orf1a;601 | 0 | 0 |
| NC_045512.2;YP_009725295.1;Wu1-SARS2_orf1a;602 | 0 | 0 |
| NC_045512.2;YP_009725295.1;Wu1-SARS2_orf1a;603 | 0 | 0 |
| NC_045512.2;YP_009725295.1;Wu1-SARS2_orf1a;604 | 0 | 0 |
| NC_045512.2;YP_009725295.1;Wu1-SARS2_orf1a;605 | 0 | 0 |
| NC_045512.2;YP_009725295.1;Wu1-SARS2_orf1a;606 | 0 | 0 |
| NC_045512.2;YP_009725295.1;Wu1-SARS2_orf1a;607 | 0 | 0 |
| NC_045512.2;YP_009725295.1;Wu1-SARS2_orf1a;608 | 0 | 0 |
| NC_045512.2;YP_009725295.1;Wu1-SARS2_orf1a;609 | 0 | 0 |
| NC_045512.2;YP_009725295.1;Wu1-SARS2_orf1a;610 | 0 | 0 |
| NC_045512.2;YP_009725295.1;Wu1-SARS2_orf1a;611 | 0 | 0 |
| NC_045512.2;YP_009725295.1;Wu1-SARS2_orf1a;612 | 0 | 0 |
| NC_045512.2;YP_009725295.1;Wu1-SARS2_orf1a;613 | 0 | 0 |
| NC_045512.2;YP_009725295.1;Wu1-SARS2_orf1a;614 | 0 | 0.05 |
| NC_045512.2;YP_009725295.1;Wu1-SARS2_orf1a;615 | 0 | 0 |
| NC_045512.2;YP_009725295.1;Wu1-SARS2_orf1a;616 | 0 | 0 |
| NC_045512.2;YP_009725295.1;Wu1-SARS2_orf1a;617 | 0.05 | 0.025 |
| NC_045512.2;YP_009725295.1;Wu1-SARS2_orf1a;618 | 0.05 | 0.05 |
| NC_045512.2;YP_009725295.1;Wu1-SARS2_orf1a;619 | 0.05 | 0.025 |
| NC_045512.2;YP_009725295.1;Wu1-SARS2_orf1a;620 | 0.05 | 0.025 |
| NC_045512.2;YP_009725295.1;Wu1-SARS2_orf1a;621 | 0.1 | 0.025 |
| NC_045512.2;YP_009725295.1;Wu1-SARS2_orf1a;622 | 0 | 0.025 |
| NC_045512.2;YP_009725295.1;Wu1-SARS2_orf1a;623 | 0 | 0.025 |
| NC_045512.2;YP_009725295.1;Wu1-SARS2_orf1a;624 | 0 | 0.05 |
| NC_045512.2;YP_009725295.1;Wu1-SARS2_orf1a;625 | 0 | 0 |
| NC_045512.2;YP_009725295.1;Wu1-SARS2_orf1a;626 | 0 | 0 |
| NC_045512.2;YP_009725295.1;Wu1-SARS2_orf1a;627 | 0 | 0 |
| NC_045512.2;YP_009725295.1;Wu1-SARS2_orf1a;628 | 0 | 0.025 |
| NC_045512.2;YP_009725295.1;Wu1-SARS2_orf1a;629 | 0 | 0.025 |
| NC_045512.2;YP_009725295.1;Wu1-SARS2_orf1a;630 | 0 | 0.025 |
| NC_045512.2;YP_009725295.1;Wu1-SARS2_orf1a;631 | 0 | 0.05 |
| NC_045512.2;YP_009725295.1;Wu1-SARS2_orf1a;632 | 0 | 0.05 |
| NC_045512.2;YP_009725295.1;Wu1-SARS2_orf1a;633 | 0 | 0.025 |

| | | |
|---|---|---|
| NC_045512.2;YP_009725295.1;Wu1-SARS2_orf1a;634 | 0 | 0 |
| NC_045512.2;YP_009725295.1;Wu1-SARS2_orf1a;635 | 0 | 0.025 |
| NC_045512.2;YP_009725295.1;Wu1-SARS2_orf1a;636 | 0 | 0.05 |
| NC_045512.2;YP_009725295.1;Wu1-SARS2_orf1a;637 | 0 | 0 |
| NC_045512.2;YP_009725295.1;Wu1-SARS2_orf1a;638 | 0.05 | 0.025 |
| NC_045512.2;YP_009725295.1;Wu1-SARS2_orf1a;639 | 0 | 0 |
| NC_045512.2;YP_009725295.1;Wu1-SARS2_orf1a;640 | 0 | 0 |
| NC_045512.2;YP_009725295.1;Wu1-SARS2_orf1a;641 | 0 | 0 |
| NC_045512.2;YP_009725295.1;Wu1-SARS2_orf1a;642 | 0 | 0 |
| NC_045512.2;YP_009725295.1;Wu1-SARS2_orf1a;643 | 0 | 0 |
| NC_045512.2;YP_009725295.1;Wu1-SARS2_orf1a;644 | 0 | 0 |
| NC_045512.2;YP_009725295.1;Wu1-SARS2_orf1a;645 | 0 | 0 |
| NC_045512.2;YP_009725295.1;Wu1-SARS2_orf1a;646 | 0 | 0 |
| NC_045512.2;YP_009725295.1;Wu1-SARS2_orf1a;647 | 0 | 0 |
| NC_045512.2;YP_009725295.1;Wu1-SARS2_orf1a;648 | 0 | 0 |
| NC_045512.2;YP_009725295.1;Wu1-SARS2_orf1a;649 | 0 | 0 |
| NC_045512.2;YP_009725295.1;Wu1-SARS2_orf1a;650 | 0 | 0 |
| NC_045512.2;YP_009725295.1;Wu1-SARS2_orf1a;651 | 0 | 0 |
| NC_045512.2;YP_009725295.1;Wu1-SARS2_orf1a;652 | 0 | 0 |
| NC_045512.2;YP_009725295.1;Wu1-SARS2_orf1a;653 | 0 | 0 |
| NC_045512.2;YP_009725295.1;Wu1-SARS2_orf1a;654 | 0 | 0 |
| NC_045512.2;YP_009725295.1;Wu1-SARS2_orf1a;655 | 0 | 0 |
| NC_045512.2;YP_009725295.1;Wu1-SARS2_orf1a;656 | 0 | 0 |
| NC_045512.2;YP_009725295.1;Wu1-SARS2_orf1a;657 | 0 | 0.025 |
| NC_045512.2;YP_009725295.1;Wu1-SARS2_orf1a;658 | 0 | 0.05 |
| NC_045512.2;YP_009725295.1;Wu1-SARS2_orf1a;659 | 0 | 0.025 |
| NC_045512.2;YP_009725295.1;Wu1-SARS2_orf1a;660 | 0 | 0.025 |
| NC_045512.2;YP_009725295.1;Wu1-SARS2_orf1a;661 | 0 | 0.025 |
| NC_045512.2;YP_009725295.1;Wu1-SARS2_orf1a;662 | 0 | 0.025 |
| NC_045512.2;YP_009725295.1;Wu1-SARS2_orf1a;663 | 0 | 0.025 |
| NC_045512.2;YP_009725295.1;Wu1-SARS2_orf1a;664 | 0 | 0.025 |
| NC_045512.2;YP_009725295.1;Wu1-SARS2_orf1a;665 | 0 | 0.025 |
| NC_045512.2;YP_009725295.1;Wu1-SARS2_orf1a;666 | 0 | 0.025 |
| NC_045512.2;YP_009725295.1;Wu1-SARS2_orf1a;667 | 0 | 0.025 |
| NC_045512.2;YP_009725295.1;Wu1-SARS2_orf1a;668 | 0 | 0 |
| NC_045512.2;YP_009725295.1;Wu1-SARS2_orf1a;669 | 0 | 0 |
| NC_045512.2;YP_009725295.1;Wu1-SARS2_orf1a;670 | 0 | 0 |
| NC_045512.2;YP_009725295.1;Wu1-SARS2_orf1a;671 | 0 | 0 |
| NC_045512.2;YP_009725295.1;Wu1-SARS2_orf1a;672 | 0 | 0 |
| NC_045512.2;YP_009725295.1;Wu1-SARS2_orf1a;673 | 0 | 0 |
| NC_045512.2;YP_009725295.1;Wu1-SARS2_orf1a;674 | 0 | 0 |
| NC_045512.2;YP_009725295.1;Wu1-SARS2_orf1a;675 | 0 | 0 |
| NC_045512.2;YP_009725295.1;Wu1-SARS2_orf1a;676 | 0 | 0 |

| | | |
|---|---|---|
| NC_045512.2;YP_009725295.1;Wu1-SARS2_orf1a;677 | 0 | 0 |
| NC_045512.2;YP_009725295.1;Wu1-SARS2_orf1a;678 | 0 | 0 |
| NC_045512.2;YP_009725295.1;Wu1-SARS2_orf1a;679 | 0 | 0 |
| NC_045512.2;YP_009725295.1;Wu1-SARS2_orf1a;680 | 0 | 0 |
| NC_045512.2;YP_009725295.1;Wu1-SARS2_orf1a;681 | 0 | 0 |
| NC_045512.2;YP_009725295.1;Wu1-SARS2_orf1a;682 | 0 | 0 |
| NC_045512.2;YP_009725295.1;Wu1-SARS2_orf1a;683 | 0 | 0 |
| NC_045512.2;YP_009725295.1;Wu1-SARS2_orf1a;684 | 0 | 0 |
| NC_045512.2;YP_009725295.1;Wu1-SARS2_orf1a;685 | 0 | 0 |
| NC_045512.2;YP_009725295.1;Wu1-SARS2_orf1a;686 | 0 | 0 |
| NC_045512.2;YP_009725295.1;Wu1-SARS2_orf1a;687 | 0 | 0 |
| NC_045512.2;YP_009725295.1;Wu1-SARS2_orf1a;688 | 0 | 0 |
| NC_045512.2;YP_009725295.1;Wu1-SARS2_orf1a;689 | 0 | 0 |
| NC_045512.2;YP_009725295.1;Wu1-SARS2_orf1a;690 | 0 | 0 |
| NC_045512.2;YP_009725295.1;Wu1-SARS2_orf1a;691 | 0 | 0 |
| NC_045512.2;YP_009725295.1;Wu1-SARS2_orf1a;692 | 0 | 0 |
| NC_045512.2;YP_009725295.1;Wu1-SARS2_orf1a;693 | 0 | 0.025 |
| NC_045512.2;YP_009725295.1;Wu1-SARS2_orf1a;694 | 0 | 0 |
| NC_045512.2;YP_009725295.1;Wu1-SARS2_orf1a;695 | 0 | 0 |
| NC_045512.2;YP_009725295.1;Wu1-SARS2_orf1a;696 | 0 | 0.025 |
| NC_045512.2;YP_009725295.1;Wu1-SARS2_orf1a;697 | 0 | 0 |
| NC_045512.2;YP_009725295.1;Wu1-SARS2_orf1a;698 | 0 | 0 |
| NC_045512.2;YP_009725295.1;Wu1-SARS2_orf1a;699 | 0 | 0 |
| NC_045512.2;YP_009725295.1;Wu1-SARS2_orf1a;700 | 0 | 0 |
| NC_045512.2;YP_009725295.1;Wu1-SARS2_orf1a;701 | 0 | 0 |
| NC_045512.2;YP_009725295.1;Wu1-SARS2_orf1a;702 | 0 | 0 |
| NC_045512.2;YP_009725295.1;Wu1-SARS2_orf1a;703 | 0 | 0 |
| NC_045512.2;YP_009725295.1;Wu1-SARS2_orf1a;704 | 0 | 0 |
| NC_045512.2;YP_009725295.1;Wu1-SARS2_orf1a;705 | 0 | 0 |
| NC_045512.2;YP_009725295.1;Wu1-SARS2_orf1a;706 | 0 | 0 |
| NC_045512.2;YP_009725295.1;Wu1-SARS2_orf1a;707 | 0 | 0 |
| NC_045512.2;YP_009725295.1;Wu1-SARS2_orf1a;708 | 0 | 0 |
| NC_045512.2;YP_009725295.1;Wu1-SARS2_orf1a;709 | 0 | 0 |
| NC_045512.2;YP_009725295.1;Wu1-SARS2_orf1a;710 | 0 | 0 |
| NC_045512.2;YP_009725295.1;Wu1-SARS2_orf1a;711 | 0 | 0 |
| NC_045512.2;YP_009725295.1;Wu1-SARS2_orf1a;712 | 0.05 | 0 |
| NC_045512.2;YP_009725295.1;Wu1-SARS2_orf1a;713 | 0 | 0 |
| NC_045512.2;YP_009725295.1;Wu1-SARS2_orf1a;714 | 0 | 0.025 |
| NC_045512.2;YP_009725295.1;Wu1-SARS2_orf1a;715 | 0 | 0 |
| NC_045512.2;YP_009725295.1;Wu1-SARS2_orf1a;716 | 0 | 0 |
| NC_045512.2;YP_009725295.1;Wu1-SARS2_orf1a;717 | 0 | 0 |
| NC_045512.2;YP_009725295.1;Wu1-SARS2_orf1a;718 | 0 | 0.025 |
| NC_045512.2;YP_009725295.1;Wu1-SARS2_orf1a;719 | 0 | 0 |

| | | |
|---|---|---|
| NC_045512.2;YP_009725295.1;Wu1-SARS2_orf1a;720 | 0 | 0 |
| NC_045512.2;YP_009725295.1;Wu1-SARS2_orf1a;721 | 0 | 0.05 |
| NC_045512.2;YP_009725295.1;Wu1-SARS2_orf1a;722 | 0 | 0.025 |
| NC_045512.2;YP_009725295.1;Wu1-SARS2_orf1a;723 | 0 | 0 |
| NC_045512.2;YP_009725295.1;Wu1-SARS2_orf1a;724 | 0 | 0 |
| NC_045512.2;YP_009725295.1;Wu1-SARS2_orf1a;725 | 0 | 0 |
| NC_045512.2;YP_009725295.1;Wu1-SARS2_orf1a;726 | 0.05 | 0 |
| NC_045512.2;YP_009725295.1;Wu1-SARS2_orf1a;727 | 0 | 0 |
| NC_045512.2;YP_009725295.1;Wu1-SARS2_orf1a;728 | 0 | 0 |
| NC_045512.2;YP_009725295.1;Wu1-SARS2_orf1a;729 | 0 | 0 |
| NC_045512.2;YP_009725295.1;Wu1-SARS2_orf1a;730 | 0 | 0 |
| NC_045512.2;YP_009725295.1;Wu1-SARS2_orf1a;731 | 0 | 0 |
| NC_045512.2;YP_009725295.1;Wu1-SARS2_orf1a;732 | 0 | 0 |
| NC_045512.2;YP_009725295.1;Wu1-SARS2_orf1a;733 | 0 | 0 |
| NC_045512.2;YP_009725295.1;Wu1-SARS2_orf1a;734 | 0 | 0 |
| NC_045512.2;YP_009725295.1;Wu1-SARS2_orf1a;735 | 0 | 0 |
| NC_045512.2;YP_009725295.1;Wu1-SARS2_orf1a;736 | 0 | 0 |
| NC_045512.2;YP_009725295.1;Wu1-SARS2_orf1a;737 | 0 | 0 |
| NC_045512.2;YP_009725295.1;Wu1-SARS2_orf1a;738 | 0 | 0 |
| NC_045512.2;YP_009725295.1;Wu1-SARS2_orf1a;739 | 0 | 0 |
| NC_045512.2;YP_009725295.1;Wu1-SARS2_orf1a;740 | 0 | 0 |
| NC_045512.2;YP_009725295.1;Wu1-SARS2_orf1a;741 | 0 | 0 |
| NC_045512.2;YP_009725295.1;Wu1-SARS2_orf1a;742 | 0 | 0 |
| NC_045512.2;YP_009725295.1;Wu1-SARS2_orf1a;743 | 0 | 0 |
| NC_045512.2;YP_009725295.1;Wu1-SARS2_orf1a;744 | 0.15 | 0.05 |
| NC_045512.2;YP_009725295.1;Wu1-SARS2_orf1a;745 | 0.1 | 0.05 |
| NC_045512.2;YP_009725295.1;Wu1-SARS2_orf1a;746 | 0.15 | 0.025 |
| NC_045512.2;YP_009725295.1;Wu1-SARS2_orf1a;747 | 0.1 | 0.025 |
| NC_045512.2;YP_009725295.1;Wu1-SARS2_orf1a;748 | 0.1 | 0.05 |
| NC_045512.2;YP_009725295.1;Wu1-SARS2_orf1a;749 | 0.15 | 0.075 |
| NC_045512.2;YP_009725295.1;Wu1-SARS2_orf1a;750 | 0.15 | 0.05 |
| NC_045512.2;YP_009725295.1;Wu1-SARS2_orf1a;751 | 0.05 | 0.05 |
| NC_045512.2;YP_009725295.1;Wu1-SARS2_orf1a;752 | 0.05 | 0.05 |
| NC_045512.2;YP_009725295.1;Wu1-SARS2_orf1a;753 | 0.05 | 0.075 |
| NC_045512.2;YP_009725295.1;Wu1-SARS2_orf1a;754 | 0.1 | 0.075 |
| NC_045512.2;YP_009725295.1;Wu1-SARS2_orf1a;755 | 0.05 | 0.05 |
| NC_045512.2;YP_009725295.1;Wu1-SARS2_orf1a;756 | 0.05 | 0.05 |
| NC_045512.2;YP_009725295.1;Wu1-SARS2_orf1a;757 | 0.05 | 0.05 |
| NC_045512.2;YP_009725295.1;Wu1-SARS2_orf1a;758 | 0.05 | 0.05 |
| NC_045512.2;YP_009725295.1;Wu1-SARS2_orf1a;759 | 0.05 | 0.075 |
| NC_045512.2;YP_009725295.1;Wu1-SARS2_orf1a;760 | 0 | 0.025 |
| NC_045512.2;YP_009725295.1;Wu1-SARS2_orf1a;761 | 0 | 0.025 |
| NC_045512.2;YP_009725295.1;Wu1-SARS2_orf1a;762 | 0 | 0.025 |

| | | |
|---|---|---|
| NC_045512.2;YP_009725295.1;Wu1-SARS2_orf1a;763 | 0 | 0.025 |
| NC_045512.2;YP_009725295.1;Wu1-SARS2_orf1a;764 | 0.05 | 0.025 |
| NC_045512.2;YP_009725295.1;Wu1-SARS2_orf1a;765 | 0.05 | 0.025 |
| NC_045512.2;YP_009725295.1;Wu1-SARS2_orf1a;766 | 0.05 | 0.075 |
| NC_045512.2;YP_009725295.1;Wu1-SARS2_orf1a;767 | 0.05 | 0.05 |
| NC_045512.2;YP_009725295.1;Wu1-SARS2_orf1a;768 | 0 | 0.075 |
| NC_045512.2;YP_009725295.1;Wu1-SARS2_orf1a;769 | 0 | 0 |
| NC_045512.2;YP_009725295.1;Wu1-SARS2_orf1a;770 | 0 | 0 |
| NC_045512.2;YP_009725295.1;Wu1-SARS2_orf1a;771 | 0 | 0 |
| NC_045512.2;YP_009725295.1;Wu1-SARS2_orf1a;772 | 0 | 0 |
| NC_045512.2;YP_009725295.1;Wu1-SARS2_orf1a;773 | 0 | 0 |
| NC_045512.2;YP_009725295.1;Wu1-SARS2_orf1a;774 | 0 | 0 |
| NC_045512.2;YP_009725295.1;Wu1-SARS2_orf1a;775 | 0.05 | 0 |
| NC_045512.2;YP_009725295.1;Wu1-SARS2_orf1a;776 | 0 | 0 |
| NC_045512.2;YP_009725295.1;Wu1-SARS2_orf1a;777 | 0 | 0 |
| NC_045512.2;YP_009725295.1;Wu1-SARS2_orf1a;778 | 0 | 0 |
| NC_045512.2;YP_009725295.1;Wu1-SARS2_orf1a;779 | 0 | 0 |
| NC_045512.2;YP_009725295.1;Wu1-SARS2_orf1a;780 | 0 | 0.025 |
| NC_045512.2;YP_009725295.1;Wu1-SARS2_orf1a;781 | 0 | 0 |
| NC_045512.2;YP_009725295.1;Wu1-SARS2_orf1a;782 | 0.05 | 0 |
| NC_045512.2;YP_009725295.1;Wu1-SARS2_orf1a;783 | 0 | 0 |
| NC_045512.2;YP_009725295.1;Wu1-SARS2_orf1a;784 | 0 | 0 |
| NC_045512.2;YP_009725295.1;Wu1-SARS2_orf1a;785 | 0 | 0 |
| NC_045512.2;YP_009725295.1;Wu1-SARS2_orf1a;786 | 0 | 0 |
| NC_045512.2;YP_009725295.1;Wu1-SARS2_orf1a;787 | 0 | 0.025 |
| NC_045512.2;YP_009725295.1;Wu1-SARS2_orf1a;788 | 0 | 0.025 |
| NC_045512.2;YP_009725295.1;Wu1-SARS2_orf1a;789 | 0.05 | 0 |
| NC_045512.2;YP_009725295.1;Wu1-SARS2_orf1a;790 | 0 | 0.025 |
| NC_045512.2;YP_009725295.1;Wu1-SARS2_orf1a;791 | 0 | 0.05 |
| NC_045512.2;YP_009725295.1;Wu1-SARS2_orf1a;792 | 0 | 0.05 |
| NC_045512.2;YP_009725295.1;Wu1-SARS2_orf1a;793 | 0 | 0.025 |
| NC_045512.2;YP_009725295.1;Wu1-SARS2_orf1a;794 | 0 | 0 |
| NC_045512.2;YP_009725295.1;Wu1-SARS2_orf1a;795 | 0 | 0 |
| NC_045512.2;YP_009725295.1;Wu1-SARS2_orf1a;796 | 0 | 0 |
| NC_045512.2;YP_009725295.1;Wu1-SARS2_orf1a;797 | 0.05 | 0.05 |
| NC_045512.2;YP_009725295.1;Wu1-SARS2_orf1a;798 | 0.05 | 0 |
| NC_045512.2;YP_009725295.1;Wu1-SARS2_orf1a;799 | 0 | 0 |
| NC_045512.2;YP_009725295.1;Wu1-SARS2_orf1a;800 | 0 | 0 |
| NC_045512.2;YP_009725295.1;Wu1-SARS2_orf1a;801 | 0.05 | 0 |
| NC_045512.2;YP_009725295.1;Wu1-SARS2_orf1a;802 | 0 | 0 |
| NC_045512.2;YP_009725295.1;Wu1-SARS2_orf1a;803 | 0.1 | 0 |
| NC_045512.2;YP_009725295.1;Wu1-SARS2_orf1a;804 | 0.05 | 0 |
| NC_045512.2;YP_009725295.1;Wu1-SARS2_orf1a;805 | 0.05 | 0 |

| | | |
|---|---|---|
| NC_045512.2;YP_009725295.1;Wu1-SARS2_orf1a;806 | 0.05 | 0 |
| NC_045512.2;YP_009725295.1;Wu1-SARS2_orf1a;807 | 0 | 0.025 |
| NC_045512.2;YP_009725295.1;Wu1-SARS2_orf1a;808 | 0 | 0 |
| NC_045512.2;YP_009725295.1;Wu1-SARS2_orf1a;809 | 0 | 0 |
| NC_045512.2;YP_009725295.1;Wu1-SARS2_orf1a;810 | 0 | 0 |
| NC_045512.2;YP_009725295.1;Wu1-SARS2_orf1a;811 | 0 | 0 |
| NC_045512.2;YP_009725295.1;Wu1-SARS2_orf1a;812 | 0 | 0 |
| NC_045512.2;YP_009725295.1;Wu1-SARS2_orf1a;813 | 0.05 | 0 |
| NC_045512.2;YP_009725295.1;Wu1-SARS2_orf1a;814 | 0 | 0.025 |
| NC_045512.2;YP_009725295.1;Wu1-SARS2_orf1a;815 | 0 | 0 |
| NC_045512.2;YP_009725295.1;Wu1-SARS2_orf1a;816 | 0 | 0 |
| NC_045512.2;YP_009725295.1;Wu1-SARS2_orf1a;817 | 0 | 0 |
| NC_045512.2;YP_009725295.1;Wu1-SARS2_orf1a;818 | 0 | 0 |
| NC_045512.2;YP_009725295.1;Wu1-SARS2_orf1a;819 | 0.05 | 0 |
| NC_045512.2;YP_009725295.1;Wu1-SARS2_orf1a;820 | 0.05 | 0 |
| NC_045512.2;YP_009725295.1;Wu1-SARS2_orf1a;821 | 0 | 0 |
| NC_045512.2;YP_009725295.1;Wu1-SARS2_orf1a;822 | 0 | 0.025 |
| NC_045512.2;YP_009725295.1;Wu1-SARS2_orf1a;823 | 0 | 0.025 |
| NC_045512.2;YP_009725295.1;Wu1-SARS2_orf1a;824 | 0 | 0.025 |
| NC_045512.2;YP_009725295.1;Wu1-SARS2_orf1a;825 | 0 | 0.025 |
| NC_045512.2;YP_009725295.1;Wu1-SARS2_orf1a;826 | 0 | 0 |
| NC_045512.2;YP_009725295.1;Wu1-SARS2_orf1a;827 | 0.1 | 0 |
| NC_045512.2;YP_009725295.1;Wu1-SARS2_orf1a;828 | 0 | 0 |
| NC_045512.2;YP_009725295.1;Wu1-SARS2_orf1a;829 | 0 | 0 |
| NC_045512.2;YP_009725295.1;Wu1-SARS2_orf1a;830 | 0 | 0 |
| NC_045512.2;YP_009725295.1;Wu1-SARS2_orf1a;831 | 0.05 | 0.05 |
| NC_045512.2;YP_009725295.1;Wu1-SARS2_orf1a;832 | 0 | 0 |
| NC_045512.2;YP_009725295.1;Wu1-SARS2_orf1a;833 | 0 | 0 |
| NC_045512.2;YP_009725295.1;Wu1-SARS2_orf1a;834 | 0 | 0 |
| NC_045512.2;YP_009725295.1;Wu1-SARS2_orf1a;835 | 0 | 0 |
| NC_045512.2;YP_009725295.1;Wu1-SARS2_orf1a;836 | 0.05 | 0 |
| NC_045512.2;YP_009725295.1;Wu1-SARS2_orf1a;837 | 0.05 | 0 |
| NC_045512.2;YP_009725295.1;Wu1-SARS2_orf1a;838 | 0 | 0 |
| NC_045512.2;YP_009725295.1;Wu1-SARS2_orf1a;839 | 0 | 0.025 |
| NC_045512.2;YP_009725295.1;Wu1-SARS2_orf1a;840 | 0.2 | 0.1 |
| NC_045512.2;YP_009725295.1;Wu1-SARS2_orf1a;841 | 0.1 | 0.15 |
| NC_045512.2;YP_009725295.1;Wu1-SARS2_orf1a;842 | 0.2 | 0.1 |
| NC_045512.2;YP_009725295.1;Wu1-SARS2_orf1a;843 | 0.25 | 0.1 |
| NC_045512.2;YP_009725295.1;Wu1-SARS2_orf1a;844 | 0.05 | 0.05 |
| NC_045512.2;YP_009725295.1;Wu1-SARS2_orf1a;845 | 0 | 0.05 |
| NC_045512.2;YP_009725295.1;Wu1-SARS2_orf1a;846 | 0 | 0.05 |
| NC_045512.2;YP_009725295.1;Wu1-SARS2_orf1a;847 | 0 | 0.025 |
| NC_045512.2;YP_009725295.1;Wu1-SARS2_orf1a;848 | 0 | 0 |

| | | |
|---|---|---|
| NC_045512.2;YP_009725295.1;Wu1-SARS2_orf1a;849 | 0 | 0 |
| NC_045512.2;YP_009725295.1;Wu1-SARS2_orf1a;850 | 0 | 0 |
| NC_045512.2;YP_009725295.1;Wu1-SARS2_orf1a;851 | 0 | 0 |
| NC_045512.2;YP_009725295.1;Wu1-SARS2_orf1a;852 | 0 | 0 |
| NC_045512.2;YP_009725295.1;Wu1-SARS2_orf1a;853 | 0 | 0 |
| NC_045512.2;YP_009725295.1;Wu1-SARS2_orf1a;854 | 0 | 0 |
| NC_045512.2;YP_009725295.1;Wu1-SARS2_orf1a;855 | 0 | 0 |
| NC_045512.2;YP_009725295.1;Wu1-SARS2_orf1a;856 | 0 | 0 |
| NC_045512.2;YP_009725295.1;Wu1-SARS2_orf1a;857 | 0 | 0 |
| NC_045512.2;YP_009725295.1;Wu1-SARS2_orf1a;858 | 0 | 0 |
| NC_045512.2;YP_009725295.1;Wu1-SARS2_orf1a;859 | 0 | 0.025 |
| NC_045512.2;YP_009725295.1;Wu1-SARS2_orf1a;860 | 0 | 0 |
| NC_045512.2;YP_009725295.1;Wu1-SARS2_orf1a;861 | 0 | 0 |
| NC_045512.2;YP_009725295.1;Wu1-SARS2_orf1a;862 | 0 | 0 |
| NC_045512.2;YP_009725295.1;Wu1-SARS2_orf1a;863 | 0 | 0 |
| NC_045512.2;YP_009725295.1;Wu1-SARS2_orf1a;864 | 0 | 0 |
| NC_045512.2;YP_009725295.1;Wu1-SARS2_orf1a;865 | 0 | 0 |
| NC_045512.2;YP_009725295.1;Wu1-SARS2_orf1a;866 | 0 | 0 |
| NC_045512.2;YP_009725295.1;Wu1-SARS2_orf1a;867 | 0 | 0 |
| NC_045512.2;YP_009725295.1;Wu1-SARS2_orf1a;868 | 0 | 0 |
| NC_045512.2;YP_009725295.1;Wu1-SARS2_orf1a;869 | 0 | 0 |
| NC_045512.2;YP_009725295.1;Wu1-SARS2_orf1a;870 | 0 | 0 |
| NC_045512.2;YP_009725295.1;Wu1-SARS2_orf1a;871 | 0 | 0 |
| NC_045512.2;YP_009725295.1;Wu1-SARS2_orf1a;872 | 0 | 0 |
| NC_045512.2;YP_009725295.1;Wu1-SARS2_orf1a;873 | 0 | 0 |
| NC_045512.2;YP_009725295.1;Wu1-SARS2_orf1a;874 | 0 | 0 |
| NC_045512.2;YP_009725295.1;Wu1-SARS2_orf1a;875 | 0 | 0.025 |
| NC_045512.2;YP_009725295.1;Wu1-SARS2_orf1a;876 | 0.05 | 0 |
| NC_045512.2;YP_009725295.1;Wu1-SARS2_orf1a;877 | 0 | 0.025 |
| NC_045512.2;YP_009725295.1;Wu1-SARS2_orf1a;878 | 0 | 0.05 |
| NC_045512.2;YP_009725295.1;Wu1-SARS2_orf1a;879 | 0 | 0.025 |
| NC_045512.2;YP_009725295.1;Wu1-SARS2_orf1a;880 | 0 | 0.025 |
| NC_045512.2;YP_009725295.1;Wu1-SARS2_orf1a;881 | 0.05 | 0.025 |
| NC_045512.2;YP_009725295.1;Wu1-SARS2_orf1a;882 | 0 | 0.025 |
| NC_045512.2;YP_009725295.1;Wu1-SARS2_orf1a;883 | 0 | 0 |
| NC_045512.2;YP_009725295.1;Wu1-SARS2_orf1a;884 | 0.05 | 0 |
| NC_045512.2;YP_009725295.1;Wu1-SARS2_orf1a;885 | 0 | 0 |
| NC_045512.2;YP_009725295.1;Wu1-SARS2_orf1a;886 | 0 | 0 |
| NC_045512.2;YP_009725295.1;Wu1-SARS2_orf1a;887 | 0 | 0 |
| NC_045512.2;YP_009725295.1;Wu1-SARS2_orf1a;888 | 0 | 0 |
| NC_045512.2;YP_009725295.1;Wu1-SARS2_orf1a;889 | 0 | 0 |
| NC_045512.2;YP_009725295.1;Wu1-SARS2_orf1a;890 | 0 | 0 |
| NC_045512.2;YP_009725295.1;Wu1-SARS2_orf1a;891 | 0 | 0 |

| | | |
|---|---|---|
| NC_045512.2;YP_009725295.1;Wu1-SARS2_orf1a;892 | 0 | 0 |
| NC_045512.2;YP_009725295.1;Wu1-SARS2_orf1a;893 | 0 | 0 |
| NC_045512.2;YP_009725295.1;Wu1-SARS2_orf1a;894 | 0 | 0 |
| NC_045512.2;YP_009725295.1;Wu1-SARS2_orf1a;895 | 0 | 0 |
| NC_045512.2;YP_009725295.1;Wu1-SARS2_orf1a;896 | 0 | 0 |
| NC_045512.2;YP_009725295.1;Wu1-SARS2_orf1a;897 | 0 | 0 |
| NC_045512.2;YP_009725295.1;Wu1-SARS2_orf1a;898 | 0 | 0 |
| NC_045512.2;YP_009725295.1;Wu1-SARS2_orf1a;899 | 0 | 0 |
| NC_045512.2;YP_009725295.1;Wu1-SARS2_orf1a;900 | 0.1 | 0 |
| NC_045512.2;YP_009725295.1;Wu1-SARS2_orf1a;901 | 0 | 0 |
| NC_045512.2;YP_009725295.1;Wu1-SARS2_orf1a;902 | 0 | 0 |
| NC_045512.2;YP_009725295.1;Wu1-SARS2_orf1a;903 | 0 | 0 |
| NC_045512.2;YP_009725295.1;Wu1-SARS2_orf1a;904 | 0 | 0 |
| NC_045512.2;YP_009725295.1;Wu1-SARS2_orf1a;905 | 0 | 0 |
| NC_045512.2;YP_009725295.1;Wu1-SARS2_orf1a;906 | 0.05 | 0 |
| NC_045512.2;YP_009725295.1;Wu1-SARS2_orf1a;907 | 0 | 0 |
| NC_045512.2;YP_009725295.1;Wu1-SARS2_orf1a;908 | 0 | 0 |
| NC_045512.2;YP_009725295.1;Wu1-SARS2_orf1a;909 | 0 | 0 |
| NC_045512.2;YP_009725295.1;Wu1-SARS2_orf1a;910 | 0 | 0 |
| NC_045512.2;YP_009725295.1;Wu1-SARS2_orf1a;911 | 0 | 0 |
| NC_045512.2;YP_009725295.1;Wu1-SARS2_orf1a;912 | 0 | 0 |
| NC_045512.2;YP_009725295.1;Wu1-SARS2_orf1a;913 | 0 | 0 |
| NC_045512.2;YP_009725295.1;Wu1-SARS2_orf1a;914 | 0 | 0 |
| NC_045512.2;YP_009725295.1;Wu1-SARS2_orf1a;915 | 0 | 0 |
| NC_045512.2;YP_009725295.1;Wu1-SARS2_orf1a;916 | 0 | 0.025 |
| NC_045512.2;YP_009725295.1;Wu1-SARS2_orf1a;917 | 0 | 0 |
| NC_045512.2;YP_009725295.1;Wu1-SARS2_orf1a;918 | 0 | 0.025 |
| NC_045512.2;YP_009725295.1;Wu1-SARS2_orf1a;919 | 0 | 0.05 |
| NC_045512.2;YP_009725295.1;Wu1-SARS2_orf1a;920 | 0 | 0.075 |
| NC_045512.2;YP_009725295.1;Wu1-SARS2_orf1a;921 | 0 | 0.1 |
| NC_045512.2;YP_009725295.1;Wu1-SARS2_orf1a;922 | 0 | 0.1 |
| NC_045512.2;YP_009725295.1;Wu1-SARS2_orf1a;923 | 0 | 0.1 |
| NC_045512.2;YP_009725295.1;Wu1-SARS2_orf1a;924 | 0 | 0.125 |
| NC_045512.2;YP_009725295.1;Wu1-SARS2_orf1a;925 | 0 | 0.1 |
| NC_045512.2;YP_009725295.1;Wu1-SARS2_orf1a;926 | 0 | 0.025 |
| NC_045512.2;YP_009725295.1;Wu1-SARS2_orf1a;927 | 0.05 | 0 |
| NC_045512.2;YP_009725295.1;Wu1-SARS2_orf1a;928 | 0.05 | 0.025 |
| NC_045512.2;YP_009725295.1;Wu1-SARS2_orf1a;929 | 0.35 | 0.25 |
| NC_045512.2;YP_009725295.1;Wu1-SARS2_orf1a;930 | 0 | 0 |
| NC_045512.2;YP_009725295.1;Wu1-SARS2_orf1a;931 | 0 | 0 |
| NC_045512.2;YP_009725295.1;Wu1-SARS2_orf1a;932 | 0 | 0 |
| NC_045512.2;YP_009725295.1;Wu1-SARS2_orf1a;933 | 0.05 | 0 |
| NC_045512.2;YP_009725295.1;Wu1-SARS2_orf1a;934 | 0 | 0 |

| | | |
|---|---|---|
| NC_045512.2;YP_009725295.1;Wu1-SARS2_orf1a;935 | 0 | 0 |
| NC_045512.2;YP_009725295.1;Wu1-SARS2_orf1a;936 | 0 | 0 |
| NC_045512.2;YP_009725295.1;Wu1-SARS2_orf1a;937 | 0 | 0 |
| NC_045512.2;YP_009725295.1;Wu1-SARS2_orf1a;938 | 0 | 0 |
| NC_045512.2;YP_009725295.1;Wu1-SARS2_orf1a;939 | 0 | 0 |
| NC_045512.2;YP_009725295.1;Wu1-SARS2_orf1a;940 | 0.05 | 0.05 |
| NC_045512.2;YP_009725295.1;Wu1-SARS2_orf1a;941 | 0 | 0.025 |
| NC_045512.2;YP_009725295.1;Wu1-SARS2_orf1a;942 | 0 | 0 |
| NC_045512.2;YP_009725295.1;Wu1-SARS2_orf1a;943 | 0 | 0 |
| NC_045512.2;YP_009725295.1;Wu1-SARS2_orf1a;944 | 0 | 0 |
| NC_045512.2;YP_009725295.1;Wu1-SARS2_orf1a;945 | 0 | 0 |
| NC_045512.2;YP_009725295.1;Wu1-SARS2_orf1a;946 | 0 | 0.025 |
| NC_045512.2;YP_009725295.1;Wu1-SARS2_orf1a;947 | 0 | 0.025 |
| NC_045512.2;YP_009725295.1;Wu1-SARS2_orf1a;948 | 0 | 0 |
| NC_045512.2;YP_009725295.1;Wu1-SARS2_orf1a;949 | 0 | 0.05 |
| NC_045512.2;YP_009725295.1;Wu1-SARS2_orf1a;950 | 0 | 0.075 |
| NC_045512.2;YP_009725295.1;Wu1-SARS2_orf1a;951 | 0 | 0 |
| NC_045512.2;YP_009725295.1;Wu1-SARS2_orf1a;952 | 0.1 | 0 |
| NC_045512.2;YP_009725295.1;Wu1-SARS2_orf1a;953 | 0.15 | 0.025 |
| NC_045512.2;YP_009725295.1;Wu1-SARS2_orf1a;954 | 0 | 0 |
| NC_045512.2;YP_009725295.1;Wu1-SARS2_orf1a;955 | 0 | 0 |
| NC_045512.2;YP_009725295.1;Wu1-SARS2_orf1a;956 | 0 | 0 |
| NC_045512.2;YP_009725295.1;Wu1-SARS2_orf1a;957 | 0 | 0 |
| NC_045512.2;YP_009725295.1;Wu1-SARS2_orf1a;958 | 0 | 0 |
| NC_045512.2;YP_009725295.1;Wu1-SARS2_orf1a;959 | 0 | 0 |
| NC_045512.2;YP_009725295.1;Wu1-SARS2_orf1a;960 | 0 | 0 |
| NC_045512.2;YP_009725295.1;Wu1-SARS2_orf1a;961 | 0 | 0 |
| NC_045512.2;YP_009725295.1;Wu1-SARS2_orf1a;962 | 0 | 0 |
| NC_045512.2;YP_009725295.1;Wu1-SARS2_orf1a;963 | 0 | 0 |
| NC_045512.2;YP_009725295.1;Wu1-SARS2_orf1a;964 | 0 | 0.025 |
| NC_045512.2;YP_009725295.1;Wu1-SARS2_orf1a;965 | 0 | 0.025 |
| NC_045512.2;YP_009725295.1;Wu1-SARS2_orf1a;966 | 0 | 0.025 |
| NC_045512.2;YP_009725295.1;Wu1-SARS2_orf1a;967 | 0 | 0.025 |
| NC_045512.2;YP_009725295.1;Wu1-SARS2_orf1a;968 | 0 | 0.025 |
| NC_045512.2;YP_009725295.1;Wu1-SARS2_orf1a;969 | 0 | 0.025 |
| NC_045512.2;YP_009725295.1;Wu1-SARS2_orf1a;970 | 0 | 0.025 |
| NC_045512.2;YP_009725295.1;Wu1-SARS2_orf1a;971 | 0 | 0.025 |
| NC_045512.2;YP_009725295.1;Wu1-SARS2_orf1a;972 | 0 | 0.05 |
| NC_045512.2;YP_009725295.1;Wu1-SARS2_orf1a;973 | 0 | 0.025 |
| NC_045512.2;YP_009725295.1;Wu1-SARS2_orf1a;974 | 0 | 0.025 |
| NC_045512.2;YP_009725295.1;Wu1-SARS2_orf1a;975 | 0 | 0.025 |
| NC_045512.2;YP_009725295.1;Wu1-SARS2_orf1a;976 | 0 | 0.075 |
| NC_045512.2;YP_009725295.1;Wu1-SARS2_orf1a;977 | 0.1 | 0.025 |

| | | |
|---|---|---|
| NC_045512.2;YP_009725295.1;Wu1-SARS2_orf1a;978 | 0 | 0.025 |
| NC_045512.2;YP_009725295.1;Wu1-SARS2_orf1a;979 | 0 | 0.025 |
| NC_045512.2;YP_009725295.1;Wu1-SARS2_orf1a;980 | 0 | 0 |
| NC_045512.2;YP_009725295.1;Wu1-SARS2_orf1a;981 | 0 | 0 |
| NC_045512.2;YP_009725295.1;Wu1-SARS2_orf1a;982 | 0 | 0 |
| NC_045512.2;YP_009725295.1;Wu1-SARS2_orf1a;983 | 0 | 0 |
| NC_045512.2;YP_009725295.1;Wu1-SARS2_orf1a;984 | 0 | 0.025 |
| NC_045512.2;YP_009725295.1;Wu1-SARS2_orf1a;985 | 0 | 0 |
| NC_045512.2;YP_009725295.1;Wu1-SARS2_orf1a;986 | 0 | 0 |
| NC_045512.2;YP_009725295.1;Wu1-SARS2_orf1a;987 | 0 | 0 |
| NC_045512.2;YP_009725295.1;Wu1-SARS2_orf1a;988 | 0 | 0 |
| NC_045512.2;YP_009725295.1;Wu1-SARS2_orf1a;989 | 0 | 0 |
| NC_045512.2;YP_009725295.1;Wu1-SARS2_orf1a;990 | 0 | 0 |
| NC_045512.2;YP_009725295.1;Wu1-SARS2_orf1a;991 | 0 | 0 |
| NC_045512.2;YP_009725295.1;Wu1-SARS2_orf1a;992 | 0 | 0 |
| NC_045512.2;YP_009725295.1;Wu1-SARS2_orf1a;993 | 0 | 0 |
| NC_045512.2;YP_009725295.1;Wu1-SARS2_orf1a;994 | 0 | 0 |
| NC_045512.2;YP_009725295.1;Wu1-SARS2_orf1a;995 | 0 | 0 |
| NC_045512.2;YP_009725295.1;Wu1-SARS2_orf1a;996 | 0 | 0.025 |
| NC_045512.2;YP_009725295.1;Wu1-SARS2_orf1a;997 | 0 | 0 |
| NC_045512.2;YP_009725295.1;Wu1-SARS2_orf1a;998 | 0 | 0 |
| NC_045512.2;YP_009725295.1;Wu1-SARS2_orf1a;999 | 0 | 0 |
| NC_045512.2;YP_009725295.1;Wu1-SARS2_orf1a;1000 | 0 | 0 |
| NC_045512.2;YP_009725295.1;Wu1-SARS2_orf1a;1001 | 0 | 0 |
| NC_045512.2;YP_009725295.1;Wu1-SARS2_orf1a;1002 | 0 | 0 |
| NC_045512.2;YP_009725295.1;Wu1-SARS2_orf1a;1003 | 0 | 0 |
| NC_045512.2;YP_009725295.1;Wu1-SARS2_orf1a;1004 | 0 | 0 |
| NC_045512.2;YP_009725295.1;Wu1-SARS2_orf1a;1005 | 0 | 0 |
| NC_045512.2;YP_009725295.1;Wu1-SARS2_orf1a;1006 | 0 | 0 |
| NC_045512.2;YP_009725295.1;Wu1-SARS2_orf1a;1007 | 0 | 0 |
| NC_045512.2;YP_009725295.1;Wu1-SARS2_orf1a;1008 | 0 | 0 |
| NC_045512.2;YP_009725295.1;Wu1-SARS2_orf1a;1009 | 0 | 0 |
| NC_045512.2;YP_009725295.1;Wu1-SARS2_orf1a;1010 | 0 | 0 |
| NC_045512.2;YP_009725295.1;Wu1-SARS2_orf1a;1011 | 0 | 0 |
| NC_045512.2;YP_009725295.1;Wu1-SARS2_orf1a;1012 | 0 | 0 |
| NC_045512.2;YP_009725295.1;Wu1-SARS2_orf1a;1013 | 0 | 0 |
| NC_045512.2;YP_009725295.1;Wu1-SARS2_orf1a;1014 | 0 | 0.05 |
| NC_045512.2;YP_009725295.1;Wu1-SARS2_orf1a;1015 | 0.1 | 0.05 |
| NC_045512.2;YP_009725295.1;Wu1-SARS2_orf1a;1016 | 0 | 0.05 |
| NC_045512.2;YP_009725295.1;Wu1-SARS2_orf1a;1017 | 0 | 0.025 |
| NC_045512.2;YP_009725295.1;Wu1-SARS2_orf1a;1018 | 0 | 0 |
| NC_045512.2;YP_009725295.1;Wu1-SARS2_orf1a;1019 | 0 | 0 |
| NC_045512.2;YP_009725295.1;Wu1-SARS2_orf1a;1020 | 0 | 0 |

| | | |
|---|---|---|
| NC_045512.2;YP_009725295.1;Wu1-SARS2_orf1a;1021 | 0 | 0 |
| NC_045512.2;YP_009725295.1;Wu1-SARS2_orf1a;1022 | 0 | 0 |
| NC_045512.2;YP_009725295.1;Wu1-SARS2_orf1a;1023 | 0.05 | 0 |
| NC_045512.2;YP_009725295.1;Wu1-SARS2_orf1a;1024 | 0.05 | 0 |
| NC_045512.2;YP_009725295.1;Wu1-SARS2_orf1a;1025 | 0.05 | 0 |
| NC_045512.2;YP_009725295.1;Wu1-SARS2_orf1a;1026 | 0.05 | 0 |
| NC_045512.2;YP_009725295.1;Wu1-SARS2_orf1a;1027 | 0.05 | 0 |
| NC_045512.2;YP_009725295.1;Wu1-SARS2_orf1a;1028 | 0.05 | 0 |
| NC_045512.2;YP_009725295.1;Wu1-SARS2_orf1a;1029 | 0.05 | 0 |
| NC_045512.2;YP_009725295.1;Wu1-SARS2_orf1a;1030 | 0.05 | 0.025 |
| NC_045512.2;YP_009725295.1;Wu1-SARS2_orf1a;1031 | 0.05 | 0 |
| NC_045512.2;YP_009725295.1;Wu1-SARS2_orf1a;1032 | 0.15 | 0.025 |
| NC_045512.2;YP_009725295.1;Wu1-SARS2_orf1a;1033 | 0.2 | 0.025 |
| NC_045512.2;YP_009725295.1;Wu1-SARS2_orf1a;1034 | 0.05 | 0.025 |
| NC_045512.2;YP_009725295.1;Wu1-SARS2_orf1a;1035 | 0 | 0.05 |
| NC_045512.2;YP_009725295.1;Wu1-SARS2_orf1a;1036 | 0 | 0.025 |
| NC_045512.2;YP_009725295.1;Wu1-SARS2_orf1a;1037 | 0 | 0 |
| NC_045512.2;YP_009725295.1;Wu1-SARS2_orf1a;1038 | 0 | 0 |
| NC_045512.2;YP_009725295.1;Wu1-SARS2_orf1a;1039 | 0 | 0 |
| NC_045512.2;YP_009725295.1;Wu1-SARS2_orf1a;1040 | 0 | 0 |
| NC_045512.2;YP_009725295.1;Wu1-SARS2_orf1a;1041 | 0 | 0 |
| NC_045512.2;YP_009725295.1;Wu1-SARS2_orf1a;1042 | 0 | 0 |
| NC_045512.2;YP_009725295.1;Wu1-SARS2_orf1a;1043 | 0 | 0.025 |
| NC_045512.2;YP_009725295.1;Wu1-SARS2_orf1a;1044 | 0 | 0 |
| NC_045512.2;YP_009725295.1;Wu1-SARS2_orf1a;1045 | 0 | 0 |
| NC_045512.2;YP_009725295.1;Wu1-SARS2_orf1a;1046 | 0 | 0.025 |
| NC_045512.2;YP_009725295.1;Wu1-SARS2_orf1a;1047 | 0 | 0.025 |
| NC_045512.2;YP_009725295.1;Wu1-SARS2_orf1a;1048 | 0.05 | 0 |
| NC_045512.2;YP_009725295.1;Wu1-SARS2_orf1a;1049 | 0 | 0 |
| NC_045512.2;YP_009725295.1;Wu1-SARS2_orf1a;1050 | 0 | 0 |
| NC_045512.2;YP_009725295.1;Wu1-SARS2_orf1a;1051 | 0 | 0 |
| NC_045512.2;YP_009725295.1;Wu1-SARS2_orf1a;1052 | 0 | 0 |
| NC_045512.2;YP_009725295.1;Wu1-SARS2_orf1a;1053 | 0 | 0 |
| NC_045512.2;YP_009725295.1;Wu1-SARS2_orf1a;1054 | 0 | 0 |
| NC_045512.2;YP_009725295.1;Wu1-SARS2_orf1a;1055 | 0.05 | 0 |
| NC_045512.2;YP_009725295.1;Wu1-SARS2_orf1a;1056 | 0 | 0 |
| NC_045512.2;YP_009725295.1;Wu1-SARS2_orf1a;1057 | 0.05 | 0.05 |
| NC_045512.2;YP_009725295.1;Wu1-SARS2_orf1a;1058 | 0 | 0 |
| NC_045512.2;YP_009725295.1;Wu1-SARS2_orf1a;1059 | 0 | 0 |
| NC_045512.2;YP_009725295.1;Wu1-SARS2_orf1a;1060 | 0 | 0 |
| NC_045512.2;YP_009725295.1;Wu1-SARS2_orf1a;1061 | 0 | 0 |
| NC_045512.2;YP_009725295.1;Wu1-SARS2_orf1a;1062 | 0 | 0 |
| NC_045512.2;YP_009725295.1;Wu1-SARS2_orf1a;1063 | 0.05 | 0 |

| | | |
|---|---|---|
| NC_045512.2;YP_009725295.1;Wu1-SARS2_orf1a;1064 | 0 | 0 |
| NC_045512.2;YP_009725295.1;Wu1-SARS2_orf1a;1065 | 0 | 0 |
| NC_045512.2;YP_009725295.1;Wu1-SARS2_orf1a;1066 | 0 | 0.025 |
| NC_045512.2;YP_009725295.1;Wu1-SARS2_orf1a;1067 | 0.1 | 0.025 |
| NC_045512.2;YP_009725295.1;Wu1-SARS2_orf1a;1068 | 0.15 | 0.025 |
| NC_045512.2;YP_009725295.1;Wu1-SARS2_orf1a;1069 | 0.05 | 0.025 |
| NC_045512.2;YP_009725295.1;Wu1-SARS2_orf1a;1070 | 0.05 | 0.025 |
| NC_045512.2;YP_009725295.1;Wu1-SARS2_orf1a;1071 | 0.15 | 0.2 |
| NC_045512.2;YP_009725295.1;Wu1-SARS2_orf1a;1072 | 0.2 | 0.2 |
| NC_045512.2;YP_009725295.1;Wu1-SARS2_orf1a;1073 | 0.2 | 0.2 |
| NC_045512.2;YP_009725295.1;Wu1-SARS2_orf1a;1074 | 0.05 | 0.15 |
| NC_045512.2;YP_009725295.1;Wu1-SARS2_orf1a;1075 | 0.05 | 0.15 |
| NC_045512.2;YP_009725295.1;Wu1-SARS2_orf1a;1076 | 0 | 0.025 |
| NC_045512.2;YP_009725295.1;Wu1-SARS2_orf1a;1077 | 0 | 0.025 |
| NC_045512.2;YP_009725295.1;Wu1-SARS2_orf1a;1078 | 0.05 | 0.025 |
| NC_045512.2;YP_009725295.1;Wu1-SARS2_orf1a;1079 | 0.05 | 0 |
| NC_045512.2;YP_009725295.1;Wu1-SARS2_orf1a;1080 | 0 | 0 |
| NC_045512.2;YP_009725295.1;Wu1-SARS2_orf1a;1081 | 0 | 0 |
| NC_045512.2;YP_009725295.1;Wu1-SARS2_orf1a;1082 | 0 | 0.025 |
| NC_045512.2;YP_009725295.1;Wu1-SARS2_orf1a;1083 | 0 | 0 |
| NC_045512.2;YP_009725295.1;Wu1-SARS2_orf1a;1084 | 0 | 0 |
| NC_045512.2;YP_009725295.1;Wu1-SARS2_orf1a;1085 | 0 | 0.025 |
| NC_045512.2;YP_009725295.1;Wu1-SARS2_orf1a;1086 | 0 | 0.05 |
| NC_045512.2;YP_009725295.1;Wu1-SARS2_orf1a;1087 | 0.15 | 0.05 |
| NC_045512.2;YP_009725295.1;Wu1-SARS2_orf1a;1088 | 0.05 | 0.05 |
| NC_045512.2;YP_009725295.1;Wu1-SARS2_orf1a;1089 | 0 | 0.025 |
| NC_045512.2;YP_009725295.1;Wu1-SARS2_orf1a;1090 | 0 | 0 |
| NC_045512.2;YP_009725295.1;Wu1-SARS2_orf1a;1091 | 0 | 0 |
| NC_045512.2;YP_009725295.1;Wu1-SARS2_orf1a;1092 | 0 | 0 |
| NC_045512.2;YP_009725295.1;Wu1-SARS2_orf1a;1093 | 0 | 0 |
| NC_045512.2;YP_009725295.1;Wu1-SARS2_orf1a;1094 | 0 | 0 |
| NC_045512.2;YP_009725295.1;Wu1-SARS2_orf1a;1095 | 0 | 0 |
| NC_045512.2;YP_009725295.1;Wu1-SARS2_orf1a;1096 | 0 | 0 |
| NC_045512.2;YP_009725295.1;Wu1-SARS2_orf1a;1097 | 0 | 0 |
| NC_045512.2;YP_009725295.1;Wu1-SARS2_orf1a;1098 | 0 | 0 |
| NC_045512.2;YP_009725295.1;Wu1-SARS2_orf1a;1099 | 0 | 0 |
| NC_045512.2;YP_009725295.1;Wu1-SARS2_orf1a;1100 | 0 | 0 |
| NC_045512.2;YP_009725295.1;Wu1-SARS2_orf1a;1101 | 0 | 0 |
| NC_045512.2;YP_009725295.1;Wu1-SARS2_orf1a;1102 | 0 | 0 |
| NC_045512.2;YP_009725295.1;Wu1-SARS2_orf1a;1103 | 0 | 0 |
| NC_045512.2;YP_009725295.1;Wu1-SARS2_orf1a;1104 | 0 | 0 |
| NC_045512.2;YP_009725295.1;Wu1-SARS2_orf1a;1105 | 0 | 0 |
| NC_045512.2;YP_009725295.1;Wu1-SARS2_orf1a;1106 | 0 | 0 |

| | | |
|---|---|---|
| NC_045512.2;YP_009725295.1;Wu1-SARS2_orf1a;1107 | 0 | 0 |
| NC_045512.2;YP_009725295.1;Wu1-SARS2_orf1a;1108 | 0 | 0 |
| NC_045512.2;YP_009725295.1;Wu1-SARS2_orf1a;1109 | 0 | 0 |
| NC_045512.2;YP_009725295.1;Wu1-SARS2_orf1a;1110 | 0 | 0 |
| NC_045512.2;YP_009725295.1;Wu1-SARS2_orf1a;1111 | 0.05 | 0 |
| NC_045512.2;YP_009725295.1;Wu1-SARS2_orf1a;1112 | 0.05 | 0 |
| NC_045512.2;YP_009725295.1;Wu1-SARS2_orf1a;1113 | 0.05 | 0.075 |
| NC_045512.2;YP_009725295.1;Wu1-SARS2_orf1a;1114 | 0.05 | 0.125 |
| NC_045512.2;YP_009725295.1;Wu1-SARS2_orf1a;1115 | 0.15 | 0.075 |
| NC_045512.2;YP_009725295.1;Wu1-SARS2_orf1a;1116 | 0 | 0.075 |
| NC_045512.2;YP_009725295.1;Wu1-SARS2_orf1a;1117 | 0.2 | 0.1 |
| NC_045512.2;YP_009725295.1;Wu1-SARS2_orf1a;1118 | 0.2 | 0.125 |
| NC_045512.2;YP_009725295.1;Wu1-SARS2_orf1a;1119 | 0.05 | 0.075 |
| NC_045512.2;YP_009725295.1;Wu1-SARS2_orf1a;1120 | 0.05 | 0.05 |
| NC_045512.2;YP_009725295.1;Wu1-SARS2_orf1a;1121 | 0.05 | 0.025 |
| NC_045512.2;YP_009725295.1;Wu1-SARS2_orf1a;1122 | 0.05 | 0.05 |
| NC_045512.2;YP_009725295.1;Wu1-SARS2_orf1a;1123 | 0 | 0.025 |
| NC_045512.2;YP_009725295.1;Wu1-SARS2_orf1a;1124 | 0 | 0.025 |
| NC_045512.2;YP_009725295.1;Wu1-SARS2_orf1a;1125 | 0.05 | 0.025 |
| NC_045512.2;YP_009725295.1;Wu1-SARS2_orf1a;1126 | 0.05 | 0.025 |
| NC_045512.2;YP_009725295.1;Wu1-SARS2_orf1a;1127 | 0 | 0 |
| NC_045512.2;YP_009725295.1;Wu1-SARS2_orf1a;1128 | 0 | 0 |
| NC_045512.2;YP_009725295.1;Wu1-SARS2_orf1a;1129 | 0 | 0 |
| NC_045512.2;YP_009725295.1;Wu1-SARS2_orf1a;1130 | 0 | 0 |
| NC_045512.2;YP_009725295.1;Wu1-SARS2_orf1a;1131 | 0 | 0.025 |
| NC_045512.2;YP_009725295.1;Wu1-SARS2_orf1a;1132 | 0 | 0.05 |
| NC_045512.2;YP_009725295.1;Wu1-SARS2_orf1a;1133 | 0 | 0 |
| NC_045512.2;YP_009725295.1;Wu1-SARS2_orf1a;1134 | 0 | 0 |
| NC_045512.2;YP_009725295.1;Wu1-SARS2_orf1a;1135 | 0 | 0 |
| NC_045512.2;YP_009725295.1;Wu1-SARS2_orf1a;1136 | 0 | 0 |
| NC_045512.2;YP_009725295.1;Wu1-SARS2_orf1a;1137 | 0 | 0 |
| NC_045512.2;YP_009725295.1;Wu1-SARS2_orf1a;1138 | 0 | 0 |
| NC_045512.2;YP_009725295.1;Wu1-SARS2_orf1a;1139 | 0 | 0 |
| NC_045512.2;YP_009725295.1;Wu1-SARS2_orf1a;1140 | 0 | 0 |
| NC_045512.2;YP_009725295.1;Wu1-SARS2_orf1a;1141 | 0 | 0 |
| NC_045512.2;YP_009725295.1;Wu1-SARS2_orf1a;1142 | 0 | 0 |
| NC_045512.2;YP_009725295.1;Wu1-SARS2_orf1a;1143 | 0 | 0 |
| NC_045512.2;YP_009725295.1;Wu1-SARS2_orf1a;1144 | 0.05 | 0.025 |
| NC_045512.2;YP_009725295.1;Wu1-SARS2_orf1a;1145 | 0 | 0 |
| NC_045512.2;YP_009725295.1;Wu1-SARS2_orf1a;1146 | 0 | 0 |
| NC_045512.2;YP_009725295.1;Wu1-SARS2_orf1a;1147 | 0 | 0.025 |
| NC_045512.2;YP_009725295.1;Wu1-SARS2_orf1a;1148 | 0 | 0 |
| NC_045512.2;YP_009725295.1;Wu1-SARS2_orf1a;1149 | 0 | 0.025 |

| | | |
|---|---|---|
| NC_045512.2;YP_009725295.1;Wu1-SARS2_orf1a;1150 | 0 | 0 |
| NC_045512.2;YP_009725295.1;Wu1-SARS2_orf1a;1151 | 0 | 0 |
| NC_045512.2;YP_009725295.1;Wu1-SARS2_orf1a;1152 | 0 | 0 |
| NC_045512.2;YP_009725295.1;Wu1-SARS2_orf1a;1153 | 0 | 0 |
| NC_045512.2;YP_009725295.1;Wu1-SARS2_orf1a;1154 | 0 | 0 |
| NC_045512.2;YP_009725295.1;Wu1-SARS2_orf1a;1155 | 0 | 0 |
| NC_045512.2;YP_009725295.1;Wu1-SARS2_orf1a;1156 | 0 | 0 |
| NC_045512.2;YP_009725295.1;Wu1-SARS2_orf1a;1157 | 0 | 0 |
| NC_045512.2;YP_009725295.1;Wu1-SARS2_orf1a;1158 | 0 | 0 |
| NC_045512.2;YP_009725295.1;Wu1-SARS2_orf1a;1159 | 0 | 0 |
| NC_045512.2;YP_009725295.1;Wu1-SARS2_orf1a;1160 | 0 | 0 |
| NC_045512.2;YP_009725295.1;Wu1-SARS2_orf1a;1161 | 0 | 0 |
| NC_045512.2;YP_009725295.1;Wu1-SARS2_orf1a;1162 | 0 | 0 |
| NC_045512.2;YP_009725295.1;Wu1-SARS2_orf1a;1163 | 0 | 0 |
| NC_045512.2;YP_009725295.1;Wu1-SARS2_orf1a;1164 | 0 | 0 |
| NC_045512.2;YP_009725295.1;Wu1-SARS2_orf1a;1165 | 0 | 0 |
| NC_045512.2;YP_009725295.1;Wu1-SARS2_orf1a;1166 | 0 | 0.05 |
| NC_045512.2;YP_009725295.1;Wu1-SARS2_orf1a;1167 | 0 | 0 |
| NC_045512.2;YP_009725295.1;Wu1-SARS2_orf1a;1168 | 0 | 0 |
| NC_045512.2;YP_009725295.1;Wu1-SARS2_orf1a;1169 | 0 | 0 |
| NC_045512.2;YP_009725295.1;Wu1-SARS2_orf1a;1170 | 0 | 0 |
| NC_045512.2;YP_009725295.1;Wu1-SARS2_orf1a;1171 | 0 | 0 |
| NC_045512.2;YP_009725295.1;Wu1-SARS2_orf1a;1172 | 0 | 0 |
| NC_045512.2;YP_009725295.1;Wu1-SARS2_orf1a;1173 | 0.05 | 0 |
| NC_045512.2;YP_009725295.1;Wu1-SARS2_orf1a;1174 | 0.05 | 0 |
| NC_045512.2;YP_009725295.1;Wu1-SARS2_orf1a;1175 | 0.05 | 0 |
| NC_045512.2;YP_009725295.1;Wu1-SARS2_orf1a;1176 | 0 | 0 |
| NC_045512.2;YP_009725295.1;Wu1-SARS2_orf1a;1177 | 0.05 | 0 |
| NC_045512.2;YP_009725295.1;Wu1-SARS2_orf1a;1178 | 0.05 | 0 |
| NC_045512.2;YP_009725295.1;Wu1-SARS2_orf1a;1179 | 0.1 | 0 |
| NC_045512.2;YP_009725295.1;Wu1-SARS2_orf1a;1180 | 0 | 0 |
| NC_045512.2;YP_009725295.1;Wu1-SARS2_orf1a;1181 | 0 | 0 |
| NC_045512.2;YP_009725295.1;Wu1-SARS2_orf1a;1182 | 0 | 0 |
| NC_045512.2;YP_009725295.1;Wu1-SARS2_orf1a;1183 | 0 | 0 |
| NC_045512.2;YP_009725295.1;Wu1-SARS2_orf1a;1184 | 0 | 0 |
| NC_045512.2;YP_009725295.1;Wu1-SARS2_orf1a;1185 | 0.05 | 0 |
| NC_045512.2;YP_009725295.1;Wu1-SARS2_orf1a;1186 | 0.05 | 0 |
| NC_045512.2;YP_009725295.1;Wu1-SARS2_orf1a;1187 | 0.1 | 0.025 |
| NC_045512.2;YP_009725295.1;Wu1-SARS2_orf1a;1188 | 0.05 | 0.025 |
| NC_045512.2;YP_009725295.1;Wu1-SARS2_orf1a;1189 | 0.05 | 0.025 |
| NC_045512.2;YP_009725295.1;Wu1-SARS2_orf1a;1190 | 0.05 | 0.075 |
| NC_045512.2;YP_009725295.1;Wu1-SARS2_orf1a;1191 | 0 | 0.05 |
| NC_045512.2;YP_009725295.1;Wu1-SARS2_orf1a;1192 | 0.05 | 0.025 |

| | | |
|---|---|---|
| NC_045512.2;YP_009725295.1;Wu1-SARS2_orf1a;1193 | 0 | 0 |
| NC_045512.2;YP_009725295.1;Wu1-SARS2_orf1a;1194 | 0.05 | 0 |
| NC_045512.2;YP_009725295.1;Wu1-SARS2_orf1a;1195 | 0.05 | 0.05 |
| NC_045512.2;YP_009725295.1;Wu1-SARS2_orf1a;1196 | 0.05 | 0.05 |
| NC_045512.2;YP_009725295.1;Wu1-SARS2_orf1a;1197 | 0.1 | 0.025 |
| NC_045512.2;YP_009725295.1;Wu1-SARS2_orf1a;1198 | 0.05 | 0.025 |
| NC_045512.2;YP_009725295.1;Wu1-SARS2_orf1a;1199 | 0.05 | 0 |
| NC_045512.2;YP_009725295.1;Wu1-SARS2_orf1a;1200 | 0.1 | 0.025 |
| NC_045512.2;YP_009725295.1;Wu1-SARS2_orf1a;1201 | 0.1 | 0.025 |
| NC_045512.2;YP_009725295.1;Wu1-SARS2_orf1a;1202 | 0 | 0.05 |
| NC_045512.2;YP_009725295.1;Wu1-SARS2_orf1a;1203 | 0 | 0.05 |
| NC_045512.2;YP_009725295.1;Wu1-SARS2_orf1a;1204 | 0.05 | 0.05 |
| NC_045512.2;YP_009725295.1;Wu1-SARS2_orf1a;1205 | 0.05 | 0.05 |
| NC_045512.2;YP_009725295.1;Wu1-SARS2_orf1a;1206 | 0.05 | 0.025 |
| NC_045512.2;YP_009725295.1;Wu1-SARS2_orf1a;1207 | 0.1 | 0.025 |
| NC_045512.2;YP_009725295.1;Wu1-SARS2_orf1a;1208 | 0.05 | 0 |
| NC_045512.2;YP_009725295.1;Wu1-SARS2_orf1a;1209 | 0.05 | 0 |
| NC_045512.2;YP_009725295.1;Wu1-SARS2_orf1a;1210 | 0.1 | 0.05 |
| NC_045512.2;YP_009725295.1;Wu1-SARS2_orf1a;1211 | 0.1 | 0.025 |
| NC_045512.2;YP_009725295.1;Wu1-SARS2_orf1a;1212 | 0.05 | 0.025 |
| NC_045512.2;YP_009725295.1;Wu1-SARS2_orf1a;1213 | 0.1 | 0.075 |
| NC_045512.2;YP_009725295.1;Wu1-SARS2_orf1a;1214 | 0.05 | 0.05 |
| NC_045512.2;YP_009725295.1;Wu1-SARS2_orf1a;1215 | 0.1 | 0.025 |
| NC_045512.2;YP_009725295.1;Wu1-SARS2_orf1a;1216 | 0.05 | 0.025 |
| NC_045512.2;YP_009725295.1;Wu1-SARS2_orf1a;1217 | 0.05 | 0.125 |
| NC_045512.2;YP_009725295.1;Wu1-SARS2_orf1a;1218 | 0.05 | 0 |
| NC_045512.2;YP_009725295.1;Wu1-SARS2_orf1a;1219 | 0 | 0 |
| NC_045512.2;YP_009725295.1;Wu1-SARS2_orf1a;1220 | 0.05 | 0 |
| NC_045512.2;YP_009725295.1;Wu1-SARS2_orf1a;1221 | 0.05 | 0 |
| NC_045512.2;YP_009725295.1;Wu1-SARS2_orf1a;1222 | 0.05 | 0 |
| NC_045512.2;YP_009725295.1;Wu1-SARS2_orf1a;1223 | 0.05 | 0 |
| NC_045512.2;YP_009725295.1;Wu1-SARS2_orf1a;1224 | 0.05 | 0 |
| NC_045512.2;YP_009725295.1;Wu1-SARS2_orf1a;1225 | 0.05 | 0.025 |
| NC_045512.2;YP_009725295.1;Wu1-SARS2_orf1a;1226 | 0.05 | 0.025 |
| NC_045512.2;YP_009725295.1;Wu1-SARS2_orf1a;1227 | 0 | 0.025 |
| NC_045512.2;YP_009725295.1;Wu1-SARS2_orf1a;1228 | 0 | 0.025 |
| NC_045512.2;YP_009725295.1;Wu1-SARS2_orf1a;1229 | 0 | 0.025 |
| NC_045512.2;YP_009725295.1;Wu1-SARS2_orf1a;1230 | 0 | 0.025 |
| NC_045512.2;YP_009725295.1;Wu1-SARS2_orf1a;1231 | 0 | 0.025 |
| NC_045512.2;YP_009725295.1;Wu1-SARS2_orf1a;1232 | 0 | 0.025 |
| NC_045512.2;YP_009725295.1;Wu1-SARS2_orf1a;1233 | 0 | 0.025 |
| NC_045512.2;YP_009725295.1;Wu1-SARS2_orf1a;1234 | 0 | 0.025 |
| NC_045512.2;YP_009725295.1;Wu1-SARS2_orf1a;1235 | 0 | 0.025 |

| | | |
|---|---|---|
| NC_045512.2;YP_009725295.1;Wu1-SARS2_orf1a;1236 | 0 | 0 |
| NC_045512.2;YP_009725295.1;Wu1-SARS2_orf1a;1237 | 0 | 0.075 |
| NC_045512.2;YP_009725295.1;Wu1-SARS2_orf1a;1238 | 0 | 0.05 |
| NC_045512.2;YP_009725295.1;Wu1-SARS2_orf1a;1239 | 0 | 0.025 |
| NC_045512.2;YP_009725295.1;Wu1-SARS2_orf1a;1240 | 0 | 0.025 |
| NC_045512.2;YP_009725295.1;Wu1-SARS2_orf1a;1241 | 0 | 0.05 |
| NC_045512.2;YP_009725295.1;Wu1-SARS2_orf1a;1242 | 0 | 0 |
| NC_045512.2;YP_009725295.1;Wu1-SARS2_orf1a;1243 | 0 | 0 |
| NC_045512.2;YP_009725295.1;Wu1-SARS2_orf1a;1244 | 0 | 0 |
| NC_045512.2;YP_009725295.1;Wu1-SARS2_orf1a;1245 | 0 | 0 |
| NC_045512.2;YP_009725295.1;Wu1-SARS2_orf1a;1246 | 0 | 0 |
| NC_045512.2;YP_009725295.1;Wu1-SARS2_orf1a;1247 | 0 | 0 |
| NC_045512.2;YP_009725295.1;Wu1-SARS2_orf1a;1248 | 0 | 0 |
| NC_045512.2;YP_009725295.1;Wu1-SARS2_orf1a;1249 | 0 | 0 |
| NC_045512.2;YP_009725295.1;Wu1-SARS2_orf1a;1250 | 0 | 0 |
| NC_045512.2;YP_009725295.1;Wu1-SARS2_orf1a;1251 | 0 | 0 |
| NC_045512.2;YP_009725295.1;Wu1-SARS2_orf1a;1252 | 0 | 0 |
| NC_045512.2;YP_009725295.1;Wu1-SARS2_orf1a;1253 | 0 | 0 |
| NC_045512.2;YP_009725295.1;Wu1-SARS2_orf1a;1254 | 0 | 0.025 |
| NC_045512.2;YP_009725295.1;Wu1-SARS2_orf1a;1255 | 0.05 | 0.075 |
| NC_045512.2;YP_009725295.1;Wu1-SARS2_orf1a;1256 | 0.1 | 0.075 |
| NC_045512.2;YP_009725295.1;Wu1-SARS2_orf1a;1257 | 0 | 0 |
| NC_045512.2;YP_009725295.1;Wu1-SARS2_orf1a;1258 | 0 | 0 |
| NC_045512.2;YP_009725295.1;Wu1-SARS2_orf1a;1259 | 0 | 0 |
| NC_045512.2;YP_009725295.1;Wu1-SARS2_orf1a;1260 | 0 | 0 |
| NC_045512.2;YP_009725295.1;Wu1-SARS2_orf1a;1261 | 0 | 0 |
| NC_045512.2;YP_009725295.1;Wu1-SARS2_orf1a;1262 | 0 | 0 |
| NC_045512.2;YP_009725295.1;Wu1-SARS2_orf1a;1263 | 0 | 0 |
| NC_045512.2;YP_009725295.1;Wu1-SARS2_orf1a;1264 | 0 | 0 |
| NC_045512.2;YP_009725295.1;Wu1-SARS2_orf1a;1265 | 0 | 0 |
| NC_045512.2;YP_009725295.1;Wu1-SARS2_orf1a;1266 | 0 | 0.025 |
| NC_045512.2;YP_009725295.1;Wu1-SARS2_orf1a;1267 | 0 | 0.025 |
| NC_045512.2;YP_009725295.1;Wu1-SARS2_orf1a;1268 | 0 | 0.025 |
| NC_045512.2;YP_009725295.1;Wu1-SARS2_orf1a;1269 | 0 | 0.05 |
| NC_045512.2;YP_009725295.1;Wu1-SARS2_orf1a;1270 | 0 | 0.075 |
| NC_045512.2;YP_009725295.1;Wu1-SARS2_orf1a;1271 | 0 | 0.075 |
| NC_045512.2;YP_009725295.1;Wu1-SARS2_orf1a;1272 | 0.05 | 0.075 |
| NC_045512.2;YP_009725295.1;Wu1-SARS2_orf1a;1273 | 0 | 0.025 |
| NC_045512.2;YP_009725295.1;Wu1-SARS2_orf1a;1274 | 0 | 0.025 |
| NC_045512.2;YP_009725295.1;Wu1-SARS2_orf1a;1275 | 0 | 0.025 |
| NC_045512.2;YP_009725295.1;Wu1-SARS2_orf1a;1276 | 0 | 0 |
| NC_045512.2;YP_009725295.1;Wu1-SARS2_orf1a;1277 | 0 | 0 |
| NC_045512.2;YP_009725295.1;Wu1-SARS2_orf1a;1278 | 0 | 0 |

| | | |
|---|---|---|
| NC_045512.2;YP_009725295.1;Wu1-SARS2_orf1a;1279 | 0 | 0 |
| NC_045512.2;YP_009725295.1;Wu1-SARS2_orf1a;1280 | 0 | 0 |
| NC_045512.2;YP_009725295.1;Wu1-SARS2_orf1a;1281 | 0 | 0 |
| NC_045512.2;YP_009725295.1;Wu1-SARS2_orf1a;1282 | 0 | 0 |
| NC_045512.2;YP_009725295.1;Wu1-SARS2_orf1a;1283 | 0 | 0 |
| NC_045512.2;YP_009725295.1;Wu1-SARS2_orf1a;1284 | 0 | 0 |
| NC_045512.2;YP_009725295.1;Wu1-SARS2_orf1a;1285 | 0 | 0 |
| NC_045512.2;YP_009725295.1;Wu1-SARS2_orf1a;1286 | 0 | 0 |
| NC_045512.2;YP_009725295.1;Wu1-SARS2_orf1a;1287 | 0 | 0 |
| NC_045512.2;YP_009725295.1;Wu1-SARS2_orf1a;1288 | 0 | 0.025 |
| NC_045512.2;YP_009725295.1;Wu1-SARS2_orf1a;1289 | 0 | 0 |
| NC_045512.2;YP_009725295.1;Wu1-SARS2_orf1a;1290 | 0 | 0 |
| NC_045512.2;YP_009725295.1;Wu1-SARS2_orf1a;1291 | 0 | 0 |
| NC_045512.2;YP_009725295.1;Wu1-SARS2_orf1a;1292 | 0 | 0 |
| NC_045512.2;YP_009725295.1;Wu1-SARS2_orf1a;1293 | 0 | 0 |
| NC_045512.2;YP_009725295.1;Wu1-SARS2_orf1a;1294 | 0 | 0 |
| NC_045512.2;YP_009725295.1;Wu1-SARS2_orf1a;1295 | 0 | 0 |
| NC_045512.2;YP_009725295.1;Wu1-SARS2_orf1a;1296 | 0.05 | 0 |
| NC_045512.2;YP_009725295.1;Wu1-SARS2_orf1a;1297 | 0.05 | 0.025 |
| NC_045512.2;YP_009725295.1;Wu1-SARS2_orf1a;1298 | 0.05 | 0.025 |
| NC_045512.2;YP_009725295.1;Wu1-SARS2_orf1a;1299 | 0.05 | 0.025 |
| NC_045512.2;YP_009725295.1;Wu1-SARS2_orf1a;1300 | 0.05 | 0 |
| NC_045512.2;YP_009725295.1;Wu1-SARS2_orf1a;1301 | 0.05 | 0 |
| NC_045512.2;YP_009725295.1;Wu1-SARS2_orf1a;1302 | 0.05 | 0 |
| NC_045512.2;YP_009725295.1;Wu1-SARS2_orf1a;1303 | 0 | 0 |
| NC_045512.2;YP_009725295.1;Wu1-SARS2_orf1a;1304 | 0 | 0 |
| NC_045512.2;YP_009725295.1;Wu1-SARS2_orf1a;1305 | 0 | 0 |
| NC_045512.2;YP_009725295.1;Wu1-SARS2_orf1a;1306 | 0 | 0 |
| NC_045512.2;YP_009725295.1;Wu1-SARS2_orf1a;1307 | 0 | 0 |
| NC_045512.2;YP_009725295.1;Wu1-SARS2_orf1a;1308 | 0 | 0 |
| NC_045512.2;YP_009725295.1;Wu1-SARS2_orf1a;1309 | 0.05 | 0 |
| NC_045512.2;YP_009725295.1;Wu1-SARS2_orf1a;1310 | 0 | 0 |
| NC_045512.2;YP_009725295.1;Wu1-SARS2_orf1a;1311 | 0 | 0 |
| NC_045512.2;YP_009725295.1;Wu1-SARS2_orf1a;1312 | 0 | 0 |
| NC_045512.2;YP_009725295.1;Wu1-SARS2_orf1a;1313 | 0 | 0 |
| NC_045512.2;YP_009725295.1;Wu1-SARS2_orf1a;1314 | 0 | 0 |
| NC_045512.2;YP_009725295.1;Wu1-SARS2_orf1a;1315 | 0 | 0 |
| NC_045512.2;YP_009725295.1;Wu1-SARS2_orf1a;1316 | 0.05 | 0 |
| NC_045512.2;YP_009725295.1;Wu1-SARS2_orf1a;1317 | 0.05 | 0 |
| NC_045512.2;YP_009725295.1;Wu1-SARS2_orf1a;1318 | 0.05 | 0.025 |
| NC_045512.2;YP_009725295.1;Wu1-SARS2_orf1a;1319 | 0 | 0 |
| NC_045512.2;YP_009725295.1;Wu1-SARS2_orf1a;1320 | 0 | 0.05 |
| NC_045512.2;YP_009725295.1;Wu1-SARS2_orf1a;1321 | 0 | 0 |

| | | |
|---|---|---|
| NC_045512.2;YP_009725295.1;Wu1-SARS2_orf1a;1322 | 0 | 0 |
| NC_045512.2;YP_009725295.1;Wu1-SARS2_orf1a;1323 | 0.05 | 0 |
| NC_045512.2;YP_009725295.1;Wu1-SARS2_orf1a;1324 | 0 | 0 |
| NC_045512.2;YP_009725295.1;Wu1-SARS2_orf1a;1325 | 0 | 0 |
| NC_045512.2;YP_009725295.1;Wu1-SARS2_orf1a;1326 | 0 | 0 |
| NC_045512.2;YP_009725295.1;Wu1-SARS2_orf1a;1327 | 0 | 0 |
| NC_045512.2;YP_009725295.1;Wu1-SARS2_orf1a;1328 | 0.05 | 0 |
| NC_045512.2;YP_009725295.1;Wu1-SARS2_orf1a;1329 | 0.05 | 0 |
| NC_045512.2;YP_009725295.1;Wu1-SARS2_orf1a;1330 | 0 | 0 |
| NC_045512.2;YP_009725295.1;Wu1-SARS2_orf1a;1331 | 0 | 0.025 |
| NC_045512.2;YP_009725295.1;Wu1-SARS2_orf1a;1332 | 0 | 0.05 |
| NC_045512.2;YP_009725295.1;Wu1-SARS2_orf1a;1333 | 0 | 0 |
| NC_045512.2;YP_009725295.1;Wu1-SARS2_orf1a;1334 | 0 | 0 |
| NC_045512.2;YP_009725295.1;Wu1-SARS2_orf1a;1335 | 0 | 0 |
| NC_045512.2;YP_009725295.1;Wu1-SARS2_orf1a;1336 | 0 | 0.025 |
| NC_045512.2;YP_009725295.1;Wu1-SARS2_orf1a;1337 | 0 | 0.05 |
| NC_045512.2;YP_009725295.1;Wu1-SARS2_orf1a;1338 | 0 | 0 |
| NC_045512.2;YP_009725295.1;Wu1-SARS2_orf1a;1339 | 0 | 0 |
| NC_045512.2;YP_009725295.1;Wu1-SARS2_orf1a;1340 | 0 | 0 |
| NC_045512.2;YP_009725295.1;Wu1-SARS2_orf1a;1341 | 0 | 0 |
| NC_045512.2;YP_009725295.1;Wu1-SARS2_orf1a;1342 | 0 | 0 |
| NC_045512.2;YP_009725295.1;Wu1-SARS2_orf1a;1343 | 0 | 0 |
| NC_045512.2;YP_009725295.1;Wu1-SARS2_orf1a;1344 | 0 | 0 |
| NC_045512.2;YP_009725295.1;Wu1-SARS2_orf1a;1345 | 0 | 0 |
| NC_045512.2;YP_009725295.1;Wu1-SARS2_orf1a;1346 | 0 | 0 |
| NC_045512.2;YP_009725295.1;Wu1-SARS2_orf1a;1347 | 0 | 0 |
| NC_045512.2;YP_009725295.1;Wu1-SARS2_orf1a;1348 | 0 | 0 |
| NC_045512.2;YP_009725295.1;Wu1-SARS2_orf1a;1349 | 0 | 0 |
| NC_045512.2;YP_009725295.1;Wu1-SARS2_orf1a;1350 | 0 | 0.025 |
| NC_045512.2;YP_009725295.1;Wu1-SARS2_orf1a;1351 | 0 | 0 |
| NC_045512.2;YP_009725295.1;Wu1-SARS2_orf1a;1352 | 0 | 0 |
| NC_045512.2;YP_009725295.1;Wu1-SARS2_orf1a;1353 | 0 | 0 |
| NC_045512.2;YP_009725295.1;Wu1-SARS2_orf1a;1354 | 0.05 | 0 |
| NC_045512.2;YP_009725295.1;Wu1-SARS2_orf1a;1355 | 0 | 0 |
| NC_045512.2;YP_009725295.1;Wu1-SARS2_orf1a;1356 | 0 | 0 |
| NC_045512.2;YP_009725295.1;Wu1-SARS2_orf1a;1357 | 0.05 | 0.025 |
| NC_045512.2;YP_009725295.1;Wu1-SARS2_orf1a;1358 | 0 | 0 |
| NC_045512.2;YP_009725295.1;Wu1-SARS2_orf1a;1359 | 0.05 | 0 |
| NC_045512.2;YP_009725295.1;Wu1-SARS2_orf1a;1360 | 0 | 0 |
| NC_045512.2;YP_009725295.1;Wu1-SARS2_orf1a;1361 | 0 | 0 |
| NC_045512.2;YP_009725295.1;Wu1-SARS2_orf1a;1362 | 0 | 0 |
| NC_045512.2;YP_009725295.1;Wu1-SARS2_orf1a;1363 | 0 | 0 |
| NC_045512.2;YP_009725295.1;Wu1-SARS2_orf1a;1364 | 0 | 0 |

| | | |
|---|---|---|
| NC_045512.2;YP_009725295.1;Wu1-SARS2_orf1a;1365 | 0.05 | 0 |
| NC_045512.2;YP_009725295.1;Wu1-SARS2_orf1a;1366 | 0 | 0 |
| NC_045512.2;YP_009725295.1;Wu1-SARS2_orf1a;1367 | 0 | 0 |
| NC_045512.2;YP_009725295.1;Wu1-SARS2_orf1a;1368 | 0 | 0 |
| NC_045512.2;YP_009725295.1;Wu1-SARS2_orf1a;1369 | 0 | 0.025 |
| NC_045512.2;YP_009725295.1;Wu1-SARS2_orf1a;1370 | 0 | 0 |
| NC_045512.2;YP_009725295.1;Wu1-SARS2_orf1a;1371 | 0 | 0 |
| NC_045512.2;YP_009725295.1;Wu1-SARS2_orf1a;1372 | 0 | 0.05 |
| NC_045512.2;YP_009725295.1;Wu1-SARS2_orf1a;1373 | 0 | 0.025 |
| NC_045512.2;YP_009725295.1;Wu1-SARS2_orf1a;1374 | 0 | 0 |
| NC_045512.2;YP_009725295.1;Wu1-SARS2_orf1a;1375 | 0 | 0.025 |
| NC_045512.2;YP_009725295.1;Wu1-SARS2_orf1a;1376 | 0 | 0 |
| NC_045512.2;YP_009725295.1;Wu1-SARS2_orf1a;1377 | 0 | 0 |
| NC_045512.2;YP_009725295.1;Wu1-SARS2_orf1a;1378 | 0 | 0 |
| NC_045512.2;YP_009725295.1;Wu1-SARS2_orf1a;1379 | 0 | 0 |
| NC_045512.2;YP_009725295.1;Wu1-SARS2_orf1a;1380 | 0 | 0 |
| NC_045512.2;YP_009725295.1;Wu1-SARS2_orf1a;1381 | 0 | 0 |
| NC_045512.2;YP_009725295.1;Wu1-SARS2_orf1a;1382 | 0.05 | 0.025 |
| NC_045512.2;YP_009725295.1;Wu1-SARS2_orf1a;1383 | 0.05 | 0.025 |
| NC_045512.2;YP_009725295.1;Wu1-SARS2_orf1a;1384 | 0 | 0 |
| NC_045512.2;YP_009725295.1;Wu1-SARS2_orf1a;1385 | 0 | 0 |
| NC_045512.2;YP_009725295.1;Wu1-SARS2_orf1a;1386 | 0 | 0 |
| NC_045512.2;YP_009725295.1;Wu1-SARS2_orf1a;1387 | 0 | 0 |
| NC_045512.2;YP_009725295.1;Wu1-SARS2_orf1a;1388 | 0 | 0 |
| NC_045512.2;YP_009725295.1;Wu1-SARS2_orf1a;1389 | 0 | 0 |
| NC_045512.2;YP_009725295.1;Wu1-SARS2_orf1a;1390 | 0 | 0 |
| NC_045512.2;YP_009725295.1;Wu1-SARS2_orf1a;1391 | 0 | 0 |
| NC_045512.2;YP_009725295.1;Wu1-SARS2_orf1a;1392 | 0 | 0 |
| NC_045512.2;YP_009725295.1;Wu1-SARS2_orf1a;1393 | 0 | 0 |
| NC_045512.2;YP_009725295.1;Wu1-SARS2_orf1a;1394 | 0 | 0 |
| NC_045512.2;YP_009725295.1;Wu1-SARS2_orf1a;1395 | 0 | 0 |
| NC_045512.2;YP_009725295.1;Wu1-SARS2_orf1a;1396 | 0.05 | 0 |
| NC_045512.2;YP_009725295.1;Wu1-SARS2_orf1a;1397 | 0.05 | 0 |
| NC_045512.2;YP_009725295.1;Wu1-SARS2_orf1a;1398 | 0.1 | 0 |
| NC_045512.2;YP_009725295.1;Wu1-SARS2_orf1a;1399 | 0.1 | 0 |
| NC_045512.2;YP_009725295.1;Wu1-SARS2_orf1a;1400 | 0.1 | 0 |
| NC_045512.2;YP_009725295.1;Wu1-SARS2_orf1a;1401 | 0.1 | 0 |
| NC_045512.2;YP_009725295.1;Wu1-SARS2_orf1a;1402 | 0.1 | 0 |
| NC_045512.2;YP_009725295.1;Wu1-SARS2_orf1a;1403 | 0.1 | 0.025 |
| NC_045512.2;YP_009725295.1;Wu1-SARS2_orf1a;1404 | 0.1 | 0.025 |
| NC_045512.2;YP_009725295.1;Wu1-SARS2_orf1a;1405 | 0.1 | 0.025 |
| NC_045512.2;YP_009725295.1;Wu1-SARS2_orf1a;1406 | 0.1 | 0.025 |
| NC_045512.2;YP_009725295.1;Wu1-SARS2_orf1a;1407 | 0 | 0.025 |

| | | |
|---|---|---|
| NC_045512.2;YP_009725295.1;Wu1-SARS2_orf1a;1408 | 0 | 0 |
| NC_045512.2;YP_009725295.1;Wu1-SARS2_orf1a;1409 | 0 | 0 |
| NC_045512.2;YP_009725295.1;Wu1-SARS2_orf1a;1410 | 0 | 0 |
| NC_045512.2;YP_009725295.1;Wu1-SARS2_orf1a;1411 | 0 | 0 |
| NC_045512.2;YP_009725295.1;Wu1-SARS2_orf1a;1412 | 0 | 0 |
| NC_045512.2;YP_009725295.1;Wu1-SARS2_orf1a;1413 | 0 | 0 |
| NC_045512.2;YP_009725295.1;Wu1-SARS2_orf1a;1414 | 0.05 | 0.1 |
| NC_045512.2;YP_009725295.1;Wu1-SARS2_orf1a;1415 | 0 | 0 |
| NC_045512.2;YP_009725295.1;Wu1-SARS2_orf1a;1416 | 0 | 0 |
| NC_045512.2;YP_009725295.1;Wu1-SARS2_orf1a;1417 | 0 | 0 |
| NC_045512.2;YP_009725295.1;Wu1-SARS2_orf1a;1418 | 0 | 0 |
| NC_045512.2;YP_009725295.1;Wu1-SARS2_orf1a;1419 | 0 | 0 |
| NC_045512.2;YP_009725295.1;Wu1-SARS2_orf1a;1420 | 0 | 0 |
| NC_045512.2;YP_009725295.1;Wu1-SARS2_orf1a;1421 | 0 | 0 |
| NC_045512.2;YP_009725295.1;Wu1-SARS2_orf1a;1422 | 0 | 0 |
| NC_045512.2;YP_009725295.1;Wu1-SARS2_orf1a;1423 | 0 | 0 |
| NC_045512.2;YP_009725295.1;Wu1-SARS2_orf1a;1424 | 0 | 0.025 |
| NC_045512.2;YP_009725295.1;Wu1-SARS2_orf1a;1425 | 0 | 0.05 |
| NC_045512.2;YP_009725295.1;Wu1-SARS2_orf1a;1426 | 0 | 0.025 |
| NC_045512.2;YP_009725295.1;Wu1-SARS2_orf1a;1427 | 0.2 | 0.1 |
| NC_045512.2;YP_009725295.1;Wu1-SARS2_orf1a;1428 | 0.25 | 0.125 |
| NC_045512.2;YP_009725295.1;Wu1-SARS2_orf1a;1429 | 0.15 | 0.25 |
| NC_045512.2;YP_009725295.1;Wu1-SARS2_orf1a;1430 | 0.2 | 0.25 |
| NC_045512.2;YP_009725295.1;Wu1-SARS2_orf1a;1431 | 0.3 | 0.35 |
| NC_045512.2;YP_009725295.1;Wu1-SARS2_orf1a;1432 | 0.2 | 0.225 |
| NC_045512.2;YP_009725295.1;Wu1-SARS2_orf1a;1433 | 0.2 | 0.175 |
| NC_045512.2;YP_009725295.1;Wu1-SARS2_orf1a;1434 | 0.2 | 0.225 |
| NC_045512.2;YP_009725295.1;Wu1-SARS2_orf1a;1435 | 0.25 | 0.2 |
| NC_045512.2;YP_009725295.1;Wu1-SARS2_orf1a;1436 | 0.2 | 0.2 |
| NC_045512.2;YP_009725295.1;Wu1-SARS2_orf1a;1437 | 0.35 | 0.3 |
| NC_045512.2;YP_009725295.1;Wu1-SARS2_orf1a;1438 | 0.15 | 0.15 |
| NC_045512.2;YP_009725295.1;Wu1-SARS2_orf1a;1439 | 0.1 | 0.1 |
| NC_045512.2;YP_009725295.1;Wu1-SARS2_orf1a;1440 | 0.1 | 0.1 |
| NC_045512.2;YP_009725295.1;Wu1-SARS2_orf1a;1441 | 0.1 | 0.1 |
| NC_045512.2;YP_009725295.1;Wu1-SARS2_orf1a;1442 | 0.05 | 0.075 |
| NC_045512.2;YP_009725295.1;Wu1-SARS2_orf1a;1443 | 0 | 0 |
| NC_045512.2;YP_009725295.1;Wu1-SARS2_orf1a;1444 | 0 | 0 |
| NC_045512.2;YP_009725295.1;Wu1-SARS2_orf1a;1445 | 0 | 0 |
| NC_045512.2;YP_009725295.1;Wu1-SARS2_orf1a;1446 | 0.05 | 0 |
| NC_045512.2;YP_009725295.1;Wu1-SARS2_orf1a;1447 | 0 | 0 |
| NC_045512.2;YP_009725295.1;Wu1-SARS2_orf1a;1448 | 0.1 | 0.05 |
| NC_045512.2;YP_009725295.1;Wu1-SARS2_orf1a;1449 | 0.05 | 0.025 |
| NC_045512.2;YP_009725295.1;Wu1-SARS2_orf1a;1450 | 0.05 | 0.025 |

| | | |
|---|---|---|
| NC_045512.2;YP_009725295.1;Wu1-SARS2_orf1a;1451 | 0.05 | 0.025 |
| NC_045512.2;YP_009725295.1;Wu1-SARS2_orf1a;1452 | 0 | 0 |
| NC_045512.2;YP_009725295.1;Wu1-SARS2_orf1a;1453 | 0 | 0 |
| NC_045512.2;YP_009725295.1;Wu1-SARS2_orf1a;1454 | 0 | 0 |
| NC_045512.2;YP_009725295.1;Wu1-SARS2_orf1a;1455 | 0 | 0 |
| NC_045512.2;YP_009725295.1;Wu1-SARS2_orf1a;1456 | 0 | 0.025 |
| NC_045512.2;YP_009725295.1;Wu1-SARS2_orf1a;1457 | 0 | 0 |
| NC_045512.2;YP_009725295.1;Wu1-SARS2_orf1a;1458 | 0 | 0 |
| NC_045512.2;YP_009725295.1;Wu1-SARS2_orf1a;1459 | 0 | 0.075 |
| NC_045512.2;YP_009725295.1;Wu1-SARS2_orf1a;1460 | 0.1 | 0.075 |
| NC_045512.2;YP_009725295.1;Wu1-SARS2_orf1a;1461 | 0.15 | 0.125 |
| NC_045512.2;YP_009725295.1;Wu1-SARS2_orf1a;1462 | 0.05 | 0 |
| NC_045512.2;YP_009725295.1;Wu1-SARS2_orf1a;1463 | 0 | 0.025 |
| NC_045512.2;YP_009725295.1;Wu1-SARS2_orf1a;1464 | 0 | 0 |
| NC_045512.2;YP_009725295.1;Wu1-SARS2_orf1a;1465 | 0 | 0 |
| NC_045512.2;YP_009725295.1;Wu1-SARS2_orf1a;1466 | 0 | 0 |
| NC_045512.2;YP_009725295.1;Wu1-SARS2_orf1a;1467 | 0 | 0 |
| NC_045512.2;YP_009725295.1;Wu1-SARS2_orf1a;1468 | 0 | 0 |
| NC_045512.2;YP_009725295.1;Wu1-SARS2_orf1a;1469 | 0 | 0 |
| NC_045512.2;YP_009725295.1;Wu1-SARS2_orf1a;1470 | 0 | 0 |
| NC_045512.2;YP_009725295.1;Wu1-SARS2_orf1a;1471 | 0 | 0 |
| NC_045512.2;YP_009725295.1;Wu1-SARS2_orf1a;1472 | 0 | 0.05 |
| NC_045512.2;YP_009725295.1;Wu1-SARS2_orf1a;1473 | 0.05 | 0.025 |
| NC_045512.2;YP_009725295.1;Wu1-SARS2_orf1a;1474 | 0 | 0.1 |
| NC_045512.2;YP_009725295.1;Wu1-SARS2_orf1a;1475 | 0 | 0.025 |
| NC_045512.2;YP_009725295.1;Wu1-SARS2_orf1a;1476 | 0 | 0 |
| NC_045512.2;YP_009725295.1;Wu1-SARS2_orf1a;1477 | 0 | 0 |
| NC_045512.2;YP_009725295.1;Wu1-SARS2_orf1a;1478 | 0 | 0 |
| NC_045512.2;YP_009725295.1;Wu1-SARS2_orf1a;1479 | 0 | 0 |
| NC_045512.2;YP_009725295.1;Wu1-SARS2_orf1a;1480 | 0.05 | 0.05 |
| NC_045512.2;YP_009725295.1;Wu1-SARS2_orf1a;1481 | 0.05 | 0.075 |
| NC_045512.2;YP_009725295.1;Wu1-SARS2_orf1a;1482 | 0.1 | 0.05 |
| NC_045512.2;YP_009725295.1;Wu1-SARS2_orf1a;1483 | 0.15 | 0.1 |
| NC_045512.2;YP_009725295.1;Wu1-SARS2_orf1a;1484 | 0.1 | 0.15 |
| NC_045512.2;YP_009725295.1;Wu1-SARS2_orf1a;1485 | 0.1 | 0.125 |
| NC_045512.2;YP_009725295.1;Wu1-SARS2_orf1a;1486 | 0.15 | 0.125 |
| NC_045512.2;YP_009725295.1;Wu1-SARS2_orf1a;1487 | 0.2 | 0.125 |
| NC_045512.2;YP_009725295.1;Wu1-SARS2_orf1a;1488 | 0.25 | 0.1 |
| NC_045512.2;YP_009725295.1;Wu1-SARS2_orf1a;1489 | 0.2 | 0.1 |
| NC_045512.2;YP_009725295.1;Wu1-SARS2_orf1a;1490 | 0.15 | 0.025 |
| NC_045512.2;YP_009725295.1;Wu1-SARS2_orf1a;1491 | 0.05 | 0 |
| NC_045512.2;YP_009725295.1;Wu1-SARS2_orf1a;1492 | 0.05 | 0 |
| NC_045512.2;YP_009725295.1;Wu1-SARS2_orf1a;1493 | 0.05 | 0 |

| | | |
|---|---|---|
| NC_045512.2;YP_009725295.1;Wu1-SARS2_orf1a;1494 | 0 | 0 |
| NC_045512.2;YP_009725295.1;Wu1-SARS2_orf1a;1495 | 0 | 0 |
| NC_045512.2;YP_009725295.1;Wu1-SARS2_orf1a;1496 | 0 | 0 |
| NC_045512.2;YP_009725295.1;Wu1-SARS2_orf1a;1497 | 0 | 0 |
| NC_045512.2;YP_009725295.1;Wu1-SARS2_orf1a;1498 | 0 | 0 |
| NC_045512.2;YP_009725295.1;Wu1-SARS2_orf1a;1499 | 0 | 0 |
| NC_045512.2;YP_009725295.1;Wu1-SARS2_orf1a;1500 | 0 | 0 |
| NC_045512.2;YP_009725295.1;Wu1-SARS2_orf1a;1501 | 0 | 0 |
| NC_045512.2;YP_009725295.1;Wu1-SARS2_orf1a;1502 | 0 | 0 |
| NC_045512.2;YP_009725295.1;Wu1-SARS2_orf1a;1503 | 0 | 0 |
| NC_045512.2;YP_009725295.1;Wu1-SARS2_orf1a;1504 | 0 | 0 |
| NC_045512.2;YP_009725295.1;Wu1-SARS2_orf1a;1505 | 0 | 0 |
| NC_045512.2;YP_009725295.1;Wu1-SARS2_orf1a;1506 | 0.05 | 0.05 |
| NC_045512.2;YP_009725295.1;Wu1-SARS2_orf1a;1507 | 0 | 0.05 |
| NC_045512.2;YP_009725295.1;Wu1-SARS2_orf1a;1508 | 0 | 0.025 |
| NC_045512.2;YP_009725295.1;Wu1-SARS2_orf1a;1509 | 0.05 | 0 |
| NC_045512.2;YP_009725295.1;Wu1-SARS2_orf1a;1510 | 0 | 0 |
| NC_045512.2;YP_009725295.1;Wu1-SARS2_orf1a;1511 | 0 | 0.025 |
| NC_045512.2;YP_009725295.1;Wu1-SARS2_orf1a;1512 | 0 | 0 |
| NC_045512.2;YP_009725295.1;Wu1-SARS2_orf1a;1513 | 0 | 0 |
| NC_045512.2;YP_009725295.1;Wu1-SARS2_orf1a;1514 | 0 | 0 |
| NC_045512.2;YP_009725295.1;Wu1-SARS2_orf1a;1515 | 0 | 0 |
| NC_045512.2;YP_009725295.1;Wu1-SARS2_orf1a;1516 | 0 | 0 |
| NC_045512.2;YP_009725295.1;Wu1-SARS2_orf1a;1517 | 0 | 0 |
| NC_045512.2;YP_009725295.1;Wu1-SARS2_orf1a;1518 | 0 | 0.075 |
| NC_045512.2;YP_009725295.1;Wu1-SARS2_orf1a;1519 | 0 | 0.075 |
| NC_045512.2;YP_009725295.1;Wu1-SARS2_orf1a;1520 | 0 | 0.075 |
| NC_045512.2;YP_009725295.1;Wu1-SARS2_orf1a;1521 | 0 | 0.075 |
| NC_045512.2;YP_009725295.1;Wu1-SARS2_orf1a;1522 | 0 | 0.05 |
| NC_045512.2;YP_009725295.1;Wu1-SARS2_orf1a;1523 | 0 | 0.05 |
| NC_045512.2;YP_009725295.1;Wu1-SARS2_orf1a;1524 | 0.05 | 0.075 |
| NC_045512.2;YP_009725295.1;Wu1-SARS2_orf1a;1525 | 0 | 0.05 |
| NC_045512.2;YP_009725295.1;Wu1-SARS2_orf1a;1526 | 0.05 | 0.075 |
| NC_045512.2;YP_009725295.1;Wu1-SARS2_orf1a;1527 | 0 | 0.025 |
| NC_045512.2;YP_009725295.1;Wu1-SARS2_orf1a;1528 | 0 | 0.025 |
| NC_045512.2;YP_009725295.1;Wu1-SARS2_orf1a;1529 | 0 | 0.025 |
| NC_045512.2;YP_009725295.1;Wu1-SARS2_orf1a;1530 | 0 | 0 |
| NC_045512.2;YP_009725295.1;Wu1-SARS2_orf1a;1531 | 0 | 0 |
| NC_045512.2;YP_009725295.1;Wu1-SARS2_orf1a;1532 | 0 | 0 |
| NC_045512.2;YP_009725295.1;Wu1-SARS2_orf1a;1533 | 0 | 0 |
| NC_045512.2;YP_009725295.1;Wu1-SARS2_orf1a;1534 | 0 | 0 |
| NC_045512.2;YP_009725295.1;Wu1-SARS2_orf1a;1535 | 0 | 0 |
| NC_045512.2;YP_009725295.1;Wu1-SARS2_orf1a;1536 | 0 | 0 |

| | | |
|---|---|---|
| NC_045512.2;YP_009725295.1;Wu1-SARS2_orf1a;1537 | 0 | 0 |
| NC_045512.2;YP_009725295.1;Wu1-SARS2_orf1a;1538 | 0 | 0 |
| NC_045512.2;YP_009725295.1;Wu1-SARS2_orf1a;1539 | 0 | 0 |
| NC_045512.2;YP_009725295.1;Wu1-SARS2_orf1a;1540 | 0 | 0 |
| NC_045512.2;YP_009725295.1;Wu1-SARS2_orf1a;1541 | 0 | 0 |
| NC_045512.2;YP_009725295.1;Wu1-SARS2_orf1a;1542 | 0 | 0.025 |
| NC_045512.2;YP_009725295.1;Wu1-SARS2_orf1a;1543 | 0 | 0.025 |
| NC_045512.2;YP_009725295.1;Wu1-SARS2_orf1a;1544 | 0 | 0.025 |
| NC_045512.2;YP_009725295.1;Wu1-SARS2_orf1a;1545 | 0 | 0.05 |
| NC_045512.2;YP_009725295.1;Wu1-SARS2_orf1a;1546 | 0 | 0.05 |
| NC_045512.2;YP_009725295.1;Wu1-SARS2_orf1a;1547 | 0 | 0.025 |
| NC_045512.2;YP_009725295.1;Wu1-SARS2_orf1a;1548 | 0 | 0 |
| NC_045512.2;YP_009725295.1;Wu1-SARS2_orf1a;1549 | 0 | 0.025 |
| NC_045512.2;YP_009725295.1;Wu1-SARS2_orf1a;1550 | 0 | 0 |
| NC_045512.2;YP_009725295.1;Wu1-SARS2_orf1a;1551 | 0 | 0.05 |
| NC_045512.2;YP_009725295.1;Wu1-SARS2_orf1a;1552 | 0 | 0 |
| NC_045512.2;YP_009725295.1;Wu1-SARS2_orf1a;1553 | 0 | 0 |
| NC_045512.2;YP_009725295.1;Wu1-SARS2_orf1a;1554 | 0 | 0 |
| NC_045512.2;YP_009725295.1;Wu1-SARS2_orf1a;1555 | 0 | 0 |
| NC_045512.2;YP_009725295.1;Wu1-SARS2_orf1a;1556 | 0 | 0 |
| NC_045512.2;YP_009725295.1;Wu1-SARS2_orf1a;1557 | 0 | 0 |
| NC_045512.2;YP_009725295.1;Wu1-SARS2_orf1a;1558 | 0 | 0 |
| NC_045512.2;YP_009725295.1;Wu1-SARS2_orf1a;1559 | 0 | 0 |
| NC_045512.2;YP_009725295.1;Wu1-SARS2_orf1a;1560 | 0 | 0 |
| NC_045512.2;YP_009725295.1;Wu1-SARS2_orf1a;1561 | 0 | 0 |
| NC_045512.2;YP_009725295.1;Wu1-SARS2_orf1a;1562 | 0 | 0 |
| NC_045512.2;YP_009725295.1;Wu1-SARS2_orf1a;1563 | 0 | 0.025 |
| NC_045512.2;YP_009725295.1;Wu1-SARS2_orf1a;1564 | 0 | 0 |
| NC_045512.2;YP_009725295.1;Wu1-SARS2_orf1a;1565 | 0 | 0 |
| NC_045512.2;YP_009725295.1;Wu1-SARS2_orf1a;1566 | 0 | 0 |
| NC_045512.2;YP_009725295.1;Wu1-SARS2_orf1a;1567 | 0 | 0.025 |
| NC_045512.2;YP_009725295.1;Wu1-SARS2_orf1a;1568 | 0 | 0.025 |
| NC_045512.2;YP_009725295.1;Wu1-SARS2_orf1a;1569 | 0 | 0.025 |
| NC_045512.2;YP_009725295.1;Wu1-SARS2_orf1a;1570 | 0 | 0.025 |
| NC_045512.2;YP_009725295.1;Wu1-SARS2_orf1a;1571 | 0 | 0.025 |
| NC_045512.2;YP_009725295.1;Wu1-SARS2_orf1a;1572 | 0 | 0.025 |
| NC_045512.2;YP_009725295.1;Wu1-SARS2_orf1a;1573 | 0.05 | 0.125 |
| NC_045512.2;YP_009725295.1;Wu1-SARS2_orf1a;1574 | 0 | 0.075 |
| NC_045512.2;YP_009725295.1;Wu1-SARS2_orf1a;1575 | 0 | 0.025 |
| NC_045512.2;YP_009725295.1;Wu1-SARS2_orf1a;1576 | 0 | 0.025 |
| NC_045512.2;YP_009725295.1;Wu1-SARS2_orf1a;1577 | 0 | 0.025 |
| NC_045512.2;YP_009725295.1;Wu1-SARS2_orf1a;1578 | 0 | 0.025 |
| NC_045512.2;YP_009725295.1;Wu1-SARS2_orf1a;1579 | 0 | 0.025 |

| | | |
|---|---|---|
| NC_045512.2;YP_009725295.1;Wu1-SARS2_orf1a;1580 | 0 | 0 |
| NC_045512.2;YP_009725295.1;Wu1-SARS2_orf1a;1581 | 0 | 0 |
| NC_045512.2;YP_009725295.1;Wu1-SARS2_orf1a;1582 | 0 | 0 |
| NC_045512.2;YP_009725295.1;Wu1-SARS2_orf1a;1583 | 0 | 0 |
| NC_045512.2;YP_009725295.1;Wu1-SARS2_orf1a;1584 | 0 | 0 |
| NC_045512.2;YP_009725295.1;Wu1-SARS2_orf1a;1585 | 0 | 0 |
| NC_045512.2;YP_009725295.1;Wu1-SARS2_orf1a;1586 | 0 | 0 |
| NC_045512.2;YP_009725295.1;Wu1-SARS2_orf1a;1587 | 0 | 0 |
| NC_045512.2;YP_009725295.1;Wu1-SARS2_orf1a;1588 | 0 | 0 |
| NC_045512.2;YP_009725295.1;Wu1-SARS2_orf1a;1589 | 0 | 0 |
| NC_045512.2;YP_009725295.1;Wu1-SARS2_orf1a;1590 | 0 | 0 |
| NC_045512.2;YP_009725295.1;Wu1-SARS2_orf1a;1591 | 0 | 0.025 |
| NC_045512.2;YP_009725295.1;Wu1-SARS2_orf1a;1592 | 0 | 0.025 |
| NC_045512.2;YP_009725295.1;Wu1-SARS2_orf1a;1593 | 0 | 0.025 |
| NC_045512.2;YP_009725295.1;Wu1-SARS2_orf1a;1594 | 0 | 0.025 |
| NC_045512.2;YP_009725295.1;Wu1-SARS2_orf1a;1595 | 0.15 | 0.125 |
| NC_045512.2;YP_009725295.1;Wu1-SARS2_orf1a;1596 | 0 | 0.025 |
| NC_045512.2;YP_009725295.1;Wu1-SARS2_orf1a;1597 | 0 | 0.025 |
| NC_045512.2;YP_009725295.1;Wu1-SARS2_orf1a;1598 | 0 | 0 |
| NC_045512.2;YP_009725295.1;Wu1-SARS2_orf1a;1599 | 0 | 0 |
| NC_045512.2;YP_009725295.1;Wu1-SARS2_orf1a;1600 | 0 | 0 |
| NC_045512.2;YP_009725295.1;Wu1-SARS2_orf1a;1601 | 0 | 0.025 |
| NC_045512.2;YP_009725295.1;Wu1-SARS2_orf1a;1602 | 0.05 | 0 |
| NC_045512.2;YP_009725295.1;Wu1-SARS2_orf1a;1603 | 0.05 | 0 |
| NC_045512.2;YP_009725295.1;Wu1-SARS2_orf1a;1604 | 0 | 0 |
| NC_045512.2;YP_009725295.1;Wu1-SARS2_orf1a;1605 | 0 | 0 |
| NC_045512.2;YP_009725295.1;Wu1-SARS2_orf1a;1606 | 0 | 0 |
| NC_045512.2;YP_009725295.1;Wu1-SARS2_orf1a;1607 | 0 | 0 |
| NC_045512.2;YP_009725295.1;Wu1-SARS2_orf1a;1608 | 0 | 0 |
| NC_045512.2;YP_009725295.1;Wu1-SARS2_orf1a;1609 | 0 | 0 |
| NC_045512.2;YP_009725295.1;Wu1-SARS2_orf1a;1610 | 0 | 0 |
| NC_045512.2;YP_009725295.1;Wu1-SARS2_orf1a;1611 | 0.05 | 0 |
| NC_045512.2;YP_009725295.1;Wu1-SARS2_orf1a;1612 | 0.05 | 0 |
| NC_045512.2;YP_009725295.1;Wu1-SARS2_orf1a;1613 | 0.05 | 0 |
| NC_045512.2;YP_009725295.1;Wu1-SARS2_orf1a;1614 | 0.05 | 0 |
| NC_045512.2;YP_009725295.1;Wu1-SARS2_orf1a;1615 | 0.05 | 0 |
| NC_045512.2;YP_009725295.1;Wu1-SARS2_orf1a;1616 | 0.05 | 0 |
| NC_045512.2;YP_009725295.1;Wu1-SARS2_orf1a;1617 | 0 | 0 |
| NC_045512.2;YP_009725295.1;Wu1-SARS2_orf1a;1618 | 0 | 0 |
| NC_045512.2;YP_009725295.1;Wu1-SARS2_orf1a;1619 | 0 | 0 |
| NC_045512.2;YP_009725295.1;Wu1-SARS2_orf1a;1620 | 0 | 0 |
| NC_045512.2;YP_009725295.1;Wu1-SARS2_orf1a;1621 | 0 | 0 |
| NC_045512.2;YP_009725295.1;Wu1-SARS2_orf1a;1622 | 0 | 0 |

| | | |
|---|---|---|
| NC_045512.2;YP_009725295.1;Wu1-SARS2_orf1a;1623 | 0 | 0 |
| NC_045512.2;YP_009725295.1;Wu1-SARS2_orf1a;1624 | 0 | 0 |
| NC_045512.2;YP_009725295.1;Wu1-SARS2_orf1a;1625 | 0 | 0 |
| NC_045512.2;YP_009725295.1;Wu1-SARS2_orf1a;1626 | 0 | 0 |
| NC_045512.2;YP_009725295.1;Wu1-SARS2_orf1a;1627 | 0 | 0 |
| NC_045512.2;YP_009725295.1;Wu1-SARS2_orf1a;1628 | 0 | 0 |
| NC_045512.2;YP_009725295.1;Wu1-SARS2_orf1a;1629 | 0 | 0 |
| NC_045512.2;YP_009725295.1;Wu1-SARS2_orf1a;1630 | 0 | 0 |
| NC_045512.2;YP_009725295.1;Wu1-SARS2_orf1a;1631 | 0.05 | 0 |
| NC_045512.2;YP_009725295.1;Wu1-SARS2_orf1a;1632 | 0.05 | 0 |
| NC_045512.2;YP_009725295.1;Wu1-SARS2_orf1a;1633 | 0.05 | 0 |
| NC_045512.2;YP_009725295.1;Wu1-SARS2_orf1a;1634 | 0.05 | 0 |
| NC_045512.2;YP_009725295.1;Wu1-SARS2_orf1a;1635 | 0 | 0 |
| NC_045512.2;YP_009725295.1;Wu1-SARS2_orf1a;1636 | 0 | 0 |
| NC_045512.2;YP_009725295.1;Wu1-SARS2_orf1a;1637 | 0 | 0 |
| NC_045512.2;YP_009725295.1;Wu1-SARS2_orf1a;1638 | 0 | 0 |
| NC_045512.2;YP_009725295.1;Wu1-SARS2_orf1a;1639 | 0.05 | 0 |
| NC_045512.2;YP_009725295.1;Wu1-SARS2_orf1a;1640 | 0 | 0 |
| NC_045512.2;YP_009725295.1;Wu1-SARS2_orf1a;1641 | 0 | 0 |
| NC_045512.2;YP_009725295.1;Wu1-SARS2_orf1a;1642 | 0 | 0 |
| NC_045512.2;YP_009725295.1;Wu1-SARS2_orf1a;1643 | 0 | 0 |
| NC_045512.2;YP_009725295.1;Wu1-SARS2_orf1a;1644 | 0 | 0 |
| NC_045512.2;YP_009725295.1;Wu1-SARS2_orf1a;1645 | 0 | 0 |
| NC_045512.2;YP_009725295.1;Wu1-SARS2_orf1a;1646 | 0 | 0 |
| NC_045512.2;YP_009725295.1;Wu1-SARS2_orf1a;1647 | 0 | 0 |
| NC_045512.2;YP_009725295.1;Wu1-SARS2_orf1a;1648 | 0 | 0 |
| NC_045512.2;YP_009725295.1;Wu1-SARS2_orf1a;1649 | 0 | 0 |
| NC_045512.2;YP_009725295.1;Wu1-SARS2_orf1a;1650 | 0 | 0 |
| NC_045512.2;YP_009725295.1;Wu1-SARS2_orf1a;1651 | 0 | 0 |
| NC_045512.2;YP_009725295.1;Wu1-SARS2_orf1a;1652 | 0 | 0 |
| NC_045512.2;YP_009725295.1;Wu1-SARS2_orf1a;1653 | 0 | 0 |
| NC_045512.2;YP_009725295.1;Wu1-SARS2_orf1a;1654 | 0 | 0 |
| NC_045512.2;YP_009725295.1;Wu1-SARS2_orf1a;1655 | 0 | 0 |
| NC_045512.2;YP_009725295.1;Wu1-SARS2_orf1a;1656 | 0 | 0 |
| NC_045512.2;YP_009725295.1;Wu1-SARS2_orf1a;1657 | 0.05 | 0.025 |
| NC_045512.2;YP_009725295.1;Wu1-SARS2_orf1a;1658 | 0.05 | 0.05 |
| NC_045512.2;YP_009725295.1;Wu1-SARS2_orf1a;1659 | 0.05 | 0.025 |
| NC_045512.2;YP_009725295.1;Wu1-SARS2_orf1a;1660 | 0.05 | 0 |
| NC_045512.2;YP_009725295.1;Wu1-SARS2_orf1a;1661 | 0.05 | 0 |
| NC_045512.2;YP_009725295.1;Wu1-SARS2_orf1a;1662 | 0.05 | 0 |
| NC_045512.2;YP_009725295.1;Wu1-SARS2_orf1a;1663 | 0.05 | 0 |
| NC_045512.2;YP_009725295.1;Wu1-SARS2_orf1a;1664 | 0.05 | 0 |
| NC_045512.2;YP_009725295.1;Wu1-SARS2_orf1a;1665 | 0 | 0 |

| | | |
|---|---|---|
| NC_045512.2;YP_009725295.1;Wu1-SARS2_orf1a;1666 | 0 | 0 |
| NC_045512.2;YP_009725295.1;Wu1-SARS2_orf1a;1667 | 0 | 0 |
| NC_045512.2;YP_009725295.1;Wu1-SARS2_orf1a;1668 | 0 | 0 |
| NC_045512.2;YP_009725295.1;Wu1-SARS2_orf1a;1669 | 0 | 0 |
| NC_045512.2;YP_009725295.1;Wu1-SARS2_orf1a;1670 | 0 | 0 |
| NC_045512.2;YP_009725295.1;Wu1-SARS2_orf1a;1671 | 0 | 0 |
| NC_045512.2;YP_009725295.1;Wu1-SARS2_orf1a;1672 | 0 | 0 |
| NC_045512.2;YP_009725295.1;Wu1-SARS2_orf1a;1673 | 0 | 0 |
| NC_045512.2;YP_009725295.1;Wu1-SARS2_orf1a;1674 | 0 | 0 |
| NC_045512.2;YP_009725295.1;Wu1-SARS2_orf1a;1675 | 0 | 0 |
| NC_045512.2;YP_009725295.1;Wu1-SARS2_orf1a;1676 | 0 | 0 |
| NC_045512.2;YP_009725295.1;Wu1-SARS2_orf1a;1677 | 0 | 0 |
| NC_045512.2;YP_009725295.1;Wu1-SARS2_orf1a;1678 | 0 | 0 |
| NC_045512.2;YP_009725295.1;Wu1-SARS2_orf1a;1679 | 0.05 | 0 |
| NC_045512.2;YP_009725295.1;Wu1-SARS2_orf1a;1680 | 0 | 0 |
| NC_045512.2;YP_009725295.1;Wu1-SARS2_orf1a;1681 | 0 | 0 |
| NC_045512.2;YP_009725295.1;Wu1-SARS2_orf1a;1682 | 0 | 0.025 |
| NC_045512.2;YP_009725295.1;Wu1-SARS2_orf1a;1683 | 0 | 0 |
| NC_045512.2;YP_009725295.1;Wu1-SARS2_orf1a;1684 | 0 | 0 |
| NC_045512.2;YP_009725295.1;Wu1-SARS2_orf1a;1685 | 0 | 0 |
| NC_045512.2;YP_009725295.1;Wu1-SARS2_orf1a;1686 | 0 | 0.05 |
| NC_045512.2;YP_009725295.1;Wu1-SARS2_orf1a;1687 | 0 | 0.025 |
| NC_045512.2;YP_009725295.1;Wu1-SARS2_orf1a;1688 | 0 | 0 |
| NC_045512.2;YP_009725295.1;Wu1-SARS2_orf1a;1689 | 0 | 0 |
| NC_045512.2;YP_009725295.1;Wu1-SARS2_orf1a;1690 | 0 | 0 |
| NC_045512.2;YP_009725295.1;Wu1-SARS2_orf1a;1691 | 0 | 0 |
| NC_045512.2;YP_009725295.1;Wu1-SARS2_orf1a;1692 | 0.05 | 0.025 |
| NC_045512.2;YP_009725295.1;Wu1-SARS2_orf1a;1693 | 0.1 | 0.075 |
| NC_045512.2;YP_009725295.1;Wu1-SARS2_orf1a;1694 | 0.05 | 0.025 |
| NC_045512.2;YP_009725295.1;Wu1-SARS2_orf1a;1695 | 0.15 | 0.025 |
| NC_045512.2;YP_009725295.1;Wu1-SARS2_orf1a;1696 | 0.05 | 0.025 |
| NC_045512.2;YP_009725295.1;Wu1-SARS2_orf1a;1697 | 0.05 | 0 |
| NC_045512.2;YP_009725295.1;Wu1-SARS2_orf1a;1698 | 0 | 0 |
| NC_045512.2;YP_009725295.1;Wu1-SARS2_orf1a;1699 | 0 | 0 |
| NC_045512.2;YP_009725295.1;Wu1-SARS2_orf1a;1700 | 0 | 0 |
| NC_045512.2;YP_009725295.1;Wu1-SARS2_orf1a;1701 | 0 | 0 |
| NC_045512.2;YP_009725295.1;Wu1-SARS2_orf1a;1702 | 0 | 0 |
| NC_045512.2;YP_009725295.1;Wu1-SARS2_orf1a;1703 | 0 | 0 |
| NC_045512.2;YP_009725295.1;Wu1-SARS2_orf1a;1704 | 0 | 0 |
| NC_045512.2;YP_009725295.1;Wu1-SARS2_orf1a;1705 | 0 | 0 |
| NC_045512.2;YP_009725295.1;Wu1-SARS2_orf1a;1706 | 0 | 0 |
| NC_045512.2;YP_009725295.1;Wu1-SARS2_orf1a;1707 | 0 | 0 |
| NC_045512.2;YP_009725295.1;Wu1-SARS2_orf1a;1708 | 0 | 0 |

| | | |
|---|---|---|
| NC_045512.2;YP_009725295.1;Wu1-SARS2_orf1a;1709 | 0 | 0 |
| NC_045512.2;YP_009725295.1;Wu1-SARS2_orf1a;1710 | 0 | 0 |
| NC_045512.2;YP_009725295.1;Wu1-SARS2_orf1a;1711 | 0 | 0.025 |
| NC_045512.2;YP_009725295.1;Wu1-SARS2_orf1a;1712 | 0 | 0.025 |
| NC_045512.2;YP_009725295.1;Wu1-SARS2_orf1a;1713 | 0 | 0.05 |
| NC_045512.2;YP_009725295.1;Wu1-SARS2_orf1a;1714 | 0 | 0.025 |
| NC_045512.2;YP_009725295.1;Wu1-SARS2_orf1a;1715 | 0 | 0 |
| NC_045512.2;YP_009725295.1;Wu1-SARS2_orf1a;1716 | 0 | 0 |
| NC_045512.2;YP_009725295.1;Wu1-SARS2_orf1a;1717 | 0 | 0 |
| NC_045512.2;YP_009725295.1;Wu1-SARS2_orf1a;1718 | 0 | 0 |
| NC_045512.2;YP_009725295.1;Wu1-SARS2_orf1a;1719 | 0 | 0.025 |
| NC_045512.2;YP_009725295.1;Wu1-SARS2_orf1a;1720 | 0 | 0.025 |
| NC_045512.2;YP_009725295.1;Wu1-SARS2_orf1a;1721 | 0 | 0 |
| NC_045512.2;YP_009725295.1;Wu1-SARS2_orf1a;1722 | 0 | 0 |
| NC_045512.2;YP_009725295.1;Wu1-SARS2_orf1a;1723 | 0.05 | 0.025 |
| NC_045512.2;YP_009725295.1;Wu1-SARS2_orf1a;1724 | 0.05 | 0 |
| NC_045512.2;YP_009725295.1;Wu1-SARS2_orf1a;1725 | 0 | 0 |
| NC_045512.2;YP_009725295.1;Wu1-SARS2_orf1a;1726 | 0.05 | 0 |
| NC_045512.2;YP_009725295.1;Wu1-SARS2_orf1a;1727 | 0 | 0 |
| NC_045512.2;YP_009725295.1;Wu1-SARS2_orf1a;1728 | 0 | 0 |
| NC_045512.2;YP_009725295.1;Wu1-SARS2_orf1a;1729 | 0 | 0 |
| NC_045512.2;YP_009725295.1;Wu1-SARS2_orf1a;1730 | 0 | 0 |
| NC_045512.2;YP_009725295.1;Wu1-SARS2_orf1a;1731 | 0 | 0 |
| NC_045512.2;YP_009725295.1;Wu1-SARS2_orf1a;1732 | 0 | 0 |
| NC_045512.2;YP_009725295.1;Wu1-SARS2_orf1a;1733 | 0 | 0 |
| NC_045512.2;YP_009725295.1;Wu1-SARS2_orf1a;1734 | 0 | 0 |
| NC_045512.2;YP_009725295.1;Wu1-SARS2_orf1a;1735 | 0 | 0 |
| NC_045512.2;YP_009725295.1;Wu1-SARS2_orf1a;1736 | 0 | 0 |
| NC_045512.2;YP_009725295.1;Wu1-SARS2_orf1a;1737 | 0 | 0 |
| NC_045512.2;YP_009725295.1;Wu1-SARS2_orf1a;1738 | 0 | 0 |
| NC_045512.2;YP_009725295.1;Wu1-SARS2_orf1a;1739 | 0 | 0 |
| NC_045512.2;YP_009725295.1;Wu1-SARS2_orf1a;1740 | 0 | 0 |
| NC_045512.2;YP_009725295.1;Wu1-SARS2_orf1a;1741 | 0 | 0 |
| NC_045512.2;YP_009725295.1;Wu1-SARS2_orf1a;1742 | 0 | 0 |
| NC_045512.2;YP_009725295.1;Wu1-SARS2_orf1a;1743 | 0 | 0 |
| NC_045512.2;YP_009725295.1;Wu1-SARS2_orf1a;1744 | 0 | 0 |
| NC_045512.2;YP_009725295.1;Wu1-SARS2_orf1a;1745 | 0 | 0.075 |
| NC_045512.2;YP_009725295.1;Wu1-SARS2_orf1a;1746 | 0 | 0.025 |
| NC_045512.2;YP_009725295.1;Wu1-SARS2_orf1a;1747 | 0 | 0.025 |
| NC_045512.2;YP_009725295.1;Wu1-SARS2_orf1a;1748 | 0.1 | 0.325 |
| NC_045512.2;YP_009725295.1;Wu1-SARS2_orf1a;1749 | 0.1 | 0.3 |
| NC_045512.2;YP_009725295.1;Wu1-SARS2_orf1a;1750 | 0.1 | 0.3 |
| NC_045512.2;YP_009725295.1;Wu1-SARS2_orf1a;1751 | 0.15 | 0.325 |

| | | |
|---|---|---|
| NC_045512.2;YP_009725295.1;Wu1-SARS2_orf1a;1752 | 0.2 | 0.325 |
| NC_045512.2;YP_009725295.1;Wu1-SARS2_orf1a;1753 | 0.1 | 0.35 |
| NC_045512.2;YP_009725295.1;Wu1-SARS2_orf1a;1754 | 0.05 | 0.275 |
| NC_045512.2;YP_009725295.1;Wu1-SARS2_orf1a;1755 | 0.05 | 0.2 |
| NC_045512.2;YP_009725295.1;Wu1-SARS2_orf1a;1756 | 0.1 | 0.25 |
| NC_045512.2;YP_009725295.1;Wu1-SARS2_orf1a;1757 | 0.05 | 0.2 |
| NC_045512.2;YP_009725295.1;Wu1-SARS2_orf1a;1758 | 0 | 0.125 |
| NC_045512.2;YP_009725295.1;Wu1-SARS2_orf1a;1759 | 0 | 0 |
| NC_045512.2;YP_009725295.1;Wu1-SARS2_orf1a;1760 | 0 | 0 |
| NC_045512.2;YP_009725295.1;Wu1-SARS2_orf1a;1761 | 0.1 | 0 |
| NC_045512.2;YP_009725295.1;Wu1-SARS2_orf1a;1762 | 0.15 | 0 |
| NC_045512.2;YP_009725295.1;Wu1-SARS2_orf1a;1763 | 0.15 | 0 |
| NC_045512.2;YP_009725295.1;Wu1-SARS2_orf1a;1764 | 0.1 | 0 |
| NC_045512.2;YP_009725295.1;Wu1-SARS2_orf1a;1765 | 0.1 | 0.05 |
| NC_045512.2;YP_009725295.1;Wu1-SARS2_orf1a;1766 | 0.1 | 0.05 |
| NC_045512.2;YP_009725295.1;Wu1-SARS2_orf1a;1767 | 0.05 | 0.025 |
| NC_045512.2;YP_009725295.1;Wu1-SARS2_orf1a;1768 | 0.05 | 0.025 |
| NC_045512.2;YP_009725295.1;Wu1-SARS2_orf1a;1769 | 0.15 | 0.05 |
| NC_045512.2;YP_009725295.1;Wu1-SARS2_orf1a;1770 | 0.05 | 0.05 |
| NC_045512.2;YP_009725295.1;Wu1-SARS2_orf1a;1771 | 0.15 | 0.05 |
| NC_045512.2;YP_009725295.1;Wu1-SARS2_orf1a;1772 | 0.15 | 0.075 |
| NC_045512.2;YP_009725295.1;Wu1-SARS2_orf1a;1773 | 0.15 | 0.05 |
| NC_045512.2;YP_009725295.1;Wu1-SARS2_orf1a;1774 | 0.15 | 0.075 |
| NC_045512.2;YP_009725295.1;Wu1-SARS2_orf1a;1775 | 0.15 | 0.1 |
| NC_045512.2;YP_009725295.1;Wu1-SARS2_orf1a;1776 | 0 | 0 |
| NC_045512.2;YP_009725295.1;Wu1-SARS2_orf1a;1777 | 0 | 0 |
| NC_045512.2;YP_009725295.1;Wu1-SARS2_orf1a;1778 | 0 | 0 |
| NC_045512.2;YP_009725295.1;Wu1-SARS2_orf1a;1779 | 0 | 0 |
| NC_045512.2;YP_009725295.1;Wu1-SARS2_orf1a;1780 | 0 | 0.025 |
| NC_045512.2;YP_009725295.1;Wu1-SARS2_orf1a;1781 | 0 | 0.025 |
| NC_045512.2;YP_009725295.1;Wu1-SARS2_orf1a;1782 | 0 | 0.025 |
| NC_045512.2;YP_009725295.1;Wu1-SARS2_orf1a;1783 | 0 | 0.025 |
| NC_045512.2;YP_009725295.1;Wu1-SARS2_orf1a;1784 | 0 | 0.025 |
| NC_045512.2;YP_009725295.1;Wu1-SARS2_orf1a;1785 | 0 | 0.025 |
| NC_045512.2;YP_009725295.1;Wu1-SARS2_orf1a;1786 | 0.05 | 0.075 |
| NC_045512.2;YP_009725295.1;Wu1-SARS2_orf1a;1787 | 0 | 0.075 |
| NC_045512.2;YP_009725295.1;Wu1-SARS2_orf1a;1788 | 0 | 0.05 |
| NC_045512.2;YP_009725295.1;Wu1-SARS2_orf1a;1789 | 0 | 0.05 |
| NC_045512.2;YP_009725295.1;Wu1-SARS2_orf1a;1790 | 0.05 | 0.05 |
| NC_045512.2;YP_009725295.1;Wu1-SARS2_orf1a;1791 | 0 | 0.025 |
| NC_045512.2;YP_009725295.1;Wu1-SARS2_orf1a;1792 | 0 | 0.025 |
| NC_045512.2;YP_009725295.1;Wu1-SARS2_orf1a;1793 | 0.05 | 0.05 |
| NC_045512.2;YP_009725295.1;Wu1-SARS2_orf1a;1794 | 0 | 0.05 |

| | | |
|---|---|---|
| NC_045512.2;YP_009725295.1;Wu1-SARS2_orf1a;1795 | 0.05 | 0.025 |
| NC_045512.2;YP_009725295.1;Wu1-SARS2_orf1a;1796 | 0 | 0 |
| NC_045512.2;YP_009725295.1;Wu1-SARS2_orf1a;1797 | 0.1 | 0 |
| NC_045512.2;YP_009725295.1;Wu1-SARS2_orf1a;1798 | 0 | 0.025 |
| NC_045512.2;YP_009725295.1;Wu1-SARS2_orf1a;1799 | 0 | 0 |
| NC_045512.2;YP_009725295.1;Wu1-SARS2_orf1a;1800 | 0 | 0 |
| NC_045512.2;YP_009725295.1;Wu1-SARS2_orf1a;1801 | 0 | 0.025 |
| NC_045512.2;YP_009725295.1;Wu1-SARS2_orf1a;1802 | 0 | 0 |
| NC_045512.2;YP_009725295.1;Wu1-SARS2_orf1a;1803 | 0 | 0 |
| NC_045512.2;YP_009725295.1;Wu1-SARS2_orf1a;1804 | 0 | 0 |
| NC_045512.2;YP_009725295.1;Wu1-SARS2_orf1a;1805 | 0 | 0 |
| NC_045512.2;YP_009725295.1;Wu1-SARS2_orf1a;1806 | 0 | 0 |
| NC_045512.2;YP_009725295.1;Wu1-SARS2_orf1a;1807 | 0 | 0 |
| NC_045512.2;YP_009725295.1;Wu1-SARS2_orf1a;1808 | 0 | 0 |
| NC_045512.2;YP_009725295.1;Wu1-SARS2_orf1a;1809 | 0 | 0 |
| NC_045512.2;YP_009725295.1;Wu1-SARS2_orf1a;1810 | 0.05 | 0.025 |
| NC_045512.2;YP_009725295.1;Wu1-SARS2_orf1a;1811 | 0 | 0 |
| NC_045512.2;YP_009725295.1;Wu1-SARS2_orf1a;1812 | 0 | 0 |
| NC_045512.2;YP_009725295.1;Wu1-SARS2_orf1a;1813 | 0 | 0 |
| NC_045512.2;YP_009725295.1;Wu1-SARS2_orf1a;1814 | 0 | 0 |
| NC_045512.2;YP_009725295.1;Wu1-SARS2_orf1a;1815 | 0 | 0 |
| NC_045512.2;YP_009725295.1;Wu1-SARS2_orf1a;1816 | 0 | 0.025 |
| NC_045512.2;YP_009725295.1;Wu1-SARS2_orf1a;1817 | 0.1 | 0.075 |
| NC_045512.2;YP_009725295.1;Wu1-SARS2_orf1a;1818 | 0 | 0 |
| NC_045512.2;YP_009725295.1;Wu1-SARS2_orf1a;1819 | 0 | 0 |
| NC_045512.2;YP_009725295.1;Wu1-SARS2_orf1a;1820 | 0 | 0 |
| NC_045512.2;YP_009725295.1;Wu1-SARS2_orf1a;1821 | 0 | 0 |
| NC_045512.2;YP_009725295.1;Wu1-SARS2_orf1a;1822 | 0 | 0 |
| NC_045512.2;YP_009725295.1;Wu1-SARS2_orf1a;1823 | 0 | 0 |
| NC_045512.2;YP_009725295.1;Wu1-SARS2_orf1a;1824 | 0 | 0 |
| NC_045512.2;YP_009725295.1;Wu1-SARS2_orf1a;1825 | 0 | 0 |
| NC_045512.2;YP_009725295.1;Wu1-SARS2_orf1a;1826 | 0 | 0.15 |
| NC_045512.2;YP_009725295.1;Wu1-SARS2_orf1a;1827 | 0 | 0 |
| NC_045512.2;YP_009725295.1;Wu1-SARS2_orf1a;1828 | 0 | 0 |
| NC_045512.2;YP_009725295.1;Wu1-SARS2_orf1a;1829 | 0 | 0 |
| NC_045512.2;YP_009725295.1;Wu1-SARS2_orf1a;1830 | 0 | 0 |
| NC_045512.2;YP_009725295.1;Wu1-SARS2_orf1a;1831 | 0.05 | 0 |
| NC_045512.2;YP_009725295.1;Wu1-SARS2_orf1a;1832 | 0.05 | 0 |
| NC_045512.2;YP_009725295.1;Wu1-SARS2_orf1a;1833 | 0 | 0 |
| NC_045512.2;YP_009725295.1;Wu1-SARS2_orf1a;1834 | 0 | 0 |
| NC_045512.2;YP_009725295.1;Wu1-SARS2_orf1a;1835 | 0 | 0 |
| NC_045512.2;YP_009725295.1;Wu1-SARS2_orf1a;1836 | 0 | 0 |
| NC_045512.2;YP_009725295.1;Wu1-SARS2_orf1a;1837 | 0 | 0 |

| | | |
|---|---|---|
| NC_045512.2;YP_009725295.1;Wu1-SARS2_orf1a;1838 | 0 | 0 |
| NC_045512.2;YP_009725295.1;Wu1-SARS2_orf1a;1839 | 0 | 0 |
| NC_045512.2;YP_009725295.1;Wu1-SARS2_orf1a;1840 | 0 | 0 |
| NC_045512.2;YP_009725295.1;Wu1-SARS2_orf1a;1841 | 0 | 0 |
| NC_045512.2;YP_009725295.1;Wu1-SARS2_orf1a;1842 | 0 | 0 |
| NC_045512.2;YP_009725295.1;Wu1-SARS2_orf1a;1843 | 0 | 0 |
| NC_045512.2;YP_009725295.1;Wu1-SARS2_orf1a;1844 | 0 | 0 |
| NC_045512.2;YP_009725295.1;Wu1-SARS2_orf1a;1845 | 0 | 0 |
| NC_045512.2;YP_009725295.1;Wu1-SARS2_orf1a;1846 | 0 | 0 |
| NC_045512.2;YP_009725295.1;Wu1-SARS2_orf1a;1847 | 0 | 0.025 |
| NC_045512.2;YP_009725295.1;Wu1-SARS2_orf1a;1848 | 0 | 0 |
| NC_045512.2;YP_009725295.1;Wu1-SARS2_orf1a;1849 | 0 | 0.025 |
| NC_045512.2;YP_009725295.1;Wu1-SARS2_orf1a;1850 | 0 | 0.075 |
| NC_045512.2;YP_009725295.1;Wu1-SARS2_orf1a;1851 | 0 | 0.05 |
| NC_045512.2;YP_009725295.1;Wu1-SARS2_orf1a;1852 | 0 | 0.025 |
| NC_045512.2;YP_009725295.1;Wu1-SARS2_orf1a;1853 | 0 | 0.025 |
| NC_045512.2;YP_009725295.1;Wu1-SARS2_orf1a;1854 | 0.05 | 0.075 |
| NC_045512.2;YP_009725295.1;Wu1-SARS2_orf1a;1855 | 0.05 | 0.1 |
| NC_045512.2;YP_009725295.1;Wu1-SARS2_orf1a;1856 | 0.1 | 0.125 |
| NC_045512.2;YP_009725295.1;Wu1-SARS2_orf1a;1857 | 0.1 | 0.1 |
| NC_045512.2;YP_009725295.1;Wu1-SARS2_orf1a;1858 | 0.1 | 0.05 |
| NC_045512.2;YP_009725295.1;Wu1-SARS2_orf1a;1859 | 0.05 | 0.075 |
| NC_045512.2;YP_009725295.1;Wu1-SARS2_orf1a;1860 | 0.05 | 0.1 |
| NC_045512.2;YP_009725295.1;Wu1-SARS2_orf1a;1861 | 0.25 | 0.15 |
| NC_045512.2;YP_009725295.1;Wu1-SARS2_orf1a;1862 | 0 | 0 |
| NC_045512.2;YP_009725295.1;Wu1-SARS2_orf1a;1863 | 0 | 0.025 |
| NC_045512.2;YP_009725295.1;Wu1-SARS2_orf1a;1864 | 0 | 0.025 |
| NC_045512.2;YP_009725295.1;Wu1-SARS2_orf1a;1865 | 0.1 | 0.025 |
| NC_045512.2;YP_009725295.1;Wu1-SARS2_orf1a;1866 | 0 | 0.025 |
| NC_045512.2;YP_009725295.1;Wu1-SARS2_orf1a;1867 | 0 | 0.025 |
| NC_045512.2;YP_009725295.1;Wu1-SARS2_orf1a;1868 | 0.05 | 0.025 |
| NC_045512.2;YP_009725295.1;Wu1-SARS2_orf1a;1869 | 0 | 0.025 |
| NC_045512.2;YP_009725295.1;Wu1-SARS2_orf1a;1870 | 0 | 0.025 |
| NC_045512.2;YP_009725295.1;Wu1-SARS2_orf1a;1871 | 0 | 0.025 |
| NC_045512.2;YP_009725295.1;Wu1-SARS2_orf1a;1872 | 0 | 0.025 |
| NC_045512.2;YP_009725295.1;Wu1-SARS2_orf1a;1873 | 0 | 0 |
| NC_045512.2;YP_009725295.1;Wu1-SARS2_orf1a;1874 | 0 | 0 |
| NC_045512.2;YP_009725295.1;Wu1-SARS2_orf1a;1875 | 0 | 0 |
| NC_045512.2;YP_009725295.1;Wu1-SARS2_orf1a;1876 | 0 | 0 |
| NC_045512.2;YP_009725295.1;Wu1-SARS2_orf1a;1877 | 0 | 0 |
| NC_045512.2;YP_009725295.1;Wu1-SARS2_orf1a;1878 | 0 | 0 |
| NC_045512.2;YP_009725295.1;Wu1-SARS2_orf1a;1879 | 0 | 0 |
| NC_045512.2;YP_009725295.1;Wu1-SARS2_orf1a;1880 | 0 | 0 |

| | | |
|---|---|---|
| NC_045512.2;YP_009725295.1;Wu1-SARS2_orf1a;1881 | 0 | 0.025 |
| NC_045512.2;YP_009725295.1;Wu1-SARS2_orf1a;1882 | 0 | 0 |
| NC_045512.2;YP_009725295.1;Wu1-SARS2_orf1a;1883 | 0 | 0 |
| NC_045512.2;YP_009725295.1;Wu1-SARS2_orf1a;1884 | 0 | 0 |
| NC_045512.2;YP_009725295.1;Wu1-SARS2_orf1a;1885 | 0 | 0 |
| NC_045512.2;YP_009725295.1;Wu1-SARS2_orf1a;1886 | 0.05 | 0.125 |
| NC_045512.2;YP_009725295.1;Wu1-SARS2_orf1a;1887 | 0 | 0.1 |
| NC_045512.2;YP_009725295.1;Wu1-SARS2_orf1a;1888 | 0 | 0.15 |
| NC_045512.2;YP_009725295.1;Wu1-SARS2_orf1a;1889 | 0 | 0.175 |
| NC_045512.2;YP_009725295.1;Wu1-SARS2_orf1a;1890 | 0.2 | 0.25 |
| NC_045512.2;YP_009725295.1;Wu1-SARS2_orf1a;1891 | 0.15 | 0.35 |
| NC_045512.2;YP_009725295.1;Wu1-SARS2_orf1a;1892 | 0.1 | 0.225 |
| NC_045512.2;YP_009725295.1;Wu1-SARS2_orf1a;1893 | 0.05 | 0.175 |
| NC_045512.2;YP_009725295.1;Wu1-SARS2_orf1a;1894 | 0.3 | 0.175 |
| NC_045512.2;YP_009725295.1;Wu1-SARS2_orf1a;1895 | 0.35 | 0.175 |
| NC_045512.2;YP_009725295.1;Wu1-SARS2_orf1a;1896 | 0.2 | 0.175 |
| NC_045512.2;YP_009725295.1;Wu1-SARS2_orf1a;1897 | 0.3 | 0.1 |
| NC_045512.2;YP_009725295.1;Wu1-SARS2_orf1a;1898 | 0.15 | 0.075 |
| NC_045512.2;YP_009725295.1;Wu1-SARS2_orf1a;1899 | 0.2 | 0.05 |
| NC_045512.2;YP_009725295.1;Wu1-SARS2_orf1a;1900 | 0.15 | 0.05 |
| NC_045512.2;YP_009725295.1;Wu1-SARS2_orf1a;1901 | 0.15 | 0.15 |
| NC_045512.2;YP_009725295.1;Wu1-SARS2_orf1a;1902 | 0 | 0.025 |
| NC_045512.2;YP_009725295.1;Wu1-SARS2_orf1a;1903 | 0.05 | 0 |
| NC_045512.2;YP_009725295.1;Wu1-SARS2_orf1a;1904 | 0.05 | 0 |
| NC_045512.2;YP_009725295.1;Wu1-SARS2_orf1a;1905 | 0.15 | 0.2 |
| NC_045512.2;YP_009725295.1;Wu1-SARS2_orf1a;1906 | 0.05 | 0 |
| NC_045512.2;YP_009725295.1;Wu1-SARS2_orf1a;1907 | 0.05 | 0 |
| NC_045512.2;YP_009725295.1;Wu1-SARS2_orf1a;1908 | 0.1 | 0 |
| NC_045512.2;YP_009725295.1;Wu1-SARS2_orf1a;1909 | 0 | 0.025 |
| NC_045512.2;YP_009725295.1;Wu1-SARS2_orf1a;1910 | 0.2 | 0.1 |
| NC_045512.2;YP_009725295.1;Wu1-SARS2_orf1a;1911 | 0 | 0.075 |
| NC_045512.2;YP_009725295.1;Wu1-SARS2_orf1a;1912 | 0 | 0.05 |
| NC_045512.2;YP_009725295.1;Wu1-SARS2_orf1a;1913 | 0 | 0.05 |
| NC_045512.2;YP_009725295.1;Wu1-SARS2_orf1a;1914 | 0.05 | 0.075 |
| NC_045512.2;YP_009725295.1;Wu1-SARS2_orf1a;1915 | 0.05 | 0.05 |
| NC_045512.2;YP_009725295.1;Wu1-SARS2_orf1a;1916 | 0.05 | 0.025 |
| NC_045512.2;YP_009725295.1;Wu1-SARS2_orf1a;1917 | 0.05 | 0.025 |
| NC_045512.2;YP_009725295.1;Wu1-SARS2_orf1a;1918 | 0.05 | 0.025 |
| NC_045512.2;YP_009725295.1;Wu1-SARS2_orf1a;1919 | 0.05 | 0.025 |
| NC_045512.2;YP_009725295.1;Wu1-SARS2_orf1a;1920 | 0.05 | 0 |
| NC_045512.2;YP_009725295.1;Wu1-SARS2_orf1a;1921 | 0.05 | 0 |
| NC_045512.2;YP_009725295.1;Wu1-SARS2_orf1a;1922 | 0 | 0 |
| NC_045512.2;YP_009725295.1;Wu1-SARS2_orf1a;1923 | 0 | 0.025 |

| | | |
|---|---|---|
| NC_045512.2;YP_009725295.1;Wu1-SARS2_orf1a;1924 | 0.05 | 0 |
| NC_045512.2;YP_009725295.1;Wu1-SARS2_orf1a;1925 | 0 | 0 |
| NC_045512.2;YP_009725295.1;Wu1-SARS2_orf1a;1926 | 0 | 0 |
| NC_045512.2;YP_009725295.1;Wu1-SARS2_orf1a;1927 | 0.2 | 0.025 |
| NC_045512.2;YP_009725295.1;Wu1-SARS2_orf1a;1928 | 0.2 | 0 |
| NC_045512.2;YP_009725295.1;Wu1-SARS2_orf1a;1929 | 0.2 | 0.05 |
| NC_045512.2;YP_009725295.1;Wu1-SARS2_orf1a;1930 | 0.3 | 0.025 |
| NC_045512.2;YP_009725295.1;Wu1-SARS2_orf1a;1931 | 0.2 | 0 |
| NC_045512.2;YP_009725295.1;Wu1-SARS2_orf1a;1932 | 0.25 | 0 |
| NC_045512.2;YP_009725295.1;Wu1-SARS2_orf1a;1933 | 0.15 | 0.075 |
| NC_045512.2;YP_009725295.1;Wu1-SARS2_orf1a;1934 | 0.1 | 0.025 |
| NC_045512.2;YP_009725295.1;Wu1-SARS2_orf1a;1935 | 0.1 | 0.05 |
| NC_045512.2;YP_009725295.1;Wu1-SARS2_orf1a;1936 | 0.1 | 0 |
| NC_045512.2;YP_009725295.1;Wu1-SARS2_orf1a;1937 | 0.05 | 0 |
| NC_045512.2;YP_009725295.1;Wu1-SARS2_orf1a;1938 | 0.1 | 0.025 |
| NC_045512.2;YP_009725295.1;Wu1-SARS2_orf1a;1939 | 0.05 | 0.05 |
| NC_045512.2;YP_009725295.1;Wu1-SARS2_orf1a;1940 | 0.05 | 0 |
| NC_045512.2;YP_009725295.1;Wu1-SARS2_orf1a;1941 | 0 | 0 |
| NC_045512.2;YP_009725295.1;Wu1-SARS2_orf1a;1942 | 0 | 0 |
| NC_045512.2;YP_009725295.1;Wu1-SARS2_orf1a;1943 | 0 | 0 |
| NC_045512.2;YP_009725295.1;Wu1-SARS2_orf1a;1944 | 0 | 0 |
| NC_045512.2;YP_009725295.1;Wu1-SARS2_orf1a;1945 | 0 | 0 |
| NC_045512.2;YP_009725295.1;Wu1-SARS2_orf1a;1946 | 0 | 0 |
| NC_045512.2;YP_009725295.1;Wu1-SARS2_orf1a;1947 | 0 | 0 |
| NC_045512.2;YP_009725295.1;Wu1-SARS2_orf1a;1948 | 0 | 0 |
| NC_045512.2;YP_009725295.1;Wu1-SARS2_orf1a;1949 | 0 | 0 |
| NC_045512.2;YP_009725295.1;Wu1-SARS2_orf1a;1950 | 0 | 0 |
| NC_045512.2;YP_009725295.1;Wu1-SARS2_orf1a;1951 | 0 | 0 |
| NC_045512.2;YP_009725295.1;Wu1-SARS2_orf1a;1952 | 0 | 0.025 |
| NC_045512.2;YP_009725295.1;Wu1-SARS2_orf1a;1953 | 0 | 0.025 |
| NC_045512.2;YP_009725295.1;Wu1-SARS2_orf1a;1954 | 0 | 0.05 |
| NC_045512.2;YP_009725295.1;Wu1-SARS2_orf1a;1955 | 0 | 0 |
| NC_045512.2;YP_009725295.1;Wu1-SARS2_orf1a;1956 | 0 | 0.025 |
| NC_045512.2;YP_009725295.1;Wu1-SARS2_orf1a;1957 | 0 | 0 |
| NC_045512.2;YP_009725295.1;Wu1-SARS2_orf1a;1958 | 0 | 0.025 |
| NC_045512.2;YP_009725295.1;Wu1-SARS2_orf1a;1959 | 0.05 | 0.025 |
| NC_045512.2;YP_009725295.1;Wu1-SARS2_orf1a;1960 | 0.05 | 0.05 |
| NC_045512.2;YP_009725295.1;Wu1-SARS2_orf1a;1961 | 0.05 | 0.075 |
| NC_045512.2;YP_009725295.1;Wu1-SARS2_orf1a;1962 | 0.05 | 0.075 |
| NC_045512.2;YP_009725295.1;Wu1-SARS2_orf1a;1963 | 0.05 | 0.025 |
| NC_045512.2;YP_009725295.1;Wu1-SARS2_orf1a;1964 | 0.05 | 0.025 |
| NC_045512.2;YP_009725295.1;Wu1-SARS2_orf1a;1965 | 0.05 | 0.075 |
| NC_045512.2;YP_009725295.1;Wu1-SARS2_orf1a;1966 | 0 | 0.025 |

| | | |
|---|---|---|
| NC_045512.2;YP_009725295.1;Wu1-SARS2_orf1a;1967 | 0 | 0.025 |
| NC_045512.2;YP_009725295.1;Wu1-SARS2_orf1a;1968 | 0 | 0.05 |
| NC_045512.2;YP_009725295.1;Wu1-SARS2_orf1a;1969 | 0 | 0.075 |
| NC_045512.2;YP_009725295.1;Wu1-SARS2_orf1a;1970 | 0 | 0 |
| NC_045512.2;YP_009725295.1;Wu1-SARS2_orf1a;1971 | 0 | 0 |
| NC_045512.2;YP_009725295.1;Wu1-SARS2_orf1a;1972 | 0 | 0 |
| NC_045512.2;YP_009725295.1;Wu1-SARS2_orf1a;1973 | 0 | 0 |
| NC_045512.2;YP_009725295.1;Wu1-SARS2_orf1a;1974 | 0 | 0 |
| NC_045512.2;YP_009725295.1;Wu1-SARS2_orf1a;1975 | 0 | 0.025 |
| NC_045512.2;YP_009725295.1;Wu1-SARS2_orf1a;1976 | 0 | 0 |
| NC_045512.2;YP_009725295.1;Wu1-SARS2_orf1a;1977 | 0 | 0 |
| NC_045512.2;YP_009725295.1;Wu1-SARS2_orf1a;1978 | 0 | 0 |
| NC_045512.2;YP_009725295.1;Wu1-SARS2_orf1a;1979 | 0 | 0 |
| NC_045512.2;YP_009725295.1;Wu1-SARS2_orf1a;1980 | 0 | 0 |
| NC_045512.2;YP_009725295.1;Wu1-SARS2_orf1a;1981 | 0 | 0 |
| NC_045512.2;YP_009725295.1;Wu1-SARS2_orf1a;1982 | 0 | 0 |
| NC_045512.2;YP_009725295.1;Wu1-SARS2_orf1a;1983 | 0 | 0 |
| NC_045512.2;YP_009725295.1;Wu1-SARS2_orf1a;1984 | 0.05 | 0 |
| NC_045512.2;YP_009725295.1;Wu1-SARS2_orf1a;1985 | 0.05 | 0 |
| NC_045512.2;YP_009725295.1;Wu1-SARS2_orf1a;1986 | 0 | 0 |
| NC_045512.2;YP_009725295.1;Wu1-SARS2_orf1a;1987 | 0.05 | 0 |
| NC_045512.2;YP_009725295.1;Wu1-SARS2_orf1a;1988 | 0.05 | 0.05 |
| NC_045512.2;YP_009725295.1;Wu1-SARS2_orf1a;1989 | 0.1 | 0.025 |
| NC_045512.2;YP_009725295.1;Wu1-SARS2_orf1a;1990 | 0.05 | 0.075 |
| NC_045512.2;YP_009725295.1;Wu1-SARS2_orf1a;1991 | 0.1 | 0.1 |
| NC_045512.2;YP_009725295.1;Wu1-SARS2_orf1a;1992 | 0.15 | 0.15 |
| NC_045512.2;YP_009725295.1;Wu1-SARS2_orf1a;1993 | 0.15 | 0.15 |
| NC_045512.2;YP_009725295.1;Wu1-SARS2_orf1a;1994 | 0.25 | 0.1 |
| NC_045512.2;YP_009725295.1;Wu1-SARS2_orf1a;1995 | 0.15 | 0.075 |
| NC_045512.2;YP_009725295.1;Wu1-SARS2_orf1a;1996 | 0.05 | 0.025 |
| NC_045512.2;YP_009725295.1;Wu1-SARS2_orf1a;1997 | 0.05 | 0.025 |
| NC_045512.2;YP_009725295.1;Wu1-SARS2_orf1a;1998 | 0 | 0 |
| NC_045512.2;YP_009725295.1;Wu1-SARS2_orf1a;1999 | 0 | 0 |
| NC_045512.2;YP_009725295.1;Wu1-SARS2_orf1a;2000 | 0 | 0 |
| NC_045512.2;YP_009725295.1;Wu1-SARS2_orf1a;2001 | 0 | 0 |
| NC_045512.2;YP_009725295.1;Wu1-SARS2_orf1a;2002 | 0 | 0 |
| NC_045512.2;YP_009725295.1;Wu1-SARS2_orf1a;2003 | 0 | 0 |
| NC_045512.2;YP_009725295.1;Wu1-SARS2_orf1a;2004 | 0 | 0 |
| NC_045512.2;YP_009725295.1;Wu1-SARS2_orf1a;2005 | 0 | 0 |
| NC_045512.2;YP_009725295.1;Wu1-SARS2_orf1a;2006 | 0 | 0 |
| NC_045512.2;YP_009725295.1;Wu1-SARS2_orf1a;2007 | 0 | 0 |
| NC_045512.2;YP_009725295.1;Wu1-SARS2_orf1a;2008 | 0 | 0 |
| NC_045512.2;YP_009725295.1;Wu1-SARS2_orf1a;2009 | 0.05 | 0 |

| | | |
|---|---|---|
| NC_045512.2;YP_009725295.1;Wu1-SARS2_orf1a;2010 | 0.1 | 0.075 |
| NC_045512.2;YP_009725295.1;Wu1-SARS2_orf1a;2011 | 0.2 | 0.1 |
| NC_045512.2;YP_009725295.1;Wu1-SARS2_orf1a;2012 | 0.1 | 0.05 |
| NC_045512.2;YP_009725295.1;Wu1-SARS2_orf1a;2013 | 0.15 | 0.075 |
| NC_045512.2;YP_009725295.1;Wu1-SARS2_orf1a;2014 | 0.35 | 0.1 |
| NC_045512.2;YP_009725295.1;Wu1-SARS2_orf1a;2015 | 0.25 | 0.1 |
| NC_045512.2;YP_009725295.1;Wu1-SARS2_orf1a;2016 | 0.1 | 0.05 |
| NC_045512.2;YP_009725295.1;Wu1-SARS2_orf1a;2017 | 0.1 | 0.05 |
| NC_045512.2;YP_009725295.1;Wu1-SARS2_orf1a;2018 | 0.15 | 0.125 |
| NC_045512.2;YP_009725295.1;Wu1-SARS2_orf1a;2019 | 0.2 | 0.05 |
| NC_045512.2;YP_009725295.1;Wu1-SARS2_orf1a;2020 | 0.05 | 0 |
| NC_045512.2;YP_009725295.1;Wu1-SARS2_orf1a;2021 | 0 | 0.025 |
| NC_045512.2;YP_009725295.1;Wu1-SARS2_orf1a;2022 | 0 | 0.025 |
| NC_045512.2;YP_009725295.1;Wu1-SARS2_orf1a;2023 | 0.05 | 0.025 |
| NC_045512.2;YP_009725295.1;Wu1-SARS2_orf1a;2024 | 0.05 | 0.025 |
| NC_045512.2;YP_009725295.1;Wu1-SARS2_orf1a;2025 | 0 | 0.025 |
| NC_045512.2;YP_009725295.1;Wu1-SARS2_orf1a;2026 | 0 | 0.025 |
| NC_045512.2;YP_009725295.1;Wu1-SARS2_orf1a;2027 | 0 | 0.025 |
| NC_045512.2;YP_009725295.1;Wu1-SARS2_orf1a;2028 | 0 | 0.025 |
| NC_045512.2;YP_009725295.1;Wu1-SARS2_orf1a;2029 | 0 | 0.025 |
| NC_045512.2;YP_009725295.1;Wu1-SARS2_orf1a;2030 | 0 | 0 |
| NC_045512.2;YP_009725295.1;Wu1-SARS2_orf1a;2031 | 0 | 0 |
| NC_045512.2;YP_009725295.1;Wu1-SARS2_orf1a;2032 | 0 | 0 |
| NC_045512.2;YP_009725295.1;Wu1-SARS2_orf1a;2033 | 0 | 0 |
| NC_045512.2;YP_009725295.1;Wu1-SARS2_orf1a;2034 | 0 | 0 |
| NC_045512.2;YP_009725295.1;Wu1-SARS2_orf1a;2035 | 0 | 0 |
| NC_045512.2;YP_009725295.1;Wu1-SARS2_orf1a;2036 | 0 | 0 |
| NC_045512.2;YP_009725295.1;Wu1-SARS2_orf1a;2037 | 0 | 0 |
| NC_045512.2;YP_009725295.1;Wu1-SARS2_orf1a;2038 | 0 | 0 |
| NC_045512.2;YP_009725295.1;Wu1-SARS2_orf1a;2039 | 0 | 0 |
| NC_045512.2;YP_009725295.1;Wu1-SARS2_orf1a;2040 | 0 | 0 |
| NC_045512.2;YP_009725295.1;Wu1-SARS2_orf1a;2041 | 0 | 0 |
| NC_045512.2;YP_009725295.1;Wu1-SARS2_orf1a;2042 | 0 | 0.025 |
| NC_045512.2;YP_009725295.1;Wu1-SARS2_orf1a;2043 | 0 | 0 |
| NC_045512.2;YP_009725295.1;Wu1-SARS2_orf1a;2044 | 0 | 0.05 |
| NC_045512.2;YP_009725295.1;Wu1-SARS2_orf1a;2045 | 0 | 0.05 |
| NC_045512.2;YP_009725295.1;Wu1-SARS2_orf1a;2046 | 0 | 0.05 |
| NC_045512.2;YP_009725295.1;Wu1-SARS2_orf1a;2047 | 0 | 0.05 |
| NC_045512.2;YP_009725295.1;Wu1-SARS2_orf1a;2048 | 0 | 0.05 |
| NC_045512.2;YP_009725295.1;Wu1-SARS2_orf1a;2049 | 0 | 0.05 |
| NC_045512.2;YP_009725295.1;Wu1-SARS2_orf1a;2050 | 0 | 0.05 |
| NC_045512.2;YP_009725295.1;Wu1-SARS2_orf1a;2051 | 0 | 0.05 |
| NC_045512.2;YP_009725295.1;Wu1-SARS2_orf1a;2052 | 0 | 0.05 |

| | | |
|---|---|---|
| NC_045512.2;YP_009725295.1;Wu1-SARS2_orf1a;2053 | 0 | 0.025 |
| NC_045512.2;YP_009725295.1;Wu1-SARS2_orf1a;2054 | 0 | 0 |
| NC_045512.2;YP_009725295.1;Wu1-SARS2_orf1a;2055 | 0 | 0 |
| NC_045512.2;YP_009725295.1;Wu1-SARS2_orf1a;2056 | 0 | 0 |
| NC_045512.2;YP_009725295.1;Wu1-SARS2_orf1a;2057 | 0 | 0 |
| NC_045512.2;YP_009725295.1;Wu1-SARS2_orf1a;2058 | 0 | 0 |
| NC_045512.2;YP_009725295.1;Wu1-SARS2_orf1a;2059 | 0 | 0 |
| NC_045512.2;YP_009725295.1;Wu1-SARS2_orf1a;2060 | 0 | 0 |
| NC_045512.2;YP_009725295.1;Wu1-SARS2_orf1a;2061 | 0 | 0 |
| NC_045512.2;YP_009725295.1;Wu1-SARS2_orf1a;2062 | 0 | 0 |
| NC_045512.2;YP_009725295.1;Wu1-SARS2_orf1a;2063 | 0 | 0 |
| NC_045512.2;YP_009725295.1;Wu1-SARS2_orf1a;2064 | 0 | 0 |
| NC_045512.2;YP_009725295.1;Wu1-SARS2_orf1a;2065 | 0 | 0 |
| NC_045512.2;YP_009725295.1;Wu1-SARS2_orf1a;2066 | 0 | 0 |
| NC_045512.2;YP_009725295.1;Wu1-SARS2_orf1a;2067 | 0 | 0 |
| NC_045512.2;YP_009725295.1;Wu1-SARS2_orf1a;2068 | 0.05 | 0.025 |
| NC_045512.2;YP_009725295.1;Wu1-SARS2_orf1a;2069 | 0.05 | 0.05 |
| NC_045512.2;YP_009725295.1;Wu1-SARS2_orf1a;2070 | 0 | 0 |
| NC_045512.2;YP_009725295.1;Wu1-SARS2_orf1a;2071 | 0 | 0 |
| NC_045512.2;YP_009725295.1;Wu1-SARS2_orf1a;2072 | 0 | 0 |
| NC_045512.2;YP_009725295.1;Wu1-SARS2_orf1a;2073 | 0 | 0.025 |
| NC_045512.2;YP_009725295.1;Wu1-SARS2_orf1a;2074 | 0 | 0 |
| NC_045512.2;YP_009725295.1;Wu1-SARS2_orf1a;2075 | 0 | 0 |
| NC_045512.2;YP_009725295.1;Wu1-SARS2_orf1a;2076 | 0 | 0 |
| NC_045512.2;YP_009725295.1;Wu1-SARS2_orf1a;2077 | 0 | 0 |
| NC_045512.2;YP_009725295.1;Wu1-SARS2_orf1a;2078 | 0 | 0 |
| NC_045512.2;YP_009725295.1;Wu1-SARS2_orf1a;2079 | 0 | 0 |
| NC_045512.2;YP_009725295.1;Wu1-SARS2_orf1a;2080 | 0 | 0 |
| NC_045512.2;YP_009725295.1;Wu1-SARS2_orf1a;2081 | 0 | 0 |
| NC_045512.2;YP_009725295.1;Wu1-SARS2_orf1a;2082 | 0 | 0 |
| NC_045512.2;YP_009725295.1;Wu1-SARS2_orf1a;2083 | 0.05 | 0.025 |
| NC_045512.2;YP_009725295.1;Wu1-SARS2_orf1a;2084 | 0 | 0 |
| NC_045512.2;YP_009725295.1;Wu1-SARS2_orf1a;2085 | 0 | 0 |
| NC_045512.2;YP_009725295.1;Wu1-SARS2_orf1a;2086 | 0 | 0 |
| NC_045512.2;YP_009725295.1;Wu1-SARS2_orf1a;2087 | 0 | 0 |
| NC_045512.2;YP_009725295.1;Wu1-SARS2_orf1a;2088 | 0 | 0 |
| NC_045512.2;YP_009725295.1;Wu1-SARS2_orf1a;2089 | 0 | 0 |
| NC_045512.2;YP_009725295.1;Wu1-SARS2_orf1a;2090 | 0 | 0 |
| NC_045512.2;YP_009725295.1;Wu1-SARS2_orf1a;2091 | 0.05 | 0.05 |
| NC_045512.2;YP_009725295.1;Wu1-SARS2_orf1a;2092 | 0 | 0.05 |
| NC_045512.2;YP_009725295.1;Wu1-SARS2_orf1a;2093 | 0 | 0.025 |
| NC_045512.2;YP_009725295.1;Wu1-SARS2_orf1a;2094 | 0 | 0 |
| NC_045512.2;YP_009725295.1;Wu1-SARS2_orf1a;2095 | 0 | 0.025 |

| | | |
|---|---|---|
| NC_045512.2;YP_009725295.1;Wu1-SARS2_orf1a;2096 | 0 | 0.025 |
| NC_045512.2;YP_009725295.1;Wu1-SARS2_orf1a;2097 | 0.05 | 0.025 |
| NC_045512.2;YP_009725295.1;Wu1-SARS2_orf1a;2098 | 0.05 | 0.05 |
| NC_045512.2;YP_009725295.1;Wu1-SARS2_orf1a;2099 | 0.25 | 0.125 |
| NC_045512.2;YP_009725295.1;Wu1-SARS2_orf1a;2100 | 0 | 0.05 |
| NC_045512.2;YP_009725295.1;Wu1-SARS2_orf1a;2101 | 0 | 0.05 |
| NC_045512.2;YP_009725295.1;Wu1-SARS2_orf1a;2102 | 0 | 0 |
| NC_045512.2;YP_009725295.1;Wu1-SARS2_orf1a;2103 | 0 | 0 |
| NC_045512.2;YP_009725295.1;Wu1-SARS2_orf1a;2104 | 0 | 0 |
| NC_045512.2;YP_009725295.1;Wu1-SARS2_orf1a;2105 | 0 | 0 |
| NC_045512.2;YP_009725295.1;Wu1-SARS2_orf1a;2106 | 0 | 0 |
| NC_045512.2;YP_009725295.1;Wu1-SARS2_orf1a;2107 | 0 | 0 |
| NC_045512.2;YP_009725295.1;Wu1-SARS2_orf1a;2108 | 0 | 0 |
| NC_045512.2;YP_009725295.1;Wu1-SARS2_orf1a;2109 | 0 | 0 |
| NC_045512.2;YP_009725295.1;Wu1-SARS2_orf1a;2110 | 0 | 0 |
| NC_045512.2;YP_009725295.1;Wu1-SARS2_orf1a;2111 | 0 | 0 |
| NC_045512.2;YP_009725295.1;Wu1-SARS2_orf1a;2112 | 0 | 0 |
| NC_045512.2;YP_009725295.1;Wu1-SARS2_orf1a;2113 | 0 | 0 |
| NC_045512.2;YP_009725295.1;Wu1-SARS2_orf1a;2114 | 0.1 | 0.025 |
| NC_045512.2;YP_009725295.1;Wu1-SARS2_orf1a;2115 | 0.05 | 0 |
| NC_045512.2;YP_009725295.1;Wu1-SARS2_orf1a;2116 | 0 | 0 |
| NC_045512.2;YP_009725295.1;Wu1-SARS2_orf1a;2117 | 0 | 0 |
| NC_045512.2;YP_009725295.1;Wu1-SARS2_orf1a;2118 | 0.05 | 0 |
| NC_045512.2;YP_009725295.1;Wu1-SARS2_orf1a;2119 | 0.05 | 0 |
| NC_045512.2;YP_009725295.1;Wu1-SARS2_orf1a;2120 | 0 | 0 |
| NC_045512.2;YP_009725295.1;Wu1-SARS2_orf1a;2121 | 0 | 0 |
| NC_045512.2;YP_009725295.1;Wu1-SARS2_orf1a;2122 | 0 | 0 |
| NC_045512.2;YP_009725295.1;Wu1-SARS2_orf1a;2123 | 0 | 0 |
| NC_045512.2;YP_009725295.1;Wu1-SARS2_orf1a;2124 | 0.05 | 0 |
| NC_045512.2;YP_009725295.1;Wu1-SARS2_orf1a;2125 | 0.05 | 0 |
| NC_045512.2;YP_009725295.1;Wu1-SARS2_orf1a;2126 | 0.05 | 0.025 |
| NC_045512.2;YP_009725295.1;Wu1-SARS2_orf1a;2127 | 0.05 | 0 |
| NC_045512.2;YP_009725295.1;Wu1-SARS2_orf1a;2128 | 0.1 | 0 |
| NC_045512.2;YP_009725295.1;Wu1-SARS2_orf1a;2129 | 0.15 | 0.025 |
| NC_045512.2;YP_009725295.1;Wu1-SARS2_orf1a;2130 | 0.1 | 0 |
| NC_045512.2;YP_009725295.1;Wu1-SARS2_orf1a;2131 | 0.05 | 0 |
| NC_045512.2;YP_009725295.1;Wu1-SARS2_orf1a;2132 | 0.1 | 0 |
| NC_045512.2;YP_009725295.1;Wu1-SARS2_orf1a;2133 | 0.1 | 0.05 |
| NC_045512.2;YP_009725295.1;Wu1-SARS2_orf1a;2134 | 0.1 | 0.025 |
| NC_045512.2;YP_009725295.1;Wu1-SARS2_orf1a;2135 | 0.1 | 0.025 |
| NC_045512.2;YP_009725295.1;Wu1-SARS2_orf1a;2136 | 0.2 | 0.075 |
| NC_045512.2;YP_009725295.1;Wu1-SARS2_orf1a;2137 | 0.05 | 0.025 |
| NC_045512.2;YP_009725295.1;Wu1-SARS2_orf1a;2138 | 0.05 | 0.05 |

| | | |
|---|---|---|
| NC_045512.2;YP_009725295.1;Wu1-SARS2_orf1a;2139 | 0.1 | 0.05 |
| NC_045512.2;YP_009725295.1;Wu1-SARS2_orf1a;2140 | 0.05 | 0.05 |
| NC_045512.2;YP_009725295.1;Wu1-SARS2_orf1a;2141 | 0.05 | 0.05 |
| NC_045512.2;YP_009725295.1;Wu1-SARS2_orf1a;2142 | 0.05 | 0.05 |
| NC_045512.2;YP_009725295.1;Wu1-SARS2_orf1a;2143 | 0.1 | 0.025 |
| NC_045512.2;YP_009725295.1;Wu1-SARS2_orf1a;2144 | 0.05 | 0 |
| NC_045512.2;YP_009725295.1;Wu1-SARS2_orf1a;2145 | 0 | 0 |
| NC_045512.2;YP_009725295.1;Wu1-SARS2_orf1a;2146 | 0 | 0 |
| NC_045512.2;YP_009725295.1;Wu1-SARS2_orf1a;2147 | 0 | 0 |
| NC_045512.2;YP_009725295.1;Wu1-SARS2_orf1a;2148 | 0 | 0 |
| NC_045512.2;YP_009725295.1;Wu1-SARS2_orf1a;2149 | 0 | 0 |
| NC_045512.2;YP_009725295.1;Wu1-SARS2_orf1a;2150 | 0 | 0 |
| NC_045512.2;YP_009725295.1;Wu1-SARS2_orf1a;2151 | 0 | 0 |
| NC_045512.2;YP_009725295.1;Wu1-SARS2_orf1a;2152 | 0 | 0 |
| NC_045512.2;YP_009725295.1;Wu1-SARS2_orf1a;2153 | 0 | 0 |
| NC_045512.2;YP_009725295.1;Wu1-SARS2_orf1a;2154 | 0 | 0 |
| NC_045512.2;YP_009725295.1;Wu1-SARS2_orf1a;2155 | 0 | 0 |
| NC_045512.2;YP_009725295.1;Wu1-SARS2_orf1a;2156 | 0 | 0 |
| NC_045512.2;YP_009725295.1;Wu1-SARS2_orf1a;2157 | 0 | 0 |
| NC_045512.2;YP_009725295.1;Wu1-SARS2_orf1a;2158 | 0 | 0 |
| NC_045512.2;YP_009725295.1;Wu1-SARS2_orf1a;2159 | 0 | 0 |
| NC_045512.2;YP_009725295.1;Wu1-SARS2_orf1a;2160 | 0 | 0 |
| NC_045512.2;YP_009725295.1;Wu1-SARS2_orf1a;2161 | 0 | 0 |
| NC_045512.2;YP_009725295.1;Wu1-SARS2_orf1a;2162 | 0 | 0 |
| NC_045512.2;YP_009725295.1;Wu1-SARS2_orf1a;2163 | 0 | 0 |
| NC_045512.2;YP_009725295.1;Wu1-SARS2_orf1a;2164 | 0 | 0 |
| NC_045512.2;YP_009725295.1;Wu1-SARS2_orf1a;2165 | 0 | 0 |
| NC_045512.2;YP_009725295.1;Wu1-SARS2_orf1a;2166 | 0 | 0 |
| NC_045512.2;YP_009725295.1;Wu1-SARS2_orf1a;2167 | 0 | 0 |
| NC_045512.2;YP_009725295.1;Wu1-SARS2_orf1a;2168 | 0 | 0 |
| NC_045512.2;YP_009725295.1;Wu1-SARS2_orf1a;2169 | 0 | 0 |
| NC_045512.2;YP_009725295.1;Wu1-SARS2_orf1a;2170 | 0 | 0 |
| NC_045512.2;YP_009725295.1;Wu1-SARS2_orf1a;2171 | 0 | 0 |
| NC_045512.2;YP_009725295.1;Wu1-SARS2_orf1a;2172 | 0 | 0 |
| NC_045512.2;YP_009725295.1;Wu1-SARS2_orf1a;2173 | 0 | 0 |
| NC_045512.2;YP_009725295.1;Wu1-SARS2_orf1a;2174 | 0 | 0 |
| NC_045512.2;YP_009725295.1;Wu1-SARS2_orf1a;2175 | 0 | 0 |
| NC_045512.2;YP_009725295.1;Wu1-SARS2_orf1a;2176 | 0 | 0 |
| NC_045512.2;YP_009725295.1;Wu1-SARS2_orf1a;2177 | 0 | 0 |
| NC_045512.2;YP_009725295.1;Wu1-SARS2_orf1a;2178 | 0 | 0 |
| NC_045512.2;YP_009725295.1;Wu1-SARS2_orf1a;2179 | 0 | 0 |
| NC_045512.2;YP_009725295.1;Wu1-SARS2_orf1a;2180 | 0.05 | 0 |
| NC_045512.2;YP_009725295.1;Wu1-SARS2_orf1a;2181 | 0.1 | 0 |

| | | |
|---|---|---|
| NC_045512.2;YP_009725295.1;Wu1-SARS2_orf1a;2182 | 0 | 0 |
| NC_045512.2;YP_009725295.1;Wu1-SARS2_orf1a;2183 | 0 | 0.025 |
| NC_045512.2;YP_009725295.1;Wu1-SARS2_orf1a;2184 | 0 | 0 |
| NC_045512.2;YP_009725295.1;Wu1-SARS2_orf1a;2185 | 0.05 | 0 |
| NC_045512.2;YP_009725295.1;Wu1-SARS2_orf1a;2186 | 0.05 | 0 |
| NC_045512.2;YP_009725295.1;Wu1-SARS2_orf1a;2187 | 0 | 0 |
| NC_045512.2;YP_009725295.1;Wu1-SARS2_orf1a;2188 | 0 | 0 |
| NC_045512.2;YP_009725295.1;Wu1-SARS2_orf1a;2189 | 0 | 0 |
| NC_045512.2;YP_009725295.1;Wu1-SARS2_orf1a;2190 | 0 | 0 |
| NC_045512.2;YP_009725295.1;Wu1-SARS2_orf1a;2191 | 0 | 0 |
| NC_045512.2;YP_009725295.1;Wu1-SARS2_orf1a;2192 | 0 | 0 |
| NC_045512.2;YP_009725295.1;Wu1-SARS2_orf1a;2193 | 0 | 0 |
| NC_045512.2;YP_009725295.1;Wu1-SARS2_orf1a;2194 | 0 | 0 |
| NC_045512.2;YP_009725295.1;Wu1-SARS2_orf1a;2195 | 0 | 0 |
| NC_045512.2;YP_009725295.1;Wu1-SARS2_orf1a;2196 | 0 | 0 |
| NC_045512.2;YP_009725295.1;Wu1-SARS2_orf1a;2197 | 0 | 0 |
| NC_045512.2;YP_009725295.1;Wu1-SARS2_orf1a;2198 | 0 | 0 |
| NC_045512.2;YP_009725295.1;Wu1-SARS2_orf1a;2199 | 0 | 0 |
| NC_045512.2;YP_009725295.1;Wu1-SARS2_orf1a;2200 | 0 | 0 |
| NC_045512.2;YP_009725295.1;Wu1-SARS2_orf1a;2201 | 0 | 0 |
| NC_045512.2;YP_009725295.1;Wu1-SARS2_orf1a;2202 | 0 | 0 |
| NC_045512.2;YP_009725295.1;Wu1-SARS2_orf1a;2203 | 0 | 0 |
| NC_045512.2;YP_009725295.1;Wu1-SARS2_orf1a;2204 | 0 | 0 |
| NC_045512.2;YP_009725295.1;Wu1-SARS2_orf1a;2205 | 0 | 0 |
| NC_045512.2;YP_009725295.1;Wu1-SARS2_orf1a;2206 | 0 | 0 |
| NC_045512.2;YP_009725295.1;Wu1-SARS2_orf1a;2207 | 0 | 0 |
| NC_045512.2;YP_009725295.1;Wu1-SARS2_orf1a;2208 | 0 | 0 |
| NC_045512.2;YP_009725295.1;Wu1-SARS2_orf1a;2209 | 0 | 0 |
| NC_045512.2;YP_009725295.1;Wu1-SARS2_orf1a;2210 | 0 | 0 |
| NC_045512.2;YP_009725295.1;Wu1-SARS2_orf1a;2211 | 0 | 0 |
| NC_045512.2;YP_009725295.1;Wu1-SARS2_orf1a;2212 | 0.05 | 0 |
| NC_045512.2;YP_009725295.1;Wu1-SARS2_orf1a;2213 | 0 | 0 |
| NC_045512.2;YP_009725295.1;Wu1-SARS2_orf1a;2214 | 0 | 0 |
| NC_045512.2;YP_009725295.1;Wu1-SARS2_orf1a;2215 | 0 | 0 |
| NC_045512.2;YP_009725295.1;Wu1-SARS2_orf1a;2216 | 0 | 0 |
| NC_045512.2;YP_009725295.1;Wu1-SARS2_orf1a;2217 | 0 | 0 |
| NC_045512.2;YP_009725295.1;Wu1-SARS2_orf1a;2218 | 0 | 0 |
| NC_045512.2;YP_009725295.1;Wu1-SARS2_orf1a;2219 | 0 | 0 |
| NC_045512.2;YP_009725295.1;Wu1-SARS2_orf1a;2220 | 0 | 0 |
| NC_045512.2;YP_009725295.1;Wu1-SARS2_orf1a;2221 | 0 | 0 |
| NC_045512.2;YP_009725295.1;Wu1-SARS2_orf1a;2222 | 0 | 0 |
| NC_045512.2;YP_009725295.1;Wu1-SARS2_orf1a;2223 | 0 | 0 |
| NC_045512.2;YP_009725295.1;Wu1-SARS2_orf1a;2224 | 0 | 0 |

| | | |
|---|---|---|
| NC_045512.2;YP_009725295.1;Wu1-SARS2_orf1a;2225 | 0 | 0 |
| NC_045512.2;YP_009725295.1;Wu1-SARS2_orf1a;2226 | 0 | 0 |
| NC_045512.2;YP_009725295.1;Wu1-SARS2_orf1a;2227 | 0 | 0 |
| NC_045512.2;YP_009725295.1;Wu1-SARS2_orf1a;2228 | 0 | 0 |
| NC_045512.2;YP_009725295.1;Wu1-SARS2_orf1a;2229 | 0 | 0 |
| NC_045512.2;YP_009725295.1;Wu1-SARS2_orf1a;2230 | 0 | 0 |
| NC_045512.2;YP_009725295.1;Wu1-SARS2_orf1a;2231 | 0 | 0 |
| NC_045512.2;YP_009725295.1;Wu1-SARS2_orf1a;2232 | 0 | 0 |
| NC_045512.2;YP_009725295.1;Wu1-SARS2_orf1a;2233 | 0 | 0 |
| NC_045512.2;YP_009725295.1;Wu1-SARS2_orf1a;2234 | 0 | 0 |
| NC_045512.2;YP_009725295.1;Wu1-SARS2_orf1a;2235 | 0 | 0 |
| NC_045512.2;YP_009725295.1;Wu1-SARS2_orf1a;2236 | 0 | 0 |
| NC_045512.2;YP_009725295.1;Wu1-SARS2_orf1a;2237 | 0 | 0 |
| NC_045512.2;YP_009725295.1;Wu1-SARS2_orf1a;2238 | 0 | 0 |
| NC_045512.2;YP_009725295.1;Wu1-SARS2_orf1a;2239 | 0 | 0 |
| NC_045512.2;YP_009725295.1;Wu1-SARS2_orf1a;2240 | 0 | 0 |
| NC_045512.2;YP_009725295.1;Wu1-SARS2_orf1a;2241 | 0 | 0 |
| NC_045512.2;YP_009725295.1;Wu1-SARS2_orf1a;2242 | 0 | 0 |
| NC_045512.2;YP_009725295.1;Wu1-SARS2_orf1a;2243 | 0 | 0 |
| NC_045512.2;YP_009725295.1;Wu1-SARS2_orf1a;2244 | 0 | 0 |
| NC_045512.2;YP_009725295.1;Wu1-SARS2_orf1a;2245 | 0 | 0 |
| NC_045512.2;YP_009725295.1;Wu1-SARS2_orf1a;2246 | 0 | 0 |
| NC_045512.2;YP_009725295.1;Wu1-SARS2_orf1a;2247 | 0 | 0 |
| NC_045512.2;YP_009725295.1;Wu1-SARS2_orf1a;2248 | 0 | 0.025 |
| NC_045512.2;YP_009725295.1;Wu1-SARS2_orf1a;2249 | 0 | 0.025 |
| NC_045512.2;YP_009725295.1;Wu1-SARS2_orf1a;2250 | 0 | 0 |
| NC_045512.2;YP_009725295.1;Wu1-SARS2_orf1a;2251 | 0 | 0 |
| NC_045512.2;YP_009725295.1;Wu1-SARS2_orf1a;2252 | 0 | 0 |
| NC_045512.2;YP_009725295.1;Wu1-SARS2_orf1a;2253 | 0 | 0 |
| NC_045512.2;YP_009725295.1;Wu1-SARS2_orf1a;2254 | 0 | 0 |
| NC_045512.2;YP_009725295.1;Wu1-SARS2_orf1a;2255 | 0 | 0 |
| NC_045512.2;YP_009725295.1;Wu1-SARS2_orf1a;2256 | 0 | 0 |
| NC_045512.2;YP_009725295.1;Wu1-SARS2_orf1a;2257 | 0 | 0 |
| NC_045512.2;YP_009725295.1;Wu1-SARS2_orf1a;2258 | 0 | 0.05 |
| NC_045512.2;YP_009725295.1;Wu1-SARS2_orf1a;2259 | 0 | 0 |
| NC_045512.2;YP_009725295.1;Wu1-SARS2_orf1a;2260 | 0.2 | 0.125 |
| NC_045512.2;YP_009725295.1;Wu1-SARS2_orf1a;2261 | 0.15 | 0.1 |
| NC_045512.2;YP_009725295.1;Wu1-SARS2_orf1a;2262 | 0.3 | 0.275 |
| NC_045512.2;YP_009725295.1;Wu1-SARS2_orf1a;2263 | 0.2 | 0.15 |
| NC_045512.2;YP_009725295.1;Wu1-SARS2_orf1a;2264 | 0.25 | 0.175 |
| NC_045512.2;YP_009725295.1;Wu1-SARS2_orf1a;2265 | 0.2 | 0.1 |
| NC_045512.2;YP_009725295.1;Wu1-SARS2_orf1a;2266 | 0.05 | 0 |
| NC_045512.2;YP_009725295.1;Wu1-SARS2_orf1a;2267 | 0.05 | 0.025 |

| | | |
|---|---|---|
| NC_045512.2;YP_009725295.1;Wu1-SARS2_orf1a;2268 | 0.05 | 0.025 |
| NC_045512.2;YP_009725295.1;Wu1-SARS2_orf1a;2269 | 0.05 | 0 |
| NC_045512.2;YP_009725295.1;Wu1-SARS2_orf1a;2270 | 0.05 | 0 |
| NC_045512.2;YP_009725295.1;Wu1-SARS2_orf1a;2271 | 0 | 0 |
| NC_045512.2;YP_009725295.1;Wu1-SARS2_orf1a;2272 | 0 | 0 |
| NC_045512.2;YP_009725295.1;Wu1-SARS2_orf1a;2273 | 0 | 0 |
| NC_045512.2;YP_009725295.1;Wu1-SARS2_orf1a;2274 | 0.05 | 0 |
| NC_045512.2;YP_009725295.1;Wu1-SARS2_orf1a;2275 | 0 | 0 |
| NC_045512.2;YP_009725295.1;Wu1-SARS2_orf1a;2276 | 0 | 0 |
| NC_045512.2;YP_009725295.1;Wu1-SARS2_orf1a;2277 | 0 | 0 |
| NC_045512.2;YP_009725295.1;Wu1-SARS2_orf1a;2278 | 0 | 0 |
| NC_045512.2;YP_009725295.1;Wu1-SARS2_orf1a;2279 | 0 | 0 |
| NC_045512.2;YP_009725295.1;Wu1-SARS2_orf1a;2280 | 0 | 0 |
| NC_045512.2;YP_009725295.1;Wu1-SARS2_orf1a;2281 | 0 | 0 |
| NC_045512.2;YP_009725295.1;Wu1-SARS2_orf1a;2282 | 0 | 0 |
| NC_045512.2;YP_009725295.1;Wu1-SARS2_orf1a;2283 | 0 | 0 |
| NC_045512.2;YP_009725295.1;Wu1-SARS2_orf1a;2284 | 0 | 0 |
| NC_045512.2;YP_009725295.1;Wu1-SARS2_orf1a;2285 | 0 | 0 |
| NC_045512.2;YP_009725295.1;Wu1-SARS2_orf1a;2286 | 0 | 0 |
| NC_045512.2;YP_009725295.1;Wu1-SARS2_orf1a;2287 | 0 | 0 |
| NC_045512.2;YP_009725295.1;Wu1-SARS2_orf1a;2288 | 0 | 0.025 |
| NC_045512.2;YP_009725295.1;Wu1-SARS2_orf1a;2289 | 0 | 0 |
| NC_045512.2;YP_009725295.1;Wu1-SARS2_orf1a;2290 | 0 | 0 |
| NC_045512.2;YP_009725295.1;Wu1-SARS2_orf1a;2291 | 0.05 | 0.025 |
| NC_045512.2;YP_009725295.1;Wu1-SARS2_orf1a;2292 | 0 | 0 |
| NC_045512.2;YP_009725295.1;Wu1-SARS2_orf1a;2293 | 0 | 0 |
| NC_045512.2;YP_009725295.1;Wu1-SARS2_orf1a;2294 | 0.05 | 0.025 |
| NC_045512.2;YP_009725295.1;Wu1-SARS2_orf1a;2295 | 0.05 | 0 |
| NC_045512.2;YP_009725295.1;Wu1-SARS2_orf1a;2296 | 0 | 0 |
| NC_045512.2;YP_009725295.1;Wu1-SARS2_orf1a;2297 | 0 | 0 |
| NC_045512.2;YP_009725295.1;Wu1-SARS2_orf1a;2298 | 0 | 0 |
| NC_045512.2;YP_009725295.1;Wu1-SARS2_orf1a;2299 | 0 | 0 |
| NC_045512.2;YP_009725295.1;Wu1-SARS2_orf1a;2300 | 0 | 0 |
| NC_045512.2;YP_009725295.1;Wu1-SARS2_orf1a;2301 | 0 | 0 |
| NC_045512.2;YP_009725295.1;Wu1-SARS2_orf1a;2302 | 0 | 0 |
| NC_045512.2;YP_009725295.1;Wu1-SARS2_orf1a;2303 | 0 | 0 |
| NC_045512.2;YP_009725295.1;Wu1-SARS2_orf1a;2304 | 0 | 0 |
| NC_045512.2;YP_009725295.1;Wu1-SARS2_orf1a;2305 | 0 | 0 |
| NC_045512.2;YP_009725295.1;Wu1-SARS2_orf1a;2306 | 0 | 0 |
| NC_045512.2;YP_009725295.1;Wu1-SARS2_orf1a;2307 | 0 | 0 |
| NC_045512.2;YP_009725295.1;Wu1-SARS2_orf1a;2308 | 0 | 0 |
| NC_045512.2;YP_009725295.1;Wu1-SARS2_orf1a;2309 | 0 | 0.075 |
| NC_045512.2;YP_009725295.1;Wu1-SARS2_orf1a;2310 | 0 | 0 |

| | | |
|---|---|---|
| NC_045512.2;YP_009725295.1;Wu1-SARS2_orf1a;2311 | 0 | 0 |
| NC_045512.2;YP_009725295.1;Wu1-SARS2_orf1a;2312 | 0 | 0 |
| NC_045512.2;YP_009725295.1;Wu1-SARS2_orf1a;2313 | 0 | 0 |
| NC_045512.2;YP_009725295.1;Wu1-SARS2_orf1a;2314 | 0 | 0 |
| NC_045512.2;YP_009725295.1;Wu1-SARS2_orf1a;2315 | 0 | 0 |
| NC_045512.2;YP_009725295.1;Wu1-SARS2_orf1a;2316 | 0 | 0 |
| NC_045512.2;YP_009725295.1;Wu1-SARS2_orf1a;2317 | 0 | 0 |
| NC_045512.2;YP_009725295.1;Wu1-SARS2_orf1a;2318 | 0 | 0 |
| NC_045512.2;YP_009725295.1;Wu1-SARS2_orf1a;2319 | 0 | 0 |
| NC_045512.2;YP_009725295.1;Wu1-SARS2_orf1a;2320 | 0 | 0 |
| NC_045512.2;YP_009725295.1;Wu1-SARS2_orf1a;2321 | 0 | 0 |
| NC_045512.2;YP_009725295.1;Wu1-SARS2_orf1a;2322 | 0 | 0 |
| NC_045512.2;YP_009725295.1;Wu1-SARS2_orf1a;2323 | 0 | 0 |
| NC_045512.2;YP_009725295.1;Wu1-SARS2_orf1a;2324 | 0 | 0 |
| NC_045512.2;YP_009725295.1;Wu1-SARS2_orf1a;2325 | 0 | 0 |
| NC_045512.2;YP_009725295.1;Wu1-SARS2_orf1a;2326 | 0 | 0 |
| NC_045512.2;YP_009725295.1;Wu1-SARS2_orf1a;2327 | 0 | 0 |
| NC_045512.2;YP_009725295.1;Wu1-SARS2_orf1a;2328 | 0 | 0 |
| NC_045512.2;YP_009725295.1;Wu1-SARS2_orf1a;2329 | 0 | 0 |
| NC_045512.2;YP_009725295.1;Wu1-SARS2_orf1a;2330 | 0 | 0 |
| NC_045512.2;YP_009725295.1;Wu1-SARS2_orf1a;2331 | 0 | 0 |
| NC_045512.2;YP_009725295.1;Wu1-SARS2_orf1a;2332 | 0 | 0 |
| NC_045512.2;YP_009725295.1;Wu1-SARS2_orf1a;2333 | 0 | 0 |
| NC_045512.2;YP_009725295.1;Wu1-SARS2_orf1a;2334 | 0 | 0 |
| NC_045512.2;YP_009725295.1;Wu1-SARS2_orf1a;2335 | 0 | 0 |
| NC_045512.2;YP_009725295.1;Wu1-SARS2_orf1a;2336 | 0 | 0 |
| NC_045512.2;YP_009725295.1;Wu1-SARS2_orf1a;2337 | 0 | 0 |
| NC_045512.2;YP_009725295.1;Wu1-SARS2_orf1a;2338 | 0 | 0 |
| NC_045512.2;YP_009725295.1;Wu1-SARS2_orf1a;2339 | 0 | 0 |
| NC_045512.2;YP_009725295.1;Wu1-SARS2_orf1a;2340 | 0 | 0 |
| NC_045512.2;YP_009725295.1;Wu1-SARS2_orf1a;2341 | 0 | 0 |
| NC_045512.2;YP_009725295.1;Wu1-SARS2_orf1a;2342 | 0 | 0 |
| NC_045512.2;YP_009725295.1;Wu1-SARS2_orf1a;2343 | 0 | 0 |
| NC_045512.2;YP_009725295.1;Wu1-SARS2_orf1a;2344 | 0 | 0 |
| NC_045512.2;YP_009725295.1;Wu1-SARS2_orf1a;2345 | 0 | 0 |
| NC_045512.2;YP_009725295.1;Wu1-SARS2_orf1a;2346 | 0 | 0 |
| NC_045512.2;YP_009725295.1;Wu1-SARS2_orf1a;2347 | 0 | 0 |
| NC_045512.2;YP_009725295.1;Wu1-SARS2_orf1a;2348 | 0 | 0 |
| NC_045512.2;YP_009725295.1;Wu1-SARS2_orf1a;2349 | 0 | 0 |
| NC_045512.2;YP_009725295.1;Wu1-SARS2_orf1a;2350 | 0 | 0 |
| NC_045512.2;YP_009725295.1;Wu1-SARS2_orf1a;2351 | 0 | 0 |
| NC_045512.2;YP_009725295.1;Wu1-SARS2_orf1a;2352 | 0 | 0 |
| NC_045512.2;YP_009725295.1;Wu1-SARS2_orf1a;2353 | 0 | 0 |

| | | |
|---|---|---|
| NC_045512.2;YP_009725295.1;Wu1-SARS2_orf1a;2354 | 0 | 0 |
| NC_045512.2;YP_009725295.1;Wu1-SARS2_orf1a;2355 | 0 | 0 |
| NC_045512.2;YP_009725295.1;Wu1-SARS2_orf1a;2356 | 0 | 0 |
| NC_045512.2;YP_009725295.1;Wu1-SARS2_orf1a;2357 | 0 | 0 |
| NC_045512.2;YP_009725295.1;Wu1-SARS2_orf1a;2358 | 0 | 0 |
| NC_045512.2;YP_009725295.1;Wu1-SARS2_orf1a;2359 | 0 | 0 |
| NC_045512.2;YP_009725295.1;Wu1-SARS2_orf1a;2360 | 0 | 0 |
| NC_045512.2;YP_009725295.1;Wu1-SARS2_orf1a;2361 | 0 | 0 |
| NC_045512.2;YP_009725295.1;Wu1-SARS2_orf1a;2362 | 0 | 0 |
| NC_045512.2;YP_009725295.1;Wu1-SARS2_orf1a;2363 | 0 | 0 |
| NC_045512.2;YP_009725295.1;Wu1-SARS2_orf1a;2364 | 0 | 0 |
| NC_045512.2;YP_009725295.1;Wu1-SARS2_orf1a;2365 | 0 | 0 |
| NC_045512.2;YP_009725295.1;Wu1-SARS2_orf1a;2366 | 0 | 0 |
| NC_045512.2;YP_009725295.1;Wu1-SARS2_orf1a;2367 | 0 | 0 |
| NC_045512.2;YP_009725295.1;Wu1-SARS2_orf1a;2368 | 0 | 0 |
| NC_045512.2;YP_009725295.1;Wu1-SARS2_orf1a;2369 | 0 | 0 |
| NC_045512.2;YP_009725295.1;Wu1-SARS2_orf1a;2370 | 0 | 0 |
| NC_045512.2;YP_009725295.1;Wu1-SARS2_orf1a;2371 | 0 | 0 |
| NC_045512.2;YP_009725295.1;Wu1-SARS2_orf1a;2372 | 0 | 0 |
| NC_045512.2;YP_009725295.1;Wu1-SARS2_orf1a;2373 | 0 | 0 |
| NC_045512.2;YP_009725295.1;Wu1-SARS2_orf1a;2374 | 0 | 0 |
| NC_045512.2;YP_009725295.1;Wu1-SARS2_orf1a;2375 | 0 | 0 |
| NC_045512.2;YP_009725295.1;Wu1-SARS2_orf1a;2376 | 0 | 0 |
| NC_045512.2;YP_009725295.1;Wu1-SARS2_orf1a;2377 | 0 | 0 |
| NC_045512.2;YP_009725295.1;Wu1-SARS2_orf1a;2378 | 0 | 0 |
| NC_045512.2;YP_009725295.1;Wu1-SARS2_orf1a;2379 | 0 | 0 |
| NC_045512.2;YP_009725295.1;Wu1-SARS2_orf1a;2380 | 0 | 0 |
| NC_045512.2;YP_009725295.1;Wu1-SARS2_orf1a;2381 | 0 | 0 |
| NC_045512.2;YP_009725295.1;Wu1-SARS2_orf1a;2382 | 0 | 0 |
| NC_045512.2;YP_009725295.1;Wu1-SARS2_orf1a;2383 | 0 | 0 |
| NC_045512.2;YP_009725295.1;Wu1-SARS2_orf1a;2384 | 0 | 0 |
| NC_045512.2;YP_009725295.1;Wu1-SARS2_orf1a;2385 | 0 | 0 |
| NC_045512.2;YP_009725295.1;Wu1-SARS2_orf1a;2386 | 0 | 0 |
| NC_045512.2;YP_009725295.1;Wu1-SARS2_orf1a;2387 | 0 | 0 |
| NC_045512.2;YP_009725295.1;Wu1-SARS2_orf1a;2388 | 0 | 0 |
| NC_045512.2;YP_009725295.1;Wu1-SARS2_orf1a;2389 | 0 | 0 |
| NC_045512.2;YP_009725295.1;Wu1-SARS2_orf1a;2390 | 0 | 0 |
| NC_045512.2;YP_009725295.1;Wu1-SARS2_orf1a;2391 | 0 | 0 |
| NC_045512.2;YP_009725295.1;Wu1-SARS2_orf1a;2392 | 0 | 0 |
| NC_045512.2;YP_009725295.1;Wu1-SARS2_orf1a;2393 | 0 | 0 |
| NC_045512.2;YP_009725295.1;Wu1-SARS2_orf1a;2394 | 0 | 0 |
| NC_045512.2;YP_009725295.1;Wu1-SARS2_orf1a;2395 | 0 | 0 |
| NC_045512.2;YP_009725295.1;Wu1-SARS2_orf1a;2396 | 0 | 0 |

| | | |
|---|---|---|
| NC_045512.2;YP_009725295.1;Wu1-SARS2_orf1a;2397 | 0 | 0 |
| NC_045512.2;YP_009725295.1;Wu1-SARS2_orf1a;2398 | 0 | 0 |
| NC_045512.2;YP_009725295.1;Wu1-SARS2_orf1a;2399 | 0 | 0 |
| NC_045512.2;YP_009725295.1;Wu1-SARS2_orf1a;2400 | 0 | 0 |
| NC_045512.2;YP_009725295.1;Wu1-SARS2_orf1a;2401 | 0 | 0 |
| NC_045512.2;YP_009725295.1;Wu1-SARS2_orf1a;2402 | 0 | 0 |
| NC_045512.2;YP_009725295.1;Wu1-SARS2_orf1a;2403 | 0 | 0 |
| NC_045512.2;YP_009725295.1;Wu1-SARS2_orf1a;2404 | 0 | 0 |
| NC_045512.2;YP_009725295.1;Wu1-SARS2_orf1a;2405 | 0 | 0 |
| NC_045512.2;YP_009725295.1;Wu1-SARS2_orf1a;2406 | 0 | 0 |
| NC_045512.2;YP_009725295.1;Wu1-SARS2_orf1a;2407 | 0 | 0 |
| NC_045512.2;YP_009725295.1;Wu1-SARS2_orf1a;2408 | 0 | 0 |
| NC_045512.2;YP_009725295.1;Wu1-SARS2_orf1a;2409 | 0 | 0 |
| NC_045512.2;YP_009725295.1;Wu1-SARS2_orf1a;2410 | 0 | 0 |
| NC_045512.2;YP_009725295.1;Wu1-SARS2_orf1a;2411 | 0 | 0 |
| NC_045512.2;YP_009725295.1;Wu1-SARS2_orf1a;2412 | 0 | 0 |
| NC_045512.2;YP_009725295.1;Wu1-SARS2_orf1a;2413 | 0 | 0 |
| NC_045512.2;YP_009725295.1;Wu1-SARS2_orf1a;2414 | 0 | 0 |
| NC_045512.2;YP_009725295.1;Wu1-SARS2_orf1a;2415 | 0 | 0 |
| NC_045512.2;YP_009725295.1;Wu1-SARS2_orf1a;2416 | 0 | 0.025 |
| NC_045512.2;YP_009725295.1;Wu1-SARS2_orf1a;2417 | 0 | 0 |
| NC_045512.2;YP_009725295.1;Wu1-SARS2_orf1a;2418 | 0 | 0 |
| NC_045512.2;YP_009725295.1;Wu1-SARS2_orf1a;2419 | 0 | 0 |
| NC_045512.2;YP_009725295.1;Wu1-SARS2_orf1a;2420 | 0 | 0 |
| NC_045512.2;YP_009725295.1;Wu1-SARS2_orf1a;2421 | 0 | 0 |
| NC_045512.2;YP_009725295.1;Wu1-SARS2_orf1a;2422 | 0 | 0 |
| NC_045512.2;YP_009725295.1;Wu1-SARS2_orf1a;2423 | 0 | 0 |
| NC_045512.2;YP_009725295.1;Wu1-SARS2_orf1a;2424 | 0 | 0 |
| NC_045512.2;YP_009725295.1;Wu1-SARS2_orf1a;2425 | 0 | 0 |
| NC_045512.2;YP_009725295.1;Wu1-SARS2_orf1a;2426 | 0 | 0 |
| NC_045512.2;YP_009725295.1;Wu1-SARS2_orf1a;2427 | 0 | 0 |
| NC_045512.2;YP_009725295.1;Wu1-SARS2_orf1a;2428 | 0 | 0 |
| NC_045512.2;YP_009725295.1;Wu1-SARS2_orf1a;2429 | 0 | 0 |
| NC_045512.2;YP_009725295.1;Wu1-SARS2_orf1a;2430 | 0 | 0 |
| NC_045512.2;YP_009725295.1;Wu1-SARS2_orf1a;2431 | 0 | 0 |
| NC_045512.2;YP_009725295.1;Wu1-SARS2_orf1a;2432 | 0 | 0 |
| NC_045512.2;YP_009725295.1;Wu1-SARS2_orf1a;2433 | 0 | 0 |
| NC_045512.2;YP_009725295.1;Wu1-SARS2_orf1a;2434 | 0 | 0 |
| NC_045512.2;YP_009725295.1;Wu1-SARS2_orf1a;2435 | 0 | 0 |
| NC_045512.2;YP_009725295.1;Wu1-SARS2_orf1a;2436 | 0 | 0 |
| NC_045512.2;YP_009725295.1;Wu1-SARS2_orf1a;2437 | 0 | 0 |
| NC_045512.2;YP_009725295.1;Wu1-SARS2_orf1a;2438 | 0 | 0 |
| NC_045512.2;YP_009725295.1;Wu1-SARS2_orf1a;2439 | 0 | 0 |

| | | |
|---|---|---|
| NC_045512.2;YP_009725295.1;Wu1-SARS2_orf1a;2440 | 0 | 0 |
| NC_045512.2;YP_009725295.1;Wu1-SARS2_orf1a;2441 | 0 | 0.025 |
| NC_045512.2;YP_009725295.1;Wu1-SARS2_orf1a;2442 | 0 | 0.025 |
| NC_045512.2;YP_009725295.1;Wu1-SARS2_orf1a;2443 | 0 | 0 |
| NC_045512.2;YP_009725295.1;Wu1-SARS2_orf1a;2444 | 0 | 0 |
| NC_045512.2;YP_009725295.1;Wu1-SARS2_orf1a;2445 | 0 | 0 |
| NC_045512.2;YP_009725295.1;Wu1-SARS2_orf1a;2446 | 0 | 0 |
| NC_045512.2;YP_009725295.1;Wu1-SARS2_orf1a;2447 | 0 | 0 |
| NC_045512.2;YP_009725295.1;Wu1-SARS2_orf1a;2448 | 0 | 0 |
| NC_045512.2;YP_009725295.1;Wu1-SARS2_orf1a;2449 | 0 | 0 |
| NC_045512.2;YP_009725295.1;Wu1-SARS2_orf1a;2450 | 0 | 0 |
| NC_045512.2;YP_009725295.1;Wu1-SARS2_orf1a;2451 | 0 | 0 |
| NC_045512.2;YP_009725295.1;Wu1-SARS2_orf1a;2452 | 0 | 0 |
| NC_045512.2;YP_009725295.1;Wu1-SARS2_orf1a;2453 | 0 | 0 |
| NC_045512.2;YP_009725295.1;Wu1-SARS2_orf1a;2454 | 0 | 0 |
| NC_045512.2;YP_009725295.1;Wu1-SARS2_orf1a;2455 | 0 | 0 |
| NC_045512.2;YP_009725295.1;Wu1-SARS2_orf1a;2456 | 0 | 0 |
| NC_045512.2;YP_009725295.1;Wu1-SARS2_orf1a;2457 | 0 | 0 |
| NC_045512.2;YP_009725295.1;Wu1-SARS2_orf1a;2458 | 0 | 0 |
| NC_045512.2;YP_009725295.1;Wu1-SARS2_orf1a;2459 | 0 | 0 |
| NC_045512.2;YP_009725295.1;Wu1-SARS2_orf1a;2460 | 0 | 0 |
| NC_045512.2;YP_009725295.1;Wu1-SARS2_orf1a;2461 | 0 | 0 |
| NC_045512.2;YP_009725295.1;Wu1-SARS2_orf1a;2462 | 0 | 0 |
| NC_045512.2;YP_009725295.1;Wu1-SARS2_orf1a;2463 | 0 | 0 |
| NC_045512.2;YP_009725295.1;Wu1-SARS2_orf1a;2464 | 0 | 0 |
| NC_045512.2;YP_009725295.1;Wu1-SARS2_orf1a;2465 | 0 | 0 |
| NC_045512.2;YP_009725295.1;Wu1-SARS2_orf1a;2466 | 0 | 0 |
| NC_045512.2;YP_009725295.1;Wu1-SARS2_orf1a;2467 | 0 | 0 |
| NC_045512.2;YP_009725295.1;Wu1-SARS2_orf1a;2468 | 0 | 0 |
| NC_045512.2;YP_009725295.1;Wu1-SARS2_orf1a;2469 | 0.05 | 0 |
| NC_045512.2;YP_009725295.1;Wu1-SARS2_orf1a;2470 | 0.05 | 0.025 |
| NC_045512.2;YP_009725295.1;Wu1-SARS2_orf1a;2471 | 0.05 | 0.025 |
| NC_045512.2;YP_009725295.1;Wu1-SARS2_orf1a;2472 | 0 | 0 |
| NC_045512.2;YP_009725295.1;Wu1-SARS2_orf1a;2473 | 0 | 0 |
| NC_045512.2;YP_009725295.1;Wu1-SARS2_orf1a;2474 | 0 | 0 |
| NC_045512.2;YP_009725295.1;Wu1-SARS2_orf1a;2475 | 0 | 0.025 |
| NC_045512.2;YP_009725295.1;Wu1-SARS2_orf1a;2476 | 0 | 0 |
| NC_045512.2;YP_009725295.1;Wu1-SARS2_orf1a;2477 | 0 | 0 |
| NC_045512.2;YP_009725295.1;Wu1-SARS2_orf1a;2478 | 0 | 0.05 |
| NC_045512.2;YP_009725295.1;Wu1-SARS2_orf1a;2479 | 0 | 0 |
| NC_045512.2;YP_009725295.1;Wu1-SARS2_orf1a;2480 | 0 | 0 |
| NC_045512.2;YP_009725295.1;Wu1-SARS2_orf1a;2481 | 0 | 0 |
| NC_045512.2;YP_009725295.1;Wu1-SARS2_orf1a;2482 | 0.05 | 0 |

| | | |
|---|---|---|
| NC_045512.2;YP_009725295.1;Wu1-SARS2_orf1a;2483 | 0 | 0 |
| NC_045512.2;YP_009725295.1;Wu1-SARS2_orf1a;2484 | 0 | 0 |
| NC_045512.2;YP_009725295.1;Wu1-SARS2_orf1a;2485 | 0 | 0 |
| NC_045512.2;YP_009725295.1;Wu1-SARS2_orf1a;2486 | 0 | 0 |
| NC_045512.2;YP_009725295.1;Wu1-SARS2_orf1a;2487 | 0 | 0 |
| NC_045512.2;YP_009725295.1;Wu1-SARS2_orf1a;2488 | 0 | 0 |
| NC_045512.2;YP_009725295.1;Wu1-SARS2_orf1a;2489 | 0 | 0 |
| NC_045512.2;YP_009725295.1;Wu1-SARS2_orf1a;2490 | 0 | 0 |
| NC_045512.2;YP_009725295.1;Wu1-SARS2_orf1a;2491 | 0 | 0 |
| NC_045512.2;YP_009725295.1;Wu1-SARS2_orf1a;2492 | 0 | 0 |
| NC_045512.2;YP_009725295.1;Wu1-SARS2_orf1a;2493 | 0 | 0 |
| NC_045512.2;YP_009725295.1;Wu1-SARS2_orf1a;2494 | 0 | 0 |
| NC_045512.2;YP_009725295.1;Wu1-SARS2_orf1a;2495 | 0 | 0 |
| NC_045512.2;YP_009725295.1;Wu1-SARS2_orf1a;2496 | 0 | 0 |
| NC_045512.2;YP_009725295.1;Wu1-SARS2_orf1a;2497 | 0 | 0 |
| NC_045512.2;YP_009725295.1;Wu1-SARS2_orf1a;2498 | 0 | 0 |
| NC_045512.2;YP_009725295.1;Wu1-SARS2_orf1a;2499 | 0 | 0 |
| NC_045512.2;YP_009725295.1;Wu1-SARS2_orf1a;2500 | 0 | 0 |
| NC_045512.2;YP_009725295.1;Wu1-SARS2_orf1a;2501 | 0 | 0 |
| NC_045512.2;YP_009725295.1;Wu1-SARS2_orf1a;2502 | 0 | 0 |
| NC_045512.2;YP_009725295.1;Wu1-SARS2_orf1a;2503 | 0 | 0 |
| NC_045512.2;YP_009725295.1;Wu1-SARS2_orf1a;2504 | 0.05 | 0.05 |
| NC_045512.2;YP_009725295.1;Wu1-SARS2_orf1a;2505 | 0 | 0 |
| NC_045512.2;YP_009725295.1;Wu1-SARS2_orf1a;2506 | 0 | 0 |
| NC_045512.2;YP_009725295.1;Wu1-SARS2_orf1a;2507 | 0 | 0 |
| NC_045512.2;YP_009725295.1;Wu1-SARS2_orf1a;2508 | 0 | 0 |
| NC_045512.2;YP_009725295.1;Wu1-SARS2_orf1a;2509 | 0 | 0 |
| NC_045512.2;YP_009725295.1;Wu1-SARS2_orf1a;2510 | 0 | 0 |
| NC_045512.2;YP_009725295.1;Wu1-SARS2_orf1a;2511 | 0 | 0 |
| NC_045512.2;YP_009725295.1;Wu1-SARS2_orf1a;2512 | 0 | 0 |
| NC_045512.2;YP_009725295.1;Wu1-SARS2_orf1a;2513 | 0 | 0 |
| NC_045512.2;YP_009725295.1;Wu1-SARS2_orf1a;2514 | 0 | 0 |
| NC_045512.2;YP_009725295.1;Wu1-SARS2_orf1a;2515 | 0 | 0 |
| NC_045512.2;YP_009725295.1;Wu1-SARS2_orf1a;2516 | 0 | 0 |
| NC_045512.2;YP_009725295.1;Wu1-SARS2_orf1a;2517 | 0 | 0 |
| NC_045512.2;YP_009725295.1;Wu1-SARS2_orf1a;2518 | 0 | 0 |
| NC_045512.2;YP_009725295.1;Wu1-SARS2_orf1a;2519 | 0 | 0 |
| NC_045512.2;YP_009725295.1;Wu1-SARS2_orf1a;2520 | 0 | 0 |
| NC_045512.2;YP_009725295.1;Wu1-SARS2_orf1a;2521 | 0 | 0.025 |
| NC_045512.2;YP_009725295.1;Wu1-SARS2_orf1a;2522 | 0.05 | 0.025 |
| NC_045512.2;YP_009725295.1;Wu1-SARS2_orf1a;2523 | 0.05 | 0.025 |
| NC_045512.2;YP_009725295.1;Wu1-SARS2_orf1a;2524 | 0.05 | 0.025 |
| NC_045512.2;YP_009725295.1;Wu1-SARS2_orf1a;2525 | 0.05 | 0.025 |

| | | |
|---|---|---|
| NC_045512.2;YP_009725295.1;Wu1-SARS2_orf1a;2526 | 0 | 0 |
| NC_045512.2;YP_009725295.1;Wu1-SARS2_orf1a;2527 | 0 | 0 |
| NC_045512.2;YP_009725295.1;Wu1-SARS2_orf1a;2528 | 0.05 | 0 |
| NC_045512.2;YP_009725295.1;Wu1-SARS2_orf1a;2529 | 0 | 0 |
| NC_045512.2;YP_009725295.1;Wu1-SARS2_orf1a;2530 | 0 | 0 |
| NC_045512.2;YP_009725295.1;Wu1-SARS2_orf1a;2531 | 0 | 0 |
| NC_045512.2;YP_009725295.1;Wu1-SARS2_orf1a;2532 | 0 | 0 |
| NC_045512.2;YP_009725295.1;Wu1-SARS2_orf1a;2533 | 0 | 0 |
| NC_045512.2;YP_009725295.1;Wu1-SARS2_orf1a;2534 | 0 | 0.025 |
| NC_045512.2;YP_009725295.1;Wu1-SARS2_orf1a;2535 | 0 | 0.025 |
| NC_045512.2;YP_009725295.1;Wu1-SARS2_orf1a;2536 | 0.05 | 0.025 |
| NC_045512.2;YP_009725295.1;Wu1-SARS2_orf1a;2537 | 0.05 | 0.025 |
| NC_045512.2;YP_009725295.1;Wu1-SARS2_orf1a;2538 | 0.05 | 0.025 |
| NC_045512.2;YP_009725295.1;Wu1-SARS2_orf1a;2539 | 0 | 0.025 |
| NC_045512.2;YP_009725295.1;Wu1-SARS2_orf1a;2540 | 0 | 0.025 |
| NC_045512.2;YP_009725295.1;Wu1-SARS2_orf1a;2541 | 0 | 0.025 |
| NC_045512.2;YP_009725295.1;Wu1-SARS2_orf1a;2542 | 0 | 0.025 |
| NC_045512.2;YP_009725295.1;Wu1-SARS2_orf1a;2543 | 0.1 | 0.05 |
| NC_045512.2;YP_009725295.1;Wu1-SARS2_orf1a;2544 | 0 | 0 |
| NC_045512.2;YP_009725295.1;Wu1-SARS2_orf1a;2545 | 0 | 0 |
| NC_045512.2;YP_009725295.1;Wu1-SARS2_orf1a;2546 | 0 | 0 |
| NC_045512.2;YP_009725295.1;Wu1-SARS2_orf1a;2547 | 0 | 0 |
| NC_045512.2;YP_009725295.1;Wu1-SARS2_orf1a;2548 | 0.05 | 0 |
| NC_045512.2;YP_009725295.1;Wu1-SARS2_orf1a;2549 | 0 | 0 |
| NC_045512.2;YP_009725295.1;Wu1-SARS2_orf1a;2550 | 0 | 0 |
| NC_045512.2;YP_009725295.1;Wu1-SARS2_orf1a;2551 | 0.05 | 0 |
| NC_045512.2;YP_009725295.1;Wu1-SARS2_orf1a;2552 | 0 | 0 |
| NC_045512.2;YP_009725295.1;Wu1-SARS2_orf1a;2553 | 0 | 0 |
| NC_045512.2;YP_009725295.1;Wu1-SARS2_orf1a;2554 | 0 | 0 |
| NC_045512.2;YP_009725295.1;Wu1-SARS2_orf1a;2555 | 0 | 0 |
| NC_045512.2;YP_009725295.1;Wu1-SARS2_orf1a;2556 | 0 | 0 |
| NC_045512.2;YP_009725295.1;Wu1-SARS2_orf1a;2557 | 0 | 0.025 |
| NC_045512.2;YP_009725295.1;Wu1-SARS2_orf1a;2558 | 0 | 0 |
| NC_045512.2;YP_009725295.1;Wu1-SARS2_orf1a;2559 | 0 | 0 |
| NC_045512.2;YP_009725295.1;Wu1-SARS2_orf1a;2560 | 0 | 0 |
| NC_045512.2;YP_009725295.1;Wu1-SARS2_orf1a;2561 | 0 | 0 |
| NC_045512.2;YP_009725295.1;Wu1-SARS2_orf1a;2562 | 0 | 0 |
| NC_045512.2;YP_009725295.1;Wu1-SARS2_orf1a;2563 | 0 | 0 |
| NC_045512.2;YP_009725295.1;Wu1-SARS2_orf1a;2564 | 0 | 0 |
| NC_045512.2;YP_009725295.1;Wu1-SARS2_orf1a;2565 | 0 | 0 |
| NC_045512.2;YP_009725295.1;Wu1-SARS2_orf1a;2566 | 0 | 0 |
| NC_045512.2;YP_009725295.1;Wu1-SARS2_orf1a;2567 | 0 | 0 |
| NC_045512.2;YP_009725295.1;Wu1-SARS2_orf1a;2568 | 0 | 0 |

| | | |
|---|---|---|
| NC_045512.2;YP_009725295.1;Wu1-SARS2_orf1a;2569 | 0 | 0.025 |
| NC_045512.2;YP_009725295.1;Wu1-SARS2_orf1a;2570 | 0 | 0.025 |
| NC_045512.2;YP_009725295.1;Wu1-SARS2_orf1a;2571 | 0.05 | 0.1 |
| NC_045512.2;YP_009725295.1;Wu1-SARS2_orf1a;2572 | 0 | 0.05 |
| NC_045512.2;YP_009725295.1;Wu1-SARS2_orf1a;2573 | 0 | 0 |
| NC_045512.2;YP_009725295.1;Wu1-SARS2_orf1a;2574 | 0 | 0 |
| NC_045512.2;YP_009725295.1;Wu1-SARS2_orf1a;2575 | 0 | 0 |
| NC_045512.2;YP_009725295.1;Wu1-SARS2_orf1a;2576 | 0 | 0 |
| NC_045512.2;YP_009725295.1;Wu1-SARS2_orf1a;2577 | 0.05 | 0 |
| NC_045512.2;YP_009725295.1;Wu1-SARS2_orf1a;2578 | 0 | 0 |
| NC_045512.2;YP_009725295.1;Wu1-SARS2_orf1a;2579 | 0.05 | 0 |
| NC_045512.2;YP_009725295.1;Wu1-SARS2_orf1a;2580 | 0 | 0 |
| NC_045512.2;YP_009725295.1;Wu1-SARS2_orf1a;2581 | 0 | 0 |
| NC_045512.2;YP_009725295.1;Wu1-SARS2_orf1a;2582 | 0 | 0 |
| NC_045512.2;YP_009725295.1;Wu1-SARS2_orf1a;2583 | 0 | 0.025 |
| NC_045512.2;YP_009725295.1;Wu1-SARS2_orf1a;2584 | 0 | 0.025 |
| NC_045512.2;YP_009725295.1;Wu1-SARS2_orf1a;2585 | 0 | 0 |
| NC_045512.2;YP_009725295.1;Wu1-SARS2_orf1a;2586 | 0 | 0 |
| NC_045512.2;YP_009725295.1;Wu1-SARS2_orf1a;2587 | 0 | 0 |
| NC_045512.2;YP_009725295.1;Wu1-SARS2_orf1a;2588 | 0 | 0 |
| NC_045512.2;YP_009725295.1;Wu1-SARS2_orf1a;2589 | 0 | 0 |
| NC_045512.2;YP_009725295.1;Wu1-SARS2_orf1a;2590 | 0 | 0.025 |
| NC_045512.2;YP_009725295.1;Wu1-SARS2_orf1a;2591 | 0 | 0 |
| NC_045512.2;YP_009725295.1;Wu1-SARS2_orf1a;2592 | 0 | 0 |
| NC_045512.2;YP_009725295.1;Wu1-SARS2_orf1a;2593 | 0.05 | 0.05 |
| NC_045512.2;YP_009725295.1;Wu1-SARS2_orf1a;2594 | 0.05 | 0.025 |
| NC_045512.2;YP_009725295.1;Wu1-SARS2_orf1a;2595 | 0.15 | 0.025 |
| NC_045512.2;YP_009725295.1;Wu1-SARS2_orf1a;2596 | 0.05 | 0 |
| NC_045512.2;YP_009725295.1;Wu1-SARS2_orf1a;2597 | 0 | 0 |
| NC_045512.2;YP_009725295.1;Wu1-SARS2_orf1a;2598 | 0 | 0 |
| NC_045512.2;YP_009725295.1;Wu1-SARS2_orf1a;2599 | 0 | 0.025 |
| NC_045512.2;YP_009725295.1;Wu1-SARS2_orf1a;2600 | 0 | 0 |
| NC_045512.2;YP_009725295.1;Wu1-SARS2_orf1a;2601 | 0 | 0 |
| NC_045512.2;YP_009725295.1;Wu1-SARS2_orf1a;2602 | 0 | 0 |
| NC_045512.2;YP_009725295.1;Wu1-SARS2_orf1a;2603 | 0 | 0 |
| NC_045512.2;YP_009725295.1;Wu1-SARS2_orf1a;2604 | 0 | 0 |
| NC_045512.2;YP_009725295.1;Wu1-SARS2_orf1a;2605 | 0 | 0 |
| NC_045512.2;YP_009725295.1;Wu1-SARS2_orf1a;2606 | 0 | 0 |
| NC_045512.2;YP_009725295.1;Wu1-SARS2_orf1a;2607 | 0 | 0.05 |
| NC_045512.2;YP_009725295.1;Wu1-SARS2_orf1a;2608 | 0 | 0 |
| NC_045512.2;YP_009725295.1;Wu1-SARS2_orf1a;2609 | 0 | 0 |
| NC_045512.2;YP_009725295.1;Wu1-SARS2_orf1a;2610 | 0 | 0 |
| NC_045512.2;YP_009725295.1;Wu1-SARS2_orf1a;2611 | 0 | 0 |

| | | |
|---|---|---|
| NC_045512.2;YP_009725295.1;Wu1-SARS2_orf1a;2612 | 0 | 0 |
| NC_045512.2;YP_009725295.1;Wu1-SARS2_orf1a;2613 | 0 | 0 |
| NC_045512.2;YP_009725295.1;Wu1-SARS2_orf1a;2614 | 0 | 0.025 |
| NC_045512.2;YP_009725295.1;Wu1-SARS2_orf1a;2615 | 0 | 0.05 |
| NC_045512.2;YP_009725295.1;Wu1-SARS2_orf1a;2616 | 0.05 | 0.025 |
| NC_045512.2;YP_009725295.1;Wu1-SARS2_orf1a;2617 | 0.1 | 0.05 |
| NC_045512.2;YP_009725295.1;Wu1-SARS2_orf1a;2618 | 0.05 | 0.025 |
| NC_045512.2;YP_009725295.1;Wu1-SARS2_orf1a;2619 | 0.05 | 0 |
| NC_045512.2;YP_009725295.1;Wu1-SARS2_orf1a;2620 | 0 | 0 |
| NC_045512.2;YP_009725295.1;Wu1-SARS2_orf1a;2621 | 0 | 0 |
| NC_045512.2;YP_009725295.1;Wu1-SARS2_orf1a;2622 | 0.05 | 0 |
| NC_045512.2;YP_009725295.1;Wu1-SARS2_orf1a;2623 | 0 | 0 |
| NC_045512.2;YP_009725295.1;Wu1-SARS2_orf1a;2624 | 0 | 0 |
| NC_045512.2;YP_009725295.1;Wu1-SARS2_orf1a;2625 | 0 | 0 |
| NC_045512.2;YP_009725295.1;Wu1-SARS2_orf1a;2626 | 0 | 0 |
| NC_045512.2;YP_009725295.1;Wu1-SARS2_orf1a;2627 | 0 | 0 |
| NC_045512.2;YP_009725295.1;Wu1-SARS2_orf1a;2628 | 0 | 0 |
| NC_045512.2;YP_009725295.1;Wu1-SARS2_orf1a;2629 | 0 | 0 |
| NC_045512.2;YP_009725295.1;Wu1-SARS2_orf1a;2630 | 0 | 0 |
| NC_045512.2;YP_009725295.1;Wu1-SARS2_orf1a;2631 | 0 | 0 |
| NC_045512.2;YP_009725295.1;Wu1-SARS2_orf1a;2632 | 0 | 0 |
| NC_045512.2;YP_009725295.1;Wu1-SARS2_orf1a;2633 | 0 | 0 |
| NC_045512.2;YP_009725295.1;Wu1-SARS2_orf1a;2634 | 0 | 0 |
| NC_045512.2;YP_009725295.1;Wu1-SARS2_orf1a;2635 | 0 | 0 |
| NC_045512.2;YP_009725295.1;Wu1-SARS2_orf1a;2636 | 0 | 0 |
| NC_045512.2;YP_009725295.1;Wu1-SARS2_orf1a;2637 | 0 | 0 |
| NC_045512.2;YP_009725295.1;Wu1-SARS2_orf1a;2638 | 0 | 0 |
| NC_045512.2;YP_009725295.1;Wu1-SARS2_orf1a;2639 | 0 | 0 |
| NC_045512.2;YP_009725295.1;Wu1-SARS2_orf1a;2640 | 0 | 0 |
| NC_045512.2;YP_009725295.1;Wu1-SARS2_orf1a;2641 | 0 | 0 |
| NC_045512.2;YP_009725295.1;Wu1-SARS2_orf1a;2642 | 0 | 0 |
| NC_045512.2;YP_009725295.1;Wu1-SARS2_orf1a;2643 | 0 | 0 |
| NC_045512.2;YP_009725295.1;Wu1-SARS2_orf1a;2644 | 0 | 0 |
| NC_045512.2;YP_009725295.1;Wu1-SARS2_orf1a;2645 | 0 | 0 |
| NC_045512.2;YP_009725295.1;Wu1-SARS2_orf1a;2646 | 0 | 0 |
| NC_045512.2;YP_009725295.1;Wu1-SARS2_orf1a;2647 | 0 | 0.025 |
| NC_045512.2;YP_009725295.1;Wu1-SARS2_orf1a;2648 | 0 | 0 |
| NC_045512.2;YP_009725295.1;Wu1-SARS2_orf1a;2649 | 0 | 0 |
| NC_045512.2;YP_009725295.1;Wu1-SARS2_orf1a;2650 | 0 | 0 |
| NC_045512.2;YP_009725295.1;Wu1-SARS2_orf1a;2651 | 0 | 0 |
| NC_045512.2;YP_009725295.1;Wu1-SARS2_orf1a;2652 | 0 | 0 |
| NC_045512.2;YP_009725295.1;Wu1-SARS2_orf1a;2653 | 0 | 0 |
| NC_045512.2;YP_009725295.1;Wu1-SARS2_orf1a;2654 | 0 | 0 |

| | | |
|---|---|---|
| NC_045512.2;YP_009725295.1;Wu1-SARS2_orf1a;2655 | 0 | 0 |
| NC_045512.2;YP_009725295.1;Wu1-SARS2_orf1a;2656 | 0 | 0 |
| NC_045512.2;YP_009725295.1;Wu1-SARS2_orf1a;2657 | 0 | 0 |
| NC_045512.2;YP_009725295.1;Wu1-SARS2_orf1a;2658 | 0 | 0 |
| NC_045512.2;YP_009725295.1;Wu1-SARS2_orf1a;2659 | 0 | 0 |
| NC_045512.2;YP_009725295.1;Wu1-SARS2_orf1a;2660 | 0 | 0 |
| NC_045512.2;YP_009725295.1;Wu1-SARS2_orf1a;2661 | 0.05 | 0 |
| NC_045512.2;YP_009725295.1;Wu1-SARS2_orf1a;2662 | 0 | 0 |
| NC_045512.2;YP_009725295.1;Wu1-SARS2_orf1a;2663 | 0.05 | 0.05 |
| NC_045512.2;YP_009725295.1;Wu1-SARS2_orf1a;2664 | 0.05 | 0.05 |
| NC_045512.2;YP_009725295.1;Wu1-SARS2_orf1a;2665 | 0 | 0 |
| NC_045512.2;YP_009725295.1;Wu1-SARS2_orf1a;2666 | 0 | 0 |
| NC_045512.2;YP_009725295.1;Wu1-SARS2_orf1a;2667 | 0 | 0.025 |
| NC_045512.2;YP_009725295.1;Wu1-SARS2_orf1a;2668 | 0.1 | 0.05 |
| NC_045512.2;YP_009725295.1;Wu1-SARS2_orf1a;2669 | 0.05 | 0 |
| NC_045512.2;YP_009725295.1;Wu1-SARS2_orf1a;2670 | 0 | 0 |
| NC_045512.2;YP_009725295.1;Wu1-SARS2_orf1a;2671 | 0 | 0 |
| NC_045512.2;YP_009725295.1;Wu1-SARS2_orf1a;2672 | 0.05 | 0 |
| NC_045512.2;YP_009725295.1;Wu1-SARS2_orf1a;2673 | 0 | 0 |
| NC_045512.2;YP_009725295.1;Wu1-SARS2_orf1a;2674 | 0 | 0.025 |
| NC_045512.2;YP_009725295.1;Wu1-SARS2_orf1a;2675 | 0.05 | 0.075 |
| NC_045512.2;YP_009725295.1;Wu1-SARS2_orf1a;2676 | 0.05 | 0.025 |
| NC_045512.2;YP_009725295.1;Wu1-SARS2_orf1a;2677 | 0.05 | 0.05 |
| NC_045512.2;YP_009725295.1;Wu1-SARS2_orf1a;2678 | 0 | 0 |
| NC_045512.2;YP_009725295.1;Wu1-SARS2_orf1a;2679 | 0 | 0 |
| NC_045512.2;YP_009725295.1;Wu1-SARS2_orf1a;2680 | 0 | 0 |
| NC_045512.2;YP_009725295.1;Wu1-SARS2_orf1a;2681 | 0 | 0 |
| NC_045512.2;YP_009725295.1;Wu1-SARS2_orf1a;2682 | 0 | 0 |
| NC_045512.2;YP_009725295.1;Wu1-SARS2_orf1a;2683 | 0 | 0 |
| NC_045512.2;YP_009725295.1;Wu1-SARS2_orf1a;2684 | 0.05 | 0 |
| NC_045512.2;YP_009725295.1;Wu1-SARS2_orf1a;2685 | 0.05 | 0 |
| NC_045512.2;YP_009725295.1;Wu1-SARS2_orf1a;2686 | 0 | 0 |
| NC_045512.2;YP_009725295.1;Wu1-SARS2_orf1a;2687 | 0 | 0 |
| NC_045512.2;YP_009725295.1;Wu1-SARS2_orf1a;2688 | 0 | 0 |
| NC_045512.2;YP_009725295.1;Wu1-SARS2_orf1a;2689 | 0.05 | 0.05 |
| NC_045512.2;YP_009725295.1;Wu1-SARS2_orf1a;2690 | 0 | 0 |
| NC_045512.2;YP_009725295.1;Wu1-SARS2_orf1a;2691 | 0.05 | 0 |
| NC_045512.2;YP_009725295.1;Wu1-SARS2_orf1a;2692 | 0 | 0 |
| NC_045512.2;YP_009725295.1;Wu1-SARS2_orf1a;2693 | 0 | 0 |
| NC_045512.2;YP_009725295.1;Wu1-SARS2_orf1a;2694 | 0 | 0 |
| NC_045512.2;YP_009725295.1;Wu1-SARS2_orf1a;2695 | 0 | 0 |
| NC_045512.2;YP_009725295.1;Wu1-SARS2_orf1a;2696 | 0 | 0 |
| NC_045512.2;YP_009725295.1;Wu1-SARS2_orf1a;2697 | 0 | 0 |

| | | |
|---|---|---|
| NC_045512.2;YP_009725295.1;Wu1-SARS2_orf1a;2698 | 0 | 0 |
| NC_045512.2;YP_009725295.1;Wu1-SARS2_orf1a;2699 | 0 | 0 |
| NC_045512.2;YP_009725295.1;Wu1-SARS2_orf1a;2700 | 0 | 0 |
| NC_045512.2;YP_009725295.1;Wu1-SARS2_orf1a;2701 | 0 | 0 |
| NC_045512.2;YP_009725295.1;Wu1-SARS2_orf1a;2702 | 0 | 0 |
| NC_045512.2;YP_009725295.1;Wu1-SARS2_orf1a;2703 | 0 | 0 |
| NC_045512.2;YP_009725295.1;Wu1-SARS2_orf1a;2704 | 0 | 0 |
| NC_045512.2;YP_009725295.1;Wu1-SARS2_orf1a;2705 | 0 | 0 |
| NC_045512.2;YP_009725295.1;Wu1-SARS2_orf1a;2706 | 0 | 0 |
| NC_045512.2;YP_009725295.1;Wu1-SARS2_orf1a;2707 | 0 | 0 |
| NC_045512.2;YP_009725295.1;Wu1-SARS2_orf1a;2708 | 0 | 0 |
| NC_045512.2;YP_009725295.1;Wu1-SARS2_orf1a;2709 | 0 | 0 |
| NC_045512.2;YP_009725295.1;Wu1-SARS2_orf1a;2710 | 0 | 0 |
| NC_045512.2;YP_009725295.1;Wu1-SARS2_orf1a;2711 | 0 | 0 |
| NC_045512.2;YP_009725295.1;Wu1-SARS2_orf1a;2712 | 0 | 0 |
| NC_045512.2;YP_009725295.1;Wu1-SARS2_orf1a;2713 | 0 | 0 |
| NC_045512.2;YP_009725295.1;Wu1-SARS2_orf1a;2714 | 0 | 0 |
| NC_045512.2;YP_009725295.1;Wu1-SARS2_orf1a;2715 | 0 | 0 |
| NC_045512.2;YP_009725295.1;Wu1-SARS2_orf1a;2716 | 0 | 0 |
| NC_045512.2;YP_009725295.1;Wu1-SARS2_orf1a;2717 | 0 | 0 |
| NC_045512.2;YP_009725295.1;Wu1-SARS2_orf1a;2718 | 0.05 | 0 |
| NC_045512.2;YP_009725295.1;Wu1-SARS2_orf1a;2719 | 0 | 0 |
| NC_045512.2;YP_009725295.1;Wu1-SARS2_orf1a;2720 | 0 | 0 |
| NC_045512.2;YP_009725295.1;Wu1-SARS2_orf1a;2721 | 0 | 0 |
| NC_045512.2;YP_009725295.1;Wu1-SARS2_orf1a;2722 | 0 | 0.025 |
| NC_045512.2;YP_009725295.1;Wu1-SARS2_orf1a;2723 | 0 | 0 |
| NC_045512.2;YP_009725295.1;Wu1-SARS2_orf1a;2724 | 0 | 0 |
| NC_045512.2;YP_009725295.1;Wu1-SARS2_orf1a;2725 | 0 | 0 |
| NC_045512.2;YP_009725295.1;Wu1-SARS2_orf1a;2726 | 0 | 0.025 |
| NC_045512.2;YP_009725295.1;Wu1-SARS2_orf1a;2727 | 0 | 0.025 |
| NC_045512.2;YP_009725295.1;Wu1-SARS2_orf1a;2728 | 0 | 0.025 |
| NC_045512.2;YP_009725295.1;Wu1-SARS2_orf1a;2729 | 0.05 | 0.025 |
| NC_045512.2;YP_009725295.1;Wu1-SARS2_orf1a;2730 | 0 | 0.025 |
| NC_045512.2;YP_009725295.1;Wu1-SARS2_orf1a;2731 | 0 | 0.025 |
| NC_045512.2;YP_009725295.1;Wu1-SARS2_orf1a;2732 | 0 | 0.025 |
| NC_045512.2;YP_009725295.1;Wu1-SARS2_orf1a;2733 | 0 | 0.025 |
| NC_045512.2;YP_009725295.1;Wu1-SARS2_orf1a;2734 | 0 | 0 |
| NC_045512.2;YP_009725295.1;Wu1-SARS2_orf1a;2735 | 0 | 0 |
| NC_045512.2;YP_009725295.1;Wu1-SARS2_orf1a;2736 | 0 | 0 |
| NC_045512.2;YP_009725295.1;Wu1-SARS2_orf1a;2737 | 0 | 0 |
| NC_045512.2;YP_009725295.1;Wu1-SARS2_orf1a;2738 | 0 | 0.025 |
| NC_045512.2;YP_009725295.1;Wu1-SARS2_orf1a;2739 | 0 | 0 |
| NC_045512.2;YP_009725295.1;Wu1-SARS2_orf1a;2740 | 0 | 0 |

| | | |
|---|---|---|
| NC_045512.2;YP_009725295.1;Wu1-SARS2_orf1a;2741 | 0 | 0.025 |
| NC_045512.2;YP_009725295.1;Wu1-SARS2_orf1a;2742 | 0 | 0 |
| NC_045512.2;YP_009725295.1;Wu1-SARS2_orf1a;2743 | 0 | 0.025 |
| NC_045512.2;YP_009725295.1;Wu1-SARS2_orf1a;2744 | 0 | 0.025 |
| NC_045512.2;YP_009725295.1;Wu1-SARS2_orf1a;2745 | 0 | 0 |
| NC_045512.2;YP_009725295.1;Wu1-SARS2_orf1a;2746 | 0 | 0 |
| NC_045512.2;YP_009725295.1;Wu1-SARS2_orf1a;2747 | 0 | 0 |
| NC_045512.2;YP_009725295.1;Wu1-SARS2_orf1a;2748 | 0 | 0 |
| NC_045512.2;YP_009725295.1;Wu1-SARS2_orf1a;2749 | 0 | 0 |
| NC_045512.2;YP_009725295.1;Wu1-SARS2_orf1a;2750 | 0 | 0 |
| NC_045512.2;YP_009725295.1;Wu1-SARS2_orf1a;2751 | 0 | 0 |
| NC_045512.2;YP_009725295.1;Wu1-SARS2_orf1a;2752 | 0 | 0 |
| NC_045512.2;YP_009725295.1;Wu1-SARS2_orf1a;2753 | 0 | 0 |
| NC_045512.2;YP_009725295.1;Wu1-SARS2_orf1a;2754 | 0 | 0 |
| NC_045512.2;YP_009725295.1;Wu1-SARS2_orf1a;2755 | 0 | 0 |
| NC_045512.2;YP_009725295.1;Wu1-SARS2_orf1a;2756 | 0 | 0 |
| NC_045512.2;YP_009725295.1;Wu1-SARS2_orf1a;2757 | 0 | 0 |
| NC_045512.2;YP_009725295.1;Wu1-SARS2_orf1a;2758 | 0 | 0 |
| NC_045512.2;YP_009725295.1;Wu1-SARS2_orf1a;2759 | 0 | 0 |
| NC_045512.2;YP_009725295.1;Wu1-SARS2_orf1a;2760 | 0 | 0 |
| NC_045512.2;YP_009725295.1;Wu1-SARS2_orf1a;2761 | 0 | 0 |
| NC_045512.2;YP_009725295.1;Wu1-SARS2_orf1a;2762 | 0 | 0 |
| NC_045512.2;YP_009725295.1;Wu1-SARS2_orf1a;2763 | 0 | 0 |
| NC_045512.2;YP_009725295.1;Wu1-SARS2_orf1a;2764 | 0 | 0 |
| NC_045512.2;YP_009725295.1;Wu1-SARS2_orf1a;2765 | 0 | 0 |
| NC_045512.2;YP_009725295.1;Wu1-SARS2_orf1a;2766 | 0 | 0 |
| NC_045512.2;YP_009725295.1;Wu1-SARS2_orf1a;2767 | 0 | 0 |
| NC_045512.2;YP_009725295.1;Wu1-SARS2_orf1a;2768 | 0 | 0 |
| NC_045512.2;YP_009725295.1;Wu1-SARS2_orf1a;2769 | 0 | 0 |
| NC_045512.2;YP_009725295.1;Wu1-SARS2_orf1a;2770 | 0 | 0 |
| NC_045512.2;YP_009725295.1;Wu1-SARS2_orf1a;2771 | 0 | 0 |
| NC_045512.2;YP_009725295.1;Wu1-SARS2_orf1a;2772 | 0 | 0 |
| NC_045512.2;YP_009725295.1;Wu1-SARS2_orf1a;2773 | 0 | 0 |
| NC_045512.2;YP_009725295.1;Wu1-SARS2_orf1a;2774 | 0 | 0 |
| NC_045512.2;YP_009725295.1;Wu1-SARS2_orf1a;2775 | 0 | 0 |
| NC_045512.2;YP_009725295.1;Wu1-SARS2_orf1a;2776 | 0 | 0 |
| NC_045512.2;YP_009725295.1;Wu1-SARS2_orf1a;2777 | 0 | 0 |
| NC_045512.2;YP_009725295.1;Wu1-SARS2_orf1a;2778 | 0 | 0 |
| NC_045512.2;YP_009725295.1;Wu1-SARS2_orf1a;2779 | 0 | 0 |
| NC_045512.2;YP_009725295.1;Wu1-SARS2_orf1a;2780 | 0 | 0 |
| NC_045512.2;YP_009725295.1;Wu1-SARS2_orf1a;2781 | 0 | 0 |
| NC_045512.2;YP_009725295.1;Wu1-SARS2_orf1a;2782 | 0 | 0 |
| NC_045512.2;YP_009725295.1;Wu1-SARS2_orf1a;2783 | 0 | 0 |

| | | |
|---|---|---|
| NC_045512.2;YP_009725295.1;Wu1-SARS2_orf1a;2784 | 0 | 0 |
| NC_045512.2;YP_009725295.1;Wu1-SARS2_orf1a;2785 | 0 | 0 |
| NC_045512.2;YP_009725295.1;Wu1-SARS2_orf1a;2786 | 0 | 0 |
| NC_045512.2;YP_009725295.1;Wu1-SARS2_orf1a;2787 | 0 | 0 |
| NC_045512.2;YP_009725295.1;Wu1-SARS2_orf1a;2788 | 0 | 0 |
| NC_045512.2;YP_009725295.1;Wu1-SARS2_orf1a;2789 | 0 | 0 |
| NC_045512.2;YP_009725295.1;Wu1-SARS2_orf1a;2790 | 0 | 0 |
| NC_045512.2;YP_009725295.1;Wu1-SARS2_orf1a;2791 | 0 | 0 |
| NC_045512.2;YP_009725295.1;Wu1-SARS2_orf1a;2792 | 0 | 0 |
| NC_045512.2;YP_009725295.1;Wu1-SARS2_orf1a;2793 | 0 | 0 |
| NC_045512.2;YP_009725295.1;Wu1-SARS2_orf1a;2794 | 0 | 0 |
| NC_045512.2;YP_009725295.1;Wu1-SARS2_orf1a;2795 | 0 | 0 |
| NC_045512.2;YP_009725295.1;Wu1-SARS2_orf1a;2796 | 0.05 | 0 |
| NC_045512.2;YP_009725295.1;Wu1-SARS2_orf1a;2797 | 0 | 0.025 |
| NC_045512.2;YP_009725295.1;Wu1-SARS2_orf1a;2798 | 0.05 | 0.025 |
| NC_045512.2;YP_009725295.1;Wu1-SARS2_orf1a;2799 | 0.05 | 0.025 |
| NC_045512.2;YP_009725295.1;Wu1-SARS2_orf1a;2800 | 0.05 | 0.025 |
| NC_045512.2;YP_009725295.1;Wu1-SARS2_orf1a;2801 | 0.05 | 0.025 |
| NC_045512.2;YP_009725295.1;Wu1-SARS2_orf1a;2802 | 0.1 | 0.025 |
| NC_045512.2;YP_009725295.1;Wu1-SARS2_orf1a;2803 | 0.05 | 0.025 |
| NC_045512.2;YP_009725295.1;Wu1-SARS2_orf1a;2804 | 0.05 | 0.075 |
| NC_045512.2;YP_009725295.1;Wu1-SARS2_orf1a;2805 | 0.05 | 0.05 |
| NC_045512.2;YP_009725295.1;Wu1-SARS2_orf1a;2806 | 0.05 | 0 |
| NC_045512.2;YP_009725295.1;Wu1-SARS2_orf1a;2807 | 0 | 0 |
| NC_045512.2;YP_009725295.1;Wu1-SARS2_orf1a;2808 | 0 | 0 |
| NC_045512.2;YP_009725295.1;Wu1-SARS2_orf1a;2809 | 0 | 0 |
| NC_045512.2;YP_009725295.1;Wu1-SARS2_orf1a;2810 | 0 | 0 |
| NC_045512.2;YP_009725295.1;Wu1-SARS2_orf1a;2811 | 0 | 0 |
| NC_045512.2;YP_009725295.1;Wu1-SARS2_orf1a;2812 | 0 | 0 |
| NC_045512.2;YP_009725295.1;Wu1-SARS2_orf1a;2813 | 0 | 0 |
| NC_045512.2;YP_009725295.1;Wu1-SARS2_orf1a;2814 | 0 | 0 |
| NC_045512.2;YP_009725295.1;Wu1-SARS2_orf1a;2815 | 0 | 0.05 |
| NC_045512.2;YP_009725295.1;Wu1-SARS2_orf1a;2816 | 0 | 0 |
| NC_045512.2;YP_009725295.1;Wu1-SARS2_orf1a;2817 | 0 | 0.025 |
| NC_045512.2;YP_009725295.1;Wu1-SARS2_orf1a;2818 | 0.05 | 0 |
| NC_045512.2;YP_009725295.1;Wu1-SARS2_orf1a;2819 | 0 | 0 |
| NC_045512.2;YP_009725295.1;Wu1-SARS2_orf1a;2820 | 0 | 0 |
| NC_045512.2;YP_009725295.1;Wu1-SARS2_orf1a;2821 | 0.05 | 0 |
| NC_045512.2;YP_009725295.1;Wu1-SARS2_orf1a;2822 | 0.1 | 0 |
| NC_045512.2;YP_009725295.1;Wu1-SARS2_orf1a;2823 | 0 | 0 |
| NC_045512.2;YP_009725295.1;Wu1-SARS2_orf1a;2824 | 0 | 0 |
| NC_045512.2;YP_009725295.1;Wu1-SARS2_orf1a;2825 | 0 | 0 |
| NC_045512.2;YP_009725295.1;Wu1-SARS2_orf1a;2826 | 0 | 0 |

| | | |
|---|---|---|
| NC_045512.2;YP_009725295.1;Wu1-SARS2_orf1a;2827 | 0 | 0 |
| NC_045512.2;YP_009725295.1;Wu1-SARS2_orf1a;2828 | 0.1 | 0.025 |
| NC_045512.2;YP_009725295.1;Wu1-SARS2_orf1a;2829 | 0 | 0 |
| NC_045512.2;YP_009725295.1;Wu1-SARS2_orf1a;2830 | 0 | 0 |
| NC_045512.2;YP_009725295.1;Wu1-SARS2_orf1a;2831 | 0 | 0 |
| NC_045512.2;YP_009725295.1;Wu1-SARS2_orf1a;2832 | 0 | 0 |
| NC_045512.2;YP_009725295.1;Wu1-SARS2_orf1a;2833 | 0 | 0 |
| NC_045512.2;YP_009725295.1;Wu1-SARS2_orf1a;2834 | 0 | 0 |
| NC_045512.2;YP_009725295.1;Wu1-SARS2_orf1a;2835 | 0.05 | 0 |
| NC_045512.2;YP_009725295.1;Wu1-SARS2_orf1a;2836 | 0 | 0.05 |
| NC_045512.2;YP_009725295.1;Wu1-SARS2_orf1a;2837 | 0 | 0 |
| NC_045512.2;YP_009725295.1;Wu1-SARS2_orf1a;2838 | 0 | 0 |
| NC_045512.2;YP_009725295.1;Wu1-SARS2_orf1a;2839 | 0.05 | 0 |
| NC_045512.2;YP_009725295.1;Wu1-SARS2_orf1a;2840 | 0 | 0.025 |
| NC_045512.2;YP_009725295.1;Wu1-SARS2_orf1a;2841 | 0.05 | 0.025 |
| NC_045512.2;YP_009725295.1;Wu1-SARS2_orf1a;2842 | 0 | 0 |
| NC_045512.2;YP_009725295.1;Wu1-SARS2_orf1a;2843 | 0 | 0 |
| NC_045512.2;YP_009725295.1;Wu1-SARS2_orf1a;2844 | 0 | 0.05 |
| NC_045512.2;YP_009725295.1;Wu1-SARS2_orf1a;2845 | 0 | 0 |
| NC_045512.2;YP_009725295.1;Wu1-SARS2_orf1a;2846 | 0 | 0 |
| NC_045512.2;YP_009725295.1;Wu1-SARS2_orf1a;2847 | 0 | 0 |
| NC_045512.2;YP_009725295.1;Wu1-SARS2_orf1a;2848 | 0 | 0 |
| NC_045512.2;YP_009725295.1;Wu1-SARS2_orf1a;2849 | 0 | 0 |
| NC_045512.2;YP_009725295.1;Wu1-SARS2_orf1a;2850 | 0 | 0 |
| NC_045512.2;YP_009725295.1;Wu1-SARS2_orf1a;2851 | 0 | 0 |
| NC_045512.2;YP_009725295.1;Wu1-SARS2_orf1a;2852 | 0 | 0 |
| NC_045512.2;YP_009725295.1;Wu1-SARS2_orf1a;2853 | 0 | 0 |
| NC_045512.2;YP_009725295.1;Wu1-SARS2_orf1a;2854 | 0 | 0 |
| NC_045512.2;YP_009725295.1;Wu1-SARS2_orf1a;2855 | 0 | 0 |
| NC_045512.2;YP_009725295.1;Wu1-SARS2_orf1a;2856 | 0 | 0 |
| NC_045512.2;YP_009725295.1;Wu1-SARS2_orf1a;2857 | 0 | 0.025 |
| NC_045512.2;YP_009725295.1;Wu1-SARS2_orf1a;2858 | 0 | 0.025 |
| NC_045512.2;YP_009725295.1;Wu1-SARS2_orf1a;2859 | 0 | 0.025 |
| NC_045512.2;YP_009725295.1;Wu1-SARS2_orf1a;2860 | 0 | 0 |
| NC_045512.2;YP_009725295.1;Wu1-SARS2_orf1a;2861 | 0 | 0 |
| NC_045512.2;YP_009725295.1;Wu1-SARS2_orf1a;2862 | 0 | 0 |
| NC_045512.2;YP_009725295.1;Wu1-SARS2_orf1a;2863 | 0 | 0 |
| NC_045512.2;YP_009725295.1;Wu1-SARS2_orf1a;2864 | 0 | 0 |
| NC_045512.2;YP_009725295.1;Wu1-SARS2_orf1a;2865 | 0 | 0 |
| NC_045512.2;YP_009725295.1;Wu1-SARS2_orf1a;2866 | 0 | 0 |
| NC_045512.2;YP_009725295.1;Wu1-SARS2_orf1a;2867 | 0 | 0 |
| NC_045512.2;YP_009725295.1;Wu1-SARS2_orf1a;2868 | 0 | 0 |
| NC_045512.2;YP_009725295.1;Wu1-SARS2_orf1a;2869 | 0 | 0 |

| | | |
|---|---|---|
| NC_045512.2;YP_009725295.1;Wu1-SARS2_orf1a;2870 | 0 | 0 |
| NC_045512.2;YP_009725295.1;Wu1-SARS2_orf1a;2871 | 0 | 0 |
| NC_045512.2;YP_009725295.1;Wu1-SARS2_orf1a;2872 | 0 | 0 |
| NC_045512.2;YP_009725295.1;Wu1-SARS2_orf1a;2873 | 0 | 0 |
| NC_045512.2;YP_009725295.1;Wu1-SARS2_orf1a;2874 | 0 | 0 |
| NC_045512.2;YP_009725295.1;Wu1-SARS2_orf1a;2875 | 0 | 0 |
| NC_045512.2;YP_009725295.1;Wu1-SARS2_orf1a;2876 | 0 | 0 |
| NC_045512.2;YP_009725295.1;Wu1-SARS2_orf1a;2877 | 0 | 0 |
| NC_045512.2;YP_009725295.1;Wu1-SARS2_orf1a;2878 | 0 | 0 |
| NC_045512.2;YP_009725295.1;Wu1-SARS2_orf1a;2879 | 0 | 0 |
| NC_045512.2;YP_009725295.1;Wu1-SARS2_orf1a;2880 | 0 | 0 |
| NC_045512.2;YP_009725295.1;Wu1-SARS2_orf1a;2881 | 0 | 0 |
| NC_045512.2;YP_009725295.1;Wu1-SARS2_orf1a;2882 | 0 | 0 |
| NC_045512.2;YP_009725295.1;Wu1-SARS2_orf1a;2883 | 0 | 0 |
| NC_045512.2;YP_009725295.1;Wu1-SARS2_orf1a;2884 | 0 | 0 |
| NC_045512.2;YP_009725295.1;Wu1-SARS2_orf1a;2885 | 0 | 0 |
| NC_045512.2;YP_009725295.1;Wu1-SARS2_orf1a;2886 | 0 | 0 |
| NC_045512.2;YP_009725295.1;Wu1-SARS2_orf1a;2887 | 0 | 0 |
| NC_045512.2;YP_009725295.1;Wu1-SARS2_orf1a;2888 | 0 | 0 |
| NC_045512.2;YP_009725295.1;Wu1-SARS2_orf1a;2889 | 0 | 0 |
| NC_045512.2;YP_009725295.1;Wu1-SARS2_orf1a;2890 | 0 | 0 |
| NC_045512.2;YP_009725295.1;Wu1-SARS2_orf1a;2891 | 0.05 | 0 |
| NC_045512.2;YP_009725295.1;Wu1-SARS2_orf1a;2892 | 0.05 | 0.025 |
| NC_045512.2;YP_009725295.1;Wu1-SARS2_orf1a;2893 | 0 | 0.025 |
| NC_045512.2;YP_009725295.1;Wu1-SARS2_orf1a;2894 | 0 | 0.025 |
| NC_045512.2;YP_009725295.1;Wu1-SARS2_orf1a;2895 | 0.05 | 0 |
| NC_045512.2;YP_009725295.1;Wu1-SARS2_orf1a;2896 | 0.05 | 0.05 |
| NC_045512.2;YP_009725295.1;Wu1-SARS2_orf1a;2897 | 0.05 | 0.025 |
| NC_045512.2;YP_009725295.1;Wu1-SARS2_orf1a;2898 | 0.05 | 0 |
| NC_045512.2;YP_009725295.1;Wu1-SARS2_orf1a;2899 | 0 | 0 |
| NC_045512.2;YP_009725295.1;Wu1-SARS2_orf1a;2900 | 0 | 0 |
| NC_045512.2;YP_009725295.1;Wu1-SARS2_orf1a;2901 | 0 | 0 |
| NC_045512.2;YP_009725295.1;Wu1-SARS2_orf1a;2902 | 0 | 0 |
| NC_045512.2;YP_009725295.1;Wu1-SARS2_orf1a;2903 | 0.05 | 0 |
| NC_045512.2;YP_009725295.1;Wu1-SARS2_orf1a;2904 | 0 | 0 |
| NC_045512.2;YP_009725295.1;Wu1-SARS2_orf1a;2905 | 0 | 0 |
| NC_045512.2;YP_009725295.1;Wu1-SARS2_orf1a;2906 | 0 | 0 |
| NC_045512.2;YP_009725295.1;Wu1-SARS2_orf1a;2907 | 0 | 0 |
| NC_045512.2;YP_009725295.1;Wu1-SARS2_orf1a;2908 | 0 | 0 |
| NC_045512.2;YP_009725295.1;Wu1-SARS2_orf1a;2909 | 0 | 0 |
| NC_045512.2;YP_009725295.1;Wu1-SARS2_orf1a;2910 | 0 | 0 |
| NC_045512.2;YP_009725295.1;Wu1-SARS2_orf1a;2911 | 0 | 0 |
| NC_045512.2;YP_009725295.1;Wu1-SARS2_orf1a;2912 | 0 | 0 |

| | | |
|---|---|---|
| NC_045512.2;YP_009725295.1;Wu1-SARS2_orf1a;2913 | 0.05 | 0.025 |
| NC_045512.2;YP_009725295.1;Wu1-SARS2_orf1a;2914 | 0.05 | 0.025 |
| NC_045512.2;YP_009725295.1;Wu1-SARS2_orf1a;2915 | 0.05 | 0.125 |
| NC_045512.2;YP_009725295.1;Wu1-SARS2_orf1a;2916 | 0.15 | 0.15 |
| NC_045512.2;YP_009725295.1;Wu1-SARS2_orf1a;2917 | 0.1 | 0.05 |
| NC_045512.2;YP_009725295.1;Wu1-SARS2_orf1a;2918 | 0.1 | 0.05 |
| NC_045512.2;YP_009725295.1;Wu1-SARS2_orf1a;2919 | 0 | 0.025 |
| NC_045512.2;YP_009725295.1;Wu1-SARS2_orf1a;2920 | 0.1 | 0.025 |
| NC_045512.2;YP_009725295.1;Wu1-SARS2_orf1a;2921 | 0.05 | 0.025 |
| NC_045512.2;YP_009725295.1;Wu1-SARS2_orf1a;2922 | 0 | 0.025 |
| NC_045512.2;YP_009725295.1;Wu1-SARS2_orf1a;2923 | 0 | 0.025 |
| NC_045512.2;YP_009725295.1;Wu1-SARS2_orf1a;2924 | 0 | 0 |
| NC_045512.2;YP_009725295.1;Wu1-SARS2_orf1a;2925 | 0 | 0 |
| NC_045512.2;YP_009725295.1;Wu1-SARS2_orf1a;2926 | 0 | 0 |
| NC_045512.2;YP_009725295.1;Wu1-SARS2_orf1a;2927 | 0 | 0 |
| NC_045512.2;YP_009725295.1;Wu1-SARS2_orf1a;2928 | 0 | 0 |
| NC_045512.2;YP_009725295.1;Wu1-SARS2_orf1a;2929 | 0 | 0.025 |
| NC_045512.2;YP_009725295.1;Wu1-SARS2_orf1a;2930 | 0 | 0 |
| NC_045512.2;YP_009725295.1;Wu1-SARS2_orf1a;2931 | 0 | 0 |
| NC_045512.2;YP_009725295.1;Wu1-SARS2_orf1a;2932 | 0 | 0 |
| NC_045512.2;YP_009725295.1;Wu1-SARS2_orf1a;2933 | 0 | 0 |
| NC_045512.2;YP_009725295.1;Wu1-SARS2_orf1a;2934 | 0 | 0 |
| NC_045512.2;YP_009725295.1;Wu1-SARS2_orf1a;2935 | 0 | 0 |
| NC_045512.2;YP_009725295.1;Wu1-SARS2_orf1a;2936 | 0.1 | 0 |
| NC_045512.2;YP_009725295.1;Wu1-SARS2_orf1a;2937 | 0 | 0.025 |
| NC_045512.2;YP_009725295.1;Wu1-SARS2_orf1a;2938 | 0 | 0.025 |
| NC_045512.2;YP_009725295.1;Wu1-SARS2_orf1a;2939 | 0.15 | 0.15 |
| NC_045512.2;YP_009725295.1;Wu1-SARS2_orf1a;2940 | 0.15 | 0.15 |
| NC_045512.2;YP_009725295.1;Wu1-SARS2_orf1a;2941 | 0.1 | 0.025 |
| NC_045512.2;YP_009725295.1;Wu1-SARS2_orf1a;2942 | 0.1 | 0.05 |
| NC_045512.2;YP_009725295.1;Wu1-SARS2_orf1a;2943 | 0.05 | 0.025 |
| NC_045512.2;YP_009725295.1;Wu1-SARS2_orf1a;2944 | 0.15 | 0.075 |
| NC_045512.2;YP_009725295.1;Wu1-SARS2_orf1a;2945 | 0.1 | 0.1 |
| NC_045512.2;YP_009725295.1;Wu1-SARS2_orf1a;2946 | 0.15 | 0.025 |
| NC_045512.2;YP_009725295.1;Wu1-SARS2_orf1a;2947 | 0.05 | 0.025 |
| NC_045512.2;YP_009725295.1;Wu1-SARS2_orf1a;2948 | 0.05 | 0.05 |
| NC_045512.2;YP_009725295.1;Wu1-SARS2_orf1a;2949 | 0.05 | 0.025 |
| NC_045512.2;YP_009725295.1;Wu1-SARS2_orf1a;2950 | 0 | 0 |
| NC_045512.2;YP_009725295.1;Wu1-SARS2_orf1a;2951 | 0 | 0 |
| NC_045512.2;YP_009725295.1;Wu1-SARS2_orf1a;2952 | 0 | 0 |
| NC_045512.2;YP_009725295.1;Wu1-SARS2_orf1a;2953 | 0 | 0 |
| NC_045512.2;YP_009725295.1;Wu1-SARS2_orf1a;2954 | 0 | 0 |
| NC_045512.2;YP_009725295.1;Wu1-SARS2_orf1a;2955 | 0 | 0 |

| | | |
|---|---|---|
| NC_045512.2;YP_009725295.1;Wu1-SARS2_orf1a;2956 | 0 | 0 |
| NC_045512.2;YP_009725295.1;Wu1-SARS2_orf1a;2957 | 0 | 0 |
| NC_045512.2;YP_009725295.1;Wu1-SARS2_orf1a;2958 | 0 | 0 |
| NC_045512.2;YP_009725295.1;Wu1-SARS2_orf1a;2959 | 0 | 0 |
| NC_045512.2;YP_009725295.1;Wu1-SARS2_orf1a;2960 | 0 | 0.025 |
| NC_045512.2;YP_009725295.1;Wu1-SARS2_orf1a;2961 | 0 | 0 |
| NC_045512.2;YP_009725295.1;Wu1-SARS2_orf1a;2962 | 0 | 0 |
| NC_045512.2;YP_009725295.1;Wu1-SARS2_orf1a;2963 | 0 | 0 |
| NC_045512.2;YP_009725295.1;Wu1-SARS2_orf1a;2964 | 0 | 0 |
| NC_045512.2;YP_009725295.1;Wu1-SARS2_orf1a;2965 | 0 | 0 |
| NC_045512.2;YP_009725295.1;Wu1-SARS2_orf1a;2966 | 0 | 0 |
| NC_045512.2;YP_009725295.1;Wu1-SARS2_orf1a;2967 | 0 | 0 |
| NC_045512.2;YP_009725295.1;Wu1-SARS2_orf1a;2968 | 0 | 0 |
| NC_045512.2;YP_009725295.1;Wu1-SARS2_orf1a;2969 | 0 | 0 |
| NC_045512.2;YP_009725295.1;Wu1-SARS2_orf1a;2970 | 0 | 0 |
| NC_045512.2;YP_009725295.1;Wu1-SARS2_orf1a;2971 | 0.05 | 0.1 |
| NC_045512.2;YP_009725295.1;Wu1-SARS2_orf1a;2972 | 0 | 0.05 |
| NC_045512.2;YP_009725295.1;Wu1-SARS2_orf1a;2973 | 0 | 0.075 |
| NC_045512.2;YP_009725295.1;Wu1-SARS2_orf1a;2974 | 0 | 0.075 |
| NC_045512.2;YP_009725295.1;Wu1-SARS2_orf1a;2975 | 0 | 0.05 |
| NC_045512.2;YP_009725295.1;Wu1-SARS2_orf1a;2976 | 0 | 0.05 |
| NC_045512.2;YP_009725295.1;Wu1-SARS2_orf1a;2977 | 0 | 0.05 |
| NC_045512.2;YP_009725295.1;Wu1-SARS2_orf1a;2978 | 0 | 0.05 |
| NC_045512.2;YP_009725295.1;Wu1-SARS2_orf1a;2979 | 0 | 0.05 |
| NC_045512.2;YP_009725295.1;Wu1-SARS2_orf1a;2980 | 0 | 0 |
| NC_045512.2;YP_009725295.1;Wu1-SARS2_orf1a;2981 | 0 | 0 |
| NC_045512.2;YP_009725295.1;Wu1-SARS2_orf1a;2982 | 0 | 0 |
| NC_045512.2;YP_009725295.1;Wu1-SARS2_orf1a;2983 | 0 | 0 |
| NC_045512.2;YP_009725295.1;Wu1-SARS2_orf1a;2984 | 0 | 0 |
| NC_045512.2;YP_009725295.1;Wu1-SARS2_orf1a;2985 | 0 | 0 |
| NC_045512.2;YP_009725295.1;Wu1-SARS2_orf1a;2986 | 0 | 0 |
| NC_045512.2;YP_009725295.1;Wu1-SARS2_orf1a;2987 | 0 | 0 |
| NC_045512.2;YP_009725295.1;Wu1-SARS2_orf1a;2988 | 0 | 0 |
| NC_045512.2;YP_009725295.1;Wu1-SARS2_orf1a;2989 | 0 | 0 |
| NC_045512.2;YP_009725295.1;Wu1-SARS2_orf1a;2990 | 0 | 0 |
| NC_045512.2;YP_009725295.1;Wu1-SARS2_orf1a;2991 | 0 | 0 |
| NC_045512.2;YP_009725295.1;Wu1-SARS2_orf1a;2992 | 0 | 0 |
| NC_045512.2;YP_009725295.1;Wu1-SARS2_orf1a;2993 | 0 | 0 |
| NC_045512.2;YP_009725295.1;Wu1-SARS2_orf1a;2994 | 0 | 0 |
| NC_045512.2;YP_009725295.1;Wu1-SARS2_orf1a;2995 | 0 | 0 |
| NC_045512.2;YP_009725295.1;Wu1-SARS2_orf1a;2996 | 0 | 0 |
| NC_045512.2;YP_009725295.1;Wu1-SARS2_orf1a;2997 | 0 | 0 |
| NC_045512.2;YP_009725295.1;Wu1-SARS2_orf1a;2998 | 0 | 0 |

| | | |
|---|---|---|
| NC_045512.2;YP_009725295.1;Wu1-SARS2_orf1a;2999 | 0 | 0 |
| NC_045512.2;YP_009725295.1;Wu1-SARS2_orf1a;3000 | 0 | 0 |
| NC_045512.2;YP_009725295.1;Wu1-SARS2_orf1a;3001 | 0 | 0 |
| NC_045512.2;YP_009725295.1;Wu1-SARS2_orf1a;3002 | 0 | 0 |
| NC_045512.2;YP_009725295.1;Wu1-SARS2_orf1a;3003 | 0 | 0 |
| NC_045512.2;YP_009725295.1;Wu1-SARS2_orf1a;3004 | 0 | 0 |
| NC_045512.2;YP_009725295.1;Wu1-SARS2_orf1a;3005 | 0 | 0 |
| NC_045512.2;YP_009725295.1;Wu1-SARS2_orf1a;3006 | 0 | 0 |
| NC_045512.2;YP_009725295.1;Wu1-SARS2_orf1a;3007 | 0 | 0 |
| NC_045512.2;YP_009725295.1;Wu1-SARS2_orf1a;3008 | 0 | 0 |
| NC_045512.2;YP_009725295.1;Wu1-SARS2_orf1a;3009 | 0 | 0 |
| NC_045512.2;YP_009725295.1;Wu1-SARS2_orf1a;3010 | 0.1 | 0.15 |
| NC_045512.2;YP_009725295.1;Wu1-SARS2_orf1a;3011 | 0.1 | 0.15 |
| NC_045512.2;YP_009725295.1;Wu1-SARS2_orf1a;3012 | 0.1 | 0.125 |
| NC_045512.2;YP_009725295.1;Wu1-SARS2_orf1a;3013 | 0.1 | 0.15 |
| NC_045512.2;YP_009725295.1;Wu1-SARS2_orf1a;3014 | 0.1 | 0.15 |
| NC_045512.2;YP_009725295.1;Wu1-SARS2_orf1a;3015 | 0.1 | 0.15 |
| NC_045512.2;YP_009725295.1;Wu1-SARS2_orf1a;3016 | 0.1 | 0.15 |
| NC_045512.2;YP_009725295.1;Wu1-SARS2_orf1a;3017 | 0.1 | 0.15 |
| NC_045512.2;YP_009725295.1;Wu1-SARS2_orf1a;3018 | 0.1 | 0.15 |
| NC_045512.2;YP_009725295.1;Wu1-SARS2_orf1a;3019 | 0.1 | 0.15 |
| NC_045512.2;YP_009725295.1;Wu1-SARS2_orf1a;3020 | 0.15 | 0.225 |
| NC_045512.2;YP_009725295.1;Wu1-SARS2_orf1a;3021 | 0.1 | 0.15 |
| NC_045512.2;YP_009725295.1;Wu1-SARS2_orf1a;3022 | 0 | 0 |
| NC_045512.2;YP_009725295.1;Wu1-SARS2_orf1a;3023 | 0 | 0 |
| NC_045512.2;YP_009725295.1;Wu1-SARS2_orf1a;3024 | 0 | 0 |
| NC_045512.2;YP_009725295.1;Wu1-SARS2_orf1a;3025 | 0.05 | 0 |
| NC_045512.2;YP_009725295.1;Wu1-SARS2_orf1a;3026 | 0 | 0 |
| NC_045512.2;YP_009725295.1;Wu1-SARS2_orf1a;3027 | 0.05 | 0 |
| NC_045512.2;YP_009725295.1;Wu1-SARS2_orf1a;3028 | 0.05 | 0 |
| NC_045512.2;YP_009725295.1;Wu1-SARS2_orf1a;3029 | 0.05 | 0 |
| NC_045512.2;YP_009725295.1;Wu1-SARS2_orf1a;3030 | 0.05 | 0 |
| NC_045512.2;YP_009725295.1;Wu1-SARS2_orf1a;3031 | 0.05 | 0 |
| NC_045512.2;YP_009725295.1;Wu1-SARS2_orf1a;3032 | 0.05 | 0 |
| NC_045512.2;YP_009725295.1;Wu1-SARS2_orf1a;3033 | 0 | 0 |
| NC_045512.2;YP_009725295.1;Wu1-SARS2_orf1a;3034 | 0 | 0 |
| NC_045512.2;YP_009725295.1;Wu1-SARS2_orf1a;3035 | 0 | 0 |
| NC_045512.2;YP_009725295.1;Wu1-SARS2_orf1a;3036 | 0.05 | 0 |
| NC_045512.2;YP_009725295.1;Wu1-SARS2_orf1a;3037 | 0 | 0 |
| NC_045512.2;YP_009725295.1;Wu1-SARS2_orf1a;3038 | 0 | 0 |
| NC_045512.2;YP_009725295.1;Wu1-SARS2_orf1a;3039 | 0 | 0.025 |
| NC_045512.2;YP_009725295.1;Wu1-SARS2_orf1a;3040 | 0 | 0 |
| NC_045512.2;YP_009725295.1;Wu1-SARS2_orf1a;3041 | 0 | 0 |

| | | |
|---|---|---|
| NC_045512.2;YP_009725295.1;Wu1-SARS2_orf1a;3042 | 0 | 0 |
| NC_045512.2;YP_009725295.1;Wu1-SARS2_orf1a;3043 | 0 | 0 |
| NC_045512.2;YP_009725295.1;Wu1-SARS2_orf1a;3044 | 0 | 0 |
| NC_045512.2;YP_009725295.1;Wu1-SARS2_orf1a;3045 | 0 | 0 |
| NC_045512.2;YP_009725295.1;Wu1-SARS2_orf1a;3046 | 0 | 0 |
| NC_045512.2;YP_009725295.1;Wu1-SARS2_orf1a;3047 | 0 | 0 |
| NC_045512.2;YP_009725295.1;Wu1-SARS2_orf1a;3048 | 0 | 0 |
| NC_045512.2;YP_009725295.1;Wu1-SARS2_orf1a;3049 | 0 | 0 |
| NC_045512.2;YP_009725295.1;Wu1-SARS2_orf1a;3050 | 0 | 0 |
| NC_045512.2;YP_009725295.1;Wu1-SARS2_orf1a;3051 | 0 | 0 |
| NC_045512.2;YP_009725295.1;Wu1-SARS2_orf1a;3052 | 0 | 0 |
| NC_045512.2;YP_009725295.1;Wu1-SARS2_orf1a;3053 | 0 | 0 |
| NC_045512.2;YP_009725295.1;Wu1-SARS2_orf1a;3054 | 0 | 0 |
| NC_045512.2;YP_009725295.1;Wu1-SARS2_orf1a;3055 | 0 | 0 |
| NC_045512.2;YP_009725295.1;Wu1-SARS2_orf1a;3056 | 0 | 0 |
| NC_045512.2;YP_009725295.1;Wu1-SARS2_orf1a;3057 | 0 | 0 |
| NC_045512.2;YP_009725295.1;Wu1-SARS2_orf1a;3058 | 0 | 0 |
| NC_045512.2;YP_009725295.1;Wu1-SARS2_orf1a;3059 | 0 | 0 |
| NC_045512.2;YP_009725295.1;Wu1-SARS2_orf1a;3060 | 0 | 0 |
| NC_045512.2;YP_009725295.1;Wu1-SARS2_orf1a;3061 | 0 | 0 |
| NC_045512.2;YP_009725295.1;Wu1-SARS2_orf1a;3062 | 0 | 0 |
| NC_045512.2;YP_009725295.1;Wu1-SARS2_orf1a;3063 | 0 | 0 |
| NC_045512.2;YP_009725295.1;Wu1-SARS2_orf1a;3064 | 0 | 0 |
| NC_045512.2;YP_009725295.1;Wu1-SARS2_orf1a;3065 | 0 | 0 |
| NC_045512.2;YP_009725295.1;Wu1-SARS2_orf1a;3066 | 0 | 0 |
| NC_045512.2;YP_009725295.1;Wu1-SARS2_orf1a;3067 | 0 | 0 |
| NC_045512.2;YP_009725295.1;Wu1-SARS2_orf1a;3068 | 0 | 0 |
| NC_045512.2;YP_009725295.1;Wu1-SARS2_orf1a;3069 | 0 | 0 |
| NC_045512.2;YP_009725295.1;Wu1-SARS2_orf1a;3070 | 0 | 0 |
| NC_045512.2;YP_009725295.1;Wu1-SARS2_orf1a;3071 | 0 | 0 |
| NC_045512.2;YP_009725295.1;Wu1-SARS2_orf1a;3072 | 0 | 0 |
| NC_045512.2;YP_009725295.1;Wu1-SARS2_orf1a;3073 | 0 | 0 |
| NC_045512.2;YP_009725295.1;Wu1-SARS2_orf1a;3074 | 0 | 0 |
| NC_045512.2;YP_009725295.1;Wu1-SARS2_orf1a;3075 | 0 | 0 |
| NC_045512.2;YP_009725295.1;Wu1-SARS2_orf1a;3076 | 0 | 0 |
| NC_045512.2;YP_009725295.1;Wu1-SARS2_orf1a;3077 | 0 | 0 |
| NC_045512.2;YP_009725295.1;Wu1-SARS2_orf1a;3078 | 0 | 0 |
| NC_045512.2;YP_009725295.1;Wu1-SARS2_orf1a;3079 | 0 | 0 |
| NC_045512.2;YP_009725295.1;Wu1-SARS2_orf1a;3080 | 0 | 0 |
| NC_045512.2;YP_009725295.1;Wu1-SARS2_orf1a;3081 | 0 | 0 |
| NC_045512.2;YP_009725295.1;Wu1-SARS2_orf1a;3082 | 0 | 0 |
| NC_045512.2;YP_009725295.1;Wu1-SARS2_orf1a;3083 | 0 | 0 |
| NC_045512.2;YP_009725295.1;Wu1-SARS2_orf1a;3084 | 0 | 0 |

| | | |
|---|---|---|
| NC_045512.2;YP_009725295.1;Wu1-SARS2_orf1a;3085 | 0 | 0 |
| NC_045512.2;YP_009725295.1;Wu1-SARS2_orf1a;3086 | 0 | 0 |
| NC_045512.2;YP_009725295.1;Wu1-SARS2_orf1a;3087 | 0 | 0 |
| NC_045512.2;YP_009725295.1;Wu1-SARS2_orf1a;3088 | 0 | 0 |
| NC_045512.2;YP_009725295.1;Wu1-SARS2_orf1a;3089 | 0 | 0 |
| NC_045512.2;YP_009725295.1;Wu1-SARS2_orf1a;3090 | 0 | 0 |
| NC_045512.2;YP_009725295.1;Wu1-SARS2_orf1a;3091 | 0 | 0 |
| NC_045512.2;YP_009725295.1;Wu1-SARS2_orf1a;3092 | 0 | 0 |
| NC_045512.2;YP_009725295.1;Wu1-SARS2_orf1a;3093 | 0 | 0 |
| NC_045512.2;YP_009725295.1;Wu1-SARS2_orf1a;3094 | 0 | 0 |
| NC_045512.2;YP_009725295.1;Wu1-SARS2_orf1a;3095 | 0 | 0 |
| NC_045512.2;YP_009725295.1;Wu1-SARS2_orf1a;3096 | 0 | 0 |
| NC_045512.2;YP_009725295.1;Wu1-SARS2_orf1a;3097 | 0 | 0 |
| NC_045512.2;YP_009725295.1;Wu1-SARS2_orf1a;3098 | 0 | 0 |
| NC_045512.2;YP_009725295.1;Wu1-SARS2_orf1a;3099 | 0 | 0 |
| NC_045512.2;YP_009725295.1;Wu1-SARS2_orf1a;3100 | 0 | 0 |
| NC_045512.2;YP_009725295.1;Wu1-SARS2_orf1a;3101 | 0 | 0 |
| NC_045512.2;YP_009725295.1;Wu1-SARS2_orf1a;3102 | 0 | 0 |
| NC_045512.2;YP_009725295.1;Wu1-SARS2_orf1a;3103 | 0 | 0 |
| NC_045512.2;YP_009725295.1;Wu1-SARS2_orf1a;3104 | 0 | 0 |
| NC_045512.2;YP_009725295.1;Wu1-SARS2_orf1a;3105 | 0 | 0 |
| NC_045512.2;YP_009725295.1;Wu1-SARS2_orf1a;3106 | 0 | 0 |
| NC_045512.2;YP_009725295.1;Wu1-SARS2_orf1a;3107 | 0 | 0 |
| NC_045512.2;YP_009725295.1;Wu1-SARS2_orf1a;3108 | 0 | 0 |
| NC_045512.2;YP_009725295.1;Wu1-SARS2_orf1a;3109 | 0 | 0 |
| NC_045512.2;YP_009725295.1;Wu1-SARS2_orf1a;3110 | 0 | 0 |
| NC_045512.2;YP_009725295.1;Wu1-SARS2_orf1a;3111 | 0 | 0 |
| NC_045512.2;YP_009725295.1;Wu1-SARS2_orf1a;3112 | 0 | 0 |
| NC_045512.2;YP_009725295.1;Wu1-SARS2_orf1a;3113 | 0 | 0 |
| NC_045512.2;YP_009725295.1;Wu1-SARS2_orf1a;3114 | 0 | 0 |
| NC_045512.2;YP_009725295.1;Wu1-SARS2_orf1a;3115 | 0 | 0 |
| NC_045512.2;YP_009725295.1;Wu1-SARS2_orf1a;3116 | 0 | 0 |
| NC_045512.2;YP_009725295.1;Wu1-SARS2_orf1a;3117 | 0 | 0 |
| NC_045512.2;YP_009725295.1;Wu1-SARS2_orf1a;3118 | 0 | 0 |
| NC_045512.2;YP_009725295.1;Wu1-SARS2_orf1a;3119 | 0 | 0 |
| NC_045512.2;YP_009725295.1;Wu1-SARS2_orf1a;3120 | 0 | 0 |
| NC_045512.2;YP_009725295.1;Wu1-SARS2_orf1a;3121 | 0 | 0 |
| NC_045512.2;YP_009725295.1;Wu1-SARS2_orf1a;3122 | 0 | 0 |
| NC_045512.2;YP_009725295.1;Wu1-SARS2_orf1a;3123 | 0 | 0 |
| NC_045512.2;YP_009725295.1;Wu1-SARS2_orf1a;3124 | 0 | 0 |
| NC_045512.2;YP_009725295.1;Wu1-SARS2_orf1a;3125 | 0 | 0 |
| NC_045512.2;YP_009725295.1;Wu1-SARS2_orf1a;3126 | 0 | 0 |
| NC_045512.2;YP_009725295.1;Wu1-SARS2_orf1a;3127 | 0 | 0 |

| | | |
|---|---|---|
| NC_045512.2;YP_009725295.1;Wu1-SARS2_orf1a;3128 | 0 | 0 |
| NC_045512.2;YP_009725295.1;Wu1-SARS2_orf1a;3129 | 0 | 0 |
| NC_045512.2;YP_009725295.1;Wu1-SARS2_orf1a;3130 | 0 | 0 |
| NC_045512.2;YP_009725295.1;Wu1-SARS2_orf1a;3131 | 0 | 0 |
| NC_045512.2;YP_009725295.1;Wu1-SARS2_orf1a;3132 | 0 | 0 |
| NC_045512.2;YP_009725295.1;Wu1-SARS2_orf1a;3133 | 0 | 0 |
| NC_045512.2;YP_009725295.1;Wu1-SARS2_orf1a;3134 | 0 | 0 |
| NC_045512.2;YP_009725295.1;Wu1-SARS2_orf1a;3135 | 0 | 0 |
| NC_045512.2;YP_009725295.1;Wu1-SARS2_orf1a;3136 | 0 | 0 |
| NC_045512.2;YP_009725295.1;Wu1-SARS2_orf1a;3137 | 0 | 0 |
| NC_045512.2;YP_009725295.1;Wu1-SARS2_orf1a;3138 | 0 | 0 |
| NC_045512.2;YP_009725295.1;Wu1-SARS2_orf1a;3139 | 0 | 0 |
| NC_045512.2;YP_009725295.1;Wu1-SARS2_orf1a;3140 | 0 | 0 |
| NC_045512.2;YP_009725295.1;Wu1-SARS2_orf1a;3141 | 0 | 0 |
| NC_045512.2;YP_009725295.1;Wu1-SARS2_orf1a;3142 | 0 | 0 |
| NC_045512.2;YP_009725295.1;Wu1-SARS2_orf1a;3143 | 0 | 0 |
| NC_045512.2;YP_009725295.1;Wu1-SARS2_orf1a;3144 | 0 | 0 |
| NC_045512.2;YP_009725295.1;Wu1-SARS2_orf1a;3145 | 0 | 0 |
| NC_045512.2;YP_009725295.1;Wu1-SARS2_orf1a;3146 | 0 | 0 |
| NC_045512.2;YP_009725295.1;Wu1-SARS2_orf1a;3147 | 0 | 0 |
| NC_045512.2;YP_009725295.1;Wu1-SARS2_orf1a;3148 | 0 | 0 |
| NC_045512.2;YP_009725295.1;Wu1-SARS2_orf1a;3149 | 0 | 0 |
| NC_045512.2;YP_009725295.1;Wu1-SARS2_orf1a;3150 | 0 | 0 |
| NC_045512.2;YP_009725295.1;Wu1-SARS2_orf1a;3151 | 0 | 0 |
| NC_045512.2;YP_009725295.1;Wu1-SARS2_orf1a;3152 | 0 | 0 |
| NC_045512.2;YP_009725295.1;Wu1-SARS2_orf1a;3153 | 0 | 0 |
| NC_045512.2;YP_009725295.1;Wu1-SARS2_orf1a;3154 | 0 | 0 |
| NC_045512.2;YP_009725295.1;Wu1-SARS2_orf1a;3155 | 0 | 0 |
| NC_045512.2;YP_009725295.1;Wu1-SARS2_orf1a;3156 | 0 | 0 |
| NC_045512.2;YP_009725295.1;Wu1-SARS2_orf1a;3157 | 0 | 0 |
| NC_045512.2;YP_009725295.1;Wu1-SARS2_orf1a;3158 | 0 | 0 |
| NC_045512.2;YP_009725295.1;Wu1-SARS2_orf1a;3159 | 0 | 0 |
| NC_045512.2;YP_009725295.1;Wu1-SARS2_orf1a;3160 | 0 | 0 |
| NC_045512.2;YP_009725295.1;Wu1-SARS2_orf1a;3161 | 0 | 0 |
| NC_045512.2;YP_009725295.1;Wu1-SARS2_orf1a;3162 | 0 | 0.075 |
| NC_045512.2;YP_009725295.1;Wu1-SARS2_orf1a;3163 | 0 | 0 |
| NC_045512.2;YP_009725295.1;Wu1-SARS2_orf1a;3164 | 0.05 | 0.025 |
| NC_045512.2;YP_009725295.1;Wu1-SARS2_orf1a;3165 | 0 | 0 |
| NC_045512.2;YP_009725295.1;Wu1-SARS2_orf1a;3166 | 0 | 0 |
| NC_045512.2;YP_009725295.1;Wu1-SARS2_orf1a;3167 | 0 | 0 |
| NC_045512.2;YP_009725295.1;Wu1-SARS2_orf1a;3168 | 0 | 0 |
| NC_045512.2;YP_009725295.1;Wu1-SARS2_orf1a;3169 | 0 | 0 |
| NC_045512.2;YP_009725295.1;Wu1-SARS2_orf1a;3170 | 0 | 0 |

| | | |
|---|---|---|
| NC_045512.2;YP_009725295.1;Wu1-SARS2_orf1a;3171 | 0 | 0 |
| NC_045512.2;YP_009725295.1;Wu1-SARS2_orf1a;3172 | 0 | 0 |
| NC_045512.2;YP_009725295.1;Wu1-SARS2_orf1a;3173 | 0 | 0 |
| NC_045512.2;YP_009725295.1;Wu1-SARS2_orf1a;3174 | 0 | 0 |
| NC_045512.2;YP_009725295.1;Wu1-SARS2_orf1a;3175 | 0 | 0 |
| NC_045512.2;YP_009725295.1;Wu1-SARS2_orf1a;3176 | 0 | 0 |
| NC_045512.2;YP_009725295.1;Wu1-SARS2_orf1a;3177 | 0 | 0 |
| NC_045512.2;YP_009725295.1;Wu1-SARS2_orf1a;3178 | 0 | 0 |
| NC_045512.2;YP_009725295.1;Wu1-SARS2_orf1a;3179 | 0 | 0 |
| NC_045512.2;YP_009725295.1;Wu1-SARS2_orf1a;3180 | 0 | 0 |
| NC_045512.2;YP_009725295.1;Wu1-SARS2_orf1a;3181 | 0 | 0 |
| NC_045512.2;YP_009725295.1;Wu1-SARS2_orf1a;3182 | 0.05 | 0 |
| NC_045512.2;YP_009725295.1;Wu1-SARS2_orf1a;3183 | 0.1 | 0 |
| NC_045512.2;YP_009725295.1;Wu1-SARS2_orf1a;3184 | 0.1 | 0 |
| NC_045512.2;YP_009725295.1;Wu1-SARS2_orf1a;3185 | 0.1 | 0 |
| NC_045512.2;YP_009725295.1;Wu1-SARS2_orf1a;3186 | 0.1 | 0 |
| NC_045512.2;YP_009725295.1;Wu1-SARS2_orf1a;3187 | 0.1 | 0 |
| NC_045512.2;YP_009725295.1;Wu1-SARS2_orf1a;3188 | 0.15 | 0 |
| NC_045512.2;YP_009725295.1;Wu1-SARS2_orf1a;3189 | 0.1 | 0 |
| NC_045512.2;YP_009725295.1;Wu1-SARS2_orf1a;3190 | 0.1 | 0 |
| NC_045512.2;YP_009725295.1;Wu1-SARS2_orf1a;3191 | 0.1 | 0.05 |
| NC_045512.2;YP_009725295.1;Wu1-SARS2_orf1a;3192 | 0.05 | 0.025 |
| NC_045512.2;YP_009725295.1;Wu1-SARS2_orf1a;3193 | 0 | 0.025 |
| NC_045512.2;YP_009725295.1;Wu1-SARS2_orf1a;3194 | 0 | 0.025 |
| NC_045512.2;YP_009725295.1;Wu1-SARS2_orf1a;3195 | 0 | 0 |
| NC_045512.2;YP_009725295.1;Wu1-SARS2_orf1a;3196 | 0 | 0 |
| NC_045512.2;YP_009725295.1;Wu1-SARS2_orf1a;3197 | 0 | 0 |
| NC_045512.2;YP_009725295.1;Wu1-SARS2_orf1a;3198 | 0 | 0 |
| NC_045512.2;YP_009725295.1;Wu1-SARS2_orf1a;3199 | 0 | 0 |
| NC_045512.2;YP_009725295.1;Wu1-SARS2_orf1a;3200 | 0 | 0 |
| NC_045512.2;YP_009725295.1;Wu1-SARS2_orf1a;3201 | 0 | 0 |
| NC_045512.2;YP_009725295.1;Wu1-SARS2_orf1a;3202 | 0 | 0 |
| NC_045512.2;YP_009725295.1;Wu1-SARS2_orf1a;3203 | 0 | 0 |
| NC_045512.2;YP_009725295.1;Wu1-SARS2_orf1a;3204 | 0 | 0 |
| NC_045512.2;YP_009725295.1;Wu1-SARS2_orf1a;3205 | 0 | 0 |
| NC_045512.2;YP_009725295.1;Wu1-SARS2_orf1a;3206 | 0 | 0 |
| NC_045512.2;YP_009725295.1;Wu1-SARS2_orf1a;3207 | 0 | 0 |
| NC_045512.2;YP_009725295.1;Wu1-SARS2_orf1a;3208 | 0 | 0 |
| NC_045512.2;YP_009725295.1;Wu1-SARS2_orf1a;3209 | 0 | 0.025 |
| NC_045512.2;YP_009725295.1;Wu1-SARS2_orf1a;3210 | 0 | 0.025 |
| NC_045512.2;YP_009725295.1;Wu1-SARS2_orf1a;3211 | 0 | 0.025 |
| NC_045512.2;YP_009725295.1;Wu1-SARS2_orf1a;3212 | 0 | 0 |
| NC_045512.2;YP_009725295.1;Wu1-SARS2_orf1a;3213 | 0 | 0 |

| | | |
|---|---|---|
| NC_045512.2;YP_009725295.1;Wu1-SARS2_orf1a;3214 | 0 | 0 |
| NC_045512.2;YP_009725295.1;Wu1-SARS2_orf1a;3215 | 0.05 | 0.075 |
| NC_045512.2;YP_009725295.1;Wu1-SARS2_orf1a;3216 | 0 | 0 |
| NC_045512.2;YP_009725295.1;Wu1-SARS2_orf1a;3217 | 0 | 0 |
| NC_045512.2;YP_009725295.1;Wu1-SARS2_orf1a;3218 | 0 | 0 |
| NC_045512.2;YP_009725295.1;Wu1-SARS2_orf1a;3219 | 0 | 0.025 |
| NC_045512.2;YP_009725295.1;Wu1-SARS2_orf1a;3220 | 0 | 0 |
| NC_045512.2;YP_009725295.1;Wu1-SARS2_orf1a;3221 | 0 | 0 |
| NC_045512.2;YP_009725295.1;Wu1-SARS2_orf1a;3222 | 0 | 0 |
| NC_045512.2;YP_009725295.1;Wu1-SARS2_orf1a;3223 | 0 | 0 |
| NC_045512.2;YP_009725295.1;Wu1-SARS2_orf1a;3224 | 0 | 0 |
| NC_045512.2;YP_009725295.1;Wu1-SARS2_orf1a;3225 | 0 | 0.075 |
| NC_045512.2;YP_009725295.1;Wu1-SARS2_orf1a;3226 | 0 | 0.025 |
| NC_045512.2;YP_009725295.1;Wu1-SARS2_orf1a;3227 | 0 | 0.025 |
| NC_045512.2;YP_009725295.1;Wu1-SARS2_orf1a;3228 | 0 | 0.025 |
| NC_045512.2;YP_009725295.1;Wu1-SARS2_orf1a;3229 | 0 | 0.025 |
| NC_045512.2;YP_009725295.1;Wu1-SARS2_orf1a;3230 | 0.05 | 0.025 |
| NC_045512.2;YP_009725295.1;Wu1-SARS2_orf1a;3231 | 0 | 0.025 |
| NC_045512.2;YP_009725295.1;Wu1-SARS2_orf1a;3232 | 0 | 0.025 |
| NC_045512.2;YP_009725295.1;Wu1-SARS2_orf1a;3233 | 0 | 0.025 |
| NC_045512.2;YP_009725295.1;Wu1-SARS2_orf1a;3234 | 0.05 | 0 |
| NC_045512.2;YP_009725295.1;Wu1-SARS2_orf1a;3235 | 0 | 0 |
| NC_045512.2;YP_009725295.1;Wu1-SARS2_orf1a;3236 | 0 | 0 |
| NC_045512.2;YP_009725295.1;Wu1-SARS2_orf1a;3237 | 0.15 | 0.05 |
| NC_045512.2;YP_009725295.1;Wu1-SARS2_orf1a;3238 | 0.3 | 0.05 |
| NC_045512.2;YP_009725295.1;Wu1-SARS2_orf1a;3239 | 0.05 | 0 |
| NC_045512.2;YP_009725295.1;Wu1-SARS2_orf1a;3240 | 0.05 | 0.05 |
| NC_045512.2;YP_009725295.1;Wu1-SARS2_orf1a;3241 | 0.1 | 0.1 |
| NC_045512.2;YP_009725295.1;Wu1-SARS2_orf1a;3242 | 0 | 0.05 |
| NC_045512.2;YP_009725295.1;Wu1-SARS2_orf1a;3243 | 0 | 0.05 |
| NC_045512.2;YP_009725295.1;Wu1-SARS2_orf1a;3244 | 0 | 0 |
| NC_045512.2;YP_009725295.1;Wu1-SARS2_orf1a;3245 | 0 | 0.025 |
| NC_045512.2;YP_009725295.1;Wu1-SARS2_orf1a;3246 | 0 | 0 |
| NC_045512.2;YP_009725295.1;Wu1-SARS2_orf1a;3247 | 0 | 0 |
| NC_045512.2;YP_009725295.1;Wu1-SARS2_orf1a;3248 | 0 | 0 |
| NC_045512.2;YP_009725295.1;Wu1-SARS2_orf1a;3249 | 0 | 0 |
| NC_045512.2;YP_009725295.1;Wu1-SARS2_orf1a;3250 | 0.05 | 0 |
| NC_045512.2;YP_009725295.1;Wu1-SARS2_orf1a;3251 | 0 | 0 |
| NC_045512.2;YP_009725295.1;Wu1-SARS2_orf1a;3252 | 0 | 0 |
| NC_045512.2;YP_009725295.1;Wu1-SARS2_orf1a;3253 | 0 | 0 |
| NC_045512.2;YP_009725295.1;Wu1-SARS2_orf1a;3254 | 0 | 0 |
| NC_045512.2;YP_009725295.1;Wu1-SARS2_orf1a;3255 | 0 | 0 |
| NC_045512.2;YP_009725295.1;Wu1-SARS2_orf1a;3256 | 0 | 0 |

| | | |
|---|---|---|
| NC_045512.2;YP_009725295.1;Wu1-SARS2_orf1a;3257 | 0 | 0 |
| NC_045512.2;YP_009725295.1;Wu1-SARS2_orf1a;3258 | 0 | 0 |
| NC_045512.2;YP_009725295.1;Wu1-SARS2_orf1a;3259 | 0 | 0 |
| NC_045512.2;YP_009725295.1;Wu1-SARS2_orf1a;3260 | 0 | 0 |
| NC_045512.2;YP_009725295.1;Wu1-SARS2_orf1a;3261 | 0 | 0 |
| NC_045512.2;YP_009725295.1;Wu1-SARS2_orf1a;3262 | 0 | 0 |
| NC_045512.2;YP_009725295.1;Wu1-SARS2_orf1a;3263 | 0 | 0 |
| NC_045512.2;YP_009725295.1;Wu1-SARS2_orf1a;3264 | 0 | 0 |
| NC_045512.2;YP_009725295.1;Wu1-SARS2_orf1a;3265 | 0 | 0 |
| NC_045512.2;YP_009725295.1;Wu1-SARS2_orf1a;3266 | 0 | 0 |
| NC_045512.2;YP_009725295.1;Wu1-SARS2_orf1a;3267 | 0 | 0 |
| NC_045512.2;YP_009725295.1;Wu1-SARS2_orf1a;3268 | 0 | 0.025 |
| NC_045512.2;YP_009725295.1;Wu1-SARS2_orf1a;3269 | 0 | 0 |
| NC_045512.2;YP_009725295.1;Wu1-SARS2_orf1a;3270 | 0.05 | 0 |
| NC_045512.2;YP_009725295.1;Wu1-SARS2_orf1a;3271 | 0 | 0 |
| NC_045512.2;YP_009725295.1;Wu1-SARS2_orf1a;3272 | 0 | 0 |
| NC_045512.2;YP_009725295.1;Wu1-SARS2_orf1a;3273 | 0 | 0 |
| NC_045512.2;YP_009725295.1;Wu1-SARS2_orf1a;3274 | 0 | 0 |
| NC_045512.2;YP_009725295.1;Wu1-SARS2_orf1a;3275 | 0 | 0 |
| NC_045512.2;YP_009725295.1;Wu1-SARS2_orf1a;3276 | 0 | 0 |
| NC_045512.2;YP_009725295.1;Wu1-SARS2_orf1a;3277 | 0 | 0 |
| NC_045512.2;YP_009725295.1;Wu1-SARS2_orf1a;3278 | 0 | 0 |
| NC_045512.2;YP_009725295.1;Wu1-SARS2_orf1a;3279 | 0 | 0 |
| NC_045512.2;YP_009725295.1;Wu1-SARS2_orf1a;3280 | 0 | 0 |
| NC_045512.2;YP_009725295.1;Wu1-SARS2_orf1a;3281 | 0 | 0 |
| NC_045512.2;YP_009725295.1;Wu1-SARS2_orf1a;3282 | 0 | 0 |
| NC_045512.2;YP_009725295.1;Wu1-SARS2_orf1a;3283 | 0 | 0 |
| NC_045512.2;YP_009725295.1;Wu1-SARS2_orf1a;3284 | 0 | 0 |
| NC_045512.2;YP_009725295.1;Wu1-SARS2_orf1a;3285 | 0 | 0 |
| NC_045512.2;YP_009725295.1;Wu1-SARS2_orf1a;3286 | 0 | 0 |
| NC_045512.2;YP_009725295.1;Wu1-SARS2_orf1a;3287 | 0 | 0 |
| NC_045512.2;YP_009725295.1;Wu1-SARS2_orf1a;3288 | 0 | 0 |
| NC_045512.2;YP_009725295.1;Wu1-SARS2_orf1a;3289 | 0 | 0 |
| NC_045512.2;YP_009725295.1;Wu1-SARS2_orf1a;3290 | 0 | 0 |
| NC_045512.2;YP_009725295.1;Wu1-SARS2_orf1a;3291 | 0 | 0 |
| NC_045512.2;YP_009725295.1;Wu1-SARS2_orf1a;3292 | 0 | 0 |
| NC_045512.2;YP_009725295.1;Wu1-SARS2_orf1a;3293 | 0 | 0 |
| NC_045512.2;YP_009725295.1;Wu1-SARS2_orf1a;3294 | 0 | 0 |
| NC_045512.2;YP_009725295.1;Wu1-SARS2_orf1a;3295 | 0 | 0 |
| NC_045512.2;YP_009725295.1;Wu1-SARS2_orf1a;3296 | 0 | 0 |
| NC_045512.2;YP_009725295.1;Wu1-SARS2_orf1a;3297 | 0 | 0 |
| NC_045512.2;YP_009725295.1;Wu1-SARS2_orf1a;3298 | 0 | 0 |
| NC_045512.2;YP_009725295.1;Wu1-SARS2_orf1a;3299 | 0 | 0.025 |

| | | |
|---|---|---|
| NC_045512.2;YP_009725295.1;Wu1-SARS2_orf1a;3300 | 0 | 0 |
| NC_045512.2;YP_009725295.1;Wu1-SARS2_orf1a;3301 | 0 | 0 |
| NC_045512.2;YP_009725295.1;Wu1-SARS2_orf1a;3302 | 0 | 0 |
| NC_045512.2;YP_009725295.1;Wu1-SARS2_orf1a;3303 | 0 | 0 |
| NC_045512.2;YP_009725295.1;Wu1-SARS2_orf1a;3304 | 0 | 0 |
| NC_045512.2;YP_009725295.1;Wu1-SARS2_orf1a;3305 | 0 | 0 |
| NC_045512.2;YP_009725295.1;Wu1-SARS2_orf1a;3306 | 0 | 0.025 |
| NC_045512.2;YP_009725295.1;Wu1-SARS2_orf1a;3307 | 0 | 0 |
| NC_045512.2;YP_009725295.1;Wu1-SARS2_orf1a;3308 | 0 | 0 |
| NC_045512.2;YP_009725295.1;Wu1-SARS2_orf1a;3309 | 0 | 0 |
| NC_045512.2;YP_009725295.1;Wu1-SARS2_orf1a;3310 | 0 | 0.025 |
| NC_045512.2;YP_009725295.1;Wu1-SARS2_orf1a;3311 | 0 | 0.025 |
| NC_045512.2;YP_009725295.1;Wu1-SARS2_orf1a;3312 | 0 | 0 |
| NC_045512.2;YP_009725295.1;Wu1-SARS2_orf1a;3313 | 0.05 | 0 |
| NC_045512.2;YP_009725295.1;Wu1-SARS2_orf1a;3314 | 0 | 0.05 |
| NC_045512.2;YP_009725295.1;Wu1-SARS2_orf1a;3315 | 0 | 0 |
| NC_045512.2;YP_009725295.1;Wu1-SARS2_orf1a;3316 | 0 | 0 |
| NC_045512.2;YP_009725295.1;Wu1-SARS2_orf1a;3317 | 0 | 0 |
| NC_045512.2;YP_009725295.1;Wu1-SARS2_orf1a;3318 | 0 | 0 |
| NC_045512.2;YP_009725295.1;Wu1-SARS2_orf1a;3319 | 0 | 0 |
| NC_045512.2;YP_009725295.1;Wu1-SARS2_orf1a;3320 | 0 | 0 |
| NC_045512.2;YP_009725295.1;Wu1-SARS2_orf1a;3321 | 0 | 0 |
| NC_045512.2;YP_009725295.1;Wu1-SARS2_orf1a;3322 | 0 | 0 |
| NC_045512.2;YP_009725295.1;Wu1-SARS2_orf1a;3323 | 0 | 0 |
| NC_045512.2;YP_009725295.1;Wu1-SARS2_orf1a;3324 | 0 | 0 |
| NC_045512.2;YP_009725295.1;Wu1-SARS2_orf1a;3325 | 0 | 0 |
| NC_045512.2;YP_009725295.1;Wu1-SARS2_orf1a;3326 | 0 | 0 |
| NC_045512.2;YP_009725295.1;Wu1-SARS2_orf1a;3327 | 0 | 0 |
| NC_045512.2;YP_009725295.1;Wu1-SARS2_orf1a;3328 | 0 | 0 |
| NC_045512.2;YP_009725295.1;Wu1-SARS2_orf1a;3329 | 0 | 0 |
| NC_045512.2;YP_009725295.1;Wu1-SARS2_orf1a;3330 | 0 | 0 |
| NC_045512.2;YP_009725295.1;Wu1-SARS2_orf1a;3331 | 0 | 0 |
| NC_045512.2;YP_009725295.1;Wu1-SARS2_orf1a;3332 | 0 | 0 |
| NC_045512.2;YP_009725295.1;Wu1-SARS2_orf1a;3333 | 0 | 0 |
| NC_045512.2;YP_009725295.1;Wu1-SARS2_orf1a;3334 | 0 | 0 |
| NC_045512.2;YP_009725295.1;Wu1-SARS2_orf1a;3335 | 0 | 0 |
| NC_045512.2;YP_009725295.1;Wu1-SARS2_orf1a;3336 | 0 | 0 |
| NC_045512.2;YP_009725295.1;Wu1-SARS2_orf1a;3337 | 0 | 0 |
| NC_045512.2;YP_009725295.1;Wu1-SARS2_orf1a;3338 | 0 | 0 |
| NC_045512.2;YP_009725295.1;Wu1-SARS2_orf1a;3339 | 0 | 0 |
| NC_045512.2;YP_009725295.1;Wu1-SARS2_orf1a;3340 | 0 | 0 |
| NC_045512.2;YP_009725295.1;Wu1-SARS2_orf1a;3341 | 0 | 0 |
| NC_045512.2;YP_009725295.1;Wu1-SARS2_orf1a;3342 | 0 | 0.025 |

| | | |
|---|---|---|
| NC_045512.2;YP_009725295.1;Wu1-SARS2_orf1a;3343 | 0.05 | 0 |
| NC_045512.2;YP_009725295.1;Wu1-SARS2_orf1a;3344 | 0 | 0 |
| NC_045512.2;YP_009725295.1;Wu1-SARS2_orf1a;3345 | 0 | 0 |
| NC_045512.2;YP_009725295.1;Wu1-SARS2_orf1a;3346 | 0 | 0 |
| NC_045512.2;YP_009725295.1;Wu1-SARS2_orf1a;3347 | 0 | 0 |
| NC_045512.2;YP_009725295.1;Wu1-SARS2_orf1a;3348 | 0 | 0 |
| NC_045512.2;YP_009725295.1;Wu1-SARS2_orf1a;3349 | 0 | 0 |
| NC_045512.2;YP_009725295.1;Wu1-SARS2_orf1a;3350 | 0 | 0 |
| NC_045512.2;YP_009725295.1;Wu1-SARS2_orf1a;3351 | 0 | 0 |
| NC_045512.2;YP_009725295.1;Wu1-SARS2_orf1a;3352 | 0 | 0.025 |
| NC_045512.2;YP_009725295.1;Wu1-SARS2_orf1a;3353 | 0 | 0 |
| NC_045512.2;YP_009725295.1;Wu1-SARS2_orf1a;3354 | 0 | 0 |
| NC_045512.2;YP_009725295.1;Wu1-SARS2_orf1a;3355 | 0 | 0 |
| NC_045512.2;YP_009725295.1;Wu1-SARS2_orf1a;3356 | 0 | 0 |
| NC_045512.2;YP_009725295.1;Wu1-SARS2_orf1a;3357 | 0 | 0 |
| NC_045512.2;YP_009725295.1;Wu1-SARS2_orf1a;3358 | 0 | 0 |
| NC_045512.2;YP_009725295.1;Wu1-SARS2_orf1a;3359 | 0 | 0 |
| NC_045512.2;YP_009725295.1;Wu1-SARS2_orf1a;3360 | 0 | 0 |
| NC_045512.2;YP_009725295.1;Wu1-SARS2_orf1a;3361 | 0 | 0 |
| NC_045512.2;YP_009725295.1;Wu1-SARS2_orf1a;3362 | 0 | 0 |
| NC_045512.2;YP_009725295.1;Wu1-SARS2_orf1a;3363 | 0 | 0 |
| NC_045512.2;YP_009725295.1;Wu1-SARS2_orf1a;3364 | 0 | 0 |
| NC_045512.2;YP_009725295.1;Wu1-SARS2_orf1a;3365 | 0 | 0 |
| NC_045512.2;YP_009725295.1;Wu1-SARS2_orf1a;3366 | 0 | 0 |
| NC_045512.2;YP_009725295.1;Wu1-SARS2_orf1a;3367 | 0 | 0 |
| NC_045512.2;YP_009725295.1;Wu1-SARS2_orf1a;3368 | 0 | 0 |
| NC_045512.2;YP_009725295.1;Wu1-SARS2_orf1a;3369 | 0 | 0 |
| NC_045512.2;YP_009725295.1;Wu1-SARS2_orf1a;3370 | 0 | 0.025 |
| NC_045512.2;YP_009725295.1;Wu1-SARS2_orf1a;3371 | 0 | 0 |
| NC_045512.2;YP_009725295.1;Wu1-SARS2_orf1a;3372 | 0 | 0 |
| NC_045512.2;YP_009725295.1;Wu1-SARS2_orf1a;3373 | 0 | 0 |
| NC_045512.2;YP_009725295.1;Wu1-SARS2_orf1a;3374 | 0 | 0 |
| NC_045512.2;YP_009725295.1;Wu1-SARS2_orf1a;3375 | 0 | 0 |
| NC_045512.2;YP_009725295.1;Wu1-SARS2_orf1a;3376 | 0 | 0 |
| NC_045512.2;YP_009725295.1;Wu1-SARS2_orf1a;3377 | 0 | 0 |
| NC_045512.2;YP_009725295.1;Wu1-SARS2_orf1a;3378 | 0 | 0 |
| NC_045512.2;YP_009725295.1;Wu1-SARS2_orf1a;3379 | 0 | 0 |
| NC_045512.2;YP_009725295.1;Wu1-SARS2_orf1a;3380 | 0 | 0 |
| NC_045512.2;YP_009725295.1;Wu1-SARS2_orf1a;3381 | 0 | 0.025 |
| NC_045512.2;YP_009725295.1;Wu1-SARS2_orf1a;3382 | 0 | 0 |
| NC_045512.2;YP_009725295.1;Wu1-SARS2_orf1a;3383 | 0 | 0 |
| NC_045512.2;YP_009725295.1;Wu1-SARS2_orf1a;3384 | 0 | 0 |
| NC_045512.2;YP_009725295.1;Wu1-SARS2_orf1a;3385 | 0 | 0 |

| | | |
|---|---|---|
| NC_045512.2;YP_009725295.1;Wu1-SARS2_orf1a;3386 | 0 | 0 |
| NC_045512.2;YP_009725295.1;Wu1-SARS2_orf1a;3387 | 0 | 0.025 |
| NC_045512.2;YP_009725295.1;Wu1-SARS2_orf1a;3388 | 0 | 0 |
| NC_045512.2;YP_009725295.1;Wu1-SARS2_orf1a;3389 | 0 | 0 |
| NC_045512.2;YP_009725295.1;Wu1-SARS2_orf1a;3390 | 0 | 0 |
| NC_045512.2;YP_009725295.1;Wu1-SARS2_orf1a;3391 | 0 | 0 |
| NC_045512.2;YP_009725295.1;Wu1-SARS2_orf1a;3392 | 0.05 | 0 |
| NC_045512.2;YP_009725295.1;Wu1-SARS2_orf1a;3393 | 0 | 0 |
| NC_045512.2;YP_009725295.1;Wu1-SARS2_orf1a;3394 | 0.05 | 0 |
| NC_045512.2;YP_009725295.1;Wu1-SARS2_orf1a;3395 | 0 | 0 |
| NC_045512.2;YP_009725295.1;Wu1-SARS2_orf1a;3396 | 0 | 0 |
| NC_045512.2;YP_009725295.1;Wu1-SARS2_orf1a;3397 | 0 | 0 |
| NC_045512.2;YP_009725295.1;Wu1-SARS2_orf1a;3398 | 0 | 0 |
| NC_045512.2;YP_009725295.1;Wu1-SARS2_orf1a;3399 | 0 | 0 |
| NC_045512.2;YP_009725295.1;Wu1-SARS2_orf1a;3400 | 0 | 0.025 |
| NC_045512.2;YP_009725295.1;Wu1-SARS2_orf1a;3401 | 0 | 0.025 |
| NC_045512.2;YP_009725295.1;Wu1-SARS2_orf1a;3402 | 0 | 0.025 |
| NC_045512.2;YP_009725295.1;Wu1-SARS2_orf1a;3403 | 0 | 0.05 |
| NC_045512.2;YP_009725295.1;Wu1-SARS2_orf1a;3404 | 0 | 0.05 |
| NC_045512.2;YP_009725295.1;Wu1-SARS2_orf1a;3405 | 0 | 0.025 |
| NC_045512.2;YP_009725295.1;Wu1-SARS2_orf1a;3406 | 0 | 0.025 |
| NC_045512.2;YP_009725295.1;Wu1-SARS2_orf1a;3407 | 0 | 0 |
| NC_045512.2;YP_009725295.1;Wu1-SARS2_orf1a;3408 | 0 | 0 |
| NC_045512.2;YP_009725295.1;Wu1-SARS2_orf1a;3409 | 0 | 0 |
| NC_045512.2;YP_009725295.1;Wu1-SARS2_orf1a;3410 | 0 | 0 |
| NC_045512.2;YP_009725295.1;Wu1-SARS2_orf1a;3411 | 0 | 0 |
| NC_045512.2;YP_009725295.1;Wu1-SARS2_orf1a;3412 | 0 | 0 |
| NC_045512.2;YP_009725295.1;Wu1-SARS2_orf1a;3413 | 0 | 0 |
| NC_045512.2;YP_009725295.1;Wu1-SARS2_orf1a;3414 | 0 | 0 |
| NC_045512.2;YP_009725295.1;Wu1-SARS2_orf1a;3415 | 0 | 0 |
| NC_045512.2;YP_009725295.1;Wu1-SARS2_orf1a;3416 | 0 | 0 |
| NC_045512.2;YP_009725295.1;Wu1-SARS2_orf1a;3417 | 0 | 0 |
| NC_045512.2;YP_009725295.1;Wu1-SARS2_orf1a;3418 | 0 | 0 |
| NC_045512.2;YP_009725295.1;Wu1-SARS2_orf1a;3419 | 0 | 0 |
| NC_045512.2;YP_009725295.1;Wu1-SARS2_orf1a;3420 | 0 | 0 |
| NC_045512.2;YP_009725295.1;Wu1-SARS2_orf1a;3421 | 0 | 0 |
| NC_045512.2;YP_009725295.1;Wu1-SARS2_orf1a;3422 | 0 | 0 |
| NC_045512.2;YP_009725295.1;Wu1-SARS2_orf1a;3423 | 0 | 0 |
| NC_045512.2;YP_009725295.1;Wu1-SARS2_orf1a;3424 | 0 | 0 |
| NC_045512.2;YP_009725295.1;Wu1-SARS2_orf1a;3425 | 0 | 0 |
| NC_045512.2;YP_009725295.1;Wu1-SARS2_orf1a;3426 | 0.15 | 0.025 |
| NC_045512.2;YP_009725295.1;Wu1-SARS2_orf1a;3427 | 0.1 | 0.05 |
| NC_045512.2;YP_009725295.1;Wu1-SARS2_orf1a;3428 | 0.05 | 0.05 |

| | | |
|---|---|---|
| NC_045512.2;YP_009725295.1;Wu1-SARS2_orf1a;3429 | 0.2 | 0.075 |
| NC_045512.2;YP_009725295.1;Wu1-SARS2_orf1a;3430 | 0.25 | 0.125 |
| NC_045512.2;YP_009725295.1;Wu1-SARS2_orf1a;3431 | 0.25 | 0.125 |
| NC_045512.2;YP_009725295.1;Wu1-SARS2_orf1a;3432 | 0.3 | 0.125 |
| NC_045512.2;YP_009725295.1;Wu1-SARS2_orf1a;3433 | 0.3 | 0.15 |
| NC_045512.2;YP_009725295.1;Wu1-SARS2_orf1a;3434 | 0.2 | 0.075 |
| NC_045512.2;YP_009725295.1;Wu1-SARS2_orf1a;3435 | 0.25 | 0.1 |
| NC_045512.2;YP_009725295.1;Wu1-SARS2_orf1a;3436 | 0.25 | 0.075 |
| NC_045512.2;YP_009725295.1;Wu1-SARS2_orf1a;3437 | 0.25 | 0.2 |
| NC_045512.2;YP_009725295.1;Wu1-SARS2_orf1a;3438 | 0.1 | 0.025 |
| NC_045512.2;YP_009725295.1;Wu1-SARS2_orf1a;3439 | 0.1 | 0.025 |
| NC_045512.2;YP_009725295.1;Wu1-SARS2_orf1a;3440 | 0.05 | 0.025 |
| NC_045512.2;YP_009725295.1;Wu1-SARS2_orf1a;3441 | 0.05 | 0 |
| NC_045512.2;YP_009725295.1;Wu1-SARS2_orf1a;3442 | 0.1 | 0.025 |
| NC_045512.2;YP_009725295.1;Wu1-SARS2_orf1a;3443 | 0.05 | 0 |
| NC_045512.2;YP_009725295.1;Wu1-SARS2_orf1a;3444 | 0.1 | 0 |
| NC_045512.2;YP_009725295.1;Wu1-SARS2_orf1a;3445 | 0 | 0 |
| NC_045512.2;YP_009725295.1;Wu1-SARS2_orf1a;3446 | 0.05 | 0 |
| NC_045512.2;YP_009725295.1;Wu1-SARS2_orf1a;3447 | 0 | 0 |
| NC_045512.2;YP_009725295.1;Wu1-SARS2_orf1a;3448 | 0.05 | 0.025 |
| NC_045512.2;YP_009725295.1;Wu1-SARS2_orf1a;3449 | 0 | 0 |
| NC_045512.2;YP_009725295.1;Wu1-SARS2_orf1a;3450 | 0 | 0 |
| NC_045512.2;YP_009725295.1;Wu1-SARS2_orf1a;3451 | 0 | 0 |
| NC_045512.2;YP_009725295.1;Wu1-SARS2_orf1a;3452 | 0 | 0 |
| NC_045512.2;YP_009725295.1;Wu1-SARS2_orf1a;3453 | 0 | 0 |
| NC_045512.2;YP_009725295.1;Wu1-SARS2_orf1a;3454 | 0 | 0 |
| NC_045512.2;YP_009725295.1;Wu1-SARS2_orf1a;3455 | 0 | 0 |
| NC_045512.2;YP_009725295.1;Wu1-SARS2_orf1a;3456 | 0 | 0 |
| NC_045512.2;YP_009725295.1;Wu1-SARS2_orf1a;3457 | 0 | 0 |
| NC_045512.2;YP_009725295.1;Wu1-SARS2_orf1a;3458 | 0 | 0 |
| NC_045512.2;YP_009725295.1;Wu1-SARS2_orf1a;3459 | 0 | 0 |
| NC_045512.2;YP_009725295.1;Wu1-SARS2_orf1a;3460 | 0 | 0 |
| NC_045512.2;YP_009725295.1;Wu1-SARS2_orf1a;3461 | 0 | 0 |
| NC_045512.2;YP_009725295.1;Wu1-SARS2_orf1a;3462 | 0 | 0 |
| NC_045512.2;YP_009725295.1;Wu1-SARS2_orf1a;3463 | 0 | 0 |
| NC_045512.2;YP_009725295.1;Wu1-SARS2_orf1a;3464 | 0 | 0 |
| NC_045512.2;YP_009725295.1;Wu1-SARS2_orf1a;3465 | 0 | 0 |
| NC_045512.2;YP_009725295.1;Wu1-SARS2_orf1a;3466 | 0 | 0 |
| NC_045512.2;YP_009725295.1;Wu1-SARS2_orf1a;3467 | 0 | 0 |
| NC_045512.2;YP_009725295.1;Wu1-SARS2_orf1a;3468 | 0 | 0 |
| NC_045512.2;YP_009725295.1;Wu1-SARS2_orf1a;3469 | 0 | 0 |
| NC_045512.2;YP_009725295.1;Wu1-SARS2_orf1a;3470 | 0 | 0 |
| NC_045512.2;YP_009725295.1;Wu1-SARS2_orf1a;3471 | 0 | 0 |

| | | |
|---|---|---|
| NC_045512.2;YP_009725295.1;Wu1-SARS2_orf1a;3472 | 0 | 0 |
| NC_045512.2;YP_009725295.1;Wu1-SARS2_orf1a;3473 | 0 | 0 |
| NC_045512.2;YP_009725295.1;Wu1-SARS2_orf1a;3474 | 0 | 0 |
| NC_045512.2;YP_009725295.1;Wu1-SARS2_orf1a;3475 | 0 | 0 |
| NC_045512.2;YP_009725295.1;Wu1-SARS2_orf1a;3476 | 0 | 0 |
| NC_045512.2;YP_009725295.1;Wu1-SARS2_orf1a;3477 | 0 | 0 |
| NC_045512.2;YP_009725295.1;Wu1-SARS2_orf1a;3478 | 0 | 0 |
| NC_045512.2;YP_009725295.1;Wu1-SARS2_orf1a;3479 | 0.05 | 0 |
| NC_045512.2;YP_009725295.1;Wu1-SARS2_orf1a;3480 | 0.05 | 0 |
| NC_045512.2;YP_009725295.1;Wu1-SARS2_orf1a;3481 | 0.05 | 0 |
| NC_045512.2;YP_009725295.1;Wu1-SARS2_orf1a;3482 | 0.05 | 0.025 |
| NC_045512.2;YP_009725295.1;Wu1-SARS2_orf1a;3483 | 0.1 | 0 |
| NC_045512.2;YP_009725295.1;Wu1-SARS2_orf1a;3484 | 0.05 | 0 |
| NC_045512.2;YP_009725295.1;Wu1-SARS2_orf1a;3485 | 0.1 | 0 |
| NC_045512.2;YP_009725295.1;Wu1-SARS2_orf1a;3486 | 0.05 | 0 |
| NC_045512.2;YP_009725295.1;Wu1-SARS2_orf1a;3487 | 0.05 | 0.025 |
| NC_045512.2;YP_009725295.1;Wu1-SARS2_orf1a;3488 | 0.05 | 0.15 |
| NC_045512.2;YP_009725295.1;Wu1-SARS2_orf1a;3489 | 0.25 | 0.1 |
| NC_045512.2;YP_009725295.1;Wu1-SARS2_orf1a;3490 | 0.05 | 0 |
| NC_045512.2;YP_009725295.1;Wu1-SARS2_orf1a;3491 | 0 | 0 |
| NC_045512.2;YP_009725295.1;Wu1-SARS2_orf1a;3492 | 0.05 | 0 |
| NC_045512.2;YP_009725295.1;Wu1-SARS2_orf1a;3493 | 0 | 0 |
| NC_045512.2;YP_009725295.1;Wu1-SARS2_orf1a;3494 | 0 | 0.025 |
| NC_045512.2;YP_009725295.1;Wu1-SARS2_orf1a;3495 | 0 | 0.025 |
| NC_045512.2;YP_009725295.1;Wu1-SARS2_orf1a;3496 | 0.05 | 0 |
| NC_045512.2;YP_009725295.1;Wu1-SARS2_orf1a;3497 | 0 | 0 |
| NC_045512.2;YP_009725295.1;Wu1-SARS2_orf1a;3498 | 0 | 0 |
| NC_045512.2;YP_009725295.1;Wu1-SARS2_orf1a;3499 | 0 | 0 |
| NC_045512.2;YP_009725295.1;Wu1-SARS2_orf1a;3500 | 0 | 0 |
| NC_045512.2;YP_009725295.1;Wu1-SARS2_orf1a;3501 | 0 | 0 |
| NC_045512.2;YP_009725295.1;Wu1-SARS2_orf1a;3502 | 0 | 0 |
| NC_045512.2;YP_009725295.1;Wu1-SARS2_orf1a;3503 | 0.05 | 0 |
| NC_045512.2;YP_009725295.1;Wu1-SARS2_orf1a;3504 | 0 | 0 |
| NC_045512.2;YP_009725295.1;Wu1-SARS2_orf1a;3505 | 0.05 | 0.025 |
| NC_045512.2;YP_009725295.1;Wu1-SARS2_orf1a;3506 | 0.15 | 0.1 |
| NC_045512.2;YP_009725295.1;Wu1-SARS2_orf1a;3507 | 0.05 | 0.05 |
| NC_045512.2;YP_009725295.1;Wu1-SARS2_orf1a;3508 | 0.1 | 0.15 |
| NC_045512.2;YP_009725295.1;Wu1-SARS2_orf1a;3509 | 0.05 | 0.025 |
| NC_045512.2;YP_009725295.1;Wu1-SARS2_orf1a;3510 | 0.05 | 0 |
| NC_045512.2;YP_009725295.1;Wu1-SARS2_orf1a;3511 | 0.05 | 0 |
| NC_045512.2;YP_009725295.1;Wu1-SARS2_orf1a;3512 | 0.05 | 0 |
| NC_045512.2;YP_009725295.1;Wu1-SARS2_orf1a;3513 | 0.05 | 0 |
| NC_045512.2;YP_009725295.1;Wu1-SARS2_orf1a;3514 | 0.05 | 0 |

| | | |
|---|---|---|
| NC_045512.2;YP_009725295.1;Wu1-SARS2_orf1a;3515 | 0.05 | 0 |
| NC_045512.2;YP_009725295.1;Wu1-SARS2_orf1a;3516 | 0 | 0 |
| NC_045512.2;YP_009725295.1;Wu1-SARS2_orf1a;3517 | 0 | 0 |
| NC_045512.2;YP_009725295.1;Wu1-SARS2_orf1a;3518 | 0 | 0 |
| NC_045512.2;YP_009725295.1;Wu1-SARS2_orf1a;3519 | 0 | 0 |
| NC_045512.2;YP_009725295.1;Wu1-SARS2_orf1a;3520 | 0 | 0 |
| NC_045512.2;YP_009725295.1;Wu1-SARS2_orf1a;3521 | 0.05 | 0 |
| NC_045512.2;YP_009725295.1;Wu1-SARS2_orf1a;3522 | 0 | 0 |
| NC_045512.2;YP_009725295.1;Wu1-SARS2_orf1a;3523 | 0 | 0 |
| NC_045512.2;YP_009725295.1;Wu1-SARS2_orf1a;3524 | 0 | 0 |
| NC_045512.2;YP_009725295.1;Wu1-SARS2_orf1a;3525 | 0 | 0 |
| NC_045512.2;YP_009725295.1;Wu1-SARS2_orf1a;3526 | 0 | 0.025 |
| NC_045512.2;YP_009725295.1;Wu1-SARS2_orf1a;3527 | 0.05 | 0 |
| NC_045512.2;YP_009725295.1;Wu1-SARS2_orf1a;3528 | 0.05 | 0 |
| NC_045512.2;YP_009725295.1;Wu1-SARS2_orf1a;3529 | 0.05 | 0.025 |
| NC_045512.2;YP_009725295.1;Wu1-SARS2_orf1a;3530 | 0 | 0.025 |
| NC_045512.2;YP_009725295.1;Wu1-SARS2_orf1a;3531 | 0 | 0.025 |
| NC_045512.2;YP_009725295.1;Wu1-SARS2_orf1a;3532 | 0 | 0.025 |
| NC_045512.2;YP_009725295.1;Wu1-SARS2_orf1a;3533 | 0 | 0 |
| NC_045512.2;YP_009725295.1;Wu1-SARS2_orf1a;3534 | 0 | 0 |
| NC_045512.2;YP_009725295.1;Wu1-SARS2_orf1a;3535 | 0 | 0 |
| NC_045512.2;YP_009725295.1;Wu1-SARS2_orf1a;3536 | 0 | 0 |
| NC_045512.2;YP_009725295.1;Wu1-SARS2_orf1a;3537 | 0 | 0 |
| NC_045512.2;YP_009725295.1;Wu1-SARS2_orf1a;3538 | 0 | 0 |
| NC_045512.2;YP_009725295.1;Wu1-SARS2_orf1a;3539 | 0 | 0.025 |
| NC_045512.2;YP_009725295.1;Wu1-SARS2_orf1a;3540 | 0 | 0 |
| NC_045512.2;YP_009725295.1;Wu1-SARS2_orf1a;3541 | 0 | 0.025 |
| NC_045512.2;YP_009725295.1;Wu1-SARS2_orf1a;3542 | 0 | 0.025 |
| NC_045512.2;YP_009725295.1;Wu1-SARS2_orf1a;3543 | 0 | 0.05 |
| NC_045512.2;YP_009725295.1;Wu1-SARS2_orf1a;3544 | 0 | 0.05 |
| NC_045512.2;YP_009725295.1;Wu1-SARS2_orf1a;3545 | 0 | 0.025 |
| NC_045512.2;YP_009725295.1;Wu1-SARS2_orf1a;3546 | 0 | 0.025 |
| NC_045512.2;YP_009725295.1;Wu1-SARS2_orf1a;3547 | 0.05 | 0.05 |
| NC_045512.2;YP_009725295.1;Wu1-SARS2_orf1a;3548 | 0.05 | 0.05 |
| NC_045512.2;YP_009725295.1;Wu1-SARS2_orf1a;3549 | 0 | 0.025 |
| NC_045512.2;YP_009725295.1;Wu1-SARS2_orf1a;3550 | 0 | 0.05 |
| NC_045512.2;YP_009725295.1;Wu1-SARS2_orf1a;3551 | 0.1 | 0.025 |
| NC_045512.2;YP_009725295.1;Wu1-SARS2_orf1a;3552 | 0 | 0 |
| NC_045512.2;YP_009725295.1;Wu1-SARS2_orf1a;3553 | 0 | 0 |
| NC_045512.2;YP_009725295.1;Wu1-SARS2_orf1a;3554 | 0 | 0 |
| NC_045512.2;YP_009725295.1;Wu1-SARS2_orf1a;3555 | 0 | 0 |
| NC_045512.2;YP_009725295.1;Wu1-SARS2_orf1a;3556 | 0 | 0 |
| NC_045512.2;YP_009725295.1;Wu1-SARS2_orf1a;3557 | 0 | 0 |

| | | |
|---|---|---|
| NC_045512.2;YP_009725295.1;Wu1-SARS2_orf1a;3558 | 0 | 0 |
| NC_045512.2;YP_009725295.1;Wu1-SARS2_orf1a;3559 | 0 | 0 |
| NC_045512.2;YP_009725295.1;Wu1-SARS2_orf1a;3560 | 0 | 0 |
| NC_045512.2;YP_009725295.1;Wu1-SARS2_orf1a;3561 | 0 | 0 |
| NC_045512.2;YP_009725295.1;Wu1-SARS2_orf1a;3562 | 0 | 0.025 |
| NC_045512.2;YP_009725295.1;Wu1-SARS2_orf1a;3563 | 0 | 0.025 |
| NC_045512.2;YP_009725295.1;Wu1-SARS2_orf1a;3564 | 0 | 0 |
| NC_045512.2;YP_009725295.1;Wu1-SARS2_orf1a;3565 | 0 | 0 |
| NC_045512.2;YP_009725295.1;Wu1-SARS2_orf1a;3566 | 0 | 0 |
| NC_045512.2;YP_009725295.1;Wu1-SARS2_orf1a;3567 | 0 | 0 |
| NC_045512.2;YP_009725295.1;Wu1-SARS2_orf1a;3568 | 0 | 0 |
| NC_045512.2;YP_009725295.1;Wu1-SARS2_orf1a;3569 | 0 | 0 |
| NC_045512.2;YP_009725295.1;Wu1-SARS2_orf1a;3570 | 0 | 0 |
| NC_045512.2;YP_009725295.1;Wu1-SARS2_orf1a;3571 | 0 | 0 |
| NC_045512.2;YP_009725295.1;Wu1-SARS2_orf1a;3572 | 0 | 0 |
| NC_045512.2;YP_009725295.1;Wu1-SARS2_orf1a;3573 | 0 | 0 |
| NC_045512.2;YP_009725295.1;Wu1-SARS2_orf1a;3574 | 0 | 0 |
| NC_045512.2;YP_009725295.1;Wu1-SARS2_orf1a;3575 | 0 | 0 |
| NC_045512.2;YP_009725295.1;Wu1-SARS2_orf1a;3576 | 0 | 0 |
| NC_045512.2;YP_009725295.1;Wu1-SARS2_orf1a;3577 | 0 | 0 |
| NC_045512.2;YP_009725295.1;Wu1-SARS2_orf1a;3578 | 0 | 0 |
| NC_045512.2;YP_009725295.1;Wu1-SARS2_orf1a;3579 | 0 | 0 |
| NC_045512.2;YP_009725295.1;Wu1-SARS2_orf1a;3580 | 0 | 0 |
| NC_045512.2;YP_009725295.1;Wu1-SARS2_orf1a;3581 | 0 | 0 |
| NC_045512.2;YP_009725295.1;Wu1-SARS2_orf1a;3582 | 0 | 0 |
| NC_045512.2;YP_009725295.1;Wu1-SARS2_orf1a;3583 | 0 | 0 |
| NC_045512.2;YP_009725295.1;Wu1-SARS2_orf1a;3584 | 0 | 0 |
| NC_045512.2;YP_009725295.1;Wu1-SARS2_orf1a;3585 | 0 | 0 |
| NC_045512.2;YP_009725295.1;Wu1-SARS2_orf1a;3586 | 0 | 0 |
| NC_045512.2;YP_009725295.1;Wu1-SARS2_orf1a;3587 | 0 | 0 |
| NC_045512.2;YP_009725295.1;Wu1-SARS2_orf1a;3588 | 0 | 0 |
| NC_045512.2;YP_009725295.1;Wu1-SARS2_orf1a;3589 | 0 | 0 |
| NC_045512.2;YP_009725295.1;Wu1-SARS2_orf1a;3590 | 0 | 0 |
| NC_045512.2;YP_009725295.1;Wu1-SARS2_orf1a;3591 | 0 | 0 |
| NC_045512.2;YP_009725295.1;Wu1-SARS2_orf1a;3592 | 0 | 0 |
| NC_045512.2;YP_009725295.1;Wu1-SARS2_orf1a;3593 | 0 | 0 |
| NC_045512.2;YP_009725295.1;Wu1-SARS2_orf1a;3594 | 0 | 0 |
| NC_045512.2;YP_009725295.1;Wu1-SARS2_orf1a;3595 | 0 | 0 |
| NC_045512.2;YP_009725295.1;Wu1-SARS2_orf1a;3596 | 0 | 0 |
| NC_045512.2;YP_009725295.1;Wu1-SARS2_orf1a;3597 | 0 | 0 |
| NC_045512.2;YP_009725295.1;Wu1-SARS2_orf1a;3598 | 0 | 0 |
| NC_045512.2;YP_009725295.1;Wu1-SARS2_orf1a;3599 | 0 | 0 |
| NC_045512.2;YP_009725295.1;Wu1-SARS2_orf1a;3600 | 0 | 0 |

| | | |
|---|---|---|
| NC_045512.2;YP_009725295.1;Wu1-SARS2_orf1a;3601 | 0 | 0 |
| NC_045512.2;YP_009725295.1;Wu1-SARS2_orf1a;3602 | 0 | 0 |
| NC_045512.2;YP_009725295.1;Wu1-SARS2_orf1a;3603 | 0 | 0 |
| NC_045512.2;YP_009725295.1;Wu1-SARS2_orf1a;3604 | 0 | 0 |
| NC_045512.2;YP_009725295.1;Wu1-SARS2_orf1a;3605 | 0 | 0 |
| NC_045512.2;YP_009725295.1;Wu1-SARS2_orf1a;3606 | 0 | 0 |
| NC_045512.2;YP_009725295.1;Wu1-SARS2_orf1a;3607 | 0 | 0 |
| NC_045512.2;YP_009725295.1;Wu1-SARS2_orf1a;3608 | 0 | 0 |
| NC_045512.2;YP_009725295.1;Wu1-SARS2_orf1a;3609 | 0 | 0 |
| NC_045512.2;YP_009725295.1;Wu1-SARS2_orf1a;3610 | 0 | 0 |
| NC_045512.2;YP_009725295.1;Wu1-SARS2_orf1a;3611 | 0 | 0 |
| NC_045512.2;YP_009725295.1;Wu1-SARS2_orf1a;3612 | 0 | 0 |
| NC_045512.2;YP_009725295.1;Wu1-SARS2_orf1a;3613 | 0 | 0 |
| NC_045512.2;YP_009725295.1;Wu1-SARS2_orf1a;3614 | 0 | 0 |
| NC_045512.2;YP_009725295.1;Wu1-SARS2_orf1a;3615 | 0 | 0 |
| NC_045512.2;YP_009725295.1;Wu1-SARS2_orf1a;3616 | 0 | 0 |
| NC_045512.2;YP_009725295.1;Wu1-SARS2_orf1a;3617 | 0 | 0 |
| NC_045512.2;YP_009725295.1;Wu1-SARS2_orf1a;3618 | 0 | 0 |
| NC_045512.2;YP_009725295.1;Wu1-SARS2_orf1a;3619 | 0 | 0 |
| NC_045512.2;YP_009725295.1;Wu1-SARS2_orf1a;3620 | 0 | 0 |
| NC_045512.2;YP_009725295.1;Wu1-SARS2_orf1a;3621 | 0 | 0 |
| NC_045512.2;YP_009725295.1;Wu1-SARS2_orf1a;3622 | 0 | 0 |
| NC_045512.2;YP_009725295.1;Wu1-SARS2_orf1a;3623 | 0 | 0 |
| NC_045512.2;YP_009725295.1;Wu1-SARS2_orf1a;3624 | 0 | 0 |
| NC_045512.2;YP_009725295.1;Wu1-SARS2_orf1a;3625 | 0 | 0 |
| NC_045512.2;YP_009725295.1;Wu1-SARS2_orf1a;3626 | 0 | 0 |
| NC_045512.2;YP_009725295.1;Wu1-SARS2_orf1a;3627 | 0 | 0 |
| NC_045512.2;YP_009725295.1;Wu1-SARS2_orf1a;3628 | 0 | 0 |
| NC_045512.2;YP_009725295.1;Wu1-SARS2_orf1a;3629 | 0 | 0 |
| NC_045512.2;YP_009725295.1;Wu1-SARS2_orf1a;3630 | 0 | 0 |
| NC_045512.2;YP_009725295.1;Wu1-SARS2_orf1a;3631 | 0 | 0 |
| NC_045512.2;YP_009725295.1;Wu1-SARS2_orf1a;3632 | 0 | 0 |
| NC_045512.2;YP_009725295.1;Wu1-SARS2_orf1a;3633 | 0 | 0 |
| NC_045512.2;YP_009725295.1;Wu1-SARS2_orf1a;3634 | 0 | 0 |
| NC_045512.2;YP_009725295.1;Wu1-SARS2_orf1a;3635 | 0 | 0 |
| NC_045512.2;YP_009725295.1;Wu1-SARS2_orf1a;3636 | 0 | 0 |
| NC_045512.2;YP_009725295.1;Wu1-SARS2_orf1a;3637 | 0 | 0 |
| NC_045512.2;YP_009725295.1;Wu1-SARS2_orf1a;3638 | 0 | 0 |
| NC_045512.2;YP_009725295.1;Wu1-SARS2_orf1a;3639 | 0 | 0 |
| NC_045512.2;YP_009725295.1;Wu1-SARS2_orf1a;3640 | 0 | 0 |
| NC_045512.2;YP_009725295.1;Wu1-SARS2_orf1a;3641 | 0 | 0 |
| NC_045512.2;YP_009725295.1;Wu1-SARS2_orf1a;3642 | 0 | 0 |
| NC_045512.2;YP_009725295.1;Wu1-SARS2_orf1a;3643 | 0 | 0 |

| | | |
|---|---|---|
| NC_045512.2;YP_009725295.1;Wu1-SARS2_orf1a;3644 | 0 | 0 |
| NC_045512.2;YP_009725295.1;Wu1-SARS2_orf1a;3645 | 0 | 0 |
| NC_045512.2;YP_009725295.1;Wu1-SARS2_orf1a;3646 | 0 | 0 |
| NC_045512.2;YP_009725295.1;Wu1-SARS2_orf1a;3647 | 0 | 0 |
| NC_045512.2;YP_009725295.1;Wu1-SARS2_orf1a;3648 | 0 | 0 |
| NC_045512.2;YP_009725295.1;Wu1-SARS2_orf1a;3649 | 0 | 0 |
| NC_045512.2;YP_009725295.1;Wu1-SARS2_orf1a;3650 | 0 | 0 |
| NC_045512.2;YP_009725295.1;Wu1-SARS2_orf1a;3651 | 0 | 0 |
| NC_045512.2;YP_009725295.1;Wu1-SARS2_orf1a;3652 | 0 | 0 |
| NC_045512.2;YP_009725295.1;Wu1-SARS2_orf1a;3653 | 0 | 0 |
| NC_045512.2;YP_009725295.1;Wu1-SARS2_orf1a;3654 | 0 | 0 |
| NC_045512.2;YP_009725295.1;Wu1-SARS2_orf1a;3655 | 0 | 0 |
| NC_045512.2;YP_009725295.1;Wu1-SARS2_orf1a;3656 | 0 | 0 |
| NC_045512.2;YP_009725295.1;Wu1-SARS2_orf1a;3657 | 0 | 0 |
| NC_045512.2;YP_009725295.1;Wu1-SARS2_orf1a;3658 | 0 | 0 |
| NC_045512.2;YP_009725295.1;Wu1-SARS2_orf1a;3659 | 0 | 0 |
| NC_045512.2;YP_009725295.1;Wu1-SARS2_orf1a;3660 | 0 | 0 |
| NC_045512.2;YP_009725295.1;Wu1-SARS2_orf1a;3661 | 0 | 0 |
| NC_045512.2;YP_009725295.1;Wu1-SARS2_orf1a;3662 | 0 | 0 |
| NC_045512.2;YP_009725295.1;Wu1-SARS2_orf1a;3663 | 0 | 0 |
| NC_045512.2;YP_009725295.1;Wu1-SARS2_orf1a;3664 | 0 | 0.025 |
| NC_045512.2;YP_009725295.1;Wu1-SARS2_orf1a;3665 | 0 | 0.025 |
| NC_045512.2;YP_009725295.1;Wu1-SARS2_orf1a;3666 | 0 | 0.05 |
| NC_045512.2;YP_009725295.1;Wu1-SARS2_orf1a;3667 | 0 | 0.025 |
| NC_045512.2;YP_009725295.1;Wu1-SARS2_orf1a;3668 | 0 | 0.025 |
| NC_045512.2;YP_009725295.1;Wu1-SARS2_orf1a;3669 | 0 | 0.025 |
| NC_045512.2;YP_009725295.1;Wu1-SARS2_orf1a;3670 | 0 | 0.025 |
| NC_045512.2;YP_009725295.1;Wu1-SARS2_orf1a;3671 | 0 | 0.025 |
| NC_045512.2;YP_009725295.1;Wu1-SARS2_orf1a;3672 | 0 | 0.025 |
| NC_045512.2;YP_009725295.1;Wu1-SARS2_orf1a;3673 | 0 | 0.025 |
| NC_045512.2;YP_009725295.1;Wu1-SARS2_orf1a;3674 | 0 | 0 |
| NC_045512.2;YP_009725295.1;Wu1-SARS2_orf1a;3675 | 0 | 0 |
| NC_045512.2;YP_009725295.1;Wu1-SARS2_orf1a;3676 | 0 | 0 |
| NC_045512.2;YP_009725295.1;Wu1-SARS2_orf1a;3677 | 0 | 0 |
| NC_045512.2;YP_009725295.1;Wu1-SARS2_orf1a;3678 | 0 | 0 |
| NC_045512.2;YP_009725295.1;Wu1-SARS2_orf1a;3679 | 0 | 0 |
| NC_045512.2;YP_009725295.1;Wu1-SARS2_orf1a;3680 | 0 | 0 |
| NC_045512.2;YP_009725295.1;Wu1-SARS2_orf1a;3681 | 0 | 0 |
| NC_045512.2;YP_009725295.1;Wu1-SARS2_orf1a;3682 | 0 | 0 |
| NC_045512.2;YP_009725295.1;Wu1-SARS2_orf1a;3683 | 0 | 0 |
| NC_045512.2;YP_009725295.1;Wu1-SARS2_orf1a;3684 | 0 | 0 |
| NC_045512.2;YP_009725295.1;Wu1-SARS2_orf1a;3685 | 0 | 0 |
| NC_045512.2;YP_009725295.1;Wu1-SARS2_orf1a;3686 | 0 | 0 |

| | | |
|---|---|---|
| NC_045512.2;YP_009725295.1;Wu1-SARS2_orf1a;3687 | 0 | 0 |
| NC_045512.2;YP_009725295.1;Wu1-SARS2_orf1a;3688 | 0 | 0 |
| NC_045512.2;YP_009725295.1;Wu1-SARS2_orf1a;3689 | 0 | 0 |
| NC_045512.2;YP_009725295.1;Wu1-SARS2_orf1a;3690 | 0 | 0 |
| NC_045512.2;YP_009725295.1;Wu1-SARS2_orf1a;3691 | 0 | 0 |
| NC_045512.2;YP_009725295.1;Wu1-SARS2_orf1a;3692 | 0 | 0 |
| NC_045512.2;YP_009725295.1;Wu1-SARS2_orf1a;3693 | 0 | 0 |
| NC_045512.2;YP_009725295.1;Wu1-SARS2_orf1a;3694 | 0 | 0 |
| NC_045512.2;YP_009725295.1;Wu1-SARS2_orf1a;3695 | 0 | 0 |
| NC_045512.2;YP_009725295.1;Wu1-SARS2_orf1a;3696 | 0 | 0 |
| NC_045512.2;YP_009725295.1;Wu1-SARS2_orf1a;3697 | 0 | 0 |
| NC_045512.2;YP_009725295.1;Wu1-SARS2_orf1a;3698 | 0 | 0.025 |
| NC_045512.2;YP_009725295.1;Wu1-SARS2_orf1a;3699 | 0 | 0 |
| NC_045512.2;YP_009725295.1;Wu1-SARS2_orf1a;3700 | 0 | 0 |
| NC_045512.2;YP_009725295.1;Wu1-SARS2_orf1a;3701 | 0 | 0 |
| NC_045512.2;YP_009725295.1;Wu1-SARS2_orf1a;3702 | 0 | 0 |
| NC_045512.2;YP_009725295.1;Wu1-SARS2_orf1a;3703 | 0 | 0 |
| NC_045512.2;YP_009725295.1;Wu1-SARS2_orf1a;3704 | 0 | 0 |
| NC_045512.2;YP_009725295.1;Wu1-SARS2_orf1a;3705 | 0 | 0 |
| NC_045512.2;YP_009725295.1;Wu1-SARS2_orf1a;3706 | 0 | 0 |
| NC_045512.2;YP_009725295.1;Wu1-SARS2_orf1a;3707 | 0 | 0 |
| NC_045512.2;YP_009725295.1;Wu1-SARS2_orf1a;3708 | 0 | 0 |
| NC_045512.2;YP_009725295.1;Wu1-SARS2_orf1a;3709 | 0 | 0 |
| NC_045512.2;YP_009725295.1;Wu1-SARS2_orf1a;3710 | 0 | 0 |
| NC_045512.2;YP_009725295.1;Wu1-SARS2_orf1a;3711 | 0 | 0 |
| NC_045512.2;YP_009725295.1;Wu1-SARS2_orf1a;3712 | 0 | 0 |
| NC_045512.2;YP_009725295.1;Wu1-SARS2_orf1a;3713 | 0 | 0 |
| NC_045512.2;YP_009725295.1;Wu1-SARS2_orf1a;3714 | 0 | 0 |
| NC_045512.2;YP_009725295.1;Wu1-SARS2_orf1a;3715 | 0 | 0 |
| NC_045512.2;YP_009725295.1;Wu1-SARS2_orf1a;3716 | 0 | 0 |
| NC_045512.2;YP_009725295.1;Wu1-SARS2_orf1a;3717 | 0 | 0 |
| NC_045512.2;YP_009725295.1;Wu1-SARS2_orf1a;3718 | 0.05 | 0 |
| NC_045512.2;YP_009725295.1;Wu1-SARS2_orf1a;3719 | 0.05 | 0 |
| NC_045512.2;YP_009725295.1;Wu1-SARS2_orf1a;3720 | 0.05 | 0 |
| NC_045512.2;YP_009725295.1;Wu1-SARS2_orf1a;3721 | 0.05 | 0.025 |
| NC_045512.2;YP_009725295.1;Wu1-SARS2_orf1a;3722 | 0.05 | 0 |
| NC_045512.2;YP_009725295.1;Wu1-SARS2_orf1a;3723 | 0 | 0 |
| NC_045512.2;YP_009725295.1;Wu1-SARS2_orf1a;3724 | 0 | 0 |
| NC_045512.2;YP_009725295.1;Wu1-SARS2_orf1a;3725 | 0 | 0 |
| NC_045512.2;YP_009725295.1;Wu1-SARS2_orf1a;3726 | 0 | 0 |
| NC_045512.2;YP_009725295.1;Wu1-SARS2_orf1a;3727 | 0 | 0 |
| NC_045512.2;YP_009725295.1;Wu1-SARS2_orf1a;3728 | 0 | 0 |
| NC_045512.2;YP_009725295.1;Wu1-SARS2_orf1a;3729 | 0 | 0 |

| | | |
|---|---|---|
| NC_045512.2;YP_009725295.1;Wu1-SARS2_orf1a;3730 | 0 | 0 |
| NC_045512.2;YP_009725295.1;Wu1-SARS2_orf1a;3731 | 0 | 0 |
| NC_045512.2;YP_009725295.1;Wu1-SARS2_orf1a;3732 | 0 | 0 |
| NC_045512.2;YP_009725295.1;Wu1-SARS2_orf1a;3733 | 0 | 0 |
| NC_045512.2;YP_009725295.1;Wu1-SARS2_orf1a;3734 | 0 | 0 |
| NC_045512.2;YP_009725295.1;Wu1-SARS2_orf1a;3735 | 0 | 0 |
| NC_045512.2;YP_009725295.1;Wu1-SARS2_orf1a;3736 | 0 | 0 |
| NC_045512.2;YP_009725295.1;Wu1-SARS2_orf1a;3737 | 0 | 0 |
| NC_045512.2;YP_009725295.1;Wu1-SARS2_orf1a;3738 | 0 | 0 |
| NC_045512.2;YP_009725295.1;Wu1-SARS2_orf1a;3739 | 0 | 0 |
| NC_045512.2;YP_009725295.1;Wu1-SARS2_orf1a;3740 | 0 | 0 |
| NC_045512.2;YP_009725295.1;Wu1-SARS2_orf1a;3741 | 0 | 0 |
| NC_045512.2;YP_009725295.1;Wu1-SARS2_orf1a;3742 | 0 | 0 |
| NC_045512.2;YP_009725295.1;Wu1-SARS2_orf1a;3743 | 0 | 0 |
| NC_045512.2;YP_009725295.1;Wu1-SARS2_orf1a;3744 | 0 | 0 |
| NC_045512.2;YP_009725295.1;Wu1-SARS2_orf1a;3745 | 0 | 0 |
| NC_045512.2;YP_009725295.1;Wu1-SARS2_orf1a;3746 | 0 | 0 |
| NC_045512.2;YP_009725295.1;Wu1-SARS2_orf1a;3747 | 0 | 0 |
| NC_045512.2;YP_009725295.1;Wu1-SARS2_orf1a;3748 | 0 | 0 |
| NC_045512.2;YP_009725295.1;Wu1-SARS2_orf1a;3749 | 0 | 0 |
| NC_045512.2;YP_009725295.1;Wu1-SARS2_orf1a;3750 | 0 | 0 |
| NC_045512.2;YP_009725295.1;Wu1-SARS2_orf1a;3751 | 0 | 0 |
| NC_045512.2;YP_009725295.1;Wu1-SARS2_orf1a;3752 | 0 | 0 |
| NC_045512.2;YP_009725295.1;Wu1-SARS2_orf1a;3753 | 0 | 0 |
| NC_045512.2;YP_009725295.1;Wu1-SARS2_orf1a;3754 | 0 | 0 |
| NC_045512.2;YP_009725295.1;Wu1-SARS2_orf1a;3755 | 0 | 0 |
| NC_045512.2;YP_009725295.1;Wu1-SARS2_orf1a;3756 | 0 | 0 |
| NC_045512.2;YP_009725295.1;Wu1-SARS2_orf1a;3757 | 0 | 0 |
| NC_045512.2;YP_009725295.1;Wu1-SARS2_orf1a;3758 | 0 | 0 |
| NC_045512.2;YP_009725295.1;Wu1-SARS2_orf1a;3759 | 0 | 0 |
| NC_045512.2;YP_009725295.1;Wu1-SARS2_orf1a;3760 | 0 | 0 |
| NC_045512.2;YP_009725295.1;Wu1-SARS2_orf1a;3761 | 0 | 0 |
| NC_045512.2;YP_009725295.1;Wu1-SARS2_orf1a;3762 | 0 | 0 |
| NC_045512.2;YP_009725295.1;Wu1-SARS2_orf1a;3763 | 0 | 0 |
| NC_045512.2;YP_009725295.1;Wu1-SARS2_orf1a;3764 | 0 | 0 |
| NC_045512.2;YP_009725295.1;Wu1-SARS2_orf1a;3765 | 0 | 0 |
| NC_045512.2;YP_009725295.1;Wu1-SARS2_orf1a;3766 | 0 | 0 |
| NC_045512.2;YP_009725295.1;Wu1-SARS2_orf1a;3767 | 0 | 0 |
| NC_045512.2;YP_009725295.1;Wu1-SARS2_orf1a;3768 | 0 | 0 |
| NC_045512.2;YP_009725295.1;Wu1-SARS2_orf1a;3769 | 0 | 0 |
| NC_045512.2;YP_009725295.1;Wu1-SARS2_orf1a;3770 | 0 | 0 |
| NC_045512.2;YP_009725295.1;Wu1-SARS2_orf1a;3771 | 0 | 0 |
| NC_045512.2;YP_009725295.1;Wu1-SARS2_orf1a;3772 | 0 | 0 |

| | | |
|---|---|---|
| NC_045512.2;YP_009725295.1;Wu1-SARS2_orf1a;3773 | 0 | 0 |
| NC_045512.2;YP_009725295.1;Wu1-SARS2_orf1a;3774 | 0 | 0 |
| NC_045512.2;YP_009725295.1;Wu1-SARS2_orf1a;3775 | 0 | 0 |
| NC_045512.2;YP_009725295.1;Wu1-SARS2_orf1a;3776 | 0 | 0 |
| NC_045512.2;YP_009725295.1;Wu1-SARS2_orf1a;3777 | 0 | 0 |
| NC_045512.2;YP_009725295.1;Wu1-SARS2_orf1a;3778 | 0 | 0 |
| NC_045512.2;YP_009725295.1;Wu1-SARS2_orf1a;3779 | 0 | 0 |
| NC_045512.2;YP_009725295.1;Wu1-SARS2_orf1a;3780 | 0 | 0 |
| NC_045512.2;YP_009725295.1;Wu1-SARS2_orf1a;3781 | 0 | 0 |
| NC_045512.2;YP_009725295.1;Wu1-SARS2_orf1a;3782 | 0 | 0 |
| NC_045512.2;YP_009725295.1;Wu1-SARS2_orf1a;3783 | 0 | 0 |
| NC_045512.2;YP_009725295.1;Wu1-SARS2_orf1a;3784 | 0 | 0 |
| NC_045512.2;YP_009725295.1;Wu1-SARS2_orf1a;3785 | 0 | 0 |
| NC_045512.2;YP_009725295.1;Wu1-SARS2_orf1a;3786 | 0 | 0 |
| NC_045512.2;YP_009725295.1;Wu1-SARS2_orf1a;3787 | 0 | 0 |
| NC_045512.2;YP_009725295.1;Wu1-SARS2_orf1a;3788 | 0 | 0 |
| NC_045512.2;YP_009725295.1;Wu1-SARS2_orf1a;3789 | 0 | 0 |
| NC_045512.2;YP_009725295.1;Wu1-SARS2_orf1a;3790 | 0 | 0 |
| NC_045512.2;YP_009725295.1;Wu1-SARS2_orf1a;3791 | 0 | 0 |
| NC_045512.2;YP_009725295.1;Wu1-SARS2_orf1a;3792 | 0 | 0 |
| NC_045512.2;YP_009725295.1;Wu1-SARS2_orf1a;3793 | 0 | 0 |
| NC_045512.2;YP_009725295.1;Wu1-SARS2_orf1a;3794 | 0 | 0 |
| NC_045512.2;YP_009725295.1;Wu1-SARS2_orf1a;3795 | 0 | 0 |
| NC_045512.2;YP_009725295.1;Wu1-SARS2_orf1a;3796 | 0 | 0 |
| NC_045512.2;YP_009725295.1;Wu1-SARS2_orf1a;3797 | 0 | 0 |
| NC_045512.2;YP_009725295.1;Wu1-SARS2_orf1a;3798 | 0 | 0 |
| NC_045512.2;YP_009725295.1;Wu1-SARS2_orf1a;3799 | 0 | 0 |
| NC_045512.2;YP_009725295.1;Wu1-SARS2_orf1a;3800 | 0 | 0 |
| NC_045512.2;YP_009725295.1;Wu1-SARS2_orf1a;3801 | 0 | 0 |
| NC_045512.2;YP_009725295.1;Wu1-SARS2_orf1a;3802 | 0 | 0 |
| NC_045512.2;YP_009725295.1;Wu1-SARS2_orf1a;3803 | 0 | 0 |
| NC_045512.2;YP_009725295.1;Wu1-SARS2_orf1a;3804 | 0 | 0 |
| NC_045512.2;YP_009725295.1;Wu1-SARS2_orf1a;3805 | 0 | 0 |
| NC_045512.2;YP_009725295.1;Wu1-SARS2_orf1a;3806 | 0 | 0 |
| NC_045512.2;YP_009725295.1;Wu1-SARS2_orf1a;3807 | 0 | 0 |
| NC_045512.2;YP_009725295.1;Wu1-SARS2_orf1a;3808 | 0 | 0 |
| NC_045512.2;YP_009725295.1;Wu1-SARS2_orf1a;3809 | 0.25 | 0.175 |
| NC_045512.2;YP_009725295.1;Wu1-SARS2_orf1a;3810 | 0 | 0 |
| NC_045512.2;YP_009725295.1;Wu1-SARS2_orf1a;3811 | 0 | 0 |
| NC_045512.2;YP_009725295.1;Wu1-SARS2_orf1a;3812 | 0 | 0 |
| NC_045512.2;YP_009725295.1;Wu1-SARS2_orf1a;3813 | 0 | 0 |
| NC_045512.2;YP_009725295.1;Wu1-SARS2_orf1a;3814 | 0 | 0 |
| NC_045512.2;YP_009725295.1;Wu1-SARS2_orf1a;3815 | 0 | 0 |

| | | |
|---|---|---|
| NC_045512.2;YP_009725295.1;Wu1-SARS2_orf1a;3816 | 0 | 0 |
| NC_045512.2;YP_009725295.1;Wu1-SARS2_orf1a;3817 | 0 | 0.05 |
| NC_045512.2;YP_009725295.1;Wu1-SARS2_orf1a;3818 | 0.05 | 0.025 |
| NC_045512.2;YP_009725295.1;Wu1-SARS2_orf1a;3819 | 0.15 | 0.075 |
| NC_045512.2;YP_009725295.1;Wu1-SARS2_orf1a;3820 | 0 | 0.025 |
| NC_045512.2;YP_009725295.1;Wu1-SARS2_orf1a;3821 | 0 | 0.025 |
| NC_045512.2;YP_009725295.1;Wu1-SARS2_orf1a;3822 | 0.05 | 0.125 |
| NC_045512.2;YP_009725295.1;Wu1-SARS2_orf1a;3823 | 0.1 | 0.05 |
| NC_045512.2;YP_009725295.1;Wu1-SARS2_orf1a;3824 | 0.3 | 0.1 |
| NC_045512.2;YP_009725295.1;Wu1-SARS2_orf1a;3825 | 0.25 | 0.1 |
| NC_045512.2;YP_009725295.1;Wu1-SARS2_orf1a;3826 | 0.25 | 0.1 |
| NC_045512.2;YP_009725295.1;Wu1-SARS2_orf1a;3827 | 0.2 | 0.1 |
| NC_045512.2;YP_009725295.1;Wu1-SARS2_orf1a;3828 | 0.15 | 0.075 |
| NC_045512.2;YP_009725295.1;Wu1-SARS2_orf1a;3829 | 0.1 | 0.075 |
| NC_045512.2;YP_009725295.1;Wu1-SARS2_orf1a;3830 | 0.05 | 0.075 |
| NC_045512.2;YP_009725295.1;Wu1-SARS2_orf1a;3831 | 0.05 | 0.05 |
| NC_045512.2;YP_009725295.1;Wu1-SARS2_orf1a;3832 | 0 | 0.05 |
| NC_045512.2;YP_009725295.1;Wu1-SARS2_orf1a;3833 | 0 | 0 |
| NC_045512.2;YP_009725295.1;Wu1-SARS2_orf1a;3834 | 0 | 0 |
| NC_045512.2;YP_009725295.1;Wu1-SARS2_orf1a;3835 | 0 | 0 |
| NC_045512.2;YP_009725295.1;Wu1-SARS2_orf1a;3836 | 0 | 0 |
| NC_045512.2;YP_009725295.1;Wu1-SARS2_orf1a;3837 | 0 | 0 |
| NC_045512.2;YP_009725295.1;Wu1-SARS2_orf1a;3838 | 0 | 0 |
| NC_045512.2;YP_009725295.1;Wu1-SARS2_orf1a;3839 | 0 | 0 |
| NC_045512.2;YP_009725295.1;Wu1-SARS2_orf1a;3840 | 0 | 0 |
| NC_045512.2;YP_009725295.1;Wu1-SARS2_orf1a;3841 | 0 | 0 |
| NC_045512.2;YP_009725295.1;Wu1-SARS2_orf1a;3842 | 0 | 0.025 |
| NC_045512.2;YP_009725295.1;Wu1-SARS2_orf1a;3843 | 0 | 0 |
| NC_045512.2;YP_009725295.1;Wu1-SARS2_orf1a;3844 | 0 | 0.025 |
| NC_045512.2;YP_009725295.1;Wu1-SARS2_orf1a;3845 | 0 | 0.025 |
| NC_045512.2;YP_009725295.1;Wu1-SARS2_orf1a;3846 | 0.05 | 0.05 |
| NC_045512.2;YP_009725295.1;Wu1-SARS2_orf1a;3847 | 0.05 | 0.05 |
| NC_045512.2;YP_009725295.1;Wu1-SARS2_orf1a;3848 | 0.05 | 0.05 |
| NC_045512.2;YP_009725295.1;Wu1-SARS2_orf1a;3849 | 0.1 | 0.1 |
| NC_045512.2;YP_009725295.1;Wu1-SARS2_orf1a;3850 | 0.05 | 0.075 |
| NC_045512.2;YP_009725295.1;Wu1-SARS2_orf1a;3851 | 0.05 | 0.05 |
| NC_045512.2;YP_009725295.1;Wu1-SARS2_orf1a;3852 | 0.05 | 0.025 |
| NC_045512.2;YP_009725295.1;Wu1-SARS2_orf1a;3853 | 0 | 0.025 |
| NC_045512.2;YP_009725295.1;Wu1-SARS2_orf1a;3854 | 0.05 | 0.025 |
| NC_045512.2;YP_009725295.1;Wu1-SARS2_orf1a;3855 | 0.05 | 0.025 |
| NC_045512.2;YP_009725295.1;Wu1-SARS2_orf1a;3856 | 0.05 | 0.025 |
| NC_045512.2;YP_009725295.1;Wu1-SARS2_orf1a;3857 | 0.05 | 0 |
| NC_045512.2;YP_009725295.1;Wu1-SARS2_orf1a;3858 | 0 | 0 |

| | | |
|---|---|---|
| NC_045512.2;YP_009725295.1;Wu1-SARS2_orf1a;3859 | 0 | 0 |
| NC_045512.2;YP_009725295.1;Wu1-SARS2_orf1a;3860 | 0 | 0 |
| NC_045512.2;YP_009725295.1;Wu1-SARS2_orf1a;3861 | 0 | 0 |
| NC_045512.2;YP_009725295.1;Wu1-SARS2_orf1a;3862 | 0 | 0 |
| NC_045512.2;YP_009725295.1;Wu1-SARS2_orf1a;3863 | 0 | 0 |
| NC_045512.2;YP_009725295.1;Wu1-SARS2_orf1a;3864 | 0 | 0 |
| NC_045512.2;YP_009725295.1;Wu1-SARS2_orf1a;3865 | 0 | 0 |
| NC_045512.2;YP_009725295.1;Wu1-SARS2_orf1a;3866 | 0 | 0 |
| NC_045512.2;YP_009725295.1;Wu1-SARS2_orf1a;3867 | 0 | 0 |
| NC_045512.2;YP_009725295.1;Wu1-SARS2_orf1a;3868 | 0 | 0 |
| NC_045512.2;YP_009725295.1;Wu1-SARS2_orf1a;3869 | 0 | 0 |
| NC_045512.2;YP_009725295.1;Wu1-SARS2_orf1a;3870 | 0 | 0 |
| NC_045512.2;YP_009725295.1;Wu1-SARS2_orf1a;3871 | 0 | 0 |
| NC_045512.2;YP_009725295.1;Wu1-SARS2_orf1a;3872 | 0 | 0 |
| NC_045512.2;YP_009725295.1;Wu1-SARS2_orf1a;3873 | 0 | 0 |
| NC_045512.2;YP_009725295.1;Wu1-SARS2_orf1a;3874 | 0 | 0 |
| NC_045512.2;YP_009725295.1;Wu1-SARS2_orf1a;3875 | 0 | 0 |
| NC_045512.2;YP_009725295.1;Wu1-SARS2_orf1a;3876 | 0 | 0 |
| NC_045512.2;YP_009725295.1;Wu1-SARS2_orf1a;3877 | 0 | 0 |
| NC_045512.2;YP_009725295.1;Wu1-SARS2_orf1a;3878 | 0 | 0 |
| NC_045512.2;YP_009725295.1;Wu1-SARS2_orf1a;3879 | 0 | 0 |
| NC_045512.2;YP_009725295.1;Wu1-SARS2_orf1a;3880 | 0 | 0 |
| NC_045512.2;YP_009725295.1;Wu1-SARS2_orf1a;3881 | 0 | 0 |
| NC_045512.2;YP_009725295.1;Wu1-SARS2_orf1a;3882 | 0 | 0.025 |
| NC_045512.2;YP_009725295.1;Wu1-SARS2_orf1a;3883 | 0 | 0 |
| NC_045512.2;YP_009725295.1;Wu1-SARS2_orf1a;3884 | 0 | 0.025 |
| NC_045512.2;YP_009725295.1;Wu1-SARS2_orf1a;3885 | 0 | 0 |
| NC_045512.2;YP_009725295.1;Wu1-SARS2_orf1a;3886 | 0 | 0 |
| NC_045512.2;YP_009725295.1;Wu1-SARS2_orf1a;3887 | 0 | 0 |
| NC_045512.2;YP_009725295.1;Wu1-SARS2_orf1a;3888 | 0.05 | 0 |
| NC_045512.2;YP_009725295.1;Wu1-SARS2_orf1a;3889 | 0.05 | 0 |
| NC_045512.2;YP_009725295.1;Wu1-SARS2_orf1a;3890 | 0.05 | 0 |
| NC_045512.2;YP_009725295.1;Wu1-SARS2_orf1a;3891 | 0 | 0 |
| NC_045512.2;YP_009725295.1;Wu1-SARS2_orf1a;3892 | 0 | 0 |
| NC_045512.2;YP_009725295.1;Wu1-SARS2_orf1a;3893 | 0 | 0 |
| NC_045512.2;YP_009725295.1;Wu1-SARS2_orf1a;3894 | 0 | 0 |
| NC_045512.2;YP_009725295.1;Wu1-SARS2_orf1a;3895 | 0 | 0 |
| NC_045512.2;YP_009725295.1;Wu1-SARS2_orf1a;3896 | 0 | 0 |
| NC_045512.2;YP_009725295.1;Wu1-SARS2_orf1a;3897 | 0 | 0 |
| NC_045512.2;YP_009725295.1;Wu1-SARS2_orf1a;3898 | 0 | 0 |
| NC_045512.2;YP_009725295.1;Wu1-SARS2_orf1a;3899 | 0 | 0 |
| NC_045512.2;YP_009725295.1;Wu1-SARS2_orf1a;3900 | 0 | 0 |
| NC_045512.2;YP_009725295.1;Wu1-SARS2_orf1a;3901 | 0 | 0 |

| | | |
|---|---|---|
| NC_045512.2;YP_009725295.1;Wu1-SARS2_orf1a;3902 | 0 | 0 |
| NC_045512.2;YP_009725295.1;Wu1-SARS2_orf1a;3903 | 0 | 0 |
| NC_045512.2;YP_009725295.1;Wu1-SARS2_orf1a;3904 | 0 | 0 |
| NC_045512.2;YP_009725295.1;Wu1-SARS2_orf1a;3905 | 0 | 0 |
| NC_045512.2;YP_009725295.1;Wu1-SARS2_orf1a;3906 | 0 | 0 |
| NC_045512.2;YP_009725295.1;Wu1-SARS2_orf1a;3907 | 0 | 0 |
| NC_045512.2;YP_009725295.1;Wu1-SARS2_orf1a;3908 | 0 | 0 |
| NC_045512.2;YP_009725295.1;Wu1-SARS2_orf1a;3909 | 0 | 0 |
| NC_045512.2;YP_009725295.1;Wu1-SARS2_orf1a;3910 | 0 | 0 |
| NC_045512.2;YP_009725295.1;Wu1-SARS2_orf1a;3911 | 0 | 0 |
| NC_045512.2;YP_009725295.1;Wu1-SARS2_orf1a;3912 | 0 | 0 |
| NC_045512.2;YP_009725295.1;Wu1-SARS2_orf1a;3913 | 0 | 0 |
| NC_045512.2;YP_009725295.1;Wu1-SARS2_orf1a;3914 | 0 | 0.05 |
| NC_045512.2;YP_009725295.1;Wu1-SARS2_orf1a;3915 | 0 | 0.025 |
| NC_045512.2;YP_009725295.1;Wu1-SARS2_orf1a;3916 | 0 | 0.025 |
| NC_045512.2;YP_009725295.1;Wu1-SARS2_orf1a;3917 | 0 | 0.025 |
| NC_045512.2;YP_009725295.1;Wu1-SARS2_orf1a;3918 | 0 | 0.025 |
| NC_045512.2;YP_009725295.1;Wu1-SARS2_orf1a;3919 | 0.05 | 0.05 |
| NC_045512.2;YP_009725295.1;Wu1-SARS2_orf1a;3920 | 0.05 | 0.05 |
| NC_045512.2;YP_009725295.1;Wu1-SARS2_orf1a;3921 | 0.05 | 0.075 |
| NC_045512.2;YP_009725295.1;Wu1-SARS2_orf1a;3922 | 0.05 | 0.05 |
| NC_045512.2;YP_009725295.1;Wu1-SARS2_orf1a;3923 | 0.1 | 0.025 |
| NC_045512.2;YP_009725295.1;Wu1-SARS2_orf1a;3924 | 0 | 0.025 |
| NC_045512.2;YP_009725295.1;Wu1-SARS2_orf1a;3925 | 0 | 0.05 |
| NC_045512.2;YP_009725295.1;Wu1-SARS2_orf1a;3926 | 0 | 0.025 |
| NC_045512.2;YP_009725295.1;Wu1-SARS2_orf1a;3927 | 0 | 0 |
| NC_045512.2;YP_009725295.1;Wu1-SARS2_orf1a;3928 | 0 | 0 |
| NC_045512.2;YP_009725295.1;Wu1-SARS2_orf1a;3929 | 0.05 | 0 |
| NC_045512.2;YP_009725295.1;Wu1-SARS2_orf1a;3930 | 0.05 | 0 |
| NC_045512.2;YP_009725295.1;Wu1-SARS2_orf1a;3931 | 0.05 | 0 |
| NC_045512.2;YP_009725295.1;Wu1-SARS2_orf1a;3932 | 0 | 0 |
| NC_045512.2;YP_009725295.1;Wu1-SARS2_orf1a;3933 | 0.05 | 0 |
| NC_045512.2;YP_009725295.1;Wu1-SARS2_orf1a;3934 | 0 | 0 |
| NC_045512.2;YP_009725295.1;Wu1-SARS2_orf1a;3935 | 0 | 0 |
| NC_045512.2;YP_009725295.1;Wu1-SARS2_orf1a;3936 | 0 | 0 |
| NC_045512.2;YP_009725295.1;Wu1-SARS2_orf1a;3937 | 0 | 0 |
| NC_045512.2;YP_009725295.1;Wu1-SARS2_orf1a;3938 | 0 | 0 |
| NC_045512.2;YP_009725295.1;Wu1-SARS2_orf1a;3939 | 0 | 0 |
| NC_045512.2;YP_009725295.1;Wu1-SARS2_orf1a;3940 | 0 | 0 |
| NC_045512.2;YP_009725295.1;Wu1-SARS2_orf1a;3941 | 0.05 | 0.05 |
| NC_045512.2;YP_009725295.1;Wu1-SARS2_orf1a;3942 | 0 | 0.025 |
| NC_045512.2;YP_009725295.1;Wu1-SARS2_orf1a;3943 | 0 | 0 |
| NC_045512.2;YP_009725295.1;Wu1-SARS2_orf1a;3944 | 0 | 0 |

| | | |
|---|---|---|
| NC_045512.2;YP_009725295.1;Wu1-SARS2_orf1a;3945 | 0 | 0 |
| NC_045512.2;YP_009725295.1;Wu1-SARS2_orf1a;3946 | 0 | 0.05 |
| NC_045512.2;YP_009725295.1;Wu1-SARS2_orf1a;3947 | 0.05 | 0 |
| NC_045512.2;YP_009725295.1;Wu1-SARS2_orf1a;3948 | 0.05 | 0 |
| NC_045512.2;YP_009725295.1;Wu1-SARS2_orf1a;3949 | 0.05 | 0 |
| NC_045512.2;YP_009725295.1;Wu1-SARS2_orf1a;3950 | 0.05 | 0.05 |
| NC_045512.2;YP_009725295.1;Wu1-SARS2_orf1a;3951 | 0.1 | 0.05 |
| NC_045512.2;YP_009725295.1;Wu1-SARS2_orf1a;3952 | 0 | 0.025 |
| NC_045512.2;YP_009725295.1;Wu1-SARS2_orf1a;3953 | 0.1 | 0.05 |
| NC_045512.2;YP_009725295.1;Wu1-SARS2_orf1a;3954 | 0.05 | 0.125 |
| NC_045512.2;YP_009725295.1;Wu1-SARS2_orf1a;3955 | 0.05 | 0.075 |
| NC_045512.2;YP_009725295.1;Wu1-SARS2_orf1a;3956 | 0.05 | 0.15 |
| NC_045512.2;YP_009725295.1;Wu1-SARS2_orf1a;3957 | 0.05 | 0.05 |
| NC_045512.2;YP_009725295.1;Wu1-SARS2_orf1a;3958 | 0 | 0.025 |
| NC_045512.2;YP_009725295.1;Wu1-SARS2_orf1a;3959 | 0 | 0 |
| NC_045512.2;YP_009725295.1;Wu1-SARS2_orf1a;3960 | 0.05 | 0 |
| NC_045512.2;YP_009725295.1;Wu1-SARS2_orf1a;3961 | 0 | 0 |
| NC_045512.2;YP_009725295.1;Wu1-SARS2_orf1a;3962 | 0 | 0 |
| NC_045512.2;YP_009725295.1;Wu1-SARS2_orf1a;3963 | 0 | 0 |
| NC_045512.2;YP_009725295.1;Wu1-SARS2_orf1a;3964 | 0 | 0 |
| NC_045512.2;YP_009725295.1;Wu1-SARS2_orf1a;3965 | 0 | 0 |
| NC_045512.2;YP_009725295.1;Wu1-SARS2_orf1a;3966 | 0 | 0 |
| NC_045512.2;YP_009725295.1;Wu1-SARS2_orf1a;3967 | 0 | 0 |
| NC_045512.2;YP_009725295.1;Wu1-SARS2_orf1a;3968 | 0 | 0 |
| NC_045512.2;YP_009725295.1;Wu1-SARS2_orf1a;3969 | 0 | 0 |
| NC_045512.2;YP_009725295.1;Wu1-SARS2_orf1a;3970 | 0 | 0 |
| NC_045512.2;YP_009725295.1;Wu1-SARS2_orf1a;3971 | 0 | 0 |
| NC_045512.2;YP_009725295.1;Wu1-SARS2_orf1a;3972 | 0 | 0 |
| NC_045512.2;YP_009725295.1;Wu1-SARS2_orf1a;3973 | 0 | 0 |
| NC_045512.2;YP_009725295.1;Wu1-SARS2_orf1a;3974 | 0 | 0 |
| NC_045512.2;YP_009725295.1;Wu1-SARS2_orf1a;3975 | 0 | 0 |
| NC_045512.2;YP_009725295.1;Wu1-SARS2_orf1a;3976 | 0 | 0 |
| NC_045512.2;YP_009725295.1;Wu1-SARS2_orf1a;3977 | 0 | 0.025 |
| NC_045512.2;YP_009725295.1;Wu1-SARS2_orf1a;3978 | 0 | 0.05 |
| NC_045512.2;YP_009725295.1;Wu1-SARS2_orf1a;3979 | 0 | 0.025 |
| NC_045512.2;YP_009725295.1;Wu1-SARS2_orf1a;3980 | 0.05 | 0.025 |
| NC_045512.2;YP_009725295.1;Wu1-SARS2_orf1a;3981 | 0.05 | 0.025 |
| NC_045512.2;YP_009725295.1;Wu1-SARS2_orf1a;3982 | 0 | 0.05 |
| NC_045512.2;YP_009725295.1;Wu1-SARS2_orf1a;3983 | 0.05 | 0.025 |
| NC_045512.2;YP_009725295.1;Wu1-SARS2_orf1a;3984 | 0 | 0.025 |
| NC_045512.2;YP_009725295.1;Wu1-SARS2_orf1a;3985 | 0 | 0 |
| NC_045512.2;YP_009725295.1;Wu1-SARS2_orf1a;3986 | 0 | 0 |
| NC_045512.2;YP_009725295.1;Wu1-SARS2_orf1a;3987 | 0 | 0 |

| | | |
|---|---|---|
| NC_045512.2;YP_009725295.1;Wu1-SARS2_orf1a;3988 | 0 | 0 |
| NC_045512.2;YP_009725295.1;Wu1-SARS2_orf1a;3989 | 0 | 0 |
| NC_045512.2;YP_009725295.1;Wu1-SARS2_orf1a;3990 | 0 | 0 |
| NC_045512.2;YP_009725295.1;Wu1-SARS2_orf1a;3991 | 0 | 0 |
| NC_045512.2;YP_009725295.1;Wu1-SARS2_orf1a;3992 | 0 | 0 |
| NC_045512.2;YP_009725295.1;Wu1-SARS2_orf1a;3993 | 0 | 0 |
| NC_045512.2;YP_009725295.1;Wu1-SARS2_orf1a;3994 | 0 | 0.025 |
| NC_045512.2;YP_009725295.1;Wu1-SARS2_orf1a;3995 | 0 | 0 |
| NC_045512.2;YP_009725295.1;Wu1-SARS2_orf1a;3996 | 0 | 0 |
| NC_045512.2;YP_009725295.1;Wu1-SARS2_orf1a;3997 | 0 | 0 |
| NC_045512.2;YP_009725295.1;Wu1-SARS2_orf1a;3998 | 0 | 0.05 |
| NC_045512.2;YP_009725295.1;Wu1-SARS2_orf1a;3999 | 0 | 0.025 |
| NC_045512.2;YP_009725295.1;Wu1-SARS2_orf1a;4000 | 0 | 0 |
| NC_045512.2;YP_009725295.1;Wu1-SARS2_orf1a;4001 | 0 | 0 |
| NC_045512.2;YP_009725295.1;Wu1-SARS2_orf1a;4002 | 0 | 0 |
| NC_045512.2;YP_009725295.1;Wu1-SARS2_orf1a;4003 | 0 | 0 |
| NC_045512.2;YP_009725295.1;Wu1-SARS2_orf1a;4004 | 0 | 0 |
| NC_045512.2;YP_009725295.1;Wu1-SARS2_orf1a;4005 | 0 | 0 |
| NC_045512.2;YP_009725295.1;Wu1-SARS2_orf1a;4006 | 0 | 0.1 |
| NC_045512.2;YP_009725295.1;Wu1-SARS2_orf1a;4007 | 0 | 0.075 |
| NC_045512.2;YP_009725295.1;Wu1-SARS2_orf1a;4008 | 0 | 0.05 |
| NC_045512.2;YP_009725295.1;Wu1-SARS2_orf1a;4009 | 0 | 0.05 |
| NC_045512.2;YP_009725295.1;Wu1-SARS2_orf1a;4010 | 0 | 0.05 |
| NC_045512.2;YP_009725295.1;Wu1-SARS2_orf1a;4011 | 0 | 0 |
| NC_045512.2;YP_009725295.1;Wu1-SARS2_orf1a;4012 | 0 | 0 |
| NC_045512.2;YP_009725295.1;Wu1-SARS2_orf1a;4013 | 0 | 0 |
| NC_045512.2;YP_009725295.1;Wu1-SARS2_orf1a;4014 | 0 | 0 |
| NC_045512.2;YP_009725295.1;Wu1-SARS2_orf1a;4015 | 0 | 0 |
| NC_045512.2;YP_009725295.1;Wu1-SARS2_orf1a;4016 | 0 | 0 |
| NC_045512.2;YP_009725295.1;Wu1-SARS2_orf1a;4017 | 0 | 0 |
| NC_045512.2;YP_009725295.1;Wu1-SARS2_orf1a;4018 | 0 | 0 |
| NC_045512.2;YP_009725295.1;Wu1-SARS2_orf1a;4019 | 0 | 0 |
| NC_045512.2;YP_009725295.1;Wu1-SARS2_orf1a;4020 | 0 | 0 |
| NC_045512.2;YP_009725295.1;Wu1-SARS2_orf1a;4021 | 0 | 0 |
| NC_045512.2;YP_009725295.1;Wu1-SARS2_orf1a;4022 | 0 | 0 |
| NC_045512.2;YP_009725295.1;Wu1-SARS2_orf1a;4023 | 0 | 0 |
| NC_045512.2;YP_009725295.1;Wu1-SARS2_orf1a;4024 | 0 | 0 |
| NC_045512.2;YP_009725295.1;Wu1-SARS2_orf1a;4025 | 0 | 0 |
| NC_045512.2;YP_009725295.1;Wu1-SARS2_orf1a;4026 | 0 | 0 |
| NC_045512.2;YP_009725295.1;Wu1-SARS2_orf1a;4027 | 0 | 0 |
| NC_045512.2;YP_009725295.1;Wu1-SARS2_orf1a;4028 | 0 | 0 |
| NC_045512.2;YP_009725295.1;Wu1-SARS2_orf1a;4029 | 0 | 0 |
| NC_045512.2;YP_009725295.1;Wu1-SARS2_orf1a;4030 | 0 | 0 |

| | | |
|---|---|---|
| NC_045512.2;YP_009725295.1;Wu1-SARS2_orf1a;4031 | 0.15 | 0.1 |
| NC_045512.2;YP_009725295.1;Wu1-SARS2_orf1a;4032 | 0.2 | 0.1 |
| NC_045512.2;YP_009725295.1;Wu1-SARS2_orf1a;4033 | 0.2 | 0.075 |
| NC_045512.2;YP_009725295.1;Wu1-SARS2_orf1a;4034 | 0.25 | 0.1 |
| NC_045512.2;YP_009725295.1;Wu1-SARS2_orf1a;4035 | 0.25 | 0.275 |
| NC_045512.2;YP_009725295.1;Wu1-SARS2_orf1a;4036 | 0.3 | 0.3 |
| NC_045512.2;YP_009725295.1;Wu1-SARS2_orf1a;4037 | 0.35 | 0.4 |
| NC_045512.2;YP_009725295.1;Wu1-SARS2_orf1a;4038 | 0.25 | 0.25 |
| NC_045512.2;YP_009725295.1;Wu1-SARS2_orf1a;4039 | 0.4 | 0.5 |
| NC_045512.2;YP_009725295.1;Wu1-SARS2_orf1a;4040 | 0.1 | 0.025 |
| NC_045512.2;YP_009725295.1;Wu1-SARS2_orf1a;4041 | 0.05 | 0.025 |
| NC_045512.2;YP_009725295.1;Wu1-SARS2_orf1a;4042 | 0 | 0 |
| NC_045512.2;YP_009725295.1;Wu1-SARS2_orf1a;4043 | 0.05 | 0.1 |
| NC_045512.2;YP_009725295.1;Wu1-SARS2_orf1a;4044 | 0.1 | 0.075 |
| NC_045512.2;YP_009725295.1;Wu1-SARS2_orf1a;4045 | 0 | 0.075 |
| NC_045512.2;YP_009725295.1;Wu1-SARS2_orf1a;4046 | 0.05 | 0.075 |
| NC_045512.2;YP_009725295.1;Wu1-SARS2_orf1a;4047 | 0.05 | 0 |
| NC_045512.2;YP_009725295.1;Wu1-SARS2_orf1a;4048 | 0 | 0.05 |
| NC_045512.2;YP_009725295.1;Wu1-SARS2_orf1a;4049 | 0 | 0 |
| NC_045512.2;YP_009725295.1;Wu1-SARS2_orf1a;4050 | 0 | 0 |
| NC_045512.2;YP_009725295.1;Wu1-SARS2_orf1a;4051 | 0 | 0 |
| NC_045512.2;YP_009725295.1;Wu1-SARS2_orf1a;4052 | 0 | 0 |
| NC_045512.2;YP_009725295.1;Wu1-SARS2_orf1a;4053 | 0.05 | 0.025 |
| NC_045512.2;YP_009725295.1;Wu1-SARS2_orf1a;4054 | 0.05 | 0.025 |
| NC_045512.2;YP_009725295.1;Wu1-SARS2_orf1a;4055 | 0 | 0 |
| NC_045512.2;YP_009725295.1;Wu1-SARS2_orf1a;4056 | 0 | 0 |
| NC_045512.2;YP_009725295.1;Wu1-SARS2_orf1a;4057 | 0 | 0 |
| NC_045512.2;YP_009725295.1;Wu1-SARS2_orf1a;4058 | 0 | 0 |
| NC_045512.2;YP_009725295.1;Wu1-SARS2_orf1a;4059 | 0 | 0 |
| NC_045512.2;YP_009725295.1;Wu1-SARS2_orf1a;4060 | 0 | 0 |
| NC_045512.2;YP_009725295.1;Wu1-SARS2_orf1a;4061 | 0 | 0 |
| NC_045512.2;YP_009725295.1;Wu1-SARS2_orf1a;4062 | 0 | 0.025 |
| NC_045512.2;YP_009725295.1;Wu1-SARS2_orf1a;4063 | 0.05 | 0 |
| NC_045512.2;YP_009725295.1;Wu1-SARS2_orf1a;4064 | 0 | 0.025 |
| NC_045512.2;YP_009725295.1;Wu1-SARS2_orf1a;4065 | 0 | 0 |
| NC_045512.2;YP_009725295.1;Wu1-SARS2_orf1a;4066 | 0 | 0 |
| NC_045512.2;YP_009725295.1;Wu1-SARS2_orf1a;4067 | 0 | 0 |
| NC_045512.2;YP_009725295.1;Wu1-SARS2_orf1a;4068 | 0 | 0.025 |
| NC_045512.2;YP_009725295.1;Wu1-SARS2_orf1a;4069 | 0 | 0 |
| NC_045512.2;YP_009725295.1;Wu1-SARS2_orf1a;4070 | 0.1 | 0 |
| NC_045512.2;YP_009725295.1;Wu1-SARS2_orf1a;4071 | 0.1 | 0.025 |
| NC_045512.2;YP_009725295.1;Wu1-SARS2_orf1a;4072 | 0.2 | 0.025 |
| NC_045512.2;YP_009725295.1;Wu1-SARS2_orf1a;4073 | 0.05 | 0.025 |

| | | |
|---|---|---|
| NC_045512.2;YP_009725295.1;Wu1-SARS2_orf1a;4074 | 0.1 | 0.05 |
| NC_045512.2;YP_009725295.1;Wu1-SARS2_orf1a;4075 | 0.05 | 0.025 |
| NC_045512.2;YP_009725295.1;Wu1-SARS2_orf1a;4076 | 0.05 | 0 |
| NC_045512.2;YP_009725295.1;Wu1-SARS2_orf1a;4077 | 0 | 0 |
| NC_045512.2;YP_009725295.1;Wu1-SARS2_orf1a;4078 | 0 | 0 |
| NC_045512.2;YP_009725295.1;Wu1-SARS2_orf1a;4079 | 0 | 0 |
| NC_045512.2;YP_009725295.1;Wu1-SARS2_orf1a;4080 | 0 | 0 |
| NC_045512.2;YP_009725295.1;Wu1-SARS2_orf1a;4081 | 0 | 0 |
| NC_045512.2;YP_009725295.1;Wu1-SARS2_orf1a;4082 | 0 | 0 |
| NC_045512.2;YP_009725295.1;Wu1-SARS2_orf1a;4083 | 0 | 0 |
| NC_045512.2;YP_009725295.1;Wu1-SARS2_orf1a;4084 | 0 | 0 |
| NC_045512.2;YP_009725295.1;Wu1-SARS2_orf1a;4085 | 0 | 0 |
| NC_045512.2;YP_009725295.1;Wu1-SARS2_orf1a;4086 | 0 | 0 |
| NC_045512.2;YP_009725295.1;Wu1-SARS2_orf1a;4087 | 0 | 0 |
| NC_045512.2;YP_009725295.1;Wu1-SARS2_orf1a;4088 | 0 | 0 |
| NC_045512.2;YP_009725295.1;Wu1-SARS2_orf1a;4089 | 0 | 0 |
| NC_045512.2;YP_009725295.1;Wu1-SARS2_orf1a;4090 | 0 | 0 |
| NC_045512.2;YP_009725295.1;Wu1-SARS2_orf1a;4091 | 0 | 0 |
| NC_045512.2;YP_009725295.1;Wu1-SARS2_orf1a;4092 | 0.05 | 0.025 |
| NC_045512.2;YP_009725295.1;Wu1-SARS2_orf1a;4093 | 0.05 | 0.05 |
| NC_045512.2;YP_009725295.1;Wu1-SARS2_orf1a;4094 | 0 | 0 |
| NC_045512.2;YP_009725295.1;Wu1-SARS2_orf1a;4095 | 0 | 0.025 |
| NC_045512.2;YP_009725295.1;Wu1-SARS2_orf1a;4096 | 0 | 0 |
| NC_045512.2;YP_009725295.1;Wu1-SARS2_orf1a;4097 | 0 | 0 |
| NC_045512.2;YP_009725295.1;Wu1-SARS2_orf1a;4098 | 0 | 0 |
| NC_045512.2;YP_009725295.1;Wu1-SARS2_orf1a;4099 | 0 | 0.025 |
| NC_045512.2;YP_009725295.1;Wu1-SARS2_orf1a;4100 | 0 | 0 |
| NC_045512.2;YP_009725295.1;Wu1-SARS2_orf1a;4101 | 0 | 0 |
| NC_045512.2;YP_009725295.1;Wu1-SARS2_orf1a;4102 | 0 | 0 |
| NC_045512.2;YP_009725295.1;Wu1-SARS2_orf1a;4103 | 0.05 | 0 |
| NC_045512.2;YP_009725295.1;Wu1-SARS2_orf1a;4104 | 0 | 0 |
| NC_045512.2;YP_009725295.1;Wu1-SARS2_orf1a;4105 | 0 | 0 |
| NC_045512.2;YP_009725295.1;Wu1-SARS2_orf1a;4106 | 0 | 0 |
| NC_045512.2;YP_009725295.1;Wu1-SARS2_orf1a;4107 | 0 | 0 |
| NC_045512.2;YP_009725295.1;Wu1-SARS2_orf1a;4108 | 0 | 0.025 |
| NC_045512.2;YP_009725295.1;Wu1-SARS2_orf1a;4109 | 0 | 0 |
| NC_045512.2;YP_009725295.1;Wu1-SARS2_orf1a;4110 | 0 | 0 |
| NC_045512.2;YP_009725295.1;Wu1-SARS2_orf1a;4111 | 0 | 0 |
| NC_045512.2;YP_009725295.1;Wu1-SARS2_orf1a;4112 | 0 | 0 |
| NC_045512.2;YP_009725295.1;Wu1-SARS2_orf1a;4113 | 0 | 0 |
| NC_045512.2;YP_009725295.1;Wu1-SARS2_orf1a;4114 | 0 | 0 |
| NC_045512.2;YP_009725295.1;Wu1-SARS2_orf1a;4115 | 0 | 0 |
| NC_045512.2;YP_009725295.1;Wu1-SARS2_orf1a;4116 | 0 | 0 |

| | | |
|---|---|---|
| NC_045512.2;YP_009725295.1;Wu1-SARS2_orf1a;4117 | 0 | 0 |
| NC_045512.2;YP_009725295.1;Wu1-SARS2_orf1a;4118 | 0 | 0 |
| NC_045512.2;YP_009725295.1;Wu1-SARS2_orf1a;4119 | 0 | 0 |
| NC_045512.2;YP_009725295.1;Wu1-SARS2_orf1a;4120 | 0 | 0 |
| NC_045512.2;YP_009725295.1;Wu1-SARS2_orf1a;4121 | 0 | 0 |
| NC_045512.2;YP_009725295.1;Wu1-SARS2_orf1a;4122 | 0 | 0 |
| NC_045512.2;YP_009725295.1;Wu1-SARS2_orf1a;4123 | 0 | 0 |
| NC_045512.2;YP_009725295.1;Wu1-SARS2_orf1a;4124 | 0 | 0 |
| NC_045512.2;YP_009725295.1;Wu1-SARS2_orf1a;4125 | 0 | 0 |
| NC_045512.2;YP_009725295.1;Wu1-SARS2_orf1a;4126 | 0 | 0 |
| NC_045512.2;YP_009725295.1;Wu1-SARS2_orf1a;4127 | 0 | 0.025 |
| NC_045512.2;YP_009725295.1;Wu1-SARS2_orf1a;4128 | 0 | 0.075 |
| NC_045512.2;YP_009725295.1;Wu1-SARS2_orf1a;4129 | 0 | 0.05 |
| NC_045512.2;YP_009725295.1;Wu1-SARS2_orf1a;4130 | 0 | 0.025 |
| NC_045512.2;YP_009725295.1;Wu1-SARS2_orf1a;4131 | 0 | 0.025 |
| NC_045512.2;YP_009725295.1;Wu1-SARS2_orf1a;4132 | 0.05 | 0.025 |
| NC_045512.2;YP_009725295.1;Wu1-SARS2_orf1a;4133 | 0.1 | 0.025 |
| NC_045512.2;YP_009725295.1;Wu1-SARS2_orf1a;4134 | 0.05 | 0.025 |
| NC_045512.2;YP_009725295.1;Wu1-SARS2_orf1a;4135 | 0 | 0.025 |
| NC_045512.2;YP_009725295.1;Wu1-SARS2_orf1a;4136 | 0 | 0 |
| NC_045512.2;YP_009725295.1;Wu1-SARS2_orf1a;4137 | 0.05 | 0.025 |
| NC_045512.2;YP_009725295.1;Wu1-SARS2_orf1a;4138 | 0 | 0 |
| NC_045512.2;YP_009725295.1;Wu1-SARS2_orf1a;4139 | 0 | 0 |
| NC_045512.2;YP_009725295.1;Wu1-SARS2_orf1a;4140 | 0 | 0.025 |
| NC_045512.2;YP_009725295.1;Wu1-SARS2_orf1a;4141 | 0.05 | 0.05 |
| NC_045512.2;YP_009725295.1;Wu1-SARS2_orf1a;4142 | 0 | 0 |
| NC_045512.2;YP_009725295.1;Wu1-SARS2_orf1a;4143 | 0 | 0 |
| NC_045512.2;YP_009725295.1;Wu1-SARS2_orf1a;4144 | 0 | 0 |
| NC_045512.2;YP_009725295.1;Wu1-SARS2_orf1a;4145 | 0 | 0 |
| NC_045512.2;YP_009725295.1;Wu1-SARS2_orf1a;4146 | 0 | 0 |
| NC_045512.2;YP_009725295.1;Wu1-SARS2_orf1a;4147 | 0 | 0 |
| NC_045512.2;YP_009725295.1;Wu1-SARS2_orf1a;4148 | 0 | 0 |
| NC_045512.2;YP_009725295.1;Wu1-SARS2_orf1a;4149 | 0 | 0 |
| NC_045512.2;YP_009725295.1;Wu1-SARS2_orf1a;4150 | 0 | 0 |
| NC_045512.2;YP_009725295.1;Wu1-SARS2_orf1a;4151 | 0 | 0 |
| NC_045512.2;YP_009725295.1;Wu1-SARS2_orf1a;4152 | 0 | 0 |
| NC_045512.2;YP_009725295.1;Wu1-SARS2_orf1a;4153 | 0 | 0 |
| NC_045512.2;YP_009725295.1;Wu1-SARS2_orf1a;4154 | 0 | 0 |
| NC_045512.2;YP_009725295.1;Wu1-SARS2_orf1a;4155 | 0 | 0 |
| NC_045512.2;YP_009725295.1;Wu1-SARS2_orf1a;4156 | 0 | 0 |
| NC_045512.2;YP_009725295.1;Wu1-SARS2_orf1a;4157 | 0 | 0 |
| NC_045512.2;YP_009725295.1;Wu1-SARS2_orf1a;4158 | 0 | 0 |
| NC_045512.2;YP_009725295.1;Wu1-SARS2_orf1a;4159 | 0 | 0 |

| | | |
|---|---|---|
| NC_045512.2;YP_009725295.1;Wu1-SARS2_orf1a;4160 | 0 | 0 |
| NC_045512.2;YP_009725295.1;Wu1-SARS2_orf1a;4161 | 0.05 | 0.05 |
| NC_045512.2;YP_009725295.1;Wu1-SARS2_orf1a;4162 | 0.1 | 0.05 |
| NC_045512.2;YP_009725295.1;Wu1-SARS2_orf1a;4163 | 0.2 | 0.05 |
| NC_045512.2;YP_009725295.1;Wu1-SARS2_orf1a;4164 | 0 | 0 |
| NC_045512.2;YP_009725295.1;Wu1-SARS2_orf1a;4165 | 0 | 0 |
| NC_045512.2;YP_009725295.1;Wu1-SARS2_orf1a;4166 | 0 | 0 |
| NC_045512.2;YP_009725295.1;Wu1-SARS2_orf1a;4167 | 0 | 0 |
| NC_045512.2;YP_009725295.1;Wu1-SARS2_orf1a;4168 | 0 | 0 |
| NC_045512.2;YP_009725295.1;Wu1-SARS2_orf1a;4169 | 0 | 0 |
| NC_045512.2;YP_009725295.1;Wu1-SARS2_orf1a;4170 | 0 | 0 |
| NC_045512.2;YP_009725295.1;Wu1-SARS2_orf1a;4171 | 0 | 0 |
| NC_045512.2;YP_009725295.1;Wu1-SARS2_orf1a;4172 | 0 | 0 |
| NC_045512.2;YP_009725295.1;Wu1-SARS2_orf1a;4173 | 0 | 0 |
| NC_045512.2;YP_009725295.1;Wu1-SARS2_orf1a;4174 | 0 | 0 |
| NC_045512.2;YP_009725295.1;Wu1-SARS2_orf1a;4175 | 0 | 0 |
| NC_045512.2;YP_009725295.1;Wu1-SARS2_orf1a;4176 | 0 | 0 |
| NC_045512.2;YP_009725295.1;Wu1-SARS2_orf1a;4177 | 0 | 0 |
| NC_045512.2;YP_009725295.1;Wu1-SARS2_orf1a;4178 | 0 | 0 |
| NC_045512.2;YP_009725295.1;Wu1-SARS2_orf1a;4179 | 0 | 0 |
| NC_045512.2;YP_009725295.1;Wu1-SARS2_orf1a;4180 | 0 | 0 |
| NC_045512.2;YP_009725295.1;Wu1-SARS2_orf1a;4181 | 0 | 0 |
| NC_045512.2;YP_009725295.1;Wu1-SARS2_orf1a;4182 | 0.05 | 0 |
| NC_045512.2;YP_009725295.1;Wu1-SARS2_orf1a;4183 | 0 | 0 |
| NC_045512.2;YP_009725295.1;Wu1-SARS2_orf1a;4184 | 0.05 | 0 |
| NC_045512.2;YP_009725295.1;Wu1-SARS2_orf1a;4185 | 0.05 | 0.05 |
| NC_045512.2;YP_009725295.1;Wu1-SARS2_orf1a;4186 | 0.15 | 0.1 |
| NC_045512.2;YP_009725295.1;Wu1-SARS2_orf1a;4187 | 0.3 | 0.25 |
| NC_045512.2;YP_009725295.1;Wu1-SARS2_orf1a;4188 | 0.15 | 0.05 |
| NC_045512.2;YP_009725295.1;Wu1-SARS2_orf1a;4189 | 0 | 0 |
| NC_045512.2;YP_009725295.1;Wu1-SARS2_orf1a;4190 | 0 | 0 |
| NC_045512.2;YP_009725295.1;Wu1-SARS2_orf1a;4191 | 0 | 0 |
| NC_045512.2;YP_009725295.1;Wu1-SARS2_orf1a;4192 | 0 | 0 |
| NC_045512.2;YP_009725295.1;Wu1-SARS2_orf1a;4193 | 0 | 0 |
| NC_045512.2;YP_009725295.1;Wu1-SARS2_orf1a;4194 | 0 | 0 |
| NC_045512.2;YP_009725295.1;Wu1-SARS2_orf1a;4195 | 0 | 0 |
| NC_045512.2;YP_009725295.1;Wu1-SARS2_orf1a;4196 | 0 | 0 |
| NC_045512.2;YP_009725295.1;Wu1-SARS2_orf1a;4197 | 0 | 0 |
| NC_045512.2;YP_009725295.1;Wu1-SARS2_orf1a;4198 | 0 | 0 |
| NC_045512.2;YP_009725295.1;Wu1-SARS2_orf1a;4199 | 0 | 0 |
| NC_045512.2;YP_009725295.1;Wu1-SARS2_orf1a;4200 | 0 | 0 |
| NC_045512.2;YP_009725295.1;Wu1-SARS2_orf1a;4201 | 0 | 0.025 |
| NC_045512.2;YP_009725295.1;Wu1-SARS2_orf1a;4202 | 0 | 0 |

| | | |
|---|---|---|
| NC_045512.2;YP_009725295.1;Wu1-SARS2_orf1a;4203 | 0.2 | 0.15 |
| NC_045512.2;YP_009725295.1;Wu1-SARS2_orf1a;4204 | 0 | 0 |
| NC_045512.2;YP_009725295.1;Wu1-SARS2_orf1a;4205 | 0.05 | 0.15 |
| NC_045512.2;YP_009725295.1;Wu1-SARS2_orf1a;4206 | 0.05 | 0.075 |
| NC_045512.2;YP_009725295.1;Wu1-SARS2_orf1a;4207 | 0.05 | 0.05 |
| NC_045512.2;YP_009725295.1;Wu1-SARS2_orf1a;4208 | 0.05 | 0.025 |
| NC_045512.2;YP_009725295.1;Wu1-SARS2_orf1a;4209 | 0 | 0 |
| NC_045512.2;YP_009725295.1;Wu1-SARS2_orf1a;4210 | 0 | 0.025 |
| NC_045512.2;YP_009725295.1;Wu1-SARS2_orf1a;4211 | 0 | 0 |
| NC_045512.2;YP_009725295.1;Wu1-SARS2_orf1a;4212 | 0 | 0 |
| NC_045512.2;YP_009725295.1;Wu1-SARS2_orf1a;4213 | 0 | 0 |
| NC_045512.2;YP_009725295.1;Wu1-SARS2_orf1a;4214 | 0 | 0 |
| NC_045512.2;YP_009725295.1;Wu1-SARS2_orf1a;4215 | 0.05 | 0 |
| NC_045512.2;YP_009725295.1;Wu1-SARS2_orf1a;4216 | 0 | 0 |
| NC_045512.2;YP_009725295.1;Wu1-SARS2_orf1a;4217 | 0 | 0 |
| NC_045512.2;YP_009725295.1;Wu1-SARS2_orf1a;4218 | 0 | 0 |
| NC_045512.2;YP_009725295.1;Wu1-SARS2_orf1a;4219 | 0 | 0 |
| NC_045512.2;YP_009725295.1;Wu1-SARS2_orf1a;4220 | 0 | 0 |
| NC_045512.2;YP_009725295.1;Wu1-SARS2_orf1a;4221 | 0 | 0 |
| NC_045512.2;YP_009725295.1;Wu1-SARS2_orf1a;4222 | 0 | 0 |
| NC_045512.2;YP_009725295.1;Wu1-SARS2_orf1a;4223 | 0 | 0 |
| NC_045512.2;YP_009725295.1;Wu1-SARS2_orf1a;4224 | 0 | 0 |
| NC_045512.2;YP_009725295.1;Wu1-SARS2_orf1a;4225 | 0 | 0 |
| NC_045512.2;YP_009725295.1;Wu1-SARS2_orf1a;4226 | 0 | 0 |
| NC_045512.2;YP_009725295.1;Wu1-SARS2_orf1a;4227 | 0 | 0 |
| NC_045512.2;YP_009725295.1;Wu1-SARS2_orf1a;4228 | 0 | 0 |
| NC_045512.2;YP_009725295.1;Wu1-SARS2_orf1a;4229 | 0.05 | 0.025 |
| NC_045512.2;YP_009725295.1;Wu1-SARS2_orf1a;4230 | 0.1 | 0.075 |
| NC_045512.2;YP_009725295.1;Wu1-SARS2_orf1a;4231 | 0.15 | 0.1 |
| NC_045512.2;YP_009725295.1;Wu1-SARS2_orf1a;4232 | 0.25 | 0.1 |
| NC_045512.2;YP_009725295.1;Wu1-SARS2_orf1a;4233 | 0.1 | 0.1 |
| NC_045512.2;YP_009725295.1;Wu1-SARS2_orf1a;4234 | 0.05 | 0.05 |
| NC_045512.2;YP_009725295.1;Wu1-SARS2_orf1a;4235 | 0.05 | 0.025 |
| NC_045512.2;YP_009725295.1;Wu1-SARS2_orf1a;4236 | 0 | 0 |
| NC_045512.2;YP_009725295.1;Wu1-SARS2_orf1a;4237 | 0 | 0 |
| NC_045512.2;YP_009725295.1;Wu1-SARS2_orf1a;4238 | 0 | 0 |
| NC_045512.2;YP_009725295.1;Wu1-SARS2_orf1a;4239 | 0 | 0 |
| NC_045512.2;YP_009725295.1;Wu1-SARS2_orf1a;4240 | 0 | 0 |
| NC_045512.2;YP_009725295.1;Wu1-SARS2_orf1a;4241 | 0 | 0 |
| NC_045512.2;YP_009725295.1;Wu1-SARS2_orf1a;4242 | 0 | 0 |
| NC_045512.2;YP_009725295.1;Wu1-SARS2_orf1a;4243 | 0 | 0 |
| NC_045512.2;YP_009725295.1;Wu1-SARS2_orf1a;4244 | 0 | 0.025 |
| NC_045512.2;YP_009725295.1;Wu1-SARS2_orf1a;4245 | 0 | 0 |

| | | |
|---|---|---|
| NC_045512.2;YP_009725295.1;Wu1-SARS2_orf1a;4246 | 0 | 0.025 |
| NC_045512.2;YP_009725295.1;Wu1-SARS2_orf1a;4247 | 0.05 | 0.025 |
| NC_045512.2;YP_009725295.1;Wu1-SARS2_orf1a;4248 | 0 | 0.025 |
| NC_045512.2;YP_009725295.1;Wu1-SARS2_orf1a;4249 | 0 | 0 |
| NC_045512.2;YP_009725295.1;Wu1-SARS2_orf1a;4250 | 0 | 0 |
| NC_045512.2;YP_009725295.1;Wu1-SARS2_orf1a;4251 | 0 | 0.025 |
| NC_045512.2;YP_009725295.1;Wu1-SARS2_orf1a;4252 | 0 | 0.025 |
| NC_045512.2;YP_009725295.1;Wu1-SARS2_orf1a;4253 | 0 | 0 |
| NC_045512.2;YP_009725295.1;Wu1-SARS2_orf1a;4254 | 0 | 0 |
| NC_045512.2;YP_009725295.1;Wu1-SARS2_orf1a;4255 | 0 | 0 |
| NC_045512.2;YP_009725295.1;Wu1-SARS2_orf1a;4256 | 0 | 0 |
| NC_045512.2;YP_009725295.1;Wu1-SARS2_orf1a;4257 | 0 | 0 |
| NC_045512.2;YP_009725295.1;Wu1-SARS2_orf1a;4258 | 0 | 0 |
| NC_045512.2;YP_009725295.1;Wu1-SARS2_orf1a;4259 | 0 | 0 |
| NC_045512.2;YP_009725295.1;Wu1-SARS2_orf1a;4260 | 0 | 0 |
| NC_045512.2;YP_009725295.1;Wu1-SARS2_orf1a;4261 | 0 | 0 |
| NC_045512.2;YP_009725295.1;Wu1-SARS2_orf1a;4262 | 0 | 0 |
| NC_045512.2;YP_009725295.1;Wu1-SARS2_orf1a;4263 | 0 | 0 |
| NC_045512.2;YP_009725295.1;Wu1-SARS2_orf1a;4264 | 0 | 0 |
| NC_045512.2;YP_009725295.1;Wu1-SARS2_orf1a;4265 | 0 | 0 |
| NC_045512.2;YP_009725295.1;Wu1-SARS2_orf1a;4266 | 0 | 0 |
| NC_045512.2;YP_009725295.1;Wu1-SARS2_orf1a;4267 | 0 | 0 |
| NC_045512.2;YP_009725295.1;Wu1-SARS2_orf1a;4268 | 0.05 | 0 |
| NC_045512.2;YP_009725295.1;Wu1-SARS2_orf1a;4269 | 0 | 0 |
| NC_045512.2;YP_009725295.1;Wu1-SARS2_orf1a;4270 | 0 | 0 |
| NC_045512.2;YP_009725295.1;Wu1-SARS2_orf1a;4271 | 0.05 | 0 |
| NC_045512.2;YP_009725295.1;Wu1-SARS2_orf1a;4272 | 0.05 | 0 |
| NC_045512.2;YP_009725295.1;Wu1-SARS2_orf1a;4273 | 0.2 | 0.05 |
| NC_045512.2;YP_009725295.1;Wu1-SARS2_orf1a;4274 | 0.15 | 0.025 |
| NC_045512.2;YP_009725295.1;Wu1-SARS2_orf1a;4275 | 0.1 | 0.025 |
| NC_045512.2;YP_009725295.1;Wu1-SARS2_orf1a;4276 | 0 | 0.025 |
| NC_045512.2;YP_009725295.1;Wu1-SARS2_orf1a;4277 | 0 | 0.05 |
| NC_045512.2;YP_009725295.1;Wu1-SARS2_orf1a;4278 | 0 | 0.05 |
| NC_045512.2;YP_009725295.1;Wu1-SARS2_orf1a;4279 | 0.15 | 0.05 |
| NC_045512.2;YP_009725295.1;Wu1-SARS2_orf1a;4280 | 0.05 | 0.05 |
| NC_045512.2;YP_009725295.1;Wu1-SARS2_orf1a;4281 | 0 | 0 |
| NC_045512.2;YP_009725295.1;Wu1-SARS2_orf1a;4282 | 0 | 0 |
| NC_045512.2;YP_009725295.1;Wu1-SARS2_orf1a;4283 | 0 | 0 |
| NC_045512.2;YP_009725295.1;Wu1-SARS2_orf1a;4284 | 0 | 0 |
| NC_045512.2;YP_009725295.1;Wu1-SARS2_orf1a;4285 | 0 | 0 |
| NC_045512.2;YP_009725295.1;Wu1-SARS2_orf1a;4286 | 0 | 0 |
| NC_045512.2;YP_009725295.1;Wu1-SARS2_orf1a;4287 | 0 | 0 |
| NC_045512.2;YP_009725295.1;Wu1-SARS2_orf1a;4288 | 0 | 0 |

| | | |
|---|---|---|
| NC_045512.2;YP_009725295.1;Wu1-SARS2_orf1a;4289 | 0 | 0.025 |
| NC_045512.2;YP_009725295.1;Wu1-SARS2_orf1a;4290 | 0 | 0 |
| NC_045512.2;YP_009725295.1;Wu1-SARS2_orf1a;4291 | 0 | 0.025 |
| NC_045512.2;YP_009725295.1;Wu1-SARS2_orf1a;4292 | 0 | 0.025 |
| NC_045512.2;YP_009725295.1;Wu1-SARS2_orf1a;4293 | 0 | 0 |
| NC_045512.2;YP_009725295.1;Wu1-SARS2_orf1a;4294 | 0 | 0 |
| NC_045512.2;YP_009725295.1;Wu1-SARS2_orf1a;4295 | 0 | 0 |
| NC_045512.2;YP_009725295.1;Wu1-SARS2_orf1a;4296 | 0 | 0 |
| NC_045512.2;YP_009725295.1;Wu1-SARS2_orf1a;4297 | 0 | 0 |
| NC_045512.2;YP_009725295.1;Wu1-SARS2_orf1a;4298 | 0 | 0 |
| NC_045512.2;YP_009725295.1;Wu1-SARS2_orf1a;4299 | 0 | 0 |
| NC_045512.2;YP_009725295.1;Wu1-SARS2_orf1a;4300 | 0 | 0 |
| NC_045512.2;YP_009725295.1;Wu1-SARS2_orf1a;4301 | 0 | 0 |
| NC_045512.2;YP_009725295.1;Wu1-SARS2_orf1a;4302 | 0 | 0.025 |
| NC_045512.2;YP_009725295.1;Wu1-SARS2_orf1a;4303 | 0 | 0.025 |
| NC_045512.2;YP_009725295.1;Wu1-SARS2_orf1a;4304 | 0 | 0.025 |
| NC_045512.2;YP_009725295.1;Wu1-SARS2_orf1a;4305 | 0 | 0.025 |
| NC_045512.2;YP_009725295.1;Wu1-SARS2_orf1a;4306 | 0 | 0.025 |
| NC_045512.2;YP_009725295.1;Wu1-SARS2_orf1a;4307 | 0 | 0.025 |
| NC_045512.2;YP_009725295.1;Wu1-SARS2_orf1a;4308 | 0.05 | 0.025 |
| NC_045512.2;YP_009725295.1;Wu1-SARS2_orf1a;4309 | 0 | 0.025 |
| NC_045512.2;YP_009725295.1;Wu1-SARS2_orf1a;4310 | 0.05 | 0.025 |
| NC_045512.2;YP_009725295.1;Wu1-SARS2_orf1a;4311 | 0 | 0 |
| NC_045512.2;YP_009725295.1;Wu1-SARS2_orf1a;4312 | 0 | 0 |
| NC_045512.2;YP_009725295.1;Wu1-SARS2_orf1a;4313 | 0 | 0 |
| NC_045512.2;YP_009725295.1;Wu1-SARS2_orf1a;4314 | 0 | 0 |
| NC_045512.2;YP_009725295.1;Wu1-SARS2_orf1a;4315 | 0 | 0 |
| NC_045512.2;YP_009725295.1;Wu1-SARS2_orf1a;4316 | 0 | 0 |
| NC_045512.2;YP_009725295.1;Wu1-SARS2_orf1a;4317 | 0 | 0 |
| NC_045512.2;YP_009725295.1;Wu1-SARS2_orf1a;4318 | 0 | 0 |
| NC_045512.2;YP_009725295.1;Wu1-SARS2_orf1a;4319 | 0 | 0 |
| NC_045512.2;YP_009725295.1;Wu1-SARS2_orf1a;4320 | 0 | 0 |
| NC_045512.2;YP_009725295.1;Wu1-SARS2_orf1a;4321 | 0 | 0 |
| NC_045512.2;YP_009725295.1;Wu1-SARS2_orf1a;4322 | 0 | 0 |
| NC_045512.2;YP_009725295.1;Wu1-SARS2_orf1a;4323 | 0 | 0 |
| NC_045512.2;YP_009725295.1;Wu1-SARS2_orf1a;4324 | 0 | 0 |
| NC_045512.2;YP_009725295.1;Wu1-SARS2_orf1a;4325 | 0 | 0 |
| NC_045512.2;YP_009725295.1;Wu1-SARS2_orf1a;4326 | 0 | 0 |
| NC_045512.2;YP_009725295.1;Wu1-SARS2_orf1a;4327 | 0 | 0 |
| NC_045512.2;YP_009725295.1;Wu1-SARS2_orf1a;4328 | 0 | 0 |
| NC_045512.2;YP_009725295.1;Wu1-SARS2_orf1a;4329 | 0 | 0.025 |
| NC_045512.2;YP_009725295.1;Wu1-SARS2_orf1a;4330 | 0 | 0.025 |
| NC_045512.2;YP_009725295.1;Wu1-SARS2_orf1a;4331 | 0 | 0.025 |

| | | |
|---|---|---|
| NC_045512.2;YP_009725295.1;Wu1-SARS2_orf1a;4332 | 0 | 0.025 |
| NC_045512.2;YP_009725295.1;Wu1-SARS2_orf1a;4333 | 0 | 0.025 |
| NC_045512.2;YP_009725295.1;Wu1-SARS2_orf1a;4334 | 0 | 0.025 |
| NC_045512.2;YP_009725295.1;Wu1-SARS2_orf1a;4335 | 0 | 0 |
| NC_045512.2;YP_009725295.1;Wu1-SARS2_orf1a;4336 | 0 | 0 |
| NC_045512.2;YP_009725295.1;Wu1-SARS2_orf1a;4337 | 0 | 0 |
| NC_045512.2;YP_009725295.1;Wu1-SARS2_orf1a;4338 | 0 | 0 |
| NC_045512.2;YP_009725295.1;Wu1-SARS2_orf1a;4339 | 0 | 0 |
| NC_045512.2;YP_009725295.1;Wu1-SARS2_orf1a;4340 | 0 | 0.05 |
| NC_045512.2;YP_009725295.1;Wu1-SARS2_orf1a;4341 | 0 | 0.05 |
| NC_045512.2;YP_009725295.1;Wu1-SARS2_orf1a;4342 | 0 | 0 |
| NC_045512.2;YP_009725295.1;Wu1-SARS2_orf1a;4343 | 0 | 0 |
| NC_045512.2;YP_009725295.1;Wu1-SARS2_orf1a;4344 | 0.05 | 0 |
| NC_045512.2;YP_009725295.1;Wu1-SARS2_orf1a;4345 | 0.05 | 0.025 |
| NC_045512.2;YP_009725295.1;Wu1-SARS2_orf1a;4346 | 0.05 | 0.05 |
| NC_045512.2;YP_009725295.1;Wu1-SARS2_orf1a;4347 | 0.05 | 0 |
| NC_045512.2;YP_009725295.1;Wu1-SARS2_orf1a;4348 | 0.05 | 0 |
| NC_045512.2;YP_009725295.1;Wu1-SARS2_orf1a;4349 | 0 | 0 |
| NC_045512.2;YP_009725295.1;Wu1-SARS2_orf1a;4350 | 0 | 0.025 |
| NC_045512.2;YP_009725295.1;Wu1-SARS2_orf1a;4351 | 0 | 0.025 |
| NC_045512.2;YP_009725295.1;Wu1-SARS2_orf1a;4352 | 0 | 0.025 |
| NC_045512.2;YP_009725295.1;Wu1-SARS2_orf1a;4353 | 0.05 | 0.05 |
| NC_045512.2;YP_009725295.1;Wu1-SARS2_orf1a;4354 | 0 | 0.025 |
| NC_045512.2;YP_009725295.1;Wu1-SARS2_orf1a;4355 | 0 | 0.025 |
| NC_045512.2;YP_009725295.1;Wu1-SARS2_orf1a;4356 | 0 | 0.025 |
| NC_045512.2;YP_009725295.1;Wu1-SARS2_orf1a;4357 | 0 | 0.025 |
| NC_045512.2;YP_009725295.1;Wu1-SARS2_orf1a;4358 | 0 | 0 |
| NC_045512.2;YP_009725295.1;Wu1-SARS2_orf1a;4359 | 0 | 0 |
| NC_045512.2;YP_009725295.1;Wu1-SARS2_orf1a;4360 | 0 | 0 |
| NC_045512.2;YP_009725295.1;Wu1-SARS2_orf1a;4361 | 0 | 0 |
| NC_045512.2;YP_009725295.1;Wu1-SARS2_orf1a;4362 | 0 | 0 |
| NC_045512.2;YP_009725295.1;Wu1-SARS2_orf1a;4363 | 0 | 0 |
| NC_045512.2;YP_009725295.1;Wu1-SARS2_orf1a;4364 | 0 | 0 |
| NC_045512.2;YP_009725295.1;Wu1-SARS2_orf1a;4365 | 0 | 0 |
| NC_045512.2;YP_009725295.1;Wu1-SARS2_orf1a;4366 | 0 | 0 |
| NC_045512.2;YP_009725295.1;Wu1-SARS2_orf1a;4367 | 0 | 0 |
| NC_045512.2;YP_009725295.1;Wu1-SARS2_orf1a;4368 | 0 | 0 |
| NC_045512.2;YP_009725295.1;Wu1-SARS2_orf1a;4369 | 0 | 0 |
| NC_045512.2;YP_009725295.1;Wu1-SARS2_orf1a;4370 | 0 | 0 |
| NC_045512.2;YP_009725295.1;Wu1-SARS2_orf1a;4371 | 0 | 0 |
| NC_045512.2;YP_009725295.1;Wu1-SARS2_orf1a;4372 | 0 | 0 |
| NC_045512.2;YP_009725295.1;Wu1-SARS2_orf1a;4373 | 0 | 0 |
| NC_045512.2;YP_009725295.1;Wu1-SARS2_orf1a;4374 | 0 | 0 |

| | | |
|---|---|---|
| NC_045512.2;YP_009725295.1;Wu1-SARS2_orf1a;4375 | 0 | 0.075 |
| NC_045512.2;YP_009725295.1;Wu1-SARS2_orf1a;4376 | 0 | 0 |
| NC_045512.2;YP_009725295.1;Wu1-SARS2_orf1a;4377 | 0 | 0.025 |
| NC_045512.2;YP_009725295.1;Wu1-SARS2_orf1a;4378 | 0 | 0 |
| NC_045512.2;YP_009725295.1;Wu1-SARS2_orf1a;4379 | 0 | 0 |
| NC_045512.2;YP_009725295.1;Wu1-SARS2_orf1a;4380 | 0 | 0 |
| NC_045512.2;YP_009725295.1;Wu1-SARS2_orf1a;4381 | 0 | 0 |
| NC_045512.2;YP_009725295.1;Wu1-SARS2_orf1a;4382 | 0 | 0 |
| NC_045512.2;YP_009725295.1;Wu1-SARS2_orf1a;4383 | 0 | 0 |
| NC_045512.2;YP_009725295.1;Wu1-SARS2_orf1a;4384 | 0 | 0 |
| NC_045512.2;YP_009725295.1;Wu1-SARS2_orf1a;4385 | 0 | 0 |
| NC_045512.2;YP_009725295.1;Wu1-SARS2_orf1a;4386 | 0.05 | 0 |
| NC_045512.2;YP_009725295.1;Wu1-SARS2_orf1a;4387 | 0.1 | 0 |
| NC_045512.2;YP_009725295.1;Wu1-SARS2_orf1a;4388 | 0.1 | 0 |
| NC_045512.2;YP_009725295.1;Wu1-SARS2_orf1a;4389 | 0 | 0 |
| NC_045512.2;YP_009725295.1;Wu1-SARS2_orf1a;4390 | 0 | 0 |
| NC_045512.2;YP_009725295.1;Wu1-SARS2_orf1a;4391 | 0 | 0 |
| NC_045512.2;YP_009725295.1;Wu1-SARS2_orf1a;4392 | 0 | 0 |
| NC_045512.2;YP_009725295.1;Wu1-SARS2_orf1a;4393 | 0 | 0 |
| NC_045512.2;YP_009725295.1;Wu1-SARS2_orf1a;4394 | 0 | 0 |
| NC_045512.2;YP_009725318.1;Wu1-SARS2_orf7b;1 | 0 | 0 |
| NC_045512.2;YP_009725318.1;Wu1-SARS2_orf7b;2 | 0 | 0 |
| NC_045512.2;YP_009725318.1;Wu1-SARS2_orf7b;3 | 0 | 0 |
| NC_045512.2;YP_009725318.1;Wu1-SARS2_orf7b;4 | 0 | 0 |
| NC_045512.2;YP_009725318.1;Wu1-SARS2_orf7b;5 | 0 | 0 |
| NC_045512.2;YP_009725318.1;Wu1-SARS2_orf7b;6 | 0 | 0 |
| NC_045512.2;YP_009725318.1;Wu1-SARS2_orf7b;7 | 0 | 0 |
| NC_045512.2;YP_009725318.1;Wu1-SARS2_orf7b;8 | 0 | 0 |
| NC_045512.2;YP_009725318.1;Wu1-SARS2_orf7b;9 | 0 | 0 |
| NC_045512.2;YP_009725318.1;Wu1-SARS2_orf7b;10 | 0 | 0 |
| NC_045512.2;YP_009725318.1;Wu1-SARS2_orf7b;11 | 0 | 0 |
| NC_045512.2;YP_009725318.1;Wu1-SARS2_orf7b;12 | 0 | 0 |
| NC_045512.2;YP_009725318.1;Wu1-SARS2_orf7b;13 | 0 | 0 |
| NC_045512.2;YP_009725318.1;Wu1-SARS2_orf7b;14 | 0 | 0 |
| NC_045512.2;YP_009725318.1;Wu1-SARS2_orf7b;15 | 0 | 0 |
| NC_045512.2;YP_009725318.1;Wu1-SARS2_orf7b;16 | 0 | 0 |
| NC_045512.2;YP_009725318.1;Wu1-SARS2_orf7b;17 | 0 | 0 |
| NC_045512.2;YP_009725318.1;Wu1-SARS2_orf7b;18 | 0 | 0 |
| NC_045512.2;YP_009725318.1;Wu1-SARS2_orf7b;19 | 0 | 0 |
| NC_045512.2;YP_009725318.1;Wu1-SARS2_orf7b;20 | 0 | 0 |
| NC_045512.2;YP_009725318.1;Wu1-SARS2_orf7b;21 | 0 | 0 |
| NC_045512.2;YP_009725318.1;Wu1-SARS2_orf7b;22 | 0 | 0 |
| NC_045512.2;YP_009725318.1;Wu1-SARS2_orf7b;23 | 0 | 0 |

| | | |
|---|---|---|
| NC_045512.2;YP_009725318.1;Wu1-SARS2_orf7b;24 | 0 | 0 |
| NC_045512.2;YP_009725318.1;Wu1-SARS2_orf7b;25 | 0 | 0 |
| NC_045512.2;YP_009725318.1;Wu1-SARS2_orf7b;26 | 0 | 0 |
| NC_045512.2;YP_009725318.1;Wu1-SARS2_orf7b;27 | 0 | 0 |
| NC_045512.2;YP_009725318.1;Wu1-SARS2_orf7b;28 | 0 | 0 |
| NC_045512.2;YP_009725318.1;Wu1-SARS2_orf7b;29 | 0 | 0 |
| NC_045512.2;YP_009725318.1;Wu1-SARS2_orf7b;30 | 0 | 0.025 |
| NC_045512.2;YP_009725318.1;Wu1-SARS2_orf7b;31 | 0 | 0 |
| NC_045512.2;YP_009725318.1;Wu1-SARS2_orf7b;32 | 0 | 0 |
| MN996532.1;QHR63299.1;BatCoV_RaTG13_orf1ab;1 | 0 | 0 |
| MN996532.1;QHR63299.1;BatCoV_RaTG13_orf1ab;2 | 0 | 0.025 |
| MN996532.1;QHR63299.1;BatCoV_RaTG13_orf1ab;3 | 0 | 0 |
| MN996532.1;QHR63299.1;BatCoV_RaTG13_orf1ab;4 | 0 | 0.025 |
| MN996532.1;QHR63299.1;BatCoV_RaTG13_orf1ab;5 | 0 | 0 |
| MN996532.1;QHR63299.1;BatCoV_RaTG13_orf1ab;6 | 0 | 0 |
| MN996532.1;QHR63299.1;BatCoV_RaTG13_orf1ab;7 | 0 | 0 |
| MN996532.1;QHR63299.1;BatCoV_RaTG13_orf1ab;8 | 0 | 0 |
| MN996532.1;QHR63299.1;BatCoV_RaTG13_orf1ab;9 | 0 | 0 |
| MN996532.1;QHR63299.1;BatCoV_RaTG13_orf1ab;10 | 0 | 0 |
| MN996532.1;QHR63299.1;BatCoV_RaTG13_orf1ab;11 | 0 | 0 |
| MN996532.1;QHR63299.1;BatCoV_RaTG13_orf1ab;12 | 0 | 0 |
| MN996532.1;QHR63299.1;BatCoV_RaTG13_orf1ab;13 | 0 | 0 |
| MN996532.1;QHR63299.1;BatCoV_RaTG13_orf1ab;14 | 0 | 0 |
| MN996532.1;QHR63299.1;BatCoV_RaTG13_orf1ab;15 | 0 | 0 |
| MN996532.1;QHR63299.1;BatCoV_RaTG13_orf1ab;16 | 0 | 0 |
| MN996532.1;QHR63299.1;BatCoV_RaTG13_orf1ab;17 | 0 | 0 |
| MN996532.1;QHR63299.1;BatCoV_RaTG13_orf1ab;18 | 0 | 0 |
| MN996532.1;QHR63299.1;BatCoV_RaTG13_orf1ab;19 | 0 | 0 |
| MN996532.1;QHR63299.1;BatCoV_RaTG13_orf1ab;20 | 0 | 0 |
| MN996532.1;QHR63299.1;BatCoV_RaTG13_orf1ab;21 | 0 | 0 |
| MN996532.1;QHR63299.1;BatCoV_RaTG13_orf1ab;22 | 0 | 0 |
| MN996532.1;QHR63299.1;BatCoV_RaTG13_orf1ab;23 | 0 | 0 |
| MN996532.1;QHR63299.1;BatCoV_RaTG13_orf1ab;24 | 0 | 0 |
| MN996532.1;QHR63299.1;BatCoV_RaTG13_orf1ab;25 | 0 | 0 |
| MN996532.1;QHR63299.1;BatCoV_RaTG13_orf1ab;26 | 0 | 0 |
| MN996532.1;QHR63299.1;BatCoV_RaTG13_orf1ab;27 | 0 | 0 |
| MN996532.1;QHR63299.1;BatCoV_RaTG13_orf1ab;28 | 0 | 0 |
| MN996532.1;QHR63299.1;BatCoV_RaTG13_orf1ab;29 | 0.1 | 0 |
| MN996532.1;QHR63299.1;BatCoV_RaTG13_orf1ab;30 | 0.3 | 0.125 |
| MN996532.1;QHR63299.1;BatCoV_RaTG13_orf1ab;31 | 0 | 0 |
| MN996532.1;QHR63299.1;BatCoV_RaTG13_orf1ab;32 | 0.05 | 0.05 |
| MN996532.1;QHR63299.1;BatCoV_RaTG13_orf1ab;33 | 0.05 | 0 |
| MN996532.1;QHR63299.1;BatCoV_RaTG13_orf1ab;34 | 0.1 | 0 |

| | | |
|---|---|---|
| MN996532.1;QHR63299.1;BatCoV_RaTG13_orf1ab;35 | 0 | 0.025 |
| MN996532.1;QHR63299.1;BatCoV_RaTG13_orf1ab;36 | 0 | 0.025 |
| MN996532.1;QHR63299.1;BatCoV_RaTG13_orf1ab;37 | 0 | 0 |
| MN996532.1;QHR63299.1;BatCoV_RaTG13_orf1ab;38 | 0 | 0 |
| MN996532.1;QHR63299.1;BatCoV_RaTG13_orf1ab;39 | 0 | 0 |
| MN996532.1;QHR63299.1;BatCoV_RaTG13_orf1ab;40 | 0 | 0 |
| MN996532.1;QHR63299.1;BatCoV_RaTG13_orf1ab;41 | 0 | 0 |
| MN996532.1;QHR63299.1;BatCoV_RaTG13_orf1ab;42 | 0 | 0 |
| MN996532.1;QHR63299.1;BatCoV_RaTG13_orf1ab;43 | 0 | 0 |
| MN996532.1;QHR63299.1;BatCoV_RaTG13_orf1ab;44 | 0.05 | 0 |
| MN996532.1;QHR63299.1;BatCoV_RaTG13_orf1ab;45 | 0 | 0 |
| MN996532.1;QHR63299.1;BatCoV_RaTG13_orf1ab;46 | 0 | 0 |
| MN996532.1;QHR63299.1;BatCoV_RaTG13_orf1ab;47 | 0 | 0 |
| MN996532.1;QHR63299.1;BatCoV_RaTG13_orf1ab;48 | 0 | 0 |
| MN996532.1;QHR63299.1;BatCoV_RaTG13_orf1ab;49 | 0 | 0 |
| MN996532.1;QHR63299.1;BatCoV_RaTG13_orf1ab;50 | 0 | 0 |
| MN996532.1;QHR63299.1;BatCoV_RaTG13_orf1ab;51 | 0 | 0 |
| MN996532.1;QHR63299.1;BatCoV_RaTG13_orf1ab;52 | 0.05 | 0.025 |
| MN996532.1;QHR63299.1;BatCoV_RaTG13_orf1ab;53 | 0 | 0.025 |
| MN996532.1;QHR63299.1;BatCoV_RaTG13_orf1ab;54 | 0.05 | 0.025 |
| MN996532.1;QHR63299.1;BatCoV_RaTG13_orf1ab;55 | 0 | 0.025 |
| MN996532.1;QHR63299.1;BatCoV_RaTG13_orf1ab;56 | 0 | 0.025 |
| MN996532.1;QHR63299.1;BatCoV_RaTG13_orf1ab;57 | 0 | 0.025 |
| MN996532.1;QHR63299.1;BatCoV_RaTG13_orf1ab;58 | 0 | 0.025 |
| MN996532.1;QHR63299.1;BatCoV_RaTG13_orf1ab;59 | 0 | 0 |
| MN996532.1;QHR63299.1;BatCoV_RaTG13_orf1ab;60 | 0 | 0 |
| MN996532.1;QHR63299.1;BatCoV_RaTG13_orf1ab;61 | 0 | 0 |
| MN996532.1;QHR63299.1;BatCoV_RaTG13_orf1ab;62 | 0 | 0 |
| MN996532.1;QHR63299.1;BatCoV_RaTG13_orf1ab;63 | 0 | 0 |
| MN996532.1;QHR63299.1;BatCoV_RaTG13_orf1ab;64 | 0 | 0 |
| MN996532.1;QHR63299.1;BatCoV_RaTG13_orf1ab;65 | 0 | 0 |
| MN996532.1;QHR63299.1;BatCoV_RaTG13_orf1ab;66 | 0 | 0 |
| MN996532.1;QHR63299.1;BatCoV_RaTG13_orf1ab;67 | 0 | 0 |
| MN996532.1;QHR63299.1;BatCoV_RaTG13_orf1ab;68 | 0 | 0 |
| MN996532.1;QHR63299.1;BatCoV_RaTG13_orf1ab;69 | 0 | 0 |
| MN996532.1;QHR63299.1;BatCoV_RaTG13_orf1ab;70 | 0 | 0 |
| MN996532.1;QHR63299.1;BatCoV_RaTG13_orf1ab;71 | 0.05 | 0 |
| MN996532.1;QHR63299.1;BatCoV_RaTG13_orf1ab;72 | 0 | 0 |
| MN996532.1;QHR63299.1;BatCoV_RaTG13_orf1ab;73 | 0 | 0 |
| MN996532.1;QHR63299.1;BatCoV_RaTG13_orf1ab;74 | 0 | 0 |
| MN996532.1;QHR63299.1;BatCoV_RaTG13_orf1ab;75 | 0 | 0.025 |
| MN996532.1;QHR63299.1;BatCoV_RaTG13_orf1ab;76 | 0 | 0 |
| MN996532.1;QHR63299.1;BatCoV_RaTG13_orf1ab;77 | 0 | 0 |

| | | |
|---|---|---|
| MN996532.1;QHR63299.1;BatCoV_RaTG13_orf1ab;78 | 0 | 0 |
| MN996532.1;QHR63299.1;BatCoV_RaTG13_orf1ab;79 | 0 | 0 |
| MN996532.1;QHR63299.1;BatCoV_RaTG13_orf1ab;80 | 0 | 0 |
| MN996532.1;QHR63299.1;BatCoV_RaTG13_orf1ab;81 | 0 | 0 |
| MN996532.1;QHR63299.1;BatCoV_RaTG13_orf1ab;82 | 0 | 0 |
| MN996532.1;QHR63299.1;BatCoV_RaTG13_orf1ab;83 | 0 | 0 |
| MN996532.1;QHR63299.1;BatCoV_RaTG13_orf1ab;84 | 0 | 0 |
| MN996532.1;QHR63299.1;BatCoV_RaTG13_orf1ab;85 | 0 | 0 |
| MN996532.1;QHR63299.1;BatCoV_RaTG13_orf1ab;86 | 0 | 0.05 |
| MN996532.1;QHR63299.1;BatCoV_RaTG13_orf1ab;87 | 0 | 0.025 |
| MN996532.1;QHR63299.1;BatCoV_RaTG13_orf1ab;88 | 0 | 0 |
| MN996532.1;QHR63299.1;BatCoV_RaTG13_orf1ab;89 | 0.05 | 0 |
| MN996532.1;QHR63299.1;BatCoV_RaTG13_orf1ab;90 | 0 | 0 |
| MN996532.1;QHR63299.1;BatCoV_RaTG13_orf1ab;91 | 0 | 0 |
| MN996532.1;QHR63299.1;BatCoV_RaTG13_orf1ab;92 | 0 | 0 |
| MN996532.1;QHR63299.1;BatCoV_RaTG13_orf1ab;93 | 0 | 0 |
| MN996532.1;QHR63299.1;BatCoV_RaTG13_orf1ab;94 | 0 | 0.025 |
| MN996532.1;QHR63299.1;BatCoV_RaTG13_orf1ab;95 | 0 | 0 |
| MN996532.1;QHR63299.1;BatCoV_RaTG13_orf1ab;96 | 0 | 0 |
| MN996532.1;QHR63299.1;BatCoV_RaTG13_orf1ab;97 | 0 | 0 |
| MN996532.1;QHR63299.1;BatCoV_RaTG13_orf1ab;98 | 0 | 0 |
| MN996532.1;QHR63299.1;BatCoV_RaTG13_orf1ab;99 | 0 | 0 |
| MN996532.1;QHR63299.1;BatCoV_RaTG13_orf1ab;100 | 0 | 0 |
| MN996532.1;QHR63299.1;BatCoV_RaTG13_orf1ab;101 | 0 | 0 |
| MN996532.1;QHR63299.1;BatCoV_RaTG13_orf1ab;102 | 0 | 0 |
| MN996532.1;QHR63299.1;BatCoV_RaTG13_orf1ab;103 | 0 | 0 |
| MN996532.1;QHR63299.1;BatCoV_RaTG13_orf1ab;104 | 0 | 0 |
| MN996532.1;QHR63299.1;BatCoV_RaTG13_orf1ab;105 | 0.05 | 0 |
| MN996532.1;QHR63299.1;BatCoV_RaTG13_orf1ab;106 | 0 | 0 |
| MN996532.1;QHR63299.1;BatCoV_RaTG13_orf1ab;107 | 0 | 0 |
| MN996532.1;QHR63299.1;BatCoV_RaTG13_orf1ab;108 | 0 | 0 |
| MN996532.1;QHR63299.1;BatCoV_RaTG13_orf1ab;109 | 0 | 0 |
| MN996532.1;QHR63299.1;BatCoV_RaTG13_orf1ab;110 | 0 | 0 |
| MN996532.1;QHR63299.1;BatCoV_RaTG13_orf1ab;111 | 0 | 0 |
| MN996532.1;QHR63299.1;BatCoV_RaTG13_orf1ab;112 | 0 | 0 |
| MN996532.1;QHR63299.1;BatCoV_RaTG13_orf1ab;113 | 0 | 0 |
| MN996532.1;QHR63299.1;BatCoV_RaTG13_orf1ab;114 | 0 | 0.025 |
| MN996532.1;QHR63299.1;BatCoV_RaTG13_orf1ab;115 | 0 | 0.025 |
| MN996532.1;QHR63299.1;BatCoV_RaTG13_orf1ab;116 | 0 | 0.025 |
| MN996532.1;QHR63299.1;BatCoV_RaTG13_orf1ab;117 | 0 | 0.025 |
| MN996532.1;QHR63299.1;BatCoV_RaTG13_orf1ab;118 | 0.05 | 0.025 |
| MN996532.1;QHR63299.1;BatCoV_RaTG13_orf1ab;119 | 0 | 0.025 |
| MN996532.1;QHR63299.1;BatCoV_RaTG13_orf1ab;120 | 0 | 0.025 |

| | | |
|---|---|---|
| MN996532.1;QHR63299.1;BatCoV_RaTG13_orf1ab;121 | 0.05 | 0.025 |
| MN996532.1;QHR63299.1;BatCoV_RaTG13_orf1ab;122 | 0 | 0.025 |
| MN996532.1;QHR63299.1;BatCoV_RaTG13_orf1ab;123 | 0 | 0.025 |
| MN996532.1;QHR63299.1;BatCoV_RaTG13_orf1ab;124 | 0 | 0.05 |
| MN996532.1;QHR63299.1;BatCoV_RaTG13_orf1ab;125 | 0.1 | 0.025 |
| MN996532.1;QHR63299.1;BatCoV_RaTG13_orf1ab;126 | 0 | 0 |
| MN996532.1;QHR63299.1;BatCoV_RaTG13_orf1ab;127 | 0 | 0 |
| MN996532.1;QHR63299.1;BatCoV_RaTG13_orf1ab;128 | 0 | 0.025 |
| MN996532.1;QHR63299.1;BatCoV_RaTG13_orf1ab;129 | 0 | 0.05 |
| MN996532.1;QHR63299.1;BatCoV_RaTG13_orf1ab;130 | 0 | 0.025 |
| MN996532.1;QHR63299.1;BatCoV_RaTG13_orf1ab;131 | 0.1 | 0.025 |
| MN996532.1;QHR63299.1;BatCoV_RaTG13_orf1ab;132 | 0.05 | 0.025 |
| MN996532.1;QHR63299.1;BatCoV_RaTG13_orf1ab;133 | 0.05 | 0 |
| MN996532.1;QHR63299.1;BatCoV_RaTG13_orf1ab;134 | 0 | 0 |
| MN996532.1;QHR63299.1;BatCoV_RaTG13_orf1ab;135 | 0.05 | 0 |
| MN996532.1;QHR63299.1;BatCoV_RaTG13_orf1ab;136 | 0 | 0.025 |
| MN996532.1;QHR63299.1;BatCoV_RaTG13_orf1ab;137 | 0 | 0 |
| MN996532.1;QHR63299.1;BatCoV_RaTG13_orf1ab;138 | 0 | 0 |
| MN996532.1;QHR63299.1;BatCoV_RaTG13_orf1ab;139 | 0.05 | 0 |
| MN996532.1;QHR63299.1;BatCoV_RaTG13_orf1ab;140 | 0.05 | 0 |
| MN996532.1;QHR63299.1;BatCoV_RaTG13_orf1ab;141 | 0 | 0 |
| MN996532.1;QHR63299.1;BatCoV_RaTG13_orf1ab;142 | 0 | 0 |
| MN996532.1;QHR63299.1;BatCoV_RaTG13_orf1ab;143 | 0.05 | 0 |
| MN996532.1;QHR63299.1;BatCoV_RaTG13_orf1ab;144 | 0 | 0.025 |
| MN996532.1;QHR63299.1;BatCoV_RaTG13_orf1ab;145 | 0.2 | 0.1 |
| MN996532.1;QHR63299.1;BatCoV_RaTG13_orf1ab;146 | 0.05 | 0 |
| MN996532.1;QHR63299.1;BatCoV_RaTG13_orf1ab;147 | 0 | 0 |
| MN996532.1;QHR63299.1;BatCoV_RaTG13_orf1ab;148 | 0.05 | 0 |
| MN996532.1;QHR63299.1;BatCoV_RaTG13_orf1ab;149 | 0.3 | 0.2 |
| MN996532.1;QHR63299.1;BatCoV_RaTG13_orf1ab;150 | 0.25 | 0.075 |
| MN996532.1;QHR63299.1;BatCoV_RaTG13_orf1ab;151 | 0.3 | 0.15 |
| MN996532.1;QHR63299.1;BatCoV_RaTG13_orf1ab;152 | 0.45 | 0.225 |
| MN996532.1;QHR63299.1;BatCoV_RaTG13_orf1ab;153 | 0.35 | 0.2 |
| MN996532.1;QHR63299.1;BatCoV_RaTG13_orf1ab;154 | 0.45 | 0.35 |
| MN996532.1;QHR63299.1;BatCoV_RaTG13_orf1ab;155 | 0.5 | 0.325 |
| MN996532.1;QHR63299.1;BatCoV_RaTG13_orf1ab;156 | 0.3 | 0.1 |
| MN996532.1;QHR63299.1;BatCoV_RaTG13_orf1ab;157 | 0.3 | 0.05 |
| MN996532.1;QHR63299.1;BatCoV_RaTG13_orf1ab;158 | 0.05 | 0.025 |
| MN996532.1;QHR63299.1;BatCoV_RaTG13_orf1ab;159 | 0 | 0 |
| MN996532.1;QHR63299.1;BatCoV_RaTG13_orf1ab;160 | 0 | 0 |
| MN996532.1;QHR63299.1;BatCoV_RaTG13_orf1ab;161 | 0 | 0 |
| MN996532.1;QHR63299.1;BatCoV_RaTG13_orf1ab;162 | 0 | 0 |
| MN996532.1;QHR63299.1;BatCoV_RaTG13_orf1ab;163 | 0 | 0.075 |

| | | |
|---|---|---|
| MN996532.1;QHR63299.1;BatCoV_RaTG13_orf1ab;164 | 0 | 0 |
| MN996532.1;QHR63299.1;BatCoV_RaTG13_orf1ab;165 | 0.05 | 0.025 |
| MN996532.1;QHR63299.1;BatCoV_RaTG13_orf1ab;166 | 0.05 | 0.025 |
| MN996532.1;QHR63299.1;BatCoV_RaTG13_orf1ab;167 | 0 | 0.05 |
| MN996532.1;QHR63299.1;BatCoV_RaTG13_orf1ab;168 | 0.15 | 0.05 |
| MN996532.1;QHR63299.1;BatCoV_RaTG13_orf1ab;169 | 0 | 0 |
| MN996532.1;QHR63299.1;BatCoV_RaTG13_orf1ab;170 | 0 | 0.025 |
| MN996532.1;QHR63299.1;BatCoV_RaTG13_orf1ab;171 | 0 | 0 |
| MN996532.1;QHR63299.1;BatCoV_RaTG13_orf1ab;172 | 0 | 0 |
| MN996532.1;QHR63299.1;BatCoV_RaTG13_orf1ab;173 | 0 | 0 |
| MN996532.1;QHR63299.1;BatCoV_RaTG13_orf1ab;174 | 0.05 | 0.025 |
| MN996532.1;QHR63299.1;BatCoV_RaTG13_orf1ab;175 | 0 | 0 |
| MN996532.1;QHR63299.1;BatCoV_RaTG13_orf1ab;176 | 0 | 0 |
| MN996532.1;QHR63299.1;BatCoV_RaTG13_orf1ab;177 | 0 | 0 |
| MN996532.1;QHR63299.1;BatCoV_RaTG13_orf1ab;178 | 0 | 0 |
| MN996532.1;QHR63299.1;BatCoV_RaTG13_orf1ab;179 | 0 | 0 |
| MN996532.1;QHR63299.1;BatCoV_RaTG13_orf1ab;180 | 0 | 0 |
| MN996532.1;QHR63299.1;BatCoV_RaTG13_orf1ab;181 | 0 | 0.025 |
| MN996532.1;QHR63299.1;BatCoV_RaTG13_orf1ab;182 | 0 | 0 |
| MN996532.1;QHR63299.1;BatCoV_RaTG13_orf1ab;183 | 0 | 0 |
| MN996532.1;QHR63299.1;BatCoV_RaTG13_orf1ab;184 | 0 | 0.025 |
| MN996532.1;QHR63299.1;BatCoV_RaTG13_orf1ab;185 | 0 | 0.025 |
| MN996532.1;QHR63299.1;BatCoV_RaTG13_orf1ab;186 | 0 | 0.025 |
| MN996532.1;QHR63299.1;BatCoV_RaTG13_orf1ab;187 | 0 | 0 |
| MN996532.1;QHR63299.1;BatCoV_RaTG13_orf1ab;188 | 0 | 0 |
| MN996532.1;QHR63299.1;BatCoV_RaTG13_orf1ab;189 | 0 | 0 |
| MN996532.1;QHR63299.1;BatCoV_RaTG13_orf1ab;190 | 0 | 0 |
| MN996532.1;QHR63299.1;BatCoV_RaTG13_orf1ab;191 | 0 | 0 |
| MN996532.1;QHR63299.1;BatCoV_RaTG13_orf1ab;192 | 0 | 0 |
| MN996532.1;QHR63299.1;BatCoV_RaTG13_orf1ab;193 | 0 | 0 |
| MN996532.1;QHR63299.1;BatCoV_RaTG13_orf1ab;194 | 0 | 0 |
| MN996532.1;QHR63299.1;BatCoV_RaTG13_orf1ab;195 | 0 | 0 |
| MN996532.1;QHR63299.1;BatCoV_RaTG13_orf1ab;196 | 0 | 0 |
| MN996532.1;QHR63299.1;BatCoV_RaTG13_orf1ab;197 | 0.05 | 0 |
| MN996532.1;QHR63299.1;BatCoV_RaTG13_orf1ab;198 | 0 | 0 |
| MN996532.1;QHR63299.1;BatCoV_RaTG13_orf1ab;199 | 0 | 0 |
| MN996532.1;QHR63299.1;BatCoV_RaTG13_orf1ab;200 | 0 | 0 |
| MN996532.1;QHR63299.1;BatCoV_RaTG13_orf1ab;201 | 0 | 0 |
| MN996532.1;QHR63299.1;BatCoV_RaTG13_orf1ab;202 | 0 | 0 |
| MN996532.1;QHR63299.1;BatCoV_RaTG13_orf1ab;203 | 0 | 0 |
| MN996532.1;QHR63299.1;BatCoV_RaTG13_orf1ab;204 | 0.05 | 0 |
| MN996532.1;QHR63299.1;BatCoV_RaTG13_orf1ab;205 | 0 | 0 |
| MN996532.1;QHR63299.1;BatCoV_RaTG13_orf1ab;206 | 0 | 0 |

| | | |
|---|---|---|
| MN996532.1;QHR63299.1;BatCoV_RaTG13_orf1ab;207 | 0 | 0 |
| MN996532.1;QHR63299.1;BatCoV_RaTG13_orf1ab;208 | 0 | 0 |
| MN996532.1;QHR63299.1;BatCoV_RaTG13_orf1ab;209 | 0 | 0 |
| MN996532.1;QHR63299.1;BatCoV_RaTG13_orf1ab;210 | 0 | 0 |
| MN996532.1;QHR63299.1;BatCoV_RaTG13_orf1ab;211 | 0.05 | 0.075 |
| MN996532.1;QHR63299.1;BatCoV_RaTG13_orf1ab;212 | 0 | 0.1 |
| MN996532.1;QHR63299.1;BatCoV_RaTG13_orf1ab;213 | 0 | 0.1 |
| MN996532.1;QHR63299.1;BatCoV_RaTG13_orf1ab;214 | 0.1 | 0.075 |
| MN996532.1;QHR63299.1;BatCoV_RaTG13_orf1ab;215 | 0.1 | 0.075 |
| MN996532.1;QHR63299.1;BatCoV_RaTG13_orf1ab;216 | 0 | 0.075 |
| MN996532.1;QHR63299.1;BatCoV_RaTG13_orf1ab;217 | 0 | 0.025 |
| MN996532.1;QHR63299.1;BatCoV_RaTG13_orf1ab;218 | 0 | 0.025 |
| MN996532.1;QHR63299.1;BatCoV_RaTG13_orf1ab;219 | 0 | 0.025 |
| MN996532.1;QHR63299.1;BatCoV_RaTG13_orf1ab;220 | 0.05 | 0.075 |
| MN996532.1;QHR63299.1;BatCoV_RaTG13_orf1ab;221 | 0 | 0.025 |
| MN996532.1;QHR63299.1;BatCoV_RaTG13_orf1ab;222 | 0 | 0.025 |
| MN996532.1;QHR63299.1;BatCoV_RaTG13_orf1ab;223 | 0 | 0.025 |
| MN996532.1;QHR63299.1;BatCoV_RaTG13_orf1ab;224 | 0 | 0 |
| MN996532.1;QHR63299.1;BatCoV_RaTG13_orf1ab;225 | 0.05 | 0 |
| MN996532.1;QHR63299.1;BatCoV_RaTG13_orf1ab;226 | 0 | 0 |
| MN996532.1;QHR63299.1;BatCoV_RaTG13_orf1ab;227 | 0 | 0 |
| MN996532.1;QHR63299.1;BatCoV_RaTG13_orf1ab;228 | 0 | 0 |
| MN996532.1;QHR63299.1;BatCoV_RaTG13_orf1ab;229 | 0 | 0 |
| MN996532.1;QHR63299.1;BatCoV_RaTG13_orf1ab;230 | 0 | 0 |
| MN996532.1;QHR63299.1;BatCoV_RaTG13_orf1ab;231 | 0 | 0 |
| MN996532.1;QHR63299.1;BatCoV_RaTG13_orf1ab;232 | 0 | 0 |
| MN996532.1;QHR63299.1;BatCoV_RaTG13_orf1ab;233 | 0 | 0 |
| MN996532.1;QHR63299.1;BatCoV_RaTG13_orf1ab;234 | 0 | 0 |
| MN996532.1;QHR63299.1;BatCoV_RaTG13_orf1ab;235 | 0 | 0 |
| MN996532.1;QHR63299.1;BatCoV_RaTG13_orf1ab;236 | 0 | 0 |
| MN996532.1;QHR63299.1;BatCoV_RaTG13_orf1ab;237 | 0 | 0 |
| MN996532.1;QHR63299.1;BatCoV_RaTG13_orf1ab;238 | 0 | 0 |
| MN996532.1;QHR63299.1;BatCoV_RaTG13_orf1ab;239 | 0 | 0 |
| MN996532.1;QHR63299.1;BatCoV_RaTG13_orf1ab;240 | 0 | 0.025 |
| MN996532.1;QHR63299.1;BatCoV_RaTG13_orf1ab;241 | 0 | 0.025 |
| MN996532.1;QHR63299.1;BatCoV_RaTG13_orf1ab;242 | 0 | 0.05 |
| MN996532.1;QHR63299.1;BatCoV_RaTG13_orf1ab;243 | 0 | 0.05 |
| MN996532.1;QHR63299.1;BatCoV_RaTG13_orf1ab;244 | 0 | 0.05 |
| MN996532.1;QHR63299.1;BatCoV_RaTG13_orf1ab;245 | 0 | 0.05 |
| MN996532.1;QHR63299.1;BatCoV_RaTG13_orf1ab;246 | 0 | 0.075 |
| MN996532.1;QHR63299.1;BatCoV_RaTG13_orf1ab;247 | 0 | 0.05 |
| MN996532.1;QHR63299.1;BatCoV_RaTG13_orf1ab;248 | 0 | 0.025 |
| MN996532.1;QHR63299.1;BatCoV_RaTG13_orf1ab;249 | 0 | 0.05 |

| | | |
|---|---|---|
| MN996532.1;QHR63299.1;BatCoV_RaTG13_orf1ab;250 | 0 | 0.025 |
| MN996532.1;QHR63299.1;BatCoV_RaTG13_orf1ab;251 | 0 | 0.025 |
| MN996532.1;QHR63299.1;BatCoV_RaTG13_orf1ab;252 | 0.05 | 0 |
| MN996532.1;QHR63299.1;BatCoV_RaTG13_orf1ab;253 | 0.05 | 0 |
| MN996532.1;QHR63299.1;BatCoV_RaTG13_orf1ab;254 | 0 | 0 |
| MN996532.1;QHR63299.1;BatCoV_RaTG13_orf1ab;255 | 0 | 0.05 |
| MN996532.1;QHR63299.1;BatCoV_RaTG13_orf1ab;256 | 0.1 | 0.1 |
| MN996532.1;QHR63299.1;BatCoV_RaTG13_orf1ab;257 | 0 | 0 |
| MN996532.1;QHR63299.1;BatCoV_RaTG13_orf1ab;258 | 0.05 | 0 |
| MN996532.1;QHR63299.1;BatCoV_RaTG13_orf1ab;259 | 0.05 | 0.05 |
| MN996532.1;QHR63299.1;BatCoV_RaTG13_orf1ab;260 | 0.05 | 0.025 |
| MN996532.1;QHR63299.1;BatCoV_RaTG13_orf1ab;261 | 0.05 | 0.025 |
| MN996532.1;QHR63299.1;BatCoV_RaTG13_orf1ab;262 | 0 | 0 |
| MN996532.1;QHR63299.1;BatCoV_RaTG13_orf1ab;263 | 0 | 0.025 |
| MN996532.1;QHR63299.1;BatCoV_RaTG13_orf1ab;264 | 0 | 0.025 |
| MN996532.1;QHR63299.1;BatCoV_RaTG13_orf1ab;265 | 0 | 0 |
| MN996532.1;QHR63299.1;BatCoV_RaTG13_orf1ab;266 | 0 | 0 |
| MN996532.1;QHR63299.1;BatCoV_RaTG13_orf1ab;267 | 0.15 | 0.025 |
| MN996532.1;QHR63299.1;BatCoV_RaTG13_orf1ab;268 | 0.15 | 0 |
| MN996532.1;QHR63299.1;BatCoV_RaTG13_orf1ab;269 | 0.15 | 0 |
| MN996532.1;QHR63299.1;BatCoV_RaTG13_orf1ab;270 | 0.2 | 0 |
| MN996532.1;QHR63299.1;BatCoV_RaTG13_orf1ab;271 | 0.15 | 0 |
| MN996532.1;QHR63299.1;BatCoV_RaTG13_orf1ab;272 | 0.1 | 0 |
| MN996532.1;QHR63299.1;BatCoV_RaTG13_orf1ab;273 | 0.1 | 0 |
| MN996532.1;QHR63299.1;BatCoV_RaTG13_orf1ab;274 | 0.15 | 0 |
| MN996532.1;QHR63299.1;BatCoV_RaTG13_orf1ab;275 | 0.15 | 0.025 |
| MN996532.1;QHR63299.1;BatCoV_RaTG13_orf1ab;276 | 0.2 | 0.05 |
| MN996532.1;QHR63299.1;BatCoV_RaTG13_orf1ab;277 | 0.2 | 0.025 |
| MN996532.1;QHR63299.1;BatCoV_RaTG13_orf1ab;278 | 0.2 | 0 |
| MN996532.1;QHR63299.1;BatCoV_RaTG13_orf1ab;279 | 0.05 | 0 |
| MN996532.1;QHR63299.1;BatCoV_RaTG13_orf1ab;280 | 0.05 | 0.025 |
| MN996532.1;QHR63299.1;BatCoV_RaTG13_orf1ab;281 | 0.1 | 0 |
| MN996532.1;QHR63299.1;BatCoV_RaTG13_orf1ab;282 | 0.05 | 0 |
| MN996532.1;QHR63299.1;BatCoV_RaTG13_orf1ab;283 | 0.1 | 0.025 |
| MN996532.1;QHR63299.1;BatCoV_RaTG13_orf1ab;284 | 0.05 | 0.025 |
| MN996532.1;QHR63299.1;BatCoV_RaTG13_orf1ab;285 | 0.05 | 0.025 |
| MN996532.1;QHR63299.1;BatCoV_RaTG13_orf1ab;286 | 0.05 | 0.025 |
| MN996532.1;QHR63299.1;BatCoV_RaTG13_orf1ab;287 | 0 | 0 |
| MN996532.1;QHR63299.1;BatCoV_RaTG13_orf1ab;288 | 0 | 0 |
| MN996532.1;QHR63299.1;BatCoV_RaTG13_orf1ab;289 | 0 | 0 |
| MN996532.1;QHR63299.1;BatCoV_RaTG13_orf1ab;290 | 0 | 0 |
| MN996532.1;QHR63299.1;BatCoV_RaTG13_orf1ab;291 | 0.05 | 0 |
| MN996532.1;QHR63299.1;BatCoV_RaTG13_orf1ab;292 | 0 | 0 |

| | | |
|---|---|---|
| MN996532.1;QHR63299.1;BatCoV_RaTG13_orf1ab;293 | 0 | 0 |
| MN996532.1;QHR63299.1;BatCoV_RaTG13_orf1ab;294 | 0 | 0 |
| MN996532.1;QHR63299.1;BatCoV_RaTG13_orf1ab;295 | 0 | 0 |
| MN996532.1;QHR63299.1;BatCoV_RaTG13_orf1ab;296 | 0.1 | 0 |
| MN996532.1;QHR63299.1;BatCoV_RaTG13_orf1ab;297 | 0.1 | 0 |
| MN996532.1;QHR63299.1;BatCoV_RaTG13_orf1ab;298 | 0.1 | 0 |
| MN996532.1;QHR63299.1;BatCoV_RaTG13_orf1ab;299 | 0.1 | 0 |
| MN996532.1;QHR63299.1;BatCoV_RaTG13_orf1ab;300 | 0.1 | 0 |
| MN996532.1;QHR63299.1;BatCoV_RaTG13_orf1ab;301 | 0.1 | 0.1 |
| MN996532.1;QHR63299.1;BatCoV_RaTG13_orf1ab;302 | 0.05 | 0 |
| MN996532.1;QHR63299.1;BatCoV_RaTG13_orf1ab;303 | 0.05 | 0 |
| MN996532.1;QHR63299.1;BatCoV_RaTG13_orf1ab;304 | 0.1 | 0 |
| MN996532.1;QHR63299.1;BatCoV_RaTG13_orf1ab;305 | 0.05 | 0 |
| MN996532.1;QHR63299.1;BatCoV_RaTG13_orf1ab;306 | 0 | 0 |
| MN996532.1;QHR63299.1;BatCoV_RaTG13_orf1ab;307 | 0 | 0 |
| MN996532.1;QHR63299.1;BatCoV_RaTG13_orf1ab;308 | 0 | 0 |
| MN996532.1;QHR63299.1;BatCoV_RaTG13_orf1ab;309 | 0 | 0 |
| MN996532.1;QHR63299.1;BatCoV_RaTG13_orf1ab;310 | 0 | 0 |
| MN996532.1;QHR63299.1;BatCoV_RaTG13_orf1ab;311 | 0 | 0 |
| MN996532.1;QHR63299.1;BatCoV_RaTG13_orf1ab;312 | 0 | 0 |
| MN996532.1;QHR63299.1;BatCoV_RaTG13_orf1ab;313 | 0 | 0 |
| MN996532.1;QHR63299.1;BatCoV_RaTG13_orf1ab;314 | 0 | 0 |
| MN996532.1;QHR63299.1;BatCoV_RaTG13_orf1ab;315 | 0 | 0 |
| MN996532.1;QHR63299.1;BatCoV_RaTG13_orf1ab;316 | 0 | 0 |
| MN996532.1;QHR63299.1;BatCoV_RaTG13_orf1ab;317 | 0 | 0 |
| MN996532.1;QHR63299.1;BatCoV_RaTG13_orf1ab;318 | 0 | 0 |
| MN996532.1;QHR63299.1;BatCoV_RaTG13_orf1ab;319 | 0 | 0 |
| MN996532.1;QHR63299.1;BatCoV_RaTG13_orf1ab;320 | 0 | 0 |
| MN996532.1;QHR63299.1;BatCoV_RaTG13_orf1ab;321 | 0 | 0 |
| MN996532.1;QHR63299.1;BatCoV_RaTG13_orf1ab;322 | 0 | 0 |
| MN996532.1;QHR63299.1;BatCoV_RaTG13_orf1ab;323 | 0 | 0 |
| MN996532.1;QHR63299.1;BatCoV_RaTG13_orf1ab;324 | 0.05 | 0 |
| MN996532.1;QHR63299.1;BatCoV_RaTG13_orf1ab;325 | 0 | 0.05 |
| MN996532.1;QHR63299.1;BatCoV_RaTG13_orf1ab;326 | 0 | 0 |
| MN996532.1;QHR63299.1;BatCoV_RaTG13_orf1ab;327 | 0 | 0 |
| MN996532.1;QHR63299.1;BatCoV_RaTG13_orf1ab;328 | 0 | 0 |
| MN996532.1;QHR63299.1;BatCoV_RaTG13_orf1ab;329 | 0 | 0 |
| MN996532.1;QHR63299.1;BatCoV_RaTG13_orf1ab;330 | 0 | 0 |
| MN996532.1;QHR63299.1;BatCoV_RaTG13_orf1ab;331 | 0 | 0 |
| MN996532.1;QHR63299.1;BatCoV_RaTG13_orf1ab;332 | 0 | 0 |
| MN996532.1;QHR63299.1;BatCoV_RaTG13_orf1ab;333 | 0 | 0 |
| MN996532.1;QHR63299.1;BatCoV_RaTG13_orf1ab;334 | 0 | 0 |
| MN996532.1;QHR63299.1;BatCoV_RaTG13_orf1ab;335 | 0 | 0 |

| | | |
|---|---|---|
| MN996532.1;QHR63299.1;BatCoV_RaTG13_orf1ab;336 | 0 | 0 |
| MN996532.1;QHR63299.1;BatCoV_RaTG13_orf1ab;337 | 0 | 0 |
| MN996532.1;QHR63299.1;BatCoV_RaTG13_orf1ab;338 | 0 | 0 |
| MN996532.1;QHR63299.1;BatCoV_RaTG13_orf1ab;339 | 0 | 0 |
| MN996532.1;QHR63299.1;BatCoV_RaTG13_orf1ab;340 | 0 | 0 |
| MN996532.1;QHR63299.1;BatCoV_RaTG13_orf1ab;341 | 0 | 0 |
| MN996532.1;QHR63299.1;BatCoV_RaTG13_orf1ab;342 | 0 | 0 |
| MN996532.1;QHR63299.1;BatCoV_RaTG13_orf1ab;343 | 0 | 0 |
| MN996532.1;QHR63299.1;BatCoV_RaTG13_orf1ab;344 | 0 | 0 |
| MN996532.1;QHR63299.1;BatCoV_RaTG13_orf1ab;345 | 0 | 0 |
| MN996532.1;QHR63299.1;BatCoV_RaTG13_orf1ab;346 | 0 | 0 |
| MN996532.1;QHR63299.1;BatCoV_RaTG13_orf1ab;347 | 0.05 | 0 |
| MN996532.1;QHR63299.1;BatCoV_RaTG13_orf1ab;348 | 0 | 0 |
| MN996532.1;QHR63299.1;BatCoV_RaTG13_orf1ab;349 | 0.05 | 0 |
| MN996532.1;QHR63299.1;BatCoV_RaTG13_orf1ab;350 | 0 | 0 |
| MN996532.1;QHR63299.1;BatCoV_RaTG13_orf1ab;351 | 0 | 0 |
| MN996532.1;QHR63299.1;BatCoV_RaTG13_orf1ab;352 | 0 | 0.025 |
| MN996532.1;QHR63299.1;BatCoV_RaTG13_orf1ab;353 | 0 | 0 |
| MN996532.1;QHR63299.1;BatCoV_RaTG13_orf1ab;354 | 0 | 0 |
| MN996532.1;QHR63299.1;BatCoV_RaTG13_orf1ab;355 | 0 | 0 |
| MN996532.1;QHR63299.1;BatCoV_RaTG13_orf1ab;356 | 0 | 0 |
| MN996532.1;QHR63299.1;BatCoV_RaTG13_orf1ab;357 | 0 | 0 |
| MN996532.1;QHR63299.1;BatCoV_RaTG13_orf1ab;358 | 0 | 0 |
| MN996532.1;QHR63299.1;BatCoV_RaTG13_orf1ab;359 | 0 | 0 |
| MN996532.1;QHR63299.1;BatCoV_RaTG13_orf1ab;360 | 0 | 0 |
| MN996532.1;QHR63299.1;BatCoV_RaTG13_orf1ab;361 | 0 | 0 |
| MN996532.1;QHR63299.1;BatCoV_RaTG13_orf1ab;362 | 0 | 0 |
| MN996532.1;QHR63299.1;BatCoV_RaTG13_orf1ab;363 | 0 | 0 |
| MN996532.1;QHR63299.1;BatCoV_RaTG13_orf1ab;364 | 0 | 0 |
| MN996532.1;QHR63299.1;BatCoV_RaTG13_orf1ab;365 | 0 | 0 |
| MN996532.1;QHR63299.1;BatCoV_RaTG13_orf1ab;366 | 0.15 | 0.05 |
| MN996532.1;QHR63299.1;BatCoV_RaTG13_orf1ab;367 | 0.05 | 0.025 |
| MN996532.1;QHR63299.1;BatCoV_RaTG13_orf1ab;368 | 0.05 | 0 |
| MN996532.1;QHR63299.1;BatCoV_RaTG13_orf1ab;369 | 0.15 | 0.05 |
| MN996532.1;QHR63299.1;BatCoV_RaTG13_orf1ab;370 | 0.1 | 0.05 |
| MN996532.1;QHR63299.1;BatCoV_RaTG13_orf1ab;371 | 0.15 | 0.05 |
| MN996532.1;QHR63299.1;BatCoV_RaTG13_orf1ab;372 | 0.05 | 0.025 |
| MN996532.1;QHR63299.1;BatCoV_RaTG13_orf1ab;373 | 0.05 | 0 |
| MN996532.1;QHR63299.1;BatCoV_RaTG13_orf1ab;374 | 0.05 | 0.025 |
| MN996532.1;QHR63299.1;BatCoV_RaTG13_orf1ab;375 | 0.05 | 0.025 |
| MN996532.1;QHR63299.1;BatCoV_RaTG13_orf1ab;376 | 0.05 | 0 |
| MN996532.1;QHR63299.1;BatCoV_RaTG13_orf1ab;377 | 0 | 0 |
| MN996532.1;QHR63299.1;BatCoV_RaTG13_orf1ab;378 | 0 | 0 |

| | | |
|---|---|---|
| MN996532.1;QHR63299.1;BatCoV_RaTG13_orf1ab;379 | 0.15 | 0.05 |
| MN996532.1;QHR63299.1;BatCoV_RaTG13_orf1ab;380 | 0 | 0.025 |
| MN996532.1;QHR63299.1;BatCoV_RaTG13_orf1ab;381 | 0 | 0.025 |
| MN996532.1;QHR63299.1;BatCoV_RaTG13_orf1ab;382 | 0 | 0.025 |
| MN996532.1;QHR63299.1;BatCoV_RaTG13_orf1ab;383 | 0 | 0.025 |
| MN996532.1;QHR63299.1;BatCoV_RaTG13_orf1ab;384 | 0 | 0.025 |
| MN996532.1;QHR63299.1;BatCoV_RaTG13_orf1ab;385 | 0 | 0.025 |
| MN996532.1;QHR63299.1;BatCoV_RaTG13_orf1ab;386 | 0.05 | 0.025 |
| MN996532.1;QHR63299.1;BatCoV_RaTG13_orf1ab;387 | 0 | 0.025 |
| MN996532.1;QHR63299.1;BatCoV_RaTG13_orf1ab;388 | 0 | 0 |
| MN996532.1;QHR63299.1;BatCoV_RaTG13_orf1ab;389 | 0 | 0 |
| MN996532.1;QHR63299.1;BatCoV_RaTG13_orf1ab;390 | 0 | 0.025 |
| MN996532.1;QHR63299.1;BatCoV_RaTG13_orf1ab;391 | 0 | 0 |
| MN996532.1;QHR63299.1;BatCoV_RaTG13_orf1ab;392 | 0 | 0 |
| MN996532.1;QHR63299.1;BatCoV_RaTG13_orf1ab;393 | 0 | 0 |
| MN996532.1;QHR63299.1;BatCoV_RaTG13_orf1ab;394 | 0 | 0 |
| MN996532.1;QHR63299.1;BatCoV_RaTG13_orf1ab;395 | 0 | 0 |
| MN996532.1;QHR63299.1;BatCoV_RaTG13_orf1ab;396 | 0 | 0 |
| MN996532.1;QHR63299.1;BatCoV_RaTG13_orf1ab;397 | 0 | 0.025 |
| MN996532.1;QHR63299.1;BatCoV_RaTG13_orf1ab;398 | 0 | 0 |
| MN996532.1;QHR63299.1;BatCoV_RaTG13_orf1ab;399 | 0 | 0 |
| MN996532.1;QHR63299.1;BatCoV_RaTG13_orf1ab;400 | 0 | 0 |
| MN996532.1;QHR63299.1;BatCoV_RaTG13_orf1ab;401 | 0 | 0 |
| MN996532.1;QHR63299.1;BatCoV_RaTG13_orf1ab;402 | 0 | 0 |
| MN996532.1;QHR63299.1;BatCoV_RaTG13_orf1ab;403 | 0 | 0 |
| MN996532.1;QHR63299.1;BatCoV_RaTG13_orf1ab;404 | 0 | 0 |
| MN996532.1;QHR63299.1;BatCoV_RaTG13_orf1ab;405 | 0 | 0 |
| MN996532.1;QHR63299.1;BatCoV_RaTG13_orf1ab;406 | 0 | 0 |
| MN996532.1;QHR63299.1;BatCoV_RaTG13_orf1ab;407 | 0 | 0 |
| MN996532.1;QHR63299.1;BatCoV_RaTG13_orf1ab;408 | 0 | 0 |
| MN996532.1;QHR63299.1;BatCoV_RaTG13_orf1ab;409 | 0 | 0 |
| MN996532.1;QHR63299.1;BatCoV_RaTG13_orf1ab;410 | 0 | 0 |
| MN996532.1;QHR63299.1;BatCoV_RaTG13_orf1ab;411 | 0 | 0 |
| MN996532.1;QHR63299.1;BatCoV_RaTG13_orf1ab;412 | 0 | 0 |
| MN996532.1;QHR63299.1;BatCoV_RaTG13_orf1ab;413 | 0 | 0 |
| MN996532.1;QHR63299.1;BatCoV_RaTG13_orf1ab;414 | 0 | 0 |
| MN996532.1;QHR63299.1;BatCoV_RaTG13_orf1ab;415 | 0 | 0 |
| MN996532.1;QHR63299.1;BatCoV_RaTG13_orf1ab;416 | 0 | 0 |
| MN996532.1;QHR63299.1;BatCoV_RaTG13_orf1ab;417 | 0 | 0 |
| MN996532.1;QHR63299.1;BatCoV_RaTG13_orf1ab;418 | 0 | 0 |
| MN996532.1;QHR63299.1;BatCoV_RaTG13_orf1ab;419 | 0 | 0 |
| MN996532.1;QHR63299.1;BatCoV_RaTG13_orf1ab;420 | 0 | 0 |
| MN996532.1;QHR63299.1;BatCoV_RaTG13_orf1ab;421 | 0 | 0 |

| | | |
|---|---|---|
| MN996532.1;QHR63299.1;BatCoV_RaTG13_orf1ab;422 | 0 | 0 |
| MN996532.1;QHR63299.1;BatCoV_RaTG13_orf1ab;423 | 0 | 0 |
| MN996532.1;QHR63299.1;BatCoV_RaTG13_orf1ab;424 | 0.05 | 0 |
| MN996532.1;QHR63299.1;BatCoV_RaTG13_orf1ab;425 | 0.05 | 0.025 |
| MN996532.1;QHR63299.1;BatCoV_RaTG13_orf1ab;426 | 0 | 0 |
| MN996532.1;QHR63299.1;BatCoV_RaTG13_orf1ab;427 | 0 | 0.025 |
| MN996532.1;QHR63299.1;BatCoV_RaTG13_orf1ab;428 | 0 | 0 |
| MN996532.1;QHR63299.1;BatCoV_RaTG13_orf1ab;429 | 0 | 0 |
| MN996532.1;QHR63299.1;BatCoV_RaTG13_orf1ab;430 | 0 | 0 |
| MN996532.1;QHR63299.1;BatCoV_RaTG13_orf1ab;431 | 0 | 0 |
| MN996532.1;QHR63299.1;BatCoV_RaTG13_orf1ab;432 | 0 | 0 |
| MN996532.1;QHR63299.1;BatCoV_RaTG13_orf1ab;433 | 0 | 0 |
| MN996532.1;QHR63299.1;BatCoV_RaTG13_orf1ab;434 | 0 | 0 |
| MN996532.1;QHR63299.1;BatCoV_RaTG13_orf1ab;435 | 0 | 0 |
| MN996532.1;QHR63299.1;BatCoV_RaTG13_orf1ab;436 | 0 | 0 |
| MN996532.1;QHR63299.1;BatCoV_RaTG13_orf1ab;437 | 0 | 0 |
| MN996532.1;QHR63299.1;BatCoV_RaTG13_orf1ab;438 | 0 | 0 |
| MN996532.1;QHR63299.1;BatCoV_RaTG13_orf1ab;439 | 0 | 0 |
| MN996532.1;QHR63299.1;BatCoV_RaTG13_orf1ab;440 | 0 | 0 |
| MN996532.1;QHR63299.1;BatCoV_RaTG13_orf1ab;441 | 0 | 0 |
| MN996532.1;QHR63299.1;BatCoV_RaTG13_orf1ab;442 | 0 | 0 |
| MN996532.1;QHR63299.1;BatCoV_RaTG13_orf1ab;443 | 0.05 | 0 |
| MN996532.1;QHR63299.1;BatCoV_RaTG13_orf1ab;444 | 0.2 | 0.025 |
| MN996532.1;QHR63299.1;BatCoV_RaTG13_orf1ab;445 | 0.5 | 0.475 |
| MN996532.1;QHR63299.1;BatCoV_RaTG13_orf1ab;446 | 0.05 | 0.05 |
| MN996532.1;QHR63299.1;BatCoV_RaTG13_orf1ab;447 | 0.05 | 0.05 |
| MN996532.1;QHR63299.1;BatCoV_RaTG13_orf1ab;448 | 0.05 | 0.05 |
| MN996532.1;QHR63299.1;BatCoV_RaTG13_orf1ab;449 | 0 | 0.025 |
| MN996532.1;QHR63299.1;BatCoV_RaTG13_orf1ab;450 | 0 | 0 |
| MN996532.1;QHR63299.1;BatCoV_RaTG13_orf1ab;451 | 0 | 0 |
| MN996532.1;QHR63299.1;BatCoV_RaTG13_orf1ab;452 | 0.05 | 0 |
| MN996532.1;QHR63299.1;BatCoV_RaTG13_orf1ab;453 | 0 | 0 |
| MN996532.1;QHR63299.1;BatCoV_RaTG13_orf1ab;454 | 0 | 0 |
| MN996532.1;QHR63299.1;BatCoV_RaTG13_orf1ab;455 | 0 | 0 |
| MN996532.1;QHR63299.1;BatCoV_RaTG13_orf1ab;456 | 0.05 | 0 |
| MN996532.1;QHR63299.1;BatCoV_RaTG13_orf1ab;457 | 0.05 | 0 |
| MN996532.1;QHR63299.1;BatCoV_RaTG13_orf1ab;458 | 0.05 | 0 |
| MN996532.1;QHR63299.1;BatCoV_RaTG13_orf1ab;459 | 0 | 0 |
| MN996532.1;QHR63299.1;BatCoV_RaTG13_orf1ab;460 | 0 | 0 |
| MN996532.1;QHR63299.1;BatCoV_RaTG13_orf1ab;461 | 0 | 0 |
| MN996532.1;QHR63299.1;BatCoV_RaTG13_orf1ab;462 | 0 | 0 |
| MN996532.1;QHR63299.1;BatCoV_RaTG13_orf1ab;463 | 0 | 0 |
| MN996532.1;QHR63299.1;BatCoV_RaTG13_orf1ab;464 | 0 | 0 |

| | | |
|---|---|---|
| MN996532.1;QHR63299.1;BatCoV_RaTG13_orf1ab;465 | 0 | 0 |
| MN996532.1;QHR63299.1;BatCoV_RaTG13_orf1ab;466 | 0 | 0 |
| MN996532.1;QHR63299.1;BatCoV_RaTG13_orf1ab;467 | 0 | 0 |
| MN996532.1;QHR63299.1;BatCoV_RaTG13_orf1ab;468 | 0 | 0 |
| MN996532.1;QHR63299.1;BatCoV_RaTG13_orf1ab;469 | 0 | 0 |
| MN996532.1;QHR63299.1;BatCoV_RaTG13_orf1ab;470 | 0 | 0 |
| MN996532.1;QHR63299.1;BatCoV_RaTG13_orf1ab;471 | 0 | 0 |
| MN996532.1;QHR63299.1;BatCoV_RaTG13_orf1ab;472 | 0 | 0 |
| MN996532.1;QHR63299.1;BatCoV_RaTG13_orf1ab;473 | 0 | 0 |
| MN996532.1;QHR63299.1;BatCoV_RaTG13_orf1ab;474 | 0 | 0 |
| MN996532.1;QHR63299.1;BatCoV_RaTG13_orf1ab;475 | 0 | 0 |
| MN996532.1;QHR63299.1;BatCoV_RaTG13_orf1ab;476 | 0 | 0 |
| MN996532.1;QHR63299.1;BatCoV_RaTG13_orf1ab;477 | 0 | 0 |
| MN996532.1;QHR63299.1;BatCoV_RaTG13_orf1ab;478 | 0 | 0 |
| MN996532.1;QHR63299.1;BatCoV_RaTG13_orf1ab;479 | 0 | 0 |
| MN996532.1;QHR63299.1;BatCoV_RaTG13_orf1ab;480 | 0 | 0.05 |
| MN996532.1;QHR63299.1;BatCoV_RaTG13_orf1ab;481 | 0 | 0.05 |
| MN996532.1;QHR63299.1;BatCoV_RaTG13_orf1ab;482 | 0 | 0.025 |
| MN996532.1;QHR63299.1;BatCoV_RaTG13_orf1ab;483 | 0 | 0.025 |
| MN996532.1;QHR63299.1;BatCoV_RaTG13_orf1ab;484 | 0 | 0 |
| MN996532.1;QHR63299.1;BatCoV_RaTG13_orf1ab;485 | 0 | 0 |
| MN996532.1;QHR63299.1;BatCoV_RaTG13_orf1ab;486 | 0 | 0.025 |
| MN996532.1;QHR63299.1;BatCoV_RaTG13_orf1ab;487 | 0 | 0 |
| MN996532.1;QHR63299.1;BatCoV_RaTG13_orf1ab;488 | 0 | 0 |
| MN996532.1;QHR63299.1;BatCoV_RaTG13_orf1ab;489 | 0 | 0.05 |
| MN996532.1;QHR63299.1;BatCoV_RaTG13_orf1ab;490 | 0 | 0.025 |
| MN996532.1;QHR63299.1;BatCoV_RaTG13_orf1ab;491 | 0 | 0 |
| MN996532.1;QHR63299.1;BatCoV_RaTG13_orf1ab;492 | 0 | 0 |
| MN996532.1;QHR63299.1;BatCoV_RaTG13_orf1ab;493 | 0 | 0.05 |
| MN996532.1;QHR63299.1;BatCoV_RaTG13_orf1ab;494 | 0 | 0.025 |
| MN996532.1;QHR63299.1;BatCoV_RaTG13_orf1ab;495 | 0 | 0 |
| MN996532.1;QHR63299.1;BatCoV_RaTG13_orf1ab;496 | 0 | 0 |
| MN996532.1;QHR63299.1;BatCoV_RaTG13_orf1ab;497 | 0.05 | 0 |
| MN996532.1;QHR63299.1;BatCoV_RaTG13_orf1ab;498 | 0 | 0 |
| MN996532.1;QHR63299.1;BatCoV_RaTG13_orf1ab;499 | 0 | 0 |
| MN996532.1;QHR63299.1;BatCoV_RaTG13_orf1ab;500 | 0 | 0 |
| MN996532.1;QHR63299.1;BatCoV_RaTG13_orf1ab;501 | 0 | 0 |
| MN996532.1;QHR63299.1;BatCoV_RaTG13_orf1ab;502 | 0 | 0 |
| MN996532.1;QHR63299.1;BatCoV_RaTG13_orf1ab;503 | 0 | 0 |
| MN996532.1;QHR63299.1;BatCoV_RaTG13_orf1ab;504 | 0 | 0 |
| MN996532.1;QHR63299.1;BatCoV_RaTG13_orf1ab;505 | 0 | 0 |
| MN996532.1;QHR63299.1;BatCoV_RaTG13_orf1ab;506 | 0 | 0 |
| MN996532.1;QHR63299.1;BatCoV_RaTG13_orf1ab;507 | 0 | 0 |

| | | |
|---|---|---|
| MN996532.1;QHR63299.1;BatCoV_RaTG13_orf1ab;508 | 0 | 0 |
| MN996532.1;QHR63299.1;BatCoV_RaTG13_orf1ab;509 | 0 | 0 |
| MN996532.1;QHR63299.1;BatCoV_RaTG13_orf1ab;510 | 0 | 0 |
| MN996532.1;QHR63299.1;BatCoV_RaTG13_orf1ab;511 | 0 | 0 |
| MN996532.1;QHR63299.1;BatCoV_RaTG13_orf1ab;512 | 0 | 0 |
| MN996532.1;QHR63299.1;BatCoV_RaTG13_orf1ab;513 | 0 | 0.025 |
| MN996532.1;QHR63299.1;BatCoV_RaTG13_orf1ab;514 | 0 | 0.025 |
| MN996532.1;QHR63299.1;BatCoV_RaTG13_orf1ab;515 | 0 | 0.025 |
| MN996532.1;QHR63299.1;BatCoV_RaTG13_orf1ab;516 | 0 | 0.025 |
| MN996532.1;QHR63299.1;BatCoV_RaTG13_orf1ab;517 | 0 | 0 |
| MN996532.1;QHR63299.1;BatCoV_RaTG13_orf1ab;518 | 0.1 | 0.025 |
| MN996532.1;QHR63299.1;BatCoV_RaTG13_orf1ab;519 | 0 | 0 |
| MN996532.1;QHR63299.1;BatCoV_RaTG13_orf1ab;520 | 0 | 0 |
| MN996532.1;QHR63299.1;BatCoV_RaTG13_orf1ab;521 | 0 | 0 |
| MN996532.1;QHR63299.1;BatCoV_RaTG13_orf1ab;522 | 0 | 0 |
| MN996532.1;QHR63299.1;BatCoV_RaTG13_orf1ab;523 | 0 | 0 |
| MN996532.1;QHR63299.1;BatCoV_RaTG13_orf1ab;524 | 0 | 0 |
| MN996532.1;QHR63299.1;BatCoV_RaTG13_orf1ab;525 | 0.05 | 0 |
| MN996532.1;QHR63299.1;BatCoV_RaTG13_orf1ab;526 | 0.05 | 0.025 |
| MN996532.1;QHR63299.1;BatCoV_RaTG13_orf1ab;527 | 0 | 0 |
| MN996532.1;QHR63299.1;BatCoV_RaTG13_orf1ab;528 | 0 | 0 |
| MN996532.1;QHR63299.1;BatCoV_RaTG13_orf1ab;529 | 0 | 0 |
| MN996532.1;QHR63299.1;BatCoV_RaTG13_orf1ab;530 | 0 | 0 |
| MN996532.1;QHR63299.1;BatCoV_RaTG13_orf1ab;531 | 0 | 0 |
| MN996532.1;QHR63299.1;BatCoV_RaTG13_orf1ab;532 | 0 | 0 |
| MN996532.1;QHR63299.1;BatCoV_RaTG13_orf1ab;533 | 0 | 0 |
| MN996532.1;QHR63299.1;BatCoV_RaTG13_orf1ab;534 | 0 | 0 |
| MN996532.1;QHR63299.1;BatCoV_RaTG13_orf1ab;535 | 0 | 0 |
| MN996532.1;QHR63299.1;BatCoV_RaTG13_orf1ab;536 | 0 | 0 |
| MN996532.1;QHR63299.1;BatCoV_RaTG13_orf1ab;537 | 0 | 0 |
| MN996532.1;QHR63299.1;BatCoV_RaTG13_orf1ab;538 | 0 | 0 |
| MN996532.1;QHR63299.1;BatCoV_RaTG13_orf1ab;539 | 0 | 0 |
| MN996532.1;QHR63299.1;BatCoV_RaTG13_orf1ab;540 | 0 | 0 |
| MN996532.1;QHR63299.1;BatCoV_RaTG13_orf1ab;541 | 0 | 0 |
| MN996532.1;QHR63299.1;BatCoV_RaTG13_orf1ab;542 | 0 | 0 |
| MN996532.1;QHR63299.1;BatCoV_RaTG13_orf1ab;543 | 0 | 0 |
| MN996532.1;QHR63299.1;BatCoV_RaTG13_orf1ab;544 | 0 | 0 |
| MN996532.1;QHR63299.1;BatCoV_RaTG13_orf1ab;545 | 0 | 0 |
| MN996532.1;QHR63299.1;BatCoV_RaTG13_orf1ab;546 | 0 | 0.025 |
| MN996532.1;QHR63299.1;BatCoV_RaTG13_orf1ab;547 | 0 | 0 |
| MN996532.1;QHR63299.1;BatCoV_RaTG13_orf1ab;548 | 0 | 0 |
| MN996532.1;QHR63299.1;BatCoV_RaTG13_orf1ab;549 | 0 | 0 |
| MN996532.1;QHR63299.1;BatCoV_RaTG13_orf1ab;550 | 0 | 0 |

| | | |
|---|---|---|
| MN996532.1;QHR63299.1;BatCoV_RaTG13_orf1ab;551 | 0.05 | 0.05 |
| MN996532.1;QHR63299.1;BatCoV_RaTG13_orf1ab;552 | 0 | 0 |
| MN996532.1;QHR63299.1;BatCoV_RaTG13_orf1ab;553 | 0.05 | 0.025 |
| MN996532.1;QHR63299.1;BatCoV_RaTG13_orf1ab;554 | 0 | 0 |
| MN996532.1;QHR63299.1;BatCoV_RaTG13_orf1ab;555 | 0.05 | 0 |
| MN996532.1;QHR63299.1;BatCoV_RaTG13_orf1ab;556 | 0 | 0 |
| MN996532.1;QHR63299.1;BatCoV_RaTG13_orf1ab;557 | 0 | 0 |
| MN996532.1;QHR63299.1;BatCoV_RaTG13_orf1ab;558 | 0 | 0 |
| MN996532.1;QHR63299.1;BatCoV_RaTG13_orf1ab;559 | 0 | 0 |
| MN996532.1;QHR63299.1;BatCoV_RaTG13_orf1ab;560 | 0 | 0 |
| MN996532.1;QHR63299.1;BatCoV_RaTG13_orf1ab;561 | 0 | 0 |
| MN996532.1;QHR63299.1;BatCoV_RaTG13_orf1ab;562 | 0 | 0 |
| MN996532.1;QHR63299.1;BatCoV_RaTG13_orf1ab;563 | 0 | 0 |
| MN996532.1;QHR63299.1;BatCoV_RaTG13_orf1ab;564 | 0 | 0 |
| MN996532.1;QHR63299.1;BatCoV_RaTG13_orf1ab;565 | 0 | 0.025 |
| MN996532.1;QHR63299.1;BatCoV_RaTG13_orf1ab;566 | 0 | 0 |
| MN996532.1;QHR63299.1;BatCoV_RaTG13_orf1ab;567 | 0 | 0 |
| MN996532.1;QHR63299.1;BatCoV_RaTG13_orf1ab;568 | 0 | 0 |
| MN996532.1;QHR63299.1;BatCoV_RaTG13_orf1ab;569 | 0 | 0.025 |
| MN996532.1;QHR63299.1;BatCoV_RaTG13_orf1ab;570 | 0 | 0.025 |
| MN996532.1;QHR63299.1;BatCoV_RaTG13_orf1ab;571 | 0 | 0.025 |
| MN996532.1;QHR63299.1;BatCoV_RaTG13_orf1ab;572 | 0.1 | 0.05 |
| MN996532.1;QHR63299.1;BatCoV_RaTG13_orf1ab;573 | 0 | 0.025 |
| MN996532.1;QHR63299.1;BatCoV_RaTG13_orf1ab;574 | 0 | 0 |
| MN996532.1;QHR63299.1;BatCoV_RaTG13_orf1ab;575 | 0 | 0 |
| MN996532.1;QHR63299.1;BatCoV_RaTG13_orf1ab;576 | 0 | 0 |
| MN996532.1;QHR63299.1;BatCoV_RaTG13_orf1ab;577 | 0 | 0 |
| MN996532.1;QHR63299.1;BatCoV_RaTG13_orf1ab;578 | 0 | 0.025 |
| MN996532.1;QHR63299.1;BatCoV_RaTG13_orf1ab;579 | 0 | 0 |
| MN996532.1;QHR63299.1;BatCoV_RaTG13_orf1ab;580 | 0 | 0 |
| MN996532.1;QHR63299.1;BatCoV_RaTG13_orf1ab;581 | 0 | 0 |
| MN996532.1;QHR63299.1;BatCoV_RaTG13_orf1ab;582 | 0 | 0 |
| MN996532.1;QHR63299.1;BatCoV_RaTG13_orf1ab;583 | 0 | 0 |
| MN996532.1;QHR63299.1;BatCoV_RaTG13_orf1ab;584 | 0 | 0 |
| MN996532.1;QHR63299.1;BatCoV_RaTG13_orf1ab;585 | 0 | 0 |
| MN996532.1;QHR63299.1;BatCoV_RaTG13_orf1ab;586 | 0 | 0 |
| MN996532.1;QHR63299.1;BatCoV_RaTG13_orf1ab;587 | 0 | 0 |
| MN996532.1;QHR63299.1;BatCoV_RaTG13_orf1ab;588 | 0 | 0 |
| MN996532.1;QHR63299.1;BatCoV_RaTG13_orf1ab;589 | 0.05 | 0 |
| MN996532.1;QHR63299.1;BatCoV_RaTG13_orf1ab;590 | 0 | 0 |
| MN996532.1;QHR63299.1;BatCoV_RaTG13_orf1ab;591 | 0 | 0 |
| MN996532.1;QHR63299.1;BatCoV_RaTG13_orf1ab;592 | 0 | 0 |
| MN996532.1;QHR63299.1;BatCoV_RaTG13_orf1ab;593 | 0 | 0 |

| | | |
|---|---|---|
| MN996532.1;QHR63299.1;BatCoV_RaTG13_orf1ab;594 | 0 | 0 |
| MN996532.1;QHR63299.1;BatCoV_RaTG13_orf1ab;595 | 0 | 0 |
| MN996532.1;QHR63299.1;BatCoV_RaTG13_orf1ab;596 | 0 | 0 |
| MN996532.1;QHR63299.1;BatCoV_RaTG13_orf1ab;597 | 0 | 0 |
| MN996532.1;QHR63299.1;BatCoV_RaTG13_orf1ab;598 | 0 | 0 |
| MN996532.1;QHR63299.1;BatCoV_RaTG13_orf1ab;599 | 0 | 0 |
| MN996532.1;QHR63299.1;BatCoV_RaTG13_orf1ab;600 | 0 | 0 |
| MN996532.1;QHR63299.1;BatCoV_RaTG13_orf1ab;601 | 0 | 0 |
| MN996532.1;QHR63299.1;BatCoV_RaTG13_orf1ab;602 | 0 | 0 |
| MN996532.1;QHR63299.1;BatCoV_RaTG13_orf1ab;603 | 0 | 0 |
| MN996532.1;QHR63299.1;BatCoV_RaTG13_orf1ab;604 | 0 | 0 |
| MN996532.1;QHR63299.1;BatCoV_RaTG13_orf1ab;605 | 0 | 0 |
| MN996532.1;QHR63299.1;BatCoV_RaTG13_orf1ab;606 | 0 | 0 |
| MN996532.1;QHR63299.1;BatCoV_RaTG13_orf1ab;607 | 0 | 0 |
| MN996532.1;QHR63299.1;BatCoV_RaTG13_orf1ab;608 | 0 | 0 |
| MN996532.1;QHR63299.1;BatCoV_RaTG13_orf1ab;609 | 0 | 0 |
| MN996532.1;QHR63299.1;BatCoV_RaTG13_orf1ab;610 | 0 | 0 |
| MN996532.1;QHR63299.1;BatCoV_RaTG13_orf1ab;611 | 0 | 0 |
| MN996532.1;QHR63299.1;BatCoV_RaTG13_orf1ab;612 | 0 | 0 |
| MN996532.1;QHR63299.1;BatCoV_RaTG13_orf1ab;613 | 0 | 0 |
| MN996532.1;QHR63299.1;BatCoV_RaTG13_orf1ab;614 | 0 | 0 |
| MN996532.1;QHR63299.1;BatCoV_RaTG13_orf1ab;615 | 0 | 0 |
| MN996532.1;QHR63299.1;BatCoV_RaTG13_orf1ab;616 | 0 | 0 |
| MN996532.1;QHR63299.1;BatCoV_RaTG13_orf1ab;617 | 0.05 | 0.025 |
| MN996532.1;QHR63299.1;BatCoV_RaTG13_orf1ab;618 | 0.1 | 0.025 |
| MN996532.1;QHR63299.1;BatCoV_RaTG13_orf1ab;619 | 0.1 | 0.025 |
| MN996532.1;QHR63299.1;BatCoV_RaTG13_orf1ab;620 | 0.1 | 0.025 |
| MN996532.1;QHR63299.1;BatCoV_RaTG13_orf1ab;621 | 0.1 | 0.05 |
| MN996532.1;QHR63299.1;BatCoV_RaTG13_orf1ab;622 | 0.1 | 0.025 |
| MN996532.1;QHR63299.1;BatCoV_RaTG13_orf1ab;623 | 0 | 0 |
| MN996532.1;QHR63299.1;BatCoV_RaTG13_orf1ab;624 | 0 | 0.05 |
| MN996532.1;QHR63299.1;BatCoV_RaTG13_orf1ab;625 | 0 | 0 |
| MN996532.1;QHR63299.1;BatCoV_RaTG13_orf1ab;626 | 0 | 0 |
| MN996532.1;QHR63299.1;BatCoV_RaTG13_orf1ab;627 | 0 | 0 |
| MN996532.1;QHR63299.1;BatCoV_RaTG13_orf1ab;628 | 0 | 0.025 |
| MN996532.1;QHR63299.1;BatCoV_RaTG13_orf1ab;629 | 0 | 0.025 |
| MN996532.1;QHR63299.1;BatCoV_RaTG13_orf1ab;630 | 0 | 0.025 |
| MN996532.1;QHR63299.1;BatCoV_RaTG13_orf1ab;631 | 0 | 0.05 |
| MN996532.1;QHR63299.1;BatCoV_RaTG13_orf1ab;632 | 0 | 0.05 |
| MN996532.1;QHR63299.1;BatCoV_RaTG13_orf1ab;633 | 0 | 0.025 |
| MN996532.1;QHR63299.1;BatCoV_RaTG13_orf1ab;634 | 0 | 0 |
| MN996532.1;QHR63299.1;BatCoV_RaTG13_orf1ab;635 | 0 | 0.025 |
| MN996532.1;QHR63299.1;BatCoV_RaTG13_orf1ab;636 | 0 | 0.05 |

| | | |
|---|---|---|
| MN996532.1;QHR63299.1;BatCoV_RaTG13_orf1ab;637 | 0 | 0 |
| MN996532.1;QHR63299.1;BatCoV_RaTG13_orf1ab;638 | 0.05 | 0.025 |
| MN996532.1;QHR63299.1;BatCoV_RaTG13_orf1ab;639 | 0 | 0 |
| MN996532.1;QHR63299.1;BatCoV_RaTG13_orf1ab;640 | 0 | 0 |
| MN996532.1;QHR63299.1;BatCoV_RaTG13_orf1ab;641 | 0 | 0 |
| MN996532.1;QHR63299.1;BatCoV_RaTG13_orf1ab;642 | 0 | 0 |
| MN996532.1;QHR63299.1;BatCoV_RaTG13_orf1ab;643 | 0 | 0 |
| MN996532.1;QHR63299.1;BatCoV_RaTG13_orf1ab;644 | 0 | 0 |
| MN996532.1;QHR63299.1;BatCoV_RaTG13_orf1ab;645 | 0 | 0 |
| MN996532.1;QHR63299.1;BatCoV_RaTG13_orf1ab;646 | 0 | 0 |
| MN996532.1;QHR63299.1;BatCoV_RaTG13_orf1ab;647 | 0 | 0 |
| MN996532.1;QHR63299.1;BatCoV_RaTG13_orf1ab;648 | 0 | 0 |
| MN996532.1;QHR63299.1;BatCoV_RaTG13_orf1ab;649 | 0 | 0 |
| MN996532.1;QHR63299.1;BatCoV_RaTG13_orf1ab;650 | 0 | 0 |
| MN996532.1;QHR63299.1;BatCoV_RaTG13_orf1ab;651 | 0 | 0 |
| MN996532.1;QHR63299.1;BatCoV_RaTG13_orf1ab;652 | 0 | 0 |
| MN996532.1;QHR63299.1;BatCoV_RaTG13_orf1ab;653 | 0 | 0 |
| MN996532.1;QHR63299.1;BatCoV_RaTG13_orf1ab;654 | 0 | 0 |
| MN996532.1;QHR63299.1;BatCoV_RaTG13_orf1ab;655 | 0 | 0 |
| MN996532.1;QHR63299.1;BatCoV_RaTG13_orf1ab;656 | 0 | 0 |
| MN996532.1;QHR63299.1;BatCoV_RaTG13_orf1ab;657 | 0 | 0.025 |
| MN996532.1;QHR63299.1;BatCoV_RaTG13_orf1ab;658 | 0 | 0.05 |
| MN996532.1;QHR63299.1;BatCoV_RaTG13_orf1ab;659 | 0 | 0.025 |
| MN996532.1;QHR63299.1;BatCoV_RaTG13_orf1ab;660 | 0 | 0.025 |
| MN996532.1;QHR63299.1;BatCoV_RaTG13_orf1ab;661 | 0 | 0.025 |
| MN996532.1;QHR63299.1;BatCoV_RaTG13_orf1ab;662 | 0 | 0.025 |
| MN996532.1;QHR63299.1;BatCoV_RaTG13_orf1ab;663 | 0 | 0.025 |
| MN996532.1;QHR63299.1;BatCoV_RaTG13_orf1ab;664 | 0 | 0.025 |
| MN996532.1;QHR63299.1;BatCoV_RaTG13_orf1ab;665 | 0 | 0.025 |
| MN996532.1;QHR63299.1;BatCoV_RaTG13_orf1ab;666 | 0 | 0.025 |
| MN996532.1;QHR63299.1;BatCoV_RaTG13_orf1ab;667 | 0 | 0.025 |
| MN996532.1;QHR63299.1;BatCoV_RaTG13_orf1ab;668 | 0 | 0 |
| MN996532.1;QHR63299.1;BatCoV_RaTG13_orf1ab;669 | 0 | 0 |
| MN996532.1;QHR63299.1;BatCoV_RaTG13_orf1ab;670 | 0 | 0 |
| MN996532.1;QHR63299.1;BatCoV_RaTG13_orf1ab;671 | 0 | 0 |
| MN996532.1;QHR63299.1;BatCoV_RaTG13_orf1ab;672 | 0 | 0 |
| MN996532.1;QHR63299.1;BatCoV_RaTG13_orf1ab;673 | 0 | 0 |
| MN996532.1;QHR63299.1;BatCoV_RaTG13_orf1ab;674 | 0 | 0 |
| MN996532.1;QHR63299.1;BatCoV_RaTG13_orf1ab;675 | 0 | 0 |
| MN996532.1;QHR63299.1;BatCoV_RaTG13_orf1ab;676 | 0 | 0 |
| MN996532.1;QHR63299.1;BatCoV_RaTG13_orf1ab;677 | 0 | 0 |
| MN996532.1;QHR63299.1;BatCoV_RaTG13_orf1ab;678 | 0 | 0 |
| MN996532.1;QHR63299.1;BatCoV_RaTG13_orf1ab;679 | 0 | 0 |

| | | |
|---|---|---|
| MN996532.1;QHR63299.1;BatCoV_RaTG13_orf1ab;680 | 0 | 0 |
| MN996532.1;QHR63299.1;BatCoV_RaTG13_orf1ab;681 | 0 | 0 |
| MN996532.1;QHR63299.1;BatCoV_RaTG13_orf1ab;682 | 0 | 0 |
| MN996532.1;QHR63299.1;BatCoV_RaTG13_orf1ab;683 | 0 | 0 |
| MN996532.1;QHR63299.1;BatCoV_RaTG13_orf1ab;684 | 0 | 0 |
| MN996532.1;QHR63299.1;BatCoV_RaTG13_orf1ab;685 | 0 | 0 |
| MN996532.1;QHR63299.1;BatCoV_RaTG13_orf1ab;686 | 0 | 0 |
| MN996532.1;QHR63299.1;BatCoV_RaTG13_orf1ab;687 | 0 | 0 |
| MN996532.1;QHR63299.1;BatCoV_RaTG13_orf1ab;688 | 0 | 0 |
| MN996532.1;QHR63299.1;BatCoV_RaTG13_orf1ab;689 | 0 | 0 |
| MN996532.1;QHR63299.1;BatCoV_RaTG13_orf1ab;690 | 0 | 0 |
| MN996532.1;QHR63299.1;BatCoV_RaTG13_orf1ab;691 | 0 | 0 |
| MN996532.1;QHR63299.1;BatCoV_RaTG13_orf1ab;692 | 0 | 0 |
| MN996532.1;QHR63299.1;BatCoV_RaTG13_orf1ab;693 | 0 | 0.025 |
| MN996532.1;QHR63299.1;BatCoV_RaTG13_orf1ab;694 | 0 | 0 |
| MN996532.1;QHR63299.1;BatCoV_RaTG13_orf1ab;695 | 0 | 0 |
| MN996532.1;QHR63299.1;BatCoV_RaTG13_orf1ab;696 | 0 | 0.025 |
| MN996532.1;QHR63299.1;BatCoV_RaTG13_orf1ab;697 | 0 | 0 |
| MN996532.1;QHR63299.1;BatCoV_RaTG13_orf1ab;698 | 0 | 0 |
| MN996532.1;QHR63299.1;BatCoV_RaTG13_orf1ab;699 | 0 | 0 |
| MN996532.1;QHR63299.1;BatCoV_RaTG13_orf1ab;700 | 0 | 0 |
| MN996532.1;QHR63299.1;BatCoV_RaTG13_orf1ab;701 | 0 | 0 |
| MN996532.1;QHR63299.1;BatCoV_RaTG13_orf1ab;702 | 0 | 0 |
| MN996532.1;QHR63299.1;BatCoV_RaTG13_orf1ab;703 | 0 | 0 |
| MN996532.1;QHR63299.1;BatCoV_RaTG13_orf1ab;704 | 0 | 0 |
| MN996532.1;QHR63299.1;BatCoV_RaTG13_orf1ab;705 | 0 | 0 |
| MN996532.1;QHR63299.1;BatCoV_RaTG13_orf1ab;706 | 0 | 0 |
| MN996532.1;QHR63299.1;BatCoV_RaTG13_orf1ab;707 | 0 | 0 |
| MN996532.1;QHR63299.1;BatCoV_RaTG13_orf1ab;708 | 0 | 0 |
| MN996532.1;QHR63299.1;BatCoV_RaTG13_orf1ab;709 | 0 | 0 |
| MN996532.1;QHR63299.1;BatCoV_RaTG13_orf1ab;710 | 0 | 0 |
| MN996532.1;QHR63299.1;BatCoV_RaTG13_orf1ab;711 | 0 | 0 |
| MN996532.1;QHR63299.1;BatCoV_RaTG13_orf1ab;712 | 0.05 | 0 |
| MN996532.1;QHR63299.1;BatCoV_RaTG13_orf1ab;713 | 0 | 0 |
| MN996532.1;QHR63299.1;BatCoV_RaTG13_orf1ab;714 | 0 | 0.025 |
| MN996532.1;QHR63299.1;BatCoV_RaTG13_orf1ab;715 | 0 | 0.075 |
| MN996532.1;QHR63299.1;BatCoV_RaTG13_orf1ab;716 | 0 | 0.05 |
| MN996532.1;QHR63299.1;BatCoV_RaTG13_orf1ab;717 | 0 | 0.05 |
| MN996532.1;QHR63299.1;BatCoV_RaTG13_orf1ab;718 | 0 | 0.075 |
| MN996532.1;QHR63299.1;BatCoV_RaTG13_orf1ab;719 | 0 | 0.05 |
| MN996532.1;QHR63299.1;BatCoV_RaTG13_orf1ab;720 | 0 | 0.05 |
| MN996532.1;QHR63299.1;BatCoV_RaTG13_orf1ab;721 | 0 | 0.05 |
| MN996532.1;QHR63299.1;BatCoV_RaTG13_orf1ab;722 | 0 | 0.05 |

| | | |
|---|---|---|
| MN996532.1;QHR63299.1;BatCoV_RaTG13_orf1ab;723 | 0 | 0.05 |
| MN996532.1;QHR63299.1;BatCoV_RaTG13_orf1ab;724 | 0 | 0.075 |
| MN996532.1;QHR63299.1;BatCoV_RaTG13_orf1ab;725 | 0 | 0 |
| MN996532.1;QHR63299.1;BatCoV_RaTG13_orf1ab;726 | 0.05 | 0 |
| MN996532.1;QHR63299.1;BatCoV_RaTG13_orf1ab;727 | 0 | 0 |
| MN996532.1;QHR63299.1;BatCoV_RaTG13_orf1ab;728 | 0 | 0 |
| MN996532.1;QHR63299.1;BatCoV_RaTG13_orf1ab;729 | 0 | 0 |
| MN996532.1;QHR63299.1;BatCoV_RaTG13_orf1ab;730 | 0 | 0 |
| MN996532.1;QHR63299.1;BatCoV_RaTG13_orf1ab;731 | 0 | 0 |
| MN996532.1;QHR63299.1;BatCoV_RaTG13_orf1ab;732 | 0 | 0 |
| MN996532.1;QHR63299.1;BatCoV_RaTG13_orf1ab;733 | 0 | 0 |
| MN996532.1;QHR63299.1;BatCoV_RaTG13_orf1ab;734 | 0 | 0 |
| MN996532.1;QHR63299.1;BatCoV_RaTG13_orf1ab;735 | 0 | 0 |
| MN996532.1;QHR63299.1;BatCoV_RaTG13_orf1ab;736 | 0 | 0 |
| MN996532.1;QHR63299.1;BatCoV_RaTG13_orf1ab;737 | 0 | 0 |
| MN996532.1;QHR63299.1;BatCoV_RaTG13_orf1ab;738 | 0 | 0 |
| MN996532.1;QHR63299.1;BatCoV_RaTG13_orf1ab;739 | 0 | 0 |
| MN996532.1;QHR63299.1;BatCoV_RaTG13_orf1ab;740 | 0 | 0 |
| MN996532.1;QHR63299.1;BatCoV_RaTG13_orf1ab;741 | 0 | 0 |
| MN996532.1;QHR63299.1;BatCoV_RaTG13_orf1ab;742 | 0 | 0 |
| MN996532.1;QHR63299.1;BatCoV_RaTG13_orf1ab;743 | 0 | 0 |
| MN996532.1;QHR63299.1;BatCoV_RaTG13_orf1ab;744 | 0.15 | 0.05 |
| MN996532.1;QHR63299.1;BatCoV_RaTG13_orf1ab;745 | 0.1 | 0.05 |
| MN996532.1;QHR63299.1;BatCoV_RaTG13_orf1ab;746 | 0.15 | 0.025 |
| MN996532.1;QHR63299.1;BatCoV_RaTG13_orf1ab;747 | 0.1 | 0.025 |
| MN996532.1;QHR63299.1;BatCoV_RaTG13_orf1ab;748 | 0.1 | 0.05 |
| MN996532.1;QHR63299.1;BatCoV_RaTG13_orf1ab;749 | 0.15 | 0.075 |
| MN996532.1;QHR63299.1;BatCoV_RaTG13_orf1ab;750 | 0.15 | 0.05 |
| MN996532.1;QHR63299.1;BatCoV_RaTG13_orf1ab;751 | 0.05 | 0.05 |
| MN996532.1;QHR63299.1;BatCoV_RaTG13_orf1ab;752 | 0.05 | 0.05 |
| MN996532.1;QHR63299.1;BatCoV_RaTG13_orf1ab;753 | 0.05 | 0.075 |
| MN996532.1;QHR63299.1;BatCoV_RaTG13_orf1ab;754 | 0.1 | 0.075 |
| MN996532.1;QHR63299.1;BatCoV_RaTG13_orf1ab;755 | 0.05 | 0.05 |
| MN996532.1;QHR63299.1;BatCoV_RaTG13_orf1ab;756 | 0.05 | 0.05 |
| MN996532.1;QHR63299.1;BatCoV_RaTG13_orf1ab;757 | 0.05 | 0.05 |
| MN996532.1;QHR63299.1;BatCoV_RaTG13_orf1ab;758 | 0.05 | 0.05 |
| MN996532.1;QHR63299.1;BatCoV_RaTG13_orf1ab;759 | 0.05 | 0.075 |
| MN996532.1;QHR63299.1;BatCoV_RaTG13_orf1ab;760 | 0 | 0.025 |
| MN996532.1;QHR63299.1;BatCoV_RaTG13_orf1ab;761 | 0 | 0.025 |
| MN996532.1;QHR63299.1;BatCoV_RaTG13_orf1ab;762 | 0 | 0.025 |
| MN996532.1;QHR63299.1;BatCoV_RaTG13_orf1ab;763 | 0 | 0.025 |
| MN996532.1;QHR63299.1;BatCoV_RaTG13_orf1ab;764 | 0.05 | 0.025 |
| MN996532.1;QHR63299.1;BatCoV_RaTG13_orf1ab;765 | 0.05 | 0.025 |

| | | |
|---|---|---|
| MN996532.1;QHR63299.1;BatCoV_RaTG13_orf1ab;766 | 0.05 | 0.075 |
| MN996532.1;QHR63299.1;BatCoV_RaTG13_orf1ab;767 | 0.05 | 0.05 |
| MN996532.1;QHR63299.1;BatCoV_RaTG13_orf1ab;768 | 0 | 0.075 |
| MN996532.1;QHR63299.1;BatCoV_RaTG13_orf1ab;769 | 0 | 0 |
| MN996532.1;QHR63299.1;BatCoV_RaTG13_orf1ab;770 | 0 | 0 |
| MN996532.1;QHR63299.1;BatCoV_RaTG13_orf1ab;771 | 0 | 0 |
| MN996532.1;QHR63299.1;BatCoV_RaTG13_orf1ab;772 | 0 | 0 |
| MN996532.1;QHR63299.1;BatCoV_RaTG13_orf1ab;773 | 0 | 0 |
| MN996532.1;QHR63299.1;BatCoV_RaTG13_orf1ab;774 | 0 | 0 |
| MN996532.1;QHR63299.1;BatCoV_RaTG13_orf1ab;775 | 0.05 | 0 |
| MN996532.1;QHR63299.1;BatCoV_RaTG13_orf1ab;776 | 0 | 0 |
| MN996532.1;QHR63299.1;BatCoV_RaTG13_orf1ab;777 | 0 | 0 |
| MN996532.1;QHR63299.1;BatCoV_RaTG13_orf1ab;778 | 0 | 0 |
| MN996532.1;QHR63299.1;BatCoV_RaTG13_orf1ab;779 | 0 | 0 |
| MN996532.1;QHR63299.1;BatCoV_RaTG13_orf1ab;780 | 0 | 0.025 |
| MN996532.1;QHR63299.1;BatCoV_RaTG13_orf1ab;781 | 0 | 0 |
| MN996532.1;QHR63299.1;BatCoV_RaTG13_orf1ab;782 | 0.05 | 0 |
| MN996532.1;QHR63299.1;BatCoV_RaTG13_orf1ab;783 | 0 | 0 |
| MN996532.1;QHR63299.1;BatCoV_RaTG13_orf1ab;784 | 0 | 0 |
| MN996532.1;QHR63299.1;BatCoV_RaTG13_orf1ab;785 | 0 | 0 |
| MN996532.1;QHR63299.1;BatCoV_RaTG13_orf1ab;786 | 0 | 0 |
| MN996532.1;QHR63299.1;BatCoV_RaTG13_orf1ab;787 | 0 | 0.025 |
| MN996532.1;QHR63299.1;BatCoV_RaTG13_orf1ab;788 | 0 | 0.025 |
| MN996532.1;QHR63299.1;BatCoV_RaTG13_orf1ab;789 | 0.05 | 0 |
| MN996532.1;QHR63299.1;BatCoV_RaTG13_orf1ab;790 | 0 | 0.025 |
| MN996532.1;QHR63299.1;BatCoV_RaTG13_orf1ab;791 | 0 | 0.05 |
| MN996532.1;QHR63299.1;BatCoV_RaTG13_orf1ab;792 | 0 | 0.05 |
| MN996532.1;QHR63299.1;BatCoV_RaTG13_orf1ab;793 | 0 | 0.025 |
| MN996532.1;QHR63299.1;BatCoV_RaTG13_orf1ab;794 | 0 | 0 |
| MN996532.1;QHR63299.1;BatCoV_RaTG13_orf1ab;795 | 0 | 0 |
| MN996532.1;QHR63299.1;BatCoV_RaTG13_orf1ab;796 | 0 | 0 |
| MN996532.1;QHR63299.1;BatCoV_RaTG13_orf1ab;797 | 0.05 | 0.05 |
| MN996532.1;QHR63299.1;BatCoV_RaTG13_orf1ab;798 | 0.05 | 0 |
| MN996532.1;QHR63299.1;BatCoV_RaTG13_orf1ab;799 | 0 | 0 |
| MN996532.1;QHR63299.1;BatCoV_RaTG13_orf1ab;800 | 0 | 0 |
| MN996532.1;QHR63299.1;BatCoV_RaTG13_orf1ab;801 | 0.05 | 0 |
| MN996532.1;QHR63299.1;BatCoV_RaTG13_orf1ab;802 | 0 | 0 |
| MN996532.1;QHR63299.1;BatCoV_RaTG13_orf1ab;803 | 0.1 | 0 |
| MN996532.1;QHR63299.1;BatCoV_RaTG13_orf1ab;804 | 0.05 | 0 |
| MN996532.1;QHR63299.1;BatCoV_RaTG13_orf1ab;805 | 0.05 | 0 |
| MN996532.1;QHR63299.1;BatCoV_RaTG13_orf1ab;806 | 0.05 | 0 |
| MN996532.1;QHR63299.1;BatCoV_RaTG13_orf1ab;807 | 0 | 0.025 |
| MN996532.1;QHR63299.1;BatCoV_RaTG13_orf1ab;808 | 0 | 0 |

| | | |
|---|---|---|
| MN996532.1;QHR63299.1;BatCoV_RaTG13_orf1ab;809 | 0 | 0 |
| MN996532.1;QHR63299.1;BatCoV_RaTG13_orf1ab;810 | 0 | 0 |
| MN996532.1;QHR63299.1;BatCoV_RaTG13_orf1ab;811 | 0 | 0 |
| MN996532.1;QHR63299.1;BatCoV_RaTG13_orf1ab;812 | 0 | 0 |
| MN996532.1;QHR63299.1;BatCoV_RaTG13_orf1ab;813 | 0.05 | 0 |
| MN996532.1;QHR63299.1;BatCoV_RaTG13_orf1ab;814 | 0 | 0.025 |
| MN996532.1;QHR63299.1;BatCoV_RaTG13_orf1ab;815 | 0 | 0 |
| MN996532.1;QHR63299.1;BatCoV_RaTG13_orf1ab;816 | 0 | 0 |
| MN996532.1;QHR63299.1;BatCoV_RaTG13_orf1ab;817 | 0 | 0 |
| MN996532.1;QHR63299.1;BatCoV_RaTG13_orf1ab;818 | 0 | 0 |
| MN996532.1;QHR63299.1;BatCoV_RaTG13_orf1ab;819 | 0.05 | 0 |
| MN996532.1;QHR63299.1;BatCoV_RaTG13_orf1ab;820 | 0.05 | 0 |
| MN996532.1;QHR63299.1;BatCoV_RaTG13_orf1ab;821 | 0 | 0 |
| MN996532.1;QHR63299.1;BatCoV_RaTG13_orf1ab;822 | 0 | 0.025 |
| MN996532.1;QHR63299.1;BatCoV_RaTG13_orf1ab;823 | 0 | 0.025 |
| MN996532.1;QHR63299.1;BatCoV_RaTG13_orf1ab;824 | 0 | 0.025 |
| MN996532.1;QHR63299.1;BatCoV_RaTG13_orf1ab;825 | 0 | 0.025 |
| MN996532.1;QHR63299.1;BatCoV_RaTG13_orf1ab;826 | 0 | 0 |
| MN996532.1;QHR63299.1;BatCoV_RaTG13_orf1ab;827 | 0.1 | 0 |
| MN996532.1;QHR63299.1;BatCoV_RaTG13_orf1ab;828 | 0 | 0 |
| MN996532.1;QHR63299.1;BatCoV_RaTG13_orf1ab;829 | 0 | 0 |
| MN996532.1;QHR63299.1;BatCoV_RaTG13_orf1ab;830 | 0 | 0 |
| MN996532.1;QHR63299.1;BatCoV_RaTG13_orf1ab;831 | 0.05 | 0.05 |
| MN996532.1;QHR63299.1;BatCoV_RaTG13_orf1ab;832 | 0 | 0 |
| MN996532.1;QHR63299.1;BatCoV_RaTG13_orf1ab;833 | 0 | 0 |
| MN996532.1;QHR63299.1;BatCoV_RaTG13_orf1ab;834 | 0 | 0 |
| MN996532.1;QHR63299.1;BatCoV_RaTG13_orf1ab;835 | 0 | 0 |
| MN996532.1;QHR63299.1;BatCoV_RaTG13_orf1ab;836 | 0.05 | 0 |
| MN996532.1;QHR63299.1;BatCoV_RaTG13_orf1ab;837 | 0.05 | 0 |
| MN996532.1;QHR63299.1;BatCoV_RaTG13_orf1ab;838 | 0 | 0 |
| MN996532.1;QHR63299.1;BatCoV_RaTG13_orf1ab;839 | 0 | 0.025 |
| MN996532.1;QHR63299.1;BatCoV_RaTG13_orf1ab;840 | 0.2 | 0.1 |
| MN996532.1;QHR63299.1;BatCoV_RaTG13_orf1ab;841 | 0.1 | 0.15 |
| MN996532.1;QHR63299.1;BatCoV_RaTG13_orf1ab;842 | 0.2 | 0.1 |
| MN996532.1;QHR63299.1;BatCoV_RaTG13_orf1ab;843 | 0.25 | 0.1 |
| MN996532.1;QHR63299.1;BatCoV_RaTG13_orf1ab;844 | 0.05 | 0.075 |
| MN996532.1;QHR63299.1;BatCoV_RaTG13_orf1ab;845 | 0 | 0 |
| MN996532.1;QHR63299.1;BatCoV_RaTG13_orf1ab;846 | 0 | 0.075 |
| MN996532.1;QHR63299.1;BatCoV_RaTG13_orf1ab;847 | 0 | 0.025 |
| MN996532.1;QHR63299.1;BatCoV_RaTG13_orf1ab;848 | 0 | 0 |
| MN996532.1;QHR63299.1;BatCoV_RaTG13_orf1ab;849 | 0 | 0 |
| MN996532.1;QHR63299.1;BatCoV_RaTG13_orf1ab;850 | 0 | 0 |
| MN996532.1;QHR63299.1;BatCoV_RaTG13_orf1ab;851 | 0 | 0 |

| | | |
|---|---|---|
| MN996532.1;QHR63299.1;BatCoV_RaTG13_orf1ab;852 | 0 | 0 |
| MN996532.1;QHR63299.1;BatCoV_RaTG13_orf1ab;853 | 0 | 0 |
| MN996532.1;QHR63299.1;BatCoV_RaTG13_orf1ab;854 | 0 | 0 |
| MN996532.1;QHR63299.1;BatCoV_RaTG13_orf1ab;855 | 0 | 0 |
| MN996532.1;QHR63299.1;BatCoV_RaTG13_orf1ab;856 | 0 | 0 |
| MN996532.1;QHR63299.1;BatCoV_RaTG13_orf1ab;857 | 0 | 0 |
| MN996532.1;QHR63299.1;BatCoV_RaTG13_orf1ab;858 | 0 | 0 |
| MN996532.1;QHR63299.1;BatCoV_RaTG13_orf1ab;859 | 0 | 0 |
| MN996532.1;QHR63299.1;BatCoV_RaTG13_orf1ab;860 | 0 | 0 |
| MN996532.1;QHR63299.1;BatCoV_RaTG13_orf1ab;861 | 0 | 0 |
| MN996532.1;QHR63299.1;BatCoV_RaTG13_orf1ab;862 | 0 | 0 |
| MN996532.1;QHR63299.1;BatCoV_RaTG13_orf1ab;863 | 0 | 0 |
| MN996532.1;QHR63299.1;BatCoV_RaTG13_orf1ab;864 | 0 | 0 |
| MN996532.1;QHR63299.1;BatCoV_RaTG13_orf1ab;865 | 0 | 0 |
| MN996532.1;QHR63299.1;BatCoV_RaTG13_orf1ab;866 | 0 | 0 |
| MN996532.1;QHR63299.1;BatCoV_RaTG13_orf1ab;867 | 0 | 0 |
| MN996532.1;QHR63299.1;BatCoV_RaTG13_orf1ab;868 | 0 | 0 |
| MN996532.1;QHR63299.1;BatCoV_RaTG13_orf1ab;869 | 0 | 0 |
| MN996532.1;QHR63299.1;BatCoV_RaTG13_orf1ab;870 | 0 | 0 |
| MN996532.1;QHR63299.1;BatCoV_RaTG13_orf1ab;871 | 0 | 0 |
| MN996532.1;QHR63299.1;BatCoV_RaTG13_orf1ab;872 | 0 | 0 |
| MN996532.1;QHR63299.1;BatCoV_RaTG13_orf1ab;873 | 0 | 0 |
| MN996532.1;QHR63299.1;BatCoV_RaTG13_orf1ab;874 | 0 | 0 |
| MN996532.1;QHR63299.1;BatCoV_RaTG13_orf1ab;875 | 0 | 0.025 |
| MN996532.1;QHR63299.1;BatCoV_RaTG13_orf1ab;876 | 0.05 | 0 |
| MN996532.1;QHR63299.1;BatCoV_RaTG13_orf1ab;877 | 0 | 0.025 |
| MN996532.1;QHR63299.1;BatCoV_RaTG13_orf1ab;878 | 0 | 0.05 |
| MN996532.1;QHR63299.1;BatCoV_RaTG13_orf1ab;879 | 0 | 0.025 |
| MN996532.1;QHR63299.1;BatCoV_RaTG13_orf1ab;880 | 0 | 0.025 |
| MN996532.1;QHR63299.1;BatCoV_RaTG13_orf1ab;881 | 0 | 0.025 |
| MN996532.1;QHR63299.1;BatCoV_RaTG13_orf1ab;882 | 0 | 0 |
| MN996532.1;QHR63299.1;BatCoV_RaTG13_orf1ab;883 | 0 | 0 |
| MN996532.1;QHR63299.1;BatCoV_RaTG13_orf1ab;884 | 0 | 0 |
| MN996532.1;QHR63299.1;BatCoV_RaTG13_orf1ab;885 | 0 | 0 |
| MN996532.1;QHR63299.1;BatCoV_RaTG13_orf1ab;886 | 0 | 0 |
| MN996532.1;QHR63299.1;BatCoV_RaTG13_orf1ab;887 | 0 | 0 |
| MN996532.1;QHR63299.1;BatCoV_RaTG13_orf1ab;888 | 0 | 0 |
| MN996532.1;QHR63299.1;BatCoV_RaTG13_orf1ab;889 | 0 | 0 |
| MN996532.1;QHR63299.1;BatCoV_RaTG13_orf1ab;890 | 0.05 | 0 |
| MN996532.1;QHR63299.1;BatCoV_RaTG13_orf1ab;891 | 0 | 0 |
| MN996532.1;QHR63299.1;BatCoV_RaTG13_orf1ab;892 | 0 | 0 |
| MN996532.1;QHR63299.1;BatCoV_RaTG13_orf1ab;893 | 0 | 0 |
| MN996532.1;QHR63299.1;BatCoV_RaTG13_orf1ab;894 | 0 | 0 |

| | | |
|---|---|---|
| MN996532.1;QHR63299.1;BatCoV_RaTG13_orf1ab;895 | 0 | 0 |
| MN996532.1;QHR63299.1;BatCoV_RaTG13_orf1ab;896 | 0 | 0 |
| MN996532.1;QHR63299.1;BatCoV_RaTG13_orf1ab;897 | 0 | 0 |
| MN996532.1;QHR63299.1;BatCoV_RaTG13_orf1ab;898 | 0 | 0 |
| MN996532.1;QHR63299.1;BatCoV_RaTG13_orf1ab;899 | 0 | 0 |
| MN996532.1;QHR63299.1;BatCoV_RaTG13_orf1ab;900 | 0.05 | 0 |
| MN996532.1;QHR63299.1;BatCoV_RaTG13_orf1ab;901 | 0 | 0 |
| MN996532.1;QHR63299.1;BatCoV_RaTG13_orf1ab;902 | 0 | 0 |
| MN996532.1;QHR63299.1;BatCoV_RaTG13_orf1ab;903 | 0 | 0 |
| MN996532.1;QHR63299.1;BatCoV_RaTG13_orf1ab;904 | 0 | 0 |
| MN996532.1;QHR63299.1;BatCoV_RaTG13_orf1ab;905 | 0 | 0 |
| MN996532.1;QHR63299.1;BatCoV_RaTG13_orf1ab;906 | 0.05 | 0 |
| MN996532.1;QHR63299.1;BatCoV_RaTG13_orf1ab;907 | 0 | 0 |
| MN996532.1;QHR63299.1;BatCoV_RaTG13_orf1ab;908 | 0 | 0 |
| MN996532.1;QHR63299.1;BatCoV_RaTG13_orf1ab;909 | 0 | 0 |
| MN996532.1;QHR63299.1;BatCoV_RaTG13_orf1ab;910 | 0 | 0 |
| MN996532.1;QHR63299.1;BatCoV_RaTG13_orf1ab;911 | 0 | 0 |
| MN996532.1;QHR63299.1;BatCoV_RaTG13_orf1ab;912 | 0 | 0 |
| MN996532.1;QHR63299.1;BatCoV_RaTG13_orf1ab;913 | 0 | 0 |
| MN996532.1;QHR63299.1;BatCoV_RaTG13_orf1ab;914 | 0 | 0 |
| MN996532.1;QHR63299.1;BatCoV_RaTG13_orf1ab;915 | 0 | 0 |
| MN996532.1;QHR63299.1;BatCoV_RaTG13_orf1ab;916 | 0 | 0.025 |
| MN996532.1;QHR63299.1;BatCoV_RaTG13_orf1ab;917 | 0 | 0 |
| MN996532.1;QHR63299.1;BatCoV_RaTG13_orf1ab;918 | 0 | 0.025 |
| MN996532.1;QHR63299.1;BatCoV_RaTG13_orf1ab;919 | 0 | 0.05 |
| MN996532.1;QHR63299.1;BatCoV_RaTG13_orf1ab;920 | 0 | 0.075 |
| MN996532.1;QHR63299.1;BatCoV_RaTG13_orf1ab;921 | 0 | 0.1 |
| MN996532.1;QHR63299.1;BatCoV_RaTG13_orf1ab;922 | 0 | 0.1 |
| MN996532.1;QHR63299.1;BatCoV_RaTG13_orf1ab;923 | 0 | 0.1 |
| MN996532.1;QHR63299.1;BatCoV_RaTG13_orf1ab;924 | 0 | 0.125 |
| MN996532.1;QHR63299.1;BatCoV_RaTG13_orf1ab;925 | 0 | 0.075 |
| MN996532.1;QHR63299.1;BatCoV_RaTG13_orf1ab;926 | 0 | 0 |
| MN996532.1;QHR63299.1;BatCoV_RaTG13_orf1ab;927 | 0.05 | 0.025 |
| MN996532.1;QHR63299.1;BatCoV_RaTG13_orf1ab;928 | 0.1 | 0.025 |
| MN996532.1;QHR63299.1;BatCoV_RaTG13_orf1ab;929 | 0.35 | 0.25 |
| MN996532.1;QHR63299.1;BatCoV_RaTG13_orf1ab;930 | 0 | 0 |
| MN996532.1;QHR63299.1;BatCoV_RaTG13_orf1ab;931 | 0 | 0 |
| MN996532.1;QHR63299.1;BatCoV_RaTG13_orf1ab;932 | 0 | 0 |
| MN996532.1;QHR63299.1;BatCoV_RaTG13_orf1ab;933 | 0 | 0 |
| MN996532.1;QHR63299.1;BatCoV_RaTG13_orf1ab;934 | 0 | 0.025 |
| MN996532.1;QHR63299.1;BatCoV_RaTG13_orf1ab;935 | 0 | 0 |
| MN996532.1;QHR63299.1;BatCoV_RaTG13_orf1ab;936 | 0 | 0.025 |
| MN996532.1;QHR63299.1;BatCoV_RaTG13_orf1ab;937 | 0 | 0 |

| | | |
|---|---|---|
| MN996532.1;QHR63299.1;BatCoV_RaTG13_orf1ab;938 | 0 | 0 |
| MN996532.1;QHR63299.1;BatCoV_RaTG13_orf1ab;939 | 0 | 0 |
| MN996532.1;QHR63299.1;BatCoV_RaTG13_orf1ab;940 | 0 | 0 |
| MN996532.1;QHR63299.1;BatCoV_RaTG13_orf1ab;941 | 0 | 0 |
| MN996532.1;QHR63299.1;BatCoV_RaTG13_orf1ab;942 | 0 | 0 |
| MN996532.1;QHR63299.1;BatCoV_RaTG13_orf1ab;943 | 0 | 0 |
| MN996532.1;QHR63299.1;BatCoV_RaTG13_orf1ab;944 | 0 | 0.025 |
| MN996532.1;QHR63299.1;BatCoV_RaTG13_orf1ab;945 | 0 | 0 |
| MN996532.1;QHR63299.1;BatCoV_RaTG13_orf1ab;946 | 0 | 0 |
| MN996532.1;QHR63299.1;BatCoV_RaTG13_orf1ab;947 | 0 | 0 |
| MN996532.1;QHR63299.1;BatCoV_RaTG13_orf1ab;948 | 0 | 0 |
| MN996532.1;QHR63299.1;BatCoV_RaTG13_orf1ab;949 | 0 | 0.025 |
| MN996532.1;QHR63299.1;BatCoV_RaTG13_orf1ab;950 | 0 | 0.075 |
| MN996532.1;QHR63299.1;BatCoV_RaTG13_orf1ab;951 | 0 | 0.025 |
| MN996532.1;QHR63299.1;BatCoV_RaTG13_orf1ab;952 | 0 | 0.025 |
| MN996532.1;QHR63299.1;BatCoV_RaTG13_orf1ab;953 | 0 | 0.025 |
| MN996532.1;QHR63299.1;BatCoV_RaTG13_orf1ab;954 | 0 | 0.025 |
| MN996532.1;QHR63299.1;BatCoV_RaTG13_orf1ab;955 | 0 | 0.025 |
| MN996532.1;QHR63299.1;BatCoV_RaTG13_orf1ab;956 | 0 | 0.025 |
| MN996532.1;QHR63299.1;BatCoV_RaTG13_orf1ab;957 | 0 | 0.025 |
| MN996532.1;QHR63299.1;BatCoV_RaTG13_orf1ab;958 | 0 | 0.025 |
| MN996532.1;QHR63299.1;BatCoV_RaTG13_orf1ab;959 | 0 | 0.025 |
| MN996532.1;QHR63299.1;BatCoV_RaTG13_orf1ab;960 | 0 | 0 |
| MN996532.1;QHR63299.1;BatCoV_RaTG13_orf1ab;961 | 0 | 0 |
| MN996532.1;QHR63299.1;BatCoV_RaTG13_orf1ab;962 | 0 | 0 |
| MN996532.1;QHR63299.1;BatCoV_RaTG13_orf1ab;963 | 0 | 0 |
| MN996532.1;QHR63299.1;BatCoV_RaTG13_orf1ab;964 | 0 | 0.025 |
| MN996532.1;QHR63299.1;BatCoV_RaTG13_orf1ab;965 | 0 | 0.025 |
| MN996532.1;QHR63299.1;BatCoV_RaTG13_orf1ab;966 | 0 | 0.025 |
| MN996532.1;QHR63299.1;BatCoV_RaTG13_orf1ab;967 | 0 | 0.025 |
| MN996532.1;QHR63299.1;BatCoV_RaTG13_orf1ab;968 | 0 | 0.025 |
| MN996532.1;QHR63299.1;BatCoV_RaTG13_orf1ab;969 | 0 | 0.025 |
| MN996532.1;QHR63299.1;BatCoV_RaTG13_orf1ab;970 | 0.1 | 0.025 |
| MN996532.1;QHR63299.1;BatCoV_RaTG13_orf1ab;971 | 0 | 0.05 |
| MN996532.1;QHR63299.1;BatCoV_RaTG13_orf1ab;972 | 0 | 0.025 |
| MN996532.1;QHR63299.1;BatCoV_RaTG13_orf1ab;973 | 0 | 0.025 |
| MN996532.1;QHR63299.1;BatCoV_RaTG13_orf1ab;974 | 0 | 0.025 |
| MN996532.1;QHR63299.1;BatCoV_RaTG13_orf1ab;975 | 0 | 0.025 |
| MN996532.1;QHR63299.1;BatCoV_RaTG13_orf1ab;976 | 0 | 0.025 |
| MN996532.1;QHR63299.1;BatCoV_RaTG13_orf1ab;977 | 0.05 | 0.025 |
| MN996532.1;QHR63299.1;BatCoV_RaTG13_orf1ab;978 | 0 | 0.025 |
| MN996532.1;QHR63299.1;BatCoV_RaTG13_orf1ab;979 | 0 | 0 |
| MN996532.1;QHR63299.1;BatCoV_RaTG13_orf1ab;980 | 0 | 0 |

| | | |
|---|---|---|
| MN996532.1;QHR63299.1;BatCoV_RaTG13_orf1ab;981 | 0 | 0.025 |
| MN996532.1;QHR63299.1;BatCoV_RaTG13_orf1ab;982 | 0 | 0.025 |
| MN996532.1;QHR63299.1;BatCoV_RaTG13_orf1ab;983 | 0 | 0.025 |
| MN996532.1;QHR63299.1;BatCoV_RaTG13_orf1ab;984 | 0 | 0.025 |
| MN996532.1;QHR63299.1;BatCoV_RaTG13_orf1ab;985 | 0 | 0.025 |
| MN996532.1;QHR63299.1;BatCoV_RaTG13_orf1ab;986 | 0 | 0.025 |
| MN996532.1;QHR63299.1;BatCoV_RaTG13_orf1ab;987 | 0.05 | 0.025 |
| MN996532.1;QHR63299.1;BatCoV_RaTG13_orf1ab;988 | 0 | 0.025 |
| MN996532.1;QHR63299.1;BatCoV_RaTG13_orf1ab;989 | 0 | 0.025 |
| MN996532.1;QHR63299.1;BatCoV_RaTG13_orf1ab;990 | 0 | 0.025 |
| MN996532.1;QHR63299.1;BatCoV_RaTG13_orf1ab;991 | 0.1 | 0.025 |
| MN996532.1;QHR63299.1;BatCoV_RaTG13_orf1ab;992 | 0 | 0 |
| MN996532.1;QHR63299.1;BatCoV_RaTG13_orf1ab;993 | 0 | 0 |
| MN996532.1;QHR63299.1;BatCoV_RaTG13_orf1ab;994 | 0 | 0 |
| MN996532.1;QHR63299.1;BatCoV_RaTG13_orf1ab;995 | 0 | 0 |
| MN996532.1;QHR63299.1;BatCoV_RaTG13_orf1ab;996 | 0 | 0 |
| MN996532.1;QHR63299.1;BatCoV_RaTG13_orf1ab;997 | 0 | 0 |
| MN996532.1;QHR63299.1;BatCoV_RaTG13_orf1ab;998 | 0 | 0 |
| MN996532.1;QHR63299.1;BatCoV_RaTG13_orf1ab;999 | 0 | 0 |
| MN996532.1;QHR63299.1;BatCoV_RaTG13_orf1ab;1000 | 0 | 0 |
| MN996532.1;QHR63299.1;BatCoV_RaTG13_orf1ab;1001 | 0 | 0 |
| MN996532.1;QHR63299.1;BatCoV_RaTG13_orf1ab;1002 | 0 | 0 |
| MN996532.1;QHR63299.1;BatCoV_RaTG13_orf1ab;1003 | 0 | 0 |
| MN996532.1;QHR63299.1;BatCoV_RaTG13_orf1ab;1004 | 0 | 0 |
| MN996532.1;QHR63299.1;BatCoV_RaTG13_orf1ab;1005 | 0 | 0 |
| MN996532.1;QHR63299.1;BatCoV_RaTG13_orf1ab;1006 | 0 | 0 |
| MN996532.1;QHR63299.1;BatCoV_RaTG13_orf1ab;1007 | 0 | 0 |
| MN996532.1;QHR63299.1;BatCoV_RaTG13_orf1ab;1008 | 0 | 0 |
| MN996532.1;QHR63299.1;BatCoV_RaTG13_orf1ab;1009 | 0 | 0 |
| MN996532.1;QHR63299.1;BatCoV_RaTG13_orf1ab;1010 | 0 | 0 |
| MN996532.1;QHR63299.1;BatCoV_RaTG13_orf1ab;1011 | 0 | 0 |
| MN996532.1;QHR63299.1;BatCoV_RaTG13_orf1ab;1012 | 0 | 0 |
| MN996532.1;QHR63299.1;BatCoV_RaTG13_orf1ab;1013 | 0 | 0 |
| MN996532.1;QHR63299.1;BatCoV_RaTG13_orf1ab;1014 | 0 | 0.025 |
| MN996532.1;QHR63299.1;BatCoV_RaTG13_orf1ab;1015 | 0 | 0.025 |
| MN996532.1;QHR63299.1;BatCoV_RaTG13_orf1ab;1016 | 0 | 0.025 |
| MN996532.1;QHR63299.1;BatCoV_RaTG13_orf1ab;1017 | 0 | 0.025 |
| MN996532.1;QHR63299.1;BatCoV_RaTG13_orf1ab;1018 | 0 | 0 |
| MN996532.1;QHR63299.1;BatCoV_RaTG13_orf1ab;1019 | 0 | 0 |
| MN996532.1;QHR63299.1;BatCoV_RaTG13_orf1ab;1020 | 0 | 0 |
| MN996532.1;QHR63299.1;BatCoV_RaTG13_orf1ab;1021 | 0 | 0 |
| MN996532.1;QHR63299.1;BatCoV_RaTG13_orf1ab;1022 | 0.05 | 0 |
| MN996532.1;QHR63299.1;BatCoV_RaTG13_orf1ab;1023 | 0.05 | 0 |

| | | |
|---|---|---|
| MN996532.1;QHR63299.1;BatCoV_RaTG13_orf1ab;1024 | 0.05 | 0 |
| MN996532.1;QHR63299.1;BatCoV_RaTG13_orf1ab;1025 | 0.05 | 0 |
| MN996532.1;QHR63299.1;BatCoV_RaTG13_orf1ab;1026 | 0.05 | 0 |
| MN996532.1;QHR63299.1;BatCoV_RaTG13_orf1ab;1027 | 0.05 | 0 |
| MN996532.1;QHR63299.1;BatCoV_RaTG13_orf1ab;1028 | 0.05 | 0 |
| MN996532.1;QHR63299.1;BatCoV_RaTG13_orf1ab;1029 | 0.05 | 0.025 |
| MN996532.1;QHR63299.1;BatCoV_RaTG13_orf1ab;1030 | 0.05 | 0 |
| MN996532.1;QHR63299.1;BatCoV_RaTG13_orf1ab;1031 | 0.15 | 0.025 |
| MN996532.1;QHR63299.1;BatCoV_RaTG13_orf1ab;1032 | 0.2 | 0.025 |
| MN996532.1;QHR63299.1;BatCoV_RaTG13_orf1ab;1033 | 0.05 | 0.025 |
| MN996532.1;QHR63299.1;BatCoV_RaTG13_orf1ab;1034 | 0 | 0.05 |
| MN996532.1;QHR63299.1;BatCoV_RaTG13_orf1ab;1035 | 0 | 0.025 |
| MN996532.1;QHR63299.1;BatCoV_RaTG13_orf1ab;1036 | 0 | 0 |
| MN996532.1;QHR63299.1;BatCoV_RaTG13_orf1ab;1037 | 0 | 0 |
| MN996532.1;QHR63299.1;BatCoV_RaTG13_orf1ab;1038 | 0 | 0 |
| MN996532.1;QHR63299.1;BatCoV_RaTG13_orf1ab;1039 | 0 | 0 |
| MN996532.1;QHR63299.1;BatCoV_RaTG13_orf1ab;1040 | 0 | 0 |
| MN996532.1;QHR63299.1;BatCoV_RaTG13_orf1ab;1041 | 0 | 0 |
| MN996532.1;QHR63299.1;BatCoV_RaTG13_orf1ab;1042 | 0 | 0.025 |
| MN996532.1;QHR63299.1;BatCoV_RaTG13_orf1ab;1043 | 0 | 0 |
| MN996532.1;QHR63299.1;BatCoV_RaTG13_orf1ab;1044 | 0 | 0 |
| MN996532.1;QHR63299.1;BatCoV_RaTG13_orf1ab;1045 | 0 | 0 |
| MN996532.1;QHR63299.1;BatCoV_RaTG13_orf1ab;1046 | 0 | 0 |
| MN996532.1;QHR63299.1;BatCoV_RaTG13_orf1ab;1047 | 0 | 0 |
| MN996532.1;QHR63299.1;BatCoV_RaTG13_orf1ab;1048 | 0 | 0 |
| MN996532.1;QHR63299.1;BatCoV_RaTG13_orf1ab;1049 | 0 | 0.025 |
| MN996532.1;QHR63299.1;BatCoV_RaTG13_orf1ab;1050 | 0 | 0 |
| MN996532.1;QHR63299.1;BatCoV_RaTG13_orf1ab;1051 | 0 | 0 |
| MN996532.1;QHR63299.1;BatCoV_RaTG13_orf1ab;1052 | 0 | 0 |
| MN996532.1;QHR63299.1;BatCoV_RaTG13_orf1ab;1053 | 0 | 0 |
| MN996532.1;QHR63299.1;BatCoV_RaTG13_orf1ab;1054 | 0.05 | 0 |
| MN996532.1;QHR63299.1;BatCoV_RaTG13_orf1ab;1055 | 0 | 0 |
| MN996532.1;QHR63299.1;BatCoV_RaTG13_orf1ab;1056 | 0.05 | 0.05 |
| MN996532.1;QHR63299.1;BatCoV_RaTG13_orf1ab;1057 | 0 | 0 |
| MN996532.1;QHR63299.1;BatCoV_RaTG13_orf1ab;1058 | 0 | 0 |
| MN996532.1;QHR63299.1;BatCoV_RaTG13_orf1ab;1059 | 0 | 0 |
| MN996532.1;QHR63299.1;BatCoV_RaTG13_orf1ab;1060 | 0 | 0 |
| MN996532.1;QHR63299.1;BatCoV_RaTG13_orf1ab;1061 | 0 | 0 |
| MN996532.1;QHR63299.1;BatCoV_RaTG13_orf1ab;1062 | 0.05 | 0 |
| MN996532.1;QHR63299.1;BatCoV_RaTG13_orf1ab;1063 | 0 | 0 |
| MN996532.1;QHR63299.1;BatCoV_RaTG13_orf1ab;1064 | 0 | 0 |
| MN996532.1;QHR63299.1;BatCoV_RaTG13_orf1ab;1065 | 0 | 0.025 |
| MN996532.1;QHR63299.1;BatCoV_RaTG13_orf1ab;1066 | 0.1 | 0.025 |

| | | |
|---|---|---|
| MN996532.1;QHR63299.1;BatCoV_RaTG13_orf1ab;1067 | 0.15 | 0.025 |
| MN996532.1;QHR63299.1;BatCoV_RaTG13_orf1ab;1068 | 0.05 | 0.025 |
| MN996532.1;QHR63299.1;BatCoV_RaTG13_orf1ab;1069 | 0.05 | 0.025 |
| MN996532.1;QHR63299.1;BatCoV_RaTG13_orf1ab;1070 | 0.15 | 0.2 |
| MN996532.1;QHR63299.1;BatCoV_RaTG13_orf1ab;1071 | 0.2 | 0.2 |
| MN996532.1;QHR63299.1;BatCoV_RaTG13_orf1ab;1072 | 0.2 | 0.2 |
| MN996532.1;QHR63299.1;BatCoV_RaTG13_orf1ab;1073 | 0.1 | 0.15 |
| MN996532.1;QHR63299.1;BatCoV_RaTG13_orf1ab;1074 | 0.05 | 0.175 |
| MN996532.1;QHR63299.1;BatCoV_RaTG13_orf1ab;1075 | 0 | 0.05 |
| MN996532.1;QHR63299.1;BatCoV_RaTG13_orf1ab;1076 | 0 | 0.05 |
| MN996532.1;QHR63299.1;BatCoV_RaTG13_orf1ab;1077 | 0.1 | 0.075 |
| MN996532.1;QHR63299.1;BatCoV_RaTG13_orf1ab;1078 | 0 | 0 |
| MN996532.1;QHR63299.1;BatCoV_RaTG13_orf1ab;1079 | 0 | 0.025 |
| MN996532.1;QHR63299.1;BatCoV_RaTG13_orf1ab;1080 | 0 | 0.025 |
| MN996532.1;QHR63299.1;BatCoV_RaTG13_orf1ab;1081 | 0 | 0 |
| MN996532.1;QHR63299.1;BatCoV_RaTG13_orf1ab;1082 | 0 | 0.025 |
| MN996532.1;QHR63299.1;BatCoV_RaTG13_orf1ab;1083 | 0 | 0.025 |
| MN996532.1;QHR63299.1;BatCoV_RaTG13_orf1ab;1084 | 0 | 0.025 |
| MN996532.1;QHR63299.1;BatCoV_RaTG13_orf1ab;1085 | 0 | 0.05 |
| MN996532.1;QHR63299.1;BatCoV_RaTG13_orf1ab;1086 | 0.05 | 0.025 |
| MN996532.1;QHR63299.1;BatCoV_RaTG13_orf1ab;1087 | 0 | 0.05 |
| MN996532.1;QHR63299.1;BatCoV_RaTG13_orf1ab;1088 | 0 | 0 |
| MN996532.1;QHR63299.1;BatCoV_RaTG13_orf1ab;1089 | 0 | 0 |
| MN996532.1;QHR63299.1;BatCoV_RaTG13_orf1ab;1090 | 0 | 0.025 |
| MN996532.1;QHR63299.1;BatCoV_RaTG13_orf1ab;1091 | 0.05 | 0.05 |
| MN996532.1;QHR63299.1;BatCoV_RaTG13_orf1ab;1092 | 0 | 0 |
| MN996532.1;QHR63299.1;BatCoV_RaTG13_orf1ab;1093 | 0 | 0 |
| MN996532.1;QHR63299.1;BatCoV_RaTG13_orf1ab;1094 | 0 | 0 |
| MN996532.1;QHR63299.1;BatCoV_RaTG13_orf1ab;1095 | 0 | 0 |
| MN996532.1;QHR63299.1;BatCoV_RaTG13_orf1ab;1096 | 0 | 0 |
| MN996532.1;QHR63299.1;BatCoV_RaTG13_orf1ab;1097 | 0 | 0 |
| MN996532.1;QHR63299.1;BatCoV_RaTG13_orf1ab;1098 | 0 | 0 |
| MN996532.1;QHR63299.1;BatCoV_RaTG13_orf1ab;1099 | 0 | 0 |
| MN996532.1;QHR63299.1;BatCoV_RaTG13_orf1ab;1100 | 0 | 0 |
| MN996532.1;QHR63299.1;BatCoV_RaTG13_orf1ab;1101 | 0 | 0 |
| MN996532.1;QHR63299.1;BatCoV_RaTG13_orf1ab;1102 | 0 | 0 |
| MN996532.1;QHR63299.1;BatCoV_RaTG13_orf1ab;1103 | 0 | 0 |
| MN996532.1;QHR63299.1;BatCoV_RaTG13_orf1ab;1104 | 0 | 0 |
| MN996532.1;QHR63299.1;BatCoV_RaTG13_orf1ab;1105 | 0 | 0 |
| MN996532.1;QHR63299.1;BatCoV_RaTG13_orf1ab;1106 | 0 | 0 |
| MN996532.1;QHR63299.1;BatCoV_RaTG13_orf1ab;1107 | 0 | 0 |
| MN996532.1;QHR63299.1;BatCoV_RaTG13_orf1ab;1108 | 0 | 0 |
| MN996532.1;QHR63299.1;BatCoV_RaTG13_orf1ab;1109 | 0 | 0 |

| | | |
|---|---|---|
| MN996532.1;QHR63299.1;BatCoV_RaTG13_orf1ab;1110 | 0.05 | 0.025 |
| MN996532.1;QHR63299.1;BatCoV_RaTG13_orf1ab;1111 | 0.05 | 0.025 |
| MN996532.1;QHR63299.1;BatCoV_RaTG13_orf1ab;1112 | 0 | 0.025 |
| MN996532.1;QHR63299.1;BatCoV_RaTG13_orf1ab;1113 | 0.05 | 0.05 |
| MN996532.1;QHR63299.1;BatCoV_RaTG13_orf1ab;1114 | 0.05 | 0.075 |
| MN996532.1;QHR63299.1;BatCoV_RaTG13_orf1ab;1115 | 0 | 0.025 |
| MN996532.1;QHR63299.1;BatCoV_RaTG13_orf1ab;1116 | 0.25 | 0.125 |
| MN996532.1;QHR63299.1;BatCoV_RaTG13_orf1ab;1117 | 0.15 | 0.15 |
| MN996532.1;QHR63299.1;BatCoV_RaTG13_orf1ab;1118 | 0.2 | 0.125 |
| MN996532.1;QHR63299.1;BatCoV_RaTG13_orf1ab;1119 | 0.05 | 0.05 |
| MN996532.1;QHR63299.1;BatCoV_RaTG13_orf1ab;1120 | 0 | 0 |
| MN996532.1;QHR63299.1;BatCoV_RaTG13_orf1ab;1121 | 0 | 0.025 |
| MN996532.1;QHR63299.1;BatCoV_RaTG13_orf1ab;1122 | 0 | 0.025 |
| MN996532.1;QHR63299.1;BatCoV_RaTG13_orf1ab;1123 | 0 | 0.05 |
| MN996532.1;QHR63299.1;BatCoV_RaTG13_orf1ab;1124 | 0.05 | 0.025 |
| MN996532.1;QHR63299.1;BatCoV_RaTG13_orf1ab;1125 | 0.05 | 0.025 |
| MN996532.1;QHR63299.1;BatCoV_RaTG13_orf1ab;1126 | 0 | 0 |
| MN996532.1;QHR63299.1;BatCoV_RaTG13_orf1ab;1127 | 0 | 0.025 |
| MN996532.1;QHR63299.1;BatCoV_RaTG13_orf1ab;1128 | 0 | 0.025 |
| MN996532.1;QHR63299.1;BatCoV_RaTG13_orf1ab;1129 | 0 | 0 |
| MN996532.1;QHR63299.1;BatCoV_RaTG13_orf1ab;1130 | 0 | 0.025 |
| MN996532.1;QHR63299.1;BatCoV_RaTG13_orf1ab;1131 | 0 | 0.05 |
| MN996532.1;QHR63299.1;BatCoV_RaTG13_orf1ab;1132 | 0 | 0 |
| MN996532.1;QHR63299.1;BatCoV_RaTG13_orf1ab;1133 | 0 | 0 |
| MN996532.1;QHR63299.1;BatCoV_RaTG13_orf1ab;1134 | 0 | 0 |
| MN996532.1;QHR63299.1;BatCoV_RaTG13_orf1ab;1135 | 0 | 0 |
| MN996532.1;QHR63299.1;BatCoV_RaTG13_orf1ab;1136 | 0 | 0 |
| MN996532.1;QHR63299.1;BatCoV_RaTG13_orf1ab;1137 | 0 | 0 |
| MN996532.1;QHR63299.1;BatCoV_RaTG13_orf1ab;1138 | 0 | 0 |
| MN996532.1;QHR63299.1;BatCoV_RaTG13_orf1ab;1139 | 0 | 0 |
| MN996532.1;QHR63299.1;BatCoV_RaTG13_orf1ab;1140 | 0 | 0 |
| MN996532.1;QHR63299.1;BatCoV_RaTG13_orf1ab;1141 | 0 | 0 |
| MN996532.1;QHR63299.1;BatCoV_RaTG13_orf1ab;1142 | 0 | 0 |
| MN996532.1;QHR63299.1;BatCoV_RaTG13_orf1ab;1143 | 0.05 | 0 |
| MN996532.1;QHR63299.1;BatCoV_RaTG13_orf1ab;1144 | 0 | 0 |
| MN996532.1;QHR63299.1;BatCoV_RaTG13_orf1ab;1145 | 0 | 0 |
| MN996532.1;QHR63299.1;BatCoV_RaTG13_orf1ab;1146 | 0 | 0 |
| MN996532.1;QHR63299.1;BatCoV_RaTG13_orf1ab;1147 | 0 | 0 |
| MN996532.1;QHR63299.1;BatCoV_RaTG13_orf1ab;1148 | 0 | 0 |
| MN996532.1;QHR63299.1;BatCoV_RaTG13_orf1ab;1149 | 0 | 0.025 |
| MN996532.1;QHR63299.1;BatCoV_RaTG13_orf1ab;1150 | 0 | 0 |
| MN996532.1;QHR63299.1;BatCoV_RaTG13_orf1ab;1151 | 0 | 0 |
| MN996532.1;QHR63299.1;BatCoV_RaTG13_orf1ab;1152 | 0 | 0 |

| | | |
|---|---|---|
| MN996532.1;QHR63299.1;BatCoV_RaTG13_orf1ab;1153 | 0 | 0 |
| MN996532.1;QHR63299.1;BatCoV_RaTG13_orf1ab;1154 | 0 | 0 |
| MN996532.1;QHR63299.1;BatCoV_RaTG13_orf1ab;1155 | 0 | 0 |
| MN996532.1;QHR63299.1;BatCoV_RaTG13_orf1ab;1156 | 0 | 0 |
| MN996532.1;QHR63299.1;BatCoV_RaTG13_orf1ab;1157 | 0 | 0 |
| MN996532.1;QHR63299.1;BatCoV_RaTG13_orf1ab;1158 | 0 | 0 |
| MN996532.1;QHR63299.1;BatCoV_RaTG13_orf1ab;1159 | 0 | 0 |
| MN996532.1;QHR63299.1;BatCoV_RaTG13_orf1ab;1160 | 0 | 0 |
| MN996532.1;QHR63299.1;BatCoV_RaTG13_orf1ab;1161 | 0 | 0 |
| MN996532.1;QHR63299.1;BatCoV_RaTG13_orf1ab;1162 | 0 | 0 |
| MN996532.1;QHR63299.1;BatCoV_RaTG13_orf1ab;1163 | 0 | 0 |
| MN996532.1;QHR63299.1;BatCoV_RaTG13_orf1ab;1164 | 0 | 0 |
| MN996532.1;QHR63299.1;BatCoV_RaTG13_orf1ab;1165 | 0 | 0.05 |
| MN996532.1;QHR63299.1;BatCoV_RaTG13_orf1ab;1166 | 0 | 0 |
| MN996532.1;QHR63299.1;BatCoV_RaTG13_orf1ab;1167 | 0 | 0 |
| MN996532.1;QHR63299.1;BatCoV_RaTG13_orf1ab;1168 | 0 | 0 |
| MN996532.1;QHR63299.1;BatCoV_RaTG13_orf1ab;1169 | 0 | 0 |
| MN996532.1;QHR63299.1;BatCoV_RaTG13_orf1ab;1170 | 0 | 0 |
| MN996532.1;QHR63299.1;BatCoV_RaTG13_orf1ab;1171 | 0 | 0 |
| MN996532.1;QHR63299.1;BatCoV_RaTG13_orf1ab;1172 | 0.05 | 0 |
| MN996532.1;QHR63299.1;BatCoV_RaTG13_orf1ab;1173 | 0.05 | 0 |
| MN996532.1;QHR63299.1;BatCoV_RaTG13_orf1ab;1174 | 0.05 | 0 |
| MN996532.1;QHR63299.1;BatCoV_RaTG13_orf1ab;1175 | 0 | 0 |
| MN996532.1;QHR63299.1;BatCoV_RaTG13_orf1ab;1176 | 0.05 | 0 |
| MN996532.1;QHR63299.1;BatCoV_RaTG13_orf1ab;1177 | 0.05 | 0 |
| MN996532.1;QHR63299.1;BatCoV_RaTG13_orf1ab;1178 | 0.1 | 0 |
| MN996532.1;QHR63299.1;BatCoV_RaTG13_orf1ab;1179 | 0 | 0 |
| MN996532.1;QHR63299.1;BatCoV_RaTG13_orf1ab;1180 | 0 | 0 |
| MN996532.1;QHR63299.1;BatCoV_RaTG13_orf1ab;1181 | 0 | 0 |
| MN996532.1;QHR63299.1;BatCoV_RaTG13_orf1ab;1182 | 0 | 0 |
| MN996532.1;QHR63299.1;BatCoV_RaTG13_orf1ab;1183 | 0 | 0 |
| MN996532.1;QHR63299.1;BatCoV_RaTG13_orf1ab;1184 | 0.05 | 0 |
| MN996532.1;QHR63299.1;BatCoV_RaTG13_orf1ab;1185 | 0.05 | 0 |
| MN996532.1;QHR63299.1;BatCoV_RaTG13_orf1ab;1186 | 0.1 | 0.025 |
| MN996532.1;QHR63299.1;BatCoV_RaTG13_orf1ab;1187 | 0.05 | 0.075 |
| MN996532.1;QHR63299.1;BatCoV_RaTG13_orf1ab;1188 | 0.05 | 0.05 |
| MN996532.1;QHR63299.1;BatCoV_RaTG13_orf1ab;1189 | 0.05 | 0.025 |
| MN996532.1;QHR63299.1;BatCoV_RaTG13_orf1ab;1190 | 0 | 0 |
| MN996532.1;QHR63299.1;BatCoV_RaTG13_orf1ab;1191 | 0.05 | 0.025 |
| MN996532.1;QHR63299.1;BatCoV_RaTG13_orf1ab;1192 | 0 | 0 |
| MN996532.1;QHR63299.1;BatCoV_RaTG13_orf1ab;1193 | 0 | 0 |
| MN996532.1;QHR63299.1;BatCoV_RaTG13_orf1ab;1194 | 0.05 | 0 |
| MN996532.1;QHR63299.1;BatCoV_RaTG13_orf1ab;1195 | 0 | 0 |

| | | |
|---|---|---|
| MN996532.1;QHR63299.1;BatCoV_RaTG13_orf1ab;1196 | 0 | 0.025 |
| MN996532.1;QHR63299.1;BatCoV_RaTG13_orf1ab;1197 | 0.05 | 0 |
| MN996532.1;QHR63299.1;BatCoV_RaTG13_orf1ab;1198 | 0 | 0 |
| MN996532.1;QHR63299.1;BatCoV_RaTG13_orf1ab;1199 | 0.05 | 0 |
| MN996532.1;QHR63299.1;BatCoV_RaTG13_orf1ab;1200 | 0 | 0 |
| MN996532.1;QHR63299.1;BatCoV_RaTG13_orf1ab;1201 | 0 | 0 |
| MN996532.1;QHR63299.1;BatCoV_RaTG13_orf1ab;1202 | 0 | 0 |
| MN996532.1;QHR63299.1;BatCoV_RaTG13_orf1ab;1203 | 0 | 0 |
| MN996532.1;QHR63299.1;BatCoV_RaTG13_orf1ab;1204 | 0 | 0 |
| MN996532.1;QHR63299.1;BatCoV_RaTG13_orf1ab;1205 | 0 | 0.025 |
| MN996532.1;QHR63299.1;BatCoV_RaTG13_orf1ab;1206 | 0 | 0.025 |
| MN996532.1;QHR63299.1;BatCoV_RaTG13_orf1ab;1207 | 0 | 0 |
| MN996532.1;QHR63299.1;BatCoV_RaTG13_orf1ab;1208 | 0 | 0 |
| MN996532.1;QHR63299.1;BatCoV_RaTG13_orf1ab;1209 | 0 | 0 |
| MN996532.1;QHR63299.1;BatCoV_RaTG13_orf1ab;1210 | 0 | 0.025 |
| MN996532.1;QHR63299.1;BatCoV_RaTG13_orf1ab;1211 | 0 | 0.025 |
| MN996532.1;QHR63299.1;BatCoV_RaTG13_orf1ab;1212 | 0 | 0 |
| MN996532.1;QHR63299.1;BatCoV_RaTG13_orf1ab;1213 | 0.05 | 0.025 |
| MN996532.1;QHR63299.1;BatCoV_RaTG13_orf1ab;1214 | 0 | 0 |
| MN996532.1;QHR63299.1;BatCoV_RaTG13_orf1ab;1215 | 0.05 | 0 |
| MN996532.1;QHR63299.1;BatCoV_RaTG13_orf1ab;1216 | 0.05 | 0.125 |
| MN996532.1;QHR63299.1;BatCoV_RaTG13_orf1ab;1217 | 0.05 | 0 |
| MN996532.1;QHR63299.1;BatCoV_RaTG13_orf1ab;1218 | 0.05 | 0 |
| MN996532.1;QHR63299.1;BatCoV_RaTG13_orf1ab;1219 | 0.05 | 0 |
| MN996532.1;QHR63299.1;BatCoV_RaTG13_orf1ab;1220 | 0.05 | 0.025 |
| MN996532.1;QHR63299.1;BatCoV_RaTG13_orf1ab;1221 | 0 | 0 |
| MN996532.1;QHR63299.1;BatCoV_RaTG13_orf1ab;1222 | 0.05 | 0 |
| MN996532.1;QHR63299.1;BatCoV_RaTG13_orf1ab;1223 | 0 | 0 |
| MN996532.1;QHR63299.1;BatCoV_RaTG13_orf1ab;1224 | 0 | 0.025 |
| MN996532.1;QHR63299.1;BatCoV_RaTG13_orf1ab;1225 | 0 | 0.05 |
| MN996532.1;QHR63299.1;BatCoV_RaTG13_orf1ab;1226 | 0 | 0.025 |
| MN996532.1;QHR63299.1;BatCoV_RaTG13_orf1ab;1227 | 0 | 0.025 |
| MN996532.1;QHR63299.1;BatCoV_RaTG13_orf1ab;1228 | 0 | 0.025 |
| MN996532.1;QHR63299.1;BatCoV_RaTG13_orf1ab;1229 | 0 | 0.025 |
| MN996532.1;QHR63299.1;BatCoV_RaTG13_orf1ab;1230 | 0 | 0.025 |
| MN996532.1;QHR63299.1;BatCoV_RaTG13_orf1ab;1231 | 0 | 0.025 |
| MN996532.1;QHR63299.1;BatCoV_RaTG13_orf1ab;1232 | 0 | 0.025 |
| MN996532.1;QHR63299.1;BatCoV_RaTG13_orf1ab;1233 | 0 | 0.025 |
| MN996532.1;QHR63299.1;BatCoV_RaTG13_orf1ab;1234 | 0 | 0.025 |
| MN996532.1;QHR63299.1;BatCoV_RaTG13_orf1ab;1235 | 0 | 0 |
| MN996532.1;QHR63299.1;BatCoV_RaTG13_orf1ab;1236 | 0 | 0.075 |
| MN996532.1;QHR63299.1;BatCoV_RaTG13_orf1ab;1237 | 0 | 0.05 |
| MN996532.1;QHR63299.1;BatCoV_RaTG13_orf1ab;1238 | 0 | 0.025 |

| | | |
|---|---|---|
| MN996532.1;QHR63299.1;BatCoV_RaTG13_orf1ab;1239 | 0 | 0.025 |
| MN996532.1;QHR63299.1;BatCoV_RaTG13_orf1ab;1240 | 0 | 0.05 |
| MN996532.1;QHR63299.1;BatCoV_RaTG13_orf1ab;1241 | 0 | 0 |
| MN996532.1;QHR63299.1;BatCoV_RaTG13_orf1ab;1242 | 0 | 0 |
| MN996532.1;QHR63299.1;BatCoV_RaTG13_orf1ab;1243 | 0 | 0 |
| MN996532.1;QHR63299.1;BatCoV_RaTG13_orf1ab;1244 | 0 | 0 |
| MN996532.1;QHR63299.1;BatCoV_RaTG13_orf1ab;1245 | 0 | 0 |
| MN996532.1;QHR63299.1;BatCoV_RaTG13_orf1ab;1246 | 0 | 0 |
| MN996532.1;QHR63299.1;BatCoV_RaTG13_orf1ab;1247 | 0 | 0 |
| MN996532.1;QHR63299.1;BatCoV_RaTG13_orf1ab;1248 | 0 | 0 |
| MN996532.1;QHR63299.1;BatCoV_RaTG13_orf1ab;1249 | 0 | 0 |
| MN996532.1;QHR63299.1;BatCoV_RaTG13_orf1ab;1250 | 0 | 0 |
| MN996532.1;QHR63299.1;BatCoV_RaTG13_orf1ab;1251 | 0 | 0 |
| MN996532.1;QHR63299.1;BatCoV_RaTG13_orf1ab;1252 | 0 | 0 |
| MN996532.1;QHR63299.1;BatCoV_RaTG13_orf1ab;1253 | 0 | 0.025 |
| MN996532.1;QHR63299.1;BatCoV_RaTG13_orf1ab;1254 | 0.05 | 0.075 |
| MN996532.1;QHR63299.1;BatCoV_RaTG13_orf1ab;1255 | 0.1 | 0.075 |
| MN996532.1;QHR63299.1;BatCoV_RaTG13_orf1ab;1256 | 0 | 0.025 |
| MN996532.1;QHR63299.1;BatCoV_RaTG13_orf1ab;1257 | 0 | 0 |
| MN996532.1;QHR63299.1;BatCoV_RaTG13_orf1ab;1258 | 0.05 | 0 |
| MN996532.1;QHR63299.1;BatCoV_RaTG13_orf1ab;1259 | 0 | 0 |
| MN996532.1;QHR63299.1;BatCoV_RaTG13_orf1ab;1260 | 0 | 0 |
| MN996532.1;QHR63299.1;BatCoV_RaTG13_orf1ab;1261 | 0 | 0 |
| MN996532.1;QHR63299.1;BatCoV_RaTG13_orf1ab;1262 | 0 | 0 |
| MN996532.1;QHR63299.1;BatCoV_RaTG13_orf1ab;1263 | 0 | 0 |
| MN996532.1;QHR63299.1;BatCoV_RaTG13_orf1ab;1264 | 0 | 0 |
| MN996532.1;QHR63299.1;BatCoV_RaTG13_orf1ab;1265 | 0 | 0.05 |
| MN996532.1;QHR63299.1;BatCoV_RaTG13_orf1ab;1266 | 0 | 0.05 |
| MN996532.1;QHR63299.1;BatCoV_RaTG13_orf1ab;1267 | 0 | 0.05 |
| MN996532.1;QHR63299.1;BatCoV_RaTG13_orf1ab;1268 | 0 | 0.05 |
| MN996532.1;QHR63299.1;BatCoV_RaTG13_orf1ab;1269 | 0 | 0.1 |
| MN996532.1;QHR63299.1;BatCoV_RaTG13_orf1ab;1270 | 0 | 0.1 |
| MN996532.1;QHR63299.1;BatCoV_RaTG13_orf1ab;1271 | 0 | 0.05 |
| MN996532.1;QHR63299.1;BatCoV_RaTG13_orf1ab;1272 | 0 | 0.025 |
| MN996532.1;QHR63299.1;BatCoV_RaTG13_orf1ab;1273 | 0 | 0.025 |
| MN996532.1;QHR63299.1;BatCoV_RaTG13_orf1ab;1274 | 0 | 0.025 |
| MN996532.1;QHR63299.1;BatCoV_RaTG13_orf1ab;1275 | 0 | 0 |
| MN996532.1;QHR63299.1;BatCoV_RaTG13_orf1ab;1276 | 0 | 0 |
| MN996532.1;QHR63299.1;BatCoV_RaTG13_orf1ab;1277 | 0 | 0 |
| MN996532.1;QHR63299.1;BatCoV_RaTG13_orf1ab;1278 | 0 | 0 |
| MN996532.1;QHR63299.1;BatCoV_RaTG13_orf1ab;1279 | 0 | 0 |
| MN996532.1;QHR63299.1;BatCoV_RaTG13_orf1ab;1280 | 0 | 0 |
| MN996532.1;QHR63299.1;BatCoV_RaTG13_orf1ab;1281 | 0 | 0 |

| | | |
|---|---|---|
| MN996532.1;QHR63299.1;BatCoV_RaTG13_orf1ab;1282 | 0 | 0 |
| MN996532.1;QHR63299.1;BatCoV_RaTG13_orf1ab;1283 | 0 | 0 |
| MN996532.1;QHR63299.1;BatCoV_RaTG13_orf1ab;1284 | 0 | 0 |
| MN996532.1;QHR63299.1;BatCoV_RaTG13_orf1ab;1285 | 0 | 0 |
| MN996532.1;QHR63299.1;BatCoV_RaTG13_orf1ab;1286 | 0 | 0 |
| MN996532.1;QHR63299.1;BatCoV_RaTG13_orf1ab;1287 | 0 | 0.025 |
| MN996532.1;QHR63299.1;BatCoV_RaTG13_orf1ab;1288 | 0 | 0 |
| MN996532.1;QHR63299.1;BatCoV_RaTG13_orf1ab;1289 | 0 | 0 |
| MN996532.1;QHR63299.1;BatCoV_RaTG13_orf1ab;1290 | 0 | 0 |
| MN996532.1;QHR63299.1;BatCoV_RaTG13_orf1ab;1291 | 0 | 0 |
| MN996532.1;QHR63299.1;BatCoV_RaTG13_orf1ab;1292 | 0 | 0 |
| MN996532.1;QHR63299.1;BatCoV_RaTG13_orf1ab;1293 | 0 | 0 |
| MN996532.1;QHR63299.1;BatCoV_RaTG13_orf1ab;1294 | 0 | 0 |
| MN996532.1;QHR63299.1;BatCoV_RaTG13_orf1ab;1295 | 0.05 | 0 |
| MN996532.1;QHR63299.1;BatCoV_RaTG13_orf1ab;1296 | 0.05 | 0.025 |
| MN996532.1;QHR63299.1;BatCoV_RaTG13_orf1ab;1297 | 0.05 | 0.025 |
| MN996532.1;QHR63299.1;BatCoV_RaTG13_orf1ab;1298 | 0.05 | 0.025 |
| MN996532.1;QHR63299.1;BatCoV_RaTG13_orf1ab;1299 | 0.05 | 0.025 |
| MN996532.1;QHR63299.1;BatCoV_RaTG13_orf1ab;1300 | 0.05 | 0 |
| MN996532.1;QHR63299.1;BatCoV_RaTG13_orf1ab;1301 | 0.05 | 0 |
| MN996532.1;QHR63299.1;BatCoV_RaTG13_orf1ab;1302 | 0 | 0 |
| MN996532.1;QHR63299.1;BatCoV_RaTG13_orf1ab;1303 | 0 | 0 |
| MN996532.1;QHR63299.1;BatCoV_RaTG13_orf1ab;1304 | 0 | 0 |
| MN996532.1;QHR63299.1;BatCoV_RaTG13_orf1ab;1305 | 0 | 0 |
| MN996532.1;QHR63299.1;BatCoV_RaTG13_orf1ab;1306 | 0 | 0 |
| MN996532.1;QHR63299.1;BatCoV_RaTG13_orf1ab;1307 | 0 | 0 |
| MN996532.1;QHR63299.1;BatCoV_RaTG13_orf1ab;1308 | 0.05 | 0 |
| MN996532.1;QHR63299.1;BatCoV_RaTG13_orf1ab;1309 | 0.05 | 0 |
| MN996532.1;QHR63299.1;BatCoV_RaTG13_orf1ab;1310 | 0 | 0.025 |
| MN996532.1;QHR63299.1;BatCoV_RaTG13_orf1ab;1311 | 0 | 0 |
| MN996532.1;QHR63299.1;BatCoV_RaTG13_orf1ab;1312 | 0 | 0.025 |
| MN996532.1;QHR63299.1;BatCoV_RaTG13_orf1ab;1313 | 0 | 0 |
| MN996532.1;QHR63299.1;BatCoV_RaTG13_orf1ab;1314 | 0 | 0 |
| MN996532.1;QHR63299.1;BatCoV_RaTG13_orf1ab;1315 | 0.05 | 0 |
| MN996532.1;QHR63299.1;BatCoV_RaTG13_orf1ab;1316 | 0.05 | 0 |
| MN996532.1;QHR63299.1;BatCoV_RaTG13_orf1ab;1317 | 0.05 | 0.025 |
| MN996532.1;QHR63299.1;BatCoV_RaTG13_orf1ab;1318 | 0 | 0 |
| MN996532.1;QHR63299.1;BatCoV_RaTG13_orf1ab;1319 | 0 | 0.05 |
| MN996532.1;QHR63299.1;BatCoV_RaTG13_orf1ab;1320 | 0 | 0 |
| MN996532.1;QHR63299.1;BatCoV_RaTG13_orf1ab;1321 | 0 | 0 |
| MN996532.1;QHR63299.1;BatCoV_RaTG13_orf1ab;1322 | 0.05 | 0 |
| MN996532.1;QHR63299.1;BatCoV_RaTG13_orf1ab;1323 | 0 | 0 |
| MN996532.1;QHR63299.1;BatCoV_RaTG13_orf1ab;1324 | 0 | 0 |

| | | |
|---|---|---|
| MN996532.1;QHR63299.1;BatCoV_RaTG13_orf1ab;1325 | 0 | 0 |
| MN996532.1;QHR63299.1;BatCoV_RaTG13_orf1ab;1326 | 0 | 0 |
| MN996532.1;QHR63299.1;BatCoV_RaTG13_orf1ab;1327 | 0 | 0.025 |
| MN996532.1;QHR63299.1;BatCoV_RaTG13_orf1ab;1328 | 0 | 0.05 |
| MN996532.1;QHR63299.1;BatCoV_RaTG13_orf1ab;1329 | 0 | 0.05 |
| MN996532.1;QHR63299.1;BatCoV_RaTG13_orf1ab;1330 | 0 | 0.05 |
| MN996532.1;QHR63299.1;BatCoV_RaTG13_orf1ab;1331 | 0 | 0.05 |
| MN996532.1;QHR63299.1;BatCoV_RaTG13_orf1ab;1332 | 0 | 0.025 |
| MN996532.1;QHR63299.1;BatCoV_RaTG13_orf1ab;1333 | 0 | 0.025 |
| MN996532.1;QHR63299.1;BatCoV_RaTG13_orf1ab;1334 | 0 | 0 |
| MN996532.1;QHR63299.1;BatCoV_RaTG13_orf1ab;1335 | 0 | 0.025 |
| MN996532.1;QHR63299.1;BatCoV_RaTG13_orf1ab;1336 | 0 | 0.05 |
| MN996532.1;QHR63299.1;BatCoV_RaTG13_orf1ab;1337 | 0 | 0 |
| MN996532.1;QHR63299.1;BatCoV_RaTG13_orf1ab;1338 | 0 | 0 |
| MN996532.1;QHR63299.1;BatCoV_RaTG13_orf1ab;1339 | 0 | 0 |
| MN996532.1;QHR63299.1;BatCoV_RaTG13_orf1ab;1340 | 0 | 0.025 |
| MN996532.1;QHR63299.1;BatCoV_RaTG13_orf1ab;1341 | 0 | 0 |
| MN996532.1;QHR63299.1;BatCoV_RaTG13_orf1ab;1342 | 0 | 0 |
| MN996532.1;QHR63299.1;BatCoV_RaTG13_orf1ab;1343 | 0 | 0 |
| MN996532.1;QHR63299.1;BatCoV_RaTG13_orf1ab;1344 | 0 | 0 |
| MN996532.1;QHR63299.1;BatCoV_RaTG13_orf1ab;1345 | 0 | 0 |
| MN996532.1;QHR63299.1;BatCoV_RaTG13_orf1ab;1346 | 0 | 0 |
| MN996532.1;QHR63299.1;BatCoV_RaTG13_orf1ab;1347 | 0 | 0 |
| MN996532.1;QHR63299.1;BatCoV_RaTG13_orf1ab;1348 | 0 | 0 |
| MN996532.1;QHR63299.1;BatCoV_RaTG13_orf1ab;1349 | 0 | 0.025 |
| MN996532.1;QHR63299.1;BatCoV_RaTG13_orf1ab;1350 | 0 | 0 |
| MN996532.1;QHR63299.1;BatCoV_RaTG13_orf1ab;1351 | 0 | 0 |
| MN996532.1;QHR63299.1;BatCoV_RaTG13_orf1ab;1352 | 0 | 0 |
| MN996532.1;QHR63299.1;BatCoV_RaTG13_orf1ab;1353 | 0.05 | 0 |
| MN996532.1;QHR63299.1;BatCoV_RaTG13_orf1ab;1354 | 0 | 0 |
| MN996532.1;QHR63299.1;BatCoV_RaTG13_orf1ab;1355 | 0 | 0 |
| MN996532.1;QHR63299.1;BatCoV_RaTG13_orf1ab;1356 | 0.05 | 0.025 |
| MN996532.1;QHR63299.1;BatCoV_RaTG13_orf1ab;1357 | 0 | 0 |
| MN996532.1;QHR63299.1;BatCoV_RaTG13_orf1ab;1358 | 0.05 | 0 |
| MN996532.1;QHR63299.1;BatCoV_RaTG13_orf1ab;1359 | 0 | 0 |
| MN996532.1;QHR63299.1;BatCoV_RaTG13_orf1ab;1360 | 0 | 0 |
| MN996532.1;QHR63299.1;BatCoV_RaTG13_orf1ab;1361 | 0 | 0 |
| MN996532.1;QHR63299.1;BatCoV_RaTG13_orf1ab;1362 | 0 | 0 |
| MN996532.1;QHR63299.1;BatCoV_RaTG13_orf1ab;1363 | 0 | 0 |
| MN996532.1;QHR63299.1;BatCoV_RaTG13_orf1ab;1364 | 0.05 | 0 |
| MN996532.1;QHR63299.1;BatCoV_RaTG13_orf1ab;1365 | 0 | 0 |
| MN996532.1;QHR63299.1;BatCoV_RaTG13_orf1ab;1366 | 0 | 0 |
| MN996532.1;QHR63299.1;BatCoV_RaTG13_orf1ab;1367 | 0 | 0 |

| | | |
|---|---|---|
| MN996532.1;QHR63299.1;BatCoV_RaTG13_orf1ab;1368 | 0 | 0.025 |
| MN996532.1;QHR63299.1;BatCoV_RaTG13_orf1ab;1369 | 0 | 0 |
| MN996532.1;QHR63299.1;BatCoV_RaTG13_orf1ab;1370 | 0 | 0 |
| MN996532.1;QHR63299.1;BatCoV_RaTG13_orf1ab;1371 | 0 | 0.05 |
| MN996532.1;QHR63299.1;BatCoV_RaTG13_orf1ab;1372 | 0 | 0.025 |
| MN996532.1;QHR63299.1;BatCoV_RaTG13_orf1ab;1373 | 0 | 0 |
| MN996532.1;QHR63299.1;BatCoV_RaTG13_orf1ab;1374 | 0 | 0.025 |
| MN996532.1;QHR63299.1;BatCoV_RaTG13_orf1ab;1375 | 0 | 0 |
| MN996532.1;QHR63299.1;BatCoV_RaTG13_orf1ab;1376 | 0 | 0 |
| MN996532.1;QHR63299.1;BatCoV_RaTG13_orf1ab;1377 | 0 | 0 |
| MN996532.1;QHR63299.1;BatCoV_RaTG13_orf1ab;1378 | 0 | 0 |
| MN996532.1;QHR63299.1;BatCoV_RaTG13_orf1ab;1379 | 0 | 0.025 |
| MN996532.1;QHR63299.1;BatCoV_RaTG13_orf1ab;1380 | 0 | 0.025 |
| MN996532.1;QHR63299.1;BatCoV_RaTG13_orf1ab;1381 | 0.05 | 0.025 |
| MN996532.1;QHR63299.1;BatCoV_RaTG13_orf1ab;1382 | 0.05 | 0.025 |
| MN996532.1;QHR63299.1;BatCoV_RaTG13_orf1ab;1383 | 0 | 0.025 |
| MN996532.1;QHR63299.1;BatCoV_RaTG13_orf1ab;1384 | 0 | 0 |
| MN996532.1;QHR63299.1;BatCoV_RaTG13_orf1ab;1385 | 0 | 0 |
| MN996532.1;QHR63299.1;BatCoV_RaTG13_orf1ab;1386 | 0 | 0 |
| MN996532.1;QHR63299.1;BatCoV_RaTG13_orf1ab;1387 | 0 | 0 |
| MN996532.1;QHR63299.1;BatCoV_RaTG13_orf1ab;1388 | 0 | 0 |
| MN996532.1;QHR63299.1;BatCoV_RaTG13_orf1ab;1389 | 0 | 0 |
| MN996532.1;QHR63299.1;BatCoV_RaTG13_orf1ab;1390 | 0 | 0 |
| MN996532.1;QHR63299.1;BatCoV_RaTG13_orf1ab;1391 | 0 | 0 |
| MN996532.1;QHR63299.1;BatCoV_RaTG13_orf1ab;1392 | 0 | 0 |
| MN996532.1;QHR63299.1;BatCoV_RaTG13_orf1ab;1393 | 0 | 0 |
| MN996532.1;QHR63299.1;BatCoV_RaTG13_orf1ab;1394 | 0 | 0 |
| MN996532.1;QHR63299.1;BatCoV_RaTG13_orf1ab;1395 | 0.05 | 0 |
| MN996532.1;QHR63299.1;BatCoV_RaTG13_orf1ab;1396 | 0.05 | 0 |
| MN996532.1;QHR63299.1;BatCoV_RaTG13_orf1ab;1397 | 0.1 | 0 |
| MN996532.1;QHR63299.1;BatCoV_RaTG13_orf1ab;1398 | 0.1 | 0 |
| MN996532.1;QHR63299.1;BatCoV_RaTG13_orf1ab;1399 | 0.1 | 0 |
| MN996532.1;QHR63299.1;BatCoV_RaTG13_orf1ab;1400 | 0.1 | 0 |
| MN996532.1;QHR63299.1;BatCoV_RaTG13_orf1ab;1401 | 0.1 | 0 |
| MN996532.1;QHR63299.1;BatCoV_RaTG13_orf1ab;1402 | 0.1 | 0.025 |
| MN996532.1;QHR63299.1;BatCoV_RaTG13_orf1ab;1403 | 0.1 | 0.025 |
| MN996532.1;QHR63299.1;BatCoV_RaTG13_orf1ab;1404 | 0.1 | 0.025 |
| MN996532.1;QHR63299.1;BatCoV_RaTG13_orf1ab;1405 | 0.1 | 0.025 |
| MN996532.1;QHR63299.1;BatCoV_RaTG13_orf1ab;1406 | 0 | 0.025 |
| MN996532.1;QHR63299.1;BatCoV_RaTG13_orf1ab;1407 | 0 | 0 |
| MN996532.1;QHR63299.1;BatCoV_RaTG13_orf1ab;1408 | 0 | 0 |
| MN996532.1;QHR63299.1;BatCoV_RaTG13_orf1ab;1409 | 0 | 0 |
| MN996532.1;QHR63299.1;BatCoV_RaTG13_orf1ab;1410 | 0 | 0 |

| | | |
|---|---|---|
| MN996532.1;QHR63299.1;BatCoV_RaTG13_orf1ab;1411 | 0 | 0 |
| MN996532.1;QHR63299.1;BatCoV_RaTG13_orf1ab;1412 | 0 | 0 |
| MN996532.1;QHR63299.1;BatCoV_RaTG13_orf1ab;1413 | 0.05 | 0.1 |
| MN996532.1;QHR63299.1;BatCoV_RaTG13_orf1ab;1414 | 0 | 0 |
| MN996532.1;QHR63299.1;BatCoV_RaTG13_orf1ab;1415 | 0 | 0 |
| MN996532.1;QHR63299.1;BatCoV_RaTG13_orf1ab;1416 | 0 | 0 |
| MN996532.1;QHR63299.1;BatCoV_RaTG13_orf1ab;1417 | 0 | 0 |
| MN996532.1;QHR63299.1;BatCoV_RaTG13_orf1ab;1418 | 0 | 0 |
| MN996532.1;QHR63299.1;BatCoV_RaTG13_orf1ab;1419 | 0 | 0 |
| MN996532.1;QHR63299.1;BatCoV_RaTG13_orf1ab;1420 | 0 | 0 |
| MN996532.1;QHR63299.1;BatCoV_RaTG13_orf1ab;1421 | 0 | 0 |
| MN996532.1;QHR63299.1;BatCoV_RaTG13_orf1ab;1422 | 0 | 0 |
| MN996532.1;QHR63299.1;BatCoV_RaTG13_orf1ab;1423 | 0 | 0.025 |
| MN996532.1;QHR63299.1;BatCoV_RaTG13_orf1ab;1424 | 0 | 0.05 |
| MN996532.1;QHR63299.1;BatCoV_RaTG13_orf1ab;1425 | 0 | 0.025 |
| MN996532.1;QHR63299.1;BatCoV_RaTG13_orf1ab;1426 | 0.2 | 0.1 |
| MN996532.1;QHR63299.1;BatCoV_RaTG13_orf1ab;1427 | 0.25 | 0.125 |
| MN996532.1;QHR63299.1;BatCoV_RaTG13_orf1ab;1428 | 0.15 | 0.25 |
| MN996532.1;QHR63299.1;BatCoV_RaTG13_orf1ab;1429 | 0.2 | 0.25 |
| MN996532.1;QHR63299.1;BatCoV_RaTG13_orf1ab;1430 | 0.3 | 0.35 |
| MN996532.1;QHR63299.1;BatCoV_RaTG13_orf1ab;1431 | 0.2 | 0.225 |
| MN996532.1;QHR63299.1;BatCoV_RaTG13_orf1ab;1432 | 0.2 | 0.175 |
| MN996532.1;QHR63299.1;BatCoV_RaTG13_orf1ab;1433 | 0.2 | 0.225 |
| MN996532.1;QHR63299.1;BatCoV_RaTG13_orf1ab;1434 | 0.25 | 0.2 |
| MN996532.1;QHR63299.1;BatCoV_RaTG13_orf1ab;1435 | 0.2 | 0.2 |
| MN996532.1;QHR63299.1;BatCoV_RaTG13_orf1ab;1436 | 0.35 | 0.3 |
| MN996532.1;QHR63299.1;BatCoV_RaTG13_orf1ab;1437 | 0.15 | 0.15 |
| MN996532.1;QHR63299.1;BatCoV_RaTG13_orf1ab;1438 | 0.1 | 0.1 |
| MN996532.1;QHR63299.1;BatCoV_RaTG13_orf1ab;1439 | 0.1 | 0.1 |
| MN996532.1;QHR63299.1;BatCoV_RaTG13_orf1ab;1440 | 0.1 | 0.1 |
| MN996532.1;QHR63299.1;BatCoV_RaTG13_orf1ab;1441 | 0.05 | 0.075 |
| MN996532.1;QHR63299.1;BatCoV_RaTG13_orf1ab;1442 | 0 | 0 |
| MN996532.1;QHR63299.1;BatCoV_RaTG13_orf1ab;1443 | 0 | 0 |
| MN996532.1;QHR63299.1;BatCoV_RaTG13_orf1ab;1444 | 0 | 0 |
| MN996532.1;QHR63299.1;BatCoV_RaTG13_orf1ab;1445 | 0.05 | 0 |
| MN996532.1;QHR63299.1;BatCoV_RaTG13_orf1ab;1446 | 0 | 0 |
| MN996532.1;QHR63299.1;BatCoV_RaTG13_orf1ab;1447 | 0.1 | 0.05 |
| MN996532.1;QHR63299.1;BatCoV_RaTG13_orf1ab;1448 | 0.05 | 0.025 |
| MN996532.1;QHR63299.1;BatCoV_RaTG13_orf1ab;1449 | 0.05 | 0.025 |
| MN996532.1;QHR63299.1;BatCoV_RaTG13_orf1ab;1450 | 0.05 | 0.025 |
| MN996532.1;QHR63299.1;BatCoV_RaTG13_orf1ab;1451 | 0 | 0 |
| MN996532.1;QHR63299.1;BatCoV_RaTG13_orf1ab;1452 | 0 | 0 |
| MN996532.1;QHR63299.1;BatCoV_RaTG13_orf1ab;1453 | 0 | 0 |

| | | |
|---|---|---|
| MN996532.1;QHR63299.1;BatCoV_RaTG13_orf1ab;1454 | 0 | 0 |
| MN996532.1;QHR63299.1;BatCoV_RaTG13_orf1ab;1455 | 0 | 0.025 |
| MN996532.1;QHR63299.1;BatCoV_RaTG13_orf1ab;1456 | 0 | 0 |
| MN996532.1;QHR63299.1;BatCoV_RaTG13_orf1ab;1457 | 0 | 0 |
| MN996532.1;QHR63299.1;BatCoV_RaTG13_orf1ab;1458 | 0 | 0.075 |
| MN996532.1;QHR63299.1;BatCoV_RaTG13_orf1ab;1459 | 0.1 | 0.075 |
| MN996532.1;QHR63299.1;BatCoV_RaTG13_orf1ab;1460 | 0.15 | 0.125 |
| MN996532.1;QHR63299.1;BatCoV_RaTG13_orf1ab;1461 | 0.05 | 0 |
| MN996532.1;QHR63299.1;BatCoV_RaTG13_orf1ab;1462 | 0 | 0.025 |
| MN996532.1;QHR63299.1;BatCoV_RaTG13_orf1ab;1463 | 0 | 0 |
| MN996532.1;QHR63299.1;BatCoV_RaTG13_orf1ab;1464 | 0 | 0 |
| MN996532.1;QHR63299.1;BatCoV_RaTG13_orf1ab;1465 | 0 | 0 |
| MN996532.1;QHR63299.1;BatCoV_RaTG13_orf1ab;1466 | 0 | 0 |
| MN996532.1;QHR63299.1;BatCoV_RaTG13_orf1ab;1467 | 0 | 0 |
| MN996532.1;QHR63299.1;BatCoV_RaTG13_orf1ab;1468 | 0 | 0 |
| MN996532.1;QHR63299.1;BatCoV_RaTG13_orf1ab;1469 | 0 | 0 |
| MN996532.1;QHR63299.1;BatCoV_RaTG13_orf1ab;1470 | 0 | 0 |
| MN996532.1;QHR63299.1;BatCoV_RaTG13_orf1ab;1471 | 0 | 0.05 |
| MN996532.1;QHR63299.1;BatCoV_RaTG13_orf1ab;1472 | 0.05 | 0.025 |
| MN996532.1;QHR63299.1;BatCoV_RaTG13_orf1ab;1473 | 0 | 0.1 |
| MN996532.1;QHR63299.1;BatCoV_RaTG13_orf1ab;1474 | 0 | 0.025 |
| MN996532.1;QHR63299.1;BatCoV_RaTG13_orf1ab;1475 | 0 | 0 |
| MN996532.1;QHR63299.1;BatCoV_RaTG13_orf1ab;1476 | 0 | 0 |
| MN996532.1;QHR63299.1;BatCoV_RaTG13_orf1ab;1477 | 0 | 0 |
| MN996532.1;QHR63299.1;BatCoV_RaTG13_orf1ab;1478 | 0 | 0 |
| MN996532.1;QHR63299.1;BatCoV_RaTG13_orf1ab;1479 | 0.05 | 0.05 |
| MN996532.1;QHR63299.1;BatCoV_RaTG13_orf1ab;1480 | 0.05 | 0.075 |
| MN996532.1;QHR63299.1;BatCoV_RaTG13_orf1ab;1481 | 0.1 | 0.05 |
| MN996532.1;QHR63299.1;BatCoV_RaTG13_orf1ab;1482 | 0.15 | 0.1 |
| MN996532.1;QHR63299.1;BatCoV_RaTG13_orf1ab;1483 | 0.1 | 0.15 |
| MN996532.1;QHR63299.1;BatCoV_RaTG13_orf1ab;1484 | 0.1 | 0.125 |
| MN996532.1;QHR63299.1;BatCoV_RaTG13_orf1ab;1485 | 0.15 | 0.125 |
| MN996532.1;QHR63299.1;BatCoV_RaTG13_orf1ab;1486 | 0.2 | 0.125 |
| MN996532.1;QHR63299.1;BatCoV_RaTG13_orf1ab;1487 | 0.25 | 0.1 |
| MN996532.1;QHR63299.1;BatCoV_RaTG13_orf1ab;1488 | 0.2 | 0.1 |
| MN996532.1;QHR63299.1;BatCoV_RaTG13_orf1ab;1489 | 0.15 | 0.025 |
| MN996532.1;QHR63299.1;BatCoV_RaTG13_orf1ab;1490 | 0.05 | 0 |
| MN996532.1;QHR63299.1;BatCoV_RaTG13_orf1ab;1491 | 0.05 | 0 |
| MN996532.1;QHR63299.1;BatCoV_RaTG13_orf1ab;1492 | 0.05 | 0 |
| MN996532.1;QHR63299.1;BatCoV_RaTG13_orf1ab;1493 | 0 | 0 |
| MN996532.1;QHR63299.1;BatCoV_RaTG13_orf1ab;1494 | 0 | 0 |
| MN996532.1;QHR63299.1;BatCoV_RaTG13_orf1ab;1495 | 0 | 0 |
| MN996532.1;QHR63299.1;BatCoV_RaTG13_orf1ab;1496 | 0 | 0 |

| | | |
|---|---|---|
| MN996532.1;QHR63299.1;BatCoV_RaTG13_orf1ab;1497 | 0 | 0 |
| MN996532.1;QHR63299.1;BatCoV_RaTG13_orf1ab;1498 | 0 | 0 |
| MN996532.1;QHR63299.1;BatCoV_RaTG13_orf1ab;1499 | 0 | 0 |
| MN996532.1;QHR63299.1;BatCoV_RaTG13_orf1ab;1500 | 0 | 0 |
| MN996532.1;QHR63299.1;BatCoV_RaTG13_orf1ab;1501 | 0 | 0 |
| MN996532.1;QHR63299.1;BatCoV_RaTG13_orf1ab;1502 | 0 | 0 |
| MN996532.1;QHR63299.1;BatCoV_RaTG13_orf1ab;1503 | 0 | 0 |
| MN996532.1;QHR63299.1;BatCoV_RaTG13_orf1ab;1504 | 0 | 0 |
| MN996532.1;QHR63299.1;BatCoV_RaTG13_orf1ab;1505 | 0.05 | 0.05 |
| MN996532.1;QHR63299.1;BatCoV_RaTG13_orf1ab;1506 | 0 | 0.05 |
| MN996532.1;QHR63299.1;BatCoV_RaTG13_orf1ab;1507 | 0 | 0.025 |
| MN996532.1;QHR63299.1;BatCoV_RaTG13_orf1ab;1508 | 0.05 | 0 |
| MN996532.1;QHR63299.1;BatCoV_RaTG13_orf1ab;1509 | 0 | 0 |
| MN996532.1;QHR63299.1;BatCoV_RaTG13_orf1ab;1510 | 0 | 0.025 |
| MN996532.1;QHR63299.1;BatCoV_RaTG13_orf1ab;1511 | 0 | 0 |
| MN996532.1;QHR63299.1;BatCoV_RaTG13_orf1ab;1512 | 0 | 0 |
| MN996532.1;QHR63299.1;BatCoV_RaTG13_orf1ab;1513 | 0 | 0 |
| MN996532.1;QHR63299.1;BatCoV_RaTG13_orf1ab;1514 | 0 | 0 |
| MN996532.1;QHR63299.1;BatCoV_RaTG13_orf1ab;1515 | 0 | 0 |
| MN996532.1;QHR63299.1;BatCoV_RaTG13_orf1ab;1516 | 0 | 0 |
| MN996532.1;QHR63299.1;BatCoV_RaTG13_orf1ab;1517 | 0 | 0.075 |
| MN996532.1;QHR63299.1;BatCoV_RaTG13_orf1ab;1518 | 0 | 0.075 |
| MN996532.1;QHR63299.1;BatCoV_RaTG13_orf1ab;1519 | 0 | 0.075 |
| MN996532.1;QHR63299.1;BatCoV_RaTG13_orf1ab;1520 | 0 | 0.075 |
| MN996532.1;QHR63299.1;BatCoV_RaTG13_orf1ab;1521 | 0 | 0.05 |
| MN996532.1;QHR63299.1;BatCoV_RaTG13_orf1ab;1522 | 0 | 0.05 |
| MN996532.1;QHR63299.1;BatCoV_RaTG13_orf1ab;1523 | 0.05 | 0.075 |
| MN996532.1;QHR63299.1;BatCoV_RaTG13_orf1ab;1524 | 0 | 0.05 |
| MN996532.1;QHR63299.1;BatCoV_RaTG13_orf1ab;1525 | 0.05 | 0.075 |
| MN996532.1;QHR63299.1;BatCoV_RaTG13_orf1ab;1526 | 0 | 0.025 |
| MN996532.1;QHR63299.1;BatCoV_RaTG13_orf1ab;1527 | 0 | 0.075 |
| MN996532.1;QHR63299.1;BatCoV_RaTG13_orf1ab;1528 | 0 | 0 |
| MN996532.1;QHR63299.1;BatCoV_RaTG13_orf1ab;1529 | 0 | 0 |
| MN996532.1;QHR63299.1;BatCoV_RaTG13_orf1ab;1530 | 0 | 0 |
| MN996532.1;QHR63299.1;BatCoV_RaTG13_orf1ab;1531 | 0 | 0 |
| MN996532.1;QHR63299.1;BatCoV_RaTG13_orf1ab;1532 | 0 | 0 |
| MN996532.1;QHR63299.1;BatCoV_RaTG13_orf1ab;1533 | 0 | 0 |
| MN996532.1;QHR63299.1;BatCoV_RaTG13_orf1ab;1534 | 0 | 0 |
| MN996532.1;QHR63299.1;BatCoV_RaTG13_orf1ab;1535 | 0 | 0 |
| MN996532.1;QHR63299.1;BatCoV_RaTG13_orf1ab;1536 | 0 | 0 |
| MN996532.1;QHR63299.1;BatCoV_RaTG13_orf1ab;1537 | 0 | 0 |
| MN996532.1;QHR63299.1;BatCoV_RaTG13_orf1ab;1538 | 0 | 0 |
| MN996532.1;QHR63299.1;BatCoV_RaTG13_orf1ab;1539 | 0 | 0.025 |

| | | |
|---|---|---|
| MN996532.1;QHR63299.1;BatCoV_RaTG13_orf1ab;1540 | 0 | 0 |
| MN996532.1;QHR63299.1;BatCoV_RaTG13_orf1ab;1541 | 0 | 0.025 |
| MN996532.1;QHR63299.1;BatCoV_RaTG13_orf1ab;1542 | 0 | 0.025 |
| MN996532.1;QHR63299.1;BatCoV_RaTG13_orf1ab;1543 | 0 | 0 |
| MN996532.1;QHR63299.1;BatCoV_RaTG13_orf1ab;1544 | 0 | 0 |
| MN996532.1;QHR63299.1;BatCoV_RaTG13_orf1ab;1545 | 0 | 0 |
| MN996532.1;QHR63299.1;BatCoV_RaTG13_orf1ab;1546 | 0 | 0 |
| MN996532.1;QHR63299.1;BatCoV_RaTG13_orf1ab;1547 | 0 | 0 |
| MN996532.1;QHR63299.1;BatCoV_RaTG13_orf1ab;1548 | 0 | 0 |
| MN996532.1;QHR63299.1;BatCoV_RaTG13_orf1ab;1549 | 0 | 0 |
| MN996532.1;QHR63299.1;BatCoV_RaTG13_orf1ab;1550 | 0 | 0 |
| MN996532.1;QHR63299.1;BatCoV_RaTG13_orf1ab;1551 | 0 | 0 |
| MN996532.1;QHR63299.1;BatCoV_RaTG13_orf1ab;1552 | 0 | 0 |
| MN996532.1;QHR63299.1;BatCoV_RaTG13_orf1ab;1553 | 0 | 0 |
| MN996532.1;QHR63299.1;BatCoV_RaTG13_orf1ab;1554 | 0 | 0 |
| MN996532.1;QHR63299.1;BatCoV_RaTG13_orf1ab;1555 | 0 | 0 |
| MN996532.1;QHR63299.1;BatCoV_RaTG13_orf1ab;1556 | 0 | 0 |
| MN996532.1;QHR63299.1;BatCoV_RaTG13_orf1ab;1557 | 0 | 0 |
| MN996532.1;QHR63299.1;BatCoV_RaTG13_orf1ab;1558 | 0 | 0 |
| MN996532.1;QHR63299.1;BatCoV_RaTG13_orf1ab;1559 | 0 | 0 |
| MN996532.1;QHR63299.1;BatCoV_RaTG13_orf1ab;1560 | 0 | 0 |
| MN996532.1;QHR63299.1;BatCoV_RaTG13_orf1ab;1561 | 0 | 0 |
| MN996532.1;QHR63299.1;BatCoV_RaTG13_orf1ab;1562 | 0 | 0.025 |
| MN996532.1;QHR63299.1;BatCoV_RaTG13_orf1ab;1563 | 0 | 0 |
| MN996532.1;QHR63299.1;BatCoV_RaTG13_orf1ab;1564 | 0 | 0 |
| MN996532.1;QHR63299.1;BatCoV_RaTG13_orf1ab;1565 | 0 | 0 |
| MN996532.1;QHR63299.1;BatCoV_RaTG13_orf1ab;1566 | 0 | 0.025 |
| MN996532.1;QHR63299.1;BatCoV_RaTG13_orf1ab;1567 | 0 | 0.025 |
| MN996532.1;QHR63299.1;BatCoV_RaTG13_orf1ab;1568 | 0 | 0.025 |
| MN996532.1;QHR63299.1;BatCoV_RaTG13_orf1ab;1569 | 0 | 0.025 |
| MN996532.1;QHR63299.1;BatCoV_RaTG13_orf1ab;1570 | 0 | 0.025 |
| MN996532.1;QHR63299.1;BatCoV_RaTG13_orf1ab;1571 | 0 | 0.025 |
| MN996532.1;QHR63299.1;BatCoV_RaTG13_orf1ab;1572 | 0.05 | 0.125 |
| MN996532.1;QHR63299.1;BatCoV_RaTG13_orf1ab;1573 | 0 | 0.075 |
| MN996532.1;QHR63299.1;BatCoV_RaTG13_orf1ab;1574 | 0 | 0.025 |
| MN996532.1;QHR63299.1;BatCoV_RaTG13_orf1ab;1575 | 0 | 0.025 |
| MN996532.1;QHR63299.1;BatCoV_RaTG13_orf1ab;1576 | 0 | 0.025 |
| MN996532.1;QHR63299.1;BatCoV_RaTG13_orf1ab;1577 | 0 | 0.025 |
| MN996532.1;QHR63299.1;BatCoV_RaTG13_orf1ab;1578 | 0 | 0.025 |
| MN996532.1;QHR63299.1;BatCoV_RaTG13_orf1ab;1579 | 0 | 0 |
| MN996532.1;QHR63299.1;BatCoV_RaTG13_orf1ab;1580 | 0 | 0 |
| MN996532.1;QHR63299.1;BatCoV_RaTG13_orf1ab;1581 | 0 | 0 |
| MN996532.1;QHR63299.1;BatCoV_RaTG13_orf1ab;1582 | 0 | 0 |

| | | |
|---|---|---|
| MN996532.1;QHR63299.1;BatCoV_RaTG13_orf1ab;1583 | 0 | 0 |
| MN996532.1;QHR63299.1;BatCoV_RaTG13_orf1ab;1584 | 0 | 0 |
| MN996532.1;QHR63299.1;BatCoV_RaTG13_orf1ab;1585 | 0 | 0 |
| MN996532.1;QHR63299.1;BatCoV_RaTG13_orf1ab;1586 | 0 | 0 |
| MN996532.1;QHR63299.1;BatCoV_RaTG13_orf1ab;1587 | 0 | 0 |
| MN996532.1;QHR63299.1;BatCoV_RaTG13_orf1ab;1588 | 0 | 0 |
| MN996532.1;QHR63299.1;BatCoV_RaTG13_orf1ab;1589 | 0 | 0 |
| MN996532.1;QHR63299.1;BatCoV_RaTG13_orf1ab;1590 | 0 | 0.025 |
| MN996532.1;QHR63299.1;BatCoV_RaTG13_orf1ab;1591 | 0 | 0.025 |
| MN996532.1;QHR63299.1;BatCoV_RaTG13_orf1ab;1592 | 0 | 0.025 |
| MN996532.1;QHR63299.1;BatCoV_RaTG13_orf1ab;1593 | 0 | 0.025 |
| MN996532.1;QHR63299.1;BatCoV_RaTG13_orf1ab;1594 | 0.15 | 0.125 |
| MN996532.1;QHR63299.1;BatCoV_RaTG13_orf1ab;1595 | 0 | 0.025 |
| MN996532.1;QHR63299.1;BatCoV_RaTG13_orf1ab;1596 | 0 | 0.025 |
| MN996532.1;QHR63299.1;BatCoV_RaTG13_orf1ab;1597 | 0 | 0 |
| MN996532.1;QHR63299.1;BatCoV_RaTG13_orf1ab;1598 | 0 | 0 |
| MN996532.1;QHR63299.1;BatCoV_RaTG13_orf1ab;1599 | 0 | 0 |
| MN996532.1;QHR63299.1;BatCoV_RaTG13_orf1ab;1600 | 0 | 0.025 |
| MN996532.1;QHR63299.1;BatCoV_RaTG13_orf1ab;1601 | 0.05 | 0 |
| MN996532.1;QHR63299.1;BatCoV_RaTG13_orf1ab;1602 | 0.05 | 0 |
| MN996532.1;QHR63299.1;BatCoV_RaTG13_orf1ab;1603 | 0 | 0 |
| MN996532.1;QHR63299.1;BatCoV_RaTG13_orf1ab;1604 | 0 | 0 |
| MN996532.1;QHR63299.1;BatCoV_RaTG13_orf1ab;1605 | 0 | 0 |
| MN996532.1;QHR63299.1;BatCoV_RaTG13_orf1ab;1606 | 0 | 0 |
| MN996532.1;QHR63299.1;BatCoV_RaTG13_orf1ab;1607 | 0 | 0 |
| MN996532.1;QHR63299.1;BatCoV_RaTG13_orf1ab;1608 | 0 | 0 |
| MN996532.1;QHR63299.1;BatCoV_RaTG13_orf1ab;1609 | 0 | 0 |
| MN996532.1;QHR63299.1;BatCoV_RaTG13_orf1ab;1610 | 0.05 | 0 |
| MN996532.1;QHR63299.1;BatCoV_RaTG13_orf1ab;1611 | 0.05 | 0 |
| MN996532.1;QHR63299.1;BatCoV_RaTG13_orf1ab;1612 | 0.05 | 0 |
| MN996532.1;QHR63299.1;BatCoV_RaTG13_orf1ab;1613 | 0.05 | 0 |
| MN996532.1;QHR63299.1;BatCoV_RaTG13_orf1ab;1614 | 0.05 | 0 |
| MN996532.1;QHR63299.1;BatCoV_RaTG13_orf1ab;1615 | 0.05 | 0 |
| MN996532.1;QHR63299.1;BatCoV_RaTG13_orf1ab;1616 | 0.05 | 0 |
| MN996532.1;QHR63299.1;BatCoV_RaTG13_orf1ab;1617 | 0 | 0 |
| MN996532.1;QHR63299.1;BatCoV_RaTG13_orf1ab;1618 | 0 | 0 |
| MN996532.1;QHR63299.1;BatCoV_RaTG13_orf1ab;1619 | 0 | 0 |
| MN996532.1;QHR63299.1;BatCoV_RaTG13_orf1ab;1620 | 0.05 | 0.025 |
| MN996532.1;QHR63299.1;BatCoV_RaTG13_orf1ab;1621 | 0 | 0 |
| MN996532.1;QHR63299.1;BatCoV_RaTG13_orf1ab;1622 | 0.05 | 0 |
| MN996532.1;QHR63299.1;BatCoV_RaTG13_orf1ab;1623 | 0 | 0 |
| MN996532.1;QHR63299.1;BatCoV_RaTG13_orf1ab;1624 | 0 | 0 |
| MN996532.1;QHR63299.1;BatCoV_RaTG13_orf1ab;1625 | 0 | 0 |

| | | |
|---|---|---|
| MN996532.1;QHR63299.1;BatCoV_RaTG13_orf1ab;1626 | 0 | 0 |
| MN996532.1;QHR63299.1;BatCoV_RaTG13_orf1ab;1627 | 0 | 0 |
| MN996532.1;QHR63299.1;BatCoV_RaTG13_orf1ab;1628 | 0 | 0.025 |
| MN996532.1;QHR63299.1;BatCoV_RaTG13_orf1ab;1629 | 0 | 0 |
| MN996532.1;QHR63299.1;BatCoV_RaTG13_orf1ab;1630 | 0.05 | 0 |
| MN996532.1;QHR63299.1;BatCoV_RaTG13_orf1ab;1631 | 0.05 | 0 |
| MN996532.1;QHR63299.1;BatCoV_RaTG13_orf1ab;1632 | 0.05 | 0 |
| MN996532.1;QHR63299.1;BatCoV_RaTG13_orf1ab;1633 | 0.05 | 0 |
| MN996532.1;QHR63299.1;BatCoV_RaTG13_orf1ab;1634 | 0 | 0 |
| MN996532.1;QHR63299.1;BatCoV_RaTG13_orf1ab;1635 | 0 | 0 |
| MN996532.1;QHR63299.1;BatCoV_RaTG13_orf1ab;1636 | 0 | 0 |
| MN996532.1;QHR63299.1;BatCoV_RaTG13_orf1ab;1637 | 0 | 0 |
| MN996532.1;QHR63299.1;BatCoV_RaTG13_orf1ab;1638 | 0.05 | 0 |
| MN996532.1;QHR63299.1;BatCoV_RaTG13_orf1ab;1639 | 0 | 0 |
| MN996532.1;QHR63299.1;BatCoV_RaTG13_orf1ab;1640 | 0 | 0 |
| MN996532.1;QHR63299.1;BatCoV_RaTG13_orf1ab;1641 | 0 | 0 |
| MN996532.1;QHR63299.1;BatCoV_RaTG13_orf1ab;1642 | 0 | 0 |
| MN996532.1;QHR63299.1;BatCoV_RaTG13_orf1ab;1643 | 0 | 0 |
| MN996532.1;QHR63299.1;BatCoV_RaTG13_orf1ab;1644 | 0 | 0 |
| MN996532.1;QHR63299.1;BatCoV_RaTG13_orf1ab;1645 | 0 | 0 |
| MN996532.1;QHR63299.1;BatCoV_RaTG13_orf1ab;1646 | 0 | 0 |
| MN996532.1;QHR63299.1;BatCoV_RaTG13_orf1ab;1647 | 0 | 0 |
| MN996532.1;QHR63299.1;BatCoV_RaTG13_orf1ab;1648 | 0 | 0 |
| MN996532.1;QHR63299.1;BatCoV_RaTG13_orf1ab;1649 | 0 | 0 |
| MN996532.1;QHR63299.1;BatCoV_RaTG13_orf1ab;1650 | 0 | 0 |
| MN996532.1;QHR63299.1;BatCoV_RaTG13_orf1ab;1651 | 0 | 0 |
| MN996532.1;QHR63299.1;BatCoV_RaTG13_orf1ab;1652 | 0 | 0 |
| MN996532.1;QHR63299.1;BatCoV_RaTG13_orf1ab;1653 | 0 | 0 |
| MN996532.1;QHR63299.1;BatCoV_RaTG13_orf1ab;1654 | 0 | 0 |
| MN996532.1;QHR63299.1;BatCoV_RaTG13_orf1ab;1655 | 0 | 0 |
| MN996532.1;QHR63299.1;BatCoV_RaTG13_orf1ab;1656 | 0.05 | 0.025 |
| MN996532.1;QHR63299.1;BatCoV_RaTG13_orf1ab;1657 | 0.05 | 0.05 |
| MN996532.1;QHR63299.1;BatCoV_RaTG13_orf1ab;1658 | 0.05 | 0.025 |
| MN996532.1;QHR63299.1;BatCoV_RaTG13_orf1ab;1659 | 0.05 | 0 |
| MN996532.1;QHR63299.1;BatCoV_RaTG13_orf1ab;1660 | 0.05 | 0 |
| MN996532.1;QHR63299.1;BatCoV_RaTG13_orf1ab;1661 | 0.05 | 0 |
| MN996532.1;QHR63299.1;BatCoV_RaTG13_orf1ab;1662 | 0.05 | 0 |
| MN996532.1;QHR63299.1;BatCoV_RaTG13_orf1ab;1663 | 0.05 | 0 |
| MN996532.1;QHR63299.1;BatCoV_RaTG13_orf1ab;1664 | 0 | 0 |
| MN996532.1;QHR63299.1;BatCoV_RaTG13_orf1ab;1665 | 0 | 0 |
| MN996532.1;QHR63299.1;BatCoV_RaTG13_orf1ab;1666 | 0 | 0 |
| MN996532.1;QHR63299.1;BatCoV_RaTG13_orf1ab;1667 | 0 | 0 |
| MN996532.1;QHR63299.1;BatCoV_RaTG13_orf1ab;1668 | 0 | 0 |

| | | |
|---|---|---|
| MN996532.1;QHR63299.1;BatCoV_RaTG13_orf1ab;1669 | 0 | 0 |
| MN996532.1;QHR63299.1;BatCoV_RaTG13_orf1ab;1670 | 0 | 0 |
| MN996532.1;QHR63299.1;BatCoV_RaTG13_orf1ab;1671 | 0 | 0 |
| MN996532.1;QHR63299.1;BatCoV_RaTG13_orf1ab;1672 | 0 | 0 |
| MN996532.1;QHR63299.1;BatCoV_RaTG13_orf1ab;1673 | 0 | 0 |
| MN996532.1;QHR63299.1;BatCoV_RaTG13_orf1ab;1674 | 0 | 0 |
| MN996532.1;QHR63299.1;BatCoV_RaTG13_orf1ab;1675 | 0 | 0 |
| MN996532.1;QHR63299.1;BatCoV_RaTG13_orf1ab;1676 | 0 | 0 |
| MN996532.1;QHR63299.1;BatCoV_RaTG13_orf1ab;1677 | 0 | 0 |
| MN996532.1;QHR63299.1;BatCoV_RaTG13_orf1ab;1678 | 0.05 | 0 |
| MN996532.1;QHR63299.1;BatCoV_RaTG13_orf1ab;1679 | 0 | 0 |
| MN996532.1;QHR63299.1;BatCoV_RaTG13_orf1ab;1680 | 0 | 0 |
| MN996532.1;QHR63299.1;BatCoV_RaTG13_orf1ab;1681 | 0 | 0.025 |
| MN996532.1;QHR63299.1;BatCoV_RaTG13_orf1ab;1682 | 0 | 0 |
| MN996532.1;QHR63299.1;BatCoV_RaTG13_orf1ab;1683 | 0 | 0 |
| MN996532.1;QHR63299.1;BatCoV_RaTG13_orf1ab;1684 | 0 | 0 |
| MN996532.1;QHR63299.1;BatCoV_RaTG13_orf1ab;1685 | 0 | 0.05 |
| MN996532.1;QHR63299.1;BatCoV_RaTG13_orf1ab;1686 | 0 | 0.025 |
| MN996532.1;QHR63299.1;BatCoV_RaTG13_orf1ab;1687 | 0 | 0 |
| MN996532.1;QHR63299.1;BatCoV_RaTG13_orf1ab;1688 | 0 | 0 |
| MN996532.1;QHR63299.1;BatCoV_RaTG13_orf1ab;1689 | 0 | 0 |
| MN996532.1;QHR63299.1;BatCoV_RaTG13_orf1ab;1690 | 0 | 0 |
| MN996532.1;QHR63299.1;BatCoV_RaTG13_orf1ab;1691 | 0.05 | 0.025 |
| MN996532.1;QHR63299.1;BatCoV_RaTG13_orf1ab;1692 | 0.1 | 0.075 |
| MN996532.1;QHR63299.1;BatCoV_RaTG13_orf1ab;1693 | 0.05 | 0.025 |
| MN996532.1;QHR63299.1;BatCoV_RaTG13_orf1ab;1694 | 0.15 | 0.025 |
| MN996532.1;QHR63299.1;BatCoV_RaTG13_orf1ab;1695 | 0.05 | 0.025 |
| MN996532.1;QHR63299.1;BatCoV_RaTG13_orf1ab;1696 | 0.05 | 0 |
| MN996532.1;QHR63299.1;BatCoV_RaTG13_orf1ab;1697 | 0 | 0 |
| MN996532.1;QHR63299.1;BatCoV_RaTG13_orf1ab;1698 | 0 | 0 |
| MN996532.1;QHR63299.1;BatCoV_RaTG13_orf1ab;1699 | 0 | 0 |
| MN996532.1;QHR63299.1;BatCoV_RaTG13_orf1ab;1700 | 0 | 0 |
| MN996532.1;QHR63299.1;BatCoV_RaTG13_orf1ab;1701 | 0 | 0 |
| MN996532.1;QHR63299.1;BatCoV_RaTG13_orf1ab;1702 | 0 | 0 |
| MN996532.1;QHR63299.1;BatCoV_RaTG13_orf1ab;1703 | 0 | 0 |
| MN996532.1;QHR63299.1;BatCoV_RaTG13_orf1ab;1704 | 0 | 0 |
| MN996532.1;QHR63299.1;BatCoV_RaTG13_orf1ab;1705 | 0 | 0 |
| MN996532.1;QHR63299.1;BatCoV_RaTG13_orf1ab;1706 | 0 | 0 |
| MN996532.1;QHR63299.1;BatCoV_RaTG13_orf1ab;1707 | 0 | 0 |
| MN996532.1;QHR63299.1;BatCoV_RaTG13_orf1ab;1708 | 0 | 0 |
| MN996532.1;QHR63299.1;BatCoV_RaTG13_orf1ab;1709 | 0 | 0 |
| MN996532.1;QHR63299.1;BatCoV_RaTG13_orf1ab;1710 | 0 | 0.025 |
| MN996532.1;QHR63299.1;BatCoV_RaTG13_orf1ab;1711 | 0 | 0.025 |

| | | |
|---|---|---|
| MN996532.1;QHR63299.1;BatCoV_RaTG13_orf1ab;1712 | 0 | 0.05 |
| MN996532.1;QHR63299.1;BatCoV_RaTG13_orf1ab;1713 | 0 | 0.025 |
| MN996532.1;QHR63299.1;BatCoV_RaTG13_orf1ab;1714 | 0 | 0 |
| MN996532.1;QHR63299.1;BatCoV_RaTG13_orf1ab;1715 | 0 | 0 |
| MN996532.1;QHR63299.1;BatCoV_RaTG13_orf1ab;1716 | 0 | 0 |
| MN996532.1;QHR63299.1;BatCoV_RaTG13_orf1ab;1717 | 0 | 0 |
| MN996532.1;QHR63299.1;BatCoV_RaTG13_orf1ab;1718 | 0 | 0.05 |
| MN996532.1;QHR63299.1;BatCoV_RaTG13_orf1ab;1719 | 0 | 0.025 |
| MN996532.1;QHR63299.1;BatCoV_RaTG13_orf1ab;1720 | 0 | 0.025 |
| MN996532.1;QHR63299.1;BatCoV_RaTG13_orf1ab;1721 | 0 | 0 |
| MN996532.1;QHR63299.1;BatCoV_RaTG13_orf1ab;1722 | 0.05 | 0.025 |
| MN996532.1;QHR63299.1;BatCoV_RaTG13_orf1ab;1723 | 0 | 0 |
| MN996532.1;QHR63299.1;BatCoV_RaTG13_orf1ab;1724 | 0 | 0 |
| MN996532.1;QHR63299.1;BatCoV_RaTG13_orf1ab;1725 | 0.05 | 0 |
| MN996532.1;QHR63299.1;BatCoV_RaTG13_orf1ab;1726 | 0.05 | 0.025 |
| MN996532.1;QHR63299.1;BatCoV_RaTG13_orf1ab;1727 | 0.05 | 0.025 |
| MN996532.1;QHR63299.1;BatCoV_RaTG13_orf1ab;1728 | 0.05 | 0.025 |
| MN996532.1;QHR63299.1;BatCoV_RaTG13_orf1ab;1729 | 0.05 | 0.025 |
| MN996532.1;QHR63299.1;BatCoV_RaTG13_orf1ab;1730 | 0 | 0 |
| MN996532.1;QHR63299.1;BatCoV_RaTG13_orf1ab;1731 | 0 | 0 |
| MN996532.1;QHR63299.1;BatCoV_RaTG13_orf1ab;1732 | 0 | 0 |
| MN996532.1;QHR63299.1;BatCoV_RaTG13_orf1ab;1733 | 0 | 0 |
| MN996532.1;QHR63299.1;BatCoV_RaTG13_orf1ab;1734 | 0 | 0 |
| MN996532.1;QHR63299.1;BatCoV_RaTG13_orf1ab;1735 | 0 | 0 |
| MN996532.1;QHR63299.1;BatCoV_RaTG13_orf1ab;1736 | 0 | 0 |
| MN996532.1;QHR63299.1;BatCoV_RaTG13_orf1ab;1737 | 0 | 0 |
| MN996532.1;QHR63299.1;BatCoV_RaTG13_orf1ab;1738 | 0 | 0 |
| MN996532.1;QHR63299.1;BatCoV_RaTG13_orf1ab;1739 | 0 | 0 |
| MN996532.1;QHR63299.1;BatCoV_RaTG13_orf1ab;1740 | 0 | 0 |
| MN996532.1;QHR63299.1;BatCoV_RaTG13_orf1ab;1741 | 0 | 0 |
| MN996532.1;QHR63299.1;BatCoV_RaTG13_orf1ab;1742 | 0 | 0 |
| MN996532.1;QHR63299.1;BatCoV_RaTG13_orf1ab;1743 | 0 | 0 |
| MN996532.1;QHR63299.1;BatCoV_RaTG13_orf1ab;1744 | 0 | 0.075 |
| MN996532.1;QHR63299.1;BatCoV_RaTG13_orf1ab;1745 | 0 | 0.025 |
| MN996532.1;QHR63299.1;BatCoV_RaTG13_orf1ab;1746 | 0 | 0.025 |
| MN996532.1;QHR63299.1;BatCoV_RaTG13_orf1ab;1747 | 0.1 | 0.325 |
| MN996532.1;QHR63299.1;BatCoV_RaTG13_orf1ab;1748 | 0.1 | 0.3 |
| MN996532.1;QHR63299.1;BatCoV_RaTG13_orf1ab;1749 | 0.1 | 0.3 |
| MN996532.1;QHR63299.1;BatCoV_RaTG13_orf1ab;1750 | 0.15 | 0.325 |
| MN996532.1;QHR63299.1;BatCoV_RaTG13_orf1ab;1751 | 0.2 | 0.325 |
| MN996532.1;QHR63299.1;BatCoV_RaTG13_orf1ab;1752 | 0.1 | 0.35 |
| MN996532.1;QHR63299.1;BatCoV_RaTG13_orf1ab;1753 | 0.05 | 0.275 |
| MN996532.1;QHR63299.1;BatCoV_RaTG13_orf1ab;1754 | 0.05 | 0.2 |

| | | |
|---|---|---|
| MN996532.1;QHR63299.1;BatCoV_RaTG13_orf1ab;1755 | 0.1 | 0.25 |
| MN996532.1;QHR63299.1;BatCoV_RaTG13_orf1ab;1756 | 0.05 | 0.2 |
| MN996532.1;QHR63299.1;BatCoV_RaTG13_orf1ab;1757 | 0 | 0.125 |
| MN996532.1;QHR63299.1;BatCoV_RaTG13_orf1ab;1758 | 0 | 0 |
| MN996532.1;QHR63299.1;BatCoV_RaTG13_orf1ab;1759 | 0 | 0 |
| MN996532.1;QHR63299.1;BatCoV_RaTG13_orf1ab;1760 | 0.1 | 0 |
| MN996532.1;QHR63299.1;BatCoV_RaTG13_orf1ab;1761 | 0.15 | 0 |
| MN996532.1;QHR63299.1;BatCoV_RaTG13_orf1ab;1762 | 0.15 | 0 |
| MN996532.1;QHR63299.1;BatCoV_RaTG13_orf1ab;1763 | 0.2 | 0 |
| MN996532.1;QHR63299.1;BatCoV_RaTG13_orf1ab;1764 | 0.1 | 0 |
| MN996532.1;QHR63299.1;BatCoV_RaTG13_orf1ab;1765 | 0.05 | 0 |
| MN996532.1;QHR63299.1;BatCoV_RaTG13_orf1ab;1766 | 0 | 0 |
| MN996532.1;QHR63299.1;BatCoV_RaTG13_orf1ab;1767 | 0.05 | 0 |
| MN996532.1;QHR63299.1;BatCoV_RaTG13_orf1ab;1768 | 0.05 | 0 |
| MN996532.1;QHR63299.1;BatCoV_RaTG13_orf1ab;1769 | 0.05 | 0 |
| MN996532.1;QHR63299.1;BatCoV_RaTG13_orf1ab;1770 | 0.05 | 0.025 |
| MN996532.1;QHR63299.1;BatCoV_RaTG13_orf1ab;1771 | 0 | 0 |
| MN996532.1;QHR63299.1;BatCoV_RaTG13_orf1ab;1772 | 0 | 0.025 |
| MN996532.1;QHR63299.1;BatCoV_RaTG13_orf1ab;1773 | 0.05 | 0.05 |
| MN996532.1;QHR63299.1;BatCoV_RaTG13_orf1ab;1774 | 0.1 | 0 |
| MN996532.1;QHR63299.1;BatCoV_RaTG13_orf1ab;1775 | 0 | 0 |
| MN996532.1;QHR63299.1;BatCoV_RaTG13_orf1ab;1776 | 0 | 0 |
| MN996532.1;QHR63299.1;BatCoV_RaTG13_orf1ab;1777 | 0 | 0 |
| MN996532.1;QHR63299.1;BatCoV_RaTG13_orf1ab;1778 | 0 | 0 |
| MN996532.1;QHR63299.1;BatCoV_RaTG13_orf1ab;1779 | 0 | 0 |
| MN996532.1;QHR63299.1;BatCoV_RaTG13_orf1ab;1780 | 0 | 0.075 |
| MN996532.1;QHR63299.1;BatCoV_RaTG13_orf1ab;1781 | 0 | 0.025 |
| MN996532.1;QHR63299.1;BatCoV_RaTG13_orf1ab;1782 | 0 | 0.025 |
| MN996532.1;QHR63299.1;BatCoV_RaTG13_orf1ab;1783 | 0 | 0.025 |
| MN996532.1;QHR63299.1;BatCoV_RaTG13_orf1ab;1784 | 0 | 0.025 |
| MN996532.1;QHR63299.1;BatCoV_RaTG13_orf1ab;1785 | 0.05 | 0.025 |
| MN996532.1;QHR63299.1;BatCoV_RaTG13_orf1ab;1786 | 0 | 0.025 |
| MN996532.1;QHR63299.1;BatCoV_RaTG13_orf1ab;1787 | 0 | 0.025 |
| MN996532.1;QHR63299.1;BatCoV_RaTG13_orf1ab;1788 | 0 | 0.025 |
| MN996532.1;QHR63299.1;BatCoV_RaTG13_orf1ab;1789 | 0 | 0.025 |
| MN996532.1;QHR63299.1;BatCoV_RaTG13_orf1ab;1790 | 0 | 0.05 |
| MN996532.1;QHR63299.1;BatCoV_RaTG13_orf1ab;1791 | 0 | 0 |
| MN996532.1;QHR63299.1;BatCoV_RaTG13_orf1ab;1792 | 0.05 | 0 |
| MN996532.1;QHR63299.1;BatCoV_RaTG13_orf1ab;1793 | 0 | 0 |
| MN996532.1;QHR63299.1;BatCoV_RaTG13_orf1ab;1794 | 0.05 | 0 |
| MN996532.1;QHR63299.1;BatCoV_RaTG13_orf1ab;1795 | 0 | 0 |
| MN996532.1;QHR63299.1;BatCoV_RaTG13_orf1ab;1796 | 0.1 | 0 |
| MN996532.1;QHR63299.1;BatCoV_RaTG13_orf1ab;1797 | 0 | 0.025 |

| | | |
|---|---|---|
| MN996532.1;QHR63299.1;BatCoV_RaTG13_orf1ab;1798 | 0 | 0 |
| MN996532.1;QHR63299.1;BatCoV_RaTG13_orf1ab;1799 | 0 | 0 |
| MN996532.1;QHR63299.1;BatCoV_RaTG13_orf1ab;1800 | 0 | 0.025 |
| MN996532.1;QHR63299.1;BatCoV_RaTG13_orf1ab;1801 | 0 | 0 |
| MN996532.1;QHR63299.1;BatCoV_RaTG13_orf1ab;1802 | 0 | 0 |
| MN996532.1;QHR63299.1;BatCoV_RaTG13_orf1ab;1803 | 0 | 0 |
| MN996532.1;QHR63299.1;BatCoV_RaTG13_orf1ab;1804 | 0 | 0 |
| MN996532.1;QHR63299.1;BatCoV_RaTG13_orf1ab;1805 | 0 | 0 |
| MN996532.1;QHR63299.1;BatCoV_RaTG13_orf1ab;1806 | 0 | 0 |
| MN996532.1;QHR63299.1;BatCoV_RaTG13_orf1ab;1807 | 0 | 0 |
| MN996532.1;QHR63299.1;BatCoV_RaTG13_orf1ab;1808 | 0 | 0 |
| MN996532.1;QHR63299.1;BatCoV_RaTG13_orf1ab;1809 | 0 | 0 |
| MN996532.1;QHR63299.1;BatCoV_RaTG13_orf1ab;1810 | 0 | 0 |
| MN996532.1;QHR63299.1;BatCoV_RaTG13_orf1ab;1811 | 0 | 0 |
| MN996532.1;QHR63299.1;BatCoV_RaTG13_orf1ab;1812 | 0 | 0 |
| MN996532.1;QHR63299.1;BatCoV_RaTG13_orf1ab;1813 | 0 | 0 |
| MN996532.1;QHR63299.1;BatCoV_RaTG13_orf1ab;1814 | 0 | 0 |
| MN996532.1;QHR63299.1;BatCoV_RaTG13_orf1ab;1815 | 0 | 0 |
| MN996532.1;QHR63299.1;BatCoV_RaTG13_orf1ab;1816 | 0 | 0 |
| MN996532.1;QHR63299.1;BatCoV_RaTG13_orf1ab;1817 | 0 | 0 |
| MN996532.1;QHR63299.1;BatCoV_RaTG13_orf1ab;1818 | 0 | 0 |
| MN996532.1;QHR63299.1;BatCoV_RaTG13_orf1ab;1819 | 0 | 0 |
| MN996532.1;QHR63299.1;BatCoV_RaTG13_orf1ab;1820 | 0 | 0 |
| MN996532.1;QHR63299.1;BatCoV_RaTG13_orf1ab;1821 | 0 | 0 |
| MN996532.1;QHR63299.1;BatCoV_RaTG13_orf1ab;1822 | 0 | 0 |
| MN996532.1;QHR63299.1;BatCoV_RaTG13_orf1ab;1823 | 0 | 0 |
| MN996532.1;QHR63299.1;BatCoV_RaTG13_orf1ab;1824 | 0 | 0 |
| MN996532.1;QHR63299.1;BatCoV_RaTG13_orf1ab;1825 | 0 | 0.15 |
| MN996532.1;QHR63299.1;BatCoV_RaTG13_orf1ab;1826 | 0 | 0 |
| MN996532.1;QHR63299.1;BatCoV_RaTG13_orf1ab;1827 | 0 | 0 |
| MN996532.1;QHR63299.1;BatCoV_RaTG13_orf1ab;1828 | 0 | 0 |
| MN996532.1;QHR63299.1;BatCoV_RaTG13_orf1ab;1829 | 0 | 0 |
| MN996532.1;QHR63299.1;BatCoV_RaTG13_orf1ab;1830 | 0.05 | 0 |
| MN996532.1;QHR63299.1;BatCoV_RaTG13_orf1ab;1831 | 0.05 | 0 |
| MN996532.1;QHR63299.1;BatCoV_RaTG13_orf1ab;1832 | 0 | 0 |
| MN996532.1;QHR63299.1;BatCoV_RaTG13_orf1ab;1833 | 0 | 0 |
| MN996532.1;QHR63299.1;BatCoV_RaTG13_orf1ab;1834 | 0 | 0 |
| MN996532.1;QHR63299.1;BatCoV_RaTG13_orf1ab;1835 | 0 | 0 |
| MN996532.1;QHR63299.1;BatCoV_RaTG13_orf1ab;1836 | 0 | 0 |
| MN996532.1;QHR63299.1;BatCoV_RaTG13_orf1ab;1837 | 0 | 0 |
| MN996532.1;QHR63299.1;BatCoV_RaTG13_orf1ab;1838 | 0 | 0 |
| MN996532.1;QHR63299.1;BatCoV_RaTG13_orf1ab;1839 | 0 | 0 |
| MN996532.1;QHR63299.1;BatCoV_RaTG13_orf1ab;1840 | 0 | 0 |

| | | |
|---|---|---|
| MN996532.1;QHR63299.1;BatCoV_RaTG13_orf1ab;1841 | 0 | 0 |
| MN996532.1;QHR63299.1;BatCoV_RaTG13_orf1ab;1842 | 0 | 0 |
| MN996532.1;QHR63299.1;BatCoV_RaTG13_orf1ab;1843 | 0 | 0 |
| MN996532.1;QHR63299.1;BatCoV_RaTG13_orf1ab;1844 | 0 | 0 |
| MN996532.1;QHR63299.1;BatCoV_RaTG13_orf1ab;1845 | 0 | 0 |
| MN996532.1;QHR63299.1;BatCoV_RaTG13_orf1ab;1846 | 0 | 0.025 |
| MN996532.1;QHR63299.1;BatCoV_RaTG13_orf1ab;1847 | 0 | 0 |
| MN996532.1;QHR63299.1;BatCoV_RaTG13_orf1ab;1848 | 0 | 0.025 |
| MN996532.1;QHR63299.1;BatCoV_RaTG13_orf1ab;1849 | 0 | 0.075 |
| MN996532.1;QHR63299.1;BatCoV_RaTG13_orf1ab;1850 | 0 | 0.05 |
| MN996532.1;QHR63299.1;BatCoV_RaTG13_orf1ab;1851 | 0 | 0 |
| MN996532.1;QHR63299.1;BatCoV_RaTG13_orf1ab;1852 | 0 | 0.025 |
| MN996532.1;QHR63299.1;BatCoV_RaTG13_orf1ab;1853 | 0 | 0 |
| MN996532.1;QHR63299.1;BatCoV_RaTG13_orf1ab;1854 | 0.05 | 0.075 |
| MN996532.1;QHR63299.1;BatCoV_RaTG13_orf1ab;1855 | 0 | 0.025 |
| MN996532.1;QHR63299.1;BatCoV_RaTG13_orf1ab;1856 | 0.05 | 0.025 |
| MN996532.1;QHR63299.1;BatCoV_RaTG13_orf1ab;1857 | 0.05 | 0.05 |
| MN996532.1;QHR63299.1;BatCoV_RaTG13_orf1ab;1858 | 0 | 0.025 |
| MN996532.1;QHR63299.1;BatCoV_RaTG13_orf1ab;1859 | 0 | 0.05 |
| MN996532.1;QHR63299.1;BatCoV_RaTG13_orf1ab;1860 | 0.2 | 0.125 |
| MN996532.1;QHR63299.1;BatCoV_RaTG13_orf1ab;1861 | 0 | 0 |
| MN996532.1;QHR63299.1;BatCoV_RaTG13_orf1ab;1862 | 0 | 0.025 |
| MN996532.1;QHR63299.1;BatCoV_RaTG13_orf1ab;1863 | 0 | 0.025 |
| MN996532.1;QHR63299.1;BatCoV_RaTG13_orf1ab;1864 | 0 | 0.025 |
| MN996532.1;QHR63299.1;BatCoV_RaTG13_orf1ab;1865 | 0 | 0.025 |
| MN996532.1;QHR63299.1;BatCoV_RaTG13_orf1ab;1866 | 0 | 0.025 |
| MN996532.1;QHR63299.1;BatCoV_RaTG13_orf1ab;1867 | 0 | 0.025 |
| MN996532.1;QHR63299.1;BatCoV_RaTG13_orf1ab;1868 | 0 | 0.025 |
| MN996532.1;QHR63299.1;BatCoV_RaTG13_orf1ab;1869 | 0 | 0.025 |
| MN996532.1;QHR63299.1;BatCoV_RaTG13_orf1ab;1870 | 0 | 0.025 |
| MN996532.1;QHR63299.1;BatCoV_RaTG13_orf1ab;1871 | 0 | 0.025 |
| MN996532.1;QHR63299.1;BatCoV_RaTG13_orf1ab;1872 | 0 | 0 |
| MN996532.1;QHR63299.1;BatCoV_RaTG13_orf1ab;1873 | 0 | 0 |
| MN996532.1;QHR63299.1;BatCoV_RaTG13_orf1ab;1874 | 0 | 0 |
| MN996532.1;QHR63299.1;BatCoV_RaTG13_orf1ab;1875 | 0 | 0 |
| MN996532.1;QHR63299.1;BatCoV_RaTG13_orf1ab;1876 | 0 | 0 |
| MN996532.1;QHR63299.1;BatCoV_RaTG13_orf1ab;1877 | 0 | 0 |
| MN996532.1;QHR63299.1;BatCoV_RaTG13_orf1ab;1878 | 0 | 0 |
| MN996532.1;QHR63299.1;BatCoV_RaTG13_orf1ab;1879 | 0 | 0 |
| MN996532.1;QHR63299.1;BatCoV_RaTG13_orf1ab;1880 | 0 | 0.025 |
| MN996532.1;QHR63299.1;BatCoV_RaTG13_orf1ab;1881 | 0 | 0 |
| MN996532.1;QHR63299.1;BatCoV_RaTG13_orf1ab;1882 | 0 | 0 |
| MN996532.1;QHR63299.1;BatCoV_RaTG13_orf1ab;1883 | 0 | 0 |

| | | |
|---|---|---|
| MN996532.1;QHR63299.1;BatCoV_RaTG13_orf1ab;1884 | 0 | 0 |
| MN996532.1;QHR63299.1;BatCoV_RaTG13_orf1ab;1885 | 0.05 | 0.125 |
| MN996532.1;QHR63299.1;BatCoV_RaTG13_orf1ab;1886 | 0 | 0.1 |
| MN996532.1;QHR63299.1;BatCoV_RaTG13_orf1ab;1887 | 0 | 0.15 |
| MN996532.1;QHR63299.1;BatCoV_RaTG13_orf1ab;1888 | 0 | 0.175 |
| MN996532.1;QHR63299.1;BatCoV_RaTG13_orf1ab;1889 | 0.2 | 0.25 |
| MN996532.1;QHR63299.1;BatCoV_RaTG13_orf1ab;1890 | 0.15 | 0.35 |
| MN996532.1;QHR63299.1;BatCoV_RaTG13_orf1ab;1891 | 0.1 | 0.225 |
| MN996532.1;QHR63299.1;BatCoV_RaTG13_orf1ab;1892 | 0.05 | 0.175 |
| MN996532.1;QHR63299.1;BatCoV_RaTG13_orf1ab;1893 | 0.3 | 0.175 |
| MN996532.1;QHR63299.1;BatCoV_RaTG13_orf1ab;1894 | 0.35 | 0.175 |
| MN996532.1;QHR63299.1;BatCoV_RaTG13_orf1ab;1895 | 0.2 | 0.175 |
| MN996532.1;QHR63299.1;BatCoV_RaTG13_orf1ab;1896 | 0.3 | 0.1 |
| MN996532.1;QHR63299.1;BatCoV_RaTG13_orf1ab;1897 | 0.15 | 0.075 |
| MN996532.1;QHR63299.1;BatCoV_RaTG13_orf1ab;1898 | 0.2 | 0.05 |
| MN996532.1;QHR63299.1;BatCoV_RaTG13_orf1ab;1899 | 0.15 | 0.05 |
| MN996532.1;QHR63299.1;BatCoV_RaTG13_orf1ab;1900 | 0.15 | 0.15 |
| MN996532.1;QHR63299.1;BatCoV_RaTG13_orf1ab;1901 | 0 | 0.025 |
| MN996532.1;QHR63299.1;BatCoV_RaTG13_orf1ab;1902 | 0.05 | 0 |
| MN996532.1;QHR63299.1;BatCoV_RaTG13_orf1ab;1903 | 0.05 | 0 |
| MN996532.1;QHR63299.1;BatCoV_RaTG13_orf1ab;1904 | 0.15 | 0.2 |
| MN996532.1;QHR63299.1;BatCoV_RaTG13_orf1ab;1905 | 0.05 | 0 |
| MN996532.1;QHR63299.1;BatCoV_RaTG13_orf1ab;1906 | 0.05 | 0 |
| MN996532.1;QHR63299.1;BatCoV_RaTG13_orf1ab;1907 | 0.1 | 0 |
| MN996532.1;QHR63299.1;BatCoV_RaTG13_orf1ab;1908 | 0 | 0.025 |
| MN996532.1;QHR63299.1;BatCoV_RaTG13_orf1ab;1909 | 0.2 | 0.1 |
| MN996532.1;QHR63299.1;BatCoV_RaTG13_orf1ab;1910 | 0 | 0.075 |
| MN996532.1;QHR63299.1;BatCoV_RaTG13_orf1ab;1911 | 0 | 0.05 |
| MN996532.1;QHR63299.1;BatCoV_RaTG13_orf1ab;1912 | 0 | 0.05 |
| MN996532.1;QHR63299.1;BatCoV_RaTG13_orf1ab;1913 | 0.05 | 0.075 |
| MN996532.1;QHR63299.1;BatCoV_RaTG13_orf1ab;1914 | 0.05 | 0.05 |
| MN996532.1;QHR63299.1;BatCoV_RaTG13_orf1ab;1915 | 0.05 | 0.025 |
| MN996532.1;QHR63299.1;BatCoV_RaTG13_orf1ab;1916 | 0.05 | 0.025 |
| MN996532.1;QHR63299.1;BatCoV_RaTG13_orf1ab;1917 | 0.05 | 0.025 |
| MN996532.1;QHR63299.1;BatCoV_RaTG13_orf1ab;1918 | 0.05 | 0.025 |
| MN996532.1;QHR63299.1;BatCoV_RaTG13_orf1ab;1919 | 0.05 | 0 |
| MN996532.1;QHR63299.1;BatCoV_RaTG13_orf1ab;1920 | 0.05 | 0 |
| MN996532.1;QHR63299.1;BatCoV_RaTG13_orf1ab;1921 | 0 | 0 |
| MN996532.1;QHR63299.1;BatCoV_RaTG13_orf1ab;1922 | 0 | 0.025 |
| MN996532.1;QHR63299.1;BatCoV_RaTG13_orf1ab;1923 | 0 | 0 |
| MN996532.1;QHR63299.1;BatCoV_RaTG13_orf1ab;1924 | 0 | 0 |
| MN996532.1;QHR63299.1;BatCoV_RaTG13_orf1ab;1925 | 0 | 0 |
| MN996532.1;QHR63299.1;BatCoV_RaTG13_orf1ab;1926 | 0.1 | 0.025 |

| | | |
|---|---|---|
| MN996532.1;QHR63299.1;BatCoV_RaTG13_orf1ab;1927 | 0.05 | 0.05 |
| MN996532.1;QHR63299.1;BatCoV_RaTG13_orf1ab;1928 | 0 | 0.075 |
| MN996532.1;QHR63299.1;BatCoV_RaTG13_orf1ab;1929 | 0 | 0.025 |
| MN996532.1;QHR63299.1;BatCoV_RaTG13_orf1ab;1930 | 0 | 0.025 |
| MN996532.1;QHR63299.1;BatCoV_RaTG13_orf1ab;1931 | 0 | 0.025 |
| MN996532.1;QHR63299.1;BatCoV_RaTG13_orf1ab;1932 | 0.05 | 0.025 |
| MN996532.1;QHR63299.1;BatCoV_RaTG13_orf1ab;1933 | 0.05 | 0 |
| MN996532.1;QHR63299.1;BatCoV_RaTG13_orf1ab;1934 | 0.1 | 0.025 |
| MN996532.1;QHR63299.1;BatCoV_RaTG13_orf1ab;1935 | 0.05 | 0 |
| MN996532.1;QHR63299.1;BatCoV_RaTG13_orf1ab;1936 | 0.05 | 0 |
| MN996532.1;QHR63299.1;BatCoV_RaTG13_orf1ab;1937 | 0.1 | 0.025 |
| MN996532.1;QHR63299.1;BatCoV_RaTG13_orf1ab;1938 | 0.05 | 0.05 |
| MN996532.1;QHR63299.1;BatCoV_RaTG13_orf1ab;1939 | 0.05 | 0.025 |
| MN996532.1;QHR63299.1;BatCoV_RaTG13_orf1ab;1940 | 0 | 0 |
| MN996532.1;QHR63299.1;BatCoV_RaTG13_orf1ab;1941 | 0 | 0 |
| MN996532.1;QHR63299.1;BatCoV_RaTG13_orf1ab;1942 | 0 | 0 |
| MN996532.1;QHR63299.1;BatCoV_RaTG13_orf1ab;1943 | 0 | 0 |
| MN996532.1;QHR63299.1;BatCoV_RaTG13_orf1ab;1944 | 0 | 0 |
| MN996532.1;QHR63299.1;BatCoV_RaTG13_orf1ab;1945 | 0 | 0 |
| MN996532.1;QHR63299.1;BatCoV_RaTG13_orf1ab;1946 | 0 | 0 |
| MN996532.1;QHR63299.1;BatCoV_RaTG13_orf1ab;1947 | 0 | 0 |
| MN996532.1;QHR63299.1;BatCoV_RaTG13_orf1ab;1948 | 0 | 0 |
| MN996532.1;QHR63299.1;BatCoV_RaTG13_orf1ab;1949 | 0 | 0 |
| MN996532.1;QHR63299.1;BatCoV_RaTG13_orf1ab;1950 | 0 | 0.025 |
| MN996532.1;QHR63299.1;BatCoV_RaTG13_orf1ab;1951 | 0 | 0 |
| MN996532.1;QHR63299.1;BatCoV_RaTG13_orf1ab;1952 | 0 | 0.025 |
| MN996532.1;QHR63299.1;BatCoV_RaTG13_orf1ab;1953 | 0.05 | 0.05 |
| MN996532.1;QHR63299.1;BatCoV_RaTG13_orf1ab;1954 | 0 | 0 |
| MN996532.1;QHR63299.1;BatCoV_RaTG13_orf1ab;1955 | 0 | 0.025 |
| MN996532.1;QHR63299.1;BatCoV_RaTG13_orf1ab;1956 | 0 | 0 |
| MN996532.1;QHR63299.1;BatCoV_RaTG13_orf1ab;1957 | 0 | 0.025 |
| MN996532.1;QHR63299.1;BatCoV_RaTG13_orf1ab;1958 | 0.05 | 0.025 |
| MN996532.1;QHR63299.1;BatCoV_RaTG13_orf1ab;1959 | 0.05 | 0.05 |
| MN996532.1;QHR63299.1;BatCoV_RaTG13_orf1ab;1960 | 0.05 | 0.075 |
| MN996532.1;QHR63299.1;BatCoV_RaTG13_orf1ab;1961 | 0.05 | 0.075 |
| MN996532.1;QHR63299.1;BatCoV_RaTG13_orf1ab;1962 | 0.05 | 0.025 |
| MN996532.1;QHR63299.1;BatCoV_RaTG13_orf1ab;1963 | 0.05 | 0.025 |
| MN996532.1;QHR63299.1;BatCoV_RaTG13_orf1ab;1964 | 0.05 | 0.075 |
| MN996532.1;QHR63299.1;BatCoV_RaTG13_orf1ab;1965 | 0 | 0.025 |
| MN996532.1;QHR63299.1;BatCoV_RaTG13_orf1ab;1966 | 0 | 0.025 |
| MN996532.1;QHR63299.1;BatCoV_RaTG13_orf1ab;1967 | 0 | 0.05 |
| MN996532.1;QHR63299.1;BatCoV_RaTG13_orf1ab;1968 | 0 | 0.075 |
| MN996532.1;QHR63299.1;BatCoV_RaTG13_orf1ab;1969 | 0 | 0 |

| | | |
|---|---|---|
| MN996532.1;QHR63299.1;BatCoV_RaTG13_orf1ab;1970 | 0 | 0 |
| MN996532.1;QHR63299.1;BatCoV_RaTG13_orf1ab;1971 | 0 | 0 |
| MN996532.1;QHR63299.1;BatCoV_RaTG13_orf1ab;1972 | 0 | 0 |
| MN996532.1;QHR63299.1;BatCoV_RaTG13_orf1ab;1973 | 0 | 0 |
| MN996532.1;QHR63299.1;BatCoV_RaTG13_orf1ab;1974 | 0 | 0.025 |
| MN996532.1;QHR63299.1;BatCoV_RaTG13_orf1ab;1975 | 0 | 0 |
| MN996532.1;QHR63299.1;BatCoV_RaTG13_orf1ab;1976 | 0 | 0 |
| MN996532.1;QHR63299.1;BatCoV_RaTG13_orf1ab;1977 | 0 | 0 |
| MN996532.1;QHR63299.1;BatCoV_RaTG13_orf1ab;1978 | 0 | 0 |
| MN996532.1;QHR63299.1;BatCoV_RaTG13_orf1ab;1979 | 0 | 0 |
| MN996532.1;QHR63299.1;BatCoV_RaTG13_orf1ab;1980 | 0 | 0 |
| MN996532.1;QHR63299.1;BatCoV_RaTG13_orf1ab;1981 | 0 | 0 |
| MN996532.1;QHR63299.1;BatCoV_RaTG13_orf1ab;1982 | 0 | 0 |
| MN996532.1;QHR63299.1;BatCoV_RaTG13_orf1ab;1983 | 0.05 | 0 |
| MN996532.1;QHR63299.1;BatCoV_RaTG13_orf1ab;1984 | 0.05 | 0 |
| MN996532.1;QHR63299.1;BatCoV_RaTG13_orf1ab;1985 | 0 | 0 |
| MN996532.1;QHR63299.1;BatCoV_RaTG13_orf1ab;1986 | 0.05 | 0 |
| MN996532.1;QHR63299.1;BatCoV_RaTG13_orf1ab;1987 | 0.05 | 0.05 |
| MN996532.1;QHR63299.1;BatCoV_RaTG13_orf1ab;1988 | 0.1 | 0.025 |
| MN996532.1;QHR63299.1;BatCoV_RaTG13_orf1ab;1989 | 0.05 | 0.075 |
| MN996532.1;QHR63299.1;BatCoV_RaTG13_orf1ab;1990 | 0.1 | 0.1 |
| MN996532.1;QHR63299.1;BatCoV_RaTG13_orf1ab;1991 | 0.15 | 0.15 |
| MN996532.1;QHR63299.1;BatCoV_RaTG13_orf1ab;1992 | 0.15 | 0.15 |
| MN996532.1;QHR63299.1;BatCoV_RaTG13_orf1ab;1993 | 0.25 | 0.1 |
| MN996532.1;QHR63299.1;BatCoV_RaTG13_orf1ab;1994 | 0.15 | 0.075 |
| MN996532.1;QHR63299.1;BatCoV_RaTG13_orf1ab;1995 | 0.05 | 0.025 |
| MN996532.1;QHR63299.1;BatCoV_RaTG13_orf1ab;1996 | 0.05 | 0.025 |
| MN996532.1;QHR63299.1;BatCoV_RaTG13_orf1ab;1997 | 0 | 0 |
| MN996532.1;QHR63299.1;BatCoV_RaTG13_orf1ab;1998 | 0 | 0 |
| MN996532.1;QHR63299.1;BatCoV_RaTG13_orf1ab;1999 | 0 | 0 |
| MN996532.1;QHR63299.1;BatCoV_RaTG13_orf1ab;2000 | 0 | 0 |
| MN996532.1;QHR63299.1;BatCoV_RaTG13_orf1ab;2001 | 0 | 0 |
| MN996532.1;QHR63299.1;BatCoV_RaTG13_orf1ab;2002 | 0 | 0 |
| MN996532.1;QHR63299.1;BatCoV_RaTG13_orf1ab;2003 | 0 | 0 |
| MN996532.1;QHR63299.1;BatCoV_RaTG13_orf1ab;2004 | 0 | 0 |
| MN996532.1;QHR63299.1;BatCoV_RaTG13_orf1ab;2005 | 0 | 0 |
| MN996532.1;QHR63299.1;BatCoV_RaTG13_orf1ab;2006 | 0 | 0 |
| MN996532.1;QHR63299.1;BatCoV_RaTG13_orf1ab;2007 | 0 | 0 |
| MN996532.1;QHR63299.1;BatCoV_RaTG13_orf1ab;2008 | 0.1 | 0.05 |
| MN996532.1;QHR63299.1;BatCoV_RaTG13_orf1ab;2009 | 0.2 | 0.075 |
| MN996532.1;QHR63299.1;BatCoV_RaTG13_orf1ab;2010 | 0.15 | 0.1 |
| MN996532.1;QHR63299.1;BatCoV_RaTG13_orf1ab;2011 | 0.25 | 0.1 |
| MN996532.1;QHR63299.1;BatCoV_RaTG13_orf1ab;2012 | 0.2 | 0.075 |

| | | |
|---|---|---|
| MN996532.1;QHR63299.1;BatCoV_RaTG13_orf1ab;2013 | 0.3 | 0.125 |
| MN996532.1;QHR63299.1;BatCoV_RaTG13_orf1ab;2014 | 0.15 | 0.1 |
| MN996532.1;QHR63299.1;BatCoV_RaTG13_orf1ab;2015 | 0.1 | 0.05 |
| MN996532.1;QHR63299.1;BatCoV_RaTG13_orf1ab;2016 | 0.1 | 0.05 |
| MN996532.1;QHR63299.1;BatCoV_RaTG13_orf1ab;2017 | 0.15 | 0.1 |
| MN996532.1;QHR63299.1;BatCoV_RaTG13_orf1ab;2018 | 0.2 | 0.075 |
| MN996532.1;QHR63299.1;BatCoV_RaTG13_orf1ab;2019 | 0.05 | 0.025 |
| MN996532.1;QHR63299.1;BatCoV_RaTG13_orf1ab;2020 | 0 | 0.025 |
| MN996532.1;QHR63299.1;BatCoV_RaTG13_orf1ab;2021 | 0 | 0.025 |
| MN996532.1;QHR63299.1;BatCoV_RaTG13_orf1ab;2022 | 0.05 | 0.025 |
| MN996532.1;QHR63299.1;BatCoV_RaTG13_orf1ab;2023 | 0.05 | 0.025 |
| MN996532.1;QHR63299.1;BatCoV_RaTG13_orf1ab;2024 | 0 | 0.025 |
| MN996532.1;QHR63299.1;BatCoV_RaTG13_orf1ab;2025 | 0 | 0.025 |
| MN996532.1;QHR63299.1;BatCoV_RaTG13_orf1ab;2026 | 0 | 0.025 |
| MN996532.1;QHR63299.1;BatCoV_RaTG13_orf1ab;2027 | 0 | 0.025 |
| MN996532.1;QHR63299.1;BatCoV_RaTG13_orf1ab;2028 | 0 | 0.025 |
| MN996532.1;QHR63299.1;BatCoV_RaTG13_orf1ab;2029 | 0 | 0 |
| MN996532.1;QHR63299.1;BatCoV_RaTG13_orf1ab;2030 | 0.15 | 0 |
| MN996532.1;QHR63299.1;BatCoV_RaTG13_orf1ab;2031 | 0 | 0 |
| MN996532.1;QHR63299.1;BatCoV_RaTG13_orf1ab;2032 | 0 | 0 |
| MN996532.1;QHR63299.1;BatCoV_RaTG13_orf1ab;2033 | 0 | 0 |
| MN996532.1;QHR63299.1;BatCoV_RaTG13_orf1ab;2034 | 0 | 0 |
| MN996532.1;QHR63299.1;BatCoV_RaTG13_orf1ab;2035 | 0 | 0 |
| MN996532.1;QHR63299.1;BatCoV_RaTG13_orf1ab;2036 | 0 | 0 |
| MN996532.1;QHR63299.1;BatCoV_RaTG13_orf1ab;2037 | 0 | 0 |
| MN996532.1;QHR63299.1;BatCoV_RaTG13_orf1ab;2038 | 0 | 0 |
| MN996532.1;QHR63299.1;BatCoV_RaTG13_orf1ab;2039 | 0 | 0 |
| MN996532.1;QHR63299.1;BatCoV_RaTG13_orf1ab;2040 | 0 | 0 |
| MN996532.1;QHR63299.1;BatCoV_RaTG13_orf1ab;2041 | 0 | 0.025 |
| MN996532.1;QHR63299.1;BatCoV_RaTG13_orf1ab;2042 | 0 | 0 |
| MN996532.1;QHR63299.1;BatCoV_RaTG13_orf1ab;2043 | 0 | 0.05 |
| MN996532.1;QHR63299.1;BatCoV_RaTG13_orf1ab;2044 | 0 | 0.05 |
| MN996532.1;QHR63299.1;BatCoV_RaTG13_orf1ab;2045 | 0 | 0.05 |
| MN996532.1;QHR63299.1;BatCoV_RaTG13_orf1ab;2046 | 0 | 0.05 |
| MN996532.1;QHR63299.1;BatCoV_RaTG13_orf1ab;2047 | 0 | 0.05 |
| MN996532.1;QHR63299.1;BatCoV_RaTG13_orf1ab;2048 | 0 | 0.025 |
| MN996532.1;QHR63299.1;BatCoV_RaTG13_orf1ab;2049 | 0 | 0.05 |
| MN996532.1;QHR63299.1;BatCoV_RaTG13_orf1ab;2050 | 0 | 0.075 |
| MN996532.1;QHR63299.1;BatCoV_RaTG13_orf1ab;2051 | 0 | 0.025 |
| MN996532.1;QHR63299.1;BatCoV_RaTG13_orf1ab;2052 | 0.05 | 0.025 |
| MN996532.1;QHR63299.1;BatCoV_RaTG13_orf1ab;2053 | 0.05 | 0 |
| MN996532.1;QHR63299.1;BatCoV_RaTG13_orf1ab;2054 | 0 | 0 |
| MN996532.1;QHR63299.1;BatCoV_RaTG13_orf1ab;2055 | 0 | 0 |

| | | |
|---|---|---|
| MN996532.1;QHR63299.1;BatCoV_RaTG13_orf1ab;2056 | 0 | 0 |
| MN996532.1;QHR63299.1;BatCoV_RaTG13_orf1ab;2057 | 0 | 0 |
| MN996532.1;QHR63299.1;BatCoV_RaTG13_orf1ab;2058 | 0 | 0 |
| MN996532.1;QHR63299.1;BatCoV_RaTG13_orf1ab;2059 | 0 | 0 |
| MN996532.1;QHR63299.1;BatCoV_RaTG13_orf1ab;2060 | 0 | 0.025 |
| MN996532.1;QHR63299.1;BatCoV_RaTG13_orf1ab;2061 | 0 | 0 |
| MN996532.1;QHR63299.1;BatCoV_RaTG13_orf1ab;2062 | 0 | 0 |
| MN996532.1;QHR63299.1;BatCoV_RaTG13_orf1ab;2063 | 0 | 0 |
| MN996532.1;QHR63299.1;BatCoV_RaTG13_orf1ab;2064 | 0 | 0 |
| MN996532.1;QHR63299.1;BatCoV_RaTG13_orf1ab;2065 | 0 | 0 |
| MN996532.1;QHR63299.1;BatCoV_RaTG13_orf1ab;2066 | 0 | 0 |
| MN996532.1;QHR63299.1;BatCoV_RaTG13_orf1ab;2067 | 0.05 | 0.025 |
| MN996532.1;QHR63299.1;BatCoV_RaTG13_orf1ab;2068 | 0.05 | 0.05 |
| MN996532.1;QHR63299.1;BatCoV_RaTG13_orf1ab;2069 | 0 | 0 |
| MN996532.1;QHR63299.1;BatCoV_RaTG13_orf1ab;2070 | 0 | 0 |
| MN996532.1;QHR63299.1;BatCoV_RaTG13_orf1ab;2071 | 0 | 0 |
| MN996532.1;QHR63299.1;BatCoV_RaTG13_orf1ab;2072 | 0 | 0.025 |
| MN996532.1;QHR63299.1;BatCoV_RaTG13_orf1ab;2073 | 0 | 0 |
| MN996532.1;QHR63299.1;BatCoV_RaTG13_orf1ab;2074 | 0 | 0 |
| MN996532.1;QHR63299.1;BatCoV_RaTG13_orf1ab;2075 | 0 | 0 |
| MN996532.1;QHR63299.1;BatCoV_RaTG13_orf1ab;2076 | 0 | 0 |
| MN996532.1;QHR63299.1;BatCoV_RaTG13_orf1ab;2077 | 0 | 0 |
| MN996532.1;QHR63299.1;BatCoV_RaTG13_orf1ab;2078 | 0 | 0 |
| MN996532.1;QHR63299.1;BatCoV_RaTG13_orf1ab;2079 | 0 | 0 |
| MN996532.1;QHR63299.1;BatCoV_RaTG13_orf1ab;2080 | 0 | 0 |
| MN996532.1;QHR63299.1;BatCoV_RaTG13_orf1ab;2081 | 0 | 0.025 |
| MN996532.1;QHR63299.1;BatCoV_RaTG13_orf1ab;2082 | 0.05 | 0.025 |
| MN996532.1;QHR63299.1;BatCoV_RaTG13_orf1ab;2083 | 0 | 0 |
| MN996532.1;QHR63299.1;BatCoV_RaTG13_orf1ab;2084 | 0 | 0 |
| MN996532.1;QHR63299.1;BatCoV_RaTG13_orf1ab;2085 | 0 | 0 |
| MN996532.1;QHR63299.1;BatCoV_RaTG13_orf1ab;2086 | 0 | 0 |
| MN996532.1;QHR63299.1;BatCoV_RaTG13_orf1ab;2087 | 0 | 0 |
| MN996532.1;QHR63299.1;BatCoV_RaTG13_orf1ab;2088 | 0 | 0 |
| MN996532.1;QHR63299.1;BatCoV_RaTG13_orf1ab;2089 | 0 | 0 |
| MN996532.1;QHR63299.1;BatCoV_RaTG13_orf1ab;2090 | 0.05 | 0.05 |
| MN996532.1;QHR63299.1;BatCoV_RaTG13_orf1ab;2091 | 0 | 0.05 |
| MN996532.1;QHR63299.1;BatCoV_RaTG13_orf1ab;2092 | 0 | 0.025 |
| MN996532.1;QHR63299.1;BatCoV_RaTG13_orf1ab;2093 | 0 | 0 |
| MN996532.1;QHR63299.1;BatCoV_RaTG13_orf1ab;2094 | 0 | 0.025 |
| MN996532.1;QHR63299.1;BatCoV_RaTG13_orf1ab;2095 | 0 | 0.025 |
| MN996532.1;QHR63299.1;BatCoV_RaTG13_orf1ab;2096 | 0.05 | 0.025 |
| MN996532.1;QHR63299.1;BatCoV_RaTG13_orf1ab;2097 | 0.05 | 0.05 |
| MN996532.1;QHR63299.1;BatCoV_RaTG13_orf1ab;2098 | 0.25 | 0.125 |

| | | |
|---|---|---|
| MN996532.1;QHR63299.1;BatCoV_RaTG13_orf1ab;2099 | 0 | 0.05 |
| MN996532.1;QHR63299.1;BatCoV_RaTG13_orf1ab;2100 | 0 | 0.05 |
| MN996532.1;QHR63299.1;BatCoV_RaTG13_orf1ab;2101 | 0 | 0 |
| MN996532.1;QHR63299.1;BatCoV_RaTG13_orf1ab;2102 | 0 | 0 |
| MN996532.1;QHR63299.1;BatCoV_RaTG13_orf1ab;2103 | 0 | 0 |
| MN996532.1;QHR63299.1;BatCoV_RaTG13_orf1ab;2104 | 0 | 0 |
| MN996532.1;QHR63299.1;BatCoV_RaTG13_orf1ab;2105 | 0 | 0 |
| MN996532.1;QHR63299.1;BatCoV_RaTG13_orf1ab;2106 | 0 | 0 |
| MN996532.1;QHR63299.1;BatCoV_RaTG13_orf1ab;2107 | 0 | 0 |
| MN996532.1;QHR63299.1;BatCoV_RaTG13_orf1ab;2108 | 0 | 0 |
| MN996532.1;QHR63299.1;BatCoV_RaTG13_orf1ab;2109 | 0 | 0 |
| MN996532.1;QHR63299.1;BatCoV_RaTG13_orf1ab;2110 | 0 | 0 |
| MN996532.1;QHR63299.1;BatCoV_RaTG13_orf1ab;2111 | 0 | 0 |
| MN996532.1;QHR63299.1;BatCoV_RaTG13_orf1ab;2112 | 0 | 0 |
| MN996532.1;QHR63299.1;BatCoV_RaTG13_orf1ab;2113 | 0 | 0 |
| MN996532.1;QHR63299.1;BatCoV_RaTG13_orf1ab;2114 | 0.05 | 0 |
| MN996532.1;QHR63299.1;BatCoV_RaTG13_orf1ab;2115 | 0 | 0 |
| MN996532.1;QHR63299.1;BatCoV_RaTG13_orf1ab;2116 | 0 | 0 |
| MN996532.1;QHR63299.1;BatCoV_RaTG13_orf1ab;2117 | 0.05 | 0 |
| MN996532.1;QHR63299.1;BatCoV_RaTG13_orf1ab;2118 | 0.05 | 0 |
| MN996532.1;QHR63299.1;BatCoV_RaTG13_orf1ab;2119 | 0 | 0 |
| MN996532.1;QHR63299.1;BatCoV_RaTG13_orf1ab;2120 | 0 | 0 |
| MN996532.1;QHR63299.1;BatCoV_RaTG13_orf1ab;2121 | 0 | 0 |
| MN996532.1;QHR63299.1;BatCoV_RaTG13_orf1ab;2122 | 0 | 0 |
| MN996532.1;QHR63299.1;BatCoV_RaTG13_orf1ab;2123 | 0.05 | 0 |
| MN996532.1;QHR63299.1;BatCoV_RaTG13_orf1ab;2124 | 0.05 | 0 |
| MN996532.1;QHR63299.1;BatCoV_RaTG13_orf1ab;2125 | 0.05 | 0.025 |
| MN996532.1;QHR63299.1;BatCoV_RaTG13_orf1ab;2126 | 0.05 | 0 |
| MN996532.1;QHR63299.1;BatCoV_RaTG13_orf1ab;2127 | 0.1 | 0 |
| MN996532.1;QHR63299.1;BatCoV_RaTG13_orf1ab;2128 | 0.15 | 0.025 |
| MN996532.1;QHR63299.1;BatCoV_RaTG13_orf1ab;2129 | 0.1 | 0 |
| MN996532.1;QHR63299.1;BatCoV_RaTG13_orf1ab;2130 | 0.05 | 0 |
| MN996532.1;QHR63299.1;BatCoV_RaTG13_orf1ab;2131 | 0.1 | 0 |
| MN996532.1;QHR63299.1;BatCoV_RaTG13_orf1ab;2132 | 0.1 | 0.05 |
| MN996532.1;QHR63299.1;BatCoV_RaTG13_orf1ab;2133 | 0.1 | 0.025 |
| MN996532.1;QHR63299.1;BatCoV_RaTG13_orf1ab;2134 | 0.1 | 0.025 |
| MN996532.1;QHR63299.1;BatCoV_RaTG13_orf1ab;2135 | 0.2 | 0.075 |
| MN996532.1;QHR63299.1;BatCoV_RaTG13_orf1ab;2136 | 0.05 | 0.025 |
| MN996532.1;QHR63299.1;BatCoV_RaTG13_orf1ab;2137 | 0.05 | 0.05 |
| MN996532.1;QHR63299.1;BatCoV_RaTG13_orf1ab;2138 | 0.1 | 0.05 |
| MN996532.1;QHR63299.1;BatCoV_RaTG13_orf1ab;2139 | 0.05 | 0.05 |
| MN996532.1;QHR63299.1;BatCoV_RaTG13_orf1ab;2140 | 0.05 | 0.05 |
| MN996532.1;QHR63299.1;BatCoV_RaTG13_orf1ab;2141 | 0.05 | 0.05 |

| | | |
|---|---|---|
| MN996532.1;QHR63299.1;BatCoV_RaTG13_orf1ab;2142 | 0.1 | 0.025 |
| MN996532.1;QHR63299.1;BatCoV_RaTG13_orf1ab;2143 | 0.05 | 0 |
| MN996532.1;QHR63299.1;BatCoV_RaTG13_orf1ab;2144 | 0 | 0 |
| MN996532.1;QHR63299.1;BatCoV_RaTG13_orf1ab;2145 | 0 | 0 |
| MN996532.1;QHR63299.1;BatCoV_RaTG13_orf1ab;2146 | 0 | 0 |
| MN996532.1;QHR63299.1;BatCoV_RaTG13_orf1ab;2147 | 0 | 0 |
| MN996532.1;QHR63299.1;BatCoV_RaTG13_orf1ab;2148 | 0 | 0 |
| MN996532.1;QHR63299.1;BatCoV_RaTG13_orf1ab;2149 | 0 | 0 |
| MN996532.1;QHR63299.1;BatCoV_RaTG13_orf1ab;2150 | 0 | 0 |
| MN996532.1;QHR63299.1;BatCoV_RaTG13_orf1ab;2151 | 0 | 0 |
| MN996532.1;QHR63299.1;BatCoV_RaTG13_orf1ab;2152 | 0 | 0 |
| MN996532.1;QHR63299.1;BatCoV_RaTG13_orf1ab;2153 | 0 | 0 |
| MN996532.1;QHR63299.1;BatCoV_RaTG13_orf1ab;2154 | 0 | 0 |
| MN996532.1;QHR63299.1;BatCoV_RaTG13_orf1ab;2155 | 0 | 0 |
| MN996532.1;QHR63299.1;BatCoV_RaTG13_orf1ab;2156 | 0 | 0 |
| MN996532.1;QHR63299.1;BatCoV_RaTG13_orf1ab;2157 | 0 | 0 |
| MN996532.1;QHR63299.1;BatCoV_RaTG13_orf1ab;2158 | 0 | 0 |
| MN996532.1;QHR63299.1;BatCoV_RaTG13_orf1ab;2159 | 0 | 0 |
| MN996532.1;QHR63299.1;BatCoV_RaTG13_orf1ab;2160 | 0 | 0 |
| MN996532.1;QHR63299.1;BatCoV_RaTG13_orf1ab;2161 | 0 | 0 |
| MN996532.1;QHR63299.1;BatCoV_RaTG13_orf1ab;2162 | 0 | 0 |
| MN996532.1;QHR63299.1;BatCoV_RaTG13_orf1ab;2163 | 0 | 0 |
| MN996532.1;QHR63299.1;BatCoV_RaTG13_orf1ab;2164 | 0 | 0 |
| MN996532.1;QHR63299.1;BatCoV_RaTG13_orf1ab;2165 | 0 | 0 |
| MN996532.1;QHR63299.1;BatCoV_RaTG13_orf1ab;2166 | 0 | 0 |
| MN996532.1;QHR63299.1;BatCoV_RaTG13_orf1ab;2167 | 0 | 0 |
| MN996532.1;QHR63299.1;BatCoV_RaTG13_orf1ab;2168 | 0 | 0 |
| MN996532.1;QHR63299.1;BatCoV_RaTG13_orf1ab;2169 | 0 | 0 |
| MN996532.1;QHR63299.1;BatCoV_RaTG13_orf1ab;2170 | 0 | 0 |
| MN996532.1;QHR63299.1;BatCoV_RaTG13_orf1ab;2171 | 0 | 0 |
| MN996532.1;QHR63299.1;BatCoV_RaTG13_orf1ab;2172 | 0 | 0 |
| MN996532.1;QHR63299.1;BatCoV_RaTG13_orf1ab;2173 | 0 | 0 |
| MN996532.1;QHR63299.1;BatCoV_RaTG13_orf1ab;2174 | 0 | 0 |
| MN996532.1;QHR63299.1;BatCoV_RaTG13_orf1ab;2175 | 0 | 0 |
| MN996532.1;QHR63299.1;BatCoV_RaTG13_orf1ab;2176 | 0 | 0 |
| MN996532.1;QHR63299.1;BatCoV_RaTG13_orf1ab;2177 | 0 | 0 |
| MN996532.1;QHR63299.1;BatCoV_RaTG13_orf1ab;2178 | 0 | 0 |
| MN996532.1;QHR63299.1;BatCoV_RaTG13_orf1ab;2179 | 0.05 | 0 |
| MN996532.1;QHR63299.1;BatCoV_RaTG13_orf1ab;2180 | 0.1 | 0 |
| MN996532.1;QHR63299.1;BatCoV_RaTG13_orf1ab;2181 | 0 | 0 |
| MN996532.1;QHR63299.1;BatCoV_RaTG13_orf1ab;2182 | 0 | 0.025 |
| MN996532.1;QHR63299.1;BatCoV_RaTG13_orf1ab;2183 | 0 | 0 |
| MN996532.1;QHR63299.1;BatCoV_RaTG13_orf1ab;2184 | 0.05 | 0 |

| | | |
|---|---|---|
| MN996532.1;QHR63299.1;BatCoV_RaTG13_orf1ab;2185 | 0.05 | 0 |
| MN996532.1;QHR63299.1;BatCoV_RaTG13_orf1ab;2186 | 0 | 0 |
| MN996532.1;QHR63299.1;BatCoV_RaTG13_orf1ab;2187 | 0 | 0 |
| MN996532.1;QHR63299.1;BatCoV_RaTG13_orf1ab;2188 | 0 | 0 |
| MN996532.1;QHR63299.1;BatCoV_RaTG13_orf1ab;2189 | 0 | 0 |
| MN996532.1;QHR63299.1;BatCoV_RaTG13_orf1ab;2190 | 0 | 0 |
| MN996532.1;QHR63299.1;BatCoV_RaTG13_orf1ab;2191 | 0 | 0 |
| MN996532.1;QHR63299.1;BatCoV_RaTG13_orf1ab;2192 | 0 | 0 |
| MN996532.1;QHR63299.1;BatCoV_RaTG13_orf1ab;2193 | 0 | 0 |
| MN996532.1;QHR63299.1;BatCoV_RaTG13_orf1ab;2194 | 0 | 0 |
| MN996532.1;QHR63299.1;BatCoV_RaTG13_orf1ab;2195 | 0 | 0 |
| MN996532.1;QHR63299.1;BatCoV_RaTG13_orf1ab;2196 | 0 | 0 |
| MN996532.1;QHR63299.1;BatCoV_RaTG13_orf1ab;2197 | 0 | 0 |
| MN996532.1;QHR63299.1;BatCoV_RaTG13_orf1ab;2198 | 0 | 0 |
| MN996532.1;QHR63299.1;BatCoV_RaTG13_orf1ab;2199 | 0 | 0 |
| MN996532.1;QHR63299.1;BatCoV_RaTG13_orf1ab;2200 | 0 | 0 |
| MN996532.1;QHR63299.1;BatCoV_RaTG13_orf1ab;2201 | 0 | 0 |
| MN996532.1;QHR63299.1;BatCoV_RaTG13_orf1ab;2202 | 0 | 0 |
| MN996532.1;QHR63299.1;BatCoV_RaTG13_orf1ab;2203 | 0 | 0 |
| MN996532.1;QHR63299.1;BatCoV_RaTG13_orf1ab;2204 | 0 | 0 |
| MN996532.1;QHR63299.1;BatCoV_RaTG13_orf1ab;2205 | 0 | 0 |
| MN996532.1;QHR63299.1;BatCoV_RaTG13_orf1ab;2206 | 0 | 0 |
| MN996532.1;QHR63299.1;BatCoV_RaTG13_orf1ab;2207 | 0 | 0 |
| MN996532.1;QHR63299.1;BatCoV_RaTG13_orf1ab;2208 | 0 | 0 |
| MN996532.1;QHR63299.1;BatCoV_RaTG13_orf1ab;2209 | 0 | 0 |
| MN996532.1;QHR63299.1;BatCoV_RaTG13_orf1ab;2210 | 0 | 0 |
| MN996532.1;QHR63299.1;BatCoV_RaTG13_orf1ab;2211 | 0 | 0 |
| MN996532.1;QHR63299.1;BatCoV_RaTG13_orf1ab;2212 | 0 | 0 |
| MN996532.1;QHR63299.1;BatCoV_RaTG13_orf1ab;2213 | 0 | 0 |
| MN996532.1;QHR63299.1;BatCoV_RaTG13_orf1ab;2214 | 0 | 0 |
| MN996532.1;QHR63299.1;BatCoV_RaTG13_orf1ab;2215 | 0 | 0 |
| MN996532.1;QHR63299.1;BatCoV_RaTG13_orf1ab;2216 | 0 | 0 |
| MN996532.1;QHR63299.1;BatCoV_RaTG13_orf1ab;2217 | 0 | 0 |
| MN996532.1;QHR63299.1;BatCoV_RaTG13_orf1ab;2218 | 0 | 0 |
| MN996532.1;QHR63299.1;BatCoV_RaTG13_orf1ab;2219 | 0 | 0 |
| MN996532.1;QHR63299.1;BatCoV_RaTG13_orf1ab;2220 | 0 | 0 |
| MN996532.1;QHR63299.1;BatCoV_RaTG13_orf1ab;2221 | 0 | 0 |
| MN996532.1;QHR63299.1;BatCoV_RaTG13_orf1ab;2222 | 0 | 0 |
| MN996532.1;QHR63299.1;BatCoV_RaTG13_orf1ab;2223 | 0 | 0 |
| MN996532.1;QHR63299.1;BatCoV_RaTG13_orf1ab;2224 | 0 | 0 |
| MN996532.1;QHR63299.1;BatCoV_RaTG13_orf1ab;2225 | 0 | 0 |
| MN996532.1;QHR63299.1;BatCoV_RaTG13_orf1ab;2226 | 0 | 0 |
| MN996532.1;QHR63299.1;BatCoV_RaTG13_orf1ab;2227 | 0 | 0 |

| | | |
|---|---|---|
| MN996532.1;QHR63299.1;BatCoV_RaTG13_orf1ab;2228 | 0 | 0 |
| MN996532.1;QHR63299.1;BatCoV_RaTG13_orf1ab;2229 | 0 | 0 |
| MN996532.1;QHR63299.1;BatCoV_RaTG13_orf1ab;2230 | 0 | 0 |
| MN996532.1;QHR63299.1;BatCoV_RaTG13_orf1ab;2231 | 0 | 0 |
| MN996532.1;QHR63299.1;BatCoV_RaTG13_orf1ab;2232 | 0 | 0 |
| MN996532.1;QHR63299.1;BatCoV_RaTG13_orf1ab;2233 | 0 | 0 |
| MN996532.1;QHR63299.1;BatCoV_RaTG13_orf1ab;2234 | 0 | 0 |
| MN996532.1;QHR63299.1;BatCoV_RaTG13_orf1ab;2235 | 0 | 0 |
| MN996532.1;QHR63299.1;BatCoV_RaTG13_orf1ab;2236 | 0 | 0 |
| MN996532.1;QHR63299.1;BatCoV_RaTG13_orf1ab;2237 | 0 | 0 |
| MN996532.1;QHR63299.1;BatCoV_RaTG13_orf1ab;2238 | 0 | 0 |
| MN996532.1;QHR63299.1;BatCoV_RaTG13_orf1ab;2239 | 0 | 0 |
| MN996532.1;QHR63299.1;BatCoV_RaTG13_orf1ab;2240 | 0 | 0 |
| MN996532.1;QHR63299.1;BatCoV_RaTG13_orf1ab;2241 | 0 | 0 |
| MN996532.1;QHR63299.1;BatCoV_RaTG13_orf1ab;2242 | 0 | 0 |
| MN996532.1;QHR63299.1;BatCoV_RaTG13_orf1ab;2243 | 0 | 0 |
| MN996532.1;QHR63299.1;BatCoV_RaTG13_orf1ab;2244 | 0 | 0 |
| MN996532.1;QHR63299.1;BatCoV_RaTG13_orf1ab;2245 | 0 | 0 |
| MN996532.1;QHR63299.1;BatCoV_RaTG13_orf1ab;2246 | 0 | 0 |
| MN996532.1;QHR63299.1;BatCoV_RaTG13_orf1ab;2247 | 0 | 0.025 |
| MN996532.1;QHR63299.1;BatCoV_RaTG13_orf1ab;2248 | 0 | 0.025 |
| MN996532.1;QHR63299.1;BatCoV_RaTG13_orf1ab;2249 | 0 | 0 |
| MN996532.1;QHR63299.1;BatCoV_RaTG13_orf1ab;2250 | 0 | 0 |
| MN996532.1;QHR63299.1;BatCoV_RaTG13_orf1ab;2251 | 0 | 0 |
| MN996532.1;QHR63299.1;BatCoV_RaTG13_orf1ab;2252 | 0 | 0 |
| MN996532.1;QHR63299.1;BatCoV_RaTG13_orf1ab;2253 | 0 | 0 |
| MN996532.1;QHR63299.1;BatCoV_RaTG13_orf1ab;2254 | 0 | 0 |
| MN996532.1;QHR63299.1;BatCoV_RaTG13_orf1ab;2255 | 0 | 0 |
| MN996532.1;QHR63299.1;BatCoV_RaTG13_orf1ab;2256 | 0 | 0 |
| MN996532.1;QHR63299.1;BatCoV_RaTG13_orf1ab;2257 | 0 | 0.05 |
| MN996532.1;QHR63299.1;BatCoV_RaTG13_orf1ab;2258 | 0 | 0 |
| MN996532.1;QHR63299.1;BatCoV_RaTG13_orf1ab;2259 | 0.2 | 0.125 |
| MN996532.1;QHR63299.1;BatCoV_RaTG13_orf1ab;2260 | 0.15 | 0.1 |
| MN996532.1;QHR63299.1;BatCoV_RaTG13_orf1ab;2261 | 0.3 | 0.275 |
| MN996532.1;QHR63299.1;BatCoV_RaTG13_orf1ab;2262 | 0.35 | 0.35 |
| MN996532.1;QHR63299.1;BatCoV_RaTG13_orf1ab;2263 | 0.35 | 0.35 |
| MN996532.1;QHR63299.1;BatCoV_RaTG13_orf1ab;2264 | 0.25 | 0.225 |
| MN996532.1;QHR63299.1;BatCoV_RaTG13_orf1ab;2265 | 0.1 | 0.025 |
| MN996532.1;QHR63299.1;BatCoV_RaTG13_orf1ab;2266 | 0.1 | 0.05 |
| MN996532.1;QHR63299.1;BatCoV_RaTG13_orf1ab;2267 | 0.05 | 0.025 |
| MN996532.1;QHR63299.1;BatCoV_RaTG13_orf1ab;2268 | 0.05 | 0 |
| MN996532.1;QHR63299.1;BatCoV_RaTG13_orf1ab;2269 | 0.05 | 0 |
| MN996532.1;QHR63299.1;BatCoV_RaTG13_orf1ab;2270 | 0.05 | 0 |

| | | |
|---|---|---|
| MN996532.1;QHR63299.1;BatCoV_RaTG13_orf1ab;2271 | 0 | 0 |
| MN996532.1;QHR63299.1;BatCoV_RaTG13_orf1ab;2272 | 0 | 0 |
| MN996532.1;QHR63299.1;BatCoV_RaTG13_orf1ab;2273 | 0.05 | 0 |
| MN996532.1;QHR63299.1;BatCoV_RaTG13_orf1ab;2274 | 0 | 0 |
| MN996532.1;QHR63299.1;BatCoV_RaTG13_orf1ab;2275 | 0 | 0 |
| MN996532.1;QHR63299.1;BatCoV_RaTG13_orf1ab;2276 | 0 | 0 |
| MN996532.1;QHR63299.1;BatCoV_RaTG13_orf1ab;2277 | 0 | 0 |
| MN996532.1;QHR63299.1;BatCoV_RaTG13_orf1ab;2278 | 0 | 0 |
| MN996532.1;QHR63299.1;BatCoV_RaTG13_orf1ab;2279 | 0 | 0 |
| MN996532.1;QHR63299.1;BatCoV_RaTG13_orf1ab;2280 | 0 | 0 |
| MN996532.1;QHR63299.1;BatCoV_RaTG13_orf1ab;2281 | 0 | 0.025 |
| MN996532.1;QHR63299.1;BatCoV_RaTG13_orf1ab;2282 | 0 | 0 |
| MN996532.1;QHR63299.1;BatCoV_RaTG13_orf1ab;2283 | 0 | 0 |
| MN996532.1;QHR63299.1;BatCoV_RaTG13_orf1ab;2284 | 0 | 0 |
| MN996532.1;QHR63299.1;BatCoV_RaTG13_orf1ab;2285 | 0 | 0 |
| MN996532.1;QHR63299.1;BatCoV_RaTG13_orf1ab;2286 | 0 | 0 |
| MN996532.1;QHR63299.1;BatCoV_RaTG13_orf1ab;2287 | 0 | 0.05 |
| MN996532.1;QHR63299.1;BatCoV_RaTG13_orf1ab;2288 | 0 | 0 |
| MN996532.1;QHR63299.1;BatCoV_RaTG13_orf1ab;2289 | 0 | 0 |
| MN996532.1;QHR63299.1;BatCoV_RaTG13_orf1ab;2290 | 0.05 | 0.025 |
| MN996532.1;QHR63299.1;BatCoV_RaTG13_orf1ab;2291 | 0 | 0 |
| MN996532.1;QHR63299.1;BatCoV_RaTG13_orf1ab;2292 | 0 | 0 |
| MN996532.1;QHR63299.1;BatCoV_RaTG13_orf1ab;2293 | 0.05 | 0.025 |
| MN996532.1;QHR63299.1;BatCoV_RaTG13_orf1ab;2294 | 0.05 | 0 |
| MN996532.1;QHR63299.1;BatCoV_RaTG13_orf1ab;2295 | 0 | 0 |
| MN996532.1;QHR63299.1;BatCoV_RaTG13_orf1ab;2296 | 0 | 0 |
| MN996532.1;QHR63299.1;BatCoV_RaTG13_orf1ab;2297 | 0 | 0 |
| MN996532.1;QHR63299.1;BatCoV_RaTG13_orf1ab;2298 | 0 | 0 |
| MN996532.1;QHR63299.1;BatCoV_RaTG13_orf1ab;2299 | 0 | 0 |
| MN996532.1;QHR63299.1;BatCoV_RaTG13_orf1ab;2300 | 0 | 0 |
| MN996532.1;QHR63299.1;BatCoV_RaTG13_orf1ab;2301 | 0 | 0 |
| MN996532.1;QHR63299.1;BatCoV_RaTG13_orf1ab;2302 | 0 | 0 |
| MN996532.1;QHR63299.1;BatCoV_RaTG13_orf1ab;2303 | 0 | 0 |
| MN996532.1;QHR63299.1;BatCoV_RaTG13_orf1ab;2304 | 0 | 0 |
| MN996532.1;QHR63299.1;BatCoV_RaTG13_orf1ab;2305 | 0 | 0 |
| MN996532.1;QHR63299.1;BatCoV_RaTG13_orf1ab;2306 | 0 | 0 |
| MN996532.1;QHR63299.1;BatCoV_RaTG13_orf1ab;2307 | 0 | 0 |
| MN996532.1;QHR63299.1;BatCoV_RaTG13_orf1ab;2308 | 0 | 0.075 |
| MN996532.1;QHR63299.1;BatCoV_RaTG13_orf1ab;2309 | 0 | 0 |
| MN996532.1;QHR63299.1;BatCoV_RaTG13_orf1ab;2310 | 0 | 0 |
| MN996532.1;QHR63299.1;BatCoV_RaTG13_orf1ab;2311 | 0 | 0 |
| MN996532.1;QHR63299.1;BatCoV_RaTG13_orf1ab;2312 | 0 | 0 |
| MN996532.1;QHR63299.1;BatCoV_RaTG13_orf1ab;2313 | 0 | 0 |

| | | |
|---|---|---|
| MN996532.1;QHR63299.1;BatCoV_RaTG13_orf1ab;2314 | 0 | 0 |
| MN996532.1;QHR63299.1;BatCoV_RaTG13_orf1ab;2315 | 0 | 0 |
| MN996532.1;QHR63299.1;BatCoV_RaTG13_orf1ab;2316 | 0 | 0 |
| MN996532.1;QHR63299.1;BatCoV_RaTG13_orf1ab;2317 | 0 | 0 |
| MN996532.1;QHR63299.1;BatCoV_RaTG13_orf1ab;2318 | 0 | 0 |
| MN996532.1;QHR63299.1;BatCoV_RaTG13_orf1ab;2319 | 0 | 0 |
| MN996532.1;QHR63299.1;BatCoV_RaTG13_orf1ab;2320 | 0 | 0 |
| MN996532.1;QHR63299.1;BatCoV_RaTG13_orf1ab;2321 | 0 | 0 |
| MN996532.1;QHR63299.1;BatCoV_RaTG13_orf1ab;2322 | 0 | 0 |
| MN996532.1;QHR63299.1;BatCoV_RaTG13_orf1ab;2323 | 0 | 0 |
| MN996532.1;QHR63299.1;BatCoV_RaTG13_orf1ab;2324 | 0 | 0 |
| MN996532.1;QHR63299.1;BatCoV_RaTG13_orf1ab;2325 | 0 | 0 |
| MN996532.1;QHR63299.1;BatCoV_RaTG13_orf1ab;2326 | 0 | 0 |
| MN996532.1;QHR63299.1;BatCoV_RaTG13_orf1ab;2327 | 0 | 0 |
| MN996532.1;QHR63299.1;BatCoV_RaTG13_orf1ab;2328 | 0 | 0 |
| MN996532.1;QHR63299.1;BatCoV_RaTG13_orf1ab;2329 | 0 | 0 |
| MN996532.1;QHR63299.1;BatCoV_RaTG13_orf1ab;2330 | 0 | 0 |
| MN996532.1;QHR63299.1;BatCoV_RaTG13_orf1ab;2331 | 0 | 0 |
| MN996532.1;QHR63299.1;BatCoV_RaTG13_orf1ab;2332 | 0 | 0 |
| MN996532.1;QHR63299.1;BatCoV_RaTG13_orf1ab;2333 | 0 | 0 |
| MN996532.1;QHR63299.1;BatCoV_RaTG13_orf1ab;2334 | 0 | 0 |
| MN996532.1;QHR63299.1;BatCoV_RaTG13_orf1ab;2335 | 0 | 0 |
| MN996532.1;QHR63299.1;BatCoV_RaTG13_orf1ab;2336 | 0 | 0 |
| MN996532.1;QHR63299.1;BatCoV_RaTG13_orf1ab;2337 | 0 | 0 |
| MN996532.1;QHR63299.1;BatCoV_RaTG13_orf1ab;2338 | 0 | 0 |
| MN996532.1;QHR63299.1;BatCoV_RaTG13_orf1ab;2339 | 0 | 0 |
| MN996532.1;QHR63299.1;BatCoV_RaTG13_orf1ab;2340 | 0 | 0 |
| MN996532.1;QHR63299.1;BatCoV_RaTG13_orf1ab;2341 | 0 | 0 |
| MN996532.1;QHR63299.1;BatCoV_RaTG13_orf1ab;2342 | 0 | 0 |
| MN996532.1;QHR63299.1;BatCoV_RaTG13_orf1ab;2343 | 0 | 0 |
| MN996532.1;QHR63299.1;BatCoV_RaTG13_orf1ab;2344 | 0 | 0 |
| MN996532.1;QHR63299.1;BatCoV_RaTG13_orf1ab;2345 | 0 | 0 |
| MN996532.1;QHR63299.1;BatCoV_RaTG13_orf1ab;2346 | 0 | 0 |
| MN996532.1;QHR63299.1;BatCoV_RaTG13_orf1ab;2347 | 0 | 0 |
| MN996532.1;QHR63299.1;BatCoV_RaTG13_orf1ab;2348 | 0 | 0 |
| MN996532.1;QHR63299.1;BatCoV_RaTG13_orf1ab;2349 | 0 | 0 |
| MN996532.1;QHR63299.1;BatCoV_RaTG13_orf1ab;2350 | 0 | 0 |
| MN996532.1;QHR63299.1;BatCoV_RaTG13_orf1ab;2351 | 0 | 0 |
| MN996532.1;QHR63299.1;BatCoV_RaTG13_orf1ab;2352 | 0 | 0 |
| MN996532.1;QHR63299.1;BatCoV_RaTG13_orf1ab;2353 | 0 | 0 |
| MN996532.1;QHR63299.1;BatCoV_RaTG13_orf1ab;2354 | 0 | 0 |
| MN996532.1;QHR63299.1;BatCoV_RaTG13_orf1ab;2355 | 0 | 0 |
| MN996532.1;QHR63299.1;BatCoV_RaTG13_orf1ab;2356 | 0 | 0 |

| | | |
|---|---|---|
| MN996532.1;QHR63299.1;BatCoV_RaTG13_orf1ab;2357 | 0 | 0 |
| MN996532.1;QHR63299.1;BatCoV_RaTG13_orf1ab;2358 | 0 | 0 |
| MN996532.1;QHR63299.1;BatCoV_RaTG13_orf1ab;2359 | 0 | 0 |
| MN996532.1;QHR63299.1;BatCoV_RaTG13_orf1ab;2360 | 0 | 0 |
| MN996532.1;QHR63299.1;BatCoV_RaTG13_orf1ab;2361 | 0 | 0 |
| MN996532.1;QHR63299.1;BatCoV_RaTG13_orf1ab;2362 | 0 | 0 |
| MN996532.1;QHR63299.1;BatCoV_RaTG13_orf1ab;2363 | 0 | 0 |
| MN996532.1;QHR63299.1;BatCoV_RaTG13_orf1ab;2364 | 0 | 0 |
| MN996532.1;QHR63299.1;BatCoV_RaTG13_orf1ab;2365 | 0 | 0 |
| MN996532.1;QHR63299.1;BatCoV_RaTG13_orf1ab;2366 | 0 | 0 |
| MN996532.1;QHR63299.1;BatCoV_RaTG13_orf1ab;2367 | 0 | 0 |
| MN996532.1;QHR63299.1;BatCoV_RaTG13_orf1ab;2368 | 0 | 0 |
| MN996532.1;QHR63299.1;BatCoV_RaTG13_orf1ab;2369 | 0 | 0 |
| MN996532.1;QHR63299.1;BatCoV_RaTG13_orf1ab;2370 | 0 | 0 |
| MN996532.1;QHR63299.1;BatCoV_RaTG13_orf1ab;2371 | 0 | 0 |
| MN996532.1;QHR63299.1;BatCoV_RaTG13_orf1ab;2372 | 0 | 0 |
| MN996532.1;QHR63299.1;BatCoV_RaTG13_orf1ab;2373 | 0 | 0 |
| MN996532.1;QHR63299.1;BatCoV_RaTG13_orf1ab;2374 | 0 | 0 |
| MN996532.1;QHR63299.1;BatCoV_RaTG13_orf1ab;2375 | 0 | 0 |
| MN996532.1;QHR63299.1;BatCoV_RaTG13_orf1ab;2376 | 0 | 0 |
| MN996532.1;QHR63299.1;BatCoV_RaTG13_orf1ab;2377 | 0 | 0 |
| MN996532.1;QHR63299.1;BatCoV_RaTG13_orf1ab;2378 | 0 | 0 |
| MN996532.1;QHR63299.1;BatCoV_RaTG13_orf1ab;2379 | 0 | 0 |
| MN996532.1;QHR63299.1;BatCoV_RaTG13_orf1ab;2380 | 0 | 0 |
| MN996532.1;QHR63299.1;BatCoV_RaTG13_orf1ab;2381 | 0 | 0 |
| MN996532.1;QHR63299.1;BatCoV_RaTG13_orf1ab;2382 | 0 | 0 |
| MN996532.1;QHR63299.1;BatCoV_RaTG13_orf1ab;2383 | 0 | 0 |
| MN996532.1;QHR63299.1;BatCoV_RaTG13_orf1ab;2384 | 0 | 0 |
| MN996532.1;QHR63299.1;BatCoV_RaTG13_orf1ab;2385 | 0 | 0 |
| MN996532.1;QHR63299.1;BatCoV_RaTG13_orf1ab;2386 | 0 | 0 |
| MN996532.1;QHR63299.1;BatCoV_RaTG13_orf1ab;2387 | 0 | 0 |
| MN996532.1;QHR63299.1;BatCoV_RaTG13_orf1ab;2388 | 0 | 0 |
| MN996532.1;QHR63299.1;BatCoV_RaTG13_orf1ab;2389 | 0 | 0 |
| MN996532.1;QHR63299.1;BatCoV_RaTG13_orf1ab;2390 | 0 | 0 |
| MN996532.1;QHR63299.1;BatCoV_RaTG13_orf1ab;2391 | 0 | 0 |
| MN996532.1;QHR63299.1;BatCoV_RaTG13_orf1ab;2392 | 0 | 0 |
| MN996532.1;QHR63299.1;BatCoV_RaTG13_orf1ab;2393 | 0 | 0 |
| MN996532.1;QHR63299.1;BatCoV_RaTG13_orf1ab;2394 | 0 | 0 |
| MN996532.1;QHR63299.1;BatCoV_RaTG13_orf1ab;2395 | 0 | 0 |
| MN996532.1;QHR63299.1;BatCoV_RaTG13_orf1ab;2396 | 0 | 0 |
| MN996532.1;QHR63299.1;BatCoV_RaTG13_orf1ab;2397 | 0 | 0 |
| MN996532.1;QHR63299.1;BatCoV_RaTG13_orf1ab;2398 | 0 | 0 |
| MN996532.1;QHR63299.1;BatCoV_RaTG13_orf1ab;2399 | 0 | 0 |

| | | |
|---|---|---|
| MN996532.1;QHR63299.1;BatCoV_RaTG13_orf1ab;2400 | 0 | 0 |
| MN996532.1;QHR63299.1;BatCoV_RaTG13_orf1ab;2401 | 0 | 0 |
| MN996532.1;QHR63299.1;BatCoV_RaTG13_orf1ab;2402 | 0 | 0 |
| MN996532.1;QHR63299.1;BatCoV_RaTG13_orf1ab;2403 | 0 | 0 |
| MN996532.1;QHR63299.1;BatCoV_RaTG13_orf1ab;2404 | 0 | 0 |
| MN996532.1;QHR63299.1;BatCoV_RaTG13_orf1ab;2405 | 0 | 0 |
| MN996532.1;QHR63299.1;BatCoV_RaTG13_orf1ab;2406 | 0 | 0 |
| MN996532.1;QHR63299.1;BatCoV_RaTG13_orf1ab;2407 | 0 | 0 |
| MN996532.1;QHR63299.1;BatCoV_RaTG13_orf1ab;2408 | 0 | 0 |
| MN996532.1;QHR63299.1;BatCoV_RaTG13_orf1ab;2409 | 0 | 0 |
| MN996532.1;QHR63299.1;BatCoV_RaTG13_orf1ab;2410 | 0 | 0 |
| MN996532.1;QHR63299.1;BatCoV_RaTG13_orf1ab;2411 | 0 | 0 |
| MN996532.1;QHR63299.1;BatCoV_RaTG13_orf1ab;2412 | 0 | 0 |
| MN996532.1;QHR63299.1;BatCoV_RaTG13_orf1ab;2413 | 0 | 0 |
| MN996532.1;QHR63299.1;BatCoV_RaTG13_orf1ab;2414 | 0 | 0 |
| MN996532.1;QHR63299.1;BatCoV_RaTG13_orf1ab;2415 | 0 | 0.025 |
| MN996532.1;QHR63299.1;BatCoV_RaTG13_orf1ab;2416 | 0 | 0 |
| MN996532.1;QHR63299.1;BatCoV_RaTG13_orf1ab;2417 | 0 | 0 |
| MN996532.1;QHR63299.1;BatCoV_RaTG13_orf1ab;2418 | 0 | 0 |
| MN996532.1;QHR63299.1;BatCoV_RaTG13_orf1ab;2419 | 0 | 0 |
| MN996532.1;QHR63299.1;BatCoV_RaTG13_orf1ab;2420 | 0 | 0 |
| MN996532.1;QHR63299.1;BatCoV_RaTG13_orf1ab;2421 | 0 | 0 |
| MN996532.1;QHR63299.1;BatCoV_RaTG13_orf1ab;2422 | 0 | 0 |
| MN996532.1;QHR63299.1;BatCoV_RaTG13_orf1ab;2423 | 0 | 0 |
| MN996532.1;QHR63299.1;BatCoV_RaTG13_orf1ab;2424 | 0 | 0 |
| MN996532.1;QHR63299.1;BatCoV_RaTG13_orf1ab;2425 | 0 | 0 |
| MN996532.1;QHR63299.1;BatCoV_RaTG13_orf1ab;2426 | 0 | 0 |
| MN996532.1;QHR63299.1;BatCoV_RaTG13_orf1ab;2427 | 0 | 0 |
| MN996532.1;QHR63299.1;BatCoV_RaTG13_orf1ab;2428 | 0 | 0 |
| MN996532.1;QHR63299.1;BatCoV_RaTG13_orf1ab;2429 | 0 | 0 |
| MN996532.1;QHR63299.1;BatCoV_RaTG13_orf1ab;2430 | 0 | 0 |
| MN996532.1;QHR63299.1;BatCoV_RaTG13_orf1ab;2431 | 0 | 0 |
| MN996532.1;QHR63299.1;BatCoV_RaTG13_orf1ab;2432 | 0 | 0 |
| MN996532.1;QHR63299.1;BatCoV_RaTG13_orf1ab;2433 | 0 | 0 |
| MN996532.1;QHR63299.1;BatCoV_RaTG13_orf1ab;2434 | 0 | 0 |
| MN996532.1;QHR63299.1;BatCoV_RaTG13_orf1ab;2435 | 0 | 0 |
| MN996532.1;QHR63299.1;BatCoV_RaTG13_orf1ab;2436 | 0 | 0 |
| MN996532.1;QHR63299.1;BatCoV_RaTG13_orf1ab;2437 | 0 | 0 |
| MN996532.1;QHR63299.1;BatCoV_RaTG13_orf1ab;2438 | 0 | 0 |
| MN996532.1;QHR63299.1;BatCoV_RaTG13_orf1ab;2439 | 0 | 0 |
| MN996532.1;QHR63299.1;BatCoV_RaTG13_orf1ab;2440 | 0 | 0.025 |
| MN996532.1;QHR63299.1;BatCoV_RaTG13_orf1ab;2441 | 0 | 0.025 |
| MN996532.1;QHR63299.1;BatCoV_RaTG13_orf1ab;2442 | 0 | 0 |

| | | |
|---|---|---|
| MN996532.1;QHR63299.1;BatCoV_RaTG13_orf1ab;2443 | 0 | 0 |
| MN996532.1;QHR63299.1;BatCoV_RaTG13_orf1ab;2444 | 0 | 0 |
| MN996532.1;QHR63299.1;BatCoV_RaTG13_orf1ab;2445 | 0 | 0 |
| MN996532.1;QHR63299.1;BatCoV_RaTG13_orf1ab;2446 | 0 | 0 |
| MN996532.1;QHR63299.1;BatCoV_RaTG13_orf1ab;2447 | 0 | 0 |
| MN996532.1;QHR63299.1;BatCoV_RaTG13_orf1ab;2448 | 0 | 0 |
| MN996532.1;QHR63299.1;BatCoV_RaTG13_orf1ab;2449 | 0 | 0 |
| MN996532.1;QHR63299.1;BatCoV_RaTG13_orf1ab;2450 | 0 | 0 |
| MN996532.1;QHR63299.1;BatCoV_RaTG13_orf1ab;2451 | 0 | 0 |
| MN996532.1;QHR63299.1;BatCoV_RaTG13_orf1ab;2452 | 0 | 0 |
| MN996532.1;QHR63299.1;BatCoV_RaTG13_orf1ab;2453 | 0 | 0 |
| MN996532.1;QHR63299.1;BatCoV_RaTG13_orf1ab;2454 | 0 | 0 |
| MN996532.1;QHR63299.1;BatCoV_RaTG13_orf1ab;2455 | 0 | 0 |
| MN996532.1;QHR63299.1;BatCoV_RaTG13_orf1ab;2456 | 0 | 0 |
| MN996532.1;QHR63299.1;BatCoV_RaTG13_orf1ab;2457 | 0 | 0 |
| MN996532.1;QHR63299.1;BatCoV_RaTG13_orf1ab;2458 | 0 | 0 |
| MN996532.1;QHR63299.1;BatCoV_RaTG13_orf1ab;2459 | 0 | 0 |
| MN996532.1;QHR63299.1;BatCoV_RaTG13_orf1ab;2460 | 0 | 0 |
| MN996532.1;QHR63299.1;BatCoV_RaTG13_orf1ab;2461 | 0 | 0 |
| MN996532.1;QHR63299.1;BatCoV_RaTG13_orf1ab;2462 | 0 | 0 |
| MN996532.1;QHR63299.1;BatCoV_RaTG13_orf1ab;2463 | 0 | 0 |
| MN996532.1;QHR63299.1;BatCoV_RaTG13_orf1ab;2464 | 0 | 0 |
| MN996532.1;QHR63299.1;BatCoV_RaTG13_orf1ab;2465 | 0 | 0 |
| MN996532.1;QHR63299.1;BatCoV_RaTG13_orf1ab;2466 | 0 | 0 |
| MN996532.1;QHR63299.1;BatCoV_RaTG13_orf1ab;2467 | 0 | 0 |
| MN996532.1;QHR63299.1;BatCoV_RaTG13_orf1ab;2468 | 0.05 | 0 |
| MN996532.1;QHR63299.1;BatCoV_RaTG13_orf1ab;2469 | 0.05 | 0.025 |
| MN996532.1;QHR63299.1;BatCoV_RaTG13_orf1ab;2470 | 0.05 | 0.025 |
| MN996532.1;QHR63299.1;BatCoV_RaTG13_orf1ab;2471 | 0 | 0 |
| MN996532.1;QHR63299.1;BatCoV_RaTG13_orf1ab;2472 | 0 | 0 |
| MN996532.1;QHR63299.1;BatCoV_RaTG13_orf1ab;2473 | 0 | 0 |
| MN996532.1;QHR63299.1;BatCoV_RaTG13_orf1ab;2474 | 0 | 0.025 |
| MN996532.1;QHR63299.1;BatCoV_RaTG13_orf1ab;2475 | 0 | 0 |
| MN996532.1;QHR63299.1;BatCoV_RaTG13_orf1ab;2476 | 0 | 0 |
| MN996532.1;QHR63299.1;BatCoV_RaTG13_orf1ab;2477 | 0 | 0.05 |
| MN996532.1;QHR63299.1;BatCoV_RaTG13_orf1ab;2478 | 0 | 0 |
| MN996532.1;QHR63299.1;BatCoV_RaTG13_orf1ab;2479 | 0 | 0 |
| MN996532.1;QHR63299.1;BatCoV_RaTG13_orf1ab;2480 | 0 | 0 |
| MN996532.1;QHR63299.1;BatCoV_RaTG13_orf1ab;2481 | 0.05 | 0 |
| MN996532.1;QHR63299.1;BatCoV_RaTG13_orf1ab;2482 | 0 | 0 |
| MN996532.1;QHR63299.1;BatCoV_RaTG13_orf1ab;2483 | 0 | 0 |
| MN996532.1;QHR63299.1;BatCoV_RaTG13_orf1ab;2484 | 0 | 0 |
| MN996532.1;QHR63299.1;BatCoV_RaTG13_orf1ab;2485 | 0 | 0 |

| | | |
|---|---|---|
| MN996532.1;QHR63299.1;BatCoV_RaTG13_orf1ab;2486 | 0 | 0 |
| MN996532.1;QHR63299.1;BatCoV_RaTG13_orf1ab;2487 | 0 | 0 |
| MN996532.1;QHR63299.1;BatCoV_RaTG13_orf1ab;2488 | 0 | 0 |
| MN996532.1;QHR63299.1;BatCoV_RaTG13_orf1ab;2489 | 0 | 0 |
| MN996532.1;QHR63299.1;BatCoV_RaTG13_orf1ab;2490 | 0 | 0 |
| MN996532.1;QHR63299.1;BatCoV_RaTG13_orf1ab;2491 | 0 | 0 |
| MN996532.1;QHR63299.1;BatCoV_RaTG13_orf1ab;2492 | 0 | 0 |
| MN996532.1;QHR63299.1;BatCoV_RaTG13_orf1ab;2493 | 0 | 0 |
| MN996532.1;QHR63299.1;BatCoV_RaTG13_orf1ab;2494 | 0 | 0 |
| MN996532.1;QHR63299.1;BatCoV_RaTG13_orf1ab;2495 | 0 | 0 |
| MN996532.1;QHR63299.1;BatCoV_RaTG13_orf1ab;2496 | 0 | 0 |
| MN996532.1;QHR63299.1;BatCoV_RaTG13_orf1ab;2497 | 0 | 0 |
| MN996532.1;QHR63299.1;BatCoV_RaTG13_orf1ab;2498 | 0 | 0 |
| MN996532.1;QHR63299.1;BatCoV_RaTG13_orf1ab;2499 | 0 | 0 |
| MN996532.1;QHR63299.1;BatCoV_RaTG13_orf1ab;2500 | 0 | 0 |
| MN996532.1;QHR63299.1;BatCoV_RaTG13_orf1ab;2501 | 0 | 0 |
| MN996532.1;QHR63299.1;BatCoV_RaTG13_orf1ab;2502 | 0 | 0 |
| MN996532.1;QHR63299.1;BatCoV_RaTG13_orf1ab;2503 | 0.05 | 0.05 |
| MN996532.1;QHR63299.1;BatCoV_RaTG13_orf1ab;2504 | 0 | 0 |
| MN996532.1;QHR63299.1;BatCoV_RaTG13_orf1ab;2505 | 0 | 0 |
| MN996532.1;QHR63299.1;BatCoV_RaTG13_orf1ab;2506 | 0 | 0 |
| MN996532.1;QHR63299.1;BatCoV_RaTG13_orf1ab;2507 | 0 | 0 |
| MN996532.1;QHR63299.1;BatCoV_RaTG13_orf1ab;2508 | 0 | 0 |
| MN996532.1;QHR63299.1;BatCoV_RaTG13_orf1ab;2509 | 0 | 0 |
| MN996532.1;QHR63299.1;BatCoV_RaTG13_orf1ab;2510 | 0 | 0 |
| MN996532.1;QHR63299.1;BatCoV_RaTG13_orf1ab;2511 | 0 | 0 |
| MN996532.1;QHR63299.1;BatCoV_RaTG13_orf1ab;2512 | 0 | 0 |
| MN996532.1;QHR63299.1;BatCoV_RaTG13_orf1ab;2513 | 0 | 0 |
| MN996532.1;QHR63299.1;BatCoV_RaTG13_orf1ab;2514 | 0 | 0 |
| MN996532.1;QHR63299.1;BatCoV_RaTG13_orf1ab;2515 | 0 | 0 |
| MN996532.1;QHR63299.1;BatCoV_RaTG13_orf1ab;2516 | 0 | 0 |
| MN996532.1;QHR63299.1;BatCoV_RaTG13_orf1ab;2517 | 0 | 0 |
| MN996532.1;QHR63299.1;BatCoV_RaTG13_orf1ab;2518 | 0 | 0 |
| MN996532.1;QHR63299.1;BatCoV_RaTG13_orf1ab;2519 | 0 | 0.025 |
| MN996532.1;QHR63299.1;BatCoV_RaTG13_orf1ab;2520 | 0.05 | 0.05 |
| MN996532.1;QHR63299.1;BatCoV_RaTG13_orf1ab;2521 | 0.1 | 0.05 |
| MN996532.1;QHR63299.1;BatCoV_RaTG13_orf1ab;2522 | 0.1 | 0.05 |
| MN996532.1;QHR63299.1;BatCoV_RaTG13_orf1ab;2523 | 0.1 | 0.075 |
| MN996532.1;QHR63299.1;BatCoV_RaTG13_orf1ab;2524 | 0.1 | 0 |
| MN996532.1;QHR63299.1;BatCoV_RaTG13_orf1ab;2525 | 0.05 | 0 |
| MN996532.1;QHR63299.1;BatCoV_RaTG13_orf1ab;2526 | 0.1 | 0.025 |
| MN996532.1;QHR63299.1;BatCoV_RaTG13_orf1ab;2527 | 0.1 | 0 |
| MN996532.1;QHR63299.1;BatCoV_RaTG13_orf1ab;2528 | 0 | 0 |

| | | |
|---|---|---|
| MN996532.1;QHR63299.1;BatCoV_RaTG13_orf1ab;2529 | 0 | 0 |
| MN996532.1;QHR63299.1;BatCoV_RaTG13_orf1ab;2530 | 0 | 0 |
| MN996532.1;QHR63299.1;BatCoV_RaTG13_orf1ab;2531 | 0 | 0 |
| MN996532.1;QHR63299.1;BatCoV_RaTG13_orf1ab;2532 | 0 | 0 |
| MN996532.1;QHR63299.1;BatCoV_RaTG13_orf1ab;2533 | 0 | 0.025 |
| MN996532.1;QHR63299.1;BatCoV_RaTG13_orf1ab;2534 | 0 | 0.025 |
| MN996532.1;QHR63299.1;BatCoV_RaTG13_orf1ab;2535 | 0.05 | 0.025 |
| MN996532.1;QHR63299.1;BatCoV_RaTG13_orf1ab;2536 | 0.05 | 0.025 |
| MN996532.1;QHR63299.1;BatCoV_RaTG13_orf1ab;2537 | 0.05 | 0.025 |
| MN996532.1;QHR63299.1;BatCoV_RaTG13_orf1ab;2538 | 0 | 0.025 |
| MN996532.1;QHR63299.1;BatCoV_RaTG13_orf1ab;2539 | 0 | 0.025 |
| MN996532.1;QHR63299.1;BatCoV_RaTG13_orf1ab;2540 | 0 | 0.025 |
| MN996532.1;QHR63299.1;BatCoV_RaTG13_orf1ab;2541 | 0 | 0.025 |
| MN996532.1;QHR63299.1;BatCoV_RaTG13_orf1ab;2542 | 0.1 | 0.05 |
| MN996532.1;QHR63299.1;BatCoV_RaTG13_orf1ab;2543 | 0 | 0 |
| MN996532.1;QHR63299.1;BatCoV_RaTG13_orf1ab;2544 | 0 | 0 |
| MN996532.1;QHR63299.1;BatCoV_RaTG13_orf1ab;2545 | 0 | 0 |
| MN996532.1;QHR63299.1;BatCoV_RaTG13_orf1ab;2546 | 0 | 0 |
| MN996532.1;QHR63299.1;BatCoV_RaTG13_orf1ab;2547 | 0.05 | 0 |
| MN996532.1;QHR63299.1;BatCoV_RaTG13_orf1ab;2548 | 0 | 0 |
| MN996532.1;QHR63299.1;BatCoV_RaTG13_orf1ab;2549 | 0 | 0 |
| MN996532.1;QHR63299.1;BatCoV_RaTG13_orf1ab;2550 | 0.05 | 0 |
| MN996532.1;QHR63299.1;BatCoV_RaTG13_orf1ab;2551 | 0 | 0 |
| MN996532.1;QHR63299.1;BatCoV_RaTG13_orf1ab;2552 | 0 | 0 |
| MN996532.1;QHR63299.1;BatCoV_RaTG13_orf1ab;2553 | 0 | 0 |
| MN996532.1;QHR63299.1;BatCoV_RaTG13_orf1ab;2554 | 0 | 0 |
| MN996532.1;QHR63299.1;BatCoV_RaTG13_orf1ab;2555 | 0 | 0 |
| MN996532.1;QHR63299.1;BatCoV_RaTG13_orf1ab;2556 | 0 | 0.025 |
| MN996532.1;QHR63299.1;BatCoV_RaTG13_orf1ab;2557 | 0 | 0 |
| MN996532.1;QHR63299.1;BatCoV_RaTG13_orf1ab;2558 | 0 | 0 |
| MN996532.1;QHR63299.1;BatCoV_RaTG13_orf1ab;2559 | 0 | 0 |
| MN996532.1;QHR63299.1;BatCoV_RaTG13_orf1ab;2560 | 0 | 0 |
| MN996532.1;QHR63299.1;BatCoV_RaTG13_orf1ab;2561 | 0 | 0 |
| MN996532.1;QHR63299.1;BatCoV_RaTG13_orf1ab;2562 | 0 | 0 |
| MN996532.1;QHR63299.1;BatCoV_RaTG13_orf1ab;2563 | 0 | 0 |
| MN996532.1;QHR63299.1;BatCoV_RaTG13_orf1ab;2564 | 0 | 0 |
| MN996532.1;QHR63299.1;BatCoV_RaTG13_orf1ab;2565 | 0 | 0 |
| MN996532.1;QHR63299.1;BatCoV_RaTG13_orf1ab;2566 | 0 | 0 |
| MN996532.1;QHR63299.1;BatCoV_RaTG13_orf1ab;2567 | 0 | 0 |
| MN996532.1;QHR63299.1;BatCoV_RaTG13_orf1ab;2568 | 0 | 0 |
| MN996532.1;QHR63299.1;BatCoV_RaTG13_orf1ab;2569 | 0 | 0.025 |
| MN996532.1;QHR63299.1;BatCoV_RaTG13_orf1ab;2570 | 0.1 | 0.1 |
| MN996532.1;QHR63299.1;BatCoV_RaTG13_orf1ab;2571 | 0 | 0.025 |

| | | |
|---|---|---|
| MN996532.1;QHR63299.1;BatCoV_RaTG13_orf1ab;2572 | 0 | 0 |
| MN996532.1;QHR63299.1;BatCoV_RaTG13_orf1ab;2573 | 0 | 0 |
| MN996532.1;QHR63299.1;BatCoV_RaTG13_orf1ab;2574 | 0 | 0 |
| MN996532.1;QHR63299.1;BatCoV_RaTG13_orf1ab;2575 | 0 | 0 |
| MN996532.1;QHR63299.1;BatCoV_RaTG13_orf1ab;2576 | 0 | 0 |
| MN996532.1;QHR63299.1;BatCoV_RaTG13_orf1ab;2577 | 0 | 0 |
| MN996532.1;QHR63299.1;BatCoV_RaTG13_orf1ab;2578 | 0 | 0 |
| MN996532.1;QHR63299.1;BatCoV_RaTG13_orf1ab;2579 | 0 | 0 |
| MN996532.1;QHR63299.1;BatCoV_RaTG13_orf1ab;2580 | 0 | 0 |
| MN996532.1;QHR63299.1;BatCoV_RaTG13_orf1ab;2581 | 0 | 0 |
| MN996532.1;QHR63299.1;BatCoV_RaTG13_orf1ab;2582 | 0 | 0 |
| MN996532.1;QHR63299.1;BatCoV_RaTG13_orf1ab;2583 | 0 | 0 |
| MN996532.1;QHR63299.1;BatCoV_RaTG13_orf1ab;2584 | 0 | 0 |
| MN996532.1;QHR63299.1;BatCoV_RaTG13_orf1ab;2585 | 0 | 0 |
| MN996532.1;QHR63299.1;BatCoV_RaTG13_orf1ab;2586 | 0 | 0 |
| MN996532.1;QHR63299.1;BatCoV_RaTG13_orf1ab;2587 | 0 | 0 |
| MN996532.1;QHR63299.1;BatCoV_RaTG13_orf1ab;2588 | 0 | 0 |
| MN996532.1;QHR63299.1;BatCoV_RaTG13_orf1ab;2589 | 0 | 0.025 |
| MN996532.1;QHR63299.1;BatCoV_RaTG13_orf1ab;2590 | 0 | 0 |
| MN996532.1;QHR63299.1;BatCoV_RaTG13_orf1ab;2591 | 0 | 0 |
| MN996532.1;QHR63299.1;BatCoV_RaTG13_orf1ab;2592 | 0.05 | 0.05 |
| MN996532.1;QHR63299.1;BatCoV_RaTG13_orf1ab;2593 | 0.05 | 0.025 |
| MN996532.1;QHR63299.1;BatCoV_RaTG13_orf1ab;2594 | 0.15 | 0.025 |
| MN996532.1;QHR63299.1;BatCoV_RaTG13_orf1ab;2595 | 0.05 | 0 |
| MN996532.1;QHR63299.1;BatCoV_RaTG13_orf1ab;2596 | 0 | 0 |
| MN996532.1;QHR63299.1;BatCoV_RaTG13_orf1ab;2597 | 0 | 0 |
| MN996532.1;QHR63299.1;BatCoV_RaTG13_orf1ab;2598 | 0 | 0.025 |
| MN996532.1;QHR63299.1;BatCoV_RaTG13_orf1ab;2599 | 0 | 0 |
| MN996532.1;QHR63299.1;BatCoV_RaTG13_orf1ab;2600 | 0 | 0 |
| MN996532.1;QHR63299.1;BatCoV_RaTG13_orf1ab;2601 | 0 | 0 |
| MN996532.1;QHR63299.1;BatCoV_RaTG13_orf1ab;2602 | 0 | 0 |
| MN996532.1;QHR63299.1;BatCoV_RaTG13_orf1ab;2603 | 0 | 0 |
| MN996532.1;QHR63299.1;BatCoV_RaTG13_orf1ab;2604 | 0 | 0 |
| MN996532.1;QHR63299.1;BatCoV_RaTG13_orf1ab;2605 | 0 | 0 |
| MN996532.1;QHR63299.1;BatCoV_RaTG13_orf1ab;2606 | 0 | 0.05 |
| MN996532.1;QHR63299.1;BatCoV_RaTG13_orf1ab;2607 | 0 | 0 |
| MN996532.1;QHR63299.1;BatCoV_RaTG13_orf1ab;2608 | 0 | 0 |
| MN996532.1;QHR63299.1;BatCoV_RaTG13_orf1ab;2609 | 0 | 0 |
| MN996532.1;QHR63299.1;BatCoV_RaTG13_orf1ab;2610 | 0 | 0 |
| MN996532.1;QHR63299.1;BatCoV_RaTG13_orf1ab;2611 | 0 | 0 |
| MN996532.1;QHR63299.1;BatCoV_RaTG13_orf1ab;2612 | 0 | 0 |
| MN996532.1;QHR63299.1;BatCoV_RaTG13_orf1ab;2613 | 0 | 0.025 |
| MN996532.1;QHR63299.1;BatCoV_RaTG13_orf1ab;2614 | 0 | 0.05 |

| | | |
|---|---|---|
| MN996532.1;QHR63299.1;BatCoV_RaTG13_orf1ab;2615 | 0.05 | 0.025 |
| MN996532.1;QHR63299.1;BatCoV_RaTG13_orf1ab;2616 | 0.1 | 0.05 |
| MN996532.1;QHR63299.1;BatCoV_RaTG13_orf1ab;2617 | 0.05 | 0.025 |
| MN996532.1;QHR63299.1;BatCoV_RaTG13_orf1ab;2618 | 0.05 | 0 |
| MN996532.1;QHR63299.1;BatCoV_RaTG13_orf1ab;2619 | 0 | 0 |
| MN996532.1;QHR63299.1;BatCoV_RaTG13_orf1ab;2620 | 0 | 0 |
| MN996532.1;QHR63299.1;BatCoV_RaTG13_orf1ab;2621 | 0.05 | 0 |
| MN996532.1;QHR63299.1;BatCoV_RaTG13_orf1ab;2622 | 0 | 0 |
| MN996532.1;QHR63299.1;BatCoV_RaTG13_orf1ab;2623 | 0 | 0 |
| MN996532.1;QHR63299.1;BatCoV_RaTG13_orf1ab;2624 | 0 | 0 |
| MN996532.1;QHR63299.1;BatCoV_RaTG13_orf1ab;2625 | 0 | 0 |
| MN996532.1;QHR63299.1;BatCoV_RaTG13_orf1ab;2626 | 0 | 0 |
| MN996532.1;QHR63299.1;BatCoV_RaTG13_orf1ab;2627 | 0 | 0 |
| MN996532.1;QHR63299.1;BatCoV_RaTG13_orf1ab;2628 | 0 | 0 |
| MN996532.1;QHR63299.1;BatCoV_RaTG13_orf1ab;2629 | 0 | 0 |
| MN996532.1;QHR63299.1;BatCoV_RaTG13_orf1ab;2630 | 0 | 0 |
| MN996532.1;QHR63299.1;BatCoV_RaTG13_orf1ab;2631 | 0 | 0 |
| MN996532.1;QHR63299.1;BatCoV_RaTG13_orf1ab;2632 | 0 | 0 |
| MN996532.1;QHR63299.1;BatCoV_RaTG13_orf1ab;2633 | 0 | 0 |
| MN996532.1;QHR63299.1;BatCoV_RaTG13_orf1ab;2634 | 0 | 0 |
| MN996532.1;QHR63299.1;BatCoV_RaTG13_orf1ab;2635 | 0 | 0 |
| MN996532.1;QHR63299.1;BatCoV_RaTG13_orf1ab;2636 | 0 | 0 |
| MN996532.1;QHR63299.1;BatCoV_RaTG13_orf1ab;2637 | 0 | 0 |
| MN996532.1;QHR63299.1;BatCoV_RaTG13_orf1ab;2638 | 0 | 0 |
| MN996532.1;QHR63299.1;BatCoV_RaTG13_orf1ab;2639 | 0 | 0 |
| MN996532.1;QHR63299.1;BatCoV_RaTG13_orf1ab;2640 | 0 | 0 |
| MN996532.1;QHR63299.1;BatCoV_RaTG13_orf1ab;2641 | 0 | 0 |
| MN996532.1;QHR63299.1;BatCoV_RaTG13_orf1ab;2642 | 0 | 0 |
| MN996532.1;QHR63299.1;BatCoV_RaTG13_orf1ab;2643 | 0 | 0 |
| MN996532.1;QHR63299.1;BatCoV_RaTG13_orf1ab;2644 | 0 | 0 |
| MN996532.1;QHR63299.1;BatCoV_RaTG13_orf1ab;2645 | 0 | 0 |
| MN996532.1;QHR63299.1;BatCoV_RaTG13_orf1ab;2646 | 0 | 0.025 |
| MN996532.1;QHR63299.1;BatCoV_RaTG13_orf1ab;2647 | 0 | 0 |
| MN996532.1;QHR63299.1;BatCoV_RaTG13_orf1ab;2648 | 0 | 0 |
| MN996532.1;QHR63299.1;BatCoV_RaTG13_orf1ab;2649 | 0 | 0 |
| MN996532.1;QHR63299.1;BatCoV_RaTG13_orf1ab;2650 | 0 | 0 |
| MN996532.1;QHR63299.1;BatCoV_RaTG13_orf1ab;2651 | 0 | 0 |
| MN996532.1;QHR63299.1;BatCoV_RaTG13_orf1ab;2652 | 0 | 0 |
| MN996532.1;QHR63299.1;BatCoV_RaTG13_orf1ab;2653 | 0 | 0 |
| MN996532.1;QHR63299.1;BatCoV_RaTG13_orf1ab;2654 | 0 | 0 |
| MN996532.1;QHR63299.1;BatCoV_RaTG13_orf1ab;2655 | 0 | 0 |
| MN996532.1;QHR63299.1;BatCoV_RaTG13_orf1ab;2656 | 0 | 0 |
| MN996532.1;QHR63299.1;BatCoV_RaTG13_orf1ab;2657 | 0 | 0 |

| | | |
|---|---|---|
| MN996532.1;QHR63299.1;BatCoV_RaTG13_orf1ab;2658 | 0 | 0 |
| MN996532.1;QHR63299.1;BatCoV_RaTG13_orf1ab;2659 | 0 | 0 |
| MN996532.1;QHR63299.1;BatCoV_RaTG13_orf1ab;2660 | 0.05 | 0 |
| MN996532.1;QHR63299.1;BatCoV_RaTG13_orf1ab;2661 | 0 | 0 |
| MN996532.1;QHR63299.1;BatCoV_RaTG13_orf1ab;2662 | 0.05 | 0.05 |
| MN996532.1;QHR63299.1;BatCoV_RaTG13_orf1ab;2663 | 0.05 | 0.05 |
| MN996532.1;QHR63299.1;BatCoV_RaTG13_orf1ab;2664 | 0 | 0 |
| MN996532.1;QHR63299.1;BatCoV_RaTG13_orf1ab;2665 | 0 | 0 |
| MN996532.1;QHR63299.1;BatCoV_RaTG13_orf1ab;2666 | 0 | 0.025 |
| MN996532.1;QHR63299.1;BatCoV_RaTG13_orf1ab;2667 | 0.1 | 0.05 |
| MN996532.1;QHR63299.1;BatCoV_RaTG13_orf1ab;2668 | 0.05 | 0 |
| MN996532.1;QHR63299.1;BatCoV_RaTG13_orf1ab;2669 | 0 | 0 |
| MN996532.1;QHR63299.1;BatCoV_RaTG13_orf1ab;2670 | 0 | 0 |
| MN996532.1;QHR63299.1;BatCoV_RaTG13_orf1ab;2671 | 0.05 | 0 |
| MN996532.1;QHR63299.1;BatCoV_RaTG13_orf1ab;2672 | 0 | 0 |
| MN996532.1;QHR63299.1;BatCoV_RaTG13_orf1ab;2673 | 0 | 0.025 |
| MN996532.1;QHR63299.1;BatCoV_RaTG13_orf1ab;2674 | 0.05 | 0.075 |
| MN996532.1;QHR63299.1;BatCoV_RaTG13_orf1ab;2675 | 0.05 | 0.025 |
| MN996532.1;QHR63299.1;BatCoV_RaTG13_orf1ab;2676 | 0.05 | 0.05 |
| MN996532.1;QHR63299.1;BatCoV_RaTG13_orf1ab;2677 | 0 | 0 |
| MN996532.1;QHR63299.1;BatCoV_RaTG13_orf1ab;2678 | 0 | 0 |
| MN996532.1;QHR63299.1;BatCoV_RaTG13_orf1ab;2679 | 0 | 0 |
| MN996532.1;QHR63299.1;BatCoV_RaTG13_orf1ab;2680 | 0 | 0 |
| MN996532.1;QHR63299.1;BatCoV_RaTG13_orf1ab;2681 | 0 | 0 |
| MN996532.1;QHR63299.1;BatCoV_RaTG13_orf1ab;2682 | 0 | 0 |
| MN996532.1;QHR63299.1;BatCoV_RaTG13_orf1ab;2683 | 0.05 | 0 |
| MN996532.1;QHR63299.1;BatCoV_RaTG13_orf1ab;2684 | 0.05 | 0 |
| MN996532.1;QHR63299.1;BatCoV_RaTG13_orf1ab;2685 | 0 | 0 |
| MN996532.1;QHR63299.1;BatCoV_RaTG13_orf1ab;2686 | 0 | 0 |
| MN996532.1;QHR63299.1;BatCoV_RaTG13_orf1ab;2687 | 0 | 0 |
| MN996532.1;QHR63299.1;BatCoV_RaTG13_orf1ab;2688 | 0.05 | 0.05 |
| MN996532.1;QHR63299.1;BatCoV_RaTG13_orf1ab;2689 | 0 | 0 |
| MN996532.1;QHR63299.1;BatCoV_RaTG13_orf1ab;2690 | 0.05 | 0 |
| MN996532.1;QHR63299.1;BatCoV_RaTG13_orf1ab;2691 | 0 | 0 |
| MN996532.1;QHR63299.1;BatCoV_RaTG13_orf1ab;2692 | 0 | 0 |
| MN996532.1;QHR63299.1;BatCoV_RaTG13_orf1ab;2693 | 0 | 0 |
| MN996532.1;QHR63299.1;BatCoV_RaTG13_orf1ab;2694 | 0 | 0 |
| MN996532.1;QHR63299.1;BatCoV_RaTG13_orf1ab;2695 | 0 | 0 |
| MN996532.1;QHR63299.1;BatCoV_RaTG13_orf1ab;2696 | 0 | 0 |
| MN996532.1;QHR63299.1;BatCoV_RaTG13_orf1ab;2697 | 0 | 0 |
| MN996532.1;QHR63299.1;BatCoV_RaTG13_orf1ab;2698 | 0 | 0 |
| MN996532.1;QHR63299.1;BatCoV_RaTG13_orf1ab;2699 | 0 | 0 |
| MN996532.1;QHR63299.1;BatCoV_RaTG13_orf1ab;2700 | 0 | 0 |

| | | |
|---|---|---|
| MN996532.1;QHR63299.1;BatCoV_RaTG13_orf1ab;2701 | 0 | 0 |
| MN996532.1;QHR63299.1;BatCoV_RaTG13_orf1ab;2702 | 0 | 0 |
| MN996532.1;QHR63299.1;BatCoV_RaTG13_orf1ab;2703 | 0 | 0 |
| MN996532.1;QHR63299.1;BatCoV_RaTG13_orf1ab;2704 | 0 | 0 |
| MN996532.1;QHR63299.1;BatCoV_RaTG13_orf1ab;2705 | 0 | 0 |
| MN996532.1;QHR63299.1;BatCoV_RaTG13_orf1ab;2706 | 0 | 0 |
| MN996532.1;QHR63299.1;BatCoV_RaTG13_orf1ab;2707 | 0 | 0 |
| MN996532.1;QHR63299.1;BatCoV_RaTG13_orf1ab;2708 | 0 | 0 |
| MN996532.1;QHR63299.1;BatCoV_RaTG13_orf1ab;2709 | 0 | 0 |
| MN996532.1;QHR63299.1;BatCoV_RaTG13_orf1ab;2710 | 0 | 0 |
| MN996532.1;QHR63299.1;BatCoV_RaTG13_orf1ab;2711 | 0 | 0 |
| MN996532.1;QHR63299.1;BatCoV_RaTG13_orf1ab;2712 | 0 | 0 |
| MN996532.1;QHR63299.1;BatCoV_RaTG13_orf1ab;2713 | 0 | 0 |
| MN996532.1;QHR63299.1;BatCoV_RaTG13_orf1ab;2714 | 0 | 0 |
| MN996532.1;QHR63299.1;BatCoV_RaTG13_orf1ab;2715 | 0 | 0 |
| MN996532.1;QHR63299.1;BatCoV_RaTG13_orf1ab;2716 | 0 | 0 |
| MN996532.1;QHR63299.1;BatCoV_RaTG13_orf1ab;2717 | 0.05 | 0 |
| MN996532.1;QHR63299.1;BatCoV_RaTG13_orf1ab;2718 | 0 | 0 |
| MN996532.1;QHR63299.1;BatCoV_RaTG13_orf1ab;2719 | 0 | 0 |
| MN996532.1;QHR63299.1;BatCoV_RaTG13_orf1ab;2720 | 0 | 0 |
| MN996532.1;QHR63299.1;BatCoV_RaTG13_orf1ab;2721 | 0 | 0.025 |
| MN996532.1;QHR63299.1;BatCoV_RaTG13_orf1ab;2722 | 0 | 0 |
| MN996532.1;QHR63299.1;BatCoV_RaTG13_orf1ab;2723 | 0 | 0 |
| MN996532.1;QHR63299.1;BatCoV_RaTG13_orf1ab;2724 | 0 | 0 |
| MN996532.1;QHR63299.1;BatCoV_RaTG13_orf1ab;2725 | 0 | 0.025 |
| MN996532.1;QHR63299.1;BatCoV_RaTG13_orf1ab;2726 | 0 | 0.025 |
| MN996532.1;QHR63299.1;BatCoV_RaTG13_orf1ab;2727 | 0 | 0.025 |
| MN996532.1;QHR63299.1;BatCoV_RaTG13_orf1ab;2728 | 0.05 | 0.025 |
| MN996532.1;QHR63299.1;BatCoV_RaTG13_orf1ab;2729 | 0 | 0.025 |
| MN996532.1;QHR63299.1;BatCoV_RaTG13_orf1ab;2730 | 0 | 0.025 |
| MN996532.1;QHR63299.1;BatCoV_RaTG13_orf1ab;2731 | 0 | 0.025 |
| MN996532.1;QHR63299.1;BatCoV_RaTG13_orf1ab;2732 | 0 | 0.025 |
| MN996532.1;QHR63299.1;BatCoV_RaTG13_orf1ab;2733 | 0 | 0 |
| MN996532.1;QHR63299.1;BatCoV_RaTG13_orf1ab;2734 | 0 | 0 |
| MN996532.1;QHR63299.1;BatCoV_RaTG13_orf1ab;2735 | 0 | 0 |
| MN996532.1;QHR63299.1;BatCoV_RaTG13_orf1ab;2736 | 0 | 0 |
| MN996532.1;QHR63299.1;BatCoV_RaTG13_orf1ab;2737 | 0 | 0.025 |
| MN996532.1;QHR63299.1;BatCoV_RaTG13_orf1ab;2738 | 0 | 0 |
| MN996532.1;QHR63299.1;BatCoV_RaTG13_orf1ab;2739 | 0 | 0 |
| MN996532.1;QHR63299.1;BatCoV_RaTG13_orf1ab;2740 | 0 | 0.025 |
| MN996532.1;QHR63299.1;BatCoV_RaTG13_orf1ab;2741 | 0 | 0 |
| MN996532.1;QHR63299.1;BatCoV_RaTG13_orf1ab;2742 | 0 | 0.025 |
| MN996532.1;QHR63299.1;BatCoV_RaTG13_orf1ab;2743 | 0 | 0.025 |

| | | |
|---|---|---|
| MN996532.1;QHR63299.1;BatCoV_RaTG13_orf1ab;2744 | 0 | 0 |
| MN996532.1;QHR63299.1;BatCoV_RaTG13_orf1ab;2745 | 0 | 0 |
| MN996532.1;QHR63299.1;BatCoV_RaTG13_orf1ab;2746 | 0 | 0 |
| MN996532.1;QHR63299.1;BatCoV_RaTG13_orf1ab;2747 | 0 | 0 |
| MN996532.1;QHR63299.1;BatCoV_RaTG13_orf1ab;2748 | 0 | 0 |
| MN996532.1;QHR63299.1;BatCoV_RaTG13_orf1ab;2749 | 0 | 0 |
| MN996532.1;QHR63299.1;BatCoV_RaTG13_orf1ab;2750 | 0 | 0 |
| MN996532.1;QHR63299.1;BatCoV_RaTG13_orf1ab;2751 | 0 | 0 |
| MN996532.1;QHR63299.1;BatCoV_RaTG13_orf1ab;2752 | 0 | 0 |
| MN996532.1;QHR63299.1;BatCoV_RaTG13_orf1ab;2753 | 0 | 0 |
| MN996532.1;QHR63299.1;BatCoV_RaTG13_orf1ab;2754 | 0 | 0 |
| MN996532.1;QHR63299.1;BatCoV_RaTG13_orf1ab;2755 | 0 | 0 |
| MN996532.1;QHR63299.1;BatCoV_RaTG13_orf1ab;2756 | 0 | 0 |
| MN996532.1;QHR63299.1;BatCoV_RaTG13_orf1ab;2757 | 0 | 0 |
| MN996532.1;QHR63299.1;BatCoV_RaTG13_orf1ab;2758 | 0 | 0 |
| MN996532.1;QHR63299.1;BatCoV_RaTG13_orf1ab;2759 | 0 | 0 |
| MN996532.1;QHR63299.1;BatCoV_RaTG13_orf1ab;2760 | 0 | 0 |
| MN996532.1;QHR63299.1;BatCoV_RaTG13_orf1ab;2761 | 0 | 0 |
| MN996532.1;QHR63299.1;BatCoV_RaTG13_orf1ab;2762 | 0 | 0 |
| MN996532.1;QHR63299.1;BatCoV_RaTG13_orf1ab;2763 | 0 | 0 |
| MN996532.1;QHR63299.1;BatCoV_RaTG13_orf1ab;2764 | 0 | 0 |
| MN996532.1;QHR63299.1;BatCoV_RaTG13_orf1ab;2765 | 0 | 0 |
| MN996532.1;QHR63299.1;BatCoV_RaTG13_orf1ab;2766 | 0 | 0 |
| MN996532.1;QHR63299.1;BatCoV_RaTG13_orf1ab;2767 | 0 | 0 |
| MN996532.1;QHR63299.1;BatCoV_RaTG13_orf1ab;2768 | 0 | 0 |
| MN996532.1;QHR63299.1;BatCoV_RaTG13_orf1ab;2769 | 0 | 0 |
| MN996532.1;QHR63299.1;BatCoV_RaTG13_orf1ab;2770 | 0 | 0 |
| MN996532.1;QHR63299.1;BatCoV_RaTG13_orf1ab;2771 | 0 | 0 |
| MN996532.1;QHR63299.1;BatCoV_RaTG13_orf1ab;2772 | 0 | 0 |
| MN996532.1;QHR63299.1;BatCoV_RaTG13_orf1ab;2773 | 0 | 0 |
| MN996532.1;QHR63299.1;BatCoV_RaTG13_orf1ab;2774 | 0 | 0 |
| MN996532.1;QHR63299.1;BatCoV_RaTG13_orf1ab;2775 | 0 | 0 |
| MN996532.1;QHR63299.1;BatCoV_RaTG13_orf1ab;2776 | 0 | 0 |
| MN996532.1;QHR63299.1;BatCoV_RaTG13_orf1ab;2777 | 0 | 0 |
| MN996532.1;QHR63299.1;BatCoV_RaTG13_orf1ab;2778 | 0 | 0 |
| MN996532.1;QHR63299.1;BatCoV_RaTG13_orf1ab;2779 | 0 | 0 |
| MN996532.1;QHR63299.1;BatCoV_RaTG13_orf1ab;2780 | 0 | 0 |
| MN996532.1;QHR63299.1;BatCoV_RaTG13_orf1ab;2781 | 0 | 0 |
| MN996532.1;QHR63299.1;BatCoV_RaTG13_orf1ab;2782 | 0 | 0 |
| MN996532.1;QHR63299.1;BatCoV_RaTG13_orf1ab;2783 | 0 | 0 |
| MN996532.1;QHR63299.1;BatCoV_RaTG13_orf1ab;2784 | 0 | 0 |
| MN996532.1;QHR63299.1;BatCoV_RaTG13_orf1ab;2785 | 0 | 0 |
| MN996532.1;QHR63299.1;BatCoV_RaTG13_orf1ab;2786 | 0 | 0 |

| | | |
|---|---|---|
| MN996532.1;QHR63299.1;BatCoV_RaTG13_orf1ab;2787 | 0 | 0 |
| MN996532.1;QHR63299.1;BatCoV_RaTG13_orf1ab;2788 | 0 | 0 |
| MN996532.1;QHR63299.1;BatCoV_RaTG13_orf1ab;2789 | 0 | 0 |
| MN996532.1;QHR63299.1;BatCoV_RaTG13_orf1ab;2790 | 0 | 0 |
| MN996532.1;QHR63299.1;BatCoV_RaTG13_orf1ab;2791 | 0 | 0 |
| MN996532.1;QHR63299.1;BatCoV_RaTG13_orf1ab;2792 | 0 | 0 |
| MN996532.1;QHR63299.1;BatCoV_RaTG13_orf1ab;2793 | 0 | 0 |
| MN996532.1;QHR63299.1;BatCoV_RaTG13_orf1ab;2794 | 0 | 0 |
| MN996532.1;QHR63299.1;BatCoV_RaTG13_orf1ab;2795 | 0.05 | 0 |
| MN996532.1;QHR63299.1;BatCoV_RaTG13_orf1ab;2796 | 0 | 0.025 |
| MN996532.1;QHR63299.1;BatCoV_RaTG13_orf1ab;2797 | 0.05 | 0.025 |
| MN996532.1;QHR63299.1;BatCoV_RaTG13_orf1ab;2798 | 0.05 | 0.025 |
| MN996532.1;QHR63299.1;BatCoV_RaTG13_orf1ab;2799 | 0.05 | 0.025 |
| MN996532.1;QHR63299.1;BatCoV_RaTG13_orf1ab;2800 | 0.05 | 0.025 |
| MN996532.1;QHR63299.1;BatCoV_RaTG13_orf1ab;2801 | 0.1 | 0.025 |
| MN996532.1;QHR63299.1;BatCoV_RaTG13_orf1ab;2802 | 0.05 | 0.025 |
| MN996532.1;QHR63299.1;BatCoV_RaTG13_orf1ab;2803 | 0.05 | 0.075 |
| MN996532.1;QHR63299.1;BatCoV_RaTG13_orf1ab;2804 | 0.05 | 0.05 |
| MN996532.1;QHR63299.1;BatCoV_RaTG13_orf1ab;2805 | 0.05 | 0 |
| MN996532.1;QHR63299.1;BatCoV_RaTG13_orf1ab;2806 | 0 | 0 |
| MN996532.1;QHR63299.1;BatCoV_RaTG13_orf1ab;2807 | 0 | 0 |
| MN996532.1;QHR63299.1;BatCoV_RaTG13_orf1ab;2808 | 0 | 0 |
| MN996532.1;QHR63299.1;BatCoV_RaTG13_orf1ab;2809 | 0 | 0 |
| MN996532.1;QHR63299.1;BatCoV_RaTG13_orf1ab;2810 | 0 | 0 |
| MN996532.1;QHR63299.1;BatCoV_RaTG13_orf1ab;2811 | 0 | 0 |
| MN996532.1;QHR63299.1;BatCoV_RaTG13_orf1ab;2812 | 0 | 0 |
| MN996532.1;QHR63299.1;BatCoV_RaTG13_orf1ab;2813 | 0 | 0 |
| MN996532.1;QHR63299.1;BatCoV_RaTG13_orf1ab;2814 | 0 | 0.05 |
| MN996532.1;QHR63299.1;BatCoV_RaTG13_orf1ab;2815 | 0 | 0 |
| MN996532.1;QHR63299.1;BatCoV_RaTG13_orf1ab;2816 | 0 | 0.025 |
| MN996532.1;QHR63299.1;BatCoV_RaTG13_orf1ab;2817 | 0.05 | 0 |
| MN996532.1;QHR63299.1;BatCoV_RaTG13_orf1ab;2818 | 0 | 0 |
| MN996532.1;QHR63299.1;BatCoV_RaTG13_orf1ab;2819 | 0 | 0 |
| MN996532.1;QHR63299.1;BatCoV_RaTG13_orf1ab;2820 | 0.05 | 0 |
| MN996532.1;QHR63299.1;BatCoV_RaTG13_orf1ab;2821 | 0.1 | 0 |
| MN996532.1;QHR63299.1;BatCoV_RaTG13_orf1ab;2822 | 0 | 0 |
| MN996532.1;QHR63299.1;BatCoV_RaTG13_orf1ab;2823 | 0 | 0 |
| MN996532.1;QHR63299.1;BatCoV_RaTG13_orf1ab;2824 | 0 | 0 |
| MN996532.1;QHR63299.1;BatCoV_RaTG13_orf1ab;2825 | 0 | 0 |
| MN996532.1;QHR63299.1;BatCoV_RaTG13_orf1ab;2826 | 0 | 0 |
| MN996532.1;QHR63299.1;BatCoV_RaTG13_orf1ab;2827 | 0.1 | 0.025 |
| MN996532.1;QHR63299.1;BatCoV_RaTG13_orf1ab;2828 | 0 | 0 |
| MN996532.1;QHR63299.1;BatCoV_RaTG13_orf1ab;2829 | 0 | 0 |

| | | |
|---|---|---|
| MN996532.1;QHR63299.1;BatCoV_RaTG13_orf1ab;2830 | 0 | 0 |
| MN996532.1;QHR63299.1;BatCoV_RaTG13_orf1ab;2831 | 0 | 0 |
| MN996532.1;QHR63299.1;BatCoV_RaTG13_orf1ab;2832 | 0 | 0 |
| MN996532.1;QHR63299.1;BatCoV_RaTG13_orf1ab;2833 | 0 | 0 |
| MN996532.1;QHR63299.1;BatCoV_RaTG13_orf1ab;2834 | 0.05 | 0 |
| MN996532.1;QHR63299.1;BatCoV_RaTG13_orf1ab;2835 | 0 | 0.05 |
| MN996532.1;QHR63299.1;BatCoV_RaTG13_orf1ab;2836 | 0 | 0 |
| MN996532.1;QHR63299.1;BatCoV_RaTG13_orf1ab;2837 | 0 | 0 |
| MN996532.1;QHR63299.1;BatCoV_RaTG13_orf1ab;2838 | 0.05 | 0 |
| MN996532.1;QHR63299.1;BatCoV_RaTG13_orf1ab;2839 | 0 | 0.025 |
| MN996532.1;QHR63299.1;BatCoV_RaTG13_orf1ab;2840 | 0.05 | 0.025 |
| MN996532.1;QHR63299.1;BatCoV_RaTG13_orf1ab;2841 | 0 | 0 |
| MN996532.1;QHR63299.1;BatCoV_RaTG13_orf1ab;2842 | 0 | 0 |
| MN996532.1;QHR63299.1;BatCoV_RaTG13_orf1ab;2843 | 0 | 0.05 |
| MN996532.1;QHR63299.1;BatCoV_RaTG13_orf1ab;2844 | 0 | 0 |
| MN996532.1;QHR63299.1;BatCoV_RaTG13_orf1ab;2845 | 0 | 0 |
| MN996532.1;QHR63299.1;BatCoV_RaTG13_orf1ab;2846 | 0 | 0 |
| MN996532.1;QHR63299.1;BatCoV_RaTG13_orf1ab;2847 | 0 | 0 |
| MN996532.1;QHR63299.1;BatCoV_RaTG13_orf1ab;2848 | 0 | 0 |
| MN996532.1;QHR63299.1;BatCoV_RaTG13_orf1ab;2849 | 0 | 0 |
| MN996532.1;QHR63299.1;BatCoV_RaTG13_orf1ab;2850 | 0 | 0 |
| MN996532.1;QHR63299.1;BatCoV_RaTG13_orf1ab;2851 | 0 | 0 |
| MN996532.1;QHR63299.1;BatCoV_RaTG13_orf1ab;2852 | 0 | 0 |
| MN996532.1;QHR63299.1;BatCoV_RaTG13_orf1ab;2853 | 0 | 0 |
| MN996532.1;QHR63299.1;BatCoV_RaTG13_orf1ab;2854 | 0 | 0 |
| MN996532.1;QHR63299.1;BatCoV_RaTG13_orf1ab;2855 | 0 | 0 |
| MN996532.1;QHR63299.1;BatCoV_RaTG13_orf1ab;2856 | 0 | 0.025 |
| MN996532.1;QHR63299.1;BatCoV_RaTG13_orf1ab;2857 | 0 | 0.025 |
| MN996532.1;QHR63299.1;BatCoV_RaTG13_orf1ab;2858 | 0 | 0.025 |
| MN996532.1;QHR63299.1;BatCoV_RaTG13_orf1ab;2859 | 0 | 0 |
| MN996532.1;QHR63299.1;BatCoV_RaTG13_orf1ab;2860 | 0 | 0 |
| MN996532.1;QHR63299.1;BatCoV_RaTG13_orf1ab;2861 | 0 | 0 |
| MN996532.1;QHR63299.1;BatCoV_RaTG13_orf1ab;2862 | 0 | 0 |
| MN996532.1;QHR63299.1;BatCoV_RaTG13_orf1ab;2863 | 0 | 0 |
| MN996532.1;QHR63299.1;BatCoV_RaTG13_orf1ab;2864 | 0 | 0 |
| MN996532.1;QHR63299.1;BatCoV_RaTG13_orf1ab;2865 | 0 | 0 |
| MN996532.1;QHR63299.1;BatCoV_RaTG13_orf1ab;2866 | 0 | 0 |
| MN996532.1;QHR63299.1;BatCoV_RaTG13_orf1ab;2867 | 0 | 0 |
| MN996532.1;QHR63299.1;BatCoV_RaTG13_orf1ab;2868 | 0 | 0 |
| MN996532.1;QHR63299.1;BatCoV_RaTG13_orf1ab;2869 | 0 | 0 |
| MN996532.1;QHR63299.1;BatCoV_RaTG13_orf1ab;2870 | 0 | 0 |
| MN996532.1;QHR63299.1;BatCoV_RaTG13_orf1ab;2871 | 0 | 0 |
| MN996532.1;QHR63299.1;BatCoV_RaTG13_orf1ab;2872 | 0 | 0 |

| | | |
|---|---|---|
| MN996532.1;QHR63299.1;BatCoV_RaTG13_orf1ab;2873 | 0 | 0 |
| MN996532.1;QHR63299.1;BatCoV_RaTG13_orf1ab;2874 | 0 | 0 |
| MN996532.1;QHR63299.1;BatCoV_RaTG13_orf1ab;2875 | 0 | 0 |
| MN996532.1;QHR63299.1;BatCoV_RaTG13_orf1ab;2876 | 0 | 0 |
| MN996532.1;QHR63299.1;BatCoV_RaTG13_orf1ab;2877 | 0 | 0 |
| MN996532.1;QHR63299.1;BatCoV_RaTG13_orf1ab;2878 | 0 | 0 |
| MN996532.1;QHR63299.1;BatCoV_RaTG13_orf1ab;2879 | 0 | 0 |
| MN996532.1;QHR63299.1;BatCoV_RaTG13_orf1ab;2880 | 0 | 0 |
| MN996532.1;QHR63299.1;BatCoV_RaTG13_orf1ab;2881 | 0 | 0 |
| MN996532.1;QHR63299.1;BatCoV_RaTG13_orf1ab;2882 | 0 | 0 |
| MN996532.1;QHR63299.1;BatCoV_RaTG13_orf1ab;2883 | 0 | 0 |
| MN996532.1;QHR63299.1;BatCoV_RaTG13_orf1ab;2884 | 0 | 0 |
| MN996532.1;QHR63299.1;BatCoV_RaTG13_orf1ab;2885 | 0 | 0 |
| MN996532.1;QHR63299.1;BatCoV_RaTG13_orf1ab;2886 | 0 | 0 |
| MN996532.1;QHR63299.1;BatCoV_RaTG13_orf1ab;2887 | 0 | 0 |
| MN996532.1;QHR63299.1;BatCoV_RaTG13_orf1ab;2888 | 0 | 0 |
| MN996532.1;QHR63299.1;BatCoV_RaTG13_orf1ab;2889 | 0 | 0 |
| MN996532.1;QHR63299.1;BatCoV_RaTG13_orf1ab;2890 | 0.05 | 0 |
| MN996532.1;QHR63299.1;BatCoV_RaTG13_orf1ab;2891 | 0.05 | 0.025 |
| MN996532.1;QHR63299.1;BatCoV_RaTG13_orf1ab;2892 | 0 | 0.025 |
| MN996532.1;QHR63299.1;BatCoV_RaTG13_orf1ab;2893 | 0 | 0.025 |
| MN996532.1;QHR63299.1;BatCoV_RaTG13_orf1ab;2894 | 0.05 | 0 |
| MN996532.1;QHR63299.1;BatCoV_RaTG13_orf1ab;2895 | 0.05 | 0.05 |
| MN996532.1;QHR63299.1;BatCoV_RaTG13_orf1ab;2896 | 0.05 | 0.025 |
| MN996532.1;QHR63299.1;BatCoV_RaTG13_orf1ab;2897 | 0.05 | 0 |
| MN996532.1;QHR63299.1;BatCoV_RaTG13_orf1ab;2898 | 0 | 0 |
| MN996532.1;QHR63299.1;BatCoV_RaTG13_orf1ab;2899 | 0 | 0 |
| MN996532.1;QHR63299.1;BatCoV_RaTG13_orf1ab;2900 | 0 | 0 |
| MN996532.1;QHR63299.1;BatCoV_RaTG13_orf1ab;2901 | 0 | 0 |
| MN996532.1;QHR63299.1;BatCoV_RaTG13_orf1ab;2902 | 0.05 | 0 |
| MN996532.1;QHR63299.1;BatCoV_RaTG13_orf1ab;2903 | 0 | 0 |
| MN996532.1;QHR63299.1;BatCoV_RaTG13_orf1ab;2904 | 0 | 0 |
| MN996532.1;QHR63299.1;BatCoV_RaTG13_orf1ab;2905 | 0 | 0 |
| MN996532.1;QHR63299.1;BatCoV_RaTG13_orf1ab;2906 | 0 | 0 |
| MN996532.1;QHR63299.1;BatCoV_RaTG13_orf1ab;2907 | 0 | 0 |
| MN996532.1;QHR63299.1;BatCoV_RaTG13_orf1ab;2908 | 0 | 0 |
| MN996532.1;QHR63299.1;BatCoV_RaTG13_orf1ab;2909 | 0 | 0 |
| MN996532.1;QHR63299.1;BatCoV_RaTG13_orf1ab;2910 | 0 | 0 |
| MN996532.1;QHR63299.1;BatCoV_RaTG13_orf1ab;2911 | 0 | 0 |
| MN996532.1;QHR63299.1;BatCoV_RaTG13_orf1ab;2912 | 0.05 | 0.025 |
| MN996532.1;QHR63299.1;BatCoV_RaTG13_orf1ab;2913 | 0.05 | 0.025 |
| MN996532.1;QHR63299.1;BatCoV_RaTG13_orf1ab;2914 | 0.05 | 0.125 |
| MN996532.1;QHR63299.1;BatCoV_RaTG13_orf1ab;2915 | 0.15 | 0.15 |

| | | |
|---|---|---|
| MN996532.1;QHR63299.1;BatCoV_RaTG13_orf1ab;2916 | 0.1 | 0.05 |
| MN996532.1;QHR63299.1;BatCoV_RaTG13_orf1ab;2917 | 0.1 | 0.05 |
| MN996532.1;QHR63299.1;BatCoV_RaTG13_orf1ab;2918 | 0 | 0.025 |
| MN996532.1;QHR63299.1;BatCoV_RaTG13_orf1ab;2919 | 0.1 | 0.025 |
| MN996532.1;QHR63299.1;BatCoV_RaTG13_orf1ab;2920 | 0.05 | 0.025 |
| MN996532.1;QHR63299.1;BatCoV_RaTG13_orf1ab;2921 | 0 | 0.025 |
| MN996532.1;QHR63299.1;BatCoV_RaTG13_orf1ab;2922 | 0 | 0.025 |
| MN996532.1;QHR63299.1;BatCoV_RaTG13_orf1ab;2923 | 0 | 0 |
| MN996532.1;QHR63299.1;BatCoV_RaTG13_orf1ab;2924 | 0 | 0 |
| MN996532.1;QHR63299.1;BatCoV_RaTG13_orf1ab;2925 | 0 | 0 |
| MN996532.1;QHR63299.1;BatCoV_RaTG13_orf1ab;2926 | 0 | 0 |
| MN996532.1;QHR63299.1;BatCoV_RaTG13_orf1ab;2927 | 0 | 0 |
| MN996532.1;QHR63299.1;BatCoV_RaTG13_orf1ab;2928 | 0 | 0.025 |
| MN996532.1;QHR63299.1;BatCoV_RaTG13_orf1ab;2929 | 0 | 0 |
| MN996532.1;QHR63299.1;BatCoV_RaTG13_orf1ab;2930 | 0 | 0 |
| MN996532.1;QHR63299.1;BatCoV_RaTG13_orf1ab;2931 | 0 | 0 |
| MN996532.1;QHR63299.1;BatCoV_RaTG13_orf1ab;2932 | 0 | 0 |
| MN996532.1;QHR63299.1;BatCoV_RaTG13_orf1ab;2933 | 0 | 0 |
| MN996532.1;QHR63299.1;BatCoV_RaTG13_orf1ab;2934 | 0 | 0 |
| MN996532.1;QHR63299.1;BatCoV_RaTG13_orf1ab;2935 | 0.1 | 0 |
| MN996532.1;QHR63299.1;BatCoV_RaTG13_orf1ab;2936 | 0 | 0.025 |
| MN996532.1;QHR63299.1;BatCoV_RaTG13_orf1ab;2937 | 0 | 0.025 |
| MN996532.1;QHR63299.1;BatCoV_RaTG13_orf1ab;2938 | 0.15 | 0.15 |
| MN996532.1;QHR63299.1;BatCoV_RaTG13_orf1ab;2939 | 0.15 | 0.15 |
| MN996532.1;QHR63299.1;BatCoV_RaTG13_orf1ab;2940 | 0.1 | 0.025 |
| MN996532.1;QHR63299.1;BatCoV_RaTG13_orf1ab;2941 | 0.1 | 0.05 |
| MN996532.1;QHR63299.1;BatCoV_RaTG13_orf1ab;2942 | 0.05 | 0.025 |
| MN996532.1;QHR63299.1;BatCoV_RaTG13_orf1ab;2943 | 0.15 | 0.075 |
| MN996532.1;QHR63299.1;BatCoV_RaTG13_orf1ab;2944 | 0.1 | 0.1 |
| MN996532.1;QHR63299.1;BatCoV_RaTG13_orf1ab;2945 | 0.15 | 0.025 |
| MN996532.1;QHR63299.1;BatCoV_RaTG13_orf1ab;2946 | 0.05 | 0.025 |
| MN996532.1;QHR63299.1;BatCoV_RaTG13_orf1ab;2947 | 0.05 | 0.05 |
| MN996532.1;QHR63299.1;BatCoV_RaTG13_orf1ab;2948 | 0.05 | 0.025 |
| MN996532.1;QHR63299.1;BatCoV_RaTG13_orf1ab;2949 | 0 | 0 |
| MN996532.1;QHR63299.1;BatCoV_RaTG13_orf1ab;2950 | 0 | 0 |
| MN996532.1;QHR63299.1;BatCoV_RaTG13_orf1ab;2951 | 0 | 0 |
| MN996532.1;QHR63299.1;BatCoV_RaTG13_orf1ab;2952 | 0 | 0 |
| MN996532.1;QHR63299.1;BatCoV_RaTG13_orf1ab;2953 | 0 | 0 |
| MN996532.1;QHR63299.1;BatCoV_RaTG13_orf1ab;2954 | 0 | 0 |
| MN996532.1;QHR63299.1;BatCoV_RaTG13_orf1ab;2955 | 0 | 0 |
| MN996532.1;QHR63299.1;BatCoV_RaTG13_orf1ab;2956 | 0 | 0 |
| MN996532.1;QHR63299.1;BatCoV_RaTG13_orf1ab;2957 | 0 | 0 |
| MN996532.1;QHR63299.1;BatCoV_RaTG13_orf1ab;2958 | 0 | 0 |

| | | |
|---|---|---|
| MN996532.1;QHR63299.1;BatCoV_RaTG13_orf1ab;2959 | 0 | 0.025 |
| MN996532.1;QHR63299.1;BatCoV_RaTG13_orf1ab;2960 | 0 | 0 |
| MN996532.1;QHR63299.1;BatCoV_RaTG13_orf1ab;2961 | 0 | 0 |
| MN996532.1;QHR63299.1;BatCoV_RaTG13_orf1ab;2962 | 0 | 0 |
| MN996532.1;QHR63299.1;BatCoV_RaTG13_orf1ab;2963 | 0 | 0 |
| MN996532.1;QHR63299.1;BatCoV_RaTG13_orf1ab;2964 | 0 | 0 |
| MN996532.1;QHR63299.1;BatCoV_RaTG13_orf1ab;2965 | 0 | 0 |
| MN996532.1;QHR63299.1;BatCoV_RaTG13_orf1ab;2966 | 0 | 0 |
| MN996532.1;QHR63299.1;BatCoV_RaTG13_orf1ab;2967 | 0 | 0 |
| MN996532.1;QHR63299.1;BatCoV_RaTG13_orf1ab;2968 | 0 | 0 |
| MN996532.1;QHR63299.1;BatCoV_RaTG13_orf1ab;2969 | 0 | 0 |
| MN996532.1;QHR63299.1;BatCoV_RaTG13_orf1ab;2970 | 0.05 | 0.1 |
| MN996532.1;QHR63299.1;BatCoV_RaTG13_orf1ab;2971 | 0 | 0.05 |
| MN996532.1;QHR63299.1;BatCoV_RaTG13_orf1ab;2972 | 0 | 0.075 |
| MN996532.1;QHR63299.1;BatCoV_RaTG13_orf1ab;2973 | 0 | 0.075 |
| MN996532.1;QHR63299.1;BatCoV_RaTG13_orf1ab;2974 | 0 | 0.05 |
| MN996532.1;QHR63299.1;BatCoV_RaTG13_orf1ab;2975 | 0 | 0.05 |
| MN996532.1;QHR63299.1;BatCoV_RaTG13_orf1ab;2976 | 0 | 0.05 |
| MN996532.1;QHR63299.1;BatCoV_RaTG13_orf1ab;2977 | 0 | 0.05 |
| MN996532.1;QHR63299.1;BatCoV_RaTG13_orf1ab;2978 | 0 | 0.05 |
| MN996532.1;QHR63299.1;BatCoV_RaTG13_orf1ab;2979 | 0 | 0 |
| MN996532.1;QHR63299.1;BatCoV_RaTG13_orf1ab;2980 | 0 | 0 |
| MN996532.1;QHR63299.1;BatCoV_RaTG13_orf1ab;2981 | 0 | 0 |
| MN996532.1;QHR63299.1;BatCoV_RaTG13_orf1ab;2982 | 0 | 0 |
| MN996532.1;QHR63299.1;BatCoV_RaTG13_orf1ab;2983 | 0 | 0 |
| MN996532.1;QHR63299.1;BatCoV_RaTG13_orf1ab;2984 | 0 | 0 |
| MN996532.1;QHR63299.1;BatCoV_RaTG13_orf1ab;2985 | 0 | 0 |
| MN996532.1;QHR63299.1;BatCoV_RaTG13_orf1ab;2986 | 0 | 0 |
| MN996532.1;QHR63299.1;BatCoV_RaTG13_orf1ab;2987 | 0 | 0 |
| MN996532.1;QHR63299.1;BatCoV_RaTG13_orf1ab;2988 | 0 | 0 |
| MN996532.1;QHR63299.1;BatCoV_RaTG13_orf1ab;2989 | 0 | 0 |
| MN996532.1;QHR63299.1;BatCoV_RaTG13_orf1ab;2990 | 0 | 0 |
| MN996532.1;QHR63299.1;BatCoV_RaTG13_orf1ab;2991 | 0 | 0 |
| MN996532.1;QHR63299.1;BatCoV_RaTG13_orf1ab;2992 | 0 | 0 |
| MN996532.1;QHR63299.1;BatCoV_RaTG13_orf1ab;2993 | 0 | 0 |
| MN996532.1;QHR63299.1;BatCoV_RaTG13_orf1ab;2994 | 0 | 0 |
| MN996532.1;QHR63299.1;BatCoV_RaTG13_orf1ab;2995 | 0 | 0 |
| MN996532.1;QHR63299.1;BatCoV_RaTG13_orf1ab;2996 | 0 | 0 |
| MN996532.1;QHR63299.1;BatCoV_RaTG13_orf1ab;2997 | 0 | 0 |
| MN996532.1;QHR63299.1;BatCoV_RaTG13_orf1ab;2998 | 0 | 0 |
| MN996532.1;QHR63299.1;BatCoV_RaTG13_orf1ab;2999 | 0 | 0 |
| MN996532.1;QHR63299.1;BatCoV_RaTG13_orf1ab;3000 | 0 | 0 |
| MN996532.1;QHR63299.1;BatCoV_RaTG13_orf1ab;3001 | 0 | 0 |

| | | |
|---|---|---|
| MN996532.1;QHR63299.1;BatCoV_RaTG13_orf1ab;3002 | 0 | 0 |
| MN996532.1;QHR63299.1;BatCoV_RaTG13_orf1ab;3003 | 0 | 0 |
| MN996532.1;QHR63299.1;BatCoV_RaTG13_orf1ab;3004 | 0 | 0 |
| MN996532.1;QHR63299.1;BatCoV_RaTG13_orf1ab;3005 | 0 | 0 |
| MN996532.1;QHR63299.1;BatCoV_RaTG13_orf1ab;3006 | 0 | 0 |
| MN996532.1;QHR63299.1;BatCoV_RaTG13_orf1ab;3007 | 0 | 0 |
| MN996532.1;QHR63299.1;BatCoV_RaTG13_orf1ab;3008 | 0 | 0 |
| MN996532.1;QHR63299.1;BatCoV_RaTG13_orf1ab;3009 | 0.1 | 0.15 |
| MN996532.1;QHR63299.1;BatCoV_RaTG13_orf1ab;3010 | 0.1 | 0.15 |
| MN996532.1;QHR63299.1;BatCoV_RaTG13_orf1ab;3011 | 0.1 | 0.125 |
| MN996532.1;QHR63299.1;BatCoV_RaTG13_orf1ab;3012 | 0.1 | 0.15 |
| MN996532.1;QHR63299.1;BatCoV_RaTG13_orf1ab;3013 | 0.1 | 0.15 |
| MN996532.1;QHR63299.1;BatCoV_RaTG13_orf1ab;3014 | 0.1 | 0.15 |
| MN996532.1;QHR63299.1;BatCoV_RaTG13_orf1ab;3015 | 0.1 | 0.15 |
| MN996532.1;QHR63299.1;BatCoV_RaTG13_orf1ab;3016 | 0.1 | 0.15 |
| MN996532.1;QHR63299.1;BatCoV_RaTG13_orf1ab;3017 | 0.1 | 0.15 |
| MN996532.1;QHR63299.1;BatCoV_RaTG13_orf1ab;3018 | 0.1 | 0.15 |
| MN996532.1;QHR63299.1;BatCoV_RaTG13_orf1ab;3019 | 0.15 | 0.225 |
| MN996532.1;QHR63299.1;BatCoV_RaTG13_orf1ab;3020 | 0.1 | 0.15 |
| MN996532.1;QHR63299.1;BatCoV_RaTG13_orf1ab;3021 | 0 | 0 |
| MN996532.1;QHR63299.1;BatCoV_RaTG13_orf1ab;3022 | 0 | 0 |
| MN996532.1;QHR63299.1;BatCoV_RaTG13_orf1ab;3023 | 0 | 0 |
| MN996532.1;QHR63299.1;BatCoV_RaTG13_orf1ab;3024 | 0.05 | 0 |
| MN996532.1;QHR63299.1;BatCoV_RaTG13_orf1ab;3025 | 0 | 0 |
| MN996532.1;QHR63299.1;BatCoV_RaTG13_orf1ab;3026 | 0.05 | 0 |
| MN996532.1;QHR63299.1;BatCoV_RaTG13_orf1ab;3027 | 0.05 | 0 |
| MN996532.1;QHR63299.1;BatCoV_RaTG13_orf1ab;3028 | 0.05 | 0 |
| MN996532.1;QHR63299.1;BatCoV_RaTG13_orf1ab;3029 | 0.05 | 0 |
| MN996532.1;QHR63299.1;BatCoV_RaTG13_orf1ab;3030 | 0.05 | 0 |
| MN996532.1;QHR63299.1;BatCoV_RaTG13_orf1ab;3031 | 0.05 | 0 |
| MN996532.1;QHR63299.1;BatCoV_RaTG13_orf1ab;3032 | 0 | 0 |
| MN996532.1;QHR63299.1;BatCoV_RaTG13_orf1ab;3033 | 0 | 0 |
| MN996532.1;QHR63299.1;BatCoV_RaTG13_orf1ab;3034 | 0 | 0 |
| MN996532.1;QHR63299.1;BatCoV_RaTG13_orf1ab;3035 | 0.05 | 0 |
| MN996532.1;QHR63299.1;BatCoV_RaTG13_orf1ab;3036 | 0 | 0 |
| MN996532.1;QHR63299.1;BatCoV_RaTG13_orf1ab;3037 | 0 | 0 |
| MN996532.1;QHR63299.1;BatCoV_RaTG13_orf1ab;3038 | 0 | 0.025 |
| MN996532.1;QHR63299.1;BatCoV_RaTG13_orf1ab;3039 | 0 | 0 |
| MN996532.1;QHR63299.1;BatCoV_RaTG13_orf1ab;3040 | 0 | 0 |
| MN996532.1;QHR63299.1;BatCoV_RaTG13_orf1ab;3041 | 0 | 0 |
| MN996532.1;QHR63299.1;BatCoV_RaTG13_orf1ab;3042 | 0 | 0 |
| MN996532.1;QHR63299.1;BatCoV_RaTG13_orf1ab;3043 | 0 | 0 |
| MN996532.1;QHR63299.1;BatCoV_RaTG13_orf1ab;3044 | 0 | 0 |

| | | |
|---|---|---|
| MN996532.1;QHR63299.1;BatCoV_RaTG13_orf1ab;3045 | 0 | 0 |
| MN996532.1;QHR63299.1;BatCoV_RaTG13_orf1ab;3046 | 0 | 0 |
| MN996532.1;QHR63299.1;BatCoV_RaTG13_orf1ab;3047 | 0 | 0 |
| MN996532.1;QHR63299.1;BatCoV_RaTG13_orf1ab;3048 | 0 | 0 |
| MN996532.1;QHR63299.1;BatCoV_RaTG13_orf1ab;3049 | 0 | 0 |
| MN996532.1;QHR63299.1;BatCoV_RaTG13_orf1ab;3050 | 0 | 0 |
| MN996532.1;QHR63299.1;BatCoV_RaTG13_orf1ab;3051 | 0 | 0 |
| MN996532.1;QHR63299.1;BatCoV_RaTG13_orf1ab;3052 | 0 | 0 |
| MN996532.1;QHR63299.1;BatCoV_RaTG13_orf1ab;3053 | 0 | 0 |
| MN996532.1;QHR63299.1;BatCoV_RaTG13_orf1ab;3054 | 0 | 0 |
| MN996532.1;QHR63299.1;BatCoV_RaTG13_orf1ab;3055 | 0 | 0 |
| MN996532.1;QHR63299.1;BatCoV_RaTG13_orf1ab;3056 | 0 | 0 |
| MN996532.1;QHR63299.1;BatCoV_RaTG13_orf1ab;3057 | 0 | 0 |
| MN996532.1;QHR63299.1;BatCoV_RaTG13_orf1ab;3058 | 0 | 0 |
| MN996532.1;QHR63299.1;BatCoV_RaTG13_orf1ab;3059 | 0 | 0 |
| MN996532.1;QHR63299.1;BatCoV_RaTG13_orf1ab;3060 | 0 | 0 |
| MN996532.1;QHR63299.1;BatCoV_RaTG13_orf1ab;3061 | 0 | 0 |
| MN996532.1;QHR63299.1;BatCoV_RaTG13_orf1ab;3062 | 0 | 0 |
| MN996532.1;QHR63299.1;BatCoV_RaTG13_orf1ab;3063 | 0 | 0 |
| MN996532.1;QHR63299.1;BatCoV_RaTG13_orf1ab;3064 | 0 | 0 |
| MN996532.1;QHR63299.1;BatCoV_RaTG13_orf1ab;3065 | 0 | 0 |
| MN996532.1;QHR63299.1;BatCoV_RaTG13_orf1ab;3066 | 0 | 0 |
| MN996532.1;QHR63299.1;BatCoV_RaTG13_orf1ab;3067 | 0 | 0 |
| MN996532.1;QHR63299.1;BatCoV_RaTG13_orf1ab;3068 | 0 | 0 |
| MN996532.1;QHR63299.1;BatCoV_RaTG13_orf1ab;3069 | 0 | 0 |
| MN996532.1;QHR63299.1;BatCoV_RaTG13_orf1ab;3070 | 0 | 0 |
| MN996532.1;QHR63299.1;BatCoV_RaTG13_orf1ab;3071 | 0 | 0 |
| MN996532.1;QHR63299.1;BatCoV_RaTG13_orf1ab;3072 | 0 | 0 |
| MN996532.1;QHR63299.1;BatCoV_RaTG13_orf1ab;3073 | 0 | 0 |
| MN996532.1;QHR63299.1;BatCoV_RaTG13_orf1ab;3074 | 0 | 0 |
| MN996532.1;QHR63299.1;BatCoV_RaTG13_orf1ab;3075 | 0 | 0 |
| MN996532.1;QHR63299.1;BatCoV_RaTG13_orf1ab;3076 | 0 | 0 |
| MN996532.1;QHR63299.1;BatCoV_RaTG13_orf1ab;3077 | 0 | 0 |
| MN996532.1;QHR63299.1;BatCoV_RaTG13_orf1ab;3078 | 0 | 0 |
| MN996532.1;QHR63299.1;BatCoV_RaTG13_orf1ab;3079 | 0 | 0 |
| MN996532.1;QHR63299.1;BatCoV_RaTG13_orf1ab;3080 | 0 | 0 |
| MN996532.1;QHR63299.1;BatCoV_RaTG13_orf1ab;3081 | 0 | 0 |
| MN996532.1;QHR63299.1;BatCoV_RaTG13_orf1ab;3082 | 0 | 0 |
| MN996532.1;QHR63299.1;BatCoV_RaTG13_orf1ab;3083 | 0 | 0 |
| MN996532.1;QHR63299.1;BatCoV_RaTG13_orf1ab;3084 | 0 | 0 |
| MN996532.1;QHR63299.1;BatCoV_RaTG13_orf1ab;3085 | 0 | 0 |
| MN996532.1;QHR63299.1;BatCoV_RaTG13_orf1ab;3086 | 0 | 0 |
| MN996532.1;QHR63299.1;BatCoV_RaTG13_orf1ab;3087 | 0 | 0 |

| | | |
|---|---|---|
| MN996532.1;QHR63299.1;BatCoV_RaTG13_orf1ab;3088 | 0 | 0 |
| MN996532.1;QHR63299.1;BatCoV_RaTG13_orf1ab;3089 | 0 | 0 |
| MN996532.1;QHR63299.1;BatCoV_RaTG13_orf1ab;3090 | 0 | 0 |
| MN996532.1;QHR63299.1;BatCoV_RaTG13_orf1ab;3091 | 0 | 0 |
| MN996532.1;QHR63299.1;BatCoV_RaTG13_orf1ab;3092 | 0 | 0 |
| MN996532.1;QHR63299.1;BatCoV_RaTG13_orf1ab;3093 | 0 | 0 |
| MN996532.1;QHR63299.1;BatCoV_RaTG13_orf1ab;3094 | 0 | 0 |
| MN996532.1;QHR63299.1;BatCoV_RaTG13_orf1ab;3095 | 0 | 0 |
| MN996532.1;QHR63299.1;BatCoV_RaTG13_orf1ab;3096 | 0 | 0 |
| MN996532.1;QHR63299.1;BatCoV_RaTG13_orf1ab;3097 | 0 | 0 |
| MN996532.1;QHR63299.1;BatCoV_RaTG13_orf1ab;3098 | 0 | 0 |
| MN996532.1;QHR63299.1;BatCoV_RaTG13_orf1ab;3099 | 0 | 0 |
| MN996532.1;QHR63299.1;BatCoV_RaTG13_orf1ab;3100 | 0 | 0 |
| MN996532.1;QHR63299.1;BatCoV_RaTG13_orf1ab;3101 | 0 | 0 |
| MN996532.1;QHR63299.1;BatCoV_RaTG13_orf1ab;3102 | 0 | 0 |
| MN996532.1;QHR63299.1;BatCoV_RaTG13_orf1ab;3103 | 0 | 0 |
| MN996532.1;QHR63299.1;BatCoV_RaTG13_orf1ab;3104 | 0 | 0 |
| MN996532.1;QHR63299.1;BatCoV_RaTG13_orf1ab;3105 | 0 | 0 |
| MN996532.1;QHR63299.1;BatCoV_RaTG13_orf1ab;3106 | 0 | 0 |
| MN996532.1;QHR63299.1;BatCoV_RaTG13_orf1ab;3107 | 0 | 0 |
| MN996532.1;QHR63299.1;BatCoV_RaTG13_orf1ab;3108 | 0 | 0 |
| MN996532.1;QHR63299.1;BatCoV_RaTG13_orf1ab;3109 | 0 | 0 |
| MN996532.1;QHR63299.1;BatCoV_RaTG13_orf1ab;3110 | 0 | 0 |
| MN996532.1;QHR63299.1;BatCoV_RaTG13_orf1ab;3111 | 0 | 0 |
| MN996532.1;QHR63299.1;BatCoV_RaTG13_orf1ab;3112 | 0 | 0 |
| MN996532.1;QHR63299.1;BatCoV_RaTG13_orf1ab;3113 | 0 | 0 |
| MN996532.1;QHR63299.1;BatCoV_RaTG13_orf1ab;3114 | 0 | 0 |
| MN996532.1;QHR63299.1;BatCoV_RaTG13_orf1ab;3115 | 0 | 0 |
| MN996532.1;QHR63299.1;BatCoV_RaTG13_orf1ab;3116 | 0 | 0 |
| MN996532.1;QHR63299.1;BatCoV_RaTG13_orf1ab;3117 | 0 | 0 |
| MN996532.1;QHR63299.1;BatCoV_RaTG13_orf1ab;3118 | 0 | 0 |
| MN996532.1;QHR63299.1;BatCoV_RaTG13_orf1ab;3119 | 0 | 0 |
| MN996532.1;QHR63299.1;BatCoV_RaTG13_orf1ab;3120 | 0 | 0 |
| MN996532.1;QHR63299.1;BatCoV_RaTG13_orf1ab;3121 | 0 | 0 |
| MN996532.1;QHR63299.1;BatCoV_RaTG13_orf1ab;3122 | 0 | 0 |
| MN996532.1;QHR63299.1;BatCoV_RaTG13_orf1ab;3123 | 0 | 0 |
| MN996532.1;QHR63299.1;BatCoV_RaTG13_orf1ab;3124 | 0 | 0 |
| MN996532.1;QHR63299.1;BatCoV_RaTG13_orf1ab;3125 | 0 | 0 |
| MN996532.1;QHR63299.1;BatCoV_RaTG13_orf1ab;3126 | 0 | 0 |
| MN996532.1;QHR63299.1;BatCoV_RaTG13_orf1ab;3127 | 0 | 0 |
| MN996532.1;QHR63299.1;BatCoV_RaTG13_orf1ab;3128 | 0 | 0 |
| MN996532.1;QHR63299.1;BatCoV_RaTG13_orf1ab;3129 | 0 | 0 |
| MN996532.1;QHR63299.1;BatCoV_RaTG13_orf1ab;3130 | 0 | 0 |

| | | |
|---|---|---|
| MN996532.1;QHR63299.1;BatCoV_RaTG13_orf1ab;3131 | 0 | 0 |
| MN996532.1;QHR63299.1;BatCoV_RaTG13_orf1ab;3132 | 0 | 0 |
| MN996532.1;QHR63299.1;BatCoV_RaTG13_orf1ab;3133 | 0 | 0 |
| MN996532.1;QHR63299.1;BatCoV_RaTG13_orf1ab;3134 | 0 | 0 |
| MN996532.1;QHR63299.1;BatCoV_RaTG13_orf1ab;3135 | 0 | 0 |
| MN996532.1;QHR63299.1;BatCoV_RaTG13_orf1ab;3136 | 0 | 0 |
| MN996532.1;QHR63299.1;BatCoV_RaTG13_orf1ab;3137 | 0 | 0 |
| MN996532.1;QHR63299.1;BatCoV_RaTG13_orf1ab;3138 | 0 | 0 |
| MN996532.1;QHR63299.1;BatCoV_RaTG13_orf1ab;3139 | 0 | 0 |
| MN996532.1;QHR63299.1;BatCoV_RaTG13_orf1ab;3140 | 0 | 0 |
| MN996532.1;QHR63299.1;BatCoV_RaTG13_orf1ab;3141 | 0 | 0 |
| MN996532.1;QHR63299.1;BatCoV_RaTG13_orf1ab;3142 | 0 | 0 |
| MN996532.1;QHR63299.1;BatCoV_RaTG13_orf1ab;3143 | 0 | 0 |
| MN996532.1;QHR63299.1;BatCoV_RaTG13_orf1ab;3144 | 0 | 0 |
| MN996532.1;QHR63299.1;BatCoV_RaTG13_orf1ab;3145 | 0 | 0 |
| MN996532.1;QHR63299.1;BatCoV_RaTG13_orf1ab;3146 | 0 | 0 |
| MN996532.1;QHR63299.1;BatCoV_RaTG13_orf1ab;3147 | 0 | 0 |
| MN996532.1;QHR63299.1;BatCoV_RaTG13_orf1ab;3148 | 0 | 0 |
| MN996532.1;QHR63299.1;BatCoV_RaTG13_orf1ab;3149 | 0 | 0 |
| MN996532.1;QHR63299.1;BatCoV_RaTG13_orf1ab;3150 | 0 | 0 |
| MN996532.1;QHR63299.1;BatCoV_RaTG13_orf1ab;3151 | 0 | 0 |
| MN996532.1;QHR63299.1;BatCoV_RaTG13_orf1ab;3152 | 0 | 0 |
| MN996532.1;QHR63299.1;BatCoV_RaTG13_orf1ab;3153 | 0 | 0 |
| MN996532.1;QHR63299.1;BatCoV_RaTG13_orf1ab;3154 | 0 | 0 |
| MN996532.1;QHR63299.1;BatCoV_RaTG13_orf1ab;3155 | 0 | 0 |
| MN996532.1;QHR63299.1;BatCoV_RaTG13_orf1ab;3156 | 0 | 0 |
| MN996532.1;QHR63299.1;BatCoV_RaTG13_orf1ab;3157 | 0 | 0 |
| MN996532.1;QHR63299.1;BatCoV_RaTG13_orf1ab;3158 | 0 | 0 |
| MN996532.1;QHR63299.1;BatCoV_RaTG13_orf1ab;3159 | 0 | 0 |
| MN996532.1;QHR63299.1;BatCoV_RaTG13_orf1ab;3160 | 0 | 0 |
| MN996532.1;QHR63299.1;BatCoV_RaTG13_orf1ab;3161 | 0 | 0.075 |
| MN996532.1;QHR63299.1;BatCoV_RaTG13_orf1ab;3162 | 0 | 0 |
| MN996532.1;QHR63299.1;BatCoV_RaTG13_orf1ab;3163 | 0.05 | 0.025 |
| MN996532.1;QHR63299.1;BatCoV_RaTG13_orf1ab;3164 | 0 | 0 |
| MN996532.1;QHR63299.1;BatCoV_RaTG13_orf1ab;3165 | 0 | 0 |
| MN996532.1;QHR63299.1;BatCoV_RaTG13_orf1ab;3166 | 0 | 0 |
| MN996532.1;QHR63299.1;BatCoV_RaTG13_orf1ab;3167 | 0 | 0 |
| MN996532.1;QHR63299.1;BatCoV_RaTG13_orf1ab;3168 | 0 | 0 |
| MN996532.1;QHR63299.1;BatCoV_RaTG13_orf1ab;3169 | 0 | 0 |
| MN996532.1;QHR63299.1;BatCoV_RaTG13_orf1ab;3170 | 0 | 0 |
| MN996532.1;QHR63299.1;BatCoV_RaTG13_orf1ab;3171 | 0 | 0 |
| MN996532.1;QHR63299.1;BatCoV_RaTG13_orf1ab;3172 | 0 | 0 |
| MN996532.1;QHR63299.1;BatCoV_RaTG13_orf1ab;3173 | 0 | 0 |

| | | |
|---|---|---|
| MN996532.1;QHR63299.1;BatCoV_RaTG13_orf1ab;3174 | 0 | 0 |
| MN996532.1;QHR63299.1;BatCoV_RaTG13_orf1ab;3175 | 0 | 0 |
| MN996532.1;QHR63299.1;BatCoV_RaTG13_orf1ab;3176 | 0 | 0 |
| MN996532.1;QHR63299.1;BatCoV_RaTG13_orf1ab;3177 | 0 | 0 |
| MN996532.1;QHR63299.1;BatCoV_RaTG13_orf1ab;3178 | 0 | 0 |
| MN996532.1;QHR63299.1;BatCoV_RaTG13_orf1ab;3179 | 0 | 0 |
| MN996532.1;QHR63299.1;BatCoV_RaTG13_orf1ab;3180 | 0 | 0 |
| MN996532.1;QHR63299.1;BatCoV_RaTG13_orf1ab;3181 | 0.05 | 0 |
| MN996532.1;QHR63299.1;BatCoV_RaTG13_orf1ab;3182 | 0.1 | 0 |
| MN996532.1;QHR63299.1;BatCoV_RaTG13_orf1ab;3183 | 0.1 | 0 |
| MN996532.1;QHR63299.1;BatCoV_RaTG13_orf1ab;3184 | 0.1 | 0 |
| MN996532.1;QHR63299.1;BatCoV_RaTG13_orf1ab;3185 | 0.1 | 0 |
| MN996532.1;QHR63299.1;BatCoV_RaTG13_orf1ab;3186 | 0.1 | 0 |
| MN996532.1;QHR63299.1;BatCoV_RaTG13_orf1ab;3187 | 0.15 | 0 |
| MN996532.1;QHR63299.1;BatCoV_RaTG13_orf1ab;3188 | 0.1 | 0 |
| MN996532.1;QHR63299.1;BatCoV_RaTG13_orf1ab;3189 | 0.1 | 0 |
| MN996532.1;QHR63299.1;BatCoV_RaTG13_orf1ab;3190 | 0.1 | 0.05 |
| MN996532.1;QHR63299.1;BatCoV_RaTG13_orf1ab;3191 | 0.05 | 0.025 |
| MN996532.1;QHR63299.1;BatCoV_RaTG13_orf1ab;3192 | 0 | 0.025 |
| MN996532.1;QHR63299.1;BatCoV_RaTG13_orf1ab;3193 | 0 | 0.025 |
| MN996532.1;QHR63299.1;BatCoV_RaTG13_orf1ab;3194 | 0 | 0 |
| MN996532.1;QHR63299.1;BatCoV_RaTG13_orf1ab;3195 | 0 | 0 |
| MN996532.1;QHR63299.1;BatCoV_RaTG13_orf1ab;3196 | 0 | 0 |
| MN996532.1;QHR63299.1;BatCoV_RaTG13_orf1ab;3197 | 0 | 0 |
| MN996532.1;QHR63299.1;BatCoV_RaTG13_orf1ab;3198 | 0 | 0 |
| MN996532.1;QHR63299.1;BatCoV_RaTG13_orf1ab;3199 | 0 | 0 |
| MN996532.1;QHR63299.1;BatCoV_RaTG13_orf1ab;3200 | 0 | 0 |
| MN996532.1;QHR63299.1;BatCoV_RaTG13_orf1ab;3201 | 0 | 0 |
| MN996532.1;QHR63299.1;BatCoV_RaTG13_orf1ab;3202 | 0 | 0 |
| MN996532.1;QHR63299.1;BatCoV_RaTG13_orf1ab;3203 | 0 | 0 |
| MN996532.1;QHR63299.1;BatCoV_RaTG13_orf1ab;3204 | 0 | 0 |
| MN996532.1;QHR63299.1;BatCoV_RaTG13_orf1ab;3205 | 0 | 0 |
| MN996532.1;QHR63299.1;BatCoV_RaTG13_orf1ab;3206 | 0 | 0 |
| MN996532.1;QHR63299.1;BatCoV_RaTG13_orf1ab;3207 | 0 | 0 |
| MN996532.1;QHR63299.1;BatCoV_RaTG13_orf1ab;3208 | 0 | 0.025 |
| MN996532.1;QHR63299.1;BatCoV_RaTG13_orf1ab;3209 | 0 | 0.025 |
| MN996532.1;QHR63299.1;BatCoV_RaTG13_orf1ab;3210 | 0 | 0.025 |
| MN996532.1;QHR63299.1;BatCoV_RaTG13_orf1ab;3211 | 0 | 0 |
| MN996532.1;QHR63299.1;BatCoV_RaTG13_orf1ab;3212 | 0 | 0 |
| MN996532.1;QHR63299.1;BatCoV_RaTG13_orf1ab;3213 | 0 | 0 |
| MN996532.1;QHR63299.1;BatCoV_RaTG13_orf1ab;3214 | 0.05 | 0.075 |
| MN996532.1;QHR63299.1;BatCoV_RaTG13_orf1ab;3215 | 0 | 0 |
| MN996532.1;QHR63299.1;BatCoV_RaTG13_orf1ab;3216 | 0 | 0 |

| | | |
|---|---|---|
| MN996532.1;QHR63299.1;BatCoV_RaTG13_orf1ab;3217 | 0 | 0 |
| MN996532.1;QHR63299.1;BatCoV_RaTG13_orf1ab;3218 | 0 | 0.025 |
| MN996532.1;QHR63299.1;BatCoV_RaTG13_orf1ab;3219 | 0 | 0 |
| MN996532.1;QHR63299.1;BatCoV_RaTG13_orf1ab;3220 | 0 | 0 |
| MN996532.1;QHR63299.1;BatCoV_RaTG13_orf1ab;3221 | 0 | 0 |
| MN996532.1;QHR63299.1;BatCoV_RaTG13_orf1ab;3222 | 0 | 0 |
| MN996532.1;QHR63299.1;BatCoV_RaTG13_orf1ab;3223 | 0 | 0 |
| MN996532.1;QHR63299.1;BatCoV_RaTG13_orf1ab;3224 | 0 | 0.075 |
| MN996532.1;QHR63299.1;BatCoV_RaTG13_orf1ab;3225 | 0 | 0.025 |
| MN996532.1;QHR63299.1;BatCoV_RaTG13_orf1ab;3226 | 0 | 0.025 |
| MN996532.1;QHR63299.1;BatCoV_RaTG13_orf1ab;3227 | 0 | 0.025 |
| MN996532.1;QHR63299.1;BatCoV_RaTG13_orf1ab;3228 | 0 | 0.025 |
| MN996532.1;QHR63299.1;BatCoV_RaTG13_orf1ab;3229 | 0.05 | 0.025 |
| MN996532.1;QHR63299.1;BatCoV_RaTG13_orf1ab;3230 | 0 | 0.025 |
| MN996532.1;QHR63299.1;BatCoV_RaTG13_orf1ab;3231 | 0 | 0.025 |
| MN996532.1;QHR63299.1;BatCoV_RaTG13_orf1ab;3232 | 0 | 0.025 |
| MN996532.1;QHR63299.1;BatCoV_RaTG13_orf1ab;3233 | 0.05 | 0 |
| MN996532.1;QHR63299.1;BatCoV_RaTG13_orf1ab;3234 | 0 | 0 |
| MN996532.1;QHR63299.1;BatCoV_RaTG13_orf1ab;3235 | 0 | 0 |
| MN996532.1;QHR63299.1;BatCoV_RaTG13_orf1ab;3236 | 0.15 | 0.05 |
| MN996532.1;QHR63299.1;BatCoV_RaTG13_orf1ab;3237 | 0.3 | 0.05 |
| MN996532.1;QHR63299.1;BatCoV_RaTG13_orf1ab;3238 | 0.05 | 0 |
| MN996532.1;QHR63299.1;BatCoV_RaTG13_orf1ab;3239 | 0.05 | 0.05 |
| MN996532.1;QHR63299.1;BatCoV_RaTG13_orf1ab;3240 | 0.1 | 0.1 |
| MN996532.1;QHR63299.1;BatCoV_RaTG13_orf1ab;3241 | 0 | 0.05 |
| MN996532.1;QHR63299.1;BatCoV_RaTG13_orf1ab;3242 | 0 | 0.05 |
| MN996532.1;QHR63299.1;BatCoV_RaTG13_orf1ab;3243 | 0 | 0 |
| MN996532.1;QHR63299.1;BatCoV_RaTG13_orf1ab;3244 | 0 | 0.025 |
| MN996532.1;QHR63299.1;BatCoV_RaTG13_orf1ab;3245 | 0 | 0 |
| MN996532.1;QHR63299.1;BatCoV_RaTG13_orf1ab;3246 | 0 | 0 |
| MN996532.1;QHR63299.1;BatCoV_RaTG13_orf1ab;3247 | 0 | 0 |
| MN996532.1;QHR63299.1;BatCoV_RaTG13_orf1ab;3248 | 0 | 0 |
| MN996532.1;QHR63299.1;BatCoV_RaTG13_orf1ab;3249 | 0.05 | 0 |
| MN996532.1;QHR63299.1;BatCoV_RaTG13_orf1ab;3250 | 0 | 0 |
| MN996532.1;QHR63299.1;BatCoV_RaTG13_orf1ab;3251 | 0 | 0 |
| MN996532.1;QHR63299.1;BatCoV_RaTG13_orf1ab;3252 | 0 | 0 |
| MN996532.1;QHR63299.1;BatCoV_RaTG13_orf1ab;3253 | 0 | 0 |
| MN996532.1;QHR63299.1;BatCoV_RaTG13_orf1ab;3254 | 0 | 0 |
| MN996532.1;QHR63299.1;BatCoV_RaTG13_orf1ab;3255 | 0 | 0 |
| MN996532.1;QHR63299.1;BatCoV_RaTG13_orf1ab;3256 | 0 | 0 |
| MN996532.1;QHR63299.1;BatCoV_RaTG13_orf1ab;3257 | 0 | 0 |
| MN996532.1;QHR63299.1;BatCoV_RaTG13_orf1ab;3258 | 0 | 0 |
| MN996532.1;QHR63299.1;BatCoV_RaTG13_orf1ab;3259 | 0 | 0 |

| | | |
|---|---|---|
| MN996532.1;QHR63299.1;BatCoV_RaTG13_orf1ab;3260 | 0 | 0 |
| MN996532.1;QHR63299.1;BatCoV_RaTG13_orf1ab;3261 | 0 | 0 |
| MN996532.1;QHR63299.1;BatCoV_RaTG13_orf1ab;3262 | 0 | 0 |
| MN996532.1;QHR63299.1;BatCoV_RaTG13_orf1ab;3263 | 0 | 0 |
| MN996532.1;QHR63299.1;BatCoV_RaTG13_orf1ab;3264 | 0 | 0 |
| MN996532.1;QHR63299.1;BatCoV_RaTG13_orf1ab;3265 | 0 | 0 |
| MN996532.1;QHR63299.1;BatCoV_RaTG13_orf1ab;3266 | 0 | 0 |
| MN996532.1;QHR63299.1;BatCoV_RaTG13_orf1ab;3267 | 0 | 0.025 |
| MN996532.1;QHR63299.1;BatCoV_RaTG13_orf1ab;3268 | 0 | 0 |
| MN996532.1;QHR63299.1;BatCoV_RaTG13_orf1ab;3269 | 0.05 | 0 |
| MN996532.1;QHR63299.1;BatCoV_RaTG13_orf1ab;3270 | 0 | 0 |
| MN996532.1;QHR63299.1;BatCoV_RaTG13_orf1ab;3271 | 0 | 0 |
| MN996532.1;QHR63299.1;BatCoV_RaTG13_orf1ab;3272 | 0 | 0 |
| MN996532.1;QHR63299.1;BatCoV_RaTG13_orf1ab;3273 | 0 | 0 |
| MN996532.1;QHR63299.1;BatCoV_RaTG13_orf1ab;3274 | 0 | 0 |
| MN996532.1;QHR63299.1;BatCoV_RaTG13_orf1ab;3275 | 0 | 0 |
| MN996532.1;QHR63299.1;BatCoV_RaTG13_orf1ab;3276 | 0 | 0 |
| MN996532.1;QHR63299.1;BatCoV_RaTG13_orf1ab;3277 | 0 | 0 |
| MN996532.1;QHR63299.1;BatCoV_RaTG13_orf1ab;3278 | 0 | 0 |
| MN996532.1;QHR63299.1;BatCoV_RaTG13_orf1ab;3279 | 0 | 0 |
| MN996532.1;QHR63299.1;BatCoV_RaTG13_orf1ab;3280 | 0 | 0 |
| MN996532.1;QHR63299.1;BatCoV_RaTG13_orf1ab;3281 | 0 | 0 |
| MN996532.1;QHR63299.1;BatCoV_RaTG13_orf1ab;3282 | 0 | 0 |
| MN996532.1;QHR63299.1;BatCoV_RaTG13_orf1ab;3283 | 0 | 0 |
| MN996532.1;QHR63299.1;BatCoV_RaTG13_orf1ab;3284 | 0 | 0 |
| MN996532.1;QHR63299.1;BatCoV_RaTG13_orf1ab;3285 | 0 | 0 |
| MN996532.1;QHR63299.1;BatCoV_RaTG13_orf1ab;3286 | 0 | 0 |
| MN996532.1;QHR63299.1;BatCoV_RaTG13_orf1ab;3287 | 0 | 0 |
| MN996532.1;QHR63299.1;BatCoV_RaTG13_orf1ab;3288 | 0 | 0 |
| MN996532.1;QHR63299.1;BatCoV_RaTG13_orf1ab;3289 | 0 | 0 |
| MN996532.1;QHR63299.1;BatCoV_RaTG13_orf1ab;3290 | 0 | 0 |
| MN996532.1;QHR63299.1;BatCoV_RaTG13_orf1ab;3291 | 0 | 0 |
| MN996532.1;QHR63299.1;BatCoV_RaTG13_orf1ab;3292 | 0 | 0 |
| MN996532.1;QHR63299.1;BatCoV_RaTG13_orf1ab;3293 | 0 | 0 |
| MN996532.1;QHR63299.1;BatCoV_RaTG13_orf1ab;3294 | 0 | 0 |
| MN996532.1;QHR63299.1;BatCoV_RaTG13_orf1ab;3295 | 0 | 0 |
| MN996532.1;QHR63299.1;BatCoV_RaTG13_orf1ab;3296 | 0 | 0 |
| MN996532.1;QHR63299.1;BatCoV_RaTG13_orf1ab;3297 | 0 | 0 |
| MN996532.1;QHR63299.1;BatCoV_RaTG13_orf1ab;3298 | 0 | 0.025 |
| MN996532.1;QHR63299.1;BatCoV_RaTG13_orf1ab;3299 | 0 | 0 |
| MN996532.1;QHR63299.1;BatCoV_RaTG13_orf1ab;3300 | 0 | 0 |
| MN996532.1;QHR63299.1;BatCoV_RaTG13_orf1ab;3301 | 0 | 0 |
| MN996532.1;QHR63299.1;BatCoV_RaTG13_orf1ab;3302 | 0 | 0 |

| | | |
|---|---|---|
| MN996532.1;QHR63299.1;BatCoV_RaTG13_orf1ab;3303 | 0 | 0 |
| MN996532.1;QHR63299.1;BatCoV_RaTG13_orf1ab;3304 | 0 | 0 |
| MN996532.1;QHR63299.1;BatCoV_RaTG13_orf1ab;3305 | 0 | 0.025 |
| MN996532.1;QHR63299.1;BatCoV_RaTG13_orf1ab;3306 | 0 | 0 |
| MN996532.1;QHR63299.1;BatCoV_RaTG13_orf1ab;3307 | 0 | 0 |
| MN996532.1;QHR63299.1;BatCoV_RaTG13_orf1ab;3308 | 0 | 0 |
| MN996532.1;QHR63299.1;BatCoV_RaTG13_orf1ab;3309 | 0 | 0.025 |
| MN996532.1;QHR63299.1;BatCoV_RaTG13_orf1ab;3310 | 0 | 0.025 |
| MN996532.1;QHR63299.1;BatCoV_RaTG13_orf1ab;3311 | 0 | 0 |
| MN996532.1;QHR63299.1;BatCoV_RaTG13_orf1ab;3312 | 0.05 | 0 |
| MN996532.1;QHR63299.1;BatCoV_RaTG13_orf1ab;3313 | 0 | 0.05 |
| MN996532.1;QHR63299.1;BatCoV_RaTG13_orf1ab;3314 | 0 | 0 |
| MN996532.1;QHR63299.1;BatCoV_RaTG13_orf1ab;3315 | 0 | 0 |
| MN996532.1;QHR63299.1;BatCoV_RaTG13_orf1ab;3316 | 0 | 0 |
| MN996532.1;QHR63299.1;BatCoV_RaTG13_orf1ab;3317 | 0 | 0 |
| MN996532.1;QHR63299.1;BatCoV_RaTG13_orf1ab;3318 | 0 | 0 |
| MN996532.1;QHR63299.1;BatCoV_RaTG13_orf1ab;3319 | 0 | 0 |
| MN996532.1;QHR63299.1;BatCoV_RaTG13_orf1ab;3320 | 0 | 0 |
| MN996532.1;QHR63299.1;BatCoV_RaTG13_orf1ab;3321 | 0 | 0 |
| MN996532.1;QHR63299.1;BatCoV_RaTG13_orf1ab;3322 | 0 | 0 |
| MN996532.1;QHR63299.1;BatCoV_RaTG13_orf1ab;3323 | 0 | 0 |
| MN996532.1;QHR63299.1;BatCoV_RaTG13_orf1ab;3324 | 0 | 0 |
| MN996532.1;QHR63299.1;BatCoV_RaTG13_orf1ab;3325 | 0 | 0 |
| MN996532.1;QHR63299.1;BatCoV_RaTG13_orf1ab;3326 | 0 | 0 |
| MN996532.1;QHR63299.1;BatCoV_RaTG13_orf1ab;3327 | 0 | 0 |
| MN996532.1;QHR63299.1;BatCoV_RaTG13_orf1ab;3328 | 0 | 0 |
| MN996532.1;QHR63299.1;BatCoV_RaTG13_orf1ab;3329 | 0 | 0 |
| MN996532.1;QHR63299.1;BatCoV_RaTG13_orf1ab;3330 | 0 | 0 |
| MN996532.1;QHR63299.1;BatCoV_RaTG13_orf1ab;3331 | 0 | 0 |
| MN996532.1;QHR63299.1;BatCoV_RaTG13_orf1ab;3332 | 0 | 0 |
| MN996532.1;QHR63299.1;BatCoV_RaTG13_orf1ab;3333 | 0 | 0 |
| MN996532.1;QHR63299.1;BatCoV_RaTG13_orf1ab;3334 | 0 | 0 |
| MN996532.1;QHR63299.1;BatCoV_RaTG13_orf1ab;3335 | 0 | 0 |
| MN996532.1;QHR63299.1;BatCoV_RaTG13_orf1ab;3336 | 0 | 0 |
| MN996532.1;QHR63299.1;BatCoV_RaTG13_orf1ab;3337 | 0 | 0 |
| MN996532.1;QHR63299.1;BatCoV_RaTG13_orf1ab;3338 | 0 | 0 |
| MN996532.1;QHR63299.1;BatCoV_RaTG13_orf1ab;3339 | 0 | 0 |
| MN996532.1;QHR63299.1;BatCoV_RaTG13_orf1ab;3340 | 0 | 0 |
| MN996532.1;QHR63299.1;BatCoV_RaTG13_orf1ab;3341 | 0 | 0.025 |
| MN996532.1;QHR63299.1;BatCoV_RaTG13_orf1ab;3342 | 0.05 | 0 |
| MN996532.1;QHR63299.1;BatCoV_RaTG13_orf1ab;3343 | 0.05 | 0 |
| MN996532.1;QHR63299.1;BatCoV_RaTG13_orf1ab;3344 | 0 | 0 |
| MN996532.1;QHR63299.1;BatCoV_RaTG13_orf1ab;3345 | 0 | 0 |

| | | |
|---|---|---|
| MN996532.1;QHR63299.1;BatCoV_RaTG13_orf1ab;3346 | 0.05 | 0 |
| MN996532.1;QHR63299.1;BatCoV_RaTG13_orf1ab;3347 | 0 | 0 |
| MN996532.1;QHR63299.1;BatCoV_RaTG13_orf1ab;3348 | 0.05 | 0 |
| MN996532.1;QHR63299.1;BatCoV_RaTG13_orf1ab;3349 | 0.05 | 0 |
| MN996532.1;QHR63299.1;BatCoV_RaTG13_orf1ab;3350 | 0.05 | 0 |
| MN996532.1;QHR63299.1;BatCoV_RaTG13_orf1ab;3351 | 0 | 0 |
| MN996532.1;QHR63299.1;BatCoV_RaTG13_orf1ab;3352 | 0.05 | 0 |
| MN996532.1;QHR63299.1;BatCoV_RaTG13_orf1ab;3353 | 0.05 | 0.05 |
| MN996532.1;QHR63299.1;BatCoV_RaTG13_orf1ab;3354 | 0 | 0 |
| MN996532.1;QHR63299.1;BatCoV_RaTG13_orf1ab;3355 | 0 | 0 |
| MN996532.1;QHR63299.1;BatCoV_RaTG13_orf1ab;3356 | 0 | 0 |
| MN996532.1;QHR63299.1;BatCoV_RaTG13_orf1ab;3357 | 0 | 0 |
| MN996532.1;QHR63299.1;BatCoV_RaTG13_orf1ab;3358 | 0 | 0 |
| MN996532.1;QHR63299.1;BatCoV_RaTG13_orf1ab;3359 | 0 | 0 |
| MN996532.1;QHR63299.1;BatCoV_RaTG13_orf1ab;3360 | 0 | 0 |
| MN996532.1;QHR63299.1;BatCoV_RaTG13_orf1ab;3361 | 0 | 0 |
| MN996532.1;QHR63299.1;BatCoV_RaTG13_orf1ab;3362 | 0 | 0 |
| MN996532.1;QHR63299.1;BatCoV_RaTG13_orf1ab;3363 | 0 | 0 |
| MN996532.1;QHR63299.1;BatCoV_RaTG13_orf1ab;3364 | 0 | 0 |
| MN996532.1;QHR63299.1;BatCoV_RaTG13_orf1ab;3365 | 0 | 0 |
| MN996532.1;QHR63299.1;BatCoV_RaTG13_orf1ab;3366 | 0 | 0 |
| MN996532.1;QHR63299.1;BatCoV_RaTG13_orf1ab;3367 | 0 | 0 |
| MN996532.1;QHR63299.1;BatCoV_RaTG13_orf1ab;3368 | 0 | 0 |
| MN996532.1;QHR63299.1;BatCoV_RaTG13_orf1ab;3369 | 0 | 0.025 |
| MN996532.1;QHR63299.1;BatCoV_RaTG13_orf1ab;3370 | 0 | 0 |
| MN996532.1;QHR63299.1;BatCoV_RaTG13_orf1ab;3371 | 0 | 0 |
| MN996532.1;QHR63299.1;BatCoV_RaTG13_orf1ab;3372 | 0 | 0 |
| MN996532.1;QHR63299.1;BatCoV_RaTG13_orf1ab;3373 | 0 | 0 |
| MN996532.1;QHR63299.1;BatCoV_RaTG13_orf1ab;3374 | 0 | 0 |
| MN996532.1;QHR63299.1;BatCoV_RaTG13_orf1ab;3375 | 0 | 0 |
| MN996532.1;QHR63299.1;BatCoV_RaTG13_orf1ab;3376 | 0 | 0 |
| MN996532.1;QHR63299.1;BatCoV_RaTG13_orf1ab;3377 | 0 | 0 |
| MN996532.1;QHR63299.1;BatCoV_RaTG13_orf1ab;3378 | 0 | 0 |
| MN996532.1;QHR63299.1;BatCoV_RaTG13_orf1ab;3379 | 0 | 0 |
| MN996532.1;QHR63299.1;BatCoV_RaTG13_orf1ab;3380 | 0 | 0.025 |
| MN996532.1;QHR63299.1;BatCoV_RaTG13_orf1ab;3381 | 0 | 0 |
| MN996532.1;QHR63299.1;BatCoV_RaTG13_orf1ab;3382 | 0 | 0 |
| MN996532.1;QHR63299.1;BatCoV_RaTG13_orf1ab;3383 | 0 | 0 |
| MN996532.1;QHR63299.1;BatCoV_RaTG13_orf1ab;3384 | 0 | 0 |
| MN996532.1;QHR63299.1;BatCoV_RaTG13_orf1ab;3385 | 0 | 0 |
| MN996532.1;QHR63299.1;BatCoV_RaTG13_orf1ab;3386 | 0 | 0.025 |
| MN996532.1;QHR63299.1;BatCoV_RaTG13_orf1ab;3387 | 0 | 0 |
| MN996532.1;QHR63299.1;BatCoV_RaTG13_orf1ab;3388 | 0 | 0 |

| | | |
|---|---|---|
| MN996532.1;QHR63299.1;BatCoV_RaTG13_orf1ab;3389 | 0 | 0 |
| MN996532.1;QHR63299.1;BatCoV_RaTG13_orf1ab;3390 | 0 | 0 |
| MN996532.1;QHR63299.1;BatCoV_RaTG13_orf1ab;3391 | 0.05 | 0 |
| MN996532.1;QHR63299.1;BatCoV_RaTG13_orf1ab;3392 | 0 | 0 |
| MN996532.1;QHR63299.1;BatCoV_RaTG13_orf1ab;3393 | 0.05 | 0 |
| MN996532.1;QHR63299.1;BatCoV_RaTG13_orf1ab;3394 | 0 | 0 |
| MN996532.1;QHR63299.1;BatCoV_RaTG13_orf1ab;3395 | 0 | 0 |
| MN996532.1;QHR63299.1;BatCoV_RaTG13_orf1ab;3396 | 0 | 0 |
| MN996532.1;QHR63299.1;BatCoV_RaTG13_orf1ab;3397 | 0 | 0 |
| MN996532.1;QHR63299.1;BatCoV_RaTG13_orf1ab;3398 | 0 | 0 |
| MN996532.1;QHR63299.1;BatCoV_RaTG13_orf1ab;3399 | 0 | 0.025 |
| MN996532.1;QHR63299.1;BatCoV_RaTG13_orf1ab;3400 | 0 | 0.025 |
| MN996532.1;QHR63299.1;BatCoV_RaTG13_orf1ab;3401 | 0 | 0.025 |
| MN996532.1;QHR63299.1;BatCoV_RaTG13_orf1ab;3402 | 0 | 0.05 |
| MN996532.1;QHR63299.1;BatCoV_RaTG13_orf1ab;3403 | 0 | 0.05 |
| MN996532.1;QHR63299.1;BatCoV_RaTG13_orf1ab;3404 | 0 | 0.025 |
| MN996532.1;QHR63299.1;BatCoV_RaTG13_orf1ab;3405 | 0 | 0.025 |
| MN996532.1;QHR63299.1;BatCoV_RaTG13_orf1ab;3406 | 0 | 0 |
| MN996532.1;QHR63299.1;BatCoV_RaTG13_orf1ab;3407 | 0 | 0 |
| MN996532.1;QHR63299.1;BatCoV_RaTG13_orf1ab;3408 | 0 | 0 |
| MN996532.1;QHR63299.1;BatCoV_RaTG13_orf1ab;3409 | 0 | 0 |
| MN996532.1;QHR63299.1;BatCoV_RaTG13_orf1ab;3410 | 0 | 0 |
| MN996532.1;QHR63299.1;BatCoV_RaTG13_orf1ab;3411 | 0 | 0 |
| MN996532.1;QHR63299.1;BatCoV_RaTG13_orf1ab;3412 | 0 | 0 |
| MN996532.1;QHR63299.1;BatCoV_RaTG13_orf1ab;3413 | 0 | 0 |
| MN996532.1;QHR63299.1;BatCoV_RaTG13_orf1ab;3414 | 0 | 0 |
| MN996532.1;QHR63299.1;BatCoV_RaTG13_orf1ab;3415 | 0 | 0 |
| MN996532.1;QHR63299.1;BatCoV_RaTG13_orf1ab;3416 | 0 | 0 |
| MN996532.1;QHR63299.1;BatCoV_RaTG13_orf1ab;3417 | 0 | 0 |
| MN996532.1;QHR63299.1;BatCoV_RaTG13_orf1ab;3418 | 0 | 0 |
| MN996532.1;QHR63299.1;BatCoV_RaTG13_orf1ab;3419 | 0 | 0 |
| MN996532.1;QHR63299.1;BatCoV_RaTG13_orf1ab;3420 | 0 | 0 |
| MN996532.1;QHR63299.1;BatCoV_RaTG13_orf1ab;3421 | 0 | 0 |
| MN996532.1;QHR63299.1;BatCoV_RaTG13_orf1ab;3422 | 0 | 0 |
| MN996532.1;QHR63299.1;BatCoV_RaTG13_orf1ab;3423 | 0 | 0 |
| MN996532.1;QHR63299.1;BatCoV_RaTG13_orf1ab;3424 | 0 | 0 |
| MN996532.1;QHR63299.1;BatCoV_RaTG13_orf1ab;3425 | 0.15 | 0.025 |
| MN996532.1;QHR63299.1;BatCoV_RaTG13_orf1ab;3426 | 0.1 | 0.05 |
| MN996532.1;QHR63299.1;BatCoV_RaTG13_orf1ab;3427 | 0.05 | 0.025 |
| MN996532.1;QHR63299.1;BatCoV_RaTG13_orf1ab;3428 | 0.2 | 0.075 |
| MN996532.1;QHR63299.1;BatCoV_RaTG13_orf1ab;3429 | 0.2 | 0.05 |
| MN996532.1;QHR63299.1;BatCoV_RaTG13_orf1ab;3430 | 0.2 | 0.05 |
| MN996532.1;QHR63299.1;BatCoV_RaTG13_orf1ab;3431 | 0.2 | 0.05 |

| | | |
|---|---|---|
| MN996532.1;QHR63299.1;BatCoV_RaTG13_orf1ab;3432 | 0.1 | 0.025 |
| MN996532.1;QHR63299.1;BatCoV_RaTG13_orf1ab;3433 | 0.05 | 0.025 |
| MN996532.1;QHR63299.1;BatCoV_RaTG13_orf1ab;3434 | 0.05 | 0.025 |
| MN996532.1;QHR63299.1;BatCoV_RaTG13_orf1ab;3435 | 0.05 | 0.05 |
| MN996532.1;QHR63299.1;BatCoV_RaTG13_orf1ab;3436 | 0.15 | 0.1 |
| MN996532.1;QHR63299.1;BatCoV_RaTG13_orf1ab;3437 | 0 | 0.025 |
| MN996532.1;QHR63299.1;BatCoV_RaTG13_orf1ab;3438 | 0 | 0 |
| MN996532.1;QHR63299.1;BatCoV_RaTG13_orf1ab;3439 | 0.05 | 0 |
| MN996532.1;QHR63299.1;BatCoV_RaTG13_orf1ab;3440 | 0.05 | 0 |
| MN996532.1;QHR63299.1;BatCoV_RaTG13_orf1ab;3441 | 0.1 | 0.025 |
| MN996532.1;QHR63299.1;BatCoV_RaTG13_orf1ab;3442 | 0.05 | 0 |
| MN996532.1;QHR63299.1;BatCoV_RaTG13_orf1ab;3443 | 0.1 | 0 |
| MN996532.1;QHR63299.1;BatCoV_RaTG13_orf1ab;3444 | 0 | 0 |
| MN996532.1;QHR63299.1;BatCoV_RaTG13_orf1ab;3445 | 0.05 | 0 |
| MN996532.1;QHR63299.1;BatCoV_RaTG13_orf1ab;3446 | 0 | 0 |
| MN996532.1;QHR63299.1;BatCoV_RaTG13_orf1ab;3447 | 0.05 | 0.025 |
| MN996532.1;QHR63299.1;BatCoV_RaTG13_orf1ab;3448 | 0 | 0 |
| MN996532.1;QHR63299.1;BatCoV_RaTG13_orf1ab;3449 | 0 | 0 |
| MN996532.1;QHR63299.1;BatCoV_RaTG13_orf1ab;3450 | 0 | 0 |
| MN996532.1;QHR63299.1;BatCoV_RaTG13_orf1ab;3451 | 0 | 0 |
| MN996532.1;QHR63299.1;BatCoV_RaTG13_orf1ab;3452 | 0 | 0 |
| MN996532.1;QHR63299.1;BatCoV_RaTG13_orf1ab;3453 | 0 | 0 |
| MN996532.1;QHR63299.1;BatCoV_RaTG13_orf1ab;3454 | 0 | 0 |
| MN996532.1;QHR63299.1;BatCoV_RaTG13_orf1ab;3455 | 0 | 0 |
| MN996532.1;QHR63299.1;BatCoV_RaTG13_orf1ab;3456 | 0 | 0 |
| MN996532.1;QHR63299.1;BatCoV_RaTG13_orf1ab;3457 | 0 | 0 |
| MN996532.1;QHR63299.1;BatCoV_RaTG13_orf1ab;3458 | 0 | 0 |
| MN996532.1;QHR63299.1;BatCoV_RaTG13_orf1ab;3459 | 0 | 0 |
| MN996532.1;QHR63299.1;BatCoV_RaTG13_orf1ab;3460 | 0 | 0 |
| MN996532.1;QHR63299.1;BatCoV_RaTG13_orf1ab;3461 | 0 | 0 |
| MN996532.1;QHR63299.1;BatCoV_RaTG13_orf1ab;3462 | 0 | 0 |
| MN996532.1;QHR63299.1;BatCoV_RaTG13_orf1ab;3463 | 0 | 0 |
| MN996532.1;QHR63299.1;BatCoV_RaTG13_orf1ab;3464 | 0 | 0 |
| MN996532.1;QHR63299.1;BatCoV_RaTG13_orf1ab;3465 | 0 | 0 |
| MN996532.1;QHR63299.1;BatCoV_RaTG13_orf1ab;3466 | 0 | 0 |
| MN996532.1;QHR63299.1;BatCoV_RaTG13_orf1ab;3467 | 0 | 0 |
| MN996532.1;QHR63299.1;BatCoV_RaTG13_orf1ab;3468 | 0 | 0 |
| MN996532.1;QHR63299.1;BatCoV_RaTG13_orf1ab;3469 | 0 | 0 |
| MN996532.1;QHR63299.1;BatCoV_RaTG13_orf1ab;3470 | 0 | 0 |
| MN996532.1;QHR63299.1;BatCoV_RaTG13_orf1ab;3471 | 0 | 0 |
| MN996532.1;QHR63299.1;BatCoV_RaTG13_orf1ab;3472 | 0 | 0 |
| MN996532.1;QHR63299.1;BatCoV_RaTG13_orf1ab;3473 | 0 | 0 |
| MN996532.1;QHR63299.1;BatCoV_RaTG13_orf1ab;3474 | 0 | 0 |

| | | |
|---|---|---|
| MN996532.1;QHR63299.1;BatCoV_RaTG13_orf1ab;3475 | 0 | 0 |
| MN996532.1;QHR63299.1;BatCoV_RaTG13_orf1ab;3476 | 0 | 0 |
| MN996532.1;QHR63299.1;BatCoV_RaTG13_orf1ab;3477 | 0 | 0 |
| MN996532.1;QHR63299.1;BatCoV_RaTG13_orf1ab;3478 | 0.05 | 0 |
| MN996532.1;QHR63299.1;BatCoV_RaTG13_orf1ab;3479 | 0.05 | 0 |
| MN996532.1;QHR63299.1;BatCoV_RaTG13_orf1ab;3480 | 0.05 | 0 |
| MN996532.1;QHR63299.1;BatCoV_RaTG13_orf1ab;3481 | 0.05 | 0.025 |
| MN996532.1;QHR63299.1;BatCoV_RaTG13_orf1ab;3482 | 0.1 | 0 |
| MN996532.1;QHR63299.1;BatCoV_RaTG13_orf1ab;3483 | 0.05 | 0 |
| MN996532.1;QHR63299.1;BatCoV_RaTG13_orf1ab;3484 | 0.1 | 0 |
| MN996532.1;QHR63299.1;BatCoV_RaTG13_orf1ab;3485 | 0.05 | 0 |
| MN996532.1;QHR63299.1;BatCoV_RaTG13_orf1ab;3486 | 0.05 | 0.025 |
| MN996532.1;QHR63299.1;BatCoV_RaTG13_orf1ab;3487 | 0.05 | 0.15 |
| MN996532.1;QHR63299.1;BatCoV_RaTG13_orf1ab;3488 | 0.25 | 0.1 |
| MN996532.1;QHR63299.1;BatCoV_RaTG13_orf1ab;3489 | 0.05 | 0 |
| MN996532.1;QHR63299.1;BatCoV_RaTG13_orf1ab;3490 | 0 | 0 |
| MN996532.1;QHR63299.1;BatCoV_RaTG13_orf1ab;3491 | 0.05 | 0 |
| MN996532.1;QHR63299.1;BatCoV_RaTG13_orf1ab;3492 | 0 | 0 |
| MN996532.1;QHR63299.1;BatCoV_RaTG13_orf1ab;3493 | 0 | 0.025 |
| MN996532.1;QHR63299.1;BatCoV_RaTG13_orf1ab;3494 | 0 | 0.025 |
| MN996532.1;QHR63299.1;BatCoV_RaTG13_orf1ab;3495 | 0.05 | 0 |
| MN996532.1;QHR63299.1;BatCoV_RaTG13_orf1ab;3496 | 0 | 0 |
| MN996532.1;QHR63299.1;BatCoV_RaTG13_orf1ab;3497 | 0 | 0 |
| MN996532.1;QHR63299.1;BatCoV_RaTG13_orf1ab;3498 | 0 | 0 |
| MN996532.1;QHR63299.1;BatCoV_RaTG13_orf1ab;3499 | 0 | 0 |
| MN996532.1;QHR63299.1;BatCoV_RaTG13_orf1ab;3500 | 0 | 0 |
| MN996532.1;QHR63299.1;BatCoV_RaTG13_orf1ab;3501 | 0 | 0 |
| MN996532.1;QHR63299.1;BatCoV_RaTG13_orf1ab;3502 | 0.05 | 0 |
| MN996532.1;QHR63299.1;BatCoV_RaTG13_orf1ab;3503 | 0 | 0 |
| MN996532.1;QHR63299.1;BatCoV_RaTG13_orf1ab;3504 | 0.05 | 0.025 |
| MN996532.1;QHR63299.1;BatCoV_RaTG13_orf1ab;3505 | 0.15 | 0.1 |
| MN996532.1;QHR63299.1;BatCoV_RaTG13_orf1ab;3506 | 0.05 | 0.05 |
| MN996532.1;QHR63299.1;BatCoV_RaTG13_orf1ab;3507 | 0.1 | 0.15 |
| MN996532.1;QHR63299.1;BatCoV_RaTG13_orf1ab;3508 | 0.05 | 0.025 |
| MN996532.1;QHR63299.1;BatCoV_RaTG13_orf1ab;3509 | 0.05 | 0 |
| MN996532.1;QHR63299.1;BatCoV_RaTG13_orf1ab;3510 | 0.05 | 0 |
| MN996532.1;QHR63299.1;BatCoV_RaTG13_orf1ab;3511 | 0.05 | 0 |
| MN996532.1;QHR63299.1;BatCoV_RaTG13_orf1ab;3512 | 0.05 | 0 |
| MN996532.1;QHR63299.1;BatCoV_RaTG13_orf1ab;3513 | 0.05 | 0 |
| MN996532.1;QHR63299.1;BatCoV_RaTG13_orf1ab;3514 | 0.05 | 0 |
| MN996532.1;QHR63299.1;BatCoV_RaTG13_orf1ab;3515 | 0 | 0 |
| MN996532.1;QHR63299.1;BatCoV_RaTG13_orf1ab;3516 | 0 | 0 |
| MN996532.1;QHR63299.1;BatCoV_RaTG13_orf1ab;3517 | 0 | 0 |

| | | |
|---|---|---|
| MN996532.1;QHR63299.1;BatCoV_RaTG13_orf1ab;3518 | 0 | 0 |
| MN996532.1;QHR63299.1;BatCoV_RaTG13_orf1ab;3519 | 0 | 0 |
| MN996532.1;QHR63299.1;BatCoV_RaTG13_orf1ab;3520 | 0.05 | 0 |
| MN996532.1;QHR63299.1;BatCoV_RaTG13_orf1ab;3521 | 0 | 0 |
| MN996532.1;QHR63299.1;BatCoV_RaTG13_orf1ab;3522 | 0 | 0 |
| MN996532.1;QHR63299.1;BatCoV_RaTG13_orf1ab;3523 | 0 | 0 |
| MN996532.1;QHR63299.1;BatCoV_RaTG13_orf1ab;3524 | 0 | 0 |
| MN996532.1;QHR63299.1;BatCoV_RaTG13_orf1ab;3525 | 0 | 0.025 |
| MN996532.1;QHR63299.1;BatCoV_RaTG13_orf1ab;3526 | 0.05 | 0 |
| MN996532.1;QHR63299.1;BatCoV_RaTG13_orf1ab;3527 | 0.05 | 0 |
| MN996532.1;QHR63299.1;BatCoV_RaTG13_orf1ab;3528 | 0.05 | 0.025 |
| MN996532.1;QHR63299.1;BatCoV_RaTG13_orf1ab;3529 | 0 | 0.025 |
| MN996532.1;QHR63299.1;BatCoV_RaTG13_orf1ab;3530 | 0 | 0.025 |
| MN996532.1;QHR63299.1;BatCoV_RaTG13_orf1ab;3531 | 0 | 0.025 |
| MN996532.1;QHR63299.1;BatCoV_RaTG13_orf1ab;3532 | 0 | 0 |
| MN996532.1;QHR63299.1;BatCoV_RaTG13_orf1ab;3533 | 0 | 0 |
| MN996532.1;QHR63299.1;BatCoV_RaTG13_orf1ab;3534 | 0 | 0 |
| MN996532.1;QHR63299.1;BatCoV_RaTG13_orf1ab;3535 | 0 | 0 |
| MN996532.1;QHR63299.1;BatCoV_RaTG13_orf1ab;3536 | 0 | 0 |
| MN996532.1;QHR63299.1;BatCoV_RaTG13_orf1ab;3537 | 0 | 0 |
| MN996532.1;QHR63299.1;BatCoV_RaTG13_orf1ab;3538 | 0 | 0.025 |
| MN996532.1;QHR63299.1;BatCoV_RaTG13_orf1ab;3539 | 0 | 0 |
| MN996532.1;QHR63299.1;BatCoV_RaTG13_orf1ab;3540 | 0 | 0.025 |
| MN996532.1;QHR63299.1;BatCoV_RaTG13_orf1ab;3541 | 0 | 0.025 |
| MN996532.1;QHR63299.1;BatCoV_RaTG13_orf1ab;3542 | 0 | 0.05 |
| MN996532.1;QHR63299.1;BatCoV_RaTG13_orf1ab;3543 | 0 | 0.05 |
| MN996532.1;QHR63299.1;BatCoV_RaTG13_orf1ab;3544 | 0 | 0.025 |
| MN996532.1;QHR63299.1;BatCoV_RaTG13_orf1ab;3545 | 0 | 0.025 |
| MN996532.1;QHR63299.1;BatCoV_RaTG13_orf1ab;3546 | 0.05 | 0.05 |
| MN996532.1;QHR63299.1;BatCoV_RaTG13_orf1ab;3547 | 0.05 | 0.05 |
| MN996532.1;QHR63299.1;BatCoV_RaTG13_orf1ab;3548 | 0 | 0.025 |
| MN996532.1;QHR63299.1;BatCoV_RaTG13_orf1ab;3549 | 0 | 0.05 |
| MN996532.1;QHR63299.1;BatCoV_RaTG13_orf1ab;3550 | 0.1 | 0.025 |
| MN996532.1;QHR63299.1;BatCoV_RaTG13_orf1ab;3551 | 0 | 0 |
| MN996532.1;QHR63299.1;BatCoV_RaTG13_orf1ab;3552 | 0 | 0 |
| MN996532.1;QHR63299.1;BatCoV_RaTG13_orf1ab;3553 | 0 | 0 |
| MN996532.1;QHR63299.1;BatCoV_RaTG13_orf1ab;3554 | 0 | 0 |
| MN996532.1;QHR63299.1;BatCoV_RaTG13_orf1ab;3555 | 0 | 0 |
| MN996532.1;QHR63299.1;BatCoV_RaTG13_orf1ab;3556 | 0 | 0 |
| MN996532.1;QHR63299.1;BatCoV_RaTG13_orf1ab;3557 | 0 | 0 |
| MN996532.1;QHR63299.1;BatCoV_RaTG13_orf1ab;3558 | 0 | 0 |
| MN996532.1;QHR63299.1;BatCoV_RaTG13_orf1ab;3559 | 0 | 0 |
| MN996532.1;QHR63299.1;BatCoV_RaTG13_orf1ab;3560 | 0 | 0 |

| | | |
|---|---|---|
| MN996532.1;QHR63299.1;BatCoV_RaTG13_orf1ab;3561 | 0 | 0.025 |
| MN996532.1;QHR63299.1;BatCoV_RaTG13_orf1ab;3562 | 0 | 0.025 |
| MN996532.1;QHR63299.1;BatCoV_RaTG13_orf1ab;3563 | 0 | 0 |
| MN996532.1;QHR63299.1;BatCoV_RaTG13_orf1ab;3564 | 0 | 0 |
| MN996532.1;QHR63299.1;BatCoV_RaTG13_orf1ab;3565 | 0 | 0 |
| MN996532.1;QHR63299.1;BatCoV_RaTG13_orf1ab;3566 | 0 | 0 |
| MN996532.1;QHR63299.1;BatCoV_RaTG13_orf1ab;3567 | 0 | 0 |
| MN996532.1;QHR63299.1;BatCoV_RaTG13_orf1ab;3568 | 0 | 0 |
| MN996532.1;QHR63299.1;BatCoV_RaTG13_orf1ab;3569 | 0 | 0 |
| MN996532.1;QHR63299.1;BatCoV_RaTG13_orf1ab;3570 | 0 | 0 |
| MN996532.1;QHR63299.1;BatCoV_RaTG13_orf1ab;3571 | 0 | 0 |
| MN996532.1;QHR63299.1;BatCoV_RaTG13_orf1ab;3572 | 0 | 0 |
| MN996532.1;QHR63299.1;BatCoV_RaTG13_orf1ab;3573 | 0 | 0 |
| MN996532.1;QHR63299.1;BatCoV_RaTG13_orf1ab;3574 | 0 | 0 |
| MN996532.1;QHR63299.1;BatCoV_RaTG13_orf1ab;3575 | 0 | 0 |
| MN996532.1;QHR63299.1;BatCoV_RaTG13_orf1ab;3576 | 0 | 0 |
| MN996532.1;QHR63299.1;BatCoV_RaTG13_orf1ab;3577 | 0 | 0 |
| MN996532.1;QHR63299.1;BatCoV_RaTG13_orf1ab;3578 | 0 | 0 |
| MN996532.1;QHR63299.1;BatCoV_RaTG13_orf1ab;3579 | 0 | 0 |
| MN996532.1;QHR63299.1;BatCoV_RaTG13_orf1ab;3580 | 0 | 0 |
| MN996532.1;QHR63299.1;BatCoV_RaTG13_orf1ab;3581 | 0 | 0 |
| MN996532.1;QHR63299.1;BatCoV_RaTG13_orf1ab;3582 | 0 | 0 |
| MN996532.1;QHR63299.1;BatCoV_RaTG13_orf1ab;3583 | 0 | 0 |
| MN996532.1;QHR63299.1;BatCoV_RaTG13_orf1ab;3584 | 0 | 0 |
| MN996532.1;QHR63299.1;BatCoV_RaTG13_orf1ab;3585 | 0 | 0 |
| MN996532.1;QHR63299.1;BatCoV_RaTG13_orf1ab;3586 | 0 | 0 |
| MN996532.1;QHR63299.1;BatCoV_RaTG13_orf1ab;3587 | 0 | 0 |
| MN996532.1;QHR63299.1;BatCoV_RaTG13_orf1ab;3588 | 0 | 0 |
| MN996532.1;QHR63299.1;BatCoV_RaTG13_orf1ab;3589 | 0 | 0 |
| MN996532.1;QHR63299.1;BatCoV_RaTG13_orf1ab;3590 | 0 | 0 |
| MN996532.1;QHR63299.1;BatCoV_RaTG13_orf1ab;3591 | 0 | 0 |
| MN996532.1;QHR63299.1;BatCoV_RaTG13_orf1ab;3592 | 0 | 0 |
| MN996532.1;QHR63299.1;BatCoV_RaTG13_orf1ab;3593 | 0 | 0 |
| MN996532.1;QHR63299.1;BatCoV_RaTG13_orf1ab;3594 | 0 | 0 |
| MN996532.1;QHR63299.1;BatCoV_RaTG13_orf1ab;3595 | 0 | 0 |
| MN996532.1;QHR63299.1;BatCoV_RaTG13_orf1ab;3596 | 0 | 0 |
| MN996532.1;QHR63299.1;BatCoV_RaTG13_orf1ab;3597 | 0 | 0 |
| MN996532.1;QHR63299.1;BatCoV_RaTG13_orf1ab;3598 | 0 | 0 |
| MN996532.1;QHR63299.1;BatCoV_RaTG13_orf1ab;3599 | 0 | 0 |
| MN996532.1;QHR63299.1;BatCoV_RaTG13_orf1ab;3600 | 0 | 0 |
| MN996532.1;QHR63299.1;BatCoV_RaTG13_orf1ab;3601 | 0 | 0 |
| MN996532.1;QHR63299.1;BatCoV_RaTG13_orf1ab;3602 | 0 | 0 |
| MN996532.1;QHR63299.1;BatCoV_RaTG13_orf1ab;3603 | 0 | 0 |

| | | |
|---|---|---|
| MN996532.1;QHR63299.1;BatCoV_RaTG13_orf1ab;3604 | 0 | 0 |
| MN996532.1;QHR63299.1;BatCoV_RaTG13_orf1ab;3605 | 0 | 0 |
| MN996532.1;QHR63299.1;BatCoV_RaTG13_orf1ab;3606 | 0 | 0 |
| MN996532.1;QHR63299.1;BatCoV_RaTG13_orf1ab;3607 | 0 | 0 |
| MN996532.1;QHR63299.1;BatCoV_RaTG13_orf1ab;3608 | 0 | 0 |
| MN996532.1;QHR63299.1;BatCoV_RaTG13_orf1ab;3609 | 0 | 0 |
| MN996532.1;QHR63299.1;BatCoV_RaTG13_orf1ab;3610 | 0 | 0 |
| MN996532.1;QHR63299.1;BatCoV_RaTG13_orf1ab;3611 | 0 | 0 |
| MN996532.1;QHR63299.1;BatCoV_RaTG13_orf1ab;3612 | 0 | 0 |
| MN996532.1;QHR63299.1;BatCoV_RaTG13_orf1ab;3613 | 0 | 0 |
| MN996532.1;QHR63299.1;BatCoV_RaTG13_orf1ab;3614 | 0 | 0 |
| MN996532.1;QHR63299.1;BatCoV_RaTG13_orf1ab;3615 | 0 | 0 |
| MN996532.1;QHR63299.1;BatCoV_RaTG13_orf1ab;3616 | 0 | 0 |
| MN996532.1;QHR63299.1;BatCoV_RaTG13_orf1ab;3617 | 0 | 0 |
| MN996532.1;QHR63299.1;BatCoV_RaTG13_orf1ab;3618 | 0 | 0 |
| MN996532.1;QHR63299.1;BatCoV_RaTG13_orf1ab;3619 | 0 | 0 |
| MN996532.1;QHR63299.1;BatCoV_RaTG13_orf1ab;3620 | 0 | 0 |
| MN996532.1;QHR63299.1;BatCoV_RaTG13_orf1ab;3621 | 0 | 0 |
| MN996532.1;QHR63299.1;BatCoV_RaTG13_orf1ab;3622 | 0 | 0 |
| MN996532.1;QHR63299.1;BatCoV_RaTG13_orf1ab;3623 | 0 | 0 |
| MN996532.1;QHR63299.1;BatCoV_RaTG13_orf1ab;3624 | 0 | 0 |
| MN996532.1;QHR63299.1;BatCoV_RaTG13_orf1ab;3625 | 0 | 0 |
| MN996532.1;QHR63299.1;BatCoV_RaTG13_orf1ab;3626 | 0 | 0 |
| MN996532.1;QHR63299.1;BatCoV_RaTG13_orf1ab;3627 | 0 | 0 |
| MN996532.1;QHR63299.1;BatCoV_RaTG13_orf1ab;3628 | 0 | 0 |
| MN996532.1;QHR63299.1;BatCoV_RaTG13_orf1ab;3629 | 0 | 0 |
| MN996532.1;QHR63299.1;BatCoV_RaTG13_orf1ab;3630 | 0 | 0 |
| MN996532.1;QHR63299.1;BatCoV_RaTG13_orf1ab;3631 | 0 | 0 |
| MN996532.1;QHR63299.1;BatCoV_RaTG13_orf1ab;3632 | 0 | 0 |
| MN996532.1;QHR63299.1;BatCoV_RaTG13_orf1ab;3633 | 0 | 0 |
| MN996532.1;QHR63299.1;BatCoV_RaTG13_orf1ab;3634 | 0 | 0 |
| MN996532.1;QHR63299.1;BatCoV_RaTG13_orf1ab;3635 | 0 | 0 |
| MN996532.1;QHR63299.1;BatCoV_RaTG13_orf1ab;3636 | 0 | 0 |
| MN996532.1;QHR63299.1;BatCoV_RaTG13_orf1ab;3637 | 0 | 0 |
| MN996532.1;QHR63299.1;BatCoV_RaTG13_orf1ab;3638 | 0 | 0 |
| MN996532.1;QHR63299.1;BatCoV_RaTG13_orf1ab;3639 | 0 | 0 |
| MN996532.1;QHR63299.1;BatCoV_RaTG13_orf1ab;3640 | 0 | 0 |
| MN996532.1;QHR63299.1;BatCoV_RaTG13_orf1ab;3641 | 0 | 0 |
| MN996532.1;QHR63299.1;BatCoV_RaTG13_orf1ab;3642 | 0 | 0 |
| MN996532.1;QHR63299.1;BatCoV_RaTG13_orf1ab;3643 | 0 | 0 |
| MN996532.1;QHR63299.1;BatCoV_RaTG13_orf1ab;3644 | 0 | 0 |
| MN996532.1;QHR63299.1;BatCoV_RaTG13_orf1ab;3645 | 0 | 0 |
| MN996532.1;QHR63299.1;BatCoV_RaTG13_orf1ab;3646 | 0 | 0 |

| | | |
|---|---|---|
| MN996532.1;QHR63299.1;BatCoV_RaTG13_orf1ab;3647 | 0 | 0 |
| MN996532.1;QHR63299.1;BatCoV_RaTG13_orf1ab;3648 | 0 | 0 |
| MN996532.1;QHR63299.1;BatCoV_RaTG13_orf1ab;3649 | 0 | 0 |
| MN996532.1;QHR63299.1;BatCoV_RaTG13_orf1ab;3650 | 0 | 0 |
| MN996532.1;QHR63299.1;BatCoV_RaTG13_orf1ab;3651 | 0 | 0 |
| MN996532.1;QHR63299.1;BatCoV_RaTG13_orf1ab;3652 | 0 | 0 |
| MN996532.1;QHR63299.1;BatCoV_RaTG13_orf1ab;3653 | 0 | 0 |
| MN996532.1;QHR63299.1;BatCoV_RaTG13_orf1ab;3654 | 0 | 0 |
| MN996532.1;QHR63299.1;BatCoV_RaTG13_orf1ab;3655 | 0 | 0 |
| MN996532.1;QHR63299.1;BatCoV_RaTG13_orf1ab;3656 | 0 | 0 |
| MN996532.1;QHR63299.1;BatCoV_RaTG13_orf1ab;3657 | 0 | 0 |
| MN996532.1;QHR63299.1;BatCoV_RaTG13_orf1ab;3658 | 0 | 0 |
| MN996532.1;QHR63299.1;BatCoV_RaTG13_orf1ab;3659 | 0 | 0 |
| MN996532.1;QHR63299.1;BatCoV_RaTG13_orf1ab;3660 | 0 | 0 |
| MN996532.1;QHR63299.1;BatCoV_RaTG13_orf1ab;3661 | 0 | 0 |
| MN996532.1;QHR63299.1;BatCoV_RaTG13_orf1ab;3662 | 0 | 0 |
| MN996532.1;QHR63299.1;BatCoV_RaTG13_orf1ab;3663 | 0 | 0.025 |
| MN996532.1;QHR63299.1;BatCoV_RaTG13_orf1ab;3664 | 0 | 0.025 |
| MN996532.1;QHR63299.1;BatCoV_RaTG13_orf1ab;3665 | 0 | 0.05 |
| MN996532.1;QHR63299.1;BatCoV_RaTG13_orf1ab;3666 | 0 | 0.025 |
| MN996532.1;QHR63299.1;BatCoV_RaTG13_orf1ab;3667 | 0 | 0.025 |
| MN996532.1;QHR63299.1;BatCoV_RaTG13_orf1ab;3668 | 0 | 0.025 |
| MN996532.1;QHR63299.1;BatCoV_RaTG13_orf1ab;3669 | 0 | 0.025 |
| MN996532.1;QHR63299.1;BatCoV_RaTG13_orf1ab;3670 | 0 | 0.025 |
| MN996532.1;QHR63299.1;BatCoV_RaTG13_orf1ab;3671 | 0 | 0.025 |
| MN996532.1;QHR63299.1;BatCoV_RaTG13_orf1ab;3672 | 0 | 0.025 |
| MN996532.1;QHR63299.1;BatCoV_RaTG13_orf1ab;3673 | 0 | 0 |
| MN996532.1;QHR63299.1;BatCoV_RaTG13_orf1ab;3674 | 0 | 0 |
| MN996532.1;QHR63299.1;BatCoV_RaTG13_orf1ab;3675 | 0 | 0 |
| MN996532.1;QHR63299.1;BatCoV_RaTG13_orf1ab;3676 | 0 | 0 |
| MN996532.1;QHR63299.1;BatCoV_RaTG13_orf1ab;3677 | 0 | 0 |
| MN996532.1;QHR63299.1;BatCoV_RaTG13_orf1ab;3678 | 0 | 0 |
| MN996532.1;QHR63299.1;BatCoV_RaTG13_orf1ab;3679 | 0 | 0 |
| MN996532.1;QHR63299.1;BatCoV_RaTG13_orf1ab;3680 | 0 | 0 |
| MN996532.1;QHR63299.1;BatCoV_RaTG13_orf1ab;3681 | 0 | 0 |
| MN996532.1;QHR63299.1;BatCoV_RaTG13_orf1ab;3682 | 0 | 0 |
| MN996532.1;QHR63299.1;BatCoV_RaTG13_orf1ab;3683 | 0 | 0 |
| MN996532.1;QHR63299.1;BatCoV_RaTG13_orf1ab;3684 | 0 | 0 |
| MN996532.1;QHR63299.1;BatCoV_RaTG13_orf1ab;3685 | 0 | 0 |
| MN996532.1;QHR63299.1;BatCoV_RaTG13_orf1ab;3686 | 0 | 0 |
| MN996532.1;QHR63299.1;BatCoV_RaTG13_orf1ab;3687 | 0 | 0 |
| MN996532.1;QHR63299.1;BatCoV_RaTG13_orf1ab;3688 | 0 | 0 |
| MN996532.1;QHR63299.1;BatCoV_RaTG13_orf1ab;3689 | 0 | 0 |

| | | |
|---|---|---|
| MN996532.1;QHR63299.1;BatCoV_RaTG13_orf1ab;3690 | 0 | 0 |
| MN996532.1;QHR63299.1;BatCoV_RaTG13_orf1ab;3691 | 0 | 0 |
| MN996532.1;QHR63299.1;BatCoV_RaTG13_orf1ab;3692 | 0 | 0 |
| MN996532.1;QHR63299.1;BatCoV_RaTG13_orf1ab;3693 | 0 | 0 |
| MN996532.1;QHR63299.1;BatCoV_RaTG13_orf1ab;3694 | 0 | 0 |
| MN996532.1;QHR63299.1;BatCoV_RaTG13_orf1ab;3695 | 0 | 0 |
| MN996532.1;QHR63299.1;BatCoV_RaTG13_orf1ab;3696 | 0 | 0 |
| MN996532.1;QHR63299.1;BatCoV_RaTG13_orf1ab;3697 | 0 | 0.025 |
| MN996532.1;QHR63299.1;BatCoV_RaTG13_orf1ab;3698 | 0 | 0 |
| MN996532.1;QHR63299.1;BatCoV_RaTG13_orf1ab;3699 | 0 | 0 |
| MN996532.1;QHR63299.1;BatCoV_RaTG13_orf1ab;3700 | 0 | 0 |
| MN996532.1;QHR63299.1;BatCoV_RaTG13_orf1ab;3701 | 0 | 0 |
| MN996532.1;QHR63299.1;BatCoV_RaTG13_orf1ab;3702 | 0 | 0 |
| MN996532.1;QHR63299.1;BatCoV_RaTG13_orf1ab;3703 | 0 | 0 |
| MN996532.1;QHR63299.1;BatCoV_RaTG13_orf1ab;3704 | 0 | 0 |
| MN996532.1;QHR63299.1;BatCoV_RaTG13_orf1ab;3705 | 0 | 0 |
| MN996532.1;QHR63299.1;BatCoV_RaTG13_orf1ab;3706 | 0 | 0 |
| MN996532.1;QHR63299.1;BatCoV_RaTG13_orf1ab;3707 | 0 | 0 |
| MN996532.1;QHR63299.1;BatCoV_RaTG13_orf1ab;3708 | 0 | 0 |
| MN996532.1;QHR63299.1;BatCoV_RaTG13_orf1ab;3709 | 0 | 0 |
| MN996532.1;QHR63299.1;BatCoV_RaTG13_orf1ab;3710 | 0 | 0 |
| MN996532.1;QHR63299.1;BatCoV_RaTG13_orf1ab;3711 | 0 | 0 |
| MN996532.1;QHR63299.1;BatCoV_RaTG13_orf1ab;3712 | 0 | 0 |
| MN996532.1;QHR63299.1;BatCoV_RaTG13_orf1ab;3713 | 0 | 0 |
| MN996532.1;QHR63299.1;BatCoV_RaTG13_orf1ab;3714 | 0 | 0 |
| MN996532.1;QHR63299.1;BatCoV_RaTG13_orf1ab;3715 | 0 | 0 |
| MN996532.1;QHR63299.1;BatCoV_RaTG13_orf1ab;3716 | 0 | 0 |
| MN996532.1;QHR63299.1;BatCoV_RaTG13_orf1ab;3717 | 0.05 | 0 |
| MN996532.1;QHR63299.1;BatCoV_RaTG13_orf1ab;3718 | 0.05 | 0 |
| MN996532.1;QHR63299.1;BatCoV_RaTG13_orf1ab;3719 | 0.05 | 0 |
| MN996532.1;QHR63299.1;BatCoV_RaTG13_orf1ab;3720 | 0.05 | 0.025 |
| MN996532.1;QHR63299.1;BatCoV_RaTG13_orf1ab;3721 | 0.05 | 0 |
| MN996532.1;QHR63299.1;BatCoV_RaTG13_orf1ab;3722 | 0 | 0 |
| MN996532.1;QHR63299.1;BatCoV_RaTG13_orf1ab;3723 | 0 | 0 |
| MN996532.1;QHR63299.1;BatCoV_RaTG13_orf1ab;3724 | 0 | 0 |
| MN996532.1;QHR63299.1;BatCoV_RaTG13_orf1ab;3725 | 0 | 0 |
| MN996532.1;QHR63299.1;BatCoV_RaTG13_orf1ab;3726 | 0 | 0 |
| MN996532.1;QHR63299.1;BatCoV_RaTG13_orf1ab;3727 | 0 | 0 |
| MN996532.1;QHR63299.1;BatCoV_RaTG13_orf1ab;3728 | 0 | 0 |
| MN996532.1;QHR63299.1;BatCoV_RaTG13_orf1ab;3729 | 0 | 0 |
| MN996532.1;QHR63299.1;BatCoV_RaTG13_orf1ab;3730 | 0 | 0 |
| MN996532.1;QHR63299.1;BatCoV_RaTG13_orf1ab;3731 | 0 | 0 |
| MN996532.1;QHR63299.1;BatCoV_RaTG13_orf1ab;3732 | 0 | 0 |

| | | |
|---|---|---|
| MN996532.1;QHR63299.1;BatCoV_RaTG13_orf1ab;3733 | 0 | 0 |
| MN996532.1;QHR63299.1;BatCoV_RaTG13_orf1ab;3734 | 0 | 0 |
| MN996532.1;QHR63299.1;BatCoV_RaTG13_orf1ab;3735 | 0 | 0 |
| MN996532.1;QHR63299.1;BatCoV_RaTG13_orf1ab;3736 | 0 | 0 |
| MN996532.1;QHR63299.1;BatCoV_RaTG13_orf1ab;3737 | 0 | 0 |
| MN996532.1;QHR63299.1;BatCoV_RaTG13_orf1ab;3738 | 0 | 0 |
| MN996532.1;QHR63299.1;BatCoV_RaTG13_orf1ab;3739 | 0 | 0 |
| MN996532.1;QHR63299.1;BatCoV_RaTG13_orf1ab;3740 | 0 | 0 |
| MN996532.1;QHR63299.1;BatCoV_RaTG13_orf1ab;3741 | 0 | 0 |
| MN996532.1;QHR63299.1;BatCoV_RaTG13_orf1ab;3742 | 0 | 0 |
| MN996532.1;QHR63299.1;BatCoV_RaTG13_orf1ab;3743 | 0 | 0 |
| MN996532.1;QHR63299.1;BatCoV_RaTG13_orf1ab;3744 | 0 | 0 |
| MN996532.1;QHR63299.1;BatCoV_RaTG13_orf1ab;3745 | 0 | 0 |
| MN996532.1;QHR63299.1;BatCoV_RaTG13_orf1ab;3746 | 0 | 0 |
| MN996532.1;QHR63299.1;BatCoV_RaTG13_orf1ab;3747 | 0 | 0 |
| MN996532.1;QHR63299.1;BatCoV_RaTG13_orf1ab;3748 | 0 | 0 |
| MN996532.1;QHR63299.1;BatCoV_RaTG13_orf1ab;3749 | 0 | 0 |
| MN996532.1;QHR63299.1;BatCoV_RaTG13_orf1ab;3750 | 0 | 0 |
| MN996532.1;QHR63299.1;BatCoV_RaTG13_orf1ab;3751 | 0 | 0 |
| MN996532.1;QHR63299.1;BatCoV_RaTG13_orf1ab;3752 | 0 | 0 |
| MN996532.1;QHR63299.1;BatCoV_RaTG13_orf1ab;3753 | 0 | 0 |
| MN996532.1;QHR63299.1;BatCoV_RaTG13_orf1ab;3754 | 0 | 0 |
| MN996532.1;QHR63299.1;BatCoV_RaTG13_orf1ab;3755 | 0 | 0 |
| MN996532.1;QHR63299.1;BatCoV_RaTG13_orf1ab;3756 | 0 | 0 |
| MN996532.1;QHR63299.1;BatCoV_RaTG13_orf1ab;3757 | 0 | 0 |
| MN996532.1;QHR63299.1;BatCoV_RaTG13_orf1ab;3758 | 0 | 0 |
| MN996532.1;QHR63299.1;BatCoV_RaTG13_orf1ab;3759 | 0 | 0 |
| MN996532.1;QHR63299.1;BatCoV_RaTG13_orf1ab;3760 | 0 | 0 |
| MN996532.1;QHR63299.1;BatCoV_RaTG13_orf1ab;3761 | 0 | 0 |
| MN996532.1;QHR63299.1;BatCoV_RaTG13_orf1ab;3762 | 0 | 0 |
| MN996532.1;QHR63299.1;BatCoV_RaTG13_orf1ab;3763 | 0 | 0 |
| MN996532.1;QHR63299.1;BatCoV_RaTG13_orf1ab;3764 | 0 | 0 |
| MN996532.1;QHR63299.1;BatCoV_RaTG13_orf1ab;3765 | 0 | 0 |
| MN996532.1;QHR63299.1;BatCoV_RaTG13_orf1ab;3766 | 0 | 0 |
| MN996532.1;QHR63299.1;BatCoV_RaTG13_orf1ab;3767 | 0 | 0 |
| MN996532.1;QHR63299.1;BatCoV_RaTG13_orf1ab;3768 | 0 | 0 |
| MN996532.1;QHR63299.1;BatCoV_RaTG13_orf1ab;3769 | 0 | 0 |
| MN996532.1;QHR63299.1;BatCoV_RaTG13_orf1ab;3770 | 0 | 0 |
| MN996532.1;QHR63299.1;BatCoV_RaTG13_orf1ab;3771 | 0 | 0 |
| MN996532.1;QHR63299.1;BatCoV_RaTG13_orf1ab;3772 | 0 | 0 |
| MN996532.1;QHR63299.1;BatCoV_RaTG13_orf1ab;3773 | 0 | 0 |
| MN996532.1;QHR63299.1;BatCoV_RaTG13_orf1ab;3774 | 0 | 0 |
| MN996532.1;QHR63299.1;BatCoV_RaTG13_orf1ab;3775 | 0 | 0 |

| | | |
|---|---|---|
| MN996532.1;QHR63299.1;BatCoV_RaTG13_orf1ab;3776 | 0 | 0 |
| MN996532.1;QHR63299.1;BatCoV_RaTG13_orf1ab;3777 | 0 | 0 |
| MN996532.1;QHR63299.1;BatCoV_RaTG13_orf1ab;3778 | 0 | 0 |
| MN996532.1;QHR63299.1;BatCoV_RaTG13_orf1ab;3779 | 0 | 0 |
| MN996532.1;QHR63299.1;BatCoV_RaTG13_orf1ab;3780 | 0 | 0 |
| MN996532.1;QHR63299.1;BatCoV_RaTG13_orf1ab;3781 | 0 | 0 |
| MN996532.1;QHR63299.1;BatCoV_RaTG13_orf1ab;3782 | 0 | 0 |
| MN996532.1;QHR63299.1;BatCoV_RaTG13_orf1ab;3783 | 0 | 0 |
| MN996532.1;QHR63299.1;BatCoV_RaTG13_orf1ab;3784 | 0 | 0 |
| MN996532.1;QHR63299.1;BatCoV_RaTG13_orf1ab;3785 | 0 | 0 |
| MN996532.1;QHR63299.1;BatCoV_RaTG13_orf1ab;3786 | 0 | 0 |
| MN996532.1;QHR63299.1;BatCoV_RaTG13_orf1ab;3787 | 0 | 0 |
| MN996532.1;QHR63299.1;BatCoV_RaTG13_orf1ab;3788 | 0 | 0 |
| MN996532.1;QHR63299.1;BatCoV_RaTG13_orf1ab;3789 | 0 | 0 |
| MN996532.1;QHR63299.1;BatCoV_RaTG13_orf1ab;3790 | 0 | 0 |
| MN996532.1;QHR63299.1;BatCoV_RaTG13_orf1ab;3791 | 0 | 0 |
| MN996532.1;QHR63299.1;BatCoV_RaTG13_orf1ab;3792 | 0 | 0 |
| MN996532.1;QHR63299.1;BatCoV_RaTG13_orf1ab;3793 | 0 | 0 |
| MN996532.1;QHR63299.1;BatCoV_RaTG13_orf1ab;3794 | 0 | 0 |
| MN996532.1;QHR63299.1;BatCoV_RaTG13_orf1ab;3795 | 0 | 0 |
| MN996532.1;QHR63299.1;BatCoV_RaTG13_orf1ab;3796 | 0 | 0 |
| MN996532.1;QHR63299.1;BatCoV_RaTG13_orf1ab;3797 | 0 | 0 |
| MN996532.1;QHR63299.1;BatCoV_RaTG13_orf1ab;3798 | 0 | 0 |
| MN996532.1;QHR63299.1;BatCoV_RaTG13_orf1ab;3799 | 0 | 0 |
| MN996532.1;QHR63299.1;BatCoV_RaTG13_orf1ab;3800 | 0 | 0 |
| MN996532.1;QHR63299.1;BatCoV_RaTG13_orf1ab;3801 | 0 | 0 |
| MN996532.1;QHR63299.1;BatCoV_RaTG13_orf1ab;3802 | 0 | 0 |
| MN996532.1;QHR63299.1;BatCoV_RaTG13_orf1ab;3803 | 0 | 0 |
| MN996532.1;QHR63299.1;BatCoV_RaTG13_orf1ab;3804 | 0 | 0 |
| MN996532.1;QHR63299.1;BatCoV_RaTG13_orf1ab;3805 | 0 | 0 |
| MN996532.1;QHR63299.1;BatCoV_RaTG13_orf1ab;3806 | 0 | 0 |
| MN996532.1;QHR63299.1;BatCoV_RaTG13_orf1ab;3807 | 0 | 0 |
| MN996532.1;QHR63299.1;BatCoV_RaTG13_orf1ab;3808 | 0.25 | 0.175 |
| MN996532.1;QHR63299.1;BatCoV_RaTG13_orf1ab;3809 | 0 | 0 |
| MN996532.1;QHR63299.1;BatCoV_RaTG13_orf1ab;3810 | 0 | 0 |
| MN996532.1;QHR63299.1;BatCoV_RaTG13_orf1ab;3811 | 0 | 0 |
| MN996532.1;QHR63299.1;BatCoV_RaTG13_orf1ab;3812 | 0 | 0 |
| MN996532.1;QHR63299.1;BatCoV_RaTG13_orf1ab;3813 | 0 | 0 |
| MN996532.1;QHR63299.1;BatCoV_RaTG13_orf1ab;3814 | 0 | 0 |
| MN996532.1;QHR63299.1;BatCoV_RaTG13_orf1ab;3815 | 0 | 0 |
| MN996532.1;QHR63299.1;BatCoV_RaTG13_orf1ab;3816 | 0 | 0.05 |
| MN996532.1;QHR63299.1;BatCoV_RaTG13_orf1ab;3817 | 0.05 | 0.025 |
| MN996532.1;QHR63299.1;BatCoV_RaTG13_orf1ab;3818 | 0.15 | 0.075 |

| | | |
|---|---|---|
| MN996532.1;QHR63299.1;BatCoV_RaTG13_orf1ab;3819 | 0 | 0.025 |
| MN996532.1;QHR63299.1;BatCoV_RaTG13_orf1ab;3820 | 0 | 0.025 |
| MN996532.1;QHR63299.1;BatCoV_RaTG13_orf1ab;3821 | 0.05 | 0.125 |
| MN996532.1;QHR63299.1;BatCoV_RaTG13_orf1ab;3822 | 0.1 | 0.05 |
| MN996532.1;QHR63299.1;BatCoV_RaTG13_orf1ab;3823 | 0.3 | 0.1 |
| MN996532.1;QHR63299.1;BatCoV_RaTG13_orf1ab;3824 | 0.25 | 0.1 |
| MN996532.1;QHR63299.1;BatCoV_RaTG13_orf1ab;3825 | 0.25 | 0.1 |
| MN996532.1;QHR63299.1;BatCoV_RaTG13_orf1ab;3826 | 0.2 | 0.1 |
| MN996532.1;QHR63299.1;BatCoV_RaTG13_orf1ab;3827 | 0.15 | 0.075 |
| MN996532.1;QHR63299.1;BatCoV_RaTG13_orf1ab;3828 | 0.1 | 0.075 |
| MN996532.1;QHR63299.1;BatCoV_RaTG13_orf1ab;3829 | 0.05 | 0.075 |
| MN996532.1;QHR63299.1;BatCoV_RaTG13_orf1ab;3830 | 0.05 | 0.05 |
| MN996532.1;QHR63299.1;BatCoV_RaTG13_orf1ab;3831 | 0 | 0.05 |
| MN996532.1;QHR63299.1;BatCoV_RaTG13_orf1ab;3832 | 0 | 0 |
| MN996532.1;QHR63299.1;BatCoV_RaTG13_orf1ab;3833 | 0 | 0 |
| MN996532.1;QHR63299.1;BatCoV_RaTG13_orf1ab;3834 | 0 | 0 |
| MN996532.1;QHR63299.1;BatCoV_RaTG13_orf1ab;3835 | 0 | 0 |
| MN996532.1;QHR63299.1;BatCoV_RaTG13_orf1ab;3836 | 0 | 0 |
| MN996532.1;QHR63299.1;BatCoV_RaTG13_orf1ab;3837 | 0 | 0 |
| MN996532.1;QHR63299.1;BatCoV_RaTG13_orf1ab;3838 | 0 | 0 |
| MN996532.1;QHR63299.1;BatCoV_RaTG13_orf1ab;3839 | 0 | 0 |
| MN996532.1;QHR63299.1;BatCoV_RaTG13_orf1ab;3840 | 0 | 0 |
| MN996532.1;QHR63299.1;BatCoV_RaTG13_orf1ab;3841 | 0 | 0.025 |
| MN996532.1;QHR63299.1;BatCoV_RaTG13_orf1ab;3842 | 0 | 0 |
| MN996532.1;QHR63299.1;BatCoV_RaTG13_orf1ab;3843 | 0 | 0.025 |
| MN996532.1;QHR63299.1;BatCoV_RaTG13_orf1ab;3844 | 0 | 0.025 |
| MN996532.1;QHR63299.1;BatCoV_RaTG13_orf1ab;3845 | 0.05 | 0.05 |
| MN996532.1;QHR63299.1;BatCoV_RaTG13_orf1ab;3846 | 0.05 | 0.05 |
| MN996532.1;QHR63299.1;BatCoV_RaTG13_orf1ab;3847 | 0.05 | 0.05 |
| MN996532.1;QHR63299.1;BatCoV_RaTG13_orf1ab;3848 | 0.1 | 0.1 |
| MN996532.1;QHR63299.1;BatCoV_RaTG13_orf1ab;3849 | 0.05 | 0.075 |
| MN996532.1;QHR63299.1;BatCoV_RaTG13_orf1ab;3850 | 0.05 | 0.05 |
| MN996532.1;QHR63299.1;BatCoV_RaTG13_orf1ab;3851 | 0.05 | 0.025 |
| MN996532.1;QHR63299.1;BatCoV_RaTG13_orf1ab;3852 | 0 | 0.025 |
| MN996532.1;QHR63299.1;BatCoV_RaTG13_orf1ab;3853 | 0.05 | 0.025 |
| MN996532.1;QHR63299.1;BatCoV_RaTG13_orf1ab;3854 | 0.05 | 0.025 |
| MN996532.1;QHR63299.1;BatCoV_RaTG13_orf1ab;3855 | 0.05 | 0.025 |
| MN996532.1;QHR63299.1;BatCoV_RaTG13_orf1ab;3856 | 0.05 | 0 |
| MN996532.1;QHR63299.1;BatCoV_RaTG13_orf1ab;3857 | 0 | 0 |
| MN996532.1;QHR63299.1;BatCoV_RaTG13_orf1ab;3858 | 0 | 0 |
| MN996532.1;QHR63299.1;BatCoV_RaTG13_orf1ab;3859 | 0 | 0 |
| MN996532.1;QHR63299.1;BatCoV_RaTG13_orf1ab;3860 | 0 | 0 |
| MN996532.1;QHR63299.1;BatCoV_RaTG13_orf1ab;3861 | 0 | 0 |

| | | |
|---|---|---|
| MN996532.1;QHR63299.1;BatCoV_RaTG13_orf1ab;3862 | 0 | 0 |
| MN996532.1;QHR63299.1;BatCoV_RaTG13_orf1ab;3863 | 0 | 0 |
| MN996532.1;QHR63299.1;BatCoV_RaTG13_orf1ab;3864 | 0 | 0 |
| MN996532.1;QHR63299.1;BatCoV_RaTG13_orf1ab;3865 | 0 | 0 |
| MN996532.1;QHR63299.1;BatCoV_RaTG13_orf1ab;3866 | 0 | 0 |
| MN996532.1;QHR63299.1;BatCoV_RaTG13_orf1ab;3867 | 0 | 0 |
| MN996532.1;QHR63299.1;BatCoV_RaTG13_orf1ab;3868 | 0 | 0 |
| MN996532.1;QHR63299.1;BatCoV_RaTG13_orf1ab;3869 | 0 | 0 |
| MN996532.1;QHR63299.1;BatCoV_RaTG13_orf1ab;3870 | 0 | 0 |
| MN996532.1;QHR63299.1;BatCoV_RaTG13_orf1ab;3871 | 0 | 0 |
| MN996532.1;QHR63299.1;BatCoV_RaTG13_orf1ab;3872 | 0 | 0 |
| MN996532.1;QHR63299.1;BatCoV_RaTG13_orf1ab;3873 | 0 | 0 |
| MN996532.1;QHR63299.1;BatCoV_RaTG13_orf1ab;3874 | 0 | 0 |
| MN996532.1;QHR63299.1;BatCoV_RaTG13_orf1ab;3875 | 0 | 0 |
| MN996532.1;QHR63299.1;BatCoV_RaTG13_orf1ab;3876 | 0 | 0 |
| MN996532.1;QHR63299.1;BatCoV_RaTG13_orf1ab;3877 | 0 | 0 |
| MN996532.1;QHR63299.1;BatCoV_RaTG13_orf1ab;3878 | 0 | 0 |
| MN996532.1;QHR63299.1;BatCoV_RaTG13_orf1ab;3879 | 0 | 0 |
| MN996532.1;QHR63299.1;BatCoV_RaTG13_orf1ab;3880 | 0 | 0 |
| MN996532.1;QHR63299.1;BatCoV_RaTG13_orf1ab;3881 | 0 | 0.025 |
| MN996532.1;QHR63299.1;BatCoV_RaTG13_orf1ab;3882 | 0 | 0 |
| MN996532.1;QHR63299.1;BatCoV_RaTG13_orf1ab;3883 | 0 | 0.025 |
| MN996532.1;QHR63299.1;BatCoV_RaTG13_orf1ab;3884 | 0 | 0 |
| MN996532.1;QHR63299.1;BatCoV_RaTG13_orf1ab;3885 | 0 | 0 |
| MN996532.1;QHR63299.1;BatCoV_RaTG13_orf1ab;3886 | 0 | 0 |
| MN996532.1;QHR63299.1;BatCoV_RaTG13_orf1ab;3887 | 0.05 | 0 |
| MN996532.1;QHR63299.1;BatCoV_RaTG13_orf1ab;3888 | 0.05 | 0 |
| MN996532.1;QHR63299.1;BatCoV_RaTG13_orf1ab;3889 | 0.05 | 0 |
| MN996532.1;QHR63299.1;BatCoV_RaTG13_orf1ab;3890 | 0 | 0 |
| MN996532.1;QHR63299.1;BatCoV_RaTG13_orf1ab;3891 | 0 | 0 |
| MN996532.1;QHR63299.1;BatCoV_RaTG13_orf1ab;3892 | 0 | 0 |
| MN996532.1;QHR63299.1;BatCoV_RaTG13_orf1ab;3893 | 0 | 0 |
| MN996532.1;QHR63299.1;BatCoV_RaTG13_orf1ab;3894 | 0 | 0 |
| MN996532.1;QHR63299.1;BatCoV_RaTG13_orf1ab;3895 | 0 | 0 |
| MN996532.1;QHR63299.1;BatCoV_RaTG13_orf1ab;3896 | 0 | 0 |
| MN996532.1;QHR63299.1;BatCoV_RaTG13_orf1ab;3897 | 0 | 0 |
| MN996532.1;QHR63299.1;BatCoV_RaTG13_orf1ab;3898 | 0 | 0 |
| MN996532.1;QHR63299.1;BatCoV_RaTG13_orf1ab;3899 | 0 | 0 |
| MN996532.1;QHR63299.1;BatCoV_RaTG13_orf1ab;3900 | 0 | 0 |
| MN996532.1;QHR63299.1;BatCoV_RaTG13_orf1ab;3901 | 0 | 0 |
| MN996532.1;QHR63299.1;BatCoV_RaTG13_orf1ab;3902 | 0 | 0 |
| MN996532.1;QHR63299.1;BatCoV_RaTG13_orf1ab;3903 | 0 | 0 |
| MN996532.1;QHR63299.1;BatCoV_RaTG13_orf1ab;3904 | 0 | 0 |

| | | |
|---|---|---|
| MN996532.1;QHR63299.1;BatCoV_RaTG13_orf1ab;3905 | 0 | 0 |
| MN996532.1;QHR63299.1;BatCoV_RaTG13_orf1ab;3906 | 0 | 0 |
| MN996532.1;QHR63299.1;BatCoV_RaTG13_orf1ab;3907 | 0 | 0 |
| MN996532.1;QHR63299.1;BatCoV_RaTG13_orf1ab;3908 | 0 | 0 |
| MN996532.1;QHR63299.1;BatCoV_RaTG13_orf1ab;3909 | 0 | 0 |
| MN996532.1;QHR63299.1;BatCoV_RaTG13_orf1ab;3910 | 0 | 0 |
| MN996532.1;QHR63299.1;BatCoV_RaTG13_orf1ab;3911 | 0 | 0 |
| MN996532.1;QHR63299.1;BatCoV_RaTG13_orf1ab;3912 | 0 | 0 |
| MN996532.1;QHR63299.1;BatCoV_RaTG13_orf1ab;3913 | 0 | 0.05 |
| MN996532.1;QHR63299.1;BatCoV_RaTG13_orf1ab;3914 | 0 | 0.025 |
| MN996532.1;QHR63299.1;BatCoV_RaTG13_orf1ab;3915 | 0 | 0.025 |
| MN996532.1;QHR63299.1;BatCoV_RaTG13_orf1ab;3916 | 0 | 0.025 |
| MN996532.1;QHR63299.1;BatCoV_RaTG13_orf1ab;3917 | 0 | 0.025 |
| MN996532.1;QHR63299.1;BatCoV_RaTG13_orf1ab;3918 | 0.05 | 0.05 |
| MN996532.1;QHR63299.1;BatCoV_RaTG13_orf1ab;3919 | 0.05 | 0.05 |
| MN996532.1;QHR63299.1;BatCoV_RaTG13_orf1ab;3920 | 0.05 | 0.075 |
| MN996532.1;QHR63299.1;BatCoV_RaTG13_orf1ab;3921 | 0.05 | 0.05 |
| MN996532.1;QHR63299.1;BatCoV_RaTG13_orf1ab;3922 | 0.1 | 0.025 |
| MN996532.1;QHR63299.1;BatCoV_RaTG13_orf1ab;3923 | 0 | 0.025 |
| MN996532.1;QHR63299.1;BatCoV_RaTG13_orf1ab;3924 | 0 | 0.05 |
| MN996532.1;QHR63299.1;BatCoV_RaTG13_orf1ab;3925 | 0 | 0.025 |
| MN996532.1;QHR63299.1;BatCoV_RaTG13_orf1ab;3926 | 0 | 0 |
| MN996532.1;QHR63299.1;BatCoV_RaTG13_orf1ab;3927 | 0 | 0 |
| MN996532.1;QHR63299.1;BatCoV_RaTG13_orf1ab;3928 | 0.05 | 0 |
| MN996532.1;QHR63299.1;BatCoV_RaTG13_orf1ab;3929 | 0.05 | 0 |
| MN996532.1;QHR63299.1;BatCoV_RaTG13_orf1ab;3930 | 0.05 | 0 |
| MN996532.1;QHR63299.1;BatCoV_RaTG13_orf1ab;3931 | 0 | 0 |
| MN996532.1;QHR63299.1;BatCoV_RaTG13_orf1ab;3932 | 0.05 | 0 |
| MN996532.1;QHR63299.1;BatCoV_RaTG13_orf1ab;3933 | 0 | 0 |
| MN996532.1;QHR63299.1;BatCoV_RaTG13_orf1ab;3934 | 0 | 0 |
| MN996532.1;QHR63299.1;BatCoV_RaTG13_orf1ab;3935 | 0 | 0 |
| MN996532.1;QHR63299.1;BatCoV_RaTG13_orf1ab;3936 | 0 | 0 |
| MN996532.1;QHR63299.1;BatCoV_RaTG13_orf1ab;3937 | 0 | 0 |
| MN996532.1;QHR63299.1;BatCoV_RaTG13_orf1ab;3938 | 0 | 0 |
| MN996532.1;QHR63299.1;BatCoV_RaTG13_orf1ab;3939 | 0 | 0 |
| MN996532.1;QHR63299.1;BatCoV_RaTG13_orf1ab;3940 | 0.05 | 0.05 |
| MN996532.1;QHR63299.1;BatCoV_RaTG13_orf1ab;3941 | 0 | 0.025 |
| MN996532.1;QHR63299.1;BatCoV_RaTG13_orf1ab;3942 | 0 | 0 |
| MN996532.1;QHR63299.1;BatCoV_RaTG13_orf1ab;3943 | 0 | 0 |
| MN996532.1;QHR63299.1;BatCoV_RaTG13_orf1ab;3944 | 0 | 0 |
| MN996532.1;QHR63299.1;BatCoV_RaTG13_orf1ab;3945 | 0 | 0.05 |
| MN996532.1;QHR63299.1;BatCoV_RaTG13_orf1ab;3946 | 0.05 | 0 |
| MN996532.1;QHR63299.1;BatCoV_RaTG13_orf1ab;3947 | 0.05 | 0 |

| | | |
|---|---|---|
| MN996532.1;QHR63299.1;BatCoV_RaTG13_orf1ab;3948 | 0.05 | 0 |
| MN996532.1;QHR63299.1;BatCoV_RaTG13_orf1ab;3949 | 0.05 | 0.05 |
| MN996532.1;QHR63299.1;BatCoV_RaTG13_orf1ab;3950 | 0.1 | 0.05 |
| MN996532.1;QHR63299.1;BatCoV_RaTG13_orf1ab;3951 | 0 | 0.025 |
| MN996532.1;QHR63299.1;BatCoV_RaTG13_orf1ab;3952 | 0.1 | 0.05 |
| MN996532.1;QHR63299.1;BatCoV_RaTG13_orf1ab;3953 | 0.05 | 0.125 |
| MN996532.1;QHR63299.1;BatCoV_RaTG13_orf1ab;3954 | 0.05 | 0.075 |
| MN996532.1;QHR63299.1;BatCoV_RaTG13_orf1ab;3955 | 0.05 | 0.15 |
| MN996532.1;QHR63299.1;BatCoV_RaTG13_orf1ab;3956 | 0.05 | 0.05 |
| MN996532.1;QHR63299.1;BatCoV_RaTG13_orf1ab;3957 | 0 | 0.025 |
| MN996532.1;QHR63299.1;BatCoV_RaTG13_orf1ab;3958 | 0 | 0 |
| MN996532.1;QHR63299.1;BatCoV_RaTG13_orf1ab;3959 | 0.05 | 0 |
| MN996532.1;QHR63299.1;BatCoV_RaTG13_orf1ab;3960 | 0 | 0 |
| MN996532.1;QHR63299.1;BatCoV_RaTG13_orf1ab;3961 | 0 | 0 |
| MN996532.1;QHR63299.1;BatCoV_RaTG13_orf1ab;3962 | 0 | 0 |
| MN996532.1;QHR63299.1;BatCoV_RaTG13_orf1ab;3963 | 0 | 0 |
| MN996532.1;QHR63299.1;BatCoV_RaTG13_orf1ab;3964 | 0 | 0 |
| MN996532.1;QHR63299.1;BatCoV_RaTG13_orf1ab;3965 | 0 | 0 |
| MN996532.1;QHR63299.1;BatCoV_RaTG13_orf1ab;3966 | 0 | 0 |
| MN996532.1;QHR63299.1;BatCoV_RaTG13_orf1ab;3967 | 0 | 0 |
| MN996532.1;QHR63299.1;BatCoV_RaTG13_orf1ab;3968 | 0 | 0 |
| MN996532.1;QHR63299.1;BatCoV_RaTG13_orf1ab;3969 | 0 | 0 |
| MN996532.1;QHR63299.1;BatCoV_RaTG13_orf1ab;3970 | 0 | 0 |
| MN996532.1;QHR63299.1;BatCoV_RaTG13_orf1ab;3971 | 0 | 0 |
| MN996532.1;QHR63299.1;BatCoV_RaTG13_orf1ab;3972 | 0 | 0 |
| MN996532.1;QHR63299.1;BatCoV_RaTG13_orf1ab;3973 | 0 | 0 |
| MN996532.1;QHR63299.1;BatCoV_RaTG13_orf1ab;3974 | 0 | 0 |
| MN996532.1;QHR63299.1;BatCoV_RaTG13_orf1ab;3975 | 0 | 0 |
| MN996532.1;QHR63299.1;BatCoV_RaTG13_orf1ab;3976 | 0 | 0.025 |
| MN996532.1;QHR63299.1;BatCoV_RaTG13_orf1ab;3977 | 0 | 0.05 |
| MN996532.1;QHR63299.1;BatCoV_RaTG13_orf1ab;3978 | 0 | 0.025 |
| MN996532.1;QHR63299.1;BatCoV_RaTG13_orf1ab;3979 | 0.05 | 0.025 |
| MN996532.1;QHR63299.1;BatCoV_RaTG13_orf1ab;3980 | 0.05 | 0.025 |
| MN996532.1;QHR63299.1;BatCoV_RaTG13_orf1ab;3981 | 0 | 0.05 |
| MN996532.1;QHR63299.1;BatCoV_RaTG13_orf1ab;3982 | 0.05 | 0.025 |
| MN996532.1;QHR63299.1;BatCoV_RaTG13_orf1ab;3983 | 0 | 0.025 |
| MN996532.1;QHR63299.1;BatCoV_RaTG13_orf1ab;3984 | 0 | 0 |
| MN996532.1;QHR63299.1;BatCoV_RaTG13_orf1ab;3985 | 0 | 0 |
| MN996532.1;QHR63299.1;BatCoV_RaTG13_orf1ab;3986 | 0 | 0 |
| MN996532.1;QHR63299.1;BatCoV_RaTG13_orf1ab;3987 | 0 | 0 |
| MN996532.1;QHR63299.1;BatCoV_RaTG13_orf1ab;3988 | 0 | 0 |
| MN996532.1;QHR63299.1;BatCoV_RaTG13_orf1ab;3989 | 0 | 0 |
| MN996532.1;QHR63299.1;BatCoV_RaTG13_orf1ab;3990 | 0 | 0 |

| | | |
|---|---|---|
| MN996532.1;QHR63299.1;BatCoV_RaTG13_orf1ab;3991 | 0 | 0 |
| MN996532.1;QHR63299.1;BatCoV_RaTG13_orf1ab;3992 | 0 | 0 |
| MN996532.1;QHR63299.1;BatCoV_RaTG13_orf1ab;3993 | 0 | 0.025 |
| MN996532.1;QHR63299.1;BatCoV_RaTG13_orf1ab;3994 | 0 | 0 |
| MN996532.1;QHR63299.1;BatCoV_RaTG13_orf1ab;3995 | 0 | 0 |
| MN996532.1;QHR63299.1;BatCoV_RaTG13_orf1ab;3996 | 0 | 0 |
| MN996532.1;QHR63299.1;BatCoV_RaTG13_orf1ab;3997 | 0 | 0.05 |
| MN996532.1;QHR63299.1;BatCoV_RaTG13_orf1ab;3998 | 0 | 0.025 |
| MN996532.1;QHR63299.1;BatCoV_RaTG13_orf1ab;3999 | 0 | 0 |
| MN996532.1;QHR63299.1;BatCoV_RaTG13_orf1ab;4000 | 0 | 0 |
| MN996532.1;QHR63299.1;BatCoV_RaTG13_orf1ab;4001 | 0 | 0 |
| MN996532.1;QHR63299.1;BatCoV_RaTG13_orf1ab;4002 | 0 | 0 |
| MN996532.1;QHR63299.1;BatCoV_RaTG13_orf1ab;4003 | 0 | 0 |
| MN996532.1;QHR63299.1;BatCoV_RaTG13_orf1ab;4004 | 0 | 0 |
| MN996532.1;QHR63299.1;BatCoV_RaTG13_orf1ab;4005 | 0 | 0.1 |
| MN996532.1;QHR63299.1;BatCoV_RaTG13_orf1ab;4006 | 0 | 0.075 |
| MN996532.1;QHR63299.1;BatCoV_RaTG13_orf1ab;4007 | 0 | 0.05 |
| MN996532.1;QHR63299.1;BatCoV_RaTG13_orf1ab;4008 | 0 | 0.05 |
| MN996532.1;QHR63299.1;BatCoV_RaTG13_orf1ab;4009 | 0 | 0.05 |
| MN996532.1;QHR63299.1;BatCoV_RaTG13_orf1ab;4010 | 0 | 0 |
| MN996532.1;QHR63299.1;BatCoV_RaTG13_orf1ab;4011 | 0 | 0 |
| MN996532.1;QHR63299.1;BatCoV_RaTG13_orf1ab;4012 | 0 | 0 |
| MN996532.1;QHR63299.1;BatCoV_RaTG13_orf1ab;4013 | 0 | 0 |
| MN996532.1;QHR63299.1;BatCoV_RaTG13_orf1ab;4014 | 0 | 0 |
| MN996532.1;QHR63299.1;BatCoV_RaTG13_orf1ab;4015 | 0 | 0 |
| MN996532.1;QHR63299.1;BatCoV_RaTG13_orf1ab;4016 | 0 | 0 |
| MN996532.1;QHR63299.1;BatCoV_RaTG13_orf1ab;4017 | 0 | 0 |
| MN996532.1;QHR63299.1;BatCoV_RaTG13_orf1ab;4018 | 0 | 0 |
| MN996532.1;QHR63299.1;BatCoV_RaTG13_orf1ab;4019 | 0 | 0 |
| MN996532.1;QHR63299.1;BatCoV_RaTG13_orf1ab;4020 | 0 | 0 |
| MN996532.1;QHR63299.1;BatCoV_RaTG13_orf1ab;4021 | 0 | 0 |
| MN996532.1;QHR63299.1;BatCoV_RaTG13_orf1ab;4022 | 0 | 0 |
| MN996532.1;QHR63299.1;BatCoV_RaTG13_orf1ab;4023 | 0 | 0 |
| MN996532.1;QHR63299.1;BatCoV_RaTG13_orf1ab;4024 | 0 | 0 |
| MN996532.1;QHR63299.1;BatCoV_RaTG13_orf1ab;4025 | 0 | 0 |
| MN996532.1;QHR63299.1;BatCoV_RaTG13_orf1ab;4026 | 0 | 0 |
| MN996532.1;QHR63299.1;BatCoV_RaTG13_orf1ab;4027 | 0 | 0 |
| MN996532.1;QHR63299.1;BatCoV_RaTG13_orf1ab;4028 | 0 | 0 |
| MN996532.1;QHR63299.1;BatCoV_RaTG13_orf1ab;4029 | 0 | 0 |
| MN996532.1;QHR63299.1;BatCoV_RaTG13_orf1ab;4030 | 0.15 | 0.1 |
| MN996532.1;QHR63299.1;BatCoV_RaTG13_orf1ab;4031 | 0.2 | 0.1 |
| MN996532.1;QHR63299.1;BatCoV_RaTG13_orf1ab;4032 | 0.2 | 0.075 |
| MN996532.1;QHR63299.1;BatCoV_RaTG13_orf1ab;4033 | 0.25 | 0.1 |

| | | |
|---|---|---|
| MN996532.1;QHR63299.1;BatCoV_RaTG13_orf1ab;4034 | 0.25 | 0.275 |
| MN996532.1;QHR63299.1;BatCoV_RaTG13_orf1ab;4035 | 0.3 | 0.3 |
| MN996532.1;QHR63299.1;BatCoV_RaTG13_orf1ab;4036 | 0.35 | 0.4 |
| MN996532.1;QHR63299.1;BatCoV_RaTG13_orf1ab;4037 | 0.25 | 0.25 |
| MN996532.1;QHR63299.1;BatCoV_RaTG13_orf1ab;4038 | 0.4 | 0.5 |
| MN996532.1;QHR63299.1;BatCoV_RaTG13_orf1ab;4039 | 0.1 | 0.025 |
| MN996532.1;QHR63299.1;BatCoV_RaTG13_orf1ab;4040 | 0.05 | 0.025 |
| MN996532.1;QHR63299.1;BatCoV_RaTG13_orf1ab;4041 | 0 | 0 |
| MN996532.1;QHR63299.1;BatCoV_RaTG13_orf1ab;4042 | 0.05 | 0.1 |
| MN996532.1;QHR63299.1;BatCoV_RaTG13_orf1ab;4043 | 0.1 | 0.075 |
| MN996532.1;QHR63299.1;BatCoV_RaTG13_orf1ab;4044 | 0 | 0.075 |
| MN996532.1;QHR63299.1;BatCoV_RaTG13_orf1ab;4045 | 0.05 | 0.075 |
| MN996532.1;QHR63299.1;BatCoV_RaTG13_orf1ab;4046 | 0.05 | 0 |
| MN996532.1;QHR63299.1;BatCoV_RaTG13_orf1ab;4047 | 0 | 0.05 |
| MN996532.1;QHR63299.1;BatCoV_RaTG13_orf1ab;4048 | 0 | 0 |
| MN996532.1;QHR63299.1;BatCoV_RaTG13_orf1ab;4049 | 0 | 0 |
| MN996532.1;QHR63299.1;BatCoV_RaTG13_orf1ab;4050 | 0 | 0 |
| MN996532.1;QHR63299.1;BatCoV_RaTG13_orf1ab;4051 | 0 | 0 |
| MN996532.1;QHR63299.1;BatCoV_RaTG13_orf1ab;4052 | 0.05 | 0.025 |
| MN996532.1;QHR63299.1;BatCoV_RaTG13_orf1ab;4053 | 0.05 | 0.025 |
| MN996532.1;QHR63299.1;BatCoV_RaTG13_orf1ab;4054 | 0 | 0 |
| MN996532.1;QHR63299.1;BatCoV_RaTG13_orf1ab;4055 | 0 | 0 |
| MN996532.1;QHR63299.1;BatCoV_RaTG13_orf1ab;4056 | 0 | 0 |
| MN996532.1;QHR63299.1;BatCoV_RaTG13_orf1ab;4057 | 0 | 0 |
| MN996532.1;QHR63299.1;BatCoV_RaTG13_orf1ab;4058 | 0 | 0 |
| MN996532.1;QHR63299.1;BatCoV_RaTG13_orf1ab;4059 | 0 | 0 |
| MN996532.1;QHR63299.1;BatCoV_RaTG13_orf1ab;4060 | 0 | 0 |
| MN996532.1;QHR63299.1;BatCoV_RaTG13_orf1ab;4061 | 0 | 0.025 |
| MN996532.1;QHR63299.1;BatCoV_RaTG13_orf1ab;4062 | 0.05 | 0 |
| MN996532.1;QHR63299.1;BatCoV_RaTG13_orf1ab;4063 | 0 | 0.025 |
| MN996532.1;QHR63299.1;BatCoV_RaTG13_orf1ab;4064 | 0 | 0 |
| MN996532.1;QHR63299.1;BatCoV_RaTG13_orf1ab;4065 | 0 | 0 |
| MN996532.1;QHR63299.1;BatCoV_RaTG13_orf1ab;4066 | 0 | 0 |
| MN996532.1;QHR63299.1;BatCoV_RaTG13_orf1ab;4067 | 0 | 0.025 |
| MN996532.1;QHR63299.1;BatCoV_RaTG13_orf1ab;4068 | 0 | 0 |
| MN996532.1;QHR63299.1;BatCoV_RaTG13_orf1ab;4069 | 0.1 | 0 |
| MN996532.1;QHR63299.1;BatCoV_RaTG13_orf1ab;4070 | 0.1 | 0.025 |
| MN996532.1;QHR63299.1;BatCoV_RaTG13_orf1ab;4071 | 0.2 | 0.025 |
| MN996532.1;QHR63299.1;BatCoV_RaTG13_orf1ab;4072 | 0.05 | 0.025 |
| MN996532.1;QHR63299.1;BatCoV_RaTG13_orf1ab;4073 | 0.1 | 0.05 |
| MN996532.1;QHR63299.1;BatCoV_RaTG13_orf1ab;4074 | 0.05 | 0.025 |
| MN996532.1;QHR63299.1;BatCoV_RaTG13_orf1ab;4075 | 0.05 | 0 |
| MN996532.1;QHR63299.1;BatCoV_RaTG13_orf1ab;4076 | 0 | 0 |

| | | |
|---|---|---|
| MN996532.1;QHR63299.1;BatCoV_RaTG13_orf1ab;4077 | 0 | 0 |
| MN996532.1;QHR63299.1;BatCoV_RaTG13_orf1ab;4078 | 0 | 0 |
| MN996532.1;QHR63299.1;BatCoV_RaTG13_orf1ab;4079 | 0 | 0 |
| MN996532.1;QHR63299.1;BatCoV_RaTG13_orf1ab;4080 | 0 | 0 |
| MN996532.1;QHR63299.1;BatCoV_RaTG13_orf1ab;4081 | 0 | 0 |
| MN996532.1;QHR63299.1;BatCoV_RaTG13_orf1ab;4082 | 0 | 0 |
| MN996532.1;QHR63299.1;BatCoV_RaTG13_orf1ab;4083 | 0 | 0 |
| MN996532.1;QHR63299.1;BatCoV_RaTG13_orf1ab;4084 | 0 | 0 |
| MN996532.1;QHR63299.1;BatCoV_RaTG13_orf1ab;4085 | 0 | 0 |
| MN996532.1;QHR63299.1;BatCoV_RaTG13_orf1ab;4086 | 0 | 0 |
| MN996532.1;QHR63299.1;BatCoV_RaTG13_orf1ab;4087 | 0 | 0 |
| MN996532.1;QHR63299.1;BatCoV_RaTG13_orf1ab;4088 | 0 | 0 |
| MN996532.1;QHR63299.1;BatCoV_RaTG13_orf1ab;4089 | 0 | 0 |
| MN996532.1;QHR63299.1;BatCoV_RaTG13_orf1ab;4090 | 0 | 0 |
| MN996532.1;QHR63299.1;BatCoV_RaTG13_orf1ab;4091 | 0.05 | 0.025 |
| MN996532.1;QHR63299.1;BatCoV_RaTG13_orf1ab;4092 | 0.05 | 0.05 |
| MN996532.1;QHR63299.1;BatCoV_RaTG13_orf1ab;4093 | 0 | 0 |
| MN996532.1;QHR63299.1;BatCoV_RaTG13_orf1ab;4094 | 0 | 0.025 |
| MN996532.1;QHR63299.1;BatCoV_RaTG13_orf1ab;4095 | 0 | 0 |
| MN996532.1;QHR63299.1;BatCoV_RaTG13_orf1ab;4096 | 0 | 0 |
| MN996532.1;QHR63299.1;BatCoV_RaTG13_orf1ab;4097 | 0 | 0 |
| MN996532.1;QHR63299.1;BatCoV_RaTG13_orf1ab;4098 | 0 | 0.025 |
| MN996532.1;QHR63299.1;BatCoV_RaTG13_orf1ab;4099 | 0 | 0 |
| MN996532.1;QHR63299.1;BatCoV_RaTG13_orf1ab;4100 | 0 | 0 |
| MN996532.1;QHR63299.1;BatCoV_RaTG13_orf1ab;4101 | 0 | 0 |
| MN996532.1;QHR63299.1;BatCoV_RaTG13_orf1ab;4102 | 0.05 | 0 |
| MN996532.1;QHR63299.1;BatCoV_RaTG13_orf1ab;4103 | 0 | 0 |
| MN996532.1;QHR63299.1;BatCoV_RaTG13_orf1ab;4104 | 0 | 0 |
| MN996532.1;QHR63299.1;BatCoV_RaTG13_orf1ab;4105 | 0 | 0 |
| MN996532.1;QHR63299.1;BatCoV_RaTG13_orf1ab;4106 | 0 | 0 |
| MN996532.1;QHR63299.1;BatCoV_RaTG13_orf1ab;4107 | 0 | 0.025 |
| MN996532.1;QHR63299.1;BatCoV_RaTG13_orf1ab;4108 | 0 | 0 |
| MN996532.1;QHR63299.1;BatCoV_RaTG13_orf1ab;4109 | 0 | 0 |
| MN996532.1;QHR63299.1;BatCoV_RaTG13_orf1ab;4110 | 0 | 0 |
| MN996532.1;QHR63299.1;BatCoV_RaTG13_orf1ab;4111 | 0 | 0 |
| MN996532.1;QHR63299.1;BatCoV_RaTG13_orf1ab;4112 | 0 | 0 |
| MN996532.1;QHR63299.1;BatCoV_RaTG13_orf1ab;4113 | 0 | 0 |
| MN996532.1;QHR63299.1;BatCoV_RaTG13_orf1ab;4114 | 0 | 0 |
| MN996532.1;QHR63299.1;BatCoV_RaTG13_orf1ab;4115 | 0 | 0 |
| MN996532.1;QHR63299.1;BatCoV_RaTG13_orf1ab;4116 | 0 | 0 |
| MN996532.1;QHR63299.1;BatCoV_RaTG13_orf1ab;4117 | 0 | 0 |
| MN996532.1;QHR63299.1;BatCoV_RaTG13_orf1ab;4118 | 0 | 0 |
| MN996532.1;QHR63299.1;BatCoV_RaTG13_orf1ab;4119 | 0 | 0 |

| | | |
|---|---|---|
| MN996532.1;QHR63299.1;BatCoV_RaTG13_orf1ab;4120 | 0 | 0 |
| MN996532.1;QHR63299.1;BatCoV_RaTG13_orf1ab;4121 | 0 | 0 |
| MN996532.1;QHR63299.1;BatCoV_RaTG13_orf1ab;4122 | 0 | 0 |
| MN996532.1;QHR63299.1;BatCoV_RaTG13_orf1ab;4123 | 0 | 0 |
| MN996532.1;QHR63299.1;BatCoV_RaTG13_orf1ab;4124 | 0 | 0 |
| MN996532.1;QHR63299.1;BatCoV_RaTG13_orf1ab;4125 | 0 | 0 |
| MN996532.1;QHR63299.1;BatCoV_RaTG13_orf1ab;4126 | 0 | 0.025 |
| MN996532.1;QHR63299.1;BatCoV_RaTG13_orf1ab;4127 | 0 | 0.075 |
| MN996532.1;QHR63299.1;BatCoV_RaTG13_orf1ab;4128 | 0 | 0.05 |
| MN996532.1;QHR63299.1;BatCoV_RaTG13_orf1ab;4129 | 0 | 0.025 |
| MN996532.1;QHR63299.1;BatCoV_RaTG13_orf1ab;4130 | 0 | 0.025 |
| MN996532.1;QHR63299.1;BatCoV_RaTG13_orf1ab;4131 | 0.05 | 0.025 |
| MN996532.1;QHR63299.1;BatCoV_RaTG13_orf1ab;4132 | 0.1 | 0.025 |
| MN996532.1;QHR63299.1;BatCoV_RaTG13_orf1ab;4133 | 0.05 | 0.025 |
| MN996532.1;QHR63299.1;BatCoV_RaTG13_orf1ab;4134 | 0 | 0.025 |
| MN996532.1;QHR63299.1;BatCoV_RaTG13_orf1ab;4135 | 0 | 0 |
| MN996532.1;QHR63299.1;BatCoV_RaTG13_orf1ab;4136 | 0.05 | 0.025 |
| MN996532.1;QHR63299.1;BatCoV_RaTG13_orf1ab;4137 | 0 | 0 |
| MN996532.1;QHR63299.1;BatCoV_RaTG13_orf1ab;4138 | 0 | 0 |
| MN996532.1;QHR63299.1;BatCoV_RaTG13_orf1ab;4139 | 0 | 0.025 |
| MN996532.1;QHR63299.1;BatCoV_RaTG13_orf1ab;4140 | 0.05 | 0.05 |
| MN996532.1;QHR63299.1;BatCoV_RaTG13_orf1ab;4141 | 0 | 0 |
| MN996532.1;QHR63299.1;BatCoV_RaTG13_orf1ab;4142 | 0 | 0 |
| MN996532.1;QHR63299.1;BatCoV_RaTG13_orf1ab;4143 | 0 | 0 |
| MN996532.1;QHR63299.1;BatCoV_RaTG13_orf1ab;4144 | 0 | 0 |
| MN996532.1;QHR63299.1;BatCoV_RaTG13_orf1ab;4145 | 0 | 0 |
| MN996532.1;QHR63299.1;BatCoV_RaTG13_orf1ab;4146 | 0 | 0 |
| MN996532.1;QHR63299.1;BatCoV_RaTG13_orf1ab;4147 | 0 | 0 |
| MN996532.1;QHR63299.1;BatCoV_RaTG13_orf1ab;4148 | 0 | 0 |
| MN996532.1;QHR63299.1;BatCoV_RaTG13_orf1ab;4149 | 0 | 0 |
| MN996532.1;QHR63299.1;BatCoV_RaTG13_orf1ab;4150 | 0 | 0 |
| MN996532.1;QHR63299.1;BatCoV_RaTG13_orf1ab;4151 | 0 | 0 |
| MN996532.1;QHR63299.1;BatCoV_RaTG13_orf1ab;4152 | 0 | 0 |
| MN996532.1;QHR63299.1;BatCoV_RaTG13_orf1ab;4153 | 0 | 0 |
| MN996532.1;QHR63299.1;BatCoV_RaTG13_orf1ab;4154 | 0 | 0 |
| MN996532.1;QHR63299.1;BatCoV_RaTG13_orf1ab;4155 | 0 | 0 |
| MN996532.1;QHR63299.1;BatCoV_RaTG13_orf1ab;4156 | 0 | 0 |
| MN996532.1;QHR63299.1;BatCoV_RaTG13_orf1ab;4157 | 0 | 0 |
| MN996532.1;QHR63299.1;BatCoV_RaTG13_orf1ab;4158 | 0 | 0 |
| MN996532.1;QHR63299.1;BatCoV_RaTG13_orf1ab;4159 | 0 | 0 |
| MN996532.1;QHR63299.1;BatCoV_RaTG13_orf1ab;4160 | 0.05 | 0.05 |
| MN996532.1;QHR63299.1;BatCoV_RaTG13_orf1ab;4161 | 0.1 | 0.05 |
| MN996532.1;QHR63299.1;BatCoV_RaTG13_orf1ab;4162 | 0.2 | 0.05 |

| | | |
|---|---|---|
| MN996532.1;QHR63299.1;BatCoV_RaTG13_orf1ab;4163 | 0 | 0 |
| MN996532.1;QHR63299.1;BatCoV_RaTG13_orf1ab;4164 | 0 | 0 |
| MN996532.1;QHR63299.1;BatCoV_RaTG13_orf1ab;4165 | 0 | 0 |
| MN996532.1;QHR63299.1;BatCoV_RaTG13_orf1ab;4166 | 0 | 0 |
| MN996532.1;QHR63299.1;BatCoV_RaTG13_orf1ab;4167 | 0 | 0 |
| MN996532.1;QHR63299.1;BatCoV_RaTG13_orf1ab;4168 | 0 | 0 |
| MN996532.1;QHR63299.1;BatCoV_RaTG13_orf1ab;4169 | 0 | 0 |
| MN996532.1;QHR63299.1;BatCoV_RaTG13_orf1ab;4170 | 0 | 0 |
| MN996532.1;QHR63299.1;BatCoV_RaTG13_orf1ab;4171 | 0 | 0 |
| MN996532.1;QHR63299.1;BatCoV_RaTG13_orf1ab;4172 | 0 | 0 |
| MN996532.1;QHR63299.1;BatCoV_RaTG13_orf1ab;4173 | 0 | 0 |
| MN996532.1;QHR63299.1;BatCoV_RaTG13_orf1ab;4174 | 0 | 0 |
| MN996532.1;QHR63299.1;BatCoV_RaTG13_orf1ab;4175 | 0 | 0 |
| MN996532.1;QHR63299.1;BatCoV_RaTG13_orf1ab;4176 | 0 | 0 |
| MN996532.1;QHR63299.1;BatCoV_RaTG13_orf1ab;4177 | 0 | 0 |
| MN996532.1;QHR63299.1;BatCoV_RaTG13_orf1ab;4178 | 0 | 0 |
| MN996532.1;QHR63299.1;BatCoV_RaTG13_orf1ab;4179 | 0 | 0 |
| MN996532.1;QHR63299.1;BatCoV_RaTG13_orf1ab;4180 | 0 | 0 |
| MN996532.1;QHR63299.1;BatCoV_RaTG13_orf1ab;4181 | 0.05 | 0 |
| MN996532.1;QHR63299.1;BatCoV_RaTG13_orf1ab;4182 | 0 | 0 |
| MN996532.1;QHR63299.1;BatCoV_RaTG13_orf1ab;4183 | 0.05 | 0 |
| MN996532.1;QHR63299.1;BatCoV_RaTG13_orf1ab;4184 | 0.05 | 0.05 |
| MN996532.1;QHR63299.1;BatCoV_RaTG13_orf1ab;4185 | 0.15 | 0.1 |
| MN996532.1;QHR63299.1;BatCoV_RaTG13_orf1ab;4186 | 0.3 | 0.25 |
| MN996532.1;QHR63299.1;BatCoV_RaTG13_orf1ab;4187 | 0.15 | 0.05 |
| MN996532.1;QHR63299.1;BatCoV_RaTG13_orf1ab;4188 | 0 | 0 |
| MN996532.1;QHR63299.1;BatCoV_RaTG13_orf1ab;4189 | 0 | 0 |
| MN996532.1;QHR63299.1;BatCoV_RaTG13_orf1ab;4190 | 0 | 0 |
| MN996532.1;QHR63299.1;BatCoV_RaTG13_orf1ab;4191 | 0 | 0 |
| MN996532.1;QHR63299.1;BatCoV_RaTG13_orf1ab;4192 | 0 | 0 |
| MN996532.1;QHR63299.1;BatCoV_RaTG13_orf1ab;4193 | 0 | 0 |
| MN996532.1;QHR63299.1;BatCoV_RaTG13_orf1ab;4194 | 0 | 0 |
| MN996532.1;QHR63299.1;BatCoV_RaTG13_orf1ab;4195 | 0 | 0 |
| MN996532.1;QHR63299.1;BatCoV_RaTG13_orf1ab;4196 | 0 | 0 |
| MN996532.1;QHR63299.1;BatCoV_RaTG13_orf1ab;4197 | 0 | 0 |
| MN996532.1;QHR63299.1;BatCoV_RaTG13_orf1ab;4198 | 0 | 0 |
| MN996532.1;QHR63299.1;BatCoV_RaTG13_orf1ab;4199 | 0 | 0 |
| MN996532.1;QHR63299.1;BatCoV_RaTG13_orf1ab;4200 | 0 | 0.025 |
| MN996532.1;QHR63299.1;BatCoV_RaTG13_orf1ab;4201 | 0 | 0 |
| MN996532.1;QHR63299.1;BatCoV_RaTG13_orf1ab;4202 | 0.2 | 0.15 |
| MN996532.1;QHR63299.1;BatCoV_RaTG13_orf1ab;4203 | 0 | 0 |
| MN996532.1;QHR63299.1;BatCoV_RaTG13_orf1ab;4204 | 0.05 | 0.15 |
| MN996532.1;QHR63299.1;BatCoV_RaTG13_orf1ab;4205 | 0.05 | 0.075 |

| | | |
|---|---|---|
| MN996532.1;QHR63299.1;BatCoV_RaTG13_orf1ab;4206 | 0.05 | 0.05 |
| MN996532.1;QHR63299.1;BatCoV_RaTG13_orf1ab;4207 | 0.05 | 0.025 |
| MN996532.1;QHR63299.1;BatCoV_RaTG13_orf1ab;4208 | 0 | 0 |
| MN996532.1;QHR63299.1;BatCoV_RaTG13_orf1ab;4209 | 0 | 0.025 |
| MN996532.1;QHR63299.1;BatCoV_RaTG13_orf1ab;4210 | 0 | 0 |
| MN996532.1;QHR63299.1;BatCoV_RaTG13_orf1ab;4211 | 0 | 0 |
| MN996532.1;QHR63299.1;BatCoV_RaTG13_orf1ab;4212 | 0 | 0 |
| MN996532.1;QHR63299.1;BatCoV_RaTG13_orf1ab;4213 | 0 | 0 |
| MN996532.1;QHR63299.1;BatCoV_RaTG13_orf1ab;4214 | 0.05 | 0 |
| MN996532.1;QHR63299.1;BatCoV_RaTG13_orf1ab;4215 | 0 | 0 |
| MN996532.1;QHR63299.1;BatCoV_RaTG13_orf1ab;4216 | 0 | 0 |
| MN996532.1;QHR63299.1;BatCoV_RaTG13_orf1ab;4217 | 0 | 0 |
| MN996532.1;QHR63299.1;BatCoV_RaTG13_orf1ab;4218 | 0 | 0 |
| MN996532.1;QHR63299.1;BatCoV_RaTG13_orf1ab;4219 | 0 | 0 |
| MN996532.1;QHR63299.1;BatCoV_RaTG13_orf1ab;4220 | 0 | 0 |
| MN996532.1;QHR63299.1;BatCoV_RaTG13_orf1ab;4221 | 0 | 0 |
| MN996532.1;QHR63299.1;BatCoV_RaTG13_orf1ab;4222 | 0 | 0 |
| MN996532.1;QHR63299.1;BatCoV_RaTG13_orf1ab;4223 | 0 | 0 |
| MN996532.1;QHR63299.1;BatCoV_RaTG13_orf1ab;4224 | 0 | 0 |
| MN996532.1;QHR63299.1;BatCoV_RaTG13_orf1ab;4225 | 0 | 0 |
| MN996532.1;QHR63299.1;BatCoV_RaTG13_orf1ab;4226 | 0 | 0 |
| MN996532.1;QHR63299.1;BatCoV_RaTG13_orf1ab;4227 | 0 | 0 |
| MN996532.1;QHR63299.1;BatCoV_RaTG13_orf1ab;4228 | 0.05 | 0.025 |
| MN996532.1;QHR63299.1;BatCoV_RaTG13_orf1ab;4229 | 0.1 | 0.075 |
| MN996532.1;QHR63299.1;BatCoV_RaTG13_orf1ab;4230 | 0.15 | 0.1 |
| MN996532.1;QHR63299.1;BatCoV_RaTG13_orf1ab;4231 | 0.25 | 0.1 |
| MN996532.1;QHR63299.1;BatCoV_RaTG13_orf1ab;4232 | 0.1 | 0.1 |
| MN996532.1;QHR63299.1;BatCoV_RaTG13_orf1ab;4233 | 0.05 | 0.05 |
| MN996532.1;QHR63299.1;BatCoV_RaTG13_orf1ab;4234 | 0.05 | 0.025 |
| MN996532.1;QHR63299.1;BatCoV_RaTG13_orf1ab;4235 | 0 | 0 |
| MN996532.1;QHR63299.1;BatCoV_RaTG13_orf1ab;4236 | 0 | 0 |
| MN996532.1;QHR63299.1;BatCoV_RaTG13_orf1ab;4237 | 0 | 0 |
| MN996532.1;QHR63299.1;BatCoV_RaTG13_orf1ab;4238 | 0 | 0 |
| MN996532.1;QHR63299.1;BatCoV_RaTG13_orf1ab;4239 | 0 | 0 |
| MN996532.1;QHR63299.1;BatCoV_RaTG13_orf1ab;4240 | 0 | 0 |
| MN996532.1;QHR63299.1;BatCoV_RaTG13_orf1ab;4241 | 0 | 0 |
| MN996532.1;QHR63299.1;BatCoV_RaTG13_orf1ab;4242 | 0 | 0 |
| MN996532.1;QHR63299.1;BatCoV_RaTG13_orf1ab;4243 | 0 | 0.025 |
| MN996532.1;QHR63299.1;BatCoV_RaTG13_orf1ab;4244 | 0 | 0 |
| MN996532.1;QHR63299.1;BatCoV_RaTG13_orf1ab;4245 | 0 | 0.025 |
| MN996532.1;QHR63299.1;BatCoV_RaTG13_orf1ab;4246 | 0.05 | 0.025 |
| MN996532.1;QHR63299.1;BatCoV_RaTG13_orf1ab;4247 | 0 | 0.025 |
| MN996532.1;QHR63299.1;BatCoV_RaTG13_orf1ab;4248 | 0 | 0 |

| | | |
|---|---|---|
| MN996532.1;QHR63299.1;BatCoV_RaTG13_orf1ab;4249 | 0 | 0 |
| MN996532.1;QHR63299.1;BatCoV_RaTG13_orf1ab;4250 | 0 | 0.025 |
| MN996532.1;QHR63299.1;BatCoV_RaTG13_orf1ab;4251 | 0 | 0.025 |
| MN996532.1;QHR63299.1;BatCoV_RaTG13_orf1ab;4252 | 0 | 0 |
| MN996532.1;QHR63299.1;BatCoV_RaTG13_orf1ab;4253 | 0 | 0 |
| MN996532.1;QHR63299.1;BatCoV_RaTG13_orf1ab;4254 | 0 | 0 |
| MN996532.1;QHR63299.1;BatCoV_RaTG13_orf1ab;4255 | 0 | 0 |
| MN996532.1;QHR63299.1;BatCoV_RaTG13_orf1ab;4256 | 0 | 0 |
| MN996532.1;QHR63299.1;BatCoV_RaTG13_orf1ab;4257 | 0 | 0 |
| MN996532.1;QHR63299.1;BatCoV_RaTG13_orf1ab;4258 | 0 | 0 |
| MN996532.1;QHR63299.1;BatCoV_RaTG13_orf1ab;4259 | 0 | 0 |
| MN996532.1;QHR63299.1;BatCoV_RaTG13_orf1ab;4260 | 0 | 0 |
| MN996532.1;QHR63299.1;BatCoV_RaTG13_orf1ab;4261 | 0 | 0 |
| MN996532.1;QHR63299.1;BatCoV_RaTG13_orf1ab;4262 | 0 | 0 |
| MN996532.1;QHR63299.1;BatCoV_RaTG13_orf1ab;4263 | 0 | 0 |
| MN996532.1;QHR63299.1;BatCoV_RaTG13_orf1ab;4264 | 0 | 0 |
| MN996532.1;QHR63299.1;BatCoV_RaTG13_orf1ab;4265 | 0 | 0 |
| MN996532.1;QHR63299.1;BatCoV_RaTG13_orf1ab;4266 | 0 | 0 |
| MN996532.1;QHR63299.1;BatCoV_RaTG13_orf1ab;4267 | 0.05 | 0 |
| MN996532.1;QHR63299.1;BatCoV_RaTG13_orf1ab;4268 | 0 | 0 |
| MN996532.1;QHR63299.1;BatCoV_RaTG13_orf1ab;4269 | 0 | 0 |
| MN996532.1;QHR63299.1;BatCoV_RaTG13_orf1ab;4270 | 0.05 | 0 |
| MN996532.1;QHR63299.1;BatCoV_RaTG13_orf1ab;4271 | 0.05 | 0 |
| MN996532.1;QHR63299.1;BatCoV_RaTG13_orf1ab;4272 | 0.2 | 0.05 |
| MN996532.1;QHR63299.1;BatCoV_RaTG13_orf1ab;4273 | 0.15 | 0.025 |
| MN996532.1;QHR63299.1;BatCoV_RaTG13_orf1ab;4274 | 0.1 | 0.025 |
| MN996532.1;QHR63299.1;BatCoV_RaTG13_orf1ab;4275 | 0 | 0.025 |
| MN996532.1;QHR63299.1;BatCoV_RaTG13_orf1ab;4276 | 0 | 0.05 |
| MN996532.1;QHR63299.1;BatCoV_RaTG13_orf1ab;4277 | 0 | 0.05 |
| MN996532.1;QHR63299.1;BatCoV_RaTG13_orf1ab;4278 | 0.15 | 0.05 |
| MN996532.1;QHR63299.1;BatCoV_RaTG13_orf1ab;4279 | 0.05 | 0.05 |
| MN996532.1;QHR63299.1;BatCoV_RaTG13_orf1ab;4280 | 0 | 0 |
| MN996532.1;QHR63299.1;BatCoV_RaTG13_orf1ab;4281 | 0 | 0 |
| MN996532.1;QHR63299.1;BatCoV_RaTG13_orf1ab;4282 | 0 | 0 |
| MN996532.1;QHR63299.1;BatCoV_RaTG13_orf1ab;4283 | 0 | 0 |
| MN996532.1;QHR63299.1;BatCoV_RaTG13_orf1ab;4284 | 0 | 0 |
| MN996532.1;QHR63299.1;BatCoV_RaTG13_orf1ab;4285 | 0 | 0 |
| MN996532.1;QHR63299.1;BatCoV_RaTG13_orf1ab;4286 | 0 | 0 |
| MN996532.1;QHR63299.1;BatCoV_RaTG13_orf1ab;4287 | 0 | 0 |
| MN996532.1;QHR63299.1;BatCoV_RaTG13_orf1ab;4288 | 0 | 0.025 |
| MN996532.1;QHR63299.1;BatCoV_RaTG13_orf1ab;4289 | 0 | 0 |
| MN996532.1;QHR63299.1;BatCoV_RaTG13_orf1ab;4290 | 0 | 0.025 |
| MN996532.1;QHR63299.1;BatCoV_RaTG13_orf1ab;4291 | 0 | 0.025 |

| | | |
|---|---|---|
| MN996532.1;QHR63299.1;BatCoV_RaTG13_orf1ab;4292 | 0 | 0 |
| MN996532.1;QHR63299.1;BatCoV_RaTG13_orf1ab;4293 | 0 | 0 |
| MN996532.1;QHR63299.1;BatCoV_RaTG13_orf1ab;4294 | 0 | 0 |
| MN996532.1;QHR63299.1;BatCoV_RaTG13_orf1ab;4295 | 0 | 0 |
| MN996532.1;QHR63299.1;BatCoV_RaTG13_orf1ab;4296 | 0 | 0 |
| MN996532.1;QHR63299.1;BatCoV_RaTG13_orf1ab;4297 | 0 | 0 |
| MN996532.1;QHR63299.1;BatCoV_RaTG13_orf1ab;4298 | 0 | 0 |
| MN996532.1;QHR63299.1;BatCoV_RaTG13_orf1ab;4299 | 0 | 0 |
| MN996532.1;QHR63299.1;BatCoV_RaTG13_orf1ab;4300 | 0 | 0 |
| MN996532.1;QHR63299.1;BatCoV_RaTG13_orf1ab;4301 | 0 | 0.025 |
| MN996532.1;QHR63299.1;BatCoV_RaTG13_orf1ab;4302 | 0 | 0.025 |
| MN996532.1;QHR63299.1;BatCoV_RaTG13_orf1ab;4303 | 0 | 0.025 |
| MN996532.1;QHR63299.1;BatCoV_RaTG13_orf1ab;4304 | 0 | 0.025 |
| MN996532.1;QHR63299.1;BatCoV_RaTG13_orf1ab;4305 | 0 | 0.025 |
| MN996532.1;QHR63299.1;BatCoV_RaTG13_orf1ab;4306 | 0 | 0.025 |
| MN996532.1;QHR63299.1;BatCoV_RaTG13_orf1ab;4307 | 0.05 | 0.025 |
| MN996532.1;QHR63299.1;BatCoV_RaTG13_orf1ab;4308 | 0 | 0.025 |
| MN996532.1;QHR63299.1;BatCoV_RaTG13_orf1ab;4309 | 0.05 | 0.025 |
| MN996532.1;QHR63299.1;BatCoV_RaTG13_orf1ab;4310 | 0 | 0 |
| MN996532.1;QHR63299.1;BatCoV_RaTG13_orf1ab;4311 | 0 | 0 |
| MN996532.1;QHR63299.1;BatCoV_RaTG13_orf1ab;4312 | 0 | 0 |
| MN996532.1;QHR63299.1;BatCoV_RaTG13_orf1ab;4313 | 0 | 0 |
| MN996532.1;QHR63299.1;BatCoV_RaTG13_orf1ab;4314 | 0 | 0 |
| MN996532.1;QHR63299.1;BatCoV_RaTG13_orf1ab;4315 | 0 | 0 |
| MN996532.1;QHR63299.1;BatCoV_RaTG13_orf1ab;4316 | 0 | 0 |
| MN996532.1;QHR63299.1;BatCoV_RaTG13_orf1ab;4317 | 0 | 0 |
| MN996532.1;QHR63299.1;BatCoV_RaTG13_orf1ab;4318 | 0 | 0 |
| MN996532.1;QHR63299.1;BatCoV_RaTG13_orf1ab;4319 | 0 | 0 |
| MN996532.1;QHR63299.1;BatCoV_RaTG13_orf1ab;4320 | 0 | 0 |
| MN996532.1;QHR63299.1;BatCoV_RaTG13_orf1ab;4321 | 0 | 0 |
| MN996532.1;QHR63299.1;BatCoV_RaTG13_orf1ab;4322 | 0 | 0 |
| MN996532.1;QHR63299.1;BatCoV_RaTG13_orf1ab;4323 | 0 | 0 |
| MN996532.1;QHR63299.1;BatCoV_RaTG13_orf1ab;4324 | 0 | 0 |
| MN996532.1;QHR63299.1;BatCoV_RaTG13_orf1ab;4325 | 0 | 0 |
| MN996532.1;QHR63299.1;BatCoV_RaTG13_orf1ab;4326 | 0 | 0 |
| MN996532.1;QHR63299.1;BatCoV_RaTG13_orf1ab;4327 | 0 | 0 |
| MN996532.1;QHR63299.1;BatCoV_RaTG13_orf1ab;4328 | 0 | 0.025 |
| MN996532.1;QHR63299.1;BatCoV_RaTG13_orf1ab;4329 | 0 | 0.025 |
| MN996532.1;QHR63299.1;BatCoV_RaTG13_orf1ab;4330 | 0 | 0.025 |
| MN996532.1;QHR63299.1;BatCoV_RaTG13_orf1ab;4331 | 0 | 0.025 |
| MN996532.1;QHR63299.1;BatCoV_RaTG13_orf1ab;4332 | 0 | 0.025 |
| MN996532.1;QHR63299.1;BatCoV_RaTG13_orf1ab;4333 | 0 | 0.025 |
| MN996532.1;QHR63299.1;BatCoV_RaTG13_orf1ab;4334 | 0 | 0 |

| | | |
|---|---|---|
| MN996532.1;QHR63299.1;BatCoV_RaTG13_orf1ab;4335 | 0 | 0 |
| MN996532.1;QHR63299.1;BatCoV_RaTG13_orf1ab;4336 | 0 | 0 |
| MN996532.1;QHR63299.1;BatCoV_RaTG13_orf1ab;4337 | 0 | 0 |
| MN996532.1;QHR63299.1;BatCoV_RaTG13_orf1ab;4338 | 0 | 0 |
| MN996532.1;QHR63299.1;BatCoV_RaTG13_orf1ab;4339 | 0 | 0.05 |
| MN996532.1;QHR63299.1;BatCoV_RaTG13_orf1ab;4340 | 0 | 0.05 |
| MN996532.1;QHR63299.1;BatCoV_RaTG13_orf1ab;4341 | 0 | 0 |
| MN996532.1;QHR63299.1;BatCoV_RaTG13_orf1ab;4342 | 0 | 0 |
| MN996532.1;QHR63299.1;BatCoV_RaTG13_orf1ab;4343 | 0.05 | 0 |
| MN996532.1;QHR63299.1;BatCoV_RaTG13_orf1ab;4344 | 0.05 | 0.025 |
| MN996532.1;QHR63299.1;BatCoV_RaTG13_orf1ab;4345 | 0.05 | 0.05 |
| MN996532.1;QHR63299.1;BatCoV_RaTG13_orf1ab;4346 | 0.05 | 0 |
| MN996532.1;QHR63299.1;BatCoV_RaTG13_orf1ab;4347 | 0.05 | 0 |
| MN996532.1;QHR63299.1;BatCoV_RaTG13_orf1ab;4348 | 0 | 0 |
| MN996532.1;QHR63299.1;BatCoV_RaTG13_orf1ab;4349 | 0 | 0.025 |
| MN996532.1;QHR63299.1;BatCoV_RaTG13_orf1ab;4350 | 0 | 0.025 |
| MN996532.1;QHR63299.1;BatCoV_RaTG13_orf1ab;4351 | 0 | 0.025 |
| MN996532.1;QHR63299.1;BatCoV_RaTG13_orf1ab;4352 | 0.05 | 0.05 |
| MN996532.1;QHR63299.1;BatCoV_RaTG13_orf1ab;4353 | 0 | 0.025 |
| MN996532.1;QHR63299.1;BatCoV_RaTG13_orf1ab;4354 | 0 | 0.025 |
| MN996532.1;QHR63299.1;BatCoV_RaTG13_orf1ab;4355 | 0 | 0.025 |
| MN996532.1;QHR63299.1;BatCoV_RaTG13_orf1ab;4356 | 0 | 0.025 |
| MN996532.1;QHR63299.1;BatCoV_RaTG13_orf1ab;4357 | 0 | 0 |
| MN996532.1;QHR63299.1;BatCoV_RaTG13_orf1ab;4358 | 0 | 0 |
| MN996532.1;QHR63299.1;BatCoV_RaTG13_orf1ab;4359 | 0 | 0 |
| MN996532.1;QHR63299.1;BatCoV_RaTG13_orf1ab;4360 | 0 | 0 |
| MN996532.1;QHR63299.1;BatCoV_RaTG13_orf1ab;4361 | 0 | 0 |
| MN996532.1;QHR63299.1;BatCoV_RaTG13_orf1ab;4362 | 0 | 0 |
| MN996532.1;QHR63299.1;BatCoV_RaTG13_orf1ab;4363 | 0 | 0 |
| MN996532.1;QHR63299.1;BatCoV_RaTG13_orf1ab;4364 | 0 | 0 |
| MN996532.1;QHR63299.1;BatCoV_RaTG13_orf1ab;4365 | 0 | 0 |
| MN996532.1;QHR63299.1;BatCoV_RaTG13_orf1ab;4366 | 0 | 0 |
| MN996532.1;QHR63299.1;BatCoV_RaTG13_orf1ab;4367 | 0 | 0 |
| MN996532.1;QHR63299.1;BatCoV_RaTG13_orf1ab;4368 | 0 | 0 |
| MN996532.1;QHR63299.1;BatCoV_RaTG13_orf1ab;4369 | 0 | 0 |
| MN996532.1;QHR63299.1;BatCoV_RaTG13_orf1ab;4370 | 0 | 0 |
| MN996532.1;QHR63299.1;BatCoV_RaTG13_orf1ab;4371 | 0 | 0 |
| MN996532.1;QHR63299.1;BatCoV_RaTG13_orf1ab;4372 | 0 | 0 |
| MN996532.1;QHR63299.1;BatCoV_RaTG13_orf1ab;4373 | 0 | 0 |
| MN996532.1;QHR63299.1;BatCoV_RaTG13_orf1ab;4374 | 0 | 0.075 |
| MN996532.1;QHR63299.1;BatCoV_RaTG13_orf1ab;4375 | 0 | 0 |
| MN996532.1;QHR63299.1;BatCoV_RaTG13_orf1ab;4376 | 0 | 0.025 |
| MN996532.1;QHR63299.1;BatCoV_RaTG13_orf1ab;4377 | 0 | 0 |

| | | |
|---|---|---|
| MN996532.1;QHR63299.1;BatCoV_RaTG13_orf1ab;4378 | 0 | 0 |
| MN996532.1;QHR63299.1;BatCoV_RaTG13_orf1ab;4379 | 0 | 0 |
| MN996532.1;QHR63299.1;BatCoV_RaTG13_orf1ab;4380 | 0 | 0 |
| MN996532.1;QHR63299.1;BatCoV_RaTG13_orf1ab;4381 | 0 | 0 |
| MN996532.1;QHR63299.1;BatCoV_RaTG13_orf1ab;4382 | 0 | 0 |
| MN996532.1;QHR63299.1;BatCoV_RaTG13_orf1ab;4383 | 0 | 0 |
| MN996532.1;QHR63299.1;BatCoV_RaTG13_orf1ab;4384 | 0 | 0 |
| MN996532.1;QHR63299.1;BatCoV_RaTG13_orf1ab;4385 | 0.05 | 0 |
| MN996532.1;QHR63299.1;BatCoV_RaTG13_orf1ab;4386 | 0.05 | 0.025 |
| MN996532.1;QHR63299.1;BatCoV_RaTG13_orf1ab;4387 | 0.15 | 0.025 |
| MN996532.1;QHR63299.1;BatCoV_RaTG13_orf1ab;4388 | 0 | 0 |
| MN996532.1;QHR63299.1;BatCoV_RaTG13_orf1ab;4389 | 0.05 | 0 |
| MN996532.1;QHR63299.1;BatCoV_RaTG13_orf1ab;4390 | 0 | 0 |
| MN996532.1;QHR63299.1;BatCoV_RaTG13_orf1ab;4391 | 0 | 0 |
| MN996532.1;QHR63299.1;BatCoV_RaTG13_orf1ab;4392 | 0 | 0 |
| MN996532.1;QHR63299.1;BatCoV_RaTG13_orf1ab;4393 | 0.05 | 0 |
| MN996532.1;QHR63299.1;BatCoV_RaTG13_orf1ab;4394 | 0 | 0 |
| MN996532.1;QHR63299.1;BatCoV_RaTG13_orf1ab;4395 | 0 | 0 |
| MN996532.1;QHR63299.1;BatCoV_RaTG13_orf1ab;4396 | 0 | 0 |
| MN996532.1;QHR63299.1;BatCoV_RaTG13_orf1ab;4397 | 0 | 0 |
| MN996532.1;QHR63299.1;BatCoV_RaTG13_orf1ab;4398 | 0 | 0 |
| MN996532.1;QHR63299.1;BatCoV_RaTG13_orf1ab;4399 | 0 | 0 |
| MN996532.1;QHR63299.1;BatCoV_RaTG13_orf1ab;4400 | 0 | 0 |
| MN996532.1;QHR63299.1;BatCoV_RaTG13_orf1ab;4401 | 0 | 0 |
| MN996532.1;QHR63299.1;BatCoV_RaTG13_orf1ab;4402 | 0 | 0 |
| MN996532.1;QHR63299.1;BatCoV_RaTG13_orf1ab;4403 | 0 | 0 |
| MN996532.1;QHR63299.1;BatCoV_RaTG13_orf1ab;4404 | 0 | 0 |
| MN996532.1;QHR63299.1;BatCoV_RaTG13_orf1ab;4405 | 0 | 0 |
| MN996532.1;QHR63299.1;BatCoV_RaTG13_orf1ab;4406 | 0 | 0.025 |
| MN996532.1;QHR63299.1;BatCoV_RaTG13_orf1ab;4407 | 0 | 0 |
| MN996532.1;QHR63299.1;BatCoV_RaTG13_orf1ab;4408 | 0 | 0 |
| MN996532.1;QHR63299.1;BatCoV_RaTG13_orf1ab;4409 | 0 | 0 |
| MN996532.1;QHR63299.1;BatCoV_RaTG13_orf1ab;4410 | 0 | 0 |
| MN996532.1;QHR63299.1;BatCoV_RaTG13_orf1ab;4411 | 0 | 0 |
| MN996532.1;QHR63299.1;BatCoV_RaTG13_orf1ab;4412 | 0 | 0 |
| MN996532.1;QHR63299.1;BatCoV_RaTG13_orf1ab;4413 | 0 | 0 |
| MN996532.1;QHR63299.1;BatCoV_RaTG13_orf1ab;4414 | 0 | 0 |
| MN996532.1;QHR63299.1;BatCoV_RaTG13_orf1ab;4415 | 0 | 0 |
| MN996532.1;QHR63299.1;BatCoV_RaTG13_orf1ab;4416 | 0 | 0 |
| MN996532.1;QHR63299.1;BatCoV_RaTG13_orf1ab;4417 | 0 | 0.025 |
| MN996532.1;QHR63299.1;BatCoV_RaTG13_orf1ab;4418 | 0.1 | 0 |
| MN996532.1;QHR63299.1;BatCoV_RaTG13_orf1ab;4419 | 0.05 | 0.05 |
| MN996532.1;QHR63299.1;BatCoV_RaTG13_orf1ab;4420 | 0.05 | 0 |

| | | |
|---|---|---|
| MN996532.1;QHR63299.1;BatCoV_RaTG13_orf1ab;4421 | 0 | 0 |
| MN996532.1;QHR63299.1;BatCoV_RaTG13_orf1ab;4422 | 0.05 | 0 |
| MN996532.1;QHR63299.1;BatCoV_RaTG13_orf1ab;4423 | 0 | 0.025 |
| MN996532.1;QHR63299.1;BatCoV_RaTG13_orf1ab;4424 | 0 | 0 |
| MN996532.1;QHR63299.1;BatCoV_RaTG13_orf1ab;4425 | 0 | 0 |
| MN996532.1;QHR63299.1;BatCoV_RaTG13_orf1ab;4426 | 0 | 0 |
| MN996532.1;QHR63299.1;BatCoV_RaTG13_orf1ab;4427 | 0 | 0 |
| MN996532.1;QHR63299.1;BatCoV_RaTG13_orf1ab;4428 | 0 | 0 |
| MN996532.1;QHR63299.1;BatCoV_RaTG13_orf1ab;4429 | 0 | 0 |
| MN996532.1;QHR63299.1;BatCoV_RaTG13_orf1ab;4430 | 0 | 0 |
| MN996532.1;QHR63299.1;BatCoV_RaTG13_orf1ab;4431 | 0 | 0 |
| MN996532.1;QHR63299.1;BatCoV_RaTG13_orf1ab;4432 | 0 | 0 |
| MN996532.1;QHR63299.1;BatCoV_RaTG13_orf1ab;4433 | 0 | 0 |
| MN996532.1;QHR63299.1;BatCoV_RaTG13_orf1ab;4434 | 0 | 0 |
| MN996532.1;QHR63299.1;BatCoV_RaTG13_orf1ab;4435 | 0 | 0 |
| MN996532.1;QHR63299.1;BatCoV_RaTG13_orf1ab;4436 | 0 | 0 |
| MN996532.1;QHR63299.1;BatCoV_RaTG13_orf1ab;4437 | 0 | 0.025 |
| MN996532.1;QHR63299.1;BatCoV_RaTG13_orf1ab;4438 | 0 | 0 |
| MN996532.1;QHR63299.1;BatCoV_RaTG13_orf1ab;4439 | 0 | 0 |
| MN996532.1;QHR63299.1;BatCoV_RaTG13_orf1ab;4440 | 0 | 0.025 |
| MN996532.1;QHR63299.1;BatCoV_RaTG13_orf1ab;4441 | 0 | 0 |
| MN996532.1;QHR63299.1;BatCoV_RaTG13_orf1ab;4442 | 0 | 0 |
| MN996532.1;QHR63299.1;BatCoV_RaTG13_orf1ab;4443 | 0 | 0.05 |
| MN996532.1;QHR63299.1;BatCoV_RaTG13_orf1ab;4444 | 0 | 0.05 |
| MN996532.1;QHR63299.1;BatCoV_RaTG13_orf1ab;4445 | 0.05 | 0.025 |
| MN996532.1;QHR63299.1;BatCoV_RaTG13_orf1ab;4446 | 0 | 0.05 |
| MN996532.1;QHR63299.1;BatCoV_RaTG13_orf1ab;4447 | 0 | 0.025 |
| MN996532.1;QHR63299.1;BatCoV_RaTG13_orf1ab;4448 | 0 | 0 |
| MN996532.1;QHR63299.1;BatCoV_RaTG13_orf1ab;4449 | 0 | 0.025 |
| MN996532.1;QHR63299.1;BatCoV_RaTG13_orf1ab;4450 | 0 | 0.025 |
| MN996532.1;QHR63299.1;BatCoV_RaTG13_orf1ab;4451 | 0 | 0.025 |
| MN996532.1;QHR63299.1;BatCoV_RaTG13_orf1ab;4452 | 0 | 0 |
| MN996532.1;QHR63299.1;BatCoV_RaTG13_orf1ab;4453 | 0 | 0 |
| MN996532.1;QHR63299.1;BatCoV_RaTG13_orf1ab;4454 | 0 | 0 |
| MN996532.1;QHR63299.1;BatCoV_RaTG13_orf1ab;4455 | 0 | 0 |
| MN996532.1;QHR63299.1;BatCoV_RaTG13_orf1ab;4456 | 0 | 0 |
| MN996532.1;QHR63299.1;BatCoV_RaTG13_orf1ab;4457 | 0 | 0 |
| MN996532.1;QHR63299.1;BatCoV_RaTG13_orf1ab;4458 | 0 | 0 |
| MN996532.1;QHR63299.1;BatCoV_RaTG13_orf1ab;4459 | 0 | 0 |
| MN996532.1;QHR63299.1;BatCoV_RaTG13_orf1ab;4460 | 0 | 0 |
| MN996532.1;QHR63299.1;BatCoV_RaTG13_orf1ab;4461 | 0 | 0 |
| MN996532.1;QHR63299.1;BatCoV_RaTG13_orf1ab;4462 | 0 | 0 |
| MN996532.1;QHR63299.1;BatCoV_RaTG13_orf1ab;4463 | 0 | 0 |

| | | |
|---|---|---|
| MN996532.1;QHR63299.1;BatCoV_RaTG13_orf1ab;4464 | 0 | 0 |
| MN996532.1;QHR63299.1;BatCoV_RaTG13_orf1ab;4465 | 0 | 0 |
| MN996532.1;QHR63299.1;BatCoV_RaTG13_orf1ab;4466 | 0 | 0 |
| MN996532.1;QHR63299.1;BatCoV_RaTG13_orf1ab;4467 | 0 | 0 |
| MN996532.1;QHR63299.1;BatCoV_RaTG13_orf1ab;4468 | 0.05 | 0.05 |
| MN996532.1;QHR63299.1;BatCoV_RaTG13_orf1ab;4469 | 0 | 0 |
| MN996532.1;QHR63299.1;BatCoV_RaTG13_orf1ab;4470 | 0 | 0 |
| MN996532.1;QHR63299.1;BatCoV_RaTG13_orf1ab;4471 | 0 | 0 |
| MN996532.1;QHR63299.1;BatCoV_RaTG13_orf1ab;4472 | 0 | 0 |
| MN996532.1;QHR63299.1;BatCoV_RaTG13_orf1ab;4473 | 0 | 0 |
| MN996532.1;QHR63299.1;BatCoV_RaTG13_orf1ab;4474 | 0 | 0 |
| MN996532.1;QHR63299.1;BatCoV_RaTG13_orf1ab;4475 | 0 | 0 |
| MN996532.1;QHR63299.1;BatCoV_RaTG13_orf1ab;4476 | 0 | 0.025 |
| MN996532.1;QHR63299.1;BatCoV_RaTG13_orf1ab;4477 | 0 | 0 |
| MN996532.1;QHR63299.1;BatCoV_RaTG13_orf1ab;4478 | 0 | 0.025 |
| MN996532.1;QHR63299.1;BatCoV_RaTG13_orf1ab;4479 | 0 | 0 |
| MN996532.1;QHR63299.1;BatCoV_RaTG13_orf1ab;4480 | 0 | 0.025 |
| MN996532.1;QHR63299.1;BatCoV_RaTG13_orf1ab;4481 | 0 | 0 |
| MN996532.1;QHR63299.1;BatCoV_RaTG13_orf1ab;4482 | 0 | 0 |
| MN996532.1;QHR63299.1;BatCoV_RaTG13_orf1ab;4483 | 0 | 0.025 |
| MN996532.1;QHR63299.1;BatCoV_RaTG13_orf1ab;4484 | 0 | 0.025 |
| MN996532.1;QHR63299.1;BatCoV_RaTG13_orf1ab;4485 | 0 | 0.05 |
| MN996532.1;QHR63299.1;BatCoV_RaTG13_orf1ab;4486 | 0 | 0.05 |
| MN996532.1;QHR63299.1;BatCoV_RaTG13_orf1ab;4487 | 0 | 0.05 |
| MN996532.1;QHR63299.1;BatCoV_RaTG13_orf1ab;4488 | 0 | 0.05 |
| MN996532.1;QHR63299.1;BatCoV_RaTG13_orf1ab;4489 | 0 | 0.025 |
| MN996532.1;QHR63299.1;BatCoV_RaTG13_orf1ab;4490 | 0 | 0.025 |
| MN996532.1;QHR63299.1;BatCoV_RaTG13_orf1ab;4491 | 0 | 0.025 |
| MN996532.1;QHR63299.1;BatCoV_RaTG13_orf1ab;4492 | 0 | 0 |
| MN996532.1;QHR63299.1;BatCoV_RaTG13_orf1ab;4493 | 0 | 0 |
| MN996532.1;QHR63299.1;BatCoV_RaTG13_orf1ab;4494 | 0 | 0 |
| MN996532.1;QHR63299.1;BatCoV_RaTG13_orf1ab;4495 | 0 | 0 |
| MN996532.1;QHR63299.1;BatCoV_RaTG13_orf1ab;4496 | 0 | 0 |
| MN996532.1;QHR63299.1;BatCoV_RaTG13_orf1ab;4497 | 0 | 0 |
| MN996532.1;QHR63299.1;BatCoV_RaTG13_orf1ab;4498 | 0 | 0 |
| MN996532.1;QHR63299.1;BatCoV_RaTG13_orf1ab;4499 | 0 | 0 |
| MN996532.1;QHR63299.1;BatCoV_RaTG13_orf1ab;4500 | 0 | 0 |
| MN996532.1;QHR63299.1;BatCoV_RaTG13_orf1ab;4501 | 0 | 0 |
| MN996532.1;QHR63299.1;BatCoV_RaTG13_orf1ab;4502 | 0 | 0 |
| MN996532.1;QHR63299.1;BatCoV_RaTG13_orf1ab;4503 | 0 | 0 |
| MN996532.1;QHR63299.1;BatCoV_RaTG13_orf1ab;4504 | 0 | 0 |
| MN996532.1;QHR63299.1;BatCoV_RaTG13_orf1ab;4505 | 0 | 0 |
| MN996532.1;QHR63299.1;BatCoV_RaTG13_orf1ab;4506 | 0 | 0 |

| | | |
|---|---|---|
| MN996532.1;QHR63299.1;BatCoV_RaTG13_orf1ab;4507 | 0 | 0 |
| MN996532.1;QHR63299.1;BatCoV_RaTG13_orf1ab;4508 | 0 | 0 |
| MN996532.1;QHR63299.1;BatCoV_RaTG13_orf1ab;4509 | 0 | 0 |
| MN996532.1;QHR63299.1;BatCoV_RaTG13_orf1ab;4510 | 0 | 0 |
| MN996532.1;QHR63299.1;BatCoV_RaTG13_orf1ab;4511 | 0 | 0 |
| MN996532.1;QHR63299.1;BatCoV_RaTG13_orf1ab;4512 | 0 | 0 |
| MN996532.1;QHR63299.1;BatCoV_RaTG13_orf1ab;4513 | 0 | 0.375 |
| MN996532.1;QHR63299.1;BatCoV_RaTG13_orf1ab;4514 | 0 | 0 |
| MN996532.1;QHR63299.1;BatCoV_RaTG13_orf1ab;4515 | 0 | 0 |
| MN996532.1;QHR63299.1;BatCoV_RaTG13_orf1ab;4516 | 0 | 0 |
| MN996532.1;QHR63299.1;BatCoV_RaTG13_orf1ab;4517 | 0 | 0 |
| MN996532.1;QHR63299.1;BatCoV_RaTG13_orf1ab;4518 | 0 | 0 |
| MN996532.1;QHR63299.1;BatCoV_RaTG13_orf1ab;4519 | 0 | 0.05 |
| MN996532.1;QHR63299.1;BatCoV_RaTG13_orf1ab;4520 | 0 | 0 |
| MN996532.1;QHR63299.1;BatCoV_RaTG13_orf1ab;4521 | 0 | 0 |
| MN996532.1;QHR63299.1;BatCoV_RaTG13_orf1ab;4522 | 0 | 0 |
| MN996532.1;QHR63299.1;BatCoV_RaTG13_orf1ab;4523 | 0 | 0 |
| MN996532.1;QHR63299.1;BatCoV_RaTG13_orf1ab;4524 | 0 | 0 |
| MN996532.1;QHR63299.1;BatCoV_RaTG13_orf1ab;4525 | 0 | 0 |
| MN996532.1;QHR63299.1;BatCoV_RaTG13_orf1ab;4526 | 0 | 0 |
| MN996532.1;QHR63299.1;BatCoV_RaTG13_orf1ab;4527 | 0 | 0 |
| MN996532.1;QHR63299.1;BatCoV_RaTG13_orf1ab;4528 | 0 | 0 |
| MN996532.1;QHR63299.1;BatCoV_RaTG13_orf1ab;4529 | 0 | 0 |
| MN996532.1;QHR63299.1;BatCoV_RaTG13_orf1ab;4530 | 0 | 0 |
| MN996532.1;QHR63299.1;BatCoV_RaTG13_orf1ab;4531 | 0 | 0 |
| MN996532.1;QHR63299.1;BatCoV_RaTG13_orf1ab;4532 | 0 | 0.025 |
| MN996532.1;QHR63299.1;BatCoV_RaTG13_orf1ab;4533 | 0 | 0 |
| MN996532.1;QHR63299.1;BatCoV_RaTG13_orf1ab;4534 | 0 | 0 |
| MN996532.1;QHR63299.1;BatCoV_RaTG13_orf1ab;4535 | 0 | 0 |
| MN996532.1;QHR63299.1;BatCoV_RaTG13_orf1ab;4536 | 0 | 0 |
| MN996532.1;QHR63299.1;BatCoV_RaTG13_orf1ab;4537 | 0 | 0.025 |
| MN996532.1;QHR63299.1;BatCoV_RaTG13_orf1ab;4538 | 0 | 0 |
| MN996532.1;QHR63299.1;BatCoV_RaTG13_orf1ab;4539 | 0 | 0 |
| MN996532.1;QHR63299.1;BatCoV_RaTG13_orf1ab;4540 | 0 | 0.025 |
| MN996532.1;QHR63299.1;BatCoV_RaTG13_orf1ab;4541 | 0 | 0.025 |
| MN996532.1;QHR63299.1;BatCoV_RaTG13_orf1ab;4542 | 0 | 0 |
| MN996532.1;QHR63299.1;BatCoV_RaTG13_orf1ab;4543 | 0 | 0 |
| MN996532.1;QHR63299.1;BatCoV_RaTG13_orf1ab;4544 | 0.2 | 0.025 |
| MN996532.1;QHR63299.1;BatCoV_RaTG13_orf1ab;4545 | 0.05 | 0 |
| MN996532.1;QHR63299.1;BatCoV_RaTG13_orf1ab;4546 | 0 | 0.025 |
| MN996532.1;QHR63299.1;BatCoV_RaTG13_orf1ab;4547 | 0 | 0.075 |
| MN996532.1;QHR63299.1;BatCoV_RaTG13_orf1ab;4548 | 0.05 | 0.075 |
| MN996532.1;QHR63299.1;BatCoV_RaTG13_orf1ab;4549 | 0.05 | 0.075 |

| | | |
|---|---|---|
| MN996532.1;QHR63299.1;BatCoV_RaTG13_orf1ab;4550 | 0.1 | 0.05 |
| MN996532.1;QHR63299.1;BatCoV_RaTG13_orf1ab;4551 | 0 | 0.025 |
| MN996532.1;QHR63299.1;BatCoV_RaTG13_orf1ab;4552 | 0 | 0 |
| MN996532.1;QHR63299.1;BatCoV_RaTG13_orf1ab;4553 | 0 | 0 |
| MN996532.1;QHR63299.1;BatCoV_RaTG13_orf1ab;4554 | 0 | 0 |
| MN996532.1;QHR63299.1;BatCoV_RaTG13_orf1ab;4555 | 0 | 0 |
| MN996532.1;QHR63299.1;BatCoV_RaTG13_orf1ab;4556 | 0.05 | 0 |
| MN996532.1;QHR63299.1;BatCoV_RaTG13_orf1ab;4557 | 0.05 | 0.025 |
| MN996532.1;QHR63299.1;BatCoV_RaTG13_orf1ab;4558 | 0.05 | 0 |
| MN996532.1;QHR63299.1;BatCoV_RaTG13_orf1ab;4559 | 0 | 0 |
| MN996532.1;QHR63299.1;BatCoV_RaTG13_orf1ab;4560 | 0 | 0 |
| MN996532.1;QHR63299.1;BatCoV_RaTG13_orf1ab;4561 | 0 | 0 |
| MN996532.1;QHR63299.1;BatCoV_RaTG13_orf1ab;4562 | 0 | 0 |
| MN996532.1;QHR63299.1;BatCoV_RaTG13_orf1ab;4563 | 0 | 0 |
| MN996532.1;QHR63299.1;BatCoV_RaTG13_orf1ab;4564 | 0.05 | 0 |
| MN996532.1;QHR63299.1;BatCoV_RaTG13_orf1ab;4565 | 0 | 0 |
| MN996532.1;QHR63299.1;BatCoV_RaTG13_orf1ab;4566 | 0 | 0 |
| MN996532.1;QHR63299.1;BatCoV_RaTG13_orf1ab;4567 | 0 | 0 |
| MN996532.1;QHR63299.1;BatCoV_RaTG13_orf1ab;4568 | 0 | 0 |
| MN996532.1;QHR63299.1;BatCoV_RaTG13_orf1ab;4569 | 0 | 0 |
| MN996532.1;QHR63299.1;BatCoV_RaTG13_orf1ab;4570 | 0 | 0 |
| MN996532.1;QHR63299.1;BatCoV_RaTG13_orf1ab;4571 | 0.05 | 0 |
| MN996532.1;QHR63299.1;BatCoV_RaTG13_orf1ab;4572 | 0 | 0 |
| MN996532.1;QHR63299.1;BatCoV_RaTG13_orf1ab;4573 | 0 | 0 |
| MN996532.1;QHR63299.1;BatCoV_RaTG13_orf1ab;4574 | 0 | 0 |
| MN996532.1;QHR63299.1;BatCoV_RaTG13_orf1ab;4575 | 0 | 0 |
| MN996532.1;QHR63299.1;BatCoV_RaTG13_orf1ab;4576 | 0 | 0 |
| MN996532.1;QHR63299.1;BatCoV_RaTG13_orf1ab;4577 | 0 | 0 |
| MN996532.1;QHR63299.1;BatCoV_RaTG13_orf1ab;4578 | 0 | 0 |
| MN996532.1;QHR63299.1;BatCoV_RaTG13_orf1ab;4579 | 0 | 0 |
| MN996532.1;QHR63299.1;BatCoV_RaTG13_orf1ab;4580 | 0 | 0 |
| MN996532.1;QHR63299.1;BatCoV_RaTG13_orf1ab;4581 | 0 | 0 |
| MN996532.1;QHR63299.1;BatCoV_RaTG13_orf1ab;4582 | 0 | 0 |
| MN996532.1;QHR63299.1;BatCoV_RaTG13_orf1ab;4583 | 0 | 0 |
| MN996532.1;QHR63299.1;BatCoV_RaTG13_orf1ab;4584 | 0 | 0 |
| MN996532.1;QHR63299.1;BatCoV_RaTG13_orf1ab;4585 | 0 | 0 |
| MN996532.1;QHR63299.1;BatCoV_RaTG13_orf1ab;4586 | 0 | 0 |
| MN996532.1;QHR63299.1;BatCoV_RaTG13_orf1ab;4587 | 0.1 | 0.025 |
| MN996532.1;QHR63299.1;BatCoV_RaTG13_orf1ab;4588 | 0.1 | 0 |
| MN996532.1;QHR63299.1;BatCoV_RaTG13_orf1ab;4589 | 0 | 0 |
| MN996532.1;QHR63299.1;BatCoV_RaTG13_orf1ab;4590 | 0.05 | 0 |
| MN996532.1;QHR63299.1;BatCoV_RaTG13_orf1ab;4591 | 0 | 0 |
| MN996532.1;QHR63299.1;BatCoV_RaTG13_orf1ab;4592 | 0 | 0 |

| | | |
|---|---|---|
| MN996532.1;QHR63299.1;BatCoV_RaTG13_orf1ab;4593 | 0 | 0 |
| MN996532.1;QHR63299.1;BatCoV_RaTG13_orf1ab;4594 | 0.05 | 0 |
| MN996532.1;QHR63299.1;BatCoV_RaTG13_orf1ab;4595 | 0 | 0 |
| MN996532.1;QHR63299.1;BatCoV_RaTG13_orf1ab;4596 | 0 | 0 |
| MN996532.1;QHR63299.1;BatCoV_RaTG13_orf1ab;4597 | 0 | 0 |
| MN996532.1;QHR63299.1;BatCoV_RaTG13_orf1ab;4598 | 0 | 0 |
| MN996532.1;QHR63299.1;BatCoV_RaTG13_orf1ab;4599 | 0 | 0 |
| MN996532.1;QHR63299.1;BatCoV_RaTG13_orf1ab;4600 | 0 | 0 |
| MN996532.1;QHR63299.1;BatCoV_RaTG13_orf1ab;4601 | 0 | 0.025 |
| MN996532.1;QHR63299.1;BatCoV_RaTG13_orf1ab;4602 | 0 | 0 |
| MN996532.1;QHR63299.1;BatCoV_RaTG13_orf1ab;4603 | 0 | 0 |
| MN996532.1;QHR63299.1;BatCoV_RaTG13_orf1ab;4604 | 0 | 0.025 |
| MN996532.1;QHR63299.1;BatCoV_RaTG13_orf1ab;4605 | 0 | 0 |
| MN996532.1;QHR63299.1;BatCoV_RaTG13_orf1ab;4606 | 0 | 0 |
| MN996532.1;QHR63299.1;BatCoV_RaTG13_orf1ab;4607 | 0 | 0 |
| MN996532.1;QHR63299.1;BatCoV_RaTG13_orf1ab;4608 | 0.1 | 0.05 |
| MN996532.1;QHR63299.1;BatCoV_RaTG13_orf1ab;4609 | 0 | 0 |
| MN996532.1;QHR63299.1;BatCoV_RaTG13_orf1ab;4610 | 0 | 0 |
| MN996532.1;QHR63299.1;BatCoV_RaTG13_orf1ab;4611 | 0 | 0 |
| MN996532.1;QHR63299.1;BatCoV_RaTG13_orf1ab;4612 | 0 | 0 |
| MN996532.1;QHR63299.1;BatCoV_RaTG13_orf1ab;4613 | 0 | 0.025 |
| MN996532.1;QHR63299.1;BatCoV_RaTG13_orf1ab;4614 | 0 | 0 |
| MN996532.1;QHR63299.1;BatCoV_RaTG13_orf1ab;4615 | 0 | 0.05 |
| MN996532.1;QHR63299.1;BatCoV_RaTG13_orf1ab;4616 | 0 | 0 |
| MN996532.1;QHR63299.1;BatCoV_RaTG13_orf1ab;4617 | 0 | 0 |
| MN996532.1;QHR63299.1;BatCoV_RaTG13_orf1ab;4618 | 0 | 0 |
| MN996532.1;QHR63299.1;BatCoV_RaTG13_orf1ab;4619 | 0 | 0 |
| MN996532.1;QHR63299.1;BatCoV_RaTG13_orf1ab;4620 | 0 | 0 |
| MN996532.1;QHR63299.1;BatCoV_RaTG13_orf1ab;4621 | 0 | 0 |
| MN996532.1;QHR63299.1;BatCoV_RaTG13_orf1ab;4622 | 0 | 0 |
| MN996532.1;QHR63299.1;BatCoV_RaTG13_orf1ab;4623 | 0 | 0 |
| MN996532.1;QHR63299.1;BatCoV_RaTG13_orf1ab;4624 | 0 | 0 |
| MN996532.1;QHR63299.1;BatCoV_RaTG13_orf1ab;4625 | 0 | 0 |
| MN996532.1;QHR63299.1;BatCoV_RaTG13_orf1ab;4626 | 0 | 0 |
| MN996532.1;QHR63299.1;BatCoV_RaTG13_orf1ab;4627 | 0 | 0 |
| MN996532.1;QHR63299.1;BatCoV_RaTG13_orf1ab;4628 | 0 | 0 |
| MN996532.1;QHR63299.1;BatCoV_RaTG13_orf1ab;4629 | 0 | 0 |
| MN996532.1;QHR63299.1;BatCoV_RaTG13_orf1ab;4630 | 0 | 0 |
| MN996532.1;QHR63299.1;BatCoV_RaTG13_orf1ab;4631 | 0 | 0 |
| MN996532.1;QHR63299.1;BatCoV_RaTG13_orf1ab;4632 | 0 | 0 |
| MN996532.1;QHR63299.1;BatCoV_RaTG13_orf1ab;4633 | 0 | 0 |
| MN996532.1;QHR63299.1;BatCoV_RaTG13_orf1ab;4634 | 0.05 | 0 |
| MN996532.1;QHR63299.1;BatCoV_RaTG13_orf1ab;4635 | 0 | 0 |

| | | |
|---|---|---|
| MN996532.1;QHR63299.1;BatCoV_RaTG13_orf1ab;4636 | 0 | 0 |
| MN996532.1;QHR63299.1;BatCoV_RaTG13_orf1ab;4637 | 0 | 0 |
| MN996532.1;QHR63299.1;BatCoV_RaTG13_orf1ab;4638 | 0 | 0 |
| MN996532.1;QHR63299.1;BatCoV_RaTG13_orf1ab;4639 | 0 | 0 |
| MN996532.1;QHR63299.1;BatCoV_RaTG13_orf1ab;4640 | 0.05 | 0 |
| MN996532.1;QHR63299.1;BatCoV_RaTG13_orf1ab;4641 | 0.05 | 0 |
| MN996532.1;QHR63299.1;BatCoV_RaTG13_orf1ab;4642 | 0.05 | 0 |
| MN996532.1;QHR63299.1;BatCoV_RaTG13_orf1ab;4643 | 0.05 | 0 |
| MN996532.1;QHR63299.1;BatCoV_RaTG13_orf1ab;4644 | 0 | 0.025 |
| MN996532.1;QHR63299.1;BatCoV_RaTG13_orf1ab;4645 | 0.05 | 0.075 |
| MN996532.1;QHR63299.1;BatCoV_RaTG13_orf1ab;4646 | 0 | 0.025 |
| MN996532.1;QHR63299.1;BatCoV_RaTG13_orf1ab;4647 | 0 | 0 |
| MN996532.1;QHR63299.1;BatCoV_RaTG13_orf1ab;4648 | 0.05 | 0.05 |
| MN996532.1;QHR63299.1;BatCoV_RaTG13_orf1ab;4649 | 0.1 | 0 |
| MN996532.1;QHR63299.1;BatCoV_RaTG13_orf1ab;4650 | 0 | 0 |
| MN996532.1;QHR63299.1;BatCoV_RaTG13_orf1ab;4651 | 0 | 0 |
| MN996532.1;QHR63299.1;BatCoV_RaTG13_orf1ab;4652 | 0 | 0 |
| MN996532.1;QHR63299.1;BatCoV_RaTG13_orf1ab;4653 | 0 | 0 |
| MN996532.1;QHR63299.1;BatCoV_RaTG13_orf1ab;4654 | 0 | 0 |
| MN996532.1;QHR63299.1;BatCoV_RaTG13_orf1ab;4655 | 0.05 | 0 |
| MN996532.1;QHR63299.1;BatCoV_RaTG13_orf1ab;4656 | 0.05 | 0 |
| MN996532.1;QHR63299.1;BatCoV_RaTG13_orf1ab;4657 | 0.1 | 0 |
| MN996532.1;QHR63299.1;BatCoV_RaTG13_orf1ab;4658 | 0.05 | 0.05 |
| MN996532.1;QHR63299.1;BatCoV_RaTG13_orf1ab;4659 | 0.1 | 0.025 |
| MN996532.1;QHR63299.1;BatCoV_RaTG13_orf1ab;4660 | 0.05 | 0.025 |
| MN996532.1;QHR63299.1;BatCoV_RaTG13_orf1ab;4661 | 0.05 | 0.05 |
| MN996532.1;QHR63299.1;BatCoV_RaTG13_orf1ab;4662 | 0.05 | 0.075 |
| MN996532.1;QHR63299.1;BatCoV_RaTG13_orf1ab;4663 | 0.05 | 0.175 |
| MN996532.1;QHR63299.1;BatCoV_RaTG13_orf1ab;4664 | 0.05 | 0.125 |
| MN996532.1;QHR63299.1;BatCoV_RaTG13_orf1ab;4665 | 0 | 0 |
| MN996532.1;QHR63299.1;BatCoV_RaTG13_orf1ab;4666 | 0 | 0 |
| MN996532.1;QHR63299.1;BatCoV_RaTG13_orf1ab;4667 | 0 | 0 |
| MN996532.1;QHR63299.1;BatCoV_RaTG13_orf1ab;4668 | 0 | 0 |
| MN996532.1;QHR63299.1;BatCoV_RaTG13_orf1ab;4669 | 0 | 0.025 |
| MN996532.1;QHR63299.1;BatCoV_RaTG13_orf1ab;4670 | 0 | 0 |
| MN996532.1;QHR63299.1;BatCoV_RaTG13_orf1ab;4671 | 0 | 0 |
| MN996532.1;QHR63299.1;BatCoV_RaTG13_orf1ab;4672 | 0 | 0 |
| MN996532.1;QHR63299.1;BatCoV_RaTG13_orf1ab;4673 | 0 | 0.025 |
| MN996532.1;QHR63299.1;BatCoV_RaTG13_orf1ab;4674 | 0 | 0 |
| MN996532.1;QHR63299.1;BatCoV_RaTG13_orf1ab;4675 | 0 | 0.025 |
| MN996532.1;QHR63299.1;BatCoV_RaTG13_orf1ab;4676 | 0 | 0.025 |
| MN996532.1;QHR63299.1;BatCoV_RaTG13_orf1ab;4677 | 0 | 0.05 |
| MN996532.1;QHR63299.1;BatCoV_RaTG13_orf1ab;4678 | 0 | 0.075 |

| | | |
|---|---|---|
| MN996532.1;QHR63299.1;BatCoV_RaTG13_orf1ab;4679 | 0 | 0.1 |
| MN996532.1;QHR63299.1;BatCoV_RaTG13_orf1ab;4680 | 0 | 0.05 |
| MN996532.1;QHR63299.1;BatCoV_RaTG13_orf1ab;4681 | 0 | 0.025 |
| MN996532.1;QHR63299.1;BatCoV_RaTG13_orf1ab;4682 | 0 | 0.05 |
| MN996532.1;QHR63299.1;BatCoV_RaTG13_orf1ab;4683 | 0 | 0.05 |
| MN996532.1;QHR63299.1;BatCoV_RaTG13_orf1ab;4684 | 0 | 0.025 |
| MN996532.1;QHR63299.1;BatCoV_RaTG13_orf1ab;4685 | 0 | 0.025 |
| MN996532.1;QHR63299.1;BatCoV_RaTG13_orf1ab;4686 | 0 | 0.025 |
| MN996532.1;QHR63299.1;BatCoV_RaTG13_orf1ab;4687 | 0 | 0 |
| MN996532.1;QHR63299.1;BatCoV_RaTG13_orf1ab;4688 | 0 | 0 |
| MN996532.1;QHR63299.1;BatCoV_RaTG13_orf1ab;4689 | 0 | 0 |
| MN996532.1;QHR63299.1;BatCoV_RaTG13_orf1ab;4690 | 0 | 0 |
| MN996532.1;QHR63299.1;BatCoV_RaTG13_orf1ab;4691 | 0 | 0 |
| MN996532.1;QHR63299.1;BatCoV_RaTG13_orf1ab;4692 | 0 | 0 |
| MN996532.1;QHR63299.1;BatCoV_RaTG13_orf1ab;4693 | 0 | 0 |
| MN996532.1;QHR63299.1;BatCoV_RaTG13_orf1ab;4694 | 0 | 0 |
| MN996532.1;QHR63299.1;BatCoV_RaTG13_orf1ab;4695 | 0 | 0 |
| MN996532.1;QHR63299.1;BatCoV_RaTG13_orf1ab;4696 | 0 | 0 |
| MN996532.1;QHR63299.1;BatCoV_RaTG13_orf1ab;4697 | 0 | 0 |
| MN996532.1;QHR63299.1;BatCoV_RaTG13_orf1ab;4698 | 0 | 0 |
| MN996532.1;QHR63299.1;BatCoV_RaTG13_orf1ab;4699 | 0 | 0 |
| MN996532.1;QHR63299.1;BatCoV_RaTG13_orf1ab;4700 | 0 | 0 |
| MN996532.1;QHR63299.1;BatCoV_RaTG13_orf1ab;4701 | 0 | 0 |
| MN996532.1;QHR63299.1;BatCoV_RaTG13_orf1ab;4702 | 0 | 0 |
| MN996532.1;QHR63299.1;BatCoV_RaTG13_orf1ab;4703 | 0 | 0 |
| MN996532.1;QHR63299.1;BatCoV_RaTG13_orf1ab;4704 | 0 | 0 |
| MN996532.1;QHR63299.1;BatCoV_RaTG13_orf1ab;4705 | 0 | 0 |
| MN996532.1;QHR63299.1;BatCoV_RaTG13_orf1ab;4706 | 0 | 0 |
| MN996532.1;QHR63299.1;BatCoV_RaTG13_orf1ab;4707 | 0 | 0 |
| MN996532.1;QHR63299.1;BatCoV_RaTG13_orf1ab;4708 | 0 | 0 |
| MN996532.1;QHR63299.1;BatCoV_RaTG13_orf1ab;4709 | 0 | 0 |
| MN996532.1;QHR63299.1;BatCoV_RaTG13_orf1ab;4710 | 0 | 0 |
| MN996532.1;QHR63299.1;BatCoV_RaTG13_orf1ab;4711 | 0 | 0 |
| MN996532.1;QHR63299.1;BatCoV_RaTG13_orf1ab;4712 | 0 | 0 |
| MN996532.1;QHR63299.1;BatCoV_RaTG13_orf1ab;4713 | 0 | 0 |
| MN996532.1;QHR63299.1;BatCoV_RaTG13_orf1ab;4714 | 0 | 0 |
| MN996532.1;QHR63299.1;BatCoV_RaTG13_orf1ab;4715 | 0 | 0 |
| MN996532.1;QHR63299.1;BatCoV_RaTG13_orf1ab;4716 | 0 | 0 |
| MN996532.1;QHR63299.1;BatCoV_RaTG13_orf1ab;4717 | 0 | 0 |
| MN996532.1;QHR63299.1;BatCoV_RaTG13_orf1ab;4718 | 0 | 0 |
| MN996532.1;QHR63299.1;BatCoV_RaTG13_orf1ab;4719 | 0 | 0 |
| MN996532.1;QHR63299.1;BatCoV_RaTG13_orf1ab;4720 | 0 | 0 |
| MN996532.1;QHR63299.1;BatCoV_RaTG13_orf1ab;4721 | 0 | 0 |

| | | |
|---|---|---|
| MN996532.1;QHR63299.1;BatCoV_RaTG13_orf1ab;4722 | 0 | 0 |
| MN996532.1;QHR63299.1;BatCoV_RaTG13_orf1ab;4723 | 0 | 0 |
| MN996532.1;QHR63299.1;BatCoV_RaTG13_orf1ab;4724 | 0 | 0 |
| MN996532.1;QHR63299.1;BatCoV_RaTG13_orf1ab;4725 | 0 | 0 |
| MN996532.1;QHR63299.1;BatCoV_RaTG13_orf1ab;4726 | 0 | 0 |
| MN996532.1;QHR63299.1;BatCoV_RaTG13_orf1ab;4727 | 0 | 0 |
| MN996532.1;QHR63299.1;BatCoV_RaTG13_orf1ab;4728 | 0 | 0 |
| MN996532.1;QHR63299.1;BatCoV_RaTG13_orf1ab;4729 | 0 | 0 |
| MN996532.1;QHR63299.1;BatCoV_RaTG13_orf1ab;4730 | 0 | 0 |
| MN996532.1;QHR63299.1;BatCoV_RaTG13_orf1ab;4731 | 0 | 0 |
| MN996532.1;QHR63299.1;BatCoV_RaTG13_orf1ab;4732 | 0 | 0 |
| MN996532.1;QHR63299.1;BatCoV_RaTG13_orf1ab;4733 | 0 | 0 |
| MN996532.1;QHR63299.1;BatCoV_RaTG13_orf1ab;4734 | 0 | 0 |
| MN996532.1;QHR63299.1;BatCoV_RaTG13_orf1ab;4735 | 0 | 0 |
| MN996532.1;QHR63299.1;BatCoV_RaTG13_orf1ab;4736 | 0 | 0 |
| MN996532.1;QHR63299.1;BatCoV_RaTG13_orf1ab;4737 | 0 | 0 |
| MN996532.1;QHR63299.1;BatCoV_RaTG13_orf1ab;4738 | 0 | 0 |
| MN996532.1;QHR63299.1;BatCoV_RaTG13_orf1ab;4739 | 0 | 0 |
| MN996532.1;QHR63299.1;BatCoV_RaTG13_orf1ab;4740 | 0 | 0 |
| MN996532.1;QHR63299.1;BatCoV_RaTG13_orf1ab;4741 | 0 | 0 |
| MN996532.1;QHR63299.1;BatCoV_RaTG13_orf1ab;4742 | 0 | 0 |
| MN996532.1;QHR63299.1;BatCoV_RaTG13_orf1ab;4743 | 0 | 0 |
| MN996532.1;QHR63299.1;BatCoV_RaTG13_orf1ab;4744 | 0 | 0 |
| MN996532.1;QHR63299.1;BatCoV_RaTG13_orf1ab;4745 | 0 | 0 |
| MN996532.1;QHR63299.1;BatCoV_RaTG13_orf1ab;4746 | 0 | 0.025 |
| MN996532.1;QHR63299.1;BatCoV_RaTG13_orf1ab;4747 | 0 | 0.075 |
| MN996532.1;QHR63299.1;BatCoV_RaTG13_orf1ab;4748 | 0.05 | 0.025 |
| MN996532.1;QHR63299.1;BatCoV_RaTG13_orf1ab;4749 | 0 | 0 |
| MN996532.1;QHR63299.1;BatCoV_RaTG13_orf1ab;4750 | 0 | 0 |
| MN996532.1;QHR63299.1;BatCoV_RaTG13_orf1ab;4751 | 0 | 0 |
| MN996532.1;QHR63299.1;BatCoV_RaTG13_orf1ab;4752 | 0 | 0 |
| MN996532.1;QHR63299.1;BatCoV_RaTG13_orf1ab;4753 | 0 | 0 |
| MN996532.1;QHR63299.1;BatCoV_RaTG13_orf1ab;4754 | 0 | 0 |
| MN996532.1;QHR63299.1;BatCoV_RaTG13_orf1ab;4755 | 0 | 0 |
| MN996532.1;QHR63299.1;BatCoV_RaTG13_orf1ab;4756 | 0.05 | 0 |
| MN996532.1;QHR63299.1;BatCoV_RaTG13_orf1ab;4757 | 0.1 | 0.025 |
| MN996532.1;QHR63299.1;BatCoV_RaTG13_orf1ab;4758 | 0.1 | 0.025 |
| MN996532.1;QHR63299.1;BatCoV_RaTG13_orf1ab;4759 | 0.05 | 0.025 |
| MN996532.1;QHR63299.1;BatCoV_RaTG13_orf1ab;4760 | 0 | 0 |
| MN996532.1;QHR63299.1;BatCoV_RaTG13_orf1ab;4761 | 0 | 0 |
| MN996532.1;QHR63299.1;BatCoV_RaTG13_orf1ab;4762 | 0 | 0 |
| MN996532.1;QHR63299.1;BatCoV_RaTG13_orf1ab;4763 | 0 | 0.025 |
| MN996532.1;QHR63299.1;BatCoV_RaTG13_orf1ab;4764 | 0 | 0.025 |

| | | |
|---|---|---|
| MN996532.1;QHR63299.1;BatCoV_RaTG13_orf1ab;4765 | 0 | 0 |
| MN996532.1;QHR63299.1;BatCoV_RaTG13_orf1ab;4766 | 0 | 0 |
| MN996532.1;QHR63299.1;BatCoV_RaTG13_orf1ab;4767 | 0 | 0 |
| MN996532.1;QHR63299.1;BatCoV_RaTG13_orf1ab;4768 | 0 | 0 |
| MN996532.1;QHR63299.1;BatCoV_RaTG13_orf1ab;4769 | 0 | 0 |
| MN996532.1;QHR63299.1;BatCoV_RaTG13_orf1ab;4770 | 0 | 0 |
| MN996532.1;QHR63299.1;BatCoV_RaTG13_orf1ab;4771 | 0 | 0 |
| MN996532.1;QHR63299.1;BatCoV_RaTG13_orf1ab;4772 | 0 | 0 |
| MN996532.1;QHR63299.1;BatCoV_RaTG13_orf1ab;4773 | 0 | 0 |
| MN996532.1;QHR63299.1;BatCoV_RaTG13_orf1ab;4774 | 0 | 0 |
| MN996532.1;QHR63299.1;BatCoV_RaTG13_orf1ab;4775 | 0 | 0 |
| MN996532.1;QHR63299.1;BatCoV_RaTG13_orf1ab;4776 | 0.05 | 0 |
| MN996532.1;QHR63299.1;BatCoV_RaTG13_orf1ab;4777 | 0 | 0 |
| MN996532.1;QHR63299.1;BatCoV_RaTG13_orf1ab;4778 | 0 | 0 |
| MN996532.1;QHR63299.1;BatCoV_RaTG13_orf1ab;4779 | 0 | 0 |
| MN996532.1;QHR63299.1;BatCoV_RaTG13_orf1ab;4780 | 0 | 0 |
| MN996532.1;QHR63299.1;BatCoV_RaTG13_orf1ab;4781 | 0 | 0 |
| MN996532.1;QHR63299.1;BatCoV_RaTG13_orf1ab;4782 | 0 | 0 |
| MN996532.1;QHR63299.1;BatCoV_RaTG13_orf1ab;4783 | 0 | 0 |
| MN996532.1;QHR63299.1;BatCoV_RaTG13_orf1ab;4784 | 0 | 0 |
| MN996532.1;QHR63299.1;BatCoV_RaTG13_orf1ab;4785 | 0 | 0 |
| MN996532.1;QHR63299.1;BatCoV_RaTG13_orf1ab;4786 | 0 | 0 |
| MN996532.1;QHR63299.1;BatCoV_RaTG13_orf1ab;4787 | 0 | 0 |
| MN996532.1;QHR63299.1;BatCoV_RaTG13_orf1ab;4788 | 0 | 0 |
| MN996532.1;QHR63299.1;BatCoV_RaTG13_orf1ab;4789 | 0 | 0 |
| MN996532.1;QHR63299.1;BatCoV_RaTG13_orf1ab;4790 | 0.05 | 0 |
| MN996532.1;QHR63299.1;BatCoV_RaTG13_orf1ab;4791 | 0.05 | 0.05 |
| MN996532.1;QHR63299.1;BatCoV_RaTG13_orf1ab;4792 | 0 | 0 |
| MN996532.1;QHR63299.1;BatCoV_RaTG13_orf1ab;4793 | 0 | 0 |
| MN996532.1;QHR63299.1;BatCoV_RaTG13_orf1ab;4794 | 0 | 0.025 |
| MN996532.1;QHR63299.1;BatCoV_RaTG13_orf1ab;4795 | 0 | 0.025 |
| MN996532.1;QHR63299.1;BatCoV_RaTG13_orf1ab;4796 | 0 | 0 |
| MN996532.1;QHR63299.1;BatCoV_RaTG13_orf1ab;4797 | 0 | 0 |
| MN996532.1;QHR63299.1;BatCoV_RaTG13_orf1ab;4798 | 0.05 | 0 |
| MN996532.1;QHR63299.1;BatCoV_RaTG13_orf1ab;4799 | 0.05 | 0.025 |
| MN996532.1;QHR63299.1;BatCoV_RaTG13_orf1ab;4800 | 0.1 | 0.025 |
| MN996532.1;QHR63299.1;BatCoV_RaTG13_orf1ab;4801 | 0.1 | 0.025 |
| MN996532.1;QHR63299.1;BatCoV_RaTG13_orf1ab;4802 | 0.1 | 0.025 |
| MN996532.1;QHR63299.1;BatCoV_RaTG13_orf1ab;4803 | 0.2 | 0.025 |
| MN996532.1;QHR63299.1;BatCoV_RaTG13_orf1ab;4804 | 0.1 | 0 |
| MN996532.1;QHR63299.1;BatCoV_RaTG13_orf1ab;4805 | 0 | 0 |
| MN996532.1;QHR63299.1;BatCoV_RaTG13_orf1ab;4806 | 0 | 0 |
| MN996532.1;QHR63299.1;BatCoV_RaTG13_orf1ab;4807 | 0.05 | 0 |

| | | |
|---|---|---|
| MN996532.1;QHR63299.1;BatCoV_RaTG13_orf1ab;4808 | 0.05 | 0 |
| MN996532.1;QHR63299.1;BatCoV_RaTG13_orf1ab;4809 | 0.05 | 0 |
| MN996532.1;QHR63299.1;BatCoV_RaTG13_orf1ab;4810 | 0 | 0 |
| MN996532.1;QHR63299.1;BatCoV_RaTG13_orf1ab;4811 | 0 | 0 |
| MN996532.1;QHR63299.1;BatCoV_RaTG13_orf1ab;4812 | 0 | 0 |
| MN996532.1;QHR63299.1;BatCoV_RaTG13_orf1ab;4813 | 0 | 0 |
| MN996532.1;QHR63299.1;BatCoV_RaTG13_orf1ab;4814 | 0 | 0 |
| MN996532.1;QHR63299.1;BatCoV_RaTG13_orf1ab;4815 | 0 | 0 |
| MN996532.1;QHR63299.1;BatCoV_RaTG13_orf1ab;4816 | 0.05 | 0 |
| MN996532.1;QHR63299.1;BatCoV_RaTG13_orf1ab;4817 | 0 | 0.025 |
| MN996532.1;QHR63299.1;BatCoV_RaTG13_orf1ab;4818 | 0 | 0 |
| MN996532.1;QHR63299.1;BatCoV_RaTG13_orf1ab;4819 | 0 | 0 |
| MN996532.1;QHR63299.1;BatCoV_RaTG13_orf1ab;4820 | 0 | 0 |
| MN996532.1;QHR63299.1;BatCoV_RaTG13_orf1ab;4821 | 0 | 0 |
| MN996532.1;QHR63299.1;BatCoV_RaTG13_orf1ab;4822 | 0 | 0 |
| MN996532.1;QHR63299.1;BatCoV_RaTG13_orf1ab;4823 | 0 | 0 |
| MN996532.1;QHR63299.1;BatCoV_RaTG13_orf1ab;4824 | 0 | 0 |
| MN996532.1;QHR63299.1;BatCoV_RaTG13_orf1ab;4825 | 0.05 | 0 |
| MN996532.1;QHR63299.1;BatCoV_RaTG13_orf1ab;4826 | 0.05 | 0 |
| MN996532.1;QHR63299.1;BatCoV_RaTG13_orf1ab;4827 | 0 | 0 |
| MN996532.1;QHR63299.1;BatCoV_RaTG13_orf1ab;4828 | 0 | 0 |
| MN996532.1;QHR63299.1;BatCoV_RaTG13_orf1ab;4829 | 0.05 | 0 |
| MN996532.1;QHR63299.1;BatCoV_RaTG13_orf1ab;4830 | 0 | 0 |
| MN996532.1;QHR63299.1;BatCoV_RaTG13_orf1ab;4831 | 0 | 0 |
| MN996532.1;QHR63299.1;BatCoV_RaTG13_orf1ab;4832 | 0 | 0 |
| MN996532.1;QHR63299.1;BatCoV_RaTG13_orf1ab;4833 | 0 | 0.025 |
| MN996532.1;QHR63299.1;BatCoV_RaTG13_orf1ab;4834 | 0 | 0 |
| MN996532.1;QHR63299.1;BatCoV_RaTG13_orf1ab;4835 | 0.1 | 0.025 |
| MN996532.1;QHR63299.1;BatCoV_RaTG13_orf1ab;4836 | 0.05 | 0.025 |
| MN996532.1;QHR63299.1;BatCoV_RaTG13_orf1ab;4837 | 0.15 | 0 |
| MN996532.1;QHR63299.1;BatCoV_RaTG13_orf1ab;4838 | 0.2 | 0 |
| MN996532.1;QHR63299.1;BatCoV_RaTG13_orf1ab;4839 | 0.15 | 0.075 |
| MN996532.1;QHR63299.1;BatCoV_RaTG13_orf1ab;4840 | 0.25 | 0.025 |
| MN996532.1;QHR63299.1;BatCoV_RaTG13_orf1ab;4841 | 0.15 | 0 |
| MN996532.1;QHR63299.1;BatCoV_RaTG13_orf1ab;4842 | 0.15 | 0 |
| MN996532.1;QHR63299.1;BatCoV_RaTG13_orf1ab;4843 | 0.15 | 0 |
| MN996532.1;QHR63299.1;BatCoV_RaTG13_orf1ab;4844 | 0 | 0 |
| MN996532.1;QHR63299.1;BatCoV_RaTG13_orf1ab;4845 | 0 | 0 |
| MN996532.1;QHR63299.1;BatCoV_RaTG13_orf1ab;4846 | 0 | 0 |
| MN996532.1;QHR63299.1;BatCoV_RaTG13_orf1ab;4847 | 0 | 0 |
| MN996532.1;QHR63299.1;BatCoV_RaTG13_orf1ab;4848 | 0 | 0 |
| MN996532.1;QHR63299.1;BatCoV_RaTG13_orf1ab;4849 | 0 | 0 |
| MN996532.1;QHR63299.1;BatCoV_RaTG13_orf1ab;4850 | 0 | 0 |

| | | |
|---|---|---|
| MN996532.1;QHR63299.1;BatCoV_RaTG13_orf1ab;4851 | 0 | 0 |
| MN996532.1;QHR63299.1;BatCoV_RaTG13_orf1ab;4852 | 0 | 0 |
| MN996532.1;QHR63299.1;BatCoV_RaTG13_orf1ab;4853 | 0 | 0 |
| MN996532.1;QHR63299.1;BatCoV_RaTG13_orf1ab;4854 | 0.05 | 0.025 |
| MN996532.1;QHR63299.1;BatCoV_RaTG13_orf1ab;4855 | 0 | 0 |
| MN996532.1;QHR63299.1;BatCoV_RaTG13_orf1ab;4856 | 0 | 0 |
| MN996532.1;QHR63299.1;BatCoV_RaTG13_orf1ab;4857 | 0 | 0.025 |
| MN996532.1;QHR63299.1;BatCoV_RaTG13_orf1ab;4858 | 0 | 0 |
| MN996532.1;QHR63299.1;BatCoV_RaTG13_orf1ab;4859 | 0 | 0 |
| MN996532.1;QHR63299.1;BatCoV_RaTG13_orf1ab;4860 | 0 | 0 |
| MN996532.1;QHR63299.1;BatCoV_RaTG13_orf1ab;4861 | 0 | 0 |
| MN996532.1;QHR63299.1;BatCoV_RaTG13_orf1ab;4862 | 0 | 0 |
| MN996532.1;QHR63299.1;BatCoV_RaTG13_orf1ab;4863 | 0 | 0.025 |
| MN996532.1;QHR63299.1;BatCoV_RaTG13_orf1ab;4864 | 0 | 0.025 |
| MN996532.1;QHR63299.1;BatCoV_RaTG13_orf1ab;4865 | 0 | 0 |
| MN996532.1;QHR63299.1;BatCoV_RaTG13_orf1ab;4866 | 0 | 0 |
| MN996532.1;QHR63299.1;BatCoV_RaTG13_orf1ab;4867 | 0.05 | 0 |
| MN996532.1;QHR63299.1;BatCoV_RaTG13_orf1ab;4868 | 0.05 | 0.075 |
| MN996532.1;QHR63299.1;BatCoV_RaTG13_orf1ab;4869 | 0.05 | 0.025 |
| MN996532.1;QHR63299.1;BatCoV_RaTG13_orf1ab;4870 | 0.05 | 0 |
| MN996532.1;QHR63299.1;BatCoV_RaTG13_orf1ab;4871 | 0 | 0.025 |
| MN996532.1;QHR63299.1;BatCoV_RaTG13_orf1ab;4872 | 0.05 | 0 |
| MN996532.1;QHR63299.1;BatCoV_RaTG13_orf1ab;4873 | 0.05 | 0 |
| MN996532.1;QHR63299.1;BatCoV_RaTG13_orf1ab;4874 | 0 | 0 |
| MN996532.1;QHR63299.1;BatCoV_RaTG13_orf1ab;4875 | 0.05 | 0.025 |
| MN996532.1;QHR63299.1;BatCoV_RaTG13_orf1ab;4876 | 0 | 0.025 |
| MN996532.1;QHR63299.1;BatCoV_RaTG13_orf1ab;4877 | 0.05 | 0.025 |
| MN996532.1;QHR63299.1;BatCoV_RaTG13_orf1ab;4878 | 0.05 | 0.025 |
| MN996532.1;QHR63299.1;BatCoV_RaTG13_orf1ab;4879 | 0.1 | 0.025 |
| MN996532.1;QHR63299.1;BatCoV_RaTG13_orf1ab;4880 | 0 | 0.025 |
| MN996532.1;QHR63299.1;BatCoV_RaTG13_orf1ab;4881 | 0.15 | 0.1 |
| MN996532.1;QHR63299.1;BatCoV_RaTG13_orf1ab;4882 | 0 | 0.05 |
| MN996532.1;QHR63299.1;BatCoV_RaTG13_orf1ab;4883 | 0.15 | 0.075 |
| MN996532.1;QHR63299.1;BatCoV_RaTG13_orf1ab;4884 | 0.05 | 0.05 |
| MN996532.1;QHR63299.1;BatCoV_RaTG13_orf1ab;4885 | 0 | 0.025 |
| MN996532.1;QHR63299.1;BatCoV_RaTG13_orf1ab;4886 | 0 | 0 |
| MN996532.1;QHR63299.1;BatCoV_RaTG13_orf1ab;4887 | 0 | 0 |
| MN996532.1;QHR63299.1;BatCoV_RaTG13_orf1ab;4888 | 0 | 0 |
| MN996532.1;QHR63299.1;BatCoV_RaTG13_orf1ab;4889 | 0 | 0 |
| MN996532.1;QHR63299.1;BatCoV_RaTG13_orf1ab;4890 | 0 | 0 |
| MN996532.1;QHR63299.1;BatCoV_RaTG13_orf1ab;4891 | 0 | 0 |
| MN996532.1;QHR63299.1;BatCoV_RaTG13_orf1ab;4892 | 0 | 0 |
| MN996532.1;QHR63299.1;BatCoV_RaTG13_orf1ab;4893 | 0 | 0 |

| | | |
|---|---|---|
| MN996532.1;QHR63299.1;BatCoV_RaTG13_orf1ab;4894 | 0 | 0 |
| MN996532.1;QHR63299.1;BatCoV_RaTG13_orf1ab;4895 | 0 | 0 |
| MN996532.1;QHR63299.1;BatCoV_RaTG13_orf1ab;4896 | 0.05 | 0 |
| MN996532.1;QHR63299.1;BatCoV_RaTG13_orf1ab;4897 | 0 | 0 |
| MN996532.1;QHR63299.1;BatCoV_RaTG13_orf1ab;4898 | 0.05 | 0 |
| MN996532.1;QHR63299.1;BatCoV_RaTG13_orf1ab;4899 | 0 | 0 |
| MN996532.1;QHR63299.1;BatCoV_RaTG13_orf1ab;4900 | 0 | 0 |
| MN996532.1;QHR63299.1;BatCoV_RaTG13_orf1ab;4901 | 0 | 0 |
| MN996532.1;QHR63299.1;BatCoV_RaTG13_orf1ab;4902 | 0 | 0 |
| MN996532.1;QHR63299.1;BatCoV_RaTG13_orf1ab;4903 | 0 | 0 |
| MN996532.1;QHR63299.1;BatCoV_RaTG13_orf1ab;4904 | 0 | 0 |
| MN996532.1;QHR63299.1;BatCoV_RaTG13_orf1ab;4905 | 0 | 0 |
| MN996532.1;QHR63299.1;BatCoV_RaTG13_orf1ab;4906 | 0 | 0 |
| MN996532.1;QHR63299.1;BatCoV_RaTG13_orf1ab;4907 | 0 | 0 |
| MN996532.1;QHR63299.1;BatCoV_RaTG13_orf1ab;4908 | 0.1 | 0.025 |
| MN996532.1;QHR63299.1;BatCoV_RaTG13_orf1ab;4909 | 0.05 | 0.025 |
| MN996532.1;QHR63299.1;BatCoV_RaTG13_orf1ab;4910 | 0.05 | 0 |
| MN996532.1;QHR63299.1;BatCoV_RaTG13_orf1ab;4911 | 0.1 | 0.05 |
| MN996532.1;QHR63299.1;BatCoV_RaTG13_orf1ab;4912 | 0.05 | 0 |
| MN996532.1;QHR63299.1;BatCoV_RaTG13_orf1ab;4913 | 0.2 | 0.025 |
| MN996532.1;QHR63299.1;BatCoV_RaTG13_orf1ab;4914 | 0.05 | 0.025 |
| MN996532.1;QHR63299.1;BatCoV_RaTG13_orf1ab;4915 | 0.05 | 0 |
| MN996532.1;QHR63299.1;BatCoV_RaTG13_orf1ab;4916 | 0.05 | 0 |
| MN996532.1;QHR63299.1;BatCoV_RaTG13_orf1ab;4917 | 0 | 0 |
| MN996532.1;QHR63299.1;BatCoV_RaTG13_orf1ab;4918 | 0 | 0 |
| MN996532.1;QHR63299.1;BatCoV_RaTG13_orf1ab;4919 | 0 | 0 |
| MN996532.1;QHR63299.1;BatCoV_RaTG13_orf1ab;4920 | 0 | 0 |
| MN996532.1;QHR63299.1;BatCoV_RaTG13_orf1ab;4921 | 0 | 0 |
| MN996532.1;QHR63299.1;BatCoV_RaTG13_orf1ab;4922 | 0.05 | 0 |
| MN996532.1;QHR63299.1;BatCoV_RaTG13_orf1ab;4923 | 0.1 | 0.15 |
| MN996532.1;QHR63299.1;BatCoV_RaTG13_orf1ab;4924 | 0 | 0 |
| MN996532.1;QHR63299.1;BatCoV_RaTG13_orf1ab;4925 | 0 | 0 |
| MN996532.1;QHR63299.1;BatCoV_RaTG13_orf1ab;4926 | 0 | 0 |
| MN996532.1;QHR63299.1;BatCoV_RaTG13_orf1ab;4927 | 0 | 0.025 |
| MN996532.1;QHR63299.1;BatCoV_RaTG13_orf1ab;4928 | 0.1 | 0.05 |
| MN996532.1;QHR63299.1;BatCoV_RaTG13_orf1ab;4929 | 0 | 0.025 |
| MN996532.1;QHR63299.1;BatCoV_RaTG13_orf1ab;4930 | 0 | 0.025 |
| MN996532.1;QHR63299.1;BatCoV_RaTG13_orf1ab;4931 | 0 | 0.025 |
| MN996532.1;QHR63299.1;BatCoV_RaTG13_orf1ab;4932 | 0 | 0.025 |
| MN996532.1;QHR63299.1;BatCoV_RaTG13_orf1ab;4933 | 0 | 0 |
| MN996532.1;QHR63299.1;BatCoV_RaTG13_orf1ab;4934 | 0 | 0 |
| MN996532.1;QHR63299.1;BatCoV_RaTG13_orf1ab;4935 | 0 | 0 |
| MN996532.1;QHR63299.1;BatCoV_RaTG13_orf1ab;4936 | 0 | 0 |

| | | |
|---|---|---|
| MN996532.1;QHR63299.1;BatCoV_RaTG13_orf1ab;4937 | 0 | 0 |
| MN996532.1;QHR63299.1;BatCoV_RaTG13_orf1ab;4938 | 0 | 0 |
| MN996532.1;QHR63299.1;BatCoV_RaTG13_orf1ab;4939 | 0 | 0 |
| MN996532.1;QHR63299.1;BatCoV_RaTG13_orf1ab;4940 | 0 | 0 |
| MN996532.1;QHR63299.1;BatCoV_RaTG13_orf1ab;4941 | 0 | 0 |
| MN996532.1;QHR63299.1;BatCoV_RaTG13_orf1ab;4942 | 0 | 0 |
| MN996532.1;QHR63299.1;BatCoV_RaTG13_orf1ab;4943 | 0 | 0 |
| MN996532.1;QHR63299.1;BatCoV_RaTG13_orf1ab;4944 | 0 | 0 |
| MN996532.1;QHR63299.1;BatCoV_RaTG13_orf1ab;4945 | 0 | 0 |
| MN996532.1;QHR63299.1;BatCoV_RaTG13_orf1ab;4946 | 0 | 0 |
| MN996532.1;QHR63299.1;BatCoV_RaTG13_orf1ab;4947 | 0 | 0 |
| MN996532.1;QHR63299.1;BatCoV_RaTG13_orf1ab;4948 | 0 | 0 |
| MN996532.1;QHR63299.1;BatCoV_RaTG13_orf1ab;4949 | 0 | 0 |
| MN996532.1;QHR63299.1;BatCoV_RaTG13_orf1ab;4950 | 0 | 0 |
| MN996532.1;QHR63299.1;BatCoV_RaTG13_orf1ab;4951 | 0 | 0 |
| MN996532.1;QHR63299.1;BatCoV_RaTG13_orf1ab;4952 | 0 | 0 |
| MN996532.1;QHR63299.1;BatCoV_RaTG13_orf1ab;4953 | 0 | 0 |
| MN996532.1;QHR63299.1;BatCoV_RaTG13_orf1ab;4954 | 0 | 0 |
| MN996532.1;QHR63299.1;BatCoV_RaTG13_orf1ab;4955 | 0 | 0 |
| MN996532.1;QHR63299.1;BatCoV_RaTG13_orf1ab;4956 | 0 | 0 |
| MN996532.1;QHR63299.1;BatCoV_RaTG13_orf1ab;4957 | 0 | 0 |
| MN996532.1;QHR63299.1;BatCoV_RaTG13_orf1ab;4958 | 0 | 0 |
| MN996532.1;QHR63299.1;BatCoV_RaTG13_orf1ab;4959 | 0 | 0 |
| MN996532.1;QHR63299.1;BatCoV_RaTG13_orf1ab;4960 | 0 | 0 |
| MN996532.1;QHR63299.1;BatCoV_RaTG13_orf1ab;4961 | 0 | 0 |
| MN996532.1;QHR63299.1;BatCoV_RaTG13_orf1ab;4962 | 0.3 | 0.175 |
| MN996532.1;QHR63299.1;BatCoV_RaTG13_orf1ab;4963 | 0.4 | 0.275 |
| MN996532.1;QHR63299.1;BatCoV_RaTG13_orf1ab;4964 | 0.35 | 0.275 |
| MN996532.1;QHR63299.1;BatCoV_RaTG13_orf1ab;4965 | 0.25 | 0.025 |
| MN996532.1;QHR63299.1;BatCoV_RaTG13_orf1ab;4966 | 0.25 | 0.075 |
| MN996532.1;QHR63299.1;BatCoV_RaTG13_orf1ab;4967 | 0.25 | 0.025 |
| MN996532.1;QHR63299.1;BatCoV_RaTG13_orf1ab;4968 | 0.25 | 0.05 |
| MN996532.1;QHR63299.1;BatCoV_RaTG13_orf1ab;4969 | 0.25 | 0.025 |
| MN996532.1;QHR63299.1;BatCoV_RaTG13_orf1ab;4970 | 0.15 | 0.025 |
| MN996532.1;QHR63299.1;BatCoV_RaTG13_orf1ab;4971 | 0.15 | 0.025 |
| MN996532.1;QHR63299.1;BatCoV_RaTG13_orf1ab;4972 | 0 | 0 |
| MN996532.1;QHR63299.1;BatCoV_RaTG13_orf1ab;4973 | 0 | 0 |
| MN996532.1;QHR63299.1;BatCoV_RaTG13_orf1ab;4974 | 0 | 0 |
| MN996532.1;QHR63299.1;BatCoV_RaTG13_orf1ab;4975 | 0 | 0 |
| MN996532.1;QHR63299.1;BatCoV_RaTG13_orf1ab;4976 | 0 | 0 |
| MN996532.1;QHR63299.1;BatCoV_RaTG13_orf1ab;4977 | 0 | 0 |
| MN996532.1;QHR63299.1;BatCoV_RaTG13_orf1ab;4978 | 0 | 0 |
| MN996532.1;QHR63299.1;BatCoV_RaTG13_orf1ab;4979 | 0 | 0 |

| | | |
|---|---|---|
| MN996532.1;QHR63299.1;BatCoV_RaTG13_orf1ab;4980 | 0 | 0 |
| MN996532.1;QHR63299.1;BatCoV_RaTG13_orf1ab;4981 | 0 | 0 |
| MN996532.1;QHR63299.1;BatCoV_RaTG13_orf1ab;4982 | 0 | 0 |
| MN996532.1;QHR63299.1;BatCoV_RaTG13_orf1ab;4983 | 0 | 0 |
| MN996532.1;QHR63299.1;BatCoV_RaTG13_orf1ab;4984 | 0 | 0.05 |
| MN996532.1;QHR63299.1;BatCoV_RaTG13_orf1ab;4985 | 0 | 0 |
| MN996532.1;QHR63299.1;BatCoV_RaTG13_orf1ab;4986 | 0 | 0 |
| MN996532.1;QHR63299.1;BatCoV_RaTG13_orf1ab;4987 | 0 | 0 |
| MN996532.1;QHR63299.1;BatCoV_RaTG13_orf1ab;4988 | 0 | 0 |
| MN996532.1;QHR63299.1;BatCoV_RaTG13_orf1ab;4989 | 0 | 0 |
| MN996532.1;QHR63299.1;BatCoV_RaTG13_orf1ab;4990 | 0 | 0 |
| MN996532.1;QHR63299.1;BatCoV_RaTG13_orf1ab;4991 | 0 | 0 |
| MN996532.1;QHR63299.1;BatCoV_RaTG13_orf1ab;4992 | 0 | 0 |
| MN996532.1;QHR63299.1;BatCoV_RaTG13_orf1ab;4993 | 0 | 0 |
| MN996532.1;QHR63299.1;BatCoV_RaTG13_orf1ab;4994 | 0 | 0 |
| MN996532.1;QHR63299.1;BatCoV_RaTG13_orf1ab;4995 | 0 | 0 |
| MN996532.1;QHR63299.1;BatCoV_RaTG13_orf1ab;4996 | 0 | 0.075 |
| MN996532.1;QHR63299.1;BatCoV_RaTG13_orf1ab;4997 | 0 | 0.025 |
| MN996532.1;QHR63299.1;BatCoV_RaTG13_orf1ab;4998 | 0 | 0.025 |
| MN996532.1;QHR63299.1;BatCoV_RaTG13_orf1ab;4999 | 0 | 0.025 |
| MN996532.1;QHR63299.1;BatCoV_RaTG13_orf1ab;5000 | 0 | 0.025 |
| MN996532.1;QHR63299.1;BatCoV_RaTG13_orf1ab;5001 | 0 | 0.025 |
| MN996532.1;QHR63299.1;BatCoV_RaTG13_orf1ab;5002 | 0 | 0.025 |
| MN996532.1;QHR63299.1;BatCoV_RaTG13_orf1ab;5003 | 0 | 0.025 |
| MN996532.1;QHR63299.1;BatCoV_RaTG13_orf1ab;5004 | 0 | 0.025 |
| MN996532.1;QHR63299.1;BatCoV_RaTG13_orf1ab;5005 | 0 | 0.025 |
| MN996532.1;QHR63299.1;BatCoV_RaTG13_orf1ab;5006 | 0 | 0.025 |
| MN996532.1;QHR63299.1;BatCoV_RaTG13_orf1ab;5007 | 0 | 0.025 |
| MN996532.1;QHR63299.1;BatCoV_RaTG13_orf1ab;5008 | 0 | 0 |
| MN996532.1;QHR63299.1;BatCoV_RaTG13_orf1ab;5009 | 0 | 0 |
| MN996532.1;QHR63299.1;BatCoV_RaTG13_orf1ab;5010 | 0 | 0 |
| MN996532.1;QHR63299.1;BatCoV_RaTG13_orf1ab;5011 | 0 | 0 |
| MN996532.1;QHR63299.1;BatCoV_RaTG13_orf1ab;5012 | 0 | 0 |
| MN996532.1;QHR63299.1;BatCoV_RaTG13_orf1ab;5013 | 0 | 0 |
| MN996532.1;QHR63299.1;BatCoV_RaTG13_orf1ab;5014 | 0 | 0 |
| MN996532.1;QHR63299.1;BatCoV_RaTG13_orf1ab;5015 | 0 | 0 |
| MN996532.1;QHR63299.1;BatCoV_RaTG13_orf1ab;5016 | 0 | 0 |
| MN996532.1;QHR63299.1;BatCoV_RaTG13_orf1ab;5017 | 0 | 0 |
| MN996532.1;QHR63299.1;BatCoV_RaTG13_orf1ab;5018 | 0 | 0 |
| MN996532.1;QHR63299.1;BatCoV_RaTG13_orf1ab;5019 | 0 | 0 |
| MN996532.1;QHR63299.1;BatCoV_RaTG13_orf1ab;5020 | 0 | 0 |
| MN996532.1;QHR63299.1;BatCoV_RaTG13_orf1ab;5021 | 0 | 0 |
| MN996532.1;QHR63299.1;BatCoV_RaTG13_orf1ab;5022 | 0 | 0 |

| | | |
|---|---|---|
| MN996532.1;QHR63299.1;BatCoV_RaTG13_orf1ab;5023 | 0 | 0 |
| MN996532.1;QHR63299.1;BatCoV_RaTG13_orf1ab;5024 | 0 | 0 |
| MN996532.1;QHR63299.1;BatCoV_RaTG13_orf1ab;5025 | 0 | 0 |
| MN996532.1;QHR63299.1;BatCoV_RaTG13_orf1ab;5026 | 0 | 0 |
| MN996532.1;QHR63299.1;BatCoV_RaTG13_orf1ab;5027 | 0 | 0 |
| MN996532.1;QHR63299.1;BatCoV_RaTG13_orf1ab;5028 | 0 | 0 |
| MN996532.1;QHR63299.1;BatCoV_RaTG13_orf1ab;5029 | 0 | 0 |
| MN996532.1;QHR63299.1;BatCoV_RaTG13_orf1ab;5030 | 0 | 0 |
| MN996532.1;QHR63299.1;BatCoV_RaTG13_orf1ab;5031 | 0 | 0 |
| MN996532.1;QHR63299.1;BatCoV_RaTG13_orf1ab;5032 | 0 | 0 |
| MN996532.1;QHR63299.1;BatCoV_RaTG13_orf1ab;5033 | 0 | 0 |
| MN996532.1;QHR63299.1;BatCoV_RaTG13_orf1ab;5034 | 0 | 0 |
| MN996532.1;QHR63299.1;BatCoV_RaTG13_orf1ab;5035 | 0 | 0 |
| MN996532.1;QHR63299.1;BatCoV_RaTG13_orf1ab;5036 | 0 | 0 |
| MN996532.1;QHR63299.1;BatCoV_RaTG13_orf1ab;5037 | 0 | 0 |
| MN996532.1;QHR63299.1;BatCoV_RaTG13_orf1ab;5038 | 0 | 0 |
| MN996532.1;QHR63299.1;BatCoV_RaTG13_orf1ab;5039 | 0 | 0 |
| MN996532.1;QHR63299.1;BatCoV_RaTG13_orf1ab;5040 | 0 | 0 |
| MN996532.1;QHR63299.1;BatCoV_RaTG13_orf1ab;5041 | 0 | 0 |
| MN996532.1;QHR63299.1;BatCoV_RaTG13_orf1ab;5042 | 0 | 0 |
| MN996532.1;QHR63299.1;BatCoV_RaTG13_orf1ab;5043 | 0 | 0 |
| MN996532.1;QHR63299.1;BatCoV_RaTG13_orf1ab;5044 | 0 | 0 |
| MN996532.1;QHR63299.1;BatCoV_RaTG13_orf1ab;5045 | 0 | 0 |
| MN996532.1;QHR63299.1;BatCoV_RaTG13_orf1ab;5046 | 0 | 0 |
| MN996532.1;QHR63299.1;BatCoV_RaTG13_orf1ab;5047 | 0 | 0 |
| MN996532.1;QHR63299.1;BatCoV_RaTG13_orf1ab;5048 | 0 | 0 |
| MN996532.1;QHR63299.1;BatCoV_RaTG13_orf1ab;5049 | 0 | 0 |
| MN996532.1;QHR63299.1;BatCoV_RaTG13_orf1ab;5050 | 0 | 0 |
| MN996532.1;QHR63299.1;BatCoV_RaTG13_orf1ab;5051 | 0 | 0 |
| MN996532.1;QHR63299.1;BatCoV_RaTG13_orf1ab;5052 | 0 | 0 |
| MN996532.1;QHR63299.1;BatCoV_RaTG13_orf1ab;5053 | 0 | 0 |
| MN996532.1;QHR63299.1;BatCoV_RaTG13_orf1ab;5054 | 0 | 0 |
| MN996532.1;QHR63299.1;BatCoV_RaTG13_orf1ab;5055 | 0 | 0 |
| MN996532.1;QHR63299.1;BatCoV_RaTG13_orf1ab;5056 | 0.05 | 0 |
| MN996532.1;QHR63299.1;BatCoV_RaTG13_orf1ab;5057 | 0 | 0 |
| MN996532.1;QHR63299.1;BatCoV_RaTG13_orf1ab;5058 | 0 | 0 |
| MN996532.1;QHR63299.1;BatCoV_RaTG13_orf1ab;5059 | 0 | 0 |
| MN996532.1;QHR63299.1;BatCoV_RaTG13_orf1ab;5060 | 0 | 0.025 |
| MN996532.1;QHR63299.1;BatCoV_RaTG13_orf1ab;5061 | 0 | 0.025 |
| MN996532.1;QHR63299.1;BatCoV_RaTG13_orf1ab;5062 | 0 | 0.025 |
| MN996532.1;QHR63299.1;BatCoV_RaTG13_orf1ab;5063 | 0 | 0.025 |
| MN996532.1;QHR63299.1;BatCoV_RaTG13_orf1ab;5064 | 0 | 0.05 |
| MN996532.1;QHR63299.1;BatCoV_RaTG13_orf1ab;5065 | 0 | 0.025 |

| | | |
|---|---|---|
| MN996532.1;QHR63299.1;BatCoV_RaTG13_orf1ab;5066 | 0 | 0.025 |
| MN996532.1;QHR63299.1;BatCoV_RaTG13_orf1ab;5067 | 0 | 0.05 |
| MN996532.1;QHR63299.1;BatCoV_RaTG13_orf1ab;5068 | 0.05 | 0.025 |
| MN996532.1;QHR63299.1;BatCoV_RaTG13_orf1ab;5069 | 0.1 | 0.05 |
| MN996532.1;QHR63299.1;BatCoV_RaTG13_orf1ab;5070 | 0 | 0 |
| MN996532.1;QHR63299.1;BatCoV_RaTG13_orf1ab;5071 | 0.05 | 0 |
| MN996532.1;QHR63299.1;BatCoV_RaTG13_orf1ab;5072 | 0 | 0 |
| MN996532.1;QHR63299.1;BatCoV_RaTG13_orf1ab;5073 | 0 | 0 |
| MN996532.1;QHR63299.1;BatCoV_RaTG13_orf1ab;5074 | 0 | 0 |
| MN996532.1;QHR63299.1;BatCoV_RaTG13_orf1ab;5075 | 0 | 0 |
| MN996532.1;QHR63299.1;BatCoV_RaTG13_orf1ab;5076 | 0 | 0 |
| MN996532.1;QHR63299.1;BatCoV_RaTG13_orf1ab;5077 | 0 | 0 |
| MN996532.1;QHR63299.1;BatCoV_RaTG13_orf1ab;5078 | 0 | 0 |
| MN996532.1;QHR63299.1;BatCoV_RaTG13_orf1ab;5079 | 0 | 0.025 |
| MN996532.1;QHR63299.1;BatCoV_RaTG13_orf1ab;5080 | 0 | 0 |
| MN996532.1;QHR63299.1;BatCoV_RaTG13_orf1ab;5081 | 0 | 0.025 |
| MN996532.1;QHR63299.1;BatCoV_RaTG13_orf1ab;5082 | 0 | 0 |
| MN996532.1;QHR63299.1;BatCoV_RaTG13_orf1ab;5083 | 0 | 0 |
| MN996532.1;QHR63299.1;BatCoV_RaTG13_orf1ab;5084 | 0 | 0 |
| MN996532.1;QHR63299.1;BatCoV_RaTG13_orf1ab;5085 | 0 | 0 |
| MN996532.1;QHR63299.1;BatCoV_RaTG13_orf1ab;5086 | 0 | 0 |
| MN996532.1;QHR63299.1;BatCoV_RaTG13_orf1ab;5087 | 0 | 0 |
| MN996532.1;QHR63299.1;BatCoV_RaTG13_orf1ab;5088 | 0 | 0.025 |
| MN996532.1;QHR63299.1;BatCoV_RaTG13_orf1ab;5089 | 0 | 0 |
| MN996532.1;QHR63299.1;BatCoV_RaTG13_orf1ab;5090 | 0 | 0.025 |
| MN996532.1;QHR63299.1;BatCoV_RaTG13_orf1ab;5091 | 0 | 0.025 |
| MN996532.1;QHR63299.1;BatCoV_RaTG13_orf1ab;5092 | 0 | 0.025 |
| MN996532.1;QHR63299.1;BatCoV_RaTG13_orf1ab;5093 | 0.05 | 0.025 |
| MN996532.1;QHR63299.1;BatCoV_RaTG13_orf1ab;5094 | 0 | 0 |
| MN996532.1;QHR63299.1;BatCoV_RaTG13_orf1ab;5095 | 0 | 0 |
| MN996532.1;QHR63299.1;BatCoV_RaTG13_orf1ab;5096 | 0.05 | 0.025 |
| MN996532.1;QHR63299.1;BatCoV_RaTG13_orf1ab;5097 | 0 | 0.05 |
| MN996532.1;QHR63299.1;BatCoV_RaTG13_orf1ab;5098 | 0 | 0.025 |
| MN996532.1;QHR63299.1;BatCoV_RaTG13_orf1ab;5099 | 0 | 0.025 |
| MN996532.1;QHR63299.1;BatCoV_RaTG13_orf1ab;5100 | 0.05 | 0 |
| MN996532.1;QHR63299.1;BatCoV_RaTG13_orf1ab;5101 | 0 | 0 |
| MN996532.1;QHR63299.1;BatCoV_RaTG13_orf1ab;5102 | 0 | 0 |
| MN996532.1;QHR63299.1;BatCoV_RaTG13_orf1ab;5103 | 0 | 0 |
| MN996532.1;QHR63299.1;BatCoV_RaTG13_orf1ab;5104 | 0 | 0 |
| MN996532.1;QHR63299.1;BatCoV_RaTG13_orf1ab;5105 | 0 | 0 |
| MN996532.1;QHR63299.1;BatCoV_RaTG13_orf1ab;5106 | 0 | 0 |
| MN996532.1;QHR63299.1;BatCoV_RaTG13_orf1ab;5107 | 0 | 0 |
| MN996532.1;QHR63299.1;BatCoV_RaTG13_orf1ab;5108 | 0 | 0 |

| | | |
|---|---|---|
| MN996532.1;QHR63299.1;BatCoV_RaTG13_orf1ab;5109 | 0 | 0 |
| MN996532.1;QHR63299.1;BatCoV_RaTG13_orf1ab;5110 | 0 | 0 |
| MN996532.1;QHR63299.1;BatCoV_RaTG13_orf1ab;5111 | 0 | 0 |
| MN996532.1;QHR63299.1;BatCoV_RaTG13_orf1ab;5112 | 0.05 | 0 |
| MN996532.1;QHR63299.1;BatCoV_RaTG13_orf1ab;5113 | 0 | 0 |
| MN996532.1;QHR63299.1;BatCoV_RaTG13_orf1ab;5114 | 0 | 0 |
| MN996532.1;QHR63299.1;BatCoV_RaTG13_orf1ab;5115 | 0 | 0 |
| MN996532.1;QHR63299.1;BatCoV_RaTG13_orf1ab;5116 | 0 | 0 |
| MN996532.1;QHR63299.1;BatCoV_RaTG13_orf1ab;5117 | 0 | 0 |
| MN996532.1;QHR63299.1;BatCoV_RaTG13_orf1ab;5118 | 0 | 0 |
| MN996532.1;QHR63299.1;BatCoV_RaTG13_orf1ab;5119 | 0 | 0 |
| MN996532.1;QHR63299.1;BatCoV_RaTG13_orf1ab;5120 | 0 | 0 |
| MN996532.1;QHR63299.1;BatCoV_RaTG13_orf1ab;5121 | 0 | 0 |
| MN996532.1;QHR63299.1;BatCoV_RaTG13_orf1ab;5122 | 0 | 0.05 |
| MN996532.1;QHR63299.1;BatCoV_RaTG13_orf1ab;5123 | 0.05 | 0.05 |
| MN996532.1;QHR63299.1;BatCoV_RaTG13_orf1ab;5124 | 0 | 0.05 |
| MN996532.1;QHR63299.1;BatCoV_RaTG13_orf1ab;5125 | 0 | 0 |
| MN996532.1;QHR63299.1;BatCoV_RaTG13_orf1ab;5126 | 0 | 0.025 |
| MN996532.1;QHR63299.1;BatCoV_RaTG13_orf1ab;5127 | 0 | 0 |
| MN996532.1;QHR63299.1;BatCoV_RaTG13_orf1ab;5128 | 0 | 0.025 |
| MN996532.1;QHR63299.1;BatCoV_RaTG13_orf1ab;5129 | 0 | 0 |
| MN996532.1;QHR63299.1;BatCoV_RaTG13_orf1ab;5130 | 0.05 | 0.025 |
| MN996532.1;QHR63299.1;BatCoV_RaTG13_orf1ab;5131 | 0 | 0 |
| MN996532.1;QHR63299.1;BatCoV_RaTG13_orf1ab;5132 | 0.05 | 0.025 |
| MN996532.1;QHR63299.1;BatCoV_RaTG13_orf1ab;5133 | 0 | 0 |
| MN996532.1;QHR63299.1;BatCoV_RaTG13_orf1ab;5134 | 0 | 0 |
| MN996532.1;QHR63299.1;BatCoV_RaTG13_orf1ab;5135 | 0 | 0 |
| MN996532.1;QHR63299.1;BatCoV_RaTG13_orf1ab;5136 | 0 | 0 |
| MN996532.1;QHR63299.1;BatCoV_RaTG13_orf1ab;5137 | 0 | 0 |
| MN996532.1;QHR63299.1;BatCoV_RaTG13_orf1ab;5138 | 0.05 | 0 |
| MN996532.1;QHR63299.1;BatCoV_RaTG13_orf1ab;5139 | 0 | 0 |
| MN996532.1;QHR63299.1;BatCoV_RaTG13_orf1ab;5140 | 0 | 0 |
| MN996532.1;QHR63299.1;BatCoV_RaTG13_orf1ab;5141 | 0 | 0 |
| MN996532.1;QHR63299.1;BatCoV_RaTG13_orf1ab;5142 | 0 | 0 |
| MN996532.1;QHR63299.1;BatCoV_RaTG13_orf1ab;5143 | 0 | 0 |
| MN996532.1;QHR63299.1;BatCoV_RaTG13_orf1ab;5144 | 0 | 0 |
| MN996532.1;QHR63299.1;BatCoV_RaTG13_orf1ab;5145 | 0 | 0 |
| MN996532.1;QHR63299.1;BatCoV_RaTG13_orf1ab;5146 | 0 | 0 |
| MN996532.1;QHR63299.1;BatCoV_RaTG13_orf1ab;5147 | 0 | 0 |
| MN996532.1;QHR63299.1;BatCoV_RaTG13_orf1ab;5148 | 0 | 0 |
| MN996532.1;QHR63299.1;BatCoV_RaTG13_orf1ab;5149 | 0 | 0 |
| MN996532.1;QHR63299.1;BatCoV_RaTG13_orf1ab;5150 | 0 | 0 |
| MN996532.1;QHR63299.1;BatCoV_RaTG13_orf1ab;5151 | 0 | 0 |

| | | |
|---|---|---|
| MN996532.1;QHR63299.1;BatCoV_RaTG13_orf1ab;5152 | 0 | 0 |
| MN996532.1;QHR63299.1;BatCoV_RaTG13_orf1ab;5153 | 0.05 | 0.05 |
| MN996532.1;QHR63299.1;BatCoV_RaTG13_orf1ab;5154 | 0 | 0 |
| MN996532.1;QHR63299.1;BatCoV_RaTG13_orf1ab;5155 | 0 | 0 |
| MN996532.1;QHR63299.1;BatCoV_RaTG13_orf1ab;5156 | 0 | 0 |
| MN996532.1;QHR63299.1;BatCoV_RaTG13_orf1ab;5157 | 0 | 0 |
| MN996532.1;QHR63299.1;BatCoV_RaTG13_orf1ab;5158 | 0 | 0 |
| MN996532.1;QHR63299.1;BatCoV_RaTG13_orf1ab;5159 | 0 | 0 |
| MN996532.1;QHR63299.1;BatCoV_RaTG13_orf1ab;5160 | 0 | 0 |
| MN996532.1;QHR63299.1;BatCoV_RaTG13_orf1ab;5161 | 0 | 0 |
| MN996532.1;QHR63299.1;BatCoV_RaTG13_orf1ab;5162 | 0 | 0 |
| MN996532.1;QHR63299.1;BatCoV_RaTG13_orf1ab;5163 | 0 | 0 |
| MN996532.1;QHR63299.1;BatCoV_RaTG13_orf1ab;5164 | 0 | 0 |
| MN996532.1;QHR63299.1;BatCoV_RaTG13_orf1ab;5165 | 0 | 0 |
| MN996532.1;QHR63299.1;BatCoV_RaTG13_orf1ab;5166 | 0 | 0 |
| MN996532.1;QHR63299.1;BatCoV_RaTG13_orf1ab;5167 | 0 | 0 |
| MN996532.1;QHR63299.1;BatCoV_RaTG13_orf1ab;5168 | 0 | 0 |
| MN996532.1;QHR63299.1;BatCoV_RaTG13_orf1ab;5169 | 0 | 0 |
| MN996532.1;QHR63299.1;BatCoV_RaTG13_orf1ab;5170 | 0 | 0 |
| MN996532.1;QHR63299.1;BatCoV_RaTG13_orf1ab;5171 | 0 | 0 |
| MN996532.1;QHR63299.1;BatCoV_RaTG13_orf1ab;5172 | 0 | 0 |
| MN996532.1;QHR63299.1;BatCoV_RaTG13_orf1ab;5173 | 0 | 0 |
| MN996532.1;QHR63299.1;BatCoV_RaTG13_orf1ab;5174 | 0.05 | 0 |
| MN996532.1;QHR63299.1;BatCoV_RaTG13_orf1ab;5175 | 0 | 0 |
| MN996532.1;QHR63299.1;BatCoV_RaTG13_orf1ab;5176 | 0 | 0 |
| MN996532.1;QHR63299.1;BatCoV_RaTG13_orf1ab;5177 | 0 | 0 |
| MN996532.1;QHR63299.1;BatCoV_RaTG13_orf1ab;5178 | 0 | 0.025 |
| MN996532.1;QHR63299.1;BatCoV_RaTG13_orf1ab;5179 | 0 | 0.025 |
| MN996532.1;QHR63299.1;BatCoV_RaTG13_orf1ab;5180 | 0 | 0.025 |
| MN996532.1;QHR63299.1;BatCoV_RaTG13_orf1ab;5181 | 0 | 0.025 |
| MN996532.1;QHR63299.1;BatCoV_RaTG13_orf1ab;5182 | 0 | 0.025 |
| MN996532.1;QHR63299.1;BatCoV_RaTG13_orf1ab;5183 | 0.05 | 0.025 |
| MN996532.1;QHR63299.1;BatCoV_RaTG13_orf1ab;5184 | 0 | 0.025 |
| MN996532.1;QHR63299.1;BatCoV_RaTG13_orf1ab;5185 | 0.05 | 0.025 |
| MN996532.1;QHR63299.1;BatCoV_RaTG13_orf1ab;5186 | 0 | 0.025 |
| MN996532.1;QHR63299.1;BatCoV_RaTG13_orf1ab;5187 | 0 | 0.025 |
| MN996532.1;QHR63299.1;BatCoV_RaTG13_orf1ab;5188 | 0.05 | 0.025 |
| MN996532.1;QHR63299.1;BatCoV_RaTG13_orf1ab;5189 | 0.05 | 0.025 |
| MN996532.1;QHR63299.1;BatCoV_RaTG13_orf1ab;5190 | 0.05 | 0.025 |
| MN996532.1;QHR63299.1;BatCoV_RaTG13_orf1ab;5191 | 0.05 | 0.025 |
| MN996532.1;QHR63299.1;BatCoV_RaTG13_orf1ab;5192 | 0 | 0 |
| MN996532.1;QHR63299.1;BatCoV_RaTG13_orf1ab;5193 | 0 | 0 |
| MN996532.1;QHR63299.1;BatCoV_RaTG13_orf1ab;5194 | 0 | 0 |

| | | |
|---|---|---|
| MN996532.1;QHR63299.1;BatCoV_RaTG13_orf1ab;5195 | 0 | 0 |
| MN996532.1;QHR63299.1;BatCoV_RaTG13_orf1ab;5196 | 0 | 0 |
| MN996532.1;QHR63299.1;BatCoV_RaTG13_orf1ab;5197 | 0 | 0 |
| MN996532.1;QHR63299.1;BatCoV_RaTG13_orf1ab;5198 | 0 | 0 |
| MN996532.1;QHR63299.1;BatCoV_RaTG13_orf1ab;5199 | 0 | 0 |
| MN996532.1;QHR63299.1;BatCoV_RaTG13_orf1ab;5200 | 0 | 0 |
| MN996532.1;QHR63299.1;BatCoV_RaTG13_orf1ab;5201 | 0 | 0 |
| MN996532.1;QHR63299.1;BatCoV_RaTG13_orf1ab;5202 | 0 | 0.025 |
| MN996532.1;QHR63299.1;BatCoV_RaTG13_orf1ab;5203 | 0 | 0.025 |
| MN996532.1;QHR63299.1;BatCoV_RaTG13_orf1ab;5204 | 0 | 0.025 |
| MN996532.1;QHR63299.1;BatCoV_RaTG13_orf1ab;5205 | 0.05 | 0 |
| MN996532.1;QHR63299.1;BatCoV_RaTG13_orf1ab;5206 | 0.05 | 0 |
| MN996532.1;QHR63299.1;BatCoV_RaTG13_orf1ab;5207 | 0 | 0 |
| MN996532.1;QHR63299.1;BatCoV_RaTG13_orf1ab;5208 | 0.05 | 0 |
| MN996532.1;QHR63299.1;BatCoV_RaTG13_orf1ab;5209 | 0 | 0 |
| MN996532.1;QHR63299.1;BatCoV_RaTG13_orf1ab;5210 | 0.1 | 0.1 |
| MN996532.1;QHR63299.1;BatCoV_RaTG13_orf1ab;5211 | 0 | 0.025 |
| MN996532.1;QHR63299.1;BatCoV_RaTG13_orf1ab;5212 | 0 | 0.025 |
| MN996532.1;QHR63299.1;BatCoV_RaTG13_orf1ab;5213 | 0 | 0 |
| MN996532.1;QHR63299.1;BatCoV_RaTG13_orf1ab;5214 | 0 | 0 |
| MN996532.1;QHR63299.1;BatCoV_RaTG13_orf1ab;5215 | 0 | 0 |
| MN996532.1;QHR63299.1;BatCoV_RaTG13_orf1ab;5216 | 0 | 0 |
| MN996532.1;QHR63299.1;BatCoV_RaTG13_orf1ab;5217 | 0 | 0 |
| MN996532.1;QHR63299.1;BatCoV_RaTG13_orf1ab;5218 | 0 | 0 |
| MN996532.1;QHR63299.1;BatCoV_RaTG13_orf1ab;5219 | 0 | 0 |
| MN996532.1;QHR63299.1;BatCoV_RaTG13_orf1ab;5220 | 0 | 0 |
| MN996532.1;QHR63299.1;BatCoV_RaTG13_orf1ab;5221 | 0 | 0 |
| MN996532.1;QHR63299.1;BatCoV_RaTG13_orf1ab;5222 | 0 | 0 |
| MN996532.1;QHR63299.1;BatCoV_RaTG13_orf1ab;5223 | 0 | 0 |
| MN996532.1;QHR63299.1;BatCoV_RaTG13_orf1ab;5224 | 0 | 0 |
| MN996532.1;QHR63299.1;BatCoV_RaTG13_orf1ab;5225 | 0 | 0 |
| MN996532.1;QHR63299.1;BatCoV_RaTG13_orf1ab;5226 | 0 | 0 |
| MN996532.1;QHR63299.1;BatCoV_RaTG13_orf1ab;5227 | 0 | 0 |
| MN996532.1;QHR63299.1;BatCoV_RaTG13_orf1ab;5228 | 0.05 | 0 |
| MN996532.1;QHR63299.1;BatCoV_RaTG13_orf1ab;5229 | 0 | 0 |
| MN996532.1;QHR63299.1;BatCoV_RaTG13_orf1ab;5230 | 0 | 0 |
| MN996532.1;QHR63299.1;BatCoV_RaTG13_orf1ab;5231 | 0 | 0 |
| MN996532.1;QHR63299.1;BatCoV_RaTG13_orf1ab;5232 | 0 | 0 |
| MN996532.1;QHR63299.1;BatCoV_RaTG13_orf1ab;5233 | 0 | 0 |
| MN996532.1;QHR63299.1;BatCoV_RaTG13_orf1ab;5234 | 0 | 0 |
| MN996532.1;QHR63299.1;BatCoV_RaTG13_orf1ab;5235 | 0 | 0 |
| MN996532.1;QHR63299.1;BatCoV_RaTG13_orf1ab;5236 | 0 | 0 |
| MN996532.1;QHR63299.1;BatCoV_RaTG13_orf1ab;5237 | 0 | 0 |

| | | |
|---|---|---|
| MN996532.1;QHR63299.1;BatCoV_RaTG13_orf1ab;5238 | 0 | 0 |
| MN996532.1;QHR63299.1;BatCoV_RaTG13_orf1ab;5239 | 0 | 0 |
| MN996532.1;QHR63299.1;BatCoV_RaTG13_orf1ab;5240 | 0 | 0 |
| MN996532.1;QHR63299.1;BatCoV_RaTG13_orf1ab;5241 | 0 | 0 |
| MN996532.1;QHR63299.1;BatCoV_RaTG13_orf1ab;5242 | 0 | 0 |
| MN996532.1;QHR63299.1;BatCoV_RaTG13_orf1ab;5243 | 0 | 0 |
| MN996532.1;QHR63299.1;BatCoV_RaTG13_orf1ab;5244 | 0 | 0 |
| MN996532.1;QHR63299.1;BatCoV_RaTG13_orf1ab;5245 | 0 | 0 |
| MN996532.1;QHR63299.1;BatCoV_RaTG13_orf1ab;5246 | 0 | 0 |
| MN996532.1;QHR63299.1;BatCoV_RaTG13_orf1ab;5247 | 0 | 0 |
| MN996532.1;QHR63299.1;BatCoV_RaTG13_orf1ab;5248 | 0 | 0 |
| MN996532.1;QHR63299.1;BatCoV_RaTG13_orf1ab;5249 | 0 | 0 |
| MN996532.1;QHR63299.1;BatCoV_RaTG13_orf1ab;5250 | 0 | 0 |
| MN996532.1;QHR63299.1;BatCoV_RaTG13_orf1ab;5251 | 0 | 0 |
| MN996532.1;QHR63299.1;BatCoV_RaTG13_orf1ab;5252 | 0 | 0 |
| MN996532.1;QHR63299.1;BatCoV_RaTG13_orf1ab;5253 | 0 | 0 |
| MN996532.1;QHR63299.1;BatCoV_RaTG13_orf1ab;5254 | 0 | 0 |
| MN996532.1;QHR63299.1;BatCoV_RaTG13_orf1ab;5255 | 0 | 0 |
| MN996532.1;QHR63299.1;BatCoV_RaTG13_orf1ab;5256 | 0 | 0 |
| MN996532.1;QHR63299.1;BatCoV_RaTG13_orf1ab;5257 | 0 | 0 |
| MN996532.1;QHR63299.1;BatCoV_RaTG13_orf1ab;5258 | 0 | 0 |
| MN996532.1;QHR63299.1;BatCoV_RaTG13_orf1ab;5259 | 0 | 0 |
| MN996532.1;QHR63299.1;BatCoV_RaTG13_orf1ab;5260 | 0 | 0 |
| MN996532.1;QHR63299.1;BatCoV_RaTG13_orf1ab;5261 | 0 | 0 |
| MN996532.1;QHR63299.1;BatCoV_RaTG13_orf1ab;5262 | 0 | 0 |
| MN996532.1;QHR63299.1;BatCoV_RaTG13_orf1ab;5263 | 0 | 0 |
| MN996532.1;QHR63299.1;BatCoV_RaTG13_orf1ab;5264 | 0 | 0 |
| MN996532.1;QHR63299.1;BatCoV_RaTG13_orf1ab;5265 | 0 | 0 |
| MN996532.1;QHR63299.1;BatCoV_RaTG13_orf1ab;5266 | 0 | 0 |
| MN996532.1;QHR63299.1;BatCoV_RaTG13_orf1ab;5267 | 0 | 0 |
| MN996532.1;QHR63299.1;BatCoV_RaTG13_orf1ab;5268 | 0 | 0 |
| MN996532.1;QHR63299.1;BatCoV_RaTG13_orf1ab;5269 | 0 | 0 |
| MN996532.1;QHR63299.1;BatCoV_RaTG13_orf1ab;5270 | 0 | 0 |
| MN996532.1;QHR63299.1;BatCoV_RaTG13_orf1ab;5271 | 0 | 0 |
| MN996532.1;QHR63299.1;BatCoV_RaTG13_orf1ab;5272 | 0 | 0 |
| MN996532.1;QHR63299.1;BatCoV_RaTG13_orf1ab;5273 | 0 | 0 |
| MN996532.1;QHR63299.1;BatCoV_RaTG13_orf1ab;5274 | 0 | 0 |
| MN996532.1;QHR63299.1;BatCoV_RaTG13_orf1ab;5275 | 0 | 0 |
| MN996532.1;QHR63299.1;BatCoV_RaTG13_orf1ab;5276 | 0 | 0 |
| MN996532.1;QHR63299.1;BatCoV_RaTG13_orf1ab;5277 | 0 | 0 |
| MN996532.1;QHR63299.1;BatCoV_RaTG13_orf1ab;5278 | 0 | 0 |
| MN996532.1;QHR63299.1;BatCoV_RaTG13_orf1ab;5279 | 0 | 0 |
| MN996532.1;QHR63299.1;BatCoV_RaTG13_orf1ab;5280 | 0 | 0.075 |

| | | |
|---|---|---|
| MN996532.1;QHR63299.1;BatCoV_RaTG13_orf1ab;5281 | 0 | 0.025 |
| MN996532.1;QHR63299.1;BatCoV_RaTG13_orf1ab;5282 | 0 | 0 |
| MN996532.1;QHR63299.1;BatCoV_RaTG13_orf1ab;5283 | 0 | 0 |
| MN996532.1;QHR63299.1;BatCoV_RaTG13_orf1ab;5284 | 0 | 0 |
| MN996532.1;QHR63299.1;BatCoV_RaTG13_orf1ab;5285 | 0 | 0 |
| MN996532.1;QHR63299.1;BatCoV_RaTG13_orf1ab;5286 | 0 | 0 |
| MN996532.1;QHR63299.1;BatCoV_RaTG13_orf1ab;5287 | 0 | 0 |
| MN996532.1;QHR63299.1;BatCoV_RaTG13_orf1ab;5288 | 0 | 0 |
| MN996532.1;QHR63299.1;BatCoV_RaTG13_orf1ab;5289 | 0 | 0 |
| MN996532.1;QHR63299.1;BatCoV_RaTG13_orf1ab;5290 | 0 | 0 |
| MN996532.1;QHR63299.1;BatCoV_RaTG13_orf1ab;5291 | 0 | 0 |
| MN996532.1;QHR63299.1;BatCoV_RaTG13_orf1ab;5292 | 0 | 0 |
| MN996532.1;QHR63299.1;BatCoV_RaTG13_orf1ab;5293 | 0 | 0 |
| MN996532.1;QHR63299.1;BatCoV_RaTG13_orf1ab;5294 | 0 | 0 |
| MN996532.1;QHR63299.1;BatCoV_RaTG13_orf1ab;5295 | 0 | 0.025 |
| MN996532.1;QHR63299.1;BatCoV_RaTG13_orf1ab;5296 | 0 | 0 |
| MN996532.1;QHR63299.1;BatCoV_RaTG13_orf1ab;5297 | 0 | 0 |
| MN996532.1;QHR63299.1;BatCoV_RaTG13_orf1ab;5298 | 0 | 0 |
| MN996532.1;QHR63299.1;BatCoV_RaTG13_orf1ab;5299 | 0 | 0 |
| MN996532.1;QHR63299.1;BatCoV_RaTG13_orf1ab;5300 | 0 | 0 |
| MN996532.1;QHR63299.1;BatCoV_RaTG13_orf1ab;5301 | 0.05 | 0 |
| MN996532.1;QHR63299.1;BatCoV_RaTG13_orf1ab;5302 | 0 | 0 |
| MN996532.1;QHR63299.1;BatCoV_RaTG13_orf1ab;5303 | 0 | 0 |
| MN996532.1;QHR63299.1;BatCoV_RaTG13_orf1ab;5304 | 0 | 0 |
| MN996532.1;QHR63299.1;BatCoV_RaTG13_orf1ab;5305 | 0 | 0 |
| MN996532.1;QHR63299.1;BatCoV_RaTG13_orf1ab;5306 | 0 | 0 |
| MN996532.1;QHR63299.1;BatCoV_RaTG13_orf1ab;5307 | 0 | 0 |
| MN996532.1;QHR63299.1;BatCoV_RaTG13_orf1ab;5308 | 0 | 0 |
| MN996532.1;QHR63299.1;BatCoV_RaTG13_orf1ab;5309 | 0 | 0 |
| MN996532.1;QHR63299.1;BatCoV_RaTG13_orf1ab;5310 | 0 | 0 |
| MN996532.1;QHR63299.1;BatCoV_RaTG13_orf1ab;5311 | 0 | 0 |
| MN996532.1;QHR63299.1;BatCoV_RaTG13_orf1ab;5312 | 0 | 0 |
| MN996532.1;QHR63299.1;BatCoV_RaTG13_orf1ab;5313 | 0 | 0 |
| MN996532.1;QHR63299.1;BatCoV_RaTG13_orf1ab;5314 | 0 | 0 |
| MN996532.1;QHR63299.1;BatCoV_RaTG13_orf1ab;5315 | 0 | 0 |
| MN996532.1;QHR63299.1;BatCoV_RaTG13_orf1ab;5316 | 0 | 0 |
| MN996532.1;QHR63299.1;BatCoV_RaTG13_orf1ab;5317 | 0 | 0 |
| MN996532.1;QHR63299.1;BatCoV_RaTG13_orf1ab;5318 | 0 | 0 |
| MN996532.1;QHR63299.1;BatCoV_RaTG13_orf1ab;5319 | 0 | 0 |
| MN996532.1;QHR63299.1;BatCoV_RaTG13_orf1ab;5320 | 0 | 0 |
| MN996532.1;QHR63299.1;BatCoV_RaTG13_orf1ab;5321 | 0 | 0 |
| MN996532.1;QHR63299.1;BatCoV_RaTG13_orf1ab;5322 | 0 | 0 |
| MN996532.1;QHR63299.1;BatCoV_RaTG13_orf1ab;5323 | 0 | 0 |

| | | |
|---|---|---|
| MN996532.1;QHR63299.1;BatCoV_RaTG13_orf1ab;5324 | 0 | 0 |
| MN996532.1;QHR63299.1;BatCoV_RaTG13_orf1ab;5325 | 0 | 0 |
| MN996532.1;QHR63299.1;BatCoV_RaTG13_orf1ab;5326 | 0 | 0 |
| MN996532.1;QHR63299.1;BatCoV_RaTG13_orf1ab;5327 | 0 | 0 |
| MN996532.1;QHR63299.1;BatCoV_RaTG13_orf1ab;5328 | 0 | 0 |
| MN996532.1;QHR63299.1;BatCoV_RaTG13_orf1ab;5329 | 0 | 0 |
| MN996532.1;QHR63299.1;BatCoV_RaTG13_orf1ab;5330 | 0 | 0 |
| MN996532.1;QHR63299.1;BatCoV_RaTG13_orf1ab;5331 | 0 | 0 |
| MN996532.1;QHR63299.1;BatCoV_RaTG13_orf1ab;5332 | 0 | 0 |
| MN996532.1;QHR63299.1;BatCoV_RaTG13_orf1ab;5333 | 0 | 0 |
| MN996532.1;QHR63299.1;BatCoV_RaTG13_orf1ab;5334 | 0 | 0 |
| MN996532.1;QHR63299.1;BatCoV_RaTG13_orf1ab;5335 | 0 | 0 |
| MN996532.1;QHR63299.1;BatCoV_RaTG13_orf1ab;5336 | 0 | 0 |
| MN996532.1;QHR63299.1;BatCoV_RaTG13_orf1ab;5337 | 0 | 0 |
| MN996532.1;QHR63299.1;BatCoV_RaTG13_orf1ab;5338 | 0 | 0 |
| MN996532.1;QHR63299.1;BatCoV_RaTG13_orf1ab;5339 | 0 | 0 |
| MN996532.1;QHR63299.1;BatCoV_RaTG13_orf1ab;5340 | 0 | 0 |
| MN996532.1;QHR63299.1;BatCoV_RaTG13_orf1ab;5341 | 0 | 0 |
| MN996532.1;QHR63299.1;BatCoV_RaTG13_orf1ab;5342 | 0 | 0 |
| MN996532.1;QHR63299.1;BatCoV_RaTG13_orf1ab;5343 | 0 | 0 |
| MN996532.1;QHR63299.1;BatCoV_RaTG13_orf1ab;5344 | 0.05 | 0 |
| MN996532.1;QHR63299.1;BatCoV_RaTG13_orf1ab;5345 | 0 | 0 |
| MN996532.1;QHR63299.1;BatCoV_RaTG13_orf1ab;5346 | 0 | 0 |
| MN996532.1;QHR63299.1;BatCoV_RaTG13_orf1ab;5347 | 0 | 0 |
| MN996532.1;QHR63299.1;BatCoV_RaTG13_orf1ab;5348 | 0 | 0 |
| MN996532.1;QHR63299.1;BatCoV_RaTG13_orf1ab;5349 | 0 | 0 |
| MN996532.1;QHR63299.1;BatCoV_RaTG13_orf1ab;5350 | 0 | 0 |
| MN996532.1;QHR63299.1;BatCoV_RaTG13_orf1ab;5351 | 0 | 0 |
| MN996532.1;QHR63299.1;BatCoV_RaTG13_orf1ab;5352 | 0 | 0 |
| MN996532.1;QHR63299.1;BatCoV_RaTG13_orf1ab;5353 | 0 | 0 |
| MN996532.1;QHR63299.1;BatCoV_RaTG13_orf1ab;5354 | 0 | 0.025 |
| MN996532.1;QHR63299.1;BatCoV_RaTG13_orf1ab;5355 | 0 | 0 |
| MN996532.1;QHR63299.1;BatCoV_RaTG13_orf1ab;5356 | 0 | 0 |
| MN996532.1;QHR63299.1;BatCoV_RaTG13_orf1ab;5357 | 0 | 0 |
| MN996532.1;QHR63299.1;BatCoV_RaTG13_orf1ab;5358 | 0 | 0 |
| MN996532.1;QHR63299.1;BatCoV_RaTG13_orf1ab;5359 | 0 | 0 |
| MN996532.1;QHR63299.1;BatCoV_RaTG13_orf1ab;5360 | 0 | 0 |
| MN996532.1;QHR63299.1;BatCoV_RaTG13_orf1ab;5361 | 0 | 0 |
| MN996532.1;QHR63299.1;BatCoV_RaTG13_orf1ab;5362 | 0 | 0 |
| MN996532.1;QHR63299.1;BatCoV_RaTG13_orf1ab;5363 | 0 | 0 |
| MN996532.1;QHR63299.1;BatCoV_RaTG13_orf1ab;5364 | 0 | 0 |
| MN996532.1;QHR63299.1;BatCoV_RaTG13_orf1ab;5365 | 0 | 0.075 |
| MN996532.1;QHR63299.1;BatCoV_RaTG13_orf1ab;5366 | 0 | 0 |

| | | |
|---|---|---|
| MN996532.1;QHR63299.1;BatCoV_RaTG13_orf1ab;5367 | 0 | 0 |
| MN996532.1;QHR63299.1;BatCoV_RaTG13_orf1ab;5368 | 0 | 0 |
| MN996532.1;QHR63299.1;BatCoV_RaTG13_orf1ab;5369 | 0.05 | 0 |
| MN996532.1;QHR63299.1;BatCoV_RaTG13_orf1ab;5370 | 0.05 | 0 |
| MN996532.1;QHR63299.1;BatCoV_RaTG13_orf1ab;5371 | 0.05 | 0 |
| MN996532.1;QHR63299.1;BatCoV_RaTG13_orf1ab;5372 | 0.05 | 0 |
| MN996532.1;QHR63299.1;BatCoV_RaTG13_orf1ab;5373 | 0 | 0 |
| MN996532.1;QHR63299.1;BatCoV_RaTG13_orf1ab;5374 | 0 | 0 |
| MN996532.1;QHR63299.1;BatCoV_RaTG13_orf1ab;5375 | 0 | 0 |
| MN996532.1;QHR63299.1;BatCoV_RaTG13_orf1ab;5376 | 0 | 0 |
| MN996532.1;QHR63299.1;BatCoV_RaTG13_orf1ab;5377 | 0.05 | 0 |
| MN996532.1;QHR63299.1;BatCoV_RaTG13_orf1ab;5378 | 0 | 0 |
| MN996532.1;QHR63299.1;BatCoV_RaTG13_orf1ab;5379 | 0 | 0 |
| MN996532.1;QHR63299.1;BatCoV_RaTG13_orf1ab;5380 | 0 | 0 |
| MN996532.1;QHR63299.1;BatCoV_RaTG13_orf1ab;5381 | 0 | 0 |
| MN996532.1;QHR63299.1;BatCoV_RaTG13_orf1ab;5382 | 0 | 0 |
| MN996532.1;QHR63299.1;BatCoV_RaTG13_orf1ab;5383 | 0 | 0 |
| MN996532.1;QHR63299.1;BatCoV_RaTG13_orf1ab;5384 | 0 | 0 |
| MN996532.1;QHR63299.1;BatCoV_RaTG13_orf1ab;5385 | 0 | 0 |
| MN996532.1;QHR63299.1;BatCoV_RaTG13_orf1ab;5386 | 0 | 0 |
| MN996532.1;QHR63299.1;BatCoV_RaTG13_orf1ab;5387 | 0 | 0 |
| MN996532.1;QHR63299.1;BatCoV_RaTG13_orf1ab;5388 | 0 | 0 |
| MN996532.1;QHR63299.1;BatCoV_RaTG13_orf1ab;5389 | 0 | 0 |
| MN996532.1;QHR63299.1;BatCoV_RaTG13_orf1ab;5390 | 0 | 0 |
| MN996532.1;QHR63299.1;BatCoV_RaTG13_orf1ab;5391 | 0 | 0 |
| MN996532.1;QHR63299.1;BatCoV_RaTG13_orf1ab;5392 | 0 | 0 |
| MN996532.1;QHR63299.1;BatCoV_RaTG13_orf1ab;5393 | 0 | 0 |
| MN996532.1;QHR63299.1;BatCoV_RaTG13_orf1ab;5394 | 0 | 0 |
| MN996532.1;QHR63299.1;BatCoV_RaTG13_orf1ab;5395 | 0 | 0 |
| MN996532.1;QHR63299.1;BatCoV_RaTG13_orf1ab;5396 | 0 | 0 |
| MN996532.1;QHR63299.1;BatCoV_RaTG13_orf1ab;5397 | 0 | 0 |
| MN996532.1;QHR63299.1;BatCoV_RaTG13_orf1ab;5398 | 0 | 0 |
| MN996532.1;QHR63299.1;BatCoV_RaTG13_orf1ab;5399 | 0 | 0 |
| MN996532.1;QHR63299.1;BatCoV_RaTG13_orf1ab;5400 | 0 | 0.025 |
| MN996532.1;QHR63299.1;BatCoV_RaTG13_orf1ab;5401 | 0 | 0 |
| MN996532.1;QHR63299.1;BatCoV_RaTG13_orf1ab;5402 | 0 | 0 |
| MN996532.1;QHR63299.1;BatCoV_RaTG13_orf1ab;5403 | 0 | 0 |
| MN996532.1;QHR63299.1;BatCoV_RaTG13_orf1ab;5404 | 0 | 0 |
| MN996532.1;QHR63299.1;BatCoV_RaTG13_orf1ab;5405 | 0 | 0 |
| MN996532.1;QHR63299.1;BatCoV_RaTG13_orf1ab;5406 | 0 | 0 |
| MN996532.1;QHR63299.1;BatCoV_RaTG13_orf1ab;5407 | 0 | 0 |
| MN996532.1;QHR63299.1;BatCoV_RaTG13_orf1ab;5408 | 0 | 0 |
| MN996532.1;QHR63299.1;BatCoV_RaTG13_orf1ab;5409 | 0 | 0 |

| | | |
|---|---|---|
| MN996532.1;QHR63299.1;BatCoV_RaTG13_orf1ab;5410 | 0 | 0 |
| MN996532.1;QHR63299.1;BatCoV_RaTG13_orf1ab;5411 | 0 | 0 |
| MN996532.1;QHR63299.1;BatCoV_RaTG13_orf1ab;5412 | 0 | 0 |
| MN996532.1;QHR63299.1;BatCoV_RaTG13_orf1ab;5413 | 0 | 0 |
| MN996532.1;QHR63299.1;BatCoV_RaTG13_orf1ab;5414 | 0.05 | 0 |
| MN996532.1;QHR63299.1;BatCoV_RaTG13_orf1ab;5415 | 0.15 | 0.125 |
| MN996532.1;QHR63299.1;BatCoV_RaTG13_orf1ab;5416 | 0.15 | 0.125 |
| MN996532.1;QHR63299.1;BatCoV_RaTG13_orf1ab;5417 | 0.2 | 0.15 |
| MN996532.1;QHR63299.1;BatCoV_RaTG13_orf1ab;5418 | 0.1 | 0.175 |
| MN996532.1;QHR63299.1;BatCoV_RaTG13_orf1ab;5419 | 0.05 | 0.1 |
| MN996532.1;QHR63299.1;BatCoV_RaTG13_orf1ab;5420 | 0.05 | 0.075 |
| MN996532.1;QHR63299.1;BatCoV_RaTG13_orf1ab;5421 | 0.05 | 0.2 |
| MN996532.1;QHR63299.1;BatCoV_RaTG13_orf1ab;5422 | 0.05 | 0.075 |
| MN996532.1;QHR63299.1;BatCoV_RaTG13_orf1ab;5423 | 0.05 | 0.025 |
| MN996532.1;QHR63299.1;BatCoV_RaTG13_orf1ab;5424 | 0.05 | 0.05 |
| MN996532.1;QHR63299.1;BatCoV_RaTG13_orf1ab;5425 | 0 | 0.025 |
| MN996532.1;QHR63299.1;BatCoV_RaTG13_orf1ab;5426 | 0 | 0 |
| MN996532.1;QHR63299.1;BatCoV_RaTG13_orf1ab;5427 | 0 | 0 |
| MN996532.1;QHR63299.1;BatCoV_RaTG13_orf1ab;5428 | 0 | 0 |
| MN996532.1;QHR63299.1;BatCoV_RaTG13_orf1ab;5429 | 0 | 0 |
| MN996532.1;QHR63299.1;BatCoV_RaTG13_orf1ab;5430 | 0 | 0 |
| MN996532.1;QHR63299.1;BatCoV_RaTG13_orf1ab;5431 | 0 | 0 |
| MN996532.1;QHR63299.1;BatCoV_RaTG13_orf1ab;5432 | 0 | 0 |
| MN996532.1;QHR63299.1;BatCoV_RaTG13_orf1ab;5433 | 0 | 0 |
| MN996532.1;QHR63299.1;BatCoV_RaTG13_orf1ab;5434 | 0 | 0 |
| MN996532.1;QHR63299.1;BatCoV_RaTG13_orf1ab;5435 | 0 | 0 |
| MN996532.1;QHR63299.1;BatCoV_RaTG13_orf1ab;5436 | 0 | 0 |
| MN996532.1;QHR63299.1;BatCoV_RaTG13_orf1ab;5437 | 0 | 0 |
| MN996532.1;QHR63299.1;BatCoV_RaTG13_orf1ab;5438 | 0 | 0 |
| MN996532.1;QHR63299.1;BatCoV_RaTG13_orf1ab;5439 | 0 | 0 |
| MN996532.1;QHR63299.1;BatCoV_RaTG13_orf1ab;5440 | 0.05 | 0 |
| MN996532.1;QHR63299.1;BatCoV_RaTG13_orf1ab;5441 | 0 | 0 |
| MN996532.1;QHR63299.1;BatCoV_RaTG13_orf1ab;5442 | 0 | 0 |
| MN996532.1;QHR63299.1;BatCoV_RaTG13_orf1ab;5443 | 0 | 0 |
| MN996532.1;QHR63299.1;BatCoV_RaTG13_orf1ab;5444 | 0 | 0 |
| MN996532.1;QHR63299.1;BatCoV_RaTG13_orf1ab;5445 | 0 | 0 |
| MN996532.1;QHR63299.1;BatCoV_RaTG13_orf1ab;5446 | 0 | 0 |
| MN996532.1;QHR63299.1;BatCoV_RaTG13_orf1ab;5447 | 0 | 0 |
| MN996532.1;QHR63299.1;BatCoV_RaTG13_orf1ab;5448 | 0 | 0.025 |
| MN996532.1;QHR63299.1;BatCoV_RaTG13_orf1ab;5449 | 0 | 0 |
| MN996532.1;QHR63299.1;BatCoV_RaTG13_orf1ab;5450 | 0 | 0 |
| MN996532.1;QHR63299.1;BatCoV_RaTG13_orf1ab;5451 | 0.05 | 0 |
| MN996532.1;QHR63299.1;BatCoV_RaTG13_orf1ab;5452 | 0.05 | 0 |

| | | |
|---|---|---|
| MN996532.1;QHR63299.1;BatCoV_RaTG13_orf1ab;5453 | 0.05 | 0 |
| MN996532.1;QHR63299.1;BatCoV_RaTG13_orf1ab;5454 | 0 | 0 |
| MN996532.1;QHR63299.1;BatCoV_RaTG13_orf1ab;5455 | 0 | 0 |
| MN996532.1;QHR63299.1;BatCoV_RaTG13_orf1ab;5456 | 0 | 0 |
| MN996532.1;QHR63299.1;BatCoV_RaTG13_orf1ab;5457 | 0 | 0 |
| MN996532.1;QHR63299.1;BatCoV_RaTG13_orf1ab;5458 | 0.05 | 0.025 |
| MN996532.1;QHR63299.1;BatCoV_RaTG13_orf1ab;5459 | 0 | 0 |
| MN996532.1;QHR63299.1;BatCoV_RaTG13_orf1ab;5460 | 0 | 0.025 |
| MN996532.1;QHR63299.1;BatCoV_RaTG13_orf1ab;5461 | 0 | 0 |
| MN996532.1;QHR63299.1;BatCoV_RaTG13_orf1ab;5462 | 0 | 0 |
| MN996532.1;QHR63299.1;BatCoV_RaTG13_orf1ab;5463 | 0 | 0 |
| MN996532.1;QHR63299.1;BatCoV_RaTG13_orf1ab;5464 | 0 | 0 |
| MN996532.1;QHR63299.1;BatCoV_RaTG13_orf1ab;5465 | 0.05 | 0 |
| MN996532.1;QHR63299.1;BatCoV_RaTG13_orf1ab;5466 | 0 | 0 |
| MN996532.1;QHR63299.1;BatCoV_RaTG13_orf1ab;5467 | 0 | 0 |
| MN996532.1;QHR63299.1;BatCoV_RaTG13_orf1ab;5468 | 0 | 0 |
| MN996532.1;QHR63299.1;BatCoV_RaTG13_orf1ab;5469 | 0.05 | 0 |
| MN996532.1;QHR63299.1;BatCoV_RaTG13_orf1ab;5470 | 0.05 | 0 |
| MN996532.1;QHR63299.1;BatCoV_RaTG13_orf1ab;5471 | 0.05 | 0 |
| MN996532.1;QHR63299.1;BatCoV_RaTG13_orf1ab;5472 | 0.05 | 0.05 |
| MN996532.1;QHR63299.1;BatCoV_RaTG13_orf1ab;5473 | 0.05 | 0.05 |
| MN996532.1;QHR63299.1;BatCoV_RaTG13_orf1ab;5474 | 0.05 | 0 |
| MN996532.1;QHR63299.1;BatCoV_RaTG13_orf1ab;5475 | 0 | 0.05 |
| MN996532.1;QHR63299.1;BatCoV_RaTG13_orf1ab;5476 | 0 | 0 |
| MN996532.1;QHR63299.1;BatCoV_RaTG13_orf1ab;5477 | 0 | 0 |
| MN996532.1;QHR63299.1;BatCoV_RaTG13_orf1ab;5478 | 0 | 0 |
| MN996532.1;QHR63299.1;BatCoV_RaTG13_orf1ab;5479 | 0 | 0.05 |
| MN996532.1;QHR63299.1;BatCoV_RaTG13_orf1ab;5480 | 0 | 0.05 |
| MN996532.1;QHR63299.1;BatCoV_RaTG13_orf1ab;5481 | 0 | 0.05 |
| MN996532.1;QHR63299.1;BatCoV_RaTG13_orf1ab;5482 | 0.05 | 0.075 |
| MN996532.1;QHR63299.1;BatCoV_RaTG13_orf1ab;5483 | 0 | 0 |
| MN996532.1;QHR63299.1;BatCoV_RaTG13_orf1ab;5484 | 0 | 0 |
| MN996532.1;QHR63299.1;BatCoV_RaTG13_orf1ab;5485 | 0 | 0 |
| MN996532.1;QHR63299.1;BatCoV_RaTG13_orf1ab;5486 | 0 | 0 |
| MN996532.1;QHR63299.1;BatCoV_RaTG13_orf1ab;5487 | 0 | 0 |
| MN996532.1;QHR63299.1;BatCoV_RaTG13_orf1ab;5488 | 0 | 0 |
| MN996532.1;QHR63299.1;BatCoV_RaTG13_orf1ab;5489 | 0 | 0 |
| MN996532.1;QHR63299.1;BatCoV_RaTG13_orf1ab;5490 | 0 | 0 |
| MN996532.1;QHR63299.1;BatCoV_RaTG13_orf1ab;5491 | 0 | 0 |
| MN996532.1;QHR63299.1;BatCoV_RaTG13_orf1ab;5492 | 0 | 0 |
| MN996532.1;QHR63299.1;BatCoV_RaTG13_orf1ab;5493 | 0 | 0 |
| MN996532.1;QHR63299.1;BatCoV_RaTG13_orf1ab;5494 | 0 | 0 |
| MN996532.1;QHR63299.1;BatCoV_RaTG13_orf1ab;5495 | 0 | 0 |

| | | |
|---|---|---|
| MN996532.1;QHR63299.1;BatCoV_RaTG13_orf1ab;5496 | 0 | 0 |
| MN996532.1;QHR63299.1;BatCoV_RaTG13_orf1ab;5497 | 0 | 0 |
| MN996532.1;QHR63299.1;BatCoV_RaTG13_orf1ab;5498 | 0 | 0 |
| MN996532.1;QHR63299.1;BatCoV_RaTG13_orf1ab;5499 | 0 | 0 |
| MN996532.1;QHR63299.1;BatCoV_RaTG13_orf1ab;5500 | 0 | 0 |
| MN996532.1;QHR63299.1;BatCoV_RaTG13_orf1ab;5501 | 0 | 0 |
| MN996532.1;QHR63299.1;BatCoV_RaTG13_orf1ab;5502 | 0 | 0 |
| MN996532.1;QHR63299.1;BatCoV_RaTG13_orf1ab;5503 | 0 | 0 |
| MN996532.1;QHR63299.1;BatCoV_RaTG13_orf1ab;5504 | 0 | 0 |
| MN996532.1;QHR63299.1;BatCoV_RaTG13_orf1ab;5505 | 0.1 | 0.025 |
| MN996532.1;QHR63299.1;BatCoV_RaTG13_orf1ab;5506 | 0.1 | 0.025 |
| MN996532.1;QHR63299.1;BatCoV_RaTG13_orf1ab;5507 | 0.1 | 0.025 |
| MN996532.1;QHR63299.1;BatCoV_RaTG13_orf1ab;5508 | 0.1 | 0 |
| MN996532.1;QHR63299.1;BatCoV_RaTG13_orf1ab;5509 | 0.05 | 0.05 |
| MN996532.1;QHR63299.1;BatCoV_RaTG13_orf1ab;5510 | 0.05 | 0 |
| MN996532.1;QHR63299.1;BatCoV_RaTG13_orf1ab;5511 | 0.1 | 0 |
| MN996532.1;QHR63299.1;BatCoV_RaTG13_orf1ab;5512 | 0.05 | 0 |
| MN996532.1;QHR63299.1;BatCoV_RaTG13_orf1ab;5513 | 0.05 | 0.025 |
| MN996532.1;QHR63299.1;BatCoV_RaTG13_orf1ab;5514 | 0.05 | 0.025 |
| MN996532.1;QHR63299.1;BatCoV_RaTG13_orf1ab;5515 | 0.05 | 0.025 |
| MN996532.1;QHR63299.1;BatCoV_RaTG13_orf1ab;5516 | 0.05 | 0.025 |
| MN996532.1;QHR63299.1;BatCoV_RaTG13_orf1ab;5517 | 0.1 | 0.025 |
| MN996532.1;QHR63299.1;BatCoV_RaTG13_orf1ab;5518 | 0.05 | 0.025 |
| MN996532.1;QHR63299.1;BatCoV_RaTG13_orf1ab;5519 | 0 | 0 |
| MN996532.1;QHR63299.1;BatCoV_RaTG13_orf1ab;5520 | 0 | 0.025 |
| MN996532.1;QHR63299.1;BatCoV_RaTG13_orf1ab;5521 | 0.1 | 0 |
| MN996532.1;QHR63299.1;BatCoV_RaTG13_orf1ab;5522 | 0 | 0.025 |
| MN996532.1;QHR63299.1;BatCoV_RaTG13_orf1ab;5523 | 0 | 0.025 |
| MN996532.1;QHR63299.1;BatCoV_RaTG13_orf1ab;5524 | 0 | 0.025 |
| MN996532.1;QHR63299.1;BatCoV_RaTG13_orf1ab;5525 | 0 | 0.025 |
| MN996532.1;QHR63299.1;BatCoV_RaTG13_orf1ab;5526 | 0.05 | 0 |
| MN996532.1;QHR63299.1;BatCoV_RaTG13_orf1ab;5527 | 0 | 0 |
| MN996532.1;QHR63299.1;BatCoV_RaTG13_orf1ab;5528 | 0 | 0 |
| MN996532.1;QHR63299.1;BatCoV_RaTG13_orf1ab;5529 | 0 | 0 |
| MN996532.1;QHR63299.1;BatCoV_RaTG13_orf1ab;5530 | 0 | 0 |
| MN996532.1;QHR63299.1;BatCoV_RaTG13_orf1ab;5531 | 0 | 0 |
| MN996532.1;QHR63299.1;BatCoV_RaTG13_orf1ab;5532 | 0 | 0 |
| MN996532.1;QHR63299.1;BatCoV_RaTG13_orf1ab;5533 | 0 | 0 |
| MN996532.1;QHR63299.1;BatCoV_RaTG13_orf1ab;5534 | 0 | 0 |
| MN996532.1;QHR63299.1;BatCoV_RaTG13_orf1ab;5535 | 0 | 0 |
| MN996532.1;QHR63299.1;BatCoV_RaTG13_orf1ab;5536 | 0 | 0 |
| MN996532.1;QHR63299.1;BatCoV_RaTG13_orf1ab;5537 | 0 | 0 |
| MN996532.1;QHR63299.1;BatCoV_RaTG13_orf1ab;5538 | 0 | 0 |

| | | |
|---|---|---|
| MN996532.1;QHR63299.1;BatCoV_RaTG13_orf1ab;5539 | 0 | 0 |
| MN996532.1;QHR63299.1;BatCoV_RaTG13_orf1ab;5540 | 0 | 0 |
| MN996532.1;QHR63299.1;BatCoV_RaTG13_orf1ab;5541 | 0 | 0 |
| MN996532.1;QHR63299.1;BatCoV_RaTG13_orf1ab;5542 | 0 | 0 |
| MN996532.1;QHR63299.1;BatCoV_RaTG13_orf1ab;5543 | 0 | 0 |
| MN996532.1;QHR63299.1;BatCoV_RaTG13_orf1ab;5544 | 0 | 0 |
| MN996532.1;QHR63299.1;BatCoV_RaTG13_orf1ab;5545 | 0 | 0 |
| MN996532.1;QHR63299.1;BatCoV_RaTG13_orf1ab;5546 | 0.05 | 0 |
| MN996532.1;QHR63299.1;BatCoV_RaTG13_orf1ab;5547 | 0 | 0 |
| MN996532.1;QHR63299.1;BatCoV_RaTG13_orf1ab;5548 | 0 | 0 |
| MN996532.1;QHR63299.1;BatCoV_RaTG13_orf1ab;5549 | 0 | 0 |
| MN996532.1;QHR63299.1;BatCoV_RaTG13_orf1ab;5550 | 0 | 0.025 |
| MN996532.1;QHR63299.1;BatCoV_RaTG13_orf1ab;5551 | 0 | 0 |
| MN996532.1;QHR63299.1;BatCoV_RaTG13_orf1ab;5552 | 0 | 0 |
| MN996532.1;QHR63299.1;BatCoV_RaTG13_orf1ab;5553 | 0 | 0 |
| MN996532.1;QHR63299.1;BatCoV_RaTG13_orf1ab;5554 | 0 | 0.025 |
| MN996532.1;QHR63299.1;BatCoV_RaTG13_orf1ab;5555 | 0 | 0.025 |
| MN996532.1;QHR63299.1;BatCoV_RaTG13_orf1ab;5556 | 0 | 0.025 |
| MN996532.1;QHR63299.1;BatCoV_RaTG13_orf1ab;5557 | 0 | 0.025 |
| MN996532.1;QHR63299.1;BatCoV_RaTG13_orf1ab;5558 | 0 | 0.025 |
| MN996532.1;QHR63299.1;BatCoV_RaTG13_orf1ab;5559 | 0 | 0.025 |
| MN996532.1;QHR63299.1;BatCoV_RaTG13_orf1ab;5560 | 0 | 0.025 |
| MN996532.1;QHR63299.1;BatCoV_RaTG13_orf1ab;5561 | 0 | 0 |
| MN996532.1;QHR63299.1;BatCoV_RaTG13_orf1ab;5562 | 0 | 0 |
| MN996532.1;QHR63299.1;BatCoV_RaTG13_orf1ab;5563 | 0 | 0.025 |
| MN996532.1;QHR63299.1;BatCoV_RaTG13_orf1ab;5564 | 0 | 0.025 |
| MN996532.1;QHR63299.1;BatCoV_RaTG13_orf1ab;5565 | 0 | 0 |
| MN996532.1;QHR63299.1;BatCoV_RaTG13_orf1ab;5566 | 0 | 0.025 |
| MN996532.1;QHR63299.1;BatCoV_RaTG13_orf1ab;5567 | 0 | 0 |
| MN996532.1;QHR63299.1;BatCoV_RaTG13_orf1ab;5568 | 0 | 0 |
| MN996532.1;QHR63299.1;BatCoV_RaTG13_orf1ab;5569 | 0 | 0 |
| MN996532.1;QHR63299.1;BatCoV_RaTG13_orf1ab;5570 | 0.05 | 0 |
| MN996532.1;QHR63299.1;BatCoV_RaTG13_orf1ab;5571 | 0.05 | 0.025 |
| MN996532.1;QHR63299.1;BatCoV_RaTG13_orf1ab;5572 | 0.05 | 0.025 |
| MN996532.1;QHR63299.1;BatCoV_RaTG13_orf1ab;5573 | 0.05 | 0.025 |
| MN996532.1;QHR63299.1;BatCoV_RaTG13_orf1ab;5574 | 0.05 | 0.025 |
| MN996532.1;QHR63299.1;BatCoV_RaTG13_orf1ab;5575 | 0.05 | 0.075 |
| MN996532.1;QHR63299.1;BatCoV_RaTG13_orf1ab;5576 | 0.05 | 0.175 |
| MN996532.1;QHR63299.1;BatCoV_RaTG13_orf1ab;5577 | 0.05 | 0.075 |
| MN996532.1;QHR63299.1;BatCoV_RaTG13_orf1ab;5578 | 0.1 | 0.125 |
| MN996532.1;QHR63299.1;BatCoV_RaTG13_orf1ab;5579 | 0.05 | 0.075 |
| MN996532.1;QHR63299.1;BatCoV_RaTG13_orf1ab;5580 | 0 | 0.05 |
| MN996532.1;QHR63299.1;BatCoV_RaTG13_orf1ab;5581 | 0.05 | 0.025 |

| | | |
|---|---|---|
| MN996532.1;QHR63299.1;BatCoV_RaTG13_orf1ab;5582 | 0 | 0 |
| MN996532.1;QHR63299.1;BatCoV_RaTG13_orf1ab;5583 | 0.1 | 0.1 |
| MN996532.1;QHR63299.1;BatCoV_RaTG13_orf1ab;5584 | 0.1 | 0.025 |
| MN996532.1;QHR63299.1;BatCoV_RaTG13_orf1ab;5585 | 0 | 0 |
| MN996532.1;QHR63299.1;BatCoV_RaTG13_orf1ab;5586 | 0.05 | 0.05 |
| MN996532.1;QHR63299.1;BatCoV_RaTG13_orf1ab;5587 | 0 | 0.025 |
| MN996532.1;QHR63299.1;BatCoV_RaTG13_orf1ab;5588 | 0 | 0.025 |
| MN996532.1;QHR63299.1;BatCoV_RaTG13_orf1ab;5589 | 0 | 0.025 |
| MN996532.1;QHR63299.1;BatCoV_RaTG13_orf1ab;5590 | 0 | 0.025 |
| MN996532.1;QHR63299.1;BatCoV_RaTG13_orf1ab;5591 | 0 | 0.025 |
| MN996532.1;QHR63299.1;BatCoV_RaTG13_orf1ab;5592 | 0 | 0.075 |
| MN996532.1;QHR63299.1;BatCoV_RaTG13_orf1ab;5593 | 0 | 0.025 |
| MN996532.1;QHR63299.1;BatCoV_RaTG13_orf1ab;5594 | 0 | 0.025 |
| MN996532.1;QHR63299.1;BatCoV_RaTG13_orf1ab;5595 | 0 | 0 |
| MN996532.1;QHR63299.1;BatCoV_RaTG13_orf1ab;5596 | 0 | 0 |
| MN996532.1;QHR63299.1;BatCoV_RaTG13_orf1ab;5597 | 0.05 | 0.025 |
| MN996532.1;QHR63299.1;BatCoV_RaTG13_orf1ab;5598 | 0 | 0 |
| MN996532.1;QHR63299.1;BatCoV_RaTG13_orf1ab;5599 | 0 | 0 |
| MN996532.1;QHR63299.1;BatCoV_RaTG13_orf1ab;5600 | 0 | 0.025 |
| MN996532.1;QHR63299.1;BatCoV_RaTG13_orf1ab;5601 | 0 | 0 |
| MN996532.1;QHR63299.1;BatCoV_RaTG13_orf1ab;5602 | 0 | 0 |
| MN996532.1;QHR63299.1;BatCoV_RaTG13_orf1ab;5603 | 0 | 0 |
| MN996532.1;QHR63299.1;BatCoV_RaTG13_orf1ab;5604 | 0 | 0 |
| MN996532.1;QHR63299.1;BatCoV_RaTG13_orf1ab;5605 | 0 | 0 |
| MN996532.1;QHR63299.1;BatCoV_RaTG13_orf1ab;5606 | 0 | 0 |
| MN996532.1;QHR63299.1;BatCoV_RaTG13_orf1ab;5607 | 0 | 0 |
| MN996532.1;QHR63299.1;BatCoV_RaTG13_orf1ab;5608 | 0 | 0 |
| MN996532.1;QHR63299.1;BatCoV_RaTG13_orf1ab;5609 | 0 | 0 |
| MN996532.1;QHR63299.1;BatCoV_RaTG13_orf1ab;5610 | 0 | 0 |
| MN996532.1;QHR63299.1;BatCoV_RaTG13_orf1ab;5611 | 0 | 0 |
| MN996532.1;QHR63299.1;BatCoV_RaTG13_orf1ab;5612 | 0 | 0 |
| MN996532.1;QHR63299.1;BatCoV_RaTG13_orf1ab;5613 | 0 | 0 |
| MN996532.1;QHR63299.1;BatCoV_RaTG13_orf1ab;5614 | 0 | 0 |
| MN996532.1;QHR63299.1;BatCoV_RaTG13_orf1ab;5615 | 0 | 0 |
| MN996532.1;QHR63299.1;BatCoV_RaTG13_orf1ab;5616 | 0 | 0 |
| MN996532.1;QHR63299.1;BatCoV_RaTG13_orf1ab;5617 | 0 | 0 |
| MN996532.1;QHR63299.1;BatCoV_RaTG13_orf1ab;5618 | 0 | 0 |
| MN996532.1;QHR63299.1;BatCoV_RaTG13_orf1ab;5619 | 0 | 0 |
| MN996532.1;QHR63299.1;BatCoV_RaTG13_orf1ab;5620 | 0 | 0 |
| MN996532.1;QHR63299.1;BatCoV_RaTG13_orf1ab;5621 | 0 | 0 |
| MN996532.1;QHR63299.1;BatCoV_RaTG13_orf1ab;5622 | 0 | 0 |
| MN996532.1;QHR63299.1;BatCoV_RaTG13_orf1ab;5623 | 0 | 0.025 |
| MN996532.1;QHR63299.1;BatCoV_RaTG13_orf1ab;5624 | 0 | 0.025 |

| | | |
|---|---|---|
| MN996532.1;QHR63299.1;BatCoV_RaTG13_orf1ab;5625 | 0 | 0.05 |
| MN996532.1;QHR63299.1;BatCoV_RaTG13_orf1ab;5626 | 0 | 0 |
| MN996532.1;QHR63299.1;BatCoV_RaTG13_orf1ab;5627 | 0 | 0 |
| MN996532.1;QHR63299.1;BatCoV_RaTG13_orf1ab;5628 | 0 | 0 |
| MN996532.1;QHR63299.1;BatCoV_RaTG13_orf1ab;5629 | 0 | 0 |
| MN996532.1;QHR63299.1;BatCoV_RaTG13_orf1ab;5630 | 0 | 0 |
| MN996532.1;QHR63299.1;BatCoV_RaTG13_orf1ab;5631 | 0 | 0 |
| MN996532.1;QHR63299.1;BatCoV_RaTG13_orf1ab;5632 | 0 | 0 |
| MN996532.1;QHR63299.1;BatCoV_RaTG13_orf1ab;5633 | 0 | 0 |
| MN996532.1;QHR63299.1;BatCoV_RaTG13_orf1ab;5634 | 0 | 0 |
| MN996532.1;QHR63299.1;BatCoV_RaTG13_orf1ab;5635 | 0 | 0 |
| MN996532.1;QHR63299.1;BatCoV_RaTG13_orf1ab;5636 | 0 | 0 |
| MN996532.1;QHR63299.1;BatCoV_RaTG13_orf1ab;5637 | 0.05 | 0 |
| MN996532.1;QHR63299.1;BatCoV_RaTG13_orf1ab;5638 | 0 | 0 |
| MN996532.1;QHR63299.1;BatCoV_RaTG13_orf1ab;5639 | 0 | 0 |
| MN996532.1;QHR63299.1;BatCoV_RaTG13_orf1ab;5640 | 0 | 0 |
| MN996532.1;QHR63299.1;BatCoV_RaTG13_orf1ab;5641 | 0 | 0 |
| MN996532.1;QHR63299.1;BatCoV_RaTG13_orf1ab;5642 | 0 | 0 |
| MN996532.1;QHR63299.1;BatCoV_RaTG13_orf1ab;5643 | 0 | 0 |
| MN996532.1;QHR63299.1;BatCoV_RaTG13_orf1ab;5644 | 0 | 0 |
| MN996532.1;QHR63299.1;BatCoV_RaTG13_orf1ab;5645 | 0 | 0 |
| MN996532.1;QHR63299.1;BatCoV_RaTG13_orf1ab;5646 | 0 | 0 |
| MN996532.1;QHR63299.1;BatCoV_RaTG13_orf1ab;5647 | 0 | 0 |
| MN996532.1;QHR63299.1;BatCoV_RaTG13_orf1ab;5648 | 0 | 0 |
| MN996532.1;QHR63299.1;BatCoV_RaTG13_orf1ab;5649 | 0 | 0 |
| MN996532.1;QHR63299.1;BatCoV_RaTG13_orf1ab;5650 | 0 | 0 |
| MN996532.1;QHR63299.1;BatCoV_RaTG13_orf1ab;5651 | 0 | 0 |
| MN996532.1;QHR63299.1;BatCoV_RaTG13_orf1ab;5652 | 0 | 0 |
| MN996532.1;QHR63299.1;BatCoV_RaTG13_orf1ab;5653 | 0 | 0 |
| MN996532.1;QHR63299.1;BatCoV_RaTG13_orf1ab;5654 | 0 | 0 |
| MN996532.1;QHR63299.1;BatCoV_RaTG13_orf1ab;5655 | 0 | 0 |
| MN996532.1;QHR63299.1;BatCoV_RaTG13_orf1ab;5656 | 0 | 0.025 |
| MN996532.1;QHR63299.1;BatCoV_RaTG13_orf1ab;5657 | 0 | 0.025 |
| MN996532.1;QHR63299.1;BatCoV_RaTG13_orf1ab;5658 | 0.05 | 0.025 |
| MN996532.1;QHR63299.1;BatCoV_RaTG13_orf1ab;5659 | 0 | 0.025 |
| MN996532.1;QHR63299.1;BatCoV_RaTG13_orf1ab;5660 | 0.05 | 0.075 |
| MN996532.1;QHR63299.1;BatCoV_RaTG13_orf1ab;5661 | 0.25 | 0.075 |
| MN996532.1;QHR63299.1;BatCoV_RaTG13_orf1ab;5662 | 0.4 | 0.125 |
| MN996532.1;QHR63299.1;BatCoV_RaTG13_orf1ab;5663 | 0.35 | 0.075 |
| MN996532.1;QHR63299.1;BatCoV_RaTG13_orf1ab;5664 | 0.3 | 0.075 |
| MN996532.1;QHR63299.1;BatCoV_RaTG13_orf1ab;5665 | 0.1 | 0.025 |
| MN996532.1;QHR63299.1;BatCoV_RaTG13_orf1ab;5666 | 0.05 | 0.025 |
| MN996532.1;QHR63299.1;BatCoV_RaTG13_orf1ab;5667 | 0 | 0.025 |

| | | |
|---|---|---|
| MN996532.1;QHR63299.1;BatCoV_RaTG13_orf1ab;5668 | 0 | 0 |
| MN996532.1;QHR63299.1;BatCoV_RaTG13_orf1ab;5669 | 0 | 0 |
| MN996532.1;QHR63299.1;BatCoV_RaTG13_orf1ab;5670 | 0 | 0 |
| MN996532.1;QHR63299.1;BatCoV_RaTG13_orf1ab;5671 | 0 | 0 |
| MN996532.1;QHR63299.1;BatCoV_RaTG13_orf1ab;5672 | 0 | 0 |
| MN996532.1;QHR63299.1;BatCoV_RaTG13_orf1ab;5673 | 0 | 0 |
| MN996532.1;QHR63299.1;BatCoV_RaTG13_orf1ab;5674 | 0 | 0 |
| MN996532.1;QHR63299.1;BatCoV_RaTG13_orf1ab;5675 | 0 | 0.025 |
| MN996532.1;QHR63299.1;BatCoV_RaTG13_orf1ab;5676 | 0 | 0 |
| MN996532.1;QHR63299.1;BatCoV_RaTG13_orf1ab;5677 | 0 | 0 |
| MN996532.1;QHR63299.1;BatCoV_RaTG13_orf1ab;5678 | 0 | 0 |
| MN996532.1;QHR63299.1;BatCoV_RaTG13_orf1ab;5679 | 0 | 0 |
| MN996532.1;QHR63299.1;BatCoV_RaTG13_orf1ab;5680 | 0 | 0 |
| MN996532.1;QHR63299.1;BatCoV_RaTG13_orf1ab;5681 | 0 | 0 |
| MN996532.1;QHR63299.1;BatCoV_RaTG13_orf1ab;5682 | 0 | 0 |
| MN996532.1;QHR63299.1;BatCoV_RaTG13_orf1ab;5683 | 0 | 0 |
| MN996532.1;QHR63299.1;BatCoV_RaTG13_orf1ab;5684 | 0 | 0 |
| MN996532.1;QHR63299.1;BatCoV_RaTG13_orf1ab;5685 | 0 | 0 |
| MN996532.1;QHR63299.1;BatCoV_RaTG13_orf1ab;5686 | 0 | 0 |
| MN996532.1;QHR63299.1;BatCoV_RaTG13_orf1ab;5687 | 0 | 0.025 |
| MN996532.1;QHR63299.1;BatCoV_RaTG13_orf1ab;5688 | 0 | 0.025 |
| MN996532.1;QHR63299.1;BatCoV_RaTG13_orf1ab;5689 | 0 | 0 |
| MN996532.1;QHR63299.1;BatCoV_RaTG13_orf1ab;5690 | 0 | 0 |
| MN996532.1;QHR63299.1;BatCoV_RaTG13_orf1ab;5691 | 0 | 0.075 |
| MN996532.1;QHR63299.1;BatCoV_RaTG13_orf1ab;5692 | 0 | 0 |
| MN996532.1;QHR63299.1;BatCoV_RaTG13_orf1ab;5693 | 0 | 0.025 |
| MN996532.1;QHR63299.1;BatCoV_RaTG13_orf1ab;5694 | 0 | 0.025 |
| MN996532.1;QHR63299.1;BatCoV_RaTG13_orf1ab;5695 | 0 | 0.05 |
| MN996532.1;QHR63299.1;BatCoV_RaTG13_orf1ab;5696 | 0 | 0.05 |
| MN996532.1;QHR63299.1;BatCoV_RaTG13_orf1ab;5697 | 0 | 0.025 |
| MN996532.1;QHR63299.1;BatCoV_RaTG13_orf1ab;5698 | 0 | 0.025 |
| MN996532.1;QHR63299.1;BatCoV_RaTG13_orf1ab;5699 | 0 | 0.025 |
| MN996532.1;QHR63299.1;BatCoV_RaTG13_orf1ab;5700 | 0 | 0.025 |
| MN996532.1;QHR63299.1;BatCoV_RaTG13_orf1ab;5701 | 0 | 0 |
| MN996532.1;QHR63299.1;BatCoV_RaTG13_orf1ab;5702 | 0 | 0 |
| MN996532.1;QHR63299.1;BatCoV_RaTG13_orf1ab;5703 | 0 | 0 |
| MN996532.1;QHR63299.1;BatCoV_RaTG13_orf1ab;5704 | 0 | 0.025 |
| MN996532.1;QHR63299.1;BatCoV_RaTG13_orf1ab;5705 | 0 | 0.025 |
| MN996532.1;QHR63299.1;BatCoV_RaTG13_orf1ab;5706 | 0 | 0.025 |
| MN996532.1;QHR63299.1;BatCoV_RaTG13_orf1ab;5707 | 0 | 0 |
| MN996532.1;QHR63299.1;BatCoV_RaTG13_orf1ab;5708 | 0 | 0 |
| MN996532.1;QHR63299.1;BatCoV_RaTG13_orf1ab;5709 | 0 | 0.025 |
| MN996532.1;QHR63299.1;BatCoV_RaTG13_orf1ab;5710 | 0 | 0.025 |

| | | |
|---|---|---|
| MN996532.1;QHR63299.1;BatCoV_RaTG13_orf1ab;5711 | 0 | 0.025 |
| MN996532.1;QHR63299.1;BatCoV_RaTG13_orf1ab;5712 | 0 | 0.025 |
| MN996532.1;QHR63299.1;BatCoV_RaTG13_orf1ab;5713 | 0 | 0.025 |
| MN996532.1;QHR63299.1;BatCoV_RaTG13_orf1ab;5714 | 0 | 0.025 |
| MN996532.1;QHR63299.1;BatCoV_RaTG13_orf1ab;5715 | 0 | 0.025 |
| MN996532.1;QHR63299.1;BatCoV_RaTG13_orf1ab;5716 | 0 | 0 |
| MN996532.1;QHR63299.1;BatCoV_RaTG13_orf1ab;5717 | 0 | 0 |
| MN996532.1;QHR63299.1;BatCoV_RaTG13_orf1ab;5718 | 0 | 0 |
| MN996532.1;QHR63299.1;BatCoV_RaTG13_orf1ab;5719 | 0 | 0 |
| MN996532.1;QHR63299.1;BatCoV_RaTG13_orf1ab;5720 | 0 | 0.025 |
| MN996532.1;QHR63299.1;BatCoV_RaTG13_orf1ab;5721 | 0 | 0 |
| MN996532.1;QHR63299.1;BatCoV_RaTG13_orf1ab;5722 | 0 | 0.025 |
| MN996532.1;QHR63299.1;BatCoV_RaTG13_orf1ab;5723 | 0 | 0.025 |
| MN996532.1;QHR63299.1;BatCoV_RaTG13_orf1ab;5724 | 0 | 0.025 |
| MN996532.1;QHR63299.1;BatCoV_RaTG13_orf1ab;5725 | 0 | 0 |
| MN996532.1;QHR63299.1;BatCoV_RaTG13_orf1ab;5726 | 0 | 0.025 |
| MN996532.1;QHR63299.1;BatCoV_RaTG13_orf1ab;5727 | 0 | 0.075 |
| MN996532.1;QHR63299.1;BatCoV_RaTG13_orf1ab;5728 | 0 | 0.075 |
| MN996532.1;QHR63299.1;BatCoV_RaTG13_orf1ab;5729 | 0.05 | 0.1 |
| MN996532.1;QHR63299.1;BatCoV_RaTG13_orf1ab;5730 | 0.1 | 0.075 |
| MN996532.1;QHR63299.1;BatCoV_RaTG13_orf1ab;5731 | 0.1 | 0.075 |
| MN996532.1;QHR63299.1;BatCoV_RaTG13_orf1ab;5732 | 0.1 | 0.05 |
| MN996532.1;QHR63299.1;BatCoV_RaTG13_orf1ab;5733 | 0.1 | 0.05 |
| MN996532.1;QHR63299.1;BatCoV_RaTG13_orf1ab;5734 | 0.1 | 0.05 |
| MN996532.1;QHR63299.1;BatCoV_RaTG13_orf1ab;5735 | 0.1 | 0.05 |
| MN996532.1;QHR63299.1;BatCoV_RaTG13_orf1ab;5736 | 0.1 | 0 |
| MN996532.1;QHR63299.1;BatCoV_RaTG13_orf1ab;5737 | 0.05 | 0.025 |
| MN996532.1;QHR63299.1;BatCoV_RaTG13_orf1ab;5738 | 0 | 0 |
| MN996532.1;QHR63299.1;BatCoV_RaTG13_orf1ab;5739 | 0 | 0 |
| MN996532.1;QHR63299.1;BatCoV_RaTG13_orf1ab;5740 | 0 | 0 |
| MN996532.1;QHR63299.1;BatCoV_RaTG13_orf1ab;5741 | 0 | 0 |
| MN996532.1;QHR63299.1;BatCoV_RaTG13_orf1ab;5742 | 0 | 0 |
| MN996532.1;QHR63299.1;BatCoV_RaTG13_orf1ab;5743 | 0 | 0 |
| MN996532.1;QHR63299.1;BatCoV_RaTG13_orf1ab;5744 | 0 | 0 |
| MN996532.1;QHR63299.1;BatCoV_RaTG13_orf1ab;5745 | 0 | 0 |
| MN996532.1;QHR63299.1;BatCoV_RaTG13_orf1ab;5746 | 0 | 0 |
| MN996532.1;QHR63299.1;BatCoV_RaTG13_orf1ab;5747 | 0 | 0 |
| MN996532.1;QHR63299.1;BatCoV_RaTG13_orf1ab;5748 | 0 | 0.025 |
| MN996532.1;QHR63299.1;BatCoV_RaTG13_orf1ab;5749 | 0 | 0 |
| MN996532.1;QHR63299.1;BatCoV_RaTG13_orf1ab;5750 | 0 | 0 |
| MN996532.1;QHR63299.1;BatCoV_RaTG13_orf1ab;5751 | 0 | 0 |
| MN996532.1;QHR63299.1;BatCoV_RaTG13_orf1ab;5752 | 0 | 0 |
| MN996532.1;QHR63299.1;BatCoV_RaTG13_orf1ab;5753 | 0 | 0 |

| | | |
|---|---|---|
| MN996532.1;QHR63299.1;BatCoV_RaTG13_orf1ab;5754 | 0 | 0 |
| MN996532.1;QHR63299.1;BatCoV_RaTG13_orf1ab;5755 | 0 | 0 |
| MN996532.1;QHR63299.1;BatCoV_RaTG13_orf1ab;5756 | 0 | 0 |
| MN996532.1;QHR63299.1;BatCoV_RaTG13_orf1ab;5757 | 0 | 0 |
| MN996532.1;QHR63299.1;BatCoV_RaTG13_orf1ab;5758 | 0 | 0 |
| MN996532.1;QHR63299.1;BatCoV_RaTG13_orf1ab;5759 | 0 | 0 |
| MN996532.1;QHR63299.1;BatCoV_RaTG13_orf1ab;5760 | 0 | 0 |
| MN996532.1;QHR63299.1;BatCoV_RaTG13_orf1ab;5761 | 0.1 | 0.05 |
| MN996532.1;QHR63299.1;BatCoV_RaTG13_orf1ab;5762 | 0.1 | 0.05 |
| MN996532.1;QHR63299.1;BatCoV_RaTG13_orf1ab;5763 | 0.15 | 0.125 |
| MN996532.1;QHR63299.1;BatCoV_RaTG13_orf1ab;5764 | 0.15 | 0.15 |
| MN996532.1;QHR63299.1;BatCoV_RaTG13_orf1ab;5765 | 0.1 | 0.05 |
| MN996532.1;QHR63299.1;BatCoV_RaTG13_orf1ab;5766 | 0.1 | 0.025 |
| MN996532.1;QHR63299.1;BatCoV_RaTG13_orf1ab;5767 | 0.1 | 0.075 |
| MN996532.1;QHR63299.1;BatCoV_RaTG13_orf1ab;5768 | 0.1 | 0.15 |
| MN996532.1;QHR63299.1;BatCoV_RaTG13_orf1ab;5769 | 0.15 | 0.225 |
| MN996532.1;QHR63299.1;BatCoV_RaTG13_orf1ab;5770 | 0.1 | 0.05 |
| MN996532.1;QHR63299.1;BatCoV_RaTG13_orf1ab;5771 | 0.1 | 0.075 |
| MN996532.1;QHR63299.1;BatCoV_RaTG13_orf1ab;5772 | 0.15 | 0.15 |
| MN996532.1;QHR63299.1;BatCoV_RaTG13_orf1ab;5773 | 0 | 0 |
| MN996532.1;QHR63299.1;BatCoV_RaTG13_orf1ab;5774 | 0 | 0 |
| MN996532.1;QHR63299.1;BatCoV_RaTG13_orf1ab;5775 | 0 | 0 |
| MN996532.1;QHR63299.1;BatCoV_RaTG13_orf1ab;5776 | 0.05 | 0 |
| MN996532.1;QHR63299.1;BatCoV_RaTG13_orf1ab;5777 | 0 | 0 |
| MN996532.1;QHR63299.1;BatCoV_RaTG13_orf1ab;5778 | 0.05 | 0.025 |
| MN996532.1;QHR63299.1;BatCoV_RaTG13_orf1ab;5779 | 0 | 0.025 |
| MN996532.1;QHR63299.1;BatCoV_RaTG13_orf1ab;5780 | 0 | 0.025 |
| MN996532.1;QHR63299.1;BatCoV_RaTG13_orf1ab;5781 | 0 | 0.025 |
| MN996532.1;QHR63299.1;BatCoV_RaTG13_orf1ab;5782 | 0 | 0.025 |
| MN996532.1;QHR63299.1;BatCoV_RaTG13_orf1ab;5783 | 0 | 0.025 |
| MN996532.1;QHR63299.1;BatCoV_RaTG13_orf1ab;5784 | 0.05 | 0 |
| MN996532.1;QHR63299.1;BatCoV_RaTG13_orf1ab;5785 | 0 | 0 |
| MN996532.1;QHR63299.1;BatCoV_RaTG13_orf1ab;5786 | 0 | 0 |
| MN996532.1;QHR63299.1;BatCoV_RaTG13_orf1ab;5787 | 0 | 0 |
| MN996532.1;QHR63299.1;BatCoV_RaTG13_orf1ab;5788 | 0 | 0 |
| MN996532.1;QHR63299.1;BatCoV_RaTG13_orf1ab;5789 | 0 | 0 |
| MN996532.1;QHR63299.1;BatCoV_RaTG13_orf1ab;5790 | 0 | 0 |
| MN996532.1;QHR63299.1;BatCoV_RaTG13_orf1ab;5791 | 0 | 0 |
| MN996532.1;QHR63299.1;BatCoV_RaTG13_orf1ab;5792 | 0 | 0 |
| MN996532.1;QHR63299.1;BatCoV_RaTG13_orf1ab;5793 | 0 | 0 |
| MN996532.1;QHR63299.1;BatCoV_RaTG13_orf1ab;5794 | 0 | 0 |
| MN996532.1;QHR63299.1;BatCoV_RaTG13_orf1ab;5795 | 0 | 0 |
| MN996532.1;QHR63299.1;BatCoV_RaTG13_orf1ab;5796 | 0 | 0 |

| | | |
|---|---|---|
| MN996532.1;QHR63299.1;BatCoV_RaTG13_orf1ab;5797 | 0 | 0 |
| MN996532.1;QHR63299.1;BatCoV_RaTG13_orf1ab;5798 | 0 | 0 |
| MN996532.1;QHR63299.1;BatCoV_RaTG13_orf1ab;5799 | 0 | 0 |
| MN996532.1;QHR63299.1;BatCoV_RaTG13_orf1ab;5800 | 0 | 0.025 |
| MN996532.1;QHR63299.1;BatCoV_RaTG13_orf1ab;5801 | 0.05 | 0 |
| MN996532.1;QHR63299.1;BatCoV_RaTG13_orf1ab;5802 | 0.05 | 0 |
| MN996532.1;QHR63299.1;BatCoV_RaTG13_orf1ab;5803 | 0.05 | 0 |
| MN996532.1;QHR63299.1;BatCoV_RaTG13_orf1ab;5804 | 0 | 0 |
| MN996532.1;QHR63299.1;BatCoV_RaTG13_orf1ab;5805 | 0 | 0 |
| MN996532.1;QHR63299.1;BatCoV_RaTG13_orf1ab;5806 | 0.05 | 0 |
| MN996532.1;QHR63299.1;BatCoV_RaTG13_orf1ab;5807 | 0 | 0 |
| MN996532.1;QHR63299.1;BatCoV_RaTG13_orf1ab;5808 | 0 | 0 |
| MN996532.1;QHR63299.1;BatCoV_RaTG13_orf1ab;5809 | 0 | 0 |
| MN996532.1;QHR63299.1;BatCoV_RaTG13_orf1ab;5810 | 0 | 0 |
| MN996532.1;QHR63299.1;BatCoV_RaTG13_orf1ab;5811 | 0 | 0 |
| MN996532.1;QHR63299.1;BatCoV_RaTG13_orf1ab;5812 | 0 | 0 |
| MN996532.1;QHR63299.1;BatCoV_RaTG13_orf1ab;5813 | 0 | 0 |
| MN996532.1;QHR63299.1;BatCoV_RaTG13_orf1ab;5814 | 0 | 0 |
| MN996532.1;QHR63299.1;BatCoV_RaTG13_orf1ab;5815 | 0 | 0 |
| MN996532.1;QHR63299.1;BatCoV_RaTG13_orf1ab;5816 | 0 | 0 |
| MN996532.1;QHR63299.1;BatCoV_RaTG13_orf1ab;5817 | 0 | 0.025 |
| MN996532.1;QHR63299.1;BatCoV_RaTG13_orf1ab;5818 | 0 | 0 |
| MN996532.1;QHR63299.1;BatCoV_RaTG13_orf1ab;5819 | 0 | 0 |
| MN996532.1;QHR63299.1;BatCoV_RaTG13_orf1ab;5820 | 0 | 0 |
| MN996532.1;QHR63299.1;BatCoV_RaTG13_orf1ab;5821 | 0 | 0 |
| MN996532.1;QHR63299.1;BatCoV_RaTG13_orf1ab;5822 | 0 | 0 |
| MN996532.1;QHR63299.1;BatCoV_RaTG13_orf1ab;5823 | 0 | 0 |
| MN996532.1;QHR63299.1;BatCoV_RaTG13_orf1ab;5824 | 0 | 0 |
| MN996532.1;QHR63299.1;BatCoV_RaTG13_orf1ab;5825 | 0 | 0.025 |
| MN996532.1;QHR63299.1;BatCoV_RaTG13_orf1ab;5826 | 0 | 0 |
| MN996532.1;QHR63299.1;BatCoV_RaTG13_orf1ab;5827 | 0.05 | 0 |
| MN996532.1;QHR63299.1;BatCoV_RaTG13_orf1ab;5828 | 0 | 0 |
| MN996532.1;QHR63299.1;BatCoV_RaTG13_orf1ab;5829 | 0 | 0 |
| MN996532.1;QHR63299.1;BatCoV_RaTG13_orf1ab;5830 | 0 | 0 |
| MN996532.1;QHR63299.1;BatCoV_RaTG13_orf1ab;5831 | 0.15 | 0.025 |
| MN996532.1;QHR63299.1;BatCoV_RaTG13_orf1ab;5832 | 0.05 | 0 |
| MN996532.1;QHR63299.1;BatCoV_RaTG13_orf1ab;5833 | 0 | 0 |
| MN996532.1;QHR63299.1;BatCoV_RaTG13_orf1ab;5834 | 0 | 0.025 |
| MN996532.1;QHR63299.1;BatCoV_RaTG13_orf1ab;5835 | 0 | 0.025 |
| MN996532.1;QHR63299.1;BatCoV_RaTG13_orf1ab;5836 | 0.1 | 0 |
| MN996532.1;QHR63299.1;BatCoV_RaTG13_orf1ab;5837 | 0.1 | 0.05 |
| MN996532.1;QHR63299.1;BatCoV_RaTG13_orf1ab;5838 | 0.05 | 0 |
| MN996532.1;QHR63299.1;BatCoV_RaTG13_orf1ab;5839 | 0 | 0 |

| | | |
|---|---|---|
| MN996532.1;QHR63299.1;BatCoV_RaTG13_orf1ab;5840 | 0 | 0 |
| MN996532.1;QHR63299.1;BatCoV_RaTG13_orf1ab;5841 | 0 | 0.05 |
| MN996532.1;QHR63299.1;BatCoV_RaTG13_orf1ab;5842 | 0 | 0 |
| MN996532.1;QHR63299.1;BatCoV_RaTG13_orf1ab;5843 | 0 | 0.025 |
| MN996532.1;QHR63299.1;BatCoV_RaTG13_orf1ab;5844 | 0 | 0 |
| MN996532.1;QHR63299.1;BatCoV_RaTG13_orf1ab;5845 | 0.05 | 0 |
| MN996532.1;QHR63299.1;BatCoV_RaTG13_orf1ab;5846 | 0 | 0 |
| MN996532.1;QHR63299.1;BatCoV_RaTG13_orf1ab;5847 | 0 | 0 |
| MN996532.1;QHR63299.1;BatCoV_RaTG13_orf1ab;5848 | 0 | 0.05 |
| MN996532.1;QHR63299.1;BatCoV_RaTG13_orf1ab;5849 | 0 | 0.025 |
| MN996532.1;QHR63299.1;BatCoV_RaTG13_orf1ab;5850 | 0 | 0.025 |
| MN996532.1;QHR63299.1;BatCoV_RaTG13_orf1ab;5851 | 0 | 0.025 |
| MN996532.1;QHR63299.1;BatCoV_RaTG13_orf1ab;5852 | 0 | 0.025 |
| MN996532.1;QHR63299.1;BatCoV_RaTG13_orf1ab;5853 | 0 | 0 |
| MN996532.1;QHR63299.1;BatCoV_RaTG13_orf1ab;5854 | 0 | 0 |
| MN996532.1;QHR63299.1;BatCoV_RaTG13_orf1ab;5855 | 0 | 0.025 |
| MN996532.1;QHR63299.1;BatCoV_RaTG13_orf1ab;5856 | 0 | 0.025 |
| MN996532.1;QHR63299.1;BatCoV_RaTG13_orf1ab;5857 | 0 | 0 |
| MN996532.1;QHR63299.1;BatCoV_RaTG13_orf1ab;5858 | 0 | 0 |
| MN996532.1;QHR63299.1;BatCoV_RaTG13_orf1ab;5859 | 0 | 0 |
| MN996532.1;QHR63299.1;BatCoV_RaTG13_orf1ab;5860 | 0 | 0 |
| MN996532.1;QHR63299.1;BatCoV_RaTG13_orf1ab;5861 | 0 | 0 |
| MN996532.1;QHR63299.1;BatCoV_RaTG13_orf1ab;5862 | 0.05 | 0 |
| MN996532.1;QHR63299.1;BatCoV_RaTG13_orf1ab;5863 | 0.05 | 0.025 |
| MN996532.1;QHR63299.1;BatCoV_RaTG13_orf1ab;5864 | 0.05 | 0 |
| MN996532.1;QHR63299.1;BatCoV_RaTG13_orf1ab;5865 | 0.05 | 0 |
| MN996532.1;QHR63299.1;BatCoV_RaTG13_orf1ab;5866 | 0.05 | 0 |
| MN996532.1;QHR63299.1;BatCoV_RaTG13_orf1ab;5867 | 0.05 | 0 |
| MN996532.1;QHR63299.1;BatCoV_RaTG13_orf1ab;5868 | 0 | 0 |
| MN996532.1;QHR63299.1;BatCoV_RaTG13_orf1ab;5869 | 0 | 0 |
| MN996532.1;QHR63299.1;BatCoV_RaTG13_orf1ab;5870 | 0.05 | 0 |
| MN996532.1;QHR63299.1;BatCoV_RaTG13_orf1ab;5871 | 0.05 | 0 |
| MN996532.1;QHR63299.1;BatCoV_RaTG13_orf1ab;5872 | 0.05 | 0.05 |
| MN996532.1;QHR63299.1;BatCoV_RaTG13_orf1ab;5873 | 0 | 0 |
| MN996532.1;QHR63299.1;BatCoV_RaTG13_orf1ab;5874 | 0 | 0 |
| MN996532.1;QHR63299.1;BatCoV_RaTG13_orf1ab;5875 | 0 | 0 |
| MN996532.1;QHR63299.1;BatCoV_RaTG13_orf1ab;5876 | 0 | 0 |
| MN996532.1;QHR63299.1;BatCoV_RaTG13_orf1ab;5877 | 0 | 0 |
| MN996532.1;QHR63299.1;BatCoV_RaTG13_orf1ab;5878 | 0 | 0 |
| MN996532.1;QHR63299.1;BatCoV_RaTG13_orf1ab;5879 | 0 | 0 |
| MN996532.1;QHR63299.1;BatCoV_RaTG13_orf1ab;5880 | 0 | 0 |
| MN996532.1;QHR63299.1;BatCoV_RaTG13_orf1ab;5881 | 0 | 0 |
| MN996532.1;QHR63299.1;BatCoV_RaTG13_orf1ab;5882 | 0 | 0 |

| | | |
|---|---|---|
| MN996532.1;QHR63299.1;BatCoV_RaTG13_orf1ab;5883 | 0 | 0 |
| MN996532.1;QHR63299.1;BatCoV_RaTG13_orf1ab;5884 | 0 | 0 |
| MN996532.1;QHR63299.1;BatCoV_RaTG13_orf1ab;5885 | 0 | 0 |
| MN996532.1;QHR63299.1;BatCoV_RaTG13_orf1ab;5886 | 0 | 0 |
| MN996532.1;QHR63299.1;BatCoV_RaTG13_orf1ab;5887 | 0 | 0 |
| MN996532.1;QHR63299.1;BatCoV_RaTG13_orf1ab;5888 | 0 | 0 |
| MN996532.1;QHR63299.1;BatCoV_RaTG13_orf1ab;5889 | 0 | 0 |
| MN996532.1;QHR63299.1;BatCoV_RaTG13_orf1ab;5890 | 0 | 0 |
| MN996532.1;QHR63299.1;BatCoV_RaTG13_orf1ab;5891 | 0 | 0 |
| MN996532.1;QHR63299.1;BatCoV_RaTG13_orf1ab;5892 | 0 | 0.025 |
| MN996532.1;QHR63299.1;BatCoV_RaTG13_orf1ab;5893 | 0 | 0 |
| MN996532.1;QHR63299.1;BatCoV_RaTG13_orf1ab;5894 | 0 | 0.05 |
| MN996532.1;QHR63299.1;BatCoV_RaTG13_orf1ab;5895 | 0 | 0.05 |
| MN996532.1;QHR63299.1;BatCoV_RaTG13_orf1ab;5896 | 0.05 | 0.05 |
| MN996532.1;QHR63299.1;BatCoV_RaTG13_orf1ab;5897 | 0.05 | 0.05 |
| MN996532.1;QHR63299.1;BatCoV_RaTG13_orf1ab;5898 | 0.1 | 0.075 |
| MN996532.1;QHR63299.1;BatCoV_RaTG13_orf1ab;5899 | 0 | 0.025 |
| MN996532.1;QHR63299.1;BatCoV_RaTG13_orf1ab;5900 | 0 | 0 |
| MN996532.1;QHR63299.1;BatCoV_RaTG13_orf1ab;5901 | 0 | 0 |
| MN996532.1;QHR63299.1;BatCoV_RaTG13_orf1ab;5902 | 0 | 0.025 |
| MN996532.1;QHR63299.1;BatCoV_RaTG13_orf1ab;5903 | 0 | 0.025 |
| MN996532.1;QHR63299.1;BatCoV_RaTG13_orf1ab;5904 | 0 | 0 |
| MN996532.1;QHR63299.1;BatCoV_RaTG13_orf1ab;5905 | 0 | 0 |
| MN996532.1;QHR63299.1;BatCoV_RaTG13_orf1ab;5906 | 0 | 0 |
| MN996532.1;QHR63299.1;BatCoV_RaTG13_orf1ab;5907 | 0 | 0.025 |
| MN996532.1;QHR63299.1;BatCoV_RaTG13_orf1ab;5908 | 0 | 0.025 |
| MN996532.1;QHR63299.1;BatCoV_RaTG13_orf1ab;5909 | 0 | 0.05 |
| MN996532.1;QHR63299.1;BatCoV_RaTG13_orf1ab;5910 | 0 | 0.05 |
| MN996532.1;QHR63299.1;BatCoV_RaTG13_orf1ab;5911 | 0.05 | 0.075 |
| MN996532.1;QHR63299.1;BatCoV_RaTG13_orf1ab;5912 | 0.1 | 0.05 |
| MN996532.1;QHR63299.1;BatCoV_RaTG13_orf1ab;5913 | 0.05 | 0.075 |
| MN996532.1;QHR63299.1;BatCoV_RaTG13_orf1ab;5914 | 0.05 | 0.05 |
| MN996532.1;QHR63299.1;BatCoV_RaTG13_orf1ab;5915 | 0.1 | 0.1 |
| MN996532.1;QHR63299.1;BatCoV_RaTG13_orf1ab;5916 | 0.15 | 0.025 |
| MN996532.1;QHR63299.1;BatCoV_RaTG13_orf1ab;5917 | 0.05 | 0 |
| MN996532.1;QHR63299.1;BatCoV_RaTG13_orf1ab;5918 | 0 | 0 |
| MN996532.1;QHR63299.1;BatCoV_RaTG13_orf1ab;5919 | 0 | 0 |
| MN996532.1;QHR63299.1;BatCoV_RaTG13_orf1ab;5920 | 0.05 | 0 |
| MN996532.1;QHR63299.1;BatCoV_RaTG13_orf1ab;5921 | 0 | 0 |
| MN996532.1;QHR63299.1;BatCoV_RaTG13_orf1ab;5922 | 0 | 0 |
| MN996532.1;QHR63299.1;BatCoV_RaTG13_orf1ab;5923 | 0 | 0 |
| MN996532.1;QHR63299.1;BatCoV_RaTG13_orf1ab;5924 | 0 | 0 |
| MN996532.1;QHR63299.1;BatCoV_RaTG13_orf1ab;5925 | 0 | 0 |

| | | |
|---|---|---|
| MN996532.1;QHR63299.1;BatCoV_RaTG13_orf1ab;5926 | 0 | 0 |
| MN996532.1;QHR63299.1;BatCoV_RaTG13_orf1ab;5927 | 0 | 0 |
| MN996532.1;QHR63299.1;BatCoV_RaTG13_orf1ab;5928 | 0 | 0 |
| MN996532.1;QHR63299.1;BatCoV_RaTG13_orf1ab;5929 | 0 | 0 |
| MN996532.1;QHR63299.1;BatCoV_RaTG13_orf1ab;5930 | 0 | 0 |
| MN996532.1;QHR63299.1;BatCoV_RaTG13_orf1ab;5931 | 0 | 0 |
| MN996532.1;QHR63299.1;BatCoV_RaTG13_orf1ab;5932 | 0 | 0 |
| MN996532.1;QHR63299.1;BatCoV_RaTG13_orf1ab;5933 | 0.05 | 0 |
| MN996532.1;QHR63299.1;BatCoV_RaTG13_orf1ab;5934 | 0 | 0 |
| MN996532.1;QHR63299.1;BatCoV_RaTG13_orf1ab;5935 | 0 | 0 |
| MN996532.1;QHR63299.1;BatCoV_RaTG13_orf1ab;5936 | 0 | 0 |
| MN996532.1;QHR63299.1;BatCoV_RaTG13_orf1ab;5937 | 0 | 0 |
| MN996532.1;QHR63299.1;BatCoV_RaTG13_orf1ab;5938 | 0 | 0 |
| MN996532.1;QHR63299.1;BatCoV_RaTG13_orf1ab;5939 | 0 | 0 |
| MN996532.1;QHR63299.1;BatCoV_RaTG13_orf1ab;5940 | 0 | 0 |
| MN996532.1;QHR63299.1;BatCoV_RaTG13_orf1ab;5941 | 0 | 0 |
| MN996532.1;QHR63299.1;BatCoV_RaTG13_orf1ab;5942 | 0 | 0 |
| MN996532.1;QHR63299.1;BatCoV_RaTG13_orf1ab;5943 | 0 | 0 |
| MN996532.1;QHR63299.1;BatCoV_RaTG13_orf1ab;5944 | 0 | 0 |
| MN996532.1;QHR63299.1;BatCoV_RaTG13_orf1ab;5945 | 0 | 0 |
| MN996532.1;QHR63299.1;BatCoV_RaTG13_orf1ab;5946 | 0 | 0 |
| MN996532.1;QHR63299.1;BatCoV_RaTG13_orf1ab;5947 | 0 | 0 |
| MN996532.1;QHR63299.1;BatCoV_RaTG13_orf1ab;5948 | 0 | 0 |
| MN996532.1;QHR63299.1;BatCoV_RaTG13_orf1ab;5949 | 0 | 0 |
| MN996532.1;QHR63299.1;BatCoV_RaTG13_orf1ab;5950 | 0 | 0 |
| MN996532.1;QHR63299.1;BatCoV_RaTG13_orf1ab;5951 | 0 | 0.025 |
| MN996532.1;QHR63299.1;BatCoV_RaTG13_orf1ab;5952 | 0 | 0 |
| MN996532.1;QHR63299.1;BatCoV_RaTG13_orf1ab;5953 | 0 | 0 |
| MN996532.1;QHR63299.1;BatCoV_RaTG13_orf1ab;5954 | 0 | 0 |
| MN996532.1;QHR63299.1;BatCoV_RaTG13_orf1ab;5955 | 0 | 0 |
| MN996532.1;QHR63299.1;BatCoV_RaTG13_orf1ab;5956 | 0 | 0 |
| MN996532.1;QHR63299.1;BatCoV_RaTG13_orf1ab;5957 | 0 | 0 |
| MN996532.1;QHR63299.1;BatCoV_RaTG13_orf1ab;5958 | 0 | 0 |
| MN996532.1;QHR63299.1;BatCoV_RaTG13_orf1ab;5959 | 0 | 0 |
| MN996532.1;QHR63299.1;BatCoV_RaTG13_orf1ab;5960 | 0 | 0 |
| MN996532.1;QHR63299.1;BatCoV_RaTG13_orf1ab;5961 | 0 | 0 |
| MN996532.1;QHR63299.1;BatCoV_RaTG13_orf1ab;5962 | 0 | 0 |
| MN996532.1;QHR63299.1;BatCoV_RaTG13_orf1ab;5963 | 0 | 0 |
| MN996532.1;QHR63299.1;BatCoV_RaTG13_orf1ab;5964 | 0 | 0 |
| MN996532.1;QHR63299.1;BatCoV_RaTG13_orf1ab;5965 | 0 | 0.025 |
| MN996532.1;QHR63299.1;BatCoV_RaTG13_orf1ab;5966 | 0 | 0 |
| MN996532.1;QHR63299.1;BatCoV_RaTG13_orf1ab;5967 | 0 | 0 |
| MN996532.1;QHR63299.1;BatCoV_RaTG13_orf1ab;5968 | 0 | 0 |

| | | |
|---|---|---|
| MN996532.1;QHR63299.1;BatCoV_RaTG13_orf1ab;5969 | 0 | 0 |
| MN996532.1;QHR63299.1;BatCoV_RaTG13_orf1ab;5970 | 0 | 0 |
| MN996532.1;QHR63299.1;BatCoV_RaTG13_orf1ab;5971 | 0 | 0 |
| MN996532.1;QHR63299.1;BatCoV_RaTG13_orf1ab;5972 | 0 | 0 |
| MN996532.1;QHR63299.1;BatCoV_RaTG13_orf1ab;5973 | 0 | 0 |
| MN996532.1;QHR63299.1;BatCoV_RaTG13_orf1ab;5974 | 0 | 0 |
| MN996532.1;QHR63299.1;BatCoV_RaTG13_orf1ab;5975 | 0 | 0 |
| MN996532.1;QHR63299.1;BatCoV_RaTG13_orf1ab;5976 | 0 | 0 |
| MN996532.1;QHR63299.1;BatCoV_RaTG13_orf1ab;5977 | 0 | 0 |
| MN996532.1;QHR63299.1;BatCoV_RaTG13_orf1ab;5978 | 0 | 0.025 |
| MN996532.1;QHR63299.1;BatCoV_RaTG13_orf1ab;5979 | 0 | 0.05 |
| MN996532.1;QHR63299.1;BatCoV_RaTG13_orf1ab;5980 | 0 | 0.05 |
| MN996532.1;QHR63299.1;BatCoV_RaTG13_orf1ab;5981 | 0.05 | 0.05 |
| MN996532.1;QHR63299.1;BatCoV_RaTG13_orf1ab;5982 | 0.1 | 0.05 |
| MN996532.1;QHR63299.1;BatCoV_RaTG13_orf1ab;5983 | 0.05 | 0.05 |
| MN996532.1;QHR63299.1;BatCoV_RaTG13_orf1ab;5984 | 0 | 0.05 |
| MN996532.1;QHR63299.1;BatCoV_RaTG13_orf1ab;5985 | 0 | 0.025 |
| MN996532.1;QHR63299.1;BatCoV_RaTG13_orf1ab;5986 | 0 | 0 |
| MN996532.1;QHR63299.1;BatCoV_RaTG13_orf1ab;5987 | 0 | 0 |
| MN996532.1;QHR63299.1;BatCoV_RaTG13_orf1ab;5988 | 0.05 | 0 |
| MN996532.1;QHR63299.1;BatCoV_RaTG13_orf1ab;5989 | 0 | 0 |
| MN996532.1;QHR63299.1;BatCoV_RaTG13_orf1ab;5990 | 0 | 0 |
| MN996532.1;QHR63299.1;BatCoV_RaTG13_orf1ab;5991 | 0 | 0 |
| MN996532.1;QHR63299.1;BatCoV_RaTG13_orf1ab;5992 | 0 | 0 |
| MN996532.1;QHR63299.1;BatCoV_RaTG13_orf1ab;5993 | 0 | 0 |
| MN996532.1;QHR63299.1;BatCoV_RaTG13_orf1ab;5994 | 0 | 0 |
| MN996532.1;QHR63299.1;BatCoV_RaTG13_orf1ab;5995 | 0 | 0 |
| MN996532.1;QHR63299.1;BatCoV_RaTG13_orf1ab;5996 | 0 | 0 |
| MN996532.1;QHR63299.1;BatCoV_RaTG13_orf1ab;5997 | 0 | 0 |
| MN996532.1;QHR63299.1;BatCoV_RaTG13_orf1ab;5998 | 0 | 0.025 |
| MN996532.1;QHR63299.1;BatCoV_RaTG13_orf1ab;5999 | 0 | 0.025 |
| MN996532.1;QHR63299.1;BatCoV_RaTG13_orf1ab;6000 | 0 | 0 |
| MN996532.1;QHR63299.1;BatCoV_RaTG13_orf1ab;6001 | 0 | 0 |
| MN996532.1;QHR63299.1;BatCoV_RaTG13_orf1ab;6002 | 0 | 0 |
| MN996532.1;QHR63299.1;BatCoV_RaTG13_orf1ab;6003 | 0 | 0 |
| MN996532.1;QHR63299.1;BatCoV_RaTG13_orf1ab;6004 | 0 | 0 |
| MN996532.1;QHR63299.1;BatCoV_RaTG13_orf1ab;6005 | 0 | 0 |
| MN996532.1;QHR63299.1;BatCoV_RaTG13_orf1ab;6006 | 0 | 0 |
| MN996532.1;QHR63299.1;BatCoV_RaTG13_orf1ab;6007 | 0 | 0 |
| MN996532.1;QHR63299.1;BatCoV_RaTG13_orf1ab;6008 | 0 | 0 |
| MN996532.1;QHR63299.1;BatCoV_RaTG13_orf1ab;6009 | 0 | 0 |
| MN996532.1;QHR63299.1;BatCoV_RaTG13_orf1ab;6010 | 0 | 0 |
| MN996532.1;QHR63299.1;BatCoV_RaTG13_orf1ab;6011 | 0.05 | 0 |

| | | |
|---|---|---|
| MN996532.1;QHR63299.1;BatCoV_RaTG13_orf1ab;6012 | 0 | 0 |
| MN996532.1;QHR63299.1;BatCoV_RaTG13_orf1ab;6013 | 0 | 0.05 |
| MN996532.1;QHR63299.1;BatCoV_RaTG13_orf1ab;6014 | 0 | 0.05 |
| MN996532.1;QHR63299.1;BatCoV_RaTG13_orf1ab;6015 | 0 | 0.05 |
| MN996532.1;QHR63299.1;BatCoV_RaTG13_orf1ab;6016 | 0 | 0.05 |
| MN996532.1;QHR63299.1;BatCoV_RaTG13_orf1ab;6017 | 0 | 0.05 |
| MN996532.1;QHR63299.1;BatCoV_RaTG13_orf1ab;6018 | 0 | 0.05 |
| MN996532.1;QHR63299.1;BatCoV_RaTG13_orf1ab;6019 | 0 | 0.05 |
| MN996532.1;QHR63299.1;BatCoV_RaTG13_orf1ab;6020 | 0 | 0.025 |
| MN996532.1;QHR63299.1;BatCoV_RaTG13_orf1ab;6021 | 0 | 0.05 |
| MN996532.1;QHR63299.1;BatCoV_RaTG13_orf1ab;6022 | 0 | 0.025 |
| MN996532.1;QHR63299.1;BatCoV_RaTG13_orf1ab;6023 | 0 | 0.025 |
| MN996532.1;QHR63299.1;BatCoV_RaTG13_orf1ab;6024 | 0 | 0.05 |
| MN996532.1;QHR63299.1;BatCoV_RaTG13_orf1ab;6025 | 0.05 | 0.05 |
| MN996532.1;QHR63299.1;BatCoV_RaTG13_orf1ab;6026 | 0 | 0 |
| MN996532.1;QHR63299.1;BatCoV_RaTG13_orf1ab;6027 | 0 | 0 |
| MN996532.1;QHR63299.1;BatCoV_RaTG13_orf1ab;6028 | 0 | 0.025 |
| MN996532.1;QHR63299.1;BatCoV_RaTG13_orf1ab;6029 | 0 | 0 |
| MN996532.1;QHR63299.1;BatCoV_RaTG13_orf1ab;6030 | 0 | 0 |
| MN996532.1;QHR63299.1;BatCoV_RaTG13_orf1ab;6031 | 0 | 0 |
| MN996532.1;QHR63299.1;BatCoV_RaTG13_orf1ab;6032 | 0 | 0 |
| MN996532.1;QHR63299.1;BatCoV_RaTG13_orf1ab;6033 | 0 | 0 |
| MN996532.1;QHR63299.1;BatCoV_RaTG13_orf1ab;6034 | 0 | 0 |
| MN996532.1;QHR63299.1;BatCoV_RaTG13_orf1ab;6035 | 0 | 0 |
| MN996532.1;QHR63299.1;BatCoV_RaTG13_orf1ab;6036 | 0 | 0 |
| MN996532.1;QHR63299.1;BatCoV_RaTG13_orf1ab;6037 | 0 | 0 |
| MN996532.1;QHR63299.1;BatCoV_RaTG13_orf1ab;6038 | 0 | 0 |
| MN996532.1;QHR63299.1;BatCoV_RaTG13_orf1ab;6039 | 0 | 0 |
| MN996532.1;QHR63299.1;BatCoV_RaTG13_orf1ab;6040 | 0 | 0.05 |
| MN996532.1;QHR63299.1;BatCoV_RaTG13_orf1ab;6041 | 0 | 0.075 |
| MN996532.1;QHR63299.1;BatCoV_RaTG13_orf1ab;6042 | 0.05 | 0.075 |
| MN996532.1;QHR63299.1;BatCoV_RaTG13_orf1ab;6043 | 0.05 | 0.025 |
| MN996532.1;QHR63299.1;BatCoV_RaTG13_orf1ab;6044 | 0.1 | 0.025 |
| MN996532.1;QHR63299.1;BatCoV_RaTG13_orf1ab;6045 | 0 | 0 |
| MN996532.1;QHR63299.1;BatCoV_RaTG13_orf1ab;6046 | 0.15 | 0.075 |
| MN996532.1;QHR63299.1;BatCoV_RaTG13_orf1ab;6047 | 0.05 | 0.025 |
| MN996532.1;QHR63299.1;BatCoV_RaTG13_orf1ab;6048 | 0 | 0.05 |
| MN996532.1;QHR63299.1;BatCoV_RaTG13_orf1ab;6049 | 0.05 | 0.075 |
| MN996532.1;QHR63299.1;BatCoV_RaTG13_orf1ab;6050 | 0 | 0.05 |
| MN996532.1;QHR63299.1;BatCoV_RaTG13_orf1ab;6051 | 0.05 | 0.025 |
| MN996532.1;QHR63299.1;BatCoV_RaTG13_orf1ab;6052 | 0 | 0.025 |
| MN996532.1;QHR63299.1;BatCoV_RaTG13_orf1ab;6053 | 0.05 | 0.075 |
| MN996532.1;QHR63299.1;BatCoV_RaTG13_orf1ab;6054 | 0 | 0.025 |

| | | |
|---|---|---|
| MN996532.1;QHR63299.1;BatCoV_RaTG13_orf1ab;6055 | 0 | 0.075 |
| MN996532.1;QHR63299.1;BatCoV_RaTG13_orf1ab;6056 | 0 | 0.1 |
| MN996532.1;QHR63299.1;BatCoV_RaTG13_orf1ab;6057 | 0 | 0.1 |
| MN996532.1;QHR63299.1;BatCoV_RaTG13_orf1ab;6058 | 0 | 0 |
| MN996532.1;QHR63299.1;BatCoV_RaTG13_orf1ab;6059 | 0 | 0 |
| MN996532.1;QHR63299.1;BatCoV_RaTG13_orf1ab;6060 | 0 | 0 |
| MN996532.1;QHR63299.1;BatCoV_RaTG13_orf1ab;6061 | 0 | 0 |
| MN996532.1;QHR63299.1;BatCoV_RaTG13_orf1ab;6062 | 0 | 0 |
| MN996532.1;QHR63299.1;BatCoV_RaTG13_orf1ab;6063 | 0 | 0 |
| MN996532.1;QHR63299.1;BatCoV_RaTG13_orf1ab;6064 | 0 | 0 |
| MN996532.1;QHR63299.1;BatCoV_RaTG13_orf1ab;6065 | 0 | 0 |
| MN996532.1;QHR63299.1;BatCoV_RaTG13_orf1ab;6066 | 0 | 0 |
| MN996532.1;QHR63299.1;BatCoV_RaTG13_orf1ab;6067 | 0 | 0 |
| MN996532.1;QHR63299.1;BatCoV_RaTG13_orf1ab;6068 | 0 | 0 |
| MN996532.1;QHR63299.1;BatCoV_RaTG13_orf1ab;6069 | 0 | 0 |
| MN996532.1;QHR63299.1;BatCoV_RaTG13_orf1ab;6070 | 0 | 0 |
| MN996532.1;QHR63299.1;BatCoV_RaTG13_orf1ab;6071 | 0 | 0 |
| MN996532.1;QHR63299.1;BatCoV_RaTG13_orf1ab;6072 | 0 | 0 |
| MN996532.1;QHR63299.1;BatCoV_RaTG13_orf1ab;6073 | 0 | 0 |
| MN996532.1;QHR63299.1;BatCoV_RaTG13_orf1ab;6074 | 0 | 0 |
| MN996532.1;QHR63299.1;BatCoV_RaTG13_orf1ab;6075 | 0 | 0 |
| MN996532.1;QHR63299.1;BatCoV_RaTG13_orf1ab;6076 | 0 | 0 |
| MN996532.1;QHR63299.1;BatCoV_RaTG13_orf1ab;6077 | 0 | 0 |
| MN996532.1;QHR63299.1;BatCoV_RaTG13_orf1ab;6078 | 0 | 0 |
| MN996532.1;QHR63299.1;BatCoV_RaTG13_orf1ab;6079 | 0 | 0 |
| MN996532.1;QHR63299.1;BatCoV_RaTG13_orf1ab;6080 | 0 | 0 |
| MN996532.1;QHR63299.1;BatCoV_RaTG13_orf1ab;6081 | 0 | 0 |
| MN996532.1;QHR63299.1;BatCoV_RaTG13_orf1ab;6082 | 0 | 0 |
| MN996532.1;QHR63299.1;BatCoV_RaTG13_orf1ab;6083 | 0 | 0 |
| MN996532.1;QHR63299.1;BatCoV_RaTG13_orf1ab;6084 | 0 | 0 |
| MN996532.1;QHR63299.1;BatCoV_RaTG13_orf1ab;6085 | 0 | 0 |
| MN996532.1;QHR63299.1;BatCoV_RaTG13_orf1ab;6086 | 0 | 0 |
| MN996532.1;QHR63299.1;BatCoV_RaTG13_orf1ab;6087 | 0 | 0 |
| MN996532.1;QHR63299.1;BatCoV_RaTG13_orf1ab;6088 | 0 | 0 |
| MN996532.1;QHR63299.1;BatCoV_RaTG13_orf1ab;6089 | 0 | 0 |
| MN996532.1;QHR63299.1;BatCoV_RaTG13_orf1ab;6090 | 0 | 0.025 |
| MN996532.1;QHR63299.1;BatCoV_RaTG13_orf1ab;6091 | 0 | 0 |
| MN996532.1;QHR63299.1;BatCoV_RaTG13_orf1ab;6092 | 0 | 0.025 |
| MN996532.1;QHR63299.1;BatCoV_RaTG13_orf1ab;6093 | 0 | 0.025 |
| MN996532.1;QHR63299.1;BatCoV_RaTG13_orf1ab;6094 | 0 | 0 |
| MN996532.1;QHR63299.1;BatCoV_RaTG13_orf1ab;6095 | 0 | 0 |
| MN996532.1;QHR63299.1;BatCoV_RaTG13_orf1ab;6096 | 0 | 0 |
| MN996532.1;QHR63299.1;BatCoV_RaTG13_orf1ab;6097 | 0 | 0 |

| | | |
|---|---|---|
| MN996532.1;QHR63299.1;BatCoV_RaTG13_orf1ab;6098 | 0 | 0 |
| MN996532.1;QHR63299.1;BatCoV_RaTG13_orf1ab;6099 | 0 | 0 |
| MN996532.1;QHR63299.1;BatCoV_RaTG13_orf1ab;6100 | 0 | 0 |
| MN996532.1;QHR63299.1;BatCoV_RaTG13_orf1ab;6101 | 0 | 0 |
| MN996532.1;QHR63299.1;BatCoV_RaTG13_orf1ab;6102 | 0 | 0 |
| MN996532.1;QHR63299.1;BatCoV_RaTG13_orf1ab;6103 | 0 | 0 |
| MN996532.1;QHR63299.1;BatCoV_RaTG13_orf1ab;6104 | 0 | 0 |
| MN996532.1;QHR63299.1;BatCoV_RaTG13_orf1ab;6105 | 0 | 0 |
| MN996532.1;QHR63299.1;BatCoV_RaTG13_orf1ab;6106 | 0 | 0 |
| MN996532.1;QHR63299.1;BatCoV_RaTG13_orf1ab;6107 | 0 | 0 |
| MN996532.1;QHR63299.1;BatCoV_RaTG13_orf1ab;6108 | 0 | 0 |
| MN996532.1;QHR63299.1;BatCoV_RaTG13_orf1ab;6109 | 0 | 0 |
| MN996532.1;QHR63299.1;BatCoV_RaTG13_orf1ab;6110 | 0 | 0 |
| MN996532.1;QHR63299.1;BatCoV_RaTG13_orf1ab;6111 | 0 | 0 |
| MN996532.1;QHR63299.1;BatCoV_RaTG13_orf1ab;6112 | 0 | 0 |
| MN996532.1;QHR63299.1;BatCoV_RaTG13_orf1ab;6113 | 0 | 0 |
| MN996532.1;QHR63299.1;BatCoV_RaTG13_orf1ab;6114 | 0 | 0 |
| MN996532.1;QHR63299.1;BatCoV_RaTG13_orf1ab;6115 | 0.05 | 0 |
| MN996532.1;QHR63299.1;BatCoV_RaTG13_orf1ab;6116 | 0 | 0 |
| MN996532.1;QHR63299.1;BatCoV_RaTG13_orf1ab;6117 | 0 | 0 |
| MN996532.1;QHR63299.1;BatCoV_RaTG13_orf1ab;6118 | 0 | 0 |
| MN996532.1;QHR63299.1;BatCoV_RaTG13_orf1ab;6119 | 0 | 0 |
| MN996532.1;QHR63299.1;BatCoV_RaTG13_orf1ab;6120 | 0 | 0 |
| MN996532.1;QHR63299.1;BatCoV_RaTG13_orf1ab;6121 | 0 | 0 |
| MN996532.1;QHR63299.1;BatCoV_RaTG13_orf1ab;6122 | 0 | 0 |
| MN996532.1;QHR63299.1;BatCoV_RaTG13_orf1ab;6123 | 0 | 0.025 |
| MN996532.1;QHR63299.1;BatCoV_RaTG13_orf1ab;6124 | 0 | 0 |
| MN996532.1;QHR63299.1;BatCoV_RaTG13_orf1ab;6125 | 0.05 | 0 |
| MN996532.1;QHR63299.1;BatCoV_RaTG13_orf1ab;6126 | 0.05 | 0 |
| MN996532.1;QHR63299.1;BatCoV_RaTG13_orf1ab;6127 | 0 | 0 |
| MN996532.1;QHR63299.1;BatCoV_RaTG13_orf1ab;6128 | 0 | 0.025 |
| MN996532.1;QHR63299.1;BatCoV_RaTG13_orf1ab;6129 | 0 | 0 |
| MN996532.1;QHR63299.1;BatCoV_RaTG13_orf1ab;6130 | 0 | 0 |
| MN996532.1;QHR63299.1;BatCoV_RaTG13_orf1ab;6131 | 0.05 | 0 |
| MN996532.1;QHR63299.1;BatCoV_RaTG13_orf1ab;6132 | 0.05 | 0.025 |
| MN996532.1;QHR63299.1;BatCoV_RaTG13_orf1ab;6133 | 0.05 | 0 |
| MN996532.1;QHR63299.1;BatCoV_RaTG13_orf1ab;6134 | 0.05 | 0.025 |
| MN996532.1;QHR63299.1;BatCoV_RaTG13_orf1ab;6135 | 0 | 0.025 |
| MN996532.1;QHR63299.1;BatCoV_RaTG13_orf1ab;6136 | 0 | 0 |
| MN996532.1;QHR63299.1;BatCoV_RaTG13_orf1ab;6137 | 0 | 0 |
| MN996532.1;QHR63299.1;BatCoV_RaTG13_orf1ab;6138 | 0 | 0 |
| MN996532.1;QHR63299.1;BatCoV_RaTG13_orf1ab;6139 | 0 | 0 |
| MN996532.1;QHR63299.1;BatCoV_RaTG13_orf1ab;6140 | 0 | 0 |

| | | |
|---|---|---|
| MN996532.1;QHR63299.1;BatCoV_RaTG13_orf1ab;6141 | 0 | 0 |
| MN996532.1;QHR63299.1;BatCoV_RaTG13_orf1ab;6142 | 0 | 0 |
| MN996532.1;QHR63299.1;BatCoV_RaTG13_orf1ab;6143 | 0 | 0 |
| MN996532.1;QHR63299.1;BatCoV_RaTG13_orf1ab;6144 | 0 | 0 |
| MN996532.1;QHR63299.1;BatCoV_RaTG13_orf1ab;6145 | 0 | 0 |
| MN996532.1;QHR63299.1;BatCoV_RaTG13_orf1ab;6146 | 0 | 0 |
| MN996532.1;QHR63299.1;BatCoV_RaTG13_orf1ab;6147 | 0 | 0 |
| MN996532.1;QHR63299.1;BatCoV_RaTG13_orf1ab;6148 | 0 | 0 |
| MN996532.1;QHR63299.1;BatCoV_RaTG13_orf1ab;6149 | 0 | 0 |
| MN996532.1;QHR63299.1;BatCoV_RaTG13_orf1ab;6150 | 0 | 0 |
| MN996532.1;QHR63299.1;BatCoV_RaTG13_orf1ab;6151 | 0 | 0 |
| MN996532.1;QHR63299.1;BatCoV_RaTG13_orf1ab;6152 | 0 | 0 |
| MN996532.1;QHR63299.1;BatCoV_RaTG13_orf1ab;6153 | 0 | 0 |
| MN996532.1;QHR63299.1;BatCoV_RaTG13_orf1ab;6154 | 0 | 0 |
| MN996532.1;QHR63299.1;BatCoV_RaTG13_orf1ab;6155 | 0 | 0 |
| MN996532.1;QHR63299.1;BatCoV_RaTG13_orf1ab;6156 | 0 | 0.05 |
| MN996532.1;QHR63299.1;BatCoV_RaTG13_orf1ab;6157 | 0 | 0 |
| MN996532.1;QHR63299.1;BatCoV_RaTG13_orf1ab;6158 | 0 | 0 |
| MN996532.1;QHR63299.1;BatCoV_RaTG13_orf1ab;6159 | 0 | 0 |
| MN996532.1;QHR63299.1;BatCoV_RaTG13_orf1ab;6160 | 0 | 0 |
| MN996532.1;QHR63299.1;BatCoV_RaTG13_orf1ab;6161 | 0 | 0 |
| MN996532.1;QHR63299.1;BatCoV_RaTG13_orf1ab;6162 | 0 | 0 |
| MN996532.1;QHR63299.1;BatCoV_RaTG13_orf1ab;6163 | 0 | 0 |
| MN996532.1;QHR63299.1;BatCoV_RaTG13_orf1ab;6164 | 0 | 0 |
| MN996532.1;QHR63299.1;BatCoV_RaTG13_orf1ab;6165 | 0 | 0 |
| MN996532.1;QHR63299.1;BatCoV_RaTG13_orf1ab;6166 | 0 | 0 |
| MN996532.1;QHR63299.1;BatCoV_RaTG13_orf1ab;6167 | 0 | 0 |
| MN996532.1;QHR63299.1;BatCoV_RaTG13_orf1ab;6168 | 0 | 0 |
| MN996532.1;QHR63299.1;BatCoV_RaTG13_orf1ab;6169 | 0 | 0 |
| MN996532.1;QHR63299.1;BatCoV_RaTG13_orf1ab;6170 | 0 | 0 |
| MN996532.1;QHR63299.1;BatCoV_RaTG13_orf1ab;6171 | 0 | 0 |
| MN996532.1;QHR63299.1;BatCoV_RaTG13_orf1ab;6172 | 0 | 0 |
| MN996532.1;QHR63299.1;BatCoV_RaTG13_orf1ab;6173 | 0 | 0 |
| MN996532.1;QHR63299.1;BatCoV_RaTG13_orf1ab;6174 | 0 | 0 |
| MN996532.1;QHR63299.1;BatCoV_RaTG13_orf1ab;6175 | 0 | 0 |
| MN996532.1;QHR63299.1;BatCoV_RaTG13_orf1ab;6176 | 0 | 0.025 |
| MN996532.1;QHR63299.1;BatCoV_RaTG13_orf1ab;6177 | 0 | 0 |
| MN996532.1;QHR63299.1;BatCoV_RaTG13_orf1ab;6178 | 0 | 0 |
| MN996532.1;QHR63299.1;BatCoV_RaTG13_orf1ab;6179 | 0.1 | 0.05 |
| MN996532.1;QHR63299.1;BatCoV_RaTG13_orf1ab;6180 | 0 | 0 |
| MN996532.1;QHR63299.1;BatCoV_RaTG13_orf1ab;6181 | 0 | 0 |
| MN996532.1;QHR63299.1;BatCoV_RaTG13_orf1ab;6182 | 0 | 0 |
| MN996532.1;QHR63299.1;BatCoV_RaTG13_orf1ab;6183 | 0 | 0 |

| | | |
|---|---|---|
| MN996532.1;QHR63299.1;BatCoV_RaTG13_orf1ab;6184 | 0 | 0 |
| MN996532.1;QHR63299.1;BatCoV_RaTG13_orf1ab;6185 | 0 | 0 |
| MN996532.1;QHR63299.1;BatCoV_RaTG13_orf1ab;6186 | 0 | 0 |
| MN996532.1;QHR63299.1;BatCoV_RaTG13_orf1ab;6187 | 0 | 0 |
| MN996532.1;QHR63299.1;BatCoV_RaTG13_orf1ab;6188 | 0 | 0 |
| MN996532.1;QHR63299.1;BatCoV_RaTG13_orf1ab;6189 | 0 | 0 |
| MN996532.1;QHR63299.1;BatCoV_RaTG13_orf1ab;6190 | 0 | 0 |
| MN996532.1;QHR63299.1;BatCoV_RaTG13_orf1ab;6191 | 0 | 0 |
| MN996532.1;QHR63299.1;BatCoV_RaTG13_orf1ab;6192 | 0 | 0 |
| MN996532.1;QHR63299.1;BatCoV_RaTG13_orf1ab;6193 | 0 | 0 |
| MN996532.1;QHR63299.1;BatCoV_RaTG13_orf1ab;6194 | 0 | 0 |
| MN996532.1;QHR63299.1;BatCoV_RaTG13_orf1ab;6195 | 0 | 0 |
| MN996532.1;QHR63299.1;BatCoV_RaTG13_orf1ab;6196 | 0 | 0 |
| MN996532.1;QHR63299.1;BatCoV_RaTG13_orf1ab;6197 | 0 | 0 |
| MN996532.1;QHR63299.1;BatCoV_RaTG13_orf1ab;6198 | 0 | 0 |
| MN996532.1;QHR63299.1;BatCoV_RaTG13_orf1ab;6199 | 0 | 0 |
| MN996532.1;QHR63299.1;BatCoV_RaTG13_orf1ab;6200 | 0 | 0 |
| MN996532.1;QHR63299.1;BatCoV_RaTG13_orf1ab;6201 | 0 | 0 |
| MN996532.1;QHR63299.1;BatCoV_RaTG13_orf1ab;6202 | 0 | 0 |
| MN996532.1;QHR63299.1;BatCoV_RaTG13_orf1ab;6203 | 0 | 0 |
| MN996532.1;QHR63299.1;BatCoV_RaTG13_orf1ab;6204 | 0 | 0 |
| MN996532.1;QHR63299.1;BatCoV_RaTG13_orf1ab;6205 | 0 | 0 |
| MN996532.1;QHR63299.1;BatCoV_RaTG13_orf1ab;6206 | 0 | 0 |
| MN996532.1;QHR63299.1;BatCoV_RaTG13_orf1ab;6207 | 0 | 0 |
| MN996532.1;QHR63299.1;BatCoV_RaTG13_orf1ab;6208 | 0 | 0 |
| MN996532.1;QHR63299.1;BatCoV_RaTG13_orf1ab;6209 | 0 | 0 |
| MN996532.1;QHR63299.1;BatCoV_RaTG13_orf1ab;6210 | 0 | 0 |
| MN996532.1;QHR63299.1;BatCoV_RaTG13_orf1ab;6211 | 0 | 0.025 |
| MN996532.1;QHR63299.1;BatCoV_RaTG13_orf1ab;6212 | 0 | 0.025 |
| MN996532.1;QHR63299.1;BatCoV_RaTG13_orf1ab;6213 | 0.1 | 0.075 |
| MN996532.1;QHR63299.1;BatCoV_RaTG13_orf1ab;6214 | 0.1 | 0.05 |
| MN996532.1;QHR63299.1;BatCoV_RaTG13_orf1ab;6215 | 0.1 | 0.05 |
| MN996532.1;QHR63299.1;BatCoV_RaTG13_orf1ab;6216 | 0.1 | 0.05 |
| MN996532.1;QHR63299.1;BatCoV_RaTG13_orf1ab;6217 | 0.1 | 0.025 |
| MN996532.1;QHR63299.1;BatCoV_RaTG13_orf1ab;6218 | 0.1 | 0.025 |
| MN996532.1;QHR63299.1;BatCoV_RaTG13_orf1ab;6219 | 0.1 | 0.05 |
| MN996532.1;QHR63299.1;BatCoV_RaTG13_orf1ab;6220 | 0.05 | 0 |
| MN996532.1;QHR63299.1;BatCoV_RaTG13_orf1ab;6221 | 0.05 | 0 |
| MN996532.1;QHR63299.1;BatCoV_RaTG13_orf1ab;6222 | 0.05 | 0 |
| MN996532.1;QHR63299.1;BatCoV_RaTG13_orf1ab;6223 | 0.1 | 0 |
| MN996532.1;QHR63299.1;BatCoV_RaTG13_orf1ab;6224 | 0 | 0 |
| MN996532.1;QHR63299.1;BatCoV_RaTG13_orf1ab;6225 | 0 | 0 |
| MN996532.1;QHR63299.1;BatCoV_RaTG13_orf1ab;6226 | 0 | 0 |

| | | |
|---|---|---|
| MN996532.1;QHR63299.1;BatCoV_RaTG13_orf1ab;6227 | 0 | 0 |
| MN996532.1;QHR63299.1;BatCoV_RaTG13_orf1ab;6228 | 0 | 0 |
| MN996532.1;QHR63299.1;BatCoV_RaTG13_orf1ab;6229 | 0 | 0 |
| MN996532.1;QHR63299.1;BatCoV_RaTG13_orf1ab;6230 | 0 | 0 |
| MN996532.1;QHR63299.1;BatCoV_RaTG13_orf1ab;6231 | 0 | 0 |
| MN996532.1;QHR63299.1;BatCoV_RaTG13_orf1ab;6232 | 0 | 0 |
| MN996532.1;QHR63299.1;BatCoV_RaTG13_orf1ab;6233 | 0 | 0 |
| MN996532.1;QHR63299.1;BatCoV_RaTG13_orf1ab;6234 | 0 | 0 |
| MN996532.1;QHR63299.1;BatCoV_RaTG13_orf1ab;6235 | 0 | 0 |
| MN996532.1;QHR63299.1;BatCoV_RaTG13_orf1ab;6236 | 0 | 0 |
| MN996532.1;QHR63299.1;BatCoV_RaTG13_orf1ab;6237 | 0 | 0 |
| MN996532.1;QHR63299.1;BatCoV_RaTG13_orf1ab;6238 | 0 | 0 |
| MN996532.1;QHR63299.1;BatCoV_RaTG13_orf1ab;6239 | 0 | 0 |
| MN996532.1;QHR63299.1;BatCoV_RaTG13_orf1ab;6240 | 0 | 0 |
| MN996532.1;QHR63299.1;BatCoV_RaTG13_orf1ab;6241 | 0.05 | 0 |
| MN996532.1;QHR63299.1;BatCoV_RaTG13_orf1ab;6242 | 0.05 | 0.025 |
| MN996532.1;QHR63299.1;BatCoV_RaTG13_orf1ab;6243 | 0 | 0 |
| MN996532.1;QHR63299.1;BatCoV_RaTG13_orf1ab;6244 | 0 | 0.025 |
| MN996532.1;QHR63299.1;BatCoV_RaTG13_orf1ab;6245 | 0 | 0 |
| MN996532.1;QHR63299.1;BatCoV_RaTG13_orf1ab;6246 | 0 | 0 |
| MN996532.1;QHR63299.1;BatCoV_RaTG13_orf1ab;6247 | 0 | 0 |
| MN996532.1;QHR63299.1;BatCoV_RaTG13_orf1ab;6248 | 0 | 0 |
| MN996532.1;QHR63299.1;BatCoV_RaTG13_orf1ab;6249 | 0 | 0 |
| MN996532.1;QHR63299.1;BatCoV_RaTG13_orf1ab;6250 | 0 | 0 |
| MN996532.1;QHR63299.1;BatCoV_RaTG13_orf1ab;6251 | 0 | 0 |
| MN996532.1;QHR63299.1;BatCoV_RaTG13_orf1ab;6252 | 0 | 0 |
| MN996532.1;QHR63299.1;BatCoV_RaTG13_orf1ab;6253 | 0 | 0 |
| MN996532.1;QHR63299.1;BatCoV_RaTG13_orf1ab;6254 | 0.05 | 0 |
| MN996532.1;QHR63299.1;BatCoV_RaTG13_orf1ab;6255 | 0 | 0 |
| MN996532.1;QHR63299.1;BatCoV_RaTG13_orf1ab;6256 | 0 | 0 |
| MN996532.1;QHR63299.1;BatCoV_RaTG13_orf1ab;6257 | 0 | 0 |
| MN996532.1;QHR63299.1;BatCoV_RaTG13_orf1ab;6258 | 0 | 0 |
| MN996532.1;QHR63299.1;BatCoV_RaTG13_orf1ab;6259 | 0 | 0 |
| MN996532.1;QHR63299.1;BatCoV_RaTG13_orf1ab;6260 | 0 | 0 |
| MN996532.1;QHR63299.1;BatCoV_RaTG13_orf1ab;6261 | 0.05 | 0.025 |
| MN996532.1;QHR63299.1;BatCoV_RaTG13_orf1ab;6262 | 0.05 | 0.025 |
| MN996532.1;QHR63299.1;BatCoV_RaTG13_orf1ab;6263 | 0.05 | 0.025 |
| MN996532.1;QHR63299.1;BatCoV_RaTG13_orf1ab;6264 | 0.05 | 0.025 |
| MN996532.1;QHR63299.1;BatCoV_RaTG13_orf1ab;6265 | 0.05 | 0.025 |
| MN996532.1;QHR63299.1;BatCoV_RaTG13_orf1ab;6266 | 0 | 0.025 |
| MN996532.1;QHR63299.1;BatCoV_RaTG13_orf1ab;6267 | 0 | 0.025 |
| MN996532.1;QHR63299.1;BatCoV_RaTG13_orf1ab;6268 | 0.1 | 0 |
| MN996532.1;QHR63299.1;BatCoV_RaTG13_orf1ab;6269 | 0.1 | 0.025 |

| | | |
|---|---|---|
| MN996532.1;QHR63299.1;BatCoV_RaTG13_orf1ab;6270 | 0.1 | 0.025 |
| MN996532.1;QHR63299.1;BatCoV_RaTG13_orf1ab;6271 | 0.05 | 0.05 |
| MN996532.1;QHR63299.1;BatCoV_RaTG13_orf1ab;6272 | 0 | 0 |
| MN996532.1;QHR63299.1;BatCoV_RaTG13_orf1ab;6273 | 0.05 | 0.025 |
| MN996532.1;QHR63299.1;BatCoV_RaTG13_orf1ab;6274 | 0 | 0 |
| MN996532.1;QHR63299.1;BatCoV_RaTG13_orf1ab;6275 | 0 | 0 |
| MN996532.1;QHR63299.1;BatCoV_RaTG13_orf1ab;6276 | 0 | 0 |
| MN996532.1;QHR63299.1;BatCoV_RaTG13_orf1ab;6277 | 0 | 0.025 |
| MN996532.1;QHR63299.1;BatCoV_RaTG13_orf1ab;6278 | 0 | 0.025 |
| MN996532.1;QHR63299.1;BatCoV_RaTG13_orf1ab;6279 | 0 | 0 |
| MN996532.1;QHR63299.1;BatCoV_RaTG13_orf1ab;6280 | 0 | 0 |
| MN996532.1;QHR63299.1;BatCoV_RaTG13_orf1ab;6281 | 0 | 0 |
| MN996532.1;QHR63299.1;BatCoV_RaTG13_orf1ab;6282 | 0 | 0 |
| MN996532.1;QHR63299.1;BatCoV_RaTG13_orf1ab;6283 | 0 | 0 |
| MN996532.1;QHR63299.1;BatCoV_RaTG13_orf1ab;6284 | 0 | 0 |
| MN996532.1;QHR63299.1;BatCoV_RaTG13_orf1ab;6285 | 0 | 0 |
| MN996532.1;QHR63299.1;BatCoV_RaTG13_orf1ab;6286 | 0 | 0 |
| MN996532.1;QHR63299.1;BatCoV_RaTG13_orf1ab;6287 | 0 | 0 |
| MN996532.1;QHR63299.1;BatCoV_RaTG13_orf1ab;6288 | 0.1 | 0.025 |
| MN996532.1;QHR63299.1;BatCoV_RaTG13_orf1ab;6289 | 0.15 | 0.025 |
| MN996532.1;QHR63299.1;BatCoV_RaTG13_orf1ab;6290 | 0.1 | 0.025 |
| MN996532.1;QHR63299.1;BatCoV_RaTG13_orf1ab;6291 | 0.2 | 0.025 |
| MN996532.1;QHR63299.1;BatCoV_RaTG13_orf1ab;6292 | 0.1 | 0.025 |
| MN996532.1;QHR63299.1;BatCoV_RaTG13_orf1ab;6293 | 0 | 0 |
| MN996532.1;QHR63299.1;BatCoV_RaTG13_orf1ab;6294 | 0 | 0 |
| MN996532.1;QHR63299.1;BatCoV_RaTG13_orf1ab;6295 | 0 | 0 |
| MN996532.1;QHR63299.1;BatCoV_RaTG13_orf1ab;6296 | 0 | 0 |
| MN996532.1;QHR63299.1;BatCoV_RaTG13_orf1ab;6297 | 0 | 0 |
| MN996532.1;QHR63299.1;BatCoV_RaTG13_orf1ab;6298 | 0 | 0 |
| MN996532.1;QHR63299.1;BatCoV_RaTG13_orf1ab;6299 | 0 | 0 |
| MN996532.1;QHR63299.1;BatCoV_RaTG13_orf1ab;6300 | 0 | 0 |
| MN996532.1;QHR63299.1;BatCoV_RaTG13_orf1ab;6301 | 0 | 0 |
| MN996532.1;QHR63299.1;BatCoV_RaTG13_orf1ab;6302 | 0 | 0 |
| MN996532.1;QHR63299.1;BatCoV_RaTG13_orf1ab;6303 | 0 | 0 |
| MN996532.1;QHR63299.1;BatCoV_RaTG13_orf1ab;6304 | 0 | 0 |
| MN996532.1;QHR63299.1;BatCoV_RaTG13_orf1ab;6305 | 0 | 0 |
| MN996532.1;QHR63299.1;BatCoV_RaTG13_orf1ab;6306 | 0 | 0 |
| MN996532.1;QHR63299.1;BatCoV_RaTG13_orf1ab;6307 | 0 | 0 |
| MN996532.1;QHR63299.1;BatCoV_RaTG13_orf1ab;6308 | 0 | 0 |
| MN996532.1;QHR63299.1;BatCoV_RaTG13_orf1ab;6309 | 0 | 0 |
| MN996532.1;QHR63299.1;BatCoV_RaTG13_orf1ab;6310 | 0 | 0 |
| MN996532.1;QHR63299.1;BatCoV_RaTG13_orf1ab;6311 | 0 | 0 |
| MN996532.1;QHR63299.1;BatCoV_RaTG13_orf1ab;6312 | 0 | 0 |

| | | |
|---|---|---|
| MN996532.1;QHR63299.1;BatCoV_RaTG13_orf1ab;6313 | 0 | 0 |
| MN996532.1;QHR63299.1;BatCoV_RaTG13_orf1ab;6314 | 0 | 0 |
| MN996532.1;QHR63299.1;BatCoV_RaTG13_orf1ab;6315 | 0 | 0 |
| MN996532.1;QHR63299.1;BatCoV_RaTG13_orf1ab;6316 | 0 | 0 |
| MN996532.1;QHR63299.1;BatCoV_RaTG13_orf1ab;6317 | 0 | 0 |
| MN996532.1;QHR63299.1;BatCoV_RaTG13_orf1ab;6318 | 0 | 0 |
| MN996532.1;QHR63299.1;BatCoV_RaTG13_orf1ab;6319 | 0 | 0 |
| MN996532.1;QHR63299.1;BatCoV_RaTG13_orf1ab;6320 | 0 | 0 |
| MN996532.1;QHR63299.1;BatCoV_RaTG13_orf1ab;6321 | 0 | 0 |
| MN996532.1;QHR63299.1;BatCoV_RaTG13_orf1ab;6322 | 0 | 0 |
| MN996532.1;QHR63299.1;BatCoV_RaTG13_orf1ab;6323 | 0 | 0 |
| MN996532.1;QHR63299.1;BatCoV_RaTG13_orf1ab;6324 | 0 | 0 |
| MN996532.1;QHR63299.1;BatCoV_RaTG13_orf1ab;6325 | 0 | 0 |
| MN996532.1;QHR63299.1;BatCoV_RaTG13_orf1ab;6326 | 0.1 | 0.025 |
| MN996532.1;QHR63299.1;BatCoV_RaTG13_orf1ab;6327 | 0.05 | 0.025 |
| MN996532.1;QHR63299.1;BatCoV_RaTG13_orf1ab;6328 | 0.05 | 0 |
| MN996532.1;QHR63299.1;BatCoV_RaTG13_orf1ab;6329 | 0.05 | 0 |
| MN996532.1;QHR63299.1;BatCoV_RaTG13_orf1ab;6330 | 0.05 | 0 |
| MN996532.1;QHR63299.1;BatCoV_RaTG13_orf1ab;6331 | 0 | 0.025 |
| MN996532.1;QHR63299.1;BatCoV_RaTG13_orf1ab;6332 | 0 | 0 |
| MN996532.1;QHR63299.1;BatCoV_RaTG13_orf1ab;6333 | 0 | 0 |
| MN996532.1;QHR63299.1;BatCoV_RaTG13_orf1ab;6334 | 0 | 0 |
| MN996532.1;QHR63299.1;BatCoV_RaTG13_orf1ab;6335 | 0 | 0 |
| MN996532.1;QHR63299.1;BatCoV_RaTG13_orf1ab;6336 | 0 | 0 |
| MN996532.1;QHR63299.1;BatCoV_RaTG13_orf1ab;6337 | 0 | 0 |
| MN996532.1;QHR63299.1;BatCoV_RaTG13_orf1ab;6338 | 0 | 0 |
| MN996532.1;QHR63299.1;BatCoV_RaTG13_orf1ab;6339 | 0 | 0 |
| MN996532.1;QHR63299.1;BatCoV_RaTG13_orf1ab;6340 | 0 | 0 |
| MN996532.1;QHR63299.1;BatCoV_RaTG13_orf1ab;6341 | 0 | 0 |
| MN996532.1;QHR63299.1;BatCoV_RaTG13_orf1ab;6342 | 0 | 0 |
| MN996532.1;QHR63299.1;BatCoV_RaTG13_orf1ab;6343 | 0 | 0 |
| MN996532.1;QHR63299.1;BatCoV_RaTG13_orf1ab;6344 | 0 | 0.025 |
| MN996532.1;QHR63299.1;BatCoV_RaTG13_orf1ab;6345 | 0 | 0.025 |
| MN996532.1;QHR63299.1;BatCoV_RaTG13_orf1ab;6346 | 0 | 0 |
| MN996532.1;QHR63299.1;BatCoV_RaTG13_orf1ab;6347 | 0 | 0.025 |
| MN996532.1;QHR63299.1;BatCoV_RaTG13_orf1ab;6348 | 0 | 0 |
| MN996532.1;QHR63299.1;BatCoV_RaTG13_orf1ab;6349 | 0 | 0.025 |
| MN996532.1;QHR63299.1;BatCoV_RaTG13_orf1ab;6350 | 0.05 | 0.025 |
| MN996532.1;QHR63299.1;BatCoV_RaTG13_orf1ab;6351 | 0.05 | 0.025 |
| MN996532.1;QHR63299.1;BatCoV_RaTG13_orf1ab;6352 | 0.05 | 0.025 |
| MN996532.1;QHR63299.1;BatCoV_RaTG13_orf1ab;6353 | 0.05 | 0.025 |
| MN996532.1;QHR63299.1;BatCoV_RaTG13_orf1ab;6354 | 0.05 | 0.025 |
| MN996532.1;QHR63299.1;BatCoV_RaTG13_orf1ab;6355 | 0.05 | 0 |

| | | |
|---|---|---|
| MN996532.1;QHR63299.1;BatCoV_RaTG13_orf1ab;6356 | 0.05 | 0.05 |
| MN996532.1;QHR63299.1;BatCoV_RaTG13_orf1ab;6357 | 0.05 | 0.025 |
| MN996532.1;QHR63299.1;BatCoV_RaTG13_orf1ab;6358 | 0 | 0 |
| MN996532.1;QHR63299.1;BatCoV_RaTG13_orf1ab;6359 | 0 | 0 |
| MN996532.1;QHR63299.1;BatCoV_RaTG13_orf1ab;6360 | 0 | 0.025 |
| MN996532.1;QHR63299.1;BatCoV_RaTG13_orf1ab;6361 | 0 | 0 |
| MN996532.1;QHR63299.1;BatCoV_RaTG13_orf1ab;6362 | 0 | 0 |
| MN996532.1;QHR63299.1;BatCoV_RaTG13_orf1ab;6363 | 0 | 0 |
| MN996532.1;QHR63299.1;BatCoV_RaTG13_orf1ab;6364 | 0 | 0 |
| MN996532.1;QHR63299.1;BatCoV_RaTG13_orf1ab;6365 | 0 | 0 |
| MN996532.1;QHR63299.1;BatCoV_RaTG13_orf1ab;6366 | 0 | 0 |
| MN996532.1;QHR63299.1;BatCoV_RaTG13_orf1ab;6367 | 0 | 0 |
| MN996532.1;QHR63299.1;BatCoV_RaTG13_orf1ab;6368 | 0 | 0 |
| MN996532.1;QHR63299.1;BatCoV_RaTG13_orf1ab;6369 | 0 | 0 |
| MN996532.1;QHR63299.1;BatCoV_RaTG13_orf1ab;6370 | 0 | 0.025 |
| MN996532.1;QHR63299.1;BatCoV_RaTG13_orf1ab;6371 | 0 | 0 |
| MN996532.1;QHR63299.1;BatCoV_RaTG13_orf1ab;6372 | 0 | 0 |
| MN996532.1;QHR63299.1;BatCoV_RaTG13_orf1ab;6373 | 0 | 0 |
| MN996532.1;QHR63299.1;BatCoV_RaTG13_orf1ab;6374 | 0 | 0 |
| MN996532.1;QHR63299.1;BatCoV_RaTG13_orf1ab;6375 | 0 | 0.05 |
| MN996532.1;QHR63299.1;BatCoV_RaTG13_orf1ab;6376 | 0 | 0.075 |
| MN996532.1;QHR63299.1;BatCoV_RaTG13_orf1ab;6377 | 0 | 0.05 |
| MN996532.1;QHR63299.1;BatCoV_RaTG13_orf1ab;6378 | 0 | 0.05 |
| MN996532.1;QHR63299.1;BatCoV_RaTG13_orf1ab;6379 | 0 | 0.075 |
| MN996532.1;QHR63299.1;BatCoV_RaTG13_orf1ab;6380 | 0 | 0.075 |
| MN996532.1;QHR63299.1;BatCoV_RaTG13_orf1ab;6381 | 0.05 | 0.05 |
| MN996532.1;QHR63299.1;BatCoV_RaTG13_orf1ab;6382 | 0 | 0.05 |
| MN996532.1;QHR63299.1;BatCoV_RaTG13_orf1ab;6383 | 0 | 0.025 |
| MN996532.1;QHR63299.1;BatCoV_RaTG13_orf1ab;6384 | 0 | 0.025 |
| MN996532.1;QHR63299.1;BatCoV_RaTG13_orf1ab;6385 | 0 | 0.025 |
| MN996532.1;QHR63299.1;BatCoV_RaTG13_orf1ab;6386 | 0 | 0 |
| MN996532.1;QHR63299.1;BatCoV_RaTG13_orf1ab;6387 | 0 | 0 |
| MN996532.1;QHR63299.1;BatCoV_RaTG13_orf1ab;6388 | 0 | 0 |
| MN996532.1;QHR63299.1;BatCoV_RaTG13_orf1ab;6389 | 0 | 0 |
| MN996532.1;QHR63299.1;BatCoV_RaTG13_orf1ab;6390 | 0 | 0 |
| MN996532.1;QHR63299.1;BatCoV_RaTG13_orf1ab;6391 | 0 | 0 |
| MN996532.1;QHR63299.1;BatCoV_RaTG13_orf1ab;6392 | 0 | 0 |
| MN996532.1;QHR63299.1;BatCoV_RaTG13_orf1ab;6393 | 0 | 0 |
| MN996532.1;QHR63299.1;BatCoV_RaTG13_orf1ab;6394 | 0 | 0 |
| MN996532.1;QHR63299.1;BatCoV_RaTG13_orf1ab;6395 | 0 | 0 |
| MN996532.1;QHR63299.1;BatCoV_RaTG13_orf1ab;6396 | 0 | 0 |
| MN996532.1;QHR63299.1;BatCoV_RaTG13_orf1ab;6397 | 0 | 0 |
| MN996532.1;QHR63299.1;BatCoV_RaTG13_orf1ab;6398 | 0 | 0 |

| | | |
|---|---|---|
| MN996532.1;QHR63299.1;BatCoV_RaTG13_orf1ab;6399 | 0 | 0 |
| MN996532.1;QHR63299.1;BatCoV_RaTG13_orf1ab;6400 | 0 | 0 |
| MN996532.1;QHR63299.1;BatCoV_RaTG13_orf1ab;6401 | 0 | 0 |
| MN996532.1;QHR63299.1;BatCoV_RaTG13_orf1ab;6402 | 0 | 0 |
| MN996532.1;QHR63299.1;BatCoV_RaTG13_orf1ab;6403 | 0 | 0 |
| MN996532.1;QHR63299.1;BatCoV_RaTG13_orf1ab;6404 | 0 | 0 |
| MN996532.1;QHR63299.1;BatCoV_RaTG13_orf1ab;6405 | 0 | 0 |
| MN996532.1;QHR63299.1;BatCoV_RaTG13_orf1ab;6406 | 0 | 0 |
| MN996532.1;QHR63299.1;BatCoV_RaTG13_orf1ab;6407 | 0 | 0 |
| MN996532.1;QHR63299.1;BatCoV_RaTG13_orf1ab;6408 | 0 | 0 |
| MN996532.1;QHR63299.1;BatCoV_RaTG13_orf1ab;6409 | 0 | 0.025 |
| MN996532.1;QHR63299.1;BatCoV_RaTG13_orf1ab;6410 | 0 | 0.05 |
| MN996532.1;QHR63299.1;BatCoV_RaTG13_orf1ab;6411 | 0 | 0.025 |
| MN996532.1;QHR63299.1;BatCoV_RaTG13_orf1ab;6412 | 0 | 0.025 |
| MN996532.1;QHR63299.1;BatCoV_RaTG13_orf1ab;6413 | 0 | 0.025 |
| MN996532.1;QHR63299.1;BatCoV_RaTG13_orf1ab;6414 | 0 | 0.025 |
| MN996532.1;QHR63299.1;BatCoV_RaTG13_orf1ab;6415 | 0 | 0.025 |
| MN996532.1;QHR63299.1;BatCoV_RaTG13_orf1ab;6416 | 0 | 0.025 |
| MN996532.1;QHR63299.1;BatCoV_RaTG13_orf1ab;6417 | 0 | 0.025 |
| MN996532.1;QHR63299.1;BatCoV_RaTG13_orf1ab;6418 | 0 | 0 |
| MN996532.1;QHR63299.1;BatCoV_RaTG13_orf1ab;6419 | 0 | 0 |
| MN996532.1;QHR63299.1;BatCoV_RaTG13_orf1ab;6420 | 0 | 0 |
| MN996532.1;QHR63299.1;BatCoV_RaTG13_orf1ab;6421 | 0 | 0 |
| MN996532.1;QHR63299.1;BatCoV_RaTG13_orf1ab;6422 | 0 | 0 |
| MN996532.1;QHR63299.1;BatCoV_RaTG13_orf1ab;6423 | 0 | 0 |
| MN996532.1;QHR63299.1;BatCoV_RaTG13_orf1ab;6424 | 0 | 0 |
| MN996532.1;QHR63299.1;BatCoV_RaTG13_orf1ab;6425 | 0 | 0 |
| MN996532.1;QHR63299.1;BatCoV_RaTG13_orf1ab;6426 | 0 | 0 |
| MN996532.1;QHR63299.1;BatCoV_RaTG13_orf1ab;6427 | 0 | 0 |
| MN996532.1;QHR63299.1;BatCoV_RaTG13_orf1ab;6428 | 0 | 0 |
| MN996532.1;QHR63299.1;BatCoV_RaTG13_orf1ab;6429 | 0 | 0 |
| MN996532.1;QHR63299.1;BatCoV_RaTG13_orf1ab;6430 | 0 | 0 |
| MN996532.1;QHR63299.1;BatCoV_RaTG13_orf1ab;6431 | 0 | 0 |
| MN996532.1;QHR63299.1;BatCoV_RaTG13_orf1ab;6432 | 0 | 0 |
| MN996532.1;QHR63299.1;BatCoV_RaTG13_orf1ab;6433 | 0 | 0 |
| MN996532.1;QHR63299.1;BatCoV_RaTG13_orf1ab;6434 | 0 | 0 |
| MN996532.1;QHR63299.1;BatCoV_RaTG13_orf1ab;6435 | 0 | 0 |
| MN996532.1;QHR63299.1;BatCoV_RaTG13_orf1ab;6436 | 0 | 0 |
| MN996532.1;QHR63299.1;BatCoV_RaTG13_orf1ab;6437 | 0 | 0 |
| MN996532.1;QHR63299.1;BatCoV_RaTG13_orf1ab;6438 | 0 | 0 |
| MN996532.1;QHR63299.1;BatCoV_RaTG13_orf1ab;6439 | 0 | 0 |
| MN996532.1;QHR63299.1;BatCoV_RaTG13_orf1ab;6440 | 0 | 0 |
| MN996532.1;QHR63299.1;BatCoV_RaTG13_orf1ab;6441 | 0 | 0 |

| | | |
|---|---|---|
| MN996532.1;QHR63299.1;BatCoV_RaTG13_orf1ab;6442 | 0 | 0 |
| MN996532.1;QHR63299.1;BatCoV_RaTG13_orf1ab;6443 | 0 | 0 |
| MN996532.1;QHR63299.1;BatCoV_RaTG13_orf1ab;6444 | 0.05 | 0 |
| MN996532.1;QHR63299.1;BatCoV_RaTG13_orf1ab;6445 | 0 | 0 |
| MN996532.1;QHR63299.1;BatCoV_RaTG13_orf1ab;6446 | 0 | 0 |
| MN996532.1;QHR63299.1;BatCoV_RaTG13_orf1ab;6447 | 0 | 0 |
| MN996532.1;QHR63299.1;BatCoV_RaTG13_orf1ab;6448 | 0 | 0 |
| MN996532.1;QHR63299.1;BatCoV_RaTG13_orf1ab;6449 | 0 | 0 |
| MN996532.1;QHR63299.1;BatCoV_RaTG13_orf1ab;6450 | 0 | 0 |
| MN996532.1;QHR63299.1;BatCoV_RaTG13_orf1ab;6451 | 0 | 0 |
| MN996532.1;QHR63299.1;BatCoV_RaTG13_orf1ab;6452 | 0 | 0 |
| MN996532.1;QHR63299.1;BatCoV_RaTG13_orf1ab;6453 | 0.05 | 0.025 |
| MN996532.1;QHR63299.1;BatCoV_RaTG13_orf1ab;6454 | 0.05 | 0.025 |
| MN996532.1;QHR63299.1;BatCoV_RaTG13_orf1ab;6455 | 0 | 0 |
| MN996532.1;QHR63299.1;BatCoV_RaTG13_orf1ab;6456 | 0.05 | 0 |
| MN996532.1;QHR63299.1;BatCoV_RaTG13_orf1ab;6457 | 0.1 | 0.025 |
| MN996532.1;QHR63299.1;BatCoV_RaTG13_orf1ab;6458 | 0.05 | 0.025 |
| MN996532.1;QHR63299.1;BatCoV_RaTG13_orf1ab;6459 | 0 | 0.025 |
| MN996532.1;QHR63299.1;BatCoV_RaTG13_orf1ab;6460 | 0.05 | 0.05 |
| MN996532.1;QHR63299.1;BatCoV_RaTG13_orf1ab;6461 | 0 | 0.025 |
| MN996532.1;QHR63299.1;BatCoV_RaTG13_orf1ab;6462 | 0 | 0.025 |
| MN996532.1;QHR63299.1;BatCoV_RaTG13_orf1ab;6463 | 0 | 0 |
| MN996532.1;QHR63299.1;BatCoV_RaTG13_orf1ab;6464 | 0 | 0 |
| MN996532.1;QHR63299.1;BatCoV_RaTG13_orf1ab;6465 | 0 | 0 |
| MN996532.1;QHR63299.1;BatCoV_RaTG13_orf1ab;6466 | 0 | 0 |
| MN996532.1;QHR63299.1;BatCoV_RaTG13_orf1ab;6467 | 0 | 0 |
| MN996532.1;QHR63299.1;BatCoV_RaTG13_orf1ab;6468 | 0 | 0 |
| MN996532.1;QHR63299.1;BatCoV_RaTG13_orf1ab;6469 | 0 | 0 |
| MN996532.1;QHR63299.1;BatCoV_RaTG13_orf1ab;6470 | 0 | 0.025 |
| MN996532.1;QHR63299.1;BatCoV_RaTG13_orf1ab;6471 | 0 | 0.025 |
| MN996532.1;QHR63299.1;BatCoV_RaTG13_orf1ab;6472 | 0 | 0.025 |
| MN996532.1;QHR63299.1;BatCoV_RaTG13_orf1ab;6473 | 0 | 0 |
| MN996532.1;QHR63299.1;BatCoV_RaTG13_orf1ab;6474 | 0.05 | 0 |
| MN996532.1;QHR63299.1;BatCoV_RaTG13_orf1ab;6475 | 0 | 0.05 |
| MN996532.1;QHR63299.1;BatCoV_RaTG13_orf1ab;6476 | 0 | 0.025 |
| MN996532.1;QHR63299.1;BatCoV_RaTG13_orf1ab;6477 | 0 | 0.025 |
| MN996532.1;QHR63299.1;BatCoV_RaTG13_orf1ab;6478 | 0 | 0.025 |
| MN996532.1;QHR63299.1;BatCoV_RaTG13_orf1ab;6479 | 0 | 0 |
| MN996532.1;QHR63299.1;BatCoV_RaTG13_orf1ab;6480 | 0 | 0 |
| MN996532.1;QHR63299.1;BatCoV_RaTG13_orf1ab;6481 | 0 | 0 |
| MN996532.1;QHR63299.1;BatCoV_RaTG13_orf1ab;6482 | 0 | 0 |
| MN996532.1;QHR63299.1;BatCoV_RaTG13_orf1ab;6483 | 0 | 0.025 |
| MN996532.1;QHR63299.1;BatCoV_RaTG13_orf1ab;6484 | 0 | 0.05 |

| | | |
|---|---|---|
| MN996532.1;QHR63299.1;BatCoV_RaTG13_orf1ab;6485 | 0 | 0.05 |
| MN996532.1;QHR63299.1;BatCoV_RaTG13_orf1ab;6486 | 0 | 0.025 |
| MN996532.1;QHR63299.1;BatCoV_RaTG13_orf1ab;6487 | 0 | 0.025 |
| MN996532.1;QHR63299.1;BatCoV_RaTG13_orf1ab;6488 | 0 | 0.05 |
| MN996532.1;QHR63299.1;BatCoV_RaTG13_orf1ab;6489 | 0 | 0.025 |
| MN996532.1;QHR63299.1;BatCoV_RaTG13_orf1ab;6490 | 0 | 0.075 |
| MN996532.1;QHR63299.1;BatCoV_RaTG13_orf1ab;6491 | 0 | 0.075 |
| MN996532.1;QHR63299.1;BatCoV_RaTG13_orf1ab;6492 | 0 | 0.05 |
| MN996532.1;QHR63299.1;BatCoV_RaTG13_orf1ab;6493 | 0 | 0.025 |
| MN996532.1;QHR63299.1;BatCoV_RaTG13_orf1ab;6494 | 0 | 0.025 |
| MN996532.1;QHR63299.1;BatCoV_RaTG13_orf1ab;6495 | 0.05 | 0.025 |
| MN996532.1;QHR63299.1;BatCoV_RaTG13_orf1ab;6496 | 0 | 0 |
| MN996532.1;QHR63299.1;BatCoV_RaTG13_orf1ab;6497 | 0 | 0 |
| MN996532.1;QHR63299.1;BatCoV_RaTG13_orf1ab;6498 | 0 | 0 |
| MN996532.1;QHR63299.1;BatCoV_RaTG13_orf1ab;6499 | 0 | 0 |
| MN996532.1;QHR63299.1;BatCoV_RaTG13_orf1ab;6500 | 0 | 0 |
| MN996532.1;QHR63299.1;BatCoV_RaTG13_orf1ab;6501 | 0 | 0 |
| MN996532.1;QHR63299.1;BatCoV_RaTG13_orf1ab;6502 | 0 | 0 |
| MN996532.1;QHR63299.1;BatCoV_RaTG13_orf1ab;6503 | 0 | 0 |
| MN996532.1;QHR63299.1;BatCoV_RaTG13_orf1ab;6504 | 0.05 | 0 |
| MN996532.1;QHR63299.1;BatCoV_RaTG13_orf1ab;6505 | 0 | 0 |
| MN996532.1;QHR63299.1;BatCoV_RaTG13_orf1ab;6506 | 0.05 | 0.1 |
| MN996532.1;QHR63299.1;BatCoV_RaTG13_orf1ab;6507 | 0.2 | 0.125 |
| MN996532.1;QHR63299.1;BatCoV_RaTG13_orf1ab;6508 | 0.05 | 0.1 |
| MN996532.1;QHR63299.1;BatCoV_RaTG13_orf1ab;6509 | 0.05 | 0.075 |
| MN996532.1;QHR63299.1;BatCoV_RaTG13_orf1ab;6510 | 0.1 | 0.075 |
| MN996532.1;QHR63299.1;BatCoV_RaTG13_orf1ab;6511 | 0.15 | 0.15 |
| MN996532.1;QHR63299.1;BatCoV_RaTG13_orf1ab;6512 | 0.15 | 0.125 |
| MN996532.1;QHR63299.1;BatCoV_RaTG13_orf1ab;6513 | 0.4 | 0.2 |
| MN996532.1;QHR63299.1;BatCoV_RaTG13_orf1ab;6514 | 0.4 | 0.2 |
| MN996532.1;QHR63299.1;BatCoV_RaTG13_orf1ab;6515 | 0.2 | 0.15 |
| MN996532.1;QHR63299.1;BatCoV_RaTG13_orf1ab;6516 | 0 | 0 |
| MN996532.1;QHR63299.1;BatCoV_RaTG13_orf1ab;6517 | 0.05 | 0 |
| MN996532.1;QHR63299.1;BatCoV_RaTG13_orf1ab;6518 | 0 | 0 |
| MN996532.1;QHR63299.1;BatCoV_RaTG13_orf1ab;6519 | 0 | 0 |
| MN996532.1;QHR63299.1;BatCoV_RaTG13_orf1ab;6520 | 0 | 0.025 |
| MN996532.1;QHR63299.1;BatCoV_RaTG13_orf1ab;6521 | 0 | 0 |
| MN996532.1;QHR63299.1;BatCoV_RaTG13_orf1ab;6522 | 0 | 0 |
| MN996532.1;QHR63299.1;BatCoV_RaTG13_orf1ab;6523 | 0 | 0 |
| MN996532.1;QHR63299.1;BatCoV_RaTG13_orf1ab;6524 | 0 | 0 |
| MN996532.1;QHR63299.1;BatCoV_RaTG13_orf1ab;6525 | 0 | 0 |
| MN996532.1;QHR63299.1;BatCoV_RaTG13_orf1ab;6526 | 0 | 0 |
| MN996532.1;QHR63299.1;BatCoV_RaTG13_orf1ab;6527 | 0.05 | 0 |

| | | |
|---|---|---|
| MN996532.1;QHR63299.1;BatCoV_RaTG13_orf1ab;6528 | 0 | 0.025 |
| MN996532.1;QHR63299.1;BatCoV_RaTG13_orf1ab;6529 | 0 | 0 |
| MN996532.1;QHR63299.1;BatCoV_RaTG13_orf1ab;6530 | 0 | 0.025 |
| MN996532.1;QHR63299.1;BatCoV_RaTG13_orf1ab;6531 | 0 | 0 |
| MN996532.1;QHR63299.1;BatCoV_RaTG13_orf1ab;6532 | 0 | 0 |
| MN996532.1;QHR63299.1;BatCoV_RaTG13_orf1ab;6533 | 0 | 0 |
| MN996532.1;QHR63299.1;BatCoV_RaTG13_orf1ab;6534 | 0 | 0 |
| MN996532.1;QHR63299.1;BatCoV_RaTG13_orf1ab;6535 | 0 | 0 |
| MN996532.1;QHR63299.1;BatCoV_RaTG13_orf1ab;6536 | 0.05 | 0 |
| MN996532.1;QHR63299.1;BatCoV_RaTG13_orf1ab;6537 | 0 | 0 |
| MN996532.1;QHR63299.1;BatCoV_RaTG13_orf1ab;6538 | 0 | 0 |
| MN996532.1;QHR63299.1;BatCoV_RaTG13_orf1ab;6539 | 0 | 0 |
| MN996532.1;QHR63299.1;BatCoV_RaTG13_orf1ab;6540 | 0.15 | 0.075 |
| MN996532.1;QHR63299.1;BatCoV_RaTG13_orf1ab;6541 | 0 | 0 |
| MN996532.1;QHR63299.1;BatCoV_RaTG13_orf1ab;6542 | 0 | 0 |
| MN996532.1;QHR63299.1;BatCoV_RaTG13_orf1ab;6543 | 0 | 0 |
| MN996532.1;QHR63299.1;BatCoV_RaTG13_orf1ab;6544 | 0 | 0 |
| MN996532.1;QHR63299.1;BatCoV_RaTG13_orf1ab;6545 | 0.05 | 0 |
| MN996532.1;QHR63299.1;BatCoV_RaTG13_orf1ab;6546 | 0.05 | 0 |
| MN996532.1;QHR63299.1;BatCoV_RaTG13_orf1ab;6547 | 0.1 | 0 |
| MN996532.1;QHR63299.1;BatCoV_RaTG13_orf1ab;6548 | 0.15 | 0.05 |
| MN996532.1;QHR63299.1;BatCoV_RaTG13_orf1ab;6549 | 0.15 | 0 |
| MN996532.1;QHR63299.1;BatCoV_RaTG13_orf1ab;6550 | 0.1 | 0 |
| MN996532.1;QHR63299.1;BatCoV_RaTG13_orf1ab;6551 | 0.1 | 0.025 |
| MN996532.1;QHR63299.1;BatCoV_RaTG13_orf1ab;6552 | 0.1 | 0.025 |
| MN996532.1;QHR63299.1;BatCoV_RaTG13_orf1ab;6553 | 0.05 | 0 |
| MN996532.1;QHR63299.1;BatCoV_RaTG13_orf1ab;6554 | 0.05 | 0.025 |
| MN996532.1;QHR63299.1;BatCoV_RaTG13_orf1ab;6555 | 0.1 | 0.05 |
| MN996532.1;QHR63299.1;BatCoV_RaTG13_orf1ab;6556 | 0.05 | 0.025 |
| MN996532.1;QHR63299.1;BatCoV_RaTG13_orf1ab;6557 | 0 | 0 |
| MN996532.1;QHR63299.1;BatCoV_RaTG13_orf1ab;6558 | 0 | 0 |
| MN996532.1;QHR63299.1;BatCoV_RaTG13_orf1ab;6559 | 0 | 0 |
| MN996532.1;QHR63299.1;BatCoV_RaTG13_orf1ab;6560 | 0 | 0 |
| MN996532.1;QHR63299.1;BatCoV_RaTG13_orf1ab;6561 | 0 | 0 |
| MN996532.1;QHR63299.1;BatCoV_RaTG13_orf1ab;6562 | 0 | 0 |
| MN996532.1;QHR63299.1;BatCoV_RaTG13_orf1ab;6563 | 0 | 0 |
| MN996532.1;QHR63299.1;BatCoV_RaTG13_orf1ab;6564 | 0 | 0 |
| MN996532.1;QHR63299.1;BatCoV_RaTG13_orf1ab;6565 | 0 | 0 |
| MN996532.1;QHR63299.1;BatCoV_RaTG13_orf1ab;6566 | 0 | 0 |
| MN996532.1;QHR63299.1;BatCoV_RaTG13_orf1ab;6567 | 0 | 0 |
| MN996532.1;QHR63299.1;BatCoV_RaTG13_orf1ab;6568 | 0.25 | 0.025 |
| MN996532.1;QHR63299.1;BatCoV_RaTG13_orf1ab;6569 | 0 | 0 |
| MN996532.1;QHR63299.1;BatCoV_RaTG13_orf1ab;6570 | 0 | 0 |

| | | |
|---|---|---|
| MN996532.1;QHR63299.1;BatCoV_RaTG13_orf1ab;6571 | 0 | 0 |
| MN996532.1;QHR63299.1;BatCoV_RaTG13_orf1ab;6572 | 0 | 0 |
| MN996532.1;QHR63299.1;BatCoV_RaTG13_orf1ab;6573 | 0 | 0 |
| MN996532.1;QHR63299.1;BatCoV_RaTG13_orf1ab;6574 | 0 | 0 |
| MN996532.1;QHR63299.1;BatCoV_RaTG13_orf1ab;6575 | 0 | 0 |
| MN996532.1;QHR63299.1;BatCoV_RaTG13_orf1ab;6576 | 0 | 0 |
| MN996532.1;QHR63299.1;BatCoV_RaTG13_orf1ab;6577 | 0 | 0 |
| MN996532.1;QHR63299.1;BatCoV_RaTG13_orf1ab;6578 | 0 | 0 |
| MN996532.1;QHR63299.1;BatCoV_RaTG13_orf1ab;6579 | 0 | 0 |
| MN996532.1;QHR63299.1;BatCoV_RaTG13_orf1ab;6580 | 0 | 0.025 |
| MN996532.1;QHR63299.1;BatCoV_RaTG13_orf1ab;6581 | 0 | 0 |
| MN996532.1;QHR63299.1;BatCoV_RaTG13_orf1ab;6582 | 0 | 0 |
| MN996532.1;QHR63299.1;BatCoV_RaTG13_orf1ab;6583 | 0 | 0 |
| MN996532.1;QHR63299.1;BatCoV_RaTG13_orf1ab;6584 | 0 | 0 |
| MN996532.1;QHR63299.1;BatCoV_RaTG13_orf1ab;6585 | 0 | 0 |
| MN996532.1;QHR63299.1;BatCoV_RaTG13_orf1ab;6586 | 0 | 0 |
| MN996532.1;QHR63299.1;BatCoV_RaTG13_orf1ab;6587 | 0 | 0 |
| MN996532.1;QHR63299.1;BatCoV_RaTG13_orf1ab;6588 | 0 | 0 |
| MN996532.1;QHR63299.1;BatCoV_RaTG13_orf1ab;6589 | 0 | 0 |
| MN996532.1;QHR63299.1;BatCoV_RaTG13_orf1ab;6590 | 0 | 0 |
| MN996532.1;QHR63299.1;BatCoV_RaTG13_orf1ab;6591 | 0.05 | 0.025 |
| MN996532.1;QHR63299.1;BatCoV_RaTG13_orf1ab;6592 | 0.05 | 0 |
| MN996532.1;QHR63299.1;BatCoV_RaTG13_orf1ab;6593 | 0.05 | 0 |
| MN996532.1;QHR63299.1;BatCoV_RaTG13_orf1ab;6594 | 0 | 0 |
| MN996532.1;QHR63299.1;BatCoV_RaTG13_orf1ab;6595 | 0.05 | 0 |
| MN996532.1;QHR63299.1;BatCoV_RaTG13_orf1ab;6596 | 0 | 0.025 |
| MN996532.1;QHR63299.1;BatCoV_RaTG13_orf1ab;6597 | 0.05 | 0.15 |
| MN996532.1;QHR63299.1;BatCoV_RaTG13_orf1ab;6598 | 0 | 0.025 |
| MN996532.1;QHR63299.1;BatCoV_RaTG13_orf1ab;6599 | 0.05 | 0 |
| MN996532.1;QHR63299.1;BatCoV_RaTG13_orf1ab;6600 | 0 | 0 |
| MN996532.1;QHR63299.1;BatCoV_RaTG13_orf1ab;6601 | 0 | 0.05 |
| MN996532.1;QHR63299.1;BatCoV_RaTG13_orf1ab;6602 | 0 | 0 |
| MN996532.1;QHR63299.1;BatCoV_RaTG13_orf1ab;6603 | 0 | 0 |
| MN996532.1;QHR63299.1;BatCoV_RaTG13_orf1ab;6604 | 0 | 0 |
| MN996532.1;QHR63299.1;BatCoV_RaTG13_orf1ab;6605 | 0 | 0 |
| MN996532.1;QHR63299.1;BatCoV_RaTG13_orf1ab;6606 | 0 | 0 |
| MN996532.1;QHR63299.1;BatCoV_RaTG13_orf1ab;6607 | 0 | 0 |
| MN996532.1;QHR63299.1;BatCoV_RaTG13_orf1ab;6608 | 0 | 0 |
| MN996532.1;QHR63299.1;BatCoV_RaTG13_orf1ab;6609 | 0 | 0 |
| MN996532.1;QHR63299.1;BatCoV_RaTG13_orf1ab;6610 | 0 | 0 |
| MN996532.1;QHR63299.1;BatCoV_RaTG13_orf1ab;6611 | 0 | 0 |
| MN996532.1;QHR63299.1;BatCoV_RaTG13_orf1ab;6612 | 0 | 0 |
| MN996532.1;QHR63299.1;BatCoV_RaTG13_orf1ab;6613 | 0 | 0 |

| | | |
|---|---|---|
| MN996532.1;QHR63299.1;BatCoV_RaTG13_orf1ab;6614 | 0 | 0 |
| MN996532.1;QHR63299.1;BatCoV_RaTG13_orf1ab;6615 | 0 | 0 |
| MN996532.1;QHR63299.1;BatCoV_RaTG13_orf1ab;6616 | 0 | 0.025 |
| MN996532.1;QHR63299.1;BatCoV_RaTG13_orf1ab;6617 | 0 | 0 |
| MN996532.1;QHR63299.1;BatCoV_RaTG13_orf1ab;6618 | 0.1 | 0 |
| MN996532.1;QHR63299.1;BatCoV_RaTG13_orf1ab;6619 | 0 | 0.025 |
| MN996532.1;QHR63299.1;BatCoV_RaTG13_orf1ab;6620 | 0 | 0.025 |
| MN996532.1;QHR63299.1;BatCoV_RaTG13_orf1ab;6621 | 0 | 0.025 |
| MN996532.1;QHR63299.1;BatCoV_RaTG13_orf1ab;6622 | 0 | 0.025 |
| MN996532.1;QHR63299.1;BatCoV_RaTG13_orf1ab;6623 | 0 | 0.025 |
| MN996532.1;QHR63299.1;BatCoV_RaTG13_orf1ab;6624 | 0 | 0.05 |
| MN996532.1;QHR63299.1;BatCoV_RaTG13_orf1ab;6625 | 0 | 0.05 |
| MN996532.1;QHR63299.1;BatCoV_RaTG13_orf1ab;6626 | 0 | 0.05 |
| MN996532.1;QHR63299.1;BatCoV_RaTG13_orf1ab;6627 | 0.1 | 0.075 |
| MN996532.1;QHR63299.1;BatCoV_RaTG13_orf1ab;6628 | 0.1 | 0.05 |
| MN996532.1;QHR63299.1;BatCoV_RaTG13_orf1ab;6629 | 0.1 | 0.05 |
| MN996532.1;QHR63299.1;BatCoV_RaTG13_orf1ab;6630 | 0.05 | 0.025 |
| MN996532.1;QHR63299.1;BatCoV_RaTG13_orf1ab;6631 | 0.05 | 0.075 |
| MN996532.1;QHR63299.1;BatCoV_RaTG13_orf1ab;6632 | 0.05 | 0.05 |
| MN996532.1;QHR63299.1;BatCoV_RaTG13_orf1ab;6633 | 0 | 0 |
| MN996532.1;QHR63299.1;BatCoV_RaTG13_orf1ab;6634 | 0 | 0 |
| MN996532.1;QHR63299.1;BatCoV_RaTG13_orf1ab;6635 | 0 | 0.025 |
| MN996532.1;QHR63299.1;BatCoV_RaTG13_orf1ab;6636 | 0 | 0 |
| MN996532.1;QHR63299.1;BatCoV_RaTG13_orf1ab;6637 | 0 | 0 |
| MN996532.1;QHR63299.1;BatCoV_RaTG13_orf1ab;6638 | 0 | 0.025 |
| MN996532.1;QHR63299.1;BatCoV_RaTG13_orf1ab;6639 | 0 | 0.025 |
| MN996532.1;QHR63299.1;BatCoV_RaTG13_orf1ab;6640 | 0 | 0 |
| MN996532.1;QHR63299.1;BatCoV_RaTG13_orf1ab;6641 | 0 | 0 |
| MN996532.1;QHR63299.1;BatCoV_RaTG13_orf1ab;6642 | 0 | 0 |
| MN996532.1;QHR63299.1;BatCoV_RaTG13_orf1ab;6643 | 0 | 0 |
| MN996532.1;QHR63299.1;BatCoV_RaTG13_orf1ab;6644 | 0 | 0 |
| MN996532.1;QHR63299.1;BatCoV_RaTG13_orf1ab;6645 | 0 | 0 |
| MN996532.1;QHR63299.1;BatCoV_RaTG13_orf1ab;6646 | 0 | 0 |
| MN996532.1;QHR63299.1;BatCoV_RaTG13_orf1ab;6647 | 0 | 0 |
| MN996532.1;QHR63299.1;BatCoV_RaTG13_orf1ab;6648 | 0 | 0 |
| MN996532.1;QHR63299.1;BatCoV_RaTG13_orf1ab;6649 | 0.05 | 0.025 |
| MN996532.1;QHR63299.1;BatCoV_RaTG13_orf1ab;6650 | 0 | 0 |
| MN996532.1;QHR63299.1;BatCoV_RaTG13_orf1ab;6651 | 0 | 0 |
| MN996532.1;QHR63299.1;BatCoV_RaTG13_orf1ab;6652 | 0 | 0 |
| MN996532.1;QHR63299.1;BatCoV_RaTG13_orf1ab;6653 | 0 | 0 |
| MN996532.1;QHR63299.1;BatCoV_RaTG13_orf1ab;6654 | 0 | 0 |
| MN996532.1;QHR63299.1;BatCoV_RaTG13_orf1ab;6655 | 0 | 0 |
| MN996532.1;QHR63299.1;BatCoV_RaTG13_orf1ab;6656 | 0 | 0 |

| | | |
|---|---|---|
| MN996532.1;QHR63299.1;BatCoV_RaTG13_orf1ab;6657 | 0 | 0.025 |
| MN996532.1;QHR63299.1;BatCoV_RaTG13_orf1ab;6658 | 0 | 0 |
| MN996532.1;QHR63299.1;BatCoV_RaTG13_orf1ab;6659 | 0 | 0 |
| MN996532.1;QHR63299.1;BatCoV_RaTG13_orf1ab;6660 | 0 | 0 |
| MN996532.1;QHR63299.1;BatCoV_RaTG13_orf1ab;6661 | 0 | 0 |
| MN996532.1;QHR63299.1;BatCoV_RaTG13_orf1ab;6662 | 0.05 | 0 |
| MN996532.1;QHR63299.1;BatCoV_RaTG13_orf1ab;6663 | 0.05 | 0 |
| MN996532.1;QHR63299.1;BatCoV_RaTG13_orf1ab;6664 | 0.05 | 0.025 |
| MN996532.1;QHR63299.1;BatCoV_RaTG13_orf1ab;6665 | 0 | 0.025 |
| MN996532.1;QHR63299.1;BatCoV_RaTG13_orf1ab;6666 | 0 | 0.025 |
| MN996532.1;QHR63299.1;BatCoV_RaTG13_orf1ab;6667 | 0.05 | 0.075 |
| MN996532.1;QHR63299.1;BatCoV_RaTG13_orf1ab;6668 | 0 | 0 |
| MN996532.1;QHR63299.1;BatCoV_RaTG13_orf1ab;6669 | 0 | 0 |
| MN996532.1;QHR63299.1;BatCoV_RaTG13_orf1ab;6670 | 0 | 0 |
| MN996532.1;QHR63299.1;BatCoV_RaTG13_orf1ab;6671 | 0 | 0 |
| MN996532.1;QHR63299.1;BatCoV_RaTG13_orf1ab;6672 | 0 | 0 |
| MN996532.1;QHR63299.1;BatCoV_RaTG13_orf1ab;6673 | 0 | 0 |
| MN996532.1;QHR63299.1;BatCoV_RaTG13_orf1ab;6674 | 0 | 0 |
| MN996532.1;QHR63299.1;BatCoV_RaTG13_orf1ab;6675 | 0 | 0 |
| MN996532.1;QHR63299.1;BatCoV_RaTG13_orf1ab;6676 | 0 | 0 |
| MN996532.1;QHR63299.1;BatCoV_RaTG13_orf1ab;6677 | 0 | 0 |
| MN996532.1;QHR63299.1;BatCoV_RaTG13_orf1ab;6678 | 0 | 0 |
| MN996532.1;QHR63299.1;BatCoV_RaTG13_orf1ab;6679 | 0 | 0 |
| MN996532.1;QHR63299.1;BatCoV_RaTG13_orf1ab;6680 | 0 | 0 |
| MN996532.1;QHR63299.1;BatCoV_RaTG13_orf1ab;6681 | 0 | 0 |
| MN996532.1;QHR63299.1;BatCoV_RaTG13_orf1ab;6682 | 0 | 0 |
| MN996532.1;QHR63299.1;BatCoV_RaTG13_orf1ab;6683 | 0 | 0 |
| MN996532.1;QHR63299.1;BatCoV_RaTG13_orf1ab;6684 | 0 | 0 |
| MN996532.1;QHR63299.1;BatCoV_RaTG13_orf1ab;6685 | 0 | 0 |
| MN996532.1;QHR63299.1;BatCoV_RaTG13_orf1ab;6686 | 0 | 0 |
| MN996532.1;QHR63299.1;BatCoV_RaTG13_orf1ab;6687 | 0 | 0 |
| MN996532.1;QHR63299.1;BatCoV_RaTG13_orf1ab;6688 | 0 | 0 |
| MN996532.1;QHR63299.1;BatCoV_RaTG13_orf1ab;6689 | 0 | 0 |
| MN996532.1;QHR63299.1;BatCoV_RaTG13_orf1ab;6690 | 0 | 0 |
| MN996532.1;QHR63299.1;BatCoV_RaTG13_orf1ab;6691 | 0 | 0 |
| MN996532.1;QHR63299.1;BatCoV_RaTG13_orf1ab;6692 | 0 | 0 |
| MN996532.1;QHR63299.1;BatCoV_RaTG13_orf1ab;6693 | 0 | 0 |
| MN996532.1;QHR63299.1;BatCoV_RaTG13_orf1ab;6694 | 0 | 0 |
| MN996532.1;QHR63299.1;BatCoV_RaTG13_orf1ab;6695 | 0 | 0 |
| MN996532.1;QHR63299.1;BatCoV_RaTG13_orf1ab;6696 | 0 | 0 |
| MN996532.1;QHR63299.1;BatCoV_RaTG13_orf1ab;6697 | 0 | 0 |
| MN996532.1;QHR63299.1;BatCoV_RaTG13_orf1ab;6698 | 0 | 0 |
| MN996532.1;QHR63299.1;BatCoV_RaTG13_orf1ab;6699 | 0 | 0 |

| | | |
|---|---|---|
| MN996532.1;QHR63299.1;BatCoV_RaTG13_orf1ab;6700 | 0.05 | 0.025 |
| MN996532.1;QHR63299.1;BatCoV_RaTG13_orf1ab;6701 | 0.05 | 0.025 |
| MN996532.1;QHR63299.1;BatCoV_RaTG13_orf1ab;6702 | 0.1 | 0.125 |
| MN996532.1;QHR63299.1;BatCoV_RaTG13_orf1ab;6703 | 0.1 | 0.075 |
| MN996532.1;QHR63299.1;BatCoV_RaTG13_orf1ab;6704 | 0.15 | 0.15 |
| MN996532.1;QHR63299.1;BatCoV_RaTG13_orf1ab;6705 | 0.1 | 0.15 |
| MN996532.1;QHR63299.1;BatCoV_RaTG13_orf1ab;6706 | 0.1 | 0.075 |
| MN996532.1;QHR63299.1;BatCoV_RaTG13_orf1ab;6707 | 0.1 | 0.075 |
| MN996532.1;QHR63299.1;BatCoV_RaTG13_orf1ab;6708 | 0.05 | 0 |
| MN996532.1;QHR63299.1;BatCoV_RaTG13_orf1ab;6709 | 0.05 | 0 |
| MN996532.1;QHR63299.1;BatCoV_RaTG13_orf1ab;6710 | 0.05 | 0 |
| MN996532.1;QHR63299.1;BatCoV_RaTG13_orf1ab;6711 | 0.05 | 0 |
| MN996532.1;QHR63299.1;BatCoV_RaTG13_orf1ab;6712 | 0 | 0 |
| MN996532.1;QHR63299.1;BatCoV_RaTG13_orf1ab;6713 | 0 | 0 |
| MN996532.1;QHR63299.1;BatCoV_RaTG13_orf1ab;6714 | 0.05 | 0 |
| MN996532.1;QHR63299.1;BatCoV_RaTG13_orf1ab;6715 | 0.05 | 0 |
| MN996532.1;QHR63299.1;BatCoV_RaTG13_orf1ab;6716 | 0.15 | 0 |
| MN996532.1;QHR63299.1;BatCoV_RaTG13_orf1ab;6717 | 0 | 0 |
| MN996532.1;QHR63299.1;BatCoV_RaTG13_orf1ab;6718 | 0.05 | 0 |
| MN996532.1;QHR63299.1;BatCoV_RaTG13_orf1ab;6719 | 0.1 | 0 |
| MN996532.1;QHR63299.1;BatCoV_RaTG13_orf1ab;6720 | 0.1 | 0.025 |
| MN996532.1;QHR63299.1;BatCoV_RaTG13_orf1ab;6721 | 0 | 0 |
| MN996532.1;QHR63299.1;BatCoV_RaTG13_orf1ab;6722 | 0.1 | 0 |
| MN996532.1;QHR63299.1;BatCoV_RaTG13_orf1ab;6723 | 0.05 | 0 |
| MN996532.1;QHR63299.1;BatCoV_RaTG13_orf1ab;6724 | 0.05 | 0 |
| MN996532.1;QHR63299.1;BatCoV_RaTG13_orf1ab;6725 | 0.1 | 0 |
| MN996532.1;QHR63299.1;BatCoV_RaTG13_orf1ab;6726 | 0.1 | 0 |
| MN996532.1;QHR63299.1;BatCoV_RaTG13_orf1ab;6727 | 0.05 | 0.025 |
| MN996532.1;QHR63299.1;BatCoV_RaTG13_orf1ab;6728 | 0 | 0 |
| MN996532.1;QHR63299.1;BatCoV_RaTG13_orf1ab;6729 | 0 | 0 |
| MN996532.1;QHR63299.1;BatCoV_RaTG13_orf1ab;6730 | 0 | 0 |
| MN996532.1;QHR63299.1;BatCoV_RaTG13_orf1ab;6731 | 0 | 0 |
| MN996532.1;QHR63299.1;BatCoV_RaTG13_orf1ab;6732 | 0 | 0 |
| MN996532.1;QHR63299.1;BatCoV_RaTG13_orf1ab;6733 | 0 | 0 |
| MN996532.1;QHR63299.1;BatCoV_RaTG13_orf1ab;6734 | 0 | 0 |
| MN996532.1;QHR63299.1;BatCoV_RaTG13_orf1ab;6735 | 0 | 0 |
| MN996532.1;QHR63299.1;BatCoV_RaTG13_orf1ab;6736 | 0 | 0 |
| MN996532.1;QHR63299.1;BatCoV_RaTG13_orf1ab;6737 | 0 | 0 |
| MN996532.1;QHR63299.1;BatCoV_RaTG13_orf1ab;6738 | 0 | 0 |
| MN996532.1;QHR63299.1;BatCoV_RaTG13_orf1ab;6739 | 0 | 0 |
| MN996532.1;QHR63299.1;BatCoV_RaTG13_orf1ab;6740 | 0 | 0 |
| MN996532.1;QHR63299.1;BatCoV_RaTG13_orf1ab;6741 | 0 | 0 |
| MN996532.1;QHR63299.1;BatCoV_RaTG13_orf1ab;6742 | 0 | 0 |

| | | |
|---|---|---|
| MN996532.1;QHR63299.1;BatCoV_RaTG13_orf1ab;6743 | 0 | 0 |
| MN996532.1;QHR63299.1;BatCoV_RaTG13_orf1ab;6744 | 0.05 | 0.025 |
| MN996532.1;QHR63299.1;BatCoV_RaTG13_orf1ab;6745 | 0.05 | 0 |
| MN996532.1;QHR63299.1;BatCoV_RaTG13_orf1ab;6746 | 0.05 | 0 |
| MN996532.1;QHR63299.1;BatCoV_RaTG13_orf1ab;6747 | 0.05 | 0 |
| MN996532.1;QHR63299.1;BatCoV_RaTG13_orf1ab;6748 | 0.1 | 0.125 |
| MN996532.1;QHR63299.1;BatCoV_RaTG13_orf1ab;6749 | 0.05 | 0 |
| MN996532.1;QHR63299.1;BatCoV_RaTG13_orf1ab;6750 | 0 | 0 |
| MN996532.1;QHR63299.1;BatCoV_RaTG13_orf1ab;6751 | 0 | 0 |
| MN996532.1;QHR63299.1;BatCoV_RaTG13_orf1ab;6752 | 0 | 0 |
| MN996532.1;QHR63299.1;BatCoV_RaTG13_orf1ab;6753 | 0 | 0 |
| MN996532.1;QHR63299.1;BatCoV_RaTG13_orf1ab;6754 | 0 | 0 |
| MN996532.1;QHR63299.1;BatCoV_RaTG13_orf1ab;6755 | 0 | 0 |
| MN996532.1;QHR63299.1;BatCoV_RaTG13_orf1ab;6756 | 0 | 0 |
| MN996532.1;QHR63299.1;BatCoV_RaTG13_orf1ab;6757 | 0 | 0 |
| MN996532.1;QHR63299.1;BatCoV_RaTG13_orf1ab;6758 | 0 | 0 |
| MN996532.1;QHR63299.1;BatCoV_RaTG13_orf1ab;6759 | 0 | 0 |
| MN996532.1;QHR63299.1;BatCoV_RaTG13_orf1ab;6760 | 0 | 0 |
| MN996532.1;QHR63299.1;BatCoV_RaTG13_orf1ab;6761 | 0 | 0 |
| MN996532.1;QHR63299.1;BatCoV_RaTG13_orf1ab;6762 | 0 | 0 |
| MN996532.1;QHR63299.1;BatCoV_RaTG13_orf1ab;6763 | 0 | 0 |
| MN996532.1;QHR63299.1;BatCoV_RaTG13_orf1ab;6764 | 0 | 0 |
| MN996532.1;QHR63299.1;BatCoV_RaTG13_orf1ab;6765 | 0 | 0 |
| MN996532.1;QHR63299.1;BatCoV_RaTG13_orf1ab;6766 | 0 | 0 |
| MN996532.1;QHR63299.1;BatCoV_RaTG13_orf1ab;6767 | 0 | 0 |
| MN996532.1;QHR63299.1;BatCoV_RaTG13_orf1ab;6768 | 0 | 0 |
| MN996532.1;QHR63299.1;BatCoV_RaTG13_orf1ab;6769 | 0 | 0 |
| MN996532.1;QHR63299.1;BatCoV_RaTG13_orf1ab;6770 | 0 | 0 |
| MN996532.1;QHR63299.1;BatCoV_RaTG13_orf1ab;6771 | 0 | 0.025 |
| MN996532.1;QHR63299.1;BatCoV_RaTG13_orf1ab;6772 | 0 | 0 |
| MN996532.1;QHR63299.1;BatCoV_RaTG13_orf1ab;6773 | 0 | 0 |
| MN996532.1;QHR63299.1;BatCoV_RaTG13_orf1ab;6774 | 0 | 0 |
| MN996532.1;QHR63299.1;BatCoV_RaTG13_orf1ab;6775 | 0 | 0 |
| MN996532.1;QHR63299.1;BatCoV_RaTG13_orf1ab;6776 | 0 | 0 |
| MN996532.1;QHR63299.1;BatCoV_RaTG13_orf1ab;6777 | 0 | 0 |
| MN996532.1;QHR63299.1;BatCoV_RaTG13_orf1ab;6778 | 0 | 0 |
| MN996532.1;QHR63299.1;BatCoV_RaTG13_orf1ab;6779 | 0 | 0 |
| MN996532.1;QHR63299.1;BatCoV_RaTG13_orf1ab;6780 | 0 | 0 |
| MN996532.1;QHR63299.1;BatCoV_RaTG13_orf1ab;6781 | 0 | 0 |
| MN996532.1;QHR63299.1;BatCoV_RaTG13_orf1ab;6782 | 0 | 0.025 |
| MN996532.1;QHR63299.1;BatCoV_RaTG13_orf1ab;6783 | 0 | 0.025 |
| MN996532.1;QHR63299.1;BatCoV_RaTG13_orf1ab;6784 | 0 | 0.025 |
| MN996532.1;QHR63299.1;BatCoV_RaTG13_orf1ab;6785 | 0 | 0.025 |

| | | |
|---|---|---|
| MN996532.1;QHR63299.1;BatCoV_RaTG13_orf1ab;6786 | 0 | 0.025 |
| MN996532.1;QHR63299.1;BatCoV_RaTG13_orf1ab;6787 | 0 | 0.025 |
| MN996532.1;QHR63299.1;BatCoV_RaTG13_orf1ab;6788 | 0 | 0.025 |
| MN996532.1;QHR63299.1;BatCoV_RaTG13_orf1ab;6789 | 0 | 0.025 |
| MN996532.1;QHR63299.1;BatCoV_RaTG13_orf1ab;6790 | 0 | 0.025 |
| MN996532.1;QHR63299.1;BatCoV_RaTG13_orf1ab;6791 | 0 | 0.025 |
| MN996532.1;QHR63299.1;BatCoV_RaTG13_orf1ab;6792 | 0.1 | 0.05 |
| MN996532.1;QHR63299.1;BatCoV_RaTG13_orf1ab;6793 | 0 | 0 |
| MN996532.1;QHR63299.1;BatCoV_RaTG13_orf1ab;6794 | 0 | 0 |
| MN996532.1;QHR63299.1;BatCoV_RaTG13_orf1ab;6795 | 0 | 0 |
| MN996532.1;QHR63299.1;BatCoV_RaTG13_orf1ab;6796 | 0 | 0 |
| MN996532.1;QHR63299.1;BatCoV_RaTG13_orf1ab;6797 | 0 | 0 |
| MN996532.1;QHR63299.1;BatCoV_RaTG13_orf1ab;6798 | 0 | 0 |
| MN996532.1;QHR63299.1;BatCoV_RaTG13_orf1ab;6799 | 0 | 0 |
| MN996532.1;QHR63299.1;BatCoV_RaTG13_orf1ab;6800 | 0 | 0 |
| MN996532.1;QHR63299.1;BatCoV_RaTG13_orf1ab;6801 | 0 | 0 |
| MN996532.1;QHR63299.1;BatCoV_RaTG13_orf1ab;6802 | 0 | 0 |
| MN996532.1;QHR63299.1;BatCoV_RaTG13_orf1ab;6803 | 0 | 0 |
| MN996532.1;QHR63299.1;BatCoV_RaTG13_orf1ab;6804 | 0 | 0 |
| MN996532.1;QHR63299.1;BatCoV_RaTG13_orf1ab;6805 | 0 | 0 |
| MN996532.1;QHR63299.1;BatCoV_RaTG13_orf1ab;6806 | 0 | 0 |
| MN996532.1;QHR63299.1;BatCoV_RaTG13_orf1ab;6807 | 0 | 0 |
| MN996532.1;QHR63299.1;BatCoV_RaTG13_orf1ab;6808 | 0 | 0.025 |
| MN996532.1;QHR63299.1;BatCoV_RaTG13_orf1ab;6809 | 0 | 0 |
| MN996532.1;QHR63299.1;BatCoV_RaTG13_orf1ab;6810 | 0 | 0 |
| MN996532.1;QHR63299.1;BatCoV_RaTG13_orf1ab;6811 | 0 | 0 |
| MN996532.1;QHR63299.1;BatCoV_RaTG13_orf1ab;6812 | 0 | 0 |
| MN996532.1;QHR63299.1;BatCoV_RaTG13_orf1ab;6813 | 0 | 0 |
| MN996532.1;QHR63299.1;BatCoV_RaTG13_orf1ab;6814 | 0 | 0 |
| MN996532.1;QHR63299.1;BatCoV_RaTG13_orf1ab;6815 | 0 | 0 |
| MN996532.1;QHR63299.1;BatCoV_RaTG13_orf1ab;6816 | 0 | 0 |
| MN996532.1;QHR63299.1;BatCoV_RaTG13_orf1ab;6817 | 0 | 0 |
| MN996532.1;QHR63299.1;BatCoV_RaTG13_orf1ab;6818 | 0 | 0 |
| MN996532.1;QHR63299.1;BatCoV_RaTG13_orf1ab;6819 | 0 | 0 |
| MN996532.1;QHR63299.1;BatCoV_RaTG13_orf1ab;6820 | 0.05 | 0.05 |
| MN996532.1;QHR63299.1;BatCoV_RaTG13_orf1ab;6821 | 0.1 | 0.075 |
| MN996532.1;QHR63299.1;BatCoV_RaTG13_orf1ab;6822 | 0.05 | 0 |
| MN996532.1;QHR63299.1;BatCoV_RaTG13_orf1ab;6823 | 0.1 | 0.025 |
| MN996532.1;QHR63299.1;BatCoV_RaTG13_orf1ab;6824 | 0.1 | 0.025 |
| MN996532.1;QHR63299.1;BatCoV_RaTG13_orf1ab;6825 | 0.1 | 0.025 |
| MN996532.1;QHR63299.1;BatCoV_RaTG13_orf1ab;6826 | 0.1 | 0.025 |
| MN996532.1;QHR63299.1;BatCoV_RaTG13_orf1ab;6827 | 0.1 | 0.025 |
| MN996532.1;QHR63299.1;BatCoV_RaTG13_orf1ab;6828 | 0 | 0 |

| | | |
|---|---|---|
| MN996532.1;QHR63299.1;BatCoV_RaTG13_orf1ab;6829 | 0.25 | 0.125 |
| MN996532.1;QHR63299.1;BatCoV_RaTG13_orf1ab;6830 | 0 | 0 |
| MN996532.1;QHR63299.1;BatCoV_RaTG13_orf1ab;6831 | 0 | 0 |
| MN996532.1;QHR63299.1;BatCoV_RaTG13_orf1ab;6832 | 0 | 0 |
| MN996532.1;QHR63299.1;BatCoV_RaTG13_orf1ab;6833 | 0 | 0 |
| MN996532.1;QHR63299.1;BatCoV_RaTG13_orf1ab;6834 | 0 | 0 |
| MN996532.1;QHR63299.1;BatCoV_RaTG13_orf1ab;6835 | 0 | 0.05 |
| MN996532.1;QHR63299.1;BatCoV_RaTG13_orf1ab;6836 | 0 | 0 |
| MN996532.1;QHR63299.1;BatCoV_RaTG13_orf1ab;6837 | 0 | 0 |
| MN996532.1;QHR63299.1;BatCoV_RaTG13_orf1ab;6838 | 0 | 0 |
| MN996532.1;QHR63299.1;BatCoV_RaTG13_orf1ab;6839 | 0 | 0 |
| MN996532.1;QHR63299.1;BatCoV_RaTG13_orf1ab;6840 | 0 | 0 |
| MN996532.1;QHR63299.1;BatCoV_RaTG13_orf1ab;6841 | 0 | 0 |
| MN996532.1;QHR63299.1;BatCoV_RaTG13_orf1ab;6842 | 0 | 0 |
| MN996532.1;QHR63299.1;BatCoV_RaTG13_orf1ab;6843 | 0 | 0 |
| MN996532.1;QHR63299.1;BatCoV_RaTG13_orf1ab;6844 | 0 | 0 |
| MN996532.1;QHR63299.1;BatCoV_RaTG13_orf1ab;6845 | 0 | 0 |
| MN996532.1;QHR63299.1;BatCoV_RaTG13_orf1ab;6846 | 0 | 0 |
| MN996532.1;QHR63299.1;BatCoV_RaTG13_orf1ab;6847 | 0 | 0 |
| MN996532.1;QHR63299.1;BatCoV_RaTG13_orf1ab;6848 | 0 | 0 |
| MN996532.1;QHR63299.1;BatCoV_RaTG13_orf1ab;6849 | 0 | 0 |
| MN996532.1;QHR63299.1;BatCoV_RaTG13_orf1ab;6850 | 0 | 0 |
| MN996532.1;QHR63299.1;BatCoV_RaTG13_orf1ab;6851 | 0 | 0 |
| MN996532.1;QHR63299.1;BatCoV_RaTG13_orf1ab;6852 | 0 | 0 |
| MN996532.1;QHR63299.1;BatCoV_RaTG13_orf1ab;6853 | 0 | 0 |
| MN996532.1;QHR63299.1;BatCoV_RaTG13_orf1ab;6854 | 0 | 0 |
| MN996532.1;QHR63299.1;BatCoV_RaTG13_orf1ab;6855 | 0 | 0 |
| MN996532.1;QHR63299.1;BatCoV_RaTG13_orf1ab;6856 | 0 | 0 |
| MN996532.1;QHR63299.1;BatCoV_RaTG13_orf1ab;6857 | 0 | 0 |
| MN996532.1;QHR63299.1;BatCoV_RaTG13_orf1ab;6858 | 0 | 0 |
| MN996532.1;QHR63299.1;BatCoV_RaTG13_orf1ab;6859 | 0 | 0 |
| MN996532.1;QHR63299.1;BatCoV_RaTG13_orf1ab;6860 | 0 | 0 |
| MN996532.1;QHR63299.1;BatCoV_RaTG13_orf1ab;6861 | 0.05 | 0.05 |
| MN996532.1;QHR63299.1;BatCoV_RaTG13_orf1ab;6862 | 0 | 0 |
| MN996532.1;QHR63299.1;BatCoV_RaTG13_orf1ab;6863 | 0 | 0 |
| MN996532.1;QHR63299.1;BatCoV_RaTG13_orf1ab;6864 | 0 | 0 |
| MN996532.1;QHR63299.1;BatCoV_RaTG13_orf1ab;6865 | 0 | 0 |
| MN996532.1;QHR63299.1;BatCoV_RaTG13_orf1ab;6866 | 0 | 0 |
| MN996532.1;QHR63299.1;BatCoV_RaTG13_orf1ab;6867 | 0 | 0 |
| MN996532.1;QHR63299.1;BatCoV_RaTG13_orf1ab;6868 | 0 | 0 |
| MN996532.1;QHR63299.1;BatCoV_RaTG13_orf1ab;6869 | 0.05 | 0 |
| MN996532.1;QHR63299.1;BatCoV_RaTG13_orf1ab;6870 | 0.05 | 0 |
| MN996532.1;QHR63299.1;BatCoV_RaTG13_orf1ab;6871 | 0 | 0 |

| | | |
|---|---|---|
| MN996532.1;QHR63299.1;BatCoV_RaTG13_orf1ab;6872 | 0 | 0 |
| MN996532.1;QHR63299.1;BatCoV_RaTG13_orf1ab;6873 | 0 | 0 |
| MN996532.1;QHR63299.1;BatCoV_RaTG13_orf1ab;6874 | 0 | 0 |
| MN996532.1;QHR63299.1;BatCoV_RaTG13_orf1ab;6875 | 0 | 0 |
| MN996532.1;QHR63299.1;BatCoV_RaTG13_orf1ab;6876 | 0 | 0 |
| MN996532.1;QHR63299.1;BatCoV_RaTG13_orf1ab;6877 | 0 | 0 |
| MN996532.1;QHR63299.1;BatCoV_RaTG13_orf1ab;6878 | 0 | 0 |
| MN996532.1;QHR63299.1;BatCoV_RaTG13_orf1ab;6879 | 0 | 0 |
| MN996532.1;QHR63299.1;BatCoV_RaTG13_orf1ab;6880 | 0 | 0 |
| MN996532.1;QHR63299.1;BatCoV_RaTG13_orf1ab;6881 | 0 | 0 |
| MN996532.1;QHR63299.1;BatCoV_RaTG13_orf1ab;6882 | 0 | 0 |
| MN996532.1;QHR63299.1;BatCoV_RaTG13_orf1ab;6883 | 0 | 0 |
| MN996532.1;QHR63299.1;BatCoV_RaTG13_orf1ab;6884 | 0 | 0 |
| MN996532.1;QHR63299.1;BatCoV_RaTG13_orf1ab;6885 | 0 | 0 |
| MN996532.1;QHR63299.1;BatCoV_RaTG13_orf1ab;6886 | 0 | 0.025 |
| MN996532.1;QHR63299.1;BatCoV_RaTG13_orf1ab;6887 | 0 | 0.025 |
| MN996532.1;QHR63299.1;BatCoV_RaTG13_orf1ab;6888 | 0 | 0.025 |
| MN996532.1;QHR63299.1;BatCoV_RaTG13_orf1ab;6889 | 0 | 0.025 |
| MN996532.1;QHR63299.1;BatCoV_RaTG13_orf1ab;6890 | 0.05 | 0.025 |
| MN996532.1;QHR63299.1;BatCoV_RaTG13_orf1ab;6891 | 0.05 | 0.05 |
| MN996532.1;QHR63299.1;BatCoV_RaTG13_orf1ab;6892 | 0.05 | 0.05 |
| MN996532.1;QHR63299.1;BatCoV_RaTG13_orf1ab;6893 | 0 | 0.025 |
| MN996532.1;QHR63299.1;BatCoV_RaTG13_orf1ab;6894 | 0 | 0.05 |
| MN996532.1;QHR63299.1;BatCoV_RaTG13_orf1ab;6895 | 0.1 | 0.05 |
| MN996532.1;QHR63299.1;BatCoV_RaTG13_orf1ab;6896 | 0 | 0.025 |
| MN996532.1;QHR63299.1;BatCoV_RaTG13_orf1ab;6897 | 0 | 0 |
| MN996532.1;QHR63299.1;BatCoV_RaTG13_orf1ab;6898 | 0.05 | 0 |
| MN996532.1;QHR63299.1;BatCoV_RaTG13_orf1ab;6899 | 0.05 | 0 |
| MN996532.1;QHR63299.1;BatCoV_RaTG13_orf1ab;6900 | 0.05 | 0 |
| MN996532.1;QHR63299.1;BatCoV_RaTG13_orf1ab;6901 | 0.05 | 0 |
| MN996532.1;QHR63299.1;BatCoV_RaTG13_orf1ab;6902 | 0.05 | 0 |
| MN996532.1;QHR63299.1;BatCoV_RaTG13_orf1ab;6903 | 0.05 | 0 |
| MN996532.1;QHR63299.1;BatCoV_RaTG13_orf1ab;6904 | 0.05 | 0 |
| MN996532.1;QHR63299.1;BatCoV_RaTG13_orf1ab;6905 | 0 | 0.025 |
| MN996532.1;QHR63299.1;BatCoV_RaTG13_orf1ab;6906 | 0 | 0 |
| MN996532.1;QHR63299.1;BatCoV_RaTG13_orf1ab;6907 | 0.05 | 0.175 |
| MN996532.1;QHR63299.1;BatCoV_RaTG13_orf1ab;6908 | 0 | 0.1 |
| MN996532.1;QHR63299.1;BatCoV_RaTG13_orf1ab;6909 | 0 | 0.025 |
| MN996532.1;QHR63299.1;BatCoV_RaTG13_orf1ab;6910 | 0 | 0 |
| MN996532.1;QHR63299.1;BatCoV_RaTG13_orf1ab;6911 | 0 | 0 |
| MN996532.1;QHR63299.1;BatCoV_RaTG13_orf1ab;6912 | 0 | 0 |
| MN996532.1;QHR63299.1;BatCoV_RaTG13_orf1ab;6913 | 0 | 0.025 |
| MN996532.1;QHR63299.1;BatCoV_RaTG13_orf1ab;6914 | 0 | 0 |

| | | |
|---|---|---|
| MN996532.1;QHR63299.1;BatCoV_RaTG13_orf1ab;6915 | 0 | 0 |
| MN996532.1;QHR63299.1;BatCoV_RaTG13_orf1ab;6916 | 0 | 0 |
| MN996532.1;QHR63299.1;BatCoV_RaTG13_orf1ab;6917 | 0 | 0 |
| MN996532.1;QHR63299.1;BatCoV_RaTG13_orf1ab;6918 | 0.15 | 0.05 |
| MN996532.1;QHR63299.1;BatCoV_RaTG13_orf1ab;6919 | 0 | 0 |
| MN996532.1;QHR63299.1;BatCoV_RaTG13_orf1ab;6920 | 0 | 0.025 |
| MN996532.1;QHR63299.1;BatCoV_RaTG13_orf1ab;6921 | 0 | 0 |
| MN996532.1;QHR63299.1;BatCoV_RaTG13_orf1ab;6922 | 0.05 | 0.025 |
| MN996532.1;QHR63299.1;BatCoV_RaTG13_orf1ab;6923 | 0.05 | 0 |
| MN996532.1;QHR63299.1;BatCoV_RaTG13_orf1ab;6924 | 0.05 | 0 |
| MN996532.1;QHR63299.1;BatCoV_RaTG13_orf1ab;6925 | 0.15 | 0.05 |
| MN996532.1;QHR63299.1;BatCoV_RaTG13_orf1ab;6926 | 0.2 | 0.075 |
| MN996532.1;QHR63299.1;BatCoV_RaTG13_orf1ab;6927 | 0.2 | 0.1 |
| MN996532.1;QHR63299.1;BatCoV_RaTG13_orf1ab;6928 | 0.1 | 0.05 |
| MN996532.1;QHR63299.1;BatCoV_RaTG13_orf1ab;6929 | 0.25 | 0.05 |
| MN996532.1;QHR63299.1;BatCoV_RaTG13_orf1ab;6930 | 0.15 | 0.05 |
| MN996532.1;QHR63299.1;BatCoV_RaTG13_orf1ab;6931 | 0.1 | 0.05 |
| MN996532.1;QHR63299.1;BatCoV_RaTG13_orf1ab;6932 | 0.05 | 0.075 |
| MN996532.1;QHR63299.1;BatCoV_RaTG13_orf1ab;6933 | 0.25 | 0.1 |
| MN996532.1;QHR63299.1;BatCoV_RaTG13_orf1ab;6934 | 0.05 | 0.025 |
| MN996532.1;QHR63299.1;BatCoV_RaTG13_orf1ab;6935 | 0.05 | 0.025 |
| MN996532.1;QHR63299.1;BatCoV_RaTG13_orf1ab;6936 | 0 | 0.025 |
| MN996532.1;QHR63299.1;BatCoV_RaTG13_orf1ab;6937 | 0 | 0.05 |
| MN996532.1;QHR63299.1;BatCoV_RaTG13_orf1ab;6938 | 0 | 0 |
| MN996532.1;QHR63299.1;BatCoV_RaTG13_orf1ab;6939 | 0 | 0 |
| MN996532.1;QHR63299.1;BatCoV_RaTG13_orf1ab;6940 | 0 | 0 |
| MN996532.1;QHR63299.1;BatCoV_RaTG13_orf1ab;6941 | 0 | 0 |
| MN996532.1;QHR63299.1;BatCoV_RaTG13_orf1ab;6942 | 0 | 0 |
| MN996532.1;QHR63299.1;BatCoV_RaTG13_orf1ab;6943 | 0 | 0 |
| MN996532.1;QHR63299.1;BatCoV_RaTG13_orf1ab;6944 | 0 | 0 |
| MN996532.1;QHR63299.1;BatCoV_RaTG13_orf1ab;6945 | 0 | 0 |
| MN996532.1;QHR63299.1;BatCoV_RaTG13_orf1ab;6946 | 0 | 0 |
| MN996532.1;QHR63299.1;BatCoV_RaTG13_orf1ab;6947 | 0 | 0 |
| MN996532.1;QHR63299.1;BatCoV_RaTG13_orf1ab;6948 | 0 | 0 |
| MN996532.1;QHR63299.1;BatCoV_RaTG13_orf1ab;6949 | 0 | 0 |
| MN996532.1;QHR63299.1;BatCoV_RaTG13_orf1ab;6950 | 0 | 0 |
| MN996532.1;QHR63299.1;BatCoV_RaTG13_orf1ab;6951 | 0 | 0 |
| MN996532.1;QHR63299.1;BatCoV_RaTG13_orf1ab;6952 | 0 | 0 |
| MN996532.1;QHR63299.1;BatCoV_RaTG13_orf1ab;6953 | 0 | 0 |
| MN996532.1;QHR63299.1;BatCoV_RaTG13_orf1ab;6954 | 0 | 0 |
| MN996532.1;QHR63299.1;BatCoV_RaTG13_orf1ab;6955 | 0 | 0 |
| MN996532.1;QHR63299.1;BatCoV_RaTG13_orf1ab;6956 | 0 | 0 |
| MN996532.1;QHR63299.1;BatCoV_RaTG13_orf1ab;6957 | 0 | 0 |

| | | |
|---|---|---|
| MN996532.1;QHR63299.1;BatCoV_RaTG13_orf1ab;6958 | 0 | 0 |
| MN996532.1;QHR63299.1;BatCoV_RaTG13_orf1ab;6959 | 0 | 0 |
| MN996532.1;QHR63299.1;BatCoV_RaTG13_orf1ab;6960 | 0 | 0 |
| MN996532.1;QHR63299.1;BatCoV_RaTG13_orf1ab;6961 | 0 | 0 |
| MN996532.1;QHR63299.1;BatCoV_RaTG13_orf1ab;6962 | 0 | 0 |
| MN996532.1;QHR63299.1;BatCoV_RaTG13_orf1ab;6963 | 0 | 0 |
| MN996532.1;QHR63299.1;BatCoV_RaTG13_orf1ab;6964 | 0 | 0.05 |
| MN996532.1;QHR63299.1;BatCoV_RaTG13_orf1ab;6965 | 0 | 0.025 |
| MN996532.1;QHR63299.1;BatCoV_RaTG13_orf1ab;6966 | 0 | 0.05 |
| MN996532.1;QHR63299.1;BatCoV_RaTG13_orf1ab;6967 | 0 | 0.025 |
| MN996532.1;QHR63299.1;BatCoV_RaTG13_orf1ab;6968 | 0 | 0.025 |
| MN996532.1;QHR63299.1;BatCoV_RaTG13_orf1ab;6969 | 0 | 0.025 |
| MN996532.1;QHR63299.1;BatCoV_RaTG13_orf1ab;6970 | 0 | 0.025 |
| MN996532.1;QHR63299.1;BatCoV_RaTG13_orf1ab;6971 | 0 | 0.025 |
| MN996532.1;QHR63299.1;BatCoV_RaTG13_orf1ab;6972 | 0 | 0.025 |
| MN996532.1;QHR63299.1;BatCoV_RaTG13_orf1ab;6973 | 0 | 0 |
| MN996532.1;QHR63299.1;BatCoV_RaTG13_orf1ab;6974 | 0 | 0 |
| MN996532.1;QHR63299.1;BatCoV_RaTG13_orf1ab;6975 | 0 | 0 |
| MN996532.1;QHR63299.1;BatCoV_RaTG13_orf1ab;6976 | 0 | 0 |
| MN996532.1;QHR63299.1;BatCoV_RaTG13_orf1ab;6977 | 0 | 0 |
| MN996532.1;QHR63299.1;BatCoV_RaTG13_orf1ab;6978 | 0 | 0 |
| MN996532.1;QHR63299.1;BatCoV_RaTG13_orf1ab;6979 | 0 | 0 |
| MN996532.1;QHR63299.1;BatCoV_RaTG13_orf1ab;6980 | 0 | 0 |
| MN996532.1;QHR63299.1;BatCoV_RaTG13_orf1ab;6981 | 0 | 0 |
| MN996532.1;QHR63299.1;BatCoV_RaTG13_orf1ab;6982 | 0 | 0 |
| MN996532.1;QHR63299.1;BatCoV_RaTG13_orf1ab;6983 | 0 | 0 |
| MN996532.1;QHR63299.1;BatCoV_RaTG13_orf1ab;6984 | 0 | 0 |
| MN996532.1;QHR63299.1;BatCoV_RaTG13_orf1ab;6985 | 0 | 0.025 |
| MN996532.1;QHR63299.1;BatCoV_RaTG13_orf1ab;6986 | 0 | 0.025 |
| MN996532.1;QHR63299.1;BatCoV_RaTG13_orf1ab;6987 | 0 | 0.05 |
| MN996532.1;QHR63299.1;BatCoV_RaTG13_orf1ab;6988 | 0 | 0 |
| MN996532.1;QHR63299.1;BatCoV_RaTG13_orf1ab;6989 | 0.05 | 0 |
| MN996532.1;QHR63299.1;BatCoV_RaTG13_orf1ab;6990 | 0 | 0 |
| MN996532.1;QHR63299.1;BatCoV_RaTG13_orf1ab;6991 | 0 | 0.05 |
| MN996532.1;QHR63299.1;BatCoV_RaTG13_orf1ab;6992 | 0 | 0 |
| MN996532.1;QHR63299.1;BatCoV_RaTG13_orf1ab;6993 | 0 | 0 |
| MN996532.1;QHR63299.1;BatCoV_RaTG13_orf1ab;6994 | 0 | 0 |
| MN996532.1;QHR63299.1;BatCoV_RaTG13_orf1ab;6995 | 0 | 0 |
| MN996532.1;QHR63299.1;BatCoV_RaTG13_orf1ab;6996 | 0 | 0 |
| MN996532.1;QHR63299.1;BatCoV_RaTG13_orf1ab;6997 | 0 | 0 |
| MN996532.1;QHR63299.1;BatCoV_RaTG13_orf1ab;6998 | 0 | 0 |
| MN996532.1;QHR63299.1;BatCoV_RaTG13_orf1ab;6999 | 0 | 0 |
| MN996532.1;QHR63299.1;BatCoV_RaTG13_orf1ab;7000 | 0 | 0 |

| | | |
|---|---|---|
| MN996532.1;QHR63299.1;BatCoV_RaTG13_orf1ab;7001 | 0 | 0 |
| MN996532.1;QHR63299.1;BatCoV_RaTG13_orf1ab;7002 | 0.05 | 0 |
| MN996532.1;QHR63299.1;BatCoV_RaTG13_orf1ab;7003 | 0.05 | 0 |
| MN996532.1;QHR63299.1;BatCoV_RaTG13_orf1ab;7004 | 0 | 0 |
| MN996532.1;QHR63299.1;BatCoV_RaTG13_orf1ab;7005 | 0.1 | 0 |
| MN996532.1;QHR63299.1;BatCoV_RaTG13_orf1ab;7006 | 0.1 | 0 |
| MN996532.1;QHR63299.1;BatCoV_RaTG13_orf1ab;7007 | 0.05 | 0 |
| MN996532.1;QHR63299.1;BatCoV_RaTG13_orf1ab;7008 | 0.1 | 0 |
| MN996532.1;QHR63299.1;BatCoV_RaTG13_orf1ab;7009 | 0.1 | 0.05 |
| MN996532.1;QHR63299.1;BatCoV_RaTG13_orf1ab;7010 | 0 | 0.05 |
| MN996532.1;QHR63299.1;BatCoV_RaTG13_orf1ab;7011 | 0 | 0.05 |
| MN996532.1;QHR63299.1;BatCoV_RaTG13_orf1ab;7012 | 0 | 0 |
| MN996532.1;QHR63299.1;BatCoV_RaTG13_orf1ab;7013 | 0 | 0 |
| MN996532.1;QHR63299.1;BatCoV_RaTG13_orf1ab;7014 | 0.05 | 0.05 |
| MN996532.1;QHR63299.1;BatCoV_RaTG13_orf1ab;7015 | 0 | 0 |
| MN996532.1;QHR63299.1;BatCoV_RaTG13_orf1ab;7016 | 0 | 0 |
| MN996532.1;QHR63299.1;BatCoV_RaTG13_orf1ab;7017 | 0 | 0 |
| MN996532.1;QHR63299.1;BatCoV_RaTG13_orf1ab;7018 | 0 | 0 |
| MN996532.1;QHR63299.1;BatCoV_RaTG13_orf1ab;7019 | 0 | 0 |
| MN996532.1;QHR63299.1;BatCoV_RaTG13_orf1ab;7020 | 0 | 0 |
| MN996532.1;QHR63299.1;BatCoV_RaTG13_orf1ab;7021 | 0 | 0 |
| MN996532.1;QHR63299.1;BatCoV_RaTG13_orf1ab;7022 | 0 | 0 |
| MN996532.1;QHR63299.1;BatCoV_RaTG13_orf1ab;7023 | 0 | 0 |
| MN996532.1;QHR63299.1;BatCoV_RaTG13_orf1ab;7024 | 0 | 0 |
| MN996532.1;QHR63299.1;BatCoV_RaTG13_orf1ab;7025 | 0 | 0 |
| MN996532.1;QHR63299.1;BatCoV_RaTG13_orf1ab;7026 | 0 | 0 |
| MN996532.1;QHR63299.1;BatCoV_RaTG13_orf1ab;7027 | 0 | 0 |
| MN996532.1;QHR63299.1;BatCoV_RaTG13_orf1ab;7028 | 0 | 0 |
| MN996532.1;QHR63299.1;BatCoV_RaTG13_orf1ab;7029 | 0 | 0 |
| MN996532.1;QHR63299.1;BatCoV_RaTG13_orf1ab;7030 | 0 | 0 |
| MN996532.1;QHR63299.1;BatCoV_RaTG13_orf1ab;7031 | 0.05 | 0.075 |
| MN996532.1;QHR63299.1;BatCoV_RaTG13_orf1ab;7032 | 0.05 | 0.075 |
| MN996532.1;QHR63299.1;BatCoV_RaTG13_orf1ab;7033 | 0.05 | 0.05 |
| MN996532.1;QHR63299.1;BatCoV_RaTG13_orf1ab;7034 | 0.05 | 0.05 |
| MN996532.1;QHR63299.1;BatCoV_RaTG13_orf1ab;7035 | 0.05 | 0.05 |
| MN996532.1;QHR63299.1;BatCoV_RaTG13_orf1ab;7036 | 0.05 | 0.075 |
| MN996532.1;QHR63299.1;BatCoV_RaTG13_orf1ab;7037 | 0.05 | 0.025 |
| MN996532.1;QHR63299.1;BatCoV_RaTG13_orf1ab;7038 | 0 | 0.025 |
| MN996532.1;QHR63299.1;BatCoV_RaTG13_orf1ab;7039 | 0 | 0 |
| MN996532.1;QHR63299.1;BatCoV_RaTG13_orf1ab;7040 | 0 | 0 |
| MN996532.1;QHR63299.1;BatCoV_RaTG13_orf1ab;7041 | 0 | 0 |
| MN996532.1;QHR63299.1;BatCoV_RaTG13_orf1ab;7042 | 0 | 0 |
| MN996532.1;QHR63299.1;BatCoV_RaTG13_orf1ab;7043 | 0 | 0 |

| | | |
|---|---|---|
| MN996532.1;QHR63299.1;BatCoV_RaTG13_orf1ab;7044 | 0 | 0 |
| MN996532.1;QHR63299.1;BatCoV_RaTG13_orf1ab;7045 | 0 | 0 |
| MN996532.1;QHR63299.1;BatCoV_RaTG13_orf1ab;7046 | 0 | 0 |
| MN996532.1;QHR63299.1;BatCoV_RaTG13_orf1ab;7047 | 0 | 0 |
| MN996532.1;QHR63299.1;BatCoV_RaTG13_orf1ab;7048 | 0 | 0 |
| MN996532.1;QHR63299.1;BatCoV_RaTG13_orf1ab;7049 | 0 | 0 |
| MN996532.1;QHR63299.1;BatCoV_RaTG13_orf1ab;7050 | 0 | 0 |
| MN996532.1;QHR63299.1;BatCoV_RaTG13_orf1ab;7051 | 0 | 0 |
| MN996532.1;QHR63299.1;BatCoV_RaTG13_orf1ab;7052 | 0.05 | 0 |
| MN996532.1;QHR63299.1;BatCoV_RaTG13_orf1ab;7053 | 0.05 | 0 |
| MN996532.1;QHR63299.1;BatCoV_RaTG13_orf1ab;7054 | 0.05 | 0 |
| MN996532.1;QHR63299.1;BatCoV_RaTG13_orf1ab;7055 | 0.1 | 0 |
| MN996532.1;QHR63299.1;BatCoV_RaTG13_orf1ab;7056 | 0.05 | 0 |
| MN996532.1;QHR63299.1;BatCoV_RaTG13_orf1ab;7057 | 0 | 0 |
| MN996532.1;QHR63299.1;BatCoV_RaTG13_orf1ab;7058 | 0 | 0 |
| MN996532.1;QHR63299.1;BatCoV_RaTG13_orf1ab;7059 | 0 | 0 |
| MN996532.1;QHR63299.1;BatCoV_RaTG13_orf1ab;7060 | 0 | 0 |
| MN996532.1;QHR63299.1;BatCoV_RaTG13_orf1ab;7061 | 0 | 0 |
| MN996532.1;QHR63299.1;BatCoV_RaTG13_orf1ab;7062 | 0 | 0 |
| MN996532.1;QHR63299.1;BatCoV_RaTG13_orf1ab;7063 | 0 | 0 |
| MN996532.1;QHR63299.1;BatCoV_RaTG13_orf1ab;7064 | 0 | 0 |
| MN996532.1;QHR63299.1;BatCoV_RaTG13_orf1ab;7065 | 0 | 0 |
| MN996532.1;QHR63299.1;BatCoV_RaTG13_orf1ab;7066 | 0 | 0 |
| MN996532.1;QHR63299.1;BatCoV_RaTG13_orf1ab;7067 | 0.05 | 0 |
| MN996532.1;QHR63299.1;BatCoV_RaTG13_orf1ab;7068 | 0 | 0 |
| MN996532.1;QHR63299.1;BatCoV_RaTG13_orf1ab;7069 | 0 | 0 |
| MN996532.1;QHR63299.1;BatCoV_RaTG13_orf1ab;7070 | 0 | 0.025 |
| MN996532.1;QHR63299.1;BatCoV_RaTG13_orf1ab;7071 | 0 | 0.025 |
| MN996532.1;QHR63299.1;BatCoV_RaTG13_orf1ab;7072 | 0 | 0.025 |
| MN996532.1;QHR63299.1;BatCoV_RaTG13_orf1ab;7073 | 0 | 0.025 |
| MN996532.1;QHR63299.1;BatCoV_RaTG13_orf1ab;7074 | 0 | 0.025 |
| MN996532.1;QHR63299.1;BatCoV_RaTG13_orf1ab;7075 | 0 | 0.025 |
| MN996532.1;QHR63299.1;BatCoV_RaTG13_orf1ab;7076 | 0 | 0.025 |
| MN996532.1;QHR63299.1;BatCoV_RaTG13_orf1ab;7077 | 0 | 0.025 |
| MN996532.1;QHR63299.1;BatCoV_RaTG13_orf1ab;7078 | 0 | 0.025 |
| MN996532.1;QHR63299.1;BatCoV_RaTG13_orf1ab;7079 | 0 | 0.025 |
| MN996532.1;QHR63299.1;BatCoV_RaTG13_orf1ab;7080 | 0 | 0.025 |
| MN996532.1;QHR63299.1;BatCoV_RaTG13_orf1ab;7081 | 0 | 0 |
| MN996532.1;QHR63299.1;BatCoV_RaTG13_orf1ab;7082 | 0 | 0 |
| MN996532.1;QHR63299.1;BatCoV_RaTG13_orf1ab;7083 | 0 | 0 |
| MN996532.1;QHR63299.1;BatCoV_RaTG13_orf1ab;7084 | 0 | 0 |
| MN996532.1;QHR63300.2;BatCoV_RaTG13_spike;1 | 0 | 0 |
| MN996532.1;QHR63300.2;BatCoV_RaTG13_spike;2 | 0 | 0 |

| | | |
|---|---|---|
| MN996532.1;QHR63300.2;BatCoV_RaTG13_spike;3 | 0 | 0 |
| MN996532.1;QHR63300.2;BatCoV_RaTG13_spike;4 | 0 | 0 |
| MN996532.1;QHR63300.2;BatCoV_RaTG13_spike;5 | 0 | 0 |
| MN996532.1;QHR63300.2;BatCoV_RaTG13_spike;6 | 0 | 0 |
| MN996532.1;QHR63300.2;BatCoV_RaTG13_spike;7 | 0 | 0 |
| MN996532.1;QHR63300.2;BatCoV_RaTG13_spike;8 | 0.05 | 0 |
| MN996532.1;QHR63300.2;BatCoV_RaTG13_spike;9 | 0 | 0 |
| MN996532.1;QHR63300.2;BatCoV_RaTG13_spike;10 | 0 | 0 |
| MN996532.1;QHR63300.2;BatCoV_RaTG13_spike;11 | 0 | 0 |
| MN996532.1;QHR63300.2;BatCoV_RaTG13_spike;12 | 0 | 0 |
| MN996532.1;QHR63300.2;BatCoV_RaTG13_spike;13 | 0 | 0 |
| MN996532.1;QHR63300.2;BatCoV_RaTG13_spike;14 | 0 | 0 |
| MN996532.1;QHR63300.2;BatCoV_RaTG13_spike;15 | 0 | 0 |
| MN996532.1;QHR63300.2;BatCoV_RaTG13_spike;16 | 0 | 0 |
| MN996532.1;QHR63300.2;BatCoV_RaTG13_spike;17 | 0.15 | 0.025 |
| MN996532.1;QHR63300.2;BatCoV_RaTG13_spike;18 | 0 | 0.025 |
| MN996532.1;QHR63300.2;BatCoV_RaTG13_spike;19 | 0 | 0 |
| MN996532.1;QHR63300.2;BatCoV_RaTG13_spike;20 | 0 | 0 |
| MN996532.1;QHR63300.2;BatCoV_RaTG13_spike;21 | 0 | 0 |
| MN996532.1;QHR63300.2;BatCoV_RaTG13_spike;22 | 0 | 0.025 |
| MN996532.1;QHR63300.2;BatCoV_RaTG13_spike;23 | 0 | 0 |
| MN996532.1;QHR63300.2;BatCoV_RaTG13_spike;24 | 0 | 0 |
| MN996532.1;QHR63300.2;BatCoV_RaTG13_spike;25 | 0 | 0.025 |
| MN996532.1;QHR63300.2;BatCoV_RaTG13_spike;26 | 0.05 | 0.025 |
| MN996532.1;QHR63300.2;BatCoV_RaTG13_spike;27 | 0.05 | 0.05 |
| MN996532.1;QHR63300.2;BatCoV_RaTG13_spike;28 | 0.05 | 0 |
| MN996532.1;QHR63300.2;BatCoV_RaTG13_spike;29 | 0 | 0 |
| MN996532.1;QHR63300.2;BatCoV_RaTG13_spike;30 | 0 | 0 |
| MN996532.1;QHR63300.2;BatCoV_RaTG13_spike;31 | 0 | 0.05 |
| MN996532.1;QHR63300.2;BatCoV_RaTG13_spike;32 | 0 | 0 |
| MN996532.1;QHR63300.2;BatCoV_RaTG13_spike;33 | 0 | 0 |
| MN996532.1;QHR63300.2;BatCoV_RaTG13_spike;34 | 0 | 0 |
| MN996532.1;QHR63300.2;BatCoV_RaTG13_spike;35 | 0 | 0 |
| MN996532.1;QHR63300.2;BatCoV_RaTG13_spike;36 | 0 | 0 |
| MN996532.1;QHR63300.2;BatCoV_RaTG13_spike;37 | 0 | 0 |
| MN996532.1;QHR63300.2;BatCoV_RaTG13_spike;38 | 0 | 0 |
| MN996532.1;QHR63300.2;BatCoV_RaTG13_spike;39 | 0 | 0 |
| MN996532.1;QHR63300.2;BatCoV_RaTG13_spike;40 | 0 | 0 |
| MN996532.1;QHR63300.2;BatCoV_RaTG13_spike;41 | 0 | 0 |
| MN996532.1;QHR63300.2;BatCoV_RaTG13_spike;42 | 0 | 0 |
| MN996532.1;QHR63300.2;BatCoV_RaTG13_spike;43 | 0 | 0 |
| MN996532.1;QHR63300.2;BatCoV_RaTG13_spike;44 | 0 | 0 |
| MN996532.1;QHR63300.2;BatCoV_RaTG13_spike;45 | 0 | 0 |

| | | |
|---|---|---|
| MN996532.1;QHR63300.2;BatCoV_RaTG13_spike;46 | 0 | 0 |
| MN996532.1;QHR63300.2;BatCoV_RaTG13_spike;47 | 0 | 0 |
| MN996532.1;QHR63300.2;BatCoV_RaTG13_spike;48 | 0 | 0 |
| MN996532.1;QHR63300.2;BatCoV_RaTG13_spike;49 | 0 | 0 |
| MN996532.1;QHR63300.2;BatCoV_RaTG13_spike;50 | 0 | 0 |
| MN996532.1;QHR63300.2;BatCoV_RaTG13_spike;51 | 0 | 0 |
| MN996532.1;QHR63300.2;BatCoV_RaTG13_spike;52 | 0 | 0 |
| MN996532.1;QHR63300.2;BatCoV_RaTG13_spike;53 | 0 | 0 |
| MN996532.1;QHR63300.2;BatCoV_RaTG13_spike;54 | 0 | 0 |
| MN996532.1;QHR63300.2;BatCoV_RaTG13_spike;55 | 0 | 0 |
| MN996532.1;QHR63300.2;BatCoV_RaTG13_spike;56 | 0 | 0 |
| MN996532.1;QHR63300.2;BatCoV_RaTG13_spike;57 | 0 | 0 |
| MN996532.1;QHR63300.2;BatCoV_RaTG13_spike;58 | 0 | 0 |
| MN996532.1;QHR63300.2;BatCoV_RaTG13_spike;59 | 0 | 0 |
| MN996532.1;QHR63300.2;BatCoV_RaTG13_spike;60 | 0 | 0 |
| MN996532.1;QHR63300.2;BatCoV_RaTG13_spike;61 | 0 | 0 |
| MN996532.1;QHR63300.2;BatCoV_RaTG13_spike;62 | 0 | 0 |
| MN996532.1;QHR63300.2;BatCoV_RaTG13_spike;63 | 0 | 0 |
| MN996532.1;QHR63300.2;BatCoV_RaTG13_spike;64 | 0 | 0 |
| MN996532.1;QHR63300.2;BatCoV_RaTG13_spike;65 | 0 | 0 |
| MN996532.1;QHR63300.2;BatCoV_RaTG13_spike;66 | 0 | 0 |
| MN996532.1;QHR63300.2;BatCoV_RaTG13_spike;67 | 0 | 0 |
| MN996532.1;QHR63300.2;BatCoV_RaTG13_spike;68 | 0 | 0 |
| MN996532.1;QHR63300.2;BatCoV_RaTG13_spike;69 | 0 | 0 |
| MN996532.1;QHR63300.2;BatCoV_RaTG13_spike;70 | 0 | 0 |
| MN996532.1;QHR63300.2;BatCoV_RaTG13_spike;71 | 0 | 0 |
| MN996532.1;QHR63300.2;BatCoV_RaTG13_spike;72 | 0 | 0 |
| MN996532.1;QHR63300.2;BatCoV_RaTG13_spike;73 | 0 | 0.025 |
| MN996532.1;QHR63300.2;BatCoV_RaTG13_spike;74 | 0 | 0 |
| MN996532.1;QHR63300.2;BatCoV_RaTG13_spike;75 | 0 | 0.025 |
| MN996532.1;QHR63300.2;BatCoV_RaTG13_spike;76 | 0 | 0.025 |
| MN996532.1;QHR63300.2;BatCoV_RaTG13_spike;77 | 0 | 0.025 |
| MN996532.1;QHR63300.2;BatCoV_RaTG13_spike;78 | 0 | 0 |
| MN996532.1;QHR63300.2;BatCoV_RaTG13_spike;79 | 0 | 0 |
| MN996532.1;QHR63300.2;BatCoV_RaTG13_spike;80 | 0 | 0 |
| MN996532.1;QHR63300.2;BatCoV_RaTG13_spike;81 | 0 | 0.025 |
| MN996532.1;QHR63300.2;BatCoV_RaTG13_spike;82 | 0.05 | 0.075 |
| MN996532.1;QHR63300.2;BatCoV_RaTG13_spike;83 | 0.05 | 0.1 |
| MN996532.1;QHR63300.2;BatCoV_RaTG13_spike;84 | 0.05 | 0.075 |
| MN996532.1;QHR63300.2;BatCoV_RaTG13_spike;85 | 0.05 | 0.05 |
| MN996532.1;QHR63300.2;BatCoV_RaTG13_spike;86 | 0.05 | 0.05 |
| MN996532.1;QHR63300.2;BatCoV_RaTG13_spike;87 | 0.05 | 0 |
| MN996532.1;QHR63300.2;BatCoV_RaTG13_spike;88 | 0.05 | 0.025 |

| | | |
|---|---|---|
| MN996532.1;QHR63300.2;BatCoV_RaTG13_spike;89 | 0.05 | 0 |
| MN996532.1;QHR63300.2;BatCoV_RaTG13_spike;90 | 0.05 | 0.025 |
| MN996532.1;QHR63300.2;BatCoV_RaTG13_spike;91 | 0.05 | 0 |
| MN996532.1;QHR63300.2;BatCoV_RaTG13_spike;92 | 0.05 | 0 |
| MN996532.1;QHR63300.2;BatCoV_RaTG13_spike;93 | 0 | 0 |
| MN996532.1;QHR63300.2;BatCoV_RaTG13_spike;94 | 0 | 0 |
| MN996532.1;QHR63300.2;BatCoV_RaTG13_spike;95 | 0 | 0 |
| MN996532.1;QHR63300.2;BatCoV_RaTG13_spike;96 | 0 | 0 |
| MN996532.1;QHR63300.2;BatCoV_RaTG13_spike;97 | 0 | 0 |
| MN996532.1;QHR63300.2;BatCoV_RaTG13_spike;98 | 0 | 0 |
| MN996532.1;QHR63300.2;BatCoV_RaTG13_spike;99 | 0 | 0 |
| MN996532.1;QHR63300.2;BatCoV_RaTG13_spike;100 | 0 | 0 |
| MN996532.1;QHR63300.2;BatCoV_RaTG13_spike;101 | 0 | 0 |
| MN996532.1;QHR63300.2;BatCoV_RaTG13_spike;102 | 0 | 0 |
| MN996532.1;QHR63300.2;BatCoV_RaTG13_spike;103 | 0 | 0 |
| MN996532.1;QHR63300.2;BatCoV_RaTG13_spike;104 | 0 | 0 |
| MN996532.1;QHR63300.2;BatCoV_RaTG13_spike;105 | 0 | 0 |
| MN996532.1;QHR63300.2;BatCoV_RaTG13_spike;106 | 0 | 0 |
| MN996532.1;QHR63300.2;BatCoV_RaTG13_spike;107 | 0 | 0 |
| MN996532.1;QHR63300.2;BatCoV_RaTG13_spike;108 | 0 | 0 |
| MN996532.1;QHR63300.2;BatCoV_RaTG13_spike;109 | 0 | 0.025 |
| MN996532.1;QHR63300.2;BatCoV_RaTG13_spike;110 | 0 | 0 |
| MN996532.1;QHR63300.2;BatCoV_RaTG13_spike;111 | 0.05 | 0.025 |
| MN996532.1;QHR63300.2;BatCoV_RaTG13_spike;112 | 0.05 | 0.025 |
| MN996532.1;QHR63300.2;BatCoV_RaTG13_spike;113 | 0.05 | 0 |
| MN996532.1;QHR63300.2;BatCoV_RaTG13_spike;114 | 0.05 | 0 |
| MN996532.1;QHR63300.2;BatCoV_RaTG13_spike;115 | 0 | 0 |
| MN996532.1;QHR63300.2;BatCoV_RaTG13_spike;116 | 0 | 0 |
| MN996532.1;QHR63300.2;BatCoV_RaTG13_spike;117 | 0 | 0 |
| MN996532.1;QHR63300.2;BatCoV_RaTG13_spike;118 | 0 | 0 |
| MN996532.1;QHR63300.2;BatCoV_RaTG13_spike;119 | 0.05 | 0 |
| MN996532.1;QHR63300.2;BatCoV_RaTG13_spike;120 | 0.05 | 0 |
| MN996532.1;QHR63300.2;BatCoV_RaTG13_spike;121 | 0 | 0 |
| MN996532.1;QHR63300.2;BatCoV_RaTG13_spike;122 | 0 | 0 |
| MN996532.1;QHR63300.2;BatCoV_RaTG13_spike;123 | 0 | 0 |
| MN996532.1;QHR63300.2;BatCoV_RaTG13_spike;124 | 0.05 | 0 |
| MN996532.1;QHR63300.2;BatCoV_RaTG13_spike;125 | 0 | 0 |
| MN996532.1;QHR63300.2;BatCoV_RaTG13_spike;126 | 0 | 0 |
| MN996532.1;QHR63300.2;BatCoV_RaTG13_spike;127 | 0 | 0 |
| MN996532.1;QHR63300.2;BatCoV_RaTG13_spike;128 | 0 | 0 |
| MN996532.1;QHR63300.2;BatCoV_RaTG13_spike;129 | 0 | 0 |
| MN996532.1;QHR63300.2;BatCoV_RaTG13_spike;130 | 0 | 0 |
| MN996532.1;QHR63300.2;BatCoV_RaTG13_spike;131 | 0 | 0 |

| | | |
|---|---|---|
| MN996532.1;QHR63300.2;BatCoV_RaTG13_spike;132 | 0 | 0 |
| MN996532.1;QHR63300.2;BatCoV_RaTG13_spike;133 | 0 | 0 |
| MN996532.1;QHR63300.2;BatCoV_RaTG13_spike;134 | 0 | 0 |
| MN996532.1;QHR63300.2;BatCoV_RaTG13_spike;135 | 0 | 0 |
| MN996532.1;QHR63300.2;BatCoV_RaTG13_spike;136 | 0 | 0 |
| MN996532.1;QHR63300.2;BatCoV_RaTG13_spike;137 | 0 | 0 |
| MN996532.1;QHR63300.2;BatCoV_RaTG13_spike;138 | 0 | 0 |
| MN996532.1;QHR63300.2;BatCoV_RaTG13_spike;139 | 0.05 | 0 |
| MN996532.1;QHR63300.2;BatCoV_RaTG13_spike;140 | 0.05 | 0.025 |
| MN996532.1;QHR63300.2;BatCoV_RaTG13_spike;141 | 0.05 | 0.075 |
| MN996532.1;QHR63300.2;BatCoV_RaTG13_spike;142 | 0.1 | 0.05 |
| MN996532.1;QHR63300.2;BatCoV_RaTG13_spike;143 | 0.05 | 0.025 |
| MN996532.1;QHR63300.2;BatCoV_RaTG13_spike;144 | 0.05 | 0 |
| MN996532.1;QHR63300.2;BatCoV_RaTG13_spike;145 | 0 | 0 |
| MN996532.1;QHR63300.2;BatCoV_RaTG13_spike;146 | 0.05 | 0 |
| MN996532.1;QHR63300.2;BatCoV_RaTG13_spike;147 | 0.05 | 0 |
| MN996532.1;QHR63300.2;BatCoV_RaTG13_spike;148 | 0.05 | 0 |
| MN996532.1;QHR63300.2;BatCoV_RaTG13_spike;149 | 0.05 | 0.025 |
| MN996532.1;QHR63300.2;BatCoV_RaTG13_spike;150 | 0 | 0.05 |
| MN996532.1;QHR63300.2;BatCoV_RaTG13_spike;151 | 0 | 0.025 |
| MN996532.1;QHR63300.2;BatCoV_RaTG13_spike;152 | 0 | 0.025 |
| MN996532.1;QHR63300.2;BatCoV_RaTG13_spike;153 | 0 | 0.025 |
| MN996532.1;QHR63300.2;BatCoV_RaTG13_spike;154 | 0 | 0.025 |
| MN996532.1;QHR63300.2;BatCoV_RaTG13_spike;155 | 0 | 0.025 |
| MN996532.1;QHR63300.2;BatCoV_RaTG13_spike;156 | 0 | 0 |
| MN996532.1;QHR63300.2;BatCoV_RaTG13_spike;157 | 0 | 0 |
| MN996532.1;QHR63300.2;BatCoV_RaTG13_spike;158 | 0.05 | 0.05 |
| MN996532.1;QHR63300.2;BatCoV_RaTG13_spike;159 | 0 | 0.1 |
| MN996532.1;QHR63300.2;BatCoV_RaTG13_spike;160 | 0 | 0 |
| MN996532.1;QHR63300.2;BatCoV_RaTG13_spike;161 | 0 | 0 |
| MN996532.1;QHR63300.2;BatCoV_RaTG13_spike;162 | 0 | 0.025 |
| MN996532.1;QHR63300.2;BatCoV_RaTG13_spike;163 | 0 | 0 |
| MN996532.1;QHR63300.2;BatCoV_RaTG13_spike;164 | 0 | 0 |
| MN996532.1;QHR63300.2;BatCoV_RaTG13_spike;165 | 0 | 0 |
| MN996532.1;QHR63300.2;BatCoV_RaTG13_spike;166 | 0 | 0 |
| MN996532.1;QHR63300.2;BatCoV_RaTG13_spike;167 | 0 | 0 |
| MN996532.1;QHR63300.2;BatCoV_RaTG13_spike;168 | 0 | 0 |
| MN996532.1;QHR63300.2;BatCoV_RaTG13_spike;169 | 0.05 | 0 |
| MN996532.1;QHR63300.2;BatCoV_RaTG13_spike;170 | 0 | 0 |
| MN996532.1;QHR63300.2;BatCoV_RaTG13_spike;171 | 0.05 | 0 |
| MN996532.1;QHR63300.2;BatCoV_RaTG13_spike;172 | 0 | 0.075 |
| MN996532.1;QHR63300.2;BatCoV_RaTG13_spike;173 | 0.05 | 0.225 |
| MN996532.1;QHR63300.2;BatCoV_RaTG13_spike;174 | 0.1 | 0.125 |

| | | |
|---|---|---|
| MN996532.1;QHR63300.2;BatCoV_RaTG13_spike;175 | 0.1 | 0.15 |
| MN996532.1;QHR63300.2;BatCoV_RaTG13_spike;176 | 0.1 | 0.175 |
| MN996532.1;QHR63300.2;BatCoV_RaTG13_spike;177 | 0.15 | 0.1 |
| MN996532.1;QHR63300.2;BatCoV_RaTG13_spike;178 | 0.05 | 0.2 |
| MN996532.1;QHR63300.2;BatCoV_RaTG13_spike;179 | 0.05 | 0.125 |
| MN996532.1;QHR63300.2;BatCoV_RaTG13_spike;180 | 0.05 | 0.05 |
| MN996532.1;QHR63300.2;BatCoV_RaTG13_spike;181 | 0 | 0.05 |
| MN996532.1;QHR63300.2;BatCoV_RaTG13_spike;182 | 0 | 0.025 |
| MN996532.1;QHR63300.2;BatCoV_RaTG13_spike;183 | 0.05 | 0.025 |
| MN996532.1;QHR63300.2;BatCoV_RaTG13_spike;184 | 0.1 | 0 |
| MN996532.1;QHR63300.2;BatCoV_RaTG13_spike;185 | 0.1 | 0 |
| MN996532.1;QHR63300.2;BatCoV_RaTG13_spike;186 | 0 | 0 |
| MN996532.1;QHR63300.2;BatCoV_RaTG13_spike;187 | 0 | 0 |
| MN996532.1;QHR63300.2;BatCoV_RaTG13_spike;188 | 0 | 0 |
| MN996532.1;QHR63300.2;BatCoV_RaTG13_spike;189 | 0 | 0 |
| MN996532.1;QHR63300.2;BatCoV_RaTG13_spike;190 | 0 | 0 |
| MN996532.1;QHR63300.2;BatCoV_RaTG13_spike;191 | 0 | 0 |
| MN996532.1;QHR63300.2;BatCoV_RaTG13_spike;192 | 0 | 0 |
| MN996532.1;QHR63300.2;BatCoV_RaTG13_spike;193 | 0 | 0 |
| MN996532.1;QHR63300.2;BatCoV_RaTG13_spike;194 | 0 | 0 |
| MN996532.1;QHR63300.2;BatCoV_RaTG13_spike;195 | 0 | 0 |
| MN996532.1;QHR63300.2;BatCoV_RaTG13_spike;196 | 0 | 0 |
| MN996532.1;QHR63300.2;BatCoV_RaTG13_spike;197 | 0 | 0 |
| MN996532.1;QHR63300.2;BatCoV_RaTG13_spike;198 | 0 | 0 |
| MN996532.1;QHR63300.2;BatCoV_RaTG13_spike;199 | 0 | 0 |
| MN996532.1;QHR63300.2;BatCoV_RaTG13_spike;200 | 0 | 0 |
| MN996532.1;QHR63300.2;BatCoV_RaTG13_spike;201 | 0 | 0 |
| MN996532.1;QHR63300.2;BatCoV_RaTG13_spike;202 | 0 | 0 |
| MN996532.1;QHR63300.2;BatCoV_RaTG13_spike;203 | 0 | 0 |
| MN996532.1;QHR63300.2;BatCoV_RaTG13_spike;204 | 0 | 0 |
| MN996532.1;QHR63300.2;BatCoV_RaTG13_spike;205 | 0 | 0 |
| MN996532.1;QHR63300.2;BatCoV_RaTG13_spike;206 | 0 | 0 |
| MN996532.1;QHR63300.2;BatCoV_RaTG13_spike;207 | 0 | 0 |
| MN996532.1;QHR63300.2;BatCoV_RaTG13_spike;208 | 0 | 0.05 |
| MN996532.1;QHR63300.2;BatCoV_RaTG13_spike;209 | 0 | 0 |
| MN996532.1;QHR63300.2;BatCoV_RaTG13_spike;210 | 0 | 0 |
| MN996532.1;QHR63300.2;BatCoV_RaTG13_spike;211 | 0 | 0 |
| MN996532.1;QHR63300.2;BatCoV_RaTG13_spike;212 | 0 | 0 |
| MN996532.1;QHR63300.2;BatCoV_RaTG13_spike;213 | 0 | 0 |
| MN996532.1;QHR63300.2;BatCoV_RaTG13_spike;214 | 0 | 0 |
| MN996532.1;QHR63300.2;BatCoV_RaTG13_spike;215 | 0 | 0 |
| MN996532.1;QHR63300.2;BatCoV_RaTG13_spike;216 | 0 | 0 |
| MN996532.1;QHR63300.2;BatCoV_RaTG13_spike;217 | 0 | 0 |

| | | |
|---|---|---|
| MN996532.1;QHR63300.2;BatCoV_RaTG13_spike;218 | 0 | 0 |
| MN996532.1;QHR63300.2;BatCoV_RaTG13_spike;219 | 0 | 0 |
| MN996532.1;QHR63300.2;BatCoV_RaTG13_spike;220 | 0 | 0 |
| MN996532.1;QHR63300.2;BatCoV_RaTG13_spike;221 | 0 | 0 |
| MN996532.1;QHR63300.2;BatCoV_RaTG13_spike;222 | 0 | 0 |
| MN996532.1;QHR63300.2;BatCoV_RaTG13_spike;223 | 0 | 0 |
| MN996532.1;QHR63300.2;BatCoV_RaTG13_spike;224 | 0.05 | 0 |
| MN996532.1;QHR63300.2;BatCoV_RaTG13_spike;225 | 0 | 0.025 |
| MN996532.1;QHR63300.2;BatCoV_RaTG13_spike;226 | 0 | 0 |
| MN996532.1;QHR63300.2;BatCoV_RaTG13_spike;227 | 0 | 0 |
| MN996532.1;QHR63300.2;BatCoV_RaTG13_spike;228 | 0 | 0 |
| MN996532.1;QHR63300.2;BatCoV_RaTG13_spike;229 | 0 | 0 |
| MN996532.1;QHR63300.2;BatCoV_RaTG13_spike;230 | 0 | 0 |
| MN996532.1;QHR63300.2;BatCoV_RaTG13_spike;231 | 0 | 0 |
| MN996532.1;QHR63300.2;BatCoV_RaTG13_spike;232 | 0 | 0 |
| MN996532.1;QHR63300.2;BatCoV_RaTG13_spike;233 | 0 | 0 |
| MN996532.1;QHR63300.2;BatCoV_RaTG13_spike;234 | 0 | 0 |
| MN996532.1;QHR63300.2;BatCoV_RaTG13_spike;235 | 0 | 0 |
| MN996532.1;QHR63300.2;BatCoV_RaTG13_spike;236 | 0 | 0 |
| MN996532.1;QHR63300.2;BatCoV_RaTG13_spike;237 | 0 | 0 |
| MN996532.1;QHR63300.2;BatCoV_RaTG13_spike;238 | 0 | 0 |
| MN996532.1;QHR63300.2;BatCoV_RaTG13_spike;239 | 0 | 0 |
| MN996532.1;QHR63300.2;BatCoV_RaTG13_spike;240 | 0 | 0.025 |
| MN996532.1;QHR63300.2;BatCoV_RaTG13_spike;241 | 0 | 0.025 |
| MN996532.1;QHR63300.2;BatCoV_RaTG13_spike;242 | 0 | 0.075 |
| MN996532.1;QHR63300.2;BatCoV_RaTG13_spike;243 | 0 | 0.025 |
| MN996532.1;QHR63300.2;BatCoV_RaTG13_spike;244 | 0 | 0.025 |
| MN996532.1;QHR63300.2;BatCoV_RaTG13_spike;245 | 0 | 0.05 |
| MN996532.1;QHR63300.2;BatCoV_RaTG13_spike;246 | 0 | 0.1 |
| MN996532.1;QHR63300.2;BatCoV_RaTG13_spike;247 | 0 | 0.1 |
| MN996532.1;QHR63300.2;BatCoV_RaTG13_spike;248 | 0.15 | 0.175 |
| MN996532.1;QHR63300.2;BatCoV_RaTG13_spike;249 | 0.1 | 0.15 |
| MN996532.1;QHR63300.2;BatCoV_RaTG13_spike;250 | 0 | 0 |
| MN996532.1;QHR63300.2;BatCoV_RaTG13_spike;251 | 0 | 0 |
| MN996532.1;QHR63300.2;BatCoV_RaTG13_spike;252 | 0 | 0.025 |
| MN996532.1;QHR63300.2;BatCoV_RaTG13_spike;253 | 0 | 0 |
| MN996532.1;QHR63300.2;BatCoV_RaTG13_spike;254 | 0 | 0 |
| MN996532.1;QHR63300.2;BatCoV_RaTG13_spike;255 | 0 | 0 |
| MN996532.1;QHR63300.2;BatCoV_RaTG13_spike;256 | 0 | 0 |
| MN996532.1;QHR63300.2;BatCoV_RaTG13_spike;257 | 0 | 0 |
| MN996532.1;QHR63300.2;BatCoV_RaTG13_spike;258 | 0 | 0 |
| MN996532.1;QHR63300.2;BatCoV_RaTG13_spike;259 | 0 | 0 |
| MN996532.1;QHR63300.2;BatCoV_RaTG13_spike;260 | 0 | 0 |

| | | |
|---|---|---|
| MN996532.1;QHR63300.2;BatCoV_RaTG13_spike;261 | 0 | 0 |
| MN996532.1;QHR63300.2;BatCoV_RaTG13_spike;262 | 0 | 0 |
| MN996532.1;QHR63300.2;BatCoV_RaTG13_spike;263 | 0 | 0 |
| MN996532.1;QHR63300.2;BatCoV_RaTG13_spike;264 | 0 | 0 |
| MN996532.1;QHR63300.2;BatCoV_RaTG13_spike;265 | 0 | 0 |
| MN996532.1;QHR63300.2;BatCoV_RaTG13_spike;266 | 0 | 0 |
| MN996532.1;QHR63300.2;BatCoV_RaTG13_spike;267 | 0 | 0 |
| MN996532.1;QHR63300.2;BatCoV_RaTG13_spike;268 | 0 | 0 |
| MN996532.1;QHR63300.2;BatCoV_RaTG13_spike;269 | 0 | 0 |
| MN996532.1;QHR63300.2;BatCoV_RaTG13_spike;270 | 0 | 0 |
| MN996532.1;QHR63300.2;BatCoV_RaTG13_spike;271 | 0 | 0 |
| MN996532.1;QHR63300.2;BatCoV_RaTG13_spike;272 | 0.05 | 0 |
| MN996532.1;QHR63300.2;BatCoV_RaTG13_spike;273 | 0.05 | 0 |
| MN996532.1;QHR63300.2;BatCoV_RaTG13_spike;274 | 0.05 | 0 |
| MN996532.1;QHR63300.2;BatCoV_RaTG13_spike;275 | 0 | 0 |
| MN996532.1;QHR63300.2;BatCoV_RaTG13_spike;276 | 0 | 0 |
| MN996532.1;QHR63300.2;BatCoV_RaTG13_spike;277 | 0.05 | 0 |
| MN996532.1;QHR63300.2;BatCoV_RaTG13_spike;278 | 0.05 | 0 |
| MN996532.1;QHR63300.2;BatCoV_RaTG13_spike;279 | 0 | 0 |
| MN996532.1;QHR63300.2;BatCoV_RaTG13_spike;280 | 0 | 0 |
| MN996532.1;QHR63300.2;BatCoV_RaTG13_spike;281 | 0 | 0 |
| MN996532.1;QHR63300.2;BatCoV_RaTG13_spike;282 | 0 | 0 |
| MN996532.1;QHR63300.2;BatCoV_RaTG13_spike;283 | 0 | 0 |
| MN996532.1;QHR63300.2;BatCoV_RaTG13_spike;284 | 0 | 0 |
| MN996532.1;QHR63300.2;BatCoV_RaTG13_spike;285 | 0 | 0 |
| MN996532.1;QHR63300.2;BatCoV_RaTG13_spike;286 | 0 | 0 |
| MN996532.1;QHR63300.2;BatCoV_RaTG13_spike;287 | 0 | 0 |
| MN996532.1;QHR63300.2;BatCoV_RaTG13_spike;288 | 0 | 0 |
| MN996532.1;QHR63300.2;BatCoV_RaTG13_spike;289 | 0 | 0 |
| MN996532.1;QHR63300.2;BatCoV_RaTG13_spike;290 | 0 | 0 |
| MN996532.1;QHR63300.2;BatCoV_RaTG13_spike;291 | 0 | 0 |
| MN996532.1;QHR63300.2;BatCoV_RaTG13_spike;292 | 0 | 0 |
| MN996532.1;QHR63300.2;BatCoV_RaTG13_spike;293 | 0 | 0 |
| MN996532.1;QHR63300.2;BatCoV_RaTG13_spike;294 | 0 | 0 |
| MN996532.1;QHR63300.2;BatCoV_RaTG13_spike;295 | 0.05 | 0 |
| MN996532.1;QHR63300.2;BatCoV_RaTG13_spike;296 | 0.1 | 0 |
| MN996532.1;QHR63300.2;BatCoV_RaTG13_spike;297 | 0 | 0 |
| MN996532.1;QHR63300.2;BatCoV_RaTG13_spike;298 | 0 | 0.025 |
| MN996532.1;QHR63300.2;BatCoV_RaTG13_spike;299 | 0 | 0 |
| MN996532.1;QHR63300.2;BatCoV_RaTG13_spike;300 | 0 | 0 |
| MN996532.1;QHR63300.2;BatCoV_RaTG13_spike;301 | 0 | 0 |
| MN996532.1;QHR63300.2;BatCoV_RaTG13_spike;302 | 0.05 | 0.075 |
| MN996532.1;QHR63300.2;BatCoV_RaTG13_spike;303 | 0 | 0.1 |

| | | |
|---|---|---|
| MN996532.1;QHR63300.2;BatCoV_RaTG13_spike;304 | 0 | 0.05 |
| MN996532.1;QHR63300.2;BatCoV_RaTG13_spike;305 | 0 | 0.05 |
| MN996532.1;QHR63300.2;BatCoV_RaTG13_spike;306 | 0 | 0.05 |
| MN996532.1;QHR63300.2;BatCoV_RaTG13_spike;307 | 0 | 0.1 |
| MN996532.1;QHR63300.2;BatCoV_RaTG13_spike;308 | 0.05 | 0.15 |
| MN996532.1;QHR63300.2;BatCoV_RaTG13_spike;309 | 0.05 | 0.275 |
| MN996532.1;QHR63300.2;BatCoV_RaTG13_spike;310 | 0.05 | 0.225 |
| MN996532.1;QHR63300.2;BatCoV_RaTG13_spike;311 | 0.1 | 0.025 |
| MN996532.1;QHR63300.2;BatCoV_RaTG13_spike;312 | 0.05 | 0.025 |
| MN996532.1;QHR63300.2;BatCoV_RaTG13_spike;313 | 0 | 0.025 |
| MN996532.1;QHR63300.2;BatCoV_RaTG13_spike;314 | 0 | 0.025 |
| MN996532.1;QHR63300.2;BatCoV_RaTG13_spike;315 | 0.05 | 0 |
| MN996532.1;QHR63300.2;BatCoV_RaTG13_spike;316 | 0 | 0 |
| MN996532.1;QHR63300.2;BatCoV_RaTG13_spike;317 | 0 | 0 |
| MN996532.1;QHR63300.2;BatCoV_RaTG13_spike;318 | 0 | 0 |
| MN996532.1;QHR63300.2;BatCoV_RaTG13_spike;319 | 0.1 | 0.025 |
| MN996532.1;QHR63300.2;BatCoV_RaTG13_spike;320 | 0.05 | 0 |
| MN996532.1;QHR63300.2;BatCoV_RaTG13_spike;321 | 0 | 0 |
| MN996532.1;QHR63300.2;BatCoV_RaTG13_spike;322 | 0 | 0 |
| MN996532.1;QHR63300.2;BatCoV_RaTG13_spike;323 | 0 | 0.025 |
| MN996532.1;QHR63300.2;BatCoV_RaTG13_spike;324 | 0 | 0 |
| MN996532.1;QHR63300.2;BatCoV_RaTG13_spike;325 | 0 | 0 |
| MN996532.1;QHR63300.2;BatCoV_RaTG13_spike;326 | 0 | 0 |
| MN996532.1;QHR63300.2;BatCoV_RaTG13_spike;327 | 0 | 0 |
| MN996532.1;QHR63300.2;BatCoV_RaTG13_spike;328 | 0 | 0 |
| MN996532.1;QHR63300.2;BatCoV_RaTG13_spike;329 | 0.05 | 0 |
| MN996532.1;QHR63300.2;BatCoV_RaTG13_spike;330 | 0.05 | 0.025 |
| MN996532.1;QHR63300.2;BatCoV_RaTG13_spike;331 | 0 | 0 |
| MN996532.1;QHR63300.2;BatCoV_RaTG13_spike;332 | 0 | 0 |
| MN996532.1;QHR63300.2;BatCoV_RaTG13_spike;333 | 0 | 0 |
| MN996532.1;QHR63300.2;BatCoV_RaTG13_spike;334 | 0 | 0 |
| MN996532.1;QHR63300.2;BatCoV_RaTG13_spike;335 | 0.05 | 0 |
| MN996532.1;QHR63300.2;BatCoV_RaTG13_spike;336 | 0 | 0 |
| MN996532.1;QHR63300.2;BatCoV_RaTG13_spike;337 | 0 | 0 |
| MN996532.1;QHR63300.2;BatCoV_RaTG13_spike;338 | 0 | 0 |
| MN996532.1;QHR63300.2;BatCoV_RaTG13_spike;339 | 0 | 0 |
| MN996532.1;QHR63300.2;BatCoV_RaTG13_spike;340 | 0 | 0 |
| MN996532.1;QHR63300.2;BatCoV_RaTG13_spike;341 | 0 | 0.025 |
| MN996532.1;QHR63300.2;BatCoV_RaTG13_spike;342 | 0 | 0 |
| MN996532.1;QHR63300.2;BatCoV_RaTG13_spike;343 | 0 | 0 |
| MN996532.1;QHR63300.2;BatCoV_RaTG13_spike;344 | 0 | 0.025 |
| MN996532.1;QHR63300.2;BatCoV_RaTG13_spike;345 | 0 | 0.025 |
| MN996532.1;QHR63300.2;BatCoV_RaTG13_spike;346 | 0 | 0.025 |

| | | |
|---|---|---|
| MN996532.1;QHR63300.2;BatCoV_RaTG13_spike;347 | 0 | 0.025 |
| MN996532.1;QHR63300.2;BatCoV_RaTG13_spike;348 | 0 | 0.025 |
| MN996532.1;QHR63300.2;BatCoV_RaTG13_spike;349 | 0.05 | 0.025 |
| MN996532.1;QHR63300.2;BatCoV_RaTG13_spike;350 | 0 | 0.025 |
| MN996532.1;QHR63300.2;BatCoV_RaTG13_spike;351 | 0 | 0.025 |
| MN996532.1;QHR63300.2;BatCoV_RaTG13_spike;352 | 0 | 0 |
| MN996532.1;QHR63300.2;BatCoV_RaTG13_spike;353 | 0 | 0 |
| MN996532.1;QHR63300.2;BatCoV_RaTG13_spike;354 | 0 | 0 |
| MN996532.1;QHR63300.2;BatCoV_RaTG13_spike;355 | 0 | 0 |
| MN996532.1;QHR63300.2;BatCoV_RaTG13_spike;356 | 0 | 0.025 |
| MN996532.1;QHR63300.2;BatCoV_RaTG13_spike;357 | 0 | 0 |
| MN996532.1;QHR63300.2;BatCoV_RaTG13_spike;358 | 0 | 0 |
| MN996532.1;QHR63300.2;BatCoV_RaTG13_spike;359 | 0 | 0 |
| MN996532.1;QHR63300.2;BatCoV_RaTG13_spike;360 | 0 | 0 |
| MN996532.1;QHR63300.2;BatCoV_RaTG13_spike;361 | 0 | 0 |
| MN996532.1;QHR63300.2;BatCoV_RaTG13_spike;362 | 0 | 0 |
| MN996532.1;QHR63300.2;BatCoV_RaTG13_spike;363 | 0.05 | 0 |
| MN996532.1;QHR63300.2;BatCoV_RaTG13_spike;364 | 0 | 0 |
| MN996532.1;QHR63300.2;BatCoV_RaTG13_spike;365 | 0 | 0 |
| MN996532.1;QHR63300.2;BatCoV_RaTG13_spike;366 | 0 | 0 |
| MN996532.1;QHR63300.2;BatCoV_RaTG13_spike;367 | 0 | 0 |
| MN996532.1;QHR63300.2;BatCoV_RaTG13_spike;368 | 0 | 0 |
| MN996532.1;QHR63300.2;BatCoV_RaTG13_spike;369 | 0 | 0 |
| MN996532.1;QHR63300.2;BatCoV_RaTG13_spike;370 | 0.05 | 0 |
| MN996532.1;QHR63300.2;BatCoV_RaTG13_spike;371 | 0 | 0 |
| MN996532.1;QHR63300.2;BatCoV_RaTG13_spike;372 | 0 | 0 |
| MN996532.1;QHR63300.2;BatCoV_RaTG13_spike;373 | 0 | 0 |
| MN996532.1;QHR63300.2;BatCoV_RaTG13_spike;374 | 0.05 | 0 |
| MN996532.1;QHR63300.2;BatCoV_RaTG13_spike;375 | 0.05 | 0 |
| MN996532.1;QHR63300.2;BatCoV_RaTG13_spike;376 | 0 | 0 |
| MN996532.1;QHR63300.2;BatCoV_RaTG13_spike;377 | 0 | 0 |
| MN996532.1;QHR63300.2;BatCoV_RaTG13_spike;378 | 0 | 0 |
| MN996532.1;QHR63300.2;BatCoV_RaTG13_spike;379 | 0 | 0 |
| MN996532.1;QHR63300.2;BatCoV_RaTG13_spike;380 | 0 | 0 |
| MN996532.1;QHR63300.2;BatCoV_RaTG13_spike;381 | 0 | 0 |
| MN996532.1;QHR63300.2;BatCoV_RaTG13_spike;382 | 0 | 0 |
| MN996532.1;QHR63300.2;BatCoV_RaTG13_spike;383 | 0 | 0.025 |
| MN996532.1;QHR63300.2;BatCoV_RaTG13_spike;384 | 0 | 0 |
| MN996532.1;QHR63300.2;BatCoV_RaTG13_spike;385 | 0 | 0.025 |
| MN996532.1;QHR63300.2;BatCoV_RaTG13_spike;386 | 0 | 0 |
| MN996532.1;QHR63300.2;BatCoV_RaTG13_spike;387 | 0 | 0 |
| MN996532.1;QHR63300.2;BatCoV_RaTG13_spike;388 | 0 | 0 |
| MN996532.1;QHR63300.2;BatCoV_RaTG13_spike;389 | 0 | 0 |

| | | |
|---|---|---|
| MN996532.1;QHR63300.2;BatCoV_RaTG13_spike;390 | 0 | 0 |
| MN996532.1;QHR63300.2;BatCoV_RaTG13_spike;391 | 0 | 0 |
| MN996532.1;QHR63300.2;BatCoV_RaTG13_spike;392 | 0 | 0.025 |
| MN996532.1;QHR63300.2;BatCoV_RaTG13_spike;393 | 0.05 | 0 |
| MN996532.1;QHR63300.2;BatCoV_RaTG13_spike;394 | 0 | 0 |
| MN996532.1;QHR63300.2;BatCoV_RaTG13_spike;395 | 0 | 0 |
| MN996532.1;QHR63300.2;BatCoV_RaTG13_spike;396 | 0 | 0 |
| MN996532.1;QHR63300.2;BatCoV_RaTG13_spike;397 | 0 | 0 |
| MN996532.1;QHR63300.2;BatCoV_RaTG13_spike;398 | 0.1 | 0.025 |
| MN996532.1;QHR63300.2;BatCoV_RaTG13_spike;399 | 0 | 0 |
| MN996532.1;QHR63300.2;BatCoV_RaTG13_spike;400 | 0 | 0 |
| MN996532.1;QHR63300.2;BatCoV_RaTG13_spike;401 | 0 | 0 |
| MN996532.1;QHR63300.2;BatCoV_RaTG13_spike;402 | 0.05 | 0.05 |
| MN996532.1;QHR63300.2;BatCoV_RaTG13_spike;403 | 0 | 0.05 |
| MN996532.1;QHR63300.2;BatCoV_RaTG13_spike;404 | 0 | 0.05 |
| MN996532.1;QHR63300.2;BatCoV_RaTG13_spike;405 | 0 | 0.05 |
| MN996532.1;QHR63300.2;BatCoV_RaTG13_spike;406 | 0 | 0.075 |
| MN996532.1;QHR63300.2;BatCoV_RaTG13_spike;407 | 0 | 0.075 |
| MN996532.1;QHR63300.2;BatCoV_RaTG13_spike;408 | 0.05 | 0.1 |
| MN996532.1;QHR63300.2;BatCoV_RaTG13_spike;409 | 0.15 | 0.125 |
| MN996532.1;QHR63300.2;BatCoV_RaTG13_spike;410 | 0 | 0.05 |
| MN996532.1;QHR63300.2;BatCoV_RaTG13_spike;411 | 0 | 0.05 |
| MN996532.1;QHR63300.2;BatCoV_RaTG13_spike;412 | 0.05 | 0.1 |
| MN996532.1;QHR63300.2;BatCoV_RaTG13_spike;413 | 0.15 | 0.075 |
| MN996532.1;QHR63300.2;BatCoV_RaTG13_spike;414 | 0.2 | 0.125 |
| MN996532.1;QHR63300.2;BatCoV_RaTG13_spike;415 | 0.1 | 0.125 |
| MN996532.1;QHR63300.2;BatCoV_RaTG13_spike;416 | 0.25 | 0.25 |
| MN996532.1;QHR63300.2;BatCoV_RaTG13_spike;417 | 0.1 | 0.05 |
| MN996532.1;QHR63300.2;BatCoV_RaTG13_spike;418 | 0 | 0.025 |
| MN996532.1;QHR63300.2;BatCoV_RaTG13_spike;419 | 0.05 | 0.025 |
| MN996532.1;QHR63300.2;BatCoV_RaTG13_spike;420 | 0 | 0.025 |
| MN996532.1;QHR63300.2;BatCoV_RaTG13_spike;421 | 0 | 0 |
| MN996532.1;QHR63300.2;BatCoV_RaTG13_spike;422 | 0 | 0 |
| MN996532.1;QHR63300.2;BatCoV_RaTG13_spike;423 | 0 | 0 |
| MN996532.1;QHR63300.2;BatCoV_RaTG13_spike;424 | 0 | 0 |
| MN996532.1;QHR63300.2;BatCoV_RaTG13_spike;425 | 0 | 0 |
| MN996532.1;QHR63300.2;BatCoV_RaTG13_spike;426 | 0 | 0 |
| MN996532.1;QHR63300.2;BatCoV_RaTG13_spike;427 | 0 | 0 |
| MN996532.1;QHR63300.2;BatCoV_RaTG13_spike;428 | 0 | 0 |
| MN996532.1;QHR63300.2;BatCoV_RaTG13_spike;429 | 0 | 0 |
| MN996532.1;QHR63300.2;BatCoV_RaTG13_spike;430 | 0 | 0 |
| MN996532.1;QHR63300.2;BatCoV_RaTG13_spike;431 | 0 | 0 |
| MN996532.1;QHR63300.2;BatCoV_RaTG13_spike;432 | 0.1 | 0.025 |

| | | |
|---|---|---|
| MN996532.1;QHR63300.2;BatCoV_RaTG13_spike;433 | 0.05 | 0.025 |
| MN996532.1;QHR63300.2;BatCoV_RaTG13_spike;434 | 0.05 | 0.075 |
| MN996532.1;QHR63300.2;BatCoV_RaTG13_spike;435 | 0.1 | 0.175 |
| MN996532.1;QHR63300.2;BatCoV_RaTG13_spike;436 | 0.1 | 0.225 |
| MN996532.1;QHR63300.2;BatCoV_RaTG13_spike;437 | 0.05 | 0.15 |
| MN996532.1;QHR63300.2;BatCoV_RaTG13_spike;438 | 0.05 | 0.025 |
| MN996532.1;QHR63300.2;BatCoV_RaTG13_spike;439 | 0.05 | 0 |
| MN996532.1;QHR63300.2;BatCoV_RaTG13_spike;440 | 0 | 0 |
| MN996532.1;QHR63300.2;BatCoV_RaTG13_spike;441 | 0 | 0 |
| MN996532.1;QHR63300.2;BatCoV_RaTG13_spike;442 | 0 | 0 |
| MN996532.1;QHR63300.2;BatCoV_RaTG13_spike;443 | 0 | 0 |
| MN996532.1;QHR63300.2;BatCoV_RaTG13_spike;444 | 0 | 0 |
| MN996532.1;QHR63300.2;BatCoV_RaTG13_spike;445 | 0.05 | 0 |
| MN996532.1;QHR63300.2;BatCoV_RaTG13_spike;446 | 0 | 0 |
| MN996532.1;QHR63300.2;BatCoV_RaTG13_spike;447 | 0 | 0 |
| MN996532.1;QHR63300.2;BatCoV_RaTG13_spike;448 | 0 | 0 |
| MN996532.1;QHR63300.2;BatCoV_RaTG13_spike;449 | 0 | 0 |
| MN996532.1;QHR63300.2;BatCoV_RaTG13_spike;450 | 0 | 0 |
| MN996532.1;QHR63300.2;BatCoV_RaTG13_spike;451 | 0 | 0 |
| MN996532.1;QHR63300.2;BatCoV_RaTG13_spike;452 | 0 | 0 |
| MN996532.1;QHR63300.2;BatCoV_RaTG13_spike;453 | 0 | 0 |
| MN996532.1;QHR63300.2;BatCoV_RaTG13_spike;454 | 0.05 | 0.025 |
| MN996532.1;QHR63300.2;BatCoV_RaTG13_spike;455 | 0.1 | 0 |
| MN996532.1;QHR63300.2;BatCoV_RaTG13_spike;456 | 0.1 | 0.025 |
| MN996532.1;QHR63300.2;BatCoV_RaTG13_spike;457 | 0.1 | 0 |
| MN996532.1;QHR63300.2;BatCoV_RaTG13_spike;458 | 0.05 | 0 |
| MN996532.1;QHR63300.2;BatCoV_RaTG13_spike;459 | 0 | 0 |
| MN996532.1;QHR63300.2;BatCoV_RaTG13_spike;460 | 0 | 0 |
| MN996532.1;QHR63300.2;BatCoV_RaTG13_spike;461 | 0 | 0 |
| MN996532.1;QHR63300.2;BatCoV_RaTG13_spike;462 | 0.05 | 0 |
| MN996532.1;QHR63300.2;BatCoV_RaTG13_spike;463 | 0 | 0 |
| MN996532.1;QHR63300.2;BatCoV_RaTG13_spike;464 | 0 | 0 |
| MN996532.1;QHR63300.2;BatCoV_RaTG13_spike;465 | 0.05 | 0 |
| MN996532.1;QHR63300.2;BatCoV_RaTG13_spike;466 | 0 | 0 |
| MN996532.1;QHR63300.2;BatCoV_RaTG13_spike;467 | 0 | 0 |
| MN996532.1;QHR63300.2;BatCoV_RaTG13_spike;468 | 0 | 0 |
| MN996532.1;QHR63300.2;BatCoV_RaTG13_spike;469 | 0 | 0 |
| MN996532.1;QHR63300.2;BatCoV_RaTG13_spike;470 | 0 | 0 |
| MN996532.1;QHR63300.2;BatCoV_RaTG13_spike;471 | 0 | 0 |
| MN996532.1;QHR63300.2;BatCoV_RaTG13_spike;472 | 0 | 0 |
| MN996532.1;QHR63300.2;BatCoV_RaTG13_spike;473 | 0 | 0 |
| MN996532.1;QHR63300.2;BatCoV_RaTG13_spike;474 | 0 | 0 |
| MN996532.1;QHR63300.2;BatCoV_RaTG13_spike;475 | 0 | 0 |

| | | |
|---|---|---|
| MN996532.1;QHR63300.2;BatCoV_RaTG13_spike;476 | 0 | 0 |
| MN996532.1;QHR63300.2;BatCoV_RaTG13_spike;477 | 0 | 0 |
| MN996532.1;QHR63300.2;BatCoV_RaTG13_spike;478 | 0 | 0 |
| MN996532.1;QHR63300.2;BatCoV_RaTG13_spike;479 | 0 | 0 |
| MN996532.1;QHR63300.2;BatCoV_RaTG13_spike;480 | 0 | 0 |
| MN996532.1;QHR63300.2;BatCoV_RaTG13_spike;481 | 0 | 0 |
| MN996532.1;QHR63300.2;BatCoV_RaTG13_spike;482 | 0 | 0 |
| MN996532.1;QHR63300.2;BatCoV_RaTG13_spike;483 | 0 | 0 |
| MN996532.1;QHR63300.2;BatCoV_RaTG13_spike;484 | 0 | 0 |
| MN996532.1;QHR63300.2;BatCoV_RaTG13_spike;485 | 0 | 0 |
| MN996532.1;QHR63300.2;BatCoV_RaTG13_spike;486 | 0 | 0 |
| MN996532.1;QHR63300.2;BatCoV_RaTG13_spike;487 | 0 | 0 |
| MN996532.1;QHR63300.2;BatCoV_RaTG13_spike;488 | 0 | 0 |
| MN996532.1;QHR63300.2;BatCoV_RaTG13_spike;489 | 0 | 0 |
| MN996532.1;QHR63300.2;BatCoV_RaTG13_spike;490 | 0 | 0 |
| MN996532.1;QHR63300.2;BatCoV_RaTG13_spike;491 | 0 | 0 |
| MN996532.1;QHR63300.2;BatCoV_RaTG13_spike;492 | 0 | 0 |
| MN996532.1;QHR63300.2;BatCoV_RaTG13_spike;493 | 0 | 0 |
| MN996532.1;QHR63300.2;BatCoV_RaTG13_spike;494 | 0 | 0 |
| MN996532.1;QHR63300.2;BatCoV_RaTG13_spike;495 | 0 | 0 |
| MN996532.1;QHR63300.2;BatCoV_RaTG13_spike;496 | 0 | 0 |
| MN996532.1;QHR63300.2;BatCoV_RaTG13_spike;497 | 0 | 0 |
| MN996532.1;QHR63300.2;BatCoV_RaTG13_spike;498 | 0 | 0 |
| MN996532.1;QHR63300.2;BatCoV_RaTG13_spike;499 | 0 | 0 |
| MN996532.1;QHR63300.2;BatCoV_RaTG13_spike;500 | 0 | 0 |
| MN996532.1;QHR63300.2;BatCoV_RaTG13_spike;501 | 0 | 0 |
| MN996532.1;QHR63300.2;BatCoV_RaTG13_spike;502 | 0 | 0 |
| MN996532.1;QHR63300.2;BatCoV_RaTG13_spike;503 | 0 | 0 |
| MN996532.1;QHR63300.2;BatCoV_RaTG13_spike;504 | 0 | 0 |
| MN996532.1;QHR63300.2;BatCoV_RaTG13_spike;505 | 0 | 0 |
| MN996532.1;QHR63300.2;BatCoV_RaTG13_spike;506 | 0 | 0 |
| MN996532.1;QHR63300.2;BatCoV_RaTG13_spike;507 | 0 | 0 |
| MN996532.1;QHR63300.2;BatCoV_RaTG13_spike;508 | 0 | 0 |
| MN996532.1;QHR63300.2;BatCoV_RaTG13_spike;509 | 0 | 0 |
| MN996532.1;QHR63300.2;BatCoV_RaTG13_spike;510 | 0 | 0 |
| MN996532.1;QHR63300.2;BatCoV_RaTG13_spike;511 | 0 | 0 |
| MN996532.1;QHR63300.2;BatCoV_RaTG13_spike;512 | 0 | 0 |
| MN996532.1;QHR63300.2;BatCoV_RaTG13_spike;513 | 0 | 0 |
| MN996532.1;QHR63300.2;BatCoV_RaTG13_spike;514 | 0 | 0 |
| MN996532.1;QHR63300.2;BatCoV_RaTG13_spike;515 | 0 | 0 |
| MN996532.1;QHR63300.2;BatCoV_RaTG13_spike;516 | 0 | 0 |
| MN996532.1;QHR63300.2;BatCoV_RaTG13_spike;517 | 0 | 0 |
| MN996532.1;QHR63300.2;BatCoV_RaTG13_spike;518 | 0 | 0 |

| | | |
|---|---|---|
| MN996532.1;QHR63300.2;BatCoV_RaTG13_spike;519 | 0 | 0 |
| MN996532.1;QHR63300.2;BatCoV_RaTG13_spike;520 | 0 | 0.025 |
| MN996532.1;QHR63300.2;BatCoV_RaTG13_spike;521 | 0 | 0.025 |
| MN996532.1;QHR63300.2;BatCoV_RaTG13_spike;522 | 0 | 0.025 |
| MN996532.1;QHR63300.2;BatCoV_RaTG13_spike;523 | 0 | 0.025 |
| MN996532.1;QHR63300.2;BatCoV_RaTG13_spike;524 | 0 | 0.025 |
| MN996532.1;QHR63300.2;BatCoV_RaTG13_spike;525 | 0 | 0.025 |
| MN996532.1;QHR63300.2;BatCoV_RaTG13_spike;526 | 0 | 0 |
| MN996532.1;QHR63300.2;BatCoV_RaTG13_spike;527 | 0 | 0.025 |
| MN996532.1;QHR63300.2;BatCoV_RaTG13_spike;528 | 0 | 0.075 |
| MN996532.1;QHR63300.2;BatCoV_RaTG13_spike;529 | 0 | 0.05 |
| MN996532.1;QHR63300.2;BatCoV_RaTG13_spike;530 | 0 | 0.075 |
| MN996532.1;QHR63300.2;BatCoV_RaTG13_spike;531 | 0 | 0 |
| MN996532.1;QHR63300.2;BatCoV_RaTG13_spike;532 | 0 | 0.025 |
| MN996532.1;QHR63300.2;BatCoV_RaTG13_spike;533 | 0 | 0.025 |
| MN996532.1;QHR63300.2;BatCoV_RaTG13_spike;534 | 0 | 0.05 |
| MN996532.1;QHR63300.2;BatCoV_RaTG13_spike;535 | 0 | 0.05 |
| MN996532.1;QHR63300.2;BatCoV_RaTG13_spike;536 | 0 | 0.025 |
| MN996532.1;QHR63300.2;BatCoV_RaTG13_spike;537 | 0 | 0.025 |
| MN996532.1;QHR63300.2;BatCoV_RaTG13_spike;538 | 0 | 0 |
| MN996532.1;QHR63300.2;BatCoV_RaTG13_spike;539 | 0 | 0 |
| MN996532.1;QHR63300.2;BatCoV_RaTG13_spike;540 | 0 | 0 |
| MN996532.1;QHR63300.2;BatCoV_RaTG13_spike;541 | 0 | 0.025 |
| MN996532.1;QHR63300.2;BatCoV_RaTG13_spike;542 | 0.05 | 0.125 |
| MN996532.1;QHR63300.2;BatCoV_RaTG13_spike;543 | 0 | 0.125 |
| MN996532.1;QHR63300.2;BatCoV_RaTG13_spike;544 | 0 | 0.125 |
| MN996532.1;QHR63300.2;BatCoV_RaTG13_spike;545 | 0 | 0.1 |
| MN996532.1;QHR63300.2;BatCoV_RaTG13_spike;546 | 0.05 | 0.125 |
| MN996532.1;QHR63300.2;BatCoV_RaTG13_spike;547 | 0 | 0.1 |
| MN996532.1;QHR63300.2;BatCoV_RaTG13_spike;548 | 0.05 | 0.125 |
| MN996532.1;QHR63300.2;BatCoV_RaTG13_spike;549 | 0 | 0.5 |
| MN996532.1;QHR63300.2;BatCoV_RaTG13_spike;550 | 0.25 | 0.475 |
| MN996532.1;QHR63300.2;BatCoV_RaTG13_spike;551 | 0.15 | 0.5 |
| MN996532.1;QHR63300.2;BatCoV_RaTG13_spike;552 | 0.5 | 0.75 |
| MN996532.1;QHR63300.2;BatCoV_RaTG13_spike;553 | 0.05 | 0.7 |
| MN996532.1;QHR63300.2;BatCoV_RaTG13_spike;554 | 0.1 | 0.775 |
| MN996532.1;QHR63300.2;BatCoV_RaTG13_spike;555 | 0.05 | 0.6 |
| MN996532.1;QHR63300.2;BatCoV_RaTG13_spike;556 | 0 | 0.7 |
| MN996532.1;QHR63300.2;BatCoV_RaTG13_spike;557 | 0 | 0.75 |
| MN996532.1;QHR63300.2;BatCoV_RaTG13_spike;558 | 0 | 0.725 |
| MN996532.1;QHR63300.2;BatCoV_RaTG13_spike;559 | 0 | 0.525 |
| MN996532.1;QHR63300.2;BatCoV_RaTG13_spike;560 | 0 | 0.525 |
| MN996532.1;QHR63300.2;BatCoV_RaTG13_spike;561 | 0 | 0.275 |

| | | |
|---|---|---|
| MN996532.1;QHR63300.2;BatCoV_RaTG13_spike;562 | 0 | 0.25 |
| MN996532.1;QHR63300.2;BatCoV_RaTG13_spike;563 | 0 | 0.025 |
| MN996532.1;QHR63300.2;BatCoV_RaTG13_spike;564 | 0.05 | 0.05 |
| MN996532.1;QHR63300.2;BatCoV_RaTG13_spike;565 | 0.05 | 0.075 |
| MN996532.1;QHR63300.2;BatCoV_RaTG13_spike;566 | 0 | 0.025 |
| MN996532.1;QHR63300.2;BatCoV_RaTG13_spike;567 | 0 | 0.05 |
| MN996532.1;QHR63300.2;BatCoV_RaTG13_spike;568 | 0 | 0.375 |
| MN996532.1;QHR63300.2;BatCoV_RaTG13_spike;569 | 0.05 | 0.6 |
| MN996532.1;QHR63300.2;BatCoV_RaTG13_spike;570 | 0 | 0.4 |
| MN996532.1;QHR63300.2;BatCoV_RaTG13_spike;571 | 0 | 0.675 |
| MN996532.1;QHR63300.2;BatCoV_RaTG13_spike;572 | 0 | 0.625 |
| MN996532.1;QHR63300.2;BatCoV_RaTG13_spike;573 | 0 | 0.475 |
| MN996532.1;QHR63300.2;BatCoV_RaTG13_spike;574 | 0 | 0.55 |
| MN996532.1;QHR63300.2;BatCoV_RaTG13_spike;575 | 0 | 0.625 |
| MN996532.1;QHR63300.2;BatCoV_RaTG13_spike;576 | 0 | 0.7 |
| MN996532.1;QHR63300.2;BatCoV_RaTG13_spike;577 | 0 | 0.325 |
| MN996532.1;QHR63300.2;BatCoV_RaTG13_spike;578 | 0 | 0 |
| MN996532.1;QHR63300.2;BatCoV_RaTG13_spike;579 | 0 | 0 |
| MN996532.1;QHR63300.2;BatCoV_RaTG13_spike;580 | 0 | 0 |
| MN996532.1;QHR63300.2;BatCoV_RaTG13_spike;581 | 0 | 0 |
| MN996532.1;QHR63300.2;BatCoV_RaTG13_spike;582 | 0 | 0.05 |
| MN996532.1;QHR63300.2;BatCoV_RaTG13_spike;583 | 0 | 0 |
| MN996532.1;QHR63300.2;BatCoV_RaTG13_spike;584 | 0 | 0 |
| MN996532.1;QHR63300.2;BatCoV_RaTG13_spike;585 | 0 | 0 |
| MN996532.1;QHR63300.2;BatCoV_RaTG13_spike;586 | 0 | 0 |
| MN996532.1;QHR63300.2;BatCoV_RaTG13_spike;587 | 0 | 0 |
| MN996532.1;QHR63300.2;BatCoV_RaTG13_spike;588 | 0 | 0 |
| MN996532.1;QHR63300.2;BatCoV_RaTG13_spike;589 | 0 | 0.05 |
| MN996532.1;QHR63300.2;BatCoV_RaTG13_spike;590 | 0.15 | 0.1 |
| MN996532.1;QHR63300.2;BatCoV_RaTG13_spike;591 | 0.1 | 0.075 |
| MN996532.1;QHR63300.2;BatCoV_RaTG13_spike;592 | 0.1 | 0.1 |
| MN996532.1;QHR63300.2;BatCoV_RaTG13_spike;593 | 0.15 | 0.1 |
| MN996532.1;QHR63300.2;BatCoV_RaTG13_spike;594 | 0.1 | 0.125 |
| MN996532.1;QHR63300.2;BatCoV_RaTG13_spike;595 | 0.15 | 0.1 |
| MN996532.1;QHR63300.2;BatCoV_RaTG13_spike;596 | 0.05 | 0.1 |
| MN996532.1;QHR63300.2;BatCoV_RaTG13_spike;597 | 0 | 0.05 |
| MN996532.1;QHR63300.2;BatCoV_RaTG13_spike;598 | 0 | 0.1 |
| MN996532.1;QHR63300.2;BatCoV_RaTG13_spike;599 | 0 | 0.075 |
| MN996532.1;QHR63300.2;BatCoV_RaTG13_spike;600 | 0 | 0.05 |
| MN996532.1;QHR63300.2;BatCoV_RaTG13_spike;601 | 0 | 0.05 |
| MN996532.1;QHR63300.2;BatCoV_RaTG13_spike;602 | 0 | 0.05 |
| MN996532.1;QHR63300.2;BatCoV_RaTG13_spike;603 | 0 | 0.025 |
| MN996532.1;QHR63300.2;BatCoV_RaTG13_spike;604 | 0 | 0.025 |

| | | |
|---|---|---|
| MN996532.1;QHR63300.2;BatCoV_RaTG13_spike;605 | 0 | 0.025 |
| MN996532.1;QHR63300.2;BatCoV_RaTG13_spike;606 | 0 | 0.025 |
| MN996532.1;QHR63300.2;BatCoV_RaTG13_spike;607 | 0 | 0.025 |
| MN996532.1;QHR63300.2;BatCoV_RaTG13_spike;608 | 0 | 0 |
| MN996532.1;QHR63300.2;BatCoV_RaTG13_spike;609 | 0 | 0 |
| MN996532.1;QHR63300.2;BatCoV_RaTG13_spike;610 | 0 | 0 |
| MN996532.1;QHR63300.2;BatCoV_RaTG13_spike;611 | 0 | 0 |
| MN996532.1;QHR63300.2;BatCoV_RaTG13_spike;612 | 0 | 0.075 |
| MN996532.1;QHR63300.2;BatCoV_RaTG13_spike;613 | 0 | 0.075 |
| MN996532.1;QHR63300.2;BatCoV_RaTG13_spike;614 | 0 | 0.1 |
| MN996532.1;QHR63300.2;BatCoV_RaTG13_spike;615 | 0 | 0.125 |
| MN996532.1;QHR63300.2;BatCoV_RaTG13_spike;616 | 0 | 0.175 |
| MN996532.1;QHR63300.2;BatCoV_RaTG13_spike;617 | 0 | 0.225 |
| MN996532.1;QHR63300.2;BatCoV_RaTG13_spike;618 | 0 | 0.125 |
| MN996532.1;QHR63300.2;BatCoV_RaTG13_spike;619 | 0 | 0.075 |
| MN996532.1;QHR63300.2;BatCoV_RaTG13_spike;620 | 0 | 0.05 |
| MN996532.1;QHR63300.2;BatCoV_RaTG13_spike;621 | 0 | 0.05 |
| MN996532.1;QHR63300.2;BatCoV_RaTG13_spike;622 | 0 | 0.075 |
| MN996532.1;QHR63300.2;BatCoV_RaTG13_spike;623 | 0.05 | 0.05 |
| MN996532.1;QHR63300.2;BatCoV_RaTG13_spike;624 | 0 | 0.025 |
| MN996532.1;QHR63300.2;BatCoV_RaTG13_spike;625 | 0 | 0.05 |
| MN996532.1;QHR63300.2;BatCoV_RaTG13_spike;626 | 0 | 0.075 |
| MN996532.1;QHR63300.2;BatCoV_RaTG13_spike;627 | 0 | 0.175 |
| MN996532.1;QHR63300.2;BatCoV_RaTG13_spike;628 | 0 | 0.05 |
| MN996532.1;QHR63300.2;BatCoV_RaTG13_spike;629 | 0 | 0.05 |
| MN996532.1;QHR63300.2;BatCoV_RaTG13_spike;630 | 0 | 0.05 |
| MN996532.1;QHR63300.2;BatCoV_RaTG13_spike;631 | 0 | 0 |
| MN996532.1;QHR63300.2;BatCoV_RaTG13_spike;632 | 0 | 0 |
| MN996532.1;QHR63300.2;BatCoV_RaTG13_spike;633 | 0 | 0.025 |
| MN996532.1;QHR63300.2;BatCoV_RaTG13_spike;634 | 0.05 | 0.025 |
| MN996532.1;QHR63300.2;BatCoV_RaTG13_spike;635 | 0 | 0.025 |
| MN996532.1;QHR63300.2;BatCoV_RaTG13_spike;636 | 0 | 0 |
| MN996532.1;QHR63300.2;BatCoV_RaTG13_spike;637 | 0 | 0 |
| MN996532.1;QHR63300.2;BatCoV_RaTG13_spike;638 | 0 | 0 |
| MN996532.1;QHR63300.2;BatCoV_RaTG13_spike;639 | 0 | 0 |
| MN996532.1;QHR63300.2;BatCoV_RaTG13_spike;640 | 0 | 0 |
| MN996532.1;QHR63300.2;BatCoV_RaTG13_spike;641 | 0 | 0 |
| MN996532.1;QHR63300.2;BatCoV_RaTG13_spike;642 | 0 | 0 |
| MN996532.1;QHR63300.2;BatCoV_RaTG13_spike;643 | 0 | 0.025 |
| MN996532.1;QHR63300.2;BatCoV_RaTG13_spike;644 | 0 | 0.025 |
| MN996532.1;QHR63300.2;BatCoV_RaTG13_spike;645 | 0 | 0.025 |
| MN996532.1;QHR63300.2;BatCoV_RaTG13_spike;646 | 0 | 0.025 |
| MN996532.1;QHR63300.2;BatCoV_RaTG13_spike;647 | 0 | 0.025 |

| | | |
|---|---|---|
| MN996532.1;QHR63300.2;BatCoV_RaTG13_spike;648 | 0 | 0.025 |
| MN996532.1;QHR63300.2;BatCoV_RaTG13_spike;649 | 0 | 0 |
| MN996532.1;QHR63300.2;BatCoV_RaTG13_spike;650 | 0 | 0.025 |
| MN996532.1;QHR63300.2;BatCoV_RaTG13_spike;651 | 0 | 0.025 |
| MN996532.1;QHR63300.2;BatCoV_RaTG13_spike;652 | 0 | 0.025 |
| MN996532.1;QHR63300.2;BatCoV_RaTG13_spike;653 | 0 | 0.075 |
| MN996532.1;QHR63300.2;BatCoV_RaTG13_spike;654 | 0 | 0.025 |
| MN996532.1;QHR63300.2;BatCoV_RaTG13_spike;655 | 0.05 | 0.05 |
| MN996532.1;QHR63300.2;BatCoV_RaTG13_spike;656 | 0.05 | 0.2 |
| MN996532.1;QHR63300.2;BatCoV_RaTG13_spike;657 | 0.05 | 0.2 |
| MN996532.1;QHR63300.2;BatCoV_RaTG13_spike;658 | 0.1 | 0.2 |
| MN996532.1;QHR63300.2;BatCoV_RaTG13_spike;659 | 0.05 | 0 |
| MN996532.1;QHR63300.2;BatCoV_RaTG13_spike;660 | 0.05 | 0 |
| MN996532.1;QHR63300.2;BatCoV_RaTG13_spike;661 | 0 | 0.025 |
| MN996532.1;QHR63300.2;BatCoV_RaTG13_spike;662 | 0 | 0.025 |
| MN996532.1;QHR63300.2;BatCoV_RaTG13_spike;663 | 0 | 0.025 |
| MN996532.1;QHR63300.2;BatCoV_RaTG13_spike;664 | 0 | 0.025 |
| MN996532.1;QHR63300.2;BatCoV_RaTG13_spike;665 | 0 | 0.025 |
| MN996532.1;QHR63300.2;BatCoV_RaTG13_spike;666 | 0 | 0.025 |
| MN996532.1;QHR63300.2;BatCoV_RaTG13_spike;667 | 0 | 0.025 |
| MN996532.1;QHR63300.2;BatCoV_RaTG13_spike;668 | 0 | 0.025 |
| MN996532.1;QHR63300.2;BatCoV_RaTG13_spike;669 | 0.05 | 0.05 |
| MN996532.1;QHR63300.2;BatCoV_RaTG13_spike;670 | 0 | 0.025 |
| MN996532.1;QHR63300.2;BatCoV_RaTG13_spike;671 | 0 | 0.025 |
| MN996532.1;QHR63300.2;BatCoV_RaTG13_spike;672 | 0.05 | 0.025 |
| MN996532.1;QHR63300.2;BatCoV_RaTG13_spike;673 | 0.1 | 0.025 |
| MN996532.1;QHR63300.2;BatCoV_RaTG13_spike;674 | 0.05 | 0.025 |
| MN996532.1;QHR63300.2;BatCoV_RaTG13_spike;675 | 0.05 | 0 |
| MN996532.1;QHR63300.2;BatCoV_RaTG13_spike;676 | 0.05 | 0 |
| MN996532.1;QHR63300.2;BatCoV_RaTG13_spike;677 | 0.05 | 0 |
| MN996532.1;QHR63300.2;BatCoV_RaTG13_spike;678 | 0.05 | 0 |
| MN996532.1;QHR63300.2;BatCoV_RaTG13_spike;679 | 0 | 0 |
| MN996532.1;QHR63300.2;BatCoV_RaTG13_spike;680 | 0 | 0.025 |
| MN996532.1;QHR63300.2;BatCoV_RaTG13_spike;681 | 0 | 0.025 |
| MN996532.1;QHR63300.2;BatCoV_RaTG13_spike;682 | 0 | 0.075 |
| MN996532.1;QHR63300.2;BatCoV_RaTG13_spike;683 | 0 | 0.125 |
| MN996532.1;QHR63300.2;BatCoV_RaTG13_spike;684 | 0 | 0.125 |
| MN996532.1;QHR63300.2;BatCoV_RaTG13_spike;685 | 0 | 0.125 |
| MN996532.1;QHR63300.2;BatCoV_RaTG13_spike;686 | 0 | 0.125 |
| MN996532.1;QHR63300.2;BatCoV_RaTG13_spike;687 | 0.1 | 0.175 |
| MN996532.1;QHR63300.2;BatCoV_RaTG13_spike;688 | 0 | 0.125 |
| MN996532.1;QHR63300.2;BatCoV_RaTG13_spike;689 | 0 | 0.125 |
| MN996532.1;QHR63300.2;BatCoV_RaTG13_spike;690 | 0 | 0.1 |

| | | |
|---|---|---|
| MN996532.1;QHR63300.2;BatCoV_RaTG13_spike;691 | 0 | 0.075 |
| MN996532.1;QHR63300.2;BatCoV_RaTG13_spike;692 | 0.05 | 0.025 |
| MN996532.1;QHR63300.2;BatCoV_RaTG13_spike;693 | 0 | 0.05 |
| MN996532.1;QHR63300.2;BatCoV_RaTG13_spike;694 | 0.05 | 0.1 |
| MN996532.1;QHR63300.2;BatCoV_RaTG13_spike;695 | 0.1 | 0.05 |
| MN996532.1;QHR63300.2;BatCoV_RaTG13_spike;696 | 0.1 | 0.1 |
| MN996532.1;QHR63300.2;BatCoV_RaTG13_spike;697 | 0.15 | 0.125 |
| MN996532.1;QHR63300.2;BatCoV_RaTG13_spike;698 | 0.15 | 0.05 |
| MN996532.1;QHR63300.2;BatCoV_RaTG13_spike;699 | 0.05 | 0.025 |
| MN996532.1;QHR63300.2;BatCoV_RaTG13_spike;700 | 0.2 | 0 |
| MN996532.1;QHR63300.2;BatCoV_RaTG13_spike;701 | 0.1 | 0.025 |
| MN996532.1;QHR63300.2;BatCoV_RaTG13_spike;702 | 0.2 | 0.125 |
| MN996532.1;QHR63300.2;BatCoV_RaTG13_spike;703 | 0.1 | 0 |
| MN996532.1;QHR63300.2;BatCoV_RaTG13_spike;704 | 0.05 | 0 |
| MN996532.1;QHR63300.2;BatCoV_RaTG13_spike;705 | 0 | 0 |
| MN996532.1;QHR63300.2;BatCoV_RaTG13_spike;706 | 0 | 0 |
| MN996532.1;QHR63300.2;BatCoV_RaTG13_spike;707 | 0 | 0.05 |
| MN996532.1;QHR63300.2;BatCoV_RaTG13_spike;708 | 0 | 0 |
| MN996532.1;QHR63300.2;BatCoV_RaTG13_spike;709 | 0 | 0.025 |
| MN996532.1;QHR63300.2;BatCoV_RaTG13_spike;710 | 0 | 0 |
| MN996532.1;QHR63300.2;BatCoV_RaTG13_spike;711 | 0 | 0 |
| MN996532.1;QHR63300.2;BatCoV_RaTG13_spike;712 | 0 | 0 |
| MN996532.1;QHR63300.2;BatCoV_RaTG13_spike;713 | 0 | 0 |
| MN996532.1;QHR63300.2;BatCoV_RaTG13_spike;714 | 0 | 0 |
| MN996532.1;QHR63300.2;BatCoV_RaTG13_spike;715 | 0 | 0 |
| MN996532.1;QHR63300.2;BatCoV_RaTG13_spike;716 | 0 | 0 |
| MN996532.1;QHR63300.2;BatCoV_RaTG13_spike;717 | 0 | 0 |
| MN996532.1;QHR63300.2;BatCoV_RaTG13_spike;718 | 0 | 0 |
| MN996532.1;QHR63300.2;BatCoV_RaTG13_spike;719 | 0 | 0 |
| MN996532.1;QHR63300.2;BatCoV_RaTG13_spike;720 | 0 | 0 |
| MN996532.1;QHR63300.2;BatCoV_RaTG13_spike;721 | 0 | 0 |
| MN996532.1;QHR63300.2;BatCoV_RaTG13_spike;722 | 0 | 0 |
| MN996532.1;QHR63300.2;BatCoV_RaTG13_spike;723 | 0 | 0 |
| MN996532.1;QHR63300.2;BatCoV_RaTG13_spike;724 | 0 | 0 |
| MN996532.1;QHR63300.2;BatCoV_RaTG13_spike;725 | 0 | 0 |
| MN996532.1;QHR63300.2;BatCoV_RaTG13_spike;726 | 0 | 0 |
| MN996532.1;QHR63300.2;BatCoV_RaTG13_spike;727 | 0 | 0 |
| MN996532.1;QHR63300.2;BatCoV_RaTG13_spike;728 | 0 | 0 |
| MN996532.1;QHR63300.2;BatCoV_RaTG13_spike;729 | 0 | 0 |
| MN996532.1;QHR63300.2;BatCoV_RaTG13_spike;730 | 0 | 0 |
| MN996532.1;QHR63300.2;BatCoV_RaTG13_spike;731 | 0 | 0 |
| MN996532.1;QHR63300.2;BatCoV_RaTG13_spike;732 | 0.05 | 0 |
| MN996532.1;QHR63300.2;BatCoV_RaTG13_spike;733 | 0.05 | 0 |

| | | |
|---|---|---|
| MN996532.1;QHR63300.2;BatCoV_RaTG13_spike;734 | 0.05 | 0 |
| MN996532.1;QHR63300.2;BatCoV_RaTG13_spike;735 | 0 | 0 |
| MN996532.1;QHR63300.2;BatCoV_RaTG13_spike;736 | 0.05 | 0 |
| MN996532.1;QHR63300.2;BatCoV_RaTG13_spike;737 | 0.05 | 0 |
| MN996532.1;QHR63300.2;BatCoV_RaTG13_spike;738 | 0.05 | 0 |
| MN996532.1;QHR63300.2;BatCoV_RaTG13_spike;739 | 0.05 | 0 |
| MN996532.1;QHR63300.2;BatCoV_RaTG13_spike;740 | 0 | 0 |
| MN996532.1;QHR63300.2;BatCoV_RaTG13_spike;741 | 0 | 0 |
| MN996532.1;QHR63300.2;BatCoV_RaTG13_spike;742 | 0 | 0 |
| MN996532.1;QHR63300.2;BatCoV_RaTG13_spike;743 | 0 | 0 |
| MN996532.1;QHR63300.2;BatCoV_RaTG13_spike;744 | 0 | 0 |
| MN996532.1;QHR63300.2;BatCoV_RaTG13_spike;745 | 0 | 0 |
| MN996532.1;QHR63300.2;BatCoV_RaTG13_spike;746 | 0 | 0 |
| MN996532.1;QHR63300.2;BatCoV_RaTG13_spike;747 | 0 | 0 |
| MN996532.1;QHR63300.2;BatCoV_RaTG13_spike;748 | 0 | 0 |
| MN996532.1;QHR63300.2;BatCoV_RaTG13_spike;749 | 0 | 0 |
| MN996532.1;QHR63300.2;BatCoV_RaTG13_spike;750 | 0 | 0 |
| MN996532.1;QHR63300.2;BatCoV_RaTG13_spike;751 | 0 | 0 |
| MN996532.1;QHR63300.2;BatCoV_RaTG13_spike;752 | 0 | 0 |
| MN996532.1;QHR63300.2;BatCoV_RaTG13_spike;753 | 0 | 0 |
| MN996532.1;QHR63300.2;BatCoV_RaTG13_spike;754 | 0 | 0 |
| MN996532.1;QHR63300.2;BatCoV_RaTG13_spike;755 | 0 | 0 |
| MN996532.1;QHR63300.2;BatCoV_RaTG13_spike;756 | 0 | 0 |
| MN996532.1;QHR63300.2;BatCoV_RaTG13_spike;757 | 0.1 | 0.275 |
| MN996532.1;QHR63300.2;BatCoV_RaTG13_spike;758 | 0.1 | 0.35 |
| MN996532.1;QHR63300.2;BatCoV_RaTG13_spike;759 | 0.15 | 0.35 |
| MN996532.1;QHR63300.2;BatCoV_RaTG13_spike;760 | 0.15 | 0.375 |
| MN996532.1;QHR63300.2;BatCoV_RaTG13_spike;761 | 0.15 | 0.4 |
| MN996532.1;QHR63300.2;BatCoV_RaTG13_spike;762 | 0.2 | 0.4 |
| MN996532.1;QHR63300.2;BatCoV_RaTG13_spike;763 | 0.15 | 0.425 |
| MN996532.1;QHR63300.2;BatCoV_RaTG13_spike;764 | 0.15 | 0.425 |
| MN996532.1;QHR63300.2;BatCoV_RaTG13_spike;765 | 0.15 | 0.4 |
| MN996532.1;QHR63300.2;BatCoV_RaTG13_spike;766 | 0 | 0.275 |
| MN996532.1;QHR63300.2;BatCoV_RaTG13_spike;767 | 0.1 | 0.225 |
| MN996532.1;QHR63300.2;BatCoV_RaTG13_spike;768 | 0.1 | 0.1 |
| MN996532.1;QHR63300.2;BatCoV_RaTG13_spike;769 | 0.1 | 0.15 |
| MN996532.1;QHR63300.2;BatCoV_RaTG13_spike;770 | 0.1 | 0.1 |
| MN996532.1;QHR63300.2;BatCoV_RaTG13_spike;771 | 0 | 0.1 |
| MN996532.1;QHR63300.2;BatCoV_RaTG13_spike;772 | 0 | 0.05 |
| MN996532.1;QHR63300.2;BatCoV_RaTG13_spike;773 | 0.05 | 0.05 |
| MN996532.1;QHR63300.2;BatCoV_RaTG13_spike;774 | 0.05 | 0.05 |
| MN996532.1;QHR63300.2;BatCoV_RaTG13_spike;775 | 0.05 | 0.075 |
| MN996532.1;QHR63300.2;BatCoV_RaTG13_spike;776 | 0 | 0 |

| | | |
|---|---|---|
| MN996532.1;QHR63300.2;BatCoV_RaTG13_spike;777 | 0 | 0 |
| MN996532.1;QHR63300.2;BatCoV_RaTG13_spike;778 | 0 | 0 |
| MN996532.1;QHR63300.2;BatCoV_RaTG13_spike;779 | 0 | 0 |
| MN996532.1;QHR63300.2;BatCoV_RaTG13_spike;780 | 0 | 0 |
| MN996532.1;QHR63300.2;BatCoV_RaTG13_spike;781 | 0 | 0.025 |
| MN996532.1;QHR63300.2;BatCoV_RaTG13_spike;782 | 0 | 0.025 |
| MN996532.1;QHR63300.2;BatCoV_RaTG13_spike;783 | 0 | 0.35 |
| MN996532.1;QHR63300.2;BatCoV_RaTG13_spike;784 | 0 | 0.475 |
| MN996532.1;QHR63300.2;BatCoV_RaTG13_spike;785 | 0 | 0.475 |
| MN996532.1;QHR63300.2;BatCoV_RaTG13_spike;786 | 0 | 0.425 |
| MN996532.1;QHR63300.2;BatCoV_RaTG13_spike;787 | 0 | 0.35 |
| MN996532.1;QHR63300.2;BatCoV_RaTG13_spike;788 | 0 | 0.45 |
| MN996532.1;QHR63300.2;BatCoV_RaTG13_spike;789 | 0 | 0.5 |
| MN996532.1;QHR63300.2;BatCoV_RaTG13_spike;790 | 0.05 | 0.55 |
| MN996532.1;QHR63300.2;BatCoV_RaTG13_spike;791 | 0 | 0.575 |
| MN996532.1;QHR63300.2;BatCoV_RaTG13_spike;792 | 0 | 0.2 |
| MN996532.1;QHR63300.2;BatCoV_RaTG13_spike;793 | 0.05 | 0.15 |
| MN996532.1;QHR63300.2;BatCoV_RaTG13_spike;794 | 0 | 0.175 |
| MN996532.1;QHR63300.2;BatCoV_RaTG13_spike;795 | 0 | 0.125 |
| MN996532.1;QHR63300.2;BatCoV_RaTG13_spike;796 | 0 | 0.05 |
| MN996532.1;QHR63300.2;BatCoV_RaTG13_spike;797 | 0 | 0.075 |
| MN996532.1;QHR63300.2;BatCoV_RaTG13_spike;798 | 0 | 0.075 |
| MN996532.1;QHR63300.2;BatCoV_RaTG13_spike;799 | 0 | 0.075 |
| MN996532.1;QHR63300.2;BatCoV_RaTG13_spike;800 | 0 | 0.15 |
| MN996532.1;QHR63300.2;BatCoV_RaTG13_spike;801 | 0 | 0.15 |
| MN996532.1;QHR63300.2;BatCoV_RaTG13_spike;802 | 0.1 | 0.25 |
| MN996532.1;QHR63300.2;BatCoV_RaTG13_spike;803 | 0.05 | 0.225 |
| MN996532.1;QHR63300.2;BatCoV_RaTG13_spike;804 | 0.05 | 0.625 |
| MN996532.1;QHR63300.2;BatCoV_RaTG13_spike;805 | 0.1 | 0.75 |
| MN996532.1;QHR63300.2;BatCoV_RaTG13_spike;806 | 0.05 | 0.85 |
| MN996532.1;QHR63300.2;BatCoV_RaTG13_spike;807 | 0.05 | 0.825 |
| MN996532.1;QHR63300.2;BatCoV_RaTG13_spike;808 | 0.05 | 0.875 |
| MN996532.1;QHR63300.2;BatCoV_RaTG13_spike;809 | 0.05 | 0.825 |
| MN996532.1;QHR63300.2;BatCoV_RaTG13_spike;810 | 0 | 0.7 |
| MN996532.1;QHR63300.2;BatCoV_RaTG13_spike;811 | 0 | 0.725 |
| MN996532.1;QHR63300.2;BatCoV_RaTG13_spike;812 | 0 | 0.35 |
| MN996532.1;QHR63300.2;BatCoV_RaTG13_spike;813 | 0.05 | 0.25 |
| MN996532.1;QHR63300.2;BatCoV_RaTG13_spike;814 | 0.1 | 0.275 |
| MN996532.1;QHR63300.2;BatCoV_RaTG13_spike;815 | 0 | 0.125 |
| MN996532.1;QHR63300.2;BatCoV_RaTG13_spike;816 | 0 | 0 |
| MN996532.1;QHR63300.2;BatCoV_RaTG13_spike;817 | 0 | 0 |
| MN996532.1;QHR63300.2;BatCoV_RaTG13_spike;818 | 0 | 0 |
| MN996532.1;QHR63300.2;BatCoV_RaTG13_spike;819 | 0 | 0.025 |

| | | |
|---|---|---|
| MN996532.1;QHR63300.2;BatCoV_RaTG13_spike;820 | 0 | 0.025 |
| MN996532.1;QHR63300.2;BatCoV_RaTG13_spike;821 | 0 | 0.025 |
| MN996532.1;QHR63300.2;BatCoV_RaTG13_spike;822 | 0 | 0.05 |
| MN996532.1;QHR63300.2;BatCoV_RaTG13_spike;823 | 0.05 | 0.025 |
| MN996532.1;QHR63300.2;BatCoV_RaTG13_spike;824 | 0.1 | 0.025 |
| MN996532.1;QHR63300.2;BatCoV_RaTG13_spike;825 | 0 | 0.025 |
| MN996532.1;QHR63300.2;BatCoV_RaTG13_spike;826 | 0 | 0.025 |
| MN996532.1;QHR63300.2;BatCoV_RaTG13_spike;827 | 0.05 | 0.025 |
| MN996532.1;QHR63300.2;BatCoV_RaTG13_spike;828 | 0 | 0.025 |
| MN996532.1;QHR63300.2;BatCoV_RaTG13_spike;829 | 0 | 0.025 |
| MN996532.1;QHR63300.2;BatCoV_RaTG13_spike;830 | 0 | 0.025 |
| MN996532.1;QHR63300.2;BatCoV_RaTG13_spike;831 | 0.05 | 0.025 |
| MN996532.1;QHR63300.2;BatCoV_RaTG13_spike;832 | 0 | 0 |
| MN996532.1;QHR63300.2;BatCoV_RaTG13_spike;833 | 0 | 0 |
| MN996532.1;QHR63300.2;BatCoV_RaTG13_spike;834 | 0 | 0 |
| MN996532.1;QHR63300.2;BatCoV_RaTG13_spike;835 | 0 | 0 |
| MN996532.1;QHR63300.2;BatCoV_RaTG13_spike;836 | 0.05 | 0.025 |
| MN996532.1;QHR63300.2;BatCoV_RaTG13_spike;837 | 0 | 0 |
| MN996532.1;QHR63300.2;BatCoV_RaTG13_spike;838 | 0.1 | 0.025 |
| MN996532.1;QHR63300.2;BatCoV_RaTG13_spike;839 | 0.05 | 0.025 |
| MN996532.1;QHR63300.2;BatCoV_RaTG13_spike;840 | 0 | 0.025 |
| MN996532.1;QHR63300.2;BatCoV_RaTG13_spike;841 | 0 | 0 |
| MN996532.1;QHR63300.2;BatCoV_RaTG13_spike;842 | 0 | 0 |
| MN996532.1;QHR63300.2;BatCoV_RaTG13_spike;843 | 0 | 0 |
| MN996532.1;QHR63300.2;BatCoV_RaTG13_spike;844 | 0.05 | 0 |
| MN996532.1;QHR63300.2;BatCoV_RaTG13_spike;845 | 0 | 0 |
| MN996532.1;QHR63300.2;BatCoV_RaTG13_spike;846 | 0 | 0 |
| MN996532.1;QHR63300.2;BatCoV_RaTG13_spike;847 | 0 | 0 |
| MN996532.1;QHR63300.2;BatCoV_RaTG13_spike;848 | 0 | 0 |
| MN996532.1;QHR63300.2;BatCoV_RaTG13_spike;849 | 0 | 0 |
| MN996532.1;QHR63300.2;BatCoV_RaTG13_spike;850 | 0 | 0 |
| MN996532.1;QHR63300.2;BatCoV_RaTG13_spike;851 | 0 | 0 |
| MN996532.1;QHR63300.2;BatCoV_RaTG13_spike;852 | 0 | 0 |
| MN996532.1;QHR63300.2;BatCoV_RaTG13_spike;853 | 0 | 0 |
| MN996532.1;QHR63300.2;BatCoV_RaTG13_spike;854 | 0 | 0 |
| MN996532.1;QHR63300.2;BatCoV_RaTG13_spike;855 | 0 | 0 |
| MN996532.1;QHR63300.2;BatCoV_RaTG13_spike;856 | 0 | 0 |
| MN996532.1;QHR63300.2;BatCoV_RaTG13_spike;857 | 0 | 0 |
| MN996532.1;QHR63300.2;BatCoV_RaTG13_spike;858 | 0 | 0 |
| MN996532.1;QHR63300.2;BatCoV_RaTG13_spike;859 | 0 | 0 |
| MN996532.1;QHR63300.2;BatCoV_RaTG13_spike;860 | 0 | 0 |
| MN996532.1;QHR63300.2;BatCoV_RaTG13_spike;861 | 0 | 0 |
| MN996532.1;QHR63300.2;BatCoV_RaTG13_spike;862 | 0 | 0 |

| | | |
|---|---|---|
| MN996532.1;QHR63300.2;BatCoV_RaTG13_spike;863 | 0 | 0 |
| MN996532.1;QHR63300.2;BatCoV_RaTG13_spike;864 | 0 | 0 |
| MN996532.1;QHR63300.2;BatCoV_RaTG13_spike;865 | 0 | 0.025 |
| MN996532.1;QHR63300.2;BatCoV_RaTG13_spike;866 | 0 | 0.025 |
| MN996532.1;QHR63300.2;BatCoV_RaTG13_spike;867 | 0 | 0 |
| MN996532.1;QHR63300.2;BatCoV_RaTG13_spike;868 | 0 | 0 |
| MN996532.1;QHR63300.2;BatCoV_RaTG13_spike;869 | 0 | 0 |
| MN996532.1;QHR63300.2;BatCoV_RaTG13_spike;870 | 0 | 0 |
| MN996532.1;QHR63300.2;BatCoV_RaTG13_spike;871 | 0 | 0 |
| MN996532.1;QHR63300.2;BatCoV_RaTG13_spike;872 | 0 | 0 |
| MN996532.1;QHR63300.2;BatCoV_RaTG13_spike;873 | 0 | 0.025 |
| MN996532.1;QHR63300.2;BatCoV_RaTG13_spike;874 | 0.1 | 0.05 |
| MN996532.1;QHR63300.2;BatCoV_RaTG13_spike;875 | 0 | 0 |
| MN996532.1;QHR63300.2;BatCoV_RaTG13_spike;876 | 0 | 0 |
| MN996532.1;QHR63300.2;BatCoV_RaTG13_spike;877 | 0 | 0 |
| MN996532.1;QHR63300.2;BatCoV_RaTG13_spike;878 | 0 | 0 |
| MN996532.1;QHR63300.2;BatCoV_RaTG13_spike;879 | 0 | 0 |
| MN996532.1;QHR63300.2;BatCoV_RaTG13_spike;880 | 0 | 0 |
| MN996532.1;QHR63300.2;BatCoV_RaTG13_spike;881 | 0 | 0 |
| MN996532.1;QHR63300.2;BatCoV_RaTG13_spike;882 | 0 | 0 |
| MN996532.1;QHR63300.2;BatCoV_RaTG13_spike;883 | 0 | 0 |
| MN996532.1;QHR63300.2;BatCoV_RaTG13_spike;884 | 0 | 0 |
| MN996532.1;QHR63300.2;BatCoV_RaTG13_spike;885 | 0 | 0.025 |
| MN996532.1;QHR63300.2;BatCoV_RaTG13_spike;886 | 0 | 0 |
| MN996532.1;QHR63300.2;BatCoV_RaTG13_spike;887 | 0 | 0 |
| MN996532.1;QHR63300.2;BatCoV_RaTG13_spike;888 | 0.05 | 0.025 |
| MN996532.1;QHR63300.2;BatCoV_RaTG13_spike;889 | 0.05 | 0.025 |
| MN996532.1;QHR63300.2;BatCoV_RaTG13_spike;890 | 0.05 | 0.025 |
| MN996532.1;QHR63300.2;BatCoV_RaTG13_spike;891 | 0.05 | 0.025 |
| MN996532.1;QHR63300.2;BatCoV_RaTG13_spike;892 | 0.05 | 0.025 |
| MN996532.1;QHR63300.2;BatCoV_RaTG13_spike;893 | 0.05 | 0.025 |
| MN996532.1;QHR63300.2;BatCoV_RaTG13_spike;894 | 0.05 | 0.025 |
| MN996532.1;QHR63300.2;BatCoV_RaTG13_spike;895 | 0.1 | 0.05 |
| MN996532.1;QHR63300.2;BatCoV_RaTG13_spike;896 | 0.15 | 0.05 |
| MN996532.1;QHR63300.2;BatCoV_RaTG13_spike;897 | 0.1 | 0.05 |
| MN996532.1;QHR63300.2;BatCoV_RaTG13_spike;898 | 0 | 0 |
| MN996532.1;QHR63300.2;BatCoV_RaTG13_spike;899 | 0 | 0 |
| MN996532.1;QHR63300.2;BatCoV_RaTG13_spike;900 | 0 | 0 |
| MN996532.1;QHR63300.2;BatCoV_RaTG13_spike;901 | 0.05 | 0.025 |
| MN996532.1;QHR63300.2;BatCoV_RaTG13_spike;902 | 0 | 0.05 |
| MN996532.1;QHR63300.2;BatCoV_RaTG13_spike;903 | 0.05 | 0.05 |
| MN996532.1;QHR63300.2;BatCoV_RaTG13_spike;904 | 0 | 0.05 |
| MN996532.1;QHR63300.2;BatCoV_RaTG13_spike;905 | 0.05 | 0.075 |

| | | |
|---|---|---|
| MN996532.1;QHR63300.2;BatCoV_RaTG13_spike;906 | 0.05 | 0.025 |
| MN996532.1;QHR63300.2;BatCoV_RaTG13_spike;907 | 0 | 0.05 |
| MN996532.1;QHR63300.2;BatCoV_RaTG13_spike;908 | 0 | 0.025 |
| MN996532.1;QHR63300.2;BatCoV_RaTG13_spike;909 | 0.05 | 0.05 |
| MN996532.1;QHR63300.2;BatCoV_RaTG13_spike;910 | 0 | 0 |
| MN996532.1;QHR63300.2;BatCoV_RaTG13_spike;911 | 0 | 0.05 |
| MN996532.1;QHR63300.2;BatCoV_RaTG13_spike;912 | 0.05 | 0.025 |
| MN996532.1;QHR63300.2;BatCoV_RaTG13_spike;913 | 0 | 0.05 |
| MN996532.1;QHR63300.2;BatCoV_RaTG13_spike;914 | 0.05 | 0.025 |
| MN996532.1;QHR63300.2;BatCoV_RaTG13_spike;915 | 0 | 0 |
| MN996532.1;QHR63300.2;BatCoV_RaTG13_spike;916 | 0.05 | 0 |
| MN996532.1;QHR63300.2;BatCoV_RaTG13_spike;917 | 0.05 | 0.025 |
| MN996532.1;QHR63300.2;BatCoV_RaTG13_spike;918 | 0.05 | 0.025 |
| MN996532.1;QHR63300.2;BatCoV_RaTG13_spike;919 | 0.05 | 0.05 |
| MN996532.1;QHR63300.2;BatCoV_RaTG13_spike;920 | 0.1 | 0.05 |
| MN996532.1;QHR63300.2;BatCoV_RaTG13_spike;921 | 0.05 | 0.025 |
| MN996532.1;QHR63300.2;BatCoV_RaTG13_spike;922 | 0.05 | 0.025 |
| MN996532.1;QHR63300.2;BatCoV_RaTG13_spike;923 | 0.05 | 0.05 |
| MN996532.1;QHR63300.2;BatCoV_RaTG13_spike;924 | 0.1 | 0.05 |
| MN996532.1;QHR63300.2;BatCoV_RaTG13_spike;925 | 0 | 0.025 |
| MN996532.1;QHR63300.2;BatCoV_RaTG13_spike;926 | 0 | 0.025 |
| MN996532.1;QHR63300.2;BatCoV_RaTG13_spike;927 | 0 | 0 |
| MN996532.1;QHR63300.2;BatCoV_RaTG13_spike;928 | 0.05 | 0.025 |
| MN996532.1;QHR63300.2;BatCoV_RaTG13_spike;929 | 0 | 0 |
| MN996532.1;QHR63300.2;BatCoV_RaTG13_spike;930 | 0 | 0 |
| MN996532.1;QHR63300.2;BatCoV_RaTG13_spike;931 | 0 | 0 |
| MN996532.1;QHR63300.2;BatCoV_RaTG13_spike;932 | 0 | 0 |
| MN996532.1;QHR63300.2;BatCoV_RaTG13_spike;933 | 0 | 0 |
| MN996532.1;QHR63300.2;BatCoV_RaTG13_spike;934 | 0 | 0 |
| MN996532.1;QHR63300.2;BatCoV_RaTG13_spike;935 | 0 | 0 |
| MN996532.1;QHR63300.2;BatCoV_RaTG13_spike;936 | 0 | 0.075 |
| MN996532.1;QHR63300.2;BatCoV_RaTG13_spike;937 | 0 | 0.025 |
| MN996532.1;QHR63300.2;BatCoV_RaTG13_spike;938 | 0 | 0.025 |
| MN996532.1;QHR63300.2;BatCoV_RaTG13_spike;939 | 0 | 0.025 |
| MN996532.1;QHR63300.2;BatCoV_RaTG13_spike;940 | 0 | 0.075 |
| MN996532.1;QHR63300.2;BatCoV_RaTG13_spike;941 | 0 | 0.125 |
| MN996532.1;QHR63300.2;BatCoV_RaTG13_spike;942 | 0.1 | 0.075 |
| MN996532.1;QHR63300.2;BatCoV_RaTG13_spike;943 | 0 | 0.1 |
| MN996532.1;QHR63300.2;BatCoV_RaTG13_spike;944 | 0 | 0.075 |
| MN996532.1;QHR63300.2;BatCoV_RaTG13_spike;945 | 0 | 0.025 |
| MN996532.1;QHR63300.2;BatCoV_RaTG13_spike;946 | 0 | 0.05 |
| MN996532.1;QHR63300.2;BatCoV_RaTG13_spike;947 | 0.05 | 0 |
| MN996532.1;QHR63300.2;BatCoV_RaTG13_spike;948 | 0 | 0 |

| | | |
|---|---|---|
| MN996532.1;QHR63300.2;BatCoV_RaTG13_spike;949 | 0 | 0.025 |
| MN996532.1;QHR63300.2;BatCoV_RaTG13_spike;950 | 0 | 0.025 |
| MN996532.1;QHR63300.2;BatCoV_RaTG13_spike;951 | 0 | 0 |
| MN996532.1;QHR63300.2;BatCoV_RaTG13_spike;952 | 0 | 0.025 |
| MN996532.1;QHR63300.2;BatCoV_RaTG13_spike;953 | 0 | 0.05 |
| MN996532.1;QHR63300.2;BatCoV_RaTG13_spike;954 | 0 | 0.025 |
| MN996532.1;QHR63300.2;BatCoV_RaTG13_spike;955 | 0 | 0 |
| MN996532.1;QHR63300.2;BatCoV_RaTG13_spike;956 | 0 | 0 |
| MN996532.1;QHR63300.2;BatCoV_RaTG13_spike;957 | 0 | 0 |
| MN996532.1;QHR63300.2;BatCoV_RaTG13_spike;958 | 0 | 0 |
| MN996532.1;QHR63300.2;BatCoV_RaTG13_spike;959 | 0 | 0 |
| MN996532.1;QHR63300.2;BatCoV_RaTG13_spike;960 | 0 | 0.025 |
| MN996532.1;QHR63300.2;BatCoV_RaTG13_spike;961 | 0 | 0 |
| MN996532.1;QHR63300.2;BatCoV_RaTG13_spike;962 | 0 | 0 |
| MN996532.1;QHR63300.2;BatCoV_RaTG13_spike;963 | 0 | 0 |
| MN996532.1;QHR63300.2;BatCoV_RaTG13_spike;964 | 0 | 0 |
| MN996532.1;QHR63300.2;BatCoV_RaTG13_spike;965 | 0 | 0 |
| MN996532.1;QHR63300.2;BatCoV_RaTG13_spike;966 | 0 | 0.025 |
| MN996532.1;QHR63300.2;BatCoV_RaTG13_spike;967 | 0 | 0.025 |
| MN996532.1;QHR63300.2;BatCoV_RaTG13_spike;968 | 0.05 | 0.05 |
| MN996532.1;QHR63300.2;BatCoV_RaTG13_spike;969 | 0 | 0.075 |
| MN996532.1;QHR63300.2;BatCoV_RaTG13_spike;970 | 0.05 | 0.075 |
| MN996532.1;QHR63300.2;BatCoV_RaTG13_spike;971 | 0.05 | 0.125 |
| MN996532.1;QHR63300.2;BatCoV_RaTG13_spike;972 | 0.05 | 0.1 |
| MN996532.1;QHR63300.2;BatCoV_RaTG13_spike;973 | 0 | 0.075 |
| MN996532.1;QHR63300.2;BatCoV_RaTG13_spike;974 | 0 | 0.05 |
| MN996532.1;QHR63300.2;BatCoV_RaTG13_spike;975 | 0.05 | 0.05 |
| MN996532.1;QHR63300.2;BatCoV_RaTG13_spike;976 | 0.05 | 0.025 |
| MN996532.1;QHR63300.2;BatCoV_RaTG13_spike;977 | 0 | 0 |
| MN996532.1;QHR63300.2;BatCoV_RaTG13_spike;978 | 0 | 0 |
| MN996532.1;QHR63300.2;BatCoV_RaTG13_spike;979 | 0 | 0 |
| MN996532.1;QHR63300.2;BatCoV_RaTG13_spike;980 | 0 | 0.025 |
| MN996532.1;QHR63300.2;BatCoV_RaTG13_spike;981 | 0 | 0 |
| MN996532.1;QHR63300.2;BatCoV_RaTG13_spike;982 | 0 | 0 |
| MN996532.1;QHR63300.2;BatCoV_RaTG13_spike;983 | 0 | 0 |
| MN996532.1;QHR63300.2;BatCoV_RaTG13_spike;984 | 0.05 | 0 |
| MN996532.1;QHR63300.2;BatCoV_RaTG13_spike;985 | 0 | 0 |
| MN996532.1;QHR63300.2;BatCoV_RaTG13_spike;986 | 0 | 0 |
| MN996532.1;QHR63300.2;BatCoV_RaTG13_spike;987 | 0 | 0 |
| MN996532.1;QHR63300.2;BatCoV_RaTG13_spike;988 | 0 | 0 |
| MN996532.1;QHR63300.2;BatCoV_RaTG13_spike;989 | 0 | 0 |
| MN996532.1;QHR63300.2;BatCoV_RaTG13_spike;990 | 0 | 0 |
| MN996532.1;QHR63300.2;BatCoV_RaTG13_spike;991 | 0 | 0 |

| | | |
|---|---|---|
| MN996532.1;QHR63300.2;BatCoV_RaTG13_spike;992 | 0 | 0 |
| MN996532.1;QHR63300.2;BatCoV_RaTG13_spike;993 | 0 | 0 |
| MN996532.1;QHR63300.2;BatCoV_RaTG13_spike;994 | 0 | 0 |
| MN996532.1;QHR63300.2;BatCoV_RaTG13_spike;995 | 0 | 0 |
| MN996532.1;QHR63300.2;BatCoV_RaTG13_spike;996 | 0 | 0 |
| MN996532.1;QHR63300.2;BatCoV_RaTG13_spike;997 | 0 | 0 |
| MN996532.1;QHR63300.2;BatCoV_RaTG13_spike;998 | 0 | 0 |
| MN996532.1;QHR63300.2;BatCoV_RaTG13_spike;999 | 0 | 0.025 |
| MN996532.1;QHR63300.2;BatCoV_RaTG13_spike;1000 | 0 | 0 |
| MN996532.1;QHR63300.2;BatCoV_RaTG13_spike;1001 | 0 | 0 |
| MN996532.1;QHR63300.2;BatCoV_RaTG13_spike;1002 | 0 | 0 |
| MN996532.1;QHR63300.2;BatCoV_RaTG13_spike;1003 | 0 | 0 |
| MN996532.1;QHR63300.2;BatCoV_RaTG13_spike;1004 | 0 | 0 |
| MN996532.1;QHR63300.2;BatCoV_RaTG13_spike;1005 | 0 | 0 |
| MN996532.1;QHR63300.2;BatCoV_RaTG13_spike;1006 | 0 | 0 |
| MN996532.1;QHR63300.2;BatCoV_RaTG13_spike;1007 | 0.05 | 0.025 |
| MN996532.1;QHR63300.2;BatCoV_RaTG13_spike;1008 | 0 | 0 |
| MN996532.1;QHR63300.2;BatCoV_RaTG13_spike;1009 | 0 | 0.025 |
| MN996532.1;QHR63300.2;BatCoV_RaTG13_spike;1010 | 0 | 0 |
| MN996532.1;QHR63300.2;BatCoV_RaTG13_spike;1011 | 0 | 0 |
| MN996532.1;QHR63300.2;BatCoV_RaTG13_spike;1012 | 0.05 | 0.025 |
| MN996532.1;QHR63300.2;BatCoV_RaTG13_spike;1013 | 0.05 | 0 |
| MN996532.1;QHR63300.2;BatCoV_RaTG13_spike;1014 | 0.05 | 0 |
| MN996532.1;QHR63300.2;BatCoV_RaTG13_spike;1015 | 0 | 0.025 |
| MN996532.1;QHR63300.2;BatCoV_RaTG13_spike;1016 | 0 | 0 |
| MN996532.1;QHR63300.2;BatCoV_RaTG13_spike;1017 | 0 | 0.025 |
| MN996532.1;QHR63300.2;BatCoV_RaTG13_spike;1018 | 0 | 0 |
| MN996532.1;QHR63300.2;BatCoV_RaTG13_spike;1019 | 0 | 0 |
| MN996532.1;QHR63300.2;BatCoV_RaTG13_spike;1020 | 0 | 0 |
| MN996532.1;QHR63300.2;BatCoV_RaTG13_spike;1021 | 0 | 0 |
| MN996532.1;QHR63300.2;BatCoV_RaTG13_spike;1022 | 0 | 0 |
| MN996532.1;QHR63300.2;BatCoV_RaTG13_spike;1023 | 0 | 0 |
| MN996532.1;QHR63300.2;BatCoV_RaTG13_spike;1024 | 0 | 0 |
| MN996532.1;QHR63300.2;BatCoV_RaTG13_spike;1025 | 0 | 0 |
| MN996532.1;QHR63300.2;BatCoV_RaTG13_spike;1026 | 0 | 0 |
| MN996532.1;QHR63300.2;BatCoV_RaTG13_spike;1027 | 0 | 0 |
| MN996532.1;QHR63300.2;BatCoV_RaTG13_spike;1028 | 0 | 0 |
| MN996532.1;QHR63300.2;BatCoV_RaTG13_spike;1029 | 0 | 0 |
| MN996532.1;QHR63300.2;BatCoV_RaTG13_spike;1030 | 0 | 0 |
| MN996532.1;QHR63300.2;BatCoV_RaTG13_spike;1031 | 0 | 0 |
| MN996532.1;QHR63300.2;BatCoV_RaTG13_spike;1032 | 0 | 0 |
| MN996532.1;QHR63300.2;BatCoV_RaTG13_spike;1033 | 0 | 0 |
| MN996532.1;QHR63300.2;BatCoV_RaTG13_spike;1034 | 0 | 0 |

| | | |
|---|---|---|
| MN996532.1;QHR63300.2;BatCoV_RaTG13_spike;1035 | 0 | 0 |
| MN996532.1;QHR63300.2;BatCoV_RaTG13_spike;1036 | 0 | 0 |
| MN996532.1;QHR63300.2;BatCoV_RaTG13_spike;1037 | 0 | 0.025 |
| MN996532.1;QHR63300.2;BatCoV_RaTG13_spike;1038 | 0.05 | 0 |
| MN996532.1;QHR63300.2;BatCoV_RaTG13_spike;1039 | 0 | 0 |
| MN996532.1;QHR63300.2;BatCoV_RaTG13_spike;1040 | 0 | 0.05 |
| MN996532.1;QHR63300.2;BatCoV_RaTG13_spike;1041 | 0 | 0.025 |
| MN996532.1;QHR63300.2;BatCoV_RaTG13_spike;1042 | 0 | 0 |
| MN996532.1;QHR63300.2;BatCoV_RaTG13_spike;1043 | 0 | 0 |
| MN996532.1;QHR63300.2;BatCoV_RaTG13_spike;1044 | 0 | 0 |
| MN996532.1;QHR63300.2;BatCoV_RaTG13_spike;1045 | 0 | 0 |
| MN996532.1;QHR63300.2;BatCoV_RaTG13_spike;1046 | 0 | 0 |
| MN996532.1;QHR63300.2;BatCoV_RaTG13_spike;1047 | 0 | 0 |
| MN996532.1;QHR63300.2;BatCoV_RaTG13_spike;1048 | 0 | 0 |
| MN996532.1;QHR63300.2;BatCoV_RaTG13_spike;1049 | 0 | 0 |
| MN996532.1;QHR63300.2;BatCoV_RaTG13_spike;1050 | 0 | 0 |
| MN996532.1;QHR63300.2;BatCoV_RaTG13_spike;1051 | 0 | 0 |
| MN996532.1;QHR63300.2;BatCoV_RaTG13_spike;1052 | 0 | 0 |
| MN996532.1;QHR63300.2;BatCoV_RaTG13_spike;1053 | 0 | 0 |
| MN996532.1;QHR63300.2;BatCoV_RaTG13_spike;1054 | 0 | 0 |
| MN996532.1;QHR63300.2;BatCoV_RaTG13_spike;1055 | 0 | 0 |
| MN996532.1;QHR63300.2;BatCoV_RaTG13_spike;1056 | 0 | 0 |
| MN996532.1;QHR63300.2;BatCoV_RaTG13_spike;1057 | 0 | 0 |
| MN996532.1;QHR63300.2;BatCoV_RaTG13_spike;1058 | 0 | 0 |
| MN996532.1;QHR63300.2;BatCoV_RaTG13_spike;1059 | 0 | 0 |
| MN996532.1;QHR63300.2;BatCoV_RaTG13_spike;1060 | 0.05 | 0 |
| MN996532.1;QHR63300.2;BatCoV_RaTG13_spike;1061 | 0.1 | 0 |
| MN996532.1;QHR63300.2;BatCoV_RaTG13_spike;1062 | 0.05 | 0 |
| MN996532.1;QHR63300.2;BatCoV_RaTG13_spike;1063 | 0.05 | 0 |
| MN996532.1;QHR63300.2;BatCoV_RaTG13_spike;1064 | 0 | 0 |
| MN996532.1;QHR63300.2;BatCoV_RaTG13_spike;1065 | 0 | 0 |
| MN996532.1;QHR63300.2;BatCoV_RaTG13_spike;1066 | 0 | 0 |
| MN996532.1;QHR63300.2;BatCoV_RaTG13_spike;1067 | 0.1 | 0 |
| MN996532.1;QHR63300.2;BatCoV_RaTG13_spike;1068 | 0 | 0 |
| MN996532.1;QHR63300.2;BatCoV_RaTG13_spike;1069 | 0 | 0.025 |
| MN996532.1;QHR63300.2;BatCoV_RaTG13_spike;1070 | 0 | 0 |
| MN996532.1;QHR63300.2;BatCoV_RaTG13_spike;1071 | 0 | 0 |
| MN996532.1;QHR63300.2;BatCoV_RaTG13_spike;1072 | 0 | 0 |
| MN996532.1;QHR63300.2;BatCoV_RaTG13_spike;1073 | 0 | 0 |
| MN996532.1;QHR63300.2;BatCoV_RaTG13_spike;1074 | 0.05 | 0 |
| MN996532.1;QHR63300.2;BatCoV_RaTG13_spike;1075 | 0.05 | 0 |
| MN996532.1;QHR63300.2;BatCoV_RaTG13_spike;1076 | 0 | 0 |
| MN996532.1;QHR63300.2;BatCoV_RaTG13_spike;1077 | 0.05 | 0.025 |

| | | |
|---|---|---|
| MN996532.1;QHR63300.2;BatCoV_RaTG13_spike;1078 | 0 | 0 |
| MN996532.1;QHR63300.2;BatCoV_RaTG13_spike;1079 | 0 | 0 |
| MN996532.1;QHR63300.2;BatCoV_RaTG13_spike;1080 | 0 | 0 |
| MN996532.1;QHR63300.2;BatCoV_RaTG13_spike;1081 | 0 | 0 |
| MN996532.1;QHR63300.2;BatCoV_RaTG13_spike;1082 | 0 | 0 |
| MN996532.1;QHR63300.2;BatCoV_RaTG13_spike;1083 | 0 | 0 |
| MN996532.1;QHR63300.2;BatCoV_RaTG13_spike;1084 | 0 | 0 |
| MN996532.1;QHR63300.2;BatCoV_RaTG13_spike;1085 | 0 | 0 |
| MN996532.1;QHR63300.2;BatCoV_RaTG13_spike;1086 | 0 | 0 |
| MN996532.1;QHR63300.2;BatCoV_RaTG13_spike;1087 | 0 | 0 |
| MN996532.1;QHR63300.2;BatCoV_RaTG13_spike;1088 | 0 | 0 |
| MN996532.1;QHR63300.2;BatCoV_RaTG13_spike;1089 | 0 | 0 |
| MN996532.1;QHR63300.2;BatCoV_RaTG13_spike;1090 | 0 | 0 |
| MN996532.1;QHR63300.2;BatCoV_RaTG13_spike;1091 | 0 | 0 |
| MN996532.1;QHR63300.2;BatCoV_RaTG13_spike;1092 | 0 | 0 |
| MN996532.1;QHR63300.2;BatCoV_RaTG13_spike;1093 | 0 | 0 |
| MN996532.1;QHR63300.2;BatCoV_RaTG13_spike;1094 | 0 | 0 |
| MN996532.1;QHR63300.2;BatCoV_RaTG13_spike;1095 | 0 | 0 |
| MN996532.1;QHR63300.2;BatCoV_RaTG13_spike;1096 | 0 | 0 |
| MN996532.1;QHR63300.2;BatCoV_RaTG13_spike;1097 | 0 | 0 |
| MN996532.1;QHR63300.2;BatCoV_RaTG13_spike;1098 | 0 | 0 |
| MN996532.1;QHR63300.2;BatCoV_RaTG13_spike;1099 | 0 | 0.025 |
| MN996532.1;QHR63300.2;BatCoV_RaTG13_spike;1100 | 0.1 | 0.025 |
| MN996532.1;QHR63300.2;BatCoV_RaTG13_spike;1101 | 0.05 | 0.025 |
| MN996532.1;QHR63300.2;BatCoV_RaTG13_spike;1102 | 0.05 | 0.025 |
| MN996532.1;QHR63300.2;BatCoV_RaTG13_spike;1103 | 0 | 0 |
| MN996532.1;QHR63300.2;BatCoV_RaTG13_spike;1104 | 0.05 | 0.025 |
| MN996532.1;QHR63300.2;BatCoV_RaTG13_spike;1105 | 0.05 | 0.025 |
| MN996532.1;QHR63300.2;BatCoV_RaTG13_spike;1106 | 0.05 | 0.05 |
| MN996532.1;QHR63300.2;BatCoV_RaTG13_spike;1107 | 0.05 | 0.025 |
| MN996532.1;QHR63300.2;BatCoV_RaTG13_spike;1108 | 0.05 | 0.025 |
| MN996532.1;QHR63300.2;BatCoV_RaTG13_spike;1109 | 0.05 | 0.025 |
| MN996532.1;QHR63300.2;BatCoV_RaTG13_spike;1110 | 0.05 | 0.025 |
| MN996532.1;QHR63300.2;BatCoV_RaTG13_spike;1111 | 0.05 | 0.025 |
| MN996532.1;QHR63300.2;BatCoV_RaTG13_spike;1112 | 0.05 | 0.025 |
| MN996532.1;QHR63300.2;BatCoV_RaTG13_spike;1113 | 0 | 0 |
| MN996532.1;QHR63300.2;BatCoV_RaTG13_spike;1114 | 0 | 0 |
| MN996532.1;QHR63300.2;BatCoV_RaTG13_spike;1115 | 0 | 0 |
| MN996532.1;QHR63300.2;BatCoV_RaTG13_spike;1116 | 0 | 0 |
| MN996532.1;QHR63300.2;BatCoV_RaTG13_spike;1117 | 0 | 0 |
| MN996532.1;QHR63300.2;BatCoV_RaTG13_spike;1118 | 0 | 0 |
| MN996532.1;QHR63300.2;BatCoV_RaTG13_spike;1119 | 0 | 0 |
| MN996532.1;QHR63300.2;BatCoV_RaTG13_spike;1120 | 0 | 0 |

| | | |
|---|---|---|
| MN996532.1;QHR63300.2;BatCoV_RaTG13_spike;1121 | 0 | 0 |
| MN996532.1;QHR63300.2;BatCoV_RaTG13_spike;1122 | 0 | 0 |
| MN996532.1;QHR63300.2;BatCoV_RaTG13_spike;1123 | 0 | 0 |
| MN996532.1;QHR63300.2;BatCoV_RaTG13_spike;1124 | 0 | 0 |
| MN996532.1;QHR63300.2;BatCoV_RaTG13_spike;1125 | 0 | 0 |
| MN996532.1;QHR63300.2;BatCoV_RaTG13_spike;1126 | 0 | 0.05 |
| MN996532.1;QHR63300.2;BatCoV_RaTG13_spike;1127 | 0 | 0 |
| MN996532.1;QHR63300.2;BatCoV_RaTG13_spike;1128 | 0 | 0 |
| MN996532.1;QHR63300.2;BatCoV_RaTG13_spike;1129 | 0 | 0.025 |
| MN996532.1;QHR63300.2;BatCoV_RaTG13_spike;1130 | 0 | 0 |
| MN996532.1;QHR63300.2;BatCoV_RaTG13_spike;1131 | 0 | 0.025 |
| MN996532.1;QHR63300.2;BatCoV_RaTG13_spike;1132 | 0 | 0 |
| MN996532.1;QHR63300.2;BatCoV_RaTG13_spike;1133 | 0 | 0 |
| MN996532.1;QHR63300.2;BatCoV_RaTG13_spike;1134 | 0 | 0 |
| MN996532.1;QHR63300.2;BatCoV_RaTG13_spike;1135 | 0 | 0.05 |
| MN996532.1;QHR63300.2;BatCoV_RaTG13_spike;1136 | 0 | 0.175 |
| MN996532.1;QHR63300.2;BatCoV_RaTG13_spike;1137 | 0.05 | 0.75 |
| MN996532.1;QHR63300.2;BatCoV_RaTG13_spike;1138 | 0.1 | 0.775 |
| MN996532.1;QHR63300.2;BatCoV_RaTG13_spike;1139 | 0.05 | 0.775 |
| MN996532.1;QHR63300.2;BatCoV_RaTG13_spike;1140 | 0.05 | 0.725 |
| MN996532.1;QHR63300.2;BatCoV_RaTG13_spike;1141 | 0.3 | 0.725 |
| MN996532.1;QHR63300.2;BatCoV_RaTG13_spike;1142 | 0.4 | 0.775 |
| MN996532.1;QHR63300.2;BatCoV_RaTG13_spike;1143 | 0.35 | 0.775 |
| MN996532.1;QHR63300.2;BatCoV_RaTG13_spike;1144 | 0.35 | 0.825 |
| MN996532.1;QHR63300.2;BatCoV_RaTG13_spike;1145 | 0.45 | 0.8 |
| MN996532.1;QHR63300.2;BatCoV_RaTG13_spike;1146 | 0.4 | 0.5 |
| MN996532.1;QHR63300.2;BatCoV_RaTG13_spike;1147 | 0.45 | 0.35 |
| MN996532.1;QHR63300.2;BatCoV_RaTG13_spike;1148 | 0.45 | 0.35 |
| MN996532.1;QHR63300.2;BatCoV_RaTG13_spike;1149 | 0.5 | 0.35 |
| MN996532.1;QHR63300.2;BatCoV_RaTG13_spike;1150 | 0.5 | 0.375 |
| MN996532.1;QHR63300.2;BatCoV_RaTG13_spike;1151 | 0 | 0.2 |
| MN996532.1;QHR63300.2;BatCoV_RaTG13_spike;1152 | 0 | 0.125 |
| MN996532.1;QHR63300.2;BatCoV_RaTG13_spike;1153 | 0 | 0.225 |
| MN996532.1;QHR63300.2;BatCoV_RaTG13_spike;1154 | 0.05 | 0.275 |
| MN996532.1;QHR63300.2;BatCoV_RaTG13_spike;1155 | 0.1 | 0.35 |
| MN996532.1;QHR63300.2;BatCoV_RaTG13_spike;1156 | 0.2 | 0.35 |
| MN996532.1;QHR63300.2;BatCoV_RaTG13_spike;1157 | 0.1 | 0.35 |
| MN996532.1;QHR63300.2;BatCoV_RaTG13_spike;1158 | 0 | 0.3 |
| MN996532.1;QHR63300.2;BatCoV_RaTG13_spike;1159 | 0 | 0.325 |
| MN996532.1;QHR63300.2;BatCoV_RaTG13_spike;1160 | 0 | 0.15 |
| MN996532.1;QHR63300.2;BatCoV_RaTG13_spike;1161 | 0 | 0.2 |
| MN996532.1;QHR63300.2;BatCoV_RaTG13_spike;1162 | 0 | 0.05 |
| MN996532.1;QHR63300.2;BatCoV_RaTG13_spike;1163 | 0.05 | 0.1 |

| | | |
|---|---|---|
| MN996532.1;QHR63300.2;BatCoV_RaTG13_spike;1164 | 0.15 | 0.05 |
| MN996532.1;QHR63300.2;BatCoV_RaTG13_spike;1165 | 0.05 | 0.125 |
| MN996532.1;QHR63300.2;BatCoV_RaTG13_spike;1166 | 0.05 | 0.05 |
| MN996532.1;QHR63300.2;BatCoV_RaTG13_spike;1167 | 0.05 | 0.1 |
| MN996532.1;QHR63300.2;BatCoV_RaTG13_spike;1168 | 0.05 | 0.175 |
| MN996532.1;QHR63300.2;BatCoV_RaTG13_spike;1169 | 0.05 | 0.225 |
| MN996532.1;QHR63300.2;BatCoV_RaTG13_spike;1170 | 0.1 | 0.225 |
| MN996532.1;QHR63300.2;BatCoV_RaTG13_spike;1171 | 0.1 | 0.275 |
| MN996532.1;QHR63300.2;BatCoV_RaTG13_spike;1172 | 0.1 | 0.275 |
| MN996532.1;QHR63300.2;BatCoV_RaTG13_spike;1173 | 0.1 | 0.3 |
| MN996532.1;QHR63300.2;BatCoV_RaTG13_spike;1174 | 0.05 | 0.3 |
| MN996532.1;QHR63300.2;BatCoV_RaTG13_spike;1175 | 0.2 | 0.325 |
| MN996532.1;QHR63300.2;BatCoV_RaTG13_spike;1176 | 0.15 | 0.3 |
| MN996532.1;QHR63300.2;BatCoV_RaTG13_spike;1177 | 0.1 | 0.175 |
| MN996532.1;QHR63300.2;BatCoV_RaTG13_spike;1178 | 0.1 | 0.15 |
| MN996532.1;QHR63300.2;BatCoV_RaTG13_spike;1179 | 0.15 | 0.125 |
| MN996532.1;QHR63300.2;BatCoV_RaTG13_spike;1180 | 0.05 | 0.125 |
| MN996532.1;QHR63300.2;BatCoV_RaTG13_spike;1181 | 0.05 | 0.125 |
| MN996532.1;QHR63300.2;BatCoV_RaTG13_spike;1182 | 0.05 | 0.1 |
| MN996532.1;QHR63300.2;BatCoV_RaTG13_spike;1183 | 0.05 | 0.125 |
| MN996532.1;QHR63300.2;BatCoV_RaTG13_spike;1184 | 0.05 | 0.1 |
| MN996532.1;QHR63300.2;BatCoV_RaTG13_spike;1185 | 0.05 | 0.125 |
| MN996532.1;QHR63300.2;BatCoV_RaTG13_spike;1186 | 0.05 | 0.125 |
| MN996532.1;QHR63300.2;BatCoV_RaTG13_spike;1187 | 0.05 | 0.075 |
| MN996532.1;QHR63300.2;BatCoV_RaTG13_spike;1188 | 0.05 | 0.125 |
| MN996532.1;QHR63300.2;BatCoV_RaTG13_spike;1189 | 0.1 | 0.15 |
| MN996532.1;QHR63300.2;BatCoV_RaTG13_spike;1190 | 0.05 | 0.125 |
| MN996532.1;QHR63300.2;BatCoV_RaTG13_spike;1191 | 0 | 0 |
| MN996532.1;QHR63300.2;BatCoV_RaTG13_spike;1192 | 0 | 0.05 |
| MN996532.1;QHR63300.2;BatCoV_RaTG13_spike;1193 | 0 | 0 |
| MN996532.1;QHR63300.2;BatCoV_RaTG13_spike;1194 | 0 | 0 |
| MN996532.1;QHR63300.2;BatCoV_RaTG13_spike;1195 | 0 | 0 |
| MN996532.1;QHR63300.2;BatCoV_RaTG13_spike;1196 | 0 | 0.025 |
| MN996532.1;QHR63300.2;BatCoV_RaTG13_spike;1197 | 0 | 0 |
| MN996532.1;QHR63300.2;BatCoV_RaTG13_spike;1198 | 0 | 0 |
| MN996532.1;QHR63300.2;BatCoV_RaTG13_spike;1199 | 0 | 0 |
| MN996532.1;QHR63300.2;BatCoV_RaTG13_spike;1200 | 0 | 0 |
| MN996532.1;QHR63300.2;BatCoV_RaTG13_spike;1201 | 0 | 0 |
| MN996532.1;QHR63300.2;BatCoV_RaTG13_spike;1202 | 0 | 0.025 |
| MN996532.1;QHR63300.2;BatCoV_RaTG13_spike;1203 | 0 | 0 |
| MN996532.1;QHR63300.2;BatCoV_RaTG13_spike;1204 | 0 | 0 |
| MN996532.1;QHR63300.2;BatCoV_RaTG13_spike;1205 | 0 | 0 |
| MN996532.1;QHR63300.2;BatCoV_RaTG13_spike;1206 | 0 | 0 |

| | | |
|---|---|---|
| MN996532.1;QHR63300.2;BatCoV_RaTG13_spike;1207 | 0 | 0 |
| MN996532.1;QHR63300.2;BatCoV_RaTG13_spike;1208 | 0 | 0 |
| MN996532.1;QHR63300.2;BatCoV_RaTG13_spike;1209 | 0 | 0 |
| MN996532.1;QHR63300.2;BatCoV_RaTG13_spike;1210 | 0 | 0 |
| MN996532.1;QHR63300.2;BatCoV_RaTG13_spike;1211 | 0 | 0 |
| MN996532.1;QHR63300.2;BatCoV_RaTG13_spike;1212 | 0 | 0 |
| MN996532.1;QHR63300.2;BatCoV_RaTG13_spike;1213 | 0 | 0 |
| MN996532.1;QHR63300.2;BatCoV_RaTG13_spike;1214 | 0 | 0 |
| MN996532.1;QHR63300.2;BatCoV_RaTG13_spike;1215 | 0 | 0 |
| MN996532.1;QHR63300.2;BatCoV_RaTG13_spike;1216 | 0 | 0 |
| MN996532.1;QHR63300.2;BatCoV_RaTG13_spike;1217 | 0 | 0 |
| MN996532.1;QHR63300.2;BatCoV_RaTG13_spike;1218 | 0 | 0 |
| MN996532.1;QHR63300.2;BatCoV_RaTG13_spike;1219 | 0 | 0 |
| MN996532.1;QHR63300.2;BatCoV_RaTG13_spike;1220 | 0 | 0 |
| MN996532.1;QHR63300.2;BatCoV_RaTG13_spike;1221 | 0 | 0 |
| MN996532.1;QHR63300.2;BatCoV_RaTG13_spike;1222 | 0 | 0 |
| MN996532.1;QHR63300.2;BatCoV_RaTG13_spike;1223 | 0 | 0 |
| MN996532.1;QHR63300.2;BatCoV_RaTG13_spike;1224 | 0 | 0 |
| MN996532.1;QHR63300.2;BatCoV_RaTG13_spike;1225 | 0 | 0 |
| MN996532.1;QHR63300.2;BatCoV_RaTG13_spike;1226 | 0 | 0 |
| MN996532.1;QHR63300.2;BatCoV_RaTG13_spike;1227 | 0 | 0 |
| MN996532.1;QHR63300.2;BatCoV_RaTG13_spike;1228 | 0 | 0 |
| MN996532.1;QHR63300.2;BatCoV_RaTG13_spike;1229 | 0 | 0 |
| MN996532.1;QHR63300.2;BatCoV_RaTG13_spike;1230 | 0 | 0 |
| MN996532.1;QHR63300.2;BatCoV_RaTG13_spike;1231 | 0 | 0 |
| MN996532.1;QHR63300.2;BatCoV_RaTG13_spike;1232 | 0 | 0 |
| MN996532.1;QHR63300.2;BatCoV_RaTG13_spike;1233 | 0 | 0 |
| MN996532.1;QHR63300.2;BatCoV_RaTG13_spike;1234 | 0 | 0 |
| MN996532.1;QHR63300.2;BatCoV_RaTG13_spike;1235 | 0 | 0 |
| MN996532.1;QHR63300.2;BatCoV_RaTG13_spike;1236 | 0 | 0 |
| MN996532.1;QHR63300.2;BatCoV_RaTG13_spike;1237 | 0 | 0 |
| MN996532.1;QHR63300.2;BatCoV_RaTG13_spike;1238 | 0 | 0 |
| MN996532.1;QHR63300.2;BatCoV_RaTG13_spike;1239 | 0 | 0 |
| MN996532.1;QHR63300.2;BatCoV_RaTG13_spike;1240 | 0 | 0 |
| MN996532.1;QHR63300.2;BatCoV_RaTG13_spike;1241 | 0 | 0.075 |
| MN996532.1;QHR63300.2;BatCoV_RaTG13_spike;1242 | 0 | 0.1 |
| MN996532.1;QHR63300.2;BatCoV_RaTG13_spike;1243 | 0 | 0.25 |
| MN996532.1;QHR63300.2;BatCoV_RaTG13_spike;1244 | 0 | 0.5 |
| MN996532.1;QHR63300.2;BatCoV_RaTG13_spike;1245 | 0 | 0.55 |
| MN996532.1;QHR63300.2;BatCoV_RaTG13_spike;1246 | 0 | 0.575 |
| MN996532.1;QHR63300.2;BatCoV_RaTG13_spike;1247 | 0 | 0.65 |
| MN996532.1;QHR63300.2;BatCoV_RaTG13_spike;1248 | 0 | 0.65 |
| MN996532.1;QHR63300.2;BatCoV_RaTG13_spike;1249 | 0 | 0.675 |

| | | |
|---|---|---|
| MN996532.1;QHR63300.2;BatCoV_RaTG13_spike;1250 | 0.1 | 0.65 |
| MN996532.1;QHR63300.2;BatCoV_RaTG13_spike;1251 | 0.05 | 0.6 |
| MN996532.1;QHR63300.2;BatCoV_RaTG13_spike;1252 | 0.05 | 0.45 |
| MN996532.1;QHR63300.2;BatCoV_RaTG13_spike;1253 | 0.05 | 0.475 |
| MN996532.1;QHR63300.2;BatCoV_RaTG13_spike;1254 | 0.1 | 0.3 |
| MN996532.1;QHR63300.2;BatCoV_RaTG13_spike;1255 | 0.05 | 0.225 |
| MN996532.1;QHR63300.2;BatCoV_RaTG13_spike;1256 | 0.05 | 0.225 |
| MN996532.1;QHR63300.2;BatCoV_RaTG13_spike;1257 | 0.05 | 0.175 |
| MN996532.1;QHR63300.2;BatCoV_RaTG13_spike;1258 | 0 | 0.175 |
| MN996532.1;QHR63301.1;BatCoV_RaTG13_orf3;1 | 0 | 0 |
| MN996532.1;QHR63301.1;BatCoV_RaTG13_orf3;2 | 0 | 0 |
| MN996532.1;QHR63301.1;BatCoV_RaTG13_orf3;3 | 0 | 0 |
| MN996532.1;QHR63301.1;BatCoV_RaTG13_orf3;4 | 0 | 0 |
| MN996532.1;QHR63301.1;BatCoV_RaTG13_orf3;5 | 0 | 0 |
| MN996532.1;QHR63301.1;BatCoV_RaTG13_orf3;6 | 0 | 0.025 |
| MN996532.1;QHR63301.1;BatCoV_RaTG13_orf3;7 | 0 | 0.125 |
| MN996532.1;QHR63301.1;BatCoV_RaTG13_orf3;8 | 0.05 | 0.1 |
| MN996532.1;QHR63301.1;BatCoV_RaTG13_orf3;9 | 0 | 0.15 |
| MN996532.1;QHR63301.1;BatCoV_RaTG13_orf3;10 | 0.05 | 0.2 |
| MN996532.1;QHR63301.1;BatCoV_RaTG13_orf3;11 | 0 | 0.175 |
| MN996532.1;QHR63301.1;BatCoV_RaTG13_orf3;12 | 0.05 | 0.175 |
| MN996532.1;QHR63301.1;BatCoV_RaTG13_orf3;13 | 0.05 | 0.2 |
| MN996532.1;QHR63301.1;BatCoV_RaTG13_orf3;14 | 0.05 | 0.175 |
| MN996532.1;QHR63301.1;BatCoV_RaTG13_orf3;15 | 0.05 | 0.225 |
| MN996532.1;QHR63301.1;BatCoV_RaTG13_orf3;16 | 0 | 0.225 |
| MN996532.1;QHR63301.1;BatCoV_RaTG13_orf3;17 | 0 | 0.05 |
| MN996532.1;QHR63301.1;BatCoV_RaTG13_orf3;18 | 0 | 0.075 |
| MN996532.1;QHR63301.1;BatCoV_RaTG13_orf3;19 | 0 | 0.05 |
| MN996532.1;QHR63301.1;BatCoV_RaTG13_orf3;20 | 0 | 0.05 |
| MN996532.1;QHR63301.1;BatCoV_RaTG13_orf3;21 | 0 | 0.05 |
| MN996532.1;QHR63301.1;BatCoV_RaTG13_orf3;22 | 0 | 0 |
| MN996532.1;QHR63301.1;BatCoV_RaTG13_orf3;23 | 0 | 0 |
| MN996532.1;QHR63301.1;BatCoV_RaTG13_orf3;24 | 0 | 0 |
| MN996532.1;QHR63301.1;BatCoV_RaTG13_orf3;25 | 0 | 0 |
| MN996532.1;QHR63301.1;BatCoV_RaTG13_orf3;26 | 0 | 0.025 |
| MN996532.1;QHR63301.1;BatCoV_RaTG13_orf3;27 | 0 | 0.05 |
| MN996532.1;QHR63301.1;BatCoV_RaTG13_orf3;28 | 0 | 0 |
| MN996532.1;QHR63301.1;BatCoV_RaTG13_orf3;29 | 0 | 0.025 |
| MN996532.1;QHR63301.1;BatCoV_RaTG13_orf3;30 | 0 | 0.025 |
| MN996532.1;QHR63301.1;BatCoV_RaTG13_orf3;31 | 0 | 0 |
| MN996532.1;QHR63301.1;BatCoV_RaTG13_orf3;32 | 0 | 0 |
| MN996532.1;QHR63301.1;BatCoV_RaTG13_orf3;33 | 0 | 0 |
| MN996532.1;QHR63301.1;BatCoV_RaTG13_orf3;34 | 0 | 0 |

| | | |
|---|---|---|
| MN996532.1;QHR63301.1;BatCoV_RaTG13_orf3;35 | 0 | 0 |
| MN996532.1;QHR63301.1;BatCoV_RaTG13_orf3;36 | 0.05 | 0 |
| MN996532.1;QHR63301.1;BatCoV_RaTG13_orf3;37 | 0 | 0 |
| MN996532.1;QHR63301.1;BatCoV_RaTG13_orf3;38 | 0 | 0 |
| MN996532.1;QHR63301.1;BatCoV_RaTG13_orf3;39 | 0 | 0 |
| MN996532.1;QHR63301.1;BatCoV_RaTG13_orf3;40 | 0 | 0 |
| MN996532.1;QHR63301.1;BatCoV_RaTG13_orf3;41 | 0 | 0 |
| MN996532.1;QHR63301.1;BatCoV_RaTG13_orf3;42 | 0 | 0 |
| MN996532.1;QHR63301.1;BatCoV_RaTG13_orf3;43 | 0 | 0 |
| MN996532.1;QHR63301.1;BatCoV_RaTG13_orf3;44 | 0 | 0 |
| MN996532.1;QHR63301.1;BatCoV_RaTG13_orf3;45 | 0 | 0 |
| MN996532.1;QHR63301.1;BatCoV_RaTG13_orf3;46 | 0 | 0 |
| MN996532.1;QHR63301.1;BatCoV_RaTG13_orf3;47 | 0 | 0 |
| MN996532.1;QHR63301.1;BatCoV_RaTG13_orf3;48 | 0 | 0 |
| MN996532.1;QHR63301.1;BatCoV_RaTG13_orf3;49 | 0 | 0 |
| MN996532.1;QHR63301.1;BatCoV_RaTG13_orf3;50 | 0 | 0 |
| MN996532.1;QHR63301.1;BatCoV_RaTG13_orf3;51 | 0 | 0 |
| MN996532.1;QHR63301.1;BatCoV_RaTG13_orf3;52 | 0 | 0 |
| MN996532.1;QHR63301.1;BatCoV_RaTG13_orf3;53 | 0 | 0 |
| MN996532.1;QHR63301.1;BatCoV_RaTG13_orf3;54 | 0 | 0 |
| MN996532.1;QHR63301.1;BatCoV_RaTG13_orf3;55 | 0 | 0 |
| MN996532.1;QHR63301.1;BatCoV_RaTG13_orf3;56 | 0 | 0 |
| MN996532.1;QHR63301.1;BatCoV_RaTG13_orf3;57 | 0 | 0 |
| MN996532.1;QHR63301.1;BatCoV_RaTG13_orf3;58 | 0 | 0 |
| MN996532.1;QHR63301.1;BatCoV_RaTG13_orf3;59 | 0 | 0 |
| MN996532.1;QHR63301.1;BatCoV_RaTG13_orf3;60 | 0 | 0 |
| MN996532.1;QHR63301.1;BatCoV_RaTG13_orf3;61 | 0 | 0 |
| MN996532.1;QHR63301.1;BatCoV_RaTG13_orf3;62 | 0 | 0 |
| MN996532.1;QHR63301.1;BatCoV_RaTG13_orf3;63 | 0 | 0 |
| MN996532.1;QHR63301.1;BatCoV_RaTG13_orf3;64 | 0 | 0 |
| MN996532.1;QHR63301.1;BatCoV_RaTG13_orf3;65 | 0 | 0 |
| MN996532.1;QHR63301.1;BatCoV_RaTG13_orf3;66 | 0 | 0 |
| MN996532.1;QHR63301.1;BatCoV_RaTG13_orf3;67 | 0 | 0 |
| MN996532.1;QHR63301.1;BatCoV_RaTG13_orf3;68 | 0 | 0 |
| MN996532.1;QHR63301.1;BatCoV_RaTG13_orf3;69 | 0 | 0 |
| MN996532.1;QHR63301.1;BatCoV_RaTG13_orf3;70 | 0 | 0 |
| MN996532.1;QHR63301.1;BatCoV_RaTG13_orf3;71 | 0 | 0 |
| MN996532.1;QHR63301.1;BatCoV_RaTG13_orf3;72 | 0 | 0 |
| MN996532.1;QHR63301.1;BatCoV_RaTG13_orf3;73 | 0 | 0 |
| MN996532.1;QHR63301.1;BatCoV_RaTG13_orf3;74 | 0 | 0 |
| MN996532.1;QHR63301.1;BatCoV_RaTG13_orf3;75 | 0 | 0 |
| MN996532.1;QHR63301.1;BatCoV_RaTG13_orf3;76 | 0 | 0 |
| MN996532.1;QHR63301.1;BatCoV_RaTG13_orf3;77 | 0 | 0 |

| | | |
|---|---|---|
| MN996532.1;QHR63301.1;BatCoV_RaTG13_orf3;78 | 0 | 0 |
| MN996532.1;QHR63301.1;BatCoV_RaTG13_orf3;79 | 0 | 0 |
| MN996532.1;QHR63301.1;BatCoV_RaTG13_orf3;80 | 0 | 0 |
| MN996532.1;QHR63301.1;BatCoV_RaTG13_orf3;81 | 0 | 0 |
| MN996532.1;QHR63301.1;BatCoV_RaTG13_orf3;82 | 0 | 0 |
| MN996532.1;QHR63301.1;BatCoV_RaTG13_orf3;83 | 0 | 0 |
| MN996532.1;QHR63301.1;BatCoV_RaTG13_orf3;84 | 0 | 0 |
| MN996532.1;QHR63301.1;BatCoV_RaTG13_orf3;85 | 0 | 0 |
| MN996532.1;QHR63301.1;BatCoV_RaTG13_orf3;86 | 0 | 0 |
| MN996532.1;QHR63301.1;BatCoV_RaTG13_orf3;87 | 0 | 0 |
| MN996532.1;QHR63301.1;BatCoV_RaTG13_orf3;88 | 0 | 0 |
| MN996532.1;QHR63301.1;BatCoV_RaTG13_orf3;89 | 0 | 0 |
| MN996532.1;QHR63301.1;BatCoV_RaTG13_orf3;90 | 0 | 0 |
| MN996532.1;QHR63301.1;BatCoV_RaTG13_orf3;91 | 0 | 0 |
| MN996532.1;QHR63301.1;BatCoV_RaTG13_orf3;92 | 0 | 0 |
| MN996532.1;QHR63301.1;BatCoV_RaTG13_orf3;93 | 0 | 0 |
| MN996532.1;QHR63301.1;BatCoV_RaTG13_orf3;94 | 0 | 0 |
| MN996532.1;QHR63301.1;BatCoV_RaTG13_orf3;95 | 0 | 0 |
| MN996532.1;QHR63301.1;BatCoV_RaTG13_orf3;96 | 0 | 0 |
| MN996532.1;QHR63301.1;BatCoV_RaTG13_orf3;97 | 0 | 0 |
| MN996532.1;QHR63301.1;BatCoV_RaTG13_orf3;98 | 0 | 0 |
| MN996532.1;QHR63301.1;BatCoV_RaTG13_orf3;99 | 0 | 0 |
| MN996532.1;QHR63301.1;BatCoV_RaTG13_orf3;100 | 0 | 0 |
| MN996532.1;QHR63301.1;BatCoV_RaTG13_orf3;101 | 0 | 0 |
| MN996532.1;QHR63301.1;BatCoV_RaTG13_orf3;102 | 0 | 0 |
| MN996532.1;QHR63301.1;BatCoV_RaTG13_orf3;103 | 0 | 0 |
| MN996532.1;QHR63301.1;BatCoV_RaTG13_orf3;104 | 0 | 0 |
| MN996532.1;QHR63301.1;BatCoV_RaTG13_orf3;105 | 0 | 0 |
| MN996532.1;QHR63301.1;BatCoV_RaTG13_orf3;106 | 0 | 0 |
| MN996532.1;QHR63301.1;BatCoV_RaTG13_orf3;107 | 0 | 0 |
| MN996532.1;QHR63301.1;BatCoV_RaTG13_orf3;108 | 0 | 0 |
| MN996532.1;QHR63301.1;BatCoV_RaTG13_orf3;109 | 0 | 0 |
| MN996532.1;QHR63301.1;BatCoV_RaTG13_orf3;110 | 0 | 0 |
| MN996532.1;QHR63301.1;BatCoV_RaTG13_orf3;111 | 0 | 0 |
| MN996532.1;QHR63301.1;BatCoV_RaTG13_orf3;112 | 0 | 0 |
| MN996532.1;QHR63301.1;BatCoV_RaTG13_orf3;113 | 0 | 0 |
| MN996532.1;QHR63301.1;BatCoV_RaTG13_orf3;114 | 0 | 0 |
| MN996532.1;QHR63301.1;BatCoV_RaTG13_orf3;115 | 0 | 0 |
| MN996532.1;QHR63301.1;BatCoV_RaTG13_orf3;116 | 0 | 0 |
| MN996532.1;QHR63301.1;BatCoV_RaTG13_orf3;117 | 0 | 0 |
| MN996532.1;QHR63301.1;BatCoV_RaTG13_orf3;118 | 0 | 0 |
| MN996532.1;QHR63301.1;BatCoV_RaTG13_orf3;119 | 0 | 0 |
| MN996532.1;QHR63301.1;BatCoV_RaTG13_orf3;120 | 0 | 0 |

| | | |
|---|---|---|
| MN996532.1;QHR63301.1;BatCoV_RaTG13_orf3;121 | 0 | 0 |
| MN996532.1;QHR63301.1;BatCoV_RaTG13_orf3;122 | 0 | 0 |
| MN996532.1;QHR63301.1;BatCoV_RaTG13_orf3;123 | 0 | 0 |
| MN996532.1;QHR63301.1;BatCoV_RaTG13_orf3;124 | 0 | 0 |
| MN996532.1;QHR63301.1;BatCoV_RaTG13_orf3;125 | 0 | 0 |
| MN996532.1;QHR63301.1;BatCoV_RaTG13_orf3;126 | 0 | 0 |
| MN996532.1;QHR63301.1;BatCoV_RaTG13_orf3;127 | 0 | 0 |
| MN996532.1;QHR63301.1;BatCoV_RaTG13_orf3;128 | 0 | 0 |
| MN996532.1;QHR63301.1;BatCoV_RaTG13_orf3;129 | 0 | 0 |
| MN996532.1;QHR63301.1;BatCoV_RaTG13_orf3;130 | 0 | 0 |
| MN996532.1;QHR63301.1;BatCoV_RaTG13_orf3;131 | 0 | 0 |
| MN996532.1;QHR63301.1;BatCoV_RaTG13_orf3;132 | 0 | 0 |
| MN996532.1;QHR63301.1;BatCoV_RaTG13_orf3;133 | 0 | 0 |
| MN996532.1;QHR63301.1;BatCoV_RaTG13_orf3;134 | 0 | 0 |
| MN996532.1;QHR63301.1;BatCoV_RaTG13_orf3;135 | 0 | 0 |
| MN996532.1;QHR63301.1;BatCoV_RaTG13_orf3;136 | 0 | 0 |
| MN996532.1;QHR63301.1;BatCoV_RaTG13_orf3;137 | 0 | 0 |
| MN996532.1;QHR63301.1;BatCoV_RaTG13_orf3;138 | 0 | 0 |
| MN996532.1;QHR63301.1;BatCoV_RaTG13_orf3;139 | 0 | 0 |
| MN996532.1;QHR63301.1;BatCoV_RaTG13_orf3;140 | 0 | 0 |
| MN996532.1;QHR63301.1;BatCoV_RaTG13_orf3;141 | 0 | 0 |
| MN996532.1;QHR63301.1;BatCoV_RaTG13_orf3;142 | 0 | 0 |
| MN996532.1;QHR63301.1;BatCoV_RaTG13_orf3;143 | 0 | 0 |
| MN996532.1;QHR63301.1;BatCoV_RaTG13_orf3;144 | 0 | 0 |
| MN996532.1;QHR63301.1;BatCoV_RaTG13_orf3;145 | 0 | 0 |
| MN996532.1;QHR63301.1;BatCoV_RaTG13_orf3;146 | 0 | 0 |
| MN996532.1;QHR63301.1;BatCoV_RaTG13_orf3;147 | 0 | 0 |
| MN996532.1;QHR63301.1;BatCoV_RaTG13_orf3;148 | 0 | 0 |
| MN996532.1;QHR63301.1;BatCoV_RaTG13_orf3;149 | 0 | 0 |
| MN996532.1;QHR63301.1;BatCoV_RaTG13_orf3;150 | 0 | 0 |
| MN996532.1;QHR63301.1;BatCoV_RaTG13_orf3;151 | 0 | 0 |
| MN996532.1;QHR63301.1;BatCoV_RaTG13_orf3;152 | 0 | 0 |
| MN996532.1;QHR63301.1;BatCoV_RaTG13_orf3;153 | 0.05 | 0 |
| MN996532.1;QHR63301.1;BatCoV_RaTG13_orf3;154 | 0 | 0 |
| MN996532.1;QHR63301.1;BatCoV_RaTG13_orf3;155 | 0 | 0 |
| MN996532.1;QHR63301.1;BatCoV_RaTG13_orf3;156 | 0 | 0 |
| MN996532.1;QHR63301.1;BatCoV_RaTG13_orf3;157 | 0 | 0 |
| MN996532.1;QHR63301.1;BatCoV_RaTG13_orf3;158 | 0 | 0 |
| MN996532.1;QHR63301.1;BatCoV_RaTG13_orf3;159 | 0 | 0 |
| MN996532.1;QHR63301.1;BatCoV_RaTG13_orf3;160 | 0 | 0 |
| MN996532.1;QHR63301.1;BatCoV_RaTG13_orf3;161 | 0 | 0 |
| MN996532.1;QHR63301.1;BatCoV_RaTG13_orf3;162 | 0 | 0.025 |
| MN996532.1;QHR63301.1;BatCoV_RaTG13_orf3;163 | 0 | 0 |

| | | |
|---|---|---|
| MN996532.1;QHR63301.1;BatCoV_RaTG13_orf3;164 | 0 | 0 |
| MN996532.1;QHR63301.1;BatCoV_RaTG13_orf3;165 | 0 | 0 |
| MN996532.1;QHR63301.1;BatCoV_RaTG13_orf3;166 | 0 | 0 |
| MN996532.1;QHR63301.1;BatCoV_RaTG13_orf3;167 | 0 | 0.025 |
| MN996532.1;QHR63301.1;BatCoV_RaTG13_orf3;168 | 0 | 0.025 |
| MN996532.1;QHR63301.1;BatCoV_RaTG13_orf3;169 | 0 | 0.025 |
| MN996532.1;QHR63301.1;BatCoV_RaTG13_orf3;170 | 0 | 0.025 |
| MN996532.1;QHR63301.1;BatCoV_RaTG13_orf3;171 | 0 | 0.025 |
| MN996532.1;QHR63301.1;BatCoV_RaTG13_orf3;172 | 0 | 0.025 |
| MN996532.1;QHR63301.1;BatCoV_RaTG13_orf3;173 | 0 | 0.075 |
| MN996532.1;QHR63301.1;BatCoV_RaTG13_orf3;174 | 0 | 0.025 |
| MN996532.1;QHR63301.1;BatCoV_RaTG13_orf3;175 | 0 | 0.025 |
| MN996532.1;QHR63301.1;BatCoV_RaTG13_orf3;176 | 0.05 | 0.025 |
| MN996532.1;QHR63301.1;BatCoV_RaTG13_orf3;177 | 0.05 | 0.1 |
| MN996532.1;QHR63301.1;BatCoV_RaTG13_orf3;178 | 0.05 | 0.075 |
| MN996532.1;QHR63301.1;BatCoV_RaTG13_orf3;179 | 0.05 | 0.05 |
| MN996532.1;QHR63301.1;BatCoV_RaTG13_orf3;180 | 0.05 | 0.05 |
| MN996532.1;QHR63301.1;BatCoV_RaTG13_orf3;181 | 0.1 | 0.05 |
| MN996532.1;QHR63301.1;BatCoV_RaTG13_orf3;182 | 0.05 | 0.05 |
| MN996532.1;QHR63301.1;BatCoV_RaTG13_orf3;183 | 0.05 | 0.15 |
| MN996532.1;QHR63301.1;BatCoV_RaTG13_orf3;184 | 0.1 | 0.075 |
| MN996532.1;QHR63301.1;BatCoV_RaTG13_orf3;185 | 0.05 | 0.05 |
| MN996532.1;QHR63301.1;BatCoV_RaTG13_orf3;186 | 0 | 0.05 |
| MN996532.1;QHR63301.1;BatCoV_RaTG13_orf3;187 | 0 | 0 |
| MN996532.1;QHR63301.1;BatCoV_RaTG13_orf3;188 | 0 | 0 |
| MN996532.1;QHR63301.1;BatCoV_RaTG13_orf3;189 | 0 | 0 |
| MN996532.1;QHR63301.1;BatCoV_RaTG13_orf3;190 | 0 | 0 |
| MN996532.1;QHR63301.1;BatCoV_RaTG13_orf3;191 | 0 | 0 |
| MN996532.1;QHR63301.1;BatCoV_RaTG13_orf3;192 | 0 | 0 |
| MN996532.1;QHR63301.1;BatCoV_RaTG13_orf3;193 | 0 | 0 |
| MN996532.1;QHR63301.1;BatCoV_RaTG13_orf3;194 | 0 | 0.025 |
| MN996532.1;QHR63301.1;BatCoV_RaTG13_orf3;195 | 0 | 0 |
| MN996532.1;QHR63301.1;BatCoV_RaTG13_orf3;196 | 0 | 0.05 |
| MN996532.1;QHR63301.1;BatCoV_RaTG13_orf3;197 | 0 | 0 |
| MN996532.1;QHR63301.1;BatCoV_RaTG13_orf3;198 | 0 | 0 |
| MN996532.1;QHR63301.1;BatCoV_RaTG13_orf3;199 | 0 | 0 |
| MN996532.1;QHR63301.1;BatCoV_RaTG13_orf3;200 | 0 | 0 |
| MN996532.1;QHR63301.1;BatCoV_RaTG13_orf3;201 | 0 | 0 |
| MN996532.1;QHR63301.1;BatCoV_RaTG13_orf3;202 | 0 | 0 |
| MN996532.1;QHR63301.1;BatCoV_RaTG13_orf3;203 | 0 | 0 |
| MN996532.1;QHR63301.1;BatCoV_RaTG13_orf3;204 | 0 | 0 |
| MN996532.1;QHR63301.1;BatCoV_RaTG13_orf3;205 | 0 | 0 |
| MN996532.1;QHR63301.1;BatCoV_RaTG13_orf3;206 | 0 | 0 |

| | | |
|---|---|---|
| MN996532.1;QHR63301.1;BatCoV_RaTG13_orf3;207 | 0 | 0.025 |
| MN996532.1;QHR63301.1;BatCoV_RaTG13_orf3;208 | 0 | 0.025 |
| MN996532.1;QHR63301.1;BatCoV_RaTG13_orf3;209 | 0.05 | 0.025 |
| MN996532.1;QHR63301.1;BatCoV_RaTG13_orf3;210 | 0 | 0.025 |
| MN996532.1;QHR63301.1;BatCoV_RaTG13_orf3;211 | 0 | 0.025 |
| MN996532.1;QHR63301.1;BatCoV_RaTG13_orf3;212 | 0 | 0.025 |
| MN996532.1;QHR63301.1;BatCoV_RaTG13_orf3;213 | 0 | 0.025 |
| MN996532.1;QHR63301.1;BatCoV_RaTG13_orf3;214 | 0 | 0.025 |
| MN996532.1;QHR63301.1;BatCoV_RaTG13_orf3;215 | 0 | 0 |
| MN996532.1;QHR63301.1;BatCoV_RaTG13_orf3;216 | 0 | 0 |
| MN996532.1;QHR63301.1;BatCoV_RaTG13_orf3;217 | 0 | 0 |
| MN996532.1;QHR63301.1;BatCoV_RaTG13_orf3;218 | 0 | 0 |
| MN996532.1;QHR63301.1;BatCoV_RaTG13_orf3;219 | 0 | 0 |
| MN996532.1;QHR63301.1;BatCoV_RaTG13_orf3;220 | 0.05 | 0 |
| MN996532.1;QHR63301.1;BatCoV_RaTG13_orf3;221 | 0 | 0 |
| MN996532.1;QHR63301.1;BatCoV_RaTG13_orf3;222 | 0 | 0 |
| MN996532.1;QHR63301.1;BatCoV_RaTG13_orf3;223 | 0 | 0 |
| MN996532.1;QHR63301.1;BatCoV_RaTG13_orf3;224 | 0 | 0 |
| MN996532.1;QHR63301.1;BatCoV_RaTG13_orf3;225 | 0 | 0 |
| MN996532.1;QHR63301.1;BatCoV_RaTG13_orf3;226 | 0 | 0 |
| MN996532.1;QHR63301.1;BatCoV_RaTG13_orf3;227 | 0 | 0 |
| MN996532.1;QHR63301.1;BatCoV_RaTG13_orf3;228 | 0 | 0 |
| MN996532.1;QHR63301.1;BatCoV_RaTG13_orf3;229 | 0 | 0 |
| MN996532.1;QHR63301.1;BatCoV_RaTG13_orf3;230 | 0 | 0.075 |
| MN996532.1;QHR63301.1;BatCoV_RaTG13_orf3;231 | 0.05 | 0.025 |
| MN996532.1;QHR63301.1;BatCoV_RaTG13_orf3;232 | 0 | 0.05 |
| MN996532.1;QHR63301.1;BatCoV_RaTG13_orf3;233 | 0 | 0.075 |
| MN996532.1;QHR63301.1;BatCoV_RaTG13_orf3;234 | 0 | 0 |
| MN996532.1;QHR63301.1;BatCoV_RaTG13_orf3;235 | 0 | 0.05 |
| MN996532.1;QHR63301.1;BatCoV_RaTG13_orf3;236 | 0 | 0.025 |
| MN996532.1;QHR63301.1;BatCoV_RaTG13_orf3;237 | 0 | 0.05 |
| MN996532.1;QHR63301.1;BatCoV_RaTG13_orf3;238 | 0 | 0.075 |
| MN996532.1;QHR63301.1;BatCoV_RaTG13_orf3;239 | 0 | 0.05 |
| MN996532.1;QHR63301.1;BatCoV_RaTG13_orf3;240 | 0 | 0.1 |
| MN996532.1;QHR63301.1;BatCoV_RaTG13_orf3;241 | 0.05 | 0.05 |
| MN996532.1;QHR63301.1;BatCoV_RaTG13_orf3;242 | 0.05 | 0.05 |
| MN996532.1;QHR63301.1;BatCoV_RaTG13_orf3;243 | 0.05 | 0.1 |
| MN996532.1;QHR63301.1;BatCoV_RaTG13_orf3;244 | 0.05 | 0.05 |
| MN996532.1;QHR63301.1;BatCoV_RaTG13_orf3;245 | 0.05 | 0.075 |
| MN996532.1;QHR63301.1;BatCoV_RaTG13_orf3;246 | 0.05 | 0.05 |
| MN996532.1;QHR63301.1;BatCoV_RaTG13_orf3;247 | 0.05 | 0.05 |
| MN996532.1;QHR63301.1;BatCoV_RaTG13_orf3;248 | 0.05 | 0.025 |
| MN996532.1;QHR63301.1;BatCoV_RaTG13_orf3;249 | 0.05 | 0.05 |

| | | |
|---|---|---|
| MN996532.1;QHR63301.1;BatCoV_RaTG13_orf3;250 | 0.05 | 0.1 |
| MN996532.1;QHR63301.1;BatCoV_RaTG13_orf3;251 | 0.05 | 0.1 |
| MN996532.1;QHR63301.1;BatCoV_RaTG13_orf3;252 | 0.05 | 0.225 |
| MN996532.1;QHR63301.1;BatCoV_RaTG13_orf3;253 | 0 | 0.275 |
| MN996532.1;QHR63301.1;BatCoV_RaTG13_orf3;254 | 0 | 0.25 |
| MN996532.1;QHR63301.1;BatCoV_RaTG13_orf3;255 | 0 | 0.275 |
| MN996532.1;QHR63301.1;BatCoV_RaTG13_orf3;256 | 0 | 0.3 |
| MN996532.1;QHR63301.1;BatCoV_RaTG13_orf3;257 | 0 | 0.375 |
| MN996532.1;QHR63301.1;BatCoV_RaTG13_orf3;258 | 0 | 0.225 |
| MN996532.1;QHR63301.1;BatCoV_RaTG13_orf3;259 | 0.05 | 0.375 |
| MN996532.1;QHR63301.1;BatCoV_RaTG13_orf3;260 | 0.05 | 0.35 |
| MN996532.1;QHR63301.1;BatCoV_RaTG13_orf3;261 | 0.05 | 0.375 |
| MN996532.1;QHR63301.1;BatCoV_RaTG13_orf3;262 | 0 | 0.075 |
| MN996532.1;QHR63301.1;BatCoV_RaTG13_orf3;263 | 0.05 | 0.1 |
| MN996532.1;QHR63301.1;BatCoV_RaTG13_orf3;264 | 0 | 0.05 |
| MN996532.1;QHR63302.1;BatCoV_RaTG13_envelop;1 | 0 | 0 |
| MN996532.1;QHR63302.1;BatCoV_RaTG13_envelop;2 | 0 | 0 |
| MN996532.1;QHR63302.1;BatCoV_RaTG13_envelop;3 | 0 | 0 |
| MN996532.1;QHR63302.1;BatCoV_RaTG13_envelop;4 | 0 | 0 |
| MN996532.1;QHR63302.1;BatCoV_RaTG13_envelop;5 | 0 | 0 |
| MN996532.1;QHR63302.1;BatCoV_RaTG13_envelop;6 | 0 | 0 |
| MN996532.1;QHR63302.1;BatCoV_RaTG13_envelop;7 | 0 | 0 |
| MN996532.1;QHR63302.1;BatCoV_RaTG13_envelop;8 | 0 | 0 |
| MN996532.1;QHR63302.1;BatCoV_RaTG13_envelop;9 | 0 | 0 |
| MN996532.1;QHR63302.1;BatCoV_RaTG13_envelop;10 | 0 | 0 |
| MN996532.1;QHR63302.1;BatCoV_RaTG13_envelop;11 | 0 | 0 |
| MN996532.1;QHR63302.1;BatCoV_RaTG13_envelop;12 | 0 | 0 |
| MN996532.1;QHR63302.1;BatCoV_RaTG13_envelop;13 | 0 | 0 |
| MN996532.1;QHR63302.1;BatCoV_RaTG13_envelop;14 | 0 | 0 |
| MN996532.1;QHR63302.1;BatCoV_RaTG13_envelop;15 | 0 | 0 |
| MN996532.1;QHR63302.1;BatCoV_RaTG13_envelop;16 | 0 | 0 |
| MN996532.1;QHR63302.1;BatCoV_RaTG13_envelop;17 | 0 | 0 |
| MN996532.1;QHR63302.1;BatCoV_RaTG13_envelop;18 | 0 | 0 |
| MN996532.1;QHR63302.1;BatCoV_RaTG13_envelop;19 | 0 | 0 |
| MN996532.1;QHR63302.1;BatCoV_RaTG13_envelop;20 | 0 | 0 |
| MN996532.1;QHR63302.1;BatCoV_RaTG13_envelop;21 | 0 | 0 |
| MN996532.1;QHR63302.1;BatCoV_RaTG13_envelop;22 | 0 | 0 |
| MN996532.1;QHR63302.1;BatCoV_RaTG13_envelop;23 | 0 | 0 |
| MN996532.1;QHR63302.1;BatCoV_RaTG13_envelop;24 | 0 | 0 |
| MN996532.1;QHR63302.1;BatCoV_RaTG13_envelop;25 | 0 | 0 |
| MN996532.1;QHR63302.1;BatCoV_RaTG13_envelop;26 | 0 | 0 |
| MN996532.1;QHR63302.1;BatCoV_RaTG13_envelop;27 | 0 | 0 |
| MN996532.1;QHR63302.1;BatCoV_RaTG13_envelop;28 | 0 | 0 |

| | | |
|---|---|---|
| MN996532.1;QHR63302.1;BatCoV_RaTG13_envelop;29 | 0 | 0 |
| MN996532.1;QHR63302.1;BatCoV_RaTG13_envelop;30 | 0 | 0 |
| MN996532.1;QHR63302.1;BatCoV_RaTG13_envelop;31 | 0 | 0 |
| MN996532.1;QHR63302.1;BatCoV_RaTG13_envelop;32 | 0 | 0 |
| MN996532.1;QHR63302.1;BatCoV_RaTG13_envelop;33 | 0 | 0 |
| MN996532.1;QHR63302.1;BatCoV_RaTG13_envelop;34 | 0 | 0 |
| MN996532.1;QHR63302.1;BatCoV_RaTG13_envelop;35 | 0 | 0 |
| MN996532.1;QHR63302.1;BatCoV_RaTG13_envelop;36 | 0 | 0 |
| MN996532.1;QHR63302.1;BatCoV_RaTG13_envelop;37 | 0 | 0 |
| MN996532.1;QHR63302.1;BatCoV_RaTG13_envelop;38 | 0 | 0 |
| MN996532.1;QHR63302.1;BatCoV_RaTG13_envelop;39 | 0 | 0 |
| MN996532.1;QHR63302.1;BatCoV_RaTG13_envelop;40 | 0 | 0.025 |
| MN996532.1;QHR63302.1;BatCoV_RaTG13_envelop;41 | 0 | 0 |
| MN996532.1;QHR63302.1;BatCoV_RaTG13_envelop;42 | 0 | 0 |
| MN996532.1;QHR63302.1;BatCoV_RaTG13_envelop;43 | 0 | 0 |
| MN996532.1;QHR63302.1;BatCoV_RaTG13_envelop;44 | 0 | 0 |
| MN996532.1;QHR63302.1;BatCoV_RaTG13_envelop;45 | 0 | 0 |
| MN996532.1;QHR63302.1;BatCoV_RaTG13_envelop;46 | 0 | 0 |
| MN996532.1;QHR63302.1;BatCoV_RaTG13_envelop;47 | 0 | 0 |
| MN996532.1;QHR63302.1;BatCoV_RaTG13_envelop;48 | 0 | 0 |
| MN996532.1;QHR63302.1;BatCoV_RaTG13_envelop;49 | 0 | 0 |
| MN996532.1;QHR63302.1;BatCoV_RaTG13_envelop;50 | 0 | 0 |
| MN996532.1;QHR63302.1;BatCoV_RaTG13_envelop;51 | 0 | 0 |
| MN996532.1;QHR63302.1;BatCoV_RaTG13_envelop;52 | 0 | 0 |
| MN996532.1;QHR63302.1;BatCoV_RaTG13_envelop;53 | 0 | 0 |
| MN996532.1;QHR63302.1;BatCoV_RaTG13_envelop;54 | 0 | 0 |
| MN996532.1;QHR63302.1;BatCoV_RaTG13_envelop;55 | 0 | 0 |
| MN996532.1;QHR63302.1;BatCoV_RaTG13_envelop;56 | 0 | 0 |
| MN996532.1;QHR63302.1;BatCoV_RaTG13_envelop;57 | 0 | 0 |
| MN996532.1;QHR63302.1;BatCoV_RaTG13_envelop;58 | 0 | 0.075 |
| MN996532.1;QHR63302.1;BatCoV_RaTG13_envelop;59 | 0 | 0 |
| MN996532.1;QHR63302.1;BatCoV_RaTG13_envelop;60 | 0.1 | 0 |
| MN996532.1;QHR63302.1;BatCoV_RaTG13_envelop;61 | 0.1 | 0 |
| MN996532.1;QHR63302.1;BatCoV_RaTG13_envelop;62 | 0.05 | 0.05 |
| MN996532.1;QHR63302.1;BatCoV_RaTG13_envelop;63 | 0.1 | 0 |
| MN996532.1;QHR63302.1;BatCoV_RaTG13_envelop;64 | 0.05 | 0.075 |
| MN996532.1;QHR63303.1;BatCoV_RaTG13_membran;1 | 0 | 0.35 |
| MN996532.1;QHR63303.1;BatCoV_RaTG13_membran;2 | 0 | 0.3 |
| MN996532.1;QHR63303.1;BatCoV_RaTG13_membran;3 | 0.05 | 0.6 |
| MN996532.1;QHR63303.1;BatCoV_RaTG13_membran;4 | 0 | 0.725 |
| MN996532.1;QHR63303.1;BatCoV_RaTG13_membran;5 | 0 | 0.65 |
| MN996532.1;QHR63303.1;BatCoV_RaTG13_membran;6 | 0 | 0.775 |
| MN996532.1;QHR63303.1;BatCoV_RaTG13_membran;7 | 0 | 0.9 |

| | | |
|---|---|---|
| MN996532.1;QHR63303.1;BatCoV_RaTG13_membran;8 | 0 | 0.4 |
| MN996532.1;QHR63303.1;BatCoV_RaTG13_membran;9 | 0 | 0.025 |
| MN996532.1;QHR63303.1;BatCoV_RaTG13_membran;10 | 0.05 | 0 |
| MN996532.1;QHR63303.1;BatCoV_RaTG13_membran;11 | 0 | 0 |
| MN996532.1;QHR63303.1;BatCoV_RaTG13_membran;12 | 0 | 0 |
| MN996532.1;QHR63303.1;BatCoV_RaTG13_membran;13 | 0 | 0 |
| MN996532.1;QHR63303.1;BatCoV_RaTG13_membran;14 | 0 | 0 |
| MN996532.1;QHR63303.1;BatCoV_RaTG13_membran;15 | 0 | 0 |
| MN996532.1;QHR63303.1;BatCoV_RaTG13_membran;16 | 0 | 0 |
| MN996532.1;QHR63303.1;BatCoV_RaTG13_membran;17 | 0 | 0 |
| MN996532.1;QHR63303.1;BatCoV_RaTG13_membran;18 | 0 | 0 |
| MN996532.1;QHR63303.1;BatCoV_RaTG13_membran;19 | 0 | 0 |
| MN996532.1;QHR63303.1;BatCoV_RaTG13_membran;20 | 0 | 0 |
| MN996532.1;QHR63303.1;BatCoV_RaTG13_membran;21 | 0 | 0 |
| MN996532.1;QHR63303.1;BatCoV_RaTG13_membran;22 | 0 | 0 |
| MN996532.1;QHR63303.1;BatCoV_RaTG13_membran;23 | 0 | 0 |
| MN996532.1;QHR63303.1;BatCoV_RaTG13_membran;24 | 0 | 0 |
| MN996532.1;QHR63303.1;BatCoV_RaTG13_membran;25 | 0 | 0 |
| MN996532.1;QHR63303.1;BatCoV_RaTG13_membran;26 | 0 | 0 |
| MN996532.1;QHR63303.1;BatCoV_RaTG13_membran;27 | 0 | 0 |
| MN996532.1;QHR63303.1;BatCoV_RaTG13_membran;28 | 0 | 0 |
| MN996532.1;QHR63303.1;BatCoV_RaTG13_membran;29 | 0 | 0 |
| MN996532.1;QHR63303.1;BatCoV_RaTG13_membran;30 | 0 | 0 |
| MN996532.1;QHR63303.1;BatCoV_RaTG13_membran;31 | 0 | 0 |
| MN996532.1;QHR63303.1;BatCoV_RaTG13_membran;32 | 0 | 0 |
| MN996532.1;QHR63303.1;BatCoV_RaTG13_membran;33 | 0 | 0 |
| MN996532.1;QHR63303.1;BatCoV_RaTG13_membran;34 | 0 | 0 |
| MN996532.1;QHR63303.1;BatCoV_RaTG13_membran;35 | 0 | 0 |
| MN996532.1;QHR63303.1;BatCoV_RaTG13_membran;36 | 0 | 0 |
| MN996532.1;QHR63303.1;BatCoV_RaTG13_membran;37 | 0 | 0 |
| MN996532.1;QHR63303.1;BatCoV_RaTG13_membran;38 | 0 | 0 |
| MN996532.1;QHR63303.1;BatCoV_RaTG13_membran;39 | 0 | 0 |
| MN996532.1;QHR63303.1;BatCoV_RaTG13_membran;40 | 0 | 0 |
| MN996532.1;QHR63303.1;BatCoV_RaTG13_membran;41 | 0 | 0 |
| MN996532.1;QHR63303.1;BatCoV_RaTG13_membran;42 | 0 | 0 |
| MN996532.1;QHR63303.1;BatCoV_RaTG13_membran;43 | 0 | 0 |
| MN996532.1;QHR63303.1;BatCoV_RaTG13_membran;44 | 0 | 0 |
| MN996532.1;QHR63303.1;BatCoV_RaTG13_membran;45 | 0 | 0 |
| MN996532.1;QHR63303.1;BatCoV_RaTG13_membran;46 | 0 | 0 |
| MN996532.1;QHR63303.1;BatCoV_RaTG13_membran;47 | 0 | 0 |
| MN996532.1;QHR63303.1;BatCoV_RaTG13_membran;48 | 0 | 0 |
| MN996532.1;QHR63303.1;BatCoV_RaTG13_membran;49 | 0 | 0 |
| MN996532.1;QHR63303.1;BatCoV_RaTG13_membran;50 | 0 | 0 |

| | | |
|---|---|---|
| MN996532.1;QHR63303.1;BatCoV_RaTG13_membran;51 | 0 | 0 |
| MN996532.1;QHR63303.1;BatCoV_RaTG13_membran;52 | 0 | 0 |
| MN996532.1;QHR63303.1;BatCoV_RaTG13_membran;53 | 0 | 0 |
| MN996532.1;QHR63303.1;BatCoV_RaTG13_membran;54 | 0 | 0 |
| MN996532.1;QHR63303.1;BatCoV_RaTG13_membran;55 | 0 | 0 |
| MN996532.1;QHR63303.1;BatCoV_RaTG13_membran;56 | 0 | 0 |
| MN996532.1;QHR63303.1;BatCoV_RaTG13_membran;57 | 0 | 0 |
| MN996532.1;QHR63303.1;BatCoV_RaTG13_membran;58 | 0 | 0 |
| MN996532.1;QHR63303.1;BatCoV_RaTG13_membran;59 | 0 | 0 |
| MN996532.1;QHR63303.1;BatCoV_RaTG13_membran;60 | 0 | 0 |
| MN996532.1;QHR63303.1;BatCoV_RaTG13_membran;61 | 0 | 0 |
| MN996532.1;QHR63303.1;BatCoV_RaTG13_membran;62 | 0 | 0 |
| MN996532.1;QHR63303.1;BatCoV_RaTG13_membran;63 | 0 | 0 |
| MN996532.1;QHR63303.1;BatCoV_RaTG13_membran;64 | 0 | 0 |
| MN996532.1;QHR63303.1;BatCoV_RaTG13_membran;65 | 0 | 0 |
| MN996532.1;QHR63303.1;BatCoV_RaTG13_membran;66 | 0 | 0 |
| MN996532.1;QHR63303.1;BatCoV_RaTG13_membran;67 | 0 | 0 |
| MN996532.1;QHR63303.1;BatCoV_RaTG13_membran;68 | 0 | 0 |
| MN996532.1;QHR63303.1;BatCoV_RaTG13_membran;69 | 0 | 0 |
| MN996532.1;QHR63303.1;BatCoV_RaTG13_membran;70 | 0 | 0 |
| MN996532.1;QHR63303.1;BatCoV_RaTG13_membran;71 | 0 | 0 |
| MN996532.1;QHR63303.1;BatCoV_RaTG13_membran;72 | 0 | 0 |
| MN996532.1;QHR63303.1;BatCoV_RaTG13_membran;73 | 0 | 0 |
| MN996532.1;QHR63303.1;BatCoV_RaTG13_membran;74 | 0 | 0 |
| MN996532.1;QHR63303.1;BatCoV_RaTG13_membran;75 | 0 | 0 |
| MN996532.1;QHR63303.1;BatCoV_RaTG13_membran;76 | 0 | 0 |
| MN996532.1;QHR63303.1;BatCoV_RaTG13_membran;77 | 0 | 0 |
| MN996532.1;QHR63303.1;BatCoV_RaTG13_membran;78 | 0 | 0 |
| MN996532.1;QHR63303.1;BatCoV_RaTG13_membran;79 | 0 | 0 |
| MN996532.1;QHR63303.1;BatCoV_RaTG13_membran;80 | 0 | 0 |
| MN996532.1;QHR63303.1;BatCoV_RaTG13_membran;81 | 0 | 0 |
| MN996532.1;QHR63303.1;BatCoV_RaTG13_membran;82 | 0 | 0 |
| MN996532.1;QHR63303.1;BatCoV_RaTG13_membran;83 | 0 | 0 |
| MN996532.1;QHR63303.1;BatCoV_RaTG13_membran;84 | 0 | 0 |
| MN996532.1;QHR63303.1;BatCoV_RaTG13_membran;85 | 0 | 0 |
| MN996532.1;QHR63303.1;BatCoV_RaTG13_membran;86 | 0 | 0 |
| MN996532.1;QHR63303.1;BatCoV_RaTG13_membran;87 | 0 | 0 |
| MN996532.1;QHR63303.1;BatCoV_RaTG13_membran;88 | 0 | 0 |
| MN996532.1;QHR63303.1;BatCoV_RaTG13_membran;89 | 0 | 0 |
| MN996532.1;QHR63303.1;BatCoV_RaTG13_membran;90 | 0 | 0 |
| MN996532.1;QHR63303.1;BatCoV_RaTG13_membran;91 | 0 | 0 |
| MN996532.1;QHR63303.1;BatCoV_RaTG13_membran;92 | 0 | 0 |
| MN996532.1;QHR63303.1;BatCoV_RaTG13_membran;93 | 0 | 0 |

| | | |
|---|---|---|
| MN996532.1;QHR63303.1;BatCoV_RaTG13_membran;94 | 0 | 0 |
| MN996532.1;QHR63303.1;BatCoV_RaTG13_membran;95 | 0 | 0 |
| MN996532.1;QHR63303.1;BatCoV_RaTG13_membran;96 | 0 | 0 |
| MN996532.1;QHR63303.1;BatCoV_RaTG13_membran;97 | 0 | 0 |
| MN996532.1;QHR63303.1;BatCoV_RaTG13_membran;98 | 0 | 0 |
| MN996532.1;QHR63303.1;BatCoV_RaTG13_membran;99 | 0 | 0 |
| MN996532.1;QHR63303.1;BatCoV_RaTG13_membran;100 | 0.05 | 0.025 |
| MN996532.1;QHR63303.1;BatCoV_RaTG13_membran;101 | 0 | 0 |
| MN996532.1;QHR63303.1;BatCoV_RaTG13_membran;102 | 0 | 0 |
| MN996532.1;QHR63303.1;BatCoV_RaTG13_membran;103 | 0 | 0 |
| MN996532.1;QHR63303.1;BatCoV_RaTG13_membran;104 | 0 | 0 |
| MN996532.1;QHR63303.1;BatCoV_RaTG13_membran;105 | 0 | 0 |
| MN996532.1;QHR63303.1;BatCoV_RaTG13_membran;106 | 0 | 0 |
| MN996532.1;QHR63303.1;BatCoV_RaTG13_membran;107 | 0 | 0.025 |
| MN996532.1;QHR63303.1;BatCoV_RaTG13_membran;108 | 0 | 0.025 |
| MN996532.1;QHR63303.1;BatCoV_RaTG13_membran;109 | 0 | 0 |
| MN996532.1;QHR63303.1;BatCoV_RaTG13_membran;110 | 0 | 0.025 |
| MN996532.1;QHR63303.1;BatCoV_RaTG13_membran;111 | 0 | 0.025 |
| MN996532.1;QHR63303.1;BatCoV_RaTG13_membran;112 | 0 | 0.025 |
| MN996532.1;QHR63303.1;BatCoV_RaTG13_membran;113 | 0 | 0 |
| MN996532.1;QHR63303.1;BatCoV_RaTG13_membran;114 | 0 | 0 |
| MN996532.1;QHR63303.1;BatCoV_RaTG13_membran;115 | 0 | 0 |
| MN996532.1;QHR63303.1;BatCoV_RaTG13_membran;116 | 0 | 0 |
| MN996532.1;QHR63303.1;BatCoV_RaTG13_membran;117 | 0 | 0 |
| MN996532.1;QHR63303.1;BatCoV_RaTG13_membran;118 | 0 | 0 |
| MN996532.1;QHR63303.1;BatCoV_RaTG13_membran;119 | 0 | 0 |
| MN996532.1;QHR63303.1;BatCoV_RaTG13_membran;120 | 0 | 0 |
| MN996532.1;QHR63303.1;BatCoV_RaTG13_membran;121 | 0 | 0 |
| MN996532.1;QHR63303.1;BatCoV_RaTG13_membran;122 | 0 | 0 |
| MN996532.1;QHR63303.1;BatCoV_RaTG13_membran;123 | 0 | 0 |
| MN996532.1;QHR63303.1;BatCoV_RaTG13_membran;124 | 0 | 0 |
| MN996532.1;QHR63303.1;BatCoV_RaTG13_membran;125 | 0 | 0 |
| MN996532.1;QHR63303.1;BatCoV_RaTG13_membran;126 | 0 | 0 |
| MN996532.1;QHR63303.1;BatCoV_RaTG13_membran;127 | 0 | 0 |
| MN996532.1;QHR63303.1;BatCoV_RaTG13_membran;128 | 0 | 0 |
| MN996532.1;QHR63303.1;BatCoV_RaTG13_membran;129 | 0 | 0 |
| MN996532.1;QHR63303.1;BatCoV_RaTG13_membran;130 | 0 | 0 |
| MN996532.1;QHR63303.1;BatCoV_RaTG13_membran;131 | 0 | 0 |
| MN996532.1;QHR63303.1;BatCoV_RaTG13_membran;132 | 0 | 0 |
| MN996532.1;QHR63303.1;BatCoV_RaTG13_membran;133 | 0 | 0 |
| MN996532.1;QHR63303.1;BatCoV_RaTG13_membran;134 | 0 | 0 |
| MN996532.1;QHR63303.1;BatCoV_RaTG13_membran;135 | 0 | 0 |
| MN996532.1;QHR63303.1;BatCoV_RaTG13_membran;136 | 0 | 0 |

| | | |
|---|---|---|
| MN996532.1;QHR63303.1;BatCoV_RaTG13_membran;137 | 0 | 0 |
| MN996532.1;QHR63303.1;BatCoV_RaTG13_membran;138 | 0 | 0 |
| MN996532.1;QHR63303.1;BatCoV_RaTG13_membran;139 | 0 | 0 |
| MN996532.1;QHR63303.1;BatCoV_RaTG13_membran;140 | 0 | 0 |
| MN996532.1;QHR63303.1;BatCoV_RaTG13_membran;141 | 0 | 0 |
| MN996532.1;QHR63303.1;BatCoV_RaTG13_membran;142 | 0 | 0 |
| MN996532.1;QHR63303.1;BatCoV_RaTG13_membran;143 | 0 | 0 |
| MN996532.1;QHR63303.1;BatCoV_RaTG13_membran;144 | 0 | 0 |
| MN996532.1;QHR63303.1;BatCoV_RaTG13_membran;145 | 0 | 0 |
| MN996532.1;QHR63303.1;BatCoV_RaTG13_membran;146 | 0 | 0 |
| MN996532.1;QHR63303.1;BatCoV_RaTG13_membran;147 | 0.05 | 0.025 |
| MN996532.1;QHR63303.1;BatCoV_RaTG13_membran;148 | 0.05 | 0.05 |
| MN996532.1;QHR63303.1;BatCoV_RaTG13_membran;149 | 0.05 | 0.05 |
| MN996532.1;QHR63303.1;BatCoV_RaTG13_membran;150 | 0.05 | 0.075 |
| MN996532.1;QHR63303.1;BatCoV_RaTG13_membran;151 | 0.05 | 0.125 |
| MN996532.1;QHR63303.1;BatCoV_RaTG13_membran;152 | 0.05 | 0.375 |
| MN996532.1;QHR63303.1;BatCoV_RaTG13_membran;153 | 0.05 | 0.45 |
| MN996532.1;QHR63303.1;BatCoV_RaTG13_membran;154 | 0.05 | 0.375 |
| MN996532.1;QHR63303.1;BatCoV_RaTG13_membran;155 | 0.1 | 0.425 |
| MN996532.1;QHR63303.1;BatCoV_RaTG13_membran;156 | 0.05 | 0.4 |
| MN996532.1;QHR63303.1;BatCoV_RaTG13_membran;157 | 0.05 | 0.35 |
| MN996532.1;QHR63303.1;BatCoV_RaTG13_membran;158 | 0 | 0.35 |
| MN996532.1;QHR63303.1;BatCoV_RaTG13_membran;159 | 0 | 0.35 |
| MN996532.1;QHR63303.1;BatCoV_RaTG13_membran;160 | 0 | 0.35 |
| MN996532.1;QHR63303.1;BatCoV_RaTG13_membran;161 | 0 | 0.35 |
| MN996532.1;QHR63303.1;BatCoV_RaTG13_membran;162 | 0 | 0.025 |
| MN996532.1;QHR63303.1;BatCoV_RaTG13_membran;163 | 0 | 0 |
| MN996532.1;QHR63303.1;BatCoV_RaTG13_membran;164 | 0 | 0 |
| MN996532.1;QHR63303.1;BatCoV_RaTG13_membran;165 | 0 | 0 |
| MN996532.1;QHR63303.1;BatCoV_RaTG13_membran;166 | 0 | 0 |
| MN996532.1;QHR63303.1;BatCoV_RaTG13_membran;167 | 0 | 0 |
| MN996532.1;QHR63303.1;BatCoV_RaTG13_membran;168 | 0 | 0 |
| MN996532.1;QHR63303.1;BatCoV_RaTG13_membran;169 | 0 | 0 |
| MN996532.1;QHR63303.1;BatCoV_RaTG13_membran;170 | 0 | 0 |
| MN996532.1;QHR63303.1;BatCoV_RaTG13_membran;171 | 0 | 0 |
| MN996532.1;QHR63303.1;BatCoV_RaTG13_membran;172 | 0 | 0 |
| MN996532.1;QHR63303.1;BatCoV_RaTG13_membran;173 | 0 | 0 |
| MN996532.1;QHR63303.1;BatCoV_RaTG13_membran;174 | 0 | 0.025 |
| MN996532.1;QHR63303.1;BatCoV_RaTG13_membran;175 | 0 | 0.075 |
| MN996532.1;QHR63303.1;BatCoV_RaTG13_membran;176 | 0 | 0.1 |
| MN996532.1;QHR63303.1;BatCoV_RaTG13_membran;177 | 0 | 0.05 |
| MN996532.1;QHR63303.1;BatCoV_RaTG13_membran;178 | 0 | 0.075 |
| MN996532.1;QHR63303.1;BatCoV_RaTG13_membran;179 | 0.05 | 0.125 |

| | | |
|---|---|---|
| MN996532.1;QHR63303.1;BatCoV_RaTG13_membran;180 | 0 | 0.35 |
| MN996532.1;QHR63303.1;BatCoV_RaTG13_membran;181 | 0 | 0.5 |
| MN996532.1;QHR63303.1;BatCoV_RaTG13_membran;182 | 0 | 0.4 |
| MN996532.1;QHR63303.1;BatCoV_RaTG13_membran;183 | 0 | 0.325 |
| MN996532.1;QHR63303.1;BatCoV_RaTG13_membran;184 | 0 | 0.275 |
| MN996532.1;QHR63303.1;BatCoV_RaTG13_membran;185 | 0 | 0.125 |
| MN996532.1;QHR63303.1;BatCoV_RaTG13_membran;186 | 0 | 0.175 |
| MN996532.1;QHR63303.1;BatCoV_RaTG13_membran;187 | 0 | 0.25 |
| MN996532.1;QHR63303.1;BatCoV_RaTG13_membran;188 | 0 | 0.225 |
| MN996532.1;QHR63303.1;BatCoV_RaTG13_membran;189 | 0 | 0.2 |
| MN996532.1;QHR63303.1;BatCoV_RaTG13_membran;190 | 0 | 0.1 |
| MN996532.1;QHR63303.1;BatCoV_RaTG13_membran;191 | 0 | 0.1 |
| MN996532.1;QHR63303.1;BatCoV_RaTG13_membran;192 | 0 | 0.075 |
| MN996532.1;QHR63303.1;BatCoV_RaTG13_membran;193 | 0 | 0.15 |
| MN996532.1;QHR63303.1;BatCoV_RaTG13_membran;194 | 0 | 0.075 |
| MN996532.1;QHR63303.1;BatCoV_RaTG13_membran;195 | 0 | 0.1 |
| MN996532.1;QHR63303.1;BatCoV_RaTG13_membran;196 | 0 | 0.125 |
| MN996532.1;QHR63303.1;BatCoV_RaTG13_membran;197 | 0 | 0.125 |
| MN996532.1;QHR63303.1;BatCoV_RaTG13_membran;198 | 0 | 0.125 |
| MN996532.1;QHR63303.1;BatCoV_RaTG13_membran;199 | 0.05 | 0.2 |
| MN996532.1;QHR63303.1;BatCoV_RaTG13_membran;200 | 0.1 | 0.125 |
| MN996532.1;QHR63303.1;BatCoV_RaTG13_membran;201 | 0.05 | 0.125 |
| MN996532.1;QHR63303.1;BatCoV_RaTG13_membran;202 | 0.05 | 0.125 |
| MN996532.1;QHR63303.1;BatCoV_RaTG13_membran;203 | 0.05 | 0.175 |
| MN996532.1;QHR63303.1;BatCoV_RaTG13_membran;204 | 0.05 | 0.225 |
| MN996532.1;QHR63303.1;BatCoV_RaTG13_membran;205 | 0 | 0.2 |
| MN996532.1;QHR63303.1;BatCoV_RaTG13_membran;206 | 0 | 0.2 |
| MN996532.1;QHR63303.1;BatCoV_RaTG13_membran;207 | 0 | 0.225 |
| MN996532.1;QHR63303.1;BatCoV_RaTG13_membran;208 | 0 | 0.175 |
| MN996532.1;QHR63303.1;BatCoV_RaTG13_membran;209 | 0 | 0.175 |
| MN996532.1;QHR63303.1;BatCoV_RaTG13_membran;210 | 0 | 0.175 |
| MN996532.1;QHR63304.1;BatCoV_RaTG13_orf6;1 | 0 | 0 |
| MN996532.1;QHR63304.1;BatCoV_RaTG13_orf6;2 | 0 | 0 |
| MN996532.1;QHR63304.1;BatCoV_RaTG13_orf6;3 | 0 | 0 |
| MN996532.1;QHR63304.1;BatCoV_RaTG13_orf6;4 | 0 | 0 |
| MN996532.1;QHR63304.1;BatCoV_RaTG13_orf6;5 | 0 | 0 |
| MN996532.1;QHR63304.1;BatCoV_RaTG13_orf6;6 | 0 | 0 |
| MN996532.1;QHR63304.1;BatCoV_RaTG13_orf6;7 | 0 | 0 |
| MN996532.1;QHR63304.1;BatCoV_RaTG13_orf6;8 | 0 | 0 |
| MN996532.1;QHR63304.1;BatCoV_RaTG13_orf6;9 | 0 | 0.025 |
| MN996532.1;QHR63304.1;BatCoV_RaTG13_orf6;10 | 0 | 0 |
| MN996532.1;QHR63304.1;BatCoV_RaTG13_orf6;11 | 0 | 0 |
| MN996532.1;QHR63304.1;BatCoV_RaTG13_orf6;12 | 0 | 0 |

| | | |
|---|---|---|
| MN996532.1;QHR63304.1;BatCoV_RaTG13_orf6;13 | 0 | 0 |
| MN996532.1;QHR63304.1;BatCoV_RaTG13_orf6;14 | 0 | 0 |
| MN996532.1;QHR63304.1;BatCoV_RaTG13_orf6;15 | 0 | 0 |
| MN996532.1;QHR63304.1;BatCoV_RaTG13_orf6;16 | 0 | 0 |
| MN996532.1;QHR63304.1;BatCoV_RaTG13_orf6;17 | 0 | 0 |
| MN996532.1;QHR63304.1;BatCoV_RaTG13_orf6;18 | 0 | 0 |
| MN996532.1;QHR63304.1;BatCoV_RaTG13_orf6;19 | 0 | 0 |
| MN996532.1;QHR63304.1;BatCoV_RaTG13_orf6;20 | 0 | 0 |
| MN996532.1;QHR63304.1;BatCoV_RaTG13_orf6;21 | 0 | 0 |
| MN996532.1;QHR63304.1;BatCoV_RaTG13_orf6;22 | 0 | 0 |
| MN996532.1;QHR63304.1;BatCoV_RaTG13_orf6;23 | 0 | 0 |
| MN996532.1;QHR63304.1;BatCoV_RaTG13_orf6;24 | 0 | 0 |
| MN996532.1;QHR63304.1;BatCoV_RaTG13_orf6;25 | 0 | 0 |
| MN996532.1;QHR63304.1;BatCoV_RaTG13_orf6;26 | 0 | 0 |
| MN996532.1;QHR63304.1;BatCoV_RaTG13_orf6;27 | 0 | 0 |
| MN996532.1;QHR63304.1;BatCoV_RaTG13_orf6;28 | 0 | 0 |
| MN996532.1;QHR63304.1;BatCoV_RaTG13_orf6;29 | 0 | 0 |
| MN996532.1;QHR63304.1;BatCoV_RaTG13_orf6;30 | 0 | 0 |
| MN996532.1;QHR63304.1;BatCoV_RaTG13_orf6;31 | 0.05 | 0 |
| MN996532.1;QHR63304.1;BatCoV_RaTG13_orf6;32 | 0.1 | 0 |
| MN996532.1;QHR63304.1;BatCoV_RaTG13_orf6;33 | 0.1 | 0 |
| MN996532.1;QHR63304.1;BatCoV_RaTG13_orf6;34 | 0.05 | 0 |
| MN996532.1;QHR63304.1;BatCoV_RaTG13_orf6;35 | 0.1 | 0 |
| MN996532.1;QHR63304.1;BatCoV_RaTG13_orf6;36 | 0.05 | 0.025 |
| MN996532.1;QHR63304.1;BatCoV_RaTG13_orf6;37 | 0.05 | 0 |
| MN996532.1;QHR63304.1;BatCoV_RaTG13_orf6;38 | 0.05 | 0.025 |
| MN996532.1;QHR63304.1;BatCoV_RaTG13_orf6;39 | 0.05 | 0.075 |
| MN996532.1;QHR63304.1;BatCoV_RaTG13_orf6;40 | 0.05 | 0.05 |
| MN996532.1;QHR63304.1;BatCoV_RaTG13_orf6;41 | 0.1 | 0.075 |
| MN996532.1;QHR63304.1;BatCoV_RaTG13_orf6;42 | 0.05 | 0.075 |
| MN996532.1;QHR63304.1;BatCoV_RaTG13_orf6;43 | 0.05 | 0.075 |
| MN996532.1;QHR63304.1;BatCoV_RaTG13_orf6;44 | 0.05 | 0.075 |
| MN996532.1;QHR63304.1;BatCoV_RaTG13_orf6;45 | 0.1 | 0.05 |
| MN996532.1;QHR63304.1;BatCoV_RaTG13_orf6;46 | 0.05 | 0.05 |
| MN996532.1;QHR63304.1;BatCoV_RaTG13_orf6;47 | 0 | 0 |
| MN996532.1;QHR63304.1;BatCoV_RaTG13_orf6;48 | 0 | 0 |
| MN996532.1;QHR63304.1;BatCoV_RaTG13_orf6;49 | 0 | 0.025 |
| MN996532.1;QHR63304.1;BatCoV_RaTG13_orf6;50 | 0 | 0 |
| MN996532.1;QHR63305.1;BatCoV_RaTG13_orf7a;1 | 0 | 0 |
| MN996532.1;QHR63305.1;BatCoV_RaTG13_orf7a;2 | 0 | 0 |
| MN996532.1;QHR63305.1;BatCoV_RaTG13_orf7a;3 | 0 | 0 |
| MN996532.1;QHR63305.1;BatCoV_RaTG13_orf7a;4 | 0 | 0 |
| MN996532.1;QHR63305.1;BatCoV_RaTG13_orf7a;5 | 0 | 0 |

| | | |
|---|---|---|
| MN996532.1;QHR63305.1;BatCoV_RaTG13_orf7a;6 | 0 | 0 |
| MN996532.1;QHR63305.1;BatCoV_RaTG13_orf7a;7 | 0 | 0 |
| MN996532.1;QHR63305.1;BatCoV_RaTG13_orf7a;8 | 0 | 0 |
| MN996532.1;QHR63305.1;BatCoV_RaTG13_orf7a;9 | 0 | 0 |
| MN996532.1;QHR63305.1;BatCoV_RaTG13_orf7a;10 | 0 | 0 |
| MN996532.1;QHR63305.1;BatCoV_RaTG13_orf7a;11 | 0 | 0 |
| MN996532.1;QHR63305.1;BatCoV_RaTG13_orf7a;12 | 0 | 0 |
| MN996532.1;QHR63305.1;BatCoV_RaTG13_orf7a;13 | 0 | 0 |
| MN996532.1;QHR63305.1;BatCoV_RaTG13_orf7a;14 | 0 | 0 |
| MN996532.1;QHR63305.1;BatCoV_RaTG13_orf7a;15 | 0 | 0 |
| MN996532.1;QHR63305.1;BatCoV_RaTG13_orf7a;16 | 0 | 0 |
| MN996532.1;QHR63305.1;BatCoV_RaTG13_orf7a;17 | 0 | 0 |
| MN996532.1;QHR63305.1;BatCoV_RaTG13_orf7a;18 | 0 | 0 |
| MN996532.1;QHR63305.1;BatCoV_RaTG13_orf7a;19 | 0 | 0 |
| MN996532.1;QHR63305.1;BatCoV_RaTG13_orf7a;20 | 0 | 0 |
| MN996532.1;QHR63305.1;BatCoV_RaTG13_orf7a;21 | 0 | 0 |
| MN996532.1;QHR63305.1;BatCoV_RaTG13_orf7a;22 | 0 | 0 |
| MN996532.1;QHR63305.1;BatCoV_RaTG13_orf7a;23 | 0.05 | 0 |
| MN996532.1;QHR63305.1;BatCoV_RaTG13_orf7a;24 | 0.05 | 0 |
| MN996532.1;QHR63305.1;BatCoV_RaTG13_orf7a;25 | 0.05 | 0 |
| MN996532.1;QHR63305.1;BatCoV_RaTG13_orf7a;26 | 0 | 0 |
| MN996532.1;QHR63305.1;BatCoV_RaTG13_orf7a;27 | 0 | 0 |
| MN996532.1;QHR63305.1;BatCoV_RaTG13_orf7a;28 | 0 | 0.025 |
| MN996532.1;QHR63305.1;BatCoV_RaTG13_orf7a;29 | 0 | 0.025 |
| MN996532.1;QHR63305.1;BatCoV_RaTG13_orf7a;30 | 0 | 0.025 |
| MN996532.1;QHR63305.1;BatCoV_RaTG13_orf7a;31 | 0 | 0 |
| MN996532.1;QHR63305.1;BatCoV_RaTG13_orf7a;32 | 0 | 0 |
| MN996532.1;QHR63305.1;BatCoV_RaTG13_orf7a;33 | 0 | 0 |
| MN996532.1;QHR63305.1;BatCoV_RaTG13_orf7a;34 | 0 | 0.025 |
| MN996532.1;QHR63305.1;BatCoV_RaTG13_orf7a;35 | 0 | 0.025 |
| MN996532.1;QHR63305.1;BatCoV_RaTG13_orf7a;36 | 0.05 | 0.025 |
| MN996532.1;QHR63305.1;BatCoV_RaTG13_orf7a;37 | 0 | 0.05 |
| MN996532.1;QHR63305.1;BatCoV_RaTG13_orf7a;38 | 0 | 0.05 |
| MN996532.1;QHR63305.1;BatCoV_RaTG13_orf7a;39 | 0 | 0.05 |
| MN996532.1;QHR63305.1;BatCoV_RaTG13_orf7a;40 | 0 | 0.05 |
| MN996532.1;QHR63305.1;BatCoV_RaTG13_orf7a;41 | 0 | 0.075 |
| MN996532.1;QHR63305.1;BatCoV_RaTG13_orf7a;42 | 0 | 0.075 |
| MN996532.1;QHR63305.1;BatCoV_RaTG13_orf7a;43 | 0 | 0.05 |
| MN996532.1;QHR63305.1;BatCoV_RaTG13_orf7a;44 | 0 | 0.025 |
| MN996532.1;QHR63305.1;BatCoV_RaTG13_orf7a;45 | 0 | 0 |
| MN996532.1;QHR63305.1;BatCoV_RaTG13_orf7a;46 | 0 | 0 |
| MN996532.1;QHR63305.1;BatCoV_RaTG13_orf7a;47 | 0 | 0 |
| MN996532.1;QHR63305.1;BatCoV_RaTG13_orf7a;48 | 0 | 0 |

| | | |
|---|---|---|
| MN996532.1;QHR63305.1;BatCoV_RaTG13_orf7a;49 | 0 | 0 |
| MN996532.1;QHR63305.1;BatCoV_RaTG13_orf7a;50 | 0 | 0 |
| MN996532.1;QHR63305.1;BatCoV_RaTG13_orf7a;51 | 0 | 0 |
| MN996532.1;QHR63305.1;BatCoV_RaTG13_orf7a;52 | 0 | 0 |
| MN996532.1;QHR63305.1;BatCoV_RaTG13_orf7a;53 | 0 | 0 |
| MN996532.1;QHR63305.1;BatCoV_RaTG13_orf7a;54 | 0 | 0 |
| MN996532.1;QHR63305.1;BatCoV_RaTG13_orf7a;55 | 0 | 0 |
| MN996532.1;QHR63305.1;BatCoV_RaTG13_orf7a;56 | 0 | 0 |
| MN996532.1;QHR63305.1;BatCoV_RaTG13_orf7a;57 | 0 | 0.025 |
| MN996532.1;QHR63305.1;BatCoV_RaTG13_orf7a;58 | 0 | 0 |
| MN996532.1;QHR63305.1;BatCoV_RaTG13_orf7a;59 | 0 | 0 |
| MN996532.1;QHR63305.1;BatCoV_RaTG13_orf7a;60 | 0 | 0.025 |
| MN996532.1;QHR63305.1;BatCoV_RaTG13_orf7a;61 | 0 | 0.025 |
| MN996532.1;QHR63305.1;BatCoV_RaTG13_orf7a;62 | 0 | 0 |
| MN996532.1;QHR63305.1;BatCoV_RaTG13_orf7a;63 | 0 | 0 |
| MN996532.1;QHR63305.1;BatCoV_RaTG13_orf7a;64 | 0 | 0 |
| MN996532.1;QHR63305.1;BatCoV_RaTG13_orf7a;65 | 0 | 0 |
| MN996532.1;QHR63305.1;BatCoV_RaTG13_orf7a;66 | 0 | 0 |
| MN996532.1;QHR63305.1;BatCoV_RaTG13_orf7a;67 | 0 | 0 |
| MN996532.1;QHR63305.1;BatCoV_RaTG13_orf7a;68 | 0 | 0 |
| MN996532.1;QHR63305.1;BatCoV_RaTG13_orf7a;69 | 0 | 0.025 |
| MN996532.1;QHR63305.1;BatCoV_RaTG13_orf7a;70 | 0 | 0 |
| MN996532.1;QHR63305.1;BatCoV_RaTG13_orf7a;71 | 0 | 0 |
| MN996532.1;QHR63305.1;BatCoV_RaTG13_orf7a;72 | 0 | 0 |
| MN996532.1;QHR63305.1;BatCoV_RaTG13_orf7a;73 | 0 | 0 |
| MN996532.1;QHR63305.1;BatCoV_RaTG13_orf7a;74 | 0 | 0 |
| MN996532.1;QHR63305.1;BatCoV_RaTG13_orf7a;75 | 0 | 0 |
| MN996532.1;QHR63305.1;BatCoV_RaTG13_orf7a;76 | 0 | 0 |
| MN996532.1;QHR63305.1;BatCoV_RaTG13_orf7a;77 | 0 | 0 |
| MN996532.1;QHR63305.1;BatCoV_RaTG13_orf7a;78 | 0 | 0 |
| MN996532.1;QHR63305.1;BatCoV_RaTG13_orf7a;79 | 0.05 | 0 |
| MN996532.1;QHR63305.1;BatCoV_RaTG13_orf7a;80 | 0.15 | 0 |
| MN996532.1;QHR63305.1;BatCoV_RaTG13_orf7a;81 | 0.05 | 0 |
| MN996532.1;QHR63305.1;BatCoV_RaTG13_orf7a;82 | 0 | 0 |
| MN996532.1;QHR63305.1;BatCoV_RaTG13_orf7a;83 | 0 | 0 |
| MN996532.1;QHR63305.1;BatCoV_RaTG13_orf7a;84 | 0.05 | 0 |
| MN996532.1;QHR63305.1;BatCoV_RaTG13_orf7a;85 | 0.05 | 0 |
| MN996532.1;QHR63305.1;BatCoV_RaTG13_orf7a;86 | 0 | 0 |
| MN996532.1;QHR63305.1;BatCoV_RaTG13_orf7a;87 | 0 | 0 |
| MN996532.1;QHR63305.1;BatCoV_RaTG13_orf7a;88 | 0 | 0 |
| MN996532.1;QHR63305.1;BatCoV_RaTG13_orf7a;89 | 0 | 0 |
| MN996532.1;QHR63305.1;BatCoV_RaTG13_orf7a;90 | 0 | 0 |
| MN996532.1;QHR63305.1;BatCoV_RaTG13_orf7a;91 | 0 | 0 |

| | | |
|---|---|---|
| MN996532.1;QHR63305.1;BatCoV_RaTG13_orf7a;92 | 0 | 0 |
| MN996532.1;QHR63305.1;BatCoV_RaTG13_orf7a;93 | 0 | 0 |
| MN996532.1;QHR63305.1;BatCoV_RaTG13_orf7a;94 | 0 | 0 |
| MN996532.1;QHR63305.1;BatCoV_RaTG13_orf7a;95 | 0 | 0 |
| MN996532.1;QHR63305.1;BatCoV_RaTG13_orf7a;96 | 0 | 0 |
| MN996532.1;QHR63305.1;BatCoV_RaTG13_orf7a;97 | 0 | 0 |
| MN996532.1;QHR63305.1;BatCoV_RaTG13_orf7a;98 | 0 | 0 |
| MN996532.1;QHR63305.1;BatCoV_RaTG13_orf7a;99 | 0 | 0 |
| MN996532.1;QHR63305.1;BatCoV_RaTG13_orf7a;100 | 0 | 0 |
| MN996532.1;QHR63305.1;BatCoV_RaTG13_orf7a;101 | 0 | 0 |
| MN996532.1;QHR63305.1;BatCoV_RaTG13_orf7a;102 | 0 | 0 |
| MN996532.1;QHR63305.1;BatCoV_RaTG13_orf7a;103 | 0 | 0 |
| MN996532.1;QHR63305.1;BatCoV_RaTG13_orf7a;104 | 0 | 0 |
| MN996532.1;QHR63305.1;BatCoV_RaTG13_orf7a;105 | 0 | 0 |
| MN996532.1;QHR63305.1;BatCoV_RaTG13_orf7a;106 | 0 | 0 |
| MN996532.1;QHR63305.1;BatCoV_RaTG13_orf7a;107 | 0 | 0 |
| MN996532.1;QHR63305.1;BatCoV_RaTG13_orf7a;108 | 0 | 0 |
| MN996532.1;QHR63305.1;BatCoV_RaTG13_orf7a;109 | 0 | 0 |
| MN996532.1;QHR63305.1;BatCoV_RaTG13_orf7a;110 | 0 | 0 |
| MN996532.1;QHR63306.1;BatCoV_RaTG13_orf7b;1 | 0 | 0 |
| MN996532.1;QHR63306.1;BatCoV_RaTG13_orf7b;2 | 0 | 0 |
| MN996532.1;QHR63306.1;BatCoV_RaTG13_orf7b;3 | 0 | 0 |
| MN996532.1;QHR63306.1;BatCoV_RaTG13_orf7b;4 | 0 | 0 |
| MN996532.1;QHR63306.1;BatCoV_RaTG13_orf7b;5 | 0 | 0 |
| MN996532.1;QHR63306.1;BatCoV_RaTG13_orf7b;6 | 0 | 0 |
| MN996532.1;QHR63306.1;BatCoV_RaTG13_orf7b;7 | 0 | 0 |
| MN996532.1;QHR63306.1;BatCoV_RaTG13_orf7b;8 | 0 | 0 |
| MN996532.1;QHR63306.1;BatCoV_RaTG13_orf7b;9 | 0 | 0 |
| MN996532.1;QHR63306.1;BatCoV_RaTG13_orf7b;10 | 0 | 0 |
| MN996532.1;QHR63306.1;BatCoV_RaTG13_orf7b;11 | 0 | 0 |
| MN996532.1;QHR63306.1;BatCoV_RaTG13_orf7b;12 | 0 | 0 |
| MN996532.1;QHR63306.1;BatCoV_RaTG13_orf7b;13 | 0 | 0 |
| MN996532.1;QHR63306.1;BatCoV_RaTG13_orf7b;14 | 0 | 0 |
| MN996532.1;QHR63306.1;BatCoV_RaTG13_orf7b;15 | 0 | 0 |
| MN996532.1;QHR63306.1;BatCoV_RaTG13_orf7b;16 | 0 | 0 |
| MN996532.1;QHR63306.1;BatCoV_RaTG13_orf7b;17 | 0 | 0 |
| MN996532.1;QHR63306.1;BatCoV_RaTG13_orf7b;18 | 0 | 0 |
| MN996532.1;QHR63306.1;BatCoV_RaTG13_orf7b;19 | 0 | 0 |
| MN996532.1;QHR63306.1;BatCoV_RaTG13_orf7b;20 | 0 | 0 |
| MN996532.1;QHR63306.1;BatCoV_RaTG13_orf7b;21 | 0 | 0 |
| MN996532.1;QHR63306.1;BatCoV_RaTG13_orf7b;22 | 0 | 0 |
| MN996532.1;QHR63306.1;BatCoV_RaTG13_orf7b;23 | 0 | 0 |
| MN996532.1;QHR63306.1;BatCoV_RaTG13_orf7b;24 | 0 | 0 |

| | | |
|---|---|---|
| MN996532.1;QHR63306.1;BatCoV_RaTG13_orf7b;25 | 0 | 0 |
| MN996532.1;QHR63306.1;BatCoV_RaTG13_orf7b;26 | 0 | 0 |
| MN996532.1;QHR63306.1;BatCoV_RaTG13_orf7b;27 | 0 | 0 |
| MN996532.1;QHR63306.1;BatCoV_RaTG13_orf7b;28 | 0 | 0 |
| MN996532.1;QHR63306.1;BatCoV_RaTG13_orf7b;29 | 0 | 0 |
| MN996532.1;QHR63306.1;BatCoV_RaTG13_orf7b;30 | 0 | 0.025 |
| MN996532.1;QHR63306.1;BatCoV_RaTG13_orf7b;31 | 0 | 0 |
| MN996532.1;QHR63306.1;BatCoV_RaTG13_orf7b;32 | 0 | 0 |
| MN996532.1;QHR63307.1;BatCoV_RaTG13_orf8;1 | 0 | 0 |
| MN996532.1;QHR63307.1;BatCoV_RaTG13_orf8;2 | 0 | 0 |
| MN996532.1;QHR63307.1;BatCoV_RaTG13_orf8;3 | 0 | 0 |
| MN996532.1;QHR63307.1;BatCoV_RaTG13_orf8;4 | 0 | 0 |
| MN996532.1;QHR63307.1;BatCoV_RaTG13_orf8;5 | 0 | 0 |
| MN996532.1;QHR63307.1;BatCoV_RaTG13_orf8;6 | 0 | 0 |
| MN996532.1;QHR63307.1;BatCoV_RaTG13_orf8;7 | 0 | 0 |
| MN996532.1;QHR63307.1;BatCoV_RaTG13_orf8;8 | 0 | 0 |
| MN996532.1;QHR63307.1;BatCoV_RaTG13_orf8;9 | 0 | 0 |
| MN996532.1;QHR63307.1;BatCoV_RaTG13_orf8;10 | 0 | 0 |
| MN996532.1;QHR63307.1;BatCoV_RaTG13_orf8;11 | 0 | 0 |
| MN996532.1;QHR63307.1;BatCoV_RaTG13_orf8;12 | 0 | 0 |
| MN996532.1;QHR63307.1;BatCoV_RaTG13_orf8;13 | 0 | 0.025 |
| MN996532.1;QHR63307.1;BatCoV_RaTG13_orf8;14 | 0 | 0.025 |
| MN996532.1;QHR63307.1;BatCoV_RaTG13_orf8;15 | 0 | 0.025 |
| MN996532.1;QHR63307.1;BatCoV_RaTG13_orf8;16 | 0 | 0.025 |
| MN996532.1;QHR63307.1;BatCoV_RaTG13_orf8;17 | 0 | 0 |
| MN996532.1;QHR63307.1;BatCoV_RaTG13_orf8;18 | 0 | 0 |
| MN996532.1;QHR63307.1;BatCoV_RaTG13_orf8;19 | 0 | 0 |
| MN996532.1;QHR63307.1;BatCoV_RaTG13_orf8;20 | 0 | 0 |
| MN996532.1;QHR63307.1;BatCoV_RaTG13_orf8;21 | 0 | 0.025 |
| MN996532.1;QHR63307.1;BatCoV_RaTG13_orf8;22 | 0 | 0 |
| MN996532.1;QHR63307.1;BatCoV_RaTG13_orf8;23 | 0 | 0 |
| MN996532.1;QHR63307.1;BatCoV_RaTG13_orf8;24 | 0.05 | 0 |
| MN996532.1;QHR63307.1;BatCoV_RaTG13_orf8;25 | 0 | 0 |
| MN996532.1;QHR63307.1;BatCoV_RaTG13_orf8;26 | 0 | 0 |
| MN996532.1;QHR63307.1;BatCoV_RaTG13_orf8;27 | 0 | 0 |
| MN996532.1;QHR63307.1;BatCoV_RaTG13_orf8;28 | 0 | 0 |
| MN996532.1;QHR63307.1;BatCoV_RaTG13_orf8;29 | 0 | 0 |
| MN996532.1;QHR63307.1;BatCoV_RaTG13_orf8;30 | 0 | 0 |
| MN996532.1;QHR63307.1;BatCoV_RaTG13_orf8;31 | 0 | 0 |
| MN996532.1;QHR63307.1;BatCoV_RaTG13_orf8;32 | 0 | 0 |
| MN996532.1;QHR63307.1;BatCoV_RaTG13_orf8;33 | 0 | 0 |
| MN996532.1;QHR63307.1;BatCoV_RaTG13_orf8;34 | 0 | 0 |
| MN996532.1;QHR63307.1;BatCoV_RaTG13_orf8;35 | 0 | 0 |

| | | |
|---|---|---|
| MN996532.1;QHR63307.1;BatCoV_RaTG13_orf8;36 | 0 | 0 |
| MN996532.1;QHR63307.1;BatCoV_RaTG13_orf8;37 | 0 | 0 |
| MN996532.1;QHR63307.1;BatCoV_RaTG13_orf8;38 | 0 | 0 |
| MN996532.1;QHR63307.1;BatCoV_RaTG13_orf8;39 | 0 | 0 |
| MN996532.1;QHR63307.1;BatCoV_RaTG13_orf8;40 | 0 | 0 |
| MN996532.1;QHR63307.1;BatCoV_RaTG13_orf8;41 | 0 | 0 |
| MN996532.1;QHR63307.1;BatCoV_RaTG13_orf8;42 | 0 | 0 |
| MN996532.1;QHR63307.1;BatCoV_RaTG13_orf8;43 | 0 | 0 |
| MN996532.1;QHR63307.1;BatCoV_RaTG13_orf8;44 | 0 | 0 |
| MN996532.1;QHR63307.1;BatCoV_RaTG13_orf8;45 | 0 | 0 |
| MN996532.1;QHR63307.1;BatCoV_RaTG13_orf8;46 | 0 | 0 |
| MN996532.1;QHR63307.1;BatCoV_RaTG13_orf8;47 | 0 | 0.025 |
| MN996532.1;QHR63307.1;BatCoV_RaTG13_orf8;48 | 0 | 0 |
| MN996532.1;QHR63307.1;BatCoV_RaTG13_orf8;49 | 0 | 0 |
| MN996532.1;QHR63307.1;BatCoV_RaTG13_orf8;50 | 0 | 0 |
| MN996532.1;QHR63307.1;BatCoV_RaTG13_orf8;51 | 0 | 0 |
| MN996532.1;QHR63307.1;BatCoV_RaTG13_orf8;52 | 0 | 0 |
| MN996532.1;QHR63307.1;BatCoV_RaTG13_orf8;53 | 0 | 0 |
| MN996532.1;QHR63307.1;BatCoV_RaTG13_orf8;54 | 0.05 | 0 |
| MN996532.1;QHR63307.1;BatCoV_RaTG13_orf8;55 | 0.05 | 0 |
| MN996532.1;QHR63307.1;BatCoV_RaTG13_orf8;56 | 0.05 | 0 |
| MN996532.1;QHR63307.1;BatCoV_RaTG13_orf8;57 | 0.05 | 0 |
| MN996532.1;QHR63307.1;BatCoV_RaTG13_orf8;58 | 0 | 0 |
| MN996532.1;QHR63307.1;BatCoV_RaTG13_orf8;59 | 0 | 0 |
| MN996532.1;QHR63307.1;BatCoV_RaTG13_orf8;60 | 0 | 0.075 |
| MN996532.1;QHR63307.1;BatCoV_RaTG13_orf8;61 | 0.05 | 0.075 |
| MN996532.1;QHR63307.1;BatCoV_RaTG13_orf8;62 | 0.1 | 0.1 |
| MN996532.1;QHR63307.1;BatCoV_RaTG13_orf8;63 | 0 | 0.125 |
| MN996532.1;QHR63307.1;BatCoV_RaTG13_orf8;64 | 0 | 0.175 |
| MN996532.1;QHR63307.1;BatCoV_RaTG13_orf8;65 | 0 | 0.15 |
| MN996532.1;QHR63307.1;BatCoV_RaTG13_orf8;66 | 0 | 0.025 |
| MN996532.1;QHR63307.1;BatCoV_RaTG13_orf8;67 | 0 | 0.075 |
| MN996532.1;QHR63307.1;BatCoV_RaTG13_orf8;68 | 0 | 0.025 |
| MN996532.1;QHR63307.1;BatCoV_RaTG13_orf8;69 | 0 | 0.05 |
| MN996532.1;QHR63307.1;BatCoV_RaTG13_orf8;70 | 0 | 0.025 |
| MN996532.1;QHR63307.1;BatCoV_RaTG13_orf8;71 | 0 | 0 |
| MN996532.1;QHR63307.1;BatCoV_RaTG13_orf8;72 | 0 | 0 |
| MN996532.1;QHR63307.1;BatCoV_RaTG13_orf8;73 | 0 | 0 |
| MN996532.1;QHR63307.1;BatCoV_RaTG13_orf8;74 | 0 | 0 |
| MN996532.1;QHR63307.1;BatCoV_RaTG13_orf8;75 | 0 | 0 |
| MN996532.1;QHR63307.1;BatCoV_RaTG13_orf8;76 | 0 | 0 |
| MN996532.1;QHR63307.1;BatCoV_RaTG13_orf8;77 | 0 | 0 |
| MN996532.1;QHR63307.1;BatCoV_RaTG13_orf8;78 | 0 | 0 |

| | | |
|---|---|---|
| MN996532.1;QHR63307.1;BatCoV_RaTG13_orf8;79 | 0 | 0 |
| MN996532.1;QHR63307.1;BatCoV_RaTG13_orf8;80 | 0 | 0 |
| MN996532.1;QHR63307.1;BatCoV_RaTG13_orf8;81 | 0 | 0 |
| MN996532.1;QHR63307.1;BatCoV_RaTG13_orf8;82 | 0 | 0 |
| MN996532.1;QHR63307.1;BatCoV_RaTG13_orf8;83 | 0.05 | 0 |
| MN996532.1;QHR63307.1;BatCoV_RaTG13_orf8;84 | 0.05 | 0.025 |
| MN996532.1;QHR63307.1;BatCoV_RaTG13_orf8;85 | 0.05 | 0.05 |
| MN996532.1;QHR63307.1;BatCoV_RaTG13_orf8;86 | 0 | 0 |
| MN996532.1;QHR63307.1;BatCoV_RaTG13_orf8;87 | 0 | 0 |
| MN996532.1;QHR63307.1;BatCoV_RaTG13_orf8;88 | 0 | 0 |
| MN996532.1;QHR63307.1;BatCoV_RaTG13_orf8;89 | 0 | 0 |
| MN996532.1;QHR63307.1;BatCoV_RaTG13_orf8;90 | 0 | 0 |
| MN996532.1;QHR63307.1;BatCoV_RaTG13_orf8;91 | 0 | 0 |
| MN996532.1;QHR63307.1;BatCoV_RaTG13_orf8;92 | 0 | 0 |
| MN996532.1;QHR63307.1;BatCoV_RaTG13_orf8;93 | 0 | 0 |
| MN996532.1;QHR63307.1;BatCoV_RaTG13_orf8;94 | 0 | 0 |
| MN996532.1;QHR63307.1;BatCoV_RaTG13_orf8;95 | 0 | 0 |
| MN996532.1;QHR63307.1;BatCoV_RaTG13_orf8;96 | 0 | 0 |
| MN996532.1;QHR63307.1;BatCoV_RaTG13_orf8;97 | 0 | 0 |
| MN996532.1;QHR63307.1;BatCoV_RaTG13_orf8;98 | 0 | 0 |
| MN996532.1;QHR63307.1;BatCoV_RaTG13_orf8;99 | 0 | 0 |
| MN996532.1;QHR63307.1;BatCoV_RaTG13_orf8;100 | 0 | 0 |
| MN996532.1;QHR63307.1;BatCoV_RaTG13_orf8;101 | 0 | 0 |
| MN996532.1;QHR63307.1;BatCoV_RaTG13_orf8;102 | 0 | 0 |
| MN996532.1;QHR63307.1;BatCoV_RaTG13_orf8;103 | 0 | 0 |
| MN996532.1;QHR63307.1;BatCoV_RaTG13_orf8;104 | 0 | 0 |
| MN996532.1;QHR63307.1;BatCoV_RaTG13_orf8;105 | 0 | 0 |
| MN996532.1;QHR63307.1;BatCoV_RaTG13_orf8;106 | 0 | 0 |
| MN996532.1;QHR63307.1;BatCoV_RaTG13_orf8;107 | 0 | 0 |
| MN996532.1;QHR63307.1;BatCoV_RaTG13_orf8;108 | 0 | 0 |
| MN996532.1;QHR63307.1;BatCoV_RaTG13_orf8;109 | 0 | 0 |
| MN996532.1;QHR63307.1;BatCoV_RaTG13_orf8;110 | 0 | 0 |
| MN996532.1;QHR63308.1;BatCoV_RaTG13_nucleoc;1 | 0.05 | 0.025 |
| MN996532.1;QHR63308.1;BatCoV_RaTG13_nucleoc;2 | 0.05 | 0.025 |
| MN996532.1;QHR63308.1;BatCoV_RaTG13_nucleoc;3 | 0.1 | 0.025 |
| MN996532.1;QHR63308.1;BatCoV_RaTG13_nucleoc;4 | 0.05 | 0.025 |
| MN996532.1;QHR63308.1;BatCoV_RaTG13_nucleoc;5 | 0.1 | 0.05 |
| MN996532.1;QHR63308.1;BatCoV_RaTG13_nucleoc;6 | 0.05 | 0.025 |
| MN996532.1;QHR63308.1;BatCoV_RaTG13_nucleoc;7 | 0.05 | 0.15 |
| MN996532.1;QHR63308.1;BatCoV_RaTG13_nucleoc;8 | 0.05 | 0.125 |
| MN996532.1;QHR63308.1;BatCoV_RaTG13_nucleoc;9 | 0.05 | 0.225 |
| MN996532.1;QHR63308.1;BatCoV_RaTG13_nucleoc;10 | 0.05 | 0.125 |
| MN996532.1;QHR63308.1;BatCoV_RaTG13_nucleoc;11 | 0.05 | 0.275 |

| | | |
|---|---|---|
| MN996532.1;QHR63308.1;BatCoV_RaTG13_nucleoc;12 | 0.05 | 0.325 |
| MN996532.1;QHR63308.1;BatCoV_RaTG13_nucleoc;13 | 0.15 | 0.2 |
| MN996532.1;QHR63308.1;BatCoV_RaTG13_nucleoc;14 | 0 | 0.3 |
| MN996532.1;QHR63308.1;BatCoV_RaTG13_nucleoc;15 | 0 | 0.15 |
| MN996532.1;QHR63308.1;BatCoV_RaTG13_nucleoc;16 | 0 | 0.1 |
| MN996532.1;QHR63308.1;BatCoV_RaTG13_nucleoc;17 | 0 | 0.1 |
| MN996532.1;QHR63308.1;BatCoV_RaTG13_nucleoc;18 | 0 | 0.1 |
| MN996532.1;QHR63308.1;BatCoV_RaTG13_nucleoc;19 | 0.05 | 0.1 |
| MN996532.1;QHR63308.1;BatCoV_RaTG13_nucleoc;20 | 0 | 0.075 |
| MN996532.1;QHR63308.1;BatCoV_RaTG13_nucleoc;21 | 0.05 | 0.125 |
| MN996532.1;QHR63308.1;BatCoV_RaTG13_nucleoc;22 | 0.05 | 0.075 |
| MN996532.1;QHR63308.1;BatCoV_RaTG13_nucleoc;23 | 0.05 | 0.2 |
| MN996532.1;QHR63308.1;BatCoV_RaTG13_nucleoc;24 | 0 | 0.2 |
| MN996532.1;QHR63308.1;BatCoV_RaTG13_nucleoc;25 | 0 | 0.15 |
| MN996532.1;QHR63308.1;BatCoV_RaTG13_nucleoc;26 | 0 | 0.15 |
| MN996532.1;QHR63308.1;BatCoV_RaTG13_nucleoc;27 | 0.05 | 0.175 |
| MN996532.1;QHR63308.1;BatCoV_RaTG13_nucleoc;28 | 0.05 | 0.125 |
| MN996532.1;QHR63308.1;BatCoV_RaTG13_nucleoc;29 | 0.1 | 0.15 |
| MN996532.1;QHR63308.1;BatCoV_RaTG13_nucleoc;30 | 0 | 0.15 |
| MN996532.1;QHR63308.1;BatCoV_RaTG13_nucleoc;31 | 0 | 0.15 |
| MN996532.1;QHR63308.1;BatCoV_RaTG13_nucleoc;32 | 0 | 0.1 |
| MN996532.1;QHR63308.1;BatCoV_RaTG13_nucleoc;33 | 0 | 0.2 |
| MN996532.1;QHR63308.1;BatCoV_RaTG13_nucleoc;34 | 0 | 0.125 |
| MN996532.1;QHR63308.1;BatCoV_RaTG13_nucleoc;35 | 0 | 0.225 |
| MN996532.1;QHR63308.1;BatCoV_RaTG13_nucleoc;36 | 0.05 | 0.15 |
| MN996532.1;QHR63308.1;BatCoV_RaTG13_nucleoc;37 | 0.05 | 0.3 |
| MN996532.1;QHR63308.1;BatCoV_RaTG13_nucleoc;38 | 0 | 0.225 |
| MN996532.1;QHR63308.1;BatCoV_RaTG13_nucleoc;39 | 0 | 0.15 |
| MN996532.1;QHR63308.1;BatCoV_RaTG13_nucleoc;40 | 0 | 0.125 |
| MN996532.1;QHR63308.1;BatCoV_RaTG13_nucleoc;41 | 0 | 0.025 |
| MN996532.1;QHR63308.1;BatCoV_RaTG13_nucleoc;42 | 0 | 0 |
| MN996532.1;QHR63308.1;BatCoV_RaTG13_nucleoc;43 | 0 | 0.025 |
| MN996532.1;QHR63308.1;BatCoV_RaTG13_nucleoc;44 | 0 | 0 |
| MN996532.1;QHR63308.1;BatCoV_RaTG13_nucleoc;45 | 0 | 0 |
| MN996532.1;QHR63308.1;BatCoV_RaTG13_nucleoc;46 | 0 | 0 |
| MN996532.1;QHR63308.1;BatCoV_RaTG13_nucleoc;47 | 0 | 0 |
| MN996532.1;QHR63308.1;BatCoV_RaTG13_nucleoc;48 | 0 | 0 |
| MN996532.1;QHR63308.1;BatCoV_RaTG13_nucleoc;49 | 0 | 0 |
| MN996532.1;QHR63308.1;BatCoV_RaTG13_nucleoc;50 | 0 | 0 |
| MN996532.1;QHR63308.1;BatCoV_RaTG13_nucleoc;51 | 0 | 0.025 |
| MN996532.1;QHR63308.1;BatCoV_RaTG13_nucleoc;52 | 0 | 0 |
| MN996532.1;QHR63308.1;BatCoV_RaTG13_nucleoc;53 | 0 | 0 |
| MN996532.1;QHR63308.1;BatCoV_RaTG13_nucleoc;54 | 0 | 0 |

| | | |
|---|---|---|
| MN996532.1;QHR63308.1;BatCoV_RaTG13_nucleoc;55 | 0 | 0 |
| MN996532.1;QHR63308.1;BatCoV_RaTG13_nucleoc;56 | 0 | 0 |
| MN996532.1;QHR63308.1;BatCoV_RaTG13_nucleoc;57 | 0 | 0 |
| MN996532.1;QHR63308.1;BatCoV_RaTG13_nucleoc;58 | 0 | 0.025 |
| MN996532.1;QHR63308.1;BatCoV_RaTG13_nucleoc;59 | 0 | 0.025 |
| MN996532.1;QHR63308.1;BatCoV_RaTG13_nucleoc;60 | 0 | 0.075 |
| MN996532.1;QHR63308.1;BatCoV_RaTG13_nucleoc;61 | 0 | 0.05 |
| MN996532.1;QHR63308.1;BatCoV_RaTG13_nucleoc;62 | 0.05 | 0.125 |
| MN996532.1;QHR63308.1;BatCoV_RaTG13_nucleoc;63 | 0 | 0.1 |
| MN996532.1;QHR63308.1;BatCoV_RaTG13_nucleoc;64 | 0.1 | 0.1 |
| MN996532.1;QHR63308.1;BatCoV_RaTG13_nucleoc;65 | 0 | 0.1 |
| MN996532.1;QHR63308.1;BatCoV_RaTG13_nucleoc;66 | 0 | 0.175 |
| MN996532.1;QHR63308.1;BatCoV_RaTG13_nucleoc;67 | 0.1 | 0.1 |
| MN996532.1;QHR63308.1;BatCoV_RaTG13_nucleoc;68 | 0 | 0.1 |
| MN996532.1;QHR63308.1;BatCoV_RaTG13_nucleoc;69 | 0 | 0.075 |
| MN996532.1;QHR63308.1;BatCoV_RaTG13_nucleoc;70 | 0 | 0.075 |
| MN996532.1;QHR63308.1;BatCoV_RaTG13_nucleoc;71 | 0 | 0.05 |
| MN996532.1;QHR63308.1;BatCoV_RaTG13_nucleoc;72 | 0 | 0.025 |
| MN996532.1;QHR63308.1;BatCoV_RaTG13_nucleoc;73 | 0.15 | 0.1 |
| MN996532.1;QHR63308.1;BatCoV_RaTG13_nucleoc;74 | 0 | 0.025 |
| MN996532.1;QHR63308.1;BatCoV_RaTG13_nucleoc;75 | 0 | 0.025 |
| MN996532.1;QHR63308.1;BatCoV_RaTG13_nucleoc;76 | 0 | 0 |
| MN996532.1;QHR63308.1;BatCoV_RaTG13_nucleoc;77 | 0 | 0 |
| MN996532.1;QHR63308.1;BatCoV_RaTG13_nucleoc;78 | 0.05 | 0 |
| MN996532.1;QHR63308.1;BatCoV_RaTG13_nucleoc;79 | 0.05 | 0 |
| MN996532.1;QHR63308.1;BatCoV_RaTG13_nucleoc;80 | 0 | 0 |
| MN996532.1;QHR63308.1;BatCoV_RaTG13_nucleoc;81 | 0 | 0 |
| MN996532.1;QHR63308.1;BatCoV_RaTG13_nucleoc;82 | 0 | 0 |
| MN996532.1;QHR63308.1;BatCoV_RaTG13_nucleoc;83 | 0 | 0 |
| MN996532.1;QHR63308.1;BatCoV_RaTG13_nucleoc;84 | 0 | 0 |
| MN996532.1;QHR63308.1;BatCoV_RaTG13_nucleoc;85 | 0.05 | 0 |
| MN996532.1;QHR63308.1;BatCoV_RaTG13_nucleoc;86 | 0 | 0 |
| MN996532.1;QHR63308.1;BatCoV_RaTG13_nucleoc;87 | 0 | 0.025 |
| MN996532.1;QHR63308.1;BatCoV_RaTG13_nucleoc;88 | 0 | 0 |
| MN996532.1;QHR63308.1;BatCoV_RaTG13_nucleoc;89 | 0 | 0.025 |
| MN996532.1;QHR63308.1;BatCoV_RaTG13_nucleoc;90 | 0 | 0.025 |
| MN996532.1;QHR63308.1;BatCoV_RaTG13_nucleoc;91 | 0 | 0.025 |
| MN996532.1;QHR63308.1;BatCoV_RaTG13_nucleoc;92 | 0 | 0.025 |
| MN996532.1;QHR63308.1;BatCoV_RaTG13_nucleoc;93 | 0 | 0.025 |
| MN996532.1;QHR63308.1;BatCoV_RaTG13_nucleoc;94 | 0 | 0.025 |
| MN996532.1;QHR63308.1;BatCoV_RaTG13_nucleoc;95 | 0 | 0.025 |
| MN996532.1;QHR63308.1;BatCoV_RaTG13_nucleoc;96 | 0 | 0.025 |
| MN996532.1;QHR63308.1;BatCoV_RaTG13_nucleoc;97 | 0 | 0.025 |

| | | |
|---|---|---|
| MN996532.1;QHR63308.1;BatCoV_RaTG13_nucleoc;98 | 0 | 0.025 |
| MN996532.1;QHR63308.1;BatCoV_RaTG13_nucleoc;99 | 0 | 0 |
| MN996532.1;QHR63308.1;BatCoV_RaTG13_nucleoc;100 | 0 | 0 |
| MN996532.1;QHR63308.1;BatCoV_RaTG13_nucleoc;101 | 0 | 0 |
| MN996532.1;QHR63308.1;BatCoV_RaTG13_nucleoc;102 | 0 | 0 |
| MN996532.1;QHR63308.1;BatCoV_RaTG13_nucleoc;103 | 0 | 0 |
| MN996532.1;QHR63308.1;BatCoV_RaTG13_nucleoc;104 | 0 | 0 |
| MN996532.1;QHR63308.1;BatCoV_RaTG13_nucleoc;105 | 0 | 0 |
| MN996532.1;QHR63308.1;BatCoV_RaTG13_nucleoc;106 | 0 | 0 |
| MN996532.1;QHR63308.1;BatCoV_RaTG13_nucleoc;107 | 0 | 0 |
| MN996532.1;QHR63308.1;BatCoV_RaTG13_nucleoc;108 | 0 | 0 |
| MN996532.1;QHR63308.1;BatCoV_RaTG13_nucleoc;109 | 0 | 0 |
| MN996532.1;QHR63308.1;BatCoV_RaTG13_nucleoc;110 | 0 | 0 |
| MN996532.1;QHR63308.1;BatCoV_RaTG13_nucleoc;111 | 0.05 | 0 |
| MN996532.1;QHR63308.1;BatCoV_RaTG13_nucleoc;112 | 0.1 | 0.125 |
| MN996532.1;QHR63308.1;BatCoV_RaTG13_nucleoc;113 | 0 | 0.1 |
| MN996532.1;QHR63308.1;BatCoV_RaTG13_nucleoc;114 | 0 | 0.1 |
| MN996532.1;QHR63308.1;BatCoV_RaTG13_nucleoc;115 | 0 | 0.1 |
| MN996532.1;QHR63308.1;BatCoV_RaTG13_nucleoc;116 | 0 | 0.15 |
| MN996532.1;QHR63308.1;BatCoV_RaTG13_nucleoc;117 | 0 | 0.15 |
| MN996532.1;QHR63308.1;BatCoV_RaTG13_nucleoc;118 | 0 | 0.05 |
| MN996532.1;QHR63308.1;BatCoV_RaTG13_nucleoc;119 | 0 | 0.05 |
| MN996532.1;QHR63308.1;BatCoV_RaTG13_nucleoc;120 | 0 | 0.05 |
| MN996532.1;QHR63308.1;BatCoV_RaTG13_nucleoc;121 | 0 | 0.025 |
| MN996532.1;QHR63308.1;BatCoV_RaTG13_nucleoc;122 | 0 | 0.025 |
| MN996532.1;QHR63308.1;BatCoV_RaTG13_nucleoc;123 | 0 | 0 |
| MN996532.1;QHR63308.1;BatCoV_RaTG13_nucleoc;124 | 0 | 0 |
| MN996532.1;QHR63308.1;BatCoV_RaTG13_nucleoc;125 | 0 | 0.025 |
| MN996532.1;QHR63308.1;BatCoV_RaTG13_nucleoc;126 | 0 | 0 |
| MN996532.1;QHR63308.1;BatCoV_RaTG13_nucleoc;127 | 0 | 0 |
| MN996532.1;QHR63308.1;BatCoV_RaTG13_nucleoc;128 | 0 | 0.05 |
| MN996532.1;QHR63308.1;BatCoV_RaTG13_nucleoc;129 | 0 | 0.075 |
| MN996532.1;QHR63308.1;BatCoV_RaTG13_nucleoc;130 | 0 | 0.05 |
| MN996532.1;QHR63308.1;BatCoV_RaTG13_nucleoc;131 | 0 | 0.075 |
| MN996532.1;QHR63308.1;BatCoV_RaTG13_nucleoc;132 | 0 | 0.075 |
| MN996532.1;QHR63308.1;BatCoV_RaTG13_nucleoc;133 | 0 | 0.05 |
| MN996532.1;QHR63308.1;BatCoV_RaTG13_nucleoc;134 | 0 | 0.025 |
| MN996532.1;QHR63308.1;BatCoV_RaTG13_nucleoc;135 | 0 | 0.025 |
| MN996532.1;QHR63308.1;BatCoV_RaTG13_nucleoc;136 | 0 | 0.025 |
| MN996532.1;QHR63308.1;BatCoV_RaTG13_nucleoc;137 | 0 | 0 |
| MN996532.1;QHR63308.1;BatCoV_RaTG13_nucleoc;138 | 0.05 | 0.05 |
| MN996532.1;QHR63308.1;BatCoV_RaTG13_nucleoc;139 | 0 | 0.05 |
| MN996532.1;QHR63308.1;BatCoV_RaTG13_nucleoc;140 | 0 | 0.075 |

| | | |
|---|---|---|
| MN996532.1;QHR63308.1;BatCoV_RaTG13_nucleoc;141 | 0.05 | 0.125 |
| MN996532.1;QHR63308.1;BatCoV_RaTG13_nucleoc;142 | 0 | 0.05 |
| MN996532.1;QHR63308.1;BatCoV_RaTG13_nucleoc;143 | 0 | 0.05 |
| MN996532.1;QHR63308.1;BatCoV_RaTG13_nucleoc;144 | 0 | 0.025 |
| MN996532.1;QHR63308.1;BatCoV_RaTG13_nucleoc;145 | 0 | 0.05 |
| MN996532.1;QHR63308.1;BatCoV_RaTG13_nucleoc;146 | 0 | 0.025 |
| MN996532.1;QHR63308.1;BatCoV_RaTG13_nucleoc;147 | 0 | 0.025 |
| MN996532.1;QHR63308.1;BatCoV_RaTG13_nucleoc;148 | 0 | 0.075 |
| MN996532.1;QHR63308.1;BatCoV_RaTG13_nucleoc;149 | 0 | 0.05 |
| MN996532.1;QHR63308.1;BatCoV_RaTG13_nucleoc;150 | 0 | 0.05 |
| MN996532.1;QHR63308.1;BatCoV_RaTG13_nucleoc;151 | 0 | 0 |
| MN996532.1;QHR63308.1;BatCoV_RaTG13_nucleoc;152 | 0 | 0.05 |
| MN996532.1;QHR63308.1;BatCoV_RaTG13_nucleoc;153 | 0 | 0.075 |
| MN996532.1;QHR63308.1;BatCoV_RaTG13_nucleoc;154 | 0.15 | 0.45 |
| MN996532.1;QHR63308.1;BatCoV_RaTG13_nucleoc;155 | 0.15 | 0.5 |
| MN996532.1;QHR63308.1;BatCoV_RaTG13_nucleoc;156 | 0.1 | 0.45 |
| MN996532.1;QHR63308.1;BatCoV_RaTG13_nucleoc;157 | 0.1 | 0.45 |
| MN996532.1;QHR63308.1;BatCoV_RaTG13_nucleoc;158 | 0.1 | 0.5 |
| MN996532.1;QHR63308.1;BatCoV_RaTG13_nucleoc;159 | 0.2 | 0.625 |
| MN996532.1;QHR63308.1;BatCoV_RaTG13_nucleoc;160 | 0.2 | 0.625 |
| MN996532.1;QHR63308.1;BatCoV_RaTG13_nucleoc;161 | 0.1 | 0.6 |
| MN996532.1;QHR63308.1;BatCoV_RaTG13_nucleoc;162 | 0.1 | 0.475 |
| MN996532.1;QHR63308.1;BatCoV_RaTG13_nucleoc;163 | 0.1 | 0.45 |
| MN996532.1;QHR63308.1;BatCoV_RaTG13_nucleoc;164 | 0.05 | 0.1 |
| MN996532.1;QHR63308.1;BatCoV_RaTG13_nucleoc;165 | 0 | 0.05 |
| MN996532.1;QHR63308.1;BatCoV_RaTG13_nucleoc;166 | 0 | 0.05 |
| MN996532.1;QHR63308.1;BatCoV_RaTG13_nucleoc;167 | 0 | 0 |
| MN996532.1;QHR63308.1;BatCoV_RaTG13_nucleoc;168 | 0.05 | 0.025 |
| MN996532.1;QHR63308.1;BatCoV_RaTG13_nucleoc;169 | 0 | 0.05 |
| MN996532.1;QHR63308.1;BatCoV_RaTG13_nucleoc;170 | 0 | 0 |
| MN996532.1;QHR63308.1;BatCoV_RaTG13_nucleoc;171 | 0 | 0 |
| MN996532.1;QHR63308.1;BatCoV_RaTG13_nucleoc;172 | 0 | 0 |
| MN996532.1;QHR63308.1;BatCoV_RaTG13_nucleoc;173 | 0 | 0.025 |
| MN996532.1;QHR63308.1;BatCoV_RaTG13_nucleoc;174 | 0 | 0.05 |
| MN996532.1;QHR63308.1;BatCoV_RaTG13_nucleoc;175 | 0 | 0.025 |
| MN996532.1;QHR63308.1;BatCoV_RaTG13_nucleoc;176 | 0 | 0.025 |
| MN996532.1;QHR63308.1;BatCoV_RaTG13_nucleoc;177 | 0 | 0.05 |
| MN996532.1;QHR63308.1;BatCoV_RaTG13_nucleoc;178 | 0 | 0.075 |
| MN996532.1;QHR63308.1;BatCoV_RaTG13_nucleoc;179 | 0 | 0.1 |
| MN996532.1;QHR63308.1;BatCoV_RaTG13_nucleoc;180 | 0 | 0.15 |
| MN996532.1;QHR63308.1;BatCoV_RaTG13_nucleoc;181 | 0.05 | 0.125 |
| MN996532.1;QHR63308.1;BatCoV_RaTG13_nucleoc;182 | 0 | 0.125 |
| MN996532.1;QHR63308.1;BatCoV_RaTG13_nucleoc;183 | 0.15 | 0.15 |

| | | |
|---|---|---|
| MN996532.1;QHR63308.1;BatCoV_RaTG13_nucleoc;184 | 0.15 | 0.125 |
| MN996532.1;QHR63308.1;BatCoV_RaTG13_nucleoc;185 | 0.15 | 0.15 |
| MN996532.1;QHR63308.1;BatCoV_RaTG13_nucleoc;186 | 0.15 | 0.2 |
| MN996532.1;QHR63308.1;BatCoV_RaTG13_nucleoc;187 | 0.15 | 0.225 |
| MN996532.1;QHR63308.1;BatCoV_RaTG13_nucleoc;188 | 0.1 | 0.2 |
| MN996532.1;QHR63308.1;BatCoV_RaTG13_nucleoc;189 | 0.1 | 0.1 |
| MN996532.1;QHR63308.1;BatCoV_RaTG13_nucleoc;190 | 0.2 | 0.225 |
| MN996532.1;QHR63308.1;BatCoV_RaTG13_nucleoc;191 | 0.05 | 0.05 |
| MN996532.1;QHR63308.1;BatCoV_RaTG13_nucleoc;192 | 0 | 0.075 |
| MN996532.1;QHR63308.1;BatCoV_RaTG13_nucleoc;193 | 0 | 0.025 |
| MN996532.1;QHR63308.1;BatCoV_RaTG13_nucleoc;194 | 0 | 0.05 |
| MN996532.1;QHR63308.1;BatCoV_RaTG13_nucleoc;195 | 0 | 0.025 |
| MN996532.1;QHR63308.1;BatCoV_RaTG13_nucleoc;196 | 0 | 0.05 |
| MN996532.1;QHR63308.1;BatCoV_RaTG13_nucleoc;197 | 0.05 | 0.05 |
| MN996532.1;QHR63308.1;BatCoV_RaTG13_nucleoc;198 | 0 | 0.025 |
| MN996532.1;QHR63308.1;BatCoV_RaTG13_nucleoc;199 | 0 | 0.025 |
| MN996532.1;QHR63308.1;BatCoV_RaTG13_nucleoc;200 | 0 | 0.075 |
| MN996532.1;QHR63308.1;BatCoV_RaTG13_nucleoc;201 | 0.05 | 0 |
| MN996532.1;QHR63308.1;BatCoV_RaTG13_nucleoc;202 | 0.05 | 0 |
| MN996532.1;QHR63308.1;BatCoV_RaTG13_nucleoc;203 | 0.05 | 0.025 |
| MN996532.1;QHR63308.1;BatCoV_RaTG13_nucleoc;204 | 0 | 0 |
| MN996532.1;QHR63308.1;BatCoV_RaTG13_nucleoc;205 | 0 | 0.025 |
| MN996532.1;QHR63308.1;BatCoV_RaTG13_nucleoc;206 | 0 | 0.025 |
| MN996532.1;QHR63308.1;BatCoV_RaTG13_nucleoc;207 | 0 | 0 |
| MN996532.1;QHR63308.1;BatCoV_RaTG13_nucleoc;208 | 0 | 0 |
| MN996532.1;QHR63308.1;BatCoV_RaTG13_nucleoc;209 | 0 | 0 |
| MN996532.1;QHR63308.1;BatCoV_RaTG13_nucleoc;210 | 0 | 0 |
| MN996532.1;QHR63308.1;BatCoV_RaTG13_nucleoc;211 | 0 | 0 |
| MN996532.1;QHR63308.1;BatCoV_RaTG13_nucleoc;212 | 0 | 0 |
| MN996532.1;QHR63308.1;BatCoV_RaTG13_nucleoc;213 | 0 | 0.075 |
| MN996532.1;QHR63308.1;BatCoV_RaTG13_nucleoc;214 | 0 | 0.1 |
| MN996532.1;QHR63308.1;BatCoV_RaTG13_nucleoc;215 | 0 | 0.075 |
| MN996532.1;QHR63308.1;BatCoV_RaTG13_nucleoc;216 | 0 | 0.05 |
| MN996532.1;QHR63308.1;BatCoV_RaTG13_nucleoc;217 | 0 | 0 |
| MN996532.1;QHR63308.1;BatCoV_RaTG13_nucleoc;218 | 0.05 | 0.05 |
| MN996532.1;QHR63308.1;BatCoV_RaTG13_nucleoc;219 | 0.05 | 0.1 |
| MN996532.1;QHR63308.1;BatCoV_RaTG13_nucleoc;220 | 0.05 | 0.225 |
| MN996532.1;QHR63308.1;BatCoV_RaTG13_nucleoc;221 | 0.05 | 0.4 |
| MN996532.1;QHR63308.1;BatCoV_RaTG13_nucleoc;222 | 0.1 | 0.325 |
| MN996532.1;QHR63308.1;BatCoV_RaTG13_nucleoc;223 | 0.05 | 0.175 |
| MN996532.1;QHR63308.1;BatCoV_RaTG13_nucleoc;224 | 0.05 | 0.075 |
| MN996532.1;QHR63308.1;BatCoV_RaTG13_nucleoc;225 | 0.05 | 0.1 |
| MN996532.1;QHR63308.1;BatCoV_RaTG13_nucleoc;226 | 0.05 | 0.175 |

| | | |
|---|---|---|
| MN996532.1;QHR63308.1;BatCoV_RaTG13_nucleoc;227 | 0 | 0.25 |
| MN996532.1;QHR63308.1;BatCoV_RaTG13_nucleoc;228 | 0 | 0.25 |
| MN996532.1;QHR63308.1;BatCoV_RaTG13_nucleoc;229 | 0.05 | 0.3 |
| MN996532.1;QHR63308.1;BatCoV_RaTG13_nucleoc;230 | 0 | 0.3 |
| MN996532.1;QHR63308.1;BatCoV_RaTG13_nucleoc;231 | 0.05 | 0.35 |
| MN996532.1;QHR63308.1;BatCoV_RaTG13_nucleoc;232 | 0 | 0.225 |
| MN996532.1;QHR63308.1;BatCoV_RaTG13_nucleoc;233 | 0 | 0.225 |
| MN996532.1;QHR63308.1;BatCoV_RaTG13_nucleoc;234 | 0 | 0 |
| MN996532.1;QHR63308.1;BatCoV_RaTG13_nucleoc;235 | 0 | 0 |
| MN996532.1;QHR63308.1;BatCoV_RaTG13_nucleoc;236 | 0 | 0 |
| MN996532.1;QHR63308.1;BatCoV_RaTG13_nucleoc;237 | 0 | 0 |
| MN996532.1;QHR63308.1;BatCoV_RaTG13_nucleoc;238 | 0 | 0.075 |
| MN996532.1;QHR63308.1;BatCoV_RaTG13_nucleoc;239 | 0 | 0.05 |
| MN996532.1;QHR63308.1;BatCoV_RaTG13_nucleoc;240 | 0.2 | 0.1 |
| MN996532.1;QHR63308.1;BatCoV_RaTG13_nucleoc;241 | 0 | 0 |
| MN996532.1;QHR63308.1;BatCoV_RaTG13_nucleoc;242 | 0 | 0.025 |
| MN996532.1;QHR63308.1;BatCoV_RaTG13_nucleoc;243 | 0 | 0.075 |
| MN996532.1;QHR63308.1;BatCoV_RaTG13_nucleoc;244 | 0 | 0.05 |
| MN996532.1;QHR63308.1;BatCoV_RaTG13_nucleoc;245 | 0 | 0.075 |
| MN996532.1;QHR63308.1;BatCoV_RaTG13_nucleoc;246 | 0 | 0.025 |
| MN996532.1;QHR63308.1;BatCoV_RaTG13_nucleoc;247 | 0.05 | 0.025 |
| MN996532.1;QHR63308.1;BatCoV_RaTG13_nucleoc;248 | 0 | 0.075 |
| MN996532.1;QHR63308.1;BatCoV_RaTG13_nucleoc;249 | 0 | 0.075 |
| MN996532.1;QHR63308.1;BatCoV_RaTG13_nucleoc;250 | 0 | 0.3 |
| MN996532.1;QHR63308.1;BatCoV_RaTG13_nucleoc;251 | 0 | 0 |
| MN996532.1;QHR63308.1;BatCoV_RaTG13_nucleoc;252 | 0.05 | 0 |
| MN996532.1;QHR63308.1;BatCoV_RaTG13_nucleoc;253 | 0 | 0 |
| MN996532.1;QHR63308.1;BatCoV_RaTG13_nucleoc;254 | 0 | 0 |
| MN996532.1;QHR63308.1;BatCoV_RaTG13_nucleoc;255 | 0 | 0 |
| MN996532.1;QHR63308.1;BatCoV_RaTG13_nucleoc;256 | 0 | 0.025 |
| MN996532.1;QHR63308.1;BatCoV_RaTG13_nucleoc;257 | 0 | 0.05 |
| MN996532.1;QHR63308.1;BatCoV_RaTG13_nucleoc;258 | 0.1 | 0.075 |
| MN996532.1;QHR63308.1;BatCoV_RaTG13_nucleoc;259 | 0.05 | 0.075 |
| MN996532.1;QHR63308.1;BatCoV_RaTG13_nucleoc;260 | 0.05 | 0.1 |
| MN996532.1;QHR63308.1;BatCoV_RaTG13_nucleoc;261 | 0.05 | 0.075 |
| MN996532.1;QHR63308.1;BatCoV_RaTG13_nucleoc;262 | 0.05 | 0.075 |
| MN996532.1;QHR63308.1;BatCoV_RaTG13_nucleoc;263 | 0.05 | 0.075 |
| MN996532.1;QHR63308.1;BatCoV_RaTG13_nucleoc;264 | 0.1 | 0.075 |
| MN996532.1;QHR63308.1;BatCoV_RaTG13_nucleoc;265 | 0.1 | 0.1 |
| MN996532.1;QHR63308.1;BatCoV_RaTG13_nucleoc;266 | 0.1 | 0.1 |
| MN996532.1;QHR63308.1;BatCoV_RaTG13_nucleoc;267 | 0.05 | 0 |
| MN996532.1;QHR63308.1;BatCoV_RaTG13_nucleoc;268 | 0.05 | 0 |
| MN996532.1;QHR63308.1;BatCoV_RaTG13_nucleoc;269 | 0.05 | 0 |

| | | |
|---|---|---|
| MN996532.1;QHR63308.1;BatCoV_RaTG13_nucleoc;270 | 0.15 | 0.05 |
| MN996532.1;QHR63308.1;BatCoV_RaTG13_nucleoc;271 | 0.15 | 0.05 |
| MN996532.1;QHR63308.1;BatCoV_RaTG13_nucleoc;272 | 0.15 | 0.05 |
| MN996532.1;QHR63308.1;BatCoV_RaTG13_nucleoc;273 | 0.1 | 0.075 |
| MN996532.1;QHR63308.1;BatCoV_RaTG13_nucleoc;274 | 0.1 | 0.05 |
| MN996532.1;QHR63308.1;BatCoV_RaTG13_nucleoc;275 | 0.05 | 0.025 |
| MN996532.1;QHR63308.1;BatCoV_RaTG13_nucleoc;276 | 0.05 | 0.05 |
| MN996532.1;QHR63308.1;BatCoV_RaTG13_nucleoc;277 | 0.05 | 0.025 |
| MN996532.1;QHR63308.1;BatCoV_RaTG13_nucleoc;278 | 0.05 | 0 |
| MN996532.1;QHR63308.1;BatCoV_RaTG13_nucleoc;279 | 0.05 | 0.05 |
| MN996532.1;QHR63308.1;BatCoV_RaTG13_nucleoc;280 | 0.05 | 0.175 |
| MN996532.1;QHR63308.1;BatCoV_RaTG13_nucleoc;281 | 0 | 0.1 |
| MN996532.1;QHR63308.1;BatCoV_RaTG13_nucleoc;282 | 0 | 0.05 |
| MN996532.1;QHR63308.1;BatCoV_RaTG13_nucleoc;283 | 0 | 0.1 |
| MN996532.1;QHR63308.1;BatCoV_RaTG13_nucleoc;284 | 0 | 0.075 |
| MN996532.1;QHR63308.1;BatCoV_RaTG13_nucleoc;285 | 0 | 0.075 |
| MN996532.1;QHR63308.1;BatCoV_RaTG13_nucleoc;286 | 0 | 0.075 |
| MN996532.1;QHR63308.1;BatCoV_RaTG13_nucleoc;287 | 0 | 0.025 |
| MN996532.1;QHR63308.1;BatCoV_RaTG13_nucleoc;288 | 0 | 0 |
| MN996532.1;QHR63308.1;BatCoV_RaTG13_nucleoc;289 | 0 | 0 |
| MN996532.1;QHR63308.1;BatCoV_RaTG13_nucleoc;290 | 0 | 0.025 |
| MN996532.1;QHR63308.1;BatCoV_RaTG13_nucleoc;291 | 0 | 0 |
| MN996532.1;QHR63308.1;BatCoV_RaTG13_nucleoc;292 | 0 | 0 |
| MN996532.1;QHR63308.1;BatCoV_RaTG13_nucleoc;293 | 0 | 0 |
| MN996532.1;QHR63308.1;BatCoV_RaTG13_nucleoc;294 | 0.05 | 0 |
| MN996532.1;QHR63308.1;BatCoV_RaTG13_nucleoc;295 | 0 | 0.05 |
| MN996532.1;QHR63308.1;BatCoV_RaTG13_nucleoc;296 | 0 | 0 |
| MN996532.1;QHR63308.1;BatCoV_RaTG13_nucleoc;297 | 0.15 | 0.025 |
| MN996532.1;QHR63308.1;BatCoV_RaTG13_nucleoc;298 | 0 | 0 |
| MN996532.1;QHR63308.1;BatCoV_RaTG13_nucleoc;299 | 0 | 0 |
| MN996532.1;QHR63308.1;BatCoV_RaTG13_nucleoc;300 | 0 | 0 |
| MN996532.1;QHR63308.1;BatCoV_RaTG13_nucleoc;301 | 0 | 0 |
| MN996532.1;QHR63308.1;BatCoV_RaTG13_nucleoc;302 | 0 | 0.025 |
| MN996532.1;QHR63308.1;BatCoV_RaTG13_nucleoc;303 | 0 | 0 |
| MN996532.1;QHR63308.1;BatCoV_RaTG13_nucleoc;304 | 0 | 0 |
| MN996532.1;QHR63308.1;BatCoV_RaTG13_nucleoc;305 | 0 | 0 |
| MN996532.1;QHR63308.1;BatCoV_RaTG13_nucleoc;306 | 0 | 0 |
| MN996532.1;QHR63308.1;BatCoV_RaTG13_nucleoc;307 | 0 | 0 |
| MN996532.1;QHR63308.1;BatCoV_RaTG13_nucleoc;308 | 0 | 0 |
| MN996532.1;QHR63308.1;BatCoV_RaTG13_nucleoc;309 | 0 | 0.025 |
| MN996532.1;QHR63308.1;BatCoV_RaTG13_nucleoc;310 | 0.05 | 0.025 |
| MN996532.1;QHR63308.1;BatCoV_RaTG13_nucleoc;311 | 0 | 0.05 |
| MN996532.1;QHR63308.1;BatCoV_RaTG13_nucleoc;312 | 0 | 0.075 |

| | | |
|---|---|---|
| MN996532.1;QHR63308.1;BatCoV_RaTG13_nucleoc;313 | 0.05 | 0.05 |
| MN996532.1;QHR63308.1;BatCoV_RaTG13_nucleoc;314 | 0.05 | 0.05 |
| MN996532.1;QHR63308.1;BatCoV_RaTG13_nucleoc;315 | 0 | 0.05 |
| MN996532.1;QHR63308.1;BatCoV_RaTG13_nucleoc;316 | 0 | 0.025 |
| MN996532.1;QHR63308.1;BatCoV_RaTG13_nucleoc;317 | 0 | 0.025 |
| MN996532.1;QHR63308.1;BatCoV_RaTG13_nucleoc;318 | 0 | 0.025 |
| MN996532.1;QHR63308.1;BatCoV_RaTG13_nucleoc;319 | 0 | 0 |
| MN996532.1;QHR63308.1;BatCoV_RaTG13_nucleoc;320 | 0 | 0 |
| MN996532.1;QHR63308.1;BatCoV_RaTG13_nucleoc;321 | 0 | 0 |
| MN996532.1;QHR63308.1;BatCoV_RaTG13_nucleoc;322 | 0 | 0 |
| MN996532.1;QHR63308.1;BatCoV_RaTG13_nucleoc;323 | 0 | 0 |
| MN996532.1;QHR63308.1;BatCoV_RaTG13_nucleoc;324 | 0 | 0 |
| MN996532.1;QHR63308.1;BatCoV_RaTG13_nucleoc;325 | 0 | 0 |
| MN996532.1;QHR63308.1;BatCoV_RaTG13_nucleoc;326 | 0 | 0 |
| MN996532.1;QHR63308.1;BatCoV_RaTG13_nucleoc;327 | 0 | 0.075 |
| MN996532.1;QHR63308.1;BatCoV_RaTG13_nucleoc;328 | 0 | 0.05 |
| MN996532.1;QHR63308.1;BatCoV_RaTG13_nucleoc;329 | 0 | 0.075 |
| MN996532.1;QHR63308.1;BatCoV_RaTG13_nucleoc;330 | 0 | 0.1 |
| MN996532.1;QHR63308.1;BatCoV_RaTG13_nucleoc;331 | 0 | 0.175 |
| MN996532.1;QHR63308.1;BatCoV_RaTG13_nucleoc;332 | 0.1 | 0.175 |
| MN996532.1;QHR63308.1;BatCoV_RaTG13_nucleoc;333 | 0.05 | 0.25 |
| MN996532.1;QHR63308.1;BatCoV_RaTG13_nucleoc;334 | 0 | 0.175 |
| MN996532.1;QHR63308.1;BatCoV_RaTG13_nucleoc;335 | 0.1 | 0.225 |
| MN996532.1;QHR63308.1;BatCoV_RaTG13_nucleoc;336 | 0 | 0.2 |
| MN996532.1;QHR63308.1;BatCoV_RaTG13_nucleoc;337 | 0.05 | 0.175 |
| MN996532.1;QHR63308.1;BatCoV_RaTG13_nucleoc;338 | 0 | 0.1 |
| MN996532.1;QHR63308.1;BatCoV_RaTG13_nucleoc;339 | 0 | 0.2 |
| MN996532.1;QHR63308.1;BatCoV_RaTG13_nucleoc;340 | 0 | 0.15 |
| MN996532.1;QHR63308.1;BatCoV_RaTG13_nucleoc;341 | 0 | 0.075 |
| MN996532.1;QHR63308.1;BatCoV_RaTG13_nucleoc;342 | 0 | 0.05 |
| MN996532.1;QHR63308.1;BatCoV_RaTG13_nucleoc;343 | 0.05 | 0.1 |
| MN996532.1;QHR63308.1;BatCoV_RaTG13_nucleoc;344 | 0.05 | 0.1 |
| MN996532.1;QHR63308.1;BatCoV_RaTG13_nucleoc;345 | 0.05 | 0.05 |
| MN996532.1;QHR63308.1;BatCoV_RaTG13_nucleoc;346 | 0 | 0.075 |
| MN996532.1;QHR63308.1;BatCoV_RaTG13_nucleoc;347 | 0 | 0.05 |
| MN996532.1;QHR63308.1;BatCoV_RaTG13_nucleoc;348 | 0 | 0 |
| MN996532.1;QHR63308.1;BatCoV_RaTG13_nucleoc;349 | 0 | 0 |
| MN996532.1;QHR63308.1;BatCoV_RaTG13_nucleoc;350 | 0 | 0 |
| MN996532.1;QHR63308.1;BatCoV_RaTG13_nucleoc;351 | 0 | 0.025 |
| MN996532.1;QHR63308.1;BatCoV_RaTG13_nucleoc;352 | 0 | 0.15 |
| MN996532.1;QHR63308.1;BatCoV_RaTG13_nucleoc;353 | 0.05 | 0.125 |
| MN996532.1;QHR63308.1;BatCoV_RaTG13_nucleoc;354 | 0 | 0.125 |
| MN996532.1;QHR63308.1;BatCoV_RaTG13_nucleoc;355 | 0 | 0.05 |

| | | |
|---|---|---|
| MN996532.1;QHR63308.1;BatCoV_RaTG13_nucleoc;356 | 0 | 0.25 |
| MN996532.1;QHR63308.1;BatCoV_RaTG13_nucleoc;357 | 0.05 | 0.125 |
| MN996532.1;QHR63308.1;BatCoV_RaTG13_nucleoc;358 | 0.05 | 0.1 |
| MN996532.1;QHR63308.1;BatCoV_RaTG13_nucleoc;359 | 0 | 0.075 |
| MN996532.1;QHR63308.1;BatCoV_RaTG13_nucleoc;360 | 0 | 0.075 |
| MN996532.1;QHR63308.1;BatCoV_RaTG13_nucleoc;361 | 0 | 0.05 |
| MN996532.1;QHR63308.1;BatCoV_RaTG13_nucleoc;362 | 0.05 | 0.1 |
| MN996532.1;QHR63308.1;BatCoV_RaTG13_nucleoc;363 | 0.05 | 0.1 |
| MN996532.1;QHR63308.1;BatCoV_RaTG13_nucleoc;364 | 0.05 | 0.025 |
| MN996532.1;QHR63308.1;BatCoV_RaTG13_nucleoc;365 | 0 | 0 |
| MN996532.1;QHR63308.1;BatCoV_RaTG13_nucleoc;366 | 0 | 0 |
| MN996532.1;QHR63308.1;BatCoV_RaTG13_nucleoc;367 | 0 | 0.025 |
| MN996532.1;QHR63308.1;BatCoV_RaTG13_nucleoc;368 | 0 | 0.05 |
| MN996532.1;QHR63308.1;BatCoV_RaTG13_nucleoc;369 | 0 | 0.075 |
| MN996532.1;QHR63308.1;BatCoV_RaTG13_nucleoc;370 | 0 | 0.075 |
| MN996532.1;QHR63308.1;BatCoV_RaTG13_nucleoc;371 | 0 | 0.05 |
| MN996532.1;QHR63308.1;BatCoV_RaTG13_nucleoc;372 | 0 | 0.1 |
| MN996532.1;QHR63308.1;BatCoV_RaTG13_nucleoc;373 | 0 | 0.1 |
| MN996532.1;QHR63308.1;BatCoV_RaTG13_nucleoc;374 | 0 | 0.125 |
| MN996532.1;QHR63308.1;BatCoV_RaTG13_nucleoc;375 | 0 | 0.1 |
| MN996532.1;QHR63308.1;BatCoV_RaTG13_nucleoc;376 | 0 | 0.125 |
| MN996532.1;QHR63308.1;BatCoV_RaTG13_nucleoc;377 | 0.05 | 0.125 |
| MN996532.1;QHR63308.1;BatCoV_RaTG13_nucleoc;378 | 0 | 0.175 |
| MN996532.1;QHR63308.1;BatCoV_RaTG13_nucleoc;379 | 0 | 0.125 |
| MN996532.1;QHR63308.1;BatCoV_RaTG13_nucleoc;380 | 0 | 0.15 |
| MN996532.1;QHR63308.1;BatCoV_RaTG13_nucleoc;381 | 0 | 0.225 |
| MN996532.1;QHR63308.1;BatCoV_RaTG13_nucleoc;382 | 0 | 0.225 |
| MN996532.1;QHR63308.1;BatCoV_RaTG13_nucleoc;383 | 0.05 | 0.125 |
| MN996532.1;QHR63308.1;BatCoV_RaTG13_nucleoc;384 | 0 | 0.125 |
| MN996532.1;QHR63308.1;BatCoV_RaTG13_nucleoc;385 | 0 | 0.15 |
| MN996532.1;QHR63308.1;BatCoV_RaTG13_nucleoc;386 | 0 | 0.4 |
| MN996532.1;QHR63308.1;BatCoV_RaTG13_nucleoc;387 | 0 | 0.5 |
| MN996532.1;QHR63308.1;BatCoV_RaTG13_nucleoc;388 | 0 | 0.575 |
| MN996532.1;QHR63308.1;BatCoV_RaTG13_nucleoc;389 | 0 | 0.3 |
| MN996532.1;QHR63308.1;BatCoV_RaTG13_nucleoc;390 | 0 | 0.7 |
| MN996532.1;QHR63308.1;BatCoV_RaTG13_nucleoc;391 | 0 | 0.75 |
| MN996532.1;QHR63308.1;BatCoV_RaTG13_nucleoc;392 | 0 | 0.775 |
| MN996532.1;QHR63308.1;BatCoV_RaTG13_nucleoc;393 | 0 | 0.8 |
| MN996532.1;QHR63308.1;BatCoV_RaTG13_nucleoc;394 | 0 | 0.825 |
| MN996532.1;QHR63308.1;BatCoV_RaTG13_nucleoc;395 | 0.05 | 0.775 |
| MN996532.1;QHR63308.1;BatCoV_RaTG13_nucleoc;396 | 0 | 0.725 |
| MN996532.1;QHR63308.1;BatCoV_RaTG13_nucleoc;397 | 0.1 | 0.7 |
| MN996532.1;QHR63308.1;BatCoV_RaTG13_nucleoc;398 | 0.25 | 0.55 |

| | | |
|---|---|---|
| MN996532.1;QHR63308.1;BatCoV_RaTG13_nucleoc;399 | 0 | 0.675 |
| MN996532.1;QHR63308.1;BatCoV_RaTG13_nucleoc;400 | 0 | 0.2 |
| MN996532.1;QHR63308.1;BatCoV_RaTG13_nucleoc;401 | 0 | 0.075 |
| MN996532.1;QHR63308.1;BatCoV_RaTG13_nucleoc;402 | 0 | 0.075 |
| MN996532.1;QHR63308.1;BatCoV_RaTG13_nucleoc;403 | 0.05 | 0.1 |
| MN996532.1;QHR63308.1;BatCoV_RaTG13_nucleoc;404 | 0 | 0.05 |
| MN996532.1;QHR63308.1;BatCoV_RaTG13_nucleoc;405 | 0 | 0.025 |
| MN996532.1;QHR63308.1;BatCoV_RaTG13_nucleoc;406 | 0 | 0.05 |
| MN996532.1;QHR63308.1;BatCoV_RaTG13_nucleoc;407 | 0 | 0.075 |
| MN996532.1;QHR63308.1;BatCoV_RaTG13_nucleoc;408 | 0 | 0.025 |
| MT072864.1;QIQ54046.1;Pangolin_CoV_orf1a;1 | 0 | 0 |
| MT072864.1;QIQ54046.1;Pangolin_CoV_orf1a;2 | 0 | 0.025 |
| MT072864.1;QIQ54046.1;Pangolin_CoV_orf1a;3 | 0 | 0 |
| MT072864.1;QIQ54046.1;Pangolin_CoV_orf1a;4 | 0 | 0.025 |
| MT072864.1;QIQ54046.1;Pangolin_CoV_orf1a;5 | 0 | 0 |
| MT072864.1;QIQ54046.1;Pangolin_CoV_orf1a;6 | 0 | 0 |
| MT072864.1;QIQ54046.1;Pangolin_CoV_orf1a;7 | 0 | 0 |
| MT072864.1;QIQ54046.1;Pangolin_CoV_orf1a;8 | 0 | 0 |
| MT072864.1;QIQ54046.1;Pangolin_CoV_orf1a;9 | 0 | 0 |
| MT072864.1;QIQ54046.1;Pangolin_CoV_orf1a;10 | 0 | 0 |
| MT072864.1;QIQ54046.1;Pangolin_CoV_orf1a;11 | 0 | 0 |
| MT072864.1;QIQ54046.1;Pangolin_CoV_orf1a;12 | 0 | 0 |
| MT072864.1;QIQ54046.1;Pangolin_CoV_orf1a;13 | 0 | 0 |
| MT072864.1;QIQ54046.1;Pangolin_CoV_orf1a;14 | 0 | 0 |
| MT072864.1;QIQ54046.1;Pangolin_CoV_orf1a;15 | 0 | 0 |
| MT072864.1;QIQ54046.1;Pangolin_CoV_orf1a;16 | 0 | 0 |
| MT072864.1;QIQ54046.1;Pangolin_CoV_orf1a;17 | 0 | 0 |
| MT072864.1;QIQ54046.1;Pangolin_CoV_orf1a;18 | 0 | 0 |
| MT072864.1;QIQ54046.1;Pangolin_CoV_orf1a;19 | 0 | 0 |
| MT072864.1;QIQ54046.1;Pangolin_CoV_orf1a;20 | 0 | 0 |
| MT072864.1;QIQ54046.1;Pangolin_CoV_orf1a;21 | 0 | 0 |
| MT072864.1;QIQ54046.1;Pangolin_CoV_orf1a;22 | 0 | 0 |
| MT072864.1;QIQ54046.1;Pangolin_CoV_orf1a;23 | 0 | 0 |
| MT072864.1;QIQ54046.1;Pangolin_CoV_orf1a;24 | 0 | 0 |
| MT072864.1;QIQ54046.1;Pangolin_CoV_orf1a;25 | 0 | 0 |
| MT072864.1;QIQ54046.1;Pangolin_CoV_orf1a;26 | 0 | 0 |
| MT072864.1;QIQ54046.1;Pangolin_CoV_orf1a;27 | 0 | 0 |
| MT072864.1;QIQ54046.1;Pangolin_CoV_orf1a;28 | 0 | 0 |
| MT072864.1;QIQ54046.1;Pangolin_CoV_orf1a;29 | 0.1 | 0 |
| MT072864.1;QIQ54046.1;Pangolin_CoV_orf1a;30 | 0.3 | 0.125 |
| MT072864.1;QIQ54046.1;Pangolin_CoV_orf1a;31 | 0 | 0 |
| MT072864.1;QIQ54046.1;Pangolin_CoV_orf1a;32 | 0.05 | 0.025 |
| MT072864.1;QIQ54046.1;Pangolin_CoV_orf1a;33 | 0.05 | 0 |

| | | |
|---|---|---|
| MT072864.1;QIQ54046.1;Pangolin_CoV_orf1a;34 | 0.1 | 0 |
| MT072864.1;QIQ54046.1;Pangolin_CoV_orf1a;35 | 0 | 0.025 |
| MT072864.1;QIQ54046.1;Pangolin_CoV_orf1a;36 | 0 | 0 |
| MT072864.1;QIQ54046.1;Pangolin_CoV_orf1a;37 | 0 | 0 |
| MT072864.1;QIQ54046.1;Pangolin_CoV_orf1a;38 | 0 | 0 |
| MT072864.1;QIQ54046.1;Pangolin_CoV_orf1a;39 | 0 | 0 |
| MT072864.1;QIQ54046.1;Pangolin_CoV_orf1a;40 | 0 | 0 |
| MT072864.1;QIQ54046.1;Pangolin_CoV_orf1a;41 | 0 | 0 |
| MT072864.1;QIQ54046.1;Pangolin_CoV_orf1a;42 | 0 | 0 |
| MT072864.1;QIQ54046.1;Pangolin_CoV_orf1a;43 | 0 | 0 |
| MT072864.1;QIQ54046.1;Pangolin_CoV_orf1a;44 | 0.05 | 0 |
| MT072864.1;QIQ54046.1;Pangolin_CoV_orf1a;45 | 0 | 0.05 |
| MT072864.1;QIQ54046.1;Pangolin_CoV_orf1a;46 | 0 | 0.075 |
| MT072864.1;QIQ54046.1;Pangolin_CoV_orf1a;47 | 0.05 | 0.075 |
| MT072864.1;QIQ54046.1;Pangolin_CoV_orf1a;48 | 0.05 | 0.05 |
| MT072864.1;QIQ54046.1;Pangolin_CoV_orf1a;49 | 0.05 | 0.05 |
| MT072864.1;QIQ54046.1;Pangolin_CoV_orf1a;50 | 0 | 0.05 |
| MT072864.1;QIQ54046.1;Pangolin_CoV_orf1a;51 | 0 | 0 |
| MT072864.1;QIQ54046.1;Pangolin_CoV_orf1a;52 | 0.1 | 0.15 |
| MT072864.1;QIQ54046.1;Pangolin_CoV_orf1a;53 | 0.1 | 0.05 |
| MT072864.1;QIQ54046.1;Pangolin_CoV_orf1a;54 | 0 | 0.025 |
| MT072864.1;QIQ54046.1;Pangolin_CoV_orf1a;55 | 0 | 0.025 |
| MT072864.1;QIQ54046.1;Pangolin_CoV_orf1a;56 | 0 | 0.025 |
| MT072864.1;QIQ54046.1;Pangolin_CoV_orf1a;57 | 0 | 0.025 |
| MT072864.1;QIQ54046.1;Pangolin_CoV_orf1a;58 | 0 | 0.025 |
| MT072864.1;QIQ54046.1;Pangolin_CoV_orf1a;59 | 0 | 0 |
| MT072864.1;QIQ54046.1;Pangolin_CoV_orf1a;60 | 0 | 0 |
| MT072864.1;QIQ54046.1;Pangolin_CoV_orf1a;61 | 0 | 0 |
| MT072864.1;QIQ54046.1;Pangolin_CoV_orf1a;62 | 0 | 0 |
| MT072864.1;QIQ54046.1;Pangolin_CoV_orf1a;63 | 0 | 0 |
| MT072864.1;QIQ54046.1;Pangolin_CoV_orf1a;64 | 0 | 0 |
| MT072864.1;QIQ54046.1;Pangolin_CoV_orf1a;65 | 0 | 0 |
| MT072864.1;QIQ54046.1;Pangolin_CoV_orf1a;66 | 0 | 0 |
| MT072864.1;QIQ54046.1;Pangolin_CoV_orf1a;67 | 0 | 0 |
| MT072864.1;QIQ54046.1;Pangolin_CoV_orf1a;68 | 0 | 0 |
| MT072864.1;QIQ54046.1;Pangolin_CoV_orf1a;69 | 0 | 0 |
| MT072864.1;QIQ54046.1;Pangolin_CoV_orf1a;70 | 0.05 | 0 |
| MT072864.1;QIQ54046.1;Pangolin_CoV_orf1a;71 | 0 | 0 |
| MT072864.1;QIQ54046.1;Pangolin_CoV_orf1a;72 | 0 | 0 |
| MT072864.1;QIQ54046.1;Pangolin_CoV_orf1a;73 | 0 | 0 |
| MT072864.1;QIQ54046.1;Pangolin_CoV_orf1a;74 | 0 | 0 |
| MT072864.1;QIQ54046.1;Pangolin_CoV_orf1a;75 | 0 | 0.025 |
| MT072864.1;QIQ54046.1;Pangolin_CoV_orf1a;76 | 0 | 0 |

| | | |
|---|---|---|
| MT072864.1;QIQ54046.1;Pangolin_CoV_orf1a;77 | 0 | 0 |
| MT072864.1;QIQ54046.1;Pangolin_CoV_orf1a;78 | 0 | 0 |
| MT072864.1;QIQ54046.1;Pangolin_CoV_orf1a;79 | 0 | 0 |
| MT072864.1;QIQ54046.1;Pangolin_CoV_orf1a;80 | 0 | 0 |
| MT072864.1;QIQ54046.1;Pangolin_CoV_orf1a;81 | 0 | 0 |
| MT072864.1;QIQ54046.1;Pangolin_CoV_orf1a;82 | 0 | 0 |
| MT072864.1;QIQ54046.1;Pangolin_CoV_orf1a;83 | 0 | 0 |
| MT072864.1;QIQ54046.1;Pangolin_CoV_orf1a;84 | 0 | 0 |
| MT072864.1;QIQ54046.1;Pangolin_CoV_orf1a;85 | 0 | 0 |
| MT072864.1;QIQ54046.1;Pangolin_CoV_orf1a;86 | 0 | 0.025 |
| MT072864.1;QIQ54046.1;Pangolin_CoV_orf1a;87 | 0 | 0 |
| MT072864.1;QIQ54046.1;Pangolin_CoV_orf1a;88 | 0 | 0 |
| MT072864.1;QIQ54046.1;Pangolin_CoV_orf1a;89 | 0 | 0 |
| MT072864.1;QIQ54046.1;Pangolin_CoV_orf1a;90 | 0.25 | 0.275 |
| MT072864.1;QIQ54046.1;Pangolin_CoV_orf1a;91 | 0.1 | 0.05 |
| MT072864.1;QIQ54046.1;Pangolin_CoV_orf1a;92 | 0 | 0 |
| MT072864.1;QIQ54046.1;Pangolin_CoV_orf1a;93 | 0 | 0 |
| MT072864.1;QIQ54046.1;Pangolin_CoV_orf1a;94 | 0 | 0.025 |
| MT072864.1;QIQ54046.1;Pangolin_CoV_orf1a;95 | 0 | 0 |
| MT072864.1;QIQ54046.1;Pangolin_CoV_orf1a;96 | 0 | 0 |
| MT072864.1;QIQ54046.1;Pangolin_CoV_orf1a;97 | 0 | 0 |
| MT072864.1;QIQ54046.1;Pangolin_CoV_orf1a;98 | 0 | 0 |
| MT072864.1;QIQ54046.1;Pangolin_CoV_orf1a;99 | 0 | 0 |
| MT072864.1;QIQ54046.1;Pangolin_CoV_orf1a;100 | 0 | 0 |
| MT072864.1;QIQ54046.1;Pangolin_CoV_orf1a;101 | 0 | 0 |
| MT072864.1;QIQ54046.1;Pangolin_CoV_orf1a;102 | 0 | 0 |
| MT072864.1;QIQ54046.1;Pangolin_CoV_orf1a;103 | 0 | 0 |
| MT072864.1;QIQ54046.1;Pangolin_CoV_orf1a;104 | 0 | 0 |
| MT072864.1;QIQ54046.1;Pangolin_CoV_orf1a;105 | 0 | 0 |
| MT072864.1;QIQ54046.1;Pangolin_CoV_orf1a;106 | 0 | 0 |
| MT072864.1;QIQ54046.1;Pangolin_CoV_orf1a;107 | 0 | 0 |
| MT072864.1;QIQ54046.1;Pangolin_CoV_orf1a;108 | 0 | 0 |
| MT072864.1;QIQ54046.1;Pangolin_CoV_orf1a;109 | 0 | 0 |
| MT072864.1;QIQ54046.1;Pangolin_CoV_orf1a;110 | 0 | 0 |
| MT072864.1;QIQ54046.1;Pangolin_CoV_orf1a;111 | 0 | 0 |
| MT072864.1;QIQ54046.1;Pangolin_CoV_orf1a;112 | 0 | 0 |
| MT072864.1;QIQ54046.1;Pangolin_CoV_orf1a;113 | 0 | 0 |
| MT072864.1;QIQ54046.1;Pangolin_CoV_orf1a;114 | 0 | 0.025 |
| MT072864.1;QIQ54046.1;Pangolin_CoV_orf1a;115 | 0 | 0.025 |
| MT072864.1;QIQ54046.1;Pangolin_CoV_orf1a;116 | 0 | 0.025 |
| MT072864.1;QIQ54046.1;Pangolin_CoV_orf1a;117 | 0 | 0.025 |
| MT072864.1;QIQ54046.1;Pangolin_CoV_orf1a;118 | 0 | 0.025 |
| MT072864.1;QIQ54046.1;Pangolin_CoV_orf1a;119 | 0 | 0.025 |

| | | |
|---|---|---|
| MT072864.1;QIQ54046.1;Pangolin_CoV_orf1a;120 | 0 | 0.025 |
| MT072864.1;QIQ54046.1;Pangolin_CoV_orf1a;121 | 0.05 | 0.05 |
| MT072864.1;QIQ54046.1;Pangolin_CoV_orf1a;122 | 0 | 0.025 |
| MT072864.1;QIQ54046.1;Pangolin_CoV_orf1a;123 | 0 | 0.025 |
| MT072864.1;QIQ54046.1;Pangolin_CoV_orf1a;124 | 0 | 0.05 |
| MT072864.1;QIQ54046.1;Pangolin_CoV_orf1a;125 | 0.1 | 0.025 |
| MT072864.1;QIQ54046.1;Pangolin_CoV_orf1a;126 | 0 | 0 |
| MT072864.1;QIQ54046.1;Pangolin_CoV_orf1a;127 | 0 | 0 |
| MT072864.1;QIQ54046.1;Pangolin_CoV_orf1a;128 | 0 | 0.025 |
| MT072864.1;QIQ54046.1;Pangolin_CoV_orf1a;129 | 0 | 0.025 |
| MT072864.1;QIQ54046.1;Pangolin_CoV_orf1a;130 | 0 | 0.025 |
| MT072864.1;QIQ54046.1;Pangolin_CoV_orf1a;131 | 0 | 0.025 |
| MT072864.1;QIQ54046.1;Pangolin_CoV_orf1a;132 | 0.05 | 0.025 |
| MT072864.1;QIQ54046.1;Pangolin_CoV_orf1a;133 | 0.05 | 0 |
| MT072864.1;QIQ54046.1;Pangolin_CoV_orf1a;134 | 0 | 0 |
| MT072864.1;QIQ54046.1;Pangolin_CoV_orf1a;135 | 0 | 0 |
| MT072864.1;QIQ54046.1;Pangolin_CoV_orf1a;136 | 0 | 0 |
| MT072864.1;QIQ54046.1;Pangolin_CoV_orf1a;137 | 0 | 0 |
| MT072864.1;QIQ54046.1;Pangolin_CoV_orf1a;138 | 0 | 0 |
| MT072864.1;QIQ54046.1;Pangolin_CoV_orf1a;139 | 0.05 | 0 |
| MT072864.1;QIQ54046.1;Pangolin_CoV_orf1a;140 | 0 | 0 |
| MT072864.1;QIQ54046.1;Pangolin_CoV_orf1a;141 | 0 | 0 |
| MT072864.1;QIQ54046.1;Pangolin_CoV_orf1a;142 | 0 | 0 |
| MT072864.1;QIQ54046.1;Pangolin_CoV_orf1a;143 | 0 | 0 |
| MT072864.1;QIQ54046.1;Pangolin_CoV_orf1a;144 | 0 | 0 |
| MT072864.1;QIQ54046.1;Pangolin_CoV_orf1a;145 | 0.1 | 0.125 |
| MT072864.1;QIQ54046.1;Pangolin_CoV_orf1a;146 | 0 | 0 |
| MT072864.1;QIQ54046.1;Pangolin_CoV_orf1a;147 | 0 | 0 |
| MT072864.1;QIQ54046.1;Pangolin_CoV_orf1a;148 | 0 | 0 |
| MT072864.1;QIQ54046.1;Pangolin_CoV_orf1a;149 | 0.3 | 0.25 |
| MT072864.1;QIQ54046.1;Pangolin_CoV_orf1a;150 | 0.25 | 0.125 |
| MT072864.1;QIQ54046.1;Pangolin_CoV_orf1a;151 | 0.3 | 0.15 |
| MT072864.1;QIQ54046.1;Pangolin_CoV_orf1a;152 | 0.35 | 0.225 |
| MT072864.1;QIQ54046.1;Pangolin_CoV_orf1a;153 | 0.25 | 0.125 |
| MT072864.1;QIQ54046.1;Pangolin_CoV_orf1a;154 | 0.35 | 0.175 |
| MT072864.1;QIQ54046.1;Pangolin_CoV_orf1a;155 | 0.3 | 0.075 |
| MT072864.1;QIQ54046.1;Pangolin_CoV_orf1a;156 | 0.2 | 0.05 |
| MT072864.1;QIQ54046.1;Pangolin_CoV_orf1a;157 | 0.1 | 0.075 |
| MT072864.1;QIQ54046.1;Pangolin_CoV_orf1a;158 | 0.1 | 0.075 |
| MT072864.1;QIQ54046.1;Pangolin_CoV_orf1a;159 | 0.05 | 0.05 |
| MT072864.1;QIQ54046.1;Pangolin_CoV_orf1a;160 | 0 | 0 |
| MT072864.1;QIQ54046.1;Pangolin_CoV_orf1a;161 | 0.05 | 0.05 |
| MT072864.1;QIQ54046.1;Pangolin_CoV_orf1a;162 | 0.05 | 0.05 |

| | | |
|---|---|---|
| MT072864.1;QIQ54046.1;Pangolin_CoV_orf1a;163 | 0 | 0.1 |
| MT072864.1;QIQ54046.1;Pangolin_CoV_orf1a;164 | 0 | 0.05 |
| MT072864.1;QIQ54046.1;Pangolin_CoV_orf1a;165 | 0.05 | 0 |
| MT072864.1;QIQ54046.1;Pangolin_CoV_orf1a;166 | 0 | 0.025 |
| MT072864.1;QIQ54046.1;Pangolin_CoV_orf1a;167 | 0 | 0.025 |
| MT072864.1;QIQ54046.1;Pangolin_CoV_orf1a;168 | 0.1 | 0.025 |
| MT072864.1;QIQ54046.1;Pangolin_CoV_orf1a;169 | 0 | 0 |
| MT072864.1;QIQ54046.1;Pangolin_CoV_orf1a;170 | 0 | 0.025 |
| MT072864.1;QIQ54046.1;Pangolin_CoV_orf1a;171 | 0 | 0 |
| MT072864.1;QIQ54046.1;Pangolin_CoV_orf1a;172 | 0 | 0 |
| MT072864.1;QIQ54046.1;Pangolin_CoV_orf1a;173 | 0 | 0 |
| MT072864.1;QIQ54046.1;Pangolin_CoV_orf1a;174 | 0.05 | 0.025 |
| MT072864.1;QIQ54046.1;Pangolin_CoV_orf1a;175 | 0 | 0 |
| MT072864.1;QIQ54046.1;Pangolin_CoV_orf1a;176 | 0 | 0 |
| MT072864.1;QIQ54046.1;Pangolin_CoV_orf1a;177 | 0 | 0 |
| MT072864.1;QIQ54046.1;Pangolin_CoV_orf1a;178 | 0 | 0 |
| MT072864.1;QIQ54046.1;Pangolin_CoV_orf1a;179 | 0 | 0 |
| MT072864.1;QIQ54046.1;Pangolin_CoV_orf1a;180 | 0 | 0 |
| MT072864.1;QIQ54046.1;Pangolin_CoV_orf1a;181 | 0 | 0.025 |
| MT072864.1;QIQ54046.1;Pangolin_CoV_orf1a;182 | 0 | 0 |
| MT072864.1;QIQ54046.1;Pangolin_CoV_orf1a;183 | 0 | 0 |
| MT072864.1;QIQ54046.1;Pangolin_CoV_orf1a;184 | 0 | 0.025 |
| MT072864.1;QIQ54046.1;Pangolin_CoV_orf1a;185 | 0 | 0.025 |
| MT072864.1;QIQ54046.1;Pangolin_CoV_orf1a;186 | 0 | 0.025 |
| MT072864.1;QIQ54046.1;Pangolin_CoV_orf1a;187 | 0 | 0 |
| MT072864.1;QIQ54046.1;Pangolin_CoV_orf1a;188 | 0 | 0 |
| MT072864.1;QIQ54046.1;Pangolin_CoV_orf1a;189 | 0 | 0 |
| MT072864.1;QIQ54046.1;Pangolin_CoV_orf1a;190 | 0 | 0 |
| MT072864.1;QIQ54046.1;Pangolin_CoV_orf1a;191 | 0 | 0 |
| MT072864.1;QIQ54046.1;Pangolin_CoV_orf1a;192 | 0 | 0 |
| MT072864.1;QIQ54046.1;Pangolin_CoV_orf1a;193 | 0 | 0 |
| MT072864.1;QIQ54046.1;Pangolin_CoV_orf1a;194 | 0 | 0 |
| MT072864.1;QIQ54046.1;Pangolin_CoV_orf1a;195 | 0 | 0 |
| MT072864.1;QIQ54046.1;Pangolin_CoV_orf1a;196 | 0 | 0 |
| MT072864.1;QIQ54046.1;Pangolin_CoV_orf1a;197 | 0 | 0 |
| MT072864.1;QIQ54046.1;Pangolin_CoV_orf1a;198 | 0 | 0 |
| MT072864.1;QIQ54046.1;Pangolin_CoV_orf1a;199 | 0 | 0 |
| MT072864.1;QIQ54046.1;Pangolin_CoV_orf1a;200 | 0 | 0 |
| MT072864.1;QIQ54046.1;Pangolin_CoV_orf1a;201 | 0 | 0 |
| MT072864.1;QIQ54046.1;Pangolin_CoV_orf1a;202 | 0 | 0 |
| MT072864.1;QIQ54046.1;Pangolin_CoV_orf1a;203 | 0 | 0 |
| MT072864.1;QIQ54046.1;Pangolin_CoV_orf1a;204 | 0.05 | 0 |
| MT072864.1;QIQ54046.1;Pangolin_CoV_orf1a;205 | 0 | 0 |

| | | |
|---|---|---|
| MT072864.1;QIQ54046.1;Pangolin_CoV_orf1a;206 | 0 | 0 |
| MT072864.1;QIQ54046.1;Pangolin_CoV_orf1a;207 | 0 | 0 |
| MT072864.1;QIQ54046.1;Pangolin_CoV_orf1a;208 | 0 | 0 |
| MT072864.1;QIQ54046.1;Pangolin_CoV_orf1a;209 | 0 | 0 |
| MT072864.1;QIQ54046.1;Pangolin_CoV_orf1a;210 | 0 | 0 |
| MT072864.1;QIQ54046.1;Pangolin_CoV_orf1a;211 | 0.1 | 0.05 |
| MT072864.1;QIQ54046.1;Pangolin_CoV_orf1a;212 | 0.05 | 0.075 |
| MT072864.1;QIQ54046.1;Pangolin_CoV_orf1a;213 | 0.05 | 0.05 |
| MT072864.1;QIQ54046.1;Pangolin_CoV_orf1a;214 | 0.15 | 0.1 |
| MT072864.1;QIQ54046.1;Pangolin_CoV_orf1a;215 | 0.1 | 0.075 |
| MT072864.1;QIQ54046.1;Pangolin_CoV_orf1a;216 | 0 | 0.075 |
| MT072864.1;QIQ54046.1;Pangolin_CoV_orf1a;217 | 0 | 0.025 |
| MT072864.1;QIQ54046.1;Pangolin_CoV_orf1a;218 | 0 | 0.025 |
| MT072864.1;QIQ54046.1;Pangolin_CoV_orf1a;219 | 0 | 0.025 |
| MT072864.1;QIQ54046.1;Pangolin_CoV_orf1a;220 | 0.05 | 0.075 |
| MT072864.1;QIQ54046.1;Pangolin_CoV_orf1a;221 | 0 | 0.025 |
| MT072864.1;QIQ54046.1;Pangolin_CoV_orf1a;222 | 0 | 0.025 |
| MT072864.1;QIQ54046.1;Pangolin_CoV_orf1a;223 | 0 | 0.025 |
| MT072864.1;QIQ54046.1;Pangolin_CoV_orf1a;224 | 0 | 0 |
| MT072864.1;QIQ54046.1;Pangolin_CoV_orf1a;225 | 0.05 | 0 |
| MT072864.1;QIQ54046.1;Pangolin_CoV_orf1a;226 | 0 | 0 |
| MT072864.1;QIQ54046.1;Pangolin_CoV_orf1a;227 | 0 | 0 |
| MT072864.1;QIQ54046.1;Pangolin_CoV_orf1a;228 | 0 | 0 |
| MT072864.1;QIQ54046.1;Pangolin_CoV_orf1a;229 | 0 | 0 |
| MT072864.1;QIQ54046.1;Pangolin_CoV_orf1a;230 | 0 | 0 |
| MT072864.1;QIQ54046.1;Pangolin_CoV_orf1a;231 | 0 | 0 |
| MT072864.1;QIQ54046.1;Pangolin_CoV_orf1a;232 | 0 | 0 |
| MT072864.1;QIQ54046.1;Pangolin_CoV_orf1a;233 | 0 | 0 |
| MT072864.1;QIQ54046.1;Pangolin_CoV_orf1a;234 | 0 | 0 |
| MT072864.1;QIQ54046.1;Pangolin_CoV_orf1a;235 | 0 | 0 |
| MT072864.1;QIQ54046.1;Pangolin_CoV_orf1a;236 | 0 | 0 |
| MT072864.1;QIQ54046.1;Pangolin_CoV_orf1a;237 | 0 | 0.025 |
| MT072864.1;QIQ54046.1;Pangolin_CoV_orf1a;238 | 0 | 0 |
| MT072864.1;QIQ54046.1;Pangolin_CoV_orf1a;239 | 0 | 0 |
| MT072864.1;QIQ54046.1;Pangolin_CoV_orf1a;240 | 0 | 0 |
| MT072864.1;QIQ54046.1;Pangolin_CoV_orf1a;241 | 0 | 0 |
| MT072864.1;QIQ54046.1;Pangolin_CoV_orf1a;242 | 0 | 0 |
| MT072864.1;QIQ54046.1;Pangolin_CoV_orf1a;243 | 0 | 0 |
| MT072864.1;QIQ54046.1;Pangolin_CoV_orf1a;244 | 0 | 0 |
| MT072864.1;QIQ54046.1;Pangolin_CoV_orf1a;245 | 0 | 0 |
| MT072864.1;QIQ54046.1;Pangolin_CoV_orf1a;246 | 0 | 0 |
| MT072864.1;QIQ54046.1;Pangolin_CoV_orf1a;247 | 0 | 0 |
| MT072864.1;QIQ54046.1;Pangolin_CoV_orf1a;248 | 0 | 0 |

| | | |
|---|---|---|
| MT072864.1;QIQ54046.1;Pangolin_CoV_orf1a;249 | 0 | 0 |
| MT072864.1;QIQ54046.1;Pangolin_CoV_orf1a;250 | 0 | 0 |
| MT072864.1;QIQ54046.1;Pangolin_CoV_orf1a;251 | 0 | 0 |
| MT072864.1;QIQ54046.1;Pangolin_CoV_orf1a;252 | 0.05 | 0.075 |
| MT072864.1;QIQ54046.1;Pangolin_CoV_orf1a;253 | 0.05 | 0.1 |
| MT072864.1;QIQ54046.1;Pangolin_CoV_orf1a;254 | 0.1 | 0.15 |
| MT072864.1;QIQ54046.1;Pangolin_CoV_orf1a;255 | 0.15 | 0.225 |
| MT072864.1;QIQ54046.1;Pangolin_CoV_orf1a;256 | 0.25 | 0.225 |
| MT072864.1;QIQ54046.1;Pangolin_CoV_orf1a;257 | 0.1 | 0.1 |
| MT072864.1;QIQ54046.1;Pangolin_CoV_orf1a;258 | 0.15 | 0.125 |
| MT072864.1;QIQ54046.1;Pangolin_CoV_orf1a;259 | 0.35 | 0.225 |
| MT072864.1;QIQ54046.1;Pangolin_CoV_orf1a;260 | 0.15 | 0.25 |
| MT072864.1;QIQ54046.1;Pangolin_CoV_orf1a;261 | 0.1 | 0.225 |
| MT072864.1;QIQ54046.1;Pangolin_CoV_orf1a;262 | 0.05 | 0.05 |
| MT072864.1;QIQ54046.1;Pangolin_CoV_orf1a;263 | 0 | 0 |
| MT072864.1;QIQ54046.1;Pangolin_CoV_orf1a;264 | 0 | 0.025 |
| MT072864.1;QIQ54046.1;Pangolin_CoV_orf1a;265 | 0 | 0 |
| MT072864.1;QIQ54046.1;Pangolin_CoV_orf1a;266 | 0 | 0 |
| MT072864.1;QIQ54046.1;Pangolin_CoV_orf1a;267 | 0.15 | 0.025 |
| MT072864.1;QIQ54046.1;Pangolin_CoV_orf1a;268 | 0.15 | 0 |
| MT072864.1;QIQ54046.1;Pangolin_CoV_orf1a;269 | 0.15 | 0 |
| MT072864.1;QIQ54046.1;Pangolin_CoV_orf1a;270 | 0.2 | 0 |
| MT072864.1;QIQ54046.1;Pangolin_CoV_orf1a;271 | 0.15 | 0 |
| MT072864.1;QIQ54046.1;Pangolin_CoV_orf1a;272 | 0.1 | 0 |
| MT072864.1;QIQ54046.1;Pangolin_CoV_orf1a;273 | 0.1 | 0 |
| MT072864.1;QIQ54046.1;Pangolin_CoV_orf1a;274 | 0.15 | 0 |
| MT072864.1;QIQ54046.1;Pangolin_CoV_orf1a;275 | 0.15 | 0.025 |
| MT072864.1;QIQ54046.1;Pangolin_CoV_orf1a;276 | 0.2 | 0.05 |
| MT072864.1;QIQ54046.1;Pangolin_CoV_orf1a;277 | 0.2 | 0.025 |
| MT072864.1;QIQ54046.1;Pangolin_CoV_orf1a;278 | 0.2 | 0 |
| MT072864.1;QIQ54046.1;Pangolin_CoV_orf1a;279 | 0.05 | 0.025 |
| MT072864.1;QIQ54046.1;Pangolin_CoV_orf1a;280 | 0.1 | 0.025 |
| MT072864.1;QIQ54046.1;Pangolin_CoV_orf1a;281 | 0.05 | 0.025 |
| MT072864.1;QIQ54046.1;Pangolin_CoV_orf1a;282 | 0.05 | 0 |
| MT072864.1;QIQ54046.1;Pangolin_CoV_orf1a;283 | 0.05 | 0 |
| MT072864.1;QIQ54046.1;Pangolin_CoV_orf1a;284 | 0.05 | 0.025 |
| MT072864.1;QIQ54046.1;Pangolin_CoV_orf1a;285 | 0.05 | 0.025 |
| MT072864.1;QIQ54046.1;Pangolin_CoV_orf1a;286 | 0.05 | 0.025 |
| MT072864.1;QIQ54046.1;Pangolin_CoV_orf1a;287 | 0 | 0 |
| MT072864.1;QIQ54046.1;Pangolin_CoV_orf1a;288 | 0 | 0 |
| MT072864.1;QIQ54046.1;Pangolin_CoV_orf1a;289 | 0 | 0 |
| MT072864.1;QIQ54046.1;Pangolin_CoV_orf1a;290 | 0 | 0 |
| MT072864.1;QIQ54046.1;Pangolin_CoV_orf1a;291 | 0 | 0 |

| | | |
|---|---|---|
| MT072864.1;QIQ54046.1;Pangolin_CoV_orf1a;292 | 0 | 0 |
| MT072864.1;QIQ54046.1;Pangolin_CoV_orf1a;293 | 0 | 0 |
| MT072864.1;QIQ54046.1;Pangolin_CoV_orf1a;294 | 0 | 0 |
| MT072864.1;QIQ54046.1;Pangolin_CoV_orf1a;295 | 0 | 0 |
| MT072864.1;QIQ54046.1;Pangolin_CoV_orf1a;296 | 0.1 | 0 |
| MT072864.1;QIQ54046.1;Pangolin_CoV_orf1a;297 | 0.1 | 0 |
| MT072864.1;QIQ54046.1;Pangolin_CoV_orf1a;298 | 0.1 | 0 |
| MT072864.1;QIQ54046.1;Pangolin_CoV_orf1a;299 | 0.1 | 0 |
| MT072864.1;QIQ54046.1;Pangolin_CoV_orf1a;300 | 0.1 | 0 |
| MT072864.1;QIQ54046.1;Pangolin_CoV_orf1a;301 | 0.1 | 0.025 |
| MT072864.1;QIQ54046.1;Pangolin_CoV_orf1a;302 | 0.05 | 0 |
| MT072864.1;QIQ54046.1;Pangolin_CoV_orf1a;303 | 0.05 | 0 |
| MT072864.1;QIQ54046.1;Pangolin_CoV_orf1a;304 | 0.05 | 0 |
| MT072864.1;QIQ54046.1;Pangolin_CoV_orf1a;305 | 0 | 0 |
| MT072864.1;QIQ54046.1;Pangolin_CoV_orf1a;306 | 0 | 0 |
| MT072864.1;QIQ54046.1;Pangolin_CoV_orf1a;307 | 0 | 0 |
| MT072864.1;QIQ54046.1;Pangolin_CoV_orf1a;308 | 0 | 0 |
| MT072864.1;QIQ54046.1;Pangolin_CoV_orf1a;309 | 0 | 0 |
| MT072864.1;QIQ54046.1;Pangolin_CoV_orf1a;310 | 0 | 0 |
| MT072864.1;QIQ54046.1;Pangolin_CoV_orf1a;311 | 0 | 0 |
| MT072864.1;QIQ54046.1;Pangolin_CoV_orf1a;312 | 0 | 0 |
| MT072864.1;QIQ54046.1;Pangolin_CoV_orf1a;313 | 0 | 0 |
| MT072864.1;QIQ54046.1;Pangolin_CoV_orf1a;314 | 0 | 0 |
| MT072864.1;QIQ54046.1;Pangolin_CoV_orf1a;315 | 0 | 0 |
| MT072864.1;QIQ54046.1;Pangolin_CoV_orf1a;316 | 0 | 0 |
| MT072864.1;QIQ54046.1;Pangolin_CoV_orf1a;317 | 0 | 0 |
| MT072864.1;QIQ54046.1;Pangolin_CoV_orf1a;318 | 0 | 0 |
| MT072864.1;QIQ54046.1;Pangolin_CoV_orf1a;319 | 0 | 0 |
| MT072864.1;QIQ54046.1;Pangolin_CoV_orf1a;320 | 0 | 0 |
| MT072864.1;QIQ54046.1;Pangolin_CoV_orf1a;321 | 0 | 0 |
| MT072864.1;QIQ54046.1;Pangolin_CoV_orf1a;322 | 0 | 0 |
| MT072864.1;QIQ54046.1;Pangolin_CoV_orf1a;323 | 0 | 0 |
| MT072864.1;QIQ54046.1;Pangolin_CoV_orf1a;324 | 0.05 | 0 |
| MT072864.1;QIQ54046.1;Pangolin_CoV_orf1a;325 | 0 | 0 |
| MT072864.1;QIQ54046.1;Pangolin_CoV_orf1a;326 | 0 | 0 |
| MT072864.1;QIQ54046.1;Pangolin_CoV_orf1a;327 | 0 | 0.025 |
| MT072864.1;QIQ54046.1;Pangolin_CoV_orf1a;328 | 0 | 0 |
| MT072864.1;QIQ54046.1;Pangolin_CoV_orf1a;329 | 0 | 0 |
| MT072864.1;QIQ54046.1;Pangolin_CoV_orf1a;330 | 0 | 0 |
| MT072864.1;QIQ54046.1;Pangolin_CoV_orf1a;331 | 0 | 0 |
| MT072864.1;QIQ54046.1;Pangolin_CoV_orf1a;332 | 0 | 0 |
| MT072864.1;QIQ54046.1;Pangolin_CoV_orf1a;333 | 0 | 0 |
| MT072864.1;QIQ54046.1;Pangolin_CoV_orf1a;334 | 0 | 0 |

| | | |
|---|---|---|
| MT072864.1;QIQ54046.1;Pangolin_CoV_orf1a;335 | 0 | 0 |
| MT072864.1;QIQ54046.1;Pangolin_CoV_orf1a;336 | 0 | 0 |
| MT072864.1;QIQ54046.1;Pangolin_CoV_orf1a;337 | 0 | 0 |
| MT072864.1;QIQ54046.1;Pangolin_CoV_orf1a;338 | 0 | 0 |
| MT072864.1;QIQ54046.1;Pangolin_CoV_orf1a;339 | 0 | 0 |
| MT072864.1;QIQ54046.1;Pangolin_CoV_orf1a;340 | 0 | 0.025 |
| MT072864.1;QIQ54046.1;Pangolin_CoV_orf1a;341 | 0 | 0.025 |
| MT072864.1;QIQ54046.1;Pangolin_CoV_orf1a;342 | 0 | 0.025 |
| MT072864.1;QIQ54046.1;Pangolin_CoV_orf1a;343 | 0 | 0.025 |
| MT072864.1;QIQ54046.1;Pangolin_CoV_orf1a;344 | 0 | 0.025 |
| MT072864.1;QIQ54046.1;Pangolin_CoV_orf1a;345 | 0 | 0.025 |
| MT072864.1;QIQ54046.1;Pangolin_CoV_orf1a;346 | 0.05 | 0 |
| MT072864.1;QIQ54046.1;Pangolin_CoV_orf1a;347 | 0 | 0 |
| MT072864.1;QIQ54046.1;Pangolin_CoV_orf1a;348 | 0 | 0 |
| MT072864.1;QIQ54046.1;Pangolin_CoV_orf1a;349 | 0.05 | 0 |
| MT072864.1;QIQ54046.1;Pangolin_CoV_orf1a;350 | 0.05 | 0 |
| MT072864.1;QIQ54046.1;Pangolin_CoV_orf1a;351 | 0.05 | 0 |
| MT072864.1;QIQ54046.1;Pangolin_CoV_orf1a;352 | 0 | 0 |
| MT072864.1;QIQ54046.1;Pangolin_CoV_orf1a;353 | 0 | 0 |
| MT072864.1;QIQ54046.1;Pangolin_CoV_orf1a;354 | 0 | 0 |
| MT072864.1;QIQ54046.1;Pangolin_CoV_orf1a;355 | 0 | 0 |
| MT072864.1;QIQ54046.1;Pangolin_CoV_orf1a;356 | 0 | 0 |
| MT072864.1;QIQ54046.1;Pangolin_CoV_orf1a;357 | 0 | 0 |
| MT072864.1;QIQ54046.1;Pangolin_CoV_orf1a;358 | 0 | 0 |
| MT072864.1;QIQ54046.1;Pangolin_CoV_orf1a;359 | 0 | 0 |
| MT072864.1;QIQ54046.1;Pangolin_CoV_orf1a;360 | 0 | 0 |
| MT072864.1;QIQ54046.1;Pangolin_CoV_orf1a;361 | 0 | 0 |
| MT072864.1;QIQ54046.1;Pangolin_CoV_orf1a;362 | 0 | 0 |
| MT072864.1;QIQ54046.1;Pangolin_CoV_orf1a;363 | 0 | 0 |
| MT072864.1;QIQ54046.1;Pangolin_CoV_orf1a;364 | 0 | 0 |
| MT072864.1;QIQ54046.1;Pangolin_CoV_orf1a;365 | 0 | 0 |
| MT072864.1;QIQ54046.1;Pangolin_CoV_orf1a;366 | 0 | 0 |
| MT072864.1;QIQ54046.1;Pangolin_CoV_orf1a;367 | 0.05 | 0 |
| MT072864.1;QIQ54046.1;Pangolin_CoV_orf1a;368 | 0.05 | 0 |
| MT072864.1;QIQ54046.1;Pangolin_CoV_orf1a;369 | 0 | 0 |
| MT072864.1;QIQ54046.1;Pangolin_CoV_orf1a;370 | 0.05 | 0 |
| MT072864.1;QIQ54046.1;Pangolin_CoV_orf1a;371 | 0.05 | 0 |
| MT072864.1;QIQ54046.1;Pangolin_CoV_orf1a;372 | 0.05 | 0 |
| MT072864.1;QIQ54046.1;Pangolin_CoV_orf1a;373 | 0 | 0 |
| MT072864.1;QIQ54046.1;Pangolin_CoV_orf1a;374 | 0 | 0 |
| MT072864.1;QIQ54046.1;Pangolin_CoV_orf1a;375 | 0.05 | 0 |
| MT072864.1;QIQ54046.1;Pangolin_CoV_orf1a;376 | 0.1 | 0 |
| MT072864.1;QIQ54046.1;Pangolin_CoV_orf1a;377 | 0 | 0 |

| | | |
|---|---|---|
| MT072864.1;QIQ54046.1;Pangolin_CoV_orf1a;378 | 0 | 0 |
| MT072864.1;QIQ54046.1;Pangolin_CoV_orf1a;379 | 0.1 | 0.075 |
| MT072864.1;QIQ54046.1;Pangolin_CoV_orf1a;380 | 0 | 0.025 |
| MT072864.1;QIQ54046.1;Pangolin_CoV_orf1a;381 | 0 | 0.025 |
| MT072864.1;QIQ54046.1;Pangolin_CoV_orf1a;382 | 0 | 0.025 |
| MT072864.1;QIQ54046.1;Pangolin_CoV_orf1a;383 | 0 | 0.025 |
| MT072864.1;QIQ54046.1;Pangolin_CoV_orf1a;384 | 0 | 0.025 |
| MT072864.1;QIQ54046.1;Pangolin_CoV_orf1a;385 | 0 | 0.025 |
| MT072864.1;QIQ54046.1;Pangolin_CoV_orf1a;386 | 0 | 0.025 |
| MT072864.1;QIQ54046.1;Pangolin_CoV_orf1a;387 | 0 | 0.025 |
| MT072864.1;QIQ54046.1;Pangolin_CoV_orf1a;388 | 0 | 0 |
| MT072864.1;QIQ54046.1;Pangolin_CoV_orf1a;389 | 0 | 0 |
| MT072864.1;QIQ54046.1;Pangolin_CoV_orf1a;390 | 0 | 0 |
| MT072864.1;QIQ54046.1;Pangolin_CoV_orf1a;391 | 0 | 0 |
| MT072864.1;QIQ54046.1;Pangolin_CoV_orf1a;392 | 0 | 0 |
| MT072864.1;QIQ54046.1;Pangolin_CoV_orf1a;393 | 0 | 0 |
| MT072864.1;QIQ54046.1;Pangolin_CoV_orf1a;394 | 0 | 0 |
| MT072864.1;QIQ54046.1;Pangolin_CoV_orf1a;395 | 0 | 0 |
| MT072864.1;QIQ54046.1;Pangolin_CoV_orf1a;396 | 0 | 0 |
| MT072864.1;QIQ54046.1;Pangolin_CoV_orf1a;397 | 0 | 0.025 |
| MT072864.1;QIQ54046.1;Pangolin_CoV_orf1a;398 | 0 | 0 |
| MT072864.1;QIQ54046.1;Pangolin_CoV_orf1a;399 | 0 | 0 |
| MT072864.1;QIQ54046.1;Pangolin_CoV_orf1a;400 | 0 | 0 |
| MT072864.1;QIQ54046.1;Pangolin_CoV_orf1a;401 | 0 | 0 |
| MT072864.1;QIQ54046.1;Pangolin_CoV_orf1a;402 | 0 | 0 |
| MT072864.1;QIQ54046.1;Pangolin_CoV_orf1a;403 | 0 | 0 |
| MT072864.1;QIQ54046.1;Pangolin_CoV_orf1a;404 | 0 | 0 |
| MT072864.1;QIQ54046.1;Pangolin_CoV_orf1a;405 | 0 | 0 |
| MT072864.1;QIQ54046.1;Pangolin_CoV_orf1a;406 | 0 | 0 |
| MT072864.1;QIQ54046.1;Pangolin_CoV_orf1a;407 | 0 | 0 |
| MT072864.1;QIQ54046.1;Pangolin_CoV_orf1a;408 | 0 | 0 |
| MT072864.1;QIQ54046.1;Pangolin_CoV_orf1a;409 | 0 | 0 |
| MT072864.1;QIQ54046.1;Pangolin_CoV_orf1a;410 | 0 | 0 |
| MT072864.1;QIQ54046.1;Pangolin_CoV_orf1a;411 | 0 | 0 |
| MT072864.1;QIQ54046.1;Pangolin_CoV_orf1a;412 | 0 | 0 |
| MT072864.1;QIQ54046.1;Pangolin_CoV_orf1a;413 | 0.05 | 0 |
| MT072864.1;QIQ54046.1;Pangolin_CoV_orf1a;414 | 0.05 | 0 |
| MT072864.1;QIQ54046.1;Pangolin_CoV_orf1a;415 | 0 | 0 |
| MT072864.1;QIQ54046.1;Pangolin_CoV_orf1a;416 | 0 | 0 |
| MT072864.1;QIQ54046.1;Pangolin_CoV_orf1a;417 | 0 | 0 |
| MT072864.1;QIQ54046.1;Pangolin_CoV_orf1a;418 | 0 | 0 |
| MT072864.1;QIQ54046.1;Pangolin_CoV_orf1a;419 | 0 | 0 |
| MT072864.1;QIQ54046.1;Pangolin_CoV_orf1a;420 | 0 | 0 |

| | | |
|---|---|---|
| MT072864.1;QIQ54046.1;Pangolin_CoV_orf1a;421 | 0 | 0 |
| MT072864.1;QIQ54046.1;Pangolin_CoV_orf1a;422 | 0 | 0 |
| MT072864.1;QIQ54046.1;Pangolin_CoV_orf1a;423 | 0 | 0 |
| MT072864.1;QIQ54046.1;Pangolin_CoV_orf1a;424 | 0.05 | 0 |
| MT072864.1;QIQ54046.1;Pangolin_CoV_orf1a;425 | 0 | 0 |
| MT072864.1;QIQ54046.1;Pangolin_CoV_orf1a;426 | 0 | 0 |
| MT072864.1;QIQ54046.1;Pangolin_CoV_orf1a;427 | 0 | 0 |
| MT072864.1;QIQ54046.1;Pangolin_CoV_orf1a;428 | 0 | 0 |
| MT072864.1;QIQ54046.1;Pangolin_CoV_orf1a;429 | 0 | 0 |
| MT072864.1;QIQ54046.1;Pangolin_CoV_orf1a;430 | 0 | 0 |
| MT072864.1;QIQ54046.1;Pangolin_CoV_orf1a;431 | 0 | 0 |
| MT072864.1;QIQ54046.1;Pangolin_CoV_orf1a;432 | 0 | 0 |
| MT072864.1;QIQ54046.1;Pangolin_CoV_orf1a;433 | 0 | 0 |
| MT072864.1;QIQ54046.1;Pangolin_CoV_orf1a;434 | 0 | 0 |
| MT072864.1;QIQ54046.1;Pangolin_CoV_orf1a;435 | 0 | 0 |
| MT072864.1;QIQ54046.1;Pangolin_CoV_orf1a;436 | 0 | 0 |
| MT072864.1;QIQ54046.1;Pangolin_CoV_orf1a;437 | 0.05 | 0.025 |
| MT072864.1;QIQ54046.1;Pangolin_CoV_orf1a;438 | 0 | 0 |
| MT072864.1;QIQ54046.1;Pangolin_CoV_orf1a;439 | 0 | 0 |
| MT072864.1;QIQ54046.1;Pangolin_CoV_orf1a;440 | 0 | 0 |
| MT072864.1;QIQ54046.1;Pangolin_CoV_orf1a;441 | 0 | 0 |
| MT072864.1;QIQ54046.1;Pangolin_CoV_orf1a;442 | 0.05 | 0 |
| MT072864.1;QIQ54046.1;Pangolin_CoV_orf1a;443 | 0 | 0 |
| MT072864.1;QIQ54046.1;Pangolin_CoV_orf1a;444 | 0.05 | 0.025 |
| MT072864.1;QIQ54046.1;Pangolin_CoV_orf1a;445 | 0.4 | 0.125 |
| MT072864.1;QIQ54046.1;Pangolin_CoV_orf1a;446 | 0 | 0 |
| MT072864.1;QIQ54046.1;Pangolin_CoV_orf1a;447 | 0.05 | 0 |
| MT072864.1;QIQ54046.1;Pangolin_CoV_orf1a;448 | 0 | 0 |
| MT072864.1;QIQ54046.1;Pangolin_CoV_orf1a;449 | 0 | 0 |
| MT072864.1;QIQ54046.1;Pangolin_CoV_orf1a;450 | 0 | 0 |
| MT072864.1;QIQ54046.1;Pangolin_CoV_orf1a;451 | 0 | 0 |
| MT072864.1;QIQ54046.1;Pangolin_CoV_orf1a;452 | 0 | 0 |
| MT072864.1;QIQ54046.1;Pangolin_CoV_orf1a;453 | 0 | 0 |
| MT072864.1;QIQ54046.1;Pangolin_CoV_orf1a;454 | 0 | 0 |
| MT072864.1;QIQ54046.1;Pangolin_CoV_orf1a;455 | 0 | 0 |
| MT072864.1;QIQ54046.1;Pangolin_CoV_orf1a;456 | 0 | 0 |
| MT072864.1;QIQ54046.1;Pangolin_CoV_orf1a;457 | 0 | 0 |
| MT072864.1;QIQ54046.1;Pangolin_CoV_orf1a;458 | 0.05 | 0 |
| MT072864.1;QIQ54046.1;Pangolin_CoV_orf1a;459 | 0 | 0 |
| MT072864.1;QIQ54046.1;Pangolin_CoV_orf1a;460 | 0 | 0 |
| MT072864.1;QIQ54046.1;Pangolin_CoV_orf1a;461 | 0 | 0 |
| MT072864.1;QIQ54046.1;Pangolin_CoV_orf1a;462 | 0 | 0 |
| MT072864.1;QIQ54046.1;Pangolin_CoV_orf1a;463 | 0 | 0 |

| | | |
|---|---|---|
| MT072864.1;QIQ54046.1;Pangolin_CoV_orf1a;464 | 0 | 0 |
| MT072864.1;QIQ54046.1;Pangolin_CoV_orf1a;465 | 0 | 0 |
| MT072864.1;QIQ54046.1;Pangolin_CoV_orf1a;466 | 0 | 0 |
| MT072864.1;QIQ54046.1;Pangolin_CoV_orf1a;467 | 0 | 0 |
| MT072864.1;QIQ54046.1;Pangolin_CoV_orf1a;468 | 0 | 0 |
| MT072864.1;QIQ54046.1;Pangolin_CoV_orf1a;469 | 0 | 0 |
| MT072864.1;QIQ54046.1;Pangolin_CoV_orf1a;470 | 0 | 0 |
| MT072864.1;QIQ54046.1;Pangolin_CoV_orf1a;471 | 0 | 0 |
| MT072864.1;QIQ54046.1;Pangolin_CoV_orf1a;472 | 0 | 0 |
| MT072864.1;QIQ54046.1;Pangolin_CoV_orf1a;473 | 0 | 0 |
| MT072864.1;QIQ54046.1;Pangolin_CoV_orf1a;474 | 0 | 0 |
| MT072864.1;QIQ54046.1;Pangolin_CoV_orf1a;475 | 0 | 0 |
| MT072864.1;QIQ54046.1;Pangolin_CoV_orf1a;476 | 0 | 0 |
| MT072864.1;QIQ54046.1;Pangolin_CoV_orf1a;477 | 0 | 0 |
| MT072864.1;QIQ54046.1;Pangolin_CoV_orf1a;478 | 0 | 0 |
| MT072864.1;QIQ54046.1;Pangolin_CoV_orf1a;479 | 0 | 0 |
| MT072864.1;QIQ54046.1;Pangolin_CoV_orf1a;480 | 0 | 0.05 |
| MT072864.1;QIQ54046.1;Pangolin_CoV_orf1a;481 | 0 | 0.05 |
| MT072864.1;QIQ54046.1;Pangolin_CoV_orf1a;482 | 0 | 0.025 |
| MT072864.1;QIQ54046.1;Pangolin_CoV_orf1a;483 | 0 | 0.025 |
| MT072864.1;QIQ54046.1;Pangolin_CoV_orf1a;484 | 0 | 0 |
| MT072864.1;QIQ54046.1;Pangolin_CoV_orf1a;485 | 0 | 0.025 |
| MT072864.1;QIQ54046.1;Pangolin_CoV_orf1a;486 | 0 | 0.025 |
| MT072864.1;QIQ54046.1;Pangolin_CoV_orf1a;487 | 0 | 0 |
| MT072864.1;QIQ54046.1;Pangolin_CoV_orf1a;488 | 0 | 0 |
| MT072864.1;QIQ54046.1;Pangolin_CoV_orf1a;489 | 0.05 | 0.075 |
| MT072864.1;QIQ54046.1;Pangolin_CoV_orf1a;490 | 0.05 | 0 |
| MT072864.1;QIQ54046.1;Pangolin_CoV_orf1a;491 | 0.05 | 0 |
| MT072864.1;QIQ54046.1;Pangolin_CoV_orf1a;492 | 0 | 0.025 |
| MT072864.1;QIQ54046.1;Pangolin_CoV_orf1a;493 | 0.1 | 0.025 |
| MT072864.1;QIQ54046.1;Pangolin_CoV_orf1a;494 | 0.1 | 0.025 |
| MT072864.1;QIQ54046.1;Pangolin_CoV_orf1a;495 | 0.05 | 0.025 |
| MT072864.1;QIQ54046.1;Pangolin_CoV_orf1a;496 | 0.05 | 0 |
| MT072864.1;QIQ54046.1;Pangolin_CoV_orf1a;497 | 0.1 | 0.075 |
| MT072864.1;QIQ54046.1;Pangolin_CoV_orf1a;498 | 0.1 | 0.025 |
| MT072864.1;QIQ54046.1;Pangolin_CoV_orf1a;499 | 0 | 0 |
| MT072864.1;QIQ54046.1;Pangolin_CoV_orf1a;500 | 0 | 0 |
| MT072864.1;QIQ54046.1;Pangolin_CoV_orf1a;501 | 0 | 0 |
| MT072864.1;QIQ54046.1;Pangolin_CoV_orf1a;502 | 0 | 0 |
| MT072864.1;QIQ54046.1;Pangolin_CoV_orf1a;503 | 0 | 0 |
| MT072864.1;QIQ54046.1;Pangolin_CoV_orf1a;504 | 0.05 | 0 |
| MT072864.1;QIQ54046.1;Pangolin_CoV_orf1a;505 | 0.05 | 0 |
| MT072864.1;QIQ54046.1;Pangolin_CoV_orf1a;506 | 0.05 | 0 |

| | | |
|---|---|---|
| MT072864.1;QIQ54046.1;Pangolin_CoV_orf1a;507 | 0.05 | 0 |
| MT072864.1;QIQ54046.1;Pangolin_CoV_orf1a;508 | 0.05 | 0 |
| MT072864.1;QIQ54046.1;Pangolin_CoV_orf1a;509 | 0.05 | 0 |
| MT072864.1;QIQ54046.1;Pangolin_CoV_orf1a;510 | 0.05 | 0 |
| MT072864.1;QIQ54046.1;Pangolin_CoV_orf1a;511 | 0 | 0.05 |
| MT072864.1;QIQ54046.1;Pangolin_CoV_orf1a;512 | 0.05 | 0.05 |
| MT072864.1;QIQ54046.1;Pangolin_CoV_orf1a;513 | 0.05 | 0.05 |
| MT072864.1;QIQ54046.1;Pangolin_CoV_orf1a;514 | 0.05 | 0.05 |
| MT072864.1;QIQ54046.1;Pangolin_CoV_orf1a;515 | 0 | 0.05 |
| MT072864.1;QIQ54046.1;Pangolin_CoV_orf1a;516 | 0 | 0.05 |
| MT072864.1;QIQ54046.1;Pangolin_CoV_orf1a;517 | 0 | 0.075 |
| MT072864.1;QIQ54046.1;Pangolin_CoV_orf1a;518 | 0.05 | 0.125 |
| MT072864.1;QIQ54046.1;Pangolin_CoV_orf1a;519 | 0 | 0 |
| MT072864.1;QIQ54046.1;Pangolin_CoV_orf1a;520 | 0 | 0 |
| MT072864.1;QIQ54046.1;Pangolin_CoV_orf1a;521 | 0 | 0 |
| MT072864.1;QIQ54046.1;Pangolin_CoV_orf1a;522 | 0 | 0 |
| MT072864.1;QIQ54046.1;Pangolin_CoV_orf1a;523 | 0 | 0.025 |
| MT072864.1;QIQ54046.1;Pangolin_CoV_orf1a;524 | 0 | 0 |
| MT072864.1;QIQ54046.1;Pangolin_CoV_orf1a;525 | 0.05 | 0 |
| MT072864.1;QIQ54046.1;Pangolin_CoV_orf1a;526 | 0.05 | 0.025 |
| MT072864.1;QIQ54046.1;Pangolin_CoV_orf1a;527 | 0 | 0 |
| MT072864.1;QIQ54046.1;Pangolin_CoV_orf1a;528 | 0 | 0 |
| MT072864.1;QIQ54046.1;Pangolin_CoV_orf1a;529 | 0 | 0 |
| MT072864.1;QIQ54046.1;Pangolin_CoV_orf1a;530 | 0 | 0 |
| MT072864.1;QIQ54046.1;Pangolin_CoV_orf1a;531 | 0 | 0 |
| MT072864.1;QIQ54046.1;Pangolin_CoV_orf1a;532 | 0 | 0 |
| MT072864.1;QIQ54046.1;Pangolin_CoV_orf1a;533 | 0 | 0 |
| MT072864.1;QIQ54046.1;Pangolin_CoV_orf1a;534 | 0 | 0 |
| MT072864.1;QIQ54046.1;Pangolin_CoV_orf1a;535 | 0 | 0 |
| MT072864.1;QIQ54046.1;Pangolin_CoV_orf1a;536 | 0 | 0 |
| MT072864.1;QIQ54046.1;Pangolin_CoV_orf1a;537 | 0 | 0 |
| MT072864.1;QIQ54046.1;Pangolin_CoV_orf1a;538 | 0 | 0 |
| MT072864.1;QIQ54046.1;Pangolin_CoV_orf1a;539 | 0 | 0 |
| MT072864.1;QIQ54046.1;Pangolin_CoV_orf1a;540 | 0 | 0 |
| MT072864.1;QIQ54046.1;Pangolin_CoV_orf1a;541 | 0 | 0 |
| MT072864.1;QIQ54046.1;Pangolin_CoV_orf1a;542 | 0 | 0 |
| MT072864.1;QIQ54046.1;Pangolin_CoV_orf1a;543 | 0 | 0 |
| MT072864.1;QIQ54046.1;Pangolin_CoV_orf1a;544 | 0 | 0 |
| MT072864.1;QIQ54046.1;Pangolin_CoV_orf1a;545 | 0 | 0 |
| MT072864.1;QIQ54046.1;Pangolin_CoV_orf1a;546 | 0 | 0.025 |
| MT072864.1;QIQ54046.1;Pangolin_CoV_orf1a;547 | 0 | 0 |
| MT072864.1;QIQ54046.1;Pangolin_CoV_orf1a;548 | 0 | 0 |
| MT072864.1;QIQ54046.1;Pangolin_CoV_orf1a;549 | 0 | 0 |

| | | |
|---|---|---|
| MT072864.1;QIQ54046.1;Pangolin_CoV_orf1a;550 | 0 | 0 |
| MT072864.1;QIQ54046.1;Pangolin_CoV_orf1a;551 | 0 | 0 |
| MT072864.1;QIQ54046.1;Pangolin_CoV_orf1a;552 | 0 | 0 |
| MT072864.1;QIQ54046.1;Pangolin_CoV_orf1a;553 | 0 | 0 |
| MT072864.1;QIQ54046.1;Pangolin_CoV_orf1a;554 | 0 | 0 |
| MT072864.1;QIQ54046.1;Pangolin_CoV_orf1a;555 | 0 | 0 |
| MT072864.1;QIQ54046.1;Pangolin_CoV_orf1a;556 | 0 | 0 |
| MT072864.1;QIQ54046.1;Pangolin_CoV_orf1a;557 | 0 | 0 |
| MT072864.1;QIQ54046.1;Pangolin_CoV_orf1a;558 | 0 | 0 |
| MT072864.1;QIQ54046.1;Pangolin_CoV_orf1a;559 | 0 | 0 |
| MT072864.1;QIQ54046.1;Pangolin_CoV_orf1a;560 | 0 | 0 |
| MT072864.1;QIQ54046.1;Pangolin_CoV_orf1a;561 | 0 | 0 |
| MT072864.1;QIQ54046.1;Pangolin_CoV_orf1a;562 | 0 | 0 |
| MT072864.1;QIQ54046.1;Pangolin_CoV_orf1a;563 | 0 | 0 |
| MT072864.1;QIQ54046.1;Pangolin_CoV_orf1a;564 | 0 | 0 |
| MT072864.1;QIQ54046.1;Pangolin_CoV_orf1a;565 | 0 | 0.025 |
| MT072864.1;QIQ54046.1;Pangolin_CoV_orf1a;566 | 0 | 0 |
| MT072864.1;QIQ54046.1;Pangolin_CoV_orf1a;567 | 0 | 0 |
| MT072864.1;QIQ54046.1;Pangolin_CoV_orf1a;568 | 0 | 0 |
| MT072864.1;QIQ54046.1;Pangolin_CoV_orf1a;569 | 0 | 0.025 |
| MT072864.1;QIQ54046.1;Pangolin_CoV_orf1a;570 | 0 | 0.025 |
| MT072864.1;QIQ54046.1;Pangolin_CoV_orf1a;571 | 0 | 0.025 |
| MT072864.1;QIQ54046.1;Pangolin_CoV_orf1a;572 | 0 | 0.025 |
| MT072864.1;QIQ54046.1;Pangolin_CoV_orf1a;573 | 0 | 0 |
| MT072864.1;QIQ54046.1;Pangolin_CoV_orf1a;574 | 0 | 0 |
| MT072864.1;QIQ54046.1;Pangolin_CoV_orf1a;575 | 0 | 0 |
| MT072864.1;QIQ54046.1;Pangolin_CoV_orf1a;576 | 0 | 0 |
| MT072864.1;QIQ54046.1;Pangolin_CoV_orf1a;577 | 0 | 0 |
| MT072864.1;QIQ54046.1;Pangolin_CoV_orf1a;578 | 0 | 0.025 |
| MT072864.1;QIQ54046.1;Pangolin_CoV_orf1a;579 | 0 | 0 |
| MT072864.1;QIQ54046.1;Pangolin_CoV_orf1a;580 | 0 | 0 |
| MT072864.1;QIQ54046.1;Pangolin_CoV_orf1a;581 | 0 | 0 |
| MT072864.1;QIQ54046.1;Pangolin_CoV_orf1a;582 | 0 | 0 |
| MT072864.1;QIQ54046.1;Pangolin_CoV_orf1a;583 | 0 | 0 |
| MT072864.1;QIQ54046.1;Pangolin_CoV_orf1a;584 | 0 | 0 |
| MT072864.1;QIQ54046.1;Pangolin_CoV_orf1a;585 | 0 | 0 |
| MT072864.1;QIQ54046.1;Pangolin_CoV_orf1a;586 | 0 | 0 |
| MT072864.1;QIQ54046.1;Pangolin_CoV_orf1a;587 | 0 | 0 |
| MT072864.1;QIQ54046.1;Pangolin_CoV_orf1a;588 | 0 | 0 |
| MT072864.1;QIQ54046.1;Pangolin_CoV_orf1a;589 | 0.05 | 0 |
| MT072864.1;QIQ54046.1;Pangolin_CoV_orf1a;590 | 0 | 0 |
| MT072864.1;QIQ54046.1;Pangolin_CoV_orf1a;591 | 0 | 0 |
| MT072864.1;QIQ54046.1;Pangolin_CoV_orf1a;592 | 0 | 0.025 |

| | | |
|---|---|---|
| MT072864.1;QIQ54046.1;Pangolin_CoV_orf1a;593 | 0 | 0 |
| MT072864.1;QIQ54046.1;Pangolin_CoV_orf1a;594 | 0 | 0 |
| MT072864.1;QIQ54046.1;Pangolin_CoV_orf1a;595 | 0 | 0 |
| MT072864.1;QIQ54046.1;Pangolin_CoV_orf1a;596 | 0 | 0 |
| MT072864.1;QIQ54046.1;Pangolin_CoV_orf1a;597 | 0 | 0.025 |
| MT072864.1;QIQ54046.1;Pangolin_CoV_orf1a;598 | 0 | 0.025 |
| MT072864.1;QIQ54046.1;Pangolin_CoV_orf1a;599 | 0 | 0.05 |
| MT072864.1;QIQ54046.1;Pangolin_CoV_orf1a;600 | 0 | 0.025 |
| MT072864.1;QIQ54046.1;Pangolin_CoV_orf1a;601 | 0 | 0.025 |
| MT072864.1;QIQ54046.1;Pangolin_CoV_orf1a;602 | 0 | 0.025 |
| MT072864.1;QIQ54046.1;Pangolin_CoV_orf1a;603 | 0 | 0.025 |
| MT072864.1;QIQ54046.1;Pangolin_CoV_orf1a;604 | 0 | 0 |
| MT072864.1;QIQ54046.1;Pangolin_CoV_orf1a;605 | 0 | 0 |
| MT072864.1;QIQ54046.1;Pangolin_CoV_orf1a;606 | 0 | 0 |
| MT072864.1;QIQ54046.1;Pangolin_CoV_orf1a;607 | 0 | 0 |
| MT072864.1;QIQ54046.1;Pangolin_CoV_orf1a;608 | 0 | 0 |
| MT072864.1;QIQ54046.1;Pangolin_CoV_orf1a;609 | 0 | 0 |
| MT072864.1;QIQ54046.1;Pangolin_CoV_orf1a;610 | 0 | 0 |
| MT072864.1;QIQ54046.1;Pangolin_CoV_orf1a;611 | 0 | 0 |
| MT072864.1;QIQ54046.1;Pangolin_CoV_orf1a;612 | 0 | 0 |
| MT072864.1;QIQ54046.1;Pangolin_CoV_orf1a;613 | 0 | 0 |
| MT072864.1;QIQ54046.1;Pangolin_CoV_orf1a;614 | 0 | 0.025 |
| MT072864.1;QIQ54046.1;Pangolin_CoV_orf1a;615 | 0 | 0 |
| MT072864.1;QIQ54046.1;Pangolin_CoV_orf1a;616 | 0 | 0 |
| MT072864.1;QIQ54046.1;Pangolin_CoV_orf1a;617 | 0.05 | 0 |
| MT072864.1;QIQ54046.1;Pangolin_CoV_orf1a;618 | 0.1 | 0.025 |
| MT072864.1;QIQ54046.1;Pangolin_CoV_orf1a;619 | 0.1 | 0.025 |
| MT072864.1;QIQ54046.1;Pangolin_CoV_orf1a;620 | 0.1 | 0.05 |
| MT072864.1;QIQ54046.1;Pangolin_CoV_orf1a;621 | 0.15 | 0.025 |
| MT072864.1;QIQ54046.1;Pangolin_CoV_orf1a;622 | 0.1 | 0.025 |
| MT072864.1;QIQ54046.1;Pangolin_CoV_orf1a;623 | 0.1 | 0.025 |
| MT072864.1;QIQ54046.1;Pangolin_CoV_orf1a;624 | 0 | 0.05 |
| MT072864.1;QIQ54046.1;Pangolin_CoV_orf1a;625 | 0 | 0 |
| MT072864.1;QIQ54046.1;Pangolin_CoV_orf1a;626 | 0 | 0 |
| MT072864.1;QIQ54046.1;Pangolin_CoV_orf1a;627 | 0 | 0.025 |
| MT072864.1;QIQ54046.1;Pangolin_CoV_orf1a;628 | 0 | 0.025 |
| MT072864.1;QIQ54046.1;Pangolin_CoV_orf1a;629 | 0 | 0.025 |
| MT072864.1;QIQ54046.1;Pangolin_CoV_orf1a;630 | 0 | 0.025 |
| MT072864.1;QIQ54046.1;Pangolin_CoV_orf1a;631 | 0 | 0.025 |
| MT072864.1;QIQ54046.1;Pangolin_CoV_orf1a;632 | 0 | 0.025 |
| MT072864.1;QIQ54046.1;Pangolin_CoV_orf1a;633 | 0 | 0.025 |
| MT072864.1;QIQ54046.1;Pangolin_CoV_orf1a;634 | 0 | 0.025 |
| MT072864.1;QIQ54046.1;Pangolin_CoV_orf1a;635 | 0 | 0.025 |

| | | |
|---|---|---|
| MT072864.1;QIQ54046.1;Pangolin_CoV_orf1a;636 | 0 | 0.025 |
| MT072864.1;QIQ54046.1;Pangolin_CoV_orf1a;637 | 0 | 0 |
| MT072864.1;QIQ54046.1;Pangolin_CoV_orf1a;638 | 0.05 | 0.05 |
| MT072864.1;QIQ54046.1;Pangolin_CoV_orf1a;639 | 0 | 0 |
| MT072864.1;QIQ54046.1;Pangolin_CoV_orf1a;640 | 0 | 0 |
| MT072864.1;QIQ54046.1;Pangolin_CoV_orf1a;641 | 0 | 0 |
| MT072864.1;QIQ54046.1;Pangolin_CoV_orf1a;642 | 0 | 0 |
| MT072864.1;QIQ54046.1;Pangolin_CoV_orf1a;643 | 0 | 0 |
| MT072864.1;QIQ54046.1;Pangolin_CoV_orf1a;644 | 0 | 0 |
| MT072864.1;QIQ54046.1;Pangolin_CoV_orf1a;645 | 0 | 0 |
| MT072864.1;QIQ54046.1;Pangolin_CoV_orf1a;646 | 0 | 0 |
| MT072864.1;QIQ54046.1;Pangolin_CoV_orf1a;647 | 0 | 0 |
| MT072864.1;QIQ54046.1;Pangolin_CoV_orf1a;648 | 0 | 0 |
| MT072864.1;QIQ54046.1;Pangolin_CoV_orf1a;649 | 0 | 0 |
| MT072864.1;QIQ54046.1;Pangolin_CoV_orf1a;650 | 0 | 0 |
| MT072864.1;QIQ54046.1;Pangolin_CoV_orf1a;651 | 0 | 0.025 |
| MT072864.1;QIQ54046.1;Pangolin_CoV_orf1a;652 | 0 | 0 |
| MT072864.1;QIQ54046.1;Pangolin_CoV_orf1a;653 | 0 | 0 |
| MT072864.1;QIQ54046.1;Pangolin_CoV_orf1a;654 | 0 | 0 |
| MT072864.1;QIQ54046.1;Pangolin_CoV_orf1a;655 | 0 | 0 |
| MT072864.1;QIQ54046.1;Pangolin_CoV_orf1a;656 | 0 | 0 |
| MT072864.1;QIQ54046.1;Pangolin_CoV_orf1a;657 | 0 | 0 |
| MT072864.1;QIQ54046.1;Pangolin_CoV_orf1a;658 | 0 | 0.025 |
| MT072864.1;QIQ54046.1;Pangolin_CoV_orf1a;659 | 0 | 0 |
| MT072864.1;QIQ54046.1;Pangolin_CoV_orf1a;660 | 0 | 0 |
| MT072864.1;QIQ54046.1;Pangolin_CoV_orf1a;661 | 0 | 0 |
| MT072864.1;QIQ54046.1;Pangolin_CoV_orf1a;662 | 0.05 | 0 |
| MT072864.1;QIQ54046.1;Pangolin_CoV_orf1a;663 | 0 | 0.025 |
| MT072864.1;QIQ54046.1;Pangolin_CoV_orf1a;664 | 0 | 0.025 |
| MT072864.1;QIQ54046.1;Pangolin_CoV_orf1a;665 | 0 | 0.025 |
| MT072864.1;QIQ54046.1;Pangolin_CoV_orf1a;666 | 0 | 0.025 |
| MT072864.1;QIQ54046.1;Pangolin_CoV_orf1a;667 | 0 | 0.025 |
| MT072864.1;QIQ54046.1;Pangolin_CoV_orf1a;668 | 0 | 0.025 |
| MT072864.1;QIQ54046.1;Pangolin_CoV_orf1a;669 | 0 | 0.025 |
| MT072864.1;QIQ54046.1;Pangolin_CoV_orf1a;670 | 0 | 0 |
| MT072864.1;QIQ54046.1;Pangolin_CoV_orf1a;671 | 0 | 0 |
| MT072864.1;QIQ54046.1;Pangolin_CoV_orf1a;672 | 0 | 0 |
| MT072864.1;QIQ54046.1;Pangolin_CoV_orf1a;673 | 0 | 0 |
| MT072864.1;QIQ54046.1;Pangolin_CoV_orf1a;674 | 0 | 0 |
| MT072864.1;QIQ54046.1;Pangolin_CoV_orf1a;675 | 0 | 0 |
| MT072864.1;QIQ54046.1;Pangolin_CoV_orf1a;676 | 0 | 0 |
| MT072864.1;QIQ54046.1;Pangolin_CoV_orf1a;677 | 0 | 0 |
| MT072864.1;QIQ54046.1;Pangolin_CoV_orf1a;678 | 0 | 0 |

| | | |
|---|---|---|
| MT072864.1;QIQ54046.1;Pangolin_CoV_orf1a;679 | 0 | 0 |
| MT072864.1;QIQ54046.1;Pangolin_CoV_orf1a;680 | 0 | 0 |
| MT072864.1;QIQ54046.1;Pangolin_CoV_orf1a;681 | 0 | 0 |
| MT072864.1;QIQ54046.1;Pangolin_CoV_orf1a;682 | 0 | 0.025 |
| MT072864.1;QIQ54046.1;Pangolin_CoV_orf1a;683 | 0 | 0 |
| MT072864.1;QIQ54046.1;Pangolin_CoV_orf1a;684 | 0.05 | 0.025 |
| MT072864.1;QIQ54046.1;Pangolin_CoV_orf1a;685 | 0.1 | 0 |
| MT072864.1;QIQ54046.1;Pangolin_CoV_orf1a;686 | 0 | 0 |
| MT072864.1;QIQ54046.1;Pangolin_CoV_orf1a;687 | 0 | 0 |
| MT072864.1;QIQ54046.1;Pangolin_CoV_orf1a;688 | 0 | 0 |
| MT072864.1;QIQ54046.1;Pangolin_CoV_orf1a;689 | 0.05 | 0.025 |
| MT072864.1;QIQ54046.1;Pangolin_CoV_orf1a;690 | 0.05 | 0 |
| MT072864.1;QIQ54046.1;Pangolin_CoV_orf1a;691 | 0.05 | 0 |
| MT072864.1;QIQ54046.1;Pangolin_CoV_orf1a;692 | 0.1 | 0.05 |
| MT072864.1;QIQ54046.1;Pangolin_CoV_orf1a;693 | 0.05 | 0.025 |
| MT072864.1;QIQ54046.1;Pangolin_CoV_orf1a;694 | 0 | 0 |
| MT072864.1;QIQ54046.1;Pangolin_CoV_orf1a;695 | 0 | 0 |
| MT072864.1;QIQ54046.1;Pangolin_CoV_orf1a;696 | 0.05 | 0.025 |
| MT072864.1;QIQ54046.1;Pangolin_CoV_orf1a;697 | 0 | 0 |
| MT072864.1;QIQ54046.1;Pangolin_CoV_orf1a;698 | 0 | 0 |
| MT072864.1;QIQ54046.1;Pangolin_CoV_orf1a;699 | 0 | 0 |
| MT072864.1;QIQ54046.1;Pangolin_CoV_orf1a;700 | 0 | 0 |
| MT072864.1;QIQ54046.1;Pangolin_CoV_orf1a;701 | 0 | 0 |
| MT072864.1;QIQ54046.1;Pangolin_CoV_orf1a;702 | 0 | 0 |
| MT072864.1;QIQ54046.1;Pangolin_CoV_orf1a;703 | 0 | 0 |
| MT072864.1;QIQ54046.1;Pangolin_CoV_orf1a;704 | 0 | 0 |
| MT072864.1;QIQ54046.1;Pangolin_CoV_orf1a;705 | 0 | 0 |
| MT072864.1;QIQ54046.1;Pangolin_CoV_orf1a;706 | 0 | 0 |
| MT072864.1;QIQ54046.1;Pangolin_CoV_orf1a;707 | 0 | 0 |
| MT072864.1;QIQ54046.1;Pangolin_CoV_orf1a;708 | 0 | 0 |
| MT072864.1;QIQ54046.1;Pangolin_CoV_orf1a;709 | 0 | 0 |
| MT072864.1;QIQ54046.1;Pangolin_CoV_orf1a;710 | 0 | 0 |
| MT072864.1;QIQ54046.1;Pangolin_CoV_orf1a;711 | 0 | 0 |
| MT072864.1;QIQ54046.1;Pangolin_CoV_orf1a;712 | 0 | 0 |
| MT072864.1;QIQ54046.1;Pangolin_CoV_orf1a;713 | 0 | 0.05 |
| MT072864.1;QIQ54046.1;Pangolin_CoV_orf1a;714 | 0.05 | 0 |
| MT072864.1;QIQ54046.1;Pangolin_CoV_orf1a;715 | 0.05 | 0 |
| MT072864.1;QIQ54046.1;Pangolin_CoV_orf1a;716 | 0.05 | 0 |
| MT072864.1;QIQ54046.1;Pangolin_CoV_orf1a;717 | 0 | 0 |
| MT072864.1;QIQ54046.1;Pangolin_CoV_orf1a;718 | 0.05 | 0.05 |
| MT072864.1;QIQ54046.1;Pangolin_CoV_orf1a;719 | 0.05 | 0 |
| MT072864.1;QIQ54046.1;Pangolin_CoV_orf1a;720 | 0.05 | 0 |
| MT072864.1;QIQ54046.1;Pangolin_CoV_orf1a;721 | 0 | 0 |

| | | |
|---|---|---|
| MT072864.1;QIQ54046.1;Pangolin_CoV_orf1a;722 | 0 | 0.025 |
| MT072864.1;QIQ54046.1;Pangolin_CoV_orf1a;723 | 0 | 0 |
| MT072864.1;QIQ54046.1;Pangolin_CoV_orf1a;724 | 0 | 0 |
| MT072864.1;QIQ54046.1;Pangolin_CoV_orf1a;725 | 0 | 0 |
| MT072864.1;QIQ54046.1;Pangolin_CoV_orf1a;726 | 0 | 0 |
| MT072864.1;QIQ54046.1;Pangolin_CoV_orf1a;727 | 0 | 0 |
| MT072864.1;QIQ54046.1;Pangolin_CoV_orf1a;728 | 0 | 0 |
| MT072864.1;QIQ54046.1;Pangolin_CoV_orf1a;729 | 0 | 0 |
| MT072864.1;QIQ54046.1;Pangolin_CoV_orf1a;730 | 0 | 0 |
| MT072864.1;QIQ54046.1;Pangolin_CoV_orf1a;731 | 0 | 0 |
| MT072864.1;QIQ54046.1;Pangolin_CoV_orf1a;732 | 0 | 0 |
| MT072864.1;QIQ54046.1;Pangolin_CoV_orf1a;733 | 0 | 0 |
| MT072864.1;QIQ54046.1;Pangolin_CoV_orf1a;734 | 0 | 0 |
| MT072864.1;QIQ54046.1;Pangolin_CoV_orf1a;735 | 0 | 0 |
| MT072864.1;QIQ54046.1;Pangolin_CoV_orf1a;736 | 0 | 0 |
| MT072864.1;QIQ54046.1;Pangolin_CoV_orf1a;737 | 0 | 0 |
| MT072864.1;QIQ54046.1;Pangolin_CoV_orf1a;738 | 0 | 0 |
| MT072864.1;QIQ54046.1;Pangolin_CoV_orf1a;739 | 0 | 0 |
| MT072864.1;QIQ54046.1;Pangolin_CoV_orf1a;740 | 0 | 0 |
| MT072864.1;QIQ54046.1;Pangolin_CoV_orf1a;741 | 0 | 0 |
| MT072864.1;QIQ54046.1;Pangolin_CoV_orf1a;742 | 0 | 0 |
| MT072864.1;QIQ54046.1;Pangolin_CoV_orf1a;743 | 0 | 0 |
| MT072864.1;QIQ54046.1;Pangolin_CoV_orf1a;744 | 0.05 | 0.125 |
| MT072864.1;QIQ54046.1;Pangolin_CoV_orf1a;745 | 0.1 | 0.075 |
| MT072864.1;QIQ54046.1;Pangolin_CoV_orf1a;746 | 0.15 | 0.075 |
| MT072864.1;QIQ54046.1;Pangolin_CoV_orf1a;747 | 0 | 0.075 |
| MT072864.1;QIQ54046.1;Pangolin_CoV_orf1a;748 | 0.05 | 0.025 |
| MT072864.1;QIQ54046.1;Pangolin_CoV_orf1a;749 | 0 | 0.05 |
| MT072864.1;QIQ54046.1;Pangolin_CoV_orf1a;750 | 0.1 | 0.075 |
| MT072864.1;QIQ54046.1;Pangolin_CoV_orf1a;751 | 0.05 | 0.075 |
| MT072864.1;QIQ54046.1;Pangolin_CoV_orf1a;752 | 0.05 | 0.05 |
| MT072864.1;QIQ54046.1;Pangolin_CoV_orf1a;753 | 0.05 | 0.025 |
| MT072864.1;QIQ54046.1;Pangolin_CoV_orf1a;754 | 0.1 | 0.05 |
| MT072864.1;QIQ54046.1;Pangolin_CoV_orf1a;755 | 0.05 | 0.025 |
| MT072864.1;QIQ54046.1;Pangolin_CoV_orf1a;756 | 0 | 0.05 |
| MT072864.1;QIQ54046.1;Pangolin_CoV_orf1a;757 | 0 | 0.025 |
| MT072864.1;QIQ54046.1;Pangolin_CoV_orf1a;758 | 0 | 0.025 |
| MT072864.1;QIQ54046.1;Pangolin_CoV_orf1a;759 | 0 | 0.025 |
| MT072864.1;QIQ54046.1;Pangolin_CoV_orf1a;760 | 0 | 0.025 |
| MT072864.1;QIQ54046.1;Pangolin_CoV_orf1a;761 | 0 | 0.05 |
| MT072864.1;QIQ54046.1;Pangolin_CoV_orf1a;762 | 0 | 0 |
| MT072864.1;QIQ54046.1;Pangolin_CoV_orf1a;763 | 0 | 0 |
| MT072864.1;QIQ54046.1;Pangolin_CoV_orf1a;764 | 0 | 0.025 |

| | | |
|---|---|---|
| MT072864.1;QIQ54046.1;Pangolin_CoV_orf1a;765 | 0 | 0.025 |
| MT072864.1;QIQ54046.1;Pangolin_CoV_orf1a;766 | 0 | 0.05 |
| MT072864.1;QIQ54046.1;Pangolin_CoV_orf1a;767 | 0 | 0 |
| MT072864.1;QIQ54046.1;Pangolin_CoV_orf1a;768 | 0 | 0 |
| MT072864.1;QIQ54046.1;Pangolin_CoV_orf1a;769 | 0 | 0 |
| MT072864.1;QIQ54046.1;Pangolin_CoV_orf1a;770 | 0 | 0 |
| MT072864.1;QIQ54046.1;Pangolin_CoV_orf1a;771 | 0 | 0 |
| MT072864.1;QIQ54046.1;Pangolin_CoV_orf1a;772 | 0 | 0 |
| MT072864.1;QIQ54046.1;Pangolin_CoV_orf1a;773 | 0 | 0 |
| MT072864.1;QIQ54046.1;Pangolin_CoV_orf1a;774 | 0 | 0 |
| MT072864.1;QIQ54046.1;Pangolin_CoV_orf1a;775 | 0 | 0 |
| MT072864.1;QIQ54046.1;Pangolin_CoV_orf1a;776 | 0 | 0 |
| MT072864.1;QIQ54046.1;Pangolin_CoV_orf1a;777 | 0 | 0 |
| MT072864.1;QIQ54046.1;Pangolin_CoV_orf1a;778 | 0 | 0 |
| MT072864.1;QIQ54046.1;Pangolin_CoV_orf1a;779 | 0 | 0 |
| MT072864.1;QIQ54046.1;Pangolin_CoV_orf1a;780 | 0 | 0.025 |
| MT072864.1;QIQ54046.1;Pangolin_CoV_orf1a;781 | 0 | 0 |
| MT072864.1;QIQ54046.1;Pangolin_CoV_orf1a;782 | 0.05 | 0 |
| MT072864.1;QIQ54046.1;Pangolin_CoV_orf1a;783 | 0 | 0 |
| MT072864.1;QIQ54046.1;Pangolin_CoV_orf1a;784 | 0 | 0 |
| MT072864.1;QIQ54046.1;Pangolin_CoV_orf1a;785 | 0 | 0 |
| MT072864.1;QIQ54046.1;Pangolin_CoV_orf1a;786 | 0 | 0 |
| MT072864.1;QIQ54046.1;Pangolin_CoV_orf1a;787 | 0 | 0.025 |
| MT072864.1;QIQ54046.1;Pangolin_CoV_orf1a;788 | 0 | 0.025 |
| MT072864.1;QIQ54046.1;Pangolin_CoV_orf1a;789 | 0.05 | 0 |
| MT072864.1;QIQ54046.1;Pangolin_CoV_orf1a;790 | 0 | 0.025 |
| MT072864.1;QIQ54046.1;Pangolin_CoV_orf1a;791 | 0 | 0.05 |
| MT072864.1;QIQ54046.1;Pangolin_CoV_orf1a;792 | 0 | 0.05 |
| MT072864.1;QIQ54046.1;Pangolin_CoV_orf1a;793 | 0 | 0.025 |
| MT072864.1;QIQ54046.1;Pangolin_CoV_orf1a;794 | 0 | 0 |
| MT072864.1;QIQ54046.1;Pangolin_CoV_orf1a;795 | 0 | 0 |
| MT072864.1;QIQ54046.1;Pangolin_CoV_orf1a;796 | 0 | 0 |
| MT072864.1;QIQ54046.1;Pangolin_CoV_orf1a;797 | 0.05 | 0.05 |
| MT072864.1;QIQ54046.1;Pangolin_CoV_orf1a;798 | 0.05 | 0 |
| MT072864.1;QIQ54046.1;Pangolin_CoV_orf1a;799 | 0 | 0 |
| MT072864.1;QIQ54046.1;Pangolin_CoV_orf1a;800 | 0 | 0 |
| MT072864.1;QIQ54046.1;Pangolin_CoV_orf1a;801 | 0.05 | 0 |
| MT072864.1;QIQ54046.1;Pangolin_CoV_orf1a;802 | 0 | 0 |
| MT072864.1;QIQ54046.1;Pangolin_CoV_orf1a;803 | 0.1 | 0 |
| MT072864.1;QIQ54046.1;Pangolin_CoV_orf1a;804 | 0.05 | 0 |
| MT072864.1;QIQ54046.1;Pangolin_CoV_orf1a;805 | 0.05 | 0 |
| MT072864.1;QIQ54046.1;Pangolin_CoV_orf1a;806 | 0.05 | 0 |
| MT072864.1;QIQ54046.1;Pangolin_CoV_orf1a;807 | 0 | 0.025 |

| | | |
|---|---|---|
| MT072864.1;QIQ54046.1;Pangolin_CoV_orf1a;808 | 0 | 0 |
| MT072864.1;QIQ54046.1;Pangolin_CoV_orf1a;809 | 0 | 0 |
| MT072864.1;QIQ54046.1;Pangolin_CoV_orf1a;810 | 0 | 0 |
| MT072864.1;QIQ54046.1;Pangolin_CoV_orf1a;811 | 0 | 0 |
| MT072864.1;QIQ54046.1;Pangolin_CoV_orf1a;812 | 0 | 0 |
| MT072864.1;QIQ54046.1;Pangolin_CoV_orf1a;813 | 0.05 | 0 |
| MT072864.1;QIQ54046.1;Pangolin_CoV_orf1a;814 | 0 | 0.025 |
| MT072864.1;QIQ54046.1;Pangolin_CoV_orf1a;815 | 0 | 0 |
| MT072864.1;QIQ54046.1;Pangolin_CoV_orf1a;816 | 0 | 0 |
| MT072864.1;QIQ54046.1;Pangolin_CoV_orf1a;817 | 0 | 0 |
| MT072864.1;QIQ54046.1;Pangolin_CoV_orf1a;818 | 0 | 0 |
| MT072864.1;QIQ54046.1;Pangolin_CoV_orf1a;819 | 0.05 | 0 |
| MT072864.1;QIQ54046.1;Pangolin_CoV_orf1a;820 | 0.05 | 0 |
| MT072864.1;QIQ54046.1;Pangolin_CoV_orf1a;821 | 0 | 0 |
| MT072864.1;QIQ54046.1;Pangolin_CoV_orf1a;822 | 0 | 0.025 |
| MT072864.1;QIQ54046.1;Pangolin_CoV_orf1a;823 | 0 | 0.025 |
| MT072864.1;QIQ54046.1;Pangolin_CoV_orf1a;824 | 0.05 | 0.025 |
| MT072864.1;QIQ54046.1;Pangolin_CoV_orf1a;825 | 0.05 | 0 |
| MT072864.1;QIQ54046.1;Pangolin_CoV_orf1a;826 | 0 | 0 |
| MT072864.1;QIQ54046.1;Pangolin_CoV_orf1a;827 | 0.05 | 0 |
| MT072864.1;QIQ54046.1;Pangolin_CoV_orf1a;828 | 0 | 0 |
| MT072864.1;QIQ54046.1;Pangolin_CoV_orf1a;829 | 0 | 0 |
| MT072864.1;QIQ54046.1;Pangolin_CoV_orf1a;830 | 0 | 0 |
| MT072864.1;QIQ54046.1;Pangolin_CoV_orf1a;831 | 0.05 | 0.05 |
| MT072864.1;QIQ54046.1;Pangolin_CoV_orf1a;832 | 0 | 0.025 |
| MT072864.1;QIQ54046.1;Pangolin_CoV_orf1a;833 | 0 | 0 |
| MT072864.1;QIQ54046.1;Pangolin_CoV_orf1a;834 | 0 | 0 |
| MT072864.1;QIQ54046.1;Pangolin_CoV_orf1a;835 | 0 | 0 |
| MT072864.1;QIQ54046.1;Pangolin_CoV_orf1a;836 | 0.05 | 0 |
| MT072864.1;QIQ54046.1;Pangolin_CoV_orf1a;837 | 0.05 | 0 |
| MT072864.1;QIQ54046.1;Pangolin_CoV_orf1a;838 | 0 | 0 |
| MT072864.1;QIQ54046.1;Pangolin_CoV_orf1a;839 | 0 | 0 |
| MT072864.1;QIQ54046.1;Pangolin_CoV_orf1a;840 | 0 | 0.025 |
| MT072864.1;QIQ54046.1;Pangolin_CoV_orf1a;841 | 0.05 | 0.05 |
| MT072864.1;QIQ54046.1;Pangolin_CoV_orf1a;842 | 0.05 | 0.025 |
| MT072864.1;QIQ54046.1;Pangolin_CoV_orf1a;843 | 0.05 | 0.025 |
| MT072864.1;QIQ54046.1;Pangolin_CoV_orf1a;844 | 0 | 0.025 |
| MT072864.1;QIQ54046.1;Pangolin_CoV_orf1a;845 | 0 | 0.025 |
| MT072864.1;QIQ54046.1;Pangolin_CoV_orf1a;846 | 0 | 0.05 |
| MT072864.1;QIQ54046.1;Pangolin_CoV_orf1a;847 | 0 | 0.025 |
| MT072864.1;QIQ54046.1;Pangolin_CoV_orf1a;848 | 0 | 0 |
| MT072864.1;QIQ54046.1;Pangolin_CoV_orf1a;849 | 0 | 0 |
| MT072864.1;QIQ54046.1;Pangolin_CoV_orf1a;850 | 0.05 | 0 |

| | | |
|---|---|---|
| MT072864.1;QIQ54046.1;Pangolin_CoV_orf1a;851 | 0 | 0 |
| MT072864.1;QIQ54046.1;Pangolin_CoV_orf1a;852 | 0 | 0 |
| MT072864.1;QIQ54046.1;Pangolin_CoV_orf1a;853 | 0 | 0 |
| MT072864.1;QIQ54046.1;Pangolin_CoV_orf1a;854 | 0 | 0 |
| MT072864.1;QIQ54046.1;Pangolin_CoV_orf1a;855 | 0 | 0.025 |
| MT072864.1;QIQ54046.1;Pangolin_CoV_orf1a;856 | 0.05 | 0 |
| MT072864.1;QIQ54046.1;Pangolin_CoV_orf1a;857 | 0.05 | 0 |
| MT072864.1;QIQ54046.1;Pangolin_CoV_orf1a;858 | 0 | 0 |
| MT072864.1;QIQ54046.1;Pangolin_CoV_orf1a;859 | 0 | 0 |
| MT072864.1;QIQ54046.1;Pangolin_CoV_orf1a;860 | 0 | 0 |
| MT072864.1;QIQ54046.1;Pangolin_CoV_orf1a;861 | 0 | 0 |
| MT072864.1;QIQ54046.1;Pangolin_CoV_orf1a;862 | 0 | 0 |
| MT072864.1;QIQ54046.1;Pangolin_CoV_orf1a;863 | 0 | 0 |
| MT072864.1;QIQ54046.1;Pangolin_CoV_orf1a;864 | 0 | 0 |
| MT072864.1;QIQ54046.1;Pangolin_CoV_orf1a;865 | 0 | 0 |
| MT072864.1;QIQ54046.1;Pangolin_CoV_orf1a;866 | 0 | 0 |
| MT072864.1;QIQ54046.1;Pangolin_CoV_orf1a;867 | 0 | 0 |
| MT072864.1;QIQ54046.1;Pangolin_CoV_orf1a;868 | 0 | 0 |
| MT072864.1;QIQ54046.1;Pangolin_CoV_orf1a;869 | 0 | 0 |
| MT072864.1;QIQ54046.1;Pangolin_CoV_orf1a;870 | 0 | 0 |
| MT072864.1;QIQ54046.1;Pangolin_CoV_orf1a;871 | 0 | 0 |
| MT072864.1;QIQ54046.1;Pangolin_CoV_orf1a;872 | 0 | 0 |
| MT072864.1;QIQ54046.1;Pangolin_CoV_orf1a;873 | 0 | 0 |
| MT072864.1;QIQ54046.1;Pangolin_CoV_orf1a;874 | 0 | 0 |
| MT072864.1;QIQ54046.1;Pangolin_CoV_orf1a;875 | 0 | 0.025 |
| MT072864.1;QIQ54046.1;Pangolin_CoV_orf1a;876 | 0 | 0.05 |
| MT072864.1;QIQ54046.1;Pangolin_CoV_orf1a;877 | 0.05 | 0.025 |
| MT072864.1;QIQ54046.1;Pangolin_CoV_orf1a;878 | 0 | 0.05 |
| MT072864.1;QIQ54046.1;Pangolin_CoV_orf1a;879 | 0 | 0.025 |
| MT072864.1;QIQ54046.1;Pangolin_CoV_orf1a;880 | 0 | 0.025 |
| MT072864.1;QIQ54046.1;Pangolin_CoV_orf1a;881 | 0.05 | 0.025 |
| MT072864.1;QIQ54046.1;Pangolin_CoV_orf1a;882 | 0 | 0.025 |
| MT072864.1;QIQ54046.1;Pangolin_CoV_orf1a;883 | 0 | 0 |
| MT072864.1;QIQ54046.1;Pangolin_CoV_orf1a;884 | 0.05 | 0 |
| MT072864.1;QIQ54046.1;Pangolin_CoV_orf1a;885 | 0 | 0 |
| MT072864.1;QIQ54046.1;Pangolin_CoV_orf1a;886 | 0 | 0 |
| MT072864.1;QIQ54046.1;Pangolin_CoV_orf1a;887 | 0 | 0 |
| MT072864.1;QIQ54046.1;Pangolin_CoV_orf1a;888 | 0 | 0.025 |
| MT072864.1;QIQ54046.1;Pangolin_CoV_orf1a;889 | 0 | 0 |
| MT072864.1;QIQ54046.1;Pangolin_CoV_orf1a;890 | 0 | 0 |
| MT072864.1;QIQ54046.1;Pangolin_CoV_orf1a;891 | 0 | 0 |
| MT072864.1;QIQ54046.1;Pangolin_CoV_orf1a;892 | 0 | 0 |
| MT072864.1;QIQ54046.1;Pangolin_CoV_orf1a;893 | 0 | 0 |

| | | |
|---|---|---|
| MT072864.1;QIQ54046.1;Pangolin_CoV_orf1a;894 | 0 | 0 |
| MT072864.1;QIQ54046.1;Pangolin_CoV_orf1a;895 | 0 | 0 |
| MT072864.1;QIQ54046.1;Pangolin_CoV_orf1a;896 | 0 | 0 |
| MT072864.1;QIQ54046.1;Pangolin_CoV_orf1a;897 | 0 | 0 |
| MT072864.1;QIQ54046.1;Pangolin_CoV_orf1a;898 | 0 | 0 |
| MT072864.1;QIQ54046.1;Pangolin_CoV_orf1a;899 | 0 | 0 |
| MT072864.1;QIQ54046.1;Pangolin_CoV_orf1a;900 | 0 | 0 |
| MT072864.1;QIQ54046.1;Pangolin_CoV_orf1a;901 | 0 | 0 |
| MT072864.1;QIQ54046.1;Pangolin_CoV_orf1a;902 | 0 | 0 |
| MT072864.1;QIQ54046.1;Pangolin_CoV_orf1a;903 | 0 | 0 |
| MT072864.1;QIQ54046.1;Pangolin_CoV_orf1a;904 | 0 | 0 |
| MT072864.1;QIQ54046.1;Pangolin_CoV_orf1a;905 | 0 | 0 |
| MT072864.1;QIQ54046.1;Pangolin_CoV_orf1a;906 | 0 | 0 |
| MT072864.1;QIQ54046.1;Pangolin_CoV_orf1a;907 | 0 | 0 |
| MT072864.1;QIQ54046.1;Pangolin_CoV_orf1a;908 | 0 | 0 |
| MT072864.1;QIQ54046.1;Pangolin_CoV_orf1a;909 | 0 | 0 |
| MT072864.1;QIQ54046.1;Pangolin_CoV_orf1a;910 | 0 | 0 |
| MT072864.1;QIQ54046.1;Pangolin_CoV_orf1a;911 | 0 | 0 |
| MT072864.1;QIQ54046.1;Pangolin_CoV_orf1a;912 | 0 | 0 |
| MT072864.1;QIQ54046.1;Pangolin_CoV_orf1a;913 | 0 | 0 |
| MT072864.1;QIQ54046.1;Pangolin_CoV_orf1a;914 | 0 | 0 |
| MT072864.1;QIQ54046.1;Pangolin_CoV_orf1a;915 | 0 | 0 |
| MT072864.1;QIQ54046.1;Pangolin_CoV_orf1a;916 | 0 | 0 |
| MT072864.1;QIQ54046.1;Pangolin_CoV_orf1a;917 | 0 | 0.025 |
| MT072864.1;QIQ54046.1;Pangolin_CoV_orf1a;918 | 0 | 0.075 |
| MT072864.1;QIQ54046.1;Pangolin_CoV_orf1a;919 | 0 | 0.1 |
| MT072864.1;QIQ54046.1;Pangolin_CoV_orf1a;920 | 0 | 0.125 |
| MT072864.1;QIQ54046.1;Pangolin_CoV_orf1a;921 | 0 | 0.1 |
| MT072864.1;QIQ54046.1;Pangolin_CoV_orf1a;922 | 0 | 0.125 |
| MT072864.1;QIQ54046.1;Pangolin_CoV_orf1a;923 | 0 | 0.125 |
| MT072864.1;QIQ54046.1;Pangolin_CoV_orf1a;924 | 0 | 0.125 |
| MT072864.1;QIQ54046.1;Pangolin_CoV_orf1a;925 | 0 | 0.05 |
| MT072864.1;QIQ54046.1;Pangolin_CoV_orf1a;926 | 0 | 0 |
| MT072864.1;QIQ54046.1;Pangolin_CoV_orf1a;927 | 0 | 0 |
| MT072864.1;QIQ54046.1;Pangolin_CoV_orf1a;928 | 0 | 0 |
| MT072864.1;QIQ54046.1;Pangolin_CoV_orf1a;929 | 0.15 | 0.175 |
| MT072864.1;QIQ54046.1;Pangolin_CoV_orf1a;930 | 0.05 | 0 |
| MT072864.1;QIQ54046.1;Pangolin_CoV_orf1a;931 | 0.1 | 0 |
| MT072864.1;QIQ54046.1;Pangolin_CoV_orf1a;932 | 0.05 | 0 |
| MT072864.1;QIQ54046.1;Pangolin_CoV_orf1a;933 | 0.1 | 0 |
| MT072864.1;QIQ54046.1;Pangolin_CoV_orf1a;934 | 0.1 | 0.025 |
| MT072864.1;QIQ54046.1;Pangolin_CoV_orf1a;935 | 0.15 | 0.025 |
| MT072864.1;QIQ54046.1;Pangolin_CoV_orf1a;936 | 0.1 | 0.025 |

| | | |
|---|---|---|
| MT072864.1;QIQ54046.1;Pangolin_CoV_orf1a;937 | 0.1 | 0.025 |
| MT072864.1;QIQ54046.1;Pangolin_CoV_orf1a;938 | 0.15 | 0.025 |
| MT072864.1;QIQ54046.1;Pangolin_CoV_orf1a;939 | 0.1 | 0.025 |
| MT072864.1;QIQ54046.1;Pangolin_CoV_orf1a;940 | 0.1 | 0.025 |
| MT072864.1;QIQ54046.1;Pangolin_CoV_orf1a;941 | 0.1 | 0.05 |
| MT072864.1;QIQ54046.1;Pangolin_CoV_orf1a;942 | 0 | 0 |
| MT072864.1;QIQ54046.1;Pangolin_CoV_orf1a;943 | 0 | 0 |
| MT072864.1;QIQ54046.1;Pangolin_CoV_orf1a;944 | 0 | 0 |
| MT072864.1;QIQ54046.1;Pangolin_CoV_orf1a;945 | 0.05 | 0.05 |
| MT072864.1;QIQ54046.1;Pangolin_CoV_orf1a;946 | 0.1 | 0.025 |
| MT072864.1;QIQ54046.1;Pangolin_CoV_orf1a;947 | 0.15 | 0.05 |
| MT072864.1;QIQ54046.1;Pangolin_CoV_orf1a;948 | 0.15 | 0.05 |
| MT072864.1;QIQ54046.1;Pangolin_CoV_orf1a;949 | 0.15 | 0.1 |
| MT072864.1;QIQ54046.1;Pangolin_CoV_orf1a;950 | 0.15 | 0.05 |
| MT072864.1;QIQ54046.1;Pangolin_CoV_orf1a;951 | 0.15 | 0.05 |
| MT072864.1;QIQ54046.1;Pangolin_CoV_orf1a;952 | 0 | 0.05 |
| MT072864.1;QIQ54046.1;Pangolin_CoV_orf1a;953 | 0 | 0.025 |
| MT072864.1;QIQ54046.1;Pangolin_CoV_orf1a;954 | 0 | 0 |
| MT072864.1;QIQ54046.1;Pangolin_CoV_orf1a;955 | 0 | 0 |
| MT072864.1;QIQ54046.1;Pangolin_CoV_orf1a;956 | 0 | 0 |
| MT072864.1;QIQ54046.1;Pangolin_CoV_orf1a;957 | 0 | 0 |
| MT072864.1;QIQ54046.1;Pangolin_CoV_orf1a;958 | 0 | 0 |
| MT072864.1;QIQ54046.1;Pangolin_CoV_orf1a;959 | 0 | 0 |
| MT072864.1;QIQ54046.1;Pangolin_CoV_orf1a;960 | 0 | 0 |
| MT072864.1;QIQ54046.1;Pangolin_CoV_orf1a;961 | 0 | 0 |
| MT072864.1;QIQ54046.1;Pangolin_CoV_orf1a;962 | 0 | 0.025 |
| MT072864.1;QIQ54046.1;Pangolin_CoV_orf1a;963 | 0 | 0.05 |
| MT072864.1;QIQ54046.1;Pangolin_CoV_orf1a;964 | 0 | 0.025 |
| MT072864.1;QIQ54046.1;Pangolin_CoV_orf1a;965 | 0 | 0.025 |
| MT072864.1;QIQ54046.1;Pangolin_CoV_orf1a;966 | 0 | 0.05 |
| MT072864.1;QIQ54046.1;Pangolin_CoV_orf1a;967 | 0 | 0.05 |
| MT072864.1;QIQ54046.1;Pangolin_CoV_orf1a;968 | 0 | 0.05 |
| MT072864.1;QIQ54046.1;Pangolin_CoV_orf1a;969 | 0 | 0.05 |
| MT072864.1;QIQ54046.1;Pangolin_CoV_orf1a;970 | 0 | 0.025 |
| MT072864.1;QIQ54046.1;Pangolin_CoV_orf1a;971 | 0 | 0.025 |
| MT072864.1;QIQ54046.1;Pangolin_CoV_orf1a;972 | 0 | 0.025 |
| MT072864.1;QIQ54046.1;Pangolin_CoV_orf1a;973 | 0 | 0.025 |
| MT072864.1;QIQ54046.1;Pangolin_CoV_orf1a;974 | 0 | 0.025 |
| MT072864.1;QIQ54046.1;Pangolin_CoV_orf1a;975 | 0 | 0 |
| MT072864.1;QIQ54046.1;Pangolin_CoV_orf1a;976 | 0 | 0 |
| MT072864.1;QIQ54046.1;Pangolin_CoV_orf1a;977 | 0 | 0 |
| MT072864.1;QIQ54046.1;Pangolin_CoV_orf1a;978 | 0 | 0 |
| MT072864.1;QIQ54046.1;Pangolin_CoV_orf1a;979 | 0 | 0 |

| | | |
|---|---|---|
| MT072864.1;QIQ54046.1;Pangolin_CoV_orf1a;980 | 0 | 0 |
| MT072864.1;QIQ54046.1;Pangolin_CoV_orf1a;981 | 0 | 0 |
| MT072864.1;QIQ54046.1;Pangolin_CoV_orf1a;982 | 0.05 | 0.1 |
| MT072864.1;QIQ54046.1;Pangolin_CoV_orf1a;983 | 0.05 | 0 |
| MT072864.1;QIQ54046.1;Pangolin_CoV_orf1a;984 | 0 | 0 |
| MT072864.1;QIQ54046.1;Pangolin_CoV_orf1a;985 | 0 | 0 |
| MT072864.1;QIQ54046.1;Pangolin_CoV_orf1a;986 | 0 | 0 |
| MT072864.1;QIQ54046.1;Pangolin_CoV_orf1a;987 | 0 | 0 |
| MT072864.1;QIQ54046.1;Pangolin_CoV_orf1a;988 | 0 | 0 |
| MT072864.1;QIQ54046.1;Pangolin_CoV_orf1a;989 | 0 | 0 |
| MT072864.1;QIQ54046.1;Pangolin_CoV_orf1a;990 | 0.05 | 0 |
| MT072864.1;QIQ54046.1;Pangolin_CoV_orf1a;991 | 0 | 0 |
| MT072864.1;QIQ54046.1;Pangolin_CoV_orf1a;992 | 0 | 0 |
| MT072864.1;QIQ54046.1;Pangolin_CoV_orf1a;993 | 0 | 0 |
| MT072864.1;QIQ54046.1;Pangolin_CoV_orf1a;994 | 0 | 0.025 |
| MT072864.1;QIQ54046.1;Pangolin_CoV_orf1a;995 | 0 | 0 |
| MT072864.1;QIQ54046.1;Pangolin_CoV_orf1a;996 | 0 | 0 |
| MT072864.1;QIQ54046.1;Pangolin_CoV_orf1a;997 | 0 | 0 |
| MT072864.1;QIQ54046.1;Pangolin_CoV_orf1a;998 | 0 | 0 |
| MT072864.1;QIQ54046.1;Pangolin_CoV_orf1a;999 | 0 | 0 |
| MT072864.1;QIQ54046.1;Pangolin_CoV_orf1a;1000 | 0 | 0 |
| MT072864.1;QIQ54046.1;Pangolin_CoV_orf1a;1001 | 0 | 0 |
| MT072864.1;QIQ54046.1;Pangolin_CoV_orf1a;1002 | 0 | 0 |
| MT072864.1;QIQ54046.1;Pangolin_CoV_orf1a;1003 | 0 | 0 |
| MT072864.1;QIQ54046.1;Pangolin_CoV_orf1a;1004 | 0 | 0 |
| MT072864.1;QIQ54046.1;Pangolin_CoV_orf1a;1005 | 0 | 0 |
| MT072864.1;QIQ54046.1;Pangolin_CoV_orf1a;1006 | 0 | 0 |
| MT072864.1;QIQ54046.1;Pangolin_CoV_orf1a;1007 | 0 | 0 |
| MT072864.1;QIQ54046.1;Pangolin_CoV_orf1a;1008 | 0 | 0 |
| MT072864.1;QIQ54046.1;Pangolin_CoV_orf1a;1009 | 0 | 0 |
| MT072864.1;QIQ54046.1;Pangolin_CoV_orf1a;1010 | 0 | 0 |
| MT072864.1;QIQ54046.1;Pangolin_CoV_orf1a;1011 | 0 | 0 |
| MT072864.1;QIQ54046.1;Pangolin_CoV_orf1a;1012 | 0 | 0 |
| MT072864.1;QIQ54046.1;Pangolin_CoV_orf1a;1013 | 0 | 0 |
| MT072864.1;QIQ54046.1;Pangolin_CoV_orf1a;1014 | 0.05 | 0 |
| MT072864.1;QIQ54046.1;Pangolin_CoV_orf1a;1015 | 0.05 | 0 |
| MT072864.1;QIQ54046.1;Pangolin_CoV_orf1a;1016 | 0.05 | 0.025 |
| MT072864.1;QIQ54046.1;Pangolin_CoV_orf1a;1017 | 0.05 | 0 |
| MT072864.1;QIQ54046.1;Pangolin_CoV_orf1a;1018 | 0.05 | 0 |
| MT072864.1;QIQ54046.1;Pangolin_CoV_orf1a;1019 | 0.05 | 0 |
| MT072864.1;QIQ54046.1;Pangolin_CoV_orf1a;1020 | 0 | 0 |
| MT072864.1;QIQ54046.1;Pangolin_CoV_orf1a;1021 | 0.05 | 0 |
| MT072864.1;QIQ54046.1;Pangolin_CoV_orf1a;1022 | 0.05 | 0 |

| | | |
|---|---|---|
| MT072864.1;QIQ54046.1;Pangolin_CoV_orf1a;1023 | 0.05 | 0 |
| MT072864.1;QIQ54046.1;Pangolin_CoV_orf1a;1024 | 0.05 | 0 |
| MT072864.1;QIQ54046.1;Pangolin_CoV_orf1a;1025 | 0.05 | 0 |
| MT072864.1;QIQ54046.1;Pangolin_CoV_orf1a;1026 | 0.05 | 0 |
| MT072864.1;QIQ54046.1;Pangolin_CoV_orf1a;1027 | 0.25 | 0.05 |
| MT072864.1;QIQ54046.1;Pangolin_CoV_orf1a;1028 | 0.1 | 0.025 |
| MT072864.1;QIQ54046.1;Pangolin_CoV_orf1a;1029 | 0.15 | 0.05 |
| MT072864.1;QIQ54046.1;Pangolin_CoV_orf1a;1030 | 0.1 | 0.025 |
| MT072864.1;QIQ54046.1;Pangolin_CoV_orf1a;1031 | 0.15 | 0.075 |
| MT072864.1;QIQ54046.1;Pangolin_CoV_orf1a;1032 | 0.3 | 0.075 |
| MT072864.1;QIQ54046.1;Pangolin_CoV_orf1a;1033 | 0.2 | 0.075 |
| MT072864.1;QIQ54046.1;Pangolin_CoV_orf1a;1034 | 0.2 | 0.075 |
| MT072864.1;QIQ54046.1;Pangolin_CoV_orf1a;1035 | 0 | 0.05 |
| MT072864.1;QIQ54046.1;Pangolin_CoV_orf1a;1036 | 0 | 0 |
| MT072864.1;QIQ54046.1;Pangolin_CoV_orf1a;1037 | 0 | 0.025 |
| MT072864.1;QIQ54046.1;Pangolin_CoV_orf1a;1038 | 0 | 0 |
| MT072864.1;QIQ54046.1;Pangolin_CoV_orf1a;1039 | 0 | 0 |
| MT072864.1;QIQ54046.1;Pangolin_CoV_orf1a;1040 | 0 | 0 |
| MT072864.1;QIQ54046.1;Pangolin_CoV_orf1a;1041 | 0 | 0 |
| MT072864.1;QIQ54046.1;Pangolin_CoV_orf1a;1042 | 0 | 0 |
| MT072864.1;QIQ54046.1;Pangolin_CoV_orf1a;1043 | 0 | 0 |
| MT072864.1;QIQ54046.1;Pangolin_CoV_orf1a;1044 | 0 | 0 |
| MT072864.1;QIQ54046.1;Pangolin_CoV_orf1a;1045 | 0 | 0 |
| MT072864.1;QIQ54046.1;Pangolin_CoV_orf1a;1046 | 0 | 0 |
| MT072864.1;QIQ54046.1;Pangolin_CoV_orf1a;1047 | 0.05 | 0 |
| MT072864.1;QIQ54046.1;Pangolin_CoV_orf1a;1048 | 0 | 0 |
| MT072864.1;QIQ54046.1;Pangolin_CoV_orf1a;1049 | 0.05 | 0.025 |
| MT072864.1;QIQ54046.1;Pangolin_CoV_orf1a;1050 | 0 | 0 |
| MT072864.1;QIQ54046.1;Pangolin_CoV_orf1a;1051 | 0 | 0 |
| MT072864.1;QIQ54046.1;Pangolin_CoV_orf1a;1052 | 0 | 0 |
| MT072864.1;QIQ54046.1;Pangolin_CoV_orf1a;1053 | 0 | 0 |
| MT072864.1;QIQ54046.1;Pangolin_CoV_orf1a;1054 | 0 | 0 |
| MT072864.1;QIQ54046.1;Pangolin_CoV_orf1a;1055 | 0 | 0 |
| MT072864.1;QIQ54046.1;Pangolin_CoV_orf1a;1056 | 0 | 0 |
| MT072864.1;QIQ54046.1;Pangolin_CoV_orf1a;1057 | 0 | 0 |
| MT072864.1;QIQ54046.1;Pangolin_CoV_orf1a;1058 | 0 | 0.025 |
| MT072864.1;QIQ54046.1;Pangolin_CoV_orf1a;1059 | 0.1 | 0.025 |
| MT072864.1;QIQ54046.1;Pangolin_CoV_orf1a;1060 | 0.15 | 0.025 |
| MT072864.1;QIQ54046.1;Pangolin_CoV_orf1a;1061 | 0.05 | 0.025 |
| MT072864.1;QIQ54046.1;Pangolin_CoV_orf1a;1062 | 0.15 | 0.025 |
| MT072864.1;QIQ54046.1;Pangolin_CoV_orf1a;1063 | 0.15 | 0.175 |
| MT072864.1;QIQ54046.1;Pangolin_CoV_orf1a;1064 | 0.25 | 0.275 |
| MT072864.1;QIQ54046.1;Pangolin_CoV_orf1a;1065 | 0.15 | 0.125 |

| | | |
|---|---|---|
| MT072864.1;QIQ54046.1;Pangolin_CoV_orf1a;1066 | 0.15 | 0.175 |
| MT072864.1;QIQ54046.1;Pangolin_CoV_orf1a;1067 | 0.05 | 0.125 |
| MT072864.1;QIQ54046.1;Pangolin_CoV_orf1a;1068 | 0 | 0.025 |
| MT072864.1;QIQ54046.1;Pangolin_CoV_orf1a;1069 | 0 | 0.025 |
| MT072864.1;QIQ54046.1;Pangolin_CoV_orf1a;1070 | 0.1 | 0.025 |
| MT072864.1;QIQ54046.1;Pangolin_CoV_orf1a;1071 | 0 | 0.025 |
| MT072864.1;QIQ54046.1;Pangolin_CoV_orf1a;1072 | 0 | 0 |
| MT072864.1;QIQ54046.1;Pangolin_CoV_orf1a;1073 | 0 | 0 |
| MT072864.1;QIQ54046.1;Pangolin_CoV_orf1a;1074 | 0 | 0.05 |
| MT072864.1;QIQ54046.1;Pangolin_CoV_orf1a;1075 | 0 | 0.025 |
| MT072864.1;QIQ54046.1;Pangolin_CoV_orf1a;1076 | 0 | 0.025 |
| MT072864.1;QIQ54046.1;Pangolin_CoV_orf1a;1077 | 0.05 | 0.025 |
| MT072864.1;QIQ54046.1;Pangolin_CoV_orf1a;1078 | 0 | 0.05 |
| MT072864.1;QIQ54046.1;Pangolin_CoV_orf1a;1079 | 0.1 | 0.1 |
| MT072864.1;QIQ54046.1;Pangolin_CoV_orf1a;1080 | 0 | 0.025 |
| MT072864.1;QIQ54046.1;Pangolin_CoV_orf1a;1081 | 0 | 0.025 |
| MT072864.1;QIQ54046.1;Pangolin_CoV_orf1a;1082 | 0 | 0 |
| MT072864.1;QIQ54046.1;Pangolin_CoV_orf1a;1083 | 0 | 0 |
| MT072864.1;QIQ54046.1;Pangolin_CoV_orf1a;1084 | 0 | 0 |
| MT072864.1;QIQ54046.1;Pangolin_CoV_orf1a;1085 | 0 | 0 |
| MT072864.1;QIQ54046.1;Pangolin_CoV_orf1a;1086 | 0 | 0 |
| MT072864.1;QIQ54046.1;Pangolin_CoV_orf1a;1087 | 0 | 0 |
| MT072864.1;QIQ54046.1;Pangolin_CoV_orf1a;1088 | 0 | 0 |
| MT072864.1;QIQ54046.1;Pangolin_CoV_orf1a;1089 | 0 | 0 |
| MT072864.1;QIQ54046.1;Pangolin_CoV_orf1a;1090 | 0 | 0 |
| MT072864.1;QIQ54046.1;Pangolin_CoV_orf1a;1091 | 0 | 0 |
| MT072864.1;QIQ54046.1;Pangolin_CoV_orf1a;1092 | 0 | 0 |
| MT072864.1;QIQ54046.1;Pangolin_CoV_orf1a;1093 | 0 | 0 |
| MT072864.1;QIQ54046.1;Pangolin_CoV_orf1a;1094 | 0 | 0 |
| MT072864.1;QIQ54046.1;Pangolin_CoV_orf1a;1095 | 0 | 0 |
| MT072864.1;QIQ54046.1;Pangolin_CoV_orf1a;1096 | 0 | 0 |
| MT072864.1;QIQ54046.1;Pangolin_CoV_orf1a;1097 | 0 | 0 |
| MT072864.1;QIQ54046.1;Pangolin_CoV_orf1a;1098 | 0 | 0 |
| MT072864.1;QIQ54046.1;Pangolin_CoV_orf1a;1099 | 0 | 0 |
| MT072864.1;QIQ54046.1;Pangolin_CoV_orf1a;1100 | 0 | 0 |
| MT072864.1;QIQ54046.1;Pangolin_CoV_orf1a;1101 | 0.05 | 0 |
| MT072864.1;QIQ54046.1;Pangolin_CoV_orf1a;1102 | 0.1 | 0 |
| MT072864.1;QIQ54046.1;Pangolin_CoV_orf1a;1103 | 0.2 | 0 |
| MT072864.1;QIQ54046.1;Pangolin_CoV_orf1a;1104 | 0.25 | 0 |
| MT072864.1;QIQ54046.1;Pangolin_CoV_orf1a;1105 | 0.15 | 0 |
| MT072864.1;QIQ54046.1;Pangolin_CoV_orf1a;1106 | 0.15 | 0.025 |
| MT072864.1;QIQ54046.1;Pangolin_CoV_orf1a;1107 | 0.25 | 0 |
| MT072864.1;QIQ54046.1;Pangolin_CoV_orf1a;1108 | 0 | 0 |

| | | |
|---|---|---|
| MT072864.1;QIQ54046.1;Pangolin_CoV_orf1a;1109 | 0.05 | 0.05 |
| MT072864.1;QIQ54046.1;Pangolin_CoV_orf1a;1110 | 0.1 | 0.05 |
| MT072864.1;QIQ54046.1;Pangolin_CoV_orf1a;1111 | 0.05 | 0.025 |
| MT072864.1;QIQ54046.1;Pangolin_CoV_orf1a;1112 | 0.05 | 0 |
| MT072864.1;QIQ54046.1;Pangolin_CoV_orf1a;1113 | 0.05 | 0.05 |
| MT072864.1;QIQ54046.1;Pangolin_CoV_orf1a;1114 | 0.05 | 0 |
| MT072864.1;QIQ54046.1;Pangolin_CoV_orf1a;1115 | 0 | 0.025 |
| MT072864.1;QIQ54046.1;Pangolin_CoV_orf1a;1116 | 0 | 0.025 |
| MT072864.1;QIQ54046.1;Pangolin_CoV_orf1a;1117 | 0 | 0 |
| MT072864.1;QIQ54046.1;Pangolin_CoV_orf1a;1118 | 0 | 0 |
| MT072864.1;QIQ54046.1;Pangolin_CoV_orf1a;1119 | 0 | 0 |
| MT072864.1;QIQ54046.1;Pangolin_CoV_orf1a;1120 | 0 | 0.025 |
| MT072864.1;QIQ54046.1;Pangolin_CoV_orf1a;1121 | 0 | 0.025 |
| MT072864.1;QIQ54046.1;Pangolin_CoV_orf1a;1122 | 0 | 0.025 |
| MT072864.1;QIQ54046.1;Pangolin_CoV_orf1a;1123 | 0 | 0.025 |
| MT072864.1;QIQ54046.1;Pangolin_CoV_orf1a;1124 | 0 | 0.05 |
| MT072864.1;QIQ54046.1;Pangolin_CoV_orf1a;1125 | 0 | 0.025 |
| MT072864.1;QIQ54046.1;Pangolin_CoV_orf1a;1126 | 0 | 0.025 |
| MT072864.1;QIQ54046.1;Pangolin_CoV_orf1a;1127 | 0 | 0.025 |
| MT072864.1;QIQ54046.1;Pangolin_CoV_orf1a;1128 | 0 | 0.025 |
| MT072864.1;QIQ54046.1;Pangolin_CoV_orf1a;1129 | 0 | 0.025 |
| MT072864.1;QIQ54046.1;Pangolin_CoV_orf1a;1130 | 0 | 0.025 |
| MT072864.1;QIQ54046.1;Pangolin_CoV_orf1a;1131 | 0 | 0.025 |
| MT072864.1;QIQ54046.1;Pangolin_CoV_orf1a;1132 | 0 | 0 |
| MT072864.1;QIQ54046.1;Pangolin_CoV_orf1a;1133 | 0 | 0 |
| MT072864.1;QIQ54046.1;Pangolin_CoV_orf1a;1134 | 0 | 0 |
| MT072864.1;QIQ54046.1;Pangolin_CoV_orf1a;1135 | 0 | 0 |
| MT072864.1;QIQ54046.1;Pangolin_CoV_orf1a;1136 | 0 | 0 |
| MT072864.1;QIQ54046.1;Pangolin_CoV_orf1a;1137 | 0 | 0 |
| MT072864.1;QIQ54046.1;Pangolin_CoV_orf1a;1138 | 0 | 0 |
| MT072864.1;QIQ54046.1;Pangolin_CoV_orf1a;1139 | 0 | 0 |
| MT072864.1;QIQ54046.1;Pangolin_CoV_orf1a;1140 | 0 | 0 |
| MT072864.1;QIQ54046.1;Pangolin_CoV_orf1a;1141 | 0 | 0 |
| MT072864.1;QIQ54046.1;Pangolin_CoV_orf1a;1142 | 0 | 0 |
| MT072864.1;QIQ54046.1;Pangolin_CoV_orf1a;1143 | 0 | 0 |
| MT072864.1;QIQ54046.1;Pangolin_CoV_orf1a;1144 | 0 | 0 |
| MT072864.1;QIQ54046.1;Pangolin_CoV_orf1a;1145 | 0 | 0 |
| MT072864.1;QIQ54046.1;Pangolin_CoV_orf1a;1146 | 0 | 0 |
| MT072864.1;QIQ54046.1;Pangolin_CoV_orf1a;1147 | 0 | 0 |
| MT072864.1;QIQ54046.1;Pangolin_CoV_orf1a;1148 | 0 | 0 |
| MT072864.1;QIQ54046.1;Pangolin_CoV_orf1a;1149 | 0.05 | 0.025 |
| MT072864.1;QIQ54046.1;Pangolin_CoV_orf1a;1150 | 0.05 | 0.05 |
| MT072864.1;QIQ54046.1;Pangolin_CoV_orf1a;1151 | 0.05 | 0.05 |

| | | |
|---|---|---|
| MT072864.1;QIQ54046.1;Pangolin_CoV_orf1a;1152 | 0 | 0 |
| MT072864.1;QIQ54046.1;Pangolin_CoV_orf1a;1153 | 0 | 0 |
| MT072864.1;QIQ54046.1;Pangolin_CoV_orf1a;1154 | 0 | 0 |
| MT072864.1;QIQ54046.1;Pangolin_CoV_orf1a;1155 | 0 | 0 |
| MT072864.1;QIQ54046.1;Pangolin_CoV_orf1a;1156 | 0 | 0 |
| MT072864.1;QIQ54046.1;Pangolin_CoV_orf1a;1157 | 0 | 0 |
| MT072864.1;QIQ54046.1;Pangolin_CoV_orf1a;1158 | 0 | 0 |
| MT072864.1;QIQ54046.1;Pangolin_CoV_orf1a;1159 | 0.05 | 0 |
| MT072864.1;QIQ54046.1;Pangolin_CoV_orf1a;1160 | 0 | 0 |
| MT072864.1;QIQ54046.1;Pangolin_CoV_orf1a;1161 | 0 | 0 |
| MT072864.1;QIQ54046.1;Pangolin_CoV_orf1a;1162 | 0 | 0 |
| MT072864.1;QIQ54046.1;Pangolin_CoV_orf1a;1163 | 0 | 0 |
| MT072864.1;QIQ54046.1;Pangolin_CoV_orf1a;1164 | 0 | 0 |
| MT072864.1;QIQ54046.1;Pangolin_CoV_orf1a;1165 | 0 | 0 |
| MT072864.1;QIQ54046.1;Pangolin_CoV_orf1a;1166 | 0 | 0 |
| MT072864.1;QIQ54046.1;Pangolin_CoV_orf1a;1167 | 0 | 0 |
| MT072864.1;QIQ54046.1;Pangolin_CoV_orf1a;1168 | 0 | 0 |
| MT072864.1;QIQ54046.1;Pangolin_CoV_orf1a;1169 | 0 | 0.025 |
| MT072864.1;QIQ54046.1;Pangolin_CoV_orf1a;1170 | 0 | 0 |
| MT072864.1;QIQ54046.1;Pangolin_CoV_orf1a;1171 | 0 | 0 |
| MT072864.1;QIQ54046.1;Pangolin_CoV_orf1a;1172 | 0 | 0 |
| MT072864.1;QIQ54046.1;Pangolin_CoV_orf1a;1173 | 0 | 0 |
| MT072864.1;QIQ54046.1;Pangolin_CoV_orf1a;1174 | 0 | 0 |
| MT072864.1;QIQ54046.1;Pangolin_CoV_orf1a;1175 | 0 | 0.1 |
| MT072864.1;QIQ54046.1;Pangolin_CoV_orf1a;1176 | 0 | 0.05 |
| MT072864.1;QIQ54046.1;Pangolin_CoV_orf1a;1177 | 0 | 0.1 |
| MT072864.1;QIQ54046.1;Pangolin_CoV_orf1a;1178 | 0 | 0.125 |
| MT072864.1;QIQ54046.1;Pangolin_CoV_orf1a;1179 | 0 | 0.1 |
| MT072864.1;QIQ54046.1;Pangolin_CoV_orf1a;1180 | 0 | 0.1 |
| MT072864.1;QIQ54046.1;Pangolin_CoV_orf1a;1181 | 0 | 0.1 |
| MT072864.1;QIQ54046.1;Pangolin_CoV_orf1a;1182 | 0 | 0.1 |
| MT072864.1;QIQ54046.1;Pangolin_CoV_orf1a;1183 | 0 | 0.075 |
| MT072864.1;QIQ54046.1;Pangolin_CoV_orf1a;1184 | 0.05 | 0.025 |
| MT072864.1;QIQ54046.1;Pangolin_CoV_orf1a;1185 | 0 | 0 |
| MT072864.1;QIQ54046.1;Pangolin_CoV_orf1a;1186 | 0 | 0.025 |
| MT072864.1;QIQ54046.1;Pangolin_CoV_orf1a;1187 | 0 | 0 |
| MT072864.1;QIQ54046.1;Pangolin_CoV_orf1a;1188 | 0 | 0.025 |
| MT072864.1;QIQ54046.1;Pangolin_CoV_orf1a;1189 | 0 | 0 |
| MT072864.1;QIQ54046.1;Pangolin_CoV_orf1a;1190 | 0 | 0 |
| MT072864.1;QIQ54046.1;Pangolin_CoV_orf1a;1191 | 0 | 0 |
| MT072864.1;QIQ54046.1;Pangolin_CoV_orf1a;1192 | 0 | 0 |
| MT072864.1;QIQ54046.1;Pangolin_CoV_orf1a;1193 | 0 | 0.025 |
| MT072864.1;QIQ54046.1;Pangolin_CoV_orf1a;1194 | 0 | 0.025 |

| | | |
|---|---|---|
| MT072864.1;QIQ54046.1;Pangolin_CoV_orf1a;1195 | 0 | 0 |
| MT072864.1;QIQ54046.1;Pangolin_CoV_orf1a;1196 | 0.05 | 0.025 |
| MT072864.1;QIQ54046.1;Pangolin_CoV_orf1a;1197 | 0 | 0.025 |
| MT072864.1;QIQ54046.1;Pangolin_CoV_orf1a;1198 | 0 | 0.025 |
| MT072864.1;QIQ54046.1;Pangolin_CoV_orf1a;1199 | 0 | 0.025 |
| MT072864.1;QIQ54046.1;Pangolin_CoV_orf1a;1200 | 0 | 0 |
| MT072864.1;QIQ54046.1;Pangolin_CoV_orf1a;1201 | 0 | 0 |
| MT072864.1;QIQ54046.1;Pangolin_CoV_orf1a;1202 | 0 | 0 |
| MT072864.1;QIQ54046.1;Pangolin_CoV_orf1a;1203 | 0 | 0 |
| MT072864.1;QIQ54046.1;Pangolin_CoV_orf1a;1204 | 0 | 0 |
| MT072864.1;QIQ54046.1;Pangolin_CoV_orf1a;1205 | 0 | 0 |
| MT072864.1;QIQ54046.1;Pangolin_CoV_orf1a;1206 | 0 | 0 |
| MT072864.1;QIQ54046.1;Pangolin_CoV_orf1a;1207 | 0 | 0.05 |
| MT072864.1;QIQ54046.1;Pangolin_CoV_orf1a;1208 | 0 | 0.1 |
| MT072864.1;QIQ54046.1;Pangolin_CoV_orf1a;1209 | 0 | 0.15 |
| MT072864.1;QIQ54046.1;Pangolin_CoV_orf1a;1210 | 0 | 0.05 |
| MT072864.1;QIQ54046.1;Pangolin_CoV_orf1a;1211 | 0 | 0.05 |
| MT072864.1;QIQ54046.1;Pangolin_CoV_orf1a;1212 | 0.05 | 0.075 |
| MT072864.1;QIQ54046.1;Pangolin_CoV_orf1a;1213 | 0.05 | 0.1 |
| MT072864.1;QIQ54046.1;Pangolin_CoV_orf1a;1214 | 0 | 0.075 |
| MT072864.1;QIQ54046.1;Pangolin_CoV_orf1a;1215 | 0.05 | 0.05 |
| MT072864.1;QIQ54046.1;Pangolin_CoV_orf1a;1216 | 0 | 0.05 |
| MT072864.1;QIQ54046.1;Pangolin_CoV_orf1a;1217 | 0 | 0.05 |
| MT072864.1;QIQ54046.1;Pangolin_CoV_orf1a;1218 | 0 | 0.025 |
| MT072864.1;QIQ54046.1;Pangolin_CoV_orf1a;1219 | 0.05 | 0.025 |
| MT072864.1;QIQ54046.1;Pangolin_CoV_orf1a;1220 | 0 | 0.025 |
| MT072864.1;QIQ54046.1;Pangolin_CoV_orf1a;1221 | 0 | 0 |
| MT072864.1;QIQ54046.1;Pangolin_CoV_orf1a;1222 | 0 | 0 |
| MT072864.1;QIQ54046.1;Pangolin_CoV_orf1a;1223 | 0 | 0 |
| MT072864.1;QIQ54046.1;Pangolin_CoV_orf1a;1224 | 0 | 0 |
| MT072864.1;QIQ54046.1;Pangolin_CoV_orf1a;1225 | 0 | 0 |
| MT072864.1;QIQ54046.1;Pangolin_CoV_orf1a;1226 | 0 | 0.025 |
| MT072864.1;QIQ54046.1;Pangolin_CoV_orf1a;1227 | 0 | 0 |
| MT072864.1;QIQ54046.1;Pangolin_CoV_orf1a;1228 | 0 | 0.075 |
| MT072864.1;QIQ54046.1;Pangolin_CoV_orf1a;1229 | 0 | 0.05 |
| MT072864.1;QIQ54046.1;Pangolin_CoV_orf1a;1230 | 0 | 0.025 |
| MT072864.1;QIQ54046.1;Pangolin_CoV_orf1a;1231 | 0 | 0.025 |
| MT072864.1;QIQ54046.1;Pangolin_CoV_orf1a;1232 | 0 | 0.05 |
| MT072864.1;QIQ54046.1;Pangolin_CoV_orf1a;1233 | 0 | 0.025 |
| MT072864.1;QIQ54046.1;Pangolin_CoV_orf1a;1234 | 0 | 0 |
| MT072864.1;QIQ54046.1;Pangolin_CoV_orf1a;1235 | 0 | 0.025 |
| MT072864.1;QIQ54046.1;Pangolin_CoV_orf1a;1236 | 0 | 0.025 |
| MT072864.1;QIQ54046.1;Pangolin_CoV_orf1a;1237 | 0 | 0 |

| | | |
|---|---|---|
| MT072864.1;QIQ54046.1;Pangolin_CoV_orf1a;1238 | 0 | 0 |
| MT072864.1;QIQ54046.1;Pangolin_CoV_orf1a;1239 | 0 | 0 |
| MT072864.1;QIQ54046.1;Pangolin_CoV_orf1a;1240 | 0 | 0 |
| MT072864.1;QIQ54046.1;Pangolin_CoV_orf1a;1241 | 0.05 | 0 |
| MT072864.1;QIQ54046.1;Pangolin_CoV_orf1a;1242 | 0 | 0 |
| MT072864.1;QIQ54046.1;Pangolin_CoV_orf1a;1243 | 0 | 0 |
| MT072864.1;QIQ54046.1;Pangolin_CoV_orf1a;1244 | 0 | 0.025 |
| MT072864.1;QIQ54046.1;Pangolin_CoV_orf1a;1245 | 0.1 | 0.125 |
| MT072864.1;QIQ54046.1;Pangolin_CoV_orf1a;1246 | 0.25 | 0.175 |
| MT072864.1;QIQ54046.1;Pangolin_CoV_orf1a;1247 | 0.15 | 0.15 |
| MT072864.1;QIQ54046.1;Pangolin_CoV_orf1a;1248 | 0 | 0.025 |
| MT072864.1;QIQ54046.1;Pangolin_CoV_orf1a;1249 | 0 | 0.025 |
| MT072864.1;QIQ54046.1;Pangolin_CoV_orf1a;1250 | 0 | 0 |
| MT072864.1;QIQ54046.1;Pangolin_CoV_orf1a;1251 | 0 | 0 |
| MT072864.1;QIQ54046.1;Pangolin_CoV_orf1a;1252 | 0 | 0 |
| MT072864.1;QIQ54046.1;Pangolin_CoV_orf1a;1253 | 0 | 0.025 |
| MT072864.1;QIQ54046.1;Pangolin_CoV_orf1a;1254 | 0 | 0.025 |
| MT072864.1;QIQ54046.1;Pangolin_CoV_orf1a;1255 | 0 | 0 |
| MT072864.1;QIQ54046.1;Pangolin_CoV_orf1a;1256 | 0 | 0.025 |
| MT072864.1;QIQ54046.1;Pangolin_CoV_orf1a;1257 | 0 | 0 |
| MT072864.1;QIQ54046.1;Pangolin_CoV_orf1a;1258 | 0 | 0.025 |
| MT072864.1;QIQ54046.1;Pangolin_CoV_orf1a;1259 | 0 | 0 |
| MT072864.1;QIQ54046.1;Pangolin_CoV_orf1a;1260 | 0 | 0 |
| MT072864.1;QIQ54046.1;Pangolin_CoV_orf1a;1261 | 0 | 0.05 |
| MT072864.1;QIQ54046.1;Pangolin_CoV_orf1a;1262 | 0 | 0.025 |
| MT072864.1;QIQ54046.1;Pangolin_CoV_orf1a;1263 | 0 | 0 |
| MT072864.1;QIQ54046.1;Pangolin_CoV_orf1a;1264 | 0 | 0.025 |
| MT072864.1;QIQ54046.1;Pangolin_CoV_orf1a;1265 | 0 | 0.025 |
| MT072864.1;QIQ54046.1;Pangolin_CoV_orf1a;1266 | 0 | 0 |
| MT072864.1;QIQ54046.1;Pangolin_CoV_orf1a;1267 | 0 | 0 |
| MT072864.1;QIQ54046.1;Pangolin_CoV_orf1a;1268 | 0 | 0 |
| MT072864.1;QIQ54046.1;Pangolin_CoV_orf1a;1269 | 0 | 0 |
| MT072864.1;QIQ54046.1;Pangolin_CoV_orf1a;1270 | 0 | 0 |
| MT072864.1;QIQ54046.1;Pangolin_CoV_orf1a;1271 | 0 | 0.025 |
| MT072864.1;QIQ54046.1;Pangolin_CoV_orf1a;1272 | 0 | 0.025 |
| MT072864.1;QIQ54046.1;Pangolin_CoV_orf1a;1273 | 0 | 0.05 |
| MT072864.1;QIQ54046.1;Pangolin_CoV_orf1a;1274 | 0 | 0.05 |
| MT072864.1;QIQ54046.1;Pangolin_CoV_orf1a;1275 | 0 | 0.025 |
| MT072864.1;QIQ54046.1;Pangolin_CoV_orf1a;1276 | 0 | 0 |
| MT072864.1;QIQ54046.1;Pangolin_CoV_orf1a;1277 | 0 | 0 |
| MT072864.1;QIQ54046.1;Pangolin_CoV_orf1a;1278 | 0 | 0 |
| MT072864.1;QIQ54046.1;Pangolin_CoV_orf1a;1279 | 0 | 0.025 |
| MT072864.1;QIQ54046.1;Pangolin_CoV_orf1a;1280 | 0 | 0.025 |

| | | |
|---|---|---|
| MT072864.1;QIQ54046.1;Pangolin_CoV_orf1a;1281 | 0.1 | 0.025 |
| MT072864.1;QIQ54046.1;Pangolin_CoV_orf1a;1282 | 0 | 0.05 |
| MT072864.1;QIQ54046.1;Pangolin_CoV_orf1a;1283 | 0 | 0 |
| MT072864.1;QIQ54046.1;Pangolin_CoV_orf1a;1284 | 0 | 0 |
| MT072864.1;QIQ54046.1;Pangolin_CoV_orf1a;1285 | 0 | 0 |
| MT072864.1;QIQ54046.1;Pangolin_CoV_orf1a;1286 | 0 | 0 |
| MT072864.1;QIQ54046.1;Pangolin_CoV_orf1a;1287 | 0.05 | 0 |
| MT072864.1;QIQ54046.1;Pangolin_CoV_orf1a;1288 | 0.05 | 0.025 |
| MT072864.1;QIQ54046.1;Pangolin_CoV_orf1a;1289 | 0.05 | 0.025 |
| MT072864.1;QIQ54046.1;Pangolin_CoV_orf1a;1290 | 0.05 | 0.025 |
| MT072864.1;QIQ54046.1;Pangolin_CoV_orf1a;1291 | 0.05 | 0 |
| MT072864.1;QIQ54046.1;Pangolin_CoV_orf1a;1292 | 0.05 | 0 |
| MT072864.1;QIQ54046.1;Pangolin_CoV_orf1a;1293 | 0.05 | 0 |
| MT072864.1;QIQ54046.1;Pangolin_CoV_orf1a;1294 | 0 | 0 |
| MT072864.1;QIQ54046.1;Pangolin_CoV_orf1a;1295 | 0 | 0 |
| MT072864.1;QIQ54046.1;Pangolin_CoV_orf1a;1296 | 0 | 0 |
| MT072864.1;QIQ54046.1;Pangolin_CoV_orf1a;1297 | 0 | 0 |
| MT072864.1;QIQ54046.1;Pangolin_CoV_orf1a;1298 | 0 | 0 |
| MT072864.1;QIQ54046.1;Pangolin_CoV_orf1a;1299 | 0 | 0 |
| MT072864.1;QIQ54046.1;Pangolin_CoV_orf1a;1300 | 0 | 0 |
| MT072864.1;QIQ54046.1;Pangolin_CoV_orf1a;1301 | 0 | 0 |
| MT072864.1;QIQ54046.1;Pangolin_CoV_orf1a;1302 | 0 | 0 |
| MT072864.1;QIQ54046.1;Pangolin_CoV_orf1a;1303 | 0 | 0 |
| MT072864.1;QIQ54046.1;Pangolin_CoV_orf1a;1304 | 0 | 0 |
| MT072864.1;QIQ54046.1;Pangolin_CoV_orf1a;1305 | 0 | 0 |
| MT072864.1;QIQ54046.1;Pangolin_CoV_orf1a;1306 | 0 | 0 |
| MT072864.1;QIQ54046.1;Pangolin_CoV_orf1a;1307 | 0 | 0 |
| MT072864.1;QIQ54046.1;Pangolin_CoV_orf1a;1308 | 0 | 0 |
| MT072864.1;QIQ54046.1;Pangolin_CoV_orf1a;1309 | 0 | 0 |
| MT072864.1;QIQ54046.1;Pangolin_CoV_orf1a;1310 | 0 | 0 |
| MT072864.1;QIQ54046.1;Pangolin_CoV_orf1a;1311 | 0 | 0 |
| MT072864.1;QIQ54046.1;Pangolin_CoV_orf1a;1312 | 0 | 0 |
| MT072864.1;QIQ54046.1;Pangolin_CoV_orf1a;1313 | 0 | 0 |
| MT072864.1;QIQ54046.1;Pangolin_CoV_orf1a;1314 | 0 | 0 |
| MT072864.1;QIQ54046.1;Pangolin_CoV_orf1a;1315 | 0 | 0 |
| MT072864.1;QIQ54046.1;Pangolin_CoV_orf1a;1316 | 0 | 0 |
| MT072864.1;QIQ54046.1;Pangolin_CoV_orf1a;1317 | 0 | 0 |
| MT072864.1;QIQ54046.1;Pangolin_CoV_orf1a;1318 | 0 | 0 |
| MT072864.1;QIQ54046.1;Pangolin_CoV_orf1a;1319 | 0.05 | 0.025 |
| MT072864.1;QIQ54046.1;Pangolin_CoV_orf1a;1320 | 0.05 | 0.025 |
| MT072864.1;QIQ54046.1;Pangolin_CoV_orf1a;1321 | 0.05 | 0.025 |
| MT072864.1;QIQ54046.1;Pangolin_CoV_orf1a;1322 | 0.05 | 0.025 |
| MT072864.1;QIQ54046.1;Pangolin_CoV_orf1a;1323 | 0.05 | 0.05 |

| | | |
|---|---|---|
| MT072864.1;QIQ54046.1;Pangolin_CoV_orf1a;1324 | 0 | 0.025 |
| MT072864.1;QIQ54046.1;Pangolin_CoV_orf1a;1325 | 0 | 0.025 |
| MT072864.1;QIQ54046.1;Pangolin_CoV_orf1a;1326 | 0 | 0.025 |
| MT072864.1;QIQ54046.1;Pangolin_CoV_orf1a;1327 | 0 | 0 |
| MT072864.1;QIQ54046.1;Pangolin_CoV_orf1a;1328 | 0 | 0 |
| MT072864.1;QIQ54046.1;Pangolin_CoV_orf1a;1329 | 0 | 0 |
| MT072864.1;QIQ54046.1;Pangolin_CoV_orf1a;1330 | 0 | 0 |
| MT072864.1;QIQ54046.1;Pangolin_CoV_orf1a;1331 | 0 | 0 |
| MT072864.1;QIQ54046.1;Pangolin_CoV_orf1a;1332 | 0 | 0 |
| MT072864.1;QIQ54046.1;Pangolin_CoV_orf1a;1333 | 0 | 0 |
| MT072864.1;QIQ54046.1;Pangolin_CoV_orf1a;1334 | 0 | 0.025 |
| MT072864.1;QIQ54046.1;Pangolin_CoV_orf1a;1335 | 0 | 0 |
| MT072864.1;QIQ54046.1;Pangolin_CoV_orf1a;1336 | 0 | 0 |
| MT072864.1;QIQ54046.1;Pangolin_CoV_orf1a;1337 | 0 | 0 |
| MT072864.1;QIQ54046.1;Pangolin_CoV_orf1a;1338 | 0 | 0.025 |
| MT072864.1;QIQ54046.1;Pangolin_CoV_orf1a;1339 | 0 | 0.025 |
| MT072864.1;QIQ54046.1;Pangolin_CoV_orf1a;1340 | 0 | 0 |
| MT072864.1;QIQ54046.1;Pangolin_CoV_orf1a;1341 | 0 | 0 |
| MT072864.1;QIQ54046.1;Pangolin_CoV_orf1a;1342 | 0 | 0 |
| MT072864.1;QIQ54046.1;Pangolin_CoV_orf1a;1343 | 0 | 0 |
| MT072864.1;QIQ54046.1;Pangolin_CoV_orf1a;1344 | 0 | 0 |
| MT072864.1;QIQ54046.1;Pangolin_CoV_orf1a;1345 | 0 | 0.025 |
| MT072864.1;QIQ54046.1;Pangolin_CoV_orf1a;1346 | 0.05 | 0.025 |
| MT072864.1;QIQ54046.1;Pangolin_CoV_orf1a;1347 | 0 | 0 |
| MT072864.1;QIQ54046.1;Pangolin_CoV_orf1a;1348 | 0.1 | 0 |
| MT072864.1;QIQ54046.1;Pangolin_CoV_orf1a;1349 | 0 | 0 |
| MT072864.1;QIQ54046.1;Pangolin_CoV_orf1a;1350 | 0.05 | 0 |
| MT072864.1;QIQ54046.1;Pangolin_CoV_orf1a;1351 | 0 | 0 |
| MT072864.1;QIQ54046.1;Pangolin_CoV_orf1a;1352 | 0 | 0 |
| MT072864.1;QIQ54046.1;Pangolin_CoV_orf1a;1353 | 0 | 0 |
| MT072864.1;QIQ54046.1;Pangolin_CoV_orf1a;1354 | 0 | 0 |
| MT072864.1;QIQ54046.1;Pangolin_CoV_orf1a;1355 | 0 | 0 |
| MT072864.1;QIQ54046.1;Pangolin_CoV_orf1a;1356 | 0.05 | 0.025 |
| MT072864.1;QIQ54046.1;Pangolin_CoV_orf1a;1357 | 0 | 0 |
| MT072864.1;QIQ54046.1;Pangolin_CoV_orf1a;1358 | 0 | 0 |
| MT072864.1;QIQ54046.1;Pangolin_CoV_orf1a;1359 | 0 | 0 |
| MT072864.1;QIQ54046.1;Pangolin_CoV_orf1a;1360 | 0 | 0.025 |
| MT072864.1;QIQ54046.1;Pangolin_CoV_orf1a;1361 | 0 | 0.025 |
| MT072864.1;QIQ54046.1;Pangolin_CoV_orf1a;1362 | 0 | 0.025 |
| MT072864.1;QIQ54046.1;Pangolin_CoV_orf1a;1363 | 0 | 0.025 |
| MT072864.1;QIQ54046.1;Pangolin_CoV_orf1a;1364 | 0 | 0 |
| MT072864.1;QIQ54046.1;Pangolin_CoV_orf1a;1365 | 0 | 0 |
| MT072864.1;QIQ54046.1;Pangolin_CoV_orf1a;1366 | 0 | 0 |

| | | |
|---|---|---|
| MT072864.1;QIQ54046.1;Pangolin_CoV_orf1a;1367 | 0 | 0 |
| MT072864.1;QIQ54046.1;Pangolin_CoV_orf1a;1368 | 0 | 0 |
| MT072864.1;QIQ54046.1;Pangolin_CoV_orf1a;1369 | 0 | 0 |
| MT072864.1;QIQ54046.1;Pangolin_CoV_orf1a;1370 | 0 | 0 |
| MT072864.1;QIQ54046.1;Pangolin_CoV_orf1a;1371 | 0 | 0 |
| MT072864.1;QIQ54046.1;Pangolin_CoV_orf1a;1372 | 0 | 0 |
| MT072864.1;QIQ54046.1;Pangolin_CoV_orf1a;1373 | 0.05 | 0.025 |
| MT072864.1;QIQ54046.1;Pangolin_CoV_orf1a;1374 | 0.05 | 0.025 |
| MT072864.1;QIQ54046.1;Pangolin_CoV_orf1a;1375 | 0 | 0 |
| MT072864.1;QIQ54046.1;Pangolin_CoV_orf1a;1376 | 0 | 0 |
| MT072864.1;QIQ54046.1;Pangolin_CoV_orf1a;1377 | 0 | 0 |
| MT072864.1;QIQ54046.1;Pangolin_CoV_orf1a;1378 | 0 | 0 |
| MT072864.1;QIQ54046.1;Pangolin_CoV_orf1a;1379 | 0 | 0 |
| MT072864.1;QIQ54046.1;Pangolin_CoV_orf1a;1380 | 0 | 0 |
| MT072864.1;QIQ54046.1;Pangolin_CoV_orf1a;1381 | 0 | 0 |
| MT072864.1;QIQ54046.1;Pangolin_CoV_orf1a;1382 | 0 | 0 |
| MT072864.1;QIQ54046.1;Pangolin_CoV_orf1a;1383 | 0 | 0 |
| MT072864.1;QIQ54046.1;Pangolin_CoV_orf1a;1384 | 0 | 0 |
| MT072864.1;QIQ54046.1;Pangolin_CoV_orf1a;1385 | 0 | 0 |
| MT072864.1;QIQ54046.1;Pangolin_CoV_orf1a;1386 | 0 | 0 |
| MT072864.1;QIQ54046.1;Pangolin_CoV_orf1a;1387 | 0.05 | 0 |
| MT072864.1;QIQ54046.1;Pangolin_CoV_orf1a;1388 | 0.05 | 0 |
| MT072864.1;QIQ54046.1;Pangolin_CoV_orf1a;1389 | 0.1 | 0 |
| MT072864.1;QIQ54046.1;Pangolin_CoV_orf1a;1390 | 0.1 | 0 |
| MT072864.1;QIQ54046.1;Pangolin_CoV_orf1a;1391 | 0.1 | 0 |
| MT072864.1;QIQ54046.1;Pangolin_CoV_orf1a;1392 | 0.1 | 0 |
| MT072864.1;QIQ54046.1;Pangolin_CoV_orf1a;1393 | 0.1 | 0 |
| MT072864.1;QIQ54046.1;Pangolin_CoV_orf1a;1394 | 0.1 | 0.025 |
| MT072864.1;QIQ54046.1;Pangolin_CoV_orf1a;1395 | 0.1 | 0.025 |
| MT072864.1;QIQ54046.1;Pangolin_CoV_orf1a;1396 | 0.1 | 0 |
| MT072864.1;QIQ54046.1;Pangolin_CoV_orf1a;1397 | 0.05 | 0.05 |
| MT072864.1;QIQ54046.1;Pangolin_CoV_orf1a;1398 | 0 | 0.025 |
| MT072864.1;QIQ54046.1;Pangolin_CoV_orf1a;1399 | 0 | 0 |
| MT072864.1;QIQ54046.1;Pangolin_CoV_orf1a;1400 | 0 | 0 |
| MT072864.1;QIQ54046.1;Pangolin_CoV_orf1a;1401 | 0 | 0 |
| MT072864.1;QIQ54046.1;Pangolin_CoV_orf1a;1402 | 0 | 0 |
| MT072864.1;QIQ54046.1;Pangolin_CoV_orf1a;1403 | 0 | 0 |
| MT072864.1;QIQ54046.1;Pangolin_CoV_orf1a;1404 | 0 | 0 |
| MT072864.1;QIQ54046.1;Pangolin_CoV_orf1a;1405 | 0 | 0.05 |
| MT072864.1;QIQ54046.1;Pangolin_CoV_orf1a;1406 | 0 | 0 |
| MT072864.1;QIQ54046.1;Pangolin_CoV_orf1a;1407 | 0 | 0 |
| MT072864.1;QIQ54046.1;Pangolin_CoV_orf1a;1408 | 0 | 0 |
| MT072864.1;QIQ54046.1;Pangolin_CoV_orf1a;1409 | 0 | 0 |

| | | |
|---|---|---|
| MT072864.1;QIQ54046.1;Pangolin_CoV_orf1a;1410 | 0 | 0 |
| MT072864.1;QIQ54046.1;Pangolin_CoV_orf1a;1411 | 0 | 0 |
| MT072864.1;QIQ54046.1;Pangolin_CoV_orf1a;1412 | 0 | 0 |
| MT072864.1;QIQ54046.1;Pangolin_CoV_orf1a;1413 | 0 | 0 |
| MT072864.1;QIQ54046.1;Pangolin_CoV_orf1a;1414 | 0 | 0 |
| MT072864.1;QIQ54046.1;Pangolin_CoV_orf1a;1415 | 0 | 0 |
| MT072864.1;QIQ54046.1;Pangolin_CoV_orf1a;1416 | 0 | 0 |
| MT072864.1;QIQ54046.1;Pangolin_CoV_orf1a;1417 | 0 | 0 |
| MT072864.1;QIQ54046.1;Pangolin_CoV_orf1a;1418 | 0 | 0 |
| MT072864.1;QIQ54046.1;Pangolin_CoV_orf1a;1419 | 0 | 0.025 |
| MT072864.1;QIQ54046.1;Pangolin_CoV_orf1a;1420 | 0.05 | 0.1 |
| MT072864.1;QIQ54046.1;Pangolin_CoV_orf1a;1421 | 0.05 | 0.075 |
| MT072864.1;QIQ54046.1;Pangolin_CoV_orf1a;1422 | 0 | 0.025 |
| MT072864.1;QIQ54046.1;Pangolin_CoV_orf1a;1423 | 0 | 0.025 |
| MT072864.1;QIQ54046.1;Pangolin_CoV_orf1a;1424 | 0 | 0.025 |
| MT072864.1;QIQ54046.1;Pangolin_CoV_orf1a;1425 | 0 | 0.025 |
| MT072864.1;QIQ54046.1;Pangolin_CoV_orf1a;1426 | 0 | 0.05 |
| MT072864.1;QIQ54046.1;Pangolin_CoV_orf1a;1427 | 0 | 0.025 |
| MT072864.1;QIQ54046.1;Pangolin_CoV_orf1a;1428 | 0 | 0.025 |
| MT072864.1;QIQ54046.1;Pangolin_CoV_orf1a;1429 | 0 | 0.025 |
| MT072864.1;QIQ54046.1;Pangolin_CoV_orf1a;1430 | 0 | 0.025 |
| MT072864.1;QIQ54046.1;Pangolin_CoV_orf1a;1431 | 0 | 0 |
| MT072864.1;QIQ54046.1;Pangolin_CoV_orf1a;1432 | 0 | 0 |
| MT072864.1;QIQ54046.1;Pangolin_CoV_orf1a;1433 | 0 | 0 |
| MT072864.1;QIQ54046.1;Pangolin_CoV_orf1a;1434 | 0 | 0 |
| MT072864.1;QIQ54046.1;Pangolin_CoV_orf1a;1435 | 0 | 0 |
| MT072864.1;QIQ54046.1;Pangolin_CoV_orf1a;1436 | 0 | 0 |
| MT072864.1;QIQ54046.1;Pangolin_CoV_orf1a;1437 | 0.05 | 0 |
| MT072864.1;QIQ54046.1;Pangolin_CoV_orf1a;1438 | 0 | 0 |
| MT072864.1;QIQ54046.1;Pangolin_CoV_orf1a;1439 | 0.1 | 0.05 |
| MT072864.1;QIQ54046.1;Pangolin_CoV_orf1a;1440 | 0.05 | 0.025 |
| MT072864.1;QIQ54046.1;Pangolin_CoV_orf1a;1441 | 0.05 | 0.025 |
| MT072864.1;QIQ54046.1;Pangolin_CoV_orf1a;1442 | 0.05 | 0.025 |
| MT072864.1;QIQ54046.1;Pangolin_CoV_orf1a;1443 | 0 | 0 |
| MT072864.1;QIQ54046.1;Pangolin_CoV_orf1a;1444 | 0 | 0 |
| MT072864.1;QIQ54046.1;Pangolin_CoV_orf1a;1445 | 0 | 0 |
| MT072864.1;QIQ54046.1;Pangolin_CoV_orf1a;1446 | 0 | 0 |
| MT072864.1;QIQ54046.1;Pangolin_CoV_orf1a;1447 | 0 | 0.025 |
| MT072864.1;QIQ54046.1;Pangolin_CoV_orf1a;1448 | 0 | 0 |
| MT072864.1;QIQ54046.1;Pangolin_CoV_orf1a;1449 | 0 | 0 |
| MT072864.1;QIQ54046.1;Pangolin_CoV_orf1a;1450 | 0 | 0.075 |
| MT072864.1;QIQ54046.1;Pangolin_CoV_orf1a;1451 | 0.1 | 0.075 |
| MT072864.1;QIQ54046.1;Pangolin_CoV_orf1a;1452 | 0.15 | 0.125 |

| | | |
|---|---|---|
| MT072864.1;QIQ54046.1;Pangolin_CoV_orf1a;1453 | 0.05 | 0 |
| MT072864.1;QIQ54046.1;Pangolin_CoV_orf1a;1454 | 0 | 0.025 |
| MT072864.1;QIQ54046.1;Pangolin_CoV_orf1a;1455 | 0 | 0 |
| MT072864.1;QIQ54046.1;Pangolin_CoV_orf1a;1456 | 0 | 0 |
| MT072864.1;QIQ54046.1;Pangolin_CoV_orf1a;1457 | 0 | 0 |
| MT072864.1;QIQ54046.1;Pangolin_CoV_orf1a;1458 | 0 | 0 |
| MT072864.1;QIQ54046.1;Pangolin_CoV_orf1a;1459 | 0 | 0 |
| MT072864.1;QIQ54046.1;Pangolin_CoV_orf1a;1460 | 0 | 0 |
| MT072864.1;QIQ54046.1;Pangolin_CoV_orf1a;1461 | 0 | 0 |
| MT072864.1;QIQ54046.1;Pangolin_CoV_orf1a;1462 | 0 | 0 |
| MT072864.1;QIQ54046.1;Pangolin_CoV_orf1a;1463 | 0 | 0.05 |
| MT072864.1;QIQ54046.1;Pangolin_CoV_orf1a;1464 | 0.05 | 0.025 |
| MT072864.1;QIQ54046.1;Pangolin_CoV_orf1a;1465 | 0 | 0.1 |
| MT072864.1;QIQ54046.1;Pangolin_CoV_orf1a;1466 | 0 | 0.025 |
| MT072864.1;QIQ54046.1;Pangolin_CoV_orf1a;1467 | 0 | 0 |
| MT072864.1;QIQ54046.1;Pangolin_CoV_orf1a;1468 | 0 | 0 |
| MT072864.1;QIQ54046.1;Pangolin_CoV_orf1a;1469 | 0 | 0 |
| MT072864.1;QIQ54046.1;Pangolin_CoV_orf1a;1470 | 0 | 0 |
| MT072864.1;QIQ54046.1;Pangolin_CoV_orf1a;1471 | 0.05 | 0.05 |
| MT072864.1;QIQ54046.1;Pangolin_CoV_orf1a;1472 | 0.05 | 0.075 |
| MT072864.1;QIQ54046.1;Pangolin_CoV_orf1a;1473 | 0.1 | 0.05 |
| MT072864.1;QIQ54046.1;Pangolin_CoV_orf1a;1474 | 0.15 | 0.1 |
| MT072864.1;QIQ54046.1;Pangolin_CoV_orf1a;1475 | 0.1 | 0.15 |
| MT072864.1;QIQ54046.1;Pangolin_CoV_orf1a;1476 | 0.1 | 0.125 |
| MT072864.1;QIQ54046.1;Pangolin_CoV_orf1a;1477 | 0.15 | 0.125 |
| MT072864.1;QIQ54046.1;Pangolin_CoV_orf1a;1478 | 0.2 | 0.125 |
| MT072864.1;QIQ54046.1;Pangolin_CoV_orf1a;1479 | 0.25 | 0.1 |
| MT072864.1;QIQ54046.1;Pangolin_CoV_orf1a;1480 | 0.2 | 0.1 |
| MT072864.1;QIQ54046.1;Pangolin_CoV_orf1a;1481 | 0.15 | 0.025 |
| MT072864.1;QIQ54046.1;Pangolin_CoV_orf1a;1482 | 0.05 | 0 |
| MT072864.1;QIQ54046.1;Pangolin_CoV_orf1a;1483 | 0.05 | 0 |
| MT072864.1;QIQ54046.1;Pangolin_CoV_orf1a;1484 | 0.05 | 0 |
| MT072864.1;QIQ54046.1;Pangolin_CoV_orf1a;1485 | 0 | 0 |
| MT072864.1;QIQ54046.1;Pangolin_CoV_orf1a;1486 | 0 | 0 |
| MT072864.1;QIQ54046.1;Pangolin_CoV_orf1a;1487 | 0 | 0 |
| MT072864.1;QIQ54046.1;Pangolin_CoV_orf1a;1488 | 0 | 0 |
| MT072864.1;QIQ54046.1;Pangolin_CoV_orf1a;1489 | 0 | 0 |
| MT072864.1;QIQ54046.1;Pangolin_CoV_orf1a;1490 | 0 | 0 |
| MT072864.1;QIQ54046.1;Pangolin_CoV_orf1a;1491 | 0 | 0 |
| MT072864.1;QIQ54046.1;Pangolin_CoV_orf1a;1492 | 0 | 0 |
| MT072864.1;QIQ54046.1;Pangolin_CoV_orf1a;1493 | 0 | 0 |
| MT072864.1;QIQ54046.1;Pangolin_CoV_orf1a;1494 | 0 | 0 |
| MT072864.1;QIQ54046.1;Pangolin_CoV_orf1a;1495 | 0 | 0 |

| | | |
|---|---|---|
| MT072864.1;QIQ54046.1;Pangolin_CoV_orf1a;1496 | 0 | 0 |
| MT072864.1;QIQ54046.1;Pangolin_CoV_orf1a;1497 | 0.05 | 0 |
| MT072864.1;QIQ54046.1;Pangolin_CoV_orf1a;1498 | 0 | 0 |
| MT072864.1;QIQ54046.1;Pangolin_CoV_orf1a;1499 | 0 | 0 |
| MT072864.1;QIQ54046.1;Pangolin_CoV_orf1a;1500 | 0.2 | 0.05 |
| MT072864.1;QIQ54046.1;Pangolin_CoV_orf1a;1501 | 0 | 0 |
| MT072864.1;QIQ54046.1;Pangolin_CoV_orf1a;1502 | 0 | 0 |
| MT072864.1;QIQ54046.1;Pangolin_CoV_orf1a;1503 | 0 | 0 |
| MT072864.1;QIQ54046.1;Pangolin_CoV_orf1a;1504 | 0 | 0 |
| MT072864.1;QIQ54046.1;Pangolin_CoV_orf1a;1505 | 0 | 0 |
| MT072864.1;QIQ54046.1;Pangolin_CoV_orf1a;1506 | 0.05 | 0 |
| MT072864.1;QIQ54046.1;Pangolin_CoV_orf1a;1507 | 0 | 0 |
| MT072864.1;QIQ54046.1;Pangolin_CoV_orf1a;1508 | 0 | 0 |
| MT072864.1;QIQ54046.1;Pangolin_CoV_orf1a;1509 | 0 | 0.075 |
| MT072864.1;QIQ54046.1;Pangolin_CoV_orf1a;1510 | 0 | 0.1 |
| MT072864.1;QIQ54046.1;Pangolin_CoV_orf1a;1511 | 0 | 0.075 |
| MT072864.1;QIQ54046.1;Pangolin_CoV_orf1a;1512 | 0 | 0.05 |
| MT072864.1;QIQ54046.1;Pangolin_CoV_orf1a;1513 | 0 | 0.05 |
| MT072864.1;QIQ54046.1;Pangolin_CoV_orf1a;1514 | 0 | 0.05 |
| MT072864.1;QIQ54046.1;Pangolin_CoV_orf1a;1515 | 0.05 | 0.075 |
| MT072864.1;QIQ54046.1;Pangolin_CoV_orf1a;1516 | 0 | 0.05 |
| MT072864.1;QIQ54046.1;Pangolin_CoV_orf1a;1517 | 0 | 0.075 |
| MT072864.1;QIQ54046.1;Pangolin_CoV_orf1a;1518 | 0 | 0.05 |
| MT072864.1;QIQ54046.1;Pangolin_CoV_orf1a;1519 | 0 | 0.05 |
| MT072864.1;QIQ54046.1;Pangolin_CoV_orf1a;1520 | 0 | 0.025 |
| MT072864.1;QIQ54046.1;Pangolin_CoV_orf1a;1521 | 0 | 0.025 |
| MT072864.1;QIQ54046.1;Pangolin_CoV_orf1a;1522 | 0 | 0.025 |
| MT072864.1;QIQ54046.1;Pangolin_CoV_orf1a;1523 | 0 | 0.025 |
| MT072864.1;QIQ54046.1;Pangolin_CoV_orf1a;1524 | 0 | 0.025 |
| MT072864.1;QIQ54046.1;Pangolin_CoV_orf1a;1525 | 0.05 | 0.025 |
| MT072864.1;QIQ54046.1;Pangolin_CoV_orf1a;1526 | 0.05 | 0.025 |
| MT072864.1;QIQ54046.1;Pangolin_CoV_orf1a;1527 | 0 | 0.025 |
| MT072864.1;QIQ54046.1;Pangolin_CoV_orf1a;1528 | 0 | 0 |
| MT072864.1;QIQ54046.1;Pangolin_CoV_orf1a;1529 | 0 | 0.05 |
| MT072864.1;QIQ54046.1;Pangolin_CoV_orf1a;1530 | 0 | 0.025 |
| MT072864.1;QIQ54046.1;Pangolin_CoV_orf1a;1531 | 0 | 0.025 |
| MT072864.1;QIQ54046.1;Pangolin_CoV_orf1a;1532 | 0 | 0.025 |
| MT072864.1;QIQ54046.1;Pangolin_CoV_orf1a;1533 | 0 | 0.025 |
| MT072864.1;QIQ54046.1;Pangolin_CoV_orf1a;1534 | 0 | 0.025 |
| MT072864.1;QIQ54046.1;Pangolin_CoV_orf1a;1535 | 0 | 0.025 |
| MT072864.1;QIQ54046.1;Pangolin_CoV_orf1a;1536 | 0 | 0.025 |
| MT072864.1;QIQ54046.1;Pangolin_CoV_orf1a;1537 | 0 | 0 |
| MT072864.1;QIQ54046.1;Pangolin_CoV_orf1a;1538 | 0 | 0 |

| | | |
|---|---|---|
| MT072864.1;QIQ54046.1;Pangolin_CoV_orf1a;1539 | 0 | 0 |
| MT072864.1;QIQ54046.1;Pangolin_CoV_orf1a;1540 | 0 | 0 |
| MT072864.1;QIQ54046.1;Pangolin_CoV_orf1a;1541 | 0 | 0 |
| MT072864.1;QIQ54046.1;Pangolin_CoV_orf1a;1542 | 0 | 0 |
| MT072864.1;QIQ54046.1;Pangolin_CoV_orf1a;1543 | 0 | 0 |
| MT072864.1;QIQ54046.1;Pangolin_CoV_orf1a;1544 | 0 | 0 |
| MT072864.1;QIQ54046.1;Pangolin_CoV_orf1a;1545 | 0 | 0.025 |
| MT072864.1;QIQ54046.1;Pangolin_CoV_orf1a;1546 | 0 | 0 |
| MT072864.1;QIQ54046.1;Pangolin_CoV_orf1a;1547 | 0 | 0 |
| MT072864.1;QIQ54046.1;Pangolin_CoV_orf1a;1548 | 0 | 0 |
| MT072864.1;QIQ54046.1;Pangolin_CoV_orf1a;1549 | 0 | 0 |
| MT072864.1;QIQ54046.1;Pangolin_CoV_orf1a;1550 | 0 | 0 |
| MT072864.1;QIQ54046.1;Pangolin_CoV_orf1a;1551 | 0 | 0 |
| MT072864.1;QIQ54046.1;Pangolin_CoV_orf1a;1552 | 0 | 0 |
| MT072864.1;QIQ54046.1;Pangolin_CoV_orf1a;1553 | 0 | 0 |
| MT072864.1;QIQ54046.1;Pangolin_CoV_orf1a;1554 | 0 | 0 |
| MT072864.1;QIQ54046.1;Pangolin_CoV_orf1a;1555 | 0 | 0 |
| MT072864.1;QIQ54046.1;Pangolin_CoV_orf1a;1556 | 0 | 0 |
| MT072864.1;QIQ54046.1;Pangolin_CoV_orf1a;1557 | 0 | 0 |
| MT072864.1;QIQ54046.1;Pangolin_CoV_orf1a;1558 | 0 | 0 |
| MT072864.1;QIQ54046.1;Pangolin_CoV_orf1a;1559 | 0 | 0.025 |
| MT072864.1;QIQ54046.1;Pangolin_CoV_orf1a;1560 | 0 | 0.025 |
| MT072864.1;QIQ54046.1;Pangolin_CoV_orf1a;1561 | 0 | 0.025 |
| MT072864.1;QIQ54046.1;Pangolin_CoV_orf1a;1562 | 0 | 0.025 |
| MT072864.1;QIQ54046.1;Pangolin_CoV_orf1a;1563 | 0 | 0.025 |
| MT072864.1;QIQ54046.1;Pangolin_CoV_orf1a;1564 | 0 | 0.025 |
| MT072864.1;QIQ54046.1;Pangolin_CoV_orf1a;1565 | 0.05 | 0.125 |
| MT072864.1;QIQ54046.1;Pangolin_CoV_orf1a;1566 | 0 | 0.075 |
| MT072864.1;QIQ54046.1;Pangolin_CoV_orf1a;1567 | 0 | 0.025 |
| MT072864.1;QIQ54046.1;Pangolin_CoV_orf1a;1568 | 0 | 0.025 |
| MT072864.1;QIQ54046.1;Pangolin_CoV_orf1a;1569 | 0 | 0.025 |
| MT072864.1;QIQ54046.1;Pangolin_CoV_orf1a;1570 | 0 | 0.025 |
| MT072864.1;QIQ54046.1;Pangolin_CoV_orf1a;1571 | 0 | 0.025 |
| MT072864.1;QIQ54046.1;Pangolin_CoV_orf1a;1572 | 0 | 0 |
| MT072864.1;QIQ54046.1;Pangolin_CoV_orf1a;1573 | 0 | 0 |
| MT072864.1;QIQ54046.1;Pangolin_CoV_orf1a;1574 | 0 | 0 |
| MT072864.1;QIQ54046.1;Pangolin_CoV_orf1a;1575 | 0 | 0 |
| MT072864.1;QIQ54046.1;Pangolin_CoV_orf1a;1576 | 0 | 0 |
| MT072864.1;QIQ54046.1;Pangolin_CoV_orf1a;1577 | 0 | 0 |
| MT072864.1;QIQ54046.1;Pangolin_CoV_orf1a;1578 | 0 | 0 |
| MT072864.1;QIQ54046.1;Pangolin_CoV_orf1a;1579 | 0 | 0 |
| MT072864.1;QIQ54046.1;Pangolin_CoV_orf1a;1580 | 0 | 0 |
| MT072864.1;QIQ54046.1;Pangolin_CoV_orf1a;1581 | 0 | 0 |

| | | |
|---|---|---|
| MT072864.1;QIQ54046.1;Pangolin_CoV_orf1a;1582 | 0 | 0 |
| MT072864.1;QIQ54046.1;Pangolin_CoV_orf1a;1583 | 0 | 0.025 |
| MT072864.1;QIQ54046.1;Pangolin_CoV_orf1a;1584 | 0 | 0.025 |
| MT072864.1;QIQ54046.1;Pangolin_CoV_orf1a;1585 | 0 | 0.025 |
| MT072864.1;QIQ54046.1;Pangolin_CoV_orf1a;1586 | 0 | 0.025 |
| MT072864.1;QIQ54046.1;Pangolin_CoV_orf1a;1587 | 0.15 | 0.125 |
| MT072864.1;QIQ54046.1;Pangolin_CoV_orf1a;1588 | 0 | 0.025 |
| MT072864.1;QIQ54046.1;Pangolin_CoV_orf1a;1589 | 0.05 | 0.05 |
| MT072864.1;QIQ54046.1;Pangolin_CoV_orf1a;1590 | 0 | 0 |
| MT072864.1;QIQ54046.1;Pangolin_CoV_orf1a;1591 | 0 | 0 |
| MT072864.1;QIQ54046.1;Pangolin_CoV_orf1a;1592 | 0 | 0 |
| MT072864.1;QIQ54046.1;Pangolin_CoV_orf1a;1593 | 0 | 0.025 |
| MT072864.1;QIQ54046.1;Pangolin_CoV_orf1a;1594 | 0 | 0 |
| MT072864.1;QIQ54046.1;Pangolin_CoV_orf1a;1595 | 0.05 | 0 |
| MT072864.1;QIQ54046.1;Pangolin_CoV_orf1a;1596 | 0 | 0 |
| MT072864.1;QIQ54046.1;Pangolin_CoV_orf1a;1597 | 0 | 0 |
| MT072864.1;QIQ54046.1;Pangolin_CoV_orf1a;1598 | 0 | 0 |
| MT072864.1;QIQ54046.1;Pangolin_CoV_orf1a;1599 | 0 | 0 |
| MT072864.1;QIQ54046.1;Pangolin_CoV_orf1a;1600 | 0 | 0 |
| MT072864.1;QIQ54046.1;Pangolin_CoV_orf1a;1601 | 0 | 0 |
| MT072864.1;QIQ54046.1;Pangolin_CoV_orf1a;1602 | 0 | 0 |
| MT072864.1;QIQ54046.1;Pangolin_CoV_orf1a;1603 | 0.05 | 0 |
| MT072864.1;QIQ54046.1;Pangolin_CoV_orf1a;1604 | 0.05 | 0 |
| MT072864.1;QIQ54046.1;Pangolin_CoV_orf1a;1605 | 0.05 | 0 |
| MT072864.1;QIQ54046.1;Pangolin_CoV_orf1a;1606 | 0.05 | 0 |
| MT072864.1;QIQ54046.1;Pangolin_CoV_orf1a;1607 | 0.1 | 0 |
| MT072864.1;QIQ54046.1;Pangolin_CoV_orf1a;1608 | 0.05 | 0 |
| MT072864.1;QIQ54046.1;Pangolin_CoV_orf1a;1609 | 0 | 0 |
| MT072864.1;QIQ54046.1;Pangolin_CoV_orf1a;1610 | 0 | 0 |
| MT072864.1;QIQ54046.1;Pangolin_CoV_orf1a;1611 | 0 | 0 |
| MT072864.1;QIQ54046.1;Pangolin_CoV_orf1a;1612 | 0 | 0 |
| MT072864.1;QIQ54046.1;Pangolin_CoV_orf1a;1613 | 0 | 0.025 |
| MT072864.1;QIQ54046.1;Pangolin_CoV_orf1a;1614 | 0 | 0 |
| MT072864.1;QIQ54046.1;Pangolin_CoV_orf1a;1615 | 0 | 0 |
| MT072864.1;QIQ54046.1;Pangolin_CoV_orf1a;1616 | 0 | 0 |
| MT072864.1;QIQ54046.1;Pangolin_CoV_orf1a;1617 | 0 | 0 |
| MT072864.1;QIQ54046.1;Pangolin_CoV_orf1a;1618 | 0 | 0 |
| MT072864.1;QIQ54046.1;Pangolin_CoV_orf1a;1619 | 0 | 0 |
| MT072864.1;QIQ54046.1;Pangolin_CoV_orf1a;1620 | 0 | 0 |
| MT072864.1;QIQ54046.1;Pangolin_CoV_orf1a;1621 | 0 | 0 |
| MT072864.1;QIQ54046.1;Pangolin_CoV_orf1a;1622 | 0 | 0 |
| MT072864.1;QIQ54046.1;Pangolin_CoV_orf1a;1623 | 0.05 | 0 |
| MT072864.1;QIQ54046.1;Pangolin_CoV_orf1a;1624 | 0.05 | 0 |

| | | |
|---|---|---|
| MT072864.1;QIQ54046.1;Pangolin_CoV_orf1a;1625 | 0.05 | 0 |
| MT072864.1;QIQ54046.1;Pangolin_CoV_orf1a;1626 | 0.05 | 0 |
| MT072864.1;QIQ54046.1;Pangolin_CoV_orf1a;1627 | 0.05 | 0 |
| MT072864.1;QIQ54046.1;Pangolin_CoV_orf1a;1628 | 0.05 | 0 |
| MT072864.1;QIQ54046.1;Pangolin_CoV_orf1a;1629 | 0.05 | 0 |
| MT072864.1;QIQ54046.1;Pangolin_CoV_orf1a;1630 | 0.05 | 0 |
| MT072864.1;QIQ54046.1;Pangolin_CoV_orf1a;1631 | 0.05 | 0 |
| MT072864.1;QIQ54046.1;Pangolin_CoV_orf1a;1632 | 0 | 0 |
| MT072864.1;QIQ54046.1;Pangolin_CoV_orf1a;1633 | 0 | 0 |
| MT072864.1;QIQ54046.1;Pangolin_CoV_orf1a;1634 | 0 | 0 |
| MT072864.1;QIQ54046.1;Pangolin_CoV_orf1a;1635 | 0 | 0 |
| MT072864.1;QIQ54046.1;Pangolin_CoV_orf1a;1636 | 0 | 0 |
| MT072864.1;QIQ54046.1;Pangolin_CoV_orf1a;1637 | 0 | 0 |
| MT072864.1;QIQ54046.1;Pangolin_CoV_orf1a;1638 | 0 | 0 |
| MT072864.1;QIQ54046.1;Pangolin_CoV_orf1a;1639 | 0 | 0 |
| MT072864.1;QIQ54046.1;Pangolin_CoV_orf1a;1640 | 0 | 0 |
| MT072864.1;QIQ54046.1;Pangolin_CoV_orf1a;1641 | 0 | 0 |
| MT072864.1;QIQ54046.1;Pangolin_CoV_orf1a;1642 | 0 | 0 |
| MT072864.1;QIQ54046.1;Pangolin_CoV_orf1a;1643 | 0 | 0 |
| MT072864.1;QIQ54046.1;Pangolin_CoV_orf1a;1644 | 0 | 0 |
| MT072864.1;QIQ54046.1;Pangolin_CoV_orf1a;1645 | 0 | 0 |
| MT072864.1;QIQ54046.1;Pangolin_CoV_orf1a;1646 | 0 | 0 |
| MT072864.1;QIQ54046.1;Pangolin_CoV_orf1a;1647 | 0 | 0 |
| MT072864.1;QIQ54046.1;Pangolin_CoV_orf1a;1648 | 0 | 0 |
| MT072864.1;QIQ54046.1;Pangolin_CoV_orf1a;1649 | 0.05 | 0.025 |
| MT072864.1;QIQ54046.1;Pangolin_CoV_orf1a;1650 | 0.05 | 0.025 |
| MT072864.1;QIQ54046.1;Pangolin_CoV_orf1a;1651 | 0.05 | 0.025 |
| MT072864.1;QIQ54046.1;Pangolin_CoV_orf1a;1652 | 0.05 | 0 |
| MT072864.1;QIQ54046.1;Pangolin_CoV_orf1a;1653 | 0.05 | 0 |
| MT072864.1;QIQ54046.1;Pangolin_CoV_orf1a;1654 | 0.05 | 0 |
| MT072864.1;QIQ54046.1;Pangolin_CoV_orf1a;1655 | 0.05 | 0 |
| MT072864.1;QIQ54046.1;Pangolin_CoV_orf1a;1656 | 0.05 | 0 |
| MT072864.1;QIQ54046.1;Pangolin_CoV_orf1a;1657 | 0 | 0 |
| MT072864.1;QIQ54046.1;Pangolin_CoV_orf1a;1658 | 0 | 0 |
| MT072864.1;QIQ54046.1;Pangolin_CoV_orf1a;1659 | 0 | 0 |
| MT072864.1;QIQ54046.1;Pangolin_CoV_orf1a;1660 | 0 | 0 |
| MT072864.1;QIQ54046.1;Pangolin_CoV_orf1a;1661 | 0 | 0 |
| MT072864.1;QIQ54046.1;Pangolin_CoV_orf1a;1662 | 0 | 0 |
| MT072864.1;QIQ54046.1;Pangolin_CoV_orf1a;1663 | 0 | 0 |
| MT072864.1;QIQ54046.1;Pangolin_CoV_orf1a;1664 | 0 | 0 |
| MT072864.1;QIQ54046.1;Pangolin_CoV_orf1a;1665 | 0 | 0 |
| MT072864.1;QIQ54046.1;Pangolin_CoV_orf1a;1666 | 0 | 0 |
| MT072864.1;QIQ54046.1;Pangolin_CoV_orf1a;1667 | 0 | 0 |

| | | |
|---|---|---|
| MT072864.1;QIQ54046.1;Pangolin_CoV_orf1a;1668 | 0 | 0 |
| MT072864.1;QIQ54046.1;Pangolin_CoV_orf1a;1669 | 0 | 0 |
| MT072864.1;QIQ54046.1;Pangolin_CoV_orf1a;1670 | 0 | 0 |
| MT072864.1;QIQ54046.1;Pangolin_CoV_orf1a;1671 | 0.05 | 0 |
| MT072864.1;QIQ54046.1;Pangolin_CoV_orf1a;1672 | 0 | 0 |
| MT072864.1;QIQ54046.1;Pangolin_CoV_orf1a;1673 | 0 | 0 |
| MT072864.1;QIQ54046.1;Pangolin_CoV_orf1a;1674 | 0 | 0.025 |
| MT072864.1;QIQ54046.1;Pangolin_CoV_orf1a;1675 | 0 | 0 |
| MT072864.1;QIQ54046.1;Pangolin_CoV_orf1a;1676 | 0 | 0 |
| MT072864.1;QIQ54046.1;Pangolin_CoV_orf1a;1677 | 0 | 0 |
| MT072864.1;QIQ54046.1;Pangolin_CoV_orf1a;1678 | 0 | 0.05 |
| MT072864.1;QIQ54046.1;Pangolin_CoV_orf1a;1679 | 0 | 0.025 |
| MT072864.1;QIQ54046.1;Pangolin_CoV_orf1a;1680 | 0 | 0 |
| MT072864.1;QIQ54046.1;Pangolin_CoV_orf1a;1681 | 0 | 0 |
| MT072864.1;QIQ54046.1;Pangolin_CoV_orf1a;1682 | 0 | 0 |
| MT072864.1;QIQ54046.1;Pangolin_CoV_orf1a;1683 | 0 | 0 |
| MT072864.1;QIQ54046.1;Pangolin_CoV_orf1a;1684 | 0.05 | 0.025 |
| MT072864.1;QIQ54046.1;Pangolin_CoV_orf1a;1685 | 0.1 | 0.075 |
| MT072864.1;QIQ54046.1;Pangolin_CoV_orf1a;1686 | 0.1 | 0.05 |
| MT072864.1;QIQ54046.1;Pangolin_CoV_orf1a;1687 | 0.05 | 0.025 |
| MT072864.1;QIQ54046.1;Pangolin_CoV_orf1a;1688 | 0.05 | 0 |
| MT072864.1;QIQ54046.1;Pangolin_CoV_orf1a;1689 | 0.05 | 0 |
| MT072864.1;QIQ54046.1;Pangolin_CoV_orf1a;1690 | 0 | 0 |
| MT072864.1;QIQ54046.1;Pangolin_CoV_orf1a;1691 | 0 | 0 |
| MT072864.1;QIQ54046.1;Pangolin_CoV_orf1a;1692 | 0 | 0 |
| MT072864.1;QIQ54046.1;Pangolin_CoV_orf1a;1693 | 0 | 0 |
| MT072864.1;QIQ54046.1;Pangolin_CoV_orf1a;1694 | 0 | 0 |
| MT072864.1;QIQ54046.1;Pangolin_CoV_orf1a;1695 | 0 | 0 |
| MT072864.1;QIQ54046.1;Pangolin_CoV_orf1a;1696 | 0 | 0 |
| MT072864.1;QIQ54046.1;Pangolin_CoV_orf1a;1697 | 0 | 0 |
| MT072864.1;QIQ54046.1;Pangolin_CoV_orf1a;1698 | 0 | 0 |
| MT072864.1;QIQ54046.1;Pangolin_CoV_orf1a;1699 | 0 | 0 |
| MT072864.1;QIQ54046.1;Pangolin_CoV_orf1a;1700 | 0 | 0 |
| MT072864.1;QIQ54046.1;Pangolin_CoV_orf1a;1701 | 0 | 0 |
| MT072864.1;QIQ54046.1;Pangolin_CoV_orf1a;1702 | 0 | 0 |
| MT072864.1;QIQ54046.1;Pangolin_CoV_orf1a;1703 | 0 | 0.025 |
| MT072864.1;QIQ54046.1;Pangolin_CoV_orf1a;1704 | 0 | 0.025 |
| MT072864.1;QIQ54046.1;Pangolin_CoV_orf1a;1705 | 0 | 0.05 |
| MT072864.1;QIQ54046.1;Pangolin_CoV_orf1a;1706 | 0 | 0.025 |
| MT072864.1;QIQ54046.1;Pangolin_CoV_orf1a;1707 | 0 | 0 |
| MT072864.1;QIQ54046.1;Pangolin_CoV_orf1a;1708 | 0 | 0 |
| MT072864.1;QIQ54046.1;Pangolin_CoV_orf1a;1709 | 0 | 0 |
| MT072864.1;QIQ54046.1;Pangolin_CoV_orf1a;1710 | 0 | 0 |

| | | |
|---|---|---|
| MT072864.1;QIQ54046.1;Pangolin_CoV_orf1a;1711 | 0 | 0 |
| MT072864.1;QIQ54046.1;Pangolin_CoV_orf1a;1712 | 0 | 0.025 |
| MT072864.1;QIQ54046.1;Pangolin_CoV_orf1a;1713 | 0 | 0 |
| MT072864.1;QIQ54046.1;Pangolin_CoV_orf1a;1714 | 0 | 0 |
| MT072864.1;QIQ54046.1;Pangolin_CoV_orf1a;1715 | 0.05 | 0.025 |
| MT072864.1;QIQ54046.1;Pangolin_CoV_orf1a;1716 | 0 | 0 |
| MT072864.1;QIQ54046.1;Pangolin_CoV_orf1a;1717 | 0 | 0 |
| MT072864.1;QIQ54046.1;Pangolin_CoV_orf1a;1718 | 0.05 | 0 |
| MT072864.1;QIQ54046.1;Pangolin_CoV_orf1a;1719 | 0.05 | 0 |
| MT072864.1;QIQ54046.1;Pangolin_CoV_orf1a;1720 | 0.05 | 0 |
| MT072864.1;QIQ54046.1;Pangolin_CoV_orf1a;1721 | 0.05 | 0 |
| MT072864.1;QIQ54046.1;Pangolin_CoV_orf1a;1722 | 0.05 | 0 |
| MT072864.1;QIQ54046.1;Pangolin_CoV_orf1a;1723 | 0 | 0 |
| MT072864.1;QIQ54046.1;Pangolin_CoV_orf1a;1724 | 0 | 0 |
| MT072864.1;QIQ54046.1;Pangolin_CoV_orf1a;1725 | 0 | 0 |
| MT072864.1;QIQ54046.1;Pangolin_CoV_orf1a;1726 | 0 | 0 |
| MT072864.1;QIQ54046.1;Pangolin_CoV_orf1a;1727 | 0 | 0 |
| MT072864.1;QIQ54046.1;Pangolin_CoV_orf1a;1728 | 0 | 0 |
| MT072864.1;QIQ54046.1;Pangolin_CoV_orf1a;1729 | 0 | 0 |
| MT072864.1;QIQ54046.1;Pangolin_CoV_orf1a;1730 | 0 | 0 |
| MT072864.1;QIQ54046.1;Pangolin_CoV_orf1a;1731 | 0 | 0 |
| MT072864.1;QIQ54046.1;Pangolin_CoV_orf1a;1732 | 0 | 0 |
| MT072864.1;QIQ54046.1;Pangolin_CoV_orf1a;1733 | 0 | 0 |
| MT072864.1;QIQ54046.1;Pangolin_CoV_orf1a;1734 | 0 | 0 |
| MT072864.1;QIQ54046.1;Pangolin_CoV_orf1a;1735 | 0 | 0 |
| MT072864.1;QIQ54046.1;Pangolin_CoV_orf1a;1736 | 0 | 0 |
| MT072864.1;QIQ54046.1;Pangolin_CoV_orf1a;1737 | 0 | 0.075 |
| MT072864.1;QIQ54046.1;Pangolin_CoV_orf1a;1738 | 0 | 0.025 |
| MT072864.1;QIQ54046.1;Pangolin_CoV_orf1a;1739 | 0 | 0.025 |
| MT072864.1;QIQ54046.1;Pangolin_CoV_orf1a;1740 | 0.1 | 0.325 |
| MT072864.1;QIQ54046.1;Pangolin_CoV_orf1a;1741 | 0.1 | 0.3 |
| MT072864.1;QIQ54046.1;Pangolin_CoV_orf1a;1742 | 0.1 | 0.3 |
| MT072864.1;QIQ54046.1;Pangolin_CoV_orf1a;1743 | 0.15 | 0.325 |
| MT072864.1;QIQ54046.1;Pangolin_CoV_orf1a;1744 | 0.2 | 0.325 |
| MT072864.1;QIQ54046.1;Pangolin_CoV_orf1a;1745 | 0.1 | 0.35 |
| MT072864.1;QIQ54046.1;Pangolin_CoV_orf1a;1746 | 0.05 | 0.275 |
| MT072864.1;QIQ54046.1;Pangolin_CoV_orf1a;1747 | 0.05 | 0.2 |
| MT072864.1;QIQ54046.1;Pangolin_CoV_orf1a;1748 | 0.1 | 0.25 |
| MT072864.1;QIQ54046.1;Pangolin_CoV_orf1a;1749 | 0.05 | 0.2 |
| MT072864.1;QIQ54046.1;Pangolin_CoV_orf1a;1750 | 0 | 0.125 |
| MT072864.1;QIQ54046.1;Pangolin_CoV_orf1a;1751 | 0 | 0 |
| MT072864.1;QIQ54046.1;Pangolin_CoV_orf1a;1752 | 0 | 0 |
| MT072864.1;QIQ54046.1;Pangolin_CoV_orf1a;1753 | 0.1 | 0 |

| | | |
|---|---|---|
| MT072864.1;QIQ54046.1;Pangolin_CoV_orf1a;1754 | 0.15 | 0 |
| MT072864.1;QIQ54046.1;Pangolin_CoV_orf1a;1755 | 0.15 | 0 |
| MT072864.1;QIQ54046.1;Pangolin_CoV_orf1a;1756 | 0.2 | 0 |
| MT072864.1;QIQ54046.1;Pangolin_CoV_orf1a;1757 | 0.1 | 0 |
| MT072864.1;QIQ54046.1;Pangolin_CoV_orf1a;1758 | 0.1 | 0 |
| MT072864.1;QIQ54046.1;Pangolin_CoV_orf1a;1759 | 0 | 0 |
| MT072864.1;QIQ54046.1;Pangolin_CoV_orf1a;1760 | 0.05 | 0 |
| MT072864.1;QIQ54046.1;Pangolin_CoV_orf1a;1761 | 0 | 0 |
| MT072864.1;QIQ54046.1;Pangolin_CoV_orf1a;1762 | 0 | 0 |
| MT072864.1;QIQ54046.1;Pangolin_CoV_orf1a;1763 | 0 | 0 |
| MT072864.1;QIQ54046.1;Pangolin_CoV_orf1a;1764 | 0 | 0 |
| MT072864.1;QIQ54046.1;Pangolin_CoV_orf1a;1765 | 0 | 0 |
| MT072864.1;QIQ54046.1;Pangolin_CoV_orf1a;1766 | 0 | 0 |
| MT072864.1;QIQ54046.1;Pangolin_CoV_orf1a;1767 | 0 | 0 |
| MT072864.1;QIQ54046.1;Pangolin_CoV_orf1a;1768 | 0 | 0 |
| MT072864.1;QIQ54046.1;Pangolin_CoV_orf1a;1769 | 0 | 0 |
| MT072864.1;QIQ54046.1;Pangolin_CoV_orf1a;1770 | 0 | 0 |
| MT072864.1;QIQ54046.1;Pangolin_CoV_orf1a;1771 | 0 | 0 |
| MT072864.1;QIQ54046.1;Pangolin_CoV_orf1a;1772 | 0.05 | 0.025 |
| MT072864.1;QIQ54046.1;Pangolin_CoV_orf1a;1773 | 0 | 0 |
| MT072864.1;QIQ54046.1;Pangolin_CoV_orf1a;1774 | 0 | 0 |
| MT072864.1;QIQ54046.1;Pangolin_CoV_orf1a;1775 | 0 | 0 |
| MT072864.1;QIQ54046.1;Pangolin_CoV_orf1a;1776 | 0 | 0 |
| MT072864.1;QIQ54046.1;Pangolin_CoV_orf1a;1777 | 0 | 0 |
| MT072864.1;QIQ54046.1;Pangolin_CoV_orf1a;1778 | 0 | 0 |
| MT072864.1;QIQ54046.1;Pangolin_CoV_orf1a;1779 | 0 | 0 |
| MT072864.1;QIQ54046.1;Pangolin_CoV_orf1a;1780 | 0 | 0 |
| MT072864.1;QIQ54046.1;Pangolin_CoV_orf1a;1781 | 0 | 0.025 |
| MT072864.1;QIQ54046.1;Pangolin_CoV_orf1a;1782 | 0 | 0.025 |
| MT072864.1;QIQ54046.1;Pangolin_CoV_orf1a;1783 | 0 | 0.025 |
| MT072864.1;QIQ54046.1;Pangolin_CoV_orf1a;1784 | 0 | 0.025 |
| MT072864.1;QIQ54046.1;Pangolin_CoV_orf1a;1785 | 0 | 0.025 |
| MT072864.1;QIQ54046.1;Pangolin_CoV_orf1a;1786 | 0 | 0 |
| MT072864.1;QIQ54046.1;Pangolin_CoV_orf1a;1787 | 0.05 | 0 |
| MT072864.1;QIQ54046.1;Pangolin_CoV_orf1a;1788 | 0 | 0.025 |
| MT072864.1;QIQ54046.1;Pangolin_CoV_orf1a;1789 | 0.1 | 0.025 |
| MT072864.1;QIQ54046.1;Pangolin_CoV_orf1a;1790 | 0 | 0 |
| MT072864.1;QIQ54046.1;Pangolin_CoV_orf1a;1791 | 0 | 0 |
| MT072864.1;QIQ54046.1;Pangolin_CoV_orf1a;1792 | 0 | 0 |
| MT072864.1;QIQ54046.1;Pangolin_CoV_orf1a;1793 | 0 | 0 |
| MT072864.1;QIQ54046.1;Pangolin_CoV_orf1a;1794 | 0 | 0 |
| MT072864.1;QIQ54046.1;Pangolin_CoV_orf1a;1795 | 0 | 0 |
| MT072864.1;QIQ54046.1;Pangolin_CoV_orf1a;1796 | 0 | 0 |

| | | |
|---|---|---|
| MT072864.1;QIQ54046.1;Pangolin_CoV_orf1a;1797 | 0 | 0 |
| MT072864.1;QIQ54046.1;Pangolin_CoV_orf1a;1798 | 0 | 0 |
| MT072864.1;QIQ54046.1;Pangolin_CoV_orf1a;1799 | 0 | 0.025 |
| MT072864.1;QIQ54046.1;Pangolin_CoV_orf1a;1800 | 0 | 0 |
| MT072864.1;QIQ54046.1;Pangolin_CoV_orf1a;1801 | 0 | 0 |
| MT072864.1;QIQ54046.1;Pangolin_CoV_orf1a;1802 | 0 | 0 |
| MT072864.1;QIQ54046.1;Pangolin_CoV_orf1a;1803 | 0 | 0 |
| MT072864.1;QIQ54046.1;Pangolin_CoV_orf1a;1804 | 0 | 0 |
| MT072864.1;QIQ54046.1;Pangolin_CoV_orf1a;1805 | 0 | 0 |
| MT072864.1;QIQ54046.1;Pangolin_CoV_orf1a;1806 | 0 | 0 |
| MT072864.1;QIQ54046.1;Pangolin_CoV_orf1a;1807 | 0 | 0 |
| MT072864.1;QIQ54046.1;Pangolin_CoV_orf1a;1808 | 0 | 0 |
| MT072864.1;QIQ54046.1;Pangolin_CoV_orf1a;1809 | 0 | 0 |
| MT072864.1;QIQ54046.1;Pangolin_CoV_orf1a;1810 | 0 | 0 |
| MT072864.1;QIQ54046.1;Pangolin_CoV_orf1a;1811 | 0 | 0 |
| MT072864.1;QIQ54046.1;Pangolin_CoV_orf1a;1812 | 0 | 0 |
| MT072864.1;QIQ54046.1;Pangolin_CoV_orf1a;1813 | 0 | 0 |
| MT072864.1;QIQ54046.1;Pangolin_CoV_orf1a;1814 | 0 | 0 |
| MT072864.1;QIQ54046.1;Pangolin_CoV_orf1a;1815 | 0 | 0 |
| MT072864.1;QIQ54046.1;Pangolin_CoV_orf1a;1816 | 0 | 0 |
| MT072864.1;QIQ54046.1;Pangolin_CoV_orf1a;1817 | 0 | 0 |
| MT072864.1;QIQ54046.1;Pangolin_CoV_orf1a;1818 | 0 | 0.15 |
| MT072864.1;QIQ54046.1;Pangolin_CoV_orf1a;1819 | 0 | 0 |
| MT072864.1;QIQ54046.1;Pangolin_CoV_orf1a;1820 | 0 | 0 |
| MT072864.1;QIQ54046.1;Pangolin_CoV_orf1a;1821 | 0 | 0 |
| MT072864.1;QIQ54046.1;Pangolin_CoV_orf1a;1822 | 0 | 0 |
| MT072864.1;QIQ54046.1;Pangolin_CoV_orf1a;1823 | 0.05 | 0 |
| MT072864.1;QIQ54046.1;Pangolin_CoV_orf1a;1824 | 0 | 0 |
| MT072864.1;QIQ54046.1;Pangolin_CoV_orf1a;1825 | 0 | 0 |
| MT072864.1;QIQ54046.1;Pangolin_CoV_orf1a;1826 | 0 | 0 |
| MT072864.1;QIQ54046.1;Pangolin_CoV_orf1a;1827 | 0 | 0 |
| MT072864.1;QIQ54046.1;Pangolin_CoV_orf1a;1828 | 0 | 0 |
| MT072864.1;QIQ54046.1;Pangolin_CoV_orf1a;1829 | 0 | 0 |
| MT072864.1;QIQ54046.1;Pangolin_CoV_orf1a;1830 | 0 | 0 |
| MT072864.1;QIQ54046.1;Pangolin_CoV_orf1a;1831 | 0 | 0 |
| MT072864.1;QIQ54046.1;Pangolin_CoV_orf1a;1832 | 0.05 | 0 |
| MT072864.1;QIQ54046.1;Pangolin_CoV_orf1a;1833 | 0 | 0 |
| MT072864.1;QIQ54046.1;Pangolin_CoV_orf1a;1834 | 0 | 0 |
| MT072864.1;QIQ54046.1;Pangolin_CoV_orf1a;1835 | 0 | 0 |
| MT072864.1;QIQ54046.1;Pangolin_CoV_orf1a;1836 | 0.05 | 0 |
| MT072864.1;QIQ54046.1;Pangolin_CoV_orf1a;1837 | 0.05 | 0.025 |
| MT072864.1;QIQ54046.1;Pangolin_CoV_orf1a;1838 | 0.05 | 0.025 |
| MT072864.1;QIQ54046.1;Pangolin_CoV_orf1a;1839 | 0.05 | 0.025 |

| | | |
|---|---|---|
| MT072864.1;QIQ54046.1;Pangolin_CoV_orf1a;1840 | 0.05 | 0.025 |
| MT072864.1;QIQ54046.1;Pangolin_CoV_orf1a;1841 | 0.05 | 0.05 |
| MT072864.1;QIQ54046.1;Pangolin_CoV_orf1a;1842 | 0.05 | 0.075 |
| MT072864.1;QIQ54046.1;Pangolin_CoV_orf1a;1843 | 0.05 | 0.05 |
| MT072864.1;QIQ54046.1;Pangolin_CoV_orf1a;1844 | 0.05 | 0.025 |
| MT072864.1;QIQ54046.1;Pangolin_CoV_orf1a;1845 | 0.05 | 0.05 |
| MT072864.1;QIQ54046.1;Pangolin_CoV_orf1a;1846 | 0 | 0.05 |
| MT072864.1;QIQ54046.1;Pangolin_CoV_orf1a;1847 | 0 | 0.1 |
| MT072864.1;QIQ54046.1;Pangolin_CoV_orf1a;1848 | 0.05 | 0.125 |
| MT072864.1;QIQ54046.1;Pangolin_CoV_orf1a;1849 | 0.05 | 0.1 |
| MT072864.1;QIQ54046.1;Pangolin_CoV_orf1a;1850 | 0.05 | 0.075 |
| MT072864.1;QIQ54046.1;Pangolin_CoV_orf1a;1851 | 0.1 | 0.1 |
| MT072864.1;QIQ54046.1;Pangolin_CoV_orf1a;1852 | 0 | 0.125 |
| MT072864.1;QIQ54046.1;Pangolin_CoV_orf1a;1853 | 0.2 | 0.125 |
| MT072864.1;QIQ54046.1;Pangolin_CoV_orf1a;1854 | 0 | 0 |
| MT072864.1;QIQ54046.1;Pangolin_CoV_orf1a;1855 | 0 | 0.025 |
| MT072864.1;QIQ54046.1;Pangolin_CoV_orf1a;1856 | 0 | 0.025 |
| MT072864.1;QIQ54046.1;Pangolin_CoV_orf1a;1857 | 0 | 0.025 |
| MT072864.1;QIQ54046.1;Pangolin_CoV_orf1a;1858 | 0 | 0.025 |
| MT072864.1;QIQ54046.1;Pangolin_CoV_orf1a;1859 | 0 | 0.025 |
| MT072864.1;QIQ54046.1;Pangolin_CoV_orf1a;1860 | 0 | 0.025 |
| MT072864.1;QIQ54046.1;Pangolin_CoV_orf1a;1861 | 0 | 0.025 |
| MT072864.1;QIQ54046.1;Pangolin_CoV_orf1a;1862 | 0.1 | 0.025 |
| MT072864.1;QIQ54046.1;Pangolin_CoV_orf1a;1863 | 0 | 0.025 |
| MT072864.1;QIQ54046.1;Pangolin_CoV_orf1a;1864 | 0 | 0.025 |
| MT072864.1;QIQ54046.1;Pangolin_CoV_orf1a;1865 | 0 | 0 |
| MT072864.1;QIQ54046.1;Pangolin_CoV_orf1a;1866 | 0 | 0 |
| MT072864.1;QIQ54046.1;Pangolin_CoV_orf1a;1867 | 0 | 0 |
| MT072864.1;QIQ54046.1;Pangolin_CoV_orf1a;1868 | 0 | 0 |
| MT072864.1;QIQ54046.1;Pangolin_CoV_orf1a;1869 | 0 | 0 |
| MT072864.1;QIQ54046.1;Pangolin_CoV_orf1a;1870 | 0 | 0.025 |
| MT072864.1;QIQ54046.1;Pangolin_CoV_orf1a;1871 | 0 | 0.025 |
| MT072864.1;QIQ54046.1;Pangolin_CoV_orf1a;1872 | 0 | 0 |
| MT072864.1;QIQ54046.1;Pangolin_CoV_orf1a;1873 | 0 | 0.025 |
| MT072864.1;QIQ54046.1;Pangolin_CoV_orf1a;1874 | 0 | 0 |
| MT072864.1;QIQ54046.1;Pangolin_CoV_orf1a;1875 | 0 | 0 |
| MT072864.1;QIQ54046.1;Pangolin_CoV_orf1a;1876 | 0 | 0 |
| MT072864.1;QIQ54046.1;Pangolin_CoV_orf1a;1877 | 0 | 0 |
| MT072864.1;QIQ54046.1;Pangolin_CoV_orf1a;1878 | 0 | 0.025 |
| MT072864.1;QIQ54046.1;Pangolin_CoV_orf1a;1879 | 0 | 0.075 |
| MT072864.1;QIQ54046.1;Pangolin_CoV_orf1a;1880 | 0 | 0.125 |
| MT072864.1;QIQ54046.1;Pangolin_CoV_orf1a;1881 | 0 | 0.15 |
| MT072864.1;QIQ54046.1;Pangolin_CoV_orf1a;1882 | 0 | 0.15 |

| | | |
|---|---|---|
| MT072864.1;QIQ54046.1;Pangolin_CoV_orf1a;1883 | 0 | 0.225 |
| MT072864.1;QIQ54046.1;Pangolin_CoV_orf1a;1884 | 0 | 0.1 |
| MT072864.1;QIQ54046.1;Pangolin_CoV_orf1a;1885 | 0 | 0.05 |
| MT072864.1;QIQ54046.1;Pangolin_CoV_orf1a;1886 | 0.1 | 0.1 |
| MT072864.1;QIQ54046.1;Pangolin_CoV_orf1a;1887 | 0.05 | 0.15 |
| MT072864.1;QIQ54046.1;Pangolin_CoV_orf1a;1888 | 0 | 0.075 |
| MT072864.1;QIQ54046.1;Pangolin_CoV_orf1a;1889 | 0.05 | 0.125 |
| MT072864.1;QIQ54046.1;Pangolin_CoV_orf1a;1890 | 0.05 | 0.025 |
| MT072864.1;QIQ54046.1;Pangolin_CoV_orf1a;1891 | 0.05 | 0.025 |
| MT072864.1;QIQ54046.1;Pangolin_CoV_orf1a;1892 | 0.05 | 0.025 |
| MT072864.1;QIQ54046.1;Pangolin_CoV_orf1a;1893 | 0.15 | 0.1 |
| MT072864.1;QIQ54046.1;Pangolin_CoV_orf1a;1894 | 0.05 | 0.025 |
| MT072864.1;QIQ54046.1;Pangolin_CoV_orf1a;1895 | 0.05 | 0 |
| MT072864.1;QIQ54046.1;Pangolin_CoV_orf1a;1896 | 0.05 | 0 |
| MT072864.1;QIQ54046.1;Pangolin_CoV_orf1a;1897 | 0.05 | 0.025 |
| MT072864.1;QIQ54046.1;Pangolin_CoV_orf1a;1898 | 0.05 | 0 |
| MT072864.1;QIQ54046.1;Pangolin_CoV_orf1a;1899 | 0.05 | 0 |
| MT072864.1;QIQ54046.1;Pangolin_CoV_orf1a;1900 | 0 | 0 |
| MT072864.1;QIQ54046.1;Pangolin_CoV_orf1a;1901 | 0 | 0 |
| MT072864.1;QIQ54046.1;Pangolin_CoV_orf1a;1902 | 0.15 | 0.075 |
| MT072864.1;QIQ54046.1;Pangolin_CoV_orf1a;1903 | 0 | 0.075 |
| MT072864.1;QIQ54046.1;Pangolin_CoV_orf1a;1904 | 0 | 0.025 |
| MT072864.1;QIQ54046.1;Pangolin_CoV_orf1a;1905 | 0 | 0.05 |
| MT072864.1;QIQ54046.1;Pangolin_CoV_orf1a;1906 | 0.15 | 0.075 |
| MT072864.1;QIQ54046.1;Pangolin_CoV_orf1a;1907 | 0.15 | 0.025 |
| MT072864.1;QIQ54046.1;Pangolin_CoV_orf1a;1908 | 0.05 | 0 |
| MT072864.1;QIQ54046.1;Pangolin_CoV_orf1a;1909 | 0.05 | 0 |
| MT072864.1;QIQ54046.1;Pangolin_CoV_orf1a;1910 | 0 | 0 |
| MT072864.1;QIQ54046.1;Pangolin_CoV_orf1a;1911 | 0.15 | 0.05 |
| MT072864.1;QIQ54046.1;Pangolin_CoV_orf1a;1912 | 0.15 | 0.05 |
| MT072864.1;QIQ54046.1;Pangolin_CoV_orf1a;1913 | 0.1 | 0.075 |
| MT072864.1;QIQ54046.1;Pangolin_CoV_orf1a;1914 | 0.1 | 0 |
| MT072864.1;QIQ54046.1;Pangolin_CoV_orf1a;1915 | 0.1 | 0.05 |
| MT072864.1;QIQ54046.1;Pangolin_CoV_orf1a;1916 | 0.1 | 0 |
| MT072864.1;QIQ54046.1;Pangolin_CoV_orf1a;1917 | 0.1 | 0 |
| MT072864.1;QIQ54046.1;Pangolin_CoV_orf1a;1918 | 0.05 | 0 |
| MT072864.1;QIQ54046.1;Pangolin_CoV_orf1a;1919 | 0.2 | 0.05 |
| MT072864.1;QIQ54046.1;Pangolin_CoV_orf1a;1920 | 0.2 | 0.025 |
| MT072864.1;QIQ54046.1;Pangolin_CoV_orf1a;1921 | 0.2 | 0.075 |
| MT072864.1;QIQ54046.1;Pangolin_CoV_orf1a;1922 | 0.3 | 0.05 |
| MT072864.1;QIQ54046.1;Pangolin_CoV_orf1a;1923 | 0.2 | 0 |
| MT072864.1;QIQ54046.1;Pangolin_CoV_orf1a;1924 | 0.15 | 0 |
| MT072864.1;QIQ54046.1;Pangolin_CoV_orf1a;1925 | 0.35 | 0.15 |

| | | |
|---|---|---|
| MT072864.1;QIQ54046.1;Pangolin_CoV_orf1a;1926 | 0.15 | 0 |
| MT072864.1;QIQ54046.1;Pangolin_CoV_orf1a;1927 | 0.1 | 0.05 |
| MT072864.1;QIQ54046.1;Pangolin_CoV_orf1a;1928 | 0.15 | 0.075 |
| MT072864.1;QIQ54046.1;Pangolin_CoV_orf1a;1929 | 0.05 | 0.025 |
| MT072864.1;QIQ54046.1;Pangolin_CoV_orf1a;1930 | 0.1 | 0.025 |
| MT072864.1;QIQ54046.1;Pangolin_CoV_orf1a;1931 | 0.1 | 0.025 |
| MT072864.1;QIQ54046.1;Pangolin_CoV_orf1a;1932 | 0 | 0 |
| MT072864.1;QIQ54046.1;Pangolin_CoV_orf1a;1933 | 0 | 0 |
| MT072864.1;QIQ54046.1;Pangolin_CoV_orf1a;1934 | 0 | 0 |
| MT072864.1;QIQ54046.1;Pangolin_CoV_orf1a;1935 | 0 | 0 |
| MT072864.1;QIQ54046.1;Pangolin_CoV_orf1a;1936 | 0 | 0 |
| MT072864.1;QIQ54046.1;Pangolin_CoV_orf1a;1937 | 0 | 0 |
| MT072864.1;QIQ54046.1;Pangolin_CoV_orf1a;1938 | 0 | 0 |
| MT072864.1;QIQ54046.1;Pangolin_CoV_orf1a;1939 | 0 | 0 |
| MT072864.1;QIQ54046.1;Pangolin_CoV_orf1a;1940 | 0 | 0 |
| MT072864.1;QIQ54046.1;Pangolin_CoV_orf1a;1941 | 0 | 0 |
| MT072864.1;QIQ54046.1;Pangolin_CoV_orf1a;1942 | 0 | 0 |
| MT072864.1;QIQ54046.1;Pangolin_CoV_orf1a;1943 | 0 | 0 |
| MT072864.1;QIQ54046.1;Pangolin_CoV_orf1a;1944 | 0 | 0 |
| MT072864.1;QIQ54046.1;Pangolin_CoV_orf1a;1945 | 0 | 0.025 |
| MT072864.1;QIQ54046.1;Pangolin_CoV_orf1a;1946 | 0 | 0.05 |
| MT072864.1;QIQ54046.1;Pangolin_CoV_orf1a;1947 | 0 | 0 |
| MT072864.1;QIQ54046.1;Pangolin_CoV_orf1a;1948 | 0 | 0.025 |
| MT072864.1;QIQ54046.1;Pangolin_CoV_orf1a;1949 | 0 | 0 |
| MT072864.1;QIQ54046.1;Pangolin_CoV_orf1a;1950 | 0 | 0.025 |
| MT072864.1;QIQ54046.1;Pangolin_CoV_orf1a;1951 | 0.05 | 0.025 |
| MT072864.1;QIQ54046.1;Pangolin_CoV_orf1a;1952 | 0.05 | 0.05 |
| MT072864.1;QIQ54046.1;Pangolin_CoV_orf1a;1953 | 0.05 | 0.075 |
| MT072864.1;QIQ54046.1;Pangolin_CoV_orf1a;1954 | 0.05 | 0.075 |
| MT072864.1;QIQ54046.1;Pangolin_CoV_orf1a;1955 | 0.05 | 0.025 |
| MT072864.1;QIQ54046.1;Pangolin_CoV_orf1a;1956 | 0.05 | 0.025 |
| MT072864.1;QIQ54046.1;Pangolin_CoV_orf1a;1957 | 0.05 | 0.075 |
| MT072864.1;QIQ54046.1;Pangolin_CoV_orf1a;1958 | 0 | 0.025 |
| MT072864.1;QIQ54046.1;Pangolin_CoV_orf1a;1959 | 0 | 0.025 |
| MT072864.1;QIQ54046.1;Pangolin_CoV_orf1a;1960 | 0 | 0.05 |
| MT072864.1;QIQ54046.1;Pangolin_CoV_orf1a;1961 | 0 | 0.075 |
| MT072864.1;QIQ54046.1;Pangolin_CoV_orf1a;1962 | 0 | 0 |
| MT072864.1;QIQ54046.1;Pangolin_CoV_orf1a;1963 | 0 | 0 |
| MT072864.1;QIQ54046.1;Pangolin_CoV_orf1a;1964 | 0 | 0 |
| MT072864.1;QIQ54046.1;Pangolin_CoV_orf1a;1965 | 0 | 0 |
| MT072864.1;QIQ54046.1;Pangolin_CoV_orf1a;1966 | 0 | 0 |
| MT072864.1;QIQ54046.1;Pangolin_CoV_orf1a;1967 | 0 | 0.025 |
| MT072864.1;QIQ54046.1;Pangolin_CoV_orf1a;1968 | 0 | 0 |

| | | |
|---|---|---|
| MT072864.1;QIQ54046.1;Pangolin_CoV_orf1a;1969 | 0 | 0 |
| MT072864.1;QIQ54046.1;Pangolin_CoV_orf1a;1970 | 0 | 0 |
| MT072864.1;QIQ54046.1;Pangolin_CoV_orf1a;1971 | 0 | 0 |
| MT072864.1;QIQ54046.1;Pangolin_CoV_orf1a;1972 | 0 | 0 |
| MT072864.1;QIQ54046.1;Pangolin_CoV_orf1a;1973 | 0 | 0 |
| MT072864.1;QIQ54046.1;Pangolin_CoV_orf1a;1974 | 0 | 0 |
| MT072864.1;QIQ54046.1;Pangolin_CoV_orf1a;1975 | 0 | 0 |
| MT072864.1;QIQ54046.1;Pangolin_CoV_orf1a;1976 | 0.1 | 0.025 |
| MT072864.1;QIQ54046.1;Pangolin_CoV_orf1a;1977 | 0.05 | 0 |
| MT072864.1;QIQ54046.1;Pangolin_CoV_orf1a;1978 | 0.1 | 0 |
| MT072864.1;QIQ54046.1;Pangolin_CoV_orf1a;1979 | 0.1 | 0.025 |
| MT072864.1;QIQ54046.1;Pangolin_CoV_orf1a;1980 | 0.1 | 0.025 |
| MT072864.1;QIQ54046.1;Pangolin_CoV_orf1a;1981 | 0.15 | 0.05 |
| MT072864.1;QIQ54046.1;Pangolin_CoV_orf1a;1982 | 0.15 | 0.075 |
| MT072864.1;QIQ54046.1;Pangolin_CoV_orf1a;1983 | 0.25 | 0.125 |
| MT072864.1;QIQ54046.1;Pangolin_CoV_orf1a;1984 | 0.25 | 0.1 |
| MT072864.1;QIQ54046.1;Pangolin_CoV_orf1a;1985 | 0.3 | 0.225 |
| MT072864.1;QIQ54046.1;Pangolin_CoV_orf1a;1986 | 0.35 | 0.1 |
| MT072864.1;QIQ54046.1;Pangolin_CoV_orf1a;1987 | 0.1 | 0.05 |
| MT072864.1;QIQ54046.1;Pangolin_CoV_orf1a;1988 | 0 | 0.025 |
| MT072864.1;QIQ54046.1;Pangolin_CoV_orf1a;1989 | 0 | 0.025 |
| MT072864.1;QIQ54046.1;Pangolin_CoV_orf1a;1990 | 0 | 0.025 |
| MT072864.1;QIQ54046.1;Pangolin_CoV_orf1a;1991 | 0 | 0 |
| MT072864.1;QIQ54046.1;Pangolin_CoV_orf1a;1992 | 0 | 0 |
| MT072864.1;QIQ54046.1;Pangolin_CoV_orf1a;1993 | 0 | 0 |
| MT072864.1;QIQ54046.1;Pangolin_CoV_orf1a;1994 | 0 | 0 |
| MT072864.1;QIQ54046.1;Pangolin_CoV_orf1a;1995 | 0 | 0 |
| MT072864.1;QIQ54046.1;Pangolin_CoV_orf1a;1996 | 0 | 0 |
| MT072864.1;QIQ54046.1;Pangolin_CoV_orf1a;1997 | 0 | 0 |
| MT072864.1;QIQ54046.1;Pangolin_CoV_orf1a;1998 | 0 | 0 |
| MT072864.1;QIQ54046.1;Pangolin_CoV_orf1a;1999 | 0 | 0 |
| MT072864.1;QIQ54046.1;Pangolin_CoV_orf1a;2000 | 0 | 0 |
| MT072864.1;QIQ54046.1;Pangolin_CoV_orf1a;2001 | 0 | 0.025 |
| MT072864.1;QIQ54046.1;Pangolin_CoV_orf1a;2002 | 0 | 0 |
| MT072864.1;QIQ54046.1;Pangolin_CoV_orf1a;2003 | 0.05 | 0.05 |
| MT072864.1;QIQ54046.1;Pangolin_CoV_orf1a;2004 | 0 | 0.025 |
| MT072864.1;QIQ54046.1;Pangolin_CoV_orf1a;2005 | 0 | 0.025 |
| MT072864.1;QIQ54046.1;Pangolin_CoV_orf1a;2006 | 0.1 | 0.1 |
| MT072864.1;QIQ54046.1;Pangolin_CoV_orf1a;2007 | 0.15 | 0.05 |
| MT072864.1;QIQ54046.1;Pangolin_CoV_orf1a;2008 | 0.2 | 0.125 |
| MT072864.1;QIQ54046.1;Pangolin_CoV_orf1a;2009 | 0.2 | 0.075 |
| MT072864.1;QIQ54046.1;Pangolin_CoV_orf1a;2010 | 0.15 | 0.05 |
| MT072864.1;QIQ54046.1;Pangolin_CoV_orf1a;2011 | 0.3 | 0.125 |

| | | |
|---|---|---|
| MT072864.1;QIQ54046.1;Pangolin_CoV_orf1a;2012 | 0 | 0.025 |
| MT072864.1;QIQ54046.1;Pangolin_CoV_orf1a;2013 | 0 | 0.025 |
| MT072864.1;QIQ54046.1;Pangolin_CoV_orf1a;2014 | 0 | 0.025 |
| MT072864.1;QIQ54046.1;Pangolin_CoV_orf1a;2015 | 0 | 0.025 |
| MT072864.1;QIQ54046.1;Pangolin_CoV_orf1a;2016 | 0 | 0 |
| MT072864.1;QIQ54046.1;Pangolin_CoV_orf1a;2017 | 0 | 0 |
| MT072864.1;QIQ54046.1;Pangolin_CoV_orf1a;2018 | 0 | 0 |
| MT072864.1;QIQ54046.1;Pangolin_CoV_orf1a;2019 | 0 | 0 |
| MT072864.1;QIQ54046.1;Pangolin_CoV_orf1a;2020 | 0 | 0 |
| MT072864.1;QIQ54046.1;Pangolin_CoV_orf1a;2021 | 0 | 0 |
| MT072864.1;QIQ54046.1;Pangolin_CoV_orf1a;2022 | 0 | 0 |
| MT072864.1;QIQ54046.1;Pangolin_CoV_orf1a;2023 | 0.05 | 0 |
| MT072864.1;QIQ54046.1;Pangolin_CoV_orf1a;2024 | 0 | 0 |
| MT072864.1;QIQ54046.1;Pangolin_CoV_orf1a;2025 | 0 | 0 |
| MT072864.1;QIQ54046.1;Pangolin_CoV_orf1a;2026 | 0 | 0.025 |
| MT072864.1;QIQ54046.1;Pangolin_CoV_orf1a;2027 | 0 | 0.025 |
| MT072864.1;QIQ54046.1;Pangolin_CoV_orf1a;2028 | 0 | 0 |
| MT072864.1;QIQ54046.1;Pangolin_CoV_orf1a;2029 | 0 | 0 |
| MT072864.1;QIQ54046.1;Pangolin_CoV_orf1a;2030 | 0 | 0 |
| MT072864.1;QIQ54046.1;Pangolin_CoV_orf1a;2031 | 0 | 0 |
| MT072864.1;QIQ54046.1;Pangolin_CoV_orf1a;2032 | 0.05 | 0 |
| MT072864.1;QIQ54046.1;Pangolin_CoV_orf1a;2033 | 0 | 0 |
| MT072864.1;QIQ54046.1;Pangolin_CoV_orf1a;2034 | 0 | 0.025 |
| MT072864.1;QIQ54046.1;Pangolin_CoV_orf1a;2035 | 0.05 | 0 |
| MT072864.1;QIQ54046.1;Pangolin_CoV_orf1a;2036 | 0.05 | 0.05 |
| MT072864.1;QIQ54046.1;Pangolin_CoV_orf1a;2037 | 0.05 | 0.025 |
| MT072864.1;QIQ54046.1;Pangolin_CoV_orf1a;2038 | 0.05 | 0.025 |
| MT072864.1;QIQ54046.1;Pangolin_CoV_orf1a;2039 | 0.05 | 0 |
| MT072864.1;QIQ54046.1;Pangolin_CoV_orf1a;2040 | 0 | 0 |
| MT072864.1;QIQ54046.1;Pangolin_CoV_orf1a;2041 | 0 | 0 |
| MT072864.1;QIQ54046.1;Pangolin_CoV_orf1a;2042 | 0.05 | 0 |
| MT072864.1;QIQ54046.1;Pangolin_CoV_orf1a;2043 | 0.05 | 0 |
| MT072864.1;QIQ54046.1;Pangolin_CoV_orf1a;2044 | 0 | 0 |
| MT072864.1;QIQ54046.1;Pangolin_CoV_orf1a;2045 | 0.05 | 0 |
| MT072864.1;QIQ54046.1;Pangolin_CoV_orf1a;2046 | 0.05 | 0 |
| MT072864.1;QIQ54046.1;Pangolin_CoV_orf1a;2047 | 0 | 0 |
| MT072864.1;QIQ54046.1;Pangolin_CoV_orf1a;2048 | 0 | 0.025 |
| MT072864.1;QIQ54046.1;Pangolin_CoV_orf1a;2049 | 0 | 0.025 |
| MT072864.1;QIQ54046.1;Pangolin_CoV_orf1a;2050 | 0 | 0.025 |
| MT072864.1;QIQ54046.1;Pangolin_CoV_orf1a;2051 | 0 | 0.025 |
| MT072864.1;QIQ54046.1;Pangolin_CoV_orf1a;2052 | 0 | 0 |
| MT072864.1;QIQ54046.1;Pangolin_CoV_orf1a;2053 | 0 | 0 |
| MT072864.1;QIQ54046.1;Pangolin_CoV_orf1a;2054 | 0 | 0 |

| | | |
|---|---|---|
| MT072864.1;QIQ54046.1;Pangolin_CoV_orf1a;2055 | 0 | 0.025 |
| MT072864.1;QIQ54046.1;Pangolin_CoV_orf1a;2056 | 0 | 0.025 |
| MT072864.1;QIQ54046.1;Pangolin_CoV_orf1a;2057 | 0 | 0 |
| MT072864.1;QIQ54046.1;Pangolin_CoV_orf1a;2058 | 0 | 0 |
| MT072864.1;QIQ54046.1;Pangolin_CoV_orf1a;2059 | 0 | 0 |
| MT072864.1;QIQ54046.1;Pangolin_CoV_orf1a;2060 | 0 | 0 |
| MT072864.1;QIQ54046.1;Pangolin_CoV_orf1a;2061 | 0.05 | 0.05 |
| MT072864.1;QIQ54046.1;Pangolin_CoV_orf1a;2062 | 0 | 0.025 |
| MT072864.1;QIQ54046.1;Pangolin_CoV_orf1a;2063 | 0 | 0.025 |
| MT072864.1;QIQ54046.1;Pangolin_CoV_orf1a;2064 | 0 | 0 |
| MT072864.1;QIQ54046.1;Pangolin_CoV_orf1a;2065 | 0 | 0.025 |
| MT072864.1;QIQ54046.1;Pangolin_CoV_orf1a;2066 | 0 | 0 |
| MT072864.1;QIQ54046.1;Pangolin_CoV_orf1a;2067 | 0 | 0 |
| MT072864.1;QIQ54046.1;Pangolin_CoV_orf1a;2068 | 0 | 0 |
| MT072864.1;QIQ54046.1;Pangolin_CoV_orf1a;2069 | 0 | 0 |
| MT072864.1;QIQ54046.1;Pangolin_CoV_orf1a;2070 | 0 | 0 |
| MT072864.1;QIQ54046.1;Pangolin_CoV_orf1a;2071 | 0 | 0 |
| MT072864.1;QIQ54046.1;Pangolin_CoV_orf1a;2072 | 0 | 0.025 |
| MT072864.1;QIQ54046.1;Pangolin_CoV_orf1a;2073 | 0 | 0.05 |
| MT072864.1;QIQ54046.1;Pangolin_CoV_orf1a;2074 | 0 | 0 |
| MT072864.1;QIQ54046.1;Pangolin_CoV_orf1a;2075 | 0 | 0.025 |
| MT072864.1;QIQ54046.1;Pangolin_CoV_orf1a;2076 | 0 | 0 |
| MT072864.1;QIQ54046.1;Pangolin_CoV_orf1a;2077 | 0 | 0 |
| MT072864.1;QIQ54046.1;Pangolin_CoV_orf1a;2078 | 0 | 0 |
| MT072864.1;QIQ54046.1;Pangolin_CoV_orf1a;2079 | 0 | 0 |
| MT072864.1;QIQ54046.1;Pangolin_CoV_orf1a;2080 | 0 | 0 |
| MT072864.1;QIQ54046.1;Pangolin_CoV_orf1a;2081 | 0 | 0 |
| MT072864.1;QIQ54046.1;Pangolin_CoV_orf1a;2082 | 0 | 0 |
| MT072864.1;QIQ54046.1;Pangolin_CoV_orf1a;2083 | 0.3 | 0.175 |
| MT072864.1;QIQ54046.1;Pangolin_CoV_orf1a;2084 | 0.05 | 0.05 |
| MT072864.1;QIQ54046.1;Pangolin_CoV_orf1a;2085 | 0 | 0.025 |
| MT072864.1;QIQ54046.1;Pangolin_CoV_orf1a;2086 | 0.05 | 0 |
| MT072864.1;QIQ54046.1;Pangolin_CoV_orf1a;2087 | 0 | 0 |
| MT072864.1;QIQ54046.1;Pangolin_CoV_orf1a;2088 | 0 | 0.025 |
| MT072864.1;QIQ54046.1;Pangolin_CoV_orf1a;2089 | 0.15 | 0.1 |
| MT072864.1;QIQ54046.1;Pangolin_CoV_orf1a;2090 | 0.2 | 0.075 |
| MT072864.1;QIQ54046.1;Pangolin_CoV_orf1a;2091 | 0.25 | 0.125 |
| MT072864.1;QIQ54046.1;Pangolin_CoV_orf1a;2092 | 0.1 | 0.025 |
| MT072864.1;QIQ54046.1;Pangolin_CoV_orf1a;2093 | 0.05 | 0 |
| MT072864.1;QIQ54046.1;Pangolin_CoV_orf1a;2094 | 0 | 0 |
| MT072864.1;QIQ54046.1;Pangolin_CoV_orf1a;2095 | 0 | 0 |
| MT072864.1;QIQ54046.1;Pangolin_CoV_orf1a;2096 | 0 | 0 |
| MT072864.1;QIQ54046.1;Pangolin_CoV_orf1a;2097 | 0 | 0 |

| | | |
|---|---|---|
| MT072864.1;QIQ54046.1;Pangolin_CoV_orf1a;2098 | 0 | 0 |
| MT072864.1;QIQ54046.1;Pangolin_CoV_orf1a;2099 | 0 | 0 |
| MT072864.1;QIQ54046.1;Pangolin_CoV_orf1a;2100 | 0 | 0 |
| MT072864.1;QIQ54046.1;Pangolin_CoV_orf1a;2101 | 0 | 0 |
| MT072864.1;QIQ54046.1;Pangolin_CoV_orf1a;2102 | 0 | 0 |
| MT072864.1;QIQ54046.1;Pangolin_CoV_orf1a;2103 | 0 | 0 |
| MT072864.1;QIQ54046.1;Pangolin_CoV_orf1a;2104 | 0 | 0 |
| MT072864.1;QIQ54046.1;Pangolin_CoV_orf1a;2105 | 0 | 0 |
| MT072864.1;QIQ54046.1;Pangolin_CoV_orf1a;2106 | 0.05 | 0 |
| MT072864.1;QIQ54046.1;Pangolin_CoV_orf1a;2107 | 0 | 0 |
| MT072864.1;QIQ54046.1;Pangolin_CoV_orf1a;2108 | 0 | 0 |
| MT072864.1;QIQ54046.1;Pangolin_CoV_orf1a;2109 | 0 | 0 |
| MT072864.1;QIQ54046.1;Pangolin_CoV_orf1a;2110 | 0.05 | 0 |
| MT072864.1;QIQ54046.1;Pangolin_CoV_orf1a;2111 | 0.05 | 0 |
| MT072864.1;QIQ54046.1;Pangolin_CoV_orf1a;2112 | 0 | 0 |
| MT072864.1;QIQ54046.1;Pangolin_CoV_orf1a;2113 | 0 | 0 |
| MT072864.1;QIQ54046.1;Pangolin_CoV_orf1a;2114 | 0 | 0 |
| MT072864.1;QIQ54046.1;Pangolin_CoV_orf1a;2115 | 0 | 0 |
| MT072864.1;QIQ54046.1;Pangolin_CoV_orf1a;2116 | 0.05 | 0 |
| MT072864.1;QIQ54046.1;Pangolin_CoV_orf1a;2117 | 0 | 0 |
| MT072864.1;QIQ54046.1;Pangolin_CoV_orf1a;2118 | 0 | 0 |
| MT072864.1;QIQ54046.1;Pangolin_CoV_orf1a;2119 | 0 | 0 |
| MT072864.1;QIQ54046.1;Pangolin_CoV_orf1a;2120 | 0 | 0 |
| MT072864.1;QIQ54046.1;Pangolin_CoV_orf1a;2121 | 0.05 | 0 |
| MT072864.1;QIQ54046.1;Pangolin_CoV_orf1a;2122 | 0 | 0 |
| MT072864.1;QIQ54046.1;Pangolin_CoV_orf1a;2123 | 0 | 0 |
| MT072864.1;QIQ54046.1;Pangolin_CoV_orf1a;2124 | 0 | 0 |
| MT072864.1;QIQ54046.1;Pangolin_CoV_orf1a;2125 | 0.05 | 0 |
| MT072864.1;QIQ54046.1;Pangolin_CoV_orf1a;2126 | 0.05 | 0 |
| MT072864.1;QIQ54046.1;Pangolin_CoV_orf1a;2127 | 0.05 | 0 |
| MT072864.1;QIQ54046.1;Pangolin_CoV_orf1a;2128 | 0.05 | 0.025 |
| MT072864.1;QIQ54046.1;Pangolin_CoV_orf1a;2129 | 0.05 | 0 |
| MT072864.1;QIQ54046.1;Pangolin_CoV_orf1a;2130 | 0.05 | 0 |
| MT072864.1;QIQ54046.1;Pangolin_CoV_orf1a;2131 | 0.1 | 0 |
| MT072864.1;QIQ54046.1;Pangolin_CoV_orf1a;2132 | 0.15 | 0.05 |
| MT072864.1;QIQ54046.1;Pangolin_CoV_orf1a;2133 | 0.1 | 0.075 |
| MT072864.1;QIQ54046.1;Pangolin_CoV_orf1a;2134 | 0.05 | 0.05 |
| MT072864.1;QIQ54046.1;Pangolin_CoV_orf1a;2135 | 0.15 | 0.075 |
| MT072864.1;QIQ54046.1;Pangolin_CoV_orf1a;2136 | 0.05 | 0 |
| MT072864.1;QIQ54046.1;Pangolin_CoV_orf1a;2137 | 0.05 | 0 |
| MT072864.1;QIQ54046.1;Pangolin_CoV_orf1a;2138 | 0 | 0 |
| MT072864.1;QIQ54046.1;Pangolin_CoV_orf1a;2139 | 0 | 0 |
| MT072864.1;QIQ54046.1;Pangolin_CoV_orf1a;2140 | 0 | 0 |

| | | |
|---|---|---|
| MT072864.1;QIQ54046.1;Pangolin_CoV_orf1a;2141 | 0 | 0 |
| MT072864.1;QIQ54046.1;Pangolin_CoV_orf1a;2142 | 0 | 0 |
| MT072864.1;QIQ54046.1;Pangolin_CoV_orf1a;2143 | 0 | 0 |
| MT072864.1;QIQ54046.1;Pangolin_CoV_orf1a;2144 | 0 | 0 |
| MT072864.1;QIQ54046.1;Pangolin_CoV_orf1a;2145 | 0 | 0 |
| MT072864.1;QIQ54046.1;Pangolin_CoV_orf1a;2146 | 0 | 0 |
| MT072864.1;QIQ54046.1;Pangolin_CoV_orf1a;2147 | 0 | 0 |
| MT072864.1;QIQ54046.1;Pangolin_CoV_orf1a;2148 | 0 | 0 |
| MT072864.1;QIQ54046.1;Pangolin_CoV_orf1a;2149 | 0 | 0 |
| MT072864.1;QIQ54046.1;Pangolin_CoV_orf1a;2150 | 0 | 0 |
| MT072864.1;QIQ54046.1;Pangolin_CoV_orf1a;2151 | 0 | 0 |
| MT072864.1;QIQ54046.1;Pangolin_CoV_orf1a;2152 | 0 | 0 |
| MT072864.1;QIQ54046.1;Pangolin_CoV_orf1a;2153 | 0 | 0 |
| MT072864.1;QIQ54046.1;Pangolin_CoV_orf1a;2154 | 0 | 0 |
| MT072864.1;QIQ54046.1;Pangolin_CoV_orf1a;2155 | 0 | 0 |
| MT072864.1;QIQ54046.1;Pangolin_CoV_orf1a;2156 | 0 | 0 |
| MT072864.1;QIQ54046.1;Pangolin_CoV_orf1a;2157 | 0 | 0 |
| MT072864.1;QIQ54046.1;Pangolin_CoV_orf1a;2158 | 0 | 0 |
| MT072864.1;QIQ54046.1;Pangolin_CoV_orf1a;2159 | 0 | 0 |
| MT072864.1;QIQ54046.1;Pangolin_CoV_orf1a;2160 | 0 | 0 |
| MT072864.1;QIQ54046.1;Pangolin_CoV_orf1a;2161 | 0 | 0 |
| MT072864.1;QIQ54046.1;Pangolin_CoV_orf1a;2162 | 0 | 0 |
| MT072864.1;QIQ54046.1;Pangolin_CoV_orf1a;2163 | 0 | 0 |
| MT072864.1;QIQ54046.1;Pangolin_CoV_orf1a;2164 | 0 | 0 |
| MT072864.1;QIQ54046.1;Pangolin_CoV_orf1a;2165 | 0 | 0 |
| MT072864.1;QIQ54046.1;Pangolin_CoV_orf1a;2166 | 0 | 0 |
| MT072864.1;QIQ54046.1;Pangolin_CoV_orf1a;2167 | 0 | 0 |
| MT072864.1;QIQ54046.1;Pangolin_CoV_orf1a;2168 | 0 | 0 |
| MT072864.1;QIQ54046.1;Pangolin_CoV_orf1a;2169 | 0 | 0 |
| MT072864.1;QIQ54046.1;Pangolin_CoV_orf1a;2170 | 0 | 0 |
| MT072864.1;QIQ54046.1;Pangolin_CoV_orf1a;2171 | 0 | 0 |
| MT072864.1;QIQ54046.1;Pangolin_CoV_orf1a;2172 | 0.05 | 0 |
| MT072864.1;QIQ54046.1;Pangolin_CoV_orf1a;2173 | 0.1 | 0 |
| MT072864.1;QIQ54046.1;Pangolin_CoV_orf1a;2174 | 0 | 0 |
| MT072864.1;QIQ54046.1;Pangolin_CoV_orf1a;2175 | 0 | 0.025 |
| MT072864.1;QIQ54046.1;Pangolin_CoV_orf1a;2176 | 0 | 0 |
| MT072864.1;QIQ54046.1;Pangolin_CoV_orf1a;2177 | 0.05 | 0 |
| MT072864.1;QIQ54046.1;Pangolin_CoV_orf1a;2178 | 0.05 | 0 |
| MT072864.1;QIQ54046.1;Pangolin_CoV_orf1a;2179 | 0 | 0 |
| MT072864.1;QIQ54046.1;Pangolin_CoV_orf1a;2180 | 0 | 0 |
| MT072864.1;QIQ54046.1;Pangolin_CoV_orf1a;2181 | 0 | 0 |
| MT072864.1;QIQ54046.1;Pangolin_CoV_orf1a;2182 | 0 | 0 |
| MT072864.1;QIQ54046.1;Pangolin_CoV_orf1a;2183 | 0 | 0 |

| | | |
|---|---|---|
| MT072864.1;QIQ54046.1;Pangolin_CoV_orf1a;2184 | 0 | 0 |
| MT072864.1;QIQ54046.1;Pangolin_CoV_orf1a;2185 | 0 | 0 |
| MT072864.1;QIQ54046.1;Pangolin_CoV_orf1a;2186 | 0 | 0 |
| MT072864.1;QIQ54046.1;Pangolin_CoV_orf1a;2187 | 0 | 0 |
| MT072864.1;QIQ54046.1;Pangolin_CoV_orf1a;2188 | 0 | 0 |
| MT072864.1;QIQ54046.1;Pangolin_CoV_orf1a;2189 | 0 | 0 |
| MT072864.1;QIQ54046.1;Pangolin_CoV_orf1a;2190 | 0 | 0 |
| MT072864.1;QIQ54046.1;Pangolin_CoV_orf1a;2191 | 0 | 0 |
| MT072864.1;QIQ54046.1;Pangolin_CoV_orf1a;2192 | 0 | 0 |
| MT072864.1;QIQ54046.1;Pangolin_CoV_orf1a;2193 | 0 | 0 |
| MT072864.1;QIQ54046.1;Pangolin_CoV_orf1a;2194 | 0 | 0 |
| MT072864.1;QIQ54046.1;Pangolin_CoV_orf1a;2195 | 0 | 0 |
| MT072864.1;QIQ54046.1;Pangolin_CoV_orf1a;2196 | 0 | 0 |
| MT072864.1;QIQ54046.1;Pangolin_CoV_orf1a;2197 | 0 | 0 |
| MT072864.1;QIQ54046.1;Pangolin_CoV_orf1a;2198 | 0 | 0 |
| MT072864.1;QIQ54046.1;Pangolin_CoV_orf1a;2199 | 0 | 0 |
| MT072864.1;QIQ54046.1;Pangolin_CoV_orf1a;2200 | 0 | 0 |
| MT072864.1;QIQ54046.1;Pangolin_CoV_orf1a;2201 | 0 | 0 |
| MT072864.1;QIQ54046.1;Pangolin_CoV_orf1a;2202 | 0 | 0 |
| MT072864.1;QIQ54046.1;Pangolin_CoV_orf1a;2203 | 0 | 0 |
| MT072864.1;QIQ54046.1;Pangolin_CoV_orf1a;2204 | 0.05 | 0 |
| MT072864.1;QIQ54046.1;Pangolin_CoV_orf1a;2205 | 0 | 0 |
| MT072864.1;QIQ54046.1;Pangolin_CoV_orf1a;2206 | 0 | 0 |
| MT072864.1;QIQ54046.1;Pangolin_CoV_orf1a;2207 | 0 | 0 |
| MT072864.1;QIQ54046.1;Pangolin_CoV_orf1a;2208 | 0 | 0 |
| MT072864.1;QIQ54046.1;Pangolin_CoV_orf1a;2209 | 0 | 0 |
| MT072864.1;QIQ54046.1;Pangolin_CoV_orf1a;2210 | 0 | 0 |
| MT072864.1;QIQ54046.1;Pangolin_CoV_orf1a;2211 | 0 | 0 |
| MT072864.1;QIQ54046.1;Pangolin_CoV_orf1a;2212 | 0 | 0 |
| MT072864.1;QIQ54046.1;Pangolin_CoV_orf1a;2213 | 0 | 0 |
| MT072864.1;QIQ54046.1;Pangolin_CoV_orf1a;2214 | 0 | 0 |
| MT072864.1;QIQ54046.1;Pangolin_CoV_orf1a;2215 | 0 | 0 |
| MT072864.1;QIQ54046.1;Pangolin_CoV_orf1a;2216 | 0 | 0 |
| MT072864.1;QIQ54046.1;Pangolin_CoV_orf1a;2217 | 0 | 0 |
| MT072864.1;QIQ54046.1;Pangolin_CoV_orf1a;2218 | 0 | 0 |
| MT072864.1;QIQ54046.1;Pangolin_CoV_orf1a;2219 | 0 | 0 |
| MT072864.1;QIQ54046.1;Pangolin_CoV_orf1a;2220 | 0 | 0 |
| MT072864.1;QIQ54046.1;Pangolin_CoV_orf1a;2221 | 0 | 0 |
| MT072864.1;QIQ54046.1;Pangolin_CoV_orf1a;2222 | 0 | 0 |
| MT072864.1;QIQ54046.1;Pangolin_CoV_orf1a;2223 | 0 | 0 |
| MT072864.1;QIQ54046.1;Pangolin_CoV_orf1a;2224 | 0 | 0 |
| MT072864.1;QIQ54046.1;Pangolin_CoV_orf1a;2225 | 0 | 0 |
| MT072864.1;QIQ54046.1;Pangolin_CoV_orf1a;2226 | 0.05 | 0 |

| | | |
|---|---|---|
| MT072864.1;QIQ54046.1;Pangolin_CoV_orf1a;2227 | 0 | 0 |
| MT072864.1;QIQ54046.1;Pangolin_CoV_orf1a;2228 | 0 | 0 |
| MT072864.1;QIQ54046.1;Pangolin_CoV_orf1a;2229 | 0 | 0 |
| MT072864.1;QIQ54046.1;Pangolin_CoV_orf1a;2230 | 0 | 0 |
| MT072864.1;QIQ54046.1;Pangolin_CoV_orf1a;2231 | 0 | 0 |
| MT072864.1;QIQ54046.1;Pangolin_CoV_orf1a;2232 | 0 | 0 |
| MT072864.1;QIQ54046.1;Pangolin_CoV_orf1a;2233 | 0 | 0 |
| MT072864.1;QIQ54046.1;Pangolin_CoV_orf1a;2234 | 0 | 0 |
| MT072864.1;QIQ54046.1;Pangolin_CoV_orf1a;2235 | 0 | 0 |
| MT072864.1;QIQ54046.1;Pangolin_CoV_orf1a;2236 | 0 | 0 |
| MT072864.1;QIQ54046.1;Pangolin_CoV_orf1a;2237 | 0.05 | 0 |
| MT072864.1;QIQ54046.1;Pangolin_CoV_orf1a;2238 | 0 | 0 |
| MT072864.1;QIQ54046.1;Pangolin_CoV_orf1a;2239 | 0 | 0 |
| MT072864.1;QIQ54046.1;Pangolin_CoV_orf1a;2240 | 0 | 0.025 |
| MT072864.1;QIQ54046.1;Pangolin_CoV_orf1a;2241 | 0 | 0.025 |
| MT072864.1;QIQ54046.1;Pangolin_CoV_orf1a;2242 | 0 | 0 |
| MT072864.1;QIQ54046.1;Pangolin_CoV_orf1a;2243 | 0 | 0 |
| MT072864.1;QIQ54046.1;Pangolin_CoV_orf1a;2244 | 0 | 0 |
| MT072864.1;QIQ54046.1;Pangolin_CoV_orf1a;2245 | 0 | 0 |
| MT072864.1;QIQ54046.1;Pangolin_CoV_orf1a;2246 | 0 | 0 |
| MT072864.1;QIQ54046.1;Pangolin_CoV_orf1a;2247 | 0 | 0 |
| MT072864.1;QIQ54046.1;Pangolin_CoV_orf1a;2248 | 0 | 0 |
| MT072864.1;QIQ54046.1;Pangolin_CoV_orf1a;2249 | 0 | 0 |
| MT072864.1;QIQ54046.1;Pangolin_CoV_orf1a;2250 | 0 | 0.05 |
| MT072864.1;QIQ54046.1;Pangolin_CoV_orf1a;2251 | 0.05 | 0 |
| MT072864.1;QIQ54046.1;Pangolin_CoV_orf1a;2252 | 0.2 | 0.05 |
| MT072864.1;QIQ54046.1;Pangolin_CoV_orf1a;2253 | 0.25 | 0.175 |
| MT072864.1;QIQ54046.1;Pangolin_CoV_orf1a;2254 | 0.3 | 0.25 |
| MT072864.1;QIQ54046.1;Pangolin_CoV_orf1a;2255 | 0.5 | 0.35 |
| MT072864.1;QIQ54046.1;Pangolin_CoV_orf1a;2256 | 0.35 | 0.35 |
| MT072864.1;QIQ54046.1;Pangolin_CoV_orf1a;2257 | 0.55 | 0.375 |
| MT072864.1;QIQ54046.1;Pangolin_CoV_orf1a;2258 | 0.4 | 0.325 |
| MT072864.1;QIQ54046.1;Pangolin_CoV_orf1a;2259 | 0.2 | 0.15 |
| MT072864.1;QIQ54046.1;Pangolin_CoV_orf1a;2260 | 0.2 | 0.2 |
| MT072864.1;QIQ54046.1;Pangolin_CoV_orf1a;2261 | 0.2 | 0.05 |
| MT072864.1;QIQ54046.1;Pangolin_CoV_orf1a;2262 | 0.2 | 0.05 |
| MT072864.1;QIQ54046.1;Pangolin_CoV_orf1a;2263 | 0.05 | 0.025 |
| MT072864.1;QIQ54046.1;Pangolin_CoV_orf1a;2264 | 0 | 0 |
| MT072864.1;QIQ54046.1;Pangolin_CoV_orf1a;2265 | 0.05 | 0 |
| MT072864.1;QIQ54046.1;Pangolin_CoV_orf1a;2266 | 0.05 | 0 |
| MT072864.1;QIQ54046.1;Pangolin_CoV_orf1a;2267 | 0 | 0 |
| MT072864.1;QIQ54046.1;Pangolin_CoV_orf1a;2268 | 0 | 0 |
| MT072864.1;QIQ54046.1;Pangolin_CoV_orf1a;2269 | 0 | 0 |

| | | |
|---|---|---|
| MT072864.1;QIQ54046.1;Pangolin_CoV_orf1a;2270 | 0 | 0 |
| MT072864.1;QIQ54046.1;Pangolin_CoV_orf1a;2271 | 0 | 0 |
| MT072864.1;QIQ54046.1;Pangolin_CoV_orf1a;2272 | 0 | 0 |
| MT072864.1;QIQ54046.1;Pangolin_CoV_orf1a;2273 | 0 | 0 |
| MT072864.1;QIQ54046.1;Pangolin_CoV_orf1a;2274 | 0 | 0 |
| MT072864.1;QIQ54046.1;Pangolin_CoV_orf1a;2275 | 0 | 0 |
| MT072864.1;QIQ54046.1;Pangolin_CoV_orf1a;2276 | 0 | 0 |
| MT072864.1;QIQ54046.1;Pangolin_CoV_orf1a;2277 | 0 | 0 |
| MT072864.1;QIQ54046.1;Pangolin_CoV_orf1a;2278 | 0 | 0.025 |
| MT072864.1;QIQ54046.1;Pangolin_CoV_orf1a;2279 | 0 | 0 |
| MT072864.1;QIQ54046.1;Pangolin_CoV_orf1a;2280 | 0 | 0.025 |
| MT072864.1;QIQ54046.1;Pangolin_CoV_orf1a;2281 | 0 | 0 |
| MT072864.1;QIQ54046.1;Pangolin_CoV_orf1a;2282 | 0 | 0 |
| MT072864.1;QIQ54046.1;Pangolin_CoV_orf1a;2283 | 0.05 | 0 |
| MT072864.1;QIQ54046.1;Pangolin_CoV_orf1a;2284 | 0 | 0 |
| MT072864.1;QIQ54046.1;Pangolin_CoV_orf1a;2285 | 0 | 0 |
| MT072864.1;QIQ54046.1;Pangolin_CoV_orf1a;2286 | 0.05 | 0 |
| MT072864.1;QIQ54046.1;Pangolin_CoV_orf1a;2287 | 0.05 | 0 |
| MT072864.1;QIQ54046.1;Pangolin_CoV_orf1a;2288 | 0 | 0 |
| MT072864.1;QIQ54046.1;Pangolin_CoV_orf1a;2289 | 0.05 | 0 |
| MT072864.1;QIQ54046.1;Pangolin_CoV_orf1a;2290 | 0 | 0 |
| MT072864.1;QIQ54046.1;Pangolin_CoV_orf1a;2291 | 0 | 0 |
| MT072864.1;QIQ54046.1;Pangolin_CoV_orf1a;2292 | 0 | 0 |
| MT072864.1;QIQ54046.1;Pangolin_CoV_orf1a;2293 | 0 | 0 |
| MT072864.1;QIQ54046.1;Pangolin_CoV_orf1a;2294 | 0 | 0 |
| MT072864.1;QIQ54046.1;Pangolin_CoV_orf1a;2295 | 0 | 0 |
| MT072864.1;QIQ54046.1;Pangolin_CoV_orf1a;2296 | 0 | 0 |
| MT072864.1;QIQ54046.1;Pangolin_CoV_orf1a;2297 | 0 | 0 |
| MT072864.1;QIQ54046.1;Pangolin_CoV_orf1a;2298 | 0 | 0 |
| MT072864.1;QIQ54046.1;Pangolin_CoV_orf1a;2299 | 0 | 0 |
| MT072864.1;QIQ54046.1;Pangolin_CoV_orf1a;2300 | 0 | 0 |
| MT072864.1;QIQ54046.1;Pangolin_CoV_orf1a;2301 | 0 | 0 |
| MT072864.1;QIQ54046.1;Pangolin_CoV_orf1a;2302 | 0 | 0 |
| MT072864.1;QIQ54046.1;Pangolin_CoV_orf1a;2303 | 0 | 0 |
| MT072864.1;QIQ54046.1;Pangolin_CoV_orf1a;2304 | 0 | 0 |
| MT072864.1;QIQ54046.1;Pangolin_CoV_orf1a;2305 | 0 | 0 |
| MT072864.1;QIQ54046.1;Pangolin_CoV_orf1a;2306 | 0 | 0 |
| MT072864.1;QIQ54046.1;Pangolin_CoV_orf1a;2307 | 0 | 0 |
| MT072864.1;QIQ54046.1;Pangolin_CoV_orf1a;2308 | 0 | 0 |
| MT072864.1;QIQ54046.1;Pangolin_CoV_orf1a;2309 | 0 | 0 |
| MT072864.1;QIQ54046.1;Pangolin_CoV_orf1a;2310 | 0 | 0 |
| MT072864.1;QIQ54046.1;Pangolin_CoV_orf1a;2311 | 0 | 0 |
| MT072864.1;QIQ54046.1;Pangolin_CoV_orf1a;2312 | 0 | 0 |

| | | |
|---|---|---|
| MT072864.1;QIQ54046.1;Pangolin_CoV_orf1a;2313 | 0 | 0 |
| MT072864.1;QIQ54046.1;Pangolin_CoV_orf1a;2314 | 0 | 0 |
| MT072864.1;QIQ54046.1;Pangolin_CoV_orf1a;2315 | 0 | 0 |
| MT072864.1;QIQ54046.1;Pangolin_CoV_orf1a;2316 | 0 | 0 |
| MT072864.1;QIQ54046.1;Pangolin_CoV_orf1a;2317 | 0 | 0 |
| MT072864.1;QIQ54046.1;Pangolin_CoV_orf1a;2318 | 0 | 0 |
| MT072864.1;QIQ54046.1;Pangolin_CoV_orf1a;2319 | 0 | 0 |
| MT072864.1;QIQ54046.1;Pangolin_CoV_orf1a;2320 | 0 | 0 |
| MT072864.1;QIQ54046.1;Pangolin_CoV_orf1a;2321 | 0 | 0 |
| MT072864.1;QIQ54046.1;Pangolin_CoV_orf1a;2322 | 0 | 0 |
| MT072864.1;QIQ54046.1;Pangolin_CoV_orf1a;2323 | 0 | 0 |
| MT072864.1;QIQ54046.1;Pangolin_CoV_orf1a;2324 | 0 | 0 |
| MT072864.1;QIQ54046.1;Pangolin_CoV_orf1a;2325 | 0 | 0 |
| MT072864.1;QIQ54046.1;Pangolin_CoV_orf1a;2326 | 0 | 0 |
| MT072864.1;QIQ54046.1;Pangolin_CoV_orf1a;2327 | 0 | 0 |
| MT072864.1;QIQ54046.1;Pangolin_CoV_orf1a;2328 | 0 | 0 |
| MT072864.1;QIQ54046.1;Pangolin_CoV_orf1a;2329 | 0 | 0 |
| MT072864.1;QIQ54046.1;Pangolin_CoV_orf1a;2330 | 0 | 0 |
| MT072864.1;QIQ54046.1;Pangolin_CoV_orf1a;2331 | 0 | 0 |
| MT072864.1;QIQ54046.1;Pangolin_CoV_orf1a;2332 | 0 | 0 |
| MT072864.1;QIQ54046.1;Pangolin_CoV_orf1a;2333 | 0 | 0 |
| MT072864.1;QIQ54046.1;Pangolin_CoV_orf1a;2334 | 0 | 0 |
| MT072864.1;QIQ54046.1;Pangolin_CoV_orf1a;2335 | 0 | 0 |
| MT072864.1;QIQ54046.1;Pangolin_CoV_orf1a;2336 | 0 | 0 |
| MT072864.1;QIQ54046.1;Pangolin_CoV_orf1a;2337 | 0 | 0 |
| MT072864.1;QIQ54046.1;Pangolin_CoV_orf1a;2338 | 0 | 0 |
| MT072864.1;QIQ54046.1;Pangolin_CoV_orf1a;2339 | 0 | 0 |
| MT072864.1;QIQ54046.1;Pangolin_CoV_orf1a;2340 | 0 | 0 |
| MT072864.1;QIQ54046.1;Pangolin_CoV_orf1a;2341 | 0 | 0 |
| MT072864.1;QIQ54046.1;Pangolin_CoV_orf1a;2342 | 0 | 0 |
| MT072864.1;QIQ54046.1;Pangolin_CoV_orf1a;2343 | 0 | 0 |
| MT072864.1;QIQ54046.1;Pangolin_CoV_orf1a;2344 | 0 | 0 |
| MT072864.1;QIQ54046.1;Pangolin_CoV_orf1a;2345 | 0 | 0 |
| MT072864.1;QIQ54046.1;Pangolin_CoV_orf1a;2346 | 0 | 0 |
| MT072864.1;QIQ54046.1;Pangolin_CoV_orf1a;2347 | 0 | 0 |
| MT072864.1;QIQ54046.1;Pangolin_CoV_orf1a;2348 | 0 | 0 |
| MT072864.1;QIQ54046.1;Pangolin_CoV_orf1a;2349 | 0 | 0 |
| MT072864.1;QIQ54046.1;Pangolin_CoV_orf1a;2350 | 0 | 0 |
| MT072864.1;QIQ54046.1;Pangolin_CoV_orf1a;2351 | 0 | 0 |
| MT072864.1;QIQ54046.1;Pangolin_CoV_orf1a;2352 | 0 | 0 |
| MT072864.1;QIQ54046.1;Pangolin_CoV_orf1a;2353 | 0 | 0 |
| MT072864.1;QIQ54046.1;Pangolin_CoV_orf1a;2354 | 0 | 0 |
| MT072864.1;QIQ54046.1;Pangolin_CoV_orf1a;2355 | 0 | 0 |

| | | |
|---|---|---|
| MT072864.1;QIQ54046.1;Pangolin_CoV_orf1a;2356 | 0 | 0 |
| MT072864.1;QIQ54046.1;Pangolin_CoV_orf1a;2357 | 0 | 0 |
| MT072864.1;QIQ54046.1;Pangolin_CoV_orf1a;2358 | 0 | 0 |
| MT072864.1;QIQ54046.1;Pangolin_CoV_orf1a;2359 | 0 | 0 |
| MT072864.1;QIQ54046.1;Pangolin_CoV_orf1a;2360 | 0 | 0 |
| MT072864.1;QIQ54046.1;Pangolin_CoV_orf1a;2361 | 0 | 0 |
| MT072864.1;QIQ54046.1;Pangolin_CoV_orf1a;2362 | 0 | 0 |
| MT072864.1;QIQ54046.1;Pangolin_CoV_orf1a;2363 | 0 | 0 |
| MT072864.1;QIQ54046.1;Pangolin_CoV_orf1a;2364 | 0 | 0 |
| MT072864.1;QIQ54046.1;Pangolin_CoV_orf1a;2365 | 0 | 0 |
| MT072864.1;QIQ54046.1;Pangolin_CoV_orf1a;2366 | 0 | 0 |
| MT072864.1;QIQ54046.1;Pangolin_CoV_orf1a;2367 | 0 | 0 |
| MT072864.1;QIQ54046.1;Pangolin_CoV_orf1a;2368 | 0 | 0 |
| MT072864.1;QIQ54046.1;Pangolin_CoV_orf1a;2369 | 0 | 0 |
| MT072864.1;QIQ54046.1;Pangolin_CoV_orf1a;2370 | 0 | 0 |
| MT072864.1;QIQ54046.1;Pangolin_CoV_orf1a;2371 | 0 | 0 |
| MT072864.1;QIQ54046.1;Pangolin_CoV_orf1a;2372 | 0 | 0 |
| MT072864.1;QIQ54046.1;Pangolin_CoV_orf1a;2373 | 0 | 0 |
| MT072864.1;QIQ54046.1;Pangolin_CoV_orf1a;2374 | 0 | 0 |
| MT072864.1;QIQ54046.1;Pangolin_CoV_orf1a;2375 | 0 | 0 |
| MT072864.1;QIQ54046.1;Pangolin_CoV_orf1a;2376 | 0 | 0 |
| MT072864.1;QIQ54046.1;Pangolin_CoV_orf1a;2377 | 0 | 0 |
| MT072864.1;QIQ54046.1;Pangolin_CoV_orf1a;2378 | 0 | 0 |
| MT072864.1;QIQ54046.1;Pangolin_CoV_orf1a;2379 | 0 | 0 |
| MT072864.1;QIQ54046.1;Pangolin_CoV_orf1a;2380 | 0 | 0 |
| MT072864.1;QIQ54046.1;Pangolin_CoV_orf1a;2381 | 0 | 0 |
| MT072864.1;QIQ54046.1;Pangolin_CoV_orf1a;2382 | 0 | 0 |
| MT072864.1;QIQ54046.1;Pangolin_CoV_orf1a;2383 | 0 | 0 |
| MT072864.1;QIQ54046.1;Pangolin_CoV_orf1a;2384 | 0 | 0 |
| MT072864.1;QIQ54046.1;Pangolin_CoV_orf1a;2385 | 0 | 0 |
| MT072864.1;QIQ54046.1;Pangolin_CoV_orf1a;2386 | 0 | 0 |
| MT072864.1;QIQ54046.1;Pangolin_CoV_orf1a;2387 | 0 | 0 |
| MT072864.1;QIQ54046.1;Pangolin_CoV_orf1a;2388 | 0 | 0.025 |
| MT072864.1;QIQ54046.1;Pangolin_CoV_orf1a;2389 | 0 | 0 |
| MT072864.1;QIQ54046.1;Pangolin_CoV_orf1a;2390 | 0 | 0 |
| MT072864.1;QIQ54046.1;Pangolin_CoV_orf1a;2391 | 0 | 0 |
| MT072864.1;QIQ54046.1;Pangolin_CoV_orf1a;2392 | 0 | 0 |
| MT072864.1;QIQ54046.1;Pangolin_CoV_orf1a;2393 | 0 | 0 |
| MT072864.1;QIQ54046.1;Pangolin_CoV_orf1a;2394 | 0 | 0 |
| MT072864.1;QIQ54046.1;Pangolin_CoV_orf1a;2395 | 0 | 0 |
| MT072864.1;QIQ54046.1;Pangolin_CoV_orf1a;2396 | 0 | 0 |
| MT072864.1;QIQ54046.1;Pangolin_CoV_orf1a;2397 | 0 | 0 |
| MT072864.1;QIQ54046.1;Pangolin_CoV_orf1a;2398 | 0 | 0 |

| | | |
|---|---|---|
| MT072864.1;QIQ54046.1;Pangolin_CoV_orf1a;2399 | 0 | 0 |
| MT072864.1;QIQ54046.1;Pangolin_CoV_orf1a;2400 | 0 | 0 |
| MT072864.1;QIQ54046.1;Pangolin_CoV_orf1a;2401 | 0 | 0 |
| MT072864.1;QIQ54046.1;Pangolin_CoV_orf1a;2402 | 0 | 0 |
| MT072864.1;QIQ54046.1;Pangolin_CoV_orf1a;2403 | 0 | 0 |
| MT072864.1;QIQ54046.1;Pangolin_CoV_orf1a;2404 | 0 | 0 |
| MT072864.1;QIQ54046.1;Pangolin_CoV_orf1a;2405 | 0 | 0 |
| MT072864.1;QIQ54046.1;Pangolin_CoV_orf1a;2406 | 0 | 0 |
| MT072864.1;QIQ54046.1;Pangolin_CoV_orf1a;2407 | 0 | 0 |
| MT072864.1;QIQ54046.1;Pangolin_CoV_orf1a;2408 | 0 | 0.025 |
| MT072864.1;QIQ54046.1;Pangolin_CoV_orf1a;2409 | 0 | 0 |
| MT072864.1;QIQ54046.1;Pangolin_CoV_orf1a;2410 | 0 | 0 |
| MT072864.1;QIQ54046.1;Pangolin_CoV_orf1a;2411 | 0 | 0 |
| MT072864.1;QIQ54046.1;Pangolin_CoV_orf1a;2412 | 0 | 0 |
| MT072864.1;QIQ54046.1;Pangolin_CoV_orf1a;2413 | 0 | 0 |
| MT072864.1;QIQ54046.1;Pangolin_CoV_orf1a;2414 | 0 | 0 |
| MT072864.1;QIQ54046.1;Pangolin_CoV_orf1a;2415 | 0 | 0 |
| MT072864.1;QIQ54046.1;Pangolin_CoV_orf1a;2416 | 0 | 0 |
| MT072864.1;QIQ54046.1;Pangolin_CoV_orf1a;2417 | 0 | 0 |
| MT072864.1;QIQ54046.1;Pangolin_CoV_orf1a;2418 | 0 | 0 |
| MT072864.1;QIQ54046.1;Pangolin_CoV_orf1a;2419 | 0 | 0 |
| MT072864.1;QIQ54046.1;Pangolin_CoV_orf1a;2420 | 0 | 0 |
| MT072864.1;QIQ54046.1;Pangolin_CoV_orf1a;2421 | 0 | 0 |
| MT072864.1;QIQ54046.1;Pangolin_CoV_orf1a;2422 | 0 | 0 |
| MT072864.1;QIQ54046.1;Pangolin_CoV_orf1a;2423 | 0 | 0 |
| MT072864.1;QIQ54046.1;Pangolin_CoV_orf1a;2424 | 0 | 0 |
| MT072864.1;QIQ54046.1;Pangolin_CoV_orf1a;2425 | 0 | 0 |
| MT072864.1;QIQ54046.1;Pangolin_CoV_orf1a;2426 | 0 | 0 |
| MT072864.1;QIQ54046.1;Pangolin_CoV_orf1a;2427 | 0 | 0 |
| MT072864.1;QIQ54046.1;Pangolin_CoV_orf1a;2428 | 0 | 0 |
| MT072864.1;QIQ54046.1;Pangolin_CoV_orf1a;2429 | 0 | 0 |
| MT072864.1;QIQ54046.1;Pangolin_CoV_orf1a;2430 | 0 | 0 |
| MT072864.1;QIQ54046.1;Pangolin_CoV_orf1a;2431 | 0 | 0 |
| MT072864.1;QIQ54046.1;Pangolin_CoV_orf1a;2432 | 0 | 0 |
| MT072864.1;QIQ54046.1;Pangolin_CoV_orf1a;2433 | 0 | 0.025 |
| MT072864.1;QIQ54046.1;Pangolin_CoV_orf1a;2434 | 0 | 0.025 |
| MT072864.1;QIQ54046.1;Pangolin_CoV_orf1a;2435 | 0 | 0 |
| MT072864.1;QIQ54046.1;Pangolin_CoV_orf1a;2436 | 0 | 0 |
| MT072864.1;QIQ54046.1;Pangolin_CoV_orf1a;2437 | 0 | 0 |
| MT072864.1;QIQ54046.1;Pangolin_CoV_orf1a;2438 | 0 | 0 |
| MT072864.1;QIQ54046.1;Pangolin_CoV_orf1a;2439 | 0 | 0 |
| MT072864.1;QIQ54046.1;Pangolin_CoV_orf1a;2440 | 0 | 0 |
| MT072864.1;QIQ54046.1;Pangolin_CoV_orf1a;2441 | 0 | 0 |

| | | |
|---|---|---|
| MT072864.1;QIQ54046.1;Pangolin_CoV_orf1a;2442 | 0 | 0 |
| MT072864.1;QIQ54046.1;Pangolin_CoV_orf1a;2443 | 0 | 0 |
| MT072864.1;QIQ54046.1;Pangolin_CoV_orf1a;2444 | 0 | 0 |
| MT072864.1;QIQ54046.1;Pangolin_CoV_orf1a;2445 | 0 | 0 |
| MT072864.1;QIQ54046.1;Pangolin_CoV_orf1a;2446 | 0 | 0 |
| MT072864.1;QIQ54046.1;Pangolin_CoV_orf1a;2447 | 0 | 0 |
| MT072864.1;QIQ54046.1;Pangolin_CoV_orf1a;2448 | 0 | 0 |
| MT072864.1;QIQ54046.1;Pangolin_CoV_orf1a;2449 | 0 | 0 |
| MT072864.1;QIQ54046.1;Pangolin_CoV_orf1a;2450 | 0 | 0 |
| MT072864.1;QIQ54046.1;Pangolin_CoV_orf1a;2451 | 0 | 0 |
| MT072864.1;QIQ54046.1;Pangolin_CoV_orf1a;2452 | 0 | 0 |
| MT072864.1;QIQ54046.1;Pangolin_CoV_orf1a;2453 | 0 | 0 |
| MT072864.1;QIQ54046.1;Pangolin_CoV_orf1a;2454 | 0 | 0 |
| MT072864.1;QIQ54046.1;Pangolin_CoV_orf1a;2455 | 0 | 0 |
| MT072864.1;QIQ54046.1;Pangolin_CoV_orf1a;2456 | 0 | 0 |
| MT072864.1;QIQ54046.1;Pangolin_CoV_orf1a;2457 | 0 | 0 |
| MT072864.1;QIQ54046.1;Pangolin_CoV_orf1a;2458 | 0 | 0 |
| MT072864.1;QIQ54046.1;Pangolin_CoV_orf1a;2459 | 0 | 0 |
| MT072864.1;QIQ54046.1;Pangolin_CoV_orf1a;2460 | 0 | 0 |
| MT072864.1;QIQ54046.1;Pangolin_CoV_orf1a;2461 | 0.05 | 0 |
| MT072864.1;QIQ54046.1;Pangolin_CoV_orf1a;2462 | 0.05 | 0.025 |
| MT072864.1;QIQ54046.1;Pangolin_CoV_orf1a;2463 | 0.05 | 0.025 |
| MT072864.1;QIQ54046.1;Pangolin_CoV_orf1a;2464 | 0 | 0 |
| MT072864.1;QIQ54046.1;Pangolin_CoV_orf1a;2465 | 0 | 0 |
| MT072864.1;QIQ54046.1;Pangolin_CoV_orf1a;2466 | 0 | 0 |
| MT072864.1;QIQ54046.1;Pangolin_CoV_orf1a;2467 | 0 | 0 |
| MT072864.1;QIQ54046.1;Pangolin_CoV_orf1a;2468 | 0 | 0 |
| MT072864.1;QIQ54046.1;Pangolin_CoV_orf1a;2469 | 0 | 0 |
| MT072864.1;QIQ54046.1;Pangolin_CoV_orf1a;2470 | 0.05 | 0.075 |
| MT072864.1;QIQ54046.1;Pangolin_CoV_orf1a;2471 | 0 | 0 |
| MT072864.1;QIQ54046.1;Pangolin_CoV_orf1a;2472 | 0 | 0 |
| MT072864.1;QIQ54046.1;Pangolin_CoV_orf1a;2473 | 0 | 0 |
| MT072864.1;QIQ54046.1;Pangolin_CoV_orf1a;2474 | 0 | 0.025 |
| MT072864.1;QIQ54046.1;Pangolin_CoV_orf1a;2475 | 0 | 0.025 |
| MT072864.1;QIQ54046.1;Pangolin_CoV_orf1a;2476 | 0 | 0.05 |
| MT072864.1;QIQ54046.1;Pangolin_CoV_orf1a;2477 | 0 | 0.025 |
| MT072864.1;QIQ54046.1;Pangolin_CoV_orf1a;2478 | 0 | 0.025 |
| MT072864.1;QIQ54046.1;Pangolin_CoV_orf1a;2479 | 0 | 0.025 |
| MT072864.1;QIQ54046.1;Pangolin_CoV_orf1a;2480 | 0 | 0.025 |
| MT072864.1;QIQ54046.1;Pangolin_CoV_orf1a;2481 | 0 | 0.025 |
| MT072864.1;QIQ54046.1;Pangolin_CoV_orf1a;2482 | 0.05 | 0.025 |
| MT072864.1;QIQ54046.1;Pangolin_CoV_orf1a;2483 | 0 | 0 |
| MT072864.1;QIQ54046.1;Pangolin_CoV_orf1a;2484 | 0 | 0 |

| | | |
|---|---|---|
| MT072864.1;QIQ54046.1;Pangolin_CoV_orf1a;2485 | 0 | 0 |
| MT072864.1;QIQ54046.1;Pangolin_CoV_orf1a;2486 | 0 | 0 |
| MT072864.1;QIQ54046.1;Pangolin_CoV_orf1a;2487 | 0 | 0 |
| MT072864.1;QIQ54046.1;Pangolin_CoV_orf1a;2488 | 0 | 0 |
| MT072864.1;QIQ54046.1;Pangolin_CoV_orf1a;2489 | 0 | 0 |
| MT072864.1;QIQ54046.1;Pangolin_CoV_orf1a;2490 | 0 | 0 |
| MT072864.1;QIQ54046.1;Pangolin_CoV_orf1a;2491 | 0 | 0 |
| MT072864.1;QIQ54046.1;Pangolin_CoV_orf1a;2492 | 0 | 0 |
| MT072864.1;QIQ54046.1;Pangolin_CoV_orf1a;2493 | 0 | 0 |
| MT072864.1;QIQ54046.1;Pangolin_CoV_orf1a;2494 | 0 | 0 |
| MT072864.1;QIQ54046.1;Pangolin_CoV_orf1a;2495 | 0 | 0 |
| MT072864.1;QIQ54046.1;Pangolin_CoV_orf1a;2496 | 0.05 | 0.05 |
| MT072864.1;QIQ54046.1;Pangolin_CoV_orf1a;2497 | 0 | 0 |
| MT072864.1;QIQ54046.1;Pangolin_CoV_orf1a;2498 | 0 | 0 |
| MT072864.1;QIQ54046.1;Pangolin_CoV_orf1a;2499 | 0 | 0 |
| MT072864.1;QIQ54046.1;Pangolin_CoV_orf1a;2500 | 0 | 0 |
| MT072864.1;QIQ54046.1;Pangolin_CoV_orf1a;2501 | 0 | 0 |
| MT072864.1;QIQ54046.1;Pangolin_CoV_orf1a;2502 | 0 | 0 |
| MT072864.1;QIQ54046.1;Pangolin_CoV_orf1a;2503 | 0 | 0 |
| MT072864.1;QIQ54046.1;Pangolin_CoV_orf1a;2504 | 0 | 0 |
| MT072864.1;QIQ54046.1;Pangolin_CoV_orf1a;2505 | 0 | 0 |
| MT072864.1;QIQ54046.1;Pangolin_CoV_orf1a;2506 | 0 | 0 |
| MT072864.1;QIQ54046.1;Pangolin_CoV_orf1a;2507 | 0 | 0 |
| MT072864.1;QIQ54046.1;Pangolin_CoV_orf1a;2508 | 0 | 0 |
| MT072864.1;QIQ54046.1;Pangolin_CoV_orf1a;2509 | 0 | 0 |
| MT072864.1;QIQ54046.1;Pangolin_CoV_orf1a;2510 | 0 | 0 |
| MT072864.1;QIQ54046.1;Pangolin_CoV_orf1a;2511 | 0 | 0 |
| MT072864.1;QIQ54046.1;Pangolin_CoV_orf1a;2512 | 0 | 0 |
| MT072864.1;QIQ54046.1;Pangolin_CoV_orf1a;2513 | 0 | 0 |
| MT072864.1;QIQ54046.1;Pangolin_CoV_orf1a;2514 | 0 | 0.025 |
| MT072864.1;QIQ54046.1;Pangolin_CoV_orf1a;2515 | 0.05 | 0.025 |
| MT072864.1;QIQ54046.1;Pangolin_CoV_orf1a;2516 | 0.05 | 0.025 |
| MT072864.1;QIQ54046.1;Pangolin_CoV_orf1a;2517 | 0.05 | 0.025 |
| MT072864.1;QIQ54046.1;Pangolin_CoV_orf1a;2518 | 0 | 0.025 |
| MT072864.1;QIQ54046.1;Pangolin_CoV_orf1a;2519 | 0 | 0 |
| MT072864.1;QIQ54046.1;Pangolin_CoV_orf1a;2520 | 0.05 | 0 |
| MT072864.1;QIQ54046.1;Pangolin_CoV_orf1a;2521 | 0 | 0 |
| MT072864.1;QIQ54046.1;Pangolin_CoV_orf1a;2522 | 0 | 0 |
| MT072864.1;QIQ54046.1;Pangolin_CoV_orf1a;2523 | 0 | 0 |
| MT072864.1;QIQ54046.1;Pangolin_CoV_orf1a;2524 | 0 | 0 |
| MT072864.1;QIQ54046.1;Pangolin_CoV_orf1a;2525 | 0 | 0.05 |
| MT072864.1;QIQ54046.1;Pangolin_CoV_orf1a;2526 | 0 | 0.025 |
| MT072864.1;QIQ54046.1;Pangolin_CoV_orf1a;2527 | 0 | 0.025 |

| | | |
|---|---|---|
| MT072864.1;QIQ54046.1;Pangolin_CoV_orf1a;2528 | 0.05 | 0.025 |
| MT072864.1;QIQ54046.1;Pangolin_CoV_orf1a;2529 | 0.05 | 0.025 |
| MT072864.1;QIQ54046.1;Pangolin_CoV_orf1a;2530 | 0.05 | 0.025 |
| MT072864.1;QIQ54046.1;Pangolin_CoV_orf1a;2531 | 0.05 | 0.025 |
| MT072864.1;QIQ54046.1;Pangolin_CoV_orf1a;2532 | 0 | 0.025 |
| MT072864.1;QIQ54046.1;Pangolin_CoV_orf1a;2533 | 0 | 0.025 |
| MT072864.1;QIQ54046.1;Pangolin_CoV_orf1a;2534 | 0.05 | 0.05 |
| MT072864.1;QIQ54046.1;Pangolin_CoV_orf1a;2535 | 0.05 | 0.025 |
| MT072864.1;QIQ54046.1;Pangolin_CoV_orf1a;2536 | 0 | 0 |
| MT072864.1;QIQ54046.1;Pangolin_CoV_orf1a;2537 | 0 | 0 |
| MT072864.1;QIQ54046.1;Pangolin_CoV_orf1a;2538 | 0 | 0 |
| MT072864.1;QIQ54046.1;Pangolin_CoV_orf1a;2539 | 0 | 0 |
| MT072864.1;QIQ54046.1;Pangolin_CoV_orf1a;2540 | 0 | 0 |
| MT072864.1;QIQ54046.1;Pangolin_CoV_orf1a;2541 | 0 | 0 |
| MT072864.1;QIQ54046.1;Pangolin_CoV_orf1a;2542 | 0 | 0 |
| MT072864.1;QIQ54046.1;Pangolin_CoV_orf1a;2543 | 0.1 | 0 |
| MT072864.1;QIQ54046.1;Pangolin_CoV_orf1a;2544 | 0 | 0 |
| MT072864.1;QIQ54046.1;Pangolin_CoV_orf1a;2545 | 0 | 0.05 |
| MT072864.1;QIQ54046.1;Pangolin_CoV_orf1a;2546 | 0 | 0 |
| MT072864.1;QIQ54046.1;Pangolin_CoV_orf1a;2547 | 0 | 0 |
| MT072864.1;QIQ54046.1;Pangolin_CoV_orf1a;2548 | 0 | 0 |
| MT072864.1;QIQ54046.1;Pangolin_CoV_orf1a;2549 | 0 | 0.025 |
| MT072864.1;QIQ54046.1;Pangolin_CoV_orf1a;2550 | 0 | 0 |
| MT072864.1;QIQ54046.1;Pangolin_CoV_orf1a;2551 | 0 | 0 |
| MT072864.1;QIQ54046.1;Pangolin_CoV_orf1a;2552 | 0 | 0 |
| MT072864.1;QIQ54046.1;Pangolin_CoV_orf1a;2553 | 0 | 0 |
| MT072864.1;QIQ54046.1;Pangolin_CoV_orf1a;2554 | 0 | 0 |
| MT072864.1;QIQ54046.1;Pangolin_CoV_orf1a;2555 | 0 | 0 |
| MT072864.1;QIQ54046.1;Pangolin_CoV_orf1a;2556 | 0 | 0 |
| MT072864.1;QIQ54046.1;Pangolin_CoV_orf1a;2557 | 0 | 0 |
| MT072864.1;QIQ54046.1;Pangolin_CoV_orf1a;2558 | 0 | 0 |
| MT072864.1;QIQ54046.1;Pangolin_CoV_orf1a;2559 | 0 | 0 |
| MT072864.1;QIQ54046.1;Pangolin_CoV_orf1a;2560 | 0 | 0 |
| MT072864.1;QIQ54046.1;Pangolin_CoV_orf1a;2561 | 0 | 0.025 |
| MT072864.1;QIQ54046.1;Pangolin_CoV_orf1a;2562 | 0 | 0.025 |
| MT072864.1;QIQ54046.1;Pangolin_CoV_orf1a;2563 | 0.05 | 0.1 |
| MT072864.1;QIQ54046.1;Pangolin_CoV_orf1a;2564 | 0 | 0.05 |
| MT072864.1;QIQ54046.1;Pangolin_CoV_orf1a;2565 | 0 | 0 |
| MT072864.1;QIQ54046.1;Pangolin_CoV_orf1a;2566 | 0 | 0 |
| MT072864.1;QIQ54046.1;Pangolin_CoV_orf1a;2567 | 0 | 0 |
| MT072864.1;QIQ54046.1;Pangolin_CoV_orf1a;2568 | 0 | 0 |
| MT072864.1;QIQ54046.1;Pangolin_CoV_orf1a;2569 | 0.05 | 0 |
| MT072864.1;QIQ54046.1;Pangolin_CoV_orf1a;2570 | 0 | 0 |

| | | |
|---|---|---|
| MT072864.1;QIQ54046.1;Pangolin_CoV_orf1a;2571 | 0.05 | 0 |
| MT072864.1;QIQ54046.1;Pangolin_CoV_orf1a;2572 | 0 | 0 |
| MT072864.1;QIQ54046.1;Pangolin_CoV_orf1a;2573 | 0 | 0 |
| MT072864.1;QIQ54046.1;Pangolin_CoV_orf1a;2574 | 0 | 0 |
| MT072864.1;QIQ54046.1;Pangolin_CoV_orf1a;2575 | 0 | 0.025 |
| MT072864.1;QIQ54046.1;Pangolin_CoV_orf1a;2576 | 0 | 0.025 |
| MT072864.1;QIQ54046.1;Pangolin_CoV_orf1a;2577 | 0 | 0 |
| MT072864.1;QIQ54046.1;Pangolin_CoV_orf1a;2578 | 0 | 0 |
| MT072864.1;QIQ54046.1;Pangolin_CoV_orf1a;2579 | 0 | 0 |
| MT072864.1;QIQ54046.1;Pangolin_CoV_orf1a;2580 | 0 | 0 |
| MT072864.1;QIQ54046.1;Pangolin_CoV_orf1a;2581 | 0 | 0 |
| MT072864.1;QIQ54046.1;Pangolin_CoV_orf1a;2582 | 0 | 0.025 |
| MT072864.1;QIQ54046.1;Pangolin_CoV_orf1a;2583 | 0 | 0 |
| MT072864.1;QIQ54046.1;Pangolin_CoV_orf1a;2584 | 0 | 0 |
| MT072864.1;QIQ54046.1;Pangolin_CoV_orf1a;2585 | 0.15 | 0.1 |
| MT072864.1;QIQ54046.1;Pangolin_CoV_orf1a;2586 | 0.25 | 0.15 |
| MT072864.1;QIQ54046.1;Pangolin_CoV_orf1a;2587 | 0.25 | 0.075 |
| MT072864.1;QIQ54046.1;Pangolin_CoV_orf1a;2588 | 0.15 | 0.075 |
| MT072864.1;QIQ54046.1;Pangolin_CoV_orf1a;2589 | 0.15 | 0.125 |
| MT072864.1;QIQ54046.1;Pangolin_CoV_orf1a;2590 | 0.1 | 0.075 |
| MT072864.1;QIQ54046.1;Pangolin_CoV_orf1a;2591 | 0.15 | 0.075 |
| MT072864.1;QIQ54046.1;Pangolin_CoV_orf1a;2592 | 0.05 | 0 |
| MT072864.1;QIQ54046.1;Pangolin_CoV_orf1a;2593 | 0 | 0.025 |
| MT072864.1;QIQ54046.1;Pangolin_CoV_orf1a;2594 | 0.05 | 0.025 |
| MT072864.1;QIQ54046.1;Pangolin_CoV_orf1a;2595 | 0 | 0 |
| MT072864.1;QIQ54046.1;Pangolin_CoV_orf1a;2596 | 0 | 0 |
| MT072864.1;QIQ54046.1;Pangolin_CoV_orf1a;2597 | 0 | 0 |
| MT072864.1;QIQ54046.1;Pangolin_CoV_orf1a;2598 | 0 | 0 |
| MT072864.1;QIQ54046.1;Pangolin_CoV_orf1a;2599 | 0 | 0 |
| MT072864.1;QIQ54046.1;Pangolin_CoV_orf1a;2600 | 0 | 0 |
| MT072864.1;QIQ54046.1;Pangolin_CoV_orf1a;2601 | 0.05 | 0 |
| MT072864.1;QIQ54046.1;Pangolin_CoV_orf1a;2602 | 0.05 | 0 |
| MT072864.1;QIQ54046.1;Pangolin_CoV_orf1a;2603 | 0.05 | 0 |
| MT072864.1;QIQ54046.1;Pangolin_CoV_orf1a;2604 | 0.05 | 0.025 |
| MT072864.1;QIQ54046.1;Pangolin_CoV_orf1a;2605 | 0.05 | 0.05 |
| MT072864.1;QIQ54046.1;Pangolin_CoV_orf1a;2606 | 0.05 | 0.075 |
| MT072864.1;QIQ54046.1;Pangolin_CoV_orf1a;2607 | 0.05 | 0.05 |
| MT072864.1;QIQ54046.1;Pangolin_CoV_orf1a;2608 | 0.1 | 0.075 |
| MT072864.1;QIQ54046.1;Pangolin_CoV_orf1a;2609 | 0.1 | 0.05 |
| MT072864.1;QIQ54046.1;Pangolin_CoV_orf1a;2610 | 0.1 | 0.05 |
| MT072864.1;QIQ54046.1;Pangolin_CoV_orf1a;2611 | 0.05 | 0 |
| MT072864.1;QIQ54046.1;Pangolin_CoV_orf1a;2612 | 0 | 0 |
| MT072864.1;QIQ54046.1;Pangolin_CoV_orf1a;2613 | 0 | 0 |

| | | |
|---|---|---|
| MT072864.1;QIQ54046.1;Pangolin_CoV_orf1a;2614 | 0.05 | 0 |
| MT072864.1;QIQ54046.1;Pangolin_CoV_orf1a;2615 | 0 | 0 |
| MT072864.1;QIQ54046.1;Pangolin_CoV_orf1a;2616 | 0 | 0 |
| MT072864.1;QIQ54046.1;Pangolin_CoV_orf1a;2617 | 0 | 0 |
| MT072864.1;QIQ54046.1;Pangolin_CoV_orf1a;2618 | 0 | 0 |
| MT072864.1;QIQ54046.1;Pangolin_CoV_orf1a;2619 | 0 | 0 |
| MT072864.1;QIQ54046.1;Pangolin_CoV_orf1a;2620 | 0 | 0 |
| MT072864.1;QIQ54046.1;Pangolin_CoV_orf1a;2621 | 0 | 0 |
| MT072864.1;QIQ54046.1;Pangolin_CoV_orf1a;2622 | 0 | 0 |
| MT072864.1;QIQ54046.1;Pangolin_CoV_orf1a;2623 | 0 | 0 |
| MT072864.1;QIQ54046.1;Pangolin_CoV_orf1a;2624 | 0 | 0 |
| MT072864.1;QIQ54046.1;Pangolin_CoV_orf1a;2625 | 0 | 0 |
| MT072864.1;QIQ54046.1;Pangolin_CoV_orf1a;2626 | 0 | 0 |
| MT072864.1;QIQ54046.1;Pangolin_CoV_orf1a;2627 | 0 | 0 |
| MT072864.1;QIQ54046.1;Pangolin_CoV_orf1a;2628 | 0 | 0 |
| MT072864.1;QIQ54046.1;Pangolin_CoV_orf1a;2629 | 0 | 0 |
| MT072864.1;QIQ54046.1;Pangolin_CoV_orf1a;2630 | 0 | 0 |
| MT072864.1;QIQ54046.1;Pangolin_CoV_orf1a;2631 | 0 | 0 |
| MT072864.1;QIQ54046.1;Pangolin_CoV_orf1a;2632 | 0 | 0 |
| MT072864.1;QIQ54046.1;Pangolin_CoV_orf1a;2633 | 0 | 0 |
| MT072864.1;QIQ54046.1;Pangolin_CoV_orf1a;2634 | 0 | 0 |
| MT072864.1;QIQ54046.1;Pangolin_CoV_orf1a;2635 | 0 | 0 |
| MT072864.1;QIQ54046.1;Pangolin_CoV_orf1a;2636 | 0 | 0 |
| MT072864.1;QIQ54046.1;Pangolin_CoV_orf1a;2637 | 0 | 0 |
| MT072864.1;QIQ54046.1;Pangolin_CoV_orf1a;2638 | 0 | 0 |
| MT072864.1;QIQ54046.1;Pangolin_CoV_orf1a;2639 | 0 | 0 |
| MT072864.1;QIQ54046.1;Pangolin_CoV_orf1a;2640 | 0 | 0 |
| MT072864.1;QIQ54046.1;Pangolin_CoV_orf1a;2641 | 0 | 0 |
| MT072864.1;QIQ54046.1;Pangolin_CoV_orf1a;2642 | 0 | 0 |
| MT072864.1;QIQ54046.1;Pangolin_CoV_orf1a;2643 | 0 | 0 |
| MT072864.1;QIQ54046.1;Pangolin_CoV_orf1a;2644 | 0 | 0 |
| MT072864.1;QIQ54046.1;Pangolin_CoV_orf1a;2645 | 0 | 0 |
| MT072864.1;QIQ54046.1;Pangolin_CoV_orf1a;2646 | 0 | 0 |
| MT072864.1;QIQ54046.1;Pangolin_CoV_orf1a;2647 | 0 | 0 |
| MT072864.1;QIQ54046.1;Pangolin_CoV_orf1a;2648 | 0 | 0 |
| MT072864.1;QIQ54046.1;Pangolin_CoV_orf1a;2649 | 0 | 0 |
| MT072864.1;QIQ54046.1;Pangolin_CoV_orf1a;2650 | 0 | 0 |
| MT072864.1;QIQ54046.1;Pangolin_CoV_orf1a;2651 | 0 | 0 |
| MT072864.1;QIQ54046.1;Pangolin_CoV_orf1a;2652 | 0 | 0 |
| MT072864.1;QIQ54046.1;Pangolin_CoV_orf1a;2653 | 0.05 | 0 |
| MT072864.1;QIQ54046.1;Pangolin_CoV_orf1a;2654 | 0 | 0 |
| MT072864.1;QIQ54046.1;Pangolin_CoV_orf1a;2655 | 0.05 | 0.05 |
| MT072864.1;QIQ54046.1;Pangolin_CoV_orf1a;2656 | 0.05 | 0.05 |

| | | |
|---|---|---|
| MT072864.1;QIQ54046.1;Pangolin_CoV_orf1a;2657 | 0 | 0 |
| MT072864.1;QIQ54046.1;Pangolin_CoV_orf1a;2658 | 0 | 0 |
| MT072864.1;QIQ54046.1;Pangolin_CoV_orf1a;2659 | 0 | 0.025 |
| MT072864.1;QIQ54046.1;Pangolin_CoV_orf1a;2660 | 0.1 | 0.05 |
| MT072864.1;QIQ54046.1;Pangolin_CoV_orf1a;2661 | 0.05 | 0 |
| MT072864.1;QIQ54046.1;Pangolin_CoV_orf1a;2662 | 0 | 0 |
| MT072864.1;QIQ54046.1;Pangolin_CoV_orf1a;2663 | 0 | 0 |
| MT072864.1;QIQ54046.1;Pangolin_CoV_orf1a;2664 | 0.05 | 0 |
| MT072864.1;QIQ54046.1;Pangolin_CoV_orf1a;2665 | 0 | 0 |
| MT072864.1;QIQ54046.1;Pangolin_CoV_orf1a;2666 | 0 | 0.025 |
| MT072864.1;QIQ54046.1;Pangolin_CoV_orf1a;2667 | 0.05 | 0.075 |
| MT072864.1;QIQ54046.1;Pangolin_CoV_orf1a;2668 | 0.05 | 0.025 |
| MT072864.1;QIQ54046.1;Pangolin_CoV_orf1a;2669 | 0.05 | 0.05 |
| MT072864.1;QIQ54046.1;Pangolin_CoV_orf1a;2670 | 0 | 0 |
| MT072864.1;QIQ54046.1;Pangolin_CoV_orf1a;2671 | 0 | 0 |
| MT072864.1;QIQ54046.1;Pangolin_CoV_orf1a;2672 | 0 | 0 |
| MT072864.1;QIQ54046.1;Pangolin_CoV_orf1a;2673 | 0 | 0 |
| MT072864.1;QIQ54046.1;Pangolin_CoV_orf1a;2674 | 0 | 0 |
| MT072864.1;QIQ54046.1;Pangolin_CoV_orf1a;2675 | 0 | 0 |
| MT072864.1;QIQ54046.1;Pangolin_CoV_orf1a;2676 | 0.05 | 0 |
| MT072864.1;QIQ54046.1;Pangolin_CoV_orf1a;2677 | 0.05 | 0 |
| MT072864.1;QIQ54046.1;Pangolin_CoV_orf1a;2678 | 0 | 0 |
| MT072864.1;QIQ54046.1;Pangolin_CoV_orf1a;2679 | 0 | 0 |
| MT072864.1;QIQ54046.1;Pangolin_CoV_orf1a;2680 | 0 | 0 |
| MT072864.1;QIQ54046.1;Pangolin_CoV_orf1a;2681 | 0.05 | 0.05 |
| MT072864.1;QIQ54046.1;Pangolin_CoV_orf1a;2682 | 0 | 0 |
| MT072864.1;QIQ54046.1;Pangolin_CoV_orf1a;2683 | 0.05 | 0 |
| MT072864.1;QIQ54046.1;Pangolin_CoV_orf1a;2684 | 0 | 0 |
| MT072864.1;QIQ54046.1;Pangolin_CoV_orf1a;2685 | 0 | 0 |
| MT072864.1;QIQ54046.1;Pangolin_CoV_orf1a;2686 | 0 | 0 |
| MT072864.1;QIQ54046.1;Pangolin_CoV_orf1a;2687 | 0 | 0 |
| MT072864.1;QIQ54046.1;Pangolin_CoV_orf1a;2688 | 0 | 0 |
| MT072864.1;QIQ54046.1;Pangolin_CoV_orf1a;2689 | 0 | 0 |
| MT072864.1;QIQ54046.1;Pangolin_CoV_orf1a;2690 | 0 | 0 |
| MT072864.1;QIQ54046.1;Pangolin_CoV_orf1a;2691 | 0 | 0 |
| MT072864.1;QIQ54046.1;Pangolin_CoV_orf1a;2692 | 0 | 0 |
| MT072864.1;QIQ54046.1;Pangolin_CoV_orf1a;2693 | 0 | 0 |
| MT072864.1;QIQ54046.1;Pangolin_CoV_orf1a;2694 | 0 | 0 |
| MT072864.1;QIQ54046.1;Pangolin_CoV_orf1a;2695 | 0 | 0 |
| MT072864.1;QIQ54046.1;Pangolin_CoV_orf1a;2696 | 0 | 0 |
| MT072864.1;QIQ54046.1;Pangolin_CoV_orf1a;2697 | 0 | 0 |
| MT072864.1;QIQ54046.1;Pangolin_CoV_orf1a;2698 | 0 | 0 |
| MT072864.1;QIQ54046.1;Pangolin_CoV_orf1a;2699 | 0 | 0 |

| | | |
|---|---|---|
| MT072864.1;QIQ54046.1;Pangolin_CoV_orf1a;2700 | 0 | 0 |
| MT072864.1;QIQ54046.1;Pangolin_CoV_orf1a;2701 | 0 | 0 |
| MT072864.1;QIQ54046.1;Pangolin_CoV_orf1a;2702 | 0 | 0 |
| MT072864.1;QIQ54046.1;Pangolin_CoV_orf1a;2703 | 0 | 0 |
| MT072864.1;QIQ54046.1;Pangolin_CoV_orf1a;2704 | 0 | 0 |
| MT072864.1;QIQ54046.1;Pangolin_CoV_orf1a;2705 | 0 | 0 |
| MT072864.1;QIQ54046.1;Pangolin_CoV_orf1a;2706 | 0 | 0 |
| MT072864.1;QIQ54046.1;Pangolin_CoV_orf1a;2707 | 0 | 0 |
| MT072864.1;QIQ54046.1;Pangolin_CoV_orf1a;2708 | 0 | 0 |
| MT072864.1;QIQ54046.1;Pangolin_CoV_orf1a;2709 | 0 | 0 |
| MT072864.1;QIQ54046.1;Pangolin_CoV_orf1a;2710 | 0.05 | 0 |
| MT072864.1;QIQ54046.1;Pangolin_CoV_orf1a;2711 | 0 | 0 |
| MT072864.1;QIQ54046.1;Pangolin_CoV_orf1a;2712 | 0 | 0 |
| MT072864.1;QIQ54046.1;Pangolin_CoV_orf1a;2713 | 0 | 0 |
| MT072864.1;QIQ54046.1;Pangolin_CoV_orf1a;2714 | 0 | 0.025 |
| MT072864.1;QIQ54046.1;Pangolin_CoV_orf1a;2715 | 0 | 0 |
| MT072864.1;QIQ54046.1;Pangolin_CoV_orf1a;2716 | 0 | 0 |
| MT072864.1;QIQ54046.1;Pangolin_CoV_orf1a;2717 | 0 | 0 |
| MT072864.1;QIQ54046.1;Pangolin_CoV_orf1a;2718 | 0 | 0.025 |
| MT072864.1;QIQ54046.1;Pangolin_CoV_orf1a;2719 | 0 | 0.025 |
| MT072864.1;QIQ54046.1;Pangolin_CoV_orf1a;2720 | 0 | 0.025 |
| MT072864.1;QIQ54046.1;Pangolin_CoV_orf1a;2721 | 0.05 | 0.025 |
| MT072864.1;QIQ54046.1;Pangolin_CoV_orf1a;2722 | 0 | 0.025 |
| MT072864.1;QIQ54046.1;Pangolin_CoV_orf1a;2723 | 0 | 0.025 |
| MT072864.1;QIQ54046.1;Pangolin_CoV_orf1a;2724 | 0 | 0.025 |
| MT072864.1;QIQ54046.1;Pangolin_CoV_orf1a;2725 | 0 | 0.025 |
| MT072864.1;QIQ54046.1;Pangolin_CoV_orf1a;2726 | 0 | 0 |
| MT072864.1;QIQ54046.1;Pangolin_CoV_orf1a;2727 | 0 | 0 |
| MT072864.1;QIQ54046.1;Pangolin_CoV_orf1a;2728 | 0 | 0 |
| MT072864.1;QIQ54046.1;Pangolin_CoV_orf1a;2729 | 0 | 0 |
| MT072864.1;QIQ54046.1;Pangolin_CoV_orf1a;2730 | 0 | 0.025 |
| MT072864.1;QIQ54046.1;Pangolin_CoV_orf1a;2731 | 0 | 0 |
| MT072864.1;QIQ54046.1;Pangolin_CoV_orf1a;2732 | 0 | 0 |
| MT072864.1;QIQ54046.1;Pangolin_CoV_orf1a;2733 | 0 | 0.025 |
| MT072864.1;QIQ54046.1;Pangolin_CoV_orf1a;2734 | 0 | 0 |
| MT072864.1;QIQ54046.1;Pangolin_CoV_orf1a;2735 | 0 | 0.025 |
| MT072864.1;QIQ54046.1;Pangolin_CoV_orf1a;2736 | 0 | 0.025 |
| MT072864.1;QIQ54046.1;Pangolin_CoV_orf1a;2737 | 0 | 0 |
| MT072864.1;QIQ54046.1;Pangolin_CoV_orf1a;2738 | 0 | 0 |
| MT072864.1;QIQ54046.1;Pangolin_CoV_orf1a;2739 | 0 | 0 |
| MT072864.1;QIQ54046.1;Pangolin_CoV_orf1a;2740 | 0 | 0 |
| MT072864.1;QIQ54046.1;Pangolin_CoV_orf1a;2741 | 0 | 0 |
| MT072864.1;QIQ54046.1;Pangolin_CoV_orf1a;2742 | 0 | 0 |

| | | |
|---|---|---|
| MT072864.1;QIQ54046.1;Pangolin_CoV_orf1a;2743 | 0 | 0 |
| MT072864.1;QIQ54046.1;Pangolin_CoV_orf1a;2744 | 0 | 0 |
| MT072864.1;QIQ54046.1;Pangolin_CoV_orf1a;2745 | 0 | 0 |
| MT072864.1;QIQ54046.1;Pangolin_CoV_orf1a;2746 | 0 | 0 |
| MT072864.1;QIQ54046.1;Pangolin_CoV_orf1a;2747 | 0 | 0 |
| MT072864.1;QIQ54046.1;Pangolin_CoV_orf1a;2748 | 0 | 0 |
| MT072864.1;QIQ54046.1;Pangolin_CoV_orf1a;2749 | 0 | 0 |
| MT072864.1;QIQ54046.1;Pangolin_CoV_orf1a;2750 | 0 | 0 |
| MT072864.1;QIQ54046.1;Pangolin_CoV_orf1a;2751 | 0 | 0 |
| MT072864.1;QIQ54046.1;Pangolin_CoV_orf1a;2752 | 0 | 0 |
| MT072864.1;QIQ54046.1;Pangolin_CoV_orf1a;2753 | 0 | 0 |
| MT072864.1;QIQ54046.1;Pangolin_CoV_orf1a;2754 | 0 | 0 |
| MT072864.1;QIQ54046.1;Pangolin_CoV_orf1a;2755 | 0 | 0 |
| MT072864.1;QIQ54046.1;Pangolin_CoV_orf1a;2756 | 0 | 0 |
| MT072864.1;QIQ54046.1;Pangolin_CoV_orf1a;2757 | 0 | 0 |
| MT072864.1;QIQ54046.1;Pangolin_CoV_orf1a;2758 | 0 | 0 |
| MT072864.1;QIQ54046.1;Pangolin_CoV_orf1a;2759 | 0 | 0 |
| MT072864.1;QIQ54046.1;Pangolin_CoV_orf1a;2760 | 0 | 0 |
| MT072864.1;QIQ54046.1;Pangolin_CoV_orf1a;2761 | 0 | 0 |
| MT072864.1;QIQ54046.1;Pangolin_CoV_orf1a;2762 | 0 | 0 |
| MT072864.1;QIQ54046.1;Pangolin_CoV_orf1a;2763 | 0 | 0 |
| MT072864.1;QIQ54046.1;Pangolin_CoV_orf1a;2764 | 0 | 0 |
| MT072864.1;QIQ54046.1;Pangolin_CoV_orf1a;2765 | 0 | 0 |
| MT072864.1;QIQ54046.1;Pangolin_CoV_orf1a;2766 | 0 | 0 |
| MT072864.1;QIQ54046.1;Pangolin_CoV_orf1a;2767 | 0 | 0 |
| MT072864.1;QIQ54046.1;Pangolin_CoV_orf1a;2768 | 0 | 0 |
| MT072864.1;QIQ54046.1;Pangolin_CoV_orf1a;2769 | 0 | 0 |
| MT072864.1;QIQ54046.1;Pangolin_CoV_orf1a;2770 | 0 | 0 |
| MT072864.1;QIQ54046.1;Pangolin_CoV_orf1a;2771 | 0 | 0 |
| MT072864.1;QIQ54046.1;Pangolin_CoV_orf1a;2772 | 0 | 0 |
| MT072864.1;QIQ54046.1;Pangolin_CoV_orf1a;2773 | 0 | 0 |
| MT072864.1;QIQ54046.1;Pangolin_CoV_orf1a;2774 | 0 | 0 |
| MT072864.1;QIQ54046.1;Pangolin_CoV_orf1a;2775 | 0 | 0 |
| MT072864.1;QIQ54046.1;Pangolin_CoV_orf1a;2776 | 0 | 0 |
| MT072864.1;QIQ54046.1;Pangolin_CoV_orf1a;2777 | 0 | 0 |
| MT072864.1;QIQ54046.1;Pangolin_CoV_orf1a;2778 | 0 | 0.025 |
| MT072864.1;QIQ54046.1;Pangolin_CoV_orf1a;2779 | 0 | 0 |
| MT072864.1;QIQ54046.1;Pangolin_CoV_orf1a;2780 | 0 | 0 |
| MT072864.1;QIQ54046.1;Pangolin_CoV_orf1a;2781 | 0 | 0 |
| MT072864.1;QIQ54046.1;Pangolin_CoV_orf1a;2782 | 0 | 0 |
| MT072864.1;QIQ54046.1;Pangolin_CoV_orf1a;2783 | 0 | 0.025 |
| MT072864.1;QIQ54046.1;Pangolin_CoV_orf1a;2784 | 0 | 0.025 |
| MT072864.1;QIQ54046.1;Pangolin_CoV_orf1a;2785 | 0 | 0.025 |

| | | |
|---|---|---|
| MT072864.1;QIQ54046.1;Pangolin_CoV_orf1a;2786 | 0 | 0.025 |
| MT072864.1;QIQ54046.1;Pangolin_CoV_orf1a;2787 | 0 | 0.025 |
| MT072864.1;QIQ54046.1;Pangolin_CoV_orf1a;2788 | 0.05 | 0.025 |
| MT072864.1;QIQ54046.1;Pangolin_CoV_orf1a;2789 | 0 | 0.05 |
| MT072864.1;QIQ54046.1;Pangolin_CoV_orf1a;2790 | 0 | 0.05 |
| MT072864.1;QIQ54046.1;Pangolin_CoV_orf1a;2791 | 0 | 0.025 |
| MT072864.1;QIQ54046.1;Pangolin_CoV_orf1a;2792 | 0 | 0.025 |
| MT072864.1;QIQ54046.1;Pangolin_CoV_orf1a;2793 | 0 | 0.025 |
| MT072864.1;QIQ54046.1;Pangolin_CoV_orf1a;2794 | 0.05 | 0.025 |
| MT072864.1;QIQ54046.1;Pangolin_CoV_orf1a;2795 | 0.05 | 0.025 |
| MT072864.1;QIQ54046.1;Pangolin_CoV_orf1a;2796 | 0.1 | 0 |
| MT072864.1;QIQ54046.1;Pangolin_CoV_orf1a;2797 | 0.15 | 0.025 |
| MT072864.1;QIQ54046.1;Pangolin_CoV_orf1a;2798 | 0.15 | 0 |
| MT072864.1;QIQ54046.1;Pangolin_CoV_orf1a;2799 | 0.1 | 0 |
| MT072864.1;QIQ54046.1;Pangolin_CoV_orf1a;2800 | 0.1 | 0 |
| MT072864.1;QIQ54046.1;Pangolin_CoV_orf1a;2801 | 0.05 | 0 |
| MT072864.1;QIQ54046.1;Pangolin_CoV_orf1a;2802 | 0.1 | 0 |
| MT072864.1;QIQ54046.1;Pangolin_CoV_orf1a;2803 | 0.3 | 0.1 |
| MT072864.1;QIQ54046.1;Pangolin_CoV_orf1a;2804 | 0.15 | 0 |
| MT072864.1;QIQ54046.1;Pangolin_CoV_orf1a;2805 | 0.05 | 0.05 |
| MT072864.1;QIQ54046.1;Pangolin_CoV_orf1a;2806 | 0 | 0 |
| MT072864.1;QIQ54046.1;Pangolin_CoV_orf1a;2807 | 0.15 | 0.05 |
| MT072864.1;QIQ54046.1;Pangolin_CoV_orf1a;2808 | 0 | 0 |
| MT072864.1;QIQ54046.1;Pangolin_CoV_orf1a;2809 | 0 | 0.025 |
| MT072864.1;QIQ54046.1;Pangolin_CoV_orf1a;2810 | 0 | 0 |
| MT072864.1;QIQ54046.1;Pangolin_CoV_orf1a;2811 | 0.05 | 0.025 |
| MT072864.1;QIQ54046.1;Pangolin_CoV_orf1a;2812 | 0 | 0.075 |
| MT072864.1;QIQ54046.1;Pangolin_CoV_orf1a;2813 | 0.15 | 0.125 |
| MT072864.1;QIQ54046.1;Pangolin_CoV_orf1a;2814 | 0 | 0.025 |
| MT072864.1;QIQ54046.1;Pangolin_CoV_orf1a;2815 | 0 | 0 |
| MT072864.1;QIQ54046.1;Pangolin_CoV_orf1a;2816 | 0 | 0 |
| MT072864.1;QIQ54046.1;Pangolin_CoV_orf1a;2817 | 0 | 0 |
| MT072864.1;QIQ54046.1;Pangolin_CoV_orf1a;2818 | 0 | 0 |
| MT072864.1;QIQ54046.1;Pangolin_CoV_orf1a;2819 | 0 | 0 |
| MT072864.1;QIQ54046.1;Pangolin_CoV_orf1a;2820 | 0.05 | 0.025 |
| MT072864.1;QIQ54046.1;Pangolin_CoV_orf1a;2821 | 0 | 0 |
| MT072864.1;QIQ54046.1;Pangolin_CoV_orf1a;2822 | 0 | 0 |
| MT072864.1;QIQ54046.1;Pangolin_CoV_orf1a;2823 | 0 | 0 |
| MT072864.1;QIQ54046.1;Pangolin_CoV_orf1a;2824 | 0 | 0 |
| MT072864.1;QIQ54046.1;Pangolin_CoV_orf1a;2825 | 0 | 0 |
| MT072864.1;QIQ54046.1;Pangolin_CoV_orf1a;2826 | 0 | 0 |
| MT072864.1;QIQ54046.1;Pangolin_CoV_orf1a;2827 | 0.15 | 0 |
| MT072864.1;QIQ54046.1;Pangolin_CoV_orf1a;2828 | 0 | 0 |

| | | |
|---|---|---|
| MT072864.1;QIQ54046.1;Pangolin_CoV_orf1a;2829 | 0 | 0 |
| MT072864.1;QIQ54046.1;Pangolin_CoV_orf1a;2830 | 0 | 0 |
| MT072864.1;QIQ54046.1;Pangolin_CoV_orf1a;2831 | 0.05 | 0 |
| MT072864.1;QIQ54046.1;Pangolin_CoV_orf1a;2832 | 0 | 0.05 |
| MT072864.1;QIQ54046.1;Pangolin_CoV_orf1a;2833 | 0.05 | 0.025 |
| MT072864.1;QIQ54046.1;Pangolin_CoV_orf1a;2834 | 0 | 0 |
| MT072864.1;QIQ54046.1;Pangolin_CoV_orf1a;2835 | 0 | 0 |
| MT072864.1;QIQ54046.1;Pangolin_CoV_orf1a;2836 | 0 | 0.05 |
| MT072864.1;QIQ54046.1;Pangolin_CoV_orf1a;2837 | 0 | 0 |
| MT072864.1;QIQ54046.1;Pangolin_CoV_orf1a;2838 | 0 | 0 |
| MT072864.1;QIQ54046.1;Pangolin_CoV_orf1a;2839 | 0 | 0 |
| MT072864.1;QIQ54046.1;Pangolin_CoV_orf1a;2840 | 0 | 0 |
| MT072864.1;QIQ54046.1;Pangolin_CoV_orf1a;2841 | 0 | 0 |
| MT072864.1;QIQ54046.1;Pangolin_CoV_orf1a;2842 | 0 | 0 |
| MT072864.1;QIQ54046.1;Pangolin_CoV_orf1a;2843 | 0 | 0 |
| MT072864.1;QIQ54046.1;Pangolin_CoV_orf1a;2844 | 0 | 0 |
| MT072864.1;QIQ54046.1;Pangolin_CoV_orf1a;2845 | 0 | 0 |
| MT072864.1;QIQ54046.1;Pangolin_CoV_orf1a;2846 | 0 | 0 |
| MT072864.1;QIQ54046.1;Pangolin_CoV_orf1a;2847 | 0 | 0.05 |
| MT072864.1;QIQ54046.1;Pangolin_CoV_orf1a;2848 | 0 | 0.125 |
| MT072864.1;QIQ54046.1;Pangolin_CoV_orf1a;2849 | 0.05 | 0.1 |
| MT072864.1;QIQ54046.1;Pangolin_CoV_orf1a;2850 | 0.05 | 0.075 |
| MT072864.1;QIQ54046.1;Pangolin_CoV_orf1a;2851 | 0.1 | 0.05 |
| MT072864.1;QIQ54046.1;Pangolin_CoV_orf1a;2852 | 0.05 | 0.025 |
| MT072864.1;QIQ54046.1;Pangolin_CoV_orf1a;2853 | 0 | 0 |
| MT072864.1;QIQ54046.1;Pangolin_CoV_orf1a;2854 | 0 | 0 |
| MT072864.1;QIQ54046.1;Pangolin_CoV_orf1a;2855 | 0 | 0 |
| MT072864.1;QIQ54046.1;Pangolin_CoV_orf1a;2856 | 0 | 0 |
| MT072864.1;QIQ54046.1;Pangolin_CoV_orf1a;2857 | 0 | 0 |
| MT072864.1;QIQ54046.1;Pangolin_CoV_orf1a;2858 | 0 | 0 |
| MT072864.1;QIQ54046.1;Pangolin_CoV_orf1a;2859 | 0 | 0 |
| MT072864.1;QIQ54046.1;Pangolin_CoV_orf1a;2860 | 0 | 0 |
| MT072864.1;QIQ54046.1;Pangolin_CoV_orf1a;2861 | 0 | 0 |
| MT072864.1;QIQ54046.1;Pangolin_CoV_orf1a;2862 | 0 | 0 |
| MT072864.1;QIQ54046.1;Pangolin_CoV_orf1a;2863 | 0 | 0 |
| MT072864.1;QIQ54046.1;Pangolin_CoV_orf1a;2864 | 0 | 0 |
| MT072864.1;QIQ54046.1;Pangolin_CoV_orf1a;2865 | 0 | 0 |
| MT072864.1;QIQ54046.1;Pangolin_CoV_orf1a;2866 | 0 | 0 |
| MT072864.1;QIQ54046.1;Pangolin_CoV_orf1a;2867 | 0 | 0 |
| MT072864.1;QIQ54046.1;Pangolin_CoV_orf1a;2868 | 0 | 0 |
| MT072864.1;QIQ54046.1;Pangolin_CoV_orf1a;2869 | 0 | 0 |
| MT072864.1;QIQ54046.1;Pangolin_CoV_orf1a;2870 | 0 | 0 |
| MT072864.1;QIQ54046.1;Pangolin_CoV_orf1a;2871 | 0 | 0 |

| | | |
|---|---|---|
| MT072864.1;QIQ54046.1;Pangolin_CoV_orf1a;2872 | 0 | 0 |
| MT072864.1;QIQ54046.1;Pangolin_CoV_orf1a;2873 | 0 | 0 |
| MT072864.1;QIQ54046.1;Pangolin_CoV_orf1a;2874 | 0 | 0 |
| MT072864.1;QIQ54046.1;Pangolin_CoV_orf1a;2875 | 0 | 0 |
| MT072864.1;QIQ54046.1;Pangolin_CoV_orf1a;2876 | 0 | 0 |
| MT072864.1;QIQ54046.1;Pangolin_CoV_orf1a;2877 | 0 | 0 |
| MT072864.1;QIQ54046.1;Pangolin_CoV_orf1a;2878 | 0 | 0 |
| MT072864.1;QIQ54046.1;Pangolin_CoV_orf1a;2879 | 0 | 0 |
| MT072864.1;QIQ54046.1;Pangolin_CoV_orf1a;2880 | 0 | 0 |
| MT072864.1;QIQ54046.1;Pangolin_CoV_orf1a;2881 | 0 | 0 |
| MT072864.1;QIQ54046.1;Pangolin_CoV_orf1a;2882 | 0 | 0 |
| MT072864.1;QIQ54046.1;Pangolin_CoV_orf1a;2883 | 0.05 | 0 |
| MT072864.1;QIQ54046.1;Pangolin_CoV_orf1a;2884 | 0.05 | 0.025 |
| MT072864.1;QIQ54046.1;Pangolin_CoV_orf1a;2885 | 0 | 0.025 |
| MT072864.1;QIQ54046.1;Pangolin_CoV_orf1a;2886 | 0 | 0.025 |
| MT072864.1;QIQ54046.1;Pangolin_CoV_orf1a;2887 | 0.05 | 0 |
| MT072864.1;QIQ54046.1;Pangolin_CoV_orf1a;2888 | 0.05 | 0.05 |
| MT072864.1;QIQ54046.1;Pangolin_CoV_orf1a;2889 | 0.05 | 0.025 |
| MT072864.1;QIQ54046.1;Pangolin_CoV_orf1a;2890 | 0.05 | 0 |
| MT072864.1;QIQ54046.1;Pangolin_CoV_orf1a;2891 | 0 | 0 |
| MT072864.1;QIQ54046.1;Pangolin_CoV_orf1a;2892 | 0 | 0 |
| MT072864.1;QIQ54046.1;Pangolin_CoV_orf1a;2893 | 0 | 0 |
| MT072864.1;QIQ54046.1;Pangolin_CoV_orf1a;2894 | 0 | 0 |
| MT072864.1;QIQ54046.1;Pangolin_CoV_orf1a;2895 | 0.05 | 0 |
| MT072864.1;QIQ54046.1;Pangolin_CoV_orf1a;2896 | 0 | 0 |
| MT072864.1;QIQ54046.1;Pangolin_CoV_orf1a;2897 | 0 | 0 |
| MT072864.1;QIQ54046.1;Pangolin_CoV_orf1a;2898 | 0 | 0 |
| MT072864.1;QIQ54046.1;Pangolin_CoV_orf1a;2899 | 0 | 0 |
| MT072864.1;QIQ54046.1;Pangolin_CoV_orf1a;2900 | 0 | 0 |
| MT072864.1;QIQ54046.1;Pangolin_CoV_orf1a;2901 | 0 | 0 |
| MT072864.1;QIQ54046.1;Pangolin_CoV_orf1a;2902 | 0 | 0 |
| MT072864.1;QIQ54046.1;Pangolin_CoV_orf1a;2903 | 0.05 | 0.025 |
| MT072864.1;QIQ54046.1;Pangolin_CoV_orf1a;2904 | 0 | 0.025 |
| MT072864.1;QIQ54046.1;Pangolin_CoV_orf1a;2905 | 0.05 | 0.05 |
| MT072864.1;QIQ54046.1;Pangolin_CoV_orf1a;2906 | 0.15 | 0.05 |
| MT072864.1;QIQ54046.1;Pangolin_CoV_orf1a;2907 | 0.3 | 0.05 |
| MT072864.1;QIQ54046.1;Pangolin_CoV_orf1a;2908 | 0.25 | 0.125 |
| MT072864.1;QIQ54046.1;Pangolin_CoV_orf1a;2909 | 0.25 | 0.05 |
| MT072864.1;QIQ54046.1;Pangolin_CoV_orf1a;2910 | 0.3 | 0.05 |
| MT072864.1;QIQ54046.1;Pangolin_CoV_orf1a;2911 | 0.05 | 0 |
| MT072864.1;QIQ54046.1;Pangolin_CoV_orf1a;2912 | 0.2 | 0.05 |
| MT072864.1;QIQ54046.1;Pangolin_CoV_orf1a;2913 | 0.3 | 0.1 |
| MT072864.1;QIQ54046.1;Pangolin_CoV_orf1a;2914 | 0.2 | 0.075 |

| | | |
|---|---|---|
| MT072864.1;QIQ54046.1;Pangolin_CoV_orf1a;2915 | 0 | 0.025 |
| MT072864.1;QIQ54046.1;Pangolin_CoV_orf1a;2916 | 0.05 | 0.05 |
| MT072864.1;QIQ54046.1;Pangolin_CoV_orf1a;2917 | 0 | 0 |
| MT072864.1;QIQ54046.1;Pangolin_CoV_orf1a;2918 | 0 | 0 |
| MT072864.1;QIQ54046.1;Pangolin_CoV_orf1a;2919 | 0 | 0 |
| MT072864.1;QIQ54046.1;Pangolin_CoV_orf1a;2920 | 0 | 0 |
| MT072864.1;QIQ54046.1;Pangolin_CoV_orf1a;2921 | 0 | 0 |
| MT072864.1;QIQ54046.1;Pangolin_CoV_orf1a;2922 | 0 | 0 |
| MT072864.1;QIQ54046.1;Pangolin_CoV_orf1a;2923 | 0 | 0 |
| MT072864.1;QIQ54046.1;Pangolin_CoV_orf1a;2924 | 0 | 0 |
| MT072864.1;QIQ54046.1;Pangolin_CoV_orf1a;2925 | 0 | 0 |
| MT072864.1;QIQ54046.1;Pangolin_CoV_orf1a;2926 | 0.05 | 0 |
| MT072864.1;QIQ54046.1;Pangolin_CoV_orf1a;2927 | 0.05 | 0 |
| MT072864.1;QIQ54046.1;Pangolin_CoV_orf1a;2928 | 0.1 | 0 |
| MT072864.1;QIQ54046.1;Pangolin_CoV_orf1a;2929 | 0 | 0.025 |
| MT072864.1;QIQ54046.1;Pangolin_CoV_orf1a;2930 | 0 | 0.05 |
| MT072864.1;QIQ54046.1;Pangolin_CoV_orf1a;2931 | 0.2 | 0.15 |
| MT072864.1;QIQ54046.1;Pangolin_CoV_orf1a;2932 | 0.15 | 0.175 |
| MT072864.1;QIQ54046.1;Pangolin_CoV_orf1a;2933 | 0.1 | 0.075 |
| MT072864.1;QIQ54046.1;Pangolin_CoV_orf1a;2934 | 0.1 | 0.025 |
| MT072864.1;QIQ54046.1;Pangolin_CoV_orf1a;2935 | 0.15 | 0.075 |
| MT072864.1;QIQ54046.1;Pangolin_CoV_orf1a;2936 | 0.05 | 0.1 |
| MT072864.1;QIQ54046.1;Pangolin_CoV_orf1a;2937 | 0.1 | 0.1 |
| MT072864.1;QIQ54046.1;Pangolin_CoV_orf1a;2938 | 0.15 | 0.025 |
| MT072864.1;QIQ54046.1;Pangolin_CoV_orf1a;2939 | 0.05 | 0.025 |
| MT072864.1;QIQ54046.1;Pangolin_CoV_orf1a;2940 | 0.05 | 0.05 |
| MT072864.1;QIQ54046.1;Pangolin_CoV_orf1a;2941 | 0.05 | 0.025 |
| MT072864.1;QIQ54046.1;Pangolin_CoV_orf1a;2942 | 0 | 0 |
| MT072864.1;QIQ54046.1;Pangolin_CoV_orf1a;2943 | 0 | 0 |
| MT072864.1;QIQ54046.1;Pangolin_CoV_orf1a;2944 | 0 | 0 |
| MT072864.1;QIQ54046.1;Pangolin_CoV_orf1a;2945 | 0 | 0 |
| MT072864.1;QIQ54046.1;Pangolin_CoV_orf1a;2946 | 0 | 0 |
| MT072864.1;QIQ54046.1;Pangolin_CoV_orf1a;2947 | 0 | 0 |
| MT072864.1;QIQ54046.1;Pangolin_CoV_orf1a;2948 | 0 | 0 |
| MT072864.1;QIQ54046.1;Pangolin_CoV_orf1a;2949 | 0 | 0 |
| MT072864.1;QIQ54046.1;Pangolin_CoV_orf1a;2950 | 0 | 0 |
| MT072864.1;QIQ54046.1;Pangolin_CoV_orf1a;2951 | 0 | 0 |
| MT072864.1;QIQ54046.1;Pangolin_CoV_orf1a;2952 | 0 | 0.025 |
| MT072864.1;QIQ54046.1;Pangolin_CoV_orf1a;2953 | 0 | 0 |
| MT072864.1;QIQ54046.1;Pangolin_CoV_orf1a;2954 | 0 | 0 |
| MT072864.1;QIQ54046.1;Pangolin_CoV_orf1a;2955 | 0 | 0 |
| MT072864.1;QIQ54046.1;Pangolin_CoV_orf1a;2956 | 0 | 0 |
| MT072864.1;QIQ54046.1;Pangolin_CoV_orf1a;2957 | 0 | 0 |

| | | |
|---|---|---|
| MT072864.1;QIQ54046.1;Pangolin_CoV_orf1a;2958 | 0 | 0 |
| MT072864.1;QIQ54046.1;Pangolin_CoV_orf1a;2959 | 0 | 0 |
| MT072864.1;QIQ54046.1;Pangolin_CoV_orf1a;2960 | 0 | 0 |
| MT072864.1;QIQ54046.1;Pangolin_CoV_orf1a;2961 | 0 | 0 |
| MT072864.1;QIQ54046.1;Pangolin_CoV_orf1a;2962 | 0 | 0 |
| MT072864.1;QIQ54046.1;Pangolin_CoV_orf1a;2963 | 0.05 | 0.1 |
| MT072864.1;QIQ54046.1;Pangolin_CoV_orf1a;2964 | 0 | 0.05 |
| MT072864.1;QIQ54046.1;Pangolin_CoV_orf1a;2965 | 0 | 0.075 |
| MT072864.1;QIQ54046.1;Pangolin_CoV_orf1a;2966 | 0 | 0.075 |
| MT072864.1;QIQ54046.1;Pangolin_CoV_orf1a;2967 | 0 | 0.05 |
| MT072864.1;QIQ54046.1;Pangolin_CoV_orf1a;2968 | 0 | 0.05 |
| MT072864.1;QIQ54046.1;Pangolin_CoV_orf1a;2969 | 0 | 0.05 |
| MT072864.1;QIQ54046.1;Pangolin_CoV_orf1a;2970 | 0 | 0.05 |
| MT072864.1;QIQ54046.1;Pangolin_CoV_orf1a;2971 | 0.05 | 0.05 |
| MT072864.1;QIQ54046.1;Pangolin_CoV_orf1a;2972 | 0 | 0 |
| MT072864.1;QIQ54046.1;Pangolin_CoV_orf1a;2973 | 0 | 0 |
| MT072864.1;QIQ54046.1;Pangolin_CoV_orf1a;2974 | 0 | 0 |
| MT072864.1;QIQ54046.1;Pangolin_CoV_orf1a;2975 | 0 | 0 |
| MT072864.1;QIQ54046.1;Pangolin_CoV_orf1a;2976 | 0 | 0 |
| MT072864.1;QIQ54046.1;Pangolin_CoV_orf1a;2977 | 0 | 0 |
| MT072864.1;QIQ54046.1;Pangolin_CoV_orf1a;2978 | 0 | 0 |
| MT072864.1;QIQ54046.1;Pangolin_CoV_orf1a;2979 | 0 | 0 |
| MT072864.1;QIQ54046.1;Pangolin_CoV_orf1a;2980 | 0 | 0 |
| MT072864.1;QIQ54046.1;Pangolin_CoV_orf1a;2981 | 0 | 0 |
| MT072864.1;QIQ54046.1;Pangolin_CoV_orf1a;2982 | 0 | 0 |
| MT072864.1;QIQ54046.1;Pangolin_CoV_orf1a;2983 | 0 | 0 |
| MT072864.1;QIQ54046.1;Pangolin_CoV_orf1a;2984 | 0 | 0 |
| MT072864.1;QIQ54046.1;Pangolin_CoV_orf1a;2985 | 0 | 0 |
| MT072864.1;QIQ54046.1;Pangolin_CoV_orf1a;2986 | 0 | 0 |
| MT072864.1;QIQ54046.1;Pangolin_CoV_orf1a;2987 | 0 | 0 |
| MT072864.1;QIQ54046.1;Pangolin_CoV_orf1a;2988 | 0 | 0 |
| MT072864.1;QIQ54046.1;Pangolin_CoV_orf1a;2989 | 0 | 0 |
| MT072864.1;QIQ54046.1;Pangolin_CoV_orf1a;2990 | 0 | 0 |
| MT072864.1;QIQ54046.1;Pangolin_CoV_orf1a;2991 | 0 | 0 |
| MT072864.1;QIQ54046.1;Pangolin_CoV_orf1a;2992 | 0 | 0 |
| MT072864.1;QIQ54046.1;Pangolin_CoV_orf1a;2993 | 0 | 0 |
| MT072864.1;QIQ54046.1;Pangolin_CoV_orf1a;2994 | 0 | 0 |
| MT072864.1;QIQ54046.1;Pangolin_CoV_orf1a;2995 | 0 | 0 |
| MT072864.1;QIQ54046.1;Pangolin_CoV_orf1a;2996 | 0 | 0 |
| MT072864.1;QIQ54046.1;Pangolin_CoV_orf1a;2997 | 0 | 0 |
| MT072864.1;QIQ54046.1;Pangolin_CoV_orf1a;2998 | 0 | 0 |
| MT072864.1;QIQ54046.1;Pangolin_CoV_orf1a;2999 | 0 | 0 |
| MT072864.1;QIQ54046.1;Pangolin_CoV_orf1a;3000 | 0 | 0 |

| | | |
|---|---|---|
| MT072864.1;QIQ54046.1;Pangolin_CoV_orf1a;3001 | 0 | 0 |
| MT072864.1;QIQ54046.1;Pangolin_CoV_orf1a;3002 | 0.15 | 0.15 |
| MT072864.1;QIQ54046.1;Pangolin_CoV_orf1a;3003 | 0.15 | 0.15 |
| MT072864.1;QIQ54046.1;Pangolin_CoV_orf1a;3004 | 0.1 | 0.125 |
| MT072864.1;QIQ54046.1;Pangolin_CoV_orf1a;3005 | 0.1 | 0.15 |
| MT072864.1;QIQ54046.1;Pangolin_CoV_orf1a;3006 | 0.1 | 0.15 |
| MT072864.1;QIQ54046.1;Pangolin_CoV_orf1a;3007 | 0.1 | 0.15 |
| MT072864.1;QIQ54046.1;Pangolin_CoV_orf1a;3008 | 0.1 | 0.125 |
| MT072864.1;QIQ54046.1;Pangolin_CoV_orf1a;3009 | 0.1 | 0.15 |
| MT072864.1;QIQ54046.1;Pangolin_CoV_orf1a;3010 | 0.1 | 0.15 |
| MT072864.1;QIQ54046.1;Pangolin_CoV_orf1a;3011 | 0.15 | 0.25 |
| MT072864.1;QIQ54046.1;Pangolin_CoV_orf1a;3012 | 0.35 | 0.325 |
| MT072864.1;QIQ54046.1;Pangolin_CoV_orf1a;3013 | 0.25 | 0.225 |
| MT072864.1;QIQ54046.1;Pangolin_CoV_orf1a;3014 | 0 | 0.025 |
| MT072864.1;QIQ54046.1;Pangolin_CoV_orf1a;3015 | 0 | 0 |
| MT072864.1;QIQ54046.1;Pangolin_CoV_orf1a;3016 | 0 | 0 |
| MT072864.1;QIQ54046.1;Pangolin_CoV_orf1a;3017 | 0.05 | 0 |
| MT072864.1;QIQ54046.1;Pangolin_CoV_orf1a;3018 | 0 | 0 |
| MT072864.1;QIQ54046.1;Pangolin_CoV_orf1a;3019 | 0.05 | 0 |
| MT072864.1;QIQ54046.1;Pangolin_CoV_orf1a;3020 | 0.05 | 0 |
| MT072864.1;QIQ54046.1;Pangolin_CoV_orf1a;3021 | 0.05 | 0 |
| MT072864.1;QIQ54046.1;Pangolin_CoV_orf1a;3022 | 0.05 | 0 |
| MT072864.1;QIQ54046.1;Pangolin_CoV_orf1a;3023 | 0.05 | 0 |
| MT072864.1;QIQ54046.1;Pangolin_CoV_orf1a;3024 | 0.05 | 0 |
| MT072864.1;QIQ54046.1;Pangolin_CoV_orf1a;3025 | 0 | 0 |
| MT072864.1;QIQ54046.1;Pangolin_CoV_orf1a;3026 | 0 | 0 |
| MT072864.1;QIQ54046.1;Pangolin_CoV_orf1a;3027 | 0 | 0 |
| MT072864.1;QIQ54046.1;Pangolin_CoV_orf1a;3028 | 0.05 | 0 |
| MT072864.1;QIQ54046.1;Pangolin_CoV_orf1a;3029 | 0 | 0 |
| MT072864.1;QIQ54046.1;Pangolin_CoV_orf1a;3030 | 0 | 0 |
| MT072864.1;QIQ54046.1;Pangolin_CoV_orf1a;3031 | 0 | 0.05 |
| MT072864.1;QIQ54046.1;Pangolin_CoV_orf1a;3032 | 0 | 0 |
| MT072864.1;QIQ54046.1;Pangolin_CoV_orf1a;3033 | 0 | 0 |
| MT072864.1;QIQ54046.1;Pangolin_CoV_orf1a;3034 | 0 | 0 |
| MT072864.1;QIQ54046.1;Pangolin_CoV_orf1a;3035 | 0 | 0 |
| MT072864.1;QIQ54046.1;Pangolin_CoV_orf1a;3036 | 0 | 0 |
| MT072864.1;QIQ54046.1;Pangolin_CoV_orf1a;3037 | 0 | 0 |
| MT072864.1;QIQ54046.1;Pangolin_CoV_orf1a;3038 | 0 | 0 |
| MT072864.1;QIQ54046.1;Pangolin_CoV_orf1a;3039 | 0 | 0 |
| MT072864.1;QIQ54046.1;Pangolin_CoV_orf1a;3040 | 0 | 0 |
| MT072864.1;QIQ54046.1;Pangolin_CoV_orf1a;3041 | 0 | 0 |
| MT072864.1;QIQ54046.1;Pangolin_CoV_orf1a;3042 | 0 | 0 |
| MT072864.1;QIQ54046.1;Pangolin_CoV_orf1a;3043 | 0 | 0 |

| | | |
|---|---|---|
| MT072864.1;QIQ54046.1;Pangolin_CoV_orf1a;3044 | 0 | 0 |
| MT072864.1;QIQ54046.1;Pangolin_CoV_orf1a;3045 | 0 | 0 |
| MT072864.1;QIQ54046.1;Pangolin_CoV_orf1a;3046 | 0 | 0 |
| MT072864.1;QIQ54046.1;Pangolin_CoV_orf1a;3047 | 0 | 0 |
| MT072864.1;QIQ54046.1;Pangolin_CoV_orf1a;3048 | 0 | 0 |
| MT072864.1;QIQ54046.1;Pangolin_CoV_orf1a;3049 | 0 | 0 |
| MT072864.1;QIQ54046.1;Pangolin_CoV_orf1a;3050 | 0 | 0 |
| MT072864.1;QIQ54046.1;Pangolin_CoV_orf1a;3051 | 0 | 0 |
| MT072864.1;QIQ54046.1;Pangolin_CoV_orf1a;3052 | 0 | 0 |
| MT072864.1;QIQ54046.1;Pangolin_CoV_orf1a;3053 | 0 | 0 |
| MT072864.1;QIQ54046.1;Pangolin_CoV_orf1a;3054 | 0 | 0 |
| MT072864.1;QIQ54046.1;Pangolin_CoV_orf1a;3055 | 0 | 0 |
| MT072864.1;QIQ54046.1;Pangolin_CoV_orf1a;3056 | 0 | 0 |
| MT072864.1;QIQ54046.1;Pangolin_CoV_orf1a;3057 | 0 | 0 |
| MT072864.1;QIQ54046.1;Pangolin_CoV_orf1a;3058 | 0 | 0 |
| MT072864.1;QIQ54046.1;Pangolin_CoV_orf1a;3059 | 0 | 0 |
| MT072864.1;QIQ54046.1;Pangolin_CoV_orf1a;3060 | 0 | 0 |
| MT072864.1;QIQ54046.1;Pangolin_CoV_orf1a;3061 | 0 | 0 |
| MT072864.1;QIQ54046.1;Pangolin_CoV_orf1a;3062 | 0 | 0 |
| MT072864.1;QIQ54046.1;Pangolin_CoV_orf1a;3063 | 0 | 0 |
| MT072864.1;QIQ54046.1;Pangolin_CoV_orf1a;3064 | 0 | 0 |
| MT072864.1;QIQ54046.1;Pangolin_CoV_orf1a;3065 | 0 | 0 |
| MT072864.1;QIQ54046.1;Pangolin_CoV_orf1a;3066 | 0 | 0 |
| MT072864.1;QIQ54046.1;Pangolin_CoV_orf1a;3067 | 0 | 0 |
| MT072864.1;QIQ54046.1;Pangolin_CoV_orf1a;3068 | 0 | 0 |
| MT072864.1;QIQ54046.1;Pangolin_CoV_orf1a;3069 | 0 | 0 |
| MT072864.1;QIQ54046.1;Pangolin_CoV_orf1a;3070 | 0 | 0 |
| MT072864.1;QIQ54046.1;Pangolin_CoV_orf1a;3071 | 0 | 0 |
| MT072864.1;QIQ54046.1;Pangolin_CoV_orf1a;3072 | 0 | 0 |
| MT072864.1;QIQ54046.1;Pangolin_CoV_orf1a;3073 | 0 | 0 |
| MT072864.1;QIQ54046.1;Pangolin_CoV_orf1a;3074 | 0 | 0 |
| MT072864.1;QIQ54046.1;Pangolin_CoV_orf1a;3075 | 0 | 0 |
| MT072864.1;QIQ54046.1;Pangolin_CoV_orf1a;3076 | 0 | 0 |
| MT072864.1;QIQ54046.1;Pangolin_CoV_orf1a;3077 | 0 | 0 |
| MT072864.1;QIQ54046.1;Pangolin_CoV_orf1a;3078 | 0 | 0 |
| MT072864.1;QIQ54046.1;Pangolin_CoV_orf1a;3079 | 0 | 0 |
| MT072864.1;QIQ54046.1;Pangolin_CoV_orf1a;3080 | 0 | 0 |
| MT072864.1;QIQ54046.1;Pangolin_CoV_orf1a;3081 | 0 | 0 |
| MT072864.1;QIQ54046.1;Pangolin_CoV_orf1a;3082 | 0 | 0 |
| MT072864.1;QIQ54046.1;Pangolin_CoV_orf1a;3083 | 0 | 0 |
| MT072864.1;QIQ54046.1;Pangolin_CoV_orf1a;3084 | 0 | 0 |
| MT072864.1;QIQ54046.1;Pangolin_CoV_orf1a;3085 | 0 | 0 |
| MT072864.1;QIQ54046.1;Pangolin_CoV_orf1a;3086 | 0 | 0 |

| | | |
|---|---|---|
| MT072864.1;QIQ54046.1;Pangolin_CoV_orf1a;3087 | 0 | 0 |
| MT072864.1;QIQ54046.1;Pangolin_CoV_orf1a;3088 | 0 | 0 |
| MT072864.1;QIQ54046.1;Pangolin_CoV_orf1a;3089 | 0 | 0 |
| MT072864.1;QIQ54046.1;Pangolin_CoV_orf1a;3090 | 0 | 0 |
| MT072864.1;QIQ54046.1;Pangolin_CoV_orf1a;3091 | 0 | 0 |
| MT072864.1;QIQ54046.1;Pangolin_CoV_orf1a;3092 | 0 | 0 |
| MT072864.1;QIQ54046.1;Pangolin_CoV_orf1a;3093 | 0 | 0 |
| MT072864.1;QIQ54046.1;Pangolin_CoV_orf1a;3094 | 0 | 0 |
| MT072864.1;QIQ54046.1;Pangolin_CoV_orf1a;3095 | 0 | 0 |
| MT072864.1;QIQ54046.1;Pangolin_CoV_orf1a;3096 | 0 | 0 |
| MT072864.1;QIQ54046.1;Pangolin_CoV_orf1a;3097 | 0 | 0 |
| MT072864.1;QIQ54046.1;Pangolin_CoV_orf1a;3098 | 0 | 0 |
| MT072864.1;QIQ54046.1;Pangolin_CoV_orf1a;3099 | 0 | 0 |
| MT072864.1;QIQ54046.1;Pangolin_CoV_orf1a;3100 | 0 | 0 |
| MT072864.1;QIQ54046.1;Pangolin_CoV_orf1a;3101 | 0 | 0 |
| MT072864.1;QIQ54046.1;Pangolin_CoV_orf1a;3102 | 0 | 0 |
| MT072864.1;QIQ54046.1;Pangolin_CoV_orf1a;3103 | 0 | 0 |
| MT072864.1;QIQ54046.1;Pangolin_CoV_orf1a;3104 | 0 | 0 |
| MT072864.1;QIQ54046.1;Pangolin_CoV_orf1a;3105 | 0 | 0 |
| MT072864.1;QIQ54046.1;Pangolin_CoV_orf1a;3106 | 0 | 0 |
| MT072864.1;QIQ54046.1;Pangolin_CoV_orf1a;3107 | 0 | 0 |
| MT072864.1;QIQ54046.1;Pangolin_CoV_orf1a;3108 | 0 | 0 |
| MT072864.1;QIQ54046.1;Pangolin_CoV_orf1a;3109 | 0 | 0 |
| MT072864.1;QIQ54046.1;Pangolin_CoV_orf1a;3110 | 0 | 0 |
| MT072864.1;QIQ54046.1;Pangolin_CoV_orf1a;3111 | 0 | 0 |
| MT072864.1;QIQ54046.1;Pangolin_CoV_orf1a;3112 | 0 | 0 |
| MT072864.1;QIQ54046.1;Pangolin_CoV_orf1a;3113 | 0 | 0 |
| MT072864.1;QIQ54046.1;Pangolin_CoV_orf1a;3114 | 0 | 0 |
| MT072864.1;QIQ54046.1;Pangolin_CoV_orf1a;3115 | 0 | 0 |
| MT072864.1;QIQ54046.1;Pangolin_CoV_orf1a;3116 | 0 | 0 |
| MT072864.1;QIQ54046.1;Pangolin_CoV_orf1a;3117 | 0 | 0 |
| MT072864.1;QIQ54046.1;Pangolin_CoV_orf1a;3118 | 0 | 0 |
| MT072864.1;QIQ54046.1;Pangolin_CoV_orf1a;3119 | 0 | 0 |
| MT072864.1;QIQ54046.1;Pangolin_CoV_orf1a;3120 | 0 | 0 |
| MT072864.1;QIQ54046.1;Pangolin_CoV_orf1a;3121 | 0 | 0 |
| MT072864.1;QIQ54046.1;Pangolin_CoV_orf1a;3122 | 0 | 0 |
| MT072864.1;QIQ54046.1;Pangolin_CoV_orf1a;3123 | 0 | 0 |
| MT072864.1;QIQ54046.1;Pangolin_CoV_orf1a;3124 | 0 | 0 |
| MT072864.1;QIQ54046.1;Pangolin_CoV_orf1a;3125 | 0 | 0 |
| MT072864.1;QIQ54046.1;Pangolin_CoV_orf1a;3126 | 0 | 0 |
| MT072864.1;QIQ54046.1;Pangolin_CoV_orf1a;3127 | 0 | 0 |
| MT072864.1;QIQ54046.1;Pangolin_CoV_orf1a;3128 | 0 | 0 |
| MT072864.1;QIQ54046.1;Pangolin_CoV_orf1a;3129 | 0 | 0 |

| | | |
|---|---|---|
| MT072864.1;QIQ54046.1;Pangolin_CoV_orf1a;3130 | 0 | 0 |
| MT072864.1;QIQ54046.1;Pangolin_CoV_orf1a;3131 | 0 | 0 |
| MT072864.1;QIQ54046.1;Pangolin_CoV_orf1a;3132 | 0 | 0 |
| MT072864.1;QIQ54046.1;Pangolin_CoV_orf1a;3133 | 0 | 0 |
| MT072864.1;QIQ54046.1;Pangolin_CoV_orf1a;3134 | 0 | 0 |
| MT072864.1;QIQ54046.1;Pangolin_CoV_orf1a;3135 | 0 | 0 |
| MT072864.1;QIQ54046.1;Pangolin_CoV_orf1a;3136 | 0 | 0 |
| MT072864.1;QIQ54046.1;Pangolin_CoV_orf1a;3137 | 0 | 0 |
| MT072864.1;QIQ54046.1;Pangolin_CoV_orf1a;3138 | 0 | 0 |
| MT072864.1;QIQ54046.1;Pangolin_CoV_orf1a;3139 | 0 | 0 |
| MT072864.1;QIQ54046.1;Pangolin_CoV_orf1a;3140 | 0 | 0 |
| MT072864.1;QIQ54046.1;Pangolin_CoV_orf1a;3141 | 0 | 0 |
| MT072864.1;QIQ54046.1;Pangolin_CoV_orf1a;3142 | 0 | 0 |
| MT072864.1;QIQ54046.1;Pangolin_CoV_orf1a;3143 | 0 | 0 |
| MT072864.1;QIQ54046.1;Pangolin_CoV_orf1a;3144 | 0 | 0 |
| MT072864.1;QIQ54046.1;Pangolin_CoV_orf1a;3145 | 0 | 0 |
| MT072864.1;QIQ54046.1;Pangolin_CoV_orf1a;3146 | 0 | 0 |
| MT072864.1;QIQ54046.1;Pangolin_CoV_orf1a;3147 | 0 | 0 |
| MT072864.1;QIQ54046.1;Pangolin_CoV_orf1a;3148 | 0 | 0 |
| MT072864.1;QIQ54046.1;Pangolin_CoV_orf1a;3149 | 0 | 0 |
| MT072864.1;QIQ54046.1;Pangolin_CoV_orf1a;3150 | 0 | 0 |
| MT072864.1;QIQ54046.1;Pangolin_CoV_orf1a;3151 | 0 | 0 |
| MT072864.1;QIQ54046.1;Pangolin_CoV_orf1a;3152 | 0 | 0 |
| MT072864.1;QIQ54046.1;Pangolin_CoV_orf1a;3153 | 0 | 0 |
| MT072864.1;QIQ54046.1;Pangolin_CoV_orf1a;3154 | 0 | 0 |
| MT072864.1;QIQ54046.1;Pangolin_CoV_orf1a;3155 | 0 | 0 |
| MT072864.1;QIQ54046.1;Pangolin_CoV_orf1a;3156 | 0.05 | 0.025 |
| MT072864.1;QIQ54046.1;Pangolin_CoV_orf1a;3157 | 0 | 0 |
| MT072864.1;QIQ54046.1;Pangolin_CoV_orf1a;3158 | 0 | 0 |
| MT072864.1;QIQ54046.1;Pangolin_CoV_orf1a;3159 | 0 | 0 |
| MT072864.1;QIQ54046.1;Pangolin_CoV_orf1a;3160 | 0 | 0 |
| MT072864.1;QIQ54046.1;Pangolin_CoV_orf1a;3161 | 0 | 0 |
| MT072864.1;QIQ54046.1;Pangolin_CoV_orf1a;3162 | 0 | 0 |
| MT072864.1;QIQ54046.1;Pangolin_CoV_orf1a;3163 | 0 | 0 |
| MT072864.1;QIQ54046.1;Pangolin_CoV_orf1a;3164 | 0 | 0 |
| MT072864.1;QIQ54046.1;Pangolin_CoV_orf1a;3165 | 0 | 0 |
| MT072864.1;QIQ54046.1;Pangolin_CoV_orf1a;3166 | 0 | 0 |
| MT072864.1;QIQ54046.1;Pangolin_CoV_orf1a;3167 | 0 | 0 |
| MT072864.1;QIQ54046.1;Pangolin_CoV_orf1a;3168 | 0 | 0 |
| MT072864.1;QIQ54046.1;Pangolin_CoV_orf1a;3169 | 0 | 0 |
| MT072864.1;QIQ54046.1;Pangolin_CoV_orf1a;3170 | 0 | 0 |
| MT072864.1;QIQ54046.1;Pangolin_CoV_orf1a;3171 | 0 | 0 |
| MT072864.1;QIQ54046.1;Pangolin_CoV_orf1a;3172 | 0 | 0 |

| | | |
|---|---|---|
| MT072864.1;QIQ54046.1;Pangolin_CoV_orf1a;3173 | 0 | 0 |
| MT072864.1;QIQ54046.1;Pangolin_CoV_orf1a;3174 | 0 | 0 |
| MT072864.1;QIQ54046.1;Pangolin_CoV_orf1a;3175 | 0 | 0 |
| MT072864.1;QIQ54046.1;Pangolin_CoV_orf1a;3176 | 0 | 0 |
| MT072864.1;QIQ54046.1;Pangolin_CoV_orf1a;3177 | 0 | 0 |
| MT072864.1;QIQ54046.1;Pangolin_CoV_orf1a;3178 | 0 | 0 |
| MT072864.1;QIQ54046.1;Pangolin_CoV_orf1a;3179 | 0 | 0 |
| MT072864.1;QIQ54046.1;Pangolin_CoV_orf1a;3180 | 0 | 0 |
| MT072864.1;QIQ54046.1;Pangolin_CoV_orf1a;3181 | 0 | 0 |
| MT072864.1;QIQ54046.1;Pangolin_CoV_orf1a;3182 | 0 | 0 |
| MT072864.1;QIQ54046.1;Pangolin_CoV_orf1a;3183 | 0 | 0 |
| MT072864.1;QIQ54046.1;Pangolin_CoV_orf1a;3184 | 0 | 0 |
| MT072864.1;QIQ54046.1;Pangolin_CoV_orf1a;3185 | 0 | 0 |
| MT072864.1;QIQ54046.1;Pangolin_CoV_orf1a;3186 | 0 | 0.025 |
| MT072864.1;QIQ54046.1;Pangolin_CoV_orf1a;3187 | 0 | 0 |
| MT072864.1;QIQ54046.1;Pangolin_CoV_orf1a;3188 | 0 | 0 |
| MT072864.1;QIQ54046.1;Pangolin_CoV_orf1a;3189 | 0 | 0 |
| MT072864.1;QIQ54046.1;Pangolin_CoV_orf1a;3190 | 0 | 0 |
| MT072864.1;QIQ54046.1;Pangolin_CoV_orf1a;3191 | 0 | 0 |
| MT072864.1;QIQ54046.1;Pangolin_CoV_orf1a;3192 | 0 | 0 |
| MT072864.1;QIQ54046.1;Pangolin_CoV_orf1a;3193 | 0 | 0 |
| MT072864.1;QIQ54046.1;Pangolin_CoV_orf1a;3194 | 0 | 0 |
| MT072864.1;QIQ54046.1;Pangolin_CoV_orf1a;3195 | 0 | 0 |
| MT072864.1;QIQ54046.1;Pangolin_CoV_orf1a;3196 | 0 | 0 |
| MT072864.1;QIQ54046.1;Pangolin_CoV_orf1a;3197 | 0 | 0 |
| MT072864.1;QIQ54046.1;Pangolin_CoV_orf1a;3198 | 0 | 0 |
| MT072864.1;QIQ54046.1;Pangolin_CoV_orf1a;3199 | 0 | 0 |
| MT072864.1;QIQ54046.1;Pangolin_CoV_orf1a;3200 | 0 | 0 |
| MT072864.1;QIQ54046.1;Pangolin_CoV_orf1a;3201 | 0 | 0.025 |
| MT072864.1;QIQ54046.1;Pangolin_CoV_orf1a;3202 | 0 | 0.025 |
| MT072864.1;QIQ54046.1;Pangolin_CoV_orf1a;3203 | 0 | 0.025 |
| MT072864.1;QIQ54046.1;Pangolin_CoV_orf1a;3204 | 0 | 0 |
| MT072864.1;QIQ54046.1;Pangolin_CoV_orf1a;3205 | 0 | 0 |
| MT072864.1;QIQ54046.1;Pangolin_CoV_orf1a;3206 | 0 | 0 |
| MT072864.1;QIQ54046.1;Pangolin_CoV_orf1a;3207 | 0.05 | 0.075 |
| MT072864.1;QIQ54046.1;Pangolin_CoV_orf1a;3208 | 0 | 0 |
| MT072864.1;QIQ54046.1;Pangolin_CoV_orf1a;3209 | 0 | 0 |
| MT072864.1;QIQ54046.1;Pangolin_CoV_orf1a;3210 | 0 | 0 |
| MT072864.1;QIQ54046.1;Pangolin_CoV_orf1a;3211 | 0 | 0.025 |
| MT072864.1;QIQ54046.1;Pangolin_CoV_orf1a;3212 | 0 | 0 |
| MT072864.1;QIQ54046.1;Pangolin_CoV_orf1a;3213 | 0 | 0 |
| MT072864.1;QIQ54046.1;Pangolin_CoV_orf1a;3214 | 0 | 0 |
| MT072864.1;QIQ54046.1;Pangolin_CoV_orf1a;3215 | 0 | 0 |

| | | |
|---|---|---|
| MT072864.1;QIQ54046.1;Pangolin_CoV_orf1a;3216 | 0 | 0 |
| MT072864.1;QIQ54046.1;Pangolin_CoV_orf1a;3217 | 0 | 0.075 |
| MT072864.1;QIQ54046.1;Pangolin_CoV_orf1a;3218 | 0 | 0.025 |
| MT072864.1;QIQ54046.1;Pangolin_CoV_orf1a;3219 | 0 | 0.025 |
| MT072864.1;QIQ54046.1;Pangolin_CoV_orf1a;3220 | 0 | 0.025 |
| MT072864.1;QIQ54046.1;Pangolin_CoV_orf1a;3221 | 0 | 0.025 |
| MT072864.1;QIQ54046.1;Pangolin_CoV_orf1a;3222 | 0.05 | 0.025 |
| MT072864.1;QIQ54046.1;Pangolin_CoV_orf1a;3223 | 0 | 0.025 |
| MT072864.1;QIQ54046.1;Pangolin_CoV_orf1a;3224 | 0 | 0.025 |
| MT072864.1;QIQ54046.1;Pangolin_CoV_orf1a;3225 | 0 | 0.025 |
| MT072864.1;QIQ54046.1;Pangolin_CoV_orf1a;3226 | 0.05 | 0 |
| MT072864.1;QIQ54046.1;Pangolin_CoV_orf1a;3227 | 0 | 0 |
| MT072864.1;QIQ54046.1;Pangolin_CoV_orf1a;3228 | 0 | 0 |
| MT072864.1;QIQ54046.1;Pangolin_CoV_orf1a;3229 | 0.15 | 0.05 |
| MT072864.1;QIQ54046.1;Pangolin_CoV_orf1a;3230 | 0.3 | 0.05 |
| MT072864.1;QIQ54046.1;Pangolin_CoV_orf1a;3231 | 0.05 | 0 |
| MT072864.1;QIQ54046.1;Pangolin_CoV_orf1a;3232 | 0.05 | 0.05 |
| MT072864.1;QIQ54046.1;Pangolin_CoV_orf1a;3233 | 0.1 | 0.1 |
| MT072864.1;QIQ54046.1;Pangolin_CoV_orf1a;3234 | 0 | 0.05 |
| MT072864.1;QIQ54046.1;Pangolin_CoV_orf1a;3235 | 0 | 0.05 |
| MT072864.1;QIQ54046.1;Pangolin_CoV_orf1a;3236 | 0 | 0 |
| MT072864.1;QIQ54046.1;Pangolin_CoV_orf1a;3237 | 0 | 0.025 |
| MT072864.1;QIQ54046.1;Pangolin_CoV_orf1a;3238 | 0 | 0 |
| MT072864.1;QIQ54046.1;Pangolin_CoV_orf1a;3239 | 0 | 0 |
| MT072864.1;QIQ54046.1;Pangolin_CoV_orf1a;3240 | 0 | 0 |
| MT072864.1;QIQ54046.1;Pangolin_CoV_orf1a;3241 | 0 | 0 |
| MT072864.1;QIQ54046.1;Pangolin_CoV_orf1a;3242 | 0.05 | 0 |
| MT072864.1;QIQ54046.1;Pangolin_CoV_orf1a;3243 | 0 | 0 |
| MT072864.1;QIQ54046.1;Pangolin_CoV_orf1a;3244 | 0 | 0 |
| MT072864.1;QIQ54046.1;Pangolin_CoV_orf1a;3245 | 0 | 0 |
| MT072864.1;QIQ54046.1;Pangolin_CoV_orf1a;3246 | 0 | 0 |
| MT072864.1;QIQ54046.1;Pangolin_CoV_orf1a;3247 | 0 | 0 |
| MT072864.1;QIQ54046.1;Pangolin_CoV_orf1a;3248 | 0 | 0 |
| MT072864.1;QIQ54046.1;Pangolin_CoV_orf1a;3249 | 0 | 0 |
| MT072864.1;QIQ54046.1;Pangolin_CoV_orf1a;3250 | 0 | 0 |
| MT072864.1;QIQ54046.1;Pangolin_CoV_orf1a;3251 | 0 | 0 |
| MT072864.1;QIQ54046.1;Pangolin_CoV_orf1a;3252 | 0 | 0 |
| MT072864.1;QIQ54046.1;Pangolin_CoV_orf1a;3253 | 0 | 0 |
| MT072864.1;QIQ54046.1;Pangolin_CoV_orf1a;3254 | 0 | 0 |
| MT072864.1;QIQ54046.1;Pangolin_CoV_orf1a;3255 | 0 | 0 |
| MT072864.1;QIQ54046.1;Pangolin_CoV_orf1a;3256 | 0 | 0 |
| MT072864.1;QIQ54046.1;Pangolin_CoV_orf1a;3257 | 0 | 0 |
| MT072864.1;QIQ54046.1;Pangolin_CoV_orf1a;3258 | 0 | 0 |

| | | |
|---|---|---|
| MT072864.1;QIQ54046.1;Pangolin_CoV_orf1a;3259 | 0 | 0 |
| MT072864.1;QIQ54046.1;Pangolin_CoV_orf1a;3260 | 0 | 0.025 |
| MT072864.1;QIQ54046.1;Pangolin_CoV_orf1a;3261 | 0 | 0 |
| MT072864.1;QIQ54046.1;Pangolin_CoV_orf1a;3262 | 0.05 | 0 |
| MT072864.1;QIQ54046.1;Pangolin_CoV_orf1a;3263 | 0 | 0 |
| MT072864.1;QIQ54046.1;Pangolin_CoV_orf1a;3264 | 0 | 0 |
| MT072864.1;QIQ54046.1;Pangolin_CoV_orf1a;3265 | 0 | 0 |
| MT072864.1;QIQ54046.1;Pangolin_CoV_orf1a;3266 | 0 | 0 |
| MT072864.1;QIQ54046.1;Pangolin_CoV_orf1a;3267 | 0 | 0 |
| MT072864.1;QIQ54046.1;Pangolin_CoV_orf1a;3268 | 0 | 0 |
| MT072864.1;QIQ54046.1;Pangolin_CoV_orf1a;3269 | 0 | 0 |
| MT072864.1;QIQ54046.1;Pangolin_CoV_orf1a;3270 | 0 | 0 |
| MT072864.1;QIQ54046.1;Pangolin_CoV_orf1a;3271 | 0 | 0 |
| MT072864.1;QIQ54046.1;Pangolin_CoV_orf1a;3272 | 0 | 0 |
| MT072864.1;QIQ54046.1;Pangolin_CoV_orf1a;3273 | 0 | 0 |
| MT072864.1;QIQ54046.1;Pangolin_CoV_orf1a;3274 | 0 | 0 |
| MT072864.1;QIQ54046.1;Pangolin_CoV_orf1a;3275 | 0 | 0 |
| MT072864.1;QIQ54046.1;Pangolin_CoV_orf1a;3276 | 0 | 0 |
| MT072864.1;QIQ54046.1;Pangolin_CoV_orf1a;3277 | 0 | 0 |
| MT072864.1;QIQ54046.1;Pangolin_CoV_orf1a;3278 | 0 | 0 |
| MT072864.1;QIQ54046.1;Pangolin_CoV_orf1a;3279 | 0 | 0 |
| MT072864.1;QIQ54046.1;Pangolin_CoV_orf1a;3280 | 0 | 0 |
| MT072864.1;QIQ54046.1;Pangolin_CoV_orf1a;3281 | 0 | 0 |
| MT072864.1;QIQ54046.1;Pangolin_CoV_orf1a;3282 | 0 | 0 |
| MT072864.1;QIQ54046.1;Pangolin_CoV_orf1a;3283 | 0 | 0 |
| MT072864.1;QIQ54046.1;Pangolin_CoV_orf1a;3284 | 0 | 0 |
| MT072864.1;QIQ54046.1;Pangolin_CoV_orf1a;3285 | 0 | 0 |
| MT072864.1;QIQ54046.1;Pangolin_CoV_orf1a;3286 | 0 | 0 |
| MT072864.1;QIQ54046.1;Pangolin_CoV_orf1a;3287 | 0 | 0 |
| MT072864.1;QIQ54046.1;Pangolin_CoV_orf1a;3288 | 0 | 0 |
| MT072864.1;QIQ54046.1;Pangolin_CoV_orf1a;3289 | 0 | 0 |
| MT072864.1;QIQ54046.1;Pangolin_CoV_orf1a;3290 | 0 | 0 |
| MT072864.1;QIQ54046.1;Pangolin_CoV_orf1a;3291 | 0 | 0.025 |
| MT072864.1;QIQ54046.1;Pangolin_CoV_orf1a;3292 | 0 | 0 |
| MT072864.1;QIQ54046.1;Pangolin_CoV_orf1a;3293 | 0 | 0 |
| MT072864.1;QIQ54046.1;Pangolin_CoV_orf1a;3294 | 0.05 | 0.025 |
| MT072864.1;QIQ54046.1;Pangolin_CoV_orf1a;3295 | 0 | 0 |
| MT072864.1;QIQ54046.1;Pangolin_CoV_orf1a;3296 | 0 | 0 |
| MT072864.1;QIQ54046.1;Pangolin_CoV_orf1a;3297 | 0 | 0 |
| MT072864.1;QIQ54046.1;Pangolin_CoV_orf1a;3298 | 0 | 0 |
| MT072864.1;QIQ54046.1;Pangolin_CoV_orf1a;3299 | 0 | 0 |
| MT072864.1;QIQ54046.1;Pangolin_CoV_orf1a;3300 | 0 | 0 |
| MT072864.1;QIQ54046.1;Pangolin_CoV_orf1a;3301 | 0 | 0.025 |

| | | |
|---|---|---|
| MT072864.1;QIQ54046.1;Pangolin_CoV_orf1a;3302 | 0 | 0.025 |
| MT072864.1;QIQ54046.1;Pangolin_CoV_orf1a;3303 | 0 | 0.025 |
| MT072864.1;QIQ54046.1;Pangolin_CoV_orf1a;3304 | 0 | 0 |
| MT072864.1;QIQ54046.1;Pangolin_CoV_orf1a;3305 | 0.05 | 0 |
| MT072864.1;QIQ54046.1;Pangolin_CoV_orf1a;3306 | 0 | 0.05 |
| MT072864.1;QIQ54046.1;Pangolin_CoV_orf1a;3307 | 0 | 0 |
| MT072864.1;QIQ54046.1;Pangolin_CoV_orf1a;3308 | 0 | 0 |
| MT072864.1;QIQ54046.1;Pangolin_CoV_orf1a;3309 | 0 | 0 |
| MT072864.1;QIQ54046.1;Pangolin_CoV_orf1a;3310 | 0 | 0 |
| MT072864.1;QIQ54046.1;Pangolin_CoV_orf1a;3311 | 0 | 0 |
| MT072864.1;QIQ54046.1;Pangolin_CoV_orf1a;3312 | 0 | 0 |
| MT072864.1;QIQ54046.1;Pangolin_CoV_orf1a;3313 | 0 | 0 |
| MT072864.1;QIQ54046.1;Pangolin_CoV_orf1a;3314 | 0 | 0 |
| MT072864.1;QIQ54046.1;Pangolin_CoV_orf1a;3315 | 0 | 0 |
| MT072864.1;QIQ54046.1;Pangolin_CoV_orf1a;3316 | 0 | 0 |
| MT072864.1;QIQ54046.1;Pangolin_CoV_orf1a;3317 | 0 | 0 |
| MT072864.1;QIQ54046.1;Pangolin_CoV_orf1a;3318 | 0 | 0 |
| MT072864.1;QIQ54046.1;Pangolin_CoV_orf1a;3319 | 0 | 0 |
| MT072864.1;QIQ54046.1;Pangolin_CoV_orf1a;3320 | 0 | 0 |
| MT072864.1;QIQ54046.1;Pangolin_CoV_orf1a;3321 | 0 | 0 |
| MT072864.1;QIQ54046.1;Pangolin_CoV_orf1a;3322 | 0 | 0 |
| MT072864.1;QIQ54046.1;Pangolin_CoV_orf1a;3323 | 0 | 0 |
| MT072864.1;QIQ54046.1;Pangolin_CoV_orf1a;3324 | 0 | 0 |
| MT072864.1;QIQ54046.1;Pangolin_CoV_orf1a;3325 | 0 | 0 |
| MT072864.1;QIQ54046.1;Pangolin_CoV_orf1a;3326 | 0 | 0 |
| MT072864.1;QIQ54046.1;Pangolin_CoV_orf1a;3327 | 0 | 0 |
| MT072864.1;QIQ54046.1;Pangolin_CoV_orf1a;3328 | 0 | 0 |
| MT072864.1;QIQ54046.1;Pangolin_CoV_orf1a;3329 | 0 | 0 |
| MT072864.1;QIQ54046.1;Pangolin_CoV_orf1a;3330 | 0 | 0 |
| MT072864.1;QIQ54046.1;Pangolin_CoV_orf1a;3331 | 0 | 0 |
| MT072864.1;QIQ54046.1;Pangolin_CoV_orf1a;3332 | 0 | 0 |
| MT072864.1;QIQ54046.1;Pangolin_CoV_orf1a;3333 | 0 | 0 |
| MT072864.1;QIQ54046.1;Pangolin_CoV_orf1a;3334 | 0.05 | 0 |
| MT072864.1;QIQ54046.1;Pangolin_CoV_orf1a;3335 | 0 | 0 |
| MT072864.1;QIQ54046.1;Pangolin_CoV_orf1a;3336 | 0 | 0 |
| MT072864.1;QIQ54046.1;Pangolin_CoV_orf1a;3337 | 0 | 0.025 |
| MT072864.1;QIQ54046.1;Pangolin_CoV_orf1a;3338 | 0 | 0 |
| MT072864.1;QIQ54046.1;Pangolin_CoV_orf1a;3339 | 0 | 0 |
| MT072864.1;QIQ54046.1;Pangolin_CoV_orf1a;3340 | 0 | 0 |
| MT072864.1;QIQ54046.1;Pangolin_CoV_orf1a;3341 | 0 | 0 |
| MT072864.1;QIQ54046.1;Pangolin_CoV_orf1a;3342 | 0 | 0 |
| MT072864.1;QIQ54046.1;Pangolin_CoV_orf1a;3343 | 0 | 0 |
| MT072864.1;QIQ54046.1;Pangolin_CoV_orf1a;3344 | 0 | 0 |

| | | |
|---|---|---|
| MT072864.1;QIQ54046.1;Pangolin_CoV_orf1a;3345 | 0 | 0 |
| MT072864.1;QIQ54046.1;Pangolin_CoV_orf1a;3346 | 0 | 0 |
| MT072864.1;QIQ54046.1;Pangolin_CoV_orf1a;3347 | 0 | 0 |
| MT072864.1;QIQ54046.1;Pangolin_CoV_orf1a;3348 | 0 | 0 |
| MT072864.1;QIQ54046.1;Pangolin_CoV_orf1a;3349 | 0 | 0 |
| MT072864.1;QIQ54046.1;Pangolin_CoV_orf1a;3350 | 0 | 0 |
| MT072864.1;QIQ54046.1;Pangolin_CoV_orf1a;3351 | 0 | 0 |
| MT072864.1;QIQ54046.1;Pangolin_CoV_orf1a;3352 | 0 | 0 |
| MT072864.1;QIQ54046.1;Pangolin_CoV_orf1a;3353 | 0 | 0 |
| MT072864.1;QIQ54046.1;Pangolin_CoV_orf1a;3354 | 0 | 0 |
| MT072864.1;QIQ54046.1;Pangolin_CoV_orf1a;3355 | 0 | 0 |
| MT072864.1;QIQ54046.1;Pangolin_CoV_orf1a;3356 | 0 | 0 |
| MT072864.1;QIQ54046.1;Pangolin_CoV_orf1a;3357 | 0 | 0 |
| MT072864.1;QIQ54046.1;Pangolin_CoV_orf1a;3358 | 0 | 0 |
| MT072864.1;QIQ54046.1;Pangolin_CoV_orf1a;3359 | 0 | 0 |
| MT072864.1;QIQ54046.1;Pangolin_CoV_orf1a;3360 | 0 | 0 |
| MT072864.1;QIQ54046.1;Pangolin_CoV_orf1a;3361 | 0 | 0 |
| MT072864.1;QIQ54046.1;Pangolin_CoV_orf1a;3362 | 0 | 0.025 |
| MT072864.1;QIQ54046.1;Pangolin_CoV_orf1a;3363 | 0 | 0 |
| MT072864.1;QIQ54046.1;Pangolin_CoV_orf1a;3364 | 0 | 0 |
| MT072864.1;QIQ54046.1;Pangolin_CoV_orf1a;3365 | 0 | 0 |
| MT072864.1;QIQ54046.1;Pangolin_CoV_orf1a;3366 | 0 | 0 |
| MT072864.1;QIQ54046.1;Pangolin_CoV_orf1a;3367 | 0 | 0 |
| MT072864.1;QIQ54046.1;Pangolin_CoV_orf1a;3368 | 0 | 0 |
| MT072864.1;QIQ54046.1;Pangolin_CoV_orf1a;3369 | 0 | 0 |
| MT072864.1;QIQ54046.1;Pangolin_CoV_orf1a;3370 | 0 | 0 |
| MT072864.1;QIQ54046.1;Pangolin_CoV_orf1a;3371 | 0 | 0 |
| MT072864.1;QIQ54046.1;Pangolin_CoV_orf1a;3372 | 0 | 0 |
| MT072864.1;QIQ54046.1;Pangolin_CoV_orf1a;3373 | 0 | 0.025 |
| MT072864.1;QIQ54046.1;Pangolin_CoV_orf1a;3374 | 0 | 0 |
| MT072864.1;QIQ54046.1;Pangolin_CoV_orf1a;3375 | 0 | 0 |
| MT072864.1;QIQ54046.1;Pangolin_CoV_orf1a;3376 | 0 | 0 |
| MT072864.1;QIQ54046.1;Pangolin_CoV_orf1a;3377 | 0 | 0 |
| MT072864.1;QIQ54046.1;Pangolin_CoV_orf1a;3378 | 0 | 0 |
| MT072864.1;QIQ54046.1;Pangolin_CoV_orf1a;3379 | 0 | 0.025 |
| MT072864.1;QIQ54046.1;Pangolin_CoV_orf1a;3380 | 0 | 0 |
| MT072864.1;QIQ54046.1;Pangolin_CoV_orf1a;3381 | 0 | 0 |
| MT072864.1;QIQ54046.1;Pangolin_CoV_orf1a;3382 | 0 | 0 |
| MT072864.1;QIQ54046.1;Pangolin_CoV_orf1a;3383 | 0 | 0 |
| MT072864.1;QIQ54046.1;Pangolin_CoV_orf1a;3384 | 0.05 | 0 |
| MT072864.1;QIQ54046.1;Pangolin_CoV_orf1a;3385 | 0 | 0 |
| MT072864.1;QIQ54046.1;Pangolin_CoV_orf1a;3386 | 0.05 | 0 |
| MT072864.1;QIQ54046.1;Pangolin_CoV_orf1a;3387 | 0 | 0 |

| | | |
|---|---|---|
| MT072864.1;QIQ54046.1;Pangolin_CoV_orf1a;3388 | 0 | 0 |
| MT072864.1;QIQ54046.1;Pangolin_CoV_orf1a;3389 | 0 | 0 |
| MT072864.1;QIQ54046.1;Pangolin_CoV_orf1a;3390 | 0 | 0 |
| MT072864.1;QIQ54046.1;Pangolin_CoV_orf1a;3391 | 0 | 0 |
| MT072864.1;QIQ54046.1;Pangolin_CoV_orf1a;3392 | 0 | 0.025 |
| MT072864.1;QIQ54046.1;Pangolin_CoV_orf1a;3393 | 0 | 0.025 |
| MT072864.1;QIQ54046.1;Pangolin_CoV_orf1a;3394 | 0 | 0.025 |
| MT072864.1;QIQ54046.1;Pangolin_CoV_orf1a;3395 | 0 | 0.05 |
| MT072864.1;QIQ54046.1;Pangolin_CoV_orf1a;3396 | 0 | 0.05 |
| MT072864.1;QIQ54046.1;Pangolin_CoV_orf1a;3397 | 0 | 0.025 |
| MT072864.1;QIQ54046.1;Pangolin_CoV_orf1a;3398 | 0 | 0.025 |
| MT072864.1;QIQ54046.1;Pangolin_CoV_orf1a;3399 | 0 | 0 |
| MT072864.1;QIQ54046.1;Pangolin_CoV_orf1a;3400 | 0 | 0 |
| MT072864.1;QIQ54046.1;Pangolin_CoV_orf1a;3401 | 0 | 0 |
| MT072864.1;QIQ54046.1;Pangolin_CoV_orf1a;3402 | 0 | 0 |
| MT072864.1;QIQ54046.1;Pangolin_CoV_orf1a;3403 | 0 | 0 |
| MT072864.1;QIQ54046.1;Pangolin_CoV_orf1a;3404 | 0 | 0 |
| MT072864.1;QIQ54046.1;Pangolin_CoV_orf1a;3405 | 0 | 0 |
| MT072864.1;QIQ54046.1;Pangolin_CoV_orf1a;3406 | 0 | 0 |
| MT072864.1;QIQ54046.1;Pangolin_CoV_orf1a;3407 | 0 | 0 |
| MT072864.1;QIQ54046.1;Pangolin_CoV_orf1a;3408 | 0 | 0 |
| MT072864.1;QIQ54046.1;Pangolin_CoV_orf1a;3409 | 0 | 0 |
| MT072864.1;QIQ54046.1;Pangolin_CoV_orf1a;3410 | 0 | 0 |
| MT072864.1;QIQ54046.1;Pangolin_CoV_orf1a;3411 | 0 | 0 |
| MT072864.1;QIQ54046.1;Pangolin_CoV_orf1a;3412 | 0 | 0 |
| MT072864.1;QIQ54046.1;Pangolin_CoV_orf1a;3413 | 0 | 0 |
| MT072864.1;QIQ54046.1;Pangolin_CoV_orf1a;3414 | 0 | 0 |
| MT072864.1;QIQ54046.1;Pangolin_CoV_orf1a;3415 | 0 | 0 |
| MT072864.1;QIQ54046.1;Pangolin_CoV_orf1a;3416 | 0 | 0 |
| MT072864.1;QIQ54046.1;Pangolin_CoV_orf1a;3417 | 0 | 0 |
| MT072864.1;QIQ54046.1;Pangolin_CoV_orf1a;3418 | 0.15 | 0.025 |
| MT072864.1;QIQ54046.1;Pangolin_CoV_orf1a;3419 | 0.1 | 0.05 |
| MT072864.1;QIQ54046.1;Pangolin_CoV_orf1a;3420 | 0.05 | 0.025 |
| MT072864.1;QIQ54046.1;Pangolin_CoV_orf1a;3421 | 0.2 | 0.075 |
| MT072864.1;QIQ54046.1;Pangolin_CoV_orf1a;3422 | 0.2 | 0.05 |
| MT072864.1;QIQ54046.1;Pangolin_CoV_orf1a;3423 | 0.2 | 0.05 |
| MT072864.1;QIQ54046.1;Pangolin_CoV_orf1a;3424 | 0.2 | 0.05 |
| MT072864.1;QIQ54046.1;Pangolin_CoV_orf1a;3425 | 0.1 | 0.025 |
| MT072864.1;QIQ54046.1;Pangolin_CoV_orf1a;3426 | 0.05 | 0.025 |
| MT072864.1;QIQ54046.1;Pangolin_CoV_orf1a;3427 | 0.05 | 0.025 |
| MT072864.1;QIQ54046.1;Pangolin_CoV_orf1a;3428 | 0.05 | 0.05 |
| MT072864.1;QIQ54046.1;Pangolin_CoV_orf1a;3429 | 0.15 | 0.1 |
| MT072864.1;QIQ54046.1;Pangolin_CoV_orf1a;3430 | 0 | 0.025 |

| | | |
|---|---|---|
| MT072864.1;QIQ54046.1;Pangolin_CoV_orf1a;3431 | 0 | 0 |
| MT072864.1;QIQ54046.1;Pangolin_CoV_orf1a;3432 | 0.05 | 0 |
| MT072864.1;QIQ54046.1;Pangolin_CoV_orf1a;3433 | 0.05 | 0 |
| MT072864.1;QIQ54046.1;Pangolin_CoV_orf1a;3434 | 0.1 | 0.025 |
| MT072864.1;QIQ54046.1;Pangolin_CoV_orf1a;3435 | 0.05 | 0 |
| MT072864.1;QIQ54046.1;Pangolin_CoV_orf1a;3436 | 0.1 | 0 |
| MT072864.1;QIQ54046.1;Pangolin_CoV_orf1a;3437 | 0 | 0 |
| MT072864.1;QIQ54046.1;Pangolin_CoV_orf1a;3438 | 0.05 | 0 |
| MT072864.1;QIQ54046.1;Pangolin_CoV_orf1a;3439 | 0 | 0 |
| MT072864.1;QIQ54046.1;Pangolin_CoV_orf1a;3440 | 0 | 0 |
| MT072864.1;QIQ54046.1;Pangolin_CoV_orf1a;3441 | 0 | 0 |
| MT072864.1;QIQ54046.1;Pangolin_CoV_orf1a;3442 | 0 | 0 |
| MT072864.1;QIQ54046.1;Pangolin_CoV_orf1a;3443 | 0 | 0 |
| MT072864.1;QIQ54046.1;Pangolin_CoV_orf1a;3444 | 0 | 0 |
| MT072864.1;QIQ54046.1;Pangolin_CoV_orf1a;3445 | 0 | 0 |
| MT072864.1;QIQ54046.1;Pangolin_CoV_orf1a;3446 | 0 | 0 |
| MT072864.1;QIQ54046.1;Pangolin_CoV_orf1a;3447 | 0 | 0 |
| MT072864.1;QIQ54046.1;Pangolin_CoV_orf1a;3448 | 0 | 0 |
| MT072864.1;QIQ54046.1;Pangolin_CoV_orf1a;3449 | 0 | 0 |
| MT072864.1;QIQ54046.1;Pangolin_CoV_orf1a;3450 | 0 | 0 |
| MT072864.1;QIQ54046.1;Pangolin_CoV_orf1a;3451 | 0 | 0 |
| MT072864.1;QIQ54046.1;Pangolin_CoV_orf1a;3452 | 0 | 0 |
| MT072864.1;QIQ54046.1;Pangolin_CoV_orf1a;3453 | 0 | 0 |
| MT072864.1;QIQ54046.1;Pangolin_CoV_orf1a;3454 | 0 | 0 |
| MT072864.1;QIQ54046.1;Pangolin_CoV_orf1a;3455 | 0 | 0 |
| MT072864.1;QIQ54046.1;Pangolin_CoV_orf1a;3456 | 0 | 0 |
| MT072864.1;QIQ54046.1;Pangolin_CoV_orf1a;3457 | 0 | 0 |
| MT072864.1;QIQ54046.1;Pangolin_CoV_orf1a;3458 | 0 | 0 |
| MT072864.1;QIQ54046.1;Pangolin_CoV_orf1a;3459 | 0 | 0 |
| MT072864.1;QIQ54046.1;Pangolin_CoV_orf1a;3460 | 0 | 0 |
| MT072864.1;QIQ54046.1;Pangolin_CoV_orf1a;3461 | 0 | 0 |
| MT072864.1;QIQ54046.1;Pangolin_CoV_orf1a;3462 | 0 | 0 |
| MT072864.1;QIQ54046.1;Pangolin_CoV_orf1a;3463 | 0 | 0 |
| MT072864.1;QIQ54046.1;Pangolin_CoV_orf1a;3464 | 0 | 0 |
| MT072864.1;QIQ54046.1;Pangolin_CoV_orf1a;3465 | 0 | 0 |
| MT072864.1;QIQ54046.1;Pangolin_CoV_orf1a;3466 | 0 | 0 |
| MT072864.1;QIQ54046.1;Pangolin_CoV_orf1a;3467 | 0 | 0 |
| MT072864.1;QIQ54046.1;Pangolin_CoV_orf1a;3468 | 0 | 0 |
| MT072864.1;QIQ54046.1;Pangolin_CoV_orf1a;3469 | 0 | 0 |
| MT072864.1;QIQ54046.1;Pangolin_CoV_orf1a;3470 | 0 | 0 |
| MT072864.1;QIQ54046.1;Pangolin_CoV_orf1a;3471 | 0.05 | 0 |
| MT072864.1;QIQ54046.1;Pangolin_CoV_orf1a;3472 | 0.05 | 0 |
| MT072864.1;QIQ54046.1;Pangolin_CoV_orf1a;3473 | 0.05 | 0 |

| | | |
|---|---|---|
| MT072864.1;QIQ54046.1;Pangolin_CoV_orf1a;3474 | 0.05 | 0.025 |
| MT072864.1;QIQ54046.1;Pangolin_CoV_orf1a;3475 | 0.05 | 0.05 |
| MT072864.1;QIQ54046.1;Pangolin_CoV_orf1a;3476 | 0.05 | 0 |
| MT072864.1;QIQ54046.1;Pangolin_CoV_orf1a;3477 | 0.1 | 0.025 |
| MT072864.1;QIQ54046.1;Pangolin_CoV_orf1a;3478 | 0.05 | 0 |
| MT072864.1;QIQ54046.1;Pangolin_CoV_orf1a;3479 | 0.05 | 0.025 |
| MT072864.1;QIQ54046.1;Pangolin_CoV_orf1a;3480 | 0.05 | 0.275 |
| MT072864.1;QIQ54046.1;Pangolin_CoV_orf1a;3481 | 0.25 | 0.075 |
| MT072864.1;QIQ54046.1;Pangolin_CoV_orf1a;3482 | 0.1 | 0 |
| MT072864.1;QIQ54046.1;Pangolin_CoV_orf1a;3483 | 0.05 | 0 |
| MT072864.1;QIQ54046.1;Pangolin_CoV_orf1a;3484 | 0.05 | 0 |
| MT072864.1;QIQ54046.1;Pangolin_CoV_orf1a;3485 | 0.05 | 0 |
| MT072864.1;QIQ54046.1;Pangolin_CoV_orf1a;3486 | 0.1 | 0.025 |
| MT072864.1;QIQ54046.1;Pangolin_CoV_orf1a;3487 | 0.1 | 0.025 |
| MT072864.1;QIQ54046.1;Pangolin_CoV_orf1a;3488 | 0.15 | 0 |
| MT072864.1;QIQ54046.1;Pangolin_CoV_orf1a;3489 | 0.1 | 0 |
| MT072864.1;QIQ54046.1;Pangolin_CoV_orf1a;3490 | 0.05 | 0 |
| MT072864.1;QIQ54046.1;Pangolin_CoV_orf1a;3491 | 0.05 | 0 |
| MT072864.1;QIQ54046.1;Pangolin_CoV_orf1a;3492 | 0 | 0.025 |
| MT072864.1;QIQ54046.1;Pangolin_CoV_orf1a;3493 | 0 | 0 |
| MT072864.1;QIQ54046.1;Pangolin_CoV_orf1a;3494 | 0 | 0 |
| MT072864.1;QIQ54046.1;Pangolin_CoV_orf1a;3495 | 0.05 | 0 |
| MT072864.1;QIQ54046.1;Pangolin_CoV_orf1a;3496 | 0 | 0 |
| MT072864.1;QIQ54046.1;Pangolin_CoV_orf1a;3497 | 0.05 | 0.025 |
| MT072864.1;QIQ54046.1;Pangolin_CoV_orf1a;3498 | 0.15 | 0.1 |
| MT072864.1;QIQ54046.1;Pangolin_CoV_orf1a;3499 | 0.05 | 0.05 |
| MT072864.1;QIQ54046.1;Pangolin_CoV_orf1a;3500 | 0.1 | 0.15 |
| MT072864.1;QIQ54046.1;Pangolin_CoV_orf1a;3501 | 0.05 | 0.025 |
| MT072864.1;QIQ54046.1;Pangolin_CoV_orf1a;3502 | 0.05 | 0 |
| MT072864.1;QIQ54046.1;Pangolin_CoV_orf1a;3503 | 0.05 | 0 |
| MT072864.1;QIQ54046.1;Pangolin_CoV_orf1a;3504 | 0.05 | 0 |
| MT072864.1;QIQ54046.1;Pangolin_CoV_orf1a;3505 | 0.05 | 0 |
| MT072864.1;QIQ54046.1;Pangolin_CoV_orf1a;3506 | 0.05 | 0 |
| MT072864.1;QIQ54046.1;Pangolin_CoV_orf1a;3507 | 0.05 | 0 |
| MT072864.1;QIQ54046.1;Pangolin_CoV_orf1a;3508 | 0 | 0 |
| MT072864.1;QIQ54046.1;Pangolin_CoV_orf1a;3509 | 0 | 0 |
| MT072864.1;QIQ54046.1;Pangolin_CoV_orf1a;3510 | 0 | 0 |
| MT072864.1;QIQ54046.1;Pangolin_CoV_orf1a;3511 | 0 | 0 |
| MT072864.1;QIQ54046.1;Pangolin_CoV_orf1a;3512 | 0 | 0 |
| MT072864.1;QIQ54046.1;Pangolin_CoV_orf1a;3513 | 0.05 | 0 |
| MT072864.1;QIQ54046.1;Pangolin_CoV_orf1a;3514 | 0 | 0 |
| MT072864.1;QIQ54046.1;Pangolin_CoV_orf1a;3515 | 0 | 0 |
| MT072864.1;QIQ54046.1;Pangolin_CoV_orf1a;3516 | 0 | 0 |

| | | |
|---|---|---|
| MT072864.1;QIQ54046.1;Pangolin_CoV_orf1a;3517 | 0.1 | 0 |
| MT072864.1;QIQ54046.1;Pangolin_CoV_orf1a;3518 | 0.05 | 0.025 |
| MT072864.1;QIQ54046.1;Pangolin_CoV_orf1a;3519 | 0.05 | 0 |
| MT072864.1;QIQ54046.1;Pangolin_CoV_orf1a;3520 | 0.1 | 0 |
| MT072864.1;QIQ54046.1;Pangolin_CoV_orf1a;3521 | 0.05 | 0.025 |
| MT072864.1;QIQ54046.1;Pangolin_CoV_orf1a;3522 | 0 | 0 |
| MT072864.1;QIQ54046.1;Pangolin_CoV_orf1a;3523 | 0 | 0.025 |
| MT072864.1;QIQ54046.1;Pangolin_CoV_orf1a;3524 | 0 | 0.025 |
| MT072864.1;QIQ54046.1;Pangolin_CoV_orf1a;3525 | 0 | 0 |
| MT072864.1;QIQ54046.1;Pangolin_CoV_orf1a;3526 | 0 | 0 |
| MT072864.1;QIQ54046.1;Pangolin_CoV_orf1a;3527 | 0 | 0 |
| MT072864.1;QIQ54046.1;Pangolin_CoV_orf1a;3528 | 0 | 0 |
| MT072864.1;QIQ54046.1;Pangolin_CoV_orf1a;3529 | 0 | 0 |
| MT072864.1;QIQ54046.1;Pangolin_CoV_orf1a;3530 | 0 | 0 |
| MT072864.1;QIQ54046.1;Pangolin_CoV_orf1a;3531 | 0 | 0 |
| MT072864.1;QIQ54046.1;Pangolin_CoV_orf1a;3532 | 0 | 0 |
| MT072864.1;QIQ54046.1;Pangolin_CoV_orf1a;3533 | 0 | 0 |
| MT072864.1;QIQ54046.1;Pangolin_CoV_orf1a;3534 | 0 | 0.025 |
| MT072864.1;QIQ54046.1;Pangolin_CoV_orf1a;3535 | 0 | 0.05 |
| MT072864.1;QIQ54046.1;Pangolin_CoV_orf1a;3536 | 0 | 0.025 |
| MT072864.1;QIQ54046.1;Pangolin_CoV_orf1a;3537 | 0 | 0.025 |
| MT072864.1;QIQ54046.1;Pangolin_CoV_orf1a;3538 | 0 | 0.025 |
| MT072864.1;QIQ54046.1;Pangolin_CoV_orf1a;3539 | 0.05 | 0.025 |
| MT072864.1;QIQ54046.1;Pangolin_CoV_orf1a;3540 | 0.05 | 0.05 |
| MT072864.1;QIQ54046.1;Pangolin_CoV_orf1a;3541 | 0 | 0.025 |
| MT072864.1;QIQ54046.1;Pangolin_CoV_orf1a;3542 | 0 | 0.05 |
| MT072864.1;QIQ54046.1;Pangolin_CoV_orf1a;3543 | 0.1 | 0.025 |
| MT072864.1;QIQ54046.1;Pangolin_CoV_orf1a;3544 | 0 | 0 |
| MT072864.1;QIQ54046.1;Pangolin_CoV_orf1a;3545 | 0 | 0 |
| MT072864.1;QIQ54046.1;Pangolin_CoV_orf1a;3546 | 0 | 0 |
| MT072864.1;QIQ54046.1;Pangolin_CoV_orf1a;3547 | 0 | 0 |
| MT072864.1;QIQ54046.1;Pangolin_CoV_orf1a;3548 | 0 | 0 |
| MT072864.1;QIQ54046.1;Pangolin_CoV_orf1a;3549 | 0 | 0 |
| MT072864.1;QIQ54046.1;Pangolin_CoV_orf1a;3550 | 0 | 0 |
| MT072864.1;QIQ54046.1;Pangolin_CoV_orf1a;3551 | 0 | 0 |
| MT072864.1;QIQ54046.1;Pangolin_CoV_orf1a;3552 | 0 | 0 |
| MT072864.1;QIQ54046.1;Pangolin_CoV_orf1a;3553 | 0 | 0 |
| MT072864.1;QIQ54046.1;Pangolin_CoV_orf1a;3554 | 0 | 0.025 |
| MT072864.1;QIQ54046.1;Pangolin_CoV_orf1a;3555 | 0 | 0 |
| MT072864.1;QIQ54046.1;Pangolin_CoV_orf1a;3556 | 0 | 0 |
| MT072864.1;QIQ54046.1;Pangolin_CoV_orf1a;3557 | 0 | 0 |
| MT072864.1;QIQ54046.1;Pangolin_CoV_orf1a;3558 | 0 | 0 |
| MT072864.1;QIQ54046.1;Pangolin_CoV_orf1a;3559 | 0 | 0 |

| | | |
|---|---|---|
| MT072864.1;QIQ54046.1;Pangolin_CoV_orf1a;3560 | 0 | 0 |
| MT072864.1;QIQ54046.1;Pangolin_CoV_orf1a;3561 | 0 | 0 |
| MT072864.1;QIQ54046.1;Pangolin_CoV_orf1a;3562 | 0 | 0 |
| MT072864.1;QIQ54046.1;Pangolin_CoV_orf1a;3563 | 0 | 0 |
| MT072864.1;QIQ54046.1;Pangolin_CoV_orf1a;3564 | 0 | 0 |
| MT072864.1;QIQ54046.1;Pangolin_CoV_orf1a;3565 | 0 | 0.025 |
| MT072864.1;QIQ54046.1;Pangolin_CoV_orf1a;3566 | 0 | 0 |
| MT072864.1;QIQ54046.1;Pangolin_CoV_orf1a;3567 | 0 | 0 |
| MT072864.1;QIQ54046.1;Pangolin_CoV_orf1a;3568 | 0 | 0 |
| MT072864.1;QIQ54046.1;Pangolin_CoV_orf1a;3569 | 0 | 0 |
| MT072864.1;QIQ54046.1;Pangolin_CoV_orf1a;3570 | 0 | 0 |
| MT072864.1;QIQ54046.1;Pangolin_CoV_orf1a;3571 | 0 | 0 |
| MT072864.1;QIQ54046.1;Pangolin_CoV_orf1a;3572 | 0 | 0 |
| MT072864.1;QIQ54046.1;Pangolin_CoV_orf1a;3573 | 0 | 0 |
| MT072864.1;QIQ54046.1;Pangolin_CoV_orf1a;3574 | 0 | 0 |
| MT072864.1;QIQ54046.1;Pangolin_CoV_orf1a;3575 | 0 | 0 |
| MT072864.1;QIQ54046.1;Pangolin_CoV_orf1a;3576 | 0 | 0 |
| MT072864.1;QIQ54046.1;Pangolin_CoV_orf1a;3577 | 0 | 0 |
| MT072864.1;QIQ54046.1;Pangolin_CoV_orf1a;3578 | 0 | 0 |
| MT072864.1;QIQ54046.1;Pangolin_CoV_orf1a;3579 | 0 | 0 |
| MT072864.1;QIQ54046.1;Pangolin_CoV_orf1a;3580 | 0 | 0 |
| MT072864.1;QIQ54046.1;Pangolin_CoV_orf1a;3581 | 0 | 0 |
| MT072864.1;QIQ54046.1;Pangolin_CoV_orf1a;3582 | 0 | 0 |
| MT072864.1;QIQ54046.1;Pangolin_CoV_orf1a;3583 | 0 | 0 |
| MT072864.1;QIQ54046.1;Pangolin_CoV_orf1a;3584 | 0 | 0 |
| MT072864.1;QIQ54046.1;Pangolin_CoV_orf1a;3585 | 0 | 0 |
| MT072864.1;QIQ54046.1;Pangolin_CoV_orf1a;3586 | 0 | 0 |
| MT072864.1;QIQ54046.1;Pangolin_CoV_orf1a;3587 | 0 | 0 |
| MT072864.1;QIQ54046.1;Pangolin_CoV_orf1a;3588 | 0 | 0 |
| MT072864.1;QIQ54046.1;Pangolin_CoV_orf1a;3589 | 0 | 0 |
| MT072864.1;QIQ54046.1;Pangolin_CoV_orf1a;3590 | 0 | 0 |
| MT072864.1;QIQ54046.1;Pangolin_CoV_orf1a;3591 | 0 | 0 |
| MT072864.1;QIQ54046.1;Pangolin_CoV_orf1a;3592 | 0 | 0 |
| MT072864.1;QIQ54046.1;Pangolin_CoV_orf1a;3593 | 0 | 0 |
| MT072864.1;QIQ54046.1;Pangolin_CoV_orf1a;3594 | 0 | 0 |
| MT072864.1;QIQ54046.1;Pangolin_CoV_orf1a;3595 | 0 | 0 |
| MT072864.1;QIQ54046.1;Pangolin_CoV_orf1a;3596 | 0 | 0 |
| MT072864.1;QIQ54046.1;Pangolin_CoV_orf1a;3597 | 0 | 0 |
| MT072864.1;QIQ54046.1;Pangolin_CoV_orf1a;3598 | 0 | 0 |
| MT072864.1;QIQ54046.1;Pangolin_CoV_orf1a;3599 | 0 | 0 |
| MT072864.1;QIQ54046.1;Pangolin_CoV_orf1a;3600 | 0 | 0 |
| MT072864.1;QIQ54046.1;Pangolin_CoV_orf1a;3601 | 0 | 0 |
| MT072864.1;QIQ54046.1;Pangolin_CoV_orf1a;3602 | 0 | 0 |

| | | |
|---|---|---|
| MT072864.1;QIQ54046.1;Pangolin_CoV_orf1a;3603 | 0 | 0 |
| MT072864.1;QIQ54046.1;Pangolin_CoV_orf1a;3604 | 0 | 0 |
| MT072864.1;QIQ54046.1;Pangolin_CoV_orf1a;3605 | 0 | 0 |
| MT072864.1;QIQ54046.1;Pangolin_CoV_orf1a;3606 | 0 | 0 |
| MT072864.1;QIQ54046.1;Pangolin_CoV_orf1a;3607 | 0 | 0 |
| MT072864.1;QIQ54046.1;Pangolin_CoV_orf1a;3608 | 0 | 0 |
| MT072864.1;QIQ54046.1;Pangolin_CoV_orf1a;3609 | 0 | 0 |
| MT072864.1;QIQ54046.1;Pangolin_CoV_orf1a;3610 | 0 | 0 |
| MT072864.1;QIQ54046.1;Pangolin_CoV_orf1a;3611 | 0 | 0 |
| MT072864.1;QIQ54046.1;Pangolin_CoV_orf1a;3612 | 0 | 0 |
| MT072864.1;QIQ54046.1;Pangolin_CoV_orf1a;3613 | 0 | 0 |
| MT072864.1;QIQ54046.1;Pangolin_CoV_orf1a;3614 | 0 | 0 |
| MT072864.1;QIQ54046.1;Pangolin_CoV_orf1a;3615 | 0 | 0 |
| MT072864.1;QIQ54046.1;Pangolin_CoV_orf1a;3616 | 0 | 0 |
| MT072864.1;QIQ54046.1;Pangolin_CoV_orf1a;3617 | 0 | 0 |
| MT072864.1;QIQ54046.1;Pangolin_CoV_orf1a;3618 | 0 | 0 |
| MT072864.1;QIQ54046.1;Pangolin_CoV_orf1a;3619 | 0 | 0 |
| MT072864.1;QIQ54046.1;Pangolin_CoV_orf1a;3620 | 0 | 0 |
| MT072864.1;QIQ54046.1;Pangolin_CoV_orf1a;3621 | 0 | 0 |
| MT072864.1;QIQ54046.1;Pangolin_CoV_orf1a;3622 | 0 | 0 |
| MT072864.1;QIQ54046.1;Pangolin_CoV_orf1a;3623 | 0 | 0 |
| MT072864.1;QIQ54046.1;Pangolin_CoV_orf1a;3624 | 0 | 0 |
| MT072864.1;QIQ54046.1;Pangolin_CoV_orf1a;3625 | 0 | 0 |
| MT072864.1;QIQ54046.1;Pangolin_CoV_orf1a;3626 | 0 | 0 |
| MT072864.1;QIQ54046.1;Pangolin_CoV_orf1a;3627 | 0 | 0 |
| MT072864.1;QIQ54046.1;Pangolin_CoV_orf1a;3628 | 0 | 0 |
| MT072864.1;QIQ54046.1;Pangolin_CoV_orf1a;3629 | 0 | 0 |
| MT072864.1;QIQ54046.1;Pangolin_CoV_orf1a;3630 | 0 | 0 |
| MT072864.1;QIQ54046.1;Pangolin_CoV_orf1a;3631 | 0 | 0 |
| MT072864.1;QIQ54046.1;Pangolin_CoV_orf1a;3632 | 0 | 0 |
| MT072864.1;QIQ54046.1;Pangolin_CoV_orf1a;3633 | 0 | 0 |
| MT072864.1;QIQ54046.1;Pangolin_CoV_orf1a;3634 | 0 | 0 |
| MT072864.1;QIQ54046.1;Pangolin_CoV_orf1a;3635 | 0 | 0 |
| MT072864.1;QIQ54046.1;Pangolin_CoV_orf1a;3636 | 0 | 0 |
| MT072864.1;QIQ54046.1;Pangolin_CoV_orf1a;3637 | 0 | 0 |
| MT072864.1;QIQ54046.1;Pangolin_CoV_orf1a;3638 | 0 | 0 |
| MT072864.1;QIQ54046.1;Pangolin_CoV_orf1a;3639 | 0 | 0 |
| MT072864.1;QIQ54046.1;Pangolin_CoV_orf1a;3640 | 0 | 0 |
| MT072864.1;QIQ54046.1;Pangolin_CoV_orf1a;3641 | 0 | 0 |
| MT072864.1;QIQ54046.1;Pangolin_CoV_orf1a;3642 | 0 | 0 |
| MT072864.1;QIQ54046.1;Pangolin_CoV_orf1a;3643 | 0 | 0 |
| MT072864.1;QIQ54046.1;Pangolin_CoV_orf1a;3644 | 0 | 0 |
| MT072864.1;QIQ54046.1;Pangolin_CoV_orf1a;3645 | 0 | 0 |

| | | |
|---|---|---|
| MT072864.1;QIQ54046.1;Pangolin_CoV_orf1a;3646 | 0 | 0 |
| MT072864.1;QIQ54046.1;Pangolin_CoV_orf1a;3647 | 0 | 0 |
| MT072864.1;QIQ54046.1;Pangolin_CoV_orf1a;3648 | 0 | 0 |
| MT072864.1;QIQ54046.1;Pangolin_CoV_orf1a;3649 | 0 | 0 |
| MT072864.1;QIQ54046.1;Pangolin_CoV_orf1a;3650 | 0 | 0 |
| MT072864.1;QIQ54046.1;Pangolin_CoV_orf1a;3651 | 0 | 0 |
| MT072864.1;QIQ54046.1;Pangolin_CoV_orf1a;3652 | 0 | 0 |
| MT072864.1;QIQ54046.1;Pangolin_CoV_orf1a;3653 | 0 | 0 |
| MT072864.1;QIQ54046.1;Pangolin_CoV_orf1a;3654 | 0 | 0 |
| MT072864.1;QIQ54046.1;Pangolin_CoV_orf1a;3655 | 0 | 0 |
| MT072864.1;QIQ54046.1;Pangolin_CoV_orf1a;3656 | 0 | 0.025 |
| MT072864.1;QIQ54046.1;Pangolin_CoV_orf1a;3657 | 0 | 0.025 |
| MT072864.1;QIQ54046.1;Pangolin_CoV_orf1a;3658 | 0 | 0.025 |
| MT072864.1;QIQ54046.1;Pangolin_CoV_orf1a;3659 | 0 | 0.025 |
| MT072864.1;QIQ54046.1;Pangolin_CoV_orf1a;3660 | 0 | 0.025 |
| MT072864.1;QIQ54046.1;Pangolin_CoV_orf1a;3661 | 0 | 0.025 |
| MT072864.1;QIQ54046.1;Pangolin_CoV_orf1a;3662 | 0 | 0.025 |
| MT072864.1;QIQ54046.1;Pangolin_CoV_orf1a;3663 | 0 | 0.025 |
| MT072864.1;QIQ54046.1;Pangolin_CoV_orf1a;3664 | 0 | 0.025 |
| MT072864.1;QIQ54046.1;Pangolin_CoV_orf1a;3665 | 0 | 0 |
| MT072864.1;QIQ54046.1;Pangolin_CoV_orf1a;3666 | 0 | 0 |
| MT072864.1;QIQ54046.1;Pangolin_CoV_orf1a;3667 | 0 | 0 |
| MT072864.1;QIQ54046.1;Pangolin_CoV_orf1a;3668 | 0 | 0 |
| MT072864.1;QIQ54046.1;Pangolin_CoV_orf1a;3669 | 0 | 0 |
| MT072864.1;QIQ54046.1;Pangolin_CoV_orf1a;3670 | 0 | 0 |
| MT072864.1;QIQ54046.1;Pangolin_CoV_orf1a;3671 | 0 | 0 |
| MT072864.1;QIQ54046.1;Pangolin_CoV_orf1a;3672 | 0 | 0 |
| MT072864.1;QIQ54046.1;Pangolin_CoV_orf1a;3673 | 0 | 0 |
| MT072864.1;QIQ54046.1;Pangolin_CoV_orf1a;3674 | 0 | 0 |
| MT072864.1;QIQ54046.1;Pangolin_CoV_orf1a;3675 | 0 | 0 |
| MT072864.1;QIQ54046.1;Pangolin_CoV_orf1a;3676 | 0 | 0 |
| MT072864.1;QIQ54046.1;Pangolin_CoV_orf1a;3677 | 0 | 0 |
| MT072864.1;QIQ54046.1;Pangolin_CoV_orf1a;3678 | 0 | 0 |
| MT072864.1;QIQ54046.1;Pangolin_CoV_orf1a;3679 | 0 | 0 |
| MT072864.1;QIQ54046.1;Pangolin_CoV_orf1a;3680 | 0 | 0 |
| MT072864.1;QIQ54046.1;Pangolin_CoV_orf1a;3681 | 0 | 0 |
| MT072864.1;QIQ54046.1;Pangolin_CoV_orf1a;3682 | 0 | 0 |
| MT072864.1;QIQ54046.1;Pangolin_CoV_orf1a;3683 | 0 | 0 |
| MT072864.1;QIQ54046.1;Pangolin_CoV_orf1a;3684 | 0 | 0 |
| MT072864.1;QIQ54046.1;Pangolin_CoV_orf1a;3685 | 0 | 0 |
| MT072864.1;QIQ54046.1;Pangolin_CoV_orf1a;3686 | 0 | 0 |
| MT072864.1;QIQ54046.1;Pangolin_CoV_orf1a;3687 | 0 | 0 |
| MT072864.1;QIQ54046.1;Pangolin_CoV_orf1a;3688 | 0 | 0 |

| | | |
|---|---|---|
| MT072864.1;QIQ54046.1;Pangolin_CoV_orf1a;3689 | 0 | 0 |
| MT072864.1;QIQ54046.1;Pangolin_CoV_orf1a;3690 | 0 | 0.025 |
| MT072864.1;QIQ54046.1;Pangolin_CoV_orf1a;3691 | 0 | 0 |
| MT072864.1;QIQ54046.1;Pangolin_CoV_orf1a;3692 | 0 | 0 |
| MT072864.1;QIQ54046.1;Pangolin_CoV_orf1a;3693 | 0 | 0 |
| MT072864.1;QIQ54046.1;Pangolin_CoV_orf1a;3694 | 0 | 0 |
| MT072864.1;QIQ54046.1;Pangolin_CoV_orf1a;3695 | 0 | 0 |
| MT072864.1;QIQ54046.1;Pangolin_CoV_orf1a;3696 | 0 | 0 |
| MT072864.1;QIQ54046.1;Pangolin_CoV_orf1a;3697 | 0 | 0 |
| MT072864.1;QIQ54046.1;Pangolin_CoV_orf1a;3698 | 0 | 0 |
| MT072864.1;QIQ54046.1;Pangolin_CoV_orf1a;3699 | 0 | 0 |
| MT072864.1;QIQ54046.1;Pangolin_CoV_orf1a;3700 | 0 | 0 |
| MT072864.1;QIQ54046.1;Pangolin_CoV_orf1a;3701 | 0 | 0 |
| MT072864.1;QIQ54046.1;Pangolin_CoV_orf1a;3702 | 0 | 0 |
| MT072864.1;QIQ54046.1;Pangolin_CoV_orf1a;3703 | 0 | 0 |
| MT072864.1;QIQ54046.1;Pangolin_CoV_orf1a;3704 | 0 | 0 |
| MT072864.1;QIQ54046.1;Pangolin_CoV_orf1a;3705 | 0 | 0 |
| MT072864.1;QIQ54046.1;Pangolin_CoV_orf1a;3706 | 0 | 0 |
| MT072864.1;QIQ54046.1;Pangolin_CoV_orf1a;3707 | 0 | 0 |
| MT072864.1;QIQ54046.1;Pangolin_CoV_orf1a;3708 | 0 | 0 |
| MT072864.1;QIQ54046.1;Pangolin_CoV_orf1a;3709 | 0 | 0 |
| MT072864.1;QIQ54046.1;Pangolin_CoV_orf1a;3710 | 0.05 | 0 |
| MT072864.1;QIQ54046.1;Pangolin_CoV_orf1a;3711 | 0.05 | 0 |
| MT072864.1;QIQ54046.1;Pangolin_CoV_orf1a;3712 | 0.05 | 0 |
| MT072864.1;QIQ54046.1;Pangolin_CoV_orf1a;3713 | 0.05 | 0.025 |
| MT072864.1;QIQ54046.1;Pangolin_CoV_orf1a;3714 | 0.05 | 0 |
| MT072864.1;QIQ54046.1;Pangolin_CoV_orf1a;3715 | 0 | 0 |
| MT072864.1;QIQ54046.1;Pangolin_CoV_orf1a;3716 | 0 | 0 |
| MT072864.1;QIQ54046.1;Pangolin_CoV_orf1a;3717 | 0 | 0 |
| MT072864.1;QIQ54046.1;Pangolin_CoV_orf1a;3718 | 0 | 0 |
| MT072864.1;QIQ54046.1;Pangolin_CoV_orf1a;3719 | 0 | 0 |
| MT072864.1;QIQ54046.1;Pangolin_CoV_orf1a;3720 | 0 | 0 |
| MT072864.1;QIQ54046.1;Pangolin_CoV_orf1a;3721 | 0 | 0 |
| MT072864.1;QIQ54046.1;Pangolin_CoV_orf1a;3722 | 0 | 0 |
| MT072864.1;QIQ54046.1;Pangolin_CoV_orf1a;3723 | 0 | 0 |
| MT072864.1;QIQ54046.1;Pangolin_CoV_orf1a;3724 | 0 | 0 |
| MT072864.1;QIQ54046.1;Pangolin_CoV_orf1a;3725 | 0 | 0 |
| MT072864.1;QIQ54046.1;Pangolin_CoV_orf1a;3726 | 0 | 0 |
| MT072864.1;QIQ54046.1;Pangolin_CoV_orf1a;3727 | 0 | 0 |
| MT072864.1;QIQ54046.1;Pangolin_CoV_orf1a;3728 | 0 | 0 |
| MT072864.1;QIQ54046.1;Pangolin_CoV_orf1a;3729 | 0 | 0 |
| MT072864.1;QIQ54046.1;Pangolin_CoV_orf1a;3730 | 0 | 0 |
| MT072864.1;QIQ54046.1;Pangolin_CoV_orf1a;3731 | 0 | 0 |

| | | |
|---|---|---|
| MT072864.1;QIQ54046.1;Pangolin_CoV_orf1a;3732 | 0 | 0 |
| MT072864.1;QIQ54046.1;Pangolin_CoV_orf1a;3733 | 0 | 0 |
| MT072864.1;QIQ54046.1;Pangolin_CoV_orf1a;3734 | 0 | 0 |
| MT072864.1;QIQ54046.1;Pangolin_CoV_orf1a;3735 | 0 | 0 |
| MT072864.1;QIQ54046.1;Pangolin_CoV_orf1a;3736 | 0 | 0 |
| MT072864.1;QIQ54046.1;Pangolin_CoV_orf1a;3737 | 0 | 0 |
| MT072864.1;QIQ54046.1;Pangolin_CoV_orf1a;3738 | 0 | 0 |
| MT072864.1;QIQ54046.1;Pangolin_CoV_orf1a;3739 | 0 | 0 |
| MT072864.1;QIQ54046.1;Pangolin_CoV_orf1a;3740 | 0 | 0 |
| MT072864.1;QIQ54046.1;Pangolin_CoV_orf1a;3741 | 0 | 0 |
| MT072864.1;QIQ54046.1;Pangolin_CoV_orf1a;3742 | 0 | 0 |
| MT072864.1;QIQ54046.1;Pangolin_CoV_orf1a;3743 | 0 | 0 |
| MT072864.1;QIQ54046.1;Pangolin_CoV_orf1a;3744 | 0 | 0 |
| MT072864.1;QIQ54046.1;Pangolin_CoV_orf1a;3745 | 0 | 0 |
| MT072864.1;QIQ54046.1;Pangolin_CoV_orf1a;3746 | 0 | 0 |
| MT072864.1;QIQ54046.1;Pangolin_CoV_orf1a;3747 | 0 | 0 |
| MT072864.1;QIQ54046.1;Pangolin_CoV_orf1a;3748 | 0 | 0 |
| MT072864.1;QIQ54046.1;Pangolin_CoV_orf1a;3749 | 0 | 0 |
| MT072864.1;QIQ54046.1;Pangolin_CoV_orf1a;3750 | 0 | 0 |
| MT072864.1;QIQ54046.1;Pangolin_CoV_orf1a;3751 | 0 | 0 |
| MT072864.1;QIQ54046.1;Pangolin_CoV_orf1a;3752 | 0 | 0 |
| MT072864.1;QIQ54046.1;Pangolin_CoV_orf1a;3753 | 0 | 0 |
| MT072864.1;QIQ54046.1;Pangolin_CoV_orf1a;3754 | 0 | 0 |
| MT072864.1;QIQ54046.1;Pangolin_CoV_orf1a;3755 | 0 | 0 |
| MT072864.1;QIQ54046.1;Pangolin_CoV_orf1a;3756 | 0 | 0 |
| MT072864.1;QIQ54046.1;Pangolin_CoV_orf1a;3757 | 0 | 0 |
| MT072864.1;QIQ54046.1;Pangolin_CoV_orf1a;3758 | 0 | 0 |
| MT072864.1;QIQ54046.1;Pangolin_CoV_orf1a;3759 | 0 | 0 |
| MT072864.1;QIQ54046.1;Pangolin_CoV_orf1a;3760 | 0 | 0 |
| MT072864.1;QIQ54046.1;Pangolin_CoV_orf1a;3761 | 0 | 0 |
| MT072864.1;QIQ54046.1;Pangolin_CoV_orf1a;3762 | 0 | 0 |
| MT072864.1;QIQ54046.1;Pangolin_CoV_orf1a;3763 | 0 | 0 |
| MT072864.1;QIQ54046.1;Pangolin_CoV_orf1a;3764 | 0 | 0 |
| MT072864.1;QIQ54046.1;Pangolin_CoV_orf1a;3765 | 0 | 0 |
| MT072864.1;QIQ54046.1;Pangolin_CoV_orf1a;3766 | 0 | 0 |
| MT072864.1;QIQ54046.1;Pangolin_CoV_orf1a;3767 | 0 | 0 |
| MT072864.1;QIQ54046.1;Pangolin_CoV_orf1a;3768 | 0 | 0 |
| MT072864.1;QIQ54046.1;Pangolin_CoV_orf1a;3769 | 0 | 0 |
| MT072864.1;QIQ54046.1;Pangolin_CoV_orf1a;3770 | 0 | 0 |
| MT072864.1;QIQ54046.1;Pangolin_CoV_orf1a;3771 | 0 | 0 |
| MT072864.1;QIQ54046.1;Pangolin_CoV_orf1a;3772 | 0 | 0 |
| MT072864.1;QIQ54046.1;Pangolin_CoV_orf1a;3773 | 0 | 0 |
| MT072864.1;QIQ54046.1;Pangolin_CoV_orf1a;3774 | 0 | 0 |

| | | |
|---|---|---|
| MT072864.1;QIQ54046.1;Pangolin_CoV_orf1a;3775 | 0 | 0 |
| MT072864.1;QIQ54046.1;Pangolin_CoV_orf1a;3776 | 0 | 0 |
| MT072864.1;QIQ54046.1;Pangolin_CoV_orf1a;3777 | 0 | 0 |
| MT072864.1;QIQ54046.1;Pangolin_CoV_orf1a;3778 | 0 | 0 |
| MT072864.1;QIQ54046.1;Pangolin_CoV_orf1a;3779 | 0 | 0 |
| MT072864.1;QIQ54046.1;Pangolin_CoV_orf1a;3780 | 0 | 0 |
| MT072864.1;QIQ54046.1;Pangolin_CoV_orf1a;3781 | 0 | 0 |
| MT072864.1;QIQ54046.1;Pangolin_CoV_orf1a;3782 | 0 | 0 |
| MT072864.1;QIQ54046.1;Pangolin_CoV_orf1a;3783 | 0 | 0 |
| MT072864.1;QIQ54046.1;Pangolin_CoV_orf1a;3784 | 0 | 0 |
| MT072864.1;QIQ54046.1;Pangolin_CoV_orf1a;3785 | 0 | 0 |
| MT072864.1;QIQ54046.1;Pangolin_CoV_orf1a;3786 | 0 | 0 |
| MT072864.1;QIQ54046.1;Pangolin_CoV_orf1a;3787 | 0 | 0 |
| MT072864.1;QIQ54046.1;Pangolin_CoV_orf1a;3788 | 0 | 0 |
| MT072864.1;QIQ54046.1;Pangolin_CoV_orf1a;3789 | 0 | 0 |
| MT072864.1;QIQ54046.1;Pangolin_CoV_orf1a;3790 | 0 | 0 |
| MT072864.1;QIQ54046.1;Pangolin_CoV_orf1a;3791 | 0 | 0 |
| MT072864.1;QIQ54046.1;Pangolin_CoV_orf1a;3792 | 0 | 0 |
| MT072864.1;QIQ54046.1;Pangolin_CoV_orf1a;3793 | 0 | 0 |
| MT072864.1;QIQ54046.1;Pangolin_CoV_orf1a;3794 | 0 | 0 |
| MT072864.1;QIQ54046.1;Pangolin_CoV_orf1a;3795 | 0 | 0 |
| MT072864.1;QIQ54046.1;Pangolin_CoV_orf1a;3796 | 0 | 0 |
| MT072864.1;QIQ54046.1;Pangolin_CoV_orf1a;3797 | 0 | 0 |
| MT072864.1;QIQ54046.1;Pangolin_CoV_orf1a;3798 | 0 | 0 |
| MT072864.1;QIQ54046.1;Pangolin_CoV_orf1a;3799 | 0 | 0 |
| MT072864.1;QIQ54046.1;Pangolin_CoV_orf1a;3800 | 0 | 0 |
| MT072864.1;QIQ54046.1;Pangolin_CoV_orf1a;3801 | 0.25 | 0.175 |
| MT072864.1;QIQ54046.1;Pangolin_CoV_orf1a;3802 | 0 | 0 |
| MT072864.1;QIQ54046.1;Pangolin_CoV_orf1a;3803 | 0 | 0 |
| MT072864.1;QIQ54046.1;Pangolin_CoV_orf1a;3804 | 0 | 0 |
| MT072864.1;QIQ54046.1;Pangolin_CoV_orf1a;3805 | 0 | 0 |
| MT072864.1;QIQ54046.1;Pangolin_CoV_orf1a;3806 | 0 | 0 |
| MT072864.1;QIQ54046.1;Pangolin_CoV_orf1a;3807 | 0 | 0 |
| MT072864.1;QIQ54046.1;Pangolin_CoV_orf1a;3808 | 0 | 0 |
| MT072864.1;QIQ54046.1;Pangolin_CoV_orf1a;3809 | 0 | 0.05 |
| MT072864.1;QIQ54046.1;Pangolin_CoV_orf1a;3810 | 0.05 | 0.025 |
| MT072864.1;QIQ54046.1;Pangolin_CoV_orf1a;3811 | 0.15 | 0.075 |
| MT072864.1;QIQ54046.1;Pangolin_CoV_orf1a;3812 | 0 | 0.025 |
| MT072864.1;QIQ54046.1;Pangolin_CoV_orf1a;3813 | 0 | 0.025 |
| MT072864.1;QIQ54046.1;Pangolin_CoV_orf1a;3814 | 0.05 | 0.125 |
| MT072864.1;QIQ54046.1;Pangolin_CoV_orf1a;3815 | 0.1 | 0.05 |
| MT072864.1;QIQ54046.1;Pangolin_CoV_orf1a;3816 | 0.3 | 0.1 |
| MT072864.1;QIQ54046.1;Pangolin_CoV_orf1a;3817 | 0.25 | 0.1 |

| | | |
|---|---|---|
| MT072864.1;QIQ54046.1;Pangolin_CoV_orf1a;3818 | 0.25 | 0.1 |
| MT072864.1;QIQ54046.1;Pangolin_CoV_orf1a;3819 | 0.2 | 0.125 |
| MT072864.1;QIQ54046.1;Pangolin_CoV_orf1a;3820 | 0.15 | 0.075 |
| MT072864.1;QIQ54046.1;Pangolin_CoV_orf1a;3821 | 0.1 | 0.075 |
| MT072864.1;QIQ54046.1;Pangolin_CoV_orf1a;3822 | 0.05 | 0.075 |
| MT072864.1;QIQ54046.1;Pangolin_CoV_orf1a;3823 | 0.05 | 0.05 |
| MT072864.1;QIQ54046.1;Pangolin_CoV_orf1a;3824 | 0 | 0.05 |
| MT072864.1;QIQ54046.1;Pangolin_CoV_orf1a;3825 | 0 | 0 |
| MT072864.1;QIQ54046.1;Pangolin_CoV_orf1a;3826 | 0 | 0 |
| MT072864.1;QIQ54046.1;Pangolin_CoV_orf1a;3827 | 0 | 0 |
| MT072864.1;QIQ54046.1;Pangolin_CoV_orf1a;3828 | 0 | 0 |
| MT072864.1;QIQ54046.1;Pangolin_CoV_orf1a;3829 | 0 | 0 |
| MT072864.1;QIQ54046.1;Pangolin_CoV_orf1a;3830 | 0 | 0 |
| MT072864.1;QIQ54046.1;Pangolin_CoV_orf1a;3831 | 0 | 0 |
| MT072864.1;QIQ54046.1;Pangolin_CoV_orf1a;3832 | 0 | 0 |
| MT072864.1;QIQ54046.1;Pangolin_CoV_orf1a;3833 | 0 | 0 |
| MT072864.1;QIQ54046.1;Pangolin_CoV_orf1a;3834 | 0 | 0 |
| MT072864.1;QIQ54046.1;Pangolin_CoV_orf1a;3835 | 0 | 0 |
| MT072864.1;QIQ54046.1;Pangolin_CoV_orf1a;3836 | 0 | 0.025 |
| MT072864.1;QIQ54046.1;Pangolin_CoV_orf1a;3837 | 0 | 0.025 |
| MT072864.1;QIQ54046.1;Pangolin_CoV_orf1a;3838 | 0.05 | 0.05 |
| MT072864.1;QIQ54046.1;Pangolin_CoV_orf1a;3839 | 0.05 | 0.05 |
| MT072864.1;QIQ54046.1;Pangolin_CoV_orf1a;3840 | 0.05 | 0.05 |
| MT072864.1;QIQ54046.1;Pangolin_CoV_orf1a;3841 | 0.1 | 0.1 |
| MT072864.1;QIQ54046.1;Pangolin_CoV_orf1a;3842 | 0.05 | 0.075 |
| MT072864.1;QIQ54046.1;Pangolin_CoV_orf1a;3843 | 0.05 | 0.05 |
| MT072864.1;QIQ54046.1;Pangolin_CoV_orf1a;3844 | 0.05 | 0.025 |
| MT072864.1;QIQ54046.1;Pangolin_CoV_orf1a;3845 | 0 | 0.025 |
| MT072864.1;QIQ54046.1;Pangolin_CoV_orf1a;3846 | 0.05 | 0.025 |
| MT072864.1;QIQ54046.1;Pangolin_CoV_orf1a;3847 | 0.05 | 0.025 |
| MT072864.1;QIQ54046.1;Pangolin_CoV_orf1a;3848 | 0.05 | 0.025 |
| MT072864.1;QIQ54046.1;Pangolin_CoV_orf1a;3849 | 0.05 | 0 |
| MT072864.1;QIQ54046.1;Pangolin_CoV_orf1a;3850 | 0 | 0 |
| MT072864.1;QIQ54046.1;Pangolin_CoV_orf1a;3851 | 0 | 0 |
| MT072864.1;QIQ54046.1;Pangolin_CoV_orf1a;3852 | 0 | 0 |
| MT072864.1;QIQ54046.1;Pangolin_CoV_orf1a;3853 | 0 | 0 |
| MT072864.1;QIQ54046.1;Pangolin_CoV_orf1a;3854 | 0 | 0 |
| MT072864.1;QIQ54046.1;Pangolin_CoV_orf1a;3855 | 0 | 0 |
| MT072864.1;QIQ54046.1;Pangolin_CoV_orf1a;3856 | 0 | 0 |
| MT072864.1;QIQ54046.1;Pangolin_CoV_orf1a;3857 | 0 | 0 |
| MT072864.1;QIQ54046.1;Pangolin_CoV_orf1a;3858 | 0 | 0 |
| MT072864.1;QIQ54046.1;Pangolin_CoV_orf1a;3859 | 0 | 0 |
| MT072864.1;QIQ54046.1;Pangolin_CoV_orf1a;3860 | 0 | 0 |

| | | |
|---|---|---|
| MT072864.1;QIQ54046.1;Pangolin_CoV_orf1a;3861 | 0 | 0 |
| MT072864.1;QIQ54046.1;Pangolin_CoV_orf1a;3862 | 0 | 0 |
| MT072864.1;QIQ54046.1;Pangolin_CoV_orf1a;3863 | 0 | 0 |
| MT072864.1;QIQ54046.1;Pangolin_CoV_orf1a;3864 | 0 | 0 |
| MT072864.1;QIQ54046.1;Pangolin_CoV_orf1a;3865 | 0 | 0 |
| MT072864.1;QIQ54046.1;Pangolin_CoV_orf1a;3866 | 0 | 0 |
| MT072864.1;QIQ54046.1;Pangolin_CoV_orf1a;3867 | 0 | 0 |
| MT072864.1;QIQ54046.1;Pangolin_CoV_orf1a;3868 | 0 | 0 |
| MT072864.1;QIQ54046.1;Pangolin_CoV_orf1a;3869 | 0 | 0 |
| MT072864.1;QIQ54046.1;Pangolin_CoV_orf1a;3870 | 0 | 0 |
| MT072864.1;QIQ54046.1;Pangolin_CoV_orf1a;3871 | 0 | 0 |
| MT072864.1;QIQ54046.1;Pangolin_CoV_orf1a;3872 | 0 | 0 |
| MT072864.1;QIQ54046.1;Pangolin_CoV_orf1a;3873 | 0 | 0 |
| MT072864.1;QIQ54046.1;Pangolin_CoV_orf1a;3874 | 0 | 0.025 |
| MT072864.1;QIQ54046.1;Pangolin_CoV_orf1a;3875 | 0 | 0 |
| MT072864.1;QIQ54046.1;Pangolin_CoV_orf1a;3876 | 0 | 0.025 |
| MT072864.1;QIQ54046.1;Pangolin_CoV_orf1a;3877 | 0 | 0 |
| MT072864.1;QIQ54046.1;Pangolin_CoV_orf1a;3878 | 0 | 0 |
| MT072864.1;QIQ54046.1;Pangolin_CoV_orf1a;3879 | 0 | 0 |
| MT072864.1;QIQ54046.1;Pangolin_CoV_orf1a;3880 | 0.05 | 0 |
| MT072864.1;QIQ54046.1;Pangolin_CoV_orf1a;3881 | 0.05 | 0 |
| MT072864.1;QIQ54046.1;Pangolin_CoV_orf1a;3882 | 0.05 | 0 |
| MT072864.1;QIQ54046.1;Pangolin_CoV_orf1a;3883 | 0 | 0 |
| MT072864.1;QIQ54046.1;Pangolin_CoV_orf1a;3884 | 0 | 0 |
| MT072864.1;QIQ54046.1;Pangolin_CoV_orf1a;3885 | 0 | 0 |
| MT072864.1;QIQ54046.1;Pangolin_CoV_orf1a;3886 | 0 | 0 |
| MT072864.1;QIQ54046.1;Pangolin_CoV_orf1a;3887 | 0 | 0 |
| MT072864.1;QIQ54046.1;Pangolin_CoV_orf1a;3888 | 0 | 0 |
| MT072864.1;QIQ54046.1;Pangolin_CoV_orf1a;3889 | 0 | 0 |
| MT072864.1;QIQ54046.1;Pangolin_CoV_orf1a;3890 | 0 | 0 |
| MT072864.1;QIQ54046.1;Pangolin_CoV_orf1a;3891 | 0 | 0 |
| MT072864.1;QIQ54046.1;Pangolin_CoV_orf1a;3892 | 0 | 0 |
| MT072864.1;QIQ54046.1;Pangolin_CoV_orf1a;3893 | 0 | 0.05 |
| MT072864.1;QIQ54046.1;Pangolin_CoV_orf1a;3894 | 0 | 0 |
| MT072864.1;QIQ54046.1;Pangolin_CoV_orf1a;3895 | 0 | 0 |
| MT072864.1;QIQ54046.1;Pangolin_CoV_orf1a;3896 | 0 | 0 |
| MT072864.1;QIQ54046.1;Pangolin_CoV_orf1a;3897 | 0 | 0 |
| MT072864.1;QIQ54046.1;Pangolin_CoV_orf1a;3898 | 0 | 0 |
| MT072864.1;QIQ54046.1;Pangolin_CoV_orf1a;3899 | 0 | 0 |
| MT072864.1;QIQ54046.1;Pangolin_CoV_orf1a;3900 | 0 | 0 |
| MT072864.1;QIQ54046.1;Pangolin_CoV_orf1a;3901 | 0 | 0 |
| MT072864.1;QIQ54046.1;Pangolin_CoV_orf1a;3902 | 0 | 0 |
| MT072864.1;QIQ54046.1;Pangolin_CoV_orf1a;3903 | 0 | 0 |

| | | |
|---|---|---|
| MT072864.1;QIQ54046.1;Pangolin_CoV_orf1a;3904 | 0 | 0 |
| MT072864.1;QIQ54046.1;Pangolin_CoV_orf1a;3905 | 0 | 0 |
| MT072864.1;QIQ54046.1;Pangolin_CoV_orf1a;3906 | 0 | 0.05 |
| MT072864.1;QIQ54046.1;Pangolin_CoV_orf1a;3907 | 0 | 0.025 |
| MT072864.1;QIQ54046.1;Pangolin_CoV_orf1a;3908 | 0 | 0.025 |
| MT072864.1;QIQ54046.1;Pangolin_CoV_orf1a;3909 | 0 | 0.025 |
| MT072864.1;QIQ54046.1;Pangolin_CoV_orf1a;3910 | 0 | 0.025 |
| MT072864.1;QIQ54046.1;Pangolin_CoV_orf1a;3911 | 0.05 | 0.05 |
| MT072864.1;QIQ54046.1;Pangolin_CoV_orf1a;3912 | 0.05 | 0.05 |
| MT072864.1;QIQ54046.1;Pangolin_CoV_orf1a;3913 | 0.05 | 0.075 |
| MT072864.1;QIQ54046.1;Pangolin_CoV_orf1a;3914 | 0.05 | 0.05 |
| MT072864.1;QIQ54046.1;Pangolin_CoV_orf1a;3915 | 0.1 | 0.025 |
| MT072864.1;QIQ54046.1;Pangolin_CoV_orf1a;3916 | 0 | 0.025 |
| MT072864.1;QIQ54046.1;Pangolin_CoV_orf1a;3917 | 0 | 0.05 |
| MT072864.1;QIQ54046.1;Pangolin_CoV_orf1a;3918 | 0 | 0.025 |
| MT072864.1;QIQ54046.1;Pangolin_CoV_orf1a;3919 | 0 | 0 |
| MT072864.1;QIQ54046.1;Pangolin_CoV_orf1a;3920 | 0 | 0 |
| MT072864.1;QIQ54046.1;Pangolin_CoV_orf1a;3921 | 0.05 | 0 |
| MT072864.1;QIQ54046.1;Pangolin_CoV_orf1a;3922 | 0.05 | 0 |
| MT072864.1;QIQ54046.1;Pangolin_CoV_orf1a;3923 | 0.05 | 0 |
| MT072864.1;QIQ54046.1;Pangolin_CoV_orf1a;3924 | 0 | 0 |
| MT072864.1;QIQ54046.1;Pangolin_CoV_orf1a;3925 | 0.05 | 0 |
| MT072864.1;QIQ54046.1;Pangolin_CoV_orf1a;3926 | 0 | 0 |
| MT072864.1;QIQ54046.1;Pangolin_CoV_orf1a;3927 | 0 | 0 |
| MT072864.1;QIQ54046.1;Pangolin_CoV_orf1a;3928 | 0 | 0 |
| MT072864.1;QIQ54046.1;Pangolin_CoV_orf1a;3929 | 0 | 0 |
| MT072864.1;QIQ54046.1;Pangolin_CoV_orf1a;3930 | 0 | 0 |
| MT072864.1;QIQ54046.1;Pangolin_CoV_orf1a;3931 | 0 | 0 |
| MT072864.1;QIQ54046.1;Pangolin_CoV_orf1a;3932 | 0 | 0 |
| MT072864.1;QIQ54046.1;Pangolin_CoV_orf1a;3933 | 0.05 | 0.05 |
| MT072864.1;QIQ54046.1;Pangolin_CoV_orf1a;3934 | 0 | 0.025 |
| MT072864.1;QIQ54046.1;Pangolin_CoV_orf1a;3935 | 0 | 0 |
| MT072864.1;QIQ54046.1;Pangolin_CoV_orf1a;3936 | 0 | 0 |
| MT072864.1;QIQ54046.1;Pangolin_CoV_orf1a;3937 | 0 | 0 |
| MT072864.1;QIQ54046.1;Pangolin_CoV_orf1a;3938 | 0 | 0.05 |
| MT072864.1;QIQ54046.1;Pangolin_CoV_orf1a;3939 | 0.05 | 0 |
| MT072864.1;QIQ54046.1;Pangolin_CoV_orf1a;3940 | 0.05 | 0 |
| MT072864.1;QIQ54046.1;Pangolin_CoV_orf1a;3941 | 0.05 | 0 |
| MT072864.1;QIQ54046.1;Pangolin_CoV_orf1a;3942 | 0.05 | 0.05 |
| MT072864.1;QIQ54046.1;Pangolin_CoV_orf1a;3943 | 0.1 | 0.05 |
| MT072864.1;QIQ54046.1;Pangolin_CoV_orf1a;3944 | 0 | 0.025 |
| MT072864.1;QIQ54046.1;Pangolin_CoV_orf1a;3945 | 0.1 | 0.05 |
| MT072864.1;QIQ54046.1;Pangolin_CoV_orf1a;3946 | 0.05 | 0.125 |

| | | |
|---|---|---|
| MT072864.1;QIQ54046.1;Pangolin_CoV_orf1a;3947 | 0.05 | 0.075 |
| MT072864.1;QIQ54046.1;Pangolin_CoV_orf1a;3948 | 0.05 | 0.15 |
| MT072864.1;QIQ54046.1;Pangolin_CoV_orf1a;3949 | 0.05 | 0.05 |
| MT072864.1;QIQ54046.1;Pangolin_CoV_orf1a;3950 | 0 | 0.025 |
| MT072864.1;QIQ54046.1;Pangolin_CoV_orf1a;3951 | 0 | 0 |
| MT072864.1;QIQ54046.1;Pangolin_CoV_orf1a;3952 | 0.05 | 0 |
| MT072864.1;QIQ54046.1;Pangolin_CoV_orf1a;3953 | 0 | 0 |
| MT072864.1;QIQ54046.1;Pangolin_CoV_orf1a;3954 | 0 | 0 |
| MT072864.1;QIQ54046.1;Pangolin_CoV_orf1a;3955 | 0 | 0 |
| MT072864.1;QIQ54046.1;Pangolin_CoV_orf1a;3956 | 0 | 0 |
| MT072864.1;QIQ54046.1;Pangolin_CoV_orf1a;3957 | 0 | 0 |
| MT072864.1;QIQ54046.1;Pangolin_CoV_orf1a;3958 | 0 | 0 |
| MT072864.1;QIQ54046.1;Pangolin_CoV_orf1a;3959 | 0 | 0 |
| MT072864.1;QIQ54046.1;Pangolin_CoV_orf1a;3960 | 0 | 0 |
| MT072864.1;QIQ54046.1;Pangolin_CoV_orf1a;3961 | 0 | 0 |
| MT072864.1;QIQ54046.1;Pangolin_CoV_orf1a;3962 | 0 | 0 |
| MT072864.1;QIQ54046.1;Pangolin_CoV_orf1a;3963 | 0 | 0 |
| MT072864.1;QIQ54046.1;Pangolin_CoV_orf1a;3964 | 0 | 0 |
| MT072864.1;QIQ54046.1;Pangolin_CoV_orf1a;3965 | 0 | 0 |
| MT072864.1;QIQ54046.1;Pangolin_CoV_orf1a;3966 | 0 | 0 |
| MT072864.1;QIQ54046.1;Pangolin_CoV_orf1a;3967 | 0 | 0 |
| MT072864.1;QIQ54046.1;Pangolin_CoV_orf1a;3968 | 0 | 0 |
| MT072864.1;QIQ54046.1;Pangolin_CoV_orf1a;3969 | 0 | 0.025 |
| MT072864.1;QIQ54046.1;Pangolin_CoV_orf1a;3970 | 0 | 0.05 |
| MT072864.1;QIQ54046.1;Pangolin_CoV_orf1a;3971 | 0 | 0.025 |
| MT072864.1;QIQ54046.1;Pangolin_CoV_orf1a;3972 | 0.05 | 0.025 |
| MT072864.1;QIQ54046.1;Pangolin_CoV_orf1a;3973 | 0.05 | 0.025 |
| MT072864.1;QIQ54046.1;Pangolin_CoV_orf1a;3974 | 0 | 0.05 |
| MT072864.1;QIQ54046.1;Pangolin_CoV_orf1a;3975 | 0.05 | 0.025 |
| MT072864.1;QIQ54046.1;Pangolin_CoV_orf1a;3976 | 0 | 0.025 |
| MT072864.1;QIQ54046.1;Pangolin_CoV_orf1a;3977 | 0 | 0 |
| MT072864.1;QIQ54046.1;Pangolin_CoV_orf1a;3978 | 0 | 0 |
| MT072864.1;QIQ54046.1;Pangolin_CoV_orf1a;3979 | 0 | 0 |
| MT072864.1;QIQ54046.1;Pangolin_CoV_orf1a;3980 | 0 | 0 |
| MT072864.1;QIQ54046.1;Pangolin_CoV_orf1a;3981 | 0 | 0 |
| MT072864.1;QIQ54046.1;Pangolin_CoV_orf1a;3982 | 0 | 0 |
| MT072864.1;QIQ54046.1;Pangolin_CoV_orf1a;3983 | 0 | 0 |
| MT072864.1;QIQ54046.1;Pangolin_CoV_orf1a;3984 | 0 | 0 |
| MT072864.1;QIQ54046.1;Pangolin_CoV_orf1a;3985 | 0 | 0 |
| MT072864.1;QIQ54046.1;Pangolin_CoV_orf1a;3986 | 0 | 0.025 |
| MT072864.1;QIQ54046.1;Pangolin_CoV_orf1a;3987 | 0 | 0 |
| MT072864.1;QIQ54046.1;Pangolin_CoV_orf1a;3988 | 0 | 0 |
| MT072864.1;QIQ54046.1;Pangolin_CoV_orf1a;3989 | 0 | 0 |

| | | |
|---|---|---|
| MT072864.1;QIQ54046.1;Pangolin_CoV_orf1a;3990 | 0 | 0.05 |
| MT072864.1;QIQ54046.1;Pangolin_CoV_orf1a;3991 | 0 | 0.025 |
| MT072864.1;QIQ54046.1;Pangolin_CoV_orf1a;3992 | 0 | 0 |
| MT072864.1;QIQ54046.1;Pangolin_CoV_orf1a;3993 | 0 | 0 |
| MT072864.1;QIQ54046.1;Pangolin_CoV_orf1a;3994 | 0 | 0 |
| MT072864.1;QIQ54046.1;Pangolin_CoV_orf1a;3995 | 0 | 0 |
| MT072864.1;QIQ54046.1;Pangolin_CoV_orf1a;3996 | 0 | 0 |
| MT072864.1;QIQ54046.1;Pangolin_CoV_orf1a;3997 | 0 | 0 |
| MT072864.1;QIQ54046.1;Pangolin_CoV_orf1a;3998 | 0 | 0.1 |
| MT072864.1;QIQ54046.1;Pangolin_CoV_orf1a;3999 | 0 | 0.075 |
| MT072864.1;QIQ54046.1;Pangolin_CoV_orf1a;4000 | 0 | 0.05 |
| MT072864.1;QIQ54046.1;Pangolin_CoV_orf1a;4001 | 0 | 0.05 |
| MT072864.1;QIQ54046.1;Pangolin_CoV_orf1a;4002 | 0 | 0.05 |
| MT072864.1;QIQ54046.1;Pangolin_CoV_orf1a;4003 | 0 | 0 |
| MT072864.1;QIQ54046.1;Pangolin_CoV_orf1a;4004 | 0 | 0 |
| MT072864.1;QIQ54046.1;Pangolin_CoV_orf1a;4005 | 0 | 0 |
| MT072864.1;QIQ54046.1;Pangolin_CoV_orf1a;4006 | 0 | 0 |
| MT072864.1;QIQ54046.1;Pangolin_CoV_orf1a;4007 | 0 | 0 |
| MT072864.1;QIQ54046.1;Pangolin_CoV_orf1a;4008 | 0 | 0 |
| MT072864.1;QIQ54046.1;Pangolin_CoV_orf1a;4009 | 0 | 0 |
| MT072864.1;QIQ54046.1;Pangolin_CoV_orf1a;4010 | 0 | 0 |
| MT072864.1;QIQ54046.1;Pangolin_CoV_orf1a;4011 | 0 | 0 |
| MT072864.1;QIQ54046.1;Pangolin_CoV_orf1a;4012 | 0 | 0 |
| MT072864.1;QIQ54046.1;Pangolin_CoV_orf1a;4013 | 0 | 0 |
| MT072864.1;QIQ54046.1;Pangolin_CoV_orf1a;4014 | 0 | 0 |
| MT072864.1;QIQ54046.1;Pangolin_CoV_orf1a;4015 | 0 | 0 |
| MT072864.1;QIQ54046.1;Pangolin_CoV_orf1a;4016 | 0 | 0 |
| MT072864.1;QIQ54046.1;Pangolin_CoV_orf1a;4017 | 0 | 0 |
| MT072864.1;QIQ54046.1;Pangolin_CoV_orf1a;4018 | 0 | 0 |
| MT072864.1;QIQ54046.1;Pangolin_CoV_orf1a;4019 | 0 | 0 |
| MT072864.1;QIQ54046.1;Pangolin_CoV_orf1a;4020 | 0 | 0 |
| MT072864.1;QIQ54046.1;Pangolin_CoV_orf1a;4021 | 0 | 0 |
| MT072864.1;QIQ54046.1;Pangolin_CoV_orf1a;4022 | 0 | 0 |
| MT072864.1;QIQ54046.1;Pangolin_CoV_orf1a;4023 | 0.15 | 0.1 |
| MT072864.1;QIQ54046.1;Pangolin_CoV_orf1a;4024 | 0.2 | 0.1 |
| MT072864.1;QIQ54046.1;Pangolin_CoV_orf1a;4025 | 0.2 | 0.075 |
| MT072864.1;QIQ54046.1;Pangolin_CoV_orf1a;4026 | 0.25 | 0.1 |
| MT072864.1;QIQ54046.1;Pangolin_CoV_orf1a;4027 | 0.25 | 0.275 |
| MT072864.1;QIQ54046.1;Pangolin_CoV_orf1a;4028 | 0.3 | 0.3 |
| MT072864.1;QIQ54046.1;Pangolin_CoV_orf1a;4029 | 0.35 | 0.4 |
| MT072864.1;QIQ54046.1;Pangolin_CoV_orf1a;4030 | 0.25 | 0.25 |
| MT072864.1;QIQ54046.1;Pangolin_CoV_orf1a;4031 | 0.4 | 0.5 |
| MT072864.1;QIQ54046.1;Pangolin_CoV_orf1a;4032 | 0.1 | 0.025 |

| | | |
|---|---|---|
| MT072864.1;QIQ54046.1;Pangolin_CoV_orf1a;4033 | 0.05 | 0.025 |
| MT072864.1;QIQ54046.1;Pangolin_CoV_orf1a;4034 | 0 | 0 |
| MT072864.1;QIQ54046.1;Pangolin_CoV_orf1a;4035 | 0.05 | 0.1 |
| MT072864.1;QIQ54046.1;Pangolin_CoV_orf1a;4036 | 0.1 | 0.075 |
| MT072864.1;QIQ54046.1;Pangolin_CoV_orf1a;4037 | 0 | 0.075 |
| MT072864.1;QIQ54046.1;Pangolin_CoV_orf1a;4038 | 0.05 | 0.075 |
| MT072864.1;QIQ54046.1;Pangolin_CoV_orf1a;4039 | 0.05 | 0 |
| MT072864.1;QIQ54046.1;Pangolin_CoV_orf1a;4040 | 0 | 0.05 |
| MT072864.1;QIQ54046.1;Pangolin_CoV_orf1a;4041 | 0 | 0 |
| MT072864.1;QIQ54046.1;Pangolin_CoV_orf1a;4042 | 0 | 0 |
| MT072864.1;QIQ54046.1;Pangolin_CoV_orf1a;4043 | 0 | 0 |
| MT072864.1;QIQ54046.1;Pangolin_CoV_orf1a;4044 | 0 | 0 |
| MT072864.1;QIQ54046.1;Pangolin_CoV_orf1a;4045 | 0.05 | 0.025 |
| MT072864.1;QIQ54046.1;Pangolin_CoV_orf1a;4046 | 0.05 | 0.025 |
| MT072864.1;QIQ54046.1;Pangolin_CoV_orf1a;4047 | 0 | 0 |
| MT072864.1;QIQ54046.1;Pangolin_CoV_orf1a;4048 | 0 | 0 |
| MT072864.1;QIQ54046.1;Pangolin_CoV_orf1a;4049 | 0 | 0 |
| MT072864.1;QIQ54046.1;Pangolin_CoV_orf1a;4050 | 0 | 0 |
| MT072864.1;QIQ54046.1;Pangolin_CoV_orf1a;4051 | 0 | 0 |
| MT072864.1;QIQ54046.1;Pangolin_CoV_orf1a;4052 | 0 | 0 |
| MT072864.1;QIQ54046.1;Pangolin_CoV_orf1a;4053 | 0 | 0 |
| MT072864.1;QIQ54046.1;Pangolin_CoV_orf1a;4054 | 0 | 0.025 |
| MT072864.1;QIQ54046.1;Pangolin_CoV_orf1a;4055 | 0.05 | 0 |
| MT072864.1;QIQ54046.1;Pangolin_CoV_orf1a;4056 | 0 | 0.025 |
| MT072864.1;QIQ54046.1;Pangolin_CoV_orf1a;4057 | 0 | 0 |
| MT072864.1;QIQ54046.1;Pangolin_CoV_orf1a;4058 | 0 | 0 |
| MT072864.1;QIQ54046.1;Pangolin_CoV_orf1a;4059 | 0 | 0.025 |
| MT072864.1;QIQ54046.1;Pangolin_CoV_orf1a;4060 | 0 | 0.05 |
| MT072864.1;QIQ54046.1;Pangolin_CoV_orf1a;4061 | 0 | 0.025 |
| MT072864.1;QIQ54046.1;Pangolin_CoV_orf1a;4062 | 0.1 | 0.025 |
| MT072864.1;QIQ54046.1;Pangolin_CoV_orf1a;4063 | 0.15 | 0.025 |
| MT072864.1;QIQ54046.1;Pangolin_CoV_orf1a;4064 | 0.2 | 0.05 |
| MT072864.1;QIQ54046.1;Pangolin_CoV_orf1a;4065 | 0.15 | 0.05 |
| MT072864.1;QIQ54046.1;Pangolin_CoV_orf1a;4066 | 0.1 | 0.05 |
| MT072864.1;QIQ54046.1;Pangolin_CoV_orf1a;4067 | 0.05 | 0.025 |
| MT072864.1;QIQ54046.1;Pangolin_CoV_orf1a;4068 | 0 | 0 |
| MT072864.1;QIQ54046.1;Pangolin_CoV_orf1a;4069 | 0 | 0 |
| MT072864.1;QIQ54046.1;Pangolin_CoV_orf1a;4070 | 0 | 0 |
| MT072864.1;QIQ54046.1;Pangolin_CoV_orf1a;4071 | 0 | 0 |
| MT072864.1;QIQ54046.1;Pangolin_CoV_orf1a;4072 | 0 | 0 |
| MT072864.1;QIQ54046.1;Pangolin_CoV_orf1a;4073 | 0 | 0 |
| MT072864.1;QIQ54046.1;Pangolin_CoV_orf1a;4074 | 0 | 0 |
| MT072864.1;QIQ54046.1;Pangolin_CoV_orf1a;4075 | 0 | 0 |

| | | |
|---|---|---|
| MT072864.1;QIQ54046.1;Pangolin_CoV_orf1a;4076 | 0 | 0 |
| MT072864.1;QIQ54046.1;Pangolin_CoV_orf1a;4077 | 0 | 0 |
| MT072864.1;QIQ54046.1;Pangolin_CoV_orf1a;4078 | 0 | 0 |
| MT072864.1;QIQ54046.1;Pangolin_CoV_orf1a;4079 | 0 | 0 |
| MT072864.1;QIQ54046.1;Pangolin_CoV_orf1a;4080 | 0 | 0 |
| MT072864.1;QIQ54046.1;Pangolin_CoV_orf1a;4081 | 0 | 0 |
| MT072864.1;QIQ54046.1;Pangolin_CoV_orf1a;4082 | 0 | 0 |
| MT072864.1;QIQ54046.1;Pangolin_CoV_orf1a;4083 | 0 | 0 |
| MT072864.1;QIQ54046.1;Pangolin_CoV_orf1a;4084 | 0.05 | 0.025 |
| MT072864.1;QIQ54046.1;Pangolin_CoV_orf1a;4085 | 0.05 | 0.05 |
| MT072864.1;QIQ54046.1;Pangolin_CoV_orf1a;4086 | 0 | 0 |
| MT072864.1;QIQ54046.1;Pangolin_CoV_orf1a;4087 | 0 | 0.025 |
| MT072864.1;QIQ54046.1;Pangolin_CoV_orf1a;4088 | 0 | 0 |
| MT072864.1;QIQ54046.1;Pangolin_CoV_orf1a;4089 | 0 | 0 |
| MT072864.1;QIQ54046.1;Pangolin_CoV_orf1a;4090 | 0 | 0 |
| MT072864.1;QIQ54046.1;Pangolin_CoV_orf1a;4091 | 0 | 0.025 |
| MT072864.1;QIQ54046.1;Pangolin_CoV_orf1a;4092 | 0 | 0 |
| MT072864.1;QIQ54046.1;Pangolin_CoV_orf1a;4093 | 0 | 0 |
| MT072864.1;QIQ54046.1;Pangolin_CoV_orf1a;4094 | 0 | 0 |
| MT072864.1;QIQ54046.1;Pangolin_CoV_orf1a;4095 | 0 | 0 |
| MT072864.1;QIQ54046.1;Pangolin_CoV_orf1a;4096 | 0 | 0 |
| MT072864.1;QIQ54046.1;Pangolin_CoV_orf1a;4097 | 0 | 0 |
| MT072864.1;QIQ54046.1;Pangolin_CoV_orf1a;4098 | 0 | 0 |
| MT072864.1;QIQ54046.1;Pangolin_CoV_orf1a;4099 | 0 | 0 |
| MT072864.1;QIQ54046.1;Pangolin_CoV_orf1a;4100 | 0 | 0.025 |
| MT072864.1;QIQ54046.1;Pangolin_CoV_orf1a;4101 | 0 | 0 |
| MT072864.1;QIQ54046.1;Pangolin_CoV_orf1a;4102 | 0 | 0 |
| MT072864.1;QIQ54046.1;Pangolin_CoV_orf1a;4103 | 0 | 0 |
| MT072864.1;QIQ54046.1;Pangolin_CoV_orf1a;4104 | 0 | 0 |
| MT072864.1;QIQ54046.1;Pangolin_CoV_orf1a;4105 | 0 | 0 |
| MT072864.1;QIQ54046.1;Pangolin_CoV_orf1a;4106 | 0 | 0 |
| MT072864.1;QIQ54046.1;Pangolin_CoV_orf1a;4107 | 0 | 0 |
| MT072864.1;QIQ54046.1;Pangolin_CoV_orf1a;4108 | 0 | 0 |
| MT072864.1;QIQ54046.1;Pangolin_CoV_orf1a;4109 | 0 | 0 |
| MT072864.1;QIQ54046.1;Pangolin_CoV_orf1a;4110 | 0 | 0 |
| MT072864.1;QIQ54046.1;Pangolin_CoV_orf1a;4111 | 0 | 0 |
| MT072864.1;QIQ54046.1;Pangolin_CoV_orf1a;4112 | 0 | 0 |
| MT072864.1;QIQ54046.1;Pangolin_CoV_orf1a;4113 | 0 | 0 |
| MT072864.1;QIQ54046.1;Pangolin_CoV_orf1a;4114 | 0 | 0 |
| MT072864.1;QIQ54046.1;Pangolin_CoV_orf1a;4115 | 0 | 0 |
| MT072864.1;QIQ54046.1;Pangolin_CoV_orf1a;4116 | 0 | 0 |
| MT072864.1;QIQ54046.1;Pangolin_CoV_orf1a;4117 | 0 | 0 |
| MT072864.1;QIQ54046.1;Pangolin_CoV_orf1a;4118 | 0 | 0 |

| | | |
|---|---|---|
| MT072864.1;QIQ54046.1;Pangolin_CoV_orf1a;4119 | 0 | 0.025 |
| MT072864.1;QIQ54046.1;Pangolin_CoV_orf1a;4120 | 0 | 0.075 |
| MT072864.1;QIQ54046.1;Pangolin_CoV_orf1a;4121 | 0 | 0.05 |
| MT072864.1;QIQ54046.1;Pangolin_CoV_orf1a;4122 | 0 | 0.025 |
| MT072864.1;QIQ54046.1;Pangolin_CoV_orf1a;4123 | 0 | 0.025 |
| MT072864.1;QIQ54046.1;Pangolin_CoV_orf1a;4124 | 0.05 | 0.025 |
| MT072864.1;QIQ54046.1;Pangolin_CoV_orf1a;4125 | 0.1 | 0.025 |
| MT072864.1;QIQ54046.1;Pangolin_CoV_orf1a;4126 | 0.05 | 0.025 |
| MT072864.1;QIQ54046.1;Pangolin_CoV_orf1a;4127 | 0 | 0.025 |
| MT072864.1;QIQ54046.1;Pangolin_CoV_orf1a;4128 | 0 | 0 |
| MT072864.1;QIQ54046.1;Pangolin_CoV_orf1a;4129 | 0.05 | 0.025 |
| MT072864.1;QIQ54046.1;Pangolin_CoV_orf1a;4130 | 0 | 0 |
| MT072864.1;QIQ54046.1;Pangolin_CoV_orf1a;4131 | 0 | 0 |
| MT072864.1;QIQ54046.1;Pangolin_CoV_orf1a;4132 | 0 | 0.025 |
| MT072864.1;QIQ54046.1;Pangolin_CoV_orf1a;4133 | 0.05 | 0.05 |
| MT072864.1;QIQ54046.1;Pangolin_CoV_orf1a;4134 | 0 | 0 |
| MT072864.1;QIQ54046.1;Pangolin_CoV_orf1a;4135 | 0 | 0 |
| MT072864.1;QIQ54046.1;Pangolin_CoV_orf1a;4136 | 0 | 0 |
| MT072864.1;QIQ54046.1;Pangolin_CoV_orf1a;4137 | 0 | 0 |
| MT072864.1;QIQ54046.1;Pangolin_CoV_orf1a;4138 | 0 | 0 |
| MT072864.1;QIQ54046.1;Pangolin_CoV_orf1a;4139 | 0 | 0 |
| MT072864.1;QIQ54046.1;Pangolin_CoV_orf1a;4140 | 0 | 0 |
| MT072864.1;QIQ54046.1;Pangolin_CoV_orf1a;4141 | 0 | 0 |
| MT072864.1;QIQ54046.1;Pangolin_CoV_orf1a;4142 | 0 | 0 |
| MT072864.1;QIQ54046.1;Pangolin_CoV_orf1a;4143 | 0 | 0 |
| MT072864.1;QIQ54046.1;Pangolin_CoV_orf1a;4144 | 0 | 0 |
| MT072864.1;QIQ54046.1;Pangolin_CoV_orf1a;4145 | 0 | 0 |
| MT072864.1;QIQ54046.1;Pangolin_CoV_orf1a;4146 | 0 | 0 |
| MT072864.1;QIQ54046.1;Pangolin_CoV_orf1a;4147 | 0 | 0 |
| MT072864.1;QIQ54046.1;Pangolin_CoV_orf1a;4148 | 0 | 0 |
| MT072864.1;QIQ54046.1;Pangolin_CoV_orf1a;4149 | 0 | 0 |
| MT072864.1;QIQ54046.1;Pangolin_CoV_orf1a;4150 | 0 | 0 |
| MT072864.1;QIQ54046.1;Pangolin_CoV_orf1a;4151 | 0 | 0 |
| MT072864.1;QIQ54046.1;Pangolin_CoV_orf1a;4152 | 0 | 0 |
| MT072864.1;QIQ54046.1;Pangolin_CoV_orf1a;4153 | 0 | 0.075 |
| MT072864.1;QIQ54046.1;Pangolin_CoV_orf1a;4154 | 0.15 | 0.025 |
| MT072864.1;QIQ54046.1;Pangolin_CoV_orf1a;4155 | 0.15 | 0.075 |
| MT072864.1;QIQ54046.1;Pangolin_CoV_orf1a;4156 | 0 | 0.025 |
| MT072864.1;QIQ54046.1;Pangolin_CoV_orf1a;4157 | 0 | 0.025 |
| MT072864.1;QIQ54046.1;Pangolin_CoV_orf1a;4158 | 0 | 0 |
| MT072864.1;QIQ54046.1;Pangolin_CoV_orf1a;4159 | 0 | 0 |
| MT072864.1;QIQ54046.1;Pangolin_CoV_orf1a;4160 | 0 | 0 |
| MT072864.1;QIQ54046.1;Pangolin_CoV_orf1a;4161 | 0 | 0 |

| | | |
|---|---|---|
| MT072864.1;QIQ54046.1;Pangolin_CoV_orf1a;4162 | 0 | 0 |
| MT072864.1;QIQ54046.1;Pangolin_CoV_orf1a;4163 | 0 | 0 |
| MT072864.1;QIQ54046.1;Pangolin_CoV_orf1a;4164 | 0 | 0 |
| MT072864.1;QIQ54046.1;Pangolin_CoV_orf1a;4165 | 0 | 0 |
| MT072864.1;QIQ54046.1;Pangolin_CoV_orf1a;4166 | 0 | 0 |
| MT072864.1;QIQ54046.1;Pangolin_CoV_orf1a;4167 | 0 | 0 |
| MT072864.1;QIQ54046.1;Pangolin_CoV_orf1a;4168 | 0 | 0 |
| MT072864.1;QIQ54046.1;Pangolin_CoV_orf1a;4169 | 0 | 0 |
| MT072864.1;QIQ54046.1;Pangolin_CoV_orf1a;4170 | 0 | 0 |
| MT072864.1;QIQ54046.1;Pangolin_CoV_orf1a;4171 | 0 | 0 |
| MT072864.1;QIQ54046.1;Pangolin_CoV_orf1a;4172 | 0 | 0 |
| MT072864.1;QIQ54046.1;Pangolin_CoV_orf1a;4173 | 0 | 0 |
| MT072864.1;QIQ54046.1;Pangolin_CoV_orf1a;4174 | 0.05 | 0 |
| MT072864.1;QIQ54046.1;Pangolin_CoV_orf1a;4175 | 0 | 0 |
| MT072864.1;QIQ54046.1;Pangolin_CoV_orf1a;4176 | 0.05 | 0 |
| MT072864.1;QIQ54046.1;Pangolin_CoV_orf1a;4177 | 0.05 | 0.05 |
| MT072864.1;QIQ54046.1;Pangolin_CoV_orf1a;4178 | 0.15 | 0.1 |
| MT072864.1;QIQ54046.1;Pangolin_CoV_orf1a;4179 | 0.3 | 0.25 |
| MT072864.1;QIQ54046.1;Pangolin_CoV_orf1a;4180 | 0.15 | 0.05 |
| MT072864.1;QIQ54046.1;Pangolin_CoV_orf1a;4181 | 0 | 0 |
| MT072864.1;QIQ54046.1;Pangolin_CoV_orf1a;4182 | 0 | 0 |
| MT072864.1;QIQ54046.1;Pangolin_CoV_orf1a;4183 | 0 | 0 |
| MT072864.1;QIQ54046.1;Pangolin_CoV_orf1a;4184 | 0 | 0 |
| MT072864.1;QIQ54046.1;Pangolin_CoV_orf1a;4185 | 0 | 0 |
| MT072864.1;QIQ54046.1;Pangolin_CoV_orf1a;4186 | 0 | 0 |
| MT072864.1;QIQ54046.1;Pangolin_CoV_orf1a;4187 | 0 | 0 |
| MT072864.1;QIQ54046.1;Pangolin_CoV_orf1a;4188 | 0 | 0 |
| MT072864.1;QIQ54046.1;Pangolin_CoV_orf1a;4189 | 0 | 0 |
| MT072864.1;QIQ54046.1;Pangolin_CoV_orf1a;4190 | 0 | 0 |
| MT072864.1;QIQ54046.1;Pangolin_CoV_orf1a;4191 | 0 | 0 |
| MT072864.1;QIQ54046.1;Pangolin_CoV_orf1a;4192 | 0 | 0 |
| MT072864.1;QIQ54046.1;Pangolin_CoV_orf1a;4193 | 0 | 0.025 |
| MT072864.1;QIQ54046.1;Pangolin_CoV_orf1a;4194 | 0 | 0 |
| MT072864.1;QIQ54046.1;Pangolin_CoV_orf1a;4195 | 0.2 | 0.15 |
| MT072864.1;QIQ54046.1;Pangolin_CoV_orf1a;4196 | 0 | 0 |
| MT072864.1;QIQ54046.1;Pangolin_CoV_orf1a;4197 | 0.05 | 0.15 |
| MT072864.1;QIQ54046.1;Pangolin_CoV_orf1a;4198 | 0.05 | 0.075 |
| MT072864.1;QIQ54046.1;Pangolin_CoV_orf1a;4199 | 0.05 | 0.05 |
| MT072864.1;QIQ54046.1;Pangolin_CoV_orf1a;4200 | 0.05 | 0.025 |
| MT072864.1;QIQ54046.1;Pangolin_CoV_orf1a;4201 | 0 | 0 |
| MT072864.1;QIQ54046.1;Pangolin_CoV_orf1a;4202 | 0 | 0.025 |
| MT072864.1;QIQ54046.1;Pangolin_CoV_orf1a;4203 | 0 | 0 |
| MT072864.1;QIQ54046.1;Pangolin_CoV_orf1a;4204 | 0 | 0 |

| | | |
|---|---|---|
| MT072864.1;QIQ54046.1;Pangolin_CoV_orf1a;4205 | 0 | 0 |
| MT072864.1;QIQ54046.1;Pangolin_CoV_orf1a;4206 | 0 | 0 |
| MT072864.1;QIQ54046.1;Pangolin_CoV_orf1a;4207 | 0.05 | 0 |
| MT072864.1;QIQ54046.1;Pangolin_CoV_orf1a;4208 | 0 | 0 |
| MT072864.1;QIQ54046.1;Pangolin_CoV_orf1a;4209 | 0 | 0 |
| MT072864.1;QIQ54046.1;Pangolin_CoV_orf1a;4210 | 0 | 0 |
| MT072864.1;QIQ54046.1;Pangolin_CoV_orf1a;4211 | 0 | 0 |
| MT072864.1;QIQ54046.1;Pangolin_CoV_orf1a;4212 | 0 | 0 |
| MT072864.1;QIQ54046.1;Pangolin_CoV_orf1a;4213 | 0 | 0 |
| MT072864.1;QIQ54046.1;Pangolin_CoV_orf1a;4214 | 0 | 0 |
| MT072864.1;QIQ54046.1;Pangolin_CoV_orf1a;4215 | 0 | 0 |
| MT072864.1;QIQ54046.1;Pangolin_CoV_orf1a;4216 | 0 | 0 |
| MT072864.1;QIQ54046.1;Pangolin_CoV_orf1a;4217 | 0 | 0 |
| MT072864.1;QIQ54046.1;Pangolin_CoV_orf1a;4218 | 0 | 0 |
| MT072864.1;QIQ54046.1;Pangolin_CoV_orf1a;4219 | 0 | 0 |
| MT072864.1;QIQ54046.1;Pangolin_CoV_orf1a;4220 | 0 | 0 |
| MT072864.1;QIQ54046.1;Pangolin_CoV_orf1a;4221 | 0.05 | 0.025 |
| MT072864.1;QIQ54046.1;Pangolin_CoV_orf1a;4222 | 0.1 | 0.075 |
| MT072864.1;QIQ54046.1;Pangolin_CoV_orf1a;4223 | 0.15 | 0.1 |
| MT072864.1;QIQ54046.1;Pangolin_CoV_orf1a;4224 | 0.25 | 0.1 |
| MT072864.1;QIQ54046.1;Pangolin_CoV_orf1a;4225 | 0.1 | 0.1 |
| MT072864.1;QIQ54046.1;Pangolin_CoV_orf1a;4226 | 0.05 | 0.05 |
| MT072864.1;QIQ54046.1;Pangolin_CoV_orf1a;4227 | 0.05 | 0.025 |
| MT072864.1;QIQ54046.1;Pangolin_CoV_orf1a;4228 | 0 | 0 |
| MT072864.1;QIQ54046.1;Pangolin_CoV_orf1a;4229 | 0 | 0 |
| MT072864.1;QIQ54046.1;Pangolin_CoV_orf1a;4230 | 0 | 0 |
| MT072864.1;QIQ54046.1;Pangolin_CoV_orf1a;4231 | 0 | 0 |
| MT072864.1;QIQ54046.1;Pangolin_CoV_orf1a;4232 | 0 | 0 |
| MT072864.1;QIQ54046.1;Pangolin_CoV_orf1a;4233 | 0 | 0 |
| MT072864.1;QIQ54046.1;Pangolin_CoV_orf1a;4234 | 0 | 0 |
| MT072864.1;QIQ54046.1;Pangolin_CoV_orf1a;4235 | 0 | 0 |
| MT072864.1;QIQ54046.1;Pangolin_CoV_orf1a;4236 | 0 | 0.025 |
| MT072864.1;QIQ54046.1;Pangolin_CoV_orf1a;4237 | 0 | 0 |
| MT072864.1;QIQ54046.1;Pangolin_CoV_orf1a;4238 | 0 | 0.025 |
| MT072864.1;QIQ54046.1;Pangolin_CoV_orf1a;4239 | 0.05 | 0.025 |
| MT072864.1;QIQ54046.1;Pangolin_CoV_orf1a;4240 | 0 | 0.025 |
| MT072864.1;QIQ54046.1;Pangolin_CoV_orf1a;4241 | 0 | 0 |
| MT072864.1;QIQ54046.1;Pangolin_CoV_orf1a;4242 | 0 | 0 |
| MT072864.1;QIQ54046.1;Pangolin_CoV_orf1a;4243 | 0 | 0.025 |
| MT072864.1;QIQ54046.1;Pangolin_CoV_orf1a;4244 | 0 | 0.025 |
| MT072864.1;QIQ54046.1;Pangolin_CoV_orf1a;4245 | 0 | 0 |
| MT072864.1;QIQ54046.1;Pangolin_CoV_orf1a;4246 | 0 | 0 |
| MT072864.1;QIQ54046.1;Pangolin_CoV_orf1a;4247 | 0 | 0 |

| | | |
|---|---|---|
| MT072864.1;QIQ54046.1;Pangolin_CoV_orf1a;4248 | 0 | 0 |
| MT072864.1;QIQ54046.1;Pangolin_CoV_orf1a;4249 | 0 | 0 |
| MT072864.1;QIQ54046.1;Pangolin_CoV_orf1a;4250 | 0 | 0 |
| MT072864.1;QIQ54046.1;Pangolin_CoV_orf1a;4251 | 0 | 0 |
| MT072864.1;QIQ54046.1;Pangolin_CoV_orf1a;4252 | 0 | 0 |
| MT072864.1;QIQ54046.1;Pangolin_CoV_orf1a;4253 | 0 | 0 |
| MT072864.1;QIQ54046.1;Pangolin_CoV_orf1a;4254 | 0 | 0 |
| MT072864.1;QIQ54046.1;Pangolin_CoV_orf1a;4255 | 0 | 0 |
| MT072864.1;QIQ54046.1;Pangolin_CoV_orf1a;4256 | 0 | 0 |
| MT072864.1;QIQ54046.1;Pangolin_CoV_orf1a;4257 | 0 | 0 |
| MT072864.1;QIQ54046.1;Pangolin_CoV_orf1a;4258 | 0 | 0 |
| MT072864.1;QIQ54046.1;Pangolin_CoV_orf1a;4259 | 0 | 0 |
| MT072864.1;QIQ54046.1;Pangolin_CoV_orf1a;4260 | 0 | 0 |
| MT072864.1;QIQ54046.1;Pangolin_CoV_orf1a;4261 | 0 | 0 |
| MT072864.1;QIQ54046.1;Pangolin_CoV_orf1a;4262 | 0.05 | 0 |
| MT072864.1;QIQ54046.1;Pangolin_CoV_orf1a;4263 | 0.1 | 0 |
| MT072864.1;QIQ54046.1;Pangolin_CoV_orf1a;4264 | 0.1 | 0 |
| MT072864.1;QIQ54046.1;Pangolin_CoV_orf1a;4265 | 0.15 | 0.075 |
| MT072864.1;QIQ54046.1;Pangolin_CoV_orf1a;4266 | 0.1 | 0.025 |
| MT072864.1;QIQ54046.1;Pangolin_CoV_orf1a;4267 | 0.35 | 0.15 |
| MT072864.1;QIQ54046.1;Pangolin_CoV_orf1a;4268 | 0.05 | 0.025 |
| MT072864.1;QIQ54046.1;Pangolin_CoV_orf1a;4269 | 0.1 | 0.025 |
| MT072864.1;QIQ54046.1;Pangolin_CoV_orf1a;4270 | 0.05 | 0.025 |
| MT072864.1;QIQ54046.1;Pangolin_CoV_orf1a;4271 | 0.1 | 0.05 |
| MT072864.1;QIQ54046.1;Pangolin_CoV_orf1a;4272 | 0.05 | 0 |
| MT072864.1;QIQ54046.1;Pangolin_CoV_orf1a;4273 | 0 | 0 |
| MT072864.1;QIQ54046.1;Pangolin_CoV_orf1a;4274 | 0 | 0 |
| MT072864.1;QIQ54046.1;Pangolin_CoV_orf1a;4275 | 0 | 0 |
| MT072864.1;QIQ54046.1;Pangolin_CoV_orf1a;4276 | 0 | 0 |
| MT072864.1;QIQ54046.1;Pangolin_CoV_orf1a;4277 | 0 | 0 |
| MT072864.1;QIQ54046.1;Pangolin_CoV_orf1a;4278 | 0 | 0 |
| MT072864.1;QIQ54046.1;Pangolin_CoV_orf1a;4279 | 0 | 0 |
| MT072864.1;QIQ54046.1;Pangolin_CoV_orf1a;4280 | 0 | 0 |
| MT072864.1;QIQ54046.1;Pangolin_CoV_orf1a;4281 | 0 | 0.025 |
| MT072864.1;QIQ54046.1;Pangolin_CoV_orf1a;4282 | 0 | 0 |
| MT072864.1;QIQ54046.1;Pangolin_CoV_orf1a;4283 | 0 | 0.025 |
| MT072864.1;QIQ54046.1;Pangolin_CoV_orf1a;4284 | 0 | 0.025 |
| MT072864.1;QIQ54046.1;Pangolin_CoV_orf1a;4285 | 0 | 0 |
| MT072864.1;QIQ54046.1;Pangolin_CoV_orf1a;4286 | 0 | 0 |
| MT072864.1;QIQ54046.1;Pangolin_CoV_orf1a;4287 | 0 | 0 |
| MT072864.1;QIQ54046.1;Pangolin_CoV_orf1a;4288 | 0 | 0 |
| MT072864.1;QIQ54046.1;Pangolin_CoV_orf1a;4289 | 0 | 0 |
| MT072864.1;QIQ54046.1;Pangolin_CoV_orf1a;4290 | 0 | 0 |

| | | |
|---|---|---|
| MT072864.1;QIQ54046.1;Pangolin_CoV_orf1a;4291 | 0 | 0 |
| MT072864.1;QIQ54046.1;Pangolin_CoV_orf1a;4292 | 0 | 0 |
| MT072864.1;QIQ54046.1;Pangolin_CoV_orf1a;4293 | 0 | 0 |
| MT072864.1;QIQ54046.1;Pangolin_CoV_orf1a;4294 | 0 | 0.025 |
| MT072864.1;QIQ54046.1;Pangolin_CoV_orf1a;4295 | 0 | 0.025 |
| MT072864.1;QIQ54046.1;Pangolin_CoV_orf1a;4296 | 0 | 0.025 |
| MT072864.1;QIQ54046.1;Pangolin_CoV_orf1a;4297 | 0 | 0.025 |
| MT072864.1;QIQ54046.1;Pangolin_CoV_orf1a;4298 | 0 | 0.025 |
| MT072864.1;QIQ54046.1;Pangolin_CoV_orf1a;4299 | 0 | 0.025 |
| MT072864.1;QIQ54046.1;Pangolin_CoV_orf1a;4300 | 0.05 | 0.025 |
| MT072864.1;QIQ54046.1;Pangolin_CoV_orf1a;4301 | 0 | 0.025 |
| MT072864.1;QIQ54046.1;Pangolin_CoV_orf1a;4302 | 0.05 | 0.025 |
| MT072864.1;QIQ54046.1;Pangolin_CoV_orf1a;4303 | 0 | 0 |
| MT072864.1;QIQ54046.1;Pangolin_CoV_orf1a;4304 | 0 | 0 |
| MT072864.1;QIQ54046.1;Pangolin_CoV_orf1a;4305 | 0 | 0 |
| MT072864.1;QIQ54046.1;Pangolin_CoV_orf1a;4306 | 0 | 0 |
| MT072864.1;QIQ54046.1;Pangolin_CoV_orf1a;4307 | 0 | 0 |
| MT072864.1;QIQ54046.1;Pangolin_CoV_orf1a;4308 | 0 | 0 |
| MT072864.1;QIQ54046.1;Pangolin_CoV_orf1a;4309 | 0 | 0 |
| MT072864.1;QIQ54046.1;Pangolin_CoV_orf1a;4310 | 0 | 0 |
| MT072864.1;QIQ54046.1;Pangolin_CoV_orf1a;4311 | 0 | 0 |
| MT072864.1;QIQ54046.1;Pangolin_CoV_orf1a;4312 | 0 | 0 |
| MT072864.1;QIQ54046.1;Pangolin_CoV_orf1a;4313 | 0 | 0 |
| MT072864.1;QIQ54046.1;Pangolin_CoV_orf1a;4314 | 0 | 0 |
| MT072864.1;QIQ54046.1;Pangolin_CoV_orf1a;4315 | 0 | 0 |
| MT072864.1;QIQ54046.1;Pangolin_CoV_orf1a;4316 | 0 | 0 |
| MT072864.1;QIQ54046.1;Pangolin_CoV_orf1a;4317 | 0 | 0 |
| MT072864.1;QIQ54046.1;Pangolin_CoV_orf1a;4318 | 0 | 0 |
| MT072864.1;QIQ54046.1;Pangolin_CoV_orf1a;4319 | 0 | 0 |
| MT072864.1;QIQ54046.1;Pangolin_CoV_orf1a;4320 | 0 | 0 |
| MT072864.1;QIQ54046.1;Pangolin_CoV_orf1a;4321 | 0 | 0.025 |
| MT072864.1;QIQ54046.1;Pangolin_CoV_orf1a;4322 | 0 | 0.025 |
| MT072864.1;QIQ54046.1;Pangolin_CoV_orf1a;4323 | 0 | 0.025 |
| MT072864.1;QIQ54046.1;Pangolin_CoV_orf1a;4324 | 0 | 0.025 |
| MT072864.1;QIQ54046.1;Pangolin_CoV_orf1a;4325 | 0 | 0 |
| MT072864.1;QIQ54046.1;Pangolin_CoV_orf1a;4326 | 0 | 0.025 |
| MT072864.1;QIQ54046.1;Pangolin_CoV_orf1a;4327 | 0 | 0 |
| MT072864.1;QIQ54046.1;Pangolin_CoV_orf1a;4328 | 0 | 0 |
| MT072864.1;QIQ54046.1;Pangolin_CoV_orf1a;4329 | 0 | 0 |
| MT072864.1;QIQ54046.1;Pangolin_CoV_orf1a;4330 | 0 | 0 |
| MT072864.1;QIQ54046.1;Pangolin_CoV_orf1a;4331 | 0 | 0 |
| MT072864.1;QIQ54046.1;Pangolin_CoV_orf1a;4332 | 0 | 0.05 |
| MT072864.1;QIQ54046.1;Pangolin_CoV_orf1a;4333 | 0 | 0 |

| | | |
|---|---|---|
| MT072864.1;QIQ54046.1;Pangolin_CoV_orf1a;4334 | 0 | 0 |
| MT072864.1;QIQ54046.1;Pangolin_CoV_orf1a;4335 | 0.05 | 0 |
| MT072864.1;QIQ54046.1;Pangolin_CoV_orf1a;4336 | 0.05 | 0 |
| MT072864.1;QIQ54046.1;Pangolin_CoV_orf1a;4337 | 0.05 | 0.025 |
| MT072864.1;QIQ54046.1;Pangolin_CoV_orf1a;4338 | 0.05 | 0.05 |
| MT072864.1;QIQ54046.1;Pangolin_CoV_orf1a;4339 | 0.05 | 0 |
| MT072864.1;QIQ54046.1;Pangolin_CoV_orf1a;4340 | 0.05 | 0 |
| MT072864.1;QIQ54046.1;Pangolin_CoV_orf1a;4341 | 0 | 0 |
| MT072864.1;QIQ54046.1;Pangolin_CoV_orf1a;4342 | 0 | 0.025 |
| MT072864.1;QIQ54046.1;Pangolin_CoV_orf1a;4343 | 0 | 0.025 |
| MT072864.1;QIQ54046.1;Pangolin_CoV_orf1a;4344 | 0 | 0.025 |
| MT072864.1;QIQ54046.1;Pangolin_CoV_orf1a;4345 | 0.05 | 0.05 |
| MT072864.1;QIQ54046.1;Pangolin_CoV_orf1a;4346 | 0 | 0.025 |
| MT072864.1;QIQ54046.1;Pangolin_CoV_orf1a;4347 | 0 | 0.025 |
| MT072864.1;QIQ54046.1;Pangolin_CoV_orf1a;4348 | 0 | 0.025 |
| MT072864.1;QIQ54046.1;Pangolin_CoV_orf1a;4349 | 0 | 0.025 |
| MT072864.1;QIQ54046.1;Pangolin_CoV_orf1a;4350 | 0 | 0 |
| MT072864.1;QIQ54046.1;Pangolin_CoV_orf1a;4351 | 0 | 0 |
| MT072864.1;QIQ54046.1;Pangolin_CoV_orf1a;4352 | 0 | 0 |
| MT072864.1;QIQ54046.1;Pangolin_CoV_orf1a;4353 | 0 | 0 |
| MT072864.1;QIQ54046.1;Pangolin_CoV_orf1a;4354 | 0 | 0 |
| MT072864.1;QIQ54046.1;Pangolin_CoV_orf1a;4355 | 0 | 0 |
| MT072864.1;QIQ54046.1;Pangolin_CoV_orf1a;4356 | 0 | 0 |
| MT072864.1;QIQ54046.1;Pangolin_CoV_orf1a;4357 | 0 | 0 |
| MT072864.1;QIQ54046.1;Pangolin_CoV_orf1a;4358 | 0 | 0 |
| MT072864.1;QIQ54046.1;Pangolin_CoV_orf1a;4359 | 0 | 0 |
| MT072864.1;QIQ54046.1;Pangolin_CoV_orf1a;4360 | 0 | 0 |
| MT072864.1;QIQ54046.1;Pangolin_CoV_orf1a;4361 | 0 | 0 |
| MT072864.1;QIQ54046.1;Pangolin_CoV_orf1a;4362 | 0 | 0 |
| MT072864.1;QIQ54046.1;Pangolin_CoV_orf1a;4363 | 0 | 0 |
| MT072864.1;QIQ54046.1;Pangolin_CoV_orf1a;4364 | 0 | 0 |
| MT072864.1;QIQ54046.1;Pangolin_CoV_orf1a;4365 | 0 | 0 |
| MT072864.1;QIQ54046.1;Pangolin_CoV_orf1a;4366 | 0 | 0 |
| MT072864.1;QIQ54046.1;Pangolin_CoV_orf1a;4367 | 0 | 0.075 |
| MT072864.1;QIQ54046.1;Pangolin_CoV_orf1a;4368 | 0 | 0 |
| MT072864.1;QIQ54046.1;Pangolin_CoV_orf1a;4369 | 0 | 0.025 |
| MT072864.1;QIQ54046.1;Pangolin_CoV_orf1a;4370 | 0 | 0 |
| MT072864.1;QIQ54046.1;Pangolin_CoV_orf1a;4371 | 0 | 0 |
| MT072864.1;QIQ54046.1;Pangolin_CoV_orf1a;4372 | 0 | 0 |
| MT072864.1;QIQ54046.1;Pangolin_CoV_orf1a;4373 | 0 | 0 |
| MT072864.1;QIQ54046.1;Pangolin_CoV_orf1a;4374 | 0 | 0 |
| MT072864.1;QIQ54046.1;Pangolin_CoV_orf1a;4375 | 0 | 0 |
| MT072864.1;QIQ54046.1;Pangolin_CoV_orf1a;4376 | 0 | 0 |

| | | |
|---|---|---|
| MT072864.1;QIQ54046.1;Pangolin_CoV_orf1a;4377 | 0 | 0 |
| MT072864.1;QIQ54046.1;Pangolin_CoV_orf1a;4378 | 0.05 | 0 |
| MT072864.1;QIQ54046.1;Pangolin_CoV_orf1a;4379 | 0.1 | 0 |
| MT072864.1;QIQ54046.1;Pangolin_CoV_orf1a;4380 | 0.1 | 0 |
| MT072864.1;QIQ54046.1;Pangolin_CoV_orf1a;4381 | 0 | 0 |
| MT072864.1;QIQ54046.1;Pangolin_CoV_orf1a;4382 | 0 | 0 |
| MT072864.1;QIQ54046.1;Pangolin_CoV_orf1a;4383 | 0 | 0 |
| MT072864.1;QIQ54046.1;Pangolin_CoV_orf1a;4384 | 0 | 0 |
| MT072864.1;QIQ54046.1;Pangolin_CoV_orf1a;4385 | 0 | 0 |
| MT072864.1;QIQ54046.1;Pangolin_CoV_orf1a;4386 | 0 | 0 |
| MT072864.1;QIQ54047.1;Pangolin_CoV_orf1ab;1 | 0 | 0 |
| MT072864.1;QIQ54047.1;Pangolin_CoV_orf1ab;2 | 0 | 0.025 |
| MT072864.1;QIQ54047.1;Pangolin_CoV_orf1ab;3 | 0 | 0 |
| MT072864.1;QIQ54047.1;Pangolin_CoV_orf1ab;4 | 0 | 0.025 |
| MT072864.1;QIQ54047.1;Pangolin_CoV_orf1ab;5 | 0 | 0 |
| MT072864.1;QIQ54047.1;Pangolin_CoV_orf1ab;6 | 0 | 0 |
| MT072864.1;QIQ54047.1;Pangolin_CoV_orf1ab;7 | 0 | 0 |
| MT072864.1;QIQ54047.1;Pangolin_CoV_orf1ab;8 | 0 | 0 |
| MT072864.1;QIQ54047.1;Pangolin_CoV_orf1ab;9 | 0 | 0 |
| MT072864.1;QIQ54047.1;Pangolin_CoV_orf1ab;10 | 0 | 0 |
| MT072864.1;QIQ54047.1;Pangolin_CoV_orf1ab;11 | 0 | 0 |
| MT072864.1;QIQ54047.1;Pangolin_CoV_orf1ab;12 | 0 | 0 |
| MT072864.1;QIQ54047.1;Pangolin_CoV_orf1ab;13 | 0 | 0 |
| MT072864.1;QIQ54047.1;Pangolin_CoV_orf1ab;14 | 0 | 0 |
| MT072864.1;QIQ54047.1;Pangolin_CoV_orf1ab;15 | 0 | 0 |
| MT072864.1;QIQ54047.1;Pangolin_CoV_orf1ab;16 | 0 | 0 |
| MT072864.1;QIQ54047.1;Pangolin_CoV_orf1ab;17 | 0 | 0 |
| MT072864.1;QIQ54047.1;Pangolin_CoV_orf1ab;18 | 0 | 0 |
| MT072864.1;QIQ54047.1;Pangolin_CoV_orf1ab;19 | 0 | 0 |
| MT072864.1;QIQ54047.1;Pangolin_CoV_orf1ab;20 | 0 | 0 |
| MT072864.1;QIQ54047.1;Pangolin_CoV_orf1ab;21 | 0 | 0 |
| MT072864.1;QIQ54047.1;Pangolin_CoV_orf1ab;22 | 0 | 0 |
| MT072864.1;QIQ54047.1;Pangolin_CoV_orf1ab;23 | 0 | 0 |
| MT072864.1;QIQ54047.1;Pangolin_CoV_orf1ab;24 | 0 | 0 |
| MT072864.1;QIQ54047.1;Pangolin_CoV_orf1ab;25 | 0 | 0 |
| MT072864.1;QIQ54047.1;Pangolin_CoV_orf1ab;26 | 0 | 0 |
| MT072864.1;QIQ54047.1;Pangolin_CoV_orf1ab;27 | 0 | 0 |
| MT072864.1;QIQ54047.1;Pangolin_CoV_orf1ab;28 | 0 | 0 |
| MT072864.1;QIQ54047.1;Pangolin_CoV_orf1ab;29 | 0.1 | 0 |
| MT072864.1;QIQ54047.1;Pangolin_CoV_orf1ab;30 | 0.3 | 0.125 |
| MT072864.1;QIQ54047.1;Pangolin_CoV_orf1ab;31 | 0 | 0 |
| MT072864.1;QIQ54047.1;Pangolin_CoV_orf1ab;32 | 0.05 | 0.025 |
| MT072864.1;QIQ54047.1;Pangolin_CoV_orf1ab;33 | 0.05 | 0 |

| | | |
|---|---|---|
| MT072864.1;QIQ54047.1;Pangolin_CoV_orf1ab;34 | 0.1 | 0 |
| MT072864.1;QIQ54047.1;Pangolin_CoV_orf1ab;35 | 0 | 0.025 |
| MT072864.1;QIQ54047.1;Pangolin_CoV_orf1ab;36 | 0 | 0 |
| MT072864.1;QIQ54047.1;Pangolin_CoV_orf1ab;37 | 0 | 0 |
| MT072864.1;QIQ54047.1;Pangolin_CoV_orf1ab;38 | 0 | 0 |
| MT072864.1;QIQ54047.1;Pangolin_CoV_orf1ab;39 | 0 | 0 |
| MT072864.1;QIQ54047.1;Pangolin_CoV_orf1ab;40 | 0 | 0 |
| MT072864.1;QIQ54047.1;Pangolin_CoV_orf1ab;41 | 0 | 0 |
| MT072864.1;QIQ54047.1;Pangolin_CoV_orf1ab;42 | 0 | 0 |
| MT072864.1;QIQ54047.1;Pangolin_CoV_orf1ab;43 | 0 | 0 |
| MT072864.1;QIQ54047.1;Pangolin_CoV_orf1ab;44 | 0.05 | 0 |
| MT072864.1;QIQ54047.1;Pangolin_CoV_orf1ab;45 | 0 | 0.05 |
| MT072864.1;QIQ54047.1;Pangolin_CoV_orf1ab;46 | 0 | 0.075 |
| MT072864.1;QIQ54047.1;Pangolin_CoV_orf1ab;47 | 0.05 | 0.075 |
| MT072864.1;QIQ54047.1;Pangolin_CoV_orf1ab;48 | 0.05 | 0.05 |
| MT072864.1;QIQ54047.1;Pangolin_CoV_orf1ab;49 | 0.05 | 0.05 |
| MT072864.1;QIQ54047.1;Pangolin_CoV_orf1ab;50 | 0 | 0.05 |
| MT072864.1;QIQ54047.1;Pangolin_CoV_orf1ab;51 | 0 | 0 |
| MT072864.1;QIQ54047.1;Pangolin_CoV_orf1ab;52 | 0.1 | 0.15 |
| MT072864.1;QIQ54047.1;Pangolin_CoV_orf1ab;53 | 0.1 | 0.05 |
| MT072864.1;QIQ54047.1;Pangolin_CoV_orf1ab;54 | 0 | 0.025 |
| MT072864.1;QIQ54047.1;Pangolin_CoV_orf1ab;55 | 0 | 0.025 |
| MT072864.1;QIQ54047.1;Pangolin_CoV_orf1ab;56 | 0 | 0.025 |
| MT072864.1;QIQ54047.1;Pangolin_CoV_orf1ab;57 | 0 | 0.025 |
| MT072864.1;QIQ54047.1;Pangolin_CoV_orf1ab;58 | 0 | 0.025 |
| MT072864.1;QIQ54047.1;Pangolin_CoV_orf1ab;59 | 0 | 0 |
| MT072864.1;QIQ54047.1;Pangolin_CoV_orf1ab;60 | 0 | 0 |
| MT072864.1;QIQ54047.1;Pangolin_CoV_orf1ab;61 | 0 | 0 |
| MT072864.1;QIQ54047.1;Pangolin_CoV_orf1ab;62 | 0 | 0 |
| MT072864.1;QIQ54047.1;Pangolin_CoV_orf1ab;63 | 0 | 0 |
| MT072864.1;QIQ54047.1;Pangolin_CoV_orf1ab;64 | 0 | 0 |
| MT072864.1;QIQ54047.1;Pangolin_CoV_orf1ab;65 | 0 | 0 |
| MT072864.1;QIQ54047.1;Pangolin_CoV_orf1ab;66 | 0 | 0 |
| MT072864.1;QIQ54047.1;Pangolin_CoV_orf1ab;67 | 0 | 0 |
| MT072864.1;QIQ54047.1;Pangolin_CoV_orf1ab;68 | 0 | 0 |
| MT072864.1;QIQ54047.1;Pangolin_CoV_orf1ab;69 | 0 | 0 |
| MT072864.1;QIQ54047.1;Pangolin_CoV_orf1ab;70 | 0.05 | 0 |
| MT072864.1;QIQ54047.1;Pangolin_CoV_orf1ab;71 | 0 | 0 |
| MT072864.1;QIQ54047.1;Pangolin_CoV_orf1ab;72 | 0 | 0 |
| MT072864.1;QIQ54047.1;Pangolin_CoV_orf1ab;73 | 0 | 0 |
| MT072864.1;QIQ54047.1;Pangolin_CoV_orf1ab;74 | 0 | 0 |
| MT072864.1;QIQ54047.1;Pangolin_CoV_orf1ab;75 | 0 | 0.025 |
| MT072864.1;QIQ54047.1;Pangolin_CoV_orf1ab;76 | 0 | 0 |

| | | |
|---|---|---|
| MT072864.1;QIQ54047.1;Pangolin_CoV_orf1ab;77 | 0 | 0 |
| MT072864.1;QIQ54047.1;Pangolin_CoV_orf1ab;78 | 0 | 0 |
| MT072864.1;QIQ54047.1;Pangolin_CoV_orf1ab;79 | 0 | 0 |
| MT072864.1;QIQ54047.1;Pangolin_CoV_orf1ab;80 | 0 | 0 |
| MT072864.1;QIQ54047.1;Pangolin_CoV_orf1ab;81 | 0 | 0 |
| MT072864.1;QIQ54047.1;Pangolin_CoV_orf1ab;82 | 0 | 0 |
| MT072864.1;QIQ54047.1;Pangolin_CoV_orf1ab;83 | 0 | 0 |
| MT072864.1;QIQ54047.1;Pangolin_CoV_orf1ab;84 | 0 | 0 |
| MT072864.1;QIQ54047.1;Pangolin_CoV_orf1ab;85 | 0 | 0 |
| MT072864.1;QIQ54047.1;Pangolin_CoV_orf1ab;86 | 0 | 0.025 |
| MT072864.1;QIQ54047.1;Pangolin_CoV_orf1ab;87 | 0 | 0 |
| MT072864.1;QIQ54047.1;Pangolin_CoV_orf1ab;88 | 0 | 0 |
| MT072864.1;QIQ54047.1;Pangolin_CoV_orf1ab;89 | 0 | 0 |
| MT072864.1;QIQ54047.1;Pangolin_CoV_orf1ab;90 | 0.25 | 0.275 |
| MT072864.1;QIQ54047.1;Pangolin_CoV_orf1ab;91 | 0.1 | 0.05 |
| MT072864.1;QIQ54047.1;Pangolin_CoV_orf1ab;92 | 0 | 0 |
| MT072864.1;QIQ54047.1;Pangolin_CoV_orf1ab;93 | 0 | 0 |
| MT072864.1;QIQ54047.1;Pangolin_CoV_orf1ab;94 | 0 | 0.025 |
| MT072864.1;QIQ54047.1;Pangolin_CoV_orf1ab;95 | 0 | 0 |
| MT072864.1;QIQ54047.1;Pangolin_CoV_orf1ab;96 | 0 | 0 |
| MT072864.1;QIQ54047.1;Pangolin_CoV_orf1ab;97 | 0 | 0 |
| MT072864.1;QIQ54047.1;Pangolin_CoV_orf1ab;98 | 0 | 0 |
| MT072864.1;QIQ54047.1;Pangolin_CoV_orf1ab;99 | 0 | 0 |
| MT072864.1;QIQ54047.1;Pangolin_CoV_orf1ab;100 | 0 | 0 |
| MT072864.1;QIQ54047.1;Pangolin_CoV_orf1ab;101 | 0 | 0 |
| MT072864.1;QIQ54047.1;Pangolin_CoV_orf1ab;102 | 0 | 0 |
| MT072864.1;QIQ54047.1;Pangolin_CoV_orf1ab;103 | 0 | 0 |
| MT072864.1;QIQ54047.1;Pangolin_CoV_orf1ab;104 | 0 | 0 |
| MT072864.1;QIQ54047.1;Pangolin_CoV_orf1ab;105 | 0 | 0 |
| MT072864.1;QIQ54047.1;Pangolin_CoV_orf1ab;106 | 0 | 0 |
| MT072864.1;QIQ54047.1;Pangolin_CoV_orf1ab;107 | 0 | 0 |
| MT072864.1;QIQ54047.1;Pangolin_CoV_orf1ab;108 | 0 | 0 |
| MT072864.1;QIQ54047.1;Pangolin_CoV_orf1ab;109 | 0 | 0 |
| MT072864.1;QIQ54047.1;Pangolin_CoV_orf1ab;110 | 0 | 0 |
| MT072864.1;QIQ54047.1;Pangolin_CoV_orf1ab;111 | 0 | 0 |
| MT072864.1;QIQ54047.1;Pangolin_CoV_orf1ab;112 | 0 | 0 |
| MT072864.1;QIQ54047.1;Pangolin_CoV_orf1ab;113 | 0 | 0 |
| MT072864.1;QIQ54047.1;Pangolin_CoV_orf1ab;114 | 0 | 0.025 |
| MT072864.1;QIQ54047.1;Pangolin_CoV_orf1ab;115 | 0 | 0.025 |
| MT072864.1;QIQ54047.1;Pangolin_CoV_orf1ab;116 | 0 | 0.025 |
| MT072864.1;QIQ54047.1;Pangolin_CoV_orf1ab;117 | 0 | 0.025 |
| MT072864.1;QIQ54047.1;Pangolin_CoV_orf1ab;118 | 0 | 0.025 |
| MT072864.1;QIQ54047.1;Pangolin_CoV_orf1ab;119 | 0 | 0.025 |

| | | |
|---|---|---|
| MT072864.1;QIQ54047.1;Pangolin_CoV_orf1ab;120 | 0 | 0.025 |
| MT072864.1;QIQ54047.1;Pangolin_CoV_orf1ab;121 | 0.05 | 0.05 |
| MT072864.1;QIQ54047.1;Pangolin_CoV_orf1ab;122 | 0 | 0.025 |
| MT072864.1;QIQ54047.1;Pangolin_CoV_orf1ab;123 | 0 | 0.025 |
| MT072864.1;QIQ54047.1;Pangolin_CoV_orf1ab;124 | 0 | 0.05 |
| MT072864.1;QIQ54047.1;Pangolin_CoV_orf1ab;125 | 0.1 | 0.025 |
| MT072864.1;QIQ54047.1;Pangolin_CoV_orf1ab;126 | 0 | 0 |
| MT072864.1;QIQ54047.1;Pangolin_CoV_orf1ab;127 | 0 | 0 |
| MT072864.1;QIQ54047.1;Pangolin_CoV_orf1ab;128 | 0 | 0.025 |
| MT072864.1;QIQ54047.1;Pangolin_CoV_orf1ab;129 | 0 | 0.025 |
| MT072864.1;QIQ54047.1;Pangolin_CoV_orf1ab;130 | 0 | 0.025 |
| MT072864.1;QIQ54047.1;Pangolin_CoV_orf1ab;131 | 0 | 0.025 |
| MT072864.1;QIQ54047.1;Pangolin_CoV_orf1ab;132 | 0.05 | 0.025 |
| MT072864.1;QIQ54047.1;Pangolin_CoV_orf1ab;133 | 0.05 | 0 |
| MT072864.1;QIQ54047.1;Pangolin_CoV_orf1ab;134 | 0 | 0 |
| MT072864.1;QIQ54047.1;Pangolin_CoV_orf1ab;135 | 0 | 0 |
| MT072864.1;QIQ54047.1;Pangolin_CoV_orf1ab;136 | 0 | 0 |
| MT072864.1;QIQ54047.1;Pangolin_CoV_orf1ab;137 | 0 | 0 |
| MT072864.1;QIQ54047.1;Pangolin_CoV_orf1ab;138 | 0 | 0 |
| MT072864.1;QIQ54047.1;Pangolin_CoV_orf1ab;139 | 0.05 | 0 |
| MT072864.1;QIQ54047.1;Pangolin_CoV_orf1ab;140 | 0 | 0 |
| MT072864.1;QIQ54047.1;Pangolin_CoV_orf1ab;141 | 0 | 0 |
| MT072864.1;QIQ54047.1;Pangolin_CoV_orf1ab;142 | 0 | 0 |
| MT072864.1;QIQ54047.1;Pangolin_CoV_orf1ab;143 | 0 | 0 |
| MT072864.1;QIQ54047.1;Pangolin_CoV_orf1ab;144 | 0 | 0 |
| MT072864.1;QIQ54047.1;Pangolin_CoV_orf1ab;145 | 0.1 | 0.125 |
| MT072864.1;QIQ54047.1;Pangolin_CoV_orf1ab;146 | 0 | 0 |
| MT072864.1;QIQ54047.1;Pangolin_CoV_orf1ab;147 | 0 | 0 |
| MT072864.1;QIQ54047.1;Pangolin_CoV_orf1ab;148 | 0 | 0 |
| MT072864.1;QIQ54047.1;Pangolin_CoV_orf1ab;149 | 0.3 | 0.25 |
| MT072864.1;QIQ54047.1;Pangolin_CoV_orf1ab;150 | 0.25 | 0.125 |
| MT072864.1;QIQ54047.1;Pangolin_CoV_orf1ab;151 | 0.3 | 0.15 |
| MT072864.1;QIQ54047.1;Pangolin_CoV_orf1ab;152 | 0.35 | 0.225 |
| MT072864.1;QIQ54047.1;Pangolin_CoV_orf1ab;153 | 0.25 | 0.125 |
| MT072864.1;QIQ54047.1;Pangolin_CoV_orf1ab;154 | 0.35 | 0.175 |
| MT072864.1;QIQ54047.1;Pangolin_CoV_orf1ab;155 | 0.3 | 0.075 |
| MT072864.1;QIQ54047.1;Pangolin_CoV_orf1ab;156 | 0.2 | 0.05 |
| MT072864.1;QIQ54047.1;Pangolin_CoV_orf1ab;157 | 0.1 | 0.075 |
| MT072864.1;QIQ54047.1;Pangolin_CoV_orf1ab;158 | 0.1 | 0.075 |
| MT072864.1;QIQ54047.1;Pangolin_CoV_orf1ab;159 | 0.05 | 0.05 |
| MT072864.1;QIQ54047.1;Pangolin_CoV_orf1ab;160 | 0 | 0 |
| MT072864.1;QIQ54047.1;Pangolin_CoV_orf1ab;161 | 0.05 | 0.05 |
| MT072864.1;QIQ54047.1;Pangolin_CoV_orf1ab;162 | 0.05 | 0.05 |

| | | |
|---|---|---|
| MT072864.1;QIQ54047.1;Pangolin_CoV_orf1ab;163 | 0 | 0.1 |
| MT072864.1;QIQ54047.1;Pangolin_CoV_orf1ab;164 | 0 | 0.05 |
| MT072864.1;QIQ54047.1;Pangolin_CoV_orf1ab;165 | 0.05 | 0 |
| MT072864.1;QIQ54047.1;Pangolin_CoV_orf1ab;166 | 0 | 0.025 |
| MT072864.1;QIQ54047.1;Pangolin_CoV_orf1ab;167 | 0 | 0.025 |
| MT072864.1;QIQ54047.1;Pangolin_CoV_orf1ab;168 | 0.1 | 0.025 |
| MT072864.1;QIQ54047.1;Pangolin_CoV_orf1ab;169 | 0 | 0 |
| MT072864.1;QIQ54047.1;Pangolin_CoV_orf1ab;170 | 0 | 0.025 |
| MT072864.1;QIQ54047.1;Pangolin_CoV_orf1ab;171 | 0 | 0 |
| MT072864.1;QIQ54047.1;Pangolin_CoV_orf1ab;172 | 0 | 0 |
| MT072864.1;QIQ54047.1;Pangolin_CoV_orf1ab;173 | 0 | 0 |
| MT072864.1;QIQ54047.1;Pangolin_CoV_orf1ab;174 | 0.05 | 0.025 |
| MT072864.1;QIQ54047.1;Pangolin_CoV_orf1ab;175 | 0 | 0 |
| MT072864.1;QIQ54047.1;Pangolin_CoV_orf1ab;176 | 0 | 0 |
| MT072864.1;QIQ54047.1;Pangolin_CoV_orf1ab;177 | 0 | 0 |
| MT072864.1;QIQ54047.1;Pangolin_CoV_orf1ab;178 | 0 | 0 |
| MT072864.1;QIQ54047.1;Pangolin_CoV_orf1ab;179 | 0 | 0 |
| MT072864.1;QIQ54047.1;Pangolin_CoV_orf1ab;180 | 0 | 0 |
| MT072864.1;QIQ54047.1;Pangolin_CoV_orf1ab;181 | 0 | 0.025 |
| MT072864.1;QIQ54047.1;Pangolin_CoV_orf1ab;182 | 0 | 0 |
| MT072864.1;QIQ54047.1;Pangolin_CoV_orf1ab;183 | 0 | 0 |
| MT072864.1;QIQ54047.1;Pangolin_CoV_orf1ab;184 | 0 | 0.025 |
| MT072864.1;QIQ54047.1;Pangolin_CoV_orf1ab;185 | 0 | 0.025 |
| MT072864.1;QIQ54047.1;Pangolin_CoV_orf1ab;186 | 0 | 0.025 |
| MT072864.1;QIQ54047.1;Pangolin_CoV_orf1ab;187 | 0 | 0 |
| MT072864.1;QIQ54047.1;Pangolin_CoV_orf1ab;188 | 0 | 0 |
| MT072864.1;QIQ54047.1;Pangolin_CoV_orf1ab;189 | 0 | 0 |
| MT072864.1;QIQ54047.1;Pangolin_CoV_orf1ab;190 | 0 | 0 |
| MT072864.1;QIQ54047.1;Pangolin_CoV_orf1ab;191 | 0 | 0 |
| MT072864.1;QIQ54047.1;Pangolin_CoV_orf1ab;192 | 0 | 0 |
| MT072864.1;QIQ54047.1;Pangolin_CoV_orf1ab;193 | 0 | 0 |
| MT072864.1;QIQ54047.1;Pangolin_CoV_orf1ab;194 | 0 | 0 |
| MT072864.1;QIQ54047.1;Pangolin_CoV_orf1ab;195 | 0 | 0 |
| MT072864.1;QIQ54047.1;Pangolin_CoV_orf1ab;196 | 0 | 0 |
| MT072864.1;QIQ54047.1;Pangolin_CoV_orf1ab;197 | 0 | 0 |
| MT072864.1;QIQ54047.1;Pangolin_CoV_orf1ab;198 | 0 | 0 |
| MT072864.1;QIQ54047.1;Pangolin_CoV_orf1ab;199 | 0 | 0 |
| MT072864.1;QIQ54047.1;Pangolin_CoV_orf1ab;200 | 0 | 0 |
| MT072864.1;QIQ54047.1;Pangolin_CoV_orf1ab;201 | 0 | 0 |
| MT072864.1;QIQ54047.1;Pangolin_CoV_orf1ab;202 | 0 | 0 |
| MT072864.1;QIQ54047.1;Pangolin_CoV_orf1ab;203 | 0 | 0 |
| MT072864.1;QIQ54047.1;Pangolin_CoV_orf1ab;204 | 0.05 | 0 |
| MT072864.1;QIQ54047.1;Pangolin_CoV_orf1ab;205 | 0 | 0 |

| | | |
|---|---|---|
| MT072864.1;QIQ54047.1;Pangolin_CoV_orf1ab;206 | 0 | 0 |
| MT072864.1;QIQ54047.1;Pangolin_CoV_orf1ab;207 | 0 | 0 |
| MT072864.1;QIQ54047.1;Pangolin_CoV_orf1ab;208 | 0 | 0 |
| MT072864.1;QIQ54047.1;Pangolin_CoV_orf1ab;209 | 0 | 0 |
| MT072864.1;QIQ54047.1;Pangolin_CoV_orf1ab;210 | 0 | 0 |
| MT072864.1;QIQ54047.1;Pangolin_CoV_orf1ab;211 | 0.1 | 0.05 |
| MT072864.1;QIQ54047.1;Pangolin_CoV_orf1ab;212 | 0.05 | 0.075 |
| MT072864.1;QIQ54047.1;Pangolin_CoV_orf1ab;213 | 0.05 | 0.05 |
| MT072864.1;QIQ54047.1;Pangolin_CoV_orf1ab;214 | 0.15 | 0.1 |
| MT072864.1;QIQ54047.1;Pangolin_CoV_orf1ab;215 | 0.1 | 0.075 |
| MT072864.1;QIQ54047.1;Pangolin_CoV_orf1ab;216 | 0 | 0.075 |
| MT072864.1;QIQ54047.1;Pangolin_CoV_orf1ab;217 | 0 | 0.025 |
| MT072864.1;QIQ54047.1;Pangolin_CoV_orf1ab;218 | 0 | 0.025 |
| MT072864.1;QIQ54047.1;Pangolin_CoV_orf1ab;219 | 0 | 0.025 |
| MT072864.1;QIQ54047.1;Pangolin_CoV_orf1ab;220 | 0.05 | 0.075 |
| MT072864.1;QIQ54047.1;Pangolin_CoV_orf1ab;221 | 0 | 0.025 |
| MT072864.1;QIQ54047.1;Pangolin_CoV_orf1ab;222 | 0 | 0.025 |
| MT072864.1;QIQ54047.1;Pangolin_CoV_orf1ab;223 | 0 | 0.025 |
| MT072864.1;QIQ54047.1;Pangolin_CoV_orf1ab;224 | 0 | 0 |
| MT072864.1;QIQ54047.1;Pangolin_CoV_orf1ab;225 | 0.05 | 0 |
| MT072864.1;QIQ54047.1;Pangolin_CoV_orf1ab;226 | 0 | 0 |
| MT072864.1;QIQ54047.1;Pangolin_CoV_orf1ab;227 | 0 | 0 |
| MT072864.1;QIQ54047.1;Pangolin_CoV_orf1ab;228 | 0 | 0 |
| MT072864.1;QIQ54047.1;Pangolin_CoV_orf1ab;229 | 0 | 0 |
| MT072864.1;QIQ54047.1;Pangolin_CoV_orf1ab;230 | 0 | 0 |
| MT072864.1;QIQ54047.1;Pangolin_CoV_orf1ab;231 | 0 | 0 |
| MT072864.1;QIQ54047.1;Pangolin_CoV_orf1ab;232 | 0 | 0 |
| MT072864.1;QIQ54047.1;Pangolin_CoV_orf1ab;233 | 0 | 0 |
| MT072864.1;QIQ54047.1;Pangolin_CoV_orf1ab;234 | 0 | 0 |
| MT072864.1;QIQ54047.1;Pangolin_CoV_orf1ab;235 | 0 | 0 |
| MT072864.1;QIQ54047.1;Pangolin_CoV_orf1ab;236 | 0 | 0 |
| MT072864.1;QIQ54047.1;Pangolin_CoV_orf1ab;237 | 0 | 0.025 |
| MT072864.1;QIQ54047.1;Pangolin_CoV_orf1ab;238 | 0 | 0 |
| MT072864.1;QIQ54047.1;Pangolin_CoV_orf1ab;239 | 0 | 0 |
| MT072864.1;QIQ54047.1;Pangolin_CoV_orf1ab;240 | 0 | 0 |
| MT072864.1;QIQ54047.1;Pangolin_CoV_orf1ab;241 | 0 | 0 |
| MT072864.1;QIQ54047.1;Pangolin_CoV_orf1ab;242 | 0 | 0 |
| MT072864.1;QIQ54047.1;Pangolin_CoV_orf1ab;243 | 0 | 0 |
| MT072864.1;QIQ54047.1;Pangolin_CoV_orf1ab;244 | 0 | 0 |
| MT072864.1;QIQ54047.1;Pangolin_CoV_orf1ab;245 | 0 | 0 |
| MT072864.1;QIQ54047.1;Pangolin_CoV_orf1ab;246 | 0 | 0 |
| MT072864.1;QIQ54047.1;Pangolin_CoV_orf1ab;247 | 0 | 0 |
| MT072864.1;QIQ54047.1;Pangolin_CoV_orf1ab;248 | 0 | 0 |

| | | |
|---|---|---|
| MT072864.1;QIQ54047.1;Pangolin_CoV_orf1ab;249 | 0 | 0 |
| MT072864.1;QIQ54047.1;Pangolin_CoV_orf1ab;250 | 0 | 0 |
| MT072864.1;QIQ54047.1;Pangolin_CoV_orf1ab;251 | 0 | 0 |
| MT072864.1;QIQ54047.1;Pangolin_CoV_orf1ab;252 | 0.05 | 0.075 |
| MT072864.1;QIQ54047.1;Pangolin_CoV_orf1ab;253 | 0.05 | 0.1 |
| MT072864.1;QIQ54047.1;Pangolin_CoV_orf1ab;254 | 0.1 | 0.15 |
| MT072864.1;QIQ54047.1;Pangolin_CoV_orf1ab;255 | 0.15 | 0.225 |
| MT072864.1;QIQ54047.1;Pangolin_CoV_orf1ab;256 | 0.25 | 0.225 |
| MT072864.1;QIQ54047.1;Pangolin_CoV_orf1ab;257 | 0.1 | 0.1 |
| MT072864.1;QIQ54047.1;Pangolin_CoV_orf1ab;258 | 0.15 | 0.125 |
| MT072864.1;QIQ54047.1;Pangolin_CoV_orf1ab;259 | 0.35 | 0.225 |
| MT072864.1;QIQ54047.1;Pangolin_CoV_orf1ab;260 | 0.15 | 0.25 |
| MT072864.1;QIQ54047.1;Pangolin_CoV_orf1ab;261 | 0.1 | 0.225 |
| MT072864.1;QIQ54047.1;Pangolin_CoV_orf1ab;262 | 0.05 | 0.05 |
| MT072864.1;QIQ54047.1;Pangolin_CoV_orf1ab;263 | 0 | 0 |
| MT072864.1;QIQ54047.1;Pangolin_CoV_orf1ab;264 | 0 | 0.025 |
| MT072864.1;QIQ54047.1;Pangolin_CoV_orf1ab;265 | 0 | 0 |
| MT072864.1;QIQ54047.1;Pangolin_CoV_orf1ab;266 | 0 | 0 |
| MT072864.1;QIQ54047.1;Pangolin_CoV_orf1ab;267 | 0.15 | 0.025 |
| MT072864.1;QIQ54047.1;Pangolin_CoV_orf1ab;268 | 0.15 | 0 |
| MT072864.1;QIQ54047.1;Pangolin_CoV_orf1ab;269 | 0.15 | 0 |
| MT072864.1;QIQ54047.1;Pangolin_CoV_orf1ab;270 | 0.2 | 0 |
| MT072864.1;QIQ54047.1;Pangolin_CoV_orf1ab;271 | 0.15 | 0 |
| MT072864.1;QIQ54047.1;Pangolin_CoV_orf1ab;272 | 0.1 | 0 |
| MT072864.1;QIQ54047.1;Pangolin_CoV_orf1ab;273 | 0.1 | 0 |
| MT072864.1;QIQ54047.1;Pangolin_CoV_orf1ab;274 | 0.15 | 0 |
| MT072864.1;QIQ54047.1;Pangolin_CoV_orf1ab;275 | 0.15 | 0.025 |
| MT072864.1;QIQ54047.1;Pangolin_CoV_orf1ab;276 | 0.2 | 0.05 |
| MT072864.1;QIQ54047.1;Pangolin_CoV_orf1ab;277 | 0.2 | 0.025 |
| MT072864.1;QIQ54047.1;Pangolin_CoV_orf1ab;278 | 0.2 | 0 |
| MT072864.1;QIQ54047.1;Pangolin_CoV_orf1ab;279 | 0.05 | 0.025 |
| MT072864.1;QIQ54047.1;Pangolin_CoV_orf1ab;280 | 0.1 | 0.025 |
| MT072864.1;QIQ54047.1;Pangolin_CoV_orf1ab;281 | 0.05 | 0.025 |
| MT072864.1;QIQ54047.1;Pangolin_CoV_orf1ab;282 | 0.05 | 0 |
| MT072864.1;QIQ54047.1;Pangolin_CoV_orf1ab;283 | 0.05 | 0 |
| MT072864.1;QIQ54047.1;Pangolin_CoV_orf1ab;284 | 0.05 | 0.025 |
| MT072864.1;QIQ54047.1;Pangolin_CoV_orf1ab;285 | 0.05 | 0.025 |
| MT072864.1;QIQ54047.1;Pangolin_CoV_orf1ab;286 | 0.05 | 0.025 |
| MT072864.1;QIQ54047.1;Pangolin_CoV_orf1ab;287 | 0 | 0 |
| MT072864.1;QIQ54047.1;Pangolin_CoV_orf1ab;288 | 0 | 0 |
| MT072864.1;QIQ54047.1;Pangolin_CoV_orf1ab;289 | 0 | 0 |
| MT072864.1;QIQ54047.1;Pangolin_CoV_orf1ab;290 | 0 | 0 |
| MT072864.1;QIQ54047.1;Pangolin_CoV_orf1ab;291 | 0 | 0 |

| | | |
|---|---|---|
| MT072864.1;QIQ54047.1;Pangolin_CoV_orf1ab;292 | 0 | 0 |
| MT072864.1;QIQ54047.1;Pangolin_CoV_orf1ab;293 | 0 | 0 |
| MT072864.1;QIQ54047.1;Pangolin_CoV_orf1ab;294 | 0 | 0 |
| MT072864.1;QIQ54047.1;Pangolin_CoV_orf1ab;295 | 0 | 0 |
| MT072864.1;QIQ54047.1;Pangolin_CoV_orf1ab;296 | 0.1 | 0 |
| MT072864.1;QIQ54047.1;Pangolin_CoV_orf1ab;297 | 0.1 | 0 |
| MT072864.1;QIQ54047.1;Pangolin_CoV_orf1ab;298 | 0.1 | 0 |
| MT072864.1;QIQ54047.1;Pangolin_CoV_orf1ab;299 | 0.1 | 0 |
| MT072864.1;QIQ54047.1;Pangolin_CoV_orf1ab;300 | 0.1 | 0 |
| MT072864.1;QIQ54047.1;Pangolin_CoV_orf1ab;301 | 0.1 | 0.025 |
| MT072864.1;QIQ54047.1;Pangolin_CoV_orf1ab;302 | 0.05 | 0 |
| MT072864.1;QIQ54047.1;Pangolin_CoV_orf1ab;303 | 0.05 | 0 |
| MT072864.1;QIQ54047.1;Pangolin_CoV_orf1ab;304 | 0.05 | 0 |
| MT072864.1;QIQ54047.1;Pangolin_CoV_orf1ab;305 | 0 | 0 |
| MT072864.1;QIQ54047.1;Pangolin_CoV_orf1ab;306 | 0 | 0 |
| MT072864.1;QIQ54047.1;Pangolin_CoV_orf1ab;307 | 0 | 0 |
| MT072864.1;QIQ54047.1;Pangolin_CoV_orf1ab;308 | 0 | 0 |
| MT072864.1;QIQ54047.1;Pangolin_CoV_orf1ab;309 | 0 | 0 |
| MT072864.1;QIQ54047.1;Pangolin_CoV_orf1ab;310 | 0 | 0 |
| MT072864.1;QIQ54047.1;Pangolin_CoV_orf1ab;311 | 0 | 0 |
| MT072864.1;QIQ54047.1;Pangolin_CoV_orf1ab;312 | 0 | 0 |
| MT072864.1;QIQ54047.1;Pangolin_CoV_orf1ab;313 | 0 | 0 |
| MT072864.1;QIQ54047.1;Pangolin_CoV_orf1ab;314 | 0 | 0 |
| MT072864.1;QIQ54047.1;Pangolin_CoV_orf1ab;315 | 0 | 0 |
| MT072864.1;QIQ54047.1;Pangolin_CoV_orf1ab;316 | 0 | 0 |
| MT072864.1;QIQ54047.1;Pangolin_CoV_orf1ab;317 | 0 | 0 |
| MT072864.1;QIQ54047.1;Pangolin_CoV_orf1ab;318 | 0 | 0 |
| MT072864.1;QIQ54047.1;Pangolin_CoV_orf1ab;319 | 0 | 0 |
| MT072864.1;QIQ54047.1;Pangolin_CoV_orf1ab;320 | 0 | 0 |
| MT072864.1;QIQ54047.1;Pangolin_CoV_orf1ab;321 | 0 | 0 |
| MT072864.1;QIQ54047.1;Pangolin_CoV_orf1ab;322 | 0 | 0 |
| MT072864.1;QIQ54047.1;Pangolin_CoV_orf1ab;323 | 0 | 0 |
| MT072864.1;QIQ54047.1;Pangolin_CoV_orf1ab;324 | 0.05 | 0 |
| MT072864.1;QIQ54047.1;Pangolin_CoV_orf1ab;325 | 0 | 0 |
| MT072864.1;QIQ54047.1;Pangolin_CoV_orf1ab;326 | 0 | 0 |
| MT072864.1;QIQ54047.1;Pangolin_CoV_orf1ab;327 | 0 | 0.025 |
| MT072864.1;QIQ54047.1;Pangolin_CoV_orf1ab;328 | 0 | 0 |
| MT072864.1;QIQ54047.1;Pangolin_CoV_orf1ab;329 | 0 | 0 |
| MT072864.1;QIQ54047.1;Pangolin_CoV_orf1ab;330 | 0 | 0 |
| MT072864.1;QIQ54047.1;Pangolin_CoV_orf1ab;331 | 0 | 0 |
| MT072864.1;QIQ54047.1;Pangolin_CoV_orf1ab;332 | 0 | 0 |
| MT072864.1;QIQ54047.1;Pangolin_CoV_orf1ab;333 | 0 | 0 |
| MT072864.1;QIQ54047.1;Pangolin_CoV_orf1ab;334 | 0 | 0 |

| | | |
|---|---|---|
| MT072864.1;QIQ54047.1;Pangolin_CoV_orf1ab;335 | 0 | 0 |
| MT072864.1;QIQ54047.1;Pangolin_CoV_orf1ab;336 | 0 | 0 |
| MT072864.1;QIQ54047.1;Pangolin_CoV_orf1ab;337 | 0 | 0 |
| MT072864.1;QIQ54047.1;Pangolin_CoV_orf1ab;338 | 0 | 0 |
| MT072864.1;QIQ54047.1;Pangolin_CoV_orf1ab;339 | 0 | 0 |
| MT072864.1;QIQ54047.1;Pangolin_CoV_orf1ab;340 | 0 | 0.025 |
| MT072864.1;QIQ54047.1;Pangolin_CoV_orf1ab;341 | 0 | 0.025 |
| MT072864.1;QIQ54047.1;Pangolin_CoV_orf1ab;342 | 0 | 0.025 |
| MT072864.1;QIQ54047.1;Pangolin_CoV_orf1ab;343 | 0 | 0.025 |
| MT072864.1;QIQ54047.1;Pangolin_CoV_orf1ab;344 | 0 | 0.025 |
| MT072864.1;QIQ54047.1;Pangolin_CoV_orf1ab;345 | 0 | 0.025 |
| MT072864.1;QIQ54047.1;Pangolin_CoV_orf1ab;346 | 0.05 | 0 |
| MT072864.1;QIQ54047.1;Pangolin_CoV_orf1ab;347 | 0 | 0 |
| MT072864.1;QIQ54047.1;Pangolin_CoV_orf1ab;348 | 0 | 0 |
| MT072864.1;QIQ54047.1;Pangolin_CoV_orf1ab;349 | 0.05 | 0 |
| MT072864.1;QIQ54047.1;Pangolin_CoV_orf1ab;350 | 0.05 | 0 |
| MT072864.1;QIQ54047.1;Pangolin_CoV_orf1ab;351 | 0.05 | 0 |
| MT072864.1;QIQ54047.1;Pangolin_CoV_orf1ab;352 | 0 | 0 |
| MT072864.1;QIQ54047.1;Pangolin_CoV_orf1ab;353 | 0 | 0 |
| MT072864.1;QIQ54047.1;Pangolin_CoV_orf1ab;354 | 0 | 0 |
| MT072864.1;QIQ54047.1;Pangolin_CoV_orf1ab;355 | 0 | 0 |
| MT072864.1;QIQ54047.1;Pangolin_CoV_orf1ab;356 | 0 | 0 |
| MT072864.1;QIQ54047.1;Pangolin_CoV_orf1ab;357 | 0 | 0 |
| MT072864.1;QIQ54047.1;Pangolin_CoV_orf1ab;358 | 0 | 0 |
| MT072864.1;QIQ54047.1;Pangolin_CoV_orf1ab;359 | 0 | 0 |
| MT072864.1;QIQ54047.1;Pangolin_CoV_orf1ab;360 | 0 | 0 |
| MT072864.1;QIQ54047.1;Pangolin_CoV_orf1ab;361 | 0 | 0 |
| MT072864.1;QIQ54047.1;Pangolin_CoV_orf1ab;362 | 0 | 0 |
| MT072864.1;QIQ54047.1;Pangolin_CoV_orf1ab;363 | 0 | 0 |
| MT072864.1;QIQ54047.1;Pangolin_CoV_orf1ab;364 | 0 | 0 |
| MT072864.1;QIQ54047.1;Pangolin_CoV_orf1ab;365 | 0 | 0 |
| MT072864.1;QIQ54047.1;Pangolin_CoV_orf1ab;366 | 0 | 0 |
| MT072864.1;QIQ54047.1;Pangolin_CoV_orf1ab;367 | 0.05 | 0 |
| MT072864.1;QIQ54047.1;Pangolin_CoV_orf1ab;368 | 0.05 | 0 |
| MT072864.1;QIQ54047.1;Pangolin_CoV_orf1ab;369 | 0 | 0 |
| MT072864.1;QIQ54047.1;Pangolin_CoV_orf1ab;370 | 0.05 | 0 |
| MT072864.1;QIQ54047.1;Pangolin_CoV_orf1ab;371 | 0.05 | 0 |
| MT072864.1;QIQ54047.1;Pangolin_CoV_orf1ab;372 | 0.05 | 0 |
| MT072864.1;QIQ54047.1;Pangolin_CoV_orf1ab;373 | 0 | 0 |
| MT072864.1;QIQ54047.1;Pangolin_CoV_orf1ab;374 | 0 | 0 |
| MT072864.1;QIQ54047.1;Pangolin_CoV_orf1ab;375 | 0.05 | 0 |
| MT072864.1;QIQ54047.1;Pangolin_CoV_orf1ab;376 | 0.1 | 0 |
| MT072864.1;QIQ54047.1;Pangolin_CoV_orf1ab;377 | 0 | 0 |

| | | |
|---|---|---|
| MT072864.1;QIQ54047.1;Pangolin_CoV_orf1ab;378 | 0 | 0 |
| MT072864.1;QIQ54047.1;Pangolin_CoV_orf1ab;379 | 0.1 | 0.075 |
| MT072864.1;QIQ54047.1;Pangolin_CoV_orf1ab;380 | 0 | 0.025 |
| MT072864.1;QIQ54047.1;Pangolin_CoV_orf1ab;381 | 0 | 0.025 |
| MT072864.1;QIQ54047.1;Pangolin_CoV_orf1ab;382 | 0 | 0.025 |
| MT072864.1;QIQ54047.1;Pangolin_CoV_orf1ab;383 | 0 | 0.025 |
| MT072864.1;QIQ54047.1;Pangolin_CoV_orf1ab;384 | 0 | 0.025 |
| MT072864.1;QIQ54047.1;Pangolin_CoV_orf1ab;385 | 0 | 0.025 |
| MT072864.1;QIQ54047.1;Pangolin_CoV_orf1ab;386 | 0 | 0.025 |
| MT072864.1;QIQ54047.1;Pangolin_CoV_orf1ab;387 | 0 | 0.025 |
| MT072864.1;QIQ54047.1;Pangolin_CoV_orf1ab;388 | 0 | 0 |
| MT072864.1;QIQ54047.1;Pangolin_CoV_orf1ab;389 | 0 | 0 |
| MT072864.1;QIQ54047.1;Pangolin_CoV_orf1ab;390 | 0 | 0 |
| MT072864.1;QIQ54047.1;Pangolin_CoV_orf1ab;391 | 0 | 0 |
| MT072864.1;QIQ54047.1;Pangolin_CoV_orf1ab;392 | 0 | 0 |
| MT072864.1;QIQ54047.1;Pangolin_CoV_orf1ab;393 | 0 | 0 |
| MT072864.1;QIQ54047.1;Pangolin_CoV_orf1ab;394 | 0 | 0 |
| MT072864.1;QIQ54047.1;Pangolin_CoV_orf1ab;395 | 0 | 0 |
| MT072864.1;QIQ54047.1;Pangolin_CoV_orf1ab;396 | 0 | 0 |
| MT072864.1;QIQ54047.1;Pangolin_CoV_orf1ab;397 | 0 | 0.025 |
| MT072864.1;QIQ54047.1;Pangolin_CoV_orf1ab;398 | 0 | 0 |
| MT072864.1;QIQ54047.1;Pangolin_CoV_orf1ab;399 | 0 | 0 |
| MT072864.1;QIQ54047.1;Pangolin_CoV_orf1ab;400 | 0 | 0 |
| MT072864.1;QIQ54047.1;Pangolin_CoV_orf1ab;401 | 0 | 0 |
| MT072864.1;QIQ54047.1;Pangolin_CoV_orf1ab;402 | 0 | 0 |
| MT072864.1;QIQ54047.1;Pangolin_CoV_orf1ab;403 | 0 | 0 |
| MT072864.1;QIQ54047.1;Pangolin_CoV_orf1ab;404 | 0 | 0 |
| MT072864.1;QIQ54047.1;Pangolin_CoV_orf1ab;405 | 0 | 0 |
| MT072864.1;QIQ54047.1;Pangolin_CoV_orf1ab;406 | 0 | 0 |
| MT072864.1;QIQ54047.1;Pangolin_CoV_orf1ab;407 | 0 | 0 |
| MT072864.1;QIQ54047.1;Pangolin_CoV_orf1ab;408 | 0 | 0 |
| MT072864.1;QIQ54047.1;Pangolin_CoV_orf1ab;409 | 0 | 0 |
| MT072864.1;QIQ54047.1;Pangolin_CoV_orf1ab;410 | 0 | 0 |
| MT072864.1;QIQ54047.1;Pangolin_CoV_orf1ab;411 | 0 | 0 |
| MT072864.1;QIQ54047.1;Pangolin_CoV_orf1ab;412 | 0 | 0 |
| MT072864.1;QIQ54047.1;Pangolin_CoV_orf1ab;413 | 0.05 | 0 |
| MT072864.1;QIQ54047.1;Pangolin_CoV_orf1ab;414 | 0.05 | 0 |
| MT072864.1;QIQ54047.1;Pangolin_CoV_orf1ab;415 | 0 | 0 |
| MT072864.1;QIQ54047.1;Pangolin_CoV_orf1ab;416 | 0 | 0 |
| MT072864.1;QIQ54047.1;Pangolin_CoV_orf1ab;417 | 0 | 0 |
| MT072864.1;QIQ54047.1;Pangolin_CoV_orf1ab;418 | 0 | 0 |
| MT072864.1;QIQ54047.1;Pangolin_CoV_orf1ab;419 | 0 | 0 |
| MT072864.1;QIQ54047.1;Pangolin_CoV_orf1ab;420 | 0 | 0 |

| | | |
|---|---|---|
| MT072864.1;QIQ54047.1;Pangolin_CoV_orf1ab;421 | 0 | 0 |
| MT072864.1;QIQ54047.1;Pangolin_CoV_orf1ab;422 | 0 | 0 |
| MT072864.1;QIQ54047.1;Pangolin_CoV_orf1ab;423 | 0 | 0 |
| MT072864.1;QIQ54047.1;Pangolin_CoV_orf1ab;424 | 0.05 | 0 |
| MT072864.1;QIQ54047.1;Pangolin_CoV_orf1ab;425 | 0 | 0 |
| MT072864.1;QIQ54047.1;Pangolin_CoV_orf1ab;426 | 0 | 0 |
| MT072864.1;QIQ54047.1;Pangolin_CoV_orf1ab;427 | 0 | 0 |
| MT072864.1;QIQ54047.1;Pangolin_CoV_orf1ab;428 | 0 | 0 |
| MT072864.1;QIQ54047.1;Pangolin_CoV_orf1ab;429 | 0 | 0 |
| MT072864.1;QIQ54047.1;Pangolin_CoV_orf1ab;430 | 0 | 0 |
| MT072864.1;QIQ54047.1;Pangolin_CoV_orf1ab;431 | 0 | 0 |
| MT072864.1;QIQ54047.1;Pangolin_CoV_orf1ab;432 | 0 | 0 |
| MT072864.1;QIQ54047.1;Pangolin_CoV_orf1ab;433 | 0 | 0 |
| MT072864.1;QIQ54047.1;Pangolin_CoV_orf1ab;434 | 0 | 0 |
| MT072864.1;QIQ54047.1;Pangolin_CoV_orf1ab;435 | 0 | 0 |
| MT072864.1;QIQ54047.1;Pangolin_CoV_orf1ab;436 | 0 | 0 |
| MT072864.1;QIQ54047.1;Pangolin_CoV_orf1ab;437 | 0.05 | 0.025 |
| MT072864.1;QIQ54047.1;Pangolin_CoV_orf1ab;438 | 0 | 0 |
| MT072864.1;QIQ54047.1;Pangolin_CoV_orf1ab;439 | 0 | 0 |
| MT072864.1;QIQ54047.1;Pangolin_CoV_orf1ab;440 | 0 | 0 |
| MT072864.1;QIQ54047.1;Pangolin_CoV_orf1ab;441 | 0 | 0 |
| MT072864.1;QIQ54047.1;Pangolin_CoV_orf1ab;442 | 0.05 | 0 |
| MT072864.1;QIQ54047.1;Pangolin_CoV_orf1ab;443 | 0 | 0 |
| MT072864.1;QIQ54047.1;Pangolin_CoV_orf1ab;444 | 0.05 | 0.025 |
| MT072864.1;QIQ54047.1;Pangolin_CoV_orf1ab;445 | 0.4 | 0.125 |
| MT072864.1;QIQ54047.1;Pangolin_CoV_orf1ab;446 | 0 | 0 |
| MT072864.1;QIQ54047.1;Pangolin_CoV_orf1ab;447 | 0.05 | 0 |
| MT072864.1;QIQ54047.1;Pangolin_CoV_orf1ab;448 | 0 | 0 |
| MT072864.1;QIQ54047.1;Pangolin_CoV_orf1ab;449 | 0 | 0 |
| MT072864.1;QIQ54047.1;Pangolin_CoV_orf1ab;450 | 0 | 0 |
| MT072864.1;QIQ54047.1;Pangolin_CoV_orf1ab;451 | 0 | 0 |
| MT072864.1;QIQ54047.1;Pangolin_CoV_orf1ab;452 | 0 | 0 |
| MT072864.1;QIQ54047.1;Pangolin_CoV_orf1ab;453 | 0 | 0 |
| MT072864.1;QIQ54047.1;Pangolin_CoV_orf1ab;454 | 0 | 0 |
| MT072864.1;QIQ54047.1;Pangolin_CoV_orf1ab;455 | 0 | 0 |
| MT072864.1;QIQ54047.1;Pangolin_CoV_orf1ab;456 | 0 | 0 |
| MT072864.1;QIQ54047.1;Pangolin_CoV_orf1ab;457 | 0 | 0 |
| MT072864.1;QIQ54047.1;Pangolin_CoV_orf1ab;458 | 0.05 | 0 |
| MT072864.1;QIQ54047.1;Pangolin_CoV_orf1ab;459 | 0 | 0 |
| MT072864.1;QIQ54047.1;Pangolin_CoV_orf1ab;460 | 0 | 0 |
| MT072864.1;QIQ54047.1;Pangolin_CoV_orf1ab;461 | 0 | 0 |
| MT072864.1;QIQ54047.1;Pangolin_CoV_orf1ab;462 | 0 | 0 |
| MT072864.1;QIQ54047.1;Pangolin_CoV_orf1ab;463 | 0 | 0 |

| | | |
|---|---|---|
| MT072864.1;QIQ54047.1;Pangolin_CoV_orf1ab;464 | 0 | 0 |
| MT072864.1;QIQ54047.1;Pangolin_CoV_orf1ab;465 | 0 | 0 |
| MT072864.1;QIQ54047.1;Pangolin_CoV_orf1ab;466 | 0 | 0 |
| MT072864.1;QIQ54047.1;Pangolin_CoV_orf1ab;467 | 0 | 0 |
| MT072864.1;QIQ54047.1;Pangolin_CoV_orf1ab;468 | 0 | 0 |
| MT072864.1;QIQ54047.1;Pangolin_CoV_orf1ab;469 | 0 | 0 |
| MT072864.1;QIQ54047.1;Pangolin_CoV_orf1ab;470 | 0 | 0 |
| MT072864.1;QIQ54047.1;Pangolin_CoV_orf1ab;471 | 0 | 0 |
| MT072864.1;QIQ54047.1;Pangolin_CoV_orf1ab;472 | 0 | 0 |
| MT072864.1;QIQ54047.1;Pangolin_CoV_orf1ab;473 | 0 | 0 |
| MT072864.1;QIQ54047.1;Pangolin_CoV_orf1ab;474 | 0 | 0 |
| MT072864.1;QIQ54047.1;Pangolin_CoV_orf1ab;475 | 0 | 0 |
| MT072864.1;QIQ54047.1;Pangolin_CoV_orf1ab;476 | 0 | 0 |
| MT072864.1;QIQ54047.1;Pangolin_CoV_orf1ab;477 | 0 | 0 |
| MT072864.1;QIQ54047.1;Pangolin_CoV_orf1ab;478 | 0 | 0 |
| MT072864.1;QIQ54047.1;Pangolin_CoV_orf1ab;479 | 0 | 0 |
| MT072864.1;QIQ54047.1;Pangolin_CoV_orf1ab;480 | 0 | 0.05 |
| MT072864.1;QIQ54047.1;Pangolin_CoV_orf1ab;481 | 0 | 0.05 |
| MT072864.1;QIQ54047.1;Pangolin_CoV_orf1ab;482 | 0 | 0.025 |
| MT072864.1;QIQ54047.1;Pangolin_CoV_orf1ab;483 | 0 | 0.025 |
| MT072864.1;QIQ54047.1;Pangolin_CoV_orf1ab;484 | 0 | 0 |
| MT072864.1;QIQ54047.1;Pangolin_CoV_orf1ab;485 | 0 | 0.025 |
| MT072864.1;QIQ54047.1;Pangolin_CoV_orf1ab;486 | 0 | 0.025 |
| MT072864.1;QIQ54047.1;Pangolin_CoV_orf1ab;487 | 0 | 0 |
| MT072864.1;QIQ54047.1;Pangolin_CoV_orf1ab;488 | 0 | 0 |
| MT072864.1;QIQ54047.1;Pangolin_CoV_orf1ab;489 | 0.05 | 0.075 |
| MT072864.1;QIQ54047.1;Pangolin_CoV_orf1ab;490 | 0.05 | 0 |
| MT072864.1;QIQ54047.1;Pangolin_CoV_orf1ab;491 | 0.05 | 0 |
| MT072864.1;QIQ54047.1;Pangolin_CoV_orf1ab;492 | 0 | 0.025 |
| MT072864.1;QIQ54047.1;Pangolin_CoV_orf1ab;493 | 0.1 | 0.025 |
| MT072864.1;QIQ54047.1;Pangolin_CoV_orf1ab;494 | 0.1 | 0.025 |
| MT072864.1;QIQ54047.1;Pangolin_CoV_orf1ab;495 | 0.05 | 0.025 |
| MT072864.1;QIQ54047.1;Pangolin_CoV_orf1ab;496 | 0.05 | 0 |
| MT072864.1;QIQ54047.1;Pangolin_CoV_orf1ab;497 | 0.1 | 0.075 |
| MT072864.1;QIQ54047.1;Pangolin_CoV_orf1ab;498 | 0.1 | 0.025 |
| MT072864.1;QIQ54047.1;Pangolin_CoV_orf1ab;499 | 0 | 0 |
| MT072864.1;QIQ54047.1;Pangolin_CoV_orf1ab;500 | 0 | 0 |
| MT072864.1;QIQ54047.1;Pangolin_CoV_orf1ab;501 | 0 | 0 |
| MT072864.1;QIQ54047.1;Pangolin_CoV_orf1ab;502 | 0 | 0 |
| MT072864.1;QIQ54047.1;Pangolin_CoV_orf1ab;503 | 0 | 0 |
| MT072864.1;QIQ54047.1;Pangolin_CoV_orf1ab;504 | 0.05 | 0 |
| MT072864.1;QIQ54047.1;Pangolin_CoV_orf1ab;505 | 0.05 | 0 |
| MT072864.1;QIQ54047.1;Pangolin_CoV_orf1ab;506 | 0.05 | 0 |

| | | |
|---|---|---|
| MT072864.1;QIQ54047.1;Pangolin_CoV_orf1ab;507 | 0.05 | 0 |
| MT072864.1;QIQ54047.1;Pangolin_CoV_orf1ab;508 | 0.05 | 0 |
| MT072864.1;QIQ54047.1;Pangolin_CoV_orf1ab;509 | 0.05 | 0 |
| MT072864.1;QIQ54047.1;Pangolin_CoV_orf1ab;510 | 0.05 | 0 |
| MT072864.1;QIQ54047.1;Pangolin_CoV_orf1ab;511 | 0 | 0.05 |
| MT072864.1;QIQ54047.1;Pangolin_CoV_orf1ab;512 | 0.05 | 0.05 |
| MT072864.1;QIQ54047.1;Pangolin_CoV_orf1ab;513 | 0.05 | 0.05 |
| MT072864.1;QIQ54047.1;Pangolin_CoV_orf1ab;514 | 0.05 | 0.05 |
| MT072864.1;QIQ54047.1;Pangolin_CoV_orf1ab;515 | 0 | 0.05 |
| MT072864.1;QIQ54047.1;Pangolin_CoV_orf1ab;516 | 0 | 0.05 |
| MT072864.1;QIQ54047.1;Pangolin_CoV_orf1ab;517 | 0 | 0.075 |
| MT072864.1;QIQ54047.1;Pangolin_CoV_orf1ab;518 | 0.05 | 0.125 |
| MT072864.1;QIQ54047.1;Pangolin_CoV_orf1ab;519 | 0 | 0 |
| MT072864.1;QIQ54047.1;Pangolin_CoV_orf1ab;520 | 0 | 0 |
| MT072864.1;QIQ54047.1;Pangolin_CoV_orf1ab;521 | 0 | 0 |
| MT072864.1;QIQ54047.1;Pangolin_CoV_orf1ab;522 | 0 | 0 |
| MT072864.1;QIQ54047.1;Pangolin_CoV_orf1ab;523 | 0 | 0.025 |
| MT072864.1;QIQ54047.1;Pangolin_CoV_orf1ab;524 | 0 | 0 |
| MT072864.1;QIQ54047.1;Pangolin_CoV_orf1ab;525 | 0.05 | 0 |
| MT072864.1;QIQ54047.1;Pangolin_CoV_orf1ab;526 | 0.05 | 0.025 |
| MT072864.1;QIQ54047.1;Pangolin_CoV_orf1ab;527 | 0 | 0 |
| MT072864.1;QIQ54047.1;Pangolin_CoV_orf1ab;528 | 0 | 0 |
| MT072864.1;QIQ54047.1;Pangolin_CoV_orf1ab;529 | 0 | 0 |
| MT072864.1;QIQ54047.1;Pangolin_CoV_orf1ab;530 | 0 | 0 |
| MT072864.1;QIQ54047.1;Pangolin_CoV_orf1ab;531 | 0 | 0 |
| MT072864.1;QIQ54047.1;Pangolin_CoV_orf1ab;532 | 0 | 0 |
| MT072864.1;QIQ54047.1;Pangolin_CoV_orf1ab;533 | 0 | 0 |
| MT072864.1;QIQ54047.1;Pangolin_CoV_orf1ab;534 | 0 | 0 |
| MT072864.1;QIQ54047.1;Pangolin_CoV_orf1ab;535 | 0 | 0 |
| MT072864.1;QIQ54047.1;Pangolin_CoV_orf1ab;536 | 0 | 0 |
| MT072864.1;QIQ54047.1;Pangolin_CoV_orf1ab;537 | 0 | 0 |
| MT072864.1;QIQ54047.1;Pangolin_CoV_orf1ab;538 | 0 | 0 |
| MT072864.1;QIQ54047.1;Pangolin_CoV_orf1ab;539 | 0 | 0 |
| MT072864.1;QIQ54047.1;Pangolin_CoV_orf1ab;540 | 0 | 0 |
| MT072864.1;QIQ54047.1;Pangolin_CoV_orf1ab;541 | 0 | 0 |
| MT072864.1;QIQ54047.1;Pangolin_CoV_orf1ab;542 | 0 | 0 |
| MT072864.1;QIQ54047.1;Pangolin_CoV_orf1ab;543 | 0 | 0 |
| MT072864.1;QIQ54047.1;Pangolin_CoV_orf1ab;544 | 0 | 0 |
| MT072864.1;QIQ54047.1;Pangolin_CoV_orf1ab;545 | 0 | 0 |
| MT072864.1;QIQ54047.1;Pangolin_CoV_orf1ab;546 | 0 | 0.025 |
| MT072864.1;QIQ54047.1;Pangolin_CoV_orf1ab;547 | 0 | 0 |
| MT072864.1;QIQ54047.1;Pangolin_CoV_orf1ab;548 | 0 | 0 |
| MT072864.1;QIQ54047.1;Pangolin_CoV_orf1ab;549 | 0 | 0 |

| | | |
|---|---|---|
| MT072864.1;QIQ54047.1;Pangolin_CoV_orf1ab;550 | 0 | 0 |
| MT072864.1;QIQ54047.1;Pangolin_CoV_orf1ab;551 | 0 | 0 |
| MT072864.1;QIQ54047.1;Pangolin_CoV_orf1ab;552 | 0 | 0 |
| MT072864.1;QIQ54047.1;Pangolin_CoV_orf1ab;553 | 0 | 0 |
| MT072864.1;QIQ54047.1;Pangolin_CoV_orf1ab;554 | 0 | 0 |
| MT072864.1;QIQ54047.1;Pangolin_CoV_orf1ab;555 | 0 | 0 |
| MT072864.1;QIQ54047.1;Pangolin_CoV_orf1ab;556 | 0 | 0 |
| MT072864.1;QIQ54047.1;Pangolin_CoV_orf1ab;557 | 0 | 0 |
| MT072864.1;QIQ54047.1;Pangolin_CoV_orf1ab;558 | 0 | 0 |
| MT072864.1;QIQ54047.1;Pangolin_CoV_orf1ab;559 | 0 | 0 |
| MT072864.1;QIQ54047.1;Pangolin_CoV_orf1ab;560 | 0 | 0 |
| MT072864.1;QIQ54047.1;Pangolin_CoV_orf1ab;561 | 0 | 0 |
| MT072864.1;QIQ54047.1;Pangolin_CoV_orf1ab;562 | 0 | 0 |
| MT072864.1;QIQ54047.1;Pangolin_CoV_orf1ab;563 | 0 | 0 |
| MT072864.1;QIQ54047.1;Pangolin_CoV_orf1ab;564 | 0 | 0 |
| MT072864.1;QIQ54047.1;Pangolin_CoV_orf1ab;565 | 0 | 0.025 |
| MT072864.1;QIQ54047.1;Pangolin_CoV_orf1ab;566 | 0 | 0 |
| MT072864.1;QIQ54047.1;Pangolin_CoV_orf1ab;567 | 0 | 0 |
| MT072864.1;QIQ54047.1;Pangolin_CoV_orf1ab;568 | 0 | 0 |
| MT072864.1;QIQ54047.1;Pangolin_CoV_orf1ab;569 | 0 | 0.025 |
| MT072864.1;QIQ54047.1;Pangolin_CoV_orf1ab;570 | 0 | 0.025 |
| MT072864.1;QIQ54047.1;Pangolin_CoV_orf1ab;571 | 0 | 0.025 |
| MT072864.1;QIQ54047.1;Pangolin_CoV_orf1ab;572 | 0 | 0.025 |
| MT072864.1;QIQ54047.1;Pangolin_CoV_orf1ab;573 | 0 | 0 |
| MT072864.1;QIQ54047.1;Pangolin_CoV_orf1ab;574 | 0 | 0 |
| MT072864.1;QIQ54047.1;Pangolin_CoV_orf1ab;575 | 0 | 0 |
| MT072864.1;QIQ54047.1;Pangolin_CoV_orf1ab;576 | 0 | 0 |
| MT072864.1;QIQ54047.1;Pangolin_CoV_orf1ab;577 | 0 | 0 |
| MT072864.1;QIQ54047.1;Pangolin_CoV_orf1ab;578 | 0 | 0.025 |
| MT072864.1;QIQ54047.1;Pangolin_CoV_orf1ab;579 | 0 | 0 |
| MT072864.1;QIQ54047.1;Pangolin_CoV_orf1ab;580 | 0 | 0 |
| MT072864.1;QIQ54047.1;Pangolin_CoV_orf1ab;581 | 0 | 0 |
| MT072864.1;QIQ54047.1;Pangolin_CoV_orf1ab;582 | 0 | 0 |
| MT072864.1;QIQ54047.1;Pangolin_CoV_orf1ab;583 | 0 | 0 |
| MT072864.1;QIQ54047.1;Pangolin_CoV_orf1ab;584 | 0 | 0 |
| MT072864.1;QIQ54047.1;Pangolin_CoV_orf1ab;585 | 0 | 0 |
| MT072864.1;QIQ54047.1;Pangolin_CoV_orf1ab;586 | 0 | 0 |
| MT072864.1;QIQ54047.1;Pangolin_CoV_orf1ab;587 | 0 | 0 |
| MT072864.1;QIQ54047.1;Pangolin_CoV_orf1ab;588 | 0 | 0 |
| MT072864.1;QIQ54047.1;Pangolin_CoV_orf1ab;589 | 0.05 | 0 |
| MT072864.1;QIQ54047.1;Pangolin_CoV_orf1ab;590 | 0 | 0 |
| MT072864.1;QIQ54047.1;Pangolin_CoV_orf1ab;591 | 0 | 0 |
| MT072864.1;QIQ54047.1;Pangolin_CoV_orf1ab;592 | 0 | 0.025 |

| | | |
|---|---|---|
| MT072864.1;QIQ54047.1;Pangolin_CoV_orf1ab;593 | 0 | 0 |
| MT072864.1;QIQ54047.1;Pangolin_CoV_orf1ab;594 | 0 | 0 |
| MT072864.1;QIQ54047.1;Pangolin_CoV_orf1ab;595 | 0 | 0 |
| MT072864.1;QIQ54047.1;Pangolin_CoV_orf1ab;596 | 0 | 0 |
| MT072864.1;QIQ54047.1;Pangolin_CoV_orf1ab;597 | 0 | 0.025 |
| MT072864.1;QIQ54047.1;Pangolin_CoV_orf1ab;598 | 0 | 0.025 |
| MT072864.1;QIQ54047.1;Pangolin_CoV_orf1ab;599 | 0 | 0.05 |
| MT072864.1;QIQ54047.1;Pangolin_CoV_orf1ab;600 | 0 | 0.025 |
| MT072864.1;QIQ54047.1;Pangolin_CoV_orf1ab;601 | 0 | 0.025 |
| MT072864.1;QIQ54047.1;Pangolin_CoV_orf1ab;602 | 0 | 0.025 |
| MT072864.1;QIQ54047.1;Pangolin_CoV_orf1ab;603 | 0 | 0.025 |
| MT072864.1;QIQ54047.1;Pangolin_CoV_orf1ab;604 | 0 | 0 |
| MT072864.1;QIQ54047.1;Pangolin_CoV_orf1ab;605 | 0 | 0 |
| MT072864.1;QIQ54047.1;Pangolin_CoV_orf1ab;606 | 0 | 0 |
| MT072864.1;QIQ54047.1;Pangolin_CoV_orf1ab;607 | 0 | 0 |
| MT072864.1;QIQ54047.1;Pangolin_CoV_orf1ab;608 | 0 | 0 |
| MT072864.1;QIQ54047.1;Pangolin_CoV_orf1ab;609 | 0 | 0 |
| MT072864.1;QIQ54047.1;Pangolin_CoV_orf1ab;610 | 0 | 0 |
| MT072864.1;QIQ54047.1;Pangolin_CoV_orf1ab;611 | 0 | 0 |
| MT072864.1;QIQ54047.1;Pangolin_CoV_orf1ab;612 | 0 | 0 |
| MT072864.1;QIQ54047.1;Pangolin_CoV_orf1ab;613 | 0 | 0 |
| MT072864.1;QIQ54047.1;Pangolin_CoV_orf1ab;614 | 0 | 0.025 |
| MT072864.1;QIQ54047.1;Pangolin_CoV_orf1ab;615 | 0 | 0 |
| MT072864.1;QIQ54047.1;Pangolin_CoV_orf1ab;616 | 0 | 0 |
| MT072864.1;QIQ54047.1;Pangolin_CoV_orf1ab;617 | 0.05 | 0 |
| MT072864.1;QIQ54047.1;Pangolin_CoV_orf1ab;618 | 0.1 | 0.025 |
| MT072864.1;QIQ54047.1;Pangolin_CoV_orf1ab;619 | 0.1 | 0.025 |
| MT072864.1;QIQ54047.1;Pangolin_CoV_orf1ab;620 | 0.1 | 0.05 |
| MT072864.1;QIQ54047.1;Pangolin_CoV_orf1ab;621 | 0.15 | 0.025 |
| MT072864.1;QIQ54047.1;Pangolin_CoV_orf1ab;622 | 0.1 | 0.025 |
| MT072864.1;QIQ54047.1;Pangolin_CoV_orf1ab;623 | 0.1 | 0.025 |
| MT072864.1;QIQ54047.1;Pangolin_CoV_orf1ab;624 | 0 | 0.05 |
| MT072864.1;QIQ54047.1;Pangolin_CoV_orf1ab;625 | 0 | 0 |
| MT072864.1;QIQ54047.1;Pangolin_CoV_orf1ab;626 | 0 | 0 |
| MT072864.1;QIQ54047.1;Pangolin_CoV_orf1ab;627 | 0 | 0.025 |
| MT072864.1;QIQ54047.1;Pangolin_CoV_orf1ab;628 | 0 | 0.025 |
| MT072864.1;QIQ54047.1;Pangolin_CoV_orf1ab;629 | 0 | 0.025 |
| MT072864.1;QIQ54047.1;Pangolin_CoV_orf1ab;630 | 0 | 0.025 |
| MT072864.1;QIQ54047.1;Pangolin_CoV_orf1ab;631 | 0 | 0.025 |
| MT072864.1;QIQ54047.1;Pangolin_CoV_orf1ab;632 | 0 | 0.025 |
| MT072864.1;QIQ54047.1;Pangolin_CoV_orf1ab;633 | 0 | 0.025 |
| MT072864.1;QIQ54047.1;Pangolin_CoV_orf1ab;634 | 0 | 0.025 |
| MT072864.1;QIQ54047.1;Pangolin_CoV_orf1ab;635 | 0 | 0.025 |

| | | |
|---|---|---|
| MT072864.1;QIQ54047.1;Pangolin_CoV_orf1ab;636 | 0 | 0.025 |
| MT072864.1;QIQ54047.1;Pangolin_CoV_orf1ab;637 | 0 | 0 |
| MT072864.1;QIQ54047.1;Pangolin_CoV_orf1ab;638 | 0.05 | 0.05 |
| MT072864.1;QIQ54047.1;Pangolin_CoV_orf1ab;639 | 0 | 0 |
| MT072864.1;QIQ54047.1;Pangolin_CoV_orf1ab;640 | 0 | 0 |
| MT072864.1;QIQ54047.1;Pangolin_CoV_orf1ab;641 | 0 | 0 |
| MT072864.1;QIQ54047.1;Pangolin_CoV_orf1ab;642 | 0 | 0 |
| MT072864.1;QIQ54047.1;Pangolin_CoV_orf1ab;643 | 0 | 0 |
| MT072864.1;QIQ54047.1;Pangolin_CoV_orf1ab;644 | 0 | 0 |
| MT072864.1;QIQ54047.1;Pangolin_CoV_orf1ab;645 | 0 | 0 |
| MT072864.1;QIQ54047.1;Pangolin_CoV_orf1ab;646 | 0 | 0 |
| MT072864.1;QIQ54047.1;Pangolin_CoV_orf1ab;647 | 0 | 0 |
| MT072864.1;QIQ54047.1;Pangolin_CoV_orf1ab;648 | 0 | 0 |
| MT072864.1;QIQ54047.1;Pangolin_CoV_orf1ab;649 | 0 | 0 |
| MT072864.1;QIQ54047.1;Pangolin_CoV_orf1ab;650 | 0 | 0 |
| MT072864.1;QIQ54047.1;Pangolin_CoV_orf1ab;651 | 0 | 0.025 |
| MT072864.1;QIQ54047.1;Pangolin_CoV_orf1ab;652 | 0 | 0 |
| MT072864.1;QIQ54047.1;Pangolin_CoV_orf1ab;653 | 0 | 0 |
| MT072864.1;QIQ54047.1;Pangolin_CoV_orf1ab;654 | 0 | 0 |
| MT072864.1;QIQ54047.1;Pangolin_CoV_orf1ab;655 | 0 | 0 |
| MT072864.1;QIQ54047.1;Pangolin_CoV_orf1ab;656 | 0 | 0 |
| MT072864.1;QIQ54047.1;Pangolin_CoV_orf1ab;657 | 0 | 0 |
| MT072864.1;QIQ54047.1;Pangolin_CoV_orf1ab;658 | 0 | 0.025 |
| MT072864.1;QIQ54047.1;Pangolin_CoV_orf1ab;659 | 0 | 0 |
| MT072864.1;QIQ54047.1;Pangolin_CoV_orf1ab;660 | 0 | 0 |
| MT072864.1;QIQ54047.1;Pangolin_CoV_orf1ab;661 | 0 | 0 |
| MT072864.1;QIQ54047.1;Pangolin_CoV_orf1ab;662 | 0.05 | 0 |
| MT072864.1;QIQ54047.1;Pangolin_CoV_orf1ab;663 | 0 | 0.025 |
| MT072864.1;QIQ54047.1;Pangolin_CoV_orf1ab;664 | 0 | 0.025 |
| MT072864.1;QIQ54047.1;Pangolin_CoV_orf1ab;665 | 0 | 0.025 |
| MT072864.1;QIQ54047.1;Pangolin_CoV_orf1ab;666 | 0 | 0.025 |
| MT072864.1;QIQ54047.1;Pangolin_CoV_orf1ab;667 | 0 | 0.025 |
| MT072864.1;QIQ54047.1;Pangolin_CoV_orf1ab;668 | 0 | 0.025 |
| MT072864.1;QIQ54047.1;Pangolin_CoV_orf1ab;669 | 0 | 0.025 |
| MT072864.1;QIQ54047.1;Pangolin_CoV_orf1ab;670 | 0 | 0 |
| MT072864.1;QIQ54047.1;Pangolin_CoV_orf1ab;671 | 0 | 0 |
| MT072864.1;QIQ54047.1;Pangolin_CoV_orf1ab;672 | 0 | 0 |
| MT072864.1;QIQ54047.1;Pangolin_CoV_orf1ab;673 | 0 | 0 |
| MT072864.1;QIQ54047.1;Pangolin_CoV_orf1ab;674 | 0 | 0 |
| MT072864.1;QIQ54047.1;Pangolin_CoV_orf1ab;675 | 0 | 0 |
| MT072864.1;QIQ54047.1;Pangolin_CoV_orf1ab;676 | 0 | 0 |
| MT072864.1;QIQ54047.1;Pangolin_CoV_orf1ab;677 | 0 | 0 |
| MT072864.1;QIQ54047.1;Pangolin_CoV_orf1ab;678 | 0 | 0 |

| | | |
|---|---|---|
| MT072864.1;QIQ54047.1;Pangolin_CoV_orf1ab;679 | 0 | 0 |
| MT072864.1;QIQ54047.1;Pangolin_CoV_orf1ab;680 | 0 | 0 |
| MT072864.1;QIQ54047.1;Pangolin_CoV_orf1ab;681 | 0 | 0 |
| MT072864.1;QIQ54047.1;Pangolin_CoV_orf1ab;682 | 0 | 0.025 |
| MT072864.1;QIQ54047.1;Pangolin_CoV_orf1ab;683 | 0 | 0 |
| MT072864.1;QIQ54047.1;Pangolin_CoV_orf1ab;684 | 0.05 | 0.025 |
| MT072864.1;QIQ54047.1;Pangolin_CoV_orf1ab;685 | 0.1 | 0 |
| MT072864.1;QIQ54047.1;Pangolin_CoV_orf1ab;686 | 0 | 0 |
| MT072864.1;QIQ54047.1;Pangolin_CoV_orf1ab;687 | 0 | 0 |
| MT072864.1;QIQ54047.1;Pangolin_CoV_orf1ab;688 | 0 | 0 |
| MT072864.1;QIQ54047.1;Pangolin_CoV_orf1ab;689 | 0.05 | 0.025 |
| MT072864.1;QIQ54047.1;Pangolin_CoV_orf1ab;690 | 0.05 | 0 |
| MT072864.1;QIQ54047.1;Pangolin_CoV_orf1ab;691 | 0.05 | 0 |
| MT072864.1;QIQ54047.1;Pangolin_CoV_orf1ab;692 | 0.1 | 0.05 |
| MT072864.1;QIQ54047.1;Pangolin_CoV_orf1ab;693 | 0.05 | 0.025 |
| MT072864.1;QIQ54047.1;Pangolin_CoV_orf1ab;694 | 0 | 0 |
| MT072864.1;QIQ54047.1;Pangolin_CoV_orf1ab;695 | 0 | 0 |
| MT072864.1;QIQ54047.1;Pangolin_CoV_orf1ab;696 | 0.05 | 0.025 |
| MT072864.1;QIQ54047.1;Pangolin_CoV_orf1ab;697 | 0 | 0 |
| MT072864.1;QIQ54047.1;Pangolin_CoV_orf1ab;698 | 0 | 0 |
| MT072864.1;QIQ54047.1;Pangolin_CoV_orf1ab;699 | 0 | 0 |
| MT072864.1;QIQ54047.1;Pangolin_CoV_orf1ab;700 | 0 | 0 |
| MT072864.1;QIQ54047.1;Pangolin_CoV_orf1ab;701 | 0 | 0 |
| MT072864.1;QIQ54047.1;Pangolin_CoV_orf1ab;702 | 0 | 0 |
| MT072864.1;QIQ54047.1;Pangolin_CoV_orf1ab;703 | 0 | 0 |
| MT072864.1;QIQ54047.1;Pangolin_CoV_orf1ab;704 | 0 | 0 |
| MT072864.1;QIQ54047.1;Pangolin_CoV_orf1ab;705 | 0 | 0 |
| MT072864.1;QIQ54047.1;Pangolin_CoV_orf1ab;706 | 0 | 0 |
| MT072864.1;QIQ54047.1;Pangolin_CoV_orf1ab;707 | 0 | 0 |
| MT072864.1;QIQ54047.1;Pangolin_CoV_orf1ab;708 | 0 | 0 |
| MT072864.1;QIQ54047.1;Pangolin_CoV_orf1ab;709 | 0 | 0 |
| MT072864.1;QIQ54047.1;Pangolin_CoV_orf1ab;710 | 0 | 0 |
| MT072864.1;QIQ54047.1;Pangolin_CoV_orf1ab;711 | 0 | 0 |
| MT072864.1;QIQ54047.1;Pangolin_CoV_orf1ab;712 | 0 | 0 |
| MT072864.1;QIQ54047.1;Pangolin_CoV_orf1ab;713 | 0 | 0.05 |
| MT072864.1;QIQ54047.1;Pangolin_CoV_orf1ab;714 | 0.05 | 0 |
| MT072864.1;QIQ54047.1;Pangolin_CoV_orf1ab;715 | 0.05 | 0 |
| MT072864.1;QIQ54047.1;Pangolin_CoV_orf1ab;716 | 0.05 | 0 |
| MT072864.1;QIQ54047.1;Pangolin_CoV_orf1ab;717 | 0 | 0 |
| MT072864.1;QIQ54047.1;Pangolin_CoV_orf1ab;718 | 0.05 | 0.05 |
| MT072864.1;QIQ54047.1;Pangolin_CoV_orf1ab;719 | 0.05 | 0 |
| MT072864.1;QIQ54047.1;Pangolin_CoV_orf1ab;720 | 0.05 | 0 |
| MT072864.1;QIQ54047.1;Pangolin_CoV_orf1ab;721 | 0 | 0 |

| | | |
|---|---|---|
| MT072864.1;QIQ54047.1;Pangolin_CoV_orf1ab;722 | 0 | 0.025 |
| MT072864.1;QIQ54047.1;Pangolin_CoV_orf1ab;723 | 0 | 0 |
| MT072864.1;QIQ54047.1;Pangolin_CoV_orf1ab;724 | 0 | 0 |
| MT072864.1;QIQ54047.1;Pangolin_CoV_orf1ab;725 | 0 | 0 |
| MT072864.1;QIQ54047.1;Pangolin_CoV_orf1ab;726 | 0 | 0 |
| MT072864.1;QIQ54047.1;Pangolin_CoV_orf1ab;727 | 0 | 0 |
| MT072864.1;QIQ54047.1;Pangolin_CoV_orf1ab;728 | 0 | 0 |
| MT072864.1;QIQ54047.1;Pangolin_CoV_orf1ab;729 | 0 | 0 |
| MT072864.1;QIQ54047.1;Pangolin_CoV_orf1ab;730 | 0 | 0 |
| MT072864.1;QIQ54047.1;Pangolin_CoV_orf1ab;731 | 0 | 0 |
| MT072864.1;QIQ54047.1;Pangolin_CoV_orf1ab;732 | 0 | 0 |
| MT072864.1;QIQ54047.1;Pangolin_CoV_orf1ab;733 | 0 | 0 |
| MT072864.1;QIQ54047.1;Pangolin_CoV_orf1ab;734 | 0 | 0 |
| MT072864.1;QIQ54047.1;Pangolin_CoV_orf1ab;735 | 0 | 0 |
| MT072864.1;QIQ54047.1;Pangolin_CoV_orf1ab;736 | 0 | 0 |
| MT072864.1;QIQ54047.1;Pangolin_CoV_orf1ab;737 | 0 | 0 |
| MT072864.1;QIQ54047.1;Pangolin_CoV_orf1ab;738 | 0 | 0 |
| MT072864.1;QIQ54047.1;Pangolin_CoV_orf1ab;739 | 0 | 0 |
| MT072864.1;QIQ54047.1;Pangolin_CoV_orf1ab;740 | 0 | 0 |
| MT072864.1;QIQ54047.1;Pangolin_CoV_orf1ab;741 | 0 | 0 |
| MT072864.1;QIQ54047.1;Pangolin_CoV_orf1ab;742 | 0 | 0 |
| MT072864.1;QIQ54047.1;Pangolin_CoV_orf1ab;743 | 0 | 0 |
| MT072864.1;QIQ54047.1;Pangolin_CoV_orf1ab;744 | 0.05 | 0.125 |
| MT072864.1;QIQ54047.1;Pangolin_CoV_orf1ab;745 | 0.1 | 0.075 |
| MT072864.1;QIQ54047.1;Pangolin_CoV_orf1ab;746 | 0.15 | 0.075 |
| MT072864.1;QIQ54047.1;Pangolin_CoV_orf1ab;747 | 0 | 0.075 |
| MT072864.1;QIQ54047.1;Pangolin_CoV_orf1ab;748 | 0.05 | 0.025 |
| MT072864.1;QIQ54047.1;Pangolin_CoV_orf1ab;749 | 0 | 0.05 |
| MT072864.1;QIQ54047.1;Pangolin_CoV_orf1ab;750 | 0.1 | 0.075 |
| MT072864.1;QIQ54047.1;Pangolin_CoV_orf1ab;751 | 0.05 | 0.075 |
| MT072864.1;QIQ54047.1;Pangolin_CoV_orf1ab;752 | 0.05 | 0.05 |
| MT072864.1;QIQ54047.1;Pangolin_CoV_orf1ab;753 | 0.05 | 0.025 |
| MT072864.1;QIQ54047.1;Pangolin_CoV_orf1ab;754 | 0.1 | 0.05 |
| MT072864.1;QIQ54047.1;Pangolin_CoV_orf1ab;755 | 0.05 | 0.025 |
| MT072864.1;QIQ54047.1;Pangolin_CoV_orf1ab;756 | 0 | 0.05 |
| MT072864.1;QIQ54047.1;Pangolin_CoV_orf1ab;757 | 0 | 0.025 |
| MT072864.1;QIQ54047.1;Pangolin_CoV_orf1ab;758 | 0 | 0.025 |
| MT072864.1;QIQ54047.1;Pangolin_CoV_orf1ab;759 | 0 | 0.025 |
| MT072864.1;QIQ54047.1;Pangolin_CoV_orf1ab;760 | 0 | 0.025 |
| MT072864.1;QIQ54047.1;Pangolin_CoV_orf1ab;761 | 0 | 0.05 |
| MT072864.1;QIQ54047.1;Pangolin_CoV_orf1ab;762 | 0 | 0 |
| MT072864.1;QIQ54047.1;Pangolin_CoV_orf1ab;763 | 0 | 0 |
| MT072864.1;QIQ54047.1;Pangolin_CoV_orf1ab;764 | 0 | 0.025 |

| | | |
|---|---|---|
| MT072864.1;QIQ54047.1;Pangolin_CoV_orf1ab;765 | 0 | 0.025 |
| MT072864.1;QIQ54047.1;Pangolin_CoV_orf1ab;766 | 0 | 0.05 |
| MT072864.1;QIQ54047.1;Pangolin_CoV_orf1ab;767 | 0 | 0 |
| MT072864.1;QIQ54047.1;Pangolin_CoV_orf1ab;768 | 0 | 0 |
| MT072864.1;QIQ54047.1;Pangolin_CoV_orf1ab;769 | 0 | 0 |
| MT072864.1;QIQ54047.1;Pangolin_CoV_orf1ab;770 | 0 | 0 |
| MT072864.1;QIQ54047.1;Pangolin_CoV_orf1ab;771 | 0 | 0 |
| MT072864.1;QIQ54047.1;Pangolin_CoV_orf1ab;772 | 0 | 0 |
| MT072864.1;QIQ54047.1;Pangolin_CoV_orf1ab;773 | 0 | 0 |
| MT072864.1;QIQ54047.1;Pangolin_CoV_orf1ab;774 | 0 | 0 |
| MT072864.1;QIQ54047.1;Pangolin_CoV_orf1ab;775 | 0 | 0 |
| MT072864.1;QIQ54047.1;Pangolin_CoV_orf1ab;776 | 0 | 0 |
| MT072864.1;QIQ54047.1;Pangolin_CoV_orf1ab;777 | 0 | 0 |
| MT072864.1;QIQ54047.1;Pangolin_CoV_orf1ab;778 | 0 | 0 |
| MT072864.1;QIQ54047.1;Pangolin_CoV_orf1ab;779 | 0 | 0 |
| MT072864.1;QIQ54047.1;Pangolin_CoV_orf1ab;780 | 0 | 0.025 |
| MT072864.1;QIQ54047.1;Pangolin_CoV_orf1ab;781 | 0 | 0 |
| MT072864.1;QIQ54047.1;Pangolin_CoV_orf1ab;782 | 0.05 | 0 |
| MT072864.1;QIQ54047.1;Pangolin_CoV_orf1ab;783 | 0 | 0 |
| MT072864.1;QIQ54047.1;Pangolin_CoV_orf1ab;784 | 0 | 0 |
| MT072864.1;QIQ54047.1;Pangolin_CoV_orf1ab;785 | 0 | 0 |
| MT072864.1;QIQ54047.1;Pangolin_CoV_orf1ab;786 | 0 | 0 |
| MT072864.1;QIQ54047.1;Pangolin_CoV_orf1ab;787 | 0 | 0.025 |
| MT072864.1;QIQ54047.1;Pangolin_CoV_orf1ab;788 | 0 | 0.025 |
| MT072864.1;QIQ54047.1;Pangolin_CoV_orf1ab;789 | 0.05 | 0 |
| MT072864.1;QIQ54047.1;Pangolin_CoV_orf1ab;790 | 0 | 0.025 |
| MT072864.1;QIQ54047.1;Pangolin_CoV_orf1ab;791 | 0 | 0.05 |
| MT072864.1;QIQ54047.1;Pangolin_CoV_orf1ab;792 | 0 | 0.05 |
| MT072864.1;QIQ54047.1;Pangolin_CoV_orf1ab;793 | 0 | 0.025 |
| MT072864.1;QIQ54047.1;Pangolin_CoV_orf1ab;794 | 0 | 0 |
| MT072864.1;QIQ54047.1;Pangolin_CoV_orf1ab;795 | 0 | 0 |
| MT072864.1;QIQ54047.1;Pangolin_CoV_orf1ab;796 | 0 | 0 |
| MT072864.1;QIQ54047.1;Pangolin_CoV_orf1ab;797 | 0.05 | 0.05 |
| MT072864.1;QIQ54047.1;Pangolin_CoV_orf1ab;798 | 0.05 | 0 |
| MT072864.1;QIQ54047.1;Pangolin_CoV_orf1ab;799 | 0 | 0 |
| MT072864.1;QIQ54047.1;Pangolin_CoV_orf1ab;800 | 0 | 0 |
| MT072864.1;QIQ54047.1;Pangolin_CoV_orf1ab;801 | 0.05 | 0 |
| MT072864.1;QIQ54047.1;Pangolin_CoV_orf1ab;802 | 0 | 0 |
| MT072864.1;QIQ54047.1;Pangolin_CoV_orf1ab;803 | 0.1 | 0 |
| MT072864.1;QIQ54047.1;Pangolin_CoV_orf1ab;804 | 0.05 | 0 |
| MT072864.1;QIQ54047.1;Pangolin_CoV_orf1ab;805 | 0.05 | 0 |
| MT072864.1;QIQ54047.1;Pangolin_CoV_orf1ab;806 | 0.05 | 0 |
| MT072864.1;QIQ54047.1;Pangolin_CoV_orf1ab;807 | 0 | 0.025 |

| | | |
|---|---|---|
| MT072864.1;QIQ54047.1;Pangolin_CoV_orf1ab;808 | 0 | 0 |
| MT072864.1;QIQ54047.1;Pangolin_CoV_orf1ab;809 | 0 | 0 |
| MT072864.1;QIQ54047.1;Pangolin_CoV_orf1ab;810 | 0 | 0 |
| MT072864.1;QIQ54047.1;Pangolin_CoV_orf1ab;811 | 0 | 0 |
| MT072864.1;QIQ54047.1;Pangolin_CoV_orf1ab;812 | 0 | 0 |
| MT072864.1;QIQ54047.1;Pangolin_CoV_orf1ab;813 | 0.05 | 0 |
| MT072864.1;QIQ54047.1;Pangolin_CoV_orf1ab;814 | 0 | 0.025 |
| MT072864.1;QIQ54047.1;Pangolin_CoV_orf1ab;815 | 0 | 0 |
| MT072864.1;QIQ54047.1;Pangolin_CoV_orf1ab;816 | 0 | 0 |
| MT072864.1;QIQ54047.1;Pangolin_CoV_orf1ab;817 | 0 | 0 |
| MT072864.1;QIQ54047.1;Pangolin_CoV_orf1ab;818 | 0 | 0 |
| MT072864.1;QIQ54047.1;Pangolin_CoV_orf1ab;819 | 0.05 | 0 |
| MT072864.1;QIQ54047.1;Pangolin_CoV_orf1ab;820 | 0.05 | 0 |
| MT072864.1;QIQ54047.1;Pangolin_CoV_orf1ab;821 | 0 | 0 |
| MT072864.1;QIQ54047.1;Pangolin_CoV_orf1ab;822 | 0 | 0.025 |
| MT072864.1;QIQ54047.1;Pangolin_CoV_orf1ab;823 | 0 | 0.025 |
| MT072864.1;QIQ54047.1;Pangolin_CoV_orf1ab;824 | 0.05 | 0.025 |
| MT072864.1;QIQ54047.1;Pangolin_CoV_orf1ab;825 | 0.05 | 0 |
| MT072864.1;QIQ54047.1;Pangolin_CoV_orf1ab;826 | 0 | 0 |
| MT072864.1;QIQ54047.1;Pangolin_CoV_orf1ab;827 | 0.05 | 0 |
| MT072864.1;QIQ54047.1;Pangolin_CoV_orf1ab;828 | 0 | 0 |
| MT072864.1;QIQ54047.1;Pangolin_CoV_orf1ab;829 | 0 | 0 |
| MT072864.1;QIQ54047.1;Pangolin_CoV_orf1ab;830 | 0 | 0 |
| MT072864.1;QIQ54047.1;Pangolin_CoV_orf1ab;831 | 0.05 | 0.05 |
| MT072864.1;QIQ54047.1;Pangolin_CoV_orf1ab;832 | 0 | 0.025 |
| MT072864.1;QIQ54047.1;Pangolin_CoV_orf1ab;833 | 0 | 0 |
| MT072864.1;QIQ54047.1;Pangolin_CoV_orf1ab;834 | 0 | 0 |
| MT072864.1;QIQ54047.1;Pangolin_CoV_orf1ab;835 | 0 | 0 |
| MT072864.1;QIQ54047.1;Pangolin_CoV_orf1ab;836 | 0.05 | 0 |
| MT072864.1;QIQ54047.1;Pangolin_CoV_orf1ab;837 | 0.05 | 0 |
| MT072864.1;QIQ54047.1;Pangolin_CoV_orf1ab;838 | 0 | 0 |
| MT072864.1;QIQ54047.1;Pangolin_CoV_orf1ab;839 | 0 | 0 |
| MT072864.1;QIQ54047.1;Pangolin_CoV_orf1ab;840 | 0 | 0.025 |
| MT072864.1;QIQ54047.1;Pangolin_CoV_orf1ab;841 | 0.05 | 0.05 |
| MT072864.1;QIQ54047.1;Pangolin_CoV_orf1ab;842 | 0.05 | 0.025 |
| MT072864.1;QIQ54047.1;Pangolin_CoV_orf1ab;843 | 0.05 | 0.025 |
| MT072864.1;QIQ54047.1;Pangolin_CoV_orf1ab;844 | 0 | 0.025 |
| MT072864.1;QIQ54047.1;Pangolin_CoV_orf1ab;845 | 0 | 0.025 |
| MT072864.1;QIQ54047.1;Pangolin_CoV_orf1ab;846 | 0 | 0.05 |
| MT072864.1;QIQ54047.1;Pangolin_CoV_orf1ab;847 | 0 | 0.025 |
| MT072864.1;QIQ54047.1;Pangolin_CoV_orf1ab;848 | 0 | 0 |
| MT072864.1;QIQ54047.1;Pangolin_CoV_orf1ab;849 | 0 | 0 |
| MT072864.1;QIQ54047.1;Pangolin_CoV_orf1ab;850 | 0.05 | 0 |

| | | |
|---|---|---|
| MT072864.1;QIQ54047.1;Pangolin_CoV_orf1ab;851 | 0 | 0 |
| MT072864.1;QIQ54047.1;Pangolin_CoV_orf1ab;852 | 0 | 0 |
| MT072864.1;QIQ54047.1;Pangolin_CoV_orf1ab;853 | 0 | 0 |
| MT072864.1;QIQ54047.1;Pangolin_CoV_orf1ab;854 | 0 | 0 |
| MT072864.1;QIQ54047.1;Pangolin_CoV_orf1ab;855 | 0 | 0.025 |
| MT072864.1;QIQ54047.1;Pangolin_CoV_orf1ab;856 | 0.05 | 0 |
| MT072864.1;QIQ54047.1;Pangolin_CoV_orf1ab;857 | 0.05 | 0 |
| MT072864.1;QIQ54047.1;Pangolin_CoV_orf1ab;858 | 0 | 0 |
| MT072864.1;QIQ54047.1;Pangolin_CoV_orf1ab;859 | 0 | 0 |
| MT072864.1;QIQ54047.1;Pangolin_CoV_orf1ab;860 | 0 | 0 |
| MT072864.1;QIQ54047.1;Pangolin_CoV_orf1ab;861 | 0 | 0 |
| MT072864.1;QIQ54047.1;Pangolin_CoV_orf1ab;862 | 0 | 0 |
| MT072864.1;QIQ54047.1;Pangolin_CoV_orf1ab;863 | 0 | 0 |
| MT072864.1;QIQ54047.1;Pangolin_CoV_orf1ab;864 | 0 | 0 |
| MT072864.1;QIQ54047.1;Pangolin_CoV_orf1ab;865 | 0 | 0 |
| MT072864.1;QIQ54047.1;Pangolin_CoV_orf1ab;866 | 0 | 0 |
| MT072864.1;QIQ54047.1;Pangolin_CoV_orf1ab;867 | 0 | 0 |
| MT072864.1;QIQ54047.1;Pangolin_CoV_orf1ab;868 | 0 | 0 |
| MT072864.1;QIQ54047.1;Pangolin_CoV_orf1ab;869 | 0 | 0 |
| MT072864.1;QIQ54047.1;Pangolin_CoV_orf1ab;870 | 0 | 0 |
| MT072864.1;QIQ54047.1;Pangolin_CoV_orf1ab;871 | 0 | 0 |
| MT072864.1;QIQ54047.1;Pangolin_CoV_orf1ab;872 | 0 | 0 |
| MT072864.1;QIQ54047.1;Pangolin_CoV_orf1ab;873 | 0 | 0 |
| MT072864.1;QIQ54047.1;Pangolin_CoV_orf1ab;874 | 0 | 0 |
| MT072864.1;QIQ54047.1;Pangolin_CoV_orf1ab;875 | 0 | 0.025 |
| MT072864.1;QIQ54047.1;Pangolin_CoV_orf1ab;876 | 0 | 0.05 |
| MT072864.1;QIQ54047.1;Pangolin_CoV_orf1ab;877 | 0.05 | 0.025 |
| MT072864.1;QIQ54047.1;Pangolin_CoV_orf1ab;878 | 0 | 0.05 |
| MT072864.1;QIQ54047.1;Pangolin_CoV_orf1ab;879 | 0 | 0.025 |
| MT072864.1;QIQ54047.1;Pangolin_CoV_orf1ab;880 | 0 | 0.025 |
| MT072864.1;QIQ54047.1;Pangolin_CoV_orf1ab;881 | 0.05 | 0.025 |
| MT072864.1;QIQ54047.1;Pangolin_CoV_orf1ab;882 | 0 | 0.025 |
| MT072864.1;QIQ54047.1;Pangolin_CoV_orf1ab;883 | 0 | 0 |
| MT072864.1;QIQ54047.1;Pangolin_CoV_orf1ab;884 | 0.05 | 0 |
| MT072864.1;QIQ54047.1;Pangolin_CoV_orf1ab;885 | 0 | 0 |
| MT072864.1;QIQ54047.1;Pangolin_CoV_orf1ab;886 | 0 | 0 |
| MT072864.1;QIQ54047.1;Pangolin_CoV_orf1ab;887 | 0 | 0 |
| MT072864.1;QIQ54047.1;Pangolin_CoV_orf1ab;888 | 0 | 0.025 |
| MT072864.1;QIQ54047.1;Pangolin_CoV_orf1ab;889 | 0 | 0 |
| MT072864.1;QIQ54047.1;Pangolin_CoV_orf1ab;890 | 0 | 0 |
| MT072864.1;QIQ54047.1;Pangolin_CoV_orf1ab;891 | 0 | 0 |
| MT072864.1;QIQ54047.1;Pangolin_CoV_orf1ab;892 | 0 | 0 |
| MT072864.1;QIQ54047.1;Pangolin_CoV_orf1ab;893 | 0 | 0 |

| | | |
|---|---|---|
| MT072864.1;QIQ54047.1;Pangolin_CoV_orf1ab;894 | 0 | 0 |
| MT072864.1;QIQ54047.1;Pangolin_CoV_orf1ab;895 | 0 | 0 |
| MT072864.1;QIQ54047.1;Pangolin_CoV_orf1ab;896 | 0 | 0 |
| MT072864.1;QIQ54047.1;Pangolin_CoV_orf1ab;897 | 0 | 0 |
| MT072864.1;QIQ54047.1;Pangolin_CoV_orf1ab;898 | 0 | 0 |
| MT072864.1;QIQ54047.1;Pangolin_CoV_orf1ab;899 | 0 | 0 |
| MT072864.1;QIQ54047.1;Pangolin_CoV_orf1ab;900 | 0 | 0 |
| MT072864.1;QIQ54047.1;Pangolin_CoV_orf1ab;901 | 0 | 0 |
| MT072864.1;QIQ54047.1;Pangolin_CoV_orf1ab;902 | 0 | 0 |
| MT072864.1;QIQ54047.1;Pangolin_CoV_orf1ab;903 | 0 | 0 |
| MT072864.1;QIQ54047.1;Pangolin_CoV_orf1ab;904 | 0 | 0 |
| MT072864.1;QIQ54047.1;Pangolin_CoV_orf1ab;905 | 0 | 0 |
| MT072864.1;QIQ54047.1;Pangolin_CoV_orf1ab;906 | 0 | 0 |
| MT072864.1;QIQ54047.1;Pangolin_CoV_orf1ab;907 | 0 | 0 |
| MT072864.1;QIQ54047.1;Pangolin_CoV_orf1ab;908 | 0 | 0 |
| MT072864.1;QIQ54047.1;Pangolin_CoV_orf1ab;909 | 0 | 0 |
| MT072864.1;QIQ54047.1;Pangolin_CoV_orf1ab;910 | 0 | 0 |
| MT072864.1;QIQ54047.1;Pangolin_CoV_orf1ab;911 | 0 | 0 |
| MT072864.1;QIQ54047.1;Pangolin_CoV_orf1ab;912 | 0 | 0 |
| MT072864.1;QIQ54047.1;Pangolin_CoV_orf1ab;913 | 0 | 0 |
| MT072864.1;QIQ54047.1;Pangolin_CoV_orf1ab;914 | 0 | 0 |
| MT072864.1;QIQ54047.1;Pangolin_CoV_orf1ab;915 | 0 | 0 |
| MT072864.1;QIQ54047.1;Pangolin_CoV_orf1ab;916 | 0 | 0 |
| MT072864.1;QIQ54047.1;Pangolin_CoV_orf1ab;917 | 0 | 0.025 |
| MT072864.1;QIQ54047.1;Pangolin_CoV_orf1ab;918 | 0 | 0.075 |
| MT072864.1;QIQ54047.1;Pangolin_CoV_orf1ab;919 | 0 | 0.1 |
| MT072864.1;QIQ54047.1;Pangolin_CoV_orf1ab;920 | 0 | 0.125 |
| MT072864.1;QIQ54047.1;Pangolin_CoV_orf1ab;921 | 0 | 0.1 |
| MT072864.1;QIQ54047.1;Pangolin_CoV_orf1ab;922 | 0 | 0.125 |
| MT072864.1;QIQ54047.1;Pangolin_CoV_orf1ab;923 | 0 | 0.125 |
| MT072864.1;QIQ54047.1;Pangolin_CoV_orf1ab;924 | 0 | 0.125 |
| MT072864.1;QIQ54047.1;Pangolin_CoV_orf1ab;925 | 0 | 0.05 |
| MT072864.1;QIQ54047.1;Pangolin_CoV_orf1ab;926 | 0 | 0 |
| MT072864.1;QIQ54047.1;Pangolin_CoV_orf1ab;927 | 0 | 0 |
| MT072864.1;QIQ54047.1;Pangolin_CoV_orf1ab;928 | 0 | 0 |
| MT072864.1;QIQ54047.1;Pangolin_CoV_orf1ab;929 | 0.15 | 0.175 |
| MT072864.1;QIQ54047.1;Pangolin_CoV_orf1ab;930 | 0.05 | 0 |
| MT072864.1;QIQ54047.1;Pangolin_CoV_orf1ab;931 | 0.1 | 0 |
| MT072864.1;QIQ54047.1;Pangolin_CoV_orf1ab;932 | 0.05 | 0 |
| MT072864.1;QIQ54047.1;Pangolin_CoV_orf1ab;933 | 0.1 | 0 |
| MT072864.1;QIQ54047.1;Pangolin_CoV_orf1ab;934 | 0.1 | 0.025 |
| MT072864.1;QIQ54047.1;Pangolin_CoV_orf1ab;935 | 0.15 | 0.025 |
| MT072864.1;QIQ54047.1;Pangolin_CoV_orf1ab;936 | 0.1 | 0.025 |

| | | |
|---|---|---|
| MT072864.1;QIQ54047.1;Pangolin_CoV_orf1ab;937 | 0.1 | 0.025 |
| MT072864.1;QIQ54047.1;Pangolin_CoV_orf1ab;938 | 0.15 | 0.025 |
| MT072864.1;QIQ54047.1;Pangolin_CoV_orf1ab;939 | 0.1 | 0.025 |
| MT072864.1;QIQ54047.1;Pangolin_CoV_orf1ab;940 | 0.1 | 0.025 |
| MT072864.1;QIQ54047.1;Pangolin_CoV_orf1ab;941 | 0.1 | 0.05 |
| MT072864.1;QIQ54047.1;Pangolin_CoV_orf1ab;942 | 0 | 0 |
| MT072864.1;QIQ54047.1;Pangolin_CoV_orf1ab;943 | 0 | 0 |
| MT072864.1;QIQ54047.1;Pangolin_CoV_orf1ab;944 | 0 | 0 |
| MT072864.1;QIQ54047.1;Pangolin_CoV_orf1ab;945 | 0.05 | 0.05 |
| MT072864.1;QIQ54047.1;Pangolin_CoV_orf1ab;946 | 0.1 | 0.025 |
| MT072864.1;QIQ54047.1;Pangolin_CoV_orf1ab;947 | 0.15 | 0.05 |
| MT072864.1;QIQ54047.1;Pangolin_CoV_orf1ab;948 | 0.15 | 0.05 |
| MT072864.1;QIQ54047.1;Pangolin_CoV_orf1ab;949 | 0.15 | 0.1 |
| MT072864.1;QIQ54047.1;Pangolin_CoV_orf1ab;950 | 0.15 | 0.05 |
| MT072864.1;QIQ54047.1;Pangolin_CoV_orf1ab;951 | 0.15 | 0.05 |
| MT072864.1;QIQ54047.1;Pangolin_CoV_orf1ab;952 | 0 | 0.05 |
| MT072864.1;QIQ54047.1;Pangolin_CoV_orf1ab;953 | 0 | 0.025 |
| MT072864.1;QIQ54047.1;Pangolin_CoV_orf1ab;954 | 0 | 0 |
| MT072864.1;QIQ54047.1;Pangolin_CoV_orf1ab;955 | 0 | 0 |
| MT072864.1;QIQ54047.1;Pangolin_CoV_orf1ab;956 | 0 | 0 |
| MT072864.1;QIQ54047.1;Pangolin_CoV_orf1ab;957 | 0 | 0 |
| MT072864.1;QIQ54047.1;Pangolin_CoV_orf1ab;958 | 0 | 0 |
| MT072864.1;QIQ54047.1;Pangolin_CoV_orf1ab;959 | 0 | 0 |
| MT072864.1;QIQ54047.1;Pangolin_CoV_orf1ab;960 | 0 | 0 |
| MT072864.1;QIQ54047.1;Pangolin_CoV_orf1ab;961 | 0 | 0 |
| MT072864.1;QIQ54047.1;Pangolin_CoV_orf1ab;962 | 0 | 0.025 |
| MT072864.1;QIQ54047.1;Pangolin_CoV_orf1ab;963 | 0 | 0.05 |
| MT072864.1;QIQ54047.1;Pangolin_CoV_orf1ab;964 | 0 | 0.025 |
| MT072864.1;QIQ54047.1;Pangolin_CoV_orf1ab;965 | 0 | 0.025 |
| MT072864.1;QIQ54047.1;Pangolin_CoV_orf1ab;966 | 0 | 0.05 |
| MT072864.1;QIQ54047.1;Pangolin_CoV_orf1ab;967 | 0 | 0.05 |
| MT072864.1;QIQ54047.1;Pangolin_CoV_orf1ab;968 | 0 | 0.05 |
| MT072864.1;QIQ54047.1;Pangolin_CoV_orf1ab;969 | 0 | 0.05 |
| MT072864.1;QIQ54047.1;Pangolin_CoV_orf1ab;970 | 0 | 0.025 |
| MT072864.1;QIQ54047.1;Pangolin_CoV_orf1ab;971 | 0 | 0.025 |
| MT072864.1;QIQ54047.1;Pangolin_CoV_orf1ab;972 | 0 | 0.025 |
| MT072864.1;QIQ54047.1;Pangolin_CoV_orf1ab;973 | 0 | 0.025 |
| MT072864.1;QIQ54047.1;Pangolin_CoV_orf1ab;974 | 0 | 0.025 |
| MT072864.1;QIQ54047.1;Pangolin_CoV_orf1ab;975 | 0 | 0 |
| MT072864.1;QIQ54047.1;Pangolin_CoV_orf1ab;976 | 0 | 0 |
| MT072864.1;QIQ54047.1;Pangolin_CoV_orf1ab;977 | 0 | 0 |
| MT072864.1;QIQ54047.1;Pangolin_CoV_orf1ab;978 | 0 | 0 |
| MT072864.1;QIQ54047.1;Pangolin_CoV_orf1ab;979 | 0 | 0 |

| | | |
|---|---|---|
| MT072864.1;QIQ54047.1;Pangolin_CoV_orf1ab;980 | 0 | 0 |
| MT072864.1;QIQ54047.1;Pangolin_CoV_orf1ab;981 | 0 | 0 |
| MT072864.1;QIQ54047.1;Pangolin_CoV_orf1ab;982 | 0.05 | 0.1 |
| MT072864.1;QIQ54047.1;Pangolin_CoV_orf1ab;983 | 0.05 | 0 |
| MT072864.1;QIQ54047.1;Pangolin_CoV_orf1ab;984 | 0 | 0 |
| MT072864.1;QIQ54047.1;Pangolin_CoV_orf1ab;985 | 0 | 0 |
| MT072864.1;QIQ54047.1;Pangolin_CoV_orf1ab;986 | 0 | 0 |
| MT072864.1;QIQ54047.1;Pangolin_CoV_orf1ab;987 | 0 | 0 |
| MT072864.1;QIQ54047.1;Pangolin_CoV_orf1ab;988 | 0 | 0 |
| MT072864.1;QIQ54047.1;Pangolin_CoV_orf1ab;989 | 0 | 0 |
| MT072864.1;QIQ54047.1;Pangolin_CoV_orf1ab;990 | 0.05 | 0 |
| MT072864.1;QIQ54047.1;Pangolin_CoV_orf1ab;991 | 0 | 0 |
| MT072864.1;QIQ54047.1;Pangolin_CoV_orf1ab;992 | 0 | 0 |
| MT072864.1;QIQ54047.1;Pangolin_CoV_orf1ab;993 | 0 | 0 |
| MT072864.1;QIQ54047.1;Pangolin_CoV_orf1ab;994 | 0 | 0.025 |
| MT072864.1;QIQ54047.1;Pangolin_CoV_orf1ab;995 | 0 | 0 |
| MT072864.1;QIQ54047.1;Pangolin_CoV_orf1ab;996 | 0 | 0 |
| MT072864.1;QIQ54047.1;Pangolin_CoV_orf1ab;997 | 0 | 0 |
| MT072864.1;QIQ54047.1;Pangolin_CoV_orf1ab;998 | 0 | 0 |
| MT072864.1;QIQ54047.1;Pangolin_CoV_orf1ab;999 | 0 | 0 |
| MT072864.1;QIQ54047.1;Pangolin_CoV_orf1ab;1000 | 0 | 0 |
| MT072864.1;QIQ54047.1;Pangolin_CoV_orf1ab;1001 | 0 | 0 |
| MT072864.1;QIQ54047.1;Pangolin_CoV_orf1ab;1002 | 0 | 0 |
| MT072864.1;QIQ54047.1;Pangolin_CoV_orf1ab;1003 | 0 | 0 |
| MT072864.1;QIQ54047.1;Pangolin_CoV_orf1ab;1004 | 0 | 0 |
| MT072864.1;QIQ54047.1;Pangolin_CoV_orf1ab;1005 | 0 | 0 |
| MT072864.1;QIQ54047.1;Pangolin_CoV_orf1ab;1006 | 0 | 0 |
| MT072864.1;QIQ54047.1;Pangolin_CoV_orf1ab;1007 | 0 | 0 |
| MT072864.1;QIQ54047.1;Pangolin_CoV_orf1ab;1008 | 0 | 0 |
| MT072864.1;QIQ54047.1;Pangolin_CoV_orf1ab;1009 | 0 | 0 |
| MT072864.1;QIQ54047.1;Pangolin_CoV_orf1ab;1010 | 0 | 0 |
| MT072864.1;QIQ54047.1;Pangolin_CoV_orf1ab;1011 | 0 | 0 |
| MT072864.1;QIQ54047.1;Pangolin_CoV_orf1ab;1012 | 0 | 0 |
| MT072864.1;QIQ54047.1;Pangolin_CoV_orf1ab;1013 | 0 | 0 |
| MT072864.1;QIQ54047.1;Pangolin_CoV_orf1ab;1014 | 0.05 | 0 |
| MT072864.1;QIQ54047.1;Pangolin_CoV_orf1ab;1015 | 0.05 | 0 |
| MT072864.1;QIQ54047.1;Pangolin_CoV_orf1ab;1016 | 0.05 | 0.025 |
| MT072864.1;QIQ54047.1;Pangolin_CoV_orf1ab;1017 | 0.05 | 0 |
| MT072864.1;QIQ54047.1;Pangolin_CoV_orf1ab;1018 | 0.05 | 0 |
| MT072864.1;QIQ54047.1;Pangolin_CoV_orf1ab;1019 | 0.05 | 0 |
| MT072864.1;QIQ54047.1;Pangolin_CoV_orf1ab;1020 | 0 | 0 |
| MT072864.1;QIQ54047.1;Pangolin_CoV_orf1ab;1021 | 0.05 | 0 |
| MT072864.1;QIQ54047.1;Pangolin_CoV_orf1ab;1022 | 0.05 | 0 |

| | | |
|---|---|---|
| MT072864.1;QIQ54047.1;Pangolin_CoV_orf1ab;1023 | 0.05 | 0 |
| MT072864.1;QIQ54047.1;Pangolin_CoV_orf1ab;1024 | 0.05 | 0 |
| MT072864.1;QIQ54047.1;Pangolin_CoV_orf1ab;1025 | 0.05 | 0 |
| MT072864.1;QIQ54047.1;Pangolin_CoV_orf1ab;1026 | 0.05 | 0 |
| MT072864.1;QIQ54047.1;Pangolin_CoV_orf1ab;1027 | 0.25 | 0.05 |
| MT072864.1;QIQ54047.1;Pangolin_CoV_orf1ab;1028 | 0.1 | 0.025 |
| MT072864.1;QIQ54047.1;Pangolin_CoV_orf1ab;1029 | 0.15 | 0.05 |
| MT072864.1;QIQ54047.1;Pangolin_CoV_orf1ab;1030 | 0.1 | 0.025 |
| MT072864.1;QIQ54047.1;Pangolin_CoV_orf1ab;1031 | 0.15 | 0.075 |
| MT072864.1;QIQ54047.1;Pangolin_CoV_orf1ab;1032 | 0.3 | 0.075 |
| MT072864.1;QIQ54047.1;Pangolin_CoV_orf1ab;1033 | 0.2 | 0.075 |
| MT072864.1;QIQ54047.1;Pangolin_CoV_orf1ab;1034 | 0.2 | 0.075 |
| MT072864.1;QIQ54047.1;Pangolin_CoV_orf1ab;1035 | 0 | 0.05 |
| MT072864.1;QIQ54047.1;Pangolin_CoV_orf1ab;1036 | 0 | 0 |
| MT072864.1;QIQ54047.1;Pangolin_CoV_orf1ab;1037 | 0 | 0.025 |
| MT072864.1;QIQ54047.1;Pangolin_CoV_orf1ab;1038 | 0 | 0 |
| MT072864.1;QIQ54047.1;Pangolin_CoV_orf1ab;1039 | 0 | 0 |
| MT072864.1;QIQ54047.1;Pangolin_CoV_orf1ab;1040 | 0 | 0 |
| MT072864.1;QIQ54047.1;Pangolin_CoV_orf1ab;1041 | 0 | 0 |
| MT072864.1;QIQ54047.1;Pangolin_CoV_orf1ab;1042 | 0 | 0 |
| MT072864.1;QIQ54047.1;Pangolin_CoV_orf1ab;1043 | 0 | 0 |
| MT072864.1;QIQ54047.1;Pangolin_CoV_orf1ab;1044 | 0 | 0 |
| MT072864.1;QIQ54047.1;Pangolin_CoV_orf1ab;1045 | 0 | 0 |
| MT072864.1;QIQ54047.1;Pangolin_CoV_orf1ab;1046 | 0 | 0 |
| MT072864.1;QIQ54047.1;Pangolin_CoV_orf1ab;1047 | 0.05 | 0 |
| MT072864.1;QIQ54047.1;Pangolin_CoV_orf1ab;1048 | 0 | 0 |
| MT072864.1;QIQ54047.1;Pangolin_CoV_orf1ab;1049 | 0.05 | 0.025 |
| MT072864.1;QIQ54047.1;Pangolin_CoV_orf1ab;1050 | 0 | 0 |
| MT072864.1;QIQ54047.1;Pangolin_CoV_orf1ab;1051 | 0 | 0 |
| MT072864.1;QIQ54047.1;Pangolin_CoV_orf1ab;1052 | 0 | 0 |
| MT072864.1;QIQ54047.1;Pangolin_CoV_orf1ab;1053 | 0 | 0 |
| MT072864.1;QIQ54047.1;Pangolin_CoV_orf1ab;1054 | 0 | 0 |
| MT072864.1;QIQ54047.1;Pangolin_CoV_orf1ab;1055 | 0 | 0 |
| MT072864.1;QIQ54047.1;Pangolin_CoV_orf1ab;1056 | 0 | 0 |
| MT072864.1;QIQ54047.1;Pangolin_CoV_orf1ab;1057 | 0 | 0 |
| MT072864.1;QIQ54047.1;Pangolin_CoV_orf1ab;1058 | 0 | 0.025 |
| MT072864.1;QIQ54047.1;Pangolin_CoV_orf1ab;1059 | 0.1 | 0.025 |
| MT072864.1;QIQ54047.1;Pangolin_CoV_orf1ab;1060 | 0.15 | 0.025 |
| MT072864.1;QIQ54047.1;Pangolin_CoV_orf1ab;1061 | 0.05 | 0.025 |
| MT072864.1;QIQ54047.1;Pangolin_CoV_orf1ab;1062 | 0.15 | 0.025 |
| MT072864.1;QIQ54047.1;Pangolin_CoV_orf1ab;1063 | 0.15 | 0.175 |
| MT072864.1;QIQ54047.1;Pangolin_CoV_orf1ab;1064 | 0.25 | 0.275 |
| MT072864.1;QIQ54047.1;Pangolin_CoV_orf1ab;1065 | 0.15 | 0.125 |

| | | |
|---|---|---|
| MT072864.1;QIQ54047.1;Pangolin_CoV_orf1ab;1066 | 0.15 | 0.175 |
| MT072864.1;QIQ54047.1;Pangolin_CoV_orf1ab;1067 | 0.05 | 0.125 |
| MT072864.1;QIQ54047.1;Pangolin_CoV_orf1ab;1068 | 0 | 0.025 |
| MT072864.1;QIQ54047.1;Pangolin_CoV_orf1ab;1069 | 0 | 0.025 |
| MT072864.1;QIQ54047.1;Pangolin_CoV_orf1ab;1070 | 0.1 | 0.025 |
| MT072864.1;QIQ54047.1;Pangolin_CoV_orf1ab;1071 | 0 | 0.025 |
| MT072864.1;QIQ54047.1;Pangolin_CoV_orf1ab;1072 | 0 | 0 |
| MT072864.1;QIQ54047.1;Pangolin_CoV_orf1ab;1073 | 0 | 0 |
| MT072864.1;QIQ54047.1;Pangolin_CoV_orf1ab;1074 | 0 | 0.05 |
| MT072864.1;QIQ54047.1;Pangolin_CoV_orf1ab;1075 | 0 | 0.025 |
| MT072864.1;QIQ54047.1;Pangolin_CoV_orf1ab;1076 | 0 | 0.025 |
| MT072864.1;QIQ54047.1;Pangolin_CoV_orf1ab;1077 | 0.05 | 0.025 |
| MT072864.1;QIQ54047.1;Pangolin_CoV_orf1ab;1078 | 0 | 0.05 |
| MT072864.1;QIQ54047.1;Pangolin_CoV_orf1ab;1079 | 0.1 | 0.1 |
| MT072864.1;QIQ54047.1;Pangolin_CoV_orf1ab;1080 | 0 | 0.025 |
| MT072864.1;QIQ54047.1;Pangolin_CoV_orf1ab;1081 | 0 | 0.025 |
| MT072864.1;QIQ54047.1;Pangolin_CoV_orf1ab;1082 | 0 | 0 |
| MT072864.1;QIQ54047.1;Pangolin_CoV_orf1ab;1083 | 0 | 0 |
| MT072864.1;QIQ54047.1;Pangolin_CoV_orf1ab;1084 | 0 | 0 |
| MT072864.1;QIQ54047.1;Pangolin_CoV_orf1ab;1085 | 0 | 0 |
| MT072864.1;QIQ54047.1;Pangolin_CoV_orf1ab;1086 | 0 | 0 |
| MT072864.1;QIQ54047.1;Pangolin_CoV_orf1ab;1087 | 0 | 0 |
| MT072864.1;QIQ54047.1;Pangolin_CoV_orf1ab;1088 | 0 | 0 |
| MT072864.1;QIQ54047.1;Pangolin_CoV_orf1ab;1089 | 0 | 0 |
| MT072864.1;QIQ54047.1;Pangolin_CoV_orf1ab;1090 | 0 | 0 |
| MT072864.1;QIQ54047.1;Pangolin_CoV_orf1ab;1091 | 0 | 0 |
| MT072864.1;QIQ54047.1;Pangolin_CoV_orf1ab;1092 | 0 | 0 |
| MT072864.1;QIQ54047.1;Pangolin_CoV_orf1ab;1093 | 0 | 0 |
| MT072864.1;QIQ54047.1;Pangolin_CoV_orf1ab;1094 | 0 | 0 |
| MT072864.1;QIQ54047.1;Pangolin_CoV_orf1ab;1095 | 0 | 0 |
| MT072864.1;QIQ54047.1;Pangolin_CoV_orf1ab;1096 | 0 | 0 |
| MT072864.1;QIQ54047.1;Pangolin_CoV_orf1ab;1097 | 0 | 0 |
| MT072864.1;QIQ54047.1;Pangolin_CoV_orf1ab;1098 | 0 | 0 |
| MT072864.1;QIQ54047.1;Pangolin_CoV_orf1ab;1099 | 0 | 0 |
| MT072864.1;QIQ54047.1;Pangolin_CoV_orf1ab;1100 | 0 | 0 |
| MT072864.1;QIQ54047.1;Pangolin_CoV_orf1ab;1101 | 0.05 | 0 |
| MT072864.1;QIQ54047.1;Pangolin_CoV_orf1ab;1102 | 0.1 | 0 |
| MT072864.1;QIQ54047.1;Pangolin_CoV_orf1ab;1103 | 0.2 | 0 |
| MT072864.1;QIQ54047.1;Pangolin_CoV_orf1ab;1104 | 0.25 | 0 |
| MT072864.1;QIQ54047.1;Pangolin_CoV_orf1ab;1105 | 0.15 | 0 |
| MT072864.1;QIQ54047.1;Pangolin_CoV_orf1ab;1106 | 0.15 | 0.025 |
| MT072864.1;QIQ54047.1;Pangolin_CoV_orf1ab;1107 | 0.25 | 0 |
| MT072864.1;QIQ54047.1;Pangolin_CoV_orf1ab;1108 | 0 | 0 |

| | | |
|---|---|---|
| MT072864.1;QIQ54047.1;Pangolin_CoV_orf1ab;1109 | 0.05 | 0.05 |
| MT072864.1;QIQ54047.1;Pangolin_CoV_orf1ab;1110 | 0.1 | 0.05 |
| MT072864.1;QIQ54047.1;Pangolin_CoV_orf1ab;1111 | 0.05 | 0.025 |
| MT072864.1;QIQ54047.1;Pangolin_CoV_orf1ab;1112 | 0.05 | 0 |
| MT072864.1;QIQ54047.1;Pangolin_CoV_orf1ab;1113 | 0.05 | 0.05 |
| MT072864.1;QIQ54047.1;Pangolin_CoV_orf1ab;1114 | 0.05 | 0 |
| MT072864.1;QIQ54047.1;Pangolin_CoV_orf1ab;1115 | 0 | 0.025 |
| MT072864.1;QIQ54047.1;Pangolin_CoV_orf1ab;1116 | 0 | 0.025 |
| MT072864.1;QIQ54047.1;Pangolin_CoV_orf1ab;1117 | 0 | 0 |
| MT072864.1;QIQ54047.1;Pangolin_CoV_orf1ab;1118 | 0 | 0 |
| MT072864.1;QIQ54047.1;Pangolin_CoV_orf1ab;1119 | 0 | 0 |
| MT072864.1;QIQ54047.1;Pangolin_CoV_orf1ab;1120 | 0 | 0.025 |
| MT072864.1;QIQ54047.1;Pangolin_CoV_orf1ab;1121 | 0 | 0.025 |
| MT072864.1;QIQ54047.1;Pangolin_CoV_orf1ab;1122 | 0 | 0.025 |
| MT072864.1;QIQ54047.1;Pangolin_CoV_orf1ab;1123 | 0 | 0.025 |
| MT072864.1;QIQ54047.1;Pangolin_CoV_orf1ab;1124 | 0 | 0.05 |
| MT072864.1;QIQ54047.1;Pangolin_CoV_orf1ab;1125 | 0 | 0.025 |
| MT072864.1;QIQ54047.1;Pangolin_CoV_orf1ab;1126 | 0 | 0.025 |
| MT072864.1;QIQ54047.1;Pangolin_CoV_orf1ab;1127 | 0 | 0.025 |
| MT072864.1;QIQ54047.1;Pangolin_CoV_orf1ab;1128 | 0 | 0.025 |
| MT072864.1;QIQ54047.1;Pangolin_CoV_orf1ab;1129 | 0 | 0.025 |
| MT072864.1;QIQ54047.1;Pangolin_CoV_orf1ab;1130 | 0 | 0.025 |
| MT072864.1;QIQ54047.1;Pangolin_CoV_orf1ab;1131 | 0 | 0.025 |
| MT072864.1;QIQ54047.1;Pangolin_CoV_orf1ab;1132 | 0 | 0 |
| MT072864.1;QIQ54047.1;Pangolin_CoV_orf1ab;1133 | 0 | 0 |
| MT072864.1;QIQ54047.1;Pangolin_CoV_orf1ab;1134 | 0 | 0 |
| MT072864.1;QIQ54047.1;Pangolin_CoV_orf1ab;1135 | 0 | 0 |
| MT072864.1;QIQ54047.1;Pangolin_CoV_orf1ab;1136 | 0 | 0 |
| MT072864.1;QIQ54047.1;Pangolin_CoV_orf1ab;1137 | 0 | 0 |
| MT072864.1;QIQ54047.1;Pangolin_CoV_orf1ab;1138 | 0 | 0 |
| MT072864.1;QIQ54047.1;Pangolin_CoV_orf1ab;1139 | 0 | 0 |
| MT072864.1;QIQ54047.1;Pangolin_CoV_orf1ab;1140 | 0 | 0 |
| MT072864.1;QIQ54047.1;Pangolin_CoV_orf1ab;1141 | 0 | 0 |
| MT072864.1;QIQ54047.1;Pangolin_CoV_orf1ab;1142 | 0 | 0 |
| MT072864.1;QIQ54047.1;Pangolin_CoV_orf1ab;1143 | 0 | 0 |
| MT072864.1;QIQ54047.1;Pangolin_CoV_orf1ab;1144 | 0 | 0 |
| MT072864.1;QIQ54047.1;Pangolin_CoV_orf1ab;1145 | 0 | 0 |
| MT072864.1;QIQ54047.1;Pangolin_CoV_orf1ab;1146 | 0 | 0 |
| MT072864.1;QIQ54047.1;Pangolin_CoV_orf1ab;1147 | 0 | 0 |
| MT072864.1;QIQ54047.1;Pangolin_CoV_orf1ab;1148 | 0 | 0 |
| MT072864.1;QIQ54047.1;Pangolin_CoV_orf1ab;1149 | 0.05 | 0.025 |
| MT072864.1;QIQ54047.1;Pangolin_CoV_orf1ab;1150 | 0.05 | 0.05 |
| MT072864.1;QIQ54047.1;Pangolin_CoV_orf1ab;1151 | 0.05 | 0.05 |

| | | |
|---|---|---|
| MT072864.1;QIQ54047.1;Pangolin_CoV_orf1ab;1152 | 0 | 0 |
| MT072864.1;QIQ54047.1;Pangolin_CoV_orf1ab;1153 | 0 | 0 |
| MT072864.1;QIQ54047.1;Pangolin_CoV_orf1ab;1154 | 0 | 0 |
| MT072864.1;QIQ54047.1;Pangolin_CoV_orf1ab;1155 | 0 | 0 |
| MT072864.1;QIQ54047.1;Pangolin_CoV_orf1ab;1156 | 0 | 0 |
| MT072864.1;QIQ54047.1;Pangolin_CoV_orf1ab;1157 | 0 | 0 |
| MT072864.1;QIQ54047.1;Pangolin_CoV_orf1ab;1158 | 0 | 0 |
| MT072864.1;QIQ54047.1;Pangolin_CoV_orf1ab;1159 | 0.05 | 0 |
| MT072864.1;QIQ54047.1;Pangolin_CoV_orf1ab;1160 | 0 | 0 |
| MT072864.1;QIQ54047.1;Pangolin_CoV_orf1ab;1161 | 0 | 0 |
| MT072864.1;QIQ54047.1;Pangolin_CoV_orf1ab;1162 | 0 | 0 |
| MT072864.1;QIQ54047.1;Pangolin_CoV_orf1ab;1163 | 0 | 0 |
| MT072864.1;QIQ54047.1;Pangolin_CoV_orf1ab;1164 | 0 | 0 |
| MT072864.1;QIQ54047.1;Pangolin_CoV_orf1ab;1165 | 0 | 0 |
| MT072864.1;QIQ54047.1;Pangolin_CoV_orf1ab;1166 | 0 | 0 |
| MT072864.1;QIQ54047.1;Pangolin_CoV_orf1ab;1167 | 0 | 0 |
| MT072864.1;QIQ54047.1;Pangolin_CoV_orf1ab;1168 | 0 | 0 |
| MT072864.1;QIQ54047.1;Pangolin_CoV_orf1ab;1169 | 0 | 0.025 |
| MT072864.1;QIQ54047.1;Pangolin_CoV_orf1ab;1170 | 0 | 0 |
| MT072864.1;QIQ54047.1;Pangolin_CoV_orf1ab;1171 | 0 | 0 |
| MT072864.1;QIQ54047.1;Pangolin_CoV_orf1ab;1172 | 0 | 0 |
| MT072864.1;QIQ54047.1;Pangolin_CoV_orf1ab;1173 | 0 | 0 |
| MT072864.1;QIQ54047.1;Pangolin_CoV_orf1ab;1174 | 0 | 0 |
| MT072864.1;QIQ54047.1;Pangolin_CoV_orf1ab;1175 | 0 | 0.1 |
| MT072864.1;QIQ54047.1;Pangolin_CoV_orf1ab;1176 | 0 | 0.05 |
| MT072864.1;QIQ54047.1;Pangolin_CoV_orf1ab;1177 | 0 | 0.1 |
| MT072864.1;QIQ54047.1;Pangolin_CoV_orf1ab;1178 | 0 | 0.125 |
| MT072864.1;QIQ54047.1;Pangolin_CoV_orf1ab;1179 | 0 | 0.1 |
| MT072864.1;QIQ54047.1;Pangolin_CoV_orf1ab;1180 | 0 | 0.1 |
| MT072864.1;QIQ54047.1;Pangolin_CoV_orf1ab;1181 | 0 | 0.1 |
| MT072864.1;QIQ54047.1;Pangolin_CoV_orf1ab;1182 | 0 | 0.1 |
| MT072864.1;QIQ54047.1;Pangolin_CoV_orf1ab;1183 | 0 | 0.075 |
| MT072864.1;QIQ54047.1;Pangolin_CoV_orf1ab;1184 | 0.05 | 0.025 |
| MT072864.1;QIQ54047.1;Pangolin_CoV_orf1ab;1185 | 0 | 0 |
| MT072864.1;QIQ54047.1;Pangolin_CoV_orf1ab;1186 | 0 | 0.025 |
| MT072864.1;QIQ54047.1;Pangolin_CoV_orf1ab;1187 | 0 | 0 |
| MT072864.1;QIQ54047.1;Pangolin_CoV_orf1ab;1188 | 0 | 0.025 |
| MT072864.1;QIQ54047.1;Pangolin_CoV_orf1ab;1189 | 0 | 0 |
| MT072864.1;QIQ54047.1;Pangolin_CoV_orf1ab;1190 | 0 | 0 |
| MT072864.1;QIQ54047.1;Pangolin_CoV_orf1ab;1191 | 0 | 0 |
| MT072864.1;QIQ54047.1;Pangolin_CoV_orf1ab;1192 | 0 | 0 |
| MT072864.1;QIQ54047.1;Pangolin_CoV_orf1ab;1193 | 0 | 0.025 |
| MT072864.1;QIQ54047.1;Pangolin_CoV_orf1ab;1194 | 0 | 0.025 |

| | | |
|---|---|---|
| MT072864.1;QIQ54047.1;Pangolin_CoV_orf1ab;1195 | 0 | 0 |
| MT072864.1;QIQ54047.1;Pangolin_CoV_orf1ab;1196 | 0.05 | 0.025 |
| MT072864.1;QIQ54047.1;Pangolin_CoV_orf1ab;1197 | 0 | 0.025 |
| MT072864.1;QIQ54047.1;Pangolin_CoV_orf1ab;1198 | 0 | 0.025 |
| MT072864.1;QIQ54047.1;Pangolin_CoV_orf1ab;1199 | 0 | 0.025 |
| MT072864.1;QIQ54047.1;Pangolin_CoV_orf1ab;1200 | 0 | 0 |
| MT072864.1;QIQ54047.1;Pangolin_CoV_orf1ab;1201 | 0 | 0 |
| MT072864.1;QIQ54047.1;Pangolin_CoV_orf1ab;1202 | 0 | 0 |
| MT072864.1;QIQ54047.1;Pangolin_CoV_orf1ab;1203 | 0 | 0 |
| MT072864.1;QIQ54047.1;Pangolin_CoV_orf1ab;1204 | 0 | 0 |
| MT072864.1;QIQ54047.1;Pangolin_CoV_orf1ab;1205 | 0 | 0 |
| MT072864.1;QIQ54047.1;Pangolin_CoV_orf1ab;1206 | 0 | 0 |
| MT072864.1;QIQ54047.1;Pangolin_CoV_orf1ab;1207 | 0 | 0.05 |
| MT072864.1;QIQ54047.1;Pangolin_CoV_orf1ab;1208 | 0 | 0.1 |
| MT072864.1;QIQ54047.1;Pangolin_CoV_orf1ab;1209 | 0 | 0.15 |
| MT072864.1;QIQ54047.1;Pangolin_CoV_orf1ab;1210 | 0 | 0.05 |
| MT072864.1;QIQ54047.1;Pangolin_CoV_orf1ab;1211 | 0 | 0.05 |
| MT072864.1;QIQ54047.1;Pangolin_CoV_orf1ab;1212 | 0.05 | 0.075 |
| MT072864.1;QIQ54047.1;Pangolin_CoV_orf1ab;1213 | 0.05 | 0.1 |
| MT072864.1;QIQ54047.1;Pangolin_CoV_orf1ab;1214 | 0 | 0.075 |
| MT072864.1;QIQ54047.1;Pangolin_CoV_orf1ab;1215 | 0.05 | 0.05 |
| MT072864.1;QIQ54047.1;Pangolin_CoV_orf1ab;1216 | 0 | 0.05 |
| MT072864.1;QIQ54047.1;Pangolin_CoV_orf1ab;1217 | 0 | 0.05 |
| MT072864.1;QIQ54047.1;Pangolin_CoV_orf1ab;1218 | 0 | 0.025 |
| MT072864.1;QIQ54047.1;Pangolin_CoV_orf1ab;1219 | 0.05 | 0.025 |
| MT072864.1;QIQ54047.1;Pangolin_CoV_orf1ab;1220 | 0 | 0.025 |
| MT072864.1;QIQ54047.1;Pangolin_CoV_orf1ab;1221 | 0 | 0 |
| MT072864.1;QIQ54047.1;Pangolin_CoV_orf1ab;1222 | 0 | 0 |
| MT072864.1;QIQ54047.1;Pangolin_CoV_orf1ab;1223 | 0 | 0 |
| MT072864.1;QIQ54047.1;Pangolin_CoV_orf1ab;1224 | 0 | 0 |
| MT072864.1;QIQ54047.1;Pangolin_CoV_orf1ab;1225 | 0 | 0 |
| MT072864.1;QIQ54047.1;Pangolin_CoV_orf1ab;1226 | 0 | 0.025 |
| MT072864.1;QIQ54047.1;Pangolin_CoV_orf1ab;1227 | 0 | 0 |
| MT072864.1;QIQ54047.1;Pangolin_CoV_orf1ab;1228 | 0 | 0.075 |
| MT072864.1;QIQ54047.1;Pangolin_CoV_orf1ab;1229 | 0 | 0.05 |
| MT072864.1;QIQ54047.1;Pangolin_CoV_orf1ab;1230 | 0 | 0.025 |
| MT072864.1;QIQ54047.1;Pangolin_CoV_orf1ab;1231 | 0 | 0.025 |
| MT072864.1;QIQ54047.1;Pangolin_CoV_orf1ab;1232 | 0 | 0.05 |
| MT072864.1;QIQ54047.1;Pangolin_CoV_orf1ab;1233 | 0 | 0.025 |
| MT072864.1;QIQ54047.1;Pangolin_CoV_orf1ab;1234 | 0 | 0 |
| MT072864.1;QIQ54047.1;Pangolin_CoV_orf1ab;1235 | 0 | 0.025 |
| MT072864.1;QIQ54047.1;Pangolin_CoV_orf1ab;1236 | 0 | 0.025 |
| MT072864.1;QIQ54047.1;Pangolin_CoV_orf1ab;1237 | 0 | 0 |

| | | |
|---|---|---|
| MT072864.1;QIQ54047.1;Pangolin_CoV_orf1ab;1238 | 0 | 0 |
| MT072864.1;QIQ54047.1;Pangolin_CoV_orf1ab;1239 | 0 | 0 |
| MT072864.1;QIQ54047.1;Pangolin_CoV_orf1ab;1240 | 0 | 0 |
| MT072864.1;QIQ54047.1;Pangolin_CoV_orf1ab;1241 | 0.05 | 0 |
| MT072864.1;QIQ54047.1;Pangolin_CoV_orf1ab;1242 | 0 | 0 |
| MT072864.1;QIQ54047.1;Pangolin_CoV_orf1ab;1243 | 0 | 0 |
| MT072864.1;QIQ54047.1;Pangolin_CoV_orf1ab;1244 | 0 | 0.025 |
| MT072864.1;QIQ54047.1;Pangolin_CoV_orf1ab;1245 | 0.1 | 0.125 |
| MT072864.1;QIQ54047.1;Pangolin_CoV_orf1ab;1246 | 0.25 | 0.175 |
| MT072864.1;QIQ54047.1;Pangolin_CoV_orf1ab;1247 | 0.15 | 0.15 |
| MT072864.1;QIQ54047.1;Pangolin_CoV_orf1ab;1248 | 0 | 0.025 |
| MT072864.1;QIQ54047.1;Pangolin_CoV_orf1ab;1249 | 0 | 0.025 |
| MT072864.1;QIQ54047.1;Pangolin_CoV_orf1ab;1250 | 0 | 0 |
| MT072864.1;QIQ54047.1;Pangolin_CoV_orf1ab;1251 | 0 | 0 |
| MT072864.1;QIQ54047.1;Pangolin_CoV_orf1ab;1252 | 0 | 0 |
| MT072864.1;QIQ54047.1;Pangolin_CoV_orf1ab;1253 | 0 | 0.025 |
| MT072864.1;QIQ54047.1;Pangolin_CoV_orf1ab;1254 | 0 | 0.025 |
| MT072864.1;QIQ54047.1;Pangolin_CoV_orf1ab;1255 | 0 | 0 |
| MT072864.1;QIQ54047.1;Pangolin_CoV_orf1ab;1256 | 0 | 0.025 |
| MT072864.1;QIQ54047.1;Pangolin_CoV_orf1ab;1257 | 0 | 0 |
| MT072864.1;QIQ54047.1;Pangolin_CoV_orf1ab;1258 | 0 | 0.025 |
| MT072864.1;QIQ54047.1;Pangolin_CoV_orf1ab;1259 | 0 | 0 |
| MT072864.1;QIQ54047.1;Pangolin_CoV_orf1ab;1260 | 0 | 0 |
| MT072864.1;QIQ54047.1;Pangolin_CoV_orf1ab;1261 | 0 | 0.05 |
| MT072864.1;QIQ54047.1;Pangolin_CoV_orf1ab;1262 | 0 | 0.025 |
| MT072864.1;QIQ54047.1;Pangolin_CoV_orf1ab;1263 | 0 | 0 |
| MT072864.1;QIQ54047.1;Pangolin_CoV_orf1ab;1264 | 0 | 0.025 |
| MT072864.1;QIQ54047.1;Pangolin_CoV_orf1ab;1265 | 0 | 0.025 |
| MT072864.1;QIQ54047.1;Pangolin_CoV_orf1ab;1266 | 0 | 0 |
| MT072864.1;QIQ54047.1;Pangolin_CoV_orf1ab;1267 | 0 | 0 |
| MT072864.1;QIQ54047.1;Pangolin_CoV_orf1ab;1268 | 0 | 0 |
| MT072864.1;QIQ54047.1;Pangolin_CoV_orf1ab;1269 | 0 | 0 |
| MT072864.1;QIQ54047.1;Pangolin_CoV_orf1ab;1270 | 0 | 0 |
| MT072864.1;QIQ54047.1;Pangolin_CoV_orf1ab;1271 | 0 | 0.025 |
| MT072864.1;QIQ54047.1;Pangolin_CoV_orf1ab;1272 | 0 | 0.025 |
| MT072864.1;QIQ54047.1;Pangolin_CoV_orf1ab;1273 | 0 | 0.05 |
| MT072864.1;QIQ54047.1;Pangolin_CoV_orf1ab;1274 | 0 | 0.05 |
| MT072864.1;QIQ54047.1;Pangolin_CoV_orf1ab;1275 | 0 | 0.025 |
| MT072864.1;QIQ54047.1;Pangolin_CoV_orf1ab;1276 | 0 | 0 |
| MT072864.1;QIQ54047.1;Pangolin_CoV_orf1ab;1277 | 0 | 0 |
| MT072864.1;QIQ54047.1;Pangolin_CoV_orf1ab;1278 | 0 | 0 |
| MT072864.1;QIQ54047.1;Pangolin_CoV_orf1ab;1279 | 0 | 0.025 |
| MT072864.1;QIQ54047.1;Pangolin_CoV_orf1ab;1280 | 0 | 0.025 |

| | | |
|---|---|---|
| MT072864.1;QIQ54047.1;Pangolin_CoV_orf1ab;1281 | 0.1 | 0.025 |
| MT072864.1;QIQ54047.1;Pangolin_CoV_orf1ab;1282 | 0 | 0.05 |
| MT072864.1;QIQ54047.1;Pangolin_CoV_orf1ab;1283 | 0 | 0 |
| MT072864.1;QIQ54047.1;Pangolin_CoV_orf1ab;1284 | 0 | 0 |
| MT072864.1;QIQ54047.1;Pangolin_CoV_orf1ab;1285 | 0 | 0 |
| MT072864.1;QIQ54047.1;Pangolin_CoV_orf1ab;1286 | 0 | 0 |
| MT072864.1;QIQ54047.1;Pangolin_CoV_orf1ab;1287 | 0.05 | 0 |
| MT072864.1;QIQ54047.1;Pangolin_CoV_orf1ab;1288 | 0.05 | 0.025 |
| MT072864.1;QIQ54047.1;Pangolin_CoV_orf1ab;1289 | 0.05 | 0.025 |
| MT072864.1;QIQ54047.1;Pangolin_CoV_orf1ab;1290 | 0.05 | 0.025 |
| MT072864.1;QIQ54047.1;Pangolin_CoV_orf1ab;1291 | 0.05 | 0 |
| MT072864.1;QIQ54047.1;Pangolin_CoV_orf1ab;1292 | 0.05 | 0 |
| MT072864.1;QIQ54047.1;Pangolin_CoV_orf1ab;1293 | 0.05 | 0 |
| MT072864.1;QIQ54047.1;Pangolin_CoV_orf1ab;1294 | 0 | 0 |
| MT072864.1;QIQ54047.1;Pangolin_CoV_orf1ab;1295 | 0 | 0 |
| MT072864.1;QIQ54047.1;Pangolin_CoV_orf1ab;1296 | 0 | 0 |
| MT072864.1;QIQ54047.1;Pangolin_CoV_orf1ab;1297 | 0 | 0 |
| MT072864.1;QIQ54047.1;Pangolin_CoV_orf1ab;1298 | 0 | 0 |
| MT072864.1;QIQ54047.1;Pangolin_CoV_orf1ab;1299 | 0 | 0 |
| MT072864.1;QIQ54047.1;Pangolin_CoV_orf1ab;1300 | 0 | 0 |
| MT072864.1;QIQ54047.1;Pangolin_CoV_orf1ab;1301 | 0 | 0 |
| MT072864.1;QIQ54047.1;Pangolin_CoV_orf1ab;1302 | 0 | 0 |
| MT072864.1;QIQ54047.1;Pangolin_CoV_orf1ab;1303 | 0 | 0 |
| MT072864.1;QIQ54047.1;Pangolin_CoV_orf1ab;1304 | 0 | 0 |
| MT072864.1;QIQ54047.1;Pangolin_CoV_orf1ab;1305 | 0 | 0 |
| MT072864.1;QIQ54047.1;Pangolin_CoV_orf1ab;1306 | 0 | 0 |
| MT072864.1;QIQ54047.1;Pangolin_CoV_orf1ab;1307 | 0 | 0 |
| MT072864.1;QIQ54047.1;Pangolin_CoV_orf1ab;1308 | 0 | 0 |
| MT072864.1;QIQ54047.1;Pangolin_CoV_orf1ab;1309 | 0 | 0 |
| MT072864.1;QIQ54047.1;Pangolin_CoV_orf1ab;1310 | 0 | 0 |
| MT072864.1;QIQ54047.1;Pangolin_CoV_orf1ab;1311 | 0 | 0 |
| MT072864.1;QIQ54047.1;Pangolin_CoV_orf1ab;1312 | 0 | 0 |
| MT072864.1;QIQ54047.1;Pangolin_CoV_orf1ab;1313 | 0 | 0 |
| MT072864.1;QIQ54047.1;Pangolin_CoV_orf1ab;1314 | 0 | 0 |
| MT072864.1;QIQ54047.1;Pangolin_CoV_orf1ab;1315 | 0 | 0 |
| MT072864.1;QIQ54047.1;Pangolin_CoV_orf1ab;1316 | 0 | 0 |
| MT072864.1;QIQ54047.1;Pangolin_CoV_orf1ab;1317 | 0 | 0 |
| MT072864.1;QIQ54047.1;Pangolin_CoV_orf1ab;1318 | 0 | 0 |
| MT072864.1;QIQ54047.1;Pangolin_CoV_orf1ab;1319 | 0.05 | 0.025 |
| MT072864.1;QIQ54047.1;Pangolin_CoV_orf1ab;1320 | 0.05 | 0.025 |
| MT072864.1;QIQ54047.1;Pangolin_CoV_orf1ab;1321 | 0.05 | 0.025 |
| MT072864.1;QIQ54047.1;Pangolin_CoV_orf1ab;1322 | 0.05 | 0.025 |
| MT072864.1;QIQ54047.1;Pangolin_CoV_orf1ab;1323 | 0.05 | 0.05 |

| | | |
|---|---|---|
| MT072864.1;QIQ54047.1;Pangolin_CoV_orf1ab;1324 | 0 | 0.025 |
| MT072864.1;QIQ54047.1;Pangolin_CoV_orf1ab;1325 | 0 | 0.025 |
| MT072864.1;QIQ54047.1;Pangolin_CoV_orf1ab;1326 | 0 | 0.025 |
| MT072864.1;QIQ54047.1;Pangolin_CoV_orf1ab;1327 | 0 | 0 |
| MT072864.1;QIQ54047.1;Pangolin_CoV_orf1ab;1328 | 0 | 0 |
| MT072864.1;QIQ54047.1;Pangolin_CoV_orf1ab;1329 | 0 | 0 |
| MT072864.1;QIQ54047.1;Pangolin_CoV_orf1ab;1330 | 0 | 0 |
| MT072864.1;QIQ54047.1;Pangolin_CoV_orf1ab;1331 | 0 | 0 |
| MT072864.1;QIQ54047.1;Pangolin_CoV_orf1ab;1332 | 0 | 0 |
| MT072864.1;QIQ54047.1;Pangolin_CoV_orf1ab;1333 | 0 | 0 |
| MT072864.1;QIQ54047.1;Pangolin_CoV_orf1ab;1334 | 0 | 0.025 |
| MT072864.1;QIQ54047.1;Pangolin_CoV_orf1ab;1335 | 0 | 0 |
| MT072864.1;QIQ54047.1;Pangolin_CoV_orf1ab;1336 | 0 | 0 |
| MT072864.1;QIQ54047.1;Pangolin_CoV_orf1ab;1337 | 0 | 0 |
| MT072864.1;QIQ54047.1;Pangolin_CoV_orf1ab;1338 | 0 | 0.025 |
| MT072864.1;QIQ54047.1;Pangolin_CoV_orf1ab;1339 | 0 | 0.025 |
| MT072864.1;QIQ54047.1;Pangolin_CoV_orf1ab;1340 | 0 | 0 |
| MT072864.1;QIQ54047.1;Pangolin_CoV_orf1ab;1341 | 0 | 0 |
| MT072864.1;QIQ54047.1;Pangolin_CoV_orf1ab;1342 | 0 | 0 |
| MT072864.1;QIQ54047.1;Pangolin_CoV_orf1ab;1343 | 0 | 0 |
| MT072864.1;QIQ54047.1;Pangolin_CoV_orf1ab;1344 | 0 | 0 |
| MT072864.1;QIQ54047.1;Pangolin_CoV_orf1ab;1345 | 0 | 0.025 |
| MT072864.1;QIQ54047.1;Pangolin_CoV_orf1ab;1346 | 0.05 | 0.025 |
| MT072864.1;QIQ54047.1;Pangolin_CoV_orf1ab;1347 | 0 | 0 |
| MT072864.1;QIQ54047.1;Pangolin_CoV_orf1ab;1348 | 0.1 | 0 |
| MT072864.1;QIQ54047.1;Pangolin_CoV_orf1ab;1349 | 0 | 0 |
| MT072864.1;QIQ54047.1;Pangolin_CoV_orf1ab;1350 | 0.05 | 0 |
| MT072864.1;QIQ54047.1;Pangolin_CoV_orf1ab;1351 | 0 | 0 |
| MT072864.1;QIQ54047.1;Pangolin_CoV_orf1ab;1352 | 0 | 0 |
| MT072864.1;QIQ54047.1;Pangolin_CoV_orf1ab;1353 | 0 | 0 |
| MT072864.1;QIQ54047.1;Pangolin_CoV_orf1ab;1354 | 0 | 0 |
| MT072864.1;QIQ54047.1;Pangolin_CoV_orf1ab;1355 | 0 | 0 |
| MT072864.1;QIQ54047.1;Pangolin_CoV_orf1ab;1356 | 0.05 | 0.025 |
| MT072864.1;QIQ54047.1;Pangolin_CoV_orf1ab;1357 | 0 | 0 |
| MT072864.1;QIQ54047.1;Pangolin_CoV_orf1ab;1358 | 0 | 0 |
| MT072864.1;QIQ54047.1;Pangolin_CoV_orf1ab;1359 | 0 | 0 |
| MT072864.1;QIQ54047.1;Pangolin_CoV_orf1ab;1360 | 0 | 0.025 |
| MT072864.1;QIQ54047.1;Pangolin_CoV_orf1ab;1361 | 0 | 0.025 |
| MT072864.1;QIQ54047.1;Pangolin_CoV_orf1ab;1362 | 0 | 0.025 |
| MT072864.1;QIQ54047.1;Pangolin_CoV_orf1ab;1363 | 0 | 0.025 |
| MT072864.1;QIQ54047.1;Pangolin_CoV_orf1ab;1364 | 0 | 0 |
| MT072864.1;QIQ54047.1;Pangolin_CoV_orf1ab;1365 | 0 | 0 |
| MT072864.1;QIQ54047.1;Pangolin_CoV_orf1ab;1366 | 0 | 0 |

| | | |
|---|---|---|
| MT072864.1;QIQ54047.1;Pangolin_CoV_orf1ab;1367 | 0 | 0 |
| MT072864.1;QIQ54047.1;Pangolin_CoV_orf1ab;1368 | 0 | 0 |
| MT072864.1;QIQ54047.1;Pangolin_CoV_orf1ab;1369 | 0 | 0 |
| MT072864.1;QIQ54047.1;Pangolin_CoV_orf1ab;1370 | 0 | 0 |
| MT072864.1;QIQ54047.1;Pangolin_CoV_orf1ab;1371 | 0 | 0 |
| MT072864.1;QIQ54047.1;Pangolin_CoV_orf1ab;1372 | 0 | 0 |
| MT072864.1;QIQ54047.1;Pangolin_CoV_orf1ab;1373 | 0.05 | 0.025 |
| MT072864.1;QIQ54047.1;Pangolin_CoV_orf1ab;1374 | 0.05 | 0.025 |
| MT072864.1;QIQ54047.1;Pangolin_CoV_orf1ab;1375 | 0 | 0 |
| MT072864.1;QIQ54047.1;Pangolin_CoV_orf1ab;1376 | 0 | 0 |
| MT072864.1;QIQ54047.1;Pangolin_CoV_orf1ab;1377 | 0 | 0 |
| MT072864.1;QIQ54047.1;Pangolin_CoV_orf1ab;1378 | 0 | 0 |
| MT072864.1;QIQ54047.1;Pangolin_CoV_orf1ab;1379 | 0 | 0 |
| MT072864.1;QIQ54047.1;Pangolin_CoV_orf1ab;1380 | 0 | 0 |
| MT072864.1;QIQ54047.1;Pangolin_CoV_orf1ab;1381 | 0 | 0 |
| MT072864.1;QIQ54047.1;Pangolin_CoV_orf1ab;1382 | 0 | 0 |
| MT072864.1;QIQ54047.1;Pangolin_CoV_orf1ab;1383 | 0 | 0 |
| MT072864.1;QIQ54047.1;Pangolin_CoV_orf1ab;1384 | 0 | 0 |
| MT072864.1;QIQ54047.1;Pangolin_CoV_orf1ab;1385 | 0 | 0 |
| MT072864.1;QIQ54047.1;Pangolin_CoV_orf1ab;1386 | 0 | 0 |
| MT072864.1;QIQ54047.1;Pangolin_CoV_orf1ab;1387 | 0.05 | 0 |
| MT072864.1;QIQ54047.1;Pangolin_CoV_orf1ab;1388 | 0.05 | 0 |
| MT072864.1;QIQ54047.1;Pangolin_CoV_orf1ab;1389 | 0.1 | 0 |
| MT072864.1;QIQ54047.1;Pangolin_CoV_orf1ab;1390 | 0.1 | 0 |
| MT072864.1;QIQ54047.1;Pangolin_CoV_orf1ab;1391 | 0.1 | 0 |
| MT072864.1;QIQ54047.1;Pangolin_CoV_orf1ab;1392 | 0.1 | 0 |
| MT072864.1;QIQ54047.1;Pangolin_CoV_orf1ab;1393 | 0.1 | 0 |
| MT072864.1;QIQ54047.1;Pangolin_CoV_orf1ab;1394 | 0.1 | 0.025 |
| MT072864.1;QIQ54047.1;Pangolin_CoV_orf1ab;1395 | 0.1 | 0.025 |
| MT072864.1;QIQ54047.1;Pangolin_CoV_orf1ab;1396 | 0.1 | 0 |
| MT072864.1;QIQ54047.1;Pangolin_CoV_orf1ab;1397 | 0.05 | 0.05 |
| MT072864.1;QIQ54047.1;Pangolin_CoV_orf1ab;1398 | 0 | 0.025 |
| MT072864.1;QIQ54047.1;Pangolin_CoV_orf1ab;1399 | 0 | 0 |
| MT072864.1;QIQ54047.1;Pangolin_CoV_orf1ab;1400 | 0 | 0 |
| MT072864.1;QIQ54047.1;Pangolin_CoV_orf1ab;1401 | 0 | 0 |
| MT072864.1;QIQ54047.1;Pangolin_CoV_orf1ab;1402 | 0 | 0 |
| MT072864.1;QIQ54047.1;Pangolin_CoV_orf1ab;1403 | 0 | 0 |
| MT072864.1;QIQ54047.1;Pangolin_CoV_orf1ab;1404 | 0 | 0 |
| MT072864.1;QIQ54047.1;Pangolin_CoV_orf1ab;1405 | 0 | 0.05 |
| MT072864.1;QIQ54047.1;Pangolin_CoV_orf1ab;1406 | 0 | 0 |
| MT072864.1;QIQ54047.1;Pangolin_CoV_orf1ab;1407 | 0 | 0 |
| MT072864.1;QIQ54047.1;Pangolin_CoV_orf1ab;1408 | 0 | 0 |
| MT072864.1;QIQ54047.1;Pangolin_CoV_orf1ab;1409 | 0 | 0 |

| | | |
|---|---|---|
| MT072864.1;QIQ54047.1;Pangolin_CoV_orf1ab;1410 | 0 | 0 |
| MT072864.1;QIQ54047.1;Pangolin_CoV_orf1ab;1411 | 0 | 0 |
| MT072864.1;QIQ54047.1;Pangolin_CoV_orf1ab;1412 | 0 | 0 |
| MT072864.1;QIQ54047.1;Pangolin_CoV_orf1ab;1413 | 0 | 0 |
| MT072864.1;QIQ54047.1;Pangolin_CoV_orf1ab;1414 | 0 | 0 |
| MT072864.1;QIQ54047.1;Pangolin_CoV_orf1ab;1415 | 0 | 0 |
| MT072864.1;QIQ54047.1;Pangolin_CoV_orf1ab;1416 | 0 | 0 |
| MT072864.1;QIQ54047.1;Pangolin_CoV_orf1ab;1417 | 0 | 0 |
| MT072864.1;QIQ54047.1;Pangolin_CoV_orf1ab;1418 | 0 | 0 |
| MT072864.1;QIQ54047.1;Pangolin_CoV_orf1ab;1419 | 0 | 0.025 |
| MT072864.1;QIQ54047.1;Pangolin_CoV_orf1ab;1420 | 0.05 | 0.1 |
| MT072864.1;QIQ54047.1;Pangolin_CoV_orf1ab;1421 | 0.05 | 0.075 |
| MT072864.1;QIQ54047.1;Pangolin_CoV_orf1ab;1422 | 0 | 0.025 |
| MT072864.1;QIQ54047.1;Pangolin_CoV_orf1ab;1423 | 0 | 0.025 |
| MT072864.1;QIQ54047.1;Pangolin_CoV_orf1ab;1424 | 0 | 0.025 |
| MT072864.1;QIQ54047.1;Pangolin_CoV_orf1ab;1425 | 0 | 0.025 |
| MT072864.1;QIQ54047.1;Pangolin_CoV_orf1ab;1426 | 0 | 0.05 |
| MT072864.1;QIQ54047.1;Pangolin_CoV_orf1ab;1427 | 0 | 0.025 |
| MT072864.1;QIQ54047.1;Pangolin_CoV_orf1ab;1428 | 0 | 0.025 |
| MT072864.1;QIQ54047.1;Pangolin_CoV_orf1ab;1429 | 0 | 0.025 |
| MT072864.1;QIQ54047.1;Pangolin_CoV_orf1ab;1430 | 0 | 0.025 |
| MT072864.1;QIQ54047.1;Pangolin_CoV_orf1ab;1431 | 0 | 0 |
| MT072864.1;QIQ54047.1;Pangolin_CoV_orf1ab;1432 | 0 | 0 |
| MT072864.1;QIQ54047.1;Pangolin_CoV_orf1ab;1433 | 0 | 0 |
| MT072864.1;QIQ54047.1;Pangolin_CoV_orf1ab;1434 | 0 | 0 |
| MT072864.1;QIQ54047.1;Pangolin_CoV_orf1ab;1435 | 0 | 0 |
| MT072864.1;QIQ54047.1;Pangolin_CoV_orf1ab;1436 | 0 | 0 |
| MT072864.1;QIQ54047.1;Pangolin_CoV_orf1ab;1437 | 0.05 | 0 |
| MT072864.1;QIQ54047.1;Pangolin_CoV_orf1ab;1438 | 0 | 0 |
| MT072864.1;QIQ54047.1;Pangolin_CoV_orf1ab;1439 | 0.1 | 0.05 |
| MT072864.1;QIQ54047.1;Pangolin_CoV_orf1ab;1440 | 0.05 | 0.025 |
| MT072864.1;QIQ54047.1;Pangolin_CoV_orf1ab;1441 | 0.05 | 0.025 |
| MT072864.1;QIQ54047.1;Pangolin_CoV_orf1ab;1442 | 0.05 | 0.025 |
| MT072864.1;QIQ54047.1;Pangolin_CoV_orf1ab;1443 | 0 | 0 |
| MT072864.1;QIQ54047.1;Pangolin_CoV_orf1ab;1444 | 0 | 0 |
| MT072864.1;QIQ54047.1;Pangolin_CoV_orf1ab;1445 | 0 | 0 |
| MT072864.1;QIQ54047.1;Pangolin_CoV_orf1ab;1446 | 0 | 0 |
| MT072864.1;QIQ54047.1;Pangolin_CoV_orf1ab;1447 | 0 | 0.025 |
| MT072864.1;QIQ54047.1;Pangolin_CoV_orf1ab;1448 | 0 | 0 |
| MT072864.1;QIQ54047.1;Pangolin_CoV_orf1ab;1449 | 0 | 0 |
| MT072864.1;QIQ54047.1;Pangolin_CoV_orf1ab;1450 | 0 | 0.075 |
| MT072864.1;QIQ54047.1;Pangolin_CoV_orf1ab;1451 | 0.1 | 0.075 |
| MT072864.1;QIQ54047.1;Pangolin_CoV_orf1ab;1452 | 0.15 | 0.125 |

| | | |
|---|---|---|
| MT072864.1;QIQ54047.1;Pangolin_CoV_orf1ab;1453 | 0.05 | 0 |
| MT072864.1;QIQ54047.1;Pangolin_CoV_orf1ab;1454 | 0 | 0.025 |
| MT072864.1;QIQ54047.1;Pangolin_CoV_orf1ab;1455 | 0 | 0 |
| MT072864.1;QIQ54047.1;Pangolin_CoV_orf1ab;1456 | 0 | 0 |
| MT072864.1;QIQ54047.1;Pangolin_CoV_orf1ab;1457 | 0 | 0 |
| MT072864.1;QIQ54047.1;Pangolin_CoV_orf1ab;1458 | 0 | 0 |
| MT072864.1;QIQ54047.1;Pangolin_CoV_orf1ab;1459 | 0 | 0 |
| MT072864.1;QIQ54047.1;Pangolin_CoV_orf1ab;1460 | 0 | 0 |
| MT072864.1;QIQ54047.1;Pangolin_CoV_orf1ab;1461 | 0 | 0 |
| MT072864.1;QIQ54047.1;Pangolin_CoV_orf1ab;1462 | 0 | 0 |
| MT072864.1;QIQ54047.1;Pangolin_CoV_orf1ab;1463 | 0 | 0.05 |
| MT072864.1;QIQ54047.1;Pangolin_CoV_orf1ab;1464 | 0.05 | 0.025 |
| MT072864.1;QIQ54047.1;Pangolin_CoV_orf1ab;1465 | 0 | 0.1 |
| MT072864.1;QIQ54047.1;Pangolin_CoV_orf1ab;1466 | 0 | 0.025 |
| MT072864.1;QIQ54047.1;Pangolin_CoV_orf1ab;1467 | 0 | 0 |
| MT072864.1;QIQ54047.1;Pangolin_CoV_orf1ab;1468 | 0 | 0 |
| MT072864.1;QIQ54047.1;Pangolin_CoV_orf1ab;1469 | 0 | 0 |
| MT072864.1;QIQ54047.1;Pangolin_CoV_orf1ab;1470 | 0 | 0 |
| MT072864.1;QIQ54047.1;Pangolin_CoV_orf1ab;1471 | 0.05 | 0.05 |
| MT072864.1;QIQ54047.1;Pangolin_CoV_orf1ab;1472 | 0.05 | 0.075 |
| MT072864.1;QIQ54047.1;Pangolin_CoV_orf1ab;1473 | 0.1 | 0.05 |
| MT072864.1;QIQ54047.1;Pangolin_CoV_orf1ab;1474 | 0.15 | 0.1 |
| MT072864.1;QIQ54047.1;Pangolin_CoV_orf1ab;1475 | 0.1 | 0.15 |
| MT072864.1;QIQ54047.1;Pangolin_CoV_orf1ab;1476 | 0.1 | 0.125 |
| MT072864.1;QIQ54047.1;Pangolin_CoV_orf1ab;1477 | 0.15 | 0.125 |
| MT072864.1;QIQ54047.1;Pangolin_CoV_orf1ab;1478 | 0.2 | 0.125 |
| MT072864.1;QIQ54047.1;Pangolin_CoV_orf1ab;1479 | 0.25 | 0.1 |
| MT072864.1;QIQ54047.1;Pangolin_CoV_orf1ab;1480 | 0.2 | 0.1 |
| MT072864.1;QIQ54047.1;Pangolin_CoV_orf1ab;1481 | 0.15 | 0.025 |
| MT072864.1;QIQ54047.1;Pangolin_CoV_orf1ab;1482 | 0.05 | 0 |
| MT072864.1;QIQ54047.1;Pangolin_CoV_orf1ab;1483 | 0.05 | 0 |
| MT072864.1;QIQ54047.1;Pangolin_CoV_orf1ab;1484 | 0.05 | 0 |
| MT072864.1;QIQ54047.1;Pangolin_CoV_orf1ab;1485 | 0 | 0 |
| MT072864.1;QIQ54047.1;Pangolin_CoV_orf1ab;1486 | 0 | 0 |
| MT072864.1;QIQ54047.1;Pangolin_CoV_orf1ab;1487 | 0 | 0 |
| MT072864.1;QIQ54047.1;Pangolin_CoV_orf1ab;1488 | 0 | 0 |
| MT072864.1;QIQ54047.1;Pangolin_CoV_orf1ab;1489 | 0 | 0 |
| MT072864.1;QIQ54047.1;Pangolin_CoV_orf1ab;1490 | 0 | 0 |
| MT072864.1;QIQ54047.1;Pangolin_CoV_orf1ab;1491 | 0 | 0 |
| MT072864.1;QIQ54047.1;Pangolin_CoV_orf1ab;1492 | 0 | 0 |
| MT072864.1;QIQ54047.1;Pangolin_CoV_orf1ab;1493 | 0 | 0 |
| MT072864.1;QIQ54047.1;Pangolin_CoV_orf1ab;1494 | 0 | 0 |
| MT072864.1;QIQ54047.1;Pangolin_CoV_orf1ab;1495 | 0 | 0 |

| | | |
|---|---|---|
| MT072864.1;QIQ54047.1;Pangolin_CoV_orf1ab;1496 | 0 | 0 |
| MT072864.1;QIQ54047.1;Pangolin_CoV_orf1ab;1497 | 0.05 | 0 |
| MT072864.1;QIQ54047.1;Pangolin_CoV_orf1ab;1498 | 0 | 0 |
| MT072864.1;QIQ54047.1;Pangolin_CoV_orf1ab;1499 | 0 | 0 |
| MT072864.1;QIQ54047.1;Pangolin_CoV_orf1ab;1500 | 0.2 | 0.05 |
| MT072864.1;QIQ54047.1;Pangolin_CoV_orf1ab;1501 | 0 | 0 |
| MT072864.1;QIQ54047.1;Pangolin_CoV_orf1ab;1502 | 0 | 0 |
| MT072864.1;QIQ54047.1;Pangolin_CoV_orf1ab;1503 | 0 | 0 |
| MT072864.1;QIQ54047.1;Pangolin_CoV_orf1ab;1504 | 0 | 0 |
| MT072864.1;QIQ54047.1;Pangolin_CoV_orf1ab;1505 | 0 | 0 |
| MT072864.1;QIQ54047.1;Pangolin_CoV_orf1ab;1506 | 0.05 | 0 |
| MT072864.1;QIQ54047.1;Pangolin_CoV_orf1ab;1507 | 0 | 0 |
| MT072864.1;QIQ54047.1;Pangolin_CoV_orf1ab;1508 | 0 | 0 |
| MT072864.1;QIQ54047.1;Pangolin_CoV_orf1ab;1509 | 0 | 0.075 |
| MT072864.1;QIQ54047.1;Pangolin_CoV_orf1ab;1510 | 0 | 0.1 |
| MT072864.1;QIQ54047.1;Pangolin_CoV_orf1ab;1511 | 0 | 0.075 |
| MT072864.1;QIQ54047.1;Pangolin_CoV_orf1ab;1512 | 0 | 0.05 |
| MT072864.1;QIQ54047.1;Pangolin_CoV_orf1ab;1513 | 0 | 0.05 |
| MT072864.1;QIQ54047.1;Pangolin_CoV_orf1ab;1514 | 0 | 0.05 |
| MT072864.1;QIQ54047.1;Pangolin_CoV_orf1ab;1515 | 0.05 | 0.075 |
| MT072864.1;QIQ54047.1;Pangolin_CoV_orf1ab;1516 | 0 | 0.05 |
| MT072864.1;QIQ54047.1;Pangolin_CoV_orf1ab;1517 | 0 | 0.075 |
| MT072864.1;QIQ54047.1;Pangolin_CoV_orf1ab;1518 | 0 | 0.05 |
| MT072864.1;QIQ54047.1;Pangolin_CoV_orf1ab;1519 | 0 | 0.05 |
| MT072864.1;QIQ54047.1;Pangolin_CoV_orf1ab;1520 | 0 | 0.025 |
| MT072864.1;QIQ54047.1;Pangolin_CoV_orf1ab;1521 | 0 | 0.025 |
| MT072864.1;QIQ54047.1;Pangolin_CoV_orf1ab;1522 | 0 | 0.025 |
| MT072864.1;QIQ54047.1;Pangolin_CoV_orf1ab;1523 | 0 | 0.025 |
| MT072864.1;QIQ54047.1;Pangolin_CoV_orf1ab;1524 | 0 | 0.025 |
| MT072864.1;QIQ54047.1;Pangolin_CoV_orf1ab;1525 | 0.05 | 0.025 |
| MT072864.1;QIQ54047.1;Pangolin_CoV_orf1ab;1526 | 0.05 | 0.025 |
| MT072864.1;QIQ54047.1;Pangolin_CoV_orf1ab;1527 | 0 | 0.025 |
| MT072864.1;QIQ54047.1;Pangolin_CoV_orf1ab;1528 | 0 | 0 |
| MT072864.1;QIQ54047.1;Pangolin_CoV_orf1ab;1529 | 0 | 0.05 |
| MT072864.1;QIQ54047.1;Pangolin_CoV_orf1ab;1530 | 0 | 0.025 |
| MT072864.1;QIQ54047.1;Pangolin_CoV_orf1ab;1531 | 0 | 0.025 |
| MT072864.1;QIQ54047.1;Pangolin_CoV_orf1ab;1532 | 0 | 0.025 |
| MT072864.1;QIQ54047.1;Pangolin_CoV_orf1ab;1533 | 0 | 0.025 |
| MT072864.1;QIQ54047.1;Pangolin_CoV_orf1ab;1534 | 0 | 0.025 |
| MT072864.1;QIQ54047.1;Pangolin_CoV_orf1ab;1535 | 0 | 0.025 |
| MT072864.1;QIQ54047.1;Pangolin_CoV_orf1ab;1536 | 0 | 0.025 |
| MT072864.1;QIQ54047.1;Pangolin_CoV_orf1ab;1537 | 0 | 0 |
| MT072864.1;QIQ54047.1;Pangolin_CoV_orf1ab;1538 | 0 | 0 |

| | | |
|---|---|---|
| MT072864.1;QIQ54047.1;Pangolin_CoV_orf1ab;1539 | 0 | 0 |
| MT072864.1;QIQ54047.1;Pangolin_CoV_orf1ab;1540 | 0 | 0 |
| MT072864.1;QIQ54047.1;Pangolin_CoV_orf1ab;1541 | 0 | 0 |
| MT072864.1;QIQ54047.1;Pangolin_CoV_orf1ab;1542 | 0 | 0 |
| MT072864.1;QIQ54047.1;Pangolin_CoV_orf1ab;1543 | 0 | 0 |
| MT072864.1;QIQ54047.1;Pangolin_CoV_orf1ab;1544 | 0 | 0 |
| MT072864.1;QIQ54047.1;Pangolin_CoV_orf1ab;1545 | 0 | 0.025 |
| MT072864.1;QIQ54047.1;Pangolin_CoV_orf1ab;1546 | 0 | 0 |
| MT072864.1;QIQ54047.1;Pangolin_CoV_orf1ab;1547 | 0 | 0 |
| MT072864.1;QIQ54047.1;Pangolin_CoV_orf1ab;1548 | 0 | 0 |
| MT072864.1;QIQ54047.1;Pangolin_CoV_orf1ab;1549 | 0 | 0 |
| MT072864.1;QIQ54047.1;Pangolin_CoV_orf1ab;1550 | 0 | 0 |
| MT072864.1;QIQ54047.1;Pangolin_CoV_orf1ab;1551 | 0 | 0 |
| MT072864.1;QIQ54047.1;Pangolin_CoV_orf1ab;1552 | 0 | 0 |
| MT072864.1;QIQ54047.1;Pangolin_CoV_orf1ab;1553 | 0 | 0 |
| MT072864.1;QIQ54047.1;Pangolin_CoV_orf1ab;1554 | 0 | 0 |
| MT072864.1;QIQ54047.1;Pangolin_CoV_orf1ab;1555 | 0 | 0 |
| MT072864.1;QIQ54047.1;Pangolin_CoV_orf1ab;1556 | 0 | 0 |
| MT072864.1;QIQ54047.1;Pangolin_CoV_orf1ab;1557 | 0 | 0 |
| MT072864.1;QIQ54047.1;Pangolin_CoV_orf1ab;1558 | 0 | 0 |
| MT072864.1;QIQ54047.1;Pangolin_CoV_orf1ab;1559 | 0 | 0.025 |
| MT072864.1;QIQ54047.1;Pangolin_CoV_orf1ab;1560 | 0 | 0.025 |
| MT072864.1;QIQ54047.1;Pangolin_CoV_orf1ab;1561 | 0 | 0.025 |
| MT072864.1;QIQ54047.1;Pangolin_CoV_orf1ab;1562 | 0 | 0.025 |
| MT072864.1;QIQ54047.1;Pangolin_CoV_orf1ab;1563 | 0 | 0.025 |
| MT072864.1;QIQ54047.1;Pangolin_CoV_orf1ab;1564 | 0 | 0.025 |
| MT072864.1;QIQ54047.1;Pangolin_CoV_orf1ab;1565 | 0.05 | 0.125 |
| MT072864.1;QIQ54047.1;Pangolin_CoV_orf1ab;1566 | 0 | 0.075 |
| MT072864.1;QIQ54047.1;Pangolin_CoV_orf1ab;1567 | 0 | 0.025 |
| MT072864.1;QIQ54047.1;Pangolin_CoV_orf1ab;1568 | 0 | 0.025 |
| MT072864.1;QIQ54047.1;Pangolin_CoV_orf1ab;1569 | 0 | 0.025 |
| MT072864.1;QIQ54047.1;Pangolin_CoV_orf1ab;1570 | 0 | 0.025 |
| MT072864.1;QIQ54047.1;Pangolin_CoV_orf1ab;1571 | 0 | 0.025 |
| MT072864.1;QIQ54047.1;Pangolin_CoV_orf1ab;1572 | 0 | 0 |
| MT072864.1;QIQ54047.1;Pangolin_CoV_orf1ab;1573 | 0 | 0 |
| MT072864.1;QIQ54047.1;Pangolin_CoV_orf1ab;1574 | 0 | 0 |
| MT072864.1;QIQ54047.1;Pangolin_CoV_orf1ab;1575 | 0 | 0 |
| MT072864.1;QIQ54047.1;Pangolin_CoV_orf1ab;1576 | 0 | 0 |
| MT072864.1;QIQ54047.1;Pangolin_CoV_orf1ab;1577 | 0 | 0 |
| MT072864.1;QIQ54047.1;Pangolin_CoV_orf1ab;1578 | 0 | 0 |
| MT072864.1;QIQ54047.1;Pangolin_CoV_orf1ab;1579 | 0 | 0 |
| MT072864.1;QIQ54047.1;Pangolin_CoV_orf1ab;1580 | 0 | 0 |
| MT072864.1;QIQ54047.1;Pangolin_CoV_orf1ab;1581 | 0 | 0 |

| | | |
|---|---|---|
| MT072864.1;QIQ54047.1;Pangolin_CoV_orf1ab;1582 | 0 | 0 |
| MT072864.1;QIQ54047.1;Pangolin_CoV_orf1ab;1583 | 0 | 0.025 |
| MT072864.1;QIQ54047.1;Pangolin_CoV_orf1ab;1584 | 0 | 0.025 |
| MT072864.1;QIQ54047.1;Pangolin_CoV_orf1ab;1585 | 0 | 0.025 |
| MT072864.1;QIQ54047.1;Pangolin_CoV_orf1ab;1586 | 0 | 0.025 |
| MT072864.1;QIQ54047.1;Pangolin_CoV_orf1ab;1587 | 0.15 | 0.125 |
| MT072864.1;QIQ54047.1;Pangolin_CoV_orf1ab;1588 | 0 | 0.025 |
| MT072864.1;QIQ54047.1;Pangolin_CoV_orf1ab;1589 | 0.05 | 0.05 |
| MT072864.1;QIQ54047.1;Pangolin_CoV_orf1ab;1590 | 0 | 0 |
| MT072864.1;QIQ54047.1;Pangolin_CoV_orf1ab;1591 | 0 | 0 |
| MT072864.1;QIQ54047.1;Pangolin_CoV_orf1ab;1592 | 0 | 0 |
| MT072864.1;QIQ54047.1;Pangolin_CoV_orf1ab;1593 | 0 | 0.025 |
| MT072864.1;QIQ54047.1;Pangolin_CoV_orf1ab;1594 | 0 | 0 |
| MT072864.1;QIQ54047.1;Pangolin_CoV_orf1ab;1595 | 0.05 | 0 |
| MT072864.1;QIQ54047.1;Pangolin_CoV_orf1ab;1596 | 0 | 0 |
| MT072864.1;QIQ54047.1;Pangolin_CoV_orf1ab;1597 | 0 | 0 |
| MT072864.1;QIQ54047.1;Pangolin_CoV_orf1ab;1598 | 0 | 0 |
| MT072864.1;QIQ54047.1;Pangolin_CoV_orf1ab;1599 | 0 | 0 |
| MT072864.1;QIQ54047.1;Pangolin_CoV_orf1ab;1600 | 0 | 0 |
| MT072864.1;QIQ54047.1;Pangolin_CoV_orf1ab;1601 | 0 | 0 |
| MT072864.1;QIQ54047.1;Pangolin_CoV_orf1ab;1602 | 0 | 0 |
| MT072864.1;QIQ54047.1;Pangolin_CoV_orf1ab;1603 | 0.05 | 0 |
| MT072864.1;QIQ54047.1;Pangolin_CoV_orf1ab;1604 | 0.05 | 0 |
| MT072864.1;QIQ54047.1;Pangolin_CoV_orf1ab;1605 | 0.05 | 0 |
| MT072864.1;QIQ54047.1;Pangolin_CoV_orf1ab;1606 | 0.05 | 0 |
| MT072864.1;QIQ54047.1;Pangolin_CoV_orf1ab;1607 | 0.1 | 0 |
| MT072864.1;QIQ54047.1;Pangolin_CoV_orf1ab;1608 | 0.05 | 0 |
| MT072864.1;QIQ54047.1;Pangolin_CoV_orf1ab;1609 | 0 | 0 |
| MT072864.1;QIQ54047.1;Pangolin_CoV_orf1ab;1610 | 0 | 0 |
| MT072864.1;QIQ54047.1;Pangolin_CoV_orf1ab;1611 | 0 | 0 |
| MT072864.1;QIQ54047.1;Pangolin_CoV_orf1ab;1612 | 0 | 0 |
| MT072864.1;QIQ54047.1;Pangolin_CoV_orf1ab;1613 | 0 | 0.025 |
| MT072864.1;QIQ54047.1;Pangolin_CoV_orf1ab;1614 | 0 | 0 |
| MT072864.1;QIQ54047.1;Pangolin_CoV_orf1ab;1615 | 0 | 0 |
| MT072864.1;QIQ54047.1;Pangolin_CoV_orf1ab;1616 | 0 | 0 |
| MT072864.1;QIQ54047.1;Pangolin_CoV_orf1ab;1617 | 0 | 0 |
| MT072864.1;QIQ54047.1;Pangolin_CoV_orf1ab;1618 | 0 | 0 |
| MT072864.1;QIQ54047.1;Pangolin_CoV_orf1ab;1619 | 0 | 0 |
| MT072864.1;QIQ54047.1;Pangolin_CoV_orf1ab;1620 | 0 | 0 |
| MT072864.1;QIQ54047.1;Pangolin_CoV_orf1ab;1621 | 0 | 0 |
| MT072864.1;QIQ54047.1;Pangolin_CoV_orf1ab;1622 | 0 | 0 |
| MT072864.1;QIQ54047.1;Pangolin_CoV_orf1ab;1623 | 0.05 | 0 |
| MT072864.1;QIQ54047.1;Pangolin_CoV_orf1ab;1624 | 0.05 | 0 |

| | | |
|---|---|---|
| MT072864.1;QIQ54047.1;Pangolin_CoV_orf1ab;1625 | 0.05 | 0 |
| MT072864.1;QIQ54047.1;Pangolin_CoV_orf1ab;1626 | 0.05 | 0 |
| MT072864.1;QIQ54047.1;Pangolin_CoV_orf1ab;1627 | 0.05 | 0 |
| MT072864.1;QIQ54047.1;Pangolin_CoV_orf1ab;1628 | 0.05 | 0 |
| MT072864.1;QIQ54047.1;Pangolin_CoV_orf1ab;1629 | 0.05 | 0 |
| MT072864.1;QIQ54047.1;Pangolin_CoV_orf1ab;1630 | 0.05 | 0 |
| MT072864.1;QIQ54047.1;Pangolin_CoV_orf1ab;1631 | 0.05 | 0 |
| MT072864.1;QIQ54047.1;Pangolin_CoV_orf1ab;1632 | 0 | 0 |
| MT072864.1;QIQ54047.1;Pangolin_CoV_orf1ab;1633 | 0 | 0 |
| MT072864.1;QIQ54047.1;Pangolin_CoV_orf1ab;1634 | 0 | 0 |
| MT072864.1;QIQ54047.1;Pangolin_CoV_orf1ab;1635 | 0 | 0 |
| MT072864.1;QIQ54047.1;Pangolin_CoV_orf1ab;1636 | 0 | 0 |
| MT072864.1;QIQ54047.1;Pangolin_CoV_orf1ab;1637 | 0 | 0 |
| MT072864.1;QIQ54047.1;Pangolin_CoV_orf1ab;1638 | 0 | 0 |
| MT072864.1;QIQ54047.1;Pangolin_CoV_orf1ab;1639 | 0 | 0 |
| MT072864.1;QIQ54047.1;Pangolin_CoV_orf1ab;1640 | 0 | 0 |
| MT072864.1;QIQ54047.1;Pangolin_CoV_orf1ab;1641 | 0 | 0 |
| MT072864.1;QIQ54047.1;Pangolin_CoV_orf1ab;1642 | 0 | 0 |
| MT072864.1;QIQ54047.1;Pangolin_CoV_orf1ab;1643 | 0 | 0 |
| MT072864.1;QIQ54047.1;Pangolin_CoV_orf1ab;1644 | 0 | 0 |
| MT072864.1;QIQ54047.1;Pangolin_CoV_orf1ab;1645 | 0 | 0 |
| MT072864.1;QIQ54047.1;Pangolin_CoV_orf1ab;1646 | 0 | 0 |
| MT072864.1;QIQ54047.1;Pangolin_CoV_orf1ab;1647 | 0 | 0 |
| MT072864.1;QIQ54047.1;Pangolin_CoV_orf1ab;1648 | 0 | 0 |
| MT072864.1;QIQ54047.1;Pangolin_CoV_orf1ab;1649 | 0.05 | 0.025 |
| MT072864.1;QIQ54047.1;Pangolin_CoV_orf1ab;1650 | 0.05 | 0.025 |
| MT072864.1;QIQ54047.1;Pangolin_CoV_orf1ab;1651 | 0.05 | 0.025 |
| MT072864.1;QIQ54047.1;Pangolin_CoV_orf1ab;1652 | 0.05 | 0 |
| MT072864.1;QIQ54047.1;Pangolin_CoV_orf1ab;1653 | 0.05 | 0 |
| MT072864.1;QIQ54047.1;Pangolin_CoV_orf1ab;1654 | 0.05 | 0 |
| MT072864.1;QIQ54047.1;Pangolin_CoV_orf1ab;1655 | 0.05 | 0 |
| MT072864.1;QIQ54047.1;Pangolin_CoV_orf1ab;1656 | 0.05 | 0 |
| MT072864.1;QIQ54047.1;Pangolin_CoV_orf1ab;1657 | 0 | 0 |
| MT072864.1;QIQ54047.1;Pangolin_CoV_orf1ab;1658 | 0 | 0 |
| MT072864.1;QIQ54047.1;Pangolin_CoV_orf1ab;1659 | 0 | 0 |
| MT072864.1;QIQ54047.1;Pangolin_CoV_orf1ab;1660 | 0 | 0 |
| MT072864.1;QIQ54047.1;Pangolin_CoV_orf1ab;1661 | 0 | 0 |
| MT072864.1;QIQ54047.1;Pangolin_CoV_orf1ab;1662 | 0 | 0 |
| MT072864.1;QIQ54047.1;Pangolin_CoV_orf1ab;1663 | 0 | 0 |
| MT072864.1;QIQ54047.1;Pangolin_CoV_orf1ab;1664 | 0 | 0 |
| MT072864.1;QIQ54047.1;Pangolin_CoV_orf1ab;1665 | 0 | 0 |
| MT072864.1;QIQ54047.1;Pangolin_CoV_orf1ab;1666 | 0 | 0 |
| MT072864.1;QIQ54047.1;Pangolin_CoV_orf1ab;1667 | 0 | 0 |

| | | |
|---|---|---|
| MT072864.1;QIQ54047.1;Pangolin_CoV_orf1ab;1668 | 0 | 0 |
| MT072864.1;QIQ54047.1;Pangolin_CoV_orf1ab;1669 | 0 | 0 |
| MT072864.1;QIQ54047.1;Pangolin_CoV_orf1ab;1670 | 0 | 0 |
| MT072864.1;QIQ54047.1;Pangolin_CoV_orf1ab;1671 | 0.05 | 0 |
| MT072864.1;QIQ54047.1;Pangolin_CoV_orf1ab;1672 | 0 | 0 |
| MT072864.1;QIQ54047.1;Pangolin_CoV_orf1ab;1673 | 0 | 0 |
| MT072864.1;QIQ54047.1;Pangolin_CoV_orf1ab;1674 | 0 | 0.025 |
| MT072864.1;QIQ54047.1;Pangolin_CoV_orf1ab;1675 | 0 | 0 |
| MT072864.1;QIQ54047.1;Pangolin_CoV_orf1ab;1676 | 0 | 0 |
| MT072864.1;QIQ54047.1;Pangolin_CoV_orf1ab;1677 | 0 | 0 |
| MT072864.1;QIQ54047.1;Pangolin_CoV_orf1ab;1678 | 0 | 0.05 |
| MT072864.1;QIQ54047.1;Pangolin_CoV_orf1ab;1679 | 0 | 0.025 |
| MT072864.1;QIQ54047.1;Pangolin_CoV_orf1ab;1680 | 0 | 0 |
| MT072864.1;QIQ54047.1;Pangolin_CoV_orf1ab;1681 | 0 | 0 |
| MT072864.1;QIQ54047.1;Pangolin_CoV_orf1ab;1682 | 0 | 0 |
| MT072864.1;QIQ54047.1;Pangolin_CoV_orf1ab;1683 | 0 | 0 |
| MT072864.1;QIQ54047.1;Pangolin_CoV_orf1ab;1684 | 0.05 | 0.025 |
| MT072864.1;QIQ54047.1;Pangolin_CoV_orf1ab;1685 | 0.1 | 0.075 |
| MT072864.1;QIQ54047.1;Pangolin_CoV_orf1ab;1686 | 0.1 | 0.05 |
| MT072864.1;QIQ54047.1;Pangolin_CoV_orf1ab;1687 | 0.05 | 0.025 |
| MT072864.1;QIQ54047.1;Pangolin_CoV_orf1ab;1688 | 0.05 | 0 |
| MT072864.1;QIQ54047.1;Pangolin_CoV_orf1ab;1689 | 0.05 | 0 |
| MT072864.1;QIQ54047.1;Pangolin_CoV_orf1ab;1690 | 0 | 0 |
| MT072864.1;QIQ54047.1;Pangolin_CoV_orf1ab;1691 | 0 | 0 |
| MT072864.1;QIQ54047.1;Pangolin_CoV_orf1ab;1692 | 0 | 0 |
| MT072864.1;QIQ54047.1;Pangolin_CoV_orf1ab;1693 | 0 | 0 |
| MT072864.1;QIQ54047.1;Pangolin_CoV_orf1ab;1694 | 0 | 0 |
| MT072864.1;QIQ54047.1;Pangolin_CoV_orf1ab;1695 | 0 | 0 |
| MT072864.1;QIQ54047.1;Pangolin_CoV_orf1ab;1696 | 0 | 0 |
| MT072864.1;QIQ54047.1;Pangolin_CoV_orf1ab;1697 | 0 | 0 |
| MT072864.1;QIQ54047.1;Pangolin_CoV_orf1ab;1698 | 0 | 0 |
| MT072864.1;QIQ54047.1;Pangolin_CoV_orf1ab;1699 | 0 | 0 |
| MT072864.1;QIQ54047.1;Pangolin_CoV_orf1ab;1700 | 0 | 0 |
| MT072864.1;QIQ54047.1;Pangolin_CoV_orf1ab;1701 | 0 | 0 |
| MT072864.1;QIQ54047.1;Pangolin_CoV_orf1ab;1702 | 0 | 0 |
| MT072864.1;QIQ54047.1;Pangolin_CoV_orf1ab;1703 | 0 | 0.025 |
| MT072864.1;QIQ54047.1;Pangolin_CoV_orf1ab;1704 | 0 | 0.025 |
| MT072864.1;QIQ54047.1;Pangolin_CoV_orf1ab;1705 | 0 | 0.05 |
| MT072864.1;QIQ54047.1;Pangolin_CoV_orf1ab;1706 | 0 | 0.025 |
| MT072864.1;QIQ54047.1;Pangolin_CoV_orf1ab;1707 | 0 | 0 |
| MT072864.1;QIQ54047.1;Pangolin_CoV_orf1ab;1708 | 0 | 0 |
| MT072864.1;QIQ54047.1;Pangolin_CoV_orf1ab;1709 | 0 | 0 |
| MT072864.1;QIQ54047.1;Pangolin_CoV_orf1ab;1710 | 0 | 0 |

| | | |
|---|---|---|
| MT072864.1;QIQ54047.1;Pangolin_CoV_orf1ab;1711 | 0 | 0 |
| MT072864.1;QIQ54047.1;Pangolin_CoV_orf1ab;1712 | 0 | 0.025 |
| MT072864.1;QIQ54047.1;Pangolin_CoV_orf1ab;1713 | 0 | 0 |
| MT072864.1;QIQ54047.1;Pangolin_CoV_orf1ab;1714 | 0 | 0 |
| MT072864.1;QIQ54047.1;Pangolin_CoV_orf1ab;1715 | 0.05 | 0.025 |
| MT072864.1;QIQ54047.1;Pangolin_CoV_orf1ab;1716 | 0 | 0 |
| MT072864.1;QIQ54047.1;Pangolin_CoV_orf1ab;1717 | 0 | 0 |
| MT072864.1;QIQ54047.1;Pangolin_CoV_orf1ab;1718 | 0.05 | 0 |
| MT072864.1;QIQ54047.1;Pangolin_CoV_orf1ab;1719 | 0.05 | 0 |
| MT072864.1;QIQ54047.1;Pangolin_CoV_orf1ab;1720 | 0.05 | 0 |
| MT072864.1;QIQ54047.1;Pangolin_CoV_orf1ab;1721 | 0.05 | 0 |
| MT072864.1;QIQ54047.1;Pangolin_CoV_orf1ab;1722 | 0.05 | 0 |
| MT072864.1;QIQ54047.1;Pangolin_CoV_orf1ab;1723 | 0 | 0 |
| MT072864.1;QIQ54047.1;Pangolin_CoV_orf1ab;1724 | 0 | 0 |
| MT072864.1;QIQ54047.1;Pangolin_CoV_orf1ab;1725 | 0 | 0 |
| MT072864.1;QIQ54047.1;Pangolin_CoV_orf1ab;1726 | 0 | 0 |
| MT072864.1;QIQ54047.1;Pangolin_CoV_orf1ab;1727 | 0 | 0 |
| MT072864.1;QIQ54047.1;Pangolin_CoV_orf1ab;1728 | 0 | 0 |
| MT072864.1;QIQ54047.1;Pangolin_CoV_orf1ab;1729 | 0 | 0 |
| MT072864.1;QIQ54047.1;Pangolin_CoV_orf1ab;1730 | 0 | 0 |
| MT072864.1;QIQ54047.1;Pangolin_CoV_orf1ab;1731 | 0 | 0 |
| MT072864.1;QIQ54047.1;Pangolin_CoV_orf1ab;1732 | 0 | 0 |
| MT072864.1;QIQ54047.1;Pangolin_CoV_orf1ab;1733 | 0 | 0 |
| MT072864.1;QIQ54047.1;Pangolin_CoV_orf1ab;1734 | 0 | 0 |
| MT072864.1;QIQ54047.1;Pangolin_CoV_orf1ab;1735 | 0 | 0 |
| MT072864.1;QIQ54047.1;Pangolin_CoV_orf1ab;1736 | 0 | 0 |
| MT072864.1;QIQ54047.1;Pangolin_CoV_orf1ab;1737 | 0 | 0.075 |
| MT072864.1;QIQ54047.1;Pangolin_CoV_orf1ab;1738 | 0 | 0.025 |
| MT072864.1;QIQ54047.1;Pangolin_CoV_orf1ab;1739 | 0 | 0.025 |
| MT072864.1;QIQ54047.1;Pangolin_CoV_orf1ab;1740 | 0.1 | 0.325 |
| MT072864.1;QIQ54047.1;Pangolin_CoV_orf1ab;1741 | 0.1 | 0.3 |
| MT072864.1;QIQ54047.1;Pangolin_CoV_orf1ab;1742 | 0.1 | 0.3 |
| MT072864.1;QIQ54047.1;Pangolin_CoV_orf1ab;1743 | 0.15 | 0.325 |
| MT072864.1;QIQ54047.1;Pangolin_CoV_orf1ab;1744 | 0.2 | 0.325 |
| MT072864.1;QIQ54047.1;Pangolin_CoV_orf1ab;1745 | 0.1 | 0.35 |
| MT072864.1;QIQ54047.1;Pangolin_CoV_orf1ab;1746 | 0.05 | 0.275 |
| MT072864.1;QIQ54047.1;Pangolin_CoV_orf1ab;1747 | 0.05 | 0.2 |
| MT072864.1;QIQ54047.1;Pangolin_CoV_orf1ab;1748 | 0.1 | 0.25 |
| MT072864.1;QIQ54047.1;Pangolin_CoV_orf1ab;1749 | 0.05 | 0.2 |
| MT072864.1;QIQ54047.1;Pangolin_CoV_orf1ab;1750 | 0 | 0.125 |
| MT072864.1;QIQ54047.1;Pangolin_CoV_orf1ab;1751 | 0 | 0 |
| MT072864.1;QIQ54047.1;Pangolin_CoV_orf1ab;1752 | 0 | 0 |
| MT072864.1;QIQ54047.1;Pangolin_CoV_orf1ab;1753 | 0.1 | 0 |

| | | |
|---|---|---|
| MT072864.1;QIQ54047.1;Pangolin_CoV_orf1ab;1754 | 0.15 | 0 |
| MT072864.1;QIQ54047.1;Pangolin_CoV_orf1ab;1755 | 0.15 | 0 |
| MT072864.1;QIQ54047.1;Pangolin_CoV_orf1ab;1756 | 0.2 | 0 |
| MT072864.1;QIQ54047.1;Pangolin_CoV_orf1ab;1757 | 0.1 | 0 |
| MT072864.1;QIQ54047.1;Pangolin_CoV_orf1ab;1758 | 0.1 | 0 |
| MT072864.1;QIQ54047.1;Pangolin_CoV_orf1ab;1759 | 0 | 0 |
| MT072864.1;QIQ54047.1;Pangolin_CoV_orf1ab;1760 | 0.05 | 0 |
| MT072864.1;QIQ54047.1;Pangolin_CoV_orf1ab;1761 | 0 | 0 |
| MT072864.1;QIQ54047.1;Pangolin_CoV_orf1ab;1762 | 0 | 0 |
| MT072864.1;QIQ54047.1;Pangolin_CoV_orf1ab;1763 | 0 | 0 |
| MT072864.1;QIQ54047.1;Pangolin_CoV_orf1ab;1764 | 0 | 0 |
| MT072864.1;QIQ54047.1;Pangolin_CoV_orf1ab;1765 | 0 | 0 |
| MT072864.1;QIQ54047.1;Pangolin_CoV_orf1ab;1766 | 0 | 0 |
| MT072864.1;QIQ54047.1;Pangolin_CoV_orf1ab;1767 | 0 | 0 |
| MT072864.1;QIQ54047.1;Pangolin_CoV_orf1ab;1768 | 0 | 0 |
| MT072864.1;QIQ54047.1;Pangolin_CoV_orf1ab;1769 | 0 | 0 |
| MT072864.1;QIQ54047.1;Pangolin_CoV_orf1ab;1770 | 0 | 0 |
| MT072864.1;QIQ54047.1;Pangolin_CoV_orf1ab;1771 | 0 | 0 |
| MT072864.1;QIQ54047.1;Pangolin_CoV_orf1ab;1772 | 0.05 | 0.025 |
| MT072864.1;QIQ54047.1;Pangolin_CoV_orf1ab;1773 | 0 | 0 |
| MT072864.1;QIQ54047.1;Pangolin_CoV_orf1ab;1774 | 0 | 0 |
| MT072864.1;QIQ54047.1;Pangolin_CoV_orf1ab;1775 | 0 | 0 |
| MT072864.1;QIQ54047.1;Pangolin_CoV_orf1ab;1776 | 0 | 0 |
| MT072864.1;QIQ54047.1;Pangolin_CoV_orf1ab;1777 | 0 | 0 |
| MT072864.1;QIQ54047.1;Pangolin_CoV_orf1ab;1778 | 0 | 0 |
| MT072864.1;QIQ54047.1;Pangolin_CoV_orf1ab;1779 | 0 | 0 |
| MT072864.1;QIQ54047.1;Pangolin_CoV_orf1ab;1780 | 0 | 0 |
| MT072864.1;QIQ54047.1;Pangolin_CoV_orf1ab;1781 | 0 | 0.025 |
| MT072864.1;QIQ54047.1;Pangolin_CoV_orf1ab;1782 | 0 | 0.025 |
| MT072864.1;QIQ54047.1;Pangolin_CoV_orf1ab;1783 | 0 | 0.025 |
| MT072864.1;QIQ54047.1;Pangolin_CoV_orf1ab;1784 | 0 | 0.025 |
| MT072864.1;QIQ54047.1;Pangolin_CoV_orf1ab;1785 | 0 | 0.025 |
| MT072864.1;QIQ54047.1;Pangolin_CoV_orf1ab;1786 | 0 | 0 |
| MT072864.1;QIQ54047.1;Pangolin_CoV_orf1ab;1787 | 0.05 | 0 |
| MT072864.1;QIQ54047.1;Pangolin_CoV_orf1ab;1788 | 0 | 0.025 |
| MT072864.1;QIQ54047.1;Pangolin_CoV_orf1ab;1789 | 0.1 | 0.025 |
| MT072864.1;QIQ54047.1;Pangolin_CoV_orf1ab;1790 | 0 | 0 |
| MT072864.1;QIQ54047.1;Pangolin_CoV_orf1ab;1791 | 0 | 0 |
| MT072864.1;QIQ54047.1;Pangolin_CoV_orf1ab;1792 | 0 | 0 |
| MT072864.1;QIQ54047.1;Pangolin_CoV_orf1ab;1793 | 0 | 0 |
| MT072864.1;QIQ54047.1;Pangolin_CoV_orf1ab;1794 | 0 | 0 |
| MT072864.1;QIQ54047.1;Pangolin_CoV_orf1ab;1795 | 0 | 0 |
| MT072864.1;QIQ54047.1;Pangolin_CoV_orf1ab;1796 | 0 | 0 |

| | | |
|---|---|---|
| MT072864.1;QIQ54047.1;Pangolin_CoV_orf1ab;1797 | 0 | 0 |
| MT072864.1;QIQ54047.1;Pangolin_CoV_orf1ab;1798 | 0 | 0 |
| MT072864.1;QIQ54047.1;Pangolin_CoV_orf1ab;1799 | 0 | 0.025 |
| MT072864.1;QIQ54047.1;Pangolin_CoV_orf1ab;1800 | 0 | 0 |
| MT072864.1;QIQ54047.1;Pangolin_CoV_orf1ab;1801 | 0 | 0 |
| MT072864.1;QIQ54047.1;Pangolin_CoV_orf1ab;1802 | 0 | 0 |
| MT072864.1;QIQ54047.1;Pangolin_CoV_orf1ab;1803 | 0 | 0 |
| MT072864.1;QIQ54047.1;Pangolin_CoV_orf1ab;1804 | 0 | 0 |
| MT072864.1;QIQ54047.1;Pangolin_CoV_orf1ab;1805 | 0 | 0 |
| MT072864.1;QIQ54047.1;Pangolin_CoV_orf1ab;1806 | 0 | 0 |
| MT072864.1;QIQ54047.1;Pangolin_CoV_orf1ab;1807 | 0 | 0 |
| MT072864.1;QIQ54047.1;Pangolin_CoV_orf1ab;1808 | 0 | 0 |
| MT072864.1;QIQ54047.1;Pangolin_CoV_orf1ab;1809 | 0 | 0 |
| MT072864.1;QIQ54047.1;Pangolin_CoV_orf1ab;1810 | 0 | 0 |
| MT072864.1;QIQ54047.1;Pangolin_CoV_orf1ab;1811 | 0 | 0 |
| MT072864.1;QIQ54047.1;Pangolin_CoV_orf1ab;1812 | 0 | 0 |
| MT072864.1;QIQ54047.1;Pangolin_CoV_orf1ab;1813 | 0 | 0 |
| MT072864.1;QIQ54047.1;Pangolin_CoV_orf1ab;1814 | 0 | 0 |
| MT072864.1;QIQ54047.1;Pangolin_CoV_orf1ab;1815 | 0 | 0 |
| MT072864.1;QIQ54047.1;Pangolin_CoV_orf1ab;1816 | 0 | 0 |
| MT072864.1;QIQ54047.1;Pangolin_CoV_orf1ab;1817 | 0 | 0 |
| MT072864.1;QIQ54047.1;Pangolin_CoV_orf1ab;1818 | 0 | 0.15 |
| MT072864.1;QIQ54047.1;Pangolin_CoV_orf1ab;1819 | 0 | 0 |
| MT072864.1;QIQ54047.1;Pangolin_CoV_orf1ab;1820 | 0 | 0 |
| MT072864.1;QIQ54047.1;Pangolin_CoV_orf1ab;1821 | 0 | 0 |
| MT072864.1;QIQ54047.1;Pangolin_CoV_orf1ab;1822 | 0 | 0 |
| MT072864.1;QIQ54047.1;Pangolin_CoV_orf1ab;1823 | 0.05 | 0 |
| MT072864.1;QIQ54047.1;Pangolin_CoV_orf1ab;1824 | 0 | 0 |
| MT072864.1;QIQ54047.1;Pangolin_CoV_orf1ab;1825 | 0 | 0 |
| MT072864.1;QIQ54047.1;Pangolin_CoV_orf1ab;1826 | 0 | 0 |
| MT072864.1;QIQ54047.1;Pangolin_CoV_orf1ab;1827 | 0 | 0 |
| MT072864.1;QIQ54047.1;Pangolin_CoV_orf1ab;1828 | 0 | 0 |
| MT072864.1;QIQ54047.1;Pangolin_CoV_orf1ab;1829 | 0 | 0 |
| MT072864.1;QIQ54047.1;Pangolin_CoV_orf1ab;1830 | 0 | 0 |
| MT072864.1;QIQ54047.1;Pangolin_CoV_orf1ab;1831 | 0 | 0 |
| MT072864.1;QIQ54047.1;Pangolin_CoV_orf1ab;1832 | 0.05 | 0 |
| MT072864.1;QIQ54047.1;Pangolin_CoV_orf1ab;1833 | 0 | 0 |
| MT072864.1;QIQ54047.1;Pangolin_CoV_orf1ab;1834 | 0 | 0 |
| MT072864.1;QIQ54047.1;Pangolin_CoV_orf1ab;1835 | 0 | 0 |
| MT072864.1;QIQ54047.1;Pangolin_CoV_orf1ab;1836 | 0.05 | 0 |
| MT072864.1;QIQ54047.1;Pangolin_CoV_orf1ab;1837 | 0.05 | 0.025 |
| MT072864.1;QIQ54047.1;Pangolin_CoV_orf1ab;1838 | 0.05 | 0.025 |
| MT072864.1;QIQ54047.1;Pangolin_CoV_orf1ab;1839 | 0.05 | 0.025 |

| | | |
|---|---|---|
| MT072864.1;QIQ54047.1;Pangolin_CoV_orf1ab;1840 | 0.05 | 0.025 |
| MT072864.1;QIQ54047.1;Pangolin_CoV_orf1ab;1841 | 0.05 | 0.05 |
| MT072864.1;QIQ54047.1;Pangolin_CoV_orf1ab;1842 | 0.05 | 0.075 |
| MT072864.1;QIQ54047.1;Pangolin_CoV_orf1ab;1843 | 0.05 | 0.05 |
| MT072864.1;QIQ54047.1;Pangolin_CoV_orf1ab;1844 | 0.05 | 0.025 |
| MT072864.1;QIQ54047.1;Pangolin_CoV_orf1ab;1845 | 0.05 | 0.05 |
| MT072864.1;QIQ54047.1;Pangolin_CoV_orf1ab;1846 | 0 | 0.05 |
| MT072864.1;QIQ54047.1;Pangolin_CoV_orf1ab;1847 | 0 | 0.1 |
| MT072864.1;QIQ54047.1;Pangolin_CoV_orf1ab;1848 | 0.05 | 0.125 |
| MT072864.1;QIQ54047.1;Pangolin_CoV_orf1ab;1849 | 0.05 | 0.1 |
| MT072864.1;QIQ54047.1;Pangolin_CoV_orf1ab;1850 | 0.05 | 0.075 |
| MT072864.1;QIQ54047.1;Pangolin_CoV_orf1ab;1851 | 0.1 | 0.1 |
| MT072864.1;QIQ54047.1;Pangolin_CoV_orf1ab;1852 | 0 | 0.125 |
| MT072864.1;QIQ54047.1;Pangolin_CoV_orf1ab;1853 | 0.2 | 0.125 |
| MT072864.1;QIQ54047.1;Pangolin_CoV_orf1ab;1854 | 0 | 0 |
| MT072864.1;QIQ54047.1;Pangolin_CoV_orf1ab;1855 | 0 | 0.025 |
| MT072864.1;QIQ54047.1;Pangolin_CoV_orf1ab;1856 | 0 | 0.025 |
| MT072864.1;QIQ54047.1;Pangolin_CoV_orf1ab;1857 | 0 | 0.025 |
| MT072864.1;QIQ54047.1;Pangolin_CoV_orf1ab;1858 | 0 | 0.025 |
| MT072864.1;QIQ54047.1;Pangolin_CoV_orf1ab;1859 | 0 | 0.025 |
| MT072864.1;QIQ54047.1;Pangolin_CoV_orf1ab;1860 | 0 | 0.025 |
| MT072864.1;QIQ54047.1;Pangolin_CoV_orf1ab;1861 | 0 | 0.025 |
| MT072864.1;QIQ54047.1;Pangolin_CoV_orf1ab;1862 | 0.1 | 0.025 |
| MT072864.1;QIQ54047.1;Pangolin_CoV_orf1ab;1863 | 0 | 0.025 |
| MT072864.1;QIQ54047.1;Pangolin_CoV_orf1ab;1864 | 0 | 0.025 |
| MT072864.1;QIQ54047.1;Pangolin_CoV_orf1ab;1865 | 0 | 0 |
| MT072864.1;QIQ54047.1;Pangolin_CoV_orf1ab;1866 | 0 | 0 |
| MT072864.1;QIQ54047.1;Pangolin_CoV_orf1ab;1867 | 0 | 0 |
| MT072864.1;QIQ54047.1;Pangolin_CoV_orf1ab;1868 | 0 | 0 |
| MT072864.1;QIQ54047.1;Pangolin_CoV_orf1ab;1869 | 0 | 0 |
| MT072864.1;QIQ54047.1;Pangolin_CoV_orf1ab;1870 | 0 | 0.025 |
| MT072864.1;QIQ54047.1;Pangolin_CoV_orf1ab;1871 | 0 | 0.025 |
| MT072864.1;QIQ54047.1;Pangolin_CoV_orf1ab;1872 | 0 | 0 |
| MT072864.1;QIQ54047.1;Pangolin_CoV_orf1ab;1873 | 0 | 0.025 |
| MT072864.1;QIQ54047.1;Pangolin_CoV_orf1ab;1874 | 0 | 0 |
| MT072864.1;QIQ54047.1;Pangolin_CoV_orf1ab;1875 | 0 | 0 |
| MT072864.1;QIQ54047.1;Pangolin_CoV_orf1ab;1876 | 0 | 0 |
| MT072864.1;QIQ54047.1;Pangolin_CoV_orf1ab;1877 | 0 | 0 |
| MT072864.1;QIQ54047.1;Pangolin_CoV_orf1ab;1878 | 0 | 0.025 |
| MT072864.1;QIQ54047.1;Pangolin_CoV_orf1ab;1879 | 0 | 0.075 |
| MT072864.1;QIQ54047.1;Pangolin_CoV_orf1ab;1880 | 0 | 0.125 |
| MT072864.1;QIQ54047.1;Pangolin_CoV_orf1ab;1881 | 0 | 0.15 |
| MT072864.1;QIQ54047.1;Pangolin_CoV_orf1ab;1882 | 0 | 0.15 |

| | | |
|---|---|---|
| MT072864.1;QIQ54047.1;Pangolin_CoV_orf1ab;1883 | 0 | 0.225 |
| MT072864.1;QIQ54047.1;Pangolin_CoV_orf1ab;1884 | 0 | 0.1 |
| MT072864.1;QIQ54047.1;Pangolin_CoV_orf1ab;1885 | 0 | 0.05 |
| MT072864.1;QIQ54047.1;Pangolin_CoV_orf1ab;1886 | 0.1 | 0.1 |
| MT072864.1;QIQ54047.1;Pangolin_CoV_orf1ab;1887 | 0.05 | 0.15 |
| MT072864.1;QIQ54047.1;Pangolin_CoV_orf1ab;1888 | 0 | 0.075 |
| MT072864.1;QIQ54047.1;Pangolin_CoV_orf1ab;1889 | 0.05 | 0.125 |
| MT072864.1;QIQ54047.1;Pangolin_CoV_orf1ab;1890 | 0.05 | 0.025 |
| MT072864.1;QIQ54047.1;Pangolin_CoV_orf1ab;1891 | 0.05 | 0.025 |
| MT072864.1;QIQ54047.1;Pangolin_CoV_orf1ab;1892 | 0.05 | 0.025 |
| MT072864.1;QIQ54047.1;Pangolin_CoV_orf1ab;1893 | 0.15 | 0.1 |
| MT072864.1;QIQ54047.1;Pangolin_CoV_orf1ab;1894 | 0.05 | 0.025 |
| MT072864.1;QIQ54047.1;Pangolin_CoV_orf1ab;1895 | 0.05 | 0 |
| MT072864.1;QIQ54047.1;Pangolin_CoV_orf1ab;1896 | 0.05 | 0 |
| MT072864.1;QIQ54047.1;Pangolin_CoV_orf1ab;1897 | 0.05 | 0.025 |
| MT072864.1;QIQ54047.1;Pangolin_CoV_orf1ab;1898 | 0.05 | 0 |
| MT072864.1;QIQ54047.1;Pangolin_CoV_orf1ab;1899 | 0.05 | 0 |
| MT072864.1;QIQ54047.1;Pangolin_CoV_orf1ab;1900 | 0 | 0 |
| MT072864.1;QIQ54047.1;Pangolin_CoV_orf1ab;1901 | 0 | 0 |
| MT072864.1;QIQ54047.1;Pangolin_CoV_orf1ab;1902 | 0.15 | 0.075 |
| MT072864.1;QIQ54047.1;Pangolin_CoV_orf1ab;1903 | 0 | 0.075 |
| MT072864.1;QIQ54047.1;Pangolin_CoV_orf1ab;1904 | 0 | 0.025 |
| MT072864.1;QIQ54047.1;Pangolin_CoV_orf1ab;1905 | 0 | 0.05 |
| MT072864.1;QIQ54047.1;Pangolin_CoV_orf1ab;1906 | 0.15 | 0.075 |
| MT072864.1;QIQ54047.1;Pangolin_CoV_orf1ab;1907 | 0.15 | 0.025 |
| MT072864.1;QIQ54047.1;Pangolin_CoV_orf1ab;1908 | 0.05 | 0 |
| MT072864.1;QIQ54047.1;Pangolin_CoV_orf1ab;1909 | 0.05 | 0 |
| MT072864.1;QIQ54047.1;Pangolin_CoV_orf1ab;1910 | 0 | 0 |
| MT072864.1;QIQ54047.1;Pangolin_CoV_orf1ab;1911 | 0.15 | 0.05 |
| MT072864.1;QIQ54047.1;Pangolin_CoV_orf1ab;1912 | 0.15 | 0.05 |
| MT072864.1;QIQ54047.1;Pangolin_CoV_orf1ab;1913 | 0.1 | 0.075 |
| MT072864.1;QIQ54047.1;Pangolin_CoV_orf1ab;1914 | 0.1 | 0 |
| MT072864.1;QIQ54047.1;Pangolin_CoV_orf1ab;1915 | 0.1 | 0.05 |
| MT072864.1;QIQ54047.1;Pangolin_CoV_orf1ab;1916 | 0.1 | 0 |
| MT072864.1;QIQ54047.1;Pangolin_CoV_orf1ab;1917 | 0.1 | 0 |
| MT072864.1;QIQ54047.1;Pangolin_CoV_orf1ab;1918 | 0.05 | 0 |
| MT072864.1;QIQ54047.1;Pangolin_CoV_orf1ab;1919 | 0.2 | 0.05 |
| MT072864.1;QIQ54047.1;Pangolin_CoV_orf1ab;1920 | 0.2 | 0.025 |
| MT072864.1;QIQ54047.1;Pangolin_CoV_orf1ab;1921 | 0.2 | 0.075 |
| MT072864.1;QIQ54047.1;Pangolin_CoV_orf1ab;1922 | 0.3 | 0.05 |
| MT072864.1;QIQ54047.1;Pangolin_CoV_orf1ab;1923 | 0.2 | 0 |
| MT072864.1;QIQ54047.1;Pangolin_CoV_orf1ab;1924 | 0.15 | 0 |
| MT072864.1;QIQ54047.1;Pangolin_CoV_orf1ab;1925 | 0.35 | 0.15 |

| | | |
|---|---|---|
| MT072864.1;QIQ54047.1;Pangolin_CoV_orf1ab;1926 | 0.15 | 0 |
| MT072864.1;QIQ54047.1;Pangolin_CoV_orf1ab;1927 | 0.1 | 0.05 |
| MT072864.1;QIQ54047.1;Pangolin_CoV_orf1ab;1928 | 0.15 | 0.075 |
| MT072864.1;QIQ54047.1;Pangolin_CoV_orf1ab;1929 | 0.05 | 0.025 |
| MT072864.1;QIQ54047.1;Pangolin_CoV_orf1ab;1930 | 0.1 | 0.025 |
| MT072864.1;QIQ54047.1;Pangolin_CoV_orf1ab;1931 | 0.1 | 0.025 |
| MT072864.1;QIQ54047.1;Pangolin_CoV_orf1ab;1932 | 0 | 0 |
| MT072864.1;QIQ54047.1;Pangolin_CoV_orf1ab;1933 | 0 | 0 |
| MT072864.1;QIQ54047.1;Pangolin_CoV_orf1ab;1934 | 0 | 0 |
| MT072864.1;QIQ54047.1;Pangolin_CoV_orf1ab;1935 | 0 | 0 |
| MT072864.1;QIQ54047.1;Pangolin_CoV_orf1ab;1936 | 0 | 0 |
| MT072864.1;QIQ54047.1;Pangolin_CoV_orf1ab;1937 | 0 | 0 |
| MT072864.1;QIQ54047.1;Pangolin_CoV_orf1ab;1938 | 0 | 0 |
| MT072864.1;QIQ54047.1;Pangolin_CoV_orf1ab;1939 | 0 | 0 |
| MT072864.1;QIQ54047.1;Pangolin_CoV_orf1ab;1940 | 0 | 0 |
| MT072864.1;QIQ54047.1;Pangolin_CoV_orf1ab;1941 | 0 | 0 |
| MT072864.1;QIQ54047.1;Pangolin_CoV_orf1ab;1942 | 0 | 0 |
| MT072864.1;QIQ54047.1;Pangolin_CoV_orf1ab;1943 | 0 | 0 |
| MT072864.1;QIQ54047.1;Pangolin_CoV_orf1ab;1944 | 0 | 0 |
| MT072864.1;QIQ54047.1;Pangolin_CoV_orf1ab;1945 | 0 | 0.025 |
| MT072864.1;QIQ54047.1;Pangolin_CoV_orf1ab;1946 | 0 | 0.05 |
| MT072864.1;QIQ54047.1;Pangolin_CoV_orf1ab;1947 | 0 | 0 |
| MT072864.1;QIQ54047.1;Pangolin_CoV_orf1ab;1948 | 0 | 0.025 |
| MT072864.1;QIQ54047.1;Pangolin_CoV_orf1ab;1949 | 0 | 0 |
| MT072864.1;QIQ54047.1;Pangolin_CoV_orf1ab;1950 | 0 | 0.025 |
| MT072864.1;QIQ54047.1;Pangolin_CoV_orf1ab;1951 | 0.05 | 0.025 |
| MT072864.1;QIQ54047.1;Pangolin_CoV_orf1ab;1952 | 0.05 | 0.05 |
| MT072864.1;QIQ54047.1;Pangolin_CoV_orf1ab;1953 | 0.05 | 0.075 |
| MT072864.1;QIQ54047.1;Pangolin_CoV_orf1ab;1954 | 0.05 | 0.075 |
| MT072864.1;QIQ54047.1;Pangolin_CoV_orf1ab;1955 | 0.05 | 0.025 |
| MT072864.1;QIQ54047.1;Pangolin_CoV_orf1ab;1956 | 0.05 | 0.025 |
| MT072864.1;QIQ54047.1;Pangolin_CoV_orf1ab;1957 | 0.05 | 0.075 |
| MT072864.1;QIQ54047.1;Pangolin_CoV_orf1ab;1958 | 0 | 0.025 |
| MT072864.1;QIQ54047.1;Pangolin_CoV_orf1ab;1959 | 0 | 0.025 |
| MT072864.1;QIQ54047.1;Pangolin_CoV_orf1ab;1960 | 0 | 0.05 |
| MT072864.1;QIQ54047.1;Pangolin_CoV_orf1ab;1961 | 0 | 0.075 |
| MT072864.1;QIQ54047.1;Pangolin_CoV_orf1ab;1962 | 0 | 0 |
| MT072864.1;QIQ54047.1;Pangolin_CoV_orf1ab;1963 | 0 | 0 |
| MT072864.1;QIQ54047.1;Pangolin_CoV_orf1ab;1964 | 0 | 0 |
| MT072864.1;QIQ54047.1;Pangolin_CoV_orf1ab;1965 | 0 | 0 |
| MT072864.1;QIQ54047.1;Pangolin_CoV_orf1ab;1966 | 0 | 0 |
| MT072864.1;QIQ54047.1;Pangolin_CoV_orf1ab;1967 | 0 | 0.025 |
| MT072864.1;QIQ54047.1;Pangolin_CoV_orf1ab;1968 | 0 | 0 |

| | | |
|---|---|---|
| MT072864.1;QIQ54047.1;Pangolin_CoV_orf1ab;1969 | 0 | 0 |
| MT072864.1;QIQ54047.1;Pangolin_CoV_orf1ab;1970 | 0 | 0 |
| MT072864.1;QIQ54047.1;Pangolin_CoV_orf1ab;1971 | 0 | 0 |
| MT072864.1;QIQ54047.1;Pangolin_CoV_orf1ab;1972 | 0 | 0 |
| MT072864.1;QIQ54047.1;Pangolin_CoV_orf1ab;1973 | 0 | 0 |
| MT072864.1;QIQ54047.1;Pangolin_CoV_orf1ab;1974 | 0 | 0 |
| MT072864.1;QIQ54047.1;Pangolin_CoV_orf1ab;1975 | 0 | 0 |
| MT072864.1;QIQ54047.1;Pangolin_CoV_orf1ab;1976 | 0.1 | 0.025 |
| MT072864.1;QIQ54047.1;Pangolin_CoV_orf1ab;1977 | 0.05 | 0 |
| MT072864.1;QIQ54047.1;Pangolin_CoV_orf1ab;1978 | 0.1 | 0 |
| MT072864.1;QIQ54047.1;Pangolin_CoV_orf1ab;1979 | 0.1 | 0.025 |
| MT072864.1;QIQ54047.1;Pangolin_CoV_orf1ab;1980 | 0.1 | 0.025 |
| MT072864.1;QIQ54047.1;Pangolin_CoV_orf1ab;1981 | 0.15 | 0.05 |
| MT072864.1;QIQ54047.1;Pangolin_CoV_orf1ab;1982 | 0.15 | 0.075 |
| MT072864.1;QIQ54047.1;Pangolin_CoV_orf1ab;1983 | 0.25 | 0.125 |
| MT072864.1;QIQ54047.1;Pangolin_CoV_orf1ab;1984 | 0.25 | 0.1 |
| MT072864.1;QIQ54047.1;Pangolin_CoV_orf1ab;1985 | 0.3 | 0.225 |
| MT072864.1;QIQ54047.1;Pangolin_CoV_orf1ab;1986 | 0.35 | 0.1 |
| MT072864.1;QIQ54047.1;Pangolin_CoV_orf1ab;1987 | 0.1 | 0.05 |
| MT072864.1;QIQ54047.1;Pangolin_CoV_orf1ab;1988 | 0 | 0.025 |
| MT072864.1;QIQ54047.1;Pangolin_CoV_orf1ab;1989 | 0 | 0.025 |
| MT072864.1;QIQ54047.1;Pangolin_CoV_orf1ab;1990 | 0 | 0.025 |
| MT072864.1;QIQ54047.1;Pangolin_CoV_orf1ab;1991 | 0 | 0 |
| MT072864.1;QIQ54047.1;Pangolin_CoV_orf1ab;1992 | 0 | 0 |
| MT072864.1;QIQ54047.1;Pangolin_CoV_orf1ab;1993 | 0 | 0 |
| MT072864.1;QIQ54047.1;Pangolin_CoV_orf1ab;1994 | 0 | 0 |
| MT072864.1;QIQ54047.1;Pangolin_CoV_orf1ab;1995 | 0 | 0 |
| MT072864.1;QIQ54047.1;Pangolin_CoV_orf1ab;1996 | 0 | 0 |
| MT072864.1;QIQ54047.1;Pangolin_CoV_orf1ab;1997 | 0 | 0 |
| MT072864.1;QIQ54047.1;Pangolin_CoV_orf1ab;1998 | 0 | 0 |
| MT072864.1;QIQ54047.1;Pangolin_CoV_orf1ab;1999 | 0 | 0 |
| MT072864.1;QIQ54047.1;Pangolin_CoV_orf1ab;2000 | 0 | 0 |
| MT072864.1;QIQ54047.1;Pangolin_CoV_orf1ab;2001 | 0 | 0.025 |
| MT072864.1;QIQ54047.1;Pangolin_CoV_orf1ab;2002 | 0 | 0 |
| MT072864.1;QIQ54047.1;Pangolin_CoV_orf1ab;2003 | 0.05 | 0.05 |
| MT072864.1;QIQ54047.1;Pangolin_CoV_orf1ab;2004 | 0 | 0.025 |
| MT072864.1;QIQ54047.1;Pangolin_CoV_orf1ab;2005 | 0 | 0.025 |
| MT072864.1;QIQ54047.1;Pangolin_CoV_orf1ab;2006 | 0.1 | 0.1 |
| MT072864.1;QIQ54047.1;Pangolin_CoV_orf1ab;2007 | 0.15 | 0.05 |
| MT072864.1;QIQ54047.1;Pangolin_CoV_orf1ab;2008 | 0.2 | 0.125 |
| MT072864.1;QIQ54047.1;Pangolin_CoV_orf1ab;2009 | 0.2 | 0.075 |
| MT072864.1;QIQ54047.1;Pangolin_CoV_orf1ab;2010 | 0.15 | 0.05 |
| MT072864.1;QIQ54047.1;Pangolin_CoV_orf1ab;2011 | 0.3 | 0.125 |

| | | |
|---|---|---|
| MT072864.1;QIQ54047.1;Pangolin_CoV_orf1ab;2012 | 0 | 0.025 |
| MT072864.1;QIQ54047.1;Pangolin_CoV_orf1ab;2013 | 0 | 0.025 |
| MT072864.1;QIQ54047.1;Pangolin_CoV_orf1ab;2014 | 0 | 0.025 |
| MT072864.1;QIQ54047.1;Pangolin_CoV_orf1ab;2015 | 0 | 0.025 |
| MT072864.1;QIQ54047.1;Pangolin_CoV_orf1ab;2016 | 0 | 0 |
| MT072864.1;QIQ54047.1;Pangolin_CoV_orf1ab;2017 | 0 | 0 |
| MT072864.1;QIQ54047.1;Pangolin_CoV_orf1ab;2018 | 0 | 0 |
| MT072864.1;QIQ54047.1;Pangolin_CoV_orf1ab;2019 | 0 | 0 |
| MT072864.1;QIQ54047.1;Pangolin_CoV_orf1ab;2020 | 0 | 0 |
| MT072864.1;QIQ54047.1;Pangolin_CoV_orf1ab;2021 | 0 | 0 |
| MT072864.1;QIQ54047.1;Pangolin_CoV_orf1ab;2022 | 0 | 0 |
| MT072864.1;QIQ54047.1;Pangolin_CoV_orf1ab;2023 | 0.05 | 0 |
| MT072864.1;QIQ54047.1;Pangolin_CoV_orf1ab;2024 | 0 | 0 |
| MT072864.1;QIQ54047.1;Pangolin_CoV_orf1ab;2025 | 0 | 0 |
| MT072864.1;QIQ54047.1;Pangolin_CoV_orf1ab;2026 | 0 | 0.025 |
| MT072864.1;QIQ54047.1;Pangolin_CoV_orf1ab;2027 | 0 | 0.025 |
| MT072864.1;QIQ54047.1;Pangolin_CoV_orf1ab;2028 | 0 | 0 |
| MT072864.1;QIQ54047.1;Pangolin_CoV_orf1ab;2029 | 0 | 0 |
| MT072864.1;QIQ54047.1;Pangolin_CoV_orf1ab;2030 | 0 | 0 |
| MT072864.1;QIQ54047.1;Pangolin_CoV_orf1ab;2031 | 0 | 0 |
| MT072864.1;QIQ54047.1;Pangolin_CoV_orf1ab;2032 | 0.05 | 0 |
| MT072864.1;QIQ54047.1;Pangolin_CoV_orf1ab;2033 | 0 | 0 |
| MT072864.1;QIQ54047.1;Pangolin_CoV_orf1ab;2034 | 0 | 0.025 |
| MT072864.1;QIQ54047.1;Pangolin_CoV_orf1ab;2035 | 0.05 | 0 |
| MT072864.1;QIQ54047.1;Pangolin_CoV_orf1ab;2036 | 0.05 | 0.05 |
| MT072864.1;QIQ54047.1;Pangolin_CoV_orf1ab;2037 | 0.05 | 0.025 |
| MT072864.1;QIQ54047.1;Pangolin_CoV_orf1ab;2038 | 0.05 | 0.025 |
| MT072864.1;QIQ54047.1;Pangolin_CoV_orf1ab;2039 | 0.05 | 0 |
| MT072864.1;QIQ54047.1;Pangolin_CoV_orf1ab;2040 | 0 | 0 |
| MT072864.1;QIQ54047.1;Pangolin_CoV_orf1ab;2041 | 0 | 0 |
| MT072864.1;QIQ54047.1;Pangolin_CoV_orf1ab;2042 | 0.05 | 0 |
| MT072864.1;QIQ54047.1;Pangolin_CoV_orf1ab;2043 | 0.05 | 0 |
| MT072864.1;QIQ54047.1;Pangolin_CoV_orf1ab;2044 | 0 | 0 |
| MT072864.1;QIQ54047.1;Pangolin_CoV_orf1ab;2045 | 0.05 | 0 |
| MT072864.1;QIQ54047.1;Pangolin_CoV_orf1ab;2046 | 0.05 | 0 |
| MT072864.1;QIQ54047.1;Pangolin_CoV_orf1ab;2047 | 0 | 0 |
| MT072864.1;QIQ54047.1;Pangolin_CoV_orf1ab;2048 | 0 | 0.025 |
| MT072864.1;QIQ54047.1;Pangolin_CoV_orf1ab;2049 | 0 | 0.025 |
| MT072864.1;QIQ54047.1;Pangolin_CoV_orf1ab;2050 | 0 | 0.025 |
| MT072864.1;QIQ54047.1;Pangolin_CoV_orf1ab;2051 | 0 | 0.025 |
| MT072864.1;QIQ54047.1;Pangolin_CoV_orf1ab;2052 | 0 | 0 |
| MT072864.1;QIQ54047.1;Pangolin_CoV_orf1ab;2053 | 0 | 0 |
| MT072864.1;QIQ54047.1;Pangolin_CoV_orf1ab;2054 | 0 | 0 |

| | | |
|---|---|---|
| MT072864.1;QIQ54047.1;Pangolin_CoV_orf1ab;2055 | 0 | 0.025 |
| MT072864.1;QIQ54047.1;Pangolin_CoV_orf1ab;2056 | 0 | 0.025 |
| MT072864.1;QIQ54047.1;Pangolin_CoV_orf1ab;2057 | 0 | 0 |
| MT072864.1;QIQ54047.1;Pangolin_CoV_orf1ab;2058 | 0 | 0 |
| MT072864.1;QIQ54047.1;Pangolin_CoV_orf1ab;2059 | 0 | 0 |
| MT072864.1;QIQ54047.1;Pangolin_CoV_orf1ab;2060 | 0 | 0 |
| MT072864.1;QIQ54047.1;Pangolin_CoV_orf1ab;2061 | 0.05 | 0.05 |
| MT072864.1;QIQ54047.1;Pangolin_CoV_orf1ab;2062 | 0 | 0.025 |
| MT072864.1;QIQ54047.1;Pangolin_CoV_orf1ab;2063 | 0 | 0.025 |
| MT072864.1;QIQ54047.1;Pangolin_CoV_orf1ab;2064 | 0 | 0 |
| MT072864.1;QIQ54047.1;Pangolin_CoV_orf1ab;2065 | 0 | 0.025 |
| MT072864.1;QIQ54047.1;Pangolin_CoV_orf1ab;2066 | 0 | 0 |
| MT072864.1;QIQ54047.1;Pangolin_CoV_orf1ab;2067 | 0 | 0 |
| MT072864.1;QIQ54047.1;Pangolin_CoV_orf1ab;2068 | 0 | 0 |
| MT072864.1;QIQ54047.1;Pangolin_CoV_orf1ab;2069 | 0 | 0 |
| MT072864.1;QIQ54047.1;Pangolin_CoV_orf1ab;2070 | 0 | 0 |
| MT072864.1;QIQ54047.1;Pangolin_CoV_orf1ab;2071 | 0 | 0 |
| MT072864.1;QIQ54047.1;Pangolin_CoV_orf1ab;2072 | 0 | 0.025 |
| MT072864.1;QIQ54047.1;Pangolin_CoV_orf1ab;2073 | 0 | 0.05 |
| MT072864.1;QIQ54047.1;Pangolin_CoV_orf1ab;2074 | 0 | 0 |
| MT072864.1;QIQ54047.1;Pangolin_CoV_orf1ab;2075 | 0 | 0.025 |
| MT072864.1;QIQ54047.1;Pangolin_CoV_orf1ab;2076 | 0 | 0 |
| MT072864.1;QIQ54047.1;Pangolin_CoV_orf1ab;2077 | 0 | 0 |
| MT072864.1;QIQ54047.1;Pangolin_CoV_orf1ab;2078 | 0 | 0 |
| MT072864.1;QIQ54047.1;Pangolin_CoV_orf1ab;2079 | 0 | 0 |
| MT072864.1;QIQ54047.1;Pangolin_CoV_orf1ab;2080 | 0 | 0 |
| MT072864.1;QIQ54047.1;Pangolin_CoV_orf1ab;2081 | 0 | 0 |
| MT072864.1;QIQ54047.1;Pangolin_CoV_orf1ab;2082 | 0 | 0 |
| MT072864.1;QIQ54047.1;Pangolin_CoV_orf1ab;2083 | 0.3 | 0.175 |
| MT072864.1;QIQ54047.1;Pangolin_CoV_orf1ab;2084 | 0.05 | 0.05 |
| MT072864.1;QIQ54047.1;Pangolin_CoV_orf1ab;2085 | 0 | 0.025 |
| MT072864.1;QIQ54047.1;Pangolin_CoV_orf1ab;2086 | 0.05 | 0 |
| MT072864.1;QIQ54047.1;Pangolin_CoV_orf1ab;2087 | 0 | 0 |
| MT072864.1;QIQ54047.1;Pangolin_CoV_orf1ab;2088 | 0 | 0.025 |
| MT072864.1;QIQ54047.1;Pangolin_CoV_orf1ab;2089 | 0.15 | 0.1 |
| MT072864.1;QIQ54047.1;Pangolin_CoV_orf1ab;2090 | 0.2 | 0.075 |
| MT072864.1;QIQ54047.1;Pangolin_CoV_orf1ab;2091 | 0.25 | 0.125 |
| MT072864.1;QIQ54047.1;Pangolin_CoV_orf1ab;2092 | 0.1 | 0.025 |
| MT072864.1;QIQ54047.1;Pangolin_CoV_orf1ab;2093 | 0.05 | 0 |
| MT072864.1;QIQ54047.1;Pangolin_CoV_orf1ab;2094 | 0 | 0 |
| MT072864.1;QIQ54047.1;Pangolin_CoV_orf1ab;2095 | 0 | 0 |
| MT072864.1;QIQ54047.1;Pangolin_CoV_orf1ab;2096 | 0 | 0 |
| MT072864.1;QIQ54047.1;Pangolin_CoV_orf1ab;2097 | 0 | 0 |

| | | |
|---|---|---|
| MT072864.1;QIQ54047.1;Pangolin_CoV_orf1ab;2098 | 0 | 0 |
| MT072864.1;QIQ54047.1;Pangolin_CoV_orf1ab;2099 | 0 | 0 |
| MT072864.1;QIQ54047.1;Pangolin_CoV_orf1ab;2100 | 0 | 0 |
| MT072864.1;QIQ54047.1;Pangolin_CoV_orf1ab;2101 | 0 | 0 |
| MT072864.1;QIQ54047.1;Pangolin_CoV_orf1ab;2102 | 0 | 0 |
| MT072864.1;QIQ54047.1;Pangolin_CoV_orf1ab;2103 | 0 | 0 |
| MT072864.1;QIQ54047.1;Pangolin_CoV_orf1ab;2104 | 0 | 0 |
| MT072864.1;QIQ54047.1;Pangolin_CoV_orf1ab;2105 | 0 | 0 |
| MT072864.1;QIQ54047.1;Pangolin_CoV_orf1ab;2106 | 0.05 | 0 |
| MT072864.1;QIQ54047.1;Pangolin_CoV_orf1ab;2107 | 0 | 0 |
| MT072864.1;QIQ54047.1;Pangolin_CoV_orf1ab;2108 | 0 | 0 |
| MT072864.1;QIQ54047.1;Pangolin_CoV_orf1ab;2109 | 0 | 0 |
| MT072864.1;QIQ54047.1;Pangolin_CoV_orf1ab;2110 | 0.05 | 0 |
| MT072864.1;QIQ54047.1;Pangolin_CoV_orf1ab;2111 | 0.05 | 0 |
| MT072864.1;QIQ54047.1;Pangolin_CoV_orf1ab;2112 | 0 | 0 |
| MT072864.1;QIQ54047.1;Pangolin_CoV_orf1ab;2113 | 0 | 0 |
| MT072864.1;QIQ54047.1;Pangolin_CoV_orf1ab;2114 | 0 | 0 |
| MT072864.1;QIQ54047.1;Pangolin_CoV_orf1ab;2115 | 0 | 0 |
| MT072864.1;QIQ54047.1;Pangolin_CoV_orf1ab;2116 | 0.05 | 0 |
| MT072864.1;QIQ54047.1;Pangolin_CoV_orf1ab;2117 | 0 | 0 |
| MT072864.1;QIQ54047.1;Pangolin_CoV_orf1ab;2118 | 0 | 0 |
| MT072864.1;QIQ54047.1;Pangolin_CoV_orf1ab;2119 | 0 | 0 |
| MT072864.1;QIQ54047.1;Pangolin_CoV_orf1ab;2120 | 0 | 0 |
| MT072864.1;QIQ54047.1;Pangolin_CoV_orf1ab;2121 | 0.05 | 0 |
| MT072864.1;QIQ54047.1;Pangolin_CoV_orf1ab;2122 | 0 | 0 |
| MT072864.1;QIQ54047.1;Pangolin_CoV_orf1ab;2123 | 0 | 0 |
| MT072864.1;QIQ54047.1;Pangolin_CoV_orf1ab;2124 | 0 | 0 |
| MT072864.1;QIQ54047.1;Pangolin_CoV_orf1ab;2125 | 0.05 | 0 |
| MT072864.1;QIQ54047.1;Pangolin_CoV_orf1ab;2126 | 0.05 | 0 |
| MT072864.1;QIQ54047.1;Pangolin_CoV_orf1ab;2127 | 0.05 | 0 |
| MT072864.1;QIQ54047.1;Pangolin_CoV_orf1ab;2128 | 0.05 | 0.025 |
| MT072864.1;QIQ54047.1;Pangolin_CoV_orf1ab;2129 | 0.05 | 0 |
| MT072864.1;QIQ54047.1;Pangolin_CoV_orf1ab;2130 | 0.05 | 0 |
| MT072864.1;QIQ54047.1;Pangolin_CoV_orf1ab;2131 | 0.1 | 0 |
| MT072864.1;QIQ54047.1;Pangolin_CoV_orf1ab;2132 | 0.15 | 0.05 |
| MT072864.1;QIQ54047.1;Pangolin_CoV_orf1ab;2133 | 0.1 | 0.075 |
| MT072864.1;QIQ54047.1;Pangolin_CoV_orf1ab;2134 | 0.05 | 0.05 |
| MT072864.1;QIQ54047.1;Pangolin_CoV_orf1ab;2135 | 0.15 | 0.075 |
| MT072864.1;QIQ54047.1;Pangolin_CoV_orf1ab;2136 | 0.05 | 0 |
| MT072864.1;QIQ54047.1;Pangolin_CoV_orf1ab;2137 | 0.05 | 0 |
| MT072864.1;QIQ54047.1;Pangolin_CoV_orf1ab;2138 | 0 | 0 |
| MT072864.1;QIQ54047.1;Pangolin_CoV_orf1ab;2139 | 0 | 0 |
| MT072864.1;QIQ54047.1;Pangolin_CoV_orf1ab;2140 | 0 | 0 |

| | | |
|---|---|---|
| MT072864.1;QIQ54047.1;Pangolin_CoV_orf1ab;2141 | 0 | 0 |
| MT072864.1;QIQ54047.1;Pangolin_CoV_orf1ab;2142 | 0 | 0 |
| MT072864.1;QIQ54047.1;Pangolin_CoV_orf1ab;2143 | 0 | 0 |
| MT072864.1;QIQ54047.1;Pangolin_CoV_orf1ab;2144 | 0 | 0 |
| MT072864.1;QIQ54047.1;Pangolin_CoV_orf1ab;2145 | 0 | 0 |
| MT072864.1;QIQ54047.1;Pangolin_CoV_orf1ab;2146 | 0 | 0 |
| MT072864.1;QIQ54047.1;Pangolin_CoV_orf1ab;2147 | 0 | 0 |
| MT072864.1;QIQ54047.1;Pangolin_CoV_orf1ab;2148 | 0 | 0 |
| MT072864.1;QIQ54047.1;Pangolin_CoV_orf1ab;2149 | 0 | 0 |
| MT072864.1;QIQ54047.1;Pangolin_CoV_orf1ab;2150 | 0 | 0 |
| MT072864.1;QIQ54047.1;Pangolin_CoV_orf1ab;2151 | 0 | 0 |
| MT072864.1;QIQ54047.1;Pangolin_CoV_orf1ab;2152 | 0 | 0 |
| MT072864.1;QIQ54047.1;Pangolin_CoV_orf1ab;2153 | 0 | 0 |
| MT072864.1;QIQ54047.1;Pangolin_CoV_orf1ab;2154 | 0 | 0 |
| MT072864.1;QIQ54047.1;Pangolin_CoV_orf1ab;2155 | 0 | 0 |
| MT072864.1;QIQ54047.1;Pangolin_CoV_orf1ab;2156 | 0 | 0 |
| MT072864.1;QIQ54047.1;Pangolin_CoV_orf1ab;2157 | 0 | 0 |
| MT072864.1;QIQ54047.1;Pangolin_CoV_orf1ab;2158 | 0 | 0 |
| MT072864.1;QIQ54047.1;Pangolin_CoV_orf1ab;2159 | 0 | 0 |
| MT072864.1;QIQ54047.1;Pangolin_CoV_orf1ab;2160 | 0 | 0 |
| MT072864.1;QIQ54047.1;Pangolin_CoV_orf1ab;2161 | 0 | 0 |
| MT072864.1;QIQ54047.1;Pangolin_CoV_orf1ab;2162 | 0 | 0 |
| MT072864.1;QIQ54047.1;Pangolin_CoV_orf1ab;2163 | 0 | 0 |
| MT072864.1;QIQ54047.1;Pangolin_CoV_orf1ab;2164 | 0 | 0 |
| MT072864.1;QIQ54047.1;Pangolin_CoV_orf1ab;2165 | 0 | 0 |
| MT072864.1;QIQ54047.1;Pangolin_CoV_orf1ab;2166 | 0 | 0 |
| MT072864.1;QIQ54047.1;Pangolin_CoV_orf1ab;2167 | 0 | 0 |
| MT072864.1;QIQ54047.1;Pangolin_CoV_orf1ab;2168 | 0 | 0 |
| MT072864.1;QIQ54047.1;Pangolin_CoV_orf1ab;2169 | 0 | 0 |
| MT072864.1;QIQ54047.1;Pangolin_CoV_orf1ab;2170 | 0 | 0 |
| MT072864.1;QIQ54047.1;Pangolin_CoV_orf1ab;2171 | 0 | 0 |
| MT072864.1;QIQ54047.1;Pangolin_CoV_orf1ab;2172 | 0.05 | 0 |
| MT072864.1;QIQ54047.1;Pangolin_CoV_orf1ab;2173 | 0.1 | 0 |
| MT072864.1;QIQ54047.1;Pangolin_CoV_orf1ab;2174 | 0 | 0 |
| MT072864.1;QIQ54047.1;Pangolin_CoV_orf1ab;2175 | 0 | 0.025 |
| MT072864.1;QIQ54047.1;Pangolin_CoV_orf1ab;2176 | 0 | 0 |
| MT072864.1;QIQ54047.1;Pangolin_CoV_orf1ab;2177 | 0.05 | 0 |
| MT072864.1;QIQ54047.1;Pangolin_CoV_orf1ab;2178 | 0.05 | 0 |
| MT072864.1;QIQ54047.1;Pangolin_CoV_orf1ab;2179 | 0 | 0 |
| MT072864.1;QIQ54047.1;Pangolin_CoV_orf1ab;2180 | 0 | 0 |
| MT072864.1;QIQ54047.1;Pangolin_CoV_orf1ab;2181 | 0 | 0 |
| MT072864.1;QIQ54047.1;Pangolin_CoV_orf1ab;2182 | 0 | 0 |
| MT072864.1;QIQ54047.1;Pangolin_CoV_orf1ab;2183 | 0 | 0 |

| | | |
|---|---|---|
| MT072864.1;QIQ54047.1;Pangolin_CoV_orf1ab;2184 | 0 | 0 |
| MT072864.1;QIQ54047.1;Pangolin_CoV_orf1ab;2185 | 0 | 0 |
| MT072864.1;QIQ54047.1;Pangolin_CoV_orf1ab;2186 | 0 | 0 |
| MT072864.1;QIQ54047.1;Pangolin_CoV_orf1ab;2187 | 0 | 0 |
| MT072864.1;QIQ54047.1;Pangolin_CoV_orf1ab;2188 | 0 | 0 |
| MT072864.1;QIQ54047.1;Pangolin_CoV_orf1ab;2189 | 0 | 0 |
| MT072864.1;QIQ54047.1;Pangolin_CoV_orf1ab;2190 | 0 | 0 |
| MT072864.1;QIQ54047.1;Pangolin_CoV_orf1ab;2191 | 0 | 0 |
| MT072864.1;QIQ54047.1;Pangolin_CoV_orf1ab;2192 | 0 | 0 |
| MT072864.1;QIQ54047.1;Pangolin_CoV_orf1ab;2193 | 0 | 0 |
| MT072864.1;QIQ54047.1;Pangolin_CoV_orf1ab;2194 | 0 | 0 |
| MT072864.1;QIQ54047.1;Pangolin_CoV_orf1ab;2195 | 0 | 0 |
| MT072864.1;QIQ54047.1;Pangolin_CoV_orf1ab;2196 | 0 | 0 |
| MT072864.1;QIQ54047.1;Pangolin_CoV_orf1ab;2197 | 0 | 0 |
| MT072864.1;QIQ54047.1;Pangolin_CoV_orf1ab;2198 | 0 | 0 |
| MT072864.1;QIQ54047.1;Pangolin_CoV_orf1ab;2199 | 0 | 0 |
| MT072864.1;QIQ54047.1;Pangolin_CoV_orf1ab;2200 | 0 | 0 |
| MT072864.1;QIQ54047.1;Pangolin_CoV_orf1ab;2201 | 0 | 0 |
| MT072864.1;QIQ54047.1;Pangolin_CoV_orf1ab;2202 | 0 | 0 |
| MT072864.1;QIQ54047.1;Pangolin_CoV_orf1ab;2203 | 0 | 0 |
| MT072864.1;QIQ54047.1;Pangolin_CoV_orf1ab;2204 | 0.05 | 0 |
| MT072864.1;QIQ54047.1;Pangolin_CoV_orf1ab;2205 | 0 | 0 |
| MT072864.1;QIQ54047.1;Pangolin_CoV_orf1ab;2206 | 0 | 0 |
| MT072864.1;QIQ54047.1;Pangolin_CoV_orf1ab;2207 | 0 | 0 |
| MT072864.1;QIQ54047.1;Pangolin_CoV_orf1ab;2208 | 0 | 0 |
| MT072864.1;QIQ54047.1;Pangolin_CoV_orf1ab;2209 | 0 | 0 |
| MT072864.1;QIQ54047.1;Pangolin_CoV_orf1ab;2210 | 0 | 0 |
| MT072864.1;QIQ54047.1;Pangolin_CoV_orf1ab;2211 | 0 | 0 |
| MT072864.1;QIQ54047.1;Pangolin_CoV_orf1ab;2212 | 0 | 0 |
| MT072864.1;QIQ54047.1;Pangolin_CoV_orf1ab;2213 | 0 | 0 |
| MT072864.1;QIQ54047.1;Pangolin_CoV_orf1ab;2214 | 0 | 0 |
| MT072864.1;QIQ54047.1;Pangolin_CoV_orf1ab;2215 | 0 | 0 |
| MT072864.1;QIQ54047.1;Pangolin_CoV_orf1ab;2216 | 0 | 0 |
| MT072864.1;QIQ54047.1;Pangolin_CoV_orf1ab;2217 | 0 | 0 |
| MT072864.1;QIQ54047.1;Pangolin_CoV_orf1ab;2218 | 0 | 0 |
| MT072864.1;QIQ54047.1;Pangolin_CoV_orf1ab;2219 | 0 | 0 |
| MT072864.1;QIQ54047.1;Pangolin_CoV_orf1ab;2220 | 0 | 0 |
| MT072864.1;QIQ54047.1;Pangolin_CoV_orf1ab;2221 | 0 | 0 |
| MT072864.1;QIQ54047.1;Pangolin_CoV_orf1ab;2222 | 0 | 0 |
| MT072864.1;QIQ54047.1;Pangolin_CoV_orf1ab;2223 | 0 | 0 |
| MT072864.1;QIQ54047.1;Pangolin_CoV_orf1ab;2224 | 0 | 0 |
| MT072864.1;QIQ54047.1;Pangolin_CoV_orf1ab;2225 | 0 | 0 |
| MT072864.1;QIQ54047.1;Pangolin_CoV_orf1ab;2226 | 0.05 | 0 |

| | | |
|---|---|---|
| MT072864.1;QIQ54047.1;Pangolin_CoV_orf1ab;2227 | 0 | 0 |
| MT072864.1;QIQ54047.1;Pangolin_CoV_orf1ab;2228 | 0 | 0 |
| MT072864.1;QIQ54047.1;Pangolin_CoV_orf1ab;2229 | 0 | 0 |
| MT072864.1;QIQ54047.1;Pangolin_CoV_orf1ab;2230 | 0 | 0 |
| MT072864.1;QIQ54047.1;Pangolin_CoV_orf1ab;2231 | 0 | 0 |
| MT072864.1;QIQ54047.1;Pangolin_CoV_orf1ab;2232 | 0 | 0 |
| MT072864.1;QIQ54047.1;Pangolin_CoV_orf1ab;2233 | 0 | 0 |
| MT072864.1;QIQ54047.1;Pangolin_CoV_orf1ab;2234 | 0 | 0 |
| MT072864.1;QIQ54047.1;Pangolin_CoV_orf1ab;2235 | 0 | 0 |
| MT072864.1;QIQ54047.1;Pangolin_CoV_orf1ab;2236 | 0 | 0 |
| MT072864.1;QIQ54047.1;Pangolin_CoV_orf1ab;2237 | 0.05 | 0 |
| MT072864.1;QIQ54047.1;Pangolin_CoV_orf1ab;2238 | 0 | 0 |
| MT072864.1;QIQ54047.1;Pangolin_CoV_orf1ab;2239 | 0 | 0 |
| MT072864.1;QIQ54047.1;Pangolin_CoV_orf1ab;2240 | 0 | 0.025 |
| MT072864.1;QIQ54047.1;Pangolin_CoV_orf1ab;2241 | 0 | 0.025 |
| MT072864.1;QIQ54047.1;Pangolin_CoV_orf1ab;2242 | 0 | 0 |
| MT072864.1;QIQ54047.1;Pangolin_CoV_orf1ab;2243 | 0 | 0 |
| MT072864.1;QIQ54047.1;Pangolin_CoV_orf1ab;2244 | 0 | 0 |
| MT072864.1;QIQ54047.1;Pangolin_CoV_orf1ab;2245 | 0 | 0 |
| MT072864.1;QIQ54047.1;Pangolin_CoV_orf1ab;2246 | 0 | 0 |
| MT072864.1;QIQ54047.1;Pangolin_CoV_orf1ab;2247 | 0 | 0 |
| MT072864.1;QIQ54047.1;Pangolin_CoV_orf1ab;2248 | 0 | 0 |
| MT072864.1;QIQ54047.1;Pangolin_CoV_orf1ab;2249 | 0 | 0 |
| MT072864.1;QIQ54047.1;Pangolin_CoV_orf1ab;2250 | 0 | 0.05 |
| MT072864.1;QIQ54047.1;Pangolin_CoV_orf1ab;2251 | 0.05 | 0 |
| MT072864.1;QIQ54047.1;Pangolin_CoV_orf1ab;2252 | 0.2 | 0.05 |
| MT072864.1;QIQ54047.1;Pangolin_CoV_orf1ab;2253 | 0.25 | 0.175 |
| MT072864.1;QIQ54047.1;Pangolin_CoV_orf1ab;2254 | 0.3 | 0.25 |
| MT072864.1;QIQ54047.1;Pangolin_CoV_orf1ab;2255 | 0.5 | 0.35 |
| MT072864.1;QIQ54047.1;Pangolin_CoV_orf1ab;2256 | 0.35 | 0.35 |
| MT072864.1;QIQ54047.1;Pangolin_CoV_orf1ab;2257 | 0.55 | 0.375 |
| MT072864.1;QIQ54047.1;Pangolin_CoV_orf1ab;2258 | 0.4 | 0.325 |
| MT072864.1;QIQ54047.1;Pangolin_CoV_orf1ab;2259 | 0.2 | 0.15 |
| MT072864.1;QIQ54047.1;Pangolin_CoV_orf1ab;2260 | 0.2 | 0.2 |
| MT072864.1;QIQ54047.1;Pangolin_CoV_orf1ab;2261 | 0.2 | 0.05 |
| MT072864.1;QIQ54047.1;Pangolin_CoV_orf1ab;2262 | 0.2 | 0.05 |
| MT072864.1;QIQ54047.1;Pangolin_CoV_orf1ab;2263 | 0.05 | 0.025 |
| MT072864.1;QIQ54047.1;Pangolin_CoV_orf1ab;2264 | 0 | 0 |
| MT072864.1;QIQ54047.1;Pangolin_CoV_orf1ab;2265 | 0.05 | 0 |
| MT072864.1;QIQ54047.1;Pangolin_CoV_orf1ab;2266 | 0.05 | 0 |
| MT072864.1;QIQ54047.1;Pangolin_CoV_orf1ab;2267 | 0 | 0 |
| MT072864.1;QIQ54047.1;Pangolin_CoV_orf1ab;2268 | 0 | 0 |
| MT072864.1;QIQ54047.1;Pangolin_CoV_orf1ab;2269 | 0 | 0 |

| | | |
|---|---|---|
| MT072864.1;QIQ54047.1;Pangolin_CoV_orf1ab;2270 | 0 | 0 |
| MT072864.1;QIQ54047.1;Pangolin_CoV_orf1ab;2271 | 0 | 0 |
| MT072864.1;QIQ54047.1;Pangolin_CoV_orf1ab;2272 | 0 | 0 |
| MT072864.1;QIQ54047.1;Pangolin_CoV_orf1ab;2273 | 0 | 0 |
| MT072864.1;QIQ54047.1;Pangolin_CoV_orf1ab;2274 | 0 | 0 |
| MT072864.1;QIQ54047.1;Pangolin_CoV_orf1ab;2275 | 0 | 0 |
| MT072864.1;QIQ54047.1;Pangolin_CoV_orf1ab;2276 | 0 | 0 |
| MT072864.1;QIQ54047.1;Pangolin_CoV_orf1ab;2277 | 0 | 0 |
| MT072864.1;QIQ54047.1;Pangolin_CoV_orf1ab;2278 | 0 | 0.025 |
| MT072864.1;QIQ54047.1;Pangolin_CoV_orf1ab;2279 | 0 | 0 |
| MT072864.1;QIQ54047.1;Pangolin_CoV_orf1ab;2280 | 0 | 0.025 |
| MT072864.1;QIQ54047.1;Pangolin_CoV_orf1ab;2281 | 0 | 0 |
| MT072864.1;QIQ54047.1;Pangolin_CoV_orf1ab;2282 | 0 | 0 |
| MT072864.1;QIQ54047.1;Pangolin_CoV_orf1ab;2283 | 0.05 | 0 |
| MT072864.1;QIQ54047.1;Pangolin_CoV_orf1ab;2284 | 0 | 0 |
| MT072864.1;QIQ54047.1;Pangolin_CoV_orf1ab;2285 | 0 | 0 |
| MT072864.1;QIQ54047.1;Pangolin_CoV_orf1ab;2286 | 0.05 | 0 |
| MT072864.1;QIQ54047.1;Pangolin_CoV_orf1ab;2287 | 0.05 | 0 |
| MT072864.1;QIQ54047.1;Pangolin_CoV_orf1ab;2288 | 0 | 0 |
| MT072864.1;QIQ54047.1;Pangolin_CoV_orf1ab;2289 | 0.05 | 0 |
| MT072864.1;QIQ54047.1;Pangolin_CoV_orf1ab;2290 | 0 | 0 |
| MT072864.1;QIQ54047.1;Pangolin_CoV_orf1ab;2291 | 0 | 0 |
| MT072864.1;QIQ54047.1;Pangolin_CoV_orf1ab;2292 | 0 | 0 |
| MT072864.1;QIQ54047.1;Pangolin_CoV_orf1ab;2293 | 0 | 0 |
| MT072864.1;QIQ54047.1;Pangolin_CoV_orf1ab;2294 | 0 | 0 |
| MT072864.1;QIQ54047.1;Pangolin_CoV_orf1ab;2295 | 0 | 0 |
| MT072864.1;QIQ54047.1;Pangolin_CoV_orf1ab;2296 | 0 | 0 |
| MT072864.1;QIQ54047.1;Pangolin_CoV_orf1ab;2297 | 0 | 0 |
| MT072864.1;QIQ54047.1;Pangolin_CoV_orf1ab;2298 | 0 | 0 |
| MT072864.1;QIQ54047.1;Pangolin_CoV_orf1ab;2299 | 0 | 0 |
| MT072864.1;QIQ54047.1;Pangolin_CoV_orf1ab;2300 | 0 | 0 |
| MT072864.1;QIQ54047.1;Pangolin_CoV_orf1ab;2301 | 0 | 0 |
| MT072864.1;QIQ54047.1;Pangolin_CoV_orf1ab;2302 | 0 | 0 |
| MT072864.1;QIQ54047.1;Pangolin_CoV_orf1ab;2303 | 0 | 0 |
| MT072864.1;QIQ54047.1;Pangolin_CoV_orf1ab;2304 | 0 | 0 |
| MT072864.1;QIQ54047.1;Pangolin_CoV_orf1ab;2305 | 0 | 0 |
| MT072864.1;QIQ54047.1;Pangolin_CoV_orf1ab;2306 | 0 | 0 |
| MT072864.1;QIQ54047.1;Pangolin_CoV_orf1ab;2307 | 0 | 0 |
| MT072864.1;QIQ54047.1;Pangolin_CoV_orf1ab;2308 | 0 | 0 |
| MT072864.1;QIQ54047.1;Pangolin_CoV_orf1ab;2309 | 0 | 0 |
| MT072864.1;QIQ54047.1;Pangolin_CoV_orf1ab;2310 | 0 | 0 |
| MT072864.1;QIQ54047.1;Pangolin_CoV_orf1ab;2311 | 0 | 0 |
| MT072864.1;QIQ54047.1;Pangolin_CoV_orf1ab;2312 | 0 | 0 |

| | | |
|---|---|---|
| MT072864.1;QIQ54047.1;Pangolin_CoV_orf1ab;2313 | 0 | 0 |
| MT072864.1;QIQ54047.1;Pangolin_CoV_orf1ab;2314 | 0 | 0 |
| MT072864.1;QIQ54047.1;Pangolin_CoV_orf1ab;2315 | 0 | 0 |
| MT072864.1;QIQ54047.1;Pangolin_CoV_orf1ab;2316 | 0 | 0 |
| MT072864.1;QIQ54047.1;Pangolin_CoV_orf1ab;2317 | 0 | 0 |
| MT072864.1;QIQ54047.1;Pangolin_CoV_orf1ab;2318 | 0 | 0 |
| MT072864.1;QIQ54047.1;Pangolin_CoV_orf1ab;2319 | 0 | 0 |
| MT072864.1;QIQ54047.1;Pangolin_CoV_orf1ab;2320 | 0 | 0 |
| MT072864.1;QIQ54047.1;Pangolin_CoV_orf1ab;2321 | 0 | 0 |
| MT072864.1;QIQ54047.1;Pangolin_CoV_orf1ab;2322 | 0 | 0 |
| MT072864.1;QIQ54047.1;Pangolin_CoV_orf1ab;2323 | 0 | 0 |
| MT072864.1;QIQ54047.1;Pangolin_CoV_orf1ab;2324 | 0 | 0 |
| MT072864.1;QIQ54047.1;Pangolin_CoV_orf1ab;2325 | 0 | 0 |
| MT072864.1;QIQ54047.1;Pangolin_CoV_orf1ab;2326 | 0 | 0 |
| MT072864.1;QIQ54047.1;Pangolin_CoV_orf1ab;2327 | 0 | 0 |
| MT072864.1;QIQ54047.1;Pangolin_CoV_orf1ab;2328 | 0 | 0 |
| MT072864.1;QIQ54047.1;Pangolin_CoV_orf1ab;2329 | 0 | 0 |
| MT072864.1;QIQ54047.1;Pangolin_CoV_orf1ab;2330 | 0 | 0 |
| MT072864.1;QIQ54047.1;Pangolin_CoV_orf1ab;2331 | 0 | 0 |
| MT072864.1;QIQ54047.1;Pangolin_CoV_orf1ab;2332 | 0 | 0 |
| MT072864.1;QIQ54047.1;Pangolin_CoV_orf1ab;2333 | 0 | 0 |
| MT072864.1;QIQ54047.1;Pangolin_CoV_orf1ab;2334 | 0 | 0 |
| MT072864.1;QIQ54047.1;Pangolin_CoV_orf1ab;2335 | 0 | 0 |
| MT072864.1;QIQ54047.1;Pangolin_CoV_orf1ab;2336 | 0 | 0 |
| MT072864.1;QIQ54047.1;Pangolin_CoV_orf1ab;2337 | 0 | 0 |
| MT072864.1;QIQ54047.1;Pangolin_CoV_orf1ab;2338 | 0 | 0 |
| MT072864.1;QIQ54047.1;Pangolin_CoV_orf1ab;2339 | 0 | 0 |
| MT072864.1;QIQ54047.1;Pangolin_CoV_orf1ab;2340 | 0 | 0 |
| MT072864.1;QIQ54047.1;Pangolin_CoV_orf1ab;2341 | 0 | 0 |
| MT072864.1;QIQ54047.1;Pangolin_CoV_orf1ab;2342 | 0 | 0 |
| MT072864.1;QIQ54047.1;Pangolin_CoV_orf1ab;2343 | 0 | 0 |
| MT072864.1;QIQ54047.1;Pangolin_CoV_orf1ab;2344 | 0 | 0 |
| MT072864.1;QIQ54047.1;Pangolin_CoV_orf1ab;2345 | 0 | 0 |
| MT072864.1;QIQ54047.1;Pangolin_CoV_orf1ab;2346 | 0 | 0 |
| MT072864.1;QIQ54047.1;Pangolin_CoV_orf1ab;2347 | 0 | 0 |
| MT072864.1;QIQ54047.1;Pangolin_CoV_orf1ab;2348 | 0 | 0 |
| MT072864.1;QIQ54047.1;Pangolin_CoV_orf1ab;2349 | 0 | 0 |
| MT072864.1;QIQ54047.1;Pangolin_CoV_orf1ab;2350 | 0 | 0 |
| MT072864.1;QIQ54047.1;Pangolin_CoV_orf1ab;2351 | 0 | 0 |
| MT072864.1;QIQ54047.1;Pangolin_CoV_orf1ab;2352 | 0 | 0 |
| MT072864.1;QIQ54047.1;Pangolin_CoV_orf1ab;2353 | 0 | 0 |
| MT072864.1;QIQ54047.1;Pangolin_CoV_orf1ab;2354 | 0 | 0 |
| MT072864.1;QIQ54047.1;Pangolin_CoV_orf1ab;2355 | 0 | 0 |

| | | |
|---|---|---|
| MT072864.1;QIQ54047.1;Pangolin_CoV_orf1ab;2356 | 0 | 0 |
| MT072864.1;QIQ54047.1;Pangolin_CoV_orf1ab;2357 | 0 | 0 |
| MT072864.1;QIQ54047.1;Pangolin_CoV_orf1ab;2358 | 0 | 0 |
| MT072864.1;QIQ54047.1;Pangolin_CoV_orf1ab;2359 | 0 | 0 |
| MT072864.1;QIQ54047.1;Pangolin_CoV_orf1ab;2360 | 0 | 0 |
| MT072864.1;QIQ54047.1;Pangolin_CoV_orf1ab;2361 | 0 | 0 |
| MT072864.1;QIQ54047.1;Pangolin_CoV_orf1ab;2362 | 0 | 0 |
| MT072864.1;QIQ54047.1;Pangolin_CoV_orf1ab;2363 | 0 | 0 |
| MT072864.1;QIQ54047.1;Pangolin_CoV_orf1ab;2364 | 0 | 0 |
| MT072864.1;QIQ54047.1;Pangolin_CoV_orf1ab;2365 | 0 | 0 |
| MT072864.1;QIQ54047.1;Pangolin_CoV_orf1ab;2366 | 0 | 0 |
| MT072864.1;QIQ54047.1;Pangolin_CoV_orf1ab;2367 | 0 | 0 |
| MT072864.1;QIQ54047.1;Pangolin_CoV_orf1ab;2368 | 0 | 0 |
| MT072864.1;QIQ54047.1;Pangolin_CoV_orf1ab;2369 | 0 | 0 |
| MT072864.1;QIQ54047.1;Pangolin_CoV_orf1ab;2370 | 0 | 0 |
| MT072864.1;QIQ54047.1;Pangolin_CoV_orf1ab;2371 | 0 | 0 |
| MT072864.1;QIQ54047.1;Pangolin_CoV_orf1ab;2372 | 0 | 0 |
| MT072864.1;QIQ54047.1;Pangolin_CoV_orf1ab;2373 | 0 | 0 |
| MT072864.1;QIQ54047.1;Pangolin_CoV_orf1ab;2374 | 0 | 0 |
| MT072864.1;QIQ54047.1;Pangolin_CoV_orf1ab;2375 | 0 | 0 |
| MT072864.1;QIQ54047.1;Pangolin_CoV_orf1ab;2376 | 0 | 0 |
| MT072864.1;QIQ54047.1;Pangolin_CoV_orf1ab;2377 | 0 | 0 |
| MT072864.1;QIQ54047.1;Pangolin_CoV_orf1ab;2378 | 0 | 0 |
| MT072864.1;QIQ54047.1;Pangolin_CoV_orf1ab;2379 | 0 | 0 |
| MT072864.1;QIQ54047.1;Pangolin_CoV_orf1ab;2380 | 0 | 0 |
| MT072864.1;QIQ54047.1;Pangolin_CoV_orf1ab;2381 | 0 | 0 |
| MT072864.1;QIQ54047.1;Pangolin_CoV_orf1ab;2382 | 0 | 0 |
| MT072864.1;QIQ54047.1;Pangolin_CoV_orf1ab;2383 | 0 | 0 |
| MT072864.1;QIQ54047.1;Pangolin_CoV_orf1ab;2384 | 0 | 0 |
| MT072864.1;QIQ54047.1;Pangolin_CoV_orf1ab;2385 | 0 | 0 |
| MT072864.1;QIQ54047.1;Pangolin_CoV_orf1ab;2386 | 0 | 0 |
| MT072864.1;QIQ54047.1;Pangolin_CoV_orf1ab;2387 | 0 | 0 |
| MT072864.1;QIQ54047.1;Pangolin_CoV_orf1ab;2388 | 0 | 0.025 |
| MT072864.1;QIQ54047.1;Pangolin_CoV_orf1ab;2389 | 0 | 0 |
| MT072864.1;QIQ54047.1;Pangolin_CoV_orf1ab;2390 | 0 | 0 |
| MT072864.1;QIQ54047.1;Pangolin_CoV_orf1ab;2391 | 0 | 0 |
| MT072864.1;QIQ54047.1;Pangolin_CoV_orf1ab;2392 | 0 | 0 |
| MT072864.1;QIQ54047.1;Pangolin_CoV_orf1ab;2393 | 0 | 0 |
| MT072864.1;QIQ54047.1;Pangolin_CoV_orf1ab;2394 | 0 | 0 |
| MT072864.1;QIQ54047.1;Pangolin_CoV_orf1ab;2395 | 0 | 0 |
| MT072864.1;QIQ54047.1;Pangolin_CoV_orf1ab;2396 | 0 | 0 |
| MT072864.1;QIQ54047.1;Pangolin_CoV_orf1ab;2397 | 0 | 0 |
| MT072864.1;QIQ54047.1;Pangolin_CoV_orf1ab;2398 | 0 | 0 |

| | | |
|---|---|---|
| MT072864.1;QIQ54047.1;Pangolin_CoV_orf1ab;2399 | 0 | 0 |
| MT072864.1;QIQ54047.1;Pangolin_CoV_orf1ab;2400 | 0 | 0 |
| MT072864.1;QIQ54047.1;Pangolin_CoV_orf1ab;2401 | 0 | 0 |
| MT072864.1;QIQ54047.1;Pangolin_CoV_orf1ab;2402 | 0 | 0 |
| MT072864.1;QIQ54047.1;Pangolin_CoV_orf1ab;2403 | 0 | 0 |
| MT072864.1;QIQ54047.1;Pangolin_CoV_orf1ab;2404 | 0 | 0 |
| MT072864.1;QIQ54047.1;Pangolin_CoV_orf1ab;2405 | 0 | 0 |
| MT072864.1;QIQ54047.1;Pangolin_CoV_orf1ab;2406 | 0 | 0 |
| MT072864.1;QIQ54047.1;Pangolin_CoV_orf1ab;2407 | 0 | 0 |
| MT072864.1;QIQ54047.1;Pangolin_CoV_orf1ab;2408 | 0 | 0.025 |
| MT072864.1;QIQ54047.1;Pangolin_CoV_orf1ab;2409 | 0 | 0 |
| MT072864.1;QIQ54047.1;Pangolin_CoV_orf1ab;2410 | 0 | 0 |
| MT072864.1;QIQ54047.1;Pangolin_CoV_orf1ab;2411 | 0 | 0 |
| MT072864.1;QIQ54047.1;Pangolin_CoV_orf1ab;2412 | 0 | 0 |
| MT072864.1;QIQ54047.1;Pangolin_CoV_orf1ab;2413 | 0 | 0 |
| MT072864.1;QIQ54047.1;Pangolin_CoV_orf1ab;2414 | 0 | 0 |
| MT072864.1;QIQ54047.1;Pangolin_CoV_orf1ab;2415 | 0 | 0 |
| MT072864.1;QIQ54047.1;Pangolin_CoV_orf1ab;2416 | 0 | 0 |
| MT072864.1;QIQ54047.1;Pangolin_CoV_orf1ab;2417 | 0 | 0 |
| MT072864.1;QIQ54047.1;Pangolin_CoV_orf1ab;2418 | 0 | 0 |
| MT072864.1;QIQ54047.1;Pangolin_CoV_orf1ab;2419 | 0 | 0 |
| MT072864.1;QIQ54047.1;Pangolin_CoV_orf1ab;2420 | 0 | 0 |
| MT072864.1;QIQ54047.1;Pangolin_CoV_orf1ab;2421 | 0 | 0 |
| MT072864.1;QIQ54047.1;Pangolin_CoV_orf1ab;2422 | 0 | 0 |
| MT072864.1;QIQ54047.1;Pangolin_CoV_orf1ab;2423 | 0 | 0 |
| MT072864.1;QIQ54047.1;Pangolin_CoV_orf1ab;2424 | 0 | 0 |
| MT072864.1;QIQ54047.1;Pangolin_CoV_orf1ab;2425 | 0 | 0 |
| MT072864.1;QIQ54047.1;Pangolin_CoV_orf1ab;2426 | 0 | 0 |
| MT072864.1;QIQ54047.1;Pangolin_CoV_orf1ab;2427 | 0 | 0 |
| MT072864.1;QIQ54047.1;Pangolin_CoV_orf1ab;2428 | 0 | 0 |
| MT072864.1;QIQ54047.1;Pangolin_CoV_orf1ab;2429 | 0 | 0 |
| MT072864.1;QIQ54047.1;Pangolin_CoV_orf1ab;2430 | 0 | 0 |
| MT072864.1;QIQ54047.1;Pangolin_CoV_orf1ab;2431 | 0 | 0 |
| MT072864.1;QIQ54047.1;Pangolin_CoV_orf1ab;2432 | 0 | 0 |
| MT072864.1;QIQ54047.1;Pangolin_CoV_orf1ab;2433 | 0 | 0.025 |
| MT072864.1;QIQ54047.1;Pangolin_CoV_orf1ab;2434 | 0 | 0.025 |
| MT072864.1;QIQ54047.1;Pangolin_CoV_orf1ab;2435 | 0 | 0 |
| MT072864.1;QIQ54047.1;Pangolin_CoV_orf1ab;2436 | 0 | 0 |
| MT072864.1;QIQ54047.1;Pangolin_CoV_orf1ab;2437 | 0 | 0 |
| MT072864.1;QIQ54047.1;Pangolin_CoV_orf1ab;2438 | 0 | 0 |
| MT072864.1;QIQ54047.1;Pangolin_CoV_orf1ab;2439 | 0 | 0 |
| MT072864.1;QIQ54047.1;Pangolin_CoV_orf1ab;2440 | 0 | 0 |
| MT072864.1;QIQ54047.1;Pangolin_CoV_orf1ab;2441 | 0 | 0 |

| | | |
|---|---|---|
| MT072864.1;QIQ54047.1;Pangolin_CoV_orf1ab;2442 | 0 | 0 |
| MT072864.1;QIQ54047.1;Pangolin_CoV_orf1ab;2443 | 0 | 0 |
| MT072864.1;QIQ54047.1;Pangolin_CoV_orf1ab;2444 | 0 | 0 |
| MT072864.1;QIQ54047.1;Pangolin_CoV_orf1ab;2445 | 0 | 0 |
| MT072864.1;QIQ54047.1;Pangolin_CoV_orf1ab;2446 | 0 | 0 |
| MT072864.1;QIQ54047.1;Pangolin_CoV_orf1ab;2447 | 0 | 0 |
| MT072864.1;QIQ54047.1;Pangolin_CoV_orf1ab;2448 | 0 | 0 |
| MT072864.1;QIQ54047.1;Pangolin_CoV_orf1ab;2449 | 0 | 0 |
| MT072864.1;QIQ54047.1;Pangolin_CoV_orf1ab;2450 | 0 | 0 |
| MT072864.1;QIQ54047.1;Pangolin_CoV_orf1ab;2451 | 0 | 0 |
| MT072864.1;QIQ54047.1;Pangolin_CoV_orf1ab;2452 | 0 | 0 |
| MT072864.1;QIQ54047.1;Pangolin_CoV_orf1ab;2453 | 0 | 0 |
| MT072864.1;QIQ54047.1;Pangolin_CoV_orf1ab;2454 | 0 | 0 |
| MT072864.1;QIQ54047.1;Pangolin_CoV_orf1ab;2455 | 0 | 0 |
| MT072864.1;QIQ54047.1;Pangolin_CoV_orf1ab;2456 | 0 | 0 |
| MT072864.1;QIQ54047.1;Pangolin_CoV_orf1ab;2457 | 0 | 0 |
| MT072864.1;QIQ54047.1;Pangolin_CoV_orf1ab;2458 | 0 | 0 |
| MT072864.1;QIQ54047.1;Pangolin_CoV_orf1ab;2459 | 0 | 0 |
| MT072864.1;QIQ54047.1;Pangolin_CoV_orf1ab;2460 | 0 | 0 |
| MT072864.1;QIQ54047.1;Pangolin_CoV_orf1ab;2461 | 0.05 | 0 |
| MT072864.1;QIQ54047.1;Pangolin_CoV_orf1ab;2462 | 0.05 | 0.025 |
| MT072864.1;QIQ54047.1;Pangolin_CoV_orf1ab;2463 | 0.05 | 0.025 |
| MT072864.1;QIQ54047.1;Pangolin_CoV_orf1ab;2464 | 0 | 0 |
| MT072864.1;QIQ54047.1;Pangolin_CoV_orf1ab;2465 | 0 | 0 |
| MT072864.1;QIQ54047.1;Pangolin_CoV_orf1ab;2466 | 0 | 0 |
| MT072864.1;QIQ54047.1;Pangolin_CoV_orf1ab;2467 | 0 | 0 |
| MT072864.1;QIQ54047.1;Pangolin_CoV_orf1ab;2468 | 0 | 0 |
| MT072864.1;QIQ54047.1;Pangolin_CoV_orf1ab;2469 | 0 | 0 |
| MT072864.1;QIQ54047.1;Pangolin_CoV_orf1ab;2470 | 0.05 | 0.075 |
| MT072864.1;QIQ54047.1;Pangolin_CoV_orf1ab;2471 | 0 | 0 |
| MT072864.1;QIQ54047.1;Pangolin_CoV_orf1ab;2472 | 0 | 0 |
| MT072864.1;QIQ54047.1;Pangolin_CoV_orf1ab;2473 | 0 | 0 |
| MT072864.1;QIQ54047.1;Pangolin_CoV_orf1ab;2474 | 0 | 0.025 |
| MT072864.1;QIQ54047.1;Pangolin_CoV_orf1ab;2475 | 0 | 0.025 |
| MT072864.1;QIQ54047.1;Pangolin_CoV_orf1ab;2476 | 0 | 0.05 |
| MT072864.1;QIQ54047.1;Pangolin_CoV_orf1ab;2477 | 0 | 0.025 |
| MT072864.1;QIQ54047.1;Pangolin_CoV_orf1ab;2478 | 0 | 0.025 |
| MT072864.1;QIQ54047.1;Pangolin_CoV_orf1ab;2479 | 0 | 0.025 |
| MT072864.1;QIQ54047.1;Pangolin_CoV_orf1ab;2480 | 0 | 0.025 |
| MT072864.1;QIQ54047.1;Pangolin_CoV_orf1ab;2481 | 0 | 0.025 |
| MT072864.1;QIQ54047.1;Pangolin_CoV_orf1ab;2482 | 0.05 | 0.025 |
| MT072864.1;QIQ54047.1;Pangolin_CoV_orf1ab;2483 | 0 | 0 |
| MT072864.1;QIQ54047.1;Pangolin_CoV_orf1ab;2484 | 0 | 0 |

| | | |
|---|---|---|
| MT072864.1;QIQ54047.1;Pangolin_CoV_orf1ab;2485 | 0 | 0 |
| MT072864.1;QIQ54047.1;Pangolin_CoV_orf1ab;2486 | 0 | 0 |
| MT072864.1;QIQ54047.1;Pangolin_CoV_orf1ab;2487 | 0 | 0 |
| MT072864.1;QIQ54047.1;Pangolin_CoV_orf1ab;2488 | 0 | 0 |
| MT072864.1;QIQ54047.1;Pangolin_CoV_orf1ab;2489 | 0 | 0 |
| MT072864.1;QIQ54047.1;Pangolin_CoV_orf1ab;2490 | 0 | 0 |
| MT072864.1;QIQ54047.1;Pangolin_CoV_orf1ab;2491 | 0 | 0 |
| MT072864.1;QIQ54047.1;Pangolin_CoV_orf1ab;2492 | 0 | 0 |
| MT072864.1;QIQ54047.1;Pangolin_CoV_orf1ab;2493 | 0 | 0 |
| MT072864.1;QIQ54047.1;Pangolin_CoV_orf1ab;2494 | 0 | 0 |
| MT072864.1;QIQ54047.1;Pangolin_CoV_orf1ab;2495 | 0 | 0 |
| MT072864.1;QIQ54047.1;Pangolin_CoV_orf1ab;2496 | 0.05 | 0.05 |
| MT072864.1;QIQ54047.1;Pangolin_CoV_orf1ab;2497 | 0 | 0 |
| MT072864.1;QIQ54047.1;Pangolin_CoV_orf1ab;2498 | 0 | 0 |
| MT072864.1;QIQ54047.1;Pangolin_CoV_orf1ab;2499 | 0 | 0 |
| MT072864.1;QIQ54047.1;Pangolin_CoV_orf1ab;2500 | 0 | 0 |
| MT072864.1;QIQ54047.1;Pangolin_CoV_orf1ab;2501 | 0 | 0 |
| MT072864.1;QIQ54047.1;Pangolin_CoV_orf1ab;2502 | 0 | 0 |
| MT072864.1;QIQ54047.1;Pangolin_CoV_orf1ab;2503 | 0 | 0 |
| MT072864.1;QIQ54047.1;Pangolin_CoV_orf1ab;2504 | 0 | 0 |
| MT072864.1;QIQ54047.1;Pangolin_CoV_orf1ab;2505 | 0 | 0 |
| MT072864.1;QIQ54047.1;Pangolin_CoV_orf1ab;2506 | 0 | 0 |
| MT072864.1;QIQ54047.1;Pangolin_CoV_orf1ab;2507 | 0 | 0 |
| MT072864.1;QIQ54047.1;Pangolin_CoV_orf1ab;2508 | 0 | 0 |
| MT072864.1;QIQ54047.1;Pangolin_CoV_orf1ab;2509 | 0 | 0 |
| MT072864.1;QIQ54047.1;Pangolin_CoV_orf1ab;2510 | 0 | 0 |
| MT072864.1;QIQ54047.1;Pangolin_CoV_orf1ab;2511 | 0 | 0 |
| MT072864.1;QIQ54047.1;Pangolin_CoV_orf1ab;2512 | 0 | 0 |
| MT072864.1;QIQ54047.1;Pangolin_CoV_orf1ab;2513 | 0 | 0 |
| MT072864.1;QIQ54047.1;Pangolin_CoV_orf1ab;2514 | 0 | 0.025 |
| MT072864.1;QIQ54047.1;Pangolin_CoV_orf1ab;2515 | 0.05 | 0.025 |
| MT072864.1;QIQ54047.1;Pangolin_CoV_orf1ab;2516 | 0.05 | 0.025 |
| MT072864.1;QIQ54047.1;Pangolin_CoV_orf1ab;2517 | 0.05 | 0.025 |
| MT072864.1;QIQ54047.1;Pangolin_CoV_orf1ab;2518 | 0 | 0.025 |
| MT072864.1;QIQ54047.1;Pangolin_CoV_orf1ab;2519 | 0 | 0 |
| MT072864.1;QIQ54047.1;Pangolin_CoV_orf1ab;2520 | 0.05 | 0 |
| MT072864.1;QIQ54047.1;Pangolin_CoV_orf1ab;2521 | 0 | 0 |
| MT072864.1;QIQ54047.1;Pangolin_CoV_orf1ab;2522 | 0 | 0 |
| MT072864.1;QIQ54047.1;Pangolin_CoV_orf1ab;2523 | 0 | 0 |
| MT072864.1;QIQ54047.1;Pangolin_CoV_orf1ab;2524 | 0 | 0 |
| MT072864.1;QIQ54047.1;Pangolin_CoV_orf1ab;2525 | 0 | 0.05 |
| MT072864.1;QIQ54047.1;Pangolin_CoV_orf1ab;2526 | 0 | 0.025 |
| MT072864.1;QIQ54047.1;Pangolin_CoV_orf1ab;2527 | 0 | 0.025 |

| | | |
|---|---|---|
| MT072864.1;QIQ54047.1;Pangolin_CoV_orf1ab;2528 | 0.05 | 0.025 |
| MT072864.1;QIQ54047.1;Pangolin_CoV_orf1ab;2529 | 0.05 | 0.025 |
| MT072864.1;QIQ54047.1;Pangolin_CoV_orf1ab;2530 | 0.05 | 0.025 |
| MT072864.1;QIQ54047.1;Pangolin_CoV_orf1ab;2531 | 0.05 | 0.025 |
| MT072864.1;QIQ54047.1;Pangolin_CoV_orf1ab;2532 | 0 | 0.025 |
| MT072864.1;QIQ54047.1;Pangolin_CoV_orf1ab;2533 | 0 | 0.025 |
| MT072864.1;QIQ54047.1;Pangolin_CoV_orf1ab;2534 | 0.05 | 0.05 |
| MT072864.1;QIQ54047.1;Pangolin_CoV_orf1ab;2535 | 0.05 | 0.025 |
| MT072864.1;QIQ54047.1;Pangolin_CoV_orf1ab;2536 | 0 | 0 |
| MT072864.1;QIQ54047.1;Pangolin_CoV_orf1ab;2537 | 0 | 0 |
| MT072864.1;QIQ54047.1;Pangolin_CoV_orf1ab;2538 | 0 | 0 |
| MT072864.1;QIQ54047.1;Pangolin_CoV_orf1ab;2539 | 0 | 0 |
| MT072864.1;QIQ54047.1;Pangolin_CoV_orf1ab;2540 | 0 | 0 |
| MT072864.1;QIQ54047.1;Pangolin_CoV_orf1ab;2541 | 0 | 0 |
| MT072864.1;QIQ54047.1;Pangolin_CoV_orf1ab;2542 | 0 | 0 |
| MT072864.1;QIQ54047.1;Pangolin_CoV_orf1ab;2543 | 0.1 | 0 |
| MT072864.1;QIQ54047.1;Pangolin_CoV_orf1ab;2544 | 0 | 0 |
| MT072864.1;QIQ54047.1;Pangolin_CoV_orf1ab;2545 | 0 | 0.05 |
| MT072864.1;QIQ54047.1;Pangolin_CoV_orf1ab;2546 | 0 | 0 |
| MT072864.1;QIQ54047.1;Pangolin_CoV_orf1ab;2547 | 0 | 0 |
| MT072864.1;QIQ54047.1;Pangolin_CoV_orf1ab;2548 | 0 | 0 |
| MT072864.1;QIQ54047.1;Pangolin_CoV_orf1ab;2549 | 0 | 0.025 |
| MT072864.1;QIQ54047.1;Pangolin_CoV_orf1ab;2550 | 0 | 0 |
| MT072864.1;QIQ54047.1;Pangolin_CoV_orf1ab;2551 | 0 | 0 |
| MT072864.1;QIQ54047.1;Pangolin_CoV_orf1ab;2552 | 0 | 0 |
| MT072864.1;QIQ54047.1;Pangolin_CoV_orf1ab;2553 | 0 | 0 |
| MT072864.1;QIQ54047.1;Pangolin_CoV_orf1ab;2554 | 0 | 0 |
| MT072864.1;QIQ54047.1;Pangolin_CoV_orf1ab;2555 | 0 | 0 |
| MT072864.1;QIQ54047.1;Pangolin_CoV_orf1ab;2556 | 0 | 0 |
| MT072864.1;QIQ54047.1;Pangolin_CoV_orf1ab;2557 | 0 | 0 |
| MT072864.1;QIQ54047.1;Pangolin_CoV_orf1ab;2558 | 0 | 0 |
| MT072864.1;QIQ54047.1;Pangolin_CoV_orf1ab;2559 | 0 | 0 |
| MT072864.1;QIQ54047.1;Pangolin_CoV_orf1ab;2560 | 0 | 0 |
| MT072864.1;QIQ54047.1;Pangolin_CoV_orf1ab;2561 | 0 | 0.025 |
| MT072864.1;QIQ54047.1;Pangolin_CoV_orf1ab;2562 | 0 | 0.025 |
| MT072864.1;QIQ54047.1;Pangolin_CoV_orf1ab;2563 | 0.05 | 0.1 |
| MT072864.1;QIQ54047.1;Pangolin_CoV_orf1ab;2564 | 0 | 0.05 |
| MT072864.1;QIQ54047.1;Pangolin_CoV_orf1ab;2565 | 0 | 0 |
| MT072864.1;QIQ54047.1;Pangolin_CoV_orf1ab;2566 | 0 | 0 |
| MT072864.1;QIQ54047.1;Pangolin_CoV_orf1ab;2567 | 0 | 0 |
| MT072864.1;QIQ54047.1;Pangolin_CoV_orf1ab;2568 | 0 | 0 |
| MT072864.1;QIQ54047.1;Pangolin_CoV_orf1ab;2569 | 0.05 | 0 |
| MT072864.1;QIQ54047.1;Pangolin_CoV_orf1ab;2570 | 0 | 0 |

| | | |
|---|---|---|
| MT072864.1;QIQ54047.1;Pangolin_CoV_orf1ab;2571 | 0.05 | 0 |
| MT072864.1;QIQ54047.1;Pangolin_CoV_orf1ab;2572 | 0 | 0 |
| MT072864.1;QIQ54047.1;Pangolin_CoV_orf1ab;2573 | 0 | 0 |
| MT072864.1;QIQ54047.1;Pangolin_CoV_orf1ab;2574 | 0 | 0 |
| MT072864.1;QIQ54047.1;Pangolin_CoV_orf1ab;2575 | 0 | 0.025 |
| MT072864.1;QIQ54047.1;Pangolin_CoV_orf1ab;2576 | 0 | 0.025 |
| MT072864.1;QIQ54047.1;Pangolin_CoV_orf1ab;2577 | 0 | 0 |
| MT072864.1;QIQ54047.1;Pangolin_CoV_orf1ab;2578 | 0 | 0 |
| MT072864.1;QIQ54047.1;Pangolin_CoV_orf1ab;2579 | 0 | 0 |
| MT072864.1;QIQ54047.1;Pangolin_CoV_orf1ab;2580 | 0 | 0 |
| MT072864.1;QIQ54047.1;Pangolin_CoV_orf1ab;2581 | 0 | 0 |
| MT072864.1;QIQ54047.1;Pangolin_CoV_orf1ab;2582 | 0 | 0.025 |
| MT072864.1;QIQ54047.1;Pangolin_CoV_orf1ab;2583 | 0 | 0 |
| MT072864.1;QIQ54047.1;Pangolin_CoV_orf1ab;2584 | 0 | 0 |
| MT072864.1;QIQ54047.1;Pangolin_CoV_orf1ab;2585 | 0.15 | 0.1 |
| MT072864.1;QIQ54047.1;Pangolin_CoV_orf1ab;2586 | 0.25 | 0.15 |
| MT072864.1;QIQ54047.1;Pangolin_CoV_orf1ab;2587 | 0.25 | 0.075 |
| MT072864.1;QIQ54047.1;Pangolin_CoV_orf1ab;2588 | 0.15 | 0.075 |
| MT072864.1;QIQ54047.1;Pangolin_CoV_orf1ab;2589 | 0.15 | 0.125 |
| MT072864.1;QIQ54047.1;Pangolin_CoV_orf1ab;2590 | 0.1 | 0.075 |
| MT072864.1;QIQ54047.1;Pangolin_CoV_orf1ab;2591 | 0.15 | 0.075 |
| MT072864.1;QIQ54047.1;Pangolin_CoV_orf1ab;2592 | 0.05 | 0 |
| MT072864.1;QIQ54047.1;Pangolin_CoV_orf1ab;2593 | 0 | 0.025 |
| MT072864.1;QIQ54047.1;Pangolin_CoV_orf1ab;2594 | 0.05 | 0.025 |
| MT072864.1;QIQ54047.1;Pangolin_CoV_orf1ab;2595 | 0 | 0 |
| MT072864.1;QIQ54047.1;Pangolin_CoV_orf1ab;2596 | 0 | 0 |
| MT072864.1;QIQ54047.1;Pangolin_CoV_orf1ab;2597 | 0 | 0 |
| MT072864.1;QIQ54047.1;Pangolin_CoV_orf1ab;2598 | 0 | 0 |
| MT072864.1;QIQ54047.1;Pangolin_CoV_orf1ab;2599 | 0 | 0 |
| MT072864.1;QIQ54047.1;Pangolin_CoV_orf1ab;2600 | 0 | 0 |
| MT072864.1;QIQ54047.1;Pangolin_CoV_orf1ab;2601 | 0.05 | 0 |
| MT072864.1;QIQ54047.1;Pangolin_CoV_orf1ab;2602 | 0.05 | 0 |
| MT072864.1;QIQ54047.1;Pangolin_CoV_orf1ab;2603 | 0.05 | 0 |
| MT072864.1;QIQ54047.1;Pangolin_CoV_orf1ab;2604 | 0.05 | 0.025 |
| MT072864.1;QIQ54047.1;Pangolin_CoV_orf1ab;2605 | 0.05 | 0.05 |
| MT072864.1;QIQ54047.1;Pangolin_CoV_orf1ab;2606 | 0.05 | 0.075 |
| MT072864.1;QIQ54047.1;Pangolin_CoV_orf1ab;2607 | 0.05 | 0.05 |
| MT072864.1;QIQ54047.1;Pangolin_CoV_orf1ab;2608 | 0.1 | 0.075 |
| MT072864.1;QIQ54047.1;Pangolin_CoV_orf1ab;2609 | 0.1 | 0.05 |
| MT072864.1;QIQ54047.1;Pangolin_CoV_orf1ab;2610 | 0.1 | 0.05 |
| MT072864.1;QIQ54047.1;Pangolin_CoV_orf1ab;2611 | 0.05 | 0 |
| MT072864.1;QIQ54047.1;Pangolin_CoV_orf1ab;2612 | 0 | 0 |
| MT072864.1;QIQ54047.1;Pangolin_CoV_orf1ab;2613 | 0 | 0 |

| | | |
|---|---|---|
| MT072864.1;QIQ54047.1;Pangolin_CoV_orf1ab;2614 | 0.05 | 0 |
| MT072864.1;QIQ54047.1;Pangolin_CoV_orf1ab;2615 | 0 | 0 |
| MT072864.1;QIQ54047.1;Pangolin_CoV_orf1ab;2616 | 0 | 0 |
| MT072864.1;QIQ54047.1;Pangolin_CoV_orf1ab;2617 | 0 | 0 |
| MT072864.1;QIQ54047.1;Pangolin_CoV_orf1ab;2618 | 0 | 0 |
| MT072864.1;QIQ54047.1;Pangolin_CoV_orf1ab;2619 | 0 | 0 |
| MT072864.1;QIQ54047.1;Pangolin_CoV_orf1ab;2620 | 0 | 0 |
| MT072864.1;QIQ54047.1;Pangolin_CoV_orf1ab;2621 | 0 | 0 |
| MT072864.1;QIQ54047.1;Pangolin_CoV_orf1ab;2622 | 0 | 0 |
| MT072864.1;QIQ54047.1;Pangolin_CoV_orf1ab;2623 | 0 | 0 |
| MT072864.1;QIQ54047.1;Pangolin_CoV_orf1ab;2624 | 0 | 0 |
| MT072864.1;QIQ54047.1;Pangolin_CoV_orf1ab;2625 | 0 | 0 |
| MT072864.1;QIQ54047.1;Pangolin_CoV_orf1ab;2626 | 0 | 0 |
| MT072864.1;QIQ54047.1;Pangolin_CoV_orf1ab;2627 | 0 | 0 |
| MT072864.1;QIQ54047.1;Pangolin_CoV_orf1ab;2628 | 0 | 0 |
| MT072864.1;QIQ54047.1;Pangolin_CoV_orf1ab;2629 | 0 | 0 |
| MT072864.1;QIQ54047.1;Pangolin_CoV_orf1ab;2630 | 0 | 0 |
| MT072864.1;QIQ54047.1;Pangolin_CoV_orf1ab;2631 | 0 | 0 |
| MT072864.1;QIQ54047.1;Pangolin_CoV_orf1ab;2632 | 0 | 0 |
| MT072864.1;QIQ54047.1;Pangolin_CoV_orf1ab;2633 | 0 | 0 |
| MT072864.1;QIQ54047.1;Pangolin_CoV_orf1ab;2634 | 0 | 0 |
| MT072864.1;QIQ54047.1;Pangolin_CoV_orf1ab;2635 | 0 | 0 |
| MT072864.1;QIQ54047.1;Pangolin_CoV_orf1ab;2636 | 0 | 0 |
| MT072864.1;QIQ54047.1;Pangolin_CoV_orf1ab;2637 | 0 | 0 |
| MT072864.1;QIQ54047.1;Pangolin_CoV_orf1ab;2638 | 0 | 0 |
| MT072864.1;QIQ54047.1;Pangolin_CoV_orf1ab;2639 | 0 | 0 |
| MT072864.1;QIQ54047.1;Pangolin_CoV_orf1ab;2640 | 0 | 0 |
| MT072864.1;QIQ54047.1;Pangolin_CoV_orf1ab;2641 | 0 | 0 |
| MT072864.1;QIQ54047.1;Pangolin_CoV_orf1ab;2642 | 0 | 0 |
| MT072864.1;QIQ54047.1;Pangolin_CoV_orf1ab;2643 | 0 | 0 |
| MT072864.1;QIQ54047.1;Pangolin_CoV_orf1ab;2644 | 0 | 0 |
| MT072864.1;QIQ54047.1;Pangolin_CoV_orf1ab;2645 | 0 | 0 |
| MT072864.1;QIQ54047.1;Pangolin_CoV_orf1ab;2646 | 0 | 0 |
| MT072864.1;QIQ54047.1;Pangolin_CoV_orf1ab;2647 | 0 | 0 |
| MT072864.1;QIQ54047.1;Pangolin_CoV_orf1ab;2648 | 0 | 0 |
| MT072864.1;QIQ54047.1;Pangolin_CoV_orf1ab;2649 | 0 | 0 |
| MT072864.1;QIQ54047.1;Pangolin_CoV_orf1ab;2650 | 0 | 0 |
| MT072864.1;QIQ54047.1;Pangolin_CoV_orf1ab;2651 | 0 | 0 |
| MT072864.1;QIQ54047.1;Pangolin_CoV_orf1ab;2652 | 0 | 0 |
| MT072864.1;QIQ54047.1;Pangolin_CoV_orf1ab;2653 | 0.05 | 0 |
| MT072864.1;QIQ54047.1;Pangolin_CoV_orf1ab;2654 | 0 | 0 |
| MT072864.1;QIQ54047.1;Pangolin_CoV_orf1ab;2655 | 0.05 | 0.05 |
| MT072864.1;QIQ54047.1;Pangolin_CoV_orf1ab;2656 | 0.05 | 0.05 |

| | | |
|---|---|---|
| MT072864.1;QIQ54047.1;Pangolin_CoV_orf1ab;2657 | 0 | 0 |
| MT072864.1;QIQ54047.1;Pangolin_CoV_orf1ab;2658 | 0 | 0 |
| MT072864.1;QIQ54047.1;Pangolin_CoV_orf1ab;2659 | 0 | 0.025 |
| MT072864.1;QIQ54047.1;Pangolin_CoV_orf1ab;2660 | 0.1 | 0.05 |
| MT072864.1;QIQ54047.1;Pangolin_CoV_orf1ab;2661 | 0.05 | 0 |
| MT072864.1;QIQ54047.1;Pangolin_CoV_orf1ab;2662 | 0 | 0 |
| MT072864.1;QIQ54047.1;Pangolin_CoV_orf1ab;2663 | 0 | 0 |
| MT072864.1;QIQ54047.1;Pangolin_CoV_orf1ab;2664 | 0.05 | 0 |
| MT072864.1;QIQ54047.1;Pangolin_CoV_orf1ab;2665 | 0 | 0 |
| MT072864.1;QIQ54047.1;Pangolin_CoV_orf1ab;2666 | 0 | 0.025 |
| MT072864.1;QIQ54047.1;Pangolin_CoV_orf1ab;2667 | 0.05 | 0.075 |
| MT072864.1;QIQ54047.1;Pangolin_CoV_orf1ab;2668 | 0.05 | 0.025 |
| MT072864.1;QIQ54047.1;Pangolin_CoV_orf1ab;2669 | 0.05 | 0.05 |
| MT072864.1;QIQ54047.1;Pangolin_CoV_orf1ab;2670 | 0 | 0 |
| MT072864.1;QIQ54047.1;Pangolin_CoV_orf1ab;2671 | 0 | 0 |
| MT072864.1;QIQ54047.1;Pangolin_CoV_orf1ab;2672 | 0 | 0 |
| MT072864.1;QIQ54047.1;Pangolin_CoV_orf1ab;2673 | 0 | 0 |
| MT072864.1;QIQ54047.1;Pangolin_CoV_orf1ab;2674 | 0 | 0 |
| MT072864.1;QIQ54047.1;Pangolin_CoV_orf1ab;2675 | 0 | 0 |
| MT072864.1;QIQ54047.1;Pangolin_CoV_orf1ab;2676 | 0.05 | 0 |
| MT072864.1;QIQ54047.1;Pangolin_CoV_orf1ab;2677 | 0.05 | 0 |
| MT072864.1;QIQ54047.1;Pangolin_CoV_orf1ab;2678 | 0 | 0 |
| MT072864.1;QIQ54047.1;Pangolin_CoV_orf1ab;2679 | 0 | 0 |
| MT072864.1;QIQ54047.1;Pangolin_CoV_orf1ab;2680 | 0 | 0 |
| MT072864.1;QIQ54047.1;Pangolin_CoV_orf1ab;2681 | 0.05 | 0.05 |
| MT072864.1;QIQ54047.1;Pangolin_CoV_orf1ab;2682 | 0 | 0 |
| MT072864.1;QIQ54047.1;Pangolin_CoV_orf1ab;2683 | 0.05 | 0 |
| MT072864.1;QIQ54047.1;Pangolin_CoV_orf1ab;2684 | 0 | 0 |
| MT072864.1;QIQ54047.1;Pangolin_CoV_orf1ab;2685 | 0 | 0 |
| MT072864.1;QIQ54047.1;Pangolin_CoV_orf1ab;2686 | 0 | 0 |
| MT072864.1;QIQ54047.1;Pangolin_CoV_orf1ab;2687 | 0 | 0 |
| MT072864.1;QIQ54047.1;Pangolin_CoV_orf1ab;2688 | 0 | 0 |
| MT072864.1;QIQ54047.1;Pangolin_CoV_orf1ab;2689 | 0 | 0 |
| MT072864.1;QIQ54047.1;Pangolin_CoV_orf1ab;2690 | 0 | 0 |
| MT072864.1;QIQ54047.1;Pangolin_CoV_orf1ab;2691 | 0 | 0 |
| MT072864.1;QIQ54047.1;Pangolin_CoV_orf1ab;2692 | 0 | 0 |
| MT072864.1;QIQ54047.1;Pangolin_CoV_orf1ab;2693 | 0 | 0 |
| MT072864.1;QIQ54047.1;Pangolin_CoV_orf1ab;2694 | 0 | 0 |
| MT072864.1;QIQ54047.1;Pangolin_CoV_orf1ab;2695 | 0 | 0 |
| MT072864.1;QIQ54047.1;Pangolin_CoV_orf1ab;2696 | 0 | 0 |
| MT072864.1;QIQ54047.1;Pangolin_CoV_orf1ab;2697 | 0 | 0 |
| MT072864.1;QIQ54047.1;Pangolin_CoV_orf1ab;2698 | 0 | 0 |
| MT072864.1;QIQ54047.1;Pangolin_CoV_orf1ab;2699 | 0 | 0 |

| | | |
|---|---|---|
| MT072864.1;QIQ54047.1;Pangolin_CoV_orf1ab;2700 | 0 | 0 |
| MT072864.1;QIQ54047.1;Pangolin_CoV_orf1ab;2701 | 0 | 0 |
| MT072864.1;QIQ54047.1;Pangolin_CoV_orf1ab;2702 | 0 | 0 |
| MT072864.1;QIQ54047.1;Pangolin_CoV_orf1ab;2703 | 0 | 0 |
| MT072864.1;QIQ54047.1;Pangolin_CoV_orf1ab;2704 | 0 | 0 |
| MT072864.1;QIQ54047.1;Pangolin_CoV_orf1ab;2705 | 0 | 0 |
| MT072864.1;QIQ54047.1;Pangolin_CoV_orf1ab;2706 | 0 | 0 |
| MT072864.1;QIQ54047.1;Pangolin_CoV_orf1ab;2707 | 0 | 0 |
| MT072864.1;QIQ54047.1;Pangolin_CoV_orf1ab;2708 | 0 | 0 |
| MT072864.1;QIQ54047.1;Pangolin_CoV_orf1ab;2709 | 0 | 0 |
| MT072864.1;QIQ54047.1;Pangolin_CoV_orf1ab;2710 | 0.05 | 0 |
| MT072864.1;QIQ54047.1;Pangolin_CoV_orf1ab;2711 | 0 | 0 |
| MT072864.1;QIQ54047.1;Pangolin_CoV_orf1ab;2712 | 0 | 0 |
| MT072864.1;QIQ54047.1;Pangolin_CoV_orf1ab;2713 | 0 | 0 |
| MT072864.1;QIQ54047.1;Pangolin_CoV_orf1ab;2714 | 0 | 0.025 |
| MT072864.1;QIQ54047.1;Pangolin_CoV_orf1ab;2715 | 0 | 0 |
| MT072864.1;QIQ54047.1;Pangolin_CoV_orf1ab;2716 | 0 | 0 |
| MT072864.1;QIQ54047.1;Pangolin_CoV_orf1ab;2717 | 0 | 0 |
| MT072864.1;QIQ54047.1;Pangolin_CoV_orf1ab;2718 | 0 | 0.025 |
| MT072864.1;QIQ54047.1;Pangolin_CoV_orf1ab;2719 | 0 | 0.025 |
| MT072864.1;QIQ54047.1;Pangolin_CoV_orf1ab;2720 | 0 | 0.025 |
| MT072864.1;QIQ54047.1;Pangolin_CoV_orf1ab;2721 | 0.05 | 0.025 |
| MT072864.1;QIQ54047.1;Pangolin_CoV_orf1ab;2722 | 0 | 0.025 |
| MT072864.1;QIQ54047.1;Pangolin_CoV_orf1ab;2723 | 0 | 0.025 |
| MT072864.1;QIQ54047.1;Pangolin_CoV_orf1ab;2724 | 0 | 0.025 |
| MT072864.1;QIQ54047.1;Pangolin_CoV_orf1ab;2725 | 0 | 0.025 |
| MT072864.1;QIQ54047.1;Pangolin_CoV_orf1ab;2726 | 0 | 0 |
| MT072864.1;QIQ54047.1;Pangolin_CoV_orf1ab;2727 | 0 | 0 |
| MT072864.1;QIQ54047.1;Pangolin_CoV_orf1ab;2728 | 0 | 0 |
| MT072864.1;QIQ54047.1;Pangolin_CoV_orf1ab;2729 | 0 | 0 |
| MT072864.1;QIQ54047.1;Pangolin_CoV_orf1ab;2730 | 0 | 0.025 |
| MT072864.1;QIQ54047.1;Pangolin_CoV_orf1ab;2731 | 0 | 0 |
| MT072864.1;QIQ54047.1;Pangolin_CoV_orf1ab;2732 | 0 | 0 |
| MT072864.1;QIQ54047.1;Pangolin_CoV_orf1ab;2733 | 0 | 0.025 |
| MT072864.1;QIQ54047.1;Pangolin_CoV_orf1ab;2734 | 0 | 0 |
| MT072864.1;QIQ54047.1;Pangolin_CoV_orf1ab;2735 | 0 | 0.025 |
| MT072864.1;QIQ54047.1;Pangolin_CoV_orf1ab;2736 | 0 | 0.025 |
| MT072864.1;QIQ54047.1;Pangolin_CoV_orf1ab;2737 | 0 | 0 |
| MT072864.1;QIQ54047.1;Pangolin_CoV_orf1ab;2738 | 0 | 0 |
| MT072864.1;QIQ54047.1;Pangolin_CoV_orf1ab;2739 | 0 | 0 |
| MT072864.1;QIQ54047.1;Pangolin_CoV_orf1ab;2740 | 0 | 0 |
| MT072864.1;QIQ54047.1;Pangolin_CoV_orf1ab;2741 | 0 | 0 |
| MT072864.1;QIQ54047.1;Pangolin_CoV_orf1ab;2742 | 0 | 0 |

| | | |
|---|---|---|
| MT072864.1;QIQ54047.1;Pangolin_CoV_orf1ab;2743 | 0 | 0 |
| MT072864.1;QIQ54047.1;Pangolin_CoV_orf1ab;2744 | 0 | 0 |
| MT072864.1;QIQ54047.1;Pangolin_CoV_orf1ab;2745 | 0 | 0 |
| MT072864.1;QIQ54047.1;Pangolin_CoV_orf1ab;2746 | 0 | 0 |
| MT072864.1;QIQ54047.1;Pangolin_CoV_orf1ab;2747 | 0 | 0 |
| MT072864.1;QIQ54047.1;Pangolin_CoV_orf1ab;2748 | 0 | 0 |
| MT072864.1;QIQ54047.1;Pangolin_CoV_orf1ab;2749 | 0 | 0 |
| MT072864.1;QIQ54047.1;Pangolin_CoV_orf1ab;2750 | 0 | 0 |
| MT072864.1;QIQ54047.1;Pangolin_CoV_orf1ab;2751 | 0 | 0 |
| MT072864.1;QIQ54047.1;Pangolin_CoV_orf1ab;2752 | 0 | 0 |
| MT072864.1;QIQ54047.1;Pangolin_CoV_orf1ab;2753 | 0 | 0 |
| MT072864.1;QIQ54047.1;Pangolin_CoV_orf1ab;2754 | 0 | 0 |
| MT072864.1;QIQ54047.1;Pangolin_CoV_orf1ab;2755 | 0 | 0 |
| MT072864.1;QIQ54047.1;Pangolin_CoV_orf1ab;2756 | 0 | 0 |
| MT072864.1;QIQ54047.1;Pangolin_CoV_orf1ab;2757 | 0 | 0 |
| MT072864.1;QIQ54047.1;Pangolin_CoV_orf1ab;2758 | 0 | 0 |
| MT072864.1;QIQ54047.1;Pangolin_CoV_orf1ab;2759 | 0 | 0 |
| MT072864.1;QIQ54047.1;Pangolin_CoV_orf1ab;2760 | 0 | 0 |
| MT072864.1;QIQ54047.1;Pangolin_CoV_orf1ab;2761 | 0 | 0 |
| MT072864.1;QIQ54047.1;Pangolin_CoV_orf1ab;2762 | 0 | 0 |
| MT072864.1;QIQ54047.1;Pangolin_CoV_orf1ab;2763 | 0 | 0 |
| MT072864.1;QIQ54047.1;Pangolin_CoV_orf1ab;2764 | 0 | 0 |
| MT072864.1;QIQ54047.1;Pangolin_CoV_orf1ab;2765 | 0 | 0 |
| MT072864.1;QIQ54047.1;Pangolin_CoV_orf1ab;2766 | 0 | 0 |
| MT072864.1;QIQ54047.1;Pangolin_CoV_orf1ab;2767 | 0 | 0 |
| MT072864.1;QIQ54047.1;Pangolin_CoV_orf1ab;2768 | 0 | 0 |
| MT072864.1;QIQ54047.1;Pangolin_CoV_orf1ab;2769 | 0 | 0 |
| MT072864.1;QIQ54047.1;Pangolin_CoV_orf1ab;2770 | 0 | 0 |
| MT072864.1;QIQ54047.1;Pangolin_CoV_orf1ab;2771 | 0 | 0 |
| MT072864.1;QIQ54047.1;Pangolin_CoV_orf1ab;2772 | 0 | 0 |
| MT072864.1;QIQ54047.1;Pangolin_CoV_orf1ab;2773 | 0 | 0 |
| MT072864.1;QIQ54047.1;Pangolin_CoV_orf1ab;2774 | 0 | 0 |
| MT072864.1;QIQ54047.1;Pangolin_CoV_orf1ab;2775 | 0 | 0 |
| MT072864.1;QIQ54047.1;Pangolin_CoV_orf1ab;2776 | 0 | 0 |
| MT072864.1;QIQ54047.1;Pangolin_CoV_orf1ab;2777 | 0 | 0 |
| MT072864.1;QIQ54047.1;Pangolin_CoV_orf1ab;2778 | 0 | 0.025 |
| MT072864.1;QIQ54047.1;Pangolin_CoV_orf1ab;2779 | 0 | 0 |
| MT072864.1;QIQ54047.1;Pangolin_CoV_orf1ab;2780 | 0 | 0 |
| MT072864.1;QIQ54047.1;Pangolin_CoV_orf1ab;2781 | 0 | 0 |
| MT072864.1;QIQ54047.1;Pangolin_CoV_orf1ab;2782 | 0 | 0 |
| MT072864.1;QIQ54047.1;Pangolin_CoV_orf1ab;2783 | 0 | 0.025 |
| MT072864.1;QIQ54047.1;Pangolin_CoV_orf1ab;2784 | 0 | 0.025 |
| MT072864.1;QIQ54047.1;Pangolin_CoV_orf1ab;2785 | 0 | 0.025 |

| | | |
|---|---|---|
| MT072864.1;QIQ54047.1;Pangolin_CoV_orf1ab;2786 | 0 | 0.025 |
| MT072864.1;QIQ54047.1;Pangolin_CoV_orf1ab;2787 | 0 | 0.025 |
| MT072864.1;QIQ54047.1;Pangolin_CoV_orf1ab;2788 | 0.05 | 0.025 |
| MT072864.1;QIQ54047.1;Pangolin_CoV_orf1ab;2789 | 0 | 0.05 |
| MT072864.1;QIQ54047.1;Pangolin_CoV_orf1ab;2790 | 0 | 0.05 |
| MT072864.1;QIQ54047.1;Pangolin_CoV_orf1ab;2791 | 0 | 0.025 |
| MT072864.1;QIQ54047.1;Pangolin_CoV_orf1ab;2792 | 0 | 0.025 |
| MT072864.1;QIQ54047.1;Pangolin_CoV_orf1ab;2793 | 0 | 0.025 |
| MT072864.1;QIQ54047.1;Pangolin_CoV_orf1ab;2794 | 0.05 | 0.025 |
| MT072864.1;QIQ54047.1;Pangolin_CoV_orf1ab;2795 | 0.05 | 0.025 |
| MT072864.1;QIQ54047.1;Pangolin_CoV_orf1ab;2796 | 0.1 | 0 |
| MT072864.1;QIQ54047.1;Pangolin_CoV_orf1ab;2797 | 0.15 | 0.025 |
| MT072864.1;QIQ54047.1;Pangolin_CoV_orf1ab;2798 | 0.15 | 0 |
| MT072864.1;QIQ54047.1;Pangolin_CoV_orf1ab;2799 | 0.1 | 0 |
| MT072864.1;QIQ54047.1;Pangolin_CoV_orf1ab;2800 | 0.1 | 0 |
| MT072864.1;QIQ54047.1;Pangolin_CoV_orf1ab;2801 | 0.05 | 0 |
| MT072864.1;QIQ54047.1;Pangolin_CoV_orf1ab;2802 | 0.1 | 0 |
| MT072864.1;QIQ54047.1;Pangolin_CoV_orf1ab;2803 | 0.3 | 0.1 |
| MT072864.1;QIQ54047.1;Pangolin_CoV_orf1ab;2804 | 0.15 | 0 |
| MT072864.1;QIQ54047.1;Pangolin_CoV_orf1ab;2805 | 0.05 | 0.05 |
| MT072864.1;QIQ54047.1;Pangolin_CoV_orf1ab;2806 | 0 | 0 |
| MT072864.1;QIQ54047.1;Pangolin_CoV_orf1ab;2807 | 0.15 | 0.05 |
| MT072864.1;QIQ54047.1;Pangolin_CoV_orf1ab;2808 | 0 | 0 |
| MT072864.1;QIQ54047.1;Pangolin_CoV_orf1ab;2809 | 0 | 0.025 |
| MT072864.1;QIQ54047.1;Pangolin_CoV_orf1ab;2810 | 0 | 0 |
| MT072864.1;QIQ54047.1;Pangolin_CoV_orf1ab;2811 | 0.05 | 0.025 |
| MT072864.1;QIQ54047.1;Pangolin_CoV_orf1ab;2812 | 0 | 0.075 |
| MT072864.1;QIQ54047.1;Pangolin_CoV_orf1ab;2813 | 0.15 | 0.125 |
| MT072864.1;QIQ54047.1;Pangolin_CoV_orf1ab;2814 | 0 | 0.025 |
| MT072864.1;QIQ54047.1;Pangolin_CoV_orf1ab;2815 | 0 | 0 |
| MT072864.1;QIQ54047.1;Pangolin_CoV_orf1ab;2816 | 0 | 0 |
| MT072864.1;QIQ54047.1;Pangolin_CoV_orf1ab;2817 | 0 | 0 |
| MT072864.1;QIQ54047.1;Pangolin_CoV_orf1ab;2818 | 0 | 0 |
| MT072864.1;QIQ54047.1;Pangolin_CoV_orf1ab;2819 | 0 | 0 |
| MT072864.1;QIQ54047.1;Pangolin_CoV_orf1ab;2820 | 0.05 | 0.025 |
| MT072864.1;QIQ54047.1;Pangolin_CoV_orf1ab;2821 | 0 | 0 |
| MT072864.1;QIQ54047.1;Pangolin_CoV_orf1ab;2822 | 0 | 0 |
| MT072864.1;QIQ54047.1;Pangolin_CoV_orf1ab;2823 | 0 | 0 |
| MT072864.1;QIQ54047.1;Pangolin_CoV_orf1ab;2824 | 0 | 0 |
| MT072864.1;QIQ54047.1;Pangolin_CoV_orf1ab;2825 | 0 | 0 |
| MT072864.1;QIQ54047.1;Pangolin_CoV_orf1ab;2826 | 0 | 0 |
| MT072864.1;QIQ54047.1;Pangolin_CoV_orf1ab;2827 | 0.15 | 0 |
| MT072864.1;QIQ54047.1;Pangolin_CoV_orf1ab;2828 | 0 | 0 |

| | | |
|---|---|---|
| MT072864.1;QIQ54047.1;Pangolin_CoV_orf1ab;2829 | 0 | 0 |
| MT072864.1;QIQ54047.1;Pangolin_CoV_orf1ab;2830 | 0 | 0 |
| MT072864.1;QIQ54047.1;Pangolin_CoV_orf1ab;2831 | 0.05 | 0 |
| MT072864.1;QIQ54047.1;Pangolin_CoV_orf1ab;2832 | 0 | 0.05 |
| MT072864.1;QIQ54047.1;Pangolin_CoV_orf1ab;2833 | 0.05 | 0.025 |
| MT072864.1;QIQ54047.1;Pangolin_CoV_orf1ab;2834 | 0 | 0 |
| MT072864.1;QIQ54047.1;Pangolin_CoV_orf1ab;2835 | 0 | 0 |
| MT072864.1;QIQ54047.1;Pangolin_CoV_orf1ab;2836 | 0 | 0.05 |
| MT072864.1;QIQ54047.1;Pangolin_CoV_orf1ab;2837 | 0 | 0 |
| MT072864.1;QIQ54047.1;Pangolin_CoV_orf1ab;2838 | 0 | 0 |
| MT072864.1;QIQ54047.1;Pangolin_CoV_orf1ab;2839 | 0 | 0 |
| MT072864.1;QIQ54047.1;Pangolin_CoV_orf1ab;2840 | 0 | 0 |
| MT072864.1;QIQ54047.1;Pangolin_CoV_orf1ab;2841 | 0 | 0 |
| MT072864.1;QIQ54047.1;Pangolin_CoV_orf1ab;2842 | 0 | 0 |
| MT072864.1;QIQ54047.1;Pangolin_CoV_orf1ab;2843 | 0 | 0 |
| MT072864.1;QIQ54047.1;Pangolin_CoV_orf1ab;2844 | 0 | 0 |
| MT072864.1;QIQ54047.1;Pangolin_CoV_orf1ab;2845 | 0 | 0 |
| MT072864.1;QIQ54047.1;Pangolin_CoV_orf1ab;2846 | 0 | 0 |
| MT072864.1;QIQ54047.1;Pangolin_CoV_orf1ab;2847 | 0 | 0.05 |
| MT072864.1;QIQ54047.1;Pangolin_CoV_orf1ab;2848 | 0 | 0.125 |
| MT072864.1;QIQ54047.1;Pangolin_CoV_orf1ab;2849 | 0.05 | 0.1 |
| MT072864.1;QIQ54047.1;Pangolin_CoV_orf1ab;2850 | 0.05 | 0.075 |
| MT072864.1;QIQ54047.1;Pangolin_CoV_orf1ab;2851 | 0.1 | 0.05 |
| MT072864.1;QIQ54047.1;Pangolin_CoV_orf1ab;2852 | 0.05 | 0.025 |
| MT072864.1;QIQ54047.1;Pangolin_CoV_orf1ab;2853 | 0 | 0 |
| MT072864.1;QIQ54047.1;Pangolin_CoV_orf1ab;2854 | 0 | 0 |
| MT072864.1;QIQ54047.1;Pangolin_CoV_orf1ab;2855 | 0 | 0 |
| MT072864.1;QIQ54047.1;Pangolin_CoV_orf1ab;2856 | 0 | 0 |
| MT072864.1;QIQ54047.1;Pangolin_CoV_orf1ab;2857 | 0 | 0 |
| MT072864.1;QIQ54047.1;Pangolin_CoV_orf1ab;2858 | 0 | 0 |
| MT072864.1;QIQ54047.1;Pangolin_CoV_orf1ab;2859 | 0 | 0 |
| MT072864.1;QIQ54047.1;Pangolin_CoV_orf1ab;2860 | 0 | 0 |
| MT072864.1;QIQ54047.1;Pangolin_CoV_orf1ab;2861 | 0 | 0 |
| MT072864.1;QIQ54047.1;Pangolin_CoV_orf1ab;2862 | 0 | 0 |
| MT072864.1;QIQ54047.1;Pangolin_CoV_orf1ab;2863 | 0 | 0 |
| MT072864.1;QIQ54047.1;Pangolin_CoV_orf1ab;2864 | 0 | 0 |
| MT072864.1;QIQ54047.1;Pangolin_CoV_orf1ab;2865 | 0 | 0 |
| MT072864.1;QIQ54047.1;Pangolin_CoV_orf1ab;2866 | 0 | 0 |
| MT072864.1;QIQ54047.1;Pangolin_CoV_orf1ab;2867 | 0 | 0 |
| MT072864.1;QIQ54047.1;Pangolin_CoV_orf1ab;2868 | 0 | 0 |
| MT072864.1;QIQ54047.1;Pangolin_CoV_orf1ab;2869 | 0 | 0 |
| MT072864.1;QIQ54047.1;Pangolin_CoV_orf1ab;2870 | 0 | 0 |
| MT072864.1;QIQ54047.1;Pangolin_CoV_orf1ab;2871 | 0 | 0 |

| | | |
|---|---|---|
| MT072864.1;QIQ54047.1;Pangolin_CoV_orf1ab;2872 | 0 | 0 |
| MT072864.1;QIQ54047.1;Pangolin_CoV_orf1ab;2873 | 0 | 0 |
| MT072864.1;QIQ54047.1;Pangolin_CoV_orf1ab;2874 | 0 | 0 |
| MT072864.1;QIQ54047.1;Pangolin_CoV_orf1ab;2875 | 0 | 0 |
| MT072864.1;QIQ54047.1;Pangolin_CoV_orf1ab;2876 | 0 | 0 |
| MT072864.1;QIQ54047.1;Pangolin_CoV_orf1ab;2877 | 0 | 0 |
| MT072864.1;QIQ54047.1;Pangolin_CoV_orf1ab;2878 | 0 | 0 |
| MT072864.1;QIQ54047.1;Pangolin_CoV_orf1ab;2879 | 0 | 0 |
| MT072864.1;QIQ54047.1;Pangolin_CoV_orf1ab;2880 | 0 | 0 |
| MT072864.1;QIQ54047.1;Pangolin_CoV_orf1ab;2881 | 0 | 0 |
| MT072864.1;QIQ54047.1;Pangolin_CoV_orf1ab;2882 | 0 | 0 |
| MT072864.1;QIQ54047.1;Pangolin_CoV_orf1ab;2883 | 0.05 | 0 |
| MT072864.1;QIQ54047.1;Pangolin_CoV_orf1ab;2884 | 0.05 | 0.025 |
| MT072864.1;QIQ54047.1;Pangolin_CoV_orf1ab;2885 | 0 | 0.025 |
| MT072864.1;QIQ54047.1;Pangolin_CoV_orf1ab;2886 | 0 | 0.025 |
| MT072864.1;QIQ54047.1;Pangolin_CoV_orf1ab;2887 | 0.05 | 0 |
| MT072864.1;QIQ54047.1;Pangolin_CoV_orf1ab;2888 | 0.05 | 0.05 |
| MT072864.1;QIQ54047.1;Pangolin_CoV_orf1ab;2889 | 0.05 | 0.025 |
| MT072864.1;QIQ54047.1;Pangolin_CoV_orf1ab;2890 | 0.05 | 0 |
| MT072864.1;QIQ54047.1;Pangolin_CoV_orf1ab;2891 | 0 | 0 |
| MT072864.1;QIQ54047.1;Pangolin_CoV_orf1ab;2892 | 0 | 0 |
| MT072864.1;QIQ54047.1;Pangolin_CoV_orf1ab;2893 | 0 | 0 |
| MT072864.1;QIQ54047.1;Pangolin_CoV_orf1ab;2894 | 0 | 0 |
| MT072864.1;QIQ54047.1;Pangolin_CoV_orf1ab;2895 | 0.05 | 0 |
| MT072864.1;QIQ54047.1;Pangolin_CoV_orf1ab;2896 | 0 | 0 |
| MT072864.1;QIQ54047.1;Pangolin_CoV_orf1ab;2897 | 0 | 0 |
| MT072864.1;QIQ54047.1;Pangolin_CoV_orf1ab;2898 | 0 | 0 |
| MT072864.1;QIQ54047.1;Pangolin_CoV_orf1ab;2899 | 0 | 0 |
| MT072864.1;QIQ54047.1;Pangolin_CoV_orf1ab;2900 | 0 | 0 |
| MT072864.1;QIQ54047.1;Pangolin_CoV_orf1ab;2901 | 0 | 0 |
| MT072864.1;QIQ54047.1;Pangolin_CoV_orf1ab;2902 | 0 | 0 |
| MT072864.1;QIQ54047.1;Pangolin_CoV_orf1ab;2903 | 0.05 | 0.025 |
| MT072864.1;QIQ54047.1;Pangolin_CoV_orf1ab;2904 | 0 | 0.025 |
| MT072864.1;QIQ54047.1;Pangolin_CoV_orf1ab;2905 | 0.05 | 0.05 |
| MT072864.1;QIQ54047.1;Pangolin_CoV_orf1ab;2906 | 0.15 | 0.05 |
| MT072864.1;QIQ54047.1;Pangolin_CoV_orf1ab;2907 | 0.3 | 0.05 |
| MT072864.1;QIQ54047.1;Pangolin_CoV_orf1ab;2908 | 0.25 | 0.125 |
| MT072864.1;QIQ54047.1;Pangolin_CoV_orf1ab;2909 | 0.25 | 0.05 |
| MT072864.1;QIQ54047.1;Pangolin_CoV_orf1ab;2910 | 0.3 | 0.05 |
| MT072864.1;QIQ54047.1;Pangolin_CoV_orf1ab;2911 | 0.05 | 0 |
| MT072864.1;QIQ54047.1;Pangolin_CoV_orf1ab;2912 | 0.2 | 0.05 |
| MT072864.1;QIQ54047.1;Pangolin_CoV_orf1ab;2913 | 0.3 | 0.1 |
| MT072864.1;QIQ54047.1;Pangolin_CoV_orf1ab;2914 | 0.2 | 0.075 |

| | | |
|---|---|---|
| MT072864.1;QIQ54047.1;Pangolin_CoV_orf1ab;2915 | 0 | 0.025 |
| MT072864.1;QIQ54047.1;Pangolin_CoV_orf1ab;2916 | 0.05 | 0.05 |
| MT072864.1;QIQ54047.1;Pangolin_CoV_orf1ab;2917 | 0 | 0 |
| MT072864.1;QIQ54047.1;Pangolin_CoV_orf1ab;2918 | 0 | 0 |
| MT072864.1;QIQ54047.1;Pangolin_CoV_orf1ab;2919 | 0 | 0 |
| MT072864.1;QIQ54047.1;Pangolin_CoV_orf1ab;2920 | 0 | 0 |
| MT072864.1;QIQ54047.1;Pangolin_CoV_orf1ab;2921 | 0 | 0 |
| MT072864.1;QIQ54047.1;Pangolin_CoV_orf1ab;2922 | 0 | 0 |
| MT072864.1;QIQ54047.1;Pangolin_CoV_orf1ab;2923 | 0 | 0 |
| MT072864.1;QIQ54047.1;Pangolin_CoV_orf1ab;2924 | 0 | 0 |
| MT072864.1;QIQ54047.1;Pangolin_CoV_orf1ab;2925 | 0 | 0 |
| MT072864.1;QIQ54047.1;Pangolin_CoV_orf1ab;2926 | 0.05 | 0 |
| MT072864.1;QIQ54047.1;Pangolin_CoV_orf1ab;2927 | 0.05 | 0 |
| MT072864.1;QIQ54047.1;Pangolin_CoV_orf1ab;2928 | 0.1 | 0 |
| MT072864.1;QIQ54047.1;Pangolin_CoV_orf1ab;2929 | 0 | 0.025 |
| MT072864.1;QIQ54047.1;Pangolin_CoV_orf1ab;2930 | 0 | 0.05 |
| MT072864.1;QIQ54047.1;Pangolin_CoV_orf1ab;2931 | 0.2 | 0.15 |
| MT072864.1;QIQ54047.1;Pangolin_CoV_orf1ab;2932 | 0.15 | 0.175 |
| MT072864.1;QIQ54047.1;Pangolin_CoV_orf1ab;2933 | 0.1 | 0.075 |
| MT072864.1;QIQ54047.1;Pangolin_CoV_orf1ab;2934 | 0.1 | 0.025 |
| MT072864.1;QIQ54047.1;Pangolin_CoV_orf1ab;2935 | 0.15 | 0.075 |
| MT072864.1;QIQ54047.1;Pangolin_CoV_orf1ab;2936 | 0.05 | 0.1 |
| MT072864.1;QIQ54047.1;Pangolin_CoV_orf1ab;2937 | 0.1 | 0.1 |
| MT072864.1;QIQ54047.1;Pangolin_CoV_orf1ab;2938 | 0.15 | 0.025 |
| MT072864.1;QIQ54047.1;Pangolin_CoV_orf1ab;2939 | 0.05 | 0.025 |
| MT072864.1;QIQ54047.1;Pangolin_CoV_orf1ab;2940 | 0.05 | 0.05 |
| MT072864.1;QIQ54047.1;Pangolin_CoV_orf1ab;2941 | 0.05 | 0.025 |
| MT072864.1;QIQ54047.1;Pangolin_CoV_orf1ab;2942 | 0 | 0 |
| MT072864.1;QIQ54047.1;Pangolin_CoV_orf1ab;2943 | 0 | 0 |
| MT072864.1;QIQ54047.1;Pangolin_CoV_orf1ab;2944 | 0 | 0 |
| MT072864.1;QIQ54047.1;Pangolin_CoV_orf1ab;2945 | 0 | 0 |
| MT072864.1;QIQ54047.1;Pangolin_CoV_orf1ab;2946 | 0 | 0 |
| MT072864.1;QIQ54047.1;Pangolin_CoV_orf1ab;2947 | 0 | 0 |
| MT072864.1;QIQ54047.1;Pangolin_CoV_orf1ab;2948 | 0 | 0 |
| MT072864.1;QIQ54047.1;Pangolin_CoV_orf1ab;2949 | 0 | 0 |
| MT072864.1;QIQ54047.1;Pangolin_CoV_orf1ab;2950 | 0 | 0 |
| MT072864.1;QIQ54047.1;Pangolin_CoV_orf1ab;2951 | 0 | 0 |
| MT072864.1;QIQ54047.1;Pangolin_CoV_orf1ab;2952 | 0 | 0.025 |
| MT072864.1;QIQ54047.1;Pangolin_CoV_orf1ab;2953 | 0 | 0 |
| MT072864.1;QIQ54047.1;Pangolin_CoV_orf1ab;2954 | 0 | 0 |
| MT072864.1;QIQ54047.1;Pangolin_CoV_orf1ab;2955 | 0 | 0 |
| MT072864.1;QIQ54047.1;Pangolin_CoV_orf1ab;2956 | 0 | 0 |
| MT072864.1;QIQ54047.1;Pangolin_CoV_orf1ab;2957 | 0 | 0 |

| | | |
|---|---|---|
| MT072864.1;QIQ54047.1;Pangolin_CoV_orf1ab;2958 | 0 | 0 |
| MT072864.1;QIQ54047.1;Pangolin_CoV_orf1ab;2959 | 0 | 0 |
| MT072864.1;QIQ54047.1;Pangolin_CoV_orf1ab;2960 | 0 | 0 |
| MT072864.1;QIQ54047.1;Pangolin_CoV_orf1ab;2961 | 0 | 0 |
| MT072864.1;QIQ54047.1;Pangolin_CoV_orf1ab;2962 | 0 | 0 |
| MT072864.1;QIQ54047.1;Pangolin_CoV_orf1ab;2963 | 0.05 | 0.1 |
| MT072864.1;QIQ54047.1;Pangolin_CoV_orf1ab;2964 | 0 | 0.05 |
| MT072864.1;QIQ54047.1;Pangolin_CoV_orf1ab;2965 | 0 | 0.075 |
| MT072864.1;QIQ54047.1;Pangolin_CoV_orf1ab;2966 | 0 | 0.075 |
| MT072864.1;QIQ54047.1;Pangolin_CoV_orf1ab;2967 | 0 | 0.05 |
| MT072864.1;QIQ54047.1;Pangolin_CoV_orf1ab;2968 | 0 | 0.05 |
| MT072864.1;QIQ54047.1;Pangolin_CoV_orf1ab;2969 | 0 | 0.05 |
| MT072864.1;QIQ54047.1;Pangolin_CoV_orf1ab;2970 | 0 | 0.05 |
| MT072864.1;QIQ54047.1;Pangolin_CoV_orf1ab;2971 | 0.05 | 0.05 |
| MT072864.1;QIQ54047.1;Pangolin_CoV_orf1ab;2972 | 0 | 0 |
| MT072864.1;QIQ54047.1;Pangolin_CoV_orf1ab;2973 | 0 | 0 |
| MT072864.1;QIQ54047.1;Pangolin_CoV_orf1ab;2974 | 0 | 0 |
| MT072864.1;QIQ54047.1;Pangolin_CoV_orf1ab;2975 | 0 | 0 |
| MT072864.1;QIQ54047.1;Pangolin_CoV_orf1ab;2976 | 0 | 0 |
| MT072864.1;QIQ54047.1;Pangolin_CoV_orf1ab;2977 | 0 | 0 |
| MT072864.1;QIQ54047.1;Pangolin_CoV_orf1ab;2978 | 0 | 0 |
| MT072864.1;QIQ54047.1;Pangolin_CoV_orf1ab;2979 | 0 | 0 |
| MT072864.1;QIQ54047.1;Pangolin_CoV_orf1ab;2980 | 0 | 0 |
| MT072864.1;QIQ54047.1;Pangolin_CoV_orf1ab;2981 | 0 | 0 |
| MT072864.1;QIQ54047.1;Pangolin_CoV_orf1ab;2982 | 0 | 0 |
| MT072864.1;QIQ54047.1;Pangolin_CoV_orf1ab;2983 | 0 | 0 |
| MT072864.1;QIQ54047.1;Pangolin_CoV_orf1ab;2984 | 0 | 0 |
| MT072864.1;QIQ54047.1;Pangolin_CoV_orf1ab;2985 | 0 | 0 |
| MT072864.1;QIQ54047.1;Pangolin_CoV_orf1ab;2986 | 0 | 0 |
| MT072864.1;QIQ54047.1;Pangolin_CoV_orf1ab;2987 | 0 | 0 |
| MT072864.1;QIQ54047.1;Pangolin_CoV_orf1ab;2988 | 0 | 0 |
| MT072864.1;QIQ54047.1;Pangolin_CoV_orf1ab;2989 | 0 | 0 |
| MT072864.1;QIQ54047.1;Pangolin_CoV_orf1ab;2990 | 0 | 0 |
| MT072864.1;QIQ54047.1;Pangolin_CoV_orf1ab;2991 | 0 | 0 |
| MT072864.1;QIQ54047.1;Pangolin_CoV_orf1ab;2992 | 0 | 0 |
| MT072864.1;QIQ54047.1;Pangolin_CoV_orf1ab;2993 | 0 | 0 |
| MT072864.1;QIQ54047.1;Pangolin_CoV_orf1ab;2994 | 0 | 0 |
| MT072864.1;QIQ54047.1;Pangolin_CoV_orf1ab;2995 | 0 | 0 |
| MT072864.1;QIQ54047.1;Pangolin_CoV_orf1ab;2996 | 0 | 0 |
| MT072864.1;QIQ54047.1;Pangolin_CoV_orf1ab;2997 | 0 | 0 |
| MT072864.1;QIQ54047.1;Pangolin_CoV_orf1ab;2998 | 0 | 0 |
| MT072864.1;QIQ54047.1;Pangolin_CoV_orf1ab;2999 | 0 | 0 |
| MT072864.1;QIQ54047.1;Pangolin_CoV_orf1ab;3000 | 0 | 0 |

| | | |
|---|---|---|
| MT072864.1;QIQ54047.1;Pangolin_CoV_orf1ab;3001 | 0 | 0 |
| MT072864.1;QIQ54047.1;Pangolin_CoV_orf1ab;3002 | 0.15 | 0.15 |
| MT072864.1;QIQ54047.1;Pangolin_CoV_orf1ab;3003 | 0.15 | 0.15 |
| MT072864.1;QIQ54047.1;Pangolin_CoV_orf1ab;3004 | 0.1 | 0.125 |
| MT072864.1;QIQ54047.1;Pangolin_CoV_orf1ab;3005 | 0.1 | 0.15 |
| MT072864.1;QIQ54047.1;Pangolin_CoV_orf1ab;3006 | 0.1 | 0.15 |
| MT072864.1;QIQ54047.1;Pangolin_CoV_orf1ab;3007 | 0.1 | 0.15 |
| MT072864.1;QIQ54047.1;Pangolin_CoV_orf1ab;3008 | 0.1 | 0.125 |
| MT072864.1;QIQ54047.1;Pangolin_CoV_orf1ab;3009 | 0.1 | 0.15 |
| MT072864.1;QIQ54047.1;Pangolin_CoV_orf1ab;3010 | 0.1 | 0.15 |
| MT072864.1;QIQ54047.1;Pangolin_CoV_orf1ab;3011 | 0.15 | 0.25 |
| MT072864.1;QIQ54047.1;Pangolin_CoV_orf1ab;3012 | 0.35 | 0.325 |
| MT072864.1;QIQ54047.1;Pangolin_CoV_orf1ab;3013 | 0.25 | 0.225 |
| MT072864.1;QIQ54047.1;Pangolin_CoV_orf1ab;3014 | 0 | 0.025 |
| MT072864.1;QIQ54047.1;Pangolin_CoV_orf1ab;3015 | 0 | 0 |
| MT072864.1;QIQ54047.1;Pangolin_CoV_orf1ab;3016 | 0 | 0 |
| MT072864.1;QIQ54047.1;Pangolin_CoV_orf1ab;3017 | 0.05 | 0 |
| MT072864.1;QIQ54047.1;Pangolin_CoV_orf1ab;3018 | 0 | 0 |
| MT072864.1;QIQ54047.1;Pangolin_CoV_orf1ab;3019 | 0.05 | 0 |
| MT072864.1;QIQ54047.1;Pangolin_CoV_orf1ab;3020 | 0.05 | 0 |
| MT072864.1;QIQ54047.1;Pangolin_CoV_orf1ab;3021 | 0.05 | 0 |
| MT072864.1;QIQ54047.1;Pangolin_CoV_orf1ab;3022 | 0.05 | 0 |
| MT072864.1;QIQ54047.1;Pangolin_CoV_orf1ab;3023 | 0.05 | 0 |
| MT072864.1;QIQ54047.1;Pangolin_CoV_orf1ab;3024 | 0.05 | 0 |
| MT072864.1;QIQ54047.1;Pangolin_CoV_orf1ab;3025 | 0 | 0 |
| MT072864.1;QIQ54047.1;Pangolin_CoV_orf1ab;3026 | 0 | 0 |
| MT072864.1;QIQ54047.1;Pangolin_CoV_orf1ab;3027 | 0 | 0 |
| MT072864.1;QIQ54047.1;Pangolin_CoV_orf1ab;3028 | 0.05 | 0 |
| MT072864.1;QIQ54047.1;Pangolin_CoV_orf1ab;3029 | 0 | 0 |
| MT072864.1;QIQ54047.1;Pangolin_CoV_orf1ab;3030 | 0 | 0 |
| MT072864.1;QIQ54047.1;Pangolin_CoV_orf1ab;3031 | 0 | 0.05 |
| MT072864.1;QIQ54047.1;Pangolin_CoV_orf1ab;3032 | 0 | 0 |
| MT072864.1;QIQ54047.1;Pangolin_CoV_orf1ab;3033 | 0 | 0 |
| MT072864.1;QIQ54047.1;Pangolin_CoV_orf1ab;3034 | 0 | 0 |
| MT072864.1;QIQ54047.1;Pangolin_CoV_orf1ab;3035 | 0 | 0 |
| MT072864.1;QIQ54047.1;Pangolin_CoV_orf1ab;3036 | 0 | 0 |
| MT072864.1;QIQ54047.1;Pangolin_CoV_orf1ab;3037 | 0 | 0 |
| MT072864.1;QIQ54047.1;Pangolin_CoV_orf1ab;3038 | 0 | 0 |
| MT072864.1;QIQ54047.1;Pangolin_CoV_orf1ab;3039 | 0 | 0 |
| MT072864.1;QIQ54047.1;Pangolin_CoV_orf1ab;3040 | 0 | 0 |
| MT072864.1;QIQ54047.1;Pangolin_CoV_orf1ab;3041 | 0 | 0 |
| MT072864.1;QIQ54047.1;Pangolin_CoV_orf1ab;3042 | 0 | 0 |
| MT072864.1;QIQ54047.1;Pangolin_CoV_orf1ab;3043 | 0 | 0 |

| | | |
|---|---|---|
| MT072864.1;QIQ54047.1;Pangolin_CoV_orf1ab;3044 | 0 | 0 |
| MT072864.1;QIQ54047.1;Pangolin_CoV_orf1ab;3045 | 0 | 0 |
| MT072864.1;QIQ54047.1;Pangolin_CoV_orf1ab;3046 | 0 | 0 |
| MT072864.1;QIQ54047.1;Pangolin_CoV_orf1ab;3047 | 0 | 0 |
| MT072864.1;QIQ54047.1;Pangolin_CoV_orf1ab;3048 | 0 | 0 |
| MT072864.1;QIQ54047.1;Pangolin_CoV_orf1ab;3049 | 0 | 0 |
| MT072864.1;QIQ54047.1;Pangolin_CoV_orf1ab;3050 | 0 | 0 |
| MT072864.1;QIQ54047.1;Pangolin_CoV_orf1ab;3051 | 0 | 0 |
| MT072864.1;QIQ54047.1;Pangolin_CoV_orf1ab;3052 | 0 | 0 |
| MT072864.1;QIQ54047.1;Pangolin_CoV_orf1ab;3053 | 0 | 0 |
| MT072864.1;QIQ54047.1;Pangolin_CoV_orf1ab;3054 | 0 | 0 |
| MT072864.1;QIQ54047.1;Pangolin_CoV_orf1ab;3055 | 0 | 0 |
| MT072864.1;QIQ54047.1;Pangolin_CoV_orf1ab;3056 | 0 | 0 |
| MT072864.1;QIQ54047.1;Pangolin_CoV_orf1ab;3057 | 0 | 0 |
| MT072864.1;QIQ54047.1;Pangolin_CoV_orf1ab;3058 | 0 | 0 |
| MT072864.1;QIQ54047.1;Pangolin_CoV_orf1ab;3059 | 0 | 0 |
| MT072864.1;QIQ54047.1;Pangolin_CoV_orf1ab;3060 | 0 | 0 |
| MT072864.1;QIQ54047.1;Pangolin_CoV_orf1ab;3061 | 0 | 0 |
| MT072864.1;QIQ54047.1;Pangolin_CoV_orf1ab;3062 | 0 | 0 |
| MT072864.1;QIQ54047.1;Pangolin_CoV_orf1ab;3063 | 0 | 0 |
| MT072864.1;QIQ54047.1;Pangolin_CoV_orf1ab;3064 | 0 | 0 |
| MT072864.1;QIQ54047.1;Pangolin_CoV_orf1ab;3065 | 0 | 0 |
| MT072864.1;QIQ54047.1;Pangolin_CoV_orf1ab;3066 | 0 | 0 |
| MT072864.1;QIQ54047.1;Pangolin_CoV_orf1ab;3067 | 0 | 0 |
| MT072864.1;QIQ54047.1;Pangolin_CoV_orf1ab;3068 | 0 | 0 |
| MT072864.1;QIQ54047.1;Pangolin_CoV_orf1ab;3069 | 0 | 0 |
| MT072864.1;QIQ54047.1;Pangolin_CoV_orf1ab;3070 | 0 | 0 |
| MT072864.1;QIQ54047.1;Pangolin_CoV_orf1ab;3071 | 0 | 0 |
| MT072864.1;QIQ54047.1;Pangolin_CoV_orf1ab;3072 | 0 | 0 |
| MT072864.1;QIQ54047.1;Pangolin_CoV_orf1ab;3073 | 0 | 0 |
| MT072864.1;QIQ54047.1;Pangolin_CoV_orf1ab;3074 | 0 | 0 |
| MT072864.1;QIQ54047.1;Pangolin_CoV_orf1ab;3075 | 0 | 0 |
| MT072864.1;QIQ54047.1;Pangolin_CoV_orf1ab;3076 | 0 | 0 |
| MT072864.1;QIQ54047.1;Pangolin_CoV_orf1ab;3077 | 0 | 0 |
| MT072864.1;QIQ54047.1;Pangolin_CoV_orf1ab;3078 | 0 | 0 |
| MT072864.1;QIQ54047.1;Pangolin_CoV_orf1ab;3079 | 0 | 0 |
| MT072864.1;QIQ54047.1;Pangolin_CoV_orf1ab;3080 | 0 | 0 |
| MT072864.1;QIQ54047.1;Pangolin_CoV_orf1ab;3081 | 0 | 0 |
| MT072864.1;QIQ54047.1;Pangolin_CoV_orf1ab;3082 | 0 | 0 |
| MT072864.1;QIQ54047.1;Pangolin_CoV_orf1ab;3083 | 0 | 0 |
| MT072864.1;QIQ54047.1;Pangolin_CoV_orf1ab;3084 | 0 | 0 |
| MT072864.1;QIQ54047.1;Pangolin_CoV_orf1ab;3085 | 0 | 0 |
| MT072864.1;QIQ54047.1;Pangolin_CoV_orf1ab;3086 | 0 | 0 |

| | | |
|---|---|---|
| MT072864.1;QIQ54047.1;Pangolin_CoV_orf1ab;3087 | 0 | 0 |
| MT072864.1;QIQ54047.1;Pangolin_CoV_orf1ab;3088 | 0 | 0 |
| MT072864.1;QIQ54047.1;Pangolin_CoV_orf1ab;3089 | 0 | 0 |
| MT072864.1;QIQ54047.1;Pangolin_CoV_orf1ab;3090 | 0 | 0 |
| MT072864.1;QIQ54047.1;Pangolin_CoV_orf1ab;3091 | 0 | 0 |
| MT072864.1;QIQ54047.1;Pangolin_CoV_orf1ab;3092 | 0 | 0 |
| MT072864.1;QIQ54047.1;Pangolin_CoV_orf1ab;3093 | 0 | 0 |
| MT072864.1;QIQ54047.1;Pangolin_CoV_orf1ab;3094 | 0 | 0 |
| MT072864.1;QIQ54047.1;Pangolin_CoV_orf1ab;3095 | 0 | 0 |
| MT072864.1;QIQ54047.1;Pangolin_CoV_orf1ab;3096 | 0 | 0 |
| MT072864.1;QIQ54047.1;Pangolin_CoV_orf1ab;3097 | 0 | 0 |
| MT072864.1;QIQ54047.1;Pangolin_CoV_orf1ab;3098 | 0 | 0 |
| MT072864.1;QIQ54047.1;Pangolin_CoV_orf1ab;3099 | 0 | 0 |
| MT072864.1;QIQ54047.1;Pangolin_CoV_orf1ab;3100 | 0 | 0 |
| MT072864.1;QIQ54047.1;Pangolin_CoV_orf1ab;3101 | 0 | 0 |
| MT072864.1;QIQ54047.1;Pangolin_CoV_orf1ab;3102 | 0 | 0 |
| MT072864.1;QIQ54047.1;Pangolin_CoV_orf1ab;3103 | 0 | 0 |
| MT072864.1;QIQ54047.1;Pangolin_CoV_orf1ab;3104 | 0 | 0 |
| MT072864.1;QIQ54047.1;Pangolin_CoV_orf1ab;3105 | 0 | 0 |
| MT072864.1;QIQ54047.1;Pangolin_CoV_orf1ab;3106 | 0 | 0 |
| MT072864.1;QIQ54047.1;Pangolin_CoV_orf1ab;3107 | 0 | 0 |
| MT072864.1;QIQ54047.1;Pangolin_CoV_orf1ab;3108 | 0 | 0 |
| MT072864.1;QIQ54047.1;Pangolin_CoV_orf1ab;3109 | 0 | 0 |
| MT072864.1;QIQ54047.1;Pangolin_CoV_orf1ab;3110 | 0 | 0 |
| MT072864.1;QIQ54047.1;Pangolin_CoV_orf1ab;3111 | 0 | 0 |
| MT072864.1;QIQ54047.1;Pangolin_CoV_orf1ab;3112 | 0 | 0 |
| MT072864.1;QIQ54047.1;Pangolin_CoV_orf1ab;3113 | 0 | 0 |
| MT072864.1;QIQ54047.1;Pangolin_CoV_orf1ab;3114 | 0 | 0 |
| MT072864.1;QIQ54047.1;Pangolin_CoV_orf1ab;3115 | 0 | 0 |
| MT072864.1;QIQ54047.1;Pangolin_CoV_orf1ab;3116 | 0 | 0 |
| MT072864.1;QIQ54047.1;Pangolin_CoV_orf1ab;3117 | 0 | 0 |
| MT072864.1;QIQ54047.1;Pangolin_CoV_orf1ab;3118 | 0 | 0 |
| MT072864.1;QIQ54047.1;Pangolin_CoV_orf1ab;3119 | 0 | 0 |
| MT072864.1;QIQ54047.1;Pangolin_CoV_orf1ab;3120 | 0 | 0 |
| MT072864.1;QIQ54047.1;Pangolin_CoV_orf1ab;3121 | 0 | 0 |
| MT072864.1;QIQ54047.1;Pangolin_CoV_orf1ab;3122 | 0 | 0 |
| MT072864.1;QIQ54047.1;Pangolin_CoV_orf1ab;3123 | 0 | 0 |
| MT072864.1;QIQ54047.1;Pangolin_CoV_orf1ab;3124 | 0 | 0 |
| MT072864.1;QIQ54047.1;Pangolin_CoV_orf1ab;3125 | 0 | 0 |
| MT072864.1;QIQ54047.1;Pangolin_CoV_orf1ab;3126 | 0 | 0 |
| MT072864.1;QIQ54047.1;Pangolin_CoV_orf1ab;3127 | 0 | 0 |
| MT072864.1;QIQ54047.1;Pangolin_CoV_orf1ab;3128 | 0 | 0 |
| MT072864.1;QIQ54047.1;Pangolin_CoV_orf1ab;3129 | 0 | 0 |

| | | |
|---|---|---|
| MT072864.1;QIQ54047.1;Pangolin_CoV_orf1ab;3130 | 0 | 0 |
| MT072864.1;QIQ54047.1;Pangolin_CoV_orf1ab;3131 | 0 | 0 |
| MT072864.1;QIQ54047.1;Pangolin_CoV_orf1ab;3132 | 0 | 0 |
| MT072864.1;QIQ54047.1;Pangolin_CoV_orf1ab;3133 | 0 | 0 |
| MT072864.1;QIQ54047.1;Pangolin_CoV_orf1ab;3134 | 0 | 0 |
| MT072864.1;QIQ54047.1;Pangolin_CoV_orf1ab;3135 | 0 | 0 |
| MT072864.1;QIQ54047.1;Pangolin_CoV_orf1ab;3136 | 0 | 0 |
| MT072864.1;QIQ54047.1;Pangolin_CoV_orf1ab;3137 | 0 | 0 |
| MT072864.1;QIQ54047.1;Pangolin_CoV_orf1ab;3138 | 0 | 0 |
| MT072864.1;QIQ54047.1;Pangolin_CoV_orf1ab;3139 | 0 | 0 |
| MT072864.1;QIQ54047.1;Pangolin_CoV_orf1ab;3140 | 0 | 0 |
| MT072864.1;QIQ54047.1;Pangolin_CoV_orf1ab;3141 | 0 | 0 |
| MT072864.1;QIQ54047.1;Pangolin_CoV_orf1ab;3142 | 0 | 0 |
| MT072864.1;QIQ54047.1;Pangolin_CoV_orf1ab;3143 | 0 | 0 |
| MT072864.1;QIQ54047.1;Pangolin_CoV_orf1ab;3144 | 0 | 0 |
| MT072864.1;QIQ54047.1;Pangolin_CoV_orf1ab;3145 | 0 | 0 |
| MT072864.1;QIQ54047.1;Pangolin_CoV_orf1ab;3146 | 0 | 0 |
| MT072864.1;QIQ54047.1;Pangolin_CoV_orf1ab;3147 | 0 | 0 |
| MT072864.1;QIQ54047.1;Pangolin_CoV_orf1ab;3148 | 0 | 0 |
| MT072864.1;QIQ54047.1;Pangolin_CoV_orf1ab;3149 | 0 | 0 |
| MT072864.1;QIQ54047.1;Pangolin_CoV_orf1ab;3150 | 0 | 0 |
| MT072864.1;QIQ54047.1;Pangolin_CoV_orf1ab;3151 | 0 | 0 |
| MT072864.1;QIQ54047.1;Pangolin_CoV_orf1ab;3152 | 0 | 0 |
| MT072864.1;QIQ54047.1;Pangolin_CoV_orf1ab;3153 | 0 | 0 |
| MT072864.1;QIQ54047.1;Pangolin_CoV_orf1ab;3154 | 0 | 0 |
| MT072864.1;QIQ54047.1;Pangolin_CoV_orf1ab;3155 | 0 | 0 |
| MT072864.1;QIQ54047.1;Pangolin_CoV_orf1ab;3156 | 0.05 | 0.025 |
| MT072864.1;QIQ54047.1;Pangolin_CoV_orf1ab;3157 | 0 | 0 |
| MT072864.1;QIQ54047.1;Pangolin_CoV_orf1ab;3158 | 0 | 0 |
| MT072864.1;QIQ54047.1;Pangolin_CoV_orf1ab;3159 | 0 | 0 |
| MT072864.1;QIQ54047.1;Pangolin_CoV_orf1ab;3160 | 0 | 0 |
| MT072864.1;QIQ54047.1;Pangolin_CoV_orf1ab;3161 | 0 | 0 |
| MT072864.1;QIQ54047.1;Pangolin_CoV_orf1ab;3162 | 0 | 0 |
| MT072864.1;QIQ54047.1;Pangolin_CoV_orf1ab;3163 | 0 | 0 |
| MT072864.1;QIQ54047.1;Pangolin_CoV_orf1ab;3164 | 0 | 0 |
| MT072864.1;QIQ54047.1;Pangolin_CoV_orf1ab;3165 | 0 | 0 |
| MT072864.1;QIQ54047.1;Pangolin_CoV_orf1ab;3166 | 0 | 0 |
| MT072864.1;QIQ54047.1;Pangolin_CoV_orf1ab;3167 | 0 | 0 |
| MT072864.1;QIQ54047.1;Pangolin_CoV_orf1ab;3168 | 0 | 0 |
| MT072864.1;QIQ54047.1;Pangolin_CoV_orf1ab;3169 | 0 | 0 |
| MT072864.1;QIQ54047.1;Pangolin_CoV_orf1ab;3170 | 0 | 0 |
| MT072864.1;QIQ54047.1;Pangolin_CoV_orf1ab;3171 | 0 | 0 |
| MT072864.1;QIQ54047.1;Pangolin_CoV_orf1ab;3172 | 0 | 0 |

| | | |
|---|---|---|
| MT072864.1;QIQ54047.1;Pangolin_CoV_orf1ab;3173 | 0 | 0 |
| MT072864.1;QIQ54047.1;Pangolin_CoV_orf1ab;3174 | 0 | 0 |
| MT072864.1;QIQ54047.1;Pangolin_CoV_orf1ab;3175 | 0 | 0 |
| MT072864.1;QIQ54047.1;Pangolin_CoV_orf1ab;3176 | 0 | 0 |
| MT072864.1;QIQ54047.1;Pangolin_CoV_orf1ab;3177 | 0 | 0 |
| MT072864.1;QIQ54047.1;Pangolin_CoV_orf1ab;3178 | 0 | 0 |
| MT072864.1;QIQ54047.1;Pangolin_CoV_orf1ab;3179 | 0 | 0 |
| MT072864.1;QIQ54047.1;Pangolin_CoV_orf1ab;3180 | 0 | 0 |
| MT072864.1;QIQ54047.1;Pangolin_CoV_orf1ab;3181 | 0 | 0 |
| MT072864.1;QIQ54047.1;Pangolin_CoV_orf1ab;3182 | 0 | 0 |
| MT072864.1;QIQ54047.1;Pangolin_CoV_orf1ab;3183 | 0 | 0 |
| MT072864.1;QIQ54047.1;Pangolin_CoV_orf1ab;3184 | 0 | 0 |
| MT072864.1;QIQ54047.1;Pangolin_CoV_orf1ab;3185 | 0 | 0 |
| MT072864.1;QIQ54047.1;Pangolin_CoV_orf1ab;3186 | 0 | 0.025 |
| MT072864.1;QIQ54047.1;Pangolin_CoV_orf1ab;3187 | 0 | 0 |
| MT072864.1;QIQ54047.1;Pangolin_CoV_orf1ab;3188 | 0 | 0 |
| MT072864.1;QIQ54047.1;Pangolin_CoV_orf1ab;3189 | 0 | 0 |
| MT072864.1;QIQ54047.1;Pangolin_CoV_orf1ab;3190 | 0 | 0 |
| MT072864.1;QIQ54047.1;Pangolin_CoV_orf1ab;3191 | 0 | 0 |
| MT072864.1;QIQ54047.1;Pangolin_CoV_orf1ab;3192 | 0 | 0 |
| MT072864.1;QIQ54047.1;Pangolin_CoV_orf1ab;3193 | 0 | 0 |
| MT072864.1;QIQ54047.1;Pangolin_CoV_orf1ab;3194 | 0 | 0 |
| MT072864.1;QIQ54047.1;Pangolin_CoV_orf1ab;3195 | 0 | 0 |
| MT072864.1;QIQ54047.1;Pangolin_CoV_orf1ab;3196 | 0 | 0 |
| MT072864.1;QIQ54047.1;Pangolin_CoV_orf1ab;3197 | 0 | 0 |
| MT072864.1;QIQ54047.1;Pangolin_CoV_orf1ab;3198 | 0 | 0 |
| MT072864.1;QIQ54047.1;Pangolin_CoV_orf1ab;3199 | 0 | 0 |
| MT072864.1;QIQ54047.1;Pangolin_CoV_orf1ab;3200 | 0 | 0 |
| MT072864.1;QIQ54047.1;Pangolin_CoV_orf1ab;3201 | 0 | 0.025 |
| MT072864.1;QIQ54047.1;Pangolin_CoV_orf1ab;3202 | 0 | 0.025 |
| MT072864.1;QIQ54047.1;Pangolin_CoV_orf1ab;3203 | 0 | 0.025 |
| MT072864.1;QIQ54047.1;Pangolin_CoV_orf1ab;3204 | 0 | 0 |
| MT072864.1;QIQ54047.1;Pangolin_CoV_orf1ab;3205 | 0 | 0 |
| MT072864.1;QIQ54047.1;Pangolin_CoV_orf1ab;3206 | 0 | 0 |
| MT072864.1;QIQ54047.1;Pangolin_CoV_orf1ab;3207 | 0.05 | 0.075 |
| MT072864.1;QIQ54047.1;Pangolin_CoV_orf1ab;3208 | 0 | 0 |
| MT072864.1;QIQ54047.1;Pangolin_CoV_orf1ab;3209 | 0 | 0 |
| MT072864.1;QIQ54047.1;Pangolin_CoV_orf1ab;3210 | 0 | 0 |
| MT072864.1;QIQ54047.1;Pangolin_CoV_orf1ab;3211 | 0 | 0.025 |
| MT072864.1;QIQ54047.1;Pangolin_CoV_orf1ab;3212 | 0 | 0 |
| MT072864.1;QIQ54047.1;Pangolin_CoV_orf1ab;3213 | 0 | 0 |
| MT072864.1;QIQ54047.1;Pangolin_CoV_orf1ab;3214 | 0 | 0 |
| MT072864.1;QIQ54047.1;Pangolin_CoV_orf1ab;3215 | 0 | 0 |

| | | |
|---|---|---|
| MT072864.1;QIQ54047.1;Pangolin_CoV_orf1ab;3216 | 0 | 0 |
| MT072864.1;QIQ54047.1;Pangolin_CoV_orf1ab;3217 | 0 | 0.075 |
| MT072864.1;QIQ54047.1;Pangolin_CoV_orf1ab;3218 | 0 | 0.025 |
| MT072864.1;QIQ54047.1;Pangolin_CoV_orf1ab;3219 | 0 | 0.025 |
| MT072864.1;QIQ54047.1;Pangolin_CoV_orf1ab;3220 | 0 | 0.025 |
| MT072864.1;QIQ54047.1;Pangolin_CoV_orf1ab;3221 | 0 | 0.025 |
| MT072864.1;QIQ54047.1;Pangolin_CoV_orf1ab;3222 | 0.05 | 0.025 |
| MT072864.1;QIQ54047.1;Pangolin_CoV_orf1ab;3223 | 0 | 0.025 |
| MT072864.1;QIQ54047.1;Pangolin_CoV_orf1ab;3224 | 0 | 0.025 |
| MT072864.1;QIQ54047.1;Pangolin_CoV_orf1ab;3225 | 0 | 0.025 |
| MT072864.1;QIQ54047.1;Pangolin_CoV_orf1ab;3226 | 0.05 | 0 |
| MT072864.1;QIQ54047.1;Pangolin_CoV_orf1ab;3227 | 0 | 0 |
| MT072864.1;QIQ54047.1;Pangolin_CoV_orf1ab;3228 | 0 | 0 |
| MT072864.1;QIQ54047.1;Pangolin_CoV_orf1ab;3229 | 0.15 | 0.05 |
| MT072864.1;QIQ54047.1;Pangolin_CoV_orf1ab;3230 | 0.3 | 0.05 |
| MT072864.1;QIQ54047.1;Pangolin_CoV_orf1ab;3231 | 0.05 | 0 |
| MT072864.1;QIQ54047.1;Pangolin_CoV_orf1ab;3232 | 0.05 | 0.05 |
| MT072864.1;QIQ54047.1;Pangolin_CoV_orf1ab;3233 | 0.1 | 0.1 |
| MT072864.1;QIQ54047.1;Pangolin_CoV_orf1ab;3234 | 0 | 0.05 |
| MT072864.1;QIQ54047.1;Pangolin_CoV_orf1ab;3235 | 0 | 0.05 |
| MT072864.1;QIQ54047.1;Pangolin_CoV_orf1ab;3236 | 0 | 0 |
| MT072864.1;QIQ54047.1;Pangolin_CoV_orf1ab;3237 | 0 | 0.025 |
| MT072864.1;QIQ54047.1;Pangolin_CoV_orf1ab;3238 | 0 | 0 |
| MT072864.1;QIQ54047.1;Pangolin_CoV_orf1ab;3239 | 0 | 0 |
| MT072864.1;QIQ54047.1;Pangolin_CoV_orf1ab;3240 | 0 | 0 |
| MT072864.1;QIQ54047.1;Pangolin_CoV_orf1ab;3241 | 0 | 0 |
| MT072864.1;QIQ54047.1;Pangolin_CoV_orf1ab;3242 | 0.05 | 0 |
| MT072864.1;QIQ54047.1;Pangolin_CoV_orf1ab;3243 | 0 | 0 |
| MT072864.1;QIQ54047.1;Pangolin_CoV_orf1ab;3244 | 0 | 0 |
| MT072864.1;QIQ54047.1;Pangolin_CoV_orf1ab;3245 | 0 | 0 |
| MT072864.1;QIQ54047.1;Pangolin_CoV_orf1ab;3246 | 0 | 0 |
| MT072864.1;QIQ54047.1;Pangolin_CoV_orf1ab;3247 | 0 | 0 |
| MT072864.1;QIQ54047.1;Pangolin_CoV_orf1ab;3248 | 0 | 0 |
| MT072864.1;QIQ54047.1;Pangolin_CoV_orf1ab;3249 | 0 | 0 |
| MT072864.1;QIQ54047.1;Pangolin_CoV_orf1ab;3250 | 0 | 0 |
| MT072864.1;QIQ54047.1;Pangolin_CoV_orf1ab;3251 | 0 | 0 |
| MT072864.1;QIQ54047.1;Pangolin_CoV_orf1ab;3252 | 0 | 0 |
| MT072864.1;QIQ54047.1;Pangolin_CoV_orf1ab;3253 | 0 | 0 |
| MT072864.1;QIQ54047.1;Pangolin_CoV_orf1ab;3254 | 0 | 0 |
| MT072864.1;QIQ54047.1;Pangolin_CoV_orf1ab;3255 | 0 | 0 |
| MT072864.1;QIQ54047.1;Pangolin_CoV_orf1ab;3256 | 0 | 0 |
| MT072864.1;QIQ54047.1;Pangolin_CoV_orf1ab;3257 | 0 | 0 |
| MT072864.1;QIQ54047.1;Pangolin_CoV_orf1ab;3258 | 0 | 0 |

| | | |
|---|---|---|
| MT072864.1;QIQ54047.1;Pangolin_CoV_orf1ab;3259 | 0 | 0 |
| MT072864.1;QIQ54047.1;Pangolin_CoV_orf1ab;3260 | 0 | 0.025 |
| MT072864.1;QIQ54047.1;Pangolin_CoV_orf1ab;3261 | 0 | 0 |
| MT072864.1;QIQ54047.1;Pangolin_CoV_orf1ab;3262 | 0.05 | 0 |
| MT072864.1;QIQ54047.1;Pangolin_CoV_orf1ab;3263 | 0 | 0 |
| MT072864.1;QIQ54047.1;Pangolin_CoV_orf1ab;3264 | 0 | 0 |
| MT072864.1;QIQ54047.1;Pangolin_CoV_orf1ab;3265 | 0 | 0 |
| MT072864.1;QIQ54047.1;Pangolin_CoV_orf1ab;3266 | 0 | 0 |
| MT072864.1;QIQ54047.1;Pangolin_CoV_orf1ab;3267 | 0 | 0 |
| MT072864.1;QIQ54047.1;Pangolin_CoV_orf1ab;3268 | 0 | 0 |
| MT072864.1;QIQ54047.1;Pangolin_CoV_orf1ab;3269 | 0 | 0 |
| MT072864.1;QIQ54047.1;Pangolin_CoV_orf1ab;3270 | 0 | 0 |
| MT072864.1;QIQ54047.1;Pangolin_CoV_orf1ab;3271 | 0 | 0 |
| MT072864.1;QIQ54047.1;Pangolin_CoV_orf1ab;3272 | 0 | 0 |
| MT072864.1;QIQ54047.1;Pangolin_CoV_orf1ab;3273 | 0 | 0 |
| MT072864.1;QIQ54047.1;Pangolin_CoV_orf1ab;3274 | 0 | 0 |
| MT072864.1;QIQ54047.1;Pangolin_CoV_orf1ab;3275 | 0 | 0 |
| MT072864.1;QIQ54047.1;Pangolin_CoV_orf1ab;3276 | 0 | 0 |
| MT072864.1;QIQ54047.1;Pangolin_CoV_orf1ab;3277 | 0 | 0 |
| MT072864.1;QIQ54047.1;Pangolin_CoV_orf1ab;3278 | 0 | 0 |
| MT072864.1;QIQ54047.1;Pangolin_CoV_orf1ab;3279 | 0 | 0 |
| MT072864.1;QIQ54047.1;Pangolin_CoV_orf1ab;3280 | 0 | 0 |
| MT072864.1;QIQ54047.1;Pangolin_CoV_orf1ab;3281 | 0 | 0 |
| MT072864.1;QIQ54047.1;Pangolin_CoV_orf1ab;3282 | 0 | 0 |
| MT072864.1;QIQ54047.1;Pangolin_CoV_orf1ab;3283 | 0 | 0 |
| MT072864.1;QIQ54047.1;Pangolin_CoV_orf1ab;3284 | 0 | 0 |
| MT072864.1;QIQ54047.1;Pangolin_CoV_orf1ab;3285 | 0 | 0 |
| MT072864.1;QIQ54047.1;Pangolin_CoV_orf1ab;3286 | 0 | 0 |
| MT072864.1;QIQ54047.1;Pangolin_CoV_orf1ab;3287 | 0 | 0 |
| MT072864.1;QIQ54047.1;Pangolin_CoV_orf1ab;3288 | 0 | 0 |
| MT072864.1;QIQ54047.1;Pangolin_CoV_orf1ab;3289 | 0 | 0 |
| MT072864.1;QIQ54047.1;Pangolin_CoV_orf1ab;3290 | 0 | 0 |
| MT072864.1;QIQ54047.1;Pangolin_CoV_orf1ab;3291 | 0 | 0.025 |
| MT072864.1;QIQ54047.1;Pangolin_CoV_orf1ab;3292 | 0 | 0 |
| MT072864.1;QIQ54047.1;Pangolin_CoV_orf1ab;3293 | 0 | 0 |
| MT072864.1;QIQ54047.1;Pangolin_CoV_orf1ab;3294 | 0.05 | 0.025 |
| MT072864.1;QIQ54047.1;Pangolin_CoV_orf1ab;3295 | 0 | 0 |
| MT072864.1;QIQ54047.1;Pangolin_CoV_orf1ab;3296 | 0 | 0 |
| MT072864.1;QIQ54047.1;Pangolin_CoV_orf1ab;3297 | 0 | 0 |
| MT072864.1;QIQ54047.1;Pangolin_CoV_orf1ab;3298 | 0 | 0 |
| MT072864.1;QIQ54047.1;Pangolin_CoV_orf1ab;3299 | 0 | 0 |
| MT072864.1;QIQ54047.1;Pangolin_CoV_orf1ab;3300 | 0 | 0 |
| MT072864.1;QIQ54047.1;Pangolin_CoV_orf1ab;3301 | 0 | 0.025 |

| | | |
|---|---|---|
| MT072864.1;QIQ54047.1;Pangolin_CoV_orf1ab;3302 | 0 | 0.025 |
| MT072864.1;QIQ54047.1;Pangolin_CoV_orf1ab;3303 | 0 | 0.025 |
| MT072864.1;QIQ54047.1;Pangolin_CoV_orf1ab;3304 | 0 | 0 |
| MT072864.1;QIQ54047.1;Pangolin_CoV_orf1ab;3305 | 0.05 | 0 |
| MT072864.1;QIQ54047.1;Pangolin_CoV_orf1ab;3306 | 0 | 0.05 |
| MT072864.1;QIQ54047.1;Pangolin_CoV_orf1ab;3307 | 0 | 0 |
| MT072864.1;QIQ54047.1;Pangolin_CoV_orf1ab;3308 | 0 | 0 |
| MT072864.1;QIQ54047.1;Pangolin_CoV_orf1ab;3309 | 0 | 0 |
| MT072864.1;QIQ54047.1;Pangolin_CoV_orf1ab;3310 | 0 | 0 |
| MT072864.1;QIQ54047.1;Pangolin_CoV_orf1ab;3311 | 0 | 0 |
| MT072864.1;QIQ54047.1;Pangolin_CoV_orf1ab;3312 | 0 | 0 |
| MT072864.1;QIQ54047.1;Pangolin_CoV_orf1ab;3313 | 0 | 0 |
| MT072864.1;QIQ54047.1;Pangolin_CoV_orf1ab;3314 | 0 | 0 |
| MT072864.1;QIQ54047.1;Pangolin_CoV_orf1ab;3315 | 0 | 0 |
| MT072864.1;QIQ54047.1;Pangolin_CoV_orf1ab;3316 | 0 | 0 |
| MT072864.1;QIQ54047.1;Pangolin_CoV_orf1ab;3317 | 0 | 0 |
| MT072864.1;QIQ54047.1;Pangolin_CoV_orf1ab;3318 | 0 | 0 |
| MT072864.1;QIQ54047.1;Pangolin_CoV_orf1ab;3319 | 0 | 0 |
| MT072864.1;QIQ54047.1;Pangolin_CoV_orf1ab;3320 | 0 | 0 |
| MT072864.1;QIQ54047.1;Pangolin_CoV_orf1ab;3321 | 0 | 0 |
| MT072864.1;QIQ54047.1;Pangolin_CoV_orf1ab;3322 | 0 | 0 |
| MT072864.1;QIQ54047.1;Pangolin_CoV_orf1ab;3323 | 0 | 0 |
| MT072864.1;QIQ54047.1;Pangolin_CoV_orf1ab;3324 | 0 | 0 |
| MT072864.1;QIQ54047.1;Pangolin_CoV_orf1ab;3325 | 0 | 0 |
| MT072864.1;QIQ54047.1;Pangolin_CoV_orf1ab;3326 | 0 | 0 |
| MT072864.1;QIQ54047.1;Pangolin_CoV_orf1ab;3327 | 0 | 0 |
| MT072864.1;QIQ54047.1;Pangolin_CoV_orf1ab;3328 | 0 | 0 |
| MT072864.1;QIQ54047.1;Pangolin_CoV_orf1ab;3329 | 0 | 0 |
| MT072864.1;QIQ54047.1;Pangolin_CoV_orf1ab;3330 | 0 | 0 |
| MT072864.1;QIQ54047.1;Pangolin_CoV_orf1ab;3331 | 0 | 0 |
| MT072864.1;QIQ54047.1;Pangolin_CoV_orf1ab;3332 | 0 | 0 |
| MT072864.1;QIQ54047.1;Pangolin_CoV_orf1ab;3333 | 0 | 0 |
| MT072864.1;QIQ54047.1;Pangolin_CoV_orf1ab;3334 | 0.05 | 0 |
| MT072864.1;QIQ54047.1;Pangolin_CoV_orf1ab;3335 | 0 | 0 |
| MT072864.1;QIQ54047.1;Pangolin_CoV_orf1ab;3336 | 0 | 0 |
| MT072864.1;QIQ54047.1;Pangolin_CoV_orf1ab;3337 | 0 | 0.025 |
| MT072864.1;QIQ54047.1;Pangolin_CoV_orf1ab;3338 | 0 | 0 |
| MT072864.1;QIQ54047.1;Pangolin_CoV_orf1ab;3339 | 0 | 0 |
| MT072864.1;QIQ54047.1;Pangolin_CoV_orf1ab;3340 | 0 | 0 |
| MT072864.1;QIQ54047.1;Pangolin_CoV_orf1ab;3341 | 0 | 0 |
| MT072864.1;QIQ54047.1;Pangolin_CoV_orf1ab;3342 | 0 | 0 |
| MT072864.1;QIQ54047.1;Pangolin_CoV_orf1ab;3343 | 0 | 0 |
| MT072864.1;QIQ54047.1;Pangolin_CoV_orf1ab;3344 | 0 | 0 |

| | | |
|---|---|---|
| MT072864.1;QIQ54047.1;Pangolin_CoV_orf1ab;3345 | 0 | 0 |
| MT072864.1;QIQ54047.1;Pangolin_CoV_orf1ab;3346 | 0 | 0 |
| MT072864.1;QIQ54047.1;Pangolin_CoV_orf1ab;3347 | 0 | 0 |
| MT072864.1;QIQ54047.1;Pangolin_CoV_orf1ab;3348 | 0 | 0 |
| MT072864.1;QIQ54047.1;Pangolin_CoV_orf1ab;3349 | 0 | 0 |
| MT072864.1;QIQ54047.1;Pangolin_CoV_orf1ab;3350 | 0 | 0 |
| MT072864.1;QIQ54047.1;Pangolin_CoV_orf1ab;3351 | 0 | 0 |
| MT072864.1;QIQ54047.1;Pangolin_CoV_orf1ab;3352 | 0 | 0 |
| MT072864.1;QIQ54047.1;Pangolin_CoV_orf1ab;3353 | 0 | 0 |
| MT072864.1;QIQ54047.1;Pangolin_CoV_orf1ab;3354 | 0 | 0 |
| MT072864.1;QIQ54047.1;Pangolin_CoV_orf1ab;3355 | 0 | 0 |
| MT072864.1;QIQ54047.1;Pangolin_CoV_orf1ab;3356 | 0 | 0 |
| MT072864.1;QIQ54047.1;Pangolin_CoV_orf1ab;3357 | 0 | 0 |
| MT072864.1;QIQ54047.1;Pangolin_CoV_orf1ab;3358 | 0 | 0 |
| MT072864.1;QIQ54047.1;Pangolin_CoV_orf1ab;3359 | 0 | 0 |
| MT072864.1;QIQ54047.1;Pangolin_CoV_orf1ab;3360 | 0 | 0 |
| MT072864.1;QIQ54047.1;Pangolin_CoV_orf1ab;3361 | 0 | 0 |
| MT072864.1;QIQ54047.1;Pangolin_CoV_orf1ab;3362 | 0 | 0.025 |
| MT072864.1;QIQ54047.1;Pangolin_CoV_orf1ab;3363 | 0 | 0 |
| MT072864.1;QIQ54047.1;Pangolin_CoV_orf1ab;3364 | 0 | 0 |
| MT072864.1;QIQ54047.1;Pangolin_CoV_orf1ab;3365 | 0 | 0 |
| MT072864.1;QIQ54047.1;Pangolin_CoV_orf1ab;3366 | 0 | 0 |
| MT072864.1;QIQ54047.1;Pangolin_CoV_orf1ab;3367 | 0 | 0 |
| MT072864.1;QIQ54047.1;Pangolin_CoV_orf1ab;3368 | 0 | 0 |
| MT072864.1;QIQ54047.1;Pangolin_CoV_orf1ab;3369 | 0 | 0 |
| MT072864.1;QIQ54047.1;Pangolin_CoV_orf1ab;3370 | 0 | 0 |
| MT072864.1;QIQ54047.1;Pangolin_CoV_orf1ab;3371 | 0 | 0 |
| MT072864.1;QIQ54047.1;Pangolin_CoV_orf1ab;3372 | 0 | 0 |
| MT072864.1;QIQ54047.1;Pangolin_CoV_orf1ab;3373 | 0 | 0.025 |
| MT072864.1;QIQ54047.1;Pangolin_CoV_orf1ab;3374 | 0 | 0 |
| MT072864.1;QIQ54047.1;Pangolin_CoV_orf1ab;3375 | 0 | 0 |
| MT072864.1;QIQ54047.1;Pangolin_CoV_orf1ab;3376 | 0 | 0 |
| MT072864.1;QIQ54047.1;Pangolin_CoV_orf1ab;3377 | 0 | 0 |
| MT072864.1;QIQ54047.1;Pangolin_CoV_orf1ab;3378 | 0 | 0 |
| MT072864.1;QIQ54047.1;Pangolin_CoV_orf1ab;3379 | 0 | 0.025 |
| MT072864.1;QIQ54047.1;Pangolin_CoV_orf1ab;3380 | 0 | 0 |
| MT072864.1;QIQ54047.1;Pangolin_CoV_orf1ab;3381 | 0 | 0 |
| MT072864.1;QIQ54047.1;Pangolin_CoV_orf1ab;3382 | 0 | 0 |
| MT072864.1;QIQ54047.1;Pangolin_CoV_orf1ab;3383 | 0 | 0 |
| MT072864.1;QIQ54047.1;Pangolin_CoV_orf1ab;3384 | 0.05 | 0 |
| MT072864.1;QIQ54047.1;Pangolin_CoV_orf1ab;3385 | 0 | 0 |
| MT072864.1;QIQ54047.1;Pangolin_CoV_orf1ab;3386 | 0.05 | 0 |
| MT072864.1;QIQ54047.1;Pangolin_CoV_orf1ab;3387 | 0 | 0 |

| | | |
|---|---|---|
| MT072864.1;QIQ54047.1;Pangolin_CoV_orf1ab;3388 | 0 | 0 |
| MT072864.1;QIQ54047.1;Pangolin_CoV_orf1ab;3389 | 0 | 0 |
| MT072864.1;QIQ54047.1;Pangolin_CoV_orf1ab;3390 | 0 | 0 |
| MT072864.1;QIQ54047.1;Pangolin_CoV_orf1ab;3391 | 0 | 0 |
| MT072864.1;QIQ54047.1;Pangolin_CoV_orf1ab;3392 | 0 | 0.025 |
| MT072864.1;QIQ54047.1;Pangolin_CoV_orf1ab;3393 | 0 | 0.025 |
| MT072864.1;QIQ54047.1;Pangolin_CoV_orf1ab;3394 | 0 | 0.025 |
| MT072864.1;QIQ54047.1;Pangolin_CoV_orf1ab;3395 | 0 | 0.05 |
| MT072864.1;QIQ54047.1;Pangolin_CoV_orf1ab;3396 | 0 | 0.05 |
| MT072864.1;QIQ54047.1;Pangolin_CoV_orf1ab;3397 | 0 | 0.025 |
| MT072864.1;QIQ54047.1;Pangolin_CoV_orf1ab;3398 | 0 | 0.025 |
| MT072864.1;QIQ54047.1;Pangolin_CoV_orf1ab;3399 | 0 | 0 |
| MT072864.1;QIQ54047.1;Pangolin_CoV_orf1ab;3400 | 0 | 0 |
| MT072864.1;QIQ54047.1;Pangolin_CoV_orf1ab;3401 | 0 | 0 |
| MT072864.1;QIQ54047.1;Pangolin_CoV_orf1ab;3402 | 0 | 0 |
| MT072864.1;QIQ54047.1;Pangolin_CoV_orf1ab;3403 | 0 | 0 |
| MT072864.1;QIQ54047.1;Pangolin_CoV_orf1ab;3404 | 0 | 0 |
| MT072864.1;QIQ54047.1;Pangolin_CoV_orf1ab;3405 | 0 | 0 |
| MT072864.1;QIQ54047.1;Pangolin_CoV_orf1ab;3406 | 0 | 0 |
| MT072864.1;QIQ54047.1;Pangolin_CoV_orf1ab;3407 | 0 | 0 |
| MT072864.1;QIQ54047.1;Pangolin_CoV_orf1ab;3408 | 0 | 0 |
| MT072864.1;QIQ54047.1;Pangolin_CoV_orf1ab;3409 | 0 | 0 |
| MT072864.1;QIQ54047.1;Pangolin_CoV_orf1ab;3410 | 0 | 0 |
| MT072864.1;QIQ54047.1;Pangolin_CoV_orf1ab;3411 | 0 | 0 |
| MT072864.1;QIQ54047.1;Pangolin_CoV_orf1ab;3412 | 0 | 0 |
| MT072864.1;QIQ54047.1;Pangolin_CoV_orf1ab;3413 | 0 | 0 |
| MT072864.1;QIQ54047.1;Pangolin_CoV_orf1ab;3414 | 0 | 0 |
| MT072864.1;QIQ54047.1;Pangolin_CoV_orf1ab;3415 | 0 | 0 |
| MT072864.1;QIQ54047.1;Pangolin_CoV_orf1ab;3416 | 0 | 0 |
| MT072864.1;QIQ54047.1;Pangolin_CoV_orf1ab;3417 | 0 | 0 |
| MT072864.1;QIQ54047.1;Pangolin_CoV_orf1ab;3418 | 0.15 | 0.025 |
| MT072864.1;QIQ54047.1;Pangolin_CoV_orf1ab;3419 | 0.1 | 0.05 |
| MT072864.1;QIQ54047.1;Pangolin_CoV_orf1ab;3420 | 0.05 | 0.025 |
| MT072864.1;QIQ54047.1;Pangolin_CoV_orf1ab;3421 | 0.2 | 0.075 |
| MT072864.1;QIQ54047.1;Pangolin_CoV_orf1ab;3422 | 0.2 | 0.05 |
| MT072864.1;QIQ54047.1;Pangolin_CoV_orf1ab;3423 | 0.2 | 0.05 |
| MT072864.1;QIQ54047.1;Pangolin_CoV_orf1ab;3424 | 0.2 | 0.05 |
| MT072864.1;QIQ54047.1;Pangolin_CoV_orf1ab;3425 | 0.1 | 0.025 |
| MT072864.1;QIQ54047.1;Pangolin_CoV_orf1ab;3426 | 0.05 | 0.025 |
| MT072864.1;QIQ54047.1;Pangolin_CoV_orf1ab;3427 | 0.05 | 0.025 |
| MT072864.1;QIQ54047.1;Pangolin_CoV_orf1ab;3428 | 0.05 | 0.05 |
| MT072864.1;QIQ54047.1;Pangolin_CoV_orf1ab;3429 | 0.15 | 0.1 |
| MT072864.1;QIQ54047.1;Pangolin_CoV_orf1ab;3430 | 0 | 0.025 |

| | | |
|---|---|---|
| MT072864.1;QIQ54047.1;Pangolin_CoV_orf1ab;3431 | 0 | 0 |
| MT072864.1;QIQ54047.1;Pangolin_CoV_orf1ab;3432 | 0.05 | 0 |
| MT072864.1;QIQ54047.1;Pangolin_CoV_orf1ab;3433 | 0.05 | 0 |
| MT072864.1;QIQ54047.1;Pangolin_CoV_orf1ab;3434 | 0.1 | 0.025 |
| MT072864.1;QIQ54047.1;Pangolin_CoV_orf1ab;3435 | 0.05 | 0 |
| MT072864.1;QIQ54047.1;Pangolin_CoV_orf1ab;3436 | 0.1 | 0 |
| MT072864.1;QIQ54047.1;Pangolin_CoV_orf1ab;3437 | 0 | 0 |
| MT072864.1;QIQ54047.1;Pangolin_CoV_orf1ab;3438 | 0.05 | 0 |
| MT072864.1;QIQ54047.1;Pangolin_CoV_orf1ab;3439 | 0 | 0 |
| MT072864.1;QIQ54047.1;Pangolin_CoV_orf1ab;3440 | 0 | 0 |
| MT072864.1;QIQ54047.1;Pangolin_CoV_orf1ab;3441 | 0 | 0 |
| MT072864.1;QIQ54047.1;Pangolin_CoV_orf1ab;3442 | 0 | 0 |
| MT072864.1;QIQ54047.1;Pangolin_CoV_orf1ab;3443 | 0 | 0 |
| MT072864.1;QIQ54047.1;Pangolin_CoV_orf1ab;3444 | 0 | 0 |
| MT072864.1;QIQ54047.1;Pangolin_CoV_orf1ab;3445 | 0 | 0 |
| MT072864.1;QIQ54047.1;Pangolin_CoV_orf1ab;3446 | 0 | 0 |
| MT072864.1;QIQ54047.1;Pangolin_CoV_orf1ab;3447 | 0 | 0 |
| MT072864.1;QIQ54047.1;Pangolin_CoV_orf1ab;3448 | 0 | 0 |
| MT072864.1;QIQ54047.1;Pangolin_CoV_orf1ab;3449 | 0 | 0 |
| MT072864.1;QIQ54047.1;Pangolin_CoV_orf1ab;3450 | 0 | 0 |
| MT072864.1;QIQ54047.1;Pangolin_CoV_orf1ab;3451 | 0 | 0 |
| MT072864.1;QIQ54047.1;Pangolin_CoV_orf1ab;3452 | 0 | 0 |
| MT072864.1;QIQ54047.1;Pangolin_CoV_orf1ab;3453 | 0 | 0 |
| MT072864.1;QIQ54047.1;Pangolin_CoV_orf1ab;3454 | 0 | 0 |
| MT072864.1;QIQ54047.1;Pangolin_CoV_orf1ab;3455 | 0 | 0 |
| MT072864.1;QIQ54047.1;Pangolin_CoV_orf1ab;3456 | 0 | 0 |
| MT072864.1;QIQ54047.1;Pangolin_CoV_orf1ab;3457 | 0 | 0 |
| MT072864.1;QIQ54047.1;Pangolin_CoV_orf1ab;3458 | 0 | 0 |
| MT072864.1;QIQ54047.1;Pangolin_CoV_orf1ab;3459 | 0 | 0 |
| MT072864.1;QIQ54047.1;Pangolin_CoV_orf1ab;3460 | 0 | 0 |
| MT072864.1;QIQ54047.1;Pangolin_CoV_orf1ab;3461 | 0 | 0 |
| MT072864.1;QIQ54047.1;Pangolin_CoV_orf1ab;3462 | 0 | 0 |
| MT072864.1;QIQ54047.1;Pangolin_CoV_orf1ab;3463 | 0 | 0 |
| MT072864.1;QIQ54047.1;Pangolin_CoV_orf1ab;3464 | 0 | 0 |
| MT072864.1;QIQ54047.1;Pangolin_CoV_orf1ab;3465 | 0 | 0 |
| MT072864.1;QIQ54047.1;Pangolin_CoV_orf1ab;3466 | 0 | 0 |
| MT072864.1;QIQ54047.1;Pangolin_CoV_orf1ab;3467 | 0 | 0 |
| MT072864.1;QIQ54047.1;Pangolin_CoV_orf1ab;3468 | 0 | 0 |
| MT072864.1;QIQ54047.1;Pangolin_CoV_orf1ab;3469 | 0 | 0 |
| MT072864.1;QIQ54047.1;Pangolin_CoV_orf1ab;3470 | 0 | 0 |
| MT072864.1;QIQ54047.1;Pangolin_CoV_orf1ab;3471 | 0.05 | 0 |
| MT072864.1;QIQ54047.1;Pangolin_CoV_orf1ab;3472 | 0.05 | 0 |
| MT072864.1;QIQ54047.1;Pangolin_CoV_orf1ab;3473 | 0.05 | 0 |

| | | |
|---|---|---|
| MT072864.1;QIQ54047.1;Pangolin_CoV_orf1ab;3474 | 0.05 | 0.025 |
| MT072864.1;QIQ54047.1;Pangolin_CoV_orf1ab;3475 | 0.05 | 0.05 |
| MT072864.1;QIQ54047.1;Pangolin_CoV_orf1ab;3476 | 0.05 | 0 |
| MT072864.1;QIQ54047.1;Pangolin_CoV_orf1ab;3477 | 0.1 | 0.025 |
| MT072864.1;QIQ54047.1;Pangolin_CoV_orf1ab;3478 | 0.05 | 0 |
| MT072864.1;QIQ54047.1;Pangolin_CoV_orf1ab;3479 | 0.05 | 0.025 |
| MT072864.1;QIQ54047.1;Pangolin_CoV_orf1ab;3480 | 0.05 | 0.275 |
| MT072864.1;QIQ54047.1;Pangolin_CoV_orf1ab;3481 | 0.25 | 0.075 |
| MT072864.1;QIQ54047.1;Pangolin_CoV_orf1ab;3482 | 0.1 | 0 |
| MT072864.1;QIQ54047.1;Pangolin_CoV_orf1ab;3483 | 0.05 | 0 |
| MT072864.1;QIQ54047.1;Pangolin_CoV_orf1ab;3484 | 0.05 | 0 |
| MT072864.1;QIQ54047.1;Pangolin_CoV_orf1ab;3485 | 0.05 | 0 |
| MT072864.1;QIQ54047.1;Pangolin_CoV_orf1ab;3486 | 0.1 | 0.025 |
| MT072864.1;QIQ54047.1;Pangolin_CoV_orf1ab;3487 | 0.1 | 0.025 |
| MT072864.1;QIQ54047.1;Pangolin_CoV_orf1ab;3488 | 0.15 | 0 |
| MT072864.1;QIQ54047.1;Pangolin_CoV_orf1ab;3489 | 0.1 | 0 |
| MT072864.1;QIQ54047.1;Pangolin_CoV_orf1ab;3490 | 0.05 | 0 |
| MT072864.1;QIQ54047.1;Pangolin_CoV_orf1ab;3491 | 0.05 | 0 |
| MT072864.1;QIQ54047.1;Pangolin_CoV_orf1ab;3492 | 0 | 0.025 |
| MT072864.1;QIQ54047.1;Pangolin_CoV_orf1ab;3493 | 0 | 0 |
| MT072864.1;QIQ54047.1;Pangolin_CoV_orf1ab;3494 | 0 | 0 |
| MT072864.1;QIQ54047.1;Pangolin_CoV_orf1ab;3495 | 0.05 | 0 |
| MT072864.1;QIQ54047.1;Pangolin_CoV_orf1ab;3496 | 0 | 0 |
| MT072864.1;QIQ54047.1;Pangolin_CoV_orf1ab;3497 | 0.05 | 0.025 |
| MT072864.1;QIQ54047.1;Pangolin_CoV_orf1ab;3498 | 0.15 | 0.1 |
| MT072864.1;QIQ54047.1;Pangolin_CoV_orf1ab;3499 | 0.05 | 0.05 |
| MT072864.1;QIQ54047.1;Pangolin_CoV_orf1ab;3500 | 0.1 | 0.15 |
| MT072864.1;QIQ54047.1;Pangolin_CoV_orf1ab;3501 | 0.05 | 0.025 |
| MT072864.1;QIQ54047.1;Pangolin_CoV_orf1ab;3502 | 0.05 | 0 |
| MT072864.1;QIQ54047.1;Pangolin_CoV_orf1ab;3503 | 0.05 | 0 |
| MT072864.1;QIQ54047.1;Pangolin_CoV_orf1ab;3504 | 0.05 | 0 |
| MT072864.1;QIQ54047.1;Pangolin_CoV_orf1ab;3505 | 0.05 | 0 |
| MT072864.1;QIQ54047.1;Pangolin_CoV_orf1ab;3506 | 0.05 | 0 |
| MT072864.1;QIQ54047.1;Pangolin_CoV_orf1ab;3507 | 0.05 | 0 |
| MT072864.1;QIQ54047.1;Pangolin_CoV_orf1ab;3508 | 0 | 0 |
| MT072864.1;QIQ54047.1;Pangolin_CoV_orf1ab;3509 | 0 | 0 |
| MT072864.1;QIQ54047.1;Pangolin_CoV_orf1ab;3510 | 0 | 0 |
| MT072864.1;QIQ54047.1;Pangolin_CoV_orf1ab;3511 | 0 | 0 |
| MT072864.1;QIQ54047.1;Pangolin_CoV_orf1ab;3512 | 0 | 0 |
| MT072864.1;QIQ54047.1;Pangolin_CoV_orf1ab;3513 | 0.05 | 0 |
| MT072864.1;QIQ54047.1;Pangolin_CoV_orf1ab;3514 | 0 | 0 |
| MT072864.1;QIQ54047.1;Pangolin_CoV_orf1ab;3515 | 0 | 0 |
| MT072864.1;QIQ54047.1;Pangolin_CoV_orf1ab;3516 | 0 | 0 |

| | | |
|---|---|---|
| MT072864.1;QIQ54047.1;Pangolin_CoV_orf1ab;3517 | 0.1 | 0 |
| MT072864.1;QIQ54047.1;Pangolin_CoV_orf1ab;3518 | 0.05 | 0.025 |
| MT072864.1;QIQ54047.1;Pangolin_CoV_orf1ab;3519 | 0.05 | 0 |
| MT072864.1;QIQ54047.1;Pangolin_CoV_orf1ab;3520 | 0.1 | 0 |
| MT072864.1;QIQ54047.1;Pangolin_CoV_orf1ab;3521 | 0.05 | 0.025 |
| MT072864.1;QIQ54047.1;Pangolin_CoV_orf1ab;3522 | 0 | 0 |
| MT072864.1;QIQ54047.1;Pangolin_CoV_orf1ab;3523 | 0 | 0.025 |
| MT072864.1;QIQ54047.1;Pangolin_CoV_orf1ab;3524 | 0 | 0.025 |
| MT072864.1;QIQ54047.1;Pangolin_CoV_orf1ab;3525 | 0 | 0 |
| MT072864.1;QIQ54047.1;Pangolin_CoV_orf1ab;3526 | 0 | 0 |
| MT072864.1;QIQ54047.1;Pangolin_CoV_orf1ab;3527 | 0 | 0 |
| MT072864.1;QIQ54047.1;Pangolin_CoV_orf1ab;3528 | 0 | 0 |
| MT072864.1;QIQ54047.1;Pangolin_CoV_orf1ab;3529 | 0 | 0 |
| MT072864.1;QIQ54047.1;Pangolin_CoV_orf1ab;3530 | 0 | 0 |
| MT072864.1;QIQ54047.1;Pangolin_CoV_orf1ab;3531 | 0 | 0 |
| MT072864.1;QIQ54047.1;Pangolin_CoV_orf1ab;3532 | 0 | 0 |
| MT072864.1;QIQ54047.1;Pangolin_CoV_orf1ab;3533 | 0 | 0 |
| MT072864.1;QIQ54047.1;Pangolin_CoV_orf1ab;3534 | 0 | 0.025 |
| MT072864.1;QIQ54047.1;Pangolin_CoV_orf1ab;3535 | 0 | 0.05 |
| MT072864.1;QIQ54047.1;Pangolin_CoV_orf1ab;3536 | 0 | 0.025 |
| MT072864.1;QIQ54047.1;Pangolin_CoV_orf1ab;3537 | 0 | 0.025 |
| MT072864.1;QIQ54047.1;Pangolin_CoV_orf1ab;3538 | 0 | 0.025 |
| MT072864.1;QIQ54047.1;Pangolin_CoV_orf1ab;3539 | 0.05 | 0.025 |
| MT072864.1;QIQ54047.1;Pangolin_CoV_orf1ab;3540 | 0.05 | 0.05 |
| MT072864.1;QIQ54047.1;Pangolin_CoV_orf1ab;3541 | 0 | 0.025 |
| MT072864.1;QIQ54047.1;Pangolin_CoV_orf1ab;3542 | 0 | 0.05 |
| MT072864.1;QIQ54047.1;Pangolin_CoV_orf1ab;3543 | 0.1 | 0.025 |
| MT072864.1;QIQ54047.1;Pangolin_CoV_orf1ab;3544 | 0 | 0 |
| MT072864.1;QIQ54047.1;Pangolin_CoV_orf1ab;3545 | 0 | 0 |
| MT072864.1;QIQ54047.1;Pangolin_CoV_orf1ab;3546 | 0 | 0 |
| MT072864.1;QIQ54047.1;Pangolin_CoV_orf1ab;3547 | 0 | 0 |
| MT072864.1;QIQ54047.1;Pangolin_CoV_orf1ab;3548 | 0 | 0 |
| MT072864.1;QIQ54047.1;Pangolin_CoV_orf1ab;3549 | 0 | 0 |
| MT072864.1;QIQ54047.1;Pangolin_CoV_orf1ab;3550 | 0 | 0 |
| MT072864.1;QIQ54047.1;Pangolin_CoV_orf1ab;3551 | 0 | 0 |
| MT072864.1;QIQ54047.1;Pangolin_CoV_orf1ab;3552 | 0 | 0 |
| MT072864.1;QIQ54047.1;Pangolin_CoV_orf1ab;3553 | 0 | 0 |
| MT072864.1;QIQ54047.1;Pangolin_CoV_orf1ab;3554 | 0 | 0.025 |
| MT072864.1;QIQ54047.1;Pangolin_CoV_orf1ab;3555 | 0 | 0 |
| MT072864.1;QIQ54047.1;Pangolin_CoV_orf1ab;3556 | 0 | 0 |
| MT072864.1;QIQ54047.1;Pangolin_CoV_orf1ab;3557 | 0 | 0 |
| MT072864.1;QIQ54047.1;Pangolin_CoV_orf1ab;3558 | 0 | 0 |
| MT072864.1;QIQ54047.1;Pangolin_CoV_orf1ab;3559 | 0 | 0 |

| | | |
|---|---|---|
| MT072864.1;QIQ54047.1;Pangolin_CoV_orf1ab;3560 | 0 | 0 |
| MT072864.1;QIQ54047.1;Pangolin_CoV_orf1ab;3561 | 0 | 0 |
| MT072864.1;QIQ54047.1;Pangolin_CoV_orf1ab;3562 | 0 | 0 |
| MT072864.1;QIQ54047.1;Pangolin_CoV_orf1ab;3563 | 0 | 0 |
| MT072864.1;QIQ54047.1;Pangolin_CoV_orf1ab;3564 | 0 | 0 |
| MT072864.1;QIQ54047.1;Pangolin_CoV_orf1ab;3565 | 0 | 0.025 |
| MT072864.1;QIQ54047.1;Pangolin_CoV_orf1ab;3566 | 0 | 0 |
| MT072864.1;QIQ54047.1;Pangolin_CoV_orf1ab;3567 | 0 | 0 |
| MT072864.1;QIQ54047.1;Pangolin_CoV_orf1ab;3568 | 0 | 0 |
| MT072864.1;QIQ54047.1;Pangolin_CoV_orf1ab;3569 | 0 | 0 |
| MT072864.1;QIQ54047.1;Pangolin_CoV_orf1ab;3570 | 0 | 0 |
| MT072864.1;QIQ54047.1;Pangolin_CoV_orf1ab;3571 | 0 | 0 |
| MT072864.1;QIQ54047.1;Pangolin_CoV_orf1ab;3572 | 0 | 0 |
| MT072864.1;QIQ54047.1;Pangolin_CoV_orf1ab;3573 | 0 | 0 |
| MT072864.1;QIQ54047.1;Pangolin_CoV_orf1ab;3574 | 0 | 0 |
| MT072864.1;QIQ54047.1;Pangolin_CoV_orf1ab;3575 | 0 | 0 |
| MT072864.1;QIQ54047.1;Pangolin_CoV_orf1ab;3576 | 0 | 0 |
| MT072864.1;QIQ54047.1;Pangolin_CoV_orf1ab;3577 | 0 | 0 |
| MT072864.1;QIQ54047.1;Pangolin_CoV_orf1ab;3578 | 0 | 0 |
| MT072864.1;QIQ54047.1;Pangolin_CoV_orf1ab;3579 | 0 | 0 |
| MT072864.1;QIQ54047.1;Pangolin_CoV_orf1ab;3580 | 0 | 0 |
| MT072864.1;QIQ54047.1;Pangolin_CoV_orf1ab;3581 | 0 | 0 |
| MT072864.1;QIQ54047.1;Pangolin_CoV_orf1ab;3582 | 0 | 0 |
| MT072864.1;QIQ54047.1;Pangolin_CoV_orf1ab;3583 | 0 | 0 |
| MT072864.1;QIQ54047.1;Pangolin_CoV_orf1ab;3584 | 0 | 0 |
| MT072864.1;QIQ54047.1;Pangolin_CoV_orf1ab;3585 | 0 | 0 |
| MT072864.1;QIQ54047.1;Pangolin_CoV_orf1ab;3586 | 0 | 0 |
| MT072864.1;QIQ54047.1;Pangolin_CoV_orf1ab;3587 | 0 | 0 |
| MT072864.1;QIQ54047.1;Pangolin_CoV_orf1ab;3588 | 0 | 0 |
| MT072864.1;QIQ54047.1;Pangolin_CoV_orf1ab;3589 | 0 | 0 |
| MT072864.1;QIQ54047.1;Pangolin_CoV_orf1ab;3590 | 0 | 0 |
| MT072864.1;QIQ54047.1;Pangolin_CoV_orf1ab;3591 | 0 | 0 |
| MT072864.1;QIQ54047.1;Pangolin_CoV_orf1ab;3592 | 0 | 0 |
| MT072864.1;QIQ54047.1;Pangolin_CoV_orf1ab;3593 | 0 | 0 |
| MT072864.1;QIQ54047.1;Pangolin_CoV_orf1ab;3594 | 0 | 0 |
| MT072864.1;QIQ54047.1;Pangolin_CoV_orf1ab;3595 | 0 | 0 |
| MT072864.1;QIQ54047.1;Pangolin_CoV_orf1ab;3596 | 0 | 0 |
| MT072864.1;QIQ54047.1;Pangolin_CoV_orf1ab;3597 | 0 | 0 |
| MT072864.1;QIQ54047.1;Pangolin_CoV_orf1ab;3598 | 0 | 0 |
| MT072864.1;QIQ54047.1;Pangolin_CoV_orf1ab;3599 | 0 | 0 |
| MT072864.1;QIQ54047.1;Pangolin_CoV_orf1ab;3600 | 0 | 0 |
| MT072864.1;QIQ54047.1;Pangolin_CoV_orf1ab;3601 | 0 | 0 |
| MT072864.1;QIQ54047.1;Pangolin_CoV_orf1ab;3602 | 0 | 0 |

| | | |
|---|---|---|
| MT072864.1;QIQ54047.1;Pangolin_CoV_orf1ab;3603 | 0 | 0 |
| MT072864.1;QIQ54047.1;Pangolin_CoV_orf1ab;3604 | 0 | 0 |
| MT072864.1;QIQ54047.1;Pangolin_CoV_orf1ab;3605 | 0 | 0 |
| MT072864.1;QIQ54047.1;Pangolin_CoV_orf1ab;3606 | 0 | 0 |
| MT072864.1;QIQ54047.1;Pangolin_CoV_orf1ab;3607 | 0 | 0 |
| MT072864.1;QIQ54047.1;Pangolin_CoV_orf1ab;3608 | 0 | 0 |
| MT072864.1;QIQ54047.1;Pangolin_CoV_orf1ab;3609 | 0 | 0 |
| MT072864.1;QIQ54047.1;Pangolin_CoV_orf1ab;3610 | 0 | 0 |
| MT072864.1;QIQ54047.1;Pangolin_CoV_orf1ab;3611 | 0 | 0 |
| MT072864.1;QIQ54047.1;Pangolin_CoV_orf1ab;3612 | 0 | 0 |
| MT072864.1;QIQ54047.1;Pangolin_CoV_orf1ab;3613 | 0 | 0 |
| MT072864.1;QIQ54047.1;Pangolin_CoV_orf1ab;3614 | 0 | 0 |
| MT072864.1;QIQ54047.1;Pangolin_CoV_orf1ab;3615 | 0 | 0 |
| MT072864.1;QIQ54047.1;Pangolin_CoV_orf1ab;3616 | 0 | 0 |
| MT072864.1;QIQ54047.1;Pangolin_CoV_orf1ab;3617 | 0 | 0 |
| MT072864.1;QIQ54047.1;Pangolin_CoV_orf1ab;3618 | 0 | 0 |
| MT072864.1;QIQ54047.1;Pangolin_CoV_orf1ab;3619 | 0 | 0 |
| MT072864.1;QIQ54047.1;Pangolin_CoV_orf1ab;3620 | 0 | 0 |
| MT072864.1;QIQ54047.1;Pangolin_CoV_orf1ab;3621 | 0 | 0 |
| MT072864.1;QIQ54047.1;Pangolin_CoV_orf1ab;3622 | 0 | 0 |
| MT072864.1;QIQ54047.1;Pangolin_CoV_orf1ab;3623 | 0 | 0 |
| MT072864.1;QIQ54047.1;Pangolin_CoV_orf1ab;3624 | 0 | 0 |
| MT072864.1;QIQ54047.1;Pangolin_CoV_orf1ab;3625 | 0 | 0 |
| MT072864.1;QIQ54047.1;Pangolin_CoV_orf1ab;3626 | 0 | 0 |
| MT072864.1;QIQ54047.1;Pangolin_CoV_orf1ab;3627 | 0 | 0 |
| MT072864.1;QIQ54047.1;Pangolin_CoV_orf1ab;3628 | 0 | 0 |
| MT072864.1;QIQ54047.1;Pangolin_CoV_orf1ab;3629 | 0 | 0 |
| MT072864.1;QIQ54047.1;Pangolin_CoV_orf1ab;3630 | 0 | 0 |
| MT072864.1;QIQ54047.1;Pangolin_CoV_orf1ab;3631 | 0 | 0 |
| MT072864.1;QIQ54047.1;Pangolin_CoV_orf1ab;3632 | 0 | 0 |
| MT072864.1;QIQ54047.1;Pangolin_CoV_orf1ab;3633 | 0 | 0 |
| MT072864.1;QIQ54047.1;Pangolin_CoV_orf1ab;3634 | 0 | 0 |
| MT072864.1;QIQ54047.1;Pangolin_CoV_orf1ab;3635 | 0 | 0 |
| MT072864.1;QIQ54047.1;Pangolin_CoV_orf1ab;3636 | 0 | 0 |
| MT072864.1;QIQ54047.1;Pangolin_CoV_orf1ab;3637 | 0 | 0 |
| MT072864.1;QIQ54047.1;Pangolin_CoV_orf1ab;3638 | 0 | 0 |
| MT072864.1;QIQ54047.1;Pangolin_CoV_orf1ab;3639 | 0 | 0 |
| MT072864.1;QIQ54047.1;Pangolin_CoV_orf1ab;3640 | 0 | 0 |
| MT072864.1;QIQ54047.1;Pangolin_CoV_orf1ab;3641 | 0 | 0 |
| MT072864.1;QIQ54047.1;Pangolin_CoV_orf1ab;3642 | 0 | 0 |
| MT072864.1;QIQ54047.1;Pangolin_CoV_orf1ab;3643 | 0 | 0 |
| MT072864.1;QIQ54047.1;Pangolin_CoV_orf1ab;3644 | 0 | 0 |
| MT072864.1;QIQ54047.1;Pangolin_CoV_orf1ab;3645 | 0 | 0 |

| | | |
|---|---|---|
| MT072864.1;QIQ54047.1;Pangolin_CoV_orf1ab;3646 | 0 | 0 |
| MT072864.1;QIQ54047.1;Pangolin_CoV_orf1ab;3647 | 0 | 0 |
| MT072864.1;QIQ54047.1;Pangolin_CoV_orf1ab;3648 | 0 | 0 |
| MT072864.1;QIQ54047.1;Pangolin_CoV_orf1ab;3649 | 0 | 0 |
| MT072864.1;QIQ54047.1;Pangolin_CoV_orf1ab;3650 | 0 | 0 |
| MT072864.1;QIQ54047.1;Pangolin_CoV_orf1ab;3651 | 0 | 0 |
| MT072864.1;QIQ54047.1;Pangolin_CoV_orf1ab;3652 | 0 | 0 |
| MT072864.1;QIQ54047.1;Pangolin_CoV_orf1ab;3653 | 0 | 0 |
| MT072864.1;QIQ54047.1;Pangolin_CoV_orf1ab;3654 | 0 | 0 |
| MT072864.1;QIQ54047.1;Pangolin_CoV_orf1ab;3655 | 0 | 0 |
| MT072864.1;QIQ54047.1;Pangolin_CoV_orf1ab;3656 | 0 | 0.025 |
| MT072864.1;QIQ54047.1;Pangolin_CoV_orf1ab;3657 | 0 | 0.025 |
| MT072864.1;QIQ54047.1;Pangolin_CoV_orf1ab;3658 | 0 | 0.025 |
| MT072864.1;QIQ54047.1;Pangolin_CoV_orf1ab;3659 | 0 | 0.025 |
| MT072864.1;QIQ54047.1;Pangolin_CoV_orf1ab;3660 | 0 | 0.025 |
| MT072864.1;QIQ54047.1;Pangolin_CoV_orf1ab;3661 | 0 | 0.025 |
| MT072864.1;QIQ54047.1;Pangolin_CoV_orf1ab;3662 | 0 | 0.025 |
| MT072864.1;QIQ54047.1;Pangolin_CoV_orf1ab;3663 | 0 | 0.025 |
| MT072864.1;QIQ54047.1;Pangolin_CoV_orf1ab;3664 | 0 | 0.025 |
| MT072864.1;QIQ54047.1;Pangolin_CoV_orf1ab;3665 | 0 | 0 |
| MT072864.1;QIQ54047.1;Pangolin_CoV_orf1ab;3666 | 0 | 0 |
| MT072864.1;QIQ54047.1;Pangolin_CoV_orf1ab;3667 | 0 | 0 |
| MT072864.1;QIQ54047.1;Pangolin_CoV_orf1ab;3668 | 0 | 0 |
| MT072864.1;QIQ54047.1;Pangolin_CoV_orf1ab;3669 | 0 | 0 |
| MT072864.1;QIQ54047.1;Pangolin_CoV_orf1ab;3670 | 0 | 0 |
| MT072864.1;QIQ54047.1;Pangolin_CoV_orf1ab;3671 | 0 | 0 |
| MT072864.1;QIQ54047.1;Pangolin_CoV_orf1ab;3672 | 0 | 0 |
| MT072864.1;QIQ54047.1;Pangolin_CoV_orf1ab;3673 | 0 | 0 |
| MT072864.1;QIQ54047.1;Pangolin_CoV_orf1ab;3674 | 0 | 0 |
| MT072864.1;QIQ54047.1;Pangolin_CoV_orf1ab;3675 | 0 | 0 |
| MT072864.1;QIQ54047.1;Pangolin_CoV_orf1ab;3676 | 0 | 0 |
| MT072864.1;QIQ54047.1;Pangolin_CoV_orf1ab;3677 | 0 | 0 |
| MT072864.1;QIQ54047.1;Pangolin_CoV_orf1ab;3678 | 0 | 0 |
| MT072864.1;QIQ54047.1;Pangolin_CoV_orf1ab;3679 | 0 | 0 |
| MT072864.1;QIQ54047.1;Pangolin_CoV_orf1ab;3680 | 0 | 0 |
| MT072864.1;QIQ54047.1;Pangolin_CoV_orf1ab;3681 | 0 | 0 |
| MT072864.1;QIQ54047.1;Pangolin_CoV_orf1ab;3682 | 0 | 0 |
| MT072864.1;QIQ54047.1;Pangolin_CoV_orf1ab;3683 | 0 | 0 |
| MT072864.1;QIQ54047.1;Pangolin_CoV_orf1ab;3684 | 0 | 0 |
| MT072864.1;QIQ54047.1;Pangolin_CoV_orf1ab;3685 | 0 | 0 |
| MT072864.1;QIQ54047.1;Pangolin_CoV_orf1ab;3686 | 0 | 0 |
| MT072864.1;QIQ54047.1;Pangolin_CoV_orf1ab;3687 | 0 | 0 |
| MT072864.1;QIQ54047.1;Pangolin_CoV_orf1ab;3688 | 0 | 0 |

| | | |
|---|---|---|
| MT072864.1;QIQ54047.1;Pangolin_CoV_orf1ab;3689 | 0 | 0 |
| MT072864.1;QIQ54047.1;Pangolin_CoV_orf1ab;3690 | 0 | 0.025 |
| MT072864.1;QIQ54047.1;Pangolin_CoV_orf1ab;3691 | 0 | 0 |
| MT072864.1;QIQ54047.1;Pangolin_CoV_orf1ab;3692 | 0 | 0 |
| MT072864.1;QIQ54047.1;Pangolin_CoV_orf1ab;3693 | 0 | 0 |
| MT072864.1;QIQ54047.1;Pangolin_CoV_orf1ab;3694 | 0 | 0 |
| MT072864.1;QIQ54047.1;Pangolin_CoV_orf1ab;3695 | 0 | 0 |
| MT072864.1;QIQ54047.1;Pangolin_CoV_orf1ab;3696 | 0 | 0 |
| MT072864.1;QIQ54047.1;Pangolin_CoV_orf1ab;3697 | 0 | 0 |
| MT072864.1;QIQ54047.1;Pangolin_CoV_orf1ab;3698 | 0 | 0 |
| MT072864.1;QIQ54047.1;Pangolin_CoV_orf1ab;3699 | 0 | 0 |
| MT072864.1;QIQ54047.1;Pangolin_CoV_orf1ab;3700 | 0 | 0 |
| MT072864.1;QIQ54047.1;Pangolin_CoV_orf1ab;3701 | 0 | 0 |
| MT072864.1;QIQ54047.1;Pangolin_CoV_orf1ab;3702 | 0 | 0 |
| MT072864.1;QIQ54047.1;Pangolin_CoV_orf1ab;3703 | 0 | 0 |
| MT072864.1;QIQ54047.1;Pangolin_CoV_orf1ab;3704 | 0 | 0 |
| MT072864.1;QIQ54047.1;Pangolin_CoV_orf1ab;3705 | 0 | 0 |
| MT072864.1;QIQ54047.1;Pangolin_CoV_orf1ab;3706 | 0 | 0 |
| MT072864.1;QIQ54047.1;Pangolin_CoV_orf1ab;3707 | 0 | 0 |
| MT072864.1;QIQ54047.1;Pangolin_CoV_orf1ab;3708 | 0 | 0 |
| MT072864.1;QIQ54047.1;Pangolin_CoV_orf1ab;3709 | 0 | 0 |
| MT072864.1;QIQ54047.1;Pangolin_CoV_orf1ab;3710 | 0.05 | 0 |
| MT072864.1;QIQ54047.1;Pangolin_CoV_orf1ab;3711 | 0.05 | 0 |
| MT072864.1;QIQ54047.1;Pangolin_CoV_orf1ab;3712 | 0.05 | 0 |
| MT072864.1;QIQ54047.1;Pangolin_CoV_orf1ab;3713 | 0.05 | 0.025 |
| MT072864.1;QIQ54047.1;Pangolin_CoV_orf1ab;3714 | 0.05 | 0 |
| MT072864.1;QIQ54047.1;Pangolin_CoV_orf1ab;3715 | 0 | 0 |
| MT072864.1;QIQ54047.1;Pangolin_CoV_orf1ab;3716 | 0 | 0 |
| MT072864.1;QIQ54047.1;Pangolin_CoV_orf1ab;3717 | 0 | 0 |
| MT072864.1;QIQ54047.1;Pangolin_CoV_orf1ab;3718 | 0 | 0 |
| MT072864.1;QIQ54047.1;Pangolin_CoV_orf1ab;3719 | 0 | 0 |
| MT072864.1;QIQ54047.1;Pangolin_CoV_orf1ab;3720 | 0 | 0 |
| MT072864.1;QIQ54047.1;Pangolin_CoV_orf1ab;3721 | 0 | 0 |
| MT072864.1;QIQ54047.1;Pangolin_CoV_orf1ab;3722 | 0 | 0 |
| MT072864.1;QIQ54047.1;Pangolin_CoV_orf1ab;3723 | 0 | 0 |
| MT072864.1;QIQ54047.1;Pangolin_CoV_orf1ab;3724 | 0 | 0 |
| MT072864.1;QIQ54047.1;Pangolin_CoV_orf1ab;3725 | 0 | 0 |
| MT072864.1;QIQ54047.1;Pangolin_CoV_orf1ab;3726 | 0 | 0 |
| MT072864.1;QIQ54047.1;Pangolin_CoV_orf1ab;3727 | 0 | 0 |
| MT072864.1;QIQ54047.1;Pangolin_CoV_orf1ab;3728 | 0 | 0 |
| MT072864.1;QIQ54047.1;Pangolin_CoV_orf1ab;3729 | 0 | 0 |
| MT072864.1;QIQ54047.1;Pangolin_CoV_orf1ab;3730 | 0 | 0 |
| MT072864.1;QIQ54047.1;Pangolin_CoV_orf1ab;3731 | 0 | 0 |

| | | |
|---|---|---|
| MT072864.1;QIQ54047.1;Pangolin_CoV_orf1ab;3732 | 0 | 0 |
| MT072864.1;QIQ54047.1;Pangolin_CoV_orf1ab;3733 | 0 | 0 |
| MT072864.1;QIQ54047.1;Pangolin_CoV_orf1ab;3734 | 0 | 0 |
| MT072864.1;QIQ54047.1;Pangolin_CoV_orf1ab;3735 | 0 | 0 |
| MT072864.1;QIQ54047.1;Pangolin_CoV_orf1ab;3736 | 0 | 0 |
| MT072864.1;QIQ54047.1;Pangolin_CoV_orf1ab;3737 | 0 | 0 |
| MT072864.1;QIQ54047.1;Pangolin_CoV_orf1ab;3738 | 0 | 0 |
| MT072864.1;QIQ54047.1;Pangolin_CoV_orf1ab;3739 | 0 | 0 |
| MT072864.1;QIQ54047.1;Pangolin_CoV_orf1ab;3740 | 0 | 0 |
| MT072864.1;QIQ54047.1;Pangolin_CoV_orf1ab;3741 | 0 | 0 |
| MT072864.1;QIQ54047.1;Pangolin_CoV_orf1ab;3742 | 0 | 0 |
| MT072864.1;QIQ54047.1;Pangolin_CoV_orf1ab;3743 | 0 | 0 |
| MT072864.1;QIQ54047.1;Pangolin_CoV_orf1ab;3744 | 0 | 0 |
| MT072864.1;QIQ54047.1;Pangolin_CoV_orf1ab;3745 | 0 | 0 |
| MT072864.1;QIQ54047.1;Pangolin_CoV_orf1ab;3746 | 0 | 0 |
| MT072864.1;QIQ54047.1;Pangolin_CoV_orf1ab;3747 | 0 | 0 |
| MT072864.1;QIQ54047.1;Pangolin_CoV_orf1ab;3748 | 0 | 0 |
| MT072864.1;QIQ54047.1;Pangolin_CoV_orf1ab;3749 | 0 | 0 |
| MT072864.1;QIQ54047.1;Pangolin_CoV_orf1ab;3750 | 0 | 0 |
| MT072864.1;QIQ54047.1;Pangolin_CoV_orf1ab;3751 | 0 | 0 |
| MT072864.1;QIQ54047.1;Pangolin_CoV_orf1ab;3752 | 0 | 0 |
| MT072864.1;QIQ54047.1;Pangolin_CoV_orf1ab;3753 | 0 | 0 |
| MT072864.1;QIQ54047.1;Pangolin_CoV_orf1ab;3754 | 0 | 0 |
| MT072864.1;QIQ54047.1;Pangolin_CoV_orf1ab;3755 | 0 | 0 |
| MT072864.1;QIQ54047.1;Pangolin_CoV_orf1ab;3756 | 0 | 0 |
| MT072864.1;QIQ54047.1;Pangolin_CoV_orf1ab;3757 | 0 | 0 |
| MT072864.1;QIQ54047.1;Pangolin_CoV_orf1ab;3758 | 0 | 0 |
| MT072864.1;QIQ54047.1;Pangolin_CoV_orf1ab;3759 | 0 | 0 |
| MT072864.1;QIQ54047.1;Pangolin_CoV_orf1ab;3760 | 0 | 0 |
| MT072864.1;QIQ54047.1;Pangolin_CoV_orf1ab;3761 | 0 | 0 |
| MT072864.1;QIQ54047.1;Pangolin_CoV_orf1ab;3762 | 0 | 0 |
| MT072864.1;QIQ54047.1;Pangolin_CoV_orf1ab;3763 | 0 | 0 |
| MT072864.1;QIQ54047.1;Pangolin_CoV_orf1ab;3764 | 0 | 0 |
| MT072864.1;QIQ54047.1;Pangolin_CoV_orf1ab;3765 | 0 | 0 |
| MT072864.1;QIQ54047.1;Pangolin_CoV_orf1ab;3766 | 0 | 0 |
| MT072864.1;QIQ54047.1;Pangolin_CoV_orf1ab;3767 | 0 | 0 |
| MT072864.1;QIQ54047.1;Pangolin_CoV_orf1ab;3768 | 0 | 0 |
| MT072864.1;QIQ54047.1;Pangolin_CoV_orf1ab;3769 | 0 | 0 |
| MT072864.1;QIQ54047.1;Pangolin_CoV_orf1ab;3770 | 0 | 0 |
| MT072864.1;QIQ54047.1;Pangolin_CoV_orf1ab;3771 | 0 | 0 |
| MT072864.1;QIQ54047.1;Pangolin_CoV_orf1ab;3772 | 0 | 0 |
| MT072864.1;QIQ54047.1;Pangolin_CoV_orf1ab;3773 | 0 | 0 |
| MT072864.1;QIQ54047.1;Pangolin_CoV_orf1ab;3774 | 0 | 0 |

| | | |
|---|---|---|
| MT072864.1;QIQ54047.1;Pangolin_CoV_orf1ab;3775 | 0 | 0 |
| MT072864.1;QIQ54047.1;Pangolin_CoV_orf1ab;3776 | 0 | 0 |
| MT072864.1;QIQ54047.1;Pangolin_CoV_orf1ab;3777 | 0 | 0 |
| MT072864.1;QIQ54047.1;Pangolin_CoV_orf1ab;3778 | 0 | 0 |
| MT072864.1;QIQ54047.1;Pangolin_CoV_orf1ab;3779 | 0 | 0 |
| MT072864.1;QIQ54047.1;Pangolin_CoV_orf1ab;3780 | 0 | 0 |
| MT072864.1;QIQ54047.1;Pangolin_CoV_orf1ab;3781 | 0 | 0 |
| MT072864.1;QIQ54047.1;Pangolin_CoV_orf1ab;3782 | 0 | 0 |
| MT072864.1;QIQ54047.1;Pangolin_CoV_orf1ab;3783 | 0 | 0 |
| MT072864.1;QIQ54047.1;Pangolin_CoV_orf1ab;3784 | 0 | 0 |
| MT072864.1;QIQ54047.1;Pangolin_CoV_orf1ab;3785 | 0 | 0 |
| MT072864.1;QIQ54047.1;Pangolin_CoV_orf1ab;3786 | 0 | 0 |
| MT072864.1;QIQ54047.1;Pangolin_CoV_orf1ab;3787 | 0 | 0 |
| MT072864.1;QIQ54047.1;Pangolin_CoV_orf1ab;3788 | 0 | 0 |
| MT072864.1;QIQ54047.1;Pangolin_CoV_orf1ab;3789 | 0 | 0 |
| MT072864.1;QIQ54047.1;Pangolin_CoV_orf1ab;3790 | 0 | 0 |
| MT072864.1;QIQ54047.1;Pangolin_CoV_orf1ab;3791 | 0 | 0 |
| MT072864.1;QIQ54047.1;Pangolin_CoV_orf1ab;3792 | 0 | 0 |
| MT072864.1;QIQ54047.1;Pangolin_CoV_orf1ab;3793 | 0 | 0 |
| MT072864.1;QIQ54047.1;Pangolin_CoV_orf1ab;3794 | 0 | 0 |
| MT072864.1;QIQ54047.1;Pangolin_CoV_orf1ab;3795 | 0 | 0 |
| MT072864.1;QIQ54047.1;Pangolin_CoV_orf1ab;3796 | 0 | 0 |
| MT072864.1;QIQ54047.1;Pangolin_CoV_orf1ab;3797 | 0 | 0 |
| MT072864.1;QIQ54047.1;Pangolin_CoV_orf1ab;3798 | 0 | 0 |
| MT072864.1;QIQ54047.1;Pangolin_CoV_orf1ab;3799 | 0 | 0 |
| MT072864.1;QIQ54047.1;Pangolin_CoV_orf1ab;3800 | 0 | 0 |
| MT072864.1;QIQ54047.1;Pangolin_CoV_orf1ab;3801 | 0.25 | 0.175 |
| MT072864.1;QIQ54047.1;Pangolin_CoV_orf1ab;3802 | 0 | 0 |
| MT072864.1;QIQ54047.1;Pangolin_CoV_orf1ab;3803 | 0 | 0 |
| MT072864.1;QIQ54047.1;Pangolin_CoV_orf1ab;3804 | 0 | 0 |
| MT072864.1;QIQ54047.1;Pangolin_CoV_orf1ab;3805 | 0 | 0 |
| MT072864.1;QIQ54047.1;Pangolin_CoV_orf1ab;3806 | 0 | 0 |
| MT072864.1;QIQ54047.1;Pangolin_CoV_orf1ab;3807 | 0 | 0 |
| MT072864.1;QIQ54047.1;Pangolin_CoV_orf1ab;3808 | 0 | 0 |
| MT072864.1;QIQ54047.1;Pangolin_CoV_orf1ab;3809 | 0 | 0.05 |
| MT072864.1;QIQ54047.1;Pangolin_CoV_orf1ab;3810 | 0.05 | 0.025 |
| MT072864.1;QIQ54047.1;Pangolin_CoV_orf1ab;3811 | 0.15 | 0.075 |
| MT072864.1;QIQ54047.1;Pangolin_CoV_orf1ab;3812 | 0 | 0.025 |
| MT072864.1;QIQ54047.1;Pangolin_CoV_orf1ab;3813 | 0 | 0.025 |
| MT072864.1;QIQ54047.1;Pangolin_CoV_orf1ab;3814 | 0.05 | 0.125 |
| MT072864.1;QIQ54047.1;Pangolin_CoV_orf1ab;3815 | 0.1 | 0.05 |
| MT072864.1;QIQ54047.1;Pangolin_CoV_orf1ab;3816 | 0.3 | 0.1 |
| MT072864.1;QIQ54047.1;Pangolin_CoV_orf1ab;3817 | 0.25 | 0.1 |

| | | |
|---|---|---|
| MT072864.1;QIQ54047.1;Pangolin_CoV_orf1ab;3818 | 0.25 | 0.1 |
| MT072864.1;QIQ54047.1;Pangolin_CoV_orf1ab;3819 | 0.2 | 0.125 |
| MT072864.1;QIQ54047.1;Pangolin_CoV_orf1ab;3820 | 0.15 | 0.075 |
| MT072864.1;QIQ54047.1;Pangolin_CoV_orf1ab;3821 | 0.1 | 0.075 |
| MT072864.1;QIQ54047.1;Pangolin_CoV_orf1ab;3822 | 0.05 | 0.075 |
| MT072864.1;QIQ54047.1;Pangolin_CoV_orf1ab;3823 | 0.05 | 0.05 |
| MT072864.1;QIQ54047.1;Pangolin_CoV_orf1ab;3824 | 0 | 0.05 |
| MT072864.1;QIQ54047.1;Pangolin_CoV_orf1ab;3825 | 0 | 0 |
| MT072864.1;QIQ54047.1;Pangolin_CoV_orf1ab;3826 | 0 | 0 |
| MT072864.1;QIQ54047.1;Pangolin_CoV_orf1ab;3827 | 0 | 0 |
| MT072864.1;QIQ54047.1;Pangolin_CoV_orf1ab;3828 | 0 | 0 |
| MT072864.1;QIQ54047.1;Pangolin_CoV_orf1ab;3829 | 0 | 0 |
| MT072864.1;QIQ54047.1;Pangolin_CoV_orf1ab;3830 | 0 | 0 |
| MT072864.1;QIQ54047.1;Pangolin_CoV_orf1ab;3831 | 0 | 0 |
| MT072864.1;QIQ54047.1;Pangolin_CoV_orf1ab;3832 | 0 | 0 |
| MT072864.1;QIQ54047.1;Pangolin_CoV_orf1ab;3833 | 0 | 0 |
| MT072864.1;QIQ54047.1;Pangolin_CoV_orf1ab;3834 | 0 | 0 |
| MT072864.1;QIQ54047.1;Pangolin_CoV_orf1ab;3835 | 0 | 0 |
| MT072864.1;QIQ54047.1;Pangolin_CoV_orf1ab;3836 | 0 | 0.025 |
| MT072864.1;QIQ54047.1;Pangolin_CoV_orf1ab;3837 | 0 | 0.025 |
| MT072864.1;QIQ54047.1;Pangolin_CoV_orf1ab;3838 | 0.05 | 0.05 |
| MT072864.1;QIQ54047.1;Pangolin_CoV_orf1ab;3839 | 0.05 | 0.05 |
| MT072864.1;QIQ54047.1;Pangolin_CoV_orf1ab;3840 | 0.05 | 0.05 |
| MT072864.1;QIQ54047.1;Pangolin_CoV_orf1ab;3841 | 0.1 | 0.1 |
| MT072864.1;QIQ54047.1;Pangolin_CoV_orf1ab;3842 | 0.05 | 0.075 |
| MT072864.1;QIQ54047.1;Pangolin_CoV_orf1ab;3843 | 0.05 | 0.05 |
| MT072864.1;QIQ54047.1;Pangolin_CoV_orf1ab;3844 | 0.05 | 0.025 |
| MT072864.1;QIQ54047.1;Pangolin_CoV_orf1ab;3845 | 0 | 0.025 |
| MT072864.1;QIQ54047.1;Pangolin_CoV_orf1ab;3846 | 0.05 | 0.025 |
| MT072864.1;QIQ54047.1;Pangolin_CoV_orf1ab;3847 | 0.05 | 0.025 |
| MT072864.1;QIQ54047.1;Pangolin_CoV_orf1ab;3848 | 0.05 | 0.025 |
| MT072864.1;QIQ54047.1;Pangolin_CoV_orf1ab;3849 | 0.05 | 0 |
| MT072864.1;QIQ54047.1;Pangolin_CoV_orf1ab;3850 | 0 | 0 |
| MT072864.1;QIQ54047.1;Pangolin_CoV_orf1ab;3851 | 0 | 0 |
| MT072864.1;QIQ54047.1;Pangolin_CoV_orf1ab;3852 | 0 | 0 |
| MT072864.1;QIQ54047.1;Pangolin_CoV_orf1ab;3853 | 0 | 0 |
| MT072864.1;QIQ54047.1;Pangolin_CoV_orf1ab;3854 | 0 | 0 |
| MT072864.1;QIQ54047.1;Pangolin_CoV_orf1ab;3855 | 0 | 0 |
| MT072864.1;QIQ54047.1;Pangolin_CoV_orf1ab;3856 | 0 | 0 |
| MT072864.1;QIQ54047.1;Pangolin_CoV_orf1ab;3857 | 0 | 0 |
| MT072864.1;QIQ54047.1;Pangolin_CoV_orf1ab;3858 | 0 | 0 |
| MT072864.1;QIQ54047.1;Pangolin_CoV_orf1ab;3859 | 0 | 0 |
| MT072864.1;QIQ54047.1;Pangolin_CoV_orf1ab;3860 | 0 | 0 |

| | | |
|---|---|---|
| MT072864.1;QIQ54047.1;Pangolin_CoV_orf1ab;3861 | 0 | 0 |
| MT072864.1;QIQ54047.1;Pangolin_CoV_orf1ab;3862 | 0 | 0 |
| MT072864.1;QIQ54047.1;Pangolin_CoV_orf1ab;3863 | 0 | 0 |
| MT072864.1;QIQ54047.1;Pangolin_CoV_orf1ab;3864 | 0 | 0 |
| MT072864.1;QIQ54047.1;Pangolin_CoV_orf1ab;3865 | 0 | 0 |
| MT072864.1;QIQ54047.1;Pangolin_CoV_orf1ab;3866 | 0 | 0 |
| MT072864.1;QIQ54047.1;Pangolin_CoV_orf1ab;3867 | 0 | 0 |
| MT072864.1;QIQ54047.1;Pangolin_CoV_orf1ab;3868 | 0 | 0 |
| MT072864.1;QIQ54047.1;Pangolin_CoV_orf1ab;3869 | 0 | 0 |
| MT072864.1;QIQ54047.1;Pangolin_CoV_orf1ab;3870 | 0 | 0 |
| MT072864.1;QIQ54047.1;Pangolin_CoV_orf1ab;3871 | 0 | 0 |
| MT072864.1;QIQ54047.1;Pangolin_CoV_orf1ab;3872 | 0 | 0 |
| MT072864.1;QIQ54047.1;Pangolin_CoV_orf1ab;3873 | 0 | 0 |
| MT072864.1;QIQ54047.1;Pangolin_CoV_orf1ab;3874 | 0 | 0.025 |
| MT072864.1;QIQ54047.1;Pangolin_CoV_orf1ab;3875 | 0 | 0 |
| MT072864.1;QIQ54047.1;Pangolin_CoV_orf1ab;3876 | 0 | 0.025 |
| MT072864.1;QIQ54047.1;Pangolin_CoV_orf1ab;3877 | 0 | 0 |
| MT072864.1;QIQ54047.1;Pangolin_CoV_orf1ab;3878 | 0 | 0 |
| MT072864.1;QIQ54047.1;Pangolin_CoV_orf1ab;3879 | 0 | 0 |
| MT072864.1;QIQ54047.1;Pangolin_CoV_orf1ab;3880 | 0.05 | 0 |
| MT072864.1;QIQ54047.1;Pangolin_CoV_orf1ab;3881 | 0.05 | 0 |
| MT072864.1;QIQ54047.1;Pangolin_CoV_orf1ab;3882 | 0.05 | 0 |
| MT072864.1;QIQ54047.1;Pangolin_CoV_orf1ab;3883 | 0 | 0 |
| MT072864.1;QIQ54047.1;Pangolin_CoV_orf1ab;3884 | 0 | 0 |
| MT072864.1;QIQ54047.1;Pangolin_CoV_orf1ab;3885 | 0 | 0 |
| MT072864.1;QIQ54047.1;Pangolin_CoV_orf1ab;3886 | 0 | 0 |
| MT072864.1;QIQ54047.1;Pangolin_CoV_orf1ab;3887 | 0 | 0 |
| MT072864.1;QIQ54047.1;Pangolin_CoV_orf1ab;3888 | 0 | 0 |
| MT072864.1;QIQ54047.1;Pangolin_CoV_orf1ab;3889 | 0 | 0 |
| MT072864.1;QIQ54047.1;Pangolin_CoV_orf1ab;3890 | 0 | 0 |
| MT072864.1;QIQ54047.1;Pangolin_CoV_orf1ab;3891 | 0 | 0 |
| MT072864.1;QIQ54047.1;Pangolin_CoV_orf1ab;3892 | 0 | 0 |
| MT072864.1;QIQ54047.1;Pangolin_CoV_orf1ab;3893 | 0 | 0.05 |
| MT072864.1;QIQ54047.1;Pangolin_CoV_orf1ab;3894 | 0 | 0 |
| MT072864.1;QIQ54047.1;Pangolin_CoV_orf1ab;3895 | 0 | 0 |
| MT072864.1;QIQ54047.1;Pangolin_CoV_orf1ab;3896 | 0 | 0 |
| MT072864.1;QIQ54047.1;Pangolin_CoV_orf1ab;3897 | 0 | 0 |
| MT072864.1;QIQ54047.1;Pangolin_CoV_orf1ab;3898 | 0 | 0 |
| MT072864.1;QIQ54047.1;Pangolin_CoV_orf1ab;3899 | 0 | 0 |
| MT072864.1;QIQ54047.1;Pangolin_CoV_orf1ab;3900 | 0 | 0 |
| MT072864.1;QIQ54047.1;Pangolin_CoV_orf1ab;3901 | 0 | 0 |
| MT072864.1;QIQ54047.1;Pangolin_CoV_orf1ab;3902 | 0 | 0 |
| MT072864.1;QIQ54047.1;Pangolin_CoV_orf1ab;3903 | 0 | 0 |

| | | |
|---|---|---|
| MT072864.1;QIQ54047.1;Pangolin_CoV_orf1ab;3904 | 0 | 0 |
| MT072864.1;QIQ54047.1;Pangolin_CoV_orf1ab;3905 | 0 | 0 |
| MT072864.1;QIQ54047.1;Pangolin_CoV_orf1ab;3906 | 0 | 0.05 |
| MT072864.1;QIQ54047.1;Pangolin_CoV_orf1ab;3907 | 0 | 0.025 |
| MT072864.1;QIQ54047.1;Pangolin_CoV_orf1ab;3908 | 0 | 0.025 |
| MT072864.1;QIQ54047.1;Pangolin_CoV_orf1ab;3909 | 0 | 0.025 |
| MT072864.1;QIQ54047.1;Pangolin_CoV_orf1ab;3910 | 0 | 0.025 |
| MT072864.1;QIQ54047.1;Pangolin_CoV_orf1ab;3911 | 0.05 | 0.05 |
| MT072864.1;QIQ54047.1;Pangolin_CoV_orf1ab;3912 | 0.05 | 0.05 |
| MT072864.1;QIQ54047.1;Pangolin_CoV_orf1ab;3913 | 0.05 | 0.075 |
| MT072864.1;QIQ54047.1;Pangolin_CoV_orf1ab;3914 | 0.05 | 0.05 |
| MT072864.1;QIQ54047.1;Pangolin_CoV_orf1ab;3915 | 0.1 | 0.025 |
| MT072864.1;QIQ54047.1;Pangolin_CoV_orf1ab;3916 | 0 | 0.025 |
| MT072864.1;QIQ54047.1;Pangolin_CoV_orf1ab;3917 | 0 | 0.05 |
| MT072864.1;QIQ54047.1;Pangolin_CoV_orf1ab;3918 | 0 | 0.025 |
| MT072864.1;QIQ54047.1;Pangolin_CoV_orf1ab;3919 | 0 | 0 |
| MT072864.1;QIQ54047.1;Pangolin_CoV_orf1ab;3920 | 0 | 0 |
| MT072864.1;QIQ54047.1;Pangolin_CoV_orf1ab;3921 | 0.05 | 0 |
| MT072864.1;QIQ54047.1;Pangolin_CoV_orf1ab;3922 | 0.05 | 0 |
| MT072864.1;QIQ54047.1;Pangolin_CoV_orf1ab;3923 | 0.05 | 0 |
| MT072864.1;QIQ54047.1;Pangolin_CoV_orf1ab;3924 | 0 | 0 |
| MT072864.1;QIQ54047.1;Pangolin_CoV_orf1ab;3925 | 0.05 | 0 |
| MT072864.1;QIQ54047.1;Pangolin_CoV_orf1ab;3926 | 0 | 0 |
| MT072864.1;QIQ54047.1;Pangolin_CoV_orf1ab;3927 | 0 | 0 |
| MT072864.1;QIQ54047.1;Pangolin_CoV_orf1ab;3928 | 0 | 0 |
| MT072864.1;QIQ54047.1;Pangolin_CoV_orf1ab;3929 | 0 | 0 |
| MT072864.1;QIQ54047.1;Pangolin_CoV_orf1ab;3930 | 0 | 0 |
| MT072864.1;QIQ54047.1;Pangolin_CoV_orf1ab;3931 | 0 | 0 |
| MT072864.1;QIQ54047.1;Pangolin_CoV_orf1ab;3932 | 0 | 0 |
| MT072864.1;QIQ54047.1;Pangolin_CoV_orf1ab;3933 | 0.05 | 0.05 |
| MT072864.1;QIQ54047.1;Pangolin_CoV_orf1ab;3934 | 0 | 0.025 |
| MT072864.1;QIQ54047.1;Pangolin_CoV_orf1ab;3935 | 0 | 0 |
| MT072864.1;QIQ54047.1;Pangolin_CoV_orf1ab;3936 | 0 | 0 |
| MT072864.1;QIQ54047.1;Pangolin_CoV_orf1ab;3937 | 0 | 0 |
| MT072864.1;QIQ54047.1;Pangolin_CoV_orf1ab;3938 | 0 | 0.05 |
| MT072864.1;QIQ54047.1;Pangolin_CoV_orf1ab;3939 | 0.05 | 0 |
| MT072864.1;QIQ54047.1;Pangolin_CoV_orf1ab;3940 | 0.05 | 0 |
| MT072864.1;QIQ54047.1;Pangolin_CoV_orf1ab;3941 | 0.05 | 0 |
| MT072864.1;QIQ54047.1;Pangolin_CoV_orf1ab;3942 | 0.05 | 0.05 |
| MT072864.1;QIQ54047.1;Pangolin_CoV_orf1ab;3943 | 0.1 | 0.05 |
| MT072864.1;QIQ54047.1;Pangolin_CoV_orf1ab;3944 | 0 | 0.025 |
| MT072864.1;QIQ54047.1;Pangolin_CoV_orf1ab;3945 | 0.1 | 0.05 |
| MT072864.1;QIQ54047.1;Pangolin_CoV_orf1ab;3946 | 0.05 | 0.125 |

| | | |
|---|---|---|
| MT072864.1;QIQ54047.1;Pangolin_CoV_orf1ab;3947 | 0.05 | 0.075 |
| MT072864.1;QIQ54047.1;Pangolin_CoV_orf1ab;3948 | 0.05 | 0.15 |
| MT072864.1;QIQ54047.1;Pangolin_CoV_orf1ab;3949 | 0.05 | 0.05 |
| MT072864.1;QIQ54047.1;Pangolin_CoV_orf1ab;3950 | 0 | 0.025 |
| MT072864.1;QIQ54047.1;Pangolin_CoV_orf1ab;3951 | 0 | 0 |
| MT072864.1;QIQ54047.1;Pangolin_CoV_orf1ab;3952 | 0.05 | 0 |
| MT072864.1;QIQ54047.1;Pangolin_CoV_orf1ab;3953 | 0 | 0 |
| MT072864.1;QIQ54047.1;Pangolin_CoV_orf1ab;3954 | 0 | 0 |
| MT072864.1;QIQ54047.1;Pangolin_CoV_orf1ab;3955 | 0 | 0 |
| MT072864.1;QIQ54047.1;Pangolin_CoV_orf1ab;3956 | 0 | 0 |
| MT072864.1;QIQ54047.1;Pangolin_CoV_orf1ab;3957 | 0 | 0 |
| MT072864.1;QIQ54047.1;Pangolin_CoV_orf1ab;3958 | 0 | 0 |
| MT072864.1;QIQ54047.1;Pangolin_CoV_orf1ab;3959 | 0 | 0 |
| MT072864.1;QIQ54047.1;Pangolin_CoV_orf1ab;3960 | 0 | 0 |
| MT072864.1;QIQ54047.1;Pangolin_CoV_orf1ab;3961 | 0 | 0 |
| MT072864.1;QIQ54047.1;Pangolin_CoV_orf1ab;3962 | 0 | 0 |
| MT072864.1;QIQ54047.1;Pangolin_CoV_orf1ab;3963 | 0 | 0 |
| MT072864.1;QIQ54047.1;Pangolin_CoV_orf1ab;3964 | 0 | 0 |
| MT072864.1;QIQ54047.1;Pangolin_CoV_orf1ab;3965 | 0 | 0 |
| MT072864.1;QIQ54047.1;Pangolin_CoV_orf1ab;3966 | 0 | 0 |
| MT072864.1;QIQ54047.1;Pangolin_CoV_orf1ab;3967 | 0 | 0 |
| MT072864.1;QIQ54047.1;Pangolin_CoV_orf1ab;3968 | 0 | 0 |
| MT072864.1;QIQ54047.1;Pangolin_CoV_orf1ab;3969 | 0 | 0.025 |
| MT072864.1;QIQ54047.1;Pangolin_CoV_orf1ab;3970 | 0 | 0.05 |
| MT072864.1;QIQ54047.1;Pangolin_CoV_orf1ab;3971 | 0 | 0.025 |
| MT072864.1;QIQ54047.1;Pangolin_CoV_orf1ab;3972 | 0.05 | 0.025 |
| MT072864.1;QIQ54047.1;Pangolin_CoV_orf1ab;3973 | 0.05 | 0.025 |
| MT072864.1;QIQ54047.1;Pangolin_CoV_orf1ab;3974 | 0 | 0.05 |
| MT072864.1;QIQ54047.1;Pangolin_CoV_orf1ab;3975 | 0.05 | 0.025 |
| MT072864.1;QIQ54047.1;Pangolin_CoV_orf1ab;3976 | 0 | 0.025 |
| MT072864.1;QIQ54047.1;Pangolin_CoV_orf1ab;3977 | 0 | 0 |
| MT072864.1;QIQ54047.1;Pangolin_CoV_orf1ab;3978 | 0 | 0 |
| MT072864.1;QIQ54047.1;Pangolin_CoV_orf1ab;3979 | 0 | 0 |
| MT072864.1;QIQ54047.1;Pangolin_CoV_orf1ab;3980 | 0 | 0 |
| MT072864.1;QIQ54047.1;Pangolin_CoV_orf1ab;3981 | 0 | 0 |
| MT072864.1;QIQ54047.1;Pangolin_CoV_orf1ab;3982 | 0 | 0 |
| MT072864.1;QIQ54047.1;Pangolin_CoV_orf1ab;3983 | 0 | 0 |
| MT072864.1;QIQ54047.1;Pangolin_CoV_orf1ab;3984 | 0 | 0 |
| MT072864.1;QIQ54047.1;Pangolin_CoV_orf1ab;3985 | 0 | 0 |
| MT072864.1;QIQ54047.1;Pangolin_CoV_orf1ab;3986 | 0 | 0.025 |
| MT072864.1;QIQ54047.1;Pangolin_CoV_orf1ab;3987 | 0 | 0 |
| MT072864.1;QIQ54047.1;Pangolin_CoV_orf1ab;3988 | 0 | 0 |
| MT072864.1;QIQ54047.1;Pangolin_CoV_orf1ab;3989 | 0 | 0 |

| | | |
|---|---|---|
| MT072864.1;QIQ54047.1;Pangolin_CoV_orf1ab;3990 | 0 | 0.05 |
| MT072864.1;QIQ54047.1;Pangolin_CoV_orf1ab;3991 | 0 | 0.025 |
| MT072864.1;QIQ54047.1;Pangolin_CoV_orf1ab;3992 | 0 | 0 |
| MT072864.1;QIQ54047.1;Pangolin_CoV_orf1ab;3993 | 0 | 0 |
| MT072864.1;QIQ54047.1;Pangolin_CoV_orf1ab;3994 | 0 | 0 |
| MT072864.1;QIQ54047.1;Pangolin_CoV_orf1ab;3995 | 0 | 0 |
| MT072864.1;QIQ54047.1;Pangolin_CoV_orf1ab;3996 | 0 | 0 |
| MT072864.1;QIQ54047.1;Pangolin_CoV_orf1ab;3997 | 0 | 0 |
| MT072864.1;QIQ54047.1;Pangolin_CoV_orf1ab;3998 | 0 | 0.1 |
| MT072864.1;QIQ54047.1;Pangolin_CoV_orf1ab;3999 | 0 | 0.075 |
| MT072864.1;QIQ54047.1;Pangolin_CoV_orf1ab;4000 | 0 | 0.05 |
| MT072864.1;QIQ54047.1;Pangolin_CoV_orf1ab;4001 | 0 | 0.05 |
| MT072864.1;QIQ54047.1;Pangolin_CoV_orf1ab;4002 | 0 | 0.05 |
| MT072864.1;QIQ54047.1;Pangolin_CoV_orf1ab;4003 | 0 | 0 |
| MT072864.1;QIQ54047.1;Pangolin_CoV_orf1ab;4004 | 0 | 0 |
| MT072864.1;QIQ54047.1;Pangolin_CoV_orf1ab;4005 | 0 | 0 |
| MT072864.1;QIQ54047.1;Pangolin_CoV_orf1ab;4006 | 0 | 0 |
| MT072864.1;QIQ54047.1;Pangolin_CoV_orf1ab;4007 | 0 | 0 |
| MT072864.1;QIQ54047.1;Pangolin_CoV_orf1ab;4008 | 0 | 0 |
| MT072864.1;QIQ54047.1;Pangolin_CoV_orf1ab;4009 | 0 | 0 |
| MT072864.1;QIQ54047.1;Pangolin_CoV_orf1ab;4010 | 0 | 0 |
| MT072864.1;QIQ54047.1;Pangolin_CoV_orf1ab;4011 | 0 | 0 |
| MT072864.1;QIQ54047.1;Pangolin_CoV_orf1ab;4012 | 0 | 0 |
| MT072864.1;QIQ54047.1;Pangolin_CoV_orf1ab;4013 | 0 | 0 |
| MT072864.1;QIQ54047.1;Pangolin_CoV_orf1ab;4014 | 0 | 0 |
| MT072864.1;QIQ54047.1;Pangolin_CoV_orf1ab;4015 | 0 | 0 |
| MT072864.1;QIQ54047.1;Pangolin_CoV_orf1ab;4016 | 0 | 0 |
| MT072864.1;QIQ54047.1;Pangolin_CoV_orf1ab;4017 | 0 | 0 |
| MT072864.1;QIQ54047.1;Pangolin_CoV_orf1ab;4018 | 0 | 0 |
| MT072864.1;QIQ54047.1;Pangolin_CoV_orf1ab;4019 | 0 | 0 |
| MT072864.1;QIQ54047.1;Pangolin_CoV_orf1ab;4020 | 0 | 0 |
| MT072864.1;QIQ54047.1;Pangolin_CoV_orf1ab;4021 | 0 | 0 |
| MT072864.1;QIQ54047.1;Pangolin_CoV_orf1ab;4022 | 0 | 0 |
| MT072864.1;QIQ54047.1;Pangolin_CoV_orf1ab;4023 | 0.15 | 0.1 |
| MT072864.1;QIQ54047.1;Pangolin_CoV_orf1ab;4024 | 0.2 | 0.1 |
| MT072864.1;QIQ54047.1;Pangolin_CoV_orf1ab;4025 | 0.2 | 0.075 |
| MT072864.1;QIQ54047.1;Pangolin_CoV_orf1ab;4026 | 0.25 | 0.1 |
| MT072864.1;QIQ54047.1;Pangolin_CoV_orf1ab;4027 | 0.25 | 0.275 |
| MT072864.1;QIQ54047.1;Pangolin_CoV_orf1ab;4028 | 0.3 | 0.3 |
| MT072864.1;QIQ54047.1;Pangolin_CoV_orf1ab;4029 | 0.35 | 0.4 |
| MT072864.1;QIQ54047.1;Pangolin_CoV_orf1ab;4030 | 0.25 | 0.25 |
| MT072864.1;QIQ54047.1;Pangolin_CoV_orf1ab;4031 | 0.4 | 0.5 |
| MT072864.1;QIQ54047.1;Pangolin_CoV_orf1ab;4032 | 0.1 | 0.025 |

| | | |
|---|---|---|
| MT072864.1;QIQ54047.1;Pangolin_CoV_orf1ab;4033 | 0.05 | 0.025 |
| MT072864.1;QIQ54047.1;Pangolin_CoV_orf1ab;4034 | 0 | 0 |
| MT072864.1;QIQ54047.1;Pangolin_CoV_orf1ab;4035 | 0.05 | 0.1 |
| MT072864.1;QIQ54047.1;Pangolin_CoV_orf1ab;4036 | 0.1 | 0.075 |
| MT072864.1;QIQ54047.1;Pangolin_CoV_orf1ab;4037 | 0 | 0.075 |
| MT072864.1;QIQ54047.1;Pangolin_CoV_orf1ab;4038 | 0.05 | 0.075 |
| MT072864.1;QIQ54047.1;Pangolin_CoV_orf1ab;4039 | 0.05 | 0 |
| MT072864.1;QIQ54047.1;Pangolin_CoV_orf1ab;4040 | 0 | 0.05 |
| MT072864.1;QIQ54047.1;Pangolin_CoV_orf1ab;4041 | 0 | 0 |
| MT072864.1;QIQ54047.1;Pangolin_CoV_orf1ab;4042 | 0 | 0 |
| MT072864.1;QIQ54047.1;Pangolin_CoV_orf1ab;4043 | 0 | 0 |
| MT072864.1;QIQ54047.1;Pangolin_CoV_orf1ab;4044 | 0 | 0 |
| MT072864.1;QIQ54047.1;Pangolin_CoV_orf1ab;4045 | 0.05 | 0.025 |
| MT072864.1;QIQ54047.1;Pangolin_CoV_orf1ab;4046 | 0.05 | 0.025 |
| MT072864.1;QIQ54047.1;Pangolin_CoV_orf1ab;4047 | 0 | 0 |
| MT072864.1;QIQ54047.1;Pangolin_CoV_orf1ab;4048 | 0 | 0 |
| MT072864.1;QIQ54047.1;Pangolin_CoV_orf1ab;4049 | 0 | 0 |
| MT072864.1;QIQ54047.1;Pangolin_CoV_orf1ab;4050 | 0 | 0 |
| MT072864.1;QIQ54047.1;Pangolin_CoV_orf1ab;4051 | 0 | 0 |
| MT072864.1;QIQ54047.1;Pangolin_CoV_orf1ab;4052 | 0 | 0 |
| MT072864.1;QIQ54047.1;Pangolin_CoV_orf1ab;4053 | 0 | 0 |
| MT072864.1;QIQ54047.1;Pangolin_CoV_orf1ab;4054 | 0 | 0.025 |
| MT072864.1;QIQ54047.1;Pangolin_CoV_orf1ab;4055 | 0.05 | 0 |
| MT072864.1;QIQ54047.1;Pangolin_CoV_orf1ab;4056 | 0 | 0.025 |
| MT072864.1;QIQ54047.1;Pangolin_CoV_orf1ab;4057 | 0 | 0 |
| MT072864.1;QIQ54047.1;Pangolin_CoV_orf1ab;4058 | 0 | 0 |
| MT072864.1;QIQ54047.1;Pangolin_CoV_orf1ab;4059 | 0 | 0.025 |
| MT072864.1;QIQ54047.1;Pangolin_CoV_orf1ab;4060 | 0 | 0.05 |
| MT072864.1;QIQ54047.1;Pangolin_CoV_orf1ab;4061 | 0 | 0.025 |
| MT072864.1;QIQ54047.1;Pangolin_CoV_orf1ab;4062 | 0.1 | 0.025 |
| MT072864.1;QIQ54047.1;Pangolin_CoV_orf1ab;4063 | 0.15 | 0.025 |
| MT072864.1;QIQ54047.1;Pangolin_CoV_orf1ab;4064 | 0.2 | 0.05 |
| MT072864.1;QIQ54047.1;Pangolin_CoV_orf1ab;4065 | 0.15 | 0.05 |
| MT072864.1;QIQ54047.1;Pangolin_CoV_orf1ab;4066 | 0.1 | 0.05 |
| MT072864.1;QIQ54047.1;Pangolin_CoV_orf1ab;4067 | 0.05 | 0.025 |
| MT072864.1;QIQ54047.1;Pangolin_CoV_orf1ab;4068 | 0 | 0 |
| MT072864.1;QIQ54047.1;Pangolin_CoV_orf1ab;4069 | 0 | 0 |
| MT072864.1;QIQ54047.1;Pangolin_CoV_orf1ab;4070 | 0 | 0 |
| MT072864.1;QIQ54047.1;Pangolin_CoV_orf1ab;4071 | 0 | 0 |
| MT072864.1;QIQ54047.1;Pangolin_CoV_orf1ab;4072 | 0 | 0 |
| MT072864.1;QIQ54047.1;Pangolin_CoV_orf1ab;4073 | 0 | 0 |
| MT072864.1;QIQ54047.1;Pangolin_CoV_orf1ab;4074 | 0 | 0 |
| MT072864.1;QIQ54047.1;Pangolin_CoV_orf1ab;4075 | 0 | 0 |

| | | |
|---|---|---|
| MT072864.1;QIQ54047.1;Pangolin_CoV_orf1ab;4076 | 0 | 0 |
| MT072864.1;QIQ54047.1;Pangolin_CoV_orf1ab;4077 | 0 | 0 |
| MT072864.1;QIQ54047.1;Pangolin_CoV_orf1ab;4078 | 0 | 0 |
| MT072864.1;QIQ54047.1;Pangolin_CoV_orf1ab;4079 | 0 | 0 |
| MT072864.1;QIQ54047.1;Pangolin_CoV_orf1ab;4080 | 0 | 0 |
| MT072864.1;QIQ54047.1;Pangolin_CoV_orf1ab;4081 | 0 | 0 |
| MT072864.1;QIQ54047.1;Pangolin_CoV_orf1ab;4082 | 0 | 0 |
| MT072864.1;QIQ54047.1;Pangolin_CoV_orf1ab;4083 | 0 | 0 |
| MT072864.1;QIQ54047.1;Pangolin_CoV_orf1ab;4084 | 0.05 | 0.025 |
| MT072864.1;QIQ54047.1;Pangolin_CoV_orf1ab;4085 | 0.05 | 0.05 |
| MT072864.1;QIQ54047.1;Pangolin_CoV_orf1ab;4086 | 0 | 0 |
| MT072864.1;QIQ54047.1;Pangolin_CoV_orf1ab;4087 | 0 | 0.025 |
| MT072864.1;QIQ54047.1;Pangolin_CoV_orf1ab;4088 | 0 | 0 |
| MT072864.1;QIQ54047.1;Pangolin_CoV_orf1ab;4089 | 0 | 0 |
| MT072864.1;QIQ54047.1;Pangolin_CoV_orf1ab;4090 | 0 | 0 |
| MT072864.1;QIQ54047.1;Pangolin_CoV_orf1ab;4091 | 0 | 0.025 |
| MT072864.1;QIQ54047.1;Pangolin_CoV_orf1ab;4092 | 0 | 0 |
| MT072864.1;QIQ54047.1;Pangolin_CoV_orf1ab;4093 | 0 | 0 |
| MT072864.1;QIQ54047.1;Pangolin_CoV_orf1ab;4094 | 0 | 0 |
| MT072864.1;QIQ54047.1;Pangolin_CoV_orf1ab;4095 | 0 | 0 |
| MT072864.1;QIQ54047.1;Pangolin_CoV_orf1ab;4096 | 0 | 0 |
| MT072864.1;QIQ54047.1;Pangolin_CoV_orf1ab;4097 | 0 | 0 |
| MT072864.1;QIQ54047.1;Pangolin_CoV_orf1ab;4098 | 0 | 0 |
| MT072864.1;QIQ54047.1;Pangolin_CoV_orf1ab;4099 | 0 | 0 |
| MT072864.1;QIQ54047.1;Pangolin_CoV_orf1ab;4100 | 0 | 0.025 |
| MT072864.1;QIQ54047.1;Pangolin_CoV_orf1ab;4101 | 0 | 0 |
| MT072864.1;QIQ54047.1;Pangolin_CoV_orf1ab;4102 | 0 | 0 |
| MT072864.1;QIQ54047.1;Pangolin_CoV_orf1ab;4103 | 0 | 0 |
| MT072864.1;QIQ54047.1;Pangolin_CoV_orf1ab;4104 | 0 | 0 |
| MT072864.1;QIQ54047.1;Pangolin_CoV_orf1ab;4105 | 0 | 0 |
| MT072864.1;QIQ54047.1;Pangolin_CoV_orf1ab;4106 | 0 | 0 |
| MT072864.1;QIQ54047.1;Pangolin_CoV_orf1ab;4107 | 0 | 0 |
| MT072864.1;QIQ54047.1;Pangolin_CoV_orf1ab;4108 | 0 | 0 |
| MT072864.1;QIQ54047.1;Pangolin_CoV_orf1ab;4109 | 0 | 0 |
| MT072864.1;QIQ54047.1;Pangolin_CoV_orf1ab;4110 | 0 | 0 |
| MT072864.1;QIQ54047.1;Pangolin_CoV_orf1ab;4111 | 0 | 0 |
| MT072864.1;QIQ54047.1;Pangolin_CoV_orf1ab;4112 | 0 | 0 |
| MT072864.1;QIQ54047.1;Pangolin_CoV_orf1ab;4113 | 0 | 0 |
| MT072864.1;QIQ54047.1;Pangolin_CoV_orf1ab;4114 | 0 | 0 |
| MT072864.1;QIQ54047.1;Pangolin_CoV_orf1ab;4115 | 0 | 0 |
| MT072864.1;QIQ54047.1;Pangolin_CoV_orf1ab;4116 | 0 | 0 |
| MT072864.1;QIQ54047.1;Pangolin_CoV_orf1ab;4117 | 0 | 0 |
| MT072864.1;QIQ54047.1;Pangolin_CoV_orf1ab;4118 | 0 | 0 |

| | | |
|---|---|---|
| MT072864.1;QIQ54047.1;Pangolin_CoV_orf1ab;4119 | 0 | 0.025 |
| MT072864.1;QIQ54047.1;Pangolin_CoV_orf1ab;4120 | 0 | 0.075 |
| MT072864.1;QIQ54047.1;Pangolin_CoV_orf1ab;4121 | 0 | 0.05 |
| MT072864.1;QIQ54047.1;Pangolin_CoV_orf1ab;4122 | 0 | 0.025 |
| MT072864.1;QIQ54047.1;Pangolin_CoV_orf1ab;4123 | 0 | 0.025 |
| MT072864.1;QIQ54047.1;Pangolin_CoV_orf1ab;4124 | 0.05 | 0.025 |
| MT072864.1;QIQ54047.1;Pangolin_CoV_orf1ab;4125 | 0.1 | 0.025 |
| MT072864.1;QIQ54047.1;Pangolin_CoV_orf1ab;4126 | 0.05 | 0.025 |
| MT072864.1;QIQ54047.1;Pangolin_CoV_orf1ab;4127 | 0 | 0.025 |
| MT072864.1;QIQ54047.1;Pangolin_CoV_orf1ab;4128 | 0 | 0 |
| MT072864.1;QIQ54047.1;Pangolin_CoV_orf1ab;4129 | 0.05 | 0.025 |
| MT072864.1;QIQ54047.1;Pangolin_CoV_orf1ab;4130 | 0 | 0 |
| MT072864.1;QIQ54047.1;Pangolin_CoV_orf1ab;4131 | 0 | 0 |
| MT072864.1;QIQ54047.1;Pangolin_CoV_orf1ab;4132 | 0 | 0.025 |
| MT072864.1;QIQ54047.1;Pangolin_CoV_orf1ab;4133 | 0.05 | 0.05 |
| MT072864.1;QIQ54047.1;Pangolin_CoV_orf1ab;4134 | 0 | 0 |
| MT072864.1;QIQ54047.1;Pangolin_CoV_orf1ab;4135 | 0 | 0 |
| MT072864.1;QIQ54047.1;Pangolin_CoV_orf1ab;4136 | 0 | 0 |
| MT072864.1;QIQ54047.1;Pangolin_CoV_orf1ab;4137 | 0 | 0 |
| MT072864.1;QIQ54047.1;Pangolin_CoV_orf1ab;4138 | 0 | 0 |
| MT072864.1;QIQ54047.1;Pangolin_CoV_orf1ab;4139 | 0 | 0 |
| MT072864.1;QIQ54047.1;Pangolin_CoV_orf1ab;4140 | 0 | 0 |
| MT072864.1;QIQ54047.1;Pangolin_CoV_orf1ab;4141 | 0 | 0 |
| MT072864.1;QIQ54047.1;Pangolin_CoV_orf1ab;4142 | 0 | 0 |
| MT072864.1;QIQ54047.1;Pangolin_CoV_orf1ab;4143 | 0 | 0 |
| MT072864.1;QIQ54047.1;Pangolin_CoV_orf1ab;4144 | 0 | 0 |
| MT072864.1;QIQ54047.1;Pangolin_CoV_orf1ab;4145 | 0 | 0 |
| MT072864.1;QIQ54047.1;Pangolin_CoV_orf1ab;4146 | 0 | 0 |
| MT072864.1;QIQ54047.1;Pangolin_CoV_orf1ab;4147 | 0 | 0 |
| MT072864.1;QIQ54047.1;Pangolin_CoV_orf1ab;4148 | 0 | 0 |
| MT072864.1;QIQ54047.1;Pangolin_CoV_orf1ab;4149 | 0 | 0 |
| MT072864.1;QIQ54047.1;Pangolin_CoV_orf1ab;4150 | 0 | 0 |
| MT072864.1;QIQ54047.1;Pangolin_CoV_orf1ab;4151 | 0 | 0 |
| MT072864.1;QIQ54047.1;Pangolin_CoV_orf1ab;4152 | 0 | 0 |
| MT072864.1;QIQ54047.1;Pangolin_CoV_orf1ab;4153 | 0 | 0.075 |
| MT072864.1;QIQ54047.1;Pangolin_CoV_orf1ab;4154 | 0.15 | 0.025 |
| MT072864.1;QIQ54047.1;Pangolin_CoV_orf1ab;4155 | 0.15 | 0.075 |
| MT072864.1;QIQ54047.1;Pangolin_CoV_orf1ab;4156 | 0 | 0.025 |
| MT072864.1;QIQ54047.1;Pangolin_CoV_orf1ab;4157 | 0 | 0.025 |
| MT072864.1;QIQ54047.1;Pangolin_CoV_orf1ab;4158 | 0 | 0 |
| MT072864.1;QIQ54047.1;Pangolin_CoV_orf1ab;4159 | 0 | 0 |
| MT072864.1;QIQ54047.1;Pangolin_CoV_orf1ab;4160 | 0 | 0 |
| MT072864.1;QIQ54047.1;Pangolin_CoV_orf1ab;4161 | 0 | 0 |

| | | |
|---|---|---|
| MT072864.1;QIQ54047.1;Pangolin_CoV_orf1ab;4162 | 0 | 0 |
| MT072864.1;QIQ54047.1;Pangolin_CoV_orf1ab;4163 | 0 | 0 |
| MT072864.1;QIQ54047.1;Pangolin_CoV_orf1ab;4164 | 0 | 0 |
| MT072864.1;QIQ54047.1;Pangolin_CoV_orf1ab;4165 | 0 | 0 |
| MT072864.1;QIQ54047.1;Pangolin_CoV_orf1ab;4166 | 0 | 0 |
| MT072864.1;QIQ54047.1;Pangolin_CoV_orf1ab;4167 | 0 | 0 |
| MT072864.1;QIQ54047.1;Pangolin_CoV_orf1ab;4168 | 0 | 0 |
| MT072864.1;QIQ54047.1;Pangolin_CoV_orf1ab;4169 | 0 | 0 |
| MT072864.1;QIQ54047.1;Pangolin_CoV_orf1ab;4170 | 0 | 0 |
| MT072864.1;QIQ54047.1;Pangolin_CoV_orf1ab;4171 | 0 | 0 |
| MT072864.1;QIQ54047.1;Pangolin_CoV_orf1ab;4172 | 0 | 0 |
| MT072864.1;QIQ54047.1;Pangolin_CoV_orf1ab;4173 | 0 | 0 |
| MT072864.1;QIQ54047.1;Pangolin_CoV_orf1ab;4174 | 0.05 | 0 |
| MT072864.1;QIQ54047.1;Pangolin_CoV_orf1ab;4175 | 0 | 0 |
| MT072864.1;QIQ54047.1;Pangolin_CoV_orf1ab;4176 | 0.05 | 0 |
| MT072864.1;QIQ54047.1;Pangolin_CoV_orf1ab;4177 | 0.05 | 0.05 |
| MT072864.1;QIQ54047.1;Pangolin_CoV_orf1ab;4178 | 0.15 | 0.1 |
| MT072864.1;QIQ54047.1;Pangolin_CoV_orf1ab;4179 | 0.3 | 0.25 |
| MT072864.1;QIQ54047.1;Pangolin_CoV_orf1ab;4180 | 0.15 | 0.05 |
| MT072864.1;QIQ54047.1;Pangolin_CoV_orf1ab;4181 | 0 | 0 |
| MT072864.1;QIQ54047.1;Pangolin_CoV_orf1ab;4182 | 0 | 0 |
| MT072864.1;QIQ54047.1;Pangolin_CoV_orf1ab;4183 | 0 | 0 |
| MT072864.1;QIQ54047.1;Pangolin_CoV_orf1ab;4184 | 0 | 0 |
| MT072864.1;QIQ54047.1;Pangolin_CoV_orf1ab;4185 | 0 | 0 |
| MT072864.1;QIQ54047.1;Pangolin_CoV_orf1ab;4186 | 0 | 0 |
| MT072864.1;QIQ54047.1;Pangolin_CoV_orf1ab;4187 | 0 | 0 |
| MT072864.1;QIQ54047.1;Pangolin_CoV_orf1ab;4188 | 0 | 0 |
| MT072864.1;QIQ54047.1;Pangolin_CoV_orf1ab;4189 | 0 | 0 |
| MT072864.1;QIQ54047.1;Pangolin_CoV_orf1ab;4190 | 0 | 0 |
| MT072864.1;QIQ54047.1;Pangolin_CoV_orf1ab;4191 | 0 | 0 |
| MT072864.1;QIQ54047.1;Pangolin_CoV_orf1ab;4192 | 0 | 0 |
| MT072864.1;QIQ54047.1;Pangolin_CoV_orf1ab;4193 | 0 | 0.025 |
| MT072864.1;QIQ54047.1;Pangolin_CoV_orf1ab;4194 | 0 | 0 |
| MT072864.1;QIQ54047.1;Pangolin_CoV_orf1ab;4195 | 0.2 | 0.15 |
| MT072864.1;QIQ54047.1;Pangolin_CoV_orf1ab;4196 | 0 | 0 |
| MT072864.1;QIQ54047.1;Pangolin_CoV_orf1ab;4197 | 0.05 | 0.15 |
| MT072864.1;QIQ54047.1;Pangolin_CoV_orf1ab;4198 | 0.05 | 0.075 |
| MT072864.1;QIQ54047.1;Pangolin_CoV_orf1ab;4199 | 0.05 | 0.05 |
| MT072864.1;QIQ54047.1;Pangolin_CoV_orf1ab;4200 | 0.05 | 0.025 |
| MT072864.1;QIQ54047.1;Pangolin_CoV_orf1ab;4201 | 0 | 0 |
| MT072864.1;QIQ54047.1;Pangolin_CoV_orf1ab;4202 | 0 | 0.025 |
| MT072864.1;QIQ54047.1;Pangolin_CoV_orf1ab;4203 | 0 | 0 |
| MT072864.1;QIQ54047.1;Pangolin_CoV_orf1ab;4204 | 0 | 0 |

| | | |
|---|---|---|
| MT072864.1;QIQ54047.1;Pangolin_CoV_orf1ab;4205 | 0 | 0 |
| MT072864.1;QIQ54047.1;Pangolin_CoV_orf1ab;4206 | 0 | 0 |
| MT072864.1;QIQ54047.1;Pangolin_CoV_orf1ab;4207 | 0.05 | 0 |
| MT072864.1;QIQ54047.1;Pangolin_CoV_orf1ab;4208 | 0 | 0 |
| MT072864.1;QIQ54047.1;Pangolin_CoV_orf1ab;4209 | 0 | 0 |
| MT072864.1;QIQ54047.1;Pangolin_CoV_orf1ab;4210 | 0 | 0 |
| MT072864.1;QIQ54047.1;Pangolin_CoV_orf1ab;4211 | 0 | 0 |
| MT072864.1;QIQ54047.1;Pangolin_CoV_orf1ab;4212 | 0 | 0 |
| MT072864.1;QIQ54047.1;Pangolin_CoV_orf1ab;4213 | 0 | 0 |
| MT072864.1;QIQ54047.1;Pangolin_CoV_orf1ab;4214 | 0 | 0 |
| MT072864.1;QIQ54047.1;Pangolin_CoV_orf1ab;4215 | 0 | 0 |
| MT072864.1;QIQ54047.1;Pangolin_CoV_orf1ab;4216 | 0 | 0 |
| MT072864.1;QIQ54047.1;Pangolin_CoV_orf1ab;4217 | 0 | 0 |
| MT072864.1;QIQ54047.1;Pangolin_CoV_orf1ab;4218 | 0 | 0 |
| MT072864.1;QIQ54047.1;Pangolin_CoV_orf1ab;4219 | 0 | 0 |
| MT072864.1;QIQ54047.1;Pangolin_CoV_orf1ab;4220 | 0 | 0 |
| MT072864.1;QIQ54047.1;Pangolin_CoV_orf1ab;4221 | 0.05 | 0.025 |
| MT072864.1;QIQ54047.1;Pangolin_CoV_orf1ab;4222 | 0.1 | 0.075 |
| MT072864.1;QIQ54047.1;Pangolin_CoV_orf1ab;4223 | 0.15 | 0.1 |
| MT072864.1;QIQ54047.1;Pangolin_CoV_orf1ab;4224 | 0.25 | 0.1 |
| MT072864.1;QIQ54047.1;Pangolin_CoV_orf1ab;4225 | 0.1 | 0.1 |
| MT072864.1;QIQ54047.1;Pangolin_CoV_orf1ab;4226 | 0.05 | 0.05 |
| MT072864.1;QIQ54047.1;Pangolin_CoV_orf1ab;4227 | 0.05 | 0.025 |
| MT072864.1;QIQ54047.1;Pangolin_CoV_orf1ab;4228 | 0 | 0 |
| MT072864.1;QIQ54047.1;Pangolin_CoV_orf1ab;4229 | 0 | 0 |
| MT072864.1;QIQ54047.1;Pangolin_CoV_orf1ab;4230 | 0 | 0 |
| MT072864.1;QIQ54047.1;Pangolin_CoV_orf1ab;4231 | 0 | 0 |
| MT072864.1;QIQ54047.1;Pangolin_CoV_orf1ab;4232 | 0 | 0 |
| MT072864.1;QIQ54047.1;Pangolin_CoV_orf1ab;4233 | 0 | 0 |
| MT072864.1;QIQ54047.1;Pangolin_CoV_orf1ab;4234 | 0 | 0 |
| MT072864.1;QIQ54047.1;Pangolin_CoV_orf1ab;4235 | 0 | 0 |
| MT072864.1;QIQ54047.1;Pangolin_CoV_orf1ab;4236 | 0 | 0.025 |
| MT072864.1;QIQ54047.1;Pangolin_CoV_orf1ab;4237 | 0 | 0 |
| MT072864.1;QIQ54047.1;Pangolin_CoV_orf1ab;4238 | 0 | 0.025 |
| MT072864.1;QIQ54047.1;Pangolin_CoV_orf1ab;4239 | 0.05 | 0.025 |
| MT072864.1;QIQ54047.1;Pangolin_CoV_orf1ab;4240 | 0 | 0.025 |
| MT072864.1;QIQ54047.1;Pangolin_CoV_orf1ab;4241 | 0 | 0 |
| MT072864.1;QIQ54047.1;Pangolin_CoV_orf1ab;4242 | 0 | 0 |
| MT072864.1;QIQ54047.1;Pangolin_CoV_orf1ab;4243 | 0 | 0.025 |
| MT072864.1;QIQ54047.1;Pangolin_CoV_orf1ab;4244 | 0 | 0.025 |
| MT072864.1;QIQ54047.1;Pangolin_CoV_orf1ab;4245 | 0 | 0 |
| MT072864.1;QIQ54047.1;Pangolin_CoV_orf1ab;4246 | 0 | 0 |
| MT072864.1;QIQ54047.1;Pangolin_CoV_orf1ab;4247 | 0 | 0 |

| | | |
|---|---|---|
| MT072864.1;QIQ54047.1;Pangolin_CoV_orf1ab;4248 | 0 | 0 |
| MT072864.1;QIQ54047.1;Pangolin_CoV_orf1ab;4249 | 0 | 0 |
| MT072864.1;QIQ54047.1;Pangolin_CoV_orf1ab;4250 | 0 | 0 |
| MT072864.1;QIQ54047.1;Pangolin_CoV_orf1ab;4251 | 0 | 0 |
| MT072864.1;QIQ54047.1;Pangolin_CoV_orf1ab;4252 | 0 | 0 |
| MT072864.1;QIQ54047.1;Pangolin_CoV_orf1ab;4253 | 0 | 0 |
| MT072864.1;QIQ54047.1;Pangolin_CoV_orf1ab;4254 | 0 | 0 |
| MT072864.1;QIQ54047.1;Pangolin_CoV_orf1ab;4255 | 0 | 0 |
| MT072864.1;QIQ54047.1;Pangolin_CoV_orf1ab;4256 | 0 | 0 |
| MT072864.1;QIQ54047.1;Pangolin_CoV_orf1ab;4257 | 0 | 0 |
| MT072864.1;QIQ54047.1;Pangolin_CoV_orf1ab;4258 | 0 | 0 |
| MT072864.1;QIQ54047.1;Pangolin_CoV_orf1ab;4259 | 0 | 0 |
| MT072864.1;QIQ54047.1;Pangolin_CoV_orf1ab;4260 | 0 | 0 |
| MT072864.1;QIQ54047.1;Pangolin_CoV_orf1ab;4261 | 0 | 0 |
| MT072864.1;QIQ54047.1;Pangolin_CoV_orf1ab;4262 | 0.05 | 0 |
| MT072864.1;QIQ54047.1;Pangolin_CoV_orf1ab;4263 | 0.1 | 0 |
| MT072864.1;QIQ54047.1;Pangolin_CoV_orf1ab;4264 | 0.1 | 0 |
| MT072864.1;QIQ54047.1;Pangolin_CoV_orf1ab;4265 | 0.15 | 0.075 |
| MT072864.1;QIQ54047.1;Pangolin_CoV_orf1ab;4266 | 0.1 | 0.025 |
| MT072864.1;QIQ54047.1;Pangolin_CoV_orf1ab;4267 | 0.35 | 0.15 |
| MT072864.1;QIQ54047.1;Pangolin_CoV_orf1ab;4268 | 0.05 | 0.025 |
| MT072864.1;QIQ54047.1;Pangolin_CoV_orf1ab;4269 | 0.1 | 0.025 |
| MT072864.1;QIQ54047.1;Pangolin_CoV_orf1ab;4270 | 0.05 | 0.025 |
| MT072864.1;QIQ54047.1;Pangolin_CoV_orf1ab;4271 | 0.1 | 0.05 |
| MT072864.1;QIQ54047.1;Pangolin_CoV_orf1ab;4272 | 0.05 | 0 |
| MT072864.1;QIQ54047.1;Pangolin_CoV_orf1ab;4273 | 0 | 0 |
| MT072864.1;QIQ54047.1;Pangolin_CoV_orf1ab;4274 | 0 | 0 |
| MT072864.1;QIQ54047.1;Pangolin_CoV_orf1ab;4275 | 0 | 0 |
| MT072864.1;QIQ54047.1;Pangolin_CoV_orf1ab;4276 | 0 | 0 |
| MT072864.1;QIQ54047.1;Pangolin_CoV_orf1ab;4277 | 0 | 0 |
| MT072864.1;QIQ54047.1;Pangolin_CoV_orf1ab;4278 | 0 | 0 |
| MT072864.1;QIQ54047.1;Pangolin_CoV_orf1ab;4279 | 0 | 0 |
| MT072864.1;QIQ54047.1;Pangolin_CoV_orf1ab;4280 | 0 | 0 |
| MT072864.1;QIQ54047.1;Pangolin_CoV_orf1ab;4281 | 0 | 0.025 |
| MT072864.1;QIQ54047.1;Pangolin_CoV_orf1ab;4282 | 0 | 0 |
| MT072864.1;QIQ54047.1;Pangolin_CoV_orf1ab;4283 | 0 | 0.025 |
| MT072864.1;QIQ54047.1;Pangolin_CoV_orf1ab;4284 | 0 | 0.025 |
| MT072864.1;QIQ54047.1;Pangolin_CoV_orf1ab;4285 | 0 | 0 |
| MT072864.1;QIQ54047.1;Pangolin_CoV_orf1ab;4286 | 0 | 0 |
| MT072864.1;QIQ54047.1;Pangolin_CoV_orf1ab;4287 | 0 | 0 |
| MT072864.1;QIQ54047.1;Pangolin_CoV_orf1ab;4288 | 0 | 0 |
| MT072864.1;QIQ54047.1;Pangolin_CoV_orf1ab;4289 | 0 | 0 |
| MT072864.1;QIQ54047.1;Pangolin_CoV_orf1ab;4290 | 0 | 0 |

| | | |
|---|---|---|
| MT072864.1;QIQ54047.1;Pangolin_CoV_orf1ab;4291 | 0 | 0 |
| MT072864.1;QIQ54047.1;Pangolin_CoV_orf1ab;4292 | 0 | 0 |
| MT072864.1;QIQ54047.1;Pangolin_CoV_orf1ab;4293 | 0 | 0 |
| MT072864.1;QIQ54047.1;Pangolin_CoV_orf1ab;4294 | 0 | 0.025 |
| MT072864.1;QIQ54047.1;Pangolin_CoV_orf1ab;4295 | 0 | 0.025 |
| MT072864.1;QIQ54047.1;Pangolin_CoV_orf1ab;4296 | 0 | 0.025 |
| MT072864.1;QIQ54047.1;Pangolin_CoV_orf1ab;4297 | 0 | 0.025 |
| MT072864.1;QIQ54047.1;Pangolin_CoV_orf1ab;4298 | 0 | 0.025 |
| MT072864.1;QIQ54047.1;Pangolin_CoV_orf1ab;4299 | 0 | 0.025 |
| MT072864.1;QIQ54047.1;Pangolin_CoV_orf1ab;4300 | 0.05 | 0.025 |
| MT072864.1;QIQ54047.1;Pangolin_CoV_orf1ab;4301 | 0 | 0.025 |
| MT072864.1;QIQ54047.1;Pangolin_CoV_orf1ab;4302 | 0.05 | 0.025 |
| MT072864.1;QIQ54047.1;Pangolin_CoV_orf1ab;4303 | 0 | 0 |
| MT072864.1;QIQ54047.1;Pangolin_CoV_orf1ab;4304 | 0 | 0 |
| MT072864.1;QIQ54047.1;Pangolin_CoV_orf1ab;4305 | 0 | 0 |
| MT072864.1;QIQ54047.1;Pangolin_CoV_orf1ab;4306 | 0 | 0 |
| MT072864.1;QIQ54047.1;Pangolin_CoV_orf1ab;4307 | 0 | 0 |
| MT072864.1;QIQ54047.1;Pangolin_CoV_orf1ab;4308 | 0 | 0 |
| MT072864.1;QIQ54047.1;Pangolin_CoV_orf1ab;4309 | 0 | 0 |
| MT072864.1;QIQ54047.1;Pangolin_CoV_orf1ab;4310 | 0 | 0 |
| MT072864.1;QIQ54047.1;Pangolin_CoV_orf1ab;4311 | 0 | 0 |
| MT072864.1;QIQ54047.1;Pangolin_CoV_orf1ab;4312 | 0 | 0 |
| MT072864.1;QIQ54047.1;Pangolin_CoV_orf1ab;4313 | 0 | 0 |
| MT072864.1;QIQ54047.1;Pangolin_CoV_orf1ab;4314 | 0 | 0 |
| MT072864.1;QIQ54047.1;Pangolin_CoV_orf1ab;4315 | 0 | 0 |
| MT072864.1;QIQ54047.1;Pangolin_CoV_orf1ab;4316 | 0 | 0 |
| MT072864.1;QIQ54047.1;Pangolin_CoV_orf1ab;4317 | 0 | 0 |
| MT072864.1;QIQ54047.1;Pangolin_CoV_orf1ab;4318 | 0 | 0 |
| MT072864.1;QIQ54047.1;Pangolin_CoV_orf1ab;4319 | 0 | 0 |
| MT072864.1;QIQ54047.1;Pangolin_CoV_orf1ab;4320 | 0 | 0 |
| MT072864.1;QIQ54047.1;Pangolin_CoV_orf1ab;4321 | 0 | 0.025 |
| MT072864.1;QIQ54047.1;Pangolin_CoV_orf1ab;4322 | 0 | 0.025 |
| MT072864.1;QIQ54047.1;Pangolin_CoV_orf1ab;4323 | 0 | 0.025 |
| MT072864.1;QIQ54047.1;Pangolin_CoV_orf1ab;4324 | 0 | 0.025 |
| MT072864.1;QIQ54047.1;Pangolin_CoV_orf1ab;4325 | 0 | 0 |
| MT072864.1;QIQ54047.1;Pangolin_CoV_orf1ab;4326 | 0 | 0.025 |
| MT072864.1;QIQ54047.1;Pangolin_CoV_orf1ab;4327 | 0 | 0 |
| MT072864.1;QIQ54047.1;Pangolin_CoV_orf1ab;4328 | 0 | 0 |
| MT072864.1;QIQ54047.1;Pangolin_CoV_orf1ab;4329 | 0 | 0 |
| MT072864.1;QIQ54047.1;Pangolin_CoV_orf1ab;4330 | 0 | 0 |
| MT072864.1;QIQ54047.1;Pangolin_CoV_orf1ab;4331 | 0 | 0 |
| MT072864.1;QIQ54047.1;Pangolin_CoV_orf1ab;4332 | 0 | 0.05 |
| MT072864.1;QIQ54047.1;Pangolin_CoV_orf1ab;4333 | 0 | 0 |

| | | |
|---|---|---|
| MT072864.1;QIQ54047.1;Pangolin_CoV_orf1ab;4334 | 0 | 0 |
| MT072864.1;QIQ54047.1;Pangolin_CoV_orf1ab;4335 | 0.05 | 0 |
| MT072864.1;QIQ54047.1;Pangolin_CoV_orf1ab;4336 | 0.05 | 0 |
| MT072864.1;QIQ54047.1;Pangolin_CoV_orf1ab;4337 | 0.05 | 0.025 |
| MT072864.1;QIQ54047.1;Pangolin_CoV_orf1ab;4338 | 0.05 | 0.05 |
| MT072864.1;QIQ54047.1;Pangolin_CoV_orf1ab;4339 | 0.05 | 0 |
| MT072864.1;QIQ54047.1;Pangolin_CoV_orf1ab;4340 | 0.05 | 0 |
| MT072864.1;QIQ54047.1;Pangolin_CoV_orf1ab;4341 | 0 | 0 |
| MT072864.1;QIQ54047.1;Pangolin_CoV_orf1ab;4342 | 0 | 0.025 |
| MT072864.1;QIQ54047.1;Pangolin_CoV_orf1ab;4343 | 0 | 0.025 |
| MT072864.1;QIQ54047.1;Pangolin_CoV_orf1ab;4344 | 0 | 0.025 |
| MT072864.1;QIQ54047.1;Pangolin_CoV_orf1ab;4345 | 0.05 | 0.05 |
| MT072864.1;QIQ54047.1;Pangolin_CoV_orf1ab;4346 | 0 | 0.025 |
| MT072864.1;QIQ54047.1;Pangolin_CoV_orf1ab;4347 | 0 | 0.025 |
| MT072864.1;QIQ54047.1;Pangolin_CoV_orf1ab;4348 | 0 | 0.025 |
| MT072864.1;QIQ54047.1;Pangolin_CoV_orf1ab;4349 | 0 | 0.025 |
| MT072864.1;QIQ54047.1;Pangolin_CoV_orf1ab;4350 | 0 | 0 |
| MT072864.1;QIQ54047.1;Pangolin_CoV_orf1ab;4351 | 0 | 0 |
| MT072864.1;QIQ54047.1;Pangolin_CoV_orf1ab;4352 | 0 | 0 |
| MT072864.1;QIQ54047.1;Pangolin_CoV_orf1ab;4353 | 0 | 0 |
| MT072864.1;QIQ54047.1;Pangolin_CoV_orf1ab;4354 | 0 | 0 |
| MT072864.1;QIQ54047.1;Pangolin_CoV_orf1ab;4355 | 0 | 0 |
| MT072864.1;QIQ54047.1;Pangolin_CoV_orf1ab;4356 | 0 | 0 |
| MT072864.1;QIQ54047.1;Pangolin_CoV_orf1ab;4357 | 0 | 0 |
| MT072864.1;QIQ54047.1;Pangolin_CoV_orf1ab;4358 | 0 | 0 |
| MT072864.1;QIQ54047.1;Pangolin_CoV_orf1ab;4359 | 0 | 0 |
| MT072864.1;QIQ54047.1;Pangolin_CoV_orf1ab;4360 | 0 | 0 |
| MT072864.1;QIQ54047.1;Pangolin_CoV_orf1ab;4361 | 0 | 0 |
| MT072864.1;QIQ54047.1;Pangolin_CoV_orf1ab;4362 | 0 | 0 |
| MT072864.1;QIQ54047.1;Pangolin_CoV_orf1ab;4363 | 0 | 0 |
| MT072864.1;QIQ54047.1;Pangolin_CoV_orf1ab;4364 | 0 | 0 |
| MT072864.1;QIQ54047.1;Pangolin_CoV_orf1ab;4365 | 0 | 0 |
| MT072864.1;QIQ54047.1;Pangolin_CoV_orf1ab;4366 | 0 | 0 |
| MT072864.1;QIQ54047.1;Pangolin_CoV_orf1ab;4367 | 0 | 0.075 |
| MT072864.1;QIQ54047.1;Pangolin_CoV_orf1ab;4368 | 0 | 0 |
| MT072864.1;QIQ54047.1;Pangolin_CoV_orf1ab;4369 | 0 | 0.025 |
| MT072864.1;QIQ54047.1;Pangolin_CoV_orf1ab;4370 | 0 | 0 |
| MT072864.1;QIQ54047.1;Pangolin_CoV_orf1ab;4371 | 0 | 0 |
| MT072864.1;QIQ54047.1;Pangolin_CoV_orf1ab;4372 | 0 | 0 |
| MT072864.1;QIQ54047.1;Pangolin_CoV_orf1ab;4373 | 0 | 0 |
| MT072864.1;QIQ54047.1;Pangolin_CoV_orf1ab;4374 | 0 | 0 |
| MT072864.1;QIQ54047.1;Pangolin_CoV_orf1ab;4375 | 0 | 0 |
| MT072864.1;QIQ54047.1;Pangolin_CoV_orf1ab;4376 | 0 | 0 |

| | | |
|---|---|---|
| MT072864.1;QIQ54047.1;Pangolin_CoV_orf1ab;4377 | 0 | 0 |
| MT072864.1;QIQ54047.1;Pangolin_CoV_orf1ab;4378 | 0.05 | 0 |
| MT072864.1;QIQ54047.1;Pangolin_CoV_orf1ab;4379 | 0.05 | 0.025 |
| MT072864.1;QIQ54047.1;Pangolin_CoV_orf1ab;4380 | 0.15 | 0.025 |
| MT072864.1;QIQ54047.1;Pangolin_CoV_orf1ab;4381 | 0 | 0 |
| MT072864.1;QIQ54047.1;Pangolin_CoV_orf1ab;4382 | 0.05 | 0 |
| MT072864.1;QIQ54047.1;Pangolin_CoV_orf1ab;4383 | 0 | 0 |
| MT072864.1;QIQ54047.1;Pangolin_CoV_orf1ab;4384 | 0 | 0 |
| MT072864.1;QIQ54047.1;Pangolin_CoV_orf1ab;4385 | 0 | 0 |
| MT072864.1;QIQ54047.1;Pangolin_CoV_orf1ab;4386 | 0.05 | 0 |
| MT072864.1;QIQ54047.1;Pangolin_CoV_orf1ab;4387 | 0 | 0 |
| MT072864.1;QIQ54047.1;Pangolin_CoV_orf1ab;4388 | 0 | 0 |
| MT072864.1;QIQ54047.1;Pangolin_CoV_orf1ab;4389 | 0 | 0 |
| MT072864.1;QIQ54047.1;Pangolin_CoV_orf1ab;4390 | 0 | 0 |
| MT072864.1;QIQ54047.1;Pangolin_CoV_orf1ab;4391 | 0 | 0 |
| MT072864.1;QIQ54047.1;Pangolin_CoV_orf1ab;4392 | 0 | 0 |
| MT072864.1;QIQ54047.1;Pangolin_CoV_orf1ab;4393 | 0 | 0 |
| MT072864.1;QIQ54047.1;Pangolin_CoV_orf1ab;4394 | 0 | 0 |
| MT072864.1;QIQ54047.1;Pangolin_CoV_orf1ab;4395 | 0 | 0 |
| MT072864.1;QIQ54047.1;Pangolin_CoV_orf1ab;4396 | 0 | 0 |
| MT072864.1;QIQ54047.1;Pangolin_CoV_orf1ab;4397 | 0 | 0 |
| MT072864.1;QIQ54047.1;Pangolin_CoV_orf1ab;4398 | 0 | 0 |
| MT072864.1;QIQ54047.1;Pangolin_CoV_orf1ab;4399 | 0 | 0.025 |
| MT072864.1;QIQ54047.1;Pangolin_CoV_orf1ab;4400 | 0 | 0 |
| MT072864.1;QIQ54047.1;Pangolin_CoV_orf1ab;4401 | 0 | 0 |
| MT072864.1;QIQ54047.1;Pangolin_CoV_orf1ab;4402 | 0 | 0 |
| MT072864.1;QIQ54047.1;Pangolin_CoV_orf1ab;4403 | 0 | 0 |
| MT072864.1;QIQ54047.1;Pangolin_CoV_orf1ab;4404 | 0 | 0 |
| MT072864.1;QIQ54047.1;Pangolin_CoV_orf1ab;4405 | 0 | 0 |
| MT072864.1;QIQ54047.1;Pangolin_CoV_orf1ab;4406 | 0 | 0 |
| MT072864.1;QIQ54047.1;Pangolin_CoV_orf1ab;4407 | 0 | 0 |
| MT072864.1;QIQ54047.1;Pangolin_CoV_orf1ab;4408 | 0 | 0 |
| MT072864.1;QIQ54047.1;Pangolin_CoV_orf1ab;4409 | 0 | 0 |
| MT072864.1;QIQ54047.1;Pangolin_CoV_orf1ab;4410 | 0.1 | 0.025 |
| MT072864.1;QIQ54047.1;Pangolin_CoV_orf1ab;4411 | 0.1 | 0 |
| MT072864.1;QIQ54047.1;Pangolin_CoV_orf1ab;4412 | 0.05 | 0.025 |
| MT072864.1;QIQ54047.1;Pangolin_CoV_orf1ab;4413 | 0 | 0 |
| MT072864.1;QIQ54047.1;Pangolin_CoV_orf1ab;4414 | 0 | 0 |
| MT072864.1;QIQ54047.1;Pangolin_CoV_orf1ab;4415 | 0 | 0 |
| MT072864.1;QIQ54047.1;Pangolin_CoV_orf1ab;4416 | 0 | 0.025 |
| MT072864.1;QIQ54047.1;Pangolin_CoV_orf1ab;4417 | 0 | 0 |
| MT072864.1;QIQ54047.1;Pangolin_CoV_orf1ab;4418 | 0 | 0 |
| MT072864.1;QIQ54047.1;Pangolin_CoV_orf1ab;4419 | 0 | 0 |

| | | |
|---|---|---|
| MT072864.1;QIQ54047.1;Pangolin_CoV_orf1ab;4420 | 0 | 0 |
| MT072864.1;QIQ54047.1;Pangolin_CoV_orf1ab;4421 | 0 | 0 |
| MT072864.1;QIQ54047.1;Pangolin_CoV_orf1ab;4422 | 0 | 0 |
| MT072864.1;QIQ54047.1;Pangolin_CoV_orf1ab;4423 | 0 | 0 |
| MT072864.1;QIQ54047.1;Pangolin_CoV_orf1ab;4424 | 0 | 0 |
| MT072864.1;QIQ54047.1;Pangolin_CoV_orf1ab;4425 | 0 | 0 |
| MT072864.1;QIQ54047.1;Pangolin_CoV_orf1ab;4426 | 0 | 0 |
| MT072864.1;QIQ54047.1;Pangolin_CoV_orf1ab;4427 | 0 | 0 |
| MT072864.1;QIQ54047.1;Pangolin_CoV_orf1ab;4428 | 0 | 0 |
| MT072864.1;QIQ54047.1;Pangolin_CoV_orf1ab;4429 | 0 | 0 |
| MT072864.1;QIQ54047.1;Pangolin_CoV_orf1ab;4430 | 0 | 0.025 |
| MT072864.1;QIQ54047.1;Pangolin_CoV_orf1ab;4431 | 0 | 0 |
| MT072864.1;QIQ54047.1;Pangolin_CoV_orf1ab;4432 | 0.05 | 0.025 |
| MT072864.1;QIQ54047.1;Pangolin_CoV_orf1ab;4433 | 0 | 0.025 |
| MT072864.1;QIQ54047.1;Pangolin_CoV_orf1ab;4434 | 0.15 | 0.075 |
| MT072864.1;QIQ54047.1;Pangolin_CoV_orf1ab;4435 | 0.2 | 0.075 |
| MT072864.1;QIQ54047.1;Pangolin_CoV_orf1ab;4436 | 0.25 | 0.1 |
| MT072864.1;QIQ54047.1;Pangolin_CoV_orf1ab;4437 | 0.3 | 0.125 |
| MT072864.1;QIQ54047.1;Pangolin_CoV_orf1ab;4438 | 0.3 | 0.2 |
| MT072864.1;QIQ54047.1;Pangolin_CoV_orf1ab;4439 | 0.3 | 0.225 |
| MT072864.1;QIQ54047.1;Pangolin_CoV_orf1ab;4440 | 0.15 | 0.075 |
| MT072864.1;QIQ54047.1;Pangolin_CoV_orf1ab;4441 | 0.05 | 0.025 |
| MT072864.1;QIQ54047.1;Pangolin_CoV_orf1ab;4442 | 0.1 | 0.05 |
| MT072864.1;QIQ54047.1;Pangolin_CoV_orf1ab;4443 | 0.05 | 0 |
| MT072864.1;QIQ54047.1;Pangolin_CoV_orf1ab;4444 | 0 | 0 |
| MT072864.1;QIQ54047.1;Pangolin_CoV_orf1ab;4445 | 0 | 0 |
| MT072864.1;QIQ54047.1;Pangolin_CoV_orf1ab;4446 | 0 | 0 |
| MT072864.1;QIQ54047.1;Pangolin_CoV_orf1ab;4447 | 0 | 0 |
| MT072864.1;QIQ54047.1;Pangolin_CoV_orf1ab;4448 | 0 | 0 |
| MT072864.1;QIQ54047.1;Pangolin_CoV_orf1ab;4449 | 0 | 0 |
| MT072864.1;QIQ54047.1;Pangolin_CoV_orf1ab;4450 | 0 | 0 |
| MT072864.1;QIQ54047.1;Pangolin_CoV_orf1ab;4451 | 0 | 0 |
| MT072864.1;QIQ54047.1;Pangolin_CoV_orf1ab;4452 | 0 | 0 |
| MT072864.1;QIQ54047.1;Pangolin_CoV_orf1ab;4453 | 0 | 0 |
| MT072864.1;QIQ54047.1;Pangolin_CoV_orf1ab;4454 | 0 | 0 |
| MT072864.1;QIQ54047.1;Pangolin_CoV_orf1ab;4455 | 0 | 0 |
| MT072864.1;QIQ54047.1;Pangolin_CoV_orf1ab;4456 | 0 | 0 |
| MT072864.1;QIQ54047.1;Pangolin_CoV_orf1ab;4457 | 0 | 0 |
| MT072864.1;QIQ54047.1;Pangolin_CoV_orf1ab;4458 | 0 | 0 |
| MT072864.1;QIQ54047.1;Pangolin_CoV_orf1ab;4459 | 0 | 0 |
| MT072864.1;QIQ54047.1;Pangolin_CoV_orf1ab;4460 | 0 | 0 |
| MT072864.1;QIQ54047.1;Pangolin_CoV_orf1ab;4461 | 0.1 | 0.05 |
| MT072864.1;QIQ54047.1;Pangolin_CoV_orf1ab;4462 | 0.05 | 0 |

| | | |
|---|---|---|
| MT072864.1;QIQ54047.1;Pangolin_CoV_orf1ab;4463 | 0 | 0 |
| MT072864.1;QIQ54047.1;Pangolin_CoV_orf1ab;4464 | 0 | 0 |
| MT072864.1;QIQ54047.1;Pangolin_CoV_orf1ab;4465 | 0 | 0 |
| MT072864.1;QIQ54047.1;Pangolin_CoV_orf1ab;4466 | 0 | 0 |
| MT072864.1;QIQ54047.1;Pangolin_CoV_orf1ab;4467 | 0 | 0 |
| MT072864.1;QIQ54047.1;Pangolin_CoV_orf1ab;4468 | 0 | 0 |
| MT072864.1;QIQ54047.1;Pangolin_CoV_orf1ab;4469 | 0.1 | 0.025 |
| MT072864.1;QIQ54047.1;Pangolin_CoV_orf1ab;4470 | 0.05 | 0 |
| MT072864.1;QIQ54047.1;Pangolin_CoV_orf1ab;4471 | 0.05 | 0 |
| MT072864.1;QIQ54047.1;Pangolin_CoV_orf1ab;4472 | 0 | 0 |
| MT072864.1;QIQ54047.1;Pangolin_CoV_orf1ab;4473 | 0 | 0 |
| MT072864.1;QIQ54047.1;Pangolin_CoV_orf1ab;4474 | 0 | 0 |
| MT072864.1;QIQ54047.1;Pangolin_CoV_orf1ab;4475 | 0 | 0 |
| MT072864.1;QIQ54047.1;Pangolin_CoV_orf1ab;4476 | 0 | 0 |
| MT072864.1;QIQ54047.1;Pangolin_CoV_orf1ab;4477 | 0 | 0 |
| MT072864.1;QIQ54047.1;Pangolin_CoV_orf1ab;4478 | 0 | 0.025 |
| MT072864.1;QIQ54047.1;Pangolin_CoV_orf1ab;4479 | 0 | 0.025 |
| MT072864.1;QIQ54047.1;Pangolin_CoV_orf1ab;4480 | 0 | 0.025 |
| MT072864.1;QIQ54047.1;Pangolin_CoV_orf1ab;4481 | 0 | 0.025 |
| MT072864.1;QIQ54047.1;Pangolin_CoV_orf1ab;4482 | 0 | 0.025 |
| MT072864.1;QIQ54047.1;Pangolin_CoV_orf1ab;4483 | 0 | 0.025 |
| MT072864.1;QIQ54047.1;Pangolin_CoV_orf1ab;4484 | 0 | 0.025 |
| MT072864.1;QIQ54047.1;Pangolin_CoV_orf1ab;4485 | 0 | 0.025 |
| MT072864.1;QIQ54047.1;Pangolin_CoV_orf1ab;4486 | 0 | 0 |
| MT072864.1;QIQ54047.1;Pangolin_CoV_orf1ab;4487 | 0 | 0 |
| MT072864.1;QIQ54047.1;Pangolin_CoV_orf1ab;4488 | 0 | 0 |
| MT072864.1;QIQ54047.1;Pangolin_CoV_orf1ab;4489 | 0 | 0 |
| MT072864.1;QIQ54047.1;Pangolin_CoV_orf1ab;4490 | 0 | 0 |
| MT072864.1;QIQ54047.1;Pangolin_CoV_orf1ab;4491 | 0 | 0 |
| MT072864.1;QIQ54047.1;Pangolin_CoV_orf1ab;4492 | 0 | 0 |
| MT072864.1;QIQ54047.1;Pangolin_CoV_orf1ab;4493 | 0 | 0 |
| MT072864.1;QIQ54047.1;Pangolin_CoV_orf1ab;4494 | 0 | 0 |
| MT072864.1;QIQ54047.1;Pangolin_CoV_orf1ab;4495 | 0 | 0 |
| MT072864.1;QIQ54047.1;Pangolin_CoV_orf1ab;4496 | 0 | 0 |
| MT072864.1;QIQ54047.1;Pangolin_CoV_orf1ab;4497 | 0 | 0 |
| MT072864.1;QIQ54047.1;Pangolin_CoV_orf1ab;4498 | 0 | 0 |
| MT072864.1;QIQ54047.1;Pangolin_CoV_orf1ab;4499 | 0 | 0 |
| MT072864.1;QIQ54047.1;Pangolin_CoV_orf1ab;4500 | 0 | 0 |
| MT072864.1;QIQ54047.1;Pangolin_CoV_orf1ab;4501 | 0 | 0 |
| MT072864.1;QIQ54047.1;Pangolin_CoV_orf1ab;4502 | 0 | 0 |
| MT072864.1;QIQ54047.1;Pangolin_CoV_orf1ab;4503 | 0 | 0 |
| MT072864.1;QIQ54047.1;Pangolin_CoV_orf1ab;4504 | 0 | 0 |
| MT072864.1;QIQ54047.1;Pangolin_CoV_orf1ab;4505 | 0 | 0 |

| | | |
|---|---|---|
| MT072864.1;QIQ54047.1;Pangolin_CoV_orf1ab;4506 | 0 | 0.375 |
| MT072864.1;QIQ54047.1;Pangolin_CoV_orf1ab;4507 | 0 | 0 |
| MT072864.1;QIQ54047.1;Pangolin_CoV_orf1ab;4508 | 0 | 0 |
| MT072864.1;QIQ54047.1;Pangolin_CoV_orf1ab;4509 | 0 | 0 |
| MT072864.1;QIQ54047.1;Pangolin_CoV_orf1ab;4510 | 0 | 0 |
| MT072864.1;QIQ54047.1;Pangolin_CoV_orf1ab;4511 | 0 | 0 |
| MT072864.1;QIQ54047.1;Pangolin_CoV_orf1ab;4512 | 0.05 | 0.1 |
| MT072864.1;QIQ54047.1;Pangolin_CoV_orf1ab;4513 | 0 | 0.025 |
| MT072864.1;QIQ54047.1;Pangolin_CoV_orf1ab;4514 | 0 | 0 |
| MT072864.1;QIQ54047.1;Pangolin_CoV_orf1ab;4515 | 0 | 0 |
| MT072864.1;QIQ54047.1;Pangolin_CoV_orf1ab;4516 | 0 | 0 |
| MT072864.1;QIQ54047.1;Pangolin_CoV_orf1ab;4517 | 0 | 0 |
| MT072864.1;QIQ54047.1;Pangolin_CoV_orf1ab;4518 | 0 | 0.025 |
| MT072864.1;QIQ54047.1;Pangolin_CoV_orf1ab;4519 | 0 | 0 |
| MT072864.1;QIQ54047.1;Pangolin_CoV_orf1ab;4520 | 0 | 0 |
| MT072864.1;QIQ54047.1;Pangolin_CoV_orf1ab;4521 | 0 | 0 |
| MT072864.1;QIQ54047.1;Pangolin_CoV_orf1ab;4522 | 0 | 0 |
| MT072864.1;QIQ54047.1;Pangolin_CoV_orf1ab;4523 | 0 | 0 |
| MT072864.1;QIQ54047.1;Pangolin_CoV_orf1ab;4524 | 0 | 0 |
| MT072864.1;QIQ54047.1;Pangolin_CoV_orf1ab;4525 | 0 | 0 |
| MT072864.1;QIQ54047.1;Pangolin_CoV_orf1ab;4526 | 0 | 0 |
| MT072864.1;QIQ54047.1;Pangolin_CoV_orf1ab;4527 | 0 | 0 |
| MT072864.1;QIQ54047.1;Pangolin_CoV_orf1ab;4528 | 0 | 0 |
| MT072864.1;QIQ54047.1;Pangolin_CoV_orf1ab;4529 | 0 | 0 |
| MT072864.1;QIQ54047.1;Pangolin_CoV_orf1ab;4530 | 0 | 0.025 |
| MT072864.1;QIQ54047.1;Pangolin_CoV_orf1ab;4531 | 0 | 0 |
| MT072864.1;QIQ54047.1;Pangolin_CoV_orf1ab;4532 | 0 | 0 |
| MT072864.1;QIQ54047.1;Pangolin_CoV_orf1ab;4533 | 0 | 0.025 |
| MT072864.1;QIQ54047.1;Pangolin_CoV_orf1ab;4534 | 0 | 0.05 |
| MT072864.1;QIQ54047.1;Pangolin_CoV_orf1ab;4535 | 0 | 0 |
| MT072864.1;QIQ54047.1;Pangolin_CoV_orf1ab;4536 | 0 | 0.025 |
| MT072864.1;QIQ54047.1;Pangolin_CoV_orf1ab;4537 | 0.2 | 0.025 |
| MT072864.1;QIQ54047.1;Pangolin_CoV_orf1ab;4538 | 0.05 | 0 |
| MT072864.1;QIQ54047.1;Pangolin_CoV_orf1ab;4539 | 0 | 0.025 |
| MT072864.1;QIQ54047.1;Pangolin_CoV_orf1ab;4540 | 0 | 0.075 |
| MT072864.1;QIQ54047.1;Pangolin_CoV_orf1ab;4541 | 0.05 | 0.075 |
| MT072864.1;QIQ54047.1;Pangolin_CoV_orf1ab;4542 | 0.05 | 0.075 |
| MT072864.1;QIQ54047.1;Pangolin_CoV_orf1ab;4543 | 0.1 | 0.05 |
| MT072864.1;QIQ54047.1;Pangolin_CoV_orf1ab;4544 | 0 | 0.025 |
| MT072864.1;QIQ54047.1;Pangolin_CoV_orf1ab;4545 | 0 | 0 |
| MT072864.1;QIQ54047.1;Pangolin_CoV_orf1ab;4546 | 0 | 0 |
| MT072864.1;QIQ54047.1;Pangolin_CoV_orf1ab;4547 | 0 | 0 |
| MT072864.1;QIQ54047.1;Pangolin_CoV_orf1ab;4548 | 0 | 0 |

| | | |
|---|---|---|
| MT072864.1;QIQ54047.1;Pangolin_CoV_orf1ab;4549 | 0.05 | 0 |
| MT072864.1;QIQ54047.1;Pangolin_CoV_orf1ab;4550 | 0.05 | 0.025 |
| MT072864.1;QIQ54047.1;Pangolin_CoV_orf1ab;4551 | 0.05 | 0 |
| MT072864.1;QIQ54047.1;Pangolin_CoV_orf1ab;4552 | 0 | 0 |
| MT072864.1;QIQ54047.1;Pangolin_CoV_orf1ab;4553 | 0 | 0 |
| MT072864.1;QIQ54047.1;Pangolin_CoV_orf1ab;4554 | 0 | 0 |
| MT072864.1;QIQ54047.1;Pangolin_CoV_orf1ab;4555 | 0 | 0 |
| MT072864.1;QIQ54047.1;Pangolin_CoV_orf1ab;4556 | 0 | 0 |
| MT072864.1;QIQ54047.1;Pangolin_CoV_orf1ab;4557 | 0.05 | 0 |
| MT072864.1;QIQ54047.1;Pangolin_CoV_orf1ab;4558 | 0 | 0 |
| MT072864.1;QIQ54047.1;Pangolin_CoV_orf1ab;4559 | 0 | 0 |
| MT072864.1;QIQ54047.1;Pangolin_CoV_orf1ab;4560 | 0 | 0 |
| MT072864.1;QIQ54047.1;Pangolin_CoV_orf1ab;4561 | 0 | 0 |
| MT072864.1;QIQ54047.1;Pangolin_CoV_orf1ab;4562 | 0 | 0 |
| MT072864.1;QIQ54047.1;Pangolin_CoV_orf1ab;4563 | 0 | 0 |
| MT072864.1;QIQ54047.1;Pangolin_CoV_orf1ab;4564 | 0.05 | 0 |
| MT072864.1;QIQ54047.1;Pangolin_CoV_orf1ab;4565 | 0 | 0 |
| MT072864.1;QIQ54047.1;Pangolin_CoV_orf1ab;4566 | 0 | 0 |
| MT072864.1;QIQ54047.1;Pangolin_CoV_orf1ab;4567 | 0 | 0 |
| MT072864.1;QIQ54047.1;Pangolin_CoV_orf1ab;4568 | 0 | 0 |
| MT072864.1;QIQ54047.1;Pangolin_CoV_orf1ab;4569 | 0 | 0 |
| MT072864.1;QIQ54047.1;Pangolin_CoV_orf1ab;4570 | 0 | 0 |
| MT072864.1;QIQ54047.1;Pangolin_CoV_orf1ab;4571 | 0 | 0 |
| MT072864.1;QIQ54047.1;Pangolin_CoV_orf1ab;4572 | 0 | 0 |
| MT072864.1;QIQ54047.1;Pangolin_CoV_orf1ab;4573 | 0 | 0 |
| MT072864.1;QIQ54047.1;Pangolin_CoV_orf1ab;4574 | 0 | 0 |
| MT072864.1;QIQ54047.1;Pangolin_CoV_orf1ab;4575 | 0 | 0 |
| MT072864.1;QIQ54047.1;Pangolin_CoV_orf1ab;4576 | 0 | 0 |
| MT072864.1;QIQ54047.1;Pangolin_CoV_orf1ab;4577 | 0 | 0 |
| MT072864.1;QIQ54047.1;Pangolin_CoV_orf1ab;4578 | 0 | 0 |
| MT072864.1;QIQ54047.1;Pangolin_CoV_orf1ab;4579 | 0 | 0 |
| MT072864.1;QIQ54047.1;Pangolin_CoV_orf1ab;4580 | 0.1 | 0.025 |
| MT072864.1;QIQ54047.1;Pangolin_CoV_orf1ab;4581 | 0.1 | 0 |
| MT072864.1;QIQ54047.1;Pangolin_CoV_orf1ab;4582 | 0 | 0 |
| MT072864.1;QIQ54047.1;Pangolin_CoV_orf1ab;4583 | 0.05 | 0 |
| MT072864.1;QIQ54047.1;Pangolin_CoV_orf1ab;4584 | 0 | 0 |
| MT072864.1;QIQ54047.1;Pangolin_CoV_orf1ab;4585 | 0 | 0 |
| MT072864.1;QIQ54047.1;Pangolin_CoV_orf1ab;4586 | 0 | 0 |
| MT072864.1;QIQ54047.1;Pangolin_CoV_orf1ab;4587 | 0.05 | 0 |
| MT072864.1;QIQ54047.1;Pangolin_CoV_orf1ab;4588 | 0 | 0 |
| MT072864.1;QIQ54047.1;Pangolin_CoV_orf1ab;4589 | 0 | 0 |
| MT072864.1;QIQ54047.1;Pangolin_CoV_orf1ab;4590 | 0 | 0 |
| MT072864.1;QIQ54047.1;Pangolin_CoV_orf1ab;4591 | 0 | 0 |

| | | |
|---|---|---|
| MT072864.1;QIQ54047.1;Pangolin_CoV_orf1ab;4592 | 0 | 0 |
| MT072864.1;QIQ54047.1;Pangolin_CoV_orf1ab;4593 | 0 | 0 |
| MT072864.1;QIQ54047.1;Pangolin_CoV_orf1ab;4594 | 0 | 0.025 |
| MT072864.1;QIQ54047.1;Pangolin_CoV_orf1ab;4595 | 0 | 0 |
| MT072864.1;QIQ54047.1;Pangolin_CoV_orf1ab;4596 | 0 | 0 |
| MT072864.1;QIQ54047.1;Pangolin_CoV_orf1ab;4597 | 0 | 0.025 |
| MT072864.1;QIQ54047.1;Pangolin_CoV_orf1ab;4598 | 0 | 0 |
| MT072864.1;QIQ54047.1;Pangolin_CoV_orf1ab;4599 | 0 | 0 |
| MT072864.1;QIQ54047.1;Pangolin_CoV_orf1ab;4600 | 0 | 0 |
| MT072864.1;QIQ54047.1;Pangolin_CoV_orf1ab;4601 | 0.1 | 0.05 |
| MT072864.1;QIQ54047.1;Pangolin_CoV_orf1ab;4602 | 0 | 0 |
| MT072864.1;QIQ54047.1;Pangolin_CoV_orf1ab;4603 | 0 | 0 |
| MT072864.1;QIQ54047.1;Pangolin_CoV_orf1ab;4604 | 0 | 0 |
| MT072864.1;QIQ54047.1;Pangolin_CoV_orf1ab;4605 | 0 | 0 |
| MT072864.1;QIQ54047.1;Pangolin_CoV_orf1ab;4606 | 0 | 0.025 |
| MT072864.1;QIQ54047.1;Pangolin_CoV_orf1ab;4607 | 0 | 0 |
| MT072864.1;QIQ54047.1;Pangolin_CoV_orf1ab;4608 | 0 | 0.05 |
| MT072864.1;QIQ54047.1;Pangolin_CoV_orf1ab;4609 | 0 | 0 |
| MT072864.1;QIQ54047.1;Pangolin_CoV_orf1ab;4610 | 0 | 0 |
| MT072864.1;QIQ54047.1;Pangolin_CoV_orf1ab;4611 | 0 | 0 |
| MT072864.1;QIQ54047.1;Pangolin_CoV_orf1ab;4612 | 0 | 0 |
| MT072864.1;QIQ54047.1;Pangolin_CoV_orf1ab;4613 | 0 | 0 |
| MT072864.1;QIQ54047.1;Pangolin_CoV_orf1ab;4614 | 0 | 0 |
| MT072864.1;QIQ54047.1;Pangolin_CoV_orf1ab;4615 | 0 | 0 |
| MT072864.1;QIQ54047.1;Pangolin_CoV_orf1ab;4616 | 0 | 0 |
| MT072864.1;QIQ54047.1;Pangolin_CoV_orf1ab;4617 | 0 | 0 |
| MT072864.1;QIQ54047.1;Pangolin_CoV_orf1ab;4618 | 0 | 0 |
| MT072864.1;QIQ54047.1;Pangolin_CoV_orf1ab;4619 | 0 | 0 |
| MT072864.1;QIQ54047.1;Pangolin_CoV_orf1ab;4620 | 0 | 0 |
| MT072864.1;QIQ54047.1;Pangolin_CoV_orf1ab;4621 | 0 | 0 |
| MT072864.1;QIQ54047.1;Pangolin_CoV_orf1ab;4622 | 0 | 0 |
| MT072864.1;QIQ54047.1;Pangolin_CoV_orf1ab;4623 | 0 | 0 |
| MT072864.1;QIQ54047.1;Pangolin_CoV_orf1ab;4624 | 0 | 0 |
| MT072864.1;QIQ54047.1;Pangolin_CoV_orf1ab;4625 | 0 | 0 |
| MT072864.1;QIQ54047.1;Pangolin_CoV_orf1ab;4626 | 0 | 0 |
| MT072864.1;QIQ54047.1;Pangolin_CoV_orf1ab;4627 | 0.05 | 0 |
| MT072864.1;QIQ54047.1;Pangolin_CoV_orf1ab;4628 | 0 | 0 |
| MT072864.1;QIQ54047.1;Pangolin_CoV_orf1ab;4629 | 0 | 0 |
| MT072864.1;QIQ54047.1;Pangolin_CoV_orf1ab;4630 | 0 | 0 |
| MT072864.1;QIQ54047.1;Pangolin_CoV_orf1ab;4631 | 0 | 0 |
| MT072864.1;QIQ54047.1;Pangolin_CoV_orf1ab;4632 | 0 | 0 |
| MT072864.1;QIQ54047.1;Pangolin_CoV_orf1ab;4633 | 0 | 0.025 |
| MT072864.1;QIQ54047.1;Pangolin_CoV_orf1ab;4634 | 0 | 0 |

| | | |
|---|---|---|
| MT072864.1;QIQ54047.1;Pangolin_CoV_orf1ab;4635 | 0 | 0 |
| MT072864.1;QIQ54047.1;Pangolin_CoV_orf1ab;4636 | 0.05 | 0.025 |
| MT072864.1;QIQ54047.1;Pangolin_CoV_orf1ab;4637 | 0 | 0 |
| MT072864.1;QIQ54047.1;Pangolin_CoV_orf1ab;4638 | 0 | 0.05 |
| MT072864.1;QIQ54047.1;Pangolin_CoV_orf1ab;4639 | 0.05 | 0.025 |
| MT072864.1;QIQ54047.1;Pangolin_CoV_orf1ab;4640 | 0 | 0 |
| MT072864.1;QIQ54047.1;Pangolin_CoV_orf1ab;4641 | 0 | 0.025 |
| MT072864.1;QIQ54047.1;Pangolin_CoV_orf1ab;4642 | 0.1 | 0 |
| MT072864.1;QIQ54047.1;Pangolin_CoV_orf1ab;4643 | 0.05 | 0.025 |
| MT072864.1;QIQ54047.1;Pangolin_CoV_orf1ab;4644 | 0 | 0 |
| MT072864.1;QIQ54047.1;Pangolin_CoV_orf1ab;4645 | 0 | 0 |
| MT072864.1;QIQ54047.1;Pangolin_CoV_orf1ab;4646 | 0 | 0 |
| MT072864.1;QIQ54047.1;Pangolin_CoV_orf1ab;4647 | 0 | 0 |
| MT072864.1;QIQ54047.1;Pangolin_CoV_orf1ab;4648 | 0.05 | 0 |
| MT072864.1;QIQ54047.1;Pangolin_CoV_orf1ab;4649 | 0.05 | 0 |
| MT072864.1;QIQ54047.1;Pangolin_CoV_orf1ab;4650 | 0.15 | 0 |
| MT072864.1;QIQ54047.1;Pangolin_CoV_orf1ab;4651 | 0.15 | 0.025 |
| MT072864.1;QIQ54047.1;Pangolin_CoV_orf1ab;4652 | 0.1 | 0 |
| MT072864.1;QIQ54047.1;Pangolin_CoV_orf1ab;4653 | 0.15 | 0 |
| MT072864.1;QIQ54047.1;Pangolin_CoV_orf1ab;4654 | 0.1 | 0.025 |
| MT072864.1;QIQ54047.1;Pangolin_CoV_orf1ab;4655 | 0.15 | 0.025 |
| MT072864.1;QIQ54047.1;Pangolin_CoV_orf1ab;4656 | 0.15 | 0 |
| MT072864.1;QIQ54047.1;Pangolin_CoV_orf1ab;4657 | 0.1 | 0 |
| MT072864.1;QIQ54047.1;Pangolin_CoV_orf1ab;4658 | 0.1 | 0 |
| MT072864.1;QIQ54047.1;Pangolin_CoV_orf1ab;4659 | 0 | 0 |
| MT072864.1;QIQ54047.1;Pangolin_CoV_orf1ab;4660 | 0.15 | 0 |
| MT072864.1;QIQ54047.1;Pangolin_CoV_orf1ab;4661 | 0.15 | 0 |
| MT072864.1;QIQ54047.1;Pangolin_CoV_orf1ab;4662 | 0.1 | 0.025 |
| MT072864.1;QIQ54047.1;Pangolin_CoV_orf1ab;4663 | 0 | 0 |
| MT072864.1;QIQ54047.1;Pangolin_CoV_orf1ab;4664 | 0 | 0 |
| MT072864.1;QIQ54047.1;Pangolin_CoV_orf1ab;4665 | 0 | 0 |
| MT072864.1;QIQ54047.1;Pangolin_CoV_orf1ab;4666 | 0 | 0 |
| MT072864.1;QIQ54047.1;Pangolin_CoV_orf1ab;4667 | 0 | 0 |
| MT072864.1;QIQ54047.1;Pangolin_CoV_orf1ab;4668 | 0 | 0 |
| MT072864.1;QIQ54047.1;Pangolin_CoV_orf1ab;4669 | 0 | 0.025 |
| MT072864.1;QIQ54047.1;Pangolin_CoV_orf1ab;4670 | 0 | 0.05 |
| MT072864.1;QIQ54047.1;Pangolin_CoV_orf1ab;4671 | 0 | 0.075 |
| MT072864.1;QIQ54047.1;Pangolin_CoV_orf1ab;4672 | 0 | 0.1 |
| MT072864.1;QIQ54047.1;Pangolin_CoV_orf1ab;4673 | 0 | 0.05 |
| MT072864.1;QIQ54047.1;Pangolin_CoV_orf1ab;4674 | 0 | 0.025 |
| MT072864.1;QIQ54047.1;Pangolin_CoV_orf1ab;4675 | 0 | 0.05 |
| MT072864.1;QIQ54047.1;Pangolin_CoV_orf1ab;4676 | 0 | 0.05 |
| MT072864.1;QIQ54047.1;Pangolin_CoV_orf1ab;4677 | 0 | 0.025 |

| | | |
|---|---|---|
| MT072864.1;QIQ54047.1;Pangolin_CoV_orf1ab;4678 | 0 | 0.025 |
| MT072864.1;QIQ54047.1;Pangolin_CoV_orf1ab;4679 | 0 | 0.025 |
| MT072864.1;QIQ54047.1;Pangolin_CoV_orf1ab;4680 | 0 | 0 |
| MT072864.1;QIQ54047.1;Pangolin_CoV_orf1ab;4681 | 0 | 0 |
| MT072864.1;QIQ54047.1;Pangolin_CoV_orf1ab;4682 | 0 | 0 |
| MT072864.1;QIQ54047.1;Pangolin_CoV_orf1ab;4683 | 0 | 0 |
| MT072864.1;QIQ54047.1;Pangolin_CoV_orf1ab;4684 | 0 | 0 |
| MT072864.1;QIQ54047.1;Pangolin_CoV_orf1ab;4685 | 0 | 0 |
| MT072864.1;QIQ54047.1;Pangolin_CoV_orf1ab;4686 | 0 | 0 |
| MT072864.1;QIQ54047.1;Pangolin_CoV_orf1ab;4687 | 0 | 0 |
| MT072864.1;QIQ54047.1;Pangolin_CoV_orf1ab;4688 | 0 | 0 |
| MT072864.1;QIQ54047.1;Pangolin_CoV_orf1ab;4689 | 0 | 0 |
| MT072864.1;QIQ54047.1;Pangolin_CoV_orf1ab;4690 | 0 | 0 |
| MT072864.1;QIQ54047.1;Pangolin_CoV_orf1ab;4691 | 0 | 0 |
| MT072864.1;QIQ54047.1;Pangolin_CoV_orf1ab;4692 | 0 | 0 |
| MT072864.1;QIQ54047.1;Pangolin_CoV_orf1ab;4693 | 0 | 0 |
| MT072864.1;QIQ54047.1;Pangolin_CoV_orf1ab;4694 | 0 | 0 |
| MT072864.1;QIQ54047.1;Pangolin_CoV_orf1ab;4695 | 0 | 0 |
| MT072864.1;QIQ54047.1;Pangolin_CoV_orf1ab;4696 | 0 | 0 |
| MT072864.1;QIQ54047.1;Pangolin_CoV_orf1ab;4697 | 0 | 0 |
| MT072864.1;QIQ54047.1;Pangolin_CoV_orf1ab;4698 | 0 | 0 |
| MT072864.1;QIQ54047.1;Pangolin_CoV_orf1ab;4699 | 0 | 0 |
| MT072864.1;QIQ54047.1;Pangolin_CoV_orf1ab;4700 | 0 | 0 |
| MT072864.1;QIQ54047.1;Pangolin_CoV_orf1ab;4701 | 0 | 0 |
| MT072864.1;QIQ54047.1;Pangolin_CoV_orf1ab;4702 | 0 | 0 |
| MT072864.1;QIQ54047.1;Pangolin_CoV_orf1ab;4703 | 0 | 0 |
| MT072864.1;QIQ54047.1;Pangolin_CoV_orf1ab;4704 | 0 | 0 |
| MT072864.1;QIQ54047.1;Pangolin_CoV_orf1ab;4705 | 0 | 0 |
| MT072864.1;QIQ54047.1;Pangolin_CoV_orf1ab;4706 | 0 | 0 |
| MT072864.1;QIQ54047.1;Pangolin_CoV_orf1ab;4707 | 0 | 0 |
| MT072864.1;QIQ54047.1;Pangolin_CoV_orf1ab;4708 | 0 | 0 |
| MT072864.1;QIQ54047.1;Pangolin_CoV_orf1ab;4709 | 0 | 0 |
| MT072864.1;QIQ54047.1;Pangolin_CoV_orf1ab;4710 | 0 | 0 |
| MT072864.1;QIQ54047.1;Pangolin_CoV_orf1ab;4711 | 0 | 0 |
| MT072864.1;QIQ54047.1;Pangolin_CoV_orf1ab;4712 | 0 | 0 |
| MT072864.1;QIQ54047.1;Pangolin_CoV_orf1ab;4713 | 0 | 0 |
| MT072864.1;QIQ54047.1;Pangolin_CoV_orf1ab;4714 | 0 | 0 |
| MT072864.1;QIQ54047.1;Pangolin_CoV_orf1ab;4715 | 0 | 0 |
| MT072864.1;QIQ54047.1;Pangolin_CoV_orf1ab;4716 | 0 | 0 |
| MT072864.1;QIQ54047.1;Pangolin_CoV_orf1ab;4717 | 0 | 0 |
| MT072864.1;QIQ54047.1;Pangolin_CoV_orf1ab;4718 | 0 | 0 |
| MT072864.1;QIQ54047.1;Pangolin_CoV_orf1ab;4719 | 0 | 0 |
| MT072864.1;QIQ54047.1;Pangolin_CoV_orf1ab;4720 | 0 | 0 |

| | | |
|---|---|---|
| MT072864.1;QIQ54047.1;Pangolin_CoV_orf1ab;4721 | 0 | 0 |
| MT072864.1;QIQ54047.1;Pangolin_CoV_orf1ab;4722 | 0 | 0 |
| MT072864.1;QIQ54047.1;Pangolin_CoV_orf1ab;4723 | 0 | 0 |
| MT072864.1;QIQ54047.1;Pangolin_CoV_orf1ab;4724 | 0 | 0 |
| MT072864.1;QIQ54047.1;Pangolin_CoV_orf1ab;4725 | 0 | 0 |
| MT072864.1;QIQ54047.1;Pangolin_CoV_orf1ab;4726 | 0 | 0 |
| MT072864.1;QIQ54047.1;Pangolin_CoV_orf1ab;4727 | 0 | 0 |
| MT072864.1;QIQ54047.1;Pangolin_CoV_orf1ab;4728 | 0 | 0 |
| MT072864.1;QIQ54047.1;Pangolin_CoV_orf1ab;4729 | 0 | 0 |
| MT072864.1;QIQ54047.1;Pangolin_CoV_orf1ab;4730 | 0 | 0 |
| MT072864.1;QIQ54047.1;Pangolin_CoV_orf1ab;4731 | 0 | 0 |
| MT072864.1;QIQ54047.1;Pangolin_CoV_orf1ab;4732 | 0 | 0 |
| MT072864.1;QIQ54047.1;Pangolin_CoV_orf1ab;4733 | 0 | 0 |
| MT072864.1;QIQ54047.1;Pangolin_CoV_orf1ab;4734 | 0 | 0 |
| MT072864.1;QIQ54047.1;Pangolin_CoV_orf1ab;4735 | 0 | 0 |
| MT072864.1;QIQ54047.1;Pangolin_CoV_orf1ab;4736 | 0 | 0 |
| MT072864.1;QIQ54047.1;Pangolin_CoV_orf1ab;4737 | 0 | 0 |
| MT072864.1;QIQ54047.1;Pangolin_CoV_orf1ab;4738 | 0 | 0 |
| MT072864.1;QIQ54047.1;Pangolin_CoV_orf1ab;4739 | 0 | 0 |
| MT072864.1;QIQ54047.1;Pangolin_CoV_orf1ab;4740 | 0 | 0.025 |
| MT072864.1;QIQ54047.1;Pangolin_CoV_orf1ab;4741 | 0.05 | 0 |
| MT072864.1;QIQ54047.1;Pangolin_CoV_orf1ab;4742 | 0 | 0 |
| MT072864.1;QIQ54047.1;Pangolin_CoV_orf1ab;4743 | 0 | 0 |
| MT072864.1;QIQ54047.1;Pangolin_CoV_orf1ab;4744 | 0 | 0 |
| MT072864.1;QIQ54047.1;Pangolin_CoV_orf1ab;4745 | 0 | 0 |
| MT072864.1;QIQ54047.1;Pangolin_CoV_orf1ab;4746 | 0 | 0 |
| MT072864.1;QIQ54047.1;Pangolin_CoV_orf1ab;4747 | 0 | 0 |
| MT072864.1;QIQ54047.1;Pangolin_CoV_orf1ab;4748 | 0 | 0 |
| MT072864.1;QIQ54047.1;Pangolin_CoV_orf1ab;4749 | 0.05 | 0 |
| MT072864.1;QIQ54047.1;Pangolin_CoV_orf1ab;4750 | 0.1 | 0.025 |
| MT072864.1;QIQ54047.1;Pangolin_CoV_orf1ab;4751 | 0.1 | 0.025 |
| MT072864.1;QIQ54047.1;Pangolin_CoV_orf1ab;4752 | 0.05 | 0.025 |
| MT072864.1;QIQ54047.1;Pangolin_CoV_orf1ab;4753 | 0 | 0 |
| MT072864.1;QIQ54047.1;Pangolin_CoV_orf1ab;4754 | 0 | 0 |
| MT072864.1;QIQ54047.1;Pangolin_CoV_orf1ab;4755 | 0 | 0 |
| MT072864.1;QIQ54047.1;Pangolin_CoV_orf1ab;4756 | 0 | 0.025 |
| MT072864.1;QIQ54047.1;Pangolin_CoV_orf1ab;4757 | 0 | 0.025 |
| MT072864.1;QIQ54047.1;Pangolin_CoV_orf1ab;4758 | 0 | 0 |
| MT072864.1;QIQ54047.1;Pangolin_CoV_orf1ab;4759 | 0 | 0 |
| MT072864.1;QIQ54047.1;Pangolin_CoV_orf1ab;4760 | 0 | 0 |
| MT072864.1;QIQ54047.1;Pangolin_CoV_orf1ab;4761 | 0 | 0 |
| MT072864.1;QIQ54047.1;Pangolin_CoV_orf1ab;4762 | 0 | 0 |
| MT072864.1;QIQ54047.1;Pangolin_CoV_orf1ab;4763 | 0 | 0 |

| | | |
|---|---|---|
| MT072864.1;QIQ54047.1;Pangolin_CoV_orf1ab;4764 | 0 | 0 |
| MT072864.1;QIQ54047.1;Pangolin_CoV_orf1ab;4765 | 0 | 0 |
| MT072864.1;QIQ54047.1;Pangolin_CoV_orf1ab;4766 | 0 | 0 |
| MT072864.1;QIQ54047.1;Pangolin_CoV_orf1ab;4767 | 0 | 0 |
| MT072864.1;QIQ54047.1;Pangolin_CoV_orf1ab;4768 | 0 | 0 |
| MT072864.1;QIQ54047.1;Pangolin_CoV_orf1ab;4769 | 0.05 | 0 |
| MT072864.1;QIQ54047.1;Pangolin_CoV_orf1ab;4770 | 0 | 0 |
| MT072864.1;QIQ54047.1;Pangolin_CoV_orf1ab;4771 | 0 | 0 |
| MT072864.1;QIQ54047.1;Pangolin_CoV_orf1ab;4772 | 0 | 0 |
| MT072864.1;QIQ54047.1;Pangolin_CoV_orf1ab;4773 | 0 | 0 |
| MT072864.1;QIQ54047.1;Pangolin_CoV_orf1ab;4774 | 0 | 0 |
| MT072864.1;QIQ54047.1;Pangolin_CoV_orf1ab;4775 | 0 | 0 |
| MT072864.1;QIQ54047.1;Pangolin_CoV_orf1ab;4776 | 0 | 0 |
| MT072864.1;QIQ54047.1;Pangolin_CoV_orf1ab;4777 | 0 | 0 |
| MT072864.1;QIQ54047.1;Pangolin_CoV_orf1ab;4778 | 0 | 0 |
| MT072864.1;QIQ54047.1;Pangolin_CoV_orf1ab;4779 | 0 | 0 |
| MT072864.1;QIQ54047.1;Pangolin_CoV_orf1ab;4780 | 0 | 0 |
| MT072864.1;QIQ54047.1;Pangolin_CoV_orf1ab;4781 | 0 | 0 |
| MT072864.1;QIQ54047.1;Pangolin_CoV_orf1ab;4782 | 0 | 0 |
| MT072864.1;QIQ54047.1;Pangolin_CoV_orf1ab;4783 | 0.05 | 0 |
| MT072864.1;QIQ54047.1;Pangolin_CoV_orf1ab;4784 | 0.05 | 0.05 |
| MT072864.1;QIQ54047.1;Pangolin_CoV_orf1ab;4785 | 0 | 0 |
| MT072864.1;QIQ54047.1;Pangolin_CoV_orf1ab;4786 | 0 | 0 |
| MT072864.1;QIQ54047.1;Pangolin_CoV_orf1ab;4787 | 0 | 0.025 |
| MT072864.1;QIQ54047.1;Pangolin_CoV_orf1ab;4788 | 0 | 0.025 |
| MT072864.1;QIQ54047.1;Pangolin_CoV_orf1ab;4789 | 0 | 0 |
| MT072864.1;QIQ54047.1;Pangolin_CoV_orf1ab;4790 | 0 | 0 |
| MT072864.1;QIQ54047.1;Pangolin_CoV_orf1ab;4791 | 0.05 | 0 |
| MT072864.1;QIQ54047.1;Pangolin_CoV_orf1ab;4792 | 0.05 | 0.025 |
| MT072864.1;QIQ54047.1;Pangolin_CoV_orf1ab;4793 | 0.1 | 0.025 |
| MT072864.1;QIQ54047.1;Pangolin_CoV_orf1ab;4794 | 0.1 | 0.025 |
| MT072864.1;QIQ54047.1;Pangolin_CoV_orf1ab;4795 | 0.1 | 0.025 |
| MT072864.1;QIQ54047.1;Pangolin_CoV_orf1ab;4796 | 0.2 | 0.025 |
| MT072864.1;QIQ54047.1;Pangolin_CoV_orf1ab;4797 | 0.1 | 0 |
| MT072864.1;QIQ54047.1;Pangolin_CoV_orf1ab;4798 | 0 | 0 |
| MT072864.1;QIQ54047.1;Pangolin_CoV_orf1ab;4799 | 0 | 0 |
| MT072864.1;QIQ54047.1;Pangolin_CoV_orf1ab;4800 | 0.05 | 0 |
| MT072864.1;QIQ54047.1;Pangolin_CoV_orf1ab;4801 | 0.05 | 0 |
| MT072864.1;QIQ54047.1;Pangolin_CoV_orf1ab;4802 | 0.05 | 0 |
| MT072864.1;QIQ54047.1;Pangolin_CoV_orf1ab;4803 | 0 | 0 |
| MT072864.1;QIQ54047.1;Pangolin_CoV_orf1ab;4804 | 0 | 0 |
| MT072864.1;QIQ54047.1;Pangolin_CoV_orf1ab;4805 | 0 | 0 |
| MT072864.1;QIQ54047.1;Pangolin_CoV_orf1ab;4806 | 0 | 0 |

| | | |
|---|---|---|
| MT072864.1;QIQ54047.1;Pangolin_CoV_orf1ab;4807 | 0 | 0 |
| MT072864.1;QIQ54047.1;Pangolin_CoV_orf1ab;4808 | 0 | 0 |
| MT072864.1;QIQ54047.1;Pangolin_CoV_orf1ab;4809 | 0.05 | 0 |
| MT072864.1;QIQ54047.1;Pangolin_CoV_orf1ab;4810 | 0 | 0.025 |
| MT072864.1;QIQ54047.1;Pangolin_CoV_orf1ab;4811 | 0 | 0 |
| MT072864.1;QIQ54047.1;Pangolin_CoV_orf1ab;4812 | 0 | 0 |
| MT072864.1;QIQ54047.1;Pangolin_CoV_orf1ab;4813 | 0 | 0 |
| MT072864.1;QIQ54047.1;Pangolin_CoV_orf1ab;4814 | 0 | 0 |
| MT072864.1;QIQ54047.1;Pangolin_CoV_orf1ab;4815 | 0 | 0 |
| MT072864.1;QIQ54047.1;Pangolin_CoV_orf1ab;4816 | 0 | 0 |
| MT072864.1;QIQ54047.1;Pangolin_CoV_orf1ab;4817 | 0 | 0 |
| MT072864.1;QIQ54047.1;Pangolin_CoV_orf1ab;4818 | 0.05 | 0 |
| MT072864.1;QIQ54047.1;Pangolin_CoV_orf1ab;4819 | 0.05 | 0 |
| MT072864.1;QIQ54047.1;Pangolin_CoV_orf1ab;4820 | 0 | 0 |
| MT072864.1;QIQ54047.1;Pangolin_CoV_orf1ab;4821 | 0 | 0 |
| MT072864.1;QIQ54047.1;Pangolin_CoV_orf1ab;4822 | 0.05 | 0 |
| MT072864.1;QIQ54047.1;Pangolin_CoV_orf1ab;4823 | 0 | 0 |
| MT072864.1;QIQ54047.1;Pangolin_CoV_orf1ab;4824 | 0 | 0 |
| MT072864.1;QIQ54047.1;Pangolin_CoV_orf1ab;4825 | 0 | 0 |
| MT072864.1;QIQ54047.1;Pangolin_CoV_orf1ab;4826 | 0 | 0.025 |
| MT072864.1;QIQ54047.1;Pangolin_CoV_orf1ab;4827 | 0 | 0 |
| MT072864.1;QIQ54047.1;Pangolin_CoV_orf1ab;4828 | 0.1 | 0.025 |
| MT072864.1;QIQ54047.1;Pangolin_CoV_orf1ab;4829 | 0.05 | 0.025 |
| MT072864.1;QIQ54047.1;Pangolin_CoV_orf1ab;4830 | 0.15 | 0 |
| MT072864.1;QIQ54047.1;Pangolin_CoV_orf1ab;4831 | 0.2 | 0 |
| MT072864.1;QIQ54047.1;Pangolin_CoV_orf1ab;4832 | 0.15 | 0.075 |
| MT072864.1;QIQ54047.1;Pangolin_CoV_orf1ab;4833 | 0.25 | 0.025 |
| MT072864.1;QIQ54047.1;Pangolin_CoV_orf1ab;4834 | 0.15 | 0 |
| MT072864.1;QIQ54047.1;Pangolin_CoV_orf1ab;4835 | 0.15 | 0 |
| MT072864.1;QIQ54047.1;Pangolin_CoV_orf1ab;4836 | 0.15 | 0 |
| MT072864.1;QIQ54047.1;Pangolin_CoV_orf1ab;4837 | 0 | 0 |
| MT072864.1;QIQ54047.1;Pangolin_CoV_orf1ab;4838 | 0 | 0 |
| MT072864.1;QIQ54047.1;Pangolin_CoV_orf1ab;4839 | 0 | 0 |
| MT072864.1;QIQ54047.1;Pangolin_CoV_orf1ab;4840 | 0 | 0 |
| MT072864.1;QIQ54047.1;Pangolin_CoV_orf1ab;4841 | 0 | 0 |
| MT072864.1;QIQ54047.1;Pangolin_CoV_orf1ab;4842 | 0 | 0 |
| MT072864.1;QIQ54047.1;Pangolin_CoV_orf1ab;4843 | 0 | 0 |
| MT072864.1;QIQ54047.1;Pangolin_CoV_orf1ab;4844 | 0 | 0 |
| MT072864.1;QIQ54047.1;Pangolin_CoV_orf1ab;4845 | 0 | 0 |
| MT072864.1;QIQ54047.1;Pangolin_CoV_orf1ab;4846 | 0 | 0 |
| MT072864.1;QIQ54047.1;Pangolin_CoV_orf1ab;4847 | 0.05 | 0.025 |
| MT072864.1;QIQ54047.1;Pangolin_CoV_orf1ab;4848 | 0 | 0 |
| MT072864.1;QIQ54047.1;Pangolin_CoV_orf1ab;4849 | 0 | 0 |

| | | |
|---|---|---|
| MT072864.1;QIQ54047.1;Pangolin_CoV_orf1ab;4850 | 0 | 0.025 |
| MT072864.1;QIQ54047.1;Pangolin_CoV_orf1ab;4851 | 0 | 0 |
| MT072864.1;QIQ54047.1;Pangolin_CoV_orf1ab;4852 | 0 | 0 |
| MT072864.1;QIQ54047.1;Pangolin_CoV_orf1ab;4853 | 0 | 0 |
| MT072864.1;QIQ54047.1;Pangolin_CoV_orf1ab;4854 | 0 | 0 |
| MT072864.1;QIQ54047.1;Pangolin_CoV_orf1ab;4855 | 0 | 0 |
| MT072864.1;QIQ54047.1;Pangolin_CoV_orf1ab;4856 | 0 | 0.025 |
| MT072864.1;QIQ54047.1;Pangolin_CoV_orf1ab;4857 | 0 | 0.025 |
| MT072864.1;QIQ54047.1;Pangolin_CoV_orf1ab;4858 | 0 | 0 |
| MT072864.1;QIQ54047.1;Pangolin_CoV_orf1ab;4859 | 0 | 0 |
| MT072864.1;QIQ54047.1;Pangolin_CoV_orf1ab;4860 | 0.05 | 0 |
| MT072864.1;QIQ54047.1;Pangolin_CoV_orf1ab;4861 | 0.05 | 0.075 |
| MT072864.1;QIQ54047.1;Pangolin_CoV_orf1ab;4862 | 0.05 | 0.025 |
| MT072864.1;QIQ54047.1;Pangolin_CoV_orf1ab;4863 | 0.05 | 0 |
| MT072864.1;QIQ54047.1;Pangolin_CoV_orf1ab;4864 | 0 | 0.025 |
| MT072864.1;QIQ54047.1;Pangolin_CoV_orf1ab;4865 | 0.05 | 0 |
| MT072864.1;QIQ54047.1;Pangolin_CoV_orf1ab;4866 | 0.05 | 0 |
| MT072864.1;QIQ54047.1;Pangolin_CoV_orf1ab;4867 | 0 | 0 |
| MT072864.1;QIQ54047.1;Pangolin_CoV_orf1ab;4868 | 0.05 | 0.025 |
| MT072864.1;QIQ54047.1;Pangolin_CoV_orf1ab;4869 | 0 | 0.025 |
| MT072864.1;QIQ54047.1;Pangolin_CoV_orf1ab;4870 | 0.05 | 0.025 |
| MT072864.1;QIQ54047.1;Pangolin_CoV_orf1ab;4871 | 0.05 | 0.025 |
| MT072864.1;QIQ54047.1;Pangolin_CoV_orf1ab;4872 | 0.1 | 0.025 |
| MT072864.1;QIQ54047.1;Pangolin_CoV_orf1ab;4873 | 0 | 0.025 |
| MT072864.1;QIQ54047.1;Pangolin_CoV_orf1ab;4874 | 0.15 | 0.1 |
| MT072864.1;QIQ54047.1;Pangolin_CoV_orf1ab;4875 | 0 | 0.05 |
| MT072864.1;QIQ54047.1;Pangolin_CoV_orf1ab;4876 | 0.15 | 0.075 |
| MT072864.1;QIQ54047.1;Pangolin_CoV_orf1ab;4877 | 0.05 | 0.05 |
| MT072864.1;QIQ54047.1;Pangolin_CoV_orf1ab;4878 | 0 | 0.025 |
| MT072864.1;QIQ54047.1;Pangolin_CoV_orf1ab;4879 | 0 | 0 |
| MT072864.1;QIQ54047.1;Pangolin_CoV_orf1ab;4880 | 0 | 0 |
| MT072864.1;QIQ54047.1;Pangolin_CoV_orf1ab;4881 | 0 | 0 |
| MT072864.1;QIQ54047.1;Pangolin_CoV_orf1ab;4882 | 0 | 0 |
| MT072864.1;QIQ54047.1;Pangolin_CoV_orf1ab;4883 | 0 | 0 |
| MT072864.1;QIQ54047.1;Pangolin_CoV_orf1ab;4884 | 0 | 0 |
| MT072864.1;QIQ54047.1;Pangolin_CoV_orf1ab;4885 | 0 | 0 |
| MT072864.1;QIQ54047.1;Pangolin_CoV_orf1ab;4886 | 0 | 0 |
| MT072864.1;QIQ54047.1;Pangolin_CoV_orf1ab;4887 | 0 | 0 |
| MT072864.1;QIQ54047.1;Pangolin_CoV_orf1ab;4888 | 0 | 0 |
| MT072864.1;QIQ54047.1;Pangolin_CoV_orf1ab;4889 | 0.05 | 0 |
| MT072864.1;QIQ54047.1;Pangolin_CoV_orf1ab;4890 | 0 | 0 |
| MT072864.1;QIQ54047.1;Pangolin_CoV_orf1ab;4891 | 0.05 | 0 |
| MT072864.1;QIQ54047.1;Pangolin_CoV_orf1ab;4892 | 0 | 0 |

| | | |
|---|---|---|
| MT072864.1;QIQ54047.1;Pangolin_CoV_orf1ab;4893 | 0 | 0 |
| MT072864.1;QIQ54047.1;Pangolin_CoV_orf1ab;4894 | 0 | 0 |
| MT072864.1;QIQ54047.1;Pangolin_CoV_orf1ab;4895 | 0 | 0 |
| MT072864.1;QIQ54047.1;Pangolin_CoV_orf1ab;4896 | 0 | 0 |
| MT072864.1;QIQ54047.1;Pangolin_CoV_orf1ab;4897 | 0 | 0 |
| MT072864.1;QIQ54047.1;Pangolin_CoV_orf1ab;4898 | 0 | 0 |
| MT072864.1;QIQ54047.1;Pangolin_CoV_orf1ab;4899 | 0 | 0 |
| MT072864.1;QIQ54047.1;Pangolin_CoV_orf1ab;4900 | 0 | 0 |
| MT072864.1;QIQ54047.1;Pangolin_CoV_orf1ab;4901 | 0.1 | 0.025 |
| MT072864.1;QIQ54047.1;Pangolin_CoV_orf1ab;4902 | 0.05 | 0.025 |
| MT072864.1;QIQ54047.1;Pangolin_CoV_orf1ab;4903 | 0.05 | 0 |
| MT072864.1;QIQ54047.1;Pangolin_CoV_orf1ab;4904 | 0.1 | 0.05 |
| MT072864.1;QIQ54047.1;Pangolin_CoV_orf1ab;4905 | 0.05 | 0 |
| MT072864.1;QIQ54047.1;Pangolin_CoV_orf1ab;4906 | 0.2 | 0.025 |
| MT072864.1;QIQ54047.1;Pangolin_CoV_orf1ab;4907 | 0.05 | 0.025 |
| MT072864.1;QIQ54047.1;Pangolin_CoV_orf1ab;4908 | 0.05 | 0 |
| MT072864.1;QIQ54047.1;Pangolin_CoV_orf1ab;4909 | 0.05 | 0 |
| MT072864.1;QIQ54047.1;Pangolin_CoV_orf1ab;4910 | 0 | 0 |
| MT072864.1;QIQ54047.1;Pangolin_CoV_orf1ab;4911 | 0 | 0 |
| MT072864.1;QIQ54047.1;Pangolin_CoV_orf1ab;4912 | 0 | 0 |
| MT072864.1;QIQ54047.1;Pangolin_CoV_orf1ab;4913 | 0 | 0 |
| MT072864.1;QIQ54047.1;Pangolin_CoV_orf1ab;4914 | 0 | 0 |
| MT072864.1;QIQ54047.1;Pangolin_CoV_orf1ab;4915 | 0.05 | 0 |
| MT072864.1;QIQ54047.1;Pangolin_CoV_orf1ab;4916 | 0.1 | 0.15 |
| MT072864.1;QIQ54047.1;Pangolin_CoV_orf1ab;4917 | 0 | 0 |
| MT072864.1;QIQ54047.1;Pangolin_CoV_orf1ab;4918 | 0 | 0 |
| MT072864.1;QIQ54047.1;Pangolin_CoV_orf1ab;4919 | 0 | 0 |
| MT072864.1;QIQ54047.1;Pangolin_CoV_orf1ab;4920 | 0 | 0.025 |
| MT072864.1;QIQ54047.1;Pangolin_CoV_orf1ab;4921 | 0.1 | 0.05 |
| MT072864.1;QIQ54047.1;Pangolin_CoV_orf1ab;4922 | 0 | 0.025 |
| MT072864.1;QIQ54047.1;Pangolin_CoV_orf1ab;4923 | 0 | 0.025 |
| MT072864.1;QIQ54047.1;Pangolin_CoV_orf1ab;4924 | 0 | 0.025 |
| MT072864.1;QIQ54047.1;Pangolin_CoV_orf1ab;4925 | 0 | 0.025 |
| MT072864.1;QIQ54047.1;Pangolin_CoV_orf1ab;4926 | 0 | 0 |
| MT072864.1;QIQ54047.1;Pangolin_CoV_orf1ab;4927 | 0 | 0 |
| MT072864.1;QIQ54047.1;Pangolin_CoV_orf1ab;4928 | 0 | 0 |
| MT072864.1;QIQ54047.1;Pangolin_CoV_orf1ab;4929 | 0 | 0 |
| MT072864.1;QIQ54047.1;Pangolin_CoV_orf1ab;4930 | 0 | 0 |
| MT072864.1;QIQ54047.1;Pangolin_CoV_orf1ab;4931 | 0 | 0 |
| MT072864.1;QIQ54047.1;Pangolin_CoV_orf1ab;4932 | 0 | 0 |
| MT072864.1;QIQ54047.1;Pangolin_CoV_orf1ab;4933 | 0 | 0 |
| MT072864.1;QIQ54047.1;Pangolin_CoV_orf1ab;4934 | 0 | 0 |
| MT072864.1;QIQ54047.1;Pangolin_CoV_orf1ab;4935 | 0 | 0 |

| | | |
|---|---|---|
| MT072864.1;QIQ54047.1;Pangolin_CoV_orf1ab;4936 | 0 | 0 |
| MT072864.1;QIQ54047.1;Pangolin_CoV_orf1ab;4937 | 0 | 0 |
| MT072864.1;QIQ54047.1;Pangolin_CoV_orf1ab;4938 | 0 | 0 |
| MT072864.1;QIQ54047.1;Pangolin_CoV_orf1ab;4939 | 0 | 0 |
| MT072864.1;QIQ54047.1;Pangolin_CoV_orf1ab;4940 | 0 | 0 |
| MT072864.1;QIQ54047.1;Pangolin_CoV_orf1ab;4941 | 0 | 0 |
| MT072864.1;QIQ54047.1;Pangolin_CoV_orf1ab;4942 | 0 | 0 |
| MT072864.1;QIQ54047.1;Pangolin_CoV_orf1ab;4943 | 0 | 0 |
| MT072864.1;QIQ54047.1;Pangolin_CoV_orf1ab;4944 | 0 | 0 |
| MT072864.1;QIQ54047.1;Pangolin_CoV_orf1ab;4945 | 0 | 0 |
| MT072864.1;QIQ54047.1;Pangolin_CoV_orf1ab;4946 | 0 | 0 |
| MT072864.1;QIQ54047.1;Pangolin_CoV_orf1ab;4947 | 0 | 0 |
| MT072864.1;QIQ54047.1;Pangolin_CoV_orf1ab;4948 | 0 | 0 |
| MT072864.1;QIQ54047.1;Pangolin_CoV_orf1ab;4949 | 0 | 0 |
| MT072864.1;QIQ54047.1;Pangolin_CoV_orf1ab;4950 | 0 | 0 |
| MT072864.1;QIQ54047.1;Pangolin_CoV_orf1ab;4951 | 0 | 0 |
| MT072864.1;QIQ54047.1;Pangolin_CoV_orf1ab;4952 | 0 | 0 |
| MT072864.1;QIQ54047.1;Pangolin_CoV_orf1ab;4953 | 0 | 0 |
| MT072864.1;QIQ54047.1;Pangolin_CoV_orf1ab;4954 | 0 | 0 |
| MT072864.1;QIQ54047.1;Pangolin_CoV_orf1ab;4955 | 0.3 | 0.175 |
| MT072864.1;QIQ54047.1;Pangolin_CoV_orf1ab;4956 | 0.4 | 0.275 |
| MT072864.1;QIQ54047.1;Pangolin_CoV_orf1ab;4957 | 0.35 | 0.275 |
| MT072864.1;QIQ54047.1;Pangolin_CoV_orf1ab;4958 | 0.25 | 0.025 |
| MT072864.1;QIQ54047.1;Pangolin_CoV_orf1ab;4959 | 0.25 | 0.075 |
| MT072864.1;QIQ54047.1;Pangolin_CoV_orf1ab;4960 | 0.25 | 0.025 |
| MT072864.1;QIQ54047.1;Pangolin_CoV_orf1ab;4961 | 0.25 | 0.05 |
| MT072864.1;QIQ54047.1;Pangolin_CoV_orf1ab;4962 | 0.25 | 0.025 |
| MT072864.1;QIQ54047.1;Pangolin_CoV_orf1ab;4963 | 0.15 | 0.025 |
| MT072864.1;QIQ54047.1;Pangolin_CoV_orf1ab;4964 | 0.15 | 0.025 |
| MT072864.1;QIQ54047.1;Pangolin_CoV_orf1ab;4965 | 0 | 0 |
| MT072864.1;QIQ54047.1;Pangolin_CoV_orf1ab;4966 | 0 | 0 |
| MT072864.1;QIQ54047.1;Pangolin_CoV_orf1ab;4967 | 0 | 0 |
| MT072864.1;QIQ54047.1;Pangolin_CoV_orf1ab;4968 | 0 | 0 |
| MT072864.1;QIQ54047.1;Pangolin_CoV_orf1ab;4969 | 0 | 0 |
| MT072864.1;QIQ54047.1;Pangolin_CoV_orf1ab;4970 | 0 | 0 |
| MT072864.1;QIQ54047.1;Pangolin_CoV_orf1ab;4971 | 0 | 0 |
| MT072864.1;QIQ54047.1;Pangolin_CoV_orf1ab;4972 | 0 | 0 |
| MT072864.1;QIQ54047.1;Pangolin_CoV_orf1ab;4973 | 0 | 0 |
| MT072864.1;QIQ54047.1;Pangolin_CoV_orf1ab;4974 | 0 | 0 |
| MT072864.1;QIQ54047.1;Pangolin_CoV_orf1ab;4975 | 0 | 0 |
| MT072864.1;QIQ54047.1;Pangolin_CoV_orf1ab;4976 | 0 | 0 |
| MT072864.1;QIQ54047.1;Pangolin_CoV_orf1ab;4977 | 0 | 0.025 |
| MT072864.1;QIQ54047.1;Pangolin_CoV_orf1ab;4978 | 0 | 0 |

| | | |
|---|---|---|
| MT072864.1;QIQ54047.1;Pangolin_CoV_orf1ab;4979 | 0 | 0 |
| MT072864.1;QIQ54047.1;Pangolin_CoV_orf1ab;4980 | 0 | 0 |
| MT072864.1;QIQ54047.1;Pangolin_CoV_orf1ab;4981 | 0 | 0 |
| MT072864.1;QIQ54047.1;Pangolin_CoV_orf1ab;4982 | 0 | 0 |
| MT072864.1;QIQ54047.1;Pangolin_CoV_orf1ab;4983 | 0 | 0 |
| MT072864.1;QIQ54047.1;Pangolin_CoV_orf1ab;4984 | 0 | 0 |
| MT072864.1;QIQ54047.1;Pangolin_CoV_orf1ab;4985 | 0 | 0 |
| MT072864.1;QIQ54047.1;Pangolin_CoV_orf1ab;4986 | 0 | 0 |
| MT072864.1;QIQ54047.1;Pangolin_CoV_orf1ab;4987 | 0 | 0 |
| MT072864.1;QIQ54047.1;Pangolin_CoV_orf1ab;4988 | 0 | 0 |
| MT072864.1;QIQ54047.1;Pangolin_CoV_orf1ab;4989 | 0 | 0.075 |
| MT072864.1;QIQ54047.1;Pangolin_CoV_orf1ab;4990 | 0 | 0.025 |
| MT072864.1;QIQ54047.1;Pangolin_CoV_orf1ab;4991 | 0 | 0.025 |
| MT072864.1;QIQ54047.1;Pangolin_CoV_orf1ab;4992 | 0 | 0.025 |
| MT072864.1;QIQ54047.1;Pangolin_CoV_orf1ab;4993 | 0 | 0.025 |
| MT072864.1;QIQ54047.1;Pangolin_CoV_orf1ab;4994 | 0 | 0.025 |
| MT072864.1;QIQ54047.1;Pangolin_CoV_orf1ab;4995 | 0 | 0.025 |
| MT072864.1;QIQ54047.1;Pangolin_CoV_orf1ab;4996 | 0 | 0.025 |
| MT072864.1;QIQ54047.1;Pangolin_CoV_orf1ab;4997 | 0 | 0.025 |
| MT072864.1;QIQ54047.1;Pangolin_CoV_orf1ab;4998 | 0 | 0.025 |
| MT072864.1;QIQ54047.1;Pangolin_CoV_orf1ab;4999 | 0 | 0.025 |
| MT072864.1;QIQ54047.1;Pangolin_CoV_orf1ab;5000 | 0 | 0.025 |
| MT072864.1;QIQ54047.1;Pangolin_CoV_orf1ab;5001 | 0 | 0 |
| MT072864.1;QIQ54047.1;Pangolin_CoV_orf1ab;5002 | 0 | 0 |
| MT072864.1;QIQ54047.1;Pangolin_CoV_orf1ab;5003 | 0 | 0 |
| MT072864.1;QIQ54047.1;Pangolin_CoV_orf1ab;5004 | 0 | 0 |
| MT072864.1;QIQ54047.1;Pangolin_CoV_orf1ab;5005 | 0 | 0 |
| MT072864.1;QIQ54047.1;Pangolin_CoV_orf1ab;5006 | 0 | 0 |
| MT072864.1;QIQ54047.1;Pangolin_CoV_orf1ab;5007 | 0 | 0 |
| MT072864.1;QIQ54047.1;Pangolin_CoV_orf1ab;5008 | 0 | 0 |
| MT072864.1;QIQ54047.1;Pangolin_CoV_orf1ab;5009 | 0 | 0 |
| MT072864.1;QIQ54047.1;Pangolin_CoV_orf1ab;5010 | 0 | 0 |
| MT072864.1;QIQ54047.1;Pangolin_CoV_orf1ab;5011 | 0 | 0 |
| MT072864.1;QIQ54047.1;Pangolin_CoV_orf1ab;5012 | 0 | 0 |
| MT072864.1;QIQ54047.1;Pangolin_CoV_orf1ab;5013 | 0 | 0 |
| MT072864.1;QIQ54047.1;Pangolin_CoV_orf1ab;5014 | 0 | 0 |
| MT072864.1;QIQ54047.1;Pangolin_CoV_orf1ab;5015 | 0 | 0 |
| MT072864.1;QIQ54047.1;Pangolin_CoV_orf1ab;5016 | 0 | 0 |
| MT072864.1;QIQ54047.1;Pangolin_CoV_orf1ab;5017 | 0 | 0 |
| MT072864.1;QIQ54047.1;Pangolin_CoV_orf1ab;5018 | 0 | 0 |
| MT072864.1;QIQ54047.1;Pangolin_CoV_orf1ab;5019 | 0 | 0 |
| MT072864.1;QIQ54047.1;Pangolin_CoV_orf1ab;5020 | 0 | 0 |
| MT072864.1;QIQ54047.1;Pangolin_CoV_orf1ab;5021 | 0 | 0 |

| | | |
|---|---|---|
| MT072864.1;QIQ54047.1;Pangolin_CoV_orf1ab;5022 | 0 | 0 |
| MT072864.1;QIQ54047.1;Pangolin_CoV_orf1ab;5023 | 0 | 0 |
| MT072864.1;QIQ54047.1;Pangolin_CoV_orf1ab;5024 | 0 | 0 |
| MT072864.1;QIQ54047.1;Pangolin_CoV_orf1ab;5025 | 0 | 0 |
| MT072864.1;QIQ54047.1;Pangolin_CoV_orf1ab;5026 | 0 | 0 |
| MT072864.1;QIQ54047.1;Pangolin_CoV_orf1ab;5027 | 0 | 0 |
| MT072864.1;QIQ54047.1;Pangolin_CoV_orf1ab;5028 | 0 | 0 |
| MT072864.1;QIQ54047.1;Pangolin_CoV_orf1ab;5029 | 0 | 0 |
| MT072864.1;QIQ54047.1;Pangolin_CoV_orf1ab;5030 | 0 | 0 |
| MT072864.1;QIQ54047.1;Pangolin_CoV_orf1ab;5031 | 0 | 0 |
| MT072864.1;QIQ54047.1;Pangolin_CoV_orf1ab;5032 | 0 | 0 |
| MT072864.1;QIQ54047.1;Pangolin_CoV_orf1ab;5033 | 0 | 0 |
| MT072864.1;QIQ54047.1;Pangolin_CoV_orf1ab;5034 | 0 | 0 |
| MT072864.1;QIQ54047.1;Pangolin_CoV_orf1ab;5035 | 0 | 0 |
| MT072864.1;QIQ54047.1;Pangolin_CoV_orf1ab;5036 | 0 | 0 |
| MT072864.1;QIQ54047.1;Pangolin_CoV_orf1ab;5037 | 0 | 0 |
| MT072864.1;QIQ54047.1;Pangolin_CoV_orf1ab;5038 | 0 | 0 |
| MT072864.1;QIQ54047.1;Pangolin_CoV_orf1ab;5039 | 0 | 0 |
| MT072864.1;QIQ54047.1;Pangolin_CoV_orf1ab;5040 | 0 | 0 |
| MT072864.1;QIQ54047.1;Pangolin_CoV_orf1ab;5041 | 0 | 0 |
| MT072864.1;QIQ54047.1;Pangolin_CoV_orf1ab;5042 | 0 | 0 |
| MT072864.1;QIQ54047.1;Pangolin_CoV_orf1ab;5043 | 0 | 0 |
| MT072864.1;QIQ54047.1;Pangolin_CoV_orf1ab;5044 | 0 | 0 |
| MT072864.1;QIQ54047.1;Pangolin_CoV_orf1ab;5045 | 0 | 0 |
| MT072864.1;QIQ54047.1;Pangolin_CoV_orf1ab;5046 | 0 | 0 |
| MT072864.1;QIQ54047.1;Pangolin_CoV_orf1ab;5047 | 0 | 0 |
| MT072864.1;QIQ54047.1;Pangolin_CoV_orf1ab;5048 | 0 | 0 |
| MT072864.1;QIQ54047.1;Pangolin_CoV_orf1ab;5049 | 0.05 | 0 |
| MT072864.1;QIQ54047.1;Pangolin_CoV_orf1ab;5050 | 0 | 0 |
| MT072864.1;QIQ54047.1;Pangolin_CoV_orf1ab;5051 | 0 | 0 |
| MT072864.1;QIQ54047.1;Pangolin_CoV_orf1ab;5052 | 0 | 0 |
| MT072864.1;QIQ54047.1;Pangolin_CoV_orf1ab;5053 | 0 | 0.025 |
| MT072864.1;QIQ54047.1;Pangolin_CoV_orf1ab;5054 | 0 | 0.025 |
| MT072864.1;QIQ54047.1;Pangolin_CoV_orf1ab;5055 | 0 | 0.025 |
| MT072864.1;QIQ54047.1;Pangolin_CoV_orf1ab;5056 | 0 | 0.025 |
| MT072864.1;QIQ54047.1;Pangolin_CoV_orf1ab;5057 | 0 | 0.05 |
| MT072864.1;QIQ54047.1;Pangolin_CoV_orf1ab;5058 | 0 | 0.025 |
| MT072864.1;QIQ54047.1;Pangolin_CoV_orf1ab;5059 | 0 | 0.025 |
| MT072864.1;QIQ54047.1;Pangolin_CoV_orf1ab;5060 | 0 | 0.05 |
| MT072864.1;QIQ54047.1;Pangolin_CoV_orf1ab;5061 | 0.05 | 0.025 |
| MT072864.1;QIQ54047.1;Pangolin_CoV_orf1ab;5062 | 0.1 | 0.05 |
| MT072864.1;QIQ54047.1;Pangolin_CoV_orf1ab;5063 | 0 | 0 |
| MT072864.1;QIQ54047.1;Pangolin_CoV_orf1ab;5064 | 0.05 | 0 |

| | | |
|---|---|---|
| MT072864.1;QIQ54047.1;Pangolin_CoV_orf1ab;5065 | 0 | 0 |
| MT072864.1;QIQ54047.1;Pangolin_CoV_orf1ab;5066 | 0 | 0 |
| MT072864.1;QIQ54047.1;Pangolin_CoV_orf1ab;5067 | 0 | 0 |
| MT072864.1;QIQ54047.1;Pangolin_CoV_orf1ab;5068 | 0 | 0 |
| MT072864.1;QIQ54047.1;Pangolin_CoV_orf1ab;5069 | 0 | 0 |
| MT072864.1;QIQ54047.1;Pangolin_CoV_orf1ab;5070 | 0 | 0 |
| MT072864.1;QIQ54047.1;Pangolin_CoV_orf1ab;5071 | 0 | 0 |
| MT072864.1;QIQ54047.1;Pangolin_CoV_orf1ab;5072 | 0 | 0.025 |
| MT072864.1;QIQ54047.1;Pangolin_CoV_orf1ab;5073 | 0 | 0 |
| MT072864.1;QIQ54047.1;Pangolin_CoV_orf1ab;5074 | 0 | 0.025 |
| MT072864.1;QIQ54047.1;Pangolin_CoV_orf1ab;5075 | 0 | 0 |
| MT072864.1;QIQ54047.1;Pangolin_CoV_orf1ab;5076 | 0 | 0 |
| MT072864.1;QIQ54047.1;Pangolin_CoV_orf1ab;5077 | 0 | 0 |
| MT072864.1;QIQ54047.1;Pangolin_CoV_orf1ab;5078 | 0 | 0 |
| MT072864.1;QIQ54047.1;Pangolin_CoV_orf1ab;5079 | 0 | 0 |
| MT072864.1;QIQ54047.1;Pangolin_CoV_orf1ab;5080 | 0 | 0 |
| MT072864.1;QIQ54047.1;Pangolin_CoV_orf1ab;5081 | 0 | 0.025 |
| MT072864.1;QIQ54047.1;Pangolin_CoV_orf1ab;5082 | 0 | 0 |
| MT072864.1;QIQ54047.1;Pangolin_CoV_orf1ab;5083 | 0 | 0.025 |
| MT072864.1;QIQ54047.1;Pangolin_CoV_orf1ab;5084 | 0 | 0.025 |
| MT072864.1;QIQ54047.1;Pangolin_CoV_orf1ab;5085 | 0 | 0.025 |
| MT072864.1;QIQ54047.1;Pangolin_CoV_orf1ab;5086 | 0 | 0.05 |
| MT072864.1;QIQ54047.1;Pangolin_CoV_orf1ab;5087 | 0 | 0 |
| MT072864.1;QIQ54047.1;Pangolin_CoV_orf1ab;5088 | 0.1 | 0.025 |
| MT072864.1;QIQ54047.1;Pangolin_CoV_orf1ab;5089 | 0.1 | 0.025 |
| MT072864.1;QIQ54047.1;Pangolin_CoV_orf1ab;5090 | 0 | 0.025 |
| MT072864.1;QIQ54047.1;Pangolin_CoV_orf1ab;5091 | 0.05 | 0.05 |
| MT072864.1;QIQ54047.1;Pangolin_CoV_orf1ab;5092 | 0.05 | 0.05 |
| MT072864.1;QIQ54047.1;Pangolin_CoV_orf1ab;5093 | 0.1 | 0.075 |
| MT072864.1;QIQ54047.1;Pangolin_CoV_orf1ab;5094 | 0 | 0 |
| MT072864.1;QIQ54047.1;Pangolin_CoV_orf1ab;5095 | 0 | 0 |
| MT072864.1;QIQ54047.1;Pangolin_CoV_orf1ab;5096 | 0.05 | 0 |
| MT072864.1;QIQ54047.1;Pangolin_CoV_orf1ab;5097 | 0 | 0 |
| MT072864.1;QIQ54047.1;Pangolin_CoV_orf1ab;5098 | 0 | 0 |
| MT072864.1;QIQ54047.1;Pangolin_CoV_orf1ab;5099 | 0 | 0 |
| MT072864.1;QIQ54047.1;Pangolin_CoV_orf1ab;5100 | 0 | 0 |
| MT072864.1;QIQ54047.1;Pangolin_CoV_orf1ab;5101 | 0 | 0 |
| MT072864.1;QIQ54047.1;Pangolin_CoV_orf1ab;5102 | 0 | 0 |
| MT072864.1;QIQ54047.1;Pangolin_CoV_orf1ab;5103 | 0 | 0 |
| MT072864.1;QIQ54047.1;Pangolin_CoV_orf1ab;5104 | 0 | 0 |
| MT072864.1;QIQ54047.1;Pangolin_CoV_orf1ab;5105 | 0.05 | 0 |
| MT072864.1;QIQ54047.1;Pangolin_CoV_orf1ab;5106 | 0 | 0 |
| MT072864.1;QIQ54047.1;Pangolin_CoV_orf1ab;5107 | 0 | 0 |

| | | |
|---|---|---|
| MT072864.1;QIQ54047.1;Pangolin_CoV_orf1ab;5108 | 0 | 0 |
| MT072864.1;QIQ54047.1;Pangolin_CoV_orf1ab;5109 | 0 | 0 |
| MT072864.1;QIQ54047.1;Pangolin_CoV_orf1ab;5110 | 0 | 0 |
| MT072864.1;QIQ54047.1;Pangolin_CoV_orf1ab;5111 | 0 | 0 |
| MT072864.1;QIQ54047.1;Pangolin_CoV_orf1ab;5112 | 0 | 0 |
| MT072864.1;QIQ54047.1;Pangolin_CoV_orf1ab;5113 | 0 | 0 |
| MT072864.1;QIQ54047.1;Pangolin_CoV_orf1ab;5114 | 0 | 0 |
| MT072864.1;QIQ54047.1;Pangolin_CoV_orf1ab;5115 | 0 | 0.05 |
| MT072864.1;QIQ54047.1;Pangolin_CoV_orf1ab;5116 | 0.05 | 0.05 |
| MT072864.1;QIQ54047.1;Pangolin_CoV_orf1ab;5117 | 0 | 0.05 |
| MT072864.1;QIQ54047.1;Pangolin_CoV_orf1ab;5118 | 0 | 0 |
| MT072864.1;QIQ54047.1;Pangolin_CoV_orf1ab;5119 | 0 | 0.025 |
| MT072864.1;QIQ54047.1;Pangolin_CoV_orf1ab;5120 | 0 | 0 |
| MT072864.1;QIQ54047.1;Pangolin_CoV_orf1ab;5121 | 0 | 0.025 |
| MT072864.1;QIQ54047.1;Pangolin_CoV_orf1ab;5122 | 0 | 0 |
| MT072864.1;QIQ54047.1;Pangolin_CoV_orf1ab;5123 | 0.05 | 0.025 |
| MT072864.1;QIQ54047.1;Pangolin_CoV_orf1ab;5124 | 0 | 0 |
| MT072864.1;QIQ54047.1;Pangolin_CoV_orf1ab;5125 | 0.05 | 0.025 |
| MT072864.1;QIQ54047.1;Pangolin_CoV_orf1ab;5126 | 0 | 0 |
| MT072864.1;QIQ54047.1;Pangolin_CoV_orf1ab;5127 | 0 | 0 |
| MT072864.1;QIQ54047.1;Pangolin_CoV_orf1ab;5128 | 0 | 0 |
| MT072864.1;QIQ54047.1;Pangolin_CoV_orf1ab;5129 | 0 | 0 |
| MT072864.1;QIQ54047.1;Pangolin_CoV_orf1ab;5130 | 0 | 0 |
| MT072864.1;QIQ54047.1;Pangolin_CoV_orf1ab;5131 | 0.05 | 0 |
| MT072864.1;QIQ54047.1;Pangolin_CoV_orf1ab;5132 | 0 | 0 |
| MT072864.1;QIQ54047.1;Pangolin_CoV_orf1ab;5133 | 0 | 0 |
| MT072864.1;QIQ54047.1;Pangolin_CoV_orf1ab;5134 | 0 | 0 |
| MT072864.1;QIQ54047.1;Pangolin_CoV_orf1ab;5135 | 0 | 0 |
| MT072864.1;QIQ54047.1;Pangolin_CoV_orf1ab;5136 | 0 | 0 |
| MT072864.1;QIQ54047.1;Pangolin_CoV_orf1ab;5137 | 0 | 0 |
| MT072864.1;QIQ54047.1;Pangolin_CoV_orf1ab;5138 | 0 | 0 |
| MT072864.1;QIQ54047.1;Pangolin_CoV_orf1ab;5139 | 0 | 0 |
| MT072864.1;QIQ54047.1;Pangolin_CoV_orf1ab;5140 | 0 | 0 |
| MT072864.1;QIQ54047.1;Pangolin_CoV_orf1ab;5141 | 0 | 0 |
| MT072864.1;QIQ54047.1;Pangolin_CoV_orf1ab;5142 | 0 | 0 |
| MT072864.1;QIQ54047.1;Pangolin_CoV_orf1ab;5143 | 0 | 0 |
| MT072864.1;QIQ54047.1;Pangolin_CoV_orf1ab;5144 | 0 | 0 |
| MT072864.1;QIQ54047.1;Pangolin_CoV_orf1ab;5145 | 0 | 0 |
| MT072864.1;QIQ54047.1;Pangolin_CoV_orf1ab;5146 | 0.05 | 0.05 |
| MT072864.1;QIQ54047.1;Pangolin_CoV_orf1ab;5147 | 0 | 0 |
| MT072864.1;QIQ54047.1;Pangolin_CoV_orf1ab;5148 | 0 | 0 |
| MT072864.1;QIQ54047.1;Pangolin_CoV_orf1ab;5149 | 0 | 0 |
| MT072864.1;QIQ54047.1;Pangolin_CoV_orf1ab;5150 | 0 | 0 |

| | | |
|---|---|---|
| MT072864.1;QIQ54047.1;Pangolin_CoV_orf1ab;5151 | 0 | 0 |
| MT072864.1;QIQ54047.1;Pangolin_CoV_orf1ab;5152 | 0 | 0 |
| MT072864.1;QIQ54047.1;Pangolin_CoV_orf1ab;5153 | 0 | 0 |
| MT072864.1;QIQ54047.1;Pangolin_CoV_orf1ab;5154 | 0 | 0 |
| MT072864.1;QIQ54047.1;Pangolin_CoV_orf1ab;5155 | 0 | 0 |
| MT072864.1;QIQ54047.1;Pangolin_CoV_orf1ab;5156 | 0 | 0 |
| MT072864.1;QIQ54047.1;Pangolin_CoV_orf1ab;5157 | 0 | 0 |
| MT072864.1;QIQ54047.1;Pangolin_CoV_orf1ab;5158 | 0 | 0 |
| MT072864.1;QIQ54047.1;Pangolin_CoV_orf1ab;5159 | 0 | 0 |
| MT072864.1;QIQ54047.1;Pangolin_CoV_orf1ab;5160 | 0 | 0 |
| MT072864.1;QIQ54047.1;Pangolin_CoV_orf1ab;5161 | 0 | 0 |
| MT072864.1;QIQ54047.1;Pangolin_CoV_orf1ab;5162 | 0 | 0 |
| MT072864.1;QIQ54047.1;Pangolin_CoV_orf1ab;5163 | 0 | 0 |
| MT072864.1;QIQ54047.1;Pangolin_CoV_orf1ab;5164 | 0 | 0 |
| MT072864.1;QIQ54047.1;Pangolin_CoV_orf1ab;5165 | 0 | 0 |
| MT072864.1;QIQ54047.1;Pangolin_CoV_orf1ab;5166 | 0 | 0 |
| MT072864.1;QIQ54047.1;Pangolin_CoV_orf1ab;5167 | 0.05 | 0 |
| MT072864.1;QIQ54047.1;Pangolin_CoV_orf1ab;5168 | 0 | 0 |
| MT072864.1;QIQ54047.1;Pangolin_CoV_orf1ab;5169 | 0 | 0 |
| MT072864.1;QIQ54047.1;Pangolin_CoV_orf1ab;5170 | 0 | 0 |
| MT072864.1;QIQ54047.1;Pangolin_CoV_orf1ab;5171 | 0 | 0.025 |
| MT072864.1;QIQ54047.1;Pangolin_CoV_orf1ab;5172 | 0 | 0.025 |
| MT072864.1;QIQ54047.1;Pangolin_CoV_orf1ab;5173 | 0 | 0.025 |
| MT072864.1;QIQ54047.1;Pangolin_CoV_orf1ab;5174 | 0 | 0.025 |
| MT072864.1;QIQ54047.1;Pangolin_CoV_orf1ab;5175 | 0 | 0.025 |
| MT072864.1;QIQ54047.1;Pangolin_CoV_orf1ab;5176 | 0.05 | 0.025 |
| MT072864.1;QIQ54047.1;Pangolin_CoV_orf1ab;5177 | 0 | 0.025 |
| MT072864.1;QIQ54047.1;Pangolin_CoV_orf1ab;5178 | 0.05 | 0.025 |
| MT072864.1;QIQ54047.1;Pangolin_CoV_orf1ab;5179 | 0 | 0.025 |
| MT072864.1;QIQ54047.1;Pangolin_CoV_orf1ab;5180 | 0 | 0.025 |
| MT072864.1;QIQ54047.1;Pangolin_CoV_orf1ab;5181 | 0.05 | 0.025 |
| MT072864.1;QIQ54047.1;Pangolin_CoV_orf1ab;5182 | 0.05 | 0.025 |
| MT072864.1;QIQ54047.1;Pangolin_CoV_orf1ab;5183 | 0.05 | 0.025 |
| MT072864.1;QIQ54047.1;Pangolin_CoV_orf1ab;5184 | 0.05 | 0.025 |
| MT072864.1;QIQ54047.1;Pangolin_CoV_orf1ab;5185 | 0 | 0 |
| MT072864.1;QIQ54047.1;Pangolin_CoV_orf1ab;5186 | 0 | 0 |
| MT072864.1;QIQ54047.1;Pangolin_CoV_orf1ab;5187 | 0 | 0 |
| MT072864.1;QIQ54047.1;Pangolin_CoV_orf1ab;5188 | 0 | 0 |
| MT072864.1;QIQ54047.1;Pangolin_CoV_orf1ab;5189 | 0 | 0 |
| MT072864.1;QIQ54047.1;Pangolin_CoV_orf1ab;5190 | 0 | 0 |
| MT072864.1;QIQ54047.1;Pangolin_CoV_orf1ab;5191 | 0 | 0 |
| MT072864.1;QIQ54047.1;Pangolin_CoV_orf1ab;5192 | 0 | 0 |
| MT072864.1;QIQ54047.1;Pangolin_CoV_orf1ab;5193 | 0 | 0 |

| | | |
|---|---|---|
| MT072864.1;QIQ54047.1;Pangolin_CoV_orf1ab;5194 | 0 | 0 |
| MT072864.1;QIQ54047.1;Pangolin_CoV_orf1ab;5195 | 0 | 0.025 |
| MT072864.1;QIQ54047.1;Pangolin_CoV_orf1ab;5196 | 0 | 0.025 |
| MT072864.1;QIQ54047.1;Pangolin_CoV_orf1ab;5197 | 0 | 0.025 |
| MT072864.1;QIQ54047.1;Pangolin_CoV_orf1ab;5198 | 0.05 | 0 |
| MT072864.1;QIQ54047.1;Pangolin_CoV_orf1ab;5199 | 0.05 | 0 |
| MT072864.1;QIQ54047.1;Pangolin_CoV_orf1ab;5200 | 0 | 0 |
| MT072864.1;QIQ54047.1;Pangolin_CoV_orf1ab;5201 | 0.05 | 0 |
| MT072864.1;QIQ54047.1;Pangolin_CoV_orf1ab;5202 | 0 | 0 |
| MT072864.1;QIQ54047.1;Pangolin_CoV_orf1ab;5203 | 0.1 | 0.1 |
| MT072864.1;QIQ54047.1;Pangolin_CoV_orf1ab;5204 | 0 | 0.025 |
| MT072864.1;QIQ54047.1;Pangolin_CoV_orf1ab;5205 | 0 | 0.025 |
| MT072864.1;QIQ54047.1;Pangolin_CoV_orf1ab;5206 | 0 | 0 |
| MT072864.1;QIQ54047.1;Pangolin_CoV_orf1ab;5207 | 0 | 0 |
| MT072864.1;QIQ54047.1;Pangolin_CoV_orf1ab;5208 | 0 | 0 |
| MT072864.1;QIQ54047.1;Pangolin_CoV_orf1ab;5209 | 0 | 0 |
| MT072864.1;QIQ54047.1;Pangolin_CoV_orf1ab;5210 | 0 | 0 |
| MT072864.1;QIQ54047.1;Pangolin_CoV_orf1ab;5211 | 0 | 0 |
| MT072864.1;QIQ54047.1;Pangolin_CoV_orf1ab;5212 | 0 | 0 |
| MT072864.1;QIQ54047.1;Pangolin_CoV_orf1ab;5213 | 0 | 0 |
| MT072864.1;QIQ54047.1;Pangolin_CoV_orf1ab;5214 | 0 | 0 |
| MT072864.1;QIQ54047.1;Pangolin_CoV_orf1ab;5215 | 0 | 0 |
| MT072864.1;QIQ54047.1;Pangolin_CoV_orf1ab;5216 | 0 | 0 |
| MT072864.1;QIQ54047.1;Pangolin_CoV_orf1ab;5217 | 0 | 0 |
| MT072864.1;QIQ54047.1;Pangolin_CoV_orf1ab;5218 | 0 | 0 |
| MT072864.1;QIQ54047.1;Pangolin_CoV_orf1ab;5219 | 0 | 0 |
| MT072864.1;QIQ54047.1;Pangolin_CoV_orf1ab;5220 | 0 | 0 |
| MT072864.1;QIQ54047.1;Pangolin_CoV_orf1ab;5221 | 0.05 | 0 |
| MT072864.1;QIQ54047.1;Pangolin_CoV_orf1ab;5222 | 0 | 0 |
| MT072864.1;QIQ54047.1;Pangolin_CoV_orf1ab;5223 | 0 | 0 |
| MT072864.1;QIQ54047.1;Pangolin_CoV_orf1ab;5224 | 0 | 0 |
| MT072864.1;QIQ54047.1;Pangolin_CoV_orf1ab;5225 | 0 | 0 |
| MT072864.1;QIQ54047.1;Pangolin_CoV_orf1ab;5226 | 0 | 0 |
| MT072864.1;QIQ54047.1;Pangolin_CoV_orf1ab;5227 | 0 | 0 |
| MT072864.1;QIQ54047.1;Pangolin_CoV_orf1ab;5228 | 0 | 0 |
| MT072864.1;QIQ54047.1;Pangolin_CoV_orf1ab;5229 | 0 | 0 |
| MT072864.1;QIQ54047.1;Pangolin_CoV_orf1ab;5230 | 0 | 0 |
| MT072864.1;QIQ54047.1;Pangolin_CoV_orf1ab;5231 | 0 | 0 |
| MT072864.1;QIQ54047.1;Pangolin_CoV_orf1ab;5232 | 0 | 0 |
| MT072864.1;QIQ54047.1;Pangolin_CoV_orf1ab;5233 | 0 | 0 |
| MT072864.1;QIQ54047.1;Pangolin_CoV_orf1ab;5234 | 0 | 0 |
| MT072864.1;QIQ54047.1;Pangolin_CoV_orf1ab;5235 | 0 | 0 |
| MT072864.1;QIQ54047.1;Pangolin_CoV_orf1ab;5236 | 0 | 0 |

| | | |
|---|---|---|
| MT072864.1;QIQ54047.1;Pangolin_CoV_orf1ab;5237 | 0 | 0 |
| MT072864.1;QIQ54047.1;Pangolin_CoV_orf1ab;5238 | 0 | 0 |
| MT072864.1;QIQ54047.1;Pangolin_CoV_orf1ab;5239 | 0 | 0 |
| MT072864.1;QIQ54047.1;Pangolin_CoV_orf1ab;5240 | 0 | 0 |
| MT072864.1;QIQ54047.1;Pangolin_CoV_orf1ab;5241 | 0 | 0 |
| MT072864.1;QIQ54047.1;Pangolin_CoV_orf1ab;5242 | 0 | 0 |
| MT072864.1;QIQ54047.1;Pangolin_CoV_orf1ab;5243 | 0 | 0 |
| MT072864.1;QIQ54047.1;Pangolin_CoV_orf1ab;5244 | 0 | 0 |
| MT072864.1;QIQ54047.1;Pangolin_CoV_orf1ab;5245 | 0 | 0 |
| MT072864.1;QIQ54047.1;Pangolin_CoV_orf1ab;5246 | 0 | 0 |
| MT072864.1;QIQ54047.1;Pangolin_CoV_orf1ab;5247 | 0 | 0 |
| MT072864.1;QIQ54047.1;Pangolin_CoV_orf1ab;5248 | 0 | 0 |
| MT072864.1;QIQ54047.1;Pangolin_CoV_orf1ab;5249 | 0 | 0 |
| MT072864.1;QIQ54047.1;Pangolin_CoV_orf1ab;5250 | 0 | 0 |
| MT072864.1;QIQ54047.1;Pangolin_CoV_orf1ab;5251 | 0 | 0 |
| MT072864.1;QIQ54047.1;Pangolin_CoV_orf1ab;5252 | 0 | 0 |
| MT072864.1;QIQ54047.1;Pangolin_CoV_orf1ab;5253 | 0 | 0 |
| MT072864.1;QIQ54047.1;Pangolin_CoV_orf1ab;5254 | 0 | 0 |
| MT072864.1;QIQ54047.1;Pangolin_CoV_orf1ab;5255 | 0 | 0 |
| MT072864.1;QIQ54047.1;Pangolin_CoV_orf1ab;5256 | 0 | 0 |
| MT072864.1;QIQ54047.1;Pangolin_CoV_orf1ab;5257 | 0 | 0 |
| MT072864.1;QIQ54047.1;Pangolin_CoV_orf1ab;5258 | 0 | 0 |
| MT072864.1;QIQ54047.1;Pangolin_CoV_orf1ab;5259 | 0 | 0 |
| MT072864.1;QIQ54047.1;Pangolin_CoV_orf1ab;5260 | 0 | 0 |
| MT072864.1;QIQ54047.1;Pangolin_CoV_orf1ab;5261 | 0 | 0 |
| MT072864.1;QIQ54047.1;Pangolin_CoV_orf1ab;5262 | 0 | 0 |
| MT072864.1;QIQ54047.1;Pangolin_CoV_orf1ab;5263 | 0 | 0 |
| MT072864.1;QIQ54047.1;Pangolin_CoV_orf1ab;5264 | 0 | 0 |
| MT072864.1;QIQ54047.1;Pangolin_CoV_orf1ab;5265 | 0 | 0 |
| MT072864.1;QIQ54047.1;Pangolin_CoV_orf1ab;5266 | 0 | 0 |
| MT072864.1;QIQ54047.1;Pangolin_CoV_orf1ab;5267 | 0 | 0 |
| MT072864.1;QIQ54047.1;Pangolin_CoV_orf1ab;5268 | 0 | 0 |
| MT072864.1;QIQ54047.1;Pangolin_CoV_orf1ab;5269 | 0 | 0 |
| MT072864.1;QIQ54047.1;Pangolin_CoV_orf1ab;5270 | 0 | 0 |
| MT072864.1;QIQ54047.1;Pangolin_CoV_orf1ab;5271 | 0 | 0 |
| MT072864.1;QIQ54047.1;Pangolin_CoV_orf1ab;5272 | 0 | 0 |
| MT072864.1;QIQ54047.1;Pangolin_CoV_orf1ab;5273 | 0 | 0.075 |
| MT072864.1;QIQ54047.1;Pangolin_CoV_orf1ab;5274 | 0 | 0.025 |
| MT072864.1;QIQ54047.1;Pangolin_CoV_orf1ab;5275 | 0 | 0 |
| MT072864.1;QIQ54047.1;Pangolin_CoV_orf1ab;5276 | 0 | 0 |
| MT072864.1;QIQ54047.1;Pangolin_CoV_orf1ab;5277 | 0 | 0 |
| MT072864.1;QIQ54047.1;Pangolin_CoV_orf1ab;5278 | 0 | 0 |
| MT072864.1;QIQ54047.1;Pangolin_CoV_orf1ab;5279 | 0 | 0 |

| | | |
|---|---|---|
| MT072864.1;QIQ54047.1;Pangolin_CoV_orf1ab;5280 | 0 | 0 |
| MT072864.1;QIQ54047.1;Pangolin_CoV_orf1ab;5281 | 0 | 0 |
| MT072864.1;QIQ54047.1;Pangolin_CoV_orf1ab;5282 | 0 | 0 |
| MT072864.1;QIQ54047.1;Pangolin_CoV_orf1ab;5283 | 0 | 0 |
| MT072864.1;QIQ54047.1;Pangolin_CoV_orf1ab;5284 | 0 | 0 |
| MT072864.1;QIQ54047.1;Pangolin_CoV_orf1ab;5285 | 0 | 0 |
| MT072864.1;QIQ54047.1;Pangolin_CoV_orf1ab;5286 | 0 | 0 |
| MT072864.1;QIQ54047.1;Pangolin_CoV_orf1ab;5287 | 0 | 0 |
| MT072864.1;QIQ54047.1;Pangolin_CoV_orf1ab;5288 | 0 | 0 |
| MT072864.1;QIQ54047.1;Pangolin_CoV_orf1ab;5289 | 0 | 0 |
| MT072864.1;QIQ54047.1;Pangolin_CoV_orf1ab;5290 | 0 | 0 |
| MT072864.1;QIQ54047.1;Pangolin_CoV_orf1ab;5291 | 0 | 0 |
| MT072864.1;QIQ54047.1;Pangolin_CoV_orf1ab;5292 | 0 | 0 |
| MT072864.1;QIQ54047.1;Pangolin_CoV_orf1ab;5293 | 0 | 0 |
| MT072864.1;QIQ54047.1;Pangolin_CoV_orf1ab;5294 | 0.05 | 0 |
| MT072864.1;QIQ54047.1;Pangolin_CoV_orf1ab;5295 | 0 | 0 |
| MT072864.1;QIQ54047.1;Pangolin_CoV_orf1ab;5296 | 0 | 0 |
| MT072864.1;QIQ54047.1;Pangolin_CoV_orf1ab;5297 | 0 | 0 |
| MT072864.1;QIQ54047.1;Pangolin_CoV_orf1ab;5298 | 0 | 0 |
| MT072864.1;QIQ54047.1;Pangolin_CoV_orf1ab;5299 | 0 | 0 |
| MT072864.1;QIQ54047.1;Pangolin_CoV_orf1ab;5300 | 0 | 0 |
| MT072864.1;QIQ54047.1;Pangolin_CoV_orf1ab;5301 | 0 | 0 |
| MT072864.1;QIQ54047.1;Pangolin_CoV_orf1ab;5302 | 0 | 0 |
| MT072864.1;QIQ54047.1;Pangolin_CoV_orf1ab;5303 | 0 | 0 |
| MT072864.1;QIQ54047.1;Pangolin_CoV_orf1ab;5304 | 0 | 0 |
| MT072864.1;QIQ54047.1;Pangolin_CoV_orf1ab;5305 | 0 | 0 |
| MT072864.1;QIQ54047.1;Pangolin_CoV_orf1ab;5306 | 0 | 0 |
| MT072864.1;QIQ54047.1;Pangolin_CoV_orf1ab;5307 | 0 | 0 |
| MT072864.1;QIQ54047.1;Pangolin_CoV_orf1ab;5308 | 0 | 0 |
| MT072864.1;QIQ54047.1;Pangolin_CoV_orf1ab;5309 | 0 | 0 |
| MT072864.1;QIQ54047.1;Pangolin_CoV_orf1ab;5310 | 0 | 0 |
| MT072864.1;QIQ54047.1;Pangolin_CoV_orf1ab;5311 | 0 | 0 |
| MT072864.1;QIQ54047.1;Pangolin_CoV_orf1ab;5312 | 0 | 0 |
| MT072864.1;QIQ54047.1;Pangolin_CoV_orf1ab;5313 | 0 | 0 |
| MT072864.1;QIQ54047.1;Pangolin_CoV_orf1ab;5314 | 0 | 0 |
| MT072864.1;QIQ54047.1;Pangolin_CoV_orf1ab;5315 | 0 | 0 |
| MT072864.1;QIQ54047.1;Pangolin_CoV_orf1ab;5316 | 0 | 0 |
| MT072864.1;QIQ54047.1;Pangolin_CoV_orf1ab;5317 | 0 | 0 |
| MT072864.1;QIQ54047.1;Pangolin_CoV_orf1ab;5318 | 0 | 0 |
| MT072864.1;QIQ54047.1;Pangolin_CoV_orf1ab;5319 | 0 | 0 |
| MT072864.1;QIQ54047.1;Pangolin_CoV_orf1ab;5320 | 0 | 0 |
| MT072864.1;QIQ54047.1;Pangolin_CoV_orf1ab;5321 | 0 | 0 |
| MT072864.1;QIQ54047.1;Pangolin_CoV_orf1ab;5322 | 0 | 0 |

| | | |
|---|---|---|
| MT072864.1;QIQ54047.1;Pangolin_CoV_orf1ab;5323 | 0 | 0 |
| MT072864.1;QIQ54047.1;Pangolin_CoV_orf1ab;5324 | 0 | 0 |
| MT072864.1;QIQ54047.1;Pangolin_CoV_orf1ab;5325 | 0 | 0 |
| MT072864.1;QIQ54047.1;Pangolin_CoV_orf1ab;5326 | 0 | 0 |
| MT072864.1;QIQ54047.1;Pangolin_CoV_orf1ab;5327 | 0 | 0 |
| MT072864.1;QIQ54047.1;Pangolin_CoV_orf1ab;5328 | 0 | 0 |
| MT072864.1;QIQ54047.1;Pangolin_CoV_orf1ab;5329 | 0 | 0 |
| MT072864.1;QIQ54047.1;Pangolin_CoV_orf1ab;5330 | 0 | 0 |
| MT072864.1;QIQ54047.1;Pangolin_CoV_orf1ab;5331 | 0 | 0 |
| MT072864.1;QIQ54047.1;Pangolin_CoV_orf1ab;5332 | 0 | 0 |
| MT072864.1;QIQ54047.1;Pangolin_CoV_orf1ab;5333 | 0 | 0 |
| MT072864.1;QIQ54047.1;Pangolin_CoV_orf1ab;5334 | 0 | 0 |
| MT072864.1;QIQ54047.1;Pangolin_CoV_orf1ab;5335 | 0 | 0 |
| MT072864.1;QIQ54047.1;Pangolin_CoV_orf1ab;5336 | 0 | 0 |
| MT072864.1;QIQ54047.1;Pangolin_CoV_orf1ab;5337 | 0.05 | 0 |
| MT072864.1;QIQ54047.1;Pangolin_CoV_orf1ab;5338 | 0 | 0 |
| MT072864.1;QIQ54047.1;Pangolin_CoV_orf1ab;5339 | 0 | 0 |
| MT072864.1;QIQ54047.1;Pangolin_CoV_orf1ab;5340 | 0 | 0 |
| MT072864.1;QIQ54047.1;Pangolin_CoV_orf1ab;5341 | 0 | 0 |
| MT072864.1;QIQ54047.1;Pangolin_CoV_orf1ab;5342 | 0 | 0 |
| MT072864.1;QIQ54047.1;Pangolin_CoV_orf1ab;5343 | 0 | 0 |
| MT072864.1;QIQ54047.1;Pangolin_CoV_orf1ab;5344 | 0 | 0 |
| MT072864.1;QIQ54047.1;Pangolin_CoV_orf1ab;5345 | 0 | 0 |
| MT072864.1;QIQ54047.1;Pangolin_CoV_orf1ab;5346 | 0 | 0 |
| MT072864.1;QIQ54047.1;Pangolin_CoV_orf1ab;5347 | 0 | 0.025 |
| MT072864.1;QIQ54047.1;Pangolin_CoV_orf1ab;5348 | 0 | 0 |
| MT072864.1;QIQ54047.1;Pangolin_CoV_orf1ab;5349 | 0 | 0 |
| MT072864.1;QIQ54047.1;Pangolin_CoV_orf1ab;5350 | 0 | 0 |
| MT072864.1;QIQ54047.1;Pangolin_CoV_orf1ab;5351 | 0 | 0 |
| MT072864.1;QIQ54047.1;Pangolin_CoV_orf1ab;5352 | 0 | 0 |
| MT072864.1;QIQ54047.1;Pangolin_CoV_orf1ab;5353 | 0 | 0 |
| MT072864.1;QIQ54047.1;Pangolin_CoV_orf1ab;5354 | 0 | 0 |
| MT072864.1;QIQ54047.1;Pangolin_CoV_orf1ab;5355 | 0 | 0 |
| MT072864.1;QIQ54047.1;Pangolin_CoV_orf1ab;5356 | 0 | 0 |
| MT072864.1;QIQ54047.1;Pangolin_CoV_orf1ab;5357 | 0 | 0 |
| MT072864.1;QIQ54047.1;Pangolin_CoV_orf1ab;5358 | 0 | 0.075 |
| MT072864.1;QIQ54047.1;Pangolin_CoV_orf1ab;5359 | 0 | 0 |
| MT072864.1;QIQ54047.1;Pangolin_CoV_orf1ab;5360 | 0 | 0 |
| MT072864.1;QIQ54047.1;Pangolin_CoV_orf1ab;5361 | 0 | 0 |
| MT072864.1;QIQ54047.1;Pangolin_CoV_orf1ab;5362 | 0.05 | 0 |
| MT072864.1;QIQ54047.1;Pangolin_CoV_orf1ab;5363 | 0.05 | 0 |
| MT072864.1;QIQ54047.1;Pangolin_CoV_orf1ab;5364 | 0.05 | 0 |
| MT072864.1;QIQ54047.1;Pangolin_CoV_orf1ab;5365 | 0.05 | 0 |

| | | |
|---|---|---|
| MT072864.1;QIQ54047.1;Pangolin_CoV_orf1ab;5366 | 0 | 0 |
| MT072864.1;QIQ54047.1;Pangolin_CoV_orf1ab;5367 | 0 | 0.025 |
| MT072864.1;QIQ54047.1;Pangolin_CoV_orf1ab;5368 | 0 | 0 |
| MT072864.1;QIQ54047.1;Pangolin_CoV_orf1ab;5369 | 0.05 | 0 |
| MT072864.1;QIQ54047.1;Pangolin_CoV_orf1ab;5370 | 0.05 | 0 |
| MT072864.1;QIQ54047.1;Pangolin_CoV_orf1ab;5371 | 0 | 0 |
| MT072864.1;QIQ54047.1;Pangolin_CoV_orf1ab;5372 | 0 | 0 |
| MT072864.1;QIQ54047.1;Pangolin_CoV_orf1ab;5373 | 0 | 0 |
| MT072864.1;QIQ54047.1;Pangolin_CoV_orf1ab;5374 | 0 | 0 |
| MT072864.1;QIQ54047.1;Pangolin_CoV_orf1ab;5375 | 0 | 0 |
| MT072864.1;QIQ54047.1;Pangolin_CoV_orf1ab;5376 | 0 | 0 |
| MT072864.1;QIQ54047.1;Pangolin_CoV_orf1ab;5377 | 0 | 0 |
| MT072864.1;QIQ54047.1;Pangolin_CoV_orf1ab;5378 | 0 | 0 |
| MT072864.1;QIQ54047.1;Pangolin_CoV_orf1ab;5379 | 0 | 0 |
| MT072864.1;QIQ54047.1;Pangolin_CoV_orf1ab;5380 | 0 | 0 |
| MT072864.1;QIQ54047.1;Pangolin_CoV_orf1ab;5381 | 0 | 0 |
| MT072864.1;QIQ54047.1;Pangolin_CoV_orf1ab;5382 | 0 | 0 |
| MT072864.1;QIQ54047.1;Pangolin_CoV_orf1ab;5383 | 0 | 0 |
| MT072864.1;QIQ54047.1;Pangolin_CoV_orf1ab;5384 | 0 | 0 |
| MT072864.1;QIQ54047.1;Pangolin_CoV_orf1ab;5385 | 0 | 0 |
| MT072864.1;QIQ54047.1;Pangolin_CoV_orf1ab;5386 | 0 | 0 |
| MT072864.1;QIQ54047.1;Pangolin_CoV_orf1ab;5387 | 0 | 0 |
| MT072864.1;QIQ54047.1;Pangolin_CoV_orf1ab;5388 | 0 | 0 |
| MT072864.1;QIQ54047.1;Pangolin_CoV_orf1ab;5389 | 0 | 0 |
| MT072864.1;QIQ54047.1;Pangolin_CoV_orf1ab;5390 | 0 | 0 |
| MT072864.1;QIQ54047.1;Pangolin_CoV_orf1ab;5391 | 0 | 0 |
| MT072864.1;QIQ54047.1;Pangolin_CoV_orf1ab;5392 | 0 | 0 |
| MT072864.1;QIQ54047.1;Pangolin_CoV_orf1ab;5393 | 0 | 0.025 |
| MT072864.1;QIQ54047.1;Pangolin_CoV_orf1ab;5394 | 0 | 0 |
| MT072864.1;QIQ54047.1;Pangolin_CoV_orf1ab;5395 | 0 | 0 |
| MT072864.1;QIQ54047.1;Pangolin_CoV_orf1ab;5396 | 0 | 0 |
| MT072864.1;QIQ54047.1;Pangolin_CoV_orf1ab;5397 | 0 | 0 |
| MT072864.1;QIQ54047.1;Pangolin_CoV_orf1ab;5398 | 0 | 0 |
| MT072864.1;QIQ54047.1;Pangolin_CoV_orf1ab;5399 | 0 | 0 |
| MT072864.1;QIQ54047.1;Pangolin_CoV_orf1ab;5400 | 0 | 0 |
| MT072864.1;QIQ54047.1;Pangolin_CoV_orf1ab;5401 | 0 | 0 |
| MT072864.1;QIQ54047.1;Pangolin_CoV_orf1ab;5402 | 0 | 0 |
| MT072864.1;QIQ54047.1;Pangolin_CoV_orf1ab;5403 | 0 | 0 |
| MT072864.1;QIQ54047.1;Pangolin_CoV_orf1ab;5404 | 0 | 0 |
| MT072864.1;QIQ54047.1;Pangolin_CoV_orf1ab;5405 | 0.05 | 0 |
| MT072864.1;QIQ54047.1;Pangolin_CoV_orf1ab;5406 | 0.05 | 0 |
| MT072864.1;QIQ54047.1;Pangolin_CoV_orf1ab;5407 | 0.1 | 0 |
| MT072864.1;QIQ54047.1;Pangolin_CoV_orf1ab;5408 | 0.2 | 0.075 |

| | | |
|---|---|---|
| MT072864.1;QIQ54047.1;Pangolin_CoV_orf1ab;5409 | 0.15 | 0.05 |
| MT072864.1;QIQ54047.1;Pangolin_CoV_orf1ab;5410 | 0.15 | 0.125 |
| MT072864.1;QIQ54047.1;Pangolin_CoV_orf1ab;5411 | 0.15 | 0.15 |
| MT072864.1;QIQ54047.1;Pangolin_CoV_orf1ab;5412 | 0.05 | 0.1 |
| MT072864.1;QIQ54047.1;Pangolin_CoV_orf1ab;5413 | 0.05 | 0.075 |
| MT072864.1;QIQ54047.1;Pangolin_CoV_orf1ab;5414 | 0.05 | 0.2 |
| MT072864.1;QIQ54047.1;Pangolin_CoV_orf1ab;5415 | 0.05 | 0.075 |
| MT072864.1;QIQ54047.1;Pangolin_CoV_orf1ab;5416 | 0.05 | 0.025 |
| MT072864.1;QIQ54047.1;Pangolin_CoV_orf1ab;5417 | 0.05 | 0.05 |
| MT072864.1;QIQ54047.1;Pangolin_CoV_orf1ab;5418 | 0 | 0.025 |
| MT072864.1;QIQ54047.1;Pangolin_CoV_orf1ab;5419 | 0 | 0 |
| MT072864.1;QIQ54047.1;Pangolin_CoV_orf1ab;5420 | 0 | 0 |
| MT072864.1;QIQ54047.1;Pangolin_CoV_orf1ab;5421 | 0 | 0 |
| MT072864.1;QIQ54047.1;Pangolin_CoV_orf1ab;5422 | 0 | 0 |
| MT072864.1;QIQ54047.1;Pangolin_CoV_orf1ab;5423 | 0 | 0 |
| MT072864.1;QIQ54047.1;Pangolin_CoV_orf1ab;5424 | 0 | 0 |
| MT072864.1;QIQ54047.1;Pangolin_CoV_orf1ab;5425 | 0 | 0 |
| MT072864.1;QIQ54047.1;Pangolin_CoV_orf1ab;5426 | 0 | 0 |
| MT072864.1;QIQ54047.1;Pangolin_CoV_orf1ab;5427 | 0 | 0 |
| MT072864.1;QIQ54047.1;Pangolin_CoV_orf1ab;5428 | 0 | 0 |
| MT072864.1;QIQ54047.1;Pangolin_CoV_orf1ab;5429 | 0 | 0 |
| MT072864.1;QIQ54047.1;Pangolin_CoV_orf1ab;5430 | 0 | 0 |
| MT072864.1;QIQ54047.1;Pangolin_CoV_orf1ab;5431 | 0 | 0 |
| MT072864.1;QIQ54047.1;Pangolin_CoV_orf1ab;5432 | 0 | 0 |
| MT072864.1;QIQ54047.1;Pangolin_CoV_orf1ab;5433 | 0.05 | 0 |
| MT072864.1;QIQ54047.1;Pangolin_CoV_orf1ab;5434 | 0 | 0 |
| MT072864.1;QIQ54047.1;Pangolin_CoV_orf1ab;5435 | 0 | 0 |
| MT072864.1;QIQ54047.1;Pangolin_CoV_orf1ab;5436 | 0 | 0 |
| MT072864.1;QIQ54047.1;Pangolin_CoV_orf1ab;5437 | 0 | 0 |
| MT072864.1;QIQ54047.1;Pangolin_CoV_orf1ab;5438 | 0 | 0 |
| MT072864.1;QIQ54047.1;Pangolin_CoV_orf1ab;5439 | 0 | 0 |
| MT072864.1;QIQ54047.1;Pangolin_CoV_orf1ab;5440 | 0 | 0 |
| MT072864.1;QIQ54047.1;Pangolin_CoV_orf1ab;5441 | 0 | 0.025 |
| MT072864.1;QIQ54047.1;Pangolin_CoV_orf1ab;5442 | 0 | 0 |
| MT072864.1;QIQ54047.1;Pangolin_CoV_orf1ab;5443 | 0 | 0 |
| MT072864.1;QIQ54047.1;Pangolin_CoV_orf1ab;5444 | 0.05 | 0 |
| MT072864.1;QIQ54047.1;Pangolin_CoV_orf1ab;5445 | 0.05 | 0 |
| MT072864.1;QIQ54047.1;Pangolin_CoV_orf1ab;5446 | 0.05 | 0 |
| MT072864.1;QIQ54047.1;Pangolin_CoV_orf1ab;5447 | 0 | 0 |
| MT072864.1;QIQ54047.1;Pangolin_CoV_orf1ab;5448 | 0 | 0 |
| MT072864.1;QIQ54047.1;Pangolin_CoV_orf1ab;5449 | 0 | 0 |
| MT072864.1;QIQ54047.1;Pangolin_CoV_orf1ab;5450 | 0 | 0 |
| MT072864.1;QIQ54047.1;Pangolin_CoV_orf1ab;5451 | 0.05 | 0.025 |

| | | |
|---|---|---|
| MT072864.1;QIQ54047.1;Pangolin_CoV_orf1ab;5452 | 0 | 0 |
| MT072864.1;QIQ54047.1;Pangolin_CoV_orf1ab;5453 | 0 | 0.025 |
| MT072864.1;QIQ54047.1;Pangolin_CoV_orf1ab;5454 | 0 | 0 |
| MT072864.1;QIQ54047.1;Pangolin_CoV_orf1ab;5455 | 0 | 0 |
| MT072864.1;QIQ54047.1;Pangolin_CoV_orf1ab;5456 | 0 | 0 |
| MT072864.1;QIQ54047.1;Pangolin_CoV_orf1ab;5457 | 0 | 0 |
| MT072864.1;QIQ54047.1;Pangolin_CoV_orf1ab;5458 | 0.05 | 0 |
| MT072864.1;QIQ54047.1;Pangolin_CoV_orf1ab;5459 | 0 | 0 |
| MT072864.1;QIQ54047.1;Pangolin_CoV_orf1ab;5460 | 0 | 0 |
| MT072864.1;QIQ54047.1;Pangolin_CoV_orf1ab;5461 | 0 | 0 |
| MT072864.1;QIQ54047.1;Pangolin_CoV_orf1ab;5462 | 0.05 | 0 |
| MT072864.1;QIQ54047.1;Pangolin_CoV_orf1ab;5463 | 0.05 | 0 |
| MT072864.1;QIQ54047.1;Pangolin_CoV_orf1ab;5464 | 0.05 | 0 |
| MT072864.1;QIQ54047.1;Pangolin_CoV_orf1ab;5465 | 0.05 | 0 |
| MT072864.1;QIQ54047.1;Pangolin_CoV_orf1ab;5466 | 0 | 0.05 |
| MT072864.1;QIQ54047.1;Pangolin_CoV_orf1ab;5467 | 0 | 0 |
| MT072864.1;QIQ54047.1;Pangolin_CoV_orf1ab;5468 | 0 | 0.025 |
| MT072864.1;QIQ54047.1;Pangolin_CoV_orf1ab;5469 | 0 | 0 |
| MT072864.1;QIQ54047.1;Pangolin_CoV_orf1ab;5470 | 0 | 0 |
| MT072864.1;QIQ54047.1;Pangolin_CoV_orf1ab;5471 | 0 | 0 |
| MT072864.1;QIQ54047.1;Pangolin_CoV_orf1ab;5472 | 0 | 0.025 |
| MT072864.1;QIQ54047.1;Pangolin_CoV_orf1ab;5473 | 0 | 0 |
| MT072864.1;QIQ54047.1;Pangolin_CoV_orf1ab;5474 | 0 | 0 |
| MT072864.1;QIQ54047.1;Pangolin_CoV_orf1ab;5475 | 0 | 0 |
| MT072864.1;QIQ54047.1;Pangolin_CoV_orf1ab;5476 | 0 | 0 |
| MT072864.1;QIQ54047.1;Pangolin_CoV_orf1ab;5477 | 0 | 0 |
| MT072864.1;QIQ54047.1;Pangolin_CoV_orf1ab;5478 | 0 | 0 |
| MT072864.1;QIQ54047.1;Pangolin_CoV_orf1ab;5479 | 0 | 0 |
| MT072864.1;QIQ54047.1;Pangolin_CoV_orf1ab;5480 | 0 | 0 |
| MT072864.1;QIQ54047.1;Pangolin_CoV_orf1ab;5481 | 0 | 0 |
| MT072864.1;QIQ54047.1;Pangolin_CoV_orf1ab;5482 | 0 | 0 |
| MT072864.1;QIQ54047.1;Pangolin_CoV_orf1ab;5483 | 0 | 0 |
| MT072864.1;QIQ54047.1;Pangolin_CoV_orf1ab;5484 | 0 | 0 |
| MT072864.1;QIQ54047.1;Pangolin_CoV_orf1ab;5485 | 0 | 0 |
| MT072864.1;QIQ54047.1;Pangolin_CoV_orf1ab;5486 | 0 | 0 |
| MT072864.1;QIQ54047.1;Pangolin_CoV_orf1ab;5487 | 0 | 0 |
| MT072864.1;QIQ54047.1;Pangolin_CoV_orf1ab;5488 | 0 | 0 |
| MT072864.1;QIQ54047.1;Pangolin_CoV_orf1ab;5489 | 0 | 0 |
| MT072864.1;QIQ54047.1;Pangolin_CoV_orf1ab;5490 | 0 | 0 |
| MT072864.1;QIQ54047.1;Pangolin_CoV_orf1ab;5491 | 0 | 0 |
| MT072864.1;QIQ54047.1;Pangolin_CoV_orf1ab;5492 | 0 | 0 |
| MT072864.1;QIQ54047.1;Pangolin_CoV_orf1ab;5493 | 0 | 0 |
| MT072864.1;QIQ54047.1;Pangolin_CoV_orf1ab;5494 | 0 | 0 |

| | | |
|---|---|---|
| MT072864.1;QIQ54047.1;Pangolin_CoV_orf1ab;5495 | 0 | 0 |
| MT072864.1;QIQ54047.1;Pangolin_CoV_orf1ab;5496 | 0 | 0 |
| MT072864.1;QIQ54047.1;Pangolin_CoV_orf1ab;5497 | 0 | 0 |
| MT072864.1;QIQ54047.1;Pangolin_CoV_orf1ab;5498 | 0.05 | 0.025 |
| MT072864.1;QIQ54047.1;Pangolin_CoV_orf1ab;5499 | 0.05 | 0.025 |
| MT072864.1;QIQ54047.1;Pangolin_CoV_orf1ab;5500 | 0.05 | 0.025 |
| MT072864.1;QIQ54047.1;Pangolin_CoV_orf1ab;5501 | 0.05 | 0 |
| MT072864.1;QIQ54047.1;Pangolin_CoV_orf1ab;5502 | 0 | 0.025 |
| MT072864.1;QIQ54047.1;Pangolin_CoV_orf1ab;5503 | 0.05 | 0 |
| MT072864.1;QIQ54047.1;Pangolin_CoV_orf1ab;5504 | 0.1 | 0 |
| MT072864.1;QIQ54047.1;Pangolin_CoV_orf1ab;5505 | 0 | 0 |
| MT072864.1;QIQ54047.1;Pangolin_CoV_orf1ab;5506 | 0.05 | 0 |
| MT072864.1;QIQ54047.1;Pangolin_CoV_orf1ab;5507 | 0.05 | 0.025 |
| MT072864.1;QIQ54047.1;Pangolin_CoV_orf1ab;5508 | 0 | 0.025 |
| MT072864.1;QIQ54047.1;Pangolin_CoV_orf1ab;5509 | 0.05 | 0 |
| MT072864.1;QIQ54047.1;Pangolin_CoV_orf1ab;5510 | 0.1 | 0.025 |
| MT072864.1;QIQ54047.1;Pangolin_CoV_orf1ab;5511 | 0.05 | 0.025 |
| MT072864.1;QIQ54047.1;Pangolin_CoV_orf1ab;5512 | 0 | 0 |
| MT072864.1;QIQ54047.1;Pangolin_CoV_orf1ab;5513 | 0 | 0.025 |
| MT072864.1;QIQ54047.1;Pangolin_CoV_orf1ab;5514 | 0.1 | 0 |
| MT072864.1;QIQ54047.1;Pangolin_CoV_orf1ab;5515 | 0 | 0.025 |
| MT072864.1;QIQ54047.1;Pangolin_CoV_orf1ab;5516 | 0 | 0.025 |
| MT072864.1;QIQ54047.1;Pangolin_CoV_orf1ab;5517 | 0 | 0.025 |
| MT072864.1;QIQ54047.1;Pangolin_CoV_orf1ab;5518 | 0 | 0.025 |
| MT072864.1;QIQ54047.1;Pangolin_CoV_orf1ab;5519 | 0.05 | 0 |
| MT072864.1;QIQ54047.1;Pangolin_CoV_orf1ab;5520 | 0 | 0 |
| MT072864.1;QIQ54047.1;Pangolin_CoV_orf1ab;5521 | 0 | 0 |
| MT072864.1;QIQ54047.1;Pangolin_CoV_orf1ab;5522 | 0 | 0 |
| MT072864.1;QIQ54047.1;Pangolin_CoV_orf1ab;5523 | 0 | 0 |
| MT072864.1;QIQ54047.1;Pangolin_CoV_orf1ab;5524 | 0 | 0 |
| MT072864.1;QIQ54047.1;Pangolin_CoV_orf1ab;5525 | 0 | 0 |
| MT072864.1;QIQ54047.1;Pangolin_CoV_orf1ab;5526 | 0 | 0 |
| MT072864.1;QIQ54047.1;Pangolin_CoV_orf1ab;5527 | 0 | 0 |
| MT072864.1;QIQ54047.1;Pangolin_CoV_orf1ab;5528 | 0 | 0 |
| MT072864.1;QIQ54047.1;Pangolin_CoV_orf1ab;5529 | 0 | 0 |
| MT072864.1;QIQ54047.1;Pangolin_CoV_orf1ab;5530 | 0 | 0 |
| MT072864.1;QIQ54047.1;Pangolin_CoV_orf1ab;5531 | 0 | 0 |
| MT072864.1;QIQ54047.1;Pangolin_CoV_orf1ab;5532 | 0 | 0 |
| MT072864.1;QIQ54047.1;Pangolin_CoV_orf1ab;5533 | 0 | 0 |
| MT072864.1;QIQ54047.1;Pangolin_CoV_orf1ab;5534 | 0 | 0 |
| MT072864.1;QIQ54047.1;Pangolin_CoV_orf1ab;5535 | 0 | 0 |
| MT072864.1;QIQ54047.1;Pangolin_CoV_orf1ab;5536 | 0 | 0 |
| MT072864.1;QIQ54047.1;Pangolin_CoV_orf1ab;5537 | 0 | 0 |

| | | |
|---|---|---|
| MT072864.1;QIQ54047.1;Pangolin_CoV_orf1ab;5538 | 0 | 0 |
| MT072864.1;QIQ54047.1;Pangolin_CoV_orf1ab;5539 | 0.05 | 0 |
| MT072864.1;QIQ54047.1;Pangolin_CoV_orf1ab;5540 | 0 | 0 |
| MT072864.1;QIQ54047.1;Pangolin_CoV_orf1ab;5541 | 0 | 0 |
| MT072864.1;QIQ54047.1;Pangolin_CoV_orf1ab;5542 | 0 | 0 |
| MT072864.1;QIQ54047.1;Pangolin_CoV_orf1ab;5543 | 0 | 0.025 |
| MT072864.1;QIQ54047.1;Pangolin_CoV_orf1ab;5544 | 0 | 0 |
| MT072864.1;QIQ54047.1;Pangolin_CoV_orf1ab;5545 | 0 | 0 |
| MT072864.1;QIQ54047.1;Pangolin_CoV_orf1ab;5546 | 0 | 0 |
| MT072864.1;QIQ54047.1;Pangolin_CoV_orf1ab;5547 | 0 | 0.025 |
| MT072864.1;QIQ54047.1;Pangolin_CoV_orf1ab;5548 | 0 | 0.025 |
| MT072864.1;QIQ54047.1;Pangolin_CoV_orf1ab;5549 | 0 | 0.025 |
| MT072864.1;QIQ54047.1;Pangolin_CoV_orf1ab;5550 | 0 | 0.025 |
| MT072864.1;QIQ54047.1;Pangolin_CoV_orf1ab;5551 | 0 | 0.025 |
| MT072864.1;QIQ54047.1;Pangolin_CoV_orf1ab;5552 | 0 | 0.025 |
| MT072864.1;QIQ54047.1;Pangolin_CoV_orf1ab;5553 | 0 | 0.025 |
| MT072864.1;QIQ54047.1;Pangolin_CoV_orf1ab;5554 | 0 | 0 |
| MT072864.1;QIQ54047.1;Pangolin_CoV_orf1ab;5555 | 0 | 0 |
| MT072864.1;QIQ54047.1;Pangolin_CoV_orf1ab;5556 | 0 | 0.025 |
| MT072864.1;QIQ54047.1;Pangolin_CoV_orf1ab;5557 | 0 | 0.025 |
| MT072864.1;QIQ54047.1;Pangolin_CoV_orf1ab;5558 | 0 | 0 |
| MT072864.1;QIQ54047.1;Pangolin_CoV_orf1ab;5559 | 0 | 0.025 |
| MT072864.1;QIQ54047.1;Pangolin_CoV_orf1ab;5560 | 0 | 0 |
| MT072864.1;QIQ54047.1;Pangolin_CoV_orf1ab;5561 | 0 | 0 |
| MT072864.1;QIQ54047.1;Pangolin_CoV_orf1ab;5562 | 0 | 0 |
| MT072864.1;QIQ54047.1;Pangolin_CoV_orf1ab;5563 | 0.05 | 0 |
| MT072864.1;QIQ54047.1;Pangolin_CoV_orf1ab;5564 | 0.05 | 0.025 |
| MT072864.1;QIQ54047.1;Pangolin_CoV_orf1ab;5565 | 0.05 | 0.025 |
| MT072864.1;QIQ54047.1;Pangolin_CoV_orf1ab;5566 | 0.05 | 0.025 |
| MT072864.1;QIQ54047.1;Pangolin_CoV_orf1ab;5567 | 0.05 | 0.025 |
| MT072864.1;QIQ54047.1;Pangolin_CoV_orf1ab;5568 | 0.05 | 0.075 |
| MT072864.1;QIQ54047.1;Pangolin_CoV_orf1ab;5569 | 0.05 | 0.175 |
| MT072864.1;QIQ54047.1;Pangolin_CoV_orf1ab;5570 | 0.05 | 0.075 |
| MT072864.1;QIQ54047.1;Pangolin_CoV_orf1ab;5571 | 0.1 | 0.125 |
| MT072864.1;QIQ54047.1;Pangolin_CoV_orf1ab;5572 | 0.05 | 0.075 |
| MT072864.1;QIQ54047.1;Pangolin_CoV_orf1ab;5573 | 0 | 0.05 |
| MT072864.1;QIQ54047.1;Pangolin_CoV_orf1ab;5574 | 0.05 | 0.025 |
| MT072864.1;QIQ54047.1;Pangolin_CoV_orf1ab;5575 | 0 | 0 |
| MT072864.1;QIQ54047.1;Pangolin_CoV_orf1ab;5576 | 0.1 | 0.1 |
| MT072864.1;QIQ54047.1;Pangolin_CoV_orf1ab;5577 | 0.05 | 0 |
| MT072864.1;QIQ54047.1;Pangolin_CoV_orf1ab;5578 | 0.05 | 0 |
| MT072864.1;QIQ54047.1;Pangolin_CoV_orf1ab;5579 | 0.05 | 0.05 |
| MT072864.1;QIQ54047.1;Pangolin_CoV_orf1ab;5580 | 0 | 0.025 |

| | | |
|---|---|---|
| MT072864.1;QIQ54047.1;Pangolin_CoV_orf1ab;5581 | 0 | 0.05 |
| MT072864.1;QIQ54047.1;Pangolin_CoV_orf1ab;5582 | 0 | 0.05 |
| MT072864.1;QIQ54047.1;Pangolin_CoV_orf1ab;5583 | 0 | 0.05 |
| MT072864.1;QIQ54047.1;Pangolin_CoV_orf1ab;5584 | 0 | 0.05 |
| MT072864.1;QIQ54047.1;Pangolin_CoV_orf1ab;5585 | 0.05 | 0.05 |
| MT072864.1;QIQ54047.1;Pangolin_CoV_orf1ab;5586 | 0.1 | 0.05 |
| MT072864.1;QIQ54047.1;Pangolin_CoV_orf1ab;5587 | 0.05 | 0.05 |
| MT072864.1;QIQ54047.1;Pangolin_CoV_orf1ab;5588 | 0 | 0 |
| MT072864.1;QIQ54047.1;Pangolin_CoV_orf1ab;5589 | 0 | 0.025 |
| MT072864.1;QIQ54047.1;Pangolin_CoV_orf1ab;5590 | 0 | 0.025 |
| MT072864.1;QIQ54047.1;Pangolin_CoV_orf1ab;5591 | 0 | 0.025 |
| MT072864.1;QIQ54047.1;Pangolin_CoV_orf1ab;5592 | 0 | 0.075 |
| MT072864.1;QIQ54047.1;Pangolin_CoV_orf1ab;5593 | 0 | 0.025 |
| MT072864.1;QIQ54047.1;Pangolin_CoV_orf1ab;5594 | 0 | 0 |
| MT072864.1;QIQ54047.1;Pangolin_CoV_orf1ab;5595 | 0 | 0 |
| MT072864.1;QIQ54047.1;Pangolin_CoV_orf1ab;5596 | 0 | 0 |
| MT072864.1;QIQ54047.1;Pangolin_CoV_orf1ab;5597 | 0 | 0 |
| MT072864.1;QIQ54047.1;Pangolin_CoV_orf1ab;5598 | 0 | 0 |
| MT072864.1;QIQ54047.1;Pangolin_CoV_orf1ab;5599 | 0 | 0 |
| MT072864.1;QIQ54047.1;Pangolin_CoV_orf1ab;5600 | 0 | 0 |
| MT072864.1;QIQ54047.1;Pangolin_CoV_orf1ab;5601 | 0 | 0 |
| MT072864.1;QIQ54047.1;Pangolin_CoV_orf1ab;5602 | 0 | 0 |
| MT072864.1;QIQ54047.1;Pangolin_CoV_orf1ab;5603 | 0 | 0 |
| MT072864.1;QIQ54047.1;Pangolin_CoV_orf1ab;5604 | 0 | 0 |
| MT072864.1;QIQ54047.1;Pangolin_CoV_orf1ab;5605 | 0 | 0 |
| MT072864.1;QIQ54047.1;Pangolin_CoV_orf1ab;5606 | 0 | 0 |
| MT072864.1;QIQ54047.1;Pangolin_CoV_orf1ab;5607 | 0 | 0 |
| MT072864.1;QIQ54047.1;Pangolin_CoV_orf1ab;5608 | 0 | 0 |
| MT072864.1;QIQ54047.1;Pangolin_CoV_orf1ab;5609 | 0 | 0 |
| MT072864.1;QIQ54047.1;Pangolin_CoV_orf1ab;5610 | 0 | 0 |
| MT072864.1;QIQ54047.1;Pangolin_CoV_orf1ab;5611 | 0 | 0 |
| MT072864.1;QIQ54047.1;Pangolin_CoV_orf1ab;5612 | 0 | 0 |
| MT072864.1;QIQ54047.1;Pangolin_CoV_orf1ab;5613 | 0 | 0 |
| MT072864.1;QIQ54047.1;Pangolin_CoV_orf1ab;5614 | 0 | 0 |
| MT072864.1;QIQ54047.1;Pangolin_CoV_orf1ab;5615 | 0 | 0 |
| MT072864.1;QIQ54047.1;Pangolin_CoV_orf1ab;5616 | 0 | 0.025 |
| MT072864.1;QIQ54047.1;Pangolin_CoV_orf1ab;5617 | 0 | 0.025 |
| MT072864.1;QIQ54047.1;Pangolin_CoV_orf1ab;5618 | 0 | 0.05 |
| MT072864.1;QIQ54047.1;Pangolin_CoV_orf1ab;5619 | 0 | 0 |
| MT072864.1;QIQ54047.1;Pangolin_CoV_orf1ab;5620 | 0 | 0 |
| MT072864.1;QIQ54047.1;Pangolin_CoV_orf1ab;5621 | 0 | 0 |
| MT072864.1;QIQ54047.1;Pangolin_CoV_orf1ab;5622 | 0 | 0 |
| MT072864.1;QIQ54047.1;Pangolin_CoV_orf1ab;5623 | 0 | 0 |

| | | |
|---|---|---|
| MT072864.1;QIQ54047.1;Pangolin_CoV_orf1ab;5624 | 0 | 0 |
| MT072864.1;QIQ54047.1;Pangolin_CoV_orf1ab;5625 | 0 | 0 |
| MT072864.1;QIQ54047.1;Pangolin_CoV_orf1ab;5626 | 0 | 0 |
| MT072864.1;QIQ54047.1;Pangolin_CoV_orf1ab;5627 | 0 | 0 |
| MT072864.1;QIQ54047.1;Pangolin_CoV_orf1ab;5628 | 0 | 0 |
| MT072864.1;QIQ54047.1;Pangolin_CoV_orf1ab;5629 | 0 | 0 |
| MT072864.1;QIQ54047.1;Pangolin_CoV_orf1ab;5630 | 0.05 | 0 |
| MT072864.1;QIQ54047.1;Pangolin_CoV_orf1ab;5631 | 0 | 0 |
| MT072864.1;QIQ54047.1;Pangolin_CoV_orf1ab;5632 | 0 | 0 |
| MT072864.1;QIQ54047.1;Pangolin_CoV_orf1ab;5633 | 0 | 0 |
| MT072864.1;QIQ54047.1;Pangolin_CoV_orf1ab;5634 | 0 | 0 |
| MT072864.1;QIQ54047.1;Pangolin_CoV_orf1ab;5635 | 0 | 0 |
| MT072864.1;QIQ54047.1;Pangolin_CoV_orf1ab;5636 | 0 | 0 |
| MT072864.1;QIQ54047.1;Pangolin_CoV_orf1ab;5637 | 0 | 0 |
| MT072864.1;QIQ54047.1;Pangolin_CoV_orf1ab;5638 | 0 | 0 |
| MT072864.1;QIQ54047.1;Pangolin_CoV_orf1ab;5639 | 0 | 0 |
| MT072864.1;QIQ54047.1;Pangolin_CoV_orf1ab;5640 | 0 | 0 |
| MT072864.1;QIQ54047.1;Pangolin_CoV_orf1ab;5641 | 0 | 0 |
| MT072864.1;QIQ54047.1;Pangolin_CoV_orf1ab;5642 | 0 | 0 |
| MT072864.1;QIQ54047.1;Pangolin_CoV_orf1ab;5643 | 0 | 0 |
| MT072864.1;QIQ54047.1;Pangolin_CoV_orf1ab;5644 | 0 | 0 |
| MT072864.1;QIQ54047.1;Pangolin_CoV_orf1ab;5645 | 0 | 0 |
| MT072864.1;QIQ54047.1;Pangolin_CoV_orf1ab;5646 | 0 | 0 |
| MT072864.1;QIQ54047.1;Pangolin_CoV_orf1ab;5647 | 0 | 0 |
| MT072864.1;QIQ54047.1;Pangolin_CoV_orf1ab;5648 | 0 | 0 |
| MT072864.1;QIQ54047.1;Pangolin_CoV_orf1ab;5649 | 0 | 0.025 |
| MT072864.1;QIQ54047.1;Pangolin_CoV_orf1ab;5650 | 0 | 0.025 |
| MT072864.1;QIQ54047.1;Pangolin_CoV_orf1ab;5651 | 0.05 | 0.025 |
| MT072864.1;QIQ54047.1;Pangolin_CoV_orf1ab;5652 | 0 | 0.025 |
| MT072864.1;QIQ54047.1;Pangolin_CoV_orf1ab;5653 | 0.05 | 0.075 |
| MT072864.1;QIQ54047.1;Pangolin_CoV_orf1ab;5654 | 0.25 | 0.075 |
| MT072864.1;QIQ54047.1;Pangolin_CoV_orf1ab;5655 | 0.4 | 0.125 |
| MT072864.1;QIQ54047.1;Pangolin_CoV_orf1ab;5656 | 0.35 | 0.075 |
| MT072864.1;QIQ54047.1;Pangolin_CoV_orf1ab;5657 | 0.3 | 0.075 |
| MT072864.1;QIQ54047.1;Pangolin_CoV_orf1ab;5658 | 0.1 | 0.025 |
| MT072864.1;QIQ54047.1;Pangolin_CoV_orf1ab;5659 | 0.05 | 0.025 |
| MT072864.1;QIQ54047.1;Pangolin_CoV_orf1ab;5660 | 0 | 0.025 |
| MT072864.1;QIQ54047.1;Pangolin_CoV_orf1ab;5661 | 0 | 0 |
| MT072864.1;QIQ54047.1;Pangolin_CoV_orf1ab;5662 | 0 | 0 |
| MT072864.1;QIQ54047.1;Pangolin_CoV_orf1ab;5663 | 0 | 0 |
| MT072864.1;QIQ54047.1;Pangolin_CoV_orf1ab;5664 | 0 | 0 |
| MT072864.1;QIQ54047.1;Pangolin_CoV_orf1ab;5665 | 0 | 0 |
| MT072864.1;QIQ54047.1;Pangolin_CoV_orf1ab;5666 | 0 | 0 |

| | | |
|---|---|---|
| MT072864.1;QIQ54047.1;Pangolin_CoV_orf1ab;5667 | 0 | 0 |
| MT072864.1;QIQ54047.1;Pangolin_CoV_orf1ab;5668 | 0 | 0.025 |
| MT072864.1;QIQ54047.1;Pangolin_CoV_orf1ab;5669 | 0 | 0 |
| MT072864.1;QIQ54047.1;Pangolin_CoV_orf1ab;5670 | 0 | 0 |
| MT072864.1;QIQ54047.1;Pangolin_CoV_orf1ab;5671 | 0 | 0 |
| MT072864.1;QIQ54047.1;Pangolin_CoV_orf1ab;5672 | 0 | 0 |
| MT072864.1;QIQ54047.1;Pangolin_CoV_orf1ab;5673 | 0 | 0 |
| MT072864.1;QIQ54047.1;Pangolin_CoV_orf1ab;5674 | 0 | 0 |
| MT072864.1;QIQ54047.1;Pangolin_CoV_orf1ab;5675 | 0 | 0 |
| MT072864.1;QIQ54047.1;Pangolin_CoV_orf1ab;5676 | 0 | 0 |
| MT072864.1;QIQ54047.1;Pangolin_CoV_orf1ab;5677 | 0 | 0 |
| MT072864.1;QIQ54047.1;Pangolin_CoV_orf1ab;5678 | 0 | 0 |
| MT072864.1;QIQ54047.1;Pangolin_CoV_orf1ab;5679 | 0 | 0 |
| MT072864.1;QIQ54047.1;Pangolin_CoV_orf1ab;5680 | 0 | 0 |
| MT072864.1;QIQ54047.1;Pangolin_CoV_orf1ab;5681 | 0 | 0 |
| MT072864.1;QIQ54047.1;Pangolin_CoV_orf1ab;5682 | 0 | 0 |
| MT072864.1;QIQ54047.1;Pangolin_CoV_orf1ab;5683 | 0 | 0 |
| MT072864.1;QIQ54047.1;Pangolin_CoV_orf1ab;5684 | 0 | 0.075 |
| MT072864.1;QIQ54047.1;Pangolin_CoV_orf1ab;5685 | 0 | 0 |
| MT072864.1;QIQ54047.1;Pangolin_CoV_orf1ab;5686 | 0 | 0.025 |
| MT072864.1;QIQ54047.1;Pangolin_CoV_orf1ab;5687 | 0 | 0.025 |
| MT072864.1;QIQ54047.1;Pangolin_CoV_orf1ab;5688 | 0 | 0.05 |
| MT072864.1;QIQ54047.1;Pangolin_CoV_orf1ab;5689 | 0 | 0.05 |
| MT072864.1;QIQ54047.1;Pangolin_CoV_orf1ab;5690 | 0 | 0.025 |
| MT072864.1;QIQ54047.1;Pangolin_CoV_orf1ab;5691 | 0 | 0.025 |
| MT072864.1;QIQ54047.1;Pangolin_CoV_orf1ab;5692 | 0 | 0.025 |
| MT072864.1;QIQ54047.1;Pangolin_CoV_orf1ab;5693 | 0 | 0.025 |
| MT072864.1;QIQ54047.1;Pangolin_CoV_orf1ab;5694 | 0 | 0 |
| MT072864.1;QIQ54047.1;Pangolin_CoV_orf1ab;5695 | 0 | 0 |
| MT072864.1;QIQ54047.1;Pangolin_CoV_orf1ab;5696 | 0 | 0 |
| MT072864.1;QIQ54047.1;Pangolin_CoV_orf1ab;5697 | 0 | 0.025 |
| MT072864.1;QIQ54047.1;Pangolin_CoV_orf1ab;5698 | 0 | 0.025 |
| MT072864.1;QIQ54047.1;Pangolin_CoV_orf1ab;5699 | 0 | 0.025 |
| MT072864.1;QIQ54047.1;Pangolin_CoV_orf1ab;5700 | 0 | 0 |
| MT072864.1;QIQ54047.1;Pangolin_CoV_orf1ab;5701 | 0 | 0 |
| MT072864.1;QIQ54047.1;Pangolin_CoV_orf1ab;5702 | 0 | 0.025 |
| MT072864.1;QIQ54047.1;Pangolin_CoV_orf1ab;5703 | 0 | 0.025 |
| MT072864.1;QIQ54047.1;Pangolin_CoV_orf1ab;5704 | 0 | 0.025 |
| MT072864.1;QIQ54047.1;Pangolin_CoV_orf1ab;5705 | 0 | 0.025 |
| MT072864.1;QIQ54047.1;Pangolin_CoV_orf1ab;5706 | 0 | 0.025 |
| MT072864.1;QIQ54047.1;Pangolin_CoV_orf1ab;5707 | 0 | 0.025 |
| MT072864.1;QIQ54047.1;Pangolin_CoV_orf1ab;5708 | 0 | 0.025 |
| MT072864.1;QIQ54047.1;Pangolin_CoV_orf1ab;5709 | 0 | 0 |

| | | |
|---|---|---|
| MT072864.1;QIQ54047.1;Pangolin_CoV_orf1ab;5710 | 0 | 0 |
| MT072864.1;QIQ54047.1;Pangolin_CoV_orf1ab;5711 | 0 | 0 |
| MT072864.1;QIQ54047.1;Pangolin_CoV_orf1ab;5712 | 0 | 0 |
| MT072864.1;QIQ54047.1;Pangolin_CoV_orf1ab;5713 | 0 | 0.025 |
| MT072864.1;QIQ54047.1;Pangolin_CoV_orf1ab;5714 | 0 | 0 |
| MT072864.1;QIQ54047.1;Pangolin_CoV_orf1ab;5715 | 0 | 0.025 |
| MT072864.1;QIQ54047.1;Pangolin_CoV_orf1ab;5716 | 0 | 0.025 |
| MT072864.1;QIQ54047.1;Pangolin_CoV_orf1ab;5717 | 0 | 0.025 |
| MT072864.1;QIQ54047.1;Pangolin_CoV_orf1ab;5718 | 0 | 0 |
| MT072864.1;QIQ54047.1;Pangolin_CoV_orf1ab;5719 | 0 | 0.025 |
| MT072864.1;QIQ54047.1;Pangolin_CoV_orf1ab;5720 | 0 | 0.075 |
| MT072864.1;QIQ54047.1;Pangolin_CoV_orf1ab;5721 | 0 | 0.075 |
| MT072864.1;QIQ54047.1;Pangolin_CoV_orf1ab;5722 | 0.05 | 0.1 |
| MT072864.1;QIQ54047.1;Pangolin_CoV_orf1ab;5723 | 0.1 | 0.075 |
| MT072864.1;QIQ54047.1;Pangolin_CoV_orf1ab;5724 | 0.1 | 0.075 |
| MT072864.1;QIQ54047.1;Pangolin_CoV_orf1ab;5725 | 0.1 | 0.05 |
| MT072864.1;QIQ54047.1;Pangolin_CoV_orf1ab;5726 | 0.1 | 0.05 |
| MT072864.1;QIQ54047.1;Pangolin_CoV_orf1ab;5727 | 0.1 | 0.05 |
| MT072864.1;QIQ54047.1;Pangolin_CoV_orf1ab;5728 | 0.1 | 0.05 |
| MT072864.1;QIQ54047.1;Pangolin_CoV_orf1ab;5729 | 0.1 | 0 |
| MT072864.1;QIQ54047.1;Pangolin_CoV_orf1ab;5730 | 0.05 | 0.025 |
| MT072864.1;QIQ54047.1;Pangolin_CoV_orf1ab;5731 | 0 | 0 |
| MT072864.1;QIQ54047.1;Pangolin_CoV_orf1ab;5732 | 0 | 0 |
| MT072864.1;QIQ54047.1;Pangolin_CoV_orf1ab;5733 | 0 | 0 |
| MT072864.1;QIQ54047.1;Pangolin_CoV_orf1ab;5734 | 0 | 0 |
| MT072864.1;QIQ54047.1;Pangolin_CoV_orf1ab;5735 | 0 | 0 |
| MT072864.1;QIQ54047.1;Pangolin_CoV_orf1ab;5736 | 0 | 0 |
| MT072864.1;QIQ54047.1;Pangolin_CoV_orf1ab;5737 | 0 | 0 |
| MT072864.1;QIQ54047.1;Pangolin_CoV_orf1ab;5738 | 0 | 0 |
| MT072864.1;QIQ54047.1;Pangolin_CoV_orf1ab;5739 | 0 | 0 |
| MT072864.1;QIQ54047.1;Pangolin_CoV_orf1ab;5740 | 0 | 0 |
| MT072864.1;QIQ54047.1;Pangolin_CoV_orf1ab;5741 | 0 | 0.025 |
| MT072864.1;QIQ54047.1;Pangolin_CoV_orf1ab;5742 | 0 | 0 |
| MT072864.1;QIQ54047.1;Pangolin_CoV_orf1ab;5743 | 0 | 0 |
| MT072864.1;QIQ54047.1;Pangolin_CoV_orf1ab;5744 | 0 | 0 |
| MT072864.1;QIQ54047.1;Pangolin_CoV_orf1ab;5745 | 0 | 0 |
| MT072864.1;QIQ54047.1;Pangolin_CoV_orf1ab;5746 | 0 | 0 |
| MT072864.1;QIQ54047.1;Pangolin_CoV_orf1ab;5747 | 0 | 0 |
| MT072864.1;QIQ54047.1;Pangolin_CoV_orf1ab;5748 | 0 | 0 |
| MT072864.1;QIQ54047.1;Pangolin_CoV_orf1ab;5749 | 0 | 0 |
| MT072864.1;QIQ54047.1;Pangolin_CoV_orf1ab;5750 | 0 | 0 |
| MT072864.1;QIQ54047.1;Pangolin_CoV_orf1ab;5751 | 0 | 0 |
| MT072864.1;QIQ54047.1;Pangolin_CoV_orf1ab;5752 | 0 | 0 |

| | | |
|---|---|---|
| MT072864.1;QIQ54047.1;Pangolin_CoV_orf1ab;5753 | 0 | 0 |
| MT072864.1;QIQ54047.1;Pangolin_CoV_orf1ab;5754 | 0.1 | 0.05 |
| MT072864.1;QIQ54047.1;Pangolin_CoV_orf1ab;5755 | 0.1 | 0.05 |
| MT072864.1;QIQ54047.1;Pangolin_CoV_orf1ab;5756 | 0.15 | 0.125 |
| MT072864.1;QIQ54047.1;Pangolin_CoV_orf1ab;5757 | 0.15 | 0.15 |
| MT072864.1;QIQ54047.1;Pangolin_CoV_orf1ab;5758 | 0.1 | 0.05 |
| MT072864.1;QIQ54047.1;Pangolin_CoV_orf1ab;5759 | 0.1 | 0.025 |
| MT072864.1;QIQ54047.1;Pangolin_CoV_orf1ab;5760 | 0.1 | 0.075 |
| MT072864.1;QIQ54047.1;Pangolin_CoV_orf1ab;5761 | 0.1 | 0.15 |
| MT072864.1;QIQ54047.1;Pangolin_CoV_orf1ab;5762 | 0.15 | 0.225 |
| MT072864.1;QIQ54047.1;Pangolin_CoV_orf1ab;5763 | 0.15 | 0.05 |
| MT072864.1;QIQ54047.1;Pangolin_CoV_orf1ab;5764 | 0.1 | 0.05 |
| MT072864.1;QIQ54047.1;Pangolin_CoV_orf1ab;5765 | 0.15 | 0.125 |
| MT072864.1;QIQ54047.1;Pangolin_CoV_orf1ab;5766 | 0 | 0 |
| MT072864.1;QIQ54047.1;Pangolin_CoV_orf1ab;5767 | 0.05 | 0 |
| MT072864.1;QIQ54047.1;Pangolin_CoV_orf1ab;5768 | 0 | 0 |
| MT072864.1;QIQ54047.1;Pangolin_CoV_orf1ab;5769 | 0.05 | 0.025 |
| MT072864.1;QIQ54047.1;Pangolin_CoV_orf1ab;5770 | 0.05 | 0.025 |
| MT072864.1;QIQ54047.1;Pangolin_CoV_orf1ab;5771 | 0 | 0.025 |
| MT072864.1;QIQ54047.1;Pangolin_CoV_orf1ab;5772 | 0 | 0.025 |
| MT072864.1;QIQ54047.1;Pangolin_CoV_orf1ab;5773 | 0 | 0.025 |
| MT072864.1;QIQ54047.1;Pangolin_CoV_orf1ab;5774 | 0 | 0 |
| MT072864.1;QIQ54047.1;Pangolin_CoV_orf1ab;5775 | 0 | 0 |
| MT072864.1;QIQ54047.1;Pangolin_CoV_orf1ab;5776 | 0 | 0.025 |
| MT072864.1;QIQ54047.1;Pangolin_CoV_orf1ab;5777 | 0.05 | 0 |
| MT072864.1;QIQ54047.1;Pangolin_CoV_orf1ab;5778 | 0 | 0 |
| MT072864.1;QIQ54047.1;Pangolin_CoV_orf1ab;5779 | 0 | 0 |
| MT072864.1;QIQ54047.1;Pangolin_CoV_orf1ab;5780 | 0 | 0 |
| MT072864.1;QIQ54047.1;Pangolin_CoV_orf1ab;5781 | 0 | 0 |
| MT072864.1;QIQ54047.1;Pangolin_CoV_orf1ab;5782 | 0 | 0 |
| MT072864.1;QIQ54047.1;Pangolin_CoV_orf1ab;5783 | 0 | 0 |
| MT072864.1;QIQ54047.1;Pangolin_CoV_orf1ab;5784 | 0 | 0 |
| MT072864.1;QIQ54047.1;Pangolin_CoV_orf1ab;5785 | 0 | 0 |
| MT072864.1;QIQ54047.1;Pangolin_CoV_orf1ab;5786 | 0 | 0 |
| MT072864.1;QIQ54047.1;Pangolin_CoV_orf1ab;5787 | 0 | 0 |
| MT072864.1;QIQ54047.1;Pangolin_CoV_orf1ab;5788 | 0 | 0 |
| MT072864.1;QIQ54047.1;Pangolin_CoV_orf1ab;5789 | 0 | 0 |
| MT072864.1;QIQ54047.1;Pangolin_CoV_orf1ab;5790 | 0 | 0 |
| MT072864.1;QIQ54047.1;Pangolin_CoV_orf1ab;5791 | 0 | 0 |
| MT072864.1;QIQ54047.1;Pangolin_CoV_orf1ab;5792 | 0 | 0 |
| MT072864.1;QIQ54047.1;Pangolin_CoV_orf1ab;5793 | 0 | 0.025 |
| MT072864.1;QIQ54047.1;Pangolin_CoV_orf1ab;5794 | 0.05 | 0 |
| MT072864.1;QIQ54047.1;Pangolin_CoV_orf1ab;5795 | 0.05 | 0 |

| | | |
|---|---|---|
| MT072864.1;QIQ54047.1;Pangolin_CoV_orf1ab;5796 | 0.05 | 0 |
| MT072864.1;QIQ54047.1;Pangolin_CoV_orf1ab;5797 | 0 | 0 |
| MT072864.1;QIQ54047.1;Pangolin_CoV_orf1ab;5798 | 0 | 0 |
| MT072864.1;QIQ54047.1;Pangolin_CoV_orf1ab;5799 | 0.05 | 0 |
| MT072864.1;QIQ54047.1;Pangolin_CoV_orf1ab;5800 | 0 | 0 |
| MT072864.1;QIQ54047.1;Pangolin_CoV_orf1ab;5801 | 0 | 0 |
| MT072864.1;QIQ54047.1;Pangolin_CoV_orf1ab;5802 | 0 | 0 |
| MT072864.1;QIQ54047.1;Pangolin_CoV_orf1ab;5803 | 0 | 0 |
| MT072864.1;QIQ54047.1;Pangolin_CoV_orf1ab;5804 | 0 | 0 |
| MT072864.1;QIQ54047.1;Pangolin_CoV_orf1ab;5805 | 0 | 0 |
| MT072864.1;QIQ54047.1;Pangolin_CoV_orf1ab;5806 | 0 | 0 |
| MT072864.1;QIQ54047.1;Pangolin_CoV_orf1ab;5807 | 0 | 0 |
| MT072864.1;QIQ54047.1;Pangolin_CoV_orf1ab;5808 | 0 | 0 |
| MT072864.1;QIQ54047.1;Pangolin_CoV_orf1ab;5809 | 0 | 0 |
| MT072864.1;QIQ54047.1;Pangolin_CoV_orf1ab;5810 | 0 | 0.025 |
| MT072864.1;QIQ54047.1;Pangolin_CoV_orf1ab;5811 | 0 | 0 |
| MT072864.1;QIQ54047.1;Pangolin_CoV_orf1ab;5812 | 0 | 0 |
| MT072864.1;QIQ54047.1;Pangolin_CoV_orf1ab;5813 | 0 | 0 |
| MT072864.1;QIQ54047.1;Pangolin_CoV_orf1ab;5814 | 0 | 0 |
| MT072864.1;QIQ54047.1;Pangolin_CoV_orf1ab;5815 | 0 | 0 |
| MT072864.1;QIQ54047.1;Pangolin_CoV_orf1ab;5816 | 0 | 0 |
| MT072864.1;QIQ54047.1;Pangolin_CoV_orf1ab;5817 | 0 | 0 |
| MT072864.1;QIQ54047.1;Pangolin_CoV_orf1ab;5818 | 0 | 0.025 |
| MT072864.1;QIQ54047.1;Pangolin_CoV_orf1ab;5819 | 0 | 0 |
| MT072864.1;QIQ54047.1;Pangolin_CoV_orf1ab;5820 | 0.05 | 0 |
| MT072864.1;QIQ54047.1;Pangolin_CoV_orf1ab;5821 | 0 | 0 |
| MT072864.1;QIQ54047.1;Pangolin_CoV_orf1ab;5822 | 0 | 0 |
| MT072864.1;QIQ54047.1;Pangolin_CoV_orf1ab;5823 | 0 | 0 |
| MT072864.1;QIQ54047.1;Pangolin_CoV_orf1ab;5824 | 0.1 | 0.05 |
| MT072864.1;QIQ54047.1;Pangolin_CoV_orf1ab;5825 | 0.15 | 0.025 |
| MT072864.1;QIQ54047.1;Pangolin_CoV_orf1ab;5826 | 0.05 | 0.025 |
| MT072864.1;QIQ54047.1;Pangolin_CoV_orf1ab;5827 | 0 | 0.025 |
| MT072864.1;QIQ54047.1;Pangolin_CoV_orf1ab;5828 | 0.2 | 0.025 |
| MT072864.1;QIQ54047.1;Pangolin_CoV_orf1ab;5829 | 0.15 | 0.05 |
| MT072864.1;QIQ54047.1;Pangolin_CoV_orf1ab;5830 | 0.1 | 0.05 |
| MT072864.1;QIQ54047.1;Pangolin_CoV_orf1ab;5831 | 0 | 0.025 |
| MT072864.1;QIQ54047.1;Pangolin_CoV_orf1ab;5832 | 0 | 0.025 |
| MT072864.1;QIQ54047.1;Pangolin_CoV_orf1ab;5833 | 0 | 0.025 |
| MT072864.1;QIQ54047.1;Pangolin_CoV_orf1ab;5834 | 0 | 0.025 |
| MT072864.1;QIQ54047.1;Pangolin_CoV_orf1ab;5835 | 0 | 0 |
| MT072864.1;QIQ54047.1;Pangolin_CoV_orf1ab;5836 | 0 | 0 |
| MT072864.1;QIQ54047.1;Pangolin_CoV_orf1ab;5837 | 0 | 0 |
| MT072864.1;QIQ54047.1;Pangolin_CoV_orf1ab;5838 | 0 | 0 |

| | | |
|---|---|---|
| MT072864.1;QIQ54047.1;Pangolin_CoV_orf1ab;5839 | 0 | 0 |
| MT072864.1;QIQ54047.1;Pangolin_CoV_orf1ab;5840 | 0 | 0 |
| MT072864.1;QIQ54047.1;Pangolin_CoV_orf1ab;5841 | 0 | 0.05 |
| MT072864.1;QIQ54047.1;Pangolin_CoV_orf1ab;5842 | 0 | 0.025 |
| MT072864.1;QIQ54047.1;Pangolin_CoV_orf1ab;5843 | 0 | 0.025 |
| MT072864.1;QIQ54047.1;Pangolin_CoV_orf1ab;5844 | 0 | 0.025 |
| MT072864.1;QIQ54047.1;Pangolin_CoV_orf1ab;5845 | 0 | 0.025 |
| MT072864.1;QIQ54047.1;Pangolin_CoV_orf1ab;5846 | 0 | 0 |
| MT072864.1;QIQ54047.1;Pangolin_CoV_orf1ab;5847 | 0 | 0 |
| MT072864.1;QIQ54047.1;Pangolin_CoV_orf1ab;5848 | 0 | 0.025 |
| MT072864.1;QIQ54047.1;Pangolin_CoV_orf1ab;5849 | 0 | 0.025 |
| MT072864.1;QIQ54047.1;Pangolin_CoV_orf1ab;5850 | 0 | 0 |
| MT072864.1;QIQ54047.1;Pangolin_CoV_orf1ab;5851 | 0 | 0 |
| MT072864.1;QIQ54047.1;Pangolin_CoV_orf1ab;5852 | 0 | 0 |
| MT072864.1;QIQ54047.1;Pangolin_CoV_orf1ab;5853 | 0 | 0 |
| MT072864.1;QIQ54047.1;Pangolin_CoV_orf1ab;5854 | 0 | 0 |
| MT072864.1;QIQ54047.1;Pangolin_CoV_orf1ab;5855 | 0.05 | 0 |
| MT072864.1;QIQ54047.1;Pangolin_CoV_orf1ab;5856 | 0.05 | 0.025 |
| MT072864.1;QIQ54047.1;Pangolin_CoV_orf1ab;5857 | 0.05 | 0 |
| MT072864.1;QIQ54047.1;Pangolin_CoV_orf1ab;5858 | 0.05 | 0 |
| MT072864.1;QIQ54047.1;Pangolin_CoV_orf1ab;5859 | 0.05 | 0 |
| MT072864.1;QIQ54047.1;Pangolin_CoV_orf1ab;5860 | 0.05 | 0 |
| MT072864.1;QIQ54047.1;Pangolin_CoV_orf1ab;5861 | 0 | 0 |
| MT072864.1;QIQ54047.1;Pangolin_CoV_orf1ab;5862 | 0 | 0 |
| MT072864.1;QIQ54047.1;Pangolin_CoV_orf1ab;5863 | 0.05 | 0 |
| MT072864.1;QIQ54047.1;Pangolin_CoV_orf1ab;5864 | 0.05 | 0 |
| MT072864.1;QIQ54047.1;Pangolin_CoV_orf1ab;5865 | 0.05 | 0.05 |
| MT072864.1;QIQ54047.1;Pangolin_CoV_orf1ab;5866 | 0 | 0 |
| MT072864.1;QIQ54047.1;Pangolin_CoV_orf1ab;5867 | 0 | 0 |
| MT072864.1;QIQ54047.1;Pangolin_CoV_orf1ab;5868 | 0 | 0 |
| MT072864.1;QIQ54047.1;Pangolin_CoV_orf1ab;5869 | 0 | 0 |
| MT072864.1;QIQ54047.1;Pangolin_CoV_orf1ab;5870 | 0 | 0 |
| MT072864.1;QIQ54047.1;Pangolin_CoV_orf1ab;5871 | 0 | 0 |
| MT072864.1;QIQ54047.1;Pangolin_CoV_orf1ab;5872 | 0 | 0 |
| MT072864.1;QIQ54047.1;Pangolin_CoV_orf1ab;5873 | 0 | 0 |
| MT072864.1;QIQ54047.1;Pangolin_CoV_orf1ab;5874 | 0 | 0 |
| MT072864.1;QIQ54047.1;Pangolin_CoV_orf1ab;5875 | 0 | 0 |
| MT072864.1;QIQ54047.1;Pangolin_CoV_orf1ab;5876 | 0 | 0 |
| MT072864.1;QIQ54047.1;Pangolin_CoV_orf1ab;5877 | 0 | 0 |
| MT072864.1;QIQ54047.1;Pangolin_CoV_orf1ab;5878 | 0 | 0 |
| MT072864.1;QIQ54047.1;Pangolin_CoV_orf1ab;5879 | 0 | 0 |
| MT072864.1;QIQ54047.1;Pangolin_CoV_orf1ab;5880 | 0 | 0 |
| MT072864.1;QIQ54047.1;Pangolin_CoV_orf1ab;5881 | 0 | 0 |

| | | |
|---|---|---|
| MT072864.1;QIQ54047.1;Pangolin_CoV_orf1ab;5882 | 0 | 0 |
| MT072864.1;QIQ54047.1;Pangolin_CoV_orf1ab;5883 | 0 | 0 |
| MT072864.1;QIQ54047.1;Pangolin_CoV_orf1ab;5884 | 0 | 0.025 |
| MT072864.1;QIQ54047.1;Pangolin_CoV_orf1ab;5885 | 0 | 0 |
| MT072864.1;QIQ54047.1;Pangolin_CoV_orf1ab;5886 | 0 | 0 |
| MT072864.1;QIQ54047.1;Pangolin_CoV_orf1ab;5887 | 0 | 0.05 |
| MT072864.1;QIQ54047.1;Pangolin_CoV_orf1ab;5888 | 0 | 0.05 |
| MT072864.1;QIQ54047.1;Pangolin_CoV_orf1ab;5889 | 0.05 | 0.05 |
| MT072864.1;QIQ54047.1;Pangolin_CoV_orf1ab;5890 | 0.05 | 0.05 |
| MT072864.1;QIQ54047.1;Pangolin_CoV_orf1ab;5891 | 0.1 | 0.075 |
| MT072864.1;QIQ54047.1;Pangolin_CoV_orf1ab;5892 | 0 | 0.025 |
| MT072864.1;QIQ54047.1;Pangolin_CoV_orf1ab;5893 | 0 | 0 |
| MT072864.1;QIQ54047.1;Pangolin_CoV_orf1ab;5894 | 0 | 0 |
| MT072864.1;QIQ54047.1;Pangolin_CoV_orf1ab;5895 | 0 | 0.05 |
| MT072864.1;QIQ54047.1;Pangolin_CoV_orf1ab;5896 | 0 | 0.05 |
| MT072864.1;QIQ54047.1;Pangolin_CoV_orf1ab;5897 | 0 | 0 |
| MT072864.1;QIQ54047.1;Pangolin_CoV_orf1ab;5898 | 0 | 0 |
| MT072864.1;QIQ54047.1;Pangolin_CoV_orf1ab;5899 | 0 | 0 |
| MT072864.1;QIQ54047.1;Pangolin_CoV_orf1ab;5900 | 0 | 0.025 |
| MT072864.1;QIQ54047.1;Pangolin_CoV_orf1ab;5901 | 0 | 0.025 |
| MT072864.1;QIQ54047.1;Pangolin_CoV_orf1ab;5902 | 0 | 0.025 |
| MT072864.1;QIQ54047.1;Pangolin_CoV_orf1ab;5903 | 0.05 | 0.025 |
| MT072864.1;QIQ54047.1;Pangolin_CoV_orf1ab;5904 | 0.05 | 0.05 |
| MT072864.1;QIQ54047.1;Pangolin_CoV_orf1ab;5905 | 0.05 | 0.025 |
| MT072864.1;QIQ54047.1;Pangolin_CoV_orf1ab;5906 | 0.05 | 0.025 |
| MT072864.1;QIQ54047.1;Pangolin_CoV_orf1ab;5907 | 0.05 | 0.025 |
| MT072864.1;QIQ54047.1;Pangolin_CoV_orf1ab;5908 | 0.1 | 0.025 |
| MT072864.1;QIQ54047.1;Pangolin_CoV_orf1ab;5909 | 0.15 | 0.025 |
| MT072864.1;QIQ54047.1;Pangolin_CoV_orf1ab;5910 | 0.05 | 0 |
| MT072864.1;QIQ54047.1;Pangolin_CoV_orf1ab;5911 | 0 | 0 |
| MT072864.1;QIQ54047.1;Pangolin_CoV_orf1ab;5912 | 0 | 0 |
| MT072864.1;QIQ54047.1;Pangolin_CoV_orf1ab;5913 | 0.05 | 0 |
| MT072864.1;QIQ54047.1;Pangolin_CoV_orf1ab;5914 | 0 | 0 |
| MT072864.1;QIQ54047.1;Pangolin_CoV_orf1ab;5915 | 0 | 0 |
| MT072864.1;QIQ54047.1;Pangolin_CoV_orf1ab;5916 | 0 | 0 |
| MT072864.1;QIQ54047.1;Pangolin_CoV_orf1ab;5917 | 0 | 0 |
| MT072864.1;QIQ54047.1;Pangolin_CoV_orf1ab;5918 | 0 | 0 |
| MT072864.1;QIQ54047.1;Pangolin_CoV_orf1ab;5919 | 0 | 0 |
| MT072864.1;QIQ54047.1;Pangolin_CoV_orf1ab;5920 | 0 | 0 |
| MT072864.1;QIQ54047.1;Pangolin_CoV_orf1ab;5921 | 0 | 0 |
| MT072864.1;QIQ54047.1;Pangolin_CoV_orf1ab;5922 | 0 | 0 |
| MT072864.1;QIQ54047.1;Pangolin_CoV_orf1ab;5923 | 0 | 0 |
| MT072864.1;QIQ54047.1;Pangolin_CoV_orf1ab;5924 | 0 | 0 |

| | | |
|---|---|---|
| MT072864.1;QIQ54047.1;Pangolin_CoV_orf1ab;5925 | 0 | 0 |
| MT072864.1;QIQ54047.1;Pangolin_CoV_orf1ab;5926 | 0.05 | 0 |
| MT072864.1;QIQ54047.1;Pangolin_CoV_orf1ab;5927 | 0 | 0 |
| MT072864.1;QIQ54047.1;Pangolin_CoV_orf1ab;5928 | 0.05 | 0 |
| MT072864.1;QIQ54047.1;Pangolin_CoV_orf1ab;5929 | 0 | 0.025 |
| MT072864.1;QIQ54047.1;Pangolin_CoV_orf1ab;5930 | 0 | 0 |
| MT072864.1;QIQ54047.1;Pangolin_CoV_orf1ab;5931 | 0 | 0 |
| MT072864.1;QIQ54047.1;Pangolin_CoV_orf1ab;5932 | 0 | 0 |
| MT072864.1;QIQ54047.1;Pangolin_CoV_orf1ab;5933 | 0 | 0 |
| MT072864.1;QIQ54047.1;Pangolin_CoV_orf1ab;5934 | 0 | 0 |
| MT072864.1;QIQ54047.1;Pangolin_CoV_orf1ab;5935 | 0 | 0 |
| MT072864.1;QIQ54047.1;Pangolin_CoV_orf1ab;5936 | 0 | 0 |
| MT072864.1;QIQ54047.1;Pangolin_CoV_orf1ab;5937 | 0 | 0 |
| MT072864.1;QIQ54047.1;Pangolin_CoV_orf1ab;5938 | 0 | 0 |
| MT072864.1;QIQ54047.1;Pangolin_CoV_orf1ab;5939 | 0 | 0 |
| MT072864.1;QIQ54047.1;Pangolin_CoV_orf1ab;5940 | 0 | 0 |
| MT072864.1;QIQ54047.1;Pangolin_CoV_orf1ab;5941 | 0 | 0 |
| MT072864.1;QIQ54047.1;Pangolin_CoV_orf1ab;5942 | 0 | 0 |
| MT072864.1;QIQ54047.1;Pangolin_CoV_orf1ab;5943 | 0 | 0 |
| MT072864.1;QIQ54047.1;Pangolin_CoV_orf1ab;5944 | 0 | 0.025 |
| MT072864.1;QIQ54047.1;Pangolin_CoV_orf1ab;5945 | 0 | 0 |
| MT072864.1;QIQ54047.1;Pangolin_CoV_orf1ab;5946 | 0 | 0 |
| MT072864.1;QIQ54047.1;Pangolin_CoV_orf1ab;5947 | 0 | 0 |
| MT072864.1;QIQ54047.1;Pangolin_CoV_orf1ab;5948 | 0 | 0 |
| MT072864.1;QIQ54047.1;Pangolin_CoV_orf1ab;5949 | 0 | 0 |
| MT072864.1;QIQ54047.1;Pangolin_CoV_orf1ab;5950 | 0 | 0 |
| MT072864.1;QIQ54047.1;Pangolin_CoV_orf1ab;5951 | 0 | 0 |
| MT072864.1;QIQ54047.1;Pangolin_CoV_orf1ab;5952 | 0 | 0 |
| MT072864.1;QIQ54047.1;Pangolin_CoV_orf1ab;5953 | 0 | 0 |
| MT072864.1;QIQ54047.1;Pangolin_CoV_orf1ab;5954 | 0 | 0 |
| MT072864.1;QIQ54047.1;Pangolin_CoV_orf1ab;5955 | 0 | 0 |
| MT072864.1;QIQ54047.1;Pangolin_CoV_orf1ab;5956 | 0 | 0 |
| MT072864.1;QIQ54047.1;Pangolin_CoV_orf1ab;5957 | 0 | 0 |
| MT072864.1;QIQ54047.1;Pangolin_CoV_orf1ab;5958 | 0 | 0.025 |
| MT072864.1;QIQ54047.1;Pangolin_CoV_orf1ab;5959 | 0 | 0 |
| MT072864.1;QIQ54047.1;Pangolin_CoV_orf1ab;5960 | 0 | 0 |
| MT072864.1;QIQ54047.1;Pangolin_CoV_orf1ab;5961 | 0 | 0 |
| MT072864.1;QIQ54047.1;Pangolin_CoV_orf1ab;5962 | 0 | 0 |
| MT072864.1;QIQ54047.1;Pangolin_CoV_orf1ab;5963 | 0 | 0 |
| MT072864.1;QIQ54047.1;Pangolin_CoV_orf1ab;5964 | 0 | 0 |
| MT072864.1;QIQ54047.1;Pangolin_CoV_orf1ab;5965 | 0 | 0 |
| MT072864.1;QIQ54047.1;Pangolin_CoV_orf1ab;5966 | 0 | 0 |
| MT072864.1;QIQ54047.1;Pangolin_CoV_orf1ab;5967 | 0 | 0 |

| | | |
|---|---|---|
| MT072864.1;QIQ54047.1;Pangolin_CoV_orf1ab;5968 | 0 | 0 |
| MT072864.1;QIQ54047.1;Pangolin_CoV_orf1ab;5969 | 0 | 0 |
| MT072864.1;QIQ54047.1;Pangolin_CoV_orf1ab;5970 | 0 | 0 |
| MT072864.1;QIQ54047.1;Pangolin_CoV_orf1ab;5971 | 0 | 0.025 |
| MT072864.1;QIQ54047.1;Pangolin_CoV_orf1ab;5972 | 0 | 0.05 |
| MT072864.1;QIQ54047.1;Pangolin_CoV_orf1ab;5973 | 0 | 0.05 |
| MT072864.1;QIQ54047.1;Pangolin_CoV_orf1ab;5974 | 0.05 | 0.05 |
| MT072864.1;QIQ54047.1;Pangolin_CoV_orf1ab;5975 | 0.1 | 0.05 |
| MT072864.1;QIQ54047.1;Pangolin_CoV_orf1ab;5976 | 0.05 | 0.05 |
| MT072864.1;QIQ54047.1;Pangolin_CoV_orf1ab;5977 | 0 | 0.05 |
| MT072864.1;QIQ54047.1;Pangolin_CoV_orf1ab;5978 | 0 | 0.025 |
| MT072864.1;QIQ54047.1;Pangolin_CoV_orf1ab;5979 | 0 | 0 |
| MT072864.1;QIQ54047.1;Pangolin_CoV_orf1ab;5980 | 0 | 0 |
| MT072864.1;QIQ54047.1;Pangolin_CoV_orf1ab;5981 | 0.05 | 0 |
| MT072864.1;QIQ54047.1;Pangolin_CoV_orf1ab;5982 | 0 | 0 |
| MT072864.1;QIQ54047.1;Pangolin_CoV_orf1ab;5983 | 0.05 | 0 |
| MT072864.1;QIQ54047.1;Pangolin_CoV_orf1ab;5984 | 0 | 0 |
| MT072864.1;QIQ54047.1;Pangolin_CoV_orf1ab;5985 | 0 | 0 |
| MT072864.1;QIQ54047.1;Pangolin_CoV_orf1ab;5986 | 0 | 0 |
| MT072864.1;QIQ54047.1;Pangolin_CoV_orf1ab;5987 | 0 | 0.025 |
| MT072864.1;QIQ54047.1;Pangolin_CoV_orf1ab;5988 | 0 | 0.025 |
| MT072864.1;QIQ54047.1;Pangolin_CoV_orf1ab;5989 | 0 | 0.025 |
| MT072864.1;QIQ54047.1;Pangolin_CoV_orf1ab;5990 | 0 | 0.025 |
| MT072864.1;QIQ54047.1;Pangolin_CoV_orf1ab;5991 | 0 | 0.075 |
| MT072864.1;QIQ54047.1;Pangolin_CoV_orf1ab;5992 | 0 | 0.025 |
| MT072864.1;QIQ54047.1;Pangolin_CoV_orf1ab;5993 | 0 | 0 |
| MT072864.1;QIQ54047.1;Pangolin_CoV_orf1ab;5994 | 0 | 0 |
| MT072864.1;QIQ54047.1;Pangolin_CoV_orf1ab;5995 | 0 | 0 |
| MT072864.1;QIQ54047.1;Pangolin_CoV_orf1ab;5996 | 0 | 0 |
| MT072864.1;QIQ54047.1;Pangolin_CoV_orf1ab;5997 | 0 | 0 |
| MT072864.1;QIQ54047.1;Pangolin_CoV_orf1ab;5998 | 0 | 0 |
| MT072864.1;QIQ54047.1;Pangolin_CoV_orf1ab;5999 | 0 | 0 |
| MT072864.1;QIQ54047.1;Pangolin_CoV_orf1ab;6000 | 0 | 0 |
| MT072864.1;QIQ54047.1;Pangolin_CoV_orf1ab;6001 | 0 | 0 |
| MT072864.1;QIQ54047.1;Pangolin_CoV_orf1ab;6002 | 0 | 0 |
| MT072864.1;QIQ54047.1;Pangolin_CoV_orf1ab;6003 | 0 | 0 |
| MT072864.1;QIQ54047.1;Pangolin_CoV_orf1ab;6004 | 0 | 0 |
| MT072864.1;QIQ54047.1;Pangolin_CoV_orf1ab;6005 | 0 | 0 |
| MT072864.1;QIQ54047.1;Pangolin_CoV_orf1ab;6006 | 0 | 0 |
| MT072864.1;QIQ54047.1;Pangolin_CoV_orf1ab;6007 | 0 | 0 |
| MT072864.1;QIQ54047.1;Pangolin_CoV_orf1ab;6008 | 0 | 0 |
| MT072864.1;QIQ54047.1;Pangolin_CoV_orf1ab;6009 | 0 | 0.025 |
| MT072864.1;QIQ54047.1;Pangolin_CoV_orf1ab;6010 | 0 | 0.025 |

| | | |
|---|---|---|
| MT072864.1;QIQ54047.1;Pangolin_CoV_orf1ab;6011 | 0 | 0 |
| MT072864.1;QIQ54047.1;Pangolin_CoV_orf1ab;6012 | 0 | 0 |
| MT072864.1;QIQ54047.1;Pangolin_CoV_orf1ab;6013 | 0 | 0 |
| MT072864.1;QIQ54047.1;Pangolin_CoV_orf1ab;6014 | 0 | 0.025 |
| MT072864.1;QIQ54047.1;Pangolin_CoV_orf1ab;6015 | 0 | 0 |
| MT072864.1;QIQ54047.1;Pangolin_CoV_orf1ab;6016 | 0 | 0 |
| MT072864.1;QIQ54047.1;Pangolin_CoV_orf1ab;6017 | 0 | 0.025 |
| MT072864.1;QIQ54047.1;Pangolin_CoV_orf1ab;6018 | 0 | 0 |
| MT072864.1;QIQ54047.1;Pangolin_CoV_orf1ab;6019 | 0 | 0 |
| MT072864.1;QIQ54047.1;Pangolin_CoV_orf1ab;6020 | 0 | 0 |
| MT072864.1;QIQ54047.1;Pangolin_CoV_orf1ab;6021 | 0 | 0.025 |
| MT072864.1;QIQ54047.1;Pangolin_CoV_orf1ab;6022 | 0 | 0 |
| MT072864.1;QIQ54047.1;Pangolin_CoV_orf1ab;6023 | 0 | 0 |
| MT072864.1;QIQ54047.1;Pangolin_CoV_orf1ab;6024 | 0 | 0 |
| MT072864.1;QIQ54047.1;Pangolin_CoV_orf1ab;6025 | 0 | 0 |
| MT072864.1;QIQ54047.1;Pangolin_CoV_orf1ab;6026 | 0 | 0 |
| MT072864.1;QIQ54047.1;Pangolin_CoV_orf1ab;6027 | 0 | 0 |
| MT072864.1;QIQ54047.1;Pangolin_CoV_orf1ab;6028 | 0 | 0 |
| MT072864.1;QIQ54047.1;Pangolin_CoV_orf1ab;6029 | 0 | 0 |
| MT072864.1;QIQ54047.1;Pangolin_CoV_orf1ab;6030 | 0 | 0 |
| MT072864.1;QIQ54047.1;Pangolin_CoV_orf1ab;6031 | 0 | 0 |
| MT072864.1;QIQ54047.1;Pangolin_CoV_orf1ab;6032 | 0 | 0 |
| MT072864.1;QIQ54047.1;Pangolin_CoV_orf1ab;6033 | 0 | 0 |
| MT072864.1;QIQ54047.1;Pangolin_CoV_orf1ab;6034 | 0 | 0 |
| MT072864.1;QIQ54047.1;Pangolin_CoV_orf1ab;6035 | 0 | 0 |
| MT072864.1;QIQ54047.1;Pangolin_CoV_orf1ab;6036 | 0 | 0 |
| MT072864.1;QIQ54047.1;Pangolin_CoV_orf1ab;6037 | 0.05 | 0 |
| MT072864.1;QIQ54047.1;Pangolin_CoV_orf1ab;6038 | 0 | 0 |
| MT072864.1;QIQ54047.1;Pangolin_CoV_orf1ab;6039 | 0 | 0 |
| MT072864.1;QIQ54047.1;Pangolin_CoV_orf1ab;6040 | 0 | 0 |
| MT072864.1;QIQ54047.1;Pangolin_CoV_orf1ab;6041 | 0 | 0 |
| MT072864.1;QIQ54047.1;Pangolin_CoV_orf1ab;6042 | 0.05 | 0 |
| MT072864.1;QIQ54047.1;Pangolin_CoV_orf1ab;6043 | 0 | 0.025 |
| MT072864.1;QIQ54047.1;Pangolin_CoV_orf1ab;6044 | 0.05 | 0.05 |
| MT072864.1;QIQ54047.1;Pangolin_CoV_orf1ab;6045 | 0 | 0 |
| MT072864.1;QIQ54047.1;Pangolin_CoV_orf1ab;6046 | 0.05 | 0.05 |
| MT072864.1;QIQ54047.1;Pangolin_CoV_orf1ab;6047 | 0.05 | 0.05 |
| MT072864.1;QIQ54047.1;Pangolin_CoV_orf1ab;6048 | 0.05 | 0.075 |
| MT072864.1;QIQ54047.1;Pangolin_CoV_orf1ab;6049 | 0.05 | 0.075 |
| MT072864.1;QIQ54047.1;Pangolin_CoV_orf1ab;6050 | 0 | 0.1 |
| MT072864.1;QIQ54047.1;Pangolin_CoV_orf1ab;6051 | 0 | 0 |
| MT072864.1;QIQ54047.1;Pangolin_CoV_orf1ab;6052 | 0 | 0 |
| MT072864.1;QIQ54047.1;Pangolin_CoV_orf1ab;6053 | 0 | 0 |

| | | |
|---|---|---|
| MT072864.1;QIQ54047.1;Pangolin_CoV_orf1ab;6054 | 0 | 0 |
| MT072864.1;QIQ54047.1;Pangolin_CoV_orf1ab;6055 | 0 | 0 |
| MT072864.1;QIQ54047.1;Pangolin_CoV_orf1ab;6056 | 0 | 0 |
| MT072864.1;QIQ54047.1;Pangolin_CoV_orf1ab;6057 | 0 | 0 |
| MT072864.1;QIQ54047.1;Pangolin_CoV_orf1ab;6058 | 0 | 0 |
| MT072864.1;QIQ54047.1;Pangolin_CoV_orf1ab;6059 | 0 | 0 |
| MT072864.1;QIQ54047.1;Pangolin_CoV_orf1ab;6060 | 0 | 0 |
| MT072864.1;QIQ54047.1;Pangolin_CoV_orf1ab;6061 | 0 | 0 |
| MT072864.1;QIQ54047.1;Pangolin_CoV_orf1ab;6062 | 0 | 0 |
| MT072864.1;QIQ54047.1;Pangolin_CoV_orf1ab;6063 | 0 | 0 |
| MT072864.1;QIQ54047.1;Pangolin_CoV_orf1ab;6064 | 0 | 0 |
| MT072864.1;QIQ54047.1;Pangolin_CoV_orf1ab;6065 | 0 | 0 |
| MT072864.1;QIQ54047.1;Pangolin_CoV_orf1ab;6066 | 0 | 0 |
| MT072864.1;QIQ54047.1;Pangolin_CoV_orf1ab;6067 | 0 | 0 |
| MT072864.1;QIQ54047.1;Pangolin_CoV_orf1ab;6068 | 0 | 0 |
| MT072864.1;QIQ54047.1;Pangolin_CoV_orf1ab;6069 | 0 | 0 |
| MT072864.1;QIQ54047.1;Pangolin_CoV_orf1ab;6070 | 0 | 0 |
| MT072864.1;QIQ54047.1;Pangolin_CoV_orf1ab;6071 | 0 | 0 |
| MT072864.1;QIQ54047.1;Pangolin_CoV_orf1ab;6072 | 0 | 0 |
| MT072864.1;QIQ54047.1;Pangolin_CoV_orf1ab;6073 | 0 | 0 |
| MT072864.1;QIQ54047.1;Pangolin_CoV_orf1ab;6074 | 0 | 0 |
| MT072864.1;QIQ54047.1;Pangolin_CoV_orf1ab;6075 | 0 | 0 |
| MT072864.1;QIQ54047.1;Pangolin_CoV_orf1ab;6076 | 0 | 0 |
| MT072864.1;QIQ54047.1;Pangolin_CoV_orf1ab;6077 | 0 | 0 |
| MT072864.1;QIQ54047.1;Pangolin_CoV_orf1ab;6078 | 0 | 0 |
| MT072864.1;QIQ54047.1;Pangolin_CoV_orf1ab;6079 | 0 | 0 |
| MT072864.1;QIQ54047.1;Pangolin_CoV_orf1ab;6080 | 0 | 0 |
| MT072864.1;QIQ54047.1;Pangolin_CoV_orf1ab;6081 | 0 | 0 |
| MT072864.1;QIQ54047.1;Pangolin_CoV_orf1ab;6082 | 0 | 0 |
| MT072864.1;QIQ54047.1;Pangolin_CoV_orf1ab;6083 | 0 | 0.025 |
| MT072864.1;QIQ54047.1;Pangolin_CoV_orf1ab;6084 | 0 | 0 |
| MT072864.1;QIQ54047.1;Pangolin_CoV_orf1ab;6085 | 0 | 0.025 |
| MT072864.1;QIQ54047.1;Pangolin_CoV_orf1ab;6086 | 0 | 0.025 |
| MT072864.1;QIQ54047.1;Pangolin_CoV_orf1ab;6087 | 0 | 0 |
| MT072864.1;QIQ54047.1;Pangolin_CoV_orf1ab;6088 | 0 | 0 |
| MT072864.1;QIQ54047.1;Pangolin_CoV_orf1ab;6089 | 0 | 0 |
| MT072864.1;QIQ54047.1;Pangolin_CoV_orf1ab;6090 | 0 | 0 |
| MT072864.1;QIQ54047.1;Pangolin_CoV_orf1ab;6091 | 0 | 0 |
| MT072864.1;QIQ54047.1;Pangolin_CoV_orf1ab;6092 | 0 | 0 |
| MT072864.1;QIQ54047.1;Pangolin_CoV_orf1ab;6093 | 0 | 0 |
| MT072864.1;QIQ54047.1;Pangolin_CoV_orf1ab;6094 | 0 | 0 |
| MT072864.1;QIQ54047.1;Pangolin_CoV_orf1ab;6095 | 0 | 0 |
| MT072864.1;QIQ54047.1;Pangolin_CoV_orf1ab;6096 | 0 | 0 |

| | | |
|---|---|---|
| MT072864.1;QIQ54047.1;Pangolin_CoV_orf1ab;6097 | 0 | 0 |
| MT072864.1;QIQ54047.1;Pangolin_CoV_orf1ab;6098 | 0 | 0 |
| MT072864.1;QIQ54047.1;Pangolin_CoV_orf1ab;6099 | 0 | 0 |
| MT072864.1;QIQ54047.1;Pangolin_CoV_orf1ab;6100 | 0 | 0 |
| MT072864.1;QIQ54047.1;Pangolin_CoV_orf1ab;6101 | 0 | 0 |
| MT072864.1;QIQ54047.1;Pangolin_CoV_orf1ab;6102 | 0 | 0 |
| MT072864.1;QIQ54047.1;Pangolin_CoV_orf1ab;6103 | 0 | 0 |
| MT072864.1;QIQ54047.1;Pangolin_CoV_orf1ab;6104 | 0 | 0 |
| MT072864.1;QIQ54047.1;Pangolin_CoV_orf1ab;6105 | 0 | 0 |
| MT072864.1;QIQ54047.1;Pangolin_CoV_orf1ab;6106 | 0 | 0 |
| MT072864.1;QIQ54047.1;Pangolin_CoV_orf1ab;6107 | 0 | 0 |
| MT072864.1;QIQ54047.1;Pangolin_CoV_orf1ab;6108 | 0.05 | 0 |
| MT072864.1;QIQ54047.1;Pangolin_CoV_orf1ab;6109 | 0 | 0 |
| MT072864.1;QIQ54047.1;Pangolin_CoV_orf1ab;6110 | 0 | 0 |
| MT072864.1;QIQ54047.1;Pangolin_CoV_orf1ab;6111 | 0 | 0 |
| MT072864.1;QIQ54047.1;Pangolin_CoV_orf1ab;6112 | 0 | 0 |
| MT072864.1;QIQ54047.1;Pangolin_CoV_orf1ab;6113 | 0 | 0 |
| MT072864.1;QIQ54047.1;Pangolin_CoV_orf1ab;6114 | 0 | 0 |
| MT072864.1;QIQ54047.1;Pangolin_CoV_orf1ab;6115 | 0 | 0 |
| MT072864.1;QIQ54047.1;Pangolin_CoV_orf1ab;6116 | 0 | 0.025 |
| MT072864.1;QIQ54047.1;Pangolin_CoV_orf1ab;6117 | 0 | 0 |
| MT072864.1;QIQ54047.1;Pangolin_CoV_orf1ab;6118 | 0.05 | 0 |
| MT072864.1;QIQ54047.1;Pangolin_CoV_orf1ab;6119 | 0.05 | 0 |
| MT072864.1;QIQ54047.1;Pangolin_CoV_orf1ab;6120 | 0 | 0 |
| MT072864.1;QIQ54047.1;Pangolin_CoV_orf1ab;6121 | 0 | 0 |
| MT072864.1;QIQ54047.1;Pangolin_CoV_orf1ab;6122 | 0 | 0 |
| MT072864.1;QIQ54047.1;Pangolin_CoV_orf1ab;6123 | 0 | 0 |
| MT072864.1;QIQ54047.1;Pangolin_CoV_orf1ab;6124 | 0 | 0 |
| MT072864.1;QIQ54047.1;Pangolin_CoV_orf1ab;6125 | 0 | 0 |
| MT072864.1;QIQ54047.1;Pangolin_CoV_orf1ab;6126 | 0 | 0 |
| MT072864.1;QIQ54047.1;Pangolin_CoV_orf1ab;6127 | 0 | 0 |
| MT072864.1;QIQ54047.1;Pangolin_CoV_orf1ab;6128 | 0 | 0 |
| MT072864.1;QIQ54047.1;Pangolin_CoV_orf1ab;6129 | 0 | 0 |
| MT072864.1;QIQ54047.1;Pangolin_CoV_orf1ab;6130 | 0 | 0 |
| MT072864.1;QIQ54047.1;Pangolin_CoV_orf1ab;6131 | 0 | 0 |
| MT072864.1;QIQ54047.1;Pangolin_CoV_orf1ab;6132 | 0 | 0 |
| MT072864.1;QIQ54047.1;Pangolin_CoV_orf1ab;6133 | 0 | 0 |
| MT072864.1;QIQ54047.1;Pangolin_CoV_orf1ab;6134 | 0 | 0 |
| MT072864.1;QIQ54047.1;Pangolin_CoV_orf1ab;6135 | 0 | 0 |
| MT072864.1;QIQ54047.1;Pangolin_CoV_orf1ab;6136 | 0 | 0 |
| MT072864.1;QIQ54047.1;Pangolin_CoV_orf1ab;6137 | 0 | 0 |
| MT072864.1;QIQ54047.1;Pangolin_CoV_orf1ab;6138 | 0 | 0 |
| MT072864.1;QIQ54047.1;Pangolin_CoV_orf1ab;6139 | 0 | 0 |

| | | |
|---|---|---|
| MT072864.1;QIQ54047.1;Pangolin_CoV_orf1ab;6140 | 0 | 0 |
| MT072864.1;QIQ54047.1;Pangolin_CoV_orf1ab;6141 | 0 | 0 |
| MT072864.1;QIQ54047.1;Pangolin_CoV_orf1ab;6142 | 0 | 0 |
| MT072864.1;QIQ54047.1;Pangolin_CoV_orf1ab;6143 | 0 | 0 |
| MT072864.1;QIQ54047.1;Pangolin_CoV_orf1ab;6144 | 0 | 0 |
| MT072864.1;QIQ54047.1;Pangolin_CoV_orf1ab;6145 | 0 | 0 |
| MT072864.1;QIQ54047.1;Pangolin_CoV_orf1ab;6146 | 0 | 0 |
| MT072864.1;QIQ54047.1;Pangolin_CoV_orf1ab;6147 | 0 | 0 |
| MT072864.1;QIQ54047.1;Pangolin_CoV_orf1ab;6148 | 0 | 0 |
| MT072864.1;QIQ54047.1;Pangolin_CoV_orf1ab;6149 | 0 | 0.05 |
| MT072864.1;QIQ54047.1;Pangolin_CoV_orf1ab;6150 | 0 | 0 |
| MT072864.1;QIQ54047.1;Pangolin_CoV_orf1ab;6151 | 0 | 0 |
| MT072864.1;QIQ54047.1;Pangolin_CoV_orf1ab;6152 | 0 | 0 |
| MT072864.1;QIQ54047.1;Pangolin_CoV_orf1ab;6153 | 0 | 0 |
| MT072864.1;QIQ54047.1;Pangolin_CoV_orf1ab;6154 | 0 | 0 |
| MT072864.1;QIQ54047.1;Pangolin_CoV_orf1ab;6155 | 0 | 0 |
| MT072864.1;QIQ54047.1;Pangolin_CoV_orf1ab;6156 | 0 | 0 |
| MT072864.1;QIQ54047.1;Pangolin_CoV_orf1ab;6157 | 0 | 0 |
| MT072864.1;QIQ54047.1;Pangolin_CoV_orf1ab;6158 | 0 | 0 |
| MT072864.1;QIQ54047.1;Pangolin_CoV_orf1ab;6159 | 0 | 0 |
| MT072864.1;QIQ54047.1;Pangolin_CoV_orf1ab;6160 | 0 | 0 |
| MT072864.1;QIQ54047.1;Pangolin_CoV_orf1ab;6161 | 0 | 0 |
| MT072864.1;QIQ54047.1;Pangolin_CoV_orf1ab;6162 | 0 | 0 |
| MT072864.1;QIQ54047.1;Pangolin_CoV_orf1ab;6163 | 0 | 0 |
| MT072864.1;QIQ54047.1;Pangolin_CoV_orf1ab;6164 | 0 | 0 |
| MT072864.1;QIQ54047.1;Pangolin_CoV_orf1ab;6165 | 0 | 0 |
| MT072864.1;QIQ54047.1;Pangolin_CoV_orf1ab;6166 | 0 | 0 |
| MT072864.1;QIQ54047.1;Pangolin_CoV_orf1ab;6167 | 0 | 0 |
| MT072864.1;QIQ54047.1;Pangolin_CoV_orf1ab;6168 | 0 | 0 |
| MT072864.1;QIQ54047.1;Pangolin_CoV_orf1ab;6169 | 0 | 0.025 |
| MT072864.1;QIQ54047.1;Pangolin_CoV_orf1ab;6170 | 0 | 0 |
| MT072864.1;QIQ54047.1;Pangolin_CoV_orf1ab;6171 | 0.05 | 0 |
| MT072864.1;QIQ54047.1;Pangolin_CoV_orf1ab;6172 | 0.1 | 0.05 |
| MT072864.1;QIQ54047.1;Pangolin_CoV_orf1ab;6173 | 0 | 0 |
| MT072864.1;QIQ54047.1;Pangolin_CoV_orf1ab;6174 | 0 | 0 |
| MT072864.1;QIQ54047.1;Pangolin_CoV_orf1ab;6175 | 0 | 0 |
| MT072864.1;QIQ54047.1;Pangolin_CoV_orf1ab;6176 | 0 | 0 |
| MT072864.1;QIQ54047.1;Pangolin_CoV_orf1ab;6177 | 0 | 0 |
| MT072864.1;QIQ54047.1;Pangolin_CoV_orf1ab;6178 | 0 | 0 |
| MT072864.1;QIQ54047.1;Pangolin_CoV_orf1ab;6179 | 0 | 0 |
| MT072864.1;QIQ54047.1;Pangolin_CoV_orf1ab;6180 | 0 | 0 |
| MT072864.1;QIQ54047.1;Pangolin_CoV_orf1ab;6181 | 0 | 0 |
| MT072864.1;QIQ54047.1;Pangolin_CoV_orf1ab;6182 | 0 | 0 |

| | | |
|---|---|---|
| MT072864.1;QIQ54047.1;Pangolin_CoV_orf1ab;6183 | 0 | 0 |
| MT072864.1;QIQ54047.1;Pangolin_CoV_orf1ab;6184 | 0 | 0 |
| MT072864.1;QIQ54047.1;Pangolin_CoV_orf1ab;6185 | 0 | 0 |
| MT072864.1;QIQ54047.1;Pangolin_CoV_orf1ab;6186 | 0 | 0 |
| MT072864.1;QIQ54047.1;Pangolin_CoV_orf1ab;6187 | 0 | 0 |
| MT072864.1;QIQ54047.1;Pangolin_CoV_orf1ab;6188 | 0 | 0 |
| MT072864.1;QIQ54047.1;Pangolin_CoV_orf1ab;6189 | 0 | 0 |
| MT072864.1;QIQ54047.1;Pangolin_CoV_orf1ab;6190 | 0 | 0 |
| MT072864.1;QIQ54047.1;Pangolin_CoV_orf1ab;6191 | 0 | 0 |
| MT072864.1;QIQ54047.1;Pangolin_CoV_orf1ab;6192 | 0 | 0 |
| MT072864.1;QIQ54047.1;Pangolin_CoV_orf1ab;6193 | 0 | 0 |
| MT072864.1;QIQ54047.1;Pangolin_CoV_orf1ab;6194 | 0 | 0 |
| MT072864.1;QIQ54047.1;Pangolin_CoV_orf1ab;6195 | 0 | 0 |
| MT072864.1;QIQ54047.1;Pangolin_CoV_orf1ab;6196 | 0 | 0 |
| MT072864.1;QIQ54047.1;Pangolin_CoV_orf1ab;6197 | 0 | 0 |
| MT072864.1;QIQ54047.1;Pangolin_CoV_orf1ab;6198 | 0 | 0 |
| MT072864.1;QIQ54047.1;Pangolin_CoV_orf1ab;6199 | 0 | 0 |
| MT072864.1;QIQ54047.1;Pangolin_CoV_orf1ab;6200 | 0 | 0 |
| MT072864.1;QIQ54047.1;Pangolin_CoV_orf1ab;6201 | 0 | 0 |
| MT072864.1;QIQ54047.1;Pangolin_CoV_orf1ab;6202 | 0 | 0 |
| MT072864.1;QIQ54047.1;Pangolin_CoV_orf1ab;6203 | 0 | 0 |
| MT072864.1;QIQ54047.1;Pangolin_CoV_orf1ab;6204 | 0 | 0.025 |
| MT072864.1;QIQ54047.1;Pangolin_CoV_orf1ab;6205 | 0 | 0.025 |
| MT072864.1;QIQ54047.1;Pangolin_CoV_orf1ab;6206 | 0.1 | 0.075 |
| MT072864.1;QIQ54047.1;Pangolin_CoV_orf1ab;6207 | 0.1 | 0.05 |
| MT072864.1;QIQ54047.1;Pangolin_CoV_orf1ab;6208 | 0.1 | 0.05 |
| MT072864.1;QIQ54047.1;Pangolin_CoV_orf1ab;6209 | 0.1 | 0.05 |
| MT072864.1;QIQ54047.1;Pangolin_CoV_orf1ab;6210 | 0.1 | 0.025 |
| MT072864.1;QIQ54047.1;Pangolin_CoV_orf1ab;6211 | 0.1 | 0.025 |
| MT072864.1;QIQ54047.1;Pangolin_CoV_orf1ab;6212 | 0.1 | 0.05 |
| MT072864.1;QIQ54047.1;Pangolin_CoV_orf1ab;6213 | 0.05 | 0 |
| MT072864.1;QIQ54047.1;Pangolin_CoV_orf1ab;6214 | 0.05 | 0 |
| MT072864.1;QIQ54047.1;Pangolin_CoV_orf1ab;6215 | 0.05 | 0 |
| MT072864.1;QIQ54047.1;Pangolin_CoV_orf1ab;6216 | 0.1 | 0 |
| MT072864.1;QIQ54047.1;Pangolin_CoV_orf1ab;6217 | 0 | 0 |
| MT072864.1;QIQ54047.1;Pangolin_CoV_orf1ab;6218 | 0 | 0 |
| MT072864.1;QIQ54047.1;Pangolin_CoV_orf1ab;6219 | 0 | 0 |
| MT072864.1;QIQ54047.1;Pangolin_CoV_orf1ab;6220 | 0 | 0 |
| MT072864.1;QIQ54047.1;Pangolin_CoV_orf1ab;6221 | 0 | 0 |
| MT072864.1;QIQ54047.1;Pangolin_CoV_orf1ab;6222 | 0 | 0 |
| MT072864.1;QIQ54047.1;Pangolin_CoV_orf1ab;6223 | 0 | 0 |
| MT072864.1;QIQ54047.1;Pangolin_CoV_orf1ab;6224 | 0 | 0 |
| MT072864.1;QIQ54047.1;Pangolin_CoV_orf1ab;6225 | 0 | 0 |

| | | |
|---|---|---|
| MT072864.1;QIQ54047.1;Pangolin_CoV_orf1ab;6226 | 0 | 0 |
| MT072864.1;QIQ54047.1;Pangolin_CoV_orf1ab;6227 | 0 | 0 |
| MT072864.1;QIQ54047.1;Pangolin_CoV_orf1ab;6228 | 0 | 0 |
| MT072864.1;QIQ54047.1;Pangolin_CoV_orf1ab;6229 | 0 | 0 |
| MT072864.1;QIQ54047.1;Pangolin_CoV_orf1ab;6230 | 0 | 0 |
| MT072864.1;QIQ54047.1;Pangolin_CoV_orf1ab;6231 | 0 | 0 |
| MT072864.1;QIQ54047.1;Pangolin_CoV_orf1ab;6232 | 0 | 0 |
| MT072864.1;QIQ54047.1;Pangolin_CoV_orf1ab;6233 | 0 | 0 |
| MT072864.1;QIQ54047.1;Pangolin_CoV_orf1ab;6234 | 0.05 | 0 |
| MT072864.1;QIQ54047.1;Pangolin_CoV_orf1ab;6235 | 0.05 | 0.025 |
| MT072864.1;QIQ54047.1;Pangolin_CoV_orf1ab;6236 | 0 | 0 |
| MT072864.1;QIQ54047.1;Pangolin_CoV_orf1ab;6237 | 0 | 0.025 |
| MT072864.1;QIQ54047.1;Pangolin_CoV_orf1ab;6238 | 0 | 0 |
| MT072864.1;QIQ54047.1;Pangolin_CoV_orf1ab;6239 | 0 | 0 |
| MT072864.1;QIQ54047.1;Pangolin_CoV_orf1ab;6240 | 0 | 0 |
| MT072864.1;QIQ54047.1;Pangolin_CoV_orf1ab;6241 | 0 | 0 |
| MT072864.1;QIQ54047.1;Pangolin_CoV_orf1ab;6242 | 0 | 0 |
| MT072864.1;QIQ54047.1;Pangolin_CoV_orf1ab;6243 | 0 | 0 |
| MT072864.1;QIQ54047.1;Pangolin_CoV_orf1ab;6244 | 0 | 0 |
| MT072864.1;QIQ54047.1;Pangolin_CoV_orf1ab;6245 | 0 | 0 |
| MT072864.1;QIQ54047.1;Pangolin_CoV_orf1ab;6246 | 0 | 0 |
| MT072864.1;QIQ54047.1;Pangolin_CoV_orf1ab;6247 | 0.05 | 0 |
| MT072864.1;QIQ54047.1;Pangolin_CoV_orf1ab;6248 | 0.1 | 0 |
| MT072864.1;QIQ54047.1;Pangolin_CoV_orf1ab;6249 | 0 | 0 |
| MT072864.1;QIQ54047.1;Pangolin_CoV_orf1ab;6250 | 0 | 0 |
| MT072864.1;QIQ54047.1;Pangolin_CoV_orf1ab;6251 | 0 | 0 |
| MT072864.1;QIQ54047.1;Pangolin_CoV_orf1ab;6252 | 0 | 0 |
| MT072864.1;QIQ54047.1;Pangolin_CoV_orf1ab;6253 | 0 | 0 |
| MT072864.1;QIQ54047.1;Pangolin_CoV_orf1ab;6254 | 0.05 | 0 |
| MT072864.1;QIQ54047.1;Pangolin_CoV_orf1ab;6255 | 0.05 | 0 |
| MT072864.1;QIQ54047.1;Pangolin_CoV_orf1ab;6256 | 0.05 | 0.025 |
| MT072864.1;QIQ54047.1;Pangolin_CoV_orf1ab;6257 | 0.05 | 0.025 |
| MT072864.1;QIQ54047.1;Pangolin_CoV_orf1ab;6258 | 0.05 | 0.025 |
| MT072864.1;QIQ54047.1;Pangolin_CoV_orf1ab;6259 | 0 | 0.025 |
| MT072864.1;QIQ54047.1;Pangolin_CoV_orf1ab;6260 | 0 | 0 |
| MT072864.1;QIQ54047.1;Pangolin_CoV_orf1ab;6261 | 0.1 | 0.025 |
| MT072864.1;QIQ54047.1;Pangolin_CoV_orf1ab;6262 | 0.05 | 0 |
| MT072864.1;QIQ54047.1;Pangolin_CoV_orf1ab;6263 | 0.1 | 0 |
| MT072864.1;QIQ54047.1;Pangolin_CoV_orf1ab;6264 | 0.05 | 0.025 |
| MT072864.1;QIQ54047.1;Pangolin_CoV_orf1ab;6265 | 0 | 0 |
| MT072864.1;QIQ54047.1;Pangolin_CoV_orf1ab;6266 | 0.05 | 0.025 |
| MT072864.1;QIQ54047.1;Pangolin_CoV_orf1ab;6267 | 0 | 0 |
| MT072864.1;QIQ54047.1;Pangolin_CoV_orf1ab;6268 | 0 | 0 |

| | | |
|---|---|---|
| MT072864.1;QIQ54047.1;Pangolin_CoV_orf1ab;6269 | 0 | 0 |
| MT072864.1;QIQ54047.1;Pangolin_CoV_orf1ab;6270 | 0 | 0.025 |
| MT072864.1;QIQ54047.1;Pangolin_CoV_orf1ab;6271 | 0 | 0.025 |
| MT072864.1;QIQ54047.1;Pangolin_CoV_orf1ab;6272 | 0 | 0 |
| MT072864.1;QIQ54047.1;Pangolin_CoV_orf1ab;6273 | 0 | 0 |
| MT072864.1;QIQ54047.1;Pangolin_CoV_orf1ab;6274 | 0 | 0 |
| MT072864.1;QIQ54047.1;Pangolin_CoV_orf1ab;6275 | 0 | 0 |
| MT072864.1;QIQ54047.1;Pangolin_CoV_orf1ab;6276 | 0 | 0 |
| MT072864.1;QIQ54047.1;Pangolin_CoV_orf1ab;6277 | 0 | 0 |
| MT072864.1;QIQ54047.1;Pangolin_CoV_orf1ab;6278 | 0 | 0 |
| MT072864.1;QIQ54047.1;Pangolin_CoV_orf1ab;6279 | 0 | 0 |
| MT072864.1;QIQ54047.1;Pangolin_CoV_orf1ab;6280 | 0 | 0.025 |
| MT072864.1;QIQ54047.1;Pangolin_CoV_orf1ab;6281 | 0.05 | 0 |
| MT072864.1;QIQ54047.1;Pangolin_CoV_orf1ab;6282 | 0 | 0.025 |
| MT072864.1;QIQ54047.1;Pangolin_CoV_orf1ab;6283 | 0 | 0.025 |
| MT072864.1;QIQ54047.1;Pangolin_CoV_orf1ab;6284 | 0.25 | 0.125 |
| MT072864.1;QIQ54047.1;Pangolin_CoV_orf1ab;6285 | 0.1 | 0.1 |
| MT072864.1;QIQ54047.1;Pangolin_CoV_orf1ab;6286 | 0 | 0.075 |
| MT072864.1;QIQ54047.1;Pangolin_CoV_orf1ab;6287 | 0 | 0 |
| MT072864.1;QIQ54047.1;Pangolin_CoV_orf1ab;6288 | 0 | 0 |
| MT072864.1;QIQ54047.1;Pangolin_CoV_orf1ab;6289 | 0 | 0.05 |
| MT072864.1;QIQ54047.1;Pangolin_CoV_orf1ab;6290 | 0.05 | 0.1 |
| MT072864.1;QIQ54047.1;Pangolin_CoV_orf1ab;6291 | 0 | 0.05 |
| MT072864.1;QIQ54047.1;Pangolin_CoV_orf1ab;6292 | 0 | 0 |
| MT072864.1;QIQ54047.1;Pangolin_CoV_orf1ab;6293 | 0 | 0 |
| MT072864.1;QIQ54047.1;Pangolin_CoV_orf1ab;6294 | 0 | 0 |
| MT072864.1;QIQ54047.1;Pangolin_CoV_orf1ab;6295 | 0 | 0 |
| MT072864.1;QIQ54047.1;Pangolin_CoV_orf1ab;6296 | 0 | 0 |
| MT072864.1;QIQ54047.1;Pangolin_CoV_orf1ab;6297 | 0 | 0 |
| MT072864.1;QIQ54047.1;Pangolin_CoV_orf1ab;6298 | 0.05 | 0 |
| MT072864.1;QIQ54047.1;Pangolin_CoV_orf1ab;6299 | 0 | 0 |
| MT072864.1;QIQ54047.1;Pangolin_CoV_orf1ab;6300 | 0 | 0 |
| MT072864.1;QIQ54047.1;Pangolin_CoV_orf1ab;6301 | 0 | 0 |
| MT072864.1;QIQ54047.1;Pangolin_CoV_orf1ab;6302 | 0 | 0 |
| MT072864.1;QIQ54047.1;Pangolin_CoV_orf1ab;6303 | 0 | 0 |
| MT072864.1;QIQ54047.1;Pangolin_CoV_orf1ab;6304 | 0 | 0 |
| MT072864.1;QIQ54047.1;Pangolin_CoV_orf1ab;6305 | 0 | 0 |
| MT072864.1;QIQ54047.1;Pangolin_CoV_orf1ab;6306 | 0 | 0 |
| MT072864.1;QIQ54047.1;Pangolin_CoV_orf1ab;6307 | 0 | 0 |
| MT072864.1;QIQ54047.1;Pangolin_CoV_orf1ab;6308 | 0 | 0 |
| MT072864.1;QIQ54047.1;Pangolin_CoV_orf1ab;6309 | 0 | 0 |
| MT072864.1;QIQ54047.1;Pangolin_CoV_orf1ab;6310 | 0 | 0 |
| MT072864.1;QIQ54047.1;Pangolin_CoV_orf1ab;6311 | 0 | 0 |

| | | |
|---|---|---|
| MT072864.1;QIQ54047.1;Pangolin_CoV_orf1ab;6312 | 0 | 0 |
| MT072864.1;QIQ54047.1;Pangolin_CoV_orf1ab;6313 | 0 | 0 |
| MT072864.1;QIQ54047.1;Pangolin_CoV_orf1ab;6314 | 0 | 0 |
| MT072864.1;QIQ54047.1;Pangolin_CoV_orf1ab;6315 | 0 | 0 |
| MT072864.1;QIQ54047.1;Pangolin_CoV_orf1ab;6316 | 0 | 0 |
| MT072864.1;QIQ54047.1;Pangolin_CoV_orf1ab;6317 | 0 | 0 |
| MT072864.1;QIQ54047.1;Pangolin_CoV_orf1ab;6318 | 0 | 0 |
| MT072864.1;QIQ54047.1;Pangolin_CoV_orf1ab;6319 | 0.1 | 0.025 |
| MT072864.1;QIQ54047.1;Pangolin_CoV_orf1ab;6320 | 0.05 | 0.025 |
| MT072864.1;QIQ54047.1;Pangolin_CoV_orf1ab;6321 | 0.05 | 0 |
| MT072864.1;QIQ54047.1;Pangolin_CoV_orf1ab;6322 | 0.05 | 0 |
| MT072864.1;QIQ54047.1;Pangolin_CoV_orf1ab;6323 | 0.05 | 0 |
| MT072864.1;QIQ54047.1;Pangolin_CoV_orf1ab;6324 | 0 | 0.025 |
| MT072864.1;QIQ54047.1;Pangolin_CoV_orf1ab;6325 | 0 | 0 |
| MT072864.1;QIQ54047.1;Pangolin_CoV_orf1ab;6326 | 0 | 0 |
| MT072864.1;QIQ54047.1;Pangolin_CoV_orf1ab;6327 | 0 | 0 |
| MT072864.1;QIQ54047.1;Pangolin_CoV_orf1ab;6328 | 0 | 0 |
| MT072864.1;QIQ54047.1;Pangolin_CoV_orf1ab;6329 | 0 | 0 |
| MT072864.1;QIQ54047.1;Pangolin_CoV_orf1ab;6330 | 0 | 0 |
| MT072864.1;QIQ54047.1;Pangolin_CoV_orf1ab;6331 | 0 | 0 |
| MT072864.1;QIQ54047.1;Pangolin_CoV_orf1ab;6332 | 0 | 0 |
| MT072864.1;QIQ54047.1;Pangolin_CoV_orf1ab;6333 | 0 | 0 |
| MT072864.1;QIQ54047.1;Pangolin_CoV_orf1ab;6334 | 0 | 0 |
| MT072864.1;QIQ54047.1;Pangolin_CoV_orf1ab;6335 | 0 | 0 |
| MT072864.1;QIQ54047.1;Pangolin_CoV_orf1ab;6336 | 0 | 0 |
| MT072864.1;QIQ54047.1;Pangolin_CoV_orf1ab;6337 | 0 | 0.025 |
| MT072864.1;QIQ54047.1;Pangolin_CoV_orf1ab;6338 | 0 | 0.025 |
| MT072864.1;QIQ54047.1;Pangolin_CoV_orf1ab;6339 | 0 | 0 |
| MT072864.1;QIQ54047.1;Pangolin_CoV_orf1ab;6340 | 0 | 0.025 |
| MT072864.1;QIQ54047.1;Pangolin_CoV_orf1ab;6341 | 0 | 0 |
| MT072864.1;QIQ54047.1;Pangolin_CoV_orf1ab;6342 | 0 | 0.025 |
| MT072864.1;QIQ54047.1;Pangolin_CoV_orf1ab;6343 | 0.05 | 0.025 |
| MT072864.1;QIQ54047.1;Pangolin_CoV_orf1ab;6344 | 0.05 | 0.025 |
| MT072864.1;QIQ54047.1;Pangolin_CoV_orf1ab;6345 | 0.05 | 0.025 |
| MT072864.1;QIQ54047.1;Pangolin_CoV_orf1ab;6346 | 0.05 | 0.025 |
| MT072864.1;QIQ54047.1;Pangolin_CoV_orf1ab;6347 | 0.05 | 0.025 |
| MT072864.1;QIQ54047.1;Pangolin_CoV_orf1ab;6348 | 0.05 | 0 |
| MT072864.1;QIQ54047.1;Pangolin_CoV_orf1ab;6349 | 0.05 | 0.05 |
| MT072864.1;QIQ54047.1;Pangolin_CoV_orf1ab;6350 | 0.05 | 0.025 |
| MT072864.1;QIQ54047.1;Pangolin_CoV_orf1ab;6351 | 0 | 0 |
| MT072864.1;QIQ54047.1;Pangolin_CoV_orf1ab;6352 | 0 | 0 |
| MT072864.1;QIQ54047.1;Pangolin_CoV_orf1ab;6353 | 0 | 0.025 |
| MT072864.1;QIQ54047.1;Pangolin_CoV_orf1ab;6354 | 0 | 0 |

| | | |
|---|---|---|
| MT072864.1;QIQ54047.1;Pangolin_CoV_orf1ab;6355 | 0 | 0 |
| MT072864.1;QIQ54047.1;Pangolin_CoV_orf1ab;6356 | 0 | 0 |
| MT072864.1;QIQ54047.1;Pangolin_CoV_orf1ab;6357 | 0 | 0 |
| MT072864.1;QIQ54047.1;Pangolin_CoV_orf1ab;6358 | 0 | 0 |
| MT072864.1;QIQ54047.1;Pangolin_CoV_orf1ab;6359 | 0 | 0 |
| MT072864.1;QIQ54047.1;Pangolin_CoV_orf1ab;6360 | 0 | 0 |
| MT072864.1;QIQ54047.1;Pangolin_CoV_orf1ab;6361 | 0 | 0 |
| MT072864.1;QIQ54047.1;Pangolin_CoV_orf1ab;6362 | 0 | 0 |
| MT072864.1;QIQ54047.1;Pangolin_CoV_orf1ab;6363 | 0 | 0.025 |
| MT072864.1;QIQ54047.1;Pangolin_CoV_orf1ab;6364 | 0 | 0 |
| MT072864.1;QIQ54047.1;Pangolin_CoV_orf1ab;6365 | 0 | 0 |
| MT072864.1;QIQ54047.1;Pangolin_CoV_orf1ab;6366 | 0 | 0 |
| MT072864.1;QIQ54047.1;Pangolin_CoV_orf1ab;6367 | 0 | 0 |
| MT072864.1;QIQ54047.1;Pangolin_CoV_orf1ab;6368 | 0 | 0.05 |
| MT072864.1;QIQ54047.1;Pangolin_CoV_orf1ab;6369 | 0 | 0.075 |
| MT072864.1;QIQ54047.1;Pangolin_CoV_orf1ab;6370 | 0 | 0.05 |
| MT072864.1;QIQ54047.1;Pangolin_CoV_orf1ab;6371 | 0 | 0.05 |
| MT072864.1;QIQ54047.1;Pangolin_CoV_orf1ab;6372 | 0 | 0.075 |
| MT072864.1;QIQ54047.1;Pangolin_CoV_orf1ab;6373 | 0 | 0.075 |
| MT072864.1;QIQ54047.1;Pangolin_CoV_orf1ab;6374 | 0.05 | 0.05 |
| MT072864.1;QIQ54047.1;Pangolin_CoV_orf1ab;6375 | 0 | 0.05 |
| MT072864.1;QIQ54047.1;Pangolin_CoV_orf1ab;6376 | 0 | 0.025 |
| MT072864.1;QIQ54047.1;Pangolin_CoV_orf1ab;6377 | 0 | 0.025 |
| MT072864.1;QIQ54047.1;Pangolin_CoV_orf1ab;6378 | 0 | 0.025 |
| MT072864.1;QIQ54047.1;Pangolin_CoV_orf1ab;6379 | 0 | 0 |
| MT072864.1;QIQ54047.1;Pangolin_CoV_orf1ab;6380 | 0 | 0 |
| MT072864.1;QIQ54047.1;Pangolin_CoV_orf1ab;6381 | 0 | 0 |
| MT072864.1;QIQ54047.1;Pangolin_CoV_orf1ab;6382 | 0 | 0 |
| MT072864.1;QIQ54047.1;Pangolin_CoV_orf1ab;6383 | 0 | 0 |
| MT072864.1;QIQ54047.1;Pangolin_CoV_orf1ab;6384 | 0 | 0 |
| MT072864.1;QIQ54047.1;Pangolin_CoV_orf1ab;6385 | 0 | 0 |
| MT072864.1;QIQ54047.1;Pangolin_CoV_orf1ab;6386 | 0 | 0 |
| MT072864.1;QIQ54047.1;Pangolin_CoV_orf1ab;6387 | 0 | 0 |
| MT072864.1;QIQ54047.1;Pangolin_CoV_orf1ab;6388 | 0 | 0 |
| MT072864.1;QIQ54047.1;Pangolin_CoV_orf1ab;6389 | 0 | 0 |
| MT072864.1;QIQ54047.1;Pangolin_CoV_orf1ab;6390 | 0 | 0 |
| MT072864.1;QIQ54047.1;Pangolin_CoV_orf1ab;6391 | 0 | 0 |
| MT072864.1;QIQ54047.1;Pangolin_CoV_orf1ab;6392 | 0 | 0 |
| MT072864.1;QIQ54047.1;Pangolin_CoV_orf1ab;6393 | 0 | 0 |
| MT072864.1;QIQ54047.1;Pangolin_CoV_orf1ab;6394 | 0 | 0 |
| MT072864.1;QIQ54047.1;Pangolin_CoV_orf1ab;6395 | 0 | 0 |
| MT072864.1;QIQ54047.1;Pangolin_CoV_orf1ab;6396 | 0 | 0 |
| MT072864.1;QIQ54047.1;Pangolin_CoV_orf1ab;6397 | 0 | 0 |

| | | |
|---|---|---|
| MT072864.1;QIQ54047.1;Pangolin_CoV_orf1ab;6398 | 0 | 0 |
| MT072864.1;QIQ54047.1;Pangolin_CoV_orf1ab;6399 | 0 | 0 |
| MT072864.1;QIQ54047.1;Pangolin_CoV_orf1ab;6400 | 0 | 0 |
| MT072864.1;QIQ54047.1;Pangolin_CoV_orf1ab;6401 | 0 | 0 |
| MT072864.1;QIQ54047.1;Pangolin_CoV_orf1ab;6402 | 0 | 0.025 |
| MT072864.1;QIQ54047.1;Pangolin_CoV_orf1ab;6403 | 0 | 0.05 |
| MT072864.1;QIQ54047.1;Pangolin_CoV_orf1ab;6404 | 0 | 0.025 |
| MT072864.1;QIQ54047.1;Pangolin_CoV_orf1ab;6405 | 0 | 0.025 |
| MT072864.1;QIQ54047.1;Pangolin_CoV_orf1ab;6406 | 0 | 0.025 |
| MT072864.1;QIQ54047.1;Pangolin_CoV_orf1ab;6407 | 0 | 0.025 |
| MT072864.1;QIQ54047.1;Pangolin_CoV_orf1ab;6408 | 0 | 0.025 |
| MT072864.1;QIQ54047.1;Pangolin_CoV_orf1ab;6409 | 0 | 0.025 |
| MT072864.1;QIQ54047.1;Pangolin_CoV_orf1ab;6410 | 0 | 0.025 |
| MT072864.1;QIQ54047.1;Pangolin_CoV_orf1ab;6411 | 0 | 0 |
| MT072864.1;QIQ54047.1;Pangolin_CoV_orf1ab;6412 | 0 | 0 |
| MT072864.1;QIQ54047.1;Pangolin_CoV_orf1ab;6413 | 0 | 0 |
| MT072864.1;QIQ54047.1;Pangolin_CoV_orf1ab;6414 | 0 | 0 |
| MT072864.1;QIQ54047.1;Pangolin_CoV_orf1ab;6415 | 0 | 0 |
| MT072864.1;QIQ54047.1;Pangolin_CoV_orf1ab;6416 | 0 | 0 |
| MT072864.1;QIQ54047.1;Pangolin_CoV_orf1ab;6417 | 0 | 0 |
| MT072864.1;QIQ54047.1;Pangolin_CoV_orf1ab;6418 | 0 | 0 |
| MT072864.1;QIQ54047.1;Pangolin_CoV_orf1ab;6419 | 0 | 0 |
| MT072864.1;QIQ54047.1;Pangolin_CoV_orf1ab;6420 | 0 | 0 |
| MT072864.1;QIQ54047.1;Pangolin_CoV_orf1ab;6421 | 0 | 0 |
| MT072864.1;QIQ54047.1;Pangolin_CoV_orf1ab;6422 | 0 | 0 |
| MT072864.1;QIQ54047.1;Pangolin_CoV_orf1ab;6423 | 0 | 0 |
| MT072864.1;QIQ54047.1;Pangolin_CoV_orf1ab;6424 | 0 | 0 |
| MT072864.1;QIQ54047.1;Pangolin_CoV_orf1ab;6425 | 0 | 0 |
| MT072864.1;QIQ54047.1;Pangolin_CoV_orf1ab;6426 | 0 | 0 |
| MT072864.1;QIQ54047.1;Pangolin_CoV_orf1ab;6427 | 0 | 0 |
| MT072864.1;QIQ54047.1;Pangolin_CoV_orf1ab;6428 | 0 | 0 |
| MT072864.1;QIQ54047.1;Pangolin_CoV_orf1ab;6429 | 0 | 0 |
| MT072864.1;QIQ54047.1;Pangolin_CoV_orf1ab;6430 | 0 | 0 |
| MT072864.1;QIQ54047.1;Pangolin_CoV_orf1ab;6431 | 0 | 0 |
| MT072864.1;QIQ54047.1;Pangolin_CoV_orf1ab;6432 | 0 | 0 |
| MT072864.1;QIQ54047.1;Pangolin_CoV_orf1ab;6433 | 0 | 0 |
| MT072864.1;QIQ54047.1;Pangolin_CoV_orf1ab;6434 | 0 | 0 |
| MT072864.1;QIQ54047.1;Pangolin_CoV_orf1ab;6435 | 0 | 0 |
| MT072864.1;QIQ54047.1;Pangolin_CoV_orf1ab;6436 | 0 | 0 |
| MT072864.1;QIQ54047.1;Pangolin_CoV_orf1ab;6437 | 0.05 | 0 |
| MT072864.1;QIQ54047.1;Pangolin_CoV_orf1ab;6438 | 0 | 0 |
| MT072864.1;QIQ54047.1;Pangolin_CoV_orf1ab;6439 | 0 | 0 |
| MT072864.1;QIQ54047.1;Pangolin_CoV_orf1ab;6440 | 0 | 0 |

| | | |
|---|---|---|
| MT072864.1;QIQ54047.1;Pangolin_CoV_orf1ab;6441 | 0 | 0 |
| MT072864.1;QIQ54047.1;Pangolin_CoV_orf1ab;6442 | 0 | 0 |
| MT072864.1;QIQ54047.1;Pangolin_CoV_orf1ab;6443 | 0.05 | 0 |
| MT072864.1;QIQ54047.1;Pangolin_CoV_orf1ab;6444 | 0.05 | 0 |
| MT072864.1;QIQ54047.1;Pangolin_CoV_orf1ab;6445 | 0.1 | 0 |
| MT072864.1;QIQ54047.1;Pangolin_CoV_orf1ab;6446 | 0.1 | 0 |
| MT072864.1;QIQ54047.1;Pangolin_CoV_orf1ab;6447 | 0.1 | 0 |
| MT072864.1;QIQ54047.1;Pangolin_CoV_orf1ab;6448 | 0.05 | 0 |
| MT072864.1;QIQ54047.1;Pangolin_CoV_orf1ab;6449 | 0.05 | 0 |
| MT072864.1;QIQ54047.1;Pangolin_CoV_orf1ab;6450 | 0.1 | 0.05 |
| MT072864.1;QIQ54047.1;Pangolin_CoV_orf1ab;6451 | 0.05 | 0.025 |
| MT072864.1;QIQ54047.1;Pangolin_CoV_orf1ab;6452 | 0.05 | 0 |
| MT072864.1;QIQ54047.1;Pangolin_CoV_orf1ab;6453 | 0 | 0.025 |
| MT072864.1;QIQ54047.1;Pangolin_CoV_orf1ab;6454 | 0 | 0.05 |
| MT072864.1;QIQ54047.1;Pangolin_CoV_orf1ab;6455 | 0.05 | 0.025 |
| MT072864.1;QIQ54047.1;Pangolin_CoV_orf1ab;6456 | 0.05 | 0.025 |
| MT072864.1;QIQ54047.1;Pangolin_CoV_orf1ab;6457 | 0 | 0 |
| MT072864.1;QIQ54047.1;Pangolin_CoV_orf1ab;6458 | 0.05 | 0 |
| MT072864.1;QIQ54047.1;Pangolin_CoV_orf1ab;6459 | 0.05 | 0 |
| MT072864.1;QIQ54047.1;Pangolin_CoV_orf1ab;6460 | 0.05 | 0 |
| MT072864.1;QIQ54047.1;Pangolin_CoV_orf1ab;6461 | 0.05 | 0 |
| MT072864.1;QIQ54047.1;Pangolin_CoV_orf1ab;6462 | 0.05 | 0 |
| MT072864.1;QIQ54047.1;Pangolin_CoV_orf1ab;6463 | 0 | 0.025 |
| MT072864.1;QIQ54047.1;Pangolin_CoV_orf1ab;6464 | 0 | 0.025 |
| MT072864.1;QIQ54047.1;Pangolin_CoV_orf1ab;6465 | 0 | 0.025 |
| MT072864.1;QIQ54047.1;Pangolin_CoV_orf1ab;6466 | 0 | 0 |
| MT072864.1;QIQ54047.1;Pangolin_CoV_orf1ab;6467 | 0.05 | 0 |
| MT072864.1;QIQ54047.1;Pangolin_CoV_orf1ab;6468 | 0 | 0.05 |
| MT072864.1;QIQ54047.1;Pangolin_CoV_orf1ab;6469 | 0 | 0.05 |
| MT072864.1;QIQ54047.1;Pangolin_CoV_orf1ab;6470 | 0 | 0.025 |
| MT072864.1;QIQ54047.1;Pangolin_CoV_orf1ab;6471 | 0 | 0 |
| MT072864.1;QIQ54047.1;Pangolin_CoV_orf1ab;6472 | 0 | 0 |
| MT072864.1;QIQ54047.1;Pangolin_CoV_orf1ab;6473 | 0 | 0 |
| MT072864.1;QIQ54047.1;Pangolin_CoV_orf1ab;6474 | 0 | 0 |
| MT072864.1;QIQ54047.1;Pangolin_CoV_orf1ab;6475 | 0 | 0 |
| MT072864.1;QIQ54047.1;Pangolin_CoV_orf1ab;6476 | 0 | 0.025 |
| MT072864.1;QIQ54047.1;Pangolin_CoV_orf1ab;6477 | 0 | 0.075 |
| MT072864.1;QIQ54047.1;Pangolin_CoV_orf1ab;6478 | 0 | 0.15 |
| MT072864.1;QIQ54047.1;Pangolin_CoV_orf1ab;6479 | 0 | 0.05 |
| MT072864.1;QIQ54047.1;Pangolin_CoV_orf1ab;6480 | 0 | 0.1 |
| MT072864.1;QIQ54047.1;Pangolin_CoV_orf1ab;6481 | 0 | 0.125 |
| MT072864.1;QIQ54047.1;Pangolin_CoV_orf1ab;6482 | 0 | 0.05 |
| MT072864.1;QIQ54047.1;Pangolin_CoV_orf1ab;6483 | 0 | 0.15 |

| | | |
|---|---|---|
| MT072864.1;QIQ54047.1;Pangolin_CoV_orf1ab;6484 | 0 | 0.125 |
| MT072864.1;QIQ54047.1;Pangolin_CoV_orf1ab;6485 | 0 | 0.075 |
| MT072864.1;QIQ54047.1;Pangolin_CoV_orf1ab;6486 | 0 | 0.025 |
| MT072864.1;QIQ54047.1;Pangolin_CoV_orf1ab;6487 | 0 | 0.025 |
| MT072864.1;QIQ54047.1;Pangolin_CoV_orf1ab;6488 | 0.05 | 0.025 |
| MT072864.1;QIQ54047.1;Pangolin_CoV_orf1ab;6489 | 0 | 0 |
| MT072864.1;QIQ54047.1;Pangolin_CoV_orf1ab;6490 | 0 | 0 |
| MT072864.1;QIQ54047.1;Pangolin_CoV_orf1ab;6491 | 0 | 0 |
| MT072864.1;QIQ54047.1;Pangolin_CoV_orf1ab;6492 | 0 | 0 |
| MT072864.1;QIQ54047.1;Pangolin_CoV_orf1ab;6493 | 0 | 0 |
| MT072864.1;QIQ54047.1;Pangolin_CoV_orf1ab;6494 | 0 | 0 |
| MT072864.1;QIQ54047.1;Pangolin_CoV_orf1ab;6495 | 0 | 0 |
| MT072864.1;QIQ54047.1;Pangolin_CoV_orf1ab;6496 | 0 | 0 |
| MT072864.1;QIQ54047.1;Pangolin_CoV_orf1ab;6497 | 0.05 | 0 |
| MT072864.1;QIQ54047.1;Pangolin_CoV_orf1ab;6498 | 0 | 0 |
| MT072864.1;QIQ54047.1;Pangolin_CoV_orf1ab;6499 | 0.05 | 0.1 |
| MT072864.1;QIQ54047.1;Pangolin_CoV_orf1ab;6500 | 0.2 | 0.125 |
| MT072864.1;QIQ54047.1;Pangolin_CoV_orf1ab;6501 | 0.05 | 0.1 |
| MT072864.1;QIQ54047.1;Pangolin_CoV_orf1ab;6502 | 0.05 | 0.075 |
| MT072864.1;QIQ54047.1;Pangolin_CoV_orf1ab;6503 | 0.1 | 0.075 |
| MT072864.1;QIQ54047.1;Pangolin_CoV_orf1ab;6504 | 0.15 | 0.15 |
| MT072864.1;QIQ54047.1;Pangolin_CoV_orf1ab;6505 | 0.15 | 0.125 |
| MT072864.1;QIQ54047.1;Pangolin_CoV_orf1ab;6506 | 0.4 | 0.2 |
| MT072864.1;QIQ54047.1;Pangolin_CoV_orf1ab;6507 | 0.4 | 0.2 |
| MT072864.1;QIQ54047.1;Pangolin_CoV_orf1ab;6508 | 0.2 | 0.15 |
| MT072864.1;QIQ54047.1;Pangolin_CoV_orf1ab;6509 | 0 | 0 |
| MT072864.1;QIQ54047.1;Pangolin_CoV_orf1ab;6510 | 0.05 | 0 |
| MT072864.1;QIQ54047.1;Pangolin_CoV_orf1ab;6511 | 0 | 0 |
| MT072864.1;QIQ54047.1;Pangolin_CoV_orf1ab;6512 | 0 | 0 |
| MT072864.1;QIQ54047.1;Pangolin_CoV_orf1ab;6513 | 0 | 0.025 |
| MT072864.1;QIQ54047.1;Pangolin_CoV_orf1ab;6514 | 0 | 0 |
| MT072864.1;QIQ54047.1;Pangolin_CoV_orf1ab;6515 | 0 | 0 |
| MT072864.1;QIQ54047.1;Pangolin_CoV_orf1ab;6516 | 0 | 0 |
| MT072864.1;QIQ54047.1;Pangolin_CoV_orf1ab;6517 | 0 | 0 |
| MT072864.1;QIQ54047.1;Pangolin_CoV_orf1ab;6518 | 0 | 0 |
| MT072864.1;QIQ54047.1;Pangolin_CoV_orf1ab;6519 | 0 | 0 |
| MT072864.1;QIQ54047.1;Pangolin_CoV_orf1ab;6520 | 0.05 | 0.025 |
| MT072864.1;QIQ54047.1;Pangolin_CoV_orf1ab;6521 | 0 | 0 |
| MT072864.1;QIQ54047.1;Pangolin_CoV_orf1ab;6522 | 0 | 0 |
| MT072864.1;QIQ54047.1;Pangolin_CoV_orf1ab;6523 | 0 | 0.025 |
| MT072864.1;QIQ54047.1;Pangolin_CoV_orf1ab;6524 | 0 | 0 |
| MT072864.1;QIQ54047.1;Pangolin_CoV_orf1ab;6525 | 0 | 0 |
| MT072864.1;QIQ54047.1;Pangolin_CoV_orf1ab;6526 | 0 | 0 |

| | | |
|---|---|---|
| MT072864.1;QIQ54047.1;Pangolin_CoV_orf1ab;6527 | 0 | 0 |
| MT072864.1;QIQ54047.1;Pangolin_CoV_orf1ab;6528 | 0.05 | 0 |
| MT072864.1;QIQ54047.1;Pangolin_CoV_orf1ab;6529 | 0.05 | 0 |
| MT072864.1;QIQ54047.1;Pangolin_CoV_orf1ab;6530 | 0 | 0 |
| MT072864.1;QIQ54047.1;Pangolin_CoV_orf1ab;6531 | 0 | 0 |
| MT072864.1;QIQ54047.1;Pangolin_CoV_orf1ab;6532 | 0 | 0 |
| MT072864.1;QIQ54047.1;Pangolin_CoV_orf1ab;6533 | 0 | 0 |
| MT072864.1;QIQ54047.1;Pangolin_CoV_orf1ab;6534 | 0 | 0 |
| MT072864.1;QIQ54047.1;Pangolin_CoV_orf1ab;6535 | 0 | 0 |
| MT072864.1;QIQ54047.1;Pangolin_CoV_orf1ab;6536 | 0 | 0 |
| MT072864.1;QIQ54047.1;Pangolin_CoV_orf1ab;6537 | 0 | 0 |
| MT072864.1;QIQ54047.1;Pangolin_CoV_orf1ab;6538 | 0 | 0 |
| MT072864.1;QIQ54047.1;Pangolin_CoV_orf1ab;6539 | 0.05 | 0 |
| MT072864.1;QIQ54047.1;Pangolin_CoV_orf1ab;6540 | 0.05 | 0 |
| MT072864.1;QIQ54047.1;Pangolin_CoV_orf1ab;6541 | 0.05 | 0.025 |
| MT072864.1;QIQ54047.1;Pangolin_CoV_orf1ab;6542 | 0.05 | 0 |
| MT072864.1;QIQ54047.1;Pangolin_CoV_orf1ab;6543 | 0.05 | 0 |
| MT072864.1;QIQ54047.1;Pangolin_CoV_orf1ab;6544 | 0.15 | 0 |
| MT072864.1;QIQ54047.1;Pangolin_CoV_orf1ab;6545 | 0.05 | 0 |
| MT072864.1;QIQ54047.1;Pangolin_CoV_orf1ab;6546 | 0 | 0 |
| MT072864.1;QIQ54047.1;Pangolin_CoV_orf1ab;6547 | 0 | 0 |
| MT072864.1;QIQ54047.1;Pangolin_CoV_orf1ab;6548 | 0.05 | 0 |
| MT072864.1;QIQ54047.1;Pangolin_CoV_orf1ab;6549 | 0.05 | 0 |
| MT072864.1;QIQ54047.1;Pangolin_CoV_orf1ab;6550 | 0 | 0 |
| MT072864.1;QIQ54047.1;Pangolin_CoV_orf1ab;6551 | 0 | 0 |
| MT072864.1;QIQ54047.1;Pangolin_CoV_orf1ab;6552 | 0 | 0 |
| MT072864.1;QIQ54047.1;Pangolin_CoV_orf1ab;6553 | 0 | 0 |
| MT072864.1;QIQ54047.1;Pangolin_CoV_orf1ab;6554 | 0 | 0 |
| MT072864.1;QIQ54047.1;Pangolin_CoV_orf1ab;6555 | 0 | 0 |
| MT072864.1;QIQ54047.1;Pangolin_CoV_orf1ab;6556 | 0 | 0 |
| MT072864.1;QIQ54047.1;Pangolin_CoV_orf1ab;6557 | 0 | 0 |
| MT072864.1;QIQ54047.1;Pangolin_CoV_orf1ab;6558 | 0 | 0 |
| MT072864.1;QIQ54047.1;Pangolin_CoV_orf1ab;6559 | 0 | 0 |
| MT072864.1;QIQ54047.1;Pangolin_CoV_orf1ab;6560 | 0 | 0 |
| MT072864.1;QIQ54047.1;Pangolin_CoV_orf1ab;6561 | 0 | 0 |
| MT072864.1;QIQ54047.1;Pangolin_CoV_orf1ab;6562 | 0 | 0 |
| MT072864.1;QIQ54047.1;Pangolin_CoV_orf1ab;6563 | 0 | 0 |
| MT072864.1;QIQ54047.1;Pangolin_CoV_orf1ab;6564 | 0 | 0 |
| MT072864.1;QIQ54047.1;Pangolin_CoV_orf1ab;6565 | 0 | 0 |
| MT072864.1;QIQ54047.1;Pangolin_CoV_orf1ab;6566 | 0 | 0 |
| MT072864.1;QIQ54047.1;Pangolin_CoV_orf1ab;6567 | 0 | 0 |
| MT072864.1;QIQ54047.1;Pangolin_CoV_orf1ab;6568 | 0.05 | 0.025 |
| MT072864.1;QIQ54047.1;Pangolin_CoV_orf1ab;6569 | 0 | 0 |

| | | |
|---|---|---|
| MT072864.1;QIQ54047.1;Pangolin_CoV_orf1ab;6570 | 0 | 0 |
| MT072864.1;QIQ54047.1;Pangolin_CoV_orf1ab;6571 | 0 | 0 |
| MT072864.1;QIQ54047.1;Pangolin_CoV_orf1ab;6572 | 0 | 0 |
| MT072864.1;QIQ54047.1;Pangolin_CoV_orf1ab;6573 | 0 | 0.025 |
| MT072864.1;QIQ54047.1;Pangolin_CoV_orf1ab;6574 | 0 | 0 |
| MT072864.1;QIQ54047.1;Pangolin_CoV_orf1ab;6575 | 0 | 0 |
| MT072864.1;QIQ54047.1;Pangolin_CoV_orf1ab;6576 | 0 | 0 |
| MT072864.1;QIQ54047.1;Pangolin_CoV_orf1ab;6577 | 0 | 0 |
| MT072864.1;QIQ54047.1;Pangolin_CoV_orf1ab;6578 | 0 | 0 |
| MT072864.1;QIQ54047.1;Pangolin_CoV_orf1ab;6579 | 0 | 0 |
| MT072864.1;QIQ54047.1;Pangolin_CoV_orf1ab;6580 | 0 | 0 |
| MT072864.1;QIQ54047.1;Pangolin_CoV_orf1ab;6581 | 0 | 0 |
| MT072864.1;QIQ54047.1;Pangolin_CoV_orf1ab;6582 | 0 | 0 |
| MT072864.1;QIQ54047.1;Pangolin_CoV_orf1ab;6583 | 0 | 0.025 |
| MT072864.1;QIQ54047.1;Pangolin_CoV_orf1ab;6584 | 0.1 | 0.05 |
| MT072864.1;QIQ54047.1;Pangolin_CoV_orf1ab;6585 | 0.1 | 0.025 |
| MT072864.1;QIQ54047.1;Pangolin_CoV_orf1ab;6586 | 0 | 0.025 |
| MT072864.1;QIQ54047.1;Pangolin_CoV_orf1ab;6587 | 0 | 0 |
| MT072864.1;QIQ54047.1;Pangolin_CoV_orf1ab;6588 | 0 | 0 |
| MT072864.1;QIQ54047.1;Pangolin_CoV_orf1ab;6589 | 0.05 | 0 |
| MT072864.1;QIQ54047.1;Pangolin_CoV_orf1ab;6590 | 0 | 0.025 |
| MT072864.1;QIQ54047.1;Pangolin_CoV_orf1ab;6591 | 0 | 0 |
| MT072864.1;QIQ54047.1;Pangolin_CoV_orf1ab;6592 | 0.05 | 0 |
| MT072864.1;QIQ54047.1;Pangolin_CoV_orf1ab;6593 | 0 | 0 |
| MT072864.1;QIQ54047.1;Pangolin_CoV_orf1ab;6594 | 0 | 0.05 |
| MT072864.1;QIQ54047.1;Pangolin_CoV_orf1ab;6595 | 0 | 0 |
| MT072864.1;QIQ54047.1;Pangolin_CoV_orf1ab;6596 | 0 | 0 |
| MT072864.1;QIQ54047.1;Pangolin_CoV_orf1ab;6597 | 0 | 0 |
| MT072864.1;QIQ54047.1;Pangolin_CoV_orf1ab;6598 | 0 | 0 |
| MT072864.1;QIQ54047.1;Pangolin_CoV_orf1ab;6599 | 0 | 0 |
| MT072864.1;QIQ54047.1;Pangolin_CoV_orf1ab;6600 | 0 | 0 |
| MT072864.1;QIQ54047.1;Pangolin_CoV_orf1ab;6601 | 0 | 0 |
| MT072864.1;QIQ54047.1;Pangolin_CoV_orf1ab;6602 | 0 | 0 |
| MT072864.1;QIQ54047.1;Pangolin_CoV_orf1ab;6603 | 0 | 0.025 |
| MT072864.1;QIQ54047.1;Pangolin_CoV_orf1ab;6604 | 0 | 0.025 |
| MT072864.1;QIQ54047.1;Pangolin_CoV_orf1ab;6605 | 0 | 0.025 |
| MT072864.1;QIQ54047.1;Pangolin_CoV_orf1ab;6606 | 0 | 0.025 |
| MT072864.1;QIQ54047.1;Pangolin_CoV_orf1ab;6607 | 0 | 0.025 |
| MT072864.1;QIQ54047.1;Pangolin_CoV_orf1ab;6608 | 0 | 0 |
| MT072864.1;QIQ54047.1;Pangolin_CoV_orf1ab;6609 | 0 | 0 |
| MT072864.1;QIQ54047.1;Pangolin_CoV_orf1ab;6610 | 0 | 0 |
| MT072864.1;QIQ54047.1;Pangolin_CoV_orf1ab;6611 | 0.05 | 0 |
| MT072864.1;QIQ54047.1;Pangolin_CoV_orf1ab;6612 | 0 | 0.05 |

| | | |
|---|---|---|
| MT072864.1;QIQ54047.1;Pangolin_CoV_orf1ab;6613 | 0.05 | 0.05 |
| MT072864.1;QIQ54047.1;Pangolin_CoV_orf1ab;6614 | 0 | 0.075 |
| MT072864.1;QIQ54047.1;Pangolin_CoV_orf1ab;6615 | 0 | 0.025 |
| MT072864.1;QIQ54047.1;Pangolin_CoV_orf1ab;6616 | 0 | 0.025 |
| MT072864.1;QIQ54047.1;Pangolin_CoV_orf1ab;6617 | 0.05 | 0.05 |
| MT072864.1;QIQ54047.1;Pangolin_CoV_orf1ab;6618 | 0.05 | 0.05 |
| MT072864.1;QIQ54047.1;Pangolin_CoV_orf1ab;6619 | 0.1 | 0.075 |
| MT072864.1;QIQ54047.1;Pangolin_CoV_orf1ab;6620 | 0 | 0.075 |
| MT072864.1;QIQ54047.1;Pangolin_CoV_orf1ab;6621 | 0.1 | 0.075 |
| MT072864.1;QIQ54047.1;Pangolin_CoV_orf1ab;6622 | 0.05 | 0.1 |
| MT072864.1;QIQ54047.1;Pangolin_CoV_orf1ab;6623 | 0.1 | 0.075 |
| MT072864.1;QIQ54047.1;Pangolin_CoV_orf1ab;6624 | 0.05 | 0.025 |
| MT072864.1;QIQ54047.1;Pangolin_CoV_orf1ab;6625 | 0.05 | 0.025 |
| MT072864.1;QIQ54047.1;Pangolin_CoV_orf1ab;6626 | 0 | 0 |
| MT072864.1;QIQ54047.1;Pangolin_CoV_orf1ab;6627 | 0 | 0 |
| MT072864.1;QIQ54047.1;Pangolin_CoV_orf1ab;6628 | 0 | 0.025 |
| MT072864.1;QIQ54047.1;Pangolin_CoV_orf1ab;6629 | 0 | 0 |
| MT072864.1;QIQ54047.1;Pangolin_CoV_orf1ab;6630 | 0 | 0 |
| MT072864.1;QIQ54047.1;Pangolin_CoV_orf1ab;6631 | 0 | 0.025 |
| MT072864.1;QIQ54047.1;Pangolin_CoV_orf1ab;6632 | 0 | 0.05 |
| MT072864.1;QIQ54047.1;Pangolin_CoV_orf1ab;6633 | 0 | 0.025 |
| MT072864.1;QIQ54047.1;Pangolin_CoV_orf1ab;6634 | 0.05 | 0 |
| MT072864.1;QIQ54047.1;Pangolin_CoV_orf1ab;6635 | 0 | 0 |
| MT072864.1;QIQ54047.1;Pangolin_CoV_orf1ab;6636 | 0 | 0 |
| MT072864.1;QIQ54047.1;Pangolin_CoV_orf1ab;6637 | 0 | 0 |
| MT072864.1;QIQ54047.1;Pangolin_CoV_orf1ab;6638 | 0 | 0 |
| MT072864.1;QIQ54047.1;Pangolin_CoV_orf1ab;6639 | 0 | 0 |
| MT072864.1;QIQ54047.1;Pangolin_CoV_orf1ab;6640 | 0 | 0 |
| MT072864.1;QIQ54047.1;Pangolin_CoV_orf1ab;6641 | 0 | 0 |
| MT072864.1;QIQ54047.1;Pangolin_CoV_orf1ab;6642 | 0 | 0 |
| MT072864.1;QIQ54047.1;Pangolin_CoV_orf1ab;6643 | 0 | 0 |
| MT072864.1;QIQ54047.1;Pangolin_CoV_orf1ab;6644 | 0 | 0 |
| MT072864.1;QIQ54047.1;Pangolin_CoV_orf1ab;6645 | 0 | 0 |
| MT072864.1;QIQ54047.1;Pangolin_CoV_orf1ab;6646 | 0 | 0 |
| MT072864.1;QIQ54047.1;Pangolin_CoV_orf1ab;6647 | 0 | 0 |
| MT072864.1;QIQ54047.1;Pangolin_CoV_orf1ab;6648 | 0 | 0 |
| MT072864.1;QIQ54047.1;Pangolin_CoV_orf1ab;6649 | 0 | 0 |
| MT072864.1;QIQ54047.1;Pangolin_CoV_orf1ab;6650 | 0 | 0.025 |
| MT072864.1;QIQ54047.1;Pangolin_CoV_orf1ab;6651 | 0 | 0 |
| MT072864.1;QIQ54047.1;Pangolin_CoV_orf1ab;6652 | 0 | 0 |
| MT072864.1;QIQ54047.1;Pangolin_CoV_orf1ab;6653 | 0 | 0 |
| MT072864.1;QIQ54047.1;Pangolin_CoV_orf1ab;6654 | 0 | 0 |
| MT072864.1;QIQ54047.1;Pangolin_CoV_orf1ab;6655 | 0.05 | 0 |

| | | |
|---|---|---|
| MT072864.1;QIQ54047.1;Pangolin_CoV_orf1ab;6656 | 0.05 | 0 |
| MT072864.1;QIQ54047.1;Pangolin_CoV_orf1ab;6657 | 0.05 | 0.025 |
| MT072864.1;QIQ54047.1;Pangolin_CoV_orf1ab;6658 | 0 | 0.025 |
| MT072864.1;QIQ54047.1;Pangolin_CoV_orf1ab;6659 | 0 | 0.025 |
| MT072864.1;QIQ54047.1;Pangolin_CoV_orf1ab;6660 | 0.05 | 0.075 |
| MT072864.1;QIQ54047.1;Pangolin_CoV_orf1ab;6661 | 0 | 0 |
| MT072864.1;QIQ54047.1;Pangolin_CoV_orf1ab;6662 | 0 | 0 |
| MT072864.1;QIQ54047.1;Pangolin_CoV_orf1ab;6663 | 0 | 0 |
| MT072864.1;QIQ54047.1;Pangolin_CoV_orf1ab;6664 | 0 | 0 |
| MT072864.1;QIQ54047.1;Pangolin_CoV_orf1ab;6665 | 0 | 0 |
| MT072864.1;QIQ54047.1;Pangolin_CoV_orf1ab;6666 | 0 | 0 |
| MT072864.1;QIQ54047.1;Pangolin_CoV_orf1ab;6667 | 0 | 0 |
| MT072864.1;QIQ54047.1;Pangolin_CoV_orf1ab;6668 | 0 | 0 |
| MT072864.1;QIQ54047.1;Pangolin_CoV_orf1ab;6669 | 0 | 0 |
| MT072864.1;QIQ54047.1;Pangolin_CoV_orf1ab;6670 | 0 | 0 |
| MT072864.1;QIQ54047.1;Pangolin_CoV_orf1ab;6671 | 0 | 0 |
| MT072864.1;QIQ54047.1;Pangolin_CoV_orf1ab;6672 | 0 | 0 |
| MT072864.1;QIQ54047.1;Pangolin_CoV_orf1ab;6673 | 0 | 0 |
| MT072864.1;QIQ54047.1;Pangolin_CoV_orf1ab;6674 | 0 | 0 |
| MT072864.1;QIQ54047.1;Pangolin_CoV_orf1ab;6675 | 0 | 0 |
| MT072864.1;QIQ54047.1;Pangolin_CoV_orf1ab;6676 | 0 | 0 |
| MT072864.1;QIQ54047.1;Pangolin_CoV_orf1ab;6677 | 0 | 0 |
| MT072864.1;QIQ54047.1;Pangolin_CoV_orf1ab;6678 | 0 | 0 |
| MT072864.1;QIQ54047.1;Pangolin_CoV_orf1ab;6679 | 0 | 0 |
| MT072864.1;QIQ54047.1;Pangolin_CoV_orf1ab;6680 | 0 | 0 |
| MT072864.1;QIQ54047.1;Pangolin_CoV_orf1ab;6681 | 0 | 0 |
| MT072864.1;QIQ54047.1;Pangolin_CoV_orf1ab;6682 | 0 | 0 |
| MT072864.1;QIQ54047.1;Pangolin_CoV_orf1ab;6683 | 0 | 0 |
| MT072864.1;QIQ54047.1;Pangolin_CoV_orf1ab;6684 | 0 | 0 |
| MT072864.1;QIQ54047.1;Pangolin_CoV_orf1ab;6685 | 0 | 0 |
| MT072864.1;QIQ54047.1;Pangolin_CoV_orf1ab;6686 | 0 | 0 |
| MT072864.1;QIQ54047.1;Pangolin_CoV_orf1ab;6687 | 0 | 0 |
| MT072864.1;QIQ54047.1;Pangolin_CoV_orf1ab;6688 | 0 | 0 |
| MT072864.1;QIQ54047.1;Pangolin_CoV_orf1ab;6689 | 0 | 0 |
| MT072864.1;QIQ54047.1;Pangolin_CoV_orf1ab;6690 | 0 | 0 |
| MT072864.1;QIQ54047.1;Pangolin_CoV_orf1ab;6691 | 0 | 0 |
| MT072864.1;QIQ54047.1;Pangolin_CoV_orf1ab;6692 | 0 | 0 |
| MT072864.1;QIQ54047.1;Pangolin_CoV_orf1ab;6693 | 0.1 | 0 |
| MT072864.1;QIQ54047.1;Pangolin_CoV_orf1ab;6694 | 0.1 | 0.05 |
| MT072864.1;QIQ54047.1;Pangolin_CoV_orf1ab;6695 | 0.15 | 0.05 |
| MT072864.1;QIQ54047.1;Pangolin_CoV_orf1ab;6696 | 0.1 | 0.05 |
| MT072864.1;QIQ54047.1;Pangolin_CoV_orf1ab;6697 | 0.1 | 0.05 |
| MT072864.1;QIQ54047.1;Pangolin_CoV_orf1ab;6698 | 0.05 | 0.05 |

| | | |
|---|---|---|
| MT072864.1;QIQ54047.1;Pangolin_CoV_orf1ab;6699 | 0.05 | 0.025 |
| MT072864.1;QIQ54047.1;Pangolin_CoV_orf1ab;6700 | 0.05 | 0.025 |
| MT072864.1;QIQ54047.1;Pangolin_CoV_orf1ab;6701 | 0.05 | 0.025 |
| MT072864.1;QIQ54047.1;Pangolin_CoV_orf1ab;6702 | 0.05 | 0.025 |
| MT072864.1;QIQ54047.1;Pangolin_CoV_orf1ab;6703 | 0.05 | 0 |
| MT072864.1;QIQ54047.1;Pangolin_CoV_orf1ab;6704 | 0 | 0 |
| MT072864.1;QIQ54047.1;Pangolin_CoV_orf1ab;6705 | 0 | 0 |
| MT072864.1;QIQ54047.1;Pangolin_CoV_orf1ab;6706 | 0 | 0 |
| MT072864.1;QIQ54047.1;Pangolin_CoV_orf1ab;6707 | 0.05 | 0 |
| MT072864.1;QIQ54047.1;Pangolin_CoV_orf1ab;6708 | 0.05 | 0 |
| MT072864.1;QIQ54047.1;Pangolin_CoV_orf1ab;6709 | 0.15 | 0 |
| MT072864.1;QIQ54047.1;Pangolin_CoV_orf1ab;6710 | 0 | 0 |
| MT072864.1;QIQ54047.1;Pangolin_CoV_orf1ab;6711 | 0.05 | 0 |
| MT072864.1;QIQ54047.1;Pangolin_CoV_orf1ab;6712 | 0.1 | 0 |
| MT072864.1;QIQ54047.1;Pangolin_CoV_orf1ab;6713 | 0.1 | 0.025 |
| MT072864.1;QIQ54047.1;Pangolin_CoV_orf1ab;6714 | 0 | 0 |
| MT072864.1;QIQ54047.1;Pangolin_CoV_orf1ab;6715 | 0.1 | 0 |
| MT072864.1;QIQ54047.1;Pangolin_CoV_orf1ab;6716 | 0.05 | 0 |
| MT072864.1;QIQ54047.1;Pangolin_CoV_orf1ab;6717 | 0.05 | 0 |
| MT072864.1;QIQ54047.1;Pangolin_CoV_orf1ab;6718 | 0.1 | 0 |
| MT072864.1;QIQ54047.1;Pangolin_CoV_orf1ab;6719 | 0.1 | 0 |
| MT072864.1;QIQ54047.1;Pangolin_CoV_orf1ab;6720 | 0.05 | 0.025 |
| MT072864.1;QIQ54047.1;Pangolin_CoV_orf1ab;6721 | 0 | 0 |
| MT072864.1;QIQ54047.1;Pangolin_CoV_orf1ab;6722 | 0 | 0 |
| MT072864.1;QIQ54047.1;Pangolin_CoV_orf1ab;6723 | 0 | 0 |
| MT072864.1;QIQ54047.1;Pangolin_CoV_orf1ab;6724 | 0 | 0 |
| MT072864.1;QIQ54047.1;Pangolin_CoV_orf1ab;6725 | 0 | 0 |
| MT072864.1;QIQ54047.1;Pangolin_CoV_orf1ab;6726 | 0 | 0 |
| MT072864.1;QIQ54047.1;Pangolin_CoV_orf1ab;6727 | 0 | 0 |
| MT072864.1;QIQ54047.1;Pangolin_CoV_orf1ab;6728 | 0 | 0 |
| MT072864.1;QIQ54047.1;Pangolin_CoV_orf1ab;6729 | 0 | 0 |
| MT072864.1;QIQ54047.1;Pangolin_CoV_orf1ab;6730 | 0 | 0 |
| MT072864.1;QIQ54047.1;Pangolin_CoV_orf1ab;6731 | 0 | 0 |
| MT072864.1;QIQ54047.1;Pangolin_CoV_orf1ab;6732 | 0 | 0 |
| MT072864.1;QIQ54047.1;Pangolin_CoV_orf1ab;6733 | 0 | 0 |
| MT072864.1;QIQ54047.1;Pangolin_CoV_orf1ab;6734 | 0 | 0 |
| MT072864.1;QIQ54047.1;Pangolin_CoV_orf1ab;6735 | 0 | 0 |
| MT072864.1;QIQ54047.1;Pangolin_CoV_orf1ab;6736 | 0 | 0 |
| MT072864.1;QIQ54047.1;Pangolin_CoV_orf1ab;6737 | 0.05 | 0.025 |
| MT072864.1;QIQ54047.1;Pangolin_CoV_orf1ab;6738 | 0.05 | 0 |
| MT072864.1;QIQ54047.1;Pangolin_CoV_orf1ab;6739 | 0.05 | 0 |
| MT072864.1;QIQ54047.1;Pangolin_CoV_orf1ab;6740 | 0.05 | 0 |
| MT072864.1;QIQ54047.1;Pangolin_CoV_orf1ab;6741 | 0.1 | 0.125 |

| | | |
|---|---|---|
| MT072864.1;QIQ54047.1;Pangolin_CoV_orf1ab;6742 | 0.05 | 0 |
| MT072864.1;QIQ54047.1;Pangolin_CoV_orf1ab;6743 | 0 | 0 |
| MT072864.1;QIQ54047.1;Pangolin_CoV_orf1ab;6744 | 0 | 0 |
| MT072864.1;QIQ54047.1;Pangolin_CoV_orf1ab;6745 | 0 | 0 |
| MT072864.1;QIQ54047.1;Pangolin_CoV_orf1ab;6746 | 0 | 0 |
| MT072864.1;QIQ54047.1;Pangolin_CoV_orf1ab;6747 | 0 | 0 |
| MT072864.1;QIQ54047.1;Pangolin_CoV_orf1ab;6748 | 0 | 0 |
| MT072864.1;QIQ54047.1;Pangolin_CoV_orf1ab;6749 | 0 | 0 |
| MT072864.1;QIQ54047.1;Pangolin_CoV_orf1ab;6750 | 0 | 0 |
| MT072864.1;QIQ54047.1;Pangolin_CoV_orf1ab;6751 | 0 | 0 |
| MT072864.1;QIQ54047.1;Pangolin_CoV_orf1ab;6752 | 0 | 0 |
| MT072864.1;QIQ54047.1;Pangolin_CoV_orf1ab;6753 | 0 | 0 |
| MT072864.1;QIQ54047.1;Pangolin_CoV_orf1ab;6754 | 0 | 0 |
| MT072864.1;QIQ54047.1;Pangolin_CoV_orf1ab;6755 | 0 | 0 |
| MT072864.1;QIQ54047.1;Pangolin_CoV_orf1ab;6756 | 0 | 0 |
| MT072864.1;QIQ54047.1;Pangolin_CoV_orf1ab;6757 | 0 | 0 |
| MT072864.1;QIQ54047.1;Pangolin_CoV_orf1ab;6758 | 0 | 0 |
| MT072864.1;QIQ54047.1;Pangolin_CoV_orf1ab;6759 | 0 | 0 |
| MT072864.1;QIQ54047.1;Pangolin_CoV_orf1ab;6760 | 0 | 0 |
| MT072864.1;QIQ54047.1;Pangolin_CoV_orf1ab;6761 | 0 | 0 |
| MT072864.1;QIQ54047.1;Pangolin_CoV_orf1ab;6762 | 0 | 0 |
| MT072864.1;QIQ54047.1;Pangolin_CoV_orf1ab;6763 | 0 | 0 |
| MT072864.1;QIQ54047.1;Pangolin_CoV_orf1ab;6764 | 0 | 0.05 |
| MT072864.1;QIQ54047.1;Pangolin_CoV_orf1ab;6765 | 0 | 0 |
| MT072864.1;QIQ54047.1;Pangolin_CoV_orf1ab;6766 | 0 | 0 |
| MT072864.1;QIQ54047.1;Pangolin_CoV_orf1ab;6767 | 0 | 0 |
| MT072864.1;QIQ54047.1;Pangolin_CoV_orf1ab;6768 | 0 | 0 |
| MT072864.1;QIQ54047.1;Pangolin_CoV_orf1ab;6769 | 0 | 0.05 |
| MT072864.1;QIQ54047.1;Pangolin_CoV_orf1ab;6770 | 0 | 0 |
| MT072864.1;QIQ54047.1;Pangolin_CoV_orf1ab;6771 | 0 | 0 |
| MT072864.1;QIQ54047.1;Pangolin_CoV_orf1ab;6772 | 0 | 0 |
| MT072864.1;QIQ54047.1;Pangolin_CoV_orf1ab;6773 | 0 | 0 |
| MT072864.1;QIQ54047.1;Pangolin_CoV_orf1ab;6774 | 0 | 0 |
| MT072864.1;QIQ54047.1;Pangolin_CoV_orf1ab;6775 | 0 | 0.025 |
| MT072864.1;QIQ54047.1;Pangolin_CoV_orf1ab;6776 | 0 | 0.025 |
| MT072864.1;QIQ54047.1;Pangolin_CoV_orf1ab;6777 | 0 | 0.025 |
| MT072864.1;QIQ54047.1;Pangolin_CoV_orf1ab;6778 | 0 | 0.025 |
| MT072864.1;QIQ54047.1;Pangolin_CoV_orf1ab;6779 | 0 | 0.025 |
| MT072864.1;QIQ54047.1;Pangolin_CoV_orf1ab;6780 | 0 | 0.025 |
| MT072864.1;QIQ54047.1;Pangolin_CoV_orf1ab;6781 | 0 | 0.025 |
| MT072864.1;QIQ54047.1;Pangolin_CoV_orf1ab;6782 | 0 | 0.025 |
| MT072864.1;QIQ54047.1;Pangolin_CoV_orf1ab;6783 | 0 | 0.025 |
| MT072864.1;QIQ54047.1;Pangolin_CoV_orf1ab;6784 | 0 | 0.025 |

| | | |
|---|---|---|
| MT072864.1;QIQ54047.1;Pangolin_CoV_orf1ab;6785 | 0.1 | 0.05 |
| MT072864.1;QIQ54047.1;Pangolin_CoV_orf1ab;6786 | 0 | 0 |
| MT072864.1;QIQ54047.1;Pangolin_CoV_orf1ab;6787 | 0 | 0 |
| MT072864.1;QIQ54047.1;Pangolin_CoV_orf1ab;6788 | 0 | 0 |
| MT072864.1;QIQ54047.1;Pangolin_CoV_orf1ab;6789 | 0 | 0 |
| MT072864.1;QIQ54047.1;Pangolin_CoV_orf1ab;6790 | 0 | 0 |
| MT072864.1;QIQ54047.1;Pangolin_CoV_orf1ab;6791 | 0 | 0 |
| MT072864.1;QIQ54047.1;Pangolin_CoV_orf1ab;6792 | 0 | 0 |
| MT072864.1;QIQ54047.1;Pangolin_CoV_orf1ab;6793 | 0 | 0 |
| MT072864.1;QIQ54047.1;Pangolin_CoV_orf1ab;6794 | 0 | 0 |
| MT072864.1;QIQ54047.1;Pangolin_CoV_orf1ab;6795 | 0 | 0 |
| MT072864.1;QIQ54047.1;Pangolin_CoV_orf1ab;6796 | 0 | 0 |
| MT072864.1;QIQ54047.1;Pangolin_CoV_orf1ab;6797 | 0 | 0 |
| MT072864.1;QIQ54047.1;Pangolin_CoV_orf1ab;6798 | 0 | 0 |
| MT072864.1;QIQ54047.1;Pangolin_CoV_orf1ab;6799 | 0 | 0 |
| MT072864.1;QIQ54047.1;Pangolin_CoV_orf1ab;6800 | 0 | 0 |
| MT072864.1;QIQ54047.1;Pangolin_CoV_orf1ab;6801 | 0.05 | 0.025 |
| MT072864.1;QIQ54047.1;Pangolin_CoV_orf1ab;6802 | 0.05 | 0.025 |
| MT072864.1;QIQ54047.1;Pangolin_CoV_orf1ab;6803 | 0.05 | 0.025 |
| MT072864.1;QIQ54047.1;Pangolin_CoV_orf1ab;6804 | 0.05 | 0.05 |
| MT072864.1;QIQ54047.1;Pangolin_CoV_orf1ab;6805 | 0.1 | 0.075 |
| MT072864.1;QIQ54047.1;Pangolin_CoV_orf1ab;6806 | 0.1 | 0.075 |
| MT072864.1;QIQ54047.1;Pangolin_CoV_orf1ab;6807 | 0.05 | 0.025 |
| MT072864.1;QIQ54047.1;Pangolin_CoV_orf1ab;6808 | 0.05 | 0.025 |
| MT072864.1;QIQ54047.1;Pangolin_CoV_orf1ab;6809 | 0.1 | 0.075 |
| MT072864.1;QIQ54047.1;Pangolin_CoV_orf1ab;6810 | 0 | 0 |
| MT072864.1;QIQ54047.1;Pangolin_CoV_orf1ab;6811 | 0 | 0 |
| MT072864.1;QIQ54047.1;Pangolin_CoV_orf1ab;6812 | 0 | 0 |
| MT072864.1;QIQ54047.1;Pangolin_CoV_orf1ab;6813 | 0 | 0 |
| MT072864.1;QIQ54047.1;Pangolin_CoV_orf1ab;6814 | 0.05 | 0 |
| MT072864.1;QIQ54047.1;Pangolin_CoV_orf1ab;6815 | 0 | 0 |
| MT072864.1;QIQ54047.1;Pangolin_CoV_orf1ab;6816 | 0 | 0 |
| MT072864.1;QIQ54047.1;Pangolin_CoV_orf1ab;6817 | 0.05 | 0 |
| MT072864.1;QIQ54047.1;Pangolin_CoV_orf1ab;6818 | 0.05 | 0 |
| MT072864.1;QIQ54047.1;Pangolin_CoV_orf1ab;6819 | 0.05 | 0 |
| MT072864.1;QIQ54047.1;Pangolin_CoV_orf1ab;6820 | 0.05 | 0 |
| MT072864.1;QIQ54047.1;Pangolin_CoV_orf1ab;6821 | 0 | 0 |
| MT072864.1;QIQ54047.1;Pangolin_CoV_orf1ab;6822 | 0 | 0 |
| MT072864.1;QIQ54047.1;Pangolin_CoV_orf1ab;6823 | 0 | 0.025 |
| MT072864.1;QIQ54047.1;Pangolin_CoV_orf1ab;6824 | 0 | 0 |
| MT072864.1;QIQ54047.1;Pangolin_CoV_orf1ab;6825 | 0 | 0 |
| MT072864.1;QIQ54047.1;Pangolin_CoV_orf1ab;6826 | 0 | 0 |
| MT072864.1;QIQ54047.1;Pangolin_CoV_orf1ab;6827 | 0 | 0 |

| | | |
|---|---|---|
| MT072864.1;QIQ54047.1;Pangolin_CoV_orf1ab;6828 | 0 | 0.05 |
| MT072864.1;QIQ54047.1;Pangolin_CoV_orf1ab;6829 | 0 | 0 |
| MT072864.1;QIQ54047.1;Pangolin_CoV_orf1ab;6830 | 0 | 0 |
| MT072864.1;QIQ54047.1;Pangolin_CoV_orf1ab;6831 | 0 | 0 |
| MT072864.1;QIQ54047.1;Pangolin_CoV_orf1ab;6832 | 0 | 0 |
| MT072864.1;QIQ54047.1;Pangolin_CoV_orf1ab;6833 | 0 | 0 |
| MT072864.1;QIQ54047.1;Pangolin_CoV_orf1ab;6834 | 0 | 0 |
| MT072864.1;QIQ54047.1;Pangolin_CoV_orf1ab;6835 | 0 | 0 |
| MT072864.1;QIQ54047.1;Pangolin_CoV_orf1ab;6836 | 0 | 0 |
| MT072864.1;QIQ54047.1;Pangolin_CoV_orf1ab;6837 | 0 | 0 |
| MT072864.1;QIQ54047.1;Pangolin_CoV_orf1ab;6838 | 0 | 0 |
| MT072864.1;QIQ54047.1;Pangolin_CoV_orf1ab;6839 | 0 | 0 |
| MT072864.1;QIQ54047.1;Pangolin_CoV_orf1ab;6840 | 0 | 0 |
| MT072864.1;QIQ54047.1;Pangolin_CoV_orf1ab;6841 | 0 | 0 |
| MT072864.1;QIQ54047.1;Pangolin_CoV_orf1ab;6842 | 0 | 0 |
| MT072864.1;QIQ54047.1;Pangolin_CoV_orf1ab;6843 | 0 | 0 |
| MT072864.1;QIQ54047.1;Pangolin_CoV_orf1ab;6844 | 0 | 0 |
| MT072864.1;QIQ54047.1;Pangolin_CoV_orf1ab;6845 | 0 | 0 |
| MT072864.1;QIQ54047.1;Pangolin_CoV_orf1ab;6846 | 0 | 0 |
| MT072864.1;QIQ54047.1;Pangolin_CoV_orf1ab;6847 | 0 | 0 |
| MT072864.1;QIQ54047.1;Pangolin_CoV_orf1ab;6848 | 0 | 0 |
| MT072864.1;QIQ54047.1;Pangolin_CoV_orf1ab;6849 | 0 | 0 |
| MT072864.1;QIQ54047.1;Pangolin_CoV_orf1ab;6850 | 0 | 0 |
| MT072864.1;QIQ54047.1;Pangolin_CoV_orf1ab;6851 | 0 | 0 |
| MT072864.1;QIQ54047.1;Pangolin_CoV_orf1ab;6852 | 0 | 0 |
| MT072864.1;QIQ54047.1;Pangolin_CoV_orf1ab;6853 | 0 | 0 |
| MT072864.1;QIQ54047.1;Pangolin_CoV_orf1ab;6854 | 0.05 | 0.05 |
| MT072864.1;QIQ54047.1;Pangolin_CoV_orf1ab;6855 | 0 | 0 |
| MT072864.1;QIQ54047.1;Pangolin_CoV_orf1ab;6856 | 0 | 0 |
| MT072864.1;QIQ54047.1;Pangolin_CoV_orf1ab;6857 | 0 | 0 |
| MT072864.1;QIQ54047.1;Pangolin_CoV_orf1ab;6858 | 0 | 0 |
| MT072864.1;QIQ54047.1;Pangolin_CoV_orf1ab;6859 | 0 | 0 |
| MT072864.1;QIQ54047.1;Pangolin_CoV_orf1ab;6860 | 0 | 0 |
| MT072864.1;QIQ54047.1;Pangolin_CoV_orf1ab;6861 | 0 | 0 |
| MT072864.1;QIQ54047.1;Pangolin_CoV_orf1ab;6862 | 0.05 | 0 |
| MT072864.1;QIQ54047.1;Pangolin_CoV_orf1ab;6863 | 0.05 | 0 |
| MT072864.1;QIQ54047.1;Pangolin_CoV_orf1ab;6864 | 0 | 0 |
| MT072864.1;QIQ54047.1;Pangolin_CoV_orf1ab;6865 | 0 | 0 |
| MT072864.1;QIQ54047.1;Pangolin_CoV_orf1ab;6866 | 0 | 0 |
| MT072864.1;QIQ54047.1;Pangolin_CoV_orf1ab;6867 | 0 | 0 |
| MT072864.1;QIQ54047.1;Pangolin_CoV_orf1ab;6868 | 0 | 0 |
| MT072864.1;QIQ54047.1;Pangolin_CoV_orf1ab;6869 | 0 | 0 |
| MT072864.1;QIQ54047.1;Pangolin_CoV_orf1ab;6870 | 0 | 0 |

| | | |
|---|---|---|
| MT072864.1;QIQ54047.1;Pangolin_CoV_orf1ab;6871 | 0 | 0 |
| MT072864.1;QIQ54047.1;Pangolin_CoV_orf1ab;6872 | 0 | 0 |
| MT072864.1;QIQ54047.1;Pangolin_CoV_orf1ab;6873 | 0 | 0 |
| MT072864.1;QIQ54047.1;Pangolin_CoV_orf1ab;6874 | 0 | 0 |
| MT072864.1;QIQ54047.1;Pangolin_CoV_orf1ab;6875 | 0 | 0 |
| MT072864.1;QIQ54047.1;Pangolin_CoV_orf1ab;6876 | 0 | 0 |
| MT072864.1;QIQ54047.1;Pangolin_CoV_orf1ab;6877 | 0 | 0 |
| MT072864.1;QIQ54047.1;Pangolin_CoV_orf1ab;6878 | 0 | 0 |
| MT072864.1;QIQ54047.1;Pangolin_CoV_orf1ab;6879 | 0 | 0.025 |
| MT072864.1;QIQ54047.1;Pangolin_CoV_orf1ab;6880 | 0 | 0.025 |
| MT072864.1;QIQ54047.1;Pangolin_CoV_orf1ab;6881 | 0 | 0.025 |
| MT072864.1;QIQ54047.1;Pangolin_CoV_orf1ab;6882 | 0 | 0.025 |
| MT072864.1;QIQ54047.1;Pangolin_CoV_orf1ab;6883 | 0.05 | 0.025 |
| MT072864.1;QIQ54047.1;Pangolin_CoV_orf1ab;6884 | 0.05 | 0.05 |
| MT072864.1;QIQ54047.1;Pangolin_CoV_orf1ab;6885 | 0.05 | 0.05 |
| MT072864.1;QIQ54047.1;Pangolin_CoV_orf1ab;6886 | 0 | 0.025 |
| MT072864.1;QIQ54047.1;Pangolin_CoV_orf1ab;6887 | 0 | 0.05 |
| MT072864.1;QIQ54047.1;Pangolin_CoV_orf1ab;6888 | 0.1 | 0.05 |
| MT072864.1;QIQ54047.1;Pangolin_CoV_orf1ab;6889 | 0 | 0.025 |
| MT072864.1;QIQ54047.1;Pangolin_CoV_orf1ab;6890 | 0 | 0 |
| MT072864.1;QIQ54047.1;Pangolin_CoV_orf1ab;6891 | 0.05 | 0 |
| MT072864.1;QIQ54047.1;Pangolin_CoV_orf1ab;6892 | 0.05 | 0 |
| MT072864.1;QIQ54047.1;Pangolin_CoV_orf1ab;6893 | 0.05 | 0 |
| MT072864.1;QIQ54047.1;Pangolin_CoV_orf1ab;6894 | 0.05 | 0 |
| MT072864.1;QIQ54047.1;Pangolin_CoV_orf1ab;6895 | 0.05 | 0 |
| MT072864.1;QIQ54047.1;Pangolin_CoV_orf1ab;6896 | 0.05 | 0 |
| MT072864.1;QIQ54047.1;Pangolin_CoV_orf1ab;6897 | 0.05 | 0 |
| MT072864.1;QIQ54047.1;Pangolin_CoV_orf1ab;6898 | 0 | 0.025 |
| MT072864.1;QIQ54047.1;Pangolin_CoV_orf1ab;6899 | 0 | 0 |
| MT072864.1;QIQ54047.1;Pangolin_CoV_orf1ab;6900 | 0.05 | 0.175 |
| MT072864.1;QIQ54047.1;Pangolin_CoV_orf1ab;6901 | 0 | 0.1 |
| MT072864.1;QIQ54047.1;Pangolin_CoV_orf1ab;6902 | 0 | 0.025 |
| MT072864.1;QIQ54047.1;Pangolin_CoV_orf1ab;6903 | 0 | 0 |
| MT072864.1;QIQ54047.1;Pangolin_CoV_orf1ab;6904 | 0 | 0 |
| MT072864.1;QIQ54047.1;Pangolin_CoV_orf1ab;6905 | 0 | 0 |
| MT072864.1;QIQ54047.1;Pangolin_CoV_orf1ab;6906 | 0 | 0.025 |
| MT072864.1;QIQ54047.1;Pangolin_CoV_orf1ab;6907 | 0 | 0 |
| MT072864.1;QIQ54047.1;Pangolin_CoV_orf1ab;6908 | 0 | 0 |
| MT072864.1;QIQ54047.1;Pangolin_CoV_orf1ab;6909 | 0 | 0 |
| MT072864.1;QIQ54047.1;Pangolin_CoV_orf1ab;6910 | 0 | 0 |
| MT072864.1;QIQ54047.1;Pangolin_CoV_orf1ab;6911 | 0.15 | 0.05 |
| MT072864.1;QIQ54047.1;Pangolin_CoV_orf1ab;6912 | 0 | 0 |
| MT072864.1;QIQ54047.1;Pangolin_CoV_orf1ab;6913 | 0 | 0.025 |

| | | |
|---|---|---|
| MT072864.1;QIQ54047.1;Pangolin_CoV_orf1ab;6914 | 0 | 0 |
| MT072864.1;QIQ54047.1;Pangolin_CoV_orf1ab;6915 | 0 | 0 |
| MT072864.1;QIQ54047.1;Pangolin_CoV_orf1ab;6916 | 0 | 0 |
| MT072864.1;QIQ54047.1;Pangolin_CoV_orf1ab;6917 | 0 | 0 |
| MT072864.1;QIQ54047.1;Pangolin_CoV_orf1ab;6918 | 0 | 0 |
| MT072864.1;QIQ54047.1;Pangolin_CoV_orf1ab;6919 | 0.05 | 0 |
| MT072864.1;QIQ54047.1;Pangolin_CoV_orf1ab;6920 | 0.05 | 0 |
| MT072864.1;QIQ54047.1;Pangolin_CoV_orf1ab;6921 | 0 | 0 |
| MT072864.1;QIQ54047.1;Pangolin_CoV_orf1ab;6922 | 0 | 0.025 |
| MT072864.1;QIQ54047.1;Pangolin_CoV_orf1ab;6923 | 0 | 0 |
| MT072864.1;QIQ54047.1;Pangolin_CoV_orf1ab;6924 | 0 | 0 |
| MT072864.1;QIQ54047.1;Pangolin_CoV_orf1ab;6925 | 0.05 | 0 |
| MT072864.1;QIQ54047.1;Pangolin_CoV_orf1ab;6926 | 0.05 | 0.025 |
| MT072864.1;QIQ54047.1;Pangolin_CoV_orf1ab;6927 | 0.05 | 0 |
| MT072864.1;QIQ54047.1;Pangolin_CoV_orf1ab;6928 | 0 | 0 |
| MT072864.1;QIQ54047.1;Pangolin_CoV_orf1ab;6929 | 0 | 0 |
| MT072864.1;QIQ54047.1;Pangolin_CoV_orf1ab;6930 | 0 | 0.05 |
| MT072864.1;QIQ54047.1;Pangolin_CoV_orf1ab;6931 | 0 | 0 |
| MT072864.1;QIQ54047.1;Pangolin_CoV_orf1ab;6932 | 0 | 0 |
| MT072864.1;QIQ54047.1;Pangolin_CoV_orf1ab;6933 | 0 | 0 |
| MT072864.1;QIQ54047.1;Pangolin_CoV_orf1ab;6934 | 0 | 0 |
| MT072864.1;QIQ54047.1;Pangolin_CoV_orf1ab;6935 | 0 | 0 |
| MT072864.1;QIQ54047.1;Pangolin_CoV_orf1ab;6936 | 0 | 0 |
| MT072864.1;QIQ54047.1;Pangolin_CoV_orf1ab;6937 | 0 | 0 |
| MT072864.1;QIQ54047.1;Pangolin_CoV_orf1ab;6938 | 0 | 0 |
| MT072864.1;QIQ54047.1;Pangolin_CoV_orf1ab;6939 | 0 | 0 |
| MT072864.1;QIQ54047.1;Pangolin_CoV_orf1ab;6940 | 0 | 0 |
| MT072864.1;QIQ54047.1;Pangolin_CoV_orf1ab;6941 | 0 | 0 |
| MT072864.1;QIQ54047.1;Pangolin_CoV_orf1ab;6942 | 0 | 0 |
| MT072864.1;QIQ54047.1;Pangolin_CoV_orf1ab;6943 | 0 | 0 |
| MT072864.1;QIQ54047.1;Pangolin_CoV_orf1ab;6944 | 0 | 0 |
| MT072864.1;QIQ54047.1;Pangolin_CoV_orf1ab;6945 | 0 | 0 |
| MT072864.1;QIQ54047.1;Pangolin_CoV_orf1ab;6946 | 0 | 0 |
| MT072864.1;QIQ54047.1;Pangolin_CoV_orf1ab;6947 | 0 | 0 |
| MT072864.1;QIQ54047.1;Pangolin_CoV_orf1ab;6948 | 0 | 0 |
| MT072864.1;QIQ54047.1;Pangolin_CoV_orf1ab;6949 | 0 | 0 |
| MT072864.1;QIQ54047.1;Pangolin_CoV_orf1ab;6950 | 0 | 0 |
| MT072864.1;QIQ54047.1;Pangolin_CoV_orf1ab;6951 | 0 | 0 |
| MT072864.1;QIQ54047.1;Pangolin_CoV_orf1ab;6952 | 0 | 0 |
| MT072864.1;QIQ54047.1;Pangolin_CoV_orf1ab;6953 | 0 | 0 |
| MT072864.1;QIQ54047.1;Pangolin_CoV_orf1ab;6954 | 0 | 0 |
| MT072864.1;QIQ54047.1;Pangolin_CoV_orf1ab;6955 | 0 | 0 |
| MT072864.1;QIQ54047.1;Pangolin_CoV_orf1ab;6956 | 0 | 0 |

| | | |
|---|---|---|
| MT072864.1;QIQ54047.1;Pangolin_CoV_orf1ab;6957 | 0 | 0.05 |
| MT072864.1;QIQ54047.1;Pangolin_CoV_orf1ab;6958 | 0 | 0.025 |
| MT072864.1;QIQ54047.1;Pangolin_CoV_orf1ab;6959 | 0 | 0.05 |
| MT072864.1;QIQ54047.1;Pangolin_CoV_orf1ab;6960 | 0 | 0.025 |
| MT072864.1;QIQ54047.1;Pangolin_CoV_orf1ab;6961 | 0 | 0.025 |
| MT072864.1;QIQ54047.1;Pangolin_CoV_orf1ab;6962 | 0 | 0.025 |
| MT072864.1;QIQ54047.1;Pangolin_CoV_orf1ab;6963 | 0 | 0.025 |
| MT072864.1;QIQ54047.1;Pangolin_CoV_orf1ab;6964 | 0 | 0.025 |
| MT072864.1;QIQ54047.1;Pangolin_CoV_orf1ab;6965 | 0 | 0.025 |
| MT072864.1;QIQ54047.1;Pangolin_CoV_orf1ab;6966 | 0 | 0 |
| MT072864.1;QIQ54047.1;Pangolin_CoV_orf1ab;6967 | 0 | 0 |
| MT072864.1;QIQ54047.1;Pangolin_CoV_orf1ab;6968 | 0 | 0 |
| MT072864.1;QIQ54047.1;Pangolin_CoV_orf1ab;6969 | 0 | 0 |
| MT072864.1;QIQ54047.1;Pangolin_CoV_orf1ab;6970 | 0 | 0 |
| MT072864.1;QIQ54047.1;Pangolin_CoV_orf1ab;6971 | 0 | 0 |
| MT072864.1;QIQ54047.1;Pangolin_CoV_orf1ab;6972 | 0 | 0 |
| MT072864.1;QIQ54047.1;Pangolin_CoV_orf1ab;6973 | 0 | 0 |
| MT072864.1;QIQ54047.1;Pangolin_CoV_orf1ab;6974 | 0 | 0 |
| MT072864.1;QIQ54047.1;Pangolin_CoV_orf1ab;6975 | 0 | 0 |
| MT072864.1;QIQ54047.1;Pangolin_CoV_orf1ab;6976 | 0 | 0 |
| MT072864.1;QIQ54047.1;Pangolin_CoV_orf1ab;6977 | 0 | 0 |
| MT072864.1;QIQ54047.1;Pangolin_CoV_orf1ab;6978 | 0 | 0.025 |
| MT072864.1;QIQ54047.1;Pangolin_CoV_orf1ab;6979 | 0 | 0.025 |
| MT072864.1;QIQ54047.1;Pangolin_CoV_orf1ab;6980 | 0 | 0.05 |
| MT072864.1;QIQ54047.1;Pangolin_CoV_orf1ab;6981 | 0 | 0 |
| MT072864.1;QIQ54047.1;Pangolin_CoV_orf1ab;6982 | 0.05 | 0 |
| MT072864.1;QIQ54047.1;Pangolin_CoV_orf1ab;6983 | 0 | 0 |
| MT072864.1;QIQ54047.1;Pangolin_CoV_orf1ab;6984 | 0 | 0.05 |
| MT072864.1;QIQ54047.1;Pangolin_CoV_orf1ab;6985 | 0 | 0 |
| MT072864.1;QIQ54047.1;Pangolin_CoV_orf1ab;6986 | 0 | 0 |
| MT072864.1;QIQ54047.1;Pangolin_CoV_orf1ab;6987 | 0 | 0 |
| MT072864.1;QIQ54047.1;Pangolin_CoV_orf1ab;6988 | 0 | 0 |
| MT072864.1;QIQ54047.1;Pangolin_CoV_orf1ab;6989 | 0 | 0 |
| MT072864.1;QIQ54047.1;Pangolin_CoV_orf1ab;6990 | 0 | 0 |
| MT072864.1;QIQ54047.1;Pangolin_CoV_orf1ab;6991 | 0 | 0 |
| MT072864.1;QIQ54047.1;Pangolin_CoV_orf1ab;6992 | 0 | 0 |
| MT072864.1;QIQ54047.1;Pangolin_CoV_orf1ab;6993 | 0 | 0 |
| MT072864.1;QIQ54047.1;Pangolin_CoV_orf1ab;6994 | 0 | 0 |
| MT072864.1;QIQ54047.1;Pangolin_CoV_orf1ab;6995 | 0.05 | 0 |
| MT072864.1;QIQ54047.1;Pangolin_CoV_orf1ab;6996 | 0.05 | 0 |
| MT072864.1;QIQ54047.1;Pangolin_CoV_orf1ab;6997 | 0 | 0 |
| MT072864.1;QIQ54047.1;Pangolin_CoV_orf1ab;6998 | 0.1 | 0 |
| MT072864.1;QIQ54047.1;Pangolin_CoV_orf1ab;6999 | 0.1 | 0 |

| | | |
|---|---|---|
| MT072864.1;QIQ54047.1;Pangolin_CoV_orf1ab;7000 | 0.05 | 0 |
| MT072864.1;QIQ54047.1;Pangolin_CoV_orf1ab;7001 | 0.1 | 0 |
| MT072864.1;QIQ54047.1;Pangolin_CoV_orf1ab;7002 | 0.1 | 0.05 |
| MT072864.1;QIQ54047.1;Pangolin_CoV_orf1ab;7003 | 0 | 0.05 |
| MT072864.1;QIQ54047.1;Pangolin_CoV_orf1ab;7004 | 0 | 0.05 |
| MT072864.1;QIQ54047.1;Pangolin_CoV_orf1ab;7005 | 0 | 0 |
| MT072864.1;QIQ54047.1;Pangolin_CoV_orf1ab;7006 | 0 | 0 |
| MT072864.1;QIQ54047.1;Pangolin_CoV_orf1ab;7007 | 0.05 | 0.05 |
| MT072864.1;QIQ54047.1;Pangolin_CoV_orf1ab;7008 | 0 | 0 |
| MT072864.1;QIQ54047.1;Pangolin_CoV_orf1ab;7009 | 0 | 0 |
| MT072864.1;QIQ54047.1;Pangolin_CoV_orf1ab;7010 | 0 | 0 |
| MT072864.1;QIQ54047.1;Pangolin_CoV_orf1ab;7011 | 0 | 0 |
| MT072864.1;QIQ54047.1;Pangolin_CoV_orf1ab;7012 | 0 | 0 |
| MT072864.1;QIQ54047.1;Pangolin_CoV_orf1ab;7013 | 0 | 0 |
| MT072864.1;QIQ54047.1;Pangolin_CoV_orf1ab;7014 | 0 | 0 |
| MT072864.1;QIQ54047.1;Pangolin_CoV_orf1ab;7015 | 0 | 0 |
| MT072864.1;QIQ54047.1;Pangolin_CoV_orf1ab;7016 | 0 | 0 |
| MT072864.1;QIQ54047.1;Pangolin_CoV_orf1ab;7017 | 0 | 0 |
| MT072864.1;QIQ54047.1;Pangolin_CoV_orf1ab;7018 | 0 | 0 |
| MT072864.1;QIQ54047.1;Pangolin_CoV_orf1ab;7019 | 0 | 0 |
| MT072864.1;QIQ54047.1;Pangolin_CoV_orf1ab;7020 | 0 | 0 |
| MT072864.1;QIQ54047.1;Pangolin_CoV_orf1ab;7021 | 0 | 0 |
| MT072864.1;QIQ54047.1;Pangolin_CoV_orf1ab;7022 | 0 | 0 |
| MT072864.1;QIQ54047.1;Pangolin_CoV_orf1ab;7023 | 0 | 0 |
| MT072864.1;QIQ54047.1;Pangolin_CoV_orf1ab;7024 | 0.05 | 0.075 |
| MT072864.1;QIQ54047.1;Pangolin_CoV_orf1ab;7025 | 0.05 | 0.075 |
| MT072864.1;QIQ54047.1;Pangolin_CoV_orf1ab;7026 | 0.05 | 0.05 |
| MT072864.1;QIQ54047.1;Pangolin_CoV_orf1ab;7027 | 0.05 | 0.05 |
| MT072864.1;QIQ54047.1;Pangolin_CoV_orf1ab;7028 | 0.05 | 0.05 |
| MT072864.1;QIQ54047.1;Pangolin_CoV_orf1ab;7029 | 0.05 | 0.075 |
| MT072864.1;QIQ54047.1;Pangolin_CoV_orf1ab;7030 | 0.05 | 0.025 |
| MT072864.1;QIQ54047.1;Pangolin_CoV_orf1ab;7031 | 0 | 0.025 |
| MT072864.1;QIQ54047.1;Pangolin_CoV_orf1ab;7032 | 0 | 0 |
| MT072864.1;QIQ54047.1;Pangolin_CoV_orf1ab;7033 | 0 | 0 |
| MT072864.1;QIQ54047.1;Pangolin_CoV_orf1ab;7034 | 0 | 0 |
| MT072864.1;QIQ54047.1;Pangolin_CoV_orf1ab;7035 | 0 | 0 |
| MT072864.1;QIQ54047.1;Pangolin_CoV_orf1ab;7036 | 0 | 0 |
| MT072864.1;QIQ54047.1;Pangolin_CoV_orf1ab;7037 | 0 | 0 |
| MT072864.1;QIQ54047.1;Pangolin_CoV_orf1ab;7038 | 0 | 0 |
| MT072864.1;QIQ54047.1;Pangolin_CoV_orf1ab;7039 | 0 | 0 |
| MT072864.1;QIQ54047.1;Pangolin_CoV_orf1ab;7040 | 0 | 0 |
| MT072864.1;QIQ54047.1;Pangolin_CoV_orf1ab;7041 | 0 | 0 |
| MT072864.1;QIQ54047.1;Pangolin_CoV_orf1ab;7042 | 0 | 0 |

| | | |
|---|---|---|
| MT072864.1;QIQ54047.1;Pangolin_CoV_orf1ab;7043 | 0 | 0 |
| MT072864.1;QIQ54047.1;Pangolin_CoV_orf1ab;7044 | 0 | 0 |
| MT072864.1;QIQ54047.1;Pangolin_CoV_orf1ab;7045 | 0.05 | 0 |
| MT072864.1;QIQ54047.1;Pangolin_CoV_orf1ab;7046 | 0.05 | 0 |
| MT072864.1;QIQ54047.1;Pangolin_CoV_orf1ab;7047 | 0.05 | 0 |
| MT072864.1;QIQ54047.1;Pangolin_CoV_orf1ab;7048 | 0.1 | 0 |
| MT072864.1;QIQ54047.1;Pangolin_CoV_orf1ab;7049 | 0.05 | 0 |
| MT072864.1;QIQ54047.1;Pangolin_CoV_orf1ab;7050 | 0 | 0 |
| MT072864.1;QIQ54047.1;Pangolin_CoV_orf1ab;7051 | 0 | 0 |
| MT072864.1;QIQ54047.1;Pangolin_CoV_orf1ab;7052 | 0 | 0 |
| MT072864.1;QIQ54047.1;Pangolin_CoV_orf1ab;7053 | 0 | 0 |
| MT072864.1;QIQ54047.1;Pangolin_CoV_orf1ab;7054 | 0 | 0 |
| MT072864.1;QIQ54047.1;Pangolin_CoV_orf1ab;7055 | 0 | 0 |
| MT072864.1;QIQ54047.1;Pangolin_CoV_orf1ab;7056 | 0 | 0 |
| MT072864.1;QIQ54047.1;Pangolin_CoV_orf1ab;7057 | 0 | 0 |
| MT072864.1;QIQ54047.1;Pangolin_CoV_orf1ab;7058 | 0 | 0 |
| MT072864.1;QIQ54047.1;Pangolin_CoV_orf1ab;7059 | 0 | 0 |
| MT072864.1;QIQ54047.1;Pangolin_CoV_orf1ab;7060 | 0.05 | 0 |
| MT072864.1;QIQ54047.1;Pangolin_CoV_orf1ab;7061 | 0 | 0 |
| MT072864.1;QIQ54047.1;Pangolin_CoV_orf1ab;7062 | 0 | 0 |
| MT072864.1;QIQ54047.1;Pangolin_CoV_orf1ab;7063 | 0 | 0 |
| MT072864.1;QIQ54047.1;Pangolin_CoV_orf1ab;7064 | 0 | 0 |
| MT072864.1;QIQ54047.1;Pangolin_CoV_orf1ab;7065 | 0 | 0 |
| MT072864.1;QIQ54047.1;Pangolin_CoV_orf1ab;7066 | 0 | 0 |
| MT072864.1;QIQ54047.1;Pangolin_CoV_orf1ab;7067 | 0 | 0.025 |
| MT072864.1;QIQ54047.1;Pangolin_CoV_orf1ab;7068 | 0 | 0 |
| MT072864.1;QIQ54047.1;Pangolin_CoV_orf1ab;7069 | 0 | 0 |
| MT072864.1;QIQ54047.1;Pangolin_CoV_orf1ab;7070 | 0 | 0 |
| MT072864.1;QIQ54047.1;Pangolin_CoV_orf1ab;7071 | 0 | 0 |
| MT072864.1;QIQ54047.1;Pangolin_CoV_orf1ab;7072 | 0 | 0 |
| MT072864.1;QIQ54047.1;Pangolin_CoV_orf1ab;7073 | 0.05 | 0 |
| MT072864.1;QIQ54047.1;Pangolin_CoV_orf1ab;7074 | 0 | 0 |
| MT072864.1;QIQ54047.1;Pangolin_CoV_orf1ab;7075 | 0 | 0 |
| MT072864.1;QIQ54047.1;Pangolin_CoV_orf1ab;7076 | 0 | 0 |
| MT072864.1;QIQ54047.1;Pangolin_CoV_orf1ab;7077 | 0 | 0 |
| MT072864.1;QIQ54048.1;Pangolin_CoV_spike;1 | 0 | 0 |
| MT072864.1;QIQ54048.1;Pangolin_CoV_spike;2 | 0 | 0 |
| MT072864.1;QIQ54048.1;Pangolin_CoV_spike;3 | 0 | 0 |
| MT072864.1;QIQ54048.1;Pangolin_CoV_spike;4 | 0 | 0 |
| MT072864.1;QIQ54048.1;Pangolin_CoV_spike;5 | 0 | 0 |
| MT072864.1;QIQ54048.1;Pangolin_CoV_spike;6 | 0 | 0 |
| MT072864.1;QIQ54048.1;Pangolin_CoV_spike;7 | 0 | 0 |
| MT072864.1;QIQ54048.1;Pangolin_CoV_spike;8 | 0.05 | 0 |

| | | |
|---|---|---|
| MT072864.1;QIQ54048.1;Pangolin_CoV_spike;9 | 0 | 0 |
| MT072864.1;QIQ54048.1;Pangolin_CoV_spike;10 | 0 | 0 |
| MT072864.1;QIQ54048.1;Pangolin_CoV_spike;11 | 0 | 0 |
| MT072864.1;QIQ54048.1;Pangolin_CoV_spike;12 | 0 | 0 |
| MT072864.1;QIQ54048.1;Pangolin_CoV_spike;13 | 0 | 0 |
| MT072864.1;QIQ54048.1;Pangolin_CoV_spike;14 | 0 | 0 |
| MT072864.1;QIQ54048.1;Pangolin_CoV_spike;15 | 0 | 0 |
| MT072864.1;QIQ54048.1;Pangolin_CoV_spike;16 | 0 | 0 |
| MT072864.1;QIQ54048.1;Pangolin_CoV_spike;17 | 0.1 | 0.025 |
| MT072864.1;QIQ54048.1;Pangolin_CoV_spike;18 | 0.05 | 0.025 |
| MT072864.1;QIQ54048.1;Pangolin_CoV_spike;19 | 0 | 0.025 |
| MT072864.1;QIQ54048.1;Pangolin_CoV_spike;20 | 0.05 | 0.025 |
| MT072864.1;QIQ54048.1;Pangolin_CoV_spike;21 | 0.05 | 0.025 |
| MT072864.1;QIQ54048.1;Pangolin_CoV_spike;22 | 0 | 0 |
| MT072864.1;QIQ54048.1;Pangolin_CoV_spike;23 | 0 | 0 |
| MT072864.1;QIQ54048.1;Pangolin_CoV_spike;24 | 0 | 0 |
| MT072864.1;QIQ54048.1;Pangolin_CoV_spike;25 | 0.1 | 0.025 |
| MT072864.1;QIQ54048.1;Pangolin_CoV_spike;26 | 0.1 | 0.025 |
| MT072864.1;QIQ54048.1;Pangolin_CoV_spike;27 | 0.1 | 0.025 |
| MT072864.1;QIQ54048.1;Pangolin_CoV_spike;28 | 0.05 | 0 |
| MT072864.1;QIQ54048.1;Pangolin_CoV_spike;29 | 0 | 0 |
| MT072864.1;QIQ54048.1;Pangolin_CoV_spike;30 | 0 | 0 |
| MT072864.1;QIQ54048.1;Pangolin_CoV_spike;31 | 0 | 0.05 |
| MT072864.1;QIQ54048.1;Pangolin_CoV_spike;32 | 0 | 0 |
| MT072864.1;QIQ54048.1;Pangolin_CoV_spike;33 | 0 | 0 |
| MT072864.1;QIQ54048.1;Pangolin_CoV_spike;34 | 0 | 0 |
| MT072864.1;QIQ54048.1;Pangolin_CoV_spike;35 | 0 | 0 |
| MT072864.1;QIQ54048.1;Pangolin_CoV_spike;36 | 0 | 0 |
| MT072864.1;QIQ54048.1;Pangolin_CoV_spike;37 | 0 | 0 |
| MT072864.1;QIQ54048.1;Pangolin_CoV_spike;38 | 0 | 0 |
| MT072864.1;QIQ54048.1;Pangolin_CoV_spike;39 | 0 | 0 |
| MT072864.1;QIQ54048.1;Pangolin_CoV_spike;40 | 0 | 0 |
| MT072864.1;QIQ54048.1;Pangolin_CoV_spike;41 | 0 | 0 |
| MT072864.1;QIQ54048.1;Pangolin_CoV_spike;42 | 0 | 0 |
| MT072864.1;QIQ54048.1;Pangolin_CoV_spike;43 | 0 | 0 |
| MT072864.1;QIQ54048.1;Pangolin_CoV_spike;44 | 0 | 0 |
| MT072864.1;QIQ54048.1;Pangolin_CoV_spike;45 | 0 | 0 |
| MT072864.1;QIQ54048.1;Pangolin_CoV_spike;46 | 0 | 0 |
| MT072864.1;QIQ54048.1;Pangolin_CoV_spike;47 | 0 | 0 |
| MT072864.1;QIQ54048.1;Pangolin_CoV_spike;48 | 0 | 0 |
| MT072864.1;QIQ54048.1;Pangolin_CoV_spike;49 | 0 | 0 |
| MT072864.1;QIQ54048.1;Pangolin_CoV_spike;50 | 0 | 0 |
| MT072864.1;QIQ54048.1;Pangolin_CoV_spike;51 | 0 | 0 |

| | | |
|---|---|---|
| MT072864.1;QIQ54048.1;Pangolin_CoV_spike;52 | 0 | 0 |
| MT072864.1;QIQ54048.1;Pangolin_CoV_spike;53 | 0 | 0 |
| MT072864.1;QIQ54048.1;Pangolin_CoV_spike;54 | 0 | 0 |
| MT072864.1;QIQ54048.1;Pangolin_CoV_spike;55 | 0 | 0 |
| MT072864.1;QIQ54048.1;Pangolin_CoV_spike;56 | 0 | 0 |
| MT072864.1;QIQ54048.1;Pangolin_CoV_spike;57 | 0 | 0 |
| MT072864.1;QIQ54048.1;Pangolin_CoV_spike;58 | 0 | 0 |
| MT072864.1;QIQ54048.1;Pangolin_CoV_spike;59 | 0 | 0 |
| MT072864.1;QIQ54048.1;Pangolin_CoV_spike;60 | 0.05 | 0 |
| MT072864.1;QIQ54048.1;Pangolin_CoV_spike;61 | 0 | 0 |
| MT072864.1;QIQ54048.1;Pangolin_CoV_spike;62 | 0 | 0 |
| MT072864.1;QIQ54048.1;Pangolin_CoV_spike;63 | 0 | 0 |
| MT072864.1;QIQ54048.1;Pangolin_CoV_spike;64 | 0 | 0 |
| MT072864.1;QIQ54048.1;Pangolin_CoV_spike;65 | 0 | 0 |
| MT072864.1;QIQ54048.1;Pangolin_CoV_spike;66 | 0 | 0 |
| MT072864.1;QIQ54048.1;Pangolin_CoV_spike;67 | 0 | 0 |
| MT072864.1;QIQ54048.1;Pangolin_CoV_spike;68 | 0 | 0 |
| MT072864.1;QIQ54048.1;Pangolin_CoV_spike;69 | 0 | 0 |
| MT072864.1;QIQ54048.1;Pangolin_CoV_spike;70 | 0 | 0 |
| MT072864.1;QIQ54048.1;Pangolin_CoV_spike;71 | 0 | 0 |
| MT072864.1;QIQ54048.1;Pangolin_CoV_spike;72 | 0.05 | 0 |
| MT072864.1;QIQ54048.1;Pangolin_CoV_spike;73 | 0.05 | 0.025 |
| MT072864.1;QIQ54048.1;Pangolin_CoV_spike;74 | 0 | 0 |
| MT072864.1;QIQ54048.1;Pangolin_CoV_spike;75 | 0 | 0 |
| MT072864.1;QIQ54048.1;Pangolin_CoV_spike;76 | 0 | 0 |
| MT072864.1;QIQ54048.1;Pangolin_CoV_spike;77 | 0.05 | 0.025 |
| MT072864.1;QIQ54048.1;Pangolin_CoV_spike;78 | 0 | 0.025 |
| MT072864.1;QIQ54048.1;Pangolin_CoV_spike;79 | 0 | 0 |
| MT072864.1;QIQ54048.1;Pangolin_CoV_spike;80 | 0 | 0 |
| MT072864.1;QIQ54048.1;Pangolin_CoV_spike;81 | 0 | 0.025 |
| MT072864.1;QIQ54048.1;Pangolin_CoV_spike;82 | 0.05 | 0.05 |
| MT072864.1;QIQ54048.1;Pangolin_CoV_spike;83 | 0.05 | 0.05 |
| MT072864.1;QIQ54048.1;Pangolin_CoV_spike;84 | 0.05 | 0.05 |
| MT072864.1;QIQ54048.1;Pangolin_CoV_spike;85 | 0.05 | 0.05 |
| MT072864.1;QIQ54048.1;Pangolin_CoV_spike;86 | 0.05 | 0.025 |
| MT072864.1;QIQ54048.1;Pangolin_CoV_spike;87 | 0.05 | 0 |
| MT072864.1;QIQ54048.1;Pangolin_CoV_spike;88 | 0.05 | 0.025 |
| MT072864.1;QIQ54048.1;Pangolin_CoV_spike;89 | 0.05 | 0 |
| MT072864.1;QIQ54048.1;Pangolin_CoV_spike;90 | 0.05 | 0.025 |
| MT072864.1;QIQ54048.1;Pangolin_CoV_spike;91 | 0.05 | 0 |
| MT072864.1;QIQ54048.1;Pangolin_CoV_spike;92 | 0.05 | 0 |
| MT072864.1;QIQ54048.1;Pangolin_CoV_spike;93 | 0 | 0 |
| MT072864.1;QIQ54048.1;Pangolin_CoV_spike;94 | 0 | 0 |

| | | |
|---|---|---|
| MT072864.1;QIQ54048.1;Pangolin_CoV_spike;95 | 0 | 0 |
| MT072864.1;QIQ54048.1;Pangolin_CoV_spike;96 | 0 | 0 |
| MT072864.1;QIQ54048.1;Pangolin_CoV_spike;97 | 0 | 0 |
| MT072864.1;QIQ54048.1;Pangolin_CoV_spike;98 | 0 | 0 |
| MT072864.1;QIQ54048.1;Pangolin_CoV_spike;99 | 0 | 0 |
| MT072864.1;QIQ54048.1;Pangolin_CoV_spike;100 | 0 | 0 |
| MT072864.1;QIQ54048.1;Pangolin_CoV_spike;101 | 0 | 0 |
| MT072864.1;QIQ54048.1;Pangolin_CoV_spike;102 | 0 | 0 |
| MT072864.1;QIQ54048.1;Pangolin_CoV_spike;103 | 0 | 0 |
| MT072864.1;QIQ54048.1;Pangolin_CoV_spike;104 | 0 | 0 |
| MT072864.1;QIQ54048.1;Pangolin_CoV_spike;105 | 0 | 0 |
| MT072864.1;QIQ54048.1;Pangolin_CoV_spike;106 | 0 | 0 |
| MT072864.1;QIQ54048.1;Pangolin_CoV_spike;107 | 0 | 0 |
| MT072864.1;QIQ54048.1;Pangolin_CoV_spike;108 | 0 | 0 |
| MT072864.1;QIQ54048.1;Pangolin_CoV_spike;109 | 0 | 0 |
| MT072864.1;QIQ54048.1;Pangolin_CoV_spike;110 | 0 | 0 |
| MT072864.1;QIQ54048.1;Pangolin_CoV_spike;111 | 0.05 | 0 |
| MT072864.1;QIQ54048.1;Pangolin_CoV_spike;112 | 0.05 | 0 |
| MT072864.1;QIQ54048.1;Pangolin_CoV_spike;113 | 0.05 | 0 |
| MT072864.1;QIQ54048.1;Pangolin_CoV_spike;114 | 0.05 | 0 |
| MT072864.1;QIQ54048.1;Pangolin_CoV_spike;115 | 0 | 0 |
| MT072864.1;QIQ54048.1;Pangolin_CoV_spike;116 | 0 | 0 |
| MT072864.1;QIQ54048.1;Pangolin_CoV_spike;117 | 0 | 0 |
| MT072864.1;QIQ54048.1;Pangolin_CoV_spike;118 | 0 | 0 |
| MT072864.1;QIQ54048.1;Pangolin_CoV_spike;119 | 0.05 | 0 |
| MT072864.1;QIQ54048.1;Pangolin_CoV_spike;120 | 0.05 | 0 |
| MT072864.1;QIQ54048.1;Pangolin_CoV_spike;121 | 0 | 0 |
| MT072864.1;QIQ54048.1;Pangolin_CoV_spike;122 | 0 | 0 |
| MT072864.1;QIQ54048.1;Pangolin_CoV_spike;123 | 0 | 0 |
| MT072864.1;QIQ54048.1;Pangolin_CoV_spike;124 | 0.05 | 0 |
| MT072864.1;QIQ54048.1;Pangolin_CoV_spike;125 | 0 | 0 |
| MT072864.1;QIQ54048.1;Pangolin_CoV_spike;126 | 0 | 0 |
| MT072864.1;QIQ54048.1;Pangolin_CoV_spike;127 | 0 | 0 |
| MT072864.1;QIQ54048.1;Pangolin_CoV_spike;128 | 0 | 0 |
| MT072864.1;QIQ54048.1;Pangolin_CoV_spike;129 | 0 | 0 |
| MT072864.1;QIQ54048.1;Pangolin_CoV_spike;130 | 0 | 0 |
| MT072864.1;QIQ54048.1;Pangolin_CoV_spike;131 | 0 | 0 |
| MT072864.1;QIQ54048.1;Pangolin_CoV_spike;132 | 0 | 0 |
| MT072864.1;QIQ54048.1;Pangolin_CoV_spike;133 | 0 | 0 |
| MT072864.1;QIQ54048.1;Pangolin_CoV_spike;134 | 0 | 0 |
| MT072864.1;QIQ54048.1;Pangolin_CoV_spike;135 | 0 | 0 |
| MT072864.1;QIQ54048.1;Pangolin_CoV_spike;136 | 0.05 | 0 |
| MT072864.1;QIQ54048.1;Pangolin_CoV_spike;137 | 0.05 | 0 |

| | | |
|---|---|---|
| MT072864.1;QIQ54048.1;Pangolin_CoV_spike;138 | 0 | 0 |
| MT072864.1;QIQ54048.1;Pangolin_CoV_spike;139 | 0.05 | 0 |
| MT072864.1;QIQ54048.1;Pangolin_CoV_spike;140 | 0.05 | 0 |
| MT072864.1;QIQ54048.1;Pangolin_CoV_spike;141 | 0.15 | 0.025 |
| MT072864.1;QIQ54048.1;Pangolin_CoV_spike;142 | 0.25 | 0.075 |
| MT072864.1;QIQ54048.1;Pangolin_CoV_spike;143 | 0.2 | 0.025 |
| MT072864.1;QIQ54048.1;Pangolin_CoV_spike;144 | 0.05 | 0 |
| MT072864.1;QIQ54048.1;Pangolin_CoV_spike;145 | 0.05 | 0 |
| MT072864.1;QIQ54048.1;Pangolin_CoV_spike;146 | 0.1 | 0 |
| MT072864.1;QIQ54048.1;Pangolin_CoV_spike;147 | 0.05 | 0 |
| MT072864.1;QIQ54048.1;Pangolin_CoV_spike;148 | 0.05 | 0 |
| MT072864.1;QIQ54048.1;Pangolin_CoV_spike;149 | 0.05 | 0.025 |
| MT072864.1;QIQ54048.1;Pangolin_CoV_spike;150 | 0 | 0.05 |
| MT072864.1;QIQ54048.1;Pangolin_CoV_spike;151 | 0 | 0.025 |
| MT072864.1;QIQ54048.1;Pangolin_CoV_spike;152 | 0 | 0.025 |
| MT072864.1;QIQ54048.1;Pangolin_CoV_spike;153 | 0.05 | 0.05 |
| MT072864.1;QIQ54048.1;Pangolin_CoV_spike;154 | 0 | 0.025 |
| MT072864.1;QIQ54048.1;Pangolin_CoV_spike;155 | 0 | 0.025 |
| MT072864.1;QIQ54048.1;Pangolin_CoV_spike;156 | 0 | 0.025 |
| MT072864.1;QIQ54048.1;Pangolin_CoV_spike;157 | 0 | 0 |
| MT072864.1;QIQ54048.1;Pangolin_CoV_spike;158 | 0.05 | 0.025 |
| MT072864.1;QIQ54048.1;Pangolin_CoV_spike;159 | 0 | 0.075 |
| MT072864.1;QIQ54048.1;Pangolin_CoV_spike;160 | 0 | 0 |
| MT072864.1;QIQ54048.1;Pangolin_CoV_spike;161 | 0 | 0 |
| MT072864.1;QIQ54048.1;Pangolin_CoV_spike;162 | 0 | 0.025 |
| MT072864.1;QIQ54048.1;Pangolin_CoV_spike;163 | 0 | 0 |
| MT072864.1;QIQ54048.1;Pangolin_CoV_spike;164 | 0 | 0 |
| MT072864.1;QIQ54048.1;Pangolin_CoV_spike;165 | 0 | 0 |
| MT072864.1;QIQ54048.1;Pangolin_CoV_spike;166 | 0 | 0 |
| MT072864.1;QIQ54048.1;Pangolin_CoV_spike;167 | 0 | 0 |
| MT072864.1;QIQ54048.1;Pangolin_CoV_spike;168 | 0 | 0 |
| MT072864.1;QIQ54048.1;Pangolin_CoV_spike;169 | 0 | 0 |
| MT072864.1;QIQ54048.1;Pangolin_CoV_spike;170 | 0 | 0 |
| MT072864.1;QIQ54048.1;Pangolin_CoV_spike;171 | 0 | 0.025 |
| MT072864.1;QIQ54048.1;Pangolin_CoV_spike;172 | 0 | 0.075 |
| MT072864.1;QIQ54048.1;Pangolin_CoV_spike;173 | 0.05 | 0.225 |
| MT072864.1;QIQ54048.1;Pangolin_CoV_spike;174 | 0.1 | 0.125 |
| MT072864.1;QIQ54048.1;Pangolin_CoV_spike;175 | 0.1 | 0.15 |
| MT072864.1;QIQ54048.1;Pangolin_CoV_spike;176 | 0.1 | 0.175 |
| MT072864.1;QIQ54048.1;Pangolin_CoV_spike;177 | 0.15 | 0.1 |
| MT072864.1;QIQ54048.1;Pangolin_CoV_spike;178 | 0.05 | 0.2 |
| MT072864.1;QIQ54048.1;Pangolin_CoV_spike;179 | 0.05 | 0.125 |
| MT072864.1;QIQ54048.1;Pangolin_CoV_spike;180 | 0.05 | 0.05 |

| | | |
|---|---|---|
| MT072864.1;QIQ54048.1;Pangolin_CoV_spike;181 | 0 | 0.05 |
| MT072864.1;QIQ54048.1;Pangolin_CoV_spike;182 | 0 | 0.025 |
| MT072864.1;QIQ54048.1;Pangolin_CoV_spike;183 | 0 | 0.025 |
| MT072864.1;QIQ54048.1;Pangolin_CoV_spike;184 | 0 | 0 |
| MT072864.1;QIQ54048.1;Pangolin_CoV_spike;185 | 0.05 | 0 |
| MT072864.1;QIQ54048.1;Pangolin_CoV_spike;186 | 0.05 | 0 |
| MT072864.1;QIQ54048.1;Pangolin_CoV_spike;187 | 0 | 0 |
| MT072864.1;QIQ54048.1;Pangolin_CoV_spike;188 | 0 | 0 |
| MT072864.1;QIQ54048.1;Pangolin_CoV_spike;189 | 0 | 0 |
| MT072864.1;QIQ54048.1;Pangolin_CoV_spike;190 | 0 | 0 |
| MT072864.1;QIQ54048.1;Pangolin_CoV_spike;191 | 0 | 0 |
| MT072864.1;QIQ54048.1;Pangolin_CoV_spike;192 | 0 | 0 |
| MT072864.1;QIQ54048.1;Pangolin_CoV_spike;193 | 0 | 0 |
| MT072864.1;QIQ54048.1;Pangolin_CoV_spike;194 | 0 | 0 |
| MT072864.1;QIQ54048.1;Pangolin_CoV_spike;195 | 0 | 0 |
| MT072864.1;QIQ54048.1;Pangolin_CoV_spike;196 | 0 | 0 |
| MT072864.1;QIQ54048.1;Pangolin_CoV_spike;197 | 0 | 0 |
| MT072864.1;QIQ54048.1;Pangolin_CoV_spike;198 | 0 | 0 |
| MT072864.1;QIQ54048.1;Pangolin_CoV_spike;199 | 0 | 0 |
| MT072864.1;QIQ54048.1;Pangolin_CoV_spike;200 | 0 | 0 |
| MT072864.1;QIQ54048.1;Pangolin_CoV_spike;201 | 0 | 0 |
| MT072864.1;QIQ54048.1;Pangolin_CoV_spike;202 | 0 | 0 |
| MT072864.1;QIQ54048.1;Pangolin_CoV_spike;203 | 0 | 0 |
| MT072864.1;QIQ54048.1;Pangolin_CoV_spike;204 | 0.05 | 0.025 |
| MT072864.1;QIQ54048.1;Pangolin_CoV_spike;205 | 0 | 0 |
| MT072864.1;QIQ54048.1;Pangolin_CoV_spike;206 | 0 | 0 |
| MT072864.1;QIQ54048.1;Pangolin_CoV_spike;207 | 0.05 | 0.025 |
| MT072864.1;QIQ54048.1;Pangolin_CoV_spike;208 | 0 | 0.025 |
| MT072864.1;QIQ54048.1;Pangolin_CoV_spike;209 | 0 | 0 |
| MT072864.1;QIQ54048.1;Pangolin_CoV_spike;210 | 0 | 0 |
| MT072864.1;QIQ54048.1;Pangolin_CoV_spike;211 | 0.05 | 0 |
| MT072864.1;QIQ54048.1;Pangolin_CoV_spike;212 | 0 | 0 |
| MT072864.1;QIQ54048.1;Pangolin_CoV_spike;213 | 0 | 0 |
| MT072864.1;QIQ54048.1;Pangolin_CoV_spike;214 | 0 | 0 |
| MT072864.1;QIQ54048.1;Pangolin_CoV_spike;215 | 0 | 0 |
| MT072864.1;QIQ54048.1;Pangolin_CoV_spike;216 | 0 | 0 |
| MT072864.1;QIQ54048.1;Pangolin_CoV_spike;217 | 0 | 0 |
| MT072864.1;QIQ54048.1;Pangolin_CoV_spike;218 | 0 | 0 |
| MT072864.1;QIQ54048.1;Pangolin_CoV_spike;219 | 0 | 0 |
| MT072864.1;QIQ54048.1;Pangolin_CoV_spike;220 | 0 | 0 |
| MT072864.1;QIQ54048.1;Pangolin_CoV_spike;221 | 0 | 0 |
| MT072864.1;QIQ54048.1;Pangolin_CoV_spike;222 | 0 | 0 |
| MT072864.1;QIQ54048.1;Pangolin_CoV_spike;223 | 0 | 0 |

| | | |
|---|---|---|
| MT072864.1;QIQ54048.1;Pangolin_CoV_spike;224 | 0.05 | 0 |
| MT072864.1;QIQ54048.1;Pangolin_CoV_spike;225 | 0 | 0.025 |
| MT072864.1;QIQ54048.1;Pangolin_CoV_spike;226 | 0 | 0 |
| MT072864.1;QIQ54048.1;Pangolin_CoV_spike;227 | 0 | 0 |
| MT072864.1;QIQ54048.1;Pangolin_CoV_spike;228 | 0 | 0 |
| MT072864.1;QIQ54048.1;Pangolin_CoV_spike;229 | 0 | 0 |
| MT072864.1;QIQ54048.1;Pangolin_CoV_spike;230 | 0 | 0 |
| MT072864.1;QIQ54048.1;Pangolin_CoV_spike;231 | 0 | 0 |
| MT072864.1;QIQ54048.1;Pangolin_CoV_spike;232 | 0 | 0 |
| MT072864.1;QIQ54048.1;Pangolin_CoV_spike;233 | 0 | 0 |
| MT072864.1;QIQ54048.1;Pangolin_CoV_spike;234 | 0 | 0 |
| MT072864.1;QIQ54048.1;Pangolin_CoV_spike;235 | 0 | 0 |
| MT072864.1;QIQ54048.1;Pangolin_CoV_spike;236 | 0 | 0 |
| MT072864.1;QIQ54048.1;Pangolin_CoV_spike;237 | 0 | 0 |
| MT072864.1;QIQ54048.1;Pangolin_CoV_spike;238 | 0 | 0 |
| MT072864.1;QIQ54048.1;Pangolin_CoV_spike;239 | 0 | 0 |
| MT072864.1;QIQ54048.1;Pangolin_CoV_spike;240 | 0 | 0 |
| MT072864.1;QIQ54048.1;Pangolin_CoV_spike;241 | 0 | 0 |
| MT072864.1;QIQ54048.1;Pangolin_CoV_spike;242 | 0 | 0.025 |
| MT072864.1;QIQ54048.1;Pangolin_CoV_spike;243 | 0 | 0 |
| MT072864.1;QIQ54048.1;Pangolin_CoV_spike;244 | 0 | 0.025 |
| MT072864.1;QIQ54048.1;Pangolin_CoV_spike;245 | 0 | 0.025 |
| MT072864.1;QIQ54048.1;Pangolin_CoV_spike;246 | 0 | 0.025 |
| MT072864.1;QIQ54048.1;Pangolin_CoV_spike;247 | 0 | 0.025 |
| MT072864.1;QIQ54048.1;Pangolin_CoV_spike;248 | 0.15 | 0.15 |
| MT072864.1;QIQ54048.1;Pangolin_CoV_spike;249 | 0.15 | 0.175 |
| MT072864.1;QIQ54048.1;Pangolin_CoV_spike;250 | 0 | 0 |
| MT072864.1;QIQ54048.1;Pangolin_CoV_spike;251 | 0 | 0 |
| MT072864.1;QIQ54048.1;Pangolin_CoV_spike;252 | 0 | 0 |
| MT072864.1;QIQ54048.1;Pangolin_CoV_spike;253 | 0 | 0 |
| MT072864.1;QIQ54048.1;Pangolin_CoV_spike;254 | 0 | 0 |
| MT072864.1;QIQ54048.1;Pangolin_CoV_spike;255 | 0 | 0 |
| MT072864.1;QIQ54048.1;Pangolin_CoV_spike;256 | 0 | 0 |
| MT072864.1;QIQ54048.1;Pangolin_CoV_spike;257 | 0 | 0.025 |
| MT072864.1;QIQ54048.1;Pangolin_CoV_spike;258 | 0 | 0 |
| MT072864.1;QIQ54048.1;Pangolin_CoV_spike;259 | 0 | 0 |
| MT072864.1;QIQ54048.1;Pangolin_CoV_spike;260 | 0 | 0 |
| MT072864.1;QIQ54048.1;Pangolin_CoV_spike;261 | 0 | 0 |
| MT072864.1;QIQ54048.1;Pangolin_CoV_spike;262 | 0 | 0 |
| MT072864.1;QIQ54048.1;Pangolin_CoV_spike;263 | 0 | 0 |
| MT072864.1;QIQ54048.1;Pangolin_CoV_spike;264 | 0 | 0 |
| MT072864.1;QIQ54048.1;Pangolin_CoV_spike;265 | 0 | 0 |
| MT072864.1;QIQ54048.1;Pangolin_CoV_spike;266 | 0 | 0 |

| | | |
|---|---|---|
| MT072864.1;QIQ54048.1;Pangolin_CoV_spike;267 | 0 | 0 |
| MT072864.1;QIQ54048.1;Pangolin_CoV_spike;268 | 0 | 0.05 |
| MT072864.1;QIQ54048.1;Pangolin_CoV_spike;269 | 0 | 0.075 |
| MT072864.1;QIQ54048.1;Pangolin_CoV_spike;270 | 0 | 0.075 |
| MT072864.1;QIQ54048.1;Pangolin_CoV_spike;271 | 0 | 0.075 |
| MT072864.1;QIQ54048.1;Pangolin_CoV_spike;272 | 0 | 0.075 |
| MT072864.1;QIQ54048.1;Pangolin_CoV_spike;273 | 0 | 0.075 |
| MT072864.1;QIQ54048.1;Pangolin_CoV_spike;274 | 0 | 0.075 |
| MT072864.1;QIQ54048.1;Pangolin_CoV_spike;275 | 0 | 0.075 |
| MT072864.1;QIQ54048.1;Pangolin_CoV_spike;276 | 0 | 0.075 |
| MT072864.1;QIQ54048.1;Pangolin_CoV_spike;277 | 0 | 0.075 |
| MT072864.1;QIQ54048.1;Pangolin_CoV_spike;278 | 0 | 0.075 |
| MT072864.1;QIQ54048.1;Pangolin_CoV_spike;279 | 0 | 0.075 |
| MT072864.1;QIQ54048.1;Pangolin_CoV_spike;280 | 0 | 0.05 |
| MT072864.1;QIQ54048.1;Pangolin_CoV_spike;281 | 0 | 0 |
| MT072864.1;QIQ54048.1;Pangolin_CoV_spike;282 | 0 | 0 |
| MT072864.1;QIQ54048.1;Pangolin_CoV_spike;283 | 0 | 0 |
| MT072864.1;QIQ54048.1;Pangolin_CoV_spike;284 | 0 | 0 |
| MT072864.1;QIQ54048.1;Pangolin_CoV_spike;285 | 0 | 0 |
| MT072864.1;QIQ54048.1;Pangolin_CoV_spike;286 | 0 | 0 |
| MT072864.1;QIQ54048.1;Pangolin_CoV_spike;287 | 0 | 0 |
| MT072864.1;QIQ54048.1;Pangolin_CoV_spike;288 | 0 | 0 |
| MT072864.1;QIQ54048.1;Pangolin_CoV_spike;289 | 0 | 0.025 |
| MT072864.1;QIQ54048.1;Pangolin_CoV_spike;290 | 0 | 0 |
| MT072864.1;QIQ54048.1;Pangolin_CoV_spike;291 | 0 | 0 |
| MT072864.1;QIQ54048.1;Pangolin_CoV_spike;292 | 0 | 0 |
| MT072864.1;QIQ54048.1;Pangolin_CoV_spike;293 | 0 | 0 |
| MT072864.1;QIQ54048.1;Pangolin_CoV_spike;294 | 0 | 0 |
| MT072864.1;QIQ54048.1;Pangolin_CoV_spike;295 | 0 | 0 |
| MT072864.1;QIQ54048.1;Pangolin_CoV_spike;296 | 0 | 0 |
| MT072864.1;QIQ54048.1;Pangolin_CoV_spike;297 | 0 | 0 |
| MT072864.1;QIQ54048.1;Pangolin_CoV_spike;298 | 0 | 0.025 |
| MT072864.1;QIQ54048.1;Pangolin_CoV_spike;299 | 0 | 0 |
| MT072864.1;QIQ54048.1;Pangolin_CoV_spike;300 | 0 | 0 |
| MT072864.1;QIQ54048.1;Pangolin_CoV_spike;301 | 0 | 0 |
| MT072864.1;QIQ54048.1;Pangolin_CoV_spike;302 | 0 | 0.075 |
| MT072864.1;QIQ54048.1;Pangolin_CoV_spike;303 | 0 | 0.1 |
| MT072864.1;QIQ54048.1;Pangolin_CoV_spike;304 | 0 | 0.05 |
| MT072864.1;QIQ54048.1;Pangolin_CoV_spike;305 | 0 | 0.05 |
| MT072864.1;QIQ54048.1;Pangolin_CoV_spike;306 | 0 | 0.05 |
| MT072864.1;QIQ54048.1;Pangolin_CoV_spike;307 | 0 | 0.1 |
| MT072864.1;QIQ54048.1;Pangolin_CoV_spike;308 | 0.05 | 0.15 |
| MT072864.1;QIQ54048.1;Pangolin_CoV_spike;309 | 0.05 | 0.175 |

| | | |
|---|---|---|
| MT072864.1;QIQ54048.1;Pangolin_CoV_spike;310 | 0 | 0.175 |
| MT072864.1;QIQ54048.1;Pangolin_CoV_spike;311 | 0 | 0.025 |
| MT072864.1;QIQ54048.1;Pangolin_CoV_spike;312 | 0 | 0.025 |
| MT072864.1;QIQ54048.1;Pangolin_CoV_spike;313 | 0 | 0.025 |
| MT072864.1;QIQ54048.1;Pangolin_CoV_spike;314 | 0 | 0.025 |
| MT072864.1;QIQ54048.1;Pangolin_CoV_spike;315 | 0 | 0 |
| MT072864.1;QIQ54048.1;Pangolin_CoV_spike;316 | 0 | 0 |
| MT072864.1;QIQ54048.1;Pangolin_CoV_spike;317 | 0 | 0 |
| MT072864.1;QIQ54048.1;Pangolin_CoV_spike;318 | 0 | 0 |
| MT072864.1;QIQ54048.1;Pangolin_CoV_spike;319 | 0 | 0 |
| MT072864.1;QIQ54048.1;Pangolin_CoV_spike;320 | 0 | 0 |
| MT072864.1;QIQ54048.1;Pangolin_CoV_spike;321 | 0 | 0 |
| MT072864.1;QIQ54048.1;Pangolin_CoV_spike;322 | 0 | 0 |
| MT072864.1;QIQ54048.1;Pangolin_CoV_spike;323 | 0 | 0 |
| MT072864.1;QIQ54048.1;Pangolin_CoV_spike;324 | 0 | 0 |
| MT072864.1;QIQ54048.1;Pangolin_CoV_spike;325 | 0 | 0 |
| MT072864.1;QIQ54048.1;Pangolin_CoV_spike;326 | 0 | 0 |
| MT072864.1;QIQ54048.1;Pangolin_CoV_spike;327 | 0 | 0 |
| MT072864.1;QIQ54048.1;Pangolin_CoV_spike;328 | 0 | 0 |
| MT072864.1;QIQ54048.1;Pangolin_CoV_spike;329 | 0.05 | 0 |
| MT072864.1;QIQ54048.1;Pangolin_CoV_spike;330 | 0.05 | 0.025 |
| MT072864.1;QIQ54048.1;Pangolin_CoV_spike;331 | 0.05 | 0.025 |
| MT072864.1;QIQ54048.1;Pangolin_CoV_spike;332 | 0.05 | 0.025 |
| MT072864.1;QIQ54048.1;Pangolin_CoV_spike;333 | 0.1 | 0.05 |
| MT072864.1;QIQ54048.1;Pangolin_CoV_spike;334 | 0.1 | 0.05 |
| MT072864.1;QIQ54048.1;Pangolin_CoV_spike;335 | 0.05 | 0.025 |
| MT072864.1;QIQ54048.1;Pangolin_CoV_spike;336 | 0 | 0.025 |
| MT072864.1;QIQ54048.1;Pangolin_CoV_spike;337 | 0.05 | 0.025 |
| MT072864.1;QIQ54048.1;Pangolin_CoV_spike;338 | 0 | 0 |
| MT072864.1;QIQ54048.1;Pangolin_CoV_spike;339 | 0 | 0 |
| MT072864.1;QIQ54048.1;Pangolin_CoV_spike;340 | 0 | 0 |
| MT072864.1;QIQ54048.1;Pangolin_CoV_spike;341 | 0 | 0 |
| MT072864.1;QIQ54048.1;Pangolin_CoV_spike;342 | 0 | 0 |
| MT072864.1;QIQ54048.1;Pangolin_CoV_spike;343 | 0 | 0.025 |
| MT072864.1;QIQ54048.1;Pangolin_CoV_spike;344 | 0 | 0.025 |
| MT072864.1;QIQ54048.1;Pangolin_CoV_spike;345 | 0 | 0.025 |
| MT072864.1;QIQ54048.1;Pangolin_CoV_spike;346 | 0 | 0.025 |
| MT072864.1;QIQ54048.1;Pangolin_CoV_spike;347 | 0 | 0.025 |
| MT072864.1;QIQ54048.1;Pangolin_CoV_spike;348 | 0 | 0.025 |
| MT072864.1;QIQ54048.1;Pangolin_CoV_spike;349 | 0.05 | 0.025 |
| MT072864.1;QIQ54048.1;Pangolin_CoV_spike;350 | 0 | 0.025 |
| MT072864.1;QIQ54048.1;Pangolin_CoV_spike;351 | 0 | 0.025 |
| MT072864.1;QIQ54048.1;Pangolin_CoV_spike;352 | 0 | 0 |

| | | |
|---|---|---|
| MT072864.1;QIQ54048.1;Pangolin_CoV_spike;353 | 0 | 0 |
| MT072864.1;QIQ54048.1;Pangolin_CoV_spike;354 | 0 | 0 |
| MT072864.1;QIQ54048.1;Pangolin_CoV_spike;355 | 0 | 0 |
| MT072864.1;QIQ54048.1;Pangolin_CoV_spike;356 | 0 | 0.025 |
| MT072864.1;QIQ54048.1;Pangolin_CoV_spike;357 | 0 | 0 |
| MT072864.1;QIQ54048.1;Pangolin_CoV_spike;358 | 0 | 0 |
| MT072864.1;QIQ54048.1;Pangolin_CoV_spike;359 | 0 | 0 |
| MT072864.1;QIQ54048.1;Pangolin_CoV_spike;360 | 0 | 0 |
| MT072864.1;QIQ54048.1;Pangolin_CoV_spike;361 | 0 | 0 |
| MT072864.1;QIQ54048.1;Pangolin_CoV_spike;362 | 0 | 0 |
| MT072864.1;QIQ54048.1;Pangolin_CoV_spike;363 | 0.05 | 0 |
| MT072864.1;QIQ54048.1;Pangolin_CoV_spike;364 | 0 | 0 |
| MT072864.1;QIQ54048.1;Pangolin_CoV_spike;365 | 0 | 0 |
| MT072864.1;QIQ54048.1;Pangolin_CoV_spike;366 | 0 | 0 |
| MT072864.1;QIQ54048.1;Pangolin_CoV_spike;367 | 0 | 0 |
| MT072864.1;QIQ54048.1;Pangolin_CoV_spike;368 | 0 | 0 |
| MT072864.1;QIQ54048.1;Pangolin_CoV_spike;369 | 0 | 0 |
| MT072864.1;QIQ54048.1;Pangolin_CoV_spike;370 | 0.05 | 0 |
| MT072864.1;QIQ54048.1;Pangolin_CoV_spike;371 | 0 | 0 |
| MT072864.1;QIQ54048.1;Pangolin_CoV_spike;372 | 0 | 0 |
| MT072864.1;QIQ54048.1;Pangolin_CoV_spike;373 | 0 | 0 |
| MT072864.1;QIQ54048.1;Pangolin_CoV_spike;374 | 0.05 | 0 |
| MT072864.1;QIQ54048.1;Pangolin_CoV_spike;375 | 0.05 | 0 |
| MT072864.1;QIQ54048.1;Pangolin_CoV_spike;376 | 0 | 0 |
| MT072864.1;QIQ54048.1;Pangolin_CoV_spike;377 | 0 | 0 |
| MT072864.1;QIQ54048.1;Pangolin_CoV_spike;378 | 0 | 0 |
| MT072864.1;QIQ54048.1;Pangolin_CoV_spike;379 | 0 | 0 |
| MT072864.1;QIQ54048.1;Pangolin_CoV_spike;380 | 0 | 0 |
| MT072864.1;QIQ54048.1;Pangolin_CoV_spike;381 | 0 | 0 |
| MT072864.1;QIQ54048.1;Pangolin_CoV_spike;382 | 0 | 0 |
| MT072864.1;QIQ54048.1;Pangolin_CoV_spike;383 | 0 | 0.025 |
| MT072864.1;QIQ54048.1;Pangolin_CoV_spike;384 | 0 | 0 |
| MT072864.1;QIQ54048.1;Pangolin_CoV_spike;385 | 0 | 0.025 |
| MT072864.1;QIQ54048.1;Pangolin_CoV_spike;386 | 0 | 0 |
| MT072864.1;QIQ54048.1;Pangolin_CoV_spike;387 | 0 | 0 |
| MT072864.1;QIQ54048.1;Pangolin_CoV_spike;388 | 0 | 0 |
| MT072864.1;QIQ54048.1;Pangolin_CoV_spike;389 | 0 | 0 |
| MT072864.1;QIQ54048.1;Pangolin_CoV_spike;390 | 0 | 0 |
| MT072864.1;QIQ54048.1;Pangolin_CoV_spike;391 | 0.05 | 0 |
| MT072864.1;QIQ54048.1;Pangolin_CoV_spike;392 | 0 | 0.025 |
| MT072864.1;QIQ54048.1;Pangolin_CoV_spike;393 | 0.1 | 0 |
| MT072864.1;QIQ54048.1;Pangolin_CoV_spike;394 | 0 | 0 |
| MT072864.1;QIQ54048.1;Pangolin_CoV_spike;395 | 0 | 0.025 |

| | | |
|---|---|---|
| MT072864.1;QIQ54048.1;Pangolin_CoV_spike;396 | 0 | 0.025 |
| MT072864.1;QIQ54048.1;Pangolin_CoV_spike;397 | 0.1 | 0 |
| MT072864.1;QIQ54048.1;Pangolin_CoV_spike;398 | 0.1 | 0.025 |
| MT072864.1;QIQ54048.1;Pangolin_CoV_spike;399 | 0 | 0 |
| MT072864.1;QIQ54048.1;Pangolin_CoV_spike;400 | 0 | 0 |
| MT072864.1;QIQ54048.1;Pangolin_CoV_spike;401 | 0.05 | 0 |
| MT072864.1;QIQ54048.1;Pangolin_CoV_spike;402 | 0.05 | 0 |
| MT072864.1;QIQ54048.1;Pangolin_CoV_spike;403 | 0 | 0.025 |
| MT072864.1;QIQ54048.1;Pangolin_CoV_spike;404 | 0 | 0.025 |
| MT072864.1;QIQ54048.1;Pangolin_CoV_spike;405 | 0 | 0 |
| MT072864.1;QIQ54048.1;Pangolin_CoV_spike;406 | 0 | 0 |
| MT072864.1;QIQ54048.1;Pangolin_CoV_spike;407 | 0 | 0 |
| MT072864.1;QIQ54048.1;Pangolin_CoV_spike;408 | 0 | 0.025 |
| MT072864.1;QIQ54048.1;Pangolin_CoV_spike;409 | 0.05 | 0.05 |
| MT072864.1;QIQ54048.1;Pangolin_CoV_spike;410 | 0.05 | 0.025 |
| MT072864.1;QIQ54048.1;Pangolin_CoV_spike;411 | 0 | 0 |
| MT072864.1;QIQ54048.1;Pangolin_CoV_spike;412 | 0 | 0.05 |
| MT072864.1;QIQ54048.1;Pangolin_CoV_spike;413 | 0.05 | 0.05 |
| MT072864.1;QIQ54048.1;Pangolin_CoV_spike;414 | 0.05 | 0.05 |
| MT072864.1;QIQ54048.1;Pangolin_CoV_spike;415 | 0 | 0.05 |
| MT072864.1;QIQ54048.1;Pangolin_CoV_spike;416 | 0.2 | 0.3 |
| MT072864.1;QIQ54048.1;Pangolin_CoV_spike;417 | 0.05 | 0.05 |
| MT072864.1;QIQ54048.1;Pangolin_CoV_spike;418 | 0 | 0.025 |
| MT072864.1;QIQ54048.1;Pangolin_CoV_spike;419 | 0.05 | 0.025 |
| MT072864.1;QIQ54048.1;Pangolin_CoV_spike;420 | 0 | 0.025 |
| MT072864.1;QIQ54048.1;Pangolin_CoV_spike;421 | 0 | 0 |
| MT072864.1;QIQ54048.1;Pangolin_CoV_spike;422 | 0 | 0 |
| MT072864.1;QIQ54048.1;Pangolin_CoV_spike;423 | 0 | 0 |
| MT072864.1;QIQ54048.1;Pangolin_CoV_spike;424 | 0 | 0 |
| MT072864.1;QIQ54048.1;Pangolin_CoV_spike;425 | 0 | 0 |
| MT072864.1;QIQ54048.1;Pangolin_CoV_spike;426 | 0 | 0 |
| MT072864.1;QIQ54048.1;Pangolin_CoV_spike;427 | 0 | 0 |
| MT072864.1;QIQ54048.1;Pangolin_CoV_spike;428 | 0 | 0 |
| MT072864.1;QIQ54048.1;Pangolin_CoV_spike;429 | 0.05 | 0 |
| MT072864.1;QIQ54048.1;Pangolin_CoV_spike;430 | 0.05 | 0.025 |
| MT072864.1;QIQ54048.1;Pangolin_CoV_spike;431 | 0.05 | 0.025 |
| MT072864.1;QIQ54048.1;Pangolin_CoV_spike;432 | 0.05 | 0.05 |
| MT072864.1;QIQ54048.1;Pangolin_CoV_spike;433 | 0.05 | 0.025 |
| MT072864.1;QIQ54048.1;Pangolin_CoV_spike;434 | 0.05 | 0.05 |
| MT072864.1;QIQ54048.1;Pangolin_CoV_spike;435 | 0 | 0.025 |
| MT072864.1;QIQ54048.1;Pangolin_CoV_spike;436 | 0 | 0.025 |
| MT072864.1;QIQ54048.1;Pangolin_CoV_spike;437 | 0 | 0.025 |
| MT072864.1;QIQ54048.1;Pangolin_CoV_spike;438 | 0 | 0.025 |

| | | |
|---|---|---|
| MT072864.1;QIQ54048.1;Pangolin_CoV_spike;439 | 0 | 0.025 |
| MT072864.1;QIQ54048.1;Pangolin_CoV_spike;440 | 0 | 0.025 |
| MT072864.1;QIQ54048.1;Pangolin_CoV_spike;441 | 0 | 0 |
| MT072864.1;QIQ54048.1;Pangolin_CoV_spike;442 | 0 | 0 |
| MT072864.1;QIQ54048.1;Pangolin_CoV_spike;443 | 0 | 0 |
| MT072864.1;QIQ54048.1;Pangolin_CoV_spike;444 | 0 | 0 |
| MT072864.1;QIQ54048.1;Pangolin_CoV_spike;445 | 0 | 0 |
| MT072864.1;QIQ54048.1;Pangolin_CoV_spike;446 | 0 | 0 |
| MT072864.1;QIQ54048.1;Pangolin_CoV_spike;447 | 0 | 0 |
| MT072864.1;QIQ54048.1;Pangolin_CoV_spike;448 | 0 | 0 |
| MT072864.1;QIQ54048.1;Pangolin_CoV_spike;449 | 0 | 0 |
| MT072864.1;QIQ54048.1;Pangolin_CoV_spike;450 | 0 | 0 |
| MT072864.1;QIQ54048.1;Pangolin_CoV_spike;451 | 0 | 0 |
| MT072864.1;QIQ54048.1;Pangolin_CoV_spike;452 | 0 | 0 |
| MT072864.1;QIQ54048.1;Pangolin_CoV_spike;453 | 0 | 0 |
| MT072864.1;QIQ54048.1;Pangolin_CoV_spike;454 | 0 | 0 |
| MT072864.1;QIQ54048.1;Pangolin_CoV_spike;455 | 0 | 0 |
| MT072864.1;QIQ54048.1;Pangolin_CoV_spike;456 | 0 | 0.025 |
| MT072864.1;QIQ54048.1;Pangolin_CoV_spike;457 | 0 | 0.025 |
| MT072864.1;QIQ54048.1;Pangolin_CoV_spike;458 | 0 | 0 |
| MT072864.1;QIQ54048.1;Pangolin_CoV_spike;459 | 0 | 0 |
| MT072864.1;QIQ54048.1;Pangolin_CoV_spike;460 | 0 | 0.025 |
| MT072864.1;QIQ54048.1;Pangolin_CoV_spike;461 | 0 | 0 |
| MT072864.1;QIQ54048.1;Pangolin_CoV_spike;462 | 0.05 | 0 |
| MT072864.1;QIQ54048.1;Pangolin_CoV_spike;463 | 0 | 0 |
| MT072864.1;QIQ54048.1;Pangolin_CoV_spike;464 | 0 | 0 |
| MT072864.1;QIQ54048.1;Pangolin_CoV_spike;465 | 0.05 | 0 |
| MT072864.1;QIQ54048.1;Pangolin_CoV_spike;466 | 0 | 0 |
| MT072864.1;QIQ54048.1;Pangolin_CoV_spike;467 | 0 | 0 |
| MT072864.1;QIQ54048.1;Pangolin_CoV_spike;468 | 0 | 0 |
| MT072864.1;QIQ54048.1;Pangolin_CoV_spike;469 | 0 | 0 |
| MT072864.1;QIQ54048.1;Pangolin_CoV_spike;470 | 0 | 0 |
| MT072864.1;QIQ54048.1;Pangolin_CoV_spike;471 | 0 | 0 |
| MT072864.1;QIQ54048.1;Pangolin_CoV_spike;472 | 0 | 0 |
| MT072864.1;QIQ54048.1;Pangolin_CoV_spike;473 | 0 | 0 |
| MT072864.1;QIQ54048.1;Pangolin_CoV_spike;474 | 0 | 0 |
| MT072864.1;QIQ54048.1;Pangolin_CoV_spike;475 | 0 | 0 |
| MT072864.1;QIQ54048.1;Pangolin_CoV_spike;476 | 0 | 0 |
| MT072864.1;QIQ54048.1;Pangolin_CoV_spike;477 | 0 | 0 |
| MT072864.1;QIQ54048.1;Pangolin_CoV_spike;478 | 0 | 0 |
| MT072864.1;QIQ54048.1;Pangolin_CoV_spike;479 | 0 | 0 |
| MT072864.1;QIQ54048.1;Pangolin_CoV_spike;480 | 0 | 0 |
| MT072864.1;QIQ54048.1;Pangolin_CoV_spike;481 | 0 | 0 |

| | | |
|---|---|---|
| MT072864.1;QIQ54048.1;Pangolin_CoV_spike;482 | 0 | 0 |
| MT072864.1;QIQ54048.1;Pangolin_CoV_spike;483 | 0 | 0 |
| MT072864.1;QIQ54048.1;Pangolin_CoV_spike;484 | 0 | 0 |
| MT072864.1;QIQ54048.1;Pangolin_CoV_spike;485 | 0 | 0 |
| MT072864.1;QIQ54048.1;Pangolin_CoV_spike;486 | 0 | 0 |
| MT072864.1;QIQ54048.1;Pangolin_CoV_spike;487 | 0 | 0 |
| MT072864.1;QIQ54048.1;Pangolin_CoV_spike;488 | 0 | 0 |
| MT072864.1;QIQ54048.1;Pangolin_CoV_spike;489 | 0 | 0 |
| MT072864.1;QIQ54048.1;Pangolin_CoV_spike;490 | 0 | 0 |
| MT072864.1;QIQ54048.1;Pangolin_CoV_spike;491 | 0 | 0.05 |
| MT072864.1;QIQ54048.1;Pangolin_CoV_spike;492 | 0.05 | 0.025 |
| MT072864.1;QIQ54048.1;Pangolin_CoV_spike;493 | 0.05 | 0 |
| MT072864.1;QIQ54048.1;Pangolin_CoV_spike;494 | 0 | 0 |
| MT072864.1;QIQ54048.1;Pangolin_CoV_spike;495 | 0 | 0 |
| MT072864.1;QIQ54048.1;Pangolin_CoV_spike;496 | 0 | 0 |
| MT072864.1;QIQ54048.1;Pangolin_CoV_spike;497 | 0 | 0 |
| MT072864.1;QIQ54048.1;Pangolin_CoV_spike;498 | 0 | 0 |
| MT072864.1;QIQ54048.1;Pangolin_CoV_spike;499 | 0 | 0 |
| MT072864.1;QIQ54048.1;Pangolin_CoV_spike;500 | 0 | 0 |
| MT072864.1;QIQ54048.1;Pangolin_CoV_spike;501 | 0 | 0 |
| MT072864.1;QIQ54048.1;Pangolin_CoV_spike;502 | 0 | 0 |
| MT072864.1;QIQ54048.1;Pangolin_CoV_spike;503 | 0 | 0 |
| MT072864.1;QIQ54048.1;Pangolin_CoV_spike;504 | 0 | 0 |
| MT072864.1;QIQ54048.1;Pangolin_CoV_spike;505 | 0 | 0 |
| MT072864.1;QIQ54048.1;Pangolin_CoV_spike;506 | 0 | 0 |
| MT072864.1;QIQ54048.1;Pangolin_CoV_spike;507 | 0 | 0 |
| MT072864.1;QIQ54048.1;Pangolin_CoV_spike;508 | 0 | 0 |
| MT072864.1;QIQ54048.1;Pangolin_CoV_spike;509 | 0 | 0 |
| MT072864.1;QIQ54048.1;Pangolin_CoV_spike;510 | 0 | 0 |
| MT072864.1;QIQ54048.1;Pangolin_CoV_spike;511 | 0 | 0.025 |
| MT072864.1;QIQ54048.1;Pangolin_CoV_spike;512 | 0 | 0.025 |
| MT072864.1;QIQ54048.1;Pangolin_CoV_spike;513 | 0 | 0.025 |
| MT072864.1;QIQ54048.1;Pangolin_CoV_spike;514 | 0 | 0 |
| MT072864.1;QIQ54048.1;Pangolin_CoV_spike;515 | 0 | 0.025 |
| MT072864.1;QIQ54048.1;Pangolin_CoV_spike;516 | 0 | 0 |
| MT072864.1;QIQ54048.1;Pangolin_CoV_spike;517 | 0 | 0 |
| MT072864.1;QIQ54048.1;Pangolin_CoV_spike;518 | 0 | 0 |
| MT072864.1;QIQ54048.1;Pangolin_CoV_spike;519 | 0 | 0 |
| MT072864.1;QIQ54048.1;Pangolin_CoV_spike;520 | 0 | 0 |
| MT072864.1;QIQ54048.1;Pangolin_CoV_spike;521 | 0 | 0 |
| MT072864.1;QIQ54048.1;Pangolin_CoV_spike;522 | 0 | 0 |
| MT072864.1;QIQ54048.1;Pangolin_CoV_spike;523 | 0 | 0 |
| MT072864.1;QIQ54048.1;Pangolin_CoV_spike;524 | 0 | 0 |

| | | |
|---|---|---|
| MT072864.1;QIQ54048.1;Pangolin_CoV_spike;525 | 0 | 0 |
| MT072864.1;QIQ54048.1;Pangolin_CoV_spike;526 | 0 | 0 |
| MT072864.1;QIQ54048.1;Pangolin_CoV_spike;527 | 0 | 0 |
| MT072864.1;QIQ54048.1;Pangolin_CoV_spike;528 | 0 | 0 |
| MT072864.1;QIQ54048.1;Pangolin_CoV_spike;529 | 0 | 0.025 |
| MT072864.1;QIQ54048.1;Pangolin_CoV_spike;530 | 0 | 0.025 |
| MT072864.1;QIQ54048.1;Pangolin_CoV_spike;531 | 0 | 0.025 |
| MT072864.1;QIQ54048.1;Pangolin_CoV_spike;532 | 0 | 0.025 |
| MT072864.1;QIQ54048.1;Pangolin_CoV_spike;533 | 0 | 0.025 |
| MT072864.1;QIQ54048.1;Pangolin_CoV_spike;534 | 0 | 0.05 |
| MT072864.1;QIQ54048.1;Pangolin_CoV_spike;535 | 0 | 0.075 |
| MT072864.1;QIQ54048.1;Pangolin_CoV_spike;536 | 0 | 0.025 |
| MT072864.1;QIQ54048.1;Pangolin_CoV_spike;537 | 0 | 0.025 |
| MT072864.1;QIQ54048.1;Pangolin_CoV_spike;538 | 0 | 0 |
| MT072864.1;QIQ54048.1;Pangolin_CoV_spike;539 | 0 | 0 |
| MT072864.1;QIQ54048.1;Pangolin_CoV_spike;540 | 0 | 0 |
| MT072864.1;QIQ54048.1;Pangolin_CoV_spike;541 | 0.05 | 0 |
| MT072864.1;QIQ54048.1;Pangolin_CoV_spike;542 | 0 | 0 |
| MT072864.1;QIQ54048.1;Pangolin_CoV_spike;543 | 0 | 0 |
| MT072864.1;QIQ54048.1;Pangolin_CoV_spike;544 | 0 | 0 |
| MT072864.1;QIQ54048.1;Pangolin_CoV_spike;545 | 0 | 0 |
| MT072864.1;QIQ54048.1;Pangolin_CoV_spike;546 | 0 | 0 |
| MT072864.1;QIQ54048.1;Pangolin_CoV_spike;547 | 0 | 0.025 |
| MT072864.1;QIQ54048.1;Pangolin_CoV_spike;548 | 0 | 0.05 |
| MT072864.1;QIQ54048.1;Pangolin_CoV_spike;549 | 0 | 0.1 |
| MT072864.1;QIQ54048.1;Pangolin_CoV_spike;550 | 0 | 0.075 |
| MT072864.1;QIQ54048.1;Pangolin_CoV_spike;551 | 0.05 | 0.1 |
| MT072864.1;QIQ54048.1;Pangolin_CoV_spike;552 | 0 | 0.125 |
| MT072864.1;QIQ54048.1;Pangolin_CoV_spike;553 | 0 | 0.425 |
| MT072864.1;QIQ54048.1;Pangolin_CoV_spike;554 | 0 | 0.45 |
| MT072864.1;QIQ54048.1;Pangolin_CoV_spike;555 | 0 | 0.425 |
| MT072864.1;QIQ54048.1;Pangolin_CoV_spike;556 | 0 | 0.675 |
| MT072864.1;QIQ54048.1;Pangolin_CoV_spike;557 | 0 | 0.65 |
| MT072864.1;QIQ54048.1;Pangolin_CoV_spike;558 | 0 | 0.5 |
| MT072864.1;QIQ54048.1;Pangolin_CoV_spike;559 | 0 | 0.475 |
| MT072864.1;QIQ54048.1;Pangolin_CoV_spike;560 | 0 | 0.55 |
| MT072864.1;QIQ54048.1;Pangolin_CoV_spike;561 | 0 | 0.25 |
| MT072864.1;QIQ54048.1;Pangolin_CoV_spike;562 | 0 | 0.25 |
| MT072864.1;QIQ54048.1;Pangolin_CoV_spike;563 | 0 | 0 |
| MT072864.1;QIQ54048.1;Pangolin_CoV_spike;564 | 0.05 | 0.075 |
| MT072864.1;QIQ54048.1;Pangolin_CoV_spike;565 | 0.05 | 0.05 |
| MT072864.1;QIQ54048.1;Pangolin_CoV_spike;566 | 0 | 0.025 |
| MT072864.1;QIQ54048.1;Pangolin_CoV_spike;567 | 0.05 | 0.05 |

| | | |
|---|---|---|
| MT072864.1;QIQ54048.1;Pangolin_CoV_spike;568 | 0.05 | 0.375 |
| MT072864.1;QIQ54048.1;Pangolin_CoV_spike;569 | 0.05 | 0.65 |
| MT072864.1;QIQ54048.1;Pangolin_CoV_spike;570 | 0 | 0.55 |
| MT072864.1;QIQ54048.1;Pangolin_CoV_spike;571 | 0 | 0.675 |
| MT072864.1;QIQ54048.1;Pangolin_CoV_spike;572 | 0 | 0.625 |
| MT072864.1;QIQ54048.1;Pangolin_CoV_spike;573 | 0 | 0.475 |
| MT072864.1;QIQ54048.1;Pangolin_CoV_spike;574 | 0 | 0.55 |
| MT072864.1;QIQ54048.1;Pangolin_CoV_spike;575 | 0 | 0.625 |
| MT072864.1;QIQ54048.1;Pangolin_CoV_spike;576 | 0 | 0.7 |
| MT072864.1;QIQ54048.1;Pangolin_CoV_spike;577 | 0 | 0.325 |
| MT072864.1;QIQ54048.1;Pangolin_CoV_spike;578 | 0 | 0 |
| MT072864.1;QIQ54048.1;Pangolin_CoV_spike;579 | 0 | 0 |
| MT072864.1;QIQ54048.1;Pangolin_CoV_spike;580 | 0 | 0 |
| MT072864.1;QIQ54048.1;Pangolin_CoV_spike;581 | 0 | 0 |
| MT072864.1;QIQ54048.1;Pangolin_CoV_spike;582 | 0 | 0.05 |
| MT072864.1;QIQ54048.1;Pangolin_CoV_spike;583 | 0 | 0 |
| MT072864.1;QIQ54048.1;Pangolin_CoV_spike;584 | 0 | 0 |
| MT072864.1;QIQ54048.1;Pangolin_CoV_spike;585 | 0 | 0 |
| MT072864.1;QIQ54048.1;Pangolin_CoV_spike;586 | 0 | 0 |
| MT072864.1;QIQ54048.1;Pangolin_CoV_spike;587 | 0 | 0 |
| MT072864.1;QIQ54048.1;Pangolin_CoV_spike;588 | 0 | 0 |
| MT072864.1;QIQ54048.1;Pangolin_CoV_spike;589 | 0 | 0.025 |
| MT072864.1;QIQ54048.1;Pangolin_CoV_spike;590 | 0.1 | 0.075 |
| MT072864.1;QIQ54048.1;Pangolin_CoV_spike;591 | 0.25 | 0.125 |
| MT072864.1;QIQ54048.1;Pangolin_CoV_spike;592 | 0.25 | 0.125 |
| MT072864.1;QIQ54048.1;Pangolin_CoV_spike;593 | 0.4 | 0.25 |
| MT072864.1;QIQ54048.1;Pangolin_CoV_spike;594 | 0.4 | 0.35 |
| MT072864.1;QIQ54048.1;Pangolin_CoV_spike;595 | 0.4 | 0.375 |
| MT072864.1;QIQ54048.1;Pangolin_CoV_spike;596 | 0.35 | 0.175 |
| MT072864.1;QIQ54048.1;Pangolin_CoV_spike;597 | 0.15 | 0.075 |
| MT072864.1;QIQ54048.1;Pangolin_CoV_spike;598 | 0.2 | 0.075 |
| MT072864.1;QIQ54048.1;Pangolin_CoV_spike;599 | 0.2 | 0.15 |
| MT072864.1;QIQ54048.1;Pangolin_CoV_spike;600 | 0.15 | 0.1 |
| MT072864.1;QIQ54048.1;Pangolin_CoV_spike;601 | 0.25 | 0.175 |
| MT072864.1;QIQ54048.1;Pangolin_CoV_spike;602 | 0.1 | 0.075 |
| MT072864.1;QIQ54048.1;Pangolin_CoV_spike;603 | 0 | 0.025 |
| MT072864.1;QIQ54048.1;Pangolin_CoV_spike;604 | 0 | 0.025 |
| MT072864.1;QIQ54048.1;Pangolin_CoV_spike;605 | 0 | 0.025 |
| MT072864.1;QIQ54048.1;Pangolin_CoV_spike;606 | 0 | 0.025 |
| MT072864.1;QIQ54048.1;Pangolin_CoV_spike;607 | 0 | 0.025 |
| MT072864.1;QIQ54048.1;Pangolin_CoV_spike;608 | 0 | 0 |
| MT072864.1;QIQ54048.1;Pangolin_CoV_spike;609 | 0 | 0 |
| MT072864.1;QIQ54048.1;Pangolin_CoV_spike;610 | 0 | 0 |

| | | |
|---|---|---|
| MT072864.1;QIQ54048.1;Pangolin_CoV_spike;611 | 0 | 0 |
| MT072864.1;QIQ54048.1;Pangolin_CoV_spike;612 | 0 | 0 |
| MT072864.1;QIQ54048.1;Pangolin_CoV_spike;613 | 0 | 0 |
| MT072864.1;QIQ54048.1;Pangolin_CoV_spike;614 | 0 | 0 |
| MT072864.1;QIQ54048.1;Pangolin_CoV_spike;615 | 0 | 0 |
| MT072864.1;QIQ54048.1;Pangolin_CoV_spike;616 | 0 | 0 |
| MT072864.1;QIQ54048.1;Pangolin_CoV_spike;617 | 0 | 0 |
| MT072864.1;QIQ54048.1;Pangolin_CoV_spike;618 | 0 | 0.025 |
| MT072864.1;QIQ54048.1;Pangolin_CoV_spike;619 | 0 | 0 |
| MT072864.1;QIQ54048.1;Pangolin_CoV_spike;620 | 0 | 0.05 |
| MT072864.1;QIQ54048.1;Pangolin_CoV_spike;621 | 0 | 0 |
| MT072864.1;QIQ54048.1;Pangolin_CoV_spike;622 | 0 | 0.05 |
| MT072864.1;QIQ54048.1;Pangolin_CoV_spike;623 | 0.05 | 0.025 |
| MT072864.1;QIQ54048.1;Pangolin_CoV_spike;624 | 0 | 0.025 |
| MT072864.1;QIQ54048.1;Pangolin_CoV_spike;625 | 0 | 0.05 |
| MT072864.1;QIQ54048.1;Pangolin_CoV_spike;626 | 0 | 0.2 |
| MT072864.1;QIQ54048.1;Pangolin_CoV_spike;627 | 0 | 0.175 |
| MT072864.1;QIQ54048.1;Pangolin_CoV_spike;628 | 0 | 0 |
| MT072864.1;QIQ54048.1;Pangolin_CoV_spike;629 | 0 | 0.025 |
| MT072864.1;QIQ54048.1;Pangolin_CoV_spike;630 | 0 | 0.05 |
| MT072864.1;QIQ54048.1;Pangolin_CoV_spike;631 | 0 | 0 |
| MT072864.1;QIQ54048.1;Pangolin_CoV_spike;632 | 0 | 0 |
| MT072864.1;QIQ54048.1;Pangolin_CoV_spike;633 | 0 | 0 |
| MT072864.1;QIQ54048.1;Pangolin_CoV_spike;634 | 0.05 | 0 |
| MT072864.1;QIQ54048.1;Pangolin_CoV_spike;635 | 0 | 0 |
| MT072864.1;QIQ54048.1;Pangolin_CoV_spike;636 | 0 | 0 |
| MT072864.1;QIQ54048.1;Pangolin_CoV_spike;637 | 0 | 0 |
| MT072864.1;QIQ54048.1;Pangolin_CoV_spike;638 | 0 | 0 |
| MT072864.1;QIQ54048.1;Pangolin_CoV_spike;639 | 0 | 0 |
| MT072864.1;QIQ54048.1;Pangolin_CoV_spike;640 | 0 | 0 |
| MT072864.1;QIQ54048.1;Pangolin_CoV_spike;641 | 0 | 0 |
| MT072864.1;QIQ54048.1;Pangolin_CoV_spike;642 | 0 | 0 |
| MT072864.1;QIQ54048.1;Pangolin_CoV_spike;643 | 0 | 0.025 |
| MT072864.1;QIQ54048.1;Pangolin_CoV_spike;644 | 0 | 0 |
| MT072864.1;QIQ54048.1;Pangolin_CoV_spike;645 | 0 | 0.025 |
| MT072864.1;QIQ54048.1;Pangolin_CoV_spike;646 | 0 | 0.025 |
| MT072864.1;QIQ54048.1;Pangolin_CoV_spike;647 | 0 | 0 |
| MT072864.1;QIQ54048.1;Pangolin_CoV_spike;648 | 0 | 0.025 |
| MT072864.1;QIQ54048.1;Pangolin_CoV_spike;649 | 0 | 0.025 |
| MT072864.1;QIQ54048.1;Pangolin_CoV_spike;650 | 0 | 0 |
| MT072864.1;QIQ54048.1;Pangolin_CoV_spike;651 | 0 | 0.025 |
| MT072864.1;QIQ54048.1;Pangolin_CoV_spike;652 | 0 | 0.025 |
| MT072864.1;QIQ54048.1;Pangolin_CoV_spike;653 | 0 | 0.075 |

| | | |
|---|---|---|
| MT072864.1;QIQ54048.1;Pangolin_CoV_spike;654 | 0 | 0.025 |
| MT072864.1;QIQ54048.1;Pangolin_CoV_spike;655 | 0 | 0.025 |
| MT072864.1;QIQ54048.1;Pangolin_CoV_spike;656 | 0.05 | 0.075 |
| MT072864.1;QIQ54048.1;Pangolin_CoV_spike;657 | 0 | 0 |
| MT072864.1;QIQ54048.1;Pangolin_CoV_spike;658 | 0.05 | 0 |
| MT072864.1;QIQ54048.1;Pangolin_CoV_spike;659 | 0 | 0 |
| MT072864.1;QIQ54048.1;Pangolin_CoV_spike;660 | 0 | 0 |
| MT072864.1;QIQ54048.1;Pangolin_CoV_spike;661 | 0 | 0 |
| MT072864.1;QIQ54048.1;Pangolin_CoV_spike;662 | 0 | 0 |
| MT072864.1;QIQ54048.1;Pangolin_CoV_spike;663 | 0 | 0 |
| MT072864.1;QIQ54048.1;Pangolin_CoV_spike;664 | 0 | 0 |
| MT072864.1;QIQ54048.1;Pangolin_CoV_spike;665 | 0 | 0 |
| MT072864.1;QIQ54048.1;Pangolin_CoV_spike;666 | 0 | 0 |
| MT072864.1;QIQ54048.1;Pangolin_CoV_spike;667 | 0 | 0 |
| MT072864.1;QIQ54048.1;Pangolin_CoV_spike;668 | 0 | 0 |
| MT072864.1;QIQ54048.1;Pangolin_CoV_spike;669 | 0 | 0 |
| MT072864.1;QIQ54048.1;Pangolin_CoV_spike;670 | 0 | 0 |
| MT072864.1;QIQ54048.1;Pangolin_CoV_spike;671 | 0 | 0 |
| MT072864.1;QIQ54048.1;Pangolin_CoV_spike;672 | 0 | 0 |
| MT072864.1;QIQ54048.1;Pangolin_CoV_spike;673 | 0 | 0 |
| MT072864.1;QIQ54048.1;Pangolin_CoV_spike;674 | 0 | 0 |
| MT072864.1;QIQ54048.1;Pangolin_CoV_spike;675 | 0 | 0 |
| MT072864.1;QIQ54048.1;Pangolin_CoV_spike;676 | 0 | 0.025 |
| MT072864.1;QIQ54048.1;Pangolin_CoV_spike;677 | 0 | 0 |
| MT072864.1;QIQ54048.1;Pangolin_CoV_spike;678 | 0 | 0 |
| MT072864.1;QIQ54048.1;Pangolin_CoV_spike;679 | 0 | 0 |
| MT072864.1;QIQ54048.1;Pangolin_CoV_spike;680 | 0 | 0.025 |
| MT072864.1;QIQ54048.1;Pangolin_CoV_spike;681 | 0 | 0.025 |
| MT072864.1;QIQ54048.1;Pangolin_CoV_spike;682 | 0 | 0.05 |
| MT072864.1;QIQ54048.1;Pangolin_CoV_spike;683 | 0 | 0.15 |
| MT072864.1;QIQ54048.1;Pangolin_CoV_spike;684 | 0 | 0.15 |
| MT072864.1;QIQ54048.1;Pangolin_CoV_spike;685 | 0 | 0.15 |
| MT072864.1;QIQ54048.1;Pangolin_CoV_spike;686 | 0 | 0.125 |
| MT072864.1;QIQ54048.1;Pangolin_CoV_spike;687 | 0.1 | 0.175 |
| MT072864.1;QIQ54048.1;Pangolin_CoV_spike;688 | 0 | 0.125 |
| MT072864.1;QIQ54048.1;Pangolin_CoV_spike;689 | 0 | 0.125 |
| MT072864.1;QIQ54048.1;Pangolin_CoV_spike;690 | 0 | 0.1 |
| MT072864.1;QIQ54048.1;Pangolin_CoV_spike;691 | 0 | 0.075 |
| MT072864.1;QIQ54048.1;Pangolin_CoV_spike;692 | 0.05 | 0.025 |
| MT072864.1;QIQ54048.1;Pangolin_CoV_spike;693 | 0 | 0.05 |
| MT072864.1;QIQ54048.1;Pangolin_CoV_spike;694 | 0.05 | 0.1 |
| MT072864.1;QIQ54048.1;Pangolin_CoV_spike;695 | 0.1 | 0.05 |
| MT072864.1;QIQ54048.1;Pangolin_CoV_spike;696 | 0.1 | 0.1 |

| | | |
|---|---|---|
| MT072864.1;QIQ54048.1;Pangolin_CoV_spike;697 | 0.15 | 0.125 |
| MT072864.1;QIQ54048.1;Pangolin_CoV_spike;698 | 0.15 | 0.05 |
| MT072864.1;QIQ54048.1;Pangolin_CoV_spike;699 | 0.05 | 0.025 |
| MT072864.1;QIQ54048.1;Pangolin_CoV_spike;700 | 0.2 | 0 |
| MT072864.1;QIQ54048.1;Pangolin_CoV_spike;701 | 0.1 | 0.025 |
| MT072864.1;QIQ54048.1;Pangolin_CoV_spike;702 | 0.2 | 0.125 |
| MT072864.1;QIQ54048.1;Pangolin_CoV_spike;703 | 0.1 | 0 |
| MT072864.1;QIQ54048.1;Pangolin_CoV_spike;704 | 0.05 | 0 |
| MT072864.1;QIQ54048.1;Pangolin_CoV_spike;705 | 0 | 0 |
| MT072864.1;QIQ54048.1;Pangolin_CoV_spike;706 | 0 | 0 |
| MT072864.1;QIQ54048.1;Pangolin_CoV_spike;707 | 0 | 0.05 |
| MT072864.1;QIQ54048.1;Pangolin_CoV_spike;708 | 0 | 0 |
| MT072864.1;QIQ54048.1;Pangolin_CoV_spike;709 | 0 | 0.025 |
| MT072864.1;QIQ54048.1;Pangolin_CoV_spike;710 | 0 | 0 |
| MT072864.1;QIQ54048.1;Pangolin_CoV_spike;711 | 0 | 0 |
| MT072864.1;QIQ54048.1;Pangolin_CoV_spike;712 | 0 | 0 |
| MT072864.1;QIQ54048.1;Pangolin_CoV_spike;713 | 0 | 0 |
| MT072864.1;QIQ54048.1;Pangolin_CoV_spike;714 | 0 | 0 |
| MT072864.1;QIQ54048.1;Pangolin_CoV_spike;715 | 0 | 0 |
| MT072864.1;QIQ54048.1;Pangolin_CoV_spike;716 | 0 | 0 |
| MT072864.1;QIQ54048.1;Pangolin_CoV_spike;717 | 0 | 0 |
| MT072864.1;QIQ54048.1;Pangolin_CoV_spike;718 | 0 | 0 |
| MT072864.1;QIQ54048.1;Pangolin_CoV_spike;719 | 0 | 0 |
| MT072864.1;QIQ54048.1;Pangolin_CoV_spike;720 | 0 | 0 |
| MT072864.1;QIQ54048.1;Pangolin_CoV_spike;721 | 0 | 0 |
| MT072864.1;QIQ54048.1;Pangolin_CoV_spike;722 | 0 | 0 |
| MT072864.1;QIQ54048.1;Pangolin_CoV_spike;723 | 0 | 0 |
| MT072864.1;QIQ54048.1;Pangolin_CoV_spike;724 | 0 | 0 |
| MT072864.1;QIQ54048.1;Pangolin_CoV_spike;725 | 0 | 0 |
| MT072864.1;QIQ54048.1;Pangolin_CoV_spike;726 | 0 | 0 |
| MT072864.1;QIQ54048.1;Pangolin_CoV_spike;727 | 0 | 0 |
| MT072864.1;QIQ54048.1;Pangolin_CoV_spike;728 | 0 | 0 |
| MT072864.1;QIQ54048.1;Pangolin_CoV_spike;729 | 0 | 0 |
| MT072864.1;QIQ54048.1;Pangolin_CoV_spike;730 | 0 | 0 |
| MT072864.1;QIQ54048.1;Pangolin_CoV_spike;731 | 0 | 0 |
| MT072864.1;QIQ54048.1;Pangolin_CoV_spike;732 | 0 | 0 |
| MT072864.1;QIQ54048.1;Pangolin_CoV_spike;733 | 0 | 0 |
| MT072864.1;QIQ54048.1;Pangolin_CoV_spike;734 | 0 | 0 |
| MT072864.1;QIQ54048.1;Pangolin_CoV_spike;735 | 0 | 0 |
| MT072864.1;QIQ54048.1;Pangolin_CoV_spike;736 | 0 | 0 |
| MT072864.1;QIQ54048.1;Pangolin_CoV_spike;737 | 0 | 0 |
| MT072864.1;QIQ54048.1;Pangolin_CoV_spike;738 | 0 | 0 |
| MT072864.1;QIQ54048.1;Pangolin_CoV_spike;739 | 0.05 | 0.025 |

| | | |
|---|---|---|
| MT072864.1;QIQ54048.1;Pangolin_CoV_spike;740 | 0 | 0 |
| MT072864.1;QIQ54048.1;Pangolin_CoV_spike;741 | 0 | 0 |
| MT072864.1;QIQ54048.1;Pangolin_CoV_spike;742 | 0 | 0 |
| MT072864.1;QIQ54048.1;Pangolin_CoV_spike;743 | 0 | 0 |
| MT072864.1;QIQ54048.1;Pangolin_CoV_spike;744 | 0 | 0 |
| MT072864.1;QIQ54048.1;Pangolin_CoV_spike;745 | 0 | 0 |
| MT072864.1;QIQ54048.1;Pangolin_CoV_spike;746 | 0 | 0 |
| MT072864.1;QIQ54048.1;Pangolin_CoV_spike;747 | 0 | 0 |
| MT072864.1;QIQ54048.1;Pangolin_CoV_spike;748 | 0 | 0 |
| MT072864.1;QIQ54048.1;Pangolin_CoV_spike;749 | 0 | 0 |
| MT072864.1;QIQ54048.1;Pangolin_CoV_spike;750 | 0 | 0 |
| MT072864.1;QIQ54048.1;Pangolin_CoV_spike;751 | 0 | 0 |
| MT072864.1;QIQ54048.1;Pangolin_CoV_spike;752 | 0 | 0 |
| MT072864.1;QIQ54048.1;Pangolin_CoV_spike;753 | 0 | 0 |
| MT072864.1;QIQ54048.1;Pangolin_CoV_spike;754 | 0 | 0 |
| MT072864.1;QIQ54048.1;Pangolin_CoV_spike;755 | 0 | 0 |
| MT072864.1;QIQ54048.1;Pangolin_CoV_spike;756 | 0 | 0 |
| MT072864.1;QIQ54048.1;Pangolin_CoV_spike;757 | 0.1 | 0.275 |
| MT072864.1;QIQ54048.1;Pangolin_CoV_spike;758 | 0.1 | 0.35 |
| MT072864.1;QIQ54048.1;Pangolin_CoV_spike;759 | 0.15 | 0.35 |
| MT072864.1;QIQ54048.1;Pangolin_CoV_spike;760 | 0.15 | 0.375 |
| MT072864.1;QIQ54048.1;Pangolin_CoV_spike;761 | 0.15 | 0.4 |
| MT072864.1;QIQ54048.1;Pangolin_CoV_spike;762 | 0.2 | 0.4 |
| MT072864.1;QIQ54048.1;Pangolin_CoV_spike;763 | 0.15 | 0.425 |
| MT072864.1;QIQ54048.1;Pangolin_CoV_spike;764 | 0.15 | 0.425 |
| MT072864.1;QIQ54048.1;Pangolin_CoV_spike;765 | 0.15 | 0.4 |
| MT072864.1;QIQ54048.1;Pangolin_CoV_spike;766 | 0 | 0.275 |
| MT072864.1;QIQ54048.1;Pangolin_CoV_spike;767 | 0.1 | 0.225 |
| MT072864.1;QIQ54048.1;Pangolin_CoV_spike;768 | 0.1 | 0.1 |
| MT072864.1;QIQ54048.1;Pangolin_CoV_spike;769 | 0.1 | 0.15 |
| MT072864.1;QIQ54048.1;Pangolin_CoV_spike;770 | 0.1 | 0.1 |
| MT072864.1;QIQ54048.1;Pangolin_CoV_spike;771 | 0 | 0.1 |
| MT072864.1;QIQ54048.1;Pangolin_CoV_spike;772 | 0 | 0.05 |
| MT072864.1;QIQ54048.1;Pangolin_CoV_spike;773 | 0.05 | 0.05 |
| MT072864.1;QIQ54048.1;Pangolin_CoV_spike;774 | 0.05 | 0.05 |
| MT072864.1;QIQ54048.1;Pangolin_CoV_spike;775 | 0.05 | 0.075 |
| MT072864.1;QIQ54048.1;Pangolin_CoV_spike;776 | 0 | 0 |
| MT072864.1;QIQ54048.1;Pangolin_CoV_spike;777 | 0 | 0 |
| MT072864.1;QIQ54048.1;Pangolin_CoV_spike;778 | 0 | 0 |
| MT072864.1;QIQ54048.1;Pangolin_CoV_spike;779 | 0 | 0 |
| MT072864.1;QIQ54048.1;Pangolin_CoV_spike;780 | 0 | 0 |
| MT072864.1;QIQ54048.1;Pangolin_CoV_spike;781 | 0 | 0.025 |
| MT072864.1;QIQ54048.1;Pangolin_CoV_spike;782 | 0 | 0.025 |

| | | |
|---|---|---|
| MT072864.1;QIQ54048.1;Pangolin_CoV_spike;783 | 0 | 0.35 |
| MT072864.1;QIQ54048.1;Pangolin_CoV_spike;784 | 0 | 0.3 |
| MT072864.1;QIQ54048.1;Pangolin_CoV_spike;785 | 0 | 0.325 |
| MT072864.1;QIQ54048.1;Pangolin_CoV_spike;786 | 0 | 0.25 |
| MT072864.1;QIQ54048.1;Pangolin_CoV_spike;787 | 0 | 0.075 |
| MT072864.1;QIQ54048.1;Pangolin_CoV_spike;788 | 0 | 0.15 |
| MT072864.1;QIQ54048.1;Pangolin_CoV_spike;789 | 0 | 0.225 |
| MT072864.1;QIQ54048.1;Pangolin_CoV_spike;790 | 0 | 0.3 |
| MT072864.1;QIQ54048.1;Pangolin_CoV_spike;791 | 0 | 0.3 |
| MT072864.1;QIQ54048.1;Pangolin_CoV_spike;792 | 0 | 0.075 |
| MT072864.1;QIQ54048.1;Pangolin_CoV_spike;793 | 0 | 0.05 |
| MT072864.1;QIQ54048.1;Pangolin_CoV_spike;794 | 0 | 0.075 |
| MT072864.1;QIQ54048.1;Pangolin_CoV_spike;795 | 0 | 0.075 |
| MT072864.1;QIQ54048.1;Pangolin_CoV_spike;796 | 0 | 0.025 |
| MT072864.1;QIQ54048.1;Pangolin_CoV_spike;797 | 0 | 0.025 |
| MT072864.1;QIQ54048.1;Pangolin_CoV_spike;798 | 0 | 0.025 |
| MT072864.1;QIQ54048.1;Pangolin_CoV_spike;799 | 0 | 0.05 |
| MT072864.1;QIQ54048.1;Pangolin_CoV_spike;800 | 0 | 0.15 |
| MT072864.1;QIQ54048.1;Pangolin_CoV_spike;801 | 0 | 0.15 |
| MT072864.1;QIQ54048.1;Pangolin_CoV_spike;802 | 0.1 | 0.25 |
| MT072864.1;QIQ54048.1;Pangolin_CoV_spike;803 | 0.05 | 0.225 |
| MT072864.1;QIQ54048.1;Pangolin_CoV_spike;804 | 0.05 | 0.625 |
| MT072864.1;QIQ54048.1;Pangolin_CoV_spike;805 | 0.1 | 0.75 |
| MT072864.1;QIQ54048.1;Pangolin_CoV_spike;806 | 0.05 | 0.85 |
| MT072864.1;QIQ54048.1;Pangolin_CoV_spike;807 | 0.05 | 0.825 |
| MT072864.1;QIQ54048.1;Pangolin_CoV_spike;808 | 0.05 | 0.875 |
| MT072864.1;QIQ54048.1;Pangolin_CoV_spike;809 | 0.05 | 0.825 |
| MT072864.1;QIQ54048.1;Pangolin_CoV_spike;810 | 0 | 0.7 |
| MT072864.1;QIQ54048.1;Pangolin_CoV_spike;811 | 0 | 0.725 |
| MT072864.1;QIQ54048.1;Pangolin_CoV_spike;812 | 0 | 0.35 |
| MT072864.1;QIQ54048.1;Pangolin_CoV_spike;813 | 0.05 | 0.25 |
| MT072864.1;QIQ54048.1;Pangolin_CoV_spike;814 | 0.1 | 0.275 |
| MT072864.1;QIQ54048.1;Pangolin_CoV_spike;815 | 0 | 0.125 |
| MT072864.1;QIQ54048.1;Pangolin_CoV_spike;816 | 0 | 0 |
| MT072864.1;QIQ54048.1;Pangolin_CoV_spike;817 | 0 | 0 |
| MT072864.1;QIQ54048.1;Pangolin_CoV_spike;818 | 0 | 0 |
| MT072864.1;QIQ54048.1;Pangolin_CoV_spike;819 | 0 | 0.025 |
| MT072864.1;QIQ54048.1;Pangolin_CoV_spike;820 | 0 | 0.025 |
| MT072864.1;QIQ54048.1;Pangolin_CoV_spike;821 | 0 | 0.025 |
| MT072864.1;QIQ54048.1;Pangolin_CoV_spike;822 | 0 | 0.05 |
| MT072864.1;QIQ54048.1;Pangolin_CoV_spike;823 | 0.05 | 0.025 |
| MT072864.1;QIQ54048.1;Pangolin_CoV_spike;824 | 0.1 | 0.025 |
| MT072864.1;QIQ54048.1;Pangolin_CoV_spike;825 | 0 | 0.025 |

| | | |
|---|---|---|
| MT072864.1;QIQ54048.1;Pangolin_CoV_spike;826 | 0 | 0.025 |
| MT072864.1;QIQ54048.1;Pangolin_CoV_spike;827 | 0.05 | 0.025 |
| MT072864.1;QIQ54048.1;Pangolin_CoV_spike;828 | 0 | 0.025 |
| MT072864.1;QIQ54048.1;Pangolin_CoV_spike;829 | 0 | 0.025 |
| MT072864.1;QIQ54048.1;Pangolin_CoV_spike;830 | 0 | 0.025 |
| MT072864.1;QIQ54048.1;Pangolin_CoV_spike;831 | 0.05 | 0.025 |
| MT072864.1;QIQ54048.1;Pangolin_CoV_spike;832 | 0 | 0 |
| MT072864.1;QIQ54048.1;Pangolin_CoV_spike;833 | 0 | 0 |
| MT072864.1;QIQ54048.1;Pangolin_CoV_spike;834 | 0 | 0 |
| MT072864.1;QIQ54048.1;Pangolin_CoV_spike;835 | 0 | 0 |
| MT072864.1;QIQ54048.1;Pangolin_CoV_spike;836 | 0.05 | 0.025 |
| MT072864.1;QIQ54048.1;Pangolin_CoV_spike;837 | 0 | 0 |
| MT072864.1;QIQ54048.1;Pangolin_CoV_spike;838 | 0.1 | 0.025 |
| MT072864.1;QIQ54048.1;Pangolin_CoV_spike;839 | 0.05 | 0.025 |
| MT072864.1;QIQ54048.1;Pangolin_CoV_spike;840 | 0 | 0.025 |
| MT072864.1;QIQ54048.1;Pangolin_CoV_spike;841 | 0 | 0 |
| MT072864.1;QIQ54048.1;Pangolin_CoV_spike;842 | 0 | 0 |
| MT072864.1;QIQ54048.1;Pangolin_CoV_spike;843 | 0 | 0 |
| MT072864.1;QIQ54048.1;Pangolin_CoV_spike;844 | 0.05 | 0 |
| MT072864.1;QIQ54048.1;Pangolin_CoV_spike;845 | 0 | 0 |
| MT072864.1;QIQ54048.1;Pangolin_CoV_spike;846 | 0 | 0 |
| MT072864.1;QIQ54048.1;Pangolin_CoV_spike;847 | 0 | 0 |
| MT072864.1;QIQ54048.1;Pangolin_CoV_spike;848 | 0 | 0 |
| MT072864.1;QIQ54048.1;Pangolin_CoV_spike;849 | 0 | 0 |
| MT072864.1;QIQ54048.1;Pangolin_CoV_spike;850 | 0 | 0 |
| MT072864.1;QIQ54048.1;Pangolin_CoV_spike;851 | 0 | 0 |
| MT072864.1;QIQ54048.1;Pangolin_CoV_spike;852 | 0 | 0 |
| MT072864.1;QIQ54048.1;Pangolin_CoV_spike;853 | 0 | 0 |
| MT072864.1;QIQ54048.1;Pangolin_CoV_spike;854 | 0 | 0 |
| MT072864.1;QIQ54048.1;Pangolin_CoV_spike;855 | 0 | 0 |
| MT072864.1;QIQ54048.1;Pangolin_CoV_spike;856 | 0 | 0 |
| MT072864.1;QIQ54048.1;Pangolin_CoV_spike;857 | 0 | 0 |
| MT072864.1;QIQ54048.1;Pangolin_CoV_spike;858 | 0 | 0 |
| MT072864.1;QIQ54048.1;Pangolin_CoV_spike;859 | 0 | 0 |
| MT072864.1;QIQ54048.1;Pangolin_CoV_spike;860 | 0 | 0 |
| MT072864.1;QIQ54048.1;Pangolin_CoV_spike;861 | 0 | 0 |
| MT072864.1;QIQ54048.1;Pangolin_CoV_spike;862 | 0 | 0 |
| MT072864.1;QIQ54048.1;Pangolin_CoV_spike;863 | 0 | 0 |
| MT072864.1;QIQ54048.1;Pangolin_CoV_spike;864 | 0 | 0 |
| MT072864.1;QIQ54048.1;Pangolin_CoV_spike;865 | 0 | 0.025 |
| MT072864.1;QIQ54048.1;Pangolin_CoV_spike;866 | 0 | 0.025 |
| MT072864.1;QIQ54048.1;Pangolin_CoV_spike;867 | 0 | 0 |
| MT072864.1;QIQ54048.1;Pangolin_CoV_spike;868 | 0 | 0 |

| | | |
|---|---|---|
| MT072864.1;QIQ54048.1;Pangolin_CoV_spike;869 | 0 | 0 |
| MT072864.1;QIQ54048.1;Pangolin_CoV_spike;870 | 0 | 0 |
| MT072864.1;QIQ54048.1;Pangolin_CoV_spike;871 | 0 | 0 |
| MT072864.1;QIQ54048.1;Pangolin_CoV_spike;872 | 0 | 0 |
| MT072864.1;QIQ54048.1;Pangolin_CoV_spike;873 | 0 | 0.025 |
| MT072864.1;QIQ54048.1;Pangolin_CoV_spike;874 | 0.1 | 0.05 |
| MT072864.1;QIQ54048.1;Pangolin_CoV_spike;875 | 0 | 0 |
| MT072864.1;QIQ54048.1;Pangolin_CoV_spike;876 | 0 | 0 |
| MT072864.1;QIQ54048.1;Pangolin_CoV_spike;877 | 0 | 0 |
| MT072864.1;QIQ54048.1;Pangolin_CoV_spike;878 | 0 | 0 |
| MT072864.1;QIQ54048.1;Pangolin_CoV_spike;879 | 0 | 0 |
| MT072864.1;QIQ54048.1;Pangolin_CoV_spike;880 | 0 | 0 |
| MT072864.1;QIQ54048.1;Pangolin_CoV_spike;881 | 0 | 0 |
| MT072864.1;QIQ54048.1;Pangolin_CoV_spike;882 | 0 | 0 |
| MT072864.1;QIQ54048.1;Pangolin_CoV_spike;883 | 0 | 0 |
| MT072864.1;QIQ54048.1;Pangolin_CoV_spike;884 | 0 | 0 |
| MT072864.1;QIQ54048.1;Pangolin_CoV_spike;885 | 0 | 0.025 |
| MT072864.1;QIQ54048.1;Pangolin_CoV_spike;886 | 0 | 0 |
| MT072864.1;QIQ54048.1;Pangolin_CoV_spike;887 | 0 | 0 |
| MT072864.1;QIQ54048.1;Pangolin_CoV_spike;888 | 0.05 | 0.025 |
| MT072864.1;QIQ54048.1;Pangolin_CoV_spike;889 | 0.05 | 0.025 |
| MT072864.1;QIQ54048.1;Pangolin_CoV_spike;890 | 0.05 | 0.025 |
| MT072864.1;QIQ54048.1;Pangolin_CoV_spike;891 | 0.05 | 0.025 |
| MT072864.1;QIQ54048.1;Pangolin_CoV_spike;892 | 0.05 | 0.025 |
| MT072864.1;QIQ54048.1;Pangolin_CoV_spike;893 | 0.05 | 0.025 |
| MT072864.1;QIQ54048.1;Pangolin_CoV_spike;894 | 0.05 | 0.025 |
| MT072864.1;QIQ54048.1;Pangolin_CoV_spike;895 | 0.1 | 0.05 |
| MT072864.1;QIQ54048.1;Pangolin_CoV_spike;896 | 0.15 | 0.05 |
| MT072864.1;QIQ54048.1;Pangolin_CoV_spike;897 | 0.1 | 0.05 |
| MT072864.1;QIQ54048.1;Pangolin_CoV_spike;898 | 0 | 0 |
| MT072864.1;QIQ54048.1;Pangolin_CoV_spike;899 | 0 | 0 |
| MT072864.1;QIQ54048.1;Pangolin_CoV_spike;900 | 0 | 0 |
| MT072864.1;QIQ54048.1;Pangolin_CoV_spike;901 | 0.05 | 0.025 |
| MT072864.1;QIQ54048.1;Pangolin_CoV_spike;902 | 0 | 0.05 |
| MT072864.1;QIQ54048.1;Pangolin_CoV_spike;903 | 0.05 | 0.05 |
| MT072864.1;QIQ54048.1;Pangolin_CoV_spike;904 | 0 | 0.05 |
| MT072864.1;QIQ54048.1;Pangolin_CoV_spike;905 | 0.05 | 0.075 |
| MT072864.1;QIQ54048.1;Pangolin_CoV_spike;906 | 0.05 | 0.025 |
| MT072864.1;QIQ54048.1;Pangolin_CoV_spike;907 | 0 | 0.05 |
| MT072864.1;QIQ54048.1;Pangolin_CoV_spike;908 | 0 | 0.025 |
| MT072864.1;QIQ54048.1;Pangolin_CoV_spike;909 | 0.05 | 0.05 |
| MT072864.1;QIQ54048.1;Pangolin_CoV_spike;910 | 0 | 0 |
| MT072864.1;QIQ54048.1;Pangolin_CoV_spike;911 | 0 | 0.05 |

| | | |
|---|---|---|
| MT072864.1;QIQ54048.1;Pangolin_CoV_spike;912 | 0.05 | 0.025 |
| MT072864.1;QIQ54048.1;Pangolin_CoV_spike;913 | 0 | 0.05 |
| MT072864.1;QIQ54048.1;Pangolin_CoV_spike;914 | 0.05 | 0.025 |
| MT072864.1;QIQ54048.1;Pangolin_CoV_spike;915 | 0 | 0 |
| MT072864.1;QIQ54048.1;Pangolin_CoV_spike;916 | 0.05 | 0 |
| MT072864.1;QIQ54048.1;Pangolin_CoV_spike;917 | 0.05 | 0.025 |
| MT072864.1;QIQ54048.1;Pangolin_CoV_spike;918 | 0.05 | 0.025 |
| MT072864.1;QIQ54048.1;Pangolin_CoV_spike;919 | 0.05 | 0.05 |
| MT072864.1;QIQ54048.1;Pangolin_CoV_spike;920 | 0.1 | 0.05 |
| MT072864.1;QIQ54048.1;Pangolin_CoV_spike;921 | 0.05 | 0.025 |
| MT072864.1;QIQ54048.1;Pangolin_CoV_spike;922 | 0.05 | 0.025 |
| MT072864.1;QIQ54048.1;Pangolin_CoV_spike;923 | 0.05 | 0.05 |
| MT072864.1;QIQ54048.1;Pangolin_CoV_spike;924 | 0.1 | 0.05 |
| MT072864.1;QIQ54048.1;Pangolin_CoV_spike;925 | 0 | 0.025 |
| MT072864.1;QIQ54048.1;Pangolin_CoV_spike;926 | 0 | 0.025 |
| MT072864.1;QIQ54048.1;Pangolin_CoV_spike;927 | 0 | 0 |
| MT072864.1;QIQ54048.1;Pangolin_CoV_spike;928 | 0.05 | 0.025 |
| MT072864.1;QIQ54048.1;Pangolin_CoV_spike;929 | 0 | 0 |
| MT072864.1;QIQ54048.1;Pangolin_CoV_spike;930 | 0 | 0 |
| MT072864.1;QIQ54048.1;Pangolin_CoV_spike;931 | 0 | 0 |
| MT072864.1;QIQ54048.1;Pangolin_CoV_spike;932 | 0 | 0 |
| MT072864.1;QIQ54048.1;Pangolin_CoV_spike;933 | 0 | 0 |
| MT072864.1;QIQ54048.1;Pangolin_CoV_spike;934 | 0 | 0 |
| MT072864.1;QIQ54048.1;Pangolin_CoV_spike;935 | 0 | 0 |
| MT072864.1;QIQ54048.1;Pangolin_CoV_spike;936 | 0 | 0.075 |
| MT072864.1;QIQ54048.1;Pangolin_CoV_spike;937 | 0 | 0.025 |
| MT072864.1;QIQ54048.1;Pangolin_CoV_spike;938 | 0 | 0.025 |
| MT072864.1;QIQ54048.1;Pangolin_CoV_spike;939 | 0 | 0.025 |
| MT072864.1;QIQ54048.1;Pangolin_CoV_spike;940 | 0 | 0.075 |
| MT072864.1;QIQ54048.1;Pangolin_CoV_spike;941 | 0 | 0.125 |
| MT072864.1;QIQ54048.1;Pangolin_CoV_spike;942 | 0.1 | 0.075 |
| MT072864.1;QIQ54048.1;Pangolin_CoV_spike;943 | 0 | 0.1 |
| MT072864.1;QIQ54048.1;Pangolin_CoV_spike;944 | 0 | 0.075 |
| MT072864.1;QIQ54048.1;Pangolin_CoV_spike;945 | 0 | 0.025 |
| MT072864.1;QIQ54048.1;Pangolin_CoV_spike;946 | 0 | 0.05 |
| MT072864.1;QIQ54048.1;Pangolin_CoV_spike;947 | 0.05 | 0 |
| MT072864.1;QIQ54048.1;Pangolin_CoV_spike;948 | 0 | 0 |
| MT072864.1;QIQ54048.1;Pangolin_CoV_spike;949 | 0 | 0.025 |
| MT072864.1;QIQ54048.1;Pangolin_CoV_spike;950 | 0 | 0.025 |
| MT072864.1;QIQ54048.1;Pangolin_CoV_spike;951 | 0 | 0 |
| MT072864.1;QIQ54048.1;Pangolin_CoV_spike;952 | 0 | 0.025 |
| MT072864.1;QIQ54048.1;Pangolin_CoV_spike;953 | 0 | 0.05 |
| MT072864.1;QIQ54048.1;Pangolin_CoV_spike;954 | 0 | 0.025 |

| | | |
|---|---|---|
| MT072864.1;QIQ54048.1;Pangolin_CoV_spike;955 | 0 | 0 |
| MT072864.1;QIQ54048.1;Pangolin_CoV_spike;956 | 0 | 0 |
| MT072864.1;QIQ54048.1;Pangolin_CoV_spike;957 | 0 | 0 |
| MT072864.1;QIQ54048.1;Pangolin_CoV_spike;958 | 0 | 0 |
| MT072864.1;QIQ54048.1;Pangolin_CoV_spike;959 | 0 | 0 |
| MT072864.1;QIQ54048.1;Pangolin_CoV_spike;960 | 0 | 0.025 |
| MT072864.1;QIQ54048.1;Pangolin_CoV_spike;961 | 0 | 0 |
| MT072864.1;QIQ54048.1;Pangolin_CoV_spike;962 | 0 | 0 |
| MT072864.1;QIQ54048.1;Pangolin_CoV_spike;963 | 0 | 0 |
| MT072864.1;QIQ54048.1;Pangolin_CoV_spike;964 | 0 | 0 |
| MT072864.1;QIQ54048.1;Pangolin_CoV_spike;965 | 0 | 0 |
| MT072864.1;QIQ54048.1;Pangolin_CoV_spike;966 | 0 | 0.025 |
| MT072864.1;QIQ54048.1;Pangolin_CoV_spike;967 | 0 | 0.025 |
| MT072864.1;QIQ54048.1;Pangolin_CoV_spike;968 | 0.05 | 0.05 |
| MT072864.1;QIQ54048.1;Pangolin_CoV_spike;969 | 0 | 0.075 |
| MT072864.1;QIQ54048.1;Pangolin_CoV_spike;970 | 0.05 | 0.075 |
| MT072864.1;QIQ54048.1;Pangolin_CoV_spike;971 | 0.05 | 0.125 |
| MT072864.1;QIQ54048.1;Pangolin_CoV_spike;972 | 0.05 | 0.1 |
| MT072864.1;QIQ54048.1;Pangolin_CoV_spike;973 | 0 | 0.075 |
| MT072864.1;QIQ54048.1;Pangolin_CoV_spike;974 | 0 | 0.05 |
| MT072864.1;QIQ54048.1;Pangolin_CoV_spike;975 | 0.05 | 0.05 |
| MT072864.1;QIQ54048.1;Pangolin_CoV_spike;976 | 0.05 | 0.025 |
| MT072864.1;QIQ54048.1;Pangolin_CoV_spike;977 | 0 | 0 |
| MT072864.1;QIQ54048.1;Pangolin_CoV_spike;978 | 0 | 0 |
| MT072864.1;QIQ54048.1;Pangolin_CoV_spike;979 | 0 | 0 |
| MT072864.1;QIQ54048.1;Pangolin_CoV_spike;980 | 0 | 0.025 |
| MT072864.1;QIQ54048.1;Pangolin_CoV_spike;981 | 0 | 0 |
| MT072864.1;QIQ54048.1;Pangolin_CoV_spike;982 | 0 | 0 |
| MT072864.1;QIQ54048.1;Pangolin_CoV_spike;983 | 0 | 0 |
| MT072864.1;QIQ54048.1;Pangolin_CoV_spike;984 | 0.05 | 0 |
| MT072864.1;QIQ54048.1;Pangolin_CoV_spike;985 | 0 | 0 |
| MT072864.1;QIQ54048.1;Pangolin_CoV_spike;986 | 0 | 0 |
| MT072864.1;QIQ54048.1;Pangolin_CoV_spike;987 | 0 | 0 |
| MT072864.1;QIQ54048.1;Pangolin_CoV_spike;988 | 0 | 0 |
| MT072864.1;QIQ54048.1;Pangolin_CoV_spike;989 | 0 | 0 |
| MT072864.1;QIQ54048.1;Pangolin_CoV_spike;990 | 0 | 0 |
| MT072864.1;QIQ54048.1;Pangolin_CoV_spike;991 | 0 | 0 |
| MT072864.1;QIQ54048.1;Pangolin_CoV_spike;992 | 0 | 0 |
| MT072864.1;QIQ54048.1;Pangolin_CoV_spike;993 | 0 | 0 |
| MT072864.1;QIQ54048.1;Pangolin_CoV_spike;994 | 0 | 0 |
| MT072864.1;QIQ54048.1;Pangolin_CoV_spike;995 | 0 | 0 |
| MT072864.1;QIQ54048.1;Pangolin_CoV_spike;996 | 0 | 0 |
| MT072864.1;QIQ54048.1;Pangolin_CoV_spike;997 | 0 | 0 |

| | | |
|---|---|---|
| MT072864.1;QIQ54048.1;Pangolin_CoV_spike;998 | 0 | 0 |
| MT072864.1;QIQ54048.1;Pangolin_CoV_spike;999 | 0 | 0.025 |
| MT072864.1;QIQ54048.1;Pangolin_CoV_spike;1000 | 0 | 0 |
| MT072864.1;QIQ54048.1;Pangolin_CoV_spike;1001 | 0 | 0 |
| MT072864.1;QIQ54048.1;Pangolin_CoV_spike;1002 | 0 | 0 |
| MT072864.1;QIQ54048.1;Pangolin_CoV_spike;1003 | 0 | 0 |
| MT072864.1;QIQ54048.1;Pangolin_CoV_spike;1004 | 0 | 0 |
| MT072864.1;QIQ54048.1;Pangolin_CoV_spike;1005 | 0 | 0 |
| MT072864.1;QIQ54048.1;Pangolin_CoV_spike;1006 | 0 | 0 |
| MT072864.1;QIQ54048.1;Pangolin_CoV_spike;1007 | 0.05 | 0.025 |
| MT072864.1;QIQ54048.1;Pangolin_CoV_spike;1008 | 0 | 0 |
| MT072864.1;QIQ54048.1;Pangolin_CoV_spike;1009 | 0 | 0.025 |
| MT072864.1;QIQ54048.1;Pangolin_CoV_spike;1010 | 0 | 0 |
| MT072864.1;QIQ54048.1;Pangolin_CoV_spike;1011 | 0 | 0 |
| MT072864.1;QIQ54048.1;Pangolin_CoV_spike;1012 | 0.05 | 0.025 |
| MT072864.1;QIQ54048.1;Pangolin_CoV_spike;1013 | 0.05 | 0 |
| MT072864.1;QIQ54048.1;Pangolin_CoV_spike;1014 | 0.05 | 0 |
| MT072864.1;QIQ54048.1;Pangolin_CoV_spike;1015 | 0 | 0.025 |
| MT072864.1;QIQ54048.1;Pangolin_CoV_spike;1016 | 0 | 0 |
| MT072864.1;QIQ54048.1;Pangolin_CoV_spike;1017 | 0 | 0.025 |
| MT072864.1;QIQ54048.1;Pangolin_CoV_spike;1018 | 0 | 0 |
| MT072864.1;QIQ54048.1;Pangolin_CoV_spike;1019 | 0 | 0 |
| MT072864.1;QIQ54048.1;Pangolin_CoV_spike;1020 | 0 | 0 |
| MT072864.1;QIQ54048.1;Pangolin_CoV_spike;1021 | 0 | 0 |
| MT072864.1;QIQ54048.1;Pangolin_CoV_spike;1022 | 0 | 0 |
| MT072864.1;QIQ54048.1;Pangolin_CoV_spike;1023 | 0 | 0 |
| MT072864.1;QIQ54048.1;Pangolin_CoV_spike;1024 | 0 | 0 |
| MT072864.1;QIQ54048.1;Pangolin_CoV_spike;1025 | 0 | 0 |
| MT072864.1;QIQ54048.1;Pangolin_CoV_spike;1026 | 0 | 0 |
| MT072864.1;QIQ54048.1;Pangolin_CoV_spike;1027 | 0 | 0 |
| MT072864.1;QIQ54048.1;Pangolin_CoV_spike;1028 | 0 | 0 |
| MT072864.1;QIQ54048.1;Pangolin_CoV_spike;1029 | 0 | 0 |
| MT072864.1;QIQ54048.1;Pangolin_CoV_spike;1030 | 0 | 0 |
| MT072864.1;QIQ54048.1;Pangolin_CoV_spike;1031 | 0 | 0 |
| MT072864.1;QIQ54048.1;Pangolin_CoV_spike;1032 | 0 | 0 |
| MT072864.1;QIQ54048.1;Pangolin_CoV_spike;1033 | 0 | 0 |
| MT072864.1;QIQ54048.1;Pangolin_CoV_spike;1034 | 0 | 0 |
| MT072864.1;QIQ54048.1;Pangolin_CoV_spike;1035 | 0 | 0 |
| MT072864.1;QIQ54048.1;Pangolin_CoV_spike;1036 | 0 | 0 |
| MT072864.1;QIQ54048.1;Pangolin_CoV_spike;1037 | 0 | 0.025 |
| MT072864.1;QIQ54048.1;Pangolin_CoV_spike;1038 | 0.05 | 0 |
| MT072864.1;QIQ54048.1;Pangolin_CoV_spike;1039 | 0 | 0 |
| MT072864.1;QIQ54048.1;Pangolin_CoV_spike;1040 | 0 | 0.05 |

| | | |
|---|---|---|
| MT072864.1;QIQ54048.1;Pangolin_CoV_spike;1041 | 0 | 0.025 |
| MT072864.1;QIQ54048.1;Pangolin_CoV_spike;1042 | 0 | 0 |
| MT072864.1;QIQ54048.1;Pangolin_CoV_spike;1043 | 0 | 0 |
| MT072864.1;QIQ54048.1;Pangolin_CoV_spike;1044 | 0 | 0 |
| MT072864.1;QIQ54048.1;Pangolin_CoV_spike;1045 | 0 | 0 |
| MT072864.1;QIQ54048.1;Pangolin_CoV_spike;1046 | 0 | 0 |
| MT072864.1;QIQ54048.1;Pangolin_CoV_spike;1047 | 0 | 0 |
| MT072864.1;QIQ54048.1;Pangolin_CoV_spike;1048 | 0 | 0 |
| MT072864.1;QIQ54048.1;Pangolin_CoV_spike;1049 | 0 | 0 |
| MT072864.1;QIQ54048.1;Pangolin_CoV_spike;1050 | 0 | 0 |
| MT072864.1;QIQ54048.1;Pangolin_CoV_spike;1051 | 0 | 0 |
| MT072864.1;QIQ54048.1;Pangolin_CoV_spike;1052 | 0 | 0 |
| MT072864.1;QIQ54048.1;Pangolin_CoV_spike;1053 | 0 | 0 |
| MT072864.1;QIQ54048.1;Pangolin_CoV_spike;1054 | 0 | 0 |
| MT072864.1;QIQ54048.1;Pangolin_CoV_spike;1055 | 0 | 0 |
| MT072864.1;QIQ54048.1;Pangolin_CoV_spike;1056 | 0 | 0 |
| MT072864.1;QIQ54048.1;Pangolin_CoV_spike;1057 | 0 | 0 |
| MT072864.1;QIQ54048.1;Pangolin_CoV_spike;1058 | 0 | 0 |
| MT072864.1;QIQ54048.1;Pangolin_CoV_spike;1059 | 0 | 0 |
| MT072864.1;QIQ54048.1;Pangolin_CoV_spike;1060 | 0.05 | 0 |
| MT072864.1;QIQ54048.1;Pangolin_CoV_spike;1061 | 0.1 | 0 |
| MT072864.1;QIQ54048.1;Pangolin_CoV_spike;1062 | 0.05 | 0 |
| MT072864.1;QIQ54048.1;Pangolin_CoV_spike;1063 | 0.05 | 0 |
| MT072864.1;QIQ54048.1;Pangolin_CoV_spike;1064 | 0 | 0 |
| MT072864.1;QIQ54048.1;Pangolin_CoV_spike;1065 | 0 | 0 |
| MT072864.1;QIQ54048.1;Pangolin_CoV_spike;1066 | 0.05 | 0 |
| MT072864.1;QIQ54048.1;Pangolin_CoV_spike;1067 | 0 | 0 |
| MT072864.1;QIQ54048.1;Pangolin_CoV_spike;1068 | 0.05 | 0 |
| MT072864.1;QIQ54048.1;Pangolin_CoV_spike;1069 | 0 | 0.025 |
| MT072864.1;QIQ54048.1;Pangolin_CoV_spike;1070 | 0 | 0 |
| MT072864.1;QIQ54048.1;Pangolin_CoV_spike;1071 | 0 | 0 |
| MT072864.1;QIQ54048.1;Pangolin_CoV_spike;1072 | 0 | 0 |
| MT072864.1;QIQ54048.1;Pangolin_CoV_spike;1073 | 0 | 0 |
| MT072864.1;QIQ54048.1;Pangolin_CoV_spike;1074 | 0.05 | 0 |
| MT072864.1;QIQ54048.1;Pangolin_CoV_spike;1075 | 0.05 | 0 |
| MT072864.1;QIQ54048.1;Pangolin_CoV_spike;1076 | 0.05 | 0.025 |
| MT072864.1;QIQ54048.1;Pangolin_CoV_spike;1077 | 0.05 | 0 |
| MT072864.1;QIQ54048.1;Pangolin_CoV_spike;1078 | 0 | 0 |
| MT072864.1;QIQ54048.1;Pangolin_CoV_spike;1079 | 0 | 0 |
| MT072864.1;QIQ54048.1;Pangolin_CoV_spike;1080 | 0.05 | 0 |
| MT072864.1;QIQ54048.1;Pangolin_CoV_spike;1081 | 0 | 0 |
| MT072864.1;QIQ54048.1;Pangolin_CoV_spike;1082 | 0 | 0 |
| MT072864.1;QIQ54048.1;Pangolin_CoV_spike;1083 | 0 | 0 |

| | | |
|---|---|---|
| MT072864.1;QIQ54048.1;Pangolin_CoV_spike;1084 | 0 | 0 |
| MT072864.1;QIQ54048.1;Pangolin_CoV_spike;1085 | 0 | 0 |
| MT072864.1;QIQ54048.1;Pangolin_CoV_spike;1086 | 0 | 0 |
| MT072864.1;QIQ54048.1;Pangolin_CoV_spike;1087 | 0 | 0 |
| MT072864.1;QIQ54048.1;Pangolin_CoV_spike;1088 | 0 | 0 |
| MT072864.1;QIQ54048.1;Pangolin_CoV_spike;1089 | 0 | 0 |
| MT072864.1;QIQ54048.1;Pangolin_CoV_spike;1090 | 0 | 0 |
| MT072864.1;QIQ54048.1;Pangolin_CoV_spike;1091 | 0 | 0 |
| MT072864.1;QIQ54048.1;Pangolin_CoV_spike;1092 | 0 | 0 |
| MT072864.1;QIQ54048.1;Pangolin_CoV_spike;1093 | 0 | 0 |
| MT072864.1;QIQ54048.1;Pangolin_CoV_spike;1094 | 0 | 0 |
| MT072864.1;QIQ54048.1;Pangolin_CoV_spike;1095 | 0 | 0 |
| MT072864.1;QIQ54048.1;Pangolin_CoV_spike;1096 | 0 | 0 |
| MT072864.1;QIQ54048.1;Pangolin_CoV_spike;1097 | 0 | 0 |
| MT072864.1;QIQ54048.1;Pangolin_CoV_spike;1098 | 0 | 0 |
| MT072864.1;QIQ54048.1;Pangolin_CoV_spike;1099 | 0 | 0.025 |
| MT072864.1;QIQ54048.1;Pangolin_CoV_spike;1100 | 0.1 | 0.05 |
| MT072864.1;QIQ54048.1;Pangolin_CoV_spike;1101 | 0.05 | 0.025 |
| MT072864.1;QIQ54048.1;Pangolin_CoV_spike;1102 | 0.05 | 0.025 |
| MT072864.1;QIQ54048.1;Pangolin_CoV_spike;1103 | 0 | 0 |
| MT072864.1;QIQ54048.1;Pangolin_CoV_spike;1104 | 0.05 | 0.025 |
| MT072864.1;QIQ54048.1;Pangolin_CoV_spike;1105 | 0.05 | 0.025 |
| MT072864.1;QIQ54048.1;Pangolin_CoV_spike;1106 | 0.05 | 0.05 |
| MT072864.1;QIQ54048.1;Pangolin_CoV_spike;1107 | 0.05 | 0.025 |
| MT072864.1;QIQ54048.1;Pangolin_CoV_spike;1108 | 0.05 | 0.025 |
| MT072864.1;QIQ54048.1;Pangolin_CoV_spike;1109 | 0.05 | 0.025 |
| MT072864.1;QIQ54048.1;Pangolin_CoV_spike;1110 | 0.05 | 0.025 |
| MT072864.1;QIQ54048.1;Pangolin_CoV_spike;1111 | 0.05 | 0.025 |
| MT072864.1;QIQ54048.1;Pangolin_CoV_spike;1112 | 0.05 | 0.025 |
| MT072864.1;QIQ54048.1;Pangolin_CoV_spike;1113 | 0 | 0 |
| MT072864.1;QIQ54048.1;Pangolin_CoV_spike;1114 | 0 | 0 |
| MT072864.1;QIQ54048.1;Pangolin_CoV_spike;1115 | 0 | 0 |
| MT072864.1;QIQ54048.1;Pangolin_CoV_spike;1116 | 0 | 0 |
| MT072864.1;QIQ54048.1;Pangolin_CoV_spike;1117 | 0 | 0 |
| MT072864.1;QIQ54048.1;Pangolin_CoV_spike;1118 | 0 | 0 |
| MT072864.1;QIQ54048.1;Pangolin_CoV_spike;1119 | 0 | 0 |
| MT072864.1;QIQ54048.1;Pangolin_CoV_spike;1120 | 0 | 0 |
| MT072864.1;QIQ54048.1;Pangolin_CoV_spike;1121 | 0 | 0 |
| MT072864.1;QIQ54048.1;Pangolin_CoV_spike;1122 | 0 | 0 |
| MT072864.1;QIQ54048.1;Pangolin_CoV_spike;1123 | 0 | 0 |
| MT072864.1;QIQ54048.1;Pangolin_CoV_spike;1124 | 0 | 0 |
| MT072864.1;QIQ54048.1;Pangolin_CoV_spike;1125 | 0 | 0 |
| MT072864.1;QIQ54048.1;Pangolin_CoV_spike;1126 | 0 | 0.05 |

| | | |
|---|---|---|
| MT072864.1;QIQ54048.1;Pangolin_CoV_spike;1127 | 0 | 0 |
| MT072864.1;QIQ54048.1;Pangolin_CoV_spike;1128 | 0 | 0 |
| MT072864.1;QIQ54048.1;Pangolin_CoV_spike;1129 | 0 | 0.025 |
| MT072864.1;QIQ54048.1;Pangolin_CoV_spike;1130 | 0 | 0 |
| MT072864.1;QIQ54048.1;Pangolin_CoV_spike;1131 | 0 | 0.025 |
| MT072864.1;QIQ54048.1;Pangolin_CoV_spike;1132 | 0 | 0 |
| MT072864.1;QIQ54048.1;Pangolin_CoV_spike;1133 | 0 | 0 |
| MT072864.1;QIQ54048.1;Pangolin_CoV_spike;1134 | 0 | 0 |
| MT072864.1;QIQ54048.1;Pangolin_CoV_spike;1135 | 0 | 0.05 |
| MT072864.1;QIQ54048.1;Pangolin_CoV_spike;1136 | 0 | 0.175 |
| MT072864.1;QIQ54048.1;Pangolin_CoV_spike;1137 | 0.05 | 0.75 |
| MT072864.1;QIQ54048.1;Pangolin_CoV_spike;1138 | 0.1 | 0.775 |
| MT072864.1;QIQ54048.1;Pangolin_CoV_spike;1139 | 0.05 | 0.775 |
| MT072864.1;QIQ54048.1;Pangolin_CoV_spike;1140 | 0.05 | 0.725 |
| MT072864.1;QIQ54048.1;Pangolin_CoV_spike;1141 | 0.3 | 0.725 |
| MT072864.1;QIQ54048.1;Pangolin_CoV_spike;1142 | 0.4 | 0.775 |
| MT072864.1;QIQ54048.1;Pangolin_CoV_spike;1143 | 0.35 | 0.775 |
| MT072864.1;QIQ54048.1;Pangolin_CoV_spike;1144 | 0.35 | 0.825 |
| MT072864.1;QIQ54048.1;Pangolin_CoV_spike;1145 | 0.45 | 0.8 |
| MT072864.1;QIQ54048.1;Pangolin_CoV_spike;1146 | 0.4 | 0.5 |
| MT072864.1;QIQ54048.1;Pangolin_CoV_spike;1147 | 0.45 | 0.35 |
| MT072864.1;QIQ54048.1;Pangolin_CoV_spike;1148 | 0.45 | 0.35 |
| MT072864.1;QIQ54048.1;Pangolin_CoV_spike;1149 | 0.5 | 0.35 |
| MT072864.1;QIQ54048.1;Pangolin_CoV_spike;1150 | 0.5 | 0.375 |
| MT072864.1;QIQ54048.1;Pangolin_CoV_spike;1151 | 0 | 0.2 |
| MT072864.1;QIQ54048.1;Pangolin_CoV_spike;1152 | 0 | 0.125 |
| MT072864.1;QIQ54048.1;Pangolin_CoV_spike;1153 | 0 | 0.225 |
| MT072864.1;QIQ54048.1;Pangolin_CoV_spike;1154 | 0.05 | 0.275 |
| MT072864.1;QIQ54048.1;Pangolin_CoV_spike;1155 | 0.1 | 0.35 |
| MT072864.1;QIQ54048.1;Pangolin_CoV_spike;1156 | 0.2 | 0.35 |
| MT072864.1;QIQ54048.1;Pangolin_CoV_spike;1157 | 0.1 | 0.35 |
| MT072864.1;QIQ54048.1;Pangolin_CoV_spike;1158 | 0 | 0.3 |
| MT072864.1;QIQ54048.1;Pangolin_CoV_spike;1159 | 0 | 0.325 |
| MT072864.1;QIQ54048.1;Pangolin_CoV_spike;1160 | 0 | 0.15 |
| MT072864.1;QIQ54048.1;Pangolin_CoV_spike;1161 | 0 | 0.2 |
| MT072864.1;QIQ54048.1;Pangolin_CoV_spike;1162 | 0 | 0.05 |
| MT072864.1;QIQ54048.1;Pangolin_CoV_spike;1163 | 0.05 | 0.1 |
| MT072864.1;QIQ54048.1;Pangolin_CoV_spike;1164 | 0.15 | 0.05 |
| MT072864.1;QIQ54048.1;Pangolin_CoV_spike;1165 | 0.05 | 0.125 |
| MT072864.1;QIQ54048.1;Pangolin_CoV_spike;1166 | 0.05 | 0.05 |
| MT072864.1;QIQ54048.1;Pangolin_CoV_spike;1167 | 0.05 | 0.1 |
| MT072864.1;QIQ54048.1;Pangolin_CoV_spike;1168 | 0.05 | 0.175 |
| MT072864.1;QIQ54048.1;Pangolin_CoV_spike;1169 | 0.05 | 0.225 |

| | | |
|---|---|---|
| MT072864.1;QIQ54048.1;Pangolin_CoV_spike;1170 | 0.1 | 0.225 |
| MT072864.1;QIQ54048.1;Pangolin_CoV_spike;1171 | 0.1 | 0.275 |
| MT072864.1;QIQ54048.1;Pangolin_CoV_spike;1172 | 0.1 | 0.275 |
| MT072864.1;QIQ54048.1;Pangolin_CoV_spike;1173 | 0.1 | 0.3 |
| MT072864.1;QIQ54048.1;Pangolin_CoV_spike;1174 | 0.05 | 0.3 |
| MT072864.1;QIQ54048.1;Pangolin_CoV_spike;1175 | 0.2 | 0.325 |
| MT072864.1;QIQ54048.1;Pangolin_CoV_spike;1176 | 0.15 | 0.3 |
| MT072864.1;QIQ54048.1;Pangolin_CoV_spike;1177 | 0.1 | 0.175 |
| MT072864.1;QIQ54048.1;Pangolin_CoV_spike;1178 | 0.1 | 0.15 |
| MT072864.1;QIQ54048.1;Pangolin_CoV_spike;1179 | 0.15 | 0.125 |
| MT072864.1;QIQ54048.1;Pangolin_CoV_spike;1180 | 0.05 | 0.125 |
| MT072864.1;QIQ54048.1;Pangolin_CoV_spike;1181 | 0.05 | 0.125 |
| MT072864.1;QIQ54048.1;Pangolin_CoV_spike;1182 | 0.05 | 0.1 |
| MT072864.1;QIQ54048.1;Pangolin_CoV_spike;1183 | 0.05 | 0.125 |
| MT072864.1;QIQ54048.1;Pangolin_CoV_spike;1184 | 0.05 | 0.1 |
| MT072864.1;QIQ54048.1;Pangolin_CoV_spike;1185 | 0.05 | 0.125 |
| MT072864.1;QIQ54048.1;Pangolin_CoV_spike;1186 | 0.05 | 0.125 |
| MT072864.1;QIQ54048.1;Pangolin_CoV_spike;1187 | 0.05 | 0.075 |
| MT072864.1;QIQ54048.1;Pangolin_CoV_spike;1188 | 0.05 | 0.125 |
| MT072864.1;QIQ54048.1;Pangolin_CoV_spike;1189 | 0.1 | 0.15 |
| MT072864.1;QIQ54048.1;Pangolin_CoV_spike;1190 | 0.05 | 0.125 |
| MT072864.1;QIQ54048.1;Pangolin_CoV_spike;1191 | 0 | 0 |
| MT072864.1;QIQ54048.1;Pangolin_CoV_spike;1192 | 0 | 0.05 |
| MT072864.1;QIQ54048.1;Pangolin_CoV_spike;1193 | 0 | 0 |
| MT072864.1;QIQ54048.1;Pangolin_CoV_spike;1194 | 0 | 0 |
| MT072864.1;QIQ54048.1;Pangolin_CoV_spike;1195 | 0 | 0 |
| MT072864.1;QIQ54048.1;Pangolin_CoV_spike;1196 | 0 | 0.025 |
| MT072864.1;QIQ54048.1;Pangolin_CoV_spike;1197 | 0 | 0 |
| MT072864.1;QIQ54048.1;Pangolin_CoV_spike;1198 | 0 | 0 |
| MT072864.1;QIQ54048.1;Pangolin_CoV_spike;1199 | 0 | 0 |
| MT072864.1;QIQ54048.1;Pangolin_CoV_spike;1200 | 0 | 0 |
| MT072864.1;QIQ54048.1;Pangolin_CoV_spike;1201 | 0 | 0 |
| MT072864.1;QIQ54048.1;Pangolin_CoV_spike;1202 | 0 | 0.025 |
| MT072864.1;QIQ54048.1;Pangolin_CoV_spike;1203 | 0 | 0 |
| MT072864.1;QIQ54048.1;Pangolin_CoV_spike;1204 | 0 | 0 |
| MT072864.1;QIQ54048.1;Pangolin_CoV_spike;1205 | 0 | 0 |
| MT072864.1;QIQ54048.1;Pangolin_CoV_spike;1206 | 0 | 0 |
| MT072864.1;QIQ54048.1;Pangolin_CoV_spike;1207 | 0 | 0 |
| MT072864.1;QIQ54048.1;Pangolin_CoV_spike;1208 | 0 | 0 |
| MT072864.1;QIQ54048.1;Pangolin_CoV_spike;1209 | 0 | 0 |
| MT072864.1;QIQ54048.1;Pangolin_CoV_spike;1210 | 0 | 0 |
| MT072864.1;QIQ54048.1;Pangolin_CoV_spike;1211 | 0 | 0 |
| MT072864.1;QIQ54048.1;Pangolin_CoV_spike;1212 | 0 | 0 |

| | | |
|---|---|---|
| MT072864.1;QIQ54048.1;Pangolin_CoV_spike;1213 | 0 | 0 |
| MT072864.1;QIQ54048.1;Pangolin_CoV_spike;1214 | 0 | 0 |
| MT072864.1;QIQ54048.1;Pangolin_CoV_spike;1215 | 0 | 0 |
| MT072864.1;QIQ54048.1;Pangolin_CoV_spike;1216 | 0 | 0 |
| MT072864.1;QIQ54048.1;Pangolin_CoV_spike;1217 | 0 | 0 |
| MT072864.1;QIQ54048.1;Pangolin_CoV_spike;1218 | 0 | 0 |
| MT072864.1;QIQ54048.1;Pangolin_CoV_spike;1219 | 0 | 0 |
| MT072864.1;QIQ54048.1;Pangolin_CoV_spike;1220 | 0 | 0 |
| MT072864.1;QIQ54048.1;Pangolin_CoV_spike;1221 | 0 | 0 |
| MT072864.1;QIQ54048.1;Pangolin_CoV_spike;1222 | 0 | 0 |
| MT072864.1;QIQ54048.1;Pangolin_CoV_spike;1223 | 0 | 0 |
| MT072864.1;QIQ54048.1;Pangolin_CoV_spike;1224 | 0 | 0 |
| MT072864.1;QIQ54048.1;Pangolin_CoV_spike;1225 | 0 | 0 |
| MT072864.1;QIQ54048.1;Pangolin_CoV_spike;1226 | 0 | 0 |
| MT072864.1;QIQ54048.1;Pangolin_CoV_spike;1227 | 0 | 0 |
| MT072864.1;QIQ54048.1;Pangolin_CoV_spike;1228 | 0 | 0 |
| MT072864.1;QIQ54048.1;Pangolin_CoV_spike;1229 | 0 | 0 |
| MT072864.1;QIQ54048.1;Pangolin_CoV_spike;1230 | 0 | 0 |
| MT072864.1;QIQ54048.1;Pangolin_CoV_spike;1231 | 0 | 0 |
| MT072864.1;QIQ54048.1;Pangolin_CoV_spike;1232 | 0 | 0 |
| MT072864.1;QIQ54048.1;Pangolin_CoV_spike;1233 | 0 | 0 |
| MT072864.1;QIQ54048.1;Pangolin_CoV_spike;1234 | 0 | 0 |
| MT072864.1;QIQ54048.1;Pangolin_CoV_spike;1235 | 0 | 0 |
| MT072864.1;QIQ54048.1;Pangolin_CoV_spike;1236 | 0 | 0 |
| MT072864.1;QIQ54048.1;Pangolin_CoV_spike;1237 | 0 | 0 |
| MT072864.1;QIQ54048.1;Pangolin_CoV_spike;1238 | 0 | 0 |
| MT072864.1;QIQ54048.1;Pangolin_CoV_spike;1239 | 0 | 0 |
| MT072864.1;QIQ54048.1;Pangolin_CoV_spike;1240 | 0 | 0 |
| MT072864.1;QIQ54048.1;Pangolin_CoV_spike;1241 | 0 | 0.075 |
| MT072864.1;QIQ54048.1;Pangolin_CoV_spike;1242 | 0 | 0.1 |
| MT072864.1;QIQ54048.1;Pangolin_CoV_spike;1243 | 0 | 0.25 |
| MT072864.1;QIQ54048.1;Pangolin_CoV_spike;1244 | 0 | 0.5 |
| MT072864.1;QIQ54048.1;Pangolin_CoV_spike;1245 | 0 | 0.55 |
| MT072864.1;QIQ54048.1;Pangolin_CoV_spike;1246 | 0 | 0.575 |
| MT072864.1;QIQ54048.1;Pangolin_CoV_spike;1247 | 0 | 0.65 |
| MT072864.1;QIQ54048.1;Pangolin_CoV_spike;1248 | 0 | 0.65 |
| MT072864.1;QIQ54048.1;Pangolin_CoV_spike;1249 | 0 | 0.675 |
| MT072864.1;QIQ54048.1;Pangolin_CoV_spike;1250 | 0.1 | 0.65 |
| MT072864.1;QIQ54048.1;Pangolin_CoV_spike;1251 | 0.05 | 0.6 |
| MT072864.1;QIQ54048.1;Pangolin_CoV_spike;1252 | 0.05 | 0.45 |
| MT072864.1;QIQ54048.1;Pangolin_CoV_spike;1253 | 0.05 | 0.475 |
| MT072864.1;QIQ54048.1;Pangolin_CoV_spike;1254 | 0.1 | 0.3 |
| MT072864.1;QIQ54048.1;Pangolin_CoV_spike;1255 | 0.05 | 0.225 |

| | | |
|---|---|---|
| MT072864.1;QIQ54048.1;Pangolin_CoV_spike;1256 | 0.05 | 0.225 |
| MT072864.1;QIQ54048.1;Pangolin_CoV_spike;1257 | 0.05 | 0.175 |
| MT072864.1;QIQ54048.1;Pangolin_CoV_spike;1258 | 0 | 0.175 |
| MT072864.1;QIQ54049.1;Pangolin_CoV_orf3;1 | 0 | 0 |
| MT072864.1;QIQ54049.1;Pangolin_CoV_orf3;2 | 0 | 0 |
| MT072864.1;QIQ54049.1;Pangolin_CoV_orf3;3 | 0 | 0 |
| MT072864.1;QIQ54049.1;Pangolin_CoV_orf3;4 | 0 | 0 |
| MT072864.1;QIQ54049.1;Pangolin_CoV_orf3;5 | 0 | 0 |
| MT072864.1;QIQ54049.1;Pangolin_CoV_orf3;6 | 0 | 0 |
| MT072864.1;QIQ54049.1;Pangolin_CoV_orf3;7 | 0 | 0 |
| MT072864.1;QIQ54049.1;Pangolin_CoV_orf3;8 | 0 | 0 |
| MT072864.1;QIQ54049.1;Pangolin_CoV_orf3;9 | 0 | 0 |
| MT072864.1;QIQ54049.1;Pangolin_CoV_orf3;10 | 0 | 0 |
| MT072864.1;QIQ54049.1;Pangolin_CoV_orf3;11 | 0 | 0 |
| MT072864.1;QIQ54049.1;Pangolin_CoV_orf3;12 | 0 | 0 |
| MT072864.1;QIQ54049.1;Pangolin_CoV_orf3;13 | 0 | 0 |
| MT072864.1;QIQ54049.1;Pangolin_CoV_orf3;14 | 0 | 0 |
| MT072864.1;QIQ54049.1;Pangolin_CoV_orf3;15 | 0 | 0 |
| MT072864.1;QIQ54049.1;Pangolin_CoV_orf3;16 | 0 | 0 |
| MT072864.1;QIQ54049.1;Pangolin_CoV_orf3;17 | 0 | 0 |
| MT072864.1;QIQ54049.1;Pangolin_CoV_orf3;18 | 0 | 0 |
| MT072864.1;QIQ54049.1;Pangolin_CoV_orf3;19 | 0 | 0 |
| MT072864.1;QIQ54049.1;Pangolin_CoV_orf3;20 | 0 | 0 |
| MT072864.1;QIQ54049.1;Pangolin_CoV_orf3;21 | 0 | 0 |
| MT072864.1;QIQ54049.1;Pangolin_CoV_orf3;22 | 0 | 0 |
| MT072864.1;QIQ54049.1;Pangolin_CoV_orf3;23 | 0 | 0 |
| MT072864.1;QIQ54049.1;Pangolin_CoV_orf3;24 | 0 | 0 |
| MT072864.1;QIQ54049.1;Pangolin_CoV_orf3;25 | 0 | 0 |
| MT072864.1;QIQ54049.1;Pangolin_CoV_orf3;26 | 0 | 0.025 |
| MT072864.1;QIQ54049.1;Pangolin_CoV_orf3;27 | 0 | 0.025 |
| MT072864.1;QIQ54049.1;Pangolin_CoV_orf3;28 | 0 | 0.025 |
| MT072864.1;QIQ54049.1;Pangolin_CoV_orf3;29 | 0 | 0 |
| MT072864.1;QIQ54049.1;Pangolin_CoV_orf3;30 | 0 | 0.025 |
| MT072864.1;QIQ54049.1;Pangolin_CoV_orf3;31 | 0 | 0 |
| MT072864.1;QIQ54049.1;Pangolin_CoV_orf3;32 | 0 | 0 |
| MT072864.1;QIQ54049.1;Pangolin_CoV_orf3;33 | 0 | 0 |
| MT072864.1;QIQ54049.1;Pangolin_CoV_orf3;34 | 0 | 0 |
| MT072864.1;QIQ54049.1;Pangolin_CoV_orf3;35 | 0 | 0 |
| MT072864.1;QIQ54049.1;Pangolin_CoV_orf3;36 | 0.05 | 0 |
| MT072864.1;QIQ54049.1;Pangolin_CoV_orf3;37 | 0 | 0 |
| MT072864.1;QIQ54049.1;Pangolin_CoV_orf3;38 | 0 | 0 |
| MT072864.1;QIQ54049.1;Pangolin_CoV_orf3;39 | 0 | 0 |
| MT072864.1;QIQ54049.1;Pangolin_CoV_orf3;40 | 0 | 0 |

| | | |
|---|---|---|
| MT072864.1;QIQ54049.1;Pangolin_CoV_orf3;41 | 0 | 0 |
| MT072864.1;QIQ54049.1;Pangolin_CoV_orf3;42 | 0 | 0 |
| MT072864.1;QIQ54049.1;Pangolin_CoV_orf3;43 | 0 | 0 |
| MT072864.1;QIQ54049.1;Pangolin_CoV_orf3;44 | 0 | 0 |
| MT072864.1;QIQ54049.1;Pangolin_CoV_orf3;45 | 0 | 0 |
| MT072864.1;QIQ54049.1;Pangolin_CoV_orf3;46 | 0 | 0 |
| MT072864.1;QIQ54049.1;Pangolin_CoV_orf3;47 | 0 | 0 |
| MT072864.1;QIQ54049.1;Pangolin_CoV_orf3;48 | 0 | 0 |
| MT072864.1;QIQ54049.1;Pangolin_CoV_orf3;49 | 0 | 0 |
| MT072864.1;QIQ54049.1;Pangolin_CoV_orf3;50 | 0 | 0 |
| MT072864.1;QIQ54049.1;Pangolin_CoV_orf3;51 | 0 | 0 |
| MT072864.1;QIQ54049.1;Pangolin_CoV_orf3;52 | 0 | 0 |
| MT072864.1;QIQ54049.1;Pangolin_CoV_orf3;53 | 0 | 0 |
| MT072864.1;QIQ54049.1;Pangolin_CoV_orf3;54 | 0 | 0 |
| MT072864.1;QIQ54049.1;Pangolin_CoV_orf3;55 | 0 | 0 |
| MT072864.1;QIQ54049.1;Pangolin_CoV_orf3;56 | 0 | 0 |
| MT072864.1;QIQ54049.1;Pangolin_CoV_orf3;57 | 0 | 0 |
| MT072864.1;QIQ54049.1;Pangolin_CoV_orf3;58 | 0 | 0 |
| MT072864.1;QIQ54049.1;Pangolin_CoV_orf3;59 | 0 | 0 |
| MT072864.1;QIQ54049.1;Pangolin_CoV_orf3;60 | 0 | 0 |
| MT072864.1;QIQ54049.1;Pangolin_CoV_orf3;61 | 0 | 0 |
| MT072864.1;QIQ54049.1;Pangolin_CoV_orf3;62 | 0 | 0 |
| MT072864.1;QIQ54049.1;Pangolin_CoV_orf3;63 | 0 | 0 |
| MT072864.1;QIQ54049.1;Pangolin_CoV_orf3;64 | 0 | 0 |
| MT072864.1;QIQ54049.1;Pangolin_CoV_orf3;65 | 0 | 0 |
| MT072864.1;QIQ54049.1;Pangolin_CoV_orf3;66 | 0 | 0 |
| MT072864.1;QIQ54049.1;Pangolin_CoV_orf3;67 | 0 | 0 |
| MT072864.1;QIQ54049.1;Pangolin_CoV_orf3;68 | 0 | 0 |
| MT072864.1;QIQ54049.1;Pangolin_CoV_orf3;69 | 0 | 0 |
| MT072864.1;QIQ54049.1;Pangolin_CoV_orf3;70 | 0 | 0 |
| MT072864.1;QIQ54049.1;Pangolin_CoV_orf3;71 | 0 | 0 |
| MT072864.1;QIQ54049.1;Pangolin_CoV_orf3;72 | 0 | 0 |
| MT072864.1;QIQ54049.1;Pangolin_CoV_orf3;73 | 0 | 0 |
| MT072864.1;QIQ54049.1;Pangolin_CoV_orf3;74 | 0 | 0 |
| MT072864.1;QIQ54049.1;Pangolin_CoV_orf3;75 | 0 | 0 |
| MT072864.1;QIQ54049.1;Pangolin_CoV_orf3;76 | 0 | 0 |
| MT072864.1;QIQ54049.1;Pangolin_CoV_orf3;77 | 0 | 0 |
| MT072864.1;QIQ54049.1;Pangolin_CoV_orf3;78 | 0 | 0 |
| MT072864.1;QIQ54049.1;Pangolin_CoV_orf3;79 | 0 | 0 |
| MT072864.1;QIQ54049.1;Pangolin_CoV_orf3;80 | 0 | 0 |
| MT072864.1;QIQ54049.1;Pangolin_CoV_orf3;81 | 0 | 0 |
| MT072864.1;QIQ54049.1;Pangolin_CoV_orf3;82 | 0 | 0 |
| MT072864.1;QIQ54049.1;Pangolin_CoV_orf3;83 | 0 | 0 |

| | | |
|---|---|---|
| MT072864.1;QIQ54049.1;Pangolin_CoV_orf3;84 | 0 | 0 |
| MT072864.1;QIQ54049.1;Pangolin_CoV_orf3;85 | 0 | 0 |
| MT072864.1;QIQ54049.1;Pangolin_CoV_orf3;86 | 0 | 0 |
| MT072864.1;QIQ54049.1;Pangolin_CoV_orf3;87 | 0 | 0 |
| MT072864.1;QIQ54049.1;Pangolin_CoV_orf3;88 | 0 | 0 |
| MT072864.1;QIQ54049.1;Pangolin_CoV_orf3;89 | 0 | 0 |
| MT072864.1;QIQ54049.1;Pangolin_CoV_orf3;90 | 0 | 0 |
| MT072864.1;QIQ54049.1;Pangolin_CoV_orf3;91 | 0 | 0 |
| MT072864.1;QIQ54049.1;Pangolin_CoV_orf3;92 | 0 | 0 |
| MT072864.1;QIQ54049.1;Pangolin_CoV_orf3;93 | 0 | 0 |
| MT072864.1;QIQ54049.1;Pangolin_CoV_orf3;94 | 0 | 0 |
| MT072864.1;QIQ54049.1;Pangolin_CoV_orf3;95 | 0 | 0 |
| MT072864.1;QIQ54049.1;Pangolin_CoV_orf3;96 | 0 | 0 |
| MT072864.1;QIQ54049.1;Pangolin_CoV_orf3;97 | 0 | 0 |
| MT072864.1;QIQ54049.1;Pangolin_CoV_orf3;98 | 0 | 0 |
| MT072864.1;QIQ54049.1;Pangolin_CoV_orf3;99 | 0 | 0 |
| MT072864.1;QIQ54049.1;Pangolin_CoV_orf3;100 | 0 | 0 |
| MT072864.1;QIQ54049.1;Pangolin_CoV_orf3;101 | 0 | 0 |
| MT072864.1;QIQ54049.1;Pangolin_CoV_orf3;102 | 0 | 0 |
| MT072864.1;QIQ54049.1;Pangolin_CoV_orf3;103 | 0 | 0 |
| MT072864.1;QIQ54049.1;Pangolin_CoV_orf3;104 | 0 | 0 |
| MT072864.1;QIQ54049.1;Pangolin_CoV_orf3;105 | 0 | 0 |
| MT072864.1;QIQ54049.1;Pangolin_CoV_orf3;106 | 0 | 0 |
| MT072864.1;QIQ54049.1;Pangolin_CoV_orf3;107 | 0 | 0 |
| MT072864.1;QIQ54049.1;Pangolin_CoV_orf3;108 | 0 | 0 |
| MT072864.1;QIQ54049.1;Pangolin_CoV_orf3;109 | 0 | 0 |
| MT072864.1;QIQ54049.1;Pangolin_CoV_orf3;110 | 0 | 0 |
| MT072864.1;QIQ54049.1;Pangolin_CoV_orf3;111 | 0 | 0 |
| MT072864.1;QIQ54049.1;Pangolin_CoV_orf3;112 | 0 | 0 |
| MT072864.1;QIQ54049.1;Pangolin_CoV_orf3;113 | 0 | 0 |
| MT072864.1;QIQ54049.1;Pangolin_CoV_orf3;114 | 0 | 0 |
| MT072864.1;QIQ54049.1;Pangolin_CoV_orf3;115 | 0 | 0 |
| MT072864.1;QIQ54049.1;Pangolin_CoV_orf3;116 | 0 | 0 |
| MT072864.1;QIQ54049.1;Pangolin_CoV_orf3;117 | 0 | 0 |
| MT072864.1;QIQ54049.1;Pangolin_CoV_orf3;118 | 0 | 0 |
| MT072864.1;QIQ54049.1;Pangolin_CoV_orf3;119 | 0 | 0 |
| MT072864.1;QIQ54049.1;Pangolin_CoV_orf3;120 | 0 | 0 |
| MT072864.1;QIQ54049.1;Pangolin_CoV_orf3;121 | 0 | 0 |
| MT072864.1;QIQ54049.1;Pangolin_CoV_orf3;122 | 0 | 0 |
| MT072864.1;QIQ54049.1;Pangolin_CoV_orf3;123 | 0 | 0 |
| MT072864.1;QIQ54049.1;Pangolin_CoV_orf3;124 | 0 | 0 |
| MT072864.1;QIQ54049.1;Pangolin_CoV_orf3;125 | 0 | 0 |
| MT072864.1;QIQ54049.1;Pangolin_CoV_orf3;126 | 0 | 0 |

| | | |
|---|---|---|
| MT072864.1;QIQ54049.1;Pangolin_CoV_orf3;127 | 0 | 0 |
| MT072864.1;QIQ54049.1;Pangolin_CoV_orf3;128 | 0 | 0 |
| MT072864.1;QIQ54049.1;Pangolin_CoV_orf3;129 | 0 | 0 |
| MT072864.1;QIQ54049.1;Pangolin_CoV_orf3;130 | 0 | 0 |
| MT072864.1;QIQ54049.1;Pangolin_CoV_orf3;131 | 0 | 0 |
| MT072864.1;QIQ54049.1;Pangolin_CoV_orf3;132 | 0 | 0 |
| MT072864.1;QIQ54049.1;Pangolin_CoV_orf3;133 | 0 | 0 |
| MT072864.1;QIQ54049.1;Pangolin_CoV_orf3;134 | 0 | 0 |
| MT072864.1;QIQ54049.1;Pangolin_CoV_orf3;135 | 0 | 0 |
| MT072864.1;QIQ54049.1;Pangolin_CoV_orf3;136 | 0 | 0 |
| MT072864.1;QIQ54049.1;Pangolin_CoV_orf3;137 | 0 | 0 |
| MT072864.1;QIQ54049.1;Pangolin_CoV_orf3;138 | 0 | 0 |
| MT072864.1;QIQ54049.1;Pangolin_CoV_orf3;139 | 0 | 0 |
| MT072864.1;QIQ54049.1;Pangolin_CoV_orf3;140 | 0 | 0 |
| MT072864.1;QIQ54049.1;Pangolin_CoV_orf3;141 | 0 | 0 |
| MT072864.1;QIQ54049.1;Pangolin_CoV_orf3;142 | 0 | 0 |
| MT072864.1;QIQ54049.1;Pangolin_CoV_orf3;143 | 0 | 0 |
| MT072864.1;QIQ54049.1;Pangolin_CoV_orf3;144 | 0 | 0 |
| MT072864.1;QIQ54049.1;Pangolin_CoV_orf3;145 | 0 | 0 |
| MT072864.1;QIQ54049.1;Pangolin_CoV_orf3;146 | 0 | 0 |
| MT072864.1;QIQ54049.1;Pangolin_CoV_orf3;147 | 0 | 0 |
| MT072864.1;QIQ54049.1;Pangolin_CoV_orf3;148 | 0 | 0 |
| MT072864.1;QIQ54049.1;Pangolin_CoV_orf3;149 | 0 | 0 |
| MT072864.1;QIQ54049.1;Pangolin_CoV_orf3;150 | 0 | 0 |
| MT072864.1;QIQ54049.1;Pangolin_CoV_orf3;151 | 0 | 0 |
| MT072864.1;QIQ54049.1;Pangolin_CoV_orf3;152 | 0 | 0 |
| MT072864.1;QIQ54049.1;Pangolin_CoV_orf3;153 | 0 | 0 |
| MT072864.1;QIQ54049.1;Pangolin_CoV_orf3;154 | 0 | 0 |
| MT072864.1;QIQ54049.1;Pangolin_CoV_orf3;155 | 0 | 0 |
| MT072864.1;QIQ54049.1;Pangolin_CoV_orf3;156 | 0 | 0 |
| MT072864.1;QIQ54049.1;Pangolin_CoV_orf3;157 | 0 | 0 |
| MT072864.1;QIQ54049.1;Pangolin_CoV_orf3;158 | 0 | 0 |
| MT072864.1;QIQ54049.1;Pangolin_CoV_orf3;159 | 0 | 0 |
| MT072864.1;QIQ54049.1;Pangolin_CoV_orf3;160 | 0 | 0 |
| MT072864.1;QIQ54049.1;Pangolin_CoV_orf3;161 | 0.05 | 0 |
| MT072864.1;QIQ54049.1;Pangolin_CoV_orf3;162 | 0 | 0 |
| MT072864.1;QIQ54049.1;Pangolin_CoV_orf3;163 | 0.05 | 0 |
| MT072864.1;QIQ54049.1;Pangolin_CoV_orf3;164 | 0.05 | 0 |
| MT072864.1;QIQ54049.1;Pangolin_CoV_orf3;165 | 0.05 | 0.025 |
| MT072864.1;QIQ54049.1;Pangolin_CoV_orf3;166 | 0 | 0 |
| MT072864.1;QIQ54049.1;Pangolin_CoV_orf3;167 | 0 | 0 |
| MT072864.1;QIQ54049.1;Pangolin_CoV_orf3;168 | 0 | 0 |
| MT072864.1;QIQ54049.1;Pangolin_CoV_orf3;169 | 0 | 0 |

| | | |
|---|---|---|
| MT072864.1;QIQ54049.1;Pangolin_CoV_orf3;170 | 0 | 0 |
| MT072864.1;QIQ54049.1;Pangolin_CoV_orf3;171 | 0 | 0 |
| MT072864.1;QIQ54049.1;Pangolin_CoV_orf3;172 | 0 | 0 |
| MT072864.1;QIQ54049.1;Pangolin_CoV_orf3;173 | 0 | 0 |
| MT072864.1;QIQ54049.1;Pangolin_CoV_orf3;174 | 0 | 0 |
| MT072864.1;QIQ54049.1;Pangolin_CoV_orf3;175 | 0 | 0 |
| MT072864.1;QIQ54049.1;Pangolin_CoV_orf3;176 | 0.05 | 0 |
| MT072864.1;QIQ54049.1;Pangolin_CoV_orf3;177 | 0.05 | 0.1 |
| MT072864.1;QIQ54049.1;Pangolin_CoV_orf3;178 | 0.05 | 0.05 |
| MT072864.1;QIQ54049.1;Pangolin_CoV_orf3;179 | 0.05 | 0.05 |
| MT072864.1;QIQ54049.1;Pangolin_CoV_orf3;180 | 0 | 0.05 |
| MT072864.1;QIQ54049.1;Pangolin_CoV_orf3;181 | 0.1 | 0.05 |
| MT072864.1;QIQ54049.1;Pangolin_CoV_orf3;182 | 0.05 | 0.05 |
| MT072864.1;QIQ54049.1;Pangolin_CoV_orf3;183 | 0.05 | 0.15 |
| MT072864.1;QIQ54049.1;Pangolin_CoV_orf3;184 | 0.1 | 0.075 |
| MT072864.1;QIQ54049.1;Pangolin_CoV_orf3;185 | 0.05 | 0.05 |
| MT072864.1;QIQ54049.1;Pangolin_CoV_orf3;186 | 0 | 0.05 |
| MT072864.1;QIQ54049.1;Pangolin_CoV_orf3;187 | 0 | 0 |
| MT072864.1;QIQ54049.1;Pangolin_CoV_orf3;188 | 0 | 0 |
| MT072864.1;QIQ54049.1;Pangolin_CoV_orf3;189 | 0 | 0 |
| MT072864.1;QIQ54049.1;Pangolin_CoV_orf3;190 | 0 | 0 |
| MT072864.1;QIQ54049.1;Pangolin_CoV_orf3;191 | 0 | 0 |
| MT072864.1;QIQ54049.1;Pangolin_CoV_orf3;192 | 0 | 0 |
| MT072864.1;QIQ54049.1;Pangolin_CoV_orf3;193 | 0 | 0 |
| MT072864.1;QIQ54049.1;Pangolin_CoV_orf3;194 | 0 | 0 |
| MT072864.1;QIQ54049.1;Pangolin_CoV_orf3;195 | 0 | 0 |
| MT072864.1;QIQ54049.1;Pangolin_CoV_orf3;196 | 0 | 0 |
| MT072864.1;QIQ54049.1;Pangolin_CoV_orf3;197 | 0 | 0 |
| MT072864.1;QIQ54049.1;Pangolin_CoV_orf3;198 | 0 | 0 |
| MT072864.1;QIQ54049.1;Pangolin_CoV_orf3;199 | 0 | 0 |
| MT072864.1;QIQ54049.1;Pangolin_CoV_orf3;200 | 0 | 0 |
| MT072864.1;QIQ54049.1;Pangolin_CoV_orf3;201 | 0 | 0 |
| MT072864.1;QIQ54049.1;Pangolin_CoV_orf3;202 | 0 | 0 |
| MT072864.1;QIQ54049.1;Pangolin_CoV_orf3;203 | 0 | 0 |
| MT072864.1;QIQ54049.1;Pangolin_CoV_orf3;204 | 0 | 0 |
| MT072864.1;QIQ54049.1;Pangolin_CoV_orf3;205 | 0 | 0 |
| MT072864.1;QIQ54049.1;Pangolin_CoV_orf3;206 | 0 | 0 |
| MT072864.1;QIQ54049.1;Pangolin_CoV_orf3;207 | 0 | 0.025 |
| MT072864.1;QIQ54049.1;Pangolin_CoV_orf3;208 | 0 | 0.025 |
| MT072864.1;QIQ54049.1;Pangolin_CoV_orf3;209 | 0.05 | 0.05 |
| MT072864.1;QIQ54049.1;Pangolin_CoV_orf3;210 | 0 | 0.025 |
| MT072864.1;QIQ54049.1;Pangolin_CoV_orf3;211 | 0.05 | 0.025 |
| MT072864.1;QIQ54049.1;Pangolin_CoV_orf3;212 | 0 | 0.025 |

| | | |
|---|---|---|
| MT072864.1;QIQ54049.1;Pangolin_CoV_orf3;213 | 0 | 0.025 |
| MT072864.1;QIQ54049.1;Pangolin_CoV_orf3;214 | 0 | 0.025 |
| MT072864.1;QIQ54049.1;Pangolin_CoV_orf3;215 | 0 | 0 |
| MT072864.1;QIQ54049.1;Pangolin_CoV_orf3;216 | 0 | 0 |
| MT072864.1;QIQ54049.1;Pangolin_CoV_orf3;217 | 0 | 0 |
| MT072864.1;QIQ54049.1;Pangolin_CoV_orf3;218 | 0 | 0 |
| MT072864.1;QIQ54049.1;Pangolin_CoV_orf3;219 | 0 | 0 |
| MT072864.1;QIQ54049.1;Pangolin_CoV_orf3;220 | 0.05 | 0 |
| MT072864.1;QIQ54049.1;Pangolin_CoV_orf3;221 | 0 | 0 |
| MT072864.1;QIQ54049.1;Pangolin_CoV_orf3;222 | 0 | 0 |
| MT072864.1;QIQ54049.1;Pangolin_CoV_orf3;223 | 0 | 0 |
| MT072864.1;QIQ54049.1;Pangolin_CoV_orf3;224 | 0 | 0 |
| MT072864.1;QIQ54049.1;Pangolin_CoV_orf3;225 | 0 | 0 |
| MT072864.1;QIQ54049.1;Pangolin_CoV_orf3;226 | 0 | 0 |
| MT072864.1;QIQ54049.1;Pangolin_CoV_orf3;227 | 0 | 0 |
| MT072864.1;QIQ54049.1;Pangolin_CoV_orf3;228 | 0 | 0 |
| MT072864.1;QIQ54049.1;Pangolin_CoV_orf3;229 | 0 | 0.025 |
| MT072864.1;QIQ54049.1;Pangolin_CoV_orf3;230 | 0 | 0.025 |
| MT072864.1;QIQ54049.1;Pangolin_CoV_orf3;231 | 0 | 0.05 |
| MT072864.1;QIQ54049.1;Pangolin_CoV_orf3;232 | 0 | 0.075 |
| MT072864.1;QIQ54049.1;Pangolin_CoV_orf3;233 | 0 | 0.1 |
| MT072864.1;QIQ54049.1;Pangolin_CoV_orf3;234 | 0 | 0.15 |
| MT072864.1;QIQ54049.1;Pangolin_CoV_orf3;235 | 0 | 0.1 |
| MT072864.1;QIQ54049.1;Pangolin_CoV_orf3;236 | 0 | 0.075 |
| MT072864.1;QIQ54049.1;Pangolin_CoV_orf3;237 | 0 | 0.075 |
| MT072864.1;QIQ54049.1;Pangolin_CoV_orf3;238 | 0 | 0.075 |
| MT072864.1;QIQ54049.1;Pangolin_CoV_orf3;239 | 0 | 0.075 |
| MT072864.1;QIQ54049.1;Pangolin_CoV_orf3;240 | 0 | 0.075 |
| MT072864.1;QIQ54049.1;Pangolin_CoV_orf3;241 | 0.05 | 0.075 |
| MT072864.1;QIQ54049.1;Pangolin_CoV_orf3;242 | 0.05 | 0.025 |
| MT072864.1;QIQ54049.1;Pangolin_CoV_orf3;243 | 0.05 | 0.1 |
| MT072864.1;QIQ54049.1;Pangolin_CoV_orf3;244 | 0.05 | 0.05 |
| MT072864.1;QIQ54049.1;Pangolin_CoV_orf3;245 | 0.05 | 0.075 |
| MT072864.1;QIQ54049.1;Pangolin_CoV_orf3;246 | 0.05 | 0.075 |
| MT072864.1;QIQ54049.1;Pangolin_CoV_orf3;247 | 0.05 | 0.05 |
| MT072864.1;QIQ54049.1;Pangolin_CoV_orf3;248 | 0.05 | 0.05 |
| MT072864.1;QIQ54049.1;Pangolin_CoV_orf3;249 | 0.05 | 0.025 |
| MT072864.1;QIQ54049.1;Pangolin_CoV_orf3;250 | 0.05 | 0.05 |
| MT072864.1;QIQ54049.1;Pangolin_CoV_orf3;251 | 0.05 | 0.05 |
| MT072864.1;QIQ54049.1;Pangolin_CoV_orf3;252 | 0.05 | 0.15 |
| MT072864.1;QIQ54049.1;Pangolin_CoV_orf3;253 | 0 | 0.125 |
| MT072864.1;QIQ54049.1;Pangolin_CoV_orf3;254 | 0 | 0.15 |
| MT072864.1;QIQ54049.1;Pangolin_CoV_orf3;255 | 0 | 0.125 |

| | | |
|---|---|---|
| MT072864.1;QIQ54049.1;Pangolin_CoV_orf3;256 | 0 | 0.125 |
| MT072864.1;QIQ54049.1;Pangolin_CoV_orf3;257 | 0 | 0.125 |
| MT072864.1;QIQ54049.1;Pangolin_CoV_orf3;258 | 0.05 | 0.125 |
| MT072864.1;QIQ54049.1;Pangolin_CoV_orf3;259 | 0.05 | 0.1 |
| MT072864.1;QIQ54049.1;Pangolin_CoV_orf3;260 | 0.05 | 0.175 |
| MT072864.1;QIQ54049.1;Pangolin_CoV_orf3;261 | 0.05 | 0.15 |
| MT072864.1;QIQ54049.1;Pangolin_CoV_orf3;262 | 0 | 0.075 |
| MT072864.1;QIQ54049.1;Pangolin_CoV_orf3;263 | 0.05 | 0.1 |
| MT072864.1;QIQ54049.1;Pangolin_CoV_orf3;264 | 0 | 0.05 |
| MT072864.1;QIQ54050.1;Pangolin_CoV_envelope;1 | 0 | 0 |
| MT072864.1;QIQ54050.1;Pangolin_CoV_envelope;2 | 0 | 0 |
| MT072864.1;QIQ54050.1;Pangolin_CoV_envelope;3 | 0 | 0 |
| MT072864.1;QIQ54050.1;Pangolin_CoV_envelope;4 | 0 | 0 |
| MT072864.1;QIQ54050.1;Pangolin_CoV_envelope;5 | 0 | 0 |
| MT072864.1;QIQ54050.1;Pangolin_CoV_envelope;6 | 0 | 0 |
| MT072864.1;QIQ54050.1;Pangolin_CoV_envelope;7 | 0 | 0 |
| MT072864.1;QIQ54050.1;Pangolin_CoV_envelope;8 | 0 | 0 |
| MT072864.1;QIQ54050.1;Pangolin_CoV_envelope;9 | 0 | 0 |
| MT072864.1;QIQ54050.1;Pangolin_CoV_envelope;10 | 0 | 0 |
| MT072864.1;QIQ54050.1;Pangolin_CoV_envelope;11 | 0 | 0 |
| MT072864.1;QIQ54050.1;Pangolin_CoV_envelope;12 | 0 | 0 |
| MT072864.1;QIQ54050.1;Pangolin_CoV_envelope;13 | 0 | 0 |
| MT072864.1;QIQ54050.1;Pangolin_CoV_envelope;14 | 0 | 0 |
| MT072864.1;QIQ54050.1;Pangolin_CoV_envelope;15 | 0 | 0 |
| MT072864.1;QIQ54050.1;Pangolin_CoV_envelope;16 | 0 | 0 |
| MT072864.1;QIQ54050.1;Pangolin_CoV_envelope;17 | 0 | 0 |
| MT072864.1;QIQ54050.1;Pangolin_CoV_envelope;18 | 0 | 0 |
| MT072864.1;QIQ54050.1;Pangolin_CoV_envelope;19 | 0 | 0 |
| MT072864.1;QIQ54050.1;Pangolin_CoV_envelope;20 | 0 | 0 |
| MT072864.1;QIQ54050.1;Pangolin_CoV_envelope;21 | 0 | 0 |
| MT072864.1;QIQ54050.1;Pangolin_CoV_envelope;22 | 0 | 0 |
| MT072864.1;QIQ54050.1;Pangolin_CoV_envelope;23 | 0 | 0 |
| MT072864.1;QIQ54050.1;Pangolin_CoV_envelope;24 | 0 | 0 |
| MT072864.1;QIQ54050.1;Pangolin_CoV_envelope;25 | 0 | 0 |
| MT072864.1;QIQ54050.1;Pangolin_CoV_envelope;26 | 0 | 0 |
| MT072864.1;QIQ54050.1;Pangolin_CoV_envelope;27 | 0 | 0 |
| MT072864.1;QIQ54050.1;Pangolin_CoV_envelope;28 | 0 | 0 |
| MT072864.1;QIQ54050.1;Pangolin_CoV_envelope;29 | 0 | 0 |
| MT072864.1;QIQ54050.1;Pangolin_CoV_envelope;30 | 0 | 0 |
| MT072864.1;QIQ54050.1;Pangolin_CoV_envelope;31 | 0 | 0 |
| MT072864.1;QIQ54050.1;Pangolin_CoV_envelope;32 | 0 | 0 |
| MT072864.1;QIQ54050.1;Pangolin_CoV_envelope;33 | 0 | 0 |
| MT072864.1;QIQ54050.1;Pangolin_CoV_envelope;34 | 0 | 0 |

| | | |
|---|---|---|
| MT072864.1;QIQ54050.1;Pangolin_CoV_envelope;35 | 0 | 0 |
| MT072864.1;QIQ54050.1;Pangolin_CoV_envelope;36 | 0 | 0 |
| MT072864.1;QIQ54050.1;Pangolin_CoV_envelope;37 | 0 | 0 |
| MT072864.1;QIQ54050.1;Pangolin_CoV_envelope;38 | 0 | 0 |
| MT072864.1;QIQ54050.1;Pangolin_CoV_envelope;39 | 0 | 0 |
| MT072864.1;QIQ54050.1;Pangolin_CoV_envelope;40 | 0 | 0.025 |
| MT072864.1;QIQ54050.1;Pangolin_CoV_envelope;41 | 0 | 0 |
| MT072864.1;QIQ54050.1;Pangolin_CoV_envelope;42 | 0 | 0 |
| MT072864.1;QIQ54050.1;Pangolin_CoV_envelope;43 | 0 | 0 |
| MT072864.1;QIQ54050.1;Pangolin_CoV_envelope;44 | 0 | 0 |
| MT072864.1;QIQ54050.1;Pangolin_CoV_envelope;45 | 0 | 0 |
| MT072864.1;QIQ54050.1;Pangolin_CoV_envelope;46 | 0 | 0 |
| MT072864.1;QIQ54050.1;Pangolin_CoV_envelope;47 | 0 | 0 |
| MT072864.1;QIQ54050.1;Pangolin_CoV_envelope;48 | 0 | 0 |
| MT072864.1;QIQ54050.1;Pangolin_CoV_envelope;49 | 0 | 0 |
| MT072864.1;QIQ54050.1;Pangolin_CoV_envelope;50 | 0 | 0 |
| MT072864.1;QIQ54050.1;Pangolin_CoV_envelope;51 | 0 | 0 |
| MT072864.1;QIQ54050.1;Pangolin_CoV_envelope;52 | 0 | 0 |
| MT072864.1;QIQ54050.1;Pangolin_CoV_envelope;53 | 0 | 0 |
| MT072864.1;QIQ54050.1;Pangolin_CoV_envelope;54 | 0 | 0 |
| MT072864.1;QIQ54050.1;Pangolin_CoV_envelope;55 | 0 | 0 |
| MT072864.1;QIQ54050.1;Pangolin_CoV_envelope;56 | 0 | 0 |
| MT072864.1;QIQ54050.1;Pangolin_CoV_envelope;57 | 0 | 0 |
| MT072864.1;QIQ54050.1;Pangolin_CoV_envelope;58 | 0 | 0.075 |
| MT072864.1;QIQ54050.1;Pangolin_CoV_envelope;59 | 0 | 0 |
| MT072864.1;QIQ54050.1;Pangolin_CoV_envelope;60 | 0.1 | 0 |
| MT072864.1;QIQ54050.1;Pangolin_CoV_envelope;61 | 0.1 | 0 |
| MT072864.1;QIQ54050.1;Pangolin_CoV_envelope;62 | 0.05 | 0.05 |
| MT072864.1;QIQ54050.1;Pangolin_CoV_envelope;63 | 0.1 | 0 |
| MT072864.1;QIQ54050.1;Pangolin_CoV_envelope;64 | 0.05 | 0.075 |
| MT072864.1;QIQ54051.1;Pangolin_CoV_membrane;1 | 0 | 0.225 |
| MT072864.1;QIQ54051.1;Pangolin_CoV_membrane;2 | 0 | 0.525 |
| MT072864.1;QIQ54051.1;Pangolin_CoV_membrane;3 | 0 | 0.4 |
| MT072864.1;QIQ54051.1;Pangolin_CoV_membrane;4 | 0 | 0.6 |
| MT072864.1;QIQ54051.1;Pangolin_CoV_membrane;5 | 0 | 0.725 |
| MT072864.1;QIQ54051.1;Pangolin_CoV_membrane;6 | 0 | 0.65 |
| MT072864.1;QIQ54051.1;Pangolin_CoV_membrane;7 | 0 | 0.775 |
| MT072864.1;QIQ54051.1;Pangolin_CoV_membrane;8 | 0 | 0.9 |
| MT072864.1;QIQ54051.1;Pangolin_CoV_membrane;9 | 0 | 0.4 |
| MT072864.1;QIQ54051.1;Pangolin_CoV_membrane;10 | 0 | 0.025 |
| MT072864.1;QIQ54051.1;Pangolin_CoV_membrane;11 | 0.05 | 0 |
| MT072864.1;QIQ54051.1;Pangolin_CoV_membrane;12 | 0 | 0 |
| MT072864.1;QIQ54051.1;Pangolin_CoV_membrane;13 | 0 | 0 |

| | | |
|---|---|---|
| MT072864.1;QIQ54051.1;Pangolin_CoV_membrane;14 | 0 | 0 |
| MT072864.1;QIQ54051.1;Pangolin_CoV_membrane;15 | 0 | 0 |
| MT072864.1;QIQ54051.1;Pangolin_CoV_membrane;16 | 0 | 0 |
| MT072864.1;QIQ54051.1;Pangolin_CoV_membrane;17 | 0 | 0 |
| MT072864.1;QIQ54051.1;Pangolin_CoV_membrane;18 | 0 | 0 |
| MT072864.1;QIQ54051.1;Pangolin_CoV_membrane;19 | 0 | 0 |
| MT072864.1;QIQ54051.1;Pangolin_CoV_membrane;20 | 0 | 0 |
| MT072864.1;QIQ54051.1;Pangolin_CoV_membrane;21 | 0 | 0 |
| MT072864.1;QIQ54051.1;Pangolin_CoV_membrane;22 | 0 | 0 |
| MT072864.1;QIQ54051.1;Pangolin_CoV_membrane;23 | 0 | 0 |
| MT072864.1;QIQ54051.1;Pangolin_CoV_membrane;24 | 0 | 0 |
| MT072864.1;QIQ54051.1;Pangolin_CoV_membrane;25 | 0 | 0 |
| MT072864.1;QIQ54051.1;Pangolin_CoV_membrane;26 | 0 | 0 |
| MT072864.1;QIQ54051.1;Pangolin_CoV_membrane;27 | 0 | 0 |
| MT072864.1;QIQ54051.1;Pangolin_CoV_membrane;28 | 0 | 0 |
| MT072864.1;QIQ54051.1;Pangolin_CoV_membrane;29 | 0 | 0 |
| MT072864.1;QIQ54051.1;Pangolin_CoV_membrane;30 | 0 | 0 |
| MT072864.1;QIQ54051.1;Pangolin_CoV_membrane;31 | 0 | 0 |
| MT072864.1;QIQ54051.1;Pangolin_CoV_membrane;32 | 0 | 0 |
| MT072864.1;QIQ54051.1;Pangolin_CoV_membrane;33 | 0 | 0 |
| MT072864.1;QIQ54051.1;Pangolin_CoV_membrane;34 | 0 | 0 |
| MT072864.1;QIQ54051.1;Pangolin_CoV_membrane;35 | 0 | 0 |
| MT072864.1;QIQ54051.1;Pangolin_CoV_membrane;36 | 0 | 0 |
| MT072864.1;QIQ54051.1;Pangolin_CoV_membrane;37 | 0 | 0 |
| MT072864.1;QIQ54051.1;Pangolin_CoV_membrane;38 | 0 | 0 |
| MT072864.1;QIQ54051.1;Pangolin_CoV_membrane;39 | 0 | 0 |
| MT072864.1;QIQ54051.1;Pangolin_CoV_membrane;40 | 0 | 0 |
| MT072864.1;QIQ54051.1;Pangolin_CoV_membrane;41 | 0 | 0 |
| MT072864.1;QIQ54051.1;Pangolin_CoV_membrane;42 | 0 | 0 |
| MT072864.1;QIQ54051.1;Pangolin_CoV_membrane;43 | 0 | 0 |
| MT072864.1;QIQ54051.1;Pangolin_CoV_membrane;44 | 0 | 0 |
| MT072864.1;QIQ54051.1;Pangolin_CoV_membrane;45 | 0 | 0 |
| MT072864.1;QIQ54051.1;Pangolin_CoV_membrane;46 | 0 | 0 |
| MT072864.1;QIQ54051.1;Pangolin_CoV_membrane;47 | 0 | 0 |
| MT072864.1;QIQ54051.1;Pangolin_CoV_membrane;48 | 0 | 0 |
| MT072864.1;QIQ54051.1;Pangolin_CoV_membrane;49 | 0 | 0 |
| MT072864.1;QIQ54051.1;Pangolin_CoV_membrane;50 | 0 | 0 |
| MT072864.1;QIQ54051.1;Pangolin_CoV_membrane;51 | 0 | 0 |
| MT072864.1;QIQ54051.1;Pangolin_CoV_membrane;52 | 0 | 0 |
| MT072864.1;QIQ54051.1;Pangolin_CoV_membrane;53 | 0 | 0 |
| MT072864.1;QIQ54051.1;Pangolin_CoV_membrane;54 | 0 | 0 |
| MT072864.1;QIQ54051.1;Pangolin_CoV_membrane;55 | 0 | 0 |
| MT072864.1;QIQ54051.1;Pangolin_CoV_membrane;56 | 0 | 0 |

| | | |
|---|---|---|
| MT072864.1;QIQ54051.1;Pangolin_CoV_membrane;57 | 0 | 0 |
| MT072864.1;QIQ54051.1;Pangolin_CoV_membrane;58 | 0 | 0 |
| MT072864.1;QIQ54051.1;Pangolin_CoV_membrane;59 | 0 | 0 |
| MT072864.1;QIQ54051.1;Pangolin_CoV_membrane;60 | 0 | 0 |
| MT072864.1;QIQ54051.1;Pangolin_CoV_membrane;61 | 0 | 0 |
| MT072864.1;QIQ54051.1;Pangolin_CoV_membrane;62 | 0 | 0 |
| MT072864.1;QIQ54051.1;Pangolin_CoV_membrane;63 | 0 | 0 |
| MT072864.1;QIQ54051.1;Pangolin_CoV_membrane;64 | 0 | 0 |
| MT072864.1;QIQ54051.1;Pangolin_CoV_membrane;65 | 0 | 0 |
| MT072864.1;QIQ54051.1;Pangolin_CoV_membrane;66 | 0 | 0 |
| MT072864.1;QIQ54051.1;Pangolin_CoV_membrane;67 | 0 | 0 |
| MT072864.1;QIQ54051.1;Pangolin_CoV_membrane;68 | 0 | 0 |
| MT072864.1;QIQ54051.1;Pangolin_CoV_membrane;69 | 0 | 0 |
| MT072864.1;QIQ54051.1;Pangolin_CoV_membrane;70 | 0 | 0 |
| MT072864.1;QIQ54051.1;Pangolin_CoV_membrane;71 | 0 | 0 |
| MT072864.1;QIQ54051.1;Pangolin_CoV_membrane;72 | 0 | 0 |
| MT072864.1;QIQ54051.1;Pangolin_CoV_membrane;73 | 0 | 0 |
| MT072864.1;QIQ54051.1;Pangolin_CoV_membrane;74 | 0 | 0 |
| MT072864.1;QIQ54051.1;Pangolin_CoV_membrane;75 | 0 | 0 |
| MT072864.1;QIQ54051.1;Pangolin_CoV_membrane;76 | 0 | 0 |
| MT072864.1;QIQ54051.1;Pangolin_CoV_membrane;77 | 0 | 0 |
| MT072864.1;QIQ54051.1;Pangolin_CoV_membrane;78 | 0 | 0 |
| MT072864.1;QIQ54051.1;Pangolin_CoV_membrane;79 | 0 | 0 |
| MT072864.1;QIQ54051.1;Pangolin_CoV_membrane;80 | 0 | 0 |
| MT072864.1;QIQ54051.1;Pangolin_CoV_membrane;81 | 0 | 0 |
| MT072864.1;QIQ54051.1;Pangolin_CoV_membrane;82 | 0 | 0 |
| MT072864.1;QIQ54051.1;Pangolin_CoV_membrane;83 | 0 | 0 |
| MT072864.1;QIQ54051.1;Pangolin_CoV_membrane;84 | 0 | 0 |
| MT072864.1;QIQ54051.1;Pangolin_CoV_membrane;85 | 0 | 0 |
| MT072864.1;QIQ54051.1;Pangolin_CoV_membrane;86 | 0 | 0 |
| MT072864.1;QIQ54051.1;Pangolin_CoV_membrane;87 | 0 | 0 |
| MT072864.1;QIQ54051.1;Pangolin_CoV_membrane;88 | 0 | 0 |
| MT072864.1;QIQ54051.1;Pangolin_CoV_membrane;89 | 0 | 0 |
| MT072864.1;QIQ54051.1;Pangolin_CoV_membrane;90 | 0 | 0 |
| MT072864.1;QIQ54051.1;Pangolin_CoV_membrane;91 | 0 | 0 |
| MT072864.1;QIQ54051.1;Pangolin_CoV_membrane;92 | 0 | 0 |
| MT072864.1;QIQ54051.1;Pangolin_CoV_membrane;93 | 0 | 0 |
| MT072864.1;QIQ54051.1;Pangolin_CoV_membrane;94 | 0 | 0 |
| MT072864.1;QIQ54051.1;Pangolin_CoV_membrane;95 | 0 | 0 |
| MT072864.1;QIQ54051.1;Pangolin_CoV_membrane;96 | 0 | 0 |
| MT072864.1;QIQ54051.1;Pangolin_CoV_membrane;97 | 0 | 0 |
| MT072864.1;QIQ54051.1;Pangolin_CoV_membrane;98 | 0 | 0 |
| MT072864.1;QIQ54051.1;Pangolin_CoV_membrane;99 | 0 | 0 |

| | | |
|---|---|---|
| MT072864.1;QIQ54051.1;Pangolin_CoV_membrane;100 | 0 | 0 |
| MT072864.1;QIQ54051.1;Pangolin_CoV_membrane;101 | 0.05 | 0.025 |
| MT072864.1;QIQ54051.1;Pangolin_CoV_membrane;102 | 0 | 0 |
| MT072864.1;QIQ54051.1;Pangolin_CoV_membrane;103 | 0 | 0 |
| MT072864.1;QIQ54051.1;Pangolin_CoV_membrane;104 | 0 | 0 |
| MT072864.1;QIQ54051.1;Pangolin_CoV_membrane;105 | 0 | 0 |
| MT072864.1;QIQ54051.1;Pangolin_CoV_membrane;106 | 0 | 0 |
| MT072864.1;QIQ54051.1;Pangolin_CoV_membrane;107 | 0 | 0 |
| MT072864.1;QIQ54051.1;Pangolin_CoV_membrane;108 | 0 | 0.025 |
| MT072864.1;QIQ54051.1;Pangolin_CoV_membrane;109 | 0 | 0.025 |
| MT072864.1;QIQ54051.1;Pangolin_CoV_membrane;110 | 0 | 0 |
| MT072864.1;QIQ54051.1;Pangolin_CoV_membrane;111 | 0 | 0.025 |
| MT072864.1;QIQ54051.1;Pangolin_CoV_membrane;112 | 0 | 0.025 |
| MT072864.1;QIQ54051.1;Pangolin_CoV_membrane;113 | 0 | 0.025 |
| MT072864.1;QIQ54051.1;Pangolin_CoV_membrane;114 | 0 | 0 |
| MT072864.1;QIQ54051.1;Pangolin_CoV_membrane;115 | 0 | 0 |
| MT072864.1;QIQ54051.1;Pangolin_CoV_membrane;116 | 0 | 0 |
| MT072864.1;QIQ54051.1;Pangolin_CoV_membrane;117 | 0 | 0 |
| MT072864.1;QIQ54051.1;Pangolin_CoV_membrane;118 | 0 | 0 |
| MT072864.1;QIQ54051.1;Pangolin_CoV_membrane;119 | 0 | 0 |
| MT072864.1;QIQ54051.1;Pangolin_CoV_membrane;120 | 0 | 0 |
| MT072864.1;QIQ54051.1;Pangolin_CoV_membrane;121 | 0 | 0 |
| MT072864.1;QIQ54051.1;Pangolin_CoV_membrane;122 | 0 | 0 |
| MT072864.1;QIQ54051.1;Pangolin_CoV_membrane;123 | 0 | 0 |
| MT072864.1;QIQ54051.1;Pangolin_CoV_membrane;124 | 0 | 0 |
| MT072864.1;QIQ54051.1;Pangolin_CoV_membrane;125 | 0 | 0 |
| MT072864.1;QIQ54051.1;Pangolin_CoV_membrane;126 | 0 | 0 |
| MT072864.1;QIQ54051.1;Pangolin_CoV_membrane;127 | 0 | 0 |
| MT072864.1;QIQ54051.1;Pangolin_CoV_membrane;128 | 0 | 0 |
| MT072864.1;QIQ54051.1;Pangolin_CoV_membrane;129 | 0 | 0 |
| MT072864.1;QIQ54051.1;Pangolin_CoV_membrane;130 | 0 | 0 |
| MT072864.1;QIQ54051.1;Pangolin_CoV_membrane;131 | 0 | 0 |
| MT072864.1;QIQ54051.1;Pangolin_CoV_membrane;132 | 0 | 0 |
| MT072864.1;QIQ54051.1;Pangolin_CoV_membrane;133 | 0 | 0 |
| MT072864.1;QIQ54051.1;Pangolin_CoV_membrane;134 | 0 | 0 |
| MT072864.1;QIQ54051.1;Pangolin_CoV_membrane;135 | 0 | 0 |
| MT072864.1;QIQ54051.1;Pangolin_CoV_membrane;136 | 0 | 0 |
| MT072864.1;QIQ54051.1;Pangolin_CoV_membrane;137 | 0 | 0 |
| MT072864.1;QIQ54051.1;Pangolin_CoV_membrane;138 | 0 | 0 |
| MT072864.1;QIQ54051.1;Pangolin_CoV_membrane;139 | 0 | 0 |
| MT072864.1;QIQ54051.1;Pangolin_CoV_membrane;140 | 0 | 0 |
| MT072864.1;QIQ54051.1;Pangolin_CoV_membrane;141 | 0 | 0 |
| MT072864.1;QIQ54051.1;Pangolin_CoV_membrane;142 | 0 | 0 |

| | | |
|---|---|---|
| MT072864.1;QIQ54051.1;Pangolin_CoV_membrane;143 | 0 | 0 |
| MT072864.1;QIQ54051.1;Pangolin_CoV_membrane;144 | 0 | 0 |
| MT072864.1;QIQ54051.1;Pangolin_CoV_membrane;145 | 0 | 0 |
| MT072864.1;QIQ54051.1;Pangolin_CoV_membrane;146 | 0 | 0 |
| MT072864.1;QIQ54051.1;Pangolin_CoV_membrane;147 | 0 | 0 |
| MT072864.1;QIQ54051.1;Pangolin_CoV_membrane;148 | 0.05 | 0.025 |
| MT072864.1;QIQ54051.1;Pangolin_CoV_membrane;149 | 0.05 | 0.05 |
| MT072864.1;QIQ54051.1;Pangolin_CoV_membrane;150 | 0.05 | 0.05 |
| MT072864.1;QIQ54051.1;Pangolin_CoV_membrane;151 | 0.05 | 0.075 |
| MT072864.1;QIQ54051.1;Pangolin_CoV_membrane;152 | 0.05 | 0.125 |
| MT072864.1;QIQ54051.1;Pangolin_CoV_membrane;153 | 0.05 | 0.375 |
| MT072864.1;QIQ54051.1;Pangolin_CoV_membrane;154 | 0.05 | 0.45 |
| MT072864.1;QIQ54051.1;Pangolin_CoV_membrane;155 | 0.05 | 0.375 |
| MT072864.1;QIQ54051.1;Pangolin_CoV_membrane;156 | 0.1 | 0.425 |
| MT072864.1;QIQ54051.1;Pangolin_CoV_membrane;157 | 0.05 | 0.4 |
| MT072864.1;QIQ54051.1;Pangolin_CoV_membrane;158 | 0.05 | 0.35 |
| MT072864.1;QIQ54051.1;Pangolin_CoV_membrane;159 | 0 | 0.35 |
| MT072864.1;QIQ54051.1;Pangolin_CoV_membrane;160 | 0 | 0.35 |
| MT072864.1;QIQ54051.1;Pangolin_CoV_membrane;161 | 0 | 0.35 |
| MT072864.1;QIQ54051.1;Pangolin_CoV_membrane;162 | 0 | 0.35 |
| MT072864.1;QIQ54051.1;Pangolin_CoV_membrane;163 | 0 | 0.025 |
| MT072864.1;QIQ54051.1;Pangolin_CoV_membrane;164 | 0 | 0 |
| MT072864.1;QIQ54051.1;Pangolin_CoV_membrane;165 | 0 | 0 |
| MT072864.1;QIQ54051.1;Pangolin_CoV_membrane;166 | 0 | 0 |
| MT072864.1;QIQ54051.1;Pangolin_CoV_membrane;167 | 0 | 0 |
| MT072864.1;QIQ54051.1;Pangolin_CoV_membrane;168 | 0 | 0 |
| MT072864.1;QIQ54051.1;Pangolin_CoV_membrane;169 | 0 | 0 |
| MT072864.1;QIQ54051.1;Pangolin_CoV_membrane;170 | 0 | 0 |
| MT072864.1;QIQ54051.1;Pangolin_CoV_membrane;171 | 0 | 0 |
| MT072864.1;QIQ54051.1;Pangolin_CoV_membrane;172 | 0 | 0 |
| MT072864.1;QIQ54051.1;Pangolin_CoV_membrane;173 | 0 | 0 |
| MT072864.1;QIQ54051.1;Pangolin_CoV_membrane;174 | 0 | 0 |
| MT072864.1;QIQ54051.1;Pangolin_CoV_membrane;175 | 0 | 0.025 |
| MT072864.1;QIQ54051.1;Pangolin_CoV_membrane;176 | 0 | 0.075 |
| MT072864.1;QIQ54051.1;Pangolin_CoV_membrane;177 | 0 | 0.1 |
| MT072864.1;QIQ54051.1;Pangolin_CoV_membrane;178 | 0 | 0.05 |
| MT072864.1;QIQ54051.1;Pangolin_CoV_membrane;179 | 0 | 0.075 |
| MT072864.1;QIQ54051.1;Pangolin_CoV_membrane;180 | 0.05 | 0.125 |
| MT072864.1;QIQ54051.1;Pangolin_CoV_membrane;181 | 0 | 0.35 |
| MT072864.1;QIQ54051.1;Pangolin_CoV_membrane;182 | 0 | 0.5 |
| MT072864.1;QIQ54051.1;Pangolin_CoV_membrane;183 | 0 | 0.4 |
| MT072864.1;QIQ54051.1;Pangolin_CoV_membrane;184 | 0 | 0.325 |
| MT072864.1;QIQ54051.1;Pangolin_CoV_membrane;185 | 0 | 0.275 |

| | | |
|---|---|---|
| MT072864.1;QIQ54051.1;Pangolin_CoV_membrane;186 | 0 | 0.125 |
| MT072864.1;QIQ54051.1;Pangolin_CoV_membrane;187 | 0 | 0.175 |
| MT072864.1;QIQ54051.1;Pangolin_CoV_membrane;188 | 0 | 0.25 |
| MT072864.1;QIQ54051.1;Pangolin_CoV_membrane;189 | 0 | 0.225 |
| MT072864.1;QIQ54051.1;Pangolin_CoV_membrane;190 | 0 | 0.2 |
| MT072864.1;QIQ54051.1;Pangolin_CoV_membrane;191 | 0 | 0.1 |
| MT072864.1;QIQ54051.1;Pangolin_CoV_membrane;192 | 0 | 0.1 |
| MT072864.1;QIQ54051.1;Pangolin_CoV_membrane;193 | 0 | 0.075 |
| MT072864.1;QIQ54051.1;Pangolin_CoV_membrane;194 | 0 | 0.15 |
| MT072864.1;QIQ54051.1;Pangolin_CoV_membrane;195 | 0 | 0.075 |
| MT072864.1;QIQ54051.1;Pangolin_CoV_membrane;196 | 0 | 0.1 |
| MT072864.1;QIQ54051.1;Pangolin_CoV_membrane;197 | 0 | 0.125 |
| MT072864.1;QIQ54051.1;Pangolin_CoV_membrane;198 | 0 | 0.1 |
| MT072864.1;QIQ54051.1;Pangolin_CoV_membrane;199 | 0 | 0.125 |
| MT072864.1;QIQ54051.1;Pangolin_CoV_membrane;200 | 0.1 | 0.225 |
| MT072864.1;QIQ54051.1;Pangolin_CoV_membrane;201 | 0.3 | 0.375 |
| MT072864.1;QIQ54051.1;Pangolin_CoV_membrane;202 | 0.3 | 0.4 |
| MT072864.1;QIQ54051.1;Pangolin_CoV_membrane;203 | 0.3 | 0.35 |
| MT072864.1;QIQ54051.1;Pangolin_CoV_membrane;204 | 0.25 | 0.45 |
| MT072864.1;QIQ54051.1;Pangolin_CoV_membrane;205 | 0.2 | 0.475 |
| MT072864.1;QIQ54051.1;Pangolin_CoV_membrane;206 | 0.15 | 0.4 |
| MT072864.1;QIQ54051.1;Pangolin_CoV_membrane;207 | 0.1 | 0.35 |
| MT072864.1;QIQ54051.1;Pangolin_CoV_membrane;208 | 0.1 | 0.35 |
| MT072864.1;QIQ54051.1;Pangolin_CoV_membrane;209 | 0.05 | 0.325 |
| MT072864.1;QIQ54051.1;Pangolin_CoV_membrane;210 | 0.1 | 0.275 |
| MT072864.1;QIQ54051.1;Pangolin_CoV_membrane;211 | 0.15 | 0.3 |
| MT072864.1;QIQ54052.1;Pangolin_CoV_orf6;1 | 0 | 0 |
| MT072864.1;QIQ54052.1;Pangolin_CoV_orf6;2 | 0 | 0 |
| MT072864.1;QIQ54052.1;Pangolin_CoV_orf6;3 | 0 | 0 |
| MT072864.1;QIQ54052.1;Pangolin_CoV_orf6;4 | 0 | 0 |
| MT072864.1;QIQ54052.1;Pangolin_CoV_orf6;5 | 0 | 0 |
| MT072864.1;QIQ54052.1;Pangolin_CoV_orf6;6 | 0 | 0 |
| MT072864.1;QIQ54052.1;Pangolin_CoV_orf6;7 | 0 | 0 |
| MT072864.1;QIQ54052.1;Pangolin_CoV_orf6;8 | 0 | 0 |
| MT072864.1;QIQ54052.1;Pangolin_CoV_orf6;9 | 0 | 0 |
| MT072864.1;QIQ54052.1;Pangolin_CoV_orf6;10 | 0 | 0 |
| MT072864.1;QIQ54052.1;Pangolin_CoV_orf6;11 | 0 | 0 |
| MT072864.1;QIQ54052.1;Pangolin_CoV_orf6;12 | 0 | 0 |
| MT072864.1;QIQ54052.1;Pangolin_CoV_orf6;13 | 0 | 0 |
| MT072864.1;QIQ54052.1;Pangolin_CoV_orf6;14 | 0 | 0 |
| MT072864.1;QIQ54052.1;Pangolin_CoV_orf6;15 | 0 | 0 |
| MT072864.1;QIQ54052.1;Pangolin_CoV_orf6;16 | 0 | 0 |
| MT072864.1;QIQ54052.1;Pangolin_CoV_orf6;17 | 0 | 0 |

| | | |
|---|---|---|
| MT072864.1;QIQ54052.1;Pangolin_CoV_orf6;18 | 0 | 0 |
| MT072864.1;QIQ54052.1;Pangolin_CoV_orf6;19 | 0 | 0 |
| MT072864.1;QIQ54052.1;Pangolin_CoV_orf6;20 | 0 | 0 |
| MT072864.1;QIQ54052.1;Pangolin_CoV_orf6;21 | 0 | 0 |
| MT072864.1;QIQ54052.1;Pangolin_CoV_orf6;22 | 0 | 0 |
| MT072864.1;QIQ54052.1;Pangolin_CoV_orf6;23 | 0 | 0 |
| MT072864.1;QIQ54052.1;Pangolin_CoV_orf6;24 | 0 | 0 |
| MT072864.1;QIQ54052.1;Pangolin_CoV_orf6;25 | 0 | 0 |
| MT072864.1;QIQ54052.1;Pangolin_CoV_orf6;26 | 0 | 0 |
| MT072864.1;QIQ54052.1;Pangolin_CoV_orf6;27 | 0 | 0 |
| MT072864.1;QIQ54052.1;Pangolin_CoV_orf6;28 | 0 | 0 |
| MT072864.1;QIQ54052.1;Pangolin_CoV_orf6;29 | 0 | 0 |
| MT072864.1;QIQ54052.1;Pangolin_CoV_orf6;30 | 0 | 0 |
| MT072864.1;QIQ54052.1;Pangolin_CoV_orf6;31 | 0 | 0 |
| MT072864.1;QIQ54052.1;Pangolin_CoV_orf6;32 | 0.05 | 0 |
| MT072864.1;QIQ54052.1;Pangolin_CoV_orf6;33 | 0 | 0 |
| MT072864.1;QIQ54052.1;Pangolin_CoV_orf6;34 | 0 | 0.025 |
| MT072864.1;QIQ54052.1;Pangolin_CoV_orf6;35 | 0 | 0.025 |
| MT072864.1;QIQ54052.1;Pangolin_CoV_orf6;36 | 0 | 0.05 |
| MT072864.1;QIQ54052.1;Pangolin_CoV_orf6;37 | 0 | 0.025 |
| MT072864.1;QIQ54052.1;Pangolin_CoV_orf6;38 | 0.05 | 0.025 |
| MT072864.1;QIQ54052.1;Pangolin_CoV_orf6;39 | 0.15 | 0.1 |
| MT072864.1;QIQ54052.1;Pangolin_CoV_orf6;40 | 0.1 | 0.075 |
| MT072864.1;QIQ54052.1;Pangolin_CoV_orf6;41 | 0.1 | 0.1 |
| MT072864.1;QIQ54052.1;Pangolin_CoV_orf6;42 | 0.05 | 0.1 |
| MT072864.1;QIQ54052.1;Pangolin_CoV_orf6;43 | 0.05 | 0.075 |
| MT072864.1;QIQ54052.1;Pangolin_CoV_orf6;44 | 0.05 | 0.075 |
| MT072864.1;QIQ54052.1;Pangolin_CoV_orf6;45 | 0.1 | 0.05 |
| MT072864.1;QIQ54052.1;Pangolin_CoV_orf6;46 | 0.05 | 0.05 |
| MT072864.1;QIQ54052.1;Pangolin_CoV_orf6;47 | 0 | 0 |
| MT072864.1;QIQ54052.1;Pangolin_CoV_orf6;48 | 0 | 0 |
| MT072864.1;QIQ54052.1;Pangolin_CoV_orf6;49 | 0 | 0.025 |
| MT072864.1;QIQ54052.1;Pangolin_CoV_orf6;50 | 0 | 0 |
| MT072864.1;QIQ54053.1;Pangolin_CoV_orf7a;1 | 0 | 0 |
| MT072864.1;QIQ54053.1;Pangolin_CoV_orf7a;2 | 0 | 0 |
| MT072864.1;QIQ54053.1;Pangolin_CoV_orf7a;3 | 0 | 0 |
| MT072864.1;QIQ54053.1;Pangolin_CoV_orf7a;4 | 0 | 0 |
| MT072864.1;QIQ54053.1;Pangolin_CoV_orf7a;5 | 0 | 0 |
| MT072864.1;QIQ54053.1;Pangolin_CoV_orf7a;6 | 0 | 0 |
| MT072864.1;QIQ54053.1;Pangolin_CoV_orf7a;7 | 0 | 0 |
| MT072864.1;QIQ54053.1;Pangolin_CoV_orf7a;8 | 0 | 0 |
| MT072864.1;QIQ54053.1;Pangolin_CoV_orf7a;9 | 0 | 0 |
| MT072864.1;QIQ54053.1;Pangolin_CoV_orf7a;10 | 0 | 0 |

| | | |
|---|---|---|
| MT072864.1;QIQ54053.1;Pangolin_CoV_orf7a;11 | 0 | 0 |
| MT072864.1;QIQ54053.1;Pangolin_CoV_orf7a;12 | 0 | 0 |
| MT072864.1;QIQ54053.1;Pangolin_CoV_orf7a;13 | 0 | 0 |
| MT072864.1;QIQ54053.1;Pangolin_CoV_orf7a;14 | 0 | 0 |
| MT072864.1;QIQ54053.1;Pangolin_CoV_orf7a;15 | 0 | 0 |
| MT072864.1;QIQ54053.1;Pangolin_CoV_orf7a;16 | 0 | 0 |
| MT072864.1;QIQ54053.1;Pangolin_CoV_orf7a;17 | 0 | 0 |
| MT072864.1;QIQ54053.1;Pangolin_CoV_orf7a;18 | 0 | 0 |
| MT072864.1;QIQ54053.1;Pangolin_CoV_orf7a;19 | 0 | 0 |
| MT072864.1;QIQ54053.1;Pangolin_CoV_orf7a;20 | 0 | 0 |
| MT072864.1;QIQ54053.1;Pangolin_CoV_orf7a;21 | 0 | 0 |
| MT072864.1;QIQ54053.1;Pangolin_CoV_orf7a;22 | 0 | 0 |
| MT072864.1;QIQ54053.1;Pangolin_CoV_orf7a;23 | 0.05 | 0 |
| MT072864.1;QIQ54053.1;Pangolin_CoV_orf7a;24 | 0 | 0 |
| MT072864.1;QIQ54053.1;Pangolin_CoV_orf7a;25 | 0.05 | 0 |
| MT072864.1;QIQ54053.1;Pangolin_CoV_orf7a;26 | 0 | 0 |
| MT072864.1;QIQ54053.1;Pangolin_CoV_orf7a;27 | 0 | 0 |
| MT072864.1;QIQ54053.1;Pangolin_CoV_orf7a;28 | 0 | 0.025 |
| MT072864.1;QIQ54053.1;Pangolin_CoV_orf7a;29 | 0 | 0.025 |
| MT072864.1;QIQ54053.1;Pangolin_CoV_orf7a;30 | 0 | 0.025 |
| MT072864.1;QIQ54053.1;Pangolin_CoV_orf7a;31 | 0 | 0.025 |
| MT072864.1;QIQ54053.1;Pangolin_CoV_orf7a;32 | 0 | 0 |
| MT072864.1;QIQ54053.1;Pangolin_CoV_orf7a;33 | 0 | 0 |
| MT072864.1;QIQ54053.1;Pangolin_CoV_orf7a;34 | 0 | 0.025 |
| MT072864.1;QIQ54053.1;Pangolin_CoV_orf7a;35 | 0 | 0.025 |
| MT072864.1;QIQ54053.1;Pangolin_CoV_orf7a;36 | 0.05 | 0.025 |
| MT072864.1;QIQ54053.1;Pangolin_CoV_orf7a;37 | 0 | 0.05 |
| MT072864.1;QIQ54053.1;Pangolin_CoV_orf7a;38 | 0 | 0.05 |
| MT072864.1;QIQ54053.1;Pangolin_CoV_orf7a;39 | 0 | 0.05 |
| MT072864.1;QIQ54053.1;Pangolin_CoV_orf7a;40 | 0 | 0.05 |
| MT072864.1;QIQ54053.1;Pangolin_CoV_orf7a;41 | 0 | 0.075 |
| MT072864.1;QIQ54053.1;Pangolin_CoV_orf7a;42 | 0 | 0.075 |
| MT072864.1;QIQ54053.1;Pangolin_CoV_orf7a;43 | 0 | 0.05 |
| MT072864.1;QIQ54053.1;Pangolin_CoV_orf7a;44 | 0 | 0.05 |
| MT072864.1;QIQ54053.1;Pangolin_CoV_orf7a;45 | 0 | 0 |
| MT072864.1;QIQ54053.1;Pangolin_CoV_orf7a;46 | 0 | 0 |
| MT072864.1;QIQ54053.1;Pangolin_CoV_orf7a;47 | 0 | 0 |
| MT072864.1;QIQ54053.1;Pangolin_CoV_orf7a;48 | 0 | 0 |
| MT072864.1;QIQ54053.1;Pangolin_CoV_orf7a;49 | 0 | 0 |
| MT072864.1;QIQ54053.1;Pangolin_CoV_orf7a;50 | 0 | 0 |
| MT072864.1;QIQ54053.1;Pangolin_CoV_orf7a;51 | 0 | 0 |
| MT072864.1;QIQ54053.1;Pangolin_CoV_orf7a;52 | 0 | 0 |
| MT072864.1;QIQ54053.1;Pangolin_CoV_orf7a;53 | 0 | 0 |

| | | |
|---|---|---|
| MT072864.1;QIQ54053.1;Pangolin_CoV_orf7a;54 | 0 | 0 |
| MT072864.1;QIQ54053.1;Pangolin_CoV_orf7a;55 | 0 | 0 |
| MT072864.1;QIQ54053.1;Pangolin_CoV_orf7a;56 | 0 | 0 |
| MT072864.1;QIQ54053.1;Pangolin_CoV_orf7a;57 | 0.05 | 0 |
| MT072864.1;QIQ54053.1;Pangolin_CoV_orf7a;58 | 0.05 | 0 |
| MT072864.1;QIQ54053.1;Pangolin_CoV_orf7a;59 | 0.05 | 0 |
| MT072864.1;QIQ54053.1;Pangolin_CoV_orf7a;60 | 0.05 | 0 |
| MT072864.1;QIQ54053.1;Pangolin_CoV_orf7a;61 | 0.05 | 0 |
| MT072864.1;QIQ54053.1;Pangolin_CoV_orf7a;62 | 0 | 0 |
| MT072864.1;QIQ54053.1;Pangolin_CoV_orf7a;63 | 0 | 0 |
| MT072864.1;QIQ54053.1;Pangolin_CoV_orf7a;64 | 0 | 0.025 |
| MT072864.1;QIQ54053.1;Pangolin_CoV_orf7a;65 | 0 | 0 |
| MT072864.1;QIQ54053.1;Pangolin_CoV_orf7a;66 | 0 | 0 |
| MT072864.1;QIQ54053.1;Pangolin_CoV_orf7a;67 | 0 | 0 |
| MT072864.1;QIQ54053.1;Pangolin_CoV_orf7a;68 | 0 | 0 |
| MT072864.1;QIQ54053.1;Pangolin_CoV_orf7a;69 | 0 | 0 |
| MT072864.1;QIQ54053.1;Pangolin_CoV_orf7a;70 | 0 | 0 |
| MT072864.1;QIQ54053.1;Pangolin_CoV_orf7a;71 | 0 | 0 |
| MT072864.1;QIQ54053.1;Pangolin_CoV_orf7a;72 | 0 | 0 |
| MT072864.1;QIQ54053.1;Pangolin_CoV_orf7a;73 | 0 | 0 |
| MT072864.1;QIQ54053.1;Pangolin_CoV_orf7a;74 | 0 | 0 |
| MT072864.1;QIQ54053.1;Pangolin_CoV_orf7a;75 | 0 | 0 |
| MT072864.1;QIQ54053.1;Pangolin_CoV_orf7a;76 | 0 | 0 |
| MT072864.1;QIQ54053.1;Pangolin_CoV_orf7a;77 | 0 | 0 |
| MT072864.1;QIQ54053.1;Pangolin_CoV_orf7a;78 | 0 | 0 |
| MT072864.1;QIQ54053.1;Pangolin_CoV_orf7a;79 | 0.05 | 0 |
| MT072864.1;QIQ54053.1;Pangolin_CoV_orf7a;80 | 0.15 | 0 |
| MT072864.1;QIQ54053.1;Pangolin_CoV_orf7a;81 | 0.05 | 0 |
| MT072864.1;QIQ54053.1;Pangolin_CoV_orf7a;82 | 0 | 0 |
| MT072864.1;QIQ54053.1;Pangolin_CoV_orf7a;83 | 0 | 0 |
| MT072864.1;QIQ54053.1;Pangolin_CoV_orf7a;84 | 0.05 | 0 |
| MT072864.1;QIQ54053.1;Pangolin_CoV_orf7a;85 | 0 | 0 |
| MT072864.1;QIQ54053.1;Pangolin_CoV_orf7a;86 | 0 | 0 |
| MT072864.1;QIQ54053.1;Pangolin_CoV_orf7a;87 | 0 | 0.025 |
| MT072864.1;QIQ54053.1;Pangolin_CoV_orf7a;88 | 0 | 0 |
| MT072864.1;QIQ54053.1;Pangolin_CoV_orf7a;89 | 0 | 0 |
| MT072864.1;QIQ54053.1;Pangolin_CoV_orf7a;90 | 0 | 0 |
| MT072864.1;QIQ54053.1;Pangolin_CoV_orf7a;91 | 0 | 0 |
| MT072864.1;QIQ54053.1;Pangolin_CoV_orf7a;92 | 0 | 0 |
| MT072864.1;QIQ54053.1;Pangolin_CoV_orf7a;93 | 0 | 0 |
| MT072864.1;QIQ54053.1;Pangolin_CoV_orf7a;94 | 0 | 0 |
| MT072864.1;QIQ54053.1;Pangolin_CoV_orf7a;95 | 0 | 0 |
| MT072864.1;QIQ54053.1;Pangolin_CoV_orf7a;96 | 0 | 0 |

| | | |
|---|---|---|
| MT072864.1;QIQ54053.1;Pangolin_CoV_orf7a;97 | 0 | 0 |
| MT072864.1;QIQ54053.1;Pangolin_CoV_orf7a;98 | 0 | 0 |
| MT072864.1;QIQ54053.1;Pangolin_CoV_orf7a;99 | 0 | 0 |
| MT072864.1;QIQ54053.1;Pangolin_CoV_orf7a;100 | 0 | 0 |
| MT072864.1;QIQ54053.1;Pangolin_CoV_orf7a;101 | 0 | 0 |
| MT072864.1;QIQ54053.1;Pangolin_CoV_orf7a;102 | 0 | 0 |
| MT072864.1;QIQ54053.1;Pangolin_CoV_orf7a;103 | 0 | 0 |
| MT072864.1;QIQ54053.1;Pangolin_CoV_orf7a;104 | 0 | 0 |
| MT072864.1;QIQ54053.1;Pangolin_CoV_orf7a;105 | 0 | 0 |
| MT072864.1;QIQ54053.1;Pangolin_CoV_orf7a;106 | 0 | 0 |
| MT072864.1;QIQ54053.1;Pangolin_CoV_orf7a;107 | 0 | 0 |
| MT072864.1;QIQ54053.1;Pangolin_CoV_orf7a;108 | 0 | 0 |
| MT072864.1;QIQ54053.1;Pangolin_CoV_orf7a;109 | 0 | 0 |
| MT072864.1;QIQ54053.1;Pangolin_CoV_orf7a;110 | 0 | 0 |
| MT072864.1;QIQ54054.1;Pangolin_CoV_orf7b;1 | 0 | 0 |
| MT072864.1;QIQ54054.1;Pangolin_CoV_orf7b;2 | 0 | 0 |
| MT072864.1;QIQ54054.1;Pangolin_CoV_orf7b;3 | 0 | 0 |
| MT072864.1;QIQ54054.1;Pangolin_CoV_orf7b;4 | 0 | 0 |
| MT072864.1;QIQ54054.1;Pangolin_CoV_orf7b;5 | 0 | 0 |
| MT072864.1;QIQ54054.1;Pangolin_CoV_orf7b;6 | 0 | 0 |
| MT072864.1;QIQ54054.1;Pangolin_CoV_orf7b;7 | 0 | 0 |
| MT072864.1;QIQ54054.1;Pangolin_CoV_orf7b;8 | 0 | 0 |
| MT072864.1;QIQ54054.1;Pangolin_CoV_orf7b;9 | 0 | 0 |
| MT072864.1;QIQ54054.1;Pangolin_CoV_orf7b;10 | 0 | 0 |
| MT072864.1;QIQ54054.1;Pangolin_CoV_orf7b;11 | 0 | 0 |
| MT072864.1;QIQ54054.1;Pangolin_CoV_orf7b;12 | 0 | 0 |
| MT072864.1;QIQ54054.1;Pangolin_CoV_orf7b;13 | 0 | 0 |
| MT072864.1;QIQ54054.1;Pangolin_CoV_orf7b;14 | 0 | 0 |
| MT072864.1;QIQ54054.1;Pangolin_CoV_orf7b;15 | 0 | 0 |
| MT072864.1;QIQ54054.1;Pangolin_CoV_orf7b;16 | 0 | 0 |
| MT072864.1;QIQ54054.1;Pangolin_CoV_orf7b;17 | 0 | 0 |
| MT072864.1;QIQ54054.1;Pangolin_CoV_orf7b;18 | 0 | 0 |
| MT072864.1;QIQ54054.1;Pangolin_CoV_orf7b;19 | 0 | 0 |
| MT072864.1;QIQ54054.1;Pangolin_CoV_orf7b;20 | 0 | 0 |
| MT072864.1;QIQ54054.1;Pangolin_CoV_orf7b;21 | 0 | 0 |
| MT072864.1;QIQ54054.1;Pangolin_CoV_orf7b;22 | 0 | 0 |
| MT072864.1;QIQ54054.1;Pangolin_CoV_orf7b;23 | 0 | 0 |
| MT072864.1;QIQ54054.1;Pangolin_CoV_orf7b;24 | 0 | 0 |
| MT072864.1;QIQ54054.1;Pangolin_CoV_orf7b;25 | 0 | 0 |
| MT072864.1;QIQ54054.1;Pangolin_CoV_orf7b;26 | 0 | 0.025 |
| MT072864.1;QIQ54054.1;Pangolin_CoV_orf7b;27 | 0 | 0 |
| MT072864.1;QIQ54054.1;Pangolin_CoV_orf7b;28 | 0 | 0 |
| MT072864.1;QIQ54054.1;Pangolin_CoV_orf7b;29 | 0 | 0.025 |

| | | |
|---|---|---|
| MT072864.1;QIQ54054.1;Pangolin_CoV_orf7b;30 | 0 | 0 |
| MT072864.1;QIQ54054.1;Pangolin_CoV_orf7b;31 | 0.05 | 0 |
| MT072864.1;QIQ54054.1;Pangolin_CoV_orf7b;32 | 0 | 0 |
| MT072864.1;QIQ54055.1;Pangolin_CoV_orf8;1 | 0 | 0 |
| MT072864.1;QIQ54055.1;Pangolin_CoV_orf8;2 | 0 | 0 |
| MT072864.1;QIQ54055.1;Pangolin_CoV_orf8;3 | 0 | 0 |
| MT072864.1;QIQ54055.1;Pangolin_CoV_orf8;4 | 0 | 0 |
| MT072864.1;QIQ54055.1;Pangolin_CoV_orf8;5 | 0 | 0 |
| MT072864.1;QIQ54055.1;Pangolin_CoV_orf8;6 | 0 | 0 |
| MT072864.1;QIQ54055.1;Pangolin_CoV_orf8;7 | 0 | 0 |
| MT072864.1;QIQ54055.1;Pangolin_CoV_orf8;8 | 0 | 0 |
| MT072864.1;QIQ54055.1;Pangolin_CoV_orf8;9 | 0 | 0 |
| MT072864.1;QIQ54055.1;Pangolin_CoV_orf8;10 | 0 | 0 |
| MT072864.1;QIQ54055.1;Pangolin_CoV_orf8;11 | 0 | 0 |
| MT072864.1;QIQ54055.1;Pangolin_CoV_orf8;12 | 0 | 0 |
| MT072864.1;QIQ54055.1;Pangolin_CoV_orf8;13 | 0 | 0.025 |
| MT072864.1;QIQ54055.1;Pangolin_CoV_orf8;14 | 0 | 0.025 |
| MT072864.1;QIQ54055.1;Pangolin_CoV_orf8;15 | 0 | 0.025 |
| MT072864.1;QIQ54055.1;Pangolin_CoV_orf8;16 | 0 | 0.025 |
| MT072864.1;QIQ54055.1;Pangolin_CoV_orf8;17 | 0 | 0 |
| MT072864.1;QIQ54055.1;Pangolin_CoV_orf8;18 | 0 | 0 |
| MT072864.1;QIQ54055.1;Pangolin_CoV_orf8;19 | 0 | 0 |
| MT072864.1;QIQ54055.1;Pangolin_CoV_orf8;20 | 0 | 0 |
| MT072864.1;QIQ54055.1;Pangolin_CoV_orf8;21 | 0 | 0 |
| MT072864.1;QIQ54055.1;Pangolin_CoV_orf8;22 | 0 | 0 |
| MT072864.1;QIQ54055.1;Pangolin_CoV_orf8;23 | 0 | 0 |
| MT072864.1;QIQ54055.1;Pangolin_CoV_orf8;24 | 0 | 0 |
| MT072864.1;QIQ54055.1;Pangolin_CoV_orf8;25 | 0 | 0 |
| MT072864.1;QIQ54055.1;Pangolin_CoV_orf8;26 | 0 | 0 |
| MT072864.1;QIQ54055.1;Pangolin_CoV_orf8;27 | 0 | 0 |
| MT072864.1;QIQ54055.1;Pangolin_CoV_orf8;28 | 0 | 0 |
| MT072864.1;QIQ54055.1;Pangolin_CoV_orf8;29 | 0 | 0.025 |
| MT072864.1;QIQ54055.1;Pangolin_CoV_orf8;30 | 0 | 0 |
| MT072864.1;QIQ54055.1;Pangolin_CoV_orf8;31 | 0 | 0 |
| MT072864.1;QIQ54055.1;Pangolin_CoV_orf8;32 | 0 | 0 |
| MT072864.1;QIQ54055.1;Pangolin_CoV_orf8;33 | 0 | 0 |
| MT072864.1;QIQ54055.1;Pangolin_CoV_orf8;34 | 0 | 0 |
| MT072864.1;QIQ54055.1;Pangolin_CoV_orf8;35 | 0 | 0 |
| MT072864.1;QIQ54055.1;Pangolin_CoV_orf8;36 | 0 | 0 |
| MT072864.1;QIQ54055.1;Pangolin_CoV_orf8;37 | 0 | 0 |
| MT072864.1;QIQ54055.1;Pangolin_CoV_orf8;38 | 0 | 0 |
| MT072864.1;QIQ54055.1;Pangolin_CoV_orf8;39 | 0 | 0 |
| MT072864.1;QIQ54055.1;Pangolin_CoV_orf8;40 | 0 | 0 |

| | | |
|---|---|---|
| MT072864.1;QIQ54055.1;Pangolin_CoV_orf8;41 | 0 | 0 |
| MT072864.1;QIQ54055.1;Pangolin_CoV_orf8;42 | 0 | 0 |
| MT072864.1;QIQ54055.1;Pangolin_CoV_orf8;43 | 0 | 0 |
| MT072864.1;QIQ54055.1;Pangolin_CoV_orf8;44 | 0 | 0 |
| MT072864.1;QIQ54055.1;Pangolin_CoV_orf8;45 | 0 | 0 |
| MT072864.1;QIQ54055.1;Pangolin_CoV_orf8;46 | 0 | 0 |
| MT072864.1;QIQ54055.1;Pangolin_CoV_orf8;47 | 0 | 0.025 |
| MT072864.1;QIQ54055.1;Pangolin_CoV_orf8;48 | 0 | 0 |
| MT072864.1;QIQ54055.1;Pangolin_CoV_orf8;49 | 0 | 0 |
| MT072864.1;QIQ54055.1;Pangolin_CoV_orf8;50 | 0 | 0 |
| MT072864.1;QIQ54055.1;Pangolin_CoV_orf8;51 | 0 | 0 |
| MT072864.1;QIQ54055.1;Pangolin_CoV_orf8;52 | 0 | 0 |
| MT072864.1;QIQ54055.1;Pangolin_CoV_orf8;53 | 0 | 0 |
| MT072864.1;QIQ54055.1;Pangolin_CoV_orf8;54 | 0.05 | 0 |
| MT072864.1;QIQ54055.1;Pangolin_CoV_orf8;55 | 0.05 | 0 |
| MT072864.1;QIQ54055.1;Pangolin_CoV_orf8;56 | 0.05 | 0 |
| MT072864.1;QIQ54055.1;Pangolin_CoV_orf8;57 | 0 | 0.025 |
| MT072864.1;QIQ54055.1;Pangolin_CoV_orf8;58 | 0.05 | 0.025 |
| MT072864.1;QIQ54055.1;Pangolin_CoV_orf8;59 | 0.05 | 0.025 |
| MT072864.1;QIQ54055.1;Pangolin_CoV_orf8;60 | 0 | 0.025 |
| MT072864.1;QIQ54055.1;Pangolin_CoV_orf8;61 | 0.05 | 0.025 |
| MT072864.1;QIQ54055.1;Pangolin_CoV_orf8;62 | 0.05 | 0.025 |
| MT072864.1;QIQ54055.1;Pangolin_CoV_orf8;63 | 0.05 | 0.075 |
| MT072864.1;QIQ54055.1;Pangolin_CoV_orf8;64 | 0.05 | 0.075 |
| MT072864.1;QIQ54055.1;Pangolin_CoV_orf8;65 | 0 | 0.025 |
| MT072864.1;QIQ54055.1;Pangolin_CoV_orf8;66 | 0 | 0 |
| MT072864.1;QIQ54055.1;Pangolin_CoV_orf8;67 | 0 | 0.025 |
| MT072864.1;QIQ54055.1;Pangolin_CoV_orf8;68 | 0 | 0 |
| MT072864.1;QIQ54055.1;Pangolin_CoV_orf8;69 | 0 | 0.05 |
| MT072864.1;QIQ54055.1;Pangolin_CoV_orf8;70 | 0 | 0.025 |
| MT072864.1;QIQ54055.1;Pangolin_CoV_orf8;71 | 0 | 0 |
| MT072864.1;QIQ54055.1;Pangolin_CoV_orf8;72 | 0 | 0 |
| MT072864.1;QIQ54055.1;Pangolin_CoV_orf8;73 | 0 | 0 |
| MT072864.1;QIQ54055.1;Pangolin_CoV_orf8;74 | 0 | 0 |
| MT072864.1;QIQ54055.1;Pangolin_CoV_orf8;75 | 0 | 0 |
| MT072864.1;QIQ54055.1;Pangolin_CoV_orf8;76 | 0 | 0 |
| MT072864.1;QIQ54055.1;Pangolin_CoV_orf8;77 | 0 | 0 |
| MT072864.1;QIQ54055.1;Pangolin_CoV_orf8;78 | 0 | 0 |
| MT072864.1;QIQ54055.1;Pangolin_CoV_orf8;79 | 0 | 0 |
| MT072864.1;QIQ54055.1;Pangolin_CoV_orf8;80 | 0 | 0 |
| MT072864.1;QIQ54055.1;Pangolin_CoV_orf8;81 | 0 | 0 |
| MT072864.1;QIQ54055.1;Pangolin_CoV_orf8;82 | 0 | 0 |
| MT072864.1;QIQ54055.1;Pangolin_CoV_orf8;83 | 0.05 | 0 |

| | | |
|---|---|---|
| MT072864.1;QIQ54055.1;Pangolin_CoV_orf8;84 | 0.05 | 0.025 |
| MT072864.1;QIQ54055.1;Pangolin_CoV_orf8;85 | 0.05 | 0.05 |
| MT072864.1;QIQ54055.1;Pangolin_CoV_orf8;86 | 0 | 0 |
| MT072864.1;QIQ54055.1;Pangolin_CoV_orf8;87 | 0 | 0 |
| MT072864.1;QIQ54055.1;Pangolin_CoV_orf8;88 | 0 | 0 |
| MT072864.1;QIQ54055.1;Pangolin_CoV_orf8;89 | 0 | 0 |
| MT072864.1;QIQ54055.1;Pangolin_CoV_orf8;90 | 0 | 0 |
| MT072864.1;QIQ54055.1;Pangolin_CoV_orf8;91 | 0 | 0 |
| MT072864.1;QIQ54055.1;Pangolin_CoV_orf8;92 | 0 | 0 |
| MT072864.1;QIQ54055.1;Pangolin_CoV_orf8;93 | 0 | 0 |
| MT072864.1;QIQ54055.1;Pangolin_CoV_orf8;94 | 0 | 0 |
| MT072864.1;QIQ54055.1;Pangolin_CoV_orf8;95 | 0 | 0 |
| MT072864.1;QIQ54055.1;Pangolin_CoV_orf8;96 | 0 | 0 |
| MT072864.1;QIQ54055.1;Pangolin_CoV_orf8;97 | 0 | 0 |
| MT072864.1;QIQ54055.1;Pangolin_CoV_orf8;98 | 0 | 0 |
| MT072864.1;QIQ54055.1;Pangolin_CoV_orf8;99 | 0 | 0 |
| MT072864.1;QIQ54055.1;Pangolin_CoV_orf8;100 | 0 | 0 |
| MT072864.1;QIQ54055.1;Pangolin_CoV_orf8;101 | 0 | 0 |
| MT072864.1;QIQ54055.1;Pangolin_CoV_orf8;102 | 0 | 0 |
| MT072864.1;QIQ54055.1;Pangolin_CoV_orf8;103 | 0 | 0 |
| MT072864.1;QIQ54055.1;Pangolin_CoV_orf8;104 | 0 | 0 |
| MT072864.1;QIQ54055.1;Pangolin_CoV_orf8;105 | 0 | 0 |
| MT072864.1;QIQ54055.1;Pangolin_CoV_orf8;106 | 0 | 0 |
| MT072864.1;QIQ54055.1;Pangolin_CoV_orf8;107 | 0 | 0 |
| MT072864.1;QIQ54055.1;Pangolin_CoV_orf8;108 | 0 | 0 |
| MT072864.1;QIQ54055.1;Pangolin_CoV_orf8;109 | 0 | 0 |
| MT072864.1;QIQ54055.1;Pangolin_CoV_orf8;110 | 0 | 0 |
| MT072864.1;QIQ54056.1;Pangolin_CoV_nucleoca;1 | 0 | 0 |
| MT072864.1;QIQ54056.1;Pangolin_CoV_nucleoca;2 | 0 | 0 |
| MT072864.1;QIQ54056.1;Pangolin_CoV_nucleoca;3 | 0.05 | 0 |
| MT072864.1;QIQ54056.1;Pangolin_CoV_nucleoca;4 | 0.05 | 0 |
| MT072864.1;QIQ54056.1;Pangolin_CoV_nucleoca;5 | 0.05 | 0 |
| MT072864.1;QIQ54056.1;Pangolin_CoV_nucleoca;6 | 0.05 | 0.1 |
| MT072864.1;QIQ54056.1;Pangolin_CoV_nucleoca;7 | 0.05 | 0.2 |
| MT072864.1;QIQ54056.1;Pangolin_CoV_nucleoca;8 | 0.1 | 0.3 |
| MT072864.1;QIQ54056.1;Pangolin_CoV_nucleoca;9 | 0.05 | 0.375 |
| MT072864.1;QIQ54056.1;Pangolin_CoV_nucleoca;10 | 0.05 | 0.35 |
| MT072864.1;QIQ54056.1;Pangolin_CoV_nucleoca;11 | 0.15 | 0.325 |
| MT072864.1;QIQ54056.1;Pangolin_CoV_nucleoca;12 | 0.05 | 0.3 |
| MT072864.1;QIQ54056.1;Pangolin_CoV_nucleoca;13 | 0.05 | 0.125 |
| MT072864.1;QIQ54056.1;Pangolin_CoV_nucleoca;14 | 0 | 0.075 |
| MT072864.1;QIQ54056.1;Pangolin_CoV_nucleoca;15 | 0 | 0.05 |
| MT072864.1;QIQ54056.1;Pangolin_CoV_nucleoca;16 | 0 | 0.1 |

| | | |
|---|---|---|
| MT072864.1;QIQ54056.1;Pangolin_CoV_nucleoca;17 | 0.05 | 0.1 |
| MT072864.1;QIQ54056.1;Pangolin_CoV_nucleoca;18 | 0 | 0.05 |
| MT072864.1;QIQ54056.1;Pangolin_CoV_nucleoca;19 | 0.05 | 0.125 |
| MT072864.1;QIQ54056.1;Pangolin_CoV_nucleoca;20 | 0.1 | 0.1 |
| MT072864.1;QIQ54056.1;Pangolin_CoV_nucleoca;21 | 0.1 | 0.175 |
| MT072864.1;QIQ54056.1;Pangolin_CoV_nucleoca;22 | 0.1 | 0.175 |
| MT072864.1;QIQ54056.1;Pangolin_CoV_nucleoca;23 | 0.05 | 0.125 |
| MT072864.1;QIQ54056.1;Pangolin_CoV_nucleoca;24 | 0.05 | 0.1 |
| MT072864.1;QIQ54056.1;Pangolin_CoV_nucleoca;25 | 0.05 | 0.15 |
| MT072864.1;QIQ54056.1;Pangolin_CoV_nucleoca;26 | 0.05 | 0.125 |
| MT072864.1;QIQ54056.1;Pangolin_CoV_nucleoca;27 | 0.05 | 0.1 |
| MT072864.1;QIQ54056.1;Pangolin_CoV_nucleoca;28 | 0.05 | 0.15 |
| MT072864.1;QIQ54056.1;Pangolin_CoV_nucleoca;29 | 0.05 | 0.175 |
| MT072864.1;QIQ54056.1;Pangolin_CoV_nucleoca;30 | 0 | 0.1 |
| MT072864.1;QIQ54056.1;Pangolin_CoV_nucleoca;31 | 0 | 0.2 |
| MT072864.1;QIQ54056.1;Pangolin_CoV_nucleoca;32 | 0 | 0.125 |
| MT072864.1;QIQ54056.1;Pangolin_CoV_nucleoca;33 | 0 | 0.225 |
| MT072864.1;QIQ54056.1;Pangolin_CoV_nucleoca;34 | 0.05 | 0.15 |
| MT072864.1;QIQ54056.1;Pangolin_CoV_nucleoca;35 | 0.05 | 0.3 |
| MT072864.1;QIQ54056.1;Pangolin_CoV_nucleoca;36 | 0 | 0.225 |
| MT072864.1;QIQ54056.1;Pangolin_CoV_nucleoca;37 | 0 | 0.15 |
| MT072864.1;QIQ54056.1;Pangolin_CoV_nucleoca;38 | 0 | 0.125 |
| MT072864.1;QIQ54056.1;Pangolin_CoV_nucleoca;39 | 0 | 0.025 |
| MT072864.1;QIQ54056.1;Pangolin_CoV_nucleoca;40 | 0 | 0 |
| MT072864.1;QIQ54056.1;Pangolin_CoV_nucleoca;41 | 0 | 0.025 |
| MT072864.1;QIQ54056.1;Pangolin_CoV_nucleoca;42 | 0 | 0 |
| MT072864.1;QIQ54056.1;Pangolin_CoV_nucleoca;43 | 0 | 0 |
| MT072864.1;QIQ54056.1;Pangolin_CoV_nucleoca;44 | 0 | 0 |
| MT072864.1;QIQ54056.1;Pangolin_CoV_nucleoca;45 | 0 | 0 |
| MT072864.1;QIQ54056.1;Pangolin_CoV_nucleoca;46 | 0 | 0 |
| MT072864.1;QIQ54056.1;Pangolin_CoV_nucleoca;47 | 0 | 0 |
| MT072864.1;QIQ54056.1;Pangolin_CoV_nucleoca;48 | 0 | 0 |
| MT072864.1;QIQ54056.1;Pangolin_CoV_nucleoca;49 | 0 | 0.025 |
| MT072864.1;QIQ54056.1;Pangolin_CoV_nucleoca;50 | 0 | 0 |
| MT072864.1;QIQ54056.1;Pangolin_CoV_nucleoca;51 | 0 | 0 |
| MT072864.1;QIQ54056.1;Pangolin_CoV_nucleoca;52 | 0 | 0 |
| MT072864.1;QIQ54056.1;Pangolin_CoV_nucleoca;53 | 0 | 0 |
| MT072864.1;QIQ54056.1;Pangolin_CoV_nucleoca;54 | 0 | 0 |
| MT072864.1;QIQ54056.1;Pangolin_CoV_nucleoca;55 | 0 | 0 |
| MT072864.1;QIQ54056.1;Pangolin_CoV_nucleoca;56 | 0 | 0.025 |
| MT072864.1;QIQ54056.1;Pangolin_CoV_nucleoca;57 | 0 | 0 |
| MT072864.1;QIQ54056.1;Pangolin_CoV_nucleoca;58 | 0 | 0.075 |
| MT072864.1;QIQ54056.1;Pangolin_CoV_nucleoca;59 | 0 | 0.025 |

| | | |
|---|---|---|
| MT072864.1;QIQ54056.1;Pangolin_CoV_nucleoca;60 | 0.05 | 0.05 |
| MT072864.1;QIQ54056.1;Pangolin_CoV_nucleoca;61 | 0.1 | 0.05 |
| MT072864.1;QIQ54056.1;Pangolin_CoV_nucleoca;62 | 0 | 0.075 |
| MT072864.1;QIQ54056.1;Pangolin_CoV_nucleoca;63 | 0.05 | 0.05 |
| MT072864.1;QIQ54056.1;Pangolin_CoV_nucleoca;64 | 0.05 | 0.1 |
| MT072864.1;QIQ54056.1;Pangolin_CoV_nucleoca;65 | 0.05 | 0.075 |
| MT072864.1;QIQ54056.1;Pangolin_CoV_nucleoca;66 | 0.1 | 0.1 |
| MT072864.1;QIQ54056.1;Pangolin_CoV_nucleoca;67 | 0 | 0 |
| MT072864.1;QIQ54056.1;Pangolin_CoV_nucleoca;68 | 0 | 0.05 |
| MT072864.1;QIQ54056.1;Pangolin_CoV_nucleoca;69 | 0.1 | 0.025 |
| MT072864.1;QIQ54056.1;Pangolin_CoV_nucleoca;70 | 0.15 | 0.075 |
| MT072864.1;QIQ54056.1;Pangolin_CoV_nucleoca;71 | 0 | 0.05 |
| MT072864.1;QIQ54056.1;Pangolin_CoV_nucleoca;72 | 0 | 0.05 |
| MT072864.1;QIQ54056.1;Pangolin_CoV_nucleoca;73 | 0.05 | 0.025 |
| MT072864.1;QIQ54056.1;Pangolin_CoV_nucleoca;74 | 0.05 | 0 |
| MT072864.1;QIQ54056.1;Pangolin_CoV_nucleoca;75 | 0.05 | 0 |
| MT072864.1;QIQ54056.1;Pangolin_CoV_nucleoca;76 | 0.05 | 0 |
| MT072864.1;QIQ54056.1;Pangolin_CoV_nucleoca;77 | 0.05 | 0 |
| MT072864.1;QIQ54056.1;Pangolin_CoV_nucleoca;78 | 0.05 | 0 |
| MT072864.1;QIQ54056.1;Pangolin_CoV_nucleoca;79 | 0 | 0 |
| MT072864.1;QIQ54056.1;Pangolin_CoV_nucleoca;80 | 0 | 0 |
| MT072864.1;QIQ54056.1;Pangolin_CoV_nucleoca;81 | 0.05 | 0 |
| MT072864.1;QIQ54056.1;Pangolin_CoV_nucleoca;82 | 0.05 | 0 |
| MT072864.1;QIQ54056.1;Pangolin_CoV_nucleoca;83 | 0.05 | 0 |
| MT072864.1;QIQ54056.1;Pangolin_CoV_nucleoca;84 | 0 | 0 |
| MT072864.1;QIQ54056.1;Pangolin_CoV_nucleoca;85 | 0 | 0.025 |
| MT072864.1;QIQ54056.1;Pangolin_CoV_nucleoca;86 | 0 | 0 |
| MT072864.1;QIQ54056.1;Pangolin_CoV_nucleoca;87 | 0 | 0.025 |
| MT072864.1;QIQ54056.1;Pangolin_CoV_nucleoca;88 | 0 | 0.025 |
| MT072864.1;QIQ54056.1;Pangolin_CoV_nucleoca;89 | 0 | 0.025 |
| MT072864.1;QIQ54056.1;Pangolin_CoV_nucleoca;90 | 0 | 0.05 |
| MT072864.1;QIQ54056.1;Pangolin_CoV_nucleoca;91 | 0 | 0.075 |
| MT072864.1;QIQ54056.1;Pangolin_CoV_nucleoca;92 | 0 | 0.05 |
| MT072864.1;QIQ54056.1;Pangolin_CoV_nucleoca;93 | 0 | 0.025 |
| MT072864.1;QIQ54056.1;Pangolin_CoV_nucleoca;94 | 0 | 0.025 |
| MT072864.1;QIQ54056.1;Pangolin_CoV_nucleoca;95 | 0 | 0.025 |
| MT072864.1;QIQ54056.1;Pangolin_CoV_nucleoca;96 | 0 | 0.025 |
| MT072864.1;QIQ54056.1;Pangolin_CoV_nucleoca;97 | 0 | 0 |
| MT072864.1;QIQ54056.1;Pangolin_CoV_nucleoca;98 | 0 | 0 |
| MT072864.1;QIQ54056.1;Pangolin_CoV_nucleoca;99 | 0 | 0 |
| MT072864.1;QIQ54056.1;Pangolin_CoV_nucleoca;100 | 0 | 0 |
| MT072864.1;QIQ54056.1;Pangolin_CoV_nucleoca;101 | 0 | 0 |
| MT072864.1;QIQ54056.1;Pangolin_CoV_nucleoca;102 | 0 | 0 |

| | | |
|---|---|---|
| MT072864.1;QIQ54056.1;Pangolin_CoV_nucleoca;103 | 0 | 0 |
| MT072864.1;QIQ54056.1;Pangolin_CoV_nucleoca;104 | 0 | 0 |
| MT072864.1;QIQ54056.1;Pangolin_CoV_nucleoca;105 | 0 | 0 |
| MT072864.1;QIQ54056.1;Pangolin_CoV_nucleoca;106 | 0 | 0 |
| MT072864.1;QIQ54056.1;Pangolin_CoV_nucleoca;107 | 0 | 0 |
| MT072864.1;QIQ54056.1;Pangolin_CoV_nucleoca;108 | 0 | 0 |
| MT072864.1;QIQ54056.1;Pangolin_CoV_nucleoca;109 | 0.05 | 0 |
| MT072864.1;QIQ54056.1;Pangolin_CoV_nucleoca;110 | 0.1 | 0.125 |
| MT072864.1;QIQ54056.1;Pangolin_CoV_nucleoca;111 | 0.05 | 0.05 |
| MT072864.1;QIQ54056.1;Pangolin_CoV_nucleoca;112 | 0 | 0.05 |
| MT072864.1;QIQ54056.1;Pangolin_CoV_nucleoca;113 | 0 | 0.075 |
| MT072864.1;QIQ54056.1;Pangolin_CoV_nucleoca;114 | 0 | 0.075 |
| MT072864.1;QIQ54056.1;Pangolin_CoV_nucleoca;115 | 0 | 0.075 |
| MT072864.1;QIQ54056.1;Pangolin_CoV_nucleoca;116 | 0 | 0.025 |
| MT072864.1;QIQ54056.1;Pangolin_CoV_nucleoca;117 | 0 | 0.025 |
| MT072864.1;QIQ54056.1;Pangolin_CoV_nucleoca;118 | 0 | 0.025 |
| MT072864.1;QIQ54056.1;Pangolin_CoV_nucleoca;119 | 0 | 0 |
| MT072864.1;QIQ54056.1;Pangolin_CoV_nucleoca;120 | 0 | 0 |
| MT072864.1;QIQ54056.1;Pangolin_CoV_nucleoca;121 | 0 | 0 |
| MT072864.1;QIQ54056.1;Pangolin_CoV_nucleoca;122 | 0 | 0 |
| MT072864.1;QIQ54056.1;Pangolin_CoV_nucleoca;123 | 0 | 0.025 |
| MT072864.1;QIQ54056.1;Pangolin_CoV_nucleoca;124 | 0 | 0 |
| MT072864.1;QIQ54056.1;Pangolin_CoV_nucleoca;125 | 0 | 0 |
| MT072864.1;QIQ54056.1;Pangolin_CoV_nucleoca;126 | 0 | 0.05 |
| MT072864.1;QIQ54056.1;Pangolin_CoV_nucleoca;127 | 0 | 0.075 |
| MT072864.1;QIQ54056.1;Pangolin_CoV_nucleoca;128 | 0 | 0.05 |
| MT072864.1;QIQ54056.1;Pangolin_CoV_nucleoca;129 | 0 | 0.075 |
| MT072864.1;QIQ54056.1;Pangolin_CoV_nucleoca;130 | 0 | 0.075 |
| MT072864.1;QIQ54056.1;Pangolin_CoV_nucleoca;131 | 0 | 0.05 |
| MT072864.1;QIQ54056.1;Pangolin_CoV_nucleoca;132 | 0 | 0.025 |
| MT072864.1;QIQ54056.1;Pangolin_CoV_nucleoca;133 | 0 | 0.025 |
| MT072864.1;QIQ54056.1;Pangolin_CoV_nucleoca;134 | 0 | 0.025 |
| MT072864.1;QIQ54056.1;Pangolin_CoV_nucleoca;135 | 0 | 0 |
| MT072864.1;QIQ54056.1;Pangolin_CoV_nucleoca;136 | 0.05 | 0.025 |
| MT072864.1;QIQ54056.1;Pangolin_CoV_nucleoca;137 | 0.1 | 0.025 |
| MT072864.1;QIQ54056.1;Pangolin_CoV_nucleoca;138 | 0.1 | 0 |
| MT072864.1;QIQ54056.1;Pangolin_CoV_nucleoca;139 | 0.15 | 0.1 |
| MT072864.1;QIQ54056.1;Pangolin_CoV_nucleoca;140 | 0.05 | 0.025 |
| MT072864.1;QIQ54056.1;Pangolin_CoV_nucleoca;141 | 0.05 | 0.05 |
| MT072864.1;QIQ54056.1;Pangolin_CoV_nucleoca;142 | 0 | 0.025 |
| MT072864.1;QIQ54056.1;Pangolin_CoV_nucleoca;143 | 0 | 0.025 |
| MT072864.1;QIQ54056.1;Pangolin_CoV_nucleoca;144 | 0 | 0 |
| MT072864.1;QIQ54056.1;Pangolin_CoV_nucleoca;145 | 0 | 0.025 |

| | | |
|---|---|---|
| MT072864.1;QIQ54056.1;Pangolin_CoV_nucleoca;146 | 0 | 0.05 |
| MT072864.1;QIQ54056.1;Pangolin_CoV_nucleoca;147 | 0.05 | 0 |
| MT072864.1;QIQ54056.1;Pangolin_CoV_nucleoca;148 | 0.1 | 0.05 |
| MT072864.1;QIQ54056.1;Pangolin_CoV_nucleoca;149 | 0 | 0 |
| MT072864.1;QIQ54056.1;Pangolin_CoV_nucleoca;150 | 0 | 0 |
| MT072864.1;QIQ54056.1;Pangolin_CoV_nucleoca;151 | 0 | 0.05 |
| MT072864.1;QIQ54056.1;Pangolin_CoV_nucleoca;152 | 0 | 0.175 |
| MT072864.1;QIQ54056.1;Pangolin_CoV_nucleoca;153 | 0 | 0.2 |
| MT072864.1;QIQ54056.1;Pangolin_CoV_nucleoca;154 | 0 | 0.175 |
| MT072864.1;QIQ54056.1;Pangolin_CoV_nucleoca;155 | 0 | 0.125 |
| MT072864.1;QIQ54056.1;Pangolin_CoV_nucleoca;156 | 0 | 0.3 |
| MT072864.1;QIQ54056.1;Pangolin_CoV_nucleoca;157 | 0 | 0.425 |
| MT072864.1;QIQ54056.1;Pangolin_CoV_nucleoca;158 | 0 | 0.475 |
| MT072864.1;QIQ54056.1;Pangolin_CoV_nucleoca;159 | 0 | 0.4 |
| MT072864.1;QIQ54056.1;Pangolin_CoV_nucleoca;160 | 0 | 0.15 |
| MT072864.1;QIQ54056.1;Pangolin_CoV_nucleoca;161 | 0 | 0.2 |
| MT072864.1;QIQ54056.1;Pangolin_CoV_nucleoca;162 | 0.05 | 0.025 |
| MT072864.1;QIQ54056.1;Pangolin_CoV_nucleoca;163 | 0 | 0.025 |
| MT072864.1;QIQ54056.1;Pangolin_CoV_nucleoca;164 | 0 | 0.05 |
| MT072864.1;QIQ54056.1;Pangolin_CoV_nucleoca;165 | 0 | 0 |
| MT072864.1;QIQ54056.1;Pangolin_CoV_nucleoca;166 | 0.05 | 0.025 |
| MT072864.1;QIQ54056.1;Pangolin_CoV_nucleoca;167 | 0 | 0.05 |
| MT072864.1;QIQ54056.1;Pangolin_CoV_nucleoca;168 | 0 | 0 |
| MT072864.1;QIQ54056.1;Pangolin_CoV_nucleoca;169 | 0 | 0 |
| MT072864.1;QIQ54056.1;Pangolin_CoV_nucleoca;170 | 0 | 0 |
| MT072864.1;QIQ54056.1;Pangolin_CoV_nucleoca;171 | 0 | 0.025 |
| MT072864.1;QIQ54056.1;Pangolin_CoV_nucleoca;172 | 0 | 0.05 |
| MT072864.1;QIQ54056.1;Pangolin_CoV_nucleoca;173 | 0 | 0.025 |
| MT072864.1;QIQ54056.1;Pangolin_CoV_nucleoca;174 | 0 | 0.025 |
| MT072864.1;QIQ54056.1;Pangolin_CoV_nucleoca;175 | 0 | 0.05 |
| MT072864.1;QIQ54056.1;Pangolin_CoV_nucleoca;176 | 0 | 0.075 |
| MT072864.1;QIQ54056.1;Pangolin_CoV_nucleoca;177 | 0 | 0.1 |
| MT072864.1;QIQ54056.1;Pangolin_CoV_nucleoca;178 | 0 | 0.15 |
| MT072864.1;QIQ54056.1;Pangolin_CoV_nucleoca;179 | 0.05 | 0.125 |
| MT072864.1;QIQ54056.1;Pangolin_CoV_nucleoca;180 | 0 | 0.125 |
| MT072864.1;QIQ54056.1;Pangolin_CoV_nucleoca;181 | 0.15 | 0.15 |
| MT072864.1;QIQ54056.1;Pangolin_CoV_nucleoca;182 | 0.15 | 0.125 |
| MT072864.1;QIQ54056.1;Pangolin_CoV_nucleoca;183 | 0.15 | 0.15 |
| MT072864.1;QIQ54056.1;Pangolin_CoV_nucleoca;184 | 0.15 | 0.2 |
| MT072864.1;QIQ54056.1;Pangolin_CoV_nucleoca;185 | 0.15 | 0.225 |
| MT072864.1;QIQ54056.1;Pangolin_CoV_nucleoca;186 | 0.1 | 0.2 |
| MT072864.1;QIQ54056.1;Pangolin_CoV_nucleoca;187 | 0.1 | 0.1 |
| MT072864.1;QIQ54056.1;Pangolin_CoV_nucleoca;188 | 0.2 | 0.225 |

| | | |
|---|---|---|
| MT072864.1;QIQ54056.1;Pangolin_CoV_nucleoca;189 | 0.05 | 0.05 |
| MT072864.1;QIQ54056.1;Pangolin_CoV_nucleoca;190 | 0 | 0.075 |
| MT072864.1;QIQ54056.1;Pangolin_CoV_nucleoca;191 | 0 | 0.025 |
| MT072864.1;QIQ54056.1;Pangolin_CoV_nucleoca;192 | 0 | 0.05 |
| MT072864.1;QIQ54056.1;Pangolin_CoV_nucleoca;193 | 0 | 0 |
| MT072864.1;QIQ54056.1;Pangolin_CoV_nucleoca;194 | 0 | 0.025 |
| MT072864.1;QIQ54056.1;Pangolin_CoV_nucleoca;195 | 0.05 | 0.025 |
| MT072864.1;QIQ54056.1;Pangolin_CoV_nucleoca;196 | 0 | 0 |
| MT072864.1;QIQ54056.1;Pangolin_CoV_nucleoca;197 | 0 | 0.025 |
| MT072864.1;QIQ54056.1;Pangolin_CoV_nucleoca;198 | 0.05 | 0.1 |
| MT072864.1;QIQ54056.1;Pangolin_CoV_nucleoca;199 | 0.05 | 0.025 |
| MT072864.1;QIQ54056.1;Pangolin_CoV_nucleoca;200 | 0.05 | 0.025 |
| MT072864.1;QIQ54056.1;Pangolin_CoV_nucleoca;201 | 0.05 | 0.025 |
| MT072864.1;QIQ54056.1;Pangolin_CoV_nucleoca;202 | 0 | 0.025 |
| MT072864.1;QIQ54056.1;Pangolin_CoV_nucleoca;203 | 0 | 0.025 |
| MT072864.1;QIQ54056.1;Pangolin_CoV_nucleoca;204 | 0 | 0.05 |
| MT072864.1;QIQ54056.1;Pangolin_CoV_nucleoca;205 | 0 | 0.025 |
| MT072864.1;QIQ54056.1;Pangolin_CoV_nucleoca;206 | 0 | 0.075 |
| MT072864.1;QIQ54056.1;Pangolin_CoV_nucleoca;207 | 0 | 0 |
| MT072864.1;QIQ54056.1;Pangolin_CoV_nucleoca;208 | 0 | 0.05 |
| MT072864.1;QIQ54056.1;Pangolin_CoV_nucleoca;209 | 0 | 0.1 |
| MT072864.1;QIQ54056.1;Pangolin_CoV_nucleoca;210 | 0 | 0.125 |
| MT072864.1;QIQ54056.1;Pangolin_CoV_nucleoca;211 | 0 | 0.125 |
| MT072864.1;QIQ54056.1;Pangolin_CoV_nucleoca;212 | 0 | 0.175 |
| MT072864.1;QIQ54056.1;Pangolin_CoV_nucleoca;213 | 0 | 0.05 |
| MT072864.1;QIQ54056.1;Pangolin_CoV_nucleoca;214 | 0 | 0.025 |
| MT072864.1;QIQ54056.1;Pangolin_CoV_nucleoca;215 | 0 | 0 |
| MT072864.1;QIQ54056.1;Pangolin_CoV_nucleoca;216 | 0.05 | 0.15 |
| MT072864.1;QIQ54056.1;Pangolin_CoV_nucleoca;217 | 0.05 | 0.15 |
| MT072864.1;QIQ54056.1;Pangolin_CoV_nucleoca;218 | 0.05 | 0.25 |
| MT072864.1;QIQ54056.1;Pangolin_CoV_nucleoca;219 | 0.05 | 0.425 |
| MT072864.1;QIQ54056.1;Pangolin_CoV_nucleoca;220 | 0.05 | 0.4 |
| MT072864.1;QIQ54056.1;Pangolin_CoV_nucleoca;221 | 0.05 | 0.275 |
| MT072864.1;QIQ54056.1;Pangolin_CoV_nucleoca;222 | 0.05 | 0.125 |
| MT072864.1;QIQ54056.1;Pangolin_CoV_nucleoca;223 | 0.05 | 0.05 |
| MT072864.1;QIQ54056.1;Pangolin_CoV_nucleoca;224 | 0 | 0.15 |
| MT072864.1;QIQ54056.1;Pangolin_CoV_nucleoca;225 | 0 | 0.225 |
| MT072864.1;QIQ54056.1;Pangolin_CoV_nucleoca;226 | 0 | 0.225 |
| MT072864.1;QIQ54056.1;Pangolin_CoV_nucleoca;227 | 0 | 0.225 |
| MT072864.1;QIQ54056.1;Pangolin_CoV_nucleoca;228 | 0 | 0.275 |
| MT072864.1;QIQ54056.1;Pangolin_CoV_nucleoca;229 | 0.05 | 0.275 |
| MT072864.1;QIQ54056.1;Pangolin_CoV_nucleoca;230 | 0 | 0.2 |
| MT072864.1;QIQ54056.1;Pangolin_CoV_nucleoca;231 | 0 | 0.15 |

| | | |
|---|---|---|
| MT072864.1;QIQ54056.1;Pangolin_CoV_nucleoca;232 | 0 | 0 |
| MT072864.1;QIQ54056.1;Pangolin_CoV_nucleoca;233 | 0 | 0 |
| MT072864.1;QIQ54056.1;Pangolin_CoV_nucleoca;234 | 0 | 0 |
| MT072864.1;QIQ54056.1;Pangolin_CoV_nucleoca;235 | 0 | 0 |
| MT072864.1;QIQ54056.1;Pangolin_CoV_nucleoca;236 | 0 | 0.05 |
| MT072864.1;QIQ54056.1;Pangolin_CoV_nucleoca;237 | 0 | 0.05 |
| MT072864.1;QIQ54056.1;Pangolin_CoV_nucleoca;238 | 0.2 | 0.1 |
| MT072864.1;QIQ54056.1;Pangolin_CoV_nucleoca;239 | 0 | 0 |
| MT072864.1;QIQ54056.1;Pangolin_CoV_nucleoca;240 | 0 | 0.025 |
| MT072864.1;QIQ54056.1;Pangolin_CoV_nucleoca;241 | 0 | 0.075 |
| MT072864.1;QIQ54056.1;Pangolin_CoV_nucleoca;242 | 0 | 0.05 |
| MT072864.1;QIQ54056.1;Pangolin_CoV_nucleoca;243 | 0 | 0.075 |
| MT072864.1;QIQ54056.1;Pangolin_CoV_nucleoca;244 | 0 | 0.025 |
| MT072864.1;QIQ54056.1;Pangolin_CoV_nucleoca;245 | 0.05 | 0.025 |
| MT072864.1;QIQ54056.1;Pangolin_CoV_nucleoca;246 | 0 | 0.075 |
| MT072864.1;QIQ54056.1;Pangolin_CoV_nucleoca;247 | 0 | 0.075 |
| MT072864.1;QIQ54056.1;Pangolin_CoV_nucleoca;248 | 0 | 0.3 |
| MT072864.1;QIQ54056.1;Pangolin_CoV_nucleoca;249 | 0 | 0 |
| MT072864.1;QIQ54056.1;Pangolin_CoV_nucleoca;250 | 0.05 | 0 |
| MT072864.1;QIQ54056.1;Pangolin_CoV_nucleoca;251 | 0 | 0 |
| MT072864.1;QIQ54056.1;Pangolin_CoV_nucleoca;252 | 0 | 0 |
| MT072864.1;QIQ54056.1;Pangolin_CoV_nucleoca;253 | 0 | 0 |
| MT072864.1;QIQ54056.1;Pangolin_CoV_nucleoca;254 | 0 | 0.025 |
| MT072864.1;QIQ54056.1;Pangolin_CoV_nucleoca;255 | 0 | 0.05 |
| MT072864.1;QIQ54056.1;Pangolin_CoV_nucleoca;256 | 0.1 | 0.075 |
| MT072864.1;QIQ54056.1;Pangolin_CoV_nucleoca;257 | 0.05 | 0.075 |
| MT072864.1;QIQ54056.1;Pangolin_CoV_nucleoca;258 | 0.05 | 0.1 |
| MT072864.1;QIQ54056.1;Pangolin_CoV_nucleoca;259 | 0.05 | 0.075 |
| MT072864.1;QIQ54056.1;Pangolin_CoV_nucleoca;260 | 0.05 | 0.075 |
| MT072864.1;QIQ54056.1;Pangolin_CoV_nucleoca;261 | 0.05 | 0.075 |
| MT072864.1;QIQ54056.1;Pangolin_CoV_nucleoca;262 | 0.1 | 0.075 |
| MT072864.1;QIQ54056.1;Pangolin_CoV_nucleoca;263 | 0.1 | 0.1 |
| MT072864.1;QIQ54056.1;Pangolin_CoV_nucleoca;264 | 0.1 | 0.1 |
| MT072864.1;QIQ54056.1;Pangolin_CoV_nucleoca;265 | 0.05 | 0 |
| MT072864.1;QIQ54056.1;Pangolin_CoV_nucleoca;266 | 0.05 | 0 |
| MT072864.1;QIQ54056.1;Pangolin_CoV_nucleoca;267 | 0.05 | 0 |
| MT072864.1;QIQ54056.1;Pangolin_CoV_nucleoca;268 | 0.15 | 0.05 |
| MT072864.1;QIQ54056.1;Pangolin_CoV_nucleoca;269 | 0.15 | 0.05 |
| MT072864.1;QIQ54056.1;Pangolin_CoV_nucleoca;270 | 0.15 | 0.05 |
| MT072864.1;QIQ54056.1;Pangolin_CoV_nucleoca;271 | 0.1 | 0.075 |
| MT072864.1;QIQ54056.1;Pangolin_CoV_nucleoca;272 | 0.1 | 0.05 |
| MT072864.1;QIQ54056.1;Pangolin_CoV_nucleoca;273 | 0.05 | 0.025 |
| MT072864.1;QIQ54056.1;Pangolin_CoV_nucleoca;274 | 0.05 | 0.05 |

| | | |
|---|---|---|
| MT072864.1;QIQ54056.1;Pangolin_CoV_nucleoca;275 | 0.05 | 0.025 |
| MT072864.1;QIQ54056.1;Pangolin_CoV_nucleoca;276 | 0.05 | 0 |
| MT072864.1;QIQ54056.1;Pangolin_CoV_nucleoca;277 | 0.05 | 0.05 |
| MT072864.1;QIQ54056.1;Pangolin_CoV_nucleoca;278 | 0.05 | 0.175 |
| MT072864.1;QIQ54056.1;Pangolin_CoV_nucleoca;279 | 0 | 0.1 |
| MT072864.1;QIQ54056.1;Pangolin_CoV_nucleoca;280 | 0 | 0.1 |
| MT072864.1;QIQ54056.1;Pangolin_CoV_nucleoca;281 | 0 | 0.1 |
| MT072864.1;QIQ54056.1;Pangolin_CoV_nucleoca;282 | 0 | 0.075 |
| MT072864.1;QIQ54056.1;Pangolin_CoV_nucleoca;283 | 0 | 0.075 |
| MT072864.1;QIQ54056.1;Pangolin_CoV_nucleoca;284 | 0 | 0.075 |
| MT072864.1;QIQ54056.1;Pangolin_CoV_nucleoca;285 | 0 | 0.025 |
| MT072864.1;QIQ54056.1;Pangolin_CoV_nucleoca;286 | 0 | 0 |
| MT072864.1;QIQ54056.1;Pangolin_CoV_nucleoca;287 | 0 | 0 |
| MT072864.1;QIQ54056.1;Pangolin_CoV_nucleoca;288 | 0.05 | 0 |
| MT072864.1;QIQ54056.1;Pangolin_CoV_nucleoca;289 | 0 | 0 |
| MT072864.1;QIQ54056.1;Pangolin_CoV_nucleoca;290 | 0 | 0 |
| MT072864.1;QIQ54056.1;Pangolin_CoV_nucleoca;291 | 0 | 0 |
| MT072864.1;QIQ54056.1;Pangolin_CoV_nucleoca;292 | 0 | 0 |
| MT072864.1;QIQ54056.1;Pangolin_CoV_nucleoca;293 | 0.1 | 0 |
| MT072864.1;QIQ54056.1;Pangolin_CoV_nucleoca;294 | 0 | 0 |
| MT072864.1;QIQ54056.1;Pangolin_CoV_nucleoca;295 | 0.15 | 0.025 |
| MT072864.1;QIQ54056.1;Pangolin_CoV_nucleoca;296 | 0 | 0 |
| MT072864.1;QIQ54056.1;Pangolin_CoV_nucleoca;297 | 0 | 0 |
| MT072864.1;QIQ54056.1;Pangolin_CoV_nucleoca;298 | 0 | 0 |
| MT072864.1;QIQ54056.1;Pangolin_CoV_nucleoca;299 | 0 | 0 |
| MT072864.1;QIQ54056.1;Pangolin_CoV_nucleoca;300 | 0 | 0.025 |
| MT072864.1;QIQ54056.1;Pangolin_CoV_nucleoca;301 | 0 | 0 |
| MT072864.1;QIQ54056.1;Pangolin_CoV_nucleoca;302 | 0 | 0 |
| MT072864.1;QIQ54056.1;Pangolin_CoV_nucleoca;303 | 0 | 0 |
| MT072864.1;QIQ54056.1;Pangolin_CoV_nucleoca;304 | 0 | 0 |
| MT072864.1;QIQ54056.1;Pangolin_CoV_nucleoca;305 | 0 | 0 |
| MT072864.1;QIQ54056.1;Pangolin_CoV_nucleoca;306 | 0 | 0 |
| MT072864.1;QIQ54056.1;Pangolin_CoV_nucleoca;307 | 0 | 0.025 |
| MT072864.1;QIQ54056.1;Pangolin_CoV_nucleoca;308 | 0.05 | 0.025 |
| MT072864.1;QIQ54056.1;Pangolin_CoV_nucleoca;309 | 0 | 0.05 |
| MT072864.1;QIQ54056.1;Pangolin_CoV_nucleoca;310 | 0 | 0.075 |
| MT072864.1;QIQ54056.1;Pangolin_CoV_nucleoca;311 | 0.05 | 0.05 |
| MT072864.1;QIQ54056.1;Pangolin_CoV_nucleoca;312 | 0.05 | 0.05 |
| MT072864.1;QIQ54056.1;Pangolin_CoV_nucleoca;313 | 0 | 0.05 |
| MT072864.1;QIQ54056.1;Pangolin_CoV_nucleoca;314 | 0 | 0.025 |
| MT072864.1;QIQ54056.1;Pangolin_CoV_nucleoca;315 | 0 | 0.025 |
| MT072864.1;QIQ54056.1;Pangolin_CoV_nucleoca;316 | 0 | 0.025 |
| MT072864.1;QIQ54056.1;Pangolin_CoV_nucleoca;317 | 0 | 0 |

| | | |
|---|---|---|
| MT072864.1;QIQ54056.1;Pangolin_CoV_nucleoca;318 | 0 | 0 |
| MT072864.1;QIQ54056.1;Pangolin_CoV_nucleoca;319 | 0 | 0 |
| MT072864.1;QIQ54056.1;Pangolin_CoV_nucleoca;320 | 0 | 0 |
| MT072864.1;QIQ54056.1;Pangolin_CoV_nucleoca;321 | 0 | 0 |
| MT072864.1;QIQ54056.1;Pangolin_CoV_nucleoca;322 | 0 | 0 |
| MT072864.1;QIQ54056.1;Pangolin_CoV_nucleoca;323 | 0 | 0 |
| MT072864.1;QIQ54056.1;Pangolin_CoV_nucleoca;324 | 0 | 0 |
| MT072864.1;QIQ54056.1;Pangolin_CoV_nucleoca;325 | 0 | 0.075 |
| MT072864.1;QIQ54056.1;Pangolin_CoV_nucleoca;326 | 0 | 0.05 |
| MT072864.1;QIQ54056.1;Pangolin_CoV_nucleoca;327 | 0 | 0.075 |
| MT072864.1;QIQ54056.1;Pangolin_CoV_nucleoca;328 | 0 | 0.075 |
| MT072864.1;QIQ54056.1;Pangolin_CoV_nucleoca;329 | 0 | 0.075 |
| MT072864.1;QIQ54056.1;Pangolin_CoV_nucleoca;330 | 0.05 | 0.1 |
| MT072864.1;QIQ54056.1;Pangolin_CoV_nucleoca;331 | 0.05 | 0.15 |
| MT072864.1;QIQ54056.1;Pangolin_CoV_nucleoca;332 | 0.1 | 0.2 |
| MT072864.1;QIQ54056.1;Pangolin_CoV_nucleoca;333 | 0 | 0.2 |
| MT072864.1;QIQ54056.1;Pangolin_CoV_nucleoca;334 | 0 | 0.125 |
| MT072864.1;QIQ54056.1;Pangolin_CoV_nucleoca;335 | 0.05 | 0.125 |
| MT072864.1;QIQ54056.1;Pangolin_CoV_nucleoca;336 | 0 | 0.05 |
| MT072864.1;QIQ54056.1;Pangolin_CoV_nucleoca;337 | 0 | 0.075 |
| MT072864.1;QIQ54056.1;Pangolin_CoV_nucleoca;338 | 0 | 0.075 |
| MT072864.1;QIQ54056.1;Pangolin_CoV_nucleoca;339 | 0 | 0.05 |
| MT072864.1;QIQ54056.1;Pangolin_CoV_nucleoca;340 | 0 | 0.05 |
| MT072864.1;QIQ54056.1;Pangolin_CoV_nucleoca;341 | 0 | 0.075 |
| MT072864.1;QIQ54056.1;Pangolin_CoV_nucleoca;342 | 0.05 | 0.075 |
| MT072864.1;QIQ54056.1;Pangolin_CoV_nucleoca;343 | 0.05 | 0.075 |
| MT072864.1;QIQ54056.1;Pangolin_CoV_nucleoca;344 | 0 | 0.1 |
| MT072864.1;QIQ54056.1;Pangolin_CoV_nucleoca;345 | 0 | 0.05 |
| MT072864.1;QIQ54056.1;Pangolin_CoV_nucleoca;346 | 0 | 0 |
| MT072864.1;QIQ54056.1;Pangolin_CoV_nucleoca;347 | 0 | 0 |
| MT072864.1;QIQ54056.1;Pangolin_CoV_nucleoca;348 | 0 | 0 |
| MT072864.1;QIQ54056.1;Pangolin_CoV_nucleoca;349 | 0 | 0.025 |
| MT072864.1;QIQ54056.1;Pangolin_CoV_nucleoca;350 | 0 | 0.15 |
| MT072864.1;QIQ54056.1;Pangolin_CoV_nucleoca;351 | 0.05 | 0.125 |
| MT072864.1;QIQ54056.1;Pangolin_CoV_nucleoca;352 | 0 | 0.125 |
| MT072864.1;QIQ54056.1;Pangolin_CoV_nucleoca;353 | 0 | 0.05 |
| MT072864.1;QIQ54056.1;Pangolin_CoV_nucleoca;354 | 0 | 0.25 |
| MT072864.1;QIQ54056.1;Pangolin_CoV_nucleoca;355 | 0.05 | 0.125 |
| MT072864.1;QIQ54056.1;Pangolin_CoV_nucleoca;356 | 0.05 | 0.1 |
| MT072864.1;QIQ54056.1;Pangolin_CoV_nucleoca;357 | 0 | 0.075 |
| MT072864.1;QIQ54056.1;Pangolin_CoV_nucleoca;358 | 0 | 0.075 |
| MT072864.1;QIQ54056.1;Pangolin_CoV_nucleoca;359 | 0 | 0.075 |
| MT072864.1;QIQ54056.1;Pangolin_CoV_nucleoca;360 | 0.05 | 0.125 |

| | | |
|---|---|---|
| MT072864.1;QIQ54056.1;Pangolin_CoV_nucleoca;361 | 0.05 | 0.1 |
| MT072864.1;QIQ54056.1;Pangolin_CoV_nucleoca;362 | 0.05 | 0.025 |
| MT072864.1;QIQ54056.1;Pangolin_CoV_nucleoca;363 | 0 | 0 |
| MT072864.1;QIQ54056.1;Pangolin_CoV_nucleoca;364 | 0 | 0 |
| MT072864.1;QIQ54056.1;Pangolin_CoV_nucleoca;365 | 0 | 0 |
| MT072864.1;QIQ54056.1;Pangolin_CoV_nucleoca;366 | 0.05 | 0.025 |
| MT072864.1;QIQ54056.1;Pangolin_CoV_nucleoca;367 | 0.05 | 0.025 |
| MT072864.1;QIQ54056.1;Pangolin_CoV_nucleoca;368 | 0 | 0.025 |
| MT072864.1;QIQ54056.1;Pangolin_CoV_nucleoca;369 | 0.05 | 0 |
| MT072864.1;QIQ54056.1;Pangolin_CoV_nucleoca;370 | 0.05 | 0.025 |
| MT072864.1;QIQ54056.1;Pangolin_CoV_nucleoca;371 | 0 | 0.075 |
| MT072864.1;QIQ54056.1;Pangolin_CoV_nucleoca;372 | 0 | 0.075 |
| MT072864.1;QIQ54056.1;Pangolin_CoV_nucleoca;373 | 0 | 0.075 |
| MT072864.1;QIQ54056.1;Pangolin_CoV_nucleoca;374 | 0 | 0.075 |
| MT072864.1;QIQ54056.1;Pangolin_CoV_nucleoca;375 | 0.05 | 0.075 |
| MT072864.1;QIQ54056.1;Pangolin_CoV_nucleoca;376 | 0 | 0.125 |
| MT072864.1;QIQ54056.1;Pangolin_CoV_nucleoca;377 | 0 | 0.15 |
| MT072864.1;QIQ54056.1;Pangolin_CoV_nucleoca;378 | 0 | 0.15 |
| MT072864.1;QIQ54056.1;Pangolin_CoV_nucleoca;379 | 0 | 0.2 |
| MT072864.1;QIQ54056.1;Pangolin_CoV_nucleoca;380 | 0 | 0.225 |
| MT072864.1;QIQ54056.1;Pangolin_CoV_nucleoca;381 | 0.05 | 0.125 |
| MT072864.1;QIQ54056.1;Pangolin_CoV_nucleoca;382 | 0 | 0.125 |
| MT072864.1;QIQ54056.1;Pangolin_CoV_nucleoca;383 | 0 | 0.15 |
| MT072864.1;QIQ54056.1;Pangolin_CoV_nucleoca;384 | 0 | 0.4 |
| MT072864.1;QIQ54056.1;Pangolin_CoV_nucleoca;385 | 0 | 0.5 |
| MT072864.1;QIQ54056.1;Pangolin_CoV_nucleoca;386 | 0 | 0.575 |
| MT072864.1;QIQ54056.1;Pangolin_CoV_nucleoca;387 | 0 | 0.3 |
| MT072864.1;QIQ54056.1;Pangolin_CoV_nucleoca;388 | 0 | 0.7 |
| MT072864.1;QIQ54056.1;Pangolin_CoV_nucleoca;389 | 0 | 0.75 |
| MT072864.1;QIQ54056.1;Pangolin_CoV_nucleoca;390 | 0 | 0.775 |
| MT072864.1;QIQ54056.1;Pangolin_CoV_nucleoca;391 | 0 | 0.8 |
| MT072864.1;QIQ54056.1;Pangolin_CoV_nucleoca;392 | 0 | 0.825 |
| MT072864.1;QIQ54056.1;Pangolin_CoV_nucleoca;393 | 0.05 | 0.775 |
| MT072864.1;QIQ54056.1;Pangolin_CoV_nucleoca;394 | 0 | 0.725 |
| MT072864.1;QIQ54056.1;Pangolin_CoV_nucleoca;395 | 0.1 | 0.7 |
| MT072864.1;QIQ54056.1;Pangolin_CoV_nucleoca;396 | 0.25 | 0.55 |
| MT072864.1;QIQ54056.1;Pangolin_CoV_nucleoca;397 | 0 | 0.675 |
| MT072864.1;QIQ54056.1;Pangolin_CoV_nucleoca;398 | 0 | 0.2 |
| MT072864.1;QIQ54056.1;Pangolin_CoV_nucleoca;399 | 0 | 0.075 |
| MT072864.1;QIQ54056.1;Pangolin_CoV_nucleoca;400 | 0 | 0.075 |
| MT072864.1;QIQ54056.1;Pangolin_CoV_nucleoca;401 | 0.05 | 0.1 |
| MT072864.1;QIQ54056.1;Pangolin_CoV_nucleoca;402 | 0 | 0.05 |
| MT072864.1;QIQ54056.1;Pangolin_CoV_nucleoca;403 | 0 | 0.025 |

| | | |
|---|---|---|
| MT072864.1;QIQ54056.1;Pangolin_CoV_nucleoca;404 | 0 | 0.05 |
| MT072864.1;QIQ54056.1;Pangolin_CoV_nucleoca;405 | 0 | 0.075 |
| MT072864.1;QIQ54056.1;Pangolin_CoV_nucleoca;406 | 0 | 0.025 |
| NC_004718.3;NP_828849.2;SARS-CoV_orf1ab;1 | 0.05 | 0 |
| NC_004718.3;NP_828849.2;SARS-CoV_orf1ab;2 | 0 | 0 |
| NC_004718.3;NP_828849.2;SARS-CoV_orf1ab;3 | 0 | 0 |
| NC_004718.3;NP_828849.2;SARS-CoV_orf1ab;4 | 0.1 | 0.025 |
| NC_004718.3;NP_828849.2;SARS-CoV_orf1ab;5 | 0.05 | 0.05 |
| NC_004718.3;NP_828849.2;SARS-CoV_orf1ab;6 | 0.05 | 0 |
| NC_004718.3;NP_828849.2;SARS-CoV_orf1ab;7 | 0 | 0 |
| NC_004718.3;NP_828849.2;SARS-CoV_orf1ab;8 | 0 | 0 |
| NC_004718.3;NP_828849.2;SARS-CoV_orf1ab;9 | 0 | 0 |
| NC_004718.3;NP_828849.2;SARS-CoV_orf1ab;10 | 0 | 0 |
| NC_004718.3;NP_828849.2;SARS-CoV_orf1ab;11 | 0 | 0 |
| NC_004718.3;NP_828849.2;SARS-CoV_orf1ab;12 | 0 | 0 |
| NC_004718.3;NP_828849.2;SARS-CoV_orf1ab;13 | 0 | 0 |
| NC_004718.3;NP_828849.2;SARS-CoV_orf1ab;14 | 0 | 0 |
| NC_004718.3;NP_828849.2;SARS-CoV_orf1ab;15 | 0 | 0 |
| NC_004718.3;NP_828849.2;SARS-CoV_orf1ab;16 | 0 | 0 |
| NC_004718.3;NP_828849.2;SARS-CoV_orf1ab;17 | 0 | 0 |
| NC_004718.3;NP_828849.2;SARS-CoV_orf1ab;18 | 0 | 0 |
| NC_004718.3;NP_828849.2;SARS-CoV_orf1ab;19 | 0 | 0 |
| NC_004718.3;NP_828849.2;SARS-CoV_orf1ab;20 | 0 | 0 |
| NC_004718.3;NP_828849.2;SARS-CoV_orf1ab;21 | 0 | 0 |
| NC_004718.3;NP_828849.2;SARS-CoV_orf1ab;22 | 0 | 0 |
| NC_004718.3;NP_828849.2;SARS-CoV_orf1ab;23 | 0 | 0 |
| NC_004718.3;NP_828849.2;SARS-CoV_orf1ab;24 | 0 | 0 |
| NC_004718.3;NP_828849.2;SARS-CoV_orf1ab;25 | 0 | 0 |
| NC_004718.3;NP_828849.2;SARS-CoV_orf1ab;26 | 0 | 0 |
| NC_004718.3;NP_828849.2;SARS-CoV_orf1ab;27 | 0 | 0 |
| NC_004718.3;NP_828849.2;SARS-CoV_orf1ab;28 | 0 | 0 |
| NC_004718.3;NP_828849.2;SARS-CoV_orf1ab;29 | 0.1 | 0 |
| NC_004718.3;NP_828849.2;SARS-CoV_orf1ab;30 | 0.25 | 0.125 |
| NC_004718.3;NP_828849.2;SARS-CoV_orf1ab;31 | 0 | 0 |
| NC_004718.3;NP_828849.2;SARS-CoV_orf1ab;32 | 0.05 | 0.025 |
| NC_004718.3;NP_828849.2;SARS-CoV_orf1ab;33 | 0.05 | 0 |
| NC_004718.3;NP_828849.2;SARS-CoV_orf1ab;34 | 0.05 | 0.025 |
| NC_004718.3;NP_828849.2;SARS-CoV_orf1ab;35 | 0 | 0.025 |
| NC_004718.3;NP_828849.2;SARS-CoV_orf1ab;36 | 0 | 0.025 |
| NC_004718.3;NP_828849.2;SARS-CoV_orf1ab;37 | 0 | 0 |
| NC_004718.3;NP_828849.2;SARS-CoV_orf1ab;38 | 0 | 0.025 |
| NC_004718.3;NP_828849.2;SARS-CoV_orf1ab;39 | 0 | 0.025 |
| NC_004718.3;NP_828849.2;SARS-CoV_orf1ab;40 | 0 | 0 |

| | | |
|---|---|---|
| NC_004718.3;NP_828849.2;SARS-CoV_orf1ab;41 | 0 | 0 |
| NC_004718.3;NP_828849.2;SARS-CoV_orf1ab;42 | 0 | 0 |
| NC_004718.3;NP_828849.2;SARS-CoV_orf1ab;43 | 0 | 0 |
| NC_004718.3;NP_828849.2;SARS-CoV_orf1ab;44 | 0.05 | 0 |
| NC_004718.3;NP_828849.2;SARS-CoV_orf1ab;45 | 0 | 0 |
| NC_004718.3;NP_828849.2;SARS-CoV_orf1ab;46 | 0 | 0 |
| NC_004718.3;NP_828849.2;SARS-CoV_orf1ab;47 | 0 | 0.025 |
| NC_004718.3;NP_828849.2;SARS-CoV_orf1ab;48 | 0 | 0 |
| NC_004718.3;NP_828849.2;SARS-CoV_orf1ab;49 | 0 | 0 |
| NC_004718.3;NP_828849.2;SARS-CoV_orf1ab;50 | 0 | 0.025 |
| NC_004718.3;NP_828849.2;SARS-CoV_orf1ab;51 | 0 | 0.05 |
| NC_004718.3;NP_828849.2;SARS-CoV_orf1ab;52 | 0.05 | 0.075 |
| NC_004718.3;NP_828849.2;SARS-CoV_orf1ab;53 | 0 | 0 |
| NC_004718.3;NP_828849.2;SARS-CoV_orf1ab;54 | 0 | 0 |
| NC_004718.3;NP_828849.2;SARS-CoV_orf1ab;55 | 0 | 0.05 |
| NC_004718.3;NP_828849.2;SARS-CoV_orf1ab;56 | 0 | 0 |
| NC_004718.3;NP_828849.2;SARS-CoV_orf1ab;57 | 0 | 0.025 |
| NC_004718.3;NP_828849.2;SARS-CoV_orf1ab;58 | 0 | 0.025 |
| NC_004718.3;NP_828849.2;SARS-CoV_orf1ab;59 | 0 | 0 |
| NC_004718.3;NP_828849.2;SARS-CoV_orf1ab;60 | 0 | 0 |
| NC_004718.3;NP_828849.2;SARS-CoV_orf1ab;61 | 0 | 0 |
| NC_004718.3;NP_828849.2;SARS-CoV_orf1ab;62 | 0 | 0 |
| NC_004718.3;NP_828849.2;SARS-CoV_orf1ab;63 | 0 | 0 |
| NC_004718.3;NP_828849.2;SARS-CoV_orf1ab;64 | 0 | 0 |
| NC_004718.3;NP_828849.2;SARS-CoV_orf1ab;65 | 0.05 | 0 |
| NC_004718.3;NP_828849.2;SARS-CoV_orf1ab;66 | 0 | 0 |
| NC_004718.3;NP_828849.2;SARS-CoV_orf1ab;67 | 0 | 0 |
| NC_004718.3;NP_828849.2;SARS-CoV_orf1ab;68 | 0 | 0 |
| NC_004718.3;NP_828849.2;SARS-CoV_orf1ab;69 | 0 | 0 |
| NC_004718.3;NP_828849.2;SARS-CoV_orf1ab;70 | 0 | 0 |
| NC_004718.3;NP_828849.2;SARS-CoV_orf1ab;71 | 0.05 | 0.025 |
| NC_004718.3;NP_828849.2;SARS-CoV_orf1ab;72 | 0.05 | 0.025 |
| NC_004718.3;NP_828849.2;SARS-CoV_orf1ab;73 | 0 | 0 |
| NC_004718.3;NP_828849.2;SARS-CoV_orf1ab;74 | 0 | 0 |
| NC_004718.3;NP_828849.2;SARS-CoV_orf1ab;75 | 0 | 0.025 |
| NC_004718.3;NP_828849.2;SARS-CoV_orf1ab;76 | 0 | 0 |
| NC_004718.3;NP_828849.2;SARS-CoV_orf1ab;77 | 0 | 0 |
| NC_004718.3;NP_828849.2;SARS-CoV_orf1ab;78 | 0 | 0 |
| NC_004718.3;NP_828849.2;SARS-CoV_orf1ab;79 | 0 | 0 |
| NC_004718.3;NP_828849.2;SARS-CoV_orf1ab;80 | 0 | 0 |
| NC_004718.3;NP_828849.2;SARS-CoV_orf1ab;81 | 0 | 0 |
| NC_004718.3;NP_828849.2;SARS-CoV_orf1ab;82 | 0 | 0 |
| NC_004718.3;NP_828849.2;SARS-CoV_orf1ab;83 | 0 | 0 |

| | | |
|---|---|---|
| NC_004718.3;NP_828849.2;SARS-CoV_orf1ab;84 | 0 | 0 |
| NC_004718.3;NP_828849.2;SARS-CoV_orf1ab;85 | 0 | 0 |
| NC_004718.3;NP_828849.2;SARS-CoV_orf1ab;86 | 0 | 0.025 |
| NC_004718.3;NP_828849.2;SARS-CoV_orf1ab;87 | 0 | 0 |
| NC_004718.3;NP_828849.2;SARS-CoV_orf1ab;88 | 0 | 0 |
| NC_004718.3;NP_828849.2;SARS-CoV_orf1ab;89 | 0 | 0 |
| NC_004718.3;NP_828849.2;SARS-CoV_orf1ab;90 | 0.2 | 0.125 |
| NC_004718.3;NP_828849.2;SARS-CoV_orf1ab;91 | 0.05 | 0.025 |
| NC_004718.3;NP_828849.2;SARS-CoV_orf1ab;92 | 0 | 0 |
| NC_004718.3;NP_828849.2;SARS-CoV_orf1ab;93 | 0 | 0 |
| NC_004718.3;NP_828849.2;SARS-CoV_orf1ab;94 | 0 | 0 |
| NC_004718.3;NP_828849.2;SARS-CoV_orf1ab;95 | 0 | 0 |
| NC_004718.3;NP_828849.2;SARS-CoV_orf1ab;96 | 0 | 0 |
| NC_004718.3;NP_828849.2;SARS-CoV_orf1ab;97 | 0 | 0 |
| NC_004718.3;NP_828849.2;SARS-CoV_orf1ab;98 | 0 | 0 |
| NC_004718.3;NP_828849.2;SARS-CoV_orf1ab;99 | 0 | 0 |
| NC_004718.3;NP_828849.2;SARS-CoV_orf1ab;100 | 0 | 0 |
| NC_004718.3;NP_828849.2;SARS-CoV_orf1ab;101 | 0 | 0 |
| NC_004718.3;NP_828849.2;SARS-CoV_orf1ab;102 | 0 | 0 |
| NC_004718.3;NP_828849.2;SARS-CoV_orf1ab;103 | 0 | 0 |
| NC_004718.3;NP_828849.2;SARS-CoV_orf1ab;104 | 0 | 0 |
| NC_004718.3;NP_828849.2;SARS-CoV_orf1ab;105 | 0 | 0 |
| NC_004718.3;NP_828849.2;SARS-CoV_orf1ab;106 | 0 | 0 |
| NC_004718.3;NP_828849.2;SARS-CoV_orf1ab;107 | 0 | 0 |
| NC_004718.3;NP_828849.2;SARS-CoV_orf1ab;108 | 0 | 0 |
| NC_004718.3;NP_828849.2;SARS-CoV_orf1ab;109 | 0 | 0 |
| NC_004718.3;NP_828849.2;SARS-CoV_orf1ab;110 | 0 | 0 |
| NC_004718.3;NP_828849.2;SARS-CoV_orf1ab;111 | 0 | 0 |
| NC_004718.3;NP_828849.2;SARS-CoV_orf1ab;112 | 0 | 0 |
| NC_004718.3;NP_828849.2;SARS-CoV_orf1ab;113 | 0 | 0 |
| NC_004718.3;NP_828849.2;SARS-CoV_orf1ab;114 | 0 | 0.05 |
| NC_004718.3;NP_828849.2;SARS-CoV_orf1ab;115 | 0 | 0.025 |
| NC_004718.3;NP_828849.2;SARS-CoV_orf1ab;116 | 0.05 | 0.025 |
| NC_004718.3;NP_828849.2;SARS-CoV_orf1ab;117 | 0.05 | 0.025 |
| NC_004718.3;NP_828849.2;SARS-CoV_orf1ab;118 | 0.1 | 0.025 |
| NC_004718.3;NP_828849.2;SARS-CoV_orf1ab;119 | 0.05 | 0.025 |
| NC_004718.3;NP_828849.2;SARS-CoV_orf1ab;120 | 0.05 | 0.025 |
| NC_004718.3;NP_828849.2;SARS-CoV_orf1ab;121 | 0.05 | 0.025 |
| NC_004718.3;NP_828849.2;SARS-CoV_orf1ab;122 | 0 | 0.025 |
| NC_004718.3;NP_828849.2;SARS-CoV_orf1ab;123 | 0 | 0.025 |
| NC_004718.3;NP_828849.2;SARS-CoV_orf1ab;124 | 0.05 | 0.05 |
| NC_004718.3;NP_828849.2;SARS-CoV_orf1ab;125 | 0.15 | 0.05 |
| NC_004718.3;NP_828849.2;SARS-CoV_orf1ab;126 | 0 | 0 |

| | | |
|---|---|---|
| NC_004718.3;NP_828849.2;SARS-CoV_orf1ab;127 | 0 | 0 |
| NC_004718.3;NP_828849.2;SARS-CoV_orf1ab;128 | 0 | 0.025 |
| NC_004718.3;NP_828849.2;SARS-CoV_orf1ab;129 | 0 | 0.025 |
| NC_004718.3;NP_828849.2;SARS-CoV_orf1ab;130 | 0 | 0.025 |
| NC_004718.3;NP_828849.2;SARS-CoV_orf1ab;131 | 0.1 | 0.025 |
| NC_004718.3;NP_828849.2;SARS-CoV_orf1ab;132 | 0.05 | 0.025 |
| NC_004718.3;NP_828849.2;SARS-CoV_orf1ab;133 | 0.05 | 0 |
| NC_004718.3;NP_828849.2;SARS-CoV_orf1ab;134 | 0 | 0 |
| NC_004718.3;NP_828849.2;SARS-CoV_orf1ab;135 | 0 | 0 |
| NC_004718.3;NP_828849.2;SARS-CoV_orf1ab;136 | 0 | 0 |
| NC_004718.3;NP_828849.2;SARS-CoV_orf1ab;137 | 0 | 0 |
| NC_004718.3;NP_828849.2;SARS-CoV_orf1ab;138 | 0 | 0 |
| NC_004718.3;NP_828849.2;SARS-CoV_orf1ab;139 | 0.05 | 0 |
| NC_004718.3;NP_828849.2;SARS-CoV_orf1ab;140 | 0.05 | 0 |
| NC_004718.3;NP_828849.2;SARS-CoV_orf1ab;141 | 0 | 0 |
| NC_004718.3;NP_828849.2;SARS-CoV_orf1ab;142 | 0 | 0 |
| NC_004718.3;NP_828849.2;SARS-CoV_orf1ab;143 | 0 | 0 |
| NC_004718.3;NP_828849.2;SARS-CoV_orf1ab;144 | 0 | 0 |
| NC_004718.3;NP_828849.2;SARS-CoV_orf1ab;145 | 0.2 | 0.15 |
| NC_004718.3;NP_828849.2;SARS-CoV_orf1ab;146 | 0 | 0 |
| NC_004718.3;NP_828849.2;SARS-CoV_orf1ab;147 | 0 | 0 |
| NC_004718.3;NP_828849.2;SARS-CoV_orf1ab;148 | 0 | 0 |
| NC_004718.3;NP_828849.2;SARS-CoV_orf1ab;149 | 0.2 | 0.125 |
| NC_004718.3;NP_828849.2;SARS-CoV_orf1ab;150 | 0.05 | 0.05 |
| NC_004718.3;NP_828849.2;SARS-CoV_orf1ab;151 | 0.05 | 0.025 |
| NC_004718.3;NP_828849.2;SARS-CoV_orf1ab;152 | 0.25 | 0.175 |
| NC_004718.3;NP_828849.2;SARS-CoV_orf1ab;153 | 0.25 | 0.25 |
| NC_004718.3;NP_828849.2;SARS-CoV_orf1ab;154 | 0.2 | 0.2 |
| NC_004718.3;NP_828849.2;SARS-CoV_orf1ab;155 | 0.1 | 0.1 |
| NC_004718.3;NP_828849.2;SARS-CoV_orf1ab;156 | 0.05 | 0.075 |
| NC_004718.3;NP_828849.2;SARS-CoV_orf1ab;157 | 0.05 | 0.05 |
| NC_004718.3;NP_828849.2;SARS-CoV_orf1ab;158 | 0 | 0 |
| NC_004718.3;NP_828849.2;SARS-CoV_orf1ab;159 | 0 | 0 |
| NC_004718.3;NP_828849.2;SARS-CoV_orf1ab;160 | 0 | 0 |
| NC_004718.3;NP_828849.2;SARS-CoV_orf1ab;161 | 0 | 0 |
| NC_004718.3;NP_828849.2;SARS-CoV_orf1ab;162 | 0 | 0 |
| NC_004718.3;NP_828849.2;SARS-CoV_orf1ab;163 | 0 | 0.025 |
| NC_004718.3;NP_828849.2;SARS-CoV_orf1ab;164 | 0 | 0 |
| NC_004718.3;NP_828849.2;SARS-CoV_orf1ab;165 | 0.05 | 0.025 |
| NC_004718.3;NP_828849.2;SARS-CoV_orf1ab;166 | 0 | 0.025 |
| NC_004718.3;NP_828849.2;SARS-CoV_orf1ab;167 | 0 | 0.025 |
| NC_004718.3;NP_828849.2;SARS-CoV_orf1ab;168 | 0.2 | 0.2 |
| NC_004718.3;NP_828849.2;SARS-CoV_orf1ab;169 | 0 | 0 |

| | | |
|---|---|---|
| NC_004718.3;NP_828849.2;SARS-CoV_orf1ab;170 | 0 | 0 |
| NC_004718.3;NP_828849.2;SARS-CoV_orf1ab;171 | 0 | 0 |
| NC_004718.3;NP_828849.2;SARS-CoV_orf1ab;172 | 0 | 0.025 |
| NC_004718.3;NP_828849.2;SARS-CoV_orf1ab;173 | 0.05 | 0.05 |
| NC_004718.3;NP_828849.2;SARS-CoV_orf1ab;174 | 0 | 0.075 |
| NC_004718.3;NP_828849.2;SARS-CoV_orf1ab;175 | 0 | 0.075 |
| NC_004718.3;NP_828849.2;SARS-CoV_orf1ab;176 | 0 | 0.025 |
| NC_004718.3;NP_828849.2;SARS-CoV_orf1ab;177 | 0 | 0.025 |
| NC_004718.3;NP_828849.2;SARS-CoV_orf1ab;178 | 0 | 0.025 |
| NC_004718.3;NP_828849.2;SARS-CoV_orf1ab;179 | 0.05 | 0.075 |
| NC_004718.3;NP_828849.2;SARS-CoV_orf1ab;180 | 0.15 | 0.15 |
| NC_004718.3;NP_828849.2;SARS-CoV_orf1ab;181 | 0 | 0.025 |
| NC_004718.3;NP_828849.2;SARS-CoV_orf1ab;182 | 0 | 0 |
| NC_004718.3;NP_828849.2;SARS-CoV_orf1ab;183 | 0 | 0 |
| NC_004718.3;NP_828849.2;SARS-CoV_orf1ab;184 | 0 | 0 |
| NC_004718.3;NP_828849.2;SARS-CoV_orf1ab;185 | 0 | 0.05 |
| NC_004718.3;NP_828849.2;SARS-CoV_orf1ab;186 | 0 | 0.025 |
| NC_004718.3;NP_828849.2;SARS-CoV_orf1ab;187 | 0 | 0 |
| NC_004718.3;NP_828849.2;SARS-CoV_orf1ab;188 | 0 | 0 |
| NC_004718.3;NP_828849.2;SARS-CoV_orf1ab;189 | 0.05 | 0 |
| NC_004718.3;NP_828849.2;SARS-CoV_orf1ab;190 | 0 | 0 |
| NC_004718.3;NP_828849.2;SARS-CoV_orf1ab;191 | 0 | 0 |
| NC_004718.3;NP_828849.2;SARS-CoV_orf1ab;192 | 0 | 0 |
| NC_004718.3;NP_828849.2;SARS-CoV_orf1ab;193 | 0 | 0 |
| NC_004718.3;NP_828849.2;SARS-CoV_orf1ab;194 | 0 | 0 |
| NC_004718.3;NP_828849.2;SARS-CoV_orf1ab;195 | 0 | 0 |
| NC_004718.3;NP_828849.2;SARS-CoV_orf1ab;196 | 0 | 0 |
| NC_004718.3;NP_828849.2;SARS-CoV_orf1ab;197 | 0 | 0 |
| NC_004718.3;NP_828849.2;SARS-CoV_orf1ab;198 | 0 | 0 |
| NC_004718.3;NP_828849.2;SARS-CoV_orf1ab;199 | 0 | 0 |
| NC_004718.3;NP_828849.2;SARS-CoV_orf1ab;200 | 0 | 0 |
| NC_004718.3;NP_828849.2;SARS-CoV_orf1ab;201 | 0 | 0 |
| NC_004718.3;NP_828849.2;SARS-CoV_orf1ab;202 | 0 | 0 |
| NC_004718.3;NP_828849.2;SARS-CoV_orf1ab;203 | 0 | 0.025 |
| NC_004718.3;NP_828849.2;SARS-CoV_orf1ab;204 | 0.05 | 0 |
| NC_004718.3;NP_828849.2;SARS-CoV_orf1ab;205 | 0 | 0 |
| NC_004718.3;NP_828849.2;SARS-CoV_orf1ab;206 | 0 | 0 |
| NC_004718.3;NP_828849.2;SARS-CoV_orf1ab;207 | 0 | 0 |
| NC_004718.3;NP_828849.2;SARS-CoV_orf1ab;208 | 0 | 0 |
| NC_004718.3;NP_828849.2;SARS-CoV_orf1ab;209 | 0 | 0 |
| NC_004718.3;NP_828849.2;SARS-CoV_orf1ab;210 | 0 | 0 |
| NC_004718.3;NP_828849.2;SARS-CoV_orf1ab;211 | 0 | 0.025 |
| NC_004718.3;NP_828849.2;SARS-CoV_orf1ab;212 | 0 | 0.05 |

| | | |
|---|---|---|
| NC_004718.3;NP_828849.2;SARS-CoV_orf1ab;213 | 0 | 0.025 |
| NC_004718.3;NP_828849.2;SARS-CoV_orf1ab;214 | 0.15 | 0.05 |
| NC_004718.3;NP_828849.2;SARS-CoV_orf1ab;215 | 0.15 | 0.075 |
| NC_004718.3;NP_828849.2;SARS-CoV_orf1ab;216 | 0.1 | 0.05 |
| NC_004718.3;NP_828849.2;SARS-CoV_orf1ab;217 | 0.05 | 0.025 |
| NC_004718.3;NP_828849.2;SARS-CoV_orf1ab;218 | 0.05 | 0.05 |
| NC_004718.3;NP_828849.2;SARS-CoV_orf1ab;219 | 0 | 0.025 |
| NC_004718.3;NP_828849.2;SARS-CoV_orf1ab;220 | 0.05 | 0.025 |
| NC_004718.3;NP_828849.2;SARS-CoV_orf1ab;221 | 0.1 | 0.025 |
| NC_004718.3;NP_828849.2;SARS-CoV_orf1ab;222 | 0.1 | 0.025 |
| NC_004718.3;NP_828849.2;SARS-CoV_orf1ab;223 | 0.15 | 0.05 |
| NC_004718.3;NP_828849.2;SARS-CoV_orf1ab;224 | 0 | 0.025 |
| NC_004718.3;NP_828849.2;SARS-CoV_orf1ab;225 | 0.05 | 0.025 |
| NC_004718.3;NP_828849.2;SARS-CoV_orf1ab;226 | 0 | 0 |
| NC_004718.3;NP_828849.2;SARS-CoV_orf1ab;227 | 0 | 0 |
| NC_004718.3;NP_828849.2;SARS-CoV_orf1ab;228 | 0 | 0 |
| NC_004718.3;NP_828849.2;SARS-CoV_orf1ab;229 | 0 | 0 |
| NC_004718.3;NP_828849.2;SARS-CoV_orf1ab;230 | 0 | 0 |
| NC_004718.3;NP_828849.2;SARS-CoV_orf1ab;231 | 0 | 0 |
| NC_004718.3;NP_828849.2;SARS-CoV_orf1ab;232 | 0 | 0 |
| NC_004718.3;NP_828849.2;SARS-CoV_orf1ab;233 | 0 | 0.05 |
| NC_004718.3;NP_828849.2;SARS-CoV_orf1ab;234 | 0 | 0 |
| NC_004718.3;NP_828849.2;SARS-CoV_orf1ab;235 | 0 | 0 |
| NC_004718.3;NP_828849.2;SARS-CoV_orf1ab;236 | 0 | 0 |
| NC_004718.3;NP_828849.2;SARS-CoV_orf1ab;237 | 0 | 0 |
| NC_004718.3;NP_828849.2;SARS-CoV_orf1ab;238 | 0 | 0.05 |
| NC_004718.3;NP_828849.2;SARS-CoV_orf1ab;239 | 0 | 0.025 |
| NC_004718.3;NP_828849.2;SARS-CoV_orf1ab;240 | 0.05 | 0.025 |
| NC_004718.3;NP_828849.2;SARS-CoV_orf1ab;241 | 0.05 | 0 |
| NC_004718.3;NP_828849.2;SARS-CoV_orf1ab;242 | 0 | 0.05 |
| NC_004718.3;NP_828849.2;SARS-CoV_orf1ab;243 | 0 | 0 |
| NC_004718.3;NP_828849.2;SARS-CoV_orf1ab;244 | 0 | 0.025 |
| NC_004718.3;NP_828849.2;SARS-CoV_orf1ab;245 | 0 | 0 |
| NC_004718.3;NP_828849.2;SARS-CoV_orf1ab;246 | 0 | 0.05 |
| NC_004718.3;NP_828849.2;SARS-CoV_orf1ab;247 | 0 | 0 |
| NC_004718.3;NP_828849.2;SARS-CoV_orf1ab;248 | 0 | 0.025 |
| NC_004718.3;NP_828849.2;SARS-CoV_orf1ab;249 | 0 | 0.025 |
| NC_004718.3;NP_828849.2;SARS-CoV_orf1ab;250 | 0 | 0 |
| NC_004718.3;NP_828849.2;SARS-CoV_orf1ab;251 | 0 | 0.025 |
| NC_004718.3;NP_828849.2;SARS-CoV_orf1ab;252 | 0 | 0 |
| NC_004718.3;NP_828849.2;SARS-CoV_orf1ab;253 | 0 | 0 |
| NC_004718.3;NP_828849.2;SARS-CoV_orf1ab;254 | 0.05 | 0 |
| NC_004718.3;NP_828849.2;SARS-CoV_orf1ab;255 | 0.05 | 0 |

| | | |
|---|---|---|
| NC_004718.3;NP_828849.2;SARS-CoV_orf1ab;256 | 0.05 | 0 |
| NC_004718.3;NP_828849.2;SARS-CoV_orf1ab;257 | 0 | 0 |
| NC_004718.3;NP_828849.2;SARS-CoV_orf1ab;258 | 0 | 0.025 |
| NC_004718.3;NP_828849.2;SARS-CoV_orf1ab;259 | 0 | 0 |
| NC_004718.3;NP_828849.2;SARS-CoV_orf1ab;260 | 0 | 0 |
| NC_004718.3;NP_828849.2;SARS-CoV_orf1ab;261 | 0 | 0 |
| NC_004718.3;NP_828849.2;SARS-CoV_orf1ab;262 | 0.05 | 0 |
| NC_004718.3;NP_828849.2;SARS-CoV_orf1ab;263 | 0.1 | 0 |
| NC_004718.3;NP_828849.2;SARS-CoV_orf1ab;264 | 0 | 0.025 |
| NC_004718.3;NP_828849.2;SARS-CoV_orf1ab;265 | 0 | 0 |
| NC_004718.3;NP_828849.2;SARS-CoV_orf1ab;266 | 0 | 0 |
| NC_004718.3;NP_828849.2;SARS-CoV_orf1ab;267 | 0 | 0.025 |
| NC_004718.3;NP_828849.2;SARS-CoV_orf1ab;268 | 0 | 0.025 |
| NC_004718.3;NP_828849.2;SARS-CoV_orf1ab;269 | 0 | 0.025 |
| NC_004718.3;NP_828849.2;SARS-CoV_orf1ab;270 | 0 | 0.025 |
| NC_004718.3;NP_828849.2;SARS-CoV_orf1ab;271 | 0.05 | 0.025 |
| NC_004718.3;NP_828849.2;SARS-CoV_orf1ab;272 | 0 | 0.025 |
| NC_004718.3;NP_828849.2;SARS-CoV_orf1ab;273 | 0 | 0.025 |
| NC_004718.3;NP_828849.2;SARS-CoV_orf1ab;274 | 0 | 0.025 |
| NC_004718.3;NP_828849.2;SARS-CoV_orf1ab;275 | 0 | 0.05 |
| NC_004718.3;NP_828849.2;SARS-CoV_orf1ab;276 | 0.05 | 0.05 |
| NC_004718.3;NP_828849.2;SARS-CoV_orf1ab;277 | 0.05 | 0.025 |
| NC_004718.3;NP_828849.2;SARS-CoV_orf1ab;278 | 0.05 | 0.025 |
| NC_004718.3;NP_828849.2;SARS-CoV_orf1ab;279 | 0.05 | 0.025 |
| NC_004718.3;NP_828849.2;SARS-CoV_orf1ab;280 | 0.05 | 0.025 |
| NC_004718.3;NP_828849.2;SARS-CoV_orf1ab;281 | 0.05 | 0.025 |
| NC_004718.3;NP_828849.2;SARS-CoV_orf1ab;282 | 0.05 | 0.025 |
| NC_004718.3;NP_828849.2;SARS-CoV_orf1ab;283 | 0.15 | 0 |
| NC_004718.3;NP_828849.2;SARS-CoV_orf1ab;284 | 0.05 | 0.025 |
| NC_004718.3;NP_828849.2;SARS-CoV_orf1ab;285 | 0.05 | 0.025 |
| NC_004718.3;NP_828849.2;SARS-CoV_orf1ab;286 | 0 | 0.025 |
| NC_004718.3;NP_828849.2;SARS-CoV_orf1ab;287 | 0 | 0.025 |
| NC_004718.3;NP_828849.2;SARS-CoV_orf1ab;288 | 0 | 0.025 |
| NC_004718.3;NP_828849.2;SARS-CoV_orf1ab;289 | 0 | 0.025 |
| NC_004718.3;NP_828849.2;SARS-CoV_orf1ab;290 | 0 | 0.025 |
| NC_004718.3;NP_828849.2;SARS-CoV_orf1ab;291 | 0 | 0.025 |
| NC_004718.3;NP_828849.2;SARS-CoV_orf1ab;292 | 0 | 0.025 |
| NC_004718.3;NP_828849.2;SARS-CoV_orf1ab;293 | 0 | 0 |
| NC_004718.3;NP_828849.2;SARS-CoV_orf1ab;294 | 0 | 0 |
| NC_004718.3;NP_828849.2;SARS-CoV_orf1ab;295 | 0 | 0 |
| NC_004718.3;NP_828849.2;SARS-CoV_orf1ab;296 | 0.05 | 0.025 |
| NC_004718.3;NP_828849.2;SARS-CoV_orf1ab;297 | 0.05 | 0 |
| NC_004718.3;NP_828849.2;SARS-CoV_orf1ab;298 | 0.05 | 0 |

| | | |
|---|---|---|
| NC_004718.3;NP_828849.2;SARS-CoV_orf1ab;299 | 0.05 | 0.025 |
| NC_004718.3;NP_828849.2;SARS-CoV_orf1ab;300 | 0.05 | 0 |
| NC_004718.3;NP_828849.2;SARS-CoV_orf1ab;301 | 0.05 | 0.075 |
| NC_004718.3;NP_828849.2;SARS-CoV_orf1ab;302 | 0 | 0 |
| NC_004718.3;NP_828849.2;SARS-CoV_orf1ab;303 | 0 | 0 |
| NC_004718.3;NP_828849.2;SARS-CoV_orf1ab;304 | 0 | 0 |
| NC_004718.3;NP_828849.2;SARS-CoV_orf1ab;305 | 0 | 0 |
| NC_004718.3;NP_828849.2;SARS-CoV_orf1ab;306 | 0 | 0 |
| NC_004718.3;NP_828849.2;SARS-CoV_orf1ab;307 | 0 | 0 |
| NC_004718.3;NP_828849.2;SARS-CoV_orf1ab;308 | 0 | 0 |
| NC_004718.3;NP_828849.2;SARS-CoV_orf1ab;309 | 0 | 0 |
| NC_004718.3;NP_828849.2;SARS-CoV_orf1ab;310 | 0 | 0 |
| NC_004718.3;NP_828849.2;SARS-CoV_orf1ab;311 | 0 | 0 |
| NC_004718.3;NP_828849.2;SARS-CoV_orf1ab;312 | 0 | 0 |
| NC_004718.3;NP_828849.2;SARS-CoV_orf1ab;313 | 0 | 0 |
| NC_004718.3;NP_828849.2;SARS-CoV_orf1ab;314 | 0 | 0 |
| NC_004718.3;NP_828849.2;SARS-CoV_orf1ab;315 | 0.1 | 0.025 |
| NC_004718.3;NP_828849.2;SARS-CoV_orf1ab;316 | 0 | 0 |
| NC_004718.3;NP_828849.2;SARS-CoV_orf1ab;317 | 0 | 0 |
| NC_004718.3;NP_828849.2;SARS-CoV_orf1ab;318 | 0 | 0 |
| NC_004718.3;NP_828849.2;SARS-CoV_orf1ab;319 | 0 | 0 |
| NC_004718.3;NP_828849.2;SARS-CoV_orf1ab;320 | 0 | 0 |
| NC_004718.3;NP_828849.2;SARS-CoV_orf1ab;321 | 0 | 0 |
| NC_004718.3;NP_828849.2;SARS-CoV_orf1ab;322 | 0 | 0 |
| NC_004718.3;NP_828849.2;SARS-CoV_orf1ab;323 | 0 | 0 |
| NC_004718.3;NP_828849.2;SARS-CoV_orf1ab;324 | 0.05 | 0 |
| NC_004718.3;NP_828849.2;SARS-CoV_orf1ab;325 | 0 | 0 |
| NC_004718.3;NP_828849.2;SARS-CoV_orf1ab;326 | 0 | 0.025 |
| NC_004718.3;NP_828849.2;SARS-CoV_orf1ab;327 | 0 | 0 |
| NC_004718.3;NP_828849.2;SARS-CoV_orf1ab;328 | 0 | 0 |
| NC_004718.3;NP_828849.2;SARS-CoV_orf1ab;329 | 0 | 0 |
| NC_004718.3;NP_828849.2;SARS-CoV_orf1ab;330 | 0 | 0 |
| NC_004718.3;NP_828849.2;SARS-CoV_orf1ab;331 | 0 | 0 |
| NC_004718.3;NP_828849.2;SARS-CoV_orf1ab;332 | 0 | 0 |
| NC_004718.3;NP_828849.2;SARS-CoV_orf1ab;333 | 0 | 0 |
| NC_004718.3;NP_828849.2;SARS-CoV_orf1ab;334 | 0 | 0 |
| NC_004718.3;NP_828849.2;SARS-CoV_orf1ab;335 | 0 | 0 |
| NC_004718.3;NP_828849.2;SARS-CoV_orf1ab;336 | 0 | 0 |
| NC_004718.3;NP_828849.2;SARS-CoV_orf1ab;337 | 0 | 0 |
| NC_004718.3;NP_828849.2;SARS-CoV_orf1ab;338 | 0 | 0 |
| NC_004718.3;NP_828849.2;SARS-CoV_orf1ab;339 | 0 | 0 |
| NC_004718.3;NP_828849.2;SARS-CoV_orf1ab;340 | 0 | 0 |
| NC_004718.3;NP_828849.2;SARS-CoV_orf1ab;341 | 0 | 0 |

| | | |
|---|---|---|
| NC_004718.3;NP_828849.2;SARS-CoV_orf1ab;342 | 0 | 0 |
| NC_004718.3;NP_828849.2;SARS-CoV_orf1ab;343 | 0 | 0 |
| NC_004718.3;NP_828849.2;SARS-CoV_orf1ab;344 | 0 | 0 |
| NC_004718.3;NP_828849.2;SARS-CoV_orf1ab;345 | 0 | 0 |
| NC_004718.3;NP_828849.2;SARS-CoV_orf1ab;346 | 0 | 0 |
| NC_004718.3;NP_828849.2;SARS-CoV_orf1ab;347 | 0 | 0 |
| NC_004718.3;NP_828849.2;SARS-CoV_orf1ab;348 | 0 | 0 |
| NC_004718.3;NP_828849.2;SARS-CoV_orf1ab;349 | 0 | 0 |
| NC_004718.3;NP_828849.2;SARS-CoV_orf1ab;350 | 0 | 0 |
| NC_004718.3;NP_828849.2;SARS-CoV_orf1ab;351 | 0 | 0 |
| NC_004718.3;NP_828849.2;SARS-CoV_orf1ab;352 | 0 | 0 |
| NC_004718.3;NP_828849.2;SARS-CoV_orf1ab;353 | 0 | 0 |
| NC_004718.3;NP_828849.2;SARS-CoV_orf1ab;354 | 0 | 0 |
| NC_004718.3;NP_828849.2;SARS-CoV_orf1ab;355 | 0 | 0 |
| NC_004718.3;NP_828849.2;SARS-CoV_orf1ab;356 | 0.05 | 0 |
| NC_004718.3;NP_828849.2;SARS-CoV_orf1ab;357 | 0.05 | 0 |
| NC_004718.3;NP_828849.2;SARS-CoV_orf1ab;358 | 0.05 | 0 |
| NC_004718.3;NP_828849.2;SARS-CoV_orf1ab;359 | 0 | 0 |
| NC_004718.3;NP_828849.2;SARS-CoV_orf1ab;360 | 0 | 0 |
| NC_004718.3;NP_828849.2;SARS-CoV_orf1ab;361 | 0 | 0 |
| NC_004718.3;NP_828849.2;SARS-CoV_orf1ab;362 | 0.05 | 0 |
| NC_004718.3;NP_828849.2;SARS-CoV_orf1ab;363 | 0 | 0 |
| NC_004718.3;NP_828849.2;SARS-CoV_orf1ab;364 | 0 | 0 |
| NC_004718.3;NP_828849.2;SARS-CoV_orf1ab;365 | 0 | 0 |
| NC_004718.3;NP_828849.2;SARS-CoV_orf1ab;366 | 0.05 | 0.025 |
| NC_004718.3;NP_828849.2;SARS-CoV_orf1ab;367 | 0 | 0 |
| NC_004718.3;NP_828849.2;SARS-CoV_orf1ab;368 | 0.05 | 0.05 |
| NC_004718.3;NP_828849.2;SARS-CoV_orf1ab;369 | 0.05 | 0.025 |
| NC_004718.3;NP_828849.2;SARS-CoV_orf1ab;370 | 0.05 | 0.025 |
| NC_004718.3;NP_828849.2;SARS-CoV_orf1ab;371 | 0.05 | 0.025 |
| NC_004718.3;NP_828849.2;SARS-CoV_orf1ab;372 | 0.05 | 0.025 |
| NC_004718.3;NP_828849.2;SARS-CoV_orf1ab;373 | 0.05 | 0 |
| NC_004718.3;NP_828849.2;SARS-CoV_orf1ab;374 | 0.05 | 0.025 |
| NC_004718.3;NP_828849.2;SARS-CoV_orf1ab;375 | 0.1 | 0 |
| NC_004718.3;NP_828849.2;SARS-CoV_orf1ab;376 | 0 | 0 |
| NC_004718.3;NP_828849.2;SARS-CoV_orf1ab;377 | 0.1 | 0 |
| NC_004718.3;NP_828849.2;SARS-CoV_orf1ab;378 | 0 | 0 |
| NC_004718.3;NP_828849.2;SARS-CoV_orf1ab;379 | 0 | 0 |
| NC_004718.3;NP_828849.2;SARS-CoV_orf1ab;380 | 0 | 0 |
| NC_004718.3;NP_828849.2;SARS-CoV_orf1ab;381 | 0 | 0 |
| NC_004718.3;NP_828849.2;SARS-CoV_orf1ab;382 | 0 | 0 |
| NC_004718.3;NP_828849.2;SARS-CoV_orf1ab;383 | 0.05 | 0.025 |
| NC_004718.3;NP_828849.2;SARS-CoV_orf1ab;384 | 0 | 0 |

| | | |
|---|---|---|
| NC_004718.3;NP_828849.2;SARS-CoV_orf1ab;385 | 0 | 0 |
| NC_004718.3;NP_828849.2;SARS-CoV_orf1ab;386 | 0 | 0 |
| NC_004718.3;NP_828849.2;SARS-CoV_orf1ab;387 | 0 | 0 |
| NC_004718.3;NP_828849.2;SARS-CoV_orf1ab;388 | 0 | 0 |
| NC_004718.3;NP_828849.2;SARS-CoV_orf1ab;389 | 0 | 0 |
| NC_004718.3;NP_828849.2;SARS-CoV_orf1ab;390 | 0 | 0 |
| NC_004718.3;NP_828849.2;SARS-CoV_orf1ab;391 | 0 | 0 |
| NC_004718.3;NP_828849.2;SARS-CoV_orf1ab;392 | 0 | 0 |
| NC_004718.3;NP_828849.2;SARS-CoV_orf1ab;393 | 0 | 0 |
| NC_004718.3;NP_828849.2;SARS-CoV_orf1ab;394 | 0 | 0 |
| NC_004718.3;NP_828849.2;SARS-CoV_orf1ab;395 | 0 | 0 |
| NC_004718.3;NP_828849.2;SARS-CoV_orf1ab;396 | 0 | 0 |
| NC_004718.3;NP_828849.2;SARS-CoV_orf1ab;397 | 0 | 0.025 |
| NC_004718.3;NP_828849.2;SARS-CoV_orf1ab;398 | 0 | 0 |
| NC_004718.3;NP_828849.2;SARS-CoV_orf1ab;399 | 0 | 0 |
| NC_004718.3;NP_828849.2;SARS-CoV_orf1ab;400 | 0 | 0 |
| NC_004718.3;NP_828849.2;SARS-CoV_orf1ab;401 | 0 | 0 |
| NC_004718.3;NP_828849.2;SARS-CoV_orf1ab;402 | 0 | 0 |
| NC_004718.3;NP_828849.2;SARS-CoV_orf1ab;403 | 0 | 0 |
| NC_004718.3;NP_828849.2;SARS-CoV_orf1ab;404 | 0 | 0 |
| NC_004718.3;NP_828849.2;SARS-CoV_orf1ab;405 | 0 | 0 |
| NC_004718.3;NP_828849.2;SARS-CoV_orf1ab;406 | 0 | 0 |
| NC_004718.3;NP_828849.2;SARS-CoV_orf1ab;407 | 0 | 0 |
| NC_004718.3;NP_828849.2;SARS-CoV_orf1ab;408 | 0 | 0 |
| NC_004718.3;NP_828849.2;SARS-CoV_orf1ab;409 | 0 | 0 |
| NC_004718.3;NP_828849.2;SARS-CoV_orf1ab;410 | 0 | 0 |
| NC_004718.3;NP_828849.2;SARS-CoV_orf1ab;411 | 0 | 0 |
| NC_004718.3;NP_828849.2;SARS-CoV_orf1ab;412 | 0 | 0 |
| NC_004718.3;NP_828849.2;SARS-CoV_orf1ab;413 | 0 | 0 |
| NC_004718.3;NP_828849.2;SARS-CoV_orf1ab;414 | 0 | 0 |
| NC_004718.3;NP_828849.2;SARS-CoV_orf1ab;415 | 0 | 0 |
| NC_004718.3;NP_828849.2;SARS-CoV_orf1ab;416 | 0 | 0 |
| NC_004718.3;NP_828849.2;SARS-CoV_orf1ab;417 | 0 | 0 |
| NC_004718.3;NP_828849.2;SARS-CoV_orf1ab;418 | 0 | 0 |
| NC_004718.3;NP_828849.2;SARS-CoV_orf1ab;419 | 0 | 0 |
| NC_004718.3;NP_828849.2;SARS-CoV_orf1ab;420 | 0 | 0 |
| NC_004718.3;NP_828849.2;SARS-CoV_orf1ab;421 | 0 | 0 |
| NC_004718.3;NP_828849.2;SARS-CoV_orf1ab;422 | 0 | 0 |
| NC_004718.3;NP_828849.2;SARS-CoV_orf1ab;423 | 0 | 0 |
| NC_004718.3;NP_828849.2;SARS-CoV_orf1ab;424 | 0 | 0 |
| NC_004718.3;NP_828849.2;SARS-CoV_orf1ab;425 | 0.05 | 0 |
| NC_004718.3;NP_828849.2;SARS-CoV_orf1ab;426 | 0 | 0 |
| NC_004718.3;NP_828849.2;SARS-CoV_orf1ab;427 | 0 | 0 |

| | | |
|---|---|---|
| NC_004718.3;NP_828849.2;SARS-CoV_orf1ab;428 | 0.05 | 0.025 |
| NC_004718.3;NP_828849.2;SARS-CoV_orf1ab;429 | 0 | 0 |
| NC_004718.3;NP_828849.2;SARS-CoV_orf1ab;430 | 0.05 | 0 |
| NC_004718.3;NP_828849.2;SARS-CoV_orf1ab;431 | 0 | 0 |
| NC_004718.3;NP_828849.2;SARS-CoV_orf1ab;432 | 0 | 0 |
| NC_004718.3;NP_828849.2;SARS-CoV_orf1ab;433 | 0 | 0 |
| NC_004718.3;NP_828849.2;SARS-CoV_orf1ab;434 | 0 | 0 |
| NC_004718.3;NP_828849.2;SARS-CoV_orf1ab;435 | 0 | 0 |
| NC_004718.3;NP_828849.2;SARS-CoV_orf1ab;436 | 0 | 0 |
| NC_004718.3;NP_828849.2;SARS-CoV_orf1ab;437 | 0 | 0.025 |
| NC_004718.3;NP_828849.2;SARS-CoV_orf1ab;438 | 0.05 | 0 |
| NC_004718.3;NP_828849.2;SARS-CoV_orf1ab;439 | 0 | 0 |
| NC_004718.3;NP_828849.2;SARS-CoV_orf1ab;440 | 0 | 0 |
| NC_004718.3;NP_828849.2;SARS-CoV_orf1ab;441 | 0 | 0 |
| NC_004718.3;NP_828849.2;SARS-CoV_orf1ab;442 | 0 | 0 |
| NC_004718.3;NP_828849.2;SARS-CoV_orf1ab;443 | 0 | 0.025 |
| NC_004718.3;NP_828849.2;SARS-CoV_orf1ab;444 | 0.05 | 0 |
| NC_004718.3;NP_828849.2;SARS-CoV_orf1ab;445 | 0.1 | 0.05 |
| NC_004718.3;NP_828849.2;SARS-CoV_orf1ab;446 | 0.05 | 0 |
| NC_004718.3;NP_828849.2;SARS-CoV_orf1ab;447 | 0.05 | 0.075 |
| NC_004718.3;NP_828849.2;SARS-CoV_orf1ab;448 | 0.05 | 0.025 |
| NC_004718.3;NP_828849.2;SARS-CoV_orf1ab;449 | 0 | 0 |
| NC_004718.3;NP_828849.2;SARS-CoV_orf1ab;450 | 0 | 0 |
| NC_004718.3;NP_828849.2;SARS-CoV_orf1ab;451 | 0 | 0.05 |
| NC_004718.3;NP_828849.2;SARS-CoV_orf1ab;452 | 0 | 0 |
| NC_004718.3;NP_828849.2;SARS-CoV_orf1ab;453 | 0.05 | 0 |
| NC_004718.3;NP_828849.2;SARS-CoV_orf1ab;454 | 0 | 0 |
| NC_004718.3;NP_828849.2;SARS-CoV_orf1ab;455 | 0 | 0.05 |
| NC_004718.3;NP_828849.2;SARS-CoV_orf1ab;456 | 0 | 0.025 |
| NC_004718.3;NP_828849.2;SARS-CoV_orf1ab;457 | 0.05 | 0 |
| NC_004718.3;NP_828849.2;SARS-CoV_orf1ab;458 | 0.05 | 0 |
| NC_004718.3;NP_828849.2;SARS-CoV_orf1ab;459 | 0 | 0 |
| NC_004718.3;NP_828849.2;SARS-CoV_orf1ab;460 | 0 | 0 |
| NC_004718.3;NP_828849.2;SARS-CoV_orf1ab;461 | 0 | 0 |
| NC_004718.3;NP_828849.2;SARS-CoV_orf1ab;462 | 0 | 0 |
| NC_004718.3;NP_828849.2;SARS-CoV_orf1ab;463 | 0 | 0 |
| NC_004718.3;NP_828849.2;SARS-CoV_orf1ab;464 | 0 | 0 |
| NC_004718.3;NP_828849.2;SARS-CoV_orf1ab;465 | 0 | 0 |
| NC_004718.3;NP_828849.2;SARS-CoV_orf1ab;466 | 0 | 0 |
| NC_004718.3;NP_828849.2;SARS-CoV_orf1ab;467 | 0 | 0 |
| NC_004718.3;NP_828849.2;SARS-CoV_orf1ab;468 | 0 | 0 |
| NC_004718.3;NP_828849.2;SARS-CoV_orf1ab;469 | 0 | 0 |
| NC_004718.3;NP_828849.2;SARS-CoV_orf1ab;470 | 0 | 0 |

| | | |
|---|---|---|
| NC_004718.3;NP_828849.2;SARS-CoV_orf1ab;471 | 0 | 0 |
| NC_004718.3;NP_828849.2;SARS-CoV_orf1ab;472 | 0 | 0 |
| NC_004718.3;NP_828849.2;SARS-CoV_orf1ab;473 | 0 | 0 |
| NC_004718.3;NP_828849.2;SARS-CoV_orf1ab;474 | 0 | 0 |
| NC_004718.3;NP_828849.2;SARS-CoV_orf1ab;475 | 0 | 0 |
| NC_004718.3;NP_828849.2;SARS-CoV_orf1ab;476 | 0 | 0 |
| NC_004718.3;NP_828849.2;SARS-CoV_orf1ab;477 | 0.05 | 0 |
| NC_004718.3;NP_828849.2;SARS-CoV_orf1ab;478 | 0.05 | 0 |
| NC_004718.3;NP_828849.2;SARS-CoV_orf1ab;479 | 0.05 | 0 |
| NC_004718.3;NP_828849.2;SARS-CoV_orf1ab;480 | 0.1 | 0 |
| NC_004718.3;NP_828849.2;SARS-CoV_orf1ab;481 | 0.05 | 0 |
| NC_004718.3;NP_828849.2;SARS-CoV_orf1ab;482 | 0.05 | 0 |
| NC_004718.3;NP_828849.2;SARS-CoV_orf1ab;483 | 0.1 | 0 |
| NC_004718.3;NP_828849.2;SARS-CoV_orf1ab;484 | 0.05 | 0 |
| NC_004718.3;NP_828849.2;SARS-CoV_orf1ab;485 | 0 | 0 |
| NC_004718.3;NP_828849.2;SARS-CoV_orf1ab;486 | 0 | 0 |
| NC_004718.3;NP_828849.2;SARS-CoV_orf1ab;487 | 0 | 0 |
| NC_004718.3;NP_828849.2;SARS-CoV_orf1ab;488 | 0 | 0 |
| NC_004718.3;NP_828849.2;SARS-CoV_orf1ab;489 | 0 | 0 |
| NC_004718.3;NP_828849.2;SARS-CoV_orf1ab;490 | 0 | 0 |
| NC_004718.3;NP_828849.2;SARS-CoV_orf1ab;491 | 0 | 0 |
| NC_004718.3;NP_828849.2;SARS-CoV_orf1ab;492 | 0 | 0 |
| NC_004718.3;NP_828849.2;SARS-CoV_orf1ab;493 | 0 | 0 |
| NC_004718.3;NP_828849.2;SARS-CoV_orf1ab;494 | 0 | 0 |
| NC_004718.3;NP_828849.2;SARS-CoV_orf1ab;495 | 0 | 0 |
| NC_004718.3;NP_828849.2;SARS-CoV_orf1ab;496 | 0 | 0 |
| NC_004718.3;NP_828849.2;SARS-CoV_orf1ab;497 | 0.05 | 0.05 |
| NC_004718.3;NP_828849.2;SARS-CoV_orf1ab;498 | 0 | 0 |
| NC_004718.3;NP_828849.2;SARS-CoV_orf1ab;499 | 0 | 0 |
| NC_004718.3;NP_828849.2;SARS-CoV_orf1ab;500 | 0 | 0 |
| NC_004718.3;NP_828849.2;SARS-CoV_orf1ab;501 | 0 | 0 |
| NC_004718.3;NP_828849.2;SARS-CoV_orf1ab;502 | 0 | 0 |
| NC_004718.3;NP_828849.2;SARS-CoV_orf1ab;503 | 0 | 0 |
| NC_004718.3;NP_828849.2;SARS-CoV_orf1ab;504 | 0 | 0 |
| NC_004718.3;NP_828849.2;SARS-CoV_orf1ab;505 | 0 | 0 |
| NC_004718.3;NP_828849.2;SARS-CoV_orf1ab;506 | 0 | 0 |
| NC_004718.3;NP_828849.2;SARS-CoV_orf1ab;507 | 0 | 0 |
| NC_004718.3;NP_828849.2;SARS-CoV_orf1ab;508 | 0 | 0 |
| NC_004718.3;NP_828849.2;SARS-CoV_orf1ab;509 | 0 | 0 |
| NC_004718.3;NP_828849.2;SARS-CoV_orf1ab;510 | 0 | 0 |
| NC_004718.3;NP_828849.2;SARS-CoV_orf1ab;511 | 0 | 0 |
| NC_004718.3;NP_828849.2;SARS-CoV_orf1ab;512 | 0 | 0 |
| NC_004718.3;NP_828849.2;SARS-CoV_orf1ab;513 | 0 | 0 |

| | | |
|---|---|---|
| NC_004718.3;NP_828849.2;SARS-CoV_orf1ab;514 | 0 | 0 |
| NC_004718.3;NP_828849.2;SARS-CoV_orf1ab;515 | 0 | 0 |
| NC_004718.3;NP_828849.2;SARS-CoV_orf1ab;516 | 0 | 0 |
| NC_004718.3;NP_828849.2;SARS-CoV_orf1ab;517 | 0 | 0 |
| NC_004718.3;NP_828849.2;SARS-CoV_orf1ab;518 | 0 | 0 |
| NC_004718.3;NP_828849.2;SARS-CoV_orf1ab;519 | 0 | 0 |
| NC_004718.3;NP_828849.2;SARS-CoV_orf1ab;520 | 0 | 0 |
| NC_004718.3;NP_828849.2;SARS-CoV_orf1ab;521 | 0 | 0 |
| NC_004718.3;NP_828849.2;SARS-CoV_orf1ab;522 | 0 | 0 |
| NC_004718.3;NP_828849.2;SARS-CoV_orf1ab;523 | 0 | 0 |
| NC_004718.3;NP_828849.2;SARS-CoV_orf1ab;524 | 0 | 0 |
| NC_004718.3;NP_828849.2;SARS-CoV_orf1ab;525 | 0 | 0 |
| NC_004718.3;NP_828849.2;SARS-CoV_orf1ab;526 | 0.05 | 0.025 |
| NC_004718.3;NP_828849.2;SARS-CoV_orf1ab;527 | 0 | 0 |
| NC_004718.3;NP_828849.2;SARS-CoV_orf1ab;528 | 0 | 0 |
| NC_004718.3;NP_828849.2;SARS-CoV_orf1ab;529 | 0 | 0 |
| NC_004718.3;NP_828849.2;SARS-CoV_orf1ab;530 | 0 | 0 |
| NC_004718.3;NP_828849.2;SARS-CoV_orf1ab;531 | 0.05 | 0 |
| NC_004718.3;NP_828849.2;SARS-CoV_orf1ab;532 | 0.05 | 0 |
| NC_004718.3;NP_828849.2;SARS-CoV_orf1ab;533 | 0 | 0 |
| NC_004718.3;NP_828849.2;SARS-CoV_orf1ab;534 | 0 | 0 |
| NC_004718.3;NP_828849.2;SARS-CoV_orf1ab;535 | 0 | 0 |
| NC_004718.3;NP_828849.2;SARS-CoV_orf1ab;536 | 0 | 0 |
| NC_004718.3;NP_828849.2;SARS-CoV_orf1ab;537 | 0 | 0 |
| NC_004718.3;NP_828849.2;SARS-CoV_orf1ab;538 | 0 | 0 |
| NC_004718.3;NP_828849.2;SARS-CoV_orf1ab;539 | 0 | 0 |
| NC_004718.3;NP_828849.2;SARS-CoV_orf1ab;540 | 0 | 0 |
| NC_004718.3;NP_828849.2;SARS-CoV_orf1ab;541 | 0 | 0 |
| NC_004718.3;NP_828849.2;SARS-CoV_orf1ab;542 | 0 | 0 |
| NC_004718.3;NP_828849.2;SARS-CoV_orf1ab;543 | 0 | 0 |
| NC_004718.3;NP_828849.2;SARS-CoV_orf1ab;544 | 0 | 0 |
| NC_004718.3;NP_828849.2;SARS-CoV_orf1ab;545 | 0.05 | 0 |
| NC_004718.3;NP_828849.2;SARS-CoV_orf1ab;546 | 0.05 | 0 |
| NC_004718.3;NP_828849.2;SARS-CoV_orf1ab;547 | 0.05 | 0 |
| NC_004718.3;NP_828849.2;SARS-CoV_orf1ab;548 | 0.05 | 0 |
| NC_004718.3;NP_828849.2;SARS-CoV_orf1ab;549 | 0.1 | 0 |
| NC_004718.3;NP_828849.2;SARS-CoV_orf1ab;550 | 0.05 | 0 |
| NC_004718.3;NP_828849.2;SARS-CoV_orf1ab;551 | 0.05 | 0.05 |
| NC_004718.3;NP_828849.2;SARS-CoV_orf1ab;552 | 0 | 0.025 |
| NC_004718.3;NP_828849.2;SARS-CoV_orf1ab;553 | 0.05 | 0.025 |
| NC_004718.3;NP_828849.2;SARS-CoV_orf1ab;554 | 0.05 | 0 |
| NC_004718.3;NP_828849.2;SARS-CoV_orf1ab;555 | 0 | 0 |
| NC_004718.3;NP_828849.2;SARS-CoV_orf1ab;556 | 0 | 0 |

| | | |
|---|---|---|
| NC_004718.3;NP_828849.2;SARS-CoV_orf1ab;557 | 0 | 0 |
| NC_004718.3;NP_828849.2;SARS-CoV_orf1ab;558 | 0 | 0 |
| NC_004718.3;NP_828849.2;SARS-CoV_orf1ab;559 | 0 | 0 |
| NC_004718.3;NP_828849.2;SARS-CoV_orf1ab;560 | 0 | 0 |
| NC_004718.3;NP_828849.2;SARS-CoV_orf1ab;561 | 0 | 0 |
| NC_004718.3;NP_828849.2;SARS-CoV_orf1ab;562 | 0 | 0 |
| NC_004718.3;NP_828849.2;SARS-CoV_orf1ab;563 | 0 | 0 |
| NC_004718.3;NP_828849.2;SARS-CoV_orf1ab;564 | 0 | 0 |
| NC_004718.3;NP_828849.2;SARS-CoV_orf1ab;565 | 0 | 0.025 |
| NC_004718.3;NP_828849.2;SARS-CoV_orf1ab;566 | 0 | 0 |
| NC_004718.3;NP_828849.2;SARS-CoV_orf1ab;567 | 0 | 0 |
| NC_004718.3;NP_828849.2;SARS-CoV_orf1ab;568 | 0 | 0 |
| NC_004718.3;NP_828849.2;SARS-CoV_orf1ab;569 | 0 | 0 |
| NC_004718.3;NP_828849.2;SARS-CoV_orf1ab;570 | 0 | 0 |
| NC_004718.3;NP_828849.2;SARS-CoV_orf1ab;571 | 0 | 0 |
| NC_004718.3;NP_828849.2;SARS-CoV_orf1ab;572 | 0.05 | 0.025 |
| NC_004718.3;NP_828849.2;SARS-CoV_orf1ab;573 | 0 | 0 |
| NC_004718.3;NP_828849.2;SARS-CoV_orf1ab;574 | 0 | 0.025 |
| NC_004718.3;NP_828849.2;SARS-CoV_orf1ab;575 | 0 | 0 |
| NC_004718.3;NP_828849.2;SARS-CoV_orf1ab;576 | 0 | 0 |
| NC_004718.3;NP_828849.2;SARS-CoV_orf1ab;577 | 0 | 0 |
| NC_004718.3;NP_828849.2;SARS-CoV_orf1ab;578 | 0 | 0.025 |
| NC_004718.3;NP_828849.2;SARS-CoV_orf1ab;579 | 0 | 0.025 |
| NC_004718.3;NP_828849.2;SARS-CoV_orf1ab;580 | 0 | 0 |
| NC_004718.3;NP_828849.2;SARS-CoV_orf1ab;581 | 0 | 0 |
| NC_004718.3;NP_828849.2;SARS-CoV_orf1ab;582 | 0 | 0 |
| NC_004718.3;NP_828849.2;SARS-CoV_orf1ab;583 | 0 | 0 |
| NC_004718.3;NP_828849.2;SARS-CoV_orf1ab;584 | 0 | 0 |
| NC_004718.3;NP_828849.2;SARS-CoV_orf1ab;585 | 0 | 0.025 |
| NC_004718.3;NP_828849.2;SARS-CoV_orf1ab;586 | 0 | 0 |
| NC_004718.3;NP_828849.2;SARS-CoV_orf1ab;587 | 0 | 0 |
| NC_004718.3;NP_828849.2;SARS-CoV_orf1ab;588 | 0 | 0 |
| NC_004718.3;NP_828849.2;SARS-CoV_orf1ab;589 | 0.05 | 0 |
| NC_004718.3;NP_828849.2;SARS-CoV_orf1ab;590 | 0 | 0 |
| NC_004718.3;NP_828849.2;SARS-CoV_orf1ab;591 | 0 | 0 |
| NC_004718.3;NP_828849.2;SARS-CoV_orf1ab;592 | 0 | 0 |
| NC_004718.3;NP_828849.2;SARS-CoV_orf1ab;593 | 0 | 0 |
| NC_004718.3;NP_828849.2;SARS-CoV_orf1ab;594 | 0 | 0 |
| NC_004718.3;NP_828849.2;SARS-CoV_orf1ab;595 | 0 | 0 |
| NC_004718.3;NP_828849.2;SARS-CoV_orf1ab;596 | 0 | 0 |
| NC_004718.3;NP_828849.2;SARS-CoV_orf1ab;597 | 0 | 0 |
| NC_004718.3;NP_828849.2;SARS-CoV_orf1ab;598 | 0 | 0 |
| NC_004718.3;NP_828849.2;SARS-CoV_orf1ab;599 | 0 | 0 |

| | | |
|---|---|---|
| NC_004718.3;NP_828849.2;SARS-CoV_orf1ab;600 | 0 | 0 |
| NC_004718.3;NP_828849.2;SARS-CoV_orf1ab;601 | 0 | 0 |
| NC_004718.3;NP_828849.2;SARS-CoV_orf1ab;602 | 0 | 0.025 |
| NC_004718.3;NP_828849.2;SARS-CoV_orf1ab;603 | 0 | 0 |
| NC_004718.3;NP_828849.2;SARS-CoV_orf1ab;604 | 0 | 0 |
| NC_004718.3;NP_828849.2;SARS-CoV_orf1ab;605 | 0.1 | 0.05 |
| NC_004718.3;NP_828849.2;SARS-CoV_orf1ab;606 | 0 | 0 |
| NC_004718.3;NP_828849.2;SARS-CoV_orf1ab;607 | 0 | 0 |
| NC_004718.3;NP_828849.2;SARS-CoV_orf1ab;608 | 0 | 0 |
| NC_004718.3;NP_828849.2;SARS-CoV_orf1ab;609 | 0 | 0 |
| NC_004718.3;NP_828849.2;SARS-CoV_orf1ab;610 | 0 | 0 |
| NC_004718.3;NP_828849.2;SARS-CoV_orf1ab;611 | 0 | 0 |
| NC_004718.3;NP_828849.2;SARS-CoV_orf1ab;612 | 0 | 0 |
| NC_004718.3;NP_828849.2;SARS-CoV_orf1ab;613 | 0 | 0 |
| NC_004718.3;NP_828849.2;SARS-CoV_orf1ab;614 | 0 | 0 |
| NC_004718.3;NP_828849.2;SARS-CoV_orf1ab;615 | 0 | 0 |
| NC_004718.3;NP_828849.2;SARS-CoV_orf1ab;616 | 0 | 0 |
| NC_004718.3;NP_828849.2;SARS-CoV_orf1ab;617 | 0 | 0 |
| NC_004718.3;NP_828849.2;SARS-CoV_orf1ab;618 | 0 | 0 |
| NC_004718.3;NP_828849.2;SARS-CoV_orf1ab;619 | 0.05 | 0 |
| NC_004718.3;NP_828849.2;SARS-CoV_orf1ab;620 | 0.05 | 0 |
| NC_004718.3;NP_828849.2;SARS-CoV_orf1ab;621 | 0.05 | 0 |
| NC_004718.3;NP_828849.2;SARS-CoV_orf1ab;622 | 0.05 | 0 |
| NC_004718.3;NP_828849.2;SARS-CoV_orf1ab;623 | 0 | 0 |
| NC_004718.3;NP_828849.2;SARS-CoV_orf1ab;624 | 0 | 0 |
| NC_004718.3;NP_828849.2;SARS-CoV_orf1ab;625 | 0 | 0 |
| NC_004718.3;NP_828849.2;SARS-CoV_orf1ab;626 | 0 | 0 |
| NC_004718.3;NP_828849.2;SARS-CoV_orf1ab;627 | 0 | 0 |
| NC_004718.3;NP_828849.2;SARS-CoV_orf1ab;628 | 0 | 0 |
| NC_004718.3;NP_828849.2;SARS-CoV_orf1ab;629 | 0 | 0 |
| NC_004718.3;NP_828849.2;SARS-CoV_orf1ab;630 | 0.05 | 0 |
| NC_004718.3;NP_828849.2;SARS-CoV_orf1ab;631 | 0 | 0 |
| NC_004718.3;NP_828849.2;SARS-CoV_orf1ab;632 | 0 | 0 |
| NC_004718.3;NP_828849.2;SARS-CoV_orf1ab;633 | 0.05 | 0.025 |
| NC_004718.3;NP_828849.2;SARS-CoV_orf1ab;634 | 0.05 | 0.025 |
| NC_004718.3;NP_828849.2;SARS-CoV_orf1ab;635 | 0.05 | 0.025 |
| NC_004718.3;NP_828849.2;SARS-CoV_orf1ab;636 | 0 | 0.025 |
| NC_004718.3;NP_828849.2;SARS-CoV_orf1ab;637 | 0 | 0 |
| NC_004718.3;NP_828849.2;SARS-CoV_orf1ab;638 | 0 | 0 |
| NC_004718.3;NP_828849.2;SARS-CoV_orf1ab;639 | 0 | 0 |
| NC_004718.3;NP_828849.2;SARS-CoV_orf1ab;640 | 0 | 0.025 |
| NC_004718.3;NP_828849.2;SARS-CoV_orf1ab;641 | 0 | 0 |
| NC_004718.3;NP_828849.2;SARS-CoV_orf1ab;642 | 0 | 0 |

| | | |
|---|---|---|
| NC_004718.3;NP_828849.2;SARS-CoV_orf1ab;643 | 0 | 0 |
| NC_004718.3;NP_828849.2;SARS-CoV_orf1ab;644 | 0 | 0 |
| NC_004718.3;NP_828849.2;SARS-CoV_orf1ab;645 | 0 | 0 |
| NC_004718.3;NP_828849.2;SARS-CoV_orf1ab;646 | 0 | 0 |
| NC_004718.3;NP_828849.2;SARS-CoV_orf1ab;647 | 0 | 0 |
| NC_004718.3;NP_828849.2;SARS-CoV_orf1ab;648 | 0 | 0 |
| NC_004718.3;NP_828849.2;SARS-CoV_orf1ab;649 | 0 | 0 |
| NC_004718.3;NP_828849.2;SARS-CoV_orf1ab;650 | 0 | 0 |
| NC_004718.3;NP_828849.2;SARS-CoV_orf1ab;651 | 0 | 0 |
| NC_004718.3;NP_828849.2;SARS-CoV_orf1ab;652 | 0 | 0.025 |
| NC_004718.3;NP_828849.2;SARS-CoV_orf1ab;653 | 0 | 0 |
| NC_004718.3;NP_828849.2;SARS-CoV_orf1ab;654 | 0 | 0.025 |
| NC_004718.3;NP_828849.2;SARS-CoV_orf1ab;655 | 0.05 | 0.05 |
| NC_004718.3;NP_828849.2;SARS-CoV_orf1ab;656 | 0 | 0.05 |
| NC_004718.3;NP_828849.2;SARS-CoV_orf1ab;657 | 0 | 0.025 |
| NC_004718.3;NP_828849.2;SARS-CoV_orf1ab;658 | 0 | 0.025 |
| NC_004718.3;NP_828849.2;SARS-CoV_orf1ab;659 | 0 | 0.05 |
| NC_004718.3;NP_828849.2;SARS-CoV_orf1ab;660 | 0.05 | 0.05 |
| NC_004718.3;NP_828849.2;SARS-CoV_orf1ab;661 | 0 | 0.05 |
| NC_004718.3;NP_828849.2;SARS-CoV_orf1ab;662 | 0 | 0 |
| NC_004718.3;NP_828849.2;SARS-CoV_orf1ab;663 | 0 | 0 |
| NC_004718.3;NP_828849.2;SARS-CoV_orf1ab;664 | 0 | 0 |
| NC_004718.3;NP_828849.2;SARS-CoV_orf1ab;665 | 0 | 0 |
| NC_004718.3;NP_828849.2;SARS-CoV_orf1ab;666 | 0.05 | 0 |
| NC_004718.3;NP_828849.2;SARS-CoV_orf1ab;667 | 0 | 0 |
| NC_004718.3;NP_828849.2;SARS-CoV_orf1ab;668 | 0 | 0 |
| NC_004718.3;NP_828849.2;SARS-CoV_orf1ab;669 | 0 | 0 |
| NC_004718.3;NP_828849.2;SARS-CoV_orf1ab;670 | 0 | 0 |
| NC_004718.3;NP_828849.2;SARS-CoV_orf1ab;671 | 0 | 0 |
| NC_004718.3;NP_828849.2;SARS-CoV_orf1ab;672 | 0 | 0 |
| NC_004718.3;NP_828849.2;SARS-CoV_orf1ab;673 | 0 | 0 |
| NC_004718.3;NP_828849.2;SARS-CoV_orf1ab;674 | 0 | 0 |
| NC_004718.3;NP_828849.2;SARS-CoV_orf1ab;675 | 0 | 0 |
| NC_004718.3;NP_828849.2;SARS-CoV_orf1ab;676 | 0 | 0 |
| NC_004718.3;NP_828849.2;SARS-CoV_orf1ab;677 | 0 | 0 |
| NC_004718.3;NP_828849.2;SARS-CoV_orf1ab;678 | 0 | 0.025 |
| NC_004718.3;NP_828849.2;SARS-CoV_orf1ab;679 | 0 | 0.1 |
| NC_004718.3;NP_828849.2;SARS-CoV_orf1ab;680 | 0 | 0 |
| NC_004718.3;NP_828849.2;SARS-CoV_orf1ab;681 | 0 | 0 |
| NC_004718.3;NP_828849.2;SARS-CoV_orf1ab;682 | 0.05 | 0 |
| NC_004718.3;NP_828849.2;SARS-CoV_orf1ab;683 | 0.05 | 0 |
| NC_004718.3;NP_828849.2;SARS-CoV_orf1ab;684 | 0.05 | 0.025 |
| NC_004718.3;NP_828849.2;SARS-CoV_orf1ab;685 | 0 | 0 |

| | | |
|---|---|---|
| NC_004718.3;NP_828849.2;SARS-CoV_orf1ab;686 | 0 | 0 |
| NC_004718.3;NP_828849.2;SARS-CoV_orf1ab;687 | 0 | 0 |
| NC_004718.3;NP_828849.2;SARS-CoV_orf1ab;688 | 0 | 0 |
| NC_004718.3;NP_828849.2;SARS-CoV_orf1ab;689 | 0 | 0 |
| NC_004718.3;NP_828849.2;SARS-CoV_orf1ab;690 | 0 | 0.025 |
| NC_004718.3;NP_828849.2;SARS-CoV_orf1ab;691 | 0 | 0 |
| NC_004718.3;NP_828849.2;SARS-CoV_orf1ab;692 | 0 | 0.025 |
| NC_004718.3;NP_828849.2;SARS-CoV_orf1ab;693 | 0 | 0.025 |
| NC_004718.3;NP_828849.2;SARS-CoV_orf1ab;694 | 0.05 | 0 |
| NC_004718.3;NP_828849.2;SARS-CoV_orf1ab;695 | 0 | 0 |
| NC_004718.3;NP_828849.2;SARS-CoV_orf1ab;696 | 0 | 0 |
| NC_004718.3;NP_828849.2;SARS-CoV_orf1ab;697 | 0 | 0 |
| NC_004718.3;NP_828849.2;SARS-CoV_orf1ab;698 | 0 | 0 |
| NC_004718.3;NP_828849.2;SARS-CoV_orf1ab;699 | 0 | 0 |
| NC_004718.3;NP_828849.2;SARS-CoV_orf1ab;700 | 0.05 | 0 |
| NC_004718.3;NP_828849.2;SARS-CoV_orf1ab;701 | 0.05 | 0 |
| NC_004718.3;NP_828849.2;SARS-CoV_orf1ab;702 | 0 | 0 |
| NC_004718.3;NP_828849.2;SARS-CoV_orf1ab;703 | 0 | 0 |
| NC_004718.3;NP_828849.2;SARS-CoV_orf1ab;704 | 0 | 0 |
| NC_004718.3;NP_828849.2;SARS-CoV_orf1ab;705 | 0 | 0 |
| NC_004718.3;NP_828849.2;SARS-CoV_orf1ab;706 | 0 | 0 |
| NC_004718.3;NP_828849.2;SARS-CoV_orf1ab;707 | 0 | 0 |
| NC_004718.3;NP_828849.2;SARS-CoV_orf1ab;708 | 0 | 0 |
| NC_004718.3;NP_828849.2;SARS-CoV_orf1ab;709 | 0 | 0.025 |
| NC_004718.3;NP_828849.2;SARS-CoV_orf1ab;710 | 0 | 0 |
| NC_004718.3;NP_828849.2;SARS-CoV_orf1ab;711 | 0 | 0 |
| NC_004718.3;NP_828849.2;SARS-CoV_orf1ab;712 | 0 | 0 |
| NC_004718.3;NP_828849.2;SARS-CoV_orf1ab;713 | 0 | 0 |
| NC_004718.3;NP_828849.2;SARS-CoV_orf1ab;714 | 0 | 0.025 |
| NC_004718.3;NP_828849.2;SARS-CoV_orf1ab;715 | 0 | 0 |
| NC_004718.3;NP_828849.2;SARS-CoV_orf1ab;716 | 0 | 0 |
| NC_004718.3;NP_828849.2;SARS-CoV_orf1ab;717 | 0 | 0 |
| NC_004718.3;NP_828849.2;SARS-CoV_orf1ab;718 | 0 | 0 |
| NC_004718.3;NP_828849.2;SARS-CoV_orf1ab;719 | 0 | 0 |
| NC_004718.3;NP_828849.2;SARS-CoV_orf1ab;720 | 0 | 0 |
| NC_004718.3;NP_828849.2;SARS-CoV_orf1ab;721 | 0 | 0 |
| NC_004718.3;NP_828849.2;SARS-CoV_orf1ab;722 | 0 | 0 |
| NC_004718.3;NP_828849.2;SARS-CoV_orf1ab;723 | 0 | 0 |
| NC_004718.3;NP_828849.2;SARS-CoV_orf1ab;724 | 0 | 0 |
| NC_004718.3;NP_828849.2;SARS-CoV_orf1ab;725 | 0 | 0 |
| NC_004718.3;NP_828849.2;SARS-CoV_orf1ab;726 | 0 | 0 |
| NC_004718.3;NP_828849.2;SARS-CoV_orf1ab;727 | 0 | 0 |
| NC_004718.3;NP_828849.2;SARS-CoV_orf1ab;728 | 0 | 0.025 |

| | | |
|---|---|---|
| NC_004718.3;NP_828849.2;SARS-CoV_orf1ab;729 | 0 | 0.025 |
| NC_004718.3;NP_828849.2;SARS-CoV_orf1ab;730 | 0 | 0.025 |
| NC_004718.3;NP_828849.2;SARS-CoV_orf1ab;731 | 0 | 0.025 |
| NC_004718.3;NP_828849.2;SARS-CoV_orf1ab;732 | 0 | 0.025 |
| NC_004718.3;NP_828849.2;SARS-CoV_orf1ab;733 | 0 | 0.025 |
| NC_004718.3;NP_828849.2;SARS-CoV_orf1ab;734 | 0 | 0.025 |
| NC_004718.3;NP_828849.2;SARS-CoV_orf1ab;735 | 0 | 0.025 |
| NC_004718.3;NP_828849.2;SARS-CoV_orf1ab;736 | 0 | 0.05 |
| NC_004718.3;NP_828849.2;SARS-CoV_orf1ab;737 | 0 | 0.025 |
| NC_004718.3;NP_828849.2;SARS-CoV_orf1ab;738 | 0 | 0.025 |
| NC_004718.3;NP_828849.2;SARS-CoV_orf1ab;739 | 0 | 0 |
| NC_004718.3;NP_828849.2;SARS-CoV_orf1ab;740 | 0 | 0 |
| NC_004718.3;NP_828849.2;SARS-CoV_orf1ab;741 | 0.05 | 0 |
| NC_004718.3;NP_828849.2;SARS-CoV_orf1ab;742 | 0 | 0 |
| NC_004718.3;NP_828849.2;SARS-CoV_orf1ab;743 | 0 | 0 |
| NC_004718.3;NP_828849.2;SARS-CoV_orf1ab;744 | 0.05 | 0.025 |
| NC_004718.3;NP_828849.2;SARS-CoV_orf1ab;745 | 0.05 | 0.05 |
| NC_004718.3;NP_828849.2;SARS-CoV_orf1ab;746 | 0.2 | 0.15 |
| NC_004718.3;NP_828849.2;SARS-CoV_orf1ab;747 | 0 | 0.05 |
| NC_004718.3;NP_828849.2;SARS-CoV_orf1ab;748 | 0.05 | 0.025 |
| NC_004718.3;NP_828849.2;SARS-CoV_orf1ab;749 | 0.05 | 0.05 |
| NC_004718.3;NP_828849.2;SARS-CoV_orf1ab;750 | 0.05 | 0.05 |
| NC_004718.3;NP_828849.2;SARS-CoV_orf1ab;751 | 0.05 | 0.025 |
| NC_004718.3;NP_828849.2;SARS-CoV_orf1ab;752 | 0.05 | 0.05 |
| NC_004718.3;NP_828849.2;SARS-CoV_orf1ab;753 | 0.05 | 0 |
| NC_004718.3;NP_828849.2;SARS-CoV_orf1ab;754 | 0.1 | 0 |
| NC_004718.3;NP_828849.2;SARS-CoV_orf1ab;755 | 0.05 | 0 |
| NC_004718.3;NP_828849.2;SARS-CoV_orf1ab;756 | 0 | 0 |
| NC_004718.3;NP_828849.2;SARS-CoV_orf1ab;757 | 0 | 0 |
| NC_004718.3;NP_828849.2;SARS-CoV_orf1ab;758 | 0 | 0 |
| NC_004718.3;NP_828849.2;SARS-CoV_orf1ab;759 | 0 | 0 |
| NC_004718.3;NP_828849.2;SARS-CoV_orf1ab;760 | 0 | 0 |
| NC_004718.3;NP_828849.2;SARS-CoV_orf1ab;761 | 0 | 0 |
| NC_004718.3;NP_828849.2;SARS-CoV_orf1ab;762 | 0 | 0 |
| NC_004718.3;NP_828849.2;SARS-CoV_orf1ab;763 | 0 | 0.025 |
| NC_004718.3;NP_828849.2;SARS-CoV_orf1ab;764 | 0 | 0 |
| NC_004718.3;NP_828849.2;SARS-CoV_orf1ab;765 | 0 | 0 |
| NC_004718.3;NP_828849.2;SARS-CoV_orf1ab;766 | 0 | 0 |
| NC_004718.3;NP_828849.2;SARS-CoV_orf1ab;767 | 0 | 0 |
| NC_004718.3;NP_828849.2;SARS-CoV_orf1ab;768 | 0 | 0 |
| NC_004718.3;NP_828849.2;SARS-CoV_orf1ab;769 | 0 | 0 |
| NC_004718.3;NP_828849.2;SARS-CoV_orf1ab;770 | 0.05 | 0.025 |
| NC_004718.3;NP_828849.2;SARS-CoV_orf1ab;771 | 0.1 | 0.025 |

| | | |
|---|---|---|
| NC_004718.3;NP_828849.2;SARS-CoV_orf1ab;772 | 0 | 0 |
| NC_004718.3;NP_828849.2;SARS-CoV_orf1ab;773 | 0 | 0 |
| NC_004718.3;NP_828849.2;SARS-CoV_orf1ab;774 | 0 | 0 |
| NC_004718.3;NP_828849.2;SARS-CoV_orf1ab;775 | 0 | 0 |
| NC_004718.3;NP_828849.2;SARS-CoV_orf1ab;776 | 0 | 0 |
| NC_004718.3;NP_828849.2;SARS-CoV_orf1ab;777 | 0 | 0 |
| NC_004718.3;NP_828849.2;SARS-CoV_orf1ab;778 | 0 | 0 |
| NC_004718.3;NP_828849.2;SARS-CoV_orf1ab;779 | 0 | 0 |
| NC_004718.3;NP_828849.2;SARS-CoV_orf1ab;780 | 0 | 0.025 |
| NC_004718.3;NP_828849.2;SARS-CoV_orf1ab;781 | 0 | 0 |
| NC_004718.3;NP_828849.2;SARS-CoV_orf1ab;782 | 0.05 | 0 |
| NC_004718.3;NP_828849.2;SARS-CoV_orf1ab;783 | 0 | 0 |
| NC_004718.3;NP_828849.2;SARS-CoV_orf1ab;784 | 0 | 0 |
| NC_004718.3;NP_828849.2;SARS-CoV_orf1ab;785 | 0 | 0 |
| NC_004718.3;NP_828849.2;SARS-CoV_orf1ab;786 | 0 | 0 |
| NC_004718.3;NP_828849.2;SARS-CoV_orf1ab;787 | 0 | 0.025 |
| NC_004718.3;NP_828849.2;SARS-CoV_orf1ab;788 | 0 | 0 |
| NC_004718.3;NP_828849.2;SARS-CoV_orf1ab;789 | 0.05 | 0 |
| NC_004718.3;NP_828849.2;SARS-CoV_orf1ab;790 | 0 | 0 |
| NC_004718.3;NP_828849.2;SARS-CoV_orf1ab;791 | 0 | 0.025 |
| NC_004718.3;NP_828849.2;SARS-CoV_orf1ab;792 | 0 | 0 |
| NC_004718.3;NP_828849.2;SARS-CoV_orf1ab;793 | 0 | 0 |
| NC_004718.3;NP_828849.2;SARS-CoV_orf1ab;794 | 0 | 0 |
| NC_004718.3;NP_828849.2;SARS-CoV_orf1ab;795 | 0 | 0 |
| NC_004718.3;NP_828849.2;SARS-CoV_orf1ab;796 | 0 | 0 |
| NC_004718.3;NP_828849.2;SARS-CoV_orf1ab;797 | 0 | 0 |
| NC_004718.3;NP_828849.2;SARS-CoV_orf1ab;798 | 0 | 0 |
| NC_004718.3;NP_828849.2;SARS-CoV_orf1ab;799 | 0 | 0 |
| NC_004718.3;NP_828849.2;SARS-CoV_orf1ab;800 | 0 | 0 |
| NC_004718.3;NP_828849.2;SARS-CoV_orf1ab;801 | 0.05 | 0 |
| NC_004718.3;NP_828849.2;SARS-CoV_orf1ab;802 | 0 | 0 |
| NC_004718.3;NP_828849.2;SARS-CoV_orf1ab;803 | 0 | 0 |
| NC_004718.3;NP_828849.2;SARS-CoV_orf1ab;804 | 0 | 0 |
| NC_004718.3;NP_828849.2;SARS-CoV_orf1ab;805 | 0 | 0 |
| NC_004718.3;NP_828849.2;SARS-CoV_orf1ab;806 | 0 | 0 |
| NC_004718.3;NP_828849.2;SARS-CoV_orf1ab;807 | 0 | 0 |
| NC_004718.3;NP_828849.2;SARS-CoV_orf1ab;808 | 0 | 0 |
| NC_004718.3;NP_828849.2;SARS-CoV_orf1ab;809 | 0 | 0 |
| NC_004718.3;NP_828849.2;SARS-CoV_orf1ab;810 | 0 | 0 |
| NC_004718.3;NP_828849.2;SARS-CoV_orf1ab;811 | 0 | 0 |
| NC_004718.3;NP_828849.2;SARS-CoV_orf1ab;812 | 0 | 0 |
| NC_004718.3;NP_828849.2;SARS-CoV_orf1ab;813 | 0 | 0 |
| NC_004718.3;NP_828849.2;SARS-CoV_orf1ab;814 | 0 | 0.025 |

| | | |
|---|---|---|
| NC_004718.3;NP_828849.2;SARS-CoV_orf1ab;815 | 0 | 0.025 |
| NC_004718.3;NP_828849.2;SARS-CoV_orf1ab;816 | 0.05 | 0.05 |
| NC_004718.3;NP_828849.2;SARS-CoV_orf1ab;817 | 0 | 0.05 |
| NC_004718.3;NP_828849.2;SARS-CoV_orf1ab;818 | 0 | 0.075 |
| NC_004718.3;NP_828849.2;SARS-CoV_orf1ab;819 | 0 | 0.025 |
| NC_004718.3;NP_828849.2;SARS-CoV_orf1ab;820 | 0 | 0.025 |
| NC_004718.3;NP_828849.2;SARS-CoV_orf1ab;821 | 0 | 0.075 |
| NC_004718.3;NP_828849.2;SARS-CoV_orf1ab;822 | 0 | 0 |
| NC_004718.3;NP_828849.2;SARS-CoV_orf1ab;823 | 0 | 0.075 |
| NC_004718.3;NP_828849.2;SARS-CoV_orf1ab;824 | 0 | 0.1 |
| NC_004718.3;NP_828849.2;SARS-CoV_orf1ab;825 | 0 | 0.075 |
| NC_004718.3;NP_828849.2;SARS-CoV_orf1ab;826 | 0.05 | 0.025 |
| NC_004718.3;NP_828849.2;SARS-CoV_orf1ab;827 | 0 | 0 |
| NC_004718.3;NP_828849.2;SARS-CoV_orf1ab;828 | 0.1 | 0.025 |
| NC_004718.3;NP_828849.2;SARS-CoV_orf1ab;829 | 0 | 0 |
| NC_004718.3;NP_828849.2;SARS-CoV_orf1ab;830 | 0 | 0 |
| NC_004718.3;NP_828849.2;SARS-CoV_orf1ab;831 | 0 | 0 |
| NC_004718.3;NP_828849.2;SARS-CoV_orf1ab;832 | 0.05 | 0 |
| NC_004718.3;NP_828849.2;SARS-CoV_orf1ab;833 | 0 | 0 |
| NC_004718.3;NP_828849.2;SARS-CoV_orf1ab;834 | 0 | 0 |
| NC_004718.3;NP_828849.2;SARS-CoV_orf1ab;835 | 0 | 0 |
| NC_004718.3;NP_828849.2;SARS-CoV_orf1ab;836 | 0 | 0 |
| NC_004718.3;NP_828849.2;SARS-CoV_orf1ab;837 | 0.05 | 0.025 |
| NC_004718.3;NP_828849.2;SARS-CoV_orf1ab;838 | 0 | 0 |
| NC_004718.3;NP_828849.2;SARS-CoV_orf1ab;839 | 0 | 0 |
| NC_004718.3;NP_828849.2;SARS-CoV_orf1ab;840 | 0 | 0 |
| NC_004718.3;NP_828849.2;SARS-CoV_orf1ab;841 | 0 | 0.025 |
| NC_004718.3;NP_828849.2;SARS-CoV_orf1ab;842 | 0.05 | 0.05 |
| NC_004718.3;NP_828849.2;SARS-CoV_orf1ab;843 | 0.05 | 0.025 |
| NC_004718.3;NP_828849.2;SARS-CoV_orf1ab;844 | 0.05 | 0.025 |
| NC_004718.3;NP_828849.2;SARS-CoV_orf1ab;845 | 0 | 0.025 |
| NC_004718.3;NP_828849.2;SARS-CoV_orf1ab;846 | 0 | 0.025 |
| NC_004718.3;NP_828849.2;SARS-CoV_orf1ab;847 | 0 | 0.05 |
| NC_004718.3;NP_828849.2;SARS-CoV_orf1ab;848 | 0 | 0 |
| NC_004718.3;NP_828849.2;SARS-CoV_orf1ab;849 | 0 | 0 |
| NC_004718.3;NP_828849.2;SARS-CoV_orf1ab;850 | 0.1 | 0.075 |
| NC_004718.3;NP_828849.2;SARS-CoV_orf1ab;851 | 0.15 | 0.1 |
| NC_004718.3;NP_828849.2;SARS-CoV_orf1ab;852 | 0.15 | 0.1 |
| NC_004718.3;NP_828849.2;SARS-CoV_orf1ab;853 | 0.15 | 0.1 |
| NC_004718.3;NP_828849.2;SARS-CoV_orf1ab;854 | 0.1 | 0.1 |
| NC_004718.3;NP_828849.2;SARS-CoV_orf1ab;855 | 0.05 | 0.05 |
| NC_004718.3;NP_828849.2;SARS-CoV_orf1ab;856 | 0 | 0.05 |
| NC_004718.3;NP_828849.2;SARS-CoV_orf1ab;857 | 0 | 0.05 |

| | | |
|---|---|---|
| NC_004718.3;NP_828849.2;SARS-CoV_orf1ab;858 | 0 | 0.05 |
| NC_004718.3;NP_828849.2;SARS-CoV_orf1ab;859 | 0 | 0 |
| NC_004718.3;NP_828849.2;SARS-CoV_orf1ab;860 | 0 | 0 |
| NC_004718.3;NP_828849.2;SARS-CoV_orf1ab;861 | 0 | 0 |
| NC_004718.3;NP_828849.2;SARS-CoV_orf1ab;862 | 0 | 0 |
| NC_004718.3;NP_828849.2;SARS-CoV_orf1ab;863 | 0 | 0 |
| NC_004718.3;NP_828849.2;SARS-CoV_orf1ab;864 | 0.05 | 0 |
| NC_004718.3;NP_828849.2;SARS-CoV_orf1ab;865 | 0 | 0 |
| NC_004718.3;NP_828849.2;SARS-CoV_orf1ab;866 | 0 | 0 |
| NC_004718.3;NP_828849.2;SARS-CoV_orf1ab;867 | 0 | 0 |
| NC_004718.3;NP_828849.2;SARS-CoV_orf1ab;868 | 0 | 0 |
| NC_004718.3;NP_828849.2;SARS-CoV_orf1ab;869 | 0 | 0 |
| NC_004718.3;NP_828849.2;SARS-CoV_orf1ab;870 | 0 | 0 |
| NC_004718.3;NP_828849.2;SARS-CoV_orf1ab;871 | 0 | 0 |
| NC_004718.3;NP_828849.2;SARS-CoV_orf1ab;872 | 0 | 0 |
| NC_004718.3;NP_828849.2;SARS-CoV_orf1ab;873 | 0 | 0 |
| NC_004718.3;NP_828849.2;SARS-CoV_orf1ab;874 | 0 | 0 |
| NC_004718.3;NP_828849.2;SARS-CoV_orf1ab;875 | 0 | 0 |
| NC_004718.3;NP_828849.2;SARS-CoV_orf1ab;876 | 0 | 0 |
| NC_004718.3;NP_828849.2;SARS-CoV_orf1ab;877 | 0 | 0.05 |
| NC_004718.3;NP_828849.2;SARS-CoV_orf1ab;878 | 0.05 | 0 |
| NC_004718.3;NP_828849.2;SARS-CoV_orf1ab;879 | 0 | 0.025 |
| NC_004718.3;NP_828849.2;SARS-CoV_orf1ab;880 | 0 | 0 |
| NC_004718.3;NP_828849.2;SARS-CoV_orf1ab;881 | 0 | 0 |
| NC_004718.3;NP_828849.2;SARS-CoV_orf1ab;882 | 0 | 0 |
| NC_004718.3;NP_828849.2;SARS-CoV_orf1ab;883 | 0 | 0 |
| NC_004718.3;NP_828849.2;SARS-CoV_orf1ab;884 | 0 | 0 |
| NC_004718.3;NP_828849.2;SARS-CoV_orf1ab;885 | 0.15 | 0.05 |
| NC_004718.3;NP_828849.2;SARS-CoV_orf1ab;886 | 0 | 0 |
| NC_004718.3;NP_828849.2;SARS-CoV_orf1ab;887 | 0 | 0 |
| NC_004718.3;NP_828849.2;SARS-CoV_orf1ab;888 | 0 | 0 |
| NC_004718.3;NP_828849.2;SARS-CoV_orf1ab;889 | 0.05 | 0 |
| NC_004718.3;NP_828849.2;SARS-CoV_orf1ab;890 | 0 | 0.025 |
| NC_004718.3;NP_828849.2;SARS-CoV_orf1ab;891 | 0 | 0 |
| NC_004718.3;NP_828849.2;SARS-CoV_orf1ab;892 | 0 | 0 |
| NC_004718.3;NP_828849.2;SARS-CoV_orf1ab;893 | 0 | 0 |
| NC_004718.3;NP_828849.2;SARS-CoV_orf1ab;894 | 0 | 0 |
| NC_004718.3;NP_828849.2;SARS-CoV_orf1ab;895 | 0 | 0 |
| NC_004718.3;NP_828849.2;SARS-CoV_orf1ab;896 | 0 | 0 |
| NC_004718.3;NP_828849.2;SARS-CoV_orf1ab;897 | 0 | 0 |
| NC_004718.3;NP_828849.2;SARS-CoV_orf1ab;898 | 0 | 0 |
| NC_004718.3;NP_828849.2;SARS-CoV_orf1ab;899 | 0 | 0.025 |
| NC_004718.3;NP_828849.2;SARS-CoV_orf1ab;900 | 0 | 0 |

| | | |
|---|---|---|
| NC_004718.3;NP_828849.2;SARS-CoV_orf1ab;901 | 0 | 0 |
| NC_004718.3;NP_828849.2;SARS-CoV_orf1ab;902 | 0 | 0 |
| NC_004718.3;NP_828849.2;SARS-CoV_orf1ab;903 | 0 | 0 |
| NC_004718.3;NP_828849.2;SARS-CoV_orf1ab;904 | 0 | 0 |
| NC_004718.3;NP_828849.2;SARS-CoV_orf1ab;905 | 0 | 0.025 |
| NC_004718.3;NP_828849.2;SARS-CoV_orf1ab;906 | 0 | 0.025 |
| NC_004718.3;NP_828849.2;SARS-CoV_orf1ab;907 | 0 | 0.05 |
| NC_004718.3;NP_828849.2;SARS-CoV_orf1ab;908 | 0 | 0 |
| NC_004718.3;NP_828849.2;SARS-CoV_orf1ab;909 | 0 | 0 |
| NC_004718.3;NP_828849.2;SARS-CoV_orf1ab;910 | 0 | 0 |
| NC_004718.3;NP_828849.2;SARS-CoV_orf1ab;911 | 0 | 0 |
| NC_004718.3;NP_828849.2;SARS-CoV_orf1ab;912 | 0 | 0 |
| NC_004718.3;NP_828849.2;SARS-CoV_orf1ab;913 | 0 | 0 |
| NC_004718.3;NP_828849.2;SARS-CoV_orf1ab;914 | 0 | 0 |
| NC_004718.3;NP_828849.2;SARS-CoV_orf1ab;915 | 0 | 0 |
| NC_004718.3;NP_828849.2;SARS-CoV_orf1ab;916 | 0 | 0 |
| NC_004718.3;NP_828849.2;SARS-CoV_orf1ab;917 | 0 | 0.025 |
| NC_004718.3;NP_828849.2;SARS-CoV_orf1ab;918 | 0 | 0.05 |
| NC_004718.3;NP_828849.2;SARS-CoV_orf1ab;919 | 0 | 0.05 |
| NC_004718.3;NP_828849.2;SARS-CoV_orf1ab;920 | 0 | 0.05 |
| NC_004718.3;NP_828849.2;SARS-CoV_orf1ab;921 | 0 | 0.05 |
| NC_004718.3;NP_828849.2;SARS-CoV_orf1ab;922 | 0 | 0.1 |
| NC_004718.3;NP_828849.2;SARS-CoV_orf1ab;923 | 0 | 0.05 |
| NC_004718.3;NP_828849.2;SARS-CoV_orf1ab;924 | 0 | 0.05 |
| NC_004718.3;NP_828849.2;SARS-CoV_orf1ab;925 | 0 | 0.075 |
| NC_004718.3;NP_828849.2;SARS-CoV_orf1ab;926 | 0 | 0.025 |
| NC_004718.3;NP_828849.2;SARS-CoV_orf1ab;927 | 0 | 0 |
| NC_004718.3;NP_828849.2;SARS-CoV_orf1ab;928 | 0 | 0 |
| NC_004718.3;NP_828849.2;SARS-CoV_orf1ab;929 | 0 | 0 |
| NC_004718.3;NP_828849.2;SARS-CoV_orf1ab;930 | 0 | 0 |
| NC_004718.3;NP_828849.2;SARS-CoV_orf1ab;931 | 0 | 0 |
| NC_004718.3;NP_828849.2;SARS-CoV_orf1ab;932 | 0 | 0.025 |
| NC_004718.3;NP_828849.2;SARS-CoV_orf1ab;933 | 0.25 | 0.05 |
| NC_004718.3;NP_828849.2;SARS-CoV_orf1ab;934 | 0 | 0 |
| NC_004718.3;NP_828849.2;SARS-CoV_orf1ab;935 | 0 | 0 |
| NC_004718.3;NP_828849.2;SARS-CoV_orf1ab;936 | 0 | 0 |
| NC_004718.3;NP_828849.2;SARS-CoV_orf1ab;937 | 0 | 0.025 |
| NC_004718.3;NP_828849.2;SARS-CoV_orf1ab;938 | 0 | 0 |
| NC_004718.3;NP_828849.2;SARS-CoV_orf1ab;939 | 0 | 0 |
| NC_004718.3;NP_828849.2;SARS-CoV_orf1ab;940 | 0 | 0 |
| NC_004718.3;NP_828849.2;SARS-CoV_orf1ab;941 | 0 | 0.025 |
| NC_004718.3;NP_828849.2;SARS-CoV_orf1ab;942 | 0 | 0 |
| NC_004718.3;NP_828849.2;SARS-CoV_orf1ab;943 | 0 | 0 |

| | | |
|---|---|---|
| NC_004718.3;NP_828849.2;SARS-CoV_orf1ab;944 | 0 | 0 |
| NC_004718.3;NP_828849.2;SARS-CoV_orf1ab;945 | 0 | 0 |
| NC_004718.3;NP_828849.2;SARS-CoV_orf1ab;946 | 0 | 0 |
| NC_004718.3;NP_828849.2;SARS-CoV_orf1ab;947 | 0 | 0 |
| NC_004718.3;NP_828849.2;SARS-CoV_orf1ab;948 | 0 | 0 |
| NC_004718.3;NP_828849.2;SARS-CoV_orf1ab;949 | 0 | 0 |
| NC_004718.3;NP_828849.2;SARS-CoV_orf1ab;950 | 0.05 | 0 |
| NC_004718.3;NP_828849.2;SARS-CoV_orf1ab;951 | 0.05 | 0 |
| NC_004718.3;NP_828849.2;SARS-CoV_orf1ab;952 | 0.05 | 0.075 |
| NC_004718.3;NP_828849.2;SARS-CoV_orf1ab;953 | 0.05 | 0 |
| NC_004718.3;NP_828849.2;SARS-CoV_orf1ab;954 | 0.15 | 0 |
| NC_004718.3;NP_828849.2;SARS-CoV_orf1ab;955 | 0.05 | 0 |
| NC_004718.3;NP_828849.2;SARS-CoV_orf1ab;956 | 0 | 0 |
| NC_004718.3;NP_828849.2;SARS-CoV_orf1ab;957 | 0 | 0 |
| NC_004718.3;NP_828849.2;SARS-CoV_orf1ab;958 | 0 | 0 |
| NC_004718.3;NP_828849.2;SARS-CoV_orf1ab;959 | 0 | 0 |
| NC_004718.3;NP_828849.2;SARS-CoV_orf1ab;960 | 0 | 0 |
| NC_004718.3;NP_828849.2;SARS-CoV_orf1ab;961 | 0 | 0 |
| NC_004718.3;NP_828849.2;SARS-CoV_orf1ab;962 | 0 | 0 |
| NC_004718.3;NP_828849.2;SARS-CoV_orf1ab;963 | 0 | 0 |
| NC_004718.3;NP_828849.2;SARS-CoV_orf1ab;964 | 0 | 0 |
| NC_004718.3;NP_828849.2;SARS-CoV_orf1ab;965 | 0 | 0 |
| NC_004718.3;NP_828849.2;SARS-CoV_orf1ab;966 | 0 | 0.025 |
| NC_004718.3;NP_828849.2;SARS-CoV_orf1ab;967 | 0 | 0.025 |
| NC_004718.3;NP_828849.2;SARS-CoV_orf1ab;968 | 0 | 0.075 |
| NC_004718.3;NP_828849.2;SARS-CoV_orf1ab;969 | 0 | 0.025 |
| NC_004718.3;NP_828849.2;SARS-CoV_orf1ab;970 | 0 | 0.025 |
| NC_004718.3;NP_828849.2;SARS-CoV_orf1ab;971 | 0 | 0.025 |
| NC_004718.3;NP_828849.2;SARS-CoV_orf1ab;972 | 0 | 0.025 |
| NC_004718.3;NP_828849.2;SARS-CoV_orf1ab;973 | 0 | 0.025 |
| NC_004718.3;NP_828849.2;SARS-CoV_orf1ab;974 | 0 | 0.05 |
| NC_004718.3;NP_828849.2;SARS-CoV_orf1ab;975 | 0 | 0.025 |
| NC_004718.3;NP_828849.2;SARS-CoV_orf1ab;976 | 0 | 0.025 |
| NC_004718.3;NP_828849.2;SARS-CoV_orf1ab;977 | 0 | 0.025 |
| NC_004718.3;NP_828849.2;SARS-CoV_orf1ab;978 | 0 | 0.025 |
| NC_004718.3;NP_828849.2;SARS-CoV_orf1ab;979 | 0.05 | 0.025 |
| NC_004718.3;NP_828849.2;SARS-CoV_orf1ab;980 | 0 | 0 |
| NC_004718.3;NP_828849.2;SARS-CoV_orf1ab;981 | 0 | 0 |
| NC_004718.3;NP_828849.2;SARS-CoV_orf1ab;982 | 0 | 0 |
| NC_004718.3;NP_828849.2;SARS-CoV_orf1ab;983 | 0 | 0 |
| NC_004718.3;NP_828849.2;SARS-CoV_orf1ab;984 | 0 | 0 |
| NC_004718.3;NP_828849.2;SARS-CoV_orf1ab;985 | 0 | 0 |
| NC_004718.3;NP_828849.2;SARS-CoV_orf1ab;986 | 0 | 0.025 |

| | | |
|---|---|---|
| NC_004718.3;NP_828849.2;SARS-CoV_orf1ab;987 | 0 | 0.025 |
| NC_004718.3;NP_828849.2;SARS-CoV_orf1ab;988 | 0 | 0.05 |
| NC_004718.3;NP_828849.2;SARS-CoV_orf1ab;989 | 0 | 0.025 |
| NC_004718.3;NP_828849.2;SARS-CoV_orf1ab;990 | 0 | 0.05 |
| NC_004718.3;NP_828849.2;SARS-CoV_orf1ab;991 | 0 | 0.025 |
| NC_004718.3;NP_828849.2;SARS-CoV_orf1ab;992 | 0.05 | 0.025 |
| NC_004718.3;NP_828849.2;SARS-CoV_orf1ab;993 | 0.05 | 0.025 |
| NC_004718.3;NP_828849.2;SARS-CoV_orf1ab;994 | 0.05 | 0.025 |
| NC_004718.3;NP_828849.2;SARS-CoV_orf1ab;995 | 0 | 0 |
| NC_004718.3;NP_828849.2;SARS-CoV_orf1ab;996 | 0.05 | 0 |
| NC_004718.3;NP_828849.2;SARS-CoV_orf1ab;997 | 0 | 0.025 |
| NC_004718.3;NP_828849.2;SARS-CoV_orf1ab;998 | 0 | 0 |
| NC_004718.3;NP_828849.2;SARS-CoV_orf1ab;999 | 0 | 0 |
| NC_004718.3;NP_828849.2;SARS-CoV_orf1ab;1000 | 0 | 0 |
| NC_004718.3;NP_828849.2;SARS-CoV_orf1ab;1001 | 0.05 | 0 |
| NC_004718.3;NP_828849.2;SARS-CoV_orf1ab;1002 | 0.05 | 0 |
| NC_004718.3;NP_828849.2;SARS-CoV_orf1ab;1003 | 0.05 | 0 |
| NC_004718.3;NP_828849.2;SARS-CoV_orf1ab;1004 | 0.05 | 0 |
| NC_004718.3;NP_828849.2;SARS-CoV_orf1ab;1005 | 0.05 | 0 |
| NC_004718.3;NP_828849.2;SARS-CoV_orf1ab;1006 | 0.05 | 0 |
| NC_004718.3;NP_828849.2;SARS-CoV_orf1ab;1007 | 0.05 | 0 |
| NC_004718.3;NP_828849.2;SARS-CoV_orf1ab;1008 | 0.05 | 0 |
| NC_004718.3;NP_828849.2;SARS-CoV_orf1ab;1009 | 0 | 0 |
| NC_004718.3;NP_828849.2;SARS-CoV_orf1ab;1010 | 0.05 | 0 |
| NC_004718.3;NP_828849.2;SARS-CoV_orf1ab;1011 | 0.05 | 0 |
| NC_004718.3;NP_828849.2;SARS-CoV_orf1ab;1012 | 0 | 0 |
| NC_004718.3;NP_828849.2;SARS-CoV_orf1ab;1013 | 0 | 0 |
| NC_004718.3;NP_828849.2;SARS-CoV_orf1ab;1014 | 0 | 0 |
| NC_004718.3;NP_828849.2;SARS-CoV_orf1ab;1015 | 0 | 0 |
| NC_004718.3;NP_828849.2;SARS-CoV_orf1ab;1016 | 0 | 0.075 |
| NC_004718.3;NP_828849.2;SARS-CoV_orf1ab;1017 | 0 | 0.05 |
| NC_004718.3;NP_828849.2;SARS-CoV_orf1ab;1018 | 0 | 0.075 |
| NC_004718.3;NP_828849.2;SARS-CoV_orf1ab;1019 | 0 | 0.075 |
| NC_004718.3;NP_828849.2;SARS-CoV_orf1ab;1020 | 0 | 0.075 |
| NC_004718.3;NP_828849.2;SARS-CoV_orf1ab;1021 | 0 | 0.1 |
| NC_004718.3;NP_828849.2;SARS-CoV_orf1ab;1022 | 0 | 0.075 |
| NC_004718.3;NP_828849.2;SARS-CoV_orf1ab;1023 | 0 | 0.075 |
| NC_004718.3;NP_828849.2;SARS-CoV_orf1ab;1024 | 0.05 | 0.05 |
| NC_004718.3;NP_828849.2;SARS-CoV_orf1ab;1025 | 0.05 | 0.05 |
| NC_004718.3;NP_828849.2;SARS-CoV_orf1ab;1026 | 0 | 0 |
| NC_004718.3;NP_828849.2;SARS-CoV_orf1ab;1027 | 0 | 0 |
| NC_004718.3;NP_828849.2;SARS-CoV_orf1ab;1028 | 0 | 0 |
| NC_004718.3;NP_828849.2;SARS-CoV_orf1ab;1029 | 0 | 0 |

| | | |
|---|---|---|
| NC_004718.3;NP_828849.2;SARS-CoV_orf1ab;1030 | 0 | 0 |
| NC_004718.3;NP_828849.2;SARS-CoV_orf1ab;1031 | 0 | 0 |
| NC_004718.3;NP_828849.2;SARS-CoV_orf1ab;1032 | 0 | 0 |
| NC_004718.3;NP_828849.2;SARS-CoV_orf1ab;1033 | 0.05 | 0 |
| NC_004718.3;NP_828849.2;SARS-CoV_orf1ab;1034 | 0 | 0 |
| NC_004718.3;NP_828849.2;SARS-CoV_orf1ab;1035 | 0.1 | 0.05 |
| NC_004718.3;NP_828849.2;SARS-CoV_orf1ab;1036 | 0 | 0 |
| NC_004718.3;NP_828849.2;SARS-CoV_orf1ab;1037 | 0 | 0 |
| NC_004718.3;NP_828849.2;SARS-CoV_orf1ab;1038 | 0 | 0 |
| NC_004718.3;NP_828849.2;SARS-CoV_orf1ab;1039 | 0 | 0 |
| NC_004718.3;NP_828849.2;SARS-CoV_orf1ab;1040 | 0 | 0 |
| NC_004718.3;NP_828849.2;SARS-CoV_orf1ab;1041 | 0 | 0 |
| NC_004718.3;NP_828849.2;SARS-CoV_orf1ab;1042 | 0 | 0 |
| NC_004718.3;NP_828849.2;SARS-CoV_orf1ab;1043 | 0 | 0 |
| NC_004718.3;NP_828849.2;SARS-CoV_orf1ab;1044 | 0 | 0 |
| NC_004718.3;NP_828849.2;SARS-CoV_orf1ab;1045 | 0.05 | 0 |
| NC_004718.3;NP_828849.2;SARS-CoV_orf1ab;1046 | 0 | 0.025 |
| NC_004718.3;NP_828849.2;SARS-CoV_orf1ab;1047 | 0.05 | 0.025 |
| NC_004718.3;NP_828849.2;SARS-CoV_orf1ab;1048 | 0 | 0 |
| NC_004718.3;NP_828849.2;SARS-CoV_orf1ab;1049 | 0.15 | 0.05 |
| NC_004718.3;NP_828849.2;SARS-CoV_orf1ab;1050 | 0.15 | 0.075 |
| NC_004718.3;NP_828849.2;SARS-CoV_orf1ab;1051 | 0.15 | 0.075 |
| NC_004718.3;NP_828849.2;SARS-CoV_orf1ab;1052 | 0.1 | 0.025 |
| NC_004718.3;NP_828849.2;SARS-CoV_orf1ab;1053 | 0.2 | 0.025 |
| NC_004718.3;NP_828849.2;SARS-CoV_orf1ab;1054 | 0.15 | 0.025 |
| NC_004718.3;NP_828849.2;SARS-CoV_orf1ab;1055 | 0.15 | 0 |
| NC_004718.3;NP_828849.2;SARS-CoV_orf1ab;1056 | 0.2 | 0.05 |
| NC_004718.3;NP_828849.2;SARS-CoV_orf1ab;1057 | 0.05 | 0 |
| NC_004718.3;NP_828849.2;SARS-CoV_orf1ab;1058 | 0.15 | 0 |
| NC_004718.3;NP_828849.2;SARS-CoV_orf1ab;1059 | 0.15 | 0.025 |
| NC_004718.3;NP_828849.2;SARS-CoV_orf1ab;1060 | 0.2 | 0.025 |
| NC_004718.3;NP_828849.2;SARS-CoV_orf1ab;1061 | 0.1 | 0.025 |
| NC_004718.3;NP_828849.2;SARS-CoV_orf1ab;1062 | 0.1 | 0.05 |
| NC_004718.3;NP_828849.2;SARS-CoV_orf1ab;1063 | 0.15 | 0.025 |
| NC_004718.3;NP_828849.2;SARS-CoV_orf1ab;1064 | 0.15 | 0.05 |
| NC_004718.3;NP_828849.2;SARS-CoV_orf1ab;1065 | 0.2 | 0.05 |
| NC_004718.3;NP_828849.2;SARS-CoV_orf1ab;1066 | 0 | 0 |
| NC_004718.3;NP_828849.2;SARS-CoV_orf1ab;1067 | 0 | 0 |
| NC_004718.3;NP_828849.2;SARS-CoV_orf1ab;1068 | 0 | 0 |
| NC_004718.3;NP_828849.2;SARS-CoV_orf1ab;1069 | 0 | 0.025 |
| NC_004718.3;NP_828849.2;SARS-CoV_orf1ab;1070 | 0 | 0 |
| NC_004718.3;NP_828849.2;SARS-CoV_orf1ab;1071 | 0 | 0 |
| NC_004718.3;NP_828849.2;SARS-CoV_orf1ab;1072 | 0 | 0 |

| | | |
|---|---|---|
| NC_004718.3;NP_828849.2;SARS-CoV_orf1ab;1073 | 0 | 0 |
| NC_004718.3;NP_828849.2;SARS-CoV_orf1ab;1074 | 0 | 0 |
| NC_004718.3;NP_828849.2;SARS-CoV_orf1ab;1075 | 0 | 0 |
| NC_004718.3;NP_828849.2;SARS-CoV_orf1ab;1076 | 0 | 0 |
| NC_004718.3;NP_828849.2;SARS-CoV_orf1ab;1077 | 0 | 0 |
| NC_004718.3;NP_828849.2;SARS-CoV_orf1ab;1078 | 0 | 0 |
| NC_004718.3;NP_828849.2;SARS-CoV_orf1ab;1079 | 0 | 0 |
| NC_004718.3;NP_828849.2;SARS-CoV_orf1ab;1080 | 0 | 0 |
| NC_004718.3;NP_828849.2;SARS-CoV_orf1ab;1081 | 0 | 0 |
| NC_004718.3;NP_828849.2;SARS-CoV_orf1ab;1082 | 0 | 0 |
| NC_004718.3;NP_828849.2;SARS-CoV_orf1ab;1083 | 0 | 0 |
| NC_004718.3;NP_828849.2;SARS-CoV_orf1ab;1084 | 0 | 0 |
| NC_004718.3;NP_828849.2;SARS-CoV_orf1ab;1085 | 0 | 0 |
| NC_004718.3;NP_828849.2;SARS-CoV_orf1ab;1086 | 0 | 0 |
| NC_004718.3;NP_828849.2;SARS-CoV_orf1ab;1087 | 0 | 0 |
| NC_004718.3;NP_828849.2;SARS-CoV_orf1ab;1088 | 0 | 0 |
| NC_004718.3;NP_828849.2;SARS-CoV_orf1ab;1089 | 0 | 0.05 |
| NC_004718.3;NP_828849.2;SARS-CoV_orf1ab;1090 | 0.05 | 0.025 |
| NC_004718.3;NP_828849.2;SARS-CoV_orf1ab;1091 | 0.05 | 0.05 |
| NC_004718.3;NP_828849.2;SARS-CoV_orf1ab;1092 | 0 | 0.05 |
| NC_004718.3;NP_828849.2;SARS-CoV_orf1ab;1093 | 0.05 | 0.025 |
| NC_004718.3;NP_828849.2;SARS-CoV_orf1ab;1094 | 0 | 0.05 |
| NC_004718.3;NP_828849.2;SARS-CoV_orf1ab;1095 | 0.05 | 0.05 |
| NC_004718.3;NP_828849.2;SARS-CoV_orf1ab;1096 | 0.05 | 0.05 |
| NC_004718.3;NP_828849.2;SARS-CoV_orf1ab;1097 | 0.05 | 0.075 |
| NC_004718.3;NP_828849.2;SARS-CoV_orf1ab;1098 | 0 | 0.025 |
| NC_004718.3;NP_828849.2;SARS-CoV_orf1ab;1099 | 0 | 0.05 |
| NC_004718.3;NP_828849.2;SARS-CoV_orf1ab;1100 | 0 | 0.025 |
| NC_004718.3;NP_828849.2;SARS-CoV_orf1ab;1101 | 0 | 0.075 |
| NC_004718.3;NP_828849.2;SARS-CoV_orf1ab;1102 | 0.05 | 0.075 |
| NC_004718.3;NP_828849.2;SARS-CoV_orf1ab;1103 | 0.05 | 0.075 |
| NC_004718.3;NP_828849.2;SARS-CoV_orf1ab;1104 | 0.1 | 0.1 |
| NC_004718.3;NP_828849.2;SARS-CoV_orf1ab;1105 | 0 | 0.05 |
| NC_004718.3;NP_828849.2;SARS-CoV_orf1ab;1106 | 0 | 0.05 |
| NC_004718.3;NP_828849.2;SARS-CoV_orf1ab;1107 | 0 | 0.075 |
| NC_004718.3;NP_828849.2;SARS-CoV_orf1ab;1108 | 0 | 0.025 |
| NC_004718.3;NP_828849.2;SARS-CoV_orf1ab;1109 | 0 | 0.025 |
| NC_004718.3;NP_828849.2;SARS-CoV_orf1ab;1110 | 0.1 | 0.05 |
| NC_004718.3;NP_828849.2;SARS-CoV_orf1ab;1111 | 0 | 0 |
| NC_004718.3;NP_828849.2;SARS-CoV_orf1ab;1112 | 0 | 0.025 |
| NC_004718.3;NP_828849.2;SARS-CoV_orf1ab;1113 | 0 | 0 |
| NC_004718.3;NP_828849.2;SARS-CoV_orf1ab;1114 | 0 | 0 |
| NC_004718.3;NP_828849.2;SARS-CoV_orf1ab;1115 | 0 | 0 |

| | | |
|---|---|---|
| NC_004718.3;NP_828849.2;SARS-CoV_orf1ab;1116 | 0 | 0 |
| NC_004718.3;NP_828849.2;SARS-CoV_orf1ab;1117 | 0 | 0 |
| NC_004718.3;NP_828849.2;SARS-CoV_orf1ab;1118 | 0 | 0 |
| NC_004718.3;NP_828849.2;SARS-CoV_orf1ab;1119 | 0 | 0 |
| NC_004718.3;NP_828849.2;SARS-CoV_orf1ab;1120 | 0 | 0 |
| NC_004718.3;NP_828849.2;SARS-CoV_orf1ab;1121 | 0 | 0 |
| NC_004718.3;NP_828849.2;SARS-CoV_orf1ab;1122 | 0 | 0 |
| NC_004718.3;NP_828849.2;SARS-CoV_orf1ab;1123 | 0 | 0 |
| NC_004718.3;NP_828849.2;SARS-CoV_orf1ab;1124 | 0 | 0 |
| NC_004718.3;NP_828849.2;SARS-CoV_orf1ab;1125 | 0 | 0 |
| NC_004718.3;NP_828849.2;SARS-CoV_orf1ab;1126 | 0 | 0 |
| NC_004718.3;NP_828849.2;SARS-CoV_orf1ab;1127 | 0 | 0 |
| NC_004718.3;NP_828849.2;SARS-CoV_orf1ab;1128 | 0 | 0 |
| NC_004718.3;NP_828849.2;SARS-CoV_orf1ab;1129 | 0 | 0 |
| NC_004718.3;NP_828849.2;SARS-CoV_orf1ab;1130 | 0 | 0 |
| NC_004718.3;NP_828849.2;SARS-CoV_orf1ab;1131 | 0 | 0 |
| NC_004718.3;NP_828849.2;SARS-CoV_orf1ab;1132 | 0 | 0 |
| NC_004718.3;NP_828849.2;SARS-CoV_orf1ab;1133 | 0 | 0 |
| NC_004718.3;NP_828849.2;SARS-CoV_orf1ab;1134 | 0 | 0 |
| NC_004718.3;NP_828849.2;SARS-CoV_orf1ab;1135 | 0 | 0 |
| NC_004718.3;NP_828849.2;SARS-CoV_orf1ab;1136 | 0 | 0 |
| NC_004718.3;NP_828849.2;SARS-CoV_orf1ab;1137 | 0 | 0 |
| NC_004718.3;NP_828849.2;SARS-CoV_orf1ab;1138 | 0 | 0 |
| NC_004718.3;NP_828849.2;SARS-CoV_orf1ab;1139 | 0 | 0 |
| NC_004718.3;NP_828849.2;SARS-CoV_orf1ab;1140 | 0 | 0 |
| NC_004718.3;NP_828849.2;SARS-CoV_orf1ab;1141 | 0 | 0 |
| NC_004718.3;NP_828849.2;SARS-CoV_orf1ab;1142 | 0 | 0 |
| NC_004718.3;NP_828849.2;SARS-CoV_orf1ab;1143 | 0 | 0 |
| NC_004718.3;NP_828849.2;SARS-CoV_orf1ab;1144 | 0 | 0 |
| NC_004718.3;NP_828849.2;SARS-CoV_orf1ab;1145 | 0 | 0 |
| NC_004718.3;NP_828849.2;SARS-CoV_orf1ab;1146 | 0 | 0 |
| NC_004718.3;NP_828849.2;SARS-CoV_orf1ab;1147 | 0 | 0 |
| NC_004718.3;NP_828849.2;SARS-CoV_orf1ab;1148 | 0 | 0 |
| NC_004718.3;NP_828849.2;SARS-CoV_orf1ab;1149 | 0 | 0 |
| NC_004718.3;NP_828849.2;SARS-CoV_orf1ab;1150 | 0 | 0.025 |
| NC_004718.3;NP_828849.2;SARS-CoV_orf1ab;1151 | 0 | 0 |
| NC_004718.3;NP_828849.2;SARS-CoV_orf1ab;1152 | 0 | 0 |
| NC_004718.3;NP_828849.2;SARS-CoV_orf1ab;1153 | 0 | 0 |
| NC_004718.3;NP_828849.2;SARS-CoV_orf1ab;1154 | 0 | 0 |
| NC_004718.3;NP_828849.2;SARS-CoV_orf1ab;1155 | 0 | 0 |
| NC_004718.3;NP_828849.2;SARS-CoV_orf1ab;1156 | 0 | 0 |
| NC_004718.3;NP_828849.2;SARS-CoV_orf1ab;1157 | 0 | 0 |
| NC_004718.3;NP_828849.2;SARS-CoV_orf1ab;1158 | 0.05 | 0.05 |

| | | |
|---|---|---|
| NC_004718.3;NP_828849.2;SARS-CoV_orf1ab;1159 | 0 | 0.025 |
| NC_004718.3;NP_828849.2;SARS-CoV_orf1ab;1160 | 0 | 0.075 |
| NC_004718.3;NP_828849.2;SARS-CoV_orf1ab;1161 | 0 | 0.075 |
| NC_004718.3;NP_828849.2;SARS-CoV_orf1ab;1162 | 0 | 0.075 |
| NC_004718.3;NP_828849.2;SARS-CoV_orf1ab;1163 | 0 | 0.075 |
| NC_004718.3;NP_828849.2;SARS-CoV_orf1ab;1164 | 0.05 | 0.15 |
| NC_004718.3;NP_828849.2;SARS-CoV_orf1ab;1165 | 0.35 | 0.475 |
| NC_004718.3;NP_828849.2;SARS-CoV_orf1ab;1166 | 0.4 | 0.45 |
| NC_004718.3;NP_828849.2;SARS-CoV_orf1ab;1167 | 0.45 | 0.4 |
| NC_004718.3;NP_828849.2;SARS-CoV_orf1ab;1168 | 0.4 | 0.4 |
| NC_004718.3;NP_828849.2;SARS-CoV_orf1ab;1169 | 0.4 | 0.25 |
| NC_004718.3;NP_828849.2;SARS-CoV_orf1ab;1170 | 0.45 | 0.375 |
| NC_004718.3;NP_828849.2;SARS-CoV_orf1ab;1171 | 0.4 | 0.35 |
| NC_004718.3;NP_828849.2;SARS-CoV_orf1ab;1172 | 0.4 | 0.4 |
| NC_004718.3;NP_828849.2;SARS-CoV_orf1ab;1173 | 0.4 | 0.325 |
| NC_004718.3;NP_828849.2;SARS-CoV_orf1ab;1174 | 0 | 0.025 |
| NC_004718.3;NP_828849.2;SARS-CoV_orf1ab;1175 | 0 | 0.025 |
| NC_004718.3;NP_828849.2;SARS-CoV_orf1ab;1176 | 0 | 0.025 |
| NC_004718.3;NP_828849.2;SARS-CoV_orf1ab;1177 | 0 | 0 |
| NC_004718.3;NP_828849.2;SARS-CoV_orf1ab;1178 | 0 | 0 |
| NC_004718.3;NP_828849.2;SARS-CoV_orf1ab;1179 | 0 | 0 |
| NC_004718.3;NP_828849.2;SARS-CoV_orf1ab;1180 | 0 | 0 |
| NC_004718.3;NP_828849.2;SARS-CoV_orf1ab;1181 | 0 | 0 |
| NC_004718.3;NP_828849.2;SARS-CoV_orf1ab;1182 | 0 | 0 |
| NC_004718.3;NP_828849.2;SARS-CoV_orf1ab;1183 | 0 | 0 |
| NC_004718.3;NP_828849.2;SARS-CoV_orf1ab;1184 | 0 | 0 |
| NC_004718.3;NP_828849.2;SARS-CoV_orf1ab;1185 | 0 | 0 |
| NC_004718.3;NP_828849.2;SARS-CoV_orf1ab;1186 | 0 | 0 |
| NC_004718.3;NP_828849.2;SARS-CoV_orf1ab;1187 | 0 | 0 |
| NC_004718.3;NP_828849.2;SARS-CoV_orf1ab;1188 | 0 | 0 |
| NC_004718.3;NP_828849.2;SARS-CoV_orf1ab;1189 | 0 | 0 |
| NC_004718.3;NP_828849.2;SARS-CoV_orf1ab;1190 | 0.05 | 0 |
| NC_004718.3;NP_828849.2;SARS-CoV_orf1ab;1191 | 0.05 | 0 |
| NC_004718.3;NP_828849.2;SARS-CoV_orf1ab;1192 | 0.1 | 0.05 |
| NC_004718.3;NP_828849.2;SARS-CoV_orf1ab;1193 | 0 | 0.05 |
| NC_004718.3;NP_828849.2;SARS-CoV_orf1ab;1194 | 0 | 0.025 |
| NC_004718.3;NP_828849.2;SARS-CoV_orf1ab;1195 | 0 | 0.025 |
| NC_004718.3;NP_828849.2;SARS-CoV_orf1ab;1196 | 0.05 | 0.025 |
| NC_004718.3;NP_828849.2;SARS-CoV_orf1ab;1197 | 0.05 | 0.025 |
| NC_004718.3;NP_828849.2;SARS-CoV_orf1ab;1198 | 0.05 | 0.025 |
| NC_004718.3;NP_828849.2;SARS-CoV_orf1ab;1199 | 0.15 | 0.075 |
| NC_004718.3;NP_828849.2;SARS-CoV_orf1ab;1200 | 0.15 | 0.175 |
| NC_004718.3;NP_828849.2;SARS-CoV_orf1ab;1201 | 0.1 | 0.2 |

| | | |
|---|---|---|
| NC_004718.3;NP_828849.2;SARS-CoV_orf1ab;1202 | 0 | 0 |
| NC_004718.3;NP_828849.2;SARS-CoV_orf1ab;1203 | 0 | 0.025 |
| NC_004718.3;NP_828849.2;SARS-CoV_orf1ab;1204 | 0 | 0.025 |
| NC_004718.3;NP_828849.2;SARS-CoV_orf1ab;1205 | 0 | 0 |
| NC_004718.3;NP_828849.2;SARS-CoV_orf1ab;1206 | 0 | 0 |
| NC_004718.3;NP_828849.2;SARS-CoV_orf1ab;1207 | 0 | 0 |
| NC_004718.3;NP_828849.2;SARS-CoV_orf1ab;1208 | 0 | 0 |
| NC_004718.3;NP_828849.2;SARS-CoV_orf1ab;1209 | 0 | 0 |
| NC_004718.3;NP_828849.2;SARS-CoV_orf1ab;1210 | 0 | 0 |
| NC_004718.3;NP_828849.2;SARS-CoV_orf1ab;1211 | 0 | 0 |
| NC_004718.3;NP_828849.2;SARS-CoV_orf1ab;1212 | 0 | 0 |
| NC_004718.3;NP_828849.2;SARS-CoV_orf1ab;1213 | 0.05 | 0.025 |
| NC_004718.3;NP_828849.2;SARS-CoV_orf1ab;1214 | 0 | 0.025 |
| NC_004718.3;NP_828849.2;SARS-CoV_orf1ab;1215 | 0 | 0.025 |
| NC_004718.3;NP_828849.2;SARS-CoV_orf1ab;1216 | 0 | 0.025 |
| NC_004718.3;NP_828849.2;SARS-CoV_orf1ab;1217 | 0 | 0.025 |
| NC_004718.3;NP_828849.2;SARS-CoV_orf1ab;1218 | 0 | 0 |
| NC_004718.3;NP_828849.2;SARS-CoV_orf1ab;1219 | 0 | 0 |
| NC_004718.3;NP_828849.2;SARS-CoV_orf1ab;1220 | 0 | 0 |
| NC_004718.3;NP_828849.2;SARS-CoV_orf1ab;1221 | 0 | 0.025 |
| NC_004718.3;NP_828849.2;SARS-CoV_orf1ab;1222 | 0 | 0 |
| NC_004718.3;NP_828849.2;SARS-CoV_orf1ab;1223 | 0 | 0 |
| NC_004718.3;NP_828849.2;SARS-CoV_orf1ab;1224 | 0 | 0 |
| NC_004718.3;NP_828849.2;SARS-CoV_orf1ab;1225 | 0 | 0 |
| NC_004718.3;NP_828849.2;SARS-CoV_orf1ab;1226 | 0 | 0 |
| NC_004718.3;NP_828849.2;SARS-CoV_orf1ab;1227 | 0 | 0 |
| NC_004718.3;NP_828849.2;SARS-CoV_orf1ab;1228 | 0 | 0 |
| NC_004718.3;NP_828849.2;SARS-CoV_orf1ab;1229 | 0 | 0 |
| NC_004718.3;NP_828849.2;SARS-CoV_orf1ab;1230 | 0 | 0.05 |
| NC_004718.3;NP_828849.2;SARS-CoV_orf1ab;1231 | 0.15 | 0.05 |
| NC_004718.3;NP_828849.2;SARS-CoV_orf1ab;1232 | 0.1 | 0.075 |
| NC_004718.3;NP_828849.2;SARS-CoV_orf1ab;1233 | 0.05 | 0.05 |
| NC_004718.3;NP_828849.2;SARS-CoV_orf1ab;1234 | 0.05 | 0.025 |
| NC_004718.3;NP_828849.2;SARS-CoV_orf1ab;1235 | 0 | 0 |
| NC_004718.3;NP_828849.2;SARS-CoV_orf1ab;1236 | 0 | 0.025 |
| NC_004718.3;NP_828849.2;SARS-CoV_orf1ab;1237 | 0 | 0 |
| NC_004718.3;NP_828849.2;SARS-CoV_orf1ab;1238 | 0 | 0 |
| NC_004718.3;NP_828849.2;SARS-CoV_orf1ab;1239 | 0 | 0 |
| NC_004718.3;NP_828849.2;SARS-CoV_orf1ab;1240 | 0.05 | 0 |
| NC_004718.3;NP_828849.2;SARS-CoV_orf1ab;1241 | 0 | 0 |
| NC_004718.3;NP_828849.2;SARS-CoV_orf1ab;1242 | 0 | 0 |
| NC_004718.3;NP_828849.2;SARS-CoV_orf1ab;1243 | 0 | 0 |
| NC_004718.3;NP_828849.2;SARS-CoV_orf1ab;1244 | 0 | 0 |

| | | |
|---|---|---|
| NC_004718.3;NP_828849.2;SARS-CoV_orf1ab;1245 | 0 | 0 |
| NC_004718.3;NP_828849.2;SARS-CoV_orf1ab;1246 | 0 | 0.025 |
| NC_004718.3;NP_828849.2;SARS-CoV_orf1ab;1247 | 0 | 0.025 |
| NC_004718.3;NP_828849.2;SARS-CoV_orf1ab;1248 | 0 | 0 |
| NC_004718.3;NP_828849.2;SARS-CoV_orf1ab;1249 | 0 | 0 |
| NC_004718.3;NP_828849.2;SARS-CoV_orf1ab;1250 | 0.05 | 0 |
| NC_004718.3;NP_828849.2;SARS-CoV_orf1ab;1251 | 0 | 0 |
| NC_004718.3;NP_828849.2;SARS-CoV_orf1ab;1252 | 0 | 0 |
| NC_004718.3;NP_828849.2;SARS-CoV_orf1ab;1253 | 0 | 0 |
| NC_004718.3;NP_828849.2;SARS-CoV_orf1ab;1254 | 0 | 0 |
| NC_004718.3;NP_828849.2;SARS-CoV_orf1ab;1255 | 0 | 0 |
| NC_004718.3;NP_828849.2;SARS-CoV_orf1ab;1256 | 0 | 0 |
| NC_004718.3;NP_828849.2;SARS-CoV_orf1ab;1257 | 0 | 0 |
| NC_004718.3;NP_828849.2;SARS-CoV_orf1ab;1258 | 0 | 0 |
| NC_004718.3;NP_828849.2;SARS-CoV_orf1ab;1259 | 0 | 0 |
| NC_004718.3;NP_828849.2;SARS-CoV_orf1ab;1260 | 0 | 0 |
| NC_004718.3;NP_828849.2;SARS-CoV_orf1ab;1261 | 0 | 0 |
| NC_004718.3;NP_828849.2;SARS-CoV_orf1ab;1262 | 0 | 0 |
| NC_004718.3;NP_828849.2;SARS-CoV_orf1ab;1263 | 0 | 0 |
| NC_004718.3;NP_828849.2;SARS-CoV_orf1ab;1264 | 0 | 0.025 |
| NC_004718.3;NP_828849.2;SARS-CoV_orf1ab;1265 | 0 | 0 |
| NC_004718.3;NP_828849.2;SARS-CoV_orf1ab;1266 | 0 | 0 |
| NC_004718.3;NP_828849.2;SARS-CoV_orf1ab;1267 | 0 | 0 |
| NC_004718.3;NP_828849.2;SARS-CoV_orf1ab;1268 | 0 | 0 |
| NC_004718.3;NP_828849.2;SARS-CoV_orf1ab;1269 | 0 | 0 |
| NC_004718.3;NP_828849.2;SARS-CoV_orf1ab;1270 | 0 | 0 |
| NC_004718.3;NP_828849.2;SARS-CoV_orf1ab;1271 | 0 | 0 |
| NC_004718.3;NP_828849.2;SARS-CoV_orf1ab;1272 | 0.05 | 0 |
| NC_004718.3;NP_828849.2;SARS-CoV_orf1ab;1273 | 0.05 | 0.025 |
| NC_004718.3;NP_828849.2;SARS-CoV_orf1ab;1274 | 0.05 | 0.025 |
| NC_004718.3;NP_828849.2;SARS-CoV_orf1ab;1275 | 0.05 | 0.05 |
| NC_004718.3;NP_828849.2;SARS-CoV_orf1ab;1276 | 0.05 | 0.025 |
| NC_004718.3;NP_828849.2;SARS-CoV_orf1ab;1277 | 0.05 | 0.025 |
| NC_004718.3;NP_828849.2;SARS-CoV_orf1ab;1278 | 0.05 | 0 |
| NC_004718.3;NP_828849.2;SARS-CoV_orf1ab;1279 | 0 | 0 |
| NC_004718.3;NP_828849.2;SARS-CoV_orf1ab;1280 | 0 | 0 |
| NC_004718.3;NP_828849.2;SARS-CoV_orf1ab;1281 | 0 | 0 |
| NC_004718.3;NP_828849.2;SARS-CoV_orf1ab;1282 | 0 | 0 |
| NC_004718.3;NP_828849.2;SARS-CoV_orf1ab;1283 | 0 | 0 |
| NC_004718.3;NP_828849.2;SARS-CoV_orf1ab;1284 | 0 | 0 |
| NC_004718.3;NP_828849.2;SARS-CoV_orf1ab;1285 | 0 | 0 |
| NC_004718.3;NP_828849.2;SARS-CoV_orf1ab;1286 | 0 | 0.025 |
| NC_004718.3;NP_828849.2;SARS-CoV_orf1ab;1287 | 0.05 | 0.025 |

| | | |
|---|---|---|
| NC_004718.3;NP_828849.2;SARS-CoV_orf1ab;1288 | 0 | 0.025 |
| NC_004718.3;NP_828849.2;SARS-CoV_orf1ab;1289 | 0 | 0.025 |
| NC_004718.3;NP_828849.2;SARS-CoV_orf1ab;1290 | 0 | 0 |
| NC_004718.3;NP_828849.2;SARS-CoV_orf1ab;1291 | 0 | 0 |
| NC_004718.3;NP_828849.2;SARS-CoV_orf1ab;1292 | 0 | 0 |
| NC_004718.3;NP_828849.2;SARS-CoV_orf1ab;1293 | 0 | 0 |
| NC_004718.3;NP_828849.2;SARS-CoV_orf1ab;1294 | 0 | 0.025 |
| NC_004718.3;NP_828849.2;SARS-CoV_orf1ab;1295 | 0.05 | 0.025 |
| NC_004718.3;NP_828849.2;SARS-CoV_orf1ab;1296 | 0.1 | 0.025 |
| NC_004718.3;NP_828849.2;SARS-CoV_orf1ab;1297 | 0.05 | 0.025 |
| NC_004718.3;NP_828849.2;SARS-CoV_orf1ab;1298 | 0 | 0 |
| NC_004718.3;NP_828849.2;SARS-CoV_orf1ab;1299 | 0 | 0 |
| NC_004718.3;NP_828849.2;SARS-CoV_orf1ab;1300 | 0.05 | 0 |
| NC_004718.3;NP_828849.2;SARS-CoV_orf1ab;1301 | 0 | 0 |
| NC_004718.3;NP_828849.2;SARS-CoV_orf1ab;1302 | 0 | 0 |
| NC_004718.3;NP_828849.2;SARS-CoV_orf1ab;1303 | 0 | 0 |
| NC_004718.3;NP_828849.2;SARS-CoV_orf1ab;1304 | 0 | 0 |
| NC_004718.3;NP_828849.2;SARS-CoV_orf1ab;1305 | 0 | 0 |
| NC_004718.3;NP_828849.2;SARS-CoV_orf1ab;1306 | 0 | 0 |
| NC_004718.3;NP_828849.2;SARS-CoV_orf1ab;1307 | 0 | 0.025 |
| NC_004718.3;NP_828849.2;SARS-CoV_orf1ab;1308 | 0 | 0.025 |
| NC_004718.3;NP_828849.2;SARS-CoV_orf1ab;1309 | 0 | 0 |
| NC_004718.3;NP_828849.2;SARS-CoV_orf1ab;1310 | 0 | 0 |
| NC_004718.3;NP_828849.2;SARS-CoV_orf1ab;1311 | 0 | 0 |
| NC_004718.3;NP_828849.2;SARS-CoV_orf1ab;1312 | 0 | 0 |
| NC_004718.3;NP_828849.2;SARS-CoV_orf1ab;1313 | 0 | 0.025 |
| NC_004718.3;NP_828849.2;SARS-CoV_orf1ab;1314 | 0 | 0 |
| NC_004718.3;NP_828849.2;SARS-CoV_orf1ab;1315 | 0 | 0 |
| NC_004718.3;NP_828849.2;SARS-CoV_orf1ab;1316 | 0 | 0 |
| NC_004718.3;NP_828849.2;SARS-CoV_orf1ab;1317 | 0 | 0 |
| NC_004718.3;NP_828849.2;SARS-CoV_orf1ab;1318 | 0 | 0 |
| NC_004718.3;NP_828849.2;SARS-CoV_orf1ab;1319 | 0 | 0.025 |
| NC_004718.3;NP_828849.2;SARS-CoV_orf1ab;1320 | 0 | 0 |
| NC_004718.3;NP_828849.2;SARS-CoV_orf1ab;1321 | 0 | 0.025 |
| NC_004718.3;NP_828849.2;SARS-CoV_orf1ab;1322 | 0 | 0.025 |
| NC_004718.3;NP_828849.2;SARS-CoV_orf1ab;1323 | 0 | 0.05 |
| NC_004718.3;NP_828849.2;SARS-CoV_orf1ab;1324 | 0 | 0 |
| NC_004718.3;NP_828849.2;SARS-CoV_orf1ab;1325 | 0 | 0 |
| NC_004718.3;NP_828849.2;SARS-CoV_orf1ab;1326 | 0 | 0 |
| NC_004718.3;NP_828849.2;SARS-CoV_orf1ab;1327 | 0 | 0 |
| NC_004718.3;NP_828849.2;SARS-CoV_orf1ab;1328 | 0 | 0 |
| NC_004718.3;NP_828849.2;SARS-CoV_orf1ab;1329 | 0 | 0 |
| NC_004718.3;NP_828849.2;SARS-CoV_orf1ab;1330 | 0 | 0 |

| | | |
|---|---|---|
| NC_004718.3;NP_828849.2;SARS-CoV_orf1ab;1331 | 0 | 0 |
| NC_004718.3;NP_828849.2;SARS-CoV_orf1ab;1332 | 0 | 0 |
| NC_004718.3;NP_828849.2;SARS-CoV_orf1ab;1333 | 0 | 0 |
| NC_004718.3;NP_828849.2;SARS-CoV_orf1ab;1334 | 0 | 0 |
| NC_004718.3;NP_828849.2;SARS-CoV_orf1ab;1335 | 0.05 | 0 |
| NC_004718.3;NP_828849.2;SARS-CoV_orf1ab;1336 | 0 | 0 |
| NC_004718.3;NP_828849.2;SARS-CoV_orf1ab;1337 | 0 | 0 |
| NC_004718.3;NP_828849.2;SARS-CoV_orf1ab;1338 | 0 | 0 |
| NC_004718.3;NP_828849.2;SARS-CoV_orf1ab;1339 | 0 | 0 |
| NC_004718.3;NP_828849.2;SARS-CoV_orf1ab;1340 | 0 | 0 |
| NC_004718.3;NP_828849.2;SARS-CoV_orf1ab;1341 | 0.05 | 0.025 |
| NC_004718.3;NP_828849.2;SARS-CoV_orf1ab;1342 | 0 | 0 |
| NC_004718.3;NP_828849.2;SARS-CoV_orf1ab;1343 | 0 | 0 |
| NC_004718.3;NP_828849.2;SARS-CoV_orf1ab;1344 | 0 | 0 |
| NC_004718.3;NP_828849.2;SARS-CoV_orf1ab;1345 | 0 | 0.025 |
| NC_004718.3;NP_828849.2;SARS-CoV_orf1ab;1346 | 0 | 0 |
| NC_004718.3;NP_828849.2;SARS-CoV_orf1ab;1347 | 0 | 0 |
| NC_004718.3;NP_828849.2;SARS-CoV_orf1ab;1348 | 0 | 0.05 |
| NC_004718.3;NP_828849.2;SARS-CoV_orf1ab;1349 | 0 | 0.025 |
| NC_004718.3;NP_828849.2;SARS-CoV_orf1ab;1350 | 0 | 0 |
| NC_004718.3;NP_828849.2;SARS-CoV_orf1ab;1351 | 0 | 0.025 |
| NC_004718.3;NP_828849.2;SARS-CoV_orf1ab;1352 | 0 | 0 |
| NC_004718.3;NP_828849.2;SARS-CoV_orf1ab;1353 | 0 | 0 |
| NC_004718.3;NP_828849.2;SARS-CoV_orf1ab;1354 | 0 | 0 |
| NC_004718.3;NP_828849.2;SARS-CoV_orf1ab;1355 | 0 | 0 |
| NC_004718.3;NP_828849.2;SARS-CoV_orf1ab;1356 | 0 | 0.025 |
| NC_004718.3;NP_828849.2;SARS-CoV_orf1ab;1357 | 0 | 0 |
| NC_004718.3;NP_828849.2;SARS-CoV_orf1ab;1358 | 0.05 | 0.025 |
| NC_004718.3;NP_828849.2;SARS-CoV_orf1ab;1359 | 0.05 | 0.025 |
| NC_004718.3;NP_828849.2;SARS-CoV_orf1ab;1360 | 0 | 0 |
| NC_004718.3;NP_828849.2;SARS-CoV_orf1ab;1361 | 0 | 0 |
| NC_004718.3;NP_828849.2;SARS-CoV_orf1ab;1362 | 0 | 0 |
| NC_004718.3;NP_828849.2;SARS-CoV_orf1ab;1363 | 0 | 0 |
| NC_004718.3;NP_828849.2;SARS-CoV_orf1ab;1364 | 0 | 0 |
| NC_004718.3;NP_828849.2;SARS-CoV_orf1ab;1365 | 0 | 0 |
| NC_004718.3;NP_828849.2;SARS-CoV_orf1ab;1366 | 0 | 0 |
| NC_004718.3;NP_828849.2;SARS-CoV_orf1ab;1367 | 0 | 0 |
| NC_004718.3;NP_828849.2;SARS-CoV_orf1ab;1368 | 0 | 0 |
| NC_004718.3;NP_828849.2;SARS-CoV_orf1ab;1369 | 0 | 0 |
| NC_004718.3;NP_828849.2;SARS-CoV_orf1ab;1370 | 0 | 0 |
| NC_004718.3;NP_828849.2;SARS-CoV_orf1ab;1371 | 0 | 0 |
| NC_004718.3;NP_828849.2;SARS-CoV_orf1ab;1372 | 0.05 | 0 |
| NC_004718.3;NP_828849.2;SARS-CoV_orf1ab;1373 | 0.05 | 0 |

| | | |
|---|---|---|
| NC_004718.3;NP_828849.2;SARS-CoV_orf1ab;1374 | 0.1 | 0 |
| NC_004718.3;NP_828849.2;SARS-CoV_orf1ab;1375 | 0.1 | 0 |
| NC_004718.3;NP_828849.2;SARS-CoV_orf1ab;1376 | 0.15 | 0.025 |
| NC_004718.3;NP_828849.2;SARS-CoV_orf1ab;1377 | 0.1 | 0 |
| NC_004718.3;NP_828849.2;SARS-CoV_orf1ab;1378 | 0.1 | 0 |
| NC_004718.3;NP_828849.2;SARS-CoV_orf1ab;1379 | 0.1 | 0 |
| NC_004718.3;NP_828849.2;SARS-CoV_orf1ab;1380 | 0.1 | 0.025 |
| NC_004718.3;NP_828849.2;SARS-CoV_orf1ab;1381 | 0.1 | 0 |
| NC_004718.3;NP_828849.2;SARS-CoV_orf1ab;1382 | 0.05 | 0.025 |
| NC_004718.3;NP_828849.2;SARS-CoV_orf1ab;1383 | 0 | 0.025 |
| NC_004718.3;NP_828849.2;SARS-CoV_orf1ab;1384 | 0 | 0 |
| NC_004718.3;NP_828849.2;SARS-CoV_orf1ab;1385 | 0 | 0 |
| NC_004718.3;NP_828849.2;SARS-CoV_orf1ab;1386 | 0 | 0 |
| NC_004718.3;NP_828849.2;SARS-CoV_orf1ab;1387 | 0 | 0 |
| NC_004718.3;NP_828849.2;SARS-CoV_orf1ab;1388 | 0 | 0 |
| NC_004718.3;NP_828849.2;SARS-CoV_orf1ab;1389 | 0 | 0 |
| NC_004718.3;NP_828849.2;SARS-CoV_orf1ab;1390 | 0.05 | 0.075 |
| NC_004718.3;NP_828849.2;SARS-CoV_orf1ab;1391 | 0 | 0 |
| NC_004718.3;NP_828849.2;SARS-CoV_orf1ab;1392 | 0 | 0 |
| NC_004718.3;NP_828849.2;SARS-CoV_orf1ab;1393 | 0 | 0 |
| NC_004718.3;NP_828849.2;SARS-CoV_orf1ab;1394 | 0 | 0 |
| NC_004718.3;NP_828849.2;SARS-CoV_orf1ab;1395 | 0 | 0 |
| NC_004718.3;NP_828849.2;SARS-CoV_orf1ab;1396 | 0 | 0 |
| NC_004718.3;NP_828849.2;SARS-CoV_orf1ab;1397 | 0 | 0 |
| NC_004718.3;NP_828849.2;SARS-CoV_orf1ab;1398 | 0 | 0 |
| NC_004718.3;NP_828849.2;SARS-CoV_orf1ab;1399 | 0 | 0 |
| NC_004718.3;NP_828849.2;SARS-CoV_orf1ab;1400 | 0 | 0 |
| NC_004718.3;NP_828849.2;SARS-CoV_orf1ab;1401 | 0 | 0 |
| NC_004718.3;NP_828849.2;SARS-CoV_orf1ab;1402 | 0 | 0 |
| NC_004718.3;NP_828849.2;SARS-CoV_orf1ab;1403 | 0 | 0.025 |
| NC_004718.3;NP_828849.2;SARS-CoV_orf1ab;1404 | 0 | 0 |
| NC_004718.3;NP_828849.2;SARS-CoV_orf1ab;1405 | 0 | 0.025 |
| NC_004718.3;NP_828849.2;SARS-CoV_orf1ab;1406 | 0.05 | 0.025 |
| NC_004718.3;NP_828849.2;SARS-CoV_orf1ab;1407 | 0 | 0.025 |
| NC_004718.3;NP_828849.2;SARS-CoV_orf1ab;1408 | 0 | 0.025 |
| NC_004718.3;NP_828849.2;SARS-CoV_orf1ab;1409 | 0.05 | 0 |
| NC_004718.3;NP_828849.2;SARS-CoV_orf1ab;1410 | 0 | 0 |
| NC_004718.3;NP_828849.2;SARS-CoV_orf1ab;1411 | 0.1 | 0 |
| NC_004718.3;NP_828849.2;SARS-CoV_orf1ab;1412 | 0.15 | 0 |
| NC_004718.3;NP_828849.2;SARS-CoV_orf1ab;1413 | 0.05 | 0 |
| NC_004718.3;NP_828849.2;SARS-CoV_orf1ab;1414 | 0 | 0.025 |
| NC_004718.3;NP_828849.2;SARS-CoV_orf1ab;1415 | 0 | 0 |
| NC_004718.3;NP_828849.2;SARS-CoV_orf1ab;1416 | 0.1 | 0.025 |

| | | |
|---|---|---|
| NC_004718.3;NP_828849.2;SARS-CoV_orf1ab;1417 | 0 | 0 |
| NC_004718.3;NP_828849.2;SARS-CoV_orf1ab;1418 | 0 | 0 |
| NC_004718.3;NP_828849.2;SARS-CoV_orf1ab;1419 | 0 | 0 |
| NC_004718.3;NP_828849.2;SARS-CoV_orf1ab;1420 | 0 | 0 |
| NC_004718.3;NP_828849.2;SARS-CoV_orf1ab;1421 | 0 | 0 |
| NC_004718.3;NP_828849.2;SARS-CoV_orf1ab;1422 | 0 | 0.025 |
| NC_004718.3;NP_828849.2;SARS-CoV_orf1ab;1423 | 0 | 0.025 |
| NC_004718.3;NP_828849.2;SARS-CoV_orf1ab;1424 | 0.05 | 0.1 |
| NC_004718.3;NP_828849.2;SARS-CoV_orf1ab;1425 | 0 | 0.05 |
| NC_004718.3;NP_828849.2;SARS-CoV_orf1ab;1426 | 0 | 0.05 |
| NC_004718.3;NP_828849.2;SARS-CoV_orf1ab;1427 | 0 | 0.05 |
| NC_004718.3;NP_828849.2;SARS-CoV_orf1ab;1428 | 0 | 0.05 |
| NC_004718.3;NP_828849.2;SARS-CoV_orf1ab;1429 | 0 | 0.05 |
| NC_004718.3;NP_828849.2;SARS-CoV_orf1ab;1430 | 0 | 0.05 |
| NC_004718.3;NP_828849.2;SARS-CoV_orf1ab;1431 | 0 | 0.025 |
| NC_004718.3;NP_828849.2;SARS-CoV_orf1ab;1432 | 0 | 0.025 |
| NC_004718.3;NP_828849.2;SARS-CoV_orf1ab;1433 | 0 | 0 |
| NC_004718.3;NP_828849.2;SARS-CoV_orf1ab;1434 | 0.05 | 0 |
| NC_004718.3;NP_828849.2;SARS-CoV_orf1ab;1435 | 0 | 0 |
| NC_004718.3;NP_828849.2;SARS-CoV_orf1ab;1436 | 0 | 0 |
| NC_004718.3;NP_828849.2;SARS-CoV_orf1ab;1437 | 0 | 0.025 |
| NC_004718.3;NP_828849.2;SARS-CoV_orf1ab;1438 | 0 | 0 |
| NC_004718.3;NP_828849.2;SARS-CoV_orf1ab;1439 | 0 | 0 |
| NC_004718.3;NP_828849.2;SARS-CoV_orf1ab;1440 | 0 | 0 |
| NC_004718.3;NP_828849.2;SARS-CoV_orf1ab;1441 | 0 | 0 |
| NC_004718.3;NP_828849.2;SARS-CoV_orf1ab;1442 | 0 | 0 |
| NC_004718.3;NP_828849.2;SARS-CoV_orf1ab;1443 | 0 | 0.025 |
| NC_004718.3;NP_828849.2;SARS-CoV_orf1ab;1444 | 0 | 0 |
| NC_004718.3;NP_828849.2;SARS-CoV_orf1ab;1445 | 0 | 0 |
| NC_004718.3;NP_828849.2;SARS-CoV_orf1ab;1446 | 0 | 0 |
| NC_004718.3;NP_828849.2;SARS-CoV_orf1ab;1447 | 0 | 0 |
| NC_004718.3;NP_828849.2;SARS-CoV_orf1ab;1448 | 0.05 | 0.05 |
| NC_004718.3;NP_828849.2;SARS-CoV_orf1ab;1449 | 0 | 0.025 |
| NC_004718.3;NP_828849.2;SARS-CoV_orf1ab;1450 | 0 | 0.075 |
| NC_004718.3;NP_828849.2;SARS-CoV_orf1ab;1451 | 0 | 0.075 |
| NC_004718.3;NP_828849.2;SARS-CoV_orf1ab;1452 | 0 | 0.025 |
| NC_004718.3;NP_828849.2;SARS-CoV_orf1ab;1453 | 0 | 0 |
| NC_004718.3;NP_828849.2;SARS-CoV_orf1ab;1454 | 0 | 0 |
| NC_004718.3;NP_828849.2;SARS-CoV_orf1ab;1455 | 0 | 0 |
| NC_004718.3;NP_828849.2;SARS-CoV_orf1ab;1456 | 0.05 | 0.075 |
| NC_004718.3;NP_828849.2;SARS-CoV_orf1ab;1457 | 0.1 | 0.05 |
| NC_004718.3;NP_828849.2;SARS-CoV_orf1ab;1458 | 0.05 | 0.05 |
| NC_004718.3;NP_828849.2;SARS-CoV_orf1ab;1459 | 0.1 | 0.15 |

| | | |
|---|---|---|
| NC_004718.3;NP_828849.2;SARS-CoV_orf1ab;1460 | 0.05 | 0.1 |
| NC_004718.3;NP_828849.2;SARS-CoV_orf1ab;1461 | 0.05 | 0.025 |
| NC_004718.3;NP_828849.2;SARS-CoV_orf1ab;1462 | 0.05 | 0.1 |
| NC_004718.3;NP_828849.2;SARS-CoV_orf1ab;1463 | 0.05 | 0.075 |
| NC_004718.3;NP_828849.2;SARS-CoV_orf1ab;1464 | 0.05 | 0.025 |
| NC_004718.3;NP_828849.2;SARS-CoV_orf1ab;1465 | 0.05 | 0.025 |
| NC_004718.3;NP_828849.2;SARS-CoV_orf1ab;1466 | 0.05 | 0.025 |
| NC_004718.3;NP_828849.2;SARS-CoV_orf1ab;1467 | 0 | 0 |
| NC_004718.3;NP_828849.2;SARS-CoV_orf1ab;1468 | 0.05 | 0.025 |
| NC_004718.3;NP_828849.2;SARS-CoV_orf1ab;1469 | 0.05 | 0.05 |
| NC_004718.3;NP_828849.2;SARS-CoV_orf1ab;1470 | 0.05 | 0 |
| NC_004718.3;NP_828849.2;SARS-CoV_orf1ab;1471 | 0.05 | 0 |
| NC_004718.3;NP_828849.2;SARS-CoV_orf1ab;1472 | 0 | 0 |
| NC_004718.3;NP_828849.2;SARS-CoV_orf1ab;1473 | 0 | 0 |
| NC_004718.3;NP_828849.2;SARS-CoV_orf1ab;1474 | 0 | 0 |
| NC_004718.3;NP_828849.2;SARS-CoV_orf1ab;1475 | 0 | 0 |
| NC_004718.3;NP_828849.2;SARS-CoV_orf1ab;1476 | 0 | 0 |
| NC_004718.3;NP_828849.2;SARS-CoV_orf1ab;1477 | 0 | 0 |
| NC_004718.3;NP_828849.2;SARS-CoV_orf1ab;1478 | 0 | 0 |
| NC_004718.3;NP_828849.2;SARS-CoV_orf1ab;1479 | 0 | 0 |
| NC_004718.3;NP_828849.2;SARS-CoV_orf1ab;1480 | 0 | 0 |
| NC_004718.3;NP_828849.2;SARS-CoV_orf1ab;1481 | 0 | 0 |
| NC_004718.3;NP_828849.2;SARS-CoV_orf1ab;1482 | 0 | 0 |
| NC_004718.3;NP_828849.2;SARS-CoV_orf1ab;1483 | 0 | 0 |
| NC_004718.3;NP_828849.2;SARS-CoV_orf1ab;1484 | 0 | 0 |
| NC_004718.3;NP_828849.2;SARS-CoV_orf1ab;1485 | 0.05 | 0.025 |
| NC_004718.3;NP_828849.2;SARS-CoV_orf1ab;1486 | 0 | 0 |
| NC_004718.3;NP_828849.2;SARS-CoV_orf1ab;1487 | 0 | 0.025 |
| NC_004718.3;NP_828849.2;SARS-CoV_orf1ab;1488 | 0 | 0 |
| NC_004718.3;NP_828849.2;SARS-CoV_orf1ab;1489 | 0 | 0 |
| NC_004718.3;NP_828849.2;SARS-CoV_orf1ab;1490 | 0 | 0 |
| NC_004718.3;NP_828849.2;SARS-CoV_orf1ab;1491 | 0 | 0 |
| NC_004718.3;NP_828849.2;SARS-CoV_orf1ab;1492 | 0 | 0 |
| NC_004718.3;NP_828849.2;SARS-CoV_orf1ab;1493 | 0 | 0 |
| NC_004718.3;NP_828849.2;SARS-CoV_orf1ab;1494 | 0 | 0.075 |
| NC_004718.3;NP_828849.2;SARS-CoV_orf1ab;1495 | 0 | 0.075 |
| NC_004718.3;NP_828849.2;SARS-CoV_orf1ab;1496 | 0 | 0.075 |
| NC_004718.3;NP_828849.2;SARS-CoV_orf1ab;1497 | 0 | 0.05 |
| NC_004718.3;NP_828849.2;SARS-CoV_orf1ab;1498 | 0 | 0.05 |
| NC_004718.3;NP_828849.2;SARS-CoV_orf1ab;1499 | 0 | 0.05 |
| NC_004718.3;NP_828849.2;SARS-CoV_orf1ab;1500 | 0 | 0.05 |
| NC_004718.3;NP_828849.2;SARS-CoV_orf1ab;1501 | 0 | 0.075 |
| NC_004718.3;NP_828849.2;SARS-CoV_orf1ab;1502 | 0 | 0.05 |

| | | |
|---|---|---|
| NC_004718.3;NP_828849.2;SARS-CoV_orf1ab;1503 | 0.05 | 0.05 |
| NC_004718.3;NP_828849.2;SARS-CoV_orf1ab;1504 | 0.05 | 0.025 |
| NC_004718.3;NP_828849.2;SARS-CoV_orf1ab;1505 | 0.05 | 0 |
| NC_004718.3;NP_828849.2;SARS-CoV_orf1ab;1506 | 0.05 | 0 |
| NC_004718.3;NP_828849.2;SARS-CoV_orf1ab;1507 | 0 | 0 |
| NC_004718.3;NP_828849.2;SARS-CoV_orf1ab;1508 | 0.05 | 0 |
| NC_004718.3;NP_828849.2;SARS-CoV_orf1ab;1509 | 0.05 | 0.025 |
| NC_004718.3;NP_828849.2;SARS-CoV_orf1ab;1510 | 0 | 0.025 |
| NC_004718.3;NP_828849.2;SARS-CoV_orf1ab;1511 | 0 | 0 |
| NC_004718.3;NP_828849.2;SARS-CoV_orf1ab;1512 | 0 | 0 |
| NC_004718.3;NP_828849.2;SARS-CoV_orf1ab;1513 | 0 | 0 |
| NC_004718.3;NP_828849.2;SARS-CoV_orf1ab;1514 | 0 | 0 |
| NC_004718.3;NP_828849.2;SARS-CoV_orf1ab;1515 | 0 | 0 |
| NC_004718.3;NP_828849.2;SARS-CoV_orf1ab;1516 | 0.05 | 0 |
| NC_004718.3;NP_828849.2;SARS-CoV_orf1ab;1517 | 0.45 | 0.35 |
| NC_004718.3;NP_828849.2;SARS-CoV_orf1ab;1518 | 0.15 | 0.125 |
| NC_004718.3;NP_828849.2;SARS-CoV_orf1ab;1519 | 0 | 0.025 |
| NC_004718.3;NP_828849.2;SARS-CoV_orf1ab;1520 | 0 | 0.025 |
| NC_004718.3;NP_828849.2;SARS-CoV_orf1ab;1521 | 0 | 0.025 |
| NC_004718.3;NP_828849.2;SARS-CoV_orf1ab;1522 | 0 | 0.025 |
| NC_004718.3;NP_828849.2;SARS-CoV_orf1ab;1523 | 0 | 0.025 |
| NC_004718.3;NP_828849.2;SARS-CoV_orf1ab;1524 | 0 | 0 |
| NC_004718.3;NP_828849.2;SARS-CoV_orf1ab;1525 | 0 | 0 |
| NC_004718.3;NP_828849.2;SARS-CoV_orf1ab;1526 | 0 | 0 |
| NC_004718.3;NP_828849.2;SARS-CoV_orf1ab;1527 | 0 | 0 |
| NC_004718.3;NP_828849.2;SARS-CoV_orf1ab;1528 | 0 | 0 |
| NC_004718.3;NP_828849.2;SARS-CoV_orf1ab;1529 | 0 | 0 |
| NC_004718.3;NP_828849.2;SARS-CoV_orf1ab;1530 | 0 | 0 |
| NC_004718.3;NP_828849.2;SARS-CoV_orf1ab;1531 | 0 | 0 |
| NC_004718.3;NP_828849.2;SARS-CoV_orf1ab;1532 | 0 | 0 |
| NC_004718.3;NP_828849.2;SARS-CoV_orf1ab;1533 | 0 | 0 |
| NC_004718.3;NP_828849.2;SARS-CoV_orf1ab;1534 | 0 | 0 |
| NC_004718.3;NP_828849.2;SARS-CoV_orf1ab;1535 | 0 | 0 |
| NC_004718.3;NP_828849.2;SARS-CoV_orf1ab;1536 | 0 | 0 |
| NC_004718.3;NP_828849.2;SARS-CoV_orf1ab;1537 | 0 | 0 |
| NC_004718.3;NP_828849.2;SARS-CoV_orf1ab;1538 | 0 | 0 |
| NC_004718.3;NP_828849.2;SARS-CoV_orf1ab;1539 | 0 | 0 |
| NC_004718.3;NP_828849.2;SARS-CoV_orf1ab;1540 | 0 | 0.025 |
| NC_004718.3;NP_828849.2;SARS-CoV_orf1ab;1541 | 0 | 0.025 |
| NC_004718.3;NP_828849.2;SARS-CoV_orf1ab;1542 | 0.05 | 0 |
| NC_004718.3;NP_828849.2;SARS-CoV_orf1ab;1543 | 0 | 0 |
| NC_004718.3;NP_828849.2;SARS-CoV_orf1ab;1544 | 0 | 0.025 |
| NC_004718.3;NP_828849.2;SARS-CoV_orf1ab;1545 | 0.05 | 0.025 |

| | | |
|---|---|---|
| NC_004718.3;NP_828849.2;SARS-CoV_orf1ab;1546 | 0.05 | 0.025 |
| NC_004718.3;NP_828849.2;SARS-CoV_orf1ab;1547 | 0 | 0.075 |
| NC_004718.3;NP_828849.2;SARS-CoV_orf1ab;1548 | 0 | 0.075 |
| NC_004718.3;NP_828849.2;SARS-CoV_orf1ab;1549 | 0 | 0.075 |
| NC_004718.3;NP_828849.2;SARS-CoV_orf1ab;1550 | 0 | 0.1 |
| NC_004718.3;NP_828849.2;SARS-CoV_orf1ab;1551 | 0 | 0.1 |
| NC_004718.3;NP_828849.2;SARS-CoV_orf1ab;1552 | 0 | 0.05 |
| NC_004718.3;NP_828849.2;SARS-CoV_orf1ab;1553 | 0 | 0.025 |
| NC_004718.3;NP_828849.2;SARS-CoV_orf1ab;1554 | 0 | 0.025 |
| NC_004718.3;NP_828849.2;SARS-CoV_orf1ab;1555 | 0 | 0.025 |
| NC_004718.3;NP_828849.2;SARS-CoV_orf1ab;1556 | 0 | 0.025 |
| NC_004718.3;NP_828849.2;SARS-CoV_orf1ab;1557 | 0 | 0 |
| NC_004718.3;NP_828849.2;SARS-CoV_orf1ab;1558 | 0 | 0 |
| NC_004718.3;NP_828849.2;SARS-CoV_orf1ab;1559 | 0 | 0 |
| NC_004718.3;NP_828849.2;SARS-CoV_orf1ab;1560 | 0.2 | 0.125 |
| NC_004718.3;NP_828849.2;SARS-CoV_orf1ab;1561 | 0 | 0 |
| NC_004718.3;NP_828849.2;SARS-CoV_orf1ab;1562 | 0 | 0 |
| NC_004718.3;NP_828849.2;SARS-CoV_orf1ab;1563 | 0 | 0 |
| NC_004718.3;NP_828849.2;SARS-CoV_orf1ab;1564 | 0 | 0 |
| NC_004718.3;NP_828849.2;SARS-CoV_orf1ab;1565 | 0 | 0 |
| NC_004718.3;NP_828849.2;SARS-CoV_orf1ab;1566 | 0 | 0 |
| NC_004718.3;NP_828849.2;SARS-CoV_orf1ab;1567 | 0 | 0 |
| NC_004718.3;NP_828849.2;SARS-CoV_orf1ab;1568 | 0 | 0.025 |
| NC_004718.3;NP_828849.2;SARS-CoV_orf1ab;1569 | 0 | 0.025 |
| NC_004718.3;NP_828849.2;SARS-CoV_orf1ab;1570 | 0 | 0.025 |
| NC_004718.3;NP_828849.2;SARS-CoV_orf1ab;1571 | 0 | 0.025 |
| NC_004718.3;NP_828849.2;SARS-CoV_orf1ab;1572 | 0.15 | 0.125 |
| NC_004718.3;NP_828849.2;SARS-CoV_orf1ab;1573 | 0 | 0.025 |
| NC_004718.3;NP_828849.2;SARS-CoV_orf1ab;1574 | 0.05 | 0 |
| NC_004718.3;NP_828849.2;SARS-CoV_orf1ab;1575 | 0 | 0 |
| NC_004718.3;NP_828849.2;SARS-CoV_orf1ab;1576 | 0 | 0 |
| NC_004718.3;NP_828849.2;SARS-CoV_orf1ab;1577 | 0 | 0 |
| NC_004718.3;NP_828849.2;SARS-CoV_orf1ab;1578 | 0 | 0.025 |
| NC_004718.3;NP_828849.2;SARS-CoV_orf1ab;1579 | 0.05 | 0 |
| NC_004718.3;NP_828849.2;SARS-CoV_orf1ab;1580 | 0 | 0 |
| NC_004718.3;NP_828849.2;SARS-CoV_orf1ab;1581 | 0 | 0 |
| NC_004718.3;NP_828849.2;SARS-CoV_orf1ab;1582 | 0 | 0 |
| NC_004718.3;NP_828849.2;SARS-CoV_orf1ab;1583 | 0 | 0 |
| NC_004718.3;NP_828849.2;SARS-CoV_orf1ab;1584 | 0 | 0 |
| NC_004718.3;NP_828849.2;SARS-CoV_orf1ab;1585 | 0 | 0 |
| NC_004718.3;NP_828849.2;SARS-CoV_orf1ab;1586 | 0 | 0 |
| NC_004718.3;NP_828849.2;SARS-CoV_orf1ab;1587 | 0 | 0 |
| NC_004718.3;NP_828849.2;SARS-CoV_orf1ab;1588 | 0 | 0 |

| | | |
|---|---|---|
| NC_004718.3;NP_828849.2;SARS-CoV_orf1ab;1589 | 0 | 0 |
| NC_004718.3;NP_828849.2;SARS-CoV_orf1ab;1590 | 0 | 0 |
| NC_004718.3;NP_828849.2;SARS-CoV_orf1ab;1591 | 0 | 0 |
| NC_004718.3;NP_828849.2;SARS-CoV_orf1ab;1592 | 0 | 0 |
| NC_004718.3;NP_828849.2;SARS-CoV_orf1ab;1593 | 0 | 0 |
| NC_004718.3;NP_828849.2;SARS-CoV_orf1ab;1594 | 0 | 0 |
| NC_004718.3;NP_828849.2;SARS-CoV_orf1ab;1595 | 0 | 0 |
| NC_004718.3;NP_828849.2;SARS-CoV_orf1ab;1596 | 0 | 0 |
| NC_004718.3;NP_828849.2;SARS-CoV_orf1ab;1597 | 0 | 0 |
| NC_004718.3;NP_828849.2;SARS-CoV_orf1ab;1598 | 0 | 0 |
| NC_004718.3;NP_828849.2;SARS-CoV_orf1ab;1599 | 0 | 0 |
| NC_004718.3;NP_828849.2;SARS-CoV_orf1ab;1600 | 0 | 0 |
| NC_004718.3;NP_828849.2;SARS-CoV_orf1ab;1601 | 0 | 0 |
| NC_004718.3;NP_828849.2;SARS-CoV_orf1ab;1602 | 0 | 0 |
| NC_004718.3;NP_828849.2;SARS-CoV_orf1ab;1603 | 0 | 0 |
| NC_004718.3;NP_828849.2;SARS-CoV_orf1ab;1604 | 0 | 0 |
| NC_004718.3;NP_828849.2;SARS-CoV_orf1ab;1605 | 0 | 0.025 |
| NC_004718.3;NP_828849.2;SARS-CoV_orf1ab;1606 | 0 | 0.025 |
| NC_004718.3;NP_828849.2;SARS-CoV_orf1ab;1607 | 0 | 0.025 |
| NC_004718.3;NP_828849.2;SARS-CoV_orf1ab;1608 | 0.05 | 0.025 |
| NC_004718.3;NP_828849.2;SARS-CoV_orf1ab;1609 | 0.05 | 0.025 |
| NC_004718.3;NP_828849.2;SARS-CoV_orf1ab;1610 | 0.05 | 0.025 |
| NC_004718.3;NP_828849.2;SARS-CoV_orf1ab;1611 | 0.05 | 0.025 |
| NC_004718.3;NP_828849.2;SARS-CoV_orf1ab;1612 | 0.05 | 0 |
| NC_004718.3;NP_828849.2;SARS-CoV_orf1ab;1613 | 0.05 | 0.025 |
| NC_004718.3;NP_828849.2;SARS-CoV_orf1ab;1614 | 0.05 | 0.05 |
| NC_004718.3;NP_828849.2;SARS-CoV_orf1ab;1615 | 0.05 | 0 |
| NC_004718.3;NP_828849.2;SARS-CoV_orf1ab;1616 | 0.05 | 0 |
| NC_004718.3;NP_828849.2;SARS-CoV_orf1ab;1617 | 0 | 0 |
| NC_004718.3;NP_828849.2;SARS-CoV_orf1ab;1618 | 0 | 0 |
| NC_004718.3;NP_828849.2;SARS-CoV_orf1ab;1619 | 0 | 0 |
| NC_004718.3;NP_828849.2;SARS-CoV_orf1ab;1620 | 0 | 0 |
| NC_004718.3;NP_828849.2;SARS-CoV_orf1ab;1621 | 0 | 0 |
| NC_004718.3;NP_828849.2;SARS-CoV_orf1ab;1622 | 0 | 0 |
| NC_004718.3;NP_828849.2;SARS-CoV_orf1ab;1623 | 0 | 0 |
| NC_004718.3;NP_828849.2;SARS-CoV_orf1ab;1624 | 0 | 0 |
| NC_004718.3;NP_828849.2;SARS-CoV_orf1ab;1625 | 0.05 | 0 |
| NC_004718.3;NP_828849.2;SARS-CoV_orf1ab;1626 | 0 | 0 |
| NC_004718.3;NP_828849.2;SARS-CoV_orf1ab;1627 | 0 | 0 |
| NC_004718.3;NP_828849.2;SARS-CoV_orf1ab;1628 | 0 | 0 |
| NC_004718.3;NP_828849.2;SARS-CoV_orf1ab;1629 | 0 | 0 |
| NC_004718.3;NP_828849.2;SARS-CoV_orf1ab;1630 | 0 | 0 |
| NC_004718.3;NP_828849.2;SARS-CoV_orf1ab;1631 | 0 | 0 |

| | | |
|---|---|---|
| NC_004718.3;NP_828849.2;SARS-CoV_orf1ab;1632 | 0 | 0 |
| NC_004718.3;NP_828849.2;SARS-CoV_orf1ab;1633 | 0 | 0 |
| NC_004718.3;NP_828849.2;SARS-CoV_orf1ab;1634 | 0.05 | 0 |
| NC_004718.3;NP_828849.2;SARS-CoV_orf1ab;1635 | 0.05 | 0 |
| NC_004718.3;NP_828849.2;SARS-CoV_orf1ab;1636 | 0.05 | 0 |
| NC_004718.3;NP_828849.2;SARS-CoV_orf1ab;1637 | 0.05 | 0 |
| NC_004718.3;NP_828849.2;SARS-CoV_orf1ab;1638 | 0.05 | 0 |
| NC_004718.3;NP_828849.2;SARS-CoV_orf1ab;1639 | 0.05 | 0 |
| NC_004718.3;NP_828849.2;SARS-CoV_orf1ab;1640 | 0.05 | 0 |
| NC_004718.3;NP_828849.2;SARS-CoV_orf1ab;1641 | 0.05 | 0 |
| NC_004718.3;NP_828849.2;SARS-CoV_orf1ab;1642 | 0 | 0 |
| NC_004718.3;NP_828849.2;SARS-CoV_orf1ab;1643 | 0 | 0 |
| NC_004718.3;NP_828849.2;SARS-CoV_orf1ab;1644 | 0 | 0 |
| NC_004718.3;NP_828849.2;SARS-CoV_orf1ab;1645 | 0 | 0 |
| NC_004718.3;NP_828849.2;SARS-CoV_orf1ab;1646 | 0 | 0 |
| NC_004718.3;NP_828849.2;SARS-CoV_orf1ab;1647 | 0 | 0 |
| NC_004718.3;NP_828849.2;SARS-CoV_orf1ab;1648 | 0 | 0 |
| NC_004718.3;NP_828849.2;SARS-CoV_orf1ab;1649 | 0 | 0 |
| NC_004718.3;NP_828849.2;SARS-CoV_orf1ab;1650 | 0 | 0 |
| NC_004718.3;NP_828849.2;SARS-CoV_orf1ab;1651 | 0 | 0 |
| NC_004718.3;NP_828849.2;SARS-CoV_orf1ab;1652 | 0 | 0 |
| NC_004718.3;NP_828849.2;SARS-CoV_orf1ab;1653 | 0 | 0 |
| NC_004718.3;NP_828849.2;SARS-CoV_orf1ab;1654 | 0 | 0 |
| NC_004718.3;NP_828849.2;SARS-CoV_orf1ab;1655 | 0 | 0 |
| NC_004718.3;NP_828849.2;SARS-CoV_orf1ab;1656 | 0 | 0.025 |
| NC_004718.3;NP_828849.2;SARS-CoV_orf1ab;1657 | 0 | 0 |
| NC_004718.3;NP_828849.2;SARS-CoV_orf1ab;1658 | 0 | 0.025 |
| NC_004718.3;NP_828849.2;SARS-CoV_orf1ab;1659 | 0 | 0 |
| NC_004718.3;NP_828849.2;SARS-CoV_orf1ab;1660 | 0 | 0 |
| NC_004718.3;NP_828849.2;SARS-CoV_orf1ab;1661 | 0.05 | 0 |
| NC_004718.3;NP_828849.2;SARS-CoV_orf1ab;1662 | 0.1 | 0.05 |
| NC_004718.3;NP_828849.2;SARS-CoV_orf1ab;1663 | 0 | 0.075 |
| NC_004718.3;NP_828849.2;SARS-CoV_orf1ab;1664 | 0.1 | 0.05 |
| NC_004718.3;NP_828849.2;SARS-CoV_orf1ab;1665 | 0 | 0 |
| NC_004718.3;NP_828849.2;SARS-CoV_orf1ab;1666 | 0 | 0 |
| NC_004718.3;NP_828849.2;SARS-CoV_orf1ab;1667 | 0 | 0.025 |
| NC_004718.3;NP_828849.2;SARS-CoV_orf1ab;1668 | 0.05 | 0 |
| NC_004718.3;NP_828849.2;SARS-CoV_orf1ab;1669 | 0.05 | 0.025 |
| NC_004718.3;NP_828849.2;SARS-CoV_orf1ab;1670 | 0.1 | 0.075 |
| NC_004718.3;NP_828849.2;SARS-CoV_orf1ab;1671 | 0.05 | 0.05 |
| NC_004718.3;NP_828849.2;SARS-CoV_orf1ab;1672 | 0.05 | 0.025 |
| NC_004718.3;NP_828849.2;SARS-CoV_orf1ab;1673 | 0 | 0.025 |
| NC_004718.3;NP_828849.2;SARS-CoV_orf1ab;1674 | 0 | 0.025 |

| | | |
|---|---|---|
| NC_004718.3;NP_828849.2;SARS-CoV_orf1ab;1675 | 0 | 0 |
| NC_004718.3;NP_828849.2;SARS-CoV_orf1ab;1676 | 0 | 0 |
| NC_004718.3;NP_828849.2;SARS-CoV_orf1ab;1677 | 0 | 0 |
| NC_004718.3;NP_828849.2;SARS-CoV_orf1ab;1678 | 0 | 0 |
| NC_004718.3;NP_828849.2;SARS-CoV_orf1ab;1679 | 0 | 0 |
| NC_004718.3;NP_828849.2;SARS-CoV_orf1ab;1680 | 0 | 0 |
| NC_004718.3;NP_828849.2;SARS-CoV_orf1ab;1681 | 0 | 0 |
| NC_004718.3;NP_828849.2;SARS-CoV_orf1ab;1682 | 0 | 0 |
| NC_004718.3;NP_828849.2;SARS-CoV_orf1ab;1683 | 0 | 0 |
| NC_004718.3;NP_828849.2;SARS-CoV_orf1ab;1684 | 0 | 0 |
| NC_004718.3;NP_828849.2;SARS-CoV_orf1ab;1685 | 0 | 0 |
| NC_004718.3;NP_828849.2;SARS-CoV_orf1ab;1686 | 0 | 0 |
| NC_004718.3;NP_828849.2;SARS-CoV_orf1ab;1687 | 0 | 0 |
| NC_004718.3;NP_828849.2;SARS-CoV_orf1ab;1688 | 0 | 0 |
| NC_004718.3;NP_828849.2;SARS-CoV_orf1ab;1689 | 0 | 0 |
| NC_004718.3;NP_828849.2;SARS-CoV_orf1ab;1690 | 0.1 | 0 |
| NC_004718.3;NP_828849.2;SARS-CoV_orf1ab;1691 | 0.15 | 0.025 |
| NC_004718.3;NP_828849.2;SARS-CoV_orf1ab;1692 | 0.05 | 0 |
| NC_004718.3;NP_828849.2;SARS-CoV_orf1ab;1693 | 0 | 0.025 |
| NC_004718.3;NP_828849.2;SARS-CoV_orf1ab;1694 | 0.05 | 0.025 |
| NC_004718.3;NP_828849.2;SARS-CoV_orf1ab;1695 | 0 | 0 |
| NC_004718.3;NP_828849.2;SARS-CoV_orf1ab;1696 | 0 | 0 |
| NC_004718.3;NP_828849.2;SARS-CoV_orf1ab;1697 | 0 | 0.025 |
| NC_004718.3;NP_828849.2;SARS-CoV_orf1ab;1698 | 0 | 0 |
| NC_004718.3;NP_828849.2;SARS-CoV_orf1ab;1699 | 0 | 0 |
| NC_004718.3;NP_828849.2;SARS-CoV_orf1ab;1700 | 0.05 | 0.025 |
| NC_004718.3;NP_828849.2;SARS-CoV_orf1ab;1701 | 0 | 0 |
| NC_004718.3;NP_828849.2;SARS-CoV_orf1ab;1702 | 0 | 0 |
| NC_004718.3;NP_828849.2;SARS-CoV_orf1ab;1703 | 0.05 | 0 |
| NC_004718.3;NP_828849.2;SARS-CoV_orf1ab;1704 | 0 | 0 |
| NC_004718.3;NP_828849.2;SARS-CoV_orf1ab;1705 | 0 | 0 |
| NC_004718.3;NP_828849.2;SARS-CoV_orf1ab;1706 | 0 | 0 |
| NC_004718.3;NP_828849.2;SARS-CoV_orf1ab;1707 | 0 | 0 |
| NC_004718.3;NP_828849.2;SARS-CoV_orf1ab;1708 | 0 | 0 |
| NC_004718.3;NP_828849.2;SARS-CoV_orf1ab;1709 | 0 | 0 |
| NC_004718.3;NP_828849.2;SARS-CoV_orf1ab;1710 | 0 | 0 |
| NC_004718.3;NP_828849.2;SARS-CoV_orf1ab;1711 | 0 | 0 |
| NC_004718.3;NP_828849.2;SARS-CoV_orf1ab;1712 | 0 | 0 |
| NC_004718.3;NP_828849.2;SARS-CoV_orf1ab;1713 | 0 | 0 |
| NC_004718.3;NP_828849.2;SARS-CoV_orf1ab;1714 | 0 | 0 |
| NC_004718.3;NP_828849.2;SARS-CoV_orf1ab;1715 | 0 | 0 |
| NC_004718.3;NP_828849.2;SARS-CoV_orf1ab;1716 | 0 | 0 |
| NC_004718.3;NP_828849.2;SARS-CoV_orf1ab;1717 | 0 | 0 |

| | | |
|---|---|---|
| NC_004718.3;NP_828849.2;SARS-CoV_orf1ab;1718 | 0 | 0 |
| NC_004718.3;NP_828849.2;SARS-CoV_orf1ab;1719 | 0.1 | 0 |
| NC_004718.3;NP_828849.2;SARS-CoV_orf1ab;1720 | 0 | 0 |
| NC_004718.3;NP_828849.2;SARS-CoV_orf1ab;1721 | 0 | 0 |
| NC_004718.3;NP_828849.2;SARS-CoV_orf1ab;1722 | 0.05 | 0 |
| NC_004718.3;NP_828849.2;SARS-CoV_orf1ab;1723 | 0.05 | 0 |
| NC_004718.3;NP_828849.2;SARS-CoV_orf1ab;1724 | 0.05 | 0 |
| NC_004718.3;NP_828849.2;SARS-CoV_orf1ab;1725 | 0.05 | 0 |
| NC_004718.3;NP_828849.2;SARS-CoV_orf1ab;1726 | 0.1 | 0.05 |
| NC_004718.3;NP_828849.2;SARS-CoV_orf1ab;1727 | 0.05 | 0.05 |
| NC_004718.3;NP_828849.2;SARS-CoV_orf1ab;1728 | 0.1 | 0.05 |
| NC_004718.3;NP_828849.2;SARS-CoV_orf1ab;1729 | 0.1 | 0.05 |
| NC_004718.3;NP_828849.2;SARS-CoV_orf1ab;1730 | 0 | 0 |
| NC_004718.3;NP_828849.2;SARS-CoV_orf1ab;1731 | 0 | 0 |
| NC_004718.3;NP_828849.2;SARS-CoV_orf1ab;1732 | 0 | 0.025 |
| NC_004718.3;NP_828849.2;SARS-CoV_orf1ab;1733 | 0.05 | 0.05 |
| NC_004718.3;NP_828849.2;SARS-CoV_orf1ab;1734 | 0.05 | 0.05 |
| NC_004718.3;NP_828849.2;SARS-CoV_orf1ab;1735 | 0 | 0.05 |
| NC_004718.3;NP_828849.2;SARS-CoV_orf1ab;1736 | 0 | 0 |
| NC_004718.3;NP_828849.2;SARS-CoV_orf1ab;1737 | 0 | 0 |
| NC_004718.3;NP_828849.2;SARS-CoV_orf1ab;1738 | 0.1 | 0 |
| NC_004718.3;NP_828849.2;SARS-CoV_orf1ab;1739 | 0.15 | 0.025 |
| NC_004718.3;NP_828849.2;SARS-CoV_orf1ab;1740 | 0.15 | 0.025 |
| NC_004718.3;NP_828849.2;SARS-CoV_orf1ab;1741 | 0.2 | 0.025 |
| NC_004718.3;NP_828849.2;SARS-CoV_orf1ab;1742 | 0.2 | 0.1 |
| NC_004718.3;NP_828849.2;SARS-CoV_orf1ab;1743 | 0.2 | 0.175 |
| NC_004718.3;NP_828849.2;SARS-CoV_orf1ab;1744 | 0.15 | 0.025 |
| NC_004718.3;NP_828849.2;SARS-CoV_orf1ab;1745 | 0.15 | 0 |
| NC_004718.3;NP_828849.2;SARS-CoV_orf1ab;1746 | 0.15 | 0.05 |
| NC_004718.3;NP_828849.2;SARS-CoV_orf1ab;1747 | 0.05 | 0.025 |
| NC_004718.3;NP_828849.2;SARS-CoV_orf1ab;1748 | 0.05 | 0.025 |
| NC_004718.3;NP_828849.2;SARS-CoV_orf1ab;1749 | 0.05 | 0 |
| NC_004718.3;NP_828849.2;SARS-CoV_orf1ab;1750 | 0.05 | 0 |
| NC_004718.3;NP_828849.2;SARS-CoV_orf1ab;1751 | 0.05 | 0 |
| NC_004718.3;NP_828849.2;SARS-CoV_orf1ab;1752 | 0 | 0.05 |
| NC_004718.3;NP_828849.2;SARS-CoV_orf1ab;1753 | 0 | 0 |
| NC_004718.3;NP_828849.2;SARS-CoV_orf1ab;1754 | 0 | 0 |
| NC_004718.3;NP_828849.2;SARS-CoV_orf1ab;1755 | 0 | 0 |
| NC_004718.3;NP_828849.2;SARS-CoV_orf1ab;1756 | 0 | 0 |
| NC_004718.3;NP_828849.2;SARS-CoV_orf1ab;1757 | 0 | 0 |
| NC_004718.3;NP_828849.2;SARS-CoV_orf1ab;1758 | 0 | 0 |
| NC_004718.3;NP_828849.2;SARS-CoV_orf1ab;1759 | 0 | 0 |
| NC_004718.3;NP_828849.2;SARS-CoV_orf1ab;1760 | 0 | 0 |

| | | |
|---|---|---|
| NC_004718.3;NP_828849.2;SARS-CoV_orf1ab;1761 | 0 | 0 |
| NC_004718.3;NP_828849.2;SARS-CoV_orf1ab;1762 | 0 | 0 |
| NC_004718.3;NP_828849.2;SARS-CoV_orf1ab;1763 | 0 | 0 |
| NC_004718.3;NP_828849.2;SARS-CoV_orf1ab;1764 | 0.05 | 0 |
| NC_004718.3;NP_828849.2;SARS-CoV_orf1ab;1765 | 0 | 0 |
| NC_004718.3;NP_828849.2;SARS-CoV_orf1ab;1766 | 0 | 0 |
| NC_004718.3;NP_828849.2;SARS-CoV_orf1ab;1767 | 0 | 0.025 |
| NC_004718.3;NP_828849.2;SARS-CoV_orf1ab;1768 | 0 | 0.025 |
| NC_004718.3;NP_828849.2;SARS-CoV_orf1ab;1769 | 0 | 0 |
| NC_004718.3;NP_828849.2;SARS-CoV_orf1ab;1770 | 0 | 0.025 |
| NC_004718.3;NP_828849.2;SARS-CoV_orf1ab;1771 | 0 | 0 |
| NC_004718.3;NP_828849.2;SARS-CoV_orf1ab;1772 | 0.05 | 0 |
| NC_004718.3;NP_828849.2;SARS-CoV_orf1ab;1773 | 0 | 0.025 |
| NC_004718.3;NP_828849.2;SARS-CoV_orf1ab;1774 | 0.1 | 0.025 |
| NC_004718.3;NP_828849.2;SARS-CoV_orf1ab;1775 | 0 | 0 |
| NC_004718.3;NP_828849.2;SARS-CoV_orf1ab;1776 | 0 | 0 |
| NC_004718.3;NP_828849.2;SARS-CoV_orf1ab;1777 | 0 | 0 |
| NC_004718.3;NP_828849.2;SARS-CoV_orf1ab;1778 | 0 | 0 |
| NC_004718.3;NP_828849.2;SARS-CoV_orf1ab;1779 | 0 | 0 |
| NC_004718.3;NP_828849.2;SARS-CoV_orf1ab;1780 | 0 | 0 |
| NC_004718.3;NP_828849.2;SARS-CoV_orf1ab;1781 | 0.05 | 0 |
| NC_004718.3;NP_828849.2;SARS-CoV_orf1ab;1782 | 0 | 0 |
| NC_004718.3;NP_828849.2;SARS-CoV_orf1ab;1783 | 0 | 0 |
| NC_004718.3;NP_828849.2;SARS-CoV_orf1ab;1784 | 0 | 0 |
| NC_004718.3;NP_828849.2;SARS-CoV_orf1ab;1785 | 0 | 0 |
| NC_004718.3;NP_828849.2;SARS-CoV_orf1ab;1786 | 0 | 0 |
| NC_004718.3;NP_828849.2;SARS-CoV_orf1ab;1787 | 0 | 0 |
| NC_004718.3;NP_828849.2;SARS-CoV_orf1ab;1788 | 0 | 0 |
| NC_004718.3;NP_828849.2;SARS-CoV_orf1ab;1789 | 0 | 0 |
| NC_004718.3;NP_828849.2;SARS-CoV_orf1ab;1790 | 0 | 0 |
| NC_004718.3;NP_828849.2;SARS-CoV_orf1ab;1791 | 0 | 0 |
| NC_004718.3;NP_828849.2;SARS-CoV_orf1ab;1792 | 0 | 0 |
| NC_004718.3;NP_828849.2;SARS-CoV_orf1ab;1793 | 0 | 0.025 |
| NC_004718.3;NP_828849.2;SARS-CoV_orf1ab;1794 | 0 | 0.025 |
| NC_004718.3;NP_828849.2;SARS-CoV_orf1ab;1795 | 0 | 0 |
| NC_004718.3;NP_828849.2;SARS-CoV_orf1ab;1796 | 0 | 0 |
| NC_004718.3;NP_828849.2;SARS-CoV_orf1ab;1797 | 0 | 0 |
| NC_004718.3;NP_828849.2;SARS-CoV_orf1ab;1798 | 0 | 0 |
| NC_004718.3;NP_828849.2;SARS-CoV_orf1ab;1799 | 0 | 0 |
| NC_004718.3;NP_828849.2;SARS-CoV_orf1ab;1800 | 0 | 0 |
| NC_004718.3;NP_828849.2;SARS-CoV_orf1ab;1801 | 0 | 0 |
| NC_004718.3;NP_828849.2;SARS-CoV_orf1ab;1802 | 0 | 0 |
| NC_004718.3;NP_828849.2;SARS-CoV_orf1ab;1803 | 0 | 0.15 |

| | | |
|---|---|---|
| NC_004718.3;NP_828849.2;SARS-CoV_orf1ab;1804 | 0 | 0 |
| NC_004718.3;NP_828849.2;SARS-CoV_orf1ab;1805 | 0 | 0.025 |
| NC_004718.3;NP_828849.2;SARS-CoV_orf1ab;1806 | 0.1 | 0 |
| NC_004718.3;NP_828849.2;SARS-CoV_orf1ab;1807 | 0 | 0 |
| NC_004718.3;NP_828849.2;SARS-CoV_orf1ab;1808 | 0.05 | 0 |
| NC_004718.3;NP_828849.2;SARS-CoV_orf1ab;1809 | 0 | 0 |
| NC_004718.3;NP_828849.2;SARS-CoV_orf1ab;1810 | 0 | 0 |
| NC_004718.3;NP_828849.2;SARS-CoV_orf1ab;1811 | 0.05 | 0 |
| NC_004718.3;NP_828849.2;SARS-CoV_orf1ab;1812 | 0.05 | 0 |
| NC_004718.3;NP_828849.2;SARS-CoV_orf1ab;1813 | 0.05 | 0 |
| NC_004718.3;NP_828849.2;SARS-CoV_orf1ab;1814 | 0.05 | 0 |
| NC_004718.3;NP_828849.2;SARS-CoV_orf1ab;1815 | 0.05 | 0 |
| NC_004718.3;NP_828849.2;SARS-CoV_orf1ab;1816 | 0.05 | 0.05 |
| NC_004718.3;NP_828849.2;SARS-CoV_orf1ab;1817 | 0 | 0 |
| NC_004718.3;NP_828849.2;SARS-CoV_orf1ab;1818 | 0.05 | 0 |
| NC_004718.3;NP_828849.2;SARS-CoV_orf1ab;1819 | 0 | 0 |
| NC_004718.3;NP_828849.2;SARS-CoV_orf1ab;1820 | 0 | 0 |
| NC_004718.3;NP_828849.2;SARS-CoV_orf1ab;1821 | 0 | 0 |
| NC_004718.3;NP_828849.2;SARS-CoV_orf1ab;1822 | 0 | 0 |
| NC_004718.3;NP_828849.2;SARS-CoV_orf1ab;1823 | 0 | 0 |
| NC_004718.3;NP_828849.2;SARS-CoV_orf1ab;1824 | 0 | 0 |
| NC_004718.3;NP_828849.2;SARS-CoV_orf1ab;1825 | 0 | 0 |
| NC_004718.3;NP_828849.2;SARS-CoV_orf1ab;1826 | 0 | 0 |
| NC_004718.3;NP_828849.2;SARS-CoV_orf1ab;1827 | 0 | 0.05 |
| NC_004718.3;NP_828849.2;SARS-CoV_orf1ab;1828 | 0 | 0 |
| NC_004718.3;NP_828849.2;SARS-CoV_orf1ab;1829 | 0 | 0 |
| NC_004718.3;NP_828849.2;SARS-CoV_orf1ab;1830 | 0 | 0.025 |
| NC_004718.3;NP_828849.2;SARS-CoV_orf1ab;1831 | 0 | 0.025 |
| NC_004718.3;NP_828849.2;SARS-CoV_orf1ab;1832 | 0 | 0.075 |
| NC_004718.3;NP_828849.2;SARS-CoV_orf1ab;1833 | 0 | 0.1 |
| NC_004718.3;NP_828849.2;SARS-CoV_orf1ab;1834 | 0 | 0.075 |
| NC_004718.3;NP_828849.2;SARS-CoV_orf1ab;1835 | 0 | 0.075 |
| NC_004718.3;NP_828849.2;SARS-CoV_orf1ab;1836 | 0 | 0.025 |
| NC_004718.3;NP_828849.2;SARS-CoV_orf1ab;1837 | 0 | 0.025 |
| NC_004718.3;NP_828849.2;SARS-CoV_orf1ab;1838 | 0.05 | 0.05 |
| NC_004718.3;NP_828849.2;SARS-CoV_orf1ab;1839 | 0 | 0 |
| NC_004718.3;NP_828849.2;SARS-CoV_orf1ab;1840 | 0 | 0.025 |
| NC_004718.3;NP_828849.2;SARS-CoV_orf1ab;1841 | 0 | 0.025 |
| NC_004718.3;NP_828849.2;SARS-CoV_orf1ab;1842 | 0 | 0.025 |
| NC_004718.3;NP_828849.2;SARS-CoV_orf1ab;1843 | 0.05 | 0.025 |
| NC_004718.3;NP_828849.2;SARS-CoV_orf1ab;1844 | 0.05 | 0.05 |
| NC_004718.3;NP_828849.2;SARS-CoV_orf1ab;1845 | 0 | 0.025 |
| NC_004718.3;NP_828849.2;SARS-CoV_orf1ab;1846 | 0.05 | 0.025 |

| | | |
|---|---|---|
| NC_004718.3;NP_828849.2;SARS-CoV_orf1ab;1847 | 0 | 0.025 |
| NC_004718.3;NP_828849.2;SARS-CoV_orf1ab;1848 | 0 | 0.025 |
| NC_004718.3;NP_828849.2;SARS-CoV_orf1ab;1849 | 0 | 0.025 |
| NC_004718.3;NP_828849.2;SARS-CoV_orf1ab;1850 | 0.05 | 0.05 |
| NC_004718.3;NP_828849.2;SARS-CoV_orf1ab;1851 | 0 | 0.05 |
| NC_004718.3;NP_828849.2;SARS-CoV_orf1ab;1852 | 0 | 0.075 |
| NC_004718.3;NP_828849.2;SARS-CoV_orf1ab;1853 | 0 | 0.025 |
| NC_004718.3;NP_828849.2;SARS-CoV_orf1ab;1854 | 0 | 0.025 |
| NC_004718.3;NP_828849.2;SARS-CoV_orf1ab;1855 | 0 | 0 |
| NC_004718.3;NP_828849.2;SARS-CoV_orf1ab;1856 | 0 | 0 |
| NC_004718.3;NP_828849.2;SARS-CoV_orf1ab;1857 | 0.05 | 0.025 |
| NC_004718.3;NP_828849.2;SARS-CoV_orf1ab;1858 | 0.05 | 0.075 |
| NC_004718.3;NP_828849.2;SARS-CoV_orf1ab;1859 | 0.05 | 0.025 |
| NC_004718.3;NP_828849.2;SARS-CoV_orf1ab;1860 | 0.05 | 0.1 |
| NC_004718.3;NP_828849.2;SARS-CoV_orf1ab;1861 | 0 | 0 |
| NC_004718.3;NP_828849.2;SARS-CoV_orf1ab;1862 | 0 | 0 |
| NC_004718.3;NP_828849.2;SARS-CoV_orf1ab;1863 | 0.2 | 0.275 |
| NC_004718.3;NP_828849.2;SARS-CoV_orf1ab;1864 | 0 | 0.075 |
| NC_004718.3;NP_828849.2;SARS-CoV_orf1ab;1865 | 0 | 0.05 |
| NC_004718.3;NP_828849.2;SARS-CoV_orf1ab;1866 | 0 | 0.075 |
| NC_004718.3;NP_828849.2;SARS-CoV_orf1ab;1867 | 0 | 0.075 |
| NC_004718.3;NP_828849.2;SARS-CoV_orf1ab;1868 | 0.05 | 0.075 |
| NC_004718.3;NP_828849.2;SARS-CoV_orf1ab;1869 | 0 | 0.075 |
| NC_004718.3;NP_828849.2;SARS-CoV_orf1ab;1870 | 0.05 | 0.075 |
| NC_004718.3;NP_828849.2;SARS-CoV_orf1ab;1871 | 0.05 | 0.075 |
| NC_004718.3;NP_828849.2;SARS-CoV_orf1ab;1872 | 0.15 | 0.1 |
| NC_004718.3;NP_828849.2;SARS-CoV_orf1ab;1873 | 0.1 | 0.075 |
| NC_004718.3;NP_828849.2;SARS-CoV_orf1ab;1874 | 0.15 | 0.075 |
| NC_004718.3;NP_828849.2;SARS-CoV_orf1ab;1875 | 0.05 | 0 |
| NC_004718.3;NP_828849.2;SARS-CoV_orf1ab;1876 | 0.15 | 0.05 |
| NC_004718.3;NP_828849.2;SARS-CoV_orf1ab;1877 | 0.1 | 0.05 |
| NC_004718.3;NP_828849.2;SARS-CoV_orf1ab;1878 | 0.15 | 0.125 |
| NC_004718.3;NP_828849.2;SARS-CoV_orf1ab;1879 | 0.05 | 0.025 |
| NC_004718.3;NP_828849.2;SARS-CoV_orf1ab;1880 | 0.05 | 0 |
| NC_004718.3;NP_828849.2;SARS-CoV_orf1ab;1881 | 0.05 | 0 |
| NC_004718.3;NP_828849.2;SARS-CoV_orf1ab;1882 | 0.05 | 0.05 |
| NC_004718.3;NP_828849.2;SARS-CoV_orf1ab;1883 | 0.05 | 0 |
| NC_004718.3;NP_828849.2;SARS-CoV_orf1ab;1884 | 0.05 | 0 |
| NC_004718.3;NP_828849.2;SARS-CoV_orf1ab;1885 | 0.1 | 0.05 |
| NC_004718.3;NP_828849.2;SARS-CoV_orf1ab;1886 | 0.1 | 0.025 |
| NC_004718.3;NP_828849.2;SARS-CoV_orf1ab;1887 | 0.1 | 0.075 |
| NC_004718.3;NP_828849.2;SARS-CoV_orf1ab;1888 | 0.05 | 0.025 |
| NC_004718.3;NP_828849.2;SARS-CoV_orf1ab;1889 | 0.05 | 0.025 |

| | | |
|---|---|---|
| NC_004718.3;NP_828849.2;SARS-CoV_orf1ab;1890 | 0.05 | 0.075 |
| NC_004718.3;NP_828849.2;SARS-CoV_orf1ab;1891 | 0.05 | 0.1 |
| NC_004718.3;NP_828849.2;SARS-CoV_orf1ab;1892 | 0.05 | 0.05 |
| NC_004718.3;NP_828849.2;SARS-CoV_orf1ab;1893 | 0.05 | 0.075 |
| NC_004718.3;NP_828849.2;SARS-CoV_orf1ab;1894 | 0.05 | 0.05 |
| NC_004718.3;NP_828849.2;SARS-CoV_orf1ab;1895 | 0.05 | 0.025 |
| NC_004718.3;NP_828849.2;SARS-CoV_orf1ab;1896 | 0.05 | 0.025 |
| NC_004718.3;NP_828849.2;SARS-CoV_orf1ab;1897 | 0.05 | 0 |
| NC_004718.3;NP_828849.2;SARS-CoV_orf1ab;1898 | 0.05 | 0 |
| NC_004718.3;NP_828849.2;SARS-CoV_orf1ab;1899 | 0 | 0 |
| NC_004718.3;NP_828849.2;SARS-CoV_orf1ab;1900 | 0 | 0.025 |
| NC_004718.3;NP_828849.2;SARS-CoV_orf1ab;1901 | 0 | 0.025 |
| NC_004718.3;NP_828849.2;SARS-CoV_orf1ab;1902 | 0 | 0 |
| NC_004718.3;NP_828849.2;SARS-CoV_orf1ab;1903 | 0 | 0 |
| NC_004718.3;NP_828849.2;SARS-CoV_orf1ab;1904 | 0.15 | 0.05 |
| NC_004718.3;NP_828849.2;SARS-CoV_orf1ab;1905 | 0.15 | 0.025 |
| NC_004718.3;NP_828849.2;SARS-CoV_orf1ab;1906 | 0.2 | 0.1 |
| NC_004718.3;NP_828849.2;SARS-CoV_orf1ab;1907 | 0.2 | 0.075 |
| NC_004718.3;NP_828849.2;SARS-CoV_orf1ab;1908 | 0.15 | 0.1 |
| NC_004718.3;NP_828849.2;SARS-CoV_orf1ab;1909 | 0.25 | 0.1 |
| NC_004718.3;NP_828849.2;SARS-CoV_orf1ab;1910 | 0.3 | 0.1 |
| NC_004718.3;NP_828849.2;SARS-CoV_orf1ab;1911 | 0.15 | 0.05 |
| NC_004718.3;NP_828849.2;SARS-CoV_orf1ab;1912 | 0.25 | 0.075 |
| NC_004718.3;NP_828849.2;SARS-CoV_orf1ab;1913 | 0.4 | 0.1 |
| NC_004718.3;NP_828849.2;SARS-CoV_orf1ab;1914 | 0.05 | 0 |
| NC_004718.3;NP_828849.2;SARS-CoV_orf1ab;1915 | 0.1 | 0.025 |
| NC_004718.3;NP_828849.2;SARS-CoV_orf1ab;1916 | 0 | 0.05 |
| NC_004718.3;NP_828849.2;SARS-CoV_orf1ab;1917 | 0 | 0 |
| NC_004718.3;NP_828849.2;SARS-CoV_orf1ab;1918 | 0 | 0 |
| NC_004718.3;NP_828849.2;SARS-CoV_orf1ab;1919 | 0 | 0 |
| NC_004718.3;NP_828849.2;SARS-CoV_orf1ab;1920 | 0 | 0 |
| NC_004718.3;NP_828849.2;SARS-CoV_orf1ab;1921 | 0 | 0 |
| NC_004718.3;NP_828849.2;SARS-CoV_orf1ab;1922 | 0 | 0 |
| NC_004718.3;NP_828849.2;SARS-CoV_orf1ab;1923 | 0 | 0 |
| NC_004718.3;NP_828849.2;SARS-CoV_orf1ab;1924 | 0 | 0 |
| NC_004718.3;NP_828849.2;SARS-CoV_orf1ab;1925 | 0 | 0 |
| NC_004718.3;NP_828849.2;SARS-CoV_orf1ab;1926 | 0 | 0 |
| NC_004718.3;NP_828849.2;SARS-CoV_orf1ab;1927 | 0 | 0 |
| NC_004718.3;NP_828849.2;SARS-CoV_orf1ab;1928 | 0 | 0 |
| NC_004718.3;NP_828849.2;SARS-CoV_orf1ab;1929 | 0 | 0 |
| NC_004718.3;NP_828849.2;SARS-CoV_orf1ab;1930 | 0 | 0.025 |
| NC_004718.3;NP_828849.2;SARS-CoV_orf1ab;1931 | 0 | 0.025 |
| NC_004718.3;NP_828849.2;SARS-CoV_orf1ab;1932 | 0 | 0 |

| | | |
|---|---|---|
| NC_004718.3;NP_828849.2;SARS-CoV_orf1ab;1933 | 0 | 0.025 |
| NC_004718.3;NP_828849.2;SARS-CoV_orf1ab;1934 | 0 | 0 |
| NC_004718.3;NP_828849.2;SARS-CoV_orf1ab;1935 | 0 | 0.025 |
| NC_004718.3;NP_828849.2;SARS-CoV_orf1ab;1936 | 0 | 0.025 |
| NC_004718.3;NP_828849.2;SARS-CoV_orf1ab;1937 | 0 | 0.025 |
| NC_004718.3;NP_828849.2;SARS-CoV_orf1ab;1938 | 0 | 0.075 |
| NC_004718.3;NP_828849.2;SARS-CoV_orf1ab;1939 | 0 | 0.025 |
| NC_004718.3;NP_828849.2;SARS-CoV_orf1ab;1940 | 0 | 0 |
| NC_004718.3;NP_828849.2;SARS-CoV_orf1ab;1941 | 0 | 0 |
| NC_004718.3;NP_828849.2;SARS-CoV_orf1ab;1942 | 0.05 | 0.05 |
| NC_004718.3;NP_828849.2;SARS-CoV_orf1ab;1943 | 0 | 0 |
| NC_004718.3;NP_828849.2;SARS-CoV_orf1ab;1944 | 0 | 0 |
| NC_004718.3;NP_828849.2;SARS-CoV_orf1ab;1945 | 0 | 0 |
| NC_004718.3;NP_828849.2;SARS-CoV_orf1ab;1946 | 0 | 0.025 |
| NC_004718.3;NP_828849.2;SARS-CoV_orf1ab;1947 | 0 | 0 |
| NC_004718.3;NP_828849.2;SARS-CoV_orf1ab;1948 | 0 | 0 |
| NC_004718.3;NP_828849.2;SARS-CoV_orf1ab;1949 | 0 | 0 |
| NC_004718.3;NP_828849.2;SARS-CoV_orf1ab;1950 | 0 | 0 |
| NC_004718.3;NP_828849.2;SARS-CoV_orf1ab;1951 | 0.05 | 0 |
| NC_004718.3;NP_828849.2;SARS-CoV_orf1ab;1952 | 0 | 0 |
| NC_004718.3;NP_828849.2;SARS-CoV_orf1ab;1953 | 0 | 0 |
| NC_004718.3;NP_828849.2;SARS-CoV_orf1ab;1954 | 0 | 0 |
| NC_004718.3;NP_828849.2;SARS-CoV_orf1ab;1955 | 0 | 0 |
| NC_004718.3;NP_828849.2;SARS-CoV_orf1ab;1956 | 0 | 0 |
| NC_004718.3;NP_828849.2;SARS-CoV_orf1ab;1957 | 0 | 0 |
| NC_004718.3;NP_828849.2;SARS-CoV_orf1ab;1958 | 0 | 0 |
| NC_004718.3;NP_828849.2;SARS-CoV_orf1ab;1959 | 0 | 0 |
| NC_004718.3;NP_828849.2;SARS-CoV_orf1ab;1960 | 0 | 0 |
| NC_004718.3;NP_828849.2;SARS-CoV_orf1ab;1961 | 0 | 0 |
| NC_004718.3;NP_828849.2;SARS-CoV_orf1ab;1962 | 0 | 0 |
| NC_004718.3;NP_828849.2;SARS-CoV_orf1ab;1963 | 0 | 0 |
| NC_004718.3;NP_828849.2;SARS-CoV_orf1ab;1964 | 0 | 0 |
| NC_004718.3;NP_828849.2;SARS-CoV_orf1ab;1965 | 0 | 0 |
| NC_004718.3;NP_828849.2;SARS-CoV_orf1ab;1966 | 0 | 0 |
| NC_004718.3;NP_828849.2;SARS-CoV_orf1ab;1967 | 0.05 | 0.025 |
| NC_004718.3;NP_828849.2;SARS-CoV_orf1ab;1968 | 0.05 | 0.025 |
| NC_004718.3;NP_828849.2;SARS-CoV_orf1ab;1969 | 0.05 | 0.025 |
| NC_004718.3;NP_828849.2;SARS-CoV_orf1ab;1970 | 0.1 | 0.025 |
| NC_004718.3;NP_828849.2;SARS-CoV_orf1ab;1971 | 0.1 | 0.025 |
| NC_004718.3;NP_828849.2;SARS-CoV_orf1ab;1972 | 0.05 | 0.05 |
| NC_004718.3;NP_828849.2;SARS-CoV_orf1ab;1973 | 0.1 | 0.025 |
| NC_004718.3;NP_828849.2;SARS-CoV_orf1ab;1974 | 0.05 | 0 |
| NC_004718.3;NP_828849.2;SARS-CoV_orf1ab;1975 | 0.05 | 0 |

| | | |
|---|---|---|
| NC_004718.3;NP_828849.2;SARS-CoV_orf1ab;1976 | 0 | 0 |
| NC_004718.3;NP_828849.2;SARS-CoV_orf1ab;1977 | 0 | 0 |
| NC_004718.3;NP_828849.2;SARS-CoV_orf1ab;1978 | 0 | 0 |
| NC_004718.3;NP_828849.2;SARS-CoV_orf1ab;1979 | 0 | 0 |
| NC_004718.3;NP_828849.2;SARS-CoV_orf1ab;1980 | 0 | 0 |
| NC_004718.3;NP_828849.2;SARS-CoV_orf1ab;1981 | 0 | 0 |
| NC_004718.3;NP_828849.2;SARS-CoV_orf1ab;1982 | 0 | 0 |
| NC_004718.3;NP_828849.2;SARS-CoV_orf1ab;1983 | 0 | 0 |
| NC_004718.3;NP_828849.2;SARS-CoV_orf1ab;1984 | 0 | 0 |
| NC_004718.3;NP_828849.2;SARS-CoV_orf1ab;1985 | 0 | 0 |
| NC_004718.3;NP_828849.2;SARS-CoV_orf1ab;1986 | 0.05 | 0 |
| NC_004718.3;NP_828849.2;SARS-CoV_orf1ab;1987 | 0.1 | 0.025 |
| NC_004718.3;NP_828849.2;SARS-CoV_orf1ab;1988 | 0.1 | 0.1 |
| NC_004718.3;NP_828849.2;SARS-CoV_orf1ab;1989 | 0.15 | 0.125 |
| NC_004718.3;NP_828849.2;SARS-CoV_orf1ab;1990 | 0.1 | 0.075 |
| NC_004718.3;NP_828849.2;SARS-CoV_orf1ab;1991 | 0.05 | 0.075 |
| NC_004718.3;NP_828849.2;SARS-CoV_orf1ab;1992 | 0 | 0.025 |
| NC_004718.3;NP_828849.2;SARS-CoV_orf1ab;1993 | 0 | 0.025 |
| NC_004718.3;NP_828849.2;SARS-CoV_orf1ab;1994 | 0.05 | 0.025 |
| NC_004718.3;NP_828849.2;SARS-CoV_orf1ab;1995 | 0.05 | 0.025 |
| NC_004718.3;NP_828849.2;SARS-CoV_orf1ab;1996 | 0.15 | 0.125 |
| NC_004718.3;NP_828849.2;SARS-CoV_orf1ab;1997 | 0 | 0.05 |
| NC_004718.3;NP_828849.2;SARS-CoV_orf1ab;1998 | 0 | 0.025 |
| NC_004718.3;NP_828849.2;SARS-CoV_orf1ab;1999 | 0 | 0 |
| NC_004718.3;NP_828849.2;SARS-CoV_orf1ab;2000 | 0.05 | 0 |
| NC_004718.3;NP_828849.2;SARS-CoV_orf1ab;2001 | 0 | 0.025 |
| NC_004718.3;NP_828849.2;SARS-CoV_orf1ab;2002 | 0 | 0.025 |
| NC_004718.3;NP_828849.2;SARS-CoV_orf1ab;2003 | 0 | 0 |
| NC_004718.3;NP_828849.2;SARS-CoV_orf1ab;2004 | 0 | 0 |
| NC_004718.3;NP_828849.2;SARS-CoV_orf1ab;2005 | 0.05 | 0 |
| NC_004718.3;NP_828849.2;SARS-CoV_orf1ab;2006 | 0.1 | 0.075 |
| NC_004718.3;NP_828849.2;SARS-CoV_orf1ab;2007 | 0 | 0 |
| NC_004718.3;NP_828849.2;SARS-CoV_orf1ab;2008 | 0.15 | 0 |
| NC_004718.3;NP_828849.2;SARS-CoV_orf1ab;2009 | 0 | 0 |
| NC_004718.3;NP_828849.2;SARS-CoV_orf1ab;2010 | 0 | 0 |
| NC_004718.3;NP_828849.2;SARS-CoV_orf1ab;2011 | 0 | 0 |
| NC_004718.3;NP_828849.2;SARS-CoV_orf1ab;2012 | 0 | 0 |
| NC_004718.3;NP_828849.2;SARS-CoV_orf1ab;2013 | 0 | 0 |
| NC_004718.3;NP_828849.2;SARS-CoV_orf1ab;2014 | 0 | 0 |
| NC_004718.3;NP_828849.2;SARS-CoV_orf1ab;2015 | 0 | 0 |
| NC_004718.3;NP_828849.2;SARS-CoV_orf1ab;2016 | 0 | 0 |
| NC_004718.3;NP_828849.2;SARS-CoV_orf1ab;2017 | 0 | 0 |
| NC_004718.3;NP_828849.2;SARS-CoV_orf1ab;2018 | 0 | 0 |

| | | |
|---|---|---|
| NC_004718.3;NP_828849.2;SARS-CoV_orf1ab;2019 | 0 | 0 |
| NC_004718.3;NP_828849.2;SARS-CoV_orf1ab;2020 | 0 | 0 |
| NC_004718.3;NP_828849.2;SARS-CoV_orf1ab;2021 | 0 | 0.05 |
| NC_004718.3;NP_828849.2;SARS-CoV_orf1ab;2022 | 0 | 0.05 |
| NC_004718.3;NP_828849.2;SARS-CoV_orf1ab;2023 | 0 | 0.05 |
| NC_004718.3;NP_828849.2;SARS-CoV_orf1ab;2024 | 0 | 0.05 |
| NC_004718.3;NP_828849.2;SARS-CoV_orf1ab;2025 | 0 | 0.05 |
| NC_004718.3;NP_828849.2;SARS-CoV_orf1ab;2026 | 0 | 0.05 |
| NC_004718.3;NP_828849.2;SARS-CoV_orf1ab;2027 | 0 | 0.05 |
| NC_004718.3;NP_828849.2;SARS-CoV_orf1ab;2028 | 0 | 0.05 |
| NC_004718.3;NP_828849.2;SARS-CoV_orf1ab;2029 | 0 | 0 |
| NC_004718.3;NP_828849.2;SARS-CoV_orf1ab;2030 | 0 | 0 |
| NC_004718.3;NP_828849.2;SARS-CoV_orf1ab;2031 | 0.05 | 0 |
| NC_004718.3;NP_828849.2;SARS-CoV_orf1ab;2032 | 0 | 0.025 |
| NC_004718.3;NP_828849.2;SARS-CoV_orf1ab;2033 | 0 | 0.025 |
| NC_004718.3;NP_828849.2;SARS-CoV_orf1ab;2034 | 0 | 0.025 |
| NC_004718.3;NP_828849.2;SARS-CoV_orf1ab;2035 | 0 | 0.025 |
| NC_004718.3;NP_828849.2;SARS-CoV_orf1ab;2036 | 0 | 0.025 |
| NC_004718.3;NP_828849.2;SARS-CoV_orf1ab;2037 | 0 | 0.025 |
| NC_004718.3;NP_828849.2;SARS-CoV_orf1ab;2038 | 0 | 0 |
| NC_004718.3;NP_828849.2;SARS-CoV_orf1ab;2039 | 0 | 0 |
| NC_004718.3;NP_828849.2;SARS-CoV_orf1ab;2040 | 0 | 0 |
| NC_004718.3;NP_828849.2;SARS-CoV_orf1ab;2041 | 0 | 0 |
| NC_004718.3;NP_828849.2;SARS-CoV_orf1ab;2042 | 0 | 0.025 |
| NC_004718.3;NP_828849.2;SARS-CoV_orf1ab;2043 | 0 | 0 |
| NC_004718.3;NP_828849.2;SARS-CoV_orf1ab;2044 | 0 | 0 |
| NC_004718.3;NP_828849.2;SARS-CoV_orf1ab;2045 | 0 | 0 |
| NC_004718.3;NP_828849.2;SARS-CoV_orf1ab;2046 | 0 | 0 |
| NC_004718.3;NP_828849.2;SARS-CoV_orf1ab;2047 | 0 | 0 |
| NC_004718.3;NP_828849.2;SARS-CoV_orf1ab;2048 | 0 | 0 |
| NC_004718.3;NP_828849.2;SARS-CoV_orf1ab;2049 | 0 | 0 |
| NC_004718.3;NP_828849.2;SARS-CoV_orf1ab;2050 | 0 | 0.05 |
| NC_004718.3;NP_828849.2;SARS-CoV_orf1ab;2051 | 0 | 0 |
| NC_004718.3;NP_828849.2;SARS-CoV_orf1ab;2052 | 0 | 0 |
| NC_004718.3;NP_828849.2;SARS-CoV_orf1ab;2053 | 0 | 0 |
| NC_004718.3;NP_828849.2;SARS-CoV_orf1ab;2054 | 0 | 0 |
| NC_004718.3;NP_828849.2;SARS-CoV_orf1ab;2055 | 0 | 0 |
| NC_004718.3;NP_828849.2;SARS-CoV_orf1ab;2056 | 0 | 0 |
| NC_004718.3;NP_828849.2;SARS-CoV_orf1ab;2057 | 0 | 0 |
| NC_004718.3;NP_828849.2;SARS-CoV_orf1ab;2058 | 0 | 0 |
| NC_004718.3;NP_828849.2;SARS-CoV_orf1ab;2059 | 0 | 0.025 |
| NC_004718.3;NP_828849.2;SARS-CoV_orf1ab;2060 | 0.05 | 0.025 |
| NC_004718.3;NP_828849.2;SARS-CoV_orf1ab;2061 | 0 | 0 |

| | | |
|---|---|---|
| NC_004718.3;NP_828849.2;SARS-CoV_orf1ab;2062 | 0 | 0 |
| NC_004718.3;NP_828849.2;SARS-CoV_orf1ab;2063 | 0 | 0 |
| NC_004718.3;NP_828849.2;SARS-CoV_orf1ab;2064 | 0 | 0 |
| NC_004718.3;NP_828849.2;SARS-CoV_orf1ab;2065 | 0 | 0 |
| NC_004718.3;NP_828849.2;SARS-CoV_orf1ab;2066 | 0 | 0 |
| NC_004718.3;NP_828849.2;SARS-CoV_orf1ab;2067 | 0 | 0 |
| NC_004718.3;NP_828849.2;SARS-CoV_orf1ab;2068 | 0.25 | 0.125 |
| NC_004718.3;NP_828849.2;SARS-CoV_orf1ab;2069 | 0 | 0 |
| NC_004718.3;NP_828849.2;SARS-CoV_orf1ab;2070 | 0 | 0 |
| NC_004718.3;NP_828849.2;SARS-CoV_orf1ab;2071 | 0 | 0 |
| NC_004718.3;NP_828849.2;SARS-CoV_orf1ab;2072 | 0 | 0 |
| NC_004718.3;NP_828849.2;SARS-CoV_orf1ab;2073 | 0 | 0 |
| NC_004718.3;NP_828849.2;SARS-CoV_orf1ab;2074 | 0.05 | 0 |
| NC_004718.3;NP_828849.2;SARS-CoV_orf1ab;2075 | 0.05 | 0.025 |
| NC_004718.3;NP_828849.2;SARS-CoV_orf1ab;2076 | 0.15 | 0 |
| NC_004718.3;NP_828849.2;SARS-CoV_orf1ab;2077 | 0.15 | 0.05 |
| NC_004718.3;NP_828849.2;SARS-CoV_orf1ab;2078 | 0.05 | 0.025 |
| NC_004718.3;NP_828849.2;SARS-CoV_orf1ab;2079 | 0 | 0 |
| NC_004718.3;NP_828849.2;SARS-CoV_orf1ab;2080 | 0 | 0 |
| NC_004718.3;NP_828849.2;SARS-CoV_orf1ab;2081 | 0 | 0 |
| NC_004718.3;NP_828849.2;SARS-CoV_orf1ab;2082 | 0 | 0 |
| NC_004718.3;NP_828849.2;SARS-CoV_orf1ab;2083 | 0 | 0 |
| NC_004718.3;NP_828849.2;SARS-CoV_orf1ab;2084 | 0 | 0 |
| NC_004718.3;NP_828849.2;SARS-CoV_orf1ab;2085 | 0 | 0 |
| NC_004718.3;NP_828849.2;SARS-CoV_orf1ab;2086 | 0 | 0 |
| NC_004718.3;NP_828849.2;SARS-CoV_orf1ab;2087 | 0 | 0 |
| NC_004718.3;NP_828849.2;SARS-CoV_orf1ab;2088 | 0 | 0 |
| NC_004718.3;NP_828849.2;SARS-CoV_orf1ab;2089 | 0 | 0 |
| NC_004718.3;NP_828849.2;SARS-CoV_orf1ab;2090 | 0 | 0 |
| NC_004718.3;NP_828849.2;SARS-CoV_orf1ab;2091 | 0.05 | 0 |
| NC_004718.3;NP_828849.2;SARS-CoV_orf1ab;2092 | 0 | 0 |
| NC_004718.3;NP_828849.2;SARS-CoV_orf1ab;2093 | 0 | 0 |
| NC_004718.3;NP_828849.2;SARS-CoV_orf1ab;2094 | 0 | 0 |
| NC_004718.3;NP_828849.2;SARS-CoV_orf1ab;2095 | 0 | 0 |
| NC_004718.3;NP_828849.2;SARS-CoV_orf1ab;2096 | 0 | 0 |
| NC_004718.3;NP_828849.2;SARS-CoV_orf1ab;2097 | 0 | 0 |
| NC_004718.3;NP_828849.2;SARS-CoV_orf1ab;2098 | 0 | 0 |
| NC_004718.3;NP_828849.2;SARS-CoV_orf1ab;2099 | 0 | 0 |
| NC_004718.3;NP_828849.2;SARS-CoV_orf1ab;2100 | 0.05 | 0.025 |
| NC_004718.3;NP_828849.2;SARS-CoV_orf1ab;2101 | 0.05 | 0.05 |
| NC_004718.3;NP_828849.2;SARS-CoV_orf1ab;2102 | 0.05 | 0.05 |
| NC_004718.3;NP_828849.2;SARS-CoV_orf1ab;2103 | 0.05 | 0.05 |
| NC_004718.3;NP_828849.2;SARS-CoV_orf1ab;2104 | 0.05 | 0 |

| | | |
|---|---|---|
| NC_004718.3;NP_828849.2;SARS-CoV_orf1ab;2105 | 0.05 | 0 |
| NC_004718.3;NP_828849.2;SARS-CoV_orf1ab;2106 | 0.05 | 0 |
| NC_004718.3;NP_828849.2;SARS-CoV_orf1ab;2107 | 0.05 | 0 |
| NC_004718.3;NP_828849.2;SARS-CoV_orf1ab;2108 | 0 | 0 |
| NC_004718.3;NP_828849.2;SARS-CoV_orf1ab;2109 | 0.05 | 0 |
| NC_004718.3;NP_828849.2;SARS-CoV_orf1ab;2110 | 0.05 | 0 |
| NC_004718.3;NP_828849.2;SARS-CoV_orf1ab;2111 | 0 | 0 |
| NC_004718.3;NP_828849.2;SARS-CoV_orf1ab;2112 | 0 | 0 |
| NC_004718.3;NP_828849.2;SARS-CoV_orf1ab;2113 | 0 | 0 |
| NC_004718.3;NP_828849.2;SARS-CoV_orf1ab;2114 | 0 | 0 |
| NC_004718.3;NP_828849.2;SARS-CoV_orf1ab;2115 | 0 | 0 |
| NC_004718.3;NP_828849.2;SARS-CoV_orf1ab;2116 | 0 | 0 |
| NC_004718.3;NP_828849.2;SARS-CoV_orf1ab;2117 | 0.05 | 0 |
| NC_004718.3;NP_828849.2;SARS-CoV_orf1ab;2118 | 0 | 0 |
| NC_004718.3;NP_828849.2;SARS-CoV_orf1ab;2119 | 0.05 | 0 |
| NC_004718.3;NP_828849.2;SARS-CoV_orf1ab;2120 | 0.1 | 0.05 |
| NC_004718.3;NP_828849.2;SARS-CoV_orf1ab;2121 | 0 | 0 |
| NC_004718.3;NP_828849.2;SARS-CoV_orf1ab;2122 | 0 | 0 |
| NC_004718.3;NP_828849.2;SARS-CoV_orf1ab;2123 | 0 | 0 |
| NC_004718.3;NP_828849.2;SARS-CoV_orf1ab;2124 | 0 | 0 |
| NC_004718.3;NP_828849.2;SARS-CoV_orf1ab;2125 | 0 | 0 |
| NC_004718.3;NP_828849.2;SARS-CoV_orf1ab;2126 | 0 | 0.025 |
| NC_004718.3;NP_828849.2;SARS-CoV_orf1ab;2127 | 0.1 | 0 |
| NC_004718.3;NP_828849.2;SARS-CoV_orf1ab;2128 | 0 | 0 |
| NC_004718.3;NP_828849.2;SARS-CoV_orf1ab;2129 | 0 | 0 |
| NC_004718.3;NP_828849.2;SARS-CoV_orf1ab;2130 | 0 | 0 |
| NC_004718.3;NP_828849.2;SARS-CoV_orf1ab;2131 | 0 | 0 |
| NC_004718.3;NP_828849.2;SARS-CoV_orf1ab;2132 | 0 | 0 |
| NC_004718.3;NP_828849.2;SARS-CoV_orf1ab;2133 | 0 | 0 |
| NC_004718.3;NP_828849.2;SARS-CoV_orf1ab;2134 | 0 | 0 |
| NC_004718.3;NP_828849.2;SARS-CoV_orf1ab;2135 | 0 | 0 |
| NC_004718.3;NP_828849.2;SARS-CoV_orf1ab;2136 | 0 | 0 |
| NC_004718.3;NP_828849.2;SARS-CoV_orf1ab;2137 | 0 | 0 |
| NC_004718.3;NP_828849.2;SARS-CoV_orf1ab;2138 | 0 | 0 |
| NC_004718.3;NP_828849.2;SARS-CoV_orf1ab;2139 | 0 | 0 |
| NC_004718.3;NP_828849.2;SARS-CoV_orf1ab;2140 | 0 | 0 |
| NC_004718.3;NP_828849.2;SARS-CoV_orf1ab;2141 | 0 | 0 |
| NC_004718.3;NP_828849.2;SARS-CoV_orf1ab;2142 | 0 | 0 |
| NC_004718.3;NP_828849.2;SARS-CoV_orf1ab;2143 | 0 | 0 |
| NC_004718.3;NP_828849.2;SARS-CoV_orf1ab;2144 | 0 | 0 |
| NC_004718.3;NP_828849.2;SARS-CoV_orf1ab;2145 | 0 | 0 |
| NC_004718.3;NP_828849.2;SARS-CoV_orf1ab;2146 | 0 | 0 |
| NC_004718.3;NP_828849.2;SARS-CoV_orf1ab;2147 | 0 | 0 |

| | | |
|---|---|---|
| NC_004718.3;NP_828849.2;SARS-CoV_orf1ab;2148 | 0 | 0 |
| NC_004718.3;NP_828849.2;SARS-CoV_orf1ab;2149 | 0 | 0 |
| NC_004718.3;NP_828849.2;SARS-CoV_orf1ab;2150 | 0 | 0 |
| NC_004718.3;NP_828849.2;SARS-CoV_orf1ab;2151 | 0 | 0 |
| NC_004718.3;NP_828849.2;SARS-CoV_orf1ab;2152 | 0 | 0 |
| NC_004718.3;NP_828849.2;SARS-CoV_orf1ab;2153 | 0 | 0 |
| NC_004718.3;NP_828849.2;SARS-CoV_orf1ab;2154 | 0 | 0 |
| NC_004718.3;NP_828849.2;SARS-CoV_orf1ab;2155 | 0 | 0.025 |
| NC_004718.3;NP_828849.2;SARS-CoV_orf1ab;2156 | 0 | 0.025 |
| NC_004718.3;NP_828849.2;SARS-CoV_orf1ab;2157 | 0 | 0.025 |
| NC_004718.3;NP_828849.2;SARS-CoV_orf1ab;2158 | 0.05 | 0.025 |
| NC_004718.3;NP_828849.2;SARS-CoV_orf1ab;2159 | 0 | 0.025 |
| NC_004718.3;NP_828849.2;SARS-CoV_orf1ab;2160 | 0 | 0.05 |
| NC_004718.3;NP_828849.2;SARS-CoV_orf1ab;2161 | 0 | 0.025 |
| NC_004718.3;NP_828849.2;SARS-CoV_orf1ab;2162 | 0 | 0 |
| NC_004718.3;NP_828849.2;SARS-CoV_orf1ab;2163 | 0 | 0 |
| NC_004718.3;NP_828849.2;SARS-CoV_orf1ab;2164 | 0 | 0.025 |
| NC_004718.3;NP_828849.2;SARS-CoV_orf1ab;2165 | 0.05 | 0 |
| NC_004718.3;NP_828849.2;SARS-CoV_orf1ab;2166 | 0 | 0 |
| NC_004718.3;NP_828849.2;SARS-CoV_orf1ab;2167 | 0 | 0 |
| NC_004718.3;NP_828849.2;SARS-CoV_orf1ab;2168 | 0 | 0 |
| NC_004718.3;NP_828849.2;SARS-CoV_orf1ab;2169 | 0 | 0 |
| NC_004718.3;NP_828849.2;SARS-CoV_orf1ab;2170 | 0 | 0 |
| NC_004718.3;NP_828849.2;SARS-CoV_orf1ab;2171 | 0 | 0 |
| NC_004718.3;NP_828849.2;SARS-CoV_orf1ab;2172 | 0 | 0 |
| NC_004718.3;NP_828849.2;SARS-CoV_orf1ab;2173 | 0 | 0 |
| NC_004718.3;NP_828849.2;SARS-CoV_orf1ab;2174 | 0 | 0 |
| NC_004718.3;NP_828849.2;SARS-CoV_orf1ab;2175 | 0 | 0 |
| NC_004718.3;NP_828849.2;SARS-CoV_orf1ab;2176 | 0 | 0 |
| NC_004718.3;NP_828849.2;SARS-CoV_orf1ab;2177 | 0 | 0 |
| NC_004718.3;NP_828849.2;SARS-CoV_orf1ab;2178 | 0.05 | 0 |
| NC_004718.3;NP_828849.2;SARS-CoV_orf1ab;2179 | 0.05 | 0 |
| NC_004718.3;NP_828849.2;SARS-CoV_orf1ab;2180 | 0.05 | 0 |
| NC_004718.3;NP_828849.2;SARS-CoV_orf1ab;2181 | 0.05 | 0 |
| NC_004718.3;NP_828849.2;SARS-CoV_orf1ab;2182 | 0.05 | 0 |
| NC_004718.3;NP_828849.2;SARS-CoV_orf1ab;2183 | 0.05 | 0 |
| NC_004718.3;NP_828849.2;SARS-CoV_orf1ab;2184 | 0.05 | 0 |
| NC_004718.3;NP_828849.2;SARS-CoV_orf1ab;2185 | 0.05 | 0 |
| NC_004718.3;NP_828849.2;SARS-CoV_orf1ab;2186 | 0 | 0 |
| NC_004718.3;NP_828849.2;SARS-CoV_orf1ab;2187 | 0.1 | 0 |
| NC_004718.3;NP_828849.2;SARS-CoV_orf1ab;2188 | 0.05 | 0 |
| NC_004718.3;NP_828849.2;SARS-CoV_orf1ab;2189 | 0.05 | 0 |
| NC_004718.3;NP_828849.2;SARS-CoV_orf1ab;2190 | 0 | 0.025 |

| | | |
|---|---|---|
| NC_004718.3;NP_828849.2;SARS-CoV_orf1ab;2191 | 0 | 0 |
| NC_004718.3;NP_828849.2;SARS-CoV_orf1ab;2192 | 0 | 0 |
| NC_004718.3;NP_828849.2;SARS-CoV_orf1ab;2193 | 0 | 0 |
| NC_004718.3;NP_828849.2;SARS-CoV_orf1ab;2194 | 0 | 0 |
| NC_004718.3;NP_828849.2;SARS-CoV_orf1ab;2195 | 0 | 0 |
| NC_004718.3;NP_828849.2;SARS-CoV_orf1ab;2196 | 0 | 0 |
| NC_004718.3;NP_828849.2;SARS-CoV_orf1ab;2197 | 0 | 0 |
| NC_004718.3;NP_828849.2;SARS-CoV_orf1ab;2198 | 0 | 0 |
| NC_004718.3;NP_828849.2;SARS-CoV_orf1ab;2199 | 0 | 0 |
| NC_004718.3;NP_828849.2;SARS-CoV_orf1ab;2200 | 0 | 0 |
| NC_004718.3;NP_828849.2;SARS-CoV_orf1ab;2201 | 0 | 0 |
| NC_004718.3;NP_828849.2;SARS-CoV_orf1ab;2202 | 0 | 0 |
| NC_004718.3;NP_828849.2;SARS-CoV_orf1ab;2203 | 0 | 0 |
| NC_004718.3;NP_828849.2;SARS-CoV_orf1ab;2204 | 0 | 0 |
| NC_004718.3;NP_828849.2;SARS-CoV_orf1ab;2205 | 0 | 0 |
| NC_004718.3;NP_828849.2;SARS-CoV_orf1ab;2206 | 0 | 0 |
| NC_004718.3;NP_828849.2;SARS-CoV_orf1ab;2207 | 0 | 0 |
| NC_004718.3;NP_828849.2;SARS-CoV_orf1ab;2208 | 0 | 0 |
| NC_004718.3;NP_828849.2;SARS-CoV_orf1ab;2209 | 0 | 0 |
| NC_004718.3;NP_828849.2;SARS-CoV_orf1ab;2210 | 0 | 0 |
| NC_004718.3;NP_828849.2;SARS-CoV_orf1ab;2211 | 0 | 0 |
| NC_004718.3;NP_828849.2;SARS-CoV_orf1ab;2212 | 0 | 0 |
| NC_004718.3;NP_828849.2;SARS-CoV_orf1ab;2213 | 0 | 0 |
| NC_004718.3;NP_828849.2;SARS-CoV_orf1ab;2214 | 0 | 0 |
| NC_004718.3;NP_828849.2;SARS-CoV_orf1ab;2215 | 0 | 0 |
| NC_004718.3;NP_828849.2;SARS-CoV_orf1ab;2216 | 0 | 0 |
| NC_004718.3;NP_828849.2;SARS-CoV_orf1ab;2217 | 0 | 0 |
| NC_004718.3;NP_828849.2;SARS-CoV_orf1ab;2218 | 0 | 0 |
| NC_004718.3;NP_828849.2;SARS-CoV_orf1ab;2219 | 0 | 0 |
| NC_004718.3;NP_828849.2;SARS-CoV_orf1ab;2220 | 0 | 0 |
| NC_004718.3;NP_828849.2;SARS-CoV_orf1ab;2221 | 0 | 0 |
| NC_004718.3;NP_828849.2;SARS-CoV_orf1ab;2222 | 0 | 0 |
| NC_004718.3;NP_828849.2;SARS-CoV_orf1ab;2223 | 0 | 0 |
| NC_004718.3;NP_828849.2;SARS-CoV_orf1ab;2224 | 0 | 0 |
| NC_004718.3;NP_828849.2;SARS-CoV_orf1ab;2225 | 0 | 0 |
| NC_004718.3;NP_828849.2;SARS-CoV_orf1ab;2226 | 0 | 0 |
| NC_004718.3;NP_828849.2;SARS-CoV_orf1ab;2227 | 0 | 0 |
| NC_004718.3;NP_828849.2;SARS-CoV_orf1ab;2228 | 0 | 0 |
| NC_004718.3;NP_828849.2;SARS-CoV_orf1ab;2229 | 0 | 0 |
| NC_004718.3;NP_828849.2;SARS-CoV_orf1ab;2230 | 0 | 0.025 |
| NC_004718.3;NP_828849.2;SARS-CoV_orf1ab;2231 | 0.05 | 0 |
| NC_004718.3;NP_828849.2;SARS-CoV_orf1ab;2232 | 0 | 0 |
| NC_004718.3;NP_828849.2;SARS-CoV_orf1ab;2233 | 0 | 0 |

| | | |
|---|---|---|
| NC_004718.3;NP_828849.2;SARS-CoV_orf1ab;2234 | 0 | 0.025 |
| NC_004718.3;NP_828849.2;SARS-CoV_orf1ab;2235 | 0 | 0.025 |
| NC_004718.3;NP_828849.2;SARS-CoV_orf1ab;2236 | 0 | 0.075 |
| NC_004718.3;NP_828849.2;SARS-CoV_orf1ab;2237 | 0 | 0.025 |
| NC_004718.3;NP_828849.2;SARS-CoV_orf1ab;2238 | 0.05 | 0.075 |
| NC_004718.3;NP_828849.2;SARS-CoV_orf1ab;2239 | 0.25 | 0.225 |
| NC_004718.3;NP_828849.2;SARS-CoV_orf1ab;2240 | 0.3 | 0.325 |
| NC_004718.3;NP_828849.2;SARS-CoV_orf1ab;2241 | 0.3 | 0.225 |
| NC_004718.3;NP_828849.2;SARS-CoV_orf1ab;2242 | 0.45 | 0.45 |
| NC_004718.3;NP_828849.2;SARS-CoV_orf1ab;2243 | 0.35 | 0.35 |
| NC_004718.3;NP_828849.2;SARS-CoV_orf1ab;2244 | 0.25 | 0.275 |
| NC_004718.3;NP_828849.2;SARS-CoV_orf1ab;2245 | 0.35 | 0.3 |
| NC_004718.3;NP_828849.2;SARS-CoV_orf1ab;2246 | 0.15 | 0.125 |
| NC_004718.3;NP_828849.2;SARS-CoV_orf1ab;2247 | 0.2 | 0.125 |
| NC_004718.3;NP_828849.2;SARS-CoV_orf1ab;2248 | 0.05 | 0.05 |
| NC_004718.3;NP_828849.2;SARS-CoV_orf1ab;2249 | 0 | 0 |
| NC_004718.3;NP_828849.2;SARS-CoV_orf1ab;2250 | 0 | 0 |
| NC_004718.3;NP_828849.2;SARS-CoV_orf1ab;2251 | 0.05 | 0 |
| NC_004718.3;NP_828849.2;SARS-CoV_orf1ab;2252 | 0 | 0 |
| NC_004718.3;NP_828849.2;SARS-CoV_orf1ab;2253 | 0 | 0 |
| NC_004718.3;NP_828849.2;SARS-CoV_orf1ab;2254 | 0 | 0 |
| NC_004718.3;NP_828849.2;SARS-CoV_orf1ab;2255 | 0 | 0 |
| NC_004718.3;NP_828849.2;SARS-CoV_orf1ab;2256 | 0 | 0 |
| NC_004718.3;NP_828849.2;SARS-CoV_orf1ab;2257 | 0 | 0 |
| NC_004718.3;NP_828849.2;SARS-CoV_orf1ab;2258 | 0 | 0 |
| NC_004718.3;NP_828849.2;SARS-CoV_orf1ab;2259 | 0 | 0 |
| NC_004718.3;NP_828849.2;SARS-CoV_orf1ab;2260 | 0 | 0.025 |
| NC_004718.3;NP_828849.2;SARS-CoV_orf1ab;2261 | 0 | 0 |
| NC_004718.3;NP_828849.2;SARS-CoV_orf1ab;2262 | 0 | 0 |
| NC_004718.3;NP_828849.2;SARS-CoV_orf1ab;2263 | 0 | 0 |
| NC_004718.3;NP_828849.2;SARS-CoV_orf1ab;2264 | 0 | 0 |
| NC_004718.3;NP_828849.2;SARS-CoV_orf1ab;2265 | 0 | 0.025 |
| NC_004718.3;NP_828849.2;SARS-CoV_orf1ab;2266 | 0 | 0 |
| NC_004718.3;NP_828849.2;SARS-CoV_orf1ab;2267 | 0 | 0 |
| NC_004718.3;NP_828849.2;SARS-CoV_orf1ab;2268 | 0.05 | 0.025 |
| NC_004718.3;NP_828849.2;SARS-CoV_orf1ab;2269 | 0 | 0 |
| NC_004718.3;NP_828849.2;SARS-CoV_orf1ab;2270 | 0.05 | 0 |
| NC_004718.3;NP_828849.2;SARS-CoV_orf1ab;2271 | 0.05 | 0.025 |
| NC_004718.3;NP_828849.2;SARS-CoV_orf1ab;2272 | 0.05 | 0 |
| NC_004718.3;NP_828849.2;SARS-CoV_orf1ab;2273 | 0.05 | 0 |
| NC_004718.3;NP_828849.2;SARS-CoV_orf1ab;2274 | 0 | 0 |
| NC_004718.3;NP_828849.2;SARS-CoV_orf1ab;2275 | 0 | 0 |
| NC_004718.3;NP_828849.2;SARS-CoV_orf1ab;2276 | 0 | 0 |

| | | |
|---|---|---|
| NC_004718.3;NP_828849.2;SARS-CoV_orf1ab;2277 | 0 | 0 |
| NC_004718.3;NP_828849.2;SARS-CoV_orf1ab;2278 | 0 | 0 |
| NC_004718.3;NP_828849.2;SARS-CoV_orf1ab;2279 | 0 | 0 |
| NC_004718.3;NP_828849.2;SARS-CoV_orf1ab;2280 | 0 | 0 |
| NC_004718.3;NP_828849.2;SARS-CoV_orf1ab;2281 | 0 | 0 |
| NC_004718.3;NP_828849.2;SARS-CoV_orf1ab;2282 | 0 | 0 |
| NC_004718.3;NP_828849.2;SARS-CoV_orf1ab;2283 | 0 | 0 |
| NC_004718.3;NP_828849.2;SARS-CoV_orf1ab;2284 | 0 | 0 |
| NC_004718.3;NP_828849.2;SARS-CoV_orf1ab;2285 | 0 | 0 |
| NC_004718.3;NP_828849.2;SARS-CoV_orf1ab;2286 | 0 | 0 |
| NC_004718.3;NP_828849.2;SARS-CoV_orf1ab;2287 | 0.05 | 0 |
| NC_004718.3;NP_828849.2;SARS-CoV_orf1ab;2288 | 0 | 0 |
| NC_004718.3;NP_828849.2;SARS-CoV_orf1ab;2289 | 0 | 0 |
| NC_004718.3;NP_828849.2;SARS-CoV_orf1ab;2290 | 0 | 0 |
| NC_004718.3;NP_828849.2;SARS-CoV_orf1ab;2291 | 0 | 0 |
| NC_004718.3;NP_828849.2;SARS-CoV_orf1ab;2292 | 0 | 0 |
| NC_004718.3;NP_828849.2;SARS-CoV_orf1ab;2293 | 0 | 0 |
| NC_004718.3;NP_828849.2;SARS-CoV_orf1ab;2294 | 0 | 0 |
| NC_004718.3;NP_828849.2;SARS-CoV_orf1ab;2295 | 0 | 0 |
| NC_004718.3;NP_828849.2;SARS-CoV_orf1ab;2296 | 0 | 0 |
| NC_004718.3;NP_828849.2;SARS-CoV_orf1ab;2297 | 0 | 0 |
| NC_004718.3;NP_828849.2;SARS-CoV_orf1ab;2298 | 0 | 0 |
| NC_004718.3;NP_828849.2;SARS-CoV_orf1ab;2299 | 0 | 0 |
| NC_004718.3;NP_828849.2;SARS-CoV_orf1ab;2300 | 0 | 0 |
| NC_004718.3;NP_828849.2;SARS-CoV_orf1ab;2301 | 0 | 0 |
| NC_004718.3;NP_828849.2;SARS-CoV_orf1ab;2302 | 0 | 0 |
| NC_004718.3;NP_828849.2;SARS-CoV_orf1ab;2303 | 0 | 0 |
| NC_004718.3;NP_828849.2;SARS-CoV_orf1ab;2304 | 0 | 0 |
| NC_004718.3;NP_828849.2;SARS-CoV_orf1ab;2305 | 0 | 0 |
| NC_004718.3;NP_828849.2;SARS-CoV_orf1ab;2306 | 0 | 0 |
| NC_004718.3;NP_828849.2;SARS-CoV_orf1ab;2307 | 0 | 0 |
| NC_004718.3;NP_828849.2;SARS-CoV_orf1ab;2308 | 0 | 0 |
| NC_004718.3;NP_828849.2;SARS-CoV_orf1ab;2309 | 0 | 0 |
| NC_004718.3;NP_828849.2;SARS-CoV_orf1ab;2310 | 0 | 0 |
| NC_004718.3;NP_828849.2;SARS-CoV_orf1ab;2311 | 0 | 0 |
| NC_004718.3;NP_828849.2;SARS-CoV_orf1ab;2312 | 0 | 0 |
| NC_004718.3;NP_828849.2;SARS-CoV_orf1ab;2313 | 0 | 0 |
| NC_004718.3;NP_828849.2;SARS-CoV_orf1ab;2314 | 0 | 0 |
| NC_004718.3;NP_828849.2;SARS-CoV_orf1ab;2315 | 0 | 0 |
| NC_004718.3;NP_828849.2;SARS-CoV_orf1ab;2316 | 0 | 0 |
| NC_004718.3;NP_828849.2;SARS-CoV_orf1ab;2317 | 0 | 0 |
| NC_004718.3;NP_828849.2;SARS-CoV_orf1ab;2318 | 0 | 0 |
| NC_004718.3;NP_828849.2;SARS-CoV_orf1ab;2319 | 0 | 0 |

| | | |
|---|---|---|
| NC_004718.3;NP_828849.2;SARS-CoV_orf1ab;2320 | 0 | 0 |
| NC_004718.3;NP_828849.2;SARS-CoV_orf1ab;2321 | 0 | 0 |
| NC_004718.3;NP_828849.2;SARS-CoV_orf1ab;2322 | 0 | 0 |
| NC_004718.3;NP_828849.2;SARS-CoV_orf1ab;2323 | 0 | 0 |
| NC_004718.3;NP_828849.2;SARS-CoV_orf1ab;2324 | 0 | 0 |
| NC_004718.3;NP_828849.2;SARS-CoV_orf1ab;2325 | 0 | 0 |
| NC_004718.3;NP_828849.2;SARS-CoV_orf1ab;2326 | 0 | 0 |
| NC_004718.3;NP_828849.2;SARS-CoV_orf1ab;2327 | 0 | 0 |
| NC_004718.3;NP_828849.2;SARS-CoV_orf1ab;2328 | 0 | 0 |
| NC_004718.3;NP_828849.2;SARS-CoV_orf1ab;2329 | 0 | 0 |
| NC_004718.3;NP_828849.2;SARS-CoV_orf1ab;2330 | 0 | 0 |
| NC_004718.3;NP_828849.2;SARS-CoV_orf1ab;2331 | 0 | 0 |
| NC_004718.3;NP_828849.2;SARS-CoV_orf1ab;2332 | 0 | 0 |
| NC_004718.3;NP_828849.2;SARS-CoV_orf1ab;2333 | 0 | 0 |
| NC_004718.3;NP_828849.2;SARS-CoV_orf1ab;2334 | 0 | 0 |
| NC_004718.3;NP_828849.2;SARS-CoV_orf1ab;2335 | 0 | 0 |
| NC_004718.3;NP_828849.2;SARS-CoV_orf1ab;2336 | 0 | 0 |
| NC_004718.3;NP_828849.2;SARS-CoV_orf1ab;2337 | 0 | 0 |
| NC_004718.3;NP_828849.2;SARS-CoV_orf1ab;2338 | 0 | 0 |
| NC_004718.3;NP_828849.2;SARS-CoV_orf1ab;2339 | 0 | 0 |
| NC_004718.3;NP_828849.2;SARS-CoV_orf1ab;2340 | 0 | 0 |
| NC_004718.3;NP_828849.2;SARS-CoV_orf1ab;2341 | 0 | 0 |
| NC_004718.3;NP_828849.2;SARS-CoV_orf1ab;2342 | 0 | 0 |
| NC_004718.3;NP_828849.2;SARS-CoV_orf1ab;2343 | 0 | 0 |
| NC_004718.3;NP_828849.2;SARS-CoV_orf1ab;2344 | 0 | 0 |
| NC_004718.3;NP_828849.2;SARS-CoV_orf1ab;2345 | 0 | 0 |
| NC_004718.3;NP_828849.2;SARS-CoV_orf1ab;2346 | 0 | 0 |
| NC_004718.3;NP_828849.2;SARS-CoV_orf1ab;2347 | 0 | 0 |
| NC_004718.3;NP_828849.2;SARS-CoV_orf1ab;2348 | 0 | 0 |
| NC_004718.3;NP_828849.2;SARS-CoV_orf1ab;2349 | 0 | 0 |
| NC_004718.3;NP_828849.2;SARS-CoV_orf1ab;2350 | 0 | 0 |
| NC_004718.3;NP_828849.2;SARS-CoV_orf1ab;2351 | 0 | 0 |
| NC_004718.3;NP_828849.2;SARS-CoV_orf1ab;2352 | 0 | 0 |
| NC_004718.3;NP_828849.2;SARS-CoV_orf1ab;2353 | 0 | 0 |
| NC_004718.3;NP_828849.2;SARS-CoV_orf1ab;2354 | 0 | 0 |
| NC_004718.3;NP_828849.2;SARS-CoV_orf1ab;2355 | 0 | 0 |
| NC_004718.3;NP_828849.2;SARS-CoV_orf1ab;2356 | 0 | 0 |
| NC_004718.3;NP_828849.2;SARS-CoV_orf1ab;2357 | 0 | 0 |
| NC_004718.3;NP_828849.2;SARS-CoV_orf1ab;2358 | 0 | 0 |
| NC_004718.3;NP_828849.2;SARS-CoV_orf1ab;2359 | 0 | 0 |
| NC_004718.3;NP_828849.2;SARS-CoV_orf1ab;2360 | 0 | 0 |
| NC_004718.3;NP_828849.2;SARS-CoV_orf1ab;2361 | 0 | 0 |
| NC_004718.3;NP_828849.2;SARS-CoV_orf1ab;2362 | 0 | 0 |

| | | |
|---|---|---|
| NC_004718.3;NP_828849.2;SARS-CoV_orf1ab;2363 | 0 | 0 |
| NC_004718.3;NP_828849.2;SARS-CoV_orf1ab;2364 | 0 | 0 |
| NC_004718.3;NP_828849.2;SARS-CoV_orf1ab;2365 | 0 | 0 |
| NC_004718.3;NP_828849.2;SARS-CoV_orf1ab;2366 | 0 | 0 |
| NC_004718.3;NP_828849.2;SARS-CoV_orf1ab;2367 | 0 | 0 |
| NC_004718.3;NP_828849.2;SARS-CoV_orf1ab;2368 | 0 | 0 |
| NC_004718.3;NP_828849.2;SARS-CoV_orf1ab;2369 | 0 | 0 |
| NC_004718.3;NP_828849.2;SARS-CoV_orf1ab;2370 | 0 | 0 |
| NC_004718.3;NP_828849.2;SARS-CoV_orf1ab;2371 | 0 | 0 |
| NC_004718.3;NP_828849.2;SARS-CoV_orf1ab;2372 | 0 | 0 |
| NC_004718.3;NP_828849.2;SARS-CoV_orf1ab;2373 | 0 | 0 |
| NC_004718.3;NP_828849.2;SARS-CoV_orf1ab;2374 | 0 | 0 |
| NC_004718.3;NP_828849.2;SARS-CoV_orf1ab;2375 | 0 | 0 |
| NC_004718.3;NP_828849.2;SARS-CoV_orf1ab;2376 | 0 | 0 |
| NC_004718.3;NP_828849.2;SARS-CoV_orf1ab;2377 | 0 | 0 |
| NC_004718.3;NP_828849.2;SARS-CoV_orf1ab;2378 | 0 | 0 |
| NC_004718.3;NP_828849.2;SARS-CoV_orf1ab;2379 | 0 | 0 |
| NC_004718.3;NP_828849.2;SARS-CoV_orf1ab;2380 | 0 | 0 |
| NC_004718.3;NP_828849.2;SARS-CoV_orf1ab;2381 | 0 | 0 |
| NC_004718.3;NP_828849.2;SARS-CoV_orf1ab;2382 | 0 | 0 |
| NC_004718.3;NP_828849.2;SARS-CoV_orf1ab;2383 | 0 | 0 |
| NC_004718.3;NP_828849.2;SARS-CoV_orf1ab;2384 | 0 | 0 |
| NC_004718.3;NP_828849.2;SARS-CoV_orf1ab;2385 | 0 | 0 |
| NC_004718.3;NP_828849.2;SARS-CoV_orf1ab;2386 | 0 | 0 |
| NC_004718.3;NP_828849.2;SARS-CoV_orf1ab;2387 | 0 | 0 |
| NC_004718.3;NP_828849.2;SARS-CoV_orf1ab;2388 | 0 | 0 |
| NC_004718.3;NP_828849.2;SARS-CoV_orf1ab;2389 | 0 | 0 |
| NC_004718.3;NP_828849.2;SARS-CoV_orf1ab;2390 | 0 | 0 |
| NC_004718.3;NP_828849.2;SARS-CoV_orf1ab;2391 | 0 | 0 |
| NC_004718.3;NP_828849.2;SARS-CoV_orf1ab;2392 | 0 | 0 |
| NC_004718.3;NP_828849.2;SARS-CoV_orf1ab;2393 | 0 | 0 |
| NC_004718.3;NP_828849.2;SARS-CoV_orf1ab;2394 | 0 | 0 |
| NC_004718.3;NP_828849.2;SARS-CoV_orf1ab;2395 | 0 | 0 |
| NC_004718.3;NP_828849.2;SARS-CoV_orf1ab;2396 | 0 | 0 |
| NC_004718.3;NP_828849.2;SARS-CoV_orf1ab;2397 | 0 | 0 |
| NC_004718.3;NP_828849.2;SARS-CoV_orf1ab;2398 | 0 | 0 |
| NC_004718.3;NP_828849.2;SARS-CoV_orf1ab;2399 | 0 | 0 |
| NC_004718.3;NP_828849.2;SARS-CoV_orf1ab;2400 | 0 | 0 |
| NC_004718.3;NP_828849.2;SARS-CoV_orf1ab;2401 | 0 | 0 |
| NC_004718.3;NP_828849.2;SARS-CoV_orf1ab;2402 | 0 | 0 |
| NC_004718.3;NP_828849.2;SARS-CoV_orf1ab;2403 | 0 | 0 |
| NC_004718.3;NP_828849.2;SARS-CoV_orf1ab;2404 | 0 | 0 |
| NC_004718.3;NP_828849.2;SARS-CoV_orf1ab;2405 | 0 | 0 |

| | | |
|---|---|---|
| NC_004718.3;NP_828849.2;SARS-CoV_orf1ab;2406 | 0 | 0 |
| NC_004718.3;NP_828849.2;SARS-CoV_orf1ab;2407 | 0 | 0 |
| NC_004718.3;NP_828849.2;SARS-CoV_orf1ab;2408 | 0 | 0 |
| NC_004718.3;NP_828849.2;SARS-CoV_orf1ab;2409 | 0 | 0 |
| NC_004718.3;NP_828849.2;SARS-CoV_orf1ab;2410 | 0 | 0 |
| NC_004718.3;NP_828849.2;SARS-CoV_orf1ab;2411 | 0 | 0 |
| NC_004718.3;NP_828849.2;SARS-CoV_orf1ab;2412 | 0 | 0 |
| NC_004718.3;NP_828849.2;SARS-CoV_orf1ab;2413 | 0 | 0 |
| NC_004718.3;NP_828849.2;SARS-CoV_orf1ab;2414 | 0 | 0 |
| NC_004718.3;NP_828849.2;SARS-CoV_orf1ab;2415 | 0 | 0 |
| NC_004718.3;NP_828849.2;SARS-CoV_orf1ab;2416 | 0 | 0 |
| NC_004718.3;NP_828849.2;SARS-CoV_orf1ab;2417 | 0 | 0 |
| NC_004718.3;NP_828849.2;SARS-CoV_orf1ab;2418 | 0 | 0 |
| NC_004718.3;NP_828849.2;SARS-CoV_orf1ab;2419 | 0 | 0 |
| NC_004718.3;NP_828849.2;SARS-CoV_orf1ab;2420 | 0 | 0 |
| NC_004718.3;NP_828849.2;SARS-CoV_orf1ab;2421 | 0 | 0 |
| NC_004718.3;NP_828849.2;SARS-CoV_orf1ab;2422 | 0 | 0 |
| NC_004718.3;NP_828849.2;SARS-CoV_orf1ab;2423 | 0 | 0 |
| NC_004718.3;NP_828849.2;SARS-CoV_orf1ab;2424 | 0 | 0 |
| NC_004718.3;NP_828849.2;SARS-CoV_orf1ab;2425 | 0 | 0 |
| NC_004718.3;NP_828849.2;SARS-CoV_orf1ab;2426 | 0 | 0 |
| NC_004718.3;NP_828849.2;SARS-CoV_orf1ab;2427 | 0 | 0 |
| NC_004718.3;NP_828849.2;SARS-CoV_orf1ab;2428 | 0 | 0 |
| NC_004718.3;NP_828849.2;SARS-CoV_orf1ab;2429 | 0 | 0 |
| NC_004718.3;NP_828849.2;SARS-CoV_orf1ab;2430 | 0 | 0 |
| NC_004718.3;NP_828849.2;SARS-CoV_orf1ab;2431 | 0 | 0 |
| NC_004718.3;NP_828849.2;SARS-CoV_orf1ab;2432 | 0 | 0 |
| NC_004718.3;NP_828849.2;SARS-CoV_orf1ab;2433 | 0 | 0 |
| NC_004718.3;NP_828849.2;SARS-CoV_orf1ab;2434 | 0 | 0 |
| NC_004718.3;NP_828849.2;SARS-CoV_orf1ab;2435 | 0 | 0 |
| NC_004718.3;NP_828849.2;SARS-CoV_orf1ab;2436 | 0 | 0 |
| NC_004718.3;NP_828849.2;SARS-CoV_orf1ab;2437 | 0 | 0 |
| NC_004718.3;NP_828849.2;SARS-CoV_orf1ab;2438 | 0 | 0 |
| NC_004718.3;NP_828849.2;SARS-CoV_orf1ab;2439 | 0 | 0 |
| NC_004718.3;NP_828849.2;SARS-CoV_orf1ab;2440 | 0 | 0 |
| NC_004718.3;NP_828849.2;SARS-CoV_orf1ab;2441 | 0 | 0 |
| NC_004718.3;NP_828849.2;SARS-CoV_orf1ab;2442 | 0 | 0 |
| NC_004718.3;NP_828849.2;SARS-CoV_orf1ab;2443 | 0 | 0 |
| NC_004718.3;NP_828849.2;SARS-CoV_orf1ab;2444 | 0 | 0 |
| NC_004718.3;NP_828849.2;SARS-CoV_orf1ab;2445 | 0 | 0 |
| NC_004718.3;NP_828849.2;SARS-CoV_orf1ab;2446 | 0.05 | 0 |
| NC_004718.3;NP_828849.2;SARS-CoV_orf1ab;2447 | 0.05 | 0.025 |
| NC_004718.3;NP_828849.2;SARS-CoV_orf1ab;2448 | 0.05 | 0.025 |

| | | |
|---|---|---|
| NC_004718.3;NP_828849.2;SARS-CoV_orf1ab;2449 | 0 | 0 |
| NC_004718.3;NP_828849.2;SARS-CoV_orf1ab;2450 | 0 | 0 |
| NC_004718.3;NP_828849.2;SARS-CoV_orf1ab;2451 | 0 | 0 |
| NC_004718.3;NP_828849.2;SARS-CoV_orf1ab;2452 | 0 | 0.025 |
| NC_004718.3;NP_828849.2;SARS-CoV_orf1ab;2453 | 0 | 0 |
| NC_004718.3;NP_828849.2;SARS-CoV_orf1ab;2454 | 0 | 0 |
| NC_004718.3;NP_828849.2;SARS-CoV_orf1ab;2455 | 0 | 0.05 |
| NC_004718.3;NP_828849.2;SARS-CoV_orf1ab;2456 | 0 | 0 |
| NC_004718.3;NP_828849.2;SARS-CoV_orf1ab;2457 | 0 | 0 |
| NC_004718.3;NP_828849.2;SARS-CoV_orf1ab;2458 | 0 | 0 |
| NC_004718.3;NP_828849.2;SARS-CoV_orf1ab;2459 | 0 | 0.025 |
| NC_004718.3;NP_828849.2;SARS-CoV_orf1ab;2460 | 0 | 0 |
| NC_004718.3;NP_828849.2;SARS-CoV_orf1ab;2461 | 0.05 | 0.025 |
| NC_004718.3;NP_828849.2;SARS-CoV_orf1ab;2462 | 0 | 0.025 |
| NC_004718.3;NP_828849.2;SARS-CoV_orf1ab;2463 | 0 | 0.025 |
| NC_004718.3;NP_828849.2;SARS-CoV_orf1ab;2464 | 0 | 0.025 |
| NC_004718.3;NP_828849.2;SARS-CoV_orf1ab;2465 | 0 | 0.025 |
| NC_004718.3;NP_828849.2;SARS-CoV_orf1ab;2466 | 0 | 0.025 |
| NC_004718.3;NP_828849.2;SARS-CoV_orf1ab;2467 | 0 | 0 |
| NC_004718.3;NP_828849.2;SARS-CoV_orf1ab;2468 | 0 | 0 |
| NC_004718.3;NP_828849.2;SARS-CoV_orf1ab;2469 | 0 | 0 |
| NC_004718.3;NP_828849.2;SARS-CoV_orf1ab;2470 | 0 | 0 |
| NC_004718.3;NP_828849.2;SARS-CoV_orf1ab;2471 | 0 | 0 |
| NC_004718.3;NP_828849.2;SARS-CoV_orf1ab;2472 | 0 | 0 |
| NC_004718.3;NP_828849.2;SARS-CoV_orf1ab;2473 | 0 | 0 |
| NC_004718.3;NP_828849.2;SARS-CoV_orf1ab;2474 | 0 | 0 |
| NC_004718.3;NP_828849.2;SARS-CoV_orf1ab;2475 | 0 | 0 |
| NC_004718.3;NP_828849.2;SARS-CoV_orf1ab;2476 | 0 | 0 |
| NC_004718.3;NP_828849.2;SARS-CoV_orf1ab;2477 | 0 | 0 |
| NC_004718.3;NP_828849.2;SARS-CoV_orf1ab;2478 | 0 | 0 |
| NC_004718.3;NP_828849.2;SARS-CoV_orf1ab;2479 | 0 | 0 |
| NC_004718.3;NP_828849.2;SARS-CoV_orf1ab;2480 | 0 | 0 |
| NC_004718.3;NP_828849.2;SARS-CoV_orf1ab;2481 | 0 | 0.025 |
| NC_004718.3;NP_828849.2;SARS-CoV_orf1ab;2482 | 0 | 0 |
| NC_004718.3;NP_828849.2;SARS-CoV_orf1ab;2483 | 0 | 0 |
| NC_004718.3;NP_828849.2;SARS-CoV_orf1ab;2484 | 0 | 0 |
| NC_004718.3;NP_828849.2;SARS-CoV_orf1ab;2485 | 0 | 0 |
| NC_004718.3;NP_828849.2;SARS-CoV_orf1ab;2486 | 0 | 0 |
| NC_004718.3;NP_828849.2;SARS-CoV_orf1ab;2487 | 0 | 0 |
| NC_004718.3;NP_828849.2;SARS-CoV_orf1ab;2488 | 0 | 0 |
| NC_004718.3;NP_828849.2;SARS-CoV_orf1ab;2489 | 0 | 0 |
| NC_004718.3;NP_828849.2;SARS-CoV_orf1ab;2490 | 0 | 0 |
| NC_004718.3;NP_828849.2;SARS-CoV_orf1ab;2491 | 0 | 0 |

| | | |
|---|---|---|
| NC_004718.3;NP_828849.2;SARS-CoV_orf1ab;2492 | 0 | 0 |
| NC_004718.3;NP_828849.2;SARS-CoV_orf1ab;2493 | 0 | 0 |
| NC_004718.3;NP_828849.2;SARS-CoV_orf1ab;2494 | 0 | 0 |
| NC_004718.3;NP_828849.2;SARS-CoV_orf1ab;2495 | 0 | 0 |
| NC_004718.3;NP_828849.2;SARS-CoV_orf1ab;2496 | 0 | 0 |
| NC_004718.3;NP_828849.2;SARS-CoV_orf1ab;2497 | 0 | 0 |
| NC_004718.3;NP_828849.2;SARS-CoV_orf1ab;2498 | 0 | 0.025 |
| NC_004718.3;NP_828849.2;SARS-CoV_orf1ab;2499 | 0.05 | 0.025 |
| NC_004718.3;NP_828849.2;SARS-CoV_orf1ab;2500 | 0.05 | 0.025 |
| NC_004718.3;NP_828849.2;SARS-CoV_orf1ab;2501 | 0.05 | 0.025 |
| NC_004718.3;NP_828849.2;SARS-CoV_orf1ab;2502 | 0.05 | 0.025 |
| NC_004718.3;NP_828849.2;SARS-CoV_orf1ab;2503 | 0 | 0 |
| NC_004718.3;NP_828849.2;SARS-CoV_orf1ab;2504 | 0 | 0 |
| NC_004718.3;NP_828849.2;SARS-CoV_orf1ab;2505 | 0.05 | 0 |
| NC_004718.3;NP_828849.2;SARS-CoV_orf1ab;2506 | 0 | 0 |
| NC_004718.3;NP_828849.2;SARS-CoV_orf1ab;2507 | 0 | 0 |
| NC_004718.3;NP_828849.2;SARS-CoV_orf1ab;2508 | 0 | 0 |
| NC_004718.3;NP_828849.2;SARS-CoV_orf1ab;2509 | 0 | 0 |
| NC_004718.3;NP_828849.2;SARS-CoV_orf1ab;2510 | 0 | 0 |
| NC_004718.3;NP_828849.2;SARS-CoV_orf1ab;2511 | 0 | 0.025 |
| NC_004718.3;NP_828849.2;SARS-CoV_orf1ab;2512 | 0 | 0.025 |
| NC_004718.3;NP_828849.2;SARS-CoV_orf1ab;2513 | 0.05 | 0.025 |
| NC_004718.3;NP_828849.2;SARS-CoV_orf1ab;2514 | 0.05 | 0.025 |
| NC_004718.3;NP_828849.2;SARS-CoV_orf1ab;2515 | 0.05 | 0.025 |
| NC_004718.3;NP_828849.2;SARS-CoV_orf1ab;2516 | 0.05 | 0.025 |
| NC_004718.3;NP_828849.2;SARS-CoV_orf1ab;2517 | 0 | 0.025 |
| NC_004718.3;NP_828849.2;SARS-CoV_orf1ab;2518 | 0.05 | 0.025 |
| NC_004718.3;NP_828849.2;SARS-CoV_orf1ab;2519 | 0.05 | 0.025 |
| NC_004718.3;NP_828849.2;SARS-CoV_orf1ab;2520 | 0.15 | 0.075 |
| NC_004718.3;NP_828849.2;SARS-CoV_orf1ab;2521 | 0 | 0 |
| NC_004718.3;NP_828849.2;SARS-CoV_orf1ab;2522 | 0 | 0.025 |
| NC_004718.3;NP_828849.2;SARS-CoV_orf1ab;2523 | 0 | 0 |
| NC_004718.3;NP_828849.2;SARS-CoV_orf1ab;2524 | 0 | 0 |
| NC_004718.3;NP_828849.2;SARS-CoV_orf1ab;2525 | 0.05 | 0 |
| NC_004718.3;NP_828849.2;SARS-CoV_orf1ab;2526 | 0 | 0 |
| NC_004718.3;NP_828849.2;SARS-CoV_orf1ab;2527 | 0.05 | 0.025 |
| NC_004718.3;NP_828849.2;SARS-CoV_orf1ab;2528 | 0.05 | 0.075 |
| NC_004718.3;NP_828849.2;SARS-CoV_orf1ab;2529 | 0 | 0 |
| NC_004718.3;NP_828849.2;SARS-CoV_orf1ab;2530 | 0 | 0.025 |
| NC_004718.3;NP_828849.2;SARS-CoV_orf1ab;2531 | 0 | 0 |
| NC_004718.3;NP_828849.2;SARS-CoV_orf1ab;2532 | 0 | 0 |
| NC_004718.3;NP_828849.2;SARS-CoV_orf1ab;2533 | 0 | 0 |
| NC_004718.3;NP_828849.2;SARS-CoV_orf1ab;2534 | 0 | 0.025 |

| | | |
|---|---|---|
| NC_004718.3;NP_828849.2;SARS-CoV_orf1ab;2535 | 0 | 0 |
| NC_004718.3;NP_828849.2;SARS-CoV_orf1ab;2536 | 0 | 0 |
| NC_004718.3;NP_828849.2;SARS-CoV_orf1ab;2537 | 0 | 0 |
| NC_004718.3;NP_828849.2;SARS-CoV_orf1ab;2538 | 0 | 0 |
| NC_004718.3;NP_828849.2;SARS-CoV_orf1ab;2539 | 0 | 0 |
| NC_004718.3;NP_828849.2;SARS-CoV_orf1ab;2540 | 0 | 0 |
| NC_004718.3;NP_828849.2;SARS-CoV_orf1ab;2541 | 0 | 0 |
| NC_004718.3;NP_828849.2;SARS-CoV_orf1ab;2542 | 0 | 0 |
| NC_004718.3;NP_828849.2;SARS-CoV_orf1ab;2543 | 0 | 0 |
| NC_004718.3;NP_828849.2;SARS-CoV_orf1ab;2544 | 0 | 0 |
| NC_004718.3;NP_828849.2;SARS-CoV_orf1ab;2545 | 0 | 0 |
| NC_004718.3;NP_828849.2;SARS-CoV_orf1ab;2546 | 0 | 0 |
| NC_004718.3;NP_828849.2;SARS-CoV_orf1ab;2547 | 0 | 0.025 |
| NC_004718.3;NP_828849.2;SARS-CoV_orf1ab;2548 | 0.1 | 0.1 |
| NC_004718.3;NP_828849.2;SARS-CoV_orf1ab;2549 | 0 | 0.05 |
| NC_004718.3;NP_828849.2;SARS-CoV_orf1ab;2550 | 0 | 0 |
| NC_004718.3;NP_828849.2;SARS-CoV_orf1ab;2551 | 0 | 0 |
| NC_004718.3;NP_828849.2;SARS-CoV_orf1ab;2552 | 0 | 0 |
| NC_004718.3;NP_828849.2;SARS-CoV_orf1ab;2553 | 0 | 0 |
| NC_004718.3;NP_828849.2;SARS-CoV_orf1ab;2554 | 0 | 0 |
| NC_004718.3;NP_828849.2;SARS-CoV_orf1ab;2555 | 0 | 0 |
| NC_004718.3;NP_828849.2;SARS-CoV_orf1ab;2556 | 0 | 0 |
| NC_004718.3;NP_828849.2;SARS-CoV_orf1ab;2557 | 0 | 0 |
| NC_004718.3;NP_828849.2;SARS-CoV_orf1ab;2558 | 0 | 0 |
| NC_004718.3;NP_828849.2;SARS-CoV_orf1ab;2559 | 0 | 0 |
| NC_004718.3;NP_828849.2;SARS-CoV_orf1ab;2560 | 0 | 0 |
| NC_004718.3;NP_828849.2;SARS-CoV_orf1ab;2561 | 0 | 0 |
| NC_004718.3;NP_828849.2;SARS-CoV_orf1ab;2562 | 0 | 0 |
| NC_004718.3;NP_828849.2;SARS-CoV_orf1ab;2563 | 0.05 | 0.025 |
| NC_004718.3;NP_828849.2;SARS-CoV_orf1ab;2564 | 0.05 | 0 |
| NC_004718.3;NP_828849.2;SARS-CoV_orf1ab;2565 | 0 | 0 |
| NC_004718.3;NP_828849.2;SARS-CoV_orf1ab;2566 | 0 | 0 |
| NC_004718.3;NP_828849.2;SARS-CoV_orf1ab;2567 | 0 | 0.025 |
| NC_004718.3;NP_828849.2;SARS-CoV_orf1ab;2568 | 0 | 0.025 |
| NC_004718.3;NP_828849.2;SARS-CoV_orf1ab;2569 | 0 | 0 |
| NC_004718.3;NP_828849.2;SARS-CoV_orf1ab;2570 | 0 | 0.05 |
| NC_004718.3;NP_828849.2;SARS-CoV_orf1ab;2571 | 0 | 0.025 |
| NC_004718.3;NP_828849.2;SARS-CoV_orf1ab;2572 | 0.05 | 0.025 |
| NC_004718.3;NP_828849.2;SARS-CoV_orf1ab;2573 | 0.1 | 0.05 |
| NC_004718.3;NP_828849.2;SARS-CoV_orf1ab;2574 | 0 | 0.025 |
| NC_004718.3;NP_828849.2;SARS-CoV_orf1ab;2575 | 0 | 0 |
| NC_004718.3;NP_828849.2;SARS-CoV_orf1ab;2576 | 0 | 0 |
| NC_004718.3;NP_828849.2;SARS-CoV_orf1ab;2577 | 0 | 0 |

| | | |
|---|---|---|
| NC_004718.3;NP_828849.2;SARS-CoV_orf1ab;2578 | 0 | 0 |
| NC_004718.3;NP_828849.2;SARS-CoV_orf1ab;2579 | 0 | 0 |
| NC_004718.3;NP_828849.2;SARS-CoV_orf1ab;2580 | 0 | 0 |
| NC_004718.3;NP_828849.2;SARS-CoV_orf1ab;2581 | 0 | 0 |
| NC_004718.3;NP_828849.2;SARS-CoV_orf1ab;2582 | 0 | 0 |
| NC_004718.3;NP_828849.2;SARS-CoV_orf1ab;2583 | 0 | 0 |
| NC_004718.3;NP_828849.2;SARS-CoV_orf1ab;2584 | 0 | 0.025 |
| NC_004718.3;NP_828849.2;SARS-CoV_orf1ab;2585 | 0 | 0 |
| NC_004718.3;NP_828849.2;SARS-CoV_orf1ab;2586 | 0.1 | 0.05 |
| NC_004718.3;NP_828849.2;SARS-CoV_orf1ab;2587 | 0.1 | 0.1 |
| NC_004718.3;NP_828849.2;SARS-CoV_orf1ab;2588 | 0.15 | 0.1 |
| NC_004718.3;NP_828849.2;SARS-CoV_orf1ab;2589 | 0.25 | 0.125 |
| NC_004718.3;NP_828849.2;SARS-CoV_orf1ab;2590 | 0.25 | 0.125 |
| NC_004718.3;NP_828849.2;SARS-CoV_orf1ab;2591 | 0.2 | 0.1 |
| NC_004718.3;NP_828849.2;SARS-CoV_orf1ab;2592 | 0.25 | 0.1 |
| NC_004718.3;NP_828849.2;SARS-CoV_orf1ab;2593 | 0.2 | 0.075 |
| NC_004718.3;NP_828849.2;SARS-CoV_orf1ab;2594 | 0.2 | 0.075 |
| NC_004718.3;NP_828849.2;SARS-CoV_orf1ab;2595 | 0.15 | 0.05 |
| NC_004718.3;NP_828849.2;SARS-CoV_orf1ab;2596 | 0.1 | 0.05 |
| NC_004718.3;NP_828849.2;SARS-CoV_orf1ab;2597 | 0 | 0 |
| NC_004718.3;NP_828849.2;SARS-CoV_orf1ab;2598 | 0 | 0 |
| NC_004718.3;NP_828849.2;SARS-CoV_orf1ab;2599 | 0.05 | 0 |
| NC_004718.3;NP_828849.2;SARS-CoV_orf1ab;2600 | 0 | 0 |
| NC_004718.3;NP_828849.2;SARS-CoV_orf1ab;2601 | 0 | 0 |
| NC_004718.3;NP_828849.2;SARS-CoV_orf1ab;2602 | 0 | 0 |
| NC_004718.3;NP_828849.2;SARS-CoV_orf1ab;2603 | 0 | 0 |
| NC_004718.3;NP_828849.2;SARS-CoV_orf1ab;2604 | 0 | 0 |
| NC_004718.3;NP_828849.2;SARS-CoV_orf1ab;2605 | 0.05 | 0 |
| NC_004718.3;NP_828849.2;SARS-CoV_orf1ab;2606 | 0 | 0 |
| NC_004718.3;NP_828849.2;SARS-CoV_orf1ab;2607 | 0 | 0 |
| NC_004718.3;NP_828849.2;SARS-CoV_orf1ab;2608 | 0 | 0 |
| NC_004718.3;NP_828849.2;SARS-CoV_orf1ab;2609 | 0 | 0.025 |
| NC_004718.3;NP_828849.2;SARS-CoV_orf1ab;2610 | 0 | 0 |
| NC_004718.3;NP_828849.2;SARS-CoV_orf1ab;2611 | 0 | 0 |
| NC_004718.3;NP_828849.2;SARS-CoV_orf1ab;2612 | 0 | 0 |
| NC_004718.3;NP_828849.2;SARS-CoV_orf1ab;2613 | 0 | 0.025 |
| NC_004718.3;NP_828849.2;SARS-CoV_orf1ab;2614 | 0 | 0 |
| NC_004718.3;NP_828849.2;SARS-CoV_orf1ab;2615 | 0.05 | 0 |
| NC_004718.3;NP_828849.2;SARS-CoV_orf1ab;2616 | 0.05 | 0 |
| NC_004718.3;NP_828849.2;SARS-CoV_orf1ab;2617 | 0 | 0 |
| NC_004718.3;NP_828849.2;SARS-CoV_orf1ab;2618 | 0 | 0 |
| NC_004718.3;NP_828849.2;SARS-CoV_orf1ab;2619 | 0 | 0 |
| NC_004718.3;NP_828849.2;SARS-CoV_orf1ab;2620 | 0 | 0 |

| | | |
|---|---|---|
| NC_004718.3;NP_828849.2;SARS-CoV_orf1ab;2621 | 0 | 0 |
| NC_004718.3;NP_828849.2;SARS-CoV_orf1ab;2622 | 0 | 0 |
| NC_004718.3;NP_828849.2;SARS-CoV_orf1ab;2623 | 0 | 0 |
| NC_004718.3;NP_828849.2;SARS-CoV_orf1ab;2624 | 0 | 0 |
| NC_004718.3;NP_828849.2;SARS-CoV_orf1ab;2625 | 0 | 0.025 |
| NC_004718.3;NP_828849.2;SARS-CoV_orf1ab;2626 | 0 | 0 |
| NC_004718.3;NP_828849.2;SARS-CoV_orf1ab;2627 | 0 | 0 |
| NC_004718.3;NP_828849.2;SARS-CoV_orf1ab;2628 | 0 | 0 |
| NC_004718.3;NP_828849.2;SARS-CoV_orf1ab;2629 | 0 | 0 |
| NC_004718.3;NP_828849.2;SARS-CoV_orf1ab;2630 | 0 | 0 |
| NC_004718.3;NP_828849.2;SARS-CoV_orf1ab;2631 | 0 | 0 |
| NC_004718.3;NP_828849.2;SARS-CoV_orf1ab;2632 | 0 | 0 |
| NC_004718.3;NP_828849.2;SARS-CoV_orf1ab;2633 | 0 | 0 |
| NC_004718.3;NP_828849.2;SARS-CoV_orf1ab;2634 | 0 | 0 |
| NC_004718.3;NP_828849.2;SARS-CoV_orf1ab;2635 | 0 | 0 |
| NC_004718.3;NP_828849.2;SARS-CoV_orf1ab;2636 | 0 | 0 |
| NC_004718.3;NP_828849.2;SARS-CoV_orf1ab;2637 | 0 | 0 |
| NC_004718.3;NP_828849.2;SARS-CoV_orf1ab;2638 | 0.05 | 0 |
| NC_004718.3;NP_828849.2;SARS-CoV_orf1ab;2639 | 0 | 0 |
| NC_004718.3;NP_828849.2;SARS-CoV_orf1ab;2640 | 0.05 | 0.05 |
| NC_004718.3;NP_828849.2;SARS-CoV_orf1ab;2641 | 0.05 | 0.05 |
| NC_004718.3;NP_828849.2;SARS-CoV_orf1ab;2642 | 0 | 0 |
| NC_004718.3;NP_828849.2;SARS-CoV_orf1ab;2643 | 0 | 0 |
| NC_004718.3;NP_828849.2;SARS-CoV_orf1ab;2644 | 0 | 0 |
| NC_004718.3;NP_828849.2;SARS-CoV_orf1ab;2645 | 0 | 0 |
| NC_004718.3;NP_828849.2;SARS-CoV_orf1ab;2646 | 0 | 0 |
| NC_004718.3;NP_828849.2;SARS-CoV_orf1ab;2647 | 0 | 0 |
| NC_004718.3;NP_828849.2;SARS-CoV_orf1ab;2648 | 0 | 0 |
| NC_004718.3;NP_828849.2;SARS-CoV_orf1ab;2649 | 0 | 0 |
| NC_004718.3;NP_828849.2;SARS-CoV_orf1ab;2650 | 0 | 0 |
| NC_004718.3;NP_828849.2;SARS-CoV_orf1ab;2651 | 0 | 0.025 |
| NC_004718.3;NP_828849.2;SARS-CoV_orf1ab;2652 | 0.05 | 0.075 |
| NC_004718.3;NP_828849.2;SARS-CoV_orf1ab;2653 | 0.05 | 0.025 |
| NC_004718.3;NP_828849.2;SARS-CoV_orf1ab;2654 | 0.05 | 0.05 |
| NC_004718.3;NP_828849.2;SARS-CoV_orf1ab;2655 | 0 | 0 |
| NC_004718.3;NP_828849.2;SARS-CoV_orf1ab;2656 | 0 | 0 |
| NC_004718.3;NP_828849.2;SARS-CoV_orf1ab;2657 | 0 | 0 |
| NC_004718.3;NP_828849.2;SARS-CoV_orf1ab;2658 | 0 | 0.025 |
| NC_004718.3;NP_828849.2;SARS-CoV_orf1ab;2659 | 0 | 0 |
| NC_004718.3;NP_828849.2;SARS-CoV_orf1ab;2660 | 0 | 0 |
| NC_004718.3;NP_828849.2;SARS-CoV_orf1ab;2661 | 0.05 | 0 |
| NC_004718.3;NP_828849.2;SARS-CoV_orf1ab;2662 | 0.05 | 0 |
| NC_004718.3;NP_828849.2;SARS-CoV_orf1ab;2663 | 0 | 0 |

| | | |
|---|---|---|
| NC_004718.3;NP_828849.2;SARS-CoV_orf1ab;2664 | 0 | 0 |
| NC_004718.3;NP_828849.2;SARS-CoV_orf1ab;2665 | 0 | 0 |
| NC_004718.3;NP_828849.2;SARS-CoV_orf1ab;2666 | 0.05 | 0.05 |
| NC_004718.3;NP_828849.2;SARS-CoV_orf1ab;2667 | 0 | 0 |
| NC_004718.3;NP_828849.2;SARS-CoV_orf1ab;2668 | 0.05 | 0 |
| NC_004718.3;NP_828849.2;SARS-CoV_orf1ab;2669 | 0 | 0 |
| NC_004718.3;NP_828849.2;SARS-CoV_orf1ab;2670 | 0.05 | 0 |
| NC_004718.3;NP_828849.2;SARS-CoV_orf1ab;2671 | 0 | 0 |
| NC_004718.3;NP_828849.2;SARS-CoV_orf1ab;2672 | 0 | 0 |
| NC_004718.3;NP_828849.2;SARS-CoV_orf1ab;2673 | 0 | 0 |
| NC_004718.3;NP_828849.2;SARS-CoV_orf1ab;2674 | 0 | 0 |
| NC_004718.3;NP_828849.2;SARS-CoV_orf1ab;2675 | 0 | 0 |
| NC_004718.3;NP_828849.2;SARS-CoV_orf1ab;2676 | 0 | 0 |
| NC_004718.3;NP_828849.2;SARS-CoV_orf1ab;2677 | 0 | 0 |
| NC_004718.3;NP_828849.2;SARS-CoV_orf1ab;2678 | 0 | 0 |
| NC_004718.3;NP_828849.2;SARS-CoV_orf1ab;2679 | 0.05 | 0 |
| NC_004718.3;NP_828849.2;SARS-CoV_orf1ab;2680 | 0 | 0 |
| NC_004718.3;NP_828849.2;SARS-CoV_orf1ab;2681 | 0 | 0 |
| NC_004718.3;NP_828849.2;SARS-CoV_orf1ab;2682 | 0 | 0 |
| NC_004718.3;NP_828849.2;SARS-CoV_orf1ab;2683 | 0 | 0 |
| NC_004718.3;NP_828849.2;SARS-CoV_orf1ab;2684 | 0 | 0 |
| NC_004718.3;NP_828849.2;SARS-CoV_orf1ab;2685 | 0 | 0 |
| NC_004718.3;NP_828849.2;SARS-CoV_orf1ab;2686 | 0 | 0.025 |
| NC_004718.3;NP_828849.2;SARS-CoV_orf1ab;2687 | 0 | 0 |
| NC_004718.3;NP_828849.2;SARS-CoV_orf1ab;2688 | 0 | 0 |
| NC_004718.3;NP_828849.2;SARS-CoV_orf1ab;2689 | 0 | 0.025 |
| NC_004718.3;NP_828849.2;SARS-CoV_orf1ab;2690 | 0 | 0 |
| NC_004718.3;NP_828849.2;SARS-CoV_orf1ab;2691 | 0 | 0.025 |
| NC_004718.3;NP_828849.2;SARS-CoV_orf1ab;2692 | 0.05 | 0.05 |
| NC_004718.3;NP_828849.2;SARS-CoV_orf1ab;2693 | 0 | 0 |
| NC_004718.3;NP_828849.2;SARS-CoV_orf1ab;2694 | 0 | 0 |
| NC_004718.3;NP_828849.2;SARS-CoV_orf1ab;2695 | 0.05 | 0 |
| NC_004718.3;NP_828849.2;SARS-CoV_orf1ab;2696 | 0 | 0 |
| NC_004718.3;NP_828849.2;SARS-CoV_orf1ab;2697 | 0 | 0 |
| NC_004718.3;NP_828849.2;SARS-CoV_orf1ab;2698 | 0 | 0 |
| NC_004718.3;NP_828849.2;SARS-CoV_orf1ab;2699 | 0 | 0.025 |
| NC_004718.3;NP_828849.2;SARS-CoV_orf1ab;2700 | 0 | 0 |
| NC_004718.3;NP_828849.2;SARS-CoV_orf1ab;2701 | 0 | 0 |
| NC_004718.3;NP_828849.2;SARS-CoV_orf1ab;2702 | 0 | 0 |
| NC_004718.3;NP_828849.2;SARS-CoV_orf1ab;2703 | 0 | 0.025 |
| NC_004718.3;NP_828849.2;SARS-CoV_orf1ab;2704 | 0 | 0 |
| NC_004718.3;NP_828849.2;SARS-CoV_orf1ab;2705 | 0.05 | 0 |
| NC_004718.3;NP_828849.2;SARS-CoV_orf1ab;2706 | 0.05 | 0 |

| | | |
|---|---|---|
| NC_004718.3;NP_828849.2;SARS-CoV_orf1ab;2707 | 0 | 0 |
| NC_004718.3;NP_828849.2;SARS-CoV_orf1ab;2708 | 0 | 0 |
| NC_004718.3;NP_828849.2;SARS-CoV_orf1ab;2709 | 0 | 0 |
| NC_004718.3;NP_828849.2;SARS-CoV_orf1ab;2710 | 0 | 0 |
| NC_004718.3;NP_828849.2;SARS-CoV_orf1ab;2711 | 0 | 0 |
| NC_004718.3;NP_828849.2;SARS-CoV_orf1ab;2712 | 0 | 0 |
| NC_004718.3;NP_828849.2;SARS-CoV_orf1ab;2713 | 0 | 0 |
| NC_004718.3;NP_828849.2;SARS-CoV_orf1ab;2714 | 0 | 0 |
| NC_004718.3;NP_828849.2;SARS-CoV_orf1ab;2715 | 0 | 0.025 |
| NC_004718.3;NP_828849.2;SARS-CoV_orf1ab;2716 | 0 | 0 |
| NC_004718.3;NP_828849.2;SARS-CoV_orf1ab;2717 | 0 | 0 |
| NC_004718.3;NP_828849.2;SARS-CoV_orf1ab;2718 | 0 | 0 |
| NC_004718.3;NP_828849.2;SARS-CoV_orf1ab;2719 | 0 | 0 |
| NC_004718.3;NP_828849.2;SARS-CoV_orf1ab;2720 | 0 | 0 |
| NC_004718.3;NP_828849.2;SARS-CoV_orf1ab;2721 | 0 | 0 |
| NC_004718.3;NP_828849.2;SARS-CoV_orf1ab;2722 | 0 | 0 |
| NC_004718.3;NP_828849.2;SARS-CoV_orf1ab;2723 | 0 | 0 |
| NC_004718.3;NP_828849.2;SARS-CoV_orf1ab;2724 | 0 | 0 |
| NC_004718.3;NP_828849.2;SARS-CoV_orf1ab;2725 | 0 | 0 |
| NC_004718.3;NP_828849.2;SARS-CoV_orf1ab;2726 | 0 | 0 |
| NC_004718.3;NP_828849.2;SARS-CoV_orf1ab;2727 | 0 | 0 |
| NC_004718.3;NP_828849.2;SARS-CoV_orf1ab;2728 | 0 | 0 |
| NC_004718.3;NP_828849.2;SARS-CoV_orf1ab;2729 | 0 | 0 |
| NC_004718.3;NP_828849.2;SARS-CoV_orf1ab;2730 | 0 | 0 |
| NC_004718.3;NP_828849.2;SARS-CoV_orf1ab;2731 | 0 | 0 |
| NC_004718.3;NP_828849.2;SARS-CoV_orf1ab;2732 | 0 | 0 |
| NC_004718.3;NP_828849.2;SARS-CoV_orf1ab;2733 | 0 | 0 |
| NC_004718.3;NP_828849.2;SARS-CoV_orf1ab;2734 | 0 | 0 |
| NC_004718.3;NP_828849.2;SARS-CoV_orf1ab;2735 | 0 | 0 |
| NC_004718.3;NP_828849.2;SARS-CoV_orf1ab;2736 | 0 | 0 |
| NC_004718.3;NP_828849.2;SARS-CoV_orf1ab;2737 | 0 | 0 |
| NC_004718.3;NP_828849.2;SARS-CoV_orf1ab;2738 | 0 | 0 |
| NC_004718.3;NP_828849.2;SARS-CoV_orf1ab;2739 | 0 | 0 |
| NC_004718.3;NP_828849.2;SARS-CoV_orf1ab;2740 | 0 | 0 |
| NC_004718.3;NP_828849.2;SARS-CoV_orf1ab;2741 | 0 | 0 |
| NC_004718.3;NP_828849.2;SARS-CoV_orf1ab;2742 | 0 | 0 |
| NC_004718.3;NP_828849.2;SARS-CoV_orf1ab;2743 | 0 | 0 |
| NC_004718.3;NP_828849.2;SARS-CoV_orf1ab;2744 | 0 | 0 |
| NC_004718.3;NP_828849.2;SARS-CoV_orf1ab;2745 | 0 | 0 |
| NC_004718.3;NP_828849.2;SARS-CoV_orf1ab;2746 | 0 | 0 |
| NC_004718.3;NP_828849.2;SARS-CoV_orf1ab;2747 | 0 | 0 |
| NC_004718.3;NP_828849.2;SARS-CoV_orf1ab;2748 | 0 | 0 |
| NC_004718.3;NP_828849.2;SARS-CoV_orf1ab;2749 | 0 | 0 |

| | | |
|---|---|---|
| NC_004718.3;NP_828849.2;SARS-CoV_orf1ab;2750 | 0 | 0 |
| NC_004718.3;NP_828849.2;SARS-CoV_orf1ab;2751 | 0 | 0 |
| NC_004718.3;NP_828849.2;SARS-CoV_orf1ab;2752 | 0 | 0 |
| NC_004718.3;NP_828849.2;SARS-CoV_orf1ab;2753 | 0 | 0 |
| NC_004718.3;NP_828849.2;SARS-CoV_orf1ab;2754 | 0 | 0 |
| NC_004718.3;NP_828849.2;SARS-CoV_orf1ab;2755 | 0 | 0 |
| NC_004718.3;NP_828849.2;SARS-CoV_orf1ab;2756 | 0 | 0 |
| NC_004718.3;NP_828849.2;SARS-CoV_orf1ab;2757 | 0 | 0 |
| NC_004718.3;NP_828849.2;SARS-CoV_orf1ab;2758 | 0 | 0 |
| NC_004718.3;NP_828849.2;SARS-CoV_orf1ab;2759 | 0 | 0 |
| NC_004718.3;NP_828849.2;SARS-CoV_orf1ab;2760 | 0 | 0 |
| NC_004718.3;NP_828849.2;SARS-CoV_orf1ab;2761 | 0 | 0 |
| NC_004718.3;NP_828849.2;SARS-CoV_orf1ab;2762 | 0 | 0 |
| NC_004718.3;NP_828849.2;SARS-CoV_orf1ab;2763 | 0 | 0 |
| NC_004718.3;NP_828849.2;SARS-CoV_orf1ab;2764 | 0 | 0 |
| NC_004718.3;NP_828849.2;SARS-CoV_orf1ab;2765 | 0 | 0 |
| NC_004718.3;NP_828849.2;SARS-CoV_orf1ab;2766 | 0 | 0 |
| NC_004718.3;NP_828849.2;SARS-CoV_orf1ab;2767 | 0.05 | 0 |
| NC_004718.3;NP_828849.2;SARS-CoV_orf1ab;2768 | 0.05 | 0 |
| NC_004718.3;NP_828849.2;SARS-CoV_orf1ab;2769 | 0 | 0 |
| NC_004718.3;NP_828849.2;SARS-CoV_orf1ab;2770 | 0.05 | 0 |
| NC_004718.3;NP_828849.2;SARS-CoV_orf1ab;2771 | 0 | 0 |
| NC_004718.3;NP_828849.2;SARS-CoV_orf1ab;2772 | 0 | 0 |
| NC_004718.3;NP_828849.2;SARS-CoV_orf1ab;2773 | 0.1 | 0 |
| NC_004718.3;NP_828849.2;SARS-CoV_orf1ab;2774 | 0.05 | 0.05 |
| NC_004718.3;NP_828849.2;SARS-CoV_orf1ab;2775 | 0 | 0.05 |
| NC_004718.3;NP_828849.2;SARS-CoV_orf1ab;2776 | 0.05 | 0.05 |
| NC_004718.3;NP_828849.2;SARS-CoV_orf1ab;2777 | 0.1 | 0.05 |
| NC_004718.3;NP_828849.2;SARS-CoV_orf1ab;2778 | 0.1 | 0.075 |
| NC_004718.3;NP_828849.2;SARS-CoV_orf1ab;2779 | 0.15 | 0.05 |
| NC_004718.3;NP_828849.2;SARS-CoV_orf1ab;2780 | 0.15 | 0.05 |
| NC_004718.3;NP_828849.2;SARS-CoV_orf1ab;2781 | 0.1 | 0.1 |
| NC_004718.3;NP_828849.2;SARS-CoV_orf1ab;2782 | 0.1 | 0.075 |
| NC_004718.3;NP_828849.2;SARS-CoV_orf1ab;2783 | 0.05 | 0 |
| NC_004718.3;NP_828849.2;SARS-CoV_orf1ab;2784 | 0.05 | 0 |
| NC_004718.3;NP_828849.2;SARS-CoV_orf1ab;2785 | 0 | 0 |
| NC_004718.3;NP_828849.2;SARS-CoV_orf1ab;2786 | 0 | 0 |
| NC_004718.3;NP_828849.2;SARS-CoV_orf1ab;2787 | 0 | 0 |
| NC_004718.3;NP_828849.2;SARS-CoV_orf1ab;2788 | 0.1 | 0 |
| NC_004718.3;NP_828849.2;SARS-CoV_orf1ab;2789 | 0 | 0 |
| NC_004718.3;NP_828849.2;SARS-CoV_orf1ab;2790 | 0 | 0 |
| NC_004718.3;NP_828849.2;SARS-CoV_orf1ab;2791 | 0 | 0 |
| NC_004718.3;NP_828849.2;SARS-CoV_orf1ab;2792 | 0.15 | 0.075 |

| | | |
|---|---|---|
| NC_004718.3;NP_828849.2;SARS-CoV_orf1ab;2793 | 0 | 0 |
| NC_004718.3;NP_828849.2;SARS-CoV_orf1ab;2794 | 0 | 0.025 |
| NC_004718.3;NP_828849.2;SARS-CoV_orf1ab;2795 | 0 | 0 |
| NC_004718.3;NP_828849.2;SARS-CoV_orf1ab;2796 | 0 | 0 |
| NC_004718.3;NP_828849.2;SARS-CoV_orf1ab;2797 | 0 | 0 |
| NC_004718.3;NP_828849.2;SARS-CoV_orf1ab;2798 | 0.05 | 0.025 |
| NC_004718.3;NP_828849.2;SARS-CoV_orf1ab;2799 | 0.05 | 0.025 |
| NC_004718.3;NP_828849.2;SARS-CoV_orf1ab;2800 | 0 | 0 |
| NC_004718.3;NP_828849.2;SARS-CoV_orf1ab;2801 | 0 | 0 |
| NC_004718.3;NP_828849.2;SARS-CoV_orf1ab;2802 | 0 | 0 |
| NC_004718.3;NP_828849.2;SARS-CoV_orf1ab;2803 | 0 | 0 |
| NC_004718.3;NP_828849.2;SARS-CoV_orf1ab;2804 | 0 | 0 |
| NC_004718.3;NP_828849.2;SARS-CoV_orf1ab;2805 | 0.05 | 0 |
| NC_004718.3;NP_828849.2;SARS-CoV_orf1ab;2806 | 0 | 0 |
| NC_004718.3;NP_828849.2;SARS-CoV_orf1ab;2807 | 0 | 0 |
| NC_004718.3;NP_828849.2;SARS-CoV_orf1ab;2808 | 0 | 0 |
| NC_004718.3;NP_828849.2;SARS-CoV_orf1ab;2809 | 0 | 0 |
| NC_004718.3;NP_828849.2;SARS-CoV_orf1ab;2810 | 0 | 0.025 |
| NC_004718.3;NP_828849.2;SARS-CoV_orf1ab;2811 | 0.05 | 0 |
| NC_004718.3;NP_828849.2;SARS-CoV_orf1ab;2812 | 0.05 | 0 |
| NC_004718.3;NP_828849.2;SARS-CoV_orf1ab;2813 | 0 | 0 |
| NC_004718.3;NP_828849.2;SARS-CoV_orf1ab;2814 | 0.05 | 0 |
| NC_004718.3;NP_828849.2;SARS-CoV_orf1ab;2815 | 0.05 | 0 |
| NC_004718.3;NP_828849.2;SARS-CoV_orf1ab;2816 | 0 | 0 |
| NC_004718.3;NP_828849.2;SARS-CoV_orf1ab;2817 | 0 | 0 |
| NC_004718.3;NP_828849.2;SARS-CoV_orf1ab;2818 | 0.05 | 0.025 |
| NC_004718.3;NP_828849.2;SARS-CoV_orf1ab;2819 | 0.1 | 0 |
| NC_004718.3;NP_828849.2;SARS-CoV_orf1ab;2820 | 0.1 | 0 |
| NC_004718.3;NP_828849.2;SARS-CoV_orf1ab;2821 | 0.05 | 0.05 |
| NC_004718.3;NP_828849.2;SARS-CoV_orf1ab;2822 | 0 | 0 |
| NC_004718.3;NP_828849.2;SARS-CoV_orf1ab;2823 | 0 | 0 |
| NC_004718.3;NP_828849.2;SARS-CoV_orf1ab;2824 | 0 | 0 |
| NC_004718.3;NP_828849.2;SARS-CoV_orf1ab;2825 | 0 | 0 |
| NC_004718.3;NP_828849.2;SARS-CoV_orf1ab;2826 | 0 | 0 |
| NC_004718.3;NP_828849.2;SARS-CoV_orf1ab;2827 | 0 | 0 |
| NC_004718.3;NP_828849.2;SARS-CoV_orf1ab;2828 | 0 | 0 |
| NC_004718.3;NP_828849.2;SARS-CoV_orf1ab;2829 | 0 | 0 |
| NC_004718.3;NP_828849.2;SARS-CoV_orf1ab;2830 | 0 | 0 |
| NC_004718.3;NP_828849.2;SARS-CoV_orf1ab;2831 | 0 | 0 |
| NC_004718.3;NP_828849.2;SARS-CoV_orf1ab;2832 | 0 | 0.025 |
| NC_004718.3;NP_828849.2;SARS-CoV_orf1ab;2833 | 0 | 0 |
| NC_004718.3;NP_828849.2;SARS-CoV_orf1ab;2834 | 0 | 0 |
| NC_004718.3;NP_828849.2;SARS-CoV_orf1ab;2835 | 0 | 0.025 |

| | | |
|---|---|---|
| NC_004718.3;NP_828849.2;SARS-CoV_orf1ab;2836 | 0 | 0.025 |
| NC_004718.3;NP_828849.2;SARS-CoV_orf1ab;2837 | 0 | 0 |
| NC_004718.3;NP_828849.2;SARS-CoV_orf1ab;2838 | 0 | 0 |
| NC_004718.3;NP_828849.2;SARS-CoV_orf1ab;2839 | 0 | 0 |
| NC_004718.3;NP_828849.2;SARS-CoV_orf1ab;2840 | 0 | 0 |
| NC_004718.3;NP_828849.2;SARS-CoV_orf1ab;2841 | 0 | 0 |
| NC_004718.3;NP_828849.2;SARS-CoV_orf1ab;2842 | 0.05 | 0 |
| NC_004718.3;NP_828849.2;SARS-CoV_orf1ab;2843 | 0.05 | 0 |
| NC_004718.3;NP_828849.2;SARS-CoV_orf1ab;2844 | 0 | 0 |
| NC_004718.3;NP_828849.2;SARS-CoV_orf1ab;2845 | 0 | 0 |
| NC_004718.3;NP_828849.2;SARS-CoV_orf1ab;2846 | 0 | 0 |
| NC_004718.3;NP_828849.2;SARS-CoV_orf1ab;2847 | 0 | 0 |
| NC_004718.3;NP_828849.2;SARS-CoV_orf1ab;2848 | 0 | 0 |
| NC_004718.3;NP_828849.2;SARS-CoV_orf1ab;2849 | 0 | 0 |
| NC_004718.3;NP_828849.2;SARS-CoV_orf1ab;2850 | 0.05 | 0 |
| NC_004718.3;NP_828849.2;SARS-CoV_orf1ab;2851 | 0 | 0 |
| NC_004718.3;NP_828849.2;SARS-CoV_orf1ab;2852 | 0 | 0 |
| NC_004718.3;NP_828849.2;SARS-CoV_orf1ab;2853 | 0 | 0 |
| NC_004718.3;NP_828849.2;SARS-CoV_orf1ab;2854 | 0 | 0 |
| NC_004718.3;NP_828849.2;SARS-CoV_orf1ab;2855 | 0 | 0 |
| NC_004718.3;NP_828849.2;SARS-CoV_orf1ab;2856 | 0 | 0 |
| NC_004718.3;NP_828849.2;SARS-CoV_orf1ab;2857 | 0 | 0 |
| NC_004718.3;NP_828849.2;SARS-CoV_orf1ab;2858 | 0 | 0 |
| NC_004718.3;NP_828849.2;SARS-CoV_orf1ab;2859 | 0 | 0 |
| NC_004718.3;NP_828849.2;SARS-CoV_orf1ab;2860 | 0 | 0 |
| NC_004718.3;NP_828849.2;SARS-CoV_orf1ab;2861 | 0 | 0 |
| NC_004718.3;NP_828849.2;SARS-CoV_orf1ab;2862 | 0 | 0 |
| NC_004718.3;NP_828849.2;SARS-CoV_orf1ab;2863 | 0 | 0 |
| NC_004718.3;NP_828849.2;SARS-CoV_orf1ab;2864 | 0 | 0 |
| NC_004718.3;NP_828849.2;SARS-CoV_orf1ab;2865 | 0 | 0 |
| NC_004718.3;NP_828849.2;SARS-CoV_orf1ab;2866 | 0 | 0 |
| NC_004718.3;NP_828849.2;SARS-CoV_orf1ab;2867 | 0 | 0 |
| NC_004718.3;NP_828849.2;SARS-CoV_orf1ab;2868 | 0 | 0.025 |
| NC_004718.3;NP_828849.2;SARS-CoV_orf1ab;2869 | 0 | 0.025 |
| NC_004718.3;NP_828849.2;SARS-CoV_orf1ab;2870 | 0 | 0.05 |
| NC_004718.3;NP_828849.2;SARS-CoV_orf1ab;2871 | 0 | 0 |
| NC_004718.3;NP_828849.2;SARS-CoV_orf1ab;2872 | 0.05 | 0 |
| NC_004718.3;NP_828849.2;SARS-CoV_orf1ab;2873 | 0.05 | 0.025 |
| NC_004718.3;NP_828849.2;SARS-CoV_orf1ab;2874 | 0.05 | 0.025 |
| NC_004718.3;NP_828849.2;SARS-CoV_orf1ab;2875 | 0.05 | 0 |
| NC_004718.3;NP_828849.2;SARS-CoV_orf1ab;2876 | 0.05 | 0 |
| NC_004718.3;NP_828849.2;SARS-CoV_orf1ab;2877 | 0 | 0 |
| NC_004718.3;NP_828849.2;SARS-CoV_orf1ab;2878 | 0.05 | 0 |

| | | |
|---|---|---|
| NC_004718.3;NP_828849.2;SARS-CoV_orf1ab;2879 | 0.05 | 0 |
| NC_004718.3;NP_828849.2;SARS-CoV_orf1ab;2880 | 0.05 | 0 |
| NC_004718.3;NP_828849.2;SARS-CoV_orf1ab;2881 | 0.05 | 0 |
| NC_004718.3;NP_828849.2;SARS-CoV_orf1ab;2882 | 0 | 0 |
| NC_004718.3;NP_828849.2;SARS-CoV_orf1ab;2883 | 0 | 0 |
| NC_004718.3;NP_828849.2;SARS-CoV_orf1ab;2884 | 0 | 0 |
| NC_004718.3;NP_828849.2;SARS-CoV_orf1ab;2885 | 0 | 0 |
| NC_004718.3;NP_828849.2;SARS-CoV_orf1ab;2886 | 0 | 0 |
| NC_004718.3;NP_828849.2;SARS-CoV_orf1ab;2887 | 0 | 0 |
| NC_004718.3;NP_828849.2;SARS-CoV_orf1ab;2888 | 0 | 0 |
| NC_004718.3;NP_828849.2;SARS-CoV_orf1ab;2889 | 0 | 0 |
| NC_004718.3;NP_828849.2;SARS-CoV_orf1ab;2890 | 0 | 0 |
| NC_004718.3;NP_828849.2;SARS-CoV_orf1ab;2891 | 0 | 0.025 |
| NC_004718.3;NP_828849.2;SARS-CoV_orf1ab;2892 | 0 | 0.025 |
| NC_004718.3;NP_828849.2;SARS-CoV_orf1ab;2893 | 0 | 0 |
| NC_004718.3;NP_828849.2;SARS-CoV_orf1ab;2894 | 0 | 0.025 |
| NC_004718.3;NP_828849.2;SARS-CoV_orf1ab;2895 | 0 | 0.025 |
| NC_004718.3;NP_828849.2;SARS-CoV_orf1ab;2896 | 0 | 0 |
| NC_004718.3;NP_828849.2;SARS-CoV_orf1ab;2897 | 0 | 0.025 |
| NC_004718.3;NP_828849.2;SARS-CoV_orf1ab;2898 | 0 | 0.025 |
| NC_004718.3;NP_828849.2;SARS-CoV_orf1ab;2899 | 0 | 0.025 |
| NC_004718.3;NP_828849.2;SARS-CoV_orf1ab;2900 | 0 | 0 |
| NC_004718.3;NP_828849.2;SARS-CoV_orf1ab;2901 | 0 | 0 |
| NC_004718.3;NP_828849.2;SARS-CoV_orf1ab;2902 | 0 | 0 |
| NC_004718.3;NP_828849.2;SARS-CoV_orf1ab;2903 | 0 | 0 |
| NC_004718.3;NP_828849.2;SARS-CoV_orf1ab;2904 | 0 | 0 |
| NC_004718.3;NP_828849.2;SARS-CoV_orf1ab;2905 | 0 | 0 |
| NC_004718.3;NP_828849.2;SARS-CoV_orf1ab;2906 | 0 | 0 |
| NC_004718.3;NP_828849.2;SARS-CoV_orf1ab;2907 | 0 | 0 |
| NC_004718.3;NP_828849.2;SARS-CoV_orf1ab;2908 | 0 | 0 |
| NC_004718.3;NP_828849.2;SARS-CoV_orf1ab;2909 | 0 | 0 |
| NC_004718.3;NP_828849.2;SARS-CoV_orf1ab;2910 | 0 | 0 |
| NC_004718.3;NP_828849.2;SARS-CoV_orf1ab;2911 | 0 | 0 |
| NC_004718.3;NP_828849.2;SARS-CoV_orf1ab;2912 | 0 | 0 |
| NC_004718.3;NP_828849.2;SARS-CoV_orf1ab;2913 | 0 | 0 |
| NC_004718.3;NP_828849.2;SARS-CoV_orf1ab;2914 | 0 | 0.025 |
| NC_004718.3;NP_828849.2;SARS-CoV_orf1ab;2915 | 0 | 0.05 |
| NC_004718.3;NP_828849.2;SARS-CoV_orf1ab;2916 | 0.25 | 0.15 |
| NC_004718.3;NP_828849.2;SARS-CoV_orf1ab;2917 | 0.15 | 0.175 |
| NC_004718.3;NP_828849.2;SARS-CoV_orf1ab;2918 | 0.1 | 0.025 |
| NC_004718.3;NP_828849.2;SARS-CoV_orf1ab;2919 | 0.1 | 0.05 |
| NC_004718.3;NP_828849.2;SARS-CoV_orf1ab;2920 | 0.1 | 0.025 |
| NC_004718.3;NP_828849.2;SARS-CoV_orf1ab;2921 | 0.15 | 0.05 |

| | | |
|---|---|---|
| NC_004718.3;NP_828849.2;SARS-CoV_orf1ab;2922 | 0.1 | 0.125 |
| NC_004718.3;NP_828849.2;SARS-CoV_orf1ab;2923 | 0.1 | 0.025 |
| NC_004718.3;NP_828849.2;SARS-CoV_orf1ab;2924 | 0.05 | 0.025 |
| NC_004718.3;NP_828849.2;SARS-CoV_orf1ab;2925 | 0.05 | 0.05 |
| NC_004718.3;NP_828849.2;SARS-CoV_orf1ab;2926 | 0.05 | 0.025 |
| NC_004718.3;NP_828849.2;SARS-CoV_orf1ab;2927 | 0 | 0 |
| NC_004718.3;NP_828849.2;SARS-CoV_orf1ab;2928 | 0 | 0 |
| NC_004718.3;NP_828849.2;SARS-CoV_orf1ab;2929 | 0 | 0 |
| NC_004718.3;NP_828849.2;SARS-CoV_orf1ab;2930 | 0 | 0 |
| NC_004718.3;NP_828849.2;SARS-CoV_orf1ab;2931 | 0 | 0 |
| NC_004718.3;NP_828849.2;SARS-CoV_orf1ab;2932 | 0 | 0 |
| NC_004718.3;NP_828849.2;SARS-CoV_orf1ab;2933 | 0 | 0 |
| NC_004718.3;NP_828849.2;SARS-CoV_orf1ab;2934 | 0 | 0 |
| NC_004718.3;NP_828849.2;SARS-CoV_orf1ab;2935 | 0 | 0 |
| NC_004718.3;NP_828849.2;SARS-CoV_orf1ab;2936 | 0 | 0 |
| NC_004718.3;NP_828849.2;SARS-CoV_orf1ab;2937 | 0 | 0.025 |
| NC_004718.3;NP_828849.2;SARS-CoV_orf1ab;2938 | 0 | 0 |
| NC_004718.3;NP_828849.2;SARS-CoV_orf1ab;2939 | 0 | 0 |
| NC_004718.3;NP_828849.2;SARS-CoV_orf1ab;2940 | 0 | 0 |
| NC_004718.3;NP_828849.2;SARS-CoV_orf1ab;2941 | 0 | 0 |
| NC_004718.3;NP_828849.2;SARS-CoV_orf1ab;2942 | 0 | 0 |
| NC_004718.3;NP_828849.2;SARS-CoV_orf1ab;2943 | 0 | 0 |
| NC_004718.3;NP_828849.2;SARS-CoV_orf1ab;2944 | 0 | 0 |
| NC_004718.3;NP_828849.2;SARS-CoV_orf1ab;2945 | 0 | 0.025 |
| NC_004718.3;NP_828849.2;SARS-CoV_orf1ab;2946 | 0 | 0.025 |
| NC_004718.3;NP_828849.2;SARS-CoV_orf1ab;2947 | 0 | 0.025 |
| NC_004718.3;NP_828849.2;SARS-CoV_orf1ab;2948 | 0.05 | 0.075 |
| NC_004718.3;NP_828849.2;SARS-CoV_orf1ab;2949 | 0 | 0.025 |
| NC_004718.3;NP_828849.2;SARS-CoV_orf1ab;2950 | 0 | 0.025 |
| NC_004718.3;NP_828849.2;SARS-CoV_orf1ab;2951 | 0 | 0.025 |
| NC_004718.3;NP_828849.2;SARS-CoV_orf1ab;2952 | 0 | 0.025 |
| NC_004718.3;NP_828849.2;SARS-CoV_orf1ab;2953 | 0 | 0 |
| NC_004718.3;NP_828849.2;SARS-CoV_orf1ab;2954 | 0 | 0 |
| NC_004718.3;NP_828849.2;SARS-CoV_orf1ab;2955 | 0 | 0 |
| NC_004718.3;NP_828849.2;SARS-CoV_orf1ab;2956 | 0 | 0 |
| NC_004718.3;NP_828849.2;SARS-CoV_orf1ab;2957 | 0 | 0 |
| NC_004718.3;NP_828849.2;SARS-CoV_orf1ab;2958 | 0 | 0 |
| NC_004718.3;NP_828849.2;SARS-CoV_orf1ab;2959 | 0 | 0 |
| NC_004718.3;NP_828849.2;SARS-CoV_orf1ab;2960 | 0 | 0 |
| NC_004718.3;NP_828849.2;SARS-CoV_orf1ab;2961 | 0 | 0 |
| NC_004718.3;NP_828849.2;SARS-CoV_orf1ab;2962 | 0 | 0 |
| NC_004718.3;NP_828849.2;SARS-CoV_orf1ab;2963 | 0 | 0 |
| NC_004718.3;NP_828849.2;SARS-CoV_orf1ab;2964 | 0 | 0 |

| | | |
|---|---|---|
| NC_004718.3;NP_828849.2;SARS-CoV_orf1ab;2965 | 0 | 0 |
| NC_004718.3;NP_828849.2;SARS-CoV_orf1ab;2966 | 0 | 0 |
| NC_004718.3;NP_828849.2;SARS-CoV_orf1ab;2967 | 0 | 0 |
| NC_004718.3;NP_828849.2;SARS-CoV_orf1ab;2968 | 0 | 0 |
| NC_004718.3;NP_828849.2;SARS-CoV_orf1ab;2969 | 0 | 0 |
| NC_004718.3;NP_828849.2;SARS-CoV_orf1ab;2970 | 0 | 0 |
| NC_004718.3;NP_828849.2;SARS-CoV_orf1ab;2971 | 0 | 0 |
| NC_004718.3;NP_828849.2;SARS-CoV_orf1ab;2972 | 0 | 0 |
| NC_004718.3;NP_828849.2;SARS-CoV_orf1ab;2973 | 0 | 0 |
| NC_004718.3;NP_828849.2;SARS-CoV_orf1ab;2974 | 0 | 0 |
| NC_004718.3;NP_828849.2;SARS-CoV_orf1ab;2975 | 0 | 0 |
| NC_004718.3;NP_828849.2;SARS-CoV_orf1ab;2976 | 0 | 0 |
| NC_004718.3;NP_828849.2;SARS-CoV_orf1ab;2977 | 0 | 0 |
| NC_004718.3;NP_828849.2;SARS-CoV_orf1ab;2978 | 0 | 0 |
| NC_004718.3;NP_828849.2;SARS-CoV_orf1ab;2979 | 0 | 0 |
| NC_004718.3;NP_828849.2;SARS-CoV_orf1ab;2980 | 0 | 0 |
| NC_004718.3;NP_828849.2;SARS-CoV_orf1ab;2981 | 0 | 0 |
| NC_004718.3;NP_828849.2;SARS-CoV_orf1ab;2982 | 0 | 0 |
| NC_004718.3;NP_828849.2;SARS-CoV_orf1ab;2983 | 0 | 0 |
| NC_004718.3;NP_828849.2;SARS-CoV_orf1ab;2984 | 0 | 0 |
| NC_004718.3;NP_828849.2;SARS-CoV_orf1ab;2985 | 0 | 0 |
| NC_004718.3;NP_828849.2;SARS-CoV_orf1ab;2986 | 0 | 0 |
| NC_004718.3;NP_828849.2;SARS-CoV_orf1ab;2987 | 0 | 0.025 |
| NC_004718.3;NP_828849.2;SARS-CoV_orf1ab;2988 | 0 | 0.025 |
| NC_004718.3;NP_828849.2;SARS-CoV_orf1ab;2989 | 0 | 0.025 |
| NC_004718.3;NP_828849.2;SARS-CoV_orf1ab;2990 | 0 | 0.05 |
| NC_004718.3;NP_828849.2;SARS-CoV_orf1ab;2991 | 0 | 0.05 |
| NC_004718.3;NP_828849.2;SARS-CoV_orf1ab;2992 | 0.05 | 0.075 |
| NC_004718.3;NP_828849.2;SARS-CoV_orf1ab;2993 | 0 | 0 |
| NC_004718.3;NP_828849.2;SARS-CoV_orf1ab;2994 | 0 | 0 |
| NC_004718.3;NP_828849.2;SARS-CoV_orf1ab;2995 | 0 | 0 |
| NC_004718.3;NP_828849.2;SARS-CoV_orf1ab;2996 | 0 | 0 |
| NC_004718.3;NP_828849.2;SARS-CoV_orf1ab;2997 | 0.05 | 0.05 |
| NC_004718.3;NP_828849.2;SARS-CoV_orf1ab;2998 | 0.05 | 0 |
| NC_004718.3;NP_828849.2;SARS-CoV_orf1ab;2999 | 0 | 0 |
| NC_004718.3;NP_828849.2;SARS-CoV_orf1ab;3000 | 0 | 0 |
| NC_004718.3;NP_828849.2;SARS-CoV_orf1ab;3001 | 0 | 0 |
| NC_004718.3;NP_828849.2;SARS-CoV_orf1ab;3002 | 0 | 0 |
| NC_004718.3;NP_828849.2;SARS-CoV_orf1ab;3003 | 0 | 0 |
| NC_004718.3;NP_828849.2;SARS-CoV_orf1ab;3004 | 0 | 0 |
| NC_004718.3;NP_828849.2;SARS-CoV_orf1ab;3005 | 0 | 0 |
| NC_004718.3;NP_828849.2;SARS-CoV_orf1ab;3006 | 0 | 0 |
| NC_004718.3;NP_828849.2;SARS-CoV_orf1ab;3007 | 0 | 0 |

| | | |
|---|---|---|
| NC_004718.3;NP_828849.2;SARS-CoV_orf1ab;3008 | 0 | 0 |
| NC_004718.3;NP_828849.2;SARS-CoV_orf1ab;3009 | 0 | 0 |
| NC_004718.3;NP_828849.2;SARS-CoV_orf1ab;3010 | 0 | 0 |
| NC_004718.3;NP_828849.2;SARS-CoV_orf1ab;3011 | 0 | 0 |
| NC_004718.3;NP_828849.2;SARS-CoV_orf1ab;3012 | 0 | 0 |
| NC_004718.3;NP_828849.2;SARS-CoV_orf1ab;3013 | 0.05 | 0 |
| NC_004718.3;NP_828849.2;SARS-CoV_orf1ab;3014 | 0 | 0 |
| NC_004718.3;NP_828849.2;SARS-CoV_orf1ab;3015 | 0 | 0 |
| NC_004718.3;NP_828849.2;SARS-CoV_orf1ab;3016 | 0 | 0 |
| NC_004718.3;NP_828849.2;SARS-CoV_orf1ab;3017 | 0 | 0 |
| NC_004718.3;NP_828849.2;SARS-CoV_orf1ab;3018 | 0 | 0 |
| NC_004718.3;NP_828849.2;SARS-CoV_orf1ab;3019 | 0 | 0 |
| NC_004718.3;NP_828849.2;SARS-CoV_orf1ab;3020 | 0 | 0 |
| NC_004718.3;NP_828849.2;SARS-CoV_orf1ab;3021 | 0 | 0 |
| NC_004718.3;NP_828849.2;SARS-CoV_orf1ab;3022 | 0 | 0 |
| NC_004718.3;NP_828849.2;SARS-CoV_orf1ab;3023 | 0 | 0 |
| NC_004718.3;NP_828849.2;SARS-CoV_orf1ab;3024 | 0 | 0 |
| NC_004718.3;NP_828849.2;SARS-CoV_orf1ab;3025 | 0 | 0 |
| NC_004718.3;NP_828849.2;SARS-CoV_orf1ab;3026 | 0 | 0 |
| NC_004718.3;NP_828849.2;SARS-CoV_orf1ab;3027 | 0 | 0 |
| NC_004718.3;NP_828849.2;SARS-CoV_orf1ab;3028 | 0 | 0 |
| NC_004718.3;NP_828849.2;SARS-CoV_orf1ab;3029 | 0 | 0 |
| NC_004718.3;NP_828849.2;SARS-CoV_orf1ab;3030 | 0 | 0 |
| NC_004718.3;NP_828849.2;SARS-CoV_orf1ab;3031 | 0 | 0 |
| NC_004718.3;NP_828849.2;SARS-CoV_orf1ab;3032 | 0 | 0 |
| NC_004718.3;NP_828849.2;SARS-CoV_orf1ab;3033 | 0 | 0 |
| NC_004718.3;NP_828849.2;SARS-CoV_orf1ab;3034 | 0 | 0 |
| NC_004718.3;NP_828849.2;SARS-CoV_orf1ab;3035 | 0 | 0 |
| NC_004718.3;NP_828849.2;SARS-CoV_orf1ab;3036 | 0 | 0 |
| NC_004718.3;NP_828849.2;SARS-CoV_orf1ab;3037 | 0 | 0 |
| NC_004718.3;NP_828849.2;SARS-CoV_orf1ab;3038 | 0 | 0 |
| NC_004718.3;NP_828849.2;SARS-CoV_orf1ab;3039 | 0 | 0 |
| NC_004718.3;NP_828849.2;SARS-CoV_orf1ab;3040 | 0 | 0 |
| NC_004718.3;NP_828849.2;SARS-CoV_orf1ab;3041 | 0 | 0 |
| NC_004718.3;NP_828849.2;SARS-CoV_orf1ab;3042 | 0.05 | 0 |
| NC_004718.3;NP_828849.2;SARS-CoV_orf1ab;3043 | 0 | 0.025 |
| NC_004718.3;NP_828849.2;SARS-CoV_orf1ab;3044 | 0 | 0 |
| NC_004718.3;NP_828849.2;SARS-CoV_orf1ab;3045 | 0 | 0 |
| NC_004718.3;NP_828849.2;SARS-CoV_orf1ab;3046 | 0 | 0 |
| NC_004718.3;NP_828849.2;SARS-CoV_orf1ab;3047 | 0 | 0 |
| NC_004718.3;NP_828849.2;SARS-CoV_orf1ab;3048 | 0 | 0 |
| NC_004718.3;NP_828849.2;SARS-CoV_orf1ab;3049 | 0 | 0 |
| NC_004718.3;NP_828849.2;SARS-CoV_orf1ab;3050 | 0 | 0 |

| | | |
|---|---|---|
| NC_004718.3;NP_828849.2;SARS-CoV_orf1ab;3051 | 0 | 0 |
| NC_004718.3;NP_828849.2;SARS-CoV_orf1ab;3052 | 0 | 0 |
| NC_004718.3;NP_828849.2;SARS-CoV_orf1ab;3053 | 0 | 0 |
| NC_004718.3;NP_828849.2;SARS-CoV_orf1ab;3054 | 0 | 0 |
| NC_004718.3;NP_828849.2;SARS-CoV_orf1ab;3055 | 0 | 0 |
| NC_004718.3;NP_828849.2;SARS-CoV_orf1ab;3056 | 0 | 0 |
| NC_004718.3;NP_828849.2;SARS-CoV_orf1ab;3057 | 0 | 0 |
| NC_004718.3;NP_828849.2;SARS-CoV_orf1ab;3058 | 0 | 0 |
| NC_004718.3;NP_828849.2;SARS-CoV_orf1ab;3059 | 0 | 0 |
| NC_004718.3;NP_828849.2;SARS-CoV_orf1ab;3060 | 0 | 0 |
| NC_004718.3;NP_828849.2;SARS-CoV_orf1ab;3061 | 0 | 0 |
| NC_004718.3;NP_828849.2;SARS-CoV_orf1ab;3062 | 0 | 0 |
| NC_004718.3;NP_828849.2;SARS-CoV_orf1ab;3063 | 0 | 0 |
| NC_004718.3;NP_828849.2;SARS-CoV_orf1ab;3064 | 0 | 0 |
| NC_004718.3;NP_828849.2;SARS-CoV_orf1ab;3065 | 0 | 0 |
| NC_004718.3;NP_828849.2;SARS-CoV_orf1ab;3066 | 0 | 0 |
| NC_004718.3;NP_828849.2;SARS-CoV_orf1ab;3067 | 0 | 0 |
| NC_004718.3;NP_828849.2;SARS-CoV_orf1ab;3068 | 0 | 0 |
| NC_004718.3;NP_828849.2;SARS-CoV_orf1ab;3069 | 0 | 0 |
| NC_004718.3;NP_828849.2;SARS-CoV_orf1ab;3070 | 0 | 0 |
| NC_004718.3;NP_828849.2;SARS-CoV_orf1ab;3071 | 0 | 0 |
| NC_004718.3;NP_828849.2;SARS-CoV_orf1ab;3072 | 0 | 0 |
| NC_004718.3;NP_828849.2;SARS-CoV_orf1ab;3073 | 0 | 0 |
| NC_004718.3;NP_828849.2;SARS-CoV_orf1ab;3074 | 0 | 0 |
| NC_004718.3;NP_828849.2;SARS-CoV_orf1ab;3075 | 0 | 0 |
| NC_004718.3;NP_828849.2;SARS-CoV_orf1ab;3076 | 0 | 0 |
| NC_004718.3;NP_828849.2;SARS-CoV_orf1ab;3077 | 0 | 0 |
| NC_004718.3;NP_828849.2;SARS-CoV_orf1ab;3078 | 0 | 0 |
| NC_004718.3;NP_828849.2;SARS-CoV_orf1ab;3079 | 0 | 0 |
| NC_004718.3;NP_828849.2;SARS-CoV_orf1ab;3080 | 0 | 0 |
| NC_004718.3;NP_828849.2;SARS-CoV_orf1ab;3081 | 0 | 0 |
| NC_004718.3;NP_828849.2;SARS-CoV_orf1ab;3082 | 0 | 0 |
| NC_004718.3;NP_828849.2;SARS-CoV_orf1ab;3083 | 0 | 0 |
| NC_004718.3;NP_828849.2;SARS-CoV_orf1ab;3084 | 0 | 0 |
| NC_004718.3;NP_828849.2;SARS-CoV_orf1ab;3085 | 0 | 0 |
| NC_004718.3;NP_828849.2;SARS-CoV_orf1ab;3086 | 0 | 0 |
| NC_004718.3;NP_828849.2;SARS-CoV_orf1ab;3087 | 0 | 0 |
| NC_004718.3;NP_828849.2;SARS-CoV_orf1ab;3088 | 0 | 0 |
| NC_004718.3;NP_828849.2;SARS-CoV_orf1ab;3089 | 0 | 0 |
| NC_004718.3;NP_828849.2;SARS-CoV_orf1ab;3090 | 0 | 0 |
| NC_004718.3;NP_828849.2;SARS-CoV_orf1ab;3091 | 0 | 0 |
| NC_004718.3;NP_828849.2;SARS-CoV_orf1ab;3092 | 0 | 0 |
| NC_004718.3;NP_828849.2;SARS-CoV_orf1ab;3093 | 0 | 0.025 |

| | | |
|---|---|---|
| NC_004718.3;NP_828849.2;SARS-CoV_orf1ab;3094 | 0.05 | 0 |
| NC_004718.3;NP_828849.2;SARS-CoV_orf1ab;3095 | 0 | 0 |
| NC_004718.3;NP_828849.2;SARS-CoV_orf1ab;3096 | 0 | 0 |
| NC_004718.3;NP_828849.2;SARS-CoV_orf1ab;3097 | 0 | 0 |
| NC_004718.3;NP_828849.2;SARS-CoV_orf1ab;3098 | 0 | 0 |
| NC_004718.3;NP_828849.2;SARS-CoV_orf1ab;3099 | 0 | 0 |
| NC_004718.3;NP_828849.2;SARS-CoV_orf1ab;3100 | 0 | 0 |
| NC_004718.3;NP_828849.2;SARS-CoV_orf1ab;3101 | 0 | 0 |
| NC_004718.3;NP_828849.2;SARS-CoV_orf1ab;3102 | 0 | 0 |
| NC_004718.3;NP_828849.2;SARS-CoV_orf1ab;3103 | 0 | 0 |
| NC_004718.3;NP_828849.2;SARS-CoV_orf1ab;3104 | 0 | 0 |
| NC_004718.3;NP_828849.2;SARS-CoV_orf1ab;3105 | 0 | 0 |
| NC_004718.3;NP_828849.2;SARS-CoV_orf1ab;3106 | 0 | 0 |
| NC_004718.3;NP_828849.2;SARS-CoV_orf1ab;3107 | 0 | 0 |
| NC_004718.3;NP_828849.2;SARS-CoV_orf1ab;3108 | 0 | 0 |
| NC_004718.3;NP_828849.2;SARS-CoV_orf1ab;3109 | 0 | 0 |
| NC_004718.3;NP_828849.2;SARS-CoV_orf1ab;3110 | 0 | 0 |
| NC_004718.3;NP_828849.2;SARS-CoV_orf1ab;3111 | 0 | 0 |
| NC_004718.3;NP_828849.2;SARS-CoV_orf1ab;3112 | 0 | 0 |
| NC_004718.3;NP_828849.2;SARS-CoV_orf1ab;3113 | 0 | 0 |
| NC_004718.3;NP_828849.2;SARS-CoV_orf1ab;3114 | 0 | 0 |
| NC_004718.3;NP_828849.2;SARS-CoV_orf1ab;3115 | 0 | 0 |
| NC_004718.3;NP_828849.2;SARS-CoV_orf1ab;3116 | 0 | 0 |
| NC_004718.3;NP_828849.2;SARS-CoV_orf1ab;3117 | 0 | 0 |
| NC_004718.3;NP_828849.2;SARS-CoV_orf1ab;3118 | 0 | 0 |
| NC_004718.3;NP_828849.2;SARS-CoV_orf1ab;3119 | 0 | 0 |
| NC_004718.3;NP_828849.2;SARS-CoV_orf1ab;3120 | 0 | 0 |
| NC_004718.3;NP_828849.2;SARS-CoV_orf1ab;3121 | 0 | 0 |
| NC_004718.3;NP_828849.2;SARS-CoV_orf1ab;3122 | 0 | 0 |
| NC_004718.3;NP_828849.2;SARS-CoV_orf1ab;3123 | 0 | 0 |
| NC_004718.3;NP_828849.2;SARS-CoV_orf1ab;3124 | 0 | 0 |
| NC_004718.3;NP_828849.2;SARS-CoV_orf1ab;3125 | 0 | 0 |
| NC_004718.3;NP_828849.2;SARS-CoV_orf1ab;3126 | 0 | 0 |
| NC_004718.3;NP_828849.2;SARS-CoV_orf1ab;3127 | 0 | 0 |
| NC_004718.3;NP_828849.2;SARS-CoV_orf1ab;3128 | 0 | 0 |
| NC_004718.3;NP_828849.2;SARS-CoV_orf1ab;3129 | 0 | 0 |
| NC_004718.3;NP_828849.2;SARS-CoV_orf1ab;3130 | 0 | 0 |
| NC_004718.3;NP_828849.2;SARS-CoV_orf1ab;3131 | 0 | 0 |
| NC_004718.3;NP_828849.2;SARS-CoV_orf1ab;3132 | 0 | 0 |
| NC_004718.3;NP_828849.2;SARS-CoV_orf1ab;3133 | 0 | 0 |
| NC_004718.3;NP_828849.2;SARS-CoV_orf1ab;3134 | 0 | 0 |
| NC_004718.3;NP_828849.2;SARS-CoV_orf1ab;3135 | 0 | 0 |
| NC_004718.3;NP_828849.2;SARS-CoV_orf1ab;3136 | 0 | 0 |

| | | |
|---|---|---|
| NC_004718.3;NP_828849.2;SARS-CoV_orf1ab;3137 | 0.05 | 0 |
| NC_004718.3;NP_828849.2;SARS-CoV_orf1ab;3138 | 0 | 0 |
| NC_004718.3;NP_828849.2;SARS-CoV_orf1ab;3139 | 0 | 0 |
| NC_004718.3;NP_828849.2;SARS-CoV_orf1ab;3140 | 0 | 0 |
| NC_004718.3;NP_828849.2;SARS-CoV_orf1ab;3141 | 0.05 | 0.025 |
| NC_004718.3;NP_828849.2;SARS-CoV_orf1ab;3142 | 0 | 0 |
| NC_004718.3;NP_828849.2;SARS-CoV_orf1ab;3143 | 0 | 0 |
| NC_004718.3;NP_828849.2;SARS-CoV_orf1ab;3144 | 0 | 0 |
| NC_004718.3;NP_828849.2;SARS-CoV_orf1ab;3145 | 0 | 0 |
| NC_004718.3;NP_828849.2;SARS-CoV_orf1ab;3146 | 0 | 0 |
| NC_004718.3;NP_828849.2;SARS-CoV_orf1ab;3147 | 0 | 0 |
| NC_004718.3;NP_828849.2;SARS-CoV_orf1ab;3148 | 0 | 0 |
| NC_004718.3;NP_828849.2;SARS-CoV_orf1ab;3149 | 0 | 0 |
| NC_004718.3;NP_828849.2;SARS-CoV_orf1ab;3150 | 0 | 0 |
| NC_004718.3;NP_828849.2;SARS-CoV_orf1ab;3151 | 0 | 0 |
| NC_004718.3;NP_828849.2;SARS-CoV_orf1ab;3152 | 0 | 0 |
| NC_004718.3;NP_828849.2;SARS-CoV_orf1ab;3153 | 0 | 0 |
| NC_004718.3;NP_828849.2;SARS-CoV_orf1ab;3154 | 0 | 0 |
| NC_004718.3;NP_828849.2;SARS-CoV_orf1ab;3155 | 0 | 0 |
| NC_004718.3;NP_828849.2;SARS-CoV_orf1ab;3156 | 0 | 0 |
| NC_004718.3;NP_828849.2;SARS-CoV_orf1ab;3157 | 0 | 0 |
| NC_004718.3;NP_828849.2;SARS-CoV_orf1ab;3158 | 0 | 0 |
| NC_004718.3;NP_828849.2;SARS-CoV_orf1ab;3159 | 0 | 0 |
| NC_004718.3;NP_828849.2;SARS-CoV_orf1ab;3160 | 0 | 0 |
| NC_004718.3;NP_828849.2;SARS-CoV_orf1ab;3161 | 0 | 0 |
| NC_004718.3;NP_828849.2;SARS-CoV_orf1ab;3162 | 0 | 0 |
| NC_004718.3;NP_828849.2;SARS-CoV_orf1ab;3163 | 0 | 0 |
| NC_004718.3;NP_828849.2;SARS-CoV_orf1ab;3164 | 0 | 0 |
| NC_004718.3;NP_828849.2;SARS-CoV_orf1ab;3165 | 0 | 0 |
| NC_004718.3;NP_828849.2;SARS-CoV_orf1ab;3166 | 0 | 0 |
| NC_004718.3;NP_828849.2;SARS-CoV_orf1ab;3167 | 0 | 0 |
| NC_004718.3;NP_828849.2;SARS-CoV_orf1ab;3168 | 0 | 0 |
| NC_004718.3;NP_828849.2;SARS-CoV_orf1ab;3169 | 0 | 0 |
| NC_004718.3;NP_828849.2;SARS-CoV_orf1ab;3170 | 0 | 0 |
| NC_004718.3;NP_828849.2;SARS-CoV_orf1ab;3171 | 0 | 0.025 |
| NC_004718.3;NP_828849.2;SARS-CoV_orf1ab;3172 | 0 | 0 |
| NC_004718.3;NP_828849.2;SARS-CoV_orf1ab;3173 | 0 | 0 |
| NC_004718.3;NP_828849.2;SARS-CoV_orf1ab;3174 | 0 | 0 |
| NC_004718.3;NP_828849.2;SARS-CoV_orf1ab;3175 | 0 | 0 |
| NC_004718.3;NP_828849.2;SARS-CoV_orf1ab;3176 | 0 | 0 |
| NC_004718.3;NP_828849.2;SARS-CoV_orf1ab;3177 | 0 | 0 |
| NC_004718.3;NP_828849.2;SARS-CoV_orf1ab;3178 | 0 | 0 |
| NC_004718.3;NP_828849.2;SARS-CoV_orf1ab;3179 | 0 | 0 |

| | | |
|---|---|---|
| NC_004718.3;NP_828849.2;SARS-CoV_orf1ab;3180 | 0 | 0 |
| NC_004718.3;NP_828849.2;SARS-CoV_orf1ab;3181 | 0 | 0 |
| NC_004718.3;NP_828849.2;SARS-CoV_orf1ab;3182 | 0 | 0 |
| NC_004718.3;NP_828849.2;SARS-CoV_orf1ab;3183 | 0 | 0 |
| NC_004718.3;NP_828849.2;SARS-CoV_orf1ab;3184 | 0 | 0 |
| NC_004718.3;NP_828849.2;SARS-CoV_orf1ab;3185 | 0 | 0 |
| NC_004718.3;NP_828849.2;SARS-CoV_orf1ab;3186 | 0 | 0 |
| NC_004718.3;NP_828849.2;SARS-CoV_orf1ab;3187 | 0 | 0.025 |
| NC_004718.3;NP_828849.2;SARS-CoV_orf1ab;3188 | 0 | 0 |
| NC_004718.3;NP_828849.2;SARS-CoV_orf1ab;3189 | 0 | 0 |
| NC_004718.3;NP_828849.2;SARS-CoV_orf1ab;3190 | 0 | 0.025 |
| NC_004718.3;NP_828849.2;SARS-CoV_orf1ab;3191 | 0 | 0 |
| NC_004718.3;NP_828849.2;SARS-CoV_orf1ab;3192 | 0.05 | 0.025 |
| NC_004718.3;NP_828849.2;SARS-CoV_orf1ab;3193 | 0 | 0 |
| NC_004718.3;NP_828849.2;SARS-CoV_orf1ab;3194 | 0 | 0 |
| NC_004718.3;NP_828849.2;SARS-CoV_orf1ab;3195 | 0 | 0 |
| NC_004718.3;NP_828849.2;SARS-CoV_orf1ab;3196 | 0 | 0.05 |
| NC_004718.3;NP_828849.2;SARS-CoV_orf1ab;3197 | 0 | 0 |
| NC_004718.3;NP_828849.2;SARS-CoV_orf1ab;3198 | 0 | 0 |
| NC_004718.3;NP_828849.2;SARS-CoV_orf1ab;3199 | 0 | 0 |
| NC_004718.3;NP_828849.2;SARS-CoV_orf1ab;3200 | 0 | 0 |
| NC_004718.3;NP_828849.2;SARS-CoV_orf1ab;3201 | 0 | 0 |
| NC_004718.3;NP_828849.2;SARS-CoV_orf1ab;3202 | 0 | 0.075 |
| NC_004718.3;NP_828849.2;SARS-CoV_orf1ab;3203 | 0 | 0.025 |
| NC_004718.3;NP_828849.2;SARS-CoV_orf1ab;3204 | 0 | 0.025 |
| NC_004718.3;NP_828849.2;SARS-CoV_orf1ab;3205 | 0 | 0.025 |
| NC_004718.3;NP_828849.2;SARS-CoV_orf1ab;3206 | 0 | 0.025 |
| NC_004718.3;NP_828849.2;SARS-CoV_orf1ab;3207 | 0.05 | 0.025 |
| NC_004718.3;NP_828849.2;SARS-CoV_orf1ab;3208 | 0 | 0.025 |
| NC_004718.3;NP_828849.2;SARS-CoV_orf1ab;3209 | 0 | 0.05 |
| NC_004718.3;NP_828849.2;SARS-CoV_orf1ab;3210 | 0 | 0.025 |
| NC_004718.3;NP_828849.2;SARS-CoV_orf1ab;3211 | 0 | 0 |
| NC_004718.3;NP_828849.2;SARS-CoV_orf1ab;3212 | 0 | 0 |
| NC_004718.3;NP_828849.2;SARS-CoV_orf1ab;3213 | 0 | 0 |
| NC_004718.3;NP_828849.2;SARS-CoV_orf1ab;3214 | 0.15 | 0.025 |
| NC_004718.3;NP_828849.2;SARS-CoV_orf1ab;3215 | 0.3 | 0.05 |
| NC_004718.3;NP_828849.2;SARS-CoV_orf1ab;3216 | 0.05 | 0.025 |
| NC_004718.3;NP_828849.2;SARS-CoV_orf1ab;3217 | 0 | 0.025 |
| NC_004718.3;NP_828849.2;SARS-CoV_orf1ab;3218 | 0.1 | 0.1 |
| NC_004718.3;NP_828849.2;SARS-CoV_orf1ab;3219 | 0 | 0.025 |
| NC_004718.3;NP_828849.2;SARS-CoV_orf1ab;3220 | 0 | 0 |
| NC_004718.3;NP_828849.2;SARS-CoV_orf1ab;3221 | 0 | 0 |
| NC_004718.3;NP_828849.2;SARS-CoV_orf1ab;3222 | 0 | 0.025 |

| | | |
|---|---|---|
| NC_004718.3;NP_828849.2;SARS-CoV_orf1ab;3223 | 0 | 0 |
| NC_004718.3;NP_828849.2;SARS-CoV_orf1ab;3224 | 0 | 0 |
| NC_004718.3;NP_828849.2;SARS-CoV_orf1ab;3225 | 0 | 0 |
| NC_004718.3;NP_828849.2;SARS-CoV_orf1ab;3226 | 0 | 0 |
| NC_004718.3;NP_828849.2;SARS-CoV_orf1ab;3227 | 0.05 | 0 |
| NC_004718.3;NP_828849.2;SARS-CoV_orf1ab;3228 | 0 | 0 |
| NC_004718.3;NP_828849.2;SARS-CoV_orf1ab;3229 | 0 | 0 |
| NC_004718.3;NP_828849.2;SARS-CoV_orf1ab;3230 | 0 | 0 |
| NC_004718.3;NP_828849.2;SARS-CoV_orf1ab;3231 | 0 | 0 |
| NC_004718.3;NP_828849.2;SARS-CoV_orf1ab;3232 | 0 | 0 |
| NC_004718.3;NP_828849.2;SARS-CoV_orf1ab;3233 | 0 | 0 |
| NC_004718.3;NP_828849.2;SARS-CoV_orf1ab;3234 | 0 | 0 |
| NC_004718.3;NP_828849.2;SARS-CoV_orf1ab;3235 | 0 | 0 |
| NC_004718.3;NP_828849.2;SARS-CoV_orf1ab;3236 | 0 | 0 |
| NC_004718.3;NP_828849.2;SARS-CoV_orf1ab;3237 | 0 | 0 |
| NC_004718.3;NP_828849.2;SARS-CoV_orf1ab;3238 | 0 | 0 |
| NC_004718.3;NP_828849.2;SARS-CoV_orf1ab;3239 | 0 | 0 |
| NC_004718.3;NP_828849.2;SARS-CoV_orf1ab;3240 | 0 | 0 |
| NC_004718.3;NP_828849.2;SARS-CoV_orf1ab;3241 | 0 | 0 |
| NC_004718.3;NP_828849.2;SARS-CoV_orf1ab;3242 | 0 | 0 |
| NC_004718.3;NP_828849.2;SARS-CoV_orf1ab;3243 | 0 | 0 |
| NC_004718.3;NP_828849.2;SARS-CoV_orf1ab;3244 | 0 | 0 |
| NC_004718.3;NP_828849.2;SARS-CoV_orf1ab;3245 | 0 | 0.025 |
| NC_004718.3;NP_828849.2;SARS-CoV_orf1ab;3246 | 0 | 0 |
| NC_004718.3;NP_828849.2;SARS-CoV_orf1ab;3247 | 0.05 | 0 |
| NC_004718.3;NP_828849.2;SARS-CoV_orf1ab;3248 | 0 | 0 |
| NC_004718.3;NP_828849.2;SARS-CoV_orf1ab;3249 | 0 | 0 |
| NC_004718.3;NP_828849.2;SARS-CoV_orf1ab;3250 | 0 | 0 |
| NC_004718.3;NP_828849.2;SARS-CoV_orf1ab;3251 | 0 | 0 |
| NC_004718.3;NP_828849.2;SARS-CoV_orf1ab;3252 | 0 | 0 |
| NC_004718.3;NP_828849.2;SARS-CoV_orf1ab;3253 | 0 | 0 |
| NC_004718.3;NP_828849.2;SARS-CoV_orf1ab;3254 | 0 | 0 |
| NC_004718.3;NP_828849.2;SARS-CoV_orf1ab;3255 | 0 | 0 |
| NC_004718.3;NP_828849.2;SARS-CoV_orf1ab;3256 | 0 | 0 |
| NC_004718.3;NP_828849.2;SARS-CoV_orf1ab;3257 | 0 | 0 |
| NC_004718.3;NP_828849.2;SARS-CoV_orf1ab;3258 | 0 | 0 |
| NC_004718.3;NP_828849.2;SARS-CoV_orf1ab;3259 | 0 | 0 |
| NC_004718.3;NP_828849.2;SARS-CoV_orf1ab;3260 | 0 | 0 |
| NC_004718.3;NP_828849.2;SARS-CoV_orf1ab;3261 | 0 | 0 |
| NC_004718.3;NP_828849.2;SARS-CoV_orf1ab;3262 | 0 | 0 |
| NC_004718.3;NP_828849.2;SARS-CoV_orf1ab;3263 | 0 | 0 |
| NC_004718.3;NP_828849.2;SARS-CoV_orf1ab;3264 | 0 | 0 |
| NC_004718.3;NP_828849.2;SARS-CoV_orf1ab;3265 | 0 | 0 |

| | | |
|---|---|---|
| NC_004718.3;NP_828849.2;SARS-CoV_orf1ab;3266 | 0 | 0 |
| NC_004718.3;NP_828849.2;SARS-CoV_orf1ab;3267 | 0 | 0 |
| NC_004718.3;NP_828849.2;SARS-CoV_orf1ab;3268 | 0 | 0 |
| NC_004718.3;NP_828849.2;SARS-CoV_orf1ab;3269 | 0 | 0 |
| NC_004718.3;NP_828849.2;SARS-CoV_orf1ab;3270 | 0 | 0 |
| NC_004718.3;NP_828849.2;SARS-CoV_orf1ab;3271 | 0 | 0 |
| NC_004718.3;NP_828849.2;SARS-CoV_orf1ab;3272 | 0 | 0 |
| NC_004718.3;NP_828849.2;SARS-CoV_orf1ab;3273 | 0 | 0 |
| NC_004718.3;NP_828849.2;SARS-CoV_orf1ab;3274 | 0 | 0 |
| NC_004718.3;NP_828849.2;SARS-CoV_orf1ab;3275 | 0 | 0 |
| NC_004718.3;NP_828849.2;SARS-CoV_orf1ab;3276 | 0 | 0.025 |
| NC_004718.3;NP_828849.2;SARS-CoV_orf1ab;3277 | 0 | 0 |
| NC_004718.3;NP_828849.2;SARS-CoV_orf1ab;3278 | 0 | 0 |
| NC_004718.3;NP_828849.2;SARS-CoV_orf1ab;3279 | 0.05 | 0.025 |
| NC_004718.3;NP_828849.2;SARS-CoV_orf1ab;3280 | 0 | 0 |
| NC_004718.3;NP_828849.2;SARS-CoV_orf1ab;3281 | 0 | 0 |
| NC_004718.3;NP_828849.2;SARS-CoV_orf1ab;3282 | 0 | 0 |
| NC_004718.3;NP_828849.2;SARS-CoV_orf1ab;3283 | 0 | 0 |
| NC_004718.3;NP_828849.2;SARS-CoV_orf1ab;3284 | 0 | 0 |
| NC_004718.3;NP_828849.2;SARS-CoV_orf1ab;3285 | 0 | 0 |
| NC_004718.3;NP_828849.2;SARS-CoV_orf1ab;3286 | 0 | 0.025 |
| NC_004718.3;NP_828849.2;SARS-CoV_orf1ab;3287 | 0 | 0.025 |
| NC_004718.3;NP_828849.2;SARS-CoV_orf1ab;3288 | 0 | 0.025 |
| NC_004718.3;NP_828849.2;SARS-CoV_orf1ab;3289 | 0 | 0 |
| NC_004718.3;NP_828849.2;SARS-CoV_orf1ab;3290 | 0 | 0 |
| NC_004718.3;NP_828849.2;SARS-CoV_orf1ab;3291 | 0 | 0.025 |
| NC_004718.3;NP_828849.2;SARS-CoV_orf1ab;3292 | 0 | 0 |
| NC_004718.3;NP_828849.2;SARS-CoV_orf1ab;3293 | 0 | 0 |
| NC_004718.3;NP_828849.2;SARS-CoV_orf1ab;3294 | 0 | 0 |
| NC_004718.3;NP_828849.2;SARS-CoV_orf1ab;3295 | 0 | 0 |
| NC_004718.3;NP_828849.2;SARS-CoV_orf1ab;3296 | 0 | 0 |
| NC_004718.3;NP_828849.2;SARS-CoV_orf1ab;3297 | 0 | 0 |
| NC_004718.3;NP_828849.2;SARS-CoV_orf1ab;3298 | 0 | 0 |
| NC_004718.3;NP_828849.2;SARS-CoV_orf1ab;3299 | 0 | 0 |
| NC_004718.3;NP_828849.2;SARS-CoV_orf1ab;3300 | 0 | 0 |
| NC_004718.3;NP_828849.2;SARS-CoV_orf1ab;3301 | 0 | 0 |
| NC_004718.3;NP_828849.2;SARS-CoV_orf1ab;3302 | 0 | 0 |
| NC_004718.3;NP_828849.2;SARS-CoV_orf1ab;3303 | 0 | 0 |
| NC_004718.3;NP_828849.2;SARS-CoV_orf1ab;3304 | 0 | 0 |
| NC_004718.3;NP_828849.2;SARS-CoV_orf1ab;3305 | 0 | 0 |
| NC_004718.3;NP_828849.2;SARS-CoV_orf1ab;3306 | 0 | 0 |
| NC_004718.3;NP_828849.2;SARS-CoV_orf1ab;3307 | 0 | 0 |
| NC_004718.3;NP_828849.2;SARS-CoV_orf1ab;3308 | 0 | 0 |

| | | |
|---|---|---|
| NC_004718.3;NP_828849.2;SARS-CoV_orf1ab;3309 | 0 | 0 |
| NC_004718.3;NP_828849.2;SARS-CoV_orf1ab;3310 | 0 | 0 |
| NC_004718.3;NP_828849.2;SARS-CoV_orf1ab;3311 | 0 | 0 |
| NC_004718.3;NP_828849.2;SARS-CoV_orf1ab;3312 | 0 | 0 |
| NC_004718.3;NP_828849.2;SARS-CoV_orf1ab;3313 | 0 | 0 |
| NC_004718.3;NP_828849.2;SARS-CoV_orf1ab;3314 | 0 | 0 |
| NC_004718.3;NP_828849.2;SARS-CoV_orf1ab;3315 | 0 | 0 |
| NC_004718.3;NP_828849.2;SARS-CoV_orf1ab;3316 | 0 | 0 |
| NC_004718.3;NP_828849.2;SARS-CoV_orf1ab;3317 | 0 | 0 |
| NC_004718.3;NP_828849.2;SARS-CoV_orf1ab;3318 | 0 | 0 |
| NC_004718.3;NP_828849.2;SARS-CoV_orf1ab;3319 | 0 | 0.025 |
| NC_004718.3;NP_828849.2;SARS-CoV_orf1ab;3320 | 0 | 0 |
| NC_004718.3;NP_828849.2;SARS-CoV_orf1ab;3321 | 0 | 0 |
| NC_004718.3;NP_828849.2;SARS-CoV_orf1ab;3322 | 0 | 0.025 |
| NC_004718.3;NP_828849.2;SARS-CoV_orf1ab;3323 | 0 | 0 |
| NC_004718.3;NP_828849.2;SARS-CoV_orf1ab;3324 | 0 | 0 |
| NC_004718.3;NP_828849.2;SARS-CoV_orf1ab;3325 | 0 | 0 |
| NC_004718.3;NP_828849.2;SARS-CoV_orf1ab;3326 | 0 | 0 |
| NC_004718.3;NP_828849.2;SARS-CoV_orf1ab;3327 | 0 | 0 |
| NC_004718.3;NP_828849.2;SARS-CoV_orf1ab;3328 | 0 | 0 |
| NC_004718.3;NP_828849.2;SARS-CoV_orf1ab;3329 | 0 | 0 |
| NC_004718.3;NP_828849.2;SARS-CoV_orf1ab;3330 | 0.15 | 0.025 |
| NC_004718.3;NP_828849.2;SARS-CoV_orf1ab;3331 | 0 | 0 |
| NC_004718.3;NP_828849.2;SARS-CoV_orf1ab;3332 | 0 | 0 |
| NC_004718.3;NP_828849.2;SARS-CoV_orf1ab;3333 | 0 | 0 |
| NC_004718.3;NP_828849.2;SARS-CoV_orf1ab;3334 | 0 | 0 |
| NC_004718.3;NP_828849.2;SARS-CoV_orf1ab;3335 | 0 | 0 |
| NC_004718.3;NP_828849.2;SARS-CoV_orf1ab;3336 | 0 | 0 |
| NC_004718.3;NP_828849.2;SARS-CoV_orf1ab;3337 | 0 | 0 |
| NC_004718.3;NP_828849.2;SARS-CoV_orf1ab;3338 | 0 | 0 |
| NC_004718.3;NP_828849.2;SARS-CoV_orf1ab;3339 | 0 | 0 |
| NC_004718.3;NP_828849.2;SARS-CoV_orf1ab;3340 | 0 | 0 |
| NC_004718.3;NP_828849.2;SARS-CoV_orf1ab;3341 | 0 | 0 |
| NC_004718.3;NP_828849.2;SARS-CoV_orf1ab;3342 | 0 | 0 |
| NC_004718.3;NP_828849.2;SARS-CoV_orf1ab;3343 | 0 | 0 |
| NC_004718.3;NP_828849.2;SARS-CoV_orf1ab;3344 | 0 | 0 |
| NC_004718.3;NP_828849.2;SARS-CoV_orf1ab;3345 | 0 | 0 |
| NC_004718.3;NP_828849.2;SARS-CoV_orf1ab;3346 | 0 | 0 |
| NC_004718.3;NP_828849.2;SARS-CoV_orf1ab;3347 | 0 | 0.025 |
| NC_004718.3;NP_828849.2;SARS-CoV_orf1ab;3348 | 0 | 0 |
| NC_004718.3;NP_828849.2;SARS-CoV_orf1ab;3349 | 0 | 0 |
| NC_004718.3;NP_828849.2;SARS-CoV_orf1ab;3350 | 0 | 0 |
| NC_004718.3;NP_828849.2;SARS-CoV_orf1ab;3351 | 0 | 0 |

| | | |
|---|---|---|
| NC_004718.3;NP_828849.2;SARS-CoV_orf1ab;3352 | 0 | 0 |
| NC_004718.3;NP_828849.2;SARS-CoV_orf1ab;3353 | 0 | 0 |
| NC_004718.3;NP_828849.2;SARS-CoV_orf1ab;3354 | 0 | 0 |
| NC_004718.3;NP_828849.2;SARS-CoV_orf1ab;3355 | 0 | 0 |
| NC_004718.3;NP_828849.2;SARS-CoV_orf1ab;3356 | 0 | 0 |
| NC_004718.3;NP_828849.2;SARS-CoV_orf1ab;3357 | 0 | 0 |
| NC_004718.3;NP_828849.2;SARS-CoV_orf1ab;3358 | 0 | 0.025 |
| NC_004718.3;NP_828849.2;SARS-CoV_orf1ab;3359 | 0 | 0 |
| NC_004718.3;NP_828849.2;SARS-CoV_orf1ab;3360 | 0 | 0 |
| NC_004718.3;NP_828849.2;SARS-CoV_orf1ab;3361 | 0 | 0 |
| NC_004718.3;NP_828849.2;SARS-CoV_orf1ab;3362 | 0 | 0.05 |
| NC_004718.3;NP_828849.2;SARS-CoV_orf1ab;3363 | 0 | 0.075 |
| NC_004718.3;NP_828849.2;SARS-CoV_orf1ab;3364 | 0 | 0.1 |
| NC_004718.3;NP_828849.2;SARS-CoV_orf1ab;3365 | 0 | 0.075 |
| NC_004718.3;NP_828849.2;SARS-CoV_orf1ab;3366 | 0 | 0.05 |
| NC_004718.3;NP_828849.2;SARS-CoV_orf1ab;3367 | 0 | 0.05 |
| NC_004718.3;NP_828849.2;SARS-CoV_orf1ab;3368 | 0 | 0.05 |
| NC_004718.3;NP_828849.2;SARS-CoV_orf1ab;3369 | 0 | 0.05 |
| NC_004718.3;NP_828849.2;SARS-CoV_orf1ab;3370 | 0 | 0.05 |
| NC_004718.3;NP_828849.2;SARS-CoV_orf1ab;3371 | 0.05 | 0.05 |
| NC_004718.3;NP_828849.2;SARS-CoV_orf1ab;3372 | 0 | 0.075 |
| NC_004718.3;NP_828849.2;SARS-CoV_orf1ab;3373 | 0 | 0 |
| NC_004718.3;NP_828849.2;SARS-CoV_orf1ab;3374 | 0 | 0 |
| NC_004718.3;NP_828849.2;SARS-CoV_orf1ab;3375 | 0 | 0 |
| NC_004718.3;NP_828849.2;SARS-CoV_orf1ab;3376 | 0 | 0 |
| NC_004718.3;NP_828849.2;SARS-CoV_orf1ab;3377 | 0 | 0.025 |
| NC_004718.3;NP_828849.2;SARS-CoV_orf1ab;3378 | 0 | 0.025 |
| NC_004718.3;NP_828849.2;SARS-CoV_orf1ab;3379 | 0 | 0.025 |
| NC_004718.3;NP_828849.2;SARS-CoV_orf1ab;3380 | 0 | 0.05 |
| NC_004718.3;NP_828849.2;SARS-CoV_orf1ab;3381 | 0 | 0.05 |
| NC_004718.3;NP_828849.2;SARS-CoV_orf1ab;3382 | 0 | 0.025 |
| NC_004718.3;NP_828849.2;SARS-CoV_orf1ab;3383 | 0 | 0.025 |
| NC_004718.3;NP_828849.2;SARS-CoV_orf1ab;3384 | 0 | 0 |
| NC_004718.3;NP_828849.2;SARS-CoV_orf1ab;3385 | 0 | 0 |
| NC_004718.3;NP_828849.2;SARS-CoV_orf1ab;3386 | 0 | 0 |
| NC_004718.3;NP_828849.2;SARS-CoV_orf1ab;3387 | 0 | 0 |
| NC_004718.3;NP_828849.2;SARS-CoV_orf1ab;3388 | 0 | 0 |
| NC_004718.3;NP_828849.2;SARS-CoV_orf1ab;3389 | 0 | 0 |
| NC_004718.3;NP_828849.2;SARS-CoV_orf1ab;3390 | 0 | 0 |
| NC_004718.3;NP_828849.2;SARS-CoV_orf1ab;3391 | 0 | 0 |
| NC_004718.3;NP_828849.2;SARS-CoV_orf1ab;3392 | 0 | 0 |
| NC_004718.3;NP_828849.2;SARS-CoV_orf1ab;3393 | 0 | 0 |
| NC_004718.3;NP_828849.2;SARS-CoV_orf1ab;3394 | 0 | 0 |

| | | |
|---|---|---|
| NC_004718.3;NP_828849.2;SARS-CoV_orf1ab;3395 | 0 | 0 |
| NC_004718.3;NP_828849.2;SARS-CoV_orf1ab;3396 | 0 | 0 |
| NC_004718.3;NP_828849.2;SARS-CoV_orf1ab;3397 | 0 | 0 |
| NC_004718.3;NP_828849.2;SARS-CoV_orf1ab;3398 | 0 | 0 |
| NC_004718.3;NP_828849.2;SARS-CoV_orf1ab;3399 | 0 | 0 |
| NC_004718.3;NP_828849.2;SARS-CoV_orf1ab;3400 | 0 | 0 |
| NC_004718.3;NP_828849.2;SARS-CoV_orf1ab;3401 | 0 | 0 |
| NC_004718.3;NP_828849.2;SARS-CoV_orf1ab;3402 | 0 | 0 |
| NC_004718.3;NP_828849.2;SARS-CoV_orf1ab;3403 | 0.15 | 0.025 |
| NC_004718.3;NP_828849.2;SARS-CoV_orf1ab;3404 | 0.1 | 0.05 |
| NC_004718.3;NP_828849.2;SARS-CoV_orf1ab;3405 | 0.05 | 0.025 |
| NC_004718.3;NP_828849.2;SARS-CoV_orf1ab;3406 | 0.2 | 0.05 |
| NC_004718.3;NP_828849.2;SARS-CoV_orf1ab;3407 | 0.2 | 0.075 |
| NC_004718.3;NP_828849.2;SARS-CoV_orf1ab;3408 | 0.15 | 0.075 |
| NC_004718.3;NP_828849.2;SARS-CoV_orf1ab;3409 | 0.2 | 0.075 |
| NC_004718.3;NP_828849.2;SARS-CoV_orf1ab;3410 | 0.25 | 0.075 |
| NC_004718.3;NP_828849.2;SARS-CoV_orf1ab;3411 | 0.2 | 0.05 |
| NC_004718.3;NP_828849.2;SARS-CoV_orf1ab;3412 | 0.2 | 0.1 |
| NC_004718.3;NP_828849.2;SARS-CoV_orf1ab;3413 | 0.15 | 0.05 |
| NC_004718.3;NP_828849.2;SARS-CoV_orf1ab;3414 | 0.25 | 0.1 |
| NC_004718.3;NP_828849.2;SARS-CoV_orf1ab;3415 | 0 | 0.025 |
| NC_004718.3;NP_828849.2;SARS-CoV_orf1ab;3416 | 0 | 0 |
| NC_004718.3;NP_828849.2;SARS-CoV_orf1ab;3417 | 0.05 | 0 |
| NC_004718.3;NP_828849.2;SARS-CoV_orf1ab;3418 | 0.05 | 0.05 |
| NC_004718.3;NP_828849.2;SARS-CoV_orf1ab;3419 | 0.1 | 0.025 |
| NC_004718.3;NP_828849.2;SARS-CoV_orf1ab;3420 | 0.05 | 0 |
| NC_004718.3;NP_828849.2;SARS-CoV_orf1ab;3421 | 0.1 | 0 |
| NC_004718.3;NP_828849.2;SARS-CoV_orf1ab;3422 | 0 | 0 |
| NC_004718.3;NP_828849.2;SARS-CoV_orf1ab;3423 | 0.05 | 0 |
| NC_004718.3;NP_828849.2;SARS-CoV_orf1ab;3424 | 0 | 0 |
| NC_004718.3;NP_828849.2;SARS-CoV_orf1ab;3425 | 0.05 | 0.025 |
| NC_004718.3;NP_828849.2;SARS-CoV_orf1ab;3426 | 0 | 0 |
| NC_004718.3;NP_828849.2;SARS-CoV_orf1ab;3427 | 0 | 0 |
| NC_004718.3;NP_828849.2;SARS-CoV_orf1ab;3428 | 0 | 0 |
| NC_004718.3;NP_828849.2;SARS-CoV_orf1ab;3429 | 0 | 0 |
| NC_004718.3;NP_828849.2;SARS-CoV_orf1ab;3430 | 0 | 0 |
| NC_004718.3;NP_828849.2;SARS-CoV_orf1ab;3431 | 0 | 0 |
| NC_004718.3;NP_828849.2;SARS-CoV_orf1ab;3432 | 0 | 0 |
| NC_004718.3;NP_828849.2;SARS-CoV_orf1ab;3433 | 0 | 0 |
| NC_004718.3;NP_828849.2;SARS-CoV_orf1ab;3434 | 0 | 0 |
| NC_004718.3;NP_828849.2;SARS-CoV_orf1ab;3435 | 0 | 0 |
| NC_004718.3;NP_828849.2;SARS-CoV_orf1ab;3436 | 0 | 0 |
| NC_004718.3;NP_828849.2;SARS-CoV_orf1ab;3437 | 0 | 0 |

| | | |
|---|---|---|
| NC_004718.3;NP_828849.2;SARS-CoV_orf1ab;3438 | 0 | 0 |
| NC_004718.3;NP_828849.2;SARS-CoV_orf1ab;3439 | 0 | 0 |
| NC_004718.3;NP_828849.2;SARS-CoV_orf1ab;3440 | 0 | 0 |
| NC_004718.3;NP_828849.2;SARS-CoV_orf1ab;3441 | 0 | 0 |
| NC_004718.3;NP_828849.2;SARS-CoV_orf1ab;3442 | 0 | 0 |
| NC_004718.3;NP_828849.2;SARS-CoV_orf1ab;3443 | 0 | 0 |
| NC_004718.3;NP_828849.2;SARS-CoV_orf1ab;3444 | 0 | 0 |
| NC_004718.3;NP_828849.2;SARS-CoV_orf1ab;3445 | 0 | 0 |
| NC_004718.3;NP_828849.2;SARS-CoV_orf1ab;3446 | 0 | 0 |
| NC_004718.3;NP_828849.2;SARS-CoV_orf1ab;3447 | 0 | 0 |
| NC_004718.3;NP_828849.2;SARS-CoV_orf1ab;3448 | 0 | 0 |
| NC_004718.3;NP_828849.2;SARS-CoV_orf1ab;3449 | 0 | 0 |
| NC_004718.3;NP_828849.2;SARS-CoV_orf1ab;3450 | 0 | 0 |
| NC_004718.3;NP_828849.2;SARS-CoV_orf1ab;3451 | 0 | 0 |
| NC_004718.3;NP_828849.2;SARS-CoV_orf1ab;3452 | 0 | 0 |
| NC_004718.3;NP_828849.2;SARS-CoV_orf1ab;3453 | 0 | 0 |
| NC_004718.3;NP_828849.2;SARS-CoV_orf1ab;3454 | 0 | 0 |
| NC_004718.3;NP_828849.2;SARS-CoV_orf1ab;3455 | 0 | 0 |
| NC_004718.3;NP_828849.2;SARS-CoV_orf1ab;3456 | 0.05 | 0 |
| NC_004718.3;NP_828849.2;SARS-CoV_orf1ab;3457 | 0.05 | 0 |
| NC_004718.3;NP_828849.2;SARS-CoV_orf1ab;3458 | 0.05 | 0 |
| NC_004718.3;NP_828849.2;SARS-CoV_orf1ab;3459 | 0.05 | 0.025 |
| NC_004718.3;NP_828849.2;SARS-CoV_orf1ab;3460 | 0.1 | 0 |
| NC_004718.3;NP_828849.2;SARS-CoV_orf1ab;3461 | 0.05 | 0 |
| NC_004718.3;NP_828849.2;SARS-CoV_orf1ab;3462 | 0.1 | 0 |
| NC_004718.3;NP_828849.2;SARS-CoV_orf1ab;3463 | 0.05 | 0 |
| NC_004718.3;NP_828849.2;SARS-CoV_orf1ab;3464 | 0.05 | 0.025 |
| NC_004718.3;NP_828849.2;SARS-CoV_orf1ab;3465 | 0.05 | 0.15 |
| NC_004718.3;NP_828849.2;SARS-CoV_orf1ab;3466 | 0.25 | 0.1 |
| NC_004718.3;NP_828849.2;SARS-CoV_orf1ab;3467 | 0.05 | 0 |
| NC_004718.3;NP_828849.2;SARS-CoV_orf1ab;3468 | 0 | 0 |
| NC_004718.3;NP_828849.2;SARS-CoV_orf1ab;3469 | 0.05 | 0 |
| NC_004718.3;NP_828849.2;SARS-CoV_orf1ab;3470 | 0 | 0 |
| NC_004718.3;NP_828849.2;SARS-CoV_orf1ab;3471 | 0 | 0.025 |
| NC_004718.3;NP_828849.2;SARS-CoV_orf1ab;3472 | 0 | 0.025 |
| NC_004718.3;NP_828849.2;SARS-CoV_orf1ab;3473 | 0.05 | 0 |
| NC_004718.3;NP_828849.2;SARS-CoV_orf1ab;3474 | 0 | 0 |
| NC_004718.3;NP_828849.2;SARS-CoV_orf1ab;3475 | 0 | 0 |
| NC_004718.3;NP_828849.2;SARS-CoV_orf1ab;3476 | 0 | 0 |
| NC_004718.3;NP_828849.2;SARS-CoV_orf1ab;3477 | 0 | 0 |
| NC_004718.3;NP_828849.2;SARS-CoV_orf1ab;3478 | 0 | 0 |
| NC_004718.3;NP_828849.2;SARS-CoV_orf1ab;3479 | 0 | 0 |
| NC_004718.3;NP_828849.2;SARS-CoV_orf1ab;3480 | 0.05 | 0 |

| | | |
|---|---|---|
| NC_004718.3;NP_828849.2;SARS-CoV_orf1ab;3481 | 0 | 0 |
| NC_004718.3;NP_828849.2;SARS-CoV_orf1ab;3482 | 0.05 | 0.025 |
| NC_004718.3;NP_828849.2;SARS-CoV_orf1ab;3483 | 0.15 | 0.1 |
| NC_004718.3;NP_828849.2;SARS-CoV_orf1ab;3484 | 0.05 | 0.05 |
| NC_004718.3;NP_828849.2;SARS-CoV_orf1ab;3485 | 0.1 | 0.15 |
| NC_004718.3;NP_828849.2;SARS-CoV_orf1ab;3486 | 0.05 | 0.025 |
| NC_004718.3;NP_828849.2;SARS-CoV_orf1ab;3487 | 0.05 | 0 |
| NC_004718.3;NP_828849.2;SARS-CoV_orf1ab;3488 | 0.05 | 0 |
| NC_004718.3;NP_828849.2;SARS-CoV_orf1ab;3489 | 0.05 | 0 |
| NC_004718.3;NP_828849.2;SARS-CoV_orf1ab;3490 | 0.05 | 0 |
| NC_004718.3;NP_828849.2;SARS-CoV_orf1ab;3491 | 0.05 | 0 |
| NC_004718.3;NP_828849.2;SARS-CoV_orf1ab;3492 | 0.1 | 0.025 |
| NC_004718.3;NP_828849.2;SARS-CoV_orf1ab;3493 | 0 | 0 |
| NC_004718.3;NP_828849.2;SARS-CoV_orf1ab;3494 | 0 | 0 |
| NC_004718.3;NP_828849.2;SARS-CoV_orf1ab;3495 | 0 | 0 |
| NC_004718.3;NP_828849.2;SARS-CoV_orf1ab;3496 | 0 | 0 |
| NC_004718.3;NP_828849.2;SARS-CoV_orf1ab;3497 | 0 | 0 |
| NC_004718.3;NP_828849.2;SARS-CoV_orf1ab;3498 | 0.05 | 0 |
| NC_004718.3;NP_828849.2;SARS-CoV_orf1ab;3499 | 0 | 0 |
| NC_004718.3;NP_828849.2;SARS-CoV_orf1ab;3500 | 0 | 0.025 |
| NC_004718.3;NP_828849.2;SARS-CoV_orf1ab;3501 | 0 | 0 |
| NC_004718.3;NP_828849.2;SARS-CoV_orf1ab;3502 | 0 | 0.025 |
| NC_004718.3;NP_828849.2;SARS-CoV_orf1ab;3503 | 0 | 0.025 |
| NC_004718.3;NP_828849.2;SARS-CoV_orf1ab;3504 | 0 | 0 |
| NC_004718.3;NP_828849.2;SARS-CoV_orf1ab;3505 | 0 | 0 |
| NC_004718.3;NP_828849.2;SARS-CoV_orf1ab;3506 | 0 | 0 |
| NC_004718.3;NP_828849.2;SARS-CoV_orf1ab;3507 | 0 | 0 |
| NC_004718.3;NP_828849.2;SARS-CoV_orf1ab;3508 | 0 | 0.025 |
| NC_004718.3;NP_828849.2;SARS-CoV_orf1ab;3509 | 0 | 0.025 |
| NC_004718.3;NP_828849.2;SARS-CoV_orf1ab;3510 | 0 | 0 |
| NC_004718.3;NP_828849.2;SARS-CoV_orf1ab;3511 | 0 | 0 |
| NC_004718.3;NP_828849.2;SARS-CoV_orf1ab;3512 | 0 | 0 |
| NC_004718.3;NP_828849.2;SARS-CoV_orf1ab;3513 | 0 | 0 |
| NC_004718.3;NP_828849.2;SARS-CoV_orf1ab;3514 | 0 | 0 |
| NC_004718.3;NP_828849.2;SARS-CoV_orf1ab;3515 | 0 | 0 |
| NC_004718.3;NP_828849.2;SARS-CoV_orf1ab;3516 | 0 | 0 |
| NC_004718.3;NP_828849.2;SARS-CoV_orf1ab;3517 | 0 | 0 |
| NC_004718.3;NP_828849.2;SARS-CoV_orf1ab;3518 | 0 | 0.025 |
| NC_004718.3;NP_828849.2;SARS-CoV_orf1ab;3519 | 0 | 0.025 |
| NC_004718.3;NP_828849.2;SARS-CoV_orf1ab;3520 | 0 | 0.05 |
| NC_004718.3;NP_828849.2;SARS-CoV_orf1ab;3521 | 0 | 0.025 |
| NC_004718.3;NP_828849.2;SARS-CoV_orf1ab;3522 | 0 | 0.025 |
| NC_004718.3;NP_828849.2;SARS-CoV_orf1ab;3523 | 0 | 0.025 |

| | | |
|---|---|---|
| NC_004718.3;NP_828849.2;SARS-CoV_orf1ab;3524 | 0 | 0.025 |
| NC_004718.3;NP_828849.2;SARS-CoV_orf1ab;3525 | 0 | 0.05 |
| NC_004718.3;NP_828849.2;SARS-CoV_orf1ab;3526 | 0 | 0.025 |
| NC_004718.3;NP_828849.2;SARS-CoV_orf1ab;3527 | 0 | 0.05 |
| NC_004718.3;NP_828849.2;SARS-CoV_orf1ab;3528 | 0.1 | 0.025 |
| NC_004718.3;NP_828849.2;SARS-CoV_orf1ab;3529 | 0 | 0 |
| NC_004718.3;NP_828849.2;SARS-CoV_orf1ab;3530 | 0 | 0 |
| NC_004718.3;NP_828849.2;SARS-CoV_orf1ab;3531 | 0 | 0 |
| NC_004718.3;NP_828849.2;SARS-CoV_orf1ab;3532 | 0 | 0 |
| NC_004718.3;NP_828849.2;SARS-CoV_orf1ab;3533 | 0 | 0 |
| NC_004718.3;NP_828849.2;SARS-CoV_orf1ab;3534 | 0 | 0 |
| NC_004718.3;NP_828849.2;SARS-CoV_orf1ab;3535 | 0 | 0 |
| NC_004718.3;NP_828849.2;SARS-CoV_orf1ab;3536 | 0.05 | 0 |
| NC_004718.3;NP_828849.2;SARS-CoV_orf1ab;3537 | 0.05 | 0 |
| NC_004718.3;NP_828849.2;SARS-CoV_orf1ab;3538 | 0.05 | 0 |
| NC_004718.3;NP_828849.2;SARS-CoV_orf1ab;3539 | 0.05 | 0 |
| NC_004718.3;NP_828849.2;SARS-CoV_orf1ab;3540 | 0.05 | 0 |
| NC_004718.3;NP_828849.2;SARS-CoV_orf1ab;3541 | 0.05 | 0 |
| NC_004718.3;NP_828849.2;SARS-CoV_orf1ab;3542 | 0.05 | 0 |
| NC_004718.3;NP_828849.2;SARS-CoV_orf1ab;3543 | 0.05 | 0 |
| NC_004718.3;NP_828849.2;SARS-CoV_orf1ab;3544 | 0 | 0 |
| NC_004718.3;NP_828849.2;SARS-CoV_orf1ab;3545 | 0.05 | 0 |
| NC_004718.3;NP_828849.2;SARS-CoV_orf1ab;3546 | 0.05 | 0 |
| NC_004718.3;NP_828849.2;SARS-CoV_orf1ab;3547 | 0.05 | 0 |
| NC_004718.3;NP_828849.2;SARS-CoV_orf1ab;3548 | 0 | 0 |
| NC_004718.3;NP_828849.2;SARS-CoV_orf1ab;3549 | 0 | 0 |
| NC_004718.3;NP_828849.2;SARS-CoV_orf1ab;3550 | 0 | 0 |
| NC_004718.3;NP_828849.2;SARS-CoV_orf1ab;3551 | 0 | 0 |
| NC_004718.3;NP_828849.2;SARS-CoV_orf1ab;3552 | 0 | 0 |
| NC_004718.3;NP_828849.2;SARS-CoV_orf1ab;3553 | 0 | 0 |
| NC_004718.3;NP_828849.2;SARS-CoV_orf1ab;3554 | 0 | 0 |
| NC_004718.3;NP_828849.2;SARS-CoV_orf1ab;3555 | 0 | 0 |
| NC_004718.3;NP_828849.2;SARS-CoV_orf1ab;3556 | 0 | 0 |
| NC_004718.3;NP_828849.2;SARS-CoV_orf1ab;3557 | 0 | 0 |
| NC_004718.3;NP_828849.2;SARS-CoV_orf1ab;3558 | 0 | 0 |
| NC_004718.3;NP_828849.2;SARS-CoV_orf1ab;3559 | 0 | 0 |
| NC_004718.3;NP_828849.2;SARS-CoV_orf1ab;3560 | 0 | 0 |
| NC_004718.3;NP_828849.2;SARS-CoV_orf1ab;3561 | 0 | 0 |
| NC_004718.3;NP_828849.2;SARS-CoV_orf1ab;3562 | 0 | 0 |
| NC_004718.3;NP_828849.2;SARS-CoV_orf1ab;3563 | 0 | 0 |
| NC_004718.3;NP_828849.2;SARS-CoV_orf1ab;3564 | 0 | 0 |
| NC_004718.3;NP_828849.2;SARS-CoV_orf1ab;3565 | 0 | 0 |
| NC_004718.3;NP_828849.2;SARS-CoV_orf1ab;3566 | 0 | 0 |

| | | |
|---|---|---|
| NC_004718.3;NP_828849.2;SARS-CoV_orf1ab;3567 | 0 | 0 |
| NC_004718.3;NP_828849.2;SARS-CoV_orf1ab;3568 | 0 | 0 |
| NC_004718.3;NP_828849.2;SARS-CoV_orf1ab;3569 | 0 | 0 |
| NC_004718.3;NP_828849.2;SARS-CoV_orf1ab;3570 | 0 | 0 |
| NC_004718.3;NP_828849.2;SARS-CoV_orf1ab;3571 | 0 | 0 |
| NC_004718.3;NP_828849.2;SARS-CoV_orf1ab;3572 | 0 | 0 |
| NC_004718.3;NP_828849.2;SARS-CoV_orf1ab;3573 | 0 | 0 |
| NC_004718.3;NP_828849.2;SARS-CoV_orf1ab;3574 | 0 | 0 |
| NC_004718.3;NP_828849.2;SARS-CoV_orf1ab;3575 | 0 | 0 |
| NC_004718.3;NP_828849.2;SARS-CoV_orf1ab;3576 | 0 | 0 |
| NC_004718.3;NP_828849.2;SARS-CoV_orf1ab;3577 | 0 | 0 |
| NC_004718.3;NP_828849.2;SARS-CoV_orf1ab;3578 | 0 | 0 |
| NC_004718.3;NP_828849.2;SARS-CoV_orf1ab;3579 | 0 | 0 |
| NC_004718.3;NP_828849.2;SARS-CoV_orf1ab;3580 | 0 | 0 |
| NC_004718.3;NP_828849.2;SARS-CoV_orf1ab;3581 | 0 | 0 |
| NC_004718.3;NP_828849.2;SARS-CoV_orf1ab;3582 | 0 | 0 |
| NC_004718.3;NP_828849.2;SARS-CoV_orf1ab;3583 | 0 | 0 |
| NC_004718.3;NP_828849.2;SARS-CoV_orf1ab;3584 | 0 | 0 |
| NC_004718.3;NP_828849.2;SARS-CoV_orf1ab;3585 | 0 | 0 |
| NC_004718.3;NP_828849.2;SARS-CoV_orf1ab;3586 | 0 | 0 |
| NC_004718.3;NP_828849.2;SARS-CoV_orf1ab;3587 | 0 | 0 |
| NC_004718.3;NP_828849.2;SARS-CoV_orf1ab;3588 | 0 | 0 |
| NC_004718.3;NP_828849.2;SARS-CoV_orf1ab;3589 | 0 | 0 |
| NC_004718.3;NP_828849.2;SARS-CoV_orf1ab;3590 | 0 | 0 |
| NC_004718.3;NP_828849.2;SARS-CoV_orf1ab;3591 | 0 | 0 |
| NC_004718.3;NP_828849.2;SARS-CoV_orf1ab;3592 | 0 | 0 |
| NC_004718.3;NP_828849.2;SARS-CoV_orf1ab;3593 | 0 | 0 |
| NC_004718.3;NP_828849.2;SARS-CoV_orf1ab;3594 | 0 | 0 |
| NC_004718.3;NP_828849.2;SARS-CoV_orf1ab;3595 | 0 | 0 |
| NC_004718.3;NP_828849.2;SARS-CoV_orf1ab;3596 | 0 | 0 |
| NC_004718.3;NP_828849.2;SARS-CoV_orf1ab;3597 | 0 | 0 |
| NC_004718.3;NP_828849.2;SARS-CoV_orf1ab;3598 | 0 | 0 |
| NC_004718.3;NP_828849.2;SARS-CoV_orf1ab;3599 | 0 | 0 |
| NC_004718.3;NP_828849.2;SARS-CoV_orf1ab;3600 | 0 | 0 |
| NC_004718.3;NP_828849.2;SARS-CoV_orf1ab;3601 | 0 | 0 |
| NC_004718.3;NP_828849.2;SARS-CoV_orf1ab;3602 | 0 | 0 |
| NC_004718.3;NP_828849.2;SARS-CoV_orf1ab;3603 | 0 | 0 |
| NC_004718.3;NP_828849.2;SARS-CoV_orf1ab;3604 | 0 | 0 |
| NC_004718.3;NP_828849.2;SARS-CoV_orf1ab;3605 | 0 | 0 |
| NC_004718.3;NP_828849.2;SARS-CoV_orf1ab;3606 | 0 | 0 |
| NC_004718.3;NP_828849.2;SARS-CoV_orf1ab;3607 | 0 | 0 |
| NC_004718.3;NP_828849.2;SARS-CoV_orf1ab;3608 | 0 | 0 |
| NC_004718.3;NP_828849.2;SARS-CoV_orf1ab;3609 | 0 | 0 |

| | | |
|---|---|---|
| NC_004718.3;NP_828849.2;SARS-CoV_orf1ab;3610 | 0 | 0 |
| NC_004718.3;NP_828849.2;SARS-CoV_orf1ab;3611 | 0 | 0 |
| NC_004718.3;NP_828849.2;SARS-CoV_orf1ab;3612 | 0 | 0 |
| NC_004718.3;NP_828849.2;SARS-CoV_orf1ab;3613 | 0 | 0 |
| NC_004718.3;NP_828849.2;SARS-CoV_orf1ab;3614 | 0 | 0 |
| NC_004718.3;NP_828849.2;SARS-CoV_orf1ab;3615 | 0 | 0 |
| NC_004718.3;NP_828849.2;SARS-CoV_orf1ab;3616 | 0 | 0 |
| NC_004718.3;NP_828849.2;SARS-CoV_orf1ab;3617 | 0 | 0 |
| NC_004718.3;NP_828849.2;SARS-CoV_orf1ab;3618 | 0 | 0 |
| NC_004718.3;NP_828849.2;SARS-CoV_orf1ab;3619 | 0 | 0 |
| NC_004718.3;NP_828849.2;SARS-CoV_orf1ab;3620 | 0 | 0 |
| NC_004718.3;NP_828849.2;SARS-CoV_orf1ab;3621 | 0 | 0 |
| NC_004718.3;NP_828849.2;SARS-CoV_orf1ab;3622 | 0 | 0 |
| NC_004718.3;NP_828849.2;SARS-CoV_orf1ab;3623 | 0 | 0 |
| NC_004718.3;NP_828849.2;SARS-CoV_orf1ab;3624 | 0 | 0 |
| NC_004718.3;NP_828849.2;SARS-CoV_orf1ab;3625 | 0 | 0 |
| NC_004718.3;NP_828849.2;SARS-CoV_orf1ab;3626 | 0 | 0 |
| NC_004718.3;NP_828849.2;SARS-CoV_orf1ab;3627 | 0 | 0 |
| NC_004718.3;NP_828849.2;SARS-CoV_orf1ab;3628 | 0 | 0 |
| NC_004718.3;NP_828849.2;SARS-CoV_orf1ab;3629 | 0 | 0 |
| NC_004718.3;NP_828849.2;SARS-CoV_orf1ab;3630 | 0 | 0 |
| NC_004718.3;NP_828849.2;SARS-CoV_orf1ab;3631 | 0 | 0 |
| NC_004718.3;NP_828849.2;SARS-CoV_orf1ab;3632 | 0 | 0 |
| NC_004718.3;NP_828849.2;SARS-CoV_orf1ab;3633 | 0 | 0 |
| NC_004718.3;NP_828849.2;SARS-CoV_orf1ab;3634 | 0 | 0 |
| NC_004718.3;NP_828849.2;SARS-CoV_orf1ab;3635 | 0 | 0 |
| NC_004718.3;NP_828849.2;SARS-CoV_orf1ab;3636 | 0 | 0 |
| NC_004718.3;NP_828849.2;SARS-CoV_orf1ab;3637 | 0 | 0 |
| NC_004718.3;NP_828849.2;SARS-CoV_orf1ab;3638 | 0 | 0 |
| NC_004718.3;NP_828849.2;SARS-CoV_orf1ab;3639 | 0 | 0 |
| NC_004718.3;NP_828849.2;SARS-CoV_orf1ab;3640 | 0 | 0 |
| NC_004718.3;NP_828849.2;SARS-CoV_orf1ab;3641 | 0 | 0 |
| NC_004718.3;NP_828849.2;SARS-CoV_orf1ab;3642 | 0 | 0 |
| NC_004718.3;NP_828849.2;SARS-CoV_orf1ab;3643 | 0 | 0.025 |
| NC_004718.3;NP_828849.2;SARS-CoV_orf1ab;3644 | 0 | 0.025 |
| NC_004718.3;NP_828849.2;SARS-CoV_orf1ab;3645 | 0 | 0 |
| NC_004718.3;NP_828849.2;SARS-CoV_orf1ab;3646 | 0 | 0 |
| NC_004718.3;NP_828849.2;SARS-CoV_orf1ab;3647 | 0 | 0.025 |
| NC_004718.3;NP_828849.2;SARS-CoV_orf1ab;3648 | 0 | 0 |
| NC_004718.3;NP_828849.2;SARS-CoV_orf1ab;3649 | 0 | 0 |
| NC_004718.3;NP_828849.2;SARS-CoV_orf1ab;3650 | 0 | 0 |
| NC_004718.3;NP_828849.2;SARS-CoV_orf1ab;3651 | 0 | 0 |
| NC_004718.3;NP_828849.2;SARS-CoV_orf1ab;3652 | 0 | 0 |

| | | |
|---|---|---|
| NC_004718.3;NP_828849.2;SARS-CoV_orf1ab;3653 | 0 | 0 |
| NC_004718.3;NP_828849.2;SARS-CoV_orf1ab;3654 | 0 | 0 |
| NC_004718.3;NP_828849.2;SARS-CoV_orf1ab;3655 | 0 | 0 |
| NC_004718.3;NP_828849.2;SARS-CoV_orf1ab;3656 | 0 | 0 |
| NC_004718.3;NP_828849.2;SARS-CoV_orf1ab;3657 | 0 | 0 |
| NC_004718.3;NP_828849.2;SARS-CoV_orf1ab;3658 | 0 | 0 |
| NC_004718.3;NP_828849.2;SARS-CoV_orf1ab;3659 | 0 | 0 |
| NC_004718.3;NP_828849.2;SARS-CoV_orf1ab;3660 | 0 | 0 |
| NC_004718.3;NP_828849.2;SARS-CoV_orf1ab;3661 | 0 | 0 |
| NC_004718.3;NP_828849.2;SARS-CoV_orf1ab;3662 | 0 | 0 |
| NC_004718.3;NP_828849.2;SARS-CoV_orf1ab;3663 | 0 | 0 |
| NC_004718.3;NP_828849.2;SARS-CoV_orf1ab;3664 | 0 | 0 |
| NC_004718.3;NP_828849.2;SARS-CoV_orf1ab;3665 | 0 | 0 |
| NC_004718.3;NP_828849.2;SARS-CoV_orf1ab;3666 | 0 | 0 |
| NC_004718.3;NP_828849.2;SARS-CoV_orf1ab;3667 | 0 | 0 |
| NC_004718.3;NP_828849.2;SARS-CoV_orf1ab;3668 | 0 | 0 |
| NC_004718.3;NP_828849.2;SARS-CoV_orf1ab;3669 | 0 | 0 |
| NC_004718.3;NP_828849.2;SARS-CoV_orf1ab;3670 | 0 | 0 |
| NC_004718.3;NP_828849.2;SARS-CoV_orf1ab;3671 | 0 | 0 |
| NC_004718.3;NP_828849.2;SARS-CoV_orf1ab;3672 | 0 | 0 |
| NC_004718.3;NP_828849.2;SARS-CoV_orf1ab;3673 | 0 | 0 |
| NC_004718.3;NP_828849.2;SARS-CoV_orf1ab;3674 | 0 | 0 |
| NC_004718.3;NP_828849.2;SARS-CoV_orf1ab;3675 | 0 | 0.025 |
| NC_004718.3;NP_828849.2;SARS-CoV_orf1ab;3676 | 0 | 0 |
| NC_004718.3;NP_828849.2;SARS-CoV_orf1ab;3677 | 0 | 0 |
| NC_004718.3;NP_828849.2;SARS-CoV_orf1ab;3678 | 0 | 0 |
| NC_004718.3;NP_828849.2;SARS-CoV_orf1ab;3679 | 0 | 0.025 |
| NC_004718.3;NP_828849.2;SARS-CoV_orf1ab;3680 | 0 | 0 |
| NC_004718.3;NP_828849.2;SARS-CoV_orf1ab;3681 | 0 | 0 |
| NC_004718.3;NP_828849.2;SARS-CoV_orf1ab;3682 | 0 | 0 |
| NC_004718.3;NP_828849.2;SARS-CoV_orf1ab;3683 | 0 | 0 |
| NC_004718.3;NP_828849.2;SARS-CoV_orf1ab;3684 | 0 | 0 |
| NC_004718.3;NP_828849.2;SARS-CoV_orf1ab;3685 | 0 | 0 |
| NC_004718.3;NP_828849.2;SARS-CoV_orf1ab;3686 | 0 | 0 |
| NC_004718.3;NP_828849.2;SARS-CoV_orf1ab;3687 | 0 | 0 |
| NC_004718.3;NP_828849.2;SARS-CoV_orf1ab;3688 | 0 | 0 |
| NC_004718.3;NP_828849.2;SARS-CoV_orf1ab;3689 | 0 | 0 |
| NC_004718.3;NP_828849.2;SARS-CoV_orf1ab;3690 | 0 | 0 |
| NC_004718.3;NP_828849.2;SARS-CoV_orf1ab;3691 | 0 | 0 |
| NC_004718.3;NP_828849.2;SARS-CoV_orf1ab;3692 | 0 | 0 |
| NC_004718.3;NP_828849.2;SARS-CoV_orf1ab;3693 | 0 | 0 |
| NC_004718.3;NP_828849.2;SARS-CoV_orf1ab;3694 | 0 | 0 |
| NC_004718.3;NP_828849.2;SARS-CoV_orf1ab;3695 | 0.05 | 0 |

| | | |
|---|---|---|
| NC_004718.3;NP_828849.2;SARS-CoV_orf1ab;3696 | 0.05 | 0 |
| NC_004718.3;NP_828849.2;SARS-CoV_orf1ab;3697 | 0.05 | 0 |
| NC_004718.3;NP_828849.2;SARS-CoV_orf1ab;3698 | 0.05 | 0.025 |
| NC_004718.3;NP_828849.2;SARS-CoV_orf1ab;3699 | 0.05 | 0 |
| NC_004718.3;NP_828849.2;SARS-CoV_orf1ab;3700 | 0 | 0 |
| NC_004718.3;NP_828849.2;SARS-CoV_orf1ab;3701 | 0 | 0 |
| NC_004718.3;NP_828849.2;SARS-CoV_orf1ab;3702 | 0 | 0 |
| NC_004718.3;NP_828849.2;SARS-CoV_orf1ab;3703 | 0 | 0 |
| NC_004718.3;NP_828849.2;SARS-CoV_orf1ab;3704 | 0 | 0 |
| NC_004718.3;NP_828849.2;SARS-CoV_orf1ab;3705 | 0 | 0 |
| NC_004718.3;NP_828849.2;SARS-CoV_orf1ab;3706 | 0 | 0 |
| NC_004718.3;NP_828849.2;SARS-CoV_orf1ab;3707 | 0 | 0 |
| NC_004718.3;NP_828849.2;SARS-CoV_orf1ab;3708 | 0 | 0 |
| NC_004718.3;NP_828849.2;SARS-CoV_orf1ab;3709 | 0 | 0 |
| NC_004718.3;NP_828849.2;SARS-CoV_orf1ab;3710 | 0 | 0 |
| NC_004718.3;NP_828849.2;SARS-CoV_orf1ab;3711 | 0 | 0 |
| NC_004718.3;NP_828849.2;SARS-CoV_orf1ab;3712 | 0 | 0 |
| NC_004718.3;NP_828849.2;SARS-CoV_orf1ab;3713 | 0 | 0 |
| NC_004718.3;NP_828849.2;SARS-CoV_orf1ab;3714 | 0 | 0 |
| NC_004718.3;NP_828849.2;SARS-CoV_orf1ab;3715 | 0 | 0 |
| NC_004718.3;NP_828849.2;SARS-CoV_orf1ab;3716 | 0 | 0 |
| NC_004718.3;NP_828849.2;SARS-CoV_orf1ab;3717 | 0 | 0 |
| NC_004718.3;NP_828849.2;SARS-CoV_orf1ab;3718 | 0 | 0 |
| NC_004718.3;NP_828849.2;SARS-CoV_orf1ab;3719 | 0 | 0 |
| NC_004718.3;NP_828849.2;SARS-CoV_orf1ab;3720 | 0 | 0 |
| NC_004718.3;NP_828849.2;SARS-CoV_orf1ab;3721 | 0 | 0 |
| NC_004718.3;NP_828849.2;SARS-CoV_orf1ab;3722 | 0 | 0 |
| NC_004718.3;NP_828849.2;SARS-CoV_orf1ab;3723 | 0 | 0 |
| NC_004718.3;NP_828849.2;SARS-CoV_orf1ab;3724 | 0 | 0 |
| NC_004718.3;NP_828849.2;SARS-CoV_orf1ab;3725 | 0 | 0 |
| NC_004718.3;NP_828849.2;SARS-CoV_orf1ab;3726 | 0 | 0 |
| NC_004718.3;NP_828849.2;SARS-CoV_orf1ab;3727 | 0 | 0 |
| NC_004718.3;NP_828849.2;SARS-CoV_orf1ab;3728 | 0 | 0 |
| NC_004718.3;NP_828849.2;SARS-CoV_orf1ab;3729 | 0 | 0 |
| NC_004718.3;NP_828849.2;SARS-CoV_orf1ab;3730 | 0 | 0 |
| NC_004718.3;NP_828849.2;SARS-CoV_orf1ab;3731 | 0 | 0 |
| NC_004718.3;NP_828849.2;SARS-CoV_orf1ab;3732 | 0 | 0 |
| NC_004718.3;NP_828849.2;SARS-CoV_orf1ab;3733 | 0 | 0 |
| NC_004718.3;NP_828849.2;SARS-CoV_orf1ab;3734 | 0 | 0 |
| NC_004718.3;NP_828849.2;SARS-CoV_orf1ab;3735 | 0 | 0 |
| NC_004718.3;NP_828849.2;SARS-CoV_orf1ab;3736 | 0 | 0 |
| NC_004718.3;NP_828849.2;SARS-CoV_orf1ab;3737 | 0 | 0 |
| NC_004718.3;NP_828849.2;SARS-CoV_orf1ab;3738 | 0 | 0 |

| | | |
|---|---|---|
| NC_004718.3;NP_828849.2;SARS-CoV_orf1ab;3739 | 0 | 0 |
| NC_004718.3;NP_828849.2;SARS-CoV_orf1ab;3740 | 0 | 0 |
| NC_004718.3;NP_828849.2;SARS-CoV_orf1ab;3741 | 0 | 0 |
| NC_004718.3;NP_828849.2;SARS-CoV_orf1ab;3742 | 0 | 0 |
| NC_004718.3;NP_828849.2;SARS-CoV_orf1ab;3743 | 0 | 0 |
| NC_004718.3;NP_828849.2;SARS-CoV_orf1ab;3744 | 0 | 0 |
| NC_004718.3;NP_828849.2;SARS-CoV_orf1ab;3745 | 0 | 0 |
| NC_004718.3;NP_828849.2;SARS-CoV_orf1ab;3746 | 0 | 0 |
| NC_004718.3;NP_828849.2;SARS-CoV_orf1ab;3747 | 0 | 0 |
| NC_004718.3;NP_828849.2;SARS-CoV_orf1ab;3748 | 0 | 0 |
| NC_004718.3;NP_828849.2;SARS-CoV_orf1ab;3749 | 0 | 0 |
| NC_004718.3;NP_828849.2;SARS-CoV_orf1ab;3750 | 0 | 0 |
| NC_004718.3;NP_828849.2;SARS-CoV_orf1ab;3751 | 0 | 0 |
| NC_004718.3;NP_828849.2;SARS-CoV_orf1ab;3752 | 0 | 0 |
| NC_004718.3;NP_828849.2;SARS-CoV_orf1ab;3753 | 0 | 0 |
| NC_004718.3;NP_828849.2;SARS-CoV_orf1ab;3754 | 0 | 0 |
| NC_004718.3;NP_828849.2;SARS-CoV_orf1ab;3755 | 0 | 0 |
| NC_004718.3;NP_828849.2;SARS-CoV_orf1ab;3756 | 0 | 0 |
| NC_004718.3;NP_828849.2;SARS-CoV_orf1ab;3757 | 0 | 0 |
| NC_004718.3;NP_828849.2;SARS-CoV_orf1ab;3758 | 0 | 0 |
| NC_004718.3;NP_828849.2;SARS-CoV_orf1ab;3759 | 0 | 0 |
| NC_004718.3;NP_828849.2;SARS-CoV_orf1ab;3760 | 0 | 0 |
| NC_004718.3;NP_828849.2;SARS-CoV_orf1ab;3761 | 0 | 0 |
| NC_004718.3;NP_828849.2;SARS-CoV_orf1ab;3762 | 0 | 0 |
| NC_004718.3;NP_828849.2;SARS-CoV_orf1ab;3763 | 0 | 0 |
| NC_004718.3;NP_828849.2;SARS-CoV_orf1ab;3764 | 0 | 0 |
| NC_004718.3;NP_828849.2;SARS-CoV_orf1ab;3765 | 0 | 0 |
| NC_004718.3;NP_828849.2;SARS-CoV_orf1ab;3766 | 0 | 0 |
| NC_004718.3;NP_828849.2;SARS-CoV_orf1ab;3767 | 0 | 0 |
| NC_004718.3;NP_828849.2;SARS-CoV_orf1ab;3768 | 0 | 0 |
| NC_004718.3;NP_828849.2;SARS-CoV_orf1ab;3769 | 0 | 0 |
| NC_004718.3;NP_828849.2;SARS-CoV_orf1ab;3770 | 0 | 0 |
| NC_004718.3;NP_828849.2;SARS-CoV_orf1ab;3771 | 0 | 0 |
| NC_004718.3;NP_828849.2;SARS-CoV_orf1ab;3772 | 0 | 0 |
| NC_004718.3;NP_828849.2;SARS-CoV_orf1ab;3773 | 0 | 0 |
| NC_004718.3;NP_828849.2;SARS-CoV_orf1ab;3774 | 0 | 0 |
| NC_004718.3;NP_828849.2;SARS-CoV_orf1ab;3775 | 0 | 0 |
| NC_004718.3;NP_828849.2;SARS-CoV_orf1ab;3776 | 0 | 0 |
| NC_004718.3;NP_828849.2;SARS-CoV_orf1ab;3777 | 0 | 0 |
| NC_004718.3;NP_828849.2;SARS-CoV_orf1ab;3778 | 0 | 0 |
| NC_004718.3;NP_828849.2;SARS-CoV_orf1ab;3779 | 0 | 0 |
| NC_004718.3;NP_828849.2;SARS-CoV_orf1ab;3780 | 0 | 0 |
| NC_004718.3;NP_828849.2;SARS-CoV_orf1ab;3781 | 0 | 0 |

| | | |
|---|---|---|
| NC_004718.3;NP_828849.2;SARS-CoV_orf1ab;3782 | 0 | 0 |
| NC_004718.3;NP_828849.2;SARS-CoV_orf1ab;3783 | 0 | 0 |
| NC_004718.3;NP_828849.2;SARS-CoV_orf1ab;3784 | 0 | 0 |
| NC_004718.3;NP_828849.2;SARS-CoV_orf1ab;3785 | 0 | 0 |
| NC_004718.3;NP_828849.2;SARS-CoV_orf1ab;3786 | 0.25 | 0.175 |
| NC_004718.3;NP_828849.2;SARS-CoV_orf1ab;3787 | 0 | 0 |
| NC_004718.3;NP_828849.2;SARS-CoV_orf1ab;3788 | 0 | 0 |
| NC_004718.3;NP_828849.2;SARS-CoV_orf1ab;3789 | 0 | 0 |
| NC_004718.3;NP_828849.2;SARS-CoV_orf1ab;3790 | 0 | 0 |
| NC_004718.3;NP_828849.2;SARS-CoV_orf1ab;3791 | 0 | 0 |
| NC_004718.3;NP_828849.2;SARS-CoV_orf1ab;3792 | 0 | 0 |
| NC_004718.3;NP_828849.2;SARS-CoV_orf1ab;3793 | 0 | 0 |
| NC_004718.3;NP_828849.2;SARS-CoV_orf1ab;3794 | 0 | 0.05 |
| NC_004718.3;NP_828849.2;SARS-CoV_orf1ab;3795 | 0.05 | 0.025 |
| NC_004718.3;NP_828849.2;SARS-CoV_orf1ab;3796 | 0.05 | 0.025 |
| NC_004718.3;NP_828849.2;SARS-CoV_orf1ab;3797 | 0 | 0.025 |
| NC_004718.3;NP_828849.2;SARS-CoV_orf1ab;3798 | 0 | 0.025 |
| NC_004718.3;NP_828849.2;SARS-CoV_orf1ab;3799 | 0 | 0.1 |
| NC_004718.3;NP_828849.2;SARS-CoV_orf1ab;3800 | 0 | 0.025 |
| NC_004718.3;NP_828849.2;SARS-CoV_orf1ab;3801 | 0.2 | 0.15 |
| NC_004718.3;NP_828849.2;SARS-CoV_orf1ab;3802 | 0.15 | 0.075 |
| NC_004718.3;NP_828849.2;SARS-CoV_orf1ab;3803 | 0.15 | 0.125 |
| NC_004718.3;NP_828849.2;SARS-CoV_orf1ab;3804 | 0.1 | 0.075 |
| NC_004718.3;NP_828849.2;SARS-CoV_orf1ab;3805 | 0.1 | 0.05 |
| NC_004718.3;NP_828849.2;SARS-CoV_orf1ab;3806 | 0.05 | 0.05 |
| NC_004718.3;NP_828849.2;SARS-CoV_orf1ab;3807 | 0 | 0.05 |
| NC_004718.3;NP_828849.2;SARS-CoV_orf1ab;3808 | 0 | 0.05 |
| NC_004718.3;NP_828849.2;SARS-CoV_orf1ab;3809 | 0 | 0.025 |
| NC_004718.3;NP_828849.2;SARS-CoV_orf1ab;3810 | 0 | 0 |
| NC_004718.3;NP_828849.2;SARS-CoV_orf1ab;3811 | 0 | 0 |
| NC_004718.3;NP_828849.2;SARS-CoV_orf1ab;3812 | 0 | 0 |
| NC_004718.3;NP_828849.2;SARS-CoV_orf1ab;3813 | 0 | 0 |
| NC_004718.3;NP_828849.2;SARS-CoV_orf1ab;3814 | 0 | 0 |
| NC_004718.3;NP_828849.2;SARS-CoV_orf1ab;3815 | 0 | 0 |
| NC_004718.3;NP_828849.2;SARS-CoV_orf1ab;3816 | 0 | 0 |
| NC_004718.3;NP_828849.2;SARS-CoV_orf1ab;3817 | 0 | 0 |
| NC_004718.3;NP_828849.2;SARS-CoV_orf1ab;3818 | 0 | 0 |
| NC_004718.3;NP_828849.2;SARS-CoV_orf1ab;3819 | 0 | 0.025 |
| NC_004718.3;NP_828849.2;SARS-CoV_orf1ab;3820 | 0 | 0 |
| NC_004718.3;NP_828849.2;SARS-CoV_orf1ab;3821 | 0 | 0.025 |
| NC_004718.3;NP_828849.2;SARS-CoV_orf1ab;3822 | 0 | 0.025 |
| NC_004718.3;NP_828849.2;SARS-CoV_orf1ab;3823 | 0.05 | 0.05 |
| NC_004718.3;NP_828849.2;SARS-CoV_orf1ab;3824 | 0.05 | 0.05 |

| | | |
|---|---|---|
| NC_004718.3;NP_828849.2;SARS-CoV_orf1ab;3825 | 0.05 | 0.05 |
| NC_004718.3;NP_828849.2;SARS-CoV_orf1ab;3826 | 0.1 | 0.1 |
| NC_004718.3;NP_828849.2;SARS-CoV_orf1ab;3827 | 0.05 | 0.075 |
| NC_004718.3;NP_828849.2;SARS-CoV_orf1ab;3828 | 0.05 | 0.05 |
| NC_004718.3;NP_828849.2;SARS-CoV_orf1ab;3829 | 0.05 | 0.025 |
| NC_004718.3;NP_828849.2;SARS-CoV_orf1ab;3830 | 0 | 0.025 |
| NC_004718.3;NP_828849.2;SARS-CoV_orf1ab;3831 | 0.05 | 0.025 |
| NC_004718.3;NP_828849.2;SARS-CoV_orf1ab;3832 | 0.05 | 0.025 |
| NC_004718.3;NP_828849.2;SARS-CoV_orf1ab;3833 | 0.05 | 0.025 |
| NC_004718.3;NP_828849.2;SARS-CoV_orf1ab;3834 | 0.05 | 0 |
| NC_004718.3;NP_828849.2;SARS-CoV_orf1ab;3835 | 0 | 0 |
| NC_004718.3;NP_828849.2;SARS-CoV_orf1ab;3836 | 0 | 0 |
| NC_004718.3;NP_828849.2;SARS-CoV_orf1ab;3837 | 0 | 0 |
| NC_004718.3;NP_828849.2;SARS-CoV_orf1ab;3838 | 0 | 0 |
| NC_004718.3;NP_828849.2;SARS-CoV_orf1ab;3839 | 0 | 0 |
| NC_004718.3;NP_828849.2;SARS-CoV_orf1ab;3840 | 0 | 0 |
| NC_004718.3;NP_828849.2;SARS-CoV_orf1ab;3841 | 0 | 0 |
| NC_004718.3;NP_828849.2;SARS-CoV_orf1ab;3842 | 0 | 0 |
| NC_004718.3;NP_828849.2;SARS-CoV_orf1ab;3843 | 0 | 0 |
| NC_004718.3;NP_828849.2;SARS-CoV_orf1ab;3844 | 0 | 0 |
| NC_004718.3;NP_828849.2;SARS-CoV_orf1ab;3845 | 0 | 0 |
| NC_004718.3;NP_828849.2;SARS-CoV_orf1ab;3846 | 0 | 0 |
| NC_004718.3;NP_828849.2;SARS-CoV_orf1ab;3847 | 0 | 0 |
| NC_004718.3;NP_828849.2;SARS-CoV_orf1ab;3848 | 0 | 0 |
| NC_004718.3;NP_828849.2;SARS-CoV_orf1ab;3849 | 0 | 0 |
| NC_004718.3;NP_828849.2;SARS-CoV_orf1ab;3850 | 0 | 0 |
| NC_004718.3;NP_828849.2;SARS-CoV_orf1ab;3851 | 0 | 0 |
| NC_004718.3;NP_828849.2;SARS-CoV_orf1ab;3852 | 0 | 0 |
| NC_004718.3;NP_828849.2;SARS-CoV_orf1ab;3853 | 0 | 0 |
| NC_004718.3;NP_828849.2;SARS-CoV_orf1ab;3854 | 0 | 0 |
| NC_004718.3;NP_828849.2;SARS-CoV_orf1ab;3855 | 0 | 0 |
| NC_004718.3;NP_828849.2;SARS-CoV_orf1ab;3856 | 0 | 0 |
| NC_004718.3;NP_828849.2;SARS-CoV_orf1ab;3857 | 0 | 0 |
| NC_004718.3;NP_828849.2;SARS-CoV_orf1ab;3858 | 0 | 0 |
| NC_004718.3;NP_828849.2;SARS-CoV_orf1ab;3859 | 0 | 0.025 |
| NC_004718.3;NP_828849.2;SARS-CoV_orf1ab;3860 | 0 | 0 |
| NC_004718.3;NP_828849.2;SARS-CoV_orf1ab;3861 | 0 | 0.025 |
| NC_004718.3;NP_828849.2;SARS-CoV_orf1ab;3862 | 0 | 0 |
| NC_004718.3;NP_828849.2;SARS-CoV_orf1ab;3863 | 0 | 0 |
| NC_004718.3;NP_828849.2;SARS-CoV_orf1ab;3864 | 0 | 0 |
| NC_004718.3;NP_828849.2;SARS-CoV_orf1ab;3865 | 0.05 | 0 |
| NC_004718.3;NP_828849.2;SARS-CoV_orf1ab;3866 | 0.05 | 0 |
| NC_004718.3;NP_828849.2;SARS-CoV_orf1ab;3867 | 0.05 | 0 |

| | | |
|---|---|---|
| NC_004718.3;NP_828849.2;SARS-CoV_orf1ab;3868 | 0 | 0 |
| NC_004718.3;NP_828849.2;SARS-CoV_orf1ab;3869 | 0 | 0 |
| NC_004718.3;NP_828849.2;SARS-CoV_orf1ab;3870 | 0 | 0 |
| NC_004718.3;NP_828849.2;SARS-CoV_orf1ab;3871 | 0 | 0 |
| NC_004718.3;NP_828849.2;SARS-CoV_orf1ab;3872 | 0 | 0 |
| NC_004718.3;NP_828849.2;SARS-CoV_orf1ab;3873 | 0 | 0 |
| NC_004718.3;NP_828849.2;SARS-CoV_orf1ab;3874 | 0 | 0 |
| NC_004718.3;NP_828849.2;SARS-CoV_orf1ab;3875 | 0 | 0 |
| NC_004718.3;NP_828849.2;SARS-CoV_orf1ab;3876 | 0 | 0 |
| NC_004718.3;NP_828849.2;SARS-CoV_orf1ab;3877 | 0 | 0 |
| NC_004718.3;NP_828849.2;SARS-CoV_orf1ab;3878 | 0 | 0 |
| NC_004718.3;NP_828849.2;SARS-CoV_orf1ab;3879 | 0 | 0 |
| NC_004718.3;NP_828849.2;SARS-CoV_orf1ab;3880 | 0 | 0 |
| NC_004718.3;NP_828849.2;SARS-CoV_orf1ab;3881 | 0 | 0 |
| NC_004718.3;NP_828849.2;SARS-CoV_orf1ab;3882 | 0 | 0 |
| NC_004718.3;NP_828849.2;SARS-CoV_orf1ab;3883 | 0 | 0 |
| NC_004718.3;NP_828849.2;SARS-CoV_orf1ab;3884 | 0 | 0 |
| NC_004718.3;NP_828849.2;SARS-CoV_orf1ab;3885 | 0 | 0 |
| NC_004718.3;NP_828849.2;SARS-CoV_orf1ab;3886 | 0 | 0 |
| NC_004718.3;NP_828849.2;SARS-CoV_orf1ab;3887 | 0 | 0 |
| NC_004718.3;NP_828849.2;SARS-CoV_orf1ab;3888 | 0 | 0 |
| NC_004718.3;NP_828849.2;SARS-CoV_orf1ab;3889 | 0 | 0 |
| NC_004718.3;NP_828849.2;SARS-CoV_orf1ab;3890 | 0 | 0 |
| NC_004718.3;NP_828849.2;SARS-CoV_orf1ab;3891 | 0 | 0.025 |
| NC_004718.3;NP_828849.2;SARS-CoV_orf1ab;3892 | 0 | 0.025 |
| NC_004718.3;NP_828849.2;SARS-CoV_orf1ab;3893 | 0 | 0.025 |
| NC_004718.3;NP_828849.2;SARS-CoV_orf1ab;3894 | 0.05 | 0.025 |
| NC_004718.3;NP_828849.2;SARS-CoV_orf1ab;3895 | 0.05 | 0.025 |
| NC_004718.3;NP_828849.2;SARS-CoV_orf1ab;3896 | 0.05 | 0.125 |
| NC_004718.3;NP_828849.2;SARS-CoV_orf1ab;3897 | 0.1 | 0.05 |
| NC_004718.3;NP_828849.2;SARS-CoV_orf1ab;3898 | 0.1 | 0.15 |
| NC_004718.3;NP_828849.2;SARS-CoV_orf1ab;3899 | 0.1 | 0.1 |
| NC_004718.3;NP_828849.2;SARS-CoV_orf1ab;3900 | 0.1 | 0.1 |
| NC_004718.3;NP_828849.2;SARS-CoV_orf1ab;3901 | 0 | 0.025 |
| NC_004718.3;NP_828849.2;SARS-CoV_orf1ab;3902 | 0 | 0.075 |
| NC_004718.3;NP_828849.2;SARS-CoV_orf1ab;3903 | 0 | 0.025 |
| NC_004718.3;NP_828849.2;SARS-CoV_orf1ab;3904 | 0 | 0 |
| NC_004718.3;NP_828849.2;SARS-CoV_orf1ab;3905 | 0 | 0 |
| NC_004718.3;NP_828849.2;SARS-CoV_orf1ab;3906 | 0.05 | 0 |
| NC_004718.3;NP_828849.2;SARS-CoV_orf1ab;3907 | 0.05 | 0 |
| NC_004718.3;NP_828849.2;SARS-CoV_orf1ab;3908 | 0.05 | 0 |
| NC_004718.3;NP_828849.2;SARS-CoV_orf1ab;3909 | 0 | 0 |
| NC_004718.3;NP_828849.2;SARS-CoV_orf1ab;3910 | 0.05 | 0 |

| | | |
|---|---|---|
| NC_004718.3;NP_828849.2;SARS-CoV_orf1ab;3911 | 0 | 0 |
| NC_004718.3;NP_828849.2;SARS-CoV_orf1ab;3912 | 0 | 0 |
| NC_004718.3;NP_828849.2;SARS-CoV_orf1ab;3913 | 0 | 0 |
| NC_004718.3;NP_828849.2;SARS-CoV_orf1ab;3914 | 0 | 0 |
| NC_004718.3;NP_828849.2;SARS-CoV_orf1ab;3915 | 0 | 0 |
| NC_004718.3;NP_828849.2;SARS-CoV_orf1ab;3916 | 0 | 0 |
| NC_004718.3;NP_828849.2;SARS-CoV_orf1ab;3917 | 0 | 0 |
| NC_004718.3;NP_828849.2;SARS-CoV_orf1ab;3918 | 0.05 | 0.05 |
| NC_004718.3;NP_828849.2;SARS-CoV_orf1ab;3919 | 0 | 0.025 |
| NC_004718.3;NP_828849.2;SARS-CoV_orf1ab;3920 | 0 | 0 |
| NC_004718.3;NP_828849.2;SARS-CoV_orf1ab;3921 | 0 | 0 |
| NC_004718.3;NP_828849.2;SARS-CoV_orf1ab;3922 | 0.05 | 0 |
| NC_004718.3;NP_828849.2;SARS-CoV_orf1ab;3923 | 0 | 0.05 |
| NC_004718.3;NP_828849.2;SARS-CoV_orf1ab;3924 | 0 | 0 |
| NC_004718.3;NP_828849.2;SARS-CoV_orf1ab;3925 | 0 | 0 |
| NC_004718.3;NP_828849.2;SARS-CoV_orf1ab;3926 | 0 | 0.025 |
| NC_004718.3;NP_828849.2;SARS-CoV_orf1ab;3927 | 0 | 0.05 |
| NC_004718.3;NP_828849.2;SARS-CoV_orf1ab;3928 | 0.05 | 0.05 |
| NC_004718.3;NP_828849.2;SARS-CoV_orf1ab;3929 | 0 | 0.025 |
| NC_004718.3;NP_828849.2;SARS-CoV_orf1ab;3930 | 0.05 | 0.075 |
| NC_004718.3;NP_828849.2;SARS-CoV_orf1ab;3931 | 0.05 | 0.15 |
| NC_004718.3;NP_828849.2;SARS-CoV_orf1ab;3932 | 0.05 | 0.15 |
| NC_004718.3;NP_828849.2;SARS-CoV_orf1ab;3933 | 0.1 | 0.2 |
| NC_004718.3;NP_828849.2;SARS-CoV_orf1ab;3934 | 0.05 | 0.1 |
| NC_004718.3;NP_828849.2;SARS-CoV_orf1ab;3935 | 0 | 0.025 |
| NC_004718.3;NP_828849.2;SARS-CoV_orf1ab;3936 | 0 | 0 |
| NC_004718.3;NP_828849.2;SARS-CoV_orf1ab;3937 | 0.05 | 0 |
| NC_004718.3;NP_828849.2;SARS-CoV_orf1ab;3938 | 0 | 0 |
| NC_004718.3;NP_828849.2;SARS-CoV_orf1ab;3939 | 0 | 0 |
| NC_004718.3;NP_828849.2;SARS-CoV_orf1ab;3940 | 0 | 0 |
| NC_004718.3;NP_828849.2;SARS-CoV_orf1ab;3941 | 0 | 0 |
| NC_004718.3;NP_828849.2;SARS-CoV_orf1ab;3942 | 0 | 0 |
| NC_004718.3;NP_828849.2;SARS-CoV_orf1ab;3943 | 0 | 0 |
| NC_004718.3;NP_828849.2;SARS-CoV_orf1ab;3944 | 0 | 0 |
| NC_004718.3;NP_828849.2;SARS-CoV_orf1ab;3945 | 0 | 0 |
| NC_004718.3;NP_828849.2;SARS-CoV_orf1ab;3946 | 0 | 0 |
| NC_004718.3;NP_828849.2;SARS-CoV_orf1ab;3947 | 0 | 0 |
| NC_004718.3;NP_828849.2;SARS-CoV_orf1ab;3948 | 0 | 0 |
| NC_004718.3;NP_828849.2;SARS-CoV_orf1ab;3949 | 0 | 0 |
| NC_004718.3;NP_828849.2;SARS-CoV_orf1ab;3950 | 0 | 0 |
| NC_004718.3;NP_828849.2;SARS-CoV_orf1ab;3951 | 0 | 0 |
| NC_004718.3;NP_828849.2;SARS-CoV_orf1ab;3952 | 0 | 0 |
| NC_004718.3;NP_828849.2;SARS-CoV_orf1ab;3953 | 0 | 0 |

| | | |
|---|---|---|
| NC_004718.3;NP_828849.2;SARS-CoV_orf1ab;3954 | 0 | 0.025 |
| NC_004718.3;NP_828849.2;SARS-CoV_orf1ab;3955 | 0 | 0.05 |
| NC_004718.3;NP_828849.2;SARS-CoV_orf1ab;3956 | 0 | 0.025 |
| NC_004718.3;NP_828849.2;SARS-CoV_orf1ab;3957 | 0.05 | 0.025 |
| NC_004718.3;NP_828849.2;SARS-CoV_orf1ab;3958 | 0.05 | 0.025 |
| NC_004718.3;NP_828849.2;SARS-CoV_orf1ab;3959 | 0 | 0.05 |
| NC_004718.3;NP_828849.2;SARS-CoV_orf1ab;3960 | 0.05 | 0.025 |
| NC_004718.3;NP_828849.2;SARS-CoV_orf1ab;3961 | 0 | 0.025 |
| NC_004718.3;NP_828849.2;SARS-CoV_orf1ab;3962 | 0 | 0 |
| NC_004718.3;NP_828849.2;SARS-CoV_orf1ab;3963 | 0 | 0 |
| NC_004718.3;NP_828849.2;SARS-CoV_orf1ab;3964 | 0 | 0 |
| NC_004718.3;NP_828849.2;SARS-CoV_orf1ab;3965 | 0 | 0 |
| NC_004718.3;NP_828849.2;SARS-CoV_orf1ab;3966 | 0 | 0 |
| NC_004718.3;NP_828849.2;SARS-CoV_orf1ab;3967 | 0 | 0 |
| NC_004718.3;NP_828849.2;SARS-CoV_orf1ab;3968 | 0 | 0 |
| NC_004718.3;NP_828849.2;SARS-CoV_orf1ab;3969 | 0 | 0 |
| NC_004718.3;NP_828849.2;SARS-CoV_orf1ab;3970 | 0 | 0 |
| NC_004718.3;NP_828849.2;SARS-CoV_orf1ab;3971 | 0 | 0.025 |
| NC_004718.3;NP_828849.2;SARS-CoV_orf1ab;3972 | 0 | 0 |
| NC_004718.3;NP_828849.2;SARS-CoV_orf1ab;3973 | 0 | 0 |
| NC_004718.3;NP_828849.2;SARS-CoV_orf1ab;3974 | 0 | 0 |
| NC_004718.3;NP_828849.2;SARS-CoV_orf1ab;3975 | 0 | 0.05 |
| NC_004718.3;NP_828849.2;SARS-CoV_orf1ab;3976 | 0 | 0.025 |
| NC_004718.3;NP_828849.2;SARS-CoV_orf1ab;3977 | 0 | 0 |
| NC_004718.3;NP_828849.2;SARS-CoV_orf1ab;3978 | 0 | 0 |
| NC_004718.3;NP_828849.2;SARS-CoV_orf1ab;3979 | 0 | 0 |
| NC_004718.3;NP_828849.2;SARS-CoV_orf1ab;3980 | 0 | 0 |
| NC_004718.3;NP_828849.2;SARS-CoV_orf1ab;3981 | 0 | 0 |
| NC_004718.3;NP_828849.2;SARS-CoV_orf1ab;3982 | 0 | 0 |
| NC_004718.3;NP_828849.2;SARS-CoV_orf1ab;3983 | 0 | 0.1 |
| NC_004718.3;NP_828849.2;SARS-CoV_orf1ab;3984 | 0 | 0.075 |
| NC_004718.3;NP_828849.2;SARS-CoV_orf1ab;3985 | 0 | 0.05 |
| NC_004718.3;NP_828849.2;SARS-CoV_orf1ab;3986 | 0 | 0.05 |
| NC_004718.3;NP_828849.2;SARS-CoV_orf1ab;3987 | 0 | 0.05 |
| NC_004718.3;NP_828849.2;SARS-CoV_orf1ab;3988 | 0 | 0 |
| NC_004718.3;NP_828849.2;SARS-CoV_orf1ab;3989 | 0 | 0 |
| NC_004718.3;NP_828849.2;SARS-CoV_orf1ab;3990 | 0 | 0 |
| NC_004718.3;NP_828849.2;SARS-CoV_orf1ab;3991 | 0 | 0 |
| NC_004718.3;NP_828849.2;SARS-CoV_orf1ab;3992 | 0 | 0 |
| NC_004718.3;NP_828849.2;SARS-CoV_orf1ab;3993 | 0 | 0 |
| NC_004718.3;NP_828849.2;SARS-CoV_orf1ab;3994 | 0 | 0 |
| NC_004718.3;NP_828849.2;SARS-CoV_orf1ab;3995 | 0 | 0 |
| NC_004718.3;NP_828849.2;SARS-CoV_orf1ab;3996 | 0 | 0 |

| | | |
|---|---|---|
| NC_004718.3;NP_828849.2;SARS-CoV_orf1ab;3997 | 0 | 0 |
| NC_004718.3;NP_828849.2;SARS-CoV_orf1ab;3998 | 0 | 0 |
| NC_004718.3;NP_828849.2;SARS-CoV_orf1ab;3999 | 0 | 0 |
| NC_004718.3;NP_828849.2;SARS-CoV_orf1ab;4000 | 0 | 0 |
| NC_004718.3;NP_828849.2;SARS-CoV_orf1ab;4001 | 0 | 0 |
| NC_004718.3;NP_828849.2;SARS-CoV_orf1ab;4002 | 0 | 0 |
| NC_004718.3;NP_828849.2;SARS-CoV_orf1ab;4003 | 0 | 0 |
| NC_004718.3;NP_828849.2;SARS-CoV_orf1ab;4004 | 0 | 0 |
| NC_004718.3;NP_828849.2;SARS-CoV_orf1ab;4005 | 0 | 0 |
| NC_004718.3;NP_828849.2;SARS-CoV_orf1ab;4006 | 0 | 0 |
| NC_004718.3;NP_828849.2;SARS-CoV_orf1ab;4007 | 0 | 0 |
| NC_004718.3;NP_828849.2;SARS-CoV_orf1ab;4008 | 0.15 | 0.1 |
| NC_004718.3;NP_828849.2;SARS-CoV_orf1ab;4009 | 0.2 | 0.1 |
| NC_004718.3;NP_828849.2;SARS-CoV_orf1ab;4010 | 0.2 | 0.075 |
| NC_004718.3;NP_828849.2;SARS-CoV_orf1ab;4011 | 0.25 | 0.1 |
| NC_004718.3;NP_828849.2;SARS-CoV_orf1ab;4012 | 0.25 | 0.275 |
| NC_004718.3;NP_828849.2;SARS-CoV_orf1ab;4013 | 0.3 | 0.3 |
| NC_004718.3;NP_828849.2;SARS-CoV_orf1ab;4014 | 0.35 | 0.4 |
| NC_004718.3;NP_828849.2;SARS-CoV_orf1ab;4015 | 0.25 | 0.25 |
| NC_004718.3;NP_828849.2;SARS-CoV_orf1ab;4016 | 0.4 | 0.5 |
| NC_004718.3;NP_828849.2;SARS-CoV_orf1ab;4017 | 0.1 | 0.025 |
| NC_004718.3;NP_828849.2;SARS-CoV_orf1ab;4018 | 0.05 | 0.025 |
| NC_004718.3;NP_828849.2;SARS-CoV_orf1ab;4019 | 0 | 0 |
| NC_004718.3;NP_828849.2;SARS-CoV_orf1ab;4020 | 0.05 | 0.1 |
| NC_004718.3;NP_828849.2;SARS-CoV_orf1ab;4021 | 0.1 | 0.075 |
| NC_004718.3;NP_828849.2;SARS-CoV_orf1ab;4022 | 0 | 0.075 |
| NC_004718.3;NP_828849.2;SARS-CoV_orf1ab;4023 | 0.05 | 0.075 |
| NC_004718.3;NP_828849.2;SARS-CoV_orf1ab;4024 | 0.05 | 0 |
| NC_004718.3;NP_828849.2;SARS-CoV_orf1ab;4025 | 0 | 0.05 |
| NC_004718.3;NP_828849.2;SARS-CoV_orf1ab;4026 | 0 | 0 |
| NC_004718.3;NP_828849.2;SARS-CoV_orf1ab;4027 | 0 | 0 |
| NC_004718.3;NP_828849.2;SARS-CoV_orf1ab;4028 | 0 | 0 |
| NC_004718.3;NP_828849.2;SARS-CoV_orf1ab;4029 | 0 | 0 |
| NC_004718.3;NP_828849.2;SARS-CoV_orf1ab;4030 | 0.05 | 0.025 |
| NC_004718.3;NP_828849.2;SARS-CoV_orf1ab;4031 | 0.05 | 0.025 |
| NC_004718.3;NP_828849.2;SARS-CoV_orf1ab;4032 | 0 | 0 |
| NC_004718.3;NP_828849.2;SARS-CoV_orf1ab;4033 | 0 | 0 |
| NC_004718.3;NP_828849.2;SARS-CoV_orf1ab;4034 | 0 | 0 |
| NC_004718.3;NP_828849.2;SARS-CoV_orf1ab;4035 | 0 | 0 |
| NC_004718.3;NP_828849.2;SARS-CoV_orf1ab;4036 | 0 | 0 |
| NC_004718.3;NP_828849.2;SARS-CoV_orf1ab;4037 | 0 | 0 |
| NC_004718.3;NP_828849.2;SARS-CoV_orf1ab;4038 | 0 | 0 |
| NC_004718.3;NP_828849.2;SARS-CoV_orf1ab;4039 | 0 | 0.025 |

| | | |
|---|---|---|
| NC_004718.3;NP_828849.2;SARS-CoV_orf1ab;4040 | 0.05 | 0 |
| NC_004718.3;NP_828849.2;SARS-CoV_orf1ab;4041 | 0.05 | 0.05 |
| NC_004718.3;NP_828849.2;SARS-CoV_orf1ab;4042 | 0.05 | 0 |
| NC_004718.3;NP_828849.2;SARS-CoV_orf1ab;4043 | 0.05 | 0 |
| NC_004718.3;NP_828849.2;SARS-CoV_orf1ab;4044 | 0.05 | 0.025 |
| NC_004718.3;NP_828849.2;SARS-CoV_orf1ab;4045 | 0.05 | 0.025 |
| NC_004718.3;NP_828849.2;SARS-CoV_orf1ab;4046 | 0.05 | 0.025 |
| NC_004718.3;NP_828849.2;SARS-CoV_orf1ab;4047 | 0.05 | 0.025 |
| NC_004718.3;NP_828849.2;SARS-CoV_orf1ab;4048 | 0.05 | 0.025 |
| NC_004718.3;NP_828849.2;SARS-CoV_orf1ab;4049 | 0.1 | 0.025 |
| NC_004718.3;NP_828849.2;SARS-CoV_orf1ab;4050 | 0.05 | 0 |
| NC_004718.3;NP_828849.2;SARS-CoV_orf1ab;4051 | 0 | 0 |
| NC_004718.3;NP_828849.2;SARS-CoV_orf1ab;4052 | 0.05 | 0 |
| NC_004718.3;NP_828849.2;SARS-CoV_orf1ab;4053 | 0 | 0 |
| NC_004718.3;NP_828849.2;SARS-CoV_orf1ab;4054 | 0 | 0 |
| NC_004718.3;NP_828849.2;SARS-CoV_orf1ab;4055 | 0 | 0 |
| NC_004718.3;NP_828849.2;SARS-CoV_orf1ab;4056 | 0 | 0 |
| NC_004718.3;NP_828849.2;SARS-CoV_orf1ab;4057 | 0 | 0 |
| NC_004718.3;NP_828849.2;SARS-CoV_orf1ab;4058 | 0 | 0 |
| NC_004718.3;NP_828849.2;SARS-CoV_orf1ab;4059 | 0 | 0 |
| NC_004718.3;NP_828849.2;SARS-CoV_orf1ab;4060 | 0 | 0 |
| NC_004718.3;NP_828849.2;SARS-CoV_orf1ab;4061 | 0 | 0 |
| NC_004718.3;NP_828849.2;SARS-CoV_orf1ab;4062 | 0 | 0 |
| NC_004718.3;NP_828849.2;SARS-CoV_orf1ab;4063 | 0 | 0 |
| NC_004718.3;NP_828849.2;SARS-CoV_orf1ab;4064 | 0 | 0 |
| NC_004718.3;NP_828849.2;SARS-CoV_orf1ab;4065 | 0 | 0 |
| NC_004718.3;NP_828849.2;SARS-CoV_orf1ab;4066 | 0 | 0 |
| NC_004718.3;NP_828849.2;SARS-CoV_orf1ab;4067 | 0 | 0 |
| NC_004718.3;NP_828849.2;SARS-CoV_orf1ab;4068 | 0 | 0 |
| NC_004718.3;NP_828849.2;SARS-CoV_orf1ab;4069 | 0.05 | 0.025 |
| NC_004718.3;NP_828849.2;SARS-CoV_orf1ab;4070 | 0.05 | 0.05 |
| NC_004718.3;NP_828849.2;SARS-CoV_orf1ab;4071 | 0 | 0 |
| NC_004718.3;NP_828849.2;SARS-CoV_orf1ab;4072 | 0 | 0.025 |
| NC_004718.3;NP_828849.2;SARS-CoV_orf1ab;4073 | 0 | 0 |
| NC_004718.3;NP_828849.2;SARS-CoV_orf1ab;4074 | 0 | 0 |
| NC_004718.3;NP_828849.2;SARS-CoV_orf1ab;4075 | 0 | 0 |
| NC_004718.3;NP_828849.2;SARS-CoV_orf1ab;4076 | 0 | 0.025 |
| NC_004718.3;NP_828849.2;SARS-CoV_orf1ab;4077 | 0 | 0 |
| NC_004718.3;NP_828849.2;SARS-CoV_orf1ab;4078 | 0 | 0 |
| NC_004718.3;NP_828849.2;SARS-CoV_orf1ab;4079 | 0 | 0 |
| NC_004718.3;NP_828849.2;SARS-CoV_orf1ab;4080 | 0.15 | 0 |
| NC_004718.3;NP_828849.2;SARS-CoV_orf1ab;4081 | 0 | 0 |
| NC_004718.3;NP_828849.2;SARS-CoV_orf1ab;4082 | 0 | 0 |

| | | |
|---|---|---|
| NC_004718.3;NP_828849.2;SARS-CoV_orf1ab;4083 | 0 | 0.025 |
| NC_004718.3;NP_828849.2;SARS-CoV_orf1ab;4084 | 0 | 0 |
| NC_004718.3;NP_828849.2;SARS-CoV_orf1ab;4085 | 0 | 0.025 |
| NC_004718.3;NP_828849.2;SARS-CoV_orf1ab;4086 | 0 | 0 |
| NC_004718.3;NP_828849.2;SARS-CoV_orf1ab;4087 | 0 | 0 |
| NC_004718.3;NP_828849.2;SARS-CoV_orf1ab;4088 | 0 | 0 |
| NC_004718.3;NP_828849.2;SARS-CoV_orf1ab;4089 | 0 | 0 |
| NC_004718.3;NP_828849.2;SARS-CoV_orf1ab;4090 | 0 | 0 |
| NC_004718.3;NP_828849.2;SARS-CoV_orf1ab;4091 | 0 | 0 |
| NC_004718.3;NP_828849.2;SARS-CoV_orf1ab;4092 | 0 | 0 |
| NC_004718.3;NP_828849.2;SARS-CoV_orf1ab;4093 | 0 | 0 |
| NC_004718.3;NP_828849.2;SARS-CoV_orf1ab;4094 | 0 | 0 |
| NC_004718.3;NP_828849.2;SARS-CoV_orf1ab;4095 | 0 | 0 |
| NC_004718.3;NP_828849.2;SARS-CoV_orf1ab;4096 | 0 | 0 |
| NC_004718.3;NP_828849.2;SARS-CoV_orf1ab;4097 | 0 | 0 |
| NC_004718.3;NP_828849.2;SARS-CoV_orf1ab;4098 | 0 | 0 |
| NC_004718.3;NP_828849.2;SARS-CoV_orf1ab;4099 | 0 | 0 |
| NC_004718.3;NP_828849.2;SARS-CoV_orf1ab;4100 | 0 | 0 |
| NC_004718.3;NP_828849.2;SARS-CoV_orf1ab;4101 | 0 | 0 |
| NC_004718.3;NP_828849.2;SARS-CoV_orf1ab;4102 | 0 | 0 |
| NC_004718.3;NP_828849.2;SARS-CoV_orf1ab;4103 | 0 | 0 |
| NC_004718.3;NP_828849.2;SARS-CoV_orf1ab;4104 | 0 | 0.025 |
| NC_004718.3;NP_828849.2;SARS-CoV_orf1ab;4105 | 0 | 0.075 |
| NC_004718.3;NP_828849.2;SARS-CoV_orf1ab;4106 | 0 | 0.05 |
| NC_004718.3;NP_828849.2;SARS-CoV_orf1ab;4107 | 0 | 0.025 |
| NC_004718.3;NP_828849.2;SARS-CoV_orf1ab;4108 | 0 | 0.025 |
| NC_004718.3;NP_828849.2;SARS-CoV_orf1ab;4109 | 0.05 | 0.025 |
| NC_004718.3;NP_828849.2;SARS-CoV_orf1ab;4110 | 0.1 | 0.025 |
| NC_004718.3;NP_828849.2;SARS-CoV_orf1ab;4111 | 0.05 | 0.025 |
| NC_004718.3;NP_828849.2;SARS-CoV_orf1ab;4112 | 0 | 0.025 |
| NC_004718.3;NP_828849.2;SARS-CoV_orf1ab;4113 | 0 | 0 |
| NC_004718.3;NP_828849.2;SARS-CoV_orf1ab;4114 | 0.05 | 0.025 |
| NC_004718.3;NP_828849.2;SARS-CoV_orf1ab;4115 | 0 | 0 |
| NC_004718.3;NP_828849.2;SARS-CoV_orf1ab;4116 | 0 | 0 |
| NC_004718.3;NP_828849.2;SARS-CoV_orf1ab;4117 | 0 | 0.025 |
| NC_004718.3;NP_828849.2;SARS-CoV_orf1ab;4118 | 0.05 | 0.05 |
| NC_004718.3;NP_828849.2;SARS-CoV_orf1ab;4119 | 0 | 0 |
| NC_004718.3;NP_828849.2;SARS-CoV_orf1ab;4120 | 0 | 0 |
| NC_004718.3;NP_828849.2;SARS-CoV_orf1ab;4121 | 0 | 0 |
| NC_004718.3;NP_828849.2;SARS-CoV_orf1ab;4122 | 0 | 0 |
| NC_004718.3;NP_828849.2;SARS-CoV_orf1ab;4123 | 0 | 0 |
| NC_004718.3;NP_828849.2;SARS-CoV_orf1ab;4124 | 0 | 0 |
| NC_004718.3;NP_828849.2;SARS-CoV_orf1ab;4125 | 0 | 0 |

| | | |
|---|---|---|
| NC_004718.3;NP_828849.2;SARS-CoV_orf1ab;4126 | 0 | 0 |
| NC_004718.3;NP_828849.2;SARS-CoV_orf1ab;4127 | 0 | 0 |
| NC_004718.3;NP_828849.2;SARS-CoV_orf1ab;4128 | 0 | 0 |
| NC_004718.3;NP_828849.2;SARS-CoV_orf1ab;4129 | 0 | 0 |
| NC_004718.3;NP_828849.2;SARS-CoV_orf1ab;4130 | 0 | 0 |
| NC_004718.3;NP_828849.2;SARS-CoV_orf1ab;4131 | 0 | 0 |
| NC_004718.3;NP_828849.2;SARS-CoV_orf1ab;4132 | 0 | 0 |
| NC_004718.3;NP_828849.2;SARS-CoV_orf1ab;4133 | 0 | 0 |
| NC_004718.3;NP_828849.2;SARS-CoV_orf1ab;4134 | 0 | 0 |
| NC_004718.3;NP_828849.2;SARS-CoV_orf1ab;4135 | 0 | 0 |
| NC_004718.3;NP_828849.2;SARS-CoV_orf1ab;4136 | 0 | 0 |
| NC_004718.3;NP_828849.2;SARS-CoV_orf1ab;4137 | 0 | 0 |
| NC_004718.3;NP_828849.2;SARS-CoV_orf1ab;4138 | 0.15 | 0.125 |
| NC_004718.3;NP_828849.2;SARS-CoV_orf1ab;4139 | 0.15 | 0.05 |
| NC_004718.3;NP_828849.2;SARS-CoV_orf1ab;4140 | 0.25 | 0.125 |
| NC_004718.3;NP_828849.2;SARS-CoV_orf1ab;4141 | 0.05 | 0.075 |
| NC_004718.3;NP_828849.2;SARS-CoV_orf1ab;4142 | 0 | 0 |
| NC_004718.3;NP_828849.2;SARS-CoV_orf1ab;4143 | 0 | 0 |
| NC_004718.3;NP_828849.2;SARS-CoV_orf1ab;4144 | 0 | 0 |
| NC_004718.3;NP_828849.2;SARS-CoV_orf1ab;4145 | 0 | 0 |
| NC_004718.3;NP_828849.2;SARS-CoV_orf1ab;4146 | 0 | 0 |
| NC_004718.3;NP_828849.2;SARS-CoV_orf1ab;4147 | 0 | 0 |
| NC_004718.3;NP_828849.2;SARS-CoV_orf1ab;4148 | 0 | 0 |
| NC_004718.3;NP_828849.2;SARS-CoV_orf1ab;4149 | 0 | 0 |
| NC_004718.3;NP_828849.2;SARS-CoV_orf1ab;4150 | 0 | 0 |
| NC_004718.3;NP_828849.2;SARS-CoV_orf1ab;4151 | 0 | 0 |
| NC_004718.3;NP_828849.2;SARS-CoV_orf1ab;4152 | 0 | 0 |
| NC_004718.3;NP_828849.2;SARS-CoV_orf1ab;4153 | 0 | 0 |
| NC_004718.3;NP_828849.2;SARS-CoV_orf1ab;4154 | 0 | 0 |
| NC_004718.3;NP_828849.2;SARS-CoV_orf1ab;4155 | 0 | 0 |
| NC_004718.3;NP_828849.2;SARS-CoV_orf1ab;4156 | 0 | 0 |
| NC_004718.3;NP_828849.2;SARS-CoV_orf1ab;4157 | 0 | 0 |
| NC_004718.3;NP_828849.2;SARS-CoV_orf1ab;4158 | 0 | 0 |
| NC_004718.3;NP_828849.2;SARS-CoV_orf1ab;4159 | 0 | 0 |
| NC_004718.3;NP_828849.2;SARS-CoV_orf1ab;4160 | 0 | 0 |
| NC_004718.3;NP_828849.2;SARS-CoV_orf1ab;4161 | 0 | 0 |
| NC_004718.3;NP_828849.2;SARS-CoV_orf1ab;4162 | 0 | 0 |
| NC_004718.3;NP_828849.2;SARS-CoV_orf1ab;4163 | 0 | 0 |
| NC_004718.3;NP_828849.2;SARS-CoV_orf1ab;4164 | 0 | 0.025 |
| NC_004718.3;NP_828849.2;SARS-CoV_orf1ab;4165 | 0 | 0.025 |
| NC_004718.3;NP_828849.2;SARS-CoV_orf1ab;4166 | 0 | 0 |
| NC_004718.3;NP_828849.2;SARS-CoV_orf1ab;4167 | 0 | 0 |
| NC_004718.3;NP_828849.2;SARS-CoV_orf1ab;4168 | 0 | 0 |

| | | |
|---|---|---|
| NC_004718.3;NP_828849.2;SARS-CoV_orf1ab;4169 | 0 | 0 |
| NC_004718.3;NP_828849.2;SARS-CoV_orf1ab;4170 | 0 | 0 |
| NC_004718.3;NP_828849.2;SARS-CoV_orf1ab;4171 | 0 | 0 |
| NC_004718.3;NP_828849.2;SARS-CoV_orf1ab;4172 | 0 | 0 |
| NC_004718.3;NP_828849.2;SARS-CoV_orf1ab;4173 | 0 | 0 |
| NC_004718.3;NP_828849.2;SARS-CoV_orf1ab;4174 | 0 | 0 |
| NC_004718.3;NP_828849.2;SARS-CoV_orf1ab;4175 | 0 | 0 |
| NC_004718.3;NP_828849.2;SARS-CoV_orf1ab;4176 | 0 | 0 |
| NC_004718.3;NP_828849.2;SARS-CoV_orf1ab;4177 | 0 | 0 |
| NC_004718.3;NP_828849.2;SARS-CoV_orf1ab;4178 | 0 | 0.025 |
| NC_004718.3;NP_828849.2;SARS-CoV_orf1ab;4179 | 0 | 0 |
| NC_004718.3;NP_828849.2;SARS-CoV_orf1ab;4180 | 0.2 | 0.15 |
| NC_004718.3;NP_828849.2;SARS-CoV_orf1ab;4181 | 0 | 0 |
| NC_004718.3;NP_828849.2;SARS-CoV_orf1ab;4182 | 0.05 | 0.15 |
| NC_004718.3;NP_828849.2;SARS-CoV_orf1ab;4183 | 0.05 | 0.075 |
| NC_004718.3;NP_828849.2;SARS-CoV_orf1ab;4184 | 0.05 | 0.05 |
| NC_004718.3;NP_828849.2;SARS-CoV_orf1ab;4185 | 0.05 | 0.025 |
| NC_004718.3;NP_828849.2;SARS-CoV_orf1ab;4186 | 0 | 0 |
| NC_004718.3;NP_828849.2;SARS-CoV_orf1ab;4187 | 0 | 0.025 |
| NC_004718.3;NP_828849.2;SARS-CoV_orf1ab;4188 | 0 | 0 |
| NC_004718.3;NP_828849.2;SARS-CoV_orf1ab;4189 | 0 | 0 |
| NC_004718.3;NP_828849.2;SARS-CoV_orf1ab;4190 | 0 | 0 |
| NC_004718.3;NP_828849.2;SARS-CoV_orf1ab;4191 | 0 | 0 |
| NC_004718.3;NP_828849.2;SARS-CoV_orf1ab;4192 | 0.05 | 0 |
| NC_004718.3;NP_828849.2;SARS-CoV_orf1ab;4193 | 0 | 0 |
| NC_004718.3;NP_828849.2;SARS-CoV_orf1ab;4194 | 0 | 0 |
| NC_004718.3;NP_828849.2;SARS-CoV_orf1ab;4195 | 0 | 0 |
| NC_004718.3;NP_828849.2;SARS-CoV_orf1ab;4196 | 0 | 0 |
| NC_004718.3;NP_828849.2;SARS-CoV_orf1ab;4197 | 0 | 0 |
| NC_004718.3;NP_828849.2;SARS-CoV_orf1ab;4198 | 0 | 0 |
| NC_004718.3;NP_828849.2;SARS-CoV_orf1ab;4199 | 0 | 0 |
| NC_004718.3;NP_828849.2;SARS-CoV_orf1ab;4200 | 0 | 0 |
| NC_004718.3;NP_828849.2;SARS-CoV_orf1ab;4201 | 0 | 0 |
| NC_004718.3;NP_828849.2;SARS-CoV_orf1ab;4202 | 0 | 0 |
| NC_004718.3;NP_828849.2;SARS-CoV_orf1ab;4203 | 0 | 0 |
| NC_004718.3;NP_828849.2;SARS-CoV_orf1ab;4204 | 0 | 0 |
| NC_004718.3;NP_828849.2;SARS-CoV_orf1ab;4205 | 0 | 0 |
| NC_004718.3;NP_828849.2;SARS-CoV_orf1ab;4206 | 0.05 | 0.025 |
| NC_004718.3;NP_828849.2;SARS-CoV_orf1ab;4207 | 0.1 | 0.075 |
| NC_004718.3;NP_828849.2;SARS-CoV_orf1ab;4208 | 0.15 | 0.1 |
| NC_004718.3;NP_828849.2;SARS-CoV_orf1ab;4209 | 0.25 | 0.1 |
| NC_004718.3;NP_828849.2;SARS-CoV_orf1ab;4210 | 0.1 | 0.1 |
| NC_004718.3;NP_828849.2;SARS-CoV_orf1ab;4211 | 0.05 | 0.05 |

| | | |
|---|---|---|
| NC_004718.3;NP_828849.2;SARS-CoV_orf1ab;4212 | 0.05 | 0.025 |
| NC_004718.3;NP_828849.2;SARS-CoV_orf1ab;4213 | 0 | 0 |
| NC_004718.3;NP_828849.2;SARS-CoV_orf1ab;4214 | 0 | 0 |
| NC_004718.3;NP_828849.2;SARS-CoV_orf1ab;4215 | 0 | 0 |
| NC_004718.3;NP_828849.2;SARS-CoV_orf1ab;4216 | 0 | 0 |
| NC_004718.3;NP_828849.2;SARS-CoV_orf1ab;4217 | 0 | 0 |
| NC_004718.3;NP_828849.2;SARS-CoV_orf1ab;4218 | 0 | 0 |
| NC_004718.3;NP_828849.2;SARS-CoV_orf1ab;4219 | 0 | 0 |
| NC_004718.3;NP_828849.2;SARS-CoV_orf1ab;4220 | 0 | 0 |
| NC_004718.3;NP_828849.2;SARS-CoV_orf1ab;4221 | 0 | 0.025 |
| NC_004718.3;NP_828849.2;SARS-CoV_orf1ab;4222 | 0 | 0 |
| NC_004718.3;NP_828849.2;SARS-CoV_orf1ab;4223 | 0 | 0.025 |
| NC_004718.3;NP_828849.2;SARS-CoV_orf1ab;4224 | 0.05 | 0.025 |
| NC_004718.3;NP_828849.2;SARS-CoV_orf1ab;4225 | 0 | 0.025 |
| NC_004718.3;NP_828849.2;SARS-CoV_orf1ab;4226 | 0 | 0 |
| NC_004718.3;NP_828849.2;SARS-CoV_orf1ab;4227 | 0 | 0 |
| NC_004718.3;NP_828849.2;SARS-CoV_orf1ab;4228 | 0 | 0.025 |
| NC_004718.3;NP_828849.2;SARS-CoV_orf1ab;4229 | 0 | 0.025 |
| NC_004718.3;NP_828849.2;SARS-CoV_orf1ab;4230 | 0 | 0 |
| NC_004718.3;NP_828849.2;SARS-CoV_orf1ab;4231 | 0 | 0 |
| NC_004718.3;NP_828849.2;SARS-CoV_orf1ab;4232 | 0 | 0 |
| NC_004718.3;NP_828849.2;SARS-CoV_orf1ab;4233 | 0 | 0 |
| NC_004718.3;NP_828849.2;SARS-CoV_orf1ab;4234 | 0 | 0 |
| NC_004718.3;NP_828849.2;SARS-CoV_orf1ab;4235 | 0 | 0 |
| NC_004718.3;NP_828849.2;SARS-CoV_orf1ab;4236 | 0 | 0 |
| NC_004718.3;NP_828849.2;SARS-CoV_orf1ab;4237 | 0 | 0 |
| NC_004718.3;NP_828849.2;SARS-CoV_orf1ab;4238 | 0 | 0 |
| NC_004718.3;NP_828849.2;SARS-CoV_orf1ab;4239 | 0 | 0 |
| NC_004718.3;NP_828849.2;SARS-CoV_orf1ab;4240 | 0 | 0 |
| NC_004718.3;NP_828849.2;SARS-CoV_orf1ab;4241 | 0 | 0 |
| NC_004718.3;NP_828849.2;SARS-CoV_orf1ab;4242 | 0 | 0 |
| NC_004718.3;NP_828849.2;SARS-CoV_orf1ab;4243 | 0 | 0 |
| NC_004718.3;NP_828849.2;SARS-CoV_orf1ab;4244 | 0.05 | 0 |
| NC_004718.3;NP_828849.2;SARS-CoV_orf1ab;4245 | 0.05 | 0 |
| NC_004718.3;NP_828849.2;SARS-CoV_orf1ab;4246 | 0.05 | 0 |
| NC_004718.3;NP_828849.2;SARS-CoV_orf1ab;4247 | 0.05 | 0 |
| NC_004718.3;NP_828849.2;SARS-CoV_orf1ab;4248 | 0.1 | 0 |
| NC_004718.3;NP_828849.2;SARS-CoV_orf1ab;4249 | 0.05 | 0.025 |
| NC_004718.3;NP_828849.2;SARS-CoV_orf1ab;4250 | 0.15 | 0.05 |
| NC_004718.3;NP_828849.2;SARS-CoV_orf1ab;4251 | 0.15 | 0.025 |
| NC_004718.3;NP_828849.2;SARS-CoV_orf1ab;4252 | 0 | 0.025 |
| NC_004718.3;NP_828849.2;SARS-CoV_orf1ab;4253 | 0 | 0.025 |
| NC_004718.3;NP_828849.2;SARS-CoV_orf1ab;4254 | 0 | 0.05 |

| | | |
|---|---|---|
| NC_004718.3;NP_828849.2;SARS-CoV_orf1ab;4255 | 0 | 0.05 |
| NC_004718.3;NP_828849.2;SARS-CoV_orf1ab;4256 | 0.15 | 0.05 |
| NC_004718.3;NP_828849.2;SARS-CoV_orf1ab;4257 | 0.05 | 0.05 |
| NC_004718.3;NP_828849.2;SARS-CoV_orf1ab;4258 | 0 | 0 |
| NC_004718.3;NP_828849.2;SARS-CoV_orf1ab;4259 | 0 | 0 |
| NC_004718.3;NP_828849.2;SARS-CoV_orf1ab;4260 | 0 | 0 |
| NC_004718.3;NP_828849.2;SARS-CoV_orf1ab;4261 | 0 | 0 |
| NC_004718.3;NP_828849.2;SARS-CoV_orf1ab;4262 | 0 | 0 |
| NC_004718.3;NP_828849.2;SARS-CoV_orf1ab;4263 | 0 | 0 |
| NC_004718.3;NP_828849.2;SARS-CoV_orf1ab;4264 | 0 | 0 |
| NC_004718.3;NP_828849.2;SARS-CoV_orf1ab;4265 | 0 | 0 |
| NC_004718.3;NP_828849.2;SARS-CoV_orf1ab;4266 | 0 | 0.025 |
| NC_004718.3;NP_828849.2;SARS-CoV_orf1ab;4267 | 0 | 0 |
| NC_004718.3;NP_828849.2;SARS-CoV_orf1ab;4268 | 0 | 0.025 |
| NC_004718.3;NP_828849.2;SARS-CoV_orf1ab;4269 | 0 | 0.025 |
| NC_004718.3;NP_828849.2;SARS-CoV_orf1ab;4270 | 0 | 0 |
| NC_004718.3;NP_828849.2;SARS-CoV_orf1ab;4271 | 0 | 0 |
| NC_004718.3;NP_828849.2;SARS-CoV_orf1ab;4272 | 0 | 0 |
| NC_004718.3;NP_828849.2;SARS-CoV_orf1ab;4273 | 0 | 0 |
| NC_004718.3;NP_828849.2;SARS-CoV_orf1ab;4274 | 0 | 0 |
| NC_004718.3;NP_828849.2;SARS-CoV_orf1ab;4275 | 0 | 0 |
| NC_004718.3;NP_828849.2;SARS-CoV_orf1ab;4276 | 0 | 0 |
| NC_004718.3;NP_828849.2;SARS-CoV_orf1ab;4277 | 0 | 0 |
| NC_004718.3;NP_828849.2;SARS-CoV_orf1ab;4278 | 0 | 0 |
| NC_004718.3;NP_828849.2;SARS-CoV_orf1ab;4279 | 0 | 0.025 |
| NC_004718.3;NP_828849.2;SARS-CoV_orf1ab;4280 | 0 | 0.025 |
| NC_004718.3;NP_828849.2;SARS-CoV_orf1ab;4281 | 0 | 0.025 |
| NC_004718.3;NP_828849.2;SARS-CoV_orf1ab;4282 | 0 | 0.025 |
| NC_004718.3;NP_828849.2;SARS-CoV_orf1ab;4283 | 0 | 0.025 |
| NC_004718.3;NP_828849.2;SARS-CoV_orf1ab;4284 | 0 | 0.025 |
| NC_004718.3;NP_828849.2;SARS-CoV_orf1ab;4285 | 0.05 | 0.025 |
| NC_004718.3;NP_828849.2;SARS-CoV_orf1ab;4286 | 0 | 0.025 |
| NC_004718.3;NP_828849.2;SARS-CoV_orf1ab;4287 | 0.05 | 0.025 |
| NC_004718.3;NP_828849.2;SARS-CoV_orf1ab;4288 | 0 | 0 |
| NC_004718.3;NP_828849.2;SARS-CoV_orf1ab;4289 | 0 | 0 |
| NC_004718.3;NP_828849.2;SARS-CoV_orf1ab;4290 | 0 | 0 |
| NC_004718.3;NP_828849.2;SARS-CoV_orf1ab;4291 | 0 | 0 |
| NC_004718.3;NP_828849.2;SARS-CoV_orf1ab;4292 | 0 | 0 |
| NC_004718.3;NP_828849.2;SARS-CoV_orf1ab;4293 | 0 | 0 |
| NC_004718.3;NP_828849.2;SARS-CoV_orf1ab;4294 | 0 | 0 |
| NC_004718.3;NP_828849.2;SARS-CoV_orf1ab;4295 | 0 | 0 |
| NC_004718.3;NP_828849.2;SARS-CoV_orf1ab;4296 | 0 | 0 |
| NC_004718.3;NP_828849.2;SARS-CoV_orf1ab;4297 | 0 | 0 |

| | | |
|---|---|---|
| NC_004718.3;NP_828849.2;SARS-CoV_orf1ab;4298 | 0 | 0 |
| NC_004718.3;NP_828849.2;SARS-CoV_orf1ab;4299 | 0 | 0 |
| NC_004718.3;NP_828849.2;SARS-CoV_orf1ab;4300 | 0 | 0 |
| NC_004718.3;NP_828849.2;SARS-CoV_orf1ab;4301 | 0 | 0 |
| NC_004718.3;NP_828849.2;SARS-CoV_orf1ab;4302 | 0 | 0 |
| NC_004718.3;NP_828849.2;SARS-CoV_orf1ab;4303 | 0 | 0 |
| NC_004718.3;NP_828849.2;SARS-CoV_orf1ab;4304 | 0 | 0 |
| NC_004718.3;NP_828849.2;SARS-CoV_orf1ab;4305 | 0 | 0 |
| NC_004718.3;NP_828849.2;SARS-CoV_orf1ab;4306 | 0 | 0.025 |
| NC_004718.3;NP_828849.2;SARS-CoV_orf1ab;4307 | 0 | 0.025 |
| NC_004718.3;NP_828849.2;SARS-CoV_orf1ab;4308 | 0 | 0.025 |
| NC_004718.3;NP_828849.2;SARS-CoV_orf1ab;4309 | 0 | 0.025 |
| NC_004718.3;NP_828849.2;SARS-CoV_orf1ab;4310 | 0 | 0.025 |
| NC_004718.3;NP_828849.2;SARS-CoV_orf1ab;4311 | 0 | 0.025 |
| NC_004718.3;NP_828849.2;SARS-CoV_orf1ab;4312 | 0 | 0 |
| NC_004718.3;NP_828849.2;SARS-CoV_orf1ab;4313 | 0 | 0 |
| NC_004718.3;NP_828849.2;SARS-CoV_orf1ab;4314 | 0 | 0 |
| NC_004718.3;NP_828849.2;SARS-CoV_orf1ab;4315 | 0 | 0 |
| NC_004718.3;NP_828849.2;SARS-CoV_orf1ab;4316 | 0 | 0 |
| NC_004718.3;NP_828849.2;SARS-CoV_orf1ab;4317 | 0 | 0.05 |
| NC_004718.3;NP_828849.2;SARS-CoV_orf1ab;4318 | 0 | 0.05 |
| NC_004718.3;NP_828849.2;SARS-CoV_orf1ab;4319 | 0 | 0 |
| NC_004718.3;NP_828849.2;SARS-CoV_orf1ab;4320 | 0 | 0 |
| NC_004718.3;NP_828849.2;SARS-CoV_orf1ab;4321 | 0.05 | 0 |
| NC_004718.3;NP_828849.2;SARS-CoV_orf1ab;4322 | 0.05 | 0.025 |
| NC_004718.3;NP_828849.2;SARS-CoV_orf1ab;4323 | 0.05 | 0.05 |
| NC_004718.3;NP_828849.2;SARS-CoV_orf1ab;4324 | 0.05 | 0 |
| NC_004718.3;NP_828849.2;SARS-CoV_orf1ab;4325 | 0.05 | 0 |
| NC_004718.3;NP_828849.2;SARS-CoV_orf1ab;4326 | 0 | 0 |
| NC_004718.3;NP_828849.2;SARS-CoV_orf1ab;4327 | 0 | 0.025 |
| NC_004718.3;NP_828849.2;SARS-CoV_orf1ab;4328 | 0 | 0.025 |
| NC_004718.3;NP_828849.2;SARS-CoV_orf1ab;4329 | 0 | 0.025 |
| NC_004718.3;NP_828849.2;SARS-CoV_orf1ab;4330 | 0 | 0.05 |
| NC_004718.3;NP_828849.2;SARS-CoV_orf1ab;4331 | 0 | 0.025 |
| NC_004718.3;NP_828849.2;SARS-CoV_orf1ab;4332 | 0 | 0.025 |
| NC_004718.3;NP_828849.2;SARS-CoV_orf1ab;4333 | 0 | 0.025 |
| NC_004718.3;NP_828849.2;SARS-CoV_orf1ab;4334 | 0 | 0.025 |
| NC_004718.3;NP_828849.2;SARS-CoV_orf1ab;4335 | 0 | 0 |
| NC_004718.3;NP_828849.2;SARS-CoV_orf1ab;4336 | 0 | 0.025 |
| NC_004718.3;NP_828849.2;SARS-CoV_orf1ab;4337 | 0 | 0 |
| NC_004718.3;NP_828849.2;SARS-CoV_orf1ab;4338 | 0 | 0 |
| NC_004718.3;NP_828849.2;SARS-CoV_orf1ab;4339 | 0 | 0 |
| NC_004718.3;NP_828849.2;SARS-CoV_orf1ab;4340 | 0 | 0 |

| | | |
|---|---|---|
| NC_004718.3;NP_828849.2;SARS-CoV_orf1ab;4341 | 0 | 0 |
| NC_004718.3;NP_828849.2;SARS-CoV_orf1ab;4342 | 0 | 0 |
| NC_004718.3;NP_828849.2;SARS-CoV_orf1ab;4343 | 0 | 0 |
| NC_004718.3;NP_828849.2;SARS-CoV_orf1ab;4344 | 0 | 0 |
| NC_004718.3;NP_828849.2;SARS-CoV_orf1ab;4345 | 0 | 0 |
| NC_004718.3;NP_828849.2;SARS-CoV_orf1ab;4346 | 0 | 0 |
| NC_004718.3;NP_828849.2;SARS-CoV_orf1ab;4347 | 0 | 0 |
| NC_004718.3;NP_828849.2;SARS-CoV_orf1ab;4348 | 0 | 0 |
| NC_004718.3;NP_828849.2;SARS-CoV_orf1ab;4349 | 0 | 0 |
| NC_004718.3;NP_828849.2;SARS-CoV_orf1ab;4350 | 0 | 0 |
| NC_004718.3;NP_828849.2;SARS-CoV_orf1ab;4351 | 0 | 0 |
| NC_004718.3;NP_828849.2;SARS-CoV_orf1ab;4352 | 0 | 0.075 |
| NC_004718.3;NP_828849.2;SARS-CoV_orf1ab;4353 | 0 | 0 |
| NC_004718.3;NP_828849.2;SARS-CoV_orf1ab;4354 | 0 | 0.025 |
| NC_004718.3;NP_828849.2;SARS-CoV_orf1ab;4355 | 0 | 0 |
| NC_004718.3;NP_828849.2;SARS-CoV_orf1ab;4356 | 0 | 0 |
| NC_004718.3;NP_828849.2;SARS-CoV_orf1ab;4357 | 0 | 0 |
| NC_004718.3;NP_828849.2;SARS-CoV_orf1ab;4358 | 0 | 0 |
| NC_004718.3;NP_828849.2;SARS-CoV_orf1ab;4359 | 0 | 0 |
| NC_004718.3;NP_828849.2;SARS-CoV_orf1ab;4360 | 0 | 0 |
| NC_004718.3;NP_828849.2;SARS-CoV_orf1ab;4361 | 0 | 0 |
| NC_004718.3;NP_828849.2;SARS-CoV_orf1ab;4362 | 0 | 0 |
| NC_004718.3;NP_828849.2;SARS-CoV_orf1ab;4363 | 0.05 | 0 |
| NC_004718.3;NP_828849.2;SARS-CoV_orf1ab;4364 | 0 | 0 |
| NC_004718.3;NP_828849.2;SARS-CoV_orf1ab;4365 | 0.05 | 0 |
| NC_004718.3;NP_828849.2;SARS-CoV_orf1ab;4366 | 0 | 0 |
| NC_004718.3;NP_828849.2;SARS-CoV_orf1ab;4367 | 0 | 0 |
| NC_004718.3;NP_828849.2;SARS-CoV_orf1ab;4368 | 0 | 0 |
| NC_004718.3;NP_828849.2;SARS-CoV_orf1ab;4369 | 0 | 0 |
| NC_004718.3;NP_828849.2;SARS-CoV_orf1ab;4370 | 0 | 0.025 |
| NC_004718.3;NP_828849.2;SARS-CoV_orf1ab;4371 | 0.05 | 0 |
| NC_004718.3;NP_828849.2;SARS-CoV_orf1ab;4372 | 0 | 0 |
| NC_004718.3;NP_828849.2;SARS-CoV_orf1ab;4373 | 0 | 0 |
| NC_004718.3;NP_828849.2;SARS-CoV_orf1ab;4374 | 0 | 0 |
| NC_004718.3;NP_828849.2;SARS-CoV_orf1ab;4375 | 0 | 0 |
| NC_004718.3;NP_828849.2;SARS-CoV_orf1ab;4376 | 0 | 0 |
| NC_004718.3;NP_828849.2;SARS-CoV_orf1ab;4377 | 0 | 0 |
| NC_004718.3;NP_828849.2;SARS-CoV_orf1ab;4378 | 0 | 0 |
| NC_004718.3;NP_828849.2;SARS-CoV_orf1ab;4379 | 0 | 0 |
| NC_004718.3;NP_828849.2;SARS-CoV_orf1ab;4380 | 0 | 0 |
| NC_004718.3;NP_828849.2;SARS-CoV_orf1ab;4381 | 0 | 0 |
| NC_004718.3;NP_828849.2;SARS-CoV_orf1ab;4382 | 0 | 0 |
| NC_004718.3;NP_828849.2;SARS-CoV_orf1ab;4383 | 0 | 0 |

| | | |
|---|---|---|
| NC_004718.3;NP_828849.2;SARS-CoV_orf1ab;4384 | 0 | 0.025 |
| NC_004718.3;NP_828849.2;SARS-CoV_orf1ab;4385 | 0 | 0 |
| NC_004718.3;NP_828849.2;SARS-CoV_orf1ab;4386 | 0 | 0 |
| NC_004718.3;NP_828849.2;SARS-CoV_orf1ab;4387 | 0 | 0 |
| NC_004718.3;NP_828849.2;SARS-CoV_orf1ab;4388 | 0 | 0 |
| NC_004718.3;NP_828849.2;SARS-CoV_orf1ab;4389 | 0 | 0 |
| NC_004718.3;NP_828849.2;SARS-CoV_orf1ab;4390 | 0 | 0 |
| NC_004718.3;NP_828849.2;SARS-CoV_orf1ab;4391 | 0 | 0 |
| NC_004718.3;NP_828849.2;SARS-CoV_orf1ab;4392 | 0 | 0 |
| NC_004718.3;NP_828849.2;SARS-CoV_orf1ab;4393 | 0 | 0 |
| NC_004718.3;NP_828849.2;SARS-CoV_orf1ab;4394 | 0 | 0 |
| NC_004718.3;NP_828849.2;SARS-CoV_orf1ab;4395 | 0.1 | 0.025 |
| NC_004718.3;NP_828849.2;SARS-CoV_orf1ab;4396 | 0.1 | 0 |
| NC_004718.3;NP_828849.2;SARS-CoV_orf1ab;4397 | 0.05 | 0.025 |
| NC_004718.3;NP_828849.2;SARS-CoV_orf1ab;4398 | 0 | 0 |
| NC_004718.3;NP_828849.2;SARS-CoV_orf1ab;4399 | 0 | 0 |
| NC_004718.3;NP_828849.2;SARS-CoV_orf1ab;4400 | 0 | 0 |
| NC_004718.3;NP_828849.2;SARS-CoV_orf1ab;4401 | 0 | 0.025 |
| NC_004718.3;NP_828849.2;SARS-CoV_orf1ab;4402 | 0 | 0 |
| NC_004718.3;NP_828849.2;SARS-CoV_orf1ab;4403 | 0 | 0 |
| NC_004718.3;NP_828849.2;SARS-CoV_orf1ab;4404 | 0 | 0 |
| NC_004718.3;NP_828849.2;SARS-CoV_orf1ab;4405 | 0 | 0 |
| NC_004718.3;NP_828849.2;SARS-CoV_orf1ab;4406 | 0 | 0 |
| NC_004718.3;NP_828849.2;SARS-CoV_orf1ab;4407 | 0 | 0 |
| NC_004718.3;NP_828849.2;SARS-CoV_orf1ab;4408 | 0 | 0 |
| NC_004718.3;NP_828849.2;SARS-CoV_orf1ab;4409 | 0 | 0 |
| NC_004718.3;NP_828849.2;SARS-CoV_orf1ab;4410 | 0 | 0 |
| NC_004718.3;NP_828849.2;SARS-CoV_orf1ab;4411 | 0 | 0 |
| NC_004718.3;NP_828849.2;SARS-CoV_orf1ab;4412 | 0 | 0 |
| NC_004718.3;NP_828849.2;SARS-CoV_orf1ab;4413 | 0 | 0 |
| NC_004718.3;NP_828849.2;SARS-CoV_orf1ab;4414 | 0 | 0 |
| NC_004718.3;NP_828849.2;SARS-CoV_orf1ab;4415 | 0 | 0.025 |
| NC_004718.3;NP_828849.2;SARS-CoV_orf1ab;4416 | 0 | 0.025 |
| NC_004718.3;NP_828849.2;SARS-CoV_orf1ab;4417 | 0 | 0.025 |
| NC_004718.3;NP_828849.2;SARS-CoV_orf1ab;4418 | 0.05 | 0.025 |
| NC_004718.3;NP_828849.2;SARS-CoV_orf1ab;4419 | 0.05 | 0.025 |
| NC_004718.3;NP_828849.2;SARS-CoV_orf1ab;4420 | 0.1 | 0.05 |
| NC_004718.3;NP_828849.2;SARS-CoV_orf1ab;4421 | 0.05 | 0 |
| NC_004718.3;NP_828849.2;SARS-CoV_orf1ab;4422 | 0.1 | 0.025 |
| NC_004718.3;NP_828849.2;SARS-CoV_orf1ab;4423 | 0.1 | 0.05 |
| NC_004718.3;NP_828849.2;SARS-CoV_orf1ab;4424 | 0.1 | 0.05 |
| NC_004718.3;NP_828849.2;SARS-CoV_orf1ab;4425 | 0.05 | 0 |
| NC_004718.3;NP_828849.2;SARS-CoV_orf1ab;4426 | 0 | 0 |

| | | |
|---|---|---|
| NC_004718.3;NP_828849.2;SARS-CoV_orf1ab;4427 | 0.05 | 0 |
| NC_004718.3;NP_828849.2;SARS-CoV_orf1ab;4428 | 0 | 0 |
| NC_004718.3;NP_828849.2;SARS-CoV_orf1ab;4429 | 0 | 0 |
| NC_004718.3;NP_828849.2;SARS-CoV_orf1ab;4430 | 0 | 0 |
| NC_004718.3;NP_828849.2;SARS-CoV_orf1ab;4431 | 0 | 0 |
| NC_004718.3;NP_828849.2;SARS-CoV_orf1ab;4432 | 0 | 0 |
| NC_004718.3;NP_828849.2;SARS-CoV_orf1ab;4433 | 0 | 0 |
| NC_004718.3;NP_828849.2;SARS-CoV_orf1ab;4434 | 0 | 0 |
| NC_004718.3;NP_828849.2;SARS-CoV_orf1ab;4435 | 0 | 0 |
| NC_004718.3;NP_828849.2;SARS-CoV_orf1ab;4436 | 0 | 0 |
| NC_004718.3;NP_828849.2;SARS-CoV_orf1ab;4437 | 0 | 0 |
| NC_004718.3;NP_828849.2;SARS-CoV_orf1ab;4438 | 0 | 0 |
| NC_004718.3;NP_828849.2;SARS-CoV_orf1ab;4439 | 0 | 0 |
| NC_004718.3;NP_828849.2;SARS-CoV_orf1ab;4440 | 0 | 0 |
| NC_004718.3;NP_828849.2;SARS-CoV_orf1ab;4441 | 0 | 0 |
| NC_004718.3;NP_828849.2;SARS-CoV_orf1ab;4442 | 0 | 0 |
| NC_004718.3;NP_828849.2;SARS-CoV_orf1ab;4443 | 0 | 0 |
| NC_004718.3;NP_828849.2;SARS-CoV_orf1ab;4444 | 0 | 0 |
| NC_004718.3;NP_828849.2;SARS-CoV_orf1ab;4445 | 0 | 0 |
| NC_004718.3;NP_828849.2;SARS-CoV_orf1ab;4446 | 0 | 0.025 |
| NC_004718.3;NP_828849.2;SARS-CoV_orf1ab;4447 | 0.05 | 0 |
| NC_004718.3;NP_828849.2;SARS-CoV_orf1ab;4448 | 0 | 0 |
| NC_004718.3;NP_828849.2;SARS-CoV_orf1ab;4449 | 0 | 0 |
| NC_004718.3;NP_828849.2;SARS-CoV_orf1ab;4450 | 0 | 0.025 |
| NC_004718.3;NP_828849.2;SARS-CoV_orf1ab;4451 | 0 | 0.025 |
| NC_004718.3;NP_828849.2;SARS-CoV_orf1ab;4452 | 0 | 0.025 |
| NC_004718.3;NP_828849.2;SARS-CoV_orf1ab;4453 | 0 | 0 |
| NC_004718.3;NP_828849.2;SARS-CoV_orf1ab;4454 | 0.05 | 0 |
| NC_004718.3;NP_828849.2;SARS-CoV_orf1ab;4455 | 0.05 | 0 |
| NC_004718.3;NP_828849.2;SARS-CoV_orf1ab;4456 | 0.05 | 0 |
| NC_004718.3;NP_828849.2;SARS-CoV_orf1ab;4457 | 0 | 0 |
| NC_004718.3;NP_828849.2;SARS-CoV_orf1ab;4458 | 0 | 0 |
| NC_004718.3;NP_828849.2;SARS-CoV_orf1ab;4459 | 0 | 0 |
| NC_004718.3;NP_828849.2;SARS-CoV_orf1ab;4460 | 0 | 0 |
| NC_004718.3;NP_828849.2;SARS-CoV_orf1ab;4461 | 0 | 0 |
| NC_004718.3;NP_828849.2;SARS-CoV_orf1ab;4462 | 0 | 0 |
| NC_004718.3;NP_828849.2;SARS-CoV_orf1ab;4463 | 0 | 0.025 |
| NC_004718.3;NP_828849.2;SARS-CoV_orf1ab;4464 | 0 | 0.025 |
| NC_004718.3;NP_828849.2;SARS-CoV_orf1ab;4465 | 0 | 0.025 |
| NC_004718.3;NP_828849.2;SARS-CoV_orf1ab;4466 | 0 | 0.025 |
| NC_004718.3;NP_828849.2;SARS-CoV_orf1ab;4467 | 0 | 0.025 |
| NC_004718.3;NP_828849.2;SARS-CoV_orf1ab;4468 | 0 | 0.025 |
| NC_004718.3;NP_828849.2;SARS-CoV_orf1ab;4469 | 0 | 0.025 |

| | | |
|---|---|---|
| NC_004718.3;NP_828849.2;SARS-CoV_orf1ab;4470 | 0 | 0.025 |
| NC_004718.3;NP_828849.2;SARS-CoV_orf1ab;4471 | 0 | 0 |
| NC_004718.3;NP_828849.2;SARS-CoV_orf1ab;4472 | 0 | 0 |
| NC_004718.3;NP_828849.2;SARS-CoV_orf1ab;4473 | 0 | 0 |
| NC_004718.3;NP_828849.2;SARS-CoV_orf1ab;4474 | 0 | 0 |
| NC_004718.3;NP_828849.2;SARS-CoV_orf1ab;4475 | 0 | 0 |
| NC_004718.3;NP_828849.2;SARS-CoV_orf1ab;4476 | 0 | 0 |
| NC_004718.3;NP_828849.2;SARS-CoV_orf1ab;4477 | 0 | 0 |
| NC_004718.3;NP_828849.2;SARS-CoV_orf1ab;4478 | 0 | 0 |
| NC_004718.3;NP_828849.2;SARS-CoV_orf1ab;4479 | 0 | 0 |
| NC_004718.3;NP_828849.2;SARS-CoV_orf1ab;4480 | 0 | 0 |
| NC_004718.3;NP_828849.2;SARS-CoV_orf1ab;4481 | 0 | 0 |
| NC_004718.3;NP_828849.2;SARS-CoV_orf1ab;4482 | 0 | 0 |
| NC_004718.3;NP_828849.2;SARS-CoV_orf1ab;4483 | 0 | 0 |
| NC_004718.3;NP_828849.2;SARS-CoV_orf1ab;4484 | 0 | 0 |
| NC_004718.3;NP_828849.2;SARS-CoV_orf1ab;4485 | 0 | 0 |
| NC_004718.3;NP_828849.2;SARS-CoV_orf1ab;4486 | 0 | 0 |
| NC_004718.3;NP_828849.2;SARS-CoV_orf1ab;4487 | 0 | 0 |
| NC_004718.3;NP_828849.2;SARS-CoV_orf1ab;4488 | 0 | 0 |
| NC_004718.3;NP_828849.2;SARS-CoV_orf1ab;4489 | 0 | 0 |
| NC_004718.3;NP_828849.2;SARS-CoV_orf1ab;4490 | 0 | 0 |
| NC_004718.3;NP_828849.2;SARS-CoV_orf1ab;4491 | 0 | 0.375 |
| NC_004718.3;NP_828849.2;SARS-CoV_orf1ab;4492 | 0 | 0 |
| NC_004718.3;NP_828849.2;SARS-CoV_orf1ab;4493 | 0 | 0 |
| NC_004718.3;NP_828849.2;SARS-CoV_orf1ab;4494 | 0 | 0 |
| NC_004718.3;NP_828849.2;SARS-CoV_orf1ab;4495 | 0 | 0 |
| NC_004718.3;NP_828849.2;SARS-CoV_orf1ab;4496 | 0 | 0 |
| NC_004718.3;NP_828849.2;SARS-CoV_orf1ab;4497 | 0.05 | 0.1 |
| NC_004718.3;NP_828849.2;SARS-CoV_orf1ab;4498 | 0 | 0.025 |
| NC_004718.3;NP_828849.2;SARS-CoV_orf1ab;4499 | 0 | 0 |
| NC_004718.3;NP_828849.2;SARS-CoV_orf1ab;4500 | 0 | 0 |
| NC_004718.3;NP_828849.2;SARS-CoV_orf1ab;4501 | 0 | 0 |
| NC_004718.3;NP_828849.2;SARS-CoV_orf1ab;4502 | 0 | 0 |
| NC_004718.3;NP_828849.2;SARS-CoV_orf1ab;4503 | 0 | 0.025 |
| NC_004718.3;NP_828849.2;SARS-CoV_orf1ab;4504 | 0 | 0 |
| NC_004718.3;NP_828849.2;SARS-CoV_orf1ab;4505 | 0 | 0 |
| NC_004718.3;NP_828849.2;SARS-CoV_orf1ab;4506 | 0 | 0 |
| NC_004718.3;NP_828849.2;SARS-CoV_orf1ab;4507 | 0 | 0 |
| NC_004718.3;NP_828849.2;SARS-CoV_orf1ab;4508 | 0 | 0 |
| NC_004718.3;NP_828849.2;SARS-CoV_orf1ab;4509 | 0 | 0 |
| NC_004718.3;NP_828849.2;SARS-CoV_orf1ab;4510 | 0 | 0.025 |
| NC_004718.3;NP_828849.2;SARS-CoV_orf1ab;4511 | 0 | 0 |
| NC_004718.3;NP_828849.2;SARS-CoV_orf1ab;4512 | 0 | 0 |

| | | |
|---|---|---|
| NC_004718.3;NP_828849.2;SARS-CoV_orf1ab;4513 | 0 | 0 |
| NC_004718.3;NP_828849.2;SARS-CoV_orf1ab;4514 | 0 | 0 |
| NC_004718.3;NP_828849.2;SARS-CoV_orf1ab;4515 | 0 | 0.025 |
| NC_004718.3;NP_828849.2;SARS-CoV_orf1ab;4516 | 0 | 0 |
| NC_004718.3;NP_828849.2;SARS-CoV_orf1ab;4517 | 0 | 0 |
| NC_004718.3;NP_828849.2;SARS-CoV_orf1ab;4518 | 0 | 0.025 |
| NC_004718.3;NP_828849.2;SARS-CoV_orf1ab;4519 | 0 | 0.025 |
| NC_004718.3;NP_828849.2;SARS-CoV_orf1ab;4520 | 0 | 0 |
| NC_004718.3;NP_828849.2;SARS-CoV_orf1ab;4521 | 0 | 0 |
| NC_004718.3;NP_828849.2;SARS-CoV_orf1ab;4522 | 0.2 | 0.025 |
| NC_004718.3;NP_828849.2;SARS-CoV_orf1ab;4523 | 0.05 | 0 |
| NC_004718.3;NP_828849.2;SARS-CoV_orf1ab;4524 | 0 | 0.025 |
| NC_004718.3;NP_828849.2;SARS-CoV_orf1ab;4525 | 0 | 0.075 |
| NC_004718.3;NP_828849.2;SARS-CoV_orf1ab;4526 | 0.05 | 0.075 |
| NC_004718.3;NP_828849.2;SARS-CoV_orf1ab;4527 | 0.05 | 0.075 |
| NC_004718.3;NP_828849.2;SARS-CoV_orf1ab;4528 | 0.1 | 0.05 |
| NC_004718.3;NP_828849.2;SARS-CoV_orf1ab;4529 | 0 | 0.025 |
| NC_004718.3;NP_828849.2;SARS-CoV_orf1ab;4530 | 0 | 0 |
| NC_004718.3;NP_828849.2;SARS-CoV_orf1ab;4531 | 0 | 0 |
| NC_004718.3;NP_828849.2;SARS-CoV_orf1ab;4532 | 0 | 0 |
| NC_004718.3;NP_828849.2;SARS-CoV_orf1ab;4533 | 0 | 0 |
| NC_004718.3;NP_828849.2;SARS-CoV_orf1ab;4534 | 0.05 | 0 |
| NC_004718.3;NP_828849.2;SARS-CoV_orf1ab;4535 | 0.05 | 0.025 |
| NC_004718.3;NP_828849.2;SARS-CoV_orf1ab;4536 | 0.05 | 0 |
| NC_004718.3;NP_828849.2;SARS-CoV_orf1ab;4537 | 0 | 0 |
| NC_004718.3;NP_828849.2;SARS-CoV_orf1ab;4538 | 0 | 0 |
| NC_004718.3;NP_828849.2;SARS-CoV_orf1ab;4539 | 0 | 0 |
| NC_004718.3;NP_828849.2;SARS-CoV_orf1ab;4540 | 0 | 0 |
| NC_004718.3;NP_828849.2;SARS-CoV_orf1ab;4541 | 0 | 0 |
| NC_004718.3;NP_828849.2;SARS-CoV_orf1ab;4542 | 0.05 | 0 |
| NC_004718.3;NP_828849.2;SARS-CoV_orf1ab;4543 | 0 | 0 |
| NC_004718.3;NP_828849.2;SARS-CoV_orf1ab;4544 | 0 | 0 |
| NC_004718.3;NP_828849.2;SARS-CoV_orf1ab;4545 | 0 | 0 |
| NC_004718.3;NP_828849.2;SARS-CoV_orf1ab;4546 | 0 | 0 |
| NC_004718.3;NP_828849.2;SARS-CoV_orf1ab;4547 | 0 | 0 |
| NC_004718.3;NP_828849.2;SARS-CoV_orf1ab;4548 | 0 | 0 |
| NC_004718.3;NP_828849.2;SARS-CoV_orf1ab;4549 | 0.05 | 0 |
| NC_004718.3;NP_828849.2;SARS-CoV_orf1ab;4550 | 0 | 0 |
| NC_004718.3;NP_828849.2;SARS-CoV_orf1ab;4551 | 0 | 0 |
| NC_004718.3;NP_828849.2;SARS-CoV_orf1ab;4552 | 0 | 0 |
| NC_004718.3;NP_828849.2;SARS-CoV_orf1ab;4553 | 0 | 0 |
| NC_004718.3;NP_828849.2;SARS-CoV_orf1ab;4554 | 0 | 0 |
| NC_004718.3;NP_828849.2;SARS-CoV_orf1ab;4555 | 0 | 0 |

| | | |
|---|---|---|
| NC_004718.3;NP_828849.2;SARS-CoV_orf1ab;4556 | 0 | 0 |
| NC_004718.3;NP_828849.2;SARS-CoV_orf1ab;4557 | 0 | 0 |
| NC_004718.3;NP_828849.2;SARS-CoV_orf1ab;4558 | 0 | 0 |
| NC_004718.3;NP_828849.2;SARS-CoV_orf1ab;4559 | 0 | 0 |
| NC_004718.3;NP_828849.2;SARS-CoV_orf1ab;4560 | 0 | 0 |
| NC_004718.3;NP_828849.2;SARS-CoV_orf1ab;4561 | 0 | 0 |
| NC_004718.3;NP_828849.2;SARS-CoV_orf1ab;4562 | 0 | 0 |
| NC_004718.3;NP_828849.2;SARS-CoV_orf1ab;4563 | 0 | 0 |
| NC_004718.3;NP_828849.2;SARS-CoV_orf1ab;4564 | 0 | 0 |
| NC_004718.3;NP_828849.2;SARS-CoV_orf1ab;4565 | 0.1 | 0.025 |
| NC_004718.3;NP_828849.2;SARS-CoV_orf1ab;4566 | 0.1 | 0 |
| NC_004718.3;NP_828849.2;SARS-CoV_orf1ab;4567 | 0 | 0 |
| NC_004718.3;NP_828849.2;SARS-CoV_orf1ab;4568 | 0.05 | 0 |
| NC_004718.3;NP_828849.2;SARS-CoV_orf1ab;4569 | 0 | 0 |
| NC_004718.3;NP_828849.2;SARS-CoV_orf1ab;4570 | 0 | 0 |
| NC_004718.3;NP_828849.2;SARS-CoV_orf1ab;4571 | 0 | 0 |
| NC_004718.3;NP_828849.2;SARS-CoV_orf1ab;4572 | 0.05 | 0 |
| NC_004718.3;NP_828849.2;SARS-CoV_orf1ab;4573 | 0 | 0 |
| NC_004718.3;NP_828849.2;SARS-CoV_orf1ab;4574 | 0 | 0 |
| NC_004718.3;NP_828849.2;SARS-CoV_orf1ab;4575 | 0 | 0 |
| NC_004718.3;NP_828849.2;SARS-CoV_orf1ab;4576 | 0 | 0 |
| NC_004718.3;NP_828849.2;SARS-CoV_orf1ab;4577 | 0 | 0 |
| NC_004718.3;NP_828849.2;SARS-CoV_orf1ab;4578 | 0 | 0 |
| NC_004718.3;NP_828849.2;SARS-CoV_orf1ab;4579 | 0 | 0.05 |
| NC_004718.3;NP_828849.2;SARS-CoV_orf1ab;4580 | 0 | 0 |
| NC_004718.3;NP_828849.2;SARS-CoV_orf1ab;4581 | 0 | 0 |
| NC_004718.3;NP_828849.2;SARS-CoV_orf1ab;4582 | 0.05 | 0 |
| NC_004718.3;NP_828849.2;SARS-CoV_orf1ab;4583 | 0 | 0 |
| NC_004718.3;NP_828849.2;SARS-CoV_orf1ab;4584 | 0 | 0 |
| NC_004718.3;NP_828849.2;SARS-CoV_orf1ab;4585 | 0 | 0 |
| NC_004718.3;NP_828849.2;SARS-CoV_orf1ab;4586 | 0.1 | 0.05 |
| NC_004718.3;NP_828849.2;SARS-CoV_orf1ab;4587 | 0.05 | 0 |
| NC_004718.3;NP_828849.2;SARS-CoV_orf1ab;4588 | 0 | 0 |
| NC_004718.3;NP_828849.2;SARS-CoV_orf1ab;4589 | 0 | 0 |
| NC_004718.3;NP_828849.2;SARS-CoV_orf1ab;4590 | 0 | 0 |
| NC_004718.3;NP_828849.2;SARS-CoV_orf1ab;4591 | 0 | 0.025 |
| NC_004718.3;NP_828849.2;SARS-CoV_orf1ab;4592 | 0 | 0 |
| NC_004718.3;NP_828849.2;SARS-CoV_orf1ab;4593 | 0 | 0 |
| NC_004718.3;NP_828849.2;SARS-CoV_orf1ab;4594 | 0 | 0 |
| NC_004718.3;NP_828849.2;SARS-CoV_orf1ab;4595 | 0 | 0 |
| NC_004718.3;NP_828849.2;SARS-CoV_orf1ab;4596 | 0 | 0 |
| NC_004718.3;NP_828849.2;SARS-CoV_orf1ab;4597 | 0 | 0 |
| NC_004718.3;NP_828849.2;SARS-CoV_orf1ab;4598 | 0 | 0 |

| | | |
|---|---|---|
| NC_004718.3;NP_828849.2;SARS-CoV_orf1ab;4599 | 0 | 0 |
| NC_004718.3;NP_828849.2;SARS-CoV_orf1ab;4600 | 0 | 0 |
| NC_004718.3;NP_828849.2;SARS-CoV_orf1ab;4601 | 0 | 0 |
| NC_004718.3;NP_828849.2;SARS-CoV_orf1ab;4602 | 0 | 0 |
| NC_004718.3;NP_828849.2;SARS-CoV_orf1ab;4603 | 0 | 0 |
| NC_004718.3;NP_828849.2;SARS-CoV_orf1ab;4604 | 0 | 0 |
| NC_004718.3;NP_828849.2;SARS-CoV_orf1ab;4605 | 0 | 0 |
| NC_004718.3;NP_828849.2;SARS-CoV_orf1ab;4606 | 0 | 0 |
| NC_004718.3;NP_828849.2;SARS-CoV_orf1ab;4607 | 0 | 0 |
| NC_004718.3;NP_828849.2;SARS-CoV_orf1ab;4608 | 0 | 0 |
| NC_004718.3;NP_828849.2;SARS-CoV_orf1ab;4609 | 0 | 0 |
| NC_004718.3;NP_828849.2;SARS-CoV_orf1ab;4610 | 0 | 0 |
| NC_004718.3;NP_828849.2;SARS-CoV_orf1ab;4611 | 0 | 0 |
| NC_004718.3;NP_828849.2;SARS-CoV_orf1ab;4612 | 0.05 | 0 |
| NC_004718.3;NP_828849.2;SARS-CoV_orf1ab;4613 | 0 | 0 |
| NC_004718.3;NP_828849.2;SARS-CoV_orf1ab;4614 | 0 | 0 |
| NC_004718.3;NP_828849.2;SARS-CoV_orf1ab;4615 | 0 | 0 |
| NC_004718.3;NP_828849.2;SARS-CoV_orf1ab;4616 | 0 | 0 |
| NC_004718.3;NP_828849.2;SARS-CoV_orf1ab;4617 | 0 | 0 |
| NC_004718.3;NP_828849.2;SARS-CoV_orf1ab;4618 | 0.05 | 0.025 |
| NC_004718.3;NP_828849.2;SARS-CoV_orf1ab;4619 | 0 | 0 |
| NC_004718.3;NP_828849.2;SARS-CoV_orf1ab;4620 | 0 | 0 |
| NC_004718.3;NP_828849.2;SARS-CoV_orf1ab;4621 | 0.05 | 0 |
| NC_004718.3;NP_828849.2;SARS-CoV_orf1ab;4622 | 0.05 | 0 |
| NC_004718.3;NP_828849.2;SARS-CoV_orf1ab;4623 | 0.05 | 0 |
| NC_004718.3;NP_828849.2;SARS-CoV_orf1ab;4624 | 0.1 | 0 |
| NC_004718.3;NP_828849.2;SARS-CoV_orf1ab;4625 | 0.05 | 0 |
| NC_004718.3;NP_828849.2;SARS-CoV_orf1ab;4626 | 0.05 | 0.025 |
| NC_004718.3;NP_828849.2;SARS-CoV_orf1ab;4627 | 0.05 | 0.025 |
| NC_004718.3;NP_828849.2;SARS-CoV_orf1ab;4628 | 0.15 | 0 |
| NC_004718.3;NP_828849.2;SARS-CoV_orf1ab;4629 | 0.05 | 0 |
| NC_004718.3;NP_828849.2;SARS-CoV_orf1ab;4630 | 0 | 0 |
| NC_004718.3;NP_828849.2;SARS-CoV_orf1ab;4631 | 0.05 | 0 |
| NC_004718.3;NP_828849.2;SARS-CoV_orf1ab;4632 | 0.05 | 0 |
| NC_004718.3;NP_828849.2;SARS-CoV_orf1ab;4633 | 0.05 | 0 |
| NC_004718.3;NP_828849.2;SARS-CoV_orf1ab;4634 | 0.05 | 0 |
| NC_004718.3;NP_828849.2;SARS-CoV_orf1ab;4635 | 0.1 | 0 |
| NC_004718.3;NP_828849.2;SARS-CoV_orf1ab;4636 | 0.05 | 0.025 |
| NC_004718.3;NP_828849.2;SARS-CoV_orf1ab;4637 | 0 | 0 |
| NC_004718.3;NP_828849.2;SARS-CoV_orf1ab;4638 | 0.05 | 0 |
| NC_004718.3;NP_828849.2;SARS-CoV_orf1ab;4639 | 0.05 | 0.025 |
| NC_004718.3;NP_828849.2;SARS-CoV_orf1ab;4640 | 0.05 | 0 |
| NC_004718.3;NP_828849.2;SARS-CoV_orf1ab;4641 | 0.05 | 0 |

| | | |
|---|---|---|
| NC_004718.3;NP_828849.2;SARS-CoV_orf1ab;4642 | 0 | 0 |
| NC_004718.3;NP_828849.2;SARS-CoV_orf1ab;4643 | 0 | 0 |
| NC_004718.3;NP_828849.2;SARS-CoV_orf1ab;4644 | 0 | 0 |
| NC_004718.3;NP_828849.2;SARS-CoV_orf1ab;4645 | 0 | 0 |
| NC_004718.3;NP_828849.2;SARS-CoV_orf1ab;4646 | 0 | 0 |
| NC_004718.3;NP_828849.2;SARS-CoV_orf1ab;4647 | 0.05 | 0.025 |
| NC_004718.3;NP_828849.2;SARS-CoV_orf1ab;4648 | 0 | 0 |
| NC_004718.3;NP_828849.2;SARS-CoV_orf1ab;4649 | 0 | 0 |
| NC_004718.3;NP_828849.2;SARS-CoV_orf1ab;4650 | 0 | 0 |
| NC_004718.3;NP_828849.2;SARS-CoV_orf1ab;4651 | 0 | 0.025 |
| NC_004718.3;NP_828849.2;SARS-CoV_orf1ab;4652 | 0 | 0 |
| NC_004718.3;NP_828849.2;SARS-CoV_orf1ab;4653 | 0 | 0.025 |
| NC_004718.3;NP_828849.2;SARS-CoV_orf1ab;4654 | 0 | 0.05 |
| NC_004718.3;NP_828849.2;SARS-CoV_orf1ab;4655 | 0 | 0.05 |
| NC_004718.3;NP_828849.2;SARS-CoV_orf1ab;4656 | 0 | 0.1 |
| NC_004718.3;NP_828849.2;SARS-CoV_orf1ab;4657 | 0 | 0.075 |
| NC_004718.3;NP_828849.2;SARS-CoV_orf1ab;4658 | 0 | 0.075 |
| NC_004718.3;NP_828849.2;SARS-CoV_orf1ab;4659 | 0 | 0.05 |
| NC_004718.3;NP_828849.2;SARS-CoV_orf1ab;4660 | 0 | 0.05 |
| NC_004718.3;NP_828849.2;SARS-CoV_orf1ab;4661 | 0 | 0.05 |
| NC_004718.3;NP_828849.2;SARS-CoV_orf1ab;4662 | 0 | 0.05 |
| NC_004718.3;NP_828849.2;SARS-CoV_orf1ab;4663 | 0 | 0.05 |
| NC_004718.3;NP_828849.2;SARS-CoV_orf1ab;4664 | 0 | 0.05 |
| NC_004718.3;NP_828849.2;SARS-CoV_orf1ab;4665 | 0 | 0.025 |
| NC_004718.3;NP_828849.2;SARS-CoV_orf1ab;4666 | 0 | 0 |
| NC_004718.3;NP_828849.2;SARS-CoV_orf1ab;4667 | 0 | 0 |
| NC_004718.3;NP_828849.2;SARS-CoV_orf1ab;4668 | 0 | 0 |
| NC_004718.3;NP_828849.2;SARS-CoV_orf1ab;4669 | 0 | 0 |
| NC_004718.3;NP_828849.2;SARS-CoV_orf1ab;4670 | 0 | 0 |
| NC_004718.3;NP_828849.2;SARS-CoV_orf1ab;4671 | 0 | 0 |
| NC_004718.3;NP_828849.2;SARS-CoV_orf1ab;4672 | 0 | 0 |
| NC_004718.3;NP_828849.2;SARS-CoV_orf1ab;4673 | 0 | 0 |
| NC_004718.3;NP_828849.2;SARS-CoV_orf1ab;4674 | 0 | 0 |
| NC_004718.3;NP_828849.2;SARS-CoV_orf1ab;4675 | 0 | 0 |
| NC_004718.3;NP_828849.2;SARS-CoV_orf1ab;4676 | 0 | 0 |
| NC_004718.3;NP_828849.2;SARS-CoV_orf1ab;4677 | 0 | 0 |
| NC_004718.3;NP_828849.2;SARS-CoV_orf1ab;4678 | 0 | 0 |
| NC_004718.3;NP_828849.2;SARS-CoV_orf1ab;4679 | 0 | 0 |
| NC_004718.3;NP_828849.2;SARS-CoV_orf1ab;4680 | 0 | 0 |
| NC_004718.3;NP_828849.2;SARS-CoV_orf1ab;4681 | 0 | 0 |
| NC_004718.3;NP_828849.2;SARS-CoV_orf1ab;4682 | 0 | 0 |
| NC_004718.3;NP_828849.2;SARS-CoV_orf1ab;4683 | 0 | 0 |
| NC_004718.3;NP_828849.2;SARS-CoV_orf1ab;4684 | 0 | 0 |

| | | |
|---|---|---|
| NC_004718.3;NP_828849.2;SARS-CoV_orf1ab;4685 | 0 | 0 |
| NC_004718.3;NP_828849.2;SARS-CoV_orf1ab;4686 | 0 | 0 |
| NC_004718.3;NP_828849.2;SARS-CoV_orf1ab;4687 | 0 | 0 |
| NC_004718.3;NP_828849.2;SARS-CoV_orf1ab;4688 | 0 | 0 |
| NC_004718.3;NP_828849.2;SARS-CoV_orf1ab;4689 | 0 | 0 |
| NC_004718.3;NP_828849.2;SARS-CoV_orf1ab;4690 | 0 | 0 |
| NC_004718.3;NP_828849.2;SARS-CoV_orf1ab;4691 | 0 | 0 |
| NC_004718.3;NP_828849.2;SARS-CoV_orf1ab;4692 | 0 | 0 |
| NC_004718.3;NP_828849.2;SARS-CoV_orf1ab;4693 | 0 | 0 |
| NC_004718.3;NP_828849.2;SARS-CoV_orf1ab;4694 | 0 | 0 |
| NC_004718.3;NP_828849.2;SARS-CoV_orf1ab;4695 | 0 | 0 |
| NC_004718.3;NP_828849.2;SARS-CoV_orf1ab;4696 | 0 | 0 |
| NC_004718.3;NP_828849.2;SARS-CoV_orf1ab;4697 | 0 | 0 |
| NC_004718.3;NP_828849.2;SARS-CoV_orf1ab;4698 | 0 | 0 |
| NC_004718.3;NP_828849.2;SARS-CoV_orf1ab;4699 | 0 | 0 |
| NC_004718.3;NP_828849.2;SARS-CoV_orf1ab;4700 | 0 | 0 |
| NC_004718.3;NP_828849.2;SARS-CoV_orf1ab;4701 | 0 | 0 |
| NC_004718.3;NP_828849.2;SARS-CoV_orf1ab;4702 | 0 | 0 |
| NC_004718.3;NP_828849.2;SARS-CoV_orf1ab;4703 | 0 | 0 |
| NC_004718.3;NP_828849.2;SARS-CoV_orf1ab;4704 | 0 | 0 |
| NC_004718.3;NP_828849.2;SARS-CoV_orf1ab;4705 | 0 | 0 |
| NC_004718.3;NP_828849.2;SARS-CoV_orf1ab;4706 | 0 | 0 |
| NC_004718.3;NP_828849.2;SARS-CoV_orf1ab;4707 | 0 | 0 |
| NC_004718.3;NP_828849.2;SARS-CoV_orf1ab;4708 | 0 | 0 |
| NC_004718.3;NP_828849.2;SARS-CoV_orf1ab;4709 | 0 | 0 |
| NC_004718.3;NP_828849.2;SARS-CoV_orf1ab;4710 | 0 | 0 |
| NC_004718.3;NP_828849.2;SARS-CoV_orf1ab;4711 | 0 | 0 |
| NC_004718.3;NP_828849.2;SARS-CoV_orf1ab;4712 | 0 | 0 |
| NC_004718.3;NP_828849.2;SARS-CoV_orf1ab;4713 | 0 | 0 |
| NC_004718.3;NP_828849.2;SARS-CoV_orf1ab;4714 | 0 | 0 |
| NC_004718.3;NP_828849.2;SARS-CoV_orf1ab;4715 | 0 | 0 |
| NC_004718.3;NP_828849.2;SARS-CoV_orf1ab;4716 | 0 | 0 |
| NC_004718.3;NP_828849.2;SARS-CoV_orf1ab;4717 | 0 | 0 |
| NC_004718.3;NP_828849.2;SARS-CoV_orf1ab;4718 | 0 | 0 |
| NC_004718.3;NP_828849.2;SARS-CoV_orf1ab;4719 | 0 | 0 |
| NC_004718.3;NP_828849.2;SARS-CoV_orf1ab;4720 | 0 | 0 |
| NC_004718.3;NP_828849.2;SARS-CoV_orf1ab;4721 | 0 | 0 |
| NC_004718.3;NP_828849.2;SARS-CoV_orf1ab;4722 | 0 | 0 |
| NC_004718.3;NP_828849.2;SARS-CoV_orf1ab;4723 | 0 | 0 |
| NC_004718.3;NP_828849.2;SARS-CoV_orf1ab;4724 | 0 | 0.025 |
| NC_004718.3;NP_828849.2;SARS-CoV_orf1ab;4725 | 0 | 0.075 |
| NC_004718.3;NP_828849.2;SARS-CoV_orf1ab;4726 | 0.05 | 0.025 |
| NC_004718.3;NP_828849.2;SARS-CoV_orf1ab;4727 | 0 | 0 |

| | | |
|---|---|---|
| NC_004718.3;NP_828849.2;SARS-CoV_orf1ab;4728 | 0 | 0 |
| NC_004718.3;NP_828849.2;SARS-CoV_orf1ab;4729 | 0 | 0 |
| NC_004718.3;NP_828849.2;SARS-CoV_orf1ab;4730 | 0 | 0 |
| NC_004718.3;NP_828849.2;SARS-CoV_orf1ab;4731 | 0 | 0 |
| NC_004718.3;NP_828849.2;SARS-CoV_orf1ab;4732 | 0 | 0 |
| NC_004718.3;NP_828849.2;SARS-CoV_orf1ab;4733 | 0 | 0 |
| NC_004718.3;NP_828849.2;SARS-CoV_orf1ab;4734 | 0.05 | 0 |
| NC_004718.3;NP_828849.2;SARS-CoV_orf1ab;4735 | 0.1 | 0.025 |
| NC_004718.3;NP_828849.2;SARS-CoV_orf1ab;4736 | 0.1 | 0.025 |
| NC_004718.3;NP_828849.2;SARS-CoV_orf1ab;4737 | 0.05 | 0.025 |
| NC_004718.3;NP_828849.2;SARS-CoV_orf1ab;4738 | 0 | 0 |
| NC_004718.3;NP_828849.2;SARS-CoV_orf1ab;4739 | 0 | 0 |
| NC_004718.3;NP_828849.2;SARS-CoV_orf1ab;4740 | 0 | 0 |
| NC_004718.3;NP_828849.2;SARS-CoV_orf1ab;4741 | 0 | 0.025 |
| NC_004718.3;NP_828849.2;SARS-CoV_orf1ab;4742 | 0 | 0.025 |
| NC_004718.3;NP_828849.2;SARS-CoV_orf1ab;4743 | 0 | 0 |
| NC_004718.3;NP_828849.2;SARS-CoV_orf1ab;4744 | 0 | 0 |
| NC_004718.3;NP_828849.2;SARS-CoV_orf1ab;4745 | 0 | 0 |
| NC_004718.3;NP_828849.2;SARS-CoV_orf1ab;4746 | 0 | 0 |
| NC_004718.3;NP_828849.2;SARS-CoV_orf1ab;4747 | 0 | 0 |
| NC_004718.3;NP_828849.2;SARS-CoV_orf1ab;4748 | 0 | 0 |
| NC_004718.3;NP_828849.2;SARS-CoV_orf1ab;4749 | 0 | 0 |
| NC_004718.3;NP_828849.2;SARS-CoV_orf1ab;4750 | 0 | 0 |
| NC_004718.3;NP_828849.2;SARS-CoV_orf1ab;4751 | 0 | 0 |
| NC_004718.3;NP_828849.2;SARS-CoV_orf1ab;4752 | 0 | 0 |
| NC_004718.3;NP_828849.2;SARS-CoV_orf1ab;4753 | 0 | 0 |
| NC_004718.3;NP_828849.2;SARS-CoV_orf1ab;4754 | 0.05 | 0 |
| NC_004718.3;NP_828849.2;SARS-CoV_orf1ab;4755 | 0 | 0 |
| NC_004718.3;NP_828849.2;SARS-CoV_orf1ab;4756 | 0 | 0 |
| NC_004718.3;NP_828849.2;SARS-CoV_orf1ab;4757 | 0 | 0 |
| NC_004718.3;NP_828849.2;SARS-CoV_orf1ab;4758 | 0 | 0 |
| NC_004718.3;NP_828849.2;SARS-CoV_orf1ab;4759 | 0 | 0 |
| NC_004718.3;NP_828849.2;SARS-CoV_orf1ab;4760 | 0 | 0 |
| NC_004718.3;NP_828849.2;SARS-CoV_orf1ab;4761 | 0 | 0 |
| NC_004718.3;NP_828849.2;SARS-CoV_orf1ab;4762 | 0 | 0 |
| NC_004718.3;NP_828849.2;SARS-CoV_orf1ab;4763 | 0 | 0 |
| NC_004718.3;NP_828849.2;SARS-CoV_orf1ab;4764 | 0 | 0 |
| NC_004718.3;NP_828849.2;SARS-CoV_orf1ab;4765 | 0 | 0 |
| NC_004718.3;NP_828849.2;SARS-CoV_orf1ab;4766 | 0 | 0 |
| NC_004718.3;NP_828849.2;SARS-CoV_orf1ab;4767 | 0 | 0 |
| NC_004718.3;NP_828849.2;SARS-CoV_orf1ab;4768 | 0.05 | 0 |
| NC_004718.3;NP_828849.2;SARS-CoV_orf1ab;4769 | 0.05 | 0.05 |
| NC_004718.3;NP_828849.2;SARS-CoV_orf1ab;4770 | 0 | 0 |

| | | |
|---|---|---|
| NC_004718.3;NP_828849.2;SARS-CoV_orf1ab;4771 | 0 | 0 |
| NC_004718.3;NP_828849.2;SARS-CoV_orf1ab;4772 | 0 | 0.025 |
| NC_004718.3;NP_828849.2;SARS-CoV_orf1ab;4773 | 0 | 0.025 |
| NC_004718.3;NP_828849.2;SARS-CoV_orf1ab;4774 | 0 | 0 |
| NC_004718.3;NP_828849.2;SARS-CoV_orf1ab;4775 | 0 | 0 |
| NC_004718.3;NP_828849.2;SARS-CoV_orf1ab;4776 | 0.05 | 0 |
| NC_004718.3;NP_828849.2;SARS-CoV_orf1ab;4777 | 0.05 | 0.025 |
| NC_004718.3;NP_828849.2;SARS-CoV_orf1ab;4778 | 0.1 | 0.025 |
| NC_004718.3;NP_828849.2;SARS-CoV_orf1ab;4779 | 0.1 | 0.025 |
| NC_004718.3;NP_828849.2;SARS-CoV_orf1ab;4780 | 0.1 | 0.025 |
| NC_004718.3;NP_828849.2;SARS-CoV_orf1ab;4781 | 0.2 | 0.025 |
| NC_004718.3;NP_828849.2;SARS-CoV_orf1ab;4782 | 0.1 | 0 |
| NC_004718.3;NP_828849.2;SARS-CoV_orf1ab;4783 | 0 | 0 |
| NC_004718.3;NP_828849.2;SARS-CoV_orf1ab;4784 | 0 | 0 |
| NC_004718.3;NP_828849.2;SARS-CoV_orf1ab;4785 | 0.05 | 0 |
| NC_004718.3;NP_828849.2;SARS-CoV_orf1ab;4786 | 0.05 | 0 |
| NC_004718.3;NP_828849.2;SARS-CoV_orf1ab;4787 | 0.05 | 0 |
| NC_004718.3;NP_828849.2;SARS-CoV_orf1ab;4788 | 0 | 0 |
| NC_004718.3;NP_828849.2;SARS-CoV_orf1ab;4789 | 0 | 0 |
| NC_004718.3;NP_828849.2;SARS-CoV_orf1ab;4790 | 0 | 0 |
| NC_004718.3;NP_828849.2;SARS-CoV_orf1ab;4791 | 0 | 0 |
| NC_004718.3;NP_828849.2;SARS-CoV_orf1ab;4792 | 0 | 0 |
| NC_004718.3;NP_828849.2;SARS-CoV_orf1ab;4793 | 0 | 0 |
| NC_004718.3;NP_828849.2;SARS-CoV_orf1ab;4794 | 0.05 | 0 |
| NC_004718.3;NP_828849.2;SARS-CoV_orf1ab;4795 | 0 | 0.025 |
| NC_004718.3;NP_828849.2;SARS-CoV_orf1ab;4796 | 0 | 0 |
| NC_004718.3;NP_828849.2;SARS-CoV_orf1ab;4797 | 0 | 0 |
| NC_004718.3;NP_828849.2;SARS-CoV_orf1ab;4798 | 0 | 0 |
| NC_004718.3;NP_828849.2;SARS-CoV_orf1ab;4799 | 0 | 0 |
| NC_004718.3;NP_828849.2;SARS-CoV_orf1ab;4800 | 0 | 0 |
| NC_004718.3;NP_828849.2;SARS-CoV_orf1ab;4801 | 0 | 0 |
| NC_004718.3;NP_828849.2;SARS-CoV_orf1ab;4802 | 0 | 0 |
| NC_004718.3;NP_828849.2;SARS-CoV_orf1ab;4803 | 0.05 | 0 |
| NC_004718.3;NP_828849.2;SARS-CoV_orf1ab;4804 | 0.05 | 0 |
| NC_004718.3;NP_828849.2;SARS-CoV_orf1ab;4805 | 0 | 0 |
| NC_004718.3;NP_828849.2;SARS-CoV_orf1ab;4806 | 0 | 0 |
| NC_004718.3;NP_828849.2;SARS-CoV_orf1ab;4807 | 0.05 | 0 |
| NC_004718.3;NP_828849.2;SARS-CoV_orf1ab;4808 | 0 | 0 |
| NC_004718.3;NP_828849.2;SARS-CoV_orf1ab;4809 | 0 | 0 |
| NC_004718.3;NP_828849.2;SARS-CoV_orf1ab;4810 | 0 | 0 |
| NC_004718.3;NP_828849.2;SARS-CoV_orf1ab;4811 | 0 | 0.025 |
| NC_004718.3;NP_828849.2;SARS-CoV_orf1ab;4812 | 0 | 0 |
| NC_004718.3;NP_828849.2;SARS-CoV_orf1ab;4813 | 0.1 | 0.025 |

| | | |
|---|---|---|
| NC_004718.3;NP_828849.2;SARS-CoV_orf1ab;4814 | 0.05 | 0.025 |
| NC_004718.3;NP_828849.2;SARS-CoV_orf1ab;4815 | 0.15 | 0 |
| NC_004718.3;NP_828849.2;SARS-CoV_orf1ab;4816 | 0.2 | 0 |
| NC_004718.3;NP_828849.2;SARS-CoV_orf1ab;4817 | 0.15 | 0.075 |
| NC_004718.3;NP_828849.2;SARS-CoV_orf1ab;4818 | 0.25 | 0.025 |
| NC_004718.3;NP_828849.2;SARS-CoV_orf1ab;4819 | 0.15 | 0 |
| NC_004718.3;NP_828849.2;SARS-CoV_orf1ab;4820 | 0.15 | 0 |
| NC_004718.3;NP_828849.2;SARS-CoV_orf1ab;4821 | 0.15 | 0 |
| NC_004718.3;NP_828849.2;SARS-CoV_orf1ab;4822 | 0 | 0 |
| NC_004718.3;NP_828849.2;SARS-CoV_orf1ab;4823 | 0 | 0 |
| NC_004718.3;NP_828849.2;SARS-CoV_orf1ab;4824 | 0 | 0 |
| NC_004718.3;NP_828849.2;SARS-CoV_orf1ab;4825 | 0 | 0 |
| NC_004718.3;NP_828849.2;SARS-CoV_orf1ab;4826 | 0 | 0 |
| NC_004718.3;NP_828849.2;SARS-CoV_orf1ab;4827 | 0 | 0 |
| NC_004718.3;NP_828849.2;SARS-CoV_orf1ab;4828 | 0 | 0 |
| NC_004718.3;NP_828849.2;SARS-CoV_orf1ab;4829 | 0 | 0 |
| NC_004718.3;NP_828849.2;SARS-CoV_orf1ab;4830 | 0 | 0 |
| NC_004718.3;NP_828849.2;SARS-CoV_orf1ab;4831 | 0 | 0 |
| NC_004718.3;NP_828849.2;SARS-CoV_orf1ab;4832 | 0.05 | 0.025 |
| NC_004718.3;NP_828849.2;SARS-CoV_orf1ab;4833 | 0 | 0 |
| NC_004718.3;NP_828849.2;SARS-CoV_orf1ab;4834 | 0 | 0 |
| NC_004718.3;NP_828849.2;SARS-CoV_orf1ab;4835 | 0 | 0.025 |
| NC_004718.3;NP_828849.2;SARS-CoV_orf1ab;4836 | 0 | 0 |
| NC_004718.3;NP_828849.2;SARS-CoV_orf1ab;4837 | 0 | 0 |
| NC_004718.3;NP_828849.2;SARS-CoV_orf1ab;4838 | 0 | 0 |
| NC_004718.3;NP_828849.2;SARS-CoV_orf1ab;4839 | 0 | 0 |
| NC_004718.3;NP_828849.2;SARS-CoV_orf1ab;4840 | 0 | 0 |
| NC_004718.3;NP_828849.2;SARS-CoV_orf1ab;4841 | 0 | 0.025 |
| NC_004718.3;NP_828849.2;SARS-CoV_orf1ab;4842 | 0 | 0.025 |
| NC_004718.3;NP_828849.2;SARS-CoV_orf1ab;4843 | 0 | 0 |
| NC_004718.3;NP_828849.2;SARS-CoV_orf1ab;4844 | 0 | 0 |
| NC_004718.3;NP_828849.2;SARS-CoV_orf1ab;4845 | 0.05 | 0 |
| NC_004718.3;NP_828849.2;SARS-CoV_orf1ab;4846 | 0.05 | 0.075 |
| NC_004718.3;NP_828849.2;SARS-CoV_orf1ab;4847 | 0.05 | 0.025 |
| NC_004718.3;NP_828849.2;SARS-CoV_orf1ab;4848 | 0.05 | 0 |
| NC_004718.3;NP_828849.2;SARS-CoV_orf1ab;4849 | 0 | 0.025 |
| NC_004718.3;NP_828849.2;SARS-CoV_orf1ab;4850 | 0.05 | 0 |
| NC_004718.3;NP_828849.2;SARS-CoV_orf1ab;4851 | 0.05 | 0 |
| NC_004718.3;NP_828849.2;SARS-CoV_orf1ab;4852 | 0 | 0 |
| NC_004718.3;NP_828849.2;SARS-CoV_orf1ab;4853 | 0.05 | 0.025 |
| NC_004718.3;NP_828849.2;SARS-CoV_orf1ab;4854 | 0 | 0.025 |
| NC_004718.3;NP_828849.2;SARS-CoV_orf1ab;4855 | 0.05 | 0.025 |
| NC_004718.3;NP_828849.2;SARS-CoV_orf1ab;4856 | 0.05 | 0.025 |

| | | |
|---|---|---|
| NC_004718.3;NP_828849.2;SARS-CoV_orf1ab;4857 | 0.1 | 0.025 |
| NC_004718.3;NP_828849.2;SARS-CoV_orf1ab;4858 | 0 | 0.025 |
| NC_004718.3;NP_828849.2;SARS-CoV_orf1ab;4859 | 0.15 | 0.1 |
| NC_004718.3;NP_828849.2;SARS-CoV_orf1ab;4860 | 0 | 0.05 |
| NC_004718.3;NP_828849.2;SARS-CoV_orf1ab;4861 | 0.15 | 0.075 |
| NC_004718.3;NP_828849.2;SARS-CoV_orf1ab;4862 | 0.05 | 0.05 |
| NC_004718.3;NP_828849.2;SARS-CoV_orf1ab;4863 | 0 | 0.025 |
| NC_004718.3;NP_828849.2;SARS-CoV_orf1ab;4864 | 0 | 0 |
| NC_004718.3;NP_828849.2;SARS-CoV_orf1ab;4865 | 0 | 0 |
| NC_004718.3;NP_828849.2;SARS-CoV_orf1ab;4866 | 0 | 0 |
| NC_004718.3;NP_828849.2;SARS-CoV_orf1ab;4867 | 0 | 0 |
| NC_004718.3;NP_828849.2;SARS-CoV_orf1ab;4868 | 0 | 0 |
| NC_004718.3;NP_828849.2;SARS-CoV_orf1ab;4869 | 0 | 0 |
| NC_004718.3;NP_828849.2;SARS-CoV_orf1ab;4870 | 0 | 0 |
| NC_004718.3;NP_828849.2;SARS-CoV_orf1ab;4871 | 0 | 0 |
| NC_004718.3;NP_828849.2;SARS-CoV_orf1ab;4872 | 0 | 0 |
| NC_004718.3;NP_828849.2;SARS-CoV_orf1ab;4873 | 0 | 0 |
| NC_004718.3;NP_828849.2;SARS-CoV_orf1ab;4874 | 0.05 | 0 |
| NC_004718.3;NP_828849.2;SARS-CoV_orf1ab;4875 | 0 | 0 |
| NC_004718.3;NP_828849.2;SARS-CoV_orf1ab;4876 | 0.05 | 0 |
| NC_004718.3;NP_828849.2;SARS-CoV_orf1ab;4877 | 0 | 0 |
| NC_004718.3;NP_828849.2;SARS-CoV_orf1ab;4878 | 0 | 0 |
| NC_004718.3;NP_828849.2;SARS-CoV_orf1ab;4879 | 0 | 0 |
| NC_004718.3;NP_828849.2;SARS-CoV_orf1ab;4880 | 0 | 0 |
| NC_004718.3;NP_828849.2;SARS-CoV_orf1ab;4881 | 0 | 0 |
| NC_004718.3;NP_828849.2;SARS-CoV_orf1ab;4882 | 0 | 0 |
| NC_004718.3;NP_828849.2;SARS-CoV_orf1ab;4883 | 0 | 0 |
| NC_004718.3;NP_828849.2;SARS-CoV_orf1ab;4884 | 0 | 0 |
| NC_004718.3;NP_828849.2;SARS-CoV_orf1ab;4885 | 0 | 0 |
| NC_004718.3;NP_828849.2;SARS-CoV_orf1ab;4886 | 0.1 | 0.025 |
| NC_004718.3;NP_828849.2;SARS-CoV_orf1ab;4887 | 0.05 | 0.025 |
| NC_004718.3;NP_828849.2;SARS-CoV_orf1ab;4888 | 0.05 | 0 |
| NC_004718.3;NP_828849.2;SARS-CoV_orf1ab;4889 | 0.1 | 0.05 |
| NC_004718.3;NP_828849.2;SARS-CoV_orf1ab;4890 | 0.05 | 0 |
| NC_004718.3;NP_828849.2;SARS-CoV_orf1ab;4891 | 0.2 | 0.025 |
| NC_004718.3;NP_828849.2;SARS-CoV_orf1ab;4892 | 0.05 | 0.025 |
| NC_004718.3;NP_828849.2;SARS-CoV_orf1ab;4893 | 0.05 | 0 |
| NC_004718.3;NP_828849.2;SARS-CoV_orf1ab;4894 | 0.05 | 0 |
| NC_004718.3;NP_828849.2;SARS-CoV_orf1ab;4895 | 0 | 0 |
| NC_004718.3;NP_828849.2;SARS-CoV_orf1ab;4896 | 0 | 0 |
| NC_004718.3;NP_828849.2;SARS-CoV_orf1ab;4897 | 0 | 0 |
| NC_004718.3;NP_828849.2;SARS-CoV_orf1ab;4898 | 0 | 0 |
| NC_004718.3;NP_828849.2;SARS-CoV_orf1ab;4899 | 0 | 0 |

| | | |
|---|---|---|
| NC_004718.3;NP_828849.2;SARS-CoV_orf1ab;4900 | 0.05 | 0 |
| NC_004718.3;NP_828849.2;SARS-CoV_orf1ab;4901 | 0.1 | 0.15 |
| NC_004718.3;NP_828849.2;SARS-CoV_orf1ab;4902 | 0 | 0 |
| NC_004718.3;NP_828849.2;SARS-CoV_orf1ab;4903 | 0 | 0 |
| NC_004718.3;NP_828849.2;SARS-CoV_orf1ab;4904 | 0 | 0 |
| NC_004718.3;NP_828849.2;SARS-CoV_orf1ab;4905 | 0 | 0.025 |
| NC_004718.3;NP_828849.2;SARS-CoV_orf1ab;4906 | 0.1 | 0.05 |
| NC_004718.3;NP_828849.2;SARS-CoV_orf1ab;4907 | 0 | 0.025 |
| NC_004718.3;NP_828849.2;SARS-CoV_orf1ab;4908 | 0 | 0.025 |
| NC_004718.3;NP_828849.2;SARS-CoV_orf1ab;4909 | 0 | 0.025 |
| NC_004718.3;NP_828849.2;SARS-CoV_orf1ab;4910 | 0 | 0.025 |
| NC_004718.3;NP_828849.2;SARS-CoV_orf1ab;4911 | 0 | 0 |
| NC_004718.3;NP_828849.2;SARS-CoV_orf1ab;4912 | 0 | 0 |
| NC_004718.3;NP_828849.2;SARS-CoV_orf1ab;4913 | 0 | 0 |
| NC_004718.3;NP_828849.2;SARS-CoV_orf1ab;4914 | 0 | 0 |
| NC_004718.3;NP_828849.2;SARS-CoV_orf1ab;4915 | 0 | 0 |
| NC_004718.3;NP_828849.2;SARS-CoV_orf1ab;4916 | 0 | 0 |
| NC_004718.3;NP_828849.2;SARS-CoV_orf1ab;4917 | 0 | 0 |
| NC_004718.3;NP_828849.2;SARS-CoV_orf1ab;4918 | 0 | 0 |
| NC_004718.3;NP_828849.2;SARS-CoV_orf1ab;4919 | 0 | 0 |
| NC_004718.3;NP_828849.2;SARS-CoV_orf1ab;4920 | 0 | 0 |
| NC_004718.3;NP_828849.2;SARS-CoV_orf1ab;4921 | 0 | 0 |
| NC_004718.3;NP_828849.2;SARS-CoV_orf1ab;4922 | 0 | 0 |
| NC_004718.3;NP_828849.2;SARS-CoV_orf1ab;4923 | 0 | 0 |
| NC_004718.3;NP_828849.2;SARS-CoV_orf1ab;4924 | 0 | 0 |
| NC_004718.3;NP_828849.2;SARS-CoV_orf1ab;4925 | 0 | 0 |
| NC_004718.3;NP_828849.2;SARS-CoV_orf1ab;4926 | 0 | 0 |
| NC_004718.3;NP_828849.2;SARS-CoV_orf1ab;4927 | 0 | 0 |
| NC_004718.3;NP_828849.2;SARS-CoV_orf1ab;4928 | 0 | 0 |
| NC_004718.3;NP_828849.2;SARS-CoV_orf1ab;4929 | 0 | 0 |
| NC_004718.3;NP_828849.2;SARS-CoV_orf1ab;4930 | 0 | 0 |
| NC_004718.3;NP_828849.2;SARS-CoV_orf1ab;4931 | 0 | 0 |
| NC_004718.3;NP_828849.2;SARS-CoV_orf1ab;4932 | 0 | 0 |
| NC_004718.3;NP_828849.2;SARS-CoV_orf1ab;4933 | 0 | 0 |
| NC_004718.3;NP_828849.2;SARS-CoV_orf1ab;4934 | 0 | 0 |
| NC_004718.3;NP_828849.2;SARS-CoV_orf1ab;4935 | 0 | 0 |
| NC_004718.3;NP_828849.2;SARS-CoV_orf1ab;4936 | 0 | 0 |
| NC_004718.3;NP_828849.2;SARS-CoV_orf1ab;4937 | 0 | 0 |
| NC_004718.3;NP_828849.2;SARS-CoV_orf1ab;4938 | 0 | 0 |
| NC_004718.3;NP_828849.2;SARS-CoV_orf1ab;4939 | 0 | 0 |
| NC_004718.3;NP_828849.2;SARS-CoV_orf1ab;4940 | 0.3 | 0.175 |
| NC_004718.3;NP_828849.2;SARS-CoV_orf1ab;4941 | 0.4 | 0.275 |
| NC_004718.3;NP_828849.2;SARS-CoV_orf1ab;4942 | 0.35 | 0.275 |

| | | |
|---|---|---|
| NC_004718.3;NP_828849.2;SARS-CoV_orf1ab;4943 | 0.25 | 0.025 |
| NC_004718.3;NP_828849.2;SARS-CoV_orf1ab;4944 | 0.25 | 0.075 |
| NC_004718.3;NP_828849.2;SARS-CoV_orf1ab;4945 | 0.25 | 0.025 |
| NC_004718.3;NP_828849.2;SARS-CoV_orf1ab;4946 | 0.25 | 0.05 |
| NC_004718.3;NP_828849.2;SARS-CoV_orf1ab;4947 | 0.25 | 0.025 |
| NC_004718.3;NP_828849.2;SARS-CoV_orf1ab;4948 | 0.15 | 0.025 |
| NC_004718.3;NP_828849.2;SARS-CoV_orf1ab;4949 | 0.15 | 0.025 |
| NC_004718.3;NP_828849.2;SARS-CoV_orf1ab;4950 | 0 | 0 |
| NC_004718.3;NP_828849.2;SARS-CoV_orf1ab;4951 | 0 | 0 |
| NC_004718.3;NP_828849.2;SARS-CoV_orf1ab;4952 | 0 | 0 |
| NC_004718.3;NP_828849.2;SARS-CoV_orf1ab;4953 | 0 | 0 |
| NC_004718.3;NP_828849.2;SARS-CoV_orf1ab;4954 | 0 | 0 |
| NC_004718.3;NP_828849.2;SARS-CoV_orf1ab;4955 | 0 | 0 |
| NC_004718.3;NP_828849.2;SARS-CoV_orf1ab;4956 | 0 | 0 |
| NC_004718.3;NP_828849.2;SARS-CoV_orf1ab;4957 | 0 | 0 |
| NC_004718.3;NP_828849.2;SARS-CoV_orf1ab;4958 | 0 | 0 |
| NC_004718.3;NP_828849.2;SARS-CoV_orf1ab;4959 | 0 | 0 |
| NC_004718.3;NP_828849.2;SARS-CoV_orf1ab;4960 | 0 | 0 |
| NC_004718.3;NP_828849.2;SARS-CoV_orf1ab;4961 | 0 | 0 |
| NC_004718.3;NP_828849.2;SARS-CoV_orf1ab;4962 | 0 | 0.05 |
| NC_004718.3;NP_828849.2;SARS-CoV_orf1ab;4963 | 0 | 0 |
| NC_004718.3;NP_828849.2;SARS-CoV_orf1ab;4964 | 0 | 0 |
| NC_004718.3;NP_828849.2;SARS-CoV_orf1ab;4965 | 0 | 0 |
| NC_004718.3;NP_828849.2;SARS-CoV_orf1ab;4966 | 0 | 0 |
| NC_004718.3;NP_828849.2;SARS-CoV_orf1ab;4967 | 0 | 0 |
| NC_004718.3;NP_828849.2;SARS-CoV_orf1ab;4968 | 0 | 0 |
| NC_004718.3;NP_828849.2;SARS-CoV_orf1ab;4969 | 0 | 0 |
| NC_004718.3;NP_828849.2;SARS-CoV_orf1ab;4970 | 0 | 0 |
| NC_004718.3;NP_828849.2;SARS-CoV_orf1ab;4971 | 0 | 0 |
| NC_004718.3;NP_828849.2;SARS-CoV_orf1ab;4972 | 0 | 0 |
| NC_004718.3;NP_828849.2;SARS-CoV_orf1ab;4973 | 0 | 0 |
| NC_004718.3;NP_828849.2;SARS-CoV_orf1ab;4974 | 0 | 0.075 |
| NC_004718.3;NP_828849.2;SARS-CoV_orf1ab;4975 | 0 | 0.025 |
| NC_004718.3;NP_828849.2;SARS-CoV_orf1ab;4976 | 0 | 0.025 |
| NC_004718.3;NP_828849.2;SARS-CoV_orf1ab;4977 | 0 | 0.025 |
| NC_004718.3;NP_828849.2;SARS-CoV_orf1ab;4978 | 0 | 0.025 |
| NC_004718.3;NP_828849.2;SARS-CoV_orf1ab;4979 | 0 | 0.025 |
| NC_004718.3;NP_828849.2;SARS-CoV_orf1ab;4980 | 0 | 0.025 |
| NC_004718.3;NP_828849.2;SARS-CoV_orf1ab;4981 | 0 | 0.025 |
| NC_004718.3;NP_828849.2;SARS-CoV_orf1ab;4982 | 0 | 0.025 |
| NC_004718.3;NP_828849.2;SARS-CoV_orf1ab;4983 | 0 | 0.025 |
| NC_004718.3;NP_828849.2;SARS-CoV_orf1ab;4984 | 0 | 0.025 |
| NC_004718.3;NP_828849.2;SARS-CoV_orf1ab;4985 | 0 | 0.025 |

| | | |
|---|---|---|
| NC_004718.3;NP_828849.2;SARS-CoV_orf1ab;4986 | 0 | 0 |
| NC_004718.3;NP_828849.2;SARS-CoV_orf1ab;4987 | 0 | 0 |
| NC_004718.3;NP_828849.2;SARS-CoV_orf1ab;4988 | 0 | 0 |
| NC_004718.3;NP_828849.2;SARS-CoV_orf1ab;4989 | 0 | 0 |
| NC_004718.3;NP_828849.2;SARS-CoV_orf1ab;4990 | 0 | 0 |
| NC_004718.3;NP_828849.2;SARS-CoV_orf1ab;4991 | 0 | 0 |
| NC_004718.3;NP_828849.2;SARS-CoV_orf1ab;4992 | 0 | 0 |
| NC_004718.3;NP_828849.2;SARS-CoV_orf1ab;4993 | 0 | 0 |
| NC_004718.3;NP_828849.2;SARS-CoV_orf1ab;4994 | 0 | 0 |
| NC_004718.3;NP_828849.2;SARS-CoV_orf1ab;4995 | 0 | 0 |
| NC_004718.3;NP_828849.2;SARS-CoV_orf1ab;4996 | 0 | 0 |
| NC_004718.3;NP_828849.2;SARS-CoV_orf1ab;4997 | 0 | 0 |
| NC_004718.3;NP_828849.2;SARS-CoV_orf1ab;4998 | 0 | 0 |
| NC_004718.3;NP_828849.2;SARS-CoV_orf1ab;4999 | 0 | 0 |
| NC_004718.3;NP_828849.2;SARS-CoV_orf1ab;5000 | 0 | 0 |
| NC_004718.3;NP_828849.2;SARS-CoV_orf1ab;5001 | 0 | 0 |
| NC_004718.3;NP_828849.2;SARS-CoV_orf1ab;5002 | 0 | 0 |
| NC_004718.3;NP_828849.2;SARS-CoV_orf1ab;5003 | 0 | 0 |
| NC_004718.3;NP_828849.2;SARS-CoV_orf1ab;5004 | 0 | 0 |
| NC_004718.3;NP_828849.2;SARS-CoV_orf1ab;5005 | 0 | 0 |
| NC_004718.3;NP_828849.2;SARS-CoV_orf1ab;5006 | 0 | 0 |
| NC_004718.3;NP_828849.2;SARS-CoV_orf1ab;5007 | 0 | 0 |
| NC_004718.3;NP_828849.2;SARS-CoV_orf1ab;5008 | 0 | 0 |
| NC_004718.3;NP_828849.2;SARS-CoV_orf1ab;5009 | 0 | 0 |
| NC_004718.3;NP_828849.2;SARS-CoV_orf1ab;5010 | 0 | 0 |
| NC_004718.3;NP_828849.2;SARS-CoV_orf1ab;5011 | 0 | 0 |
| NC_004718.3;NP_828849.2;SARS-CoV_orf1ab;5012 | 0 | 0 |
| NC_004718.3;NP_828849.2;SARS-CoV_orf1ab;5013 | 0 | 0 |
| NC_004718.3;NP_828849.2;SARS-CoV_orf1ab;5014 | 0 | 0 |
| NC_004718.3;NP_828849.2;SARS-CoV_orf1ab;5015 | 0 | 0 |
| NC_004718.3;NP_828849.2;SARS-CoV_orf1ab;5016 | 0 | 0 |
| NC_004718.3;NP_828849.2;SARS-CoV_orf1ab;5017 | 0 | 0 |
| NC_004718.3;NP_828849.2;SARS-CoV_orf1ab;5018 | 0 | 0 |
| NC_004718.3;NP_828849.2;SARS-CoV_orf1ab;5019 | 0 | 0 |
| NC_004718.3;NP_828849.2;SARS-CoV_orf1ab;5020 | 0 | 0 |
| NC_004718.3;NP_828849.2;SARS-CoV_orf1ab;5021 | 0 | 0 |
| NC_004718.3;NP_828849.2;SARS-CoV_orf1ab;5022 | 0 | 0 |
| NC_004718.3;NP_828849.2;SARS-CoV_orf1ab;5023 | 0 | 0 |
| NC_004718.3;NP_828849.2;SARS-CoV_orf1ab;5024 | 0 | 0 |
| NC_004718.3;NP_828849.2;SARS-CoV_orf1ab;5025 | 0 | 0 |
| NC_004718.3;NP_828849.2;SARS-CoV_orf1ab;5026 | 0 | 0 |
| NC_004718.3;NP_828849.2;SARS-CoV_orf1ab;5027 | 0 | 0 |
| NC_004718.3;NP_828849.2;SARS-CoV_orf1ab;5028 | 0 | 0 |

| | | |
|---|---|---|
| NC_004718.3;NP_828849.2;SARS-CoV_orf1ab;5029 | 0 | 0 |
| NC_004718.3;NP_828849.2;SARS-CoV_orf1ab;5030 | 0 | 0 |
| NC_004718.3;NP_828849.2;SARS-CoV_orf1ab;5031 | 0 | 0 |
| NC_004718.3;NP_828849.2;SARS-CoV_orf1ab;5032 | 0 | 0 |
| NC_004718.3;NP_828849.2;SARS-CoV_orf1ab;5033 | 0 | 0 |
| NC_004718.3;NP_828849.2;SARS-CoV_orf1ab;5034 | 0.05 | 0 |
| NC_004718.3;NP_828849.2;SARS-CoV_orf1ab;5035 | 0 | 0 |
| NC_004718.3;NP_828849.2;SARS-CoV_orf1ab;5036 | 0 | 0 |
| NC_004718.3;NP_828849.2;SARS-CoV_orf1ab;5037 | 0 | 0 |
| NC_004718.3;NP_828849.2;SARS-CoV_orf1ab;5038 | 0 | 0.025 |
| NC_004718.3;NP_828849.2;SARS-CoV_orf1ab;5039 | 0 | 0.025 |
| NC_004718.3;NP_828849.2;SARS-CoV_orf1ab;5040 | 0 | 0.025 |
| NC_004718.3;NP_828849.2;SARS-CoV_orf1ab;5041 | 0 | 0.025 |
| NC_004718.3;NP_828849.2;SARS-CoV_orf1ab;5042 | 0 | 0.05 |
| NC_004718.3;NP_828849.2;SARS-CoV_orf1ab;5043 | 0 | 0.025 |
| NC_004718.3;NP_828849.2;SARS-CoV_orf1ab;5044 | 0 | 0.025 |
| NC_004718.3;NP_828849.2;SARS-CoV_orf1ab;5045 | 0 | 0.05 |
| NC_004718.3;NP_828849.2;SARS-CoV_orf1ab;5046 | 0.05 | 0.025 |
| NC_004718.3;NP_828849.2;SARS-CoV_orf1ab;5047 | 0.1 | 0.05 |
| NC_004718.3;NP_828849.2;SARS-CoV_orf1ab;5048 | 0 | 0 |
| NC_004718.3;NP_828849.2;SARS-CoV_orf1ab;5049 | 0.05 | 0 |
| NC_004718.3;NP_828849.2;SARS-CoV_orf1ab;5050 | 0 | 0 |
| NC_004718.3;NP_828849.2;SARS-CoV_orf1ab;5051 | 0 | 0 |
| NC_004718.3;NP_828849.2;SARS-CoV_orf1ab;5052 | 0 | 0 |
| NC_004718.3;NP_828849.2;SARS-CoV_orf1ab;5053 | 0 | 0 |
| NC_004718.3;NP_828849.2;SARS-CoV_orf1ab;5054 | 0 | 0 |
| NC_004718.3;NP_828849.2;SARS-CoV_orf1ab;5055 | 0 | 0 |
| NC_004718.3;NP_828849.2;SARS-CoV_orf1ab;5056 | 0 | 0 |
| NC_004718.3;NP_828849.2;SARS-CoV_orf1ab;5057 | 0 | 0.025 |
| NC_004718.3;NP_828849.2;SARS-CoV_orf1ab;5058 | 0 | 0 |
| NC_004718.3;NP_828849.2;SARS-CoV_orf1ab;5059 | 0 | 0.025 |
| NC_004718.3;NP_828849.2;SARS-CoV_orf1ab;5060 | 0 | 0 |
| NC_004718.3;NP_828849.2;SARS-CoV_orf1ab;5061 | 0 | 0 |
| NC_004718.3;NP_828849.2;SARS-CoV_orf1ab;5062 | 0 | 0 |
| NC_004718.3;NP_828849.2;SARS-CoV_orf1ab;5063 | 0 | 0 |
| NC_004718.3;NP_828849.2;SARS-CoV_orf1ab;5064 | 0 | 0 |
| NC_004718.3;NP_828849.2;SARS-CoV_orf1ab;5065 | 0 | 0 |
| NC_004718.3;NP_828849.2;SARS-CoV_orf1ab;5066 | 0 | 0.025 |
| NC_004718.3;NP_828849.2;SARS-CoV_orf1ab;5067 | 0 | 0 |
| NC_004718.3;NP_828849.2;SARS-CoV_orf1ab;5068 | 0 | 0.025 |
| NC_004718.3;NP_828849.2;SARS-CoV_orf1ab;5069 | 0 | 0.025 |
| NC_004718.3;NP_828849.2;SARS-CoV_orf1ab;5070 | 0 | 0.025 |
| NC_004718.3;NP_828849.2;SARS-CoV_orf1ab;5071 | 0.05 | 0.025 |

| | | |
|---|---|---|
| NC_004718.3;NP_828849.2;SARS-CoV_orf1ab;5072 | 0 | 0 |
| NC_004718.3;NP_828849.2;SARS-CoV_orf1ab;5073 | 0 | 0.075 |
| NC_004718.3;NP_828849.2;SARS-CoV_orf1ab;5074 | 0.05 | 0.05 |
| NC_004718.3;NP_828849.2;SARS-CoV_orf1ab;5075 | 0 | 0.025 |
| NC_004718.3;NP_828849.2;SARS-CoV_orf1ab;5076 | 0 | 0.05 |
| NC_004718.3;NP_828849.2;SARS-CoV_orf1ab;5077 | 0 | 0.05 |
| NC_004718.3;NP_828849.2;SARS-CoV_orf1ab;5078 | 0.05 | 0.025 |
| NC_004718.3;NP_828849.2;SARS-CoV_orf1ab;5079 | 0 | 0 |
| NC_004718.3;NP_828849.2;SARS-CoV_orf1ab;5080 | 0 | 0 |
| NC_004718.3;NP_828849.2;SARS-CoV_orf1ab;5081 | 0 | 0 |
| NC_004718.3;NP_828849.2;SARS-CoV_orf1ab;5082 | 0 | 0 |
| NC_004718.3;NP_828849.2;SARS-CoV_orf1ab;5083 | 0 | 0 |
| NC_004718.3;NP_828849.2;SARS-CoV_orf1ab;5084 | 0 | 0 |
| NC_004718.3;NP_828849.2;SARS-CoV_orf1ab;5085 | 0 | 0 |
| NC_004718.3;NP_828849.2;SARS-CoV_orf1ab;5086 | 0 | 0 |
| NC_004718.3;NP_828849.2;SARS-CoV_orf1ab;5087 | 0 | 0 |
| NC_004718.3;NP_828849.2;SARS-CoV_orf1ab;5088 | 0 | 0 |
| NC_004718.3;NP_828849.2;SARS-CoV_orf1ab;5089 | 0 | 0 |
| NC_004718.3;NP_828849.2;SARS-CoV_orf1ab;5090 | 0.05 | 0 |
| NC_004718.3;NP_828849.2;SARS-CoV_orf1ab;5091 | 0 | 0 |
| NC_004718.3;NP_828849.2;SARS-CoV_orf1ab;5092 | 0 | 0 |
| NC_004718.3;NP_828849.2;SARS-CoV_orf1ab;5093 | 0 | 0 |
| NC_004718.3;NP_828849.2;SARS-CoV_orf1ab;5094 | 0 | 0.025 |
| NC_004718.3;NP_828849.2;SARS-CoV_orf1ab;5095 | 0 | 0 |
| NC_004718.3;NP_828849.2;SARS-CoV_orf1ab;5096 | 0 | 0 |
| NC_004718.3;NP_828849.2;SARS-CoV_orf1ab;5097 | 0 | 0 |
| NC_004718.3;NP_828849.2;SARS-CoV_orf1ab;5098 | 0 | 0 |
| NC_004718.3;NP_828849.2;SARS-CoV_orf1ab;5099 | 0 | 0 |
| NC_004718.3;NP_828849.2;SARS-CoV_orf1ab;5100 | 0 | 0 |
| NC_004718.3;NP_828849.2;SARS-CoV_orf1ab;5101 | 0.05 | 0 |
| NC_004718.3;NP_828849.2;SARS-CoV_orf1ab;5102 | 0 | 0 |
| NC_004718.3;NP_828849.2;SARS-CoV_orf1ab;5103 | 0 | 0 |
| NC_004718.3;NP_828849.2;SARS-CoV_orf1ab;5104 | 0 | 0 |
| NC_004718.3;NP_828849.2;SARS-CoV_orf1ab;5105 | 0 | 0 |
| NC_004718.3;NP_828849.2;SARS-CoV_orf1ab;5106 | 0 | 0 |
| NC_004718.3;NP_828849.2;SARS-CoV_orf1ab;5107 | 0 | 0 |
| NC_004718.3;NP_828849.2;SARS-CoV_orf1ab;5108 | 0.05 | 0 |
| NC_004718.3;NP_828849.2;SARS-CoV_orf1ab;5109 | 0 | 0 |
| NC_004718.3;NP_828849.2;SARS-CoV_orf1ab;5110 | 0 | 0 |
| NC_004718.3;NP_828849.2;SARS-CoV_orf1ab;5111 | 0 | 0 |
| NC_004718.3;NP_828849.2;SARS-CoV_orf1ab;5112 | 0 | 0 |
| NC_004718.3;NP_828849.2;SARS-CoV_orf1ab;5113 | 0 | 0 |
| NC_004718.3;NP_828849.2;SARS-CoV_orf1ab;5114 | 0 | 0 |

| | | |
|---|---|---|
| NC_004718.3;NP_828849.2;SARS-CoV_orf1ab;5115 | 0 | 0 |
| NC_004718.3;NP_828849.2;SARS-CoV_orf1ab;5116 | 0.05 | 0 |
| NC_004718.3;NP_828849.2;SARS-CoV_orf1ab;5117 | 0 | 0 |
| NC_004718.3;NP_828849.2;SARS-CoV_orf1ab;5118 | 0 | 0 |
| NC_004718.3;NP_828849.2;SARS-CoV_orf1ab;5119 | 0 | 0 |
| NC_004718.3;NP_828849.2;SARS-CoV_orf1ab;5120 | 0 | 0 |
| NC_004718.3;NP_828849.2;SARS-CoV_orf1ab;5121 | 0 | 0 |
| NC_004718.3;NP_828849.2;SARS-CoV_orf1ab;5122 | 0 | 0 |
| NC_004718.3;NP_828849.2;SARS-CoV_orf1ab;5123 | 0 | 0 |
| NC_004718.3;NP_828849.2;SARS-CoV_orf1ab;5124 | 0 | 0 |
| NC_004718.3;NP_828849.2;SARS-CoV_orf1ab;5125 | 0 | 0 |
| NC_004718.3;NP_828849.2;SARS-CoV_orf1ab;5126 | 0.05 | 0.025 |
| NC_004718.3;NP_828849.2;SARS-CoV_orf1ab;5127 | 0.05 | 0 |
| NC_004718.3;NP_828849.2;SARS-CoV_orf1ab;5128 | 0 | 0.025 |
| NC_004718.3;NP_828849.2;SARS-CoV_orf1ab;5129 | 0 | 0.025 |
| NC_004718.3;NP_828849.2;SARS-CoV_orf1ab;5130 | 0 | 0.025 |
| NC_004718.3;NP_828849.2;SARS-CoV_orf1ab;5131 | 0.1 | 0.075 |
| NC_004718.3;NP_828849.2;SARS-CoV_orf1ab;5132 | 0.05 | 0 |
| NC_004718.3;NP_828849.2;SARS-CoV_orf1ab;5133 | 0 | 0 |
| NC_004718.3;NP_828849.2;SARS-CoV_orf1ab;5134 | 0 | 0 |
| NC_004718.3;NP_828849.2;SARS-CoV_orf1ab;5135 | 0 | 0 |
| NC_004718.3;NP_828849.2;SARS-CoV_orf1ab;5136 | 0 | 0 |
| NC_004718.3;NP_828849.2;SARS-CoV_orf1ab;5137 | 0 | 0 |
| NC_004718.3;NP_828849.2;SARS-CoV_orf1ab;5138 | 0 | 0 |
| NC_004718.3;NP_828849.2;SARS-CoV_orf1ab;5139 | 0 | 0 |
| NC_004718.3;NP_828849.2;SARS-CoV_orf1ab;5140 | 0 | 0 |
| NC_004718.3;NP_828849.2;SARS-CoV_orf1ab;5141 | 0 | 0 |
| NC_004718.3;NP_828849.2;SARS-CoV_orf1ab;5142 | 0 | 0 |
| NC_004718.3;NP_828849.2;SARS-CoV_orf1ab;5143 | 0 | 0 |
| NC_004718.3;NP_828849.2;SARS-CoV_orf1ab;5144 | 0 | 0 |
| NC_004718.3;NP_828849.2;SARS-CoV_orf1ab;5145 | 0 | 0 |
| NC_004718.3;NP_828849.2;SARS-CoV_orf1ab;5146 | 0 | 0 |
| NC_004718.3;NP_828849.2;SARS-CoV_orf1ab;5147 | 0 | 0 |
| NC_004718.3;NP_828849.2;SARS-CoV_orf1ab;5148 | 0 | 0 |
| NC_004718.3;NP_828849.2;SARS-CoV_orf1ab;5149 | 0 | 0 |
| NC_004718.3;NP_828849.2;SARS-CoV_orf1ab;5150 | 0 | 0 |
| NC_004718.3;NP_828849.2;SARS-CoV_orf1ab;5151 | 0 | 0 |
| NC_004718.3;NP_828849.2;SARS-CoV_orf1ab;5152 | 0.05 | 0 |
| NC_004718.3;NP_828849.2;SARS-CoV_orf1ab;5153 | 0 | 0 |
| NC_004718.3;NP_828849.2;SARS-CoV_orf1ab;5154 | 0 | 0 |
| NC_004718.3;NP_828849.2;SARS-CoV_orf1ab;5155 | 0 | 0 |
| NC_004718.3;NP_828849.2;SARS-CoV_orf1ab;5156 | 0 | 0.025 |
| NC_004718.3;NP_828849.2;SARS-CoV_orf1ab;5157 | 0 | 0.025 |

| | | |
|---|---|---|
| NC_004718.3;NP_828849.2;SARS-CoV_orf1ab;5158 | 0 | 0.025 |
| NC_004718.3;NP_828849.2;SARS-CoV_orf1ab;5159 | 0 | 0.025 |
| NC_004718.3;NP_828849.2;SARS-CoV_orf1ab;5160 | 0 | 0.025 |
| NC_004718.3;NP_828849.2;SARS-CoV_orf1ab;5161 | 0.05 | 0.025 |
| NC_004718.3;NP_828849.2;SARS-CoV_orf1ab;5162 | 0 | 0.025 |
| NC_004718.3;NP_828849.2;SARS-CoV_orf1ab;5163 | 0.05 | 0.025 |
| NC_004718.3;NP_828849.2;SARS-CoV_orf1ab;5164 | 0 | 0.025 |
| NC_004718.3;NP_828849.2;SARS-CoV_orf1ab;5165 | 0 | 0.025 |
| NC_004718.3;NP_828849.2;SARS-CoV_orf1ab;5166 | 0.05 | 0.025 |
| NC_004718.3;NP_828849.2;SARS-CoV_orf1ab;5167 | 0.05 | 0.025 |
| NC_004718.3;NP_828849.2;SARS-CoV_orf1ab;5168 | 0.05 | 0.025 |
| NC_004718.3;NP_828849.2;SARS-CoV_orf1ab;5169 | 0.05 | 0.025 |
| NC_004718.3;NP_828849.2;SARS-CoV_orf1ab;5170 | 0 | 0 |
| NC_004718.3;NP_828849.2;SARS-CoV_orf1ab;5171 | 0 | 0 |
| NC_004718.3;NP_828849.2;SARS-CoV_orf1ab;5172 | 0 | 0 |
| NC_004718.3;NP_828849.2;SARS-CoV_orf1ab;5173 | 0 | 0 |
| NC_004718.3;NP_828849.2;SARS-CoV_orf1ab;5174 | 0 | 0 |
| NC_004718.3;NP_828849.2;SARS-CoV_orf1ab;5175 | 0 | 0 |
| NC_004718.3;NP_828849.2;SARS-CoV_orf1ab;5176 | 0 | 0 |
| NC_004718.3;NP_828849.2;SARS-CoV_orf1ab;5177 | 0 | 0 |
| NC_004718.3;NP_828849.2;SARS-CoV_orf1ab;5178 | 0 | 0 |
| NC_004718.3;NP_828849.2;SARS-CoV_orf1ab;5179 | 0 | 0 |
| NC_004718.3;NP_828849.2;SARS-CoV_orf1ab;5180 | 0 | 0.025 |
| NC_004718.3;NP_828849.2;SARS-CoV_orf1ab;5181 | 0 | 0.025 |
| NC_004718.3;NP_828849.2;SARS-CoV_orf1ab;5182 | 0 | 0.025 |
| NC_004718.3;NP_828849.2;SARS-CoV_orf1ab;5183 | 0.05 | 0 |
| NC_004718.3;NP_828849.2;SARS-CoV_orf1ab;5184 | 0.05 | 0 |
| NC_004718.3;NP_828849.2;SARS-CoV_orf1ab;5185 | 0 | 0 |
| NC_004718.3;NP_828849.2;SARS-CoV_orf1ab;5186 | 0.05 | 0 |
| NC_004718.3;NP_828849.2;SARS-CoV_orf1ab;5187 | 0 | 0 |
| NC_004718.3;NP_828849.2;SARS-CoV_orf1ab;5188 | 0.1 | 0.1 |
| NC_004718.3;NP_828849.2;SARS-CoV_orf1ab;5189 | 0 | 0.025 |
| NC_004718.3;NP_828849.2;SARS-CoV_orf1ab;5190 | 0 | 0.025 |
| NC_004718.3;NP_828849.2;SARS-CoV_orf1ab;5191 | 0 | 0 |
| NC_004718.3;NP_828849.2;SARS-CoV_orf1ab;5192 | 0 | 0 |
| NC_004718.3;NP_828849.2;SARS-CoV_orf1ab;5193 | 0 | 0 |
| NC_004718.3;NP_828849.2;SARS-CoV_orf1ab;5194 | 0 | 0 |
| NC_004718.3;NP_828849.2;SARS-CoV_orf1ab;5195 | 0 | 0 |
| NC_004718.3;NP_828849.2;SARS-CoV_orf1ab;5196 | 0 | 0 |
| NC_004718.3;NP_828849.2;SARS-CoV_orf1ab;5197 | 0 | 0 |
| NC_004718.3;NP_828849.2;SARS-CoV_orf1ab;5198 | 0 | 0 |
| NC_004718.3;NP_828849.2;SARS-CoV_orf1ab;5199 | 0 | 0 |
| NC_004718.3;NP_828849.2;SARS-CoV_orf1ab;5200 | 0 | 0 |

| | | |
|---|---|---|
| NC_004718.3;NP_828849.2;SARS-CoV_orf1ab;5201 | 0 | 0 |
| NC_004718.3;NP_828849.2;SARS-CoV_orf1ab;5202 | 0 | 0 |
| NC_004718.3;NP_828849.2;SARS-CoV_orf1ab;5203 | 0 | 0 |
| NC_004718.3;NP_828849.2;SARS-CoV_orf1ab;5204 | 0 | 0 |
| NC_004718.3;NP_828849.2;SARS-CoV_orf1ab;5205 | 0 | 0 |
| NC_004718.3;NP_828849.2;SARS-CoV_orf1ab;5206 | 0.05 | 0 |
| NC_004718.3;NP_828849.2;SARS-CoV_orf1ab;5207 | 0 | 0 |
| NC_004718.3;NP_828849.2;SARS-CoV_orf1ab;5208 | 0 | 0 |
| NC_004718.3;NP_828849.2;SARS-CoV_orf1ab;5209 | 0 | 0 |
| NC_004718.3;NP_828849.2;SARS-CoV_orf1ab;5210 | 0 | 0 |
| NC_004718.3;NP_828849.2;SARS-CoV_orf1ab;5211 | 0 | 0 |
| NC_004718.3;NP_828849.2;SARS-CoV_orf1ab;5212 | 0 | 0 |
| NC_004718.3;NP_828849.2;SARS-CoV_orf1ab;5213 | 0 | 0 |
| NC_004718.3;NP_828849.2;SARS-CoV_orf1ab;5214 | 0 | 0 |
| NC_004718.3;NP_828849.2;SARS-CoV_orf1ab;5215 | 0 | 0 |
| NC_004718.3;NP_828849.2;SARS-CoV_orf1ab;5216 | 0 | 0 |
| NC_004718.3;NP_828849.2;SARS-CoV_orf1ab;5217 | 0 | 0 |
| NC_004718.3;NP_828849.2;SARS-CoV_orf1ab;5218 | 0 | 0 |
| NC_004718.3;NP_828849.2;SARS-CoV_orf1ab;5219 | 0 | 0 |
| NC_004718.3;NP_828849.2;SARS-CoV_orf1ab;5220 | 0 | 0 |
| NC_004718.3;NP_828849.2;SARS-CoV_orf1ab;5221 | 0 | 0 |
| NC_004718.3;NP_828849.2;SARS-CoV_orf1ab;5222 | 0 | 0 |
| NC_004718.3;NP_828849.2;SARS-CoV_orf1ab;5223 | 0 | 0 |
| NC_004718.3;NP_828849.2;SARS-CoV_orf1ab;5224 | 0 | 0 |
| NC_004718.3;NP_828849.2;SARS-CoV_orf1ab;5225 | 0 | 0 |
| NC_004718.3;NP_828849.2;SARS-CoV_orf1ab;5226 | 0 | 0 |
| NC_004718.3;NP_828849.2;SARS-CoV_orf1ab;5227 | 0 | 0 |
| NC_004718.3;NP_828849.2;SARS-CoV_orf1ab;5228 | 0 | 0 |
| NC_004718.3;NP_828849.2;SARS-CoV_orf1ab;5229 | 0 | 0 |
| NC_004718.3;NP_828849.2;SARS-CoV_orf1ab;5230 | 0 | 0 |
| NC_004718.3;NP_828849.2;SARS-CoV_orf1ab;5231 | 0 | 0 |
| NC_004718.3;NP_828849.2;SARS-CoV_orf1ab;5232 | 0 | 0 |
| NC_004718.3;NP_828849.2;SARS-CoV_orf1ab;5233 | 0 | 0 |
| NC_004718.3;NP_828849.2;SARS-CoV_orf1ab;5234 | 0 | 0 |
| NC_004718.3;NP_828849.2;SARS-CoV_orf1ab;5235 | 0 | 0 |
| NC_004718.3;NP_828849.2;SARS-CoV_orf1ab;5236 | 0 | 0 |
| NC_004718.3;NP_828849.2;SARS-CoV_orf1ab;5237 | 0 | 0 |
| NC_004718.3;NP_828849.2;SARS-CoV_orf1ab;5238 | 0 | 0 |
| NC_004718.3;NP_828849.2;SARS-CoV_orf1ab;5239 | 0 | 0 |
| NC_004718.3;NP_828849.2;SARS-CoV_orf1ab;5240 | 0 | 0 |
| NC_004718.3;NP_828849.2;SARS-CoV_orf1ab;5241 | 0 | 0 |
| NC_004718.3;NP_828849.2;SARS-CoV_orf1ab;5242 | 0 | 0 |
| NC_004718.3;NP_828849.2;SARS-CoV_orf1ab;5243 | 0 | 0 |

| | | |
|---|---|---|
| NC_004718.3;NP_828849.2;SARS-CoV_orf1ab;5244 | 0 | 0 |
| NC_004718.3;NP_828849.2;SARS-CoV_orf1ab;5245 | 0 | 0 |
| NC_004718.3;NP_828849.2;SARS-CoV_orf1ab;5246 | 0 | 0 |
| NC_004718.3;NP_828849.2;SARS-CoV_orf1ab;5247 | 0 | 0 |
| NC_004718.3;NP_828849.2;SARS-CoV_orf1ab;5248 | 0 | 0 |
| NC_004718.3;NP_828849.2;SARS-CoV_orf1ab;5249 | 0 | 0 |
| NC_004718.3;NP_828849.2;SARS-CoV_orf1ab;5250 | 0 | 0 |
| NC_004718.3;NP_828849.2;SARS-CoV_orf1ab;5251 | 0 | 0 |
| NC_004718.3;NP_828849.2;SARS-CoV_orf1ab;5252 | 0 | 0 |
| NC_004718.3;NP_828849.2;SARS-CoV_orf1ab;5253 | 0 | 0 |
| NC_004718.3;NP_828849.2;SARS-CoV_orf1ab;5254 | 0 | 0 |
| NC_004718.3;NP_828849.2;SARS-CoV_orf1ab;5255 | 0 | 0 |
| NC_004718.3;NP_828849.2;SARS-CoV_orf1ab;5256 | 0 | 0 |
| NC_004718.3;NP_828849.2;SARS-CoV_orf1ab;5257 | 0 | 0 |
| NC_004718.3;NP_828849.2;SARS-CoV_orf1ab;5258 | 0 | 0.075 |
| NC_004718.3;NP_828849.2;SARS-CoV_orf1ab;5259 | 0 | 0.025 |
| NC_004718.3;NP_828849.2;SARS-CoV_orf1ab;5260 | 0 | 0 |
| NC_004718.3;NP_828849.2;SARS-CoV_orf1ab;5261 | 0 | 0 |
| NC_004718.3;NP_828849.2;SARS-CoV_orf1ab;5262 | 0 | 0 |
| NC_004718.3;NP_828849.2;SARS-CoV_orf1ab;5263 | 0 | 0 |
| NC_004718.3;NP_828849.2;SARS-CoV_orf1ab;5264 | 0 | 0 |
| NC_004718.3;NP_828849.2;SARS-CoV_orf1ab;5265 | 0 | 0 |
| NC_004718.3;NP_828849.2;SARS-CoV_orf1ab;5266 | 0 | 0 |
| NC_004718.3;NP_828849.2;SARS-CoV_orf1ab;5267 | 0 | 0 |
| NC_004718.3;NP_828849.2;SARS-CoV_orf1ab;5268 | 0 | 0 |
| NC_004718.3;NP_828849.2;SARS-CoV_orf1ab;5269 | 0 | 0 |
| NC_004718.3;NP_828849.2;SARS-CoV_orf1ab;5270 | 0 | 0 |
| NC_004718.3;NP_828849.2;SARS-CoV_orf1ab;5271 | 0 | 0 |
| NC_004718.3;NP_828849.2;SARS-CoV_orf1ab;5272 | 0 | 0 |
| NC_004718.3;NP_828849.2;SARS-CoV_orf1ab;5273 | 0 | 0.025 |
| NC_004718.3;NP_828849.2;SARS-CoV_orf1ab;5274 | 0 | 0 |
| NC_004718.3;NP_828849.2;SARS-CoV_orf1ab;5275 | 0 | 0 |
| NC_004718.3;NP_828849.2;SARS-CoV_orf1ab;5276 | 0 | 0 |
| NC_004718.3;NP_828849.2;SARS-CoV_orf1ab;5277 | 0 | 0 |
| NC_004718.3;NP_828849.2;SARS-CoV_orf1ab;5278 | 0 | 0 |
| NC_004718.3;NP_828849.2;SARS-CoV_orf1ab;5279 | 0.05 | 0 |
| NC_004718.3;NP_828849.2;SARS-CoV_orf1ab;5280 | 0 | 0 |
| NC_004718.3;NP_828849.2;SARS-CoV_orf1ab;5281 | 0 | 0 |
| NC_004718.3;NP_828849.2;SARS-CoV_orf1ab;5282 | 0 | 0 |
| NC_004718.3;NP_828849.2;SARS-CoV_orf1ab;5283 | 0 | 0 |
| NC_004718.3;NP_828849.2;SARS-CoV_orf1ab;5284 | 0 | 0 |
| NC_004718.3;NP_828849.2;SARS-CoV_orf1ab;5285 | 0 | 0 |
| NC_004718.3;NP_828849.2;SARS-CoV_orf1ab;5286 | 0 | 0 |

| | | |
|---|---|---|
| NC_004718.3;NP_828849.2;SARS-CoV_orf1ab;5287 | 0 | 0 |
| NC_004718.3;NP_828849.2;SARS-CoV_orf1ab;5288 | 0 | 0 |
| NC_004718.3;NP_828849.2;SARS-CoV_orf1ab;5289 | 0 | 0 |
| NC_004718.3;NP_828849.2;SARS-CoV_orf1ab;5290 | 0 | 0 |
| NC_004718.3;NP_828849.2;SARS-CoV_orf1ab;5291 | 0 | 0 |
| NC_004718.3;NP_828849.2;SARS-CoV_orf1ab;5292 | 0 | 0 |
| NC_004718.3;NP_828849.2;SARS-CoV_orf1ab;5293 | 0 | 0 |
| NC_004718.3;NP_828849.2;SARS-CoV_orf1ab;5294 | 0 | 0 |
| NC_004718.3;NP_828849.2;SARS-CoV_orf1ab;5295 | 0 | 0 |
| NC_004718.3;NP_828849.2;SARS-CoV_orf1ab;5296 | 0 | 0 |
| NC_004718.3;NP_828849.2;SARS-CoV_orf1ab;5297 | 0 | 0 |
| NC_004718.3;NP_828849.2;SARS-CoV_orf1ab;5298 | 0 | 0 |
| NC_004718.3;NP_828849.2;SARS-CoV_orf1ab;5299 | 0 | 0 |
| NC_004718.3;NP_828849.2;SARS-CoV_orf1ab;5300 | 0 | 0 |
| NC_004718.3;NP_828849.2;SARS-CoV_orf1ab;5301 | 0 | 0 |
| NC_004718.3;NP_828849.2;SARS-CoV_orf1ab;5302 | 0 | 0 |
| NC_004718.3;NP_828849.2;SARS-CoV_orf1ab;5303 | 0 | 0 |
| NC_004718.3;NP_828849.2;SARS-CoV_orf1ab;5304 | 0 | 0 |
| NC_004718.3;NP_828849.2;SARS-CoV_orf1ab;5305 | 0 | 0 |
| NC_004718.3;NP_828849.2;SARS-CoV_orf1ab;5306 | 0 | 0 |
| NC_004718.3;NP_828849.2;SARS-CoV_orf1ab;5307 | 0 | 0 |
| NC_004718.3;NP_828849.2;SARS-CoV_orf1ab;5308 | 0 | 0 |
| NC_004718.3;NP_828849.2;SARS-CoV_orf1ab;5309 | 0 | 0 |
| NC_004718.3;NP_828849.2;SARS-CoV_orf1ab;5310 | 0 | 0 |
| NC_004718.3;NP_828849.2;SARS-CoV_orf1ab;5311 | 0 | 0 |
| NC_004718.3;NP_828849.2;SARS-CoV_orf1ab;5312 | 0 | 0 |
| NC_004718.3;NP_828849.2;SARS-CoV_orf1ab;5313 | 0 | 0 |
| NC_004718.3;NP_828849.2;SARS-CoV_orf1ab;5314 | 0 | 0 |
| NC_004718.3;NP_828849.2;SARS-CoV_orf1ab;5315 | 0 | 0 |
| NC_004718.3;NP_828849.2;SARS-CoV_orf1ab;5316 | 0 | 0 |
| NC_004718.3;NP_828849.2;SARS-CoV_orf1ab;5317 | 0 | 0 |
| NC_004718.3;NP_828849.2;SARS-CoV_orf1ab;5318 | 0 | 0 |
| NC_004718.3;NP_828849.2;SARS-CoV_orf1ab;5319 | 0 | 0 |
| NC_004718.3;NP_828849.2;SARS-CoV_orf1ab;5320 | 0 | 0 |
| NC_004718.3;NP_828849.2;SARS-CoV_orf1ab;5321 | 0 | 0 |
| NC_004718.3;NP_828849.2;SARS-CoV_orf1ab;5322 | 0.05 | 0 |
| NC_004718.3;NP_828849.2;SARS-CoV_orf1ab;5323 | 0 | 0 |
| NC_004718.3;NP_828849.2;SARS-CoV_orf1ab;5324 | 0 | 0 |
| NC_004718.3;NP_828849.2;SARS-CoV_orf1ab;5325 | 0 | 0 |
| NC_004718.3;NP_828849.2;SARS-CoV_orf1ab;5326 | 0 | 0 |
| NC_004718.3;NP_828849.2;SARS-CoV_orf1ab;5327 | 0 | 0 |
| NC_004718.3;NP_828849.2;SARS-CoV_orf1ab;5328 | 0 | 0 |
| NC_004718.3;NP_828849.2;SARS-CoV_orf1ab;5329 | 0 | 0 |

| | | |
|---|---|---|
| NC_004718.3;NP_828849.2;SARS-CoV_orf1ab;5330 | 0 | 0 |
| NC_004718.3;NP_828849.2;SARS-CoV_orf1ab;5331 | 0 | 0 |
| NC_004718.3;NP_828849.2;SARS-CoV_orf1ab;5332 | 0 | 0.025 |
| NC_004718.3;NP_828849.2;SARS-CoV_orf1ab;5333 | 0 | 0 |
| NC_004718.3;NP_828849.2;SARS-CoV_orf1ab;5334 | 0 | 0 |
| NC_004718.3;NP_828849.2;SARS-CoV_orf1ab;5335 | 0 | 0 |
| NC_004718.3;NP_828849.2;SARS-CoV_orf1ab;5336 | 0 | 0 |
| NC_004718.3;NP_828849.2;SARS-CoV_orf1ab;5337 | 0 | 0 |
| NC_004718.3;NP_828849.2;SARS-CoV_orf1ab;5338 | 0 | 0 |
| NC_004718.3;NP_828849.2;SARS-CoV_orf1ab;5339 | 0 | 0 |
| NC_004718.3;NP_828849.2;SARS-CoV_orf1ab;5340 | 0 | 0 |
| NC_004718.3;NP_828849.2;SARS-CoV_orf1ab;5341 | 0 | 0 |
| NC_004718.3;NP_828849.2;SARS-CoV_orf1ab;5342 | 0 | 0 |
| NC_004718.3;NP_828849.2;SARS-CoV_orf1ab;5343 | 0 | 0.075 |
| NC_004718.3;NP_828849.2;SARS-CoV_orf1ab;5344 | 0 | 0 |
| NC_004718.3;NP_828849.2;SARS-CoV_orf1ab;5345 | 0 | 0 |
| NC_004718.3;NP_828849.2;SARS-CoV_orf1ab;5346 | 0 | 0 |
| NC_004718.3;NP_828849.2;SARS-CoV_orf1ab;5347 | 0.05 | 0 |
| NC_004718.3;NP_828849.2;SARS-CoV_orf1ab;5348 | 0.05 | 0 |
| NC_004718.3;NP_828849.2;SARS-CoV_orf1ab;5349 | 0.05 | 0 |
| NC_004718.3;NP_828849.2;SARS-CoV_orf1ab;5350 | 0.05 | 0 |
| NC_004718.3;NP_828849.2;SARS-CoV_orf1ab;5351 | 0 | 0 |
| NC_004718.3;NP_828849.2;SARS-CoV_orf1ab;5352 | 0 | 0 |
| NC_004718.3;NP_828849.2;SARS-CoV_orf1ab;5353 | 0 | 0 |
| NC_004718.3;NP_828849.2;SARS-CoV_orf1ab;5354 | 0 | 0 |
| NC_004718.3;NP_828849.2;SARS-CoV_orf1ab;5355 | 0.05 | 0 |
| NC_004718.3;NP_828849.2;SARS-CoV_orf1ab;5356 | 0 | 0 |
| NC_004718.3;NP_828849.2;SARS-CoV_orf1ab;5357 | 0 | 0 |
| NC_004718.3;NP_828849.2;SARS-CoV_orf1ab;5358 | 0 | 0 |
| NC_004718.3;NP_828849.2;SARS-CoV_orf1ab;5359 | 0 | 0 |
| NC_004718.3;NP_828849.2;SARS-CoV_orf1ab;5360 | 0 | 0 |
| NC_004718.3;NP_828849.2;SARS-CoV_orf1ab;5361 | 0 | 0 |
| NC_004718.3;NP_828849.2;SARS-CoV_orf1ab;5362 | 0 | 0 |
| NC_004718.3;NP_828849.2;SARS-CoV_orf1ab;5363 | 0 | 0 |
| NC_004718.3;NP_828849.2;SARS-CoV_orf1ab;5364 | 0 | 0 |
| NC_004718.3;NP_828849.2;SARS-CoV_orf1ab;5365 | 0 | 0 |
| NC_004718.3;NP_828849.2;SARS-CoV_orf1ab;5366 | 0 | 0 |
| NC_004718.3;NP_828849.2;SARS-CoV_orf1ab;5367 | 0 | 0 |
| NC_004718.3;NP_828849.2;SARS-CoV_orf1ab;5368 | 0 | 0 |
| NC_004718.3;NP_828849.2;SARS-CoV_orf1ab;5369 | 0 | 0 |
| NC_004718.3;NP_828849.2;SARS-CoV_orf1ab;5370 | 0 | 0 |
| NC_004718.3;NP_828849.2;SARS-CoV_orf1ab;5371 | 0 | 0 |
| NC_004718.3;NP_828849.2;SARS-CoV_orf1ab;5372 | 0 | 0 |

| | | |
|---|---|---|
| NC_004718.3;NP_828849.2;SARS-CoV_orf1ab;5373 | 0 | 0 |
| NC_004718.3;NP_828849.2;SARS-CoV_orf1ab;5374 | 0 | 0 |
| NC_004718.3;NP_828849.2;SARS-CoV_orf1ab;5375 | 0 | 0 |
| NC_004718.3;NP_828849.2;SARS-CoV_orf1ab;5376 | 0 | 0 |
| NC_004718.3;NP_828849.2;SARS-CoV_orf1ab;5377 | 0 | 0 |
| NC_004718.3;NP_828849.2;SARS-CoV_orf1ab;5378 | 0 | 0.025 |
| NC_004718.3;NP_828849.2;SARS-CoV_orf1ab;5379 | 0 | 0 |
| NC_004718.3;NP_828849.2;SARS-CoV_orf1ab;5380 | 0 | 0 |
| NC_004718.3;NP_828849.2;SARS-CoV_orf1ab;5381 | 0 | 0 |
| NC_004718.3;NP_828849.2;SARS-CoV_orf1ab;5382 | 0 | 0 |
| NC_004718.3;NP_828849.2;SARS-CoV_orf1ab;5383 | 0 | 0 |
| NC_004718.3;NP_828849.2;SARS-CoV_orf1ab;5384 | 0 | 0 |
| NC_004718.3;NP_828849.2;SARS-CoV_orf1ab;5385 | 0 | 0 |
| NC_004718.3;NP_828849.2;SARS-CoV_orf1ab;5386 | 0 | 0 |
| NC_004718.3;NP_828849.2;SARS-CoV_orf1ab;5387 | 0 | 0 |
| NC_004718.3;NP_828849.2;SARS-CoV_orf1ab;5388 | 0 | 0 |
| NC_004718.3;NP_828849.2;SARS-CoV_orf1ab;5389 | 0 | 0 |
| NC_004718.3;NP_828849.2;SARS-CoV_orf1ab;5390 | 0 | 0 |
| NC_004718.3;NP_828849.2;SARS-CoV_orf1ab;5391 | 0 | 0 |
| NC_004718.3;NP_828849.2;SARS-CoV_orf1ab;5392 | 0.05 | 0 |
| NC_004718.3;NP_828849.2;SARS-CoV_orf1ab;5393 | 0.15 | 0.125 |
| NC_004718.3;NP_828849.2;SARS-CoV_orf1ab;5394 | 0.15 | 0.125 |
| NC_004718.3;NP_828849.2;SARS-CoV_orf1ab;5395 | 0.2 | 0.15 |
| NC_004718.3;NP_828849.2;SARS-CoV_orf1ab;5396 | 0.1 | 0.175 |
| NC_004718.3;NP_828849.2;SARS-CoV_orf1ab;5397 | 0.05 | 0.1 |
| NC_004718.3;NP_828849.2;SARS-CoV_orf1ab;5398 | 0.05 | 0.075 |
| NC_004718.3;NP_828849.2;SARS-CoV_orf1ab;5399 | 0.05 | 0.2 |
| NC_004718.3;NP_828849.2;SARS-CoV_orf1ab;5400 | 0.05 | 0.075 |
| NC_004718.3;NP_828849.2;SARS-CoV_orf1ab;5401 | 0.05 | 0.025 |
| NC_004718.3;NP_828849.2;SARS-CoV_orf1ab;5402 | 0.05 | 0.05 |
| NC_004718.3;NP_828849.2;SARS-CoV_orf1ab;5403 | 0 | 0.025 |
| NC_004718.3;NP_828849.2;SARS-CoV_orf1ab;5404 | 0 | 0 |
| NC_004718.3;NP_828849.2;SARS-CoV_orf1ab;5405 | 0 | 0 |
| NC_004718.3;NP_828849.2;SARS-CoV_orf1ab;5406 | 0 | 0 |
| NC_004718.3;NP_828849.2;SARS-CoV_orf1ab;5407 | 0 | 0 |
| NC_004718.3;NP_828849.2;SARS-CoV_orf1ab;5408 | 0 | 0 |
| NC_004718.3;NP_828849.2;SARS-CoV_orf1ab;5409 | 0 | 0 |
| NC_004718.3;NP_828849.2;SARS-CoV_orf1ab;5410 | 0 | 0 |
| NC_004718.3;NP_828849.2;SARS-CoV_orf1ab;5411 | 0 | 0 |
| NC_004718.3;NP_828849.2;SARS-CoV_orf1ab;5412 | 0 | 0 |
| NC_004718.3;NP_828849.2;SARS-CoV_orf1ab;5413 | 0 | 0 |
| NC_004718.3;NP_828849.2;SARS-CoV_orf1ab;5414 | 0 | 0 |
| NC_004718.3;NP_828849.2;SARS-CoV_orf1ab;5415 | 0 | 0 |

| | | |
|---|---|---|
| NC_004718.3;NP_828849.2;SARS-CoV_orf1ab;5416 | 0 | 0 |
| NC_004718.3;NP_828849.2;SARS-CoV_orf1ab;5417 | 0 | 0 |
| NC_004718.3;NP_828849.2;SARS-CoV_orf1ab;5418 | 0.05 | 0 |
| NC_004718.3;NP_828849.2;SARS-CoV_orf1ab;5419 | 0 | 0 |
| NC_004718.3;NP_828849.2;SARS-CoV_orf1ab;5420 | 0 | 0 |
| NC_004718.3;NP_828849.2;SARS-CoV_orf1ab;5421 | 0 | 0 |
| NC_004718.3;NP_828849.2;SARS-CoV_orf1ab;5422 | 0 | 0 |
| NC_004718.3;NP_828849.2;SARS-CoV_orf1ab;5423 | 0 | 0 |
| NC_004718.3;NP_828849.2;SARS-CoV_orf1ab;5424 | 0 | 0 |
| NC_004718.3;NP_828849.2;SARS-CoV_orf1ab;5425 | 0 | 0 |
| NC_004718.3;NP_828849.2;SARS-CoV_orf1ab;5426 | 0 | 0.025 |
| NC_004718.3;NP_828849.2;SARS-CoV_orf1ab;5427 | 0 | 0 |
| NC_004718.3;NP_828849.2;SARS-CoV_orf1ab;5428 | 0 | 0 |
| NC_004718.3;NP_828849.2;SARS-CoV_orf1ab;5429 | 0.05 | 0 |
| NC_004718.3;NP_828849.2;SARS-CoV_orf1ab;5430 | 0.05 | 0 |
| NC_004718.3;NP_828849.2;SARS-CoV_orf1ab;5431 | 0.05 | 0 |
| NC_004718.3;NP_828849.2;SARS-CoV_orf1ab;5432 | 0 | 0 |
| NC_004718.3;NP_828849.2;SARS-CoV_orf1ab;5433 | 0 | 0 |
| NC_004718.3;NP_828849.2;SARS-CoV_orf1ab;5434 | 0 | 0 |
| NC_004718.3;NP_828849.2;SARS-CoV_orf1ab;5435 | 0 | 0 |
| NC_004718.3;NP_828849.2;SARS-CoV_orf1ab;5436 | 0.05 | 0.025 |
| NC_004718.3;NP_828849.2;SARS-CoV_orf1ab;5437 | 0 | 0 |
| NC_004718.3;NP_828849.2;SARS-CoV_orf1ab;5438 | 0 | 0.025 |
| NC_004718.3;NP_828849.2;SARS-CoV_orf1ab;5439 | 0 | 0 |
| NC_004718.3;NP_828849.2;SARS-CoV_orf1ab;5440 | 0 | 0 |
| NC_004718.3;NP_828849.2;SARS-CoV_orf1ab;5441 | 0 | 0 |
| NC_004718.3;NP_828849.2;SARS-CoV_orf1ab;5442 | 0 | 0 |
| NC_004718.3;NP_828849.2;SARS-CoV_orf1ab;5443 | 0.05 | 0 |
| NC_004718.3;NP_828849.2;SARS-CoV_orf1ab;5444 | 0 | 0 |
| NC_004718.3;NP_828849.2;SARS-CoV_orf1ab;5445 | 0 | 0 |
| NC_004718.3;NP_828849.2;SARS-CoV_orf1ab;5446 | 0 | 0 |
| NC_004718.3;NP_828849.2;SARS-CoV_orf1ab;5447 | 0.05 | 0 |
| NC_004718.3;NP_828849.2;SARS-CoV_orf1ab;5448 | 0.05 | 0 |
| NC_004718.3;NP_828849.2;SARS-CoV_orf1ab;5449 | 0.05 | 0 |
| NC_004718.3;NP_828849.2;SARS-CoV_orf1ab;5450 | 0.05 | 0.05 |
| NC_004718.3;NP_828849.2;SARS-CoV_orf1ab;5451 | 0.05 | 0.05 |
| NC_004718.3;NP_828849.2;SARS-CoV_orf1ab;5452 | 0.05 | 0 |
| NC_004718.3;NP_828849.2;SARS-CoV_orf1ab;5453 | 0 | 0.05 |
| NC_004718.3;NP_828849.2;SARS-CoV_orf1ab;5454 | 0 | 0 |
| NC_004718.3;NP_828849.2;SARS-CoV_orf1ab;5455 | 0 | 0 |
| NC_004718.3;NP_828849.2;SARS-CoV_orf1ab;5456 | 0 | 0 |
| NC_004718.3;NP_828849.2;SARS-CoV_orf1ab;5457 | 0 | 0.05 |
| NC_004718.3;NP_828849.2;SARS-CoV_orf1ab;5458 | 0 | 0.05 |

| | | |
|---|---|---|
| NC_004718.3;NP_828849.2;SARS-CoV_orf1ab;5459 | 0 | 0.05 |
| NC_004718.3;NP_828849.2;SARS-CoV_orf1ab;5460 | 0.05 | 0.075 |
| NC_004718.3;NP_828849.2;SARS-CoV_orf1ab;5461 | 0 | 0 |
| NC_004718.3;NP_828849.2;SARS-CoV_orf1ab;5462 | 0 | 0 |
| NC_004718.3;NP_828849.2;SARS-CoV_orf1ab;5463 | 0 | 0 |
| NC_004718.3;NP_828849.2;SARS-CoV_orf1ab;5464 | 0 | 0 |
| NC_004718.3;NP_828849.2;SARS-CoV_orf1ab;5465 | 0 | 0 |
| NC_004718.3;NP_828849.2;SARS-CoV_orf1ab;5466 | 0 | 0 |
| NC_004718.3;NP_828849.2;SARS-CoV_orf1ab;5467 | 0 | 0 |
| NC_004718.3;NP_828849.2;SARS-CoV_orf1ab;5468 | 0 | 0 |
| NC_004718.3;NP_828849.2;SARS-CoV_orf1ab;5469 | 0 | 0 |
| NC_004718.3;NP_828849.2;SARS-CoV_orf1ab;5470 | 0 | 0 |
| NC_004718.3;NP_828849.2;SARS-CoV_orf1ab;5471 | 0 | 0 |
| NC_004718.3;NP_828849.2;SARS-CoV_orf1ab;5472 | 0 | 0 |
| NC_004718.3;NP_828849.2;SARS-CoV_orf1ab;5473 | 0 | 0 |
| NC_004718.3;NP_828849.2;SARS-CoV_orf1ab;5474 | 0 | 0 |
| NC_004718.3;NP_828849.2;SARS-CoV_orf1ab;5475 | 0 | 0 |
| NC_004718.3;NP_828849.2;SARS-CoV_orf1ab;5476 | 0 | 0 |
| NC_004718.3;NP_828849.2;SARS-CoV_orf1ab;5477 | 0 | 0 |
| NC_004718.3;NP_828849.2;SARS-CoV_orf1ab;5478 | 0 | 0 |
| NC_004718.3;NP_828849.2;SARS-CoV_orf1ab;5479 | 0 | 0 |
| NC_004718.3;NP_828849.2;SARS-CoV_orf1ab;5480 | 0 | 0 |
| NC_004718.3;NP_828849.2;SARS-CoV_orf1ab;5481 | 0 | 0 |
| NC_004718.3;NP_828849.2;SARS-CoV_orf1ab;5482 | 0 | 0 |
| NC_004718.3;NP_828849.2;SARS-CoV_orf1ab;5483 | 0.1 | 0.025 |
| NC_004718.3;NP_828849.2;SARS-CoV_orf1ab;5484 | 0.1 | 0.025 |
| NC_004718.3;NP_828849.2;SARS-CoV_orf1ab;5485 | 0.1 | 0.025 |
| NC_004718.3;NP_828849.2;SARS-CoV_orf1ab;5486 | 0.1 | 0 |
| NC_004718.3;NP_828849.2;SARS-CoV_orf1ab;5487 | 0.05 | 0.05 |
| NC_004718.3;NP_828849.2;SARS-CoV_orf1ab;5488 | 0.05 | 0 |
| NC_004718.3;NP_828849.2;SARS-CoV_orf1ab;5489 | 0.1 | 0 |
| NC_004718.3;NP_828849.2;SARS-CoV_orf1ab;5490 | 0.05 | 0 |
| NC_004718.3;NP_828849.2;SARS-CoV_orf1ab;5491 | 0.05 | 0.025 |
| NC_004718.3;NP_828849.2;SARS-CoV_orf1ab;5492 | 0.05 | 0.025 |
| NC_004718.3;NP_828849.2;SARS-CoV_orf1ab;5493 | 0.05 | 0.025 |
| NC_004718.3;NP_828849.2;SARS-CoV_orf1ab;5494 | 0.05 | 0.025 |
| NC_004718.3;NP_828849.2;SARS-CoV_orf1ab;5495 | 0.1 | 0.025 |
| NC_004718.3;NP_828849.2;SARS-CoV_orf1ab;5496 | 0.05 | 0.025 |
| NC_004718.3;NP_828849.2;SARS-CoV_orf1ab;5497 | 0 | 0 |
| NC_004718.3;NP_828849.2;SARS-CoV_orf1ab;5498 | 0 | 0.025 |
| NC_004718.3;NP_828849.2;SARS-CoV_orf1ab;5499 | 0.1 | 0 |
| NC_004718.3;NP_828849.2;SARS-CoV_orf1ab;5500 | 0 | 0.025 |
| NC_004718.3;NP_828849.2;SARS-CoV_orf1ab;5501 | 0 | 0.025 |

| | | |
|---|---|---|
| NC_004718.3;NP_828849.2;SARS-CoV_orf1ab;5502 | 0 | 0.025 |
| NC_004718.3;NP_828849.2;SARS-CoV_orf1ab;5503 | 0 | 0.025 |
| NC_004718.3;NP_828849.2;SARS-CoV_orf1ab;5504 | 0.05 | 0 |
| NC_004718.3;NP_828849.2;SARS-CoV_orf1ab;5505 | 0 | 0 |
| NC_004718.3;NP_828849.2;SARS-CoV_orf1ab;5506 | 0 | 0 |
| NC_004718.3;NP_828849.2;SARS-CoV_orf1ab;5507 | 0 | 0 |
| NC_004718.3;NP_828849.2;SARS-CoV_orf1ab;5508 | 0 | 0 |
| NC_004718.3;NP_828849.2;SARS-CoV_orf1ab;5509 | 0 | 0 |
| NC_004718.3;NP_828849.2;SARS-CoV_orf1ab;5510 | 0 | 0 |
| NC_004718.3;NP_828849.2;SARS-CoV_orf1ab;5511 | 0 | 0 |
| NC_004718.3;NP_828849.2;SARS-CoV_orf1ab;5512 | 0 | 0 |
| NC_004718.3;NP_828849.2;SARS-CoV_orf1ab;5513 | 0 | 0 |
| NC_004718.3;NP_828849.2;SARS-CoV_orf1ab;5514 | 0 | 0 |
| NC_004718.3;NP_828849.2;SARS-CoV_orf1ab;5515 | 0 | 0 |
| NC_004718.3;NP_828849.2;SARS-CoV_orf1ab;5516 | 0 | 0 |
| NC_004718.3;NP_828849.2;SARS-CoV_orf1ab;5517 | 0 | 0 |
| NC_004718.3;NP_828849.2;SARS-CoV_orf1ab;5518 | 0 | 0 |
| NC_004718.3;NP_828849.2;SARS-CoV_orf1ab;5519 | 0 | 0 |
| NC_004718.3;NP_828849.2;SARS-CoV_orf1ab;5520 | 0 | 0 |
| NC_004718.3;NP_828849.2;SARS-CoV_orf1ab;5521 | 0 | 0 |
| NC_004718.3;NP_828849.2;SARS-CoV_orf1ab;5522 | 0 | 0 |
| NC_004718.3;NP_828849.2;SARS-CoV_orf1ab;5523 | 0 | 0 |
| NC_004718.3;NP_828849.2;SARS-CoV_orf1ab;5524 | 0.05 | 0 |
| NC_004718.3;NP_828849.2;SARS-CoV_orf1ab;5525 | 0 | 0 |
| NC_004718.3;NP_828849.2;SARS-CoV_orf1ab;5526 | 0 | 0 |
| NC_004718.3;NP_828849.2;SARS-CoV_orf1ab;5527 | 0 | 0 |
| NC_004718.3;NP_828849.2;SARS-CoV_orf1ab;5528 | 0 | 0.025 |
| NC_004718.3;NP_828849.2;SARS-CoV_orf1ab;5529 | 0 | 0 |
| NC_004718.3;NP_828849.2;SARS-CoV_orf1ab;5530 | 0 | 0 |
| NC_004718.3;NP_828849.2;SARS-CoV_orf1ab;5531 | 0 | 0 |
| NC_004718.3;NP_828849.2;SARS-CoV_orf1ab;5532 | 0 | 0.025 |
| NC_004718.3;NP_828849.2;SARS-CoV_orf1ab;5533 | 0 | 0.025 |
| NC_004718.3;NP_828849.2;SARS-CoV_orf1ab;5534 | 0 | 0.025 |
| NC_004718.3;NP_828849.2;SARS-CoV_orf1ab;5535 | 0 | 0.025 |
| NC_004718.3;NP_828849.2;SARS-CoV_orf1ab;5536 | 0 | 0.025 |
| NC_004718.3;NP_828849.2;SARS-CoV_orf1ab;5537 | 0 | 0.025 |
| NC_004718.3;NP_828849.2;SARS-CoV_orf1ab;5538 | 0 | 0.025 |
| NC_004718.3;NP_828849.2;SARS-CoV_orf1ab;5539 | 0 | 0 |
| NC_004718.3;NP_828849.2;SARS-CoV_orf1ab;5540 | 0 | 0 |
| NC_004718.3;NP_828849.2;SARS-CoV_orf1ab;5541 | 0 | 0.025 |
| NC_004718.3;NP_828849.2;SARS-CoV_orf1ab;5542 | 0 | 0.025 |
| NC_004718.3;NP_828849.2;SARS-CoV_orf1ab;5543 | 0 | 0 |
| NC_004718.3;NP_828849.2;SARS-CoV_orf1ab;5544 | 0 | 0.025 |

| | | |
|---|---|---|
| NC_004718.3;NP_828849.2;SARS-CoV_orf1ab;5545 | 0 | 0 |
| NC_004718.3;NP_828849.2;SARS-CoV_orf1ab;5546 | 0 | 0 |
| NC_004718.3;NP_828849.2;SARS-CoV_orf1ab;5547 | 0 | 0 |
| NC_004718.3;NP_828849.2;SARS-CoV_orf1ab;5548 | 0.05 | 0 |
| NC_004718.3;NP_828849.2;SARS-CoV_orf1ab;5549 | 0.05 | 0.025 |
| NC_004718.3;NP_828849.2;SARS-CoV_orf1ab;5550 | 0.05 | 0.025 |
| NC_004718.3;NP_828849.2;SARS-CoV_orf1ab;5551 | 0.05 | 0.025 |
| NC_004718.3;NP_828849.2;SARS-CoV_orf1ab;5552 | 0.05 | 0.025 |
| NC_004718.3;NP_828849.2;SARS-CoV_orf1ab;5553 | 0.05 | 0.075 |
| NC_004718.3;NP_828849.2;SARS-CoV_orf1ab;5554 | 0.05 | 0.175 |
| NC_004718.3;NP_828849.2;SARS-CoV_orf1ab;5555 | 0.05 | 0.075 |
| NC_004718.3;NP_828849.2;SARS-CoV_orf1ab;5556 | 0.1 | 0.125 |
| NC_004718.3;NP_828849.2;SARS-CoV_orf1ab;5557 | 0.05 | 0.075 |
| NC_004718.3;NP_828849.2;SARS-CoV_orf1ab;5558 | 0 | 0.05 |
| NC_004718.3;NP_828849.2;SARS-CoV_orf1ab;5559 | 0.05 | 0.025 |
| NC_004718.3;NP_828849.2;SARS-CoV_orf1ab;5560 | 0 | 0 |
| NC_004718.3;NP_828849.2;SARS-CoV_orf1ab;5561 | 0.1 | 0.1 |
| NC_004718.3;NP_828849.2;SARS-CoV_orf1ab;5562 | 0.1 | 0.025 |
| NC_004718.3;NP_828849.2;SARS-CoV_orf1ab;5563 | 0 | 0 |
| NC_004718.3;NP_828849.2;SARS-CoV_orf1ab;5564 | 0.05 | 0.05 |
| NC_004718.3;NP_828849.2;SARS-CoV_orf1ab;5565 | 0 | 0.025 |
| NC_004718.3;NP_828849.2;SARS-CoV_orf1ab;5566 | 0 | 0.025 |
| NC_004718.3;NP_828849.2;SARS-CoV_orf1ab;5567 | 0 | 0.025 |
| NC_004718.3;NP_828849.2;SARS-CoV_orf1ab;5568 | 0 | 0.025 |
| NC_004718.3;NP_828849.2;SARS-CoV_orf1ab;5569 | 0 | 0.025 |
| NC_004718.3;NP_828849.2;SARS-CoV_orf1ab;5570 | 0 | 0.075 |
| NC_004718.3;NP_828849.2;SARS-CoV_orf1ab;5571 | 0 | 0.025 |
| NC_004718.3;NP_828849.2;SARS-CoV_orf1ab;5572 | 0 | 0.025 |
| NC_004718.3;NP_828849.2;SARS-CoV_orf1ab;5573 | 0 | 0 |
| NC_004718.3;NP_828849.2;SARS-CoV_orf1ab;5574 | 0 | 0 |
| NC_004718.3;NP_828849.2;SARS-CoV_orf1ab;5575 | 0.05 | 0.025 |
| NC_004718.3;NP_828849.2;SARS-CoV_orf1ab;5576 | 0 | 0 |
| NC_004718.3;NP_828849.2;SARS-CoV_orf1ab;5577 | 0 | 0 |
| NC_004718.3;NP_828849.2;SARS-CoV_orf1ab;5578 | 0 | 0.025 |
| NC_004718.3;NP_828849.2;SARS-CoV_orf1ab;5579 | 0 | 0 |
| NC_004718.3;NP_828849.2;SARS-CoV_orf1ab;5580 | 0 | 0 |
| NC_004718.3;NP_828849.2;SARS-CoV_orf1ab;5581 | 0 | 0 |
| NC_004718.3;NP_828849.2;SARS-CoV_orf1ab;5582 | 0 | 0 |
| NC_004718.3;NP_828849.2;SARS-CoV_orf1ab;5583 | 0 | 0 |
| NC_004718.3;NP_828849.2;SARS-CoV_orf1ab;5584 | 0 | 0 |
| NC_004718.3;NP_828849.2;SARS-CoV_orf1ab;5585 | 0 | 0 |
| NC_004718.3;NP_828849.2;SARS-CoV_orf1ab;5586 | 0 | 0 |
| NC_004718.3;NP_828849.2;SARS-CoV_orf1ab;5587 | 0 | 0 |

| | | |
|---|---|---|
| NC_004718.3;NP_828849.2;SARS-CoV_orf1ab;5588 | 0 | 0 |
| NC_004718.3;NP_828849.2;SARS-CoV_orf1ab;5589 | 0 | 0 |
| NC_004718.3;NP_828849.2;SARS-CoV_orf1ab;5590 | 0 | 0 |
| NC_004718.3;NP_828849.2;SARS-CoV_orf1ab;5591 | 0 | 0 |
| NC_004718.3;NP_828849.2;SARS-CoV_orf1ab;5592 | 0 | 0 |
| NC_004718.3;NP_828849.2;SARS-CoV_orf1ab;5593 | 0 | 0 |
| NC_004718.3;NP_828849.2;SARS-CoV_orf1ab;5594 | 0 | 0 |
| NC_004718.3;NP_828849.2;SARS-CoV_orf1ab;5595 | 0 | 0 |
| NC_004718.3;NP_828849.2;SARS-CoV_orf1ab;5596 | 0 | 0 |
| NC_004718.3;NP_828849.2;SARS-CoV_orf1ab;5597 | 0 | 0 |
| NC_004718.3;NP_828849.2;SARS-CoV_orf1ab;5598 | 0 | 0 |
| NC_004718.3;NP_828849.2;SARS-CoV_orf1ab;5599 | 0 | 0 |
| NC_004718.3;NP_828849.2;SARS-CoV_orf1ab;5600 | 0 | 0 |
| NC_004718.3;NP_828849.2;SARS-CoV_orf1ab;5601 | 0 | 0.025 |
| NC_004718.3;NP_828849.2;SARS-CoV_orf1ab;5602 | 0 | 0.025 |
| NC_004718.3;NP_828849.2;SARS-CoV_orf1ab;5603 | 0 | 0.05 |
| NC_004718.3;NP_828849.2;SARS-CoV_orf1ab;5604 | 0 | 0 |
| NC_004718.3;NP_828849.2;SARS-CoV_orf1ab;5605 | 0 | 0 |
| NC_004718.3;NP_828849.2;SARS-CoV_orf1ab;5606 | 0 | 0 |
| NC_004718.3;NP_828849.2;SARS-CoV_orf1ab;5607 | 0 | 0 |
| NC_004718.3;NP_828849.2;SARS-CoV_orf1ab;5608 | 0 | 0 |
| NC_004718.3;NP_828849.2;SARS-CoV_orf1ab;5609 | 0 | 0 |
| NC_004718.3;NP_828849.2;SARS-CoV_orf1ab;5610 | 0 | 0 |
| NC_004718.3;NP_828849.2;SARS-CoV_orf1ab;5611 | 0 | 0 |
| NC_004718.3;NP_828849.2;SARS-CoV_orf1ab;5612 | 0 | 0 |
| NC_004718.3;NP_828849.2;SARS-CoV_orf1ab;5613 | 0 | 0 |
| NC_004718.3;NP_828849.2;SARS-CoV_orf1ab;5614 | 0 | 0 |
| NC_004718.3;NP_828849.2;SARS-CoV_orf1ab;5615 | 0.05 | 0 |
| NC_004718.3;NP_828849.2;SARS-CoV_orf1ab;5616 | 0 | 0 |
| NC_004718.3;NP_828849.2;SARS-CoV_orf1ab;5617 | 0 | 0 |
| NC_004718.3;NP_828849.2;SARS-CoV_orf1ab;5618 | 0 | 0 |
| NC_004718.3;NP_828849.2;SARS-CoV_orf1ab;5619 | 0 | 0 |
| NC_004718.3;NP_828849.2;SARS-CoV_orf1ab;5620 | 0 | 0 |
| NC_004718.3;NP_828849.2;SARS-CoV_orf1ab;5621 | 0 | 0 |
| NC_004718.3;NP_828849.2;SARS-CoV_orf1ab;5622 | 0 | 0 |
| NC_004718.3;NP_828849.2;SARS-CoV_orf1ab;5623 | 0 | 0 |
| NC_004718.3;NP_828849.2;SARS-CoV_orf1ab;5624 | 0 | 0 |
| NC_004718.3;NP_828849.2;SARS-CoV_orf1ab;5625 | 0 | 0 |
| NC_004718.3;NP_828849.2;SARS-CoV_orf1ab;5626 | 0 | 0 |
| NC_004718.3;NP_828849.2;SARS-CoV_orf1ab;5627 | 0 | 0 |
| NC_004718.3;NP_828849.2;SARS-CoV_orf1ab;5628 | 0 | 0 |
| NC_004718.3;NP_828849.2;SARS-CoV_orf1ab;5629 | 0 | 0 |
| NC_004718.3;NP_828849.2;SARS-CoV_orf1ab;5630 | 0 | 0 |

| | | |
|---|---|---|
| NC_004718.3;NP_828849.2;SARS-CoV_orf1ab;5631 | 0 | 0 |
| NC_004718.3;NP_828849.2;SARS-CoV_orf1ab;5632 | 0 | 0 |
| NC_004718.3;NP_828849.2;SARS-CoV_orf1ab;5633 | 0 | 0 |
| NC_004718.3;NP_828849.2;SARS-CoV_orf1ab;5634 | 0 | 0.025 |
| NC_004718.3;NP_828849.2;SARS-CoV_orf1ab;5635 | 0 | 0.025 |
| NC_004718.3;NP_828849.2;SARS-CoV_orf1ab;5636 | 0.05 | 0.025 |
| NC_004718.3;NP_828849.2;SARS-CoV_orf1ab;5637 | 0 | 0.025 |
| NC_004718.3;NP_828849.2;SARS-CoV_orf1ab;5638 | 0.05 | 0.075 |
| NC_004718.3;NP_828849.2;SARS-CoV_orf1ab;5639 | 0.25 | 0.075 |
| NC_004718.3;NP_828849.2;SARS-CoV_orf1ab;5640 | 0.4 | 0.125 |
| NC_004718.3;NP_828849.2;SARS-CoV_orf1ab;5641 | 0.35 | 0.075 |
| NC_004718.3;NP_828849.2;SARS-CoV_orf1ab;5642 | 0.3 | 0.075 |
| NC_004718.3;NP_828849.2;SARS-CoV_orf1ab;5643 | 0.1 | 0.025 |
| NC_004718.3;NP_828849.2;SARS-CoV_orf1ab;5644 | 0.05 | 0.025 |
| NC_004718.3;NP_828849.2;SARS-CoV_orf1ab;5645 | 0 | 0.025 |
| NC_004718.3;NP_828849.2;SARS-CoV_orf1ab;5646 | 0 | 0 |
| NC_004718.3;NP_828849.2;SARS-CoV_orf1ab;5647 | 0 | 0 |
| NC_004718.3;NP_828849.2;SARS-CoV_orf1ab;5648 | 0 | 0 |
| NC_004718.3;NP_828849.2;SARS-CoV_orf1ab;5649 | 0 | 0 |
| NC_004718.3;NP_828849.2;SARS-CoV_orf1ab;5650 | 0 | 0 |
| NC_004718.3;NP_828849.2;SARS-CoV_orf1ab;5651 | 0 | 0 |
| NC_004718.3;NP_828849.2;SARS-CoV_orf1ab;5652 | 0 | 0 |
| NC_004718.3;NP_828849.2;SARS-CoV_orf1ab;5653 | 0 | 0.025 |
| NC_004718.3;NP_828849.2;SARS-CoV_orf1ab;5654 | 0 | 0 |
| NC_004718.3;NP_828849.2;SARS-CoV_orf1ab;5655 | 0 | 0 |
| NC_004718.3;NP_828849.2;SARS-CoV_orf1ab;5656 | 0 | 0 |
| NC_004718.3;NP_828849.2;SARS-CoV_orf1ab;5657 | 0 | 0 |
| NC_004718.3;NP_828849.2;SARS-CoV_orf1ab;5658 | 0 | 0 |
| NC_004718.3;NP_828849.2;SARS-CoV_orf1ab;5659 | 0 | 0 |
| NC_004718.3;NP_828849.2;SARS-CoV_orf1ab;5660 | 0 | 0 |
| NC_004718.3;NP_828849.2;SARS-CoV_orf1ab;5661 | 0 | 0 |
| NC_004718.3;NP_828849.2;SARS-CoV_orf1ab;5662 | 0 | 0 |
| NC_004718.3;NP_828849.2;SARS-CoV_orf1ab;5663 | 0 | 0 |
| NC_004718.3;NP_828849.2;SARS-CoV_orf1ab;5664 | 0 | 0 |
| NC_004718.3;NP_828849.2;SARS-CoV_orf1ab;5665 | 0 | 0.025 |
| NC_004718.3;NP_828849.2;SARS-CoV_orf1ab;5666 | 0 | 0.025 |
| NC_004718.3;NP_828849.2;SARS-CoV_orf1ab;5667 | 0 | 0 |
| NC_004718.3;NP_828849.2;SARS-CoV_orf1ab;5668 | 0 | 0 |
| NC_004718.3;NP_828849.2;SARS-CoV_orf1ab;5669 | 0 | 0.075 |
| NC_004718.3;NP_828849.2;SARS-CoV_orf1ab;5670 | 0 | 0 |
| NC_004718.3;NP_828849.2;SARS-CoV_orf1ab;5671 | 0 | 0.025 |
| NC_004718.3;NP_828849.2;SARS-CoV_orf1ab;5672 | 0 | 0.025 |
| NC_004718.3;NP_828849.2;SARS-CoV_orf1ab;5673 | 0 | 0.05 |

| | | |
|---|---|---|
| NC_004718.3;NP_828849.2;SARS-CoV_orf1ab;5674 | 0 | 0.05 |
| NC_004718.3;NP_828849.2;SARS-CoV_orf1ab;5675 | 0 | 0.025 |
| NC_004718.3;NP_828849.2;SARS-CoV_orf1ab;5676 | 0 | 0.025 |
| NC_004718.3;NP_828849.2;SARS-CoV_orf1ab;5677 | 0 | 0.025 |
| NC_004718.3;NP_828849.2;SARS-CoV_orf1ab;5678 | 0 | 0.025 |
| NC_004718.3;NP_828849.2;SARS-CoV_orf1ab;5679 | 0 | 0 |
| NC_004718.3;NP_828849.2;SARS-CoV_orf1ab;5680 | 0 | 0 |
| NC_004718.3;NP_828849.2;SARS-CoV_orf1ab;5681 | 0 | 0 |
| NC_004718.3;NP_828849.2;SARS-CoV_orf1ab;5682 | 0 | 0.025 |
| NC_004718.3;NP_828849.2;SARS-CoV_orf1ab;5683 | 0 | 0.025 |
| NC_004718.3;NP_828849.2;SARS-CoV_orf1ab;5684 | 0 | 0.025 |
| NC_004718.3;NP_828849.2;SARS-CoV_orf1ab;5685 | 0 | 0 |
| NC_004718.3;NP_828849.2;SARS-CoV_orf1ab;5686 | 0 | 0 |
| NC_004718.3;NP_828849.2;SARS-CoV_orf1ab;5687 | 0 | 0.025 |
| NC_004718.3;NP_828849.2;SARS-CoV_orf1ab;5688 | 0 | 0.025 |
| NC_004718.3;NP_828849.2;SARS-CoV_orf1ab;5689 | 0 | 0.025 |
| NC_004718.3;NP_828849.2;SARS-CoV_orf1ab;5690 | 0 | 0.025 |
| NC_004718.3;NP_828849.2;SARS-CoV_orf1ab;5691 | 0 | 0.025 |
| NC_004718.3;NP_828849.2;SARS-CoV_orf1ab;5692 | 0 | 0.025 |
| NC_004718.3;NP_828849.2;SARS-CoV_orf1ab;5693 | 0 | 0.025 |
| NC_004718.3;NP_828849.2;SARS-CoV_orf1ab;5694 | 0 | 0 |
| NC_004718.3;NP_828849.2;SARS-CoV_orf1ab;5695 | 0 | 0 |
| NC_004718.3;NP_828849.2;SARS-CoV_orf1ab;5696 | 0 | 0 |
| NC_004718.3;NP_828849.2;SARS-CoV_orf1ab;5697 | 0 | 0 |
| NC_004718.3;NP_828849.2;SARS-CoV_orf1ab;5698 | 0 | 0.025 |
| NC_004718.3;NP_828849.2;SARS-CoV_orf1ab;5699 | 0 | 0 |
| NC_004718.3;NP_828849.2;SARS-CoV_orf1ab;5700 | 0 | 0.025 |
| NC_004718.3;NP_828849.2;SARS-CoV_orf1ab;5701 | 0 | 0.025 |
| NC_004718.3;NP_828849.2;SARS-CoV_orf1ab;5702 | 0 | 0.025 |
| NC_004718.3;NP_828849.2;SARS-CoV_orf1ab;5703 | 0 | 0 |
| NC_004718.3;NP_828849.2;SARS-CoV_orf1ab;5704 | 0 | 0.025 |
| NC_004718.3;NP_828849.2;SARS-CoV_orf1ab;5705 | 0 | 0.075 |
| NC_004718.3;NP_828849.2;SARS-CoV_orf1ab;5706 | 0 | 0.075 |
| NC_004718.3;NP_828849.2;SARS-CoV_orf1ab;5707 | 0.05 | 0.1 |
| NC_004718.3;NP_828849.2;SARS-CoV_orf1ab;5708 | 0.1 | 0.075 |
| NC_004718.3;NP_828849.2;SARS-CoV_orf1ab;5709 | 0.1 | 0.075 |
| NC_004718.3;NP_828849.2;SARS-CoV_orf1ab;5710 | 0.1 | 0.05 |
| NC_004718.3;NP_828849.2;SARS-CoV_orf1ab;5711 | 0.1 | 0.05 |
| NC_004718.3;NP_828849.2;SARS-CoV_orf1ab;5712 | 0.1 | 0.05 |
| NC_004718.3;NP_828849.2;SARS-CoV_orf1ab;5713 | 0.1 | 0.05 |
| NC_004718.3;NP_828849.2;SARS-CoV_orf1ab;5714 | 0.1 | 0 |
| NC_004718.3;NP_828849.2;SARS-CoV_orf1ab;5715 | 0.05 | 0.025 |
| NC_004718.3;NP_828849.2;SARS-CoV_orf1ab;5716 | 0 | 0 |

| | | |
|---|---|---|
| NC_004718.3;NP_828849.2;SARS-CoV_orf1ab;5717 | 0 | 0 |
| NC_004718.3;NP_828849.2;SARS-CoV_orf1ab;5718 | 0 | 0 |
| NC_004718.3;NP_828849.2;SARS-CoV_orf1ab;5719 | 0 | 0 |
| NC_004718.3;NP_828849.2;SARS-CoV_orf1ab;5720 | 0 | 0 |
| NC_004718.3;NP_828849.2;SARS-CoV_orf1ab;5721 | 0 | 0 |
| NC_004718.3;NP_828849.2;SARS-CoV_orf1ab;5722 | 0 | 0 |
| NC_004718.3;NP_828849.2;SARS-CoV_orf1ab;5723 | 0 | 0 |
| NC_004718.3;NP_828849.2;SARS-CoV_orf1ab;5724 | 0 | 0 |
| NC_004718.3;NP_828849.2;SARS-CoV_orf1ab;5725 | 0 | 0 |
| NC_004718.3;NP_828849.2;SARS-CoV_orf1ab;5726 | 0 | 0.025 |
| NC_004718.3;NP_828849.2;SARS-CoV_orf1ab;5727 | 0 | 0 |
| NC_004718.3;NP_828849.2;SARS-CoV_orf1ab;5728 | 0 | 0 |
| NC_004718.3;NP_828849.2;SARS-CoV_orf1ab;5729 | 0 | 0 |
| NC_004718.3;NP_828849.2;SARS-CoV_orf1ab;5730 | 0 | 0 |
| NC_004718.3;NP_828849.2;SARS-CoV_orf1ab;5731 | 0 | 0 |
| NC_004718.3;NP_828849.2;SARS-CoV_orf1ab;5732 | 0 | 0 |
| NC_004718.3;NP_828849.2;SARS-CoV_orf1ab;5733 | 0 | 0 |
| NC_004718.3;NP_828849.2;SARS-CoV_orf1ab;5734 | 0 | 0 |
| NC_004718.3;NP_828849.2;SARS-CoV_orf1ab;5735 | 0 | 0 |
| NC_004718.3;NP_828849.2;SARS-CoV_orf1ab;5736 | 0 | 0 |
| NC_004718.3;NP_828849.2;SARS-CoV_orf1ab;5737 | 0 | 0 |
| NC_004718.3;NP_828849.2;SARS-CoV_orf1ab;5738 | 0 | 0 |
| NC_004718.3;NP_828849.2;SARS-CoV_orf1ab;5739 | 0.1 | 0.05 |
| NC_004718.3;NP_828849.2;SARS-CoV_orf1ab;5740 | 0.1 | 0.05 |
| NC_004718.3;NP_828849.2;SARS-CoV_orf1ab;5741 | 0.15 | 0.125 |
| NC_004718.3;NP_828849.2;SARS-CoV_orf1ab;5742 | 0.15 | 0.15 |
| NC_004718.3;NP_828849.2;SARS-CoV_orf1ab;5743 | 0.1 | 0.05 |
| NC_004718.3;NP_828849.2;SARS-CoV_orf1ab;5744 | 0.1 | 0.025 |
| NC_004718.3;NP_828849.2;SARS-CoV_orf1ab;5745 | 0.1 | 0.075 |
| NC_004718.3;NP_828849.2;SARS-CoV_orf1ab;5746 | 0.1 | 0.15 |
| NC_004718.3;NP_828849.2;SARS-CoV_orf1ab;5747 | 0.15 | 0.225 |
| NC_004718.3;NP_828849.2;SARS-CoV_orf1ab;5748 | 0.1 | 0.05 |
| NC_004718.3;NP_828849.2;SARS-CoV_orf1ab;5749 | 0.1 | 0.075 |
| NC_004718.3;NP_828849.2;SARS-CoV_orf1ab;5750 | 0.15 | 0.15 |
| NC_004718.3;NP_828849.2;SARS-CoV_orf1ab;5751 | 0 | 0 |
| NC_004718.3;NP_828849.2;SARS-CoV_orf1ab;5752 | 0 | 0 |
| NC_004718.3;NP_828849.2;SARS-CoV_orf1ab;5753 | 0 | 0 |
| NC_004718.3;NP_828849.2;SARS-CoV_orf1ab;5754 | 0.05 | 0 |
| NC_004718.3;NP_828849.2;SARS-CoV_orf1ab;5755 | 0 | 0 |
| NC_004718.3;NP_828849.2;SARS-CoV_orf1ab;5756 | 0.05 | 0.025 |
| NC_004718.3;NP_828849.2;SARS-CoV_orf1ab;5757 | 0 | 0.025 |
| NC_004718.3;NP_828849.2;SARS-CoV_orf1ab;5758 | 0 | 0.025 |
| NC_004718.3;NP_828849.2;SARS-CoV_orf1ab;5759 | 0 | 0.025 |

| | | |
|---|---|---|
| NC_004718.3;NP_828849.2;SARS-CoV_orf1ab;5760 | 0 | 0.025 |
| NC_004718.3;NP_828849.2;SARS-CoV_orf1ab;5761 | 0 | 0.025 |
| NC_004718.3;NP_828849.2;SARS-CoV_orf1ab;5762 | 0.05 | 0 |
| NC_004718.3;NP_828849.2;SARS-CoV_orf1ab;5763 | 0 | 0 |
| NC_004718.3;NP_828849.2;SARS-CoV_orf1ab;5764 | 0 | 0 |
| NC_004718.3;NP_828849.2;SARS-CoV_orf1ab;5765 | 0 | 0 |
| NC_004718.3;NP_828849.2;SARS-CoV_orf1ab;5766 | 0 | 0 |
| NC_004718.3;NP_828849.2;SARS-CoV_orf1ab;5767 | 0 | 0 |
| NC_004718.3;NP_828849.2;SARS-CoV_orf1ab;5768 | 0 | 0 |
| NC_004718.3;NP_828849.2;SARS-CoV_orf1ab;5769 | 0 | 0 |
| NC_004718.3;NP_828849.2;SARS-CoV_orf1ab;5770 | 0 | 0 |
| NC_004718.3;NP_828849.2;SARS-CoV_orf1ab;5771 | 0 | 0 |
| NC_004718.3;NP_828849.2;SARS-CoV_orf1ab;5772 | 0 | 0 |
| NC_004718.3;NP_828849.2;SARS-CoV_orf1ab;5773 | 0 | 0 |
| NC_004718.3;NP_828849.2;SARS-CoV_orf1ab;5774 | 0 | 0 |
| NC_004718.3;NP_828849.2;SARS-CoV_orf1ab;5775 | 0 | 0 |
| NC_004718.3;NP_828849.2;SARS-CoV_orf1ab;5776 | 0 | 0 |
| NC_004718.3;NP_828849.2;SARS-CoV_orf1ab;5777 | 0 | 0 |
| NC_004718.3;NP_828849.2;SARS-CoV_orf1ab;5778 | 0 | 0.025 |
| NC_004718.3;NP_828849.2;SARS-CoV_orf1ab;5779 | 0.05 | 0 |
| NC_004718.3;NP_828849.2;SARS-CoV_orf1ab;5780 | 0.05 | 0 |
| NC_004718.3;NP_828849.2;SARS-CoV_orf1ab;5781 | 0.05 | 0 |
| NC_004718.3;NP_828849.2;SARS-CoV_orf1ab;5782 | 0 | 0 |
| NC_004718.3;NP_828849.2;SARS-CoV_orf1ab;5783 | 0 | 0 |
| NC_004718.3;NP_828849.2;SARS-CoV_orf1ab;5784 | 0.05 | 0 |
| NC_004718.3;NP_828849.2;SARS-CoV_orf1ab;5785 | 0 | 0 |
| NC_004718.3;NP_828849.2;SARS-CoV_orf1ab;5786 | 0 | 0 |
| NC_004718.3;NP_828849.2;SARS-CoV_orf1ab;5787 | 0 | 0 |
| NC_004718.3;NP_828849.2;SARS-CoV_orf1ab;5788 | 0 | 0 |
| NC_004718.3;NP_828849.2;SARS-CoV_orf1ab;5789 | 0 | 0 |
| NC_004718.3;NP_828849.2;SARS-CoV_orf1ab;5790 | 0 | 0 |
| NC_004718.3;NP_828849.2;SARS-CoV_orf1ab;5791 | 0 | 0 |
| NC_004718.3;NP_828849.2;SARS-CoV_orf1ab;5792 | 0 | 0 |
| NC_004718.3;NP_828849.2;SARS-CoV_orf1ab;5793 | 0 | 0 |
| NC_004718.3;NP_828849.2;SARS-CoV_orf1ab;5794 | 0 | 0 |
| NC_004718.3;NP_828849.2;SARS-CoV_orf1ab;5795 | 0 | 0.025 |
| NC_004718.3;NP_828849.2;SARS-CoV_orf1ab;5796 | 0 | 0 |
| NC_004718.3;NP_828849.2;SARS-CoV_orf1ab;5797 | 0 | 0 |
| NC_004718.3;NP_828849.2;SARS-CoV_orf1ab;5798 | 0 | 0 |
| NC_004718.3;NP_828849.2;SARS-CoV_orf1ab;5799 | 0 | 0 |
| NC_004718.3;NP_828849.2;SARS-CoV_orf1ab;5800 | 0 | 0 |
| NC_004718.3;NP_828849.2;SARS-CoV_orf1ab;5801 | 0 | 0 |
| NC_004718.3;NP_828849.2;SARS-CoV_orf1ab;5802 | 0 | 0 |

| | | |
|---|---|---|
| NC_004718.3;NP_828849.2;SARS-CoV_orf1ab;5803 | 0 | 0.05 |
| NC_004718.3;NP_828849.2;SARS-CoV_orf1ab;5804 | 0 | 0 |
| NC_004718.3;NP_828849.2;SARS-CoV_orf1ab;5805 | 0.05 | 0 |
| NC_004718.3;NP_828849.2;SARS-CoV_orf1ab;5806 | 0 | 0 |
| NC_004718.3;NP_828849.2;SARS-CoV_orf1ab;5807 | 0 | 0 |
| NC_004718.3;NP_828849.2;SARS-CoV_orf1ab;5808 | 0 | 0 |
| NC_004718.3;NP_828849.2;SARS-CoV_orf1ab;5809 | 0.15 | 0.025 |
| NC_004718.3;NP_828849.2;SARS-CoV_orf1ab;5810 | 0.05 | 0 |
| NC_004718.3;NP_828849.2;SARS-CoV_orf1ab;5811 | 0 | 0 |
| NC_004718.3;NP_828849.2;SARS-CoV_orf1ab;5812 | 0 | 0.025 |
| NC_004718.3;NP_828849.2;SARS-CoV_orf1ab;5813 | 0 | 0.025 |
| NC_004718.3;NP_828849.2;SARS-CoV_orf1ab;5814 | 0.1 | 0 |
| NC_004718.3;NP_828849.2;SARS-CoV_orf1ab;5815 | 0.1 | 0.05 |
| NC_004718.3;NP_828849.2;SARS-CoV_orf1ab;5816 | 0.05 | 0 |
| NC_004718.3;NP_828849.2;SARS-CoV_orf1ab;5817 | 0 | 0 |
| NC_004718.3;NP_828849.2;SARS-CoV_orf1ab;5818 | 0 | 0 |
| NC_004718.3;NP_828849.2;SARS-CoV_orf1ab;5819 | 0 | 0.05 |
| NC_004718.3;NP_828849.2;SARS-CoV_orf1ab;5820 | 0 | 0 |
| NC_004718.3;NP_828849.2;SARS-CoV_orf1ab;5821 | 0 | 0.025 |
| NC_004718.3;NP_828849.2;SARS-CoV_orf1ab;5822 | 0 | 0 |
| NC_004718.3;NP_828849.2;SARS-CoV_orf1ab;5823 | 0.05 | 0 |
| NC_004718.3;NP_828849.2;SARS-CoV_orf1ab;5824 | 0 | 0 |
| NC_004718.3;NP_828849.2;SARS-CoV_orf1ab;5825 | 0 | 0 |
| NC_004718.3;NP_828849.2;SARS-CoV_orf1ab;5826 | 0 | 0.05 |
| NC_004718.3;NP_828849.2;SARS-CoV_orf1ab;5827 | 0 | 0.025 |
| NC_004718.3;NP_828849.2;SARS-CoV_orf1ab;5828 | 0 | 0.025 |
| NC_004718.3;NP_828849.2;SARS-CoV_orf1ab;5829 | 0 | 0.025 |
| NC_004718.3;NP_828849.2;SARS-CoV_orf1ab;5830 | 0 | 0.025 |
| NC_004718.3;NP_828849.2;SARS-CoV_orf1ab;5831 | 0 | 0 |
| NC_004718.3;NP_828849.2;SARS-CoV_orf1ab;5832 | 0 | 0 |
| NC_004718.3;NP_828849.2;SARS-CoV_orf1ab;5833 | 0 | 0.025 |
| NC_004718.3;NP_828849.2;SARS-CoV_orf1ab;5834 | 0 | 0.025 |
| NC_004718.3;NP_828849.2;SARS-CoV_orf1ab;5835 | 0 | 0 |
| NC_004718.3;NP_828849.2;SARS-CoV_orf1ab;5836 | 0 | 0 |
| NC_004718.3;NP_828849.2;SARS-CoV_orf1ab;5837 | 0 | 0 |
| NC_004718.3;NP_828849.2;SARS-CoV_orf1ab;5838 | 0 | 0 |
| NC_004718.3;NP_828849.2;SARS-CoV_orf1ab;5839 | 0 | 0 |
| NC_004718.3;NP_828849.2;SARS-CoV_orf1ab;5840 | 0.05 | 0 |
| NC_004718.3;NP_828849.2;SARS-CoV_orf1ab;5841 | 0.05 | 0.025 |
| NC_004718.3;NP_828849.2;SARS-CoV_orf1ab;5842 | 0.05 | 0 |
| NC_004718.3;NP_828849.2;SARS-CoV_orf1ab;5843 | 0.05 | 0 |
| NC_004718.3;NP_828849.2;SARS-CoV_orf1ab;5844 | 0.05 | 0 |
| NC_004718.3;NP_828849.2;SARS-CoV_orf1ab;5845 | 0.05 | 0 |

| | | |
|---|---|---|
| NC_004718.3;NP_828849.2;SARS-CoV_orf1ab;5846 | 0 | 0 |
| NC_004718.3;NP_828849.2;SARS-CoV_orf1ab;5847 | 0 | 0 |
| NC_004718.3;NP_828849.2;SARS-CoV_orf1ab;5848 | 0.05 | 0 |
| NC_004718.3;NP_828849.2;SARS-CoV_orf1ab;5849 | 0.05 | 0 |
| NC_004718.3;NP_828849.2;SARS-CoV_orf1ab;5850 | 0.05 | 0.05 |
| NC_004718.3;NP_828849.2;SARS-CoV_orf1ab;5851 | 0 | 0 |
| NC_004718.3;NP_828849.2;SARS-CoV_orf1ab;5852 | 0 | 0 |
| NC_004718.3;NP_828849.2;SARS-CoV_orf1ab;5853 | 0 | 0 |
| NC_004718.3;NP_828849.2;SARS-CoV_orf1ab;5854 | 0 | 0 |
| NC_004718.3;NP_828849.2;SARS-CoV_orf1ab;5855 | 0 | 0 |
| NC_004718.3;NP_828849.2;SARS-CoV_orf1ab;5856 | 0 | 0 |
| NC_004718.3;NP_828849.2;SARS-CoV_orf1ab;5857 | 0 | 0 |
| NC_004718.3;NP_828849.2;SARS-CoV_orf1ab;5858 | 0 | 0 |
| NC_004718.3;NP_828849.2;SARS-CoV_orf1ab;5859 | 0 | 0 |
| NC_004718.3;NP_828849.2;SARS-CoV_orf1ab;5860 | 0 | 0 |
| NC_004718.3;NP_828849.2;SARS-CoV_orf1ab;5861 | 0 | 0 |
| NC_004718.3;NP_828849.2;SARS-CoV_orf1ab;5862 | 0 | 0 |
| NC_004718.3;NP_828849.2;SARS-CoV_orf1ab;5863 | 0 | 0 |
| NC_004718.3;NP_828849.2;SARS-CoV_orf1ab;5864 | 0 | 0 |
| NC_004718.3;NP_828849.2;SARS-CoV_orf1ab;5865 | 0 | 0 |
| NC_004718.3;NP_828849.2;SARS-CoV_orf1ab;5866 | 0 | 0 |
| NC_004718.3;NP_828849.2;SARS-CoV_orf1ab;5867 | 0 | 0 |
| NC_004718.3;NP_828849.2;SARS-CoV_orf1ab;5868 | 0 | 0 |
| NC_004718.3;NP_828849.2;SARS-CoV_orf1ab;5869 | 0 | 0.025 |
| NC_004718.3;NP_828849.2;SARS-CoV_orf1ab;5870 | 0 | 0 |
| NC_004718.3;NP_828849.2;SARS-CoV_orf1ab;5871 | 0 | 0 |
| NC_004718.3;NP_828849.2;SARS-CoV_orf1ab;5872 | 0 | 0.05 |
| NC_004718.3;NP_828849.2;SARS-CoV_orf1ab;5873 | 0 | 0.05 |
| NC_004718.3;NP_828849.2;SARS-CoV_orf1ab;5874 | 0.05 | 0.05 |
| NC_004718.3;NP_828849.2;SARS-CoV_orf1ab;5875 | 0.05 | 0.05 |
| NC_004718.3;NP_828849.2;SARS-CoV_orf1ab;5876 | 0.1 | 0.075 |
| NC_004718.3;NP_828849.2;SARS-CoV_orf1ab;5877 | 0 | 0.025 |
| NC_004718.3;NP_828849.2;SARS-CoV_orf1ab;5878 | 0 | 0 |
| NC_004718.3;NP_828849.2;SARS-CoV_orf1ab;5879 | 0 | 0 |
| NC_004718.3;NP_828849.2;SARS-CoV_orf1ab;5880 | 0 | 0.025 |
| NC_004718.3;NP_828849.2;SARS-CoV_orf1ab;5881 | 0 | 0.025 |
| NC_004718.3;NP_828849.2;SARS-CoV_orf1ab;5882 | 0 | 0 |
| NC_004718.3;NP_828849.2;SARS-CoV_orf1ab;5883 | 0 | 0 |
| NC_004718.3;NP_828849.2;SARS-CoV_orf1ab;5884 | 0 | 0 |
| NC_004718.3;NP_828849.2;SARS-CoV_orf1ab;5885 | 0 | 0.025 |
| NC_004718.3;NP_828849.2;SARS-CoV_orf1ab;5886 | 0 | 0.025 |
| NC_004718.3;NP_828849.2;SARS-CoV_orf1ab;5887 | 0 | 0.05 |
| NC_004718.3;NP_828849.2;SARS-CoV_orf1ab;5888 | 0 | 0.05 |

| | | |
|---|---|---|
| NC_004718.3;NP_828849.2;SARS-CoV_orf1ab;5889 | 0.05 | 0.075 |
| NC_004718.3;NP_828849.2;SARS-CoV_orf1ab;5890 | 0.1 | 0.05 |
| NC_004718.3;NP_828849.2;SARS-CoV_orf1ab;5891 | 0.05 | 0.075 |
| NC_004718.3;NP_828849.2;SARS-CoV_orf1ab;5892 | 0.05 | 0.05 |
| NC_004718.3;NP_828849.2;SARS-CoV_orf1ab;5893 | 0.1 | 0.1 |
| NC_004718.3;NP_828849.2;SARS-CoV_orf1ab;5894 | 0.15 | 0.025 |
| NC_004718.3;NP_828849.2;SARS-CoV_orf1ab;5895 | 0.05 | 0 |
| NC_004718.3;NP_828849.2;SARS-CoV_orf1ab;5896 | 0 | 0 |
| NC_004718.3;NP_828849.2;SARS-CoV_orf1ab;5897 | 0 | 0 |
| NC_004718.3;NP_828849.2;SARS-CoV_orf1ab;5898 | 0.05 | 0 |
| NC_004718.3;NP_828849.2;SARS-CoV_orf1ab;5899 | 0 | 0 |
| NC_004718.3;NP_828849.2;SARS-CoV_orf1ab;5900 | 0 | 0 |
| NC_004718.3;NP_828849.2;SARS-CoV_orf1ab;5901 | 0 | 0 |
| NC_004718.3;NP_828849.2;SARS-CoV_orf1ab;5902 | 0 | 0 |
| NC_004718.3;NP_828849.2;SARS-CoV_orf1ab;5903 | 0 | 0 |
| NC_004718.3;NP_828849.2;SARS-CoV_orf1ab;5904 | 0 | 0 |
| NC_004718.3;NP_828849.2;SARS-CoV_orf1ab;5905 | 0 | 0 |
| NC_004718.3;NP_828849.2;SARS-CoV_orf1ab;5906 | 0 | 0 |
| NC_004718.3;NP_828849.2;SARS-CoV_orf1ab;5907 | 0 | 0 |
| NC_004718.3;NP_828849.2;SARS-CoV_orf1ab;5908 | 0 | 0 |
| NC_004718.3;NP_828849.2;SARS-CoV_orf1ab;5909 | 0 | 0 |
| NC_004718.3;NP_828849.2;SARS-CoV_orf1ab;5910 | 0 | 0 |
| NC_004718.3;NP_828849.2;SARS-CoV_orf1ab;5911 | 0.05 | 0 |
| NC_004718.3;NP_828849.2;SARS-CoV_orf1ab;5912 | 0 | 0 |
| NC_004718.3;NP_828849.2;SARS-CoV_orf1ab;5913 | 0 | 0 |
| NC_004718.3;NP_828849.2;SARS-CoV_orf1ab;5914 | 0 | 0.025 |
| NC_004718.3;NP_828849.2;SARS-CoV_orf1ab;5915 | 0 | 0 |
| NC_004718.3;NP_828849.2;SARS-CoV_orf1ab;5916 | 0 | 0 |
| NC_004718.3;NP_828849.2;SARS-CoV_orf1ab;5917 | 0 | 0 |
| NC_004718.3;NP_828849.2;SARS-CoV_orf1ab;5918 | 0 | 0 |
| NC_004718.3;NP_828849.2;SARS-CoV_orf1ab;5919 | 0 | 0 |
| NC_004718.3;NP_828849.2;SARS-CoV_orf1ab;5920 | 0 | 0 |
| NC_004718.3;NP_828849.2;SARS-CoV_orf1ab;5921 | 0 | 0 |
| NC_004718.3;NP_828849.2;SARS-CoV_orf1ab;5922 | 0 | 0 |
| NC_004718.3;NP_828849.2;SARS-CoV_orf1ab;5923 | 0 | 0 |
| NC_004718.3;NP_828849.2;SARS-CoV_orf1ab;5924 | 0 | 0 |
| NC_004718.3;NP_828849.2;SARS-CoV_orf1ab;5925 | 0 | 0 |
| NC_004718.3;NP_828849.2;SARS-CoV_orf1ab;5926 | 0 | 0 |
| NC_004718.3;NP_828849.2;SARS-CoV_orf1ab;5927 | 0 | 0 |
| NC_004718.3;NP_828849.2;SARS-CoV_orf1ab;5928 | 0 | 0 |
| NC_004718.3;NP_828849.2;SARS-CoV_orf1ab;5929 | 0.05 | 0.05 |
| NC_004718.3;NP_828849.2;SARS-CoV_orf1ab;5930 | 0.05 | 0.05 |
| NC_004718.3;NP_828849.2;SARS-CoV_orf1ab;5931 | 0 | 0 |

| | | |
|---|---|---|
| NC_004718.3;NP_828849.2;SARS-CoV_orf1ab;5932 | 0 | 0 |
| NC_004718.3;NP_828849.2;SARS-CoV_orf1ab;5933 | 0 | 0.025 |
| NC_004718.3;NP_828849.2;SARS-CoV_orf1ab;5934 | 0 | 0 |
| NC_004718.3;NP_828849.2;SARS-CoV_orf1ab;5935 | 0 | 0 |
| NC_004718.3;NP_828849.2;SARS-CoV_orf1ab;5936 | 0 | 0 |
| NC_004718.3;NP_828849.2;SARS-CoV_orf1ab;5937 | 0 | 0 |
| NC_004718.3;NP_828849.2;SARS-CoV_orf1ab;5938 | 0 | 0 |
| NC_004718.3;NP_828849.2;SARS-CoV_orf1ab;5939 | 0 | 0 |
| NC_004718.3;NP_828849.2;SARS-CoV_orf1ab;5940 | 0 | 0 |
| NC_004718.3;NP_828849.2;SARS-CoV_orf1ab;5941 | 0 | 0 |
| NC_004718.3;NP_828849.2;SARS-CoV_orf1ab;5942 | 0 | 0 |
| NC_004718.3;NP_828849.2;SARS-CoV_orf1ab;5943 | 0 | 0.025 |
| NC_004718.3;NP_828849.2;SARS-CoV_orf1ab;5944 | 0 | 0 |
| NC_004718.3;NP_828849.2;SARS-CoV_orf1ab;5945 | 0 | 0 |
| NC_004718.3;NP_828849.2;SARS-CoV_orf1ab;5946 | 0 | 0 |
| NC_004718.3;NP_828849.2;SARS-CoV_orf1ab;5947 | 0 | 0 |
| NC_004718.3;NP_828849.2;SARS-CoV_orf1ab;5948 | 0 | 0 |
| NC_004718.3;NP_828849.2;SARS-CoV_orf1ab;5949 | 0 | 0 |
| NC_004718.3;NP_828849.2;SARS-CoV_orf1ab;5950 | 0 | 0 |
| NC_004718.3;NP_828849.2;SARS-CoV_orf1ab;5951 | 0 | 0 |
| NC_004718.3;NP_828849.2;SARS-CoV_orf1ab;5952 | 0 | 0 |
| NC_004718.3;NP_828849.2;SARS-CoV_orf1ab;5953 | 0 | 0 |
| NC_004718.3;NP_828849.2;SARS-CoV_orf1ab;5954 | 0 | 0 |
| NC_004718.3;NP_828849.2;SARS-CoV_orf1ab;5955 | 0 | 0 |
| NC_004718.3;NP_828849.2;SARS-CoV_orf1ab;5956 | 0 | 0.025 |
| NC_004718.3;NP_828849.2;SARS-CoV_orf1ab;5957 | 0 | 0.05 |
| NC_004718.3;NP_828849.2;SARS-CoV_orf1ab;5958 | 0 | 0.05 |
| NC_004718.3;NP_828849.2;SARS-CoV_orf1ab;5959 | 0.05 | 0.05 |
| NC_004718.3;NP_828849.2;SARS-CoV_orf1ab;5960 | 0.1 | 0.05 |
| NC_004718.3;NP_828849.2;SARS-CoV_orf1ab;5961 | 0.05 | 0.05 |
| NC_004718.3;NP_828849.2;SARS-CoV_orf1ab;5962 | 0 | 0.05 |
| NC_004718.3;NP_828849.2;SARS-CoV_orf1ab;5963 | 0 | 0.025 |
| NC_004718.3;NP_828849.2;SARS-CoV_orf1ab;5964 | 0 | 0 |
| NC_004718.3;NP_828849.2;SARS-CoV_orf1ab;5965 | 0 | 0 |
| NC_004718.3;NP_828849.2;SARS-CoV_orf1ab;5966 | 0.05 | 0 |
| NC_004718.3;NP_828849.2;SARS-CoV_orf1ab;5967 | 0 | 0 |
| NC_004718.3;NP_828849.2;SARS-CoV_orf1ab;5968 | 0 | 0 |
| NC_004718.3;NP_828849.2;SARS-CoV_orf1ab;5969 | 0 | 0 |
| NC_004718.3;NP_828849.2;SARS-CoV_orf1ab;5970 | 0 | 0 |
| NC_004718.3;NP_828849.2;SARS-CoV_orf1ab;5971 | 0 | 0 |
| NC_004718.3;NP_828849.2;SARS-CoV_orf1ab;5972 | 0 | 0 |
| NC_004718.3;NP_828849.2;SARS-CoV_orf1ab;5973 | 0 | 0 |
| NC_004718.3;NP_828849.2;SARS-CoV_orf1ab;5974 | 0 | 0 |

| | | |
|---|---|---|
| NC_004718.3;NP_828849.2;SARS-CoV_orf1ab;5975 | 0 | 0 |
| NC_004718.3;NP_828849.2;SARS-CoV_orf1ab;5976 | 0 | 0.025 |
| NC_004718.3;NP_828849.2;SARS-CoV_orf1ab;5977 | 0 | 0 |
| NC_004718.3;NP_828849.2;SARS-CoV_orf1ab;5978 | 0 | 0 |
| NC_004718.3;NP_828849.2;SARS-CoV_orf1ab;5979 | 0 | 0 |
| NC_004718.3;NP_828849.2;SARS-CoV_orf1ab;5980 | 0 | 0 |
| NC_004718.3;NP_828849.2;SARS-CoV_orf1ab;5981 | 0 | 0 |
| NC_004718.3;NP_828849.2;SARS-CoV_orf1ab;5982 | 0 | 0 |
| NC_004718.3;NP_828849.2;SARS-CoV_orf1ab;5983 | 0 | 0 |
| NC_004718.3;NP_828849.2;SARS-CoV_orf1ab;5984 | 0 | 0 |
| NC_004718.3;NP_828849.2;SARS-CoV_orf1ab;5985 | 0 | 0 |
| NC_004718.3;NP_828849.2;SARS-CoV_orf1ab;5986 | 0 | 0 |
| NC_004718.3;NP_828849.2;SARS-CoV_orf1ab;5987 | 0 | 0.025 |
| NC_004718.3;NP_828849.2;SARS-CoV_orf1ab;5988 | 0 | 0 |
| NC_004718.3;NP_828849.2;SARS-CoV_orf1ab;5989 | 0 | 0 |
| NC_004718.3;NP_828849.2;SARS-CoV_orf1ab;5990 | 0 | 0 |
| NC_004718.3;NP_828849.2;SARS-CoV_orf1ab;5991 | 0 | 0 |
| NC_004718.3;NP_828849.2;SARS-CoV_orf1ab;5992 | 0 | 0 |
| NC_004718.3;NP_828849.2;SARS-CoV_orf1ab;5993 | 0 | 0 |
| NC_004718.3;NP_828849.2;SARS-CoV_orf1ab;5994 | 0 | 0 |
| NC_004718.3;NP_828849.2;SARS-CoV_orf1ab;5995 | 0 | 0 |
| NC_004718.3;NP_828849.2;SARS-CoV_orf1ab;5996 | 0 | 0 |
| NC_004718.3;NP_828849.2;SARS-CoV_orf1ab;5997 | 0 | 0 |
| NC_004718.3;NP_828849.2;SARS-CoV_orf1ab;5998 | 0 | 0.025 |
| NC_004718.3;NP_828849.2;SARS-CoV_orf1ab;5999 | 0 | 0.05 |
| NC_004718.3;NP_828849.2;SARS-CoV_orf1ab;6000 | 0 | 0.025 |
| NC_004718.3;NP_828849.2;SARS-CoV_orf1ab;6001 | 0 | 0 |
| NC_004718.3;NP_828849.2;SARS-CoV_orf1ab;6002 | 0 | 0.025 |
| NC_004718.3;NP_828849.2;SARS-CoV_orf1ab;6003 | 0 | 0 |
| NC_004718.3;NP_828849.2;SARS-CoV_orf1ab;6004 | 0 | 0 |
| NC_004718.3;NP_828849.2;SARS-CoV_orf1ab;6005 | 0 | 0 |
| NC_004718.3;NP_828849.2;SARS-CoV_orf1ab;6006 | 0 | 0.025 |
| NC_004718.3;NP_828849.2;SARS-CoV_orf1ab;6007 | 0 | 0 |
| NC_004718.3;NP_828849.2;SARS-CoV_orf1ab;6008 | 0 | 0 |
| NC_004718.3;NP_828849.2;SARS-CoV_orf1ab;6009 | 0 | 0 |
| NC_004718.3;NP_828849.2;SARS-CoV_orf1ab;6010 | 0 | 0 |
| NC_004718.3;NP_828849.2;SARS-CoV_orf1ab;6011 | 0 | 0 |
| NC_004718.3;NP_828849.2;SARS-CoV_orf1ab;6012 | 0 | 0 |
| NC_004718.3;NP_828849.2;SARS-CoV_orf1ab;6013 | 0 | 0 |
| NC_004718.3;NP_828849.2;SARS-CoV_orf1ab;6014 | 0 | 0 |
| NC_004718.3;NP_828849.2;SARS-CoV_orf1ab;6015 | 0 | 0 |
| NC_004718.3;NP_828849.2;SARS-CoV_orf1ab;6016 | 0 | 0 |
| NC_004718.3;NP_828849.2;SARS-CoV_orf1ab;6017 | 0 | 0 |

| | | |
|---|---|---|
| NC_004718.3;NP_828849.2;SARS-CoV_orf1ab;6018 | 0 | 0 |
| NC_004718.3;NP_828849.2;SARS-CoV_orf1ab;6019 | 0 | 0.025 |
| NC_004718.3;NP_828849.2;SARS-CoV_orf1ab;6020 | 0 | 0.025 |
| NC_004718.3;NP_828849.2;SARS-CoV_orf1ab;6021 | 0 | 0.025 |
| NC_004718.3;NP_828849.2;SARS-CoV_orf1ab;6022 | 0 | 0.025 |
| NC_004718.3;NP_828849.2;SARS-CoV_orf1ab;6023 | 0 | 0 |
| NC_004718.3;NP_828849.2;SARS-CoV_orf1ab;6024 | 0.1 | 0.05 |
| NC_004718.3;NP_828849.2;SARS-CoV_orf1ab;6025 | 0.05 | 0.05 |
| NC_004718.3;NP_828849.2;SARS-CoV_orf1ab;6026 | 0 | 0.1 |
| NC_004718.3;NP_828849.2;SARS-CoV_orf1ab;6027 | 0.2 | 0.075 |
| NC_004718.3;NP_828849.2;SARS-CoV_orf1ab;6028 | 0 | 0.025 |
| NC_004718.3;NP_828849.2;SARS-CoV_orf1ab;6029 | 0.1 | 0.05 |
| NC_004718.3;NP_828849.2;SARS-CoV_orf1ab;6030 | 0 | 0.05 |
| NC_004718.3;NP_828849.2;SARS-CoV_orf1ab;6031 | 0 | 0.025 |
| NC_004718.3;NP_828849.2;SARS-CoV_orf1ab;6032 | 0 | 0.025 |
| NC_004718.3;NP_828849.2;SARS-CoV_orf1ab;6033 | 0.1 | 0.1 |
| NC_004718.3;NP_828849.2;SARS-CoV_orf1ab;6034 | 0.05 | 0.075 |
| NC_004718.3;NP_828849.2;SARS-CoV_orf1ab;6035 | 0 | 0 |
| NC_004718.3;NP_828849.2;SARS-CoV_orf1ab;6036 | 0.05 | 0.05 |
| NC_004718.3;NP_828849.2;SARS-CoV_orf1ab;6037 | 0.1 | 0.1 |
| NC_004718.3;NP_828849.2;SARS-CoV_orf1ab;6038 | 0 | 0 |
| NC_004718.3;NP_828849.2;SARS-CoV_orf1ab;6039 | 0 | 0 |
| NC_004718.3;NP_828849.2;SARS-CoV_orf1ab;6040 | 0 | 0 |
| NC_004718.3;NP_828849.2;SARS-CoV_orf1ab;6041 | 0 | 0 |
| NC_004718.3;NP_828849.2;SARS-CoV_orf1ab;6042 | 0 | 0 |
| NC_004718.3;NP_828849.2;SARS-CoV_orf1ab;6043 | 0 | 0 |
| NC_004718.3;NP_828849.2;SARS-CoV_orf1ab;6044 | 0 | 0 |
| NC_004718.3;NP_828849.2;SARS-CoV_orf1ab;6045 | 0 | 0 |
| NC_004718.3;NP_828849.2;SARS-CoV_orf1ab;6046 | 0 | 0 |
| NC_004718.3;NP_828849.2;SARS-CoV_orf1ab;6047 | 0 | 0 |
| NC_004718.3;NP_828849.2;SARS-CoV_orf1ab;6048 | 0 | 0 |
| NC_004718.3;NP_828849.2;SARS-CoV_orf1ab;6049 | 0 | 0 |
| NC_004718.3;NP_828849.2;SARS-CoV_orf1ab;6050 | 0 | 0 |
| NC_004718.3;NP_828849.2;SARS-CoV_orf1ab;6051 | 0 | 0 |
| NC_004718.3;NP_828849.2;SARS-CoV_orf1ab;6052 | 0 | 0 |
| NC_004718.3;NP_828849.2;SARS-CoV_orf1ab;6053 | 0 | 0 |
| NC_004718.3;NP_828849.2;SARS-CoV_orf1ab;6054 | 0 | 0 |
| NC_004718.3;NP_828849.2;SARS-CoV_orf1ab;6055 | 0 | 0 |
| NC_004718.3;NP_828849.2;SARS-CoV_orf1ab;6056 | 0 | 0 |
| NC_004718.3;NP_828849.2;SARS-CoV_orf1ab;6057 | 0 | 0 |
| NC_004718.3;NP_828849.2;SARS-CoV_orf1ab;6058 | 0 | 0 |
| NC_004718.3;NP_828849.2;SARS-CoV_orf1ab;6059 | 0 | 0 |
| NC_004718.3;NP_828849.2;SARS-CoV_orf1ab;6060 | 0 | 0 |

| | | |
|---|---|---|
| NC_004718.3;NP_828849.2;SARS-CoV_orf1ab;6061 | 0 | 0 |
| NC_004718.3;NP_828849.2;SARS-CoV_orf1ab;6062 | 0 | 0 |
| NC_004718.3;NP_828849.2;SARS-CoV_orf1ab;6063 | 0 | 0 |
| NC_004718.3;NP_828849.2;SARS-CoV_orf1ab;6064 | 0 | 0 |
| NC_004718.3;NP_828849.2;SARS-CoV_orf1ab;6065 | 0 | 0 |
| NC_004718.3;NP_828849.2;SARS-CoV_orf1ab;6066 | 0 | 0 |
| NC_004718.3;NP_828849.2;SARS-CoV_orf1ab;6067 | 0 | 0 |
| NC_004718.3;NP_828849.2;SARS-CoV_orf1ab;6068 | 0 | 0 |
| NC_004718.3;NP_828849.2;SARS-CoV_orf1ab;6069 | 0 | 0 |
| NC_004718.3;NP_828849.2;SARS-CoV_orf1ab;6070 | 0 | 0 |
| NC_004718.3;NP_828849.2;SARS-CoV_orf1ab;6071 | 0 | 0 |
| NC_004718.3;NP_828849.2;SARS-CoV_orf1ab;6072 | 0 | 0 |
| NC_004718.3;NP_828849.2;SARS-CoV_orf1ab;6073 | 0 | 0 |
| NC_004718.3;NP_828849.2;SARS-CoV_orf1ab;6074 | 0 | 0 |
| NC_004718.3;NP_828849.2;SARS-CoV_orf1ab;6075 | 0 | 0 |
| NC_004718.3;NP_828849.2;SARS-CoV_orf1ab;6076 | 0 | 0 |
| NC_004718.3;NP_828849.2;SARS-CoV_orf1ab;6077 | 0 | 0 |
| NC_004718.3;NP_828849.2;SARS-CoV_orf1ab;6078 | 0 | 0 |
| NC_004718.3;NP_828849.2;SARS-CoV_orf1ab;6079 | 0 | 0 |
| NC_004718.3;NP_828849.2;SARS-CoV_orf1ab;6080 | 0 | 0 |
| NC_004718.3;NP_828849.2;SARS-CoV_orf1ab;6081 | 0 | 0 |
| NC_004718.3;NP_828849.2;SARS-CoV_orf1ab;6082 | 0 | 0 |
| NC_004718.3;NP_828849.2;SARS-CoV_orf1ab;6083 | 0 | 0 |
| NC_004718.3;NP_828849.2;SARS-CoV_orf1ab;6084 | 0 | 0 |
| NC_004718.3;NP_828849.2;SARS-CoV_orf1ab;6085 | 0 | 0 |
| NC_004718.3;NP_828849.2;SARS-CoV_orf1ab;6086 | 0 | 0 |
| NC_004718.3;NP_828849.2;SARS-CoV_orf1ab;6087 | 0 | 0 |
| NC_004718.3;NP_828849.2;SARS-CoV_orf1ab;6088 | 0 | 0 |
| NC_004718.3;NP_828849.2;SARS-CoV_orf1ab;6089 | 0 | 0 |
| NC_004718.3;NP_828849.2;SARS-CoV_orf1ab;6090 | 0 | 0 |
| NC_004718.3;NP_828849.2;SARS-CoV_orf1ab;6091 | 0 | 0 |
| NC_004718.3;NP_828849.2;SARS-CoV_orf1ab;6092 | 0 | 0 |
| NC_004718.3;NP_828849.2;SARS-CoV_orf1ab;6093 | 0.05 | 0 |
| NC_004718.3;NP_828849.2;SARS-CoV_orf1ab;6094 | 0 | 0 |
| NC_004718.3;NP_828849.2;SARS-CoV_orf1ab;6095 | 0 | 0 |
| NC_004718.3;NP_828849.2;SARS-CoV_orf1ab;6096 | 0 | 0 |
| NC_004718.3;NP_828849.2;SARS-CoV_orf1ab;6097 | 0 | 0 |
| NC_004718.3;NP_828849.2;SARS-CoV_orf1ab;6098 | 0 | 0 |
| NC_004718.3;NP_828849.2;SARS-CoV_orf1ab;6099 | 0 | 0 |
| NC_004718.3;NP_828849.2;SARS-CoV_orf1ab;6100 | 0 | 0 |
| NC_004718.3;NP_828849.2;SARS-CoV_orf1ab;6101 | 0 | 0.025 |
| NC_004718.3;NP_828849.2;SARS-CoV_orf1ab;6102 | 0 | 0 |
| NC_004718.3;NP_828849.2;SARS-CoV_orf1ab;6103 | 0.05 | 0 |

| | | |
|---|---|---|
| NC_004718.3;NP_828849.2;SARS-CoV_orf1ab;6104 | 0.05 | 0 |
| NC_004718.3;NP_828849.2;SARS-CoV_orf1ab;6105 | 0 | 0 |
| NC_004718.3;NP_828849.2;SARS-CoV_orf1ab;6106 | 0 | 0.025 |
| NC_004718.3;NP_828849.2;SARS-CoV_orf1ab;6107 | 0 | 0 |
| NC_004718.3;NP_828849.2;SARS-CoV_orf1ab;6108 | 0 | 0 |
| NC_004718.3;NP_828849.2;SARS-CoV_orf1ab;6109 | 0 | 0 |
| NC_004718.3;NP_828849.2;SARS-CoV_orf1ab;6110 | 0.05 | 0.025 |
| NC_004718.3;NP_828849.2;SARS-CoV_orf1ab;6111 | 0 | 0 |
| NC_004718.3;NP_828849.2;SARS-CoV_orf1ab;6112 | 0.05 | 0 |
| NC_004718.3;NP_828849.2;SARS-CoV_orf1ab;6113 | 0 | 0.025 |
| NC_004718.3;NP_828849.2;SARS-CoV_orf1ab;6114 | 0 | 0 |
| NC_004718.3;NP_828849.2;SARS-CoV_orf1ab;6115 | 0 | 0 |
| NC_004718.3;NP_828849.2;SARS-CoV_orf1ab;6116 | 0 | 0 |
| NC_004718.3;NP_828849.2;SARS-CoV_orf1ab;6117 | 0 | 0 |
| NC_004718.3;NP_828849.2;SARS-CoV_orf1ab;6118 | 0 | 0 |
| NC_004718.3;NP_828849.2;SARS-CoV_orf1ab;6119 | 0 | 0 |
| NC_004718.3;NP_828849.2;SARS-CoV_orf1ab;6120 | 0 | 0.025 |
| NC_004718.3;NP_828849.2;SARS-CoV_orf1ab;6121 | 0 | 0 |
| NC_004718.3;NP_828849.2;SARS-CoV_orf1ab;6122 | 0 | 0 |
| NC_004718.3;NP_828849.2;SARS-CoV_orf1ab;6123 | 0 | 0 |
| NC_004718.3;NP_828849.2;SARS-CoV_orf1ab;6124 | 0 | 0 |
| NC_004718.3;NP_828849.2;SARS-CoV_orf1ab;6125 | 0 | 0 |
| NC_004718.3;NP_828849.2;SARS-CoV_orf1ab;6126 | 0 | 0 |
| NC_004718.3;NP_828849.2;SARS-CoV_orf1ab;6127 | 0 | 0 |
| NC_004718.3;NP_828849.2;SARS-CoV_orf1ab;6128 | 0 | 0 |
| NC_004718.3;NP_828849.2;SARS-CoV_orf1ab;6129 | 0 | 0 |
| NC_004718.3;NP_828849.2;SARS-CoV_orf1ab;6130 | 0 | 0 |
| NC_004718.3;NP_828849.2;SARS-CoV_orf1ab;6131 | 0 | 0 |
| NC_004718.3;NP_828849.2;SARS-CoV_orf1ab;6132 | 0 | 0 |
| NC_004718.3;NP_828849.2;SARS-CoV_orf1ab;6133 | 0 | 0 |
| NC_004718.3;NP_828849.2;SARS-CoV_orf1ab;6134 | 0 | 0.05 |
| NC_004718.3;NP_828849.2;SARS-CoV_orf1ab;6135 | 0 | 0 |
| NC_004718.3;NP_828849.2;SARS-CoV_orf1ab;6136 | 0 | 0 |
| NC_004718.3;NP_828849.2;SARS-CoV_orf1ab;6137 | 0 | 0 |
| NC_004718.3;NP_828849.2;SARS-CoV_orf1ab;6138 | 0 | 0 |
| NC_004718.3;NP_828849.2;SARS-CoV_orf1ab;6139 | 0 | 0 |
| NC_004718.3;NP_828849.2;SARS-CoV_orf1ab;6140 | 0 | 0 |
| NC_004718.3;NP_828849.2;SARS-CoV_orf1ab;6141 | 0 | 0 |
| NC_004718.3;NP_828849.2;SARS-CoV_orf1ab;6142 | 0 | 0 |
| NC_004718.3;NP_828849.2;SARS-CoV_orf1ab;6143 | 0 | 0 |
| NC_004718.3;NP_828849.2;SARS-CoV_orf1ab;6144 | 0 | 0 |
| NC_004718.3;NP_828849.2;SARS-CoV_orf1ab;6145 | 0 | 0 |
| NC_004718.3;NP_828849.2;SARS-CoV_orf1ab;6146 | 0 | 0 |

| | | |
|---|---|---|
| NC_004718.3;NP_828849.2;SARS-CoV_orf1ab;6147 | 0 | 0 |
| NC_004718.3;NP_828849.2;SARS-CoV_orf1ab;6148 | 0 | 0 |
| NC_004718.3;NP_828849.2;SARS-CoV_orf1ab;6149 | 0 | 0 |
| NC_004718.3;NP_828849.2;SARS-CoV_orf1ab;6150 | 0 | 0 |
| NC_004718.3;NP_828849.2;SARS-CoV_orf1ab;6151 | 0 | 0 |
| NC_004718.3;NP_828849.2;SARS-CoV_orf1ab;6152 | 0 | 0 |
| NC_004718.3;NP_828849.2;SARS-CoV_orf1ab;6153 | 0 | 0 |
| NC_004718.3;NP_828849.2;SARS-CoV_orf1ab;6154 | 0 | 0.025 |
| NC_004718.3;NP_828849.2;SARS-CoV_orf1ab;6155 | 0 | 0 |
| NC_004718.3;NP_828849.2;SARS-CoV_orf1ab;6156 | 0.05 | 0 |
| NC_004718.3;NP_828849.2;SARS-CoV_orf1ab;6157 | 0.05 | 0.05 |
| NC_004718.3;NP_828849.2;SARS-CoV_orf1ab;6158 | 0 | 0 |
| NC_004718.3;NP_828849.2;SARS-CoV_orf1ab;6159 | 0 | 0 |
| NC_004718.3;NP_828849.2;SARS-CoV_orf1ab;6160 | 0 | 0 |
| NC_004718.3;NP_828849.2;SARS-CoV_orf1ab;6161 | 0 | 0 |
| NC_004718.3;NP_828849.2;SARS-CoV_orf1ab;6162 | 0 | 0 |
| NC_004718.3;NP_828849.2;SARS-CoV_orf1ab;6163 | 0 | 0 |
| NC_004718.3;NP_828849.2;SARS-CoV_orf1ab;6164 | 0 | 0 |
| NC_004718.3;NP_828849.2;SARS-CoV_orf1ab;6165 | 0 | 0 |
| NC_004718.3;NP_828849.2;SARS-CoV_orf1ab;6166 | 0 | 0 |
| NC_004718.3;NP_828849.2;SARS-CoV_orf1ab;6167 | 0 | 0 |
| NC_004718.3;NP_828849.2;SARS-CoV_orf1ab;6168 | 0 | 0 |
| NC_004718.3;NP_828849.2;SARS-CoV_orf1ab;6169 | 0 | 0 |
| NC_004718.3;NP_828849.2;SARS-CoV_orf1ab;6170 | 0 | 0 |
| NC_004718.3;NP_828849.2;SARS-CoV_orf1ab;6171 | 0 | 0 |
| NC_004718.3;NP_828849.2;SARS-CoV_orf1ab;6172 | 0 | 0 |
| NC_004718.3;NP_828849.2;SARS-CoV_orf1ab;6173 | 0 | 0 |
| NC_004718.3;NP_828849.2;SARS-CoV_orf1ab;6174 | 0 | 0 |
| NC_004718.3;NP_828849.2;SARS-CoV_orf1ab;6175 | 0 | 0 |
| NC_004718.3;NP_828849.2;SARS-CoV_orf1ab;6176 | 0 | 0 |
| NC_004718.3;NP_828849.2;SARS-CoV_orf1ab;6177 | 0 | 0 |
| NC_004718.3;NP_828849.2;SARS-CoV_orf1ab;6178 | 0 | 0 |
| NC_004718.3;NP_828849.2;SARS-CoV_orf1ab;6179 | 0 | 0 |
| NC_004718.3;NP_828849.2;SARS-CoV_orf1ab;6180 | 0 | 0 |
| NC_004718.3;NP_828849.2;SARS-CoV_orf1ab;6181 | 0 | 0 |
| NC_004718.3;NP_828849.2;SARS-CoV_orf1ab;6182 | 0 | 0 |
| NC_004718.3;NP_828849.2;SARS-CoV_orf1ab;6183 | 0 | 0 |
| NC_004718.3;NP_828849.2;SARS-CoV_orf1ab;6184 | 0 | 0 |
| NC_004718.3;NP_828849.2;SARS-CoV_orf1ab;6185 | 0 | 0 |
| NC_004718.3;NP_828849.2;SARS-CoV_orf1ab;6186 | 0 | 0 |
| NC_004718.3;NP_828849.2;SARS-CoV_orf1ab;6187 | 0 | 0 |
| NC_004718.3;NP_828849.2;SARS-CoV_orf1ab;6188 | 0 | 0.025 |
| NC_004718.3;NP_828849.2;SARS-CoV_orf1ab;6189 | 0 | 0.025 |

| | | |
|---|---|---|
| NC_004718.3;NP_828849.2;SARS-CoV_orf1ab;6190 | 0.05 | 0.025 |
| NC_004718.3;NP_828849.2;SARS-CoV_orf1ab;6191 | 0 | 0.025 |
| NC_004718.3;NP_828849.2;SARS-CoV_orf1ab;6192 | 0 | 0.025 |
| NC_004718.3;NP_828849.2;SARS-CoV_orf1ab;6193 | 0 | 0.025 |
| NC_004718.3;NP_828849.2;SARS-CoV_orf1ab;6194 | 0 | 0.05 |
| NC_004718.3;NP_828849.2;SARS-CoV_orf1ab;6195 | 0 | 0.05 |
| NC_004718.3;NP_828849.2;SARS-CoV_orf1ab;6196 | 0 | 0.025 |
| NC_004718.3;NP_828849.2;SARS-CoV_orf1ab;6197 | 0 | 0.075 |
| NC_004718.3;NP_828849.2;SARS-CoV_orf1ab;6198 | 0 | 0 |
| NC_004718.3;NP_828849.2;SARS-CoV_orf1ab;6199 | 0 | 0 |
| NC_004718.3;NP_828849.2;SARS-CoV_orf1ab;6200 | 0 | 0 |
| NC_004718.3;NP_828849.2;SARS-CoV_orf1ab;6201 | 0.1 | 0 |
| NC_004718.3;NP_828849.2;SARS-CoV_orf1ab;6202 | 0 | 0 |
| NC_004718.3;NP_828849.2;SARS-CoV_orf1ab;6203 | 0 | 0 |
| NC_004718.3;NP_828849.2;SARS-CoV_orf1ab;6204 | 0 | 0 |
| NC_004718.3;NP_828849.2;SARS-CoV_orf1ab;6205 | 0 | 0 |
| NC_004718.3;NP_828849.2;SARS-CoV_orf1ab;6206 | 0.05 | 0 |
| NC_004718.3;NP_828849.2;SARS-CoV_orf1ab;6207 | 0 | 0 |
| NC_004718.3;NP_828849.2;SARS-CoV_orf1ab;6208 | 0 | 0 |
| NC_004718.3;NP_828849.2;SARS-CoV_orf1ab;6209 | 0 | 0 |
| NC_004718.3;NP_828849.2;SARS-CoV_orf1ab;6210 | 0 | 0 |
| NC_004718.3;NP_828849.2;SARS-CoV_orf1ab;6211 | 0 | 0 |
| NC_004718.3;NP_828849.2;SARS-CoV_orf1ab;6212 | 0 | 0 |
| NC_004718.3;NP_828849.2;SARS-CoV_orf1ab;6213 | 0 | 0 |
| NC_004718.3;NP_828849.2;SARS-CoV_orf1ab;6214 | 0 | 0 |
| NC_004718.3;NP_828849.2;SARS-CoV_orf1ab;6215 | 0 | 0 |
| NC_004718.3;NP_828849.2;SARS-CoV_orf1ab;6216 | 0 | 0 |
| NC_004718.3;NP_828849.2;SARS-CoV_orf1ab;6217 | 0 | 0 |
| NC_004718.3;NP_828849.2;SARS-CoV_orf1ab;6218 | 0 | 0 |
| NC_004718.3;NP_828849.2;SARS-CoV_orf1ab;6219 | 0 | 0.025 |
| NC_004718.3;NP_828849.2;SARS-CoV_orf1ab;6220 | 0 | 0.025 |
| NC_004718.3;NP_828849.2;SARS-CoV_orf1ab;6221 | 0 | 0.075 |
| NC_004718.3;NP_828849.2;SARS-CoV_orf1ab;6222 | 0 | 0.025 |
| NC_004718.3;NP_828849.2;SARS-CoV_orf1ab;6223 | 0 | 0 |
| NC_004718.3;NP_828849.2;SARS-CoV_orf1ab;6224 | 0 | 0 |
| NC_004718.3;NP_828849.2;SARS-CoV_orf1ab;6225 | 0 | 0 |
| NC_004718.3;NP_828849.2;SARS-CoV_orf1ab;6226 | 0 | 0 |
| NC_004718.3;NP_828849.2;SARS-CoV_orf1ab;6227 | 0 | 0 |
| NC_004718.3;NP_828849.2;SARS-CoV_orf1ab;6228 | 0 | 0 |
| NC_004718.3;NP_828849.2;SARS-CoV_orf1ab;6229 | 0 | 0 |
| NC_004718.3;NP_828849.2;SARS-CoV_orf1ab;6230 | 0 | 0 |
| NC_004718.3;NP_828849.2;SARS-CoV_orf1ab;6231 | 0 | 0 |
| NC_004718.3;NP_828849.2;SARS-CoV_orf1ab;6232 | 0.05 | 0 |

| | | |
|---|---|---|
| NC_004718.3;NP_828849.2;SARS-CoV_orf1ab;6233 | 0.05 | 0 |
| NC_004718.3;NP_828849.2;SARS-CoV_orf1ab;6234 | 0 | 0 |
| NC_004718.3;NP_828849.2;SARS-CoV_orf1ab;6235 | 0 | 0 |
| NC_004718.3;NP_828849.2;SARS-CoV_orf1ab;6236 | 0.05 | 0 |
| NC_004718.3;NP_828849.2;SARS-CoV_orf1ab;6237 | 0.05 | 0 |
| NC_004718.3;NP_828849.2;SARS-CoV_orf1ab;6238 | 0 | 0 |
| NC_004718.3;NP_828849.2;SARS-CoV_orf1ab;6239 | 0 | 0.025 |
| NC_004718.3;NP_828849.2;SARS-CoV_orf1ab;6240 | 0 | 0.025 |
| NC_004718.3;NP_828849.2;SARS-CoV_orf1ab;6241 | 0 | 0.025 |
| NC_004718.3;NP_828849.2;SARS-CoV_orf1ab;6242 | 0 | 0.025 |
| NC_004718.3;NP_828849.2;SARS-CoV_orf1ab;6243 | 0 | 0.025 |
| NC_004718.3;NP_828849.2;SARS-CoV_orf1ab;6244 | 0 | 0.025 |
| NC_004718.3;NP_828849.2;SARS-CoV_orf1ab;6245 | 0 | 0.025 |
| NC_004718.3;NP_828849.2;SARS-CoV_orf1ab;6246 | 0 | 0.025 |
| NC_004718.3;NP_828849.2;SARS-CoV_orf1ab;6247 | 0.05 | 0.05 |
| NC_004718.3;NP_828849.2;SARS-CoV_orf1ab;6248 | 0.1 | 0.025 |
| NC_004718.3;NP_828849.2;SARS-CoV_orf1ab;6249 | 0.05 | 0.05 |
| NC_004718.3;NP_828849.2;SARS-CoV_orf1ab;6250 | 0 | 0 |
| NC_004718.3;NP_828849.2;SARS-CoV_orf1ab;6251 | 0.05 | 0.025 |
| NC_004718.3;NP_828849.2;SARS-CoV_orf1ab;6252 | 0 | 0 |
| NC_004718.3;NP_828849.2;SARS-CoV_orf1ab;6253 | 0 | 0 |
| NC_004718.3;NP_828849.2;SARS-CoV_orf1ab;6254 | 0 | 0 |
| NC_004718.3;NP_828849.2;SARS-CoV_orf1ab;6255 | 0 | 0.025 |
| NC_004718.3;NP_828849.2;SARS-CoV_orf1ab;6256 | 0 | 0.025 |
| NC_004718.3;NP_828849.2;SARS-CoV_orf1ab;6257 | 0 | 0 |
| NC_004718.3;NP_828849.2;SARS-CoV_orf1ab;6258 | 0 | 0 |
| NC_004718.3;NP_828849.2;SARS-CoV_orf1ab;6259 | 0 | 0 |
| NC_004718.3;NP_828849.2;SARS-CoV_orf1ab;6260 | 0 | 0 |
| NC_004718.3;NP_828849.2;SARS-CoV_orf1ab;6261 | 0 | 0 |
| NC_004718.3;NP_828849.2;SARS-CoV_orf1ab;6262 | 0 | 0 |
| NC_004718.3;NP_828849.2;SARS-CoV_orf1ab;6263 | 0 | 0.025 |
| NC_004718.3;NP_828849.2;SARS-CoV_orf1ab;6264 | 0 | 0.025 |
| NC_004718.3;NP_828849.2;SARS-CoV_orf1ab;6265 | 0.05 | 0.05 |
| NC_004718.3;NP_828849.2;SARS-CoV_orf1ab;6266 | 0.1 | 0.025 |
| NC_004718.3;NP_828849.2;SARS-CoV_orf1ab;6267 | 0.15 | 0.025 |
| NC_004718.3;NP_828849.2;SARS-CoV_orf1ab;6268 | 0.1 | 0.025 |
| NC_004718.3;NP_828849.2;SARS-CoV_orf1ab;6269 | 0.25 | 0.025 |
| NC_004718.3;NP_828849.2;SARS-CoV_orf1ab;6270 | 0.1 | 0.05 |
| NC_004718.3;NP_828849.2;SARS-CoV_orf1ab;6271 | 0 | 0.025 |
| NC_004718.3;NP_828849.2;SARS-CoV_orf1ab;6272 | 0 | 0.025 |
| NC_004718.3;NP_828849.2;SARS-CoV_orf1ab;6273 | 0 | 0.025 |
| NC_004718.3;NP_828849.2;SARS-CoV_orf1ab;6274 | 0 | 0.025 |
| NC_004718.3;NP_828849.2;SARS-CoV_orf1ab;6275 | 0 | 0 |

| | | |
|---|---|---|
| NC_004718.3;NP_828849.2;SARS-CoV_orf1ab;6276 | 0 | 0 |
| NC_004718.3;NP_828849.2;SARS-CoV_orf1ab;6277 | 0 | 0 |
| NC_004718.3;NP_828849.2;SARS-CoV_orf1ab;6278 | 0 | 0 |
| NC_004718.3;NP_828849.2;SARS-CoV_orf1ab;6279 | 0 | 0 |
| NC_004718.3;NP_828849.2;SARS-CoV_orf1ab;6280 | 0 | 0 |
| NC_004718.3;NP_828849.2;SARS-CoV_orf1ab;6281 | 0 | 0 |
| NC_004718.3;NP_828849.2;SARS-CoV_orf1ab;6282 | 0 | 0 |
| NC_004718.3;NP_828849.2;SARS-CoV_orf1ab;6283 | 0.05 | 0 |
| NC_004718.3;NP_828849.2;SARS-CoV_orf1ab;6284 | 0 | 0 |
| NC_004718.3;NP_828849.2;SARS-CoV_orf1ab;6285 | 0 | 0 |
| NC_004718.3;NP_828849.2;SARS-CoV_orf1ab;6286 | 0 | 0 |
| NC_004718.3;NP_828849.2;SARS-CoV_orf1ab;6287 | 0 | 0 |
| NC_004718.3;NP_828849.2;SARS-CoV_orf1ab;6288 | 0 | 0 |
| NC_004718.3;NP_828849.2;SARS-CoV_orf1ab;6289 | 0 | 0 |
| NC_004718.3;NP_828849.2;SARS-CoV_orf1ab;6290 | 0 | 0 |
| NC_004718.3;NP_828849.2;SARS-CoV_orf1ab;6291 | 0 | 0 |
| NC_004718.3;NP_828849.2;SARS-CoV_orf1ab;6292 | 0 | 0 |
| NC_004718.3;NP_828849.2;SARS-CoV_orf1ab;6293 | 0 | 0 |
| NC_004718.3;NP_828849.2;SARS-CoV_orf1ab;6294 | 0 | 0 |
| NC_004718.3;NP_828849.2;SARS-CoV_orf1ab;6295 | 0 | 0 |
| NC_004718.3;NP_828849.2;SARS-CoV_orf1ab;6296 | 0 | 0 |
| NC_004718.3;NP_828849.2;SARS-CoV_orf1ab;6297 | 0 | 0 |
| NC_004718.3;NP_828849.2;SARS-CoV_orf1ab;6298 | 0 | 0 |
| NC_004718.3;NP_828849.2;SARS-CoV_orf1ab;6299 | 0 | 0 |
| NC_004718.3;NP_828849.2;SARS-CoV_orf1ab;6300 | 0 | 0 |
| NC_004718.3;NP_828849.2;SARS-CoV_orf1ab;6301 | 0 | 0 |
| NC_004718.3;NP_828849.2;SARS-CoV_orf1ab;6302 | 0 | 0 |
| NC_004718.3;NP_828849.2;SARS-CoV_orf1ab;6303 | 0 | 0 |
| NC_004718.3;NP_828849.2;SARS-CoV_orf1ab;6304 | 0.1 | 0.025 |
| NC_004718.3;NP_828849.2;SARS-CoV_orf1ab;6305 | 0.05 | 0.025 |
| NC_004718.3;NP_828849.2;SARS-CoV_orf1ab;6306 | 0.05 | 0 |
| NC_004718.3;NP_828849.2;SARS-CoV_orf1ab;6307 | 0.05 | 0 |
| NC_004718.3;NP_828849.2;SARS-CoV_orf1ab;6308 | 0.05 | 0 |
| NC_004718.3;NP_828849.2;SARS-CoV_orf1ab;6309 | 0 | 0.025 |
| NC_004718.3;NP_828849.2;SARS-CoV_orf1ab;6310 | 0 | 0 |
| NC_004718.3;NP_828849.2;SARS-CoV_orf1ab;6311 | 0 | 0 |
| NC_004718.3;NP_828849.2;SARS-CoV_orf1ab;6312 | 0 | 0 |
| NC_004718.3;NP_828849.2;SARS-CoV_orf1ab;6313 | 0 | 0 |
| NC_004718.3;NP_828849.2;SARS-CoV_orf1ab;6314 | 0 | 0 |
| NC_004718.3;NP_828849.2;SARS-CoV_orf1ab;6315 | 0 | 0 |
| NC_004718.3;NP_828849.2;SARS-CoV_orf1ab;6316 | 0 | 0 |
| NC_004718.3;NP_828849.2;SARS-CoV_orf1ab;6317 | 0 | 0 |
| NC_004718.3;NP_828849.2;SARS-CoV_orf1ab;6318 | 0 | 0 |

| | | |
|---|---|---|
| NC_004718.3;NP_828849.2;SARS-CoV_orf1ab;6319 | 0 | 0 |
| NC_004718.3;NP_828849.2;SARS-CoV_orf1ab;6320 | 0 | 0 |
| NC_004718.3;NP_828849.2;SARS-CoV_orf1ab;6321 | 0 | 0 |
| NC_004718.3;NP_828849.2;SARS-CoV_orf1ab;6322 | 0 | 0.025 |
| NC_004718.3;NP_828849.2;SARS-CoV_orf1ab;6323 | 0 | 0.025 |
| NC_004718.3;NP_828849.2;SARS-CoV_orf1ab;6324 | 0 | 0.025 |
| NC_004718.3;NP_828849.2;SARS-CoV_orf1ab;6325 | 0 | 0.025 |
| NC_004718.3;NP_828849.2;SARS-CoV_orf1ab;6326 | 0 | 0.025 |
| NC_004718.3;NP_828849.2;SARS-CoV_orf1ab;6327 | 0.05 | 0.05 |
| NC_004718.3;NP_828849.2;SARS-CoV_orf1ab;6328 | 0.05 | 0.05 |
| NC_004718.3;NP_828849.2;SARS-CoV_orf1ab;6329 | 0.1 | 0.025 |
| NC_004718.3;NP_828849.2;SARS-CoV_orf1ab;6330 | 0.05 | 0.05 |
| NC_004718.3;NP_828849.2;SARS-CoV_orf1ab;6331 | 0.05 | 0.025 |
| NC_004718.3;NP_828849.2;SARS-CoV_orf1ab;6332 | 0.05 | 0.025 |
| NC_004718.3;NP_828849.2;SARS-CoV_orf1ab;6333 | 0.05 | 0 |
| NC_004718.3;NP_828849.2;SARS-CoV_orf1ab;6334 | 0.05 | 0.05 |
| NC_004718.3;NP_828849.2;SARS-CoV_orf1ab;6335 | 0.05 | 0.05 |
| NC_004718.3;NP_828849.2;SARS-CoV_orf1ab;6336 | 0 | 0 |
| NC_004718.3;NP_828849.2;SARS-CoV_orf1ab;6337 | 0 | 0 |
| NC_004718.3;NP_828849.2;SARS-CoV_orf1ab;6338 | 0 | 0 |
| NC_004718.3;NP_828849.2;SARS-CoV_orf1ab;6339 | 0 | 0 |
| NC_004718.3;NP_828849.2;SARS-CoV_orf1ab;6340 | 0 | 0 |
| NC_004718.3;NP_828849.2;SARS-CoV_orf1ab;6341 | 0 | 0 |
| NC_004718.3;NP_828849.2;SARS-CoV_orf1ab;6342 | 0 | 0 |
| NC_004718.3;NP_828849.2;SARS-CoV_orf1ab;6343 | 0 | 0 |
| NC_004718.3;NP_828849.2;SARS-CoV_orf1ab;6344 | 0 | 0 |
| NC_004718.3;NP_828849.2;SARS-CoV_orf1ab;6345 | 0 | 0 |
| NC_004718.3;NP_828849.2;SARS-CoV_orf1ab;6346 | 0 | 0 |
| NC_004718.3;NP_828849.2;SARS-CoV_orf1ab;6347 | 0 | 0 |
| NC_004718.3;NP_828849.2;SARS-CoV_orf1ab;6348 | 0 | 0.025 |
| NC_004718.3;NP_828849.2;SARS-CoV_orf1ab;6349 | 0 | 0 |
| NC_004718.3;NP_828849.2;SARS-CoV_orf1ab;6350 | 0 | 0 |
| NC_004718.3;NP_828849.2;SARS-CoV_orf1ab;6351 | 0 | 0 |
| NC_004718.3;NP_828849.2;SARS-CoV_orf1ab;6352 | 0 | 0 |
| NC_004718.3;NP_828849.2;SARS-CoV_orf1ab;6353 | 0 | 0.05 |
| NC_004718.3;NP_828849.2;SARS-CoV_orf1ab;6354 | 0 | 0.075 |
| NC_004718.3;NP_828849.2;SARS-CoV_orf1ab;6355 | 0 | 0.05 |
| NC_004718.3;NP_828849.2;SARS-CoV_orf1ab;6356 | 0 | 0.05 |
| NC_004718.3;NP_828849.2;SARS-CoV_orf1ab;6357 | 0 | 0.075 |
| NC_004718.3;NP_828849.2;SARS-CoV_orf1ab;6358 | 0 | 0.075 |
| NC_004718.3;NP_828849.2;SARS-CoV_orf1ab;6359 | 0.05 | 0.05 |
| NC_004718.3;NP_828849.2;SARS-CoV_orf1ab;6360 | 0 | 0.05 |
| NC_004718.3;NP_828849.2;SARS-CoV_orf1ab;6361 | 0 | 0.025 |

| | | |
|---|---|---|
| NC_004718.3;NP_828849.2;SARS-CoV_orf1ab;6362 | 0 | 0.025 |
| NC_004718.3;NP_828849.2;SARS-CoV_orf1ab;6363 | 0 | 0.025 |
| NC_004718.3;NP_828849.2;SARS-CoV_orf1ab;6364 | 0 | 0 |
| NC_004718.3;NP_828849.2;SARS-CoV_orf1ab;6365 | 0 | 0 |
| NC_004718.3;NP_828849.2;SARS-CoV_orf1ab;6366 | 0 | 0 |
| NC_004718.3;NP_828849.2;SARS-CoV_orf1ab;6367 | 0 | 0 |
| NC_004718.3;NP_828849.2;SARS-CoV_orf1ab;6368 | 0 | 0 |
| NC_004718.3;NP_828849.2;SARS-CoV_orf1ab;6369 | 0 | 0 |
| NC_004718.3;NP_828849.2;SARS-CoV_orf1ab;6370 | 0 | 0 |
| NC_004718.3;NP_828849.2;SARS-CoV_orf1ab;6371 | 0 | 0 |
| NC_004718.3;NP_828849.2;SARS-CoV_orf1ab;6372 | 0 | 0 |
| NC_004718.3;NP_828849.2;SARS-CoV_orf1ab;6373 | 0 | 0 |
| NC_004718.3;NP_828849.2;SARS-CoV_orf1ab;6374 | 0 | 0 |
| NC_004718.3;NP_828849.2;SARS-CoV_orf1ab;6375 | 0 | 0 |
| NC_004718.3;NP_828849.2;SARS-CoV_orf1ab;6376 | 0 | 0 |
| NC_004718.3;NP_828849.2;SARS-CoV_orf1ab;6377 | 0 | 0 |
| NC_004718.3;NP_828849.2;SARS-CoV_orf1ab;6378 | 0 | 0 |
| NC_004718.3;NP_828849.2;SARS-CoV_orf1ab;6379 | 0 | 0 |
| NC_004718.3;NP_828849.2;SARS-CoV_orf1ab;6380 | 0 | 0 |
| NC_004718.3;NP_828849.2;SARS-CoV_orf1ab;6381 | 0 | 0 |
| NC_004718.3;NP_828849.2;SARS-CoV_orf1ab;6382 | 0 | 0 |
| NC_004718.3;NP_828849.2;SARS-CoV_orf1ab;6383 | 0 | 0 |
| NC_004718.3;NP_828849.2;SARS-CoV_orf1ab;6384 | 0 | 0 |
| NC_004718.3;NP_828849.2;SARS-CoV_orf1ab;6385 | 0 | 0.025 |
| NC_004718.3;NP_828849.2;SARS-CoV_orf1ab;6386 | 0 | 0.05 |
| NC_004718.3;NP_828849.2;SARS-CoV_orf1ab;6387 | 0 | 0.025 |
| NC_004718.3;NP_828849.2;SARS-CoV_orf1ab;6388 | 0.05 | 0.05 |
| NC_004718.3;NP_828849.2;SARS-CoV_orf1ab;6389 | 0 | 0.025 |
| NC_004718.3;NP_828849.2;SARS-CoV_orf1ab;6390 | 0 | 0.025 |
| NC_004718.3;NP_828849.2;SARS-CoV_orf1ab;6391 | 0 | 0.025 |
| NC_004718.3;NP_828849.2;SARS-CoV_orf1ab;6392 | 0 | 0.05 |
| NC_004718.3;NP_828849.2;SARS-CoV_orf1ab;6393 | 0 | 0.025 |
| NC_004718.3;NP_828849.2;SARS-CoV_orf1ab;6394 | 0 | 0.025 |
| NC_004718.3;NP_828849.2;SARS-CoV_orf1ab;6395 | 0 | 0.025 |
| NC_004718.3;NP_828849.2;SARS-CoV_orf1ab;6396 | 0 | 0 |
| NC_004718.3;NP_828849.2;SARS-CoV_orf1ab;6397 | 0 | 0 |
| NC_004718.3;NP_828849.2;SARS-CoV_orf1ab;6398 | 0 | 0 |
| NC_004718.3;NP_828849.2;SARS-CoV_orf1ab;6399 | 0 | 0 |
| NC_004718.3;NP_828849.2;SARS-CoV_orf1ab;6400 | 0 | 0 |
| NC_004718.3;NP_828849.2;SARS-CoV_orf1ab;6401 | 0 | 0 |
| NC_004718.3;NP_828849.2;SARS-CoV_orf1ab;6402 | 0 | 0 |
| NC_004718.3;NP_828849.2;SARS-CoV_orf1ab;6403 | 0 | 0 |
| NC_004718.3;NP_828849.2;SARS-CoV_orf1ab;6404 | 0 | 0 |

| | | |
|---|---|---|
| NC_004718.3;NP_828849.2;SARS-CoV_orf1ab;6405 | 0 | 0 |
| NC_004718.3;NP_828849.2;SARS-CoV_orf1ab;6406 | 0 | 0 |
| NC_004718.3;NP_828849.2;SARS-CoV_orf1ab;6407 | 0 | 0 |
| NC_004718.3;NP_828849.2;SARS-CoV_orf1ab;6408 | 0 | 0 |
| NC_004718.3;NP_828849.2;SARS-CoV_orf1ab;6409 | 0 | 0 |
| NC_004718.3;NP_828849.2;SARS-CoV_orf1ab;6410 | 0 | 0 |
| NC_004718.3;NP_828849.2;SARS-CoV_orf1ab;6411 | 0 | 0 |
| NC_004718.3;NP_828849.2;SARS-CoV_orf1ab;6412 | 0 | 0 |
| NC_004718.3;NP_828849.2;SARS-CoV_orf1ab;6413 | 0 | 0 |
| NC_004718.3;NP_828849.2;SARS-CoV_orf1ab;6414 | 0 | 0 |
| NC_004718.3;NP_828849.2;SARS-CoV_orf1ab;6415 | 0 | 0 |
| NC_004718.3;NP_828849.2;SARS-CoV_orf1ab;6416 | 0 | 0 |
| NC_004718.3;NP_828849.2;SARS-CoV_orf1ab;6417 | 0 | 0 |
| NC_004718.3;NP_828849.2;SARS-CoV_orf1ab;6418 | 0 | 0 |
| NC_004718.3;NP_828849.2;SARS-CoV_orf1ab;6419 | 0 | 0 |
| NC_004718.3;NP_828849.2;SARS-CoV_orf1ab;6420 | 0 | 0 |
| NC_004718.3;NP_828849.2;SARS-CoV_orf1ab;6421 | 0 | 0 |
| NC_004718.3;NP_828849.2;SARS-CoV_orf1ab;6422 | 0 | 0 |
| NC_004718.3;NP_828849.2;SARS-CoV_orf1ab;6423 | 0 | 0 |
| NC_004718.3;NP_828849.2;SARS-CoV_orf1ab;6424 | 0 | 0 |
| NC_004718.3;NP_828849.2;SARS-CoV_orf1ab;6425 | 0.05 | 0 |
| NC_004718.3;NP_828849.2;SARS-CoV_orf1ab;6426 | 0 | 0 |
| NC_004718.3;NP_828849.2;SARS-CoV_orf1ab;6427 | 0.05 | 0 |
| NC_004718.3;NP_828849.2;SARS-CoV_orf1ab;6428 | 0 | 0 |
| NC_004718.3;NP_828849.2;SARS-CoV_orf1ab;6429 | 0 | 0.025 |
| NC_004718.3;NP_828849.2;SARS-CoV_orf1ab;6430 | 0.05 | 0 |
| NC_004718.3;NP_828849.2;SARS-CoV_orf1ab;6431 | 0.05 | 0.025 |
| NC_004718.3;NP_828849.2;SARS-CoV_orf1ab;6432 | 0 | 0 |
| NC_004718.3;NP_828849.2;SARS-CoV_orf1ab;6433 | 0 | 0 |
| NC_004718.3;NP_828849.2;SARS-CoV_orf1ab;6434 | 0 | 0 |
| NC_004718.3;NP_828849.2;SARS-CoV_orf1ab;6435 | 0 | 0.025 |
| NC_004718.3;NP_828849.2;SARS-CoV_orf1ab;6436 | 0.05 | 0 |
| NC_004718.3;NP_828849.2;SARS-CoV_orf1ab;6437 | 0 | 0 |
| NC_004718.3;NP_828849.2;SARS-CoV_orf1ab;6438 | 0.05 | 0 |
| NC_004718.3;NP_828849.2;SARS-CoV_orf1ab;6439 | 0 | 0.025 |
| NC_004718.3;NP_828849.2;SARS-CoV_orf1ab;6440 | 0 | 0 |
| NC_004718.3;NP_828849.2;SARS-CoV_orf1ab;6441 | 0 | 0 |
| NC_004718.3;NP_828849.2;SARS-CoV_orf1ab;6442 | 0 | 0 |
| NC_004718.3;NP_828849.2;SARS-CoV_orf1ab;6443 | 0 | 0 |
| NC_004718.3;NP_828849.2;SARS-CoV_orf1ab;6444 | 0 | 0 |
| NC_004718.3;NP_828849.2;SARS-CoV_orf1ab;6445 | 0 | 0 |
| NC_004718.3;NP_828849.2;SARS-CoV_orf1ab;6446 | 0.05 | 0 |
| NC_004718.3;NP_828849.2;SARS-CoV_orf1ab;6447 | 0 | 0 |

| | | |
|---|---|---|
| NC_004718.3;NP_828849.2;SARS-CoV_orf1ab;6448 | 0 | 0.025 |
| NC_004718.3;NP_828849.2;SARS-CoV_orf1ab;6449 | 0 | 0.05 |
| NC_004718.3;NP_828849.2;SARS-CoV_orf1ab;6450 | 0 | 0.05 |
| NC_004718.3;NP_828849.2;SARS-CoV_orf1ab;6451 | 0 | 0.025 |
| NC_004718.3;NP_828849.2;SARS-CoV_orf1ab;6452 | 0.05 | 0.05 |
| NC_004718.3;NP_828849.2;SARS-CoV_orf1ab;6453 | 0 | 0.025 |
| NC_004718.3;NP_828849.2;SARS-CoV_orf1ab;6454 | 0 | 0.025 |
| NC_004718.3;NP_828849.2;SARS-CoV_orf1ab;6455 | 0.05 | 0.05 |
| NC_004718.3;NP_828849.2;SARS-CoV_orf1ab;6456 | 0 | 0.025 |
| NC_004718.3;NP_828849.2;SARS-CoV_orf1ab;6457 | 0 | 0.025 |
| NC_004718.3;NP_828849.2;SARS-CoV_orf1ab;6458 | 0 | 0.025 |
| NC_004718.3;NP_828849.2;SARS-CoV_orf1ab;6459 | 0 | 0.025 |
| NC_004718.3;NP_828849.2;SARS-CoV_orf1ab;6460 | 0 | 0.05 |
| NC_004718.3;NP_828849.2;SARS-CoV_orf1ab;6461 | 0 | 0.075 |
| NC_004718.3;NP_828849.2;SARS-CoV_orf1ab;6462 | 0 | 0.075 |
| NC_004718.3;NP_828849.2;SARS-CoV_orf1ab;6463 | 0 | 0.075 |
| NC_004718.3;NP_828849.2;SARS-CoV_orf1ab;6464 | 0 | 0.025 |
| NC_004718.3;NP_828849.2;SARS-CoV_orf1ab;6465 | 0 | 0.025 |
| NC_004718.3;NP_828849.2;SARS-CoV_orf1ab;6466 | 0.1 | 0.075 |
| NC_004718.3;NP_828849.2;SARS-CoV_orf1ab;6467 | 0 | 0.025 |
| NC_004718.3;NP_828849.2;SARS-CoV_orf1ab;6468 | 0.1 | 0.05 |
| NC_004718.3;NP_828849.2;SARS-CoV_orf1ab;6469 | 0 | 0.05 |
| NC_004718.3;NP_828849.2;SARS-CoV_orf1ab;6470 | 0 | 0.05 |
| NC_004718.3;NP_828849.2;SARS-CoV_orf1ab;6471 | 0 | 0.025 |
| NC_004718.3;NP_828849.2;SARS-CoV_orf1ab;6472 | 0 | 0.025 |
| NC_004718.3;NP_828849.2;SARS-CoV_orf1ab;6473 | 0.05 | 0.025 |
| NC_004718.3;NP_828849.2;SARS-CoV_orf1ab;6474 | 0 | 0 |
| NC_004718.3;NP_828849.2;SARS-CoV_orf1ab;6475 | 0 | 0 |
| NC_004718.3;NP_828849.2;SARS-CoV_orf1ab;6476 | 0 | 0 |
| NC_004718.3;NP_828849.2;SARS-CoV_orf1ab;6477 | 0 | 0 |
| NC_004718.3;NP_828849.2;SARS-CoV_orf1ab;6478 | 0 | 0 |
| NC_004718.3;NP_828849.2;SARS-CoV_orf1ab;6479 | 0 | 0 |
| NC_004718.3;NP_828849.2;SARS-CoV_orf1ab;6480 | 0 | 0 |
| NC_004718.3;NP_828849.2;SARS-CoV_orf1ab;6481 | 0 | 0 |
| NC_004718.3;NP_828849.2;SARS-CoV_orf1ab;6482 | 0.05 | 0 |
| NC_004718.3;NP_828849.2;SARS-CoV_orf1ab;6483 | 0 | 0 |
| NC_004718.3;NP_828849.2;SARS-CoV_orf1ab;6484 | 0 | 0.075 |
| NC_004718.3;NP_828849.2;SARS-CoV_orf1ab;6485 | 0.1 | 0.075 |
| NC_004718.3;NP_828849.2;SARS-CoV_orf1ab;6486 | 0 | 0.05 |
| NC_004718.3;NP_828849.2;SARS-CoV_orf1ab;6487 | 0 | 0.05 |
| NC_004718.3;NP_828849.2;SARS-CoV_orf1ab;6488 | 0.05 | 0.075 |
| NC_004718.3;NP_828849.2;SARS-CoV_orf1ab;6489 | 0.1 | 0.125 |
| NC_004718.3;NP_828849.2;SARS-CoV_orf1ab;6490 | 0 | 0.1 |

| | | |
|---|---|---|
| NC_004718.3;NP_828849.2;SARS-CoV_orf1ab;6491 | 0.3 | 0.15 |
| NC_004718.3;NP_828849.2;SARS-CoV_orf1ab;6492 | 0.25 | 0.2 |
| NC_004718.3;NP_828849.2;SARS-CoV_orf1ab;6493 | 0.15 | 0.1 |
| NC_004718.3;NP_828849.2;SARS-CoV_orf1ab;6494 | 0 | 0 |
| NC_004718.3;NP_828849.2;SARS-CoV_orf1ab;6495 | 0.05 | 0 |
| NC_004718.3;NP_828849.2;SARS-CoV_orf1ab;6496 | 0 | 0.05 |
| NC_004718.3;NP_828849.2;SARS-CoV_orf1ab;6497 | 0 | 0 |
| NC_004718.3;NP_828849.2;SARS-CoV_orf1ab;6498 | 0 | 0.025 |
| NC_004718.3;NP_828849.2;SARS-CoV_orf1ab;6499 | 0 | 0 |
| NC_004718.3;NP_828849.2;SARS-CoV_orf1ab;6500 | 0 | 0 |
| NC_004718.3;NP_828849.2;SARS-CoV_orf1ab;6501 | 0 | 0 |
| NC_004718.3;NP_828849.2;SARS-CoV_orf1ab;6502 | 0 | 0 |
| NC_004718.3;NP_828849.2;SARS-CoV_orf1ab;6503 | 0 | 0 |
| NC_004718.3;NP_828849.2;SARS-CoV_orf1ab;6504 | 0 | 0 |
| NC_004718.3;NP_828849.2;SARS-CoV_orf1ab;6505 | 0.05 | 0.025 |
| NC_004718.3;NP_828849.2;SARS-CoV_orf1ab;6506 | 0 | 0 |
| NC_004718.3;NP_828849.2;SARS-CoV_orf1ab;6507 | 0 | 0 |
| NC_004718.3;NP_828849.2;SARS-CoV_orf1ab;6508 | 0 | 0.025 |
| NC_004718.3;NP_828849.2;SARS-CoV_orf1ab;6509 | 0 | 0 |
| NC_004718.3;NP_828849.2;SARS-CoV_orf1ab;6510 | 0 | 0 |
| NC_004718.3;NP_828849.2;SARS-CoV_orf1ab;6511 | 0 | 0 |
| NC_004718.3;NP_828849.2;SARS-CoV_orf1ab;6512 | 0 | 0 |
| NC_004718.3;NP_828849.2;SARS-CoV_orf1ab;6513 | 0.05 | 0 |
| NC_004718.3;NP_828849.2;SARS-CoV_orf1ab;6514 | 0.05 | 0 |
| NC_004718.3;NP_828849.2;SARS-CoV_orf1ab;6515 | 0 | 0 |
| NC_004718.3;NP_828849.2;SARS-CoV_orf1ab;6516 | 0 | 0 |
| NC_004718.3;NP_828849.2;SARS-CoV_orf1ab;6517 | 0 | 0 |
| NC_004718.3;NP_828849.2;SARS-CoV_orf1ab;6518 | 0 | 0 |
| NC_004718.3;NP_828849.2;SARS-CoV_orf1ab;6519 | 0 | 0 |
| NC_004718.3;NP_828849.2;SARS-CoV_orf1ab;6520 | 0 | 0 |
| NC_004718.3;NP_828849.2;SARS-CoV_orf1ab;6521 | 0 | 0 |
| NC_004718.3;NP_828849.2;SARS-CoV_orf1ab;6522 | 0 | 0 |
| NC_004718.3;NP_828849.2;SARS-CoV_orf1ab;6523 | 0 | 0 |
| NC_004718.3;NP_828849.2;SARS-CoV_orf1ab;6524 | 0.05 | 0 |
| NC_004718.3;NP_828849.2;SARS-CoV_orf1ab;6525 | 0.05 | 0.025 |
| NC_004718.3;NP_828849.2;SARS-CoV_orf1ab;6526 | 0.05 | 0.05 |
| NC_004718.3;NP_828849.2;SARS-CoV_orf1ab;6527 | 0.1 | 0.025 |
| NC_004718.3;NP_828849.2;SARS-CoV_orf1ab;6528 | 0.05 | 0.025 |
| NC_004718.3;NP_828849.2;SARS-CoV_orf1ab;6529 | 0.1 | 0.025 |
| NC_004718.3;NP_828849.2;SARS-CoV_orf1ab;6530 | 0.1 | 0.025 |
| NC_004718.3;NP_828849.2;SARS-CoV_orf1ab;6531 | 0.05 | 0.025 |
| NC_004718.3;NP_828849.2;SARS-CoV_orf1ab;6532 | 0.05 | 0.025 |
| NC_004718.3;NP_828849.2;SARS-CoV_orf1ab;6533 | 0.05 | 0.05 |

| | | |
|---|---|---|
| NC_004718.3;NP_828849.2;SARS-CoV_orf1ab;6534 | 0.05 | 0.025 |
| NC_004718.3;NP_828849.2;SARS-CoV_orf1ab;6535 | 0 | 0 |
| NC_004718.3;NP_828849.2;SARS-CoV_orf1ab;6536 | 0 | 0 |
| NC_004718.3;NP_828849.2;SARS-CoV_orf1ab;6537 | 0 | 0 |
| NC_004718.3;NP_828849.2;SARS-CoV_orf1ab;6538 | 0 | 0 |
| NC_004718.3;NP_828849.2;SARS-CoV_orf1ab;6539 | 0 | 0 |
| NC_004718.3;NP_828849.2;SARS-CoV_orf1ab;6540 | 0 | 0 |
| NC_004718.3;NP_828849.2;SARS-CoV_orf1ab;6541 | 0 | 0 |
| NC_004718.3;NP_828849.2;SARS-CoV_orf1ab;6542 | 0 | 0 |
| NC_004718.3;NP_828849.2;SARS-CoV_orf1ab;6543 | 0 | 0 |
| NC_004718.3;NP_828849.2;SARS-CoV_orf1ab;6544 | 0 | 0 |
| NC_004718.3;NP_828849.2;SARS-CoV_orf1ab;6545 | 0 | 0 |
| NC_004718.3;NP_828849.2;SARS-CoV_orf1ab;6546 | 0 | 0 |
| NC_004718.3;NP_828849.2;SARS-CoV_orf1ab;6547 | 0 | 0 |
| NC_004718.3;NP_828849.2;SARS-CoV_orf1ab;6548 | 0 | 0 |
| NC_004718.3;NP_828849.2;SARS-CoV_orf1ab;6549 | 0 | 0 |
| NC_004718.3;NP_828849.2;SARS-CoV_orf1ab;6550 | 0 | 0 |
| NC_004718.3;NP_828849.2;SARS-CoV_orf1ab;6551 | 0 | 0 |
| NC_004718.3;NP_828849.2;SARS-CoV_orf1ab;6552 | 0 | 0 |
| NC_004718.3;NP_828849.2;SARS-CoV_orf1ab;6553 | 0.05 | 0.025 |
| NC_004718.3;NP_828849.2;SARS-CoV_orf1ab;6554 | 0 | 0 |
| NC_004718.3;NP_828849.2;SARS-CoV_orf1ab;6555 | 0 | 0 |
| NC_004718.3;NP_828849.2;SARS-CoV_orf1ab;6556 | 0 | 0 |
| NC_004718.3;NP_828849.2;SARS-CoV_orf1ab;6557 | 0 | 0 |
| NC_004718.3;NP_828849.2;SARS-CoV_orf1ab;6558 | 0 | 0.025 |
| NC_004718.3;NP_828849.2;SARS-CoV_orf1ab;6559 | 0 | 0 |
| NC_004718.3;NP_828849.2;SARS-CoV_orf1ab;6560 | 0 | 0 |
| NC_004718.3;NP_828849.2;SARS-CoV_orf1ab;6561 | 0 | 0 |
| NC_004718.3;NP_828849.2;SARS-CoV_orf1ab;6562 | 0 | 0 |
| NC_004718.3;NP_828849.2;SARS-CoV_orf1ab;6563 | 0 | 0 |
| NC_004718.3;NP_828849.2;SARS-CoV_orf1ab;6564 | 0 | 0 |
| NC_004718.3;NP_828849.2;SARS-CoV_orf1ab;6565 | 0 | 0 |
| NC_004718.3;NP_828849.2;SARS-CoV_orf1ab;6566 | 0 | 0 |
| NC_004718.3;NP_828849.2;SARS-CoV_orf1ab;6567 | 0 | 0 |
| NC_004718.3;NP_828849.2;SARS-CoV_orf1ab;6568 | 0 | 0 |
| NC_004718.3;NP_828849.2;SARS-CoV_orf1ab;6569 | 0 | 0 |
| NC_004718.3;NP_828849.2;SARS-CoV_orf1ab;6570 | 0 | 0.025 |
| NC_004718.3;NP_828849.2;SARS-CoV_orf1ab;6571 | 0 | 0.025 |
| NC_004718.3;NP_828849.2;SARS-CoV_orf1ab;6572 | 0 | 0.025 |
| NC_004718.3;NP_828849.2;SARS-CoV_orf1ab;6573 | 0 | 0.025 |
| NC_004718.3;NP_828849.2;SARS-CoV_orf1ab;6574 | 0 | 0.025 |
| NC_004718.3;NP_828849.2;SARS-CoV_orf1ab;6575 | 0.05 | 0.175 |
| NC_004718.3;NP_828849.2;SARS-CoV_orf1ab;6576 | 0 | 0.025 |

| | | |
|---|---|---|
| NC_004718.3;NP_828849.2;SARS-CoV_orf1ab;6577 | 0 | 0 |
| NC_004718.3;NP_828849.2;SARS-CoV_orf1ab;6578 | 0 | 0 |
| NC_004718.3;NP_828849.2;SARS-CoV_orf1ab;6579 | 0 | 0.025 |
| NC_004718.3;NP_828849.2;SARS-CoV_orf1ab;6580 | 0 | 0 |
| NC_004718.3;NP_828849.2;SARS-CoV_orf1ab;6581 | 0.05 | 0.025 |
| NC_004718.3;NP_828849.2;SARS-CoV_orf1ab;6582 | 0 | 0 |
| NC_004718.3;NP_828849.2;SARS-CoV_orf1ab;6583 | 0 | 0 |
| NC_004718.3;NP_828849.2;SARS-CoV_orf1ab;6584 | 0 | 0 |
| NC_004718.3;NP_828849.2;SARS-CoV_orf1ab;6585 | 0 | 0 |
| NC_004718.3;NP_828849.2;SARS-CoV_orf1ab;6586 | 0 | 0 |
| NC_004718.3;NP_828849.2;SARS-CoV_orf1ab;6587 | 0 | 0 |
| NC_004718.3;NP_828849.2;SARS-CoV_orf1ab;6588 | 0 | 0 |
| NC_004718.3;NP_828849.2;SARS-CoV_orf1ab;6589 | 0 | 0 |
| NC_004718.3;NP_828849.2;SARS-CoV_orf1ab;6590 | 0 | 0 |
| NC_004718.3;NP_828849.2;SARS-CoV_orf1ab;6591 | 0 | 0 |
| NC_004718.3;NP_828849.2;SARS-CoV_orf1ab;6592 | 0 | 0 |
| NC_004718.3;NP_828849.2;SARS-CoV_orf1ab;6593 | 0 | 0 |
| NC_004718.3;NP_828849.2;SARS-CoV_orf1ab;6594 | 0 | 0.025 |
| NC_004718.3;NP_828849.2;SARS-CoV_orf1ab;6595 | 0 | 0 |
| NC_004718.3;NP_828849.2;SARS-CoV_orf1ab;6596 | 0 | 0.025 |
| NC_004718.3;NP_828849.2;SARS-CoV_orf1ab;6597 | 0 | 0.025 |
| NC_004718.3;NP_828849.2;SARS-CoV_orf1ab;6598 | 0.1 | 0.05 |
| NC_004718.3;NP_828849.2;SARS-CoV_orf1ab;6599 | 0.1 | 0.1 |
| NC_004718.3;NP_828849.2;SARS-CoV_orf1ab;6600 | 0 | 0.025 |
| NC_004718.3;NP_828849.2;SARS-CoV_orf1ab;6601 | 0 | 0.025 |
| NC_004718.3;NP_828849.2;SARS-CoV_orf1ab;6602 | 0 | 0.025 |
| NC_004718.3;NP_828849.2;SARS-CoV_orf1ab;6603 | 0 | 0.05 |
| NC_004718.3;NP_828849.2;SARS-CoV_orf1ab;6604 | 0 | 0.075 |
| NC_004718.3;NP_828849.2;SARS-CoV_orf1ab;6605 | 0 | 0.1 |
| NC_004718.3;NP_828849.2;SARS-CoV_orf1ab;6606 | 0.1 | 0.075 |
| NC_004718.3;NP_828849.2;SARS-CoV_orf1ab;6607 | 0.1 | 0.15 |
| NC_004718.3;NP_828849.2;SARS-CoV_orf1ab;6608 | 0.05 | 0.175 |
| NC_004718.3;NP_828849.2;SARS-CoV_orf1ab;6609 | 0 | 0.125 |
| NC_004718.3;NP_828849.2;SARS-CoV_orf1ab;6610 | 0.05 | 0.175 |
| NC_004718.3;NP_828849.2;SARS-CoV_orf1ab;6611 | 0 | 0 |
| NC_004718.3;NP_828849.2;SARS-CoV_orf1ab;6612 | 0 | 0 |
| NC_004718.3;NP_828849.2;SARS-CoV_orf1ab;6613 | 0 | 0 |
| NC_004718.3;NP_828849.2;SARS-CoV_orf1ab;6614 | 0 | 0 |
| NC_004718.3;NP_828849.2;SARS-CoV_orf1ab;6615 | 0 | 0 |
| NC_004718.3;NP_828849.2;SARS-CoV_orf1ab;6616 | 0 | 0 |
| NC_004718.3;NP_828849.2;SARS-CoV_orf1ab;6617 | 0 | 0 |
| NC_004718.3;NP_828849.2;SARS-CoV_orf1ab;6618 | 0 | 0.05 |
| NC_004718.3;NP_828849.2;SARS-CoV_orf1ab;6619 | 0 | 0.025 |

| | | |
|---|---|---|
| NC_004718.3;NP_828849.2;SARS-CoV_orf1ab;6620 | 0.05 | 0.05 |
| NC_004718.3;NP_828849.2;SARS-CoV_orf1ab;6621 | 0.05 | 0.075 |
| NC_004718.3;NP_828849.2;SARS-CoV_orf1ab;6622 | 0.05 | 0.025 |
| NC_004718.3;NP_828849.2;SARS-CoV_orf1ab;6623 | 0.1 | 0.075 |
| NC_004718.3;NP_828849.2;SARS-CoV_orf1ab;6624 | 0.05 | 0.025 |
| NC_004718.3;NP_828849.2;SARS-CoV_orf1ab;6625 | 0.05 | 0.025 |
| NC_004718.3;NP_828849.2;SARS-CoV_orf1ab;6626 | 0.05 | 0.025 |
| NC_004718.3;NP_828849.2;SARS-CoV_orf1ab;6627 | 0.05 | 0.025 |
| NC_004718.3;NP_828849.2;SARS-CoV_orf1ab;6628 | 0.15 | 0.075 |
| NC_004718.3;NP_828849.2;SARS-CoV_orf1ab;6629 | 0.2 | 0.05 |
| NC_004718.3;NP_828849.2;SARS-CoV_orf1ab;6630 | 0 | 0.025 |
| NC_004718.3;NP_828849.2;SARS-CoV_orf1ab;6631 | 0 | 0.025 |
| NC_004718.3;NP_828849.2;SARS-CoV_orf1ab;6632 | 0 | 0.025 |
| NC_004718.3;NP_828849.2;SARS-CoV_orf1ab;6633 | 0 | 0.025 |
| NC_004718.3;NP_828849.2;SARS-CoV_orf1ab;6634 | 0 | 0.025 |
| NC_004718.3;NP_828849.2;SARS-CoV_orf1ab;6635 | 0 | 0.025 |
| NC_004718.3;NP_828849.2;SARS-CoV_orf1ab;6636 | 0 | 0.025 |
| NC_004718.3;NP_828849.2;SARS-CoV_orf1ab;6637 | 0 | 0.025 |
| NC_004718.3;NP_828849.2;SARS-CoV_orf1ab;6638 | 0 | 0.025 |
| NC_004718.3;NP_828849.2;SARS-CoV_orf1ab;6639 | 0 | 0.025 |
| NC_004718.3;NP_828849.2;SARS-CoV_orf1ab;6640 | 0 | 0 |
| NC_004718.3;NP_828849.2;SARS-CoV_orf1ab;6641 | 0 | 0 |
| NC_004718.3;NP_828849.2;SARS-CoV_orf1ab;6642 | 0 | 0.025 |
| NC_004718.3;NP_828849.2;SARS-CoV_orf1ab;6643 | 0 | 0.025 |
| NC_004718.3;NP_828849.2;SARS-CoV_orf1ab;6644 | 0 | 0.025 |
| NC_004718.3;NP_828849.2;SARS-CoV_orf1ab;6645 | 0 | 0.1 |
| NC_004718.3;NP_828849.2;SARS-CoV_orf1ab;6646 | 0 | 0.025 |
| NC_004718.3;NP_828849.2;SARS-CoV_orf1ab;6647 | 0 | 0 |
| NC_004718.3;NP_828849.2;SARS-CoV_orf1ab;6648 | 0 | 0 |
| NC_004718.3;NP_828849.2;SARS-CoV_orf1ab;6649 | 0 | 0 |
| NC_004718.3;NP_828849.2;SARS-CoV_orf1ab;6650 | 0 | 0 |
| NC_004718.3;NP_828849.2;SARS-CoV_orf1ab;6651 | 0 | 0 |
| NC_004718.3;NP_828849.2;SARS-CoV_orf1ab;6652 | 0 | 0 |
| NC_004718.3;NP_828849.2;SARS-CoV_orf1ab;6653 | 0 | 0 |
| NC_004718.3;NP_828849.2;SARS-CoV_orf1ab;6654 | 0 | 0 |
| NC_004718.3;NP_828849.2;SARS-CoV_orf1ab;6655 | 0 | 0 |
| NC_004718.3;NP_828849.2;SARS-CoV_orf1ab;6656 | 0 | 0 |
| NC_004718.3;NP_828849.2;SARS-CoV_orf1ab;6657 | 0 | 0 |
| NC_004718.3;NP_828849.2;SARS-CoV_orf1ab;6658 | 0 | 0 |
| NC_004718.3;NP_828849.2;SARS-CoV_orf1ab;6659 | 0 | 0 |
| NC_004718.3;NP_828849.2;SARS-CoV_orf1ab;6660 | 0 | 0 |
| NC_004718.3;NP_828849.2;SARS-CoV_orf1ab;6661 | 0 | 0 |
| NC_004718.3;NP_828849.2;SARS-CoV_orf1ab;6662 | 0 | 0 |

| | | |
|---|---|---|
| NC_004718.3;NP_828849.2;SARS-CoV_orf1ab;6663 | 0 | 0 |
| NC_004718.3;NP_828849.2;SARS-CoV_orf1ab;6664 | 0 | 0 |
| NC_004718.3;NP_828849.2;SARS-CoV_orf1ab;6665 | 0 | 0 |
| NC_004718.3;NP_828849.2;SARS-CoV_orf1ab;6666 | 0 | 0 |
| NC_004718.3;NP_828849.2;SARS-CoV_orf1ab;6667 | 0 | 0 |
| NC_004718.3;NP_828849.2;SARS-CoV_orf1ab;6668 | 0 | 0 |
| NC_004718.3;NP_828849.2;SARS-CoV_orf1ab;6669 | 0 | 0 |
| NC_004718.3;NP_828849.2;SARS-CoV_orf1ab;6670 | 0 | 0 |
| NC_004718.3;NP_828849.2;SARS-CoV_orf1ab;6671 | 0 | 0 |
| NC_004718.3;NP_828849.2;SARS-CoV_orf1ab;6672 | 0 | 0 |
| NC_004718.3;NP_828849.2;SARS-CoV_orf1ab;6673 | 0 | 0 |
| NC_004718.3;NP_828849.2;SARS-CoV_orf1ab;6674 | 0 | 0 |
| NC_004718.3;NP_828849.2;SARS-CoV_orf1ab;6675 | 0 | 0 |
| NC_004718.3;NP_828849.2;SARS-CoV_orf1ab;6676 | 0 | 0 |
| NC_004718.3;NP_828849.2;SARS-CoV_orf1ab;6677 | 0 | 0 |
| NC_004718.3;NP_828849.2;SARS-CoV_orf1ab;6678 | 0 | 0 |
| NC_004718.3;NP_828849.2;SARS-CoV_orf1ab;6679 | 0 | 0 |
| NC_004718.3;NP_828849.2;SARS-CoV_orf1ab;6680 | 0 | 0 |
| NC_004718.3;NP_828849.2;SARS-CoV_orf1ab;6681 | 0 | 0 |
| NC_004718.3;NP_828849.2;SARS-CoV_orf1ab;6682 | 0 | 0 |
| NC_004718.3;NP_828849.2;SARS-CoV_orf1ab;6683 | 0 | 0 |
| NC_004718.3;NP_828849.2;SARS-CoV_orf1ab;6684 | 0 | 0 |
| NC_004718.3;NP_828849.2;SARS-CoV_orf1ab;6685 | 0.05 | 0 |
| NC_004718.3;NP_828849.2;SARS-CoV_orf1ab;6686 | 0 | 0 |
| NC_004718.3;NP_828849.2;SARS-CoV_orf1ab;6687 | 0.05 | 0.15 |
| NC_004718.3;NP_828849.2;SARS-CoV_orf1ab;6688 | 0.05 | 0.05 |
| NC_004718.3;NP_828849.2;SARS-CoV_orf1ab;6689 | 0.05 | 0 |
| NC_004718.3;NP_828849.2;SARS-CoV_orf1ab;6690 | 0 | 0.025 |
| NC_004718.3;NP_828849.2;SARS-CoV_orf1ab;6691 | 0 | 0 |
| NC_004718.3;NP_828849.2;SARS-CoV_orf1ab;6692 | 0.05 | 0 |
| NC_004718.3;NP_828849.2;SARS-CoV_orf1ab;6693 | 0.05 | 0 |
| NC_004718.3;NP_828849.2;SARS-CoV_orf1ab;6694 | 0.15 | 0 |
| NC_004718.3;NP_828849.2;SARS-CoV_orf1ab;6695 | 0 | 0 |
| NC_004718.3;NP_828849.2;SARS-CoV_orf1ab;6696 | 0.05 | 0 |
| NC_004718.3;NP_828849.2;SARS-CoV_orf1ab;6697 | 0.1 | 0 |
| NC_004718.3;NP_828849.2;SARS-CoV_orf1ab;6698 | 0.1 | 0.025 |
| NC_004718.3;NP_828849.2;SARS-CoV_orf1ab;6699 | 0 | 0 |
| NC_004718.3;NP_828849.2;SARS-CoV_orf1ab;6700 | 0.1 | 0 |
| NC_004718.3;NP_828849.2;SARS-CoV_orf1ab;6701 | 0.05 | 0 |
| NC_004718.3;NP_828849.2;SARS-CoV_orf1ab;6702 | 0.05 | 0 |
| NC_004718.3;NP_828849.2;SARS-CoV_orf1ab;6703 | 0.1 | 0 |
| NC_004718.3;NP_828849.2;SARS-CoV_orf1ab;6704 | 0.1 | 0 |
| NC_004718.3;NP_828849.2;SARS-CoV_orf1ab;6705 | 0.05 | 0.025 |

| | | |
|---|---|---|
| NC_004718.3;NP_828849.2;SARS-CoV_orf1ab;6706 | 0 | 0 |
| NC_004718.3;NP_828849.2;SARS-CoV_orf1ab;6707 | 0 | 0 |
| NC_004718.3;NP_828849.2;SARS-CoV_orf1ab;6708 | 0 | 0 |
| NC_004718.3;NP_828849.2;SARS-CoV_orf1ab;6709 | 0 | 0 |
| NC_004718.3;NP_828849.2;SARS-CoV_orf1ab;6710 | 0 | 0 |
| NC_004718.3;NP_828849.2;SARS-CoV_orf1ab;6711 | 0 | 0 |
| NC_004718.3;NP_828849.2;SARS-CoV_orf1ab;6712 | 0 | 0 |
| NC_004718.3;NP_828849.2;SARS-CoV_orf1ab;6713 | 0 | 0 |
| NC_004718.3;NP_828849.2;SARS-CoV_orf1ab;6714 | 0 | 0 |
| NC_004718.3;NP_828849.2;SARS-CoV_orf1ab;6715 | 0 | 0 |
| NC_004718.3;NP_828849.2;SARS-CoV_orf1ab;6716 | 0 | 0 |
| NC_004718.3;NP_828849.2;SARS-CoV_orf1ab;6717 | 0 | 0 |
| NC_004718.3;NP_828849.2;SARS-CoV_orf1ab;6718 | 0 | 0 |
| NC_004718.3;NP_828849.2;SARS-CoV_orf1ab;6719 | 0 | 0 |
| NC_004718.3;NP_828849.2;SARS-CoV_orf1ab;6720 | 0 | 0 |
| NC_004718.3;NP_828849.2;SARS-CoV_orf1ab;6721 | 0 | 0 |
| NC_004718.3;NP_828849.2;SARS-CoV_orf1ab;6722 | 0.05 | 0.025 |
| NC_004718.3;NP_828849.2;SARS-CoV_orf1ab;6723 | 0.05 | 0 |
| NC_004718.3;NP_828849.2;SARS-CoV_orf1ab;6724 | 0.05 | 0 |
| NC_004718.3;NP_828849.2;SARS-CoV_orf1ab;6725 | 0.05 | 0 |
| NC_004718.3;NP_828849.2;SARS-CoV_orf1ab;6726 | 0.1 | 0.125 |
| NC_004718.3;NP_828849.2;SARS-CoV_orf1ab;6727 | 0.05 | 0 |
| NC_004718.3;NP_828849.2;SARS-CoV_orf1ab;6728 | 0 | 0 |
| NC_004718.3;NP_828849.2;SARS-CoV_orf1ab;6729 | 0 | 0 |
| NC_004718.3;NP_828849.2;SARS-CoV_orf1ab;6730 | 0 | 0 |
| NC_004718.3;NP_828849.2;SARS-CoV_orf1ab;6731 | 0 | 0 |
| NC_004718.3;NP_828849.2;SARS-CoV_orf1ab;6732 | 0.05 | 0 |
| NC_004718.3;NP_828849.2;SARS-CoV_orf1ab;6733 | 0 | 0 |
| NC_004718.3;NP_828849.2;SARS-CoV_orf1ab;6734 | 0 | 0 |
| NC_004718.3;NP_828849.2;SARS-CoV_orf1ab;6735 | 0 | 0 |
| NC_004718.3;NP_828849.2;SARS-CoV_orf1ab;6736 | 0 | 0 |
| NC_004718.3;NP_828849.2;SARS-CoV_orf1ab;6737 | 0 | 0 |
| NC_004718.3;NP_828849.2;SARS-CoV_orf1ab;6738 | 0 | 0 |
| NC_004718.3;NP_828849.2;SARS-CoV_orf1ab;6739 | 0 | 0 |
| NC_004718.3;NP_828849.2;SARS-CoV_orf1ab;6740 | 0 | 0 |
| NC_004718.3;NP_828849.2;SARS-CoV_orf1ab;6741 | 0 | 0 |
| NC_004718.3;NP_828849.2;SARS-CoV_orf1ab;6742 | 0 | 0 |
| NC_004718.3;NP_828849.2;SARS-CoV_orf1ab;6743 | 0 | 0 |
| NC_004718.3;NP_828849.2;SARS-CoV_orf1ab;6744 | 0 | 0 |
| NC_004718.3;NP_828849.2;SARS-CoV_orf1ab;6745 | 0 | 0 |
| NC_004718.3;NP_828849.2;SARS-CoV_orf1ab;6746 | 0 | 0 |
| NC_004718.3;NP_828849.2;SARS-CoV_orf1ab;6747 | 0 | 0 |
| NC_004718.3;NP_828849.2;SARS-CoV_orf1ab;6748 | 0 | 0 |

| | | |
|---|---|---|
| NC_004718.3;NP_828849.2;SARS-CoV_orf1ab;6749 | 0 | 0.025 |
| NC_004718.3;NP_828849.2;SARS-CoV_orf1ab;6750 | 0 | 0 |
| NC_004718.3;NP_828849.2;SARS-CoV_orf1ab;6751 | 0 | 0 |
| NC_004718.3;NP_828849.2;SARS-CoV_orf1ab;6752 | 0 | 0 |
| NC_004718.3;NP_828849.2;SARS-CoV_orf1ab;6753 | 0 | 0 |
| NC_004718.3;NP_828849.2;SARS-CoV_orf1ab;6754 | 0 | 0.025 |
| NC_004718.3;NP_828849.2;SARS-CoV_orf1ab;6755 | 0 | 0 |
| NC_004718.3;NP_828849.2;SARS-CoV_orf1ab;6756 | 0 | 0 |
| NC_004718.3;NP_828849.2;SARS-CoV_orf1ab;6757 | 0 | 0 |
| NC_004718.3;NP_828849.2;SARS-CoV_orf1ab;6758 | 0 | 0 |
| NC_004718.3;NP_828849.2;SARS-CoV_orf1ab;6759 | 0 | 0 |
| NC_004718.3;NP_828849.2;SARS-CoV_orf1ab;6760 | 0 | 0.025 |
| NC_004718.3;NP_828849.2;SARS-CoV_orf1ab;6761 | 0 | 0.025 |
| NC_004718.3;NP_828849.2;SARS-CoV_orf1ab;6762 | 0 | 0.025 |
| NC_004718.3;NP_828849.2;SARS-CoV_orf1ab;6763 | 0 | 0.025 |
| NC_004718.3;NP_828849.2;SARS-CoV_orf1ab;6764 | 0 | 0.05 |
| NC_004718.3;NP_828849.2;SARS-CoV_orf1ab;6765 | 0 | 0.025 |
| NC_004718.3;NP_828849.2;SARS-CoV_orf1ab;6766 | 0 | 0.05 |
| NC_004718.3;NP_828849.2;SARS-CoV_orf1ab;6767 | 0 | 0.05 |
| NC_004718.3;NP_828849.2;SARS-CoV_orf1ab;6768 | 0 | 0.05 |
| NC_004718.3;NP_828849.2;SARS-CoV_orf1ab;6769 | 0 | 0.05 |
| NC_004718.3;NP_828849.2;SARS-CoV_orf1ab;6770 | 0.1 | 0.075 |
| NC_004718.3;NP_828849.2;SARS-CoV_orf1ab;6771 | 0 | 0.025 |
| NC_004718.3;NP_828849.2;SARS-CoV_orf1ab;6772 | 0.05 | 0.05 |
| NC_004718.3;NP_828849.2;SARS-CoV_orf1ab;6773 | 0 | 0.025 |
| NC_004718.3;NP_828849.2;SARS-CoV_orf1ab;6774 | 0.05 | 0.025 |
| NC_004718.3;NP_828849.2;SARS-CoV_orf1ab;6775 | 0 | 0 |
| NC_004718.3;NP_828849.2;SARS-CoV_orf1ab;6776 | 0 | 0 |
| NC_004718.3;NP_828849.2;SARS-CoV_orf1ab;6777 | 0 | 0 |
| NC_004718.3;NP_828849.2;SARS-CoV_orf1ab;6778 | 0 | 0 |
| NC_004718.3;NP_828849.2;SARS-CoV_orf1ab;6779 | 0 | 0 |
| NC_004718.3;NP_828849.2;SARS-CoV_orf1ab;6780 | 0 | 0 |
| NC_004718.3;NP_828849.2;SARS-CoV_orf1ab;6781 | 0 | 0 |
| NC_004718.3;NP_828849.2;SARS-CoV_orf1ab;6782 | 0 | 0 |
| NC_004718.3;NP_828849.2;SARS-CoV_orf1ab;6783 | 0 | 0 |
| NC_004718.3;NP_828849.2;SARS-CoV_orf1ab;6784 | 0 | 0 |
| NC_004718.3;NP_828849.2;SARS-CoV_orf1ab;6785 | 0 | 0 |
| NC_004718.3;NP_828849.2;SARS-CoV_orf1ab;6786 | 0 | 0.025 |
| NC_004718.3;NP_828849.2;SARS-CoV_orf1ab;6787 | 0 | 0 |
| NC_004718.3;NP_828849.2;SARS-CoV_orf1ab;6788 | 0 | 0 |
| NC_004718.3;NP_828849.2;SARS-CoV_orf1ab;6789 | 0 | 0 |
| NC_004718.3;NP_828849.2;SARS-CoV_orf1ab;6790 | 0 | 0 |
| NC_004718.3;NP_828849.2;SARS-CoV_orf1ab;6791 | 0 | 0 |

| | | |
|---|---|---|
| NC_004718.3;NP_828849.2;SARS-CoV_orf1ab;6792 | 0 | 0 |
| NC_004718.3;NP_828849.2;SARS-CoV_orf1ab;6793 | 0 | 0 |
| NC_004718.3;NP_828849.2;SARS-CoV_orf1ab;6794 | 0.05 | 0 |
| NC_004718.3;NP_828849.2;SARS-CoV_orf1ab;6795 | 0 | 0 |
| NC_004718.3;NP_828849.2;SARS-CoV_orf1ab;6796 | 0 | 0 |
| NC_004718.3;NP_828849.2;SARS-CoV_orf1ab;6797 | 0.05 | 0 |
| NC_004718.3;NP_828849.2;SARS-CoV_orf1ab;6798 | 0.05 | 0 |
| NC_004718.3;NP_828849.2;SARS-CoV_orf1ab;6799 | 0.05 | 0.025 |
| NC_004718.3;NP_828849.2;SARS-CoV_orf1ab;6800 | 0 | 0.025 |
| NC_004718.3;NP_828849.2;SARS-CoV_orf1ab;6801 | 0.05 | 0.025 |
| NC_004718.3;NP_828849.2;SARS-CoV_orf1ab;6802 | 0.05 | 0 |
| NC_004718.3;NP_828849.2;SARS-CoV_orf1ab;6803 | 0.05 | 0.025 |
| NC_004718.3;NP_828849.2;SARS-CoV_orf1ab;6804 | 0.05 | 0 |
| NC_004718.3;NP_828849.2;SARS-CoV_orf1ab;6805 | 0.05 | 0.025 |
| NC_004718.3;NP_828849.2;SARS-CoV_orf1ab;6806 | 0 | 0.025 |
| NC_004718.3;NP_828849.2;SARS-CoV_orf1ab;6807 | 0 | 0 |
| NC_004718.3;NP_828849.2;SARS-CoV_orf1ab;6808 | 0 | 0 |
| NC_004718.3;NP_828849.2;SARS-CoV_orf1ab;6809 | 0 | 0 |
| NC_004718.3;NP_828849.2;SARS-CoV_orf1ab;6810 | 0 | 0 |
| NC_004718.3;NP_828849.2;SARS-CoV_orf1ab;6811 | 0 | 0 |
| NC_004718.3;NP_828849.2;SARS-CoV_orf1ab;6812 | 0 | 0 |
| NC_004718.3;NP_828849.2;SARS-CoV_orf1ab;6813 | 0 | 0.05 |
| NC_004718.3;NP_828849.2;SARS-CoV_orf1ab;6814 | 0 | 0 |
| NC_004718.3;NP_828849.2;SARS-CoV_orf1ab;6815 | 0 | 0 |
| NC_004718.3;NP_828849.2;SARS-CoV_orf1ab;6816 | 0 | 0 |
| NC_004718.3;NP_828849.2;SARS-CoV_orf1ab;6817 | 0 | 0 |
| NC_004718.3;NP_828849.2;SARS-CoV_orf1ab;6818 | 0 | 0 |
| NC_004718.3;NP_828849.2;SARS-CoV_orf1ab;6819 | 0 | 0 |
| NC_004718.3;NP_828849.2;SARS-CoV_orf1ab;6820 | 0 | 0 |
| NC_004718.3;NP_828849.2;SARS-CoV_orf1ab;6821 | 0 | 0 |
| NC_004718.3;NP_828849.2;SARS-CoV_orf1ab;6822 | 0 | 0 |
| NC_004718.3;NP_828849.2;SARS-CoV_orf1ab;6823 | 0 | 0 |
| NC_004718.3;NP_828849.2;SARS-CoV_orf1ab;6824 | 0 | 0 |
| NC_004718.3;NP_828849.2;SARS-CoV_orf1ab;6825 | 0 | 0 |
| NC_004718.3;NP_828849.2;SARS-CoV_orf1ab;6826 | 0 | 0 |
| NC_004718.3;NP_828849.2;SARS-CoV_orf1ab;6827 | 0 | 0 |
| NC_004718.3;NP_828849.2;SARS-CoV_orf1ab;6828 | 0 | 0 |
| NC_004718.3;NP_828849.2;SARS-CoV_orf1ab;6829 | 0 | 0 |
| NC_004718.3;NP_828849.2;SARS-CoV_orf1ab;6830 | 0 | 0 |
| NC_004718.3;NP_828849.2;SARS-CoV_orf1ab;6831 | 0 | 0 |
| NC_004718.3;NP_828849.2;SARS-CoV_orf1ab;6832 | 0 | 0 |
| NC_004718.3;NP_828849.2;SARS-CoV_orf1ab;6833 | 0 | 0 |
| NC_004718.3;NP_828849.2;SARS-CoV_orf1ab;6834 | 0 | 0 |

| | | |
|---|---|---|
| NC_004718.3;NP_828849.2;SARS-CoV_orf1ab;6835 | 0 | 0 |
| NC_004718.3;NP_828849.2;SARS-CoV_orf1ab;6836 | 0 | 0 |
| NC_004718.3;NP_828849.2;SARS-CoV_orf1ab;6837 | 0 | 0 |
| NC_004718.3;NP_828849.2;SARS-CoV_orf1ab;6838 | 0 | 0 |
| NC_004718.3;NP_828849.2;SARS-CoV_orf1ab;6839 | 0.05 | 0.05 |
| NC_004718.3;NP_828849.2;SARS-CoV_orf1ab;6840 | 0 | 0 |
| NC_004718.3;NP_828849.2;SARS-CoV_orf1ab;6841 | 0 | 0 |
| NC_004718.3;NP_828849.2;SARS-CoV_orf1ab;6842 | 0 | 0 |
| NC_004718.3;NP_828849.2;SARS-CoV_orf1ab;6843 | 0 | 0 |
| NC_004718.3;NP_828849.2;SARS-CoV_orf1ab;6844 | 0 | 0 |
| NC_004718.3;NP_828849.2;SARS-CoV_orf1ab;6845 | 0 | 0 |
| NC_004718.3;NP_828849.2;SARS-CoV_orf1ab;6846 | 0 | 0 |
| NC_004718.3;NP_828849.2;SARS-CoV_orf1ab;6847 | 0.05 | 0 |
| NC_004718.3;NP_828849.2;SARS-CoV_orf1ab;6848 | 0.05 | 0 |
| NC_004718.3;NP_828849.2;SARS-CoV_orf1ab;6849 | 0 | 0 |
| NC_004718.3;NP_828849.2;SARS-CoV_orf1ab;6850 | 0 | 0 |
| NC_004718.3;NP_828849.2;SARS-CoV_orf1ab;6851 | 0 | 0 |
| NC_004718.3;NP_828849.2;SARS-CoV_orf1ab;6852 | 0 | 0 |
| NC_004718.3;NP_828849.2;SARS-CoV_orf1ab;6853 | 0 | 0 |
| NC_004718.3;NP_828849.2;SARS-CoV_orf1ab;6854 | 0 | 0 |
| NC_004718.3;NP_828849.2;SARS-CoV_orf1ab;6855 | 0 | 0 |
| NC_004718.3;NP_828849.2;SARS-CoV_orf1ab;6856 | 0 | 0 |
| NC_004718.3;NP_828849.2;SARS-CoV_orf1ab;6857 | 0 | 0 |
| NC_004718.3;NP_828849.2;SARS-CoV_orf1ab;6858 | 0 | 0 |
| NC_004718.3;NP_828849.2;SARS-CoV_orf1ab;6859 | 0 | 0 |
| NC_004718.3;NP_828849.2;SARS-CoV_orf1ab;6860 | 0 | 0 |
| NC_004718.3;NP_828849.2;SARS-CoV_orf1ab;6861 | 0 | 0 |
| NC_004718.3;NP_828849.2;SARS-CoV_orf1ab;6862 | 0 | 0 |
| NC_004718.3;NP_828849.2;SARS-CoV_orf1ab;6863 | 0 | 0 |
| NC_004718.3;NP_828849.2;SARS-CoV_orf1ab;6864 | 0 | 0.025 |
| NC_004718.3;NP_828849.2;SARS-CoV_orf1ab;6865 | 0 | 0.025 |
| NC_004718.3;NP_828849.2;SARS-CoV_orf1ab;6866 | 0 | 0.025 |
| NC_004718.3;NP_828849.2;SARS-CoV_orf1ab;6867 | 0 | 0.025 |
| NC_004718.3;NP_828849.2;SARS-CoV_orf1ab;6868 | 0.05 | 0.025 |
| NC_004718.3;NP_828849.2;SARS-CoV_orf1ab;6869 | 0.05 | 0.05 |
| NC_004718.3;NP_828849.2;SARS-CoV_orf1ab;6870 | 0.05 | 0.05 |
| NC_004718.3;NP_828849.2;SARS-CoV_orf1ab;6871 | 0 | 0.025 |
| NC_004718.3;NP_828849.2;SARS-CoV_orf1ab;6872 | 0 | 0.05 |
| NC_004718.3;NP_828849.2;SARS-CoV_orf1ab;6873 | 0.1 | 0.05 |
| NC_004718.3;NP_828849.2;SARS-CoV_orf1ab;6874 | 0 | 0.025 |
| NC_004718.3;NP_828849.2;SARS-CoV_orf1ab;6875 | 0 | 0 |
| NC_004718.3;NP_828849.2;SARS-CoV_orf1ab;6876 | 0.05 | 0 |
| NC_004718.3;NP_828849.2;SARS-CoV_orf1ab;6877 | 0.05 | 0 |

| | | |
|---|---|---|
| NC_004718.3;NP_828849.2;SARS-CoV_orf1ab;6878 | 0.05 | 0 |
| NC_004718.3;NP_828849.2;SARS-CoV_orf1ab;6879 | 0.05 | 0 |
| NC_004718.3;NP_828849.2;SARS-CoV_orf1ab;6880 | 0.05 | 0 |
| NC_004718.3;NP_828849.2;SARS-CoV_orf1ab;6881 | 0.05 | 0 |
| NC_004718.3;NP_828849.2;SARS-CoV_orf1ab;6882 | 0.05 | 0 |
| NC_004718.3;NP_828849.2;SARS-CoV_orf1ab;6883 | 0 | 0.025 |
| NC_004718.3;NP_828849.2;SARS-CoV_orf1ab;6884 | 0 | 0 |
| NC_004718.3;NP_828849.2;SARS-CoV_orf1ab;6885 | 0.05 | 0.175 |
| NC_004718.3;NP_828849.2;SARS-CoV_orf1ab;6886 | 0 | 0.1 |
| NC_004718.3;NP_828849.2;SARS-CoV_orf1ab;6887 | 0 | 0.025 |
| NC_004718.3;NP_828849.2;SARS-CoV_orf1ab;6888 | 0 | 0 |
| NC_004718.3;NP_828849.2;SARS-CoV_orf1ab;6889 | 0 | 0 |
| NC_004718.3;NP_828849.2;SARS-CoV_orf1ab;6890 | 0 | 0 |
| NC_004718.3;NP_828849.2;SARS-CoV_orf1ab;6891 | 0 | 0.025 |
| NC_004718.3;NP_828849.2;SARS-CoV_orf1ab;6892 | 0 | 0 |
| NC_004718.3;NP_828849.2;SARS-CoV_orf1ab;6893 | 0 | 0 |
| NC_004718.3;NP_828849.2;SARS-CoV_orf1ab;6894 | 0 | 0 |
| NC_004718.3;NP_828849.2;SARS-CoV_orf1ab;6895 | 0 | 0 |
| NC_004718.3;NP_828849.2;SARS-CoV_orf1ab;6896 | 0.1 | 0.05 |
| NC_004718.3;NP_828849.2;SARS-CoV_orf1ab;6897 | 0 | 0 |
| NC_004718.3;NP_828849.2;SARS-CoV_orf1ab;6898 | 0 | 0.025 |
| NC_004718.3;NP_828849.2;SARS-CoV_orf1ab;6899 | 0.05 | 0 |
| NC_004718.3;NP_828849.2;SARS-CoV_orf1ab;6900 | 0.05 | 0 |
| NC_004718.3;NP_828849.2;SARS-CoV_orf1ab;6901 | 0 | 0 |
| NC_004718.3;NP_828849.2;SARS-CoV_orf1ab;6902 | 0 | 0.05 |
| NC_004718.3;NP_828849.2;SARS-CoV_orf1ab;6903 | 0.05 | 0.05 |
| NC_004718.3;NP_828849.2;SARS-CoV_orf1ab;6904 | 0.05 | 0.025 |
| NC_004718.3;NP_828849.2;SARS-CoV_orf1ab;6905 | 0.05 | 0.05 |
| NC_004718.3;NP_828849.2;SARS-CoV_orf1ab;6906 | 0 | 0.075 |
| NC_004718.3;NP_828849.2;SARS-CoV_orf1ab;6907 | 0.05 | 0.075 |
| NC_004718.3;NP_828849.2;SARS-CoV_orf1ab;6908 | 0.05 | 0.075 |
| NC_004718.3;NP_828849.2;SARS-CoV_orf1ab;6909 | 0.05 | 0.025 |
| NC_004718.3;NP_828849.2;SARS-CoV_orf1ab;6910 | 0.05 | 0.025 |
| NC_004718.3;NP_828849.2;SARS-CoV_orf1ab;6911 | 0.05 | 0.05 |
| NC_004718.3;NP_828849.2;SARS-CoV_orf1ab;6912 | 0.05 | 0.025 |
| NC_004718.3;NP_828849.2;SARS-CoV_orf1ab;6913 | 0.05 | 0.025 |
| NC_004718.3;NP_828849.2;SARS-CoV_orf1ab;6914 | 0 | 0.025 |
| NC_004718.3;NP_828849.2;SARS-CoV_orf1ab;6915 | 0 | 0.05 |
| NC_004718.3;NP_828849.2;SARS-CoV_orf1ab;6916 | 0 | 0 |
| NC_004718.3;NP_828849.2;SARS-CoV_orf1ab;6917 | 0 | 0 |
| NC_004718.3;NP_828849.2;SARS-CoV_orf1ab;6918 | 0 | 0 |
| NC_004718.3;NP_828849.2;SARS-CoV_orf1ab;6919 | 0 | 0 |
| NC_004718.3;NP_828849.2;SARS-CoV_orf1ab;6920 | 0 | 0 |

| | | |
|---|---|---|
| NC_004718.3;NP_828849.2;SARS-CoV_orf1ab;6921 | 0 | 0 |
| NC_004718.3;NP_828849.2;SARS-CoV_orf1ab;6922 | 0 | 0 |
| NC_004718.3;NP_828849.2;SARS-CoV_orf1ab;6923 | 0 | 0 |
| NC_004718.3;NP_828849.2;SARS-CoV_orf1ab;6924 | 0 | 0 |
| NC_004718.3;NP_828849.2;SARS-CoV_orf1ab;6925 | 0 | 0 |
| NC_004718.3;NP_828849.2;SARS-CoV_orf1ab;6926 | 0 | 0 |
| NC_004718.3;NP_828849.2;SARS-CoV_orf1ab;6927 | 0 | 0 |
| NC_004718.3;NP_828849.2;SARS-CoV_orf1ab;6928 | 0 | 0 |
| NC_004718.3;NP_828849.2;SARS-CoV_orf1ab;6929 | 0 | 0 |
| NC_004718.3;NP_828849.2;SARS-CoV_orf1ab;6930 | 0 | 0 |
| NC_004718.3;NP_828849.2;SARS-CoV_orf1ab;6931 | 0 | 0 |
| NC_004718.3;NP_828849.2;SARS-CoV_orf1ab;6932 | 0 | 0 |
| NC_004718.3;NP_828849.2;SARS-CoV_orf1ab;6933 | 0 | 0 |
| NC_004718.3;NP_828849.2;SARS-CoV_orf1ab;6934 | 0 | 0 |
| NC_004718.3;NP_828849.2;SARS-CoV_orf1ab;6935 | 0 | 0 |
| NC_004718.3;NP_828849.2;SARS-CoV_orf1ab;6936 | 0 | 0 |
| NC_004718.3;NP_828849.2;SARS-CoV_orf1ab;6937 | 0 | 0 |
| NC_004718.3;NP_828849.2;SARS-CoV_orf1ab;6938 | 0 | 0 |
| NC_004718.3;NP_828849.2;SARS-CoV_orf1ab;6939 | 0 | 0 |
| NC_004718.3;NP_828849.2;SARS-CoV_orf1ab;6940 | 0 | 0 |
| NC_004718.3;NP_828849.2;SARS-CoV_orf1ab;6941 | 0 | 0 |
| NC_004718.3;NP_828849.2;SARS-CoV_orf1ab;6942 | 0 | 0.025 |
| NC_004718.3;NP_828849.2;SARS-CoV_orf1ab;6943 | 0 | 0.025 |
| NC_004718.3;NP_828849.2;SARS-CoV_orf1ab;6944 | 0 | 0.05 |
| NC_004718.3;NP_828849.2;SARS-CoV_orf1ab;6945 | 0 | 0.025 |
| NC_004718.3;NP_828849.2;SARS-CoV_orf1ab;6946 | 0 | 0.025 |
| NC_004718.3;NP_828849.2;SARS-CoV_orf1ab;6947 | 0 | 0.025 |
| NC_004718.3;NP_828849.2;SARS-CoV_orf1ab;6948 | 0 | 0.025 |
| NC_004718.3;NP_828849.2;SARS-CoV_orf1ab;6949 | 0 | 0.025 |
| NC_004718.3;NP_828849.2;SARS-CoV_orf1ab;6950 | 0 | 0 |
| NC_004718.3;NP_828849.2;SARS-CoV_orf1ab;6951 | 0 | 0 |
| NC_004718.3;NP_828849.2;SARS-CoV_orf1ab;6952 | 0 | 0 |
| NC_004718.3;NP_828849.2;SARS-CoV_orf1ab;6953 | 0 | 0 |
| NC_004718.3;NP_828849.2;SARS-CoV_orf1ab;6954 | 0 | 0 |
| NC_004718.3;NP_828849.2;SARS-CoV_orf1ab;6955 | 0 | 0 |
| NC_004718.3;NP_828849.2;SARS-CoV_orf1ab;6956 | 0 | 0 |
| NC_004718.3;NP_828849.2;SARS-CoV_orf1ab;6957 | 0 | 0 |
| NC_004718.3;NP_828849.2;SARS-CoV_orf1ab;6958 | 0 | 0 |
| NC_004718.3;NP_828849.2;SARS-CoV_orf1ab;6959 | 0 | 0 |
| NC_004718.3;NP_828849.2;SARS-CoV_orf1ab;6960 | 0 | 0 |
| NC_004718.3;NP_828849.2;SARS-CoV_orf1ab;6961 | 0 | 0 |
| NC_004718.3;NP_828849.2;SARS-CoV_orf1ab;6962 | 0 | 0 |
| NC_004718.3;NP_828849.2;SARS-CoV_orf1ab;6963 | 0 | 0.025 |

| | | |
|---|---|---|
| NC_004718.3;NP_828849.2;SARS-CoV_orf1ab;6964 | 0 | 0.025 |
| NC_004718.3;NP_828849.2;SARS-CoV_orf1ab;6965 | 0 | 0.05 |
| NC_004718.3;NP_828849.2;SARS-CoV_orf1ab;6966 | 0 | 0 |
| NC_004718.3;NP_828849.2;SARS-CoV_orf1ab;6967 | 0.05 | 0 |
| NC_004718.3;NP_828849.2;SARS-CoV_orf1ab;6968 | 0 | 0 |
| NC_004718.3;NP_828849.2;SARS-CoV_orf1ab;6969 | 0 | 0.075 |
| NC_004718.3;NP_828849.2;SARS-CoV_orf1ab;6970 | 0.05 | 0.025 |
| NC_004718.3;NP_828849.2;SARS-CoV_orf1ab;6971 | 0 | 0 |
| NC_004718.3;NP_828849.2;SARS-CoV_orf1ab;6972 | 0 | 0.025 |
| NC_004718.3;NP_828849.2;SARS-CoV_orf1ab;6973 | 0 | 0.025 |
| NC_004718.3;NP_828849.2;SARS-CoV_orf1ab;6974 | 0 | 0 |
| NC_004718.3;NP_828849.2;SARS-CoV_orf1ab;6975 | 0 | 0 |
| NC_004718.3;NP_828849.2;SARS-CoV_orf1ab;6976 | 0 | 0 |
| NC_004718.3;NP_828849.2;SARS-CoV_orf1ab;6977 | 0.05 | 0 |
| NC_004718.3;NP_828849.2;SARS-CoV_orf1ab;6978 | 0.05 | 0 |
| NC_004718.3;NP_828849.2;SARS-CoV_orf1ab;6979 | 0 | 0 |
| NC_004718.3;NP_828849.2;SARS-CoV_orf1ab;6980 | 0.05 | 0 |
| NC_004718.3;NP_828849.2;SARS-CoV_orf1ab;6981 | 0.25 | 0.025 |
| NC_004718.3;NP_828849.2;SARS-CoV_orf1ab;6982 | 0.1 | 0.175 |
| NC_004718.3;NP_828849.2;SARS-CoV_orf1ab;6983 | 0.05 | 0.05 |
| NC_004718.3;NP_828849.2;SARS-CoV_orf1ab;6984 | 0.05 | 0.05 |
| NC_004718.3;NP_828849.2;SARS-CoV_orf1ab;6985 | 0.05 | 0.025 |
| NC_004718.3;NP_828849.2;SARS-CoV_orf1ab;6986 | 0.1 | 0.025 |
| NC_004718.3;NP_828849.2;SARS-CoV_orf1ab;6987 | 0 | 0.025 |
| NC_004718.3;NP_828849.2;SARS-CoV_orf1ab;6988 | 0 | 0.025 |
| NC_004718.3;NP_828849.2;SARS-CoV_orf1ab;6989 | 0 | 0.025 |
| NC_004718.3;NP_828849.2;SARS-CoV_orf1ab;6990 | 0 | 0.025 |
| NC_004718.3;NP_828849.2;SARS-CoV_orf1ab;6991 | 0 | 0 |
| NC_004718.3;NP_828849.2;SARS-CoV_orf1ab;6992 | 0.05 | 0.05 |
| NC_004718.3;NP_828849.2;SARS-CoV_orf1ab;6993 | 0 | 0 |
| NC_004718.3;NP_828849.2;SARS-CoV_orf1ab;6994 | 0 | 0 |
| NC_004718.3;NP_828849.2;SARS-CoV_orf1ab;6995 | 0 | 0 |
| NC_004718.3;NP_828849.2;SARS-CoV_orf1ab;6996 | 0 | 0 |
| NC_004718.3;NP_828849.2;SARS-CoV_orf1ab;6997 | 0 | 0 |
| NC_004718.3;NP_828849.2;SARS-CoV_orf1ab;6998 | 0 | 0 |
| NC_004718.3;NP_828849.2;SARS-CoV_orf1ab;6999 | 0 | 0 |
| NC_004718.3;NP_828849.2;SARS-CoV_orf1ab;7000 | 0 | 0 |
| NC_004718.3;NP_828849.2;SARS-CoV_orf1ab;7001 | 0 | 0 |
| NC_004718.3;NP_828849.2;SARS-CoV_orf1ab;7002 | 0 | 0 |
| NC_004718.3;NP_828849.2;SARS-CoV_orf1ab;7003 | 0 | 0 |
| NC_004718.3;NP_828849.2;SARS-CoV_orf1ab;7004 | 0 | 0 |
| NC_004718.3;NP_828849.2;SARS-CoV_orf1ab;7005 | 0 | 0 |
| NC_004718.3;NP_828849.2;SARS-CoV_orf1ab;7006 | 0 | 0 |

| | | |
|---|---|---|
| NC_004718.3;NP_828849.2;SARS-CoV_orf1ab;7007 | 0 | 0 |
| NC_004718.3;NP_828849.2;SARS-CoV_orf1ab;7008 | 0 | 0 |
| NC_004718.3;NP_828849.2;SARS-CoV_orf1ab;7009 | 0.05 | 0.075 |
| NC_004718.3;NP_828849.2;SARS-CoV_orf1ab;7010 | 0.05 | 0.075 |
| NC_004718.3;NP_828849.2;SARS-CoV_orf1ab;7011 | 0.05 | 0.05 |
| NC_004718.3;NP_828849.2;SARS-CoV_orf1ab;7012 | 0.05 | 0.05 |
| NC_004718.3;NP_828849.2;SARS-CoV_orf1ab;7013 | 0.05 | 0.05 |
| NC_004718.3;NP_828849.2;SARS-CoV_orf1ab;7014 | 0.05 | 0.075 |
| NC_004718.3;NP_828849.2;SARS-CoV_orf1ab;7015 | 0.05 | 0.025 |
| NC_004718.3;NP_828849.2;SARS-CoV_orf1ab;7016 | 0 | 0.025 |
| NC_004718.3;NP_828849.2;SARS-CoV_orf1ab;7017 | 0 | 0 |
| NC_004718.3;NP_828849.2;SARS-CoV_orf1ab;7018 | 0 | 0 |
| NC_004718.3;NP_828849.2;SARS-CoV_orf1ab;7019 | 0 | 0 |
| NC_004718.3;NP_828849.2;SARS-CoV_orf1ab;7020 | 0 | 0 |
| NC_004718.3;NP_828849.2;SARS-CoV_orf1ab;7021 | 0 | 0 |
| NC_004718.3;NP_828849.2;SARS-CoV_orf1ab;7022 | 0 | 0 |
| NC_004718.3;NP_828849.2;SARS-CoV_orf1ab;7023 | 0 | 0 |
| NC_004718.3;NP_828849.2;SARS-CoV_orf1ab;7024 | 0 | 0 |
| NC_004718.3;NP_828849.2;SARS-CoV_orf1ab;7025 | 0 | 0 |
| NC_004718.3;NP_828849.2;SARS-CoV_orf1ab;7026 | 0.05 | 0 |
| NC_004718.3;NP_828849.2;SARS-CoV_orf1ab;7027 | 0 | 0 |
| NC_004718.3;NP_828849.2;SARS-CoV_orf1ab;7028 | 0 | 0 |
| NC_004718.3;NP_828849.2;SARS-CoV_orf1ab;7029 | 0 | 0.025 |
| NC_004718.3;NP_828849.2;SARS-CoV_orf1ab;7030 | 0 | 0.025 |
| NC_004718.3;NP_828849.2;SARS-CoV_orf1ab;7031 | 0 | 0.025 |
| NC_004718.3;NP_828849.2;SARS-CoV_orf1ab;7032 | 0 | 0.025 |
| NC_004718.3;NP_828849.2;SARS-CoV_orf1ab;7033 | 0 | 0.025 |
| NC_004718.3;NP_828849.2;SARS-CoV_orf1ab;7034 | 0 | 0.05 |
| NC_004718.3;NP_828849.2;SARS-CoV_orf1ab;7035 | 0.05 | 0.025 |
| NC_004718.3;NP_828849.2;SARS-CoV_orf1ab;7036 | 0 | 0 |
| NC_004718.3;NP_828849.2;SARS-CoV_orf1ab;7037 | 0 | 0.025 |
| NC_004718.3;NP_828849.2;SARS-CoV_orf1ab;7038 | 0 | 0.025 |
| NC_004718.3;NP_828849.2;SARS-CoV_orf1ab;7039 | 0 | 0.025 |
| NC_004718.3;NP_828849.2;SARS-CoV_orf1ab;7040 | 0 | 0 |
| NC_004718.3;NP_828849.2;SARS-CoV_orf1ab;7041 | 0 | 0 |
| NC_004718.3;NP_828849.2;SARS-CoV_orf1ab;7042 | 0 | 0 |
| NC_004718.3;NP_828849.2;SARS-CoV_orf1ab;7043 | 0 | 0 |
| NC_004718.3;NP_828849.2;SARS-CoV_orf1ab;7044 | 0 | 0 |
| NC_004718.3;NP_828849.2;SARS-CoV_orf1ab;7045 | 0 | 0 |
| NC_004718.3;NP_828849.2;SARS-CoV_orf1ab;7046 | 0 | 0 |
| NC_004718.3;NP_828849.2;SARS-CoV_orf1ab;7047 | 0 | 0 |
| NC_004718.3;NP_828849.2;SARS-CoV_orf1ab;7048 | 0 | 0.025 |
| NC_004718.3;NP_828849.2;SARS-CoV_orf1ab;7049 | 0 | 0.05 |

| | | |
|---|---|---|
| NC_004718.3;NP_828849.2;SARS-CoV_orf1ab;7050 | 0 | 0.025 |
| NC_004718.3;NP_828849.2;SARS-CoV_orf1ab;7051 | 0 | 0.05 |
| NC_004718.3;NP_828849.2;SARS-CoV_orf1ab;7052 | 0 | 0.025 |
| NC_004718.3;NP_828849.2;SARS-CoV_orf1ab;7053 | 0 | 0.025 |
| NC_004718.3;NP_828849.2;SARS-CoV_orf1ab;7054 | 0 | 0.025 |
| NC_004718.3;NP_828849.2;SARS-CoV_orf1ab;7055 | 0 | 0.025 |
| NC_004718.3;NP_828849.2;SARS-CoV_orf1ab;7056 | 0 | 0.025 |
| NC_004718.3;NP_828849.2;SARS-CoV_orf1ab;7057 | 0 | 0.025 |
| NC_004718.3;NP_828849.2;SARS-CoV_orf1ab;7058 | 0 | 0.025 |
| NC_004718.3;NP_828849.2;SARS-CoV_orf1ab;7059 | 0 | 0 |
| NC_004718.3;NP_828849.2;SARS-CoV_orf1ab;7060 | 0 | 0 |
| NC_004718.3;NP_828849.2;SARS-CoV_orf1ab;7061 | 0 | 0 |
| NC_004718.3;NP_828849.2;SARS-CoV_orf1ab;7062 | 0 | 0 |
| NC_004718.3;NP_828850.1;SARS-CoV_orf1a;1 | 0.05 | 0 |
| NC_004718.3;NP_828850.1;SARS-CoV_orf1a;2 | 0 | 0 |
| NC_004718.3;NP_828850.1;SARS-CoV_orf1a;3 | 0 | 0 |
| NC_004718.3;NP_828850.1;SARS-CoV_orf1a;4 | 0.1 | 0.025 |
| NC_004718.3;NP_828850.1;SARS-CoV_orf1a;5 | 0.05 | 0.05 |
| NC_004718.3;NP_828850.1;SARS-CoV_orf1a;6 | 0.05 | 0 |
| NC_004718.3;NP_828850.1;SARS-CoV_orf1a;7 | 0 | 0 |
| NC_004718.3;NP_828850.1;SARS-CoV_orf1a;8 | 0 | 0 |
| NC_004718.3;NP_828850.1;SARS-CoV_orf1a;9 | 0 | 0 |
| NC_004718.3;NP_828850.1;SARS-CoV_orf1a;10 | 0 | 0 |
| NC_004718.3;NP_828850.1;SARS-CoV_orf1a;11 | 0 | 0 |
| NC_004718.3;NP_828850.1;SARS-CoV_orf1a;12 | 0 | 0 |
| NC_004718.3;NP_828850.1;SARS-CoV_orf1a;13 | 0 | 0 |
| NC_004718.3;NP_828850.1;SARS-CoV_orf1a;14 | 0 | 0 |
| NC_004718.3;NP_828850.1;SARS-CoV_orf1a;15 | 0 | 0 |
| NC_004718.3;NP_828850.1;SARS-CoV_orf1a;16 | 0 | 0 |
| NC_004718.3;NP_828850.1;SARS-CoV_orf1a;17 | 0 | 0 |
| NC_004718.3;NP_828850.1;SARS-CoV_orf1a;18 | 0 | 0 |
| NC_004718.3;NP_828850.1;SARS-CoV_orf1a;19 | 0 | 0 |
| NC_004718.3;NP_828850.1;SARS-CoV_orf1a;20 | 0 | 0 |
| NC_004718.3;NP_828850.1;SARS-CoV_orf1a;21 | 0 | 0 |
| NC_004718.3;NP_828850.1;SARS-CoV_orf1a;22 | 0 | 0 |
| NC_004718.3;NP_828850.1;SARS-CoV_orf1a;23 | 0 | 0 |
| NC_004718.3;NP_828850.1;SARS-CoV_orf1a;24 | 0 | 0 |
| NC_004718.3;NP_828850.1;SARS-CoV_orf1a;25 | 0 | 0 |
| NC_004718.3;NP_828850.1;SARS-CoV_orf1a;26 | 0 | 0 |
| NC_004718.3;NP_828850.1;SARS-CoV_orf1a;27 | 0 | 0 |
| NC_004718.3;NP_828850.1;SARS-CoV_orf1a;28 | 0 | 0 |
| NC_004718.3;NP_828850.1;SARS-CoV_orf1a;29 | 0.1 | 0 |
| NC_004718.3;NP_828850.1;SARS-CoV_orf1a;30 | 0.25 | 0.125 |

| | | |
|---|---|---|
| NC_004718.3;NP_828850.1;SARS-CoV_orf1a;31 | 0 | 0 |
| NC_004718.3;NP_828850.1;SARS-CoV_orf1a;32 | 0.05 | 0.025 |
| NC_004718.3;NP_828850.1;SARS-CoV_orf1a;33 | 0.05 | 0 |
| NC_004718.3;NP_828850.1;SARS-CoV_orf1a;34 | 0.05 | 0.025 |
| NC_004718.3;NP_828850.1;SARS-CoV_orf1a;35 | 0 | 0.025 |
| NC_004718.3;NP_828850.1;SARS-CoV_orf1a;36 | 0 | 0.025 |
| NC_004718.3;NP_828850.1;SARS-CoV_orf1a;37 | 0 | 0 |
| NC_004718.3;NP_828850.1;SARS-CoV_orf1a;38 | 0 | 0.025 |
| NC_004718.3;NP_828850.1;SARS-CoV_orf1a;39 | 0 | 0.025 |
| NC_004718.3;NP_828850.1;SARS-CoV_orf1a;40 | 0 | 0 |
| NC_004718.3;NP_828850.1;SARS-CoV_orf1a;41 | 0 | 0 |
| NC_004718.3;NP_828850.1;SARS-CoV_orf1a;42 | 0 | 0 |
| NC_004718.3;NP_828850.1;SARS-CoV_orf1a;43 | 0 | 0 |
| NC_004718.3;NP_828850.1;SARS-CoV_orf1a;44 | 0.05 | 0 |
| NC_004718.3;NP_828850.1;SARS-CoV_orf1a;45 | 0 | 0 |
| NC_004718.3;NP_828850.1;SARS-CoV_orf1a;46 | 0 | 0 |
| NC_004718.3;NP_828850.1;SARS-CoV_orf1a;47 | 0 | 0.025 |
| NC_004718.3;NP_828850.1;SARS-CoV_orf1a;48 | 0 | 0 |
| NC_004718.3;NP_828850.1;SARS-CoV_orf1a;49 | 0 | 0 |
| NC_004718.3;NP_828850.1;SARS-CoV_orf1a;50 | 0 | 0.025 |
| NC_004718.3;NP_828850.1;SARS-CoV_orf1a;51 | 0 | 0.05 |
| NC_004718.3;NP_828850.1;SARS-CoV_orf1a;52 | 0.05 | 0.075 |
| NC_004718.3;NP_828850.1;SARS-CoV_orf1a;53 | 0 | 0 |
| NC_004718.3;NP_828850.1;SARS-CoV_orf1a;54 | 0 | 0 |
| NC_004718.3;NP_828850.1;SARS-CoV_orf1a;55 | 0 | 0.05 |
| NC_004718.3;NP_828850.1;SARS-CoV_orf1a;56 | 0 | 0 |
| NC_004718.3;NP_828850.1;SARS-CoV_orf1a;57 | 0 | 0.025 |
| NC_004718.3;NP_828850.1;SARS-CoV_orf1a;58 | 0 | 0.025 |
| NC_004718.3;NP_828850.1;SARS-CoV_orf1a;59 | 0 | 0 |
| NC_004718.3;NP_828850.1;SARS-CoV_orf1a;60 | 0 | 0 |
| NC_004718.3;NP_828850.1;SARS-CoV_orf1a;61 | 0 | 0 |
| NC_004718.3;NP_828850.1;SARS-CoV_orf1a;62 | 0 | 0 |
| NC_004718.3;NP_828850.1;SARS-CoV_orf1a;63 | 0 | 0 |
| NC_004718.3;NP_828850.1;SARS-CoV_orf1a;64 | 0 | 0 |
| NC_004718.3;NP_828850.1;SARS-CoV_orf1a;65 | 0.05 | 0 |
| NC_004718.3;NP_828850.1;SARS-CoV_orf1a;66 | 0 | 0 |
| NC_004718.3;NP_828850.1;SARS-CoV_orf1a;67 | 0 | 0 |
| NC_004718.3;NP_828850.1;SARS-CoV_orf1a;68 | 0 | 0 |
| NC_004718.3;NP_828850.1;SARS-CoV_orf1a;69 | 0 | 0 |
| NC_004718.3;NP_828850.1;SARS-CoV_orf1a;70 | 0 | 0 |
| NC_004718.3;NP_828850.1;SARS-CoV_orf1a;71 | 0.05 | 0.025 |
| NC_004718.3;NP_828850.1;SARS-CoV_orf1a;72 | 0.05 | 0.025 |
| NC_004718.3;NP_828850.1;SARS-CoV_orf1a;73 | 0 | 0 |

| | | |
|---|---|---|
| NC_004718.3;NP_828850.1;SARS-CoV_orf1a;74 | 0 | 0 |
| NC_004718.3;NP_828850.1;SARS-CoV_orf1a;75 | 0 | 0.025 |
| NC_004718.3;NP_828850.1;SARS-CoV_orf1a;76 | 0 | 0 |
| NC_004718.3;NP_828850.1;SARS-CoV_orf1a;77 | 0 | 0 |
| NC_004718.3;NP_828850.1;SARS-CoV_orf1a;78 | 0 | 0 |
| NC_004718.3;NP_828850.1;SARS-CoV_orf1a;79 | 0 | 0 |
| NC_004718.3;NP_828850.1;SARS-CoV_orf1a;80 | 0 | 0 |
| NC_004718.3;NP_828850.1;SARS-CoV_orf1a;81 | 0 | 0 |
| NC_004718.3;NP_828850.1;SARS-CoV_orf1a;82 | 0 | 0 |
| NC_004718.3;NP_828850.1;SARS-CoV_orf1a;83 | 0 | 0 |
| NC_004718.3;NP_828850.1;SARS-CoV_orf1a;84 | 0 | 0 |
| NC_004718.3;NP_828850.1;SARS-CoV_orf1a;85 | 0 | 0 |
| NC_004718.3;NP_828850.1;SARS-CoV_orf1a;86 | 0 | 0.025 |
| NC_004718.3;NP_828850.1;SARS-CoV_orf1a;87 | 0 | 0 |
| NC_004718.3;NP_828850.1;SARS-CoV_orf1a;88 | 0 | 0 |
| NC_004718.3;NP_828850.1;SARS-CoV_orf1a;89 | 0 | 0 |
| NC_004718.3;NP_828850.1;SARS-CoV_orf1a;90 | 0.2 | 0.125 |
| NC_004718.3;NP_828850.1;SARS-CoV_orf1a;91 | 0.05 | 0.025 |
| NC_004718.3;NP_828850.1;SARS-CoV_orf1a;92 | 0 | 0 |
| NC_004718.3;NP_828850.1;SARS-CoV_orf1a;93 | 0 | 0 |
| NC_004718.3;NP_828850.1;SARS-CoV_orf1a;94 | 0 | 0 |
| NC_004718.3;NP_828850.1;SARS-CoV_orf1a;95 | 0 | 0 |
| NC_004718.3;NP_828850.1;SARS-CoV_orf1a;96 | 0 | 0 |
| NC_004718.3;NP_828850.1;SARS-CoV_orf1a;97 | 0 | 0 |
| NC_004718.3;NP_828850.1;SARS-CoV_orf1a;98 | 0 | 0 |
| NC_004718.3;NP_828850.1;SARS-CoV_orf1a;99 | 0 | 0 |
| NC_004718.3;NP_828850.1;SARS-CoV_orf1a;100 | 0 | 0 |
| NC_004718.3;NP_828850.1;SARS-CoV_orf1a;101 | 0 | 0 |
| NC_004718.3;NP_828850.1;SARS-CoV_orf1a;102 | 0 | 0 |
| NC_004718.3;NP_828850.1;SARS-CoV_orf1a;103 | 0 | 0 |
| NC_004718.3;NP_828850.1;SARS-CoV_orf1a;104 | 0 | 0 |
| NC_004718.3;NP_828850.1;SARS-CoV_orf1a;105 | 0 | 0 |
| NC_004718.3;NP_828850.1;SARS-CoV_orf1a;106 | 0 | 0 |
| NC_004718.3;NP_828850.1;SARS-CoV_orf1a;107 | 0 | 0 |
| NC_004718.3;NP_828850.1;SARS-CoV_orf1a;108 | 0 | 0 |
| NC_004718.3;NP_828850.1;SARS-CoV_orf1a;109 | 0 | 0 |
| NC_004718.3;NP_828850.1;SARS-CoV_orf1a;110 | 0 | 0 |
| NC_004718.3;NP_828850.1;SARS-CoV_orf1a;111 | 0 | 0 |
| NC_004718.3;NP_828850.1;SARS-CoV_orf1a;112 | 0 | 0 |
| NC_004718.3;NP_828850.1;SARS-CoV_orf1a;113 | 0 | 0 |
| NC_004718.3;NP_828850.1;SARS-CoV_orf1a;114 | 0 | 0.05 |
| NC_004718.3;NP_828850.1;SARS-CoV_orf1a;115 | 0 | 0.025 |
| NC_004718.3;NP_828850.1;SARS-CoV_orf1a;116 | 0.05 | 0.025 |

| | | |
|---|---|---|
| NC_004718.3;NP_828850.1;SARS-CoV_orf1a;117 | 0.05 | 0.025 |
| NC_004718.3;NP_828850.1;SARS-CoV_orf1a;118 | 0.1 | 0.025 |
| NC_004718.3;NP_828850.1;SARS-CoV_orf1a;119 | 0.05 | 0.025 |
| NC_004718.3;NP_828850.1;SARS-CoV_orf1a;120 | 0.05 | 0.025 |
| NC_004718.3;NP_828850.1;SARS-CoV_orf1a;121 | 0.05 | 0.025 |
| NC_004718.3;NP_828850.1;SARS-CoV_orf1a;122 | 0 | 0.025 |
| NC_004718.3;NP_828850.1;SARS-CoV_orf1a;123 | 0 | 0.025 |
| NC_004718.3;NP_828850.1;SARS-CoV_orf1a;124 | 0.05 | 0.05 |
| NC_004718.3;NP_828850.1;SARS-CoV_orf1a;125 | 0.15 | 0.05 |
| NC_004718.3;NP_828850.1;SARS-CoV_orf1a;126 | 0 | 0 |
| NC_004718.3;NP_828850.1;SARS-CoV_orf1a;127 | 0 | 0 |
| NC_004718.3;NP_828850.1;SARS-CoV_orf1a;128 | 0 | 0.025 |
| NC_004718.3;NP_828850.1;SARS-CoV_orf1a;129 | 0 | 0.025 |
| NC_004718.3;NP_828850.1;SARS-CoV_orf1a;130 | 0 | 0.025 |
| NC_004718.3;NP_828850.1;SARS-CoV_orf1a;131 | 0.1 | 0.025 |
| NC_004718.3;NP_828850.1;SARS-CoV_orf1a;132 | 0.05 | 0.025 |
| NC_004718.3;NP_828850.1;SARS-CoV_orf1a;133 | 0.05 | 0 |
| NC_004718.3;NP_828850.1;SARS-CoV_orf1a;134 | 0 | 0 |
| NC_004718.3;NP_828850.1;SARS-CoV_orf1a;135 | 0 | 0 |
| NC_004718.3;NP_828850.1;SARS-CoV_orf1a;136 | 0 | 0 |
| NC_004718.3;NP_828850.1;SARS-CoV_orf1a;137 | 0 | 0 |
| NC_004718.3;NP_828850.1;SARS-CoV_orf1a;138 | 0 | 0 |
| NC_004718.3;NP_828850.1;SARS-CoV_orf1a;139 | 0.05 | 0 |
| NC_004718.3;NP_828850.1;SARS-CoV_orf1a;140 | 0.05 | 0 |
| NC_004718.3;NP_828850.1;SARS-CoV_orf1a;141 | 0 | 0 |
| NC_004718.3;NP_828850.1;SARS-CoV_orf1a;142 | 0 | 0 |
| NC_004718.3;NP_828850.1;SARS-CoV_orf1a;143 | 0 | 0 |
| NC_004718.3;NP_828850.1;SARS-CoV_orf1a;144 | 0 | 0 |
| NC_004718.3;NP_828850.1;SARS-CoV_orf1a;145 | 0.2 | 0.15 |
| NC_004718.3;NP_828850.1;SARS-CoV_orf1a;146 | 0 | 0 |
| NC_004718.3;NP_828850.1;SARS-CoV_orf1a;147 | 0 | 0 |
| NC_004718.3;NP_828850.1;SARS-CoV_orf1a;148 | 0 | 0 |
| NC_004718.3;NP_828850.1;SARS-CoV_orf1a;149 | 0.2 | 0.125 |
| NC_004718.3;NP_828850.1;SARS-CoV_orf1a;150 | 0.05 | 0.05 |
| NC_004718.3;NP_828850.1;SARS-CoV_orf1a;151 | 0.05 | 0.025 |
| NC_004718.3;NP_828850.1;SARS-CoV_orf1a;152 | 0.25 | 0.175 |
| NC_004718.3;NP_828850.1;SARS-CoV_orf1a;153 | 0.25 | 0.25 |
| NC_004718.3;NP_828850.1;SARS-CoV_orf1a;154 | 0.2 | 0.2 |
| NC_004718.3;NP_828850.1;SARS-CoV_orf1a;155 | 0.1 | 0.1 |
| NC_004718.3;NP_828850.1;SARS-CoV_orf1a;156 | 0.05 | 0.075 |
| NC_004718.3;NP_828850.1;SARS-CoV_orf1a;157 | 0.05 | 0.05 |
| NC_004718.3;NP_828850.1;SARS-CoV_orf1a;158 | 0 | 0 |
| NC_004718.3;NP_828850.1;SARS-CoV_orf1a;159 | 0 | 0 |

| | | |
|---|---|---|
| NC_004718.3;NP_828850.1;SARS-CoV_orf1a;160 | 0 | 0 |
| NC_004718.3;NP_828850.1;SARS-CoV_orf1a;161 | 0 | 0 |
| NC_004718.3;NP_828850.1;SARS-CoV_orf1a;162 | 0 | 0 |
| NC_004718.3;NP_828850.1;SARS-CoV_orf1a;163 | 0 | 0.025 |
| NC_004718.3;NP_828850.1;SARS-CoV_orf1a;164 | 0 | 0 |
| NC_004718.3;NP_828850.1;SARS-CoV_orf1a;165 | 0.05 | 0.025 |
| NC_004718.3;NP_828850.1;SARS-CoV_orf1a;166 | 0 | 0.025 |
| NC_004718.3;NP_828850.1;SARS-CoV_orf1a;167 | 0 | 0.025 |
| NC_004718.3;NP_828850.1;SARS-CoV_orf1a;168 | 0.2 | 0.2 |
| NC_004718.3;NP_828850.1;SARS-CoV_orf1a;169 | 0 | 0 |
| NC_004718.3;NP_828850.1;SARS-CoV_orf1a;170 | 0 | 0 |
| NC_004718.3;NP_828850.1;SARS-CoV_orf1a;171 | 0 | 0 |
| NC_004718.3;NP_828850.1;SARS-CoV_orf1a;172 | 0 | 0.025 |
| NC_004718.3;NP_828850.1;SARS-CoV_orf1a;173 | 0.05 | 0.05 |
| NC_004718.3;NP_828850.1;SARS-CoV_orf1a;174 | 0 | 0.075 |
| NC_004718.3;NP_828850.1;SARS-CoV_orf1a;175 | 0 | 0.075 |
| NC_004718.3;NP_828850.1;SARS-CoV_orf1a;176 | 0 | 0.025 |
| NC_004718.3;NP_828850.1;SARS-CoV_orf1a;177 | 0 | 0.025 |
| NC_004718.3;NP_828850.1;SARS-CoV_orf1a;178 | 0 | 0.025 |
| NC_004718.3;NP_828850.1;SARS-CoV_orf1a;179 | 0.05 | 0.075 |
| NC_004718.3;NP_828850.1;SARS-CoV_orf1a;180 | 0.15 | 0.15 |
| NC_004718.3;NP_828850.1;SARS-CoV_orf1a;181 | 0 | 0.025 |
| NC_004718.3;NP_828850.1;SARS-CoV_orf1a;182 | 0 | 0 |
| NC_004718.3;NP_828850.1;SARS-CoV_orf1a;183 | 0 | 0 |
| NC_004718.3;NP_828850.1;SARS-CoV_orf1a;184 | 0 | 0 |
| NC_004718.3;NP_828850.1;SARS-CoV_orf1a;185 | 0 | 0.05 |
| NC_004718.3;NP_828850.1;SARS-CoV_orf1a;186 | 0 | 0.025 |
| NC_004718.3;NP_828850.1;SARS-CoV_orf1a;187 | 0 | 0 |
| NC_004718.3;NP_828850.1;SARS-CoV_orf1a;188 | 0 | 0 |
| NC_004718.3;NP_828850.1;SARS-CoV_orf1a;189 | 0.05 | 0 |
| NC_004718.3;NP_828850.1;SARS-CoV_orf1a;190 | 0 | 0 |
| NC_004718.3;NP_828850.1;SARS-CoV_orf1a;191 | 0 | 0 |
| NC_004718.3;NP_828850.1;SARS-CoV_orf1a;192 | 0 | 0 |
| NC_004718.3;NP_828850.1;SARS-CoV_orf1a;193 | 0 | 0 |
| NC_004718.3;NP_828850.1;SARS-CoV_orf1a;194 | 0 | 0 |
| NC_004718.3;NP_828850.1;SARS-CoV_orf1a;195 | 0 | 0 |
| NC_004718.3;NP_828850.1;SARS-CoV_orf1a;196 | 0 | 0 |
| NC_004718.3;NP_828850.1;SARS-CoV_orf1a;197 | 0 | 0 |
| NC_004718.3;NP_828850.1;SARS-CoV_orf1a;198 | 0 | 0 |
| NC_004718.3;NP_828850.1;SARS-CoV_orf1a;199 | 0 | 0 |
| NC_004718.3;NP_828850.1;SARS-CoV_orf1a;200 | 0 | 0 |
| NC_004718.3;NP_828850.1;SARS-CoV_orf1a;201 | 0 | 0 |
| NC_004718.3;NP_828850.1;SARS-CoV_orf1a;202 | 0 | 0 |

| | | |
|---|---|---|
| NC_004718.3;NP_828850.1;SARS-CoV_orf1a;203 | 0 | 0.025 |
| NC_004718.3;NP_828850.1;SARS-CoV_orf1a;204 | 0.05 | 0 |
| NC_004718.3;NP_828850.1;SARS-CoV_orf1a;205 | 0 | 0 |
| NC_004718.3;NP_828850.1;SARS-CoV_orf1a;206 | 0 | 0 |
| NC_004718.3;NP_828850.1;SARS-CoV_orf1a;207 | 0 | 0 |
| NC_004718.3;NP_828850.1;SARS-CoV_orf1a;208 | 0 | 0 |
| NC_004718.3;NP_828850.1;SARS-CoV_orf1a;209 | 0 | 0 |
| NC_004718.3;NP_828850.1;SARS-CoV_orf1a;210 | 0 | 0 |
| NC_004718.3;NP_828850.1;SARS-CoV_orf1a;211 | 0 | 0.025 |
| NC_004718.3;NP_828850.1;SARS-CoV_orf1a;212 | 0 | 0.05 |
| NC_004718.3;NP_828850.1;SARS-CoV_orf1a;213 | 0 | 0.025 |
| NC_004718.3;NP_828850.1;SARS-CoV_orf1a;214 | 0.15 | 0.05 |
| NC_004718.3;NP_828850.1;SARS-CoV_orf1a;215 | 0.15 | 0.075 |
| NC_004718.3;NP_828850.1;SARS-CoV_orf1a;216 | 0.1 | 0.05 |
| NC_004718.3;NP_828850.1;SARS-CoV_orf1a;217 | 0.05 | 0.025 |
| NC_004718.3;NP_828850.1;SARS-CoV_orf1a;218 | 0.05 | 0.05 |
| NC_004718.3;NP_828850.1;SARS-CoV_orf1a;219 | 0 | 0.025 |
| NC_004718.3;NP_828850.1;SARS-CoV_orf1a;220 | 0.05 | 0.025 |
| NC_004718.3;NP_828850.1;SARS-CoV_orf1a;221 | 0.1 | 0.025 |
| NC_004718.3;NP_828850.1;SARS-CoV_orf1a;222 | 0.1 | 0.025 |
| NC_004718.3;NP_828850.1;SARS-CoV_orf1a;223 | 0.15 | 0.05 |
| NC_004718.3;NP_828850.1;SARS-CoV_orf1a;224 | 0 | 0.025 |
| NC_004718.3;NP_828850.1;SARS-CoV_orf1a;225 | 0.05 | 0.025 |
| NC_004718.3;NP_828850.1;SARS-CoV_orf1a;226 | 0 | 0 |
| NC_004718.3;NP_828850.1;SARS-CoV_orf1a;227 | 0 | 0 |
| NC_004718.3;NP_828850.1;SARS-CoV_orf1a;228 | 0 | 0 |
| NC_004718.3;NP_828850.1;SARS-CoV_orf1a;229 | 0 | 0 |
| NC_004718.3;NP_828850.1;SARS-CoV_orf1a;230 | 0 | 0 |
| NC_004718.3;NP_828850.1;SARS-CoV_orf1a;231 | 0 | 0 |
| NC_004718.3;NP_828850.1;SARS-CoV_orf1a;232 | 0 | 0 |
| NC_004718.3;NP_828850.1;SARS-CoV_orf1a;233 | 0 | 0.05 |
| NC_004718.3;NP_828850.1;SARS-CoV_orf1a;234 | 0 | 0 |
| NC_004718.3;NP_828850.1;SARS-CoV_orf1a;235 | 0 | 0 |
| NC_004718.3;NP_828850.1;SARS-CoV_orf1a;236 | 0 | 0 |
| NC_004718.3;NP_828850.1;SARS-CoV_orf1a;237 | 0 | 0 |
| NC_004718.3;NP_828850.1;SARS-CoV_orf1a;238 | 0 | 0.05 |
| NC_004718.3;NP_828850.1;SARS-CoV_orf1a;239 | 0 | 0.025 |
| NC_004718.3;NP_828850.1;SARS-CoV_orf1a;240 | 0.05 | 0.025 |
| NC_004718.3;NP_828850.1;SARS-CoV_orf1a;241 | 0.05 | 0 |
| NC_004718.3;NP_828850.1;SARS-CoV_orf1a;242 | 0 | 0.05 |
| NC_004718.3;NP_828850.1;SARS-CoV_orf1a;243 | 0 | 0 |
| NC_004718.3;NP_828850.1;SARS-CoV_orf1a;244 | 0 | 0.025 |
| NC_004718.3;NP_828850.1;SARS-CoV_orf1a;245 | 0 | 0 |

| | | |
|---|---|---|
| NC_004718.3;NP_828850.1;SARS-CoV_orf1a;246 | 0 | 0.05 |
| NC_004718.3;NP_828850.1;SARS-CoV_orf1a;247 | 0 | 0 |
| NC_004718.3;NP_828850.1;SARS-CoV_orf1a;248 | 0 | 0.025 |
| NC_004718.3;NP_828850.1;SARS-CoV_orf1a;249 | 0 | 0.025 |
| NC_004718.3;NP_828850.1;SARS-CoV_orf1a;250 | 0 | 0 |
| NC_004718.3;NP_828850.1;SARS-CoV_orf1a;251 | 0 | 0.025 |
| NC_004718.3;NP_828850.1;SARS-CoV_orf1a;252 | 0 | 0 |
| NC_004718.3;NP_828850.1;SARS-CoV_orf1a;253 | 0 | 0 |
| NC_004718.3;NP_828850.1;SARS-CoV_orf1a;254 | 0.05 | 0 |
| NC_004718.3;NP_828850.1;SARS-CoV_orf1a;255 | 0.05 | 0 |
| NC_004718.3;NP_828850.1;SARS-CoV_orf1a;256 | 0.05 | 0 |
| NC_004718.3;NP_828850.1;SARS-CoV_orf1a;257 | 0 | 0 |
| NC_004718.3;NP_828850.1;SARS-CoV_orf1a;258 | 0 | 0.025 |
| NC_004718.3;NP_828850.1;SARS-CoV_orf1a;259 | 0 | 0 |
| NC_004718.3;NP_828850.1;SARS-CoV_orf1a;260 | 0 | 0 |
| NC_004718.3;NP_828850.1;SARS-CoV_orf1a;261 | 0 | 0 |
| NC_004718.3;NP_828850.1;SARS-CoV_orf1a;262 | 0.05 | 0 |
| NC_004718.3;NP_828850.1;SARS-CoV_orf1a;263 | 0.1 | 0 |
| NC_004718.3;NP_828850.1;SARS-CoV_orf1a;264 | 0 | 0.025 |
| NC_004718.3;NP_828850.1;SARS-CoV_orf1a;265 | 0 | 0 |
| NC_004718.3;NP_828850.1;SARS-CoV_orf1a;266 | 0 | 0 |
| NC_004718.3;NP_828850.1;SARS-CoV_orf1a;267 | 0 | 0.025 |
| NC_004718.3;NP_828850.1;SARS-CoV_orf1a;268 | 0 | 0.025 |
| NC_004718.3;NP_828850.1;SARS-CoV_orf1a;269 | 0 | 0.025 |
| NC_004718.3;NP_828850.1;SARS-CoV_orf1a;270 | 0 | 0.025 |
| NC_004718.3;NP_828850.1;SARS-CoV_orf1a;271 | 0.05 | 0.025 |
| NC_004718.3;NP_828850.1;SARS-CoV_orf1a;272 | 0 | 0.025 |
| NC_004718.3;NP_828850.1;SARS-CoV_orf1a;273 | 0 | 0.025 |
| NC_004718.3;NP_828850.1;SARS-CoV_orf1a;274 | 0 | 0.025 |
| NC_004718.3;NP_828850.1;SARS-CoV_orf1a;275 | 0 | 0.05 |
| NC_004718.3;NP_828850.1;SARS-CoV_orf1a;276 | 0.05 | 0.05 |
| NC_004718.3;NP_828850.1;SARS-CoV_orf1a;277 | 0.05 | 0.025 |
| NC_004718.3;NP_828850.1;SARS-CoV_orf1a;278 | 0.05 | 0.025 |
| NC_004718.3;NP_828850.1;SARS-CoV_orf1a;279 | 0.05 | 0.025 |
| NC_004718.3;NP_828850.1;SARS-CoV_orf1a;280 | 0.05 | 0.025 |
| NC_004718.3;NP_828850.1;SARS-CoV_orf1a;281 | 0.05 | 0.025 |
| NC_004718.3;NP_828850.1;SARS-CoV_orf1a;282 | 0.05 | 0.025 |
| NC_004718.3;NP_828850.1;SARS-CoV_orf1a;283 | 0.15 | 0 |
| NC_004718.3;NP_828850.1;SARS-CoV_orf1a;284 | 0.05 | 0.025 |
| NC_004718.3;NP_828850.1;SARS-CoV_orf1a;285 | 0.05 | 0.025 |
| NC_004718.3;NP_828850.1;SARS-CoV_orf1a;286 | 0 | 0.025 |
| NC_004718.3;NP_828850.1;SARS-CoV_orf1a;287 | 0 | 0.025 |
| NC_004718.3;NP_828850.1;SARS-CoV_orf1a;288 | 0 | 0.025 |

| | | |
|---|---|---|
| NC_004718.3;NP_828850.1;SARS-CoV_orf1a;289 | 0 | 0.025 |
| NC_004718.3;NP_828850.1;SARS-CoV_orf1a;290 | 0 | 0.025 |
| NC_004718.3;NP_828850.1;SARS-CoV_orf1a;291 | 0 | 0.025 |
| NC_004718.3;NP_828850.1;SARS-CoV_orf1a;292 | 0 | 0.025 |
| NC_004718.3;NP_828850.1;SARS-CoV_orf1a;293 | 0 | 0 |
| NC_004718.3;NP_828850.1;SARS-CoV_orf1a;294 | 0 | 0 |
| NC_004718.3;NP_828850.1;SARS-CoV_orf1a;295 | 0 | 0 |
| NC_004718.3;NP_828850.1;SARS-CoV_orf1a;296 | 0.05 | 0.025 |
| NC_004718.3;NP_828850.1;SARS-CoV_orf1a;297 | 0.05 | 0 |
| NC_004718.3;NP_828850.1;SARS-CoV_orf1a;298 | 0.05 | 0 |
| NC_004718.3;NP_828850.1;SARS-CoV_orf1a;299 | 0.05 | 0.025 |
| NC_004718.3;NP_828850.1;SARS-CoV_orf1a;300 | 0.05 | 0 |
| NC_004718.3;NP_828850.1;SARS-CoV_orf1a;301 | 0.05 | 0.075 |
| NC_004718.3;NP_828850.1;SARS-CoV_orf1a;302 | 0 | 0 |
| NC_004718.3;NP_828850.1;SARS-CoV_orf1a;303 | 0 | 0 |
| NC_004718.3;NP_828850.1;SARS-CoV_orf1a;304 | 0 | 0 |
| NC_004718.3;NP_828850.1;SARS-CoV_orf1a;305 | 0 | 0 |
| NC_004718.3;NP_828850.1;SARS-CoV_orf1a;306 | 0 | 0 |
| NC_004718.3;NP_828850.1;SARS-CoV_orf1a;307 | 0 | 0 |
| NC_004718.3;NP_828850.1;SARS-CoV_orf1a;308 | 0 | 0 |
| NC_004718.3;NP_828850.1;SARS-CoV_orf1a;309 | 0 | 0 |
| NC_004718.3;NP_828850.1;SARS-CoV_orf1a;310 | 0 | 0 |
| NC_004718.3;NP_828850.1;SARS-CoV_orf1a;311 | 0 | 0 |
| NC_004718.3;NP_828850.1;SARS-CoV_orf1a;312 | 0 | 0 |
| NC_004718.3;NP_828850.1;SARS-CoV_orf1a;313 | 0 | 0 |
| NC_004718.3;NP_828850.1;SARS-CoV_orf1a;314 | 0 | 0 |
| NC_004718.3;NP_828850.1;SARS-CoV_orf1a;315 | 0.1 | 0.025 |
| NC_004718.3;NP_828850.1;SARS-CoV_orf1a;316 | 0 | 0 |
| NC_004718.3;NP_828850.1;SARS-CoV_orf1a;317 | 0 | 0 |
| NC_004718.3;NP_828850.1;SARS-CoV_orf1a;318 | 0 | 0 |
| NC_004718.3;NP_828850.1;SARS-CoV_orf1a;319 | 0 | 0 |
| NC_004718.3;NP_828850.1;SARS-CoV_orf1a;320 | 0 | 0 |
| NC_004718.3;NP_828850.1;SARS-CoV_orf1a;321 | 0 | 0 |
| NC_004718.3;NP_828850.1;SARS-CoV_orf1a;322 | 0 | 0 |
| NC_004718.3;NP_828850.1;SARS-CoV_orf1a;323 | 0 | 0 |
| NC_004718.3;NP_828850.1;SARS-CoV_orf1a;324 | 0.05 | 0 |
| NC_004718.3;NP_828850.1;SARS-CoV_orf1a;325 | 0 | 0 |
| NC_004718.3;NP_828850.1;SARS-CoV_orf1a;326 | 0 | 0.025 |
| NC_004718.3;NP_828850.1;SARS-CoV_orf1a;327 | 0 | 0 |
| NC_004718.3;NP_828850.1;SARS-CoV_orf1a;328 | 0 | 0 |
| NC_004718.3;NP_828850.1;SARS-CoV_orf1a;329 | 0 | 0 |
| NC_004718.3;NP_828850.1;SARS-CoV_orf1a;330 | 0 | 0 |
| NC_004718.3;NP_828850.1;SARS-CoV_orf1a;331 | 0 | 0 |

| | | |
|---|---|---|
| NC_004718.3;NP_828850.1;SARS-CoV_orf1a;332 | 0 | 0 |
| NC_004718.3;NP_828850.1;SARS-CoV_orf1a;333 | 0 | 0 |
| NC_004718.3;NP_828850.1;SARS-CoV_orf1a;334 | 0 | 0 |
| NC_004718.3;NP_828850.1;SARS-CoV_orf1a;335 | 0 | 0 |
| NC_004718.3;NP_828850.1;SARS-CoV_orf1a;336 | 0 | 0 |
| NC_004718.3;NP_828850.1;SARS-CoV_orf1a;337 | 0 | 0 |
| NC_004718.3;NP_828850.1;SARS-CoV_orf1a;338 | 0 | 0 |
| NC_004718.3;NP_828850.1;SARS-CoV_orf1a;339 | 0 | 0 |
| NC_004718.3;NP_828850.1;SARS-CoV_orf1a;340 | 0 | 0 |
| NC_004718.3;NP_828850.1;SARS-CoV_orf1a;341 | 0 | 0 |
| NC_004718.3;NP_828850.1;SARS-CoV_orf1a;342 | 0 | 0 |
| NC_004718.3;NP_828850.1;SARS-CoV_orf1a;343 | 0 | 0 |
| NC_004718.3;NP_828850.1;SARS-CoV_orf1a;344 | 0 | 0 |
| NC_004718.3;NP_828850.1;SARS-CoV_orf1a;345 | 0 | 0 |
| NC_004718.3;NP_828850.1;SARS-CoV_orf1a;346 | 0 | 0 |
| NC_004718.3;NP_828850.1;SARS-CoV_orf1a;347 | 0 | 0 |
| NC_004718.3;NP_828850.1;SARS-CoV_orf1a;348 | 0 | 0 |
| NC_004718.3;NP_828850.1;SARS-CoV_orf1a;349 | 0 | 0 |
| NC_004718.3;NP_828850.1;SARS-CoV_orf1a;350 | 0 | 0 |
| NC_004718.3;NP_828850.1;SARS-CoV_orf1a;351 | 0 | 0 |
| NC_004718.3;NP_828850.1;SARS-CoV_orf1a;352 | 0 | 0 |
| NC_004718.3;NP_828850.1;SARS-CoV_orf1a;353 | 0 | 0 |
| NC_004718.3;NP_828850.1;SARS-CoV_orf1a;354 | 0 | 0 |
| NC_004718.3;NP_828850.1;SARS-CoV_orf1a;355 | 0 | 0 |
| NC_004718.3;NP_828850.1;SARS-CoV_orf1a;356 | 0.05 | 0 |
| NC_004718.3;NP_828850.1;SARS-CoV_orf1a;357 | 0.05 | 0 |
| NC_004718.3;NP_828850.1;SARS-CoV_orf1a;358 | 0.05 | 0 |
| NC_004718.3;NP_828850.1;SARS-CoV_orf1a;359 | 0 | 0 |
| NC_004718.3;NP_828850.1;SARS-CoV_orf1a;360 | 0 | 0 |
| NC_004718.3;NP_828850.1;SARS-CoV_orf1a;361 | 0 | 0 |
| NC_004718.3;NP_828850.1;SARS-CoV_orf1a;362 | 0.05 | 0 |
| NC_004718.3;NP_828850.1;SARS-CoV_orf1a;363 | 0 | 0 |
| NC_004718.3;NP_828850.1;SARS-CoV_orf1a;364 | 0 | 0 |
| NC_004718.3;NP_828850.1;SARS-CoV_orf1a;365 | 0 | 0 |
| NC_004718.3;NP_828850.1;SARS-CoV_orf1a;366 | 0.05 | 0.025 |
| NC_004718.3;NP_828850.1;SARS-CoV_orf1a;367 | 0 | 0 |
| NC_004718.3;NP_828850.1;SARS-CoV_orf1a;368 | 0.05 | 0.05 |
| NC_004718.3;NP_828850.1;SARS-CoV_orf1a;369 | 0.05 | 0.025 |
| NC_004718.3;NP_828850.1;SARS-CoV_orf1a;370 | 0.05 | 0.025 |
| NC_004718.3;NP_828850.1;SARS-CoV_orf1a;371 | 0.05 | 0.025 |
| NC_004718.3;NP_828850.1;SARS-CoV_orf1a;372 | 0.05 | 0.025 |
| NC_004718.3;NP_828850.1;SARS-CoV_orf1a;373 | 0.05 | 0 |
| NC_004718.3;NP_828850.1;SARS-CoV_orf1a;374 | 0.05 | 0.025 |

| | | |
|---|---|---|
| NC_004718.3;NP_828850.1;SARS-CoV_orf1a;375 | 0.1 | 0 |
| NC_004718.3;NP_828850.1;SARS-CoV_orf1a;376 | 0 | 0 |
| NC_004718.3;NP_828850.1;SARS-CoV_orf1a;377 | 0.1 | 0 |
| NC_004718.3;NP_828850.1;SARS-CoV_orf1a;378 | 0 | 0 |
| NC_004718.3;NP_828850.1;SARS-CoV_orf1a;379 | 0 | 0 |
| NC_004718.3;NP_828850.1;SARS-CoV_orf1a;380 | 0 | 0 |
| NC_004718.3;NP_828850.1;SARS-CoV_orf1a;381 | 0 | 0 |
| NC_004718.3;NP_828850.1;SARS-CoV_orf1a;382 | 0 | 0 |
| NC_004718.3;NP_828850.1;SARS-CoV_orf1a;383 | 0.05 | 0.025 |
| NC_004718.3;NP_828850.1;SARS-CoV_orf1a;384 | 0 | 0 |
| NC_004718.3;NP_828850.1;SARS-CoV_orf1a;385 | 0 | 0 |
| NC_004718.3;NP_828850.1;SARS-CoV_orf1a;386 | 0 | 0 |
| NC_004718.3;NP_828850.1;SARS-CoV_orf1a;387 | 0 | 0 |
| NC_004718.3;NP_828850.1;SARS-CoV_orf1a;388 | 0 | 0 |
| NC_004718.3;NP_828850.1;SARS-CoV_orf1a;389 | 0 | 0 |
| NC_004718.3;NP_828850.1;SARS-CoV_orf1a;390 | 0 | 0 |
| NC_004718.3;NP_828850.1;SARS-CoV_orf1a;391 | 0 | 0 |
| NC_004718.3;NP_828850.1;SARS-CoV_orf1a;392 | 0 | 0 |
| NC_004718.3;NP_828850.1;SARS-CoV_orf1a;393 | 0 | 0 |
| NC_004718.3;NP_828850.1;SARS-CoV_orf1a;394 | 0 | 0 |
| NC_004718.3;NP_828850.1;SARS-CoV_orf1a;395 | 0 | 0 |
| NC_004718.3;NP_828850.1;SARS-CoV_orf1a;396 | 0 | 0 |
| NC_004718.3;NP_828850.1;SARS-CoV_orf1a;397 | 0 | 0.025 |
| NC_004718.3;NP_828850.1;SARS-CoV_orf1a;398 | 0 | 0 |
| NC_004718.3;NP_828850.1;SARS-CoV_orf1a;399 | 0 | 0 |
| NC_004718.3;NP_828850.1;SARS-CoV_orf1a;400 | 0 | 0 |
| NC_004718.3;NP_828850.1;SARS-CoV_orf1a;401 | 0 | 0 |
| NC_004718.3;NP_828850.1;SARS-CoV_orf1a;402 | 0 | 0 |
| NC_004718.3;NP_828850.1;SARS-CoV_orf1a;403 | 0 | 0 |
| NC_004718.3;NP_828850.1;SARS-CoV_orf1a;404 | 0 | 0 |
| NC_004718.3;NP_828850.1;SARS-CoV_orf1a;405 | 0 | 0 |
| NC_004718.3;NP_828850.1;SARS-CoV_orf1a;406 | 0 | 0 |
| NC_004718.3;NP_828850.1;SARS-CoV_orf1a;407 | 0 | 0 |
| NC_004718.3;NP_828850.1;SARS-CoV_orf1a;408 | 0 | 0 |
| NC_004718.3;NP_828850.1;SARS-CoV_orf1a;409 | 0 | 0 |
| NC_004718.3;NP_828850.1;SARS-CoV_orf1a;410 | 0 | 0 |
| NC_004718.3;NP_828850.1;SARS-CoV_orf1a;411 | 0 | 0 |
| NC_004718.3;NP_828850.1;SARS-CoV_orf1a;412 | 0 | 0 |
| NC_004718.3;NP_828850.1;SARS-CoV_orf1a;413 | 0 | 0 |
| NC_004718.3;NP_828850.1;SARS-CoV_orf1a;414 | 0 | 0 |
| NC_004718.3;NP_828850.1;SARS-CoV_orf1a;415 | 0 | 0 |
| NC_004718.3;NP_828850.1;SARS-CoV_orf1a;416 | 0 | 0 |
| NC_004718.3;NP_828850.1;SARS-CoV_orf1a;417 | 0 | 0 |

| | | |
|---|---|---|
| NC_004718.3;NP_828850.1;SARS-CoV_orf1a;418 | 0 | 0 |
| NC_004718.3;NP_828850.1;SARS-CoV_orf1a;419 | 0 | 0 |
| NC_004718.3;NP_828850.1;SARS-CoV_orf1a;420 | 0 | 0 |
| NC_004718.3;NP_828850.1;SARS-CoV_orf1a;421 | 0 | 0 |
| NC_004718.3;NP_828850.1;SARS-CoV_orf1a;422 | 0 | 0 |
| NC_004718.3;NP_828850.1;SARS-CoV_orf1a;423 | 0 | 0 |
| NC_004718.3;NP_828850.1;SARS-CoV_orf1a;424 | 0 | 0 |
| NC_004718.3;NP_828850.1;SARS-CoV_orf1a;425 | 0.05 | 0 |
| NC_004718.3;NP_828850.1;SARS-CoV_orf1a;426 | 0 | 0 |
| NC_004718.3;NP_828850.1;SARS-CoV_orf1a;427 | 0 | 0 |
| NC_004718.3;NP_828850.1;SARS-CoV_orf1a;428 | 0.05 | 0.025 |
| NC_004718.3;NP_828850.1;SARS-CoV_orf1a;429 | 0 | 0 |
| NC_004718.3;NP_828850.1;SARS-CoV_orf1a;430 | 0.05 | 0 |
| NC_004718.3;NP_828850.1;SARS-CoV_orf1a;431 | 0 | 0 |
| NC_004718.3;NP_828850.1;SARS-CoV_orf1a;432 | 0 | 0 |
| NC_004718.3;NP_828850.1;SARS-CoV_orf1a;433 | 0 | 0 |
| NC_004718.3;NP_828850.1;SARS-CoV_orf1a;434 | 0 | 0 |
| NC_004718.3;NP_828850.1;SARS-CoV_orf1a;435 | 0 | 0 |
| NC_004718.3;NP_828850.1;SARS-CoV_orf1a;436 | 0 | 0 |
| NC_004718.3;NP_828850.1;SARS-CoV_orf1a;437 | 0 | 0.025 |
| NC_004718.3;NP_828850.1;SARS-CoV_orf1a;438 | 0.05 | 0 |
| NC_004718.3;NP_828850.1;SARS-CoV_orf1a;439 | 0 | 0 |
| NC_004718.3;NP_828850.1;SARS-CoV_orf1a;440 | 0 | 0 |
| NC_004718.3;NP_828850.1;SARS-CoV_orf1a;441 | 0 | 0 |
| NC_004718.3;NP_828850.1;SARS-CoV_orf1a;442 | 0 | 0 |
| NC_004718.3;NP_828850.1;SARS-CoV_orf1a;443 | 0 | 0.025 |
| NC_004718.3;NP_828850.1;SARS-CoV_orf1a;444 | 0.05 | 0 |
| NC_004718.3;NP_828850.1;SARS-CoV_orf1a;445 | 0.1 | 0.05 |
| NC_004718.3;NP_828850.1;SARS-CoV_orf1a;446 | 0.05 | 0 |
| NC_004718.3;NP_828850.1;SARS-CoV_orf1a;447 | 0.05 | 0.075 |
| NC_004718.3;NP_828850.1;SARS-CoV_orf1a;448 | 0.05 | 0.025 |
| NC_004718.3;NP_828850.1;SARS-CoV_orf1a;449 | 0 | 0 |
| NC_004718.3;NP_828850.1;SARS-CoV_orf1a;450 | 0 | 0 |
| NC_004718.3;NP_828850.1;SARS-CoV_orf1a;451 | 0 | 0.05 |
| NC_004718.3;NP_828850.1;SARS-CoV_orf1a;452 | 0 | 0 |
| NC_004718.3;NP_828850.1;SARS-CoV_orf1a;453 | 0.05 | 0 |
| NC_004718.3;NP_828850.1;SARS-CoV_orf1a;454 | 0 | 0 |
| NC_004718.3;NP_828850.1;SARS-CoV_orf1a;455 | 0 | 0.05 |
| NC_004718.3;NP_828850.1;SARS-CoV_orf1a;456 | 0 | 0.025 |
| NC_004718.3;NP_828850.1;SARS-CoV_orf1a;457 | 0.05 | 0 |
| NC_004718.3;NP_828850.1;SARS-CoV_orf1a;458 | 0.05 | 0 |
| NC_004718.3;NP_828850.1;SARS-CoV_orf1a;459 | 0 | 0 |
| NC_004718.3;NP_828850.1;SARS-CoV_orf1a;460 | 0 | 0 |

| | | |
|---|---|---|
| NC_004718.3;NP_828850.1;SARS-CoV_orf1a;461 | 0 | 0 |
| NC_004718.3;NP_828850.1;SARS-CoV_orf1a;462 | 0 | 0 |
| NC_004718.3;NP_828850.1;SARS-CoV_orf1a;463 | 0 | 0 |
| NC_004718.3;NP_828850.1;SARS-CoV_orf1a;464 | 0 | 0 |
| NC_004718.3;NP_828850.1;SARS-CoV_orf1a;465 | 0 | 0 |
| NC_004718.3;NP_828850.1;SARS-CoV_orf1a;466 | 0 | 0 |
| NC_004718.3;NP_828850.1;SARS-CoV_orf1a;467 | 0 | 0 |
| NC_004718.3;NP_828850.1;SARS-CoV_orf1a;468 | 0 | 0 |
| NC_004718.3;NP_828850.1;SARS-CoV_orf1a;469 | 0 | 0 |
| NC_004718.3;NP_828850.1;SARS-CoV_orf1a;470 | 0 | 0 |
| NC_004718.3;NP_828850.1;SARS-CoV_orf1a;471 | 0 | 0 |
| NC_004718.3;NP_828850.1;SARS-CoV_orf1a;472 | 0 | 0 |
| NC_004718.3;NP_828850.1;SARS-CoV_orf1a;473 | 0 | 0 |
| NC_004718.3;NP_828850.1;SARS-CoV_orf1a;474 | 0 | 0 |
| NC_004718.3;NP_828850.1;SARS-CoV_orf1a;475 | 0 | 0 |
| NC_004718.3;NP_828850.1;SARS-CoV_orf1a;476 | 0 | 0 |
| NC_004718.3;NP_828850.1;SARS-CoV_orf1a;477 | 0.05 | 0 |
| NC_004718.3;NP_828850.1;SARS-CoV_orf1a;478 | 0.05 | 0 |
| NC_004718.3;NP_828850.1;SARS-CoV_orf1a;479 | 0.05 | 0 |
| NC_004718.3;NP_828850.1;SARS-CoV_orf1a;480 | 0.1 | 0 |
| NC_004718.3;NP_828850.1;SARS-CoV_orf1a;481 | 0.05 | 0 |
| NC_004718.3;NP_828850.1;SARS-CoV_orf1a;482 | 0.05 | 0 |
| NC_004718.3;NP_828850.1;SARS-CoV_orf1a;483 | 0.1 | 0 |
| NC_004718.3;NP_828850.1;SARS-CoV_orf1a;484 | 0.05 | 0 |
| NC_004718.3;NP_828850.1;SARS-CoV_orf1a;485 | 0 | 0 |
| NC_004718.3;NP_828850.1;SARS-CoV_orf1a;486 | 0 | 0 |
| NC_004718.3;NP_828850.1;SARS-CoV_orf1a;487 | 0 | 0 |
| NC_004718.3;NP_828850.1;SARS-CoV_orf1a;488 | 0 | 0 |
| NC_004718.3;NP_828850.1;SARS-CoV_orf1a;489 | 0 | 0 |
| NC_004718.3;NP_828850.1;SARS-CoV_orf1a;490 | 0 | 0 |
| NC_004718.3;NP_828850.1;SARS-CoV_orf1a;491 | 0 | 0 |
| NC_004718.3;NP_828850.1;SARS-CoV_orf1a;492 | 0 | 0 |
| NC_004718.3;NP_828850.1;SARS-CoV_orf1a;493 | 0 | 0 |
| NC_004718.3;NP_828850.1;SARS-CoV_orf1a;494 | 0 | 0 |
| NC_004718.3;NP_828850.1;SARS-CoV_orf1a;495 | 0 | 0 |
| NC_004718.3;NP_828850.1;SARS-CoV_orf1a;496 | 0 | 0 |
| NC_004718.3;NP_828850.1;SARS-CoV_orf1a;497 | 0.05 | 0.05 |
| NC_004718.3;NP_828850.1;SARS-CoV_orf1a;498 | 0 | 0 |
| NC_004718.3;NP_828850.1;SARS-CoV_orf1a;499 | 0 | 0 |
| NC_004718.3;NP_828850.1;SARS-CoV_orf1a;500 | 0 | 0 |
| NC_004718.3;NP_828850.1;SARS-CoV_orf1a;501 | 0 | 0 |
| NC_004718.3;NP_828850.1;SARS-CoV_orf1a;502 | 0 | 0 |
| NC_004718.3;NP_828850.1;SARS-CoV_orf1a;503 | 0 | 0 |

| | | |
|---|---|---|
| NC_004718.3;NP_828850.1;SARS-CoV_orf1a;504 | 0 | 0 |
| NC_004718.3;NP_828850.1;SARS-CoV_orf1a;505 | 0 | 0 |
| NC_004718.3;NP_828850.1;SARS-CoV_orf1a;506 | 0 | 0 |
| NC_004718.3;NP_828850.1;SARS-CoV_orf1a;507 | 0 | 0 |
| NC_004718.3;NP_828850.1;SARS-CoV_orf1a;508 | 0 | 0 |
| NC_004718.3;NP_828850.1;SARS-CoV_orf1a;509 | 0 | 0 |
| NC_004718.3;NP_828850.1;SARS-CoV_orf1a;510 | 0 | 0 |
| NC_004718.3;NP_828850.1;SARS-CoV_orf1a;511 | 0 | 0 |
| NC_004718.3;NP_828850.1;SARS-CoV_orf1a;512 | 0 | 0 |
| NC_004718.3;NP_828850.1;SARS-CoV_orf1a;513 | 0 | 0 |
| NC_004718.3;NP_828850.1;SARS-CoV_orf1a;514 | 0 | 0 |
| NC_004718.3;NP_828850.1;SARS-CoV_orf1a;515 | 0 | 0 |
| NC_004718.3;NP_828850.1;SARS-CoV_orf1a;516 | 0 | 0 |
| NC_004718.3;NP_828850.1;SARS-CoV_orf1a;517 | 0 | 0 |
| NC_004718.3;NP_828850.1;SARS-CoV_orf1a;518 | 0 | 0 |
| NC_004718.3;NP_828850.1;SARS-CoV_orf1a;519 | 0 | 0 |
| NC_004718.3;NP_828850.1;SARS-CoV_orf1a;520 | 0 | 0 |
| NC_004718.3;NP_828850.1;SARS-CoV_orf1a;521 | 0 | 0 |
| NC_004718.3;NP_828850.1;SARS-CoV_orf1a;522 | 0 | 0 |
| NC_004718.3;NP_828850.1;SARS-CoV_orf1a;523 | 0 | 0 |
| NC_004718.3;NP_828850.1;SARS-CoV_orf1a;524 | 0 | 0 |
| NC_004718.3;NP_828850.1;SARS-CoV_orf1a;525 | 0 | 0 |
| NC_004718.3;NP_828850.1;SARS-CoV_orf1a;526 | 0.05 | 0.025 |
| NC_004718.3;NP_828850.1;SARS-CoV_orf1a;527 | 0 | 0 |
| NC_004718.3;NP_828850.1;SARS-CoV_orf1a;528 | 0 | 0 |
| NC_004718.3;NP_828850.1;SARS-CoV_orf1a;529 | 0 | 0 |
| NC_004718.3;NP_828850.1;SARS-CoV_orf1a;530 | 0 | 0 |
| NC_004718.3;NP_828850.1;SARS-CoV_orf1a;531 | 0.05 | 0 |
| NC_004718.3;NP_828850.1;SARS-CoV_orf1a;532 | 0.05 | 0 |
| NC_004718.3;NP_828850.1;SARS-CoV_orf1a;533 | 0 | 0 |
| NC_004718.3;NP_828850.1;SARS-CoV_orf1a;534 | 0 | 0 |
| NC_004718.3;NP_828850.1;SARS-CoV_orf1a;535 | 0 | 0 |
| NC_004718.3;NP_828850.1;SARS-CoV_orf1a;536 | 0 | 0 |
| NC_004718.3;NP_828850.1;SARS-CoV_orf1a;537 | 0 | 0 |
| NC_004718.3;NP_828850.1;SARS-CoV_orf1a;538 | 0 | 0 |
| NC_004718.3;NP_828850.1;SARS-CoV_orf1a;539 | 0 | 0 |
| NC_004718.3;NP_828850.1;SARS-CoV_orf1a;540 | 0 | 0 |
| NC_004718.3;NP_828850.1;SARS-CoV_orf1a;541 | 0 | 0 |
| NC_004718.3;NP_828850.1;SARS-CoV_orf1a;542 | 0 | 0 |
| NC_004718.3;NP_828850.1;SARS-CoV_orf1a;543 | 0 | 0 |
| NC_004718.3;NP_828850.1;SARS-CoV_orf1a;544 | 0 | 0 |
| NC_004718.3;NP_828850.1;SARS-CoV_orf1a;545 | 0.05 | 0 |
| NC_004718.3;NP_828850.1;SARS-CoV_orf1a;546 | 0.05 | 0 |

| | | |
|---|---|---|
| NC_004718.3;NP_828850.1;SARS-CoV_orf1a;547 | 0.05 | 0 |
| NC_004718.3;NP_828850.1;SARS-CoV_orf1a;548 | 0.05 | 0 |
| NC_004718.3;NP_828850.1;SARS-CoV_orf1a;549 | 0.1 | 0 |
| NC_004718.3;NP_828850.1;SARS-CoV_orf1a;550 | 0.05 | 0 |
| NC_004718.3;NP_828850.1;SARS-CoV_orf1a;551 | 0.05 | 0.05 |
| NC_004718.3;NP_828850.1;SARS-CoV_orf1a;552 | 0 | 0.025 |
| NC_004718.3;NP_828850.1;SARS-CoV_orf1a;553 | 0.05 | 0.025 |
| NC_004718.3;NP_828850.1;SARS-CoV_orf1a;554 | 0.05 | 0 |
| NC_004718.3;NP_828850.1;SARS-CoV_orf1a;555 | 0 | 0 |
| NC_004718.3;NP_828850.1;SARS-CoV_orf1a;556 | 0 | 0 |
| NC_004718.3;NP_828850.1;SARS-CoV_orf1a;557 | 0 | 0 |
| NC_004718.3;NP_828850.1;SARS-CoV_orf1a;558 | 0 | 0 |
| NC_004718.3;NP_828850.1;SARS-CoV_orf1a;559 | 0 | 0 |
| NC_004718.3;NP_828850.1;SARS-CoV_orf1a;560 | 0 | 0 |
| NC_004718.3;NP_828850.1;SARS-CoV_orf1a;561 | 0 | 0 |
| NC_004718.3;NP_828850.1;SARS-CoV_orf1a;562 | 0 | 0 |
| NC_004718.3;NP_828850.1;SARS-CoV_orf1a;563 | 0 | 0 |
| NC_004718.3;NP_828850.1;SARS-CoV_orf1a;564 | 0 | 0 |
| NC_004718.3;NP_828850.1;SARS-CoV_orf1a;565 | 0 | 0.025 |
| NC_004718.3;NP_828850.1;SARS-CoV_orf1a;566 | 0 | 0 |
| NC_004718.3;NP_828850.1;SARS-CoV_orf1a;567 | 0 | 0 |
| NC_004718.3;NP_828850.1;SARS-CoV_orf1a;568 | 0 | 0 |
| NC_004718.3;NP_828850.1;SARS-CoV_orf1a;569 | 0 | 0 |
| NC_004718.3;NP_828850.1;SARS-CoV_orf1a;570 | 0 | 0 |
| NC_004718.3;NP_828850.1;SARS-CoV_orf1a;571 | 0 | 0 |
| NC_004718.3;NP_828850.1;SARS-CoV_orf1a;572 | 0.05 | 0.025 |
| NC_004718.3;NP_828850.1;SARS-CoV_orf1a;573 | 0 | 0 |
| NC_004718.3;NP_828850.1;SARS-CoV_orf1a;574 | 0 | 0.025 |
| NC_004718.3;NP_828850.1;SARS-CoV_orf1a;575 | 0 | 0 |
| NC_004718.3;NP_828850.1;SARS-CoV_orf1a;576 | 0 | 0 |
| NC_004718.3;NP_828850.1;SARS-CoV_orf1a;577 | 0 | 0 |
| NC_004718.3;NP_828850.1;SARS-CoV_orf1a;578 | 0 | 0.025 |
| NC_004718.3;NP_828850.1;SARS-CoV_orf1a;579 | 0 | 0.025 |
| NC_004718.3;NP_828850.1;SARS-CoV_orf1a;580 | 0 | 0 |
| NC_004718.3;NP_828850.1;SARS-CoV_orf1a;581 | 0 | 0 |
| NC_004718.3;NP_828850.1;SARS-CoV_orf1a;582 | 0 | 0 |
| NC_004718.3;NP_828850.1;SARS-CoV_orf1a;583 | 0 | 0 |
| NC_004718.3;NP_828850.1;SARS-CoV_orf1a;584 | 0 | 0 |
| NC_004718.3;NP_828850.1;SARS-CoV_orf1a;585 | 0 | 0.025 |
| NC_004718.3;NP_828850.1;SARS-CoV_orf1a;586 | 0 | 0 |
| NC_004718.3;NP_828850.1;SARS-CoV_orf1a;587 | 0 | 0 |
| NC_004718.3;NP_828850.1;SARS-CoV_orf1a;588 | 0 | 0 |
| NC_004718.3;NP_828850.1;SARS-CoV_orf1a;589 | 0.05 | 0 |

| | | |
|---|---|---|
| NC_004718.3;NP_828850.1;SARS-CoV_orf1a;590 | 0 | 0 |
| NC_004718.3;NP_828850.1;SARS-CoV_orf1a;591 | 0 | 0 |
| NC_004718.3;NP_828850.1;SARS-CoV_orf1a;592 | 0 | 0 |
| NC_004718.3;NP_828850.1;SARS-CoV_orf1a;593 | 0 | 0 |
| NC_004718.3;NP_828850.1;SARS-CoV_orf1a;594 | 0 | 0 |
| NC_004718.3;NP_828850.1;SARS-CoV_orf1a;595 | 0 | 0 |
| NC_004718.3;NP_828850.1;SARS-CoV_orf1a;596 | 0 | 0 |
| NC_004718.3;NP_828850.1;SARS-CoV_orf1a;597 | 0 | 0 |
| NC_004718.3;NP_828850.1;SARS-CoV_orf1a;598 | 0 | 0 |
| NC_004718.3;NP_828850.1;SARS-CoV_orf1a;599 | 0 | 0 |
| NC_004718.3;NP_828850.1;SARS-CoV_orf1a;600 | 0 | 0 |
| NC_004718.3;NP_828850.1;SARS-CoV_orf1a;601 | 0 | 0 |
| NC_004718.3;NP_828850.1;SARS-CoV_orf1a;602 | 0 | 0.025 |
| NC_004718.3;NP_828850.1;SARS-CoV_orf1a;603 | 0 | 0 |
| NC_004718.3;NP_828850.1;SARS-CoV_orf1a;604 | 0 | 0 |
| NC_004718.3;NP_828850.1;SARS-CoV_orf1a;605 | 0.1 | 0.05 |
| NC_004718.3;NP_828850.1;SARS-CoV_orf1a;606 | 0 | 0 |
| NC_004718.3;NP_828850.1;SARS-CoV_orf1a;607 | 0 | 0 |
| NC_004718.3;NP_828850.1;SARS-CoV_orf1a;608 | 0 | 0 |
| NC_004718.3;NP_828850.1;SARS-CoV_orf1a;609 | 0 | 0 |
| NC_004718.3;NP_828850.1;SARS-CoV_orf1a;610 | 0 | 0 |
| NC_004718.3;NP_828850.1;SARS-CoV_orf1a;611 | 0 | 0 |
| NC_004718.3;NP_828850.1;SARS-CoV_orf1a;612 | 0 | 0 |
| NC_004718.3;NP_828850.1;SARS-CoV_orf1a;613 | 0 | 0 |
| NC_004718.3;NP_828850.1;SARS-CoV_orf1a;614 | 0 | 0 |
| NC_004718.3;NP_828850.1;SARS-CoV_orf1a;615 | 0 | 0 |
| NC_004718.3;NP_828850.1;SARS-CoV_orf1a;616 | 0 | 0 |
| NC_004718.3;NP_828850.1;SARS-CoV_orf1a;617 | 0 | 0 |
| NC_004718.3;NP_828850.1;SARS-CoV_orf1a;618 | 0 | 0 |
| NC_004718.3;NP_828850.1;SARS-CoV_orf1a;619 | 0.05 | 0 |
| NC_004718.3;NP_828850.1;SARS-CoV_orf1a;620 | 0.05 | 0 |
| NC_004718.3;NP_828850.1;SARS-CoV_orf1a;621 | 0.05 | 0 |
| NC_004718.3;NP_828850.1;SARS-CoV_orf1a;622 | 0.05 | 0 |
| NC_004718.3;NP_828850.1;SARS-CoV_orf1a;623 | 0 | 0 |
| NC_004718.3;NP_828850.1;SARS-CoV_orf1a;624 | 0 | 0 |
| NC_004718.3;NP_828850.1;SARS-CoV_orf1a;625 | 0 | 0 |
| NC_004718.3;NP_828850.1;SARS-CoV_orf1a;626 | 0 | 0 |
| NC_004718.3;NP_828850.1;SARS-CoV_orf1a;627 | 0 | 0 |
| NC_004718.3;NP_828850.1;SARS-CoV_orf1a;628 | 0 | 0 |
| NC_004718.3;NP_828850.1;SARS-CoV_orf1a;629 | 0 | 0 |
| NC_004718.3;NP_828850.1;SARS-CoV_orf1a;630 | 0.05 | 0 |
| NC_004718.3;NP_828850.1;SARS-CoV_orf1a;631 | 0 | 0 |
| NC_004718.3;NP_828850.1;SARS-CoV_orf1a;632 | 0 | 0 |

| | | |
|---|---|---|
| NC_004718.3;NP_828850.1;SARS-CoV_orf1a;633 | 0.05 | 0.025 |
| NC_004718.3;NP_828850.1;SARS-CoV_orf1a;634 | 0.05 | 0.025 |
| NC_004718.3;NP_828850.1;SARS-CoV_orf1a;635 | 0.05 | 0.025 |
| NC_004718.3;NP_828850.1;SARS-CoV_orf1a;636 | 0 | 0.025 |
| NC_004718.3;NP_828850.1;SARS-CoV_orf1a;637 | 0 | 0 |
| NC_004718.3;NP_828850.1;SARS-CoV_orf1a;638 | 0 | 0 |
| NC_004718.3;NP_828850.1;SARS-CoV_orf1a;639 | 0 | 0 |
| NC_004718.3;NP_828850.1;SARS-CoV_orf1a;640 | 0 | 0.025 |
| NC_004718.3;NP_828850.1;SARS-CoV_orf1a;641 | 0 | 0 |
| NC_004718.3;NP_828850.1;SARS-CoV_orf1a;642 | 0 | 0 |
| NC_004718.3;NP_828850.1;SARS-CoV_orf1a;643 | 0 | 0 |
| NC_004718.3;NP_828850.1;SARS-CoV_orf1a;644 | 0 | 0 |
| NC_004718.3;NP_828850.1;SARS-CoV_orf1a;645 | 0 | 0 |
| NC_004718.3;NP_828850.1;SARS-CoV_orf1a;646 | 0 | 0 |
| NC_004718.3;NP_828850.1;SARS-CoV_orf1a;647 | 0 | 0 |
| NC_004718.3;NP_828850.1;SARS-CoV_orf1a;648 | 0 | 0 |
| NC_004718.3;NP_828850.1;SARS-CoV_orf1a;649 | 0 | 0 |
| NC_004718.3;NP_828850.1;SARS-CoV_orf1a;650 | 0 | 0 |
| NC_004718.3;NP_828850.1;SARS-CoV_orf1a;651 | 0 | 0 |
| NC_004718.3;NP_828850.1;SARS-CoV_orf1a;652 | 0 | 0.025 |
| NC_004718.3;NP_828850.1;SARS-CoV_orf1a;653 | 0 | 0 |
| NC_004718.3;NP_828850.1;SARS-CoV_orf1a;654 | 0 | 0.025 |
| NC_004718.3;NP_828850.1;SARS-CoV_orf1a;655 | 0.05 | 0.05 |
| NC_004718.3;NP_828850.1;SARS-CoV_orf1a;656 | 0 | 0.05 |
| NC_004718.3;NP_828850.1;SARS-CoV_orf1a;657 | 0 | 0.025 |
| NC_004718.3;NP_828850.1;SARS-CoV_orf1a;658 | 0 | 0.025 |
| NC_004718.3;NP_828850.1;SARS-CoV_orf1a;659 | 0 | 0.05 |
| NC_004718.3;NP_828850.1;SARS-CoV_orf1a;660 | 0.05 | 0.05 |
| NC_004718.3;NP_828850.1;SARS-CoV_orf1a;661 | 0 | 0.05 |
| NC_004718.3;NP_828850.1;SARS-CoV_orf1a;662 | 0 | 0 |
| NC_004718.3;NP_828850.1;SARS-CoV_orf1a;663 | 0 | 0 |
| NC_004718.3;NP_828850.1;SARS-CoV_orf1a;664 | 0 | 0 |
| NC_004718.3;NP_828850.1;SARS-CoV_orf1a;665 | 0 | 0 |
| NC_004718.3;NP_828850.1;SARS-CoV_orf1a;666 | 0.05 | 0 |
| NC_004718.3;NP_828850.1;SARS-CoV_orf1a;667 | 0 | 0 |
| NC_004718.3;NP_828850.1;SARS-CoV_orf1a;668 | 0 | 0 |
| NC_004718.3;NP_828850.1;SARS-CoV_orf1a;669 | 0 | 0 |
| NC_004718.3;NP_828850.1;SARS-CoV_orf1a;670 | 0 | 0 |
| NC_004718.3;NP_828850.1;SARS-CoV_orf1a;671 | 0 | 0 |
| NC_004718.3;NP_828850.1;SARS-CoV_orf1a;672 | 0 | 0 |
| NC_004718.3;NP_828850.1;SARS-CoV_orf1a;673 | 0 | 0 |
| NC_004718.3;NP_828850.1;SARS-CoV_orf1a;674 | 0 | 0 |
| NC_004718.3;NP_828850.1;SARS-CoV_orf1a;675 | 0 | 0 |

| | | |
|---|---|---|
| NC_004718.3;NP_828850.1;SARS-CoV_orf1a;676 | 0 | 0 |
| NC_004718.3;NP_828850.1;SARS-CoV_orf1a;677 | 0 | 0 |
| NC_004718.3;NP_828850.1;SARS-CoV_orf1a;678 | 0 | 0.025 |
| NC_004718.3;NP_828850.1;SARS-CoV_orf1a;679 | 0 | 0.1 |
| NC_004718.3;NP_828850.1;SARS-CoV_orf1a;680 | 0 | 0 |
| NC_004718.3;NP_828850.1;SARS-CoV_orf1a;681 | 0 | 0 |
| NC_004718.3;NP_828850.1;SARS-CoV_orf1a;682 | 0.05 | 0 |
| NC_004718.3;NP_828850.1;SARS-CoV_orf1a;683 | 0.05 | 0 |
| NC_004718.3;NP_828850.1;SARS-CoV_orf1a;684 | 0.05 | 0.025 |
| NC_004718.3;NP_828850.1;SARS-CoV_orf1a;685 | 0 | 0 |
| NC_004718.3;NP_828850.1;SARS-CoV_orf1a;686 | 0 | 0 |
| NC_004718.3;NP_828850.1;SARS-CoV_orf1a;687 | 0 | 0 |
| NC_004718.3;NP_828850.1;SARS-CoV_orf1a;688 | 0 | 0 |
| NC_004718.3;NP_828850.1;SARS-CoV_orf1a;689 | 0 | 0 |
| NC_004718.3;NP_828850.1;SARS-CoV_orf1a;690 | 0 | 0.025 |
| NC_004718.3;NP_828850.1;SARS-CoV_orf1a;691 | 0 | 0 |
| NC_004718.3;NP_828850.1;SARS-CoV_orf1a;692 | 0 | 0.025 |
| NC_004718.3;NP_828850.1;SARS-CoV_orf1a;693 | 0 | 0.025 |
| NC_004718.3;NP_828850.1;SARS-CoV_orf1a;694 | 0.05 | 0 |
| NC_004718.3;NP_828850.1;SARS-CoV_orf1a;695 | 0 | 0 |
| NC_004718.3;NP_828850.1;SARS-CoV_orf1a;696 | 0 | 0 |
| NC_004718.3;NP_828850.1;SARS-CoV_orf1a;697 | 0 | 0 |
| NC_004718.3;NP_828850.1;SARS-CoV_orf1a;698 | 0 | 0 |
| NC_004718.3;NP_828850.1;SARS-CoV_orf1a;699 | 0 | 0 |
| NC_004718.3;NP_828850.1;SARS-CoV_orf1a;700 | 0.05 | 0 |
| NC_004718.3;NP_828850.1;SARS-CoV_orf1a;701 | 0.05 | 0 |
| NC_004718.3;NP_828850.1;SARS-CoV_orf1a;702 | 0 | 0 |
| NC_004718.3;NP_828850.1;SARS-CoV_orf1a;703 | 0 | 0 |
| NC_004718.3;NP_828850.1;SARS-CoV_orf1a;704 | 0 | 0 |
| NC_004718.3;NP_828850.1;SARS-CoV_orf1a;705 | 0 | 0 |
| NC_004718.3;NP_828850.1;SARS-CoV_orf1a;706 | 0 | 0 |
| NC_004718.3;NP_828850.1;SARS-CoV_orf1a;707 | 0 | 0 |
| NC_004718.3;NP_828850.1;SARS-CoV_orf1a;708 | 0 | 0 |
| NC_004718.3;NP_828850.1;SARS-CoV_orf1a;709 | 0 | 0.025 |
| NC_004718.3;NP_828850.1;SARS-CoV_orf1a;710 | 0 | 0 |
| NC_004718.3;NP_828850.1;SARS-CoV_orf1a;711 | 0 | 0 |
| NC_004718.3;NP_828850.1;SARS-CoV_orf1a;712 | 0 | 0 |
| NC_004718.3;NP_828850.1;SARS-CoV_orf1a;713 | 0 | 0 |
| NC_004718.3;NP_828850.1;SARS-CoV_orf1a;714 | 0 | 0.025 |
| NC_004718.3;NP_828850.1;SARS-CoV_orf1a;715 | 0 | 0 |
| NC_004718.3;NP_828850.1;SARS-CoV_orf1a;716 | 0 | 0 |
| NC_004718.3;NP_828850.1;SARS-CoV_orf1a;717 | 0 | 0 |
| NC_004718.3;NP_828850.1;SARS-CoV_orf1a;718 | 0 | 0 |

| | | |
|---|---|---|
| NC_004718.3;NP_828850.1;SARS-CoV_orf1a;719 | 0 | 0 |
| NC_004718.3;NP_828850.1;SARS-CoV_orf1a;720 | 0 | 0 |
| NC_004718.3;NP_828850.1;SARS-CoV_orf1a;721 | 0 | 0 |
| NC_004718.3;NP_828850.1;SARS-CoV_orf1a;722 | 0 | 0 |
| NC_004718.3;NP_828850.1;SARS-CoV_orf1a;723 | 0 | 0 |
| NC_004718.3;NP_828850.1;SARS-CoV_orf1a;724 | 0 | 0 |
| NC_004718.3;NP_828850.1;SARS-CoV_orf1a;725 | 0 | 0 |
| NC_004718.3;NP_828850.1;SARS-CoV_orf1a;726 | 0 | 0 |
| NC_004718.3;NP_828850.1;SARS-CoV_orf1a;727 | 0 | 0 |
| NC_004718.3;NP_828850.1;SARS-CoV_orf1a;728 | 0 | 0.025 |
| NC_004718.3;NP_828850.1;SARS-CoV_orf1a;729 | 0 | 0.025 |
| NC_004718.3;NP_828850.1;SARS-CoV_orf1a;730 | 0 | 0.025 |
| NC_004718.3;NP_828850.1;SARS-CoV_orf1a;731 | 0 | 0.025 |
| NC_004718.3;NP_828850.1;SARS-CoV_orf1a;732 | 0 | 0.025 |
| NC_004718.3;NP_828850.1;SARS-CoV_orf1a;733 | 0 | 0.025 |
| NC_004718.3;NP_828850.1;SARS-CoV_orf1a;734 | 0 | 0.025 |
| NC_004718.3;NP_828850.1;SARS-CoV_orf1a;735 | 0 | 0.025 |
| NC_004718.3;NP_828850.1;SARS-CoV_orf1a;736 | 0 | 0.05 |
| NC_004718.3;NP_828850.1;SARS-CoV_orf1a;737 | 0 | 0.025 |
| NC_004718.3;NP_828850.1;SARS-CoV_orf1a;738 | 0 | 0.025 |
| NC_004718.3;NP_828850.1;SARS-CoV_orf1a;739 | 0 | 0 |
| NC_004718.3;NP_828850.1;SARS-CoV_orf1a;740 | 0 | 0 |
| NC_004718.3;NP_828850.1;SARS-CoV_orf1a;741 | 0.05 | 0 |
| NC_004718.3;NP_828850.1;SARS-CoV_orf1a;742 | 0 | 0 |
| NC_004718.3;NP_828850.1;SARS-CoV_orf1a;743 | 0 | 0 |
| NC_004718.3;NP_828850.1;SARS-CoV_orf1a;744 | 0.05 | 0.025 |
| NC_004718.3;NP_828850.1;SARS-CoV_orf1a;745 | 0.05 | 0.05 |
| NC_004718.3;NP_828850.1;SARS-CoV_orf1a;746 | 0.2 | 0.15 |
| NC_004718.3;NP_828850.1;SARS-CoV_orf1a;747 | 0 | 0.05 |
| NC_004718.3;NP_828850.1;SARS-CoV_orf1a;748 | 0.05 | 0.025 |
| NC_004718.3;NP_828850.1;SARS-CoV_orf1a;749 | 0.05 | 0.05 |
| NC_004718.3;NP_828850.1;SARS-CoV_orf1a;750 | 0.05 | 0.05 |
| NC_004718.3;NP_828850.1;SARS-CoV_orf1a;751 | 0.05 | 0.025 |
| NC_004718.3;NP_828850.1;SARS-CoV_orf1a;752 | 0.05 | 0.05 |
| NC_004718.3;NP_828850.1;SARS-CoV_orf1a;753 | 0.05 | 0 |
| NC_004718.3;NP_828850.1;SARS-CoV_orf1a;754 | 0.1 | 0 |
| NC_004718.3;NP_828850.1;SARS-CoV_orf1a;755 | 0.05 | 0 |
| NC_004718.3;NP_828850.1;SARS-CoV_orf1a;756 | 0 | 0 |
| NC_004718.3;NP_828850.1;SARS-CoV_orf1a;757 | 0 | 0 |
| NC_004718.3;NP_828850.1;SARS-CoV_orf1a;758 | 0 | 0 |
| NC_004718.3;NP_828850.1;SARS-CoV_orf1a;759 | 0 | 0 |
| NC_004718.3;NP_828850.1;SARS-CoV_orf1a;760 | 0 | 0 |
| NC_004718.3;NP_828850.1;SARS-CoV_orf1a;761 | 0 | 0 |

| | | |
|---|---|---|
| NC_004718.3;NP_828850.1;SARS-CoV_orf1a;762 | 0 | 0 |
| NC_004718.3;NP_828850.1;SARS-CoV_orf1a;763 | 0 | 0.025 |
| NC_004718.3;NP_828850.1;SARS-CoV_orf1a;764 | 0 | 0 |
| NC_004718.3;NP_828850.1;SARS-CoV_orf1a;765 | 0 | 0 |
| NC_004718.3;NP_828850.1;SARS-CoV_orf1a;766 | 0 | 0 |
| NC_004718.3;NP_828850.1;SARS-CoV_orf1a;767 | 0 | 0 |
| NC_004718.3;NP_828850.1;SARS-CoV_orf1a;768 | 0 | 0 |
| NC_004718.3;NP_828850.1;SARS-CoV_orf1a;769 | 0 | 0 |
| NC_004718.3;NP_828850.1;SARS-CoV_orf1a;770 | 0.05 | 0.025 |
| NC_004718.3;NP_828850.1;SARS-CoV_orf1a;771 | 0.1 | 0.025 |
| NC_004718.3;NP_828850.1;SARS-CoV_orf1a;772 | 0 | 0 |
| NC_004718.3;NP_828850.1;SARS-CoV_orf1a;773 | 0 | 0 |
| NC_004718.3;NP_828850.1;SARS-CoV_orf1a;774 | 0 | 0 |
| NC_004718.3;NP_828850.1;SARS-CoV_orf1a;775 | 0 | 0 |
| NC_004718.3;NP_828850.1;SARS-CoV_orf1a;776 | 0 | 0 |
| NC_004718.3;NP_828850.1;SARS-CoV_orf1a;777 | 0 | 0 |
| NC_004718.3;NP_828850.1;SARS-CoV_orf1a;778 | 0 | 0 |
| NC_004718.3;NP_828850.1;SARS-CoV_orf1a;779 | 0 | 0 |
| NC_004718.3;NP_828850.1;SARS-CoV_orf1a;780 | 0 | 0.025 |
| NC_004718.3;NP_828850.1;SARS-CoV_orf1a;781 | 0 | 0 |
| NC_004718.3;NP_828850.1;SARS-CoV_orf1a;782 | 0.05 | 0 |
| NC_004718.3;NP_828850.1;SARS-CoV_orf1a;783 | 0 | 0 |
| NC_004718.3;NP_828850.1;SARS-CoV_orf1a;784 | 0 | 0 |
| NC_004718.3;NP_828850.1;SARS-CoV_orf1a;785 | 0 | 0 |
| NC_004718.3;NP_828850.1;SARS-CoV_orf1a;786 | 0 | 0 |
| NC_004718.3;NP_828850.1;SARS-CoV_orf1a;787 | 0 | 0.025 |
| NC_004718.3;NP_828850.1;SARS-CoV_orf1a;788 | 0 | 0 |
| NC_004718.3;NP_828850.1;SARS-CoV_orf1a;789 | 0.05 | 0 |
| NC_004718.3;NP_828850.1;SARS-CoV_orf1a;790 | 0 | 0 |
| NC_004718.3;NP_828850.1;SARS-CoV_orf1a;791 | 0 | 0.025 |
| NC_004718.3;NP_828850.1;SARS-CoV_orf1a;792 | 0 | 0 |
| NC_004718.3;NP_828850.1;SARS-CoV_orf1a;793 | 0 | 0 |
| NC_004718.3;NP_828850.1;SARS-CoV_orf1a;794 | 0 | 0 |
| NC_004718.3;NP_828850.1;SARS-CoV_orf1a;795 | 0 | 0 |
| NC_004718.3;NP_828850.1;SARS-CoV_orf1a;796 | 0 | 0 |
| NC_004718.3;NP_828850.1;SARS-CoV_orf1a;797 | 0 | 0 |
| NC_004718.3;NP_828850.1;SARS-CoV_orf1a;798 | 0 | 0 |
| NC_004718.3;NP_828850.1;SARS-CoV_orf1a;799 | 0 | 0 |
| NC_004718.3;NP_828850.1;SARS-CoV_orf1a;800 | 0 | 0 |
| NC_004718.3;NP_828850.1;SARS-CoV_orf1a;801 | 0.05 | 0 |
| NC_004718.3;NP_828850.1;SARS-CoV_orf1a;802 | 0 | 0 |
| NC_004718.3;NP_828850.1;SARS-CoV_orf1a;803 | 0 | 0 |
| NC_004718.3;NP_828850.1;SARS-CoV_orf1a;804 | 0 | 0 |

| | | |
|---|---|---|
| NC_004718.3;NP_828850.1;SARS-CoV_orf1a;805 | 0 | 0 |
| NC_004718.3;NP_828850.1;SARS-CoV_orf1a;806 | 0 | 0 |
| NC_004718.3;NP_828850.1;SARS-CoV_orf1a;807 | 0 | 0 |
| NC_004718.3;NP_828850.1;SARS-CoV_orf1a;808 | 0 | 0 |
| NC_004718.3;NP_828850.1;SARS-CoV_orf1a;809 | 0 | 0 |
| NC_004718.3;NP_828850.1;SARS-CoV_orf1a;810 | 0 | 0 |
| NC_004718.3;NP_828850.1;SARS-CoV_orf1a;811 | 0 | 0 |
| NC_004718.3;NP_828850.1;SARS-CoV_orf1a;812 | 0 | 0 |
| NC_004718.3;NP_828850.1;SARS-CoV_orf1a;813 | 0 | 0 |
| NC_004718.3;NP_828850.1;SARS-CoV_orf1a;814 | 0 | 0.025 |
| NC_004718.3;NP_828850.1;SARS-CoV_orf1a;815 | 0 | 0.025 |
| NC_004718.3;NP_828850.1;SARS-CoV_orf1a;816 | 0.05 | 0.05 |
| NC_004718.3;NP_828850.1;SARS-CoV_orf1a;817 | 0 | 0.05 |
| NC_004718.3;NP_828850.1;SARS-CoV_orf1a;818 | 0 | 0.075 |
| NC_004718.3;NP_828850.1;SARS-CoV_orf1a;819 | 0 | 0.025 |
| NC_004718.3;NP_828850.1;SARS-CoV_orf1a;820 | 0 | 0.025 |
| NC_004718.3;NP_828850.1;SARS-CoV_orf1a;821 | 0 | 0.075 |
| NC_004718.3;NP_828850.1;SARS-CoV_orf1a;822 | 0 | 0 |
| NC_004718.3;NP_828850.1;SARS-CoV_orf1a;823 | 0 | 0.075 |
| NC_004718.3;NP_828850.1;SARS-CoV_orf1a;824 | 0 | 0.1 |
| NC_004718.3;NP_828850.1;SARS-CoV_orf1a;825 | 0 | 0.075 |
| NC_004718.3;NP_828850.1;SARS-CoV_orf1a;826 | 0.05 | 0.025 |
| NC_004718.3;NP_828850.1;SARS-CoV_orf1a;827 | 0 | 0 |
| NC_004718.3;NP_828850.1;SARS-CoV_orf1a;828 | 0.1 | 0.025 |
| NC_004718.3;NP_828850.1;SARS-CoV_orf1a;829 | 0 | 0 |
| NC_004718.3;NP_828850.1;SARS-CoV_orf1a;830 | 0 | 0 |
| NC_004718.3;NP_828850.1;SARS-CoV_orf1a;831 | 0 | 0 |
| NC_004718.3;NP_828850.1;SARS-CoV_orf1a;832 | 0.05 | 0 |
| NC_004718.3;NP_828850.1;SARS-CoV_orf1a;833 | 0 | 0 |
| NC_004718.3;NP_828850.1;SARS-CoV_orf1a;834 | 0 | 0 |
| NC_004718.3;NP_828850.1;SARS-CoV_orf1a;835 | 0 | 0 |
| NC_004718.3;NP_828850.1;SARS-CoV_orf1a;836 | 0 | 0 |
| NC_004718.3;NP_828850.1;SARS-CoV_orf1a;837 | 0.05 | 0.025 |
| NC_004718.3;NP_828850.1;SARS-CoV_orf1a;838 | 0 | 0 |
| NC_004718.3;NP_828850.1;SARS-CoV_orf1a;839 | 0 | 0 |
| NC_004718.3;NP_828850.1;SARS-CoV_orf1a;840 | 0 | 0 |
| NC_004718.3;NP_828850.1;SARS-CoV_orf1a;841 | 0 | 0.025 |
| NC_004718.3;NP_828850.1;SARS-CoV_orf1a;842 | 0.05 | 0.05 |
| NC_004718.3;NP_828850.1;SARS-CoV_orf1a;843 | 0.05 | 0.025 |
| NC_004718.3;NP_828850.1;SARS-CoV_orf1a;844 | 0.05 | 0.025 |
| NC_004718.3;NP_828850.1;SARS-CoV_orf1a;845 | 0 | 0.025 |
| NC_004718.3;NP_828850.1;SARS-CoV_orf1a;846 | 0 | 0.025 |
| NC_004718.3;NP_828850.1;SARS-CoV_orf1a;847 | 0 | 0.05 |

| | | |
|---|---|---|
| NC_004718.3;NP_828850.1;SARS-CoV_orf1a;848 | 0 | 0 |
| NC_004718.3;NP_828850.1;SARS-CoV_orf1a;849 | 0 | 0 |
| NC_004718.3;NP_828850.1;SARS-CoV_orf1a;850 | 0.1 | 0.075 |
| NC_004718.3;NP_828850.1;SARS-CoV_orf1a;851 | 0.15 | 0.1 |
| NC_004718.3;NP_828850.1;SARS-CoV_orf1a;852 | 0.15 | 0.1 |
| NC_004718.3;NP_828850.1;SARS-CoV_orf1a;853 | 0.15 | 0.1 |
| NC_004718.3;NP_828850.1;SARS-CoV_orf1a;854 | 0.1 | 0.1 |
| NC_004718.3;NP_828850.1;SARS-CoV_orf1a;855 | 0.05 | 0.05 |
| NC_004718.3;NP_828850.1;SARS-CoV_orf1a;856 | 0 | 0.05 |
| NC_004718.3;NP_828850.1;SARS-CoV_orf1a;857 | 0 | 0.05 |
| NC_004718.3;NP_828850.1;SARS-CoV_orf1a;858 | 0 | 0.05 |
| NC_004718.3;NP_828850.1;SARS-CoV_orf1a;859 | 0 | 0 |
| NC_004718.3;NP_828850.1;SARS-CoV_orf1a;860 | 0 | 0 |
| NC_004718.3;NP_828850.1;SARS-CoV_orf1a;861 | 0 | 0 |
| NC_004718.3;NP_828850.1;SARS-CoV_orf1a;862 | 0 | 0 |
| NC_004718.3;NP_828850.1;SARS-CoV_orf1a;863 | 0 | 0 |
| NC_004718.3;NP_828850.1;SARS-CoV_orf1a;864 | 0.05 | 0 |
| NC_004718.3;NP_828850.1;SARS-CoV_orf1a;865 | 0 | 0 |
| NC_004718.3;NP_828850.1;SARS-CoV_orf1a;866 | 0 | 0 |
| NC_004718.3;NP_828850.1;SARS-CoV_orf1a;867 | 0 | 0 |
| NC_004718.3;NP_828850.1;SARS-CoV_orf1a;868 | 0 | 0 |
| NC_004718.3;NP_828850.1;SARS-CoV_orf1a;869 | 0 | 0 |
| NC_004718.3;NP_828850.1;SARS-CoV_orf1a;870 | 0 | 0 |
| NC_004718.3;NP_828850.1;SARS-CoV_orf1a;871 | 0 | 0 |
| NC_004718.3;NP_828850.1;SARS-CoV_orf1a;872 | 0 | 0 |
| NC_004718.3;NP_828850.1;SARS-CoV_orf1a;873 | 0 | 0 |
| NC_004718.3;NP_828850.1;SARS-CoV_orf1a;874 | 0 | 0 |
| NC_004718.3;NP_828850.1;SARS-CoV_orf1a;875 | 0 | 0 |
| NC_004718.3;NP_828850.1;SARS-CoV_orf1a;876 | 0 | 0 |
| NC_004718.3;NP_828850.1;SARS-CoV_orf1a;877 | 0 | 0.05 |
| NC_004718.3;NP_828850.1;SARS-CoV_orf1a;878 | 0.05 | 0 |
| NC_004718.3;NP_828850.1;SARS-CoV_orf1a;879 | 0 | 0.025 |
| NC_004718.3;NP_828850.1;SARS-CoV_orf1a;880 | 0 | 0 |
| NC_004718.3;NP_828850.1;SARS-CoV_orf1a;881 | 0 | 0 |
| NC_004718.3;NP_828850.1;SARS-CoV_orf1a;882 | 0 | 0 |
| NC_004718.3;NP_828850.1;SARS-CoV_orf1a;883 | 0 | 0 |
| NC_004718.3;NP_828850.1;SARS-CoV_orf1a;884 | 0 | 0 |
| NC_004718.3;NP_828850.1;SARS-CoV_orf1a;885 | 0.15 | 0.05 |
| NC_004718.3;NP_828850.1;SARS-CoV_orf1a;886 | 0 | 0 |
| NC_004718.3;NP_828850.1;SARS-CoV_orf1a;887 | 0 | 0 |
| NC_004718.3;NP_828850.1;SARS-CoV_orf1a;888 | 0 | 0 |
| NC_004718.3;NP_828850.1;SARS-CoV_orf1a;889 | 0.05 | 0 |
| NC_004718.3;NP_828850.1;SARS-CoV_orf1a;890 | 0 | 0.025 |

| | | |
|---|---|---|
| NC_004718.3;NP_828850.1;SARS-CoV_orf1a;891 | 0 | 0 |
| NC_004718.3;NP_828850.1;SARS-CoV_orf1a;892 | 0 | 0 |
| NC_004718.3;NP_828850.1;SARS-CoV_orf1a;893 | 0 | 0 |
| NC_004718.3;NP_828850.1;SARS-CoV_orf1a;894 | 0 | 0 |
| NC_004718.3;NP_828850.1;SARS-CoV_orf1a;895 | 0 | 0 |
| NC_004718.3;NP_828850.1;SARS-CoV_orf1a;896 | 0 | 0 |
| NC_004718.3;NP_828850.1;SARS-CoV_orf1a;897 | 0 | 0 |
| NC_004718.3;NP_828850.1;SARS-CoV_orf1a;898 | 0 | 0 |
| NC_004718.3;NP_828850.1;SARS-CoV_orf1a;899 | 0 | 0.025 |
| NC_004718.3;NP_828850.1;SARS-CoV_orf1a;900 | 0 | 0 |
| NC_004718.3;NP_828850.1;SARS-CoV_orf1a;901 | 0 | 0 |
| NC_004718.3;NP_828850.1;SARS-CoV_orf1a;902 | 0 | 0 |
| NC_004718.3;NP_828850.1;SARS-CoV_orf1a;903 | 0 | 0 |
| NC_004718.3;NP_828850.1;SARS-CoV_orf1a;904 | 0 | 0 |
| NC_004718.3;NP_828850.1;SARS-CoV_orf1a;905 | 0 | 0.025 |
| NC_004718.3;NP_828850.1;SARS-CoV_orf1a;906 | 0 | 0.025 |
| NC_004718.3;NP_828850.1;SARS-CoV_orf1a;907 | 0 | 0.05 |
| NC_004718.3;NP_828850.1;SARS-CoV_orf1a;908 | 0 | 0 |
| NC_004718.3;NP_828850.1;SARS-CoV_orf1a;909 | 0 | 0 |
| NC_004718.3;NP_828850.1;SARS-CoV_orf1a;910 | 0 | 0 |
| NC_004718.3;NP_828850.1;SARS-CoV_orf1a;911 | 0 | 0 |
| NC_004718.3;NP_828850.1;SARS-CoV_orf1a;912 | 0 | 0 |
| NC_004718.3;NP_828850.1;SARS-CoV_orf1a;913 | 0 | 0 |
| NC_004718.3;NP_828850.1;SARS-CoV_orf1a;914 | 0 | 0 |
| NC_004718.3;NP_828850.1;SARS-CoV_orf1a;915 | 0 | 0 |
| NC_004718.3;NP_828850.1;SARS-CoV_orf1a;916 | 0 | 0 |
| NC_004718.3;NP_828850.1;SARS-CoV_orf1a;917 | 0 | 0.025 |
| NC_004718.3;NP_828850.1;SARS-CoV_orf1a;918 | 0 | 0.05 |
| NC_004718.3;NP_828850.1;SARS-CoV_orf1a;919 | 0 | 0.05 |
| NC_004718.3;NP_828850.1;SARS-CoV_orf1a;920 | 0 | 0.05 |
| NC_004718.3;NP_828850.1;SARS-CoV_orf1a;921 | 0 | 0.05 |
| NC_004718.3;NP_828850.1;SARS-CoV_orf1a;922 | 0 | 0.1 |
| NC_004718.3;NP_828850.1;SARS-CoV_orf1a;923 | 0 | 0.05 |
| NC_004718.3;NP_828850.1;SARS-CoV_orf1a;924 | 0 | 0.05 |
| NC_004718.3;NP_828850.1;SARS-CoV_orf1a;925 | 0 | 0.075 |
| NC_004718.3;NP_828850.1;SARS-CoV_orf1a;926 | 0 | 0.025 |
| NC_004718.3;NP_828850.1;SARS-CoV_orf1a;927 | 0 | 0 |
| NC_004718.3;NP_828850.1;SARS-CoV_orf1a;928 | 0 | 0 |
| NC_004718.3;NP_828850.1;SARS-CoV_orf1a;929 | 0 | 0 |
| NC_004718.3;NP_828850.1;SARS-CoV_orf1a;930 | 0 | 0 |
| NC_004718.3;NP_828850.1;SARS-CoV_orf1a;931 | 0 | 0 |
| NC_004718.3;NP_828850.1;SARS-CoV_orf1a;932 | 0 | 0.025 |
| NC_004718.3;NP_828850.1;SARS-CoV_orf1a;933 | 0.25 | 0.05 |

| | | |
|---|---|---|
| NC_004718.3;NP_828850.1;SARS-CoV_orf1a;934 | 0 | 0 |
| NC_004718.3;NP_828850.1;SARS-CoV_orf1a;935 | 0 | 0 |
| NC_004718.3;NP_828850.1;SARS-CoV_orf1a;936 | 0 | 0 |
| NC_004718.3;NP_828850.1;SARS-CoV_orf1a;937 | 0 | 0.025 |
| NC_004718.3;NP_828850.1;SARS-CoV_orf1a;938 | 0 | 0 |
| NC_004718.3;NP_828850.1;SARS-CoV_orf1a;939 | 0 | 0 |
| NC_004718.3;NP_828850.1;SARS-CoV_orf1a;940 | 0 | 0 |
| NC_004718.3;NP_828850.1;SARS-CoV_orf1a;941 | 0 | 0.025 |
| NC_004718.3;NP_828850.1;SARS-CoV_orf1a;942 | 0 | 0 |
| NC_004718.3;NP_828850.1;SARS-CoV_orf1a;943 | 0 | 0 |
| NC_004718.3;NP_828850.1;SARS-CoV_orf1a;944 | 0 | 0 |
| NC_004718.3;NP_828850.1;SARS-CoV_orf1a;945 | 0 | 0 |
| NC_004718.3;NP_828850.1;SARS-CoV_orf1a;946 | 0 | 0 |
| NC_004718.3;NP_828850.1;SARS-CoV_orf1a;947 | 0 | 0 |
| NC_004718.3;NP_828850.1;SARS-CoV_orf1a;948 | 0 | 0 |
| NC_004718.3;NP_828850.1;SARS-CoV_orf1a;949 | 0 | 0 |
| NC_004718.3;NP_828850.1;SARS-CoV_orf1a;950 | 0.05 | 0 |
| NC_004718.3;NP_828850.1;SARS-CoV_orf1a;951 | 0.05 | 0 |
| NC_004718.3;NP_828850.1;SARS-CoV_orf1a;952 | 0.05 | 0.075 |
| NC_004718.3;NP_828850.1;SARS-CoV_orf1a;953 | 0.05 | 0 |
| NC_004718.3;NP_828850.1;SARS-CoV_orf1a;954 | 0.15 | 0 |
| NC_004718.3;NP_828850.1;SARS-CoV_orf1a;955 | 0.05 | 0 |
| NC_004718.3;NP_828850.1;SARS-CoV_orf1a;956 | 0 | 0 |
| NC_004718.3;NP_828850.1;SARS-CoV_orf1a;957 | 0 | 0 |
| NC_004718.3;NP_828850.1;SARS-CoV_orf1a;958 | 0 | 0 |
| NC_004718.3;NP_828850.1;SARS-CoV_orf1a;959 | 0 | 0 |
| NC_004718.3;NP_828850.1;SARS-CoV_orf1a;960 | 0 | 0 |
| NC_004718.3;NP_828850.1;SARS-CoV_orf1a;961 | 0 | 0 |
| NC_004718.3;NP_828850.1;SARS-CoV_orf1a;962 | 0 | 0 |
| NC_004718.3;NP_828850.1;SARS-CoV_orf1a;963 | 0 | 0 |
| NC_004718.3;NP_828850.1;SARS-CoV_orf1a;964 | 0 | 0 |
| NC_004718.3;NP_828850.1;SARS-CoV_orf1a;965 | 0 | 0 |
| NC_004718.3;NP_828850.1;SARS-CoV_orf1a;966 | 0 | 0.025 |
| NC_004718.3;NP_828850.1;SARS-CoV_orf1a;967 | 0 | 0.025 |
| NC_004718.3;NP_828850.1;SARS-CoV_orf1a;968 | 0 | 0.075 |
| NC_004718.3;NP_828850.1;SARS-CoV_orf1a;969 | 0 | 0.025 |
| NC_004718.3;NP_828850.1;SARS-CoV_orf1a;970 | 0 | 0.025 |
| NC_004718.3;NP_828850.1;SARS-CoV_orf1a;971 | 0 | 0.025 |
| NC_004718.3;NP_828850.1;SARS-CoV_orf1a;972 | 0 | 0.025 |
| NC_004718.3;NP_828850.1;SARS-CoV_orf1a;973 | 0 | 0.025 |
| NC_004718.3;NP_828850.1;SARS-CoV_orf1a;974 | 0 | 0.05 |
| NC_004718.3;NP_828850.1;SARS-CoV_orf1a;975 | 0 | 0.025 |
| NC_004718.3;NP_828850.1;SARS-CoV_orf1a;976 | 0 | 0.025 |

| | | |
|---|---|---|
| NC_004718.3;NP_828850.1;SARS-CoV_orf1a;977 | 0 | 0.025 |
| NC_004718.3;NP_828850.1;SARS-CoV_orf1a;978 | 0 | 0.025 |
| NC_004718.3;NP_828850.1;SARS-CoV_orf1a;979 | 0.05 | 0.025 |
| NC_004718.3;NP_828850.1;SARS-CoV_orf1a;980 | 0 | 0 |
| NC_004718.3;NP_828850.1;SARS-CoV_orf1a;981 | 0 | 0 |
| NC_004718.3;NP_828850.1;SARS-CoV_orf1a;982 | 0 | 0 |
| NC_004718.3;NP_828850.1;SARS-CoV_orf1a;983 | 0 | 0 |
| NC_004718.3;NP_828850.1;SARS-CoV_orf1a;984 | 0 | 0 |
| NC_004718.3;NP_828850.1;SARS-CoV_orf1a;985 | 0 | 0 |
| NC_004718.3;NP_828850.1;SARS-CoV_orf1a;986 | 0 | 0.025 |
| NC_004718.3;NP_828850.1;SARS-CoV_orf1a;987 | 0 | 0.025 |
| NC_004718.3;NP_828850.1;SARS-CoV_orf1a;988 | 0 | 0.05 |
| NC_004718.3;NP_828850.1;SARS-CoV_orf1a;989 | 0 | 0.025 |
| NC_004718.3;NP_828850.1;SARS-CoV_orf1a;990 | 0 | 0.05 |
| NC_004718.3;NP_828850.1;SARS-CoV_orf1a;991 | 0 | 0.025 |
| NC_004718.3;NP_828850.1;SARS-CoV_orf1a;992 | 0.05 | 0.025 |
| NC_004718.3;NP_828850.1;SARS-CoV_orf1a;993 | 0.05 | 0.025 |
| NC_004718.3;NP_828850.1;SARS-CoV_orf1a;994 | 0.05 | 0.025 |
| NC_004718.3;NP_828850.1;SARS-CoV_orf1a;995 | 0 | 0 |
| NC_004718.3;NP_828850.1;SARS-CoV_orf1a;996 | 0.05 | 0 |
| NC_004718.3;NP_828850.1;SARS-CoV_orf1a;997 | 0 | 0.025 |
| NC_004718.3;NP_828850.1;SARS-CoV_orf1a;998 | 0 | 0 |
| NC_004718.3;NP_828850.1;SARS-CoV_orf1a;999 | 0 | 0 |
| NC_004718.3;NP_828850.1;SARS-CoV_orf1a;1000 | 0 | 0 |
| NC_004718.3;NP_828850.1;SARS-CoV_orf1a;1001 | 0.05 | 0 |
| NC_004718.3;NP_828850.1;SARS-CoV_orf1a;1002 | 0.05 | 0 |
| NC_004718.3;NP_828850.1;SARS-CoV_orf1a;1003 | 0.05 | 0 |
| NC_004718.3;NP_828850.1;SARS-CoV_orf1a;1004 | 0.05 | 0 |
| NC_004718.3;NP_828850.1;SARS-CoV_orf1a;1005 | 0.05 | 0 |
| NC_004718.3;NP_828850.1;SARS-CoV_orf1a;1006 | 0.05 | 0 |
| NC_004718.3;NP_828850.1;SARS-CoV_orf1a;1007 | 0.05 | 0 |
| NC_004718.3;NP_828850.1;SARS-CoV_orf1a;1008 | 0.05 | 0 |
| NC_004718.3;NP_828850.1;SARS-CoV_orf1a;1009 | 0 | 0 |
| NC_004718.3;NP_828850.1;SARS-CoV_orf1a;1010 | 0.05 | 0 |
| NC_004718.3;NP_828850.1;SARS-CoV_orf1a;1011 | 0.05 | 0 |
| NC_004718.3;NP_828850.1;SARS-CoV_orf1a;1012 | 0 | 0 |
| NC_004718.3;NP_828850.1;SARS-CoV_orf1a;1013 | 0 | 0 |
| NC_004718.3;NP_828850.1;SARS-CoV_orf1a;1014 | 0 | 0 |
| NC_004718.3;NP_828850.1;SARS-CoV_orf1a;1015 | 0 | 0 |
| NC_004718.3;NP_828850.1;SARS-CoV_orf1a;1016 | 0 | 0.075 |
| NC_004718.3;NP_828850.1;SARS-CoV_orf1a;1017 | 0 | 0.05 |
| NC_004718.3;NP_828850.1;SARS-CoV_orf1a;1018 | 0 | 0.075 |
| NC_004718.3;NP_828850.1;SARS-CoV_orf1a;1019 | 0 | 0.075 |

| | | |
|---|---|---|
| NC_004718.3;NP_828850.1;SARS-CoV_orf1a;1020 | 0 | 0.075 |
| NC_004718.3;NP_828850.1;SARS-CoV_orf1a;1021 | 0 | 0.1 |
| NC_004718.3;NP_828850.1;SARS-CoV_orf1a;1022 | 0 | 0.075 |
| NC_004718.3;NP_828850.1;SARS-CoV_orf1a;1023 | 0 | 0.075 |
| NC_004718.3;NP_828850.1;SARS-CoV_orf1a;1024 | 0.05 | 0.05 |
| NC_004718.3;NP_828850.1;SARS-CoV_orf1a;1025 | 0.05 | 0.05 |
| NC_004718.3;NP_828850.1;SARS-CoV_orf1a;1026 | 0 | 0 |
| NC_004718.3;NP_828850.1;SARS-CoV_orf1a;1027 | 0 | 0 |
| NC_004718.3;NP_828850.1;SARS-CoV_orf1a;1028 | 0 | 0 |
| NC_004718.3;NP_828850.1;SARS-CoV_orf1a;1029 | 0 | 0 |
| NC_004718.3;NP_828850.1;SARS-CoV_orf1a;1030 | 0 | 0 |
| NC_004718.3;NP_828850.1;SARS-CoV_orf1a;1031 | 0 | 0 |
| NC_004718.3;NP_828850.1;SARS-CoV_orf1a;1032 | 0 | 0 |
| NC_004718.3;NP_828850.1;SARS-CoV_orf1a;1033 | 0.05 | 0 |
| NC_004718.3;NP_828850.1;SARS-CoV_orf1a;1034 | 0 | 0 |
| NC_004718.3;NP_828850.1;SARS-CoV_orf1a;1035 | 0.1 | 0.05 |
| NC_004718.3;NP_828850.1;SARS-CoV_orf1a;1036 | 0 | 0 |
| NC_004718.3;NP_828850.1;SARS-CoV_orf1a;1037 | 0 | 0 |
| NC_004718.3;NP_828850.1;SARS-CoV_orf1a;1038 | 0 | 0 |
| NC_004718.3;NP_828850.1;SARS-CoV_orf1a;1039 | 0 | 0 |
| NC_004718.3;NP_828850.1;SARS-CoV_orf1a;1040 | 0 | 0 |
| NC_004718.3;NP_828850.1;SARS-CoV_orf1a;1041 | 0 | 0 |
| NC_004718.3;NP_828850.1;SARS-CoV_orf1a;1042 | 0 | 0 |
| NC_004718.3;NP_828850.1;SARS-CoV_orf1a;1043 | 0 | 0 |
| NC_004718.3;NP_828850.1;SARS-CoV_orf1a;1044 | 0 | 0 |
| NC_004718.3;NP_828850.1;SARS-CoV_orf1a;1045 | 0.05 | 0 |
| NC_004718.3;NP_828850.1;SARS-CoV_orf1a;1046 | 0 | 0.025 |
| NC_004718.3;NP_828850.1;SARS-CoV_orf1a;1047 | 0.05 | 0.025 |
| NC_004718.3;NP_828850.1;SARS-CoV_orf1a;1048 | 0 | 0 |
| NC_004718.3;NP_828850.1;SARS-CoV_orf1a;1049 | 0.15 | 0.05 |
| NC_004718.3;NP_828850.1;SARS-CoV_orf1a;1050 | 0.15 | 0.075 |
| NC_004718.3;NP_828850.1;SARS-CoV_orf1a;1051 | 0.15 | 0.075 |
| NC_004718.3;NP_828850.1;SARS-CoV_orf1a;1052 | 0.1 | 0.025 |
| NC_004718.3;NP_828850.1;SARS-CoV_orf1a;1053 | 0.2 | 0.025 |
| NC_004718.3;NP_828850.1;SARS-CoV_orf1a;1054 | 0.15 | 0.025 |
| NC_004718.3;NP_828850.1;SARS-CoV_orf1a;1055 | 0.15 | 0 |
| NC_004718.3;NP_828850.1;SARS-CoV_orf1a;1056 | 0.2 | 0.05 |
| NC_004718.3;NP_828850.1;SARS-CoV_orf1a;1057 | 0.05 | 0 |
| NC_004718.3;NP_828850.1;SARS-CoV_orf1a;1058 | 0.15 | 0 |
| NC_004718.3;NP_828850.1;SARS-CoV_orf1a;1059 | 0.15 | 0.025 |
| NC_004718.3;NP_828850.1;SARS-CoV_orf1a;1060 | 0.2 | 0.025 |
| NC_004718.3;NP_828850.1;SARS-CoV_orf1a;1061 | 0.1 | 0.025 |
| NC_004718.3;NP_828850.1;SARS-CoV_orf1a;1062 | 0.1 | 0.05 |

| | | |
|---|---|---|
| NC_004718.3;NP_828850.1;SARS-CoV_orf1a;1063 | 0.15 | 0.025 |
| NC_004718.3;NP_828850.1;SARS-CoV_orf1a;1064 | 0.15 | 0.05 |
| NC_004718.3;NP_828850.1;SARS-CoV_orf1a;1065 | 0.2 | 0.05 |
| NC_004718.3;NP_828850.1;SARS-CoV_orf1a;1066 | 0 | 0 |
| NC_004718.3;NP_828850.1;SARS-CoV_orf1a;1067 | 0 | 0 |
| NC_004718.3;NP_828850.1;SARS-CoV_orf1a;1068 | 0 | 0 |
| NC_004718.3;NP_828850.1;SARS-CoV_orf1a;1069 | 0 | 0.025 |
| NC_004718.3;NP_828850.1;SARS-CoV_orf1a;1070 | 0 | 0 |
| NC_004718.3;NP_828850.1;SARS-CoV_orf1a;1071 | 0 | 0 |
| NC_004718.3;NP_828850.1;SARS-CoV_orf1a;1072 | 0 | 0 |
| NC_004718.3;NP_828850.1;SARS-CoV_orf1a;1073 | 0 | 0 |
| NC_004718.3;NP_828850.1;SARS-CoV_orf1a;1074 | 0 | 0 |
| NC_004718.3;NP_828850.1;SARS-CoV_orf1a;1075 | 0 | 0 |
| NC_004718.3;NP_828850.1;SARS-CoV_orf1a;1076 | 0 | 0 |
| NC_004718.3;NP_828850.1;SARS-CoV_orf1a;1077 | 0 | 0 |
| NC_004718.3;NP_828850.1;SARS-CoV_orf1a;1078 | 0 | 0 |
| NC_004718.3;NP_828850.1;SARS-CoV_orf1a;1079 | 0 | 0 |
| NC_004718.3;NP_828850.1;SARS-CoV_orf1a;1080 | 0 | 0 |
| NC_004718.3;NP_828850.1;SARS-CoV_orf1a;1081 | 0 | 0 |
| NC_004718.3;NP_828850.1;SARS-CoV_orf1a;1082 | 0 | 0 |
| NC_004718.3;NP_828850.1;SARS-CoV_orf1a;1083 | 0 | 0 |
| NC_004718.3;NP_828850.1;SARS-CoV_orf1a;1084 | 0 | 0 |
| NC_004718.3;NP_828850.1;SARS-CoV_orf1a;1085 | 0 | 0 |
| NC_004718.3;NP_828850.1;SARS-CoV_orf1a;1086 | 0 | 0 |
| NC_004718.3;NP_828850.1;SARS-CoV_orf1a;1087 | 0 | 0 |
| NC_004718.3;NP_828850.1;SARS-CoV_orf1a;1088 | 0 | 0 |
| NC_004718.3;NP_828850.1;SARS-CoV_orf1a;1089 | 0 | 0.05 |
| NC_004718.3;NP_828850.1;SARS-CoV_orf1a;1090 | 0.05 | 0.025 |
| NC_004718.3;NP_828850.1;SARS-CoV_orf1a;1091 | 0.05 | 0.05 |
| NC_004718.3;NP_828850.1;SARS-CoV_orf1a;1092 | 0 | 0.05 |
| NC_004718.3;NP_828850.1;SARS-CoV_orf1a;1093 | 0.05 | 0.025 |
| NC_004718.3;NP_828850.1;SARS-CoV_orf1a;1094 | 0 | 0.05 |
| NC_004718.3;NP_828850.1;SARS-CoV_orf1a;1095 | 0.05 | 0.05 |
| NC_004718.3;NP_828850.1;SARS-CoV_orf1a;1096 | 0.05 | 0.05 |
| NC_004718.3;NP_828850.1;SARS-CoV_orf1a;1097 | 0.05 | 0.075 |
| NC_004718.3;NP_828850.1;SARS-CoV_orf1a;1098 | 0 | 0.025 |
| NC_004718.3;NP_828850.1;SARS-CoV_orf1a;1099 | 0 | 0.05 |
| NC_004718.3;NP_828850.1;SARS-CoV_orf1a;1100 | 0 | 0.025 |
| NC_004718.3;NP_828850.1;SARS-CoV_orf1a;1101 | 0 | 0.075 |
| NC_004718.3;NP_828850.1;SARS-CoV_orf1a;1102 | 0.05 | 0.075 |
| NC_004718.3;NP_828850.1;SARS-CoV_orf1a;1103 | 0.05 | 0.075 |
| NC_004718.3;NP_828850.1;SARS-CoV_orf1a;1104 | 0.1 | 0.1 |
| NC_004718.3;NP_828850.1;SARS-CoV_orf1a;1105 | 0 | 0.05 |

| | | |
|---|---|---|
| NC_004718.3;NP_828850.1;SARS-CoV_orf1a;1106 | 0 | 0.05 |
| NC_004718.3;NP_828850.1;SARS-CoV_orf1a;1107 | 0 | 0.075 |
| NC_004718.3;NP_828850.1;SARS-CoV_orf1a;1108 | 0 | 0.025 |
| NC_004718.3;NP_828850.1;SARS-CoV_orf1a;1109 | 0 | 0.025 |
| NC_004718.3;NP_828850.1;SARS-CoV_orf1a;1110 | 0.1 | 0.05 |
| NC_004718.3;NP_828850.1;SARS-CoV_orf1a;1111 | 0 | 0 |
| NC_004718.3;NP_828850.1;SARS-CoV_orf1a;1112 | 0 | 0.025 |
| NC_004718.3;NP_828850.1;SARS-CoV_orf1a;1113 | 0 | 0 |
| NC_004718.3;NP_828850.1;SARS-CoV_orf1a;1114 | 0 | 0 |
| NC_004718.3;NP_828850.1;SARS-CoV_orf1a;1115 | 0 | 0 |
| NC_004718.3;NP_828850.1;SARS-CoV_orf1a;1116 | 0 | 0 |
| NC_004718.3;NP_828850.1;SARS-CoV_orf1a;1117 | 0 | 0 |
| NC_004718.3;NP_828850.1;SARS-CoV_orf1a;1118 | 0 | 0 |
| NC_004718.3;NP_828850.1;SARS-CoV_orf1a;1119 | 0 | 0 |
| NC_004718.3;NP_828850.1;SARS-CoV_orf1a;1120 | 0 | 0 |
| NC_004718.3;NP_828850.1;SARS-CoV_orf1a;1121 | 0 | 0 |
| NC_004718.3;NP_828850.1;SARS-CoV_orf1a;1122 | 0 | 0 |
| NC_004718.3;NP_828850.1;SARS-CoV_orf1a;1123 | 0 | 0 |
| NC_004718.3;NP_828850.1;SARS-CoV_orf1a;1124 | 0 | 0 |
| NC_004718.3;NP_828850.1;SARS-CoV_orf1a;1125 | 0 | 0 |
| NC_004718.3;NP_828850.1;SARS-CoV_orf1a;1126 | 0 | 0 |
| NC_004718.3;NP_828850.1;SARS-CoV_orf1a;1127 | 0 | 0 |
| NC_004718.3;NP_828850.1;SARS-CoV_orf1a;1128 | 0 | 0 |
| NC_004718.3;NP_828850.1;SARS-CoV_orf1a;1129 | 0 | 0 |
| NC_004718.3;NP_828850.1;SARS-CoV_orf1a;1130 | 0 | 0 |
| NC_004718.3;NP_828850.1;SARS-CoV_orf1a;1131 | 0 | 0 |
| NC_004718.3;NP_828850.1;SARS-CoV_orf1a;1132 | 0 | 0 |
| NC_004718.3;NP_828850.1;SARS-CoV_orf1a;1133 | 0 | 0 |
| NC_004718.3;NP_828850.1;SARS-CoV_orf1a;1134 | 0 | 0 |
| NC_004718.3;NP_828850.1;SARS-CoV_orf1a;1135 | 0 | 0 |
| NC_004718.3;NP_828850.1;SARS-CoV_orf1a;1136 | 0 | 0 |
| NC_004718.3;NP_828850.1;SARS-CoV_orf1a;1137 | 0 | 0 |
| NC_004718.3;NP_828850.1;SARS-CoV_orf1a;1138 | 0 | 0 |
| NC_004718.3;NP_828850.1;SARS-CoV_orf1a;1139 | 0 | 0 |
| NC_004718.3;NP_828850.1;SARS-CoV_orf1a;1140 | 0 | 0 |
| NC_004718.3;NP_828850.1;SARS-CoV_orf1a;1141 | 0 | 0 |
| NC_004718.3;NP_828850.1;SARS-CoV_orf1a;1142 | 0 | 0 |
| NC_004718.3;NP_828850.1;SARS-CoV_orf1a;1143 | 0 | 0 |
| NC_004718.3;NP_828850.1;SARS-CoV_orf1a;1144 | 0 | 0 |
| NC_004718.3;NP_828850.1;SARS-CoV_orf1a;1145 | 0 | 0 |
| NC_004718.3;NP_828850.1;SARS-CoV_orf1a;1146 | 0 | 0 |
| NC_004718.3;NP_828850.1;SARS-CoV_orf1a;1147 | 0 | 0 |
| NC_004718.3;NP_828850.1;SARS-CoV_orf1a;1148 | 0 | 0 |

| | | |
|---|---|---|
| NC_004718.3;NP_828850.1;SARS-CoV_orf1a;1149 | 0 | 0 |
| NC_004718.3;NP_828850.1;SARS-CoV_orf1a;1150 | 0 | 0.025 |
| NC_004718.3;NP_828850.1;SARS-CoV_orf1a;1151 | 0 | 0 |
| NC_004718.3;NP_828850.1;SARS-CoV_orf1a;1152 | 0 | 0 |
| NC_004718.3;NP_828850.1;SARS-CoV_orf1a;1153 | 0 | 0 |
| NC_004718.3;NP_828850.1;SARS-CoV_orf1a;1154 | 0 | 0 |
| NC_004718.3;NP_828850.1;SARS-CoV_orf1a;1155 | 0 | 0 |
| NC_004718.3;NP_828850.1;SARS-CoV_orf1a;1156 | 0 | 0 |
| NC_004718.3;NP_828850.1;SARS-CoV_orf1a;1157 | 0 | 0 |
| NC_004718.3;NP_828850.1;SARS-CoV_orf1a;1158 | 0.05 | 0.05 |
| NC_004718.3;NP_828850.1;SARS-CoV_orf1a;1159 | 0 | 0.025 |
| NC_004718.3;NP_828850.1;SARS-CoV_orf1a;1160 | 0 | 0.075 |
| NC_004718.3;NP_828850.1;SARS-CoV_orf1a;1161 | 0 | 0.075 |
| NC_004718.3;NP_828850.1;SARS-CoV_orf1a;1162 | 0 | 0.075 |
| NC_004718.3;NP_828850.1;SARS-CoV_orf1a;1163 | 0 | 0.075 |
| NC_004718.3;NP_828850.1;SARS-CoV_orf1a;1164 | 0.05 | 0.15 |
| NC_004718.3;NP_828850.1;SARS-CoV_orf1a;1165 | 0.35 | 0.475 |
| NC_004718.3;NP_828850.1;SARS-CoV_orf1a;1166 | 0.4 | 0.45 |
| NC_004718.3;NP_828850.1;SARS-CoV_orf1a;1167 | 0.45 | 0.4 |
| NC_004718.3;NP_828850.1;SARS-CoV_orf1a;1168 | 0.4 | 0.4 |
| NC_004718.3;NP_828850.1;SARS-CoV_orf1a;1169 | 0.4 | 0.25 |
| NC_004718.3;NP_828850.1;SARS-CoV_orf1a;1170 | 0.45 | 0.375 |
| NC_004718.3;NP_828850.1;SARS-CoV_orf1a;1171 | 0.4 | 0.35 |
| NC_004718.3;NP_828850.1;SARS-CoV_orf1a;1172 | 0.4 | 0.4 |
| NC_004718.3;NP_828850.1;SARS-CoV_orf1a;1173 | 0.4 | 0.325 |
| NC_004718.3;NP_828850.1;SARS-CoV_orf1a;1174 | 0 | 0.025 |
| NC_004718.3;NP_828850.1;SARS-CoV_orf1a;1175 | 0 | 0.025 |
| NC_004718.3;NP_828850.1;SARS-CoV_orf1a;1176 | 0 | 0.025 |
| NC_004718.3;NP_828850.1;SARS-CoV_orf1a;1177 | 0 | 0 |
| NC_004718.3;NP_828850.1;SARS-CoV_orf1a;1178 | 0 | 0 |
| NC_004718.3;NP_828850.1;SARS-CoV_orf1a;1179 | 0 | 0 |
| NC_004718.3;NP_828850.1;SARS-CoV_orf1a;1180 | 0 | 0 |
| NC_004718.3;NP_828850.1;SARS-CoV_orf1a;1181 | 0 | 0 |
| NC_004718.3;NP_828850.1;SARS-CoV_orf1a;1182 | 0 | 0 |
| NC_004718.3;NP_828850.1;SARS-CoV_orf1a;1183 | 0 | 0 |
| NC_004718.3;NP_828850.1;SARS-CoV_orf1a;1184 | 0 | 0 |
| NC_004718.3;NP_828850.1;SARS-CoV_orf1a;1185 | 0 | 0 |
| NC_004718.3;NP_828850.1;SARS-CoV_orf1a;1186 | 0 | 0 |
| NC_004718.3;NP_828850.1;SARS-CoV_orf1a;1187 | 0 | 0 |
| NC_004718.3;NP_828850.1;SARS-CoV_orf1a;1188 | 0 | 0 |
| NC_004718.3;NP_828850.1;SARS-CoV_orf1a;1189 | 0 | 0 |
| NC_004718.3;NP_828850.1;SARS-CoV_orf1a;1190 | 0.05 | 0 |
| NC_004718.3;NP_828850.1;SARS-CoV_orf1a;1191 | 0.05 | 0 |

| | | |
|---|---|---|
| NC_004718.3;NP_828850.1;SARS-CoV_orf1a;1192 | 0.1 | 0.05 |
| NC_004718.3;NP_828850.1;SARS-CoV_orf1a;1193 | 0 | 0.05 |
| NC_004718.3;NP_828850.1;SARS-CoV_orf1a;1194 | 0 | 0.025 |
| NC_004718.3;NP_828850.1;SARS-CoV_orf1a;1195 | 0 | 0.025 |
| NC_004718.3;NP_828850.1;SARS-CoV_orf1a;1196 | 0.05 | 0.025 |
| NC_004718.3;NP_828850.1;SARS-CoV_orf1a;1197 | 0.05 | 0.025 |
| NC_004718.3;NP_828850.1;SARS-CoV_orf1a;1198 | 0.05 | 0.025 |
| NC_004718.3;NP_828850.1;SARS-CoV_orf1a;1199 | 0.15 | 0.075 |
| NC_004718.3;NP_828850.1;SARS-CoV_orf1a;1200 | 0.15 | 0.175 |
| NC_004718.3;NP_828850.1;SARS-CoV_orf1a;1201 | 0.1 | 0.2 |
| NC_004718.3;NP_828850.1;SARS-CoV_orf1a;1202 | 0 | 0 |
| NC_004718.3;NP_828850.1;SARS-CoV_orf1a;1203 | 0 | 0.025 |
| NC_004718.3;NP_828850.1;SARS-CoV_orf1a;1204 | 0 | 0.025 |
| NC_004718.3;NP_828850.1;SARS-CoV_orf1a;1205 | 0 | 0 |
| NC_004718.3;NP_828850.1;SARS-CoV_orf1a;1206 | 0 | 0 |
| NC_004718.3;NP_828850.1;SARS-CoV_orf1a;1207 | 0 | 0 |
| NC_004718.3;NP_828850.1;SARS-CoV_orf1a;1208 | 0 | 0 |
| NC_004718.3;NP_828850.1;SARS-CoV_orf1a;1209 | 0 | 0 |
| NC_004718.3;NP_828850.1;SARS-CoV_orf1a;1210 | 0 | 0 |
| NC_004718.3;NP_828850.1;SARS-CoV_orf1a;1211 | 0 | 0 |
| NC_004718.3;NP_828850.1;SARS-CoV_orf1a;1212 | 0 | 0 |
| NC_004718.3;NP_828850.1;SARS-CoV_orf1a;1213 | 0.05 | 0.025 |
| NC_004718.3;NP_828850.1;SARS-CoV_orf1a;1214 | 0 | 0.025 |
| NC_004718.3;NP_828850.1;SARS-CoV_orf1a;1215 | 0 | 0.025 |
| NC_004718.3;NP_828850.1;SARS-CoV_orf1a;1216 | 0 | 0.025 |
| NC_004718.3;NP_828850.1;SARS-CoV_orf1a;1217 | 0 | 0.025 |
| NC_004718.3;NP_828850.1;SARS-CoV_orf1a;1218 | 0 | 0 |
| NC_004718.3;NP_828850.1;SARS-CoV_orf1a;1219 | 0 | 0 |
| NC_004718.3;NP_828850.1;SARS-CoV_orf1a;1220 | 0 | 0 |
| NC_004718.3;NP_828850.1;SARS-CoV_orf1a;1221 | 0 | 0.025 |
| NC_004718.3;NP_828850.1;SARS-CoV_orf1a;1222 | 0 | 0 |
| NC_004718.3;NP_828850.1;SARS-CoV_orf1a;1223 | 0 | 0 |
| NC_004718.3;NP_828850.1;SARS-CoV_orf1a;1224 | 0 | 0 |
| NC_004718.3;NP_828850.1;SARS-CoV_orf1a;1225 | 0 | 0 |
| NC_004718.3;NP_828850.1;SARS-CoV_orf1a;1226 | 0 | 0 |
| NC_004718.3;NP_828850.1;SARS-CoV_orf1a;1227 | 0 | 0 |
| NC_004718.3;NP_828850.1;SARS-CoV_orf1a;1228 | 0 | 0 |
| NC_004718.3;NP_828850.1;SARS-CoV_orf1a;1229 | 0 | 0 |
| NC_004718.3;NP_828850.1;SARS-CoV_orf1a;1230 | 0 | 0.05 |
| NC_004718.3;NP_828850.1;SARS-CoV_orf1a;1231 | 0.15 | 0.05 |
| NC_004718.3;NP_828850.1;SARS-CoV_orf1a;1232 | 0.1 | 0.075 |
| NC_004718.3;NP_828850.1;SARS-CoV_orf1a;1233 | 0.05 | 0.05 |
| NC_004718.3;NP_828850.1;SARS-CoV_orf1a;1234 | 0.05 | 0.025 |

| | | |
|---|---|---|
| NC_004718.3;NP_828850.1;SARS-CoV_orf1a;1235 | 0 | 0 |
| NC_004718.3;NP_828850.1;SARS-CoV_orf1a;1236 | 0 | 0.025 |
| NC_004718.3;NP_828850.1;SARS-CoV_orf1a;1237 | 0 | 0 |
| NC_004718.3;NP_828850.1;SARS-CoV_orf1a;1238 | 0 | 0 |
| NC_004718.3;NP_828850.1;SARS-CoV_orf1a;1239 | 0 | 0 |
| NC_004718.3;NP_828850.1;SARS-CoV_orf1a;1240 | 0.05 | 0 |
| NC_004718.3;NP_828850.1;SARS-CoV_orf1a;1241 | 0 | 0 |
| NC_004718.3;NP_828850.1;SARS-CoV_orf1a;1242 | 0 | 0 |
| NC_004718.3;NP_828850.1;SARS-CoV_orf1a;1243 | 0 | 0 |
| NC_004718.3;NP_828850.1;SARS-CoV_orf1a;1244 | 0 | 0 |
| NC_004718.3;NP_828850.1;SARS-CoV_orf1a;1245 | 0 | 0 |
| NC_004718.3;NP_828850.1;SARS-CoV_orf1a;1246 | 0 | 0.025 |
| NC_004718.3;NP_828850.1;SARS-CoV_orf1a;1247 | 0 | 0.025 |
| NC_004718.3;NP_828850.1;SARS-CoV_orf1a;1248 | 0 | 0 |
| NC_004718.3;NP_828850.1;SARS-CoV_orf1a;1249 | 0 | 0 |
| NC_004718.3;NP_828850.1;SARS-CoV_orf1a;1250 | 0.05 | 0 |
| NC_004718.3;NP_828850.1;SARS-CoV_orf1a;1251 | 0 | 0 |
| NC_004718.3;NP_828850.1;SARS-CoV_orf1a;1252 | 0 | 0 |
| NC_004718.3;NP_828850.1;SARS-CoV_orf1a;1253 | 0 | 0 |
| NC_004718.3;NP_828850.1;SARS-CoV_orf1a;1254 | 0 | 0 |
| NC_004718.3;NP_828850.1;SARS-CoV_orf1a;1255 | 0 | 0 |
| NC_004718.3;NP_828850.1;SARS-CoV_orf1a;1256 | 0 | 0 |
| NC_004718.3;NP_828850.1;SARS-CoV_orf1a;1257 | 0 | 0 |
| NC_004718.3;NP_828850.1;SARS-CoV_orf1a;1258 | 0 | 0 |
| NC_004718.3;NP_828850.1;SARS-CoV_orf1a;1259 | 0 | 0 |
| NC_004718.3;NP_828850.1;SARS-CoV_orf1a;1260 | 0 | 0 |
| NC_004718.3;NP_828850.1;SARS-CoV_orf1a;1261 | 0 | 0 |
| NC_004718.3;NP_828850.1;SARS-CoV_orf1a;1262 | 0 | 0 |
| NC_004718.3;NP_828850.1;SARS-CoV_orf1a;1263 | 0 | 0 |
| NC_004718.3;NP_828850.1;SARS-CoV_orf1a;1264 | 0 | 0.025 |
| NC_004718.3;NP_828850.1;SARS-CoV_orf1a;1265 | 0 | 0 |
| NC_004718.3;NP_828850.1;SARS-CoV_orf1a;1266 | 0 | 0 |
| NC_004718.3;NP_828850.1;SARS-CoV_orf1a;1267 | 0 | 0 |
| NC_004718.3;NP_828850.1;SARS-CoV_orf1a;1268 | 0 | 0 |
| NC_004718.3;NP_828850.1;SARS-CoV_orf1a;1269 | 0 | 0 |
| NC_004718.3;NP_828850.1;SARS-CoV_orf1a;1270 | 0 | 0 |
| NC_004718.3;NP_828850.1;SARS-CoV_orf1a;1271 | 0 | 0 |
| NC_004718.3;NP_828850.1;SARS-CoV_orf1a;1272 | 0.05 | 0 |
| NC_004718.3;NP_828850.1;SARS-CoV_orf1a;1273 | 0.05 | 0.025 |
| NC_004718.3;NP_828850.1;SARS-CoV_orf1a;1274 | 0.05 | 0.025 |
| NC_004718.3;NP_828850.1;SARS-CoV_orf1a;1275 | 0.05 | 0.05 |
| NC_004718.3;NP_828850.1;SARS-CoV_orf1a;1276 | 0.05 | 0.025 |
| NC_004718.3;NP_828850.1;SARS-CoV_orf1a;1277 | 0.05 | 0.025 |

| | | |
|---|---|---|
| NC_004718.3;NP_828850.1;SARS-CoV_orf1a;1278 | 0.05 | 0 |
| NC_004718.3;NP_828850.1;SARS-CoV_orf1a;1279 | 0 | 0 |
| NC_004718.3;NP_828850.1;SARS-CoV_orf1a;1280 | 0 | 0 |
| NC_004718.3;NP_828850.1;SARS-CoV_orf1a;1281 | 0 | 0 |
| NC_004718.3;NP_828850.1;SARS-CoV_orf1a;1282 | 0 | 0 |
| NC_004718.3;NP_828850.1;SARS-CoV_orf1a;1283 | 0 | 0 |
| NC_004718.3;NP_828850.1;SARS-CoV_orf1a;1284 | 0 | 0 |
| NC_004718.3;NP_828850.1;SARS-CoV_orf1a;1285 | 0 | 0 |
| NC_004718.3;NP_828850.1;SARS-CoV_orf1a;1286 | 0 | 0.025 |
| NC_004718.3;NP_828850.1;SARS-CoV_orf1a;1287 | 0.05 | 0.025 |
| NC_004718.3;NP_828850.1;SARS-CoV_orf1a;1288 | 0 | 0.025 |
| NC_004718.3;NP_828850.1;SARS-CoV_orf1a;1289 | 0 | 0.025 |
| NC_004718.3;NP_828850.1;SARS-CoV_orf1a;1290 | 0 | 0 |
| NC_004718.3;NP_828850.1;SARS-CoV_orf1a;1291 | 0 | 0 |
| NC_004718.3;NP_828850.1;SARS-CoV_orf1a;1292 | 0 | 0 |
| NC_004718.3;NP_828850.1;SARS-CoV_orf1a;1293 | 0 | 0 |
| NC_004718.3;NP_828850.1;SARS-CoV_orf1a;1294 | 0 | 0.025 |
| NC_004718.3;NP_828850.1;SARS-CoV_orf1a;1295 | 0.05 | 0.025 |
| NC_004718.3;NP_828850.1;SARS-CoV_orf1a;1296 | 0.1 | 0.025 |
| NC_004718.3;NP_828850.1;SARS-CoV_orf1a;1297 | 0.05 | 0.025 |
| NC_004718.3;NP_828850.1;SARS-CoV_orf1a;1298 | 0 | 0 |
| NC_004718.3;NP_828850.1;SARS-CoV_orf1a;1299 | 0 | 0 |
| NC_004718.3;NP_828850.1;SARS-CoV_orf1a;1300 | 0.05 | 0 |
| NC_004718.3;NP_828850.1;SARS-CoV_orf1a;1301 | 0 | 0 |
| NC_004718.3;NP_828850.1;SARS-CoV_orf1a;1302 | 0 | 0 |
| NC_004718.3;NP_828850.1;SARS-CoV_orf1a;1303 | 0 | 0 |
| NC_004718.3;NP_828850.1;SARS-CoV_orf1a;1304 | 0 | 0 |
| NC_004718.3;NP_828850.1;SARS-CoV_orf1a;1305 | 0 | 0 |
| NC_004718.3;NP_828850.1;SARS-CoV_orf1a;1306 | 0 | 0 |
| NC_004718.3;NP_828850.1;SARS-CoV_orf1a;1307 | 0 | 0.025 |
| NC_004718.3;NP_828850.1;SARS-CoV_orf1a;1308 | 0 | 0.025 |
| NC_004718.3;NP_828850.1;SARS-CoV_orf1a;1309 | 0 | 0 |
| NC_004718.3;NP_828850.1;SARS-CoV_orf1a;1310 | 0 | 0 |
| NC_004718.3;NP_828850.1;SARS-CoV_orf1a;1311 | 0 | 0 |
| NC_004718.3;NP_828850.1;SARS-CoV_orf1a;1312 | 0 | 0 |
| NC_004718.3;NP_828850.1;SARS-CoV_orf1a;1313 | 0 | 0.025 |
| NC_004718.3;NP_828850.1;SARS-CoV_orf1a;1314 | 0 | 0 |
| NC_004718.3;NP_828850.1;SARS-CoV_orf1a;1315 | 0 | 0 |
| NC_004718.3;NP_828850.1;SARS-CoV_orf1a;1316 | 0 | 0 |
| NC_004718.3;NP_828850.1;SARS-CoV_orf1a;1317 | 0 | 0 |
| NC_004718.3;NP_828850.1;SARS-CoV_orf1a;1318 | 0 | 0 |
| NC_004718.3;NP_828850.1;SARS-CoV_orf1a;1319 | 0 | 0.025 |
| NC_004718.3;NP_828850.1;SARS-CoV_orf1a;1320 | 0 | 0 |

| | | |
|---|---|---|
| NC_004718.3;NP_828850.1;SARS-CoV_orf1a;1321 | 0 | 0.025 |
| NC_004718.3;NP_828850.1;SARS-CoV_orf1a;1322 | 0 | 0.025 |
| NC_004718.3;NP_828850.1;SARS-CoV_orf1a;1323 | 0 | 0.05 |
| NC_004718.3;NP_828850.1;SARS-CoV_orf1a;1324 | 0 | 0 |
| NC_004718.3;NP_828850.1;SARS-CoV_orf1a;1325 | 0 | 0 |
| NC_004718.3;NP_828850.1;SARS-CoV_orf1a;1326 | 0 | 0 |
| NC_004718.3;NP_828850.1;SARS-CoV_orf1a;1327 | 0 | 0 |
| NC_004718.3;NP_828850.1;SARS-CoV_orf1a;1328 | 0 | 0 |
| NC_004718.3;NP_828850.1;SARS-CoV_orf1a;1329 | 0 | 0 |
| NC_004718.3;NP_828850.1;SARS-CoV_orf1a;1330 | 0 | 0 |
| NC_004718.3;NP_828850.1;SARS-CoV_orf1a;1331 | 0 | 0 |
| NC_004718.3;NP_828850.1;SARS-CoV_orf1a;1332 | 0 | 0 |
| NC_004718.3;NP_828850.1;SARS-CoV_orf1a;1333 | 0 | 0 |
| NC_004718.3;NP_828850.1;SARS-CoV_orf1a;1334 | 0 | 0 |
| NC_004718.3;NP_828850.1;SARS-CoV_orf1a;1335 | 0.05 | 0 |
| NC_004718.3;NP_828850.1;SARS-CoV_orf1a;1336 | 0 | 0 |
| NC_004718.3;NP_828850.1;SARS-CoV_orf1a;1337 | 0 | 0 |
| NC_004718.3;NP_828850.1;SARS-CoV_orf1a;1338 | 0 | 0 |
| NC_004718.3;NP_828850.1;SARS-CoV_orf1a;1339 | 0 | 0 |
| NC_004718.3;NP_828850.1;SARS-CoV_orf1a;1340 | 0 | 0 |
| NC_004718.3;NP_828850.1;SARS-CoV_orf1a;1341 | 0.05 | 0.025 |
| NC_004718.3;NP_828850.1;SARS-CoV_orf1a;1342 | 0 | 0 |
| NC_004718.3;NP_828850.1;SARS-CoV_orf1a;1343 | 0 | 0 |
| NC_004718.3;NP_828850.1;SARS-CoV_orf1a;1344 | 0 | 0 |
| NC_004718.3;NP_828850.1;SARS-CoV_orf1a;1345 | 0 | 0.025 |
| NC_004718.3;NP_828850.1;SARS-CoV_orf1a;1346 | 0 | 0 |
| NC_004718.3;NP_828850.1;SARS-CoV_orf1a;1347 | 0 | 0 |
| NC_004718.3;NP_828850.1;SARS-CoV_orf1a;1348 | 0 | 0.05 |
| NC_004718.3;NP_828850.1;SARS-CoV_orf1a;1349 | 0 | 0.025 |
| NC_004718.3;NP_828850.1;SARS-CoV_orf1a;1350 | 0 | 0 |
| NC_004718.3;NP_828850.1;SARS-CoV_orf1a;1351 | 0 | 0.025 |
| NC_004718.3;NP_828850.1;SARS-CoV_orf1a;1352 | 0 | 0 |
| NC_004718.3;NP_828850.1;SARS-CoV_orf1a;1353 | 0 | 0 |
| NC_004718.3;NP_828850.1;SARS-CoV_orf1a;1354 | 0 | 0 |
| NC_004718.3;NP_828850.1;SARS-CoV_orf1a;1355 | 0 | 0 |
| NC_004718.3;NP_828850.1;SARS-CoV_orf1a;1356 | 0 | 0.025 |
| NC_004718.3;NP_828850.1;SARS-CoV_orf1a;1357 | 0 | 0 |
| NC_004718.3;NP_828850.1;SARS-CoV_orf1a;1358 | 0.05 | 0.025 |
| NC_004718.3;NP_828850.1;SARS-CoV_orf1a;1359 | 0.05 | 0.025 |
| NC_004718.3;NP_828850.1;SARS-CoV_orf1a;1360 | 0 | 0 |
| NC_004718.3;NP_828850.1;SARS-CoV_orf1a;1361 | 0 | 0 |
| NC_004718.3;NP_828850.1;SARS-CoV_orf1a;1362 | 0 | 0 |
| NC_004718.3;NP_828850.1;SARS-CoV_orf1a;1363 | 0 | 0 |

| | | |
|---|---|---|
| NC_004718.3;NP_828850.1;SARS-CoV_orf1a;1364 | 0 | 0 |
| NC_004718.3;NP_828850.1;SARS-CoV_orf1a;1365 | 0 | 0 |
| NC_004718.3;NP_828850.1;SARS-CoV_orf1a;1366 | 0 | 0 |
| NC_004718.3;NP_828850.1;SARS-CoV_orf1a;1367 | 0 | 0 |
| NC_004718.3;NP_828850.1;SARS-CoV_orf1a;1368 | 0 | 0 |
| NC_004718.3;NP_828850.1;SARS-CoV_orf1a;1369 | 0 | 0 |
| NC_004718.3;NP_828850.1;SARS-CoV_orf1a;1370 | 0 | 0 |
| NC_004718.3;NP_828850.1;SARS-CoV_orf1a;1371 | 0 | 0 |
| NC_004718.3;NP_828850.1;SARS-CoV_orf1a;1372 | 0.05 | 0 |
| NC_004718.3;NP_828850.1;SARS-CoV_orf1a;1373 | 0.05 | 0 |
| NC_004718.3;NP_828850.1;SARS-CoV_orf1a;1374 | 0.1 | 0 |
| NC_004718.3;NP_828850.1;SARS-CoV_orf1a;1375 | 0.1 | 0 |
| NC_004718.3;NP_828850.1;SARS-CoV_orf1a;1376 | 0.15 | 0.025 |
| NC_004718.3;NP_828850.1;SARS-CoV_orf1a;1377 | 0.1 | 0 |
| NC_004718.3;NP_828850.1;SARS-CoV_orf1a;1378 | 0.1 | 0 |
| NC_004718.3;NP_828850.1;SARS-CoV_orf1a;1379 | 0.1 | 0 |
| NC_004718.3;NP_828850.1;SARS-CoV_orf1a;1380 | 0.1 | 0.025 |
| NC_004718.3;NP_828850.1;SARS-CoV_orf1a;1381 | 0.1 | 0 |
| NC_004718.3;NP_828850.1;SARS-CoV_orf1a;1382 | 0.05 | 0.025 |
| NC_004718.3;NP_828850.1;SARS-CoV_orf1a;1383 | 0 | 0.025 |
| NC_004718.3;NP_828850.1;SARS-CoV_orf1a;1384 | 0 | 0 |
| NC_004718.3;NP_828850.1;SARS-CoV_orf1a;1385 | 0 | 0 |
| NC_004718.3;NP_828850.1;SARS-CoV_orf1a;1386 | 0 | 0 |
| NC_004718.3;NP_828850.1;SARS-CoV_orf1a;1387 | 0 | 0 |
| NC_004718.3;NP_828850.1;SARS-CoV_orf1a;1388 | 0 | 0 |
| NC_004718.3;NP_828850.1;SARS-CoV_orf1a;1389 | 0 | 0 |
| NC_004718.3;NP_828850.1;SARS-CoV_orf1a;1390 | 0.05 | 0.075 |
| NC_004718.3;NP_828850.1;SARS-CoV_orf1a;1391 | 0 | 0 |
| NC_004718.3;NP_828850.1;SARS-CoV_orf1a;1392 | 0 | 0 |
| NC_004718.3;NP_828850.1;SARS-CoV_orf1a;1393 | 0 | 0 |
| NC_004718.3;NP_828850.1;SARS-CoV_orf1a;1394 | 0 | 0 |
| NC_004718.3;NP_828850.1;SARS-CoV_orf1a;1395 | 0 | 0 |
| NC_004718.3;NP_828850.1;SARS-CoV_orf1a;1396 | 0 | 0 |
| NC_004718.3;NP_828850.1;SARS-CoV_orf1a;1397 | 0 | 0 |
| NC_004718.3;NP_828850.1;SARS-CoV_orf1a;1398 | 0 | 0 |
| NC_004718.3;NP_828850.1;SARS-CoV_orf1a;1399 | 0 | 0 |
| NC_004718.3;NP_828850.1;SARS-CoV_orf1a;1400 | 0 | 0 |
| NC_004718.3;NP_828850.1;SARS-CoV_orf1a;1401 | 0 | 0 |
| NC_004718.3;NP_828850.1;SARS-CoV_orf1a;1402 | 0 | 0 |
| NC_004718.3;NP_828850.1;SARS-CoV_orf1a;1403 | 0 | 0.025 |
| NC_004718.3;NP_828850.1;SARS-CoV_orf1a;1404 | 0 | 0 |
| NC_004718.3;NP_828850.1;SARS-CoV_orf1a;1405 | 0 | 0.025 |
| NC_004718.3;NP_828850.1;SARS-CoV_orf1a;1406 | 0.05 | 0.025 |

| | | |
|---|---|---|
| NC_004718.3;NP_828850.1;SARS-CoV_orf1a;1407 | 0 | 0.025 |
| NC_004718.3;NP_828850.1;SARS-CoV_orf1a;1408 | 0 | 0.025 |
| NC_004718.3;NP_828850.1;SARS-CoV_orf1a;1409 | 0.05 | 0 |
| NC_004718.3;NP_828850.1;SARS-CoV_orf1a;1410 | 0 | 0 |
| NC_004718.3;NP_828850.1;SARS-CoV_orf1a;1411 | 0.1 | 0 |
| NC_004718.3;NP_828850.1;SARS-CoV_orf1a;1412 | 0.15 | 0 |
| NC_004718.3;NP_828850.1;SARS-CoV_orf1a;1413 | 0.05 | 0 |
| NC_004718.3;NP_828850.1;SARS-CoV_orf1a;1414 | 0 | 0.025 |
| NC_004718.3;NP_828850.1;SARS-CoV_orf1a;1415 | 0 | 0 |
| NC_004718.3;NP_828850.1;SARS-CoV_orf1a;1416 | 0.1 | 0.025 |
| NC_004718.3;NP_828850.1;SARS-CoV_orf1a;1417 | 0 | 0 |
| NC_004718.3;NP_828850.1;SARS-CoV_orf1a;1418 | 0 | 0 |
| NC_004718.3;NP_828850.1;SARS-CoV_orf1a;1419 | 0 | 0 |
| NC_004718.3;NP_828850.1;SARS-CoV_orf1a;1420 | 0 | 0 |
| NC_004718.3;NP_828850.1;SARS-CoV_orf1a;1421 | 0 | 0 |
| NC_004718.3;NP_828850.1;SARS-CoV_orf1a;1422 | 0 | 0.025 |
| NC_004718.3;NP_828850.1;SARS-CoV_orf1a;1423 | 0 | 0.025 |
| NC_004718.3;NP_828850.1;SARS-CoV_orf1a;1424 | 0.05 | 0.1 |
| NC_004718.3;NP_828850.1;SARS-CoV_orf1a;1425 | 0 | 0.05 |
| NC_004718.3;NP_828850.1;SARS-CoV_orf1a;1426 | 0 | 0.05 |
| NC_004718.3;NP_828850.1;SARS-CoV_orf1a;1427 | 0 | 0.05 |
| NC_004718.3;NP_828850.1;SARS-CoV_orf1a;1428 | 0 | 0.05 |
| NC_004718.3;NP_828850.1;SARS-CoV_orf1a;1429 | 0 | 0.05 |
| NC_004718.3;NP_828850.1;SARS-CoV_orf1a;1430 | 0 | 0.05 |
| NC_004718.3;NP_828850.1;SARS-CoV_orf1a;1431 | 0 | 0.025 |
| NC_004718.3;NP_828850.1;SARS-CoV_orf1a;1432 | 0 | 0.025 |
| NC_004718.3;NP_828850.1;SARS-CoV_orf1a;1433 | 0 | 0 |
| NC_004718.3;NP_828850.1;SARS-CoV_orf1a;1434 | 0.05 | 0 |
| NC_004718.3;NP_828850.1;SARS-CoV_orf1a;1435 | 0 | 0 |
| NC_004718.3;NP_828850.1;SARS-CoV_orf1a;1436 | 0 | 0 |
| NC_004718.3;NP_828850.1;SARS-CoV_orf1a;1437 | 0 | 0.025 |
| NC_004718.3;NP_828850.1;SARS-CoV_orf1a;1438 | 0 | 0 |
| NC_004718.3;NP_828850.1;SARS-CoV_orf1a;1439 | 0 | 0 |
| NC_004718.3;NP_828850.1;SARS-CoV_orf1a;1440 | 0 | 0 |
| NC_004718.3;NP_828850.1;SARS-CoV_orf1a;1441 | 0 | 0 |
| NC_004718.3;NP_828850.1;SARS-CoV_orf1a;1442 | 0 | 0 |
| NC_004718.3;NP_828850.1;SARS-CoV_orf1a;1443 | 0 | 0.025 |
| NC_004718.3;NP_828850.1;SARS-CoV_orf1a;1444 | 0 | 0 |
| NC_004718.3;NP_828850.1;SARS-CoV_orf1a;1445 | 0 | 0 |
| NC_004718.3;NP_828850.1;SARS-CoV_orf1a;1446 | 0 | 0 |
| NC_004718.3;NP_828850.1;SARS-CoV_orf1a;1447 | 0 | 0 |
| NC_004718.3;NP_828850.1;SARS-CoV_orf1a;1448 | 0.05 | 0.05 |
| NC_004718.3;NP_828850.1;SARS-CoV_orf1a;1449 | 0 | 0.025 |

| | | |
|---|---|---|
| NC_004718.3;NP_828850.1;SARS-CoV_orf1a;1450 | 0 | 0.075 |
| NC_004718.3;NP_828850.1;SARS-CoV_orf1a;1451 | 0 | 0.075 |
| NC_004718.3;NP_828850.1;SARS-CoV_orf1a;1452 | 0 | 0.025 |
| NC_004718.3;NP_828850.1;SARS-CoV_orf1a;1453 | 0 | 0 |
| NC_004718.3;NP_828850.1;SARS-CoV_orf1a;1454 | 0 | 0 |
| NC_004718.3;NP_828850.1;SARS-CoV_orf1a;1455 | 0 | 0 |
| NC_004718.3;NP_828850.1;SARS-CoV_orf1a;1456 | 0.05 | 0.075 |
| NC_004718.3;NP_828850.1;SARS-CoV_orf1a;1457 | 0.1 | 0.05 |
| NC_004718.3;NP_828850.1;SARS-CoV_orf1a;1458 | 0.05 | 0.05 |
| NC_004718.3;NP_828850.1;SARS-CoV_orf1a;1459 | 0.1 | 0.15 |
| NC_004718.3;NP_828850.1;SARS-CoV_orf1a;1460 | 0.05 | 0.1 |
| NC_004718.3;NP_828850.1;SARS-CoV_orf1a;1461 | 0.05 | 0.025 |
| NC_004718.3;NP_828850.1;SARS-CoV_orf1a;1462 | 0.05 | 0.1 |
| NC_004718.3;NP_828850.1;SARS-CoV_orf1a;1463 | 0.05 | 0.075 |
| NC_004718.3;NP_828850.1;SARS-CoV_orf1a;1464 | 0.05 | 0.025 |
| NC_004718.3;NP_828850.1;SARS-CoV_orf1a;1465 | 0.05 | 0.025 |
| NC_004718.3;NP_828850.1;SARS-CoV_orf1a;1466 | 0.05 | 0.025 |
| NC_004718.3;NP_828850.1;SARS-CoV_orf1a;1467 | 0 | 0 |
| NC_004718.3;NP_828850.1;SARS-CoV_orf1a;1468 | 0.05 | 0.025 |
| NC_004718.3;NP_828850.1;SARS-CoV_orf1a;1469 | 0.05 | 0.05 |
| NC_004718.3;NP_828850.1;SARS-CoV_orf1a;1470 | 0.05 | 0 |
| NC_004718.3;NP_828850.1;SARS-CoV_orf1a;1471 | 0.05 | 0 |
| NC_004718.3;NP_828850.1;SARS-CoV_orf1a;1472 | 0 | 0 |
| NC_004718.3;NP_828850.1;SARS-CoV_orf1a;1473 | 0 | 0 |
| NC_004718.3;NP_828850.1;SARS-CoV_orf1a;1474 | 0 | 0 |
| NC_004718.3;NP_828850.1;SARS-CoV_orf1a;1475 | 0 | 0 |
| NC_004718.3;NP_828850.1;SARS-CoV_orf1a;1476 | 0 | 0 |
| NC_004718.3;NP_828850.1;SARS-CoV_orf1a;1477 | 0 | 0 |
| NC_004718.3;NP_828850.1;SARS-CoV_orf1a;1478 | 0 | 0 |
| NC_004718.3;NP_828850.1;SARS-CoV_orf1a;1479 | 0 | 0 |
| NC_004718.3;NP_828850.1;SARS-CoV_orf1a;1480 | 0 | 0 |
| NC_004718.3;NP_828850.1;SARS-CoV_orf1a;1481 | 0 | 0 |
| NC_004718.3;NP_828850.1;SARS-CoV_orf1a;1482 | 0 | 0 |
| NC_004718.3;NP_828850.1;SARS-CoV_orf1a;1483 | 0 | 0 |
| NC_004718.3;NP_828850.1;SARS-CoV_orf1a;1484 | 0 | 0 |
| NC_004718.3;NP_828850.1;SARS-CoV_orf1a;1485 | 0.05 | 0.025 |
| NC_004718.3;NP_828850.1;SARS-CoV_orf1a;1486 | 0 | 0 |
| NC_004718.3;NP_828850.1;SARS-CoV_orf1a;1487 | 0 | 0.025 |
| NC_004718.3;NP_828850.1;SARS-CoV_orf1a;1488 | 0 | 0 |
| NC_004718.3;NP_828850.1;SARS-CoV_orf1a;1489 | 0 | 0 |
| NC_004718.3;NP_828850.1;SARS-CoV_orf1a;1490 | 0 | 0 |
| NC_004718.3;NP_828850.1;SARS-CoV_orf1a;1491 | 0 | 0 |
| NC_004718.3;NP_828850.1;SARS-CoV_orf1a;1492 | 0 | 0 |

| | | |
|---|---|---|
| NC_004718.3;NP_828850.1;SARS-CoV_orf1a;1493 | 0 | 0 |
| NC_004718.3;NP_828850.1;SARS-CoV_orf1a;1494 | 0 | 0.075 |
| NC_004718.3;NP_828850.1;SARS-CoV_orf1a;1495 | 0 | 0.075 |
| NC_004718.3;NP_828850.1;SARS-CoV_orf1a;1496 | 0 | 0.075 |
| NC_004718.3;NP_828850.1;SARS-CoV_orf1a;1497 | 0 | 0.05 |
| NC_004718.3;NP_828850.1;SARS-CoV_orf1a;1498 | 0 | 0.05 |
| NC_004718.3;NP_828850.1;SARS-CoV_orf1a;1499 | 0 | 0.05 |
| NC_004718.3;NP_828850.1;SARS-CoV_orf1a;1500 | 0 | 0.05 |
| NC_004718.3;NP_828850.1;SARS-CoV_orf1a;1501 | 0 | 0.075 |
| NC_004718.3;NP_828850.1;SARS-CoV_orf1a;1502 | 0 | 0.05 |
| NC_004718.3;NP_828850.1;SARS-CoV_orf1a;1503 | 0.05 | 0.05 |
| NC_004718.3;NP_828850.1;SARS-CoV_orf1a;1504 | 0.05 | 0.025 |
| NC_004718.3;NP_828850.1;SARS-CoV_orf1a;1505 | 0.05 | 0 |
| NC_004718.3;NP_828850.1;SARS-CoV_orf1a;1506 | 0.05 | 0 |
| NC_004718.3;NP_828850.1;SARS-CoV_orf1a;1507 | 0 | 0 |
| NC_004718.3;NP_828850.1;SARS-CoV_orf1a;1508 | 0.05 | 0 |
| NC_004718.3;NP_828850.1;SARS-CoV_orf1a;1509 | 0.05 | 0.025 |
| NC_004718.3;NP_828850.1;SARS-CoV_orf1a;1510 | 0 | 0.025 |
| NC_004718.3;NP_828850.1;SARS-CoV_orf1a;1511 | 0 | 0 |
| NC_004718.3;NP_828850.1;SARS-CoV_orf1a;1512 | 0 | 0 |
| NC_004718.3;NP_828850.1;SARS-CoV_orf1a;1513 | 0 | 0 |
| NC_004718.3;NP_828850.1;SARS-CoV_orf1a;1514 | 0 | 0 |
| NC_004718.3;NP_828850.1;SARS-CoV_orf1a;1515 | 0 | 0 |
| NC_004718.3;NP_828850.1;SARS-CoV_orf1a;1516 | 0.05 | 0 |
| NC_004718.3;NP_828850.1;SARS-CoV_orf1a;1517 | 0.45 | 0.35 |
| NC_004718.3;NP_828850.1;SARS-CoV_orf1a;1518 | 0.15 | 0.125 |
| NC_004718.3;NP_828850.1;SARS-CoV_orf1a;1519 | 0 | 0.025 |
| NC_004718.3;NP_828850.1;SARS-CoV_orf1a;1520 | 0 | 0.025 |
| NC_004718.3;NP_828850.1;SARS-CoV_orf1a;1521 | 0 | 0.025 |
| NC_004718.3;NP_828850.1;SARS-CoV_orf1a;1522 | 0 | 0.025 |
| NC_004718.3;NP_828850.1;SARS-CoV_orf1a;1523 | 0 | 0.025 |
| NC_004718.3;NP_828850.1;SARS-CoV_orf1a;1524 | 0 | 0 |
| NC_004718.3;NP_828850.1;SARS-CoV_orf1a;1525 | 0 | 0 |
| NC_004718.3;NP_828850.1;SARS-CoV_orf1a;1526 | 0 | 0 |
| NC_004718.3;NP_828850.1;SARS-CoV_orf1a;1527 | 0 | 0 |
| NC_004718.3;NP_828850.1;SARS-CoV_orf1a;1528 | 0 | 0 |
| NC_004718.3;NP_828850.1;SARS-CoV_orf1a;1529 | 0 | 0 |
| NC_004718.3;NP_828850.1;SARS-CoV_orf1a;1530 | 0 | 0 |
| NC_004718.3;NP_828850.1;SARS-CoV_orf1a;1531 | 0 | 0 |
| NC_004718.3;NP_828850.1;SARS-CoV_orf1a;1532 | 0 | 0 |
| NC_004718.3;NP_828850.1;SARS-CoV_orf1a;1533 | 0 | 0 |
| NC_004718.3;NP_828850.1;SARS-CoV_orf1a;1534 | 0 | 0 |
| NC_004718.3;NP_828850.1;SARS-CoV_orf1a;1535 | 0 | 0 |

| | | |
|---|---|---|
| NC_004718.3;NP_828850.1;SARS-CoV_orf1a;1536 | 0 | 0 |
| NC_004718.3;NP_828850.1;SARS-CoV_orf1a;1537 | 0 | 0 |
| NC_004718.3;NP_828850.1;SARS-CoV_orf1a;1538 | 0 | 0 |
| NC_004718.3;NP_828850.1;SARS-CoV_orf1a;1539 | 0 | 0 |
| NC_004718.3;NP_828850.1;SARS-CoV_orf1a;1540 | 0 | 0.025 |
| NC_004718.3;NP_828850.1;SARS-CoV_orf1a;1541 | 0 | 0.025 |
| NC_004718.3;NP_828850.1;SARS-CoV_orf1a;1542 | 0.05 | 0 |
| NC_004718.3;NP_828850.1;SARS-CoV_orf1a;1543 | 0 | 0 |
| NC_004718.3;NP_828850.1;SARS-CoV_orf1a;1544 | 0 | 0.025 |
| NC_004718.3;NP_828850.1;SARS-CoV_orf1a;1545 | 0.05 | 0.025 |
| NC_004718.3;NP_828850.1;SARS-CoV_orf1a;1546 | 0.05 | 0.025 |
| NC_004718.3;NP_828850.1;SARS-CoV_orf1a;1547 | 0 | 0.075 |
| NC_004718.3;NP_828850.1;SARS-CoV_orf1a;1548 | 0 | 0.075 |
| NC_004718.3;NP_828850.1;SARS-CoV_orf1a;1549 | 0 | 0.075 |
| NC_004718.3;NP_828850.1;SARS-CoV_orf1a;1550 | 0 | 0.1 |
| NC_004718.3;NP_828850.1;SARS-CoV_orf1a;1551 | 0 | 0.1 |
| NC_004718.3;NP_828850.1;SARS-CoV_orf1a;1552 | 0 | 0.05 |
| NC_004718.3;NP_828850.1;SARS-CoV_orf1a;1553 | 0 | 0.025 |
| NC_004718.3;NP_828850.1;SARS-CoV_orf1a;1554 | 0 | 0.025 |
| NC_004718.3;NP_828850.1;SARS-CoV_orf1a;1555 | 0 | 0.025 |
| NC_004718.3;NP_828850.1;SARS-CoV_orf1a;1556 | 0 | 0.025 |
| NC_004718.3;NP_828850.1;SARS-CoV_orf1a;1557 | 0 | 0 |
| NC_004718.3;NP_828850.1;SARS-CoV_orf1a;1558 | 0 | 0 |
| NC_004718.3;NP_828850.1;SARS-CoV_orf1a;1559 | 0 | 0 |
| NC_004718.3;NP_828850.1;SARS-CoV_orf1a;1560 | 0.2 | 0.125 |
| NC_004718.3;NP_828850.1;SARS-CoV_orf1a;1561 | 0 | 0 |
| NC_004718.3;NP_828850.1;SARS-CoV_orf1a;1562 | 0 | 0 |
| NC_004718.3;NP_828850.1;SARS-CoV_orf1a;1563 | 0 | 0 |
| NC_004718.3;NP_828850.1;SARS-CoV_orf1a;1564 | 0 | 0 |
| NC_004718.3;NP_828850.1;SARS-CoV_orf1a;1565 | 0 | 0 |
| NC_004718.3;NP_828850.1;SARS-CoV_orf1a;1566 | 0 | 0 |
| NC_004718.3;NP_828850.1;SARS-CoV_orf1a;1567 | 0 | 0 |
| NC_004718.3;NP_828850.1;SARS-CoV_orf1a;1568 | 0 | 0.025 |
| NC_004718.3;NP_828850.1;SARS-CoV_orf1a;1569 | 0 | 0.025 |
| NC_004718.3;NP_828850.1;SARS-CoV_orf1a;1570 | 0 | 0.025 |
| NC_004718.3;NP_828850.1;SARS-CoV_orf1a;1571 | 0 | 0.025 |
| NC_004718.3;NP_828850.1;SARS-CoV_orf1a;1572 | 0.15 | 0.125 |
| NC_004718.3;NP_828850.1;SARS-CoV_orf1a;1573 | 0 | 0.025 |
| NC_004718.3;NP_828850.1;SARS-CoV_orf1a;1574 | 0.05 | 0 |
| NC_004718.3;NP_828850.1;SARS-CoV_orf1a;1575 | 0 | 0 |
| NC_004718.3;NP_828850.1;SARS-CoV_orf1a;1576 | 0 | 0 |
| NC_004718.3;NP_828850.1;SARS-CoV_orf1a;1577 | 0 | 0 |
| NC_004718.3;NP_828850.1;SARS-CoV_orf1a;1578 | 0 | 0.025 |

| | | |
|---|---|---|
| NC_004718.3;NP_828850.1;SARS-CoV_orf1a;1579 | 0.05 | 0 |
| NC_004718.3;NP_828850.1;SARS-CoV_orf1a;1580 | 0 | 0 |
| NC_004718.3;NP_828850.1;SARS-CoV_orf1a;1581 | 0 | 0 |
| NC_004718.3;NP_828850.1;SARS-CoV_orf1a;1582 | 0 | 0 |
| NC_004718.3;NP_828850.1;SARS-CoV_orf1a;1583 | 0 | 0 |
| NC_004718.3;NP_828850.1;SARS-CoV_orf1a;1584 | 0 | 0 |
| NC_004718.3;NP_828850.1;SARS-CoV_orf1a;1585 | 0 | 0 |
| NC_004718.3;NP_828850.1;SARS-CoV_orf1a;1586 | 0 | 0 |
| NC_004718.3;NP_828850.1;SARS-CoV_orf1a;1587 | 0 | 0 |
| NC_004718.3;NP_828850.1;SARS-CoV_orf1a;1588 | 0 | 0 |
| NC_004718.3;NP_828850.1;SARS-CoV_orf1a;1589 | 0 | 0 |
| NC_004718.3;NP_828850.1;SARS-CoV_orf1a;1590 | 0 | 0 |
| NC_004718.3;NP_828850.1;SARS-CoV_orf1a;1591 | 0 | 0 |
| NC_004718.3;NP_828850.1;SARS-CoV_orf1a;1592 | 0 | 0 |
| NC_004718.3;NP_828850.1;SARS-CoV_orf1a;1593 | 0 | 0 |
| NC_004718.3;NP_828850.1;SARS-CoV_orf1a;1594 | 0 | 0 |
| NC_004718.3;NP_828850.1;SARS-CoV_orf1a;1595 | 0 | 0 |
| NC_004718.3;NP_828850.1;SARS-CoV_orf1a;1596 | 0 | 0 |
| NC_004718.3;NP_828850.1;SARS-CoV_orf1a;1597 | 0 | 0 |
| NC_004718.3;NP_828850.1;SARS-CoV_orf1a;1598 | 0 | 0 |
| NC_004718.3;NP_828850.1;SARS-CoV_orf1a;1599 | 0 | 0 |
| NC_004718.3;NP_828850.1;SARS-CoV_orf1a;1600 | 0 | 0 |
| NC_004718.3;NP_828850.1;SARS-CoV_orf1a;1601 | 0 | 0 |
| NC_004718.3;NP_828850.1;SARS-CoV_orf1a;1602 | 0 | 0 |
| NC_004718.3;NP_828850.1;SARS-CoV_orf1a;1603 | 0 | 0 |
| NC_004718.3;NP_828850.1;SARS-CoV_orf1a;1604 | 0 | 0 |
| NC_004718.3;NP_828850.1;SARS-CoV_orf1a;1605 | 0 | 0.025 |
| NC_004718.3;NP_828850.1;SARS-CoV_orf1a;1606 | 0 | 0.025 |
| NC_004718.3;NP_828850.1;SARS-CoV_orf1a;1607 | 0 | 0.025 |
| NC_004718.3;NP_828850.1;SARS-CoV_orf1a;1608 | 0.05 | 0.025 |
| NC_004718.3;NP_828850.1;SARS-CoV_orf1a;1609 | 0.05 | 0.025 |
| NC_004718.3;NP_828850.1;SARS-CoV_orf1a;1610 | 0.05 | 0.025 |
| NC_004718.3;NP_828850.1;SARS-CoV_orf1a;1611 | 0.05 | 0.025 |
| NC_004718.3;NP_828850.1;SARS-CoV_orf1a;1612 | 0.05 | 0 |
| NC_004718.3;NP_828850.1;SARS-CoV_orf1a;1613 | 0.05 | 0.025 |
| NC_004718.3;NP_828850.1;SARS-CoV_orf1a;1614 | 0.05 | 0.05 |
| NC_004718.3;NP_828850.1;SARS-CoV_orf1a;1615 | 0.05 | 0 |
| NC_004718.3;NP_828850.1;SARS-CoV_orf1a;1616 | 0.05 | 0 |
| NC_004718.3;NP_828850.1;SARS-CoV_orf1a;1617 | 0 | 0 |
| NC_004718.3;NP_828850.1;SARS-CoV_orf1a;1618 | 0 | 0 |
| NC_004718.3;NP_828850.1;SARS-CoV_orf1a;1619 | 0 | 0 |
| NC_004718.3;NP_828850.1;SARS-CoV_orf1a;1620 | 0 | 0 |
| NC_004718.3;NP_828850.1;SARS-CoV_orf1a;1621 | 0 | 0 |

| | | |
|---|---|---|
| NC_004718.3;NP_828850.1;SARS-CoV_orf1a;1622 | 0 | 0 |
| NC_004718.3;NP_828850.1;SARS-CoV_orf1a;1623 | 0 | 0 |
| NC_004718.3;NP_828850.1;SARS-CoV_orf1a;1624 | 0 | 0 |
| NC_004718.3;NP_828850.1;SARS-CoV_orf1a;1625 | 0.05 | 0 |
| NC_004718.3;NP_828850.1;SARS-CoV_orf1a;1626 | 0 | 0 |
| NC_004718.3;NP_828850.1;SARS-CoV_orf1a;1627 | 0 | 0 |
| NC_004718.3;NP_828850.1;SARS-CoV_orf1a;1628 | 0 | 0 |
| NC_004718.3;NP_828850.1;SARS-CoV_orf1a;1629 | 0 | 0 |
| NC_004718.3;NP_828850.1;SARS-CoV_orf1a;1630 | 0 | 0 |
| NC_004718.3;NP_828850.1;SARS-CoV_orf1a;1631 | 0 | 0 |
| NC_004718.3;NP_828850.1;SARS-CoV_orf1a;1632 | 0 | 0 |
| NC_004718.3;NP_828850.1;SARS-CoV_orf1a;1633 | 0 | 0 |
| NC_004718.3;NP_828850.1;SARS-CoV_orf1a;1634 | 0.05 | 0 |
| NC_004718.3;NP_828850.1;SARS-CoV_orf1a;1635 | 0.05 | 0 |
| NC_004718.3;NP_828850.1;SARS-CoV_orf1a;1636 | 0.05 | 0 |
| NC_004718.3;NP_828850.1;SARS-CoV_orf1a;1637 | 0.05 | 0 |
| NC_004718.3;NP_828850.1;SARS-CoV_orf1a;1638 | 0.05 | 0 |
| NC_004718.3;NP_828850.1;SARS-CoV_orf1a;1639 | 0.05 | 0 |
| NC_004718.3;NP_828850.1;SARS-CoV_orf1a;1640 | 0.05 | 0 |
| NC_004718.3;NP_828850.1;SARS-CoV_orf1a;1641 | 0.05 | 0 |
| NC_004718.3;NP_828850.1;SARS-CoV_orf1a;1642 | 0 | 0 |
| NC_004718.3;NP_828850.1;SARS-CoV_orf1a;1643 | 0 | 0 |
| NC_004718.3;NP_828850.1;SARS-CoV_orf1a;1644 | 0 | 0 |
| NC_004718.3;NP_828850.1;SARS-CoV_orf1a;1645 | 0 | 0 |
| NC_004718.3;NP_828850.1;SARS-CoV_orf1a;1646 | 0 | 0 |
| NC_004718.3;NP_828850.1;SARS-CoV_orf1a;1647 | 0 | 0 |
| NC_004718.3;NP_828850.1;SARS-CoV_orf1a;1648 | 0 | 0 |
| NC_004718.3;NP_828850.1;SARS-CoV_orf1a;1649 | 0 | 0 |
| NC_004718.3;NP_828850.1;SARS-CoV_orf1a;1650 | 0 | 0 |
| NC_004718.3;NP_828850.1;SARS-CoV_orf1a;1651 | 0 | 0 |
| NC_004718.3;NP_828850.1;SARS-CoV_orf1a;1652 | 0 | 0 |
| NC_004718.3;NP_828850.1;SARS-CoV_orf1a;1653 | 0 | 0 |
| NC_004718.3;NP_828850.1;SARS-CoV_orf1a;1654 | 0 | 0 |
| NC_004718.3;NP_828850.1;SARS-CoV_orf1a;1655 | 0 | 0 |
| NC_004718.3;NP_828850.1;SARS-CoV_orf1a;1656 | 0 | 0.025 |
| NC_004718.3;NP_828850.1;SARS-CoV_orf1a;1657 | 0 | 0 |
| NC_004718.3;NP_828850.1;SARS-CoV_orf1a;1658 | 0 | 0.025 |
| NC_004718.3;NP_828850.1;SARS-CoV_orf1a;1659 | 0 | 0 |
| NC_004718.3;NP_828850.1;SARS-CoV_orf1a;1660 | 0 | 0 |
| NC_004718.3;NP_828850.1;SARS-CoV_orf1a;1661 | 0.05 | 0 |
| NC_004718.3;NP_828850.1;SARS-CoV_orf1a;1662 | 0.1 | 0.05 |
| NC_004718.3;NP_828850.1;SARS-CoV_orf1a;1663 | 0 | 0.075 |
| NC_004718.3;NP_828850.1;SARS-CoV_orf1a;1664 | 0.1 | 0.05 |

| | | |
|---|---|---|
| NC_004718.3;NP_828850.1;SARS-CoV_orf1a;1665 | 0 | 0 |
| NC_004718.3;NP_828850.1;SARS-CoV_orf1a;1666 | 0 | 0 |
| NC_004718.3;NP_828850.1;SARS-CoV_orf1a;1667 | 0 | 0.025 |
| NC_004718.3;NP_828850.1;SARS-CoV_orf1a;1668 | 0.05 | 0 |
| NC_004718.3;NP_828850.1;SARS-CoV_orf1a;1669 | 0.05 | 0.025 |
| NC_004718.3;NP_828850.1;SARS-CoV_orf1a;1670 | 0.1 | 0.075 |
| NC_004718.3;NP_828850.1;SARS-CoV_orf1a;1671 | 0.05 | 0.05 |
| NC_004718.3;NP_828850.1;SARS-CoV_orf1a;1672 | 0.05 | 0.025 |
| NC_004718.3;NP_828850.1;SARS-CoV_orf1a;1673 | 0 | 0.025 |
| NC_004718.3;NP_828850.1;SARS-CoV_orf1a;1674 | 0 | 0.025 |
| NC_004718.3;NP_828850.1;SARS-CoV_orf1a;1675 | 0 | 0 |
| NC_004718.3;NP_828850.1;SARS-CoV_orf1a;1676 | 0 | 0 |
| NC_004718.3;NP_828850.1;SARS-CoV_orf1a;1677 | 0 | 0 |
| NC_004718.3;NP_828850.1;SARS-CoV_orf1a;1678 | 0 | 0 |
| NC_004718.3;NP_828850.1;SARS-CoV_orf1a;1679 | 0 | 0 |
| NC_004718.3;NP_828850.1;SARS-CoV_orf1a;1680 | 0 | 0 |
| NC_004718.3;NP_828850.1;SARS-CoV_orf1a;1681 | 0 | 0 |
| NC_004718.3;NP_828850.1;SARS-CoV_orf1a;1682 | 0 | 0 |
| NC_004718.3;NP_828850.1;SARS-CoV_orf1a;1683 | 0 | 0 |
| NC_004718.3;NP_828850.1;SARS-CoV_orf1a;1684 | 0 | 0 |
| NC_004718.3;NP_828850.1;SARS-CoV_orf1a;1685 | 0 | 0 |
| NC_004718.3;NP_828850.1;SARS-CoV_orf1a;1686 | 0 | 0 |
| NC_004718.3;NP_828850.1;SARS-CoV_orf1a;1687 | 0 | 0 |
| NC_004718.3;NP_828850.1;SARS-CoV_orf1a;1688 | 0 | 0 |
| NC_004718.3;NP_828850.1;SARS-CoV_orf1a;1689 | 0 | 0 |
| NC_004718.3;NP_828850.1;SARS-CoV_orf1a;1690 | 0.1 | 0 |
| NC_004718.3;NP_828850.1;SARS-CoV_orf1a;1691 | 0.15 | 0.025 |
| NC_004718.3;NP_828850.1;SARS-CoV_orf1a;1692 | 0.05 | 0 |
| NC_004718.3;NP_828850.1;SARS-CoV_orf1a;1693 | 0 | 0.025 |
| NC_004718.3;NP_828850.1;SARS-CoV_orf1a;1694 | 0.05 | 0.025 |
| NC_004718.3;NP_828850.1;SARS-CoV_orf1a;1695 | 0 | 0 |
| NC_004718.3;NP_828850.1;SARS-CoV_orf1a;1696 | 0 | 0 |
| NC_004718.3;NP_828850.1;SARS-CoV_orf1a;1697 | 0 | 0.025 |
| NC_004718.3;NP_828850.1;SARS-CoV_orf1a;1698 | 0 | 0 |
| NC_004718.3;NP_828850.1;SARS-CoV_orf1a;1699 | 0 | 0 |
| NC_004718.3;NP_828850.1;SARS-CoV_orf1a;1700 | 0.05 | 0.025 |
| NC_004718.3;NP_828850.1;SARS-CoV_orf1a;1701 | 0 | 0 |
| NC_004718.3;NP_828850.1;SARS-CoV_orf1a;1702 | 0 | 0 |
| NC_004718.3;NP_828850.1;SARS-CoV_orf1a;1703 | 0.05 | 0 |
| NC_004718.3;NP_828850.1;SARS-CoV_orf1a;1704 | 0 | 0 |
| NC_004718.3;NP_828850.1;SARS-CoV_orf1a;1705 | 0 | 0 |
| NC_004718.3;NP_828850.1;SARS-CoV_orf1a;1706 | 0 | 0 |
| NC_004718.3;NP_828850.1;SARS-CoV_orf1a;1707 | 0 | 0 |

| | | |
|---|---|---|
| NC_004718.3;NP_828850.1;SARS-CoV_orf1a;1708 | 0 | 0 |
| NC_004718.3;NP_828850.1;SARS-CoV_orf1a;1709 | 0 | 0 |
| NC_004718.3;NP_828850.1;SARS-CoV_orf1a;1710 | 0 | 0 |
| NC_004718.3;NP_828850.1;SARS-CoV_orf1a;1711 | 0 | 0 |
| NC_004718.3;NP_828850.1;SARS-CoV_orf1a;1712 | 0 | 0 |
| NC_004718.3;NP_828850.1;SARS-CoV_orf1a;1713 | 0 | 0 |
| NC_004718.3;NP_828850.1;SARS-CoV_orf1a;1714 | 0 | 0 |
| NC_004718.3;NP_828850.1;SARS-CoV_orf1a;1715 | 0 | 0 |
| NC_004718.3;NP_828850.1;SARS-CoV_orf1a;1716 | 0 | 0 |
| NC_004718.3;NP_828850.1;SARS-CoV_orf1a;1717 | 0 | 0 |
| NC_004718.3;NP_828850.1;SARS-CoV_orf1a;1718 | 0 | 0 |
| NC_004718.3;NP_828850.1;SARS-CoV_orf1a;1719 | 0.1 | 0 |
| NC_004718.3;NP_828850.1;SARS-CoV_orf1a;1720 | 0 | 0 |
| NC_004718.3;NP_828850.1;SARS-CoV_orf1a;1721 | 0 | 0 |
| NC_004718.3;NP_828850.1;SARS-CoV_orf1a;1722 | 0.05 | 0 |
| NC_004718.3;NP_828850.1;SARS-CoV_orf1a;1723 | 0.05 | 0 |
| NC_004718.3;NP_828850.1;SARS-CoV_orf1a;1724 | 0.05 | 0 |
| NC_004718.3;NP_828850.1;SARS-CoV_orf1a;1725 | 0.05 | 0 |
| NC_004718.3;NP_828850.1;SARS-CoV_orf1a;1726 | 0.1 | 0.05 |
| NC_004718.3;NP_828850.1;SARS-CoV_orf1a;1727 | 0.05 | 0.05 |
| NC_004718.3;NP_828850.1;SARS-CoV_orf1a;1728 | 0.1 | 0.05 |
| NC_004718.3;NP_828850.1;SARS-CoV_orf1a;1729 | 0.1 | 0.05 |
| NC_004718.3;NP_828850.1;SARS-CoV_orf1a;1730 | 0 | 0 |
| NC_004718.3;NP_828850.1;SARS-CoV_orf1a;1731 | 0 | 0 |
| NC_004718.3;NP_828850.1;SARS-CoV_orf1a;1732 | 0 | 0.025 |
| NC_004718.3;NP_828850.1;SARS-CoV_orf1a;1733 | 0.05 | 0.05 |
| NC_004718.3;NP_828850.1;SARS-CoV_orf1a;1734 | 0.05 | 0.05 |
| NC_004718.3;NP_828850.1;SARS-CoV_orf1a;1735 | 0 | 0.05 |
| NC_004718.3;NP_828850.1;SARS-CoV_orf1a;1736 | 0 | 0 |
| NC_004718.3;NP_828850.1;SARS-CoV_orf1a;1737 | 0 | 0 |
| NC_004718.3;NP_828850.1;SARS-CoV_orf1a;1738 | 0.1 | 0 |
| NC_004718.3;NP_828850.1;SARS-CoV_orf1a;1739 | 0.15 | 0.025 |
| NC_004718.3;NP_828850.1;SARS-CoV_orf1a;1740 | 0.15 | 0.025 |
| NC_004718.3;NP_828850.1;SARS-CoV_orf1a;1741 | 0.2 | 0.025 |
| NC_004718.3;NP_828850.1;SARS-CoV_orf1a;1742 | 0.2 | 0.1 |
| NC_004718.3;NP_828850.1;SARS-CoV_orf1a;1743 | 0.2 | 0.175 |
| NC_004718.3;NP_828850.1;SARS-CoV_orf1a;1744 | 0.15 | 0.025 |
| NC_004718.3;NP_828850.1;SARS-CoV_orf1a;1745 | 0.15 | 0 |
| NC_004718.3;NP_828850.1;SARS-CoV_orf1a;1746 | 0.15 | 0.05 |
| NC_004718.3;NP_828850.1;SARS-CoV_orf1a;1747 | 0.05 | 0.025 |
| NC_004718.3;NP_828850.1;SARS-CoV_orf1a;1748 | 0.05 | 0.025 |
| NC_004718.3;NP_828850.1;SARS-CoV_orf1a;1749 | 0.05 | 0 |
| NC_004718.3;NP_828850.1;SARS-CoV_orf1a;1750 | 0.05 | 0 |

| | | |
|---|---|---|
| NC_004718.3;NP_828850.1;SARS-CoV_orf1a;1751 | 0.05 | 0 |
| NC_004718.3;NP_828850.1;SARS-CoV_orf1a;1752 | 0 | 0.05 |
| NC_004718.3;NP_828850.1;SARS-CoV_orf1a;1753 | 0 | 0 |
| NC_004718.3;NP_828850.1;SARS-CoV_orf1a;1754 | 0 | 0 |
| NC_004718.3;NP_828850.1;SARS-CoV_orf1a;1755 | 0 | 0 |
| NC_004718.3;NP_828850.1;SARS-CoV_orf1a;1756 | 0 | 0 |
| NC_004718.3;NP_828850.1;SARS-CoV_orf1a;1757 | 0 | 0 |
| NC_004718.3;NP_828850.1;SARS-CoV_orf1a;1758 | 0 | 0 |
| NC_004718.3;NP_828850.1;SARS-CoV_orf1a;1759 | 0 | 0 |
| NC_004718.3;NP_828850.1;SARS-CoV_orf1a;1760 | 0 | 0 |
| NC_004718.3;NP_828850.1;SARS-CoV_orf1a;1761 | 0 | 0 |
| NC_004718.3;NP_828850.1;SARS-CoV_orf1a;1762 | 0 | 0 |
| NC_004718.3;NP_828850.1;SARS-CoV_orf1a;1763 | 0 | 0 |
| NC_004718.3;NP_828850.1;SARS-CoV_orf1a;1764 | 0.05 | 0 |
| NC_004718.3;NP_828850.1;SARS-CoV_orf1a;1765 | 0 | 0 |
| NC_004718.3;NP_828850.1;SARS-CoV_orf1a;1766 | 0 | 0 |
| NC_004718.3;NP_828850.1;SARS-CoV_orf1a;1767 | 0 | 0.025 |
| NC_004718.3;NP_828850.1;SARS-CoV_orf1a;1768 | 0 | 0.025 |
| NC_004718.3;NP_828850.1;SARS-CoV_orf1a;1769 | 0 | 0 |
| NC_004718.3;NP_828850.1;SARS-CoV_orf1a;1770 | 0 | 0.025 |
| NC_004718.3;NP_828850.1;SARS-CoV_orf1a;1771 | 0 | 0 |
| NC_004718.3;NP_828850.1;SARS-CoV_orf1a;1772 | 0.05 | 0 |
| NC_004718.3;NP_828850.1;SARS-CoV_orf1a;1773 | 0 | 0.025 |
| NC_004718.3;NP_828850.1;SARS-CoV_orf1a;1774 | 0.1 | 0.025 |
| NC_004718.3;NP_828850.1;SARS-CoV_orf1a;1775 | 0 | 0 |
| NC_004718.3;NP_828850.1;SARS-CoV_orf1a;1776 | 0 | 0 |
| NC_004718.3;NP_828850.1;SARS-CoV_orf1a;1777 | 0 | 0 |
| NC_004718.3;NP_828850.1;SARS-CoV_orf1a;1778 | 0 | 0 |
| NC_004718.3;NP_828850.1;SARS-CoV_orf1a;1779 | 0 | 0 |
| NC_004718.3;NP_828850.1;SARS-CoV_orf1a;1780 | 0 | 0 |
| NC_004718.3;NP_828850.1;SARS-CoV_orf1a;1781 | 0.05 | 0 |
| NC_004718.3;NP_828850.1;SARS-CoV_orf1a;1782 | 0 | 0 |
| NC_004718.3;NP_828850.1;SARS-CoV_orf1a;1783 | 0 | 0 |
| NC_004718.3;NP_828850.1;SARS-CoV_orf1a;1784 | 0 | 0 |
| NC_004718.3;NP_828850.1;SARS-CoV_orf1a;1785 | 0 | 0 |
| NC_004718.3;NP_828850.1;SARS-CoV_orf1a;1786 | 0 | 0 |
| NC_004718.3;NP_828850.1;SARS-CoV_orf1a;1787 | 0 | 0 |
| NC_004718.3;NP_828850.1;SARS-CoV_orf1a;1788 | 0 | 0 |
| NC_004718.3;NP_828850.1;SARS-CoV_orf1a;1789 | 0 | 0 |
| NC_004718.3;NP_828850.1;SARS-CoV_orf1a;1790 | 0 | 0 |
| NC_004718.3;NP_828850.1;SARS-CoV_orf1a;1791 | 0 | 0 |
| NC_004718.3;NP_828850.1;SARS-CoV_orf1a;1792 | 0 | 0 |
| NC_004718.3;NP_828850.1;SARS-CoV_orf1a;1793 | 0 | 0.025 |

| | | |
|---|---|---|
| NC_004718.3;NP_828850.1;SARS-CoV_orf1a;1794 | 0 | 0.025 |
| NC_004718.3;NP_828850.1;SARS-CoV_orf1a;1795 | 0 | 0 |
| NC_004718.3;NP_828850.1;SARS-CoV_orf1a;1796 | 0 | 0 |
| NC_004718.3;NP_828850.1;SARS-CoV_orf1a;1797 | 0 | 0 |
| NC_004718.3;NP_828850.1;SARS-CoV_orf1a;1798 | 0 | 0 |
| NC_004718.3;NP_828850.1;SARS-CoV_orf1a;1799 | 0 | 0 |
| NC_004718.3;NP_828850.1;SARS-CoV_orf1a;1800 | 0 | 0 |
| NC_004718.3;NP_828850.1;SARS-CoV_orf1a;1801 | 0 | 0 |
| NC_004718.3;NP_828850.1;SARS-CoV_orf1a;1802 | 0 | 0 |
| NC_004718.3;NP_828850.1;SARS-CoV_orf1a;1803 | 0 | 0.15 |
| NC_004718.3;NP_828850.1;SARS-CoV_orf1a;1804 | 0 | 0 |
| NC_004718.3;NP_828850.1;SARS-CoV_orf1a;1805 | 0 | 0.025 |
| NC_004718.3;NP_828850.1;SARS-CoV_orf1a;1806 | 0.1 | 0 |
| NC_004718.3;NP_828850.1;SARS-CoV_orf1a;1807 | 0 | 0 |
| NC_004718.3;NP_828850.1;SARS-CoV_orf1a;1808 | 0.05 | 0 |
| NC_004718.3;NP_828850.1;SARS-CoV_orf1a;1809 | 0 | 0 |
| NC_004718.3;NP_828850.1;SARS-CoV_orf1a;1810 | 0 | 0 |
| NC_004718.3;NP_828850.1;SARS-CoV_orf1a;1811 | 0.05 | 0 |
| NC_004718.3;NP_828850.1;SARS-CoV_orf1a;1812 | 0.05 | 0 |
| NC_004718.3;NP_828850.1;SARS-CoV_orf1a;1813 | 0.05 | 0 |
| NC_004718.3;NP_828850.1;SARS-CoV_orf1a;1814 | 0.05 | 0 |
| NC_004718.3;NP_828850.1;SARS-CoV_orf1a;1815 | 0.05 | 0 |
| NC_004718.3;NP_828850.1;SARS-CoV_orf1a;1816 | 0.05 | 0.05 |
| NC_004718.3;NP_828850.1;SARS-CoV_orf1a;1817 | 0 | 0 |
| NC_004718.3;NP_828850.1;SARS-CoV_orf1a;1818 | 0.05 | 0 |
| NC_004718.3;NP_828850.1;SARS-CoV_orf1a;1819 | 0 | 0 |
| NC_004718.3;NP_828850.1;SARS-CoV_orf1a;1820 | 0 | 0 |
| NC_004718.3;NP_828850.1;SARS-CoV_orf1a;1821 | 0 | 0 |
| NC_004718.3;NP_828850.1;SARS-CoV_orf1a;1822 | 0 | 0 |
| NC_004718.3;NP_828850.1;SARS-CoV_orf1a;1823 | 0 | 0 |
| NC_004718.3;NP_828850.1;SARS-CoV_orf1a;1824 | 0 | 0 |
| NC_004718.3;NP_828850.1;SARS-CoV_orf1a;1825 | 0 | 0 |
| NC_004718.3;NP_828850.1;SARS-CoV_orf1a;1826 | 0 | 0 |
| NC_004718.3;NP_828850.1;SARS-CoV_orf1a;1827 | 0 | 0.05 |
| NC_004718.3;NP_828850.1;SARS-CoV_orf1a;1828 | 0 | 0 |
| NC_004718.3;NP_828850.1;SARS-CoV_orf1a;1829 | 0 | 0 |
| NC_004718.3;NP_828850.1;SARS-CoV_orf1a;1830 | 0 | 0.025 |
| NC_004718.3;NP_828850.1;SARS-CoV_orf1a;1831 | 0 | 0.025 |
| NC_004718.3;NP_828850.1;SARS-CoV_orf1a;1832 | 0 | 0.075 |
| NC_004718.3;NP_828850.1;SARS-CoV_orf1a;1833 | 0 | 0.1 |
| NC_004718.3;NP_828850.1;SARS-CoV_orf1a;1834 | 0 | 0.075 |
| NC_004718.3;NP_828850.1;SARS-CoV_orf1a;1835 | 0 | 0.075 |
| NC_004718.3;NP_828850.1;SARS-CoV_orf1a;1836 | 0 | 0.025 |

| | | |
|---|---|---|
| NC_004718.3;NP_828850.1;SARS-CoV_orf1a;1837 | 0 | 0.025 |
| NC_004718.3;NP_828850.1;SARS-CoV_orf1a;1838 | 0.05 | 0.05 |
| NC_004718.3;NP_828850.1;SARS-CoV_orf1a;1839 | 0 | 0 |
| NC_004718.3;NP_828850.1;SARS-CoV_orf1a;1840 | 0 | 0.025 |
| NC_004718.3;NP_828850.1;SARS-CoV_orf1a;1841 | 0 | 0.025 |
| NC_004718.3;NP_828850.1;SARS-CoV_orf1a;1842 | 0 | 0.025 |
| NC_004718.3;NP_828850.1;SARS-CoV_orf1a;1843 | 0.05 | 0.025 |
| NC_004718.3;NP_828850.1;SARS-CoV_orf1a;1844 | 0.05 | 0.05 |
| NC_004718.3;NP_828850.1;SARS-CoV_orf1a;1845 | 0 | 0.025 |
| NC_004718.3;NP_828850.1;SARS-CoV_orf1a;1846 | 0.05 | 0.025 |
| NC_004718.3;NP_828850.1;SARS-CoV_orf1a;1847 | 0 | 0.025 |
| NC_004718.3;NP_828850.1;SARS-CoV_orf1a;1848 | 0 | 0.025 |
| NC_004718.3;NP_828850.1;SARS-CoV_orf1a;1849 | 0 | 0.025 |
| NC_004718.3;NP_828850.1;SARS-CoV_orf1a;1850 | 0.05 | 0.05 |
| NC_004718.3;NP_828850.1;SARS-CoV_orf1a;1851 | 0 | 0.05 |
| NC_004718.3;NP_828850.1;SARS-CoV_orf1a;1852 | 0 | 0.075 |
| NC_004718.3;NP_828850.1;SARS-CoV_orf1a;1853 | 0 | 0.025 |
| NC_004718.3;NP_828850.1;SARS-CoV_orf1a;1854 | 0 | 0.025 |
| NC_004718.3;NP_828850.1;SARS-CoV_orf1a;1855 | 0 | 0 |
| NC_004718.3;NP_828850.1;SARS-CoV_orf1a;1856 | 0 | 0 |
| NC_004718.3;NP_828850.1;SARS-CoV_orf1a;1857 | 0.05 | 0.025 |
| NC_004718.3;NP_828850.1;SARS-CoV_orf1a;1858 | 0.05 | 0.075 |
| NC_004718.3;NP_828850.1;SARS-CoV_orf1a;1859 | 0.05 | 0.025 |
| NC_004718.3;NP_828850.1;SARS-CoV_orf1a;1860 | 0.05 | 0.1 |
| NC_004718.3;NP_828850.1;SARS-CoV_orf1a;1861 | 0 | 0 |
| NC_004718.3;NP_828850.1;SARS-CoV_orf1a;1862 | 0 | 0 |
| NC_004718.3;NP_828850.1;SARS-CoV_orf1a;1863 | 0.2 | 0.275 |
| NC_004718.3;NP_828850.1;SARS-CoV_orf1a;1864 | 0 | 0.075 |
| NC_004718.3;NP_828850.1;SARS-CoV_orf1a;1865 | 0 | 0.05 |
| NC_004718.3;NP_828850.1;SARS-CoV_orf1a;1866 | 0 | 0.075 |
| NC_004718.3;NP_828850.1;SARS-CoV_orf1a;1867 | 0 | 0.075 |
| NC_004718.3;NP_828850.1;SARS-CoV_orf1a;1868 | 0.05 | 0.075 |
| NC_004718.3;NP_828850.1;SARS-CoV_orf1a;1869 | 0 | 0.075 |
| NC_004718.3;NP_828850.1;SARS-CoV_orf1a;1870 | 0.05 | 0.075 |
| NC_004718.3;NP_828850.1;SARS-CoV_orf1a;1871 | 0.05 | 0.075 |
| NC_004718.3;NP_828850.1;SARS-CoV_orf1a;1872 | 0.15 | 0.1 |
| NC_004718.3;NP_828850.1;SARS-CoV_orf1a;1873 | 0.1 | 0.075 |
| NC_004718.3;NP_828850.1;SARS-CoV_orf1a;1874 | 0.15 | 0.075 |
| NC_004718.3;NP_828850.1;SARS-CoV_orf1a;1875 | 0.05 | 0 |
| NC_004718.3;NP_828850.1;SARS-CoV_orf1a;1876 | 0.15 | 0.05 |
| NC_004718.3;NP_828850.1;SARS-CoV_orf1a;1877 | 0.1 | 0.05 |
| NC_004718.3;NP_828850.1;SARS-CoV_orf1a;1878 | 0.15 | 0.125 |
| NC_004718.3;NP_828850.1;SARS-CoV_orf1a;1879 | 0.05 | 0.025 |

| | | |
|---|---|---|
| NC_004718.3;NP_828850.1;SARS-CoV_orf1a;1880 | 0.05 | 0 |
| NC_004718.3;NP_828850.1;SARS-CoV_orf1a;1881 | 0.05 | 0 |
| NC_004718.3;NP_828850.1;SARS-CoV_orf1a;1882 | 0.05 | 0.05 |
| NC_004718.3;NP_828850.1;SARS-CoV_orf1a;1883 | 0.05 | 0 |
| NC_004718.3;NP_828850.1;SARS-CoV_orf1a;1884 | 0.05 | 0 |
| NC_004718.3;NP_828850.1;SARS-CoV_orf1a;1885 | 0.1 | 0.05 |
| NC_004718.3;NP_828850.1;SARS-CoV_orf1a;1886 | 0.1 | 0.025 |
| NC_004718.3;NP_828850.1;SARS-CoV_orf1a;1887 | 0.1 | 0.075 |
| NC_004718.3;NP_828850.1;SARS-CoV_orf1a;1888 | 0.05 | 0.025 |
| NC_004718.3;NP_828850.1;SARS-CoV_orf1a;1889 | 0.05 | 0.025 |
| NC_004718.3;NP_828850.1;SARS-CoV_orf1a;1890 | 0.05 | 0.075 |
| NC_004718.3;NP_828850.1;SARS-CoV_orf1a;1891 | 0.05 | 0.1 |
| NC_004718.3;NP_828850.1;SARS-CoV_orf1a;1892 | 0.05 | 0.05 |
| NC_004718.3;NP_828850.1;SARS-CoV_orf1a;1893 | 0.05 | 0.075 |
| NC_004718.3;NP_828850.1;SARS-CoV_orf1a;1894 | 0.05 | 0.05 |
| NC_004718.3;NP_828850.1;SARS-CoV_orf1a;1895 | 0.05 | 0.025 |
| NC_004718.3;NP_828850.1;SARS-CoV_orf1a;1896 | 0.05 | 0.025 |
| NC_004718.3;NP_828850.1;SARS-CoV_orf1a;1897 | 0.05 | 0 |
| NC_004718.3;NP_828850.1;SARS-CoV_orf1a;1898 | 0.05 | 0 |
| NC_004718.3;NP_828850.1;SARS-CoV_orf1a;1899 | 0 | 0 |
| NC_004718.3;NP_828850.1;SARS-CoV_orf1a;1900 | 0 | 0.025 |
| NC_004718.3;NP_828850.1;SARS-CoV_orf1a;1901 | 0 | 0.025 |
| NC_004718.3;NP_828850.1;SARS-CoV_orf1a;1902 | 0 | 0 |
| NC_004718.3;NP_828850.1;SARS-CoV_orf1a;1903 | 0 | 0 |
| NC_004718.3;NP_828850.1;SARS-CoV_orf1a;1904 | 0.15 | 0.05 |
| NC_004718.3;NP_828850.1;SARS-CoV_orf1a;1905 | 0.15 | 0.025 |
| NC_004718.3;NP_828850.1;SARS-CoV_orf1a;1906 | 0.2 | 0.1 |
| NC_004718.3;NP_828850.1;SARS-CoV_orf1a;1907 | 0.2 | 0.075 |
| NC_004718.3;NP_828850.1;SARS-CoV_orf1a;1908 | 0.15 | 0.1 |
| NC_004718.3;NP_828850.1;SARS-CoV_orf1a;1909 | 0.25 | 0.1 |
| NC_004718.3;NP_828850.1;SARS-CoV_orf1a;1910 | 0.3 | 0.1 |
| NC_004718.3;NP_828850.1;SARS-CoV_orf1a;1911 | 0.15 | 0.05 |
| NC_004718.3;NP_828850.1;SARS-CoV_orf1a;1912 | 0.25 | 0.075 |
| NC_004718.3;NP_828850.1;SARS-CoV_orf1a;1913 | 0.4 | 0.1 |
| NC_004718.3;NP_828850.1;SARS-CoV_orf1a;1914 | 0.05 | 0 |
| NC_004718.3;NP_828850.1;SARS-CoV_orf1a;1915 | 0.1 | 0.025 |
| NC_004718.3;NP_828850.1;SARS-CoV_orf1a;1916 | 0 | 0.05 |
| NC_004718.3;NP_828850.1;SARS-CoV_orf1a;1917 | 0 | 0 |
| NC_004718.3;NP_828850.1;SARS-CoV_orf1a;1918 | 0 | 0 |
| NC_004718.3;NP_828850.1;SARS-CoV_orf1a;1919 | 0 | 0 |
| NC_004718.3;NP_828850.1;SARS-CoV_orf1a;1920 | 0 | 0 |
| NC_004718.3;NP_828850.1;SARS-CoV_orf1a;1921 | 0 | 0 |
| NC_004718.3;NP_828850.1;SARS-CoV_orf1a;1922 | 0 | 0 |

| | | |
|---|---|---|
| NC_004718.3;NP_828850.1;SARS-CoV_orf1a;1923 | 0 | 0 |
| NC_004718.3;NP_828850.1;SARS-CoV_orf1a;1924 | 0 | 0 |
| NC_004718.3;NP_828850.1;SARS-CoV_orf1a;1925 | 0 | 0 |
| NC_004718.3;NP_828850.1;SARS-CoV_orf1a;1926 | 0 | 0 |
| NC_004718.3;NP_828850.1;SARS-CoV_orf1a;1927 | 0 | 0 |
| NC_004718.3;NP_828850.1;SARS-CoV_orf1a;1928 | 0 | 0 |
| NC_004718.3;NP_828850.1;SARS-CoV_orf1a;1929 | 0 | 0 |
| NC_004718.3;NP_828850.1;SARS-CoV_orf1a;1930 | 0 | 0.025 |
| NC_004718.3;NP_828850.1;SARS-CoV_orf1a;1931 | 0 | 0.025 |
| NC_004718.3;NP_828850.1;SARS-CoV_orf1a;1932 | 0 | 0 |
| NC_004718.3;NP_828850.1;SARS-CoV_orf1a;1933 | 0 | 0.025 |
| NC_004718.3;NP_828850.1;SARS-CoV_orf1a;1934 | 0 | 0 |
| NC_004718.3;NP_828850.1;SARS-CoV_orf1a;1935 | 0 | 0.025 |
| NC_004718.3;NP_828850.1;SARS-CoV_orf1a;1936 | 0 | 0.025 |
| NC_004718.3;NP_828850.1;SARS-CoV_orf1a;1937 | 0 | 0.025 |
| NC_004718.3;NP_828850.1;SARS-CoV_orf1a;1938 | 0 | 0.075 |
| NC_004718.3;NP_828850.1;SARS-CoV_orf1a;1939 | 0 | 0.025 |
| NC_004718.3;NP_828850.1;SARS-CoV_orf1a;1940 | 0 | 0 |
| NC_004718.3;NP_828850.1;SARS-CoV_orf1a;1941 | 0 | 0 |
| NC_004718.3;NP_828850.1;SARS-CoV_orf1a;1942 | 0.05 | 0.05 |
| NC_004718.3;NP_828850.1;SARS-CoV_orf1a;1943 | 0 | 0 |
| NC_004718.3;NP_828850.1;SARS-CoV_orf1a;1944 | 0 | 0 |
| NC_004718.3;NP_828850.1;SARS-CoV_orf1a;1945 | 0 | 0 |
| NC_004718.3;NP_828850.1;SARS-CoV_orf1a;1946 | 0 | 0.025 |
| NC_004718.3;NP_828850.1;SARS-CoV_orf1a;1947 | 0 | 0 |
| NC_004718.3;NP_828850.1;SARS-CoV_orf1a;1948 | 0 | 0 |
| NC_004718.3;NP_828850.1;SARS-CoV_orf1a;1949 | 0 | 0 |
| NC_004718.3;NP_828850.1;SARS-CoV_orf1a;1950 | 0 | 0 |
| NC_004718.3;NP_828850.1;SARS-CoV_orf1a;1951 | 0.05 | 0 |
| NC_004718.3;NP_828850.1;SARS-CoV_orf1a;1952 | 0 | 0 |
| NC_004718.3;NP_828850.1;SARS-CoV_orf1a;1953 | 0 | 0 |
| NC_004718.3;NP_828850.1;SARS-CoV_orf1a;1954 | 0 | 0 |
| NC_004718.3;NP_828850.1;SARS-CoV_orf1a;1955 | 0 | 0 |
| NC_004718.3;NP_828850.1;SARS-CoV_orf1a;1956 | 0 | 0 |
| NC_004718.3;NP_828850.1;SARS-CoV_orf1a;1957 | 0 | 0 |
| NC_004718.3;NP_828850.1;SARS-CoV_orf1a;1958 | 0 | 0 |
| NC_004718.3;NP_828850.1;SARS-CoV_orf1a;1959 | 0 | 0 |
| NC_004718.3;NP_828850.1;SARS-CoV_orf1a;1960 | 0 | 0 |
| NC_004718.3;NP_828850.1;SARS-CoV_orf1a;1961 | 0 | 0 |
| NC_004718.3;NP_828850.1;SARS-CoV_orf1a;1962 | 0 | 0 |
| NC_004718.3;NP_828850.1;SARS-CoV_orf1a;1963 | 0 | 0 |
| NC_004718.3;NP_828850.1;SARS-CoV_orf1a;1964 | 0 | 0 |
| NC_004718.3;NP_828850.1;SARS-CoV_orf1a;1965 | 0 | 0 |

| | | |
|---|---|---|
| NC_004718.3;NP_828850.1;SARS-CoV_orf1a;1966 | 0 | 0 |
| NC_004718.3;NP_828850.1;SARS-CoV_orf1a;1967 | 0.05 | 0.025 |
| NC_004718.3;NP_828850.1;SARS-CoV_orf1a;1968 | 0.05 | 0.025 |
| NC_004718.3;NP_828850.1;SARS-CoV_orf1a;1969 | 0.05 | 0.025 |
| NC_004718.3;NP_828850.1;SARS-CoV_orf1a;1970 | 0.1 | 0.025 |
| NC_004718.3;NP_828850.1;SARS-CoV_orf1a;1971 | 0.1 | 0.025 |
| NC_004718.3;NP_828850.1;SARS-CoV_orf1a;1972 | 0.05 | 0.05 |
| NC_004718.3;NP_828850.1;SARS-CoV_orf1a;1973 | 0.1 | 0.025 |
| NC_004718.3;NP_828850.1;SARS-CoV_orf1a;1974 | 0.05 | 0 |
| NC_004718.3;NP_828850.1;SARS-CoV_orf1a;1975 | 0.05 | 0 |
| NC_004718.3;NP_828850.1;SARS-CoV_orf1a;1976 | 0 | 0 |
| NC_004718.3;NP_828850.1;SARS-CoV_orf1a;1977 | 0 | 0 |
| NC_004718.3;NP_828850.1;SARS-CoV_orf1a;1978 | 0 | 0 |
| NC_004718.3;NP_828850.1;SARS-CoV_orf1a;1979 | 0 | 0 |
| NC_004718.3;NP_828850.1;SARS-CoV_orf1a;1980 | 0 | 0 |
| NC_004718.3;NP_828850.1;SARS-CoV_orf1a;1981 | 0 | 0 |
| NC_004718.3;NP_828850.1;SARS-CoV_orf1a;1982 | 0 | 0 |
| NC_004718.3;NP_828850.1;SARS-CoV_orf1a;1983 | 0 | 0 |
| NC_004718.3;NP_828850.1;SARS-CoV_orf1a;1984 | 0 | 0 |
| NC_004718.3;NP_828850.1;SARS-CoV_orf1a;1985 | 0 | 0 |
| NC_004718.3;NP_828850.1;SARS-CoV_orf1a;1986 | 0.05 | 0 |
| NC_004718.3;NP_828850.1;SARS-CoV_orf1a;1987 | 0.1 | 0.025 |
| NC_004718.3;NP_828850.1;SARS-CoV_orf1a;1988 | 0.1 | 0.1 |
| NC_004718.3;NP_828850.1;SARS-CoV_orf1a;1989 | 0.15 | 0.125 |
| NC_004718.3;NP_828850.1;SARS-CoV_orf1a;1990 | 0.1 | 0.075 |
| NC_004718.3;NP_828850.1;SARS-CoV_orf1a;1991 | 0.05 | 0.075 |
| NC_004718.3;NP_828850.1;SARS-CoV_orf1a;1992 | 0 | 0.025 |
| NC_004718.3;NP_828850.1;SARS-CoV_orf1a;1993 | 0 | 0.025 |
| NC_004718.3;NP_828850.1;SARS-CoV_orf1a;1994 | 0.05 | 0.025 |
| NC_004718.3;NP_828850.1;SARS-CoV_orf1a;1995 | 0.05 | 0.025 |
| NC_004718.3;NP_828850.1;SARS-CoV_orf1a;1996 | 0.15 | 0.125 |
| NC_004718.3;NP_828850.1;SARS-CoV_orf1a;1997 | 0 | 0.05 |
| NC_004718.3;NP_828850.1;SARS-CoV_orf1a;1998 | 0 | 0.025 |
| NC_004718.3;NP_828850.1;SARS-CoV_orf1a;1999 | 0 | 0 |
| NC_004718.3;NP_828850.1;SARS-CoV_orf1a;2000 | 0.05 | 0 |
| NC_004718.3;NP_828850.1;SARS-CoV_orf1a;2001 | 0 | 0.025 |
| NC_004718.3;NP_828850.1;SARS-CoV_orf1a;2002 | 0 | 0.025 |
| NC_004718.3;NP_828850.1;SARS-CoV_orf1a;2003 | 0 | 0 |
| NC_004718.3;NP_828850.1;SARS-CoV_orf1a;2004 | 0 | 0 |
| NC_004718.3;NP_828850.1;SARS-CoV_orf1a;2005 | 0.05 | 0 |
| NC_004718.3;NP_828850.1;SARS-CoV_orf1a;2006 | 0.1 | 0.075 |
| NC_004718.3;NP_828850.1;SARS-CoV_orf1a;2007 | 0 | 0 |
| NC_004718.3;NP_828850.1;SARS-CoV_orf1a;2008 | 0.15 | 0 |

| | | |
|---|---|---|
| NC_004718.3;NP_828850.1;SARS-CoV_orf1a;2009 | 0 | 0 |
| NC_004718.3;NP_828850.1;SARS-CoV_orf1a;2010 | 0 | 0 |
| NC_004718.3;NP_828850.1;SARS-CoV_orf1a;2011 | 0 | 0 |
| NC_004718.3;NP_828850.1;SARS-CoV_orf1a;2012 | 0 | 0 |
| NC_004718.3;NP_828850.1;SARS-CoV_orf1a;2013 | 0 | 0 |
| NC_004718.3;NP_828850.1;SARS-CoV_orf1a;2014 | 0 | 0 |
| NC_004718.3;NP_828850.1;SARS-CoV_orf1a;2015 | 0 | 0 |
| NC_004718.3;NP_828850.1;SARS-CoV_orf1a;2016 | 0 | 0 |
| NC_004718.3;NP_828850.1;SARS-CoV_orf1a;2017 | 0 | 0 |
| NC_004718.3;NP_828850.1;SARS-CoV_orf1a;2018 | 0 | 0 |
| NC_004718.3;NP_828850.1;SARS-CoV_orf1a;2019 | 0 | 0 |
| NC_004718.3;NP_828850.1;SARS-CoV_orf1a;2020 | 0 | 0 |
| NC_004718.3;NP_828850.1;SARS-CoV_orf1a;2021 | 0 | 0.05 |
| NC_004718.3;NP_828850.1;SARS-CoV_orf1a;2022 | 0 | 0.05 |
| NC_004718.3;NP_828850.1;SARS-CoV_orf1a;2023 | 0 | 0.05 |
| NC_004718.3;NP_828850.1;SARS-CoV_orf1a;2024 | 0 | 0.05 |
| NC_004718.3;NP_828850.1;SARS-CoV_orf1a;2025 | 0 | 0.05 |
| NC_004718.3;NP_828850.1;SARS-CoV_orf1a;2026 | 0 | 0.05 |
| NC_004718.3;NP_828850.1;SARS-CoV_orf1a;2027 | 0 | 0.05 |
| NC_004718.3;NP_828850.1;SARS-CoV_orf1a;2028 | 0 | 0.05 |
| NC_004718.3;NP_828850.1;SARS-CoV_orf1a;2029 | 0 | 0 |
| NC_004718.3;NP_828850.1;SARS-CoV_orf1a;2030 | 0 | 0 |
| NC_004718.3;NP_828850.1;SARS-CoV_orf1a;2031 | 0.05 | 0 |
| NC_004718.3;NP_828850.1;SARS-CoV_orf1a;2032 | 0 | 0.025 |
| NC_004718.3;NP_828850.1;SARS-CoV_orf1a;2033 | 0 | 0.025 |
| NC_004718.3;NP_828850.1;SARS-CoV_orf1a;2034 | 0 | 0.025 |
| NC_004718.3;NP_828850.1;SARS-CoV_orf1a;2035 | 0 | 0.025 |
| NC_004718.3;NP_828850.1;SARS-CoV_orf1a;2036 | 0 | 0.025 |
| NC_004718.3;NP_828850.1;SARS-CoV_orf1a;2037 | 0 | 0.025 |
| NC_004718.3;NP_828850.1;SARS-CoV_orf1a;2038 | 0 | 0 |
| NC_004718.3;NP_828850.1;SARS-CoV_orf1a;2039 | 0 | 0 |
| NC_004718.3;NP_828850.1;SARS-CoV_orf1a;2040 | 0 | 0 |
| NC_004718.3;NP_828850.1;SARS-CoV_orf1a;2041 | 0 | 0 |
| NC_004718.3;NP_828850.1;SARS-CoV_orf1a;2042 | 0 | 0.025 |
| NC_004718.3;NP_828850.1;SARS-CoV_orf1a;2043 | 0 | 0 |
| NC_004718.3;NP_828850.1;SARS-CoV_orf1a;2044 | 0 | 0 |
| NC_004718.3;NP_828850.1;SARS-CoV_orf1a;2045 | 0 | 0 |
| NC_004718.3;NP_828850.1;SARS-CoV_orf1a;2046 | 0 | 0 |
| NC_004718.3;NP_828850.1;SARS-CoV_orf1a;2047 | 0 | 0 |
| NC_004718.3;NP_828850.1;SARS-CoV_orf1a;2048 | 0 | 0 |
| NC_004718.3;NP_828850.1;SARS-CoV_orf1a;2049 | 0 | 0 |
| NC_004718.3;NP_828850.1;SARS-CoV_orf1a;2050 | 0 | 0.05 |
| NC_004718.3;NP_828850.1;SARS-CoV_orf1a;2051 | 0 | 0 |

| | | |
|---|---|---|
| NC_004718.3;NP_828850.1;SARS-CoV_orf1a;2052 | 0 | 0 |
| NC_004718.3;NP_828850.1;SARS-CoV_orf1a;2053 | 0 | 0 |
| NC_004718.3;NP_828850.1;SARS-CoV_orf1a;2054 | 0 | 0 |
| NC_004718.3;NP_828850.1;SARS-CoV_orf1a;2055 | 0 | 0 |
| NC_004718.3;NP_828850.1;SARS-CoV_orf1a;2056 | 0 | 0 |
| NC_004718.3;NP_828850.1;SARS-CoV_orf1a;2057 | 0 | 0 |
| NC_004718.3;NP_828850.1;SARS-CoV_orf1a;2058 | 0 | 0 |
| NC_004718.3;NP_828850.1;SARS-CoV_orf1a;2059 | 0 | 0.025 |
| NC_004718.3;NP_828850.1;SARS-CoV_orf1a;2060 | 0.05 | 0.025 |
| NC_004718.3;NP_828850.1;SARS-CoV_orf1a;2061 | 0 | 0 |
| NC_004718.3;NP_828850.1;SARS-CoV_orf1a;2062 | 0 | 0 |
| NC_004718.3;NP_828850.1;SARS-CoV_orf1a;2063 | 0 | 0 |
| NC_004718.3;NP_828850.1;SARS-CoV_orf1a;2064 | 0 | 0 |
| NC_004718.3;NP_828850.1;SARS-CoV_orf1a;2065 | 0 | 0 |
| NC_004718.3;NP_828850.1;SARS-CoV_orf1a;2066 | 0 | 0 |
| NC_004718.3;NP_828850.1;SARS-CoV_orf1a;2067 | 0 | 0 |
| NC_004718.3;NP_828850.1;SARS-CoV_orf1a;2068 | 0.25 | 0.125 |
| NC_004718.3;NP_828850.1;SARS-CoV_orf1a;2069 | 0 | 0 |
| NC_004718.3;NP_828850.1;SARS-CoV_orf1a;2070 | 0 | 0 |
| NC_004718.3;NP_828850.1;SARS-CoV_orf1a;2071 | 0 | 0 |
| NC_004718.3;NP_828850.1;SARS-CoV_orf1a;2072 | 0 | 0 |
| NC_004718.3;NP_828850.1;SARS-CoV_orf1a;2073 | 0 | 0 |
| NC_004718.3;NP_828850.1;SARS-CoV_orf1a;2074 | 0.05 | 0 |
| NC_004718.3;NP_828850.1;SARS-CoV_orf1a;2075 | 0.05 | 0.025 |
| NC_004718.3;NP_828850.1;SARS-CoV_orf1a;2076 | 0.15 | 0 |
| NC_004718.3;NP_828850.1;SARS-CoV_orf1a;2077 | 0.15 | 0.05 |
| NC_004718.3;NP_828850.1;SARS-CoV_orf1a;2078 | 0.05 | 0.025 |
| NC_004718.3;NP_828850.1;SARS-CoV_orf1a;2079 | 0 | 0 |
| NC_004718.3;NP_828850.1;SARS-CoV_orf1a;2080 | 0 | 0 |
| NC_004718.3;NP_828850.1;SARS-CoV_orf1a;2081 | 0 | 0 |
| NC_004718.3;NP_828850.1;SARS-CoV_orf1a;2082 | 0 | 0 |
| NC_004718.3;NP_828850.1;SARS-CoV_orf1a;2083 | 0 | 0 |
| NC_004718.3;NP_828850.1;SARS-CoV_orf1a;2084 | 0 | 0 |
| NC_004718.3;NP_828850.1;SARS-CoV_orf1a;2085 | 0 | 0 |
| NC_004718.3;NP_828850.1;SARS-CoV_orf1a;2086 | 0 | 0 |
| NC_004718.3;NP_828850.1;SARS-CoV_orf1a;2087 | 0 | 0 |
| NC_004718.3;NP_828850.1;SARS-CoV_orf1a;2088 | 0 | 0 |
| NC_004718.3;NP_828850.1;SARS-CoV_orf1a;2089 | 0 | 0 |
| NC_004718.3;NP_828850.1;SARS-CoV_orf1a;2090 | 0 | 0 |
| NC_004718.3;NP_828850.1;SARS-CoV_orf1a;2091 | 0.05 | 0 |
| NC_004718.3;NP_828850.1;SARS-CoV_orf1a;2092 | 0 | 0 |
| NC_004718.3;NP_828850.1;SARS-CoV_orf1a;2093 | 0 | 0 |
| NC_004718.3;NP_828850.1;SARS-CoV_orf1a;2094 | 0 | 0 |

| | | |
|---|---|---|
| NC_004718.3;NP_828850.1;SARS-CoV_orf1a;2095 | 0 | 0 |
| NC_004718.3;NP_828850.1;SARS-CoV_orf1a;2096 | 0 | 0 |
| NC_004718.3;NP_828850.1;SARS-CoV_orf1a;2097 | 0 | 0 |
| NC_004718.3;NP_828850.1;SARS-CoV_orf1a;2098 | 0 | 0 |
| NC_004718.3;NP_828850.1;SARS-CoV_orf1a;2099 | 0 | 0 |
| NC_004718.3;NP_828850.1;SARS-CoV_orf1a;2100 | 0.05 | 0.025 |
| NC_004718.3;NP_828850.1;SARS-CoV_orf1a;2101 | 0.05 | 0.05 |
| NC_004718.3;NP_828850.1;SARS-CoV_orf1a;2102 | 0.05 | 0.05 |
| NC_004718.3;NP_828850.1;SARS-CoV_orf1a;2103 | 0.05 | 0.05 |
| NC_004718.3;NP_828850.1;SARS-CoV_orf1a;2104 | 0.05 | 0 |
| NC_004718.3;NP_828850.1;SARS-CoV_orf1a;2105 | 0.05 | 0 |
| NC_004718.3;NP_828850.1;SARS-CoV_orf1a;2106 | 0.05 | 0 |
| NC_004718.3;NP_828850.1;SARS-CoV_orf1a;2107 | 0.05 | 0 |
| NC_004718.3;NP_828850.1;SARS-CoV_orf1a;2108 | 0 | 0 |
| NC_004718.3;NP_828850.1;SARS-CoV_orf1a;2109 | 0.05 | 0 |
| NC_004718.3;NP_828850.1;SARS-CoV_orf1a;2110 | 0.05 | 0 |
| NC_004718.3;NP_828850.1;SARS-CoV_orf1a;2111 | 0 | 0 |
| NC_004718.3;NP_828850.1;SARS-CoV_orf1a;2112 | 0 | 0 |
| NC_004718.3;NP_828850.1;SARS-CoV_orf1a;2113 | 0 | 0 |
| NC_004718.3;NP_828850.1;SARS-CoV_orf1a;2114 | 0 | 0 |
| NC_004718.3;NP_828850.1;SARS-CoV_orf1a;2115 | 0 | 0 |
| NC_004718.3;NP_828850.1;SARS-CoV_orf1a;2116 | 0 | 0 |
| NC_004718.3;NP_828850.1;SARS-CoV_orf1a;2117 | 0.05 | 0 |
| NC_004718.3;NP_828850.1;SARS-CoV_orf1a;2118 | 0 | 0 |
| NC_004718.3;NP_828850.1;SARS-CoV_orf1a;2119 | 0.05 | 0 |
| NC_004718.3;NP_828850.1;SARS-CoV_orf1a;2120 | 0.1 | 0.05 |
| NC_004718.3;NP_828850.1;SARS-CoV_orf1a;2121 | 0 | 0 |
| NC_004718.3;NP_828850.1;SARS-CoV_orf1a;2122 | 0 | 0 |
| NC_004718.3;NP_828850.1;SARS-CoV_orf1a;2123 | 0 | 0 |
| NC_004718.3;NP_828850.1;SARS-CoV_orf1a;2124 | 0 | 0 |
| NC_004718.3;NP_828850.1;SARS-CoV_orf1a;2125 | 0 | 0 |
| NC_004718.3;NP_828850.1;SARS-CoV_orf1a;2126 | 0 | 0.025 |
| NC_004718.3;NP_828850.1;SARS-CoV_orf1a;2127 | 0.1 | 0 |
| NC_004718.3;NP_828850.1;SARS-CoV_orf1a;2128 | 0 | 0 |
| NC_004718.3;NP_828850.1;SARS-CoV_orf1a;2129 | 0 | 0 |
| NC_004718.3;NP_828850.1;SARS-CoV_orf1a;2130 | 0 | 0 |
| NC_004718.3;NP_828850.1;SARS-CoV_orf1a;2131 | 0 | 0 |
| NC_004718.3;NP_828850.1;SARS-CoV_orf1a;2132 | 0 | 0 |
| NC_004718.3;NP_828850.1;SARS-CoV_orf1a;2133 | 0 | 0 |
| NC_004718.3;NP_828850.1;SARS-CoV_orf1a;2134 | 0 | 0 |
| NC_004718.3;NP_828850.1;SARS-CoV_orf1a;2135 | 0 | 0 |
| NC_004718.3;NP_828850.1;SARS-CoV_orf1a;2136 | 0 | 0 |
| NC_004718.3;NP_828850.1;SARS-CoV_orf1a;2137 | 0 | 0 |

| | | |
|---|---|---|
| NC_004718.3;NP_828850.1;SARS-CoV_orf1a;2138 | 0 | 0 |
| NC_004718.3;NP_828850.1;SARS-CoV_orf1a;2139 | 0 | 0 |
| NC_004718.3;NP_828850.1;SARS-CoV_orf1a;2140 | 0 | 0 |
| NC_004718.3;NP_828850.1;SARS-CoV_orf1a;2141 | 0 | 0 |
| NC_004718.3;NP_828850.1;SARS-CoV_orf1a;2142 | 0 | 0 |
| NC_004718.3;NP_828850.1;SARS-CoV_orf1a;2143 | 0 | 0 |
| NC_004718.3;NP_828850.1;SARS-CoV_orf1a;2144 | 0 | 0 |
| NC_004718.3;NP_828850.1;SARS-CoV_orf1a;2145 | 0 | 0 |
| NC_004718.3;NP_828850.1;SARS-CoV_orf1a;2146 | 0 | 0 |
| NC_004718.3;NP_828850.1;SARS-CoV_orf1a;2147 | 0 | 0 |
| NC_004718.3;NP_828850.1;SARS-CoV_orf1a;2148 | 0 | 0 |
| NC_004718.3;NP_828850.1;SARS-CoV_orf1a;2149 | 0 | 0 |
| NC_004718.3;NP_828850.1;SARS-CoV_orf1a;2150 | 0 | 0 |
| NC_004718.3;NP_828850.1;SARS-CoV_orf1a;2151 | 0 | 0 |
| NC_004718.3;NP_828850.1;SARS-CoV_orf1a;2152 | 0 | 0 |
| NC_004718.3;NP_828850.1;SARS-CoV_orf1a;2153 | 0 | 0 |
| NC_004718.3;NP_828850.1;SARS-CoV_orf1a;2154 | 0 | 0 |
| NC_004718.3;NP_828850.1;SARS-CoV_orf1a;2155 | 0 | 0.025 |
| NC_004718.3;NP_828850.1;SARS-CoV_orf1a;2156 | 0 | 0.025 |
| NC_004718.3;NP_828850.1;SARS-CoV_orf1a;2157 | 0 | 0.025 |
| NC_004718.3;NP_828850.1;SARS-CoV_orf1a;2158 | 0.05 | 0.025 |
| NC_004718.3;NP_828850.1;SARS-CoV_orf1a;2159 | 0 | 0.025 |
| NC_004718.3;NP_828850.1;SARS-CoV_orf1a;2160 | 0 | 0.05 |
| NC_004718.3;NP_828850.1;SARS-CoV_orf1a;2161 | 0 | 0.025 |
| NC_004718.3;NP_828850.1;SARS-CoV_orf1a;2162 | 0 | 0 |
| NC_004718.3;NP_828850.1;SARS-CoV_orf1a;2163 | 0 | 0 |
| NC_004718.3;NP_828850.1;SARS-CoV_orf1a;2164 | 0 | 0.025 |
| NC_004718.3;NP_828850.1;SARS-CoV_orf1a;2165 | 0.05 | 0 |
| NC_004718.3;NP_828850.1;SARS-CoV_orf1a;2166 | 0 | 0 |
| NC_004718.3;NP_828850.1;SARS-CoV_orf1a;2167 | 0 | 0 |
| NC_004718.3;NP_828850.1;SARS-CoV_orf1a;2168 | 0 | 0 |
| NC_004718.3;NP_828850.1;SARS-CoV_orf1a;2169 | 0 | 0 |
| NC_004718.3;NP_828850.1;SARS-CoV_orf1a;2170 | 0 | 0 |
| NC_004718.3;NP_828850.1;SARS-CoV_orf1a;2171 | 0 | 0 |
| NC_004718.3;NP_828850.1;SARS-CoV_orf1a;2172 | 0 | 0 |
| NC_004718.3;NP_828850.1;SARS-CoV_orf1a;2173 | 0 | 0 |
| NC_004718.3;NP_828850.1;SARS-CoV_orf1a;2174 | 0 | 0 |
| NC_004718.3;NP_828850.1;SARS-CoV_orf1a;2175 | 0 | 0 |
| NC_004718.3;NP_828850.1;SARS-CoV_orf1a;2176 | 0 | 0 |
| NC_004718.3;NP_828850.1;SARS-CoV_orf1a;2177 | 0 | 0 |
| NC_004718.3;NP_828850.1;SARS-CoV_orf1a;2178 | 0.05 | 0 |
| NC_004718.3;NP_828850.1;SARS-CoV_orf1a;2179 | 0.05 | 0 |
| NC_004718.3;NP_828850.1;SARS-CoV_orf1a;2180 | 0.05 | 0 |

| | | |
|---|---|---|
| NC_004718.3;NP_828850.1;SARS-CoV_orf1a;2181 | 0.05 | 0 |
| NC_004718.3;NP_828850.1;SARS-CoV_orf1a;2182 | 0.05 | 0 |
| NC_004718.3;NP_828850.1;SARS-CoV_orf1a;2183 | 0.05 | 0 |
| NC_004718.3;NP_828850.1;SARS-CoV_orf1a;2184 | 0.05 | 0 |
| NC_004718.3;NP_828850.1;SARS-CoV_orf1a;2185 | 0.05 | 0 |
| NC_004718.3;NP_828850.1;SARS-CoV_orf1a;2186 | 0 | 0 |
| NC_004718.3;NP_828850.1;SARS-CoV_orf1a;2187 | 0.1 | 0 |
| NC_004718.3;NP_828850.1;SARS-CoV_orf1a;2188 | 0.05 | 0 |
| NC_004718.3;NP_828850.1;SARS-CoV_orf1a;2189 | 0.05 | 0 |
| NC_004718.3;NP_828850.1;SARS-CoV_orf1a;2190 | 0 | 0.025 |
| NC_004718.3;NP_828850.1;SARS-CoV_orf1a;2191 | 0 | 0 |
| NC_004718.3;NP_828850.1;SARS-CoV_orf1a;2192 | 0 | 0 |
| NC_004718.3;NP_828850.1;SARS-CoV_orf1a;2193 | 0 | 0 |
| NC_004718.3;NP_828850.1;SARS-CoV_orf1a;2194 | 0 | 0 |
| NC_004718.3;NP_828850.1;SARS-CoV_orf1a;2195 | 0 | 0 |
| NC_004718.3;NP_828850.1;SARS-CoV_orf1a;2196 | 0 | 0 |
| NC_004718.3;NP_828850.1;SARS-CoV_orf1a;2197 | 0 | 0 |
| NC_004718.3;NP_828850.1;SARS-CoV_orf1a;2198 | 0 | 0 |
| NC_004718.3;NP_828850.1;SARS-CoV_orf1a;2199 | 0 | 0 |
| NC_004718.3;NP_828850.1;SARS-CoV_orf1a;2200 | 0 | 0 |
| NC_004718.3;NP_828850.1;SARS-CoV_orf1a;2201 | 0 | 0 |
| NC_004718.3;NP_828850.1;SARS-CoV_orf1a;2202 | 0 | 0 |
| NC_004718.3;NP_828850.1;SARS-CoV_orf1a;2203 | 0 | 0 |
| NC_004718.3;NP_828850.1;SARS-CoV_orf1a;2204 | 0 | 0 |
| NC_004718.3;NP_828850.1;SARS-CoV_orf1a;2205 | 0 | 0 |
| NC_004718.3;NP_828850.1;SARS-CoV_orf1a;2206 | 0 | 0 |
| NC_004718.3;NP_828850.1;SARS-CoV_orf1a;2207 | 0 | 0 |
| NC_004718.3;NP_828850.1;SARS-CoV_orf1a;2208 | 0 | 0 |
| NC_004718.3;NP_828850.1;SARS-CoV_orf1a;2209 | 0 | 0 |
| NC_004718.3;NP_828850.1;SARS-CoV_orf1a;2210 | 0 | 0 |
| NC_004718.3;NP_828850.1;SARS-CoV_orf1a;2211 | 0 | 0 |
| NC_004718.3;NP_828850.1;SARS-CoV_orf1a;2212 | 0 | 0 |
| NC_004718.3;NP_828850.1;SARS-CoV_orf1a;2213 | 0 | 0 |
| NC_004718.3;NP_828850.1;SARS-CoV_orf1a;2214 | 0 | 0 |
| NC_004718.3;NP_828850.1;SARS-CoV_orf1a;2215 | 0 | 0 |
| NC_004718.3;NP_828850.1;SARS-CoV_orf1a;2216 | 0 | 0 |
| NC_004718.3;NP_828850.1;SARS-CoV_orf1a;2217 | 0 | 0 |
| NC_004718.3;NP_828850.1;SARS-CoV_orf1a;2218 | 0 | 0 |
| NC_004718.3;NP_828850.1;SARS-CoV_orf1a;2219 | 0 | 0 |
| NC_004718.3;NP_828850.1;SARS-CoV_orf1a;2220 | 0 | 0 |
| NC_004718.3;NP_828850.1;SARS-CoV_orf1a;2221 | 0 | 0 |
| NC_004718.3;NP_828850.1;SARS-CoV_orf1a;2222 | 0 | 0 |
| NC_004718.3;NP_828850.1;SARS-CoV_orf1a;2223 | 0 | 0 |

| | | |
|---|---|---|
| NC_004718.3;NP_828850.1;SARS-CoV_orf1a;2224 | 0 | 0 |
| NC_004718.3;NP_828850.1;SARS-CoV_orf1a;2225 | 0 | 0 |
| NC_004718.3;NP_828850.1;SARS-CoV_orf1a;2226 | 0 | 0 |
| NC_004718.3;NP_828850.1;SARS-CoV_orf1a;2227 | 0 | 0 |
| NC_004718.3;NP_828850.1;SARS-CoV_orf1a;2228 | 0 | 0 |
| NC_004718.3;NP_828850.1;SARS-CoV_orf1a;2229 | 0 | 0 |
| NC_004718.3;NP_828850.1;SARS-CoV_orf1a;2230 | 0 | 0.025 |
| NC_004718.3;NP_828850.1;SARS-CoV_orf1a;2231 | 0.05 | 0 |
| NC_004718.3;NP_828850.1;SARS-CoV_orf1a;2232 | 0 | 0 |
| NC_004718.3;NP_828850.1;SARS-CoV_orf1a;2233 | 0 | 0 |
| NC_004718.3;NP_828850.1;SARS-CoV_orf1a;2234 | 0 | 0.025 |
| NC_004718.3;NP_828850.1;SARS-CoV_orf1a;2235 | 0 | 0.025 |
| NC_004718.3;NP_828850.1;SARS-CoV_orf1a;2236 | 0 | 0.075 |
| NC_004718.3;NP_828850.1;SARS-CoV_orf1a;2237 | 0 | 0.025 |
| NC_004718.3;NP_828850.1;SARS-CoV_orf1a;2238 | 0.05 | 0.075 |
| NC_004718.3;NP_828850.1;SARS-CoV_orf1a;2239 | 0.25 | 0.225 |
| NC_004718.3;NP_828850.1;SARS-CoV_orf1a;2240 | 0.3 | 0.325 |
| NC_004718.3;NP_828850.1;SARS-CoV_orf1a;2241 | 0.3 | 0.225 |
| NC_004718.3;NP_828850.1;SARS-CoV_orf1a;2242 | 0.45 | 0.45 |
| NC_004718.3;NP_828850.1;SARS-CoV_orf1a;2243 | 0.35 | 0.35 |
| NC_004718.3;NP_828850.1;SARS-CoV_orf1a;2244 | 0.25 | 0.275 |
| NC_004718.3;NP_828850.1;SARS-CoV_orf1a;2245 | 0.35 | 0.3 |
| NC_004718.3;NP_828850.1;SARS-CoV_orf1a;2246 | 0.15 | 0.125 |
| NC_004718.3;NP_828850.1;SARS-CoV_orf1a;2247 | 0.2 | 0.125 |
| NC_004718.3;NP_828850.1;SARS-CoV_orf1a;2248 | 0.05 | 0.05 |
| NC_004718.3;NP_828850.1;SARS-CoV_orf1a;2249 | 0 | 0 |
| NC_004718.3;NP_828850.1;SARS-CoV_orf1a;2250 | 0 | 0 |
| NC_004718.3;NP_828850.1;SARS-CoV_orf1a;2251 | 0.05 | 0 |
| NC_004718.3;NP_828850.1;SARS-CoV_orf1a;2252 | 0 | 0 |
| NC_004718.3;NP_828850.1;SARS-CoV_orf1a;2253 | 0 | 0 |
| NC_004718.3;NP_828850.1;SARS-CoV_orf1a;2254 | 0 | 0 |
| NC_004718.3;NP_828850.1;SARS-CoV_orf1a;2255 | 0 | 0 |
| NC_004718.3;NP_828850.1;SARS-CoV_orf1a;2256 | 0 | 0 |
| NC_004718.3;NP_828850.1;SARS-CoV_orf1a;2257 | 0 | 0 |
| NC_004718.3;NP_828850.1;SARS-CoV_orf1a;2258 | 0 | 0 |
| NC_004718.3;NP_828850.1;SARS-CoV_orf1a;2259 | 0 | 0 |
| NC_004718.3;NP_828850.1;SARS-CoV_orf1a;2260 | 0 | 0.025 |
| NC_004718.3;NP_828850.1;SARS-CoV_orf1a;2261 | 0 | 0 |
| NC_004718.3;NP_828850.1;SARS-CoV_orf1a;2262 | 0 | 0 |
| NC_004718.3;NP_828850.1;SARS-CoV_orf1a;2263 | 0 | 0 |
| NC_004718.3;NP_828850.1;SARS-CoV_orf1a;2264 | 0 | 0 |
| NC_004718.3;NP_828850.1;SARS-CoV_orf1a;2265 | 0 | 0.025 |
| NC_004718.3;NP_828850.1;SARS-CoV_orf1a;2266 | 0 | 0 |

| | | |
|---|---|---|
| NC_004718.3;NP_828850.1;SARS-CoV_orf1a;2267 | 0 | 0 |
| NC_004718.3;NP_828850.1;SARS-CoV_orf1a;2268 | 0.05 | 0.025 |
| NC_004718.3;NP_828850.1;SARS-CoV_orf1a;2269 | 0 | 0 |
| NC_004718.3;NP_828850.1;SARS-CoV_orf1a;2270 | 0.05 | 0 |
| NC_004718.3;NP_828850.1;SARS-CoV_orf1a;2271 | 0.05 | 0.025 |
| NC_004718.3;NP_828850.1;SARS-CoV_orf1a;2272 | 0.05 | 0 |
| NC_004718.3;NP_828850.1;SARS-CoV_orf1a;2273 | 0.05 | 0 |
| NC_004718.3;NP_828850.1;SARS-CoV_orf1a;2274 | 0 | 0 |
| NC_004718.3;NP_828850.1;SARS-CoV_orf1a;2275 | 0 | 0 |
| NC_004718.3;NP_828850.1;SARS-CoV_orf1a;2276 | 0 | 0 |
| NC_004718.3;NP_828850.1;SARS-CoV_orf1a;2277 | 0 | 0 |
| NC_004718.3;NP_828850.1;SARS-CoV_orf1a;2278 | 0 | 0 |
| NC_004718.3;NP_828850.1;SARS-CoV_orf1a;2279 | 0 | 0 |
| NC_004718.3;NP_828850.1;SARS-CoV_orf1a;2280 | 0 | 0 |
| NC_004718.3;NP_828850.1;SARS-CoV_orf1a;2281 | 0 | 0 |
| NC_004718.3;NP_828850.1;SARS-CoV_orf1a;2282 | 0 | 0 |
| NC_004718.3;NP_828850.1;SARS-CoV_orf1a;2283 | 0 | 0 |
| NC_004718.3;NP_828850.1;SARS-CoV_orf1a;2284 | 0 | 0 |
| NC_004718.3;NP_828850.1;SARS-CoV_orf1a;2285 | 0 | 0 |
| NC_004718.3;NP_828850.1;SARS-CoV_orf1a;2286 | 0 | 0 |
| NC_004718.3;NP_828850.1;SARS-CoV_orf1a;2287 | 0.05 | 0 |
| NC_004718.3;NP_828850.1;SARS-CoV_orf1a;2288 | 0 | 0 |
| NC_004718.3;NP_828850.1;SARS-CoV_orf1a;2289 | 0 | 0 |
| NC_004718.3;NP_828850.1;SARS-CoV_orf1a;2290 | 0 | 0 |
| NC_004718.3;NP_828850.1;SARS-CoV_orf1a;2291 | 0 | 0 |
| NC_004718.3;NP_828850.1;SARS-CoV_orf1a;2292 | 0 | 0 |
| NC_004718.3;NP_828850.1;SARS-CoV_orf1a;2293 | 0 | 0 |
| NC_004718.3;NP_828850.1;SARS-CoV_orf1a;2294 | 0 | 0 |
| NC_004718.3;NP_828850.1;SARS-CoV_orf1a;2295 | 0 | 0 |
| NC_004718.3;NP_828850.1;SARS-CoV_orf1a;2296 | 0 | 0 |
| NC_004718.3;NP_828850.1;SARS-CoV_orf1a;2297 | 0 | 0 |
| NC_004718.3;NP_828850.1;SARS-CoV_orf1a;2298 | 0 | 0 |
| NC_004718.3;NP_828850.1;SARS-CoV_orf1a;2299 | 0 | 0 |
| NC_004718.3;NP_828850.1;SARS-CoV_orf1a;2300 | 0 | 0 |
| NC_004718.3;NP_828850.1;SARS-CoV_orf1a;2301 | 0 | 0 |
| NC_004718.3;NP_828850.1;SARS-CoV_orf1a;2302 | 0 | 0 |
| NC_004718.3;NP_828850.1;SARS-CoV_orf1a;2303 | 0 | 0 |
| NC_004718.3;NP_828850.1;SARS-CoV_orf1a;2304 | 0 | 0 |
| NC_004718.3;NP_828850.1;SARS-CoV_orf1a;2305 | 0 | 0 |
| NC_004718.3;NP_828850.1;SARS-CoV_orf1a;2306 | 0 | 0 |
| NC_004718.3;NP_828850.1;SARS-CoV_orf1a;2307 | 0 | 0 |
| NC_004718.3;NP_828850.1;SARS-CoV_orf1a;2308 | 0 | 0 |
| NC_004718.3;NP_828850.1;SARS-CoV_orf1a;2309 | 0 | 0 |

| | | |
|---|---|---|
| NC_004718.3;NP_828850.1;SARS-CoV_orf1a;2310 | 0 | 0 |
| NC_004718.3;NP_828850.1;SARS-CoV_orf1a;2311 | 0 | 0 |
| NC_004718.3;NP_828850.1;SARS-CoV_orf1a;2312 | 0 | 0 |
| NC_004718.3;NP_828850.1;SARS-CoV_orf1a;2313 | 0 | 0 |
| NC_004718.3;NP_828850.1;SARS-CoV_orf1a;2314 | 0 | 0 |
| NC_004718.3;NP_828850.1;SARS-CoV_orf1a;2315 | 0 | 0 |
| NC_004718.3;NP_828850.1;SARS-CoV_orf1a;2316 | 0 | 0 |
| NC_004718.3;NP_828850.1;SARS-CoV_orf1a;2317 | 0 | 0 |
| NC_004718.3;NP_828850.1;SARS-CoV_orf1a;2318 | 0 | 0 |
| NC_004718.3;NP_828850.1;SARS-CoV_orf1a;2319 | 0 | 0 |
| NC_004718.3;NP_828850.1;SARS-CoV_orf1a;2320 | 0 | 0 |
| NC_004718.3;NP_828850.1;SARS-CoV_orf1a;2321 | 0 | 0 |
| NC_004718.3;NP_828850.1;SARS-CoV_orf1a;2322 | 0 | 0 |
| NC_004718.3;NP_828850.1;SARS-CoV_orf1a;2323 | 0 | 0 |
| NC_004718.3;NP_828850.1;SARS-CoV_orf1a;2324 | 0 | 0 |
| NC_004718.3;NP_828850.1;SARS-CoV_orf1a;2325 | 0 | 0 |
| NC_004718.3;NP_828850.1;SARS-CoV_orf1a;2326 | 0 | 0 |
| NC_004718.3;NP_828850.1;SARS-CoV_orf1a;2327 | 0 | 0 |
| NC_004718.3;NP_828850.1;SARS-CoV_orf1a;2328 | 0 | 0 |
| NC_004718.3;NP_828850.1;SARS-CoV_orf1a;2329 | 0 | 0 |
| NC_004718.3;NP_828850.1;SARS-CoV_orf1a;2330 | 0 | 0 |
| NC_004718.3;NP_828850.1;SARS-CoV_orf1a;2331 | 0 | 0 |
| NC_004718.3;NP_828850.1;SARS-CoV_orf1a;2332 | 0 | 0 |
| NC_004718.3;NP_828850.1;SARS-CoV_orf1a;2333 | 0 | 0 |
| NC_004718.3;NP_828850.1;SARS-CoV_orf1a;2334 | 0 | 0 |
| NC_004718.3;NP_828850.1;SARS-CoV_orf1a;2335 | 0 | 0 |
| NC_004718.3;NP_828850.1;SARS-CoV_orf1a;2336 | 0 | 0 |
| NC_004718.3;NP_828850.1;SARS-CoV_orf1a;2337 | 0 | 0 |
| NC_004718.3;NP_828850.1;SARS-CoV_orf1a;2338 | 0 | 0 |
| NC_004718.3;NP_828850.1;SARS-CoV_orf1a;2339 | 0 | 0 |
| NC_004718.3;NP_828850.1;SARS-CoV_orf1a;2340 | 0 | 0 |
| NC_004718.3;NP_828850.1;SARS-CoV_orf1a;2341 | 0 | 0 |
| NC_004718.3;NP_828850.1;SARS-CoV_orf1a;2342 | 0 | 0 |
| NC_004718.3;NP_828850.1;SARS-CoV_orf1a;2343 | 0 | 0 |
| NC_004718.3;NP_828850.1;SARS-CoV_orf1a;2344 | 0 | 0 |
| NC_004718.3;NP_828850.1;SARS-CoV_orf1a;2345 | 0 | 0 |
| NC_004718.3;NP_828850.1;SARS-CoV_orf1a;2346 | 0 | 0 |
| NC_004718.3;NP_828850.1;SARS-CoV_orf1a;2347 | 0 | 0 |
| NC_004718.3;NP_828850.1;SARS-CoV_orf1a;2348 | 0 | 0 |
| NC_004718.3;NP_828850.1;SARS-CoV_orf1a;2349 | 0 | 0 |
| NC_004718.3;NP_828850.1;SARS-CoV_orf1a;2350 | 0 | 0 |
| NC_004718.3;NP_828850.1;SARS-CoV_orf1a;2351 | 0 | 0 |
| NC_004718.3;NP_828850.1;SARS-CoV_orf1a;2352 | 0 | 0 |

| | | |
|---|---|---|
| NC_004718.3;NP_828850.1;SARS-CoV_orf1a;2353 | 0 | 0 |
| NC_004718.3;NP_828850.1;SARS-CoV_orf1a;2354 | 0 | 0 |
| NC_004718.3;NP_828850.1;SARS-CoV_orf1a;2355 | 0 | 0 |
| NC_004718.3;NP_828850.1;SARS-CoV_orf1a;2356 | 0 | 0 |
| NC_004718.3;NP_828850.1;SARS-CoV_orf1a;2357 | 0 | 0 |
| NC_004718.3;NP_828850.1;SARS-CoV_orf1a;2358 | 0 | 0 |
| NC_004718.3;NP_828850.1;SARS-CoV_orf1a;2359 | 0 | 0 |
| NC_004718.3;NP_828850.1;SARS-CoV_orf1a;2360 | 0 | 0 |
| NC_004718.3;NP_828850.1;SARS-CoV_orf1a;2361 | 0 | 0 |
| NC_004718.3;NP_828850.1;SARS-CoV_orf1a;2362 | 0 | 0 |
| NC_004718.3;NP_828850.1;SARS-CoV_orf1a;2363 | 0 | 0 |
| NC_004718.3;NP_828850.1;SARS-CoV_orf1a;2364 | 0 | 0 |
| NC_004718.3;NP_828850.1;SARS-CoV_orf1a;2365 | 0 | 0 |
| NC_004718.3;NP_828850.1;SARS-CoV_orf1a;2366 | 0 | 0 |
| NC_004718.3;NP_828850.1;SARS-CoV_orf1a;2367 | 0 | 0 |
| NC_004718.3;NP_828850.1;SARS-CoV_orf1a;2368 | 0 | 0 |
| NC_004718.3;NP_828850.1;SARS-CoV_orf1a;2369 | 0 | 0 |
| NC_004718.3;NP_828850.1;SARS-CoV_orf1a;2370 | 0 | 0 |
| NC_004718.3;NP_828850.1;SARS-CoV_orf1a;2371 | 0 | 0 |
| NC_004718.3;NP_828850.1;SARS-CoV_orf1a;2372 | 0 | 0 |
| NC_004718.3;NP_828850.1;SARS-CoV_orf1a;2373 | 0 | 0 |
| NC_004718.3;NP_828850.1;SARS-CoV_orf1a;2374 | 0 | 0 |
| NC_004718.3;NP_828850.1;SARS-CoV_orf1a;2375 | 0 | 0 |
| NC_004718.3;NP_828850.1;SARS-CoV_orf1a;2376 | 0 | 0 |
| NC_004718.3;NP_828850.1;SARS-CoV_orf1a;2377 | 0 | 0 |
| NC_004718.3;NP_828850.1;SARS-CoV_orf1a;2378 | 0 | 0 |
| NC_004718.3;NP_828850.1;SARS-CoV_orf1a;2379 | 0 | 0 |
| NC_004718.3;NP_828850.1;SARS-CoV_orf1a;2380 | 0 | 0 |
| NC_004718.3;NP_828850.1;SARS-CoV_orf1a;2381 | 0 | 0 |
| NC_004718.3;NP_828850.1;SARS-CoV_orf1a;2382 | 0 | 0 |
| NC_004718.3;NP_828850.1;SARS-CoV_orf1a;2383 | 0 | 0 |
| NC_004718.3;NP_828850.1;SARS-CoV_orf1a;2384 | 0 | 0 |
| NC_004718.3;NP_828850.1;SARS-CoV_orf1a;2385 | 0 | 0 |
| NC_004718.3;NP_828850.1;SARS-CoV_orf1a;2386 | 0 | 0 |
| NC_004718.3;NP_828850.1;SARS-CoV_orf1a;2387 | 0 | 0 |
| NC_004718.3;NP_828850.1;SARS-CoV_orf1a;2388 | 0 | 0 |
| NC_004718.3;NP_828850.1;SARS-CoV_orf1a;2389 | 0 | 0 |
| NC_004718.3;NP_828850.1;SARS-CoV_orf1a;2390 | 0 | 0 |
| NC_004718.3;NP_828850.1;SARS-CoV_orf1a;2391 | 0 | 0 |
| NC_004718.3;NP_828850.1;SARS-CoV_orf1a;2392 | 0 | 0 |
| NC_004718.3;NP_828850.1;SARS-CoV_orf1a;2393 | 0 | 0 |
| NC_004718.3;NP_828850.1;SARS-CoV_orf1a;2394 | 0 | 0 |
| NC_004718.3;NP_828850.1;SARS-CoV_orf1a;2395 | 0 | 0 |

| | | |
|---|---|---|
| NC_004718.3;NP_828850.1;SARS-CoV_orf1a;2396 | 0 | 0 |
| NC_004718.3;NP_828850.1;SARS-CoV_orf1a;2397 | 0 | 0 |
| NC_004718.3;NP_828850.1;SARS-CoV_orf1a;2398 | 0 | 0 |
| NC_004718.3;NP_828850.1;SARS-CoV_orf1a;2399 | 0 | 0 |
| NC_004718.3;NP_828850.1;SARS-CoV_orf1a;2400 | 0 | 0 |
| NC_004718.3;NP_828850.1;SARS-CoV_orf1a;2401 | 0 | 0 |
| NC_004718.3;NP_828850.1;SARS-CoV_orf1a;2402 | 0 | 0 |
| NC_004718.3;NP_828850.1;SARS-CoV_orf1a;2403 | 0 | 0 |
| NC_004718.3;NP_828850.1;SARS-CoV_orf1a;2404 | 0 | 0 |
| NC_004718.3;NP_828850.1;SARS-CoV_orf1a;2405 | 0 | 0 |
| NC_004718.3;NP_828850.1;SARS-CoV_orf1a;2406 | 0 | 0 |
| NC_004718.3;NP_828850.1;SARS-CoV_orf1a;2407 | 0 | 0 |
| NC_004718.3;NP_828850.1;SARS-CoV_orf1a;2408 | 0 | 0 |
| NC_004718.3;NP_828850.1;SARS-CoV_orf1a;2409 | 0 | 0 |
| NC_004718.3;NP_828850.1;SARS-CoV_orf1a;2410 | 0 | 0 |
| NC_004718.3;NP_828850.1;SARS-CoV_orf1a;2411 | 0 | 0 |
| NC_004718.3;NP_828850.1;SARS-CoV_orf1a;2412 | 0 | 0 |
| NC_004718.3;NP_828850.1;SARS-CoV_orf1a;2413 | 0 | 0 |
| NC_004718.3;NP_828850.1;SARS-CoV_orf1a;2414 | 0 | 0 |
| NC_004718.3;NP_828850.1;SARS-CoV_orf1a;2415 | 0 | 0 |
| NC_004718.3;NP_828850.1;SARS-CoV_orf1a;2416 | 0 | 0 |
| NC_004718.3;NP_828850.1;SARS-CoV_orf1a;2417 | 0 | 0 |
| NC_004718.3;NP_828850.1;SARS-CoV_orf1a;2418 | 0 | 0 |
| NC_004718.3;NP_828850.1;SARS-CoV_orf1a;2419 | 0 | 0 |
| NC_004718.3;NP_828850.1;SARS-CoV_orf1a;2420 | 0 | 0 |
| NC_004718.3;NP_828850.1;SARS-CoV_orf1a;2421 | 0 | 0 |
| NC_004718.3;NP_828850.1;SARS-CoV_orf1a;2422 | 0 | 0 |
| NC_004718.3;NP_828850.1;SARS-CoV_orf1a;2423 | 0 | 0 |
| NC_004718.3;NP_828850.1;SARS-CoV_orf1a;2424 | 0 | 0 |
| NC_004718.3;NP_828850.1;SARS-CoV_orf1a;2425 | 0 | 0 |
| NC_004718.3;NP_828850.1;SARS-CoV_orf1a;2426 | 0 | 0 |
| NC_004718.3;NP_828850.1;SARS-CoV_orf1a;2427 | 0 | 0 |
| NC_004718.3;NP_828850.1;SARS-CoV_orf1a;2428 | 0 | 0 |
| NC_004718.3;NP_828850.1;SARS-CoV_orf1a;2429 | 0 | 0 |
| NC_004718.3;NP_828850.1;SARS-CoV_orf1a;2430 | 0 | 0 |
| NC_004718.3;NP_828850.1;SARS-CoV_orf1a;2431 | 0 | 0 |
| NC_004718.3;NP_828850.1;SARS-CoV_orf1a;2432 | 0 | 0 |
| NC_004718.3;NP_828850.1;SARS-CoV_orf1a;2433 | 0 | 0 |
| NC_004718.3;NP_828850.1;SARS-CoV_orf1a;2434 | 0 | 0 |
| NC_004718.3;NP_828850.1;SARS-CoV_orf1a;2435 | 0 | 0 |
| NC_004718.3;NP_828850.1;SARS-CoV_orf1a;2436 | 0 | 0 |
| NC_004718.3;NP_828850.1;SARS-CoV_orf1a;2437 | 0 | 0 |
| NC_004718.3;NP_828850.1;SARS-CoV_orf1a;2438 | 0 | 0 |

| | | |
|---|---|---|
| NC_004718.3;NP_828850.1;SARS-CoV_orf1a;2439 | 0 | 0 |
| NC_004718.3;NP_828850.1;SARS-CoV_orf1a;2440 | 0 | 0 |
| NC_004718.3;NP_828850.1;SARS-CoV_orf1a;2441 | 0 | 0 |
| NC_004718.3;NP_828850.1;SARS-CoV_orf1a;2442 | 0 | 0 |
| NC_004718.3;NP_828850.1;SARS-CoV_orf1a;2443 | 0 | 0 |
| NC_004718.3;NP_828850.1;SARS-CoV_orf1a;2444 | 0 | 0 |
| NC_004718.3;NP_828850.1;SARS-CoV_orf1a;2445 | 0 | 0 |
| NC_004718.3;NP_828850.1;SARS-CoV_orf1a;2446 | 0.05 | 0 |
| NC_004718.3;NP_828850.1;SARS-CoV_orf1a;2447 | 0.05 | 0.025 |
| NC_004718.3;NP_828850.1;SARS-CoV_orf1a;2448 | 0.05 | 0.025 |
| NC_004718.3;NP_828850.1;SARS-CoV_orf1a;2449 | 0 | 0 |
| NC_004718.3;NP_828850.1;SARS-CoV_orf1a;2450 | 0 | 0 |
| NC_004718.3;NP_828850.1;SARS-CoV_orf1a;2451 | 0 | 0 |
| NC_004718.3;NP_828850.1;SARS-CoV_orf1a;2452 | 0 | 0.025 |
| NC_004718.3;NP_828850.1;SARS-CoV_orf1a;2453 | 0 | 0 |
| NC_004718.3;NP_828850.1;SARS-CoV_orf1a;2454 | 0 | 0 |
| NC_004718.3;NP_828850.1;SARS-CoV_orf1a;2455 | 0 | 0.05 |
| NC_004718.3;NP_828850.1;SARS-CoV_orf1a;2456 | 0 | 0 |
| NC_004718.3;NP_828850.1;SARS-CoV_orf1a;2457 | 0 | 0 |
| NC_004718.3;NP_828850.1;SARS-CoV_orf1a;2458 | 0 | 0 |
| NC_004718.3;NP_828850.1;SARS-CoV_orf1a;2459 | 0 | 0.025 |
| NC_004718.3;NP_828850.1;SARS-CoV_orf1a;2460 | 0 | 0 |
| NC_004718.3;NP_828850.1;SARS-CoV_orf1a;2461 | 0.05 | 0.025 |
| NC_004718.3;NP_828850.1;SARS-CoV_orf1a;2462 | 0 | 0.025 |
| NC_004718.3;NP_828850.1;SARS-CoV_orf1a;2463 | 0 | 0.025 |
| NC_004718.3;NP_828850.1;SARS-CoV_orf1a;2464 | 0 | 0.025 |
| NC_004718.3;NP_828850.1;SARS-CoV_orf1a;2465 | 0 | 0.025 |
| NC_004718.3;NP_828850.1;SARS-CoV_orf1a;2466 | 0 | 0.025 |
| NC_004718.3;NP_828850.1;SARS-CoV_orf1a;2467 | 0 | 0 |
| NC_004718.3;NP_828850.1;SARS-CoV_orf1a;2468 | 0 | 0 |
| NC_004718.3;NP_828850.1;SARS-CoV_orf1a;2469 | 0 | 0 |
| NC_004718.3;NP_828850.1;SARS-CoV_orf1a;2470 | 0 | 0 |
| NC_004718.3;NP_828850.1;SARS-CoV_orf1a;2471 | 0 | 0 |
| NC_004718.3;NP_828850.1;SARS-CoV_orf1a;2472 | 0 | 0 |
| NC_004718.3;NP_828850.1;SARS-CoV_orf1a;2473 | 0 | 0 |
| NC_004718.3;NP_828850.1;SARS-CoV_orf1a;2474 | 0 | 0 |
| NC_004718.3;NP_828850.1;SARS-CoV_orf1a;2475 | 0 | 0 |
| NC_004718.3;NP_828850.1;SARS-CoV_orf1a;2476 | 0 | 0 |
| NC_004718.3;NP_828850.1;SARS-CoV_orf1a;2477 | 0 | 0 |
| NC_004718.3;NP_828850.1;SARS-CoV_orf1a;2478 | 0 | 0 |
| NC_004718.3;NP_828850.1;SARS-CoV_orf1a;2479 | 0 | 0 |
| NC_004718.3;NP_828850.1;SARS-CoV_orf1a;2480 | 0 | 0 |
| NC_004718.3;NP_828850.1;SARS-CoV_orf1a;2481 | 0 | 0.025 |

| | | |
|---|---|---|
| NC_004718.3;NP_828850.1;SARS-CoV_orf1a;2482 | 0 | 0 |
| NC_004718.3;NP_828850.1;SARS-CoV_orf1a;2483 | 0 | 0 |
| NC_004718.3;NP_828850.1;SARS-CoV_orf1a;2484 | 0 | 0 |
| NC_004718.3;NP_828850.1;SARS-CoV_orf1a;2485 | 0 | 0 |
| NC_004718.3;NP_828850.1;SARS-CoV_orf1a;2486 | 0 | 0 |
| NC_004718.3;NP_828850.1;SARS-CoV_orf1a;2487 | 0 | 0 |
| NC_004718.3;NP_828850.1;SARS-CoV_orf1a;2488 | 0 | 0 |
| NC_004718.3;NP_828850.1;SARS-CoV_orf1a;2489 | 0 | 0 |
| NC_004718.3;NP_828850.1;SARS-CoV_orf1a;2490 | 0 | 0 |
| NC_004718.3;NP_828850.1;SARS-CoV_orf1a;2491 | 0 | 0 |
| NC_004718.3;NP_828850.1;SARS-CoV_orf1a;2492 | 0 | 0 |
| NC_004718.3;NP_828850.1;SARS-CoV_orf1a;2493 | 0 | 0 |
| NC_004718.3;NP_828850.1;SARS-CoV_orf1a;2494 | 0 | 0 |
| NC_004718.3;NP_828850.1;SARS-CoV_orf1a;2495 | 0 | 0 |
| NC_004718.3;NP_828850.1;SARS-CoV_orf1a;2496 | 0 | 0 |
| NC_004718.3;NP_828850.1;SARS-CoV_orf1a;2497 | 0 | 0 |
| NC_004718.3;NP_828850.1;SARS-CoV_orf1a;2498 | 0 | 0.025 |
| NC_004718.3;NP_828850.1;SARS-CoV_orf1a;2499 | 0.05 | 0.025 |
| NC_004718.3;NP_828850.1;SARS-CoV_orf1a;2500 | 0.05 | 0.025 |
| NC_004718.3;NP_828850.1;SARS-CoV_orf1a;2501 | 0.05 | 0.025 |
| NC_004718.3;NP_828850.1;SARS-CoV_orf1a;2502 | 0.05 | 0.025 |
| NC_004718.3;NP_828850.1;SARS-CoV_orf1a;2503 | 0 | 0 |
| NC_004718.3;NP_828850.1;SARS-CoV_orf1a;2504 | 0 | 0 |
| NC_004718.3;NP_828850.1;SARS-CoV_orf1a;2505 | 0.05 | 0 |
| NC_004718.3;NP_828850.1;SARS-CoV_orf1a;2506 | 0 | 0 |
| NC_004718.3;NP_828850.1;SARS-CoV_orf1a;2507 | 0 | 0 |
| NC_004718.3;NP_828850.1;SARS-CoV_orf1a;2508 | 0 | 0 |
| NC_004718.3;NP_828850.1;SARS-CoV_orf1a;2509 | 0 | 0 |
| NC_004718.3;NP_828850.1;SARS-CoV_orf1a;2510 | 0 | 0 |
| NC_004718.3;NP_828850.1;SARS-CoV_orf1a;2511 | 0 | 0.025 |
| NC_004718.3;NP_828850.1;SARS-CoV_orf1a;2512 | 0 | 0.025 |
| NC_004718.3;NP_828850.1;SARS-CoV_orf1a;2513 | 0.05 | 0.025 |
| NC_004718.3;NP_828850.1;SARS-CoV_orf1a;2514 | 0.05 | 0.025 |
| NC_004718.3;NP_828850.1;SARS-CoV_orf1a;2515 | 0.05 | 0.025 |
| NC_004718.3;NP_828850.1;SARS-CoV_orf1a;2516 | 0.05 | 0.025 |
| NC_004718.3;NP_828850.1;SARS-CoV_orf1a;2517 | 0 | 0.025 |
| NC_004718.3;NP_828850.1;SARS-CoV_orf1a;2518 | 0.05 | 0.025 |
| NC_004718.3;NP_828850.1;SARS-CoV_orf1a;2519 | 0.05 | 0.025 |
| NC_004718.3;NP_828850.1;SARS-CoV_orf1a;2520 | 0.15 | 0.075 |
| NC_004718.3;NP_828850.1;SARS-CoV_orf1a;2521 | 0 | 0 |
| NC_004718.3;NP_828850.1;SARS-CoV_orf1a;2522 | 0 | 0.025 |
| NC_004718.3;NP_828850.1;SARS-CoV_orf1a;2523 | 0 | 0 |
| NC_004718.3;NP_828850.1;SARS-CoV_orf1a;2524 | 0 | 0 |

| | | |
|---|---|---|
| NC_004718.3;NP_828850.1;SARS-CoV_orf1a;2525 | 0.05 | 0 |
| NC_004718.3;NP_828850.1;SARS-CoV_orf1a;2526 | 0 | 0 |
| NC_004718.3;NP_828850.1;SARS-CoV_orf1a;2527 | 0.05 | 0.025 |
| NC_004718.3;NP_828850.1;SARS-CoV_orf1a;2528 | 0.05 | 0.075 |
| NC_004718.3;NP_828850.1;SARS-CoV_orf1a;2529 | 0 | 0 |
| NC_004718.3;NP_828850.1;SARS-CoV_orf1a;2530 | 0 | 0.025 |
| NC_004718.3;NP_828850.1;SARS-CoV_orf1a;2531 | 0 | 0 |
| NC_004718.3;NP_828850.1;SARS-CoV_orf1a;2532 | 0 | 0 |
| NC_004718.3;NP_828850.1;SARS-CoV_orf1a;2533 | 0 | 0 |
| NC_004718.3;NP_828850.1;SARS-CoV_orf1a;2534 | 0 | 0.025 |
| NC_004718.3;NP_828850.1;SARS-CoV_orf1a;2535 | 0 | 0 |
| NC_004718.3;NP_828850.1;SARS-CoV_orf1a;2536 | 0 | 0 |
| NC_004718.3;NP_828850.1;SARS-CoV_orf1a;2537 | 0 | 0 |
| NC_004718.3;NP_828850.1;SARS-CoV_orf1a;2538 | 0 | 0 |
| NC_004718.3;NP_828850.1;SARS-CoV_orf1a;2539 | 0 | 0 |
| NC_004718.3;NP_828850.1;SARS-CoV_orf1a;2540 | 0 | 0 |
| NC_004718.3;NP_828850.1;SARS-CoV_orf1a;2541 | 0 | 0 |
| NC_004718.3;NP_828850.1;SARS-CoV_orf1a;2542 | 0 | 0 |
| NC_004718.3;NP_828850.1;SARS-CoV_orf1a;2543 | 0 | 0 |
| NC_004718.3;NP_828850.1;SARS-CoV_orf1a;2544 | 0 | 0 |
| NC_004718.3;NP_828850.1;SARS-CoV_orf1a;2545 | 0 | 0 |
| NC_004718.3;NP_828850.1;SARS-CoV_orf1a;2546 | 0 | 0 |
| NC_004718.3;NP_828850.1;SARS-CoV_orf1a;2547 | 0 | 0.025 |
| NC_004718.3;NP_828850.1;SARS-CoV_orf1a;2548 | 0.1 | 0.1 |
| NC_004718.3;NP_828850.1;SARS-CoV_orf1a;2549 | 0 | 0.05 |
| NC_004718.3;NP_828850.1;SARS-CoV_orf1a;2550 | 0 | 0 |
| NC_004718.3;NP_828850.1;SARS-CoV_orf1a;2551 | 0 | 0 |
| NC_004718.3;NP_828850.1;SARS-CoV_orf1a;2552 | 0 | 0 |
| NC_004718.3;NP_828850.1;SARS-CoV_orf1a;2553 | 0 | 0 |
| NC_004718.3;NP_828850.1;SARS-CoV_orf1a;2554 | 0 | 0 |
| NC_004718.3;NP_828850.1;SARS-CoV_orf1a;2555 | 0 | 0 |
| NC_004718.3;NP_828850.1;SARS-CoV_orf1a;2556 | 0 | 0 |
| NC_004718.3;NP_828850.1;SARS-CoV_orf1a;2557 | 0 | 0 |
| NC_004718.3;NP_828850.1;SARS-CoV_orf1a;2558 | 0 | 0 |
| NC_004718.3;NP_828850.1;SARS-CoV_orf1a;2559 | 0 | 0 |
| NC_004718.3;NP_828850.1;SARS-CoV_orf1a;2560 | 0 | 0 |
| NC_004718.3;NP_828850.1;SARS-CoV_orf1a;2561 | 0 | 0 |
| NC_004718.3;NP_828850.1;SARS-CoV_orf1a;2562 | 0 | 0 |
| NC_004718.3;NP_828850.1;SARS-CoV_orf1a;2563 | 0.05 | 0.025 |
| NC_004718.3;NP_828850.1;SARS-CoV_orf1a;2564 | 0.05 | 0 |
| NC_004718.3;NP_828850.1;SARS-CoV_orf1a;2565 | 0 | 0 |
| NC_004718.3;NP_828850.1;SARS-CoV_orf1a;2566 | 0 | 0 |
| NC_004718.3;NP_828850.1;SARS-CoV_orf1a;2567 | 0 | 0.025 |

| | | |
|---|---|---|
| NC_004718.3;NP_828850.1;SARS-CoV_orf1a;2568 | 0 | 0.025 |
| NC_004718.3;NP_828850.1;SARS-CoV_orf1a;2569 | 0 | 0 |
| NC_004718.3;NP_828850.1;SARS-CoV_orf1a;2570 | 0 | 0.05 |
| NC_004718.3;NP_828850.1;SARS-CoV_orf1a;2571 | 0 | 0.025 |
| NC_004718.3;NP_828850.1;SARS-CoV_orf1a;2572 | 0.05 | 0.025 |
| NC_004718.3;NP_828850.1;SARS-CoV_orf1a;2573 | 0.1 | 0.05 |
| NC_004718.3;NP_828850.1;SARS-CoV_orf1a;2574 | 0 | 0.025 |
| NC_004718.3;NP_828850.1;SARS-CoV_orf1a;2575 | 0 | 0 |
| NC_004718.3;NP_828850.1;SARS-CoV_orf1a;2576 | 0 | 0 |
| NC_004718.3;NP_828850.1;SARS-CoV_orf1a;2577 | 0 | 0 |
| NC_004718.3;NP_828850.1;SARS-CoV_orf1a;2578 | 0 | 0 |
| NC_004718.3;NP_828850.1;SARS-CoV_orf1a;2579 | 0 | 0 |
| NC_004718.3;NP_828850.1;SARS-CoV_orf1a;2580 | 0 | 0 |
| NC_004718.3;NP_828850.1;SARS-CoV_orf1a;2581 | 0 | 0 |
| NC_004718.3;NP_828850.1;SARS-CoV_orf1a;2582 | 0 | 0 |
| NC_004718.3;NP_828850.1;SARS-CoV_orf1a;2583 | 0 | 0 |
| NC_004718.3;NP_828850.1;SARS-CoV_orf1a;2584 | 0 | 0.025 |
| NC_004718.3;NP_828850.1;SARS-CoV_orf1a;2585 | 0 | 0 |
| NC_004718.3;NP_828850.1;SARS-CoV_orf1a;2586 | 0.1 | 0.05 |
| NC_004718.3;NP_828850.1;SARS-CoV_orf1a;2587 | 0.1 | 0.1 |
| NC_004718.3;NP_828850.1;SARS-CoV_orf1a;2588 | 0.15 | 0.1 |
| NC_004718.3;NP_828850.1;SARS-CoV_orf1a;2589 | 0.25 | 0.125 |
| NC_004718.3;NP_828850.1;SARS-CoV_orf1a;2590 | 0.25 | 0.125 |
| NC_004718.3;NP_828850.1;SARS-CoV_orf1a;2591 | 0.2 | 0.1 |
| NC_004718.3;NP_828850.1;SARS-CoV_orf1a;2592 | 0.25 | 0.1 |
| NC_004718.3;NP_828850.1;SARS-CoV_orf1a;2593 | 0.2 | 0.075 |
| NC_004718.3;NP_828850.1;SARS-CoV_orf1a;2594 | 0.2 | 0.075 |
| NC_004718.3;NP_828850.1;SARS-CoV_orf1a;2595 | 0.15 | 0.05 |
| NC_004718.3;NP_828850.1;SARS-CoV_orf1a;2596 | 0.1 | 0.05 |
| NC_004718.3;NP_828850.1;SARS-CoV_orf1a;2597 | 0 | 0 |
| NC_004718.3;NP_828850.1;SARS-CoV_orf1a;2598 | 0 | 0 |
| NC_004718.3;NP_828850.1;SARS-CoV_orf1a;2599 | 0.05 | 0 |
| NC_004718.3;NP_828850.1;SARS-CoV_orf1a;2600 | 0 | 0 |
| NC_004718.3;NP_828850.1;SARS-CoV_orf1a;2601 | 0 | 0 |
| NC_004718.3;NP_828850.1;SARS-CoV_orf1a;2602 | 0 | 0 |
| NC_004718.3;NP_828850.1;SARS-CoV_orf1a;2603 | 0 | 0 |
| NC_004718.3;NP_828850.1;SARS-CoV_orf1a;2604 | 0 | 0 |
| NC_004718.3;NP_828850.1;SARS-CoV_orf1a;2605 | 0.05 | 0 |
| NC_004718.3;NP_828850.1;SARS-CoV_orf1a;2606 | 0 | 0 |
| NC_004718.3;NP_828850.1;SARS-CoV_orf1a;2607 | 0 | 0 |
| NC_004718.3;NP_828850.1;SARS-CoV_orf1a;2608 | 0 | 0 |
| NC_004718.3;NP_828850.1;SARS-CoV_orf1a;2609 | 0 | 0.025 |
| NC_004718.3;NP_828850.1;SARS-CoV_orf1a;2610 | 0 | 0 |

| | | |
|---|---|---|
| NC_004718.3;NP_828850.1;SARS-CoV_orf1a;2611 | 0 | 0 |
| NC_004718.3;NP_828850.1;SARS-CoV_orf1a;2612 | 0 | 0 |
| NC_004718.3;NP_828850.1;SARS-CoV_orf1a;2613 | 0 | 0.025 |
| NC_004718.3;NP_828850.1;SARS-CoV_orf1a;2614 | 0 | 0 |
| NC_004718.3;NP_828850.1;SARS-CoV_orf1a;2615 | 0.05 | 0 |
| NC_004718.3;NP_828850.1;SARS-CoV_orf1a;2616 | 0.05 | 0 |
| NC_004718.3;NP_828850.1;SARS-CoV_orf1a;2617 | 0 | 0 |
| NC_004718.3;NP_828850.1;SARS-CoV_orf1a;2618 | 0 | 0 |
| NC_004718.3;NP_828850.1;SARS-CoV_orf1a;2619 | 0 | 0 |
| NC_004718.3;NP_828850.1;SARS-CoV_orf1a;2620 | 0 | 0 |
| NC_004718.3;NP_828850.1;SARS-CoV_orf1a;2621 | 0 | 0 |
| NC_004718.3;NP_828850.1;SARS-CoV_orf1a;2622 | 0 | 0 |
| NC_004718.3;NP_828850.1;SARS-CoV_orf1a;2623 | 0 | 0 |
| NC_004718.3;NP_828850.1;SARS-CoV_orf1a;2624 | 0 | 0 |
| NC_004718.3;NP_828850.1;SARS-CoV_orf1a;2625 | 0 | 0.025 |
| NC_004718.3;NP_828850.1;SARS-CoV_orf1a;2626 | 0 | 0 |
| NC_004718.3;NP_828850.1;SARS-CoV_orf1a;2627 | 0 | 0 |
| NC_004718.3;NP_828850.1;SARS-CoV_orf1a;2628 | 0 | 0 |
| NC_004718.3;NP_828850.1;SARS-CoV_orf1a;2629 | 0 | 0 |
| NC_004718.3;NP_828850.1;SARS-CoV_orf1a;2630 | 0 | 0 |
| NC_004718.3;NP_828850.1;SARS-CoV_orf1a;2631 | 0 | 0 |
| NC_004718.3;NP_828850.1;SARS-CoV_orf1a;2632 | 0 | 0 |
| NC_004718.3;NP_828850.1;SARS-CoV_orf1a;2633 | 0 | 0 |
| NC_004718.3;NP_828850.1;SARS-CoV_orf1a;2634 | 0 | 0 |
| NC_004718.3;NP_828850.1;SARS-CoV_orf1a;2635 | 0 | 0 |
| NC_004718.3;NP_828850.1;SARS-CoV_orf1a;2636 | 0 | 0 |
| NC_004718.3;NP_828850.1;SARS-CoV_orf1a;2637 | 0 | 0 |
| NC_004718.3;NP_828850.1;SARS-CoV_orf1a;2638 | 0.05 | 0 |
| NC_004718.3;NP_828850.1;SARS-CoV_orf1a;2639 | 0 | 0 |
| NC_004718.3;NP_828850.1;SARS-CoV_orf1a;2640 | 0.05 | 0.05 |
| NC_004718.3;NP_828850.1;SARS-CoV_orf1a;2641 | 0.05 | 0.05 |
| NC_004718.3;NP_828850.1;SARS-CoV_orf1a;2642 | 0 | 0 |
| NC_004718.3;NP_828850.1;SARS-CoV_orf1a;2643 | 0 | 0 |
| NC_004718.3;NP_828850.1;SARS-CoV_orf1a;2644 | 0 | 0 |
| NC_004718.3;NP_828850.1;SARS-CoV_orf1a;2645 | 0 | 0 |
| NC_004718.3;NP_828850.1;SARS-CoV_orf1a;2646 | 0 | 0 |
| NC_004718.3;NP_828850.1;SARS-CoV_orf1a;2647 | 0 | 0 |
| NC_004718.3;NP_828850.1;SARS-CoV_orf1a;2648 | 0 | 0 |
| NC_004718.3;NP_828850.1;SARS-CoV_orf1a;2649 | 0 | 0 |
| NC_004718.3;NP_828850.1;SARS-CoV_orf1a;2650 | 0 | 0 |
| NC_004718.3;NP_828850.1;SARS-CoV_orf1a;2651 | 0 | 0.025 |
| NC_004718.3;NP_828850.1;SARS-CoV_orf1a;2652 | 0.05 | 0.075 |
| NC_004718.3;NP_828850.1;SARS-CoV_orf1a;2653 | 0.05 | 0.025 |

| | | |
|---|---|---|
| NC_004718.3;NP_828850.1;SARS-CoV_orf1a;2654 | 0.05 | 0.05 |
| NC_004718.3;NP_828850.1;SARS-CoV_orf1a;2655 | 0 | 0 |
| NC_004718.3;NP_828850.1;SARS-CoV_orf1a;2656 | 0 | 0 |
| NC_004718.3;NP_828850.1;SARS-CoV_orf1a;2657 | 0 | 0 |
| NC_004718.3;NP_828850.1;SARS-CoV_orf1a;2658 | 0 | 0.025 |
| NC_004718.3;NP_828850.1;SARS-CoV_orf1a;2659 | 0 | 0 |
| NC_004718.3;NP_828850.1;SARS-CoV_orf1a;2660 | 0 | 0 |
| NC_004718.3;NP_828850.1;SARS-CoV_orf1a;2661 | 0.05 | 0 |
| NC_004718.3;NP_828850.1;SARS-CoV_orf1a;2662 | 0.05 | 0 |
| NC_004718.3;NP_828850.1;SARS-CoV_orf1a;2663 | 0 | 0 |
| NC_004718.3;NP_828850.1;SARS-CoV_orf1a;2664 | 0 | 0 |
| NC_004718.3;NP_828850.1;SARS-CoV_orf1a;2665 | 0 | 0 |
| NC_004718.3;NP_828850.1;SARS-CoV_orf1a;2666 | 0.05 | 0.05 |
| NC_004718.3;NP_828850.1;SARS-CoV_orf1a;2667 | 0 | 0 |
| NC_004718.3;NP_828850.1;SARS-CoV_orf1a;2668 | 0.05 | 0 |
| NC_004718.3;NP_828850.1;SARS-CoV_orf1a;2669 | 0 | 0 |
| NC_004718.3;NP_828850.1;SARS-CoV_orf1a;2670 | 0.05 | 0 |
| NC_004718.3;NP_828850.1;SARS-CoV_orf1a;2671 | 0 | 0 |
| NC_004718.3;NP_828850.1;SARS-CoV_orf1a;2672 | 0 | 0 |
| NC_004718.3;NP_828850.1;SARS-CoV_orf1a;2673 | 0 | 0 |
| NC_004718.3;NP_828850.1;SARS-CoV_orf1a;2674 | 0 | 0 |
| NC_004718.3;NP_828850.1;SARS-CoV_orf1a;2675 | 0 | 0 |
| NC_004718.3;NP_828850.1;SARS-CoV_orf1a;2676 | 0 | 0 |
| NC_004718.3;NP_828850.1;SARS-CoV_orf1a;2677 | 0 | 0 |
| NC_004718.3;NP_828850.1;SARS-CoV_orf1a;2678 | 0 | 0 |
| NC_004718.3;NP_828850.1;SARS-CoV_orf1a;2679 | 0.05 | 0 |
| NC_004718.3;NP_828850.1;SARS-CoV_orf1a;2680 | 0 | 0 |
| NC_004718.3;NP_828850.1;SARS-CoV_orf1a;2681 | 0 | 0 |
| NC_004718.3;NP_828850.1;SARS-CoV_orf1a;2682 | 0 | 0 |
| NC_004718.3;NP_828850.1;SARS-CoV_orf1a;2683 | 0 | 0 |
| NC_004718.3;NP_828850.1;SARS-CoV_orf1a;2684 | 0 | 0 |
| NC_004718.3;NP_828850.1;SARS-CoV_orf1a;2685 | 0 | 0 |
| NC_004718.3;NP_828850.1;SARS-CoV_orf1a;2686 | 0 | 0.025 |
| NC_004718.3;NP_828850.1;SARS-CoV_orf1a;2687 | 0 | 0 |
| NC_004718.3;NP_828850.1;SARS-CoV_orf1a;2688 | 0 | 0 |
| NC_004718.3;NP_828850.1;SARS-CoV_orf1a;2689 | 0 | 0.025 |
| NC_004718.3;NP_828850.1;SARS-CoV_orf1a;2690 | 0 | 0 |
| NC_004718.3;NP_828850.1;SARS-CoV_orf1a;2691 | 0 | 0.025 |
| NC_004718.3;NP_828850.1;SARS-CoV_orf1a;2692 | 0.05 | 0.05 |
| NC_004718.3;NP_828850.1;SARS-CoV_orf1a;2693 | 0 | 0 |
| NC_004718.3;NP_828850.1;SARS-CoV_orf1a;2694 | 0 | 0 |
| NC_004718.3;NP_828850.1;SARS-CoV_orf1a;2695 | 0.05 | 0 |
| NC_004718.3;NP_828850.1;SARS-CoV_orf1a;2696 | 0 | 0 |

| | | |
|---|---|---|
| NC_004718.3;NP_828850.1;SARS-CoV_orf1a;2697 | 0 | 0 |
| NC_004718.3;NP_828850.1;SARS-CoV_orf1a;2698 | 0 | 0 |
| NC_004718.3;NP_828850.1;SARS-CoV_orf1a;2699 | 0 | 0.025 |
| NC_004718.3;NP_828850.1;SARS-CoV_orf1a;2700 | 0 | 0 |
| NC_004718.3;NP_828850.1;SARS-CoV_orf1a;2701 | 0 | 0 |
| NC_004718.3;NP_828850.1;SARS-CoV_orf1a;2702 | 0 | 0 |
| NC_004718.3;NP_828850.1;SARS-CoV_orf1a;2703 | 0 | 0.025 |
| NC_004718.3;NP_828850.1;SARS-CoV_orf1a;2704 | 0 | 0 |
| NC_004718.3;NP_828850.1;SARS-CoV_orf1a;2705 | 0.05 | 0 |
| NC_004718.3;NP_828850.1;SARS-CoV_orf1a;2706 | 0.05 | 0 |
| NC_004718.3;NP_828850.1;SARS-CoV_orf1a;2707 | 0 | 0 |
| NC_004718.3;NP_828850.1;SARS-CoV_orf1a;2708 | 0 | 0 |
| NC_004718.3;NP_828850.1;SARS-CoV_orf1a;2709 | 0 | 0 |
| NC_004718.3;NP_828850.1;SARS-CoV_orf1a;2710 | 0 | 0 |
| NC_004718.3;NP_828850.1;SARS-CoV_orf1a;2711 | 0 | 0 |
| NC_004718.3;NP_828850.1;SARS-CoV_orf1a;2712 | 0 | 0 |
| NC_004718.3;NP_828850.1;SARS-CoV_orf1a;2713 | 0 | 0 |
| NC_004718.3;NP_828850.1;SARS-CoV_orf1a;2714 | 0 | 0 |
| NC_004718.3;NP_828850.1;SARS-CoV_orf1a;2715 | 0 | 0.025 |
| NC_004718.3;NP_828850.1;SARS-CoV_orf1a;2716 | 0 | 0 |
| NC_004718.3;NP_828850.1;SARS-CoV_orf1a;2717 | 0 | 0 |
| NC_004718.3;NP_828850.1;SARS-CoV_orf1a;2718 | 0 | 0 |
| NC_004718.3;NP_828850.1;SARS-CoV_orf1a;2719 | 0 | 0 |
| NC_004718.3;NP_828850.1;SARS-CoV_orf1a;2720 | 0 | 0 |
| NC_004718.3;NP_828850.1;SARS-CoV_orf1a;2721 | 0 | 0 |
| NC_004718.3;NP_828850.1;SARS-CoV_orf1a;2722 | 0 | 0 |
| NC_004718.3;NP_828850.1;SARS-CoV_orf1a;2723 | 0 | 0 |
| NC_004718.3;NP_828850.1;SARS-CoV_orf1a;2724 | 0 | 0 |
| NC_004718.3;NP_828850.1;SARS-CoV_orf1a;2725 | 0 | 0 |
| NC_004718.3;NP_828850.1;SARS-CoV_orf1a;2726 | 0 | 0 |
| NC_004718.3;NP_828850.1;SARS-CoV_orf1a;2727 | 0 | 0 |
| NC_004718.3;NP_828850.1;SARS-CoV_orf1a;2728 | 0 | 0 |
| NC_004718.3;NP_828850.1;SARS-CoV_orf1a;2729 | 0 | 0 |
| NC_004718.3;NP_828850.1;SARS-CoV_orf1a;2730 | 0 | 0 |
| NC_004718.3;NP_828850.1;SARS-CoV_orf1a;2731 | 0 | 0 |
| NC_004718.3;NP_828850.1;SARS-CoV_orf1a;2732 | 0 | 0 |
| NC_004718.3;NP_828850.1;SARS-CoV_orf1a;2733 | 0 | 0 |
| NC_004718.3;NP_828850.1;SARS-CoV_orf1a;2734 | 0 | 0 |
| NC_004718.3;NP_828850.1;SARS-CoV_orf1a;2735 | 0 | 0 |
| NC_004718.3;NP_828850.1;SARS-CoV_orf1a;2736 | 0 | 0 |
| NC_004718.3;NP_828850.1;SARS-CoV_orf1a;2737 | 0 | 0 |
| NC_004718.3;NP_828850.1;SARS-CoV_orf1a;2738 | 0 | 0 |
| NC_004718.3;NP_828850.1;SARS-CoV_orf1a;2739 | 0 | 0 |

| | | |
|---|---|---|
| NC_004718.3;NP_828850.1;SARS-CoV_orf1a;2740 | 0 | 0 |
| NC_004718.3;NP_828850.1;SARS-CoV_orf1a;2741 | 0 | 0 |
| NC_004718.3;NP_828850.1;SARS-CoV_orf1a;2742 | 0 | 0 |
| NC_004718.3;NP_828850.1;SARS-CoV_orf1a;2743 | 0 | 0 |
| NC_004718.3;NP_828850.1;SARS-CoV_orf1a;2744 | 0 | 0 |
| NC_004718.3;NP_828850.1;SARS-CoV_orf1a;2745 | 0 | 0 |
| NC_004718.3;NP_828850.1;SARS-CoV_orf1a;2746 | 0 | 0 |
| NC_004718.3;NP_828850.1;SARS-CoV_orf1a;2747 | 0 | 0 |
| NC_004718.3;NP_828850.1;SARS-CoV_orf1a;2748 | 0 | 0 |
| NC_004718.3;NP_828850.1;SARS-CoV_orf1a;2749 | 0 | 0 |
| NC_004718.3;NP_828850.1;SARS-CoV_orf1a;2750 | 0 | 0 |
| NC_004718.3;NP_828850.1;SARS-CoV_orf1a;2751 | 0 | 0 |
| NC_004718.3;NP_828850.1;SARS-CoV_orf1a;2752 | 0 | 0 |
| NC_004718.3;NP_828850.1;SARS-CoV_orf1a;2753 | 0 | 0 |
| NC_004718.3;NP_828850.1;SARS-CoV_orf1a;2754 | 0 | 0 |
| NC_004718.3;NP_828850.1;SARS-CoV_orf1a;2755 | 0 | 0 |
| NC_004718.3;NP_828850.1;SARS-CoV_orf1a;2756 | 0 | 0 |
| NC_004718.3;NP_828850.1;SARS-CoV_orf1a;2757 | 0 | 0 |
| NC_004718.3;NP_828850.1;SARS-CoV_orf1a;2758 | 0 | 0 |
| NC_004718.3;NP_828850.1;SARS-CoV_orf1a;2759 | 0 | 0 |
| NC_004718.3;NP_828850.1;SARS-CoV_orf1a;2760 | 0 | 0 |
| NC_004718.3;NP_828850.1;SARS-CoV_orf1a;2761 | 0 | 0 |
| NC_004718.3;NP_828850.1;SARS-CoV_orf1a;2762 | 0 | 0 |
| NC_004718.3;NP_828850.1;SARS-CoV_orf1a;2763 | 0 | 0 |
| NC_004718.3;NP_828850.1;SARS-CoV_orf1a;2764 | 0 | 0 |
| NC_004718.3;NP_828850.1;SARS-CoV_orf1a;2765 | 0 | 0 |
| NC_004718.3;NP_828850.1;SARS-CoV_orf1a;2766 | 0 | 0 |
| NC_004718.3;NP_828850.1;SARS-CoV_orf1a;2767 | 0.05 | 0 |
| NC_004718.3;NP_828850.1;SARS-CoV_orf1a;2768 | 0.05 | 0 |
| NC_004718.3;NP_828850.1;SARS-CoV_orf1a;2769 | 0 | 0 |
| NC_004718.3;NP_828850.1;SARS-CoV_orf1a;2770 | 0.05 | 0 |
| NC_004718.3;NP_828850.1;SARS-CoV_orf1a;2771 | 0 | 0 |
| NC_004718.3;NP_828850.1;SARS-CoV_orf1a;2772 | 0 | 0 |
| NC_004718.3;NP_828850.1;SARS-CoV_orf1a;2773 | 0.1 | 0 |
| NC_004718.3;NP_828850.1;SARS-CoV_orf1a;2774 | 0.05 | 0.05 |
| NC_004718.3;NP_828850.1;SARS-CoV_orf1a;2775 | 0 | 0.05 |
| NC_004718.3;NP_828850.1;SARS-CoV_orf1a;2776 | 0.05 | 0.05 |
| NC_004718.3;NP_828850.1;SARS-CoV_orf1a;2777 | 0.1 | 0.05 |
| NC_004718.3;NP_828850.1;SARS-CoV_orf1a;2778 | 0.1 | 0.075 |
| NC_004718.3;NP_828850.1;SARS-CoV_orf1a;2779 | 0.15 | 0.05 |
| NC_004718.3;NP_828850.1;SARS-CoV_orf1a;2780 | 0.15 | 0.05 |
| NC_004718.3;NP_828850.1;SARS-CoV_orf1a;2781 | 0.1 | 0.1 |
| NC_004718.3;NP_828850.1;SARS-CoV_orf1a;2782 | 0.1 | 0.075 |

| | | |
|---|---|---|
| NC_004718.3;NP_828850.1;SARS-CoV_orf1a;2783 | 0.05 | 0 |
| NC_004718.3;NP_828850.1;SARS-CoV_orf1a;2784 | 0.05 | 0 |
| NC_004718.3;NP_828850.1;SARS-CoV_orf1a;2785 | 0 | 0 |
| NC_004718.3;NP_828850.1;SARS-CoV_orf1a;2786 | 0 | 0 |
| NC_004718.3;NP_828850.1;SARS-CoV_orf1a;2787 | 0 | 0 |
| NC_004718.3;NP_828850.1;SARS-CoV_orf1a;2788 | 0.1 | 0 |
| NC_004718.3;NP_828850.1;SARS-CoV_orf1a;2789 | 0 | 0 |
| NC_004718.3;NP_828850.1;SARS-CoV_orf1a;2790 | 0 | 0 |
| NC_004718.3;NP_828850.1;SARS-CoV_orf1a;2791 | 0 | 0 |
| NC_004718.3;NP_828850.1;SARS-CoV_orf1a;2792 | 0.15 | 0.075 |
| NC_004718.3;NP_828850.1;SARS-CoV_orf1a;2793 | 0 | 0 |
| NC_004718.3;NP_828850.1;SARS-CoV_orf1a;2794 | 0 | 0.025 |
| NC_004718.3;NP_828850.1;SARS-CoV_orf1a;2795 | 0 | 0 |
| NC_004718.3;NP_828850.1;SARS-CoV_orf1a;2796 | 0 | 0 |
| NC_004718.3;NP_828850.1;SARS-CoV_orf1a;2797 | 0 | 0 |
| NC_004718.3;NP_828850.1;SARS-CoV_orf1a;2798 | 0.05 | 0.025 |
| NC_004718.3;NP_828850.1;SARS-CoV_orf1a;2799 | 0.05 | 0.025 |
| NC_004718.3;NP_828850.1;SARS-CoV_orf1a;2800 | 0 | 0 |
| NC_004718.3;NP_828850.1;SARS-CoV_orf1a;2801 | 0 | 0 |
| NC_004718.3;NP_828850.1;SARS-CoV_orf1a;2802 | 0 | 0 |
| NC_004718.3;NP_828850.1;SARS-CoV_orf1a;2803 | 0 | 0 |
| NC_004718.3;NP_828850.1;SARS-CoV_orf1a;2804 | 0 | 0 |
| NC_004718.3;NP_828850.1;SARS-CoV_orf1a;2805 | 0.05 | 0 |
| NC_004718.3;NP_828850.1;SARS-CoV_orf1a;2806 | 0 | 0 |
| NC_004718.3;NP_828850.1;SARS-CoV_orf1a;2807 | 0 | 0 |
| NC_004718.3;NP_828850.1;SARS-CoV_orf1a;2808 | 0 | 0 |
| NC_004718.3;NP_828850.1;SARS-CoV_orf1a;2809 | 0 | 0 |
| NC_004718.3;NP_828850.1;SARS-CoV_orf1a;2810 | 0 | 0.025 |
| NC_004718.3;NP_828850.1;SARS-CoV_orf1a;2811 | 0.05 | 0 |
| NC_004718.3;NP_828850.1;SARS-CoV_orf1a;2812 | 0.05 | 0 |
| NC_004718.3;NP_828850.1;SARS-CoV_orf1a;2813 | 0 | 0 |
| NC_004718.3;NP_828850.1;SARS-CoV_orf1a;2814 | 0.05 | 0 |
| NC_004718.3;NP_828850.1;SARS-CoV_orf1a;2815 | 0.05 | 0 |
| NC_004718.3;NP_828850.1;SARS-CoV_orf1a;2816 | 0 | 0 |
| NC_004718.3;NP_828850.1;SARS-CoV_orf1a;2817 | 0 | 0 |
| NC_004718.3;NP_828850.1;SARS-CoV_orf1a;2818 | 0.05 | 0.025 |
| NC_004718.3;NP_828850.1;SARS-CoV_orf1a;2819 | 0.1 | 0 |
| NC_004718.3;NP_828850.1;SARS-CoV_orf1a;2820 | 0.1 | 0 |
| NC_004718.3;NP_828850.1;SARS-CoV_orf1a;2821 | 0.05 | 0.05 |
| NC_004718.3;NP_828850.1;SARS-CoV_orf1a;2822 | 0 | 0 |
| NC_004718.3;NP_828850.1;SARS-CoV_orf1a;2823 | 0 | 0 |
| NC_004718.3;NP_828850.1;SARS-CoV_orf1a;2824 | 0 | 0 |
| NC_004718.3;NP_828850.1;SARS-CoV_orf1a;2825 | 0 | 0 |

| | | |
|---|---|---|
| NC_004718.3;NP_828850.1;SARS-CoV_orf1a;2826 | 0 | 0 |
| NC_004718.3;NP_828850.1;SARS-CoV_orf1a;2827 | 0 | 0 |
| NC_004718.3;NP_828850.1;SARS-CoV_orf1a;2828 | 0 | 0 |
| NC_004718.3;NP_828850.1;SARS-CoV_orf1a;2829 | 0 | 0 |
| NC_004718.3;NP_828850.1;SARS-CoV_orf1a;2830 | 0 | 0 |
| NC_004718.3;NP_828850.1;SARS-CoV_orf1a;2831 | 0 | 0 |
| NC_004718.3;NP_828850.1;SARS-CoV_orf1a;2832 | 0 | 0.025 |
| NC_004718.3;NP_828850.1;SARS-CoV_orf1a;2833 | 0 | 0 |
| NC_004718.3;NP_828850.1;SARS-CoV_orf1a;2834 | 0 | 0 |
| NC_004718.3;NP_828850.1;SARS-CoV_orf1a;2835 | 0 | 0.025 |
| NC_004718.3;NP_828850.1;SARS-CoV_orf1a;2836 | 0 | 0.025 |
| NC_004718.3;NP_828850.1;SARS-CoV_orf1a;2837 | 0 | 0 |
| NC_004718.3;NP_828850.1;SARS-CoV_orf1a;2838 | 0 | 0 |
| NC_004718.3;NP_828850.1;SARS-CoV_orf1a;2839 | 0 | 0 |
| NC_004718.3;NP_828850.1;SARS-CoV_orf1a;2840 | 0 | 0 |
| NC_004718.3;NP_828850.1;SARS-CoV_orf1a;2841 | 0 | 0 |
| NC_004718.3;NP_828850.1;SARS-CoV_orf1a;2842 | 0.05 | 0 |
| NC_004718.3;NP_828850.1;SARS-CoV_orf1a;2843 | 0.05 | 0 |
| NC_004718.3;NP_828850.1;SARS-CoV_orf1a;2844 | 0 | 0 |
| NC_004718.3;NP_828850.1;SARS-CoV_orf1a;2845 | 0 | 0 |
| NC_004718.3;NP_828850.1;SARS-CoV_orf1a;2846 | 0 | 0 |
| NC_004718.3;NP_828850.1;SARS-CoV_orf1a;2847 | 0 | 0 |
| NC_004718.3;NP_828850.1;SARS-CoV_orf1a;2848 | 0 | 0 |
| NC_004718.3;NP_828850.1;SARS-CoV_orf1a;2849 | 0 | 0 |
| NC_004718.3;NP_828850.1;SARS-CoV_orf1a;2850 | 0.05 | 0 |
| NC_004718.3;NP_828850.1;SARS-CoV_orf1a;2851 | 0 | 0 |
| NC_004718.3;NP_828850.1;SARS-CoV_orf1a;2852 | 0 | 0 |
| NC_004718.3;NP_828850.1;SARS-CoV_orf1a;2853 | 0 | 0 |
| NC_004718.3;NP_828850.1;SARS-CoV_orf1a;2854 | 0 | 0 |
| NC_004718.3;NP_828850.1;SARS-CoV_orf1a;2855 | 0 | 0 |
| NC_004718.3;NP_828850.1;SARS-CoV_orf1a;2856 | 0 | 0 |
| NC_004718.3;NP_828850.1;SARS-CoV_orf1a;2857 | 0 | 0 |
| NC_004718.3;NP_828850.1;SARS-CoV_orf1a;2858 | 0 | 0 |
| NC_004718.3;NP_828850.1;SARS-CoV_orf1a;2859 | 0 | 0 |
| NC_004718.3;NP_828850.1;SARS-CoV_orf1a;2860 | 0 | 0 |
| NC_004718.3;NP_828850.1;SARS-CoV_orf1a;2861 | 0 | 0 |
| NC_004718.3;NP_828850.1;SARS-CoV_orf1a;2862 | 0 | 0 |
| NC_004718.3;NP_828850.1;SARS-CoV_orf1a;2863 | 0 | 0 |
| NC_004718.3;NP_828850.1;SARS-CoV_orf1a;2864 | 0 | 0 |
| NC_004718.3;NP_828850.1;SARS-CoV_orf1a;2865 | 0 | 0 |
| NC_004718.3;NP_828850.1;SARS-CoV_orf1a;2866 | 0 | 0 |
| NC_004718.3;NP_828850.1;SARS-CoV_orf1a;2867 | 0 | 0 |
| NC_004718.3;NP_828850.1;SARS-CoV_orf1a;2868 | 0 | 0.025 |

| | | |
|---|---|---|
| NC_004718.3;NP_828850.1;SARS-CoV_orf1a;2869 | 0 | 0.025 |
| NC_004718.3;NP_828850.1;SARS-CoV_orf1a;2870 | 0 | 0.05 |
| NC_004718.3;NP_828850.1;SARS-CoV_orf1a;2871 | 0 | 0 |
| NC_004718.3;NP_828850.1;SARS-CoV_orf1a;2872 | 0.05 | 0 |
| NC_004718.3;NP_828850.1;SARS-CoV_orf1a;2873 | 0.05 | 0.025 |
| NC_004718.3;NP_828850.1;SARS-CoV_orf1a;2874 | 0.05 | 0.025 |
| NC_004718.3;NP_828850.1;SARS-CoV_orf1a;2875 | 0.05 | 0 |
| NC_004718.3;NP_828850.1;SARS-CoV_orf1a;2876 | 0.05 | 0 |
| NC_004718.3;NP_828850.1;SARS-CoV_orf1a;2877 | 0 | 0 |
| NC_004718.3;NP_828850.1;SARS-CoV_orf1a;2878 | 0.05 | 0 |
| NC_004718.3;NP_828850.1;SARS-CoV_orf1a;2879 | 0.05 | 0 |
| NC_004718.3;NP_828850.1;SARS-CoV_orf1a;2880 | 0.05 | 0 |
| NC_004718.3;NP_828850.1;SARS-CoV_orf1a;2881 | 0.05 | 0 |
| NC_004718.3;NP_828850.1;SARS-CoV_orf1a;2882 | 0 | 0 |
| NC_004718.3;NP_828850.1;SARS-CoV_orf1a;2883 | 0 | 0 |
| NC_004718.3;NP_828850.1;SARS-CoV_orf1a;2884 | 0 | 0 |
| NC_004718.3;NP_828850.1;SARS-CoV_orf1a;2885 | 0 | 0 |
| NC_004718.3;NP_828850.1;SARS-CoV_orf1a;2886 | 0 | 0 |
| NC_004718.3;NP_828850.1;SARS-CoV_orf1a;2887 | 0 | 0 |
| NC_004718.3;NP_828850.1;SARS-CoV_orf1a;2888 | 0 | 0 |
| NC_004718.3;NP_828850.1;SARS-CoV_orf1a;2889 | 0 | 0 |
| NC_004718.3;NP_828850.1;SARS-CoV_orf1a;2890 | 0 | 0 |
| NC_004718.3;NP_828850.1;SARS-CoV_orf1a;2891 | 0 | 0.025 |
| NC_004718.3;NP_828850.1;SARS-CoV_orf1a;2892 | 0 | 0.025 |
| NC_004718.3;NP_828850.1;SARS-CoV_orf1a;2893 | 0 | 0 |
| NC_004718.3;NP_828850.1;SARS-CoV_orf1a;2894 | 0 | 0.025 |
| NC_004718.3;NP_828850.1;SARS-CoV_orf1a;2895 | 0 | 0.025 |
| NC_004718.3;NP_828850.1;SARS-CoV_orf1a;2896 | 0 | 0 |
| NC_004718.3;NP_828850.1;SARS-CoV_orf1a;2897 | 0 | 0.025 |
| NC_004718.3;NP_828850.1;SARS-CoV_orf1a;2898 | 0 | 0.025 |
| NC_004718.3;NP_828850.1;SARS-CoV_orf1a;2899 | 0 | 0.025 |
| NC_004718.3;NP_828850.1;SARS-CoV_orf1a;2900 | 0 | 0 |
| NC_004718.3;NP_828850.1;SARS-CoV_orf1a;2901 | 0 | 0 |
| NC_004718.3;NP_828850.1;SARS-CoV_orf1a;2902 | 0 | 0 |
| NC_004718.3;NP_828850.1;SARS-CoV_orf1a;2903 | 0 | 0 |
| NC_004718.3;NP_828850.1;SARS-CoV_orf1a;2904 | 0 | 0 |
| NC_004718.3;NP_828850.1;SARS-CoV_orf1a;2905 | 0 | 0 |
| NC_004718.3;NP_828850.1;SARS-CoV_orf1a;2906 | 0 | 0 |
| NC_004718.3;NP_828850.1;SARS-CoV_orf1a;2907 | 0 | 0 |
| NC_004718.3;NP_828850.1;SARS-CoV_orf1a;2908 | 0 | 0 |
| NC_004718.3;NP_828850.1;SARS-CoV_orf1a;2909 | 0 | 0 |
| NC_004718.3;NP_828850.1;SARS-CoV_orf1a;2910 | 0 | 0 |
| NC_004718.3;NP_828850.1;SARS-CoV_orf1a;2911 | 0 | 0 |

| | | |
|---|---|---|
| NC_004718.3;NP_828850.1;SARS-CoV_orf1a;2912 | 0 | 0 |
| NC_004718.3;NP_828850.1;SARS-CoV_orf1a;2913 | 0 | 0 |
| NC_004718.3;NP_828850.1;SARS-CoV_orf1a;2914 | 0 | 0.025 |
| NC_004718.3;NP_828850.1;SARS-CoV_orf1a;2915 | 0 | 0.05 |
| NC_004718.3;NP_828850.1;SARS-CoV_orf1a;2916 | 0.25 | 0.15 |
| NC_004718.3;NP_828850.1;SARS-CoV_orf1a;2917 | 0.15 | 0.175 |
| NC_004718.3;NP_828850.1;SARS-CoV_orf1a;2918 | 0.1 | 0.025 |
| NC_004718.3;NP_828850.1;SARS-CoV_orf1a;2919 | 0.1 | 0.05 |
| NC_004718.3;NP_828850.1;SARS-CoV_orf1a;2920 | 0.1 | 0.025 |
| NC_004718.3;NP_828850.1;SARS-CoV_orf1a;2921 | 0.15 | 0.05 |
| NC_004718.3;NP_828850.1;SARS-CoV_orf1a;2922 | 0.1 | 0.125 |
| NC_004718.3;NP_828850.1;SARS-CoV_orf1a;2923 | 0.1 | 0.025 |
| NC_004718.3;NP_828850.1;SARS-CoV_orf1a;2924 | 0.05 | 0.025 |
| NC_004718.3;NP_828850.1;SARS-CoV_orf1a;2925 | 0.05 | 0.05 |
| NC_004718.3;NP_828850.1;SARS-CoV_orf1a;2926 | 0.05 | 0.025 |
| NC_004718.3;NP_828850.1;SARS-CoV_orf1a;2927 | 0 | 0 |
| NC_004718.3;NP_828850.1;SARS-CoV_orf1a;2928 | 0 | 0 |
| NC_004718.3;NP_828850.1;SARS-CoV_orf1a;2929 | 0 | 0 |
| NC_004718.3;NP_828850.1;SARS-CoV_orf1a;2930 | 0 | 0 |
| NC_004718.3;NP_828850.1;SARS-CoV_orf1a;2931 | 0 | 0 |
| NC_004718.3;NP_828850.1;SARS-CoV_orf1a;2932 | 0 | 0 |
| NC_004718.3;NP_828850.1;SARS-CoV_orf1a;2933 | 0 | 0 |
| NC_004718.3;NP_828850.1;SARS-CoV_orf1a;2934 | 0 | 0 |
| NC_004718.3;NP_828850.1;SARS-CoV_orf1a;2935 | 0 | 0 |
| NC_004718.3;NP_828850.1;SARS-CoV_orf1a;2936 | 0 | 0 |
| NC_004718.3;NP_828850.1;SARS-CoV_orf1a;2937 | 0 | 0.025 |
| NC_004718.3;NP_828850.1;SARS-CoV_orf1a;2938 | 0 | 0 |
| NC_004718.3;NP_828850.1;SARS-CoV_orf1a;2939 | 0 | 0 |
| NC_004718.3;NP_828850.1;SARS-CoV_orf1a;2940 | 0 | 0 |
| NC_004718.3;NP_828850.1;SARS-CoV_orf1a;2941 | 0 | 0 |
| NC_004718.3;NP_828850.1;SARS-CoV_orf1a;2942 | 0 | 0 |
| NC_004718.3;NP_828850.1;SARS-CoV_orf1a;2943 | 0 | 0 |
| NC_004718.3;NP_828850.1;SARS-CoV_orf1a;2944 | 0 | 0 |
| NC_004718.3;NP_828850.1;SARS-CoV_orf1a;2945 | 0 | 0.025 |
| NC_004718.3;NP_828850.1;SARS-CoV_orf1a;2946 | 0 | 0.025 |
| NC_004718.3;NP_828850.1;SARS-CoV_orf1a;2947 | 0 | 0.025 |
| NC_004718.3;NP_828850.1;SARS-CoV_orf1a;2948 | 0.05 | 0.075 |
| NC_004718.3;NP_828850.1;SARS-CoV_orf1a;2949 | 0 | 0.025 |
| NC_004718.3;NP_828850.1;SARS-CoV_orf1a;2950 | 0 | 0.025 |
| NC_004718.3;NP_828850.1;SARS-CoV_orf1a;2951 | 0 | 0.025 |
| NC_004718.3;NP_828850.1;SARS-CoV_orf1a;2952 | 0 | 0.025 |
| NC_004718.3;NP_828850.1;SARS-CoV_orf1a;2953 | 0 | 0 |
| NC_004718.3;NP_828850.1;SARS-CoV_orf1a;2954 | 0 | 0 |

| | | |
|---|---|---|
| NC_004718.3;NP_828850.1;SARS-CoV_orf1a;2955 | 0 | 0 |
| NC_004718.3;NP_828850.1;SARS-CoV_orf1a;2956 | 0 | 0 |
| NC_004718.3;NP_828850.1;SARS-CoV_orf1a;2957 | 0 | 0 |
| NC_004718.3;NP_828850.1;SARS-CoV_orf1a;2958 | 0 | 0 |
| NC_004718.3;NP_828850.1;SARS-CoV_orf1a;2959 | 0 | 0 |
| NC_004718.3;NP_828850.1;SARS-CoV_orf1a;2960 | 0 | 0 |
| NC_004718.3;NP_828850.1;SARS-CoV_orf1a;2961 | 0 | 0 |
| NC_004718.3;NP_828850.1;SARS-CoV_orf1a;2962 | 0 | 0 |
| NC_004718.3;NP_828850.1;SARS-CoV_orf1a;2963 | 0 | 0 |
| NC_004718.3;NP_828850.1;SARS-CoV_orf1a;2964 | 0 | 0 |
| NC_004718.3;NP_828850.1;SARS-CoV_orf1a;2965 | 0 | 0 |
| NC_004718.3;NP_828850.1;SARS-CoV_orf1a;2966 | 0 | 0 |
| NC_004718.3;NP_828850.1;SARS-CoV_orf1a;2967 | 0 | 0 |
| NC_004718.3;NP_828850.1;SARS-CoV_orf1a;2968 | 0 | 0 |
| NC_004718.3;NP_828850.1;SARS-CoV_orf1a;2969 | 0 | 0 |
| NC_004718.3;NP_828850.1;SARS-CoV_orf1a;2970 | 0 | 0 |
| NC_004718.3;NP_828850.1;SARS-CoV_orf1a;2971 | 0 | 0 |
| NC_004718.3;NP_828850.1;SARS-CoV_orf1a;2972 | 0 | 0 |
| NC_004718.3;NP_828850.1;SARS-CoV_orf1a;2973 | 0 | 0 |
| NC_004718.3;NP_828850.1;SARS-CoV_orf1a;2974 | 0 | 0 |
| NC_004718.3;NP_828850.1;SARS-CoV_orf1a;2975 | 0 | 0 |
| NC_004718.3;NP_828850.1;SARS-CoV_orf1a;2976 | 0 | 0 |
| NC_004718.3;NP_828850.1;SARS-CoV_orf1a;2977 | 0 | 0 |
| NC_004718.3;NP_828850.1;SARS-CoV_orf1a;2978 | 0 | 0 |
| NC_004718.3;NP_828850.1;SARS-CoV_orf1a;2979 | 0 | 0 |
| NC_004718.3;NP_828850.1;SARS-CoV_orf1a;2980 | 0 | 0 |
| NC_004718.3;NP_828850.1;SARS-CoV_orf1a;2981 | 0 | 0 |
| NC_004718.3;NP_828850.1;SARS-CoV_orf1a;2982 | 0 | 0 |
| NC_004718.3;NP_828850.1;SARS-CoV_orf1a;2983 | 0 | 0 |
| NC_004718.3;NP_828850.1;SARS-CoV_orf1a;2984 | 0 | 0 |
| NC_004718.3;NP_828850.1;SARS-CoV_orf1a;2985 | 0 | 0 |
| NC_004718.3;NP_828850.1;SARS-CoV_orf1a;2986 | 0 | 0 |
| NC_004718.3;NP_828850.1;SARS-CoV_orf1a;2987 | 0 | 0.025 |
| NC_004718.3;NP_828850.1;SARS-CoV_orf1a;2988 | 0 | 0.025 |
| NC_004718.3;NP_828850.1;SARS-CoV_orf1a;2989 | 0 | 0.025 |
| NC_004718.3;NP_828850.1;SARS-CoV_orf1a;2990 | 0 | 0.05 |
| NC_004718.3;NP_828850.1;SARS-CoV_orf1a;2991 | 0 | 0.05 |
| NC_004718.3;NP_828850.1;SARS-CoV_orf1a;2992 | 0.05 | 0.075 |
| NC_004718.3;NP_828850.1;SARS-CoV_orf1a;2993 | 0 | 0 |
| NC_004718.3;NP_828850.1;SARS-CoV_orf1a;2994 | 0 | 0 |
| NC_004718.3;NP_828850.1;SARS-CoV_orf1a;2995 | 0 | 0 |
| NC_004718.3;NP_828850.1;SARS-CoV_orf1a;2996 | 0 | 0 |
| NC_004718.3;NP_828850.1;SARS-CoV_orf1a;2997 | 0.05 | 0.05 |

| | | |
|---|---|---|
| NC_004718.3;NP_828850.1;SARS-CoV_orf1a;2998 | 0.05 | 0 |
| NC_004718.3;NP_828850.1;SARS-CoV_orf1a;2999 | 0 | 0 |
| NC_004718.3;NP_828850.1;SARS-CoV_orf1a;3000 | 0 | 0 |
| NC_004718.3;NP_828850.1;SARS-CoV_orf1a;3001 | 0 | 0 |
| NC_004718.3;NP_828850.1;SARS-CoV_orf1a;3002 | 0 | 0 |
| NC_004718.3;NP_828850.1;SARS-CoV_orf1a;3003 | 0 | 0 |
| NC_004718.3;NP_828850.1;SARS-CoV_orf1a;3004 | 0 | 0 |
| NC_004718.3;NP_828850.1;SARS-CoV_orf1a;3005 | 0 | 0 |
| NC_004718.3;NP_828850.1;SARS-CoV_orf1a;3006 | 0 | 0 |
| NC_004718.3;NP_828850.1;SARS-CoV_orf1a;3007 | 0 | 0 |
| NC_004718.3;NP_828850.1;SARS-CoV_orf1a;3008 | 0 | 0 |
| NC_004718.3;NP_828850.1;SARS-CoV_orf1a;3009 | 0 | 0 |
| NC_004718.3;NP_828850.1;SARS-CoV_orf1a;3010 | 0 | 0 |
| NC_004718.3;NP_828850.1;SARS-CoV_orf1a;3011 | 0 | 0 |
| NC_004718.3;NP_828850.1;SARS-CoV_orf1a;3012 | 0 | 0 |
| NC_004718.3;NP_828850.1;SARS-CoV_orf1a;3013 | 0.05 | 0 |
| NC_004718.3;NP_828850.1;SARS-CoV_orf1a;3014 | 0 | 0 |
| NC_004718.3;NP_828850.1;SARS-CoV_orf1a;3015 | 0 | 0 |
| NC_004718.3;NP_828850.1;SARS-CoV_orf1a;3016 | 0 | 0 |
| NC_004718.3;NP_828850.1;SARS-CoV_orf1a;3017 | 0 | 0 |
| NC_004718.3;NP_828850.1;SARS-CoV_orf1a;3018 | 0 | 0 |
| NC_004718.3;NP_828850.1;SARS-CoV_orf1a;3019 | 0 | 0 |
| NC_004718.3;NP_828850.1;SARS-CoV_orf1a;3020 | 0 | 0 |
| NC_004718.3;NP_828850.1;SARS-CoV_orf1a;3021 | 0 | 0 |
| NC_004718.3;NP_828850.1;SARS-CoV_orf1a;3022 | 0 | 0 |
| NC_004718.3;NP_828850.1;SARS-CoV_orf1a;3023 | 0 | 0 |
| NC_004718.3;NP_828850.1;SARS-CoV_orf1a;3024 | 0 | 0 |
| NC_004718.3;NP_828850.1;SARS-CoV_orf1a;3025 | 0 | 0 |
| NC_004718.3;NP_828850.1;SARS-CoV_orf1a;3026 | 0 | 0 |
| NC_004718.3;NP_828850.1;SARS-CoV_orf1a;3027 | 0 | 0 |
| NC_004718.3;NP_828850.1;SARS-CoV_orf1a;3028 | 0 | 0 |
| NC_004718.3;NP_828850.1;SARS-CoV_orf1a;3029 | 0 | 0 |
| NC_004718.3;NP_828850.1;SARS-CoV_orf1a;3030 | 0 | 0 |
| NC_004718.3;NP_828850.1;SARS-CoV_orf1a;3031 | 0 | 0 |
| NC_004718.3;NP_828850.1;SARS-CoV_orf1a;3032 | 0 | 0 |
| NC_004718.3;NP_828850.1;SARS-CoV_orf1a;3033 | 0 | 0 |
| NC_004718.3;NP_828850.1;SARS-CoV_orf1a;3034 | 0 | 0 |
| NC_004718.3;NP_828850.1;SARS-CoV_orf1a;3035 | 0 | 0 |
| NC_004718.3;NP_828850.1;SARS-CoV_orf1a;3036 | 0 | 0 |
| NC_004718.3;NP_828850.1;SARS-CoV_orf1a;3037 | 0 | 0 |
| NC_004718.3;NP_828850.1;SARS-CoV_orf1a;3038 | 0 | 0 |
| NC_004718.3;NP_828850.1;SARS-CoV_orf1a;3039 | 0 | 0 |
| NC_004718.3;NP_828850.1;SARS-CoV_orf1a;3040 | 0 | 0 |

| | | |
|---|---|---|
| NC_004718.3;NP_828850.1;SARS-CoV_orf1a;3041 | 0 | 0 |
| NC_004718.3;NP_828850.1;SARS-CoV_orf1a;3042 | 0.05 | 0 |
| NC_004718.3;NP_828850.1;SARS-CoV_orf1a;3043 | 0 | 0.025 |
| NC_004718.3;NP_828850.1;SARS-CoV_orf1a;3044 | 0 | 0 |
| NC_004718.3;NP_828850.1;SARS-CoV_orf1a;3045 | 0 | 0 |
| NC_004718.3;NP_828850.1;SARS-CoV_orf1a;3046 | 0 | 0 |
| NC_004718.3;NP_828850.1;SARS-CoV_orf1a;3047 | 0 | 0 |
| NC_004718.3;NP_828850.1;SARS-CoV_orf1a;3048 | 0 | 0 |
| NC_004718.3;NP_828850.1;SARS-CoV_orf1a;3049 | 0 | 0 |
| NC_004718.3;NP_828850.1;SARS-CoV_orf1a;3050 | 0 | 0 |
| NC_004718.3;NP_828850.1;SARS-CoV_orf1a;3051 | 0 | 0 |
| NC_004718.3;NP_828850.1;SARS-CoV_orf1a;3052 | 0 | 0 |
| NC_004718.3;NP_828850.1;SARS-CoV_orf1a;3053 | 0 | 0 |
| NC_004718.3;NP_828850.1;SARS-CoV_orf1a;3054 | 0 | 0 |
| NC_004718.3;NP_828850.1;SARS-CoV_orf1a;3055 | 0 | 0 |
| NC_004718.3;NP_828850.1;SARS-CoV_orf1a;3056 | 0 | 0 |
| NC_004718.3;NP_828850.1;SARS-CoV_orf1a;3057 | 0 | 0 |
| NC_004718.3;NP_828850.1;SARS-CoV_orf1a;3058 | 0 | 0 |
| NC_004718.3;NP_828850.1;SARS-CoV_orf1a;3059 | 0 | 0 |
| NC_004718.3;NP_828850.1;SARS-CoV_orf1a;3060 | 0 | 0 |
| NC_004718.3;NP_828850.1;SARS-CoV_orf1a;3061 | 0 | 0 |
| NC_004718.3;NP_828850.1;SARS-CoV_orf1a;3062 | 0 | 0 |
| NC_004718.3;NP_828850.1;SARS-CoV_orf1a;3063 | 0 | 0 |
| NC_004718.3;NP_828850.1;SARS-CoV_orf1a;3064 | 0 | 0 |
| NC_004718.3;NP_828850.1;SARS-CoV_orf1a;3065 | 0 | 0 |
| NC_004718.3;NP_828850.1;SARS-CoV_orf1a;3066 | 0 | 0 |
| NC_004718.3;NP_828850.1;SARS-CoV_orf1a;3067 | 0 | 0 |
| NC_004718.3;NP_828850.1;SARS-CoV_orf1a;3068 | 0 | 0 |
| NC_004718.3;NP_828850.1;SARS-CoV_orf1a;3069 | 0 | 0 |
| NC_004718.3;NP_828850.1;SARS-CoV_orf1a;3070 | 0 | 0 |
| NC_004718.3;NP_828850.1;SARS-CoV_orf1a;3071 | 0 | 0 |
| NC_004718.3;NP_828850.1;SARS-CoV_orf1a;3072 | 0 | 0 |
| NC_004718.3;NP_828850.1;SARS-CoV_orf1a;3073 | 0 | 0 |
| NC_004718.3;NP_828850.1;SARS-CoV_orf1a;3074 | 0 | 0 |
| NC_004718.3;NP_828850.1;SARS-CoV_orf1a;3075 | 0 | 0 |
| NC_004718.3;NP_828850.1;SARS-CoV_orf1a;3076 | 0 | 0 |
| NC_004718.3;NP_828850.1;SARS-CoV_orf1a;3077 | 0 | 0 |
| NC_004718.3;NP_828850.1;SARS-CoV_orf1a;3078 | 0 | 0 |
| NC_004718.3;NP_828850.1;SARS-CoV_orf1a;3079 | 0 | 0 |
| NC_004718.3;NP_828850.1;SARS-CoV_orf1a;3080 | 0 | 0 |
| NC_004718.3;NP_828850.1;SARS-CoV_orf1a;3081 | 0 | 0 |
| NC_004718.3;NP_828850.1;SARS-CoV_orf1a;3082 | 0 | 0 |
| NC_004718.3;NP_828850.1;SARS-CoV_orf1a;3083 | 0 | 0 |

| | | |
|---|---|---|
| NC_004718.3;NP_828850.1;SARS-CoV_orf1a;3084 | 0 | 0 |
| NC_004718.3;NP_828850.1;SARS-CoV_orf1a;3085 | 0 | 0 |
| NC_004718.3;NP_828850.1;SARS-CoV_orf1a;3086 | 0 | 0 |
| NC_004718.3;NP_828850.1;SARS-CoV_orf1a;3087 | 0 | 0 |
| NC_004718.3;NP_828850.1;SARS-CoV_orf1a;3088 | 0 | 0 |
| NC_004718.3;NP_828850.1;SARS-CoV_orf1a;3089 | 0 | 0 |
| NC_004718.3;NP_828850.1;SARS-CoV_orf1a;3090 | 0 | 0 |
| NC_004718.3;NP_828850.1;SARS-CoV_orf1a;3091 | 0 | 0 |
| NC_004718.3;NP_828850.1;SARS-CoV_orf1a;3092 | 0 | 0 |
| NC_004718.3;NP_828850.1;SARS-CoV_orf1a;3093 | 0 | 0.025 |
| NC_004718.3;NP_828850.1;SARS-CoV_orf1a;3094 | 0.05 | 0 |
| NC_004718.3;NP_828850.1;SARS-CoV_orf1a;3095 | 0 | 0 |
| NC_004718.3;NP_828850.1;SARS-CoV_orf1a;3096 | 0 | 0 |
| NC_004718.3;NP_828850.1;SARS-CoV_orf1a;3097 | 0 | 0 |
| NC_004718.3;NP_828850.1;SARS-CoV_orf1a;3098 | 0 | 0 |
| NC_004718.3;NP_828850.1;SARS-CoV_orf1a;3099 | 0 | 0 |
| NC_004718.3;NP_828850.1;SARS-CoV_orf1a;3100 | 0 | 0 |
| NC_004718.3;NP_828850.1;SARS-CoV_orf1a;3101 | 0 | 0 |
| NC_004718.3;NP_828850.1;SARS-CoV_orf1a;3102 | 0 | 0 |
| NC_004718.3;NP_828850.1;SARS-CoV_orf1a;3103 | 0 | 0 |
| NC_004718.3;NP_828850.1;SARS-CoV_orf1a;3104 | 0 | 0 |
| NC_004718.3;NP_828850.1;SARS-CoV_orf1a;3105 | 0 | 0 |
| NC_004718.3;NP_828850.1;SARS-CoV_orf1a;3106 | 0 | 0 |
| NC_004718.3;NP_828850.1;SARS-CoV_orf1a;3107 | 0 | 0 |
| NC_004718.3;NP_828850.1;SARS-CoV_orf1a;3108 | 0 | 0 |
| NC_004718.3;NP_828850.1;SARS-CoV_orf1a;3109 | 0 | 0 |
| NC_004718.3;NP_828850.1;SARS-CoV_orf1a;3110 | 0 | 0 |
| NC_004718.3;NP_828850.1;SARS-CoV_orf1a;3111 | 0 | 0 |
| NC_004718.3;NP_828850.1;SARS-CoV_orf1a;3112 | 0 | 0 |
| NC_004718.3;NP_828850.1;SARS-CoV_orf1a;3113 | 0 | 0 |
| NC_004718.3;NP_828850.1;SARS-CoV_orf1a;3114 | 0 | 0 |
| NC_004718.3;NP_828850.1;SARS-CoV_orf1a;3115 | 0 | 0 |
| NC_004718.3;NP_828850.1;SARS-CoV_orf1a;3116 | 0 | 0 |
| NC_004718.3;NP_828850.1;SARS-CoV_orf1a;3117 | 0 | 0 |
| NC_004718.3;NP_828850.1;SARS-CoV_orf1a;3118 | 0 | 0 |
| NC_004718.3;NP_828850.1;SARS-CoV_orf1a;3119 | 0 | 0 |
| NC_004718.3;NP_828850.1;SARS-CoV_orf1a;3120 | 0 | 0 |
| NC_004718.3;NP_828850.1;SARS-CoV_orf1a;3121 | 0 | 0 |
| NC_004718.3;NP_828850.1;SARS-CoV_orf1a;3122 | 0 | 0 |
| NC_004718.3;NP_828850.1;SARS-CoV_orf1a;3123 | 0 | 0 |
| NC_004718.3;NP_828850.1;SARS-CoV_orf1a;3124 | 0 | 0 |
| NC_004718.3;NP_828850.1;SARS-CoV_orf1a;3125 | 0 | 0 |
| NC_004718.3;NP_828850.1;SARS-CoV_orf1a;3126 | 0 | 0 |

| | | |
|---|---|---|
| NC_004718.3;NP_828850.1;SARS-CoV_orf1a;3127 | 0 | 0 |
| NC_004718.3;NP_828850.1;SARS-CoV_orf1a;3128 | 0 | 0 |
| NC_004718.3;NP_828850.1;SARS-CoV_orf1a;3129 | 0 | 0 |
| NC_004718.3;NP_828850.1;SARS-CoV_orf1a;3130 | 0 | 0 |
| NC_004718.3;NP_828850.1;SARS-CoV_orf1a;3131 | 0 | 0 |
| NC_004718.3;NP_828850.1;SARS-CoV_orf1a;3132 | 0 | 0 |
| NC_004718.3;NP_828850.1;SARS-CoV_orf1a;3133 | 0 | 0 |
| NC_004718.3;NP_828850.1;SARS-CoV_orf1a;3134 | 0 | 0 |
| NC_004718.3;NP_828850.1;SARS-CoV_orf1a;3135 | 0 | 0 |
| NC_004718.3;NP_828850.1;SARS-CoV_orf1a;3136 | 0 | 0 |
| NC_004718.3;NP_828850.1;SARS-CoV_orf1a;3137 | 0.05 | 0 |
| NC_004718.3;NP_828850.1;SARS-CoV_orf1a;3138 | 0 | 0 |
| NC_004718.3;NP_828850.1;SARS-CoV_orf1a;3139 | 0 | 0 |
| NC_004718.3;NP_828850.1;SARS-CoV_orf1a;3140 | 0 | 0 |
| NC_004718.3;NP_828850.1;SARS-CoV_orf1a;3141 | 0.05 | 0.025 |
| NC_004718.3;NP_828850.1;SARS-CoV_orf1a;3142 | 0 | 0 |
| NC_004718.3;NP_828850.1;SARS-CoV_orf1a;3143 | 0 | 0 |
| NC_004718.3;NP_828850.1;SARS-CoV_orf1a;3144 | 0 | 0 |
| NC_004718.3;NP_828850.1;SARS-CoV_orf1a;3145 | 0 | 0 |
| NC_004718.3;NP_828850.1;SARS-CoV_orf1a;3146 | 0 | 0 |
| NC_004718.3;NP_828850.1;SARS-CoV_orf1a;3147 | 0 | 0 |
| NC_004718.3;NP_828850.1;SARS-CoV_orf1a;3148 | 0 | 0 |
| NC_004718.3;NP_828850.1;SARS-CoV_orf1a;3149 | 0 | 0 |
| NC_004718.3;NP_828850.1;SARS-CoV_orf1a;3150 | 0 | 0 |
| NC_004718.3;NP_828850.1;SARS-CoV_orf1a;3151 | 0 | 0 |
| NC_004718.3;NP_828850.1;SARS-CoV_orf1a;3152 | 0 | 0 |
| NC_004718.3;NP_828850.1;SARS-CoV_orf1a;3153 | 0 | 0 |
| NC_004718.3;NP_828850.1;SARS-CoV_orf1a;3154 | 0 | 0 |
| NC_004718.3;NP_828850.1;SARS-CoV_orf1a;3155 | 0 | 0 |
| NC_004718.3;NP_828850.1;SARS-CoV_orf1a;3156 | 0 | 0 |
| NC_004718.3;NP_828850.1;SARS-CoV_orf1a;3157 | 0 | 0 |
| NC_004718.3;NP_828850.1;SARS-CoV_orf1a;3158 | 0 | 0 |
| NC_004718.3;NP_828850.1;SARS-CoV_orf1a;3159 | 0 | 0 |
| NC_004718.3;NP_828850.1;SARS-CoV_orf1a;3160 | 0 | 0 |
| NC_004718.3;NP_828850.1;SARS-CoV_orf1a;3161 | 0 | 0 |
| NC_004718.3;NP_828850.1;SARS-CoV_orf1a;3162 | 0 | 0 |
| NC_004718.3;NP_828850.1;SARS-CoV_orf1a;3163 | 0 | 0 |
| NC_004718.3;NP_828850.1;SARS-CoV_orf1a;3164 | 0 | 0 |
| NC_004718.3;NP_828850.1;SARS-CoV_orf1a;3165 | 0 | 0 |
| NC_004718.3;NP_828850.1;SARS-CoV_orf1a;3166 | 0 | 0 |
| NC_004718.3;NP_828850.1;SARS-CoV_orf1a;3167 | 0 | 0 |
| NC_004718.3;NP_828850.1;SARS-CoV_orf1a;3168 | 0 | 0 |
| NC_004718.3;NP_828850.1;SARS-CoV_orf1a;3169 | 0 | 0 |

| | | |
|---|---|---|
| NC_004718.3;NP_828850.1;SARS-CoV_orf1a;3170 | 0 | 0 |
| NC_004718.3;NP_828850.1;SARS-CoV_orf1a;3171 | 0 | 0.025 |
| NC_004718.3;NP_828850.1;SARS-CoV_orf1a;3172 | 0 | 0 |
| NC_004718.3;NP_828850.1;SARS-CoV_orf1a;3173 | 0 | 0 |
| NC_004718.3;NP_828850.1;SARS-CoV_orf1a;3174 | 0 | 0 |
| NC_004718.3;NP_828850.1;SARS-CoV_orf1a;3175 | 0 | 0 |
| NC_004718.3;NP_828850.1;SARS-CoV_orf1a;3176 | 0 | 0 |
| NC_004718.3;NP_828850.1;SARS-CoV_orf1a;3177 | 0 | 0 |
| NC_004718.3;NP_828850.1;SARS-CoV_orf1a;3178 | 0 | 0 |
| NC_004718.3;NP_828850.1;SARS-CoV_orf1a;3179 | 0 | 0 |
| NC_004718.3;NP_828850.1;SARS-CoV_orf1a;3180 | 0 | 0 |
| NC_004718.3;NP_828850.1;SARS-CoV_orf1a;3181 | 0 | 0 |
| NC_004718.3;NP_828850.1;SARS-CoV_orf1a;3182 | 0 | 0 |
| NC_004718.3;NP_828850.1;SARS-CoV_orf1a;3183 | 0 | 0 |
| NC_004718.3;NP_828850.1;SARS-CoV_orf1a;3184 | 0 | 0 |
| NC_004718.3;NP_828850.1;SARS-CoV_orf1a;3185 | 0 | 0 |
| NC_004718.3;NP_828850.1;SARS-CoV_orf1a;3186 | 0 | 0 |
| NC_004718.3;NP_828850.1;SARS-CoV_orf1a;3187 | 0 | 0.025 |
| NC_004718.3;NP_828850.1;SARS-CoV_orf1a;3188 | 0 | 0 |
| NC_004718.3;NP_828850.1;SARS-CoV_orf1a;3189 | 0 | 0 |
| NC_004718.3;NP_828850.1;SARS-CoV_orf1a;3190 | 0 | 0.025 |
| NC_004718.3;NP_828850.1;SARS-CoV_orf1a;3191 | 0 | 0 |
| NC_004718.3;NP_828850.1;SARS-CoV_orf1a;3192 | 0.05 | 0.025 |
| NC_004718.3;NP_828850.1;SARS-CoV_orf1a;3193 | 0 | 0 |
| NC_004718.3;NP_828850.1;SARS-CoV_orf1a;3194 | 0 | 0 |
| NC_004718.3;NP_828850.1;SARS-CoV_orf1a;3195 | 0 | 0 |
| NC_004718.3;NP_828850.1;SARS-CoV_orf1a;3196 | 0 | 0.05 |
| NC_004718.3;NP_828850.1;SARS-CoV_orf1a;3197 | 0 | 0 |
| NC_004718.3;NP_828850.1;SARS-CoV_orf1a;3198 | 0 | 0 |
| NC_004718.3;NP_828850.1;SARS-CoV_orf1a;3199 | 0 | 0 |
| NC_004718.3;NP_828850.1;SARS-CoV_orf1a;3200 | 0 | 0 |
| NC_004718.3;NP_828850.1;SARS-CoV_orf1a;3201 | 0 | 0 |
| NC_004718.3;NP_828850.1;SARS-CoV_orf1a;3202 | 0 | 0.075 |
| NC_004718.3;NP_828850.1;SARS-CoV_orf1a;3203 | 0 | 0.025 |
| NC_004718.3;NP_828850.1;SARS-CoV_orf1a;3204 | 0 | 0.025 |
| NC_004718.3;NP_828850.1;SARS-CoV_orf1a;3205 | 0 | 0.025 |
| NC_004718.3;NP_828850.1;SARS-CoV_orf1a;3206 | 0 | 0.025 |
| NC_004718.3;NP_828850.1;SARS-CoV_orf1a;3207 | 0.05 | 0.025 |
| NC_004718.3;NP_828850.1;SARS-CoV_orf1a;3208 | 0 | 0.025 |
| NC_004718.3;NP_828850.1;SARS-CoV_orf1a;3209 | 0 | 0.05 |
| NC_004718.3;NP_828850.1;SARS-CoV_orf1a;3210 | 0 | 0.025 |
| NC_004718.3;NP_828850.1;SARS-CoV_orf1a;3211 | 0 | 0 |
| NC_004718.3;NP_828850.1;SARS-CoV_orf1a;3212 | 0 | 0 |

| | | |
|---|---|---|
| NC_004718.3;NP_828850.1;SARS-CoV_orf1a;3213 | 0 | 0 |
| NC_004718.3;NP_828850.1;SARS-CoV_orf1a;3214 | 0.15 | 0.025 |
| NC_004718.3;NP_828850.1;SARS-CoV_orf1a;3215 | 0.3 | 0.05 |
| NC_004718.3;NP_828850.1;SARS-CoV_orf1a;3216 | 0.05 | 0.025 |
| NC_004718.3;NP_828850.1;SARS-CoV_orf1a;3217 | 0 | 0.025 |
| NC_004718.3;NP_828850.1;SARS-CoV_orf1a;3218 | 0.1 | 0.1 |
| NC_004718.3;NP_828850.1;SARS-CoV_orf1a;3219 | 0 | 0.025 |
| NC_004718.3;NP_828850.1;SARS-CoV_orf1a;3220 | 0 | 0 |
| NC_004718.3;NP_828850.1;SARS-CoV_orf1a;3221 | 0 | 0 |
| NC_004718.3;NP_828850.1;SARS-CoV_orf1a;3222 | 0 | 0.025 |
| NC_004718.3;NP_828850.1;SARS-CoV_orf1a;3223 | 0 | 0 |
| NC_004718.3;NP_828850.1;SARS-CoV_orf1a;3224 | 0 | 0 |
| NC_004718.3;NP_828850.1;SARS-CoV_orf1a;3225 | 0 | 0 |
| NC_004718.3;NP_828850.1;SARS-CoV_orf1a;3226 | 0 | 0 |
| NC_004718.3;NP_828850.1;SARS-CoV_orf1a;3227 | 0.05 | 0 |
| NC_004718.3;NP_828850.1;SARS-CoV_orf1a;3228 | 0 | 0 |
| NC_004718.3;NP_828850.1;SARS-CoV_orf1a;3229 | 0 | 0 |
| NC_004718.3;NP_828850.1;SARS-CoV_orf1a;3230 | 0 | 0 |
| NC_004718.3;NP_828850.1;SARS-CoV_orf1a;3231 | 0 | 0 |
| NC_004718.3;NP_828850.1;SARS-CoV_orf1a;3232 | 0 | 0 |
| NC_004718.3;NP_828850.1;SARS-CoV_orf1a;3233 | 0 | 0 |
| NC_004718.3;NP_828850.1;SARS-CoV_orf1a;3234 | 0 | 0 |
| NC_004718.3;NP_828850.1;SARS-CoV_orf1a;3235 | 0 | 0 |
| NC_004718.3;NP_828850.1;SARS-CoV_orf1a;3236 | 0 | 0 |
| NC_004718.3;NP_828850.1;SARS-CoV_orf1a;3237 | 0 | 0 |
| NC_004718.3;NP_828850.1;SARS-CoV_orf1a;3238 | 0 | 0 |
| NC_004718.3;NP_828850.1;SARS-CoV_orf1a;3239 | 0 | 0 |
| NC_004718.3;NP_828850.1;SARS-CoV_orf1a;3240 | 0 | 0 |
| NC_004718.3;NP_828850.1;SARS-CoV_orf1a;3241 | 0 | 0 |
| NC_004718.3;NP_828850.1;SARS-CoV_orf1a;3242 | 0 | 0 |
| NC_004718.3;NP_828850.1;SARS-CoV_orf1a;3243 | 0 | 0 |
| NC_004718.3;NP_828850.1;SARS-CoV_orf1a;3244 | 0 | 0 |
| NC_004718.3;NP_828850.1;SARS-CoV_orf1a;3245 | 0 | 0.025 |
| NC_004718.3;NP_828850.1;SARS-CoV_orf1a;3246 | 0 | 0 |
| NC_004718.3;NP_828850.1;SARS-CoV_orf1a;3247 | 0.05 | 0 |
| NC_004718.3;NP_828850.1;SARS-CoV_orf1a;3248 | 0 | 0 |
| NC_004718.3;NP_828850.1;SARS-CoV_orf1a;3249 | 0 | 0 |
| NC_004718.3;NP_828850.1;SARS-CoV_orf1a;3250 | 0 | 0 |
| NC_004718.3;NP_828850.1;SARS-CoV_orf1a;3251 | 0 | 0 |
| NC_004718.3;NP_828850.1;SARS-CoV_orf1a;3252 | 0 | 0 |
| NC_004718.3;NP_828850.1;SARS-CoV_orf1a;3253 | 0 | 0 |
| NC_004718.3;NP_828850.1;SARS-CoV_orf1a;3254 | 0 | 0 |
| NC_004718.3;NP_828850.1;SARS-CoV_orf1a;3255 | 0 | 0 |

| | | |
|---|---|---|
| NC_004718.3;NP_828850.1;SARS-CoV_orf1a;3256 | 0 | 0 |
| NC_004718.3;NP_828850.1;SARS-CoV_orf1a;3257 | 0 | 0 |
| NC_004718.3;NP_828850.1;SARS-CoV_orf1a;3258 | 0 | 0 |
| NC_004718.3;NP_828850.1;SARS-CoV_orf1a;3259 | 0 | 0 |
| NC_004718.3;NP_828850.1;SARS-CoV_orf1a;3260 | 0 | 0 |
| NC_004718.3;NP_828850.1;SARS-CoV_orf1a;3261 | 0 | 0 |
| NC_004718.3;NP_828850.1;SARS-CoV_orf1a;3262 | 0 | 0 |
| NC_004718.3;NP_828850.1;SARS-CoV_orf1a;3263 | 0 | 0 |
| NC_004718.3;NP_828850.1;SARS-CoV_orf1a;3264 | 0 | 0 |
| NC_004718.3;NP_828850.1;SARS-CoV_orf1a;3265 | 0 | 0 |
| NC_004718.3;NP_828850.1;SARS-CoV_orf1a;3266 | 0 | 0 |
| NC_004718.3;NP_828850.1;SARS-CoV_orf1a;3267 | 0 | 0 |
| NC_004718.3;NP_828850.1;SARS-CoV_orf1a;3268 | 0 | 0 |
| NC_004718.3;NP_828850.1;SARS-CoV_orf1a;3269 | 0 | 0 |
| NC_004718.3;NP_828850.1;SARS-CoV_orf1a;3270 | 0 | 0 |
| NC_004718.3;NP_828850.1;SARS-CoV_orf1a;3271 | 0 | 0 |
| NC_004718.3;NP_828850.1;SARS-CoV_orf1a;3272 | 0 | 0 |
| NC_004718.3;NP_828850.1;SARS-CoV_orf1a;3273 | 0 | 0 |
| NC_004718.3;NP_828850.1;SARS-CoV_orf1a;3274 | 0 | 0 |
| NC_004718.3;NP_828850.1;SARS-CoV_orf1a;3275 | 0 | 0 |
| NC_004718.3;NP_828850.1;SARS-CoV_orf1a;3276 | 0 | 0.025 |
| NC_004718.3;NP_828850.1;SARS-CoV_orf1a;3277 | 0 | 0 |
| NC_004718.3;NP_828850.1;SARS-CoV_orf1a;3278 | 0 | 0 |
| NC_004718.3;NP_828850.1;SARS-CoV_orf1a;3279 | 0.05 | 0.025 |
| NC_004718.3;NP_828850.1;SARS-CoV_orf1a;3280 | 0 | 0 |
| NC_004718.3;NP_828850.1;SARS-CoV_orf1a;3281 | 0 | 0 |
| NC_004718.3;NP_828850.1;SARS-CoV_orf1a;3282 | 0 | 0 |
| NC_004718.3;NP_828850.1;SARS-CoV_orf1a;3283 | 0 | 0 |
| NC_004718.3;NP_828850.1;SARS-CoV_orf1a;3284 | 0 | 0 |
| NC_004718.3;NP_828850.1;SARS-CoV_orf1a;3285 | 0 | 0 |
| NC_004718.3;NP_828850.1;SARS-CoV_orf1a;3286 | 0 | 0.025 |
| NC_004718.3;NP_828850.1;SARS-CoV_orf1a;3287 | 0 | 0.025 |
| NC_004718.3;NP_828850.1;SARS-CoV_orf1a;3288 | 0 | 0.025 |
| NC_004718.3;NP_828850.1;SARS-CoV_orf1a;3289 | 0 | 0 |
| NC_004718.3;NP_828850.1;SARS-CoV_orf1a;3290 | 0 | 0 |
| NC_004718.3;NP_828850.1;SARS-CoV_orf1a;3291 | 0 | 0.025 |
| NC_004718.3;NP_828850.1;SARS-CoV_orf1a;3292 | 0 | 0 |
| NC_004718.3;NP_828850.1;SARS-CoV_orf1a;3293 | 0 | 0 |
| NC_004718.3;NP_828850.1;SARS-CoV_orf1a;3294 | 0 | 0 |
| NC_004718.3;NP_828850.1;SARS-CoV_orf1a;3295 | 0 | 0 |
| NC_004718.3;NP_828850.1;SARS-CoV_orf1a;3296 | 0 | 0 |
| NC_004718.3;NP_828850.1;SARS-CoV_orf1a;3297 | 0 | 0 |
| NC_004718.3;NP_828850.1;SARS-CoV_orf1a;3298 | 0 | 0 |

| | | |
|---|---|---|
| NC_004718.3;NP_828850.1;SARS-CoV_orf1a;3299 | 0 | 0 |
| NC_004718.3;NP_828850.1;SARS-CoV_orf1a;3300 | 0 | 0 |
| NC_004718.3;NP_828850.1;SARS-CoV_orf1a;3301 | 0 | 0 |
| NC_004718.3;NP_828850.1;SARS-CoV_orf1a;3302 | 0 | 0 |
| NC_004718.3;NP_828850.1;SARS-CoV_orf1a;3303 | 0 | 0 |
| NC_004718.3;NP_828850.1;SARS-CoV_orf1a;3304 | 0 | 0 |
| NC_004718.3;NP_828850.1;SARS-CoV_orf1a;3305 | 0 | 0 |
| NC_004718.3;NP_828850.1;SARS-CoV_orf1a;3306 | 0 | 0 |
| NC_004718.3;NP_828850.1;SARS-CoV_orf1a;3307 | 0 | 0 |
| NC_004718.3;NP_828850.1;SARS-CoV_orf1a;3308 | 0 | 0 |
| NC_004718.3;NP_828850.1;SARS-CoV_orf1a;3309 | 0 | 0 |
| NC_004718.3;NP_828850.1;SARS-CoV_orf1a;3310 | 0 | 0 |
| NC_004718.3;NP_828850.1;SARS-CoV_orf1a;3311 | 0 | 0 |
| NC_004718.3;NP_828850.1;SARS-CoV_orf1a;3312 | 0 | 0 |
| NC_004718.3;NP_828850.1;SARS-CoV_orf1a;3313 | 0 | 0 |
| NC_004718.3;NP_828850.1;SARS-CoV_orf1a;3314 | 0 | 0 |
| NC_004718.3;NP_828850.1;SARS-CoV_orf1a;3315 | 0 | 0 |
| NC_004718.3;NP_828850.1;SARS-CoV_orf1a;3316 | 0 | 0 |
| NC_004718.3;NP_828850.1;SARS-CoV_orf1a;3317 | 0 | 0 |
| NC_004718.3;NP_828850.1;SARS-CoV_orf1a;3318 | 0 | 0 |
| NC_004718.3;NP_828850.1;SARS-CoV_orf1a;3319 | 0 | 0.025 |
| NC_004718.3;NP_828850.1;SARS-CoV_orf1a;3320 | 0 | 0 |
| NC_004718.3;NP_828850.1;SARS-CoV_orf1a;3321 | 0 | 0 |
| NC_004718.3;NP_828850.1;SARS-CoV_orf1a;3322 | 0 | 0.025 |
| NC_004718.3;NP_828850.1;SARS-CoV_orf1a;3323 | 0 | 0 |
| NC_004718.3;NP_828850.1;SARS-CoV_orf1a;3324 | 0 | 0 |
| NC_004718.3;NP_828850.1;SARS-CoV_orf1a;3325 | 0 | 0 |
| NC_004718.3;NP_828850.1;SARS-CoV_orf1a;3326 | 0 | 0 |
| NC_004718.3;NP_828850.1;SARS-CoV_orf1a;3327 | 0 | 0 |
| NC_004718.3;NP_828850.1;SARS-CoV_orf1a;3328 | 0 | 0 |
| NC_004718.3;NP_828850.1;SARS-CoV_orf1a;3329 | 0 | 0 |
| NC_004718.3;NP_828850.1;SARS-CoV_orf1a;3330 | 0.15 | 0.025 |
| NC_004718.3;NP_828850.1;SARS-CoV_orf1a;3331 | 0 | 0 |
| NC_004718.3;NP_828850.1;SARS-CoV_orf1a;3332 | 0 | 0 |
| NC_004718.3;NP_828850.1;SARS-CoV_orf1a;3333 | 0 | 0 |
| NC_004718.3;NP_828850.1;SARS-CoV_orf1a;3334 | 0 | 0 |
| NC_004718.3;NP_828850.1;SARS-CoV_orf1a;3335 | 0 | 0 |
| NC_004718.3;NP_828850.1;SARS-CoV_orf1a;3336 | 0 | 0 |
| NC_004718.3;NP_828850.1;SARS-CoV_orf1a;3337 | 0 | 0 |
| NC_004718.3;NP_828850.1;SARS-CoV_orf1a;3338 | 0 | 0 |
| NC_004718.3;NP_828850.1;SARS-CoV_orf1a;3339 | 0 | 0 |
| NC_004718.3;NP_828850.1;SARS-CoV_orf1a;3340 | 0 | 0 |
| NC_004718.3;NP_828850.1;SARS-CoV_orf1a;3341 | 0 | 0 |

| | | |
|---|---|---|
| NC_004718.3;NP_828850.1;SARS-CoV_orf1a;3342 | 0 | 0 |
| NC_004718.3;NP_828850.1;SARS-CoV_orf1a;3343 | 0 | 0 |
| NC_004718.3;NP_828850.1;SARS-CoV_orf1a;3344 | 0 | 0 |
| NC_004718.3;NP_828850.1;SARS-CoV_orf1a;3345 | 0 | 0 |
| NC_004718.3;NP_828850.1;SARS-CoV_orf1a;3346 | 0 | 0 |
| NC_004718.3;NP_828850.1;SARS-CoV_orf1a;3347 | 0 | 0.025 |
| NC_004718.3;NP_828850.1;SARS-CoV_orf1a;3348 | 0 | 0 |
| NC_004718.3;NP_828850.1;SARS-CoV_orf1a;3349 | 0 | 0 |
| NC_004718.3;NP_828850.1;SARS-CoV_orf1a;3350 | 0 | 0 |
| NC_004718.3;NP_828850.1;SARS-CoV_orf1a;3351 | 0 | 0 |
| NC_004718.3;NP_828850.1;SARS-CoV_orf1a;3352 | 0 | 0 |
| NC_004718.3;NP_828850.1;SARS-CoV_orf1a;3353 | 0 | 0 |
| NC_004718.3;NP_828850.1;SARS-CoV_orf1a;3354 | 0 | 0 |
| NC_004718.3;NP_828850.1;SARS-CoV_orf1a;3355 | 0 | 0 |
| NC_004718.3;NP_828850.1;SARS-CoV_orf1a;3356 | 0 | 0 |
| NC_004718.3;NP_828850.1;SARS-CoV_orf1a;3357 | 0 | 0 |
| NC_004718.3;NP_828850.1;SARS-CoV_orf1a;3358 | 0 | 0.025 |
| NC_004718.3;NP_828850.1;SARS-CoV_orf1a;3359 | 0 | 0 |
| NC_004718.3;NP_828850.1;SARS-CoV_orf1a;3360 | 0 | 0 |
| NC_004718.3;NP_828850.1;SARS-CoV_orf1a;3361 | 0 | 0 |
| NC_004718.3;NP_828850.1;SARS-CoV_orf1a;3362 | 0 | 0.05 |
| NC_004718.3;NP_828850.1;SARS-CoV_orf1a;3363 | 0 | 0.075 |
| NC_004718.3;NP_828850.1;SARS-CoV_orf1a;3364 | 0 | 0.1 |
| NC_004718.3;NP_828850.1;SARS-CoV_orf1a;3365 | 0 | 0.075 |
| NC_004718.3;NP_828850.1;SARS-CoV_orf1a;3366 | 0 | 0.05 |
| NC_004718.3;NP_828850.1;SARS-CoV_orf1a;3367 | 0 | 0.05 |
| NC_004718.3;NP_828850.1;SARS-CoV_orf1a;3368 | 0 | 0.05 |
| NC_004718.3;NP_828850.1;SARS-CoV_orf1a;3369 | 0 | 0.05 |
| NC_004718.3;NP_828850.1;SARS-CoV_orf1a;3370 | 0 | 0.05 |
| NC_004718.3;NP_828850.1;SARS-CoV_orf1a;3371 | 0.05 | 0.05 |
| NC_004718.3;NP_828850.1;SARS-CoV_orf1a;3372 | 0 | 0.075 |
| NC_004718.3;NP_828850.1;SARS-CoV_orf1a;3373 | 0 | 0 |
| NC_004718.3;NP_828850.1;SARS-CoV_orf1a;3374 | 0 | 0 |
| NC_004718.3;NP_828850.1;SARS-CoV_orf1a;3375 | 0 | 0 |
| NC_004718.3;NP_828850.1;SARS-CoV_orf1a;3376 | 0 | 0 |
| NC_004718.3;NP_828850.1;SARS-CoV_orf1a;3377 | 0 | 0.025 |
| NC_004718.3;NP_828850.1;SARS-CoV_orf1a;3378 | 0 | 0.025 |
| NC_004718.3;NP_828850.1;SARS-CoV_orf1a;3379 | 0 | 0.025 |
| NC_004718.3;NP_828850.1;SARS-CoV_orf1a;3380 | 0 | 0.05 |
| NC_004718.3;NP_828850.1;SARS-CoV_orf1a;3381 | 0 | 0.05 |
| NC_004718.3;NP_828850.1;SARS-CoV_orf1a;3382 | 0 | 0.025 |
| NC_004718.3;NP_828850.1;SARS-CoV_orf1a;3383 | 0 | 0.025 |
| NC_004718.3;NP_828850.1;SARS-CoV_orf1a;3384 | 0 | 0 |

| | | |
|---|---|---|
| NC_004718.3;NP_828850.1;SARS-CoV_orf1a;3385 | 0 | 0 |
| NC_004718.3;NP_828850.1;SARS-CoV_orf1a;3386 | 0 | 0 |
| NC_004718.3;NP_828850.1;SARS-CoV_orf1a;3387 | 0 | 0 |
| NC_004718.3;NP_828850.1;SARS-CoV_orf1a;3388 | 0 | 0 |
| NC_004718.3;NP_828850.1;SARS-CoV_orf1a;3389 | 0 | 0 |
| NC_004718.3;NP_828850.1;SARS-CoV_orf1a;3390 | 0 | 0 |
| NC_004718.3;NP_828850.1;SARS-CoV_orf1a;3391 | 0 | 0 |
| NC_004718.3;NP_828850.1;SARS-CoV_orf1a;3392 | 0 | 0 |
| NC_004718.3;NP_828850.1;SARS-CoV_orf1a;3393 | 0 | 0 |
| NC_004718.3;NP_828850.1;SARS-CoV_orf1a;3394 | 0 | 0 |
| NC_004718.3;NP_828850.1;SARS-CoV_orf1a;3395 | 0 | 0 |
| NC_004718.3;NP_828850.1;SARS-CoV_orf1a;3396 | 0 | 0 |
| NC_004718.3;NP_828850.1;SARS-CoV_orf1a;3397 | 0 | 0 |
| NC_004718.3;NP_828850.1;SARS-CoV_orf1a;3398 | 0 | 0 |
| NC_004718.3;NP_828850.1;SARS-CoV_orf1a;3399 | 0 | 0 |
| NC_004718.3;NP_828850.1;SARS-CoV_orf1a;3400 | 0 | 0 |
| NC_004718.3;NP_828850.1;SARS-CoV_orf1a;3401 | 0 | 0 |
| NC_004718.3;NP_828850.1;SARS-CoV_orf1a;3402 | 0 | 0 |
| NC_004718.3;NP_828850.1;SARS-CoV_orf1a;3403 | 0.15 | 0.025 |
| NC_004718.3;NP_828850.1;SARS-CoV_orf1a;3404 | 0.1 | 0.05 |
| NC_004718.3;NP_828850.1;SARS-CoV_orf1a;3405 | 0.05 | 0.025 |
| NC_004718.3;NP_828850.1;SARS-CoV_orf1a;3406 | 0.2 | 0.05 |
| NC_004718.3;NP_828850.1;SARS-CoV_orf1a;3407 | 0.2 | 0.075 |
| NC_004718.3;NP_828850.1;SARS-CoV_orf1a;3408 | 0.15 | 0.075 |
| NC_004718.3;NP_828850.1;SARS-CoV_orf1a;3409 | 0.2 | 0.075 |
| NC_004718.3;NP_828850.1;SARS-CoV_orf1a;3410 | 0.25 | 0.075 |
| NC_004718.3;NP_828850.1;SARS-CoV_orf1a;3411 | 0.2 | 0.05 |
| NC_004718.3;NP_828850.1;SARS-CoV_orf1a;3412 | 0.2 | 0.1 |
| NC_004718.3;NP_828850.1;SARS-CoV_orf1a;3413 | 0.15 | 0.05 |
| NC_004718.3;NP_828850.1;SARS-CoV_orf1a;3414 | 0.25 | 0.1 |
| NC_004718.3;NP_828850.1;SARS-CoV_orf1a;3415 | 0 | 0.025 |
| NC_004718.3;NP_828850.1;SARS-CoV_orf1a;3416 | 0 | 0 |
| NC_004718.3;NP_828850.1;SARS-CoV_orf1a;3417 | 0.05 | 0 |
| NC_004718.3;NP_828850.1;SARS-CoV_orf1a;3418 | 0.05 | 0.05 |
| NC_004718.3;NP_828850.1;SARS-CoV_orf1a;3419 | 0.1 | 0.025 |
| NC_004718.3;NP_828850.1;SARS-CoV_orf1a;3420 | 0.05 | 0 |
| NC_004718.3;NP_828850.1;SARS-CoV_orf1a;3421 | 0.1 | 0 |
| NC_004718.3;NP_828850.1;SARS-CoV_orf1a;3422 | 0 | 0 |
| NC_004718.3;NP_828850.1;SARS-CoV_orf1a;3423 | 0.05 | 0 |
| NC_004718.3;NP_828850.1;SARS-CoV_orf1a;3424 | 0 | 0 |
| NC_004718.3;NP_828850.1;SARS-CoV_orf1a;3425 | 0.05 | 0.025 |
| NC_004718.3;NP_828850.1;SARS-CoV_orf1a;3426 | 0 | 0 |
| NC_004718.3;NP_828850.1;SARS-CoV_orf1a;3427 | 0 | 0 |

| | | |
|---|---|---|
| NC_004718.3;NP_828850.1;SARS-CoV_orf1a;3428 | 0 | 0 |
| NC_004718.3;NP_828850.1;SARS-CoV_orf1a;3429 | 0 | 0 |
| NC_004718.3;NP_828850.1;SARS-CoV_orf1a;3430 | 0 | 0 |
| NC_004718.3;NP_828850.1;SARS-CoV_orf1a;3431 | 0 | 0 |
| NC_004718.3;NP_828850.1;SARS-CoV_orf1a;3432 | 0 | 0 |
| NC_004718.3;NP_828850.1;SARS-CoV_orf1a;3433 | 0 | 0 |
| NC_004718.3;NP_828850.1;SARS-CoV_orf1a;3434 | 0 | 0 |
| NC_004718.3;NP_828850.1;SARS-CoV_orf1a;3435 | 0 | 0 |
| NC_004718.3;NP_828850.1;SARS-CoV_orf1a;3436 | 0 | 0 |
| NC_004718.3;NP_828850.1;SARS-CoV_orf1a;3437 | 0 | 0 |
| NC_004718.3;NP_828850.1;SARS-CoV_orf1a;3438 | 0 | 0 |
| NC_004718.3;NP_828850.1;SARS-CoV_orf1a;3439 | 0 | 0 |
| NC_004718.3;NP_828850.1;SARS-CoV_orf1a;3440 | 0 | 0 |
| NC_004718.3;NP_828850.1;SARS-CoV_orf1a;3441 | 0 | 0 |
| NC_004718.3;NP_828850.1;SARS-CoV_orf1a;3442 | 0 | 0 |
| NC_004718.3;NP_828850.1;SARS-CoV_orf1a;3443 | 0 | 0 |
| NC_004718.3;NP_828850.1;SARS-CoV_orf1a;3444 | 0 | 0 |
| NC_004718.3;NP_828850.1;SARS-CoV_orf1a;3445 | 0 | 0 |
| NC_004718.3;NP_828850.1;SARS-CoV_orf1a;3446 | 0 | 0 |
| NC_004718.3;NP_828850.1;SARS-CoV_orf1a;3447 | 0 | 0 |
| NC_004718.3;NP_828850.1;SARS-CoV_orf1a;3448 | 0 | 0 |
| NC_004718.3;NP_828850.1;SARS-CoV_orf1a;3449 | 0 | 0 |
| NC_004718.3;NP_828850.1;SARS-CoV_orf1a;3450 | 0 | 0 |
| NC_004718.3;NP_828850.1;SARS-CoV_orf1a;3451 | 0 | 0 |
| NC_004718.3;NP_828850.1;SARS-CoV_orf1a;3452 | 0 | 0 |
| NC_004718.3;NP_828850.1;SARS-CoV_orf1a;3453 | 0 | 0 |
| NC_004718.3;NP_828850.1;SARS-CoV_orf1a;3454 | 0 | 0 |
| NC_004718.3;NP_828850.1;SARS-CoV_orf1a;3455 | 0 | 0 |
| NC_004718.3;NP_828850.1;SARS-CoV_orf1a;3456 | 0.05 | 0 |
| NC_004718.3;NP_828850.1;SARS-CoV_orf1a;3457 | 0.05 | 0 |
| NC_004718.3;NP_828850.1;SARS-CoV_orf1a;3458 | 0.05 | 0 |
| NC_004718.3;NP_828850.1;SARS-CoV_orf1a;3459 | 0.05 | 0.025 |
| NC_004718.3;NP_828850.1;SARS-CoV_orf1a;3460 | 0.1 | 0 |
| NC_004718.3;NP_828850.1;SARS-CoV_orf1a;3461 | 0.05 | 0 |
| NC_004718.3;NP_828850.1;SARS-CoV_orf1a;3462 | 0.1 | 0 |
| NC_004718.3;NP_828850.1;SARS-CoV_orf1a;3463 | 0.05 | 0 |
| NC_004718.3;NP_828850.1;SARS-CoV_orf1a;3464 | 0.05 | 0.025 |
| NC_004718.3;NP_828850.1;SARS-CoV_orf1a;3465 | 0.05 | 0.15 |
| NC_004718.3;NP_828850.1;SARS-CoV_orf1a;3466 | 0.25 | 0.1 |
| NC_004718.3;NP_828850.1;SARS-CoV_orf1a;3467 | 0.05 | 0 |
| NC_004718.3;NP_828850.1;SARS-CoV_orf1a;3468 | 0 | 0 |
| NC_004718.3;NP_828850.1;SARS-CoV_orf1a;3469 | 0.05 | 0 |
| NC_004718.3;NP_828850.1;SARS-CoV_orf1a;3470 | 0 | 0 |

| | | |
|---|---|---|
| NC_004718.3;NP_828850.1;SARS-CoV_orf1a;3471 | 0 | 0.025 |
| NC_004718.3;NP_828850.1;SARS-CoV_orf1a;3472 | 0 | 0.025 |
| NC_004718.3;NP_828850.1;SARS-CoV_orf1a;3473 | 0.05 | 0 |
| NC_004718.3;NP_828850.1;SARS-CoV_orf1a;3474 | 0 | 0 |
| NC_004718.3;NP_828850.1;SARS-CoV_orf1a;3475 | 0 | 0 |
| NC_004718.3;NP_828850.1;SARS-CoV_orf1a;3476 | 0 | 0 |
| NC_004718.3;NP_828850.1;SARS-CoV_orf1a;3477 | 0 | 0 |
| NC_004718.3;NP_828850.1;SARS-CoV_orf1a;3478 | 0 | 0 |
| NC_004718.3;NP_828850.1;SARS-CoV_orf1a;3479 | 0 | 0 |
| NC_004718.3;NP_828850.1;SARS-CoV_orf1a;3480 | 0.05 | 0 |
| NC_004718.3;NP_828850.1;SARS-CoV_orf1a;3481 | 0 | 0 |
| NC_004718.3;NP_828850.1;SARS-CoV_orf1a;3482 | 0.05 | 0.025 |
| NC_004718.3;NP_828850.1;SARS-CoV_orf1a;3483 | 0.15 | 0.1 |
| NC_004718.3;NP_828850.1;SARS-CoV_orf1a;3484 | 0.05 | 0.05 |
| NC_004718.3;NP_828850.1;SARS-CoV_orf1a;3485 | 0.1 | 0.15 |
| NC_004718.3;NP_828850.1;SARS-CoV_orf1a;3486 | 0.05 | 0.025 |
| NC_004718.3;NP_828850.1;SARS-CoV_orf1a;3487 | 0.05 | 0 |
| NC_004718.3;NP_828850.1;SARS-CoV_orf1a;3488 | 0.05 | 0 |
| NC_004718.3;NP_828850.1;SARS-CoV_orf1a;3489 | 0.05 | 0 |
| NC_004718.3;NP_828850.1;SARS-CoV_orf1a;3490 | 0.05 | 0 |
| NC_004718.3;NP_828850.1;SARS-CoV_orf1a;3491 | 0.05 | 0 |
| NC_004718.3;NP_828850.1;SARS-CoV_orf1a;3492 | 0.1 | 0.025 |
| NC_004718.3;NP_828850.1;SARS-CoV_orf1a;3493 | 0 | 0 |
| NC_004718.3;NP_828850.1;SARS-CoV_orf1a;3494 | 0 | 0 |
| NC_004718.3;NP_828850.1;SARS-CoV_orf1a;3495 | 0 | 0 |
| NC_004718.3;NP_828850.1;SARS-CoV_orf1a;3496 | 0 | 0 |
| NC_004718.3;NP_828850.1;SARS-CoV_orf1a;3497 | 0 | 0 |
| NC_004718.3;NP_828850.1;SARS-CoV_orf1a;3498 | 0.05 | 0 |
| NC_004718.3;NP_828850.1;SARS-CoV_orf1a;3499 | 0 | 0 |
| NC_004718.3;NP_828850.1;SARS-CoV_orf1a;3500 | 0 | 0.025 |
| NC_004718.3;NP_828850.1;SARS-CoV_orf1a;3501 | 0 | 0 |
| NC_004718.3;NP_828850.1;SARS-CoV_orf1a;3502 | 0 | 0.025 |
| NC_004718.3;NP_828850.1;SARS-CoV_orf1a;3503 | 0 | 0.025 |
| NC_004718.3;NP_828850.1;SARS-CoV_orf1a;3504 | 0 | 0 |
| NC_004718.3;NP_828850.1;SARS-CoV_orf1a;3505 | 0 | 0 |
| NC_004718.3;NP_828850.1;SARS-CoV_orf1a;3506 | 0 | 0 |
| NC_004718.3;NP_828850.1;SARS-CoV_orf1a;3507 | 0 | 0 |
| NC_004718.3;NP_828850.1;SARS-CoV_orf1a;3508 | 0 | 0.025 |
| NC_004718.3;NP_828850.1;SARS-CoV_orf1a;3509 | 0 | 0.025 |
| NC_004718.3;NP_828850.1;SARS-CoV_orf1a;3510 | 0 | 0 |
| NC_004718.3;NP_828850.1;SARS-CoV_orf1a;3511 | 0 | 0 |
| NC_004718.3;NP_828850.1;SARS-CoV_orf1a;3512 | 0 | 0 |
| NC_004718.3;NP_828850.1;SARS-CoV_orf1a;3513 | 0 | 0 |

| | | |
|---|---|---|
| NC_004718.3;NP_828850.1;SARS-CoV_orf1a;3514 | 0 | 0 |
| NC_004718.3;NP_828850.1;SARS-CoV_orf1a;3515 | 0 | 0 |
| NC_004718.3;NP_828850.1;SARS-CoV_orf1a;3516 | 0 | 0 |
| NC_004718.3;NP_828850.1;SARS-CoV_orf1a;3517 | 0 | 0 |
| NC_004718.3;NP_828850.1;SARS-CoV_orf1a;3518 | 0 | 0.025 |
| NC_004718.3;NP_828850.1;SARS-CoV_orf1a;3519 | 0 | 0.025 |
| NC_004718.3;NP_828850.1;SARS-CoV_orf1a;3520 | 0 | 0.05 |
| NC_004718.3;NP_828850.1;SARS-CoV_orf1a;3521 | 0 | 0.025 |
| NC_004718.3;NP_828850.1;SARS-CoV_orf1a;3522 | 0 | 0.025 |
| NC_004718.3;NP_828850.1;SARS-CoV_orf1a;3523 | 0 | 0.025 |
| NC_004718.3;NP_828850.1;SARS-CoV_orf1a;3524 | 0 | 0.025 |
| NC_004718.3;NP_828850.1;SARS-CoV_orf1a;3525 | 0 | 0.05 |
| NC_004718.3;NP_828850.1;SARS-CoV_orf1a;3526 | 0 | 0.025 |
| NC_004718.3;NP_828850.1;SARS-CoV_orf1a;3527 | 0 | 0.05 |
| NC_004718.3;NP_828850.1;SARS-CoV_orf1a;3528 | 0.1 | 0.025 |
| NC_004718.3;NP_828850.1;SARS-CoV_orf1a;3529 | 0 | 0 |
| NC_004718.3;NP_828850.1;SARS-CoV_orf1a;3530 | 0 | 0 |
| NC_004718.3;NP_828850.1;SARS-CoV_orf1a;3531 | 0 | 0 |
| NC_004718.3;NP_828850.1;SARS-CoV_orf1a;3532 | 0 | 0 |
| NC_004718.3;NP_828850.1;SARS-CoV_orf1a;3533 | 0 | 0 |
| NC_004718.3;NP_828850.1;SARS-CoV_orf1a;3534 | 0 | 0 |
| NC_004718.3;NP_828850.1;SARS-CoV_orf1a;3535 | 0 | 0 |
| NC_004718.3;NP_828850.1;SARS-CoV_orf1a;3536 | 0.05 | 0 |
| NC_004718.3;NP_828850.1;SARS-CoV_orf1a;3537 | 0.05 | 0 |
| NC_004718.3;NP_828850.1;SARS-CoV_orf1a;3538 | 0.05 | 0 |
| NC_004718.3;NP_828850.1;SARS-CoV_orf1a;3539 | 0.05 | 0 |
| NC_004718.3;NP_828850.1;SARS-CoV_orf1a;3540 | 0.05 | 0 |
| NC_004718.3;NP_828850.1;SARS-CoV_orf1a;3541 | 0.05 | 0 |
| NC_004718.3;NP_828850.1;SARS-CoV_orf1a;3542 | 0.05 | 0 |
| NC_004718.3;NP_828850.1;SARS-CoV_orf1a;3543 | 0.05 | 0 |
| NC_004718.3;NP_828850.1;SARS-CoV_orf1a;3544 | 0 | 0 |
| NC_004718.3;NP_828850.1;SARS-CoV_orf1a;3545 | 0.05 | 0 |
| NC_004718.3;NP_828850.1;SARS-CoV_orf1a;3546 | 0.05 | 0 |
| NC_004718.3;NP_828850.1;SARS-CoV_orf1a;3547 | 0.05 | 0 |
| NC_004718.3;NP_828850.1;SARS-CoV_orf1a;3548 | 0 | 0 |
| NC_004718.3;NP_828850.1;SARS-CoV_orf1a;3549 | 0 | 0 |
| NC_004718.3;NP_828850.1;SARS-CoV_orf1a;3550 | 0 | 0 |
| NC_004718.3;NP_828850.1;SARS-CoV_orf1a;3551 | 0 | 0 |
| NC_004718.3;NP_828850.1;SARS-CoV_orf1a;3552 | 0 | 0 |
| NC_004718.3;NP_828850.1;SARS-CoV_orf1a;3553 | 0 | 0 |
| NC_004718.3;NP_828850.1;SARS-CoV_orf1a;3554 | 0 | 0 |
| NC_004718.3;NP_828850.1;SARS-CoV_orf1a;3555 | 0 | 0 |
| NC_004718.3;NP_828850.1;SARS-CoV_orf1a;3556 | 0 | 0 |

| | | |
|---|---|---|
| NC_004718.3;NP_828850.1;SARS-CoV_orf1a;3557 | 0 | 0 |
| NC_004718.3;NP_828850.1;SARS-CoV_orf1a;3558 | 0 | 0 |
| NC_004718.3;NP_828850.1;SARS-CoV_orf1a;3559 | 0 | 0 |
| NC_004718.3;NP_828850.1;SARS-CoV_orf1a;3560 | 0 | 0 |
| NC_004718.3;NP_828850.1;SARS-CoV_orf1a;3561 | 0 | 0 |
| NC_004718.3;NP_828850.1;SARS-CoV_orf1a;3562 | 0 | 0 |
| NC_004718.3;NP_828850.1;SARS-CoV_orf1a;3563 | 0 | 0 |
| NC_004718.3;NP_828850.1;SARS-CoV_orf1a;3564 | 0 | 0 |
| NC_004718.3;NP_828850.1;SARS-CoV_orf1a;3565 | 0 | 0 |
| NC_004718.3;NP_828850.1;SARS-CoV_orf1a;3566 | 0 | 0 |
| NC_004718.3;NP_828850.1;SARS-CoV_orf1a;3567 | 0 | 0 |
| NC_004718.3;NP_828850.1;SARS-CoV_orf1a;3568 | 0 | 0 |
| NC_004718.3;NP_828850.1;SARS-CoV_orf1a;3569 | 0 | 0 |
| NC_004718.3;NP_828850.1;SARS-CoV_orf1a;3570 | 0 | 0 |
| NC_004718.3;NP_828850.1;SARS-CoV_orf1a;3571 | 0 | 0 |
| NC_004718.3;NP_828850.1;SARS-CoV_orf1a;3572 | 0 | 0 |
| NC_004718.3;NP_828850.1;SARS-CoV_orf1a;3573 | 0 | 0 |
| NC_004718.3;NP_828850.1;SARS-CoV_orf1a;3574 | 0 | 0 |
| NC_004718.3;NP_828850.1;SARS-CoV_orf1a;3575 | 0 | 0 |
| NC_004718.3;NP_828850.1;SARS-CoV_orf1a;3576 | 0 | 0 |
| NC_004718.3;NP_828850.1;SARS-CoV_orf1a;3577 | 0 | 0 |
| NC_004718.3;NP_828850.1;SARS-CoV_orf1a;3578 | 0 | 0 |
| NC_004718.3;NP_828850.1;SARS-CoV_orf1a;3579 | 0 | 0 |
| NC_004718.3;NP_828850.1;SARS-CoV_orf1a;3580 | 0 | 0 |
| NC_004718.3;NP_828850.1;SARS-CoV_orf1a;3581 | 0 | 0 |
| NC_004718.3;NP_828850.1;SARS-CoV_orf1a;3582 | 0 | 0 |
| NC_004718.3;NP_828850.1;SARS-CoV_orf1a;3583 | 0 | 0 |
| NC_004718.3;NP_828850.1;SARS-CoV_orf1a;3584 | 0 | 0 |
| NC_004718.3;NP_828850.1;SARS-CoV_orf1a;3585 | 0 | 0 |
| NC_004718.3;NP_828850.1;SARS-CoV_orf1a;3586 | 0 | 0 |
| NC_004718.3;NP_828850.1;SARS-CoV_orf1a;3587 | 0 | 0 |
| NC_004718.3;NP_828850.1;SARS-CoV_orf1a;3588 | 0 | 0 |
| NC_004718.3;NP_828850.1;SARS-CoV_orf1a;3589 | 0 | 0 |
| NC_004718.3;NP_828850.1;SARS-CoV_orf1a;3590 | 0 | 0 |
| NC_004718.3;NP_828850.1;SARS-CoV_orf1a;3591 | 0 | 0 |
| NC_004718.3;NP_828850.1;SARS-CoV_orf1a;3592 | 0 | 0 |
| NC_004718.3;NP_828850.1;SARS-CoV_orf1a;3593 | 0 | 0 |
| NC_004718.3;NP_828850.1;SARS-CoV_orf1a;3594 | 0 | 0 |
| NC_004718.3;NP_828850.1;SARS-CoV_orf1a;3595 | 0 | 0 |
| NC_004718.3;NP_828850.1;SARS-CoV_orf1a;3596 | 0 | 0 |
| NC_004718.3;NP_828850.1;SARS-CoV_orf1a;3597 | 0 | 0 |
| NC_004718.3;NP_828850.1;SARS-CoV_orf1a;3598 | 0 | 0 |
| NC_004718.3;NP_828850.1;SARS-CoV_orf1a;3599 | 0 | 0 |

| | | |
|---|---|---|
| NC_004718.3;NP_828850.1;SARS-CoV_orf1a;3600 | 0 | 0 |
| NC_004718.3;NP_828850.1;SARS-CoV_orf1a;3601 | 0 | 0 |
| NC_004718.3;NP_828850.1;SARS-CoV_orf1a;3602 | 0 | 0 |
| NC_004718.3;NP_828850.1;SARS-CoV_orf1a;3603 | 0 | 0 |
| NC_004718.3;NP_828850.1;SARS-CoV_orf1a;3604 | 0 | 0 |
| NC_004718.3;NP_828850.1;SARS-CoV_orf1a;3605 | 0 | 0 |
| NC_004718.3;NP_828850.1;SARS-CoV_orf1a;3606 | 0 | 0 |
| NC_004718.3;NP_828850.1;SARS-CoV_orf1a;3607 | 0 | 0 |
| NC_004718.3;NP_828850.1;SARS-CoV_orf1a;3608 | 0 | 0 |
| NC_004718.3;NP_828850.1;SARS-CoV_orf1a;3609 | 0 | 0 |
| NC_004718.3;NP_828850.1;SARS-CoV_orf1a;3610 | 0 | 0 |
| NC_004718.3;NP_828850.1;SARS-CoV_orf1a;3611 | 0 | 0 |
| NC_004718.3;NP_828850.1;SARS-CoV_orf1a;3612 | 0 | 0 |
| NC_004718.3;NP_828850.1;SARS-CoV_orf1a;3613 | 0 | 0 |
| NC_004718.3;NP_828850.1;SARS-CoV_orf1a;3614 | 0 | 0 |
| NC_004718.3;NP_828850.1;SARS-CoV_orf1a;3615 | 0 | 0 |
| NC_004718.3;NP_828850.1;SARS-CoV_orf1a;3616 | 0 | 0 |
| NC_004718.3;NP_828850.1;SARS-CoV_orf1a;3617 | 0 | 0 |
| NC_004718.3;NP_828850.1;SARS-CoV_orf1a;3618 | 0 | 0 |
| NC_004718.3;NP_828850.1;SARS-CoV_orf1a;3619 | 0 | 0 |
| NC_004718.3;NP_828850.1;SARS-CoV_orf1a;3620 | 0 | 0 |
| NC_004718.3;NP_828850.1;SARS-CoV_orf1a;3621 | 0 | 0 |
| NC_004718.3;NP_828850.1;SARS-CoV_orf1a;3622 | 0 | 0 |
| NC_004718.3;NP_828850.1;SARS-CoV_orf1a;3623 | 0 | 0 |
| NC_004718.3;NP_828850.1;SARS-CoV_orf1a;3624 | 0 | 0 |
| NC_004718.3;NP_828850.1;SARS-CoV_orf1a;3625 | 0 | 0 |
| NC_004718.3;NP_828850.1;SARS-CoV_orf1a;3626 | 0 | 0 |
| NC_004718.3;NP_828850.1;SARS-CoV_orf1a;3627 | 0 | 0 |
| NC_004718.3;NP_828850.1;SARS-CoV_orf1a;3628 | 0 | 0 |
| NC_004718.3;NP_828850.1;SARS-CoV_orf1a;3629 | 0 | 0 |
| NC_004718.3;NP_828850.1;SARS-CoV_orf1a;3630 | 0 | 0 |
| NC_004718.3;NP_828850.1;SARS-CoV_orf1a;3631 | 0 | 0 |
| NC_004718.3;NP_828850.1;SARS-CoV_orf1a;3632 | 0 | 0 |
| NC_004718.3;NP_828850.1;SARS-CoV_orf1a;3633 | 0 | 0 |
| NC_004718.3;NP_828850.1;SARS-CoV_orf1a;3634 | 0 | 0 |
| NC_004718.3;NP_828850.1;SARS-CoV_orf1a;3635 | 0 | 0 |
| NC_004718.3;NP_828850.1;SARS-CoV_orf1a;3636 | 0 | 0 |
| NC_004718.3;NP_828850.1;SARS-CoV_orf1a;3637 | 0 | 0 |
| NC_004718.3;NP_828850.1;SARS-CoV_orf1a;3638 | 0 | 0 |
| NC_004718.3;NP_828850.1;SARS-CoV_orf1a;3639 | 0 | 0 |
| NC_004718.3;NP_828850.1;SARS-CoV_orf1a;3640 | 0 | 0 |
| NC_004718.3;NP_828850.1;SARS-CoV_orf1a;3641 | 0 | 0 |
| NC_004718.3;NP_828850.1;SARS-CoV_orf1a;3642 | 0 | 0 |

| | | |
|---|---|---|
| NC_004718.3;NP_828850.1;SARS-CoV_orf1a;3643 | 0 | 0.025 |
| NC_004718.3;NP_828850.1;SARS-CoV_orf1a;3644 | 0 | 0.025 |
| NC_004718.3;NP_828850.1;SARS-CoV_orf1a;3645 | 0 | 0 |
| NC_004718.3;NP_828850.1;SARS-CoV_orf1a;3646 | 0 | 0 |
| NC_004718.3;NP_828850.1;SARS-CoV_orf1a;3647 | 0 | 0.025 |
| NC_004718.3;NP_828850.1;SARS-CoV_orf1a;3648 | 0 | 0 |
| NC_004718.3;NP_828850.1;SARS-CoV_orf1a;3649 | 0 | 0 |
| NC_004718.3;NP_828850.1;SARS-CoV_orf1a;3650 | 0 | 0 |
| NC_004718.3;NP_828850.1;SARS-CoV_orf1a;3651 | 0 | 0 |
| NC_004718.3;NP_828850.1;SARS-CoV_orf1a;3652 | 0 | 0 |
| NC_004718.3;NP_828850.1;SARS-CoV_orf1a;3653 | 0 | 0 |
| NC_004718.3;NP_828850.1;SARS-CoV_orf1a;3654 | 0 | 0 |
| NC_004718.3;NP_828850.1;SARS-CoV_orf1a;3655 | 0 | 0 |
| NC_004718.3;NP_828850.1;SARS-CoV_orf1a;3656 | 0 | 0 |
| NC_004718.3;NP_828850.1;SARS-CoV_orf1a;3657 | 0 | 0 |
| NC_004718.3;NP_828850.1;SARS-CoV_orf1a;3658 | 0 | 0 |
| NC_004718.3;NP_828850.1;SARS-CoV_orf1a;3659 | 0 | 0 |
| NC_004718.3;NP_828850.1;SARS-CoV_orf1a;3660 | 0 | 0 |
| NC_004718.3;NP_828850.1;SARS-CoV_orf1a;3661 | 0 | 0 |
| NC_004718.3;NP_828850.1;SARS-CoV_orf1a;3662 | 0 | 0 |
| NC_004718.3;NP_828850.1;SARS-CoV_orf1a;3663 | 0 | 0 |
| NC_004718.3;NP_828850.1;SARS-CoV_orf1a;3664 | 0 | 0 |
| NC_004718.3;NP_828850.1;SARS-CoV_orf1a;3665 | 0 | 0 |
| NC_004718.3;NP_828850.1;SARS-CoV_orf1a;3666 | 0 | 0 |
| NC_004718.3;NP_828850.1;SARS-CoV_orf1a;3667 | 0 | 0 |
| NC_004718.3;NP_828850.1;SARS-CoV_orf1a;3668 | 0 | 0 |
| NC_004718.3;NP_828850.1;SARS-CoV_orf1a;3669 | 0 | 0 |
| NC_004718.3;NP_828850.1;SARS-CoV_orf1a;3670 | 0 | 0 |
| NC_004718.3;NP_828850.1;SARS-CoV_orf1a;3671 | 0 | 0 |
| NC_004718.3;NP_828850.1;SARS-CoV_orf1a;3672 | 0 | 0 |
| NC_004718.3;NP_828850.1;SARS-CoV_orf1a;3673 | 0 | 0 |
| NC_004718.3;NP_828850.1;SARS-CoV_orf1a;3674 | 0 | 0 |
| NC_004718.3;NP_828850.1;SARS-CoV_orf1a;3675 | 0 | 0.025 |
| NC_004718.3;NP_828850.1;SARS-CoV_orf1a;3676 | 0 | 0 |
| NC_004718.3;NP_828850.1;SARS-CoV_orf1a;3677 | 0 | 0 |
| NC_004718.3;NP_828850.1;SARS-CoV_orf1a;3678 | 0 | 0 |
| NC_004718.3;NP_828850.1;SARS-CoV_orf1a;3679 | 0 | 0.025 |
| NC_004718.3;NP_828850.1;SARS-CoV_orf1a;3680 | 0 | 0 |
| NC_004718.3;NP_828850.1;SARS-CoV_orf1a;3681 | 0 | 0 |
| NC_004718.3;NP_828850.1;SARS-CoV_orf1a;3682 | 0 | 0 |
| NC_004718.3;NP_828850.1;SARS-CoV_orf1a;3683 | 0 | 0 |
| NC_004718.3;NP_828850.1;SARS-CoV_orf1a;3684 | 0 | 0 |
| NC_004718.3;NP_828850.1;SARS-CoV_orf1a;3685 | 0 | 0 |

| | | |
|---|---|---|
| NC_004718.3;NP_828850.1;SARS-CoV_orf1a;3686 | 0 | 0 |
| NC_004718.3;NP_828850.1;SARS-CoV_orf1a;3687 | 0 | 0 |
| NC_004718.3;NP_828850.1;SARS-CoV_orf1a;3688 | 0 | 0 |
| NC_004718.3;NP_828850.1;SARS-CoV_orf1a;3689 | 0 | 0 |
| NC_004718.3;NP_828850.1;SARS-CoV_orf1a;3690 | 0 | 0 |
| NC_004718.3;NP_828850.1;SARS-CoV_orf1a;3691 | 0 | 0 |
| NC_004718.3;NP_828850.1;SARS-CoV_orf1a;3692 | 0 | 0 |
| NC_004718.3;NP_828850.1;SARS-CoV_orf1a;3693 | 0 | 0 |
| NC_004718.3;NP_828850.1;SARS-CoV_orf1a;3694 | 0 | 0 |
| NC_004718.3;NP_828850.1;SARS-CoV_orf1a;3695 | 0.05 | 0 |
| NC_004718.3;NP_828850.1;SARS-CoV_orf1a;3696 | 0.05 | 0 |
| NC_004718.3;NP_828850.1;SARS-CoV_orf1a;3697 | 0.05 | 0 |
| NC_004718.3;NP_828850.1;SARS-CoV_orf1a;3698 | 0.05 | 0.025 |
| NC_004718.3;NP_828850.1;SARS-CoV_orf1a;3699 | 0.05 | 0 |
| NC_004718.3;NP_828850.1;SARS-CoV_orf1a;3700 | 0 | 0 |
| NC_004718.3;NP_828850.1;SARS-CoV_orf1a;3701 | 0 | 0 |
| NC_004718.3;NP_828850.1;SARS-CoV_orf1a;3702 | 0 | 0 |
| NC_004718.3;NP_828850.1;SARS-CoV_orf1a;3703 | 0 | 0 |
| NC_004718.3;NP_828850.1;SARS-CoV_orf1a;3704 | 0 | 0 |
| NC_004718.3;NP_828850.1;SARS-CoV_orf1a;3705 | 0 | 0 |
| NC_004718.3;NP_828850.1;SARS-CoV_orf1a;3706 | 0 | 0 |
| NC_004718.3;NP_828850.1;SARS-CoV_orf1a;3707 | 0 | 0 |
| NC_004718.3;NP_828850.1;SARS-CoV_orf1a;3708 | 0 | 0 |
| NC_004718.3;NP_828850.1;SARS-CoV_orf1a;3709 | 0 | 0 |
| NC_004718.3;NP_828850.1;SARS-CoV_orf1a;3710 | 0 | 0 |
| NC_004718.3;NP_828850.1;SARS-CoV_orf1a;3711 | 0 | 0 |
| NC_004718.3;NP_828850.1;SARS-CoV_orf1a;3712 | 0 | 0 |
| NC_004718.3;NP_828850.1;SARS-CoV_orf1a;3713 | 0 | 0 |
| NC_004718.3;NP_828850.1;SARS-CoV_orf1a;3714 | 0 | 0 |
| NC_004718.3;NP_828850.1;SARS-CoV_orf1a;3715 | 0 | 0 |
| NC_004718.3;NP_828850.1;SARS-CoV_orf1a;3716 | 0 | 0 |
| NC_004718.3;NP_828850.1;SARS-CoV_orf1a;3717 | 0 | 0 |
| NC_004718.3;NP_828850.1;SARS-CoV_orf1a;3718 | 0 | 0 |
| NC_004718.3;NP_828850.1;SARS-CoV_orf1a;3719 | 0 | 0 |
| NC_004718.3;NP_828850.1;SARS-CoV_orf1a;3720 | 0 | 0 |
| NC_004718.3;NP_828850.1;SARS-CoV_orf1a;3721 | 0 | 0 |
| NC_004718.3;NP_828850.1;SARS-CoV_orf1a;3722 | 0 | 0 |
| NC_004718.3;NP_828850.1;SARS-CoV_orf1a;3723 | 0 | 0 |
| NC_004718.3;NP_828850.1;SARS-CoV_orf1a;3724 | 0 | 0 |
| NC_004718.3;NP_828850.1;SARS-CoV_orf1a;3725 | 0 | 0 |
| NC_004718.3;NP_828850.1;SARS-CoV_orf1a;3726 | 0 | 0 |
| NC_004718.3;NP_828850.1;SARS-CoV_orf1a;3727 | 0 | 0 |
| NC_004718.3;NP_828850.1;SARS-CoV_orf1a;3728 | 0 | 0 |

| | | |
|---|---|---|
| NC_004718.3;NP_828850.1;SARS-CoV_orf1a;3729 | 0 | 0 |
| NC_004718.3;NP_828850.1;SARS-CoV_orf1a;3730 | 0 | 0 |
| NC_004718.3;NP_828850.1;SARS-CoV_orf1a;3731 | 0 | 0 |
| NC_004718.3;NP_828850.1;SARS-CoV_orf1a;3732 | 0 | 0 |
| NC_004718.3;NP_828850.1;SARS-CoV_orf1a;3733 | 0 | 0 |
| NC_004718.3;NP_828850.1;SARS-CoV_orf1a;3734 | 0 | 0 |
| NC_004718.3;NP_828850.1;SARS-CoV_orf1a;3735 | 0 | 0 |
| NC_004718.3;NP_828850.1;SARS-CoV_orf1a;3736 | 0 | 0 |
| NC_004718.3;NP_828850.1;SARS-CoV_orf1a;3737 | 0 | 0 |
| NC_004718.3;NP_828850.1;SARS-CoV_orf1a;3738 | 0 | 0 |
| NC_004718.3;NP_828850.1;SARS-CoV_orf1a;3739 | 0 | 0 |
| NC_004718.3;NP_828850.1;SARS-CoV_orf1a;3740 | 0 | 0 |
| NC_004718.3;NP_828850.1;SARS-CoV_orf1a;3741 | 0 | 0 |
| NC_004718.3;NP_828850.1;SARS-CoV_orf1a;3742 | 0 | 0 |
| NC_004718.3;NP_828850.1;SARS-CoV_orf1a;3743 | 0 | 0 |
| NC_004718.3;NP_828850.1;SARS-CoV_orf1a;3744 | 0 | 0 |
| NC_004718.3;NP_828850.1;SARS-CoV_orf1a;3745 | 0 | 0 |
| NC_004718.3;NP_828850.1;SARS-CoV_orf1a;3746 | 0 | 0 |
| NC_004718.3;NP_828850.1;SARS-CoV_orf1a;3747 | 0 | 0 |
| NC_004718.3;NP_828850.1;SARS-CoV_orf1a;3748 | 0 | 0 |
| NC_004718.3;NP_828850.1;SARS-CoV_orf1a;3749 | 0 | 0 |
| NC_004718.3;NP_828850.1;SARS-CoV_orf1a;3750 | 0 | 0 |
| NC_004718.3;NP_828850.1;SARS-CoV_orf1a;3751 | 0 | 0 |
| NC_004718.3;NP_828850.1;SARS-CoV_orf1a;3752 | 0 | 0 |
| NC_004718.3;NP_828850.1;SARS-CoV_orf1a;3753 | 0 | 0 |
| NC_004718.3;NP_828850.1;SARS-CoV_orf1a;3754 | 0 | 0 |
| NC_004718.3;NP_828850.1;SARS-CoV_orf1a;3755 | 0 | 0 |
| NC_004718.3;NP_828850.1;SARS-CoV_orf1a;3756 | 0 | 0 |
| NC_004718.3;NP_828850.1;SARS-CoV_orf1a;3757 | 0 | 0 |
| NC_004718.3;NP_828850.1;SARS-CoV_orf1a;3758 | 0 | 0 |
| NC_004718.3;NP_828850.1;SARS-CoV_orf1a;3759 | 0 | 0 |
| NC_004718.3;NP_828850.1;SARS-CoV_orf1a;3760 | 0 | 0 |
| NC_004718.3;NP_828850.1;SARS-CoV_orf1a;3761 | 0 | 0 |
| NC_004718.3;NP_828850.1;SARS-CoV_orf1a;3762 | 0 | 0 |
| NC_004718.3;NP_828850.1;SARS-CoV_orf1a;3763 | 0 | 0 |
| NC_004718.3;NP_828850.1;SARS-CoV_orf1a;3764 | 0 | 0 |
| NC_004718.3;NP_828850.1;SARS-CoV_orf1a;3765 | 0 | 0 |
| NC_004718.3;NP_828850.1;SARS-CoV_orf1a;3766 | 0 | 0 |
| NC_004718.3;NP_828850.1;SARS-CoV_orf1a;3767 | 0 | 0 |
| NC_004718.3;NP_828850.1;SARS-CoV_orf1a;3768 | 0 | 0 |
| NC_004718.3;NP_828850.1;SARS-CoV_orf1a;3769 | 0 | 0 |
| NC_004718.3;NP_828850.1;SARS-CoV_orf1a;3770 | 0 | 0 |
| NC_004718.3;NP_828850.1;SARS-CoV_orf1a;3771 | 0 | 0 |

| | | |
|---|---|---|
| NC_004718.3;NP_828850.1;SARS-CoV_orf1a;3772 | 0 | 0 |
| NC_004718.3;NP_828850.1;SARS-CoV_orf1a;3773 | 0 | 0 |
| NC_004718.3;NP_828850.1;SARS-CoV_orf1a;3774 | 0 | 0 |
| NC_004718.3;NP_828850.1;SARS-CoV_orf1a;3775 | 0 | 0 |
| NC_004718.3;NP_828850.1;SARS-CoV_orf1a;3776 | 0 | 0 |
| NC_004718.3;NP_828850.1;SARS-CoV_orf1a;3777 | 0 | 0 |
| NC_004718.3;NP_828850.1;SARS-CoV_orf1a;3778 | 0 | 0 |
| NC_004718.3;NP_828850.1;SARS-CoV_orf1a;3779 | 0 | 0 |
| NC_004718.3;NP_828850.1;SARS-CoV_orf1a;3780 | 0 | 0 |
| NC_004718.3;NP_828850.1;SARS-CoV_orf1a;3781 | 0 | 0 |
| NC_004718.3;NP_828850.1;SARS-CoV_orf1a;3782 | 0 | 0 |
| NC_004718.3;NP_828850.1;SARS-CoV_orf1a;3783 | 0 | 0 |
| NC_004718.3;NP_828850.1;SARS-CoV_orf1a;3784 | 0 | 0 |
| NC_004718.3;NP_828850.1;SARS-CoV_orf1a;3785 | 0 | 0 |
| NC_004718.3;NP_828850.1;SARS-CoV_orf1a;3786 | 0.25 | 0.175 |
| NC_004718.3;NP_828850.1;SARS-CoV_orf1a;3787 | 0 | 0 |
| NC_004718.3;NP_828850.1;SARS-CoV_orf1a;3788 | 0 | 0 |
| NC_004718.3;NP_828850.1;SARS-CoV_orf1a;3789 | 0 | 0 |
| NC_004718.3;NP_828850.1;SARS-CoV_orf1a;3790 | 0 | 0 |
| NC_004718.3;NP_828850.1;SARS-CoV_orf1a;3791 | 0 | 0 |
| NC_004718.3;NP_828850.1;SARS-CoV_orf1a;3792 | 0 | 0 |
| NC_004718.3;NP_828850.1;SARS-CoV_orf1a;3793 | 0 | 0 |
| NC_004718.3;NP_828850.1;SARS-CoV_orf1a;3794 | 0 | 0.05 |
| NC_004718.3;NP_828850.1;SARS-CoV_orf1a;3795 | 0.05 | 0.025 |
| NC_004718.3;NP_828850.1;SARS-CoV_orf1a;3796 | 0.05 | 0.025 |
| NC_004718.3;NP_828850.1;SARS-CoV_orf1a;3797 | 0 | 0.025 |
| NC_004718.3;NP_828850.1;SARS-CoV_orf1a;3798 | 0 | 0.025 |
| NC_004718.3;NP_828850.1;SARS-CoV_orf1a;3799 | 0 | 0.1 |
| NC_004718.3;NP_828850.1;SARS-CoV_orf1a;3800 | 0 | 0.025 |
| NC_004718.3;NP_828850.1;SARS-CoV_orf1a;3801 | 0.2 | 0.15 |
| NC_004718.3;NP_828850.1;SARS-CoV_orf1a;3802 | 0.15 | 0.075 |
| NC_004718.3;NP_828850.1;SARS-CoV_orf1a;3803 | 0.15 | 0.125 |
| NC_004718.3;NP_828850.1;SARS-CoV_orf1a;3804 | 0.1 | 0.075 |
| NC_004718.3;NP_828850.1;SARS-CoV_orf1a;3805 | 0.1 | 0.05 |
| NC_004718.3;NP_828850.1;SARS-CoV_orf1a;3806 | 0.05 | 0.05 |
| NC_004718.3;NP_828850.1;SARS-CoV_orf1a;3807 | 0 | 0.05 |
| NC_004718.3;NP_828850.1;SARS-CoV_orf1a;3808 | 0 | 0.05 |
| NC_004718.3;NP_828850.1;SARS-CoV_orf1a;3809 | 0 | 0.025 |
| NC_004718.3;NP_828850.1;SARS-CoV_orf1a;3810 | 0 | 0 |
| NC_004718.3;NP_828850.1;SARS-CoV_orf1a;3811 | 0 | 0 |
| NC_004718.3;NP_828850.1;SARS-CoV_orf1a;3812 | 0 | 0 |
| NC_004718.3;NP_828850.1;SARS-CoV_orf1a;3813 | 0 | 0 |
| NC_004718.3;NP_828850.1;SARS-CoV_orf1a;3814 | 0 | 0 |

| | | |
|---|---|---|
| NC_004718.3;NP_828850.1;SARS-CoV_orf1a;3815 | 0 | 0 |
| NC_004718.3;NP_828850.1;SARS-CoV_orf1a;3816 | 0 | 0 |
| NC_004718.3;NP_828850.1;SARS-CoV_orf1a;3817 | 0 | 0 |
| NC_004718.3;NP_828850.1;SARS-CoV_orf1a;3818 | 0 | 0 |
| NC_004718.3;NP_828850.1;SARS-CoV_orf1a;3819 | 0 | 0.025 |
| NC_004718.3;NP_828850.1;SARS-CoV_orf1a;3820 | 0 | 0 |
| NC_004718.3;NP_828850.1;SARS-CoV_orf1a;3821 | 0 | 0.025 |
| NC_004718.3;NP_828850.1;SARS-CoV_orf1a;3822 | 0 | 0.025 |
| NC_004718.3;NP_828850.1;SARS-CoV_orf1a;3823 | 0.05 | 0.05 |
| NC_004718.3;NP_828850.1;SARS-CoV_orf1a;3824 | 0.05 | 0.05 |
| NC_004718.3;NP_828850.1;SARS-CoV_orf1a;3825 | 0.05 | 0.05 |
| NC_004718.3;NP_828850.1;SARS-CoV_orf1a;3826 | 0.1 | 0.1 |
| NC_004718.3;NP_828850.1;SARS-CoV_orf1a;3827 | 0.05 | 0.075 |
| NC_004718.3;NP_828850.1;SARS-CoV_orf1a;3828 | 0.05 | 0.05 |
| NC_004718.3;NP_828850.1;SARS-CoV_orf1a;3829 | 0.05 | 0.025 |
| NC_004718.3;NP_828850.1;SARS-CoV_orf1a;3830 | 0 | 0.025 |
| NC_004718.3;NP_828850.1;SARS-CoV_orf1a;3831 | 0.05 | 0.025 |
| NC_004718.3;NP_828850.1;SARS-CoV_orf1a;3832 | 0.05 | 0.025 |
| NC_004718.3;NP_828850.1;SARS-CoV_orf1a;3833 | 0.05 | 0.025 |
| NC_004718.3;NP_828850.1;SARS-CoV_orf1a;3834 | 0.05 | 0 |
| NC_004718.3;NP_828850.1;SARS-CoV_orf1a;3835 | 0 | 0 |
| NC_004718.3;NP_828850.1;SARS-CoV_orf1a;3836 | 0 | 0 |
| NC_004718.3;NP_828850.1;SARS-CoV_orf1a;3837 | 0 | 0 |
| NC_004718.3;NP_828850.1;SARS-CoV_orf1a;3838 | 0 | 0 |
| NC_004718.3;NP_828850.1;SARS-CoV_orf1a;3839 | 0 | 0 |
| NC_004718.3;NP_828850.1;SARS-CoV_orf1a;3840 | 0 | 0 |
| NC_004718.3;NP_828850.1;SARS-CoV_orf1a;3841 | 0 | 0 |
| NC_004718.3;NP_828850.1;SARS-CoV_orf1a;3842 | 0 | 0 |
| NC_004718.3;NP_828850.1;SARS-CoV_orf1a;3843 | 0 | 0 |
| NC_004718.3;NP_828850.1;SARS-CoV_orf1a;3844 | 0 | 0 |
| NC_004718.3;NP_828850.1;SARS-CoV_orf1a;3845 | 0 | 0 |
| NC_004718.3;NP_828850.1;SARS-CoV_orf1a;3846 | 0 | 0 |
| NC_004718.3;NP_828850.1;SARS-CoV_orf1a;3847 | 0 | 0 |
| NC_004718.3;NP_828850.1;SARS-CoV_orf1a;3848 | 0 | 0 |
| NC_004718.3;NP_828850.1;SARS-CoV_orf1a;3849 | 0 | 0 |
| NC_004718.3;NP_828850.1;SARS-CoV_orf1a;3850 | 0 | 0 |
| NC_004718.3;NP_828850.1;SARS-CoV_orf1a;3851 | 0 | 0 |
| NC_004718.3;NP_828850.1;SARS-CoV_orf1a;3852 | 0 | 0 |
| NC_004718.3;NP_828850.1;SARS-CoV_orf1a;3853 | 0 | 0 |
| NC_004718.3;NP_828850.1;SARS-CoV_orf1a;3854 | 0 | 0 |
| NC_004718.3;NP_828850.1;SARS-CoV_orf1a;3855 | 0 | 0 |
| NC_004718.3;NP_828850.1;SARS-CoV_orf1a;3856 | 0 | 0 |
| NC_004718.3;NP_828850.1;SARS-CoV_orf1a;3857 | 0 | 0 |

| | | |
|---|---|---|
| NC_004718.3;NP_828850.1;SARS-CoV_orf1a;3858 | 0 | 0 |
| NC_004718.3;NP_828850.1;SARS-CoV_orf1a;3859 | 0 | 0.025 |
| NC_004718.3;NP_828850.1;SARS-CoV_orf1a;3860 | 0 | 0 |
| NC_004718.3;NP_828850.1;SARS-CoV_orf1a;3861 | 0 | 0.025 |
| NC_004718.3;NP_828850.1;SARS-CoV_orf1a;3862 | 0 | 0 |
| NC_004718.3;NP_828850.1;SARS-CoV_orf1a;3863 | 0 | 0 |
| NC_004718.3;NP_828850.1;SARS-CoV_orf1a;3864 | 0 | 0 |
| NC_004718.3;NP_828850.1;SARS-CoV_orf1a;3865 | 0.05 | 0 |
| NC_004718.3;NP_828850.1;SARS-CoV_orf1a;3866 | 0.05 | 0 |
| NC_004718.3;NP_828850.1;SARS-CoV_orf1a;3867 | 0.05 | 0 |
| NC_004718.3;NP_828850.1;SARS-CoV_orf1a;3868 | 0 | 0 |
| NC_004718.3;NP_828850.1;SARS-CoV_orf1a;3869 | 0 | 0 |
| NC_004718.3;NP_828850.1;SARS-CoV_orf1a;3870 | 0 | 0 |
| NC_004718.3;NP_828850.1;SARS-CoV_orf1a;3871 | 0 | 0 |
| NC_004718.3;NP_828850.1;SARS-CoV_orf1a;3872 | 0 | 0 |
| NC_004718.3;NP_828850.1;SARS-CoV_orf1a;3873 | 0 | 0 |
| NC_004718.3;NP_828850.1;SARS-CoV_orf1a;3874 | 0 | 0 |
| NC_004718.3;NP_828850.1;SARS-CoV_orf1a;3875 | 0 | 0 |
| NC_004718.3;NP_828850.1;SARS-CoV_orf1a;3876 | 0 | 0 |
| NC_004718.3;NP_828850.1;SARS-CoV_orf1a;3877 | 0 | 0 |
| NC_004718.3;NP_828850.1;SARS-CoV_orf1a;3878 | 0 | 0 |
| NC_004718.3;NP_828850.1;SARS-CoV_orf1a;3879 | 0 | 0 |
| NC_004718.3;NP_828850.1;SARS-CoV_orf1a;3880 | 0 | 0 |
| NC_004718.3;NP_828850.1;SARS-CoV_orf1a;3881 | 0 | 0 |
| NC_004718.3;NP_828850.1;SARS-CoV_orf1a;3882 | 0 | 0 |
| NC_004718.3;NP_828850.1;SARS-CoV_orf1a;3883 | 0 | 0 |
| NC_004718.3;NP_828850.1;SARS-CoV_orf1a;3884 | 0 | 0 |
| NC_004718.3;NP_828850.1;SARS-CoV_orf1a;3885 | 0 | 0 |
| NC_004718.3;NP_828850.1;SARS-CoV_orf1a;3886 | 0 | 0 |
| NC_004718.3;NP_828850.1;SARS-CoV_orf1a;3887 | 0 | 0 |
| NC_004718.3;NP_828850.1;SARS-CoV_orf1a;3888 | 0 | 0 |
| NC_004718.3;NP_828850.1;SARS-CoV_orf1a;3889 | 0 | 0 |
| NC_004718.3;NP_828850.1;SARS-CoV_orf1a;3890 | 0 | 0 |
| NC_004718.3;NP_828850.1;SARS-CoV_orf1a;3891 | 0 | 0.025 |
| NC_004718.3;NP_828850.1;SARS-CoV_orf1a;3892 | 0 | 0.025 |
| NC_004718.3;NP_828850.1;SARS-CoV_orf1a;3893 | 0 | 0.025 |
| NC_004718.3;NP_828850.1;SARS-CoV_orf1a;3894 | 0.05 | 0.025 |
| NC_004718.3;NP_828850.1;SARS-CoV_orf1a;3895 | 0.05 | 0.025 |
| NC_004718.3;NP_828850.1;SARS-CoV_orf1a;3896 | 0.05 | 0.125 |
| NC_004718.3;NP_828850.1;SARS-CoV_orf1a;3897 | 0.1 | 0.05 |
| NC_004718.3;NP_828850.1;SARS-CoV_orf1a;3898 | 0.1 | 0.15 |
| NC_004718.3;NP_828850.1;SARS-CoV_orf1a;3899 | 0.1 | 0.1 |
| NC_004718.3;NP_828850.1;SARS-CoV_orf1a;3900 | 0.1 | 0.1 |

| | | |
|---|---|---|
| NC_004718.3;NP_828850.1;SARS-CoV_orf1a;3901 | 0 | 0.025 |
| NC_004718.3;NP_828850.1;SARS-CoV_orf1a;3902 | 0 | 0.075 |
| NC_004718.3;NP_828850.1;SARS-CoV_orf1a;3903 | 0 | 0.025 |
| NC_004718.3;NP_828850.1;SARS-CoV_orf1a;3904 | 0 | 0 |
| NC_004718.3;NP_828850.1;SARS-CoV_orf1a;3905 | 0 | 0 |
| NC_004718.3;NP_828850.1;SARS-CoV_orf1a;3906 | 0.05 | 0 |
| NC_004718.3;NP_828850.1;SARS-CoV_orf1a;3907 | 0.05 | 0 |
| NC_004718.3;NP_828850.1;SARS-CoV_orf1a;3908 | 0.05 | 0 |
| NC_004718.3;NP_828850.1;SARS-CoV_orf1a;3909 | 0 | 0 |
| NC_004718.3;NP_828850.1;SARS-CoV_orf1a;3910 | 0.05 | 0 |
| NC_004718.3;NP_828850.1;SARS-CoV_orf1a;3911 | 0 | 0 |
| NC_004718.3;NP_828850.1;SARS-CoV_orf1a;3912 | 0 | 0 |
| NC_004718.3;NP_828850.1;SARS-CoV_orf1a;3913 | 0 | 0 |
| NC_004718.3;NP_828850.1;SARS-CoV_orf1a;3914 | 0 | 0 |
| NC_004718.3;NP_828850.1;SARS-CoV_orf1a;3915 | 0 | 0 |
| NC_004718.3;NP_828850.1;SARS-CoV_orf1a;3916 | 0 | 0 |
| NC_004718.3;NP_828850.1;SARS-CoV_orf1a;3917 | 0 | 0 |
| NC_004718.3;NP_828850.1;SARS-CoV_orf1a;3918 | 0.05 | 0.05 |
| NC_004718.3;NP_828850.1;SARS-CoV_orf1a;3919 | 0 | 0.025 |
| NC_004718.3;NP_828850.1;SARS-CoV_orf1a;3920 | 0 | 0 |
| NC_004718.3;NP_828850.1;SARS-CoV_orf1a;3921 | 0 | 0 |
| NC_004718.3;NP_828850.1;SARS-CoV_orf1a;3922 | 0.05 | 0 |
| NC_004718.3;NP_828850.1;SARS-CoV_orf1a;3923 | 0 | 0.05 |
| NC_004718.3;NP_828850.1;SARS-CoV_orf1a;3924 | 0 | 0 |
| NC_004718.3;NP_828850.1;SARS-CoV_orf1a;3925 | 0 | 0 |
| NC_004718.3;NP_828850.1;SARS-CoV_orf1a;3926 | 0 | 0.025 |
| NC_004718.3;NP_828850.1;SARS-CoV_orf1a;3927 | 0 | 0.05 |
| NC_004718.3;NP_828850.1;SARS-CoV_orf1a;3928 | 0.05 | 0.05 |
| NC_004718.3;NP_828850.1;SARS-CoV_orf1a;3929 | 0 | 0.025 |
| NC_004718.3;NP_828850.1;SARS-CoV_orf1a;3930 | 0.05 | 0.075 |
| NC_004718.3;NP_828850.1;SARS-CoV_orf1a;3931 | 0.05 | 0.15 |
| NC_004718.3;NP_828850.1;SARS-CoV_orf1a;3932 | 0.05 | 0.15 |
| NC_004718.3;NP_828850.1;SARS-CoV_orf1a;3933 | 0.1 | 0.2 |
| NC_004718.3;NP_828850.1;SARS-CoV_orf1a;3934 | 0.05 | 0.1 |
| NC_004718.3;NP_828850.1;SARS-CoV_orf1a;3935 | 0 | 0.025 |
| NC_004718.3;NP_828850.1;SARS-CoV_orf1a;3936 | 0 | 0 |
| NC_004718.3;NP_828850.1;SARS-CoV_orf1a;3937 | 0.05 | 0 |
| NC_004718.3;NP_828850.1;SARS-CoV_orf1a;3938 | 0 | 0 |
| NC_004718.3;NP_828850.1;SARS-CoV_orf1a;3939 | 0 | 0 |
| NC_004718.3;NP_828850.1;SARS-CoV_orf1a;3940 | 0 | 0 |
| NC_004718.3;NP_828850.1;SARS-CoV_orf1a;3941 | 0 | 0 |
| NC_004718.3;NP_828850.1;SARS-CoV_orf1a;3942 | 0 | 0 |
| NC_004718.3;NP_828850.1;SARS-CoV_orf1a;3943 | 0 | 0 |

| | | |
|---|---|---|
| NC_004718.3;NP_828850.1;SARS-CoV_orf1a;3944 | 0 | 0 |
| NC_004718.3;NP_828850.1;SARS-CoV_orf1a;3945 | 0 | 0 |
| NC_004718.3;NP_828850.1;SARS-CoV_orf1a;3946 | 0 | 0 |
| NC_004718.3;NP_828850.1;SARS-CoV_orf1a;3947 | 0 | 0 |
| NC_004718.3;NP_828850.1;SARS-CoV_orf1a;3948 | 0 | 0 |
| NC_004718.3;NP_828850.1;SARS-CoV_orf1a;3949 | 0 | 0 |
| NC_004718.3;NP_828850.1;SARS-CoV_orf1a;3950 | 0 | 0 |
| NC_004718.3;NP_828850.1;SARS-CoV_orf1a;3951 | 0 | 0 |
| NC_004718.3;NP_828850.1;SARS-CoV_orf1a;3952 | 0 | 0 |
| NC_004718.3;NP_828850.1;SARS-CoV_orf1a;3953 | 0 | 0 |
| NC_004718.3;NP_828850.1;SARS-CoV_orf1a;3954 | 0 | 0.025 |
| NC_004718.3;NP_828850.1;SARS-CoV_orf1a;3955 | 0 | 0.05 |
| NC_004718.3;NP_828850.1;SARS-CoV_orf1a;3956 | 0 | 0.025 |
| NC_004718.3;NP_828850.1;SARS-CoV_orf1a;3957 | 0.05 | 0.025 |
| NC_004718.3;NP_828850.1;SARS-CoV_orf1a;3958 | 0.05 | 0.025 |
| NC_004718.3;NP_828850.1;SARS-CoV_orf1a;3959 | 0 | 0.05 |
| NC_004718.3;NP_828850.1;SARS-CoV_orf1a;3960 | 0.05 | 0.025 |
| NC_004718.3;NP_828850.1;SARS-CoV_orf1a;3961 | 0 | 0.025 |
| NC_004718.3;NP_828850.1;SARS-CoV_orf1a;3962 | 0 | 0 |
| NC_004718.3;NP_828850.1;SARS-CoV_orf1a;3963 | 0 | 0 |
| NC_004718.3;NP_828850.1;SARS-CoV_orf1a;3964 | 0 | 0 |
| NC_004718.3;NP_828850.1;SARS-CoV_orf1a;3965 | 0 | 0 |
| NC_004718.3;NP_828850.1;SARS-CoV_orf1a;3966 | 0 | 0 |
| NC_004718.3;NP_828850.1;SARS-CoV_orf1a;3967 | 0 | 0 |
| NC_004718.3;NP_828850.1;SARS-CoV_orf1a;3968 | 0 | 0 |
| NC_004718.3;NP_828850.1;SARS-CoV_orf1a;3969 | 0 | 0 |
| NC_004718.3;NP_828850.1;SARS-CoV_orf1a;3970 | 0 | 0 |
| NC_004718.3;NP_828850.1;SARS-CoV_orf1a;3971 | 0 | 0.025 |
| NC_004718.3;NP_828850.1;SARS-CoV_orf1a;3972 | 0 | 0 |
| NC_004718.3;NP_828850.1;SARS-CoV_orf1a;3973 | 0 | 0 |
| NC_004718.3;NP_828850.1;SARS-CoV_orf1a;3974 | 0 | 0 |
| NC_004718.3;NP_828850.1;SARS-CoV_orf1a;3975 | 0 | 0.05 |
| NC_004718.3;NP_828850.1;SARS-CoV_orf1a;3976 | 0 | 0.025 |
| NC_004718.3;NP_828850.1;SARS-CoV_orf1a;3977 | 0 | 0 |
| NC_004718.3;NP_828850.1;SARS-CoV_orf1a;3978 | 0 | 0 |
| NC_004718.3;NP_828850.1;SARS-CoV_orf1a;3979 | 0 | 0 |
| NC_004718.3;NP_828850.1;SARS-CoV_orf1a;3980 | 0 | 0 |
| NC_004718.3;NP_828850.1;SARS-CoV_orf1a;3981 | 0 | 0 |
| NC_004718.3;NP_828850.1;SARS-CoV_orf1a;3982 | 0 | 0 |
| NC_004718.3;NP_828850.1;SARS-CoV_orf1a;3983 | 0 | 0.1 |
| NC_004718.3;NP_828850.1;SARS-CoV_orf1a;3984 | 0 | 0.075 |
| NC_004718.3;NP_828850.1;SARS-CoV_orf1a;3985 | 0 | 0.05 |
| NC_004718.3;NP_828850.1;SARS-CoV_orf1a;3986 | 0 | 0.05 |

| | | |
|---|---|---|
| NC_004718.3;NP_828850.1;SARS-CoV_orf1a;3987 | 0 | 0.05 |
| NC_004718.3;NP_828850.1;SARS-CoV_orf1a;3988 | 0 | 0 |
| NC_004718.3;NP_828850.1;SARS-CoV_orf1a;3989 | 0 | 0 |
| NC_004718.3;NP_828850.1;SARS-CoV_orf1a;3990 | 0 | 0 |
| NC_004718.3;NP_828850.1;SARS-CoV_orf1a;3991 | 0 | 0 |
| NC_004718.3;NP_828850.1;SARS-CoV_orf1a;3992 | 0 | 0 |
| NC_004718.3;NP_828850.1;SARS-CoV_orf1a;3993 | 0 | 0 |
| NC_004718.3;NP_828850.1;SARS-CoV_orf1a;3994 | 0 | 0 |
| NC_004718.3;NP_828850.1;SARS-CoV_orf1a;3995 | 0 | 0 |
| NC_004718.3;NP_828850.1;SARS-CoV_orf1a;3996 | 0 | 0 |
| NC_004718.3;NP_828850.1;SARS-CoV_orf1a;3997 | 0 | 0 |
| NC_004718.3;NP_828850.1;SARS-CoV_orf1a;3998 | 0 | 0 |
| NC_004718.3;NP_828850.1;SARS-CoV_orf1a;3999 | 0 | 0 |
| NC_004718.3;NP_828850.1;SARS-CoV_orf1a;4000 | 0 | 0 |
| NC_004718.3;NP_828850.1;SARS-CoV_orf1a;4001 | 0 | 0 |
| NC_004718.3;NP_828850.1;SARS-CoV_orf1a;4002 | 0 | 0 |
| NC_004718.3;NP_828850.1;SARS-CoV_orf1a;4003 | 0 | 0 |
| NC_004718.3;NP_828850.1;SARS-CoV_orf1a;4004 | 0 | 0 |
| NC_004718.3;NP_828850.1;SARS-CoV_orf1a;4005 | 0 | 0 |
| NC_004718.3;NP_828850.1;SARS-CoV_orf1a;4006 | 0 | 0 |
| NC_004718.3;NP_828850.1;SARS-CoV_orf1a;4007 | 0 | 0 |
| NC_004718.3;NP_828850.1;SARS-CoV_orf1a;4008 | 0.15 | 0.1 |
| NC_004718.3;NP_828850.1;SARS-CoV_orf1a;4009 | 0.2 | 0.1 |
| NC_004718.3;NP_828850.1;SARS-CoV_orf1a;4010 | 0.2 | 0.075 |
| NC_004718.3;NP_828850.1;SARS-CoV_orf1a;4011 | 0.25 | 0.1 |
| NC_004718.3;NP_828850.1;SARS-CoV_orf1a;4012 | 0.25 | 0.275 |
| NC_004718.3;NP_828850.1;SARS-CoV_orf1a;4013 | 0.3 | 0.3 |
| NC_004718.3;NP_828850.1;SARS-CoV_orf1a;4014 | 0.35 | 0.4 |
| NC_004718.3;NP_828850.1;SARS-CoV_orf1a;4015 | 0.25 | 0.25 |
| NC_004718.3;NP_828850.1;SARS-CoV_orf1a;4016 | 0.4 | 0.5 |
| NC_004718.3;NP_828850.1;SARS-CoV_orf1a;4017 | 0.1 | 0.025 |
| NC_004718.3;NP_828850.1;SARS-CoV_orf1a;4018 | 0.05 | 0.025 |
| NC_004718.3;NP_828850.1;SARS-CoV_orf1a;4019 | 0 | 0 |
| NC_004718.3;NP_828850.1;SARS-CoV_orf1a;4020 | 0.05 | 0.1 |
| NC_004718.3;NP_828850.1;SARS-CoV_orf1a;4021 | 0.1 | 0.075 |
| NC_004718.3;NP_828850.1;SARS-CoV_orf1a;4022 | 0 | 0.075 |
| NC_004718.3;NP_828850.1;SARS-CoV_orf1a;4023 | 0.05 | 0.075 |
| NC_004718.3;NP_828850.1;SARS-CoV_orf1a;4024 | 0.05 | 0 |
| NC_004718.3;NP_828850.1;SARS-CoV_orf1a;4025 | 0 | 0.05 |
| NC_004718.3;NP_828850.1;SARS-CoV_orf1a;4026 | 0 | 0 |
| NC_004718.3;NP_828850.1;SARS-CoV_orf1a;4027 | 0 | 0 |
| NC_004718.3;NP_828850.1;SARS-CoV_orf1a;4028 | 0 | 0 |
| NC_004718.3;NP_828850.1;SARS-CoV_orf1a;4029 | 0 | 0 |

| | | |
|---|---|---|
| NC_004718.3;NP_828850.1;SARS-CoV_orf1a;4030 | 0.05 | 0.025 |
| NC_004718.3;NP_828850.1;SARS-CoV_orf1a;4031 | 0.05 | 0.025 |
| NC_004718.3;NP_828850.1;SARS-CoV_orf1a;4032 | 0 | 0 |
| NC_004718.3;NP_828850.1;SARS-CoV_orf1a;4033 | 0 | 0 |
| NC_004718.3;NP_828850.1;SARS-CoV_orf1a;4034 | 0 | 0 |
| NC_004718.3;NP_828850.1;SARS-CoV_orf1a;4035 | 0 | 0 |
| NC_004718.3;NP_828850.1;SARS-CoV_orf1a;4036 | 0 | 0 |
| NC_004718.3;NP_828850.1;SARS-CoV_orf1a;4037 | 0 | 0 |
| NC_004718.3;NP_828850.1;SARS-CoV_orf1a;4038 | 0 | 0 |
| NC_004718.3;NP_828850.1;SARS-CoV_orf1a;4039 | 0 | 0.025 |
| NC_004718.3;NP_828850.1;SARS-CoV_orf1a;4040 | 0.05 | 0 |
| NC_004718.3;NP_828850.1;SARS-CoV_orf1a;4041 | 0.05 | 0.05 |
| NC_004718.3;NP_828850.1;SARS-CoV_orf1a;4042 | 0.05 | 0 |
| NC_004718.3;NP_828850.1;SARS-CoV_orf1a;4043 | 0.05 | 0 |
| NC_004718.3;NP_828850.1;SARS-CoV_orf1a;4044 | 0.05 | 0.025 |
| NC_004718.3;NP_828850.1;SARS-CoV_orf1a;4045 | 0.05 | 0.025 |
| NC_004718.3;NP_828850.1;SARS-CoV_orf1a;4046 | 0.05 | 0.025 |
| NC_004718.3;NP_828850.1;SARS-CoV_orf1a;4047 | 0.05 | 0.025 |
| NC_004718.3;NP_828850.1;SARS-CoV_orf1a;4048 | 0.05 | 0.025 |
| NC_004718.3;NP_828850.1;SARS-CoV_orf1a;4049 | 0.1 | 0.025 |
| NC_004718.3;NP_828850.1;SARS-CoV_orf1a;4050 | 0.05 | 0 |
| NC_004718.3;NP_828850.1;SARS-CoV_orf1a;4051 | 0 | 0 |
| NC_004718.3;NP_828850.1;SARS-CoV_orf1a;4052 | 0.05 | 0 |
| NC_004718.3;NP_828850.1;SARS-CoV_orf1a;4053 | 0 | 0 |
| NC_004718.3;NP_828850.1;SARS-CoV_orf1a;4054 | 0 | 0 |
| NC_004718.3;NP_828850.1;SARS-CoV_orf1a;4055 | 0 | 0 |
| NC_004718.3;NP_828850.1;SARS-CoV_orf1a;4056 | 0 | 0 |
| NC_004718.3;NP_828850.1;SARS-CoV_orf1a;4057 | 0 | 0 |
| NC_004718.3;NP_828850.1;SARS-CoV_orf1a;4058 | 0 | 0 |
| NC_004718.3;NP_828850.1;SARS-CoV_orf1a;4059 | 0 | 0 |
| NC_004718.3;NP_828850.1;SARS-CoV_orf1a;4060 | 0 | 0 |
| NC_004718.3;NP_828850.1;SARS-CoV_orf1a;4061 | 0 | 0 |
| NC_004718.3;NP_828850.1;SARS-CoV_orf1a;4062 | 0 | 0 |
| NC_004718.3;NP_828850.1;SARS-CoV_orf1a;4063 | 0 | 0 |
| NC_004718.3;NP_828850.1;SARS-CoV_orf1a;4064 | 0 | 0 |
| NC_004718.3;NP_828850.1;SARS-CoV_orf1a;4065 | 0 | 0 |
| NC_004718.3;NP_828850.1;SARS-CoV_orf1a;4066 | 0 | 0 |
| NC_004718.3;NP_828850.1;SARS-CoV_orf1a;4067 | 0 | 0 |
| NC_004718.3;NP_828850.1;SARS-CoV_orf1a;4068 | 0 | 0 |
| NC_004718.3;NP_828850.1;SARS-CoV_orf1a;4069 | 0.05 | 0.025 |
| NC_004718.3;NP_828850.1;SARS-CoV_orf1a;4070 | 0.05 | 0.05 |
| NC_004718.3;NP_828850.1;SARS-CoV_orf1a;4071 | 0 | 0 |
| NC_004718.3;NP_828850.1;SARS-CoV_orf1a;4072 | 0 | 0.025 |

| | | |
|---|---|---|
| NC_004718.3;NP_828850.1;SARS-CoV_orf1a;4073 | 0 | 0 |
| NC_004718.3;NP_828850.1;SARS-CoV_orf1a;4074 | 0 | 0 |
| NC_004718.3;NP_828850.1;SARS-CoV_orf1a;4075 | 0 | 0 |
| NC_004718.3;NP_828850.1;SARS-CoV_orf1a;4076 | 0 | 0.025 |
| NC_004718.3;NP_828850.1;SARS-CoV_orf1a;4077 | 0 | 0 |
| NC_004718.3;NP_828850.1;SARS-CoV_orf1a;4078 | 0 | 0 |
| NC_004718.3;NP_828850.1;SARS-CoV_orf1a;4079 | 0 | 0 |
| NC_004718.3;NP_828850.1;SARS-CoV_orf1a;4080 | 0.15 | 0 |
| NC_004718.3;NP_828850.1;SARS-CoV_orf1a;4081 | 0 | 0 |
| NC_004718.3;NP_828850.1;SARS-CoV_orf1a;4082 | 0 | 0 |
| NC_004718.3;NP_828850.1;SARS-CoV_orf1a;4083 | 0 | 0.025 |
| NC_004718.3;NP_828850.1;SARS-CoV_orf1a;4084 | 0 | 0 |
| NC_004718.3;NP_828850.1;SARS-CoV_orf1a;4085 | 0 | 0.025 |
| NC_004718.3;NP_828850.1;SARS-CoV_orf1a;4086 | 0 | 0 |
| NC_004718.3;NP_828850.1;SARS-CoV_orf1a;4087 | 0 | 0 |
| NC_004718.3;NP_828850.1;SARS-CoV_orf1a;4088 | 0 | 0 |
| NC_004718.3;NP_828850.1;SARS-CoV_orf1a;4089 | 0 | 0 |
| NC_004718.3;NP_828850.1;SARS-CoV_orf1a;4090 | 0 | 0 |
| NC_004718.3;NP_828850.1;SARS-CoV_orf1a;4091 | 0 | 0 |
| NC_004718.3;NP_828850.1;SARS-CoV_orf1a;4092 | 0 | 0 |
| NC_004718.3;NP_828850.1;SARS-CoV_orf1a;4093 | 0 | 0 |
| NC_004718.3;NP_828850.1;SARS-CoV_orf1a;4094 | 0 | 0 |
| NC_004718.3;NP_828850.1;SARS-CoV_orf1a;4095 | 0 | 0 |
| NC_004718.3;NP_828850.1;SARS-CoV_orf1a;4096 | 0 | 0 |
| NC_004718.3;NP_828850.1;SARS-CoV_orf1a;4097 | 0 | 0 |
| NC_004718.3;NP_828850.1;SARS-CoV_orf1a;4098 | 0 | 0 |
| NC_004718.3;NP_828850.1;SARS-CoV_orf1a;4099 | 0 | 0 |
| NC_004718.3;NP_828850.1;SARS-CoV_orf1a;4100 | 0 | 0 |
| NC_004718.3;NP_828850.1;SARS-CoV_orf1a;4101 | 0 | 0 |
| NC_004718.3;NP_828850.1;SARS-CoV_orf1a;4102 | 0 | 0 |
| NC_004718.3;NP_828850.1;SARS-CoV_orf1a;4103 | 0 | 0 |
| NC_004718.3;NP_828850.1;SARS-CoV_orf1a;4104 | 0 | 0.025 |
| NC_004718.3;NP_828850.1;SARS-CoV_orf1a;4105 | 0 | 0.075 |
| NC_004718.3;NP_828850.1;SARS-CoV_orf1a;4106 | 0 | 0.05 |
| NC_004718.3;NP_828850.1;SARS-CoV_orf1a;4107 | 0 | 0.025 |
| NC_004718.3;NP_828850.1;SARS-CoV_orf1a;4108 | 0 | 0.025 |
| NC_004718.3;NP_828850.1;SARS-CoV_orf1a;4109 | 0.05 | 0.025 |
| NC_004718.3;NP_828850.1;SARS-CoV_orf1a;4110 | 0.1 | 0.025 |
| NC_004718.3;NP_828850.1;SARS-CoV_orf1a;4111 | 0.05 | 0.025 |
| NC_004718.3;NP_828850.1;SARS-CoV_orf1a;4112 | 0 | 0.025 |
| NC_004718.3;NP_828850.1;SARS-CoV_orf1a;4113 | 0 | 0 |
| NC_004718.3;NP_828850.1;SARS-CoV_orf1a;4114 | 0.05 | 0.025 |
| NC_004718.3;NP_828850.1;SARS-CoV_orf1a;4115 | 0 | 0 |

| | | |
|---|---|---|
| NC_004718.3;NP_828850.1;SARS-CoV_orf1a;4116 | 0 | 0 |
| NC_004718.3;NP_828850.1;SARS-CoV_orf1a;4117 | 0 | 0.025 |
| NC_004718.3;NP_828850.1;SARS-CoV_orf1a;4118 | 0.05 | 0.05 |
| NC_004718.3;NP_828850.1;SARS-CoV_orf1a;4119 | 0 | 0 |
| NC_004718.3;NP_828850.1;SARS-CoV_orf1a;4120 | 0 | 0 |
| NC_004718.3;NP_828850.1;SARS-CoV_orf1a;4121 | 0 | 0 |
| NC_004718.3;NP_828850.1;SARS-CoV_orf1a;4122 | 0 | 0 |
| NC_004718.3;NP_828850.1;SARS-CoV_orf1a;4123 | 0 | 0 |
| NC_004718.3;NP_828850.1;SARS-CoV_orf1a;4124 | 0 | 0 |
| NC_004718.3;NP_828850.1;SARS-CoV_orf1a;4125 | 0 | 0 |
| NC_004718.3;NP_828850.1;SARS-CoV_orf1a;4126 | 0 | 0 |
| NC_004718.3;NP_828850.1;SARS-CoV_orf1a;4127 | 0 | 0 |
| NC_004718.3;NP_828850.1;SARS-CoV_orf1a;4128 | 0 | 0 |
| NC_004718.3;NP_828850.1;SARS-CoV_orf1a;4129 | 0 | 0 |
| NC_004718.3;NP_828850.1;SARS-CoV_orf1a;4130 | 0 | 0 |
| NC_004718.3;NP_828850.1;SARS-CoV_orf1a;4131 | 0 | 0 |
| NC_004718.3;NP_828850.1;SARS-CoV_orf1a;4132 | 0 | 0 |
| NC_004718.3;NP_828850.1;SARS-CoV_orf1a;4133 | 0 | 0 |
| NC_004718.3;NP_828850.1;SARS-CoV_orf1a;4134 | 0 | 0 |
| NC_004718.3;NP_828850.1;SARS-CoV_orf1a;4135 | 0 | 0 |
| NC_004718.3;NP_828850.1;SARS-CoV_orf1a;4136 | 0 | 0 |
| NC_004718.3;NP_828850.1;SARS-CoV_orf1a;4137 | 0 | 0 |
| NC_004718.3;NP_828850.1;SARS-CoV_orf1a;4138 | 0.15 | 0.125 |
| NC_004718.3;NP_828850.1;SARS-CoV_orf1a;4139 | 0.15 | 0.05 |
| NC_004718.3;NP_828850.1;SARS-CoV_orf1a;4140 | 0.25 | 0.125 |
| NC_004718.3;NP_828850.1;SARS-CoV_orf1a;4141 | 0.05 | 0.075 |
| NC_004718.3;NP_828850.1;SARS-CoV_orf1a;4142 | 0 | 0 |
| NC_004718.3;NP_828850.1;SARS-CoV_orf1a;4143 | 0 | 0 |
| NC_004718.3;NP_828850.1;SARS-CoV_orf1a;4144 | 0 | 0 |
| NC_004718.3;NP_828850.1;SARS-CoV_orf1a;4145 | 0 | 0 |
| NC_004718.3;NP_828850.1;SARS-CoV_orf1a;4146 | 0 | 0 |
| NC_004718.3;NP_828850.1;SARS-CoV_orf1a;4147 | 0 | 0 |
| NC_004718.3;NP_828850.1;SARS-CoV_orf1a;4148 | 0 | 0 |
| NC_004718.3;NP_828850.1;SARS-CoV_orf1a;4149 | 0 | 0 |
| NC_004718.3;NP_828850.1;SARS-CoV_orf1a;4150 | 0 | 0 |
| NC_004718.3;NP_828850.1;SARS-CoV_orf1a;4151 | 0 | 0 |
| NC_004718.3;NP_828850.1;SARS-CoV_orf1a;4152 | 0 | 0 |
| NC_004718.3;NP_828850.1;SARS-CoV_orf1a;4153 | 0 | 0 |
| NC_004718.3;NP_828850.1;SARS-CoV_orf1a;4154 | 0 | 0 |
| NC_004718.3;NP_828850.1;SARS-CoV_orf1a;4155 | 0 | 0 |
| NC_004718.3;NP_828850.1;SARS-CoV_orf1a;4156 | 0 | 0 |
| NC_004718.3;NP_828850.1;SARS-CoV_orf1a;4157 | 0 | 0 |
| NC_004718.3;NP_828850.1;SARS-CoV_orf1a;4158 | 0 | 0 |

| | | |
|---|---|---|
| NC_004718.3;NP_828850.1;SARS-CoV_orf1a;4159 | 0 | 0 |
| NC_004718.3;NP_828850.1;SARS-CoV_orf1a;4160 | 0 | 0 |
| NC_004718.3;NP_828850.1;SARS-CoV_orf1a;4161 | 0 | 0 |
| NC_004718.3;NP_828850.1;SARS-CoV_orf1a;4162 | 0 | 0 |
| NC_004718.3;NP_828850.1;SARS-CoV_orf1a;4163 | 0 | 0 |
| NC_004718.3;NP_828850.1;SARS-CoV_orf1a;4164 | 0 | 0.025 |
| NC_004718.3;NP_828850.1;SARS-CoV_orf1a;4165 | 0 | 0.025 |
| NC_004718.3;NP_828850.1;SARS-CoV_orf1a;4166 | 0 | 0 |
| NC_004718.3;NP_828850.1;SARS-CoV_orf1a;4167 | 0 | 0 |
| NC_004718.3;NP_828850.1;SARS-CoV_orf1a;4168 | 0 | 0 |
| NC_004718.3;NP_828850.1;SARS-CoV_orf1a;4169 | 0 | 0 |
| NC_004718.3;NP_828850.1;SARS-CoV_orf1a;4170 | 0 | 0 |
| NC_004718.3;NP_828850.1;SARS-CoV_orf1a;4171 | 0 | 0 |
| NC_004718.3;NP_828850.1;SARS-CoV_orf1a;4172 | 0 | 0 |
| NC_004718.3;NP_828850.1;SARS-CoV_orf1a;4173 | 0 | 0 |
| NC_004718.3;NP_828850.1;SARS-CoV_orf1a;4174 | 0 | 0 |
| NC_004718.3;NP_828850.1;SARS-CoV_orf1a;4175 | 0 | 0 |
| NC_004718.3;NP_828850.1;SARS-CoV_orf1a;4176 | 0 | 0 |
| NC_004718.3;NP_828850.1;SARS-CoV_orf1a;4177 | 0 | 0 |
| NC_004718.3;NP_828850.1;SARS-CoV_orf1a;4178 | 0 | 0.025 |
| NC_004718.3;NP_828850.1;SARS-CoV_orf1a;4179 | 0 | 0 |
| NC_004718.3;NP_828850.1;SARS-CoV_orf1a;4180 | 0.2 | 0.15 |
| NC_004718.3;NP_828850.1;SARS-CoV_orf1a;4181 | 0 | 0 |
| NC_004718.3;NP_828850.1;SARS-CoV_orf1a;4182 | 0.05 | 0.15 |
| NC_004718.3;NP_828850.1;SARS-CoV_orf1a;4183 | 0.05 | 0.075 |
| NC_004718.3;NP_828850.1;SARS-CoV_orf1a;4184 | 0.05 | 0.05 |
| NC_004718.3;NP_828850.1;SARS-CoV_orf1a;4185 | 0.05 | 0.025 |
| NC_004718.3;NP_828850.1;SARS-CoV_orf1a;4186 | 0 | 0 |
| NC_004718.3;NP_828850.1;SARS-CoV_orf1a;4187 | 0 | 0.025 |
| NC_004718.3;NP_828850.1;SARS-CoV_orf1a;4188 | 0 | 0 |
| NC_004718.3;NP_828850.1;SARS-CoV_orf1a;4189 | 0 | 0 |
| NC_004718.3;NP_828850.1;SARS-CoV_orf1a;4190 | 0 | 0 |
| NC_004718.3;NP_828850.1;SARS-CoV_orf1a;4191 | 0 | 0 |
| NC_004718.3;NP_828850.1;SARS-CoV_orf1a;4192 | 0.05 | 0 |
| NC_004718.3;NP_828850.1;SARS-CoV_orf1a;4193 | 0 | 0 |
| NC_004718.3;NP_828850.1;SARS-CoV_orf1a;4194 | 0 | 0 |
| NC_004718.3;NP_828850.1;SARS-CoV_orf1a;4195 | 0 | 0 |
| NC_004718.3;NP_828850.1;SARS-CoV_orf1a;4196 | 0 | 0 |
| NC_004718.3;NP_828850.1;SARS-CoV_orf1a;4197 | 0 | 0 |
| NC_004718.3;NP_828850.1;SARS-CoV_orf1a;4198 | 0 | 0 |
| NC_004718.3;NP_828850.1;SARS-CoV_orf1a;4199 | 0 | 0 |
| NC_004718.3;NP_828850.1;SARS-CoV_orf1a;4200 | 0 | 0 |
| NC_004718.3;NP_828850.1;SARS-CoV_orf1a;4201 | 0 | 0 |

| | | |
|---|---|---|
| NC_004718.3;NP_828850.1;SARS-CoV_orf1a;4202 | 0 | 0 |
| NC_004718.3;NP_828850.1;SARS-CoV_orf1a;4203 | 0 | 0 |
| NC_004718.3;NP_828850.1;SARS-CoV_orf1a;4204 | 0 | 0 |
| NC_004718.3;NP_828850.1;SARS-CoV_orf1a;4205 | 0 | 0 |
| NC_004718.3;NP_828850.1;SARS-CoV_orf1a;4206 | 0.05 | 0.025 |
| NC_004718.3;NP_828850.1;SARS-CoV_orf1a;4207 | 0.1 | 0.075 |
| NC_004718.3;NP_828850.1;SARS-CoV_orf1a;4208 | 0.15 | 0.1 |
| NC_004718.3;NP_828850.1;SARS-CoV_orf1a;4209 | 0.25 | 0.1 |
| NC_004718.3;NP_828850.1;SARS-CoV_orf1a;4210 | 0.1 | 0.1 |
| NC_004718.3;NP_828850.1;SARS-CoV_orf1a;4211 | 0.05 | 0.05 |
| NC_004718.3;NP_828850.1;SARS-CoV_orf1a;4212 | 0.05 | 0.025 |
| NC_004718.3;NP_828850.1;SARS-CoV_orf1a;4213 | 0 | 0 |
| NC_004718.3;NP_828850.1;SARS-CoV_orf1a;4214 | 0 | 0 |
| NC_004718.3;NP_828850.1;SARS-CoV_orf1a;4215 | 0 | 0 |
| NC_004718.3;NP_828850.1;SARS-CoV_orf1a;4216 | 0 | 0 |
| NC_004718.3;NP_828850.1;SARS-CoV_orf1a;4217 | 0 | 0 |
| NC_004718.3;NP_828850.1;SARS-CoV_orf1a;4218 | 0 | 0 |
| NC_004718.3;NP_828850.1;SARS-CoV_orf1a;4219 | 0 | 0 |
| NC_004718.3;NP_828850.1;SARS-CoV_orf1a;4220 | 0 | 0 |
| NC_004718.3;NP_828850.1;SARS-CoV_orf1a;4221 | 0 | 0.025 |
| NC_004718.3;NP_828850.1;SARS-CoV_orf1a;4222 | 0 | 0 |
| NC_004718.3;NP_828850.1;SARS-CoV_orf1a;4223 | 0 | 0.025 |
| NC_004718.3;NP_828850.1;SARS-CoV_orf1a;4224 | 0.05 | 0.025 |
| NC_004718.3;NP_828850.1;SARS-CoV_orf1a;4225 | 0 | 0.025 |
| NC_004718.3;NP_828850.1;SARS-CoV_orf1a;4226 | 0 | 0 |
| NC_004718.3;NP_828850.1;SARS-CoV_orf1a;4227 | 0 | 0 |
| NC_004718.3;NP_828850.1;SARS-CoV_orf1a;4228 | 0 | 0.025 |
| NC_004718.3;NP_828850.1;SARS-CoV_orf1a;4229 | 0 | 0.025 |
| NC_004718.3;NP_828850.1;SARS-CoV_orf1a;4230 | 0 | 0 |
| NC_004718.3;NP_828850.1;SARS-CoV_orf1a;4231 | 0 | 0 |
| NC_004718.3;NP_828850.1;SARS-CoV_orf1a;4232 | 0 | 0 |
| NC_004718.3;NP_828850.1;SARS-CoV_orf1a;4233 | 0 | 0 |
| NC_004718.3;NP_828850.1;SARS-CoV_orf1a;4234 | 0 | 0 |
| NC_004718.3;NP_828850.1;SARS-CoV_orf1a;4235 | 0 | 0 |
| NC_004718.3;NP_828850.1;SARS-CoV_orf1a;4236 | 0 | 0 |
| NC_004718.3;NP_828850.1;SARS-CoV_orf1a;4237 | 0 | 0 |
| NC_004718.3;NP_828850.1;SARS-CoV_orf1a;4238 | 0 | 0 |
| NC_004718.3;NP_828850.1;SARS-CoV_orf1a;4239 | 0 | 0 |
| NC_004718.3;NP_828850.1;SARS-CoV_orf1a;4240 | 0 | 0 |
| NC_004718.3;NP_828850.1;SARS-CoV_orf1a;4241 | 0 | 0 |
| NC_004718.3;NP_828850.1;SARS-CoV_orf1a;4242 | 0 | 0 |
| NC_004718.3;NP_828850.1;SARS-CoV_orf1a;4243 | 0 | 0 |
| NC_004718.3;NP_828850.1;SARS-CoV_orf1a;4244 | 0.05 | 0 |

| | | |
|---|---|---|
| NC_004718.3;NP_828850.1;SARS-CoV_orf1a;4245 | 0.05 | 0 |
| NC_004718.3;NP_828850.1;SARS-CoV_orf1a;4246 | 0.05 | 0 |
| NC_004718.3;NP_828850.1;SARS-CoV_orf1a;4247 | 0.05 | 0 |
| NC_004718.3;NP_828850.1;SARS-CoV_orf1a;4248 | 0.1 | 0 |
| NC_004718.3;NP_828850.1;SARS-CoV_orf1a;4249 | 0.05 | 0.025 |
| NC_004718.3;NP_828850.1;SARS-CoV_orf1a;4250 | 0.15 | 0.05 |
| NC_004718.3;NP_828850.1;SARS-CoV_orf1a;4251 | 0.15 | 0.025 |
| NC_004718.3;NP_828850.1;SARS-CoV_orf1a;4252 | 0 | 0.025 |
| NC_004718.3;NP_828850.1;SARS-CoV_orf1a;4253 | 0 | 0.025 |
| NC_004718.3;NP_828850.1;SARS-CoV_orf1a;4254 | 0 | 0.05 |
| NC_004718.3;NP_828850.1;SARS-CoV_orf1a;4255 | 0 | 0.05 |
| NC_004718.3;NP_828850.1;SARS-CoV_orf1a;4256 | 0.15 | 0.05 |
| NC_004718.3;NP_828850.1;SARS-CoV_orf1a;4257 | 0.05 | 0.05 |
| NC_004718.3;NP_828850.1;SARS-CoV_orf1a;4258 | 0 | 0 |
| NC_004718.3;NP_828850.1;SARS-CoV_orf1a;4259 | 0 | 0 |
| NC_004718.3;NP_828850.1;SARS-CoV_orf1a;4260 | 0 | 0 |
| NC_004718.3;NP_828850.1;SARS-CoV_orf1a;4261 | 0 | 0 |
| NC_004718.3;NP_828850.1;SARS-CoV_orf1a;4262 | 0 | 0 |
| NC_004718.3;NP_828850.1;SARS-CoV_orf1a;4263 | 0 | 0 |
| NC_004718.3;NP_828850.1;SARS-CoV_orf1a;4264 | 0 | 0 |
| NC_004718.3;NP_828850.1;SARS-CoV_orf1a;4265 | 0 | 0 |
| NC_004718.3;NP_828850.1;SARS-CoV_orf1a;4266 | 0 | 0.025 |
| NC_004718.3;NP_828850.1;SARS-CoV_orf1a;4267 | 0 | 0 |
| NC_004718.3;NP_828850.1;SARS-CoV_orf1a;4268 | 0 | 0.025 |
| NC_004718.3;NP_828850.1;SARS-CoV_orf1a;4269 | 0 | 0.025 |
| NC_004718.3;NP_828850.1;SARS-CoV_orf1a;4270 | 0 | 0 |
| NC_004718.3;NP_828850.1;SARS-CoV_orf1a;4271 | 0 | 0 |
| NC_004718.3;NP_828850.1;SARS-CoV_orf1a;4272 | 0 | 0 |
| NC_004718.3;NP_828850.1;SARS-CoV_orf1a;4273 | 0 | 0 |
| NC_004718.3;NP_828850.1;SARS-CoV_orf1a;4274 | 0 | 0 |
| NC_004718.3;NP_828850.1;SARS-CoV_orf1a;4275 | 0 | 0 |
| NC_004718.3;NP_828850.1;SARS-CoV_orf1a;4276 | 0 | 0 |
| NC_004718.3;NP_828850.1;SARS-CoV_orf1a;4277 | 0 | 0 |
| NC_004718.3;NP_828850.1;SARS-CoV_orf1a;4278 | 0 | 0 |
| NC_004718.3;NP_828850.1;SARS-CoV_orf1a;4279 | 0 | 0.025 |
| NC_004718.3;NP_828850.1;SARS-CoV_orf1a;4280 | 0 | 0.025 |
| NC_004718.3;NP_828850.1;SARS-CoV_orf1a;4281 | 0 | 0.025 |
| NC_004718.3;NP_828850.1;SARS-CoV_orf1a;4282 | 0 | 0.025 |
| NC_004718.3;NP_828850.1;SARS-CoV_orf1a;4283 | 0 | 0.025 |
| NC_004718.3;NP_828850.1;SARS-CoV_orf1a;4284 | 0 | 0.025 |
| NC_004718.3;NP_828850.1;SARS-CoV_orf1a;4285 | 0.05 | 0.025 |
| NC_004718.3;NP_828850.1;SARS-CoV_orf1a;4286 | 0 | 0.025 |
| NC_004718.3;NP_828850.1;SARS-CoV_orf1a;4287 | 0.05 | 0.025 |

| | | |
|---|---|---|
| NC_004718.3;NP_828850.1;SARS-CoV_orf1a;4288 | 0 | 0 |
| NC_004718.3;NP_828850.1;SARS-CoV_orf1a;4289 | 0 | 0 |
| NC_004718.3;NP_828850.1;SARS-CoV_orf1a;4290 | 0 | 0 |
| NC_004718.3;NP_828850.1;SARS-CoV_orf1a;4291 | 0 | 0 |
| NC_004718.3;NP_828850.1;SARS-CoV_orf1a;4292 | 0 | 0 |
| NC_004718.3;NP_828850.1;SARS-CoV_orf1a;4293 | 0 | 0 |
| NC_004718.3;NP_828850.1;SARS-CoV_orf1a;4294 | 0 | 0 |
| NC_004718.3;NP_828850.1;SARS-CoV_orf1a;4295 | 0 | 0 |
| NC_004718.3;NP_828850.1;SARS-CoV_orf1a;4296 | 0 | 0 |
| NC_004718.3;NP_828850.1;SARS-CoV_orf1a;4297 | 0 | 0 |
| NC_004718.3;NP_828850.1;SARS-CoV_orf1a;4298 | 0 | 0 |
| NC_004718.3;NP_828850.1;SARS-CoV_orf1a;4299 | 0 | 0 |
| NC_004718.3;NP_828850.1;SARS-CoV_orf1a;4300 | 0 | 0 |
| NC_004718.3;NP_828850.1;SARS-CoV_orf1a;4301 | 0 | 0 |
| NC_004718.3;NP_828850.1;SARS-CoV_orf1a;4302 | 0 | 0 |
| NC_004718.3;NP_828850.1;SARS-CoV_orf1a;4303 | 0 | 0 |
| NC_004718.3;NP_828850.1;SARS-CoV_orf1a;4304 | 0 | 0 |
| NC_004718.3;NP_828850.1;SARS-CoV_orf1a;4305 | 0 | 0 |
| NC_004718.3;NP_828850.1;SARS-CoV_orf1a;4306 | 0 | 0.025 |
| NC_004718.3;NP_828850.1;SARS-CoV_orf1a;4307 | 0 | 0.025 |
| NC_004718.3;NP_828850.1;SARS-CoV_orf1a;4308 | 0 | 0.025 |
| NC_004718.3;NP_828850.1;SARS-CoV_orf1a;4309 | 0 | 0.025 |
| NC_004718.3;NP_828850.1;SARS-CoV_orf1a;4310 | 0 | 0.025 |
| NC_004718.3;NP_828850.1;SARS-CoV_orf1a;4311 | 0 | 0.025 |
| NC_004718.3;NP_828850.1;SARS-CoV_orf1a;4312 | 0 | 0 |
| NC_004718.3;NP_828850.1;SARS-CoV_orf1a;4313 | 0 | 0 |
| NC_004718.3;NP_828850.1;SARS-CoV_orf1a;4314 | 0 | 0 |
| NC_004718.3;NP_828850.1;SARS-CoV_orf1a;4315 | 0 | 0 |
| NC_004718.3;NP_828850.1;SARS-CoV_orf1a;4316 | 0 | 0 |
| NC_004718.3;NP_828850.1;SARS-CoV_orf1a;4317 | 0 | 0.05 |
| NC_004718.3;NP_828850.1;SARS-CoV_orf1a;4318 | 0 | 0.05 |
| NC_004718.3;NP_828850.1;SARS-CoV_orf1a;4319 | 0 | 0 |
| NC_004718.3;NP_828850.1;SARS-CoV_orf1a;4320 | 0 | 0 |
| NC_004718.3;NP_828850.1;SARS-CoV_orf1a;4321 | 0.05 | 0 |
| NC_004718.3;NP_828850.1;SARS-CoV_orf1a;4322 | 0.05 | 0.025 |
| NC_004718.3;NP_828850.1;SARS-CoV_orf1a;4323 | 0.05 | 0.05 |
| NC_004718.3;NP_828850.1;SARS-CoV_orf1a;4324 | 0.05 | 0 |
| NC_004718.3;NP_828850.1;SARS-CoV_orf1a;4325 | 0.05 | 0 |
| NC_004718.3;NP_828850.1;SARS-CoV_orf1a;4326 | 0 | 0 |
| NC_004718.3;NP_828850.1;SARS-CoV_orf1a;4327 | 0 | 0.025 |
| NC_004718.3;NP_828850.1;SARS-CoV_orf1a;4328 | 0 | 0.025 |
| NC_004718.3;NP_828850.1;SARS-CoV_orf1a;4329 | 0 | 0.025 |
| NC_004718.3;NP_828850.1;SARS-CoV_orf1a;4330 | 0 | 0.05 |

| | | |
|---|---|---|
| NC_004718.3;NP_828850.1;SARS-CoV_orf1a;4331 | 0 | 0.025 |
| NC_004718.3;NP_828850.1;SARS-CoV_orf1a;4332 | 0 | 0.025 |
| NC_004718.3;NP_828850.1;SARS-CoV_orf1a;4333 | 0 | 0.025 |
| NC_004718.3;NP_828850.1;SARS-CoV_orf1a;4334 | 0 | 0.025 |
| NC_004718.3;NP_828850.1;SARS-CoV_orf1a;4335 | 0 | 0 |
| NC_004718.3;NP_828850.1;SARS-CoV_orf1a;4336 | 0 | 0.025 |
| NC_004718.3;NP_828850.1;SARS-CoV_orf1a;4337 | 0 | 0 |
| NC_004718.3;NP_828850.1;SARS-CoV_orf1a;4338 | 0 | 0 |
| NC_004718.3;NP_828850.1;SARS-CoV_orf1a;4339 | 0 | 0 |
| NC_004718.3;NP_828850.1;SARS-CoV_orf1a;4340 | 0 | 0 |
| NC_004718.3;NP_828850.1;SARS-CoV_orf1a;4341 | 0 | 0 |
| NC_004718.3;NP_828850.1;SARS-CoV_orf1a;4342 | 0 | 0 |
| NC_004718.3;NP_828850.1;SARS-CoV_orf1a;4343 | 0 | 0 |
| NC_004718.3;NP_828850.1;SARS-CoV_orf1a;4344 | 0 | 0 |
| NC_004718.3;NP_828850.1;SARS-CoV_orf1a;4345 | 0 | 0 |
| NC_004718.3;NP_828850.1;SARS-CoV_orf1a;4346 | 0 | 0 |
| NC_004718.3;NP_828850.1;SARS-CoV_orf1a;4347 | 0 | 0 |
| NC_004718.3;NP_828850.1;SARS-CoV_orf1a;4348 | 0 | 0 |
| NC_004718.3;NP_828850.1;SARS-CoV_orf1a;4349 | 0 | 0 |
| NC_004718.3;NP_828850.1;SARS-CoV_orf1a;4350 | 0 | 0 |
| NC_004718.3;NP_828850.1;SARS-CoV_orf1a;4351 | 0 | 0 |
| NC_004718.3;NP_828850.1;SARS-CoV_orf1a;4352 | 0 | 0.075 |
| NC_004718.3;NP_828850.1;SARS-CoV_orf1a;4353 | 0 | 0 |
| NC_004718.3;NP_828850.1;SARS-CoV_orf1a;4354 | 0 | 0.025 |
| NC_004718.3;NP_828850.1;SARS-CoV_orf1a;4355 | 0 | 0 |
| NC_004718.3;NP_828850.1;SARS-CoV_orf1a;4356 | 0 | 0 |
| NC_004718.3;NP_828850.1;SARS-CoV_orf1a;4357 | 0 | 0 |
| NC_004718.3;NP_828850.1;SARS-CoV_orf1a;4358 | 0 | 0 |
| NC_004718.3;NP_828850.1;SARS-CoV_orf1a;4359 | 0 | 0 |
| NC_004718.3;NP_828850.1;SARS-CoV_orf1a;4360 | 0 | 0 |
| NC_004718.3;NP_828850.1;SARS-CoV_orf1a;4361 | 0 | 0 |
| NC_004718.3;NP_828850.1;SARS-CoV_orf1a;4362 | 0 | 0 |
| NC_004718.3;NP_828850.1;SARS-CoV_orf1a;4363 | 0.05 | 0 |
| NC_004718.3;NP_828850.1;SARS-CoV_orf1a;4364 | 0.05 | 0 |
| NC_004718.3;NP_828850.1;SARS-CoV_orf1a;4365 | 0 | 0 |
| NC_004718.3;NP_828850.1;SARS-CoV_orf1a;4366 | 0 | 0 |
| NC_004718.3;NP_828850.1;SARS-CoV_orf1a;4367 | 0 | 0 |
| NC_004718.3;NP_828850.1;SARS-CoV_orf1a;4368 | 0 | 0 |
| NC_004718.3;NP_828850.1;SARS-CoV_orf1a;4369 | 0 | 0 |
| NC_004718.3;NP_828850.1;SARS-CoV_orf1a;4370 | 0 | 0 |
| NC_004718.3;NP_828850.1;SARS-CoV_orf1a;4371 | 0 | 0 |
| NC_004718.3;NP_828851.1;SARS-CoV_E2-glycopr;1 | 0 | 0 |
| NC_004718.3;NP_828851.1;SARS-CoV_E2-glycopr;2 | 0 | 0 |

| | | |
|---|---|---|
| NC_004718.3;NP_828851.1;SARS-CoV_E2-glycopr;3 | 0 | 0 |
| NC_004718.3;NP_828851.1;SARS-CoV_E2-glycopr;4 | 0 | 0 |
| NC_004718.3;NP_828851.1;SARS-CoV_E2-glycopr;5 | 0 | 0 |
| NC_004718.3;NP_828851.1;SARS-CoV_E2-glycopr;6 | 0 | 0.025 |
| NC_004718.3;NP_828851.1;SARS-CoV_E2-glycopr;7 | 0 | 0 |
| NC_004718.3;NP_828851.1;SARS-CoV_E2-glycopr;8 | 0.05 | 0.025 |
| NC_004718.3;NP_828851.1;SARS-CoV_E2-glycopr;9 | 0.05 | 0.025 |
| NC_004718.3;NP_828851.1;SARS-CoV_E2-glycopr;10 | 0.05 | 0.025 |
| NC_004718.3;NP_828851.1;SARS-CoV_E2-glycopr;11 | 0.05 | 0.025 |
| NC_004718.3;NP_828851.1;SARS-CoV_E2-glycopr;12 | 0 | 0 |
| NC_004718.3;NP_828851.1;SARS-CoV_E2-glycopr;13 | 0 | 0 |
| NC_004718.3;NP_828851.1;SARS-CoV_E2-glycopr;14 | 0 | 0 |
| NC_004718.3;NP_828851.1;SARS-CoV_E2-glycopr;15 | 0 | 0 |
| NC_004718.3;NP_828851.1;SARS-CoV_E2-glycopr;16 | 0 | 0 |
| NC_004718.3;NP_828851.1;SARS-CoV_E2-glycopr;17 | 0.05 | 0 |
| NC_004718.3;NP_828851.1;SARS-CoV_E2-glycopr;18 | 0.05 | 0.025 |
| NC_004718.3;NP_828851.1;SARS-CoV_E2-glycopr;19 | 0.05 | 0.025 |
| NC_004718.3;NP_828851.1;SARS-CoV_E2-glycopr;20 | 0.05 | 0 |
| NC_004718.3;NP_828851.1;SARS-CoV_E2-glycopr;21 | 0.05 | 0.025 |
| NC_004718.3;NP_828851.1;SARS-CoV_E2-glycopr;22 | 0.05 | 0.025 |
| NC_004718.3;NP_828851.1;SARS-CoV_E2-glycopr;23 | 0.05 | 0.025 |
| NC_004718.3;NP_828851.1;SARS-CoV_E2-glycopr;24 | 0 | 0 |
| NC_004718.3;NP_828851.1;SARS-CoV_E2-glycopr;25 | 0 | 0 |
| NC_004718.3;NP_828851.1;SARS-CoV_E2-glycopr;26 | 0 | 0.025 |
| NC_004718.3;NP_828851.1;SARS-CoV_E2-glycopr;27 | 0 | 0 |
| NC_004718.3;NP_828851.1;SARS-CoV_E2-glycopr;28 | 0 | 0 |
| NC_004718.3;NP_828851.1;SARS-CoV_E2-glycopr;29 | 0 | 0 |
| NC_004718.3;NP_828851.1;SARS-CoV_E2-glycopr;30 | 0.05 | 0 |
| NC_004718.3;NP_828851.1;SARS-CoV_E2-glycopr;31 | 0 | 0 |
| NC_004718.3;NP_828851.1;SARS-CoV_E2-glycopr;32 | 0 | 0 |
| NC_004718.3;NP_828851.1;SARS-CoV_E2-glycopr;33 | 0 | 0 |
| NC_004718.3;NP_828851.1;SARS-CoV_E2-glycopr;34 | 0 | 0 |
| NC_004718.3;NP_828851.1;SARS-CoV_E2-glycopr;35 | 0 | 0 |
| NC_004718.3;NP_828851.1;SARS-CoV_E2-glycopr;36 | 0 | 0 |
| NC_004718.3;NP_828851.1;SARS-CoV_E2-glycopr;37 | 0 | 0 |
| NC_004718.3;NP_828851.1;SARS-CoV_E2-glycopr;38 | 0 | 0 |
| NC_004718.3;NP_828851.1;SARS-CoV_E2-glycopr;39 | 0 | 0 |
| NC_004718.3;NP_828851.1;SARS-CoV_E2-glycopr;40 | 0 | 0 |
| NC_004718.3;NP_828851.1;SARS-CoV_E2-glycopr;41 | 0 | 0 |
| NC_004718.3;NP_828851.1;SARS-CoV_E2-glycopr;42 | 0 | 0 |
| NC_004718.3;NP_828851.1;SARS-CoV_E2-glycopr;43 | 0 | 0 |
| NC_004718.3;NP_828851.1;SARS-CoV_E2-glycopr;44 | 0 | 0 |
| NC_004718.3;NP_828851.1;SARS-CoV_E2-glycopr;45 | 0 | 0 |

| | | |
|---|---|---|
| NC_004718.3;NP_828851.1;SARS-CoV_E2-glycopr;46 | 0 | 0 |
| NC_004718.3;NP_828851.1;SARS-CoV_E2-glycopr;47 | 0 | 0 |
| NC_004718.3;NP_828851.1;SARS-CoV_E2-glycopr;48 | 0 | 0.025 |
| NC_004718.3;NP_828851.1;SARS-CoV_E2-glycopr;49 | 0 | 0 |
| NC_004718.3;NP_828851.1;SARS-CoV_E2-glycopr;50 | 0 | 0 |
| NC_004718.3;NP_828851.1;SARS-CoV_E2-glycopr;51 | 0 | 0 |
| NC_004718.3;NP_828851.1;SARS-CoV_E2-glycopr;52 | 0 | 0 |
| NC_004718.3;NP_828851.1;SARS-CoV_E2-glycopr;53 | 0 | 0 |
| NC_004718.3;NP_828851.1;SARS-CoV_E2-glycopr;54 | 0 | 0 |
| NC_004718.3;NP_828851.1;SARS-CoV_E2-glycopr;55 | 0 | 0 |
| NC_004718.3;NP_828851.1;SARS-CoV_E2-glycopr;56 | 0 | 0 |
| NC_004718.3;NP_828851.1;SARS-CoV_E2-glycopr;57 | 0 | 0 |
| NC_004718.3;NP_828851.1;SARS-CoV_E2-glycopr;58 | 0 | 0 |
| NC_004718.3;NP_828851.1;SARS-CoV_E2-glycopr;59 | 0 | 0 |
| NC_004718.3;NP_828851.1;SARS-CoV_E2-glycopr;60 | 0 | 0 |
| NC_004718.3;NP_828851.1;SARS-CoV_E2-glycopr;61 | 0 | 0 |
| NC_004718.3;NP_828851.1;SARS-CoV_E2-glycopr;62 | 0 | 0 |
| NC_004718.3;NP_828851.1;SARS-CoV_E2-glycopr;63 | 0 | 0 |
| NC_004718.3;NP_828851.1;SARS-CoV_E2-glycopr;64 | 0 | 0 |
| NC_004718.3;NP_828851.1;SARS-CoV_E2-glycopr;65 | 0 | 0 |
| NC_004718.3;NP_828851.1;SARS-CoV_E2-glycopr;66 | 0 | 0 |
| NC_004718.3;NP_828851.1;SARS-CoV_E2-glycopr;67 | 0 | 0 |
| NC_004718.3;NP_828851.1;SARS-CoV_E2-glycopr;68 | 0 | 0 |
| NC_004718.3;NP_828851.1;SARS-CoV_E2-glycopr;69 | 0 | 0 |
| NC_004718.3;NP_828851.1;SARS-CoV_E2-glycopr;70 | 0 | 0.025 |
| NC_004718.3;NP_828851.1;SARS-CoV_E2-glycopr;71 | 0 | 0.05 |
| NC_004718.3;NP_828851.1;SARS-CoV_E2-glycopr;72 | 0 | 0.025 |
| NC_004718.3;NP_828851.1;SARS-CoV_E2-glycopr;73 | 0 | 0 |
| NC_004718.3;NP_828851.1;SARS-CoV_E2-glycopr;74 | 0 | 0.025 |
| NC_004718.3;NP_828851.1;SARS-CoV_E2-glycopr;75 | 0 | 0 |
| NC_004718.3;NP_828851.1;SARS-CoV_E2-glycopr;76 | 0.05 | 0 |
| NC_004718.3;NP_828851.1;SARS-CoV_E2-glycopr;77 | 0 | 0 |
| NC_004718.3;NP_828851.1;SARS-CoV_E2-glycopr;78 | 0 | 0 |
| NC_004718.3;NP_828851.1;SARS-CoV_E2-glycopr;79 | 0 | 0.025 |
| NC_004718.3;NP_828851.1;SARS-CoV_E2-glycopr;80 | 0 | 0.025 |
| NC_004718.3;NP_828851.1;SARS-CoV_E2-glycopr;81 | 0 | 0.025 |
| NC_004718.3;NP_828851.1;SARS-CoV_E2-glycopr;82 | 0 | 0.05 |
| NC_004718.3;NP_828851.1;SARS-CoV_E2-glycopr;83 | 0.05 | 0.075 |
| NC_004718.3;NP_828851.1;SARS-CoV_E2-glycopr;84 | 0 | 0.05 |
| NC_004718.3;NP_828851.1;SARS-CoV_E2-glycopr;85 | 0 | 0.05 |
| NC_004718.3;NP_828851.1;SARS-CoV_E2-glycopr;86 | 0 | 0.025 |
| NC_004718.3;NP_828851.1;SARS-CoV_E2-glycopr;87 | 0.05 | 0.025 |
| NC_004718.3;NP_828851.1;SARS-CoV_E2-glycopr;88 | 0.05 | 0 |

| | | |
|---|---|---|
| NC_004718.3;NP_828851.1;SARS-CoV_E2-glycopr;89 | 0.05 | 0 |
| NC_004718.3;NP_828851.1;SARS-CoV_E2-glycopr;90 | 0.05 | 0 |
| NC_004718.3;NP_828851.1;SARS-CoV_E2-glycopr;91 | 0 | 0 |
| NC_004718.3;NP_828851.1;SARS-CoV_E2-glycopr;92 | 0 | 0 |
| NC_004718.3;NP_828851.1;SARS-CoV_E2-glycopr;93 | 0.05 | 0 |
| NC_004718.3;NP_828851.1;SARS-CoV_E2-glycopr;94 | 0 | 0 |
| NC_004718.3;NP_828851.1;SARS-CoV_E2-glycopr;95 | 0 | 0 |
| NC_004718.3;NP_828851.1;SARS-CoV_E2-glycopr;96 | 0 | 0 |
| NC_004718.3;NP_828851.1;SARS-CoV_E2-glycopr;97 | 0 | 0 |
| NC_004718.3;NP_828851.1;SARS-CoV_E2-glycopr;98 | 0 | 0 |
| NC_004718.3;NP_828851.1;SARS-CoV_E2-glycopr;99 | 0 | 0.025 |
| NC_004718.3;NP_828851.1;SARS-CoV_E2-glycopr;100 | 0 | 0 |
| NC_004718.3;NP_828851.1;SARS-CoV_E2-glycopr;101 | 0 | 0 |
| NC_004718.3;NP_828851.1;SARS-CoV_E2-glycopr;102 | 0 | 0 |
| NC_004718.3;NP_828851.1;SARS-CoV_E2-glycopr;103 | 0 | 0 |
| NC_004718.3;NP_828851.1;SARS-CoV_E2-glycopr;104 | 0 | 0.025 |
| NC_004718.3;NP_828851.1;SARS-CoV_E2-glycopr;105 | 0 | 0 |
| NC_004718.3;NP_828851.1;SARS-CoV_E2-glycopr;106 | 0 | 0 |
| NC_004718.3;NP_828851.1;SARS-CoV_E2-glycopr;107 | 0.1 | 0.025 |
| NC_004718.3;NP_828851.1;SARS-CoV_E2-glycopr;108 | 0.1 | 0.125 |
| NC_004718.3;NP_828851.1;SARS-CoV_E2-glycopr;109 | 0.1 | 0.125 |
| NC_004718.3;NP_828851.1;SARS-CoV_E2-glycopr;110 | 0.05 | 0.025 |
| NC_004718.3;NP_828851.1;SARS-CoV_E2-glycopr;111 | 0.05 | 0.025 |
| NC_004718.3;NP_828851.1;SARS-CoV_E2-glycopr;112 | 0.05 | 0.025 |
| NC_004718.3;NP_828851.1;SARS-CoV_E2-glycopr;113 | 0.05 | 0 |
| NC_004718.3;NP_828851.1;SARS-CoV_E2-glycopr;114 | 0.05 | 0.025 |
| NC_004718.3;NP_828851.1;SARS-CoV_E2-glycopr;115 | 0.05 | 0.025 |
| NC_004718.3;NP_828851.1;SARS-CoV_E2-glycopr;116 | 0.25 | 0.275 |
| NC_004718.3;NP_828851.1;SARS-CoV_E2-glycopr;117 | 0.2 | 0.175 |
| NC_004718.3;NP_828851.1;SARS-CoV_E2-glycopr;118 | 0.05 | 0.025 |
| NC_004718.3;NP_828851.1;SARS-CoV_E2-glycopr;119 | 0 | 0 |
| NC_004718.3;NP_828851.1;SARS-CoV_E2-glycopr;120 | 0 | 0 |
| NC_004718.3;NP_828851.1;SARS-CoV_E2-glycopr;121 | 0.05 | 0 |
| NC_004718.3;NP_828851.1;SARS-CoV_E2-glycopr;122 | 0 | 0 |
| NC_004718.3;NP_828851.1;SARS-CoV_E2-glycopr;123 | 0 | 0 |
| NC_004718.3;NP_828851.1;SARS-CoV_E2-glycopr;124 | 0 | 0 |
| NC_004718.3;NP_828851.1;SARS-CoV_E2-glycopr;125 | 0 | 0 |
| NC_004718.3;NP_828851.1;SARS-CoV_E2-glycopr;126 | 0 | 0 |
| NC_004718.3;NP_828851.1;SARS-CoV_E2-glycopr;127 | 0 | 0 |
| NC_004718.3;NP_828851.1;SARS-CoV_E2-glycopr;128 | 0 | 0 |
| NC_004718.3;NP_828851.1;SARS-CoV_E2-glycopr;129 | 0 | 0.025 |
| NC_004718.3;NP_828851.1;SARS-CoV_E2-glycopr;130 | 0 | 0 |
| NC_004718.3;NP_828851.1;SARS-CoV_E2-glycopr;131 | 0 | 0.025 |

| | | |
|---|---|---|
| NC_004718.3;NP_828851.1;SARS-CoV_E2-glycopr;132 | 0 | 0.025 |
| NC_004718.3;NP_828851.1;SARS-CoV_E2-glycopr;133 | 0 | 0.025 |
| NC_004718.3;NP_828851.1;SARS-CoV_E2-glycopr;134 | 0.05 | 0 |
| NC_004718.3;NP_828851.1;SARS-CoV_E2-glycopr;135 | 0.05 | 0 |
| NC_004718.3;NP_828851.1;SARS-CoV_E2-glycopr;136 | 0 | 0 |
| NC_004718.3;NP_828851.1;SARS-CoV_E2-glycopr;137 | 0 | 0 |
| NC_004718.3;NP_828851.1;SARS-CoV_E2-glycopr;138 | 0 | 0 |
| NC_004718.3;NP_828851.1;SARS-CoV_E2-glycopr;139 | 0 | 0 |
| NC_004718.3;NP_828851.1;SARS-CoV_E2-glycopr;140 | 0 | 0 |
| NC_004718.3;NP_828851.1;SARS-CoV_E2-glycopr;141 | 0 | 0 |
| NC_004718.3;NP_828851.1;SARS-CoV_E2-glycopr;142 | 0 | 0 |
| NC_004718.3;NP_828851.1;SARS-CoV_E2-glycopr;143 | 0 | 0 |
| NC_004718.3;NP_828851.1;SARS-CoV_E2-glycopr;144 | 0 | 0 |
| NC_004718.3;NP_828851.1;SARS-CoV_E2-glycopr;145 | 0 | 0 |
| NC_004718.3;NP_828851.1;SARS-CoV_E2-glycopr;146 | 0 | 0 |
| NC_004718.3;NP_828851.1;SARS-CoV_E2-glycopr;147 | 0 | 0 |
| NC_004718.3;NP_828851.1;SARS-CoV_E2-glycopr;148 | 0 | 0 |
| NC_004718.3;NP_828851.1;SARS-CoV_E2-glycopr;149 | 0 | 0 |
| NC_004718.3;NP_828851.1;SARS-CoV_E2-glycopr;150 | 0 | 0 |
| NC_004718.3;NP_828851.1;SARS-CoV_E2-glycopr;151 | 0 | 0 |
| NC_004718.3;NP_828851.1;SARS-CoV_E2-glycopr;152 | 0 | 0 |
| NC_004718.3;NP_828851.1;SARS-CoV_E2-glycopr;153 | 0 | 0 |
| NC_004718.3;NP_828851.1;SARS-CoV_E2-glycopr;154 | 0 | 0 |
| NC_004718.3;NP_828851.1;SARS-CoV_E2-glycopr;155 | 0 | 0 |
| NC_004718.3;NP_828851.1;SARS-CoV_E2-glycopr;156 | 0 | 0 |
| NC_004718.3;NP_828851.1;SARS-CoV_E2-glycopr;157 | 0 | 0 |
| NC_004718.3;NP_828851.1;SARS-CoV_E2-glycopr;158 | 0 | 0 |
| NC_004718.3;NP_828851.1;SARS-CoV_E2-glycopr;159 | 0 | 0 |
| NC_004718.3;NP_828851.1;SARS-CoV_E2-glycopr;160 | 0 | 0 |
| NC_004718.3;NP_828851.1;SARS-CoV_E2-glycopr;161 | 0 | 0 |
| NC_004718.3;NP_828851.1;SARS-CoV_E2-glycopr;162 | 0 | 0 |
| NC_004718.3;NP_828851.1;SARS-CoV_E2-glycopr;163 | 0 | 0 |
| NC_004718.3;NP_828851.1;SARS-CoV_E2-glycopr;164 | 0 | 0 |
| NC_004718.3;NP_828851.1;SARS-CoV_E2-glycopr;165 | 0 | 0.075 |
| NC_004718.3;NP_828851.1;SARS-CoV_E2-glycopr;166 | 0 | 0.025 |
| NC_004718.3;NP_828851.1;SARS-CoV_E2-glycopr;167 | 0 | 0.05 |
| NC_004718.3;NP_828851.1;SARS-CoV_E2-glycopr;168 | 0.05 | 0.05 |
| NC_004718.3;NP_828851.1;SARS-CoV_E2-glycopr;169 | 0.1 | 0.025 |
| NC_004718.3;NP_828851.1;SARS-CoV_E2-glycopr;170 | 0.15 | 0.05 |
| NC_004718.3;NP_828851.1;SARS-CoV_E2-glycopr;171 | 0 | 0.05 |
| NC_004718.3;NP_828851.1;SARS-CoV_E2-glycopr;172 | 0 | 0.025 |
| NC_004718.3;NP_828851.1;SARS-CoV_E2-glycopr;173 | 0 | 0.025 |
| NC_004718.3;NP_828851.1;SARS-CoV_E2-glycopr;174 | 0 | 0.025 |

| | | |
|---|---|---|
| NC_004718.3;NP_828851.1;SARS-CoV_E2-glycopr;175 | 0 | 0 |
| NC_004718.3;NP_828851.1;SARS-CoV_E2-glycopr;176 | 0.05 | 0 |
| NC_004718.3;NP_828851.1;SARS-CoV_E2-glycopr;177 | 0 | 0 |
| NC_004718.3;NP_828851.1;SARS-CoV_E2-glycopr;178 | 0 | 0 |
| NC_004718.3;NP_828851.1;SARS-CoV_E2-glycopr;179 | 0.05 | 0 |
| NC_004718.3;NP_828851.1;SARS-CoV_E2-glycopr;180 | 0.05 | 0 |
| NC_004718.3;NP_828851.1;SARS-CoV_E2-glycopr;181 | 0.05 | 0 |
| NC_004718.3;NP_828851.1;SARS-CoV_E2-glycopr;182 | 0 | 0 |
| NC_004718.3;NP_828851.1;SARS-CoV_E2-glycopr;183 | 0 | 0 |
| NC_004718.3;NP_828851.1;SARS-CoV_E2-glycopr;184 | 0 | 0 |
| NC_004718.3;NP_828851.1;SARS-CoV_E2-glycopr;185 | 0 | 0 |
| NC_004718.3;NP_828851.1;SARS-CoV_E2-glycopr;186 | 0 | 0 |
| NC_004718.3;NP_828851.1;SARS-CoV_E2-glycopr;187 | 0 | 0 |
| NC_004718.3;NP_828851.1;SARS-CoV_E2-glycopr;188 | 0 | 0 |
| NC_004718.3;NP_828851.1;SARS-CoV_E2-glycopr;189 | 0 | 0.025 |
| NC_004718.3;NP_828851.1;SARS-CoV_E2-glycopr;190 | 0 | 0.025 |
| NC_004718.3;NP_828851.1;SARS-CoV_E2-glycopr;191 | 0 | 0 |
| NC_004718.3;NP_828851.1;SARS-CoV_E2-glycopr;192 | 0 | 0 |
| NC_004718.3;NP_828851.1;SARS-CoV_E2-glycopr;193 | 0.1 | 0 |
| NC_004718.3;NP_828851.1;SARS-CoV_E2-glycopr;194 | 0.05 | 0.05 |
| NC_004718.3;NP_828851.1;SARS-CoV_E2-glycopr;195 | 0.1 | 0.075 |
| NC_004718.3;NP_828851.1;SARS-CoV_E2-glycopr;196 | 0.15 | 0.05 |
| NC_004718.3;NP_828851.1;SARS-CoV_E2-glycopr;197 | 0.1 | 0.05 |
| NC_004718.3;NP_828851.1;SARS-CoV_E2-glycopr;198 | 0.1 | 0.075 |
| NC_004718.3;NP_828851.1;SARS-CoV_E2-glycopr;199 | 0.1 | 0.075 |
| NC_004718.3;NP_828851.1;SARS-CoV_E2-glycopr;200 | 0.1 | 0.1 |
| NC_004718.3;NP_828851.1;SARS-CoV_E2-glycopr;201 | 0.2 | 0.2 |
| NC_004718.3;NP_828851.1;SARS-CoV_E2-glycopr;202 | 0 | 0 |
| NC_004718.3;NP_828851.1;SARS-CoV_E2-glycopr;203 | 0 | 0.1 |
| NC_004718.3;NP_828851.1;SARS-CoV_E2-glycopr;204 | 0.05 | 0.025 |
| NC_004718.3;NP_828851.1;SARS-CoV_E2-glycopr;205 | 0 | 0 |
| NC_004718.3;NP_828851.1;SARS-CoV_E2-glycopr;206 | 0 | 0 |
| NC_004718.3;NP_828851.1;SARS-CoV_E2-glycopr;207 | 0 | 0 |
| NC_004718.3;NP_828851.1;SARS-CoV_E2-glycopr;208 | 0 | 0 |
| NC_004718.3;NP_828851.1;SARS-CoV_E2-glycopr;209 | 0 | 0 |
| NC_004718.3;NP_828851.1;SARS-CoV_E2-glycopr;210 | 0 | 0 |
| NC_004718.3;NP_828851.1;SARS-CoV_E2-glycopr;211 | 0 | 0 |
| NC_004718.3;NP_828851.1;SARS-CoV_E2-glycopr;212 | 0 | 0 |
| NC_004718.3;NP_828851.1;SARS-CoV_E2-glycopr;213 | 0 | 0 |
| NC_004718.3;NP_828851.1;SARS-CoV_E2-glycopr;214 | 0 | 0 |
| NC_004718.3;NP_828851.1;SARS-CoV_E2-glycopr;215 | 0 | 0 |
| NC_004718.3;NP_828851.1;SARS-CoV_E2-glycopr;216 | 0 | 0 |
| NC_004718.3;NP_828851.1;SARS-CoV_E2-glycopr;217 | 0 | 0 |

| | | |
|---|---|---|
| NC_004718.3;NP_828851.1;SARS-CoV_E2-glycopr;218 | 0 | 0 |
| NC_004718.3;NP_828851.1;SARS-CoV_E2-glycopr;219 | 0 | 0 |
| NC_004718.3;NP_828851.1;SARS-CoV_E2-glycopr;220 | 0 | 0 |
| NC_004718.3;NP_828851.1;SARS-CoV_E2-glycopr;221 | 0 | 0 |
| NC_004718.3;NP_828851.1;SARS-CoV_E2-glycopr;222 | 0 | 0 |
| NC_004718.3;NP_828851.1;SARS-CoV_E2-glycopr;223 | 0 | 0 |
| NC_004718.3;NP_828851.1;SARS-CoV_E2-glycopr;224 | 0 | 0 |
| NC_004718.3;NP_828851.1;SARS-CoV_E2-glycopr;225 | 0 | 0 |
| NC_004718.3;NP_828851.1;SARS-CoV_E2-glycopr;226 | 0 | 0 |
| NC_004718.3;NP_828851.1;SARS-CoV_E2-glycopr;227 | 0 | 0 |
| NC_004718.3;NP_828851.1;SARS-CoV_E2-glycopr;228 | 0 | 0 |
| NC_004718.3;NP_828851.1;SARS-CoV_E2-glycopr;229 | 0 | 0 |
| NC_004718.3;NP_828851.1;SARS-CoV_E2-glycopr;230 | 0 | 0 |
| NC_004718.3;NP_828851.1;SARS-CoV_E2-glycopr;231 | 0 | 0 |
| NC_004718.3;NP_828851.1;SARS-CoV_E2-glycopr;232 | 0 | 0 |
| NC_004718.3;NP_828851.1;SARS-CoV_E2-glycopr;233 | 0 | 0 |
| NC_004718.3;NP_828851.1;SARS-CoV_E2-glycopr;234 | 0 | 0 |
| NC_004718.3;NP_828851.1;SARS-CoV_E2-glycopr;235 | 0.05 | 0.025 |
| NC_004718.3;NP_828851.1;SARS-CoV_E2-glycopr;236 | 0.05 | 0.05 |
| NC_004718.3;NP_828851.1;SARS-CoV_E2-glycopr;237 | 0 | 0 |
| NC_004718.3;NP_828851.1;SARS-CoV_E2-glycopr;238 | 0 | 0 |
| NC_004718.3;NP_828851.1;SARS-CoV_E2-glycopr;239 | 0 | 0 |
| NC_004718.3;NP_828851.1;SARS-CoV_E2-glycopr;240 | 0 | 0 |
| NC_004718.3;NP_828851.1;SARS-CoV_E2-glycopr;241 | 0 | 0 |
| NC_004718.3;NP_828851.1;SARS-CoV_E2-glycopr;242 | 0 | 0 |
| NC_004718.3;NP_828851.1;SARS-CoV_E2-glycopr;243 | 0 | 0 |
| NC_004718.3;NP_828851.1;SARS-CoV_E2-glycopr;244 | 0 | 0 |
| NC_004718.3;NP_828851.1;SARS-CoV_E2-glycopr;245 | 0 | 0 |
| NC_004718.3;NP_828851.1;SARS-CoV_E2-glycopr;246 | 0 | 0 |
| NC_004718.3;NP_828851.1;SARS-CoV_E2-glycopr;247 | 0 | 0 |
| NC_004718.3;NP_828851.1;SARS-CoV_E2-glycopr;248 | 0 | 0 |
| NC_004718.3;NP_828851.1;SARS-CoV_E2-glycopr;249 | 0 | 0 |
| NC_004718.3;NP_828851.1;SARS-CoV_E2-glycopr;250 | 0 | 0 |
| NC_004718.3;NP_828851.1;SARS-CoV_E2-glycopr;251 | 0 | 0 |
| NC_004718.3;NP_828851.1;SARS-CoV_E2-glycopr;252 | 0 | 0 |
| NC_004718.3;NP_828851.1;SARS-CoV_E2-glycopr;253 | 0 | 0 |
| NC_004718.3;NP_828851.1;SARS-CoV_E2-glycopr;254 | 0 | 0 |
| NC_004718.3;NP_828851.1;SARS-CoV_E2-glycopr;255 | 0.05 | 0 |
| NC_004718.3;NP_828851.1;SARS-CoV_E2-glycopr;256 | 0 | 0.025 |
| NC_004718.3;NP_828851.1;SARS-CoV_E2-glycopr;257 | 0 | 0 |
| NC_004718.3;NP_828851.1;SARS-CoV_E2-glycopr;258 | 0.05 | 0.075 |
| NC_004718.3;NP_828851.1;SARS-CoV_E2-glycopr;259 | 0.05 | 0.05 |
| NC_004718.3;NP_828851.1;SARS-CoV_E2-glycopr;260 | 0.05 | 0.025 |

| | | |
|---|---|---|
| NC_004718.3;NP_828851.1;SARS-CoV_E2-glycopr;261 | 0.05 | 0.075 |
| NC_004718.3;NP_828851.1;SARS-CoV_E2-glycopr;262 | 0 | 0.025 |
| NC_004718.3;NP_828851.1;SARS-CoV_E2-glycopr;263 | 0 | 0.05 |
| NC_004718.3;NP_828851.1;SARS-CoV_E2-glycopr;264 | 0 | 0.05 |
| NC_004718.3;NP_828851.1;SARS-CoV_E2-glycopr;265 | 0 | 0.025 |
| NC_004718.3;NP_828851.1;SARS-CoV_E2-glycopr;266 | 0 | 0.025 |
| NC_004718.3;NP_828851.1;SARS-CoV_E2-glycopr;267 | 0 | 0.025 |
| NC_004718.3;NP_828851.1;SARS-CoV_E2-glycopr;268 | 0 | 0 |
| NC_004718.3;NP_828851.1;SARS-CoV_E2-glycopr;269 | 0 | 0 |
| NC_004718.3;NP_828851.1;SARS-CoV_E2-glycopr;270 | 0 | 0 |
| NC_004718.3;NP_828851.1;SARS-CoV_E2-glycopr;271 | 0 | 0 |
| NC_004718.3;NP_828851.1;SARS-CoV_E2-glycopr;272 | 0 | 0 |
| NC_004718.3;NP_828851.1;SARS-CoV_E2-glycopr;273 | 0 | 0 |
| NC_004718.3;NP_828851.1;SARS-CoV_E2-glycopr;274 | 0 | 0 |
| NC_004718.3;NP_828851.1;SARS-CoV_E2-glycopr;275 | 0 | 0 |
| NC_004718.3;NP_828851.1;SARS-CoV_E2-glycopr;276 | 0 | 0 |
| NC_004718.3;NP_828851.1;SARS-CoV_E2-glycopr;277 | 0 | 0 |
| NC_004718.3;NP_828851.1;SARS-CoV_E2-glycopr;278 | 0 | 0 |
| NC_004718.3;NP_828851.1;SARS-CoV_E2-glycopr;279 | 0 | 0 |
| NC_004718.3;NP_828851.1;SARS-CoV_E2-glycopr;280 | 0 | 0 |
| NC_004718.3;NP_828851.1;SARS-CoV_E2-glycopr;281 | 0 | 0 |
| NC_004718.3;NP_828851.1;SARS-CoV_E2-glycopr;282 | 0 | 0.025 |
| NC_004718.3;NP_828851.1;SARS-CoV_E2-glycopr;283 | 0 | 0.025 |
| NC_004718.3;NP_828851.1;SARS-CoV_E2-glycopr;284 | 0.05 | 0.05 |
| NC_004718.3;NP_828851.1;SARS-CoV_E2-glycopr;285 | 0 | 0 |
| NC_004718.3;NP_828851.1;SARS-CoV_E2-glycopr;286 | 0.05 | 0.075 |
| NC_004718.3;NP_828851.1;SARS-CoV_E2-glycopr;287 | 0.05 | 0.075 |
| NC_004718.3;NP_828851.1;SARS-CoV_E2-glycopr;288 | 0.05 | 0.075 |
| NC_004718.3;NP_828851.1;SARS-CoV_E2-glycopr;289 | 0.15 | 0.075 |
| NC_004718.3;NP_828851.1;SARS-CoV_E2-glycopr;290 | 0.15 | 0.175 |
| NC_004718.3;NP_828851.1;SARS-CoV_E2-glycopr;291 | 0.1 | 0.125 |
| NC_004718.3;NP_828851.1;SARS-CoV_E2-glycopr;292 | 0 | 0.075 |
| NC_004718.3;NP_828851.1;SARS-CoV_E2-glycopr;293 | 0 | 0.05 |
| NC_004718.3;NP_828851.1;SARS-CoV_E2-glycopr;294 | 0.05 | 0.075 |
| NC_004718.3;NP_828851.1;SARS-CoV_E2-glycopr;295 | 0 | 0.1 |
| NC_004718.3;NP_828851.1;SARS-CoV_E2-glycopr;296 | 0 | 0.15 |
| NC_004718.3;NP_828851.1;SARS-CoV_E2-glycopr;297 | 0.05 | 0.225 |
| NC_004718.3;NP_828851.1;SARS-CoV_E2-glycopr;298 | 0.1 | 0.025 |
| NC_004718.3;NP_828851.1;SARS-CoV_E2-glycopr;299 | 0.05 | 0.025 |
| NC_004718.3;NP_828851.1;SARS-CoV_E2-glycopr;300 | 0 | 0.025 |
| NC_004718.3;NP_828851.1;SARS-CoV_E2-glycopr;301 | 0 | 0.025 |
| NC_004718.3;NP_828851.1;SARS-CoV_E2-glycopr;302 | 0.05 | 0 |
| NC_004718.3;NP_828851.1;SARS-CoV_E2-glycopr;303 | 0 | 0 |

| | | |
|---|---|---|
| NC_004718.3;NP_828851.1;SARS-CoV_E2-glycopr;304 | 0 | 0 |
| NC_004718.3;NP_828851.1;SARS-CoV_E2-glycopr;305 | 0 | 0 |
| NC_004718.3;NP_828851.1;SARS-CoV_E2-glycopr;306 | 0.1 | 0.15 |
| NC_004718.3;NP_828851.1;SARS-CoV_E2-glycopr;307 | 0 | 0 |
| NC_004718.3;NP_828851.1;SARS-CoV_E2-glycopr;308 | 0 | 0 |
| NC_004718.3;NP_828851.1;SARS-CoV_E2-glycopr;309 | 0 | 0 |
| NC_004718.3;NP_828851.1;SARS-CoV_E2-glycopr;310 | 0 | 0 |
| NC_004718.3;NP_828851.1;SARS-CoV_E2-glycopr;311 | 0 | 0 |
| NC_004718.3;NP_828851.1;SARS-CoV_E2-glycopr;312 | 0 | 0 |
| NC_004718.3;NP_828851.1;SARS-CoV_E2-glycopr;313 | 0.05 | 0 |
| NC_004718.3;NP_828851.1;SARS-CoV_E2-glycopr;314 | 0 | 0 |
| NC_004718.3;NP_828851.1;SARS-CoV_E2-glycopr;315 | 0 | 0 |
| NC_004718.3;NP_828851.1;SARS-CoV_E2-glycopr;316 | 0.05 | 0 |
| NC_004718.3;NP_828851.1;SARS-CoV_E2-glycopr;317 | 0.05 | 0.025 |
| NC_004718.3;NP_828851.1;SARS-CoV_E2-glycopr;318 | 0.05 | 0.025 |
| NC_004718.3;NP_828851.1;SARS-CoV_E2-glycopr;319 | 0.05 | 0.025 |
| NC_004718.3;NP_828851.1;SARS-CoV_E2-glycopr;320 | 0.05 | 0.05 |
| NC_004718.3;NP_828851.1;SARS-CoV_E2-glycopr;321 | 0.05 | 0.05 |
| NC_004718.3;NP_828851.1;SARS-CoV_E2-glycopr;322 | 0.05 | 0.025 |
| NC_004718.3;NP_828851.1;SARS-CoV_E2-glycopr;323 | 0.05 | 0.05 |
| NC_004718.3;NP_828851.1;SARS-CoV_E2-glycopr;324 | 0.05 | 0.025 |
| NC_004718.3;NP_828851.1;SARS-CoV_E2-glycopr;325 | 0 | 0.025 |
| NC_004718.3;NP_828851.1;SARS-CoV_E2-glycopr;326 | 0 | 0.025 |
| NC_004718.3;NP_828851.1;SARS-CoV_E2-glycopr;327 | 0 | 0.025 |
| NC_004718.3;NP_828851.1;SARS-CoV_E2-glycopr;328 | 0 | 0 |
| NC_004718.3;NP_828851.1;SARS-CoV_E2-glycopr;329 | 0 | 0 |
| NC_004718.3;NP_828851.1;SARS-CoV_E2-glycopr;330 | 0.1 | 0.075 |
| NC_004718.3;NP_828851.1;SARS-CoV_E2-glycopr;331 | 0 | 0 |
| NC_004718.3;NP_828851.1;SARS-CoV_E2-glycopr;332 | 0 | 0 |
| NC_004718.3;NP_828851.1;SARS-CoV_E2-glycopr;333 | 0 | 0 |
| NC_004718.3;NP_828851.1;SARS-CoV_E2-glycopr;334 | 0 | 0 |
| NC_004718.3;NP_828851.1;SARS-CoV_E2-glycopr;335 | 0 | 0 |
| NC_004718.3;NP_828851.1;SARS-CoV_E2-glycopr;336 | 0 | 0 |
| NC_004718.3;NP_828851.1;SARS-CoV_E2-glycopr;337 | 0 | 0 |
| NC_004718.3;NP_828851.1;SARS-CoV_E2-glycopr;338 | 0 | 0 |
| NC_004718.3;NP_828851.1;SARS-CoV_E2-glycopr;339 | 0 | 0 |
| NC_004718.3;NP_828851.1;SARS-CoV_E2-glycopr;340 | 0.05 | 0 |
| NC_004718.3;NP_828851.1;SARS-CoV_E2-glycopr;341 | 0.05 | 0 |
| NC_004718.3;NP_828851.1;SARS-CoV_E2-glycopr;342 | 0 | 0 |
| NC_004718.3;NP_828851.1;SARS-CoV_E2-glycopr;343 | 0 | 0 |
| NC_004718.3;NP_828851.1;SARS-CoV_E2-glycopr;344 | 0 | 0 |
| NC_004718.3;NP_828851.1;SARS-CoV_E2-glycopr;345 | 0 | 0 |
| NC_004718.3;NP_828851.1;SARS-CoV_E2-glycopr;346 | 0 | 0 |

| | | |
|---|---|---|
| NC_004718.3;NP_828851.1;SARS-CoV_E2-glycopr;347 | 0 | 0 |
| NC_004718.3;NP_828851.1;SARS-CoV_E2-glycopr;348 | 0 | 0 |
| NC_004718.3;NP_828851.1;SARS-CoV_E2-glycopr;349 | 0 | 0 |
| NC_004718.3;NP_828851.1;SARS-CoV_E2-glycopr;350 | 0 | 0 |
| NC_004718.3;NP_828851.1;SARS-CoV_E2-glycopr;351 | 0 | 0 |
| NC_004718.3;NP_828851.1;SARS-CoV_E2-glycopr;352 | 0 | 0 |
| NC_004718.3;NP_828851.1;SARS-CoV_E2-glycopr;353 | 0 | 0 |
| NC_004718.3;NP_828851.1;SARS-CoV_E2-glycopr;354 | 0 | 0 |
| NC_004718.3;NP_828851.1;SARS-CoV_E2-glycopr;355 | 0 | 0 |
| NC_004718.3;NP_828851.1;SARS-CoV_E2-glycopr;356 | 0 | 0 |
| NC_004718.3;NP_828851.1;SARS-CoV_E2-glycopr;357 | 0 | 0 |
| NC_004718.3;NP_828851.1;SARS-CoV_E2-glycopr;358 | 0 | 0 |
| NC_004718.3;NP_828851.1;SARS-CoV_E2-glycopr;359 | 0 | 0 |
| NC_004718.3;NP_828851.1;SARS-CoV_E2-glycopr;360 | 0 | 0 |
| NC_004718.3;NP_828851.1;SARS-CoV_E2-glycopr;361 | 0 | 0.025 |
| NC_004718.3;NP_828851.1;SARS-CoV_E2-glycopr;362 | 0 | 0.05 |
| NC_004718.3;NP_828851.1;SARS-CoV_E2-glycopr;363 | 0 | 0.025 |
| NC_004718.3;NP_828851.1;SARS-CoV_E2-glycopr;364 | 0 | 0.025 |
| NC_004718.3;NP_828851.1;SARS-CoV_E2-glycopr;365 | 0 | 0.025 |
| NC_004718.3;NP_828851.1;SARS-CoV_E2-glycopr;366 | 0 | 0.05 |
| NC_004718.3;NP_828851.1;SARS-CoV_E2-glycopr;367 | 0 | 0.05 |
| NC_004718.3;NP_828851.1;SARS-CoV_E2-glycopr;368 | 0 | 0.05 |
| NC_004718.3;NP_828851.1;SARS-CoV_E2-glycopr;369 | 0 | 0.025 |
| NC_004718.3;NP_828851.1;SARS-CoV_E2-glycopr;370 | 0 | 0.05 |
| NC_004718.3;NP_828851.1;SARS-CoV_E2-glycopr;371 | 0 | 0 |
| NC_004718.3;NP_828851.1;SARS-CoV_E2-glycopr;372 | 0 | 0.025 |
| NC_004718.3;NP_828851.1;SARS-CoV_E2-glycopr;373 | 0 | 0 |
| NC_004718.3;NP_828851.1;SARS-CoV_E2-glycopr;374 | 0 | 0 |
| NC_004718.3;NP_828851.1;SARS-CoV_E2-glycopr;375 | 0 | 0 |
| NC_004718.3;NP_828851.1;SARS-CoV_E2-glycopr;376 | 0 | 0 |
| NC_004718.3;NP_828851.1;SARS-CoV_E2-glycopr;377 | 0 | 0 |
| NC_004718.3;NP_828851.1;SARS-CoV_E2-glycopr;378 | 0.05 | 0 |
| NC_004718.3;NP_828851.1;SARS-CoV_E2-glycopr;379 | 0.05 | 0.025 |
| NC_004718.3;NP_828851.1;SARS-CoV_E2-glycopr;380 | 0.05 | 0 |
| NC_004718.3;NP_828851.1;SARS-CoV_E2-glycopr;381 | 0 | 0 |
| NC_004718.3;NP_828851.1;SARS-CoV_E2-glycopr;382 | 0 | 0 |
| NC_004718.3;NP_828851.1;SARS-CoV_E2-glycopr;383 | 0 | 0 |
| NC_004718.3;NP_828851.1;SARS-CoV_E2-glycopr;384 | 0.1 | 0 |
| NC_004718.3;NP_828851.1;SARS-CoV_E2-glycopr;385 | 0.1 | 0.025 |
| NC_004718.3;NP_828851.1;SARS-CoV_E2-glycopr;386 | 0 | 0.025 |
| NC_004718.3;NP_828851.1;SARS-CoV_E2-glycopr;387 | 0 | 0 |
| NC_004718.3;NP_828851.1;SARS-CoV_E2-glycopr;388 | 0 | 0 |
| NC_004718.3;NP_828851.1;SARS-CoV_E2-glycopr;389 | 0 | 0 |

| | | |
|---|---|---|
| NC_004718.3;NP_828851.1;SARS-CoV_E2-glycopr;390 | 0 | 0 |
| NC_004718.3;NP_828851.1;SARS-CoV_E2-glycopr;391 | 0 | 0.025 |
| NC_004718.3;NP_828851.1;SARS-CoV_E2-glycopr;392 | 0 | 0 |
| NC_004718.3;NP_828851.1;SARS-CoV_E2-glycopr;393 | 0 | 0 |
| NC_004718.3;NP_828851.1;SARS-CoV_E2-glycopr;394 | 0 | 0 |
| NC_004718.3;NP_828851.1;SARS-CoV_E2-glycopr;395 | 0 | 0.025 |
| NC_004718.3;NP_828851.1;SARS-CoV_E2-glycopr;396 | 0.05 | 0.05 |
| NC_004718.3;NP_828851.1;SARS-CoV_E2-glycopr;397 | 0.05 | 0.025 |
| NC_004718.3;NP_828851.1;SARS-CoV_E2-glycopr;398 | 0 | 0 |
| NC_004718.3;NP_828851.1;SARS-CoV_E2-glycopr;399 | 0 | 0.05 |
| NC_004718.3;NP_828851.1;SARS-CoV_E2-glycopr;400 | 0.05 | 0.05 |
| NC_004718.3;NP_828851.1;SARS-CoV_E2-glycopr;401 | 0.05 | 0.05 |
| NC_004718.3;NP_828851.1;SARS-CoV_E2-glycopr;402 | 0.05 | 0.05 |
| NC_004718.3;NP_828851.1;SARS-CoV_E2-glycopr;403 | 0.1 | 0.225 |
| NC_004718.3;NP_828851.1;SARS-CoV_E2-glycopr;404 | 0 | 0.1 |
| NC_004718.3;NP_828851.1;SARS-CoV_E2-glycopr;405 | 0 | 0.025 |
| NC_004718.3;NP_828851.1;SARS-CoV_E2-glycopr;406 | 0.05 | 0 |
| NC_004718.3;NP_828851.1;SARS-CoV_E2-glycopr;407 | 0.05 | 0 |
| NC_004718.3;NP_828851.1;SARS-CoV_E2-glycopr;408 | 0 | 0 |
| NC_004718.3;NP_828851.1;SARS-CoV_E2-glycopr;409 | 0 | 0 |
| NC_004718.3;NP_828851.1;SARS-CoV_E2-glycopr;410 | 0 | 0 |
| NC_004718.3;NP_828851.1;SARS-CoV_E2-glycopr;411 | 0 | 0 |
| NC_004718.3;NP_828851.1;SARS-CoV_E2-glycopr;412 | 0 | 0 |
| NC_004718.3;NP_828851.1;SARS-CoV_E2-glycopr;413 | 0 | 0 |
| NC_004718.3;NP_828851.1;SARS-CoV_E2-glycopr;414 | 0 | 0 |
| NC_004718.3;NP_828851.1;SARS-CoV_E2-glycopr;415 | 0 | 0 |
| NC_004718.3;NP_828851.1;SARS-CoV_E2-glycopr;416 | 0 | 0 |
| NC_004718.3;NP_828851.1;SARS-CoV_E2-glycopr;417 | 0 | 0 |
| NC_004718.3;NP_828851.1;SARS-CoV_E2-glycopr;418 | 0 | 0 |
| NC_004718.3;NP_828851.1;SARS-CoV_E2-glycopr;419 | 0.05 | 0 |
| NC_004718.3;NP_828851.1;SARS-CoV_E2-glycopr;420 | 0.1 | 0.025 |
| NC_004718.3;NP_828851.1;SARS-CoV_E2-glycopr;421 | 0.15 | 0.025 |
| NC_004718.3;NP_828851.1;SARS-CoV_E2-glycopr;422 | 0.2 | 0.075 |
| NC_004718.3;NP_828851.1;SARS-CoV_E2-glycopr;423 | 0.15 | 0.025 |
| NC_004718.3;NP_828851.1;SARS-CoV_E2-glycopr;424 | 0.25 | 0.075 |
| NC_004718.3;NP_828851.1;SARS-CoV_E2-glycopr;425 | 0.15 | 0.075 |
| NC_004718.3;NP_828851.1;SARS-CoV_E2-glycopr;426 | 0.05 | 0.05 |
| NC_004718.3;NP_828851.1;SARS-CoV_E2-glycopr;427 | 0.1 | 0.075 |
| NC_004718.3;NP_828851.1;SARS-CoV_E2-glycopr;428 | 0.05 | 0.025 |
| NC_004718.3;NP_828851.1;SARS-CoV_E2-glycopr;429 | 0 | 0 |
| NC_004718.3;NP_828851.1;SARS-CoV_E2-glycopr;430 | 0 | 0 |
| NC_004718.3;NP_828851.1;SARS-CoV_E2-glycopr;431 | 0 | 0 |
| NC_004718.3;NP_828851.1;SARS-CoV_E2-glycopr;432 | 0 | 0 |

| | | |
|---|---|---|
| NC_004718.3;NP_828851.1;SARS-CoV_E2-glycopr;433 | 0 | 0 |
| NC_004718.3;NP_828851.1;SARS-CoV_E2-glycopr;434 | 0 | 0 |
| NC_004718.3;NP_828851.1;SARS-CoV_E2-glycopr;435 | 0 | 0 |
| NC_004718.3;NP_828851.1;SARS-CoV_E2-glycopr;436 | 0 | 0 |
| NC_004718.3;NP_828851.1;SARS-CoV_E2-glycopr;437 | 0 | 0 |
| NC_004718.3;NP_828851.1;SARS-CoV_E2-glycopr;438 | 0 | 0 |
| NC_004718.3;NP_828851.1;SARS-CoV_E2-glycopr;439 | 0 | 0 |
| NC_004718.3;NP_828851.1;SARS-CoV_E2-glycopr;440 | 0 | 0 |
| NC_004718.3;NP_828851.1;SARS-CoV_E2-glycopr;441 | 0 | 0.025 |
| NC_004718.3;NP_828851.1;SARS-CoV_E2-glycopr;442 | 0 | 0.05 |
| NC_004718.3;NP_828851.1;SARS-CoV_E2-glycopr;443 | 0 | 0.025 |
| NC_004718.3;NP_828851.1;SARS-CoV_E2-glycopr;444 | 0.05 | 0.025 |
| NC_004718.3;NP_828851.1;SARS-CoV_E2-glycopr;445 | 0 | 0.05 |
| NC_004718.3;NP_828851.1;SARS-CoV_E2-glycopr;446 | 0.05 | 0.15 |
| NC_004718.3;NP_828851.1;SARS-CoV_E2-glycopr;447 | 0 | 0.15 |
| NC_004718.3;NP_828851.1;SARS-CoV_E2-glycopr;448 | 0.05 | 0.05 |
| NC_004718.3;NP_828851.1;SARS-CoV_E2-glycopr;449 | 0.1 | 0 |
| NC_004718.3;NP_828851.1;SARS-CoV_E2-glycopr;450 | 0 | 0 |
| NC_004718.3;NP_828851.1;SARS-CoV_E2-glycopr;451 | 0 | 0 |
| NC_004718.3;NP_828851.1;SARS-CoV_E2-glycopr;452 | 0.05 | 0 |
| NC_004718.3;NP_828851.1;SARS-CoV_E2-glycopr;453 | 0 | 0 |
| NC_004718.3;NP_828851.1;SARS-CoV_E2-glycopr;454 | 0 | 0 |
| NC_004718.3;NP_828851.1;SARS-CoV_E2-glycopr;455 | 0.05 | 0 |
| NC_004718.3;NP_828851.1;SARS-CoV_E2-glycopr;456 | 0.1 | 0 |
| NC_004718.3;NP_828851.1;SARS-CoV_E2-glycopr;457 | 0 | 0 |
| NC_004718.3;NP_828851.1;SARS-CoV_E2-glycopr;458 | 0 | 0 |
| NC_004718.3;NP_828851.1;SARS-CoV_E2-glycopr;459 | 0 | 0 |
| NC_004718.3;NP_828851.1;SARS-CoV_E2-glycopr;460 | 0 | 0 |
| NC_004718.3;NP_828851.1;SARS-CoV_E2-glycopr;461 | 0 | 0 |
| NC_004718.3;NP_828851.1;SARS-CoV_E2-glycopr;462 | 0 | 0 |
| NC_004718.3;NP_828851.1;SARS-CoV_E2-glycopr;463 | 0 | 0 |
| NC_004718.3;NP_828851.1;SARS-CoV_E2-glycopr;464 | 0 | 0 |
| NC_004718.3;NP_828851.1;SARS-CoV_E2-glycopr;465 | 0 | 0.025 |
| NC_004718.3;NP_828851.1;SARS-CoV_E2-glycopr;466 | 0 | 0 |
| NC_004718.3;NP_828851.1;SARS-CoV_E2-glycopr;467 | 0 | 0 |
| NC_004718.3;NP_828851.1;SARS-CoV_E2-glycopr;468 | 0 | 0 |
| NC_004718.3;NP_828851.1;SARS-CoV_E2-glycopr;469 | 0 | 0 |
| NC_004718.3;NP_828851.1;SARS-CoV_E2-glycopr;470 | 0 | 0 |
| NC_004718.3;NP_828851.1;SARS-CoV_E2-glycopr;471 | 0 | 0 |
| NC_004718.3;NP_828851.1;SARS-CoV_E2-glycopr;472 | 0 | 0 |
| NC_004718.3;NP_828851.1;SARS-CoV_E2-glycopr;473 | 0 | 0 |
| NC_004718.3;NP_828851.1;SARS-CoV_E2-glycopr;474 | 0 | 0 |
| NC_004718.3;NP_828851.1;SARS-CoV_E2-glycopr;475 | 0 | 0 |

| | | |
|---|---|---|
| NC_004718.3;NP_828851.1;SARS-CoV_E2-glycopr;476 | 0 | 0 |
| NC_004718.3;NP_828851.1;SARS-CoV_E2-glycopr;477 | 0 | 0 |
| NC_004718.3;NP_828851.1;SARS-CoV_E2-glycopr;478 | 0 | 0 |
| NC_004718.3;NP_828851.1;SARS-CoV_E2-glycopr;479 | 0 | 0 |
| NC_004718.3;NP_828851.1;SARS-CoV_E2-glycopr;480 | 0 | 0 |
| NC_004718.3;NP_828851.1;SARS-CoV_E2-glycopr;481 | 0 | 0 |
| NC_004718.3;NP_828851.1;SARS-CoV_E2-glycopr;482 | 0 | 0 |
| NC_004718.3;NP_828851.1;SARS-CoV_E2-glycopr;483 | 0 | 0 |
| NC_004718.3;NP_828851.1;SARS-CoV_E2-glycopr;484 | 0 | 0 |
| NC_004718.3;NP_828851.1;SARS-CoV_E2-glycopr;485 | 0 | 0 |
| NC_004718.3;NP_828851.1;SARS-CoV_E2-glycopr;486 | 0 | 0 |
| NC_004718.3;NP_828851.1;SARS-CoV_E2-glycopr;487 | 0 | 0 |
| NC_004718.3;NP_828851.1;SARS-CoV_E2-glycopr;488 | 0 | 0 |
| NC_004718.3;NP_828851.1;SARS-CoV_E2-glycopr;489 | 0 | 0 |
| NC_004718.3;NP_828851.1;SARS-CoV_E2-glycopr;490 | 0 | 0 |
| NC_004718.3;NP_828851.1;SARS-CoV_E2-glycopr;491 | 0 | 0 |
| NC_004718.3;NP_828851.1;SARS-CoV_E2-glycopr;492 | 0 | 0 |
| NC_004718.3;NP_828851.1;SARS-CoV_E2-glycopr;493 | 0 | 0 |
| NC_004718.3;NP_828851.1;SARS-CoV_E2-glycopr;494 | 0 | 0 |
| NC_004718.3;NP_828851.1;SARS-CoV_E2-glycopr;495 | 0 | 0 |
| NC_004718.3;NP_828851.1;SARS-CoV_E2-glycopr;496 | 0 | 0 |
| NC_004718.3;NP_828851.1;SARS-CoV_E2-glycopr;497 | 0 | 0 |
| NC_004718.3;NP_828851.1;SARS-CoV_E2-glycopr;498 | 0 | 0 |
| NC_004718.3;NP_828851.1;SARS-CoV_E2-glycopr;499 | 0 | 0 |
| NC_004718.3;NP_828851.1;SARS-CoV_E2-glycopr;500 | 0 | 0 |
| NC_004718.3;NP_828851.1;SARS-CoV_E2-glycopr;501 | 0 | 0.025 |
| NC_004718.3;NP_828851.1;SARS-CoV_E2-glycopr;502 | 0 | 0.025 |
| NC_004718.3;NP_828851.1;SARS-CoV_E2-glycopr;503 | 0 | 0 |
| NC_004718.3;NP_828851.1;SARS-CoV_E2-glycopr;504 | 0 | 0 |
| NC_004718.3;NP_828851.1;SARS-CoV_E2-glycopr;505 | 0 | 0 |
| NC_004718.3;NP_828851.1;SARS-CoV_E2-glycopr;506 | 0.15 | 0 |
| NC_004718.3;NP_828851.1;SARS-CoV_E2-glycopr;507 | 0.05 | 0.025 |
| NC_004718.3;NP_828851.1;SARS-CoV_E2-glycopr;508 | 0.05 | 0.025 |
| NC_004718.3;NP_828851.1;SARS-CoV_E2-glycopr;509 | 0.05 | 0.025 |
| NC_004718.3;NP_828851.1;SARS-CoV_E2-glycopr;510 | 0.05 | 0.025 |
| NC_004718.3;NP_828851.1;SARS-CoV_E2-glycopr;511 | 0.05 | 0.05 |
| NC_004718.3;NP_828851.1;SARS-CoV_E2-glycopr;512 | 0 | 0.025 |
| NC_004718.3;NP_828851.1;SARS-CoV_E2-glycopr;513 | 0 | 0.05 |
| NC_004718.3;NP_828851.1;SARS-CoV_E2-glycopr;514 | 0 | 0.025 |
| NC_004718.3;NP_828851.1;SARS-CoV_E2-glycopr;515 | 0 | 0 |
| NC_004718.3;NP_828851.1;SARS-CoV_E2-glycopr;516 | 0.1 | 0 |
| NC_004718.3;NP_828851.1;SARS-CoV_E2-glycopr;517 | 0 | 0 |
| NC_004718.3;NP_828851.1;SARS-CoV_E2-glycopr;518 | 0.05 | 0 |

| | | |
|---|---|---|
| NC_004718.3;NP_828851.1;SARS-CoV_E2-glycopr;519 | 0 | 0 |
| NC_004718.3;NP_828851.1;SARS-CoV_E2-glycopr;520 | 0.05 | 0 |
| NC_004718.3;NP_828851.1;SARS-CoV_E2-glycopr;521 | 0.05 | 0.025 |
| NC_004718.3;NP_828851.1;SARS-CoV_E2-glycopr;522 | 0 | 0 |
| NC_004718.3;NP_828851.1;SARS-CoV_E2-glycopr;523 | 0 | 0 |
| NC_004718.3;NP_828851.1;SARS-CoV_E2-glycopr;524 | 0 | 0 |
| NC_004718.3;NP_828851.1;SARS-CoV_E2-glycopr;525 | 0 | 0 |
| NC_004718.3;NP_828851.1;SARS-CoV_E2-glycopr;526 | 0 | 0 |
| NC_004718.3;NP_828851.1;SARS-CoV_E2-glycopr;527 | 0 | 0.025 |
| NC_004718.3;NP_828851.1;SARS-CoV_E2-glycopr;528 | 0 | 0 |
| NC_004718.3;NP_828851.1;SARS-CoV_E2-glycopr;529 | 0 | 0 |
| NC_004718.3;NP_828851.1;SARS-CoV_E2-glycopr;530 | 0 | 0 |
| NC_004718.3;NP_828851.1;SARS-CoV_E2-glycopr;531 | 0 | 0 |
| NC_004718.3;NP_828851.1;SARS-CoV_E2-glycopr;532 | 0 | 0 |
| NC_004718.3;NP_828851.1;SARS-CoV_E2-glycopr;533 | 0 | 0 |
| NC_004718.3;NP_828851.1;SARS-CoV_E2-glycopr;534 | 0 | 0 |
| NC_004718.3;NP_828851.1;SARS-CoV_E2-glycopr;535 | 0 | 0 |
| NC_004718.3;NP_828851.1;SARS-CoV_E2-glycopr;536 | 0 | 0 |
| NC_004718.3;NP_828851.1;SARS-CoV_E2-glycopr;537 | 0.05 | 0 |
| NC_004718.3;NP_828851.1;SARS-CoV_E2-glycopr;538 | 0 | 0.025 |
| NC_004718.3;NP_828851.1;SARS-CoV_E2-glycopr;539 | 0 | 0.125 |
| NC_004718.3;NP_828851.1;SARS-CoV_E2-glycopr;540 | 0 | 0.175 |
| NC_004718.3;NP_828851.1;SARS-CoV_E2-glycopr;541 | 0 | 0.2 |
| NC_004718.3;NP_828851.1;SARS-CoV_E2-glycopr;542 | 0.05 | 0.5 |
| NC_004718.3;NP_828851.1;SARS-CoV_E2-glycopr;543 | 0 | 0.4 |
| NC_004718.3;NP_828851.1;SARS-CoV_E2-glycopr;544 | 0 | 0.325 |
| NC_004718.3;NP_828851.1;SARS-CoV_E2-glycopr;545 | 0 | 0.275 |
| NC_004718.3;NP_828851.1;SARS-CoV_E2-glycopr;546 | 0 | 0.325 |
| NC_004718.3;NP_828851.1;SARS-CoV_E2-glycopr;547 | 0 | 0.175 |
| NC_004718.3;NP_828851.1;SARS-CoV_E2-glycopr;548 | 0 | 0.15 |
| NC_004718.3;NP_828851.1;SARS-CoV_E2-glycopr;549 | 0.1 | 0.025 |
| NC_004718.3;NP_828851.1;SARS-CoV_E2-glycopr;550 | 0.05 | 0 |
| NC_004718.3;NP_828851.1;SARS-CoV_E2-glycopr;551 | 0.15 | 0 |
| NC_004718.3;NP_828851.1;SARS-CoV_E2-glycopr;552 | 0.1 | 0.025 |
| NC_004718.3;NP_828851.1;SARS-CoV_E2-glycopr;553 | 0.05 | 0 |
| NC_004718.3;NP_828851.1;SARS-CoV_E2-glycopr;554 | 0.05 | 0 |
| NC_004718.3;NP_828851.1;SARS-CoV_E2-glycopr;555 | 0.05 | 0 |
| NC_004718.3;NP_828851.1;SARS-CoV_E2-glycopr;556 | 0.05 | 0 |
| NC_004718.3;NP_828851.1;SARS-CoV_E2-glycopr;557 | 0.1 | 0 |
| NC_004718.3;NP_828851.1;SARS-CoV_E2-glycopr;558 | 0 | 0 |
| NC_004718.3;NP_828851.1;SARS-CoV_E2-glycopr;559 | 0 | 0.025 |
| NC_004718.3;NP_828851.1;SARS-CoV_E2-glycopr;560 | 0 | 0.025 |
| NC_004718.3;NP_828851.1;SARS-CoV_E2-glycopr;561 | 0.05 | 0.025 |

| | | |
|---|---|---|
| NC_004718.3;NP_828851.1;SARS-CoV_E2-glycopr;562 | 0.05 | 0.025 |
| NC_004718.3;NP_828851.1;SARS-CoV_E2-glycopr;563 | 0 | 0 |
| NC_004718.3;NP_828851.1;SARS-CoV_E2-glycopr;564 | 0 | 0 |
| NC_004718.3;NP_828851.1;SARS-CoV_E2-glycopr;565 | 0 | 0 |
| NC_004718.3;NP_828851.1;SARS-CoV_E2-glycopr;566 | 0 | 0 |
| NC_004718.3;NP_828851.1;SARS-CoV_E2-glycopr;567 | 0 | 0 |
| NC_004718.3;NP_828851.1;SARS-CoV_E2-glycopr;568 | 0 | 0.025 |
| NC_004718.3;NP_828851.1;SARS-CoV_E2-glycopr;569 | 0 | 0 |
| NC_004718.3;NP_828851.1;SARS-CoV_E2-glycopr;570 | 0 | 0 |
| NC_004718.3;NP_828851.1;SARS-CoV_E2-glycopr;571 | 0 | 0 |
| NC_004718.3;NP_828851.1;SARS-CoV_E2-glycopr;572 | 0 | 0 |
| NC_004718.3;NP_828851.1;SARS-CoV_E2-glycopr;573 | 0 | 0 |
| NC_004718.3;NP_828851.1;SARS-CoV_E2-glycopr;574 | 0 | 0 |
| NC_004718.3;NP_828851.1;SARS-CoV_E2-glycopr;575 | 0 | 0.05 |
| NC_004718.3;NP_828851.1;SARS-CoV_E2-glycopr;576 | 0.1 | 0.075 |
| NC_004718.3;NP_828851.1;SARS-CoV_E2-glycopr;577 | 0.05 | 0.1 |
| NC_004718.3;NP_828851.1;SARS-CoV_E2-glycopr;578 | 0.05 | 0.025 |
| NC_004718.3;NP_828851.1;SARS-CoV_E2-glycopr;579 | 0 | 0.05 |
| NC_004718.3;NP_828851.1;SARS-CoV_E2-glycopr;580 | 0 | 0.025 |
| NC_004718.3;NP_828851.1;SARS-CoV_E2-glycopr;581 | 0 | 0.025 |
| NC_004718.3;NP_828851.1;SARS-CoV_E2-glycopr;582 | 0 | 0.025 |
| NC_004718.3;NP_828851.1;SARS-CoV_E2-glycopr;583 | 0 | 0 |
| NC_004718.3;NP_828851.1;SARS-CoV_E2-glycopr;584 | 0 | 0 |
| NC_004718.3;NP_828851.1;SARS-CoV_E2-glycopr;585 | 0 | 0 |
| NC_004718.3;NP_828851.1;SARS-CoV_E2-glycopr;586 | 0 | 0 |
| NC_004718.3;NP_828851.1;SARS-CoV_E2-glycopr;587 | 0 | 0 |
| NC_004718.3;NP_828851.1;SARS-CoV_E2-glycopr;588 | 0 | 0 |
| NC_004718.3;NP_828851.1;SARS-CoV_E2-glycopr;589 | 0 | 0 |
| NC_004718.3;NP_828851.1;SARS-CoV_E2-glycopr;590 | 0 | 0 |
| NC_004718.3;NP_828851.1;SARS-CoV_E2-glycopr;591 | 0 | 0 |
| NC_004718.3;NP_828851.1;SARS-CoV_E2-glycopr;592 | 0 | 0 |
| NC_004718.3;NP_828851.1;SARS-CoV_E2-glycopr;593 | 0 | 0 |
| NC_004718.3;NP_828851.1;SARS-CoV_E2-glycopr;594 | 0 | 0 |
| NC_004718.3;NP_828851.1;SARS-CoV_E2-glycopr;595 | 0 | 0 |
| NC_004718.3;NP_828851.1;SARS-CoV_E2-glycopr;596 | 0 | 0 |
| NC_004718.3;NP_828851.1;SARS-CoV_E2-glycopr;597 | 0 | 0 |
| NC_004718.3;NP_828851.1;SARS-CoV_E2-glycopr;598 | 0 | 0.025 |
| NC_004718.3;NP_828851.1;SARS-CoV_E2-glycopr;599 | 0 | 0.05 |
| NC_004718.3;NP_828851.1;SARS-CoV_E2-glycopr;600 | 0 | 0.05 |
| NC_004718.3;NP_828851.1;SARS-CoV_E2-glycopr;601 | 0 | 0.05 |
| NC_004718.3;NP_828851.1;SARS-CoV_E2-glycopr;602 | 0 | 0.075 |
| NC_004718.3;NP_828851.1;SARS-CoV_E2-glycopr;603 | 0 | 0.1 |
| NC_004718.3;NP_828851.1;SARS-CoV_E2-glycopr;604 | 0 | 0.1 |

| | | |
|---|---|---|
| NC_004718.3;NP_828851.1;SARS-CoV_E2-glycopr;605 | 0 | 0.075 |
| NC_004718.3;NP_828851.1;SARS-CoV_E2-glycopr;606 | 0 | 0.075 |
| NC_004718.3;NP_828851.1;SARS-CoV_E2-glycopr;607 | 0 | 0.05 |
| NC_004718.3;NP_828851.1;SARS-CoV_E2-glycopr;608 | 0 | 0.05 |
| NC_004718.3;NP_828851.1;SARS-CoV_E2-glycopr;609 | 0.05 | 0.025 |
| NC_004718.3;NP_828851.1;SARS-CoV_E2-glycopr;610 | 0 | 0 |
| NC_004718.3;NP_828851.1;SARS-CoV_E2-glycopr;611 | 0 | 0.05 |
| NC_004718.3;NP_828851.1;SARS-CoV_E2-glycopr;612 | 0 | 0.075 |
| NC_004718.3;NP_828851.1;SARS-CoV_E2-glycopr;613 | 0.05 | 0.15 |
| NC_004718.3;NP_828851.1;SARS-CoV_E2-glycopr;614 | 0.05 | 0 |
| NC_004718.3;NP_828851.1;SARS-CoV_E2-glycopr;615 | 0.05 | 0 |
| NC_004718.3;NP_828851.1;SARS-CoV_E2-glycopr;616 | 0.05 | 0.05 |
| NC_004718.3;NP_828851.1;SARS-CoV_E2-glycopr;617 | 0.05 | 0 |
| NC_004718.3;NP_828851.1;SARS-CoV_E2-glycopr;618 | 0.05 | 0.025 |
| NC_004718.3;NP_828851.1;SARS-CoV_E2-glycopr;619 | 0.05 | 0.025 |
| NC_004718.3;NP_828851.1;SARS-CoV_E2-glycopr;620 | 0.05 | 0.05 |
| NC_004718.3;NP_828851.1;SARS-CoV_E2-glycopr;621 | 0.05 | 0.1 |
| NC_004718.3;NP_828851.1;SARS-CoV_E2-glycopr;622 | 0.05 | 0.05 |
| NC_004718.3;NP_828851.1;SARS-CoV_E2-glycopr;623 | 0 | 0 |
| NC_004718.3;NP_828851.1;SARS-CoV_E2-glycopr;624 | 0 | 0 |
| NC_004718.3;NP_828851.1;SARS-CoV_E2-glycopr;625 | 0 | 0 |
| NC_004718.3;NP_828851.1;SARS-CoV_E2-glycopr;626 | 0 | 0 |
| NC_004718.3;NP_828851.1;SARS-CoV_E2-glycopr;627 | 0 | 0 |
| NC_004718.3;NP_828851.1;SARS-CoV_E2-glycopr;628 | 0 | 0 |
| NC_004718.3;NP_828851.1;SARS-CoV_E2-glycopr;629 | 0.05 | 0 |
| NC_004718.3;NP_828851.1;SARS-CoV_E2-glycopr;630 | 0 | 0 |
| NC_004718.3;NP_828851.1;SARS-CoV_E2-glycopr;631 | 0 | 0 |
| NC_004718.3;NP_828851.1;SARS-CoV_E2-glycopr;632 | 0 | 0 |
| NC_004718.3;NP_828851.1;SARS-CoV_E2-glycopr;633 | 0 | 0 |
| NC_004718.3;NP_828851.1;SARS-CoV_E2-glycopr;634 | 0 | 0 |
| NC_004718.3;NP_828851.1;SARS-CoV_E2-glycopr;635 | 0 | 0 |
| NC_004718.3;NP_828851.1;SARS-CoV_E2-glycopr;636 | 0 | 0 |
| NC_004718.3;NP_828851.1;SARS-CoV_E2-glycopr;637 | 0 | 0 |
| NC_004718.3;NP_828851.1;SARS-CoV_E2-glycopr;638 | 0 | 0 |
| NC_004718.3;NP_828851.1;SARS-CoV_E2-glycopr;639 | 0 | 0.025 |
| NC_004718.3;NP_828851.1;SARS-CoV_E2-glycopr;640 | 0 | 0 |
| NC_004718.3;NP_828851.1;SARS-CoV_E2-glycopr;641 | 0 | 0.025 |
| NC_004718.3;NP_828851.1;SARS-CoV_E2-glycopr;642 | 0.05 | 0.075 |
| NC_004718.3;NP_828851.1;SARS-CoV_E2-glycopr;643 | 0.05 | 0.125 |
| NC_004718.3;NP_828851.1;SARS-CoV_E2-glycopr;644 | 0.1 | 0.2 |
| NC_004718.3;NP_828851.1;SARS-CoV_E2-glycopr;645 | 0.05 | 0 |
| NC_004718.3;NP_828851.1;SARS-CoV_E2-glycopr;646 | 0 | 0 |
| NC_004718.3;NP_828851.1;SARS-CoV_E2-glycopr;647 | 0 | 0 |

| | | |
|---|---|---|
| NC_004718.3;NP_828851.1;SARS-CoV_E2-glycopr;648 | 0 | 0 |
| NC_004718.3;NP_828851.1;SARS-CoV_E2-glycopr;649 | 0 | 0 |
| NC_004718.3;NP_828851.1;SARS-CoV_E2-glycopr;650 | 0 | 0 |
| NC_004718.3;NP_828851.1;SARS-CoV_E2-glycopr;651 | 0 | 0 |
| NC_004718.3;NP_828851.1;SARS-CoV_E2-glycopr;652 | 0 | 0 |
| NC_004718.3;NP_828851.1;SARS-CoV_E2-glycopr;653 | 0 | 0 |
| NC_004718.3;NP_828851.1;SARS-CoV_E2-glycopr;654 | 0 | 0 |
| NC_004718.3;NP_828851.1;SARS-CoV_E2-glycopr;655 | 0 | 0 |
| NC_004718.3;NP_828851.1;SARS-CoV_E2-glycopr;656 | 0 | 0 |
| NC_004718.3;NP_828851.1;SARS-CoV_E2-glycopr;657 | 0 | 0 |
| NC_004718.3;NP_828851.1;SARS-CoV_E2-glycopr;658 | 0 | 0 |
| NC_004718.3;NP_828851.1;SARS-CoV_E2-glycopr;659 | 0 | 0.025 |
| NC_004718.3;NP_828851.1;SARS-CoV_E2-glycopr;660 | 0 | 0 |
| NC_004718.3;NP_828851.1;SARS-CoV_E2-glycopr;661 | 0 | 0 |
| NC_004718.3;NP_828851.1;SARS-CoV_E2-glycopr;662 | 0 | 0 |
| NC_004718.3;NP_828851.1;SARS-CoV_E2-glycopr;663 | 0 | 0 |
| NC_004718.3;NP_828851.1;SARS-CoV_E2-glycopr;664 | 0 | 0 |
| NC_004718.3;NP_828851.1;SARS-CoV_E2-glycopr;665 | 0 | 0 |
| NC_004718.3;NP_828851.1;SARS-CoV_E2-glycopr;666 | 0 | 0.025 |
| NC_004718.3;NP_828851.1;SARS-CoV_E2-glycopr;667 | 0 | 0.025 |
| NC_004718.3;NP_828851.1;SARS-CoV_E2-glycopr;668 | 0 | 0.05 |
| NC_004718.3;NP_828851.1;SARS-CoV_E2-glycopr;669 | 0 | 0.125 |
| NC_004718.3;NP_828851.1;SARS-CoV_E2-glycopr;670 | 0 | 0.125 |
| NC_004718.3;NP_828851.1;SARS-CoV_E2-glycopr;671 | 0 | 0.15 |
| NC_004718.3;NP_828851.1;SARS-CoV_E2-glycopr;672 | 0 | 0.1 |
| NC_004718.3;NP_828851.1;SARS-CoV_E2-glycopr;673 | 0 | 0.15 |
| NC_004718.3;NP_828851.1;SARS-CoV_E2-glycopr;674 | 0 | 0.1 |
| NC_004718.3;NP_828851.1;SARS-CoV_E2-glycopr;675 | 0 | 0.075 |
| NC_004718.3;NP_828851.1;SARS-CoV_E2-glycopr;676 | 0.05 | 0.1 |
| NC_004718.3;NP_828851.1;SARS-CoV_E2-glycopr;677 | 0.05 | 0.125 |
| NC_004718.3;NP_828851.1;SARS-CoV_E2-glycopr;678 | 0.05 | 0.075 |
| NC_004718.3;NP_828851.1;SARS-CoV_E2-glycopr;679 | 0.05 | 0.075 |
| NC_004718.3;NP_828851.1;SARS-CoV_E2-glycopr;680 | 0.15 | 0.125 |
| NC_004718.3;NP_828851.1;SARS-CoV_E2-glycopr;681 | 0.05 | 0.025 |
| NC_004718.3;NP_828851.1;SARS-CoV_E2-glycopr;682 | 0.1 | 0.075 |
| NC_004718.3;NP_828851.1;SARS-CoV_E2-glycopr;683 | 0.1 | 0.025 |
| NC_004718.3;NP_828851.1;SARS-CoV_E2-glycopr;684 | 0.2 | 0 |
| NC_004718.3;NP_828851.1;SARS-CoV_E2-glycopr;685 | 0.05 | 0.025 |
| NC_004718.3;NP_828851.1;SARS-CoV_E2-glycopr;686 | 0.05 | 0 |
| NC_004718.3;NP_828851.1;SARS-CoV_E2-glycopr;687 | 0.1 | 0 |
| NC_004718.3;NP_828851.1;SARS-CoV_E2-glycopr;688 | 0.15 | 0.1 |
| NC_004718.3;NP_828851.1;SARS-CoV_E2-glycopr;689 | 0.05 | 0 |
| NC_004718.3;NP_828851.1;SARS-CoV_E2-glycopr;690 | 0 | 0 |

| | | |
|---|---|---|
| NC_004718.3;NP_828851.1;SARS-CoV_E2-glycopr;691 | 0 | 0 |
| NC_004718.3;NP_828851.1;SARS-CoV_E2-glycopr;692 | 0 | 0 |
| NC_004718.3;NP_828851.1;SARS-CoV_E2-glycopr;693 | 0 | 0 |
| NC_004718.3;NP_828851.1;SARS-CoV_E2-glycopr;694 | 0 | 0 |
| NC_004718.3;NP_828851.1;SARS-CoV_E2-glycopr;695 | 0 | 0.025 |
| NC_004718.3;NP_828851.1;SARS-CoV_E2-glycopr;696 | 0 | 0 |
| NC_004718.3;NP_828851.1;SARS-CoV_E2-glycopr;697 | 0 | 0 |
| NC_004718.3;NP_828851.1;SARS-CoV_E2-glycopr;698 | 0 | 0 |
| NC_004718.3;NP_828851.1;SARS-CoV_E2-glycopr;699 | 0 | 0 |
| NC_004718.3;NP_828851.1;SARS-CoV_E2-glycopr;700 | 0 | 0 |
| NC_004718.3;NP_828851.1;SARS-CoV_E2-glycopr;701 | 0 | 0 |
| NC_004718.3;NP_828851.1;SARS-CoV_E2-glycopr;702 | 0 | 0 |
| NC_004718.3;NP_828851.1;SARS-CoV_E2-glycopr;703 | 0 | 0 |
| NC_004718.3;NP_828851.1;SARS-CoV_E2-glycopr;704 | 0 | 0 |
| NC_004718.3;NP_828851.1;SARS-CoV_E2-glycopr;705 | 0 | 0 |
| NC_004718.3;NP_828851.1;SARS-CoV_E2-glycopr;706 | 0 | 0 |
| NC_004718.3;NP_828851.1;SARS-CoV_E2-glycopr;707 | 0 | 0 |
| NC_004718.3;NP_828851.1;SARS-CoV_E2-glycopr;708 | 0 | 0 |
| NC_004718.3;NP_828851.1;SARS-CoV_E2-glycopr;709 | 0 | 0 |
| NC_004718.3;NP_828851.1;SARS-CoV_E2-glycopr;710 | 0 | 0 |
| NC_004718.3;NP_828851.1;SARS-CoV_E2-glycopr;711 | 0 | 0 |
| NC_004718.3;NP_828851.1;SARS-CoV_E2-glycopr;712 | 0 | 0 |
| NC_004718.3;NP_828851.1;SARS-CoV_E2-glycopr;713 | 0 | 0.025 |
| NC_004718.3;NP_828851.1;SARS-CoV_E2-glycopr;714 | 0 | 0 |
| NC_004718.3;NP_828851.1;SARS-CoV_E2-glycopr;715 | 0 | 0.025 |
| NC_004718.3;NP_828851.1;SARS-CoV_E2-glycopr;716 | 0 | 0 |
| NC_004718.3;NP_828851.1;SARS-CoV_E2-glycopr;717 | 0.05 | 0 |
| NC_004718.3;NP_828851.1;SARS-CoV_E2-glycopr;718 | 0.1 | 0 |
| NC_004718.3;NP_828851.1;SARS-CoV_E2-glycopr;719 | 0.1 | 0.025 |
| NC_004718.3;NP_828851.1;SARS-CoV_E2-glycopr;720 | 0.1 | 0 |
| NC_004718.3;NP_828851.1;SARS-CoV_E2-glycopr;721 | 0.05 | 0 |
| NC_004718.3;NP_828851.1;SARS-CoV_E2-glycopr;722 | 0.05 | 0 |
| NC_004718.3;NP_828851.1;SARS-CoV_E2-glycopr;723 | 0.05 | 0 |
| NC_004718.3;NP_828851.1;SARS-CoV_E2-glycopr;724 | 0.05 | 0 |
| NC_004718.3;NP_828851.1;SARS-CoV_E2-glycopr;725 | 0.1 | 0 |
| NC_004718.3;NP_828851.1;SARS-CoV_E2-glycopr;726 | 0 | 0 |
| NC_004718.3;NP_828851.1;SARS-CoV_E2-glycopr;727 | 0 | 0 |
| NC_004718.3;NP_828851.1;SARS-CoV_E2-glycopr;728 | 0 | 0 |
| NC_004718.3;NP_828851.1;SARS-CoV_E2-glycopr;729 | 0 | 0 |
| NC_004718.3;NP_828851.1;SARS-CoV_E2-glycopr;730 | 0 | 0 |
| NC_004718.3;NP_828851.1;SARS-CoV_E2-glycopr;731 | 0 | 0 |
| NC_004718.3;NP_828851.1;SARS-CoV_E2-glycopr;732 | 0 | 0 |
| NC_004718.3;NP_828851.1;SARS-CoV_E2-glycopr;733 | 0 | 0 |

| | | |
|---|---|---|
| NC_004718.3;NP_828851.1;SARS-CoV_E2-glycopr;734 | 0 | 0 |
| NC_004718.3;NP_828851.1;SARS-CoV_E2-glycopr;735 | 0 | 0 |
| NC_004718.3;NP_828851.1;SARS-CoV_E2-glycopr;736 | 0 | 0 |
| NC_004718.3;NP_828851.1;SARS-CoV_E2-glycopr;737 | 0 | 0 |
| NC_004718.3;NP_828851.1;SARS-CoV_E2-glycopr;738 | 0 | 0 |
| NC_004718.3;NP_828851.1;SARS-CoV_E2-glycopr;739 | 0.05 | 0.05 |
| NC_004718.3;NP_828851.1;SARS-CoV_E2-glycopr;740 | 0 | 0.025 |
| NC_004718.3;NP_828851.1;SARS-CoV_E2-glycopr;741 | 0 | 0.025 |
| NC_004718.3;NP_828851.1;SARS-CoV_E2-glycopr;742 | 0 | 0.025 |
| NC_004718.3;NP_828851.1;SARS-CoV_E2-glycopr;743 | 0.05 | 0.025 |
| NC_004718.3;NP_828851.1;SARS-CoV_E2-glycopr;744 | 0.05 | 0.025 |
| NC_004718.3;NP_828851.1;SARS-CoV_E2-glycopr;745 | 0.05 | 0.025 |
| NC_004718.3;NP_828851.1;SARS-CoV_E2-glycopr;746 | 0.05 | 0.025 |
| NC_004718.3;NP_828851.1;SARS-CoV_E2-glycopr;747 | 0.05 | 0.025 |
| NC_004718.3;NP_828851.1;SARS-CoV_E2-glycopr;748 | 0.05 | 0 |
| NC_004718.3;NP_828851.1;SARS-CoV_E2-glycopr;749 | 0.05 | 0 |
| NC_004718.3;NP_828851.1;SARS-CoV_E2-glycopr;750 | 0.05 | 0 |
| NC_004718.3;NP_828851.1;SARS-CoV_E2-glycopr;751 | 0.05 | 0.025 |
| NC_004718.3;NP_828851.1;SARS-CoV_E2-glycopr;752 | 0 | 0 |
| NC_004718.3;NP_828851.1;SARS-CoV_E2-glycopr;753 | 0.05 | 0.125 |
| NC_004718.3;NP_828851.1;SARS-CoV_E2-glycopr;754 | 0.05 | 0.15 |
| NC_004718.3;NP_828851.1;SARS-CoV_E2-glycopr;755 | 0.05 | 0.175 |
| NC_004718.3;NP_828851.1;SARS-CoV_E2-glycopr;756 | 0.05 | 0.125 |
| NC_004718.3;NP_828851.1;SARS-CoV_E2-glycopr;757 | 0.1 | 0.05 |
| NC_004718.3;NP_828851.1;SARS-CoV_E2-glycopr;758 | 0.05 | 0.025 |
| NC_004718.3;NP_828851.1;SARS-CoV_E2-glycopr;759 | 0.05 | 0.025 |
| NC_004718.3;NP_828851.1;SARS-CoV_E2-glycopr;760 | 0.05 | 0.075 |
| NC_004718.3;NP_828851.1;SARS-CoV_E2-glycopr;761 | 0.05 | 0.075 |
| NC_004718.3;NP_828851.1;SARS-CoV_E2-glycopr;762 | 0 | 0 |
| NC_004718.3;NP_828851.1;SARS-CoV_E2-glycopr;763 | 0 | 0 |
| NC_004718.3;NP_828851.1;SARS-CoV_E2-glycopr;764 | 0 | 0 |
| NC_004718.3;NP_828851.1;SARS-CoV_E2-glycopr;765 | 0 | 0 |
| NC_004718.3;NP_828851.1;SARS-CoV_E2-glycopr;766 | 0 | 0 |
| NC_004718.3;NP_828851.1;SARS-CoV_E2-glycopr;767 | 0 | 0 |
| NC_004718.3;NP_828851.1;SARS-CoV_E2-glycopr;768 | 0 | 0 |
| NC_004718.3;NP_828851.1;SARS-CoV_E2-glycopr;769 | 0 | 0 |
| NC_004718.3;NP_828851.1;SARS-CoV_E2-glycopr;770 | 0 | 0.025 |
| NC_004718.3;NP_828851.1;SARS-CoV_E2-glycopr;771 | 0 | 0.025 |
| NC_004718.3;NP_828851.1;SARS-CoV_E2-glycopr;772 | 0 | 0.025 |
| NC_004718.3;NP_828851.1;SARS-CoV_E2-glycopr;773 | 0 | 0 |
| NC_004718.3;NP_828851.1;SARS-CoV_E2-glycopr;774 | 0 | 0 |
| NC_004718.3;NP_828851.1;SARS-CoV_E2-glycopr;775 | 0 | 0.025 |
| NC_004718.3;NP_828851.1;SARS-CoV_E2-glycopr;776 | 0 | 0.075 |

| | | |
|---|---|---|
| NC_004718.3;NP_828851.1;SARS-CoV_E2-glycopr;777 | 0 | 0.075 |
| NC_004718.3;NP_828851.1;SARS-CoV_E2-glycopr;778 | 0 | 0.025 |
| NC_004718.3;NP_828851.1;SARS-CoV_E2-glycopr;779 | 0.05 | 0.1 |
| NC_004718.3;NP_828851.1;SARS-CoV_E2-glycopr;780 | 0.05 | 0.1 |
| NC_004718.3;NP_828851.1;SARS-CoV_E2-glycopr;781 | 0 | 0.05 |
| NC_004718.3;NP_828851.1;SARS-CoV_E2-glycopr;782 | 0 | 0 |
| NC_004718.3;NP_828851.1;SARS-CoV_E2-glycopr;783 | 0.05 | 0.025 |
| NC_004718.3;NP_828851.1;SARS-CoV_E2-glycopr;784 | 0.05 | 0 |
| NC_004718.3;NP_828851.1;SARS-CoV_E2-glycopr;785 | 0.05 | 0.025 |
| NC_004718.3;NP_828851.1;SARS-CoV_E2-glycopr;786 | 0.05 | 0.025 |
| NC_004718.3;NP_828851.1;SARS-CoV_E2-glycopr;787 | 0.1 | 0.025 |
| NC_004718.3;NP_828851.1;SARS-CoV_E2-glycopr;788 | 0.1 | 0.175 |
| NC_004718.3;NP_828851.1;SARS-CoV_E2-glycopr;789 | 0.05 | 0.225 |
| NC_004718.3;NP_828851.1;SARS-CoV_E2-glycopr;790 | 0.05 | 0.6 |
| NC_004718.3;NP_828851.1;SARS-CoV_E2-glycopr;791 | 0.1 | 0.725 |
| NC_004718.3;NP_828851.1;SARS-CoV_E2-glycopr;792 | 0.05 | 0.8 |
| NC_004718.3;NP_828851.1;SARS-CoV_E2-glycopr;793 | 0.05 | 0.8 |
| NC_004718.3;NP_828851.1;SARS-CoV_E2-glycopr;794 | 0.05 | 0.85 |
| NC_004718.3;NP_828851.1;SARS-CoV_E2-glycopr;795 | 0.05 | 0.8 |
| NC_004718.3;NP_828851.1;SARS-CoV_E2-glycopr;796 | 0 | 0.7 |
| NC_004718.3;NP_828851.1;SARS-CoV_E2-glycopr;797 | 0 | 0.725 |
| NC_004718.3;NP_828851.1;SARS-CoV_E2-glycopr;798 | 0 | 0.35 |
| NC_004718.3;NP_828851.1;SARS-CoV_E2-glycopr;799 | 0.05 | 0.25 |
| NC_004718.3;NP_828851.1;SARS-CoV_E2-glycopr;800 | 0.1 | 0.275 |
| NC_004718.3;NP_828851.1;SARS-CoV_E2-glycopr;801 | 0 | 0.15 |
| NC_004718.3;NP_828851.1;SARS-CoV_E2-glycopr;802 | 0 | 0 |
| NC_004718.3;NP_828851.1;SARS-CoV_E2-glycopr;803 | 0 | 0 |
| NC_004718.3;NP_828851.1;SARS-CoV_E2-glycopr;804 | 0 | 0 |
| NC_004718.3;NP_828851.1;SARS-CoV_E2-glycopr;805 | 0 | 0 |
| NC_004718.3;NP_828851.1;SARS-CoV_E2-glycopr;806 | 0 | 0.025 |
| NC_004718.3;NP_828851.1;SARS-CoV_E2-glycopr;807 | 0 | 0.025 |
| NC_004718.3;NP_828851.1;SARS-CoV_E2-glycopr;808 | 0 | 0.05 |
| NC_004718.3;NP_828851.1;SARS-CoV_E2-glycopr;809 | 0 | 0.025 |
| NC_004718.3;NP_828851.1;SARS-CoV_E2-glycopr;810 | 0.1 | 0.025 |
| NC_004718.3;NP_828851.1;SARS-CoV_E2-glycopr;811 | 0 | 0.025 |
| NC_004718.3;NP_828851.1;SARS-CoV_E2-glycopr;812 | 0 | 0.025 |
| NC_004718.3;NP_828851.1;SARS-CoV_E2-glycopr;813 | 0 | 0.025 |
| NC_004718.3;NP_828851.1;SARS-CoV_E2-glycopr;814 | 0.05 | 0.025 |
| NC_004718.3;NP_828851.1;SARS-CoV_E2-glycopr;815 | 0 | 0.025 |
| NC_004718.3;NP_828851.1;SARS-CoV_E2-glycopr;816 | 0 | 0.05 |
| NC_004718.3;NP_828851.1;SARS-CoV_E2-glycopr;817 | 0 | 0.025 |
| NC_004718.3;NP_828851.1;SARS-CoV_E2-glycopr;818 | 0 | 0 |
| NC_004718.3;NP_828851.1;SARS-CoV_E2-glycopr;819 | 0 | 0 |

| | | |
|---|---|---|
| NC_004718.3;NP_828851.1;SARS-CoV_E2-glycopr;820 | 0 | 0 |
| NC_004718.3;NP_828851.1;SARS-CoV_E2-glycopr;821 | 0 | 0 |
| NC_004718.3;NP_828851.1;SARS-CoV_E2-glycopr;822 | 0 | 0 |
| NC_004718.3;NP_828851.1;SARS-CoV_E2-glycopr;823 | 0 | 0 |
| NC_004718.3;NP_828851.1;SARS-CoV_E2-glycopr;824 | 0 | 0.025 |
| NC_004718.3;NP_828851.1;SARS-CoV_E2-glycopr;825 | 0 | 0 |
| NC_004718.3;NP_828851.1;SARS-CoV_E2-glycopr;826 | 0 | 0 |
| NC_004718.3;NP_828851.1;SARS-CoV_E2-glycopr;827 | 0 | 0 |
| NC_004718.3;NP_828851.1;SARS-CoV_E2-glycopr;828 | 0 | 0 |
| NC_004718.3;NP_828851.1;SARS-CoV_E2-glycopr;829 | 0 | 0 |
| NC_004718.3;NP_828851.1;SARS-CoV_E2-glycopr;830 | 0.05 | 0 |
| NC_004718.3;NP_828851.1;SARS-CoV_E2-glycopr;831 | 0 | 0 |
| NC_004718.3;NP_828851.1;SARS-CoV_E2-glycopr;832 | 0 | 0 |
| NC_004718.3;NP_828851.1;SARS-CoV_E2-glycopr;833 | 0 | 0 |
| NC_004718.3;NP_828851.1;SARS-CoV_E2-glycopr;834 | 0 | 0 |
| NC_004718.3;NP_828851.1;SARS-CoV_E2-glycopr;835 | 0 | 0 |
| NC_004718.3;NP_828851.1;SARS-CoV_E2-glycopr;836 | 0 | 0 |
| NC_004718.3;NP_828851.1;SARS-CoV_E2-glycopr;837 | 0 | 0 |
| NC_004718.3;NP_828851.1;SARS-CoV_E2-glycopr;838 | 0 | 0 |
| NC_004718.3;NP_828851.1;SARS-CoV_E2-glycopr;839 | 0 | 0 |
| NC_004718.3;NP_828851.1;SARS-CoV_E2-glycopr;840 | 0 | 0 |
| NC_004718.3;NP_828851.1;SARS-CoV_E2-glycopr;841 | 0 | 0 |
| NC_004718.3;NP_828851.1;SARS-CoV_E2-glycopr;842 | 0 | 0 |
| NC_004718.3;NP_828851.1;SARS-CoV_E2-glycopr;843 | 0.05 | 0 |
| NC_004718.3;NP_828851.1;SARS-CoV_E2-glycopr;844 | 0 | 0.025 |
| NC_004718.3;NP_828851.1;SARS-CoV_E2-glycopr;845 | 0 | 0 |
| NC_004718.3;NP_828851.1;SARS-CoV_E2-glycopr;846 | 0 | 0 |
| NC_004718.3;NP_828851.1;SARS-CoV_E2-glycopr;847 | 0 | 0 |
| NC_004718.3;NP_828851.1;SARS-CoV_E2-glycopr;848 | 0 | 0 |
| NC_004718.3;NP_828851.1;SARS-CoV_E2-glycopr;849 | 0 | 0 |
| NC_004718.3;NP_828851.1;SARS-CoV_E2-glycopr;850 | 0 | 0 |
| NC_004718.3;NP_828851.1;SARS-CoV_E2-glycopr;851 | 0 | 0.025 |
| NC_004718.3;NP_828851.1;SARS-CoV_E2-glycopr;852 | 0 | 0.025 |
| NC_004718.3;NP_828851.1;SARS-CoV_E2-glycopr;853 | 0 | 0 |
| NC_004718.3;NP_828851.1;SARS-CoV_E2-glycopr;854 | 0 | 0 |
| NC_004718.3;NP_828851.1;SARS-CoV_E2-glycopr;855 | 0 | 0 |
| NC_004718.3;NP_828851.1;SARS-CoV_E2-glycopr;856 | 0 | 0 |
| NC_004718.3;NP_828851.1;SARS-CoV_E2-glycopr;857 | 0 | 0 |
| NC_004718.3;NP_828851.1;SARS-CoV_E2-glycopr;858 | 0 | 0 |
| NC_004718.3;NP_828851.1;SARS-CoV_E2-glycopr;859 | 0 | 0.025 |
| NC_004718.3;NP_828851.1;SARS-CoV_E2-glycopr;860 | 0.1 | 0.075 |
| NC_004718.3;NP_828851.1;SARS-CoV_E2-glycopr;861 | 0 | 0 |
| NC_004718.3;NP_828851.1;SARS-CoV_E2-glycopr;862 | 0 | 0 |

| | | |
|---|---|---|
| NC_004718.3;NP_828851.1;SARS-CoV_E2-glycopr;863 | 0 | 0 |
| NC_004718.3;NP_828851.1;SARS-CoV_E2-glycopr;864 | 0 | 0 |
| NC_004718.3;NP_828851.1;SARS-CoV_E2-glycopr;865 | 0 | 0 |
| NC_004718.3;NP_828851.1;SARS-CoV_E2-glycopr;866 | 0 | 0 |
| NC_004718.3;NP_828851.1;SARS-CoV_E2-glycopr;867 | 0 | 0 |
| NC_004718.3;NP_828851.1;SARS-CoV_E2-glycopr;868 | 0 | 0 |
| NC_004718.3;NP_828851.1;SARS-CoV_E2-glycopr;869 | 0 | 0 |
| NC_004718.3;NP_828851.1;SARS-CoV_E2-glycopr;870 | 0 | 0 |
| NC_004718.3;NP_828851.1;SARS-CoV_E2-glycopr;871 | 0 | 0.025 |
| NC_004718.3;NP_828851.1;SARS-CoV_E2-glycopr;872 | 0 | 0 |
| NC_004718.3;NP_828851.1;SARS-CoV_E2-glycopr;873 | 0 | 0 |
| NC_004718.3;NP_828851.1;SARS-CoV_E2-glycopr;874 | 0.05 | 0.025 |
| NC_004718.3;NP_828851.1;SARS-CoV_E2-glycopr;875 | 0.05 | 0.025 |
| NC_004718.3;NP_828851.1;SARS-CoV_E2-glycopr;876 | 0.05 | 0.025 |
| NC_004718.3;NP_828851.1;SARS-CoV_E2-glycopr;877 | 0.05 | 0.025 |
| NC_004718.3;NP_828851.1;SARS-CoV_E2-glycopr;878 | 0.05 | 0.025 |
| NC_004718.3;NP_828851.1;SARS-CoV_E2-glycopr;879 | 0.05 | 0.025 |
| NC_004718.3;NP_828851.1;SARS-CoV_E2-glycopr;880 | 0.05 | 0.025 |
| NC_004718.3;NP_828851.1;SARS-CoV_E2-glycopr;881 | 0.1 | 0.05 |
| NC_004718.3;NP_828851.1;SARS-CoV_E2-glycopr;882 | 0.15 | 0.05 |
| NC_004718.3;NP_828851.1;SARS-CoV_E2-glycopr;883 | 0.1 | 0.05 |
| NC_004718.3;NP_828851.1;SARS-CoV_E2-glycopr;884 | 0 | 0 |
| NC_004718.3;NP_828851.1;SARS-CoV_E2-glycopr;885 | 0 | 0 |
| NC_004718.3;NP_828851.1;SARS-CoV_E2-glycopr;886 | 0 | 0 |
| NC_004718.3;NP_828851.1;SARS-CoV_E2-glycopr;887 | 0.05 | 0.025 |
| NC_004718.3;NP_828851.1;SARS-CoV_E2-glycopr;888 | 0 | 0.05 |
| NC_004718.3;NP_828851.1;SARS-CoV_E2-glycopr;889 | 0 | 0.05 |
| NC_004718.3;NP_828851.1;SARS-CoV_E2-glycopr;890 | 0.05 | 0.075 |
| NC_004718.3;NP_828851.1;SARS-CoV_E2-glycopr;891 | 0.05 | 0.075 |
| NC_004718.3;NP_828851.1;SARS-CoV_E2-glycopr;892 | 0.05 | 0.025 |
| NC_004718.3;NP_828851.1;SARS-CoV_E2-glycopr;893 | 0 | 0.075 |
| NC_004718.3;NP_828851.1;SARS-CoV_E2-glycopr;894 | 0.05 | 0.05 |
| NC_004718.3;NP_828851.1;SARS-CoV_E2-glycopr;895 | 0.2 | 0.2 |
| NC_004718.3;NP_828851.1;SARS-CoV_E2-glycopr;896 | 0.15 | 0.1 |
| NC_004718.3;NP_828851.1;SARS-CoV_E2-glycopr;897 | 0.2 | 0.15 |
| NC_004718.3;NP_828851.1;SARS-CoV_E2-glycopr;898 | 0.3 | 0.225 |
| NC_004718.3;NP_828851.1;SARS-CoV_E2-glycopr;899 | 0.35 | 0.3 |
| NC_004718.3;NP_828851.1;SARS-CoV_E2-glycopr;900 | 0.25 | 0.25 |
| NC_004718.3;NP_828851.1;SARS-CoV_E2-glycopr;901 | 0.2 | 0.125 |
| NC_004718.3;NP_828851.1;SARS-CoV_E2-glycopr;902 | 0.25 | 0.15 |
| NC_004718.3;NP_828851.1;SARS-CoV_E2-glycopr;903 | 0.3 | 0.225 |
| NC_004718.3;NP_828851.1;SARS-CoV_E2-glycopr;904 | 0.3 | 0.2 |
| NC_004718.3;NP_828851.1;SARS-CoV_E2-glycopr;905 | 0.05 | 0 |

| | | |
|---|---|---|
| NC_004718.3;NP_828851.1;SARS-CoV_E2-glycopr;906 | 0.05 | 0.025 |
| NC_004718.3;NP_828851.1;SARS-CoV_E2-glycopr;907 | 0 | 0 |
| NC_004718.3;NP_828851.1;SARS-CoV_E2-glycopr;908 | 0 | 0 |
| NC_004718.3;NP_828851.1;SARS-CoV_E2-glycopr;909 | 0 | 0 |
| NC_004718.3;NP_828851.1;SARS-CoV_E2-glycopr;910 | 0 | 0 |
| NC_004718.3;NP_828851.1;SARS-CoV_E2-glycopr;911 | 0 | 0 |
| NC_004718.3;NP_828851.1;SARS-CoV_E2-glycopr;912 | 0.05 | 0.05 |
| NC_004718.3;NP_828851.1;SARS-CoV_E2-glycopr;913 | 0 | 0 |
| NC_004718.3;NP_828851.1;SARS-CoV_E2-glycopr;914 | 0 | 0 |
| NC_004718.3;NP_828851.1;SARS-CoV_E2-glycopr;915 | 0 | 0 |
| NC_004718.3;NP_828851.1;SARS-CoV_E2-glycopr;916 | 0 | 0 |
| NC_004718.3;NP_828851.1;SARS-CoV_E2-glycopr;917 | 0 | 0 |
| NC_004718.3;NP_828851.1;SARS-CoV_E2-glycopr;918 | 0 | 0 |
| NC_004718.3;NP_828851.1;SARS-CoV_E2-glycopr;919 | 0 | 0 |
| NC_004718.3;NP_828851.1;SARS-CoV_E2-glycopr;920 | 0 | 0 |
| NC_004718.3;NP_828851.1;SARS-CoV_E2-glycopr;921 | 0 | 0 |
| NC_004718.3;NP_828851.1;SARS-CoV_E2-glycopr;922 | 0 | 0.075 |
| NC_004718.3;NP_828851.1;SARS-CoV_E2-glycopr;923 | 0 | 0.025 |
| NC_004718.3;NP_828851.1;SARS-CoV_E2-glycopr;924 | 0 | 0.025 |
| NC_004718.3;NP_828851.1;SARS-CoV_E2-glycopr;925 | 0 | 0.025 |
| NC_004718.3;NP_828851.1;SARS-CoV_E2-glycopr;926 | 0 | 0.075 |
| NC_004718.3;NP_828851.1;SARS-CoV_E2-glycopr;927 | 0 | 0.125 |
| NC_004718.3;NP_828851.1;SARS-CoV_E2-glycopr;928 | 0.1 | 0.075 |
| NC_004718.3;NP_828851.1;SARS-CoV_E2-glycopr;929 | 0 | 0.1 |
| NC_004718.3;NP_828851.1;SARS-CoV_E2-glycopr;930 | 0 | 0.075 |
| NC_004718.3;NP_828851.1;SARS-CoV_E2-glycopr;931 | 0 | 0.025 |
| NC_004718.3;NP_828851.1;SARS-CoV_E2-glycopr;932 | 0 | 0.05 |
| NC_004718.3;NP_828851.1;SARS-CoV_E2-glycopr;933 | 0.05 | 0 |
| NC_004718.3;NP_828851.1;SARS-CoV_E2-glycopr;934 | 0 | 0 |
| NC_004718.3;NP_828851.1;SARS-CoV_E2-glycopr;935 | 0 | 0.025 |
| NC_004718.3;NP_828851.1;SARS-CoV_E2-glycopr;936 | 0 | 0.025 |
| NC_004718.3;NP_828851.1;SARS-CoV_E2-glycopr;937 | 0 | 0 |
| NC_004718.3;NP_828851.1;SARS-CoV_E2-glycopr;938 | 0 | 0.025 |
| NC_004718.3;NP_828851.1;SARS-CoV_E2-glycopr;939 | 0 | 0.05 |
| NC_004718.3;NP_828851.1;SARS-CoV_E2-glycopr;940 | 0 | 0.025 |
| NC_004718.3;NP_828851.1;SARS-CoV_E2-glycopr;941 | 0 | 0 |
| NC_004718.3;NP_828851.1;SARS-CoV_E2-glycopr;942 | 0 | 0 |
| NC_004718.3;NP_828851.1;SARS-CoV_E2-glycopr;943 | 0 | 0 |
| NC_004718.3;NP_828851.1;SARS-CoV_E2-glycopr;944 | 0 | 0 |
| NC_004718.3;NP_828851.1;SARS-CoV_E2-glycopr;945 | 0 | 0 |
| NC_004718.3;NP_828851.1;SARS-CoV_E2-glycopr;946 | 0 | 0.025 |
| NC_004718.3;NP_828851.1;SARS-CoV_E2-glycopr;947 | 0 | 0 |
| NC_004718.3;NP_828851.1;SARS-CoV_E2-glycopr;948 | 0 | 0 |

| | | |
|---|---|---|
| NC_004718.3;NP_828851.1;SARS-CoV_E2-glycopr;949 | 0 | 0 |
| NC_004718.3;NP_828851.1;SARS-CoV_E2-glycopr;950 | 0 | 0 |
| NC_004718.3;NP_828851.1;SARS-CoV_E2-glycopr;951 | 0 | 0 |
| NC_004718.3;NP_828851.1;SARS-CoV_E2-glycopr;952 | 0 | 0.025 |
| NC_004718.3;NP_828851.1;SARS-CoV_E2-glycopr;953 | 0 | 0.025 |
| NC_004718.3;NP_828851.1;SARS-CoV_E2-glycopr;954 | 0.05 | 0.05 |
| NC_004718.3;NP_828851.1;SARS-CoV_E2-glycopr;955 | 0 | 0.075 |
| NC_004718.3;NP_828851.1;SARS-CoV_E2-glycopr;956 | 0.05 | 0.075 |
| NC_004718.3;NP_828851.1;SARS-CoV_E2-glycopr;957 | 0.05 | 0.125 |
| NC_004718.3;NP_828851.1;SARS-CoV_E2-glycopr;958 | 0.05 | 0.1 |
| NC_004718.3;NP_828851.1;SARS-CoV_E2-glycopr;959 | 0 | 0.075 |
| NC_004718.3;NP_828851.1;SARS-CoV_E2-glycopr;960 | 0 | 0.05 |
| NC_004718.3;NP_828851.1;SARS-CoV_E2-glycopr;961 | 0.05 | 0.05 |
| NC_004718.3;NP_828851.1;SARS-CoV_E2-glycopr;962 | 0.05 | 0.025 |
| NC_004718.3;NP_828851.1;SARS-CoV_E2-glycopr;963 | 0 | 0 |
| NC_004718.3;NP_828851.1;SARS-CoV_E2-glycopr;964 | 0 | 0 |
| NC_004718.3;NP_828851.1;SARS-CoV_E2-glycopr;965 | 0 | 0 |
| NC_004718.3;NP_828851.1;SARS-CoV_E2-glycopr;966 | 0 | 0.025 |
| NC_004718.3;NP_828851.1;SARS-CoV_E2-glycopr;967 | 0 | 0 |
| NC_004718.3;NP_828851.1;SARS-CoV_E2-glycopr;968 | 0 | 0 |
| NC_004718.3;NP_828851.1;SARS-CoV_E2-glycopr;969 | 0 | 0 |
| NC_004718.3;NP_828851.1;SARS-CoV_E2-glycopr;970 | 0.05 | 0 |
| NC_004718.3;NP_828851.1;SARS-CoV_E2-glycopr;971 | 0 | 0 |
| NC_004718.3;NP_828851.1;SARS-CoV_E2-glycopr;972 | 0 | 0 |
| NC_004718.3;NP_828851.1;SARS-CoV_E2-glycopr;973 | 0 | 0 |
| NC_004718.3;NP_828851.1;SARS-CoV_E2-glycopr;974 | 0 | 0 |
| NC_004718.3;NP_828851.1;SARS-CoV_E2-glycopr;975 | 0 | 0 |
| NC_004718.3;NP_828851.1;SARS-CoV_E2-glycopr;976 | 0 | 0 |
| NC_004718.3;NP_828851.1;SARS-CoV_E2-glycopr;977 | 0 | 0 |
| NC_004718.3;NP_828851.1;SARS-CoV_E2-glycopr;978 | 0 | 0 |
| NC_004718.3;NP_828851.1;SARS-CoV_E2-glycopr;979 | 0 | 0 |
| NC_004718.3;NP_828851.1;SARS-CoV_E2-glycopr;980 | 0 | 0 |
| NC_004718.3;NP_828851.1;SARS-CoV_E2-glycopr;981 | 0 | 0 |
| NC_004718.3;NP_828851.1;SARS-CoV_E2-glycopr;982 | 0 | 0 |
| NC_004718.3;NP_828851.1;SARS-CoV_E2-glycopr;983 | 0 | 0 |
| NC_004718.3;NP_828851.1;SARS-CoV_E2-glycopr;984 | 0 | 0 |
| NC_004718.3;NP_828851.1;SARS-CoV_E2-glycopr;985 | 0 | 0.025 |
| NC_004718.3;NP_828851.1;SARS-CoV_E2-glycopr;986 | 0 | 0 |
| NC_004718.3;NP_828851.1;SARS-CoV_E2-glycopr;987 | 0 | 0 |
| NC_004718.3;NP_828851.1;SARS-CoV_E2-glycopr;988 | 0 | 0 |
| NC_004718.3;NP_828851.1;SARS-CoV_E2-glycopr;989 | 0 | 0 |
| NC_004718.3;NP_828851.1;SARS-CoV_E2-glycopr;990 | 0 | 0 |
| NC_004718.3;NP_828851.1;SARS-CoV_E2-glycopr;991 | 0 | 0 |

| | | |
|---|---|---|
| NC_004718.3;NP_828851.1;SARS-CoV_E2-glycopr;992 | 0 | 0 |
| NC_004718.3;NP_828851.1;SARS-CoV_E2-glycopr;993 | 0.05 | 0.025 |
| NC_004718.3;NP_828851.1;SARS-CoV_E2-glycopr;994 | 0 | 0 |
| NC_004718.3;NP_828851.1;SARS-CoV_E2-glycopr;995 | 0 | 0.025 |
| NC_004718.3;NP_828851.1;SARS-CoV_E2-glycopr;996 | 0 | 0 |
| NC_004718.3;NP_828851.1;SARS-CoV_E2-glycopr;997 | 0 | 0 |
| NC_004718.3;NP_828851.1;SARS-CoV_E2-glycopr;998 | 0.05 | 0.025 |
| NC_004718.3;NP_828851.1;SARS-CoV_E2-glycopr;999 | 0.05 | 0 |
| NC_004718.3;NP_828851.1;SARS-CoV_E2-glycopr;1000 | 0.05 | 0 |
| NC_004718.3;NP_828851.1;SARS-CoV_E2-glycopr;1001 | 0 | 0.025 |
| NC_004718.3;NP_828851.1;SARS-CoV_E2-glycopr;1002 | 0 | 0 |
| NC_004718.3;NP_828851.1;SARS-CoV_E2-glycopr;1003 | 0 | 0.025 |
| NC_004718.3;NP_828851.1;SARS-CoV_E2-glycopr;1004 | 0 | 0 |
| NC_004718.3;NP_828851.1;SARS-CoV_E2-glycopr;1005 | 0 | 0 |
| NC_004718.3;NP_828851.1;SARS-CoV_E2-glycopr;1006 | 0 | 0 |
| NC_004718.3;NP_828851.1;SARS-CoV_E2-glycopr;1007 | 0 | 0 |
| NC_004718.3;NP_828851.1;SARS-CoV_E2-glycopr;1008 | 0 | 0 |
| NC_004718.3;NP_828851.1;SARS-CoV_E2-glycopr;1009 | 0 | 0 |
| NC_004718.3;NP_828851.1;SARS-CoV_E2-glycopr;1010 | 0 | 0 |
| NC_004718.3;NP_828851.1;SARS-CoV_E2-glycopr;1011 | 0 | 0 |
| NC_004718.3;NP_828851.1;SARS-CoV_E2-glycopr;1012 | 0 | 0 |
| NC_004718.3;NP_828851.1;SARS-CoV_E2-glycopr;1013 | 0 | 0 |
| NC_004718.3;NP_828851.1;SARS-CoV_E2-glycopr;1014 | 0 | 0 |
| NC_004718.3;NP_828851.1;SARS-CoV_E2-glycopr;1015 | 0 | 0 |
| NC_004718.3;NP_828851.1;SARS-CoV_E2-glycopr;1016 | 0 | 0 |
| NC_004718.3;NP_828851.1;SARS-CoV_E2-glycopr;1017 | 0 | 0 |
| NC_004718.3;NP_828851.1;SARS-CoV_E2-glycopr;1018 | 0 | 0 |
| NC_004718.3;NP_828851.1;SARS-CoV_E2-glycopr;1019 | 0 | 0 |
| NC_004718.3;NP_828851.1;SARS-CoV_E2-glycopr;1020 | 0 | 0 |
| NC_004718.3;NP_828851.1;SARS-CoV_E2-glycopr;1021 | 0 | 0 |
| NC_004718.3;NP_828851.1;SARS-CoV_E2-glycopr;1022 | 0 | 0 |
| NC_004718.3;NP_828851.1;SARS-CoV_E2-glycopr;1023 | 0.05 | 0.05 |
| NC_004718.3;NP_828851.1;SARS-CoV_E2-glycopr;1024 | 0 | 0 |
| NC_004718.3;NP_828851.1;SARS-CoV_E2-glycopr;1025 | 0 | 0.025 |
| NC_004718.3;NP_828851.1;SARS-CoV_E2-glycopr;1026 | 0 | 0.025 |
| NC_004718.3;NP_828851.1;SARS-CoV_E2-glycopr;1027 | 0 | 0.025 |
| NC_004718.3;NP_828851.1;SARS-CoV_E2-glycopr;1028 | 0 | 0 |
| NC_004718.3;NP_828851.1;SARS-CoV_E2-glycopr;1029 | 0 | 0 |
| NC_004718.3;NP_828851.1;SARS-CoV_E2-glycopr;1030 | 0 | 0 |
| NC_004718.3;NP_828851.1;SARS-CoV_E2-glycopr;1031 | 0 | 0 |
| NC_004718.3;NP_828851.1;SARS-CoV_E2-glycopr;1032 | 0 | 0 |
| NC_004718.3;NP_828851.1;SARS-CoV_E2-glycopr;1033 | 0 | 0 |
| NC_004718.3;NP_828851.1;SARS-CoV_E2-glycopr;1034 | 0 | 0 |

| | | |
|---|---|---|
| NC_004718.3;NP_828851.1;SARS-CoV_E2-glycopr;1035 | 0 | 0 |
| NC_004718.3;NP_828851.1;SARS-CoV_E2-glycopr;1036 | 0 | 0 |
| NC_004718.3;NP_828851.1;SARS-CoV_E2-glycopr;1037 | 0 | 0 |
| NC_004718.3;NP_828851.1;SARS-CoV_E2-glycopr;1038 | 0 | 0 |
| NC_004718.3;NP_828851.1;SARS-CoV_E2-glycopr;1039 | 0 | 0 |
| NC_004718.3;NP_828851.1;SARS-CoV_E2-glycopr;1040 | 0 | 0 |
| NC_004718.3;NP_828851.1;SARS-CoV_E2-glycopr;1041 | 0 | 0 |
| NC_004718.3;NP_828851.1;SARS-CoV_E2-glycopr;1042 | 0 | 0 |
| NC_004718.3;NP_828851.1;SARS-CoV_E2-glycopr;1043 | 0 | 0 |
| NC_004718.3;NP_828851.1;SARS-CoV_E2-glycopr;1044 | 0 | 0 |
| NC_004718.3;NP_828851.1;SARS-CoV_E2-glycopr;1045 | 0 | 0 |
| NC_004718.3;NP_828851.1;SARS-CoV_E2-glycopr;1046 | 0 | 0 |
| NC_004718.3;NP_828851.1;SARS-CoV_E2-glycopr;1047 | 0 | 0.025 |
| NC_004718.3;NP_828851.1;SARS-CoV_E2-glycopr;1048 | 0.2 | 0.025 |
| NC_004718.3;NP_828851.1;SARS-CoV_E2-glycopr;1049 | 0.1 | 0 |
| NC_004718.3;NP_828851.1;SARS-CoV_E2-glycopr;1050 | 0 | 0 |
| NC_004718.3;NP_828851.1;SARS-CoV_E2-glycopr;1051 | 0 | 0 |
| NC_004718.3;NP_828851.1;SARS-CoV_E2-glycopr;1052 | 0.2 | 0.05 |
| NC_004718.3;NP_828851.1;SARS-CoV_E2-glycopr;1053 | 0.05 | 0 |
| NC_004718.3;NP_828851.1;SARS-CoV_E2-glycopr;1054 | 0 | 0 |
| NC_004718.3;NP_828851.1;SARS-CoV_E2-glycopr;1055 | 0 | 0 |
| NC_004718.3;NP_828851.1;SARS-CoV_E2-glycopr;1056 | 0 | 0 |
| NC_004718.3;NP_828851.1;SARS-CoV_E2-glycopr;1057 | 0 | 0 |
| NC_004718.3;NP_828851.1;SARS-CoV_E2-glycopr;1058 | 0 | 0 |
| NC_004718.3;NP_828851.1;SARS-CoV_E2-glycopr;1059 | 0.05 | 0 |
| NC_004718.3;NP_828851.1;SARS-CoV_E2-glycopr;1060 | 0.05 | 0 |
| NC_004718.3;NP_828851.1;SARS-CoV_E2-glycopr;1061 | 0.05 | 0 |
| NC_004718.3;NP_828851.1;SARS-CoV_E2-glycopr;1062 | 0.05 | 0 |
| NC_004718.3;NP_828851.1;SARS-CoV_E2-glycopr;1063 | 0 | 0 |
| NC_004718.3;NP_828851.1;SARS-CoV_E2-glycopr;1064 | 0 | 0 |
| NC_004718.3;NP_828851.1;SARS-CoV_E2-glycopr;1065 | 0 | 0 |
| NC_004718.3;NP_828851.1;SARS-CoV_E2-glycopr;1066 | 0 | 0 |
| NC_004718.3;NP_828851.1;SARS-CoV_E2-glycopr;1067 | 0 | 0 |
| NC_004718.3;NP_828851.1;SARS-CoV_E2-glycopr;1068 | 0 | 0 |
| NC_004718.3;NP_828851.1;SARS-CoV_E2-glycopr;1069 | 0 | 0 |
| NC_004718.3;NP_828851.1;SARS-CoV_E2-glycopr;1070 | 0 | 0 |
| NC_004718.3;NP_828851.1;SARS-CoV_E2-glycopr;1071 | 0 | 0 |
| NC_004718.3;NP_828851.1;SARS-CoV_E2-glycopr;1072 | 0 | 0 |
| NC_004718.3;NP_828851.1;SARS-CoV_E2-glycopr;1073 | 0 | 0 |
| NC_004718.3;NP_828851.1;SARS-CoV_E2-glycopr;1074 | 0 | 0 |
| NC_004718.3;NP_828851.1;SARS-CoV_E2-glycopr;1075 | 0 | 0 |
| NC_004718.3;NP_828851.1;SARS-CoV_E2-glycopr;1076 | 0 | 0 |
| NC_004718.3;NP_828851.1;SARS-CoV_E2-glycopr;1077 | 0 | 0 |

| | | |
|---|---|---|
| NC_004718.3;NP_828851.1;SARS-CoV_E2-glycopr;1078 | 0 | 0 |
| NC_004718.3;NP_828851.1;SARS-CoV_E2-glycopr;1079 | 0 | 0 |
| NC_004718.3;NP_828851.1;SARS-CoV_E2-glycopr;1080 | 0 | 0 |
| NC_004718.3;NP_828851.1;SARS-CoV_E2-glycopr;1081 | 0 | 0 |
| NC_004718.3;NP_828851.1;SARS-CoV_E2-glycopr;1082 | 0 | 0 |
| NC_004718.3;NP_828851.1;SARS-CoV_E2-glycopr;1083 | 0 | 0 |
| NC_004718.3;NP_828851.1;SARS-CoV_E2-glycopr;1084 | 0 | 0 |
| NC_004718.3;NP_828851.1;SARS-CoV_E2-glycopr;1085 | 0 | 0 |
| NC_004718.3;NP_828851.1;SARS-CoV_E2-glycopr;1086 | 0 | 0 |
| NC_004718.3;NP_828851.1;SARS-CoV_E2-glycopr;1087 | 0 | 0 |
| NC_004718.3;NP_828851.1;SARS-CoV_E2-glycopr;1088 | 0 | 0 |
| NC_004718.3;NP_828851.1;SARS-CoV_E2-glycopr;1089 | 0 | 0 |
| NC_004718.3;NP_828851.1;SARS-CoV_E2-glycopr;1090 | 0.05 | 0.025 |
| NC_004718.3;NP_828851.1;SARS-CoV_E2-glycopr;1091 | 0.05 | 0.025 |
| NC_004718.3;NP_828851.1;SARS-CoV_E2-glycopr;1092 | 0.05 | 0.075 |
| NC_004718.3;NP_828851.1;SARS-CoV_E2-glycopr;1093 | 0.05 | 0.025 |
| NC_004718.3;NP_828851.1;SARS-CoV_E2-glycopr;1094 | 0.05 | 0.05 |
| NC_004718.3;NP_828851.1;SARS-CoV_E2-glycopr;1095 | 0.05 | 0.075 |
| NC_004718.3;NP_828851.1;SARS-CoV_E2-glycopr;1096 | 0.05 | 0.05 |
| NC_004718.3;NP_828851.1;SARS-CoV_E2-glycopr;1097 | 0.05 | 0.025 |
| NC_004718.3;NP_828851.1;SARS-CoV_E2-glycopr;1098 | 0.05 | 0.05 |
| NC_004718.3;NP_828851.1;SARS-CoV_E2-glycopr;1099 | 0 | 0.025 |
| NC_004718.3;NP_828851.1;SARS-CoV_E2-glycopr;1100 | 0 | 0 |
| NC_004718.3;NP_828851.1;SARS-CoV_E2-glycopr;1101 | 0 | 0 |
| NC_004718.3;NP_828851.1;SARS-CoV_E2-glycopr;1102 | 0 | 0 |
| NC_004718.3;NP_828851.1;SARS-CoV_E2-glycopr;1103 | 0 | 0 |
| NC_004718.3;NP_828851.1;SARS-CoV_E2-glycopr;1104 | 0 | 0 |
| NC_004718.3;NP_828851.1;SARS-CoV_E2-glycopr;1105 | 0 | 0 |
| NC_004718.3;NP_828851.1;SARS-CoV_E2-glycopr;1106 | 0 | 0 |
| NC_004718.3;NP_828851.1;SARS-CoV_E2-glycopr;1107 | 0 | 0 |
| NC_004718.3;NP_828851.1;SARS-CoV_E2-glycopr;1108 | 0 | 0 |
| NC_004718.3;NP_828851.1;SARS-CoV_E2-glycopr;1109 | 0 | 0 |
| NC_004718.3;NP_828851.1;SARS-CoV_E2-glycopr;1110 | 0 | 0 |
| NC_004718.3;NP_828851.1;SARS-CoV_E2-glycopr;1111 | 0 | 0 |
| NC_004718.3;NP_828851.1;SARS-CoV_E2-glycopr;1112 | 0 | 0.025 |
| NC_004718.3;NP_828851.1;SARS-CoV_E2-glycopr;1113 | 0 | 0 |
| NC_004718.3;NP_828851.1;SARS-CoV_E2-glycopr;1114 | 0 | 0 |
| NC_004718.3;NP_828851.1;SARS-CoV_E2-glycopr;1115 | 0 | 0.025 |
| NC_004718.3;NP_828851.1;SARS-CoV_E2-glycopr;1116 | 0 | 0 |
| NC_004718.3;NP_828851.1;SARS-CoV_E2-glycopr;1117 | 0 | 0.025 |
| NC_004718.3;NP_828851.1;SARS-CoV_E2-glycopr;1118 | 0 | 0 |
| NC_004718.3;NP_828851.1;SARS-CoV_E2-glycopr;1119 | 0 | 0 |
| NC_004718.3;NP_828851.1;SARS-CoV_E2-glycopr;1120 | 0 | 0 |

| | | |
|---|---|---|
| NC_004718.3;NP_828851.1;SARS-CoV_E2-glycopr;1121 | 0 | 0.05 |
| NC_004718.3;NP_828851.1;SARS-CoV_E2-glycopr;1122 | 0 | 0.175 |
| NC_004718.3;NP_828851.1;SARS-CoV_E2-glycopr;1123 | 0.05 | 0.75 |
| NC_004718.3;NP_828851.1;SARS-CoV_E2-glycopr;1124 | 0.1 | 0.775 |
| NC_004718.3;NP_828851.1;SARS-CoV_E2-glycopr;1125 | 0.05 | 0.775 |
| NC_004718.3;NP_828851.1;SARS-CoV_E2-glycopr;1126 | 0.05 | 0.725 |
| NC_004718.3;NP_828851.1;SARS-CoV_E2-glycopr;1127 | 0.3 | 0.725 |
| NC_004718.3;NP_828851.1;SARS-CoV_E2-glycopr;1128 | 0.4 | 0.775 |
| NC_004718.3;NP_828851.1;SARS-CoV_E2-glycopr;1129 | 0.35 | 0.775 |
| NC_004718.3;NP_828851.1;SARS-CoV_E2-glycopr;1130 | 0.35 | 0.825 |
| NC_004718.3;NP_828851.1;SARS-CoV_E2-glycopr;1131 | 0.45 | 0.8 |
| NC_004718.3;NP_828851.1;SARS-CoV_E2-glycopr;1132 | 0.4 | 0.5 |
| NC_004718.3;NP_828851.1;SARS-CoV_E2-glycopr;1133 | 0.45 | 0.35 |
| NC_004718.3;NP_828851.1;SARS-CoV_E2-glycopr;1134 | 0.45 | 0.35 |
| NC_004718.3;NP_828851.1;SARS-CoV_E2-glycopr;1135 | 0.5 | 0.35 |
| NC_004718.3;NP_828851.1;SARS-CoV_E2-glycopr;1136 | 0.5 | 0.375 |
| NC_004718.3;NP_828851.1;SARS-CoV_E2-glycopr;1137 | 0 | 0.2 |
| NC_004718.3;NP_828851.1;SARS-CoV_E2-glycopr;1138 | 0 | 0.125 |
| NC_004718.3;NP_828851.1;SARS-CoV_E2-glycopr;1139 | 0 | 0.225 |
| NC_004718.3;NP_828851.1;SARS-CoV_E2-glycopr;1140 | 0.05 | 0.275 |
| NC_004718.3;NP_828851.1;SARS-CoV_E2-glycopr;1141 | 0.1 | 0.35 |
| NC_004718.3;NP_828851.1;SARS-CoV_E2-glycopr;1142 | 0.2 | 0.35 |
| NC_004718.3;NP_828851.1;SARS-CoV_E2-glycopr;1143 | 0.1 | 0.35 |
| NC_004718.3;NP_828851.1;SARS-CoV_E2-glycopr;1144 | 0 | 0.3 |
| NC_004718.3;NP_828851.1;SARS-CoV_E2-glycopr;1145 | 0 | 0.325 |
| NC_004718.3;NP_828851.1;SARS-CoV_E2-glycopr;1146 | 0 | 0.15 |
| NC_004718.3;NP_828851.1;SARS-CoV_E2-glycopr;1147 | 0 | 0.2 |
| NC_004718.3;NP_828851.1;SARS-CoV_E2-glycopr;1148 | 0 | 0.05 |
| NC_004718.3;NP_828851.1;SARS-CoV_E2-glycopr;1149 | 0.05 | 0.1 |
| NC_004718.3;NP_828851.1;SARS-CoV_E2-glycopr;1150 | 0.15 | 0.05 |
| NC_004718.3;NP_828851.1;SARS-CoV_E2-glycopr;1151 | 0.05 | 0.125 |
| NC_004718.3;NP_828851.1;SARS-CoV_E2-glycopr;1152 | 0.05 | 0.05 |
| NC_004718.3;NP_828851.1;SARS-CoV_E2-glycopr;1153 | 0.05 | 0.1 |
| NC_004718.3;NP_828851.1;SARS-CoV_E2-glycopr;1154 | 0.05 | 0.175 |
| NC_004718.3;NP_828851.1;SARS-CoV_E2-glycopr;1155 | 0.05 | 0.225 |
| NC_004718.3;NP_828851.1;SARS-CoV_E2-glycopr;1156 | 0.1 | 0.225 |
| NC_004718.3;NP_828851.1;SARS-CoV_E2-glycopr;1157 | 0.1 | 0.275 |
| NC_004718.3;NP_828851.1;SARS-CoV_E2-glycopr;1158 | 0.1 | 0.275 |
| NC_004718.3;NP_828851.1;SARS-CoV_E2-glycopr;1159 | 0.1 | 0.3 |
| NC_004718.3;NP_828851.1;SARS-CoV_E2-glycopr;1160 | 0.05 | 0.3 |
| NC_004718.3;NP_828851.1;SARS-CoV_E2-glycopr;1161 | 0.2 | 0.325 |
| NC_004718.3;NP_828851.1;SARS-CoV_E2-glycopr;1162 | 0.15 | 0.3 |
| NC_004718.3;NP_828851.1;SARS-CoV_E2-glycopr;1163 | 0.1 | 0.175 |

| | | |
|---|---|---|
| NC_004718.3;NP_828851.1;SARS-CoV_E2-glycopr;1164 | 0.1 | 0.15 |
| NC_004718.3;NP_828851.1;SARS-CoV_E2-glycopr;1165 | 0.15 | 0.125 |
| NC_004718.3;NP_828851.1;SARS-CoV_E2-glycopr;1166 | 0.05 | 0.125 |
| NC_004718.3;NP_828851.1;SARS-CoV_E2-glycopr;1167 | 0.05 | 0.125 |
| NC_004718.3;NP_828851.1;SARS-CoV_E2-glycopr;1168 | 0.05 | 0.1 |
| NC_004718.3;NP_828851.1;SARS-CoV_E2-glycopr;1169 | 0.05 | 0.125 |
| NC_004718.3;NP_828851.1;SARS-CoV_E2-glycopr;1170 | 0.05 | 0.1 |
| NC_004718.3;NP_828851.1;SARS-CoV_E2-glycopr;1171 | 0.05 | 0.125 |
| NC_004718.3;NP_828851.1;SARS-CoV_E2-glycopr;1172 | 0.05 | 0.125 |
| NC_004718.3;NP_828851.1;SARS-CoV_E2-glycopr;1173 | 0.05 | 0.075 |
| NC_004718.3;NP_828851.1;SARS-CoV_E2-glycopr;1174 | 0.05 | 0.125 |
| NC_004718.3;NP_828851.1;SARS-CoV_E2-glycopr;1175 | 0.1 | 0.15 |
| NC_004718.3;NP_828851.1;SARS-CoV_E2-glycopr;1176 | 0.05 | 0.125 |
| NC_004718.3;NP_828851.1;SARS-CoV_E2-glycopr;1177 | 0 | 0 |
| NC_004718.3;NP_828851.1;SARS-CoV_E2-glycopr;1178 | 0 | 0.05 |
| NC_004718.3;NP_828851.1;SARS-CoV_E2-glycopr;1179 | 0 | 0 |
| NC_004718.3;NP_828851.1;SARS-CoV_E2-glycopr;1180 | 0 | 0 |
| NC_004718.3;NP_828851.1;SARS-CoV_E2-glycopr;1181 | 0 | 0 |
| NC_004718.3;NP_828851.1;SARS-CoV_E2-glycopr;1182 | 0 | 0.025 |
| NC_004718.3;NP_828851.1;SARS-CoV_E2-glycopr;1183 | 0 | 0 |
| NC_004718.3;NP_828851.1;SARS-CoV_E2-glycopr;1184 | 0 | 0 |
| NC_004718.3;NP_828851.1;SARS-CoV_E2-glycopr;1185 | 0 | 0 |
| NC_004718.3;NP_828851.1;SARS-CoV_E2-glycopr;1186 | 0 | 0 |
| NC_004718.3;NP_828851.1;SARS-CoV_E2-glycopr;1187 | 0 | 0 |
| NC_004718.3;NP_828851.1;SARS-CoV_E2-glycopr;1188 | 0 | 0.025 |
| NC_004718.3;NP_828851.1;SARS-CoV_E2-glycopr;1189 | 0 | 0 |
| NC_004718.3;NP_828851.1;SARS-CoV_E2-glycopr;1190 | 0 | 0 |
| NC_004718.3;NP_828851.1;SARS-CoV_E2-glycopr;1191 | 0 | 0 |
| NC_004718.3;NP_828851.1;SARS-CoV_E2-glycopr;1192 | 0 | 0 |
| NC_004718.3;NP_828851.1;SARS-CoV_E2-glycopr;1193 | 0 | 0 |
| NC_004718.3;NP_828851.1;SARS-CoV_E2-glycopr;1194 | 0 | 0 |
| NC_004718.3;NP_828851.1;SARS-CoV_E2-glycopr;1195 | 0 | 0 |
| NC_004718.3;NP_828851.1;SARS-CoV_E2-glycopr;1196 | 0 | 0 |
| NC_004718.3;NP_828851.1;SARS-CoV_E2-glycopr;1197 | 0 | 0 |
| NC_004718.3;NP_828851.1;SARS-CoV_E2-glycopr;1198 | 0 | 0 |
| NC_004718.3;NP_828851.1;SARS-CoV_E2-glycopr;1199 | 0 | 0 |
| NC_004718.3;NP_828851.1;SARS-CoV_E2-glycopr;1200 | 0 | 0 |
| NC_004718.3;NP_828851.1;SARS-CoV_E2-glycopr;1201 | 0 | 0 |
| NC_004718.3;NP_828851.1;SARS-CoV_E2-glycopr;1202 | 0 | 0 |
| NC_004718.3;NP_828851.1;SARS-CoV_E2-glycopr;1203 | 0 | 0 |
| NC_004718.3;NP_828851.1;SARS-CoV_E2-glycopr;1204 | 0 | 0 |
| NC_004718.3;NP_828851.1;SARS-CoV_E2-glycopr;1205 | 0 | 0 |
| NC_004718.3;NP_828851.1;SARS-CoV_E2-glycopr;1206 | 0 | 0 |

| | | |
|---|---|---|
| NC_004718.3;NP_828851.1;SARS-CoV_E2-glycopr;1207 | 0 | 0 |
| NC_004718.3;NP_828851.1;SARS-CoV_E2-glycopr;1208 | 0 | 0 |
| NC_004718.3;NP_828851.1;SARS-CoV_E2-glycopr;1209 | 0 | 0 |
| NC_004718.3;NP_828851.1;SARS-CoV_E2-glycopr;1210 | 0 | 0 |
| NC_004718.3;NP_828851.1;SARS-CoV_E2-glycopr;1211 | 0 | 0 |
| NC_004718.3;NP_828851.1;SARS-CoV_E2-glycopr;1212 | 0 | 0 |
| NC_004718.3;NP_828851.1;SARS-CoV_E2-glycopr;1213 | 0 | 0 |
| NC_004718.3;NP_828851.1;SARS-CoV_E2-glycopr;1214 | 0 | 0 |
| NC_004718.3;NP_828851.1;SARS-CoV_E2-glycopr;1215 | 0 | 0 |
| NC_004718.3;NP_828851.1;SARS-CoV_E2-glycopr;1216 | 0 | 0 |
| NC_004718.3;NP_828851.1;SARS-CoV_E2-glycopr;1217 | 0 | 0 |
| NC_004718.3;NP_828851.1;SARS-CoV_E2-glycopr;1218 | 0 | 0 |
| NC_004718.3;NP_828851.1;SARS-CoV_E2-glycopr;1219 | 0 | 0 |
| NC_004718.3;NP_828851.1;SARS-CoV_E2-glycopr;1220 | 0 | 0 |
| NC_004718.3;NP_828851.1;SARS-CoV_E2-glycopr;1221 | 0 | 0 |
| NC_004718.3;NP_828851.1;SARS-CoV_E2-glycopr;1222 | 0 | 0 |
| NC_004718.3;NP_828851.1;SARS-CoV_E2-glycopr;1223 | 0 | 0 |
| NC_004718.3;NP_828851.1;SARS-CoV_E2-glycopr;1224 | 0 | 0 |
| NC_004718.3;NP_828851.1;SARS-CoV_E2-glycopr;1225 | 0 | 0 |
| NC_004718.3;NP_828851.1;SARS-CoV_E2-glycopr;1226 | 0 | 0 |
| NC_004718.3;NP_828851.1;SARS-CoV_E2-glycopr;1227 | 0 | 0.075 |
| NC_004718.3;NP_828851.1;SARS-CoV_E2-glycopr;1228 | 0 | 0.1 |
| NC_004718.3;NP_828851.1;SARS-CoV_E2-glycopr;1229 | 0 | 0.25 |
| NC_004718.3;NP_828851.1;SARS-CoV_E2-glycopr;1230 | 0 | 0.5 |
| NC_004718.3;NP_828851.1;SARS-CoV_E2-glycopr;1231 | 0 | 0.55 |
| NC_004718.3;NP_828851.1;SARS-CoV_E2-glycopr;1232 | 0 | 0.575 |
| NC_004718.3;NP_828851.1;SARS-CoV_E2-glycopr;1233 | 0 | 0.65 |
| NC_004718.3;NP_828851.1;SARS-CoV_E2-glycopr;1234 | 0 | 0.65 |
| NC_004718.3;NP_828851.1;SARS-CoV_E2-glycopr;1235 | 0 | 0.675 |
| NC_004718.3;NP_828851.1;SARS-CoV_E2-glycopr;1236 | 0.1 | 0.65 |
| NC_004718.3;NP_828851.1;SARS-CoV_E2-glycopr;1237 | 0.05 | 0.6 |
| NC_004718.3;NP_828851.1;SARS-CoV_E2-glycopr;1238 | 0.05 | 0.45 |
| NC_004718.3;NP_828851.1;SARS-CoV_E2-glycopr;1239 | 0.05 | 0.475 |
| NC_004718.3;NP_828851.1;SARS-CoV_E2-glycopr;1240 | 0.1 | 0.3 |
| NC_004718.3;NP_828851.1;SARS-CoV_E2-glycopr;1241 | 0.05 | 0.225 |
| NC_004718.3;NP_828851.1;SARS-CoV_E2-glycopr;1242 | 0.05 | 0.225 |
| NC_004718.3;NP_828851.1;SARS-CoV_E2-glycopr;1243 | 0.05 | 0.175 |
| NC_004718.3;NP_828851.1;SARS-CoV_E2-glycopr;1244 | 0 | 0.175 |
| NC_004718.3;NP_828852.2;SARS-CoV_3a;1 | 0 | 0 |
| NC_004718.3;NP_828852.2;SARS-CoV_3a;2 | 0 | 0 |
| NC_004718.3;NP_828852.2;SARS-CoV_3a;3 | 0 | 0 |
| NC_004718.3;NP_828852.2;SARS-CoV_3a;4 | 0 | 0 |
| NC_004718.3;NP_828852.2;SARS-CoV_3a;5 | 0 | 0 |

| | | |
|---|---|---|
| NC_004718.3;NP_828852.2;SARS-CoV_3a;6 | 0 | 0 |
| NC_004718.3;NP_828852.2;SARS-CoV_3a;7 | 0 | 0 |
| NC_004718.3;NP_828852.2;SARS-CoV_3a;8 | 0.05 | 0 |
| NC_004718.3;NP_828852.2;SARS-CoV_3a;9 | 0 | 0.025 |
| NC_004718.3;NP_828852.2;SARS-CoV_3a;10 | 0 | 0.025 |
| NC_004718.3;NP_828852.2;SARS-CoV_3a;11 | 0 | 0 |
| NC_004718.3;NP_828852.2;SARS-CoV_3a;12 | 0 | 0.05 |
| NC_004718.3;NP_828852.2;SARS-CoV_3a;13 | 0.1 | 0.075 |
| NC_004718.3;NP_828852.2;SARS-CoV_3a;14 | 0.1 | 0.025 |
| NC_004718.3;NP_828852.2;SARS-CoV_3a;15 | 0.1 | 0.025 |
| NC_004718.3;NP_828852.2;SARS-CoV_3a;16 | 0.05 | 0.025 |
| NC_004718.3;NP_828852.2;SARS-CoV_3a;17 | 0.05 | 0.025 |
| NC_004718.3;NP_828852.2;SARS-CoV_3a;18 | 0.05 | 0.025 |
| NC_004718.3;NP_828852.2;SARS-CoV_3a;19 | 0.05 | 0.025 |
| NC_004718.3;NP_828852.2;SARS-CoV_3a;20 | 0 | 0 |
| NC_004718.3;NP_828852.2;SARS-CoV_3a;21 | 0 | 0.025 |
| NC_004718.3;NP_828852.2;SARS-CoV_3a;22 | 0 | 0 |
| NC_004718.3;NP_828852.2;SARS-CoV_3a;23 | 0 | 0 |
| NC_004718.3;NP_828852.2;SARS-CoV_3a;24 | 0 | 0.125 |
| NC_004718.3;NP_828852.2;SARS-CoV_3a;25 | 0 | 0.025 |
| NC_004718.3;NP_828852.2;SARS-CoV_3a;26 | 0 | 0 |
| NC_004718.3;NP_828852.2;SARS-CoV_3a;27 | 0 | 0 |
| NC_004718.3;NP_828852.2;SARS-CoV_3a;28 | 0 | 0 |
| NC_004718.3;NP_828852.2;SARS-CoV_3a;29 | 0 | 0 |
| NC_004718.3;NP_828852.2;SARS-CoV_3a;30 | 0 | 0 |
| NC_004718.3;NP_828852.2;SARS-CoV_3a;31 | 0 | 0 |
| NC_004718.3;NP_828852.2;SARS-CoV_3a;32 | 0 | 0 |
| NC_004718.3;NP_828852.2;SARS-CoV_3a;33 | 0 | 0 |
| NC_004718.3;NP_828852.2;SARS-CoV_3a;34 | 0 | 0 |
| NC_004718.3;NP_828852.2;SARS-CoV_3a;35 | 0 | 0 |
| NC_004718.3;NP_828852.2;SARS-CoV_3a;36 | 0.05 | 0.025 |
| NC_004718.3;NP_828852.2;SARS-CoV_3a;37 | 0 | 0 |
| NC_004718.3;NP_828852.2;SARS-CoV_3a;38 | 0 | 0 |
| NC_004718.3;NP_828852.2;SARS-CoV_3a;39 | 0 | 0 |
| NC_004718.3;NP_828852.2;SARS-CoV_3a;40 | 0 | 0 |
| NC_004718.3;NP_828852.2;SARS-CoV_3a;41 | 0 | 0 |
| NC_004718.3;NP_828852.2;SARS-CoV_3a;42 | 0 | 0 |
| NC_004718.3;NP_828852.2;SARS-CoV_3a;43 | 0 | 0 |
| NC_004718.3;NP_828852.2;SARS-CoV_3a;44 | 0 | 0 |
| NC_004718.3;NP_828852.2;SARS-CoV_3a;45 | 0 | 0 |
| NC_004718.3;NP_828852.2;SARS-CoV_3a;46 | 0 | 0 |
| NC_004718.3;NP_828852.2;SARS-CoV_3a;47 | 0 | 0 |
| NC_004718.3;NP_828852.2;SARS-CoV_3a;48 | 0 | 0 |

| | | |
|---|---|---|
| NC_004718.3;NP_828852.2;SARS-CoV_3a;49 | 0 | 0.025 |
| NC_004718.3;NP_828852.2;SARS-CoV_3a;50 | 0 | 0 |
| NC_004718.3;NP_828852.2;SARS-CoV_3a;51 | 0 | 0 |
| NC_004718.3;NP_828852.2;SARS-CoV_3a;52 | 0 | 0 |
| NC_004718.3;NP_828852.2;SARS-CoV_3a;53 | 0 | 0 |
| NC_004718.3;NP_828852.2;SARS-CoV_3a;54 | 0 | 0 |
| NC_004718.3;NP_828852.2;SARS-CoV_3a;55 | 0 | 0 |
| NC_004718.3;NP_828852.2;SARS-CoV_3a;56 | 0 | 0 |
| NC_004718.3;NP_828852.2;SARS-CoV_3a;57 | 0 | 0.025 |
| NC_004718.3;NP_828852.2;SARS-CoV_3a;58 | 0 | 0.025 |
| NC_004718.3;NP_828852.2;SARS-CoV_3a;59 | 0 | 0 |
| NC_004718.3;NP_828852.2;SARS-CoV_3a;60 | 0 | 0 |
| NC_004718.3;NP_828852.2;SARS-CoV_3a;61 | 0 | 0 |
| NC_004718.3;NP_828852.2;SARS-CoV_3a;62 | 0 | 0 |
| NC_004718.3;NP_828852.2;SARS-CoV_3a;63 | 0 | 0 |
| NC_004718.3;NP_828852.2;SARS-CoV_3a;64 | 0 | 0 |
| NC_004718.3;NP_828852.2;SARS-CoV_3a;65 | 0 | 0 |
| NC_004718.3;NP_828852.2;SARS-CoV_3a;66 | 0 | 0 |
| NC_004718.3;NP_828852.2;SARS-CoV_3a;67 | 0 | 0 |
| NC_004718.3;NP_828852.2;SARS-CoV_3a;68 | 0 | 0 |
| NC_004718.3;NP_828852.2;SARS-CoV_3a;69 | 0 | 0 |
| NC_004718.3;NP_828852.2;SARS-CoV_3a;70 | 0 | 0 |
| NC_004718.3;NP_828852.2;SARS-CoV_3a;71 | 0 | 0 |
| NC_004718.3;NP_828852.2;SARS-CoV_3a;72 | 0 | 0 |
| NC_004718.3;NP_828852.2;SARS-CoV_3a;73 | 0 | 0 |
| NC_004718.3;NP_828852.2;SARS-CoV_3a;74 | 0 | 0 |
| NC_004718.3;NP_828852.2;SARS-CoV_3a;75 | 0 | 0 |
| NC_004718.3;NP_828852.2;SARS-CoV_3a;76 | 0 | 0 |
| NC_004718.3;NP_828852.2;SARS-CoV_3a;77 | 0 | 0 |
| NC_004718.3;NP_828852.2;SARS-CoV_3a;78 | 0 | 0 |
| NC_004718.3;NP_828852.2;SARS-CoV_3a;79 | 0 | 0 |
| NC_004718.3;NP_828852.2;SARS-CoV_3a;80 | 0 | 0 |
| NC_004718.3;NP_828852.2;SARS-CoV_3a;81 | 0 | 0 |
| NC_004718.3;NP_828852.2;SARS-CoV_3a;82 | 0 | 0 |
| NC_004718.3;NP_828852.2;SARS-CoV_3a;83 | 0 | 0 |
| NC_004718.3;NP_828852.2;SARS-CoV_3a;84 | 0 | 0 |
| NC_004718.3;NP_828852.2;SARS-CoV_3a;85 | 0 | 0 |
| NC_004718.3;NP_828852.2;SARS-CoV_3a;86 | 0 | 0 |
| NC_004718.3;NP_828852.2;SARS-CoV_3a;87 | 0 | 0 |
| NC_004718.3;NP_828852.2;SARS-CoV_3a;88 | 0 | 0 |
| NC_004718.3;NP_828852.2;SARS-CoV_3a;89 | 0 | 0 |
| NC_004718.3;NP_828852.2;SARS-CoV_3a;90 | 0 | 0 |
| NC_004718.3;NP_828852.2;SARS-CoV_3a;91 | 0 | 0 |

| | | |
|---|---|---|
| NC_004718.3;NP_828852.2;SARS-CoV_3a;92 | 0 | 0 |
| NC_004718.3;NP_828852.2;SARS-CoV_3a;93 | 0 | 0 |
| NC_004718.3;NP_828852.2;SARS-CoV_3a;94 | 0 | 0 |
| NC_004718.3;NP_828852.2;SARS-CoV_3a;95 | 0 | 0 |
| NC_004718.3;NP_828852.2;SARS-CoV_3a;96 | 0 | 0 |
| NC_004718.3;NP_828852.2;SARS-CoV_3a;97 | 0 | 0 |
| NC_004718.3;NP_828852.2;SARS-CoV_3a;98 | 0 | 0 |
| NC_004718.3;NP_828852.2;SARS-CoV_3a;99 | 0 | 0 |
| NC_004718.3;NP_828852.2;SARS-CoV_3a;100 | 0 | 0 |
| NC_004718.3;NP_828852.2;SARS-CoV_3a;101 | 0 | 0 |
| NC_004718.3;NP_828852.2;SARS-CoV_3a;102 | 0 | 0 |
| NC_004718.3;NP_828852.2;SARS-CoV_3a;103 | 0 | 0 |
| NC_004718.3;NP_828852.2;SARS-CoV_3a;104 | 0 | 0 |
| NC_004718.3;NP_828852.2;SARS-CoV_3a;105 | 0 | 0 |
| NC_004718.3;NP_828852.2;SARS-CoV_3a;106 | 0 | 0 |
| NC_004718.3;NP_828852.2;SARS-CoV_3a;107 | 0 | 0 |
| NC_004718.3;NP_828852.2;SARS-CoV_3a;108 | 0 | 0 |
| NC_004718.3;NP_828852.2;SARS-CoV_3a;109 | 0 | 0 |
| NC_004718.3;NP_828852.2;SARS-CoV_3a;110 | 0 | 0 |
| NC_004718.3;NP_828852.2;SARS-CoV_3a;111 | 0 | 0 |
| NC_004718.3;NP_828852.2;SARS-CoV_3a;112 | 0 | 0 |
| NC_004718.3;NP_828852.2;SARS-CoV_3a;113 | 0 | 0 |
| NC_004718.3;NP_828852.2;SARS-CoV_3a;114 | 0 | 0 |
| NC_004718.3;NP_828852.2;SARS-CoV_3a;115 | 0 | 0 |
| NC_004718.3;NP_828852.2;SARS-CoV_3a;116 | 0 | 0 |
| NC_004718.3;NP_828852.2;SARS-CoV_3a;117 | 0 | 0 |
| NC_004718.3;NP_828852.2;SARS-CoV_3a;118 | 0 | 0 |
| NC_004718.3;NP_828852.2;SARS-CoV_3a;119 | 0 | 0 |
| NC_004718.3;NP_828852.2;SARS-CoV_3a;120 | 0 | 0 |
| NC_004718.3;NP_828852.2;SARS-CoV_3a;121 | 0 | 0 |
| NC_004718.3;NP_828852.2;SARS-CoV_3a;122 | 0 | 0 |
| NC_004718.3;NP_828852.2;SARS-CoV_3a;123 | 0 | 0 |
| NC_004718.3;NP_828852.2;SARS-CoV_3a;124 | 0 | 0 |
| NC_004718.3;NP_828852.2;SARS-CoV_3a;125 | 0 | 0 |
| NC_004718.3;NP_828852.2;SARS-CoV_3a;126 | 0 | 0 |
| NC_004718.3;NP_828852.2;SARS-CoV_3a;127 | 0 | 0 |
| NC_004718.3;NP_828852.2;SARS-CoV_3a;128 | 0 | 0 |
| NC_004718.3;NP_828852.2;SARS-CoV_3a;129 | 0.05 | 0 |
| NC_004718.3;NP_828852.2;SARS-CoV_3a;130 | 0 | 0.025 |
| NC_004718.3;NP_828852.2;SARS-CoV_3a;131 | 0 | 0 |
| NC_004718.3;NP_828852.2;SARS-CoV_3a;132 | 0 | 0 |
| NC_004718.3;NP_828852.2;SARS-CoV_3a;133 | 0 | 0 |
| NC_004718.3;NP_828852.2;SARS-CoV_3a;134 | 0 | 0 |

| | | |
|---|---|---|
| NC_004718.3;NP_828852.2;SARS-CoV_3a;135 | 0 | 0 |
| NC_004718.3;NP_828852.2;SARS-CoV_3a;136 | 0 | 0 |
| NC_004718.3;NP_828852.2;SARS-CoV_3a;137 | 0 | 0 |
| NC_004718.3;NP_828852.2;SARS-CoV_3a;138 | 0 | 0 |
| NC_004718.3;NP_828852.2;SARS-CoV_3a;139 | 0 | 0 |
| NC_004718.3;NP_828852.2;SARS-CoV_3a;140 | 0 | 0 |
| NC_004718.3;NP_828852.2;SARS-CoV_3a;141 | 0 | 0 |
| NC_004718.3;NP_828852.2;SARS-CoV_3a;142 | 0 | 0 |
| NC_004718.3;NP_828852.2;SARS-CoV_3a;143 | 0 | 0 |
| NC_004718.3;NP_828852.2;SARS-CoV_3a;144 | 0 | 0 |
| NC_004718.3;NP_828852.2;SARS-CoV_3a;145 | 0 | 0 |
| NC_004718.3;NP_828852.2;SARS-CoV_3a;146 | 0 | 0 |
| NC_004718.3;NP_828852.2;SARS-CoV_3a;147 | 0 | 0 |
| NC_004718.3;NP_828852.2;SARS-CoV_3a;148 | 0 | 0 |
| NC_004718.3;NP_828852.2;SARS-CoV_3a;149 | 0 | 0 |
| NC_004718.3;NP_828852.2;SARS-CoV_3a;150 | 0 | 0 |
| NC_004718.3;NP_828852.2;SARS-CoV_3a;151 | 0 | 0 |
| NC_004718.3;NP_828852.2;SARS-CoV_3a;152 | 0 | 0 |
| NC_004718.3;NP_828852.2;SARS-CoV_3a;153 | 0 | 0 |
| NC_004718.3;NP_828852.2;SARS-CoV_3a;154 | 0 | 0 |
| NC_004718.3;NP_828852.2;SARS-CoV_3a;155 | 0 | 0 |
| NC_004718.3;NP_828852.2;SARS-CoV_3a;156 | 0 | 0 |
| NC_004718.3;NP_828852.2;SARS-CoV_3a;157 | 0 | 0 |
| NC_004718.3;NP_828852.2;SARS-CoV_3a;158 | 0 | 0 |
| NC_004718.3;NP_828852.2;SARS-CoV_3a;159 | 0 | 0 |
| NC_004718.3;NP_828852.2;SARS-CoV_3a;160 | 0 | 0 |
| NC_004718.3;NP_828852.2;SARS-CoV_3a;161 | 0 | 0 |
| NC_004718.3;NP_828852.2;SARS-CoV_3a;162 | 0 | 0.025 |
| NC_004718.3;NP_828852.2;SARS-CoV_3a;163 | 0 | 0.025 |
| NC_004718.3;NP_828852.2;SARS-CoV_3a;164 | 0 | 0.025 |
| NC_004718.3;NP_828852.2;SARS-CoV_3a;165 | 0 | 0.025 |
| NC_004718.3;NP_828852.2;SARS-CoV_3a;166 | 0 | 0 |
| NC_004718.3;NP_828852.2;SARS-CoV_3a;167 | 0 | 0 |
| NC_004718.3;NP_828852.2;SARS-CoV_3a;168 | 0 | 0 |
| NC_004718.3;NP_828852.2;SARS-CoV_3a;169 | 0 | 0 |
| NC_004718.3;NP_828852.2;SARS-CoV_3a;170 | 0 | 0 |
| NC_004718.3;NP_828852.2;SARS-CoV_3a;171 | 0.1 | 0 |
| NC_004718.3;NP_828852.2;SARS-CoV_3a;172 | 0.2 | 0 |
| NC_004718.3;NP_828852.2;SARS-CoV_3a;173 | 0.15 | 0.075 |
| NC_004718.3;NP_828852.2;SARS-CoV_3a;174 | 0.15 | 0.075 |
| NC_004718.3;NP_828852.2;SARS-CoV_3a;175 | 0.25 | 0.05 |
| NC_004718.3;NP_828852.2;SARS-CoV_3a;176 | 0.25 | 0.075 |
| NC_004718.3;NP_828852.2;SARS-CoV_3a;177 | 0.2 | 0.025 |

| | | |
|---|---|---|
| NC_004718.3;NP_828852.2;SARS-CoV_3a;178 | 0.2 | 0.075 |
| NC_004718.3;NP_828852.2;SARS-CoV_3a;179 | 0.2 | 0.025 |
| NC_004718.3;NP_828852.2;SARS-CoV_3a;180 | 0.2 | 0.025 |
| NC_004718.3;NP_828852.2;SARS-CoV_3a;181 | 0.15 | 0.05 |
| NC_004718.3;NP_828852.2;SARS-CoV_3a;182 | 0.1 | 0 |
| NC_004718.3;NP_828852.2;SARS-CoV_3a;183 | 0.1 | 0.025 |
| NC_004718.3;NP_828852.2;SARS-CoV_3a;184 | 0.1 | 0 |
| NC_004718.3;NP_828852.2;SARS-CoV_3a;185 | 0.1 | 0 |
| NC_004718.3;NP_828852.2;SARS-CoV_3a;186 | 0.05 | 0 |
| NC_004718.3;NP_828852.2;SARS-CoV_3a;187 | 0.05 | 0 |
| NC_004718.3;NP_828852.2;SARS-CoV_3a;188 | 0.05 | 0 |
| NC_004718.3;NP_828852.2;SARS-CoV_3a;189 | 0.05 | 0 |
| NC_004718.3;NP_828852.2;SARS-CoV_3a;190 | 0 | 0 |
| NC_004718.3;NP_828852.2;SARS-CoV_3a;191 | 0 | 0 |
| NC_004718.3;NP_828852.2;SARS-CoV_3a;192 | 0 | 0 |
| NC_004718.3;NP_828852.2;SARS-CoV_3a;193 | 0 | 0 |
| NC_004718.3;NP_828852.2;SARS-CoV_3a;194 | 0 | 0 |
| NC_004718.3;NP_828852.2;SARS-CoV_3a;195 | 0 | 0 |
| NC_004718.3;NP_828852.2;SARS-CoV_3a;196 | 0.1 | 0.1 |
| NC_004718.3;NP_828852.2;SARS-CoV_3a;197 | 0 | 0 |
| NC_004718.3;NP_828852.2;SARS-CoV_3a;198 | 0 | 0 |
| NC_004718.3;NP_828852.2;SARS-CoV_3a;199 | 0 | 0 |
| NC_004718.3;NP_828852.2;SARS-CoV_3a;200 | 0 | 0 |
| NC_004718.3;NP_828852.2;SARS-CoV_3a;201 | 0 | 0 |
| NC_004718.3;NP_828852.2;SARS-CoV_3a;202 | 0 | 0 |
| NC_004718.3;NP_828852.2;SARS-CoV_3a;203 | 0 | 0 |
| NC_004718.3;NP_828852.2;SARS-CoV_3a;204 | 0 | 0 |
| NC_004718.3;NP_828852.2;SARS-CoV_3a;205 | 0 | 0 |
| NC_004718.3;NP_828852.2;SARS-CoV_3a;206 | 0 | 0 |
| NC_004718.3;NP_828852.2;SARS-CoV_3a;207 | 0 | 0 |
| NC_004718.3;NP_828852.2;SARS-CoV_3a;208 | 0 | 0 |
| NC_004718.3;NP_828852.2;SARS-CoV_3a;209 | 0.05 | 0 |
| NC_004718.3;NP_828852.2;SARS-CoV_3a;210 | 0.05 | 0 |
| NC_004718.3;NP_828852.2;SARS-CoV_3a;211 | 0 | 0.025 |
| NC_004718.3;NP_828852.2;SARS-CoV_3a;212 | 0.05 | 0.025 |
| NC_004718.3;NP_828852.2;SARS-CoV_3a;213 | 0.05 | 0 |
| NC_004718.3;NP_828852.2;SARS-CoV_3a;214 | 0 | 0 |
| NC_004718.3;NP_828852.2;SARS-CoV_3a;215 | 0 | 0.05 |
| NC_004718.3;NP_828852.2;SARS-CoV_3a;216 | 0 | 0 |
| NC_004718.3;NP_828852.2;SARS-CoV_3a;217 | 0 | 0 |
| NC_004718.3;NP_828852.2;SARS-CoV_3a;218 | 0 | 0 |
| NC_004718.3;NP_828852.2;SARS-CoV_3a;219 | 0 | 0 |
| NC_004718.3;NP_828852.2;SARS-CoV_3a;220 | 0 | 0 |

| | | |
|---|---|---|
| NC_004718.3;NP_828852.2;SARS-CoV_3a;221 | 0 | 0 |
| NC_004718.3;NP_828852.2;SARS-CoV_3a;222 | 0 | 0 |
| NC_004718.3;NP_828852.2;SARS-CoV_3a;223 | 0 | 0 |
| NC_004718.3;NP_828852.2;SARS-CoV_3a;224 | 0 | 0.025 |
| NC_004718.3;NP_828852.2;SARS-CoV_3a;225 | 0 | 0 |
| NC_004718.3;NP_828852.2;SARS-CoV_3a;226 | 0 | 0 |
| NC_004718.3;NP_828852.2;SARS-CoV_3a;227 | 0 | 0 |
| NC_004718.3;NP_828852.2;SARS-CoV_3a;228 | 0 | 0 |
| NC_004718.3;NP_828852.2;SARS-CoV_3a;229 | 0 | 0.025 |
| NC_004718.3;NP_828852.2;SARS-CoV_3a;230 | 0 | 0.025 |
| NC_004718.3;NP_828852.2;SARS-CoV_3a;231 | 0 | 0.025 |
| NC_004718.3;NP_828852.2;SARS-CoV_3a;232 | 0 | 0.025 |
| NC_004718.3;NP_828852.2;SARS-CoV_3a;233 | 0 | 0.025 |
| NC_004718.3;NP_828852.2;SARS-CoV_3a;234 | 0 | 0.05 |
| NC_004718.3;NP_828852.2;SARS-CoV_3a;235 | 0.05 | 0.025 |
| NC_004718.3;NP_828852.2;SARS-CoV_3a;236 | 0 | 0.075 |
| NC_004718.3;NP_828852.2;SARS-CoV_3a;237 | 0 | 0.075 |
| NC_004718.3;NP_828852.2;SARS-CoV_3a;238 | 0 | 0.05 |
| NC_004718.3;NP_828852.2;SARS-CoV_3a;239 | 0.05 | 0.05 |
| NC_004718.3;NP_828852.2;SARS-CoV_3a;240 | 0.1 | 0.125 |
| NC_004718.3;NP_828852.2;SARS-CoV_3a;241 | 0.05 | 0.075 |
| NC_004718.3;NP_828852.2;SARS-CoV_3a;242 | 0.05 | 0.075 |
| NC_004718.3;NP_828852.2;SARS-CoV_3a;243 | 0.05 | 0.1 |
| NC_004718.3;NP_828852.2;SARS-CoV_3a;244 | 0.05 | 0.075 |
| NC_004718.3;NP_828852.2;SARS-CoV_3a;245 | 0.05 | 0.075 |
| NC_004718.3;NP_828852.2;SARS-CoV_3a;246 | 0.05 | 0.075 |
| NC_004718.3;NP_828852.2;SARS-CoV_3a;247 | 0.05 | 0.025 |
| NC_004718.3;NP_828852.2;SARS-CoV_3a;248 | 0.05 | 0.025 |
| NC_004718.3;NP_828852.2;SARS-CoV_3a;249 | 0.05 | 0.05 |
| NC_004718.3;NP_828852.2;SARS-CoV_3a;250 | 0 | 0.025 |
| NC_004718.3;NP_828852.2;SARS-CoV_3a;251 | 0.05 | 0.125 |
| NC_004718.3;NP_828852.2;SARS-CoV_3a;252 | 0 | 0.075 |
| NC_004718.3;NP_828852.2;SARS-CoV_3a;253 | 0 | 0.125 |
| NC_004718.3;NP_828852.2;SARS-CoV_3a;254 | 0 | 0.1 |
| NC_004718.3;NP_828852.2;SARS-CoV_3a;255 | 0 | 0.1 |
| NC_004718.3;NP_828852.2;SARS-CoV_3a;256 | 0 | 0.125 |
| NC_004718.3;NP_828852.2;SARS-CoV_3a;257 | 0.05 | 0.125 |
| NC_004718.3;NP_828852.2;SARS-CoV_3a;258 | 0.05 | 0.1 |
| NC_004718.3;NP_828852.2;SARS-CoV_3a;259 | 0.05 | 0.175 |
| NC_004718.3;NP_828852.2;SARS-CoV_3a;260 | 0.05 | 0.15 |
| NC_004718.3;NP_828852.2;SARS-CoV_3a;261 | 0 | 0.075 |
| NC_004718.3;NP_828852.2;SARS-CoV_3a;262 | 0.05 | 0.1 |
| NC_004718.3;NP_828852.2;SARS-CoV_3a;263 | 0 | 0.05 |

| | | |
|---|---|---|
| NC_004718.3;NP_828853.1;SARS-CoV_3b;1 | 0 | 0 |
| NC_004718.3;NP_828853.1;SARS-CoV_3b;2 | 0 | 0 |
| NC_004718.3;NP_828853.1;SARS-CoV_3b;3 | 0 | 0 |
| NC_004718.3;NP_828853.1;SARS-CoV_3b;4 | 0 | 0 |
| NC_004718.3;NP_828853.1;SARS-CoV_3b;5 | 0 | 0 |
| NC_004718.3;NP_828853.1;SARS-CoV_3b;6 | 0 | 0 |
| NC_004718.3;NP_828853.1;SARS-CoV_3b;7 | 0 | 0 |
| NC_004718.3;NP_828853.1;SARS-CoV_3b;8 | 0 | 0 |
| NC_004718.3;NP_828853.1;SARS-CoV_3b;9 | 0 | 0 |
| NC_004718.3;NP_828853.1;SARS-CoV_3b;10 | 0 | 0 |
| NC_004718.3;NP_828853.1;SARS-CoV_3b;11 | 0 | 0 |
| NC_004718.3;NP_828853.1;SARS-CoV_3b;12 | 0 | 0 |
| NC_004718.3;NP_828853.1;SARS-CoV_3b;13 | 0 | 0 |
| NC_004718.3;NP_828853.1;SARS-CoV_3b;14 | 0 | 0 |
| NC_004718.3;NP_828853.1;SARS-CoV_3b;15 | 0 | 0 |
| NC_004718.3;NP_828853.1;SARS-CoV_3b;16 | 0 | 0 |
| NC_004718.3;NP_828853.1;SARS-CoV_3b;17 | 0 | 0 |
| NC_004718.3;NP_828853.1;SARS-CoV_3b;18 | 0 | 0 |
| NC_004718.3;NP_828853.1;SARS-CoV_3b;19 | 0 | 0 |
| NC_004718.3;NP_828853.1;SARS-CoV_3b;20 | 0 | 0.05 |
| NC_004718.3;NP_828853.1;SARS-CoV_3b;21 | 0 | 0 |
| NC_004718.3;NP_828853.1;SARS-CoV_3b;22 | 0 | 0 |
| NC_004718.3;NP_828853.1;SARS-CoV_3b;23 | 0 | 0 |
| NC_004718.3;NP_828853.1;SARS-CoV_3b;24 | 0 | 0 |
| NC_004718.3;NP_828853.1;SARS-CoV_3b;25 | 0 | 0 |
| NC_004718.3;NP_828853.1;SARS-CoV_3b;26 | 0 | 0 |
| NC_004718.3;NP_828853.1;SARS-CoV_3b;27 | 0 | 0 |
| NC_004718.3;NP_828853.1;SARS-CoV_3b;28 | 0 | 0 |
| NC_004718.3;NP_828853.1;SARS-CoV_3b;29 | 0 | 0 |
| NC_004718.3;NP_828853.1;SARS-CoV_3b;30 | 0 | 0 |
| NC_004718.3;NP_828853.1;SARS-CoV_3b;31 | 0 | 0 |
| NC_004718.3;NP_828853.1;SARS-CoV_3b;32 | 0 | 0 |
| NC_004718.3;NP_828853.1;SARS-CoV_3b;33 | 0 | 0 |
| NC_004718.3;NP_828853.1;SARS-CoV_3b;34 | 0 | 0 |
| NC_004718.3;NP_828853.1;SARS-CoV_3b;35 | 0 | 0 |
| NC_004718.3;NP_828853.1;SARS-CoV_3b;36 | 0 | 0 |
| NC_004718.3;NP_828853.1;SARS-CoV_3b;37 | 0 | 0 |
| NC_004718.3;NP_828853.1;SARS-CoV_3b;38 | 0 | 0 |
| NC_004718.3;NP_828853.1;SARS-CoV_3b;39 | 0 | 0 |
| NC_004718.3;NP_828853.1;SARS-CoV_3b;40 | 0 | 0 |
| NC_004718.3;NP_828853.1;SARS-CoV_3b;41 | 0 | 0 |
| NC_004718.3;NP_828853.1;SARS-CoV_3b;42 | 0 | 0 |
| NC_004718.3;NP_828853.1;SARS-CoV_3b;43 | 0 | 0 |

| | | |
|---|---|---|
| NC_004718.3;NP_828853.1;SARS-CoV_3b;44 | 0 | 0 |
| NC_004718.3;NP_828853.1;SARS-CoV_3b;45 | 0 | 0 |
| NC_004718.3;NP_828853.1;SARS-CoV_3b;46 | 0 | 0 |
| NC_004718.3;NP_828853.1;SARS-CoV_3b;47 | 0 | 0 |
| NC_004718.3;NP_828853.1;SARS-CoV_3b;48 | 0 | 0 |
| NC_004718.3;NP_828853.1;SARS-CoV_3b;49 | 0 | 0 |
| NC_004718.3;NP_828853.1;SARS-CoV_3b;50 | 0 | 0 |
| NC_004718.3;NP_828853.1;SARS-CoV_3b;51 | 0 | 0 |
| NC_004718.3;NP_828853.1;SARS-CoV_3b;52 | 0 | 0 |
| NC_004718.3;NP_828853.1;SARS-CoV_3b;53 | 0 | 0 |
| NC_004718.3;NP_828853.1;SARS-CoV_3b;54 | 0 | 0 |
| NC_004718.3;NP_828853.1;SARS-CoV_3b;55 | 0 | 0 |
| NC_004718.3;NP_828853.1;SARS-CoV_3b;56 | 0 | 0 |
| NC_004718.3;NP_828853.1;SARS-CoV_3b;57 | 0 | 0 |
| NC_004718.3;NP_828853.1;SARS-CoV_3b;58 | 0 | 0 |
| NC_004718.3;NP_828853.1;SARS-CoV_3b;59 | 0 | 0 |
| NC_004718.3;NP_828853.1;SARS-CoV_3b;60 | 0 | 0.025 |
| NC_004718.3;NP_828853.1;SARS-CoV_3b;61 | 0 | 0 |
| NC_004718.3;NP_828853.1;SARS-CoV_3b;62 | 0 | 0 |
| NC_004718.3;NP_828853.1;SARS-CoV_3b;63 | 0 | 0 |
| NC_004718.3;NP_828853.1;SARS-CoV_3b;64 | 0 | 0 |
| NC_004718.3;NP_828853.1;SARS-CoV_3b;65 | 0 | 0 |
| NC_004718.3;NP_828853.1;SARS-CoV_3b;66 | 0 | 0 |
| NC_004718.3;NP_828853.1;SARS-CoV_3b;67 | 0 | 0 |
| NC_004718.3;NP_828853.1;SARS-CoV_3b;68 | 0 | 0 |
| NC_004718.3;NP_828853.1;SARS-CoV_3b;69 | 0 | 0 |
| NC_004718.3;NP_828853.1;SARS-CoV_3b;70 | 0 | 0 |
| NC_004718.3;NP_828853.1;SARS-CoV_3b;71 | 0 | 0 |
| NC_004718.3;NP_828853.1;SARS-CoV_3b;72 | 0 | 0 |
| NC_004718.3;NP_828853.1;SARS-CoV_3b;73 | 0 | 0 |
| NC_004718.3;NP_828853.1;SARS-CoV_3b;74 | 0 | 0 |
| NC_004718.3;NP_828853.1;SARS-CoV_3b;75 | 0 | 0 |
| NC_004718.3;NP_828853.1;SARS-CoV_3b;76 | 0 | 0 |
| NC_004718.3;NP_828853.1;SARS-CoV_3b;77 | 0 | 0 |
| NC_004718.3;NP_828853.1;SARS-CoV_3b;78 | 0 | 0 |
| NC_004718.3;NP_828853.1;SARS-CoV_3b;79 | 0 | 0 |
| NC_004718.3;NP_828853.1;SARS-CoV_3b;80 | 0 | 0 |
| NC_004718.3;NP_828853.1;SARS-CoV_3b;81 | 0 | 0 |
| NC_004718.3;NP_828853.1;SARS-CoV_3b;82 | 0 | 0 |
| NC_004718.3;NP_828853.1;SARS-CoV_3b;83 | 0 | 0 |
| NC_004718.3;NP_828853.1;SARS-CoV_3b;84 | 0 | 0 |
| NC_004718.3;NP_828853.1;SARS-CoV_3b;85 | 0 | 0 |
| NC_004718.3;NP_828853.1;SARS-CoV_3b;86 | 0 | 0 |

| | | |
|---|---|---|
| NC_004718.3;NP_828853.1;SARS-CoV_3b;87 | 0 | 0 |
| NC_004718.3;NP_828853.1;SARS-CoV_3b;88 | 0 | 0 |
| NC_004718.3;NP_828853.1;SARS-CoV_3b;89 | 0 | 0 |
| NC_004718.3;NP_828853.1;SARS-CoV_3b;90 | 0 | 0 |
| NC_004718.3;NP_828853.1;SARS-CoV_3b;91 | 0 | 0 |
| NC_004718.3;NP_828853.1;SARS-CoV_3b;92 | 0 | 0 |
| NC_004718.3;NP_828853.1;SARS-CoV_3b;93 | 0 | 0 |
| NC_004718.3;NP_828853.1;SARS-CoV_3b;94 | 0 | 0 |
| NC_004718.3;NP_828853.1;SARS-CoV_3b;95 | 0 | 0 |
| NC_004718.3;NP_828853.1;SARS-CoV_3b;96 | 0 | 0 |
| NC_004718.3;NP_828853.1;SARS-CoV_3b;97 | 0 | 0 |
| NC_004718.3;NP_828853.1;SARS-CoV_3b;98 | 0 | 0 |
| NC_004718.3;NP_828853.1;SARS-CoV_3b;99 | 0 | 0 |
| NC_004718.3;NP_828853.1;SARS-CoV_3b;100 | 0 | 0 |
| NC_004718.3;NP_828853.1;SARS-CoV_3b;101 | 0 | 0 |
| NC_004718.3;NP_828853.1;SARS-CoV_3b;102 | 0 | 0 |
| NC_004718.3;NP_828853.1;SARS-CoV_3b;103 | 0 | 0 |
| NC_004718.3;NP_828853.1;SARS-CoV_3b;104 | 0 | 0 |
| NC_004718.3;NP_828853.1;SARS-CoV_3b;105 | 0 | 0 |
| NC_004718.3;NP_828853.1;SARS-CoV_3b;106 | 0 | 0 |
| NC_004718.3;NP_828853.1;SARS-CoV_3b;107 | 0 | 0 |
| NC_004718.3;NP_828853.1;SARS-CoV_3b;108 | 0 | 0 |
| NC_004718.3;NP_828853.1;SARS-CoV_3b;109 | 0 | 0 |
| NC_004718.3;NP_828853.1;SARS-CoV_3b;110 | 0 | 0 |
| NC_004718.3;NP_828853.1;SARS-CoV_3b;111 | 0 | 0 |
| NC_004718.3;NP_828853.1;SARS-CoV_3b;112 | 0 | 0 |
| NC_004718.3;NP_828853.1;SARS-CoV_3b;113 | 0 | 0 |
| NC_004718.3;NP_828853.1;SARS-CoV_3b;114 | 0 | 0 |
| NC_004718.3;NP_828853.1;SARS-CoV_3b;115 | 0 | 0 |
| NC_004718.3;NP_828853.1;SARS-CoV_3b;116 | 0 | 0 |
| NC_004718.3;NP_828853.1;SARS-CoV_3b;117 | 0 | 0 |
| NC_004718.3;NP_828853.1;SARS-CoV_3b;118 | 0 | 0 |
| NC_004718.3;NP_828853.1;SARS-CoV_3b;119 | 0 | 0 |
| NC_004718.3;NP_828853.1;SARS-CoV_3b;120 | 0 | 0 |
| NC_004718.3;NP_828853.1;SARS-CoV_3b;121 | 0 | 0 |
| NC_004718.3;NP_828853.1;SARS-CoV_3b;122 | 0 | 0 |
| NC_004718.3;NP_828853.1;SARS-CoV_3b;123 | 0 | 0 |
| NC_004718.3;NP_828853.1;SARS-CoV_3b;124 | 0 | 0 |
| NC_004718.3;NP_828853.1;SARS-CoV_3b;125 | 0 | 0 |
| NC_004718.3;NP_828853.1;SARS-CoV_3b;126 | 0 | 0 |
| NC_004718.3;NP_828853.1;SARS-CoV_3b;127 | 0 | 0 |
| NC_004718.3;NP_828853.1;SARS-CoV_3b;128 | 0 | 0 |
| NC_004718.3;NP_828853.1;SARS-CoV_3b;129 | 0 | 0 |

| | | |
|---|---|---|
| NC_004718.3;NP_828853.1;SARS-CoV_3b;130 | 0.05 | 0 |
| NC_004718.3;NP_828853.1;SARS-CoV_3b;131 | 0 | 0 |
| NC_004718.3;NP_828853.1;SARS-CoV_3b;132 | 0.05 | 0 |
| NC_004718.3;NP_828853.1;SARS-CoV_3b;133 | 0 | 0 |
| NC_004718.3;NP_828853.1;SARS-CoV_3b;134 | 0 | 0 |
| NC_004718.3;NP_828853.1;SARS-CoV_3b;135 | 0 | 0 |
| NC_004718.3;NP_828853.1;SARS-CoV_3b;136 | 0 | 0 |
| NC_004718.3;NP_828853.1;SARS-CoV_3b;137 | 0.05 | 0 |
| NC_004718.3;NP_828853.1;SARS-CoV_3b;138 | 0 | 0 |
| NC_004718.3;NP_828853.1;SARS-CoV_3b;139 | 0 | 0 |
| NC_004718.3;NP_828853.1;SARS-CoV_3b;140 | 0 | 0 |
| NC_004718.3;NP_828853.1;SARS-CoV_3b;141 | 0 | 0 |
| NC_004718.3;NP_828853.1;SARS-CoV_3b;142 | 0 | 0 |
| NC_004718.3;NP_828853.1;SARS-CoV_3b;143 | 0 | 0 |
| NC_004718.3;NP_828854.1;SARS-CoV_E;1 | 0 | 0 |
| NC_004718.3;NP_828854.1;SARS-CoV_E;2 | 0 | 0 |
| NC_004718.3;NP_828854.1;SARS-CoV_E;3 | 0 | 0 |
| NC_004718.3;NP_828854.1;SARS-CoV_E;4 | 0 | 0 |
| NC_004718.3;NP_828854.1;SARS-CoV_E;5 | 0 | 0 |
| NC_004718.3;NP_828854.1;SARS-CoV_E;6 | 0 | 0 |
| NC_004718.3;NP_828854.1;SARS-CoV_E;7 | 0 | 0 |
| NC_004718.3;NP_828854.1;SARS-CoV_E;8 | 0 | 0 |
| NC_004718.3;NP_828854.1;SARS-CoV_E;9 | 0 | 0 |
| NC_004718.3;NP_828854.1;SARS-CoV_E;10 | 0 | 0 |
| NC_004718.3;NP_828854.1;SARS-CoV_E;11 | 0 | 0 |
| NC_004718.3;NP_828854.1;SARS-CoV_E;12 | 0 | 0 |
| NC_004718.3;NP_828854.1;SARS-CoV_E;13 | 0 | 0 |
| NC_004718.3;NP_828854.1;SARS-CoV_E;14 | 0 | 0 |
| NC_004718.3;NP_828854.1;SARS-CoV_E;15 | 0 | 0 |
| NC_004718.3;NP_828854.1;SARS-CoV_E;16 | 0 | 0 |
| NC_004718.3;NP_828854.1;SARS-CoV_E;17 | 0 | 0 |
| NC_004718.3;NP_828854.1;SARS-CoV_E;18 | 0 | 0 |
| NC_004718.3;NP_828854.1;SARS-CoV_E;19 | 0 | 0 |
| NC_004718.3;NP_828854.1;SARS-CoV_E;20 | 0 | 0 |
| NC_004718.3;NP_828854.1;SARS-CoV_E;21 | 0 | 0 |
| NC_004718.3;NP_828854.1;SARS-CoV_E;22 | 0 | 0 |
| NC_004718.3;NP_828854.1;SARS-CoV_E;23 | 0 | 0 |
| NC_004718.3;NP_828854.1;SARS-CoV_E;24 | 0 | 0 |
| NC_004718.3;NP_828854.1;SARS-CoV_E;25 | 0 | 0 |
| NC_004718.3;NP_828854.1;SARS-CoV_E;26 | 0 | 0 |
| NC_004718.3;NP_828854.1;SARS-CoV_E;27 | 0 | 0 |
| NC_004718.3;NP_828854.1;SARS-CoV_E;28 | 0 | 0 |
| NC_004718.3;NP_828854.1;SARS-CoV_E;29 | 0 | 0 |

| | | |
|---|---|---|
| NC_004718.3;NP_828854.1;SARS-CoV_E;30 | 0 | 0 |
| NC_004718.3;NP_828854.1;SARS-CoV_E;31 | 0 | 0 |
| NC_004718.3;NP_828854.1;SARS-CoV_E;32 | 0 | 0 |
| NC_004718.3;NP_828854.1;SARS-CoV_E;33 | 0 | 0 |
| NC_004718.3;NP_828854.1;SARS-CoV_E;34 | 0 | 0 |
| NC_004718.3;NP_828854.1;SARS-CoV_E;35 | 0 | 0 |
| NC_004718.3;NP_828854.1;SARS-CoV_E;36 | 0 | 0 |
| NC_004718.3;NP_828854.1;SARS-CoV_E;37 | 0 | 0 |
| NC_004718.3;NP_828854.1;SARS-CoV_E;38 | 0 | 0 |
| NC_004718.3;NP_828854.1;SARS-CoV_E;39 | 0 | 0 |
| NC_004718.3;NP_828854.1;SARS-CoV_E;40 | 0 | 0 |
| NC_004718.3;NP_828854.1;SARS-CoV_E;41 | 0 | 0 |
| NC_004718.3;NP_828854.1;SARS-CoV_E;42 | 0 | 0 |
| NC_004718.3;NP_828854.1;SARS-CoV_E;43 | 0 | 0 |
| NC_004718.3;NP_828854.1;SARS-CoV_E;44 | 0 | 0 |
| NC_004718.3;NP_828854.1;SARS-CoV_E;45 | 0 | 0 |
| NC_004718.3;NP_828854.1;SARS-CoV_E;46 | 0 | 0 |
| NC_004718.3;NP_828854.1;SARS-CoV_E;47 | 0 | 0 |
| NC_004718.3;NP_828854.1;SARS-CoV_E;48 | 0 | 0 |
| NC_004718.3;NP_828854.1;SARS-CoV_E;49 | 0 | 0 |
| NC_004718.3;NP_828854.1;SARS-CoV_E;50 | 0 | 0 |
| NC_004718.3;NP_828854.1;SARS-CoV_E;51 | 0 | 0 |
| NC_004718.3;NP_828854.1;SARS-CoV_E;52 | 0 | 0 |
| NC_004718.3;NP_828854.1;SARS-CoV_E;53 | 0 | 0 |
| NC_004718.3;NP_828854.1;SARS-CoV_E;54 | 0 | 0 |
| NC_004718.3;NP_828854.1;SARS-CoV_E;55 | 0 | 0 |
| NC_004718.3;NP_828854.1;SARS-CoV_E;56 | 0 | 0 |
| NC_004718.3;NP_828854.1;SARS-CoV_E;57 | 0.05 | 0.025 |
| NC_004718.3;NP_828854.1;SARS-CoV_E;58 | 0 | 0.075 |
| NC_004718.3;NP_828854.1;SARS-CoV_E;59 | 0 | 0.075 |
| NC_004718.3;NP_828854.1;SARS-CoV_E;60 | 0.1 | 0 |
| NC_004718.3;NP_828854.1;SARS-CoV_E;61 | 0.05 | 0 |
| NC_004718.3;NP_828854.1;SARS-CoV_E;62 | 0 | 0 |
| NC_004718.3;NP_828854.1;SARS-CoV_E;63 | 0 | 0 |
| NC_004718.3;NP_828854.1;SARS-CoV_E;64 | 0 | 0 |
| NC_004718.3;NP_828854.1;SARS-CoV_E;65 | 0 | 0 |
| NC_004718.3;NP_828855.1;SARS-CoV_matrix;1 | 0 | 0.475 |
| NC_004718.3;NP_828855.1;SARS-CoV_matrix;2 | 0 | 0.4 |
| NC_004718.3;NP_828855.1;SARS-CoV_matrix;3 | 0.05 | 0.625 |
| NC_004718.3;NP_828855.1;SARS-CoV_matrix;4 | 0.05 | 0.5 |
| NC_004718.3;NP_828855.1;SARS-CoV_matrix;5 | 0 | 0.6 |
| NC_004718.3;NP_828855.1;SARS-CoV_matrix;6 | 0 | 0.725 |
| NC_004718.3;NP_828855.1;SARS-CoV_matrix;7 | 0 | 0.9 |

| | | |
|---|---|---|
| NC_004718.3;NP_828855.1;SARS-CoV_matrix;8 | 0 | 0.3 |
| NC_004718.3;NP_828855.1;SARS-CoV_matrix;9 | 0 | 0 |
| NC_004718.3;NP_828855.1;SARS-CoV_matrix;10 | 0 | 0 |
| NC_004718.3;NP_828855.1;SARS-CoV_matrix;11 | 0 | 0 |
| NC_004718.3;NP_828855.1;SARS-CoV_matrix;12 | 0 | 0 |
| NC_004718.3;NP_828855.1;SARS-CoV_matrix;13 | 0.05 | 0 |
| NC_004718.3;NP_828855.1;SARS-CoV_matrix;14 | 0 | 0 |
| NC_004718.3;NP_828855.1;SARS-CoV_matrix;15 | 0 | 0 |
| NC_004718.3;NP_828855.1;SARS-CoV_matrix;16 | 0 | 0 |
| NC_004718.3;NP_828855.1;SARS-CoV_matrix;17 | 0 | 0 |
| NC_004718.3;NP_828855.1;SARS-CoV_matrix;18 | 0 | 0 |
| NC_004718.3;NP_828855.1;SARS-CoV_matrix;19 | 0 | 0 |
| NC_004718.3;NP_828855.1;SARS-CoV_matrix;20 | 0 | 0 |
| NC_004718.3;NP_828855.1;SARS-CoV_matrix;21 | 0 | 0 |
| NC_004718.3;NP_828855.1;SARS-CoV_matrix;22 | 0 | 0 |
| NC_004718.3;NP_828855.1;SARS-CoV_matrix;23 | 0 | 0 |
| NC_004718.3;NP_828855.1;SARS-CoV_matrix;24 | 0 | 0 |
| NC_004718.3;NP_828855.1;SARS-CoV_matrix;25 | 0 | 0 |
| NC_004718.3;NP_828855.1;SARS-CoV_matrix;26 | 0 | 0 |
| NC_004718.3;NP_828855.1;SARS-CoV_matrix;27 | 0 | 0 |
| NC_004718.3;NP_828855.1;SARS-CoV_matrix;28 | 0 | 0 |
| NC_004718.3;NP_828855.1;SARS-CoV_matrix;29 | 0 | 0 |
| NC_004718.3;NP_828855.1;SARS-CoV_matrix;30 | 0 | 0 |
| NC_004718.3;NP_828855.1;SARS-CoV_matrix;31 | 0 | 0 |
| NC_004718.3;NP_828855.1;SARS-CoV_matrix;32 | 0 | 0 |
| NC_004718.3;NP_828855.1;SARS-CoV_matrix;33 | 0 | 0 |
| NC_004718.3;NP_828855.1;SARS-CoV_matrix;34 | 0 | 0 |
| NC_004718.3;NP_828855.1;SARS-CoV_matrix;35 | 0 | 0 |
| NC_004718.3;NP_828855.1;SARS-CoV_matrix;36 | 0 | 0 |
| NC_004718.3;NP_828855.1;SARS-CoV_matrix;37 | 0 | 0 |
| NC_004718.3;NP_828855.1;SARS-CoV_matrix;38 | 0.05 | 0 |
| NC_004718.3;NP_828855.1;SARS-CoV_matrix;39 | 0.05 | 0 |
| NC_004718.3;NP_828855.1;SARS-CoV_matrix;40 | 0 | 0 |
| NC_004718.3;NP_828855.1;SARS-CoV_matrix;41 | 0 | 0 |
| NC_004718.3;NP_828855.1;SARS-CoV_matrix;42 | 0 | 0 |
| NC_004718.3;NP_828855.1;SARS-CoV_matrix;43 | 0 | 0 |
| NC_004718.3;NP_828855.1;SARS-CoV_matrix;44 | 0 | 0 |
| NC_004718.3;NP_828855.1;SARS-CoV_matrix;45 | 0 | 0 |
| NC_004718.3;NP_828855.1;SARS-CoV_matrix;46 | 0 | 0 |
| NC_004718.3;NP_828855.1;SARS-CoV_matrix;47 | 0 | 0 |
| NC_004718.3;NP_828855.1;SARS-CoV_matrix;48 | 0 | 0 |
| NC_004718.3;NP_828855.1;SARS-CoV_matrix;49 | 0 | 0 |
| NC_004718.3;NP_828855.1;SARS-CoV_matrix;50 | 0 | 0 |

| | | |
|---|---|---|
| NC_004718.3;NP_828855.1;SARS-CoV_matrix;51 | 0 | 0 |
| NC_004718.3;NP_828855.1;SARS-CoV_matrix;52 | 0 | 0 |
| NC_004718.3;NP_828855.1;SARS-CoV_matrix;53 | 0 | 0 |
| NC_004718.3;NP_828855.1;SARS-CoV_matrix;54 | 0 | 0 |
| NC_004718.3;NP_828855.1;SARS-CoV_matrix;55 | 0 | 0 |
| NC_004718.3;NP_828855.1;SARS-CoV_matrix;56 | 0 | 0 |
| NC_004718.3;NP_828855.1;SARS-CoV_matrix;57 | 0 | 0 |
| NC_004718.3;NP_828855.1;SARS-CoV_matrix;58 | 0 | 0 |
| NC_004718.3;NP_828855.1;SARS-CoV_matrix;59 | 0 | 0 |
| NC_004718.3;NP_828855.1;SARS-CoV_matrix;60 | 0 | 0 |
| NC_004718.3;NP_828855.1;SARS-CoV_matrix;61 | 0 | 0 |
| NC_004718.3;NP_828855.1;SARS-CoV_matrix;62 | 0 | 0 |
| NC_004718.3;NP_828855.1;SARS-CoV_matrix;63 | 0 | 0 |
| NC_004718.3;NP_828855.1;SARS-CoV_matrix;64 | 0 | 0 |
| NC_004718.3;NP_828855.1;SARS-CoV_matrix;65 | 0 | 0 |
| NC_004718.3;NP_828855.1;SARS-CoV_matrix;66 | 0 | 0 |
| NC_004718.3;NP_828855.1;SARS-CoV_matrix;67 | 0 | 0 |
| NC_004718.3;NP_828855.1;SARS-CoV_matrix;68 | 0 | 0 |
| NC_004718.3;NP_828855.1;SARS-CoV_matrix;69 | 0 | 0 |
| NC_004718.3;NP_828855.1;SARS-CoV_matrix;70 | 0 | 0 |
| NC_004718.3;NP_828855.1;SARS-CoV_matrix;71 | 0 | 0 |
| NC_004718.3;NP_828855.1;SARS-CoV_matrix;72 | 0 | 0 |
| NC_004718.3;NP_828855.1;SARS-CoV_matrix;73 | 0 | 0 |
| NC_004718.3;NP_828855.1;SARS-CoV_matrix;74 | 0 | 0 |
| NC_004718.3;NP_828855.1;SARS-CoV_matrix;75 | 0 | 0 |
| NC_004718.3;NP_828855.1;SARS-CoV_matrix;76 | 0 | 0 |
| NC_004718.3;NP_828855.1;SARS-CoV_matrix;77 | 0 | 0 |
| NC_004718.3;NP_828855.1;SARS-CoV_matrix;78 | 0 | 0 |
| NC_004718.3;NP_828855.1;SARS-CoV_matrix;79 | 0 | 0 |
| NC_004718.3;NP_828855.1;SARS-CoV_matrix;80 | 0 | 0 |
| NC_004718.3;NP_828855.1;SARS-CoV_matrix;81 | 0 | 0 |
| NC_004718.3;NP_828855.1;SARS-CoV_matrix;82 | 0 | 0 |
| NC_004718.3;NP_828855.1;SARS-CoV_matrix;83 | 0 | 0 |
| NC_004718.3;NP_828855.1;SARS-CoV_matrix;84 | 0 | 0 |
| NC_004718.3;NP_828855.1;SARS-CoV_matrix;85 | 0 | 0 |
| NC_004718.3;NP_828855.1;SARS-CoV_matrix;86 | 0 | 0 |
| NC_004718.3;NP_828855.1;SARS-CoV_matrix;87 | 0 | 0 |
| NC_004718.3;NP_828855.1;SARS-CoV_matrix;88 | 0 | 0 |
| NC_004718.3;NP_828855.1;SARS-CoV_matrix;89 | 0 | 0 |
| NC_004718.3;NP_828855.1;SARS-CoV_matrix;90 | 0 | 0 |
| NC_004718.3;NP_828855.1;SARS-CoV_matrix;91 | 0 | 0 |
| NC_004718.3;NP_828855.1;SARS-CoV_matrix;92 | 0 | 0 |
| NC_004718.3;NP_828855.1;SARS-CoV_matrix;93 | 0 | 0 |

| | | |
|---|---|---|
| NC_004718.3;NP_828855.1;SARS-CoV_matrix;94 | 0 | 0 |
| NC_004718.3;NP_828855.1;SARS-CoV_matrix;95 | 0 | 0 |
| NC_004718.3;NP_828855.1;SARS-CoV_matrix;96 | 0 | 0 |
| NC_004718.3;NP_828855.1;SARS-CoV_matrix;97 | 0 | 0 |
| NC_004718.3;NP_828855.1;SARS-CoV_matrix;98 | 0 | 0 |
| NC_004718.3;NP_828855.1;SARS-CoV_matrix;99 | 0 | 0 |
| NC_004718.3;NP_828855.1;SARS-CoV_matrix;100 | 0.05 | 0.025 |
| NC_004718.3;NP_828855.1;SARS-CoV_matrix;101 | 0 | 0 |
| NC_004718.3;NP_828855.1;SARS-CoV_matrix;102 | 0 | 0 |
| NC_004718.3;NP_828855.1;SARS-CoV_matrix;103 | 0 | 0 |
| NC_004718.3;NP_828855.1;SARS-CoV_matrix;104 | 0 | 0 |
| NC_004718.3;NP_828855.1;SARS-CoV_matrix;105 | 0 | 0 |
| NC_004718.3;NP_828855.1;SARS-CoV_matrix;106 | 0 | 0 |
| NC_004718.3;NP_828855.1;SARS-CoV_matrix;107 | 0 | 0.025 |
| NC_004718.3;NP_828855.1;SARS-CoV_matrix;108 | 0 | 0.025 |
| NC_004718.3;NP_828855.1;SARS-CoV_matrix;109 | 0 | 0 |
| NC_004718.3;NP_828855.1;SARS-CoV_matrix;110 | 0 | 0.025 |
| NC_004718.3;NP_828855.1;SARS-CoV_matrix;111 | 0 | 0.025 |
| NC_004718.3;NP_828855.1;SARS-CoV_matrix;112 | 0 | 0.025 |
| NC_004718.3;NP_828855.1;SARS-CoV_matrix;113 | 0 | 0 |
| NC_004718.3;NP_828855.1;SARS-CoV_matrix;114 | 0 | 0 |
| NC_004718.3;NP_828855.1;SARS-CoV_matrix;115 | 0 | 0 |
| NC_004718.3;NP_828855.1;SARS-CoV_matrix;116 | 0.05 | 0 |
| NC_004718.3;NP_828855.1;SARS-CoV_matrix;117 | 0 | 0 |
| NC_004718.3;NP_828855.1;SARS-CoV_matrix;118 | 0 | 0 |
| NC_004718.3;NP_828855.1;SARS-CoV_matrix;119 | 0 | 0 |
| NC_004718.3;NP_828855.1;SARS-CoV_matrix;120 | 0 | 0 |
| NC_004718.3;NP_828855.1;SARS-CoV_matrix;121 | 0 | 0.025 |
| NC_004718.3;NP_828855.1;SARS-CoV_matrix;122 | 0 | 0.025 |
| NC_004718.3;NP_828855.1;SARS-CoV_matrix;123 | 0 | 0.025 |
| NC_004718.3;NP_828855.1;SARS-CoV_matrix;124 | 0 | 0 |
| NC_004718.3;NP_828855.1;SARS-CoV_matrix;125 | 0 | 0 |
| NC_004718.3;NP_828855.1;SARS-CoV_matrix;126 | 0 | 0 |
| NC_004718.3;NP_828855.1;SARS-CoV_matrix;127 | 0 | 0 |
| NC_004718.3;NP_828855.1;SARS-CoV_matrix;128 | 0 | 0 |
| NC_004718.3;NP_828855.1;SARS-CoV_matrix;129 | 0 | 0 |
| NC_004718.3;NP_828855.1;SARS-CoV_matrix;130 | 0 | 0 |
| NC_004718.3;NP_828855.1;SARS-CoV_matrix;131 | 0 | 0 |
| NC_004718.3;NP_828855.1;SARS-CoV_matrix;132 | 0 | 0 |
| NC_004718.3;NP_828855.1;SARS-CoV_matrix;133 | 0 | 0 |
| NC_004718.3;NP_828855.1;SARS-CoV_matrix;134 | 0 | 0 |
| NC_004718.3;NP_828855.1;SARS-CoV_matrix;135 | 0 | 0 |
| NC_004718.3;NP_828855.1;SARS-CoV_matrix;136 | 0 | 0 |

| | | |
|---|---|---|
| NC_004718.3;NP_828855.1;SARS-CoV_matrix;137 | 0 | 0 |
| NC_004718.3;NP_828855.1;SARS-CoV_matrix;138 | 0 | 0 |
| NC_004718.3;NP_828855.1;SARS-CoV_matrix;139 | 0 | 0 |
| NC_004718.3;NP_828855.1;SARS-CoV_matrix;140 | 0 | 0.025 |
| NC_004718.3;NP_828855.1;SARS-CoV_matrix;141 | 0 | 0 |
| NC_004718.3;NP_828855.1;SARS-CoV_matrix;142 | 0 | 0 |
| NC_004718.3;NP_828855.1;SARS-CoV_matrix;143 | 0 | 0 |
| NC_004718.3;NP_828855.1;SARS-CoV_matrix;144 | 0 | 0 |
| NC_004718.3;NP_828855.1;SARS-CoV_matrix;145 | 0 | 0 |
| NC_004718.3;NP_828855.1;SARS-CoV_matrix;146 | 0 | 0 |
| NC_004718.3;NP_828855.1;SARS-CoV_matrix;147 | 0.05 | 0 |
| NC_004718.3;NP_828855.1;SARS-CoV_matrix;148 | 0.05 | 0.05 |
| NC_004718.3;NP_828855.1;SARS-CoV_matrix;149 | 0.05 | 0.025 |
| NC_004718.3;NP_828855.1;SARS-CoV_matrix;150 | 0.05 | 0.05 |
| NC_004718.3;NP_828855.1;SARS-CoV_matrix;151 | 0.05 | 0.15 |
| NC_004718.3;NP_828855.1;SARS-CoV_matrix;152 | 0.05 | 0.375 |
| NC_004718.3;NP_828855.1;SARS-CoV_matrix;153 | 0.05 | 0.45 |
| NC_004718.3;NP_828855.1;SARS-CoV_matrix;154 | 0.05 | 0.425 |
| NC_004718.3;NP_828855.1;SARS-CoV_matrix;155 | 0.1 | 0.425 |
| NC_004718.3;NP_828855.1;SARS-CoV_matrix;156 | 0.05 | 0.4 |
| NC_004718.3;NP_828855.1;SARS-CoV_matrix;157 | 0.05 | 0.35 |
| NC_004718.3;NP_828855.1;SARS-CoV_matrix;158 | 0 | 0.35 |
| NC_004718.3;NP_828855.1;SARS-CoV_matrix;159 | 0 | 0.35 |
| NC_004718.3;NP_828855.1;SARS-CoV_matrix;160 | 0 | 0.35 |
| NC_004718.3;NP_828855.1;SARS-CoV_matrix;161 | 0 | 0.35 |
| NC_004718.3;NP_828855.1;SARS-CoV_matrix;162 | 0 | 0.025 |
| NC_004718.3;NP_828855.1;SARS-CoV_matrix;163 | 0 | 0 |
| NC_004718.3;NP_828855.1;SARS-CoV_matrix;164 | 0 | 0 |
| NC_004718.3;NP_828855.1;SARS-CoV_matrix;165 | 0 | 0 |
| NC_004718.3;NP_828855.1;SARS-CoV_matrix;166 | 0 | 0 |
| NC_004718.3;NP_828855.1;SARS-CoV_matrix;167 | 0 | 0 |
| NC_004718.3;NP_828855.1;SARS-CoV_matrix;168 | 0 | 0 |
| NC_004718.3;NP_828855.1;SARS-CoV_matrix;169 | 0 | 0 |
| NC_004718.3;NP_828855.1;SARS-CoV_matrix;170 | 0 | 0 |
| NC_004718.3;NP_828855.1;SARS-CoV_matrix;171 | 0 | 0 |
| NC_004718.3;NP_828855.1;SARS-CoV_matrix;172 | 0 | 0 |
| NC_004718.3;NP_828855.1;SARS-CoV_matrix;173 | 0 | 0 |
| NC_004718.3;NP_828855.1;SARS-CoV_matrix;174 | 0 | 0.075 |
| NC_004718.3;NP_828855.1;SARS-CoV_matrix;175 | 0 | 0.075 |
| NC_004718.3;NP_828855.1;SARS-CoV_matrix;176 | 0 | 0.125 |
| NC_004718.3;NP_828855.1;SARS-CoV_matrix;177 | 0 | 0.025 |
| NC_004718.3;NP_828855.1;SARS-CoV_matrix;178 | 0 | 0.075 |
| NC_004718.3;NP_828855.1;SARS-CoV_matrix;179 | 0.05 | 0.1 |

| | | |
|---|---|---|
| NC_004718.3;NP_828855.1;SARS-CoV_matrix;180 | 0 | 0.325 |
| NC_004718.3;NP_828855.1;SARS-CoV_matrix;181 | 0 | 0.425 |
| NC_004718.3;NP_828855.1;SARS-CoV_matrix;182 | 0 | 0.375 |
| NC_004718.3;NP_828855.1;SARS-CoV_matrix;183 | 0 | 0.4 |
| NC_004718.3;NP_828855.1;SARS-CoV_matrix;184 | 0 | 0.175 |
| NC_004718.3;NP_828855.1;SARS-CoV_matrix;185 | 0 | 0.125 |
| NC_004718.3;NP_828855.1;SARS-CoV_matrix;186 | 0 | 0.225 |
| NC_004718.3;NP_828855.1;SARS-CoV_matrix;187 | 0 | 0.25 |
| NC_004718.3;NP_828855.1;SARS-CoV_matrix;188 | 0 | 0.25 |
| NC_004718.3;NP_828855.1;SARS-CoV_matrix;189 | 0 | 0.225 |
| NC_004718.3;NP_828855.1;SARS-CoV_matrix;190 | 0 | 0.05 |
| NC_004718.3;NP_828855.1;SARS-CoV_matrix;191 | 0 | 0.05 |
| NC_004718.3;NP_828855.1;SARS-CoV_matrix;192 | 0 | 0.05 |
| NC_004718.3;NP_828855.1;SARS-CoV_matrix;193 | 0 | 0.15 |
| NC_004718.3;NP_828855.1;SARS-CoV_matrix;194 | 0 | 0.1 |
| NC_004718.3;NP_828855.1;SARS-CoV_matrix;195 | 0 | 0.1 |
| NC_004718.3;NP_828855.1;SARS-CoV_matrix;196 | 0 | 0.1 |
| NC_004718.3;NP_828855.1;SARS-CoV_matrix;197 | 0 | 0.1 |
| NC_004718.3;NP_828855.1;SARS-CoV_matrix;198 | 0 | 0.125 |
| NC_004718.3;NP_828855.1;SARS-CoV_matrix;199 | 0.05 | 0.2 |
| NC_004718.3;NP_828855.1;SARS-CoV_matrix;200 | 0.1 | 0.125 |
| NC_004718.3;NP_828855.1;SARS-CoV_matrix;201 | 0 | 0.1 |
| NC_004718.3;NP_828855.1;SARS-CoV_matrix;202 | 0 | 0.075 |
| NC_004718.3;NP_828855.1;SARS-CoV_matrix;203 | 0 | 0.025 |
| NC_004718.3;NP_828855.1;SARS-CoV_matrix;204 | 0 | 0.125 |
| NC_004718.3;NP_828855.1;SARS-CoV_matrix;205 | 0 | 0.225 |
| NC_004718.3;NP_828855.1;SARS-CoV_matrix;206 | 0 | 0.2 |
| NC_004718.3;NP_828855.1;SARS-CoV_matrix;207 | 0 | 0.175 |
| NC_004718.3;NP_828855.1;SARS-CoV_matrix;208 | 0 | 0.175 |
| NC_004718.3;NP_828855.1;SARS-CoV_matrix;209 | 0 | 0.175 |
| NC_004718.3;NP_828855.1;SARS-CoV_matrix;210 | 0 | 0.175 |
| NC_004718.3;NP_828856.1;SARS-CoV_6;1 | 0 | 0 |
| NC_004718.3;NP_828856.1;SARS-CoV_6;2 | 0 | 0 |
| NC_004718.3;NP_828856.1;SARS-CoV_6;3 | 0 | 0 |
| NC_004718.3;NP_828856.1;SARS-CoV_6;4 | 0 | 0 |
| NC_004718.3;NP_828856.1;SARS-CoV_6;5 | 0 | 0 |
| NC_004718.3;NP_828856.1;SARS-CoV_6;6 | 0 | 0 |
| NC_004718.3;NP_828856.1;SARS-CoV_6;7 | 0 | 0 |
| NC_004718.3;NP_828856.1;SARS-CoV_6;8 | 0 | 0 |
| NC_004718.3;NP_828856.1;SARS-CoV_6;9 | 0 | 0 |
| NC_004718.3;NP_828856.1;SARS-CoV_6;10 | 0 | 0 |
| NC_004718.3;NP_828856.1;SARS-CoV_6;11 | 0 | 0 |
| NC_004718.3;NP_828856.1;SARS-CoV_6;12 | 0 | 0 |

| | | |
|---|---|---|
| NC_004718.3;NP_828856.1;SARS-CoV_6;13 | 0 | 0 |
| NC_004718.3;NP_828856.1;SARS-CoV_6;14 | 0 | 0 |
| NC_004718.3;NP_828856.1;SARS-CoV_6;15 | 0 | 0 |
| NC_004718.3;NP_828856.1;SARS-CoV_6;16 | 0 | 0 |
| NC_004718.3;NP_828856.1;SARS-CoV_6;17 | 0 | 0 |
| NC_004718.3;NP_828856.1;SARS-CoV_6;18 | 0 | 0 |
| NC_004718.3;NP_828856.1;SARS-CoV_6;19 | 0 | 0 |
| NC_004718.3;NP_828856.1;SARS-CoV_6;20 | 0 | 0 |
| NC_004718.3;NP_828856.1;SARS-CoV_6;21 | 0 | 0 |
| NC_004718.3;NP_828856.1;SARS-CoV_6;22 | 0 | 0 |
| NC_004718.3;NP_828856.1;SARS-CoV_6;23 | 0 | 0 |
| NC_004718.3;NP_828856.1;SARS-CoV_6;24 | 0 | 0 |
| NC_004718.3;NP_828856.1;SARS-CoV_6;25 | 0 | 0 |
| NC_004718.3;NP_828856.1;SARS-CoV_6;26 | 0 | 0 |
| NC_004718.3;NP_828856.1;SARS-CoV_6;27 | 0 | 0 |
| NC_004718.3;NP_828856.1;SARS-CoV_6;28 | 0 | 0 |
| NC_004718.3;NP_828856.1;SARS-CoV_6;29 | 0 | 0 |
| NC_004718.3;NP_828856.1;SARS-CoV_6;30 | 0 | 0 |
| NC_004718.3;NP_828856.1;SARS-CoV_6;31 | 0 | 0 |
| NC_004718.3;NP_828856.1;SARS-CoV_6;32 | 0 | 0 |
| NC_004718.3;NP_828856.1;SARS-CoV_6;33 | 0 | 0 |
| NC_004718.3;NP_828856.1;SARS-CoV_6;34 | 0 | 0 |
| NC_004718.3;NP_828856.1;SARS-CoV_6;35 | 0 | 0 |
| NC_004718.3;NP_828856.1;SARS-CoV_6;36 | 0 | 0 |
| NC_004718.3;NP_828856.1;SARS-CoV_6;37 | 0 | 0 |
| NC_004718.3;NP_828856.1;SARS-CoV_6;38 | 0 | 0.025 |
| NC_004718.3;NP_828856.1;SARS-CoV_6;39 | 0.05 | 0.025 |
| NC_004718.3;NP_828856.1;SARS-CoV_6;40 | 0.05 | 0.025 |
| NC_004718.3;NP_828856.1;SARS-CoV_6;41 | 0 | 0 |
| NC_004718.3;NP_828856.1;SARS-CoV_6;42 | 0 | 0 |
| NC_004718.3;NP_828856.1;SARS-CoV_6;43 | 0 | 0.025 |
| NC_004718.3;NP_828856.1;SARS-CoV_6;44 | 0 | 0 |
| NC_004718.3;NP_828856.1;SARS-CoV_6;45 | 0 | 0 |
| NC_004718.3;NP_828856.1;SARS-CoV_6;46 | 0 | 0 |
| NC_004718.3;NP_828856.1;SARS-CoV_6;47 | 0 | 0 |
| NC_004718.3;NP_828856.1;SARS-CoV_6;48 | 0.05 | 0.025 |
| NC_004718.3;NP_828856.1;SARS-CoV_6;49 | 0.05 | 0 |
| NC_004718.3;NP_828856.1;SARS-CoV_6;50 | 0.05 | 0 |
| NC_004718.3;NP_828856.1;SARS-CoV_6;51 | 0 | 0 |
| NC_004718.3;NP_828856.1;SARS-CoV_6;52 | 0 | 0 |
| NC_004718.3;NP_828857.1;SARS-CoV_7;1 | 0 | 0 |
| NC_004718.3;NP_828857.1;SARS-CoV_7;2 | 0 | 0 |
| NC_004718.3;NP_828857.1;SARS-CoV_7;3 | 0 | 0 |

| | | |
|---|---|---|
| NC_004718.3;NP_828857.1;SARS-CoV_7;4 | 0 | 0 |
| NC_004718.3;NP_828857.1;SARS-CoV_7;5 | 0 | 0 |
| NC_004718.3;NP_828857.1;SARS-CoV_7;6 | 0 | 0 |
| NC_004718.3;NP_828857.1;SARS-CoV_7;7 | 0 | 0 |
| NC_004718.3;NP_828857.1;SARS-CoV_7;8 | 0 | 0 |
| NC_004718.3;NP_828857.1;SARS-CoV_7;9 | 0 | 0 |
| NC_004718.3;NP_828857.1;SARS-CoV_7;10 | 0 | 0 |
| NC_004718.3;NP_828857.1;SARS-CoV_7;11 | 0 | 0 |
| NC_004718.3;NP_828857.1;SARS-CoV_7;12 | 0 | 0 |
| NC_004718.3;NP_828857.1;SARS-CoV_7;13 | 0 | 0 |
| NC_004718.3;NP_828857.1;SARS-CoV_7;14 | 0 | 0 |
| NC_004718.3;NP_828857.1;SARS-CoV_7;15 | 0 | 0 |
| NC_004718.3;NP_828857.1;SARS-CoV_7;16 | 0 | 0 |
| NC_004718.3;NP_828857.1;SARS-CoV_7;17 | 0 | 0 |
| NC_004718.3;NP_828857.1;SARS-CoV_7;18 | 0 | 0 |
| NC_004718.3;NP_828857.1;SARS-CoV_7;19 | 0 | 0 |
| NC_004718.3;NP_828857.1;SARS-CoV_7;20 | 0 | 0 |
| NC_004718.3;NP_828857.1;SARS-CoV_7;21 | 0 | 0 |
| NC_004718.3;NP_828857.1;SARS-CoV_7;22 | 0 | 0 |
| NC_004718.3;NP_828857.1;SARS-CoV_7;23 | 0.05 | 0 |
| NC_004718.3;NP_828857.1;SARS-CoV_7;24 | 0.05 | 0 |
| NC_004718.3;NP_828857.1;SARS-CoV_7;25 | 0.05 | 0.025 |
| NC_004718.3;NP_828857.1;SARS-CoV_7;26 | 0 | 0.025 |
| NC_004718.3;NP_828857.1;SARS-CoV_7;27 | 0 | 0 |
| NC_004718.3;NP_828857.1;SARS-CoV_7;28 | 0 | 0 |
| NC_004718.3;NP_828857.1;SARS-CoV_7;29 | 0 | 0 |
| NC_004718.3;NP_828857.1;SARS-CoV_7;30 | 0 | 0 |
| NC_004718.3;NP_828857.1;SARS-CoV_7;31 | 0 | 0 |
| NC_004718.3;NP_828857.1;SARS-CoV_7;32 | 0 | 0 |
| NC_004718.3;NP_828857.1;SARS-CoV_7;33 | 0 | 0 |
| NC_004718.3;NP_828857.1;SARS-CoV_7;34 | 0 | 0.025 |
| NC_004718.3;NP_828857.1;SARS-CoV_7;35 | 0 | 0.025 |
| NC_004718.3;NP_828857.1;SARS-CoV_7;36 | 0 | 0.025 |
| NC_004718.3;NP_828857.1;SARS-CoV_7;37 | 0 | 0.05 |
| NC_004718.3;NP_828857.1;SARS-CoV_7;38 | 0 | 0.05 |
| NC_004718.3;NP_828857.1;SARS-CoV_7;39 | 0 | 0.05 |
| NC_004718.3;NP_828857.1;SARS-CoV_7;40 | 0 | 0.05 |
| NC_004718.3;NP_828857.1;SARS-CoV_7;41 | 0 | 0.075 |
| NC_004718.3;NP_828857.1;SARS-CoV_7;42 | 0 | 0.075 |
| NC_004718.3;NP_828857.1;SARS-CoV_7;43 | 0 | 0.05 |
| NC_004718.3;NP_828857.1;SARS-CoV_7;44 | 0 | 0.025 |
| NC_004718.3;NP_828857.1;SARS-CoV_7;45 | 0 | 0 |
| NC_004718.3;NP_828857.1;SARS-CoV_7;46 | 0 | 0 |

| | | |
|---|---|---|
| NC_004718.3;NP_828857.1;SARS-CoV_7;47 | 0 | 0 |
| NC_004718.3;NP_828857.1;SARS-CoV_7;48 | 0 | 0 |
| NC_004718.3;NP_828857.1;SARS-CoV_7;49 | 0 | 0 |
| NC_004718.3;NP_828857.1;SARS-CoV_7;50 | 0 | 0 |
| NC_004718.3;NP_828857.1;SARS-CoV_7;51 | 0 | 0 |
| NC_004718.3;NP_828857.1;SARS-CoV_7;52 | 0 | 0 |
| NC_004718.3;NP_828857.1;SARS-CoV_7;53 | 0 | 0 |
| NC_004718.3;NP_828857.1;SARS-CoV_7;54 | 0 | 0 |
| NC_004718.3;NP_828857.1;SARS-CoV_7;55 | 0 | 0 |
| NC_004718.3;NP_828857.1;SARS-CoV_7;56 | 0 | 0 |
| NC_004718.3;NP_828857.1;SARS-CoV_7;57 | 0 | 0 |
| NC_004718.3;NP_828857.1;SARS-CoV_7;58 | 0 | 0 |
| NC_004718.3;NP_828857.1;SARS-CoV_7;59 | 0 | 0 |
| NC_004718.3;NP_828857.1;SARS-CoV_7;60 | 0 | 0 |
| NC_004718.3;NP_828857.1;SARS-CoV_7;61 | 0 | 0 |
| NC_004718.3;NP_828857.1;SARS-CoV_7;62 | 0 | 0 |
| NC_004718.3;NP_828857.1;SARS-CoV_7;63 | 0 | 0 |
| NC_004718.3;NP_828857.1;SARS-CoV_7;64 | 0 | 0 |
| NC_004718.3;NP_828857.1;SARS-CoV_7;65 | 0 | 0 |
| NC_004718.3;NP_828857.1;SARS-CoV_7;66 | 0 | 0 |
| NC_004718.3;NP_828857.1;SARS-CoV_7;67 | 0 | 0 |
| NC_004718.3;NP_828857.1;SARS-CoV_7;68 | 0 | 0 |
| NC_004718.3;NP_828857.1;SARS-CoV_7;69 | 0 | 0 |
| NC_004718.3;NP_828857.1;SARS-CoV_7;70 | 0 | 0 |
| NC_004718.3;NP_828857.1;SARS-CoV_7;71 | 0 | 0 |
| NC_004718.3;NP_828857.1;SARS-CoV_7;72 | 0 | 0 |
| NC_004718.3;NP_828857.1;SARS-CoV_7;73 | 0 | 0 |
| NC_004718.3;NP_828857.1;SARS-CoV_7;74 | 0 | 0 |
| NC_004718.3;NP_828857.1;SARS-CoV_7;75 | 0 | 0 |
| NC_004718.3;NP_828857.1;SARS-CoV_7;76 | 0 | 0 |
| NC_004718.3;NP_828857.1;SARS-CoV_7;77 | 0 | 0 |
| NC_004718.3;NP_828857.1;SARS-CoV_7;78 | 0 | 0 |
| NC_004718.3;NP_828857.1;SARS-CoV_7;79 | 0.05 | 0 |
| NC_004718.3;NP_828857.1;SARS-CoV_7;80 | 0.1 | 0 |
| NC_004718.3;NP_828857.1;SARS-CoV_7;81 | 0.05 | 0 |
| NC_004718.3;NP_828857.1;SARS-CoV_7;82 | 0.1 | 0 |
| NC_004718.3;NP_828857.1;SARS-CoV_7;83 | 0.05 | 0 |
| NC_004718.3;NP_828857.1;SARS-CoV_7;84 | 0.1 | 0 |
| NC_004718.3;NP_828857.1;SARS-CoV_7;85 | 0.1 | 0 |
| NC_004718.3;NP_828857.1;SARS-CoV_7;86 | 0 | 0 |
| NC_004718.3;NP_828857.1;SARS-CoV_7;87 | 0 | 0.025 |
| NC_004718.3;NP_828857.1;SARS-CoV_7;88 | 0.05 | 0.05 |
| NC_004718.3;NP_828857.1;SARS-CoV_7;89 | 0 | 0 |

| | | |
|---|---|---|
| NC_004718.3;NP_828857.1;SARS-CoV_7;90 | 0 | 0 |
| NC_004718.3;NP_828857.1;SARS-CoV_7;91 | 0.05 | 0.025 |
| NC_004718.3;NP_828857.1;SARS-CoV_7;92 | 0 | 0 |
| NC_004718.3;NP_828857.1;SARS-CoV_7;93 | 0 | 0 |
| NC_004718.3;NP_828857.1;SARS-CoV_7;94 | 0 | 0 |
| NC_004718.3;NP_828857.1;SARS-CoV_7;95 | 0 | 0 |
| NC_004718.3;NP_828857.1;SARS-CoV_7;96 | 0 | 0 |
| NC_004718.3;NP_828857.1;SARS-CoV_7;97 | 0 | 0 |
| NC_004718.3;NP_828857.1;SARS-CoV_7;98 | 0 | 0 |
| NC_004718.3;NP_828857.1;SARS-CoV_7;99 | 0 | 0 |
| NC_004718.3;NP_828857.1;SARS-CoV_7;100 | 0 | 0 |
| NC_004718.3;NP_828857.1;SARS-CoV_7;101 | 0 | 0 |
| NC_004718.3;NP_828857.1;SARS-CoV_7;102 | 0 | 0 |
| NC_004718.3;NP_828857.1;SARS-CoV_7;103 | 0 | 0 |
| NC_004718.3;NP_828857.1;SARS-CoV_7;104 | 0 | 0 |
| NC_004718.3;NP_828857.1;SARS-CoV_7;105 | 0 | 0 |
| NC_004718.3;NP_828857.1;SARS-CoV_7;106 | 0 | 0 |
| NC_004718.3;NP_828857.1;SARS-CoV_7;107 | 0 | 0 |
| NC_004718.3;NP_828857.1;SARS-CoV_7;108 | 0 | 0 |
| NC_004718.3;NP_828857.1;SARS-CoV_7;109 | 0 | 0 |
| NC_004718.3;NP_828857.1;SARS-CoV_7;110 | 0 | 0 |
| NC_004718.3;NP_828857.1;SARS-CoV_7;111 | 0 | 0 |
| NC_004718.3;NP_828858.1;SARS-CoV_nucleocapsid;1 | 0 | 0 |
| NC_004718.3;NP_828858.1;SARS-CoV_nucleocapsid;2 | 0 | 0.025 |
| NC_004718.3;NP_828858.1;SARS-CoV_nucleocapsid;3 | 0.05 | 0 |
| NC_004718.3;NP_828858.1;SARS-CoV_nucleocapsid;4 | 0 | 0 |
| NC_004718.3;NP_828858.1;SARS-CoV_nucleocapsid;5 | 0 | 0.025 |
| NC_004718.3;NP_828858.1;SARS-CoV_nucleocapsid;6 | 0.05 | 0.025 |
| NC_004718.3;NP_828858.1;SARS-CoV_nucleocapsid;7 | 0.1 | 0 |
| NC_004718.3;NP_828858.1;SARS-CoV_nucleocapsid;8 | 0.05 | 0 |
| NC_004718.3;NP_828858.1;SARS-CoV_nucleocapsid;9 | 0.05 | 0.05 |
| NC_004718.3;NP_828858.1;SARS-CoV_nucleocapsid;10 | 0.05 | 0.075 |
| NC_004718.3;NP_828858.1;SARS-CoV_nucleocapsid;11 | 0.05 | 0.225 |
| NC_004718.3;NP_828858.1;SARS-CoV_nucleocapsid;12 | 0.05 | 0.275 |
| NC_004718.3;NP_828858.1;SARS-CoV_nucleocapsid;13 | 0.05 | 0.25 |
| NC_004718.3;NP_828858.1;SARS-CoV_nucleocapsid;14 | 0.1 | 0.2 |
| NC_004718.3;NP_828858.1;SARS-CoV_nucleocapsid;15 | 0.05 | 0.2 |
| NC_004718.3;NP_828858.1;SARS-CoV_nucleocapsid;16 | 0 | 0.1 |
| NC_004718.3;NP_828858.1;SARS-CoV_nucleocapsid;17 | 0.05 | 0.075 |
| NC_004718.3;NP_828858.1;SARS-CoV_nucleocapsid;18 | 0.05 | 0.075 |
| NC_004718.3;NP_828858.1;SARS-CoV_nucleocapsid;19 | 0.1 | 0.075 |
| NC_004718.3;NP_828858.1;SARS-CoV_nucleocapsid;20 | 0.15 | 0.075 |
| NC_004718.3;NP_828858.1;SARS-CoV_nucleocapsid;21 | 0.1 | 0.05 |

| | | |
|---|---|---|
| NC_004718.3;NP_828858.1;SARS-CoV_nucleocapsid;22 | 0.2 | 0.125 |
| NC_004718.3;NP_828858.1;SARS-CoV_nucleocapsid;23 | 0.15 | 0.1 |
| NC_004718.3;NP_828858.1;SARS-CoV_nucleocapsid;24 | 0.15 | 0.2 |
| NC_004718.3;NP_828858.1;SARS-CoV_nucleocapsid;25 | 0.1 | 0.2 |
| NC_004718.3;NP_828858.1;SARS-CoV_nucleocapsid;26 | 0.1 | 0.125 |
| NC_004718.3;NP_828858.1;SARS-CoV_nucleocapsid;27 | 0.05 | 0.1 |
| NC_004718.3;NP_828858.1;SARS-CoV_nucleocapsid;28 | 0.05 | 0.125 |
| NC_004718.3;NP_828858.1;SARS-CoV_nucleocapsid;29 | 0 | 0.1 |
| NC_004718.3;NP_828858.1;SARS-CoV_nucleocapsid;30 | 0.05 | 0.125 |
| NC_004718.3;NP_828858.1;SARS-CoV_nucleocapsid;31 | 0 | 0.1 |
| NC_004718.3;NP_828858.1;SARS-CoV_nucleocapsid;32 | 0 | 0.1 |
| NC_004718.3;NP_828858.1;SARS-CoV_nucleocapsid;33 | 0 | 0.125 |
| NC_004718.3;NP_828858.1;SARS-CoV_nucleocapsid;34 | 0 | 0.2 |
| NC_004718.3;NP_828858.1;SARS-CoV_nucleocapsid;35 | 0 | 0.125 |
| NC_004718.3;NP_828858.1;SARS-CoV_nucleocapsid;36 | 0 | 0.225 |
| NC_004718.3;NP_828858.1;SARS-CoV_nucleocapsid;37 | 0.05 | 0.15 |
| NC_004718.3;NP_828858.1;SARS-CoV_nucleocapsid;38 | 0.05 | 0.3 |
| NC_004718.3;NP_828858.1;SARS-CoV_nucleocapsid;39 | 0 | 0.225 |
| NC_004718.3;NP_828858.1;SARS-CoV_nucleocapsid;40 | 0 | 0.15 |
| NC_004718.3;NP_828858.1;SARS-CoV_nucleocapsid;41 | 0 | 0.125 |
| NC_004718.3;NP_828858.1;SARS-CoV_nucleocapsid;42 | 0 | 0.025 |
| NC_004718.3;NP_828858.1;SARS-CoV_nucleocapsid;43 | 0 | 0 |
| NC_004718.3;NP_828858.1;SARS-CoV_nucleocapsid;44 | 0 | 0.025 |
| NC_004718.3;NP_828858.1;SARS-CoV_nucleocapsid;45 | 0 | 0 |
| NC_004718.3;NP_828858.1;SARS-CoV_nucleocapsid;46 | 0 | 0 |
| NC_004718.3;NP_828858.1;SARS-CoV_nucleocapsid;47 | 0 | 0 |
| NC_004718.3;NP_828858.1;SARS-CoV_nucleocapsid;48 | 0 | 0 |
| NC_004718.3;NP_828858.1;SARS-CoV_nucleocapsid;49 | 0 | 0 |
| NC_004718.3;NP_828858.1;SARS-CoV_nucleocapsid;50 | 0 | 0 |
| NC_004718.3;NP_828858.1;SARS-CoV_nucleocapsid;51 | 0 | 0 |
| NC_004718.3;NP_828858.1;SARS-CoV_nucleocapsid;52 | 0 | 0.025 |
| NC_004718.3;NP_828858.1;SARS-CoV_nucleocapsid;53 | 0 | 0 |
| NC_004718.3;NP_828858.1;SARS-CoV_nucleocapsid;54 | 0 | 0 |
| NC_004718.3;NP_828858.1;SARS-CoV_nucleocapsid;55 | 0 | 0 |
| NC_004718.3;NP_828858.1;SARS-CoV_nucleocapsid;56 | 0 | 0 |
| NC_004718.3;NP_828858.1;SARS-CoV_nucleocapsid;57 | 0 | 0 |
| NC_004718.3;NP_828858.1;SARS-CoV_nucleocapsid;58 | 0 | 0 |
| NC_004718.3;NP_828858.1;SARS-CoV_nucleocapsid;59 | 0 | 0.025 |
| NC_004718.3;NP_828858.1;SARS-CoV_nucleocapsid;60 | 0 | 0 |
| NC_004718.3;NP_828858.1;SARS-CoV_nucleocapsid;61 | 0 | 0.025 |
| NC_004718.3;NP_828858.1;SARS-CoV_nucleocapsid;62 | 0 | 0.025 |
| NC_004718.3;NP_828858.1;SARS-CoV_nucleocapsid;63 | 0.05 | 0.05 |
| NC_004718.3;NP_828858.1;SARS-CoV_nucleocapsid;64 | 0.05 | 0.05 |

| | | |
|---|---|---|
| NC_004718.3;NP_828858.1;SARS-CoV_nucleocapsid;65 | 0.05 | 0.025 |
| NC_004718.3;NP_828858.1;SARS-CoV_nucleocapsid;66 | 0.1 | 0.125 |
| NC_004718.3;NP_828858.1;SARS-CoV_nucleocapsid;67 | 0.2 | 0.125 |
| NC_004718.3;NP_828858.1;SARS-CoV_nucleocapsid;68 | 0.05 | 0.075 |
| NC_004718.3;NP_828858.1;SARS-CoV_nucleocapsid;69 | 0.15 | 0.05 |
| NC_004718.3;NP_828858.1;SARS-CoV_nucleocapsid;70 | 0.15 | 0.1 |
| NC_004718.3;NP_828858.1;SARS-CoV_nucleocapsid;71 | 0.1 | 0.075 |
| NC_004718.3;NP_828858.1;SARS-CoV_nucleocapsid;72 | 0.1 | 0.025 |
| NC_004718.3;NP_828858.1;SARS-CoV_nucleocapsid;73 | 0.1 | 0.05 |
| NC_004718.3;NP_828858.1;SARS-CoV_nucleocapsid;74 | 0.1 | 0.05 |
| NC_004718.3;NP_828858.1;SARS-CoV_nucleocapsid;75 | 0.1 | 0.075 |
| NC_004718.3;NP_828858.1;SARS-CoV_nucleocapsid;76 | 0 | 0 |
| NC_004718.3;NP_828858.1;SARS-CoV_nucleocapsid;77 | 0 | 0 |
| NC_004718.3;NP_828858.1;SARS-CoV_nucleocapsid;78 | 0 | 0 |
| NC_004718.3;NP_828858.1;SARS-CoV_nucleocapsid;79 | 0.05 | 0 |
| NC_004718.3;NP_828858.1;SARS-CoV_nucleocapsid;80 | 0.05 | 0 |
| NC_004718.3;NP_828858.1;SARS-CoV_nucleocapsid;81 | 0.05 | 0 |
| NC_004718.3;NP_828858.1;SARS-CoV_nucleocapsid;82 | 0 | 0 |
| NC_004718.3;NP_828858.1;SARS-CoV_nucleocapsid;83 | 0 | 0 |
| NC_004718.3;NP_828858.1;SARS-CoV_nucleocapsid;84 | 0.05 | 0 |
| NC_004718.3;NP_828858.1;SARS-CoV_nucleocapsid;85 | 0.05 | 0 |
| NC_004718.3;NP_828858.1;SARS-CoV_nucleocapsid;86 | 0.05 | 0 |
| NC_004718.3;NP_828858.1;SARS-CoV_nucleocapsid;87 | 0 | 0 |
| NC_004718.3;NP_828858.1;SARS-CoV_nucleocapsid;88 | 0 | 0.025 |
| NC_004718.3;NP_828858.1;SARS-CoV_nucleocapsid;89 | 0 | 0 |
| NC_004718.3;NP_828858.1;SARS-CoV_nucleocapsid;90 | 0 | 0.025 |
| NC_004718.3;NP_828858.1;SARS-CoV_nucleocapsid;91 | 0 | 0.025 |
| NC_004718.3;NP_828858.1;SARS-CoV_nucleocapsid;92 | 0 | 0.025 |
| NC_004718.3;NP_828858.1;SARS-CoV_nucleocapsid;93 | 0 | 0.05 |
| NC_004718.3;NP_828858.1;SARS-CoV_nucleocapsid;94 | 0 | 0.05 |
| NC_004718.3;NP_828858.1;SARS-CoV_nucleocapsid;95 | 0 | 0.05 |
| NC_004718.3;NP_828858.1;SARS-CoV_nucleocapsid;96 | 0 | 0.025 |
| NC_004718.3;NP_828858.1;SARS-CoV_nucleocapsid;97 | 0 | 0.025 |
| NC_004718.3;NP_828858.1;SARS-CoV_nucleocapsid;98 | 0 | 0.025 |
| NC_004718.3;NP_828858.1;SARS-CoV_nucleocapsid;99 | 0 | 0 |
| NC_004718.3;NP_828858.1;SARS-CoV_nucleocapsid;100 | 0 | 0 |
| NC_004718.3;NP_828858.1;SARS-CoV_nucleocapsid;101 | 0 | 0 |
| NC_004718.3;NP_828858.1;SARS-CoV_nucleocapsid;102 | 0 | 0 |
| NC_004718.3;NP_828858.1;SARS-CoV_nucleocapsid;103 | 0 | 0 |
| NC_004718.3;NP_828858.1;SARS-CoV_nucleocapsid;104 | 0 | 0 |
| NC_004718.3;NP_828858.1;SARS-CoV_nucleocapsid;105 | 0 | 0 |
| NC_004718.3;NP_828858.1;SARS-CoV_nucleocapsid;106 | 0 | 0 |
| NC_004718.3;NP_828858.1;SARS-CoV_nucleocapsid;107 | 0 | 0 |

| | | |
|---|---|---|
| NC_004718.3;NP_828858.1;SARS-CoV_nucleocapsid;108 | 0 | 0 |
| NC_004718.3;NP_828858.1;SARS-CoV_nucleocapsid;109 | 0 | 0 |
| NC_004718.3;NP_828858.1;SARS-CoV_nucleocapsid;110 | 0 | 0 |
| NC_004718.3;NP_828858.1;SARS-CoV_nucleocapsid;111 | 0 | 0 |
| NC_004718.3;NP_828858.1;SARS-CoV_nucleocapsid;112 | 0 | 0 |
| NC_004718.3;NP_828858.1;SARS-CoV_nucleocapsid;113 | 0.1 | 0.025 |
| NC_004718.3;NP_828858.1;SARS-CoV_nucleocapsid;114 | 0.05 | 0.05 |
| NC_004718.3;NP_828858.1;SARS-CoV_nucleocapsid;115 | 0 | 0.025 |
| NC_004718.3;NP_828858.1;SARS-CoV_nucleocapsid;116 | 0 | 0.025 |
| NC_004718.3;NP_828858.1;SARS-CoV_nucleocapsid;117 | 0 | 0.025 |
| NC_004718.3;NP_828858.1;SARS-CoV_nucleocapsid;118 | 0 | 0.025 |
| NC_004718.3;NP_828858.1;SARS-CoV_nucleocapsid;119 | 0 | 0.025 |
| NC_004718.3;NP_828858.1;SARS-CoV_nucleocapsid;120 | 0 | 0.025 |
| NC_004718.3;NP_828858.1;SARS-CoV_nucleocapsid;121 | 0 | 0.025 |
| NC_004718.3;NP_828858.1;SARS-CoV_nucleocapsid;122 | 0 | 0 |
| NC_004718.3;NP_828858.1;SARS-CoV_nucleocapsid;123 | 0 | 0 |
| NC_004718.3;NP_828858.1;SARS-CoV_nucleocapsid;124 | 0 | 0 |
| NC_004718.3;NP_828858.1;SARS-CoV_nucleocapsid;125 | 0 | 0 |
| NC_004718.3;NP_828858.1;SARS-CoV_nucleocapsid;126 | 0 | 0.05 |
| NC_004718.3;NP_828858.1;SARS-CoV_nucleocapsid;127 | 0 | 0.025 |
| NC_004718.3;NP_828858.1;SARS-CoV_nucleocapsid;128 | 0 | 0.025 |
| NC_004718.3;NP_828858.1;SARS-CoV_nucleocapsid;129 | 0 | 0.075 |
| NC_004718.3;NP_828858.1;SARS-CoV_nucleocapsid;130 | 0 | 0.1 |
| NC_004718.3;NP_828858.1;SARS-CoV_nucleocapsid;131 | 0 | 0.05 |
| NC_004718.3;NP_828858.1;SARS-CoV_nucleocapsid;132 | 0 | 0.1 |
| NC_004718.3;NP_828858.1;SARS-CoV_nucleocapsid;133 | 0 | 0.075 |
| NC_004718.3;NP_828858.1;SARS-CoV_nucleocapsid;134 | 0 | 0.05 |
| NC_004718.3;NP_828858.1;SARS-CoV_nucleocapsid;135 | 0 | 0.025 |
| NC_004718.3;NP_828858.1;SARS-CoV_nucleocapsid;136 | 0 | 0.025 |
| NC_004718.3;NP_828858.1;SARS-CoV_nucleocapsid;137 | 0 | 0.025 |
| NC_004718.3;NP_828858.1;SARS-CoV_nucleocapsid;138 | 0 | 0 |
| NC_004718.3;NP_828858.1;SARS-CoV_nucleocapsid;139 | 0.05 | 0.025 |
| NC_004718.3;NP_828858.1;SARS-CoV_nucleocapsid;140 | 0.1 | 0.025 |
| NC_004718.3;NP_828858.1;SARS-CoV_nucleocapsid;141 | 0.1 | 0 |
| NC_004718.3;NP_828858.1;SARS-CoV_nucleocapsid;142 | 0.15 | 0.1 |
| NC_004718.3;NP_828858.1;SARS-CoV_nucleocapsid;143 | 0.05 | 0.05 |
| NC_004718.3;NP_828858.1;SARS-CoV_nucleocapsid;144 | 0.1 | 0.05 |
| NC_004718.3;NP_828858.1;SARS-CoV_nucleocapsid;145 | 0 | 0.075 |
| NC_004718.3;NP_828858.1;SARS-CoV_nucleocapsid;146 | 0 | 0.1 |
| NC_004718.3;NP_828858.1;SARS-CoV_nucleocapsid;147 | 0 | 0.075 |
| NC_004718.3;NP_828858.1;SARS-CoV_nucleocapsid;148 | 0 | 0.05 |
| NC_004718.3;NP_828858.1;SARS-CoV_nucleocapsid;149 | 0.1 | 0.125 |
| NC_004718.3;NP_828858.1;SARS-CoV_nucleocapsid;150 | 0.2 | 0.125 |

| | | |
|---|---|---|
| NC_004718.3;NP_828858.1;SARS-CoV_nucleocapsid;151 | 0.2 | 0.125 |
| NC_004718.3;NP_828858.1;SARS-CoV_nucleocapsid;152 | 0 | 0.05 |
| NC_004718.3;NP_828858.1;SARS-CoV_nucleocapsid;153 | 0 | 0.075 |
| NC_004718.3;NP_828858.1;SARS-CoV_nucleocapsid;154 | 0 | 0.25 |
| NC_004718.3;NP_828858.1;SARS-CoV_nucleocapsid;155 | 0.15 | 0.525 |
| NC_004718.3;NP_828858.1;SARS-CoV_nucleocapsid;156 | 0.15 | 0.55 |
| NC_004718.3;NP_828858.1;SARS-CoV_nucleocapsid;157 | 0.15 | 0.55 |
| NC_004718.3;NP_828858.1;SARS-CoV_nucleocapsid;158 | 0.1 | 0.525 |
| NC_004718.3;NP_828858.1;SARS-CoV_nucleocapsid;159 | 0.1 | 0.5 |
| NC_004718.3;NP_828858.1;SARS-CoV_nucleocapsid;160 | 0.2 | 0.625 |
| NC_004718.3;NP_828858.1;SARS-CoV_nucleocapsid;161 | 0.2 | 0.625 |
| NC_004718.3;NP_828858.1;SARS-CoV_nucleocapsid;162 | 0.1 | 0.6 |
| NC_004718.3;NP_828858.1;SARS-CoV_nucleocapsid;163 | 0.1 | 0.475 |
| NC_004718.3;NP_828858.1;SARS-CoV_nucleocapsid;164 | 0.1 | 0.45 |
| NC_004718.3;NP_828858.1;SARS-CoV_nucleocapsid;165 | 0.05 | 0.1 |
| NC_004718.3;NP_828858.1;SARS-CoV_nucleocapsid;166 | 0 | 0.05 |
| NC_004718.3;NP_828858.1;SARS-CoV_nucleocapsid;167 | 0 | 0.05 |
| NC_004718.3;NP_828858.1;SARS-CoV_nucleocapsid;168 | 0 | 0 |
| NC_004718.3;NP_828858.1;SARS-CoV_nucleocapsid;169 | 0.05 | 0.025 |
| NC_004718.3;NP_828858.1;SARS-CoV_nucleocapsid;170 | 0 | 0.05 |
| NC_004718.3;NP_828858.1;SARS-CoV_nucleocapsid;171 | 0 | 0 |
| NC_004718.3;NP_828858.1;SARS-CoV_nucleocapsid;172 | 0 | 0 |
| NC_004718.3;NP_828858.1;SARS-CoV_nucleocapsid;173 | 0 | 0 |
| NC_004718.3;NP_828858.1;SARS-CoV_nucleocapsid;174 | 0 | 0.025 |
| NC_004718.3;NP_828858.1;SARS-CoV_nucleocapsid;175 | 0 | 0.05 |
| NC_004718.3;NP_828858.1;SARS-CoV_nucleocapsid;176 | 0 | 0.025 |
| NC_004718.3;NP_828858.1;SARS-CoV_nucleocapsid;177 | 0 | 0.025 |
| NC_004718.3;NP_828858.1;SARS-CoV_nucleocapsid;178 | 0 | 0.025 |
| NC_004718.3;NP_828858.1;SARS-CoV_nucleocapsid;179 | 0 | 0.025 |
| NC_004718.3;NP_828858.1;SARS-CoV_nucleocapsid;180 | 0 | 0.05 |
| NC_004718.3;NP_828858.1;SARS-CoV_nucleocapsid;181 | 0 | 0.025 |
| NC_004718.3;NP_828858.1;SARS-CoV_nucleocapsid;182 | 0 | 0.05 |
| NC_004718.3;NP_828858.1;SARS-CoV_nucleocapsid;183 | 0 | 0.025 |
| NC_004718.3;NP_828858.1;SARS-CoV_nucleocapsid;184 | 0 | 0.025 |
| NC_004718.3;NP_828858.1;SARS-CoV_nucleocapsid;185 | 0 | 0.025 |
| NC_004718.3;NP_828858.1;SARS-CoV_nucleocapsid;186 | 0 | 0.025 |
| NC_004718.3;NP_828858.1;SARS-CoV_nucleocapsid;187 | 0 | 0.05 |
| NC_004718.3;NP_828858.1;SARS-CoV_nucleocapsid;188 | 0 | 0.025 |
| NC_004718.3;NP_828858.1;SARS-CoV_nucleocapsid;189 | 0 | 0 |
| NC_004718.3;NP_828858.1;SARS-CoV_nucleocapsid;190 | 0 | 0 |
| NC_004718.3;NP_828858.1;SARS-CoV_nucleocapsid;191 | 0 | 0.075 |
| NC_004718.3;NP_828858.1;SARS-CoV_nucleocapsid;192 | 0 | 0 |
| NC_004718.3;NP_828858.1;SARS-CoV_nucleocapsid;193 | 0 | 0.05 |

| | | |
|---|---|---|
| NC_004718.3;NP_828858.1;SARS-CoV_nucleocapsid;194 | 0 | 0.025 |
| NC_004718.3;NP_828858.1;SARS-CoV_nucleocapsid;195 | 0 | 0 |
| NC_004718.3;NP_828858.1;SARS-CoV_nucleocapsid;196 | 0 | 0.025 |
| NC_004718.3;NP_828858.1;SARS-CoV_nucleocapsid;197 | 0.05 | 0.05 |
| NC_004718.3;NP_828858.1;SARS-CoV_nucleocapsid;198 | 0.1 | 0.05 |
| NC_004718.3;NP_828858.1;SARS-CoV_nucleocapsid;199 | 0.05 | 0.025 |
| NC_004718.3;NP_828858.1;SARS-CoV_nucleocapsid;200 | 0.05 | 0 |
| NC_004718.3;NP_828858.1;SARS-CoV_nucleocapsid;201 | 0.1 | 0.125 |
| NC_004718.3;NP_828858.1;SARS-CoV_nucleocapsid;202 | 0.15 | 0.025 |
| NC_004718.3;NP_828858.1;SARS-CoV_nucleocapsid;203 | 0.1 | 0 |
| NC_004718.3;NP_828858.1;SARS-CoV_nucleocapsid;204 | 0 | 0 |
| NC_004718.3;NP_828858.1;SARS-CoV_nucleocapsid;205 | 0 | 0 |
| NC_004718.3;NP_828858.1;SARS-CoV_nucleocapsid;206 | 0 | 0 |
| NC_004718.3;NP_828858.1;SARS-CoV_nucleocapsid;207 | 0 | 0.025 |
| NC_004718.3;NP_828858.1;SARS-CoV_nucleocapsid;208 | 0 | 0.025 |
| NC_004718.3;NP_828858.1;SARS-CoV_nucleocapsid;209 | 0 | 0 |
| NC_004718.3;NP_828858.1;SARS-CoV_nucleocapsid;210 | 0 | 0 |
| NC_004718.3;NP_828858.1;SARS-CoV_nucleocapsid;211 | 0 | 0 |
| NC_004718.3;NP_828858.1;SARS-CoV_nucleocapsid;212 | 0 | 0 |
| NC_004718.3;NP_828858.1;SARS-CoV_nucleocapsid;213 | 0 | 0 |
| NC_004718.3;NP_828858.1;SARS-CoV_nucleocapsid;214 | 0 | 0 |
| NC_004718.3;NP_828858.1;SARS-CoV_nucleocapsid;215 | 0 | 0 |
| NC_004718.3;NP_828858.1;SARS-CoV_nucleocapsid;216 | 0 | 0 |
| NC_004718.3;NP_828858.1;SARS-CoV_nucleocapsid;217 | 0 | 0 |
| NC_004718.3;NP_828858.1;SARS-CoV_nucleocapsid;218 | 0 | 0 |
| NC_004718.3;NP_828858.1;SARS-CoV_nucleocapsid;219 | 0.05 | 0.05 |
| NC_004718.3;NP_828858.1;SARS-CoV_nucleocapsid;220 | 0.05 | 0.15 |
| NC_004718.3;NP_828858.1;SARS-CoV_nucleocapsid;221 | 0.05 | 0.2 |
| NC_004718.3;NP_828858.1;SARS-CoV_nucleocapsid;222 | 0.05 | 0.3 |
| NC_004718.3;NP_828858.1;SARS-CoV_nucleocapsid;223 | 0.1 | 0.25 |
| NC_004718.3;NP_828858.1;SARS-CoV_nucleocapsid;224 | 0.05 | 0.175 |
| NC_004718.3;NP_828858.1;SARS-CoV_nucleocapsid;225 | 0.05 | 0.1 |
| NC_004718.3;NP_828858.1;SARS-CoV_nucleocapsid;226 | 0.05 | 0.025 |
| NC_004718.3;NP_828858.1;SARS-CoV_nucleocapsid;227 | 0 | 0.1 |
| NC_004718.3;NP_828858.1;SARS-CoV_nucleocapsid;228 | 0 | 0.2 |
| NC_004718.3;NP_828858.1;SARS-CoV_nucleocapsid;229 | 0 | 0.175 |
| NC_004718.3;NP_828858.1;SARS-CoV_nucleocapsid;230 | 0.05 | 0.175 |
| NC_004718.3;NP_828858.1;SARS-CoV_nucleocapsid;231 | 0 | 0.225 |
| NC_004718.3;NP_828858.1;SARS-CoV_nucleocapsid;232 | 0.05 | 0.275 |
| NC_004718.3;NP_828858.1;SARS-CoV_nucleocapsid;233 | 0 | 0.25 |
| NC_004718.3;NP_828858.1;SARS-CoV_nucleocapsid;234 | 0 | 0.175 |
| NC_004718.3;NP_828858.1;SARS-CoV_nucleocapsid;235 | 0 | 0.05 |
| NC_004718.3;NP_828858.1;SARS-CoV_nucleocapsid;236 | 0 | 0.1 |

| | | |
|---|---|---|
| NC_004718.3;NP_828858.1;SARS-CoV_nucleocapsid;237 | 0 | 0.05 |
| NC_004718.3;NP_828858.1;SARS-CoV_nucleocapsid;238 | 0 | 0.025 |
| NC_004718.3;NP_828858.1;SARS-CoV_nucleocapsid;239 | 0.05 | 0.25 |
| NC_004718.3;NP_828858.1;SARS-CoV_nucleocapsid;240 | 0.05 | 0.175 |
| NC_004718.3;NP_828858.1;SARS-CoV_nucleocapsid;241 | 0.25 | 0.225 |
| NC_004718.3;NP_828858.1;SARS-CoV_nucleocapsid;242 | 0.05 | 0.075 |
| NC_004718.3;NP_828858.1;SARS-CoV_nucleocapsid;243 | 0 | 0.025 |
| NC_004718.3;NP_828858.1;SARS-CoV_nucleocapsid;244 | 0 | 0.075 |
| NC_004718.3;NP_828858.1;SARS-CoV_nucleocapsid;245 | 0 | 0.05 |
| NC_004718.3;NP_828858.1;SARS-CoV_nucleocapsid;246 | 0 | 0.075 |
| NC_004718.3;NP_828858.1;SARS-CoV_nucleocapsid;247 | 0 | 0.025 |
| NC_004718.3;NP_828858.1;SARS-CoV_nucleocapsid;248 | 0.05 | 0.025 |
| NC_004718.3;NP_828858.1;SARS-CoV_nucleocapsid;249 | 0 | 0.075 |
| NC_004718.3;NP_828858.1;SARS-CoV_nucleocapsid;250 | 0 | 0.075 |
| NC_004718.3;NP_828858.1;SARS-CoV_nucleocapsid;251 | 0 | 0.3 |
| NC_004718.3;NP_828858.1;SARS-CoV_nucleocapsid;252 | 0 | 0 |
| NC_004718.3;NP_828858.1;SARS-CoV_nucleocapsid;253 | 0.05 | 0 |
| NC_004718.3;NP_828858.1;SARS-CoV_nucleocapsid;254 | 0 | 0 |
| NC_004718.3;NP_828858.1;SARS-CoV_nucleocapsid;255 | 0 | 0 |
| NC_004718.3;NP_828858.1;SARS-CoV_nucleocapsid;256 | 0 | 0 |
| NC_004718.3;NP_828858.1;SARS-CoV_nucleocapsid;257 | 0 | 0.025 |
| NC_004718.3;NP_828858.1;SARS-CoV_nucleocapsid;258 | 0 | 0.05 |
| NC_004718.3;NP_828858.1;SARS-CoV_nucleocapsid;259 | 0.1 | 0.075 |
| NC_004718.3;NP_828858.1;SARS-CoV_nucleocapsid;260 | 0.05 | 0.075 |
| NC_004718.3;NP_828858.1;SARS-CoV_nucleocapsid;261 | 0.05 | 0.1 |
| NC_004718.3;NP_828858.1;SARS-CoV_nucleocapsid;262 | 0.05 | 0.075 |
| NC_004718.3;NP_828858.1;SARS-CoV_nucleocapsid;263 | 0.05 | 0.075 |
| NC_004718.3;NP_828858.1;SARS-CoV_nucleocapsid;264 | 0.05 | 0.075 |
| NC_004718.3;NP_828858.1;SARS-CoV_nucleocapsid;265 | 0.1 | 0.075 |
| NC_004718.3;NP_828858.1;SARS-CoV_nucleocapsid;266 | 0.1 | 0.1 |
| NC_004718.3;NP_828858.1;SARS-CoV_nucleocapsid;267 | 0.1 | 0.1 |
| NC_004718.3;NP_828858.1;SARS-CoV_nucleocapsid;268 | 0.05 | 0 |
| NC_004718.3;NP_828858.1;SARS-CoV_nucleocapsid;269 | 0.05 | 0 |
| NC_004718.3;NP_828858.1;SARS-CoV_nucleocapsid;270 | 0.05 | 0 |
| NC_004718.3;NP_828858.1;SARS-CoV_nucleocapsid;271 | 0.15 | 0.05 |
| NC_004718.3;NP_828858.1;SARS-CoV_nucleocapsid;272 | 0.15 | 0.05 |
| NC_004718.3;NP_828858.1;SARS-CoV_nucleocapsid;273 | 0.15 | 0.05 |
| NC_004718.3;NP_828858.1;SARS-CoV_nucleocapsid;274 | 0.1 | 0.075 |
| NC_004718.3;NP_828858.1;SARS-CoV_nucleocapsid;275 | 0.1 | 0.05 |
| NC_004718.3;NP_828858.1;SARS-CoV_nucleocapsid;276 | 0.05 | 0 |
| NC_004718.3;NP_828858.1;SARS-CoV_nucleocapsid;277 | 0.05 | 0.025 |
| NC_004718.3;NP_828858.1;SARS-CoV_nucleocapsid;278 | 0.05 | 0.025 |
| NC_004718.3;NP_828858.1;SARS-CoV_nucleocapsid;279 | 0.1 | 0.05 |

| | | |
|---|---|---|
| NC_004718.3;NP_828858.1;SARS-CoV_nucleocapsid;280 | 0.1 | 0.1 |
| NC_004718.3;NP_828858.1;SARS-CoV_nucleocapsid;281 | 0.1 | 0.175 |
| NC_004718.3;NP_828858.1;SARS-CoV_nucleocapsid;282 | 0.1 | 0.15 |
| NC_004718.3;NP_828858.1;SARS-CoV_nucleocapsid;283 | 0.1 | 0.1 |
| NC_004718.3;NP_828858.1;SARS-CoV_nucleocapsid;284 | 0.05 | 0.1 |
| NC_004718.3;NP_828858.1;SARS-CoV_nucleocapsid;285 | 0.05 | 0.125 |
| NC_004718.3;NP_828858.1;SARS-CoV_nucleocapsid;286 | 0.05 | 0.1 |
| NC_004718.3;NP_828858.1;SARS-CoV_nucleocapsid;287 | 0.05 | 0.1 |
| NC_004718.3;NP_828858.1;SARS-CoV_nucleocapsid;288 | 0.05 | 0.05 |
| NC_004718.3;NP_828858.1;SARS-CoV_nucleocapsid;289 | 0.05 | 0.025 |
| NC_004718.3;NP_828858.1;SARS-CoV_nucleocapsid;290 | 0 | 0.025 |
| NC_004718.3;NP_828858.1;SARS-CoV_nucleocapsid;291 | 0 | 0 |
| NC_004718.3;NP_828858.1;SARS-CoV_nucleocapsid;292 | 0 | 0 |
| NC_004718.3;NP_828858.1;SARS-CoV_nucleocapsid;293 | 0 | 0 |
| NC_004718.3;NP_828858.1;SARS-CoV_nucleocapsid;294 | 0 | 0 |
| NC_004718.3;NP_828858.1;SARS-CoV_nucleocapsid;295 | 0.05 | 0 |
| NC_004718.3;NP_828858.1;SARS-CoV_nucleocapsid;296 | 0 | 0.05 |
| NC_004718.3;NP_828858.1;SARS-CoV_nucleocapsid;297 | 0 | 0 |
| NC_004718.3;NP_828858.1;SARS-CoV_nucleocapsid;298 | 0.15 | 0.025 |
| NC_004718.3;NP_828858.1;SARS-CoV_nucleocapsid;299 | 0 | 0 |
| NC_004718.3;NP_828858.1;SARS-CoV_nucleocapsid;300 | 0 | 0 |
| NC_004718.3;NP_828858.1;SARS-CoV_nucleocapsid;301 | 0 | 0 |
| NC_004718.3;NP_828858.1;SARS-CoV_nucleocapsid;302 | 0 | 0 |
| NC_004718.3;NP_828858.1;SARS-CoV_nucleocapsid;303 | 0 | 0.025 |
| NC_004718.3;NP_828858.1;SARS-CoV_nucleocapsid;304 | 0 | 0 |
| NC_004718.3;NP_828858.1;SARS-CoV_nucleocapsid;305 | 0 | 0 |
| NC_004718.3;NP_828858.1;SARS-CoV_nucleocapsid;306 | 0 | 0 |
| NC_004718.3;NP_828858.1;SARS-CoV_nucleocapsid;307 | 0 | 0 |
| NC_004718.3;NP_828858.1;SARS-CoV_nucleocapsid;308 | 0 | 0 |
| NC_004718.3;NP_828858.1;SARS-CoV_nucleocapsid;309 | 0 | 0 |
| NC_004718.3;NP_828858.1;SARS-CoV_nucleocapsid;310 | 0 | 0.025 |
| NC_004718.3;NP_828858.1;SARS-CoV_nucleocapsid;311 | 0.05 | 0.025 |
| NC_004718.3;NP_828858.1;SARS-CoV_nucleocapsid;312 | 0 | 0.05 |
| NC_004718.3;NP_828858.1;SARS-CoV_nucleocapsid;313 | 0 | 0.075 |
| NC_004718.3;NP_828858.1;SARS-CoV_nucleocapsid;314 | 0.05 | 0.05 |
| NC_004718.3;NP_828858.1;SARS-CoV_nucleocapsid;315 | 0.05 | 0.05 |
| NC_004718.3;NP_828858.1;SARS-CoV_nucleocapsid;316 | 0 | 0.05 |
| NC_004718.3;NP_828858.1;SARS-CoV_nucleocapsid;317 | 0 | 0.025 |
| NC_004718.3;NP_828858.1;SARS-CoV_nucleocapsid;318 | 0 | 0.025 |
| NC_004718.3;NP_828858.1;SARS-CoV_nucleocapsid;319 | 0 | 0.025 |
| NC_004718.3;NP_828858.1;SARS-CoV_nucleocapsid;320 | 0 | 0 |
| NC_004718.3;NP_828858.1;SARS-CoV_nucleocapsid;321 | 0 | 0 |
| NC_004718.3;NP_828858.1;SARS-CoV_nucleocapsid;322 | 0 | 0 |

| | | |
|---|---|---|
| NC_004718.3;NP_828858.1;SARS-CoV_nucleocapsid;323 | 0 | 0 |
| NC_004718.3;NP_828858.1;SARS-CoV_nucleocapsid;324 | 0 | 0 |
| NC_004718.3;NP_828858.1;SARS-CoV_nucleocapsid;325 | 0 | 0 |
| NC_004718.3;NP_828858.1;SARS-CoV_nucleocapsid;326 | 0 | 0 |
| NC_004718.3;NP_828858.1;SARS-CoV_nucleocapsid;327 | 0 | 0 |
| NC_004718.3;NP_828858.1;SARS-CoV_nucleocapsid;328 | 0 | 0.025 |
| NC_004718.3;NP_828858.1;SARS-CoV_nucleocapsid;329 | 0 | 0.025 |
| NC_004718.3;NP_828858.1;SARS-CoV_nucleocapsid;330 | 0 | 0.05 |
| NC_004718.3;NP_828858.1;SARS-CoV_nucleocapsid;331 | 0 | 0.05 |
| NC_004718.3;NP_828858.1;SARS-CoV_nucleocapsid;332 | 0 | 0.05 |
| NC_004718.3;NP_828858.1;SARS-CoV_nucleocapsid;333 | 0.05 | 0.075 |
| NC_004718.3;NP_828858.1;SARS-CoV_nucleocapsid;334 | 0 | 0.075 |
| NC_004718.3;NP_828858.1;SARS-CoV_nucleocapsid;335 | 0 | 0.1 |
| NC_004718.3;NP_828858.1;SARS-CoV_nucleocapsid;336 | 0.05 | 0.175 |
| NC_004718.3;NP_828858.1;SARS-CoV_nucleocapsid;337 | 0 | 0.075 |
| NC_004718.3;NP_828858.1;SARS-CoV_nucleocapsid;338 | 0.05 | 0.075 |
| NC_004718.3;NP_828858.1;SARS-CoV_nucleocapsid;339 | 0 | 0.025 |
| NC_004718.3;NP_828858.1;SARS-CoV_nucleocapsid;340 | 0 | 0.05 |
| NC_004718.3;NP_828858.1;SARS-CoV_nucleocapsid;341 | 0 | 0.05 |
| NC_004718.3;NP_828858.1;SARS-CoV_nucleocapsid;342 | 0.05 | 0.025 |
| NC_004718.3;NP_828858.1;SARS-CoV_nucleocapsid;343 | 0 | 0.025 |
| NC_004718.3;NP_828858.1;SARS-CoV_nucleocapsid;344 | 0 | 0.075 |
| NC_004718.3;NP_828858.1;SARS-CoV_nucleocapsid;345 | 0 | 0.075 |
| NC_004718.3;NP_828858.1;SARS-CoV_nucleocapsid;346 | 0.05 | 0.05 |
| NC_004718.3;NP_828858.1;SARS-CoV_nucleocapsid;347 | 0 | 0.1 |
| NC_004718.3;NP_828858.1;SARS-CoV_nucleocapsid;348 | 0 | 0.05 |
| NC_004718.3;NP_828858.1;SARS-CoV_nucleocapsid;349 | 0 | 0 |
| NC_004718.3;NP_828858.1;SARS-CoV_nucleocapsid;350 | 0 | 0 |
| NC_004718.3;NP_828858.1;SARS-CoV_nucleocapsid;351 | 0 | 0 |
| NC_004718.3;NP_828858.1;SARS-CoV_nucleocapsid;352 | 0 | 0.025 |
| NC_004718.3;NP_828858.1;SARS-CoV_nucleocapsid;353 | 0 | 0.15 |
| NC_004718.3;NP_828858.1;SARS-CoV_nucleocapsid;354 | 0.05 | 0.125 |
| NC_004718.3;NP_828858.1;SARS-CoV_nucleocapsid;355 | 0 | 0.125 |
| NC_004718.3;NP_828858.1;SARS-CoV_nucleocapsid;356 | 0 | 0.05 |
| NC_004718.3;NP_828858.1;SARS-CoV_nucleocapsid;357 | 0 | 0.25 |
| NC_004718.3;NP_828858.1;SARS-CoV_nucleocapsid;358 | 0.05 | 0.125 |
| NC_004718.3;NP_828858.1;SARS-CoV_nucleocapsid;359 | 0.05 | 0.1 |
| NC_004718.3;NP_828858.1;SARS-CoV_nucleocapsid;360 | 0 | 0.075 |
| NC_004718.3;NP_828858.1;SARS-CoV_nucleocapsid;361 | 0 | 0.075 |
| NC_004718.3;NP_828858.1;SARS-CoV_nucleocapsid;362 | 0 | 0.075 |
| NC_004718.3;NP_828858.1;SARS-CoV_nucleocapsid;363 | 0.05 | 0.125 |
| NC_004718.3;NP_828858.1;SARS-CoV_nucleocapsid;364 | 0.05 | 0.1 |
| NC_004718.3;NP_828858.1;SARS-CoV_nucleocapsid;365 | 0.05 | 0.025 |

| | | |
|---|---|---|
| NC_004718.3;NP_828858.1;SARS-CoV_nucleocapsid;366 | 0 | 0 |
| NC_004718.3;NP_828858.1;SARS-CoV_nucleocapsid;367 | 0 | 0.05 |
| NC_004718.3;NP_828858.1;SARS-CoV_nucleocapsid;368 | 0 | 0.025 |
| NC_004718.3;NP_828858.1;SARS-CoV_nucleocapsid;369 | 0 | 0 |
| NC_004718.3;NP_828858.1;SARS-CoV_nucleocapsid;370 | 0.05 | 0.025 |
| NC_004718.3;NP_828858.1;SARS-CoV_nucleocapsid;371 | 0 | 0.025 |
| NC_004718.3;NP_828858.1;SARS-CoV_nucleocapsid;372 | 0.05 | 0 |
| NC_004718.3;NP_828858.1;SARS-CoV_nucleocapsid;373 | 0.1 | 0.025 |
| NC_004718.3;NP_828858.1;SARS-CoV_nucleocapsid;374 | 0.05 | 0.05 |
| NC_004718.3;NP_828858.1;SARS-CoV_nucleocapsid;375 | 0 | 0.075 |
| NC_004718.3;NP_828858.1;SARS-CoV_nucleocapsid;376 | 0.05 | 0.075 |
| NC_004718.3;NP_828858.1;SARS-CoV_nucleocapsid;377 | 0.05 | 0.125 |
| NC_004718.3;NP_828858.1;SARS-CoV_nucleocapsid;378 | 0.05 | 0.15 |
| NC_004718.3;NP_828858.1;SARS-CoV_nucleocapsid;379 | 0.05 | 0.1 |
| NC_004718.3;NP_828858.1;SARS-CoV_nucleocapsid;380 | 0 | 0.1 |
| NC_004718.3;NP_828858.1;SARS-CoV_nucleocapsid;381 | 0 | 0.15 |
| NC_004718.3;NP_828858.1;SARS-CoV_nucleocapsid;382 | 0 | 0.15 |
| NC_004718.3;NP_828858.1;SARS-CoV_nucleocapsid;383 | 0 | 0.1 |
| NC_004718.3;NP_828858.1;SARS-CoV_nucleocapsid;384 | 0.05 | 0 |
| NC_004718.3;NP_828858.1;SARS-CoV_nucleocapsid;385 | 0 | 0 |
| NC_004718.3;NP_828858.1;SARS-CoV_nucleocapsid;386 | 0 | 0.025 |
| NC_004718.3;NP_828858.1;SARS-CoV_nucleocapsid;387 | 0 | 0.025 |
| NC_004718.3;NP_828858.1;SARS-CoV_nucleocapsid;388 | 0 | 0.075 |
| NC_004718.3;NP_828858.1;SARS-CoV_nucleocapsid;389 | 0 | 0.075 |
| NC_004718.3;NP_828858.1;SARS-CoV_nucleocapsid;390 | 0 | 0.025 |
| NC_004718.3;NP_828858.1;SARS-CoV_nucleocapsid;391 | 0 | 0.1 |
| NC_004718.3;NP_828858.1;SARS-CoV_nucleocapsid;392 | 0 | 0.075 |
| NC_004718.3;NP_828858.1;SARS-CoV_nucleocapsid;393 | 0 | 0.1 |
| NC_004718.3;NP_828858.1;SARS-CoV_nucleocapsid;394 | 0 | 0.175 |
| NC_004718.3;NP_828858.1;SARS-CoV_nucleocapsid;395 | 0 | 0.125 |
| NC_004718.3;NP_828858.1;SARS-CoV_nucleocapsid;396 | 0 | 0.125 |
| NC_004718.3;NP_828858.1;SARS-CoV_nucleocapsid;397 | 0 | 0.025 |
| NC_004718.3;NP_828858.1;SARS-CoV_nucleocapsid;398 | 0 | 0.075 |
| NC_004718.3;NP_828858.1;SARS-CoV_nucleocapsid;399 | 0 | 0.025 |
| NC_004718.3;NP_828858.1;SARS-CoV_nucleocapsid;400 | 0 | 0.075 |
| NC_004718.3;NP_828858.1;SARS-CoV_nucleocapsid;401 | 0 | 0.1 |
| NC_004718.3;NP_828858.1;SARS-CoV_nucleocapsid;402 | 0.05 | 0.05 |
| NC_004718.3;NP_828858.1;SARS-CoV_nucleocapsid;403 | 0 | 0 |
| NC_004718.3;NP_828858.1;SARS-CoV_nucleocapsid;404 | 0 | 0.025 |
| NC_004718.3;NP_828858.1;SARS-CoV_nucleocapsid;405 | 0 | 0.025 |
| NC_004718.3;NP_828858.1;SARS-CoV_nucleocapsid;406 | 0 | 0.025 |
| NC_004718.3;NP_828858.1;SARS-CoV_nucleocapsid;407 | 0 | 0 |
| NC_004718.3;NP_828858.1;SARS-CoV_nucleocapsid;408 | 0 | 0.025 |

| | | |
|---|---|---|
| NC_004718.3;NP_828858.1;SARS-CoV_nucleocapsid;409 | 0 | 0.025 |
| NC_004718.3;NP_828858.1;SARS-CoV_nucleocapsid;410 | 0.05 | 0.025 |
| NC_004718.3;NP_828858.1;SARS-CoV_nucleocapsid;411 | 0.05 | 0 |
| NC_004718.3;NP_828859.1;SARS-CoV_9b;1 | 0.2 | 0.075 |
| NC_004718.3;NP_828859.1;SARS-CoV_9b;2 | 0.2 | 0.125 |
| NC_004718.3;NP_828859.1;SARS-CoV_9b;3 | 0.1 | 0 |
| NC_004718.3;NP_828859.1;SARS-CoV_9b;4 | 0.1 | 0.025 |
| NC_004718.3;NP_828859.1;SARS-CoV_9b;5 | 0.05 | 0.025 |
| NC_004718.3;NP_828859.1;SARS-CoV_9b;6 | 0.05 | 0.025 |
| NC_004718.3;NP_828859.1;SARS-CoV_9b;7 | 0 | 0.025 |
| NC_004718.3;NP_828859.1;SARS-CoV_9b;8 | 0 | 0.025 |
| NC_004718.3;NP_828859.1;SARS-CoV_9b;9 | 0 | 0.025 |
| NC_004718.3;NP_828859.1;SARS-CoV_9b;10 | 0 | 0.025 |
| NC_004718.3;NP_828859.1;SARS-CoV_9b;11 | 0 | 0 |
| NC_004718.3;NP_828859.1;SARS-CoV_9b;12 | 0 | 0 |
| NC_004718.3;NP_828859.1;SARS-CoV_9b;13 | 0 | 0 |
| NC_004718.3;NP_828859.1;SARS-CoV_9b;14 | 0 | 0 |
| NC_004718.3;NP_828859.1;SARS-CoV_9b;15 | 0 | 0 |
| NC_004718.3;NP_828859.1;SARS-CoV_9b;16 | 0 | 0 |
| NC_004718.3;NP_828859.1;SARS-CoV_9b;17 | 0 | 0 |
| NC_004718.3;NP_828859.1;SARS-CoV_9b;18 | 0 | 0 |
| NC_004718.3;NP_828859.1;SARS-CoV_9b;19 | 0 | 0 |
| NC_004718.3;NP_828859.1;SARS-CoV_9b;20 | 0 | 0 |
| NC_004718.3;NP_828859.1;SARS-CoV_9b;21 | 0 | 0 |
| NC_004718.3;NP_828859.1;SARS-CoV_9b;22 | 0 | 0 |
| NC_004718.3;NP_828859.1;SARS-CoV_9b;23 | 0 | 0 |
| NC_004718.3;NP_828859.1;SARS-CoV_9b;24 | 0 | 0 |
| NC_004718.3;NP_828859.1;SARS-CoV_9b;25 | 0 | 0.025 |
| NC_004718.3;NP_828859.1;SARS-CoV_9b;26 | 0 | 0.05 |
| NC_004718.3;NP_828859.1;SARS-CoV_9b;27 | 0 | 0.025 |
| NC_004718.3;NP_828859.1;SARS-CoV_9b;28 | 0 | 0.1 |
| NC_004718.3;NP_828859.1;SARS-CoV_9b;29 | 0 | 0.05 |
| NC_004718.3;NP_828859.1;SARS-CoV_9b;30 | 0 | 0.05 |
| NC_004718.3;NP_828859.1;SARS-CoV_9b;31 | 0 | 0.05 |
| NC_004718.3;NP_828859.1;SARS-CoV_9b;32 | 0 | 0.05 |
| NC_004718.3;NP_828859.1;SARS-CoV_9b;33 | 0 | 0.025 |
| NC_004718.3;NP_828859.1;SARS-CoV_9b;34 | 0 | 0.05 |
| NC_004718.3;NP_828859.1;SARS-CoV_9b;35 | 0 | 0.025 |
| NC_004718.3;NP_828859.1;SARS-CoV_9b;36 | 0 | 0 |
| NC_004718.3;NP_828859.1;SARS-CoV_9b;37 | 0 | 0 |
| NC_004718.3;NP_828859.1;SARS-CoV_9b;38 | 0 | 0 |
| NC_004718.3;NP_828859.1;SARS-CoV_9b;39 | 0 | 0 |
| NC_004718.3;NP_828859.1;SARS-CoV_9b;40 | 0 | 0 |

| | | |
|---|---|---|
| NC_004718.3;NP_828859.1;SARS-CoV_9b;41 | 0 | 0 |
| NC_004718.3;NP_828859.1;SARS-CoV_9b;42 | 0 | 0 |
| NC_004718.3;NP_828859.1;SARS-CoV_9b;43 | 0 | 0 |
| NC_004718.3;NP_828859.1;SARS-CoV_9b;44 | 0 | 0 |
| NC_004718.3;NP_828859.1;SARS-CoV_9b;45 | 0 | 0 |
| NC_004718.3;NP_828859.1;SARS-CoV_9b;46 | 0 | 0 |
| NC_004718.3;NP_828859.1;SARS-CoV_9b;47 | 0 | 0 |
| NC_004718.3;NP_828859.1;SARS-CoV_9b;48 | 0 | 0 |
| NC_004718.3;NP_828859.1;SARS-CoV_9b;49 | 0 | 0 |
| NC_004718.3;NP_828859.1;SARS-CoV_9b;50 | 0.05 | 0 |
| NC_004718.3;NP_828859.1;SARS-CoV_9b;51 | 0 | 0.025 |
| NC_004718.3;NP_828859.1;SARS-CoV_9b;52 | 0.1 | 0.05 |
| NC_004718.3;NP_828859.1;SARS-CoV_9b;53 | 0.1 | 0.025 |
| NC_004718.3;NP_828859.1;SARS-CoV_9b;54 | 0.05 | 0.075 |
| NC_004718.3;NP_828859.1;SARS-CoV_9b;55 | 0.1 | 0.075 |
| NC_004718.3;NP_828859.1;SARS-CoV_9b;56 | 0.1 | 0.075 |
| NC_004718.3;NP_828859.1;SARS-CoV_9b;57 | 0.1 | 0.1 |
| NC_004718.3;NP_828859.1;SARS-CoV_9b;58 | 0.1 | 0 |
| NC_004718.3;NP_828859.1;SARS-CoV_9b;59 | 0.1 | 0.025 |
| NC_004718.3;NP_828859.1;SARS-CoV_9b;60 | 0.05 | 0.05 |
| NC_004718.3;NP_828859.1;SARS-CoV_9b;61 | 0.05 | 0.025 |
| NC_004718.3;NP_828859.1;SARS-CoV_9b;62 | 0 | 0 |
| NC_004718.3;NP_828859.1;SARS-CoV_9b;63 | 0 | 0 |
| NC_004718.3;NP_828859.1;SARS-CoV_9b;64 | 0 | 0 |
| NC_004718.3;NP_828859.1;SARS-CoV_9b;65 | 0 | 0 |
| NC_004718.3;NP_828859.1;SARS-CoV_9b;66 | 0 | 0.025 |
| NC_004718.3;NP_828859.1;SARS-CoV_9b;67 | 0 | 0 |
| NC_004718.3;NP_828859.1;SARS-CoV_9b;68 | 0 | 0 |
| NC_004718.3;NP_828859.1;SARS-CoV_9b;69 | 0 | 0 |
| NC_004718.3;NP_828859.1;SARS-CoV_9b;70 | 0 | 0 |
| NC_004718.3;NP_828859.1;SARS-CoV_9b;71 | 0 | 0 |
| NC_004718.3;NP_828859.1;SARS-CoV_9b;72 | 0 | 0 |
| NC_004718.3;NP_828859.1;SARS-CoV_9b;73 | 0 | 0 |
| NC_004718.3;NP_828859.1;SARS-CoV_9b;74 | 0 | 0 |
| NC_004718.3;NP_828859.1;SARS-CoV_9b;75 | 0 | 0 |
| NC_004718.3;NP_828859.1;SARS-CoV_9b;76 | 0 | 0 |
| NC_004718.3;NP_828859.1;SARS-CoV_9b;77 | 0 | 0 |
| NC_004718.3;NP_828859.1;SARS-CoV_9b;78 | 0.05 | 0 |
| NC_004718.3;NP_828859.1;SARS-CoV_9b;79 | 0 | 0 |
| NC_004718.3;NP_828859.1;SARS-CoV_9b;80 | 0 | 0 |
| NC_004718.3;NP_828859.1;SARS-CoV_9b;81 | 0 | 0 |
| NC_004718.3;NP_828859.1;SARS-CoV_9b;82 | 0 | 0 |
| NC_004718.3;NP_828859.1;SARS-CoV_9b;83 | 0.05 | 0 |

| | | |
|---|---|---|
| NC_004718.3;NP_828859.1;SARS-CoV_9b;84 | 0 | 0 |
| NC_004718.3;NP_828859.1;SARS-CoV_9b;85 | 0.05 | 0 |
| NC_004718.3;NP_828859.1;SARS-CoV_9b;86 | 0.05 | 0 |
| NC_004718.3;NP_828859.1;SARS-CoV_9b;87 | 0.05 | 0 |
| NC_004718.3;NP_849175.1;SARS-CoV_7b;1 | 0 | 0 |
| NC_004718.3;NP_849175.1;SARS-CoV_7b;2 | 0 | 0 |
| NC_004718.3;NP_849175.1;SARS-CoV_7b;3 | 0 | 0 |
| NC_004718.3;NP_849175.1;SARS-CoV_7b;4 | 0 | 0 |
| NC_004718.3;NP_849175.1;SARS-CoV_7b;5 | 0 | 0 |
| NC_004718.3;NP_849175.1;SARS-CoV_7b;6 | 0 | 0 |
| NC_004718.3;NP_849175.1;SARS-CoV_7b;7 | 0 | 0 |
| NC_004718.3;NP_849175.1;SARS-CoV_7b;8 | 0 | 0 |
| NC_004718.3;NP_849175.1;SARS-CoV_7b;9 | 0 | 0 |
| NC_004718.3;NP_849175.1;SARS-CoV_7b;10 | 0 | 0 |
| NC_004718.3;NP_849175.1;SARS-CoV_7b;11 | 0 | 0 |
| NC_004718.3;NP_849175.1;SARS-CoV_7b;12 | 0 | 0 |
| NC_004718.3;NP_849175.1;SARS-CoV_7b;13 | 0 | 0 |
| NC_004718.3;NP_849175.1;SARS-CoV_7b;14 | 0 | 0 |
| NC_004718.3;NP_849175.1;SARS-CoV_7b;15 | 0 | 0 |
| NC_004718.3;NP_849175.1;SARS-CoV_7b;16 | 0 | 0 |
| NC_004718.3;NP_849175.1;SARS-CoV_7b;17 | 0 | 0 |
| NC_004718.3;NP_849175.1;SARS-CoV_7b;18 | 0 | 0 |
| NC_004718.3;NP_849175.1;SARS-CoV_7b;19 | 0 | 0 |
| NC_004718.3;NP_849175.1;SARS-CoV_7b;20 | 0 | 0 |
| NC_004718.3;NP_849175.1;SARS-CoV_7b;21 | 0 | 0 |
| NC_004718.3;NP_849175.1;SARS-CoV_7b;22 | 0 | 0 |
| NC_004718.3;NP_849175.1;SARS-CoV_7b;23 | 0 | 0 |
| NC_004718.3;NP_849175.1;SARS-CoV_7b;24 | 0 | 0 |
| NC_004718.3;NP_849175.1;SARS-CoV_7b;25 | 0 | 0 |
| NC_004718.3;NP_849175.1;SARS-CoV_7b;26 | 0 | 0 |
| NC_004718.3;NP_849175.1;SARS-CoV_7b;27 | 0 | 0 |
| NC_004718.3;NP_849175.1;SARS-CoV_7b;28 | 0 | 0 |
| NC_004718.3;NP_849175.1;SARS-CoV_7b;29 | 0 | 0.05 |
| NC_004718.3;NP_849175.1;SARS-CoV_7b;30 | 0 | 0.025 |
| NC_004718.3;NP_849175.1;SARS-CoV_7b;31 | 0 | 0.025 |
| NC_004718.3;NP_849175.1;SARS-CoV_7b;32 | 0 | 0 |
| NC_004718.3;NP_849175.1;SARS-CoV_7b;33 | 0 | 0 |
| NC_004718.3;NP_849176.1;SARS-CoV_8a;1 | 0 | 0 |
| NC_004718.3;NP_849176.1;SARS-CoV_8a;2 | 0 | 0 |
| NC_004718.3;NP_849176.1;SARS-CoV_8a;3 | 0 | 0 |
| NC_004718.3;NP_849176.1;SARS-CoV_8a;4 | 0 | 0 |
| NC_004718.3;NP_849176.1;SARS-CoV_8a;5 | 0 | 0 |
| NC_004718.3;NP_849176.1;SARS-CoV_8a;6 | 0 | 0 |

| | | |
|---|---|---|
| NC_004718.3;NP_849176.1;SARS-CoV_8a;7 | 0 | 0 |
| NC_004718.3;NP_849176.1;SARS-CoV_8a;8 | 0 | 0 |
| NC_004718.3;NP_849176.1;SARS-CoV_8a;9 | 0 | 0 |
| NC_004718.3;NP_849176.1;SARS-CoV_8a;10 | 0 | 0 |
| NC_004718.3;NP_849176.1;SARS-CoV_8a;11 | 0 | 0 |
| NC_004718.3;NP_849176.1;SARS-CoV_8a;12 | 0 | 0 |
| NC_004718.3;NP_849176.1;SARS-CoV_8a;13 | 0 | 0 |
| NC_004718.3;NP_849176.1;SARS-CoV_8a;14 | 0 | 0 |
| NC_004718.3;NP_849176.1;SARS-CoV_8a;15 | 0 | 0 |
| NC_004718.3;NP_849176.1;SARS-CoV_8a;16 | 0 | 0 |
| NC_004718.3;NP_849176.1;SARS-CoV_8a;17 | 0.05 | 0 |
| NC_004718.3;NP_849176.1;SARS-CoV_8a;18 | 0 | 0.05 |
| NC_004718.3;NP_849176.1;SARS-CoV_8a;19 | 0.1 | 0.025 |
| NC_004718.3;NP_849176.1;SARS-CoV_8a;20 | 0.1 | 0.025 |
| NC_004718.3;NP_849176.1;SARS-CoV_8a;21 | 0.1 | 0 |
| NC_004718.3;NP_849176.1;SARS-CoV_8a;22 | 0.05 | 0 |
| NC_004718.3;NP_849176.1;SARS-CoV_8a;23 | 0.05 | 0 |
| NC_004718.3;NP_849176.1;SARS-CoV_8a;24 | 0.1 | 0.025 |
| NC_004718.3;NP_849176.1;SARS-CoV_8a;25 | 0.1 | 0 |
| NC_004718.3;NP_849176.1;SARS-CoV_8a;26 | 0 | 0 |
| NC_004718.3;NP_849176.1;SARS-CoV_8a;27 | 0 | 0 |
| NC_004718.3;NP_849176.1;SARS-CoV_8a;28 | 0 | 0 |
| NC_004718.3;NP_849177.1;SARS-CoV_8b;1 | 0 | 0 |
| NC_004718.3;NP_849177.1;SARS-CoV_8b;2 | 0 | 0 |
| NC_004718.3;NP_849177.1;SARS-CoV_8b;3 | 0 | 0 |
| NC_004718.3;NP_849177.1;SARS-CoV_8b;4 | 0 | 0 |
| NC_004718.3;NP_849177.1;SARS-CoV_8b;5 | 0 | 0 |
| NC_004718.3;NP_849177.1;SARS-CoV_8b;6 | 0 | 0 |
| NC_004718.3;NP_849177.1;SARS-CoV_8b;7 | 0 | 0 |
| NC_004718.3;NP_849177.1;SARS-CoV_8b;8 | 0 | 0 |
| NC_004718.3;NP_849177.1;SARS-CoV_8b;9 | 0 | 0 |
| NC_004718.3;NP_849177.1;SARS-CoV_8b;10 | 0 | 0 |
| NC_004718.3;NP_849177.1;SARS-CoV_8b;11 | 0 | 0 |
| NC_004718.3;NP_849177.1;SARS-CoV_8b;12 | 0 | 0 |
| NC_004718.3;NP_849177.1;SARS-CoV_8b;13 | 0 | 0 |
| NC_004718.3;NP_849177.1;SARS-CoV_8b;14 | 0 | 0 |
| NC_004718.3;NP_849177.1;SARS-CoV_8b;15 | 0 | 0 |
| NC_004718.3;NP_849177.1;SARS-CoV_8b;16 | 0 | 0 |
| NC_004718.3;NP_849177.1;SARS-CoV_8b;17 | 0 | 0 |
| NC_004718.3;NP_849177.1;SARS-CoV_8b;18 | 0 | 0 |
| NC_004718.3;NP_849177.1;SARS-CoV_8b;19 | 0 | 0 |
| NC_004718.3;NP_849177.1;SARS-CoV_8b;20 | 0 | 0 |
| NC_004718.3;NP_849177.1;SARS-CoV_8b;21 | 0 | 0 |

| | | |
|---|---|---|
| NC_004718.3;NP_849177.1;SARS-CoV_8b;22 | 0 | 0 |
| NC_004718.3;NP_849177.1;SARS-CoV_8b;23 | 0 | 0 |
| NC_004718.3;NP_849177.1;SARS-CoV_8b;24 | 0 | 0 |
| NC_004718.3;NP_849177.1;SARS-CoV_8b;25 | 0 | 0 |
| NC_004718.3;NP_849177.1;SARS-CoV_8b;26 | 0 | 0 |
| NC_004718.3;NP_849177.1;SARS-CoV_8b;27 | 0 | 0 |
| NC_004718.3;NP_849177.1;SARS-CoV_8b;28 | 0 | 0 |
| NC_004718.3;NP_849177.1;SARS-CoV_8b;29 | 0 | 0 |
| NC_004718.3;NP_849177.1;SARS-CoV_8b;30 | 0 | 0 |
| NC_004718.3;NP_849177.1;SARS-CoV_8b;31 | 0 | 0 |
| NC_004718.3;NP_849177.1;SARS-CoV_8b;32 | 0.15 | 0.05 |
| NC_004718.3;NP_849177.1;SARS-CoV_8b;33 | 0.05 | 0 |
| NC_004718.3;NP_849177.1;SARS-CoV_8b;34 | 0.05 | 0 |
| NC_004718.3;NP_849177.1;SARS-CoV_8b;35 | 0.1 | 0.125 |
| NC_004718.3;NP_849177.1;SARS-CoV_8b;36 | 0.15 | 0.125 |
| NC_004718.3;NP_849177.1;SARS-CoV_8b;37 | 0.15 | 0.125 |
| NC_004718.3;NP_849177.1;SARS-CoV_8b;38 | 0.15 | 0.15 |
| NC_004718.3;NP_849177.1;SARS-CoV_8b;39 | 0.2 | 0.15 |
| NC_004718.3;NP_849177.1;SARS-CoV_8b;40 | 0.15 | 0.225 |
| NC_004718.3;NP_849177.1;SARS-CoV_8b;41 | 0.2 | 0.35 |
| NC_004718.3;NP_849177.1;SARS-CoV_8b;42 | 0.15 | 0.15 |
| NC_004718.3;NP_849177.1;SARS-CoV_8b;43 | 0.15 | 0.05 |
| NC_004718.3;NP_849177.1;SARS-CoV_8b;44 | 0 | 0 |
| NC_004718.3;NP_849177.1;SARS-CoV_8b;45 | 0 | 0 |
| NC_004718.3;NP_849177.1;SARS-CoV_8b;46 | 0 | 0 |
| NC_004718.3;NP_849177.1;SARS-CoV_8b;47 | 0 | 0 |
| NC_004718.3;NP_849177.1;SARS-CoV_8b;48 | 0 | 0 |
| NC_004718.3;NP_849177.1;SARS-CoV_8b;49 | 0 | 0 |
| NC_004718.3;NP_849177.1;SARS-CoV_8b;50 | 0 | 0 |
| NC_004718.3;NP_849177.1;SARS-CoV_8b;51 | 0 | 0 |
| NC_004718.3;NP_849177.1;SARS-CoV_8b;52 | 0 | 0 |
| NC_004718.3;NP_849177.1;SARS-CoV_8b;53 | 0 | 0 |
| NC_004718.3;NP_849177.1;SARS-CoV_8b;54 | 0 | 0 |
| NC_004718.3;NP_849177.1;SARS-CoV_8b;55 | 0 | 0 |
| NC_004718.3;NP_849177.1;SARS-CoV_8b;56 | 0.05 | 0.025 |
| NC_004718.3;NP_849177.1;SARS-CoV_8b;57 | 0 | 0 |
| NC_004718.3;NP_849177.1;SARS-CoV_8b;58 | 0 | 0 |
| NC_004718.3;NP_849177.1;SARS-CoV_8b;59 | 0 | 0.025 |
| NC_004718.3;NP_849177.1;SARS-CoV_8b;60 | 0 | 0 |
| NC_004718.3;NP_849177.1;SARS-CoV_8b;61 | 0 | 0 |
| NC_004718.3;NP_849177.1;SARS-CoV_8b;62 | 0 | 0 |
| NC_004718.3;NP_849177.1;SARS-CoV_8b;63 | 0 | 0 |
| NC_004718.3;NP_849177.1;SARS-CoV_8b;64 | 0 | 0 |

| | | |
|---|---|---|
| NC_004718.3;NP_849177.1;SARS-CoV_8b;65 | 0 | 0 |
| NC_004718.3;NP_849177.1;SARS-CoV_8b;66 | 0 | 0 |
| NC_004718.3;NP_849177.1;SARS-CoV_8b;67 | 0 | 0 |
| NC_004718.3;NP_849177.1;SARS-CoV_8b;68 | 0 | 0.025 |
| NC_004718.3;NP_849177.1;SARS-CoV_8b;69 | 0 | 0.025 |
| NC_004718.3;NP_849177.1;SARS-CoV_8b;70 | 0 | 0.025 |
| NC_004718.3;NP_849177.1;SARS-CoV_8b;71 | 0 | 0.025 |
| NC_004718.3;NP_849177.1;SARS-CoV_8b;72 | 0 | 0 |
| NC_004718.3;NP_849177.1;SARS-CoV_8b;73 | 0 | 0.025 |
| NC_019843.3;YP_009047202.1;MERS_1ab;1 | 0 | 0 |
| NC_019843.3;YP_009047202.1;MERS_1ab;2 | 0 | 0 |
| NC_019843.3;YP_009047202.1;MERS_1ab;3 | 0 | 0 |
| NC_019843.3;YP_009047202.1;MERS_1ab;4 | 0.05 | 0 |
| NC_019843.3;YP_009047202.1;MERS_1ab;5 | 0 | 0 |
| NC_019843.3;YP_009047202.1;MERS_1ab;6 | 0 | 0 |
| NC_019843.3;YP_009047202.1;MERS_1ab;7 | 0 | 0 |
| NC_019843.3;YP_009047202.1;MERS_1ab;8 | 0.05 | 0 |
| NC_019843.3;YP_009047202.1;MERS_1ab;9 | 0 | 0.025 |
| NC_019843.3;YP_009047202.1;MERS_1ab;10 | 0 | 0.025 |
| NC_019843.3;YP_009047202.1;MERS_1ab;11 | 0 | 0.025 |
| NC_019843.3;YP_009047202.1;MERS_1ab;12 | 0 | 0.025 |
| NC_019843.3;YP_009047202.1;MERS_1ab;13 | 0 | 0.05 |
| NC_019843.3;YP_009047202.1;MERS_1ab;14 | 0.05 | 0 |
| NC_019843.3;YP_009047202.1;MERS_1ab;15 | 0 | 0 |
| NC_019843.3;YP_009047202.1;MERS_1ab;16 | 0.05 | 0 |
| NC_019843.3;YP_009047202.1;MERS_1ab;17 | 0 | 0 |
| NC_019843.3;YP_009047202.1;MERS_1ab;18 | 0 | 0 |
| NC_019843.3;YP_009047202.1;MERS_1ab;19 | 0.1 | 0.025 |
| NC_019843.3;YP_009047202.1;MERS_1ab;20 | 0 | 0.025 |
| NC_019843.3;YP_009047202.1;MERS_1ab;21 | 0 | 0.025 |
| NC_019843.3;YP_009047202.1;MERS_1ab;22 | 0 | 0.025 |
| NC_019843.3;YP_009047202.1;MERS_1ab;23 | 0 | 0.025 |
| NC_019843.3;YP_009047202.1;MERS_1ab;24 | 0 | 0 |
| NC_019843.3;YP_009047202.1;MERS_1ab;25 | 0 | 0 |
| NC_019843.3;YP_009047202.1;MERS_1ab;26 | 0 | 0 |
| NC_019843.3;YP_009047202.1;MERS_1ab;27 | 0 | 0 |
| NC_019843.3;YP_009047202.1;MERS_1ab;28 | 0 | 0 |
| NC_019843.3;YP_009047202.1;MERS_1ab;29 | 0.05 | 0 |
| NC_019843.3;YP_009047202.1;MERS_1ab;30 | 0.05 | 0 |
| NC_019843.3;YP_009047202.1;MERS_1ab;31 | 0.05 | 0 |
| NC_019843.3;YP_009047202.1;MERS_1ab;32 | 0.05 | 0 |
| NC_019843.3;YP_009047202.1;MERS_1ab;33 | 0.05 | 0 |
| NC_019843.3;YP_009047202.1;MERS_1ab;34 | 0.05 | 0.025 |

| | | |
|---|---|---|
| NC_019843.3;YP_009047202.1;MERS_1ab;35 | 0.05 | 0 |
| NC_019843.3;YP_009047202.1;MERS_1ab;36 | 0.15 | 0.025 |
| NC_019843.3;YP_009047202.1;MERS_1ab;37 | 0.05 | 0.05 |
| NC_019843.3;YP_009047202.1;MERS_1ab;38 | 0.05 | 0.025 |
| NC_019843.3;YP_009047202.1;MERS_1ab;39 | 0.05 | 0.025 |
| NC_019843.3;YP_009047202.1;MERS_1ab;40 | 0 | 0.025 |
| NC_019843.3;YP_009047202.1;MERS_1ab;41 | 0 | 0.025 |
| NC_019843.3;YP_009047202.1;MERS_1ab;42 | 0 | 0.025 |
| NC_019843.3;YP_009047202.1;MERS_1ab;43 | 0 | 0.025 |
| NC_019843.3;YP_009047202.1;MERS_1ab;44 | 0 | 0.025 |
| NC_019843.3;YP_009047202.1;MERS_1ab;45 | 0 | 0.025 |
| NC_019843.3;YP_009047202.1;MERS_1ab;46 | 0.05 | 0 |
| NC_019843.3;YP_009047202.1;MERS_1ab;47 | 0 | 0.025 |
| NC_019843.3;YP_009047202.1;MERS_1ab;48 | 0 | 0.025 |
| NC_019843.3;YP_009047202.1;MERS_1ab;49 | 0 | 0 |
| NC_019843.3;YP_009047202.1;MERS_1ab;50 | 0 | 0 |
| NC_019843.3;YP_009047202.1;MERS_1ab;51 | 0 | 0 |
| NC_019843.3;YP_009047202.1;MERS_1ab;52 | 0 | 0 |
| NC_019843.3;YP_009047202.1;MERS_1ab;53 | 0 | 0 |
| NC_019843.3;YP_009047202.1;MERS_1ab;54 | 0 | 0 |
| NC_019843.3;YP_009047202.1;MERS_1ab;55 | 0 | 0 |
| NC_019843.3;YP_009047202.1;MERS_1ab;56 | 0 | 0 |
| NC_019843.3;YP_009047202.1;MERS_1ab;57 | 0 | 0 |
| NC_019843.3;YP_009047202.1;MERS_1ab;58 | 0 | 0 |
| NC_019843.3;YP_009047202.1;MERS_1ab;59 | 0 | 0 |
| NC_019843.3;YP_009047202.1;MERS_1ab;60 | 0 | 0 |
| NC_019843.3;YP_009047202.1;MERS_1ab;61 | 0 | 0 |
| NC_019843.3;YP_009047202.1;MERS_1ab;62 | 0 | 0 |
| NC_019843.3;YP_009047202.1;MERS_1ab;63 | 0 | 0 |
| NC_019843.3;YP_009047202.1;MERS_1ab;64 | 0 | 0 |
| NC_019843.3;YP_009047202.1;MERS_1ab;65 | 0 | 0 |
| NC_019843.3;YP_009047202.1;MERS_1ab;66 | 0 | 0 |
| NC_019843.3;YP_009047202.1;MERS_1ab;67 | 0 | 0 |
| NC_019843.3;YP_009047202.1;MERS_1ab;68 | 0 | 0 |
| NC_019843.3;YP_009047202.1;MERS_1ab;69 | 0 | 0 |
| NC_019843.3;YP_009047202.1;MERS_1ab;70 | 0 | 0 |
| NC_019843.3;YP_009047202.1;MERS_1ab;71 | 0 | 0 |
| NC_019843.3;YP_009047202.1;MERS_1ab;72 | 0 | 0 |
| NC_019843.3;YP_009047202.1;MERS_1ab;73 | 0 | 0 |
| NC_019843.3;YP_009047202.1;MERS_1ab;74 | 0 | 0 |
| NC_019843.3;YP_009047202.1;MERS_1ab;75 | 0 | 0 |
| NC_019843.3;YP_009047202.1;MERS_1ab;76 | 0 | 0 |
| NC_019843.3;YP_009047202.1;MERS_1ab;77 | 0 | 0 |

| | | |
|---|---|---|
| NC_019843.3;YP_009047202.1;MERS_1ab;78 | 0 | 0 |
| NC_019843.3;YP_009047202.1;MERS_1ab;79 | 0 | 0 |
| NC_019843.3;YP_009047202.1;MERS_1ab;80 | 0 | 0 |
| NC_019843.3;YP_009047202.1;MERS_1ab;81 | 0 | 0 |
| NC_019843.3;YP_009047202.1;MERS_1ab;82 | 0 | 0 |
| NC_019843.3;YP_009047202.1;MERS_1ab;83 | 0 | 0 |
| NC_019843.3;YP_009047202.1;MERS_1ab;84 | 0 | 0 |
| NC_019843.3;YP_009047202.1;MERS_1ab;85 | 0 | 0 |
| NC_019843.3;YP_009047202.1;MERS_1ab;86 | 0 | 0 |
| NC_019843.3;YP_009047202.1;MERS_1ab;87 | 0 | 0 |
| NC_019843.3;YP_009047202.1;MERS_1ab;88 | 0 | 0 |
| NC_019843.3;YP_009047202.1;MERS_1ab;89 | 0 | 0 |
| NC_019843.3;YP_009047202.1;MERS_1ab;90 | 0 | 0 |
| NC_019843.3;YP_009047202.1;MERS_1ab;91 | 0 | 0 |
| NC_019843.3;YP_009047202.1;MERS_1ab;92 | 0.1 | 0.025 |
| NC_019843.3;YP_009047202.1;MERS_1ab;93 | 0 | 0 |
| NC_019843.3;YP_009047202.1;MERS_1ab;94 | 0 | 0 |
| NC_019843.3;YP_009047202.1;MERS_1ab;95 | 0 | 0 |
| NC_019843.3;YP_009047202.1;MERS_1ab;96 | 0 | 0 |
| NC_019843.3;YP_009047202.1;MERS_1ab;97 | 0 | 0 |
| NC_019843.3;YP_009047202.1;MERS_1ab;98 | 0.05 | 0.075 |
| NC_019843.3;YP_009047202.1;MERS_1ab;99 | 0 | 0.025 |
| NC_019843.3;YP_009047202.1;MERS_1ab;100 | 0 | 0.025 |
| NC_019843.3;YP_009047202.1;MERS_1ab;101 | 0.05 | 0.025 |
| NC_019843.3;YP_009047202.1;MERS_1ab;102 | 0.05 | 0 |
| NC_019843.3;YP_009047202.1;MERS_1ab;103 | 0.05 | 0.025 |
| NC_019843.3;YP_009047202.1;MERS_1ab;104 | 0.05 | 0.05 |
| NC_019843.3;YP_009047202.1;MERS_1ab;105 | 0 | 0 |
| NC_019843.3;YP_009047202.1;MERS_1ab;106 | 0 | 0 |
| NC_019843.3;YP_009047202.1;MERS_1ab;107 | 0.05 | 0.1 |
| NC_019843.3;YP_009047202.1;MERS_1ab;108 | 0 | 0 |
| NC_019843.3;YP_009047202.1;MERS_1ab;109 | 0 | 0 |
| NC_019843.3;YP_009047202.1;MERS_1ab;110 | 0 | 0 |
| NC_019843.3;YP_009047202.1;MERS_1ab;111 | 0 | 0 |
| NC_019843.3;YP_009047202.1;MERS_1ab;112 | 0 | 0 |
| NC_019843.3;YP_009047202.1;MERS_1ab;113 | 0 | 0 |
| NC_019843.3;YP_009047202.1;MERS_1ab;114 | 0 | 0 |
| NC_019843.3;YP_009047202.1;MERS_1ab;115 | 0 | 0.025 |
| NC_019843.3;YP_009047202.1;MERS_1ab;116 | 0 | 0 |
| NC_019843.3;YP_009047202.1;MERS_1ab;117 | 0 | 0 |
| NC_019843.3;YP_009047202.1;MERS_1ab;118 | 0 | 0 |
| NC_019843.3;YP_009047202.1;MERS_1ab;119 | 0 | 0 |
| NC_019843.3;YP_009047202.1;MERS_1ab;120 | 0.05 | 0 |

| | | |
|---|---|---|
| NC_019843.3;YP_009047202.1;MERS_1ab;121 | 0 | 0 |
| NC_019843.3;YP_009047202.1;MERS_1ab;122 | 0 | 0 |
| NC_019843.3;YP_009047202.1;MERS_1ab;123 | 0 | 0 |
| NC_019843.3;YP_009047202.1;MERS_1ab;124 | 0 | 0 |
| NC_019843.3;YP_009047202.1;MERS_1ab;125 | 0 | 0.025 |
| NC_019843.3;YP_009047202.1;MERS_1ab;126 | 0.05 | 0 |
| NC_019843.3;YP_009047202.1;MERS_1ab;127 | 0.05 | 0 |
| NC_019843.3;YP_009047202.1;MERS_1ab;128 | 0.1 | 0 |
| NC_019843.3;YP_009047202.1;MERS_1ab;129 | 0 | 0 |
| NC_019843.3;YP_009047202.1;MERS_1ab;130 | 0 | 0 |
| NC_019843.3;YP_009047202.1;MERS_1ab;131 | 0 | 0 |
| NC_019843.3;YP_009047202.1;MERS_1ab;132 | 0 | 0 |
| NC_019843.3;YP_009047202.1;MERS_1ab;133 | 0 | 0 |
| NC_019843.3;YP_009047202.1;MERS_1ab;134 | 0 | 0 |
| NC_019843.3;YP_009047202.1;MERS_1ab;135 | 0 | 0 |
| NC_019843.3;YP_009047202.1;MERS_1ab;136 | 0 | 0 |
| NC_019843.3;YP_009047202.1;MERS_1ab;137 | 0 | 0 |
| NC_019843.3;YP_009047202.1;MERS_1ab;138 | 0 | 0 |
| NC_019843.3;YP_009047202.1;MERS_1ab;139 | 0 | 0 |
| NC_019843.3;YP_009047202.1;MERS_1ab;140 | 0 | 0 |
| NC_019843.3;YP_009047202.1;MERS_1ab;141 | 0 | 0 |
| NC_019843.3;YP_009047202.1;MERS_1ab;142 | 0 | 0 |
| NC_019843.3;YP_009047202.1;MERS_1ab;143 | 0 | 0 |
| NC_019843.3;YP_009047202.1;MERS_1ab;144 | 0 | 0 |
| NC_019843.3;YP_009047202.1;MERS_1ab;145 | 0 | 0 |
| NC_019843.3;YP_009047202.1;MERS_1ab;146 | 0 | 0 |
| NC_019843.3;YP_009047202.1;MERS_1ab;147 | 0 | 0 |
| NC_019843.3;YP_009047202.1;MERS_1ab;148 | 0 | 0 |
| NC_019843.3;YP_009047202.1;MERS_1ab;149 | 0 | 0.025 |
| NC_019843.3;YP_009047202.1;MERS_1ab;150 | 0 | 0.025 |
| NC_019843.3;YP_009047202.1;MERS_1ab;151 | 0 | 0.025 |
| NC_019843.3;YP_009047202.1;MERS_1ab;152 | 0 | 0.025 |
| NC_019843.3;YP_009047202.1;MERS_1ab;153 | 0 | 0.025 |
| NC_019843.3;YP_009047202.1;MERS_1ab;154 | 0 | 0.025 |
| NC_019843.3;YP_009047202.1;MERS_1ab;155 | 0 | 0.025 |
| NC_019843.3;YP_009047202.1;MERS_1ab;156 | 0 | 0.025 |
| NC_019843.3;YP_009047202.1;MERS_1ab;157 | 0 | 0.025 |
| NC_019843.3;YP_009047202.1;MERS_1ab;158 | 0 | 0.025 |
| NC_019843.3;YP_009047202.1;MERS_1ab;159 | 0 | 0 |
| NC_019843.3;YP_009047202.1;MERS_1ab;160 | 0 | 0.05 |
| NC_019843.3;YP_009047202.1;MERS_1ab;161 | 0 | 0 |
| NC_019843.3;YP_009047202.1;MERS_1ab;162 | 0 | 0 |
| NC_019843.3;YP_009047202.1;MERS_1ab;163 | 0.05 | 0.025 |

| | | |
|---|---|---|
| NC_019843.3;YP_009047202.1;MERS_1ab;164 | 0.1 | 0.025 |
| NC_019843.3;YP_009047202.1;MERS_1ab;165 | 0.1 | 0.025 |
| NC_019843.3;YP_009047202.1;MERS_1ab;166 | 0.1 | 0.05 |
| NC_019843.3;YP_009047202.1;MERS_1ab;167 | 0.1 | 0.075 |
| NC_019843.3;YP_009047202.1;MERS_1ab;168 | 0.1 | 0.05 |
| NC_019843.3;YP_009047202.1;MERS_1ab;169 | 0.1 | 0.025 |
| NC_019843.3;YP_009047202.1;MERS_1ab;170 | 0.1 | 0.05 |
| NC_019843.3;YP_009047202.1;MERS_1ab;171 | 0 | 0 |
| NC_019843.3;YP_009047202.1;MERS_1ab;172 | 0 | 0 |
| NC_019843.3;YP_009047202.1;MERS_1ab;173 | 0 | 0.075 |
| NC_019843.3;YP_009047202.1;MERS_1ab;174 | 0 | 0 |
| NC_019843.3;YP_009047202.1;MERS_1ab;175 | 0 | 0 |
| NC_019843.3;YP_009047202.1;MERS_1ab;176 | 0.05 | 0 |
| NC_019843.3;YP_009047202.1;MERS_1ab;177 | 0 | 0 |
| NC_019843.3;YP_009047202.1;MERS_1ab;178 | 0 | 0 |
| NC_019843.3;YP_009047202.1;MERS_1ab;179 | 0 | 0.05 |
| NC_019843.3;YP_009047202.1;MERS_1ab;180 | 0 | 0.025 |
| NC_019843.3;YP_009047202.1;MERS_1ab;181 | 0 | 0 |
| NC_019843.3;YP_009047202.1;MERS_1ab;182 | 0 | 0 |
| NC_019843.3;YP_009047202.1;MERS_1ab;183 | 0 | 0 |
| NC_019843.3;YP_009047202.1;MERS_1ab;184 | 0.05 | 0 |
| NC_019843.3;YP_009047202.1;MERS_1ab;185 | 0 | 0 |
| NC_019843.3;YP_009047202.1;MERS_1ab;186 | 0.1 | 0 |
| NC_019843.3;YP_009047202.1;MERS_1ab;187 | 0.1 | 0.025 |
| NC_019843.3;YP_009047202.1;MERS_1ab;188 | 0.05 | 0.025 |
| NC_019843.3;YP_009047202.1;MERS_1ab;189 | 0 | 0 |
| NC_019843.3;YP_009047202.1;MERS_1ab;190 | 0 | 0 |
| NC_019843.3;YP_009047202.1;MERS_1ab;191 | 0 | 0 |
| NC_019843.3;YP_009047202.1;MERS_1ab;192 | 0 | 0 |
| NC_019843.3;YP_009047202.1;MERS_1ab;193 | 0 | 0 |
| NC_019843.3;YP_009047202.1;MERS_1ab;194 | 0 | 0 |
| NC_019843.3;YP_009047202.1;MERS_1ab;195 | 0 | 0 |
| NC_019843.3;YP_009047202.1;MERS_1ab;196 | 0.05 | 0 |
| NC_019843.3;YP_009047202.1;MERS_1ab;197 | 0.1 | 0.025 |
| NC_019843.3;YP_009047202.1;MERS_1ab;198 | 0 | 0.025 |
| NC_019843.3;YP_009047202.1;MERS_1ab;199 | 0.05 | 0.05 |
| NC_019843.3;YP_009047202.1;MERS_1ab;200 | 0 | 0.025 |
| NC_019843.3;YP_009047202.1;MERS_1ab;201 | 0 | 0 |
| NC_019843.3;YP_009047202.1;MERS_1ab;202 | 0 | 0 |
| NC_019843.3;YP_009047202.1;MERS_1ab;203 | 0 | 0 |
| NC_019843.3;YP_009047202.1;MERS_1ab;204 | 0 | 0 |
| NC_019843.3;YP_009047202.1;MERS_1ab;205 | 0 | 0 |
| NC_019843.3;YP_009047202.1;MERS_1ab;206 | 0 | 0 |

| | | |
|---|---|---|
| NC_019843.3;YP_009047202.1;MERS_1ab;207 | 0 | 0 |
| NC_019843.3;YP_009047202.1;MERS_1ab;208 | 0 | 0 |
| NC_019843.3;YP_009047202.1;MERS_1ab;209 | 0 | 0 |
| NC_019843.3;YP_009047202.1;MERS_1ab;210 | 0 | 0 |
| NC_019843.3;YP_009047202.1;MERS_1ab;211 | 0 | 0 |
| NC_019843.3;YP_009047202.1;MERS_1ab;212 | 0 | 0 |
| NC_019843.3;YP_009047202.1;MERS_1ab;213 | 0 | 0 |
| NC_019843.3;YP_009047202.1;MERS_1ab;214 | 0 | 0 |
| NC_019843.3;YP_009047202.1;MERS_1ab;215 | 0 | 0 |
| NC_019843.3;YP_009047202.1;MERS_1ab;216 | 0 | 0 |
| NC_019843.3;YP_009047202.1;MERS_1ab;217 | 0 | 0 |
| NC_019843.3;YP_009047202.1;MERS_1ab;218 | 0 | 0 |
| NC_019843.3;YP_009047202.1;MERS_1ab;219 | 0.05 | 0 |
| NC_019843.3;YP_009047202.1;MERS_1ab;220 | 0 | 0 |
| NC_019843.3;YP_009047202.1;MERS_1ab;221 | 0 | 0 |
| NC_019843.3;YP_009047202.1;MERS_1ab;222 | 0 | 0 |
| NC_019843.3;YP_009047202.1;MERS_1ab;223 | 0 | 0 |
| NC_019843.3;YP_009047202.1;MERS_1ab;224 | 0 | 0 |
| NC_019843.3;YP_009047202.1;MERS_1ab;225 | 0 | 0 |
| NC_019843.3;YP_009047202.1;MERS_1ab;226 | 0 | 0 |
| NC_019843.3;YP_009047202.1;MERS_1ab;227 | 0 | 0 |
| NC_019843.3;YP_009047202.1;MERS_1ab;228 | 0 | 0 |
| NC_019843.3;YP_009047202.1;MERS_1ab;229 | 0 | 0 |
| NC_019843.3;YP_009047202.1;MERS_1ab;230 | 0 | 0 |
| NC_019843.3;YP_009047202.1;MERS_1ab;231 | 0 | 0 |
| NC_019843.3;YP_009047202.1;MERS_1ab;232 | 0.05 | 0 |
| NC_019843.3;YP_009047202.1;MERS_1ab;233 | 0 | 0 |
| NC_019843.3;YP_009047202.1;MERS_1ab;234 | 0 | 0 |
| NC_019843.3;YP_009047202.1;MERS_1ab;235 | 0 | 0 |
| NC_019843.3;YP_009047202.1;MERS_1ab;236 | 0 | 0 |
| NC_019843.3;YP_009047202.1;MERS_1ab;237 | 0 | 0 |
| NC_019843.3;YP_009047202.1;MERS_1ab;238 | 0 | 0 |
| NC_019843.3;YP_009047202.1;MERS_1ab;239 | 0 | 0 |
| NC_019843.3;YP_009047202.1;MERS_1ab;240 | 0 | 0 |
| NC_019843.3;YP_009047202.1;MERS_1ab;241 | 0 | 0 |
| NC_019843.3;YP_009047202.1;MERS_1ab;242 | 0 | 0 |
| NC_019843.3;YP_009047202.1;MERS_1ab;243 | 0 | 0 |
| NC_019843.3;YP_009047202.1;MERS_1ab;244 | 0 | 0.025 |
| NC_019843.3;YP_009047202.1;MERS_1ab;245 | 0 | 0 |
| NC_019843.3;YP_009047202.1;MERS_1ab;246 | 0 | 0 |
| NC_019843.3;YP_009047202.1;MERS_1ab;247 | 0 | 0.025 |
| NC_019843.3;YP_009047202.1;MERS_1ab;248 | 0 | 0 |
| NC_019843.3;YP_009047202.1;MERS_1ab;249 | 0 | 0 |

| | | |
|---|---|---|
| NC_019843.3;YP_009047202.1;MERS_1ab;250 | 0 | 0 |
| NC_019843.3;YP_009047202.1;MERS_1ab;251 | 0 | 0.05 |
| NC_019843.3;YP_009047202.1;MERS_1ab;252 | 0 | 0.05 |
| NC_019843.3;YP_009047202.1;MERS_1ab;253 | 0 | 0.025 |
| NC_019843.3;YP_009047202.1;MERS_1ab;254 | 0 | 0 |
| NC_019843.3;YP_009047202.1;MERS_1ab;255 | 0 | 0 |
| NC_019843.3;YP_009047202.1;MERS_1ab;256 | 0 | 0 |
| NC_019843.3;YP_009047202.1;MERS_1ab;257 | 0 | 0 |
| NC_019843.3;YP_009047202.1;MERS_1ab;258 | 0 | 0 |
| NC_019843.3;YP_009047202.1;MERS_1ab;259 | 0.05 | 0.025 |
| NC_019843.3;YP_009047202.1;MERS_1ab;260 | 0.05 | 0.025 |
| NC_019843.3;YP_009047202.1;MERS_1ab;261 | 0 | 0 |
| NC_019843.3;YP_009047202.1;MERS_1ab;262 | 0 | 0.025 |
| NC_019843.3;YP_009047202.1;MERS_1ab;263 | 0 | 0.025 |
| NC_019843.3;YP_009047202.1;MERS_1ab;264 | 0 | 0.025 |
| NC_019843.3;YP_009047202.1;MERS_1ab;265 | 0 | 0.025 |
| NC_019843.3;YP_009047202.1;MERS_1ab;266 | 0 | 0.025 |
| NC_019843.3;YP_009047202.1;MERS_1ab;267 | 0.05 | 0.05 |
| NC_019843.3;YP_009047202.1;MERS_1ab;268 | 0 | 0 |
| NC_019843.3;YP_009047202.1;MERS_1ab;269 | 0 | 0.025 |
| NC_019843.3;YP_009047202.1;MERS_1ab;270 | 0 | 0 |
| NC_019843.3;YP_009047202.1;MERS_1ab;271 | 0 | 0 |
| NC_019843.3;YP_009047202.1;MERS_1ab;272 | 0 | 0 |
| NC_019843.3;YP_009047202.1;MERS_1ab;273 | 0 | 0 |
| NC_019843.3;YP_009047202.1;MERS_1ab;274 | 0 | 0 |
| NC_019843.3;YP_009047202.1;MERS_1ab;275 | 0 | 0 |
| NC_019843.3;YP_009047202.1;MERS_1ab;276 | 0 | 0 |
| NC_019843.3;YP_009047202.1;MERS_1ab;277 | 0 | 0 |
| NC_019843.3;YP_009047202.1;MERS_1ab;278 | 0 | 0 |
| NC_019843.3;YP_009047202.1;MERS_1ab;279 | 0 | 0 |
| NC_019843.3;YP_009047202.1;MERS_1ab;280 | 0 | 0 |
| NC_019843.3;YP_009047202.1;MERS_1ab;281 | 0 | 0 |
| NC_019843.3;YP_009047202.1;MERS_1ab;282 | 0 | 0 |
| NC_019843.3;YP_009047202.1;MERS_1ab;283 | 0 | 0 |
| NC_019843.3;YP_009047202.1;MERS_1ab;284 | 0 | 0 |
| NC_019843.3;YP_009047202.1;MERS_1ab;285 | 0 | 0 |
| NC_019843.3;YP_009047202.1;MERS_1ab;286 | 0 | 0 |
| NC_019843.3;YP_009047202.1;MERS_1ab;287 | 0 | 0 |
| NC_019843.3;YP_009047202.1;MERS_1ab;288 | 0 | 0 |
| NC_019843.3;YP_009047202.1;MERS_1ab;289 | 0 | 0 |
| NC_019843.3;YP_009047202.1;MERS_1ab;290 | 0 | 0 |
| NC_019843.3;YP_009047202.1;MERS_1ab;291 | 0 | 0 |
| NC_019843.3;YP_009047202.1;MERS_1ab;292 | 0 | 0 |

| | | |
|---|---|---|
| NC_019843.3;YP_009047202.1;MERS_1ab;293 | 0 | 0 |
| NC_019843.3;YP_009047202.1;MERS_1ab;294 | 0 | 0 |
| NC_019843.3;YP_009047202.1;MERS_1ab;295 | 0 | 0 |
| NC_019843.3;YP_009047202.1;MERS_1ab;296 | 0 | 0 |
| NC_019843.3;YP_009047202.1;MERS_1ab;297 | 0 | 0 |
| NC_019843.3;YP_009047202.1;MERS_1ab;298 | 0 | 0 |
| NC_019843.3;YP_009047202.1;MERS_1ab;299 | 0 | 0 |
| NC_019843.3;YP_009047202.1;MERS_1ab;300 | 0 | 0 |
| NC_019843.3;YP_009047202.1;MERS_1ab;301 | 0 | 0 |
| NC_019843.3;YP_009047202.1;MERS_1ab;302 | 0 | 0 |
| NC_019843.3;YP_009047202.1;MERS_1ab;303 | 0 | 0 |
| NC_019843.3;YP_009047202.1;MERS_1ab;304 | 0.05 | 0.05 |
| NC_019843.3;YP_009047202.1;MERS_1ab;305 | 0 | 0 |
| NC_019843.3;YP_009047202.1;MERS_1ab;306 | 0 | 0 |
| NC_019843.3;YP_009047202.1;MERS_1ab;307 | 0 | 0 |
| NC_019843.3;YP_009047202.1;MERS_1ab;308 | 0 | 0 |
| NC_019843.3;YP_009047202.1;MERS_1ab;309 | 0 | 0 |
| NC_019843.3;YP_009047202.1;MERS_1ab;310 | 0 | 0 |
| NC_019843.3;YP_009047202.1;MERS_1ab;311 | 0 | 0 |
| NC_019843.3;YP_009047202.1;MERS_1ab;312 | 0 | 0 |
| NC_019843.3;YP_009047202.1;MERS_1ab;313 | 0 | 0 |
| NC_019843.3;YP_009047202.1;MERS_1ab;314 | 0 | 0 |
| NC_019843.3;YP_009047202.1;MERS_1ab;315 | 0 | 0 |
| NC_019843.3;YP_009047202.1;MERS_1ab;316 | 0 | 0 |
| NC_019843.3;YP_009047202.1;MERS_1ab;317 | 0 | 0 |
| NC_019843.3;YP_009047202.1;MERS_1ab;318 | 0 | 0 |
| NC_019843.3;YP_009047202.1;MERS_1ab;319 | 0 | 0 |
| NC_019843.3;YP_009047202.1;MERS_1ab;320 | 0 | 0 |
| NC_019843.3;YP_009047202.1;MERS_1ab;321 | 0 | 0 |
| NC_019843.3;YP_009047202.1;MERS_1ab;322 | 0 | 0 |
| NC_019843.3;YP_009047202.1;MERS_1ab;323 | 0 | 0 |
| NC_019843.3;YP_009047202.1;MERS_1ab;324 | 0 | 0 |
| NC_019843.3;YP_009047202.1;MERS_1ab;325 | 0 | 0 |
| NC_019843.3;YP_009047202.1;MERS_1ab;326 | 0 | 0 |
| NC_019843.3;YP_009047202.1;MERS_1ab;327 | 0 | 0 |
| NC_019843.3;YP_009047202.1;MERS_1ab;328 | 0 | 0 |
| NC_019843.3;YP_009047202.1;MERS_1ab;329 | 0 | 0 |
| NC_019843.3;YP_009047202.1;MERS_1ab;330 | 0 | 0 |
| NC_019843.3;YP_009047202.1;MERS_1ab;331 | 0 | 0 |
| NC_019843.3;YP_009047202.1;MERS_1ab;332 | 0 | 0 |
| NC_019843.3;YP_009047202.1;MERS_1ab;333 | 0 | 0 |
| NC_019843.3;YP_009047202.1;MERS_1ab;334 | 0.05 | 0 |
| NC_019843.3;YP_009047202.1;MERS_1ab;335 | 0 | 0 |

| | | |
|---|---|---|
| NC_019843.3;YP_009047202.1;MERS_1ab;336 | 0 | 0.025 |
| NC_019843.3;YP_009047202.1;MERS_1ab;337 | 0 | 0.025 |
| NC_019843.3;YP_009047202.1;MERS_1ab;338 | 0 | 0.025 |
| NC_019843.3;YP_009047202.1;MERS_1ab;339 | 0 | 0 |
| NC_019843.3;YP_009047202.1;MERS_1ab;340 | 0.05 | 0.025 |
| NC_019843.3;YP_009047202.1;MERS_1ab;341 | 0 | 0.05 |
| NC_019843.3;YP_009047202.1;MERS_1ab;342 | 0 | 0 |
| NC_019843.3;YP_009047202.1;MERS_1ab;343 | 0 | 0 |
| NC_019843.3;YP_009047202.1;MERS_1ab;344 | 0 | 0 |
| NC_019843.3;YP_009047202.1;MERS_1ab;345 | 0 | 0 |
| NC_019843.3;YP_009047202.1;MERS_1ab;346 | 0 | 0 |
| NC_019843.3;YP_009047202.1;MERS_1ab;347 | 0.05 | 0 |
| NC_019843.3;YP_009047202.1;MERS_1ab;348 | 0 | 0 |
| NC_019843.3;YP_009047202.1;MERS_1ab;349 | 0 | 0 |
| NC_019843.3;YP_009047202.1;MERS_1ab;350 | 0 | 0 |
| NC_019843.3;YP_009047202.1;MERS_1ab;351 | 0 | 0 |
| NC_019843.3;YP_009047202.1;MERS_1ab;352 | 0 | 0.025 |
| NC_019843.3;YP_009047202.1;MERS_1ab;353 | 0 | 0.025 |
| NC_019843.3;YP_009047202.1;MERS_1ab;354 | 0 | 0.025 |
| NC_019843.3;YP_009047202.1;MERS_1ab;355 | 0 | 0.025 |
| NC_019843.3;YP_009047202.1;MERS_1ab;356 | 0.05 | 0.05 |
| NC_019843.3;YP_009047202.1;MERS_1ab;357 | 0 | 0.05 |
| NC_019843.3;YP_009047202.1;MERS_1ab;358 | 0 | 0.05 |
| NC_019843.3;YP_009047202.1;MERS_1ab;359 | 0 | 0.025 |
| NC_019843.3;YP_009047202.1;MERS_1ab;360 | 0 | 0 |
| NC_019843.3;YP_009047202.1;MERS_1ab;361 | 0 | 0.025 |
| NC_019843.3;YP_009047202.1;MERS_1ab;362 | 0 | 0 |
| NC_019843.3;YP_009047202.1;MERS_1ab;363 | 0 | 0.025 |
| NC_019843.3;YP_009047202.1;MERS_1ab;364 | 0 | 0 |
| NC_019843.3;YP_009047202.1;MERS_1ab;365 | 0 | 0 |
| NC_019843.3;YP_009047202.1;MERS_1ab;366 | 0 | 0 |
| NC_019843.3;YP_009047202.1;MERS_1ab;367 | 0 | 0 |
| NC_019843.3;YP_009047202.1;MERS_1ab;368 | 0 | 0 |
| NC_019843.3;YP_009047202.1;MERS_1ab;369 | 0 | 0 |
| NC_019843.3;YP_009047202.1;MERS_1ab;370 | 0 | 0 |
| NC_019843.3;YP_009047202.1;MERS_1ab;371 | 0 | 0 |
| NC_019843.3;YP_009047202.1;MERS_1ab;372 | 0 | 0 |
| NC_019843.3;YP_009047202.1;MERS_1ab;373 | 0 | 0 |
| NC_019843.3;YP_009047202.1;MERS_1ab;374 | 0 | 0 |
| NC_019843.3;YP_009047202.1;MERS_1ab;375 | 0 | 0 |
| NC_019843.3;YP_009047202.1;MERS_1ab;376 | 0 | 0 |
| NC_019843.3;YP_009047202.1;MERS_1ab;377 | 0 | 0 |
| NC_019843.3;YP_009047202.1;MERS_1ab;378 | 0 | 0.025 |

| | | |
|---|---|---|
| NC_019843.3;YP_009047202.1;MERS_1ab;379 | 0 | 0 |
| NC_019843.3;YP_009047202.1;MERS_1ab;380 | 0 | 0 |
| NC_019843.3;YP_009047202.1;MERS_1ab;381 | 0 | 0.025 |
| NC_019843.3;YP_009047202.1;MERS_1ab;382 | 0 | 0.025 |
| NC_019843.3;YP_009047202.1;MERS_1ab;383 | 0 | 0.025 |
| NC_019843.3;YP_009047202.1;MERS_1ab;384 | 0 | 0.025 |
| NC_019843.3;YP_009047202.1;MERS_1ab;385 | 0 | 0 |
| NC_019843.3;YP_009047202.1;MERS_1ab;386 | 0 | 0.025 |
| NC_019843.3;YP_009047202.1;MERS_1ab;387 | 0 | 0 |
| NC_019843.3;YP_009047202.1;MERS_1ab;388 | 0.05 | 0.05 |
| NC_019843.3;YP_009047202.1;MERS_1ab;389 | 0 | 0.025 |
| NC_019843.3;YP_009047202.1;MERS_1ab;390 | 0 | 0.025 |
| NC_019843.3;YP_009047202.1;MERS_1ab;391 | 0 | 0 |
| NC_019843.3;YP_009047202.1;MERS_1ab;392 | 0.05 | 0 |
| NC_019843.3;YP_009047202.1;MERS_1ab;393 | 0 | 0.025 |
| NC_019843.3;YP_009047202.1;MERS_1ab;394 | 0 | 0 |
| NC_019843.3;YP_009047202.1;MERS_1ab;395 | 0 | 0 |
| NC_019843.3;YP_009047202.1;MERS_1ab;396 | 0 | 0 |
| NC_019843.3;YP_009047202.1;MERS_1ab;397 | 0 | 0 |
| NC_019843.3;YP_009047202.1;MERS_1ab;398 | 0 | 0 |
| NC_019843.3;YP_009047202.1;MERS_1ab;399 | 0 | 0 |
| NC_019843.3;YP_009047202.1;MERS_1ab;400 | 0 | 0 |
| NC_019843.3;YP_009047202.1;MERS_1ab;401 | 0 | 0 |
| NC_019843.3;YP_009047202.1;MERS_1ab;402 | 0 | 0 |
| NC_019843.3;YP_009047202.1;MERS_1ab;403 | 0 | 0 |
| NC_019843.3;YP_009047202.1;MERS_1ab;404 | 0 | 0 |
| NC_019843.3;YP_009047202.1;MERS_1ab;405 | 0.05 | 0 |
| NC_019843.3;YP_009047202.1;MERS_1ab;406 | 0.1 | 0.025 |
| NC_019843.3;YP_009047202.1;MERS_1ab;407 | 0.1 | 0.05 |
| NC_019843.3;YP_009047202.1;MERS_1ab;408 | 0.05 | 0.025 |
| NC_019843.3;YP_009047202.1;MERS_1ab;409 | 0 | 0.05 |
| NC_019843.3;YP_009047202.1;MERS_1ab;410 | 0 | 0 |
| NC_019843.3;YP_009047202.1;MERS_1ab;411 | 0.05 | 0 |
| NC_019843.3;YP_009047202.1;MERS_1ab;412 | 0 | 0 |
| NC_019843.3;YP_009047202.1;MERS_1ab;413 | 0 | 0 |
| NC_019843.3;YP_009047202.1;MERS_1ab;414 | 0 | 0 |
| NC_019843.3;YP_009047202.1;MERS_1ab;415 | 0 | 0 |
| NC_019843.3;YP_009047202.1;MERS_1ab;416 | 0 | 0 |
| NC_019843.3;YP_009047202.1;MERS_1ab;417 | 0 | 0 |
| NC_019843.3;YP_009047202.1;MERS_1ab;418 | 0 | 0 |
| NC_019843.3;YP_009047202.1;MERS_1ab;419 | 0 | 0 |
| NC_019843.3;YP_009047202.1;MERS_1ab;420 | 0 | 0 |
| NC_019843.3;YP_009047202.1;MERS_1ab;421 | 0 | 0 |

| | | |
|---|---|---|
| NC_019843.3;YP_009047202.1;MERS_1ab;422 | 0 | 0 |
| NC_019843.3;YP_009047202.1;MERS_1ab;423 | 0 | 0 |
| NC_019843.3;YP_009047202.1;MERS_1ab;424 | 0 | 0 |
| NC_019843.3;YP_009047202.1;MERS_1ab;425 | 0 | 0 |
| NC_019843.3;YP_009047202.1;MERS_1ab;426 | 0 | 0 |
| NC_019843.3;YP_009047202.1;MERS_1ab;427 | 0 | 0 |
| NC_019843.3;YP_009047202.1;MERS_1ab;428 | 0 | 0 |
| NC_019843.3;YP_009047202.1;MERS_1ab;429 | 0 | 0 |
| NC_019843.3;YP_009047202.1;MERS_1ab;430 | 0 | 0 |
| NC_019843.3;YP_009047202.1;MERS_1ab;431 | 0 | 0 |
| NC_019843.3;YP_009047202.1;MERS_1ab;432 | 0 | 0 |
| NC_019843.3;YP_009047202.1;MERS_1ab;433 | 0 | 0 |
| NC_019843.3;YP_009047202.1;MERS_1ab;434 | 0 | 0 |
| NC_019843.3;YP_009047202.1;MERS_1ab;435 | 0 | 0.025 |
| NC_019843.3;YP_009047202.1;MERS_1ab;436 | 0 | 0.025 |
| NC_019843.3;YP_009047202.1;MERS_1ab;437 | 0 | 0 |
| NC_019843.3;YP_009047202.1;MERS_1ab;438 | 0 | 0 |
| NC_019843.3;YP_009047202.1;MERS_1ab;439 | 0 | 0 |
| NC_019843.3;YP_009047202.1;MERS_1ab;440 | 0 | 0 |
| NC_019843.3;YP_009047202.1;MERS_1ab;441 | 0 | 0 |
| NC_019843.3;YP_009047202.1;MERS_1ab;442 | 0 | 0 |
| NC_019843.3;YP_009047202.1;MERS_1ab;443 | 0 | 0 |
| NC_019843.3;YP_009047202.1;MERS_1ab;444 | 0 | 0 |
| NC_019843.3;YP_009047202.1;MERS_1ab;445 | 0 | 0 |
| NC_019843.3;YP_009047202.1;MERS_1ab;446 | 0 | 0 |
| NC_019843.3;YP_009047202.1;MERS_1ab;447 | 0 | 0 |
| NC_019843.3;YP_009047202.1;MERS_1ab;448 | 0 | 0 |
| NC_019843.3;YP_009047202.1;MERS_1ab;449 | 0 | 0 |
| NC_019843.3;YP_009047202.1;MERS_1ab;450 | 0 | 0 |
| NC_019843.3;YP_009047202.1;MERS_1ab;451 | 0 | 0 |
| NC_019843.3;YP_009047202.1;MERS_1ab;452 | 0 | 0 |
| NC_019843.3;YP_009047202.1;MERS_1ab;453 | 0 | 0 |
| NC_019843.3;YP_009047202.1;MERS_1ab;454 | 0 | 0 |
| NC_019843.3;YP_009047202.1;MERS_1ab;455 | 0 | 0 |
| NC_019843.3;YP_009047202.1;MERS_1ab;456 | 0 | 0 |
| NC_019843.3;YP_009047202.1;MERS_1ab;457 | 0 | 0 |
| NC_019843.3;YP_009047202.1;MERS_1ab;458 | 0 | 0 |
| NC_019843.3;YP_009047202.1;MERS_1ab;459 | 0 | 0 |
| NC_019843.3;YP_009047202.1;MERS_1ab;460 | 0 | 0 |
| NC_019843.3;YP_009047202.1;MERS_1ab;461 | 0 | 0 |
| NC_019843.3;YP_009047202.1;MERS_1ab;462 | 0 | 0 |
| NC_019843.3;YP_009047202.1;MERS_1ab;463 | 0 | 0 |
| NC_019843.3;YP_009047202.1;MERS_1ab;464 | 0 | 0 |

| | | |
|---|---|---|
| NC_019843.3;YP_009047202.1;MERS_1ab;465 | 0 | 0 |
| NC_019843.3;YP_009047202.1;MERS_1ab;466 | 0 | 0.025 |
| NC_019843.3;YP_009047202.1;MERS_1ab;467 | 0.05 | 0 |
| NC_019843.3;YP_009047202.1;MERS_1ab;468 | 0 | 0 |
| NC_019843.3;YP_009047202.1;MERS_1ab;469 | 0 | 0 |
| NC_019843.3;YP_009047202.1;MERS_1ab;470 | 0 | 0 |
| NC_019843.3;YP_009047202.1;MERS_1ab;471 | 0 | 0 |
| NC_019843.3;YP_009047202.1;MERS_1ab;472 | 0 | 0.025 |
| NC_019843.3;YP_009047202.1;MERS_1ab;473 | 0 | 0 |
| NC_019843.3;YP_009047202.1;MERS_1ab;474 | 0 | 0 |
| NC_019843.3;YP_009047202.1;MERS_1ab;475 | 0 | 0 |
| NC_019843.3;YP_009047202.1;MERS_1ab;476 | 0 | 0 |
| NC_019843.3;YP_009047202.1;MERS_1ab;477 | 0 | 0 |
| NC_019843.3;YP_009047202.1;MERS_1ab;478 | 0 | 0 |
| NC_019843.3;YP_009047202.1;MERS_1ab;479 | 0 | 0 |
| NC_019843.3;YP_009047202.1;MERS_1ab;480 | 0 | 0 |
| NC_019843.3;YP_009047202.1;MERS_1ab;481 | 0 | 0 |
| NC_019843.3;YP_009047202.1;MERS_1ab;482 | 0 | 0 |
| NC_019843.3;YP_009047202.1;MERS_1ab;483 | 0 | 0 |
| NC_019843.3;YP_009047202.1;MERS_1ab;484 | 0 | 0 |
| NC_019843.3;YP_009047202.1;MERS_1ab;485 | 0 | 0 |
| NC_019843.3;YP_009047202.1;MERS_1ab;486 | 0 | 0 |
| NC_019843.3;YP_009047202.1;MERS_1ab;487 | 0 | 0 |
| NC_019843.3;YP_009047202.1;MERS_1ab;488 | 0 | 0 |
| NC_019843.3;YP_009047202.1;MERS_1ab;489 | 0.05 | 0 |
| NC_019843.3;YP_009047202.1;MERS_1ab;490 | 0 | 0 |
| NC_019843.3;YP_009047202.1;MERS_1ab;491 | 0 | 0 |
| NC_019843.3;YP_009047202.1;MERS_1ab;492 | 0 | 0 |
| NC_019843.3;YP_009047202.1;MERS_1ab;493 | 0 | 0 |
| NC_019843.3;YP_009047202.1;MERS_1ab;494 | 0 | 0 |
| NC_019843.3;YP_009047202.1;MERS_1ab;495 | 0 | 0 |
| NC_019843.3;YP_009047202.1;MERS_1ab;496 | 0 | 0 |
| NC_019843.3;YP_009047202.1;MERS_1ab;497 | 0.05 | 0 |
| NC_019843.3;YP_009047202.1;MERS_1ab;498 | 0.05 | 0 |
| NC_019843.3;YP_009047202.1;MERS_1ab;499 | 0.05 | 0 |
| NC_019843.3;YP_009047202.1;MERS_1ab;500 | 0.05 | 0 |
| NC_019843.3;YP_009047202.1;MERS_1ab;501 | 0.05 | 0 |
| NC_019843.3;YP_009047202.1;MERS_1ab;502 | 0.05 | 0 |
| NC_019843.3;YP_009047202.1;MERS_1ab;503 | 0 | 0 |
| NC_019843.3;YP_009047202.1;MERS_1ab;504 | 0 | 0 |
| NC_019843.3;YP_009047202.1;MERS_1ab;505 | 0.05 | 0 |
| NC_019843.3;YP_009047202.1;MERS_1ab;506 | 0 | 0 |
| NC_019843.3;YP_009047202.1;MERS_1ab;507 | 0 | 0 |

| | | |
|---|---|---|
| NC_019843.3;YP_009047202.1;MERS_1ab;508 | 0 | 0 |
| NC_019843.3;YP_009047202.1;MERS_1ab;509 | 0 | 0 |
| NC_019843.3;YP_009047202.1;MERS_1ab;510 | 0 | 0 |
| NC_019843.3;YP_009047202.1;MERS_1ab;511 | 0 | 0 |
| NC_019843.3;YP_009047202.1;MERS_1ab;512 | 0.05 | 0 |
| NC_019843.3;YP_009047202.1;MERS_1ab;513 | 0.1 | 0 |
| NC_019843.3;YP_009047202.1;MERS_1ab;514 | 0.1 | 0.025 |
| NC_019843.3;YP_009047202.1;MERS_1ab;515 | 0.05 | 0.05 |
| NC_019843.3;YP_009047202.1;MERS_1ab;516 | 0.1 | 0.05 |
| NC_019843.3;YP_009047202.1;MERS_1ab;517 | 0.05 | 0.3 |
| NC_019843.3;YP_009047202.1;MERS_1ab;518 | 0.05 | 0 |
| NC_019843.3;YP_009047202.1;MERS_1ab;519 | 0.05 | 0 |
| NC_019843.3;YP_009047202.1;MERS_1ab;520 | 0 | 0.05 |
| NC_019843.3;YP_009047202.1;MERS_1ab;521 | 0.05 | 0.025 |
| NC_019843.3;YP_009047202.1;MERS_1ab;522 | 0 | 0.025 |
| NC_019843.3;YP_009047202.1;MERS_1ab;523 | 0 | 0 |
| NC_019843.3;YP_009047202.1;MERS_1ab;524 | 0 | 0 |
| NC_019843.3;YP_009047202.1;MERS_1ab;525 | 0 | 0.025 |
| NC_019843.3;YP_009047202.1;MERS_1ab;526 | 0.05 | 0.025 |
| NC_019843.3;YP_009047202.1;MERS_1ab;527 | 0 | 0.025 |
| NC_019843.3;YP_009047202.1;MERS_1ab;528 | 0 | 0.025 |
| NC_019843.3;YP_009047202.1;MERS_1ab;529 | 0.05 | 0.025 |
| NC_019843.3;YP_009047202.1;MERS_1ab;530 | 0.05 | 0.025 |
| NC_019843.3;YP_009047202.1;MERS_1ab;531 | 0 | 0 |
| NC_019843.3;YP_009047202.1;MERS_1ab;532 | 0 | 0 |
| NC_019843.3;YP_009047202.1;MERS_1ab;533 | 0 | 0 |
| NC_019843.3;YP_009047202.1;MERS_1ab;534 | 0 | 0 |
| NC_019843.3;YP_009047202.1;MERS_1ab;535 | 0.05 | 0.025 |
| NC_019843.3;YP_009047202.1;MERS_1ab;536 | 0.05 | 0.025 |
| NC_019843.3;YP_009047202.1;MERS_1ab;537 | 0 | 0.025 |
| NC_019843.3;YP_009047202.1;MERS_1ab;538 | 0 | 0.025 |
| NC_019843.3;YP_009047202.1;MERS_1ab;539 | 0.1 | 0.05 |
| NC_019843.3;YP_009047202.1;MERS_1ab;540 | 0 | 0 |
| NC_019843.3;YP_009047202.1;MERS_1ab;541 | 0 | 0 |
| NC_019843.3;YP_009047202.1;MERS_1ab;542 | 0 | 0 |
| NC_019843.3;YP_009047202.1;MERS_1ab;543 | 0 | 0 |
| NC_019843.3;YP_009047202.1;MERS_1ab;544 | 0 | 0 |
| NC_019843.3;YP_009047202.1;MERS_1ab;545 | 0 | 0 |
| NC_019843.3;YP_009047202.1;MERS_1ab;546 | 0 | 0 |
| NC_019843.3;YP_009047202.1;MERS_1ab;547 | 0 | 0 |
| NC_019843.3;YP_009047202.1;MERS_1ab;548 | 0 | 0 |
| NC_019843.3;YP_009047202.1;MERS_1ab;549 | 0 | 0 |
| NC_019843.3;YP_009047202.1;MERS_1ab;550 | 0 | 0 |

| | | |
|---|---|---|
| NC_019843.3;YP_009047202.1;MERS_1ab;551 | 0 | 0 |
| NC_019843.3;YP_009047202.1;MERS_1ab;552 | 0 | 0 |
| NC_019843.3;YP_009047202.1;MERS_1ab;553 | 0 | 0.025 |
| NC_019843.3;YP_009047202.1;MERS_1ab;554 | 0 | 0 |
| NC_019843.3;YP_009047202.1;MERS_1ab;555 | 0 | 0.075 |
| NC_019843.3;YP_009047202.1;MERS_1ab;556 | 0 | 0.025 |
| NC_019843.3;YP_009047202.1;MERS_1ab;557 | 0 | 0 |
| NC_019843.3;YP_009047202.1;MERS_1ab;558 | 0 | 0 |
| NC_019843.3;YP_009047202.1;MERS_1ab;559 | 0 | 0 |
| NC_019843.3;YP_009047202.1;MERS_1ab;560 | 0 | 0 |
| NC_019843.3;YP_009047202.1;MERS_1ab;561 | 0 | 0 |
| NC_019843.3;YP_009047202.1;MERS_1ab;562 | 0 | 0 |
| NC_019843.3;YP_009047202.1;MERS_1ab;563 | 0 | 0 |
| NC_019843.3;YP_009047202.1;MERS_1ab;564 | 0.05 | 0 |
| NC_019843.3;YP_009047202.1;MERS_1ab;565 | 0 | 0 |
| NC_019843.3;YP_009047202.1;MERS_1ab;566 | 0 | 0 |
| NC_019843.3;YP_009047202.1;MERS_1ab;567 | 0.05 | 0.025 |
| NC_019843.3;YP_009047202.1;MERS_1ab;568 | 0 | 0 |
| NC_019843.3;YP_009047202.1;MERS_1ab;569 | 0 | 0 |
| NC_019843.3;YP_009047202.1;MERS_1ab;570 | 0 | 0 |
| NC_019843.3;YP_009047202.1;MERS_1ab;571 | 0 | 0 |
| NC_019843.3;YP_009047202.1;MERS_1ab;572 | 0 | 0.025 |
| NC_019843.3;YP_009047202.1;MERS_1ab;573 | 0 | 0.025 |
| NC_019843.3;YP_009047202.1;MERS_1ab;574 | 0.05 | 0 |
| NC_019843.3;YP_009047202.1;MERS_1ab;575 | 0 | 0 |
| NC_019843.3;YP_009047202.1;MERS_1ab;576 | 0 | 0.025 |
| NC_019843.3;YP_009047202.1;MERS_1ab;577 | 0 | 0 |
| NC_019843.3;YP_009047202.1;MERS_1ab;578 | 0 | 0 |
| NC_019843.3;YP_009047202.1;MERS_1ab;579 | 0 | 0 |
| NC_019843.3;YP_009047202.1;MERS_1ab;580 | 0 | 0 |
| NC_019843.3;YP_009047202.1;MERS_1ab;581 | 0 | 0 |
| NC_019843.3;YP_009047202.1;MERS_1ab;582 | 0 | 0 |
| NC_019843.3;YP_009047202.1;MERS_1ab;583 | 0 | 0 |
| NC_019843.3;YP_009047202.1;MERS_1ab;584 | 0 | 0 |
| NC_019843.3;YP_009047202.1;MERS_1ab;585 | 0 | 0 |
| NC_019843.3;YP_009047202.1;MERS_1ab;586 | 0 | 0 |
| NC_019843.3;YP_009047202.1;MERS_1ab;587 | 0 | 0 |
| NC_019843.3;YP_009047202.1;MERS_1ab;588 | 0 | 0 |
| NC_019843.3;YP_009047202.1;MERS_1ab;589 | 0 | 0 |
| NC_019843.3;YP_009047202.1;MERS_1ab;590 | 0 | 0 |
| NC_019843.3;YP_009047202.1;MERS_1ab;591 | 0 | 0 |
| NC_019843.3;YP_009047202.1;MERS_1ab;592 | 0 | 0 |
| NC_019843.3;YP_009047202.1;MERS_1ab;593 | 0 | 0 |

| | | |
|---|---|---|
| NC_019843.3;YP_009047202.1;MERS_1ab;594 | 0.05 | 0 |
| NC_019843.3;YP_009047202.1;MERS_1ab;595 | 0.05 | 0 |
| NC_019843.3;YP_009047202.1;MERS_1ab;596 | 0.05 | 0 |
| NC_019843.3;YP_009047202.1;MERS_1ab;597 | 0.15 | 0.075 |
| NC_019843.3;YP_009047202.1;MERS_1ab;598 | 0.05 | 0.025 |
| NC_019843.3;YP_009047202.1;MERS_1ab;599 | 0 | 0 |
| NC_019843.3;YP_009047202.1;MERS_1ab;600 | 0 | 0 |
| NC_019843.3;YP_009047202.1;MERS_1ab;601 | 0 | 0 |
| NC_019843.3;YP_009047202.1;MERS_1ab;602 | 0 | 0 |
| NC_019843.3;YP_009047202.1;MERS_1ab;603 | 0 | 0 |
| NC_019843.3;YP_009047202.1;MERS_1ab;604 | 0 | 0 |
| NC_019843.3;YP_009047202.1;MERS_1ab;605 | 0 | 0 |
| NC_019843.3;YP_009047202.1;MERS_1ab;606 | 0 | 0 |
| NC_019843.3;YP_009047202.1;MERS_1ab;607 | 0 | 0 |
| NC_019843.3;YP_009047202.1;MERS_1ab;608 | 0 | 0 |
| NC_019843.3;YP_009047202.1;MERS_1ab;609 | 0 | 0 |
| NC_019843.3;YP_009047202.1;MERS_1ab;610 | 0 | 0 |
| NC_019843.3;YP_009047202.1;MERS_1ab;611 | 0 | 0 |
| NC_019843.3;YP_009047202.1;MERS_1ab;612 | 0 | 0 |
| NC_019843.3;YP_009047202.1;MERS_1ab;613 | 0 | 0 |
| NC_019843.3;YP_009047202.1;MERS_1ab;614 | 0 | 0 |
| NC_019843.3;YP_009047202.1;MERS_1ab;615 | 0 | 0 |
| NC_019843.3;YP_009047202.1;MERS_1ab;616 | 0 | 0 |
| NC_019843.3;YP_009047202.1;MERS_1ab;617 | 0 | 0 |
| NC_019843.3;YP_009047202.1;MERS_1ab;618 | 0 | 0 |
| NC_019843.3;YP_009047202.1;MERS_1ab;619 | 0 | 0 |
| NC_019843.3;YP_009047202.1;MERS_1ab;620 | 0 | 0 |
| NC_019843.3;YP_009047202.1;MERS_1ab;621 | 0 | 0 |
| NC_019843.3;YP_009047202.1;MERS_1ab;622 | 0 | 0 |
| NC_019843.3;YP_009047202.1;MERS_1ab;623 | 0 | 0 |
| NC_019843.3;YP_009047202.1;MERS_1ab;624 | 0 | 0 |
| NC_019843.3;YP_009047202.1;MERS_1ab;625 | 0 | 0 |
| NC_019843.3;YP_009047202.1;MERS_1ab;626 | 0 | 0 |
| NC_019843.3;YP_009047202.1;MERS_1ab;627 | 0 | 0 |
| NC_019843.3;YP_009047202.1;MERS_1ab;628 | 0 | 0 |
| NC_019843.3;YP_009047202.1;MERS_1ab;629 | 0 | 0 |
| NC_019843.3;YP_009047202.1;MERS_1ab;630 | 0 | 0 |
| NC_019843.3;YP_009047202.1;MERS_1ab;631 | 0 | 0 |
| NC_019843.3;YP_009047202.1;MERS_1ab;632 | 0 | 0 |
| NC_019843.3;YP_009047202.1;MERS_1ab;633 | 0 | 0 |
| NC_019843.3;YP_009047202.1;MERS_1ab;634 | 0 | 0 |
| NC_019843.3;YP_009047202.1;MERS_1ab;635 | 0 | 0 |
| NC_019843.3;YP_009047202.1;MERS_1ab;636 | 0 | 0 |

| | | |
|---|---|---|
| NC_019843.3;YP_009047202.1;MERS_1ab;637 | 0 | 0 |
| NC_019843.3;YP_009047202.1;MERS_1ab;638 | 0 | 0 |
| NC_019843.3;YP_009047202.1;MERS_1ab;639 | 0 | 0 |
| NC_019843.3;YP_009047202.1;MERS_1ab;640 | 0 | 0 |
| NC_019843.3;YP_009047202.1;MERS_1ab;641 | 0 | 0 |
| NC_019843.3;YP_009047202.1;MERS_1ab;642 | 0 | 0 |
| NC_019843.3;YP_009047202.1;MERS_1ab;643 | 0 | 0 |
| NC_019843.3;YP_009047202.1;MERS_1ab;644 | 0 | 0 |
| NC_019843.3;YP_009047202.1;MERS_1ab;645 | 0 | 0 |
| NC_019843.3;YP_009047202.1;MERS_1ab;646 | 0 | 0 |
| NC_019843.3;YP_009047202.1;MERS_1ab;647 | 0 | 0 |
| NC_019843.3;YP_009047202.1;MERS_1ab;648 | 0 | 0 |
| NC_019843.3;YP_009047202.1;MERS_1ab;649 | 0 | 0 |
| NC_019843.3;YP_009047202.1;MERS_1ab;650 | 0 | 0 |
| NC_019843.3;YP_009047202.1;MERS_1ab;651 | 0 | 0 |
| NC_019843.3;YP_009047202.1;MERS_1ab;652 | 0 | 0.075 |
| NC_019843.3;YP_009047202.1;MERS_1ab;653 | 0 | 0 |
| NC_019843.3;YP_009047202.1;MERS_1ab;654 | 0 | 0 |
| NC_019843.3;YP_009047202.1;MERS_1ab;655 | 0 | 0 |
| NC_019843.3;YP_009047202.1;MERS_1ab;656 | 0 | 0 |
| NC_019843.3;YP_009047202.1;MERS_1ab;657 | 0 | 0 |
| NC_019843.3;YP_009047202.1;MERS_1ab;658 | 0 | 0 |
| NC_019843.3;YP_009047202.1;MERS_1ab;659 | 0 | 0 |
| NC_019843.3;YP_009047202.1;MERS_1ab;660 | 0 | 0 |
| NC_019843.3;YP_009047202.1;MERS_1ab;661 | 0 | 0 |
| NC_019843.3;YP_009047202.1;MERS_1ab;662 | 0 | 0 |
| NC_019843.3;YP_009047202.1;MERS_1ab;663 | 0 | 0 |
| NC_019843.3;YP_009047202.1;MERS_1ab;664 | 0 | 0 |
| NC_019843.3;YP_009047202.1;MERS_1ab;665 | 0 | 0 |
| NC_019843.3;YP_009047202.1;MERS_1ab;666 | 0 | 0 |
| NC_019843.3;YP_009047202.1;MERS_1ab;667 | 0 | 0 |
| NC_019843.3;YP_009047202.1;MERS_1ab;668 | 0 | 0 |
| NC_019843.3;YP_009047202.1;MERS_1ab;669 | 0 | 0 |
| NC_019843.3;YP_009047202.1;MERS_1ab;670 | 0 | 0 |
| NC_019843.3;YP_009047202.1;MERS_1ab;671 | 0 | 0 |
| NC_019843.3;YP_009047202.1;MERS_1ab;672 | 0 | 0 |
| NC_019843.3;YP_009047202.1;MERS_1ab;673 | 0 | 0 |
| NC_019843.3;YP_009047202.1;MERS_1ab;674 | 0 | 0 |
| NC_019843.3;YP_009047202.1;MERS_1ab;675 | 0 | 0 |
| NC_019843.3;YP_009047202.1;MERS_1ab;676 | 0 | 0 |
| NC_019843.3;YP_009047202.1;MERS_1ab;677 | 0 | 0 |
| NC_019843.3;YP_009047202.1;MERS_1ab;678 | 0 | 0 |
| NC_019843.3;YP_009047202.1;MERS_1ab;679 | 0 | 0 |

| | | |
|---|---|---|
| NC_019843.3;YP_009047202.1;MERS_1ab;680 | 0 | 0 |
| NC_019843.3;YP_009047202.1;MERS_1ab;681 | 0 | 0 |
| NC_019843.3;YP_009047202.1;MERS_1ab;682 | 0 | 0 |
| NC_019843.3;YP_009047202.1;MERS_1ab;683 | 0 | 0 |
| NC_019843.3;YP_009047202.1;MERS_1ab;684 | 0 | 0 |
| NC_019843.3;YP_009047202.1;MERS_1ab;685 | 0 | 0 |
| NC_019843.3;YP_009047202.1;MERS_1ab;686 | 0 | 0 |
| NC_019843.3;YP_009047202.1;MERS_1ab;687 | 0 | 0 |
| NC_019843.3;YP_009047202.1;MERS_1ab;688 | 0 | 0 |
| NC_019843.3;YP_009047202.1;MERS_1ab;689 | 0 | 0 |
| NC_019843.3;YP_009047202.1;MERS_1ab;690 | 0 | 0 |
| NC_019843.3;YP_009047202.1;MERS_1ab;691 | 0 | 0 |
| NC_019843.3;YP_009047202.1;MERS_1ab;692 | 0.05 | 0 |
| NC_019843.3;YP_009047202.1;MERS_1ab;693 | 0 | 0 |
| NC_019843.3;YP_009047202.1;MERS_1ab;694 | 0 | 0 |
| NC_019843.3;YP_009047202.1;MERS_1ab;695 | 0 | 0 |
| NC_019843.3;YP_009047202.1;MERS_1ab;696 | 0 | 0 |
| NC_019843.3;YP_009047202.1;MERS_1ab;697 | 0 | 0 |
| NC_019843.3;YP_009047202.1;MERS_1ab;698 | 0 | 0 |
| NC_019843.3;YP_009047202.1;MERS_1ab;699 | 0 | 0 |
| NC_019843.3;YP_009047202.1;MERS_1ab;700 | 0 | 0 |
| NC_019843.3;YP_009047202.1;MERS_1ab;701 | 0.1 | 0 |
| NC_019843.3;YP_009047202.1;MERS_1ab;702 | 0.1 | 0 |
| NC_019843.3;YP_009047202.1;MERS_1ab;703 | 0.15 | 0.025 |
| NC_019843.3;YP_009047202.1;MERS_1ab;704 | 0 | 0 |
| NC_019843.3;YP_009047202.1;MERS_1ab;705 | 0 | 0 |
| NC_019843.3;YP_009047202.1;MERS_1ab;706 | 0 | 0 |
| NC_019843.3;YP_009047202.1;MERS_1ab;707 | 0 | 0 |
| NC_019843.3;YP_009047202.1;MERS_1ab;708 | 0 | 0 |
| NC_019843.3;YP_009047202.1;MERS_1ab;709 | 0 | 0 |
| NC_019843.3;YP_009047202.1;MERS_1ab;710 | 0 | 0 |
| NC_019843.3;YP_009047202.1;MERS_1ab;711 | 0 | 0 |
| NC_019843.3;YP_009047202.1;MERS_1ab;712 | 0 | 0 |
| NC_019843.3;YP_009047202.1;MERS_1ab;713 | 0 | 0 |
| NC_019843.3;YP_009047202.1;MERS_1ab;714 | 0 | 0 |
| NC_019843.3;YP_009047202.1;MERS_1ab;715 | 0 | 0 |
| NC_019843.3;YP_009047202.1;MERS_1ab;716 | 0 | 0 |
| NC_019843.3;YP_009047202.1;MERS_1ab;717 | 0 | 0 |
| NC_019843.3;YP_009047202.1;MERS_1ab;718 | 0 | 0 |
| NC_019843.3;YP_009047202.1;MERS_1ab;719 | 0 | 0 |
| NC_019843.3;YP_009047202.1;MERS_1ab;720 | 0 | 0 |
| NC_019843.3;YP_009047202.1;MERS_1ab;721 | 0 | 0 |
| NC_019843.3;YP_009047202.1;MERS_1ab;722 | 0 | 0 |

| | | |
|---|---|---|
| NC_019843.3;YP_009047202.1;MERS_1ab;723 | 0 | 0 |
| NC_019843.3;YP_009047202.1;MERS_1ab;724 | 0 | 0 |
| NC_019843.3;YP_009047202.1;MERS_1ab;725 | 0 | 0 |
| NC_019843.3;YP_009047202.1;MERS_1ab;726 | 0 | 0 |
| NC_019843.3;YP_009047202.1;MERS_1ab;727 | 0 | 0.025 |
| NC_019843.3;YP_009047202.1;MERS_1ab;728 | 0 | 0 |
| NC_019843.3;YP_009047202.1;MERS_1ab;729 | 0.05 | 0.025 |
| NC_019843.3;YP_009047202.1;MERS_1ab;730 | 0 | 0 |
| NC_019843.3;YP_009047202.1;MERS_1ab;731 | 0 | 0 |
| NC_019843.3;YP_009047202.1;MERS_1ab;732 | 0 | 0 |
| NC_019843.3;YP_009047202.1;MERS_1ab;733 | 0 | 0 |
| NC_019843.3;YP_009047202.1;MERS_1ab;734 | 0 | 0 |
| NC_019843.3;YP_009047202.1;MERS_1ab;735 | 0 | 0 |
| NC_019843.3;YP_009047202.1;MERS_1ab;736 | 0 | 0 |
| NC_019843.3;YP_009047202.1;MERS_1ab;737 | 0 | 0 |
| NC_019843.3;YP_009047202.1;MERS_1ab;738 | 0 | 0 |
| NC_019843.3;YP_009047202.1;MERS_1ab;739 | 0 | 0 |
| NC_019843.3;YP_009047202.1;MERS_1ab;740 | 0 | 0 |
| NC_019843.3;YP_009047202.1;MERS_1ab;741 | 0 | 0 |
| NC_019843.3;YP_009047202.1;MERS_1ab;742 | 0 | 0 |
| NC_019843.3;YP_009047202.1;MERS_1ab;743 | 0 | 0.025 |
| NC_019843.3;YP_009047202.1;MERS_1ab;744 | 0 | 0.025 |
| NC_019843.3;YP_009047202.1;MERS_1ab;745 | 0.05 | 0.025 |
| NC_019843.3;YP_009047202.1;MERS_1ab;746 | 0.05 | 0.025 |
| NC_019843.3;YP_009047202.1;MERS_1ab;747 | 0 | 0 |
| NC_019843.3;YP_009047202.1;MERS_1ab;748 | 0 | 0.025 |
| NC_019843.3;YP_009047202.1;MERS_1ab;749 | 0.05 | 0 |
| NC_019843.3;YP_009047202.1;MERS_1ab;750 | 0 | 0 |
| NC_019843.3;YP_009047202.1;MERS_1ab;751 | 0.1 | 0 |
| NC_019843.3;YP_009047202.1;MERS_1ab;752 | 0 | 0 |
| NC_019843.3;YP_009047202.1;MERS_1ab;753 | 0 | 0 |
| NC_019843.3;YP_009047202.1;MERS_1ab;754 | 0 | 0 |
| NC_019843.3;YP_009047202.1;MERS_1ab;755 | 0 | 0 |
| NC_019843.3;YP_009047202.1;MERS_1ab;756 | 0 | 0 |
| NC_019843.3;YP_009047202.1;MERS_1ab;757 | 0.05 | 0 |
| NC_019843.3;YP_009047202.1;MERS_1ab;758 | 0 | 0.025 |
| NC_019843.3;YP_009047202.1;MERS_1ab;759 | 0 | 0 |
| NC_019843.3;YP_009047202.1;MERS_1ab;760 | 0 | 0 |
| NC_019843.3;YP_009047202.1;MERS_1ab;761 | 0 | 0.025 |
| NC_019843.3;YP_009047202.1;MERS_1ab;762 | 0 | 0.025 |
| NC_019843.3;YP_009047202.1;MERS_1ab;763 | 0 | 0.025 |
| NC_019843.3;YP_009047202.1;MERS_1ab;764 | 0 | 0.025 |
| NC_019843.3;YP_009047202.1;MERS_1ab;765 | 0 | 0.025 |

| | | |
|---|---|---|
| NC_019843.3;YP_009047202.1;MERS_1ab;766 | 0.05 | 0.075 |
| NC_019843.3;YP_009047202.1;MERS_1ab;767 | 0 | 0.075 |
| NC_019843.3;YP_009047202.1;MERS_1ab;768 | 0 | 0 |
| NC_019843.3;YP_009047202.1;MERS_1ab;769 | 0 | 0 |
| NC_019843.3;YP_009047202.1;MERS_1ab;770 | 0 | 0 |
| NC_019843.3;YP_009047202.1;MERS_1ab;771 | 0.05 | 0 |
| NC_019843.3;YP_009047202.1;MERS_1ab;772 | 0.05 | 0 |
| NC_019843.3;YP_009047202.1;MERS_1ab;773 | 0 | 0 |
| NC_019843.3;YP_009047202.1;MERS_1ab;774 | 0 | 0 |
| NC_019843.3;YP_009047202.1;MERS_1ab;775 | 0 | 0 |
| NC_019843.3;YP_009047202.1;MERS_1ab;776 | 0 | 0.025 |
| NC_019843.3;YP_009047202.1;MERS_1ab;777 | 0 | 0.025 |
| NC_019843.3;YP_009047202.1;MERS_1ab;778 | 0.05 | 0.05 |
| NC_019843.3;YP_009047202.1;MERS_1ab;779 | 0.05 | 0.05 |
| NC_019843.3;YP_009047202.1;MERS_1ab;780 | 0.1 | 0.025 |
| NC_019843.3;YP_009047202.1;MERS_1ab;781 | 0.1 | 0.025 |
| NC_019843.3;YP_009047202.1;MERS_1ab;782 | 0 | 0.05 |
| NC_019843.3;YP_009047202.1;MERS_1ab;783 | 0.05 | 0.05 |
| NC_019843.3;YP_009047202.1;MERS_1ab;784 | 0.05 | 0.05 |
| NC_019843.3;YP_009047202.1;MERS_1ab;785 | 0.05 | 0.05 |
| NC_019843.3;YP_009047202.1;MERS_1ab;786 | 0.05 | 0.075 |
| NC_019843.3;YP_009047202.1;MERS_1ab;787 | 0.05 | 0.05 |
| NC_019843.3;YP_009047202.1;MERS_1ab;788 | 0 | 0.025 |
| NC_019843.3;YP_009047202.1;MERS_1ab;789 | 0 | 0.05 |
| NC_019843.3;YP_009047202.1;MERS_1ab;790 | 0.05 | 0.075 |
| NC_019843.3;YP_009047202.1;MERS_1ab;791 | 0 | 0.025 |
| NC_019843.3;YP_009047202.1;MERS_1ab;792 | 0 | 0 |
| NC_019843.3;YP_009047202.1;MERS_1ab;793 | 0 | 0 |
| NC_019843.3;YP_009047202.1;MERS_1ab;794 | 0 | 0 |
| NC_019843.3;YP_009047202.1;MERS_1ab;795 | 0 | 0 |
| NC_019843.3;YP_009047202.1;MERS_1ab;796 | 0 | 0 |
| NC_019843.3;YP_009047202.1;MERS_1ab;797 | 0 | 0 |
| NC_019843.3;YP_009047202.1;MERS_1ab;798 | 0 | 0 |
| NC_019843.3;YP_009047202.1;MERS_1ab;799 | 0 | 0 |
| NC_019843.3;YP_009047202.1;MERS_1ab;800 | 0 | 0 |
| NC_019843.3;YP_009047202.1;MERS_1ab;801 | 0 | 0 |
| NC_019843.3;YP_009047202.1;MERS_1ab;802 | 0 | 0 |
| NC_019843.3;YP_009047202.1;MERS_1ab;803 | 0 | 0 |
| NC_019843.3;YP_009047202.1;MERS_1ab;804 | 0 | 0 |
| NC_019843.3;YP_009047202.1;MERS_1ab;805 | 0 | 0 |
| NC_019843.3;YP_009047202.1;MERS_1ab;806 | 0 | 0 |
| NC_019843.3;YP_009047202.1;MERS_1ab;807 | 0 | 0 |
| NC_019843.3;YP_009047202.1;MERS_1ab;808 | 0 | 0 |

| | | |
|---|---|---|
| NC_019843.3;YP_009047202.1;MERS_1ab;809 | 0 | 0 |
| NC_019843.3;YP_009047202.1;MERS_1ab;810 | 0 | 0 |
| NC_019843.3;YP_009047202.1;MERS_1ab;811 | 0 | 0 |
| NC_019843.3;YP_009047202.1;MERS_1ab;812 | 0 | 0 |
| NC_019843.3;YP_009047202.1;MERS_1ab;813 | 0 | 0 |
| NC_019843.3;YP_009047202.1;MERS_1ab;814 | 0 | 0 |
| NC_019843.3;YP_009047202.1;MERS_1ab;815 | 0 | 0 |
| NC_019843.3;YP_009047202.1;MERS_1ab;816 | 0 | 0 |
| NC_019843.3;YP_009047202.1;MERS_1ab;817 | 0.05 | 0 |
| NC_019843.3;YP_009047202.1;MERS_1ab;818 | 0.05 | 0 |
| NC_019843.3;YP_009047202.1;MERS_1ab;819 | 0 | 0 |
| NC_019843.3;YP_009047202.1;MERS_1ab;820 | 0.05 | 0 |
| NC_019843.3;YP_009047202.1;MERS_1ab;821 | 0.05 | 0 |
| NC_019843.3;YP_009047202.1;MERS_1ab;822 | 0.1 | 0 |
| NC_019843.3;YP_009047202.1;MERS_1ab;823 | 0 | 0 |
| NC_019843.3;YP_009047202.1;MERS_1ab;824 | 0.05 | 0 |
| NC_019843.3;YP_009047202.1;MERS_1ab;825 | 0.05 | 0 |
| NC_019843.3;YP_009047202.1;MERS_1ab;826 | 0.1 | 0 |
| NC_019843.3;YP_009047202.1;MERS_1ab;827 | 0.05 | 0 |
| NC_019843.3;YP_009047202.1;MERS_1ab;828 | 0.05 | 0.025 |
| NC_019843.3;YP_009047202.1;MERS_1ab;829 | 0 | 0 |
| NC_019843.3;YP_009047202.1;MERS_1ab;830 | 0 | 0 |
| NC_019843.3;YP_009047202.1;MERS_1ab;831 | 0 | 0 |
| NC_019843.3;YP_009047202.1;MERS_1ab;832 | 0 | 0 |
| NC_019843.3;YP_009047202.1;MERS_1ab;833 | 0 | 0 |
| NC_019843.3;YP_009047202.1;MERS_1ab;834 | 0 | 0 |
| NC_019843.3;YP_009047202.1;MERS_1ab;835 | 0 | 0 |
| NC_019843.3;YP_009047202.1;MERS_1ab;836 | 0 | 0 |
| NC_019843.3;YP_009047202.1;MERS_1ab;837 | 0 | 0 |
| NC_019843.3;YP_009047202.1;MERS_1ab;838 | 0 | 0 |
| NC_019843.3;YP_009047202.1;MERS_1ab;839 | 0 | 0 |
| NC_019843.3;YP_009047202.1;MERS_1ab;840 | 0 | 0 |
| NC_019843.3;YP_009047202.1;MERS_1ab;841 | 0 | 0 |
| NC_019843.3;YP_009047202.1;MERS_1ab;842 | 0 | 0 |
| NC_019843.3;YP_009047202.1;MERS_1ab;843 | 0 | 0 |
| NC_019843.3;YP_009047202.1;MERS_1ab;844 | 0 | 0 |
| NC_019843.3;YP_009047202.1;MERS_1ab;845 | 0 | 0 |
| NC_019843.3;YP_009047202.1;MERS_1ab;846 | 0 | 0 |
| NC_019843.3;YP_009047202.1;MERS_1ab;847 | 0 | 0 |
| NC_019843.3;YP_009047202.1;MERS_1ab;848 | 0 | 0 |
| NC_019843.3;YP_009047202.1;MERS_1ab;849 | 0 | 0 |
| NC_019843.3;YP_009047202.1;MERS_1ab;850 | 0 | 0.025 |
| NC_019843.3;YP_009047202.1;MERS_1ab;851 | 0 | 0.025 |

| | | |
|---|---|---|
| NC_019843.3;YP_009047202.1;MERS_1ab;852 | 0.05 | 0.025 |
| NC_019843.3;YP_009047202.1;MERS_1ab;853 | 0 | 0.05 |
| NC_019843.3;YP_009047202.1;MERS_1ab;854 | 0 | 0 |
| NC_019843.3;YP_009047202.1;MERS_1ab;855 | 0 | 0.075 |
| NC_019843.3;YP_009047202.1;MERS_1ab;856 | 0.05 | 0.025 |
| NC_019843.3;YP_009047202.1;MERS_1ab;857 | 0 | 0 |
| NC_019843.3;YP_009047202.1;MERS_1ab;858 | 0 | 0.025 |
| NC_019843.3;YP_009047202.1;MERS_1ab;859 | 0 | 0.025 |
| NC_019843.3;YP_009047202.1;MERS_1ab;860 | 0 | 0.025 |
| NC_019843.3;YP_009047202.1;MERS_1ab;861 | 0 | 0.025 |
| NC_019843.3;YP_009047202.1;MERS_1ab;862 | 0 | 0 |
| NC_019843.3;YP_009047202.1;MERS_1ab;863 | 0 | 0 |
| NC_019843.3;YP_009047202.1;MERS_1ab;864 | 0 | 0 |
| NC_019843.3;YP_009047202.1;MERS_1ab;865 | 0 | 0 |
| NC_019843.3;YP_009047202.1;MERS_1ab;866 | 0 | 0 |
| NC_019843.3;YP_009047202.1;MERS_1ab;867 | 0 | 0 |
| NC_019843.3;YP_009047202.1;MERS_1ab;868 | 0 | 0 |
| NC_019843.3;YP_009047202.1;MERS_1ab;869 | 0 | 0 |
| NC_019843.3;YP_009047202.1;MERS_1ab;870 | 0 | 0 |
| NC_019843.3;YP_009047202.1;MERS_1ab;871 | 0 | 0 |
| NC_019843.3;YP_009047202.1;MERS_1ab;872 | 0 | 0 |
| NC_019843.3;YP_009047202.1;MERS_1ab;873 | 0 | 0 |
| NC_019843.3;YP_009047202.1;MERS_1ab;874 | 0 | 0 |
| NC_019843.3;YP_009047202.1;MERS_1ab;875 | 0 | 0 |
| NC_019843.3;YP_009047202.1;MERS_1ab;876 | 0 | 0 |
| NC_019843.3;YP_009047202.1;MERS_1ab;877 | 0 | 0 |
| NC_019843.3;YP_009047202.1;MERS_1ab;878 | 0 | 0 |
| NC_019843.3;YP_009047202.1;MERS_1ab;879 | 0 | 0 |
| NC_019843.3;YP_009047202.1;MERS_1ab;880 | 0 | 0 |
| NC_019843.3;YP_009047202.1;MERS_1ab;881 | 0.05 | 0 |
| NC_019843.3;YP_009047202.1;MERS_1ab;882 | 0 | 0 |
| NC_019843.3;YP_009047202.1;MERS_1ab;883 | 0 | 0 |
| NC_019843.3;YP_009047202.1;MERS_1ab;884 | 0 | 0 |
| NC_019843.3;YP_009047202.1;MERS_1ab;885 | 0 | 0 |
| NC_019843.3;YP_009047202.1;MERS_1ab;886 | 0 | 0 |
| NC_019843.3;YP_009047202.1;MERS_1ab;887 | 0 | 0 |
| NC_019843.3;YP_009047202.1;MERS_1ab;888 | 0 | 0 |
| NC_019843.3;YP_009047202.1;MERS_1ab;889 | 0.05 | 0 |
| NC_019843.3;YP_009047202.1;MERS_1ab;890 | 0.05 | 0 |
| NC_019843.3;YP_009047202.1;MERS_1ab;891 | 0 | 0 |
| NC_019843.3;YP_009047202.1;MERS_1ab;892 | 0 | 0 |
| NC_019843.3;YP_009047202.1;MERS_1ab;893 | 0 | 0 |
| NC_019843.3;YP_009047202.1;MERS_1ab;894 | 0 | 0 |

| | | |
|---|---|---|
| NC_019843.3;YP_009047202.1;MERS_1ab;895 | 0.05 | 0.025 |
| NC_019843.3;YP_009047202.1;MERS_1ab;896 | 0 | 0.025 |
| NC_019843.3;YP_009047202.1;MERS_1ab;897 | 0 | 0.025 |
| NC_019843.3;YP_009047202.1;MERS_1ab;898 | 0 | 0 |
| NC_019843.3;YP_009047202.1;MERS_1ab;899 | 0 | 0 |
| NC_019843.3;YP_009047202.1;MERS_1ab;900 | 0 | 0 |
| NC_019843.3;YP_009047202.1;MERS_1ab;901 | 0 | 0 |
| NC_019843.3;YP_009047202.1;MERS_1ab;902 | 0 | 0.1 |
| NC_019843.3;YP_009047202.1;MERS_1ab;903 | 0 | 0.05 |
| NC_019843.3;YP_009047202.1;MERS_1ab;904 | 0 | 0 |
| NC_019843.3;YP_009047202.1;MERS_1ab;905 | 0 | 0.025 |
| NC_019843.3;YP_009047202.1;MERS_1ab;906 | 0 | 0 |
| NC_019843.3;YP_009047202.1;MERS_1ab;907 | 0 | 0 |
| NC_019843.3;YP_009047202.1;MERS_1ab;908 | 0 | 0 |
| NC_019843.3;YP_009047202.1;MERS_1ab;909 | 0 | 0.025 |
| NC_019843.3;YP_009047202.1;MERS_1ab;910 | 0 | 0.025 |
| NC_019843.3;YP_009047202.1;MERS_1ab;911 | 0 | 0 |
| NC_019843.3;YP_009047202.1;MERS_1ab;912 | 0 | 0 |
| NC_019843.3;YP_009047202.1;MERS_1ab;913 | 0 | 0 |
| NC_019843.3;YP_009047202.1;MERS_1ab;914 | 0 | 0.025 |
| NC_019843.3;YP_009047202.1;MERS_1ab;915 | 0.05 | 0 |
| NC_019843.3;YP_009047202.1;MERS_1ab;916 | 0 | 0 |
| NC_019843.3;YP_009047202.1;MERS_1ab;917 | 0 | 0 |
| NC_019843.3;YP_009047202.1;MERS_1ab;918 | 0 | 0 |
| NC_019843.3;YP_009047202.1;MERS_1ab;919 | 0 | 0 |
| NC_019843.3;YP_009047202.1;MERS_1ab;920 | 0.05 | 0 |
| NC_019843.3;YP_009047202.1;MERS_1ab;921 | 0.05 | 0 |
| NC_019843.3;YP_009047202.1;MERS_1ab;922 | 0.05 | 0 |
| NC_019843.3;YP_009047202.1;MERS_1ab;923 | 0.1 | 0 |
| NC_019843.3;YP_009047202.1;MERS_1ab;924 | 0.1 | 0.025 |
| NC_019843.3;YP_009047202.1;MERS_1ab;925 | 0.15 | 0.05 |
| NC_019843.3;YP_009047202.1;MERS_1ab;926 | 0.1 | 0.05 |
| NC_019843.3;YP_009047202.1;MERS_1ab;927 | 0.1 | 0.025 |
| NC_019843.3;YP_009047202.1;MERS_1ab;928 | 0.05 | 0.025 |
| NC_019843.3;YP_009047202.1;MERS_1ab;929 | 0 | 0.025 |
| NC_019843.3;YP_009047202.1;MERS_1ab;930 | 0.05 | 0 |
| NC_019843.3;YP_009047202.1;MERS_1ab;931 | 0.05 | 0 |
| NC_019843.3;YP_009047202.1;MERS_1ab;932 | 0 | 0 |
| NC_019843.3;YP_009047202.1;MERS_1ab;933 | 0 | 0 |
| NC_019843.3;YP_009047202.1;MERS_1ab;934 | 0 | 0 |
| NC_019843.3;YP_009047202.1;MERS_1ab;935 | 0 | 0 |
| NC_019843.3;YP_009047202.1;MERS_1ab;936 | 0 | 0 |
| NC_019843.3;YP_009047202.1;MERS_1ab;937 | 0 | 0 |

| | | |
|---|---|---|
| NC_019843.3;YP_009047202.1;MERS_1ab;938 | 0 | 0 |
| NC_019843.3;YP_009047202.1;MERS_1ab;939 | 0 | 0 |
| NC_019843.3;YP_009047202.1;MERS_1ab;940 | 0 | 0 |
| NC_019843.3;YP_009047202.1;MERS_1ab;941 | 0 | 0 |
| NC_019843.3;YP_009047202.1;MERS_1ab;942 | 0 | 0 |
| NC_019843.3;YP_009047202.1;MERS_1ab;943 | 0 | 0 |
| NC_019843.3;YP_009047202.1;MERS_1ab;944 | 0 | 0 |
| NC_019843.3;YP_009047202.1;MERS_1ab;945 | 0 | 0 |
| NC_019843.3;YP_009047202.1;MERS_1ab;946 | 0 | 0 |
| NC_019843.3;YP_009047202.1;MERS_1ab;947 | 0 | 0 |
| NC_019843.3;YP_009047202.1;MERS_1ab;948 | 0 | 0 |
| NC_019843.3;YP_009047202.1;MERS_1ab;949 | 0 | 0 |
| NC_019843.3;YP_009047202.1;MERS_1ab;950 | 0 | 0 |
| NC_019843.3;YP_009047202.1;MERS_1ab;951 | 0 | 0 |
| NC_019843.3;YP_009047202.1;MERS_1ab;952 | 0 | 0 |
| NC_019843.3;YP_009047202.1;MERS_1ab;953 | 0 | 0 |
| NC_019843.3;YP_009047202.1;MERS_1ab;954 | 0 | 0 |
| NC_019843.3;YP_009047202.1;MERS_1ab;955 | 0 | 0 |
| NC_019843.3;YP_009047202.1;MERS_1ab;956 | 0 | 0 |
| NC_019843.3;YP_009047202.1;MERS_1ab;957 | 0 | 0 |
| NC_019843.3;YP_009047202.1;MERS_1ab;958 | 0 | 0 |
| NC_019843.3;YP_009047202.1;MERS_1ab;959 | 0.05 | 0 |
| NC_019843.3;YP_009047202.1;MERS_1ab;960 | 0 | 0 |
| NC_019843.3;YP_009047202.1;MERS_1ab;961 | 0 | 0 |
| NC_019843.3;YP_009047202.1;MERS_1ab;962 | 0 | 0 |
| NC_019843.3;YP_009047202.1;MERS_1ab;963 | 0 | 0 |
| NC_019843.3;YP_009047202.1;MERS_1ab;964 | 0 | 0 |
| NC_019843.3;YP_009047202.1;MERS_1ab;965 | 0 | 0.025 |
| NC_019843.3;YP_009047202.1;MERS_1ab;966 | 0 | 0.025 |
| NC_019843.3;YP_009047202.1;MERS_1ab;967 | 0 | 0 |
| NC_019843.3;YP_009047202.1;MERS_1ab;968 | 0 | 0 |
| NC_019843.3;YP_009047202.1;MERS_1ab;969 | 0 | 0 |
| NC_019843.3;YP_009047202.1;MERS_1ab;970 | 0 | 0 |
| NC_019843.3;YP_009047202.1;MERS_1ab;971 | 0 | 0 |
| NC_019843.3;YP_009047202.1;MERS_1ab;972 | 0 | 0 |
| NC_019843.3;YP_009047202.1;MERS_1ab;973 | 0 | 0 |
| NC_019843.3;YP_009047202.1;MERS_1ab;974 | 0 | 0 |
| NC_019843.3;YP_009047202.1;MERS_1ab;975 | 0 | 0 |
| NC_019843.3;YP_009047202.1;MERS_1ab;976 | 0 | 0 |
| NC_019843.3;YP_009047202.1;MERS_1ab;977 | 0 | 0 |
| NC_019843.3;YP_009047202.1;MERS_1ab;978 | 0 | 0 |
| NC_019843.3;YP_009047202.1;MERS_1ab;979 | 0 | 0 |
| NC_019843.3;YP_009047202.1;MERS_1ab;980 | 0 | 0 |

| | | |
|---|---|---|
| NC_019843.3;YP_009047202.1;MERS_1ab;981 | 0 | 0 |
| NC_019843.3;YP_009047202.1;MERS_1ab;982 | 0.05 | 0 |
| NC_019843.3;YP_009047202.1;MERS_1ab;983 | 0 | 0 |
| NC_019843.3;YP_009047202.1;MERS_1ab;984 | 0 | 0 |
| NC_019843.3;YP_009047202.1;MERS_1ab;985 | 0 | 0 |
| NC_019843.3;YP_009047202.1;MERS_1ab;986 | 0 | 0 |
| NC_019843.3;YP_009047202.1;MERS_1ab;987 | 0 | 0 |
| NC_019843.3;YP_009047202.1;MERS_1ab;988 | 0 | 0 |
| NC_019843.3;YP_009047202.1;MERS_1ab;989 | 0 | 0 |
| NC_019843.3;YP_009047202.1;MERS_1ab;990 | 0 | 0 |
| NC_019843.3;YP_009047202.1;MERS_1ab;991 | 0 | 0 |
| NC_019843.3;YP_009047202.1;MERS_1ab;992 | 0.05 | 0 |
| NC_019843.3;YP_009047202.1;MERS_1ab;993 | 0.05 | 0 |
| NC_019843.3;YP_009047202.1;MERS_1ab;994 | 0 | 0.025 |
| NC_019843.3;YP_009047202.1;MERS_1ab;995 | 0.05 | 0 |
| NC_019843.3;YP_009047202.1;MERS_1ab;996 | 0.05 | 0 |
| NC_019843.3;YP_009047202.1;MERS_1ab;997 | 0.05 | 0 |
| NC_019843.3;YP_009047202.1;MERS_1ab;998 | 0.05 | 0 |
| NC_019843.3;YP_009047202.1;MERS_1ab;999 | 0.05 | 0 |
| NC_019843.3;YP_009047202.1;MERS_1ab;1000 | 0.1 | 0 |
| NC_019843.3;YP_009047202.1;MERS_1ab;1001 | 0.05 | 0.025 |
| NC_019843.3;YP_009047202.1;MERS_1ab;1002 | 0.1 | 0.025 |
| NC_019843.3;YP_009047202.1;MERS_1ab;1003 | 0.05 | 0.025 |
| NC_019843.3;YP_009047202.1;MERS_1ab;1004 | 0.15 | 0.075 |
| NC_019843.3;YP_009047202.1;MERS_1ab;1005 | 0.15 | 0.05 |
| NC_019843.3;YP_009047202.1;MERS_1ab;1006 | 0.15 | 0.05 |
| NC_019843.3;YP_009047202.1;MERS_1ab;1007 | 0.1 | 0.05 |
| NC_019843.3;YP_009047202.1;MERS_1ab;1008 | 0.1 | 0.025 |
| NC_019843.3;YP_009047202.1;MERS_1ab;1009 | 0.05 | 0.025 |
| NC_019843.3;YP_009047202.1;MERS_1ab;1010 | 0.05 | 0.025 |
| NC_019843.3;YP_009047202.1;MERS_1ab;1011 | 0 | 0.025 |
| NC_019843.3;YP_009047202.1;MERS_1ab;1012 | 0 | 0.025 |
| NC_019843.3;YP_009047202.1;MERS_1ab;1013 | 0 | 0 |
| NC_019843.3;YP_009047202.1;MERS_1ab;1014 | 0 | 0 |
| NC_019843.3;YP_009047202.1;MERS_1ab;1015 | 0 | 0 |
| NC_019843.3;YP_009047202.1;MERS_1ab;1016 | 0 | 0 |
| NC_019843.3;YP_009047202.1;MERS_1ab;1017 | 0 | 0 |
| NC_019843.3;YP_009047202.1;MERS_1ab;1018 | 0 | 0 |
| NC_019843.3;YP_009047202.1;MERS_1ab;1019 | 0.1 | 0.025 |
| NC_019843.3;YP_009047202.1;MERS_1ab;1020 | 0 | 0 |
| NC_019843.3;YP_009047202.1;MERS_1ab;1021 | 0 | 0 |
| NC_019843.3;YP_009047202.1;MERS_1ab;1022 | 0 | 0 |
| NC_019843.3;YP_009047202.1;MERS_1ab;1023 | 0 | 0 |

| | | |
|---|---|---|
| NC_019843.3;YP_009047202.1;MERS_1ab;1024 | 0 | 0 |
| NC_019843.3;YP_009047202.1;MERS_1ab;1025 | 0 | 0 |
| NC_019843.3;YP_009047202.1;MERS_1ab;1026 | 0 | 0 |
| NC_019843.3;YP_009047202.1;MERS_1ab;1027 | 0 | 0 |
| NC_019843.3;YP_009047202.1;MERS_1ab;1028 | 0 | 0 |
| NC_019843.3;YP_009047202.1;MERS_1ab;1029 | 0 | 0 |
| NC_019843.3;YP_009047202.1;MERS_1ab;1030 | 0 | 0 |
| NC_019843.3;YP_009047202.1;MERS_1ab;1031 | 0 | 0 |
| NC_019843.3;YP_009047202.1;MERS_1ab;1032 | 0 | 0 |
| NC_019843.3;YP_009047202.1;MERS_1ab;1033 | 0 | 0 |
| NC_019843.3;YP_009047202.1;MERS_1ab;1034 | 0.05 | 0.05 |
| NC_019843.3;YP_009047202.1;MERS_1ab;1035 | 0.05 | 0 |
| NC_019843.3;YP_009047202.1;MERS_1ab;1036 | 0 | 0.05 |
| NC_019843.3;YP_009047202.1;MERS_1ab;1037 | 0 | 0.05 |
| NC_019843.3;YP_009047202.1;MERS_1ab;1038 | 0 | 0.05 |
| NC_019843.3;YP_009047202.1;MERS_1ab;1039 | 0 | 0.05 |
| NC_019843.3;YP_009047202.1;MERS_1ab;1040 | 0.05 | 0.05 |
| NC_019843.3;YP_009047202.1;MERS_1ab;1041 | 0.05 | 0.05 |
| NC_019843.3;YP_009047202.1;MERS_1ab;1042 | 0.05 | 0.075 |
| NC_019843.3;YP_009047202.1;MERS_1ab;1043 | 0.05 | 0.075 |
| NC_019843.3;YP_009047202.1;MERS_1ab;1044 | 0.05 | 0.025 |
| NC_019843.3;YP_009047202.1;MERS_1ab;1045 | 0 | 0.025 |
| NC_019843.3;YP_009047202.1;MERS_1ab;1046 | 0 | 0.025 |
| NC_019843.3;YP_009047202.1;MERS_1ab;1047 | 0 | 0.025 |
| NC_019843.3;YP_009047202.1;MERS_1ab;1048 | 0 | 0.025 |
| NC_019843.3;YP_009047202.1;MERS_1ab;1049 | 0 | 0.025 |
| NC_019843.3;YP_009047202.1;MERS_1ab;1050 | 0 | 0.05 |
| NC_019843.3;YP_009047202.1;MERS_1ab;1051 | 0.05 | 0.025 |
| NC_019843.3;YP_009047202.1;MERS_1ab;1052 | 0 | 0.025 |
| NC_019843.3;YP_009047202.1;MERS_1ab;1053 | 0 | 0.025 |
| NC_019843.3;YP_009047202.1;MERS_1ab;1054 | 0 | 0.025 |
| NC_019843.3;YP_009047202.1;MERS_1ab;1055 | 0 | 0 |
| NC_019843.3;YP_009047202.1;MERS_1ab;1056 | 0.05 | 0 |
| NC_019843.3;YP_009047202.1;MERS_1ab;1057 | 0.1 | 0 |
| NC_019843.3;YP_009047202.1;MERS_1ab;1058 | 0.05 | 0 |
| NC_019843.3;YP_009047202.1;MERS_1ab;1059 | 0.05 | 0.025 |
| NC_019843.3;YP_009047202.1;MERS_1ab;1060 | 0.05 | 0 |
| NC_019843.3;YP_009047202.1;MERS_1ab;1061 | 0 | 0.025 |
| NC_019843.3;YP_009047202.1;MERS_1ab;1062 | 0.05 | 0.075 |
| NC_019843.3;YP_009047202.1;MERS_1ab;1063 | 0.1 | 0.05 |
| NC_019843.3;YP_009047202.1;MERS_1ab;1064 | 0 | 0.025 |
| NC_019843.3;YP_009047202.1;MERS_1ab;1065 | 0 | 0.05 |
| NC_019843.3;YP_009047202.1;MERS_1ab;1066 | 0.05 | 0.075 |

| | | |
|---|---|---|
| NC_019843.3;YP_009047202.1;MERS_1ab;1067 | 0 | 0.05 |
| NC_019843.3;YP_009047202.1;MERS_1ab;1068 | 0 | 0 |
| NC_019843.3;YP_009047202.1;MERS_1ab;1069 | 0 | 0.025 |
| NC_019843.3;YP_009047202.1;MERS_1ab;1070 | 0 | 0.025 |
| NC_019843.3;YP_009047202.1;MERS_1ab;1071 | 0 | 0 |
| NC_019843.3;YP_009047202.1;MERS_1ab;1072 | 0 | 0 |
| NC_019843.3;YP_009047202.1;MERS_1ab;1073 | 0 | 0 |
| NC_019843.3;YP_009047202.1;MERS_1ab;1074 | 0 | 0 |
| NC_019843.3;YP_009047202.1;MERS_1ab;1075 | 0 | 0 |
| NC_019843.3;YP_009047202.1;MERS_1ab;1076 | 0 | 0 |
| NC_019843.3;YP_009047202.1;MERS_1ab;1077 | 0 | 0 |
| NC_019843.3;YP_009047202.1;MERS_1ab;1078 | 0 | 0 |
| NC_019843.3;YP_009047202.1;MERS_1ab;1079 | 0 | 0 |
| NC_019843.3;YP_009047202.1;MERS_1ab;1080 | 0 | 0 |
| NC_019843.3;YP_009047202.1;MERS_1ab;1081 | 0 | 0.025 |
| NC_019843.3;YP_009047202.1;MERS_1ab;1082 | 0.2 | 0.25 |
| NC_019843.3;YP_009047202.1;MERS_1ab;1083 | 0.1 | 0.075 |
| NC_019843.3;YP_009047202.1;MERS_1ab;1084 | 0.15 | 0.125 |
| NC_019843.3;YP_009047202.1;MERS_1ab;1085 | 0.2 | 0.175 |
| NC_019843.3;YP_009047202.1;MERS_1ab;1086 | 0.15 | 0.15 |
| NC_019843.3;YP_009047202.1;MERS_1ab;1087 | 0.1 | 0.075 |
| NC_019843.3;YP_009047202.1;MERS_1ab;1088 | 0.05 | 0.075 |
| NC_019843.3;YP_009047202.1;MERS_1ab;1089 | 0.05 | 0.025 |
| NC_019843.3;YP_009047202.1;MERS_1ab;1090 | 0.05 | 0.05 |
| NC_019843.3;YP_009047202.1;MERS_1ab;1091 | 0.05 | 0 |
| NC_019843.3;YP_009047202.1;MERS_1ab;1092 | 0.1 | 0.025 |
| NC_019843.3;YP_009047202.1;MERS_1ab;1093 | 0.1 | 0.025 |
| NC_019843.3;YP_009047202.1;MERS_1ab;1094 | 0.05 | 0 |
| NC_019843.3;YP_009047202.1;MERS_1ab;1095 | 0 | 0 |
| NC_019843.3;YP_009047202.1;MERS_1ab;1096 | 0 | 0 |
| NC_019843.3;YP_009047202.1;MERS_1ab;1097 | 0 | 0.025 |
| NC_019843.3;YP_009047202.1;MERS_1ab;1098 | 0 | 0.025 |
| NC_019843.3;YP_009047202.1;MERS_1ab;1099 | 0.05 | 0.025 |
| NC_019843.3;YP_009047202.1;MERS_1ab;1100 | 0 | 0.075 |
| NC_019843.3;YP_009047202.1;MERS_1ab;1101 | 0 | 0.05 |
| NC_019843.3;YP_009047202.1;MERS_1ab;1102 | 0 | 0.025 |
| NC_019843.3;YP_009047202.1;MERS_1ab;1103 | 0 | 0.025 |
| NC_019843.3;YP_009047202.1;MERS_1ab;1104 | 0 | 0.125 |
| NC_019843.3;YP_009047202.1;MERS_1ab;1105 | 0.15 | 0.15 |
| NC_019843.3;YP_009047202.1;MERS_1ab;1106 | 0.05 | 0 |
| NC_019843.3;YP_009047202.1;MERS_1ab;1107 | 0 | 0.025 |
| NC_019843.3;YP_009047202.1;MERS_1ab;1108 | 0 | 0 |
| NC_019843.3;YP_009047202.1;MERS_1ab;1109 | 0 | 0 |

| | | |
|---|---|---|
| NC_019843.3;YP_009047202.1;MERS_1ab;1110 | 0 | 0 |
| NC_019843.3;YP_009047202.1;MERS_1ab;1111 | 0 | 0 |
| NC_019843.3;YP_009047202.1;MERS_1ab;1112 | 0 | 0 |
| NC_019843.3;YP_009047202.1;MERS_1ab;1113 | 0 | 0 |
| NC_019843.3;YP_009047202.1;MERS_1ab;1114 | 0 | 0 |
| NC_019843.3;YP_009047202.1;MERS_1ab;1115 | 0 | 0 |
| NC_019843.3;YP_009047202.1;MERS_1ab;1116 | 0 | 0 |
| NC_019843.3;YP_009047202.1;MERS_1ab;1117 | 0 | 0 |
| NC_019843.3;YP_009047202.1;MERS_1ab;1118 | 0 | 0 |
| NC_019843.3;YP_009047202.1;MERS_1ab;1119 | 0 | 0 |
| NC_019843.3;YP_009047202.1;MERS_1ab;1120 | 0 | 0 |
| NC_019843.3;YP_009047202.1;MERS_1ab;1121 | 0 | 0 |
| NC_019843.3;YP_009047202.1;MERS_1ab;1122 | 0 | 0 |
| NC_019843.3;YP_009047202.1;MERS_1ab;1123 | 0 | 0 |
| NC_019843.3;YP_009047202.1;MERS_1ab;1124 | 0 | 0 |
| NC_019843.3;YP_009047202.1;MERS_1ab;1125 | 0 | 0 |
| NC_019843.3;YP_009047202.1;MERS_1ab;1126 | 0.05 | 0 |
| NC_019843.3;YP_009047202.1;MERS_1ab;1127 | 0 | 0 |
| NC_019843.3;YP_009047202.1;MERS_1ab;1128 | 0 | 0 |
| NC_019843.3;YP_009047202.1;MERS_1ab;1129 | 0 | 0 |
| NC_019843.3;YP_009047202.1;MERS_1ab;1130 | 0 | 0 |
| NC_019843.3;YP_009047202.1;MERS_1ab;1131 | 0.05 | 0.025 |
| NC_019843.3;YP_009047202.1;MERS_1ab;1132 | 0 | 0 |
| NC_019843.3;YP_009047202.1;MERS_1ab;1133 | 0 | 0 |
| NC_019843.3;YP_009047202.1;MERS_1ab;1134 | 0 | 0 |
| NC_019843.3;YP_009047202.1;MERS_1ab;1135 | 0 | 0 |
| NC_019843.3;YP_009047202.1;MERS_1ab;1136 | 0 | 0 |
| NC_019843.3;YP_009047202.1;MERS_1ab;1137 | 0 | 0 |
| NC_019843.3;YP_009047202.1;MERS_1ab;1138 | 0 | 0 |
| NC_019843.3;YP_009047202.1;MERS_1ab;1139 | 0.05 | 0 |
| NC_019843.3;YP_009047202.1;MERS_1ab;1140 | 0.05 | 0 |
| NC_019843.3;YP_009047202.1;MERS_1ab;1141 | 0 | 0 |
| NC_019843.3;YP_009047202.1;MERS_1ab;1142 | 0.4 | 0.3 |
| NC_019843.3;YP_009047202.1;MERS_1ab;1143 | 0 | 0.025 |
| NC_019843.3;YP_009047202.1;MERS_1ab;1144 | 0 | 0.025 |
| NC_019843.3;YP_009047202.1;MERS_1ab;1145 | 0 | 0.025 |
| NC_019843.3;YP_009047202.1;MERS_1ab;1146 | 0 | 0.025 |
| NC_019843.3;YP_009047202.1;MERS_1ab;1147 | 0 | 0.025 |
| NC_019843.3;YP_009047202.1;MERS_1ab;1148 | 0.05 | 0.05 |
| NC_019843.3;YP_009047202.1;MERS_1ab;1149 | 0 | 0.025 |
| NC_019843.3;YP_009047202.1;MERS_1ab;1150 | 0 | 0 |
| NC_019843.3;YP_009047202.1;MERS_1ab;1151 | 0 | 0 |
| NC_019843.3;YP_009047202.1;MERS_1ab;1152 | 0 | 0 |

| | | |
|---|---|---|
| NC_019843.3;YP_009047202.1;MERS_1ab;1153 | 0.05 | 0.05 |
| NC_019843.3;YP_009047202.1;MERS_1ab;1154 | 0.05 | 0.025 |
| NC_019843.3;YP_009047202.1;MERS_1ab;1155 | 0.05 | 0 |
| NC_019843.3;YP_009047202.1;MERS_1ab;1156 | 0.05 | 0 |
| NC_019843.3;YP_009047202.1;MERS_1ab;1157 | 0.05 | 0.025 |
| NC_019843.3;YP_009047202.1;MERS_1ab;1158 | 0.1 | 0.05 |
| NC_019843.3;YP_009047202.1;MERS_1ab;1159 | 0.05 | 0.025 |
| NC_019843.3;YP_009047202.1;MERS_1ab;1160 | 0.05 | 0.025 |
| NC_019843.3;YP_009047202.1;MERS_1ab;1161 | 0.05 | 0 |
| NC_019843.3;YP_009047202.1;MERS_1ab;1162 | 0.05 | 0 |
| NC_019843.3;YP_009047202.1;MERS_1ab;1163 | 0 | 0.025 |
| NC_019843.3;YP_009047202.1;MERS_1ab;1164 | 0 | 0.025 |
| NC_019843.3;YP_009047202.1;MERS_1ab;1165 | 0 | 0.05 |
| NC_019843.3;YP_009047202.1;MERS_1ab;1166 | 0 | 0.025 |
| NC_019843.3;YP_009047202.1;MERS_1ab;1167 | 0.05 | 0.025 |
| NC_019843.3;YP_009047202.1;MERS_1ab;1168 | 0.05 | 0.025 |
| NC_019843.3;YP_009047202.1;MERS_1ab;1169 | 0 | 0.025 |
| NC_019843.3;YP_009047202.1;MERS_1ab;1170 | 0.05 | 0.05 |
| NC_019843.3;YP_009047202.1;MERS_1ab;1171 | 0 | 0.075 |
| NC_019843.3;YP_009047202.1;MERS_1ab;1172 | 0.1 | 0.075 |
| NC_019843.3;YP_009047202.1;MERS_1ab;1173 | 0 | 0.05 |
| NC_019843.3;YP_009047202.1;MERS_1ab;1174 | 0 | 0 |
| NC_019843.3;YP_009047202.1;MERS_1ab;1175 | 0 | 0 |
| NC_019843.3;YP_009047202.1;MERS_1ab;1176 | 0 | 0 |
| NC_019843.3;YP_009047202.1;MERS_1ab;1177 | 0 | 0 |
| NC_019843.3;YP_009047202.1;MERS_1ab;1178 | 0 | 0 |
| NC_019843.3;YP_009047202.1;MERS_1ab;1179 | 0 | 0 |
| NC_019843.3;YP_009047202.1;MERS_1ab;1180 | 0 | 0 |
| NC_019843.3;YP_009047202.1;MERS_1ab;1181 | 0 | 0 |
| NC_019843.3;YP_009047202.1;MERS_1ab;1182 | 0 | 0 |
| NC_019843.3;YP_009047202.1;MERS_1ab;1183 | 0 | 0 |
| NC_019843.3;YP_009047202.1;MERS_1ab;1184 | 0 | 0 |
| NC_019843.3;YP_009047202.1;MERS_1ab;1185 | 0 | 0 |
| NC_019843.3;YP_009047202.1;MERS_1ab;1186 | 0 | 0 |
| NC_019843.3;YP_009047202.1;MERS_1ab;1187 | 0 | 0 |
| NC_019843.3;YP_009047202.1;MERS_1ab;1188 | 0 | 0 |
| NC_019843.3;YP_009047202.1;MERS_1ab;1189 | 0 | 0 |
| NC_019843.3;YP_009047202.1;MERS_1ab;1190 | 0 | 0 |
| NC_019843.3;YP_009047202.1;MERS_1ab;1191 | 0 | 0 |
| NC_019843.3;YP_009047202.1;MERS_1ab;1192 | 0 | 0 |
| NC_019843.3;YP_009047202.1;MERS_1ab;1193 | 0 | 0 |
| NC_019843.3;YP_009047202.1;MERS_1ab;1194 | 0 | 0 |
| NC_019843.3;YP_009047202.1;MERS_1ab;1195 | 0 | 0 |

| | | |
|---|---|---|
| NC_019843.3;YP_009047202.1;MERS_1ab;1196 | 0 | 0 |
| NC_019843.3;YP_009047202.1;MERS_1ab;1197 | 0 | 0 |
| NC_019843.3;YP_009047202.1;MERS_1ab;1198 | 0 | 0.025 |
| NC_019843.3;YP_009047202.1;MERS_1ab;1199 | 0 | 0 |
| NC_019843.3;YP_009047202.1;MERS_1ab;1200 | 0 | 0.025 |
| NC_019843.3;YP_009047202.1;MERS_1ab;1201 | 0 | 0.025 |
| NC_019843.3;YP_009047202.1;MERS_1ab;1202 | 0 | 0.05 |
| NC_019843.3;YP_009047202.1;MERS_1ab;1203 | 0.05 | 0.025 |
| NC_019843.3;YP_009047202.1;MERS_1ab;1204 | 0.05 | 0 |
| NC_019843.3;YP_009047202.1;MERS_1ab;1205 | 0.05 | 0.025 |
| NC_019843.3;YP_009047202.1;MERS_1ab;1206 | 0.05 | 0 |
| NC_019843.3;YP_009047202.1;MERS_1ab;1207 | 0.05 | 0 |
| NC_019843.3;YP_009047202.1;MERS_1ab;1208 | 0.05 | 0 |
| NC_019843.3;YP_009047202.1;MERS_1ab;1209 | 0.05 | 0.025 |
| NC_019843.3;YP_009047202.1;MERS_1ab;1210 | 0.1 | 0.025 |
| NC_019843.3;YP_009047202.1;MERS_1ab;1211 | 0 | 0 |
| NC_019843.3;YP_009047202.1;MERS_1ab;1212 | 0 | 0 |
| NC_019843.3;YP_009047202.1;MERS_1ab;1213 | 0 | 0 |
| NC_019843.3;YP_009047202.1;MERS_1ab;1214 | 0 | 0 |
| NC_019843.3;YP_009047202.1;MERS_1ab;1215 | 0 | 0 |
| NC_019843.3;YP_009047202.1;MERS_1ab;1216 | 0 | 0 |
| NC_019843.3;YP_009047202.1;MERS_1ab;1217 | 0 | 0 |
| NC_019843.3;YP_009047202.1;MERS_1ab;1218 | 0 | 0 |
| NC_019843.3;YP_009047202.1;MERS_1ab;1219 | 0 | 0 |
| NC_019843.3;YP_009047202.1;MERS_1ab;1220 | 0 | 0 |
| NC_019843.3;YP_009047202.1;MERS_1ab;1221 | 0 | 0 |
| NC_019843.3;YP_009047202.1;MERS_1ab;1222 | 0 | 0 |
| NC_019843.3;YP_009047202.1;MERS_1ab;1223 | 0 | 0 |
| NC_019843.3;YP_009047202.1;MERS_1ab;1224 | 0 | 0 |
| NC_019843.3;YP_009047202.1;MERS_1ab;1225 | 0 | 0 |
| NC_019843.3;YP_009047202.1;MERS_1ab;1226 | 0 | 0 |
| NC_019843.3;YP_009047202.1;MERS_1ab;1227 | 0 | 0 |
| NC_019843.3;YP_009047202.1;MERS_1ab;1228 | 0 | 0 |
| NC_019843.3;YP_009047202.1;MERS_1ab;1229 | 0 | 0 |
| NC_019843.3;YP_009047202.1;MERS_1ab;1230 | 0 | 0 |
| NC_019843.3;YP_009047202.1;MERS_1ab;1231 | 0 | 0 |
| NC_019843.3;YP_009047202.1;MERS_1ab;1232 | 0 | 0 |
| NC_019843.3;YP_009047202.1;MERS_1ab;1233 | 0 | 0 |
| NC_019843.3;YP_009047202.1;MERS_1ab;1234 | 0 | 0 |
| NC_019843.3;YP_009047202.1;MERS_1ab;1235 | 0 | 0 |
| NC_019843.3;YP_009047202.1;MERS_1ab;1236 | 0 | 0 |
| NC_019843.3;YP_009047202.1;MERS_1ab;1237 | 0 | 0 |
| NC_019843.3;YP_009047202.1;MERS_1ab;1238 | 0 | 0 |

| | | |
|---|---|---|
| NC_019843.3;YP_009047202.1;MERS_1ab;1239 | 0 | 0 |
| NC_019843.3;YP_009047202.1;MERS_1ab;1240 | 0 | 0 |
| NC_019843.3;YP_009047202.1;MERS_1ab;1241 | 0 | 0 |
| NC_019843.3;YP_009047202.1;MERS_1ab;1242 | 0 | 0 |
| NC_019843.3;YP_009047202.1;MERS_1ab;1243 | 0 | 0 |
| NC_019843.3;YP_009047202.1;MERS_1ab;1244 | 0 | 0.025 |
| NC_019843.3;YP_009047202.1;MERS_1ab;1245 | 0 | 0 |
| NC_019843.3;YP_009047202.1;MERS_1ab;1246 | 0 | 0 |
| NC_019843.3;YP_009047202.1;MERS_1ab;1247 | 0 | 0 |
| NC_019843.3;YP_009047202.1;MERS_1ab;1248 | 0 | 0 |
| NC_019843.3;YP_009047202.1;MERS_1ab;1249 | 0 | 0 |
| NC_019843.3;YP_009047202.1;MERS_1ab;1250 | 0 | 0 |
| NC_019843.3;YP_009047202.1;MERS_1ab;1251 | 0 | 0 |
| NC_019843.3;YP_009047202.1;MERS_1ab;1252 | 0 | 0 |
| NC_019843.3;YP_009047202.1;MERS_1ab;1253 | 0 | 0 |
| NC_019843.3;YP_009047202.1;MERS_1ab;1254 | 0 | 0 |
| NC_019843.3;YP_009047202.1;MERS_1ab;1255 | 0 | 0 |
| NC_019843.3;YP_009047202.1;MERS_1ab;1256 | 0 | 0 |
| NC_019843.3;YP_009047202.1;MERS_1ab;1257 | 0 | 0 |
| NC_019843.3;YP_009047202.1;MERS_1ab;1258 | 0 | 0 |
| NC_019843.3;YP_009047202.1;MERS_1ab;1259 | 0 | 0 |
| NC_019843.3;YP_009047202.1;MERS_1ab;1260 | 0 | 0.025 |
| NC_019843.3;YP_009047202.1;MERS_1ab;1261 | 0 | 0 |
| NC_019843.3;YP_009047202.1;MERS_1ab;1262 | 0 | 0 |
| NC_019843.3;YP_009047202.1;MERS_1ab;1263 | 0 | 0 |
| NC_019843.3;YP_009047202.1;MERS_1ab;1264 | 0 | 0.025 |
| NC_019843.3;YP_009047202.1;MERS_1ab;1265 | 0 | 0.025 |
| NC_019843.3;YP_009047202.1;MERS_1ab;1266 | 0 | 0 |
| NC_019843.3;YP_009047202.1;MERS_1ab;1267 | 0 | 0 |
| NC_019843.3;YP_009047202.1;MERS_1ab;1268 | 0 | 0 |
| NC_019843.3;YP_009047202.1;MERS_1ab;1269 | 0 | 0 |
| NC_019843.3;YP_009047202.1;MERS_1ab;1270 | 0 | 0 |
| NC_019843.3;YP_009047202.1;MERS_1ab;1271 | 0 | 0.025 |
| NC_019843.3;YP_009047202.1;MERS_1ab;1272 | 0 | 0.025 |
| NC_019843.3;YP_009047202.1;MERS_1ab;1273 | 0 | 0 |
| NC_019843.3;YP_009047202.1;MERS_1ab;1274 | 0 | 0 |
| NC_019843.3;YP_009047202.1;MERS_1ab;1275 | 0 | 0 |
| NC_019843.3;YP_009047202.1;MERS_1ab;1276 | 0 | 0 |
| NC_019843.3;YP_009047202.1;MERS_1ab;1277 | 0 | 0 |
| NC_019843.3;YP_009047202.1;MERS_1ab;1278 | 0 | 0.025 |
| NC_019843.3;YP_009047202.1;MERS_1ab;1279 | 0 | 0.025 |
| NC_019843.3;YP_009047202.1;MERS_1ab;1280 | 0 | 0.025 |
| NC_019843.3;YP_009047202.1;MERS_1ab;1281 | 0 | 0.025 |

| | | |
|---|---|---|
| NC_019843.3;YP_009047202.1;MERS_1ab;1282 | 0 | 0.05 |
| NC_019843.3;YP_009047202.1;MERS_1ab;1283 | 0 | 0.075 |
| NC_019843.3;YP_009047202.1;MERS_1ab;1284 | 0 | 0.025 |
| NC_019843.3;YP_009047202.1;MERS_1ab;1285 | 0 | 0.1 |
| NC_019843.3;YP_009047202.1;MERS_1ab;1286 | 0 | 0.075 |
| NC_019843.3;YP_009047202.1;MERS_1ab;1287 | 0 | 0 |
| NC_019843.3;YP_009047202.1;MERS_1ab;1288 | 0 | 0 |
| NC_019843.3;YP_009047202.1;MERS_1ab;1289 | 0.05 | 0 |
| NC_019843.3;YP_009047202.1;MERS_1ab;1290 | 0 | 0 |
| NC_019843.3;YP_009047202.1;MERS_1ab;1291 | 0 | 0 |
| NC_019843.3;YP_009047202.1;MERS_1ab;1292 | 0 | 0 |
| NC_019843.3;YP_009047202.1;MERS_1ab;1293 | 0 | 0 |
| NC_019843.3;YP_009047202.1;MERS_1ab;1294 | 0 | 0 |
| NC_019843.3;YP_009047202.1;MERS_1ab;1295 | 0 | 0 |
| NC_019843.3;YP_009047202.1;MERS_1ab;1296 | 0 | 0 |
| NC_019843.3;YP_009047202.1;MERS_1ab;1297 | 0 | 0 |
| NC_019843.3;YP_009047202.1;MERS_1ab;1298 | 0 | 0 |
| NC_019843.3;YP_009047202.1;MERS_1ab;1299 | 0 | 0 |
| NC_019843.3;YP_009047202.1;MERS_1ab;1300 | 0 | 0 |
| NC_019843.3;YP_009047202.1;MERS_1ab;1301 | 0 | 0.05 |
| NC_019843.3;YP_009047202.1;MERS_1ab;1302 | 0 | 0.025 |
| NC_019843.3;YP_009047202.1;MERS_1ab;1303 | 0 | 0.05 |
| NC_019843.3;YP_009047202.1;MERS_1ab;1304 | 0 | 0.025 |
| NC_019843.3;YP_009047202.1;MERS_1ab;1305 | 0 | 0.1 |
| NC_019843.3;YP_009047202.1;MERS_1ab;1306 | 0.1 | 0.05 |
| NC_019843.3;YP_009047202.1;MERS_1ab;1307 | 0 | 0 |
| NC_019843.3;YP_009047202.1;MERS_1ab;1308 | 0 | 0 |
| NC_019843.3;YP_009047202.1;MERS_1ab;1309 | 0 | 0 |
| NC_019843.3;YP_009047202.1;MERS_1ab;1310 | 0 | 0 |
| NC_019843.3;YP_009047202.1;MERS_1ab;1311 | 0 | 0 |
| NC_019843.3;YP_009047202.1;MERS_1ab;1312 | 0 | 0 |
| NC_019843.3;YP_009047202.1;MERS_1ab;1313 | 0 | 0 |
| NC_019843.3;YP_009047202.1;MERS_1ab;1314 | 0 | 0 |
| NC_019843.3;YP_009047202.1;MERS_1ab;1315 | 0 | 0 |
| NC_019843.3;YP_009047202.1;MERS_1ab;1316 | 0 | 0 |
| NC_019843.3;YP_009047202.1;MERS_1ab;1317 | 0 | 0.025 |
| NC_019843.3;YP_009047202.1;MERS_1ab;1318 | 0.1 | 0.075 |
| NC_019843.3;YP_009047202.1;MERS_1ab;1319 | 0.05 | 0 |
| NC_019843.3;YP_009047202.1;MERS_1ab;1320 | 0 | 0 |
| NC_019843.3;YP_009047202.1;MERS_1ab;1321 | 0 | 0 |
| NC_019843.3;YP_009047202.1;MERS_1ab;1322 | 0 | 0 |
| NC_019843.3;YP_009047202.1;MERS_1ab;1323 | 0 | 0 |
| NC_019843.3;YP_009047202.1;MERS_1ab;1324 | 0 | 0 |

| | | |
|---|---|---|
| NC_019843.3;YP_009047202.1;MERS_1ab;1325 | 0 | 0 |
| NC_019843.3;YP_009047202.1;MERS_1ab;1326 | 0 | 0 |
| NC_019843.3;YP_009047202.1;MERS_1ab;1327 | 0 | 0 |
| NC_019843.3;YP_009047202.1;MERS_1ab;1328 | 0 | 0 |
| NC_019843.3;YP_009047202.1;MERS_1ab;1329 | 0 | 0 |
| NC_019843.3;YP_009047202.1;MERS_1ab;1330 | 0 | 0 |
| NC_019843.3;YP_009047202.1;MERS_1ab;1331 | 0 | 0 |
| NC_019843.3;YP_009047202.1;MERS_1ab;1332 | 0 | 0 |
| NC_019843.3;YP_009047202.1;MERS_1ab;1333 | 0 | 0 |
| NC_019843.3;YP_009047202.1;MERS_1ab;1334 | 0 | 0 |
| NC_019843.3;YP_009047202.1;MERS_1ab;1335 | 0 | 0 |
| NC_019843.3;YP_009047202.1;MERS_1ab;1336 | 0.15 | 0.075 |
| NC_019843.3;YP_009047202.1;MERS_1ab;1337 | 0.1 | 0.025 |
| NC_019843.3;YP_009047202.1;MERS_1ab;1338 | 0.1 | 0.025 |
| NC_019843.3;YP_009047202.1;MERS_1ab;1339 | 0.15 | 0.025 |
| NC_019843.3;YP_009047202.1;MERS_1ab;1340 | 0.15 | 0.075 |
| NC_019843.3;YP_009047202.1;MERS_1ab;1341 | 0 | 0.05 |
| NC_019843.3;YP_009047202.1;MERS_1ab;1342 | 0 | 0.025 |
| NC_019843.3;YP_009047202.1;MERS_1ab;1343 | 0.05 | 0.025 |
| NC_019843.3;YP_009047202.1;MERS_1ab;1344 | 0.05 | 0.05 |
| NC_019843.3;YP_009047202.1;MERS_1ab;1345 | 0.1 | 0.075 |
| NC_019843.3;YP_009047202.1;MERS_1ab;1346 | 0.1 | 0 |
| NC_019843.3;YP_009047202.1;MERS_1ab;1347 | 0 | 0 |
| NC_019843.3;YP_009047202.1;MERS_1ab;1348 | 0 | 0 |
| NC_019843.3;YP_009047202.1;MERS_1ab;1349 | 0 | 0.025 |
| NC_019843.3;YP_009047202.1;MERS_1ab;1350 | 0 | 0 |
| NC_019843.3;YP_009047202.1;MERS_1ab;1351 | 0 | 0 |
| NC_019843.3;YP_009047202.1;MERS_1ab;1352 | 0 | 0 |
| NC_019843.3;YP_009047202.1;MERS_1ab;1353 | 0 | 0 |
| NC_019843.3;YP_009047202.1;MERS_1ab;1354 | 0 | 0 |
| NC_019843.3;YP_009047202.1;MERS_1ab;1355 | 0 | 0 |
| NC_019843.3;YP_009047202.1;MERS_1ab;1356 | 0 | 0 |
| NC_019843.3;YP_009047202.1;MERS_1ab;1357 | 0 | 0 |
| NC_019843.3;YP_009047202.1;MERS_1ab;1358 | 0 | 0 |
| NC_019843.3;YP_009047202.1;MERS_1ab;1359 | 0 | 0 |
| NC_019843.3;YP_009047202.1;MERS_1ab;1360 | 0 | 0 |
| NC_019843.3;YP_009047202.1;MERS_1ab;1361 | 0 | 0 |
| NC_019843.3;YP_009047202.1;MERS_1ab;1362 | 0 | 0 |
| NC_019843.3;YP_009047202.1;MERS_1ab;1363 | 0 | 0 |
| NC_019843.3;YP_009047202.1;MERS_1ab;1364 | 0 | 0 |
| NC_019843.3;YP_009047202.1;MERS_1ab;1365 | 0 | 0 |
| NC_019843.3;YP_009047202.1;MERS_1ab;1366 | 0 | 0 |
| NC_019843.3;YP_009047202.1;MERS_1ab;1367 | 0 | 0 |

| | | |
|---|---|---|
| NC_019843.3;YP_009047202.1;MERS_1ab;1368 | 0 | 0 |
| NC_019843.3;YP_009047202.1;MERS_1ab;1369 | 0 | 0 |
| NC_019843.3;YP_009047202.1;MERS_1ab;1370 | 0 | 0 |
| NC_019843.3;YP_009047202.1;MERS_1ab;1371 | 0 | 0 |
| NC_019843.3;YP_009047202.1;MERS_1ab;1372 | 0 | 0 |
| NC_019843.3;YP_009047202.1;MERS_1ab;1373 | 0 | 0 |
| NC_019843.3;YP_009047202.1;MERS_1ab;1374 | 0 | 0 |
| NC_019843.3;YP_009047202.1;MERS_1ab;1375 | 0 | 0 |
| NC_019843.3;YP_009047202.1;MERS_1ab;1376 | 0 | 0.025 |
| NC_019843.3;YP_009047202.1;MERS_1ab;1377 | 0 | 0 |
| NC_019843.3;YP_009047202.1;MERS_1ab;1378 | 0.05 | 0.025 |
| NC_019843.3;YP_009047202.1;MERS_1ab;1379 | 0.05 | 0 |
| NC_019843.3;YP_009047202.1;MERS_1ab;1380 | 0.05 | 0 |
| NC_019843.3;YP_009047202.1;MERS_1ab;1381 | 0.05 | 0 |
| NC_019843.3;YP_009047202.1;MERS_1ab;1382 | 0 | 0 |
| NC_019843.3;YP_009047202.1;MERS_1ab;1383 | 0 | 0 |
| NC_019843.3;YP_009047202.1;MERS_1ab;1384 | 0 | 0 |
| NC_019843.3;YP_009047202.1;MERS_1ab;1385 | 0 | 0 |
| NC_019843.3;YP_009047202.1;MERS_1ab;1386 | 0 | 0 |
| NC_019843.3;YP_009047202.1;MERS_1ab;1387 | 0 | 0.025 |
| NC_019843.3;YP_009047202.1;MERS_1ab;1388 | 0 | 0 |
| NC_019843.3;YP_009047202.1;MERS_1ab;1389 | 0 | 0.025 |
| NC_019843.3;YP_009047202.1;MERS_1ab;1390 | 0 | 0 |
| NC_019843.3;YP_009047202.1;MERS_1ab;1391 | 0 | 0.025 |
| NC_019843.3;YP_009047202.1;MERS_1ab;1392 | 0 | 0.025 |
| NC_019843.3;YP_009047202.1;MERS_1ab;1393 | 0 | 0.05 |
| NC_019843.3;YP_009047202.1;MERS_1ab;1394 | 0.05 | 0.05 |
| NC_019843.3;YP_009047202.1;MERS_1ab;1395 | 0.05 | 0.075 |
| NC_019843.3;YP_009047202.1;MERS_1ab;1396 | 0 | 0.025 |
| NC_019843.3;YP_009047202.1;MERS_1ab;1397 | 0 | 0.025 |
| NC_019843.3;YP_009047202.1;MERS_1ab;1398 | 0.05 | 0.025 |
| NC_019843.3;YP_009047202.1;MERS_1ab;1399 | 0 | 0 |
| NC_019843.3;YP_009047202.1;MERS_1ab;1400 | 0 | 0 |
| NC_019843.3;YP_009047202.1;MERS_1ab;1401 | 0.05 | 0 |
| NC_019843.3;YP_009047202.1;MERS_1ab;1402 | 0 | 0 |
| NC_019843.3;YP_009047202.1;MERS_1ab;1403 | 0.05 | 0.025 |
| NC_019843.3;YP_009047202.1;MERS_1ab;1404 | 0.4 | 0.425 |
| NC_019843.3;YP_009047202.1;MERS_1ab;1405 | 0 | 0.025 |
| NC_019843.3;YP_009047202.1;MERS_1ab;1406 | 0 | 0.025 |
| NC_019843.3;YP_009047202.1;MERS_1ab;1407 | 0 | 0 |
| NC_019843.3;YP_009047202.1;MERS_1ab;1408 | 0 | 0 |
| NC_019843.3;YP_009047202.1;MERS_1ab;1409 | 0 | 0 |
| NC_019843.3;YP_009047202.1;MERS_1ab;1410 | 0 | 0 |

| | | |
|---|---|---|
| NC_019843.3;YP_009047202.1;MERS_1ab;1411 | 0 | 0 |
| NC_019843.3;YP_009047202.1;MERS_1ab;1412 | 0 | 0 |
| NC_019843.3;YP_009047202.1;MERS_1ab;1413 | 0 | 0.025 |
| NC_019843.3;YP_009047202.1;MERS_1ab;1414 | 0 | 0.025 |
| NC_019843.3;YP_009047202.1;MERS_1ab;1415 | 0 | 0 |
| NC_019843.3;YP_009047202.1;MERS_1ab;1416 | 0 | 0 |
| NC_019843.3;YP_009047202.1;MERS_1ab;1417 | 0 | 0 |
| NC_019843.3;YP_009047202.1;MERS_1ab;1418 | 0 | 0 |
| NC_019843.3;YP_009047202.1;MERS_1ab;1419 | 0 | 0 |
| NC_019843.3;YP_009047202.1;MERS_1ab;1420 | 0 | 0 |
| NC_019843.3;YP_009047202.1;MERS_1ab;1421 | 0 | 0 |
| NC_019843.3;YP_009047202.1;MERS_1ab;1422 | 0 | 0 |
| NC_019843.3;YP_009047202.1;MERS_1ab;1423 | 0 | 0 |
| NC_019843.3;YP_009047202.1;MERS_1ab;1424 | 0.05 | 0 |
| NC_019843.3;YP_009047202.1;MERS_1ab;1425 | 0 | 0 |
| NC_019843.3;YP_009047202.1;MERS_1ab;1426 | 0 | 0 |
| NC_019843.3;YP_009047202.1;MERS_1ab;1427 | 0 | 0 |
| NC_019843.3;YP_009047202.1;MERS_1ab;1428 | 0 | 0 |
| NC_019843.3;YP_009047202.1;MERS_1ab;1429 | 0 | 0 |
| NC_019843.3;YP_009047202.1;MERS_1ab;1430 | 0 | 0 |
| NC_019843.3;YP_009047202.1;MERS_1ab;1431 | 0 | 0 |
| NC_019843.3;YP_009047202.1;MERS_1ab;1432 | 0 | 0 |
| NC_019843.3;YP_009047202.1;MERS_1ab;1433 | 0 | 0 |
| NC_019843.3;YP_009047202.1;MERS_1ab;1434 | 0 | 0 |
| NC_019843.3;YP_009047202.1;MERS_1ab;1435 | 0 | 0 |
| NC_019843.3;YP_009047202.1;MERS_1ab;1436 | 0 | 0.025 |
| NC_019843.3;YP_009047202.1;MERS_1ab;1437 | 0.05 | 0.05 |
| NC_019843.3;YP_009047202.1;MERS_1ab;1438 | 0 | 0.025 |
| NC_019843.3;YP_009047202.1;MERS_1ab;1439 | 0 | 0 |
| NC_019843.3;YP_009047202.1;MERS_1ab;1440 | 0 | 0 |
| NC_019843.3;YP_009047202.1;MERS_1ab;1441 | 0 | 0 |
| NC_019843.3;YP_009047202.1;MERS_1ab;1442 | 0 | 0 |
| NC_019843.3;YP_009047202.1;MERS_1ab;1443 | 0 | 0 |
| NC_019843.3;YP_009047202.1;MERS_1ab;1444 | 0 | 0 |
| NC_019843.3;YP_009047202.1;MERS_1ab;1445 | 0 | 0 |
| NC_019843.3;YP_009047202.1;MERS_1ab;1446 | 0 | 0 |
| NC_019843.3;YP_009047202.1;MERS_1ab;1447 | 0 | 0 |
| NC_019843.3;YP_009047202.1;MERS_1ab;1448 | 0 | 0 |
| NC_019843.3;YP_009047202.1;MERS_1ab;1449 | 0 | 0.025 |
| NC_019843.3;YP_009047202.1;MERS_1ab;1450 | 0 | 0 |
| NC_019843.3;YP_009047202.1;MERS_1ab;1451 | 0 | 0.025 |
| NC_019843.3;YP_009047202.1;MERS_1ab;1452 | 0 | 0 |
| NC_019843.3;YP_009047202.1;MERS_1ab;1453 | 0 | 0 |

| | | |
|---|---|---|
| NC_019843.3;YP_009047202.1;MERS_1ab;1454 | 0 | 0.025 |
| NC_019843.3;YP_009047202.1;MERS_1ab;1455 | 0 | 0.025 |
| NC_019843.3;YP_009047202.1;MERS_1ab;1456 | 0 | 0.025 |
| NC_019843.3;YP_009047202.1;MERS_1ab;1457 | 0 | 0.05 |
| NC_019843.3;YP_009047202.1;MERS_1ab;1458 | 0 | 0 |
| NC_019843.3;YP_009047202.1;MERS_1ab;1459 | 0 | 0 |
| NC_019843.3;YP_009047202.1;MERS_1ab;1460 | 0 | 0 |
| NC_019843.3;YP_009047202.1;MERS_1ab;1461 | 0 | 0 |
| NC_019843.3;YP_009047202.1;MERS_1ab;1462 | 0 | 0 |
| NC_019843.3;YP_009047202.1;MERS_1ab;1463 | 0 | 0 |
| NC_019843.3;YP_009047202.1;MERS_1ab;1464 | 0 | 0 |
| NC_019843.3;YP_009047202.1;MERS_1ab;1465 | 0 | 0 |
| NC_019843.3;YP_009047202.1;MERS_1ab;1466 | 0 | 0 |
| NC_019843.3;YP_009047202.1;MERS_1ab;1467 | 0 | 0 |
| NC_019843.3;YP_009047202.1;MERS_1ab;1468 | 0 | 0 |
| NC_019843.3;YP_009047202.1;MERS_1ab;1469 | 0 | 0 |
| NC_019843.3;YP_009047202.1;MERS_1ab;1470 | 0 | 0 |
| NC_019843.3;YP_009047202.1;MERS_1ab;1471 | 0 | 0 |
| NC_019843.3;YP_009047202.1;MERS_1ab;1472 | 0 | 0 |
| NC_019843.3;YP_009047202.1;MERS_1ab;1473 | 0.05 | 0 |
| NC_019843.3;YP_009047202.1;MERS_1ab;1474 | 0 | 0 |
| NC_019843.3;YP_009047202.1;MERS_1ab;1475 | 0 | 0 |
| NC_019843.3;YP_009047202.1;MERS_1ab;1476 | 0 | 0 |
| NC_019843.3;YP_009047202.1;MERS_1ab;1477 | 0 | 0 |
| NC_019843.3;YP_009047202.1;MERS_1ab;1478 | 0 | 0 |
| NC_019843.3;YP_009047202.1;MERS_1ab;1479 | 0 | 0 |
| NC_019843.3;YP_009047202.1;MERS_1ab;1480 | 0 | 0 |
| NC_019843.3;YP_009047202.1;MERS_1ab;1481 | 0 | 0 |
| NC_019843.3;YP_009047202.1;MERS_1ab;1482 | 0 | 0 |
| NC_019843.3;YP_009047202.1;MERS_1ab;1483 | 0 | 0 |
| NC_019843.3;YP_009047202.1;MERS_1ab;1484 | 0 | 0 |
| NC_019843.3;YP_009047202.1;MERS_1ab;1485 | 0 | 0 |
| NC_019843.3;YP_009047202.1;MERS_1ab;1486 | 0 | 0 |
| NC_019843.3;YP_009047202.1;MERS_1ab;1487 | 0 | 0 |
| NC_019843.3;YP_009047202.1;MERS_1ab;1488 | 0 | 0 |
| NC_019843.3;YP_009047202.1;MERS_1ab;1489 | 0 | 0.025 |
| NC_019843.3;YP_009047202.1;MERS_1ab;1490 | 0.1 | 0.075 |
| NC_019843.3;YP_009047202.1;MERS_1ab;1491 | 0 | 0.075 |
| NC_019843.3;YP_009047202.1;MERS_1ab;1492 | 0 | 0 |
| NC_019843.3;YP_009047202.1;MERS_1ab;1493 | 0 | 0.025 |
| NC_019843.3;YP_009047202.1;MERS_1ab;1494 | 0 | 0 |
| NC_019843.3;YP_009047202.1;MERS_1ab;1495 | 0 | 0 |
| NC_019843.3;YP_009047202.1;MERS_1ab;1496 | 0 | 0 |

| | | |
|---|---|---|
| NC_019843.3;YP_009047202.1;MERS_1ab;1497 | 0 | 0 |
| NC_019843.3;YP_009047202.1;MERS_1ab;1498 | 0 | 0 |
| NC_019843.3;YP_009047202.1;MERS_1ab;1499 | 0 | 0 |
| NC_019843.3;YP_009047202.1;MERS_1ab;1500 | 0 | 0 |
| NC_019843.3;YP_009047202.1;MERS_1ab;1501 | 0 | 0 |
| NC_019843.3;YP_009047202.1;MERS_1ab;1502 | 0 | 0 |
| NC_019843.3;YP_009047202.1;MERS_1ab;1503 | 0 | 0 |
| NC_019843.3;YP_009047202.1;MERS_1ab;1504 | 0 | 0 |
| NC_019843.3;YP_009047202.1;MERS_1ab;1505 | 0 | 0 |
| NC_019843.3;YP_009047202.1;MERS_1ab;1506 | 0 | 0 |
| NC_019843.3;YP_009047202.1;MERS_1ab;1507 | 0 | 0 |
| NC_019843.3;YP_009047202.1;MERS_1ab;1508 | 0 | 0 |
| NC_019843.3;YP_009047202.1;MERS_1ab;1509 | 0 | 0 |
| NC_019843.3;YP_009047202.1;MERS_1ab;1510 | 0 | 0 |
| NC_019843.3;YP_009047202.1;MERS_1ab;1511 | 0 | 0 |
| NC_019843.3;YP_009047202.1;MERS_1ab;1512 | 0 | 0 |
| NC_019843.3;YP_009047202.1;MERS_1ab;1513 | 0 | 0 |
| NC_019843.3;YP_009047202.1;MERS_1ab;1514 | 0.05 | 0 |
| NC_019843.3;YP_009047202.1;MERS_1ab;1515 | 0.05 | 0 |
| NC_019843.3;YP_009047202.1;MERS_1ab;1516 | 0.05 | 0 |
| NC_019843.3;YP_009047202.1;MERS_1ab;1517 | 0 | 0 |
| NC_019843.3;YP_009047202.1;MERS_1ab;1518 | 0.05 | 0.025 |
| NC_019843.3;YP_009047202.1;MERS_1ab;1519 | 0 | 0 |
| NC_019843.3;YP_009047202.1;MERS_1ab;1520 | 0.05 | 0 |
| NC_019843.3;YP_009047202.1;MERS_1ab;1521 | 0.05 | 0 |
| NC_019843.3;YP_009047202.1;MERS_1ab;1522 | 0 | 0 |
| NC_019843.3;YP_009047202.1;MERS_1ab;1523 | 0.05 | 0 |
| NC_019843.3;YP_009047202.1;MERS_1ab;1524 | 0 | 0 |
| NC_019843.3;YP_009047202.1;MERS_1ab;1525 | 0 | 0 |
| NC_019843.3;YP_009047202.1;MERS_1ab;1526 | 0 | 0 |
| NC_019843.3;YP_009047202.1;MERS_1ab;1527 | 0 | 0.025 |
| NC_019843.3;YP_009047202.1;MERS_1ab;1528 | 0 | 0.025 |
| NC_019843.3;YP_009047202.1;MERS_1ab;1529 | 0 | 0.025 |
| NC_019843.3;YP_009047202.1;MERS_1ab;1530 | 0.05 | 0 |
| NC_019843.3;YP_009047202.1;MERS_1ab;1531 | 0.1 | 0 |
| NC_019843.3;YP_009047202.1;MERS_1ab;1532 | 0.05 | 0 |
| NC_019843.3;YP_009047202.1;MERS_1ab;1533 | 0.05 | 0 |
| NC_019843.3;YP_009047202.1;MERS_1ab;1534 | 0.05 | 0 |
| NC_019843.3;YP_009047202.1;MERS_1ab;1535 | 0.15 | 0 |
| NC_019843.3;YP_009047202.1;MERS_1ab;1536 | 0.1 | 0 |
| NC_019843.3;YP_009047202.1;MERS_1ab;1537 | 0.05 | 0 |
| NC_019843.3;YP_009047202.1;MERS_1ab;1538 | 0.05 | 0 |
| NC_019843.3;YP_009047202.1;MERS_1ab;1539 | 0.05 | 0 |

| | | |
|---|---|---|
| NC_019843.3;YP_009047202.1;MERS_1ab;1540 | 0.1 | 0 |
| NC_019843.3;YP_009047202.1;MERS_1ab;1541 | 0.1 | 0.05 |
| NC_019843.3;YP_009047202.1;MERS_1ab;1542 | 0.15 | 0 |
| NC_019843.3;YP_009047202.1;MERS_1ab;1543 | 0.1 | 0 |
| NC_019843.3;YP_009047202.1;MERS_1ab;1544 | 0.1 | 0 |
| NC_019843.3;YP_009047202.1;MERS_1ab;1545 | 0.1 | 0.05 |
| NC_019843.3;YP_009047202.1;MERS_1ab;1546 | 0 | 0.025 |
| NC_019843.3;YP_009047202.1;MERS_1ab;1547 | 0 | 0 |
| NC_019843.3;YP_009047202.1;MERS_1ab;1548 | 0 | 0 |
| NC_019843.3;YP_009047202.1;MERS_1ab;1549 | 0 | 0.025 |
| NC_019843.3;YP_009047202.1;MERS_1ab;1550 | 0 | 0 |
| NC_019843.3;YP_009047202.1;MERS_1ab;1551 | 0 | 0 |
| NC_019843.3;YP_009047202.1;MERS_1ab;1552 | 0 | 0 |
| NC_019843.3;YP_009047202.1;MERS_1ab;1553 | 0 | 0 |
| NC_019843.3;YP_009047202.1;MERS_1ab;1554 | 0 | 0.025 |
| NC_019843.3;YP_009047202.1;MERS_1ab;1555 | 0 | 0 |
| NC_019843.3;YP_009047202.1;MERS_1ab;1556 | 0 | 0 |
| NC_019843.3;YP_009047202.1;MERS_1ab;1557 | 0 | 0 |
| NC_019843.3;YP_009047202.1;MERS_1ab;1558 | 0 | 0 |
| NC_019843.3;YP_009047202.1;MERS_1ab;1559 | 0 | 0 |
| NC_019843.3;YP_009047202.1;MERS_1ab;1560 | 0 | 0 |
| NC_019843.3;YP_009047202.1;MERS_1ab;1561 | 0 | 0 |
| NC_019843.3;YP_009047202.1;MERS_1ab;1562 | 0 | 0 |
| NC_019843.3;YP_009047202.1;MERS_1ab;1563 | 0 | 0 |
| NC_019843.3;YP_009047202.1;MERS_1ab;1564 | 0 | 0 |
| NC_019843.3;YP_009047202.1;MERS_1ab;1565 | 0 | 0 |
| NC_019843.3;YP_009047202.1;MERS_1ab;1566 | 0 | 0 |
| NC_019843.3;YP_009047202.1;MERS_1ab;1567 | 0 | 0.025 |
| NC_019843.3;YP_009047202.1;MERS_1ab;1568 | 0 | 0 |
| NC_019843.3;YP_009047202.1;MERS_1ab;1569 | 0 | 0.025 |
| NC_019843.3;YP_009047202.1;MERS_1ab;1570 | 0 | 0 |
| NC_019843.3;YP_009047202.1;MERS_1ab;1571 | 0 | 0 |
| NC_019843.3;YP_009047202.1;MERS_1ab;1572 | 0 | 0 |
| NC_019843.3;YP_009047202.1;MERS_1ab;1573 | 0 | 0 |
| NC_019843.3;YP_009047202.1;MERS_1ab;1574 | 0 | 0 |
| NC_019843.3;YP_009047202.1;MERS_1ab;1575 | 0 | 0 |
| NC_019843.3;YP_009047202.1;MERS_1ab;1576 | 0 | 0.075 |
| NC_019843.3;YP_009047202.1;MERS_1ab;1577 | 0 | 0.025 |
| NC_019843.3;YP_009047202.1;MERS_1ab;1578 | 0 | 0 |
| NC_019843.3;YP_009047202.1;MERS_1ab;1579 | 0 | 0.025 |
| NC_019843.3;YP_009047202.1;MERS_1ab;1580 | 0 | 0.025 |
| NC_019843.3;YP_009047202.1;MERS_1ab;1581 | 0 | 0.05 |
| NC_019843.3;YP_009047202.1;MERS_1ab;1582 | 0 | 0.025 |

| | | |
|---|---|---|
| NC_019843.3;YP_009047202.1;MERS_1ab;1583 | 0 | 0 |
| NC_019843.3;YP_009047202.1;MERS_1ab;1584 | 0 | 0.025 |
| NC_019843.3;YP_009047202.1;MERS_1ab;1585 | 0 | 0.025 |
| NC_019843.3;YP_009047202.1;MERS_1ab;1586 | 0 | 0.025 |
| NC_019843.3;YP_009047202.1;MERS_1ab;1587 | 0 | 0 |
| NC_019843.3;YP_009047202.1;MERS_1ab;1588 | 0 | 0 |
| NC_019843.3;YP_009047202.1;MERS_1ab;1589 | 0 | 0 |
| NC_019843.3;YP_009047202.1;MERS_1ab;1590 | 0 | 0 |
| NC_019843.3;YP_009047202.1;MERS_1ab;1591 | 0 | 0 |
| NC_019843.3;YP_009047202.1;MERS_1ab;1592 | 0 | 0 |
| NC_019843.3;YP_009047202.1;MERS_1ab;1593 | 0 | 0 |
| NC_019843.3;YP_009047202.1;MERS_1ab;1594 | 0 | 0 |
| NC_019843.3;YP_009047202.1;MERS_1ab;1595 | 0 | 0 |
| NC_019843.3;YP_009047202.1;MERS_1ab;1596 | 0 | 0.025 |
| NC_019843.3;YP_009047202.1;MERS_1ab;1597 | 0 | 0 |
| NC_019843.3;YP_009047202.1;MERS_1ab;1598 | 0 | 0 |
| NC_019843.3;YP_009047202.1;MERS_1ab;1599 | 0 | 0.15 |
| NC_019843.3;YP_009047202.1;MERS_1ab;1600 | 0 | 0 |
| NC_019843.3;YP_009047202.1;MERS_1ab;1601 | 0 | 0 |
| NC_019843.3;YP_009047202.1;MERS_1ab;1602 | 0 | 0.025 |
| NC_019843.3;YP_009047202.1;MERS_1ab;1603 | 0.05 | 0.025 |
| NC_019843.3;YP_009047202.1;MERS_1ab;1604 | 0.15 | 0.025 |
| NC_019843.3;YP_009047202.1;MERS_1ab;1605 | 0 | 0 |
| NC_019843.3;YP_009047202.1;MERS_1ab;1606 | 0 | 0 |
| NC_019843.3;YP_009047202.1;MERS_1ab;1607 | 0 | 0 |
| NC_019843.3;YP_009047202.1;MERS_1ab;1608 | 0 | 0 |
| NC_019843.3;YP_009047202.1;MERS_1ab;1609 | 0 | 0 |
| NC_019843.3;YP_009047202.1;MERS_1ab;1610 | 0 | 0 |
| NC_019843.3;YP_009047202.1;MERS_1ab;1611 | 0 | 0 |
| NC_019843.3;YP_009047202.1;MERS_1ab;1612 | 0 | 0.025 |
| NC_019843.3;YP_009047202.1;MERS_1ab;1613 | 0.05 | 0.025 |
| NC_019843.3;YP_009047202.1;MERS_1ab;1614 | 0.05 | 0.025 |
| NC_019843.3;YP_009047202.1;MERS_1ab;1615 | 0.05 | 0.025 |
| NC_019843.3;YP_009047202.1;MERS_1ab;1616 | 0.1 | 0.025 |
| NC_019843.3;YP_009047202.1;MERS_1ab;1617 | 0.1 | 0.025 |
| NC_019843.3;YP_009047202.1;MERS_1ab;1618 | 0 | 0.025 |
| NC_019843.3;YP_009047202.1;MERS_1ab;1619 | 0 | 0 |
| NC_019843.3;YP_009047202.1;MERS_1ab;1620 | 0 | 0 |
| NC_019843.3;YP_009047202.1;MERS_1ab;1621 | 0 | 0 |
| NC_019843.3;YP_009047202.1;MERS_1ab;1622 | 0 | 0 |
| NC_019843.3;YP_009047202.1;MERS_1ab;1623 | 0 | 0 |
| NC_019843.3;YP_009047202.1;MERS_1ab;1624 | 0 | 0 |
| NC_019843.3;YP_009047202.1;MERS_1ab;1625 | 0 | 0 |

| | | |
|---|---|---|
| NC_019843.3;YP_009047202.1;MERS_1ab;1626 | 0 | 0 |
| NC_019843.3;YP_009047202.1;MERS_1ab;1627 | 0 | 0 |
| NC_019843.3;YP_009047202.1;MERS_1ab;1628 | 0 | 0 |
| NC_019843.3;YP_009047202.1;MERS_1ab;1629 | 0 | 0 |
| NC_019843.3;YP_009047202.1;MERS_1ab;1630 | 0 | 0 |
| NC_019843.3;YP_009047202.1;MERS_1ab;1631 | 0 | 0 |
| NC_019843.3;YP_009047202.1;MERS_1ab;1632 | 0 | 0.025 |
| NC_019843.3;YP_009047202.1;MERS_1ab;1633 | 0.1 | 0 |
| NC_019843.3;YP_009047202.1;MERS_1ab;1634 | 0.15 | 0 |
| NC_019843.3;YP_009047202.1;MERS_1ab;1635 | 0.15 | 0.05 |
| NC_019843.3;YP_009047202.1;MERS_1ab;1636 | 0.1 | 0 |
| NC_019843.3;YP_009047202.1;MERS_1ab;1637 | 0.05 | 0 |
| NC_019843.3;YP_009047202.1;MERS_1ab;1638 | 0.05 | 0 |
| NC_019843.3;YP_009047202.1;MERS_1ab;1639 | 0.1 | 0 |
| NC_019843.3;YP_009047202.1;MERS_1ab;1640 | 0.05 | 0 |
| NC_019843.3;YP_009047202.1;MERS_1ab;1641 | 0.05 | 0 |
| NC_019843.3;YP_009047202.1;MERS_1ab;1642 | 0 | 0 |
| NC_019843.3;YP_009047202.1;MERS_1ab;1643 | 0 | 0 |
| NC_019843.3;YP_009047202.1;MERS_1ab;1644 | 0 | 0.025 |
| NC_019843.3;YP_009047202.1;MERS_1ab;1645 | 0.15 | 0.075 |
| NC_019843.3;YP_009047202.1;MERS_1ab;1646 | 0 | 0 |
| NC_019843.3;YP_009047202.1;MERS_1ab;1647 | 0 | 0 |
| NC_019843.3;YP_009047202.1;MERS_1ab;1648 | 0 | 0 |
| NC_019843.3;YP_009047202.1;MERS_1ab;1649 | 0 | 0 |
| NC_019843.3;YP_009047202.1;MERS_1ab;1650 | 0 | 0 |
| NC_019843.3;YP_009047202.1;MERS_1ab;1651 | 0 | 0 |
| NC_019843.3;YP_009047202.1;MERS_1ab;1652 | 0 | 0 |
| NC_019843.3;YP_009047202.1;MERS_1ab;1653 | 0 | 0 |
| NC_019843.3;YP_009047202.1;MERS_1ab;1654 | 0 | 0 |
| NC_019843.3;YP_009047202.1;MERS_1ab;1655 | 0 | 0 |
| NC_019843.3;YP_009047202.1;MERS_1ab;1656 | 0 | 0 |
| NC_019843.3;YP_009047202.1;MERS_1ab;1657 | 0 | 0 |
| NC_019843.3;YP_009047202.1;MERS_1ab;1658 | 0 | 0 |
| NC_019843.3;YP_009047202.1;MERS_1ab;1659 | 0 | 0 |
| NC_019843.3;YP_009047202.1;MERS_1ab;1660 | 0 | 0 |
| NC_019843.3;YP_009047202.1;MERS_1ab;1661 | 0 | 0 |
| NC_019843.3;YP_009047202.1;MERS_1ab;1662 | 0 | 0 |
| NC_019843.3;YP_009047202.1;MERS_1ab;1663 | 0 | 0 |
| NC_019843.3;YP_009047202.1;MERS_1ab;1664 | 0 | 0 |
| NC_019843.3;YP_009047202.1;MERS_1ab;1665 | 0 | 0 |
| NC_019843.3;YP_009047202.1;MERS_1ab;1666 | 0 | 0 |
| NC_019843.3;YP_009047202.1;MERS_1ab;1667 | 0 | 0 |
| NC_019843.3;YP_009047202.1;MERS_1ab;1668 | 0 | 0 |

| | | |
|---|---|---|
| NC_019843.3;YP_009047202.1;MERS_1ab;1669 | 0 | 0 |
| NC_019843.3;YP_009047202.1;MERS_1ab;1670 | 0 | 0 |
| NC_019843.3;YP_009047202.1;MERS_1ab;1671 | 0 | 0 |
| NC_019843.3;YP_009047202.1;MERS_1ab;1672 | 0 | 0 |
| NC_019843.3;YP_009047202.1;MERS_1ab;1673 | 0 | 0 |
| NC_019843.3;YP_009047202.1;MERS_1ab;1674 | 0 | 0 |
| NC_019843.3;YP_009047202.1;MERS_1ab;1675 | 0 | 0 |
| NC_019843.3;YP_009047202.1;MERS_1ab;1676 | 0.15 | 0.125 |
| NC_019843.3;YP_009047202.1;MERS_1ab;1677 | 0 | 0 |
| NC_019843.3;YP_009047202.1;MERS_1ab;1678 | 0 | 0 |
| NC_019843.3;YP_009047202.1;MERS_1ab;1679 | 0 | 0 |
| NC_019843.3;YP_009047202.1;MERS_1ab;1680 | 0 | 0 |
| NC_019843.3;YP_009047202.1;MERS_1ab;1681 | 0 | 0 |
| NC_019843.3;YP_009047202.1;MERS_1ab;1682 | 0 | 0 |
| NC_019843.3;YP_009047202.1;MERS_1ab;1683 | 0 | 0 |
| NC_019843.3;YP_009047202.1;MERS_1ab;1684 | 0 | 0 |
| NC_019843.3;YP_009047202.1;MERS_1ab;1685 | 0 | 0 |
| NC_019843.3;YP_009047202.1;MERS_1ab;1686 | 0 | 0 |
| NC_019843.3;YP_009047202.1;MERS_1ab;1687 | 0 | 0 |
| NC_019843.3;YP_009047202.1;MERS_1ab;1688 | 0 | 0.025 |
| NC_019843.3;YP_009047202.1;MERS_1ab;1689 | 0 | 0.025 |
| NC_019843.3;YP_009047202.1;MERS_1ab;1690 | 0 | 0.025 |
| NC_019843.3;YP_009047202.1;MERS_1ab;1691 | 0 | 0 |
| NC_019843.3;YP_009047202.1;MERS_1ab;1692 | 0 | 0.025 |
| NC_019843.3;YP_009047202.1;MERS_1ab;1693 | 0 | 0.175 |
| NC_019843.3;YP_009047202.1;MERS_1ab;1694 | 0 | 0.225 |
| NC_019843.3;YP_009047202.1;MERS_1ab;1695 | 0 | 0.025 |
| NC_019843.3;YP_009047202.1;MERS_1ab;1696 | 0 | 0 |
| NC_019843.3;YP_009047202.1;MERS_1ab;1697 | 0 | 0.025 |
| NC_019843.3;YP_009047202.1;MERS_1ab;1698 | 0 | 0 |
| NC_019843.3;YP_009047202.1;MERS_1ab;1699 | 0 | 0 |
| NC_019843.3;YP_009047202.1;MERS_1ab;1700 | 0 | 0 |
| NC_019843.3;YP_009047202.1;MERS_1ab;1701 | 0 | 0 |
| NC_019843.3;YP_009047202.1;MERS_1ab;1702 | 0 | 0 |
| NC_019843.3;YP_009047202.1;MERS_1ab;1703 | 0 | 0 |
| NC_019843.3;YP_009047202.1;MERS_1ab;1704 | 0 | 0 |
| NC_019843.3;YP_009047202.1;MERS_1ab;1705 | 0 | 0 |
| NC_019843.3;YP_009047202.1;MERS_1ab;1706 | 0 | 0 |
| NC_019843.3;YP_009047202.1;MERS_1ab;1707 | 0 | 0 |
| NC_019843.3;YP_009047202.1;MERS_1ab;1708 | 0 | 0 |
| NC_019843.3;YP_009047202.1;MERS_1ab;1709 | 0 | 0 |
| NC_019843.3;YP_009047202.1;MERS_1ab;1710 | 0 | 0.025 |
| NC_019843.3;YP_009047202.1;MERS_1ab;1711 | 0 | 0 |

| | | |
|---|---|---|
| NC_019843.3;YP_009047202.1;MERS_1ab;1712 | 0 | 0 |
| NC_019843.3;YP_009047202.1;MERS_1ab;1713 | 0 | 0 |
| NC_019843.3;YP_009047202.1;MERS_1ab;1714 | 0 | 0 |
| NC_019843.3;YP_009047202.1;MERS_1ab;1715 | 0 | 0 |
| NC_019843.3;YP_009047202.1;MERS_1ab;1716 | 0 | 0 |
| NC_019843.3;YP_009047202.1;MERS_1ab;1717 | 0 | 0 |
| NC_019843.3;YP_009047202.1;MERS_1ab;1718 | 0 | 0 |
| NC_019843.3;YP_009047202.1;MERS_1ab;1719 | 0 | 0 |
| NC_019843.3;YP_009047202.1;MERS_1ab;1720 | 0 | 0 |
| NC_019843.3;YP_009047202.1;MERS_1ab;1721 | 0 | 0 |
| NC_019843.3;YP_009047202.1;MERS_1ab;1722 | 0 | 0 |
| NC_019843.3;YP_009047202.1;MERS_1ab;1723 | 0 | 0 |
| NC_019843.3;YP_009047202.1;MERS_1ab;1724 | 0 | 0 |
| NC_019843.3;YP_009047202.1;MERS_1ab;1725 | 0 | 0 |
| NC_019843.3;YP_009047202.1;MERS_1ab;1726 | 0 | 0 |
| NC_019843.3;YP_009047202.1;MERS_1ab;1727 | 0 | 0 |
| NC_019843.3;YP_009047202.1;MERS_1ab;1728 | 0 | 0 |
| NC_019843.3;YP_009047202.1;MERS_1ab;1729 | 0 | 0 |
| NC_019843.3;YP_009047202.1;MERS_1ab;1730 | 0 | 0 |
| NC_019843.3;YP_009047202.1;MERS_1ab;1731 | 0 | 0 |
| NC_019843.3;YP_009047202.1;MERS_1ab;1732 | 0 | 0 |
| NC_019843.3;YP_009047202.1;MERS_1ab;1733 | 0 | 0 |
| NC_019843.3;YP_009047202.1;MERS_1ab;1734 | 0 | 0 |
| NC_019843.3;YP_009047202.1;MERS_1ab;1735 | 0 | 0 |
| NC_019843.3;YP_009047202.1;MERS_1ab;1736 | 0 | 0 |
| NC_019843.3;YP_009047202.1;MERS_1ab;1737 | 0 | 0 |
| NC_019843.3;YP_009047202.1;MERS_1ab;1738 | 0 | 0 |
| NC_019843.3;YP_009047202.1;MERS_1ab;1739 | 0 | 0 |
| NC_019843.3;YP_009047202.1;MERS_1ab;1740 | 0 | 0 |
| NC_019843.3;YP_009047202.1;MERS_1ab;1741 | 0 | 0 |
| NC_019843.3;YP_009047202.1;MERS_1ab;1742 | 0 | 0 |
| NC_019843.3;YP_009047202.1;MERS_1ab;1743 | 0 | 0 |
| NC_019843.3;YP_009047202.1;MERS_1ab;1744 | 0 | 0 |
| NC_019843.3;YP_009047202.1;MERS_1ab;1745 | 0 | 0 |
| NC_019843.3;YP_009047202.1;MERS_1ab;1746 | 0 | 0 |
| NC_019843.3;YP_009047202.1;MERS_1ab;1747 | 0 | 0 |
| NC_019843.3;YP_009047202.1;MERS_1ab;1748 | 0 | 0 |
| NC_019843.3;YP_009047202.1;MERS_1ab;1749 | 0 | 0 |
| NC_019843.3;YP_009047202.1;MERS_1ab;1750 | 0 | 0 |
| NC_019843.3;YP_009047202.1;MERS_1ab;1751 | 0 | 0 |
| NC_019843.3;YP_009047202.1;MERS_1ab;1752 | 0.1 | 0.025 |
| NC_019843.3;YP_009047202.1;MERS_1ab;1753 | 0 | 0 |
| NC_019843.3;YP_009047202.1;MERS_1ab;1754 | 0 | 0 |

| | | |
|---|---|---|
| NC_019843.3;YP_009047202.1;MERS_1ab;1755 | 0 | 0 |
| NC_019843.3;YP_009047202.1;MERS_1ab;1756 | 0 | 0 |
| NC_019843.3;YP_009047202.1;MERS_1ab;1757 | 0 | 0 |
| NC_019843.3;YP_009047202.1;MERS_1ab;1758 | 0 | 0 |
| NC_019843.3;YP_009047202.1;MERS_1ab;1759 | 0 | 0.025 |
| NC_019843.3;YP_009047202.1;MERS_1ab;1760 | 0 | 0.025 |
| NC_019843.3;YP_009047202.1;MERS_1ab;1761 | 0 | 0.025 |
| NC_019843.3;YP_009047202.1;MERS_1ab;1762 | 0 | 0.025 |
| NC_019843.3;YP_009047202.1;MERS_1ab;1763 | 0 | 0.025 |
| NC_019843.3;YP_009047202.1;MERS_1ab;1764 | 0 | 0.025 |
| NC_019843.3;YP_009047202.1;MERS_1ab;1765 | 0 | 0.025 |
| NC_019843.3;YP_009047202.1;MERS_1ab;1766 | 0 | 0.05 |
| NC_019843.3;YP_009047202.1;MERS_1ab;1767 | 0 | 0 |
| NC_019843.3;YP_009047202.1;MERS_1ab;1768 | 0 | 0 |
| NC_019843.3;YP_009047202.1;MERS_1ab;1769 | 0 | 0 |
| NC_019843.3;YP_009047202.1;MERS_1ab;1770 | 0 | 0.025 |
| NC_019843.3;YP_009047202.1;MERS_1ab;1771 | 0 | 0.025 |
| NC_019843.3;YP_009047202.1;MERS_1ab;1772 | 0 | 0.025 |
| NC_019843.3;YP_009047202.1;MERS_1ab;1773 | 0.05 | 0.025 |
| NC_019843.3;YP_009047202.1;MERS_1ab;1774 | 0.05 | 0.025 |
| NC_019843.3;YP_009047202.1;MERS_1ab;1775 | 0 | 0.025 |
| NC_019843.3;YP_009047202.1;MERS_1ab;1776 | 0 | 0.025 |
| NC_019843.3;YP_009047202.1;MERS_1ab;1777 | 0 | 0.025 |
| NC_019843.3;YP_009047202.1;MERS_1ab;1778 | 0 | 0.025 |
| NC_019843.3;YP_009047202.1;MERS_1ab;1779 | 0 | 0.025 |
| NC_019843.3;YP_009047202.1;MERS_1ab;1780 | 0 | 0 |
| NC_019843.3;YP_009047202.1;MERS_1ab;1781 | 0 | 0 |
| NC_019843.3;YP_009047202.1;MERS_1ab;1782 | 0 | 0 |
| NC_019843.3;YP_009047202.1;MERS_1ab;1783 | 0 | 0 |
| NC_019843.3;YP_009047202.1;MERS_1ab;1784 | 0 | 0 |
| NC_019843.3;YP_009047202.1;MERS_1ab;1785 | 0 | 0.025 |
| NC_019843.3;YP_009047202.1;MERS_1ab;1786 | 0 | 0.05 |
| NC_019843.3;YP_009047202.1;MERS_1ab;1787 | 0.05 | 0.025 |
| NC_019843.3;YP_009047202.1;MERS_1ab;1788 | 0 | 0.025 |
| NC_019843.3;YP_009047202.1;MERS_1ab;1789 | 0.05 | 0.025 |
| NC_019843.3;YP_009047202.1;MERS_1ab;1790 | 0 | 0 |
| NC_019843.3;YP_009047202.1;MERS_1ab;1791 | 0 | 0 |
| NC_019843.3;YP_009047202.1;MERS_1ab;1792 | 0 | 0 |
| NC_019843.3;YP_009047202.1;MERS_1ab;1793 | 0 | 0 |
| NC_019843.3;YP_009047202.1;MERS_1ab;1794 | 0 | 0 |
| NC_019843.3;YP_009047202.1;MERS_1ab;1795 | 0 | 0 |
| NC_019843.3;YP_009047202.1;MERS_1ab;1796 | 0 | 0 |
| NC_019843.3;YP_009047202.1;MERS_1ab;1797 | 0 | 0 |

| | | |
|---|---|---|
| NC_019843.3;YP_009047202.1;MERS_1ab;1798 | 0 | 0 |
| NC_019843.3;YP_009047202.1;MERS_1ab;1799 | 0 | 0 |
| NC_019843.3;YP_009047202.1;MERS_1ab;1800 | 0 | 0 |
| NC_019843.3;YP_009047202.1;MERS_1ab;1801 | 0 | 0 |
| NC_019843.3;YP_009047202.1;MERS_1ab;1802 | 0 | 0 |
| NC_019843.3;YP_009047202.1;MERS_1ab;1803 | 0.2 | 0.025 |
| NC_019843.3;YP_009047202.1;MERS_1ab;1804 | 0 | 0 |
| NC_019843.3;YP_009047202.1;MERS_1ab;1805 | 0.05 | 0 |
| NC_019843.3;YP_009047202.1;MERS_1ab;1806 | 0.1 | 0 |
| NC_019843.3;YP_009047202.1;MERS_1ab;1807 | 0.1 | 0 |
| NC_019843.3;YP_009047202.1;MERS_1ab;1808 | 0.05 | 0 |
| NC_019843.3;YP_009047202.1;MERS_1ab;1809 | 0.05 | 0 |
| NC_019843.3;YP_009047202.1;MERS_1ab;1810 | 0.05 | 0 |
| NC_019843.3;YP_009047202.1;MERS_1ab;1811 | 0 | 0 |
| NC_019843.3;YP_009047202.1;MERS_1ab;1812 | 0 | 0 |
| NC_019843.3;YP_009047202.1;MERS_1ab;1813 | 0 | 0 |
| NC_019843.3;YP_009047202.1;MERS_1ab;1814 | 0 | 0 |
| NC_019843.3;YP_009047202.1;MERS_1ab;1815 | 0.05 | 0 |
| NC_019843.3;YP_009047202.1;MERS_1ab;1816 | 0.1 | 0.025 |
| NC_019843.3;YP_009047202.1;MERS_1ab;1817 | 0.15 | 0.05 |
| NC_019843.3;YP_009047202.1;MERS_1ab;1818 | 0.2 | 0.025 |
| NC_019843.3;YP_009047202.1;MERS_1ab;1819 | 0.15 | 0 |
| NC_019843.3;YP_009047202.1;MERS_1ab;1820 | 0.15 | 0.025 |
| NC_019843.3;YP_009047202.1;MERS_1ab;1821 | 0.2 | 0 |
| NC_019843.3;YP_009047202.1;MERS_1ab;1822 | 0.2 | 0.075 |
| NC_019843.3;YP_009047202.1;MERS_1ab;1823 | 0.3 | 0.1 |
| NC_019843.3;YP_009047202.1;MERS_1ab;1824 | 0.25 | 0.05 |
| NC_019843.3;YP_009047202.1;MERS_1ab;1825 | 0.25 | 0.05 |
| NC_019843.3;YP_009047202.1;MERS_1ab;1826 | 0 | 0.025 |
| NC_019843.3;YP_009047202.1;MERS_1ab;1827 | 0 | 0.025 |
| NC_019843.3;YP_009047202.1;MERS_1ab;1828 | 0.05 | 0 |
| NC_019843.3;YP_009047202.1;MERS_1ab;1829 | 0 | 0 |
| NC_019843.3;YP_009047202.1;MERS_1ab;1830 | 0 | 0 |
| NC_019843.3;YP_009047202.1;MERS_1ab;1831 | 0 | 0 |
| NC_019843.3;YP_009047202.1;MERS_1ab;1832 | 0 | 0.025 |
| NC_019843.3;YP_009047202.1;MERS_1ab;1833 | 0 | 0 |
| NC_019843.3;YP_009047202.1;MERS_1ab;1834 | 0 | 0 |
| NC_019843.3;YP_009047202.1;MERS_1ab;1835 | 0 | 0 |
| NC_019843.3;YP_009047202.1;MERS_1ab;1836 | 0 | 0 |
| NC_019843.3;YP_009047202.1;MERS_1ab;1837 | 0 | 0 |
| NC_019843.3;YP_009047202.1;MERS_1ab;1838 | 0 | 0 |
| NC_019843.3;YP_009047202.1;MERS_1ab;1839 | 0 | 0 |
| NC_019843.3;YP_009047202.1;MERS_1ab;1840 | 0 | 0 |

| | | |
|---|---|---|
| NC_019843.3;YP_009047202.1;MERS_1ab;1841 | 0 | 0 |
| NC_019843.3;YP_009047202.1;MERS_1ab;1842 | 0 | 0 |
| NC_019843.3;YP_009047202.1;MERS_1ab;1843 | 0 | 0 |
| NC_019843.3;YP_009047202.1;MERS_1ab;1844 | 0 | 0 |
| NC_019843.3;YP_009047202.1;MERS_1ab;1845 | 0 | 0 |
| NC_019843.3;YP_009047202.1;MERS_1ab;1846 | 0 | 0 |
| NC_019843.3;YP_009047202.1;MERS_1ab;1847 | 0 | 0 |
| NC_019843.3;YP_009047202.1;MERS_1ab;1848 | 0 | 0.025 |
| NC_019843.3;YP_009047202.1;MERS_1ab;1849 | 0 | 0.025 |
| NC_019843.3;YP_009047202.1;MERS_1ab;1850 | 0.1 | 0.025 |
| NC_019843.3;YP_009047202.1;MERS_1ab;1851 | 0 | 0.025 |
| NC_019843.3;YP_009047202.1;MERS_1ab;1852 | 0 | 0.025 |
| NC_019843.3;YP_009047202.1;MERS_1ab;1853 | 0 | 0.025 |
| NC_019843.3;YP_009047202.1;MERS_1ab;1854 | 0 | 0.025 |
| NC_019843.3;YP_009047202.1;MERS_1ab;1855 | 0 | 0.025 |
| NC_019843.3;YP_009047202.1;MERS_1ab;1856 | 0 | 0.025 |
| NC_019843.3;YP_009047202.1;MERS_1ab;1857 | 0 | 0 |
| NC_019843.3;YP_009047202.1;MERS_1ab;1858 | 0.05 | 0.05 |
| NC_019843.3;YP_009047202.1;MERS_1ab;1859 | 0 | 0 |
| NC_019843.3;YP_009047202.1;MERS_1ab;1860 | 0 | 0 |
| NC_019843.3;YP_009047202.1;MERS_1ab;1861 | 0 | 0 |
| NC_019843.3;YP_009047202.1;MERS_1ab;1862 | 0 | 0 |
| NC_019843.3;YP_009047202.1;MERS_1ab;1863 | 0.05 | 0.025 |
| NC_019843.3;YP_009047202.1;MERS_1ab;1864 | 0 | 0.025 |
| NC_019843.3;YP_009047202.1;MERS_1ab;1865 | 0 | 0 |
| NC_019843.3;YP_009047202.1;MERS_1ab;1866 | 0 | 0 |
| NC_019843.3;YP_009047202.1;MERS_1ab;1867 | 0 | 0.025 |
| NC_019843.3;YP_009047202.1;MERS_1ab;1868 | 0.05 | 0.025 |
| NC_019843.3;YP_009047202.1;MERS_1ab;1869 | 0 | 0.025 |
| NC_019843.3;YP_009047202.1;MERS_1ab;1870 | 0 | 0.05 |
| NC_019843.3;YP_009047202.1;MERS_1ab;1871 | 0.05 | 0.075 |
| NC_019843.3;YP_009047202.1;MERS_1ab;1872 | 0 | 0.025 |
| NC_019843.3;YP_009047202.1;MERS_1ab;1873 | 0.05 | 0.025 |
| NC_019843.3;YP_009047202.1;MERS_1ab;1874 | 0.05 | 0.025 |
| NC_019843.3;YP_009047202.1;MERS_1ab;1875 | 0 | 0.025 |
| NC_019843.3;YP_009047202.1;MERS_1ab;1876 | 0 | 0.025 |
| NC_019843.3;YP_009047202.1;MERS_1ab;1877 | 0 | 0.075 |
| NC_019843.3;YP_009047202.1;MERS_1ab;1878 | 0 | 0 |
| NC_019843.3;YP_009047202.1;MERS_1ab;1879 | 0 | 0 |
| NC_019843.3;YP_009047202.1;MERS_1ab;1880 | 0 | 0 |
| NC_019843.3;YP_009047202.1;MERS_1ab;1881 | 0 | 0 |
| NC_019843.3;YP_009047202.1;MERS_1ab;1882 | 0 | 0 |
| NC_019843.3;YP_009047202.1;MERS_1ab;1883 | 0 | 0.025 |

| | | |
|---|---|---|
| NC_019843.3;YP_009047202.1;MERS_1ab;1884 | 0 | 0.025 |
| NC_019843.3;YP_009047202.1;MERS_1ab;1885 | 0 | 0.025 |
| NC_019843.3;YP_009047202.1;MERS_1ab;1886 | 0 | 0.05 |
| NC_019843.3;YP_009047202.1;MERS_1ab;1887 | 0 | 0 |
| NC_019843.3;YP_009047202.1;MERS_1ab;1888 | 0.05 | 0.025 |
| NC_019843.3;YP_009047202.1;MERS_1ab;1889 | 0.05 | 0.05 |
| NC_019843.3;YP_009047202.1;MERS_1ab;1890 | 0 | 0.025 |
| NC_019843.3;YP_009047202.1;MERS_1ab;1891 | 0 | 0.025 |
| NC_019843.3;YP_009047202.1;MERS_1ab;1892 | 0 | 0.025 |
| NC_019843.3;YP_009047202.1;MERS_1ab;1893 | 0 | 0.075 |
| NC_019843.3;YP_009047202.1;MERS_1ab;1894 | 0 | 0.025 |
| NC_019843.3;YP_009047202.1;MERS_1ab;1895 | 0.05 | 0.05 |
| NC_019843.3;YP_009047202.1;MERS_1ab;1896 | 0 | 0.025 |
| NC_019843.3;YP_009047202.1;MERS_1ab;1897 | 0 | 0.025 |
| NC_019843.3;YP_009047202.1;MERS_1ab;1898 | 0 | 0.075 |
| NC_019843.3;YP_009047202.1;MERS_1ab;1899 | 0 | 0 |
| NC_019843.3;YP_009047202.1;MERS_1ab;1900 | 0 | 0 |
| NC_019843.3;YP_009047202.1;MERS_1ab;1901 | 0.05 | 0 |
| NC_019843.3;YP_009047202.1;MERS_1ab;1902 | 0 | 0.025 |
| NC_019843.3;YP_009047202.1;MERS_1ab;1903 | 0 | 0.05 |
| NC_019843.3;YP_009047202.1;MERS_1ab;1904 | 0 | 0.025 |
| NC_019843.3;YP_009047202.1;MERS_1ab;1905 | 0.05 | 0 |
| NC_019843.3;YP_009047202.1;MERS_1ab;1906 | 0 | 0 |
| NC_019843.3;YP_009047202.1;MERS_1ab;1907 | 0 | 0 |
| NC_019843.3;YP_009047202.1;MERS_1ab;1908 | 0 | 0 |
| NC_019843.3;YP_009047202.1;MERS_1ab;1909 | 0 | 0 |
| NC_019843.3;YP_009047202.1;MERS_1ab;1910 | 0 | 0.025 |
| NC_019843.3;YP_009047202.1;MERS_1ab;1911 | 0 | 0 |
| NC_019843.3;YP_009047202.1;MERS_1ab;1912 | 0 | 0 |
| NC_019843.3;YP_009047202.1;MERS_1ab;1913 | 0 | 0 |
| NC_019843.3;YP_009047202.1;MERS_1ab;1914 | 0 | 0 |
| NC_019843.3;YP_009047202.1;MERS_1ab;1915 | 0 | 0 |
| NC_019843.3;YP_009047202.1;MERS_1ab;1916 | 0 | 0 |
| NC_019843.3;YP_009047202.1;MERS_1ab;1917 | 0 | 0 |
| NC_019843.3;YP_009047202.1;MERS_1ab;1918 | 0 | 0 |
| NC_019843.3;YP_009047202.1;MERS_1ab;1919 | 0 | 0.025 |
| NC_019843.3;YP_009047202.1;MERS_1ab;1920 | 0.1 | 0.05 |
| NC_019843.3;YP_009047202.1;MERS_1ab;1921 | 0.1 | 0.1 |
| NC_019843.3;YP_009047202.1;MERS_1ab;1922 | 0.05 | 0.05 |
| NC_019843.3;YP_009047202.1;MERS_1ab;1923 | 0 | 0.025 |
| NC_019843.3;YP_009047202.1;MERS_1ab;1924 | 0.1 | 0.025 |
| NC_019843.3;YP_009047202.1;MERS_1ab;1925 | 0.1 | 0.05 |
| NC_019843.3;YP_009047202.1;MERS_1ab;1926 | 0.15 | 0.05 |

| | | |
|---|---|---|
| NC_019843.3;YP_009047202.1;MERS_1ab;1927 | 0.15 | 0.075 |
| NC_019843.3;YP_009047202.1;MERS_1ab;1928 | 0.1 | 0.1 |
| NC_019843.3;YP_009047202.1;MERS_1ab;1929 | 0.05 | 0 |
| NC_019843.3;YP_009047202.1;MERS_1ab;1930 | 0.1 | 0 |
| NC_019843.3;YP_009047202.1;MERS_1ab;1931 | 0 | 0 |
| NC_019843.3;YP_009047202.1;MERS_1ab;1932 | 0.05 | 0 |
| NC_019843.3;YP_009047202.1;MERS_1ab;1933 | 0 | 0 |
| NC_019843.3;YP_009047202.1;MERS_1ab;1934 | 0 | 0 |
| NC_019843.3;YP_009047202.1;MERS_1ab;1935 | 0 | 0.025 |
| NC_019843.3;YP_009047202.1;MERS_1ab;1936 | 0 | 0 |
| NC_019843.3;YP_009047202.1;MERS_1ab;1937 | 0 | 0 |
| NC_019843.3;YP_009047202.1;MERS_1ab;1938 | 0.05 | 0 |
| NC_019843.3;YP_009047202.1;MERS_1ab;1939 | 0 | 0 |
| NC_019843.3;YP_009047202.1;MERS_1ab;1940 | 0 | 0.025 |
| NC_019843.3;YP_009047202.1;MERS_1ab;1941 | 0.05 | 0.025 |
| NC_019843.3;YP_009047202.1;MERS_1ab;1942 | 0.1 | 0.075 |
| NC_019843.3;YP_009047202.1;MERS_1ab;1943 | 0.1 | 0.15 |
| NC_019843.3;YP_009047202.1;MERS_1ab;1944 | 0.3 | 0.175 |
| NC_019843.3;YP_009047202.1;MERS_1ab;1945 | 0.1 | 0.1 |
| NC_019843.3;YP_009047202.1;MERS_1ab;1946 | 0.05 | 0.05 |
| NC_019843.3;YP_009047202.1;MERS_1ab;1947 | 0.2 | 0.075 |
| NC_019843.3;YP_009047202.1;MERS_1ab;1948 | 0.15 | 0 |
| NC_019843.3;YP_009047202.1;MERS_1ab;1949 | 0.15 | 0.05 |
| NC_019843.3;YP_009047202.1;MERS_1ab;1950 | 0 | 0.125 |
| NC_019843.3;YP_009047202.1;MERS_1ab;1951 | 0 | 0 |
| NC_019843.3;YP_009047202.1;MERS_1ab;1952 | 0 | 0 |
| NC_019843.3;YP_009047202.1;MERS_1ab;1953 | 0 | 0 |
| NC_019843.3;YP_009047202.1;MERS_1ab;1954 | 0 | 0 |
| NC_019843.3;YP_009047202.1;MERS_1ab;1955 | 0.05 | 0 |
| NC_019843.3;YP_009047202.1;MERS_1ab;1956 | 0 | 0 |
| NC_019843.3;YP_009047202.1;MERS_1ab;1957 | 0 | 0 |
| NC_019843.3;YP_009047202.1;MERS_1ab;1958 | 0 | 0 |
| NC_019843.3;YP_009047202.1;MERS_1ab;1959 | 0 | 0 |
| NC_019843.3;YP_009047202.1;MERS_1ab;1960 | 0 | 0 |
| NC_019843.3;YP_009047202.1;MERS_1ab;1961 | 0 | 0 |
| NC_019843.3;YP_009047202.1;MERS_1ab;1962 | 0 | 0 |
| NC_019843.3;YP_009047202.1;MERS_1ab;1963 | 0 | 0 |
| NC_019843.3;YP_009047202.1;MERS_1ab;1964 | 0.05 | 0 |
| NC_019843.3;YP_009047202.1;MERS_1ab;1965 | 0.05 | 0 |
| NC_019843.3;YP_009047202.1;MERS_1ab;1966 | 0.05 | 0 |
| NC_019843.3;YP_009047202.1;MERS_1ab;1967 | 0.05 | 0 |
| NC_019843.3;YP_009047202.1;MERS_1ab;1968 | 0.05 | 0 |
| NC_019843.3;YP_009047202.1;MERS_1ab;1969 | 0.05 | 0 |

| | | |
|---|---|---|
| NC_019843.3;YP_009047202.1;MERS_1ab;1970 | 0.05 | 0 |
| NC_019843.3;YP_009047202.1;MERS_1ab;1971 | 0.05 | 0 |
| NC_019843.3;YP_009047202.1;MERS_1ab;1972 | 0.05 | 0 |
| NC_019843.3;YP_009047202.1;MERS_1ab;1973 | 0.05 | 0.025 |
| NC_019843.3;YP_009047202.1;MERS_1ab;1974 | 0 | 0 |
| NC_019843.3;YP_009047202.1;MERS_1ab;1975 | 0 | 0 |
| NC_019843.3;YP_009047202.1;MERS_1ab;1976 | 0 | 0 |
| NC_019843.3;YP_009047202.1;MERS_1ab;1977 | 0 | 0 |
| NC_019843.3;YP_009047202.1;MERS_1ab;1978 | 0 | 0.025 |
| NC_019843.3;YP_009047202.1;MERS_1ab;1979 | 0.05 | 0 |
| NC_019843.3;YP_009047202.1;MERS_1ab;1980 | 0.05 | 0 |
| NC_019843.3;YP_009047202.1;MERS_1ab;1981 | 0 | 0 |
| NC_019843.3;YP_009047202.1;MERS_1ab;1982 | 0.05 | 0.025 |
| NC_019843.3;YP_009047202.1;MERS_1ab;1983 | 0.05 | 0.025 |
| NC_019843.3;YP_009047202.1;MERS_1ab;1984 | 0.1 | 0.025 |
| NC_019843.3;YP_009047202.1;MERS_1ab;1985 | 0.1 | 0.025 |
| NC_019843.3;YP_009047202.1;MERS_1ab;1986 | 0.1 | 0.05 |
| NC_019843.3;YP_009047202.1;MERS_1ab;1987 | 0.1 | 0 |
| NC_019843.3;YP_009047202.1;MERS_1ab;1988 | 0.05 | 0 |
| NC_019843.3;YP_009047202.1;MERS_1ab;1989 | 0 | 0.025 |
| NC_019843.3;YP_009047202.1;MERS_1ab;1990 | 0 | 0 |
| NC_019843.3;YP_009047202.1;MERS_1ab;1991 | 0 | 0 |
| NC_019843.3;YP_009047202.1;MERS_1ab;1992 | 0.05 | 0 |
| NC_019843.3;YP_009047202.1;MERS_1ab;1993 | 0 | 0 |
| NC_019843.3;YP_009047202.1;MERS_1ab;1994 | 0 | 0 |
| NC_019843.3;YP_009047202.1;MERS_1ab;1995 | 0 | 0.025 |
| NC_019843.3;YP_009047202.1;MERS_1ab;1996 | 0 | 0 |
| NC_019843.3;YP_009047202.1;MERS_1ab;1997 | 0 | 0 |
| NC_019843.3;YP_009047202.1;MERS_1ab;1998 | 0 | 0 |
| NC_019843.3;YP_009047202.1;MERS_1ab;1999 | 0 | 0 |
| NC_019843.3;YP_009047202.1;MERS_1ab;2000 | 0 | 0 |
| NC_019843.3;YP_009047202.1;MERS_1ab;2001 | 0.05 | 0 |
| NC_019843.3;YP_009047202.1;MERS_1ab;2002 | 0.05 | 0 |
| NC_019843.3;YP_009047202.1;MERS_1ab;2003 | 0.05 | 0 |
| NC_019843.3;YP_009047202.1;MERS_1ab;2004 | 0.05 | 0 |
| NC_019843.3;YP_009047202.1;MERS_1ab;2005 | 0.05 | 0 |
| NC_019843.3;YP_009047202.1;MERS_1ab;2006 | 0.05 | 0.025 |
| NC_019843.3;YP_009047202.1;MERS_1ab;2007 | 0.05 | 0.025 |
| NC_019843.3;YP_009047202.1;MERS_1ab;2008 | 0.05 | 0.025 |
| NC_019843.3;YP_009047202.1;MERS_1ab;2009 | 0 | 0.025 |
| NC_019843.3;YP_009047202.1;MERS_1ab;2010 | 0 | 0.025 |
| NC_019843.3;YP_009047202.1;MERS_1ab;2011 | 0 | 0 |
| NC_019843.3;YP_009047202.1;MERS_1ab;2012 | 0 | 0 |

| | | |
|---|---|---|
| NC_019843.3;YP_009047202.1;MERS_1ab;2013 | 0.05 | 0 |
| NC_019843.3;YP_009047202.1;MERS_1ab;2014 | 0 | 0 |
| NC_019843.3;YP_009047202.1;MERS_1ab;2015 | 0.05 | 0 |
| NC_019843.3;YP_009047202.1;MERS_1ab;2016 | 0 | 0 |
| NC_019843.3;YP_009047202.1;MERS_1ab;2017 | 0 | 0 |
| NC_019843.3;YP_009047202.1;MERS_1ab;2018 | 0 | 0 |
| NC_019843.3;YP_009047202.1;MERS_1ab;2019 | 0 | 0 |
| NC_019843.3;YP_009047202.1;MERS_1ab;2020 | 0 | 0 |
| NC_019843.3;YP_009047202.1;MERS_1ab;2021 | 0 | 0 |
| NC_019843.3;YP_009047202.1;MERS_1ab;2022 | 0 | 0 |
| NC_019843.3;YP_009047202.1;MERS_1ab;2023 | 0 | 0 |
| NC_019843.3;YP_009047202.1;MERS_1ab;2024 | 0 | 0.025 |
| NC_019843.3;YP_009047202.1;MERS_1ab;2025 | 0 | 0 |
| NC_019843.3;YP_009047202.1;MERS_1ab;2026 | 0 | 0 |
| NC_019843.3;YP_009047202.1;MERS_1ab;2027 | 0 | 0 |
| NC_019843.3;YP_009047202.1;MERS_1ab;2028 | 0 | 0 |
| NC_019843.3;YP_009047202.1;MERS_1ab;2029 | 0 | 0 |
| NC_019843.3;YP_009047202.1;MERS_1ab;2030 | 0 | 0 |
| NC_019843.3;YP_009047202.1;MERS_1ab;2031 | 0 | 0 |
| NC_019843.3;YP_009047202.1;MERS_1ab;2032 | 0 | 0 |
| NC_019843.3;YP_009047202.1;MERS_1ab;2033 | 0 | 0 |
| NC_019843.3;YP_009047202.1;MERS_1ab;2034 | 0 | 0 |
| NC_019843.3;YP_009047202.1;MERS_1ab;2035 | 0 | 0 |
| NC_019843.3;YP_009047202.1;MERS_1ab;2036 | 0 | 0 |
| NC_019843.3;YP_009047202.1;MERS_1ab;2037 | 0 | 0 |
| NC_019843.3;YP_009047202.1;MERS_1ab;2038 | 0 | 0 |
| NC_019843.3;YP_009047202.1;MERS_1ab;2039 | 0 | 0 |
| NC_019843.3;YP_009047202.1;MERS_1ab;2040 | 0 | 0 |
| NC_019843.3;YP_009047202.1;MERS_1ab;2041 | 0 | 0.025 |
| NC_019843.3;YP_009047202.1;MERS_1ab;2042 | 0.05 | 0.025 |
| NC_019843.3;YP_009047202.1;MERS_1ab;2043 | 0 | 0 |
| NC_019843.3;YP_009047202.1;MERS_1ab;2044 | 0 | 0 |
| NC_019843.3;YP_009047202.1;MERS_1ab;2045 | 0 | 0 |
| NC_019843.3;YP_009047202.1;MERS_1ab;2046 | 0 | 0 |
| NC_019843.3;YP_009047202.1;MERS_1ab;2047 | 0 | 0 |
| NC_019843.3;YP_009047202.1;MERS_1ab;2048 | 0 | 0 |
| NC_019843.3;YP_009047202.1;MERS_1ab;2049 | 0 | 0 |
| NC_019843.3;YP_009047202.1;MERS_1ab;2050 | 0 | 0 |
| NC_019843.3;YP_009047202.1;MERS_1ab;2051 | 0 | 0 |
| NC_019843.3;YP_009047202.1;MERS_1ab;2052 | 0 | 0 |
| NC_019843.3;YP_009047202.1;MERS_1ab;2053 | 0 | 0 |
| NC_019843.3;YP_009047202.1;MERS_1ab;2054 | 0 | 0 |
| NC_019843.3;YP_009047202.1;MERS_1ab;2055 | 0 | 0 |

| | | |
|---|---|---|
| NC_019843.3;YP_009047202.1;MERS_1ab;2056 | 0 | 0 |
| NC_019843.3;YP_009047202.1;MERS_1ab;2057 | 0 | 0 |
| NC_019843.3;YP_009047202.1;MERS_1ab;2058 | 0 | 0 |
| NC_019843.3;YP_009047202.1;MERS_1ab;2059 | 0 | 0 |
| NC_019843.3;YP_009047202.1;MERS_1ab;2060 | 0 | 0 |
| NC_019843.3;YP_009047202.1;MERS_1ab;2061 | 0 | 0 |
| NC_019843.3;YP_009047202.1;MERS_1ab;2062 | 0 | 0 |
| NC_019843.3;YP_009047202.1;MERS_1ab;2063 | 0 | 0 |
| NC_019843.3;YP_009047202.1;MERS_1ab;2064 | 0 | 0 |
| NC_019843.3;YP_009047202.1;MERS_1ab;2065 | 0 | 0 |
| NC_019843.3;YP_009047202.1;MERS_1ab;2066 | 0 | 0 |
| NC_019843.3;YP_009047202.1;MERS_1ab;2067 | 0 | 0 |
| NC_019843.3;YP_009047202.1;MERS_1ab;2068 | 0 | 0 |
| NC_019843.3;YP_009047202.1;MERS_1ab;2069 | 0 | 0 |
| NC_019843.3;YP_009047202.1;MERS_1ab;2070 | 0 | 0 |
| NC_019843.3;YP_009047202.1;MERS_1ab;2071 | 0 | 0 |
| NC_019843.3;YP_009047202.1;MERS_1ab;2072 | 0 | 0 |
| NC_019843.3;YP_009047202.1;MERS_1ab;2073 | 0 | 0 |
| NC_019843.3;YP_009047202.1;MERS_1ab;2074 | 0 | 0 |
| NC_019843.3;YP_009047202.1;MERS_1ab;2075 | 0 | 0 |
| NC_019843.3;YP_009047202.1;MERS_1ab;2076 | 0 | 0 |
| NC_019843.3;YP_009047202.1;MERS_1ab;2077 | 0 | 0.025 |
| NC_019843.3;YP_009047202.1;MERS_1ab;2078 | 0 | 0 |
| NC_019843.3;YP_009047202.1;MERS_1ab;2079 | 0 | 0 |
| NC_019843.3;YP_009047202.1;MERS_1ab;2080 | 0 | 0 |
| NC_019843.3;YP_009047202.1;MERS_1ab;2081 | 0 | 0 |
| NC_019843.3;YP_009047202.1;MERS_1ab;2082 | 0 | 0 |
| NC_019843.3;YP_009047202.1;MERS_1ab;2083 | 0 | 0 |
| NC_019843.3;YP_009047202.1;MERS_1ab;2084 | 0 | 0 |
| NC_019843.3;YP_009047202.1;MERS_1ab;2085 | 0 | 0 |
| NC_019843.3;YP_009047202.1;MERS_1ab;2086 | 0 | 0 |
| NC_019843.3;YP_009047202.1;MERS_1ab;2087 | 0 | 0 |
| NC_019843.3;YP_009047202.1;MERS_1ab;2088 | 0 | 0 |
| NC_019843.3;YP_009047202.1;MERS_1ab;2089 | 0 | 0 |
| NC_019843.3;YP_009047202.1;MERS_1ab;2090 | 0 | 0 |
| NC_019843.3;YP_009047202.1;MERS_1ab;2091 | 0 | 0 |
| NC_019843.3;YP_009047202.1;MERS_1ab;2092 | 0 | 0 |
| NC_019843.3;YP_009047202.1;MERS_1ab;2093 | 0 | 0 |
| NC_019843.3;YP_009047202.1;MERS_1ab;2094 | 0 | 0 |
| NC_019843.3;YP_009047202.1;MERS_1ab;2095 | 0 | 0 |
| NC_019843.3;YP_009047202.1;MERS_1ab;2096 | 0 | 0 |
| NC_019843.3;YP_009047202.1;MERS_1ab;2097 | 0 | 0 |
| NC_019843.3;YP_009047202.1;MERS_1ab;2098 | 0 | 0 |

| | | |
|---|---|---|
| NC_019843.3;YP_009047202.1;MERS_1ab;2099 | 0 | 0 |
| NC_019843.3;YP_009047202.1;MERS_1ab;2100 | 0 | 0 |
| NC_019843.3;YP_009047202.1;MERS_1ab;2101 | 0 | 0 |
| NC_019843.3;YP_009047202.1;MERS_1ab;2102 | 0 | 0 |
| NC_019843.3;YP_009047202.1;MERS_1ab;2103 | 0 | 0 |
| NC_019843.3;YP_009047202.1;MERS_1ab;2104 | 0 | 0 |
| NC_019843.3;YP_009047202.1;MERS_1ab;2105 | 0 | 0 |
| NC_019843.3;YP_009047202.1;MERS_1ab;2106 | 0 | 0 |
| NC_019843.3;YP_009047202.1;MERS_1ab;2107 | 0 | 0 |
| NC_019843.3;YP_009047202.1;MERS_1ab;2108 | 0 | 0 |
| NC_019843.3;YP_009047202.1;MERS_1ab;2109 | 0 | 0 |
| NC_019843.3;YP_009047202.1;MERS_1ab;2110 | 0 | 0 |
| NC_019843.3;YP_009047202.1;MERS_1ab;2111 | 0 | 0 |
| NC_019843.3;YP_009047202.1;MERS_1ab;2112 | 0 | 0 |
| NC_019843.3;YP_009047202.1;MERS_1ab;2113 | 0 | 0 |
| NC_019843.3;YP_009047202.1;MERS_1ab;2114 | 0 | 0 |
| NC_019843.3;YP_009047202.1;MERS_1ab;2115 | 0 | 0 |
| NC_019843.3;YP_009047202.1;MERS_1ab;2116 | 0 | 0 |
| NC_019843.3;YP_009047202.1;MERS_1ab;2117 | 0 | 0 |
| NC_019843.3;YP_009047202.1;MERS_1ab;2118 | 0 | 0 |
| NC_019843.3;YP_009047202.1;MERS_1ab;2119 | 0.05 | 0.025 |
| NC_019843.3;YP_009047202.1;MERS_1ab;2120 | 0.05 | 0.025 |
| NC_019843.3;YP_009047202.1;MERS_1ab;2121 | 0.05 | 0 |
| NC_019843.3;YP_009047202.1;MERS_1ab;2122 | 0 | 0 |
| NC_019843.3;YP_009047202.1;MERS_1ab;2123 | 0 | 0 |
| NC_019843.3;YP_009047202.1;MERS_1ab;2124 | 0 | 0 |
| NC_019843.3;YP_009047202.1;MERS_1ab;2125 | 0 | 0 |
| NC_019843.3;YP_009047202.1;MERS_1ab;2126 | 0 | 0 |
| NC_019843.3;YP_009047202.1;MERS_1ab;2127 | 0 | 0 |
| NC_019843.3;YP_009047202.1;MERS_1ab;2128 | 0 | 0 |
| NC_019843.3;YP_009047202.1;MERS_1ab;2129 | 0 | 0 |
| NC_019843.3;YP_009047202.1;MERS_1ab;2130 | 0 | 0 |
| NC_019843.3;YP_009047202.1;MERS_1ab;2131 | 0 | 0 |
| NC_019843.3;YP_009047202.1;MERS_1ab;2132 | 0 | 0 |
| NC_019843.3;YP_009047202.1;MERS_1ab;2133 | 0 | 0 |
| NC_019843.3;YP_009047202.1;MERS_1ab;2134 | 0 | 0 |
| NC_019843.3;YP_009047202.1;MERS_1ab;2135 | 0 | 0 |
| NC_019843.3;YP_009047202.1;MERS_1ab;2136 | 0 | 0.025 |
| NC_019843.3;YP_009047202.1;MERS_1ab;2137 | 0 | 0.025 |
| NC_019843.3;YP_009047202.1;MERS_1ab;2138 | 0 | 0.025 |
| NC_019843.3;YP_009047202.1;MERS_1ab;2139 | 0 | 0.025 |
| NC_019843.3;YP_009047202.1;MERS_1ab;2140 | 0 | 0.025 |
| NC_019843.3;YP_009047202.1;MERS_1ab;2141 | 0 | 0.025 |

| | | |
|---|---|---|
| NC_019843.3;YP_009047202.1;MERS_1ab;2142 | 0 | 0.025 |
| NC_019843.3;YP_009047202.1;MERS_1ab;2143 | 0.05 | 0.1 |
| NC_019843.3;YP_009047202.1;MERS_1ab;2144 | 0 | 0.075 |
| NC_019843.3;YP_009047202.1;MERS_1ab;2145 | 0 | 0.05 |
| NC_019843.3;YP_009047202.1;MERS_1ab;2146 | 0 | 0 |
| NC_019843.3;YP_009047202.1;MERS_1ab;2147 | 0 | 0 |
| NC_019843.3;YP_009047202.1;MERS_1ab;2148 | 0 | 0 |
| NC_019843.3;YP_009047202.1;MERS_1ab;2149 | 0 | 0 |
| NC_019843.3;YP_009047202.1;MERS_1ab;2150 | 0 | 0 |
| NC_019843.3;YP_009047202.1;MERS_1ab;2151 | 0 | 0 |
| NC_019843.3;YP_009047202.1;MERS_1ab;2152 | 0 | 0 |
| NC_019843.3;YP_009047202.1;MERS_1ab;2153 | 0 | 0 |
| NC_019843.3;YP_009047202.1;MERS_1ab;2154 | 0 | 0 |
| NC_019843.3;YP_009047202.1;MERS_1ab;2155 | 0 | 0 |
| NC_019843.3;YP_009047202.1;MERS_1ab;2156 | 0 | 0 |
| NC_019843.3;YP_009047202.1;MERS_1ab;2157 | 0 | 0.05 |
| NC_019843.3;YP_009047202.1;MERS_1ab;2158 | 0 | 0 |
| NC_019843.3;YP_009047202.1;MERS_1ab;2159 | 0 | 0 |
| NC_019843.3;YP_009047202.1;MERS_1ab;2160 | 0 | 0 |
| NC_019843.3;YP_009047202.1;MERS_1ab;2161 | 0 | 0 |
| NC_019843.3;YP_009047202.1;MERS_1ab;2162 | 0 | 0 |
| NC_019843.3;YP_009047202.1;MERS_1ab;2163 | 0 | 0 |
| NC_019843.3;YP_009047202.1;MERS_1ab;2164 | 0 | 0 |
| NC_019843.3;YP_009047202.1;MERS_1ab;2165 | 0 | 0 |
| NC_019843.3;YP_009047202.1;MERS_1ab;2166 | 0 | 0 |
| NC_019843.3;YP_009047202.1;MERS_1ab;2167 | 0 | 0 |
| NC_019843.3;YP_009047202.1;MERS_1ab;2168 | 0 | 0 |
| NC_019843.3;YP_009047202.1;MERS_1ab;2169 | 0 | 0 |
| NC_019843.3;YP_009047202.1;MERS_1ab;2170 | 0 | 0 |
| NC_019843.3;YP_009047202.1;MERS_1ab;2171 | 0 | 0 |
| NC_019843.3;YP_009047202.1;MERS_1ab;2172 | 0 | 0 |
| NC_019843.3;YP_009047202.1;MERS_1ab;2173 | 0 | 0 |
| NC_019843.3;YP_009047202.1;MERS_1ab;2174 | 0 | 0 |
| NC_019843.3;YP_009047202.1;MERS_1ab;2175 | 0 | 0 |
| NC_019843.3;YP_009047202.1;MERS_1ab;2176 | 0 | 0 |
| NC_019843.3;YP_009047202.1;MERS_1ab;2177 | 0 | 0 |
| NC_019843.3;YP_009047202.1;MERS_1ab;2178 | 0 | 0 |
| NC_019843.3;YP_009047202.1;MERS_1ab;2179 | 0 | 0 |
| NC_019843.3;YP_009047202.1;MERS_1ab;2180 | 0 | 0 |
| NC_019843.3;YP_009047202.1;MERS_1ab;2181 | 0 | 0 |
| NC_019843.3;YP_009047202.1;MERS_1ab;2182 | 0 | 0 |
| NC_019843.3;YP_009047202.1;MERS_1ab;2183 | 0 | 0 |
| NC_019843.3;YP_009047202.1;MERS_1ab;2184 | 0 | 0 |

| | | |
|---|---|---|
| NC_019843.3;YP_009047202.1;MERS_1ab;2185 | 0 | 0 |
| NC_019843.3;YP_009047202.1;MERS_1ab;2186 | 0 | 0 |
| NC_019843.3;YP_009047202.1;MERS_1ab;2187 | 0 | 0 |
| NC_019843.3;YP_009047202.1;MERS_1ab;2188 | 0 | 0 |
| NC_019843.3;YP_009047202.1;MERS_1ab;2189 | 0 | 0 |
| NC_019843.3;YP_009047202.1;MERS_1ab;2190 | 0 | 0 |
| NC_019843.3;YP_009047202.1;MERS_1ab;2191 | 0 | 0 |
| NC_019843.3;YP_009047202.1;MERS_1ab;2192 | 0 | 0 |
| NC_019843.3;YP_009047202.1;MERS_1ab;2193 | 0 | 0 |
| NC_019843.3;YP_009047202.1;MERS_1ab;2194 | 0 | 0 |
| NC_019843.3;YP_009047202.1;MERS_1ab;2195 | 0 | 0 |
| NC_019843.3;YP_009047202.1;MERS_1ab;2196 | 0 | 0 |
| NC_019843.3;YP_009047202.1;MERS_1ab;2197 | 0 | 0 |
| NC_019843.3;YP_009047202.1;MERS_1ab;2198 | 0 | 0 |
| NC_019843.3;YP_009047202.1;MERS_1ab;2199 | 0 | 0 |
| NC_019843.3;YP_009047202.1;MERS_1ab;2200 | 0 | 0 |
| NC_019843.3;YP_009047202.1;MERS_1ab;2201 | 0 | 0 |
| NC_019843.3;YP_009047202.1;MERS_1ab;2202 | 0 | 0 |
| NC_019843.3;YP_009047202.1;MERS_1ab;2203 | 0 | 0 |
| NC_019843.3;YP_009047202.1;MERS_1ab;2204 | 0 | 0 |
| NC_019843.3;YP_009047202.1;MERS_1ab;2205 | 0 | 0 |
| NC_019843.3;YP_009047202.1;MERS_1ab;2206 | 0 | 0 |
| NC_019843.3;YP_009047202.1;MERS_1ab;2207 | 0 | 0 |
| NC_019843.3;YP_009047202.1;MERS_1ab;2208 | 0.05 | 0 |
| NC_019843.3;YP_009047202.1;MERS_1ab;2209 | 0 | 0 |
| NC_019843.3;YP_009047202.1;MERS_1ab;2210 | 0 | 0 |
| NC_019843.3;YP_009047202.1;MERS_1ab;2211 | 0 | 0 |
| NC_019843.3;YP_009047202.1;MERS_1ab;2212 | 0 | 0 |
| NC_019843.3;YP_009047202.1;MERS_1ab;2213 | 0 | 0 |
| NC_019843.3;YP_009047202.1;MERS_1ab;2214 | 0 | 0 |
| NC_019843.3;YP_009047202.1;MERS_1ab;2215 | 0 | 0 |
| NC_019843.3;YP_009047202.1;MERS_1ab;2216 | 0 | 0 |
| NC_019843.3;YP_009047202.1;MERS_1ab;2217 | 0 | 0 |
| NC_019843.3;YP_009047202.1;MERS_1ab;2218 | 0 | 0 |
| NC_019843.3;YP_009047202.1;MERS_1ab;2219 | 0 | 0 |
| NC_019843.3;YP_009047202.1;MERS_1ab;2220 | 0.05 | 0 |
| NC_019843.3;YP_009047202.1;MERS_1ab;2221 | 0 | 0 |
| NC_019843.3;YP_009047202.1;MERS_1ab;2222 | 0.05 | 0 |
| NC_019843.3;YP_009047202.1;MERS_1ab;2223 | 0 | 0 |
| NC_019843.3;YP_009047202.1;MERS_1ab;2224 | 0 | 0 |
| NC_019843.3;YP_009047202.1;MERS_1ab;2225 | 0 | 0 |
| NC_019843.3;YP_009047202.1;MERS_1ab;2226 | 0 | 0 |
| NC_019843.3;YP_009047202.1;MERS_1ab;2227 | 0 | 0 |

| | | |
|---|---|---|
| NC_019843.3;YP_009047202.1;MERS_1ab;2228 | 0.15 | 0.05 |
| NC_019843.3;YP_009047202.1;MERS_1ab;2229 | 0.05 | 0.05 |
| NC_019843.3;YP_009047202.1;MERS_1ab;2230 | 0.1 | 0.05 |
| NC_019843.3;YP_009047202.1;MERS_1ab;2231 | 0.15 | 0.05 |
| NC_019843.3;YP_009047202.1;MERS_1ab;2232 | 0.15 | 0.05 |
| NC_019843.3;YP_009047202.1;MERS_1ab;2233 | 0.15 | 0.05 |
| NC_019843.3;YP_009047202.1;MERS_1ab;2234 | 0.1 | 0.075 |
| NC_019843.3;YP_009047202.1;MERS_1ab;2235 | 0.1 | 0.05 |
| NC_019843.3;YP_009047202.1;MERS_1ab;2236 | 0 | 0 |
| NC_019843.3;YP_009047202.1;MERS_1ab;2237 | 0 | 0 |
| NC_019843.3;YP_009047202.1;MERS_1ab;2238 | 0 | 0 |
| NC_019843.3;YP_009047202.1;MERS_1ab;2239 | 0.05 | 0 |
| NC_019843.3;YP_009047202.1;MERS_1ab;2240 | 0 | 0.025 |
| NC_019843.3;YP_009047202.1;MERS_1ab;2241 | 0 | 0.025 |
| NC_019843.3;YP_009047202.1;MERS_1ab;2242 | 0 | 0 |
| NC_019843.3;YP_009047202.1;MERS_1ab;2243 | 0 | 0 |
| NC_019843.3;YP_009047202.1;MERS_1ab;2244 | 0 | 0 |
| NC_019843.3;YP_009047202.1;MERS_1ab;2245 | 0 | 0 |
| NC_019843.3;YP_009047202.1;MERS_1ab;2246 | 0 | 0 |
| NC_019843.3;YP_009047202.1;MERS_1ab;2247 | 0 | 0 |
| NC_019843.3;YP_009047202.1;MERS_1ab;2248 | 0 | 0 |
| NC_019843.3;YP_009047202.1;MERS_1ab;2249 | 0 | 0 |
| NC_019843.3;YP_009047202.1;MERS_1ab;2250 | 0 | 0 |
| NC_019843.3;YP_009047202.1;MERS_1ab;2251 | 0 | 0 |
| NC_019843.3;YP_009047202.1;MERS_1ab;2252 | 0 | 0 |
| NC_019843.3;YP_009047202.1;MERS_1ab;2253 | 0 | 0 |
| NC_019843.3;YP_009047202.1;MERS_1ab;2254 | 0 | 0 |
| NC_019843.3;YP_009047202.1;MERS_1ab;2255 | 0 | 0 |
| NC_019843.3;YP_009047202.1;MERS_1ab;2256 | 0 | 0 |
| NC_019843.3;YP_009047202.1;MERS_1ab;2257 | 0 | 0 |
| NC_019843.3;YP_009047202.1;MERS_1ab;2258 | 0 | 0 |
| NC_019843.3;YP_009047202.1;MERS_1ab;2259 | 0.15 | 0.075 |
| NC_019843.3;YP_009047202.1;MERS_1ab;2260 | 0 | 0.025 |
| NC_019843.3;YP_009047202.1;MERS_1ab;2261 | 0 | 0.15 |
| NC_019843.3;YP_009047202.1;MERS_1ab;2262 | 0 | 0 |
| NC_019843.3;YP_009047202.1;MERS_1ab;2263 | 0 | 0 |
| NC_019843.3;YP_009047202.1;MERS_1ab;2264 | 0 | 0 |
| NC_019843.3;YP_009047202.1;MERS_1ab;2265 | 0 | 0 |
| NC_019843.3;YP_009047202.1;MERS_1ab;2266 | 0 | 0 |
| NC_019843.3;YP_009047202.1;MERS_1ab;2267 | 0 | 0 |
| NC_019843.3;YP_009047202.1;MERS_1ab;2268 | 0 | 0 |
| NC_019843.3;YP_009047202.1;MERS_1ab;2269 | 0 | 0 |
| NC_019843.3;YP_009047202.1;MERS_1ab;2270 | 0 | 0 |

| | | |
|---|---|---|
| NC_019843.3;YP_009047202.1;MERS_1ab;2271 | 0 | 0 |
| NC_019843.3;YP_009047202.1;MERS_1ab;2272 | 0 | 0 |
| NC_019843.3;YP_009047202.1;MERS_1ab;2273 | 0 | 0 |
| NC_019843.3;YP_009047202.1;MERS_1ab;2274 | 0 | 0 |
| NC_019843.3;YP_009047202.1;MERS_1ab;2275 | 0 | 0 |
| NC_019843.3;YP_009047202.1;MERS_1ab;2276 | 0 | 0 |
| NC_019843.3;YP_009047202.1;MERS_1ab;2277 | 0 | 0 |
| NC_019843.3;YP_009047202.1;MERS_1ab;2278 | 0 | 0 |
| NC_019843.3;YP_009047202.1;MERS_1ab;2279 | 0 | 0 |
| NC_019843.3;YP_009047202.1;MERS_1ab;2280 | 0 | 0 |
| NC_019843.3;YP_009047202.1;MERS_1ab;2281 | 0 | 0 |
| NC_019843.3;YP_009047202.1;MERS_1ab;2282 | 0 | 0 |
| NC_019843.3;YP_009047202.1;MERS_1ab;2283 | 0 | 0 |
| NC_019843.3;YP_009047202.1;MERS_1ab;2284 | 0 | 0 |
| NC_019843.3;YP_009047202.1;MERS_1ab;2285 | 0 | 0 |
| NC_019843.3;YP_009047202.1;MERS_1ab;2286 | 0 | 0 |
| NC_019843.3;YP_009047202.1;MERS_1ab;2287 | 0 | 0 |
| NC_019843.3;YP_009047202.1;MERS_1ab;2288 | 0 | 0 |
| NC_019843.3;YP_009047202.1;MERS_1ab;2289 | 0 | 0 |
| NC_019843.3;YP_009047202.1;MERS_1ab;2290 | 0 | 0 |
| NC_019843.3;YP_009047202.1;MERS_1ab;2291 | 0 | 0 |
| NC_019843.3;YP_009047202.1;MERS_1ab;2292 | 0 | 0 |
| NC_019843.3;YP_009047202.1;MERS_1ab;2293 | 0 | 0 |
| NC_019843.3;YP_009047202.1;MERS_1ab;2294 | 0 | 0 |
| NC_019843.3;YP_009047202.1;MERS_1ab;2295 | 0 | 0 |
| NC_019843.3;YP_009047202.1;MERS_1ab;2296 | 0 | 0 |
| NC_019843.3;YP_009047202.1;MERS_1ab;2297 | 0 | 0 |
| NC_019843.3;YP_009047202.1;MERS_1ab;2298 | 0 | 0 |
| NC_019843.3;YP_009047202.1;MERS_1ab;2299 | 0 | 0 |
| NC_019843.3;YP_009047202.1;MERS_1ab;2300 | 0 | 0 |
| NC_019843.3;YP_009047202.1;MERS_1ab;2301 | 0 | 0 |
| NC_019843.3;YP_009047202.1;MERS_1ab;2302 | 0 | 0 |
| NC_019843.3;YP_009047202.1;MERS_1ab;2303 | 0 | 0 |
| NC_019843.3;YP_009047202.1;MERS_1ab;2304 | 0 | 0 |
| NC_019843.3;YP_009047202.1;MERS_1ab;2305 | 0 | 0 |
| NC_019843.3;YP_009047202.1;MERS_1ab;2306 | 0 | 0 |
| NC_019843.3;YP_009047202.1;MERS_1ab;2307 | 0 | 0 |
| NC_019843.3;YP_009047202.1;MERS_1ab;2308 | 0 | 0 |
| NC_019843.3;YP_009047202.1;MERS_1ab;2309 | 0 | 0 |
| NC_019843.3;YP_009047202.1;MERS_1ab;2310 | 0 | 0 |
| NC_019843.3;YP_009047202.1;MERS_1ab;2311 | 0 | 0 |
| NC_019843.3;YP_009047202.1;MERS_1ab;2312 | 0 | 0 |
| NC_019843.3;YP_009047202.1;MERS_1ab;2313 | 0 | 0 |

| | | |
|---|---|---|
| NC_019843.3;YP_009047202.1;MERS_1ab;2314 | 0 | 0 |
| NC_019843.3;YP_009047202.1;MERS_1ab;2315 | 0 | 0 |
| NC_019843.3;YP_009047202.1;MERS_1ab;2316 | 0 | 0 |
| NC_019843.3;YP_009047202.1;MERS_1ab;2317 | 0 | 0 |
| NC_019843.3;YP_009047202.1;MERS_1ab;2318 | 0 | 0 |
| NC_019843.3;YP_009047202.1;MERS_1ab;2319 | 0 | 0 |
| NC_019843.3;YP_009047202.1;MERS_1ab;2320 | 0 | 0 |
| NC_019843.3;YP_009047202.1;MERS_1ab;2321 | 0 | 0 |
| NC_019843.3;YP_009047202.1;MERS_1ab;2322 | 0 | 0 |
| NC_019843.3;YP_009047202.1;MERS_1ab;2323 | 0 | 0 |
| NC_019843.3;YP_009047202.1;MERS_1ab;2324 | 0 | 0 |
| NC_019843.3;YP_009047202.1;MERS_1ab;2325 | 0 | 0 |
| NC_019843.3;YP_009047202.1;MERS_1ab;2326 | 0 | 0 |
| NC_019843.3;YP_009047202.1;MERS_1ab;2327 | 0 | 0 |
| NC_019843.3;YP_009047202.1;MERS_1ab;2328 | 0 | 0 |
| NC_019843.3;YP_009047202.1;MERS_1ab;2329 | 0 | 0 |
| NC_019843.3;YP_009047202.1;MERS_1ab;2330 | 0 | 0 |
| NC_019843.3;YP_009047202.1;MERS_1ab;2331 | 0 | 0 |
| NC_019843.3;YP_009047202.1;MERS_1ab;2332 | 0 | 0 |
| NC_019843.3;YP_009047202.1;MERS_1ab;2333 | 0.05 | 0 |
| NC_019843.3;YP_009047202.1;MERS_1ab;2334 | 0 | 0.025 |
| NC_019843.3;YP_009047202.1;MERS_1ab;2335 | 0 | 0 |
| NC_019843.3;YP_009047202.1;MERS_1ab;2336 | 0 | 0 |
| NC_019843.3;YP_009047202.1;MERS_1ab;2337 | 0 | 0 |
| NC_019843.3;YP_009047202.1;MERS_1ab;2338 | 0 | 0 |
| NC_019843.3;YP_009047202.1;MERS_1ab;2339 | 0 | 0 |
| NC_019843.3;YP_009047202.1;MERS_1ab;2340 | 0 | 0 |
| NC_019843.3;YP_009047202.1;MERS_1ab;2341 | 0 | 0 |
| NC_019843.3;YP_009047202.1;MERS_1ab;2342 | 0 | 0 |
| NC_019843.3;YP_009047202.1;MERS_1ab;2343 | 0 | 0 |
| NC_019843.3;YP_009047202.1;MERS_1ab;2344 | 0 | 0 |
| NC_019843.3;YP_009047202.1;MERS_1ab;2345 | 0 | 0 |
| NC_019843.3;YP_009047202.1;MERS_1ab;2346 | 0 | 0 |
| NC_019843.3;YP_009047202.1;MERS_1ab;2347 | 0 | 0 |
| NC_019843.3;YP_009047202.1;MERS_1ab;2348 | 0 | 0 |
| NC_019843.3;YP_009047202.1;MERS_1ab;2349 | 0 | 0 |
| NC_019843.3;YP_009047202.1;MERS_1ab;2350 | 0 | 0 |
| NC_019843.3;YP_009047202.1;MERS_1ab;2351 | 0 | 0 |
| NC_019843.3;YP_009047202.1;MERS_1ab;2352 | 0 | 0 |
| NC_019843.3;YP_009047202.1;MERS_1ab;2353 | 0 | 0 |
| NC_019843.3;YP_009047202.1;MERS_1ab;2354 | 0 | 0 |
| NC_019843.3;YP_009047202.1;MERS_1ab;2355 | 0 | 0 |
| NC_019843.3;YP_009047202.1;MERS_1ab;2356 | 0 | 0 |

| | | |
|---|---|---|
| NC_019843.3;YP_009047202.1;MERS_1ab;2357 | 0 | 0 |
| NC_019843.3;YP_009047202.1;MERS_1ab;2358 | 0 | 0 |
| NC_019843.3;YP_009047202.1;MERS_1ab;2359 | 0 | 0 |
| NC_019843.3;YP_009047202.1;MERS_1ab;2360 | 0 | 0 |
| NC_019843.3;YP_009047202.1;MERS_1ab;2361 | 0 | 0 |
| NC_019843.3;YP_009047202.1;MERS_1ab;2362 | 0 | 0 |
| NC_019843.3;YP_009047202.1;MERS_1ab;2363 | 0 | 0 |
| NC_019843.3;YP_009047202.1;MERS_1ab;2364 | 0 | 0.025 |
| NC_019843.3;YP_009047202.1;MERS_1ab;2365 | 0 | 0 |
| NC_019843.3;YP_009047202.1;MERS_1ab;2366 | 0 | 0 |
| NC_019843.3;YP_009047202.1;MERS_1ab;2367 | 0 | 0 |
| NC_019843.3;YP_009047202.1;MERS_1ab;2368 | 0 | 0 |
| NC_019843.3;YP_009047202.1;MERS_1ab;2369 | 0 | 0 |
| NC_019843.3;YP_009047202.1;MERS_1ab;2370 | 0 | 0 |
| NC_019843.3;YP_009047202.1;MERS_1ab;2371 | 0 | 0 |
| NC_019843.3;YP_009047202.1;MERS_1ab;2372 | 0 | 0 |
| NC_019843.3;YP_009047202.1;MERS_1ab;2373 | 0 | 0 |
| NC_019843.3;YP_009047202.1;MERS_1ab;2374 | 0 | 0 |
| NC_019843.3;YP_009047202.1;MERS_1ab;2375 | 0 | 0 |
| NC_019843.3;YP_009047202.1;MERS_1ab;2376 | 0 | 0 |
| NC_019843.3;YP_009047202.1;MERS_1ab;2377 | 0 | 0 |
| NC_019843.3;YP_009047202.1;MERS_1ab;2378 | 0 | 0 |
| NC_019843.3;YP_009047202.1;MERS_1ab;2379 | 0 | 0 |
| NC_019843.3;YP_009047202.1;MERS_1ab;2380 | 0 | 0 |
| NC_019843.3;YP_009047202.1;MERS_1ab;2381 | 0 | 0 |
| NC_019843.3;YP_009047202.1;MERS_1ab;2382 | 0 | 0 |
| NC_019843.3;YP_009047202.1;MERS_1ab;2383 | 0 | 0 |
| NC_019843.3;YP_009047202.1;MERS_1ab;2384 | 0 | 0 |
| NC_019843.3;YP_009047202.1;MERS_1ab;2385 | 0 | 0 |
| NC_019843.3;YP_009047202.1;MERS_1ab;2386 | 0 | 0 |
| NC_019843.3;YP_009047202.1;MERS_1ab;2387 | 0 | 0 |
| NC_019843.3;YP_009047202.1;MERS_1ab;2388 | 0 | 0 |
| NC_019843.3;YP_009047202.1;MERS_1ab;2389 | 0 | 0 |
| NC_019843.3;YP_009047202.1;MERS_1ab;2390 | 0 | 0 |
| NC_019843.3;YP_009047202.1;MERS_1ab;2391 | 0 | 0 |
| NC_019843.3;YP_009047202.1;MERS_1ab;2392 | 0 | 0 |
| NC_019843.3;YP_009047202.1;MERS_1ab;2393 | 0 | 0 |
| NC_019843.3;YP_009047202.1;MERS_1ab;2394 | 0 | 0 |
| NC_019843.3;YP_009047202.1;MERS_1ab;2395 | 0 | 0 |
| NC_019843.3;YP_009047202.1;MERS_1ab;2396 | 0 | 0 |
| NC_019843.3;YP_009047202.1;MERS_1ab;2397 | 0 | 0 |
| NC_019843.3;YP_009047202.1;MERS_1ab;2398 | 0 | 0 |
| NC_019843.3;YP_009047202.1;MERS_1ab;2399 | 0 | 0 |

| | | |
|---|---|---|
| NC_019843.3;YP_009047202.1;MERS_1ab;2400 | 0 | 0 |
| NC_019843.3;YP_009047202.1;MERS_1ab;2401 | 0 | 0 |
| NC_019843.3;YP_009047202.1;MERS_1ab;2402 | 0 | 0 |
| NC_019843.3;YP_009047202.1;MERS_1ab;2403 | 0 | 0 |
| NC_019843.3;YP_009047202.1;MERS_1ab;2404 | 0 | 0 |
| NC_019843.3;YP_009047202.1;MERS_1ab;2405 | 0 | 0 |
| NC_019843.3;YP_009047202.1;MERS_1ab;2406 | 0 | 0 |
| NC_019843.3;YP_009047202.1;MERS_1ab;2407 | 0 | 0 |
| NC_019843.3;YP_009047202.1;MERS_1ab;2408 | 0 | 0 |
| NC_019843.3;YP_009047202.1;MERS_1ab;2409 | 0 | 0 |
| NC_019843.3;YP_009047202.1;MERS_1ab;2410 | 0 | 0.025 |
| NC_019843.3;YP_009047202.1;MERS_1ab;2411 | 0 | 0 |
| NC_019843.3;YP_009047202.1;MERS_1ab;2412 | 0 | 0 |
| NC_019843.3;YP_009047202.1;MERS_1ab;2413 | 0 | 0 |
| NC_019843.3;YP_009047202.1;MERS_1ab;2414 | 0 | 0 |
| NC_019843.3;YP_009047202.1;MERS_1ab;2415 | 0 | 0 |
| NC_019843.3;YP_009047202.1;MERS_1ab;2416 | 0 | 0 |
| NC_019843.3;YP_009047202.1;MERS_1ab;2417 | 0 | 0 |
| NC_019843.3;YP_009047202.1;MERS_1ab;2418 | 0 | 0 |
| NC_019843.3;YP_009047202.1;MERS_1ab;2419 | 0 | 0 |
| NC_019843.3;YP_009047202.1;MERS_1ab;2420 | 0 | 0 |
| NC_019843.3;YP_009047202.1;MERS_1ab;2421 | 0 | 0 |
| NC_019843.3;YP_009047202.1;MERS_1ab;2422 | 0 | 0 |
| NC_019843.3;YP_009047202.1;MERS_1ab;2423 | 0 | 0 |
| NC_019843.3;YP_009047202.1;MERS_1ab;2424 | 0 | 0 |
| NC_019843.3;YP_009047202.1;MERS_1ab;2425 | 0 | 0 |
| NC_019843.3;YP_009047202.1;MERS_1ab;2426 | 0 | 0 |
| NC_019843.3;YP_009047202.1;MERS_1ab;2427 | 0 | 0 |
| NC_019843.3;YP_009047202.1;MERS_1ab;2428 | 0 | 0.075 |
| NC_019843.3;YP_009047202.1;MERS_1ab;2429 | 0 | 0 |
| NC_019843.3;YP_009047202.1;MERS_1ab;2430 | 0 | 0 |
| NC_019843.3;YP_009047202.1;MERS_1ab;2431 | 0 | 0 |
| NC_019843.3;YP_009047202.1;MERS_1ab;2432 | 0 | 0 |
| NC_019843.3;YP_009047202.1;MERS_1ab;2433 | 0 | 0.025 |
| NC_019843.3;YP_009047202.1;MERS_1ab;2434 | 0 | 0 |
| NC_019843.3;YP_009047202.1;MERS_1ab;2435 | 0 | 0.025 |
| NC_019843.3;YP_009047202.1;MERS_1ab;2436 | 0 | 0 |
| NC_019843.3;YP_009047202.1;MERS_1ab;2437 | 0.05 | 0 |
| NC_019843.3;YP_009047202.1;MERS_1ab;2438 | 0 | 0 |
| NC_019843.3;YP_009047202.1;MERS_1ab;2439 | 0.05 | 0 |
| NC_019843.3;YP_009047202.1;MERS_1ab;2440 | 0.05 | 0 |
| NC_019843.3;YP_009047202.1;MERS_1ab;2441 | 0.05 | 0 |
| NC_019843.3;YP_009047202.1;MERS_1ab;2442 | 0.05 | 0 |

| | | |
|---|---|---|
| NC_019843.3;YP_009047202.1;MERS_1ab;2443 | 0.05 | 0 |
| NC_019843.3;YP_009047202.1;MERS_1ab;2444 | 0.05 | 0 |
| NC_019843.3;YP_009047202.1;MERS_1ab;2445 | 0.05 | 0 |
| NC_019843.3;YP_009047202.1;MERS_1ab;2446 | 0.1 | 0 |
| NC_019843.3;YP_009047202.1;MERS_1ab;2447 | 0.05 | 0 |
| NC_019843.3;YP_009047202.1;MERS_1ab;2448 | 0 | 0 |
| NC_019843.3;YP_009047202.1;MERS_1ab;2449 | 0 | 0.025 |
| NC_019843.3;YP_009047202.1;MERS_1ab;2450 | 0 | 0 |
| NC_019843.3;YP_009047202.1;MERS_1ab;2451 | 0 | 0 |
| NC_019843.3;YP_009047202.1;MERS_1ab;2452 | 0 | 0 |
| NC_019843.3;YP_009047202.1;MERS_1ab;2453 | 0 | 0 |
| NC_019843.3;YP_009047202.1;MERS_1ab;2454 | 0 | 0 |
| NC_019843.3;YP_009047202.1;MERS_1ab;2455 | 0 | 0 |
| NC_019843.3;YP_009047202.1;MERS_1ab;2456 | 0 | 0 |
| NC_019843.3;YP_009047202.1;MERS_1ab;2457 | 0 | 0 |
| NC_019843.3;YP_009047202.1;MERS_1ab;2458 | 0 | 0 |
| NC_019843.3;YP_009047202.1;MERS_1ab;2459 | 0 | 0 |
| NC_019843.3;YP_009047202.1;MERS_1ab;2460 | 0 | 0 |
| NC_019843.3;YP_009047202.1;MERS_1ab;2461 | 0 | 0 |
| NC_019843.3;YP_009047202.1;MERS_1ab;2462 | 0 | 0 |
| NC_019843.3;YP_009047202.1;MERS_1ab;2463 | 0 | 0.025 |
| NC_019843.3;YP_009047202.1;MERS_1ab;2464 | 0 | 0 |
| NC_019843.3;YP_009047202.1;MERS_1ab;2465 | 0 | 0 |
| NC_019843.3;YP_009047202.1;MERS_1ab;2466 | 0 | 0 |
| NC_019843.3;YP_009047202.1;MERS_1ab;2467 | 0 | 0 |
| NC_019843.3;YP_009047202.1;MERS_1ab;2468 | 0 | 0 |
| NC_019843.3;YP_009047202.1;MERS_1ab;2469 | 0 | 0 |
| NC_019843.3;YP_009047202.1;MERS_1ab;2470 | 0 | 0 |
| NC_019843.3;YP_009047202.1;MERS_1ab;2471 | 0 | 0 |
| NC_019843.3;YP_009047202.1;MERS_1ab;2472 | 0 | 0 |
| NC_019843.3;YP_009047202.1;MERS_1ab;2473 | 0 | 0 |
| NC_019843.3;YP_009047202.1;MERS_1ab;2474 | 0 | 0 |
| NC_019843.3;YP_009047202.1;MERS_1ab;2475 | 0 | 0 |
| NC_019843.3;YP_009047202.1;MERS_1ab;2476 | 0 | 0 |
| NC_019843.3;YP_009047202.1;MERS_1ab;2477 | 0 | 0 |
| NC_019843.3;YP_009047202.1;MERS_1ab;2478 | 0 | 0 |
| NC_019843.3;YP_009047202.1;MERS_1ab;2479 | 0 | 0 |
| NC_019843.3;YP_009047202.1;MERS_1ab;2480 | 0 | 0 |
| NC_019843.3;YP_009047202.1;MERS_1ab;2481 | 0 | 0 |
| NC_019843.3;YP_009047202.1;MERS_1ab;2482 | 0 | 0 |
| NC_019843.3;YP_009047202.1;MERS_1ab;2483 | 0.05 | 0 |
| NC_019843.3;YP_009047202.1;MERS_1ab;2484 | 0 | 0 |
| NC_019843.3;YP_009047202.1;MERS_1ab;2485 | 0 | 0 |

| | | |
|---|---|---|
| NC_019843.3;YP_009047202.1;MERS_1ab;2486 | 0 | 0 |
| NC_019843.3;YP_009047202.1;MERS_1ab;2487 | 0 | 0 |
| NC_019843.3;YP_009047202.1;MERS_1ab;2488 | 0 | 0 |
| NC_019843.3;YP_009047202.1;MERS_1ab;2489 | 0 | 0 |
| NC_019843.3;YP_009047202.1;MERS_1ab;2490 | 0 | 0 |
| NC_019843.3;YP_009047202.1;MERS_1ab;2491 | 0 | 0 |
| NC_019843.3;YP_009047202.1;MERS_1ab;2492 | 0 | 0 |
| NC_019843.3;YP_009047202.1;MERS_1ab;2493 | 0 | 0 |
| NC_019843.3;YP_009047202.1;MERS_1ab;2494 | 0 | 0 |
| NC_019843.3;YP_009047202.1;MERS_1ab;2495 | 0 | 0 |
| NC_019843.3;YP_009047202.1;MERS_1ab;2496 | 0 | 0 |
| NC_019843.3;YP_009047202.1;MERS_1ab;2497 | 0 | 0.025 |
| NC_019843.3;YP_009047202.1;MERS_1ab;2498 | 0.05 | 0 |
| NC_019843.3;YP_009047202.1;MERS_1ab;2499 | 0.05 | 0.025 |
| NC_019843.3;YP_009047202.1;MERS_1ab;2500 | 0 | 0 |
| NC_019843.3;YP_009047202.1;MERS_1ab;2501 | 0 | 0 |
| NC_019843.3;YP_009047202.1;MERS_1ab;2502 | 0 | 0 |
| NC_019843.3;YP_009047202.1;MERS_1ab;2503 | 0 | 0 |
| NC_019843.3;YP_009047202.1;MERS_1ab;2504 | 0 | 0 |
| NC_019843.3;YP_009047202.1;MERS_1ab;2505 | 0 | 0 |
| NC_019843.3;YP_009047202.1;MERS_1ab;2506 | 0 | 0 |
| NC_019843.3;YP_009047202.1;MERS_1ab;2507 | 0 | 0 |
| NC_019843.3;YP_009047202.1;MERS_1ab;2508 | 0 | 0 |
| NC_019843.3;YP_009047202.1;MERS_1ab;2509 | 0 | 0 |
| NC_019843.3;YP_009047202.1;MERS_1ab;2510 | 0 | 0.025 |
| NC_019843.3;YP_009047202.1;MERS_1ab;2511 | 0 | 0.025 |
| NC_019843.3;YP_009047202.1;MERS_1ab;2512 | 0 | 0.025 |
| NC_019843.3;YP_009047202.1;MERS_1ab;2513 | 0 | 0.025 |
| NC_019843.3;YP_009047202.1;MERS_1ab;2514 | 0 | 0.05 |
| NC_019843.3;YP_009047202.1;MERS_1ab;2515 | 0 | 0.05 |
| NC_019843.3;YP_009047202.1;MERS_1ab;2516 | 0 | 0 |
| NC_019843.3;YP_009047202.1;MERS_1ab;2517 | 0 | 0 |
| NC_019843.3;YP_009047202.1;MERS_1ab;2518 | 0 | 0 |
| NC_019843.3;YP_009047202.1;MERS_1ab;2519 | 0 | 0 |
| NC_019843.3;YP_009047202.1;MERS_1ab;2520 | 0 | 0 |
| NC_019843.3;YP_009047202.1;MERS_1ab;2521 | 0 | 0 |
| NC_019843.3;YP_009047202.1;MERS_1ab;2522 | 0 | 0 |
| NC_019843.3;YP_009047202.1;MERS_1ab;2523 | 0 | 0.025 |
| NC_019843.3;YP_009047202.1;MERS_1ab;2524 | 0 | 0 |
| NC_019843.3;YP_009047202.1;MERS_1ab;2525 | 0 | 0 |
| NC_019843.3;YP_009047202.1;MERS_1ab;2526 | 0 | 0 |
| NC_019843.3;YP_009047202.1;MERS_1ab;2527 | 0 | 0 |
| NC_019843.3;YP_009047202.1;MERS_1ab;2528 | 0 | 0 |

| | | |
|---|---|---|
| NC_019843.3;YP_009047202.1;MERS_1ab;2529 | 0 | 0 |
| NC_019843.3;YP_009047202.1;MERS_1ab;2530 | 0 | 0 |
| NC_019843.3;YP_009047202.1;MERS_1ab;2531 | 0 | 0 |
| NC_019843.3;YP_009047202.1;MERS_1ab;2532 | 0 | 0 |
| NC_019843.3;YP_009047202.1;MERS_1ab;2533 | 0 | 0 |
| NC_019843.3;YP_009047202.1;MERS_1ab;2534 | 0 | 0 |
| NC_019843.3;YP_009047202.1;MERS_1ab;2535 | 0 | 0 |
| NC_019843.3;YP_009047202.1;MERS_1ab;2536 | 0 | 0 |
| NC_019843.3;YP_009047202.1;MERS_1ab;2537 | 0 | 0 |
| NC_019843.3;YP_009047202.1;MERS_1ab;2538 | 0 | 0 |
| NC_019843.3;YP_009047202.1;MERS_1ab;2539 | 0 | 0 |
| NC_019843.3;YP_009047202.1;MERS_1ab;2540 | 0 | 0 |
| NC_019843.3;YP_009047202.1;MERS_1ab;2541 | 0 | 0 |
| NC_019843.3;YP_009047202.1;MERS_1ab;2542 | 0 | 0 |
| NC_019843.3;YP_009047202.1;MERS_1ab;2543 | 0 | 0 |
| NC_019843.3;YP_009047202.1;MERS_1ab;2544 | 0.25 | 0.2 |
| NC_019843.3;YP_009047202.1;MERS_1ab;2545 | 0 | 0 |
| NC_019843.3;YP_009047202.1;MERS_1ab;2546 | 0 | 0 |
| NC_019843.3;YP_009047202.1;MERS_1ab;2547 | 0 | 0 |
| NC_019843.3;YP_009047202.1;MERS_1ab;2548 | 0 | 0 |
| NC_019843.3;YP_009047202.1;MERS_1ab;2549 | 0 | 0 |
| NC_019843.3;YP_009047202.1;MERS_1ab;2550 | 0 | 0 |
| NC_019843.3;YP_009047202.1;MERS_1ab;2551 | 0 | 0 |
| NC_019843.3;YP_009047202.1;MERS_1ab;2552 | 0 | 0 |
| NC_019843.3;YP_009047202.1;MERS_1ab;2553 | 0 | 0 |
| NC_019843.3;YP_009047202.1;MERS_1ab;2554 | 0 | 0 |
| NC_019843.3;YP_009047202.1;MERS_1ab;2555 | 0 | 0 |
| NC_019843.3;YP_009047202.1;MERS_1ab;2556 | 0 | 0 |
| NC_019843.3;YP_009047202.1;MERS_1ab;2557 | 0 | 0.05 |
| NC_019843.3;YP_009047202.1;MERS_1ab;2558 | 0 | 0.025 |
| NC_019843.3;YP_009047202.1;MERS_1ab;2559 | 0 | 0.025 |
| NC_019843.3;YP_009047202.1;MERS_1ab;2560 | 0 | 0.025 |
| NC_019843.3;YP_009047202.1;MERS_1ab;2561 | 0 | 0.025 |
| NC_019843.3;YP_009047202.1;MERS_1ab;2562 | 0 | 0.025 |
| NC_019843.3;YP_009047202.1;MERS_1ab;2563 | 0 | 0.05 |
| NC_019843.3;YP_009047202.1;MERS_1ab;2564 | 0 | 0 |
| NC_019843.3;YP_009047202.1;MERS_1ab;2565 | 0.05 | 0.025 |
| NC_019843.3;YP_009047202.1;MERS_1ab;2566 | 0 | 0 |
| NC_019843.3;YP_009047202.1;MERS_1ab;2567 | 0 | 0.025 |
| NC_019843.3;YP_009047202.1;MERS_1ab;2568 | 0 | 0 |
| NC_019843.3;YP_009047202.1;MERS_1ab;2569 | 0 | 0 |
| NC_019843.3;YP_009047202.1;MERS_1ab;2570 | 0 | 0 |
| NC_019843.3;YP_009047202.1;MERS_1ab;2571 | 0 | 0 |

| | | |
|---|---|---|
| NC_019843.3;YP_009047202.1;MERS_1ab;2572 | 0 | 0 |
| NC_019843.3;YP_009047202.1;MERS_1ab;2573 | 0 | 0 |
| NC_019843.3;YP_009047202.1;MERS_1ab;2574 | 0 | 0 |
| NC_019843.3;YP_009047202.1;MERS_1ab;2575 | 0 | 0 |
| NC_019843.3;YP_009047202.1;MERS_1ab;2576 | 0 | 0 |
| NC_019843.3;YP_009047202.1;MERS_1ab;2577 | 0.05 | 0.025 |
| NC_019843.3;YP_009047202.1;MERS_1ab;2578 | 0.05 | 0.025 |
| NC_019843.3;YP_009047202.1;MERS_1ab;2579 | 0.05 | 0.025 |
| NC_019843.3;YP_009047202.1;MERS_1ab;2580 | 0.05 | 0.025 |
| NC_019843.3;YP_009047202.1;MERS_1ab;2581 | 0.05 | 0.025 |
| NC_019843.3;YP_009047202.1;MERS_1ab;2582 | 0 | 0.075 |
| NC_019843.3;YP_009047202.1;MERS_1ab;2583 | 0 | 0.1 |
| NC_019843.3;YP_009047202.1;MERS_1ab;2584 | 0 | 0.05 |
| NC_019843.3;YP_009047202.1;MERS_1ab;2585 | 0 | 0.025 |
| NC_019843.3;YP_009047202.1;MERS_1ab;2586 | 0 | 0.05 |
| NC_019843.3;YP_009047202.1;MERS_1ab;2587 | 0 | 0.075 |
| NC_019843.3;YP_009047202.1;MERS_1ab;2588 | 0.05 | 0.025 |
| NC_019843.3;YP_009047202.1;MERS_1ab;2589 | 0.05 | 0.075 |
| NC_019843.3;YP_009047202.1;MERS_1ab;2590 | 0 | 0.025 |
| NC_019843.3;YP_009047202.1;MERS_1ab;2591 | 0 | 0 |
| NC_019843.3;YP_009047202.1;MERS_1ab;2592 | 0 | 0 |
| NC_019843.3;YP_009047202.1;MERS_1ab;2593 | 0 | 0 |
| NC_019843.3;YP_009047202.1;MERS_1ab;2594 | 0 | 0 |
| NC_019843.3;YP_009047202.1;MERS_1ab;2595 | 0 | 0 |
| NC_019843.3;YP_009047202.1;MERS_1ab;2596 | 0 | 0 |
| NC_019843.3;YP_009047202.1;MERS_1ab;2597 | 0 | 0 |
| NC_019843.3;YP_009047202.1;MERS_1ab;2598 | 0 | 0 |
| NC_019843.3;YP_009047202.1;MERS_1ab;2599 | 0 | 0.025 |
| NC_019843.3;YP_009047202.1;MERS_1ab;2600 | 0 | 0.025 |
| NC_019843.3;YP_009047202.1;MERS_1ab;2601 | 0 | 0.025 |
| NC_019843.3;YP_009047202.1;MERS_1ab;2602 | 0 | 0.05 |
| NC_019843.3;YP_009047202.1;MERS_1ab;2603 | 0 | 0.025 |
| NC_019843.3;YP_009047202.1;MERS_1ab;2604 | 0 | 0.025 |
| NC_019843.3;YP_009047202.1;MERS_1ab;2605 | 0 | 0.025 |
| NC_019843.3;YP_009047202.1;MERS_1ab;2606 | 0 | 0.025 |
| NC_019843.3;YP_009047202.1;MERS_1ab;2607 | 0 | 0 |
| NC_019843.3;YP_009047202.1;MERS_1ab;2608 | 0.05 | 0 |
| NC_019843.3;YP_009047202.1;MERS_1ab;2609 | 0 | 0 |
| NC_019843.3;YP_009047202.1;MERS_1ab;2610 | 0.05 | 0 |
| NC_019843.3;YP_009047202.1;MERS_1ab;2611 | 0.05 | 0 |
| NC_019843.3;YP_009047202.1;MERS_1ab;2612 | 0.1 | 0.025 |
| NC_019843.3;YP_009047202.1;MERS_1ab;2613 | 0.05 | 0 |
| NC_019843.3;YP_009047202.1;MERS_1ab;2614 | 0.05 | 0 |

| | | |
|---|---|---|
| NC_019843.3;YP_009047202.1;MERS_1ab;2615 | 0.05 | 0.05 |
| NC_019843.3;YP_009047202.1;MERS_1ab;2616 | 0.05 | 0 |
| NC_019843.3;YP_009047202.1;MERS_1ab;2617 | 0 | 0 |
| NC_019843.3;YP_009047202.1;MERS_1ab;2618 | 0 | 0 |
| NC_019843.3;YP_009047202.1;MERS_1ab;2619 | 0 | 0 |
| NC_019843.3;YP_009047202.1;MERS_1ab;2620 | 0 | 0 |
| NC_019843.3;YP_009047202.1;MERS_1ab;2621 | 0 | 0 |
| NC_019843.3;YP_009047202.1;MERS_1ab;2622 | 0 | 0 |
| NC_019843.3;YP_009047202.1;MERS_1ab;2623 | 0 | 0 |
| NC_019843.3;YP_009047202.1;MERS_1ab;2624 | 0 | 0 |
| NC_019843.3;YP_009047202.1;MERS_1ab;2625 | 0 | 0 |
| NC_019843.3;YP_009047202.1;MERS_1ab;2626 | 0.1 | 0.05 |
| NC_019843.3;YP_009047202.1;MERS_1ab;2627 | 0.05 | 0.025 |
| NC_019843.3;YP_009047202.1;MERS_1ab;2628 | 0.1 | 0.025 |
| NC_019843.3;YP_009047202.1;MERS_1ab;2629 | 0.15 | 0.075 |
| NC_019843.3;YP_009047202.1;MERS_1ab;2630 | 0.2 | 0.075 |
| NC_019843.3;YP_009047202.1;MERS_1ab;2631 | 0.15 | 0.075 |
| NC_019843.3;YP_009047202.1;MERS_1ab;2632 | 0.15 | 0.075 |
| NC_019843.3;YP_009047202.1;MERS_1ab;2633 | 0.15 | 0.075 |
| NC_019843.3;YP_009047202.1;MERS_1ab;2634 | 0.2 | 0.05 |
| NC_019843.3;YP_009047202.1;MERS_1ab;2635 | 0 | 0.175 |
| NC_019843.3;YP_009047202.1;MERS_1ab;2636 | 0 | 0.175 |
| NC_019843.3;YP_009047202.1;MERS_1ab;2637 | 0.1 | 0.15 |
| NC_019843.3;YP_009047202.1;MERS_1ab;2638 | 0.1 | 0.025 |
| NC_019843.3;YP_009047202.1;MERS_1ab;2639 | 0.1 | 0.1 |
| NC_019843.3;YP_009047202.1;MERS_1ab;2640 | 0 | 0.075 |
| NC_019843.3;YP_009047202.1;MERS_1ab;2641 | 0.05 | 0.025 |
| NC_019843.3;YP_009047202.1;MERS_1ab;2642 | 0.1 | 0.075 |
| NC_019843.3;YP_009047202.1;MERS_1ab;2643 | 0.45 | 0.175 |
| NC_019843.3;YP_009047202.1;MERS_1ab;2644 | 0.15 | 0.125 |
| NC_019843.3;YP_009047202.1;MERS_1ab;2645 | 0 | 0.075 |
| NC_019843.3;YP_009047202.1;MERS_1ab;2646 | 0 | 0.05 |
| NC_019843.3;YP_009047202.1;MERS_1ab;2647 | 0 | 0.025 |
| NC_019843.3;YP_009047202.1;MERS_1ab;2648 | 0.05 | 0.05 |
| NC_019843.3;YP_009047202.1;MERS_1ab;2649 | 0 | 0 |
| NC_019843.3;YP_009047202.1;MERS_1ab;2650 | 0 | 0 |
| NC_019843.3;YP_009047202.1;MERS_1ab;2651 | 0 | 0 |
| NC_019843.3;YP_009047202.1;MERS_1ab;2652 | 0 | 0 |
| NC_019843.3;YP_009047202.1;MERS_1ab;2653 | 0 | 0 |
| NC_019843.3;YP_009047202.1;MERS_1ab;2654 | 0 | 0 |
| NC_019843.3;YP_009047202.1;MERS_1ab;2655 | 0 | 0 |
| NC_019843.3;YP_009047202.1;MERS_1ab;2656 | 0 | 0.025 |
| NC_019843.3;YP_009047202.1;MERS_1ab;2657 | 0.05 | 0 |

| | | |
|---|---|---|
| NC_019843.3;YP_009047202.1;MERS_1ab;2658 | 0 | 0 |
| NC_019843.3;YP_009047202.1;MERS_1ab;2659 | 0 | 0 |
| NC_019843.3;YP_009047202.1;MERS_1ab;2660 | 0 | 0 |
| NC_019843.3;YP_009047202.1;MERS_1ab;2661 | 0 | 0 |
| NC_019843.3;YP_009047202.1;MERS_1ab;2662 | 0 | 0.025 |
| NC_019843.3;YP_009047202.1;MERS_1ab;2663 | 0 | 0 |
| NC_019843.3;YP_009047202.1;MERS_1ab;2664 | 0 | 0.025 |
| NC_019843.3;YP_009047202.1;MERS_1ab;2665 | 0 | 0 |
| NC_019843.3;YP_009047202.1;MERS_1ab;2666 | 0 | 0 |
| NC_019843.3;YP_009047202.1;MERS_1ab;2667 | 0 | 0 |
| NC_019843.3;YP_009047202.1;MERS_1ab;2668 | 0 | 0.025 |
| NC_019843.3;YP_009047202.1;MERS_1ab;2669 | 0 | 0.025 |
| NC_019843.3;YP_009047202.1;MERS_1ab;2670 | 0 | 0 |
| NC_019843.3;YP_009047202.1;MERS_1ab;2671 | 0 | 0 |
| NC_019843.3;YP_009047202.1;MERS_1ab;2672 | 0 | 0 |
| NC_019843.3;YP_009047202.1;MERS_1ab;2673 | 0 | 0 |
| NC_019843.3;YP_009047202.1;MERS_1ab;2674 | 0 | 0 |
| NC_019843.3;YP_009047202.1;MERS_1ab;2675 | 0 | 0 |
| NC_019843.3;YP_009047202.1;MERS_1ab;2676 | 0.05 | 0 |
| NC_019843.3;YP_009047202.1;MERS_1ab;2677 | 0 | 0 |
| NC_019843.3;YP_009047202.1;MERS_1ab;2678 | 0 | 0 |
| NC_019843.3;YP_009047202.1;MERS_1ab;2679 | 0 | 0 |
| NC_019843.3;YP_009047202.1;MERS_1ab;2680 | 0 | 0 |
| NC_019843.3;YP_009047202.1;MERS_1ab;2681 | 0 | 0 |
| NC_019843.3;YP_009047202.1;MERS_1ab;2682 | 0.05 | 0 |
| NC_019843.3;YP_009047202.1;MERS_1ab;2683 | 0.05 | 0 |
| NC_019843.3;YP_009047202.1;MERS_1ab;2684 | 0.05 | 0 |
| NC_019843.3;YP_009047202.1;MERS_1ab;2685 | 0 | 0 |
| NC_019843.3;YP_009047202.1;MERS_1ab;2686 | 0.05 | 0 |
| NC_019843.3;YP_009047202.1;MERS_1ab;2687 | 0 | 0 |
| NC_019843.3;YP_009047202.1;MERS_1ab;2688 | 0 | 0 |
| NC_019843.3;YP_009047202.1;MERS_1ab;2689 | 0 | 0 |
| NC_019843.3;YP_009047202.1;MERS_1ab;2690 | 0 | 0 |
| NC_019843.3;YP_009047202.1;MERS_1ab;2691 | 0 | 0 |
| NC_019843.3;YP_009047202.1;MERS_1ab;2692 | 0 | 0 |
| NC_019843.3;YP_009047202.1;MERS_1ab;2693 | 0 | 0 |
| NC_019843.3;YP_009047202.1;MERS_1ab;2694 | 0 | 0 |
| NC_019843.3;YP_009047202.1;MERS_1ab;2695 | 0 | 0 |
| NC_019843.3;YP_009047202.1;MERS_1ab;2696 | 0 | 0 |
| NC_019843.3;YP_009047202.1;MERS_1ab;2697 | 0 | 0 |
| NC_019843.3;YP_009047202.1;MERS_1ab;2698 | 0 | 0 |
| NC_019843.3;YP_009047202.1;MERS_1ab;2699 | 0 | 0 |
| NC_019843.3;YP_009047202.1;MERS_1ab;2700 | 0 | 0.025 |

| | | |
|---|---|---|
| NC_019843.3;YP_009047202.1;MERS_1ab;2701 | 0 | 0 |
| NC_019843.3;YP_009047202.1;MERS_1ab;2702 | 0 | 0 |
| NC_019843.3;YP_009047202.1;MERS_1ab;2703 | 0 | 0 |
| NC_019843.3;YP_009047202.1;MERS_1ab;2704 | 0 | 0 |
| NC_019843.3;YP_009047202.1;MERS_1ab;2705 | 0 | 0 |
| NC_019843.3;YP_009047202.1;MERS_1ab;2706 | 0 | 0 |
| NC_019843.3;YP_009047202.1;MERS_1ab;2707 | 0 | 0 |
| NC_019843.3;YP_009047202.1;MERS_1ab;2708 | 0 | 0 |
| NC_019843.3;YP_009047202.1;MERS_1ab;2709 | 0 | 0 |
| NC_019843.3;YP_009047202.1;MERS_1ab;2710 | 0 | 0 |
| NC_019843.3;YP_009047202.1;MERS_1ab;2711 | 0 | 0 |
| NC_019843.3;YP_009047202.1;MERS_1ab;2712 | 0 | 0 |
| NC_019843.3;YP_009047202.1;MERS_1ab;2713 | 0 | 0 |
| NC_019843.3;YP_009047202.1;MERS_1ab;2714 | 0 | 0 |
| NC_019843.3;YP_009047202.1;MERS_1ab;2715 | 0.05 | 0 |
| NC_019843.3;YP_009047202.1;MERS_1ab;2716 | 0 | 0 |
| NC_019843.3;YP_009047202.1;MERS_1ab;2717 | 0 | 0 |
| NC_019843.3;YP_009047202.1;MERS_1ab;2718 | 0 | 0 |
| NC_019843.3;YP_009047202.1;MERS_1ab;2719 | 0 | 0 |
| NC_019843.3;YP_009047202.1;MERS_1ab;2720 | 0 | 0 |
| NC_019843.3;YP_009047202.1;MERS_1ab;2721 | 0 | 0 |
| NC_019843.3;YP_009047202.1;MERS_1ab;2722 | 0 | 0 |
| NC_019843.3;YP_009047202.1;MERS_1ab;2723 | 0 | 0 |
| NC_019843.3;YP_009047202.1;MERS_1ab;2724 | 0 | 0 |
| NC_019843.3;YP_009047202.1;MERS_1ab;2725 | 0 | 0 |
| NC_019843.3;YP_009047202.1;MERS_1ab;2726 | 0 | 0 |
| NC_019843.3;YP_009047202.1;MERS_1ab;2727 | 0 | 0 |
| NC_019843.3;YP_009047202.1;MERS_1ab;2728 | 0 | 0 |
| NC_019843.3;YP_009047202.1;MERS_1ab;2729 | 0 | 0 |
| NC_019843.3;YP_009047202.1;MERS_1ab;2730 | 0 | 0 |
| NC_019843.3;YP_009047202.1;MERS_1ab;2731 | 0.05 | 0 |
| NC_019843.3;YP_009047202.1;MERS_1ab;2732 | 0 | 0.025 |
| NC_019843.3;YP_009047202.1;MERS_1ab;2733 | 0 | 0.025 |
| NC_019843.3;YP_009047202.1;MERS_1ab;2734 | 0 | 0.025 |
| NC_019843.3;YP_009047202.1;MERS_1ab;2735 | 0 | 0.025 |
| NC_019843.3;YP_009047202.1;MERS_1ab;2736 | 0 | 0.025 |
| NC_019843.3;YP_009047202.1;MERS_1ab;2737 | 0 | 0 |
| NC_019843.3;YP_009047202.1;MERS_1ab;2738 | 0 | 0 |
| NC_019843.3;YP_009047202.1;MERS_1ab;2739 | 0 | 0 |
| NC_019843.3;YP_009047202.1;MERS_1ab;2740 | 0 | 0 |
| NC_019843.3;YP_009047202.1;MERS_1ab;2741 | 0 | 0 |
| NC_019843.3;YP_009047202.1;MERS_1ab;2742 | 0 | 0 |
| NC_019843.3;YP_009047202.1;MERS_1ab;2743 | 0 | 0 |

| | | |
|---|---|---|
| NC_019843.3;YP_009047202.1;MERS_1ab;2744 | 0 | 0 |
| NC_019843.3;YP_009047202.1;MERS_1ab;2745 | 0 | 0 |
| NC_019843.3;YP_009047202.1;MERS_1ab;2746 | 0 | 0 |
| NC_019843.3;YP_009047202.1;MERS_1ab;2747 | 0 | 0 |
| NC_019843.3;YP_009047202.1;MERS_1ab;2748 | 0 | 0 |
| NC_019843.3;YP_009047202.1;MERS_1ab;2749 | 0 | 0 |
| NC_019843.3;YP_009047202.1;MERS_1ab;2750 | 0 | 0 |
| NC_019843.3;YP_009047202.1;MERS_1ab;2751 | 0 | 0 |
| NC_019843.3;YP_009047202.1;MERS_1ab;2752 | 0 | 0 |
| NC_019843.3;YP_009047202.1;MERS_1ab;2753 | 0 | 0 |
| NC_019843.3;YP_009047202.1;MERS_1ab;2754 | 0 | 0 |
| NC_019843.3;YP_009047202.1;MERS_1ab;2755 | 0 | 0 |
| NC_019843.3;YP_009047202.1;MERS_1ab;2756 | 0 | 0 |
| NC_019843.3;YP_009047202.1;MERS_1ab;2757 | 0 | 0 |
| NC_019843.3;YP_009047202.1;MERS_1ab;2758 | 0 | 0 |
| NC_019843.3;YP_009047202.1;MERS_1ab;2759 | 0 | 0 |
| NC_019843.3;YP_009047202.1;MERS_1ab;2760 | 0 | 0 |
| NC_019843.3;YP_009047202.1;MERS_1ab;2761 | 0 | 0 |
| NC_019843.3;YP_009047202.1;MERS_1ab;2762 | 0 | 0 |
| NC_019843.3;YP_009047202.1;MERS_1ab;2763 | 0 | 0 |
| NC_019843.3;YP_009047202.1;MERS_1ab;2764 | 0 | 0 |
| NC_019843.3;YP_009047202.1;MERS_1ab;2765 | 0 | 0 |
| NC_019843.3;YP_009047202.1;MERS_1ab;2766 | 0 | 0 |
| NC_019843.3;YP_009047202.1;MERS_1ab;2767 | 0 | 0 |
| NC_019843.3;YP_009047202.1;MERS_1ab;2768 | 0 | 0 |
| NC_019843.3;YP_009047202.1;MERS_1ab;2769 | 0 | 0 |
| NC_019843.3;YP_009047202.1;MERS_1ab;2770 | 0 | 0 |
| NC_019843.3;YP_009047202.1;MERS_1ab;2771 | 0 | 0 |
| NC_019843.3;YP_009047202.1;MERS_1ab;2772 | 0 | 0 |
| NC_019843.3;YP_009047202.1;MERS_1ab;2773 | 0 | 0.025 |
| NC_019843.3;YP_009047202.1;MERS_1ab;2774 | 0 | 0.075 |
| NC_019843.3;YP_009047202.1;MERS_1ab;2775 | 0 | 0.025 |
| NC_019843.3;YP_009047202.1;MERS_1ab;2776 | 0 | 0.025 |
| NC_019843.3;YP_009047202.1;MERS_1ab;2777 | 0 | 0.05 |
| NC_019843.3;YP_009047202.1;MERS_1ab;2778 | 0 | 0.125 |
| NC_019843.3;YP_009047202.1;MERS_1ab;2779 | 0 | 0.05 |
| NC_019843.3;YP_009047202.1;MERS_1ab;2780 | 0.2 | 0.25 |
| NC_019843.3;YP_009047202.1;MERS_1ab;2781 | 0.05 | 0.05 |
| NC_019843.3;YP_009047202.1;MERS_1ab;2782 | 0 | 0.025 |
| NC_019843.3;YP_009047202.1;MERS_1ab;2783 | 0 | 0.05 |
| NC_019843.3;YP_009047202.1;MERS_1ab;2784 | 0 | 0 |
| NC_019843.3;YP_009047202.1;MERS_1ab;2785 | 0 | 0 |
| NC_019843.3;YP_009047202.1;MERS_1ab;2786 | 0 | 0 |

| | | |
|---|---|---|
| NC_019843.3;YP_009047202.1;MERS_1ab;2787 | 0 | 0 |
| NC_019843.3;YP_009047202.1;MERS_1ab;2788 | 0.05 | 0 |
| NC_019843.3;YP_009047202.1;MERS_1ab;2789 | 0.05 | 0.05 |
| NC_019843.3;YP_009047202.1;MERS_1ab;2790 | 0 | 0 |
| NC_019843.3;YP_009047202.1;MERS_1ab;2791 | 0 | 0 |
| NC_019843.3;YP_009047202.1;MERS_1ab;2792 | 0 | 0.025 |
| NC_019843.3;YP_009047202.1;MERS_1ab;2793 | 0 | 0.05 |
| NC_019843.3;YP_009047202.1;MERS_1ab;2794 | 0.05 | 0 |
| NC_019843.3;YP_009047202.1;MERS_1ab;2795 | 0 | 0 |
| NC_019843.3;YP_009047202.1;MERS_1ab;2796 | 0 | 0 |
| NC_019843.3;YP_009047202.1;MERS_1ab;2797 | 0 | 0.05 |
| NC_019843.3;YP_009047202.1;MERS_1ab;2798 | 0 | 0.075 |
| NC_019843.3;YP_009047202.1;MERS_1ab;2799 | 0 | 0.05 |
| NC_019843.3;YP_009047202.1;MERS_1ab;2800 | 0 | 0 |
| NC_019843.3;YP_009047202.1;MERS_1ab;2801 | 0 | 0 |
| NC_019843.3;YP_009047202.1;MERS_1ab;2802 | 0 | 0 |
| NC_019843.3;YP_009047202.1;MERS_1ab;2803 | 0 | 0.025 |
| NC_019843.3;YP_009047202.1;MERS_1ab;2804 | 0 | 0.025 |
| NC_019843.3;YP_009047202.1;MERS_1ab;2805 | 0 | 0 |
| NC_019843.3;YP_009047202.1;MERS_1ab;2806 | 0 | 0 |
| NC_019843.3;YP_009047202.1;MERS_1ab;2807 | 0 | 0 |
| NC_019843.3;YP_009047202.1;MERS_1ab;2808 | 0 | 0 |
| NC_019843.3;YP_009047202.1;MERS_1ab;2809 | 0 | 0 |
| NC_019843.3;YP_009047202.1;MERS_1ab;2810 | 0 | 0 |
| NC_019843.3;YP_009047202.1;MERS_1ab;2811 | 0 | 0 |
| NC_019843.3;YP_009047202.1;MERS_1ab;2812 | 0 | 0 |
| NC_019843.3;YP_009047202.1;MERS_1ab;2813 | 0 | 0 |
| NC_019843.3;YP_009047202.1;MERS_1ab;2814 | 0 | 0 |
| NC_019843.3;YP_009047202.1;MERS_1ab;2815 | 0.05 | 0 |
| NC_019843.3;YP_009047202.1;MERS_1ab;2816 | 0.05 | 0 |
| NC_019843.3;YP_009047202.1;MERS_1ab;2817 | 0.1 | 0 |
| NC_019843.3;YP_009047202.1;MERS_1ab;2818 | 0.05 | 0.025 |
| NC_019843.3;YP_009047202.1;MERS_1ab;2819 | 0.05 | 0.05 |
| NC_019843.3;YP_009047202.1;MERS_1ab;2820 | 0 | 0 |
| NC_019843.3;YP_009047202.1;MERS_1ab;2821 | 0.05 | 0 |
| NC_019843.3;YP_009047202.1;MERS_1ab;2822 | 0.05 | 0 |
| NC_019843.3;YP_009047202.1;MERS_1ab;2823 | 0.05 | 0 |
| NC_019843.3;YP_009047202.1;MERS_1ab;2824 | 0 | 0 |
| NC_019843.3;YP_009047202.1;MERS_1ab;2825 | 0 | 0.025 |
| NC_019843.3;YP_009047202.1;MERS_1ab;2826 | 0 | 0 |
| NC_019843.3;YP_009047202.1;MERS_1ab;2827 | 0 | 0 |
| NC_019843.3;YP_009047202.1;MERS_1ab;2828 | 0 | 0 |
| NC_019843.3;YP_009047202.1;MERS_1ab;2829 | 0 | 0 |

| | | |
|---|---|---|
| NC_019843.3;YP_009047202.1;MERS_1ab;2830 | 0 | 0 |
| NC_019843.3;YP_009047202.1;MERS_1ab;2831 | 0 | 0 |
| NC_019843.3;YP_009047202.1;MERS_1ab;2832 | 0 | 0 |
| NC_019843.3;YP_009047202.1;MERS_1ab;2833 | 0 | 0 |
| NC_019843.3;YP_009047202.1;MERS_1ab;2834 | 0 | 0 |
| NC_019843.3;YP_009047202.1;MERS_1ab;2835 | 0 | 0 |
| NC_019843.3;YP_009047202.1;MERS_1ab;2836 | 0 | 0 |
| NC_019843.3;YP_009047202.1;MERS_1ab;2837 | 0 | 0 |
| NC_019843.3;YP_009047202.1;MERS_1ab;2838 | 0 | 0 |
| NC_019843.3;YP_009047202.1;MERS_1ab;2839 | 0 | 0 |
| NC_019843.3;YP_009047202.1;MERS_1ab;2840 | 0.05 | 0 |
| NC_019843.3;YP_009047202.1;MERS_1ab;2841 | 0.05 | 0 |
| NC_019843.3;YP_009047202.1;MERS_1ab;2842 | 0.15 | 0.025 |
| NC_019843.3;YP_009047202.1;MERS_1ab;2843 | 0 | 0 |
| NC_019843.3;YP_009047202.1;MERS_1ab;2844 | 0.05 | 0 |
| NC_019843.3;YP_009047202.1;MERS_1ab;2845 | 0 | 0 |
| NC_019843.3;YP_009047202.1;MERS_1ab;2846 | 0 | 0 |
| NC_019843.3;YP_009047202.1;MERS_1ab;2847 | 0.05 | 0.025 |
| NC_019843.3;YP_009047202.1;MERS_1ab;2848 | 0.05 | 0 |
| NC_019843.3;YP_009047202.1;MERS_1ab;2849 | 0.1 | 0 |
| NC_019843.3;YP_009047202.1;MERS_1ab;2850 | 0 | 0 |
| NC_019843.3;YP_009047202.1;MERS_1ab;2851 | 0 | 0 |
| NC_019843.3;YP_009047202.1;MERS_1ab;2852 | 0 | 0 |
| NC_019843.3;YP_009047202.1;MERS_1ab;2853 | 0 | 0 |
| NC_019843.3;YP_009047202.1;MERS_1ab;2854 | 0 | 0 |
| NC_019843.3;YP_009047202.1;MERS_1ab;2855 | 0 | 0 |
| NC_019843.3;YP_009047202.1;MERS_1ab;2856 | 0 | 0 |
| NC_019843.3;YP_009047202.1;MERS_1ab;2857 | 0 | 0 |
| NC_019843.3;YP_009047202.1;MERS_1ab;2858 | 0 | 0 |
| NC_019843.3;YP_009047202.1;MERS_1ab;2859 | 0 | 0 |
| NC_019843.3;YP_009047202.1;MERS_1ab;2860 | 0 | 0 |
| NC_019843.3;YP_009047202.1;MERS_1ab;2861 | 0 | 0 |
| NC_019843.3;YP_009047202.1;MERS_1ab;2862 | 0 | 0 |
| NC_019843.3;YP_009047202.1;MERS_1ab;2863 | 0 | 0 |
| NC_019843.3;YP_009047202.1;MERS_1ab;2864 | 0 | 0 |
| NC_019843.3;YP_009047202.1;MERS_1ab;2865 | 0 | 0 |
| NC_019843.3;YP_009047202.1;MERS_1ab;2866 | 0 | 0 |
| NC_019843.3;YP_009047202.1;MERS_1ab;2867 | 0 | 0 |
| NC_019843.3;YP_009047202.1;MERS_1ab;2868 | 0 | 0 |
| NC_019843.3;YP_009047202.1;MERS_1ab;2869 | 0 | 0 |
| NC_019843.3;YP_009047202.1;MERS_1ab;2870 | 0.05 | 0 |
| NC_019843.3;YP_009047202.1;MERS_1ab;2871 | 0 | 0 |
| NC_019843.3;YP_009047202.1;MERS_1ab;2872 | 0 | 0.025 |

| | | |
|---|---|---|
| NC_019843.3;YP_009047202.1;MERS_1ab;2873 | 0 | 0.025 |
| NC_019843.3;YP_009047202.1;MERS_1ab;2874 | 0 | 0.025 |
| NC_019843.3;YP_009047202.1;MERS_1ab;2875 | 0 | 0 |
| NC_019843.3;YP_009047202.1;MERS_1ab;2876 | 0 | 0 |
| NC_019843.3;YP_009047202.1;MERS_1ab;2877 | 0 | 0 |
| NC_019843.3;YP_009047202.1;MERS_1ab;2878 | 0 | 0 |
| NC_019843.3;YP_009047202.1;MERS_1ab;2879 | 0.05 | 0 |
| NC_019843.3;YP_009047202.1;MERS_1ab;2880 | 0.05 | 0 |
| NC_019843.3;YP_009047202.1;MERS_1ab;2881 | 0 | 0 |
| NC_019843.3;YP_009047202.1;MERS_1ab;2882 | 0 | 0 |
| NC_019843.3;YP_009047202.1;MERS_1ab;2883 | 0 | 0 |
| NC_019843.3;YP_009047202.1;MERS_1ab;2884 | 0.05 | 0 |
| NC_019843.3;YP_009047202.1;MERS_1ab;2885 | 0 | 0.05 |
| NC_019843.3;YP_009047202.1;MERS_1ab;2886 | 0 | 0.025 |
| NC_019843.3;YP_009047202.1;MERS_1ab;2887 | 0 | 0 |
| NC_019843.3;YP_009047202.1;MERS_1ab;2888 | 0 | 0 |
| NC_019843.3;YP_009047202.1;MERS_1ab;2889 | 0 | 0 |
| NC_019843.3;YP_009047202.1;MERS_1ab;2890 | 0 | 0 |
| NC_019843.3;YP_009047202.1;MERS_1ab;2891 | 0 | 0 |
| NC_019843.3;YP_009047202.1;MERS_1ab;2892 | 0 | 0.025 |
| NC_019843.3;YP_009047202.1;MERS_1ab;2893 | 0 | 0 |
| NC_019843.3;YP_009047202.1;MERS_1ab;2894 | 0 | 0 |
| NC_019843.3;YP_009047202.1;MERS_1ab;2895 | 0 | 0 |
| NC_019843.3;YP_009047202.1;MERS_1ab;2896 | 0 | 0 |
| NC_019843.3;YP_009047202.1;MERS_1ab;2897 | 0.05 | 0 |
| NC_019843.3;YP_009047202.1;MERS_1ab;2898 | 0.05 | 0 |
| NC_019843.3;YP_009047202.1;MERS_1ab;2899 | 0.05 | 0 |
| NC_019843.3;YP_009047202.1;MERS_1ab;2900 | 0.05 | 0 |
| NC_019843.3;YP_009047202.1;MERS_1ab;2901 | 0.05 | 0 |
| NC_019843.3;YP_009047202.1;MERS_1ab;2902 | 0.05 | 0 |
| NC_019843.3;YP_009047202.1;MERS_1ab;2903 | 0.05 | 0 |
| NC_019843.3;YP_009047202.1;MERS_1ab;2904 | 0.05 | 0 |
| NC_019843.3;YP_009047202.1;MERS_1ab;2905 | 0.05 | 0.025 |
| NC_019843.3;YP_009047202.1;MERS_1ab;2906 | 0.05 | 0 |
| NC_019843.3;YP_009047202.1;MERS_1ab;2907 | 0 | 0 |
| NC_019843.3;YP_009047202.1;MERS_1ab;2908 | 0 | 0 |
| NC_019843.3;YP_009047202.1;MERS_1ab;2909 | 0 | 0 |
| NC_019843.3;YP_009047202.1;MERS_1ab;2910 | 0.05 | 0 |
| NC_019843.3;YP_009047202.1;MERS_1ab;2911 | 0 | 0 |
| NC_019843.3;YP_009047202.1;MERS_1ab;2912 | 0.05 | 0 |
| NC_019843.3;YP_009047202.1;MERS_1ab;2913 | 0 | 0 |
| NC_019843.3;YP_009047202.1;MERS_1ab;2914 | 0 | 0 |
| NC_019843.3;YP_009047202.1;MERS_1ab;2915 | 0 | 0 |

| | | |
|---|---|---|
| NC_019843.3;YP_009047202.1;MERS_1ab;2916 | 0 | 0 |
| NC_019843.3;YP_009047202.1;MERS_1ab;2917 | 0 | 0 |
| NC_019843.3;YP_009047202.1;MERS_1ab;2918 | 0.1 | 0 |
| NC_019843.3;YP_009047202.1;MERS_1ab;2919 | 0 | 0 |
| NC_019843.3;YP_009047202.1;MERS_1ab;2920 | 0 | 0 |
| NC_019843.3;YP_009047202.1;MERS_1ab;2921 | 0 | 0 |
| NC_019843.3;YP_009047202.1;MERS_1ab;2922 | 0 | 0 |
| NC_019843.3;YP_009047202.1;MERS_1ab;2923 | 0 | 0 |
| NC_019843.3;YP_009047202.1;MERS_1ab;2924 | 0.05 | 0 |
| NC_019843.3;YP_009047202.1;MERS_1ab;2925 | 0 | 0 |
| NC_019843.3;YP_009047202.1;MERS_1ab;2926 | 0 | 0 |
| NC_019843.3;YP_009047202.1;MERS_1ab;2927 | 0 | 0.075 |
| NC_019843.3;YP_009047202.1;MERS_1ab;2928 | 0 | 0.025 |
| NC_019843.3;YP_009047202.1;MERS_1ab;2929 | 0 | 0 |
| NC_019843.3;YP_009047202.1;MERS_1ab;2930 | 0.05 | 0.025 |
| NC_019843.3;YP_009047202.1;MERS_1ab;2931 | 0 | 0 |
| NC_019843.3;YP_009047202.1;MERS_1ab;2932 | 0 | 0 |
| NC_019843.3;YP_009047202.1;MERS_1ab;2933 | 0 | 0 |
| NC_019843.3;YP_009047202.1;MERS_1ab;2934 | 0 | 0 |
| NC_019843.3;YP_009047202.1;MERS_1ab;2935 | 0.05 | 0 |
| NC_019843.3;YP_009047202.1;MERS_1ab;2936 | 0.05 | 0 |
| NC_019843.3;YP_009047202.1;MERS_1ab;2937 | 0.05 | 0 |
| NC_019843.3;YP_009047202.1;MERS_1ab;2938 | 0.05 | 0 |
| NC_019843.3;YP_009047202.1;MERS_1ab;2939 | 0 | 0.025 |
| NC_019843.3;YP_009047202.1;MERS_1ab;2940 | 0 | 0.025 |
| NC_019843.3;YP_009047202.1;MERS_1ab;2941 | 0 | 0 |
| NC_019843.3;YP_009047202.1;MERS_1ab;2942 | 0 | 0 |
| NC_019843.3;YP_009047202.1;MERS_1ab;2943 | 0 | 0 |
| NC_019843.3;YP_009047202.1;MERS_1ab;2944 | 0 | 0 |
| NC_019843.3;YP_009047202.1;MERS_1ab;2945 | 0 | 0 |
| NC_019843.3;YP_009047202.1;MERS_1ab;2946 | 0 | 0 |
| NC_019843.3;YP_009047202.1;MERS_1ab;2947 | 0 | 0 |
| NC_019843.3;YP_009047202.1;MERS_1ab;2948 | 0 | 0 |
| NC_019843.3;YP_009047202.1;MERS_1ab;2949 | 0 | 0 |
| NC_019843.3;YP_009047202.1;MERS_1ab;2950 | 0 | 0 |
| NC_019843.3;YP_009047202.1;MERS_1ab;2951 | 0 | 0 |
| NC_019843.3;YP_009047202.1;MERS_1ab;2952 | 0 | 0 |
| NC_019843.3;YP_009047202.1;MERS_1ab;2953 | 0 | 0 |
| NC_019843.3;YP_009047202.1;MERS_1ab;2954 | 0 | 0 |
| NC_019843.3;YP_009047202.1;MERS_1ab;2955 | 0 | 0 |
| NC_019843.3;YP_009047202.1;MERS_1ab;2956 | 0 | 0 |
| NC_019843.3;YP_009047202.1;MERS_1ab;2957 | 0 | 0 |
| NC_019843.3;YP_009047202.1;MERS_1ab;2958 | 0 | 0 |

| | | |
|---|---|---|
| NC_019843.3;YP_009047202.1;MERS_1ab;2959 | 0 | 0 |
| NC_019843.3;YP_009047202.1;MERS_1ab;2960 | 0 | 0 |
| NC_019843.3;YP_009047202.1;MERS_1ab;2961 | 0 | 0 |
| NC_019843.3;YP_009047202.1;MERS_1ab;2962 | 0 | 0 |
| NC_019843.3;YP_009047202.1;MERS_1ab;2963 | 0 | 0 |
| NC_019843.3;YP_009047202.1;MERS_1ab;2964 | 0 | 0 |
| NC_019843.3;YP_009047202.1;MERS_1ab;2965 | 0 | 0 |
| NC_019843.3;YP_009047202.1;MERS_1ab;2966 | 0 | 0 |
| NC_019843.3;YP_009047202.1;MERS_1ab;2967 | 0 | 0 |
| NC_019843.3;YP_009047202.1;MERS_1ab;2968 | 0 | 0 |
| NC_019843.3;YP_009047202.1;MERS_1ab;2969 | 0 | 0 |
| NC_019843.3;YP_009047202.1;MERS_1ab;2970 | 0 | 0 |
| NC_019843.3;YP_009047202.1;MERS_1ab;2971 | 0 | 0 |
| NC_019843.3;YP_009047202.1;MERS_1ab;2972 | 0 | 0 |
| NC_019843.3;YP_009047202.1;MERS_1ab;2973 | 0 | 0 |
| NC_019843.3;YP_009047202.1;MERS_1ab;2974 | 0 | 0 |
| NC_019843.3;YP_009047202.1;MERS_1ab;2975 | 0 | 0 |
| NC_019843.3;YP_009047202.1;MERS_1ab;2976 | 0 | 0 |
| NC_019843.3;YP_009047202.1;MERS_1ab;2977 | 0 | 0 |
| NC_019843.3;YP_009047202.1;MERS_1ab;2978 | 0 | 0.025 |
| NC_019843.3;YP_009047202.1;MERS_1ab;2979 | 0 | 0.025 |
| NC_019843.3;YP_009047202.1;MERS_1ab;2980 | 0 | 0.025 |
| NC_019843.3;YP_009047202.1;MERS_1ab;2981 | 0 | 0.025 |
| NC_019843.3;YP_009047202.1;MERS_1ab;2982 | 0 | 0.025 |
| NC_019843.3;YP_009047202.1;MERS_1ab;2983 | 0 | 0.025 |
| NC_019843.3;YP_009047202.1;MERS_1ab;2984 | 0 | 0.025 |
| NC_019843.3;YP_009047202.1;MERS_1ab;2985 | 0 | 0.025 |
| NC_019843.3;YP_009047202.1;MERS_1ab;2986 | 0 | 0.025 |
| NC_019843.3;YP_009047202.1;MERS_1ab;2987 | 0 | 0.025 |
| NC_019843.3;YP_009047202.1;MERS_1ab;2988 | 0 | 0.025 |
| NC_019843.3;YP_009047202.1;MERS_1ab;2989 | 0 | 0.025 |
| NC_019843.3;YP_009047202.1;MERS_1ab;2990 | 0 | 0 |
| NC_019843.3;YP_009047202.1;MERS_1ab;2991 | 0 | 0 |
| NC_019843.3;YP_009047202.1;MERS_1ab;2992 | 0 | 0 |
| NC_019843.3;YP_009047202.1;MERS_1ab;2993 | 0 | 0 |
| NC_019843.3;YP_009047202.1;MERS_1ab;2994 | 0 | 0 |
| NC_019843.3;YP_009047202.1;MERS_1ab;2995 | 0 | 0 |
| NC_019843.3;YP_009047202.1;MERS_1ab;2996 | 0 | 0 |
| NC_019843.3;YP_009047202.1;MERS_1ab;2997 | 0 | 0 |
| NC_019843.3;YP_009047202.1;MERS_1ab;2998 | 0 | 0 |
| NC_019843.3;YP_009047202.1;MERS_1ab;2999 | 0 | 0 |
| NC_019843.3;YP_009047202.1;MERS_1ab;3000 | 0 | 0 |
| NC_019843.3;YP_009047202.1;MERS_1ab;3001 | 0 | 0 |

| | | |
|---|---|---|
| NC_019843.3;YP_009047202.1;MERS_1ab;3002 | 0 | 0 |
| NC_019843.3;YP_009047202.1;MERS_1ab;3003 | 0 | 0 |
| NC_019843.3;YP_009047202.1;MERS_1ab;3004 | 0 | 0 |
| NC_019843.3;YP_009047202.1;MERS_1ab;3005 | 0 | 0 |
| NC_019843.3;YP_009047202.1;MERS_1ab;3006 | 0 | 0 |
| NC_019843.3;YP_009047202.1;MERS_1ab;3007 | 0 | 0 |
| NC_019843.3;YP_009047202.1;MERS_1ab;3008 | 0 | 0 |
| NC_019843.3;YP_009047202.1;MERS_1ab;3009 | 0 | 0 |
| NC_019843.3;YP_009047202.1;MERS_1ab;3010 | 0 | 0 |
| NC_019843.3;YP_009047202.1;MERS_1ab;3011 | 0 | 0 |
| NC_019843.3;YP_009047202.1;MERS_1ab;3012 | 0 | 0 |
| NC_019843.3;YP_009047202.1;MERS_1ab;3013 | 0 | 0 |
| NC_019843.3;YP_009047202.1;MERS_1ab;3014 | 0 | 0 |
| NC_019843.3;YP_009047202.1;MERS_1ab;3015 | 0 | 0 |
| NC_019843.3;YP_009047202.1;MERS_1ab;3016 | 0 | 0 |
| NC_019843.3;YP_009047202.1;MERS_1ab;3017 | 0 | 0 |
| NC_019843.3;YP_009047202.1;MERS_1ab;3018 | 0 | 0 |
| NC_019843.3;YP_009047202.1;MERS_1ab;3019 | 0 | 0 |
| NC_019843.3;YP_009047202.1;MERS_1ab;3020 | 0 | 0 |
| NC_019843.3;YP_009047202.1;MERS_1ab;3021 | 0 | 0 |
| NC_019843.3;YP_009047202.1;MERS_1ab;3022 | 0 | 0 |
| NC_019843.3;YP_009047202.1;MERS_1ab;3023 | 0 | 0 |
| NC_019843.3;YP_009047202.1;MERS_1ab;3024 | 0 | 0 |
| NC_019843.3;YP_009047202.1;MERS_1ab;3025 | 0 | 0 |
| NC_019843.3;YP_009047202.1;MERS_1ab;3026 | 0 | 0 |
| NC_019843.3;YP_009047202.1;MERS_1ab;3027 | 0 | 0 |
| NC_019843.3;YP_009047202.1;MERS_1ab;3028 | 0 | 0 |
| NC_019843.3;YP_009047202.1;MERS_1ab;3029 | 0 | 0 |
| NC_019843.3;YP_009047202.1;MERS_1ab;3030 | 0 | 0 |
| NC_019843.3;YP_009047202.1;MERS_1ab;3031 | 0 | 0 |
| NC_019843.3;YP_009047202.1;MERS_1ab;3032 | 0 | 0 |
| NC_019843.3;YP_009047202.1;MERS_1ab;3033 | 0 | 0 |
| NC_019843.3;YP_009047202.1;MERS_1ab;3034 | 0 | 0 |
| NC_019843.3;YP_009047202.1;MERS_1ab;3035 | 0 | 0 |
| NC_019843.3;YP_009047202.1;MERS_1ab;3036 | 0 | 0 |
| NC_019843.3;YP_009047202.1;MERS_1ab;3037 | 0 | 0 |
| NC_019843.3;YP_009047202.1;MERS_1ab;3038 | 0 | 0 |
| NC_019843.3;YP_009047202.1;MERS_1ab;3039 | 0 | 0 |
| NC_019843.3;YP_009047202.1;MERS_1ab;3040 | 0 | 0 |
| NC_019843.3;YP_009047202.1;MERS_1ab;3041 | 0 | 0 |
| NC_019843.3;YP_009047202.1;MERS_1ab;3042 | 0 | 0 |
| NC_019843.3;YP_009047202.1;MERS_1ab;3043 | 0 | 0 |
| NC_019843.3;YP_009047202.1;MERS_1ab;3044 | 0 | 0 |

| | | |
|---|---|---|
| NC_019843.3;YP_009047202.1;MERS_1ab;3045 | 0 | 0 |
| NC_019843.3;YP_009047202.1;MERS_1ab;3046 | 0.05 | 0 |
| NC_019843.3;YP_009047202.1;MERS_1ab;3047 | 0.05 | 0 |
| NC_019843.3;YP_009047202.1;MERS_1ab;3048 | 0 | 0 |
| NC_019843.3;YP_009047202.1;MERS_1ab;3049 | 0 | 0 |
| NC_019843.3;YP_009047202.1;MERS_1ab;3050 | 0 | 0 |
| NC_019843.3;YP_009047202.1;MERS_1ab;3051 | 0 | 0 |
| NC_019843.3;YP_009047202.1;MERS_1ab;3052 | 0 | 0 |
| NC_019843.3;YP_009047202.1;MERS_1ab;3053 | 0 | 0 |
| NC_019843.3;YP_009047202.1;MERS_1ab;3054 | 0 | 0 |
| NC_019843.3;YP_009047202.1;MERS_1ab;3055 | 0 | 0 |
| NC_019843.3;YP_009047202.1;MERS_1ab;3056 | 0 | 0 |
| NC_019843.3;YP_009047202.1;MERS_1ab;3057 | 0 | 0 |
| NC_019843.3;YP_009047202.1;MERS_1ab;3058 | 0 | 0.025 |
| NC_019843.3;YP_009047202.1;MERS_1ab;3059 | 0 | 0 |
| NC_019843.3;YP_009047202.1;MERS_1ab;3060 | 0 | 0 |
| NC_019843.3;YP_009047202.1;MERS_1ab;3061 | 0 | 0 |
| NC_019843.3;YP_009047202.1;MERS_1ab;3062 | 0 | 0 |
| NC_019843.3;YP_009047202.1;MERS_1ab;3063 | 0 | 0 |
| NC_019843.3;YP_009047202.1;MERS_1ab;3064 | 0 | 0 |
| NC_019843.3;YP_009047202.1;MERS_1ab;3065 | 0 | 0 |
| NC_019843.3;YP_009047202.1;MERS_1ab;3066 | 0 | 0 |
| NC_019843.3;YP_009047202.1;MERS_1ab;3067 | 0 | 0 |
| NC_019843.3;YP_009047202.1;MERS_1ab;3068 | 0 | 0 |
| NC_019843.3;YP_009047202.1;MERS_1ab;3069 | 0 | 0 |
| NC_019843.3;YP_009047202.1;MERS_1ab;3070 | 0 | 0 |
| NC_019843.3;YP_009047202.1;MERS_1ab;3071 | 0 | 0 |
| NC_019843.3;YP_009047202.1;MERS_1ab;3072 | 0 | 0 |
| NC_019843.3;YP_009047202.1;MERS_1ab;3073 | 0 | 0 |
| NC_019843.3;YP_009047202.1;MERS_1ab;3074 | 0 | 0 |
| NC_019843.3;YP_009047202.1;MERS_1ab;3075 | 0 | 0 |
| NC_019843.3;YP_009047202.1;MERS_1ab;3076 | 0 | 0 |
| NC_019843.3;YP_009047202.1;MERS_1ab;3077 | 0 | 0 |
| NC_019843.3;YP_009047202.1;MERS_1ab;3078 | 0 | 0 |
| NC_019843.3;YP_009047202.1;MERS_1ab;3079 | 0 | 0 |
| NC_019843.3;YP_009047202.1;MERS_1ab;3080 | 0 | 0 |
| NC_019843.3;YP_009047202.1;MERS_1ab;3081 | 0 | 0 |
| NC_019843.3;YP_009047202.1;MERS_1ab;3082 | 0 | 0 |
| NC_019843.3;YP_009047202.1;MERS_1ab;3083 | 0 | 0 |
| NC_019843.3;YP_009047202.1;MERS_1ab;3084 | 0 | 0 |
| NC_019843.3;YP_009047202.1;MERS_1ab;3085 | 0 | 0 |
| NC_019843.3;YP_009047202.1;MERS_1ab;3086 | 0 | 0 |
| NC_019843.3;YP_009047202.1;MERS_1ab;3087 | 0 | 0 |

| | | |
|---|---|---|
| NC_019843.3;YP_009047202.1;MERS_1ab;3088 | 0 | 0 |
| NC_019843.3;YP_009047202.1;MERS_1ab;3089 | 0 | 0 |
| NC_019843.3;YP_009047202.1;MERS_1ab;3090 | 0 | 0 |
| NC_019843.3;YP_009047202.1;MERS_1ab;3091 | 0 | 0 |
| NC_019843.3;YP_009047202.1;MERS_1ab;3092 | 0 | 0 |
| NC_019843.3;YP_009047202.1;MERS_1ab;3093 | 0 | 0 |
| NC_019843.3;YP_009047202.1;MERS_1ab;3094 | 0 | 0 |
| NC_019843.3;YP_009047202.1;MERS_1ab;3095 | 0 | 0 |
| NC_019843.3;YP_009047202.1;MERS_1ab;3096 | 0 | 0 |
| NC_019843.3;YP_009047202.1;MERS_1ab;3097 | 0 | 0 |
| NC_019843.3;YP_009047202.1;MERS_1ab;3098 | 0 | 0 |
| NC_019843.3;YP_009047202.1;MERS_1ab;3099 | 0 | 0 |
| NC_019843.3;YP_009047202.1;MERS_1ab;3100 | 0 | 0.05 |
| NC_019843.3;YP_009047202.1;MERS_1ab;3101 | 0 | 0 |
| NC_019843.3;YP_009047202.1;MERS_1ab;3102 | 0 | 0 |
| NC_019843.3;YP_009047202.1;MERS_1ab;3103 | 0 | 0 |
| NC_019843.3;YP_009047202.1;MERS_1ab;3104 | 0 | 0 |
| NC_019843.3;YP_009047202.1;MERS_1ab;3105 | 0 | 0 |
| NC_019843.3;YP_009047202.1;MERS_1ab;3106 | 0 | 0 |
| NC_019843.3;YP_009047202.1;MERS_1ab;3107 | 0 | 0 |
| NC_019843.3;YP_009047202.1;MERS_1ab;3108 | 0 | 0 |
| NC_019843.3;YP_009047202.1;MERS_1ab;3109 | 0 | 0 |
| NC_019843.3;YP_009047202.1;MERS_1ab;3110 | 0 | 0 |
| NC_019843.3;YP_009047202.1;MERS_1ab;3111 | 0 | 0 |
| NC_019843.3;YP_009047202.1;MERS_1ab;3112 | 0 | 0 |
| NC_019843.3;YP_009047202.1;MERS_1ab;3113 | 0 | 0 |
| NC_019843.3;YP_009047202.1;MERS_1ab;3114 | 0 | 0 |
| NC_019843.3;YP_009047202.1;MERS_1ab;3115 | 0 | 0 |
| NC_019843.3;YP_009047202.1;MERS_1ab;3116 | 0 | 0 |
| NC_019843.3;YP_009047202.1;MERS_1ab;3117 | 0 | 0 |
| NC_019843.3;YP_009047202.1;MERS_1ab;3118 | 0 | 0 |
| NC_019843.3;YP_009047202.1;MERS_1ab;3119 | 0 | 0 |
| NC_019843.3;YP_009047202.1;MERS_1ab;3120 | 0 | 0 |
| NC_019843.3;YP_009047202.1;MERS_1ab;3121 | 0 | 0 |
| NC_019843.3;YP_009047202.1;MERS_1ab;3122 | 0 | 0 |
| NC_019843.3;YP_009047202.1;MERS_1ab;3123 | 0 | 0 |
| NC_019843.3;YP_009047202.1;MERS_1ab;3124 | 0 | 0 |
| NC_019843.3;YP_009047202.1;MERS_1ab;3125 | 0 | 0 |
| NC_019843.3;YP_009047202.1;MERS_1ab;3126 | 0 | 0 |
| NC_019843.3;YP_009047202.1;MERS_1ab;3127 | 0 | 0 |
| NC_019843.3;YP_009047202.1;MERS_1ab;3128 | 0 | 0 |
| NC_019843.3;YP_009047202.1;MERS_1ab;3129 | 0 | 0 |
| NC_019843.3;YP_009047202.1;MERS_1ab;3130 | 0 | 0 |

| | | |
|---|---|---|
| NC_019843.3;YP_009047202.1;MERS_1ab;3131 | 0 | 0 |
| NC_019843.3;YP_009047202.1;MERS_1ab;3132 | 0 | 0 |
| NC_019843.3;YP_009047202.1;MERS_1ab;3133 | 0 | 0 |
| NC_019843.3;YP_009047202.1;MERS_1ab;3134 | 0 | 0 |
| NC_019843.3;YP_009047202.1;MERS_1ab;3135 | 0 | 0 |
| NC_019843.3;YP_009047202.1;MERS_1ab;3136 | 0 | 0 |
| NC_019843.3;YP_009047202.1;MERS_1ab;3137 | 0 | 0 |
| NC_019843.3;YP_009047202.1;MERS_1ab;3138 | 0 | 0 |
| NC_019843.3;YP_009047202.1;MERS_1ab;3139 | 0 | 0 |
| NC_019843.3;YP_009047202.1;MERS_1ab;3140 | 0 | 0.025 |
| NC_019843.3;YP_009047202.1;MERS_1ab;3141 | 0 | 0.025 |
| NC_019843.3;YP_009047202.1;MERS_1ab;3142 | 0.05 | 0.05 |
| NC_019843.3;YP_009047202.1;MERS_1ab;3143 | 0.05 | 0.075 |
| NC_019843.3;YP_009047202.1;MERS_1ab;3144 | 0 | 0.05 |
| NC_019843.3;YP_009047202.1;MERS_1ab;3145 | 0 | 0.05 |
| NC_019843.3;YP_009047202.1;MERS_1ab;3146 | 0 | 0 |
| NC_019843.3;YP_009047202.1;MERS_1ab;3147 | 0 | 0 |
| NC_019843.3;YP_009047202.1;MERS_1ab;3148 | 0 | 0.05 |
| NC_019843.3;YP_009047202.1;MERS_1ab;3149 | 0 | 0.025 |
| NC_019843.3;YP_009047202.1;MERS_1ab;3150 | 0.05 | 0.05 |
| NC_019843.3;YP_009047202.1;MERS_1ab;3151 | 0 | 0.05 |
| NC_019843.3;YP_009047202.1;MERS_1ab;3152 | 0.05 | 0.05 |
| NC_019843.3;YP_009047202.1;MERS_1ab;3153 | 0.05 | 0.05 |
| NC_019843.3;YP_009047202.1;MERS_1ab;3154 | 0.05 | 0.05 |
| NC_019843.3;YP_009047202.1;MERS_1ab;3155 | 0.05 | 0.025 |
| NC_019843.3;YP_009047202.1;MERS_1ab;3156 | 0 | 0 |
| NC_019843.3;YP_009047202.1;MERS_1ab;3157 | 0 | 0.025 |
| NC_019843.3;YP_009047202.1;MERS_1ab;3158 | 0 | 0.025 |
| NC_019843.3;YP_009047202.1;MERS_1ab;3159 | 0 | 0 |
| NC_019843.3;YP_009047202.1;MERS_1ab;3160 | 0 | 0.05 |
| NC_019843.3;YP_009047202.1;MERS_1ab;3161 | 0 | 0.025 |
| NC_019843.3;YP_009047202.1;MERS_1ab;3162 | 0 | 0 |
| NC_019843.3;YP_009047202.1;MERS_1ab;3163 | 0.05 | 0.025 |
| NC_019843.3;YP_009047202.1;MERS_1ab;3164 | 0 | 0 |
| NC_019843.3;YP_009047202.1;MERS_1ab;3165 | 0 | 0 |
| NC_019843.3;YP_009047202.1;MERS_1ab;3166 | 0 | 0.025 |
| NC_019843.3;YP_009047202.1;MERS_1ab;3167 | 0 | 0.025 |
| NC_019843.3;YP_009047202.1;MERS_1ab;3168 | 0 | 0 |
| NC_019843.3;YP_009047202.1;MERS_1ab;3169 | 0 | 0 |
| NC_019843.3;YP_009047202.1;MERS_1ab;3170 | 0 | 0 |
| NC_019843.3;YP_009047202.1;MERS_1ab;3171 | 0.05 | 0 |
| NC_019843.3;YP_009047202.1;MERS_1ab;3172 | 0 | 0 |
| NC_019843.3;YP_009047202.1;MERS_1ab;3173 | 0.05 | 0 |

| | | |
|---|---|---|
| NC_019843.3;YP_009047202.1;MERS_1ab;3174 | 0 | 0 |
| NC_019843.3;YP_009047202.1;MERS_1ab;3175 | 0 | 0.025 |
| NC_019843.3;YP_009047202.1;MERS_1ab;3176 | 0 | 0 |
| NC_019843.3;YP_009047202.1;MERS_1ab;3177 | 0 | 0 |
| NC_019843.3;YP_009047202.1;MERS_1ab;3178 | 0 | 0.025 |
| NC_019843.3;YP_009047202.1;MERS_1ab;3179 | 0 | 0 |
| NC_019843.3;YP_009047202.1;MERS_1ab;3180 | 0 | 0 |
| NC_019843.3;YP_009047202.1;MERS_1ab;3181 | 0.05 | 0 |
| NC_019843.3;YP_009047202.1;MERS_1ab;3182 | 0 | 0 |
| NC_019843.3;YP_009047202.1;MERS_1ab;3183 | 0 | 0 |
| NC_019843.3;YP_009047202.1;MERS_1ab;3184 | 0 | 0 |
| NC_019843.3;YP_009047202.1;MERS_1ab;3185 | 0 | 0 |
| NC_019843.3;YP_009047202.1;MERS_1ab;3186 | 0 | 0 |
| NC_019843.3;YP_009047202.1;MERS_1ab;3187 | 0 | 0 |
| NC_019843.3;YP_009047202.1;MERS_1ab;3188 | 0 | 0 |
| NC_019843.3;YP_009047202.1;MERS_1ab;3189 | 0 | 0 |
| NC_019843.3;YP_009047202.1;MERS_1ab;3190 | 0 | 0 |
| NC_019843.3;YP_009047202.1;MERS_1ab;3191 | 0 | 0 |
| NC_019843.3;YP_009047202.1;MERS_1ab;3192 | 0 | 0 |
| NC_019843.3;YP_009047202.1;MERS_1ab;3193 | 0 | 0 |
| NC_019843.3;YP_009047202.1;MERS_1ab;3194 | 0.05 | 0.025 |
| NC_019843.3;YP_009047202.1;MERS_1ab;3195 | 0 | 0.025 |
| NC_019843.3;YP_009047202.1;MERS_1ab;3196 | 0 | 0 |
| NC_019843.3;YP_009047202.1;MERS_1ab;3197 | 0 | 0 |
| NC_019843.3;YP_009047202.1;MERS_1ab;3198 | 0 | 0 |
| NC_019843.3;YP_009047202.1;MERS_1ab;3199 | 0.05 | 0.05 |
| NC_019843.3;YP_009047202.1;MERS_1ab;3200 | 0.05 | 0.025 |
| NC_019843.3;YP_009047202.1;MERS_1ab;3201 | 0 | 0 |
| NC_019843.3;YP_009047202.1;MERS_1ab;3202 | 0 | 0 |
| NC_019843.3;YP_009047202.1;MERS_1ab;3203 | 0 | 0 |
| NC_019843.3;YP_009047202.1;MERS_1ab;3204 | 0 | 0 |
| NC_019843.3;YP_009047202.1;MERS_1ab;3205 | 0 | 0 |
| NC_019843.3;YP_009047202.1;MERS_1ab;3206 | 0 | 0 |
| NC_019843.3;YP_009047202.1;MERS_1ab;3207 | 0 | 0 |
| NC_019843.3;YP_009047202.1;MERS_1ab;3208 | 0 | 0 |
| NC_019843.3;YP_009047202.1;MERS_1ab;3209 | 0 | 0 |
| NC_019843.3;YP_009047202.1;MERS_1ab;3210 | 0 | 0 |
| NC_019843.3;YP_009047202.1;MERS_1ab;3211 | 0 | 0 |
| NC_019843.3;YP_009047202.1;MERS_1ab;3212 | 0 | 0 |
| NC_019843.3;YP_009047202.1;MERS_1ab;3213 | 0 | 0 |
| NC_019843.3;YP_009047202.1;MERS_1ab;3214 | 0.2 | 0.025 |
| NC_019843.3;YP_009047202.1;MERS_1ab;3215 | 0.1 | 0 |
| NC_019843.3;YP_009047202.1;MERS_1ab;3216 | 0.05 | 0 |

| | | |
|---|---|---|
| NC_019843.3;YP_009047202.1;MERS_1ab;3217 | 0.05 | 0 |
| NC_019843.3;YP_009047202.1;MERS_1ab;3218 | 0.05 | 0 |
| NC_019843.3;YP_009047202.1;MERS_1ab;3219 | 0.05 | 0 |
| NC_019843.3;YP_009047202.1;MERS_1ab;3220 | 0.1 | 0 |
| NC_019843.3;YP_009047202.1;MERS_1ab;3221 | 0.05 | 0 |
| NC_019843.3;YP_009047202.1;MERS_1ab;3222 | 0.05 | 0 |
| NC_019843.3;YP_009047202.1;MERS_1ab;3223 | 0.05 | 0 |
| NC_019843.3;YP_009047202.1;MERS_1ab;3224 | 0.05 | 0 |
| NC_019843.3;YP_009047202.1;MERS_1ab;3225 | 0 | 0 |
| NC_019843.3;YP_009047202.1;MERS_1ab;3226 | 0 | 0 |
| NC_019843.3;YP_009047202.1;MERS_1ab;3227 | 0 | 0.025 |
| NC_019843.3;YP_009047202.1;MERS_1ab;3228 | 0 | 0 |
| NC_019843.3;YP_009047202.1;MERS_1ab;3229 | 0.05 | 0 |
| NC_019843.3;YP_009047202.1;MERS_1ab;3230 | 0 | 0 |
| NC_019843.3;YP_009047202.1;MERS_1ab;3231 | 0 | 0 |
| NC_019843.3;YP_009047202.1;MERS_1ab;3232 | 0 | 0.025 |
| NC_019843.3;YP_009047202.1;MERS_1ab;3233 | 0 | 0 |
| NC_019843.3;YP_009047202.1;MERS_1ab;3234 | 0.05 | 0 |
| NC_019843.3;YP_009047202.1;MERS_1ab;3235 | 0 | 0 |
| NC_019843.3;YP_009047202.1;MERS_1ab;3236 | 0 | 0 |
| NC_019843.3;YP_009047202.1;MERS_1ab;3237 | 0 | 0 |
| NC_019843.3;YP_009047202.1;MERS_1ab;3238 | 0 | 0 |
| NC_019843.3;YP_009047202.1;MERS_1ab;3239 | 0 | 0 |
| NC_019843.3;YP_009047202.1;MERS_1ab;3240 | 0 | 0 |
| NC_019843.3;YP_009047202.1;MERS_1ab;3241 | 0 | 0 |
| NC_019843.3;YP_009047202.1;MERS_1ab;3242 | 0 | 0 |
| NC_019843.3;YP_009047202.1;MERS_1ab;3243 | 0 | 0.025 |
| NC_019843.3;YP_009047202.1;MERS_1ab;3244 | 0.05 | 0.025 |
| NC_019843.3;YP_009047202.1;MERS_1ab;3245 | 0.05 | 0 |
| NC_019843.3;YP_009047202.1;MERS_1ab;3246 | 0.1 | 0 |
| NC_019843.3;YP_009047202.1;MERS_1ab;3247 | 0.05 | 0.025 |
| NC_019843.3;YP_009047202.1;MERS_1ab;3248 | 0 | 0 |
| NC_019843.3;YP_009047202.1;MERS_1ab;3249 | 0.05 | 0 |
| NC_019843.3;YP_009047202.1;MERS_1ab;3250 | 0 | 0 |
| NC_019843.3;YP_009047202.1;MERS_1ab;3251 | 0 | 0 |
| NC_019843.3;YP_009047202.1;MERS_1ab;3252 | 0 | 0 |
| NC_019843.3;YP_009047202.1;MERS_1ab;3253 | 0 | 0 |
| NC_019843.3;YP_009047202.1;MERS_1ab;3254 | 0 | 0 |
| NC_019843.3;YP_009047202.1;MERS_1ab;3255 | 0 | 0 |
| NC_019843.3;YP_009047202.1;MERS_1ab;3256 | 0 | 0 |
| NC_019843.3;YP_009047202.1;MERS_1ab;3257 | 0 | 0 |
| NC_019843.3;YP_009047202.1;MERS_1ab;3258 | 0 | 0.05 |
| NC_019843.3;YP_009047202.1;MERS_1ab;3259 | 0 | 0 |

| | | |
|---|---|---|
| NC_019843.3;YP_009047202.1;MERS_1ab;3260 | 0 | 0 |
| NC_019843.3;YP_009047202.1;MERS_1ab;3261 | 0 | 0 |
| NC_019843.3;YP_009047202.1;MERS_1ab;3262 | 0 | 0 |
| NC_019843.3;YP_009047202.1;MERS_1ab;3263 | 0 | 0 |
| NC_019843.3;YP_009047202.1;MERS_1ab;3264 | 0 | 0 |
| NC_019843.3;YP_009047202.1;MERS_1ab;3265 | 0 | 0 |
| NC_019843.3;YP_009047202.1;MERS_1ab;3266 | 0 | 0 |
| NC_019843.3;YP_009047202.1;MERS_1ab;3267 | 0 | 0 |
| NC_019843.3;YP_009047202.1;MERS_1ab;3268 | 0 | 0 |
| NC_019843.3;YP_009047202.1;MERS_1ab;3269 | 0 | 0 |
| NC_019843.3;YP_009047202.1;MERS_1ab;3270 | 0 | 0 |
| NC_019843.3;YP_009047202.1;MERS_1ab;3271 | 0 | 0 |
| NC_019843.3;YP_009047202.1;MERS_1ab;3272 | 0 | 0 |
| NC_019843.3;YP_009047202.1;MERS_1ab;3273 | 0 | 0 |
| NC_019843.3;YP_009047202.1;MERS_1ab;3274 | 0 | 0 |
| NC_019843.3;YP_009047202.1;MERS_1ab;3275 | 0 | 0 |
| NC_019843.3;YP_009047202.1;MERS_1ab;3276 | 0 | 0 |
| NC_019843.3;YP_009047202.1;MERS_1ab;3277 | 0 | 0 |
| NC_019843.3;YP_009047202.1;MERS_1ab;3278 | 0 | 0 |
| NC_019843.3;YP_009047202.1;MERS_1ab;3279 | 0 | 0 |
| NC_019843.3;YP_009047202.1;MERS_1ab;3280 | 0 | 0 |
| NC_019843.3;YP_009047202.1;MERS_1ab;3281 | 0 | 0 |
| NC_019843.3;YP_009047202.1;MERS_1ab;3282 | 0 | 0 |
| NC_019843.3;YP_009047202.1;MERS_1ab;3283 | 0.05 | 0 |
| NC_019843.3;YP_009047202.1;MERS_1ab;3284 | 0 | 0 |
| NC_019843.3;YP_009047202.1;MERS_1ab;3285 | 0 | 0 |
| NC_019843.3;YP_009047202.1;MERS_1ab;3286 | 0 | 0 |
| NC_019843.3;YP_009047202.1;MERS_1ab;3287 | 0 | 0 |
| NC_019843.3;YP_009047202.1;MERS_1ab;3288 | 0 | 0 |
| NC_019843.3;YP_009047202.1;MERS_1ab;3289 | 0 | 0 |
| NC_019843.3;YP_009047202.1;MERS_1ab;3290 | 0 | 0 |
| NC_019843.3;YP_009047202.1;MERS_1ab;3291 | 0 | 0 |
| NC_019843.3;YP_009047202.1;MERS_1ab;3292 | 0 | 0 |
| NC_019843.3;YP_009047202.1;MERS_1ab;3293 | 0 | 0 |
| NC_019843.3;YP_009047202.1;MERS_1ab;3294 | 0 | 0 |
| NC_019843.3;YP_009047202.1;MERS_1ab;3295 | 0 | 0 |
| NC_019843.3;YP_009047202.1;MERS_1ab;3296 | 0 | 0 |
| NC_019843.3;YP_009047202.1;MERS_1ab;3297 | 0 | 0 |
| NC_019843.3;YP_009047202.1;MERS_1ab;3298 | 0 | 0 |
| NC_019843.3;YP_009047202.1;MERS_1ab;3299 | 0 | 0 |
| NC_019843.3;YP_009047202.1;MERS_1ab;3300 | 0 | 0 |
| NC_019843.3;YP_009047202.1;MERS_1ab;3301 | 0 | 0 |
| NC_019843.3;YP_009047202.1;MERS_1ab;3302 | 0 | 0 |

| | | |
|---|---|---|
| NC_019843.3;YP_009047202.1;MERS_1ab;3303 | 0 | 0 |
| NC_019843.3;YP_009047202.1;MERS_1ab;3304 | 0 | 0 |
| NC_019843.3;YP_009047202.1;MERS_1ab;3305 | 0 | 0 |
| NC_019843.3;YP_009047202.1;MERS_1ab;3306 | 0.05 | 0 |
| NC_019843.3;YP_009047202.1;MERS_1ab;3307 | 0.15 | 0 |
| NC_019843.3;YP_009047202.1;MERS_1ab;3308 | 0.1 | 0.025 |
| NC_019843.3;YP_009047202.1;MERS_1ab;3309 | 0.1 | 0.05 |
| NC_019843.3;YP_009047202.1;MERS_1ab;3310 | 0.1 | 0.025 |
| NC_019843.3;YP_009047202.1;MERS_1ab;3311 | 0.1 | 0.025 |
| NC_019843.3;YP_009047202.1;MERS_1ab;3312 | 0.15 | 0.025 |
| NC_019843.3;YP_009047202.1;MERS_1ab;3313 | 0.1 | 0.025 |
| NC_019843.3;YP_009047202.1;MERS_1ab;3314 | 0.1 | 0.05 |
| NC_019843.3;YP_009047202.1;MERS_1ab;3315 | 0.1 | 0.025 |
| NC_019843.3;YP_009047202.1;MERS_1ab;3316 | 0.1 | 0.025 |
| NC_019843.3;YP_009047202.1;MERS_1ab;3317 | 0.1 | 0.025 |
| NC_019843.3;YP_009047202.1;MERS_1ab;3318 | 0.15 | 0.05 |
| NC_019843.3;YP_009047202.1;MERS_1ab;3319 | 0 | 0.025 |
| NC_019843.3;YP_009047202.1;MERS_1ab;3320 | 0 | 0 |
| NC_019843.3;YP_009047202.1;MERS_1ab;3321 | 0 | 0 |
| NC_019843.3;YP_009047202.1;MERS_1ab;3322 | 0 | 0 |
| NC_019843.3;YP_009047202.1;MERS_1ab;3323 | 0 | 0.025 |
| NC_019843.3;YP_009047202.1;MERS_1ab;3324 | 0 | 0.025 |
| NC_019843.3;YP_009047202.1;MERS_1ab;3325 | 0 | 0 |
| NC_019843.3;YP_009047202.1;MERS_1ab;3326 | 0 | 0 |
| NC_019843.3;YP_009047202.1;MERS_1ab;3327 | 0 | 0 |
| NC_019843.3;YP_009047202.1;MERS_1ab;3328 | 0 | 0 |
| NC_019843.3;YP_009047202.1;MERS_1ab;3329 | 0 | 0 |
| NC_019843.3;YP_009047202.1;MERS_1ab;3330 | 0 | 0 |
| NC_019843.3;YP_009047202.1;MERS_1ab;3331 | 0 | 0 |
| NC_019843.3;YP_009047202.1;MERS_1ab;3332 | 0 | 0 |
| NC_019843.3;YP_009047202.1;MERS_1ab;3333 | 0 | 0 |
| NC_019843.3;YP_009047202.1;MERS_1ab;3334 | 0 | 0 |
| NC_019843.3;YP_009047202.1;MERS_1ab;3335 | 0 | 0 |
| NC_019843.3;YP_009047202.1;MERS_1ab;3336 | 0 | 0 |
| NC_019843.3;YP_009047202.1;MERS_1ab;3337 | 0 | 0 |
| NC_019843.3;YP_009047202.1;MERS_1ab;3338 | 0 | 0 |
| NC_019843.3;YP_009047202.1;MERS_1ab;3339 | 0 | 0.05 |
| NC_019843.3;YP_009047202.1;MERS_1ab;3340 | 0 | 0.025 |
| NC_019843.3;YP_009047202.1;MERS_1ab;3341 | 0.05 | 0.075 |
| NC_019843.3;YP_009047202.1;MERS_1ab;3342 | 0 | 0 |
| NC_019843.3;YP_009047202.1;MERS_1ab;3343 | 0 | 0.025 |
| NC_019843.3;YP_009047202.1;MERS_1ab;3344 | 0 | 0.025 |
| NC_019843.3;YP_009047202.1;MERS_1ab;3345 | 0.05 | 0.025 |

| | | |
|---|---|---|
| NC_019843.3;YP_009047202.1;MERS_1ab;3346 | 0.1 | 0.075 |
| NC_019843.3;YP_009047202.1;MERS_1ab;3347 | 0 | 0 |
| NC_019843.3;YP_009047202.1;MERS_1ab;3348 | 0 | 0 |
| NC_019843.3;YP_009047202.1;MERS_1ab;3349 | 0 | 0 |
| NC_019843.3;YP_009047202.1;MERS_1ab;3350 | 0 | 0 |
| NC_019843.3;YP_009047202.1;MERS_1ab;3351 | 0 | 0 |
| NC_019843.3;YP_009047202.1;MERS_1ab;3352 | 0 | 0 |
| NC_019843.3;YP_009047202.1;MERS_1ab;3353 | 0 | 0 |
| NC_019843.3;YP_009047202.1;MERS_1ab;3354 | 0 | 0 |
| NC_019843.3;YP_009047202.1;MERS_1ab;3355 | 0 | 0 |
| NC_019843.3;YP_009047202.1;MERS_1ab;3356 | 0 | 0 |
| NC_019843.3;YP_009047202.1;MERS_1ab;3357 | 0 | 0 |
| NC_019843.3;YP_009047202.1;MERS_1ab;3358 | 0 | 0 |
| NC_019843.3;YP_009047202.1;MERS_1ab;3359 | 0 | 0 |
| NC_019843.3;YP_009047202.1;MERS_1ab;3360 | 0 | 0 |
| NC_019843.3;YP_009047202.1;MERS_1ab;3361 | 0 | 0 |
| NC_019843.3;YP_009047202.1;MERS_1ab;3362 | 0 | 0 |
| NC_019843.3;YP_009047202.1;MERS_1ab;3363 | 0 | 0 |
| NC_019843.3;YP_009047202.1;MERS_1ab;3364 | 0 | 0 |
| NC_019843.3;YP_009047202.1;MERS_1ab;3365 | 0 | 0 |
| NC_019843.3;YP_009047202.1;MERS_1ab;3366 | 0 | 0 |
| NC_019843.3;YP_009047202.1;MERS_1ab;3367 | 0 | 0 |
| NC_019843.3;YP_009047202.1;MERS_1ab;3368 | 0 | 0 |
| NC_019843.3;YP_009047202.1;MERS_1ab;3369 | 0 | 0 |
| NC_019843.3;YP_009047202.1;MERS_1ab;3370 | 0 | 0 |
| NC_019843.3;YP_009047202.1;MERS_1ab;3371 | 0 | 0 |
| NC_019843.3;YP_009047202.1;MERS_1ab;3372 | 0 | 0.025 |
| NC_019843.3;YP_009047202.1;MERS_1ab;3373 | 0 | 0.025 |
| NC_019843.3;YP_009047202.1;MERS_1ab;3374 | 0 | 0.05 |
| NC_019843.3;YP_009047202.1;MERS_1ab;3375 | 0 | 0 |
| NC_019843.3;YP_009047202.1;MERS_1ab;3376 | 0 | 0 |
| NC_019843.3;YP_009047202.1;MERS_1ab;3377 | 0 | 0 |
| NC_019843.3;YP_009047202.1;MERS_1ab;3378 | 0 | 0 |
| NC_019843.3;YP_009047202.1;MERS_1ab;3379 | 0 | 0 |
| NC_019843.3;YP_009047202.1;MERS_1ab;3380 | 0 | 0 |
| NC_019843.3;YP_009047202.1;MERS_1ab;3381 | 0 | 0 |
| NC_019843.3;YP_009047202.1;MERS_1ab;3382 | 0 | 0 |
| NC_019843.3;YP_009047202.1;MERS_1ab;3383 | 0 | 0 |
| NC_019843.3;YP_009047202.1;MERS_1ab;3384 | 0 | 0 |
| NC_019843.3;YP_009047202.1;MERS_1ab;3385 | 0 | 0 |
| NC_019843.3;YP_009047202.1;MERS_1ab;3386 | 0 | 0 |
| NC_019843.3;YP_009047202.1;MERS_1ab;3387 | 0 | 0.025 |
| NC_019843.3;YP_009047202.1;MERS_1ab;3388 | 0 | 0 |

| | | |
|---|---|---|
| NC_019843.3;YP_009047202.1;MERS_1ab;3389 | 0.05 | 0 |
| NC_019843.3;YP_009047202.1;MERS_1ab;3390 | 0.1 | 0 |
| NC_019843.3;YP_009047202.1;MERS_1ab;3391 | 0.1 | 0 |
| NC_019843.3;YP_009047202.1;MERS_1ab;3392 | 0.05 | 0.025 |
| NC_019843.3;YP_009047202.1;MERS_1ab;3393 | 0.1 | 0.025 |
| NC_019843.3;YP_009047202.1;MERS_1ab;3394 | 0 | 0 |
| NC_019843.3;YP_009047202.1;MERS_1ab;3395 | 0 | 0 |
| NC_019843.3;YP_009047202.1;MERS_1ab;3396 | 0 | 0 |
| NC_019843.3;YP_009047202.1;MERS_1ab;3397 | 0 | 0 |
| NC_019843.3;YP_009047202.1;MERS_1ab;3398 | 0 | 0 |
| NC_019843.3;YP_009047202.1;MERS_1ab;3399 | 0 | 0 |
| NC_019843.3;YP_009047202.1;MERS_1ab;3400 | 0.05 | 0 |
| NC_019843.3;YP_009047202.1;MERS_1ab;3401 | 0 | 0 |
| NC_019843.3;YP_009047202.1;MERS_1ab;3402 | 0 | 0 |
| NC_019843.3;YP_009047202.1;MERS_1ab;3403 | 0 | 0 |
| NC_019843.3;YP_009047202.1;MERS_1ab;3404 | 0 | 0 |
| NC_019843.3;YP_009047202.1;MERS_1ab;3405 | 0 | 0 |
| NC_019843.3;YP_009047202.1;MERS_1ab;3406 | 0 | 0 |
| NC_019843.3;YP_009047202.1;MERS_1ab;3407 | 0 | 0 |
| NC_019843.3;YP_009047202.1;MERS_1ab;3408 | 0 | 0 |
| NC_019843.3;YP_009047202.1;MERS_1ab;3409 | 0 | 0 |
| NC_019843.3;YP_009047202.1;MERS_1ab;3410 | 0 | 0 |
| NC_019843.3;YP_009047202.1;MERS_1ab;3411 | 0 | 0 |
| NC_019843.3;YP_009047202.1;MERS_1ab;3412 | 0 | 0 |
| NC_019843.3;YP_009047202.1;MERS_1ab;3413 | 0.05 | 0 |
| NC_019843.3;YP_009047202.1;MERS_1ab;3414 | 0.05 | 0 |
| NC_019843.3;YP_009047202.1;MERS_1ab;3415 | 0.05 | 0 |
| NC_019843.3;YP_009047202.1;MERS_1ab;3416 | 0.05 | 0 |
| NC_019843.3;YP_009047202.1;MERS_1ab;3417 | 0.05 | 0 |
| NC_019843.3;YP_009047202.1;MERS_1ab;3418 | 0.05 | 0.05 |
| NC_019843.3;YP_009047202.1;MERS_1ab;3419 | 0.05 | 0 |
| NC_019843.3;YP_009047202.1;MERS_1ab;3420 | 0.05 | 0 |
| NC_019843.3;YP_009047202.1;MERS_1ab;3421 | 0.05 | 0 |
| NC_019843.3;YP_009047202.1;MERS_1ab;3422 | 0 | 0 |
| NC_019843.3;YP_009047202.1;MERS_1ab;3423 | 0.05 | 0 |
| NC_019843.3;YP_009047202.1;MERS_1ab;3424 | 0 | 0 |
| NC_019843.3;YP_009047202.1;MERS_1ab;3425 | 0 | 0 |
| NC_019843.3;YP_009047202.1;MERS_1ab;3426 | 0 | 0 |
| NC_019843.3;YP_009047202.1;MERS_1ab;3427 | 0 | 0 |
| NC_019843.3;YP_009047202.1;MERS_1ab;3428 | 0 | 0 |
| NC_019843.3;YP_009047202.1;MERS_1ab;3429 | 0 | 0 |
| NC_019843.3;YP_009047202.1;MERS_1ab;3430 | 0 | 0 |
| NC_019843.3;YP_009047202.1;MERS_1ab;3431 | 0 | 0 |

| | | |
|---|---|---|
| NC_019843.3;YP_009047202.1;MERS_1ab;3432 | 0 | 0.025 |
| NC_019843.3;YP_009047202.1;MERS_1ab;3433 | 0 | 0 |
| NC_019843.3;YP_009047202.1;MERS_1ab;3434 | 0 | 0.025 |
| NC_019843.3;YP_009047202.1;MERS_1ab;3435 | 0 | 0 |
| NC_019843.3;YP_009047202.1;MERS_1ab;3436 | 0 | 0 |
| NC_019843.3;YP_009047202.1;MERS_1ab;3437 | 0 | 0 |
| NC_019843.3;YP_009047202.1;MERS_1ab;3438 | 0 | 0 |
| NC_019843.3;YP_009047202.1;MERS_1ab;3439 | 0 | 0 |
| NC_019843.3;YP_009047202.1;MERS_1ab;3440 | 0 | 0 |
| NC_019843.3;YP_009047202.1;MERS_1ab;3441 | 0 | 0 |
| NC_019843.3;YP_009047202.1;MERS_1ab;3442 | 0 | 0 |
| NC_019843.3;YP_009047202.1;MERS_1ab;3443 | 0 | 0 |
| NC_019843.3;YP_009047202.1;MERS_1ab;3444 | 0 | 0 |
| NC_019843.3;YP_009047202.1;MERS_1ab;3445 | 0 | 0 |
| NC_019843.3;YP_009047202.1;MERS_1ab;3446 | 0 | 0 |
| NC_019843.3;YP_009047202.1;MERS_1ab;3447 | 0 | 0 |
| NC_019843.3;YP_009047202.1;MERS_1ab;3448 | 0 | 0 |
| NC_019843.3;YP_009047202.1;MERS_1ab;3449 | 0 | 0 |
| NC_019843.3;YP_009047202.1;MERS_1ab;3450 | 0 | 0 |
| NC_019843.3;YP_009047202.1;MERS_1ab;3451 | 0 | 0 |
| NC_019843.3;YP_009047202.1;MERS_1ab;3452 | 0 | 0 |
| NC_019843.3;YP_009047202.1;MERS_1ab;3453 | 0 | 0 |
| NC_019843.3;YP_009047202.1;MERS_1ab;3454 | 0 | 0 |
| NC_019843.3;YP_009047202.1;MERS_1ab;3455 | 0 | 0 |
| NC_019843.3;YP_009047202.1;MERS_1ab;3456 | 0 | 0 |
| NC_019843.3;YP_009047202.1;MERS_1ab;3457 | 0 | 0 |
| NC_019843.3;YP_009047202.1;MERS_1ab;3458 | 0 | 0 |
| NC_019843.3;YP_009047202.1;MERS_1ab;3459 | 0 | 0 |
| NC_019843.3;YP_009047202.1;MERS_1ab;3460 | 0 | 0.025 |
| NC_019843.3;YP_009047202.1;MERS_1ab;3461 | 0 | 0 |
| NC_019843.3;YP_009047202.1;MERS_1ab;3462 | 0 | 0 |
| NC_019843.3;YP_009047202.1;MERS_1ab;3463 | 0 | 0 |
| NC_019843.3;YP_009047202.1;MERS_1ab;3464 | 0 | 0 |
| NC_019843.3;YP_009047202.1;MERS_1ab;3465 | 0 | 0 |
| NC_019843.3;YP_009047202.1;MERS_1ab;3466 | 0 | 0 |
| NC_019843.3;YP_009047202.1;MERS_1ab;3467 | 0 | 0 |
| NC_019843.3;YP_009047202.1;MERS_1ab;3468 | 0 | 0 |
| NC_019843.3;YP_009047202.1;MERS_1ab;3469 | 0 | 0 |
| NC_019843.3;YP_009047202.1;MERS_1ab;3470 | 0 | 0.025 |
| NC_019843.3;YP_009047202.1;MERS_1ab;3471 | 0 | 0.025 |
| NC_019843.3;YP_009047202.1;MERS_1ab;3472 | 0 | 0 |
| NC_019843.3;YP_009047202.1;MERS_1ab;3473 | 0 | 0 |
| NC_019843.3;YP_009047202.1;MERS_1ab;3474 | 0 | 0 |

| | | |
|---|---|---|
| NC_019843.3;YP_009047202.1;MERS_1ab;3475 | 0 | 0 |
| NC_019843.3;YP_009047202.1;MERS_1ab;3476 | 0 | 0.025 |
| NC_019843.3;YP_009047202.1;MERS_1ab;3477 | 0 | 0.025 |
| NC_019843.3;YP_009047202.1;MERS_1ab;3478 | 0 | 0 |
| NC_019843.3;YP_009047202.1;MERS_1ab;3479 | 0 | 0.025 |
| NC_019843.3;YP_009047202.1;MERS_1ab;3480 | 0 | 0 |
| NC_019843.3;YP_009047202.1;MERS_1ab;3481 | 0 | 0 |
| NC_019843.3;YP_009047202.1;MERS_1ab;3482 | 0 | 0 |
| NC_019843.3;YP_009047202.1;MERS_1ab;3483 | 0 | 0 |
| NC_019843.3;YP_009047202.1;MERS_1ab;3484 | 0 | 0.025 |
| NC_019843.3;YP_009047202.1;MERS_1ab;3485 | 0 | 0.025 |
| NC_019843.3;YP_009047202.1;MERS_1ab;3486 | 0 | 0.05 |
| NC_019843.3;YP_009047202.1;MERS_1ab;3487 | 0 | 0.025 |
| NC_019843.3;YP_009047202.1;MERS_1ab;3488 | 0 | 0.025 |
| NC_019843.3;YP_009047202.1;MERS_1ab;3489 | 0 | 0 |
| NC_019843.3;YP_009047202.1;MERS_1ab;3490 | 0 | 0 |
| NC_019843.3;YP_009047202.1;MERS_1ab;3491 | 0.05 | 0 |
| NC_019843.3;YP_009047202.1;MERS_1ab;3492 | 0 | 0.025 |
| NC_019843.3;YP_009047202.1;MERS_1ab;3493 | 0 | 0.025 |
| NC_019843.3;YP_009047202.1;MERS_1ab;3494 | 0 | 0 |
| NC_019843.3;YP_009047202.1;MERS_1ab;3495 | 0 | 0 |
| NC_019843.3;YP_009047202.1;MERS_1ab;3496 | 0 | 0 |
| NC_019843.3;YP_009047202.1;MERS_1ab;3497 | 0 | 0 |
| NC_019843.3;YP_009047202.1;MERS_1ab;3498 | 0 | 0 |
| NC_019843.3;YP_009047202.1;MERS_1ab;3499 | 0 | 0 |
| NC_019843.3;YP_009047202.1;MERS_1ab;3500 | 0 | 0 |
| NC_019843.3;YP_009047202.1;MERS_1ab;3501 | 0 | 0 |
| NC_019843.3;YP_009047202.1;MERS_1ab;3502 | 0 | 0 |
| NC_019843.3;YP_009047202.1;MERS_1ab;3503 | 0 | 0 |
| NC_019843.3;YP_009047202.1;MERS_1ab;3504 | 0 | 0 |
| NC_019843.3;YP_009047202.1;MERS_1ab;3505 | 0 | 0 |
| NC_019843.3;YP_009047202.1;MERS_1ab;3506 | 0 | 0 |
| NC_019843.3;YP_009047202.1;MERS_1ab;3507 | 0 | 0.225 |
| NC_019843.3;YP_009047202.1;MERS_1ab;3508 | 0 | 0 |
| NC_019843.3;YP_009047202.1;MERS_1ab;3509 | 0 | 0 |
| NC_019843.3;YP_009047202.1;MERS_1ab;3510 | 0 | 0 |
| NC_019843.3;YP_009047202.1;MERS_1ab;3511 | 0 | 0 |
| NC_019843.3;YP_009047202.1;MERS_1ab;3512 | 0 | 0 |
| NC_019843.3;YP_009047202.1;MERS_1ab;3513 | 0 | 0 |
| NC_019843.3;YP_009047202.1;MERS_1ab;3514 | 0 | 0 |
| NC_019843.3;YP_009047202.1;MERS_1ab;3515 | 0 | 0 |
| NC_019843.3;YP_009047202.1;MERS_1ab;3516 | 0 | 0 |
| NC_019843.3;YP_009047202.1;MERS_1ab;3517 | 0 | 0 |

| | | |
|---|---|---|
| NC_019843.3;YP_009047202.1;MERS_1ab;3518 | 0 | 0 |
| NC_019843.3;YP_009047202.1;MERS_1ab;3519 | 0 | 0 |
| NC_019843.3;YP_009047202.1;MERS_1ab;3520 | 0 | 0 |
| NC_019843.3;YP_009047202.1;MERS_1ab;3521 | 0 | 0 |
| NC_019843.3;YP_009047202.1;MERS_1ab;3522 | 0 | 0 |
| NC_019843.3;YP_009047202.1;MERS_1ab;3523 | 0 | 0 |
| NC_019843.3;YP_009047202.1;MERS_1ab;3524 | 0 | 0 |
| NC_019843.3;YP_009047202.1;MERS_1ab;3525 | 0 | 0 |
| NC_019843.3;YP_009047202.1;MERS_1ab;3526 | 0 | 0 |
| NC_019843.3;YP_009047202.1;MERS_1ab;3527 | 0 | 0 |
| NC_019843.3;YP_009047202.1;MERS_1ab;3528 | 0 | 0 |
| NC_019843.3;YP_009047202.1;MERS_1ab;3529 | 0 | 0 |
| NC_019843.3;YP_009047202.1;MERS_1ab;3530 | 0 | 0 |
| NC_019843.3;YP_009047202.1;MERS_1ab;3531 | 0 | 0 |
| NC_019843.3;YP_009047202.1;MERS_1ab;3532 | 0 | 0 |
| NC_019843.3;YP_009047202.1;MERS_1ab;3533 | 0 | 0 |
| NC_019843.3;YP_009047202.1;MERS_1ab;3534 | 0 | 0.025 |
| NC_019843.3;YP_009047202.1;MERS_1ab;3535 | 0.1 | 0.025 |
| NC_019843.3;YP_009047202.1;MERS_1ab;3536 | 0 | 0 |
| NC_019843.3;YP_009047202.1;MERS_1ab;3537 | 0.05 | 0 |
| NC_019843.3;YP_009047202.1;MERS_1ab;3538 | 0 | 0.05 |
| NC_019843.3;YP_009047202.1;MERS_1ab;3539 | 0 | 0 |
| NC_019843.3;YP_009047202.1;MERS_1ab;3540 | 0 | 0 |
| NC_019843.3;YP_009047202.1;MERS_1ab;3541 | 0 | 0 |
| NC_019843.3;YP_009047202.1;MERS_1ab;3542 | 0 | 0.025 |
| NC_019843.3;YP_009047202.1;MERS_1ab;3543 | 0 | 0 |
| NC_019843.3;YP_009047202.1;MERS_1ab;3544 | 0 | 0 |
| NC_019843.3;YP_009047202.1;MERS_1ab;3545 | 0 | 0 |
| NC_019843.3;YP_009047202.1;MERS_1ab;3546 | 0 | 0 |
| NC_019843.3;YP_009047202.1;MERS_1ab;3547 | 0 | 0 |
| NC_019843.3;YP_009047202.1;MERS_1ab;3548 | 0 | 0 |
| NC_019843.3;YP_009047202.1;MERS_1ab;3549 | 0 | 0 |
| NC_019843.3;YP_009047202.1;MERS_1ab;3550 | 0.05 | 0 |
| NC_019843.3;YP_009047202.1;MERS_1ab;3551 | 0 | 0 |
| NC_019843.3;YP_009047202.1;MERS_1ab;3552 | 0 | 0 |
| NC_019843.3;YP_009047202.1;MERS_1ab;3553 | 0 | 0 |
| NC_019843.3;YP_009047202.1;MERS_1ab;3554 | 0 | 0 |
| NC_019843.3;YP_009047202.1;MERS_1ab;3555 | 0 | 0 |
| NC_019843.3;YP_009047202.1;MERS_1ab;3556 | 0 | 0 |
| NC_019843.3;YP_009047202.1;MERS_1ab;3557 | 0 | 0 |
| NC_019843.3;YP_009047202.1;MERS_1ab;3558 | 0 | 0 |
| NC_019843.3;YP_009047202.1;MERS_1ab;3559 | 0 | 0 |
| NC_019843.3;YP_009047202.1;MERS_1ab;3560 | 0 | 0 |

| | | |
|---|---|---|
| NC_019843.3;YP_009047202.1;MERS_1ab;3561 | 0 | 0 |
| NC_019843.3;YP_009047202.1;MERS_1ab;3562 | 0 | 0 |
| NC_019843.3;YP_009047202.1;MERS_1ab;3563 | 0 | 0 |
| NC_019843.3;YP_009047202.1;MERS_1ab;3564 | 0 | 0 |
| NC_019843.3;YP_009047202.1;MERS_1ab;3565 | 0 | 0 |
| NC_019843.3;YP_009047202.1;MERS_1ab;3566 | 0 | 0 |
| NC_019843.3;YP_009047202.1;MERS_1ab;3567 | 0 | 0 |
| NC_019843.3;YP_009047202.1;MERS_1ab;3568 | 0 | 0 |
| NC_019843.3;YP_009047202.1;MERS_1ab;3569 | 0 | 0 |
| NC_019843.3;YP_009047202.1;MERS_1ab;3570 | 0 | 0 |
| NC_019843.3;YP_009047202.1;MERS_1ab;3571 | 0 | 0 |
| NC_019843.3;YP_009047202.1;MERS_1ab;3572 | 0 | 0 |
| NC_019843.3;YP_009047202.1;MERS_1ab;3573 | 0 | 0 |
| NC_019843.3;YP_009047202.1;MERS_1ab;3574 | 0 | 0 |
| NC_019843.3;YP_009047202.1;MERS_1ab;3575 | 0 | 0 |
| NC_019843.3;YP_009047202.1;MERS_1ab;3576 | 0 | 0 |
| NC_019843.3;YP_009047202.1;MERS_1ab;3577 | 0 | 0 |
| NC_019843.3;YP_009047202.1;MERS_1ab;3578 | 0 | 0 |
| NC_019843.3;YP_009047202.1;MERS_1ab;3579 | 0 | 0 |
| NC_019843.3;YP_009047202.1;MERS_1ab;3580 | 0 | 0 |
| NC_019843.3;YP_009047202.1;MERS_1ab;3581 | 0 | 0 |
| NC_019843.3;YP_009047202.1;MERS_1ab;3582 | 0 | 0 |
| NC_019843.3;YP_009047202.1;MERS_1ab;3583 | 0 | 0 |
| NC_019843.3;YP_009047202.1;MERS_1ab;3584 | 0 | 0 |
| NC_019843.3;YP_009047202.1;MERS_1ab;3585 | 0 | 0 |
| NC_019843.3;YP_009047202.1;MERS_1ab;3586 | 0 | 0 |
| NC_019843.3;YP_009047202.1;MERS_1ab;3587 | 0 | 0 |
| NC_019843.3;YP_009047202.1;MERS_1ab;3588 | 0 | 0 |
| NC_019843.3;YP_009047202.1;MERS_1ab;3589 | 0 | 0 |
| NC_019843.3;YP_009047202.1;MERS_1ab;3590 | 0 | 0 |
| NC_019843.3;YP_009047202.1;MERS_1ab;3591 | 0 | 0 |
| NC_019843.3;YP_009047202.1;MERS_1ab;3592 | 0 | 0 |
| NC_019843.3;YP_009047202.1;MERS_1ab;3593 | 0 | 0 |
| NC_019843.3;YP_009047202.1;MERS_1ab;3594 | 0 | 0 |
| NC_019843.3;YP_009047202.1;MERS_1ab;3595 | 0 | 0 |
| NC_019843.3;YP_009047202.1;MERS_1ab;3596 | 0 | 0 |
| NC_019843.3;YP_009047202.1;MERS_1ab;3597 | 0 | 0 |
| NC_019843.3;YP_009047202.1;MERS_1ab;3598 | 0 | 0 |
| NC_019843.3;YP_009047202.1;MERS_1ab;3599 | 0 | 0 |
| NC_019843.3;YP_009047202.1;MERS_1ab;3600 | 0 | 0 |
| NC_019843.3;YP_009047202.1;MERS_1ab;3601 | 0 | 0 |
| NC_019843.3;YP_009047202.1;MERS_1ab;3602 | 0 | 0 |
| NC_019843.3;YP_009047202.1;MERS_1ab;3603 | 0 | 0 |

| | | |
|---|---|---|
| NC_019843.3;YP_009047202.1;MERS_1ab;3604 | 0 | 0 |
| NC_019843.3;YP_009047202.1;MERS_1ab;3605 | 0 | 0 |
| NC_019843.3;YP_009047202.1;MERS_1ab;3606 | 0 | 0 |
| NC_019843.3;YP_009047202.1;MERS_1ab;3607 | 0 | 0 |
| NC_019843.3;YP_009047202.1;MERS_1ab;3608 | 0 | 0 |
| NC_019843.3;YP_009047202.1;MERS_1ab;3609 | 0 | 0 |
| NC_019843.3;YP_009047202.1;MERS_1ab;3610 | 0 | 0 |
| NC_019843.3;YP_009047202.1;MERS_1ab;3611 | 0 | 0 |
| NC_019843.3;YP_009047202.1;MERS_1ab;3612 | 0 | 0 |
| NC_019843.3;YP_009047202.1;MERS_1ab;3613 | 0 | 0 |
| NC_019843.3;YP_009047202.1;MERS_1ab;3614 | 0 | 0 |
| NC_019843.3;YP_009047202.1;MERS_1ab;3615 | 0 | 0 |
| NC_019843.3;YP_009047202.1;MERS_1ab;3616 | 0 | 0 |
| NC_019843.3;YP_009047202.1;MERS_1ab;3617 | 0 | 0 |
| NC_019843.3;YP_009047202.1;MERS_1ab;3618 | 0 | 0 |
| NC_019843.3;YP_009047202.1;MERS_1ab;3619 | 0 | 0 |
| NC_019843.3;YP_009047202.1;MERS_1ab;3620 | 0 | 0 |
| NC_019843.3;YP_009047202.1;MERS_1ab;3621 | 0 | 0 |
| NC_019843.3;YP_009047202.1;MERS_1ab;3622 | 0 | 0 |
| NC_019843.3;YP_009047202.1;MERS_1ab;3623 | 0 | 0 |
| NC_019843.3;YP_009047202.1;MERS_1ab;3624 | 0 | 0 |
| NC_019843.3;YP_009047202.1;MERS_1ab;3625 | 0 | 0 |
| NC_019843.3;YP_009047202.1;MERS_1ab;3626 | 0 | 0 |
| NC_019843.3;YP_009047202.1;MERS_1ab;3627 | 0 | 0 |
| NC_019843.3;YP_009047202.1;MERS_1ab;3628 | 0 | 0 |
| NC_019843.3;YP_009047202.1;MERS_1ab;3629 | 0 | 0 |
| NC_019843.3;YP_009047202.1;MERS_1ab;3630 | 0 | 0 |
| NC_019843.3;YP_009047202.1;MERS_1ab;3631 | 0 | 0 |
| NC_019843.3;YP_009047202.1;MERS_1ab;3632 | 0 | 0 |
| NC_019843.3;YP_009047202.1;MERS_1ab;3633 | 0 | 0 |
| NC_019843.3;YP_009047202.1;MERS_1ab;3634 | 0 | 0.025 |
| NC_019843.3;YP_009047202.1;MERS_1ab;3635 | 0 | 0 |
| NC_019843.3;YP_009047202.1;MERS_1ab;3636 | 0 | 0 |
| NC_019843.3;YP_009047202.1;MERS_1ab;3637 | 0 | 0 |
| NC_019843.3;YP_009047202.1;MERS_1ab;3638 | 0 | 0 |
| NC_019843.3;YP_009047202.1;MERS_1ab;3639 | 0 | 0 |
| NC_019843.3;YP_009047202.1;MERS_1ab;3640 | 0 | 0 |
| NC_019843.3;YP_009047202.1;MERS_1ab;3641 | 0 | 0 |
| NC_019843.3;YP_009047202.1;MERS_1ab;3642 | 0 | 0 |
| NC_019843.3;YP_009047202.1;MERS_1ab;3643 | 0 | 0 |
| NC_019843.3;YP_009047202.1;MERS_1ab;3644 | 0 | 0 |
| NC_019843.3;YP_009047202.1;MERS_1ab;3645 | 0 | 0 |
| NC_019843.3;YP_009047202.1;MERS_1ab;3646 | 0.05 | 0.075 |

| | | |
|---|---|---|
| NC_019843.3;YP_009047202.1;MERS_1ab;3647 | 0.15 | 0.175 |
| NC_019843.3;YP_009047202.1;MERS_1ab;3648 | 0.15 | 0.1 |
| NC_019843.3;YP_009047202.1;MERS_1ab;3649 | 0.15 | 0.225 |
| NC_019843.3;YP_009047202.1;MERS_1ab;3650 | 0.25 | 0.3 |
| NC_019843.3;YP_009047202.1;MERS_1ab;3651 | 0.2 | 0.25 |
| NC_019843.3;YP_009047202.1;MERS_1ab;3652 | 0.2 | 0.25 |
| NC_019843.3;YP_009047202.1;MERS_1ab;3653 | 0.25 | 0.3 |
| NC_019843.3;YP_009047202.1;MERS_1ab;3654 | 0.2 | 0.225 |
| NC_019843.3;YP_009047202.1;MERS_1ab;3655 | 0.35 | 0.325 |
| NC_019843.3;YP_009047202.1;MERS_1ab;3656 | 0.3 | 0.35 |
| NC_019843.3;YP_009047202.1;MERS_1ab;3657 | 0.05 | 0.1 |
| NC_019843.3;YP_009047202.1;MERS_1ab;3658 | 0 | 0 |
| NC_019843.3;YP_009047202.1;MERS_1ab;3659 | 0 | 0 |
| NC_019843.3;YP_009047202.1;MERS_1ab;3660 | 0 | 0 |
| NC_019843.3;YP_009047202.1;MERS_1ab;3661 | 0 | 0 |
| NC_019843.3;YP_009047202.1;MERS_1ab;3662 | 0 | 0 |
| NC_019843.3;YP_009047202.1;MERS_1ab;3663 | 0 | 0 |
| NC_019843.3;YP_009047202.1;MERS_1ab;3664 | 0 | 0 |
| NC_019843.3;YP_009047202.1;MERS_1ab;3665 | 0 | 0 |
| NC_019843.3;YP_009047202.1;MERS_1ab;3666 | 0 | 0 |
| NC_019843.3;YP_009047202.1;MERS_1ab;3667 | 0 | 0 |
| NC_019843.3;YP_009047202.1;MERS_1ab;3668 | 0 | 0 |
| NC_019843.3;YP_009047202.1;MERS_1ab;3669 | 0 | 0 |
| NC_019843.3;YP_009047202.1;MERS_1ab;3670 | 0 | 0 |
| NC_019843.3;YP_009047202.1;MERS_1ab;3671 | 0 | 0 |
| NC_019843.3;YP_009047202.1;MERS_1ab;3672 | 0 | 0 |
| NC_019843.3;YP_009047202.1;MERS_1ab;3673 | 0 | 0 |
| NC_019843.3;YP_009047202.1;MERS_1ab;3674 | 0 | 0 |
| NC_019843.3;YP_009047202.1;MERS_1ab;3675 | 0 | 0 |
| NC_019843.3;YP_009047202.1;MERS_1ab;3676 | 0 | 0 |
| NC_019843.3;YP_009047202.1;MERS_1ab;3677 | 0 | 0 |
| NC_019843.3;YP_009047202.1;MERS_1ab;3678 | 0 | 0 |
| NC_019843.3;YP_009047202.1;MERS_1ab;3679 | 0 | 0 |
| NC_019843.3;YP_009047202.1;MERS_1ab;3680 | 0 | 0 |
| NC_019843.3;YP_009047202.1;MERS_1ab;3681 | 0 | 0 |
| NC_019843.3;YP_009047202.1;MERS_1ab;3682 | 0 | 0 |
| NC_019843.3;YP_009047202.1;MERS_1ab;3683 | 0 | 0 |
| NC_019843.3;YP_009047202.1;MERS_1ab;3684 | 0 | 0 |
| NC_019843.3;YP_009047202.1;MERS_1ab;3685 | 0 | 0 |
| NC_019843.3;YP_009047202.1;MERS_1ab;3686 | 0 | 0 |
| NC_019843.3;YP_009047202.1;MERS_1ab;3687 | 0 | 0 |
| NC_019843.3;YP_009047202.1;MERS_1ab;3688 | 0 | 0 |
| NC_019843.3;YP_009047202.1;MERS_1ab;3689 | 0 | 0 |

| | | |
|---|---|---|
| NC_019843.3;YP_009047202.1;MERS_1ab;3690 | 0 | 0 |
| NC_019843.3;YP_009047202.1;MERS_1ab;3691 | 0 | 0 |
| NC_019843.3;YP_009047202.1;MERS_1ab;3692 | 0 | 0 |
| NC_019843.3;YP_009047202.1;MERS_1ab;3693 | 0 | 0 |
| NC_019843.3;YP_009047202.1;MERS_1ab;3694 | 0 | 0 |
| NC_019843.3;YP_009047202.1;MERS_1ab;3695 | 0 | 0 |
| NC_019843.3;YP_009047202.1;MERS_1ab;3696 | 0 | 0 |
| NC_019843.3;YP_009047202.1;MERS_1ab;3697 | 0 | 0 |
| NC_019843.3;YP_009047202.1;MERS_1ab;3698 | 0 | 0 |
| NC_019843.3;YP_009047202.1;MERS_1ab;3699 | 0.05 | 0.025 |
| NC_019843.3;YP_009047202.1;MERS_1ab;3700 | 0.05 | 0.025 |
| NC_019843.3;YP_009047202.1;MERS_1ab;3701 | 0.05 | 0.025 |
| NC_019843.3;YP_009047202.1;MERS_1ab;3702 | 0.05 | 0.025 |
| NC_019843.3;YP_009047202.1;MERS_1ab;3703 | 0.05 | 0.025 |
| NC_019843.3;YP_009047202.1;MERS_1ab;3704 | 0.05 | 0.025 |
| NC_019843.3;YP_009047202.1;MERS_1ab;3705 | 0.1 | 0 |
| NC_019843.3;YP_009047202.1;MERS_1ab;3706 | 0 | 0 |
| NC_019843.3;YP_009047202.1;MERS_1ab;3707 | 0 | 0 |
| NC_019843.3;YP_009047202.1;MERS_1ab;3708 | 0 | 0 |
| NC_019843.3;YP_009047202.1;MERS_1ab;3709 | 0 | 0 |
| NC_019843.3;YP_009047202.1;MERS_1ab;3710 | 0 | 0 |
| NC_019843.3;YP_009047202.1;MERS_1ab;3711 | 0 | 0 |
| NC_019843.3;YP_009047202.1;MERS_1ab;3712 | 0 | 0 |
| NC_019843.3;YP_009047202.1;MERS_1ab;3713 | 0 | 0 |
| NC_019843.3;YP_009047202.1;MERS_1ab;3714 | 0 | 0 |
| NC_019843.3;YP_009047202.1;MERS_1ab;3715 | 0 | 0 |
| NC_019843.3;YP_009047202.1;MERS_1ab;3716 | 0 | 0 |
| NC_019843.3;YP_009047202.1;MERS_1ab;3717 | 0 | 0 |
| NC_019843.3;YP_009047202.1;MERS_1ab;3718 | 0 | 0 |
| NC_019843.3;YP_009047202.1;MERS_1ab;3719 | 0 | 0 |
| NC_019843.3;YP_009047202.1;MERS_1ab;3720 | 0 | 0 |
| NC_019843.3;YP_009047202.1;MERS_1ab;3721 | 0 | 0 |
| NC_019843.3;YP_009047202.1;MERS_1ab;3722 | 0 | 0 |
| NC_019843.3;YP_009047202.1;MERS_1ab;3723 | 0 | 0 |
| NC_019843.3;YP_009047202.1;MERS_1ab;3724 | 0 | 0 |
| NC_019843.3;YP_009047202.1;MERS_1ab;3725 | 0 | 0 |
| NC_019843.3;YP_009047202.1;MERS_1ab;3726 | 0 | 0 |
| NC_019843.3;YP_009047202.1;MERS_1ab;3727 | 0 | 0 |
| NC_019843.3;YP_009047202.1;MERS_1ab;3728 | 0 | 0 |
| NC_019843.3;YP_009047202.1;MERS_1ab;3729 | 0 | 0 |
| NC_019843.3;YP_009047202.1;MERS_1ab;3730 | 0 | 0 |
| NC_019843.3;YP_009047202.1;MERS_1ab;3731 | 0 | 0 |
| NC_019843.3;YP_009047202.1;MERS_1ab;3732 | 0 | 0 |

| | | |
|---|---|---|
| NC_019843.3;YP_009047202.1;MERS_1ab;3733 | 0 | 0 |
| NC_019843.3;YP_009047202.1;MERS_1ab;3734 | 0 | 0 |
| NC_019843.3;YP_009047202.1;MERS_1ab;3735 | 0 | 0 |
| NC_019843.3;YP_009047202.1;MERS_1ab;3736 | 0 | 0 |
| NC_019843.3;YP_009047202.1;MERS_1ab;3737 | 0 | 0 |
| NC_019843.3;YP_009047202.1;MERS_1ab;3738 | 0 | 0 |
| NC_019843.3;YP_009047202.1;MERS_1ab;3739 | 0 | 0 |
| NC_019843.3;YP_009047202.1;MERS_1ab;3740 | 0 | 0 |
| NC_019843.3;YP_009047202.1;MERS_1ab;3741 | 0 | 0 |
| NC_019843.3;YP_009047202.1;MERS_1ab;3742 | 0 | 0 |
| NC_019843.3;YP_009047202.1;MERS_1ab;3743 | 0 | 0 |
| NC_019843.3;YP_009047202.1;MERS_1ab;3744 | 0 | 0 |
| NC_019843.3;YP_009047202.1;MERS_1ab;3745 | 0 | 0 |
| NC_019843.3;YP_009047202.1;MERS_1ab;3746 | 0 | 0 |
| NC_019843.3;YP_009047202.1;MERS_1ab;3747 | 0 | 0 |
| NC_019843.3;YP_009047202.1;MERS_1ab;3748 | 0 | 0 |
| NC_019843.3;YP_009047202.1;MERS_1ab;3749 | 0 | 0 |
| NC_019843.3;YP_009047202.1;MERS_1ab;3750 | 0 | 0.025 |
| NC_019843.3;YP_009047202.1;MERS_1ab;3751 | 0 | 0 |
| NC_019843.3;YP_009047202.1;MERS_1ab;3752 | 0 | 0 |
| NC_019843.3;YP_009047202.1;MERS_1ab;3753 | 0 | 0 |
| NC_019843.3;YP_009047202.1;MERS_1ab;3754 | 0 | 0 |
| NC_019843.3;YP_009047202.1;MERS_1ab;3755 | 0 | 0 |
| NC_019843.3;YP_009047202.1;MERS_1ab;3756 | 0 | 0 |
| NC_019843.3;YP_009047202.1;MERS_1ab;3757 | 0 | 0 |
| NC_019843.3;YP_009047202.1;MERS_1ab;3758 | 0 | 0 |
| NC_019843.3;YP_009047202.1;MERS_1ab;3759 | 0 | 0 |
| NC_019843.3;YP_009047202.1;MERS_1ab;3760 | 0 | 0 |
| NC_019843.3;YP_009047202.1;MERS_1ab;3761 | 0 | 0 |
| NC_019843.3;YP_009047202.1;MERS_1ab;3762 | 0 | 0 |
| NC_019843.3;YP_009047202.1;MERS_1ab;3763 | 0 | 0 |
| NC_019843.3;YP_009047202.1;MERS_1ab;3764 | 0 | 0 |
| NC_019843.3;YP_009047202.1;MERS_1ab;3765 | 0 | 0 |
| NC_019843.3;YP_009047202.1;MERS_1ab;3766 | 0 | 0 |
| NC_019843.3;YP_009047202.1;MERS_1ab;3767 | 0 | 0 |
| NC_019843.3;YP_009047202.1;MERS_1ab;3768 | 0 | 0 |
| NC_019843.3;YP_009047202.1;MERS_1ab;3769 | 0 | 0 |
| NC_019843.3;YP_009047202.1;MERS_1ab;3770 | 0 | 0 |
| NC_019843.3;YP_009047202.1;MERS_1ab;3771 | 0 | 0 |
| NC_019843.3;YP_009047202.1;MERS_1ab;3772 | 0 | 0 |
| NC_019843.3;YP_009047202.1;MERS_1ab;3773 | 0 | 0 |
| NC_019843.3;YP_009047202.1;MERS_1ab;3774 | 0 | 0 |
| NC_019843.3;YP_009047202.1;MERS_1ab;3775 | 0 | 0 |

| | | |
|---|---|---|
| NC_019843.3;YP_009047202.1;MERS_1ab;3776 | 0 | 0 |
| NC_019843.3;YP_009047202.1;MERS_1ab;3777 | 0 | 0 |
| NC_019843.3;YP_009047202.1;MERS_1ab;3778 | 0 | 0 |
| NC_019843.3;YP_009047202.1;MERS_1ab;3779 | 0 | 0 |
| NC_019843.3;YP_009047202.1;MERS_1ab;3780 | 0 | 0 |
| NC_019843.3;YP_009047202.1;MERS_1ab;3781 | 0 | 0 |
| NC_019843.3;YP_009047202.1;MERS_1ab;3782 | 0 | 0 |
| NC_019843.3;YP_009047202.1;MERS_1ab;3783 | 0 | 0 |
| NC_019843.3;YP_009047202.1;MERS_1ab;3784 | 0 | 0 |
| NC_019843.3;YP_009047202.1;MERS_1ab;3785 | 0 | 0 |
| NC_019843.3;YP_009047202.1;MERS_1ab;3786 | 0 | 0 |
| NC_019843.3;YP_009047202.1;MERS_1ab;3787 | 0 | 0 |
| NC_019843.3;YP_009047202.1;MERS_1ab;3788 | 0 | 0 |
| NC_019843.3;YP_009047202.1;MERS_1ab;3789 | 0 | 0 |
| NC_019843.3;YP_009047202.1;MERS_1ab;3790 | 0 | 0 |
| NC_019843.3;YP_009047202.1;MERS_1ab;3791 | 0 | 0 |
| NC_019843.3;YP_009047202.1;MERS_1ab;3792 | 0 | 0 |
| NC_019843.3;YP_009047202.1;MERS_1ab;3793 | 0 | 0 |
| NC_019843.3;YP_009047202.1;MERS_1ab;3794 | 0 | 0 |
| NC_019843.3;YP_009047202.1;MERS_1ab;3795 | 0 | 0.075 |
| NC_019843.3;YP_009047202.1;MERS_1ab;3796 | 0 | 0 |
| NC_019843.3;YP_009047202.1;MERS_1ab;3797 | 0 | 0 |
| NC_019843.3;YP_009047202.1;MERS_1ab;3798 | 0 | 0 |
| NC_019843.3;YP_009047202.1;MERS_1ab;3799 | 0 | 0 |
| NC_019843.3;YP_009047202.1;MERS_1ab;3800 | 0 | 0.025 |
| NC_019843.3;YP_009047202.1;MERS_1ab;3801 | 0 | 0.025 |
| NC_019843.3;YP_009047202.1;MERS_1ab;3802 | 0 | 0 |
| NC_019843.3;YP_009047202.1;MERS_1ab;3803 | 0 | 0.025 |
| NC_019843.3;YP_009047202.1;MERS_1ab;3804 | 0 | 0 |
| NC_019843.3;YP_009047202.1;MERS_1ab;3805 | 0.05 | 0.025 |
| NC_019843.3;YP_009047202.1;MERS_1ab;3806 | 0 | 0 |
| NC_019843.3;YP_009047202.1;MERS_1ab;3807 | 0 | 0.025 |
| NC_019843.3;YP_009047202.1;MERS_1ab;3808 | 0 | 0 |
| NC_019843.3;YP_009047202.1;MERS_1ab;3809 | 0.05 | 0 |
| NC_019843.3;YP_009047202.1;MERS_1ab;3810 | 0 | 0 |
| NC_019843.3;YP_009047202.1;MERS_1ab;3811 | 0.15 | 0.025 |
| NC_019843.3;YP_009047202.1;MERS_1ab;3812 | 0 | 0.025 |
| NC_019843.3;YP_009047202.1;MERS_1ab;3813 | 0 | 0 |
| NC_019843.3;YP_009047202.1;MERS_1ab;3814 | 0.05 | 0 |
| NC_019843.3;YP_009047202.1;MERS_1ab;3815 | 0.05 | 0.025 |
| NC_019843.3;YP_009047202.1;MERS_1ab;3816 | 0 | 0 |
| NC_019843.3;YP_009047202.1;MERS_1ab;3817 | 0.05 | 0.05 |
| NC_019843.3;YP_009047202.1;MERS_1ab;3818 | 0 | 0 |

| | | |
|---|---|---|
| NC_019843.3;YP_009047202.1;MERS_1ab;3819 | 0 | 0 |
| NC_019843.3;YP_009047202.1;MERS_1ab;3820 | 0 | 0 |
| NC_019843.3;YP_009047202.1;MERS_1ab;3821 | 0 | 0 |
| NC_019843.3;YP_009047202.1;MERS_1ab;3822 | 0 | 0 |
| NC_019843.3;YP_009047202.1;MERS_1ab;3823 | 0 | 0 |
| NC_019843.3;YP_009047202.1;MERS_1ab;3824 | 0 | 0 |
| NC_019843.3;YP_009047202.1;MERS_1ab;3825 | 0 | 0 |
| NC_019843.3;YP_009047202.1;MERS_1ab;3826 | 0 | 0 |
| NC_019843.3;YP_009047202.1;MERS_1ab;3827 | 0 | 0 |
| NC_019843.3;YP_009047202.1;MERS_1ab;3828 | 0 | 0 |
| NC_019843.3;YP_009047202.1;MERS_1ab;3829 | 0 | 0 |
| NC_019843.3;YP_009047202.1;MERS_1ab;3830 | 0 | 0 |
| NC_019843.3;YP_009047202.1;MERS_1ab;3831 | 0 | 0 |
| NC_019843.3;YP_009047202.1;MERS_1ab;3832 | 0.05 | 0 |
| NC_019843.3;YP_009047202.1;MERS_1ab;3833 | 0.05 | 0 |
| NC_019843.3;YP_009047202.1;MERS_1ab;3834 | 0.05 | 0 |
| NC_019843.3;YP_009047202.1;MERS_1ab;3835 | 0.05 | 0 |
| NC_019843.3;YP_009047202.1;MERS_1ab;3836 | 0.1 | 0 |
| NC_019843.3;YP_009047202.1;MERS_1ab;3837 | 0.05 | 0 |
| NC_019843.3;YP_009047202.1;MERS_1ab;3838 | 0.05 | 0 |
| NC_019843.3;YP_009047202.1;MERS_1ab;3839 | 0.05 | 0 |
| NC_019843.3;YP_009047202.1;MERS_1ab;3840 | 0.05 | 0 |
| NC_019843.3;YP_009047202.1;MERS_1ab;3841 | 0 | 0 |
| NC_019843.3;YP_009047202.1;MERS_1ab;3842 | 0 | 0 |
| NC_019843.3;YP_009047202.1;MERS_1ab;3843 | 0 | 0 |
| NC_019843.3;YP_009047202.1;MERS_1ab;3844 | 0 | 0 |
| NC_019843.3;YP_009047202.1;MERS_1ab;3845 | 0 | 0 |
| NC_019843.3;YP_009047202.1;MERS_1ab;3846 | 0 | 0 |
| NC_019843.3;YP_009047202.1;MERS_1ab;3847 | 0 | 0 |
| NC_019843.3;YP_009047202.1;MERS_1ab;3848 | 0 | 0 |
| NC_019843.3;YP_009047202.1;MERS_1ab;3849 | 0 | 0 |
| NC_019843.3;YP_009047202.1;MERS_1ab;3850 | 0 | 0 |
| NC_019843.3;YP_009047202.1;MERS_1ab;3851 | 0 | 0 |
| NC_019843.3;YP_009047202.1;MERS_1ab;3852 | 0 | 0 |
| NC_019843.3;YP_009047202.1;MERS_1ab;3853 | 0 | 0 |
| NC_019843.3;YP_009047202.1;MERS_1ab;3854 | 0 | 0 |
| NC_019843.3;YP_009047202.1;MERS_1ab;3855 | 0 | 0 |
| NC_019843.3;YP_009047202.1;MERS_1ab;3856 | 0 | 0 |
| NC_019843.3;YP_009047202.1;MERS_1ab;3857 | 0 | 0 |
| NC_019843.3;YP_009047202.1;MERS_1ab;3858 | 0 | 0 |
| NC_019843.3;YP_009047202.1;MERS_1ab;3859 | 0 | 0 |
| NC_019843.3;YP_009047202.1;MERS_1ab;3860 | 0 | 0 |
| NC_019843.3;YP_009047202.1;MERS_1ab;3861 | 0 | 0 |

| | | |
|---|---|---|
| NC_019843.3;YP_009047202.1;MERS_1ab;3862 | 0 | 0 |
| NC_019843.3;YP_009047202.1;MERS_1ab;3863 | 0 | 0 |
| NC_019843.3;YP_009047202.1;MERS_1ab;3864 | 0 | 0 |
| NC_019843.3;YP_009047202.1;MERS_1ab;3865 | 0 | 0 |
| NC_019843.3;YP_009047202.1;MERS_1ab;3866 | 0 | 0 |
| NC_019843.3;YP_009047202.1;MERS_1ab;3867 | 0 | 0 |
| NC_019843.3;YP_009047202.1;MERS_1ab;3868 | 0 | 0 |
| NC_019843.3;YP_009047202.1;MERS_1ab;3869 | 0 | 0 |
| NC_019843.3;YP_009047202.1;MERS_1ab;3870 | 0 | 0 |
| NC_019843.3;YP_009047202.1;MERS_1ab;3871 | 0 | 0 |
| NC_019843.3;YP_009047202.1;MERS_1ab;3872 | 0.05 | 0.025 |
| NC_019843.3;YP_009047202.1;MERS_1ab;3873 | 0 | 0 |
| NC_019843.3;YP_009047202.1;MERS_1ab;3874 | 0 | 0 |
| NC_019843.3;YP_009047202.1;MERS_1ab;3875 | 0 | 0 |
| NC_019843.3;YP_009047202.1;MERS_1ab;3876 | 0 | 0 |
| NC_019843.3;YP_009047202.1;MERS_1ab;3877 | 0 | 0 |
| NC_019843.3;YP_009047202.1;MERS_1ab;3878 | 0 | 0 |
| NC_019843.3;YP_009047202.1;MERS_1ab;3879 | 0 | 0 |
| NC_019843.3;YP_009047202.1;MERS_1ab;3880 | 0 | 0 |
| NC_019843.3;YP_009047202.1;MERS_1ab;3881 | 0 | 0 |
| NC_019843.3;YP_009047202.1;MERS_1ab;3882 | 0 | 0 |
| NC_019843.3;YP_009047202.1;MERS_1ab;3883 | 0 | 0 |
| NC_019843.3;YP_009047202.1;MERS_1ab;3884 | 0 | 0 |
| NC_019843.3;YP_009047202.1;MERS_1ab;3885 | 0 | 0 |
| NC_019843.3;YP_009047202.1;MERS_1ab;3886 | 0 | 0 |
| NC_019843.3;YP_009047202.1;MERS_1ab;3887 | 0 | 0 |
| NC_019843.3;YP_009047202.1;MERS_1ab;3888 | 0 | 0 |
| NC_019843.3;YP_009047202.1;MERS_1ab;3889 | 0 | 0 |
| NC_019843.3;YP_009047202.1;MERS_1ab;3890 | 0 | 0 |
| NC_019843.3;YP_009047202.1;MERS_1ab;3891 | 0 | 0 |
| NC_019843.3;YP_009047202.1;MERS_1ab;3892 | 0 | 0 |
| NC_019843.3;YP_009047202.1;MERS_1ab;3893 | 0 | 0 |
| NC_019843.3;YP_009047202.1;MERS_1ab;3894 | 0 | 0 |
| NC_019843.3;YP_009047202.1;MERS_1ab;3895 | 0 | 0 |
| NC_019843.3;YP_009047202.1;MERS_1ab;3896 | 0 | 0 |
| NC_019843.3;YP_009047202.1;MERS_1ab;3897 | 0 | 0 |
| NC_019843.3;YP_009047202.1;MERS_1ab;3898 | 0 | 0 |
| NC_019843.3;YP_009047202.1;MERS_1ab;3899 | 0 | 0.025 |
| NC_019843.3;YP_009047202.1;MERS_1ab;3900 | 0 | 0.025 |
| NC_019843.3;YP_009047202.1;MERS_1ab;3901 | 0 | 0.05 |
| NC_019843.3;YP_009047202.1;MERS_1ab;3902 | 0 | 0.05 |
| NC_019843.3;YP_009047202.1;MERS_1ab;3903 | 0 | 0.075 |
| NC_019843.3;YP_009047202.1;MERS_1ab;3904 | 0 | 0.05 |

| | | |
|---|---|---|
| NC_019843.3;YP_009047202.1;MERS_1ab;3905 | 0 | 0.075 |
| NC_019843.3;YP_009047202.1;MERS_1ab;3906 | 0 | 0.1 |
| NC_019843.3;YP_009047202.1;MERS_1ab;3907 | 0 | 0.075 |
| NC_019843.3;YP_009047202.1;MERS_1ab;3908 | 0 | 0.05 |
| NC_019843.3;YP_009047202.1;MERS_1ab;3909 | 0 | 0.05 |
| NC_019843.3;YP_009047202.1;MERS_1ab;3910 | 0 | 0.025 |
| NC_019843.3;YP_009047202.1;MERS_1ab;3911 | 0 | 0 |
| NC_019843.3;YP_009047202.1;MERS_1ab;3912 | 0 | 0 |
| NC_019843.3;YP_009047202.1;MERS_1ab;3913 | 0 | 0 |
| NC_019843.3;YP_009047202.1;MERS_1ab;3914 | 0 | 0 |
| NC_019843.3;YP_009047202.1;MERS_1ab;3915 | 0 | 0.025 |
| NC_019843.3;YP_009047202.1;MERS_1ab;3916 | 0 | 0 |
| NC_019843.3;YP_009047202.1;MERS_1ab;3917 | 0.05 | 0 |
| NC_019843.3;YP_009047202.1;MERS_1ab;3918 | 0 | 0 |
| NC_019843.3;YP_009047202.1;MERS_1ab;3919 | 0 | 0 |
| NC_019843.3;YP_009047202.1;MERS_1ab;3920 | 0 | 0 |
| NC_019843.3;YP_009047202.1;MERS_1ab;3921 | 0 | 0 |
| NC_019843.3;YP_009047202.1;MERS_1ab;3922 | 0 | 0 |
| NC_019843.3;YP_009047202.1;MERS_1ab;3923 | 0 | 0 |
| NC_019843.3;YP_009047202.1;MERS_1ab;3924 | 0 | 0 |
| NC_019843.3;YP_009047202.1;MERS_1ab;3925 | 0 | 0 |
| NC_019843.3;YP_009047202.1;MERS_1ab;3926 | 0 | 0 |
| NC_019843.3;YP_009047202.1;MERS_1ab;3927 | 0 | 0 |
| NC_019843.3;YP_009047202.1;MERS_1ab;3928 | 0 | 0 |
| NC_019843.3;YP_009047202.1;MERS_1ab;3929 | 0 | 0 |
| NC_019843.3;YP_009047202.1;MERS_1ab;3930 | 0 | 0 |
| NC_019843.3;YP_009047202.1;MERS_1ab;3931 | 0 | 0.025 |
| NC_019843.3;YP_009047202.1;MERS_1ab;3932 | 0 | 0 |
| NC_019843.3;YP_009047202.1;MERS_1ab;3933 | 0 | 0.025 |
| NC_019843.3;YP_009047202.1;MERS_1ab;3934 | 0 | 0 |
| NC_019843.3;YP_009047202.1;MERS_1ab;3935 | 0 | 0.05 |
| NC_019843.3;YP_009047202.1;MERS_1ab;3936 | 0.05 | 0.05 |
| NC_019843.3;YP_009047202.1;MERS_1ab;3937 | 0 | 0.075 |
| NC_019843.3;YP_009047202.1;MERS_1ab;3938 | 0.05 | 0.15 |
| NC_019843.3;YP_009047202.1;MERS_1ab;3939 | 0.05 | 0.125 |
| NC_019843.3;YP_009047202.1;MERS_1ab;3940 | 0.05 | 0.075 |
| NC_019843.3;YP_009047202.1;MERS_1ab;3941 | 0.05 | 0 |
| NC_019843.3;YP_009047202.1;MERS_1ab;3942 | 0.05 | 0 |
| NC_019843.3;YP_009047202.1;MERS_1ab;3943 | 0 | 0 |
| NC_019843.3;YP_009047202.1;MERS_1ab;3944 | 0 | 0 |
| NC_019843.3;YP_009047202.1;MERS_1ab;3945 | 0 | 0.025 |
| NC_019843.3;YP_009047202.1;MERS_1ab;3946 | 0 | 0 |
| NC_019843.3;YP_009047202.1;MERS_1ab;3947 | 0 | 0 |

| | | |
|---|---|---|
| NC_019843.3;YP_009047202.1;MERS_1ab;3948 | 0 | 0 |
| NC_019843.3;YP_009047202.1;MERS_1ab;3949 | 0 | 0 |
| NC_019843.3;YP_009047202.1;MERS_1ab;3950 | 0 | 0 |
| NC_019843.3;YP_009047202.1;MERS_1ab;3951 | 0 | 0 |
| NC_019843.3;YP_009047202.1;MERS_1ab;3952 | 0 | 0 |
| NC_019843.3;YP_009047202.1;MERS_1ab;3953 | 0 | 0 |
| NC_019843.3;YP_009047202.1;MERS_1ab;3954 | 0 | 0 |
| NC_019843.3;YP_009047202.1;MERS_1ab;3955 | 0 | 0 |
| NC_019843.3;YP_009047202.1;MERS_1ab;3956 | 0 | 0 |
| NC_019843.3;YP_009047202.1;MERS_1ab;3957 | 0 | 0 |
| NC_019843.3;YP_009047202.1;MERS_1ab;3958 | 0 | 0.025 |
| NC_019843.3;YP_009047202.1;MERS_1ab;3959 | 0 | 0 |
| NC_019843.3;YP_009047202.1;MERS_1ab;3960 | 0 | 0 |
| NC_019843.3;YP_009047202.1;MERS_1ab;3961 | 0 | 0 |
| NC_019843.3;YP_009047202.1;MERS_1ab;3962 | 0 | 0 |
| NC_019843.3;YP_009047202.1;MERS_1ab;3963 | 0 | 0 |
| NC_019843.3;YP_009047202.1;MERS_1ab;3964 | 0 | 0 |
| NC_019843.3;YP_009047202.1;MERS_1ab;3965 | 0 | 0 |
| NC_019843.3;YP_009047202.1;MERS_1ab;3966 | 0.1 | 0 |
| NC_019843.3;YP_009047202.1;MERS_1ab;3967 | 0.05 | 0 |
| NC_019843.3;YP_009047202.1;MERS_1ab;3968 | 0.05 | 0 |
| NC_019843.3;YP_009047202.1;MERS_1ab;3969 | 0.05 | 0.025 |
| NC_019843.3;YP_009047202.1;MERS_1ab;3970 | 0.1 | 0.025 |
| NC_019843.3;YP_009047202.1;MERS_1ab;3971 | 0.05 | 0.025 |
| NC_019843.3;YP_009047202.1;MERS_1ab;3972 | 0 | 0 |
| NC_019843.3;YP_009047202.1;MERS_1ab;3973 | 0 | 0 |
| NC_019843.3;YP_009047202.1;MERS_1ab;3974 | 0 | 0.025 |
| NC_019843.3;YP_009047202.1;MERS_1ab;3975 | 0 | 0.025 |
| NC_019843.3;YP_009047202.1;MERS_1ab;3976 | 0 | 0.025 |
| NC_019843.3;YP_009047202.1;MERS_1ab;3977 | 0 | 0.05 |
| NC_019843.3;YP_009047202.1;MERS_1ab;3978 | 0 | 0.025 |
| NC_019843.3;YP_009047202.1;MERS_1ab;3979 | 0 | 0 |
| NC_019843.3;YP_009047202.1;MERS_1ab;3980 | 0 | 0 |
| NC_019843.3;YP_009047202.1;MERS_1ab;3981 | 0.05 | 0 |
| NC_019843.3;YP_009047202.1;MERS_1ab;3982 | 0 | 0.025 |
| NC_019843.3;YP_009047202.1;MERS_1ab;3983 | 0 | 0 |
| NC_019843.3;YP_009047202.1;MERS_1ab;3984 | 0 | 0 |
| NC_019843.3;YP_009047202.1;MERS_1ab;3985 | 0 | 0 |
| NC_019843.3;YP_009047202.1;MERS_1ab;3986 | 0 | 0.025 |
| NC_019843.3;YP_009047202.1;MERS_1ab;3987 | 0 | 0.025 |
| NC_019843.3;YP_009047202.1;MERS_1ab;3988 | 0 | 0 |
| NC_019843.3;YP_009047202.1;MERS_1ab;3989 | 0 | 0 |
| NC_019843.3;YP_009047202.1;MERS_1ab;3990 | 0 | 0 |

| | | |
|---|---|---|
| NC_019843.3;YP_009047202.1;MERS_1ab;3991 | 0 | 0 |
| NC_019843.3;YP_009047202.1;MERS_1ab;3992 | 0.05 | 0.025 |
| NC_019843.3;YP_009047202.1;MERS_1ab;3993 | 0.1 | 0.025 |
| NC_019843.3;YP_009047202.1;MERS_1ab;3994 | 0.05 | 0.025 |
| NC_019843.3;YP_009047202.1;MERS_1ab;3995 | 0.05 | 0.025 |
| NC_019843.3;YP_009047202.1;MERS_1ab;3996 | 0.05 | 0.025 |
| NC_019843.3;YP_009047202.1;MERS_1ab;3997 | 0 | 0.025 |
| NC_019843.3;YP_009047202.1;MERS_1ab;3998 | 0 | 0.025 |
| NC_019843.3;YP_009047202.1;MERS_1ab;3999 | 0 | 0.025 |
| NC_019843.3;YP_009047202.1;MERS_1ab;4000 | 0 | 0.025 |
| NC_019843.3;YP_009047202.1;MERS_1ab;4001 | 0 | 0.025 |
| NC_019843.3;YP_009047202.1;MERS_1ab;4002 | 0 | 0 |
| NC_019843.3;YP_009047202.1;MERS_1ab;4003 | 0 | 0.025 |
| NC_019843.3;YP_009047202.1;MERS_1ab;4004 | 0 | 0 |
| NC_019843.3;YP_009047202.1;MERS_1ab;4005 | 0 | 0 |
| NC_019843.3;YP_009047202.1;MERS_1ab;4006 | 0 | 0 |
| NC_019843.3;YP_009047202.1;MERS_1ab;4007 | 0 | 0 |
| NC_019843.3;YP_009047202.1;MERS_1ab;4008 | 0 | 0.025 |
| NC_019843.3;YP_009047202.1;MERS_1ab;4009 | 0 | 0 |
| NC_019843.3;YP_009047202.1;MERS_1ab;4010 | 0 | 0 |
| NC_019843.3;YP_009047202.1;MERS_1ab;4011 | 0 | 0 |
| NC_019843.3;YP_009047202.1;MERS_1ab;4012 | 0 | 0 |
| NC_019843.3;YP_009047202.1;MERS_1ab;4013 | 0 | 0 |
| NC_019843.3;YP_009047202.1;MERS_1ab;4014 | 0 | 0 |
| NC_019843.3;YP_009047202.1;MERS_1ab;4015 | 0 | 0 |
| NC_019843.3;YP_009047202.1;MERS_1ab;4016 | 0 | 0 |
| NC_019843.3;YP_009047202.1;MERS_1ab;4017 | 0 | 0 |
| NC_019843.3;YP_009047202.1;MERS_1ab;4018 | 0.2 | 0.125 |
| NC_019843.3;YP_009047202.1;MERS_1ab;4019 | 0.15 | 0.125 |
| NC_019843.3;YP_009047202.1;MERS_1ab;4020 | 0.2 | 0.125 |
| NC_019843.3;YP_009047202.1;MERS_1ab;4021 | 0.25 | 0.15 |
| NC_019843.3;YP_009047202.1;MERS_1ab;4022 | 0.2 | 0.175 |
| NC_019843.3;YP_009047202.1;MERS_1ab;4023 | 0.25 | 0.225 |
| NC_019843.3;YP_009047202.1;MERS_1ab;4024 | 0.3 | 0.225 |
| NC_019843.3;YP_009047202.1;MERS_1ab;4025 | 0.2 | 0.075 |
| NC_019843.3;YP_009047202.1;MERS_1ab;4026 | 0.2 | 0.175 |
| NC_019843.3;YP_009047202.1;MERS_1ab;4027 | 0.1 | 0.025 |
| NC_019843.3;YP_009047202.1;MERS_1ab;4028 | 0.05 | 0 |
| NC_019843.3;YP_009047202.1;MERS_1ab;4029 | 0.05 | 0 |
| NC_019843.3;YP_009047202.1;MERS_1ab;4030 | 0.05 | 0 |
| NC_019843.3;YP_009047202.1;MERS_1ab;4031 | 0.05 | 0 |
| NC_019843.3;YP_009047202.1;MERS_1ab;4032 | 0.05 | 0.025 |
| NC_019843.3;YP_009047202.1;MERS_1ab;4033 | 0.1 | 0.025 |

| | | |
|---|---|---|
| NC_019843.3;YP_009047202.1;MERS_1ab;4034 | 0.05 | 0.025 |
| NC_019843.3;YP_009047202.1;MERS_1ab;4035 | 0 | 0 |
| NC_019843.3;YP_009047202.1;MERS_1ab;4036 | 0.05 | 0 |
| NC_019843.3;YP_009047202.1;MERS_1ab;4037 | 0 | 0 |
| NC_019843.3;YP_009047202.1;MERS_1ab;4038 | 0 | 0 |
| NC_019843.3;YP_009047202.1;MERS_1ab;4039 | 0 | 0 |
| NC_019843.3;YP_009047202.1;MERS_1ab;4040 | 0 | 0 |
| NC_019843.3;YP_009047202.1;MERS_1ab;4041 | 0 | 0 |
| NC_019843.3;YP_009047202.1;MERS_1ab;4042 | 0 | 0 |
| NC_019843.3;YP_009047202.1;MERS_1ab;4043 | 0 | 0 |
| NC_019843.3;YP_009047202.1;MERS_1ab;4044 | 0 | 0 |
| NC_019843.3;YP_009047202.1;MERS_1ab;4045 | 0 | 0 |
| NC_019843.3;YP_009047202.1;MERS_1ab;4046 | 0 | 0 |
| NC_019843.3;YP_009047202.1;MERS_1ab;4047 | 0 | 0 |
| NC_019843.3;YP_009047202.1;MERS_1ab;4048 | 0 | 0 |
| NC_019843.3;YP_009047202.1;MERS_1ab;4049 | 0.05 | 0 |
| NC_019843.3;YP_009047202.1;MERS_1ab;4050 | 0.05 | 0.05 |
| NC_019843.3;YP_009047202.1;MERS_1ab;4051 | 0 | 0 |
| NC_019843.3;YP_009047202.1;MERS_1ab;4052 | 0 | 0 |
| NC_019843.3;YP_009047202.1;MERS_1ab;4053 | 0.05 | 0 |
| NC_019843.3;YP_009047202.1;MERS_1ab;4054 | 0 | 0 |
| NC_019843.3;YP_009047202.1;MERS_1ab;4055 | 0 | 0 |
| NC_019843.3;YP_009047202.1;MERS_1ab;4056 | 0 | 0 |
| NC_019843.3;YP_009047202.1;MERS_1ab;4057 | 0 | 0 |
| NC_019843.3;YP_009047202.1;MERS_1ab;4058 | 0 | 0 |
| NC_019843.3;YP_009047202.1;MERS_1ab;4059 | 0 | 0 |
| NC_019843.3;YP_009047202.1;MERS_1ab;4060 | 0 | 0.025 |
| NC_019843.3;YP_009047202.1;MERS_1ab;4061 | 0 | 0 |
| NC_019843.3;YP_009047202.1;MERS_1ab;4062 | 0 | 0 |
| NC_019843.3;YP_009047202.1;MERS_1ab;4063 | 0 | 0 |
| NC_019843.3;YP_009047202.1;MERS_1ab;4064 | 0 | 0 |
| NC_019843.3;YP_009047202.1;MERS_1ab;4065 | 0 | 0 |
| NC_019843.3;YP_009047202.1;MERS_1ab;4066 | 0 | 0 |
| NC_019843.3;YP_009047202.1;MERS_1ab;4067 | 0 | 0 |
| NC_019843.3;YP_009047202.1;MERS_1ab;4068 | 0 | 0 |
| NC_019843.3;YP_009047202.1;MERS_1ab;4069 | 0 | 0.025 |
| NC_019843.3;YP_009047202.1;MERS_1ab;4070 | 0 | 0.025 |
| NC_019843.3;YP_009047202.1;MERS_1ab;4071 | 0 | 0 |
| NC_019843.3;YP_009047202.1;MERS_1ab;4072 | 0 | 0 |
| NC_019843.3;YP_009047202.1;MERS_1ab;4073 | 0 | 0 |
| NC_019843.3;YP_009047202.1;MERS_1ab;4074 | 0 | 0 |
| NC_019843.3;YP_009047202.1;MERS_1ab;4075 | 0.05 | 0 |
| NC_019843.3;YP_009047202.1;MERS_1ab;4076 | 0 | 0 |

| | | |
|---|---|---|
| NC_019843.3;YP_009047202.1;MERS_1ab;4077 | 0.05 | 0 |
| NC_019843.3;YP_009047202.1;MERS_1ab;4078 | 0.05 | 0 |
| NC_019843.3;YP_009047202.1;MERS_1ab;4079 | 0.05 | 0 |
| NC_019843.3;YP_009047202.1;MERS_1ab;4080 | 0 | 0 |
| NC_019843.3;YP_009047202.1;MERS_1ab;4081 | 0 | 0 |
| NC_019843.3;YP_009047202.1;MERS_1ab;4082 | 0 | 0.05 |
| NC_019843.3;YP_009047202.1;MERS_1ab;4083 | 0 | 0.025 |
| NC_019843.3;YP_009047202.1;MERS_1ab;4084 | 0 | 0 |
| NC_019843.3;YP_009047202.1;MERS_1ab;4085 | 0 | 0 |
| NC_019843.3;YP_009047202.1;MERS_1ab;4086 | 0 | 0 |
| NC_019843.3;YP_009047202.1;MERS_1ab;4087 | 0 | 0 |
| NC_019843.3;YP_009047202.1;MERS_1ab;4088 | 0 | 0 |
| NC_019843.3;YP_009047202.1;MERS_1ab;4089 | 0 | 0 |
| NC_019843.3;YP_009047202.1;MERS_1ab;4090 | 0 | 0 |
| NC_019843.3;YP_009047202.1;MERS_1ab;4091 | 0 | 0 |
| NC_019843.3;YP_009047202.1;MERS_1ab;4092 | 0 | 0 |
| NC_019843.3;YP_009047202.1;MERS_1ab;4093 | 0 | 0 |
| NC_019843.3;YP_009047202.1;MERS_1ab;4094 | 0.05 | 0.05 |
| NC_019843.3;YP_009047202.1;MERS_1ab;4095 | 0 | 0.025 |
| NC_019843.3;YP_009047202.1;MERS_1ab;4096 | 0.1 | 0.075 |
| NC_019843.3;YP_009047202.1;MERS_1ab;4097 | 0.05 | 0.075 |
| NC_019843.3;YP_009047202.1;MERS_1ab;4098 | 0.05 | 0 |
| NC_019843.3;YP_009047202.1;MERS_1ab;4099 | 0 | 0 |
| NC_019843.3;YP_009047202.1;MERS_1ab;4100 | 0 | 0 |
| NC_019843.3;YP_009047202.1;MERS_1ab;4101 | 0 | 0 |
| NC_019843.3;YP_009047202.1;MERS_1ab;4102 | 0 | 0 |
| NC_019843.3;YP_009047202.1;MERS_1ab;4103 | 0 | 0 |
| NC_019843.3;YP_009047202.1;MERS_1ab;4104 | 0 | 0 |
| NC_019843.3;YP_009047202.1;MERS_1ab;4105 | 0 | 0 |
| NC_019843.3;YP_009047202.1;MERS_1ab;4106 | 0 | 0 |
| NC_019843.3;YP_009047202.1;MERS_1ab;4107 | 0 | 0 |
| NC_019843.3;YP_009047202.1;MERS_1ab;4108 | 0 | 0 |
| NC_019843.3;YP_009047202.1;MERS_1ab;4109 | 0 | 0 |
| NC_019843.3;YP_009047202.1;MERS_1ab;4110 | 0 | 0 |
| NC_019843.3;YP_009047202.1;MERS_1ab;4111 | 0 | 0 |
| NC_019843.3;YP_009047202.1;MERS_1ab;4112 | 0 | 0 |
| NC_019843.3;YP_009047202.1;MERS_1ab;4113 | 0 | 0 |
| NC_019843.3;YP_009047202.1;MERS_1ab;4114 | 0.05 | 0 |
| NC_019843.3;YP_009047202.1;MERS_1ab;4115 | 0.05 | 0.05 |
| NC_019843.3;YP_009047202.1;MERS_1ab;4116 | 0.05 | 0.05 |
| NC_019843.3;YP_009047202.1;MERS_1ab;4117 | 0 | 0 |
| NC_019843.3;YP_009047202.1;MERS_1ab;4118 | 0 | 0 |
| NC_019843.3;YP_009047202.1;MERS_1ab;4119 | 0 | 0.1 |

| | | |
|---|---|---|
| NC_019843.3;YP_009047202.1;MERS_1ab;4120 | 0 | 0.025 |
| NC_019843.3;YP_009047202.1;MERS_1ab;4121 | 0 | 0 |
| NC_019843.3;YP_009047202.1;MERS_1ab;4122 | 0 | 0 |
| NC_019843.3;YP_009047202.1;MERS_1ab;4123 | 0 | 0 |
| NC_019843.3;YP_009047202.1;MERS_1ab;4124 | 0 | 0 |
| NC_019843.3;YP_009047202.1;MERS_1ab;4125 | 0 | 0 |
| NC_019843.3;YP_009047202.1;MERS_1ab;4126 | 0.05 | 0.05 |
| NC_019843.3;YP_009047202.1;MERS_1ab;4127 | 0 | 0.025 |
| NC_019843.3;YP_009047202.1;MERS_1ab;4128 | 0 | 0.025 |
| NC_019843.3;YP_009047202.1;MERS_1ab;4129 | 0 | 0.025 |
| NC_019843.3;YP_009047202.1;MERS_1ab;4130 | 0 | 0 |
| NC_019843.3;YP_009047202.1;MERS_1ab;4131 | 0 | 0 |
| NC_019843.3;YP_009047202.1;MERS_1ab;4132 | 0 | 0 |
| NC_019843.3;YP_009047202.1;MERS_1ab;4133 | 0 | 0 |
| NC_019843.3;YP_009047202.1;MERS_1ab;4134 | 0 | 0 |
| NC_019843.3;YP_009047202.1;MERS_1ab;4135 | 0 | 0 |
| NC_019843.3;YP_009047202.1;MERS_1ab;4136 | 0 | 0 |
| NC_019843.3;YP_009047202.1;MERS_1ab;4137 | 0 | 0 |
| NC_019843.3;YP_009047202.1;MERS_1ab;4138 | 0 | 0 |
| NC_019843.3;YP_009047202.1;MERS_1ab;4139 | 0 | 0 |
| NC_019843.3;YP_009047202.1;MERS_1ab;4140 | 0 | 0 |
| NC_019843.3;YP_009047202.1;MERS_1ab;4141 | 0 | 0 |
| NC_019843.3;YP_009047202.1;MERS_1ab;4142 | 0 | 0 |
| NC_019843.3;YP_009047202.1;MERS_1ab;4143 | 0 | 0 |
| NC_019843.3;YP_009047202.1;MERS_1ab;4144 | 0 | 0 |
| NC_019843.3;YP_009047202.1;MERS_1ab;4145 | 0 | 0 |
| NC_019843.3;YP_009047202.1;MERS_1ab;4146 | 0 | 0.075 |
| NC_019843.3;YP_009047202.1;MERS_1ab;4147 | 0 | 0 |
| NC_019843.3;YP_009047202.1;MERS_1ab;4148 | 0 | 0 |
| NC_019843.3;YP_009047202.1;MERS_1ab;4149 | 0 | 0 |
| NC_019843.3;YP_009047202.1;MERS_1ab;4150 | 0 | 0 |
| NC_019843.3;YP_009047202.1;MERS_1ab;4151 | 0 | 0 |
| NC_019843.3;YP_009047202.1;MERS_1ab;4152 | 0 | 0 |
| NC_019843.3;YP_009047202.1;MERS_1ab;4153 | 0 | 0 |
| NC_019843.3;YP_009047202.1;MERS_1ab;4154 | 0 | 0 |
| NC_019843.3;YP_009047202.1;MERS_1ab;4155 | 0 | 0 |
| NC_019843.3;YP_009047202.1;MERS_1ab;4156 | 0 | 0.025 |
| NC_019843.3;YP_009047202.1;MERS_1ab;4157 | 0 | 0 |
| NC_019843.3;YP_009047202.1;MERS_1ab;4158 | 0 | 0 |
| NC_019843.3;YP_009047202.1;MERS_1ab;4159 | 0.05 | 0 |
| NC_019843.3;YP_009047202.1;MERS_1ab;4160 | 0.05 | 0 |
| NC_019843.3;YP_009047202.1;MERS_1ab;4161 | 0 | 0 |
| NC_019843.3;YP_009047202.1;MERS_1ab;4162 | 0 | 0 |

| | | |
|---|---|---|
| NC_019843.3;YP_009047202.1;MERS_1ab;4163 | 0.05 | 0.025 |
| NC_019843.3;YP_009047202.1;MERS_1ab;4164 | 0 | 0 |
| NC_019843.3;YP_009047202.1;MERS_1ab;4165 | 0 | 0 |
| NC_019843.3;YP_009047202.1;MERS_1ab;4166 | 0 | 0 |
| NC_019843.3;YP_009047202.1;MERS_1ab;4167 | 0 | 0 |
| NC_019843.3;YP_009047202.1;MERS_1ab;4168 | 0 | 0 |
| NC_019843.3;YP_009047202.1;MERS_1ab;4169 | 0 | 0 |
| NC_019843.3;YP_009047202.1;MERS_1ab;4170 | 0 | 0 |
| NC_019843.3;YP_009047202.1;MERS_1ab;4171 | 0 | 0.025 |
| NC_019843.3;YP_009047202.1;MERS_1ab;4172 | 0.05 | 0 |
| NC_019843.3;YP_009047202.1;MERS_1ab;4173 | 0.15 | 0.025 |
| NC_019843.3;YP_009047202.1;MERS_1ab;4174 | 0.05 | 0 |
| NC_019843.3;YP_009047202.1;MERS_1ab;4175 | 0.1 | 0 |
| NC_019843.3;YP_009047202.1;MERS_1ab;4176 | 0.05 | 0 |
| NC_019843.3;YP_009047202.1;MERS_1ab;4177 | 0.05 | 0 |
| NC_019843.3;YP_009047202.1;MERS_1ab;4178 | 0 | 0 |
| NC_019843.3;YP_009047202.1;MERS_1ab;4179 | 0 | 0 |
| NC_019843.3;YP_009047202.1;MERS_1ab;4180 | 0.05 | 0 |
| NC_019843.3;YP_009047202.1;MERS_1ab;4181 | 0.05 | 0 |
| NC_019843.3;YP_009047202.1;MERS_1ab;4182 | 0.05 | 0 |
| NC_019843.3;YP_009047202.1;MERS_1ab;4183 | 0.05 | 0 |
| NC_019843.3;YP_009047202.1;MERS_1ab;4184 | 0 | 0 |
| NC_019843.3;YP_009047202.1;MERS_1ab;4185 | 0 | 0 |
| NC_019843.3;YP_009047202.1;MERS_1ab;4186 | 0 | 0 |
| NC_019843.3;YP_009047202.1;MERS_1ab;4187 | 0 | 0 |
| NC_019843.3;YP_009047202.1;MERS_1ab;4188 | 0 | 0 |
| NC_019843.3;YP_009047202.1;MERS_1ab;4189 | 0 | 0 |
| NC_019843.3;YP_009047202.1;MERS_1ab;4190 | 0 | 0 |
| NC_019843.3;YP_009047202.1;MERS_1ab;4191 | 0 | 0 |
| NC_019843.3;YP_009047202.1;MERS_1ab;4192 | 0 | 0 |
| NC_019843.3;YP_009047202.1;MERS_1ab;4193 | 0 | 0 |
| NC_019843.3;YP_009047202.1;MERS_1ab;4194 | 0 | 0 |
| NC_019843.3;YP_009047202.1;MERS_1ab;4195 | 0 | 0 |
| NC_019843.3;YP_009047202.1;MERS_1ab;4196 | 0 | 0 |
| NC_019843.3;YP_009047202.1;MERS_1ab;4197 | 0 | 0 |
| NC_019843.3;YP_009047202.1;MERS_1ab;4198 | 0 | 0 |
| NC_019843.3;YP_009047202.1;MERS_1ab;4199 | 0 | 0 |
| NC_019843.3;YP_009047202.1;MERS_1ab;4200 | 0 | 0 |
| NC_019843.3;YP_009047202.1;MERS_1ab;4201 | 0 | 0 |
| NC_019843.3;YP_009047202.1;MERS_1ab;4202 | 0 | 0 |
| NC_019843.3;YP_009047202.1;MERS_1ab;4203 | 0 | 0 |
| NC_019843.3;YP_009047202.1;MERS_1ab;4204 | 0 | 0 |
| NC_019843.3;YP_009047202.1;MERS_1ab;4205 | 0 | 0 |

| | | |
|---|---|---|
| NC_019843.3;YP_009047202.1;MERS_1ab;4206 | 0 | 0 |
| NC_019843.3;YP_009047202.1;MERS_1ab;4207 | 0 | 0 |
| NC_019843.3;YP_009047202.1;MERS_1ab;4208 | 0 | 0 |
| NC_019843.3;YP_009047202.1;MERS_1ab;4209 | 0 | 0 |
| NC_019843.3;YP_009047202.1;MERS_1ab;4210 | 0 | 0 |
| NC_019843.3;YP_009047202.1;MERS_1ab;4211 | 0 | 0 |
| NC_019843.3;YP_009047202.1;MERS_1ab;4212 | 0 | 0 |
| NC_019843.3;YP_009047202.1;MERS_1ab;4213 | 0 | 0 |
| NC_019843.3;YP_009047202.1;MERS_1ab;4214 | 0 | 0 |
| NC_019843.3;YP_009047202.1;MERS_1ab;4215 | 0 | 0 |
| NC_019843.3;YP_009047202.1;MERS_1ab;4216 | 0 | 0 |
| NC_019843.3;YP_009047202.1;MERS_1ab;4217 | 0.1 | 0 |
| NC_019843.3;YP_009047202.1;MERS_1ab;4218 | 0 | 0 |
| NC_019843.3;YP_009047202.1;MERS_1ab;4219 | 0 | 0 |
| NC_019843.3;YP_009047202.1;MERS_1ab;4220 | 0 | 0 |
| NC_019843.3;YP_009047202.1;MERS_1ab;4221 | 0 | 0 |
| NC_019843.3;YP_009047202.1;MERS_1ab;4222 | 0 | 0 |
| NC_019843.3;YP_009047202.1;MERS_1ab;4223 | 0 | 0 |
| NC_019843.3;YP_009047202.1;MERS_1ab;4224 | 0 | 0 |
| NC_019843.3;YP_009047202.1;MERS_1ab;4225 | 0 | 0 |
| NC_019843.3;YP_009047202.1;MERS_1ab;4226 | 0 | 0 |
| NC_019843.3;YP_009047202.1;MERS_1ab;4227 | 0 | 0 |
| NC_019843.3;YP_009047202.1;MERS_1ab;4228 | 0 | 0 |
| NC_019843.3;YP_009047202.1;MERS_1ab;4229 | 0 | 0 |
| NC_019843.3;YP_009047202.1;MERS_1ab;4230 | 0 | 0 |
| NC_019843.3;YP_009047202.1;MERS_1ab;4231 | 0.1 | 0.025 |
| NC_019843.3;YP_009047202.1;MERS_1ab;4232 | 0 | 0 |
| NC_019843.3;YP_009047202.1;MERS_1ab;4233 | 0 | 0 |
| NC_019843.3;YP_009047202.1;MERS_1ab;4234 | 0 | 0.025 |
| NC_019843.3;YP_009047202.1;MERS_1ab;4235 | 0 | 0.075 |
| NC_019843.3;YP_009047202.1;MERS_1ab;4236 | 0.05 | 0.025 |
| NC_019843.3;YP_009047202.1;MERS_1ab;4237 | 0 | 0 |
| NC_019843.3;YP_009047202.1;MERS_1ab;4238 | 0 | 0 |
| NC_019843.3;YP_009047202.1;MERS_1ab;4239 | 0 | 0 |
| NC_019843.3;YP_009047202.1;MERS_1ab;4240 | 0 | 0 |
| NC_019843.3;YP_009047202.1;MERS_1ab;4241 | 0 | 0 |
| NC_019843.3;YP_009047202.1;MERS_1ab;4242 | 0 | 0 |
| NC_019843.3;YP_009047202.1;MERS_1ab;4243 | 0 | 0 |
| NC_019843.3;YP_009047202.1;MERS_1ab;4244 | 0 | 0 |
| NC_019843.3;YP_009047202.1;MERS_1ab;4245 | 0.1 | 0 |
| NC_019843.3;YP_009047202.1;MERS_1ab;4246 | 0 | 0.025 |
| NC_019843.3;YP_009047202.1;MERS_1ab;4247 | 0 | 0 |
| NC_019843.3;YP_009047202.1;MERS_1ab;4248 | 0 | 0 |

| | | |
|---|---|---|
| NC_019843.3;YP_009047202.1;MERS_1ab;4249 | 0 | 0 |
| NC_019843.3;YP_009047202.1;MERS_1ab;4250 | 0 | 0.025 |
| NC_019843.3;YP_009047202.1;MERS_1ab;4251 | 0 | 0.025 |
| NC_019843.3;YP_009047202.1;MERS_1ab;4252 | 0 | 0.025 |
| NC_019843.3;YP_009047202.1;MERS_1ab;4253 | 0 | 0.025 |
| NC_019843.3;YP_009047202.1;MERS_1ab;4254 | 0.05 | 0.025 |
| NC_019843.3;YP_009047202.1;MERS_1ab;4255 | 0.05 | 0.025 |
| NC_019843.3;YP_009047202.1;MERS_1ab;4256 | 0.15 | 0.05 |
| NC_019843.3;YP_009047202.1;MERS_1ab;4257 | 0.1 | 0.1 |
| NC_019843.3;YP_009047202.1;MERS_1ab;4258 | 0.1 | 0.075 |
| NC_019843.3;YP_009047202.1;MERS_1ab;4259 | 0.15 | 0.075 |
| NC_019843.3;YP_009047202.1;MERS_1ab;4260 | 0.1 | 0.05 |
| NC_019843.3;YP_009047202.1;MERS_1ab;4261 | 0.1 | 0.05 |
| NC_019843.3;YP_009047202.1;MERS_1ab;4262 | 0.1 | 0.05 |
| NC_019843.3;YP_009047202.1;MERS_1ab;4263 | 0.1 | 0.075 |
| NC_019843.3;YP_009047202.1;MERS_1ab;4264 | 0 | 0.025 |
| NC_019843.3;YP_009047202.1;MERS_1ab;4265 | 0 | 0.075 |
| NC_019843.3;YP_009047202.1;MERS_1ab;4266 | 0 | 0 |
| NC_019843.3;YP_009047202.1;MERS_1ab;4267 | 0 | 0.025 |
| NC_019843.3;YP_009047202.1;MERS_1ab;4268 | 0 | 0.025 |
| NC_019843.3;YP_009047202.1;MERS_1ab;4269 | 0 | 0 |
| NC_019843.3;YP_009047202.1;MERS_1ab;4270 | 0 | 0 |
| NC_019843.3;YP_009047202.1;MERS_1ab;4271 | 0 | 0 |
| NC_019843.3;YP_009047202.1;MERS_1ab;4272 | 0 | 0 |
| NC_019843.3;YP_009047202.1;MERS_1ab;4273 | 0.05 | 0 |
| NC_019843.3;YP_009047202.1;MERS_1ab;4274 | 0 | 0 |
| NC_019843.3;YP_009047202.1;MERS_1ab;4275 | 0 | 0 |
| NC_019843.3;YP_009047202.1;MERS_1ab;4276 | 0.05 | 0 |
| NC_019843.3;YP_009047202.1;MERS_1ab;4277 | 0 | 0 |
| NC_019843.3;YP_009047202.1;MERS_1ab;4278 | 0.05 | 0 |
| NC_019843.3;YP_009047202.1;MERS_1ab;4279 | 0 | 0 |
| NC_019843.3;YP_009047202.1;MERS_1ab;4280 | 0 | 0 |
| NC_019843.3;YP_009047202.1;MERS_1ab;4281 | 0 | 0 |
| NC_019843.3;YP_009047202.1;MERS_1ab;4282 | 0 | 0 |
| NC_019843.3;YP_009047202.1;MERS_1ab;4283 | 0.05 | 0 |
| NC_019843.3;YP_009047202.1;MERS_1ab;4284 | 0.05 | 0 |
| NC_019843.3;YP_009047202.1;MERS_1ab;4285 | 0.05 | 0 |
| NC_019843.3;YP_009047202.1;MERS_1ab;4286 | 0 | 0 |
| NC_019843.3;YP_009047202.1;MERS_1ab;4287 | 0 | 0 |
| NC_019843.3;YP_009047202.1;MERS_1ab;4288 | 0 | 0 |
| NC_019843.3;YP_009047202.1;MERS_1ab;4289 | 0 | 0 |
| NC_019843.3;YP_009047202.1;MERS_1ab;4290 | 0 | 0 |
| NC_019843.3;YP_009047202.1;MERS_1ab;4291 | 0 | 0 |

| | | |
|---|---|---|
| NC_019843.3;YP_009047202.1;MERS_1ab;4292 | 0 | 0 |
| NC_019843.3;YP_009047202.1;MERS_1ab;4293 | 0 | 0 |
| NC_019843.3;YP_009047202.1;MERS_1ab;4294 | 0 | 0 |
| NC_019843.3;YP_009047202.1;MERS_1ab;4295 | 0 | 0 |
| NC_019843.3;YP_009047202.1;MERS_1ab;4296 | 0 | 0 |
| NC_019843.3;YP_009047202.1;MERS_1ab;4297 | 0 | 0.075 |
| NC_019843.3;YP_009047202.1;MERS_1ab;4298 | 0 | 0 |
| NC_019843.3;YP_009047202.1;MERS_1ab;4299 | 0 | 0 |
| NC_019843.3;YP_009047202.1;MERS_1ab;4300 | 0 | 0 |
| NC_019843.3;YP_009047202.1;MERS_1ab;4301 | 0 | 0 |
| NC_019843.3;YP_009047202.1;MERS_1ab;4302 | 0 | 0 |
| NC_019843.3;YP_009047202.1;MERS_1ab;4303 | 0 | 0 |
| NC_019843.3;YP_009047202.1;MERS_1ab;4304 | 0 | 0 |
| NC_019843.3;YP_009047202.1;MERS_1ab;4305 | 0 | 0 |
| NC_019843.3;YP_009047202.1;MERS_1ab;4306 | 0 | 0 |
| NC_019843.3;YP_009047202.1;MERS_1ab;4307 | 0 | 0 |
| NC_019843.3;YP_009047202.1;MERS_1ab;4308 | 0 | 0 |
| NC_019843.3;YP_009047202.1;MERS_1ab;4309 | 0 | 0 |
| NC_019843.3;YP_009047202.1;MERS_1ab;4310 | 0 | 0 |
| NC_019843.3;YP_009047202.1;MERS_1ab;4311 | 0 | 0 |
| NC_019843.3;YP_009047202.1;MERS_1ab;4312 | 0 | 0.025 |
| NC_019843.3;YP_009047202.1;MERS_1ab;4313 | 0 | 0 |
| NC_019843.3;YP_009047202.1;MERS_1ab;4314 | 0 | 0 |
| NC_019843.3;YP_009047202.1;MERS_1ab;4315 | 0 | 0 |
| NC_019843.3;YP_009047202.1;MERS_1ab;4316 | 0 | 0 |
| NC_019843.3;YP_009047202.1;MERS_1ab;4317 | 0 | 0 |
| NC_019843.3;YP_009047202.1;MERS_1ab;4318 | 0 | 0 |
| NC_019843.3;YP_009047202.1;MERS_1ab;4319 | 0 | 0 |
| NC_019843.3;YP_009047202.1;MERS_1ab;4320 | 0 | 0 |
| NC_019843.3;YP_009047202.1;MERS_1ab;4321 | 0 | 0 |
| NC_019843.3;YP_009047202.1;MERS_1ab;4322 | 0 | 0 |
| NC_019843.3;YP_009047202.1;MERS_1ab;4323 | 0 | 0 |
| NC_019843.3;YP_009047202.1;MERS_1ab;4324 | 0 | 0 |
| NC_019843.3;YP_009047202.1;MERS_1ab;4325 | 0 | 0 |
| NC_019843.3;YP_009047202.1;MERS_1ab;4326 | 0 | 0 |
| NC_019843.3;YP_009047202.1;MERS_1ab;4327 | 0 | 0 |
| NC_019843.3;YP_009047202.1;MERS_1ab;4328 | 0 | 0 |
| NC_019843.3;YP_009047202.1;MERS_1ab;4329 | 0.05 | 0 |
| NC_019843.3;YP_009047202.1;MERS_1ab;4330 | 0 | 0.05 |
| NC_019843.3;YP_009047202.1;MERS_1ab;4331 | 0.05 | 0 |
| NC_019843.3;YP_009047202.1;MERS_1ab;4332 | 0.05 | 0 |
| NC_019843.3;YP_009047202.1;MERS_1ab;4333 | 0 | 0.025 |
| NC_019843.3;YP_009047202.1;MERS_1ab;4334 | 0 | 0 |

| | | |
|---|---|---|
| NC_019843.3;YP_009047202.1;MERS_1ab;4335 | 0 | 0 |
| NC_019843.3;YP_009047202.1;MERS_1ab;4336 | 0 | 0 |
| NC_019843.3;YP_009047202.1;MERS_1ab;4337 | 0 | 0 |
| NC_019843.3;YP_009047202.1;MERS_1ab;4338 | 0 | 0 |
| NC_019843.3;YP_009047202.1;MERS_1ab;4339 | 0 | 0 |
| NC_019843.3;YP_009047202.1;MERS_1ab;4340 | 0 | 0 |
| NC_019843.3;YP_009047202.1;MERS_1ab;4341 | 0 | 0 |
| NC_019843.3;YP_009047202.1;MERS_1ab;4342 | 0 | 0 |
| NC_019843.3;YP_009047202.1;MERS_1ab;4343 | 0.05 | 0.025 |
| NC_019843.3;YP_009047202.1;MERS_1ab;4344 | 0 | 0 |
| NC_019843.3;YP_009047202.1;MERS_1ab;4345 | 0 | 0 |
| NC_019843.3;YP_009047202.1;MERS_1ab;4346 | 0 | 0 |
| NC_019843.3;YP_009047202.1;MERS_1ab;4347 | 0 | 0 |
| NC_019843.3;YP_009047202.1;MERS_1ab;4348 | 0 | 0 |
| NC_019843.3;YP_009047202.1;MERS_1ab;4349 | 0 | 0 |
| NC_019843.3;YP_009047202.1;MERS_1ab;4350 | 0 | 0 |
| NC_019843.3;YP_009047202.1;MERS_1ab;4351 | 0 | 0 |
| NC_019843.3;YP_009047202.1;MERS_1ab;4352 | 0 | 0 |
| NC_019843.3;YP_009047202.1;MERS_1ab;4353 | 0 | 0 |
| NC_019843.3;YP_009047202.1;MERS_1ab;4354 | 0 | 0 |
| NC_019843.3;YP_009047202.1;MERS_1ab;4355 | 0 | 0 |
| NC_019843.3;YP_009047202.1;MERS_1ab;4356 | 0 | 0 |
| NC_019843.3;YP_009047202.1;MERS_1ab;4357 | 0 | 0 |
| NC_019843.3;YP_009047202.1;MERS_1ab;4358 | 0 | 0 |
| NC_019843.3;YP_009047202.1;MERS_1ab;4359 | 0.05 | 0.025 |
| NC_019843.3;YP_009047202.1;MERS_1ab;4360 | 0 | 0 |
| NC_019843.3;YP_009047202.1;MERS_1ab;4361 | 0 | 0 |
| NC_019843.3;YP_009047202.1;MERS_1ab;4362 | 0 | 0 |
| NC_019843.3;YP_009047202.1;MERS_1ab;4363 | 0 | 0 |
| NC_019843.3;YP_009047202.1;MERS_1ab;4364 | 0 | 0 |
| NC_019843.3;YP_009047202.1;MERS_1ab;4365 | 0 | 0 |
| NC_019843.3;YP_009047202.1;MERS_1ab;4366 | 0 | 0 |
| NC_019843.3;YP_009047202.1;MERS_1ab;4367 | 0 | 0 |
| NC_019843.3;YP_009047202.1;MERS_1ab;4368 | 0.1 | 0.05 |
| NC_019843.3;YP_009047202.1;MERS_1ab;4369 | 0 | 0 |
| NC_019843.3;YP_009047202.1;MERS_1ab;4370 | 0.05 | 0.075 |
| NC_019843.3;YP_009047202.1;MERS_1ab;4371 | 0.05 | 0.075 |
| NC_019843.3;YP_009047202.1;MERS_1ab;4372 | 0.15 | 0.05 |
| NC_019843.3;YP_009047202.1;MERS_1ab;4373 | 0.1 | 0.05 |
| NC_019843.3;YP_009047202.1;MERS_1ab;4374 | 0.1 | 0.05 |
| NC_019843.3;YP_009047202.1;MERS_1ab;4375 | 0.15 | 0.05 |
| NC_019843.3;YP_009047202.1;MERS_1ab;4376 | 0.1 | 0.05 |
| NC_019843.3;YP_009047202.1;MERS_1ab;4377 | 0.1 | 0.05 |

| | | |
|---|---|---|
| NC_019843.3;YP_009047202.1;MERS_1ab;4378 | 0.1 | 0.05 |
| NC_019843.3;YP_009047202.1;MERS_1ab;4379 | 0.1 | 0.05 |
| NC_019843.3;YP_009047202.1;MERS_1ab;4380 | 0.05 | 0 |
| NC_019843.3;YP_009047202.1;MERS_1ab;4381 | 0 | 0 |
| NC_019843.3;YP_009047202.1;MERS_1ab;4382 | 0 | 0 |
| NC_019843.3;YP_009047202.1;MERS_1ab;4383 | 0 | 0 |
| NC_019843.3;YP_009047202.1;MERS_1ab;4384 | 0 | 0 |
| NC_019843.3;YP_009047202.1;MERS_1ab;4385 | 0 | 0 |
| NC_019843.3;YP_009047202.1;MERS_1ab;4386 | 0 | 0.025 |
| NC_019843.3;YP_009047202.1;MERS_1ab;4387 | 0 | 0.025 |
| NC_019843.3;YP_009047202.1;MERS_1ab;4388 | 0 | 0.05 |
| NC_019843.3;YP_009047202.1;MERS_1ab;4389 | 0 | 0 |
| NC_019843.3;YP_009047202.1;MERS_1ab;4390 | 0 | 0 |
| NC_019843.3;YP_009047202.1;MERS_1ab;4391 | 0 | 0.05 |
| NC_019843.3;YP_009047202.1;MERS_1ab;4392 | 0 | 0 |
| NC_019843.3;YP_009047202.1;MERS_1ab;4393 | 0 | 0 |
| NC_019843.3;YP_009047202.1;MERS_1ab;4394 | 0 | 0 |
| NC_019843.3;YP_009047202.1;MERS_1ab;4395 | 0 | 0 |
| NC_019843.3;YP_009047202.1;MERS_1ab;4396 | 0 | 0 |
| NC_019843.3;YP_009047202.1;MERS_1ab;4397 | 0 | 0 |
| NC_019843.3;YP_009047202.1;MERS_1ab;4398 | 0 | 0 |
| NC_019843.3;YP_009047202.1;MERS_1ab;4399 | 0 | 0 |
| NC_019843.3;YP_009047202.1;MERS_1ab;4400 | 0.05 | 0 |
| NC_019843.3;YP_009047202.1;MERS_1ab;4401 | 0 | 0 |
| NC_019843.3;YP_009047202.1;MERS_1ab;4402 | 0 | 0 |
| NC_019843.3;YP_009047202.1;MERS_1ab;4403 | 0 | 0 |
| NC_019843.3;YP_009047202.1;MERS_1ab;4404 | 0 | 0 |
| NC_019843.3;YP_009047202.1;MERS_1ab;4405 | 0 | 0 |
| NC_019843.3;YP_009047202.1;MERS_1ab;4406 | 0 | 0 |
| NC_019843.3;YP_009047202.1;MERS_1ab;4407 | 0 | 0 |
| NC_019843.3;YP_009047202.1;MERS_1ab;4408 | 0 | 0.025 |
| NC_019843.3;YP_009047202.1;MERS_1ab;4409 | 0 | 0 |
| NC_019843.3;YP_009047202.1;MERS_1ab;4410 | 0 | 0 |
| NC_019843.3;YP_009047202.1;MERS_1ab;4411 | 0 | 0 |
| NC_019843.3;YP_009047202.1;MERS_1ab;4412 | 0 | 0 |
| NC_019843.3;YP_009047202.1;MERS_1ab;4413 | 0 | 0 |
| NC_019843.3;YP_009047202.1;MERS_1ab;4414 | 0 | 0 |
| NC_019843.3;YP_009047202.1;MERS_1ab;4415 | 0 | 0 |
| NC_019843.3;YP_009047202.1;MERS_1ab;4416 | 0 | 0 |
| NC_019843.3;YP_009047202.1;MERS_1ab;4417 | 0 | 0 |
| NC_019843.3;YP_009047202.1;MERS_1ab;4418 | 0 | 0 |
| NC_019843.3;YP_009047202.1;MERS_1ab;4419 | 0 | 0 |
| NC_019843.3;YP_009047202.1;MERS_1ab;4420 | 0 | 0 |

| | | |
|---|---|---|
| NC_019843.3;YP_009047202.1;MERS_1ab;4421 | 0 | 0 |
| NC_019843.3;YP_009047202.1;MERS_1ab;4422 | 0 | 0 |
| NC_019843.3;YP_009047202.1;MERS_1ab;4423 | 0 | 0 |
| NC_019843.3;YP_009047202.1;MERS_1ab;4424 | 0 | 0 |
| NC_019843.3;YP_009047202.1;MERS_1ab;4425 | 0 | 0 |
| NC_019843.3;YP_009047202.1;MERS_1ab;4426 | 0 | 0 |
| NC_019843.3;YP_009047202.1;MERS_1ab;4427 | 0.05 | 0 |
| NC_019843.3;YP_009047202.1;MERS_1ab;4428 | 0.1 | 0 |
| NC_019843.3;YP_009047202.1;MERS_1ab;4429 | 0.05 | 0 |
| NC_019843.3;YP_009047202.1;MERS_1ab;4430 | 0.05 | 0 |
| NC_019843.3;YP_009047202.1;MERS_1ab;4431 | 0.05 | 0 |
| NC_019843.3;YP_009047202.1;MERS_1ab;4432 | 0.05 | 0 |
| NC_019843.3;YP_009047202.1;MERS_1ab;4433 | 0.15 | 0 |
| NC_019843.3;YP_009047202.1;MERS_1ab;4434 | 0 | 0 |
| NC_019843.3;YP_009047202.1;MERS_1ab;4435 | 0 | 0 |
| NC_019843.3;YP_009047202.1;MERS_1ab;4436 | 0 | 0 |
| NC_019843.3;YP_009047202.1;MERS_1ab;4437 | 0 | 0 |
| NC_019843.3;YP_009047202.1;MERS_1ab;4438 | 0 | 0 |
| NC_019843.3;YP_009047202.1;MERS_1ab;4439 | 0 | 0 |
| NC_019843.3;YP_009047202.1;MERS_1ab;4440 | 0 | 0 |
| NC_019843.3;YP_009047202.1;MERS_1ab;4441 | 0 | 0 |
| NC_019843.3;YP_009047202.1;MERS_1ab;4442 | 0 | 0 |
| NC_019843.3;YP_009047202.1;MERS_1ab;4443 | 0 | 0 |
| NC_019843.3;YP_009047202.1;MERS_1ab;4444 | 0 | 0 |
| NC_019843.3;YP_009047202.1;MERS_1ab;4445 | 0 | 0 |
| NC_019843.3;YP_009047202.1;MERS_1ab;4446 | 0.05 | 0 |
| NC_019843.3;YP_009047202.1;MERS_1ab;4447 | 0 | 0 |
| NC_019843.3;YP_009047202.1;MERS_1ab;4448 | 0 | 0 |
| NC_019843.3;YP_009047202.1;MERS_1ab;4449 | 0 | 0 |
| NC_019843.3;YP_009047202.1;MERS_1ab;4450 | 0 | 0 |
| NC_019843.3;YP_009047202.1;MERS_1ab;4451 | 0 | 0 |
| NC_019843.3;YP_009047202.1;MERS_1ab;4452 | 0 | 0 |
| NC_019843.3;YP_009047202.1;MERS_1ab;4453 | 0 | 0 |
| NC_019843.3;YP_009047202.1;MERS_1ab;4454 | 0 | 0 |
| NC_019843.3;YP_009047202.1;MERS_1ab;4455 | 0 | 0 |
| NC_019843.3;YP_009047202.1;MERS_1ab;4456 | 0 | 0 |
| NC_019843.3;YP_009047202.1;MERS_1ab;4457 | 0 | 0 |
| NC_019843.3;YP_009047202.1;MERS_1ab;4458 | 0 | 0 |
| NC_019843.3;YP_009047202.1;MERS_1ab;4459 | 0 | 0 |
| NC_019843.3;YP_009047202.1;MERS_1ab;4460 | 0.05 | 0 |
| NC_019843.3;YP_009047202.1;MERS_1ab;4461 | 0 | 0 |
| NC_019843.3;YP_009047202.1;MERS_1ab;4462 | 0 | 0 |
| NC_019843.3;YP_009047202.1;MERS_1ab;4463 | 0 | 0 |

| | | |
|---|---|---|
| NC_019843.3;YP_009047202.1;MERS_1ab;4464 | 0 | 0 |
| NC_019843.3;YP_009047202.1;MERS_1ab;4465 | 0 | 0 |
| NC_019843.3;YP_009047202.1;MERS_1ab;4466 | 0 | 0 |
| NC_019843.3;YP_009047202.1;MERS_1ab;4467 | 0 | 0 |
| NC_019843.3;YP_009047202.1;MERS_1ab;4468 | 0 | 0 |
| NC_019843.3;YP_009047202.1;MERS_1ab;4469 | 0 | 0 |
| NC_019843.3;YP_009047202.1;MERS_1ab;4470 | 0 | 0 |
| NC_019843.3;YP_009047202.1;MERS_1ab;4471 | 0 | 0 |
| NC_019843.3;YP_009047202.1;MERS_1ab;4472 | 0 | 0 |
| NC_019843.3;YP_009047202.1;MERS_1ab;4473 | 0 | 0 |
| NC_019843.3;YP_009047202.1;MERS_1ab;4474 | 0 | 0 |
| NC_019843.3;YP_009047202.1;MERS_1ab;4475 | 0 | 0 |
| NC_019843.3;YP_009047202.1;MERS_1ab;4476 | 0 | 0 |
| NC_019843.3;YP_009047202.1;MERS_1ab;4477 | 0 | 0 |
| NC_019843.3;YP_009047202.1;MERS_1ab;4478 | 0 | 0 |
| NC_019843.3;YP_009047202.1;MERS_1ab;4479 | 0 | 0 |
| NC_019843.3;YP_009047202.1;MERS_1ab;4480 | 0 | 0 |
| NC_019843.3;YP_009047202.1;MERS_1ab;4481 | 0 | 0 |
| NC_019843.3;YP_009047202.1;MERS_1ab;4482 | 0 | 0 |
| NC_019843.3;YP_009047202.1;MERS_1ab;4483 | 0 | 0 |
| NC_019843.3;YP_009047202.1;MERS_1ab;4484 | 0 | 0 |
| NC_019843.3;YP_009047202.1;MERS_1ab;4485 | 0 | 0 |
| NC_019843.3;YP_009047202.1;MERS_1ab;4486 | 0 | 0 |
| NC_019843.3;YP_009047202.1;MERS_1ab;4487 | 0 | 0 |
| NC_019843.3;YP_009047202.1;MERS_1ab;4488 | 0 | 0 |
| NC_019843.3;YP_009047202.1;MERS_1ab;4489 | 0 | 0 |
| NC_019843.3;YP_009047202.1;MERS_1ab;4490 | 0 | 0 |
| NC_019843.3;YP_009047202.1;MERS_1ab;4491 | 0 | 0 |
| NC_019843.3;YP_009047202.1;MERS_1ab;4492 | 0 | 0 |
| NC_019843.3;YP_009047202.1;MERS_1ab;4493 | 0 | 0 |
| NC_019843.3;YP_009047202.1;MERS_1ab;4494 | 0 | 0 |
| NC_019843.3;YP_009047202.1;MERS_1ab;4495 | 0 | 0 |
| NC_019843.3;YP_009047202.1;MERS_1ab;4496 | 0 | 0 |
| NC_019843.3;YP_009047202.1;MERS_1ab;4497 | 0 | 0 |
| NC_019843.3;YP_009047202.1;MERS_1ab;4498 | 0 | 0 |
| NC_019843.3;YP_009047202.1;MERS_1ab;4499 | 0 | 0 |
| NC_019843.3;YP_009047202.1;MERS_1ab;4500 | 0 | 0 |
| NC_019843.3;YP_009047202.1;MERS_1ab;4501 | 0 | 0.225 |
| NC_019843.3;YP_009047202.1;MERS_1ab;4502 | 0 | 0 |
| NC_019843.3;YP_009047202.1;MERS_1ab;4503 | 0 | 0 |
| NC_019843.3;YP_009047202.1;MERS_1ab;4504 | 0 | 0.025 |
| NC_019843.3;YP_009047202.1;MERS_1ab;4505 | 0 | 0 |
| NC_019843.3;YP_009047202.1;MERS_1ab;4506 | 0 | 0.025 |

| | | |
|---|---|---|
| NC_019843.3;YP_009047202.1;MERS_1ab;4507 | 0 | 0.05 |
| NC_019843.3;YP_009047202.1;MERS_1ab;4508 | 0 | 0.025 |
| NC_019843.3;YP_009047202.1;MERS_1ab;4509 | 0 | 0 |
| NC_019843.3;YP_009047202.1;MERS_1ab;4510 | 0 | 0 |
| NC_019843.3;YP_009047202.1;MERS_1ab;4511 | 0 | 0 |
| NC_019843.3;YP_009047202.1;MERS_1ab;4512 | 0 | 0 |
| NC_019843.3;YP_009047202.1;MERS_1ab;4513 | 0 | 0 |
| NC_019843.3;YP_009047202.1;MERS_1ab;4514 | 0 | 0 |
| NC_019843.3;YP_009047202.1;MERS_1ab;4515 | 0 | 0 |
| NC_019843.3;YP_009047202.1;MERS_1ab;4516 | 0 | 0 |
| NC_019843.3;YP_009047202.1;MERS_1ab;4517 | 0 | 0 |
| NC_019843.3;YP_009047202.1;MERS_1ab;4518 | 0 | 0 |
| NC_019843.3;YP_009047202.1;MERS_1ab;4519 | 0 | 0 |
| NC_019843.3;YP_009047202.1;MERS_1ab;4520 | 0 | 0 |
| NC_019843.3;YP_009047202.1;MERS_1ab;4521 | 0 | 0 |
| NC_019843.3;YP_009047202.1;MERS_1ab;4522 | 0 | 0 |
| NC_019843.3;YP_009047202.1;MERS_1ab;4523 | 0.05 | 0 |
| NC_019843.3;YP_009047202.1;MERS_1ab;4524 | 0.2 | 0 |
| NC_019843.3;YP_009047202.1;MERS_1ab;4525 | 0.05 | 0 |
| NC_019843.3;YP_009047202.1;MERS_1ab;4526 | 0 | 0 |
| NC_019843.3;YP_009047202.1;MERS_1ab;4527 | 0 | 0 |
| NC_019843.3;YP_009047202.1;MERS_1ab;4528 | 0 | 0 |
| NC_019843.3;YP_009047202.1;MERS_1ab;4529 | 0 | 0 |
| NC_019843.3;YP_009047202.1;MERS_1ab;4530 | 0 | 0 |
| NC_019843.3;YP_009047202.1;MERS_1ab;4531 | 0.05 | 0 |
| NC_019843.3;YP_009047202.1;MERS_1ab;4532 | 0 | 0 |
| NC_019843.3;YP_009047202.1;MERS_1ab;4533 | 0 | 0 |
| NC_019843.3;YP_009047202.1;MERS_1ab;4534 | 0 | 0 |
| NC_019843.3;YP_009047202.1;MERS_1ab;4535 | 0.05 | 0 |
| NC_019843.3;YP_009047202.1;MERS_1ab;4536 | 0 | 0 |
| NC_019843.3;YP_009047202.1;MERS_1ab;4537 | 0 | 0 |
| NC_019843.3;YP_009047202.1;MERS_1ab;4538 | 0 | 0 |
| NC_019843.3;YP_009047202.1;MERS_1ab;4539 | 0 | 0 |
| NC_019843.3;YP_009047202.1;MERS_1ab;4540 | 0 | 0 |
| NC_019843.3;YP_009047202.1;MERS_1ab;4541 | 0 | 0 |
| NC_019843.3;YP_009047202.1;MERS_1ab;4542 | 0 | 0 |
| NC_019843.3;YP_009047202.1;MERS_1ab;4543 | 0 | 0 |
| NC_019843.3;YP_009047202.1;MERS_1ab;4544 | 0 | 0 |
| NC_019843.3;YP_009047202.1;MERS_1ab;4545 | 0 | 0 |
| NC_019843.3;YP_009047202.1;MERS_1ab;4546 | 0 | 0 |
| NC_019843.3;YP_009047202.1;MERS_1ab;4547 | 0 | 0 |
| NC_019843.3;YP_009047202.1;MERS_1ab;4548 | 0 | 0 |
| NC_019843.3;YP_009047202.1;MERS_1ab;4549 | 0 | 0 |

| | | |
|---|---|---|
| NC_019843.3;YP_009047202.1;MERS_1ab;4550 | 0 | 0 |
| NC_019843.3;YP_009047202.1;MERS_1ab;4551 | 0 | 0 |
| NC_019843.3;YP_009047202.1;MERS_1ab;4552 | 0 | 0 |
| NC_019843.3;YP_009047202.1;MERS_1ab;4553 | 0 | 0 |
| NC_019843.3;YP_009047202.1;MERS_1ab;4554 | 0 | 0 |
| NC_019843.3;YP_009047202.1;MERS_1ab;4555 | 0 | 0 |
| NC_019843.3;YP_009047202.1;MERS_1ab;4556 | 0 | 0 |
| NC_019843.3;YP_009047202.1;MERS_1ab;4557 | 0 | 0.025 |
| NC_019843.3;YP_009047202.1;MERS_1ab;4558 | 0.05 | 0 |
| NC_019843.3;YP_009047202.1;MERS_1ab;4559 | 0 | 0 |
| NC_019843.3;YP_009047202.1;MERS_1ab;4560 | 0 | 0 |
| NC_019843.3;YP_009047202.1;MERS_1ab;4561 | 0 | 0 |
| NC_019843.3;YP_009047202.1;MERS_1ab;4562 | 0 | 0 |
| NC_019843.3;YP_009047202.1;MERS_1ab;4563 | 0 | 0.025 |
| NC_019843.3;YP_009047202.1;MERS_1ab;4564 | 0 | 0.025 |
| NC_019843.3;YP_009047202.1;MERS_1ab;4565 | 0 | 0 |
| NC_019843.3;YP_009047202.1;MERS_1ab;4566 | 0 | 0 |
| NC_019843.3;YP_009047202.1;MERS_1ab;4567 | 0 | 0 |
| NC_019843.3;YP_009047202.1;MERS_1ab;4568 | 0 | 0 |
| NC_019843.3;YP_009047202.1;MERS_1ab;4569 | 0 | 0 |
| NC_019843.3;YP_009047202.1;MERS_1ab;4570 | 0 | 0 |
| NC_019843.3;YP_009047202.1;MERS_1ab;4571 | 0 | 0 |
| NC_019843.3;YP_009047202.1;MERS_1ab;4572 | 0 | 0 |
| NC_019843.3;YP_009047202.1;MERS_1ab;4573 | 0 | 0 |
| NC_019843.3;YP_009047202.1;MERS_1ab;4574 | 0 | 0 |
| NC_019843.3;YP_009047202.1;MERS_1ab;4575 | 0 | 0 |
| NC_019843.3;YP_009047202.1;MERS_1ab;4576 | 0 | 0 |
| NC_019843.3;YP_009047202.1;MERS_1ab;4577 | 0.05 | 0.025 |
| NC_019843.3;YP_009047202.1;MERS_1ab;4578 | 0 | 0.025 |
| NC_019843.3;YP_009047202.1;MERS_1ab;4579 | 0 | 0.025 |
| NC_019843.3;YP_009047202.1;MERS_1ab;4580 | 0 | 0 |
| NC_019843.3;YP_009047202.1;MERS_1ab;4581 | 0.15 | 0.05 |
| NC_019843.3;YP_009047202.1;MERS_1ab;4582 | 0 | 0.025 |
| NC_019843.3;YP_009047202.1;MERS_1ab;4583 | 0.05 | 0.025 |
| NC_019843.3;YP_009047202.1;MERS_1ab;4584 | 0 | 0.025 |
| NC_019843.3;YP_009047202.1;MERS_1ab;4585 | 0 | 0.025 |
| NC_019843.3;YP_009047202.1;MERS_1ab;4586 | 0 | 0.025 |
| NC_019843.3;YP_009047202.1;MERS_1ab;4587 | 0 | 0 |
| NC_019843.3;YP_009047202.1;MERS_1ab;4588 | 0 | 0.025 |
| NC_019843.3;YP_009047202.1;MERS_1ab;4589 | 0 | 0 |
| NC_019843.3;YP_009047202.1;MERS_1ab;4590 | 0 | 0 |
| NC_019843.3;YP_009047202.1;MERS_1ab;4591 | 0 | 0 |
| NC_019843.3;YP_009047202.1;MERS_1ab;4592 | 0 | 0 |

| | | |
|---|---|---|
| NC_019843.3;YP_009047202.1;MERS_1ab;4593 | 0 | 0 |
| NC_019843.3;YP_009047202.1;MERS_1ab;4594 | 0 | 0 |
| NC_019843.3;YP_009047202.1;MERS_1ab;4595 | 0.1 | 0.05 |
| NC_019843.3;YP_009047202.1;MERS_1ab;4596 | 0 | 0 |
| NC_019843.3;YP_009047202.1;MERS_1ab;4597 | 0 | 0 |
| NC_019843.3;YP_009047202.1;MERS_1ab;4598 | 0 | 0 |
| NC_019843.3;YP_009047202.1;MERS_1ab;4599 | 0 | 0 |
| NC_019843.3;YP_009047202.1;MERS_1ab;4600 | 0 | 0 |
| NC_019843.3;YP_009047202.1;MERS_1ab;4601 | 0 | 0 |
| NC_019843.3;YP_009047202.1;MERS_1ab;4602 | 0 | 0 |
| NC_019843.3;YP_009047202.1;MERS_1ab;4603 | 0 | 0 |
| NC_019843.3;YP_009047202.1;MERS_1ab;4604 | 0 | 0 |
| NC_019843.3;YP_009047202.1;MERS_1ab;4605 | 0 | 0 |
| NC_019843.3;YP_009047202.1;MERS_1ab;4606 | 0 | 0.025 |
| NC_019843.3;YP_009047202.1;MERS_1ab;4607 | 0 | 0 |
| NC_019843.3;YP_009047202.1;MERS_1ab;4608 | 0 | 0 |
| NC_019843.3;YP_009047202.1;MERS_1ab;4609 | 0 | 0 |
| NC_019843.3;YP_009047202.1;MERS_1ab;4610 | 0 | 0 |
| NC_019843.3;YP_009047202.1;MERS_1ab;4611 | 0 | 0 |
| NC_019843.3;YP_009047202.1;MERS_1ab;4612 | 0 | 0 |
| NC_019843.3;YP_009047202.1;MERS_1ab;4613 | 0 | 0 |
| NC_019843.3;YP_009047202.1;MERS_1ab;4614 | 0 | 0 |
| NC_019843.3;YP_009047202.1;MERS_1ab;4615 | 0 | 0 |
| NC_019843.3;YP_009047202.1;MERS_1ab;4616 | 0.05 | 0.025 |
| NC_019843.3;YP_009047202.1;MERS_1ab;4617 | 0 | 0 |
| NC_019843.3;YP_009047202.1;MERS_1ab;4618 | 0 | 0 |
| NC_019843.3;YP_009047202.1;MERS_1ab;4619 | 0 | 0 |
| NC_019843.3;YP_009047202.1;MERS_1ab;4620 | 0 | 0 |
| NC_019843.3;YP_009047202.1;MERS_1ab;4621 | 0 | 0 |
| NC_019843.3;YP_009047202.1;MERS_1ab;4622 | 0 | 0.025 |
| NC_019843.3;YP_009047202.1;MERS_1ab;4623 | 0 | 0 |
| NC_019843.3;YP_009047202.1;MERS_1ab;4624 | 0 | 0 |
| NC_019843.3;YP_009047202.1;MERS_1ab;4625 | 0 | 0 |
| NC_019843.3;YP_009047202.1;MERS_1ab;4626 | 0 | 0.025 |
| NC_019843.3;YP_009047202.1;MERS_1ab;4627 | 0 | 0.025 |
| NC_019843.3;YP_009047202.1;MERS_1ab;4628 | 0 | 0 |
| NC_019843.3;YP_009047202.1;MERS_1ab;4629 | 0 | 0 |
| NC_019843.3;YP_009047202.1;MERS_1ab;4630 | 0 | 0.025 |
| NC_019843.3;YP_009047202.1;MERS_1ab;4631 | 0 | 0.025 |
| NC_019843.3;YP_009047202.1;MERS_1ab;4632 | 0 | 0.075 |
| NC_019843.3;YP_009047202.1;MERS_1ab;4633 | 0 | 0.025 |
| NC_019843.3;YP_009047202.1;MERS_1ab;4634 | 0 | 0.05 |
| NC_019843.3;YP_009047202.1;MERS_1ab;4635 | 0 | 0.05 |

| | | |
|---|---|---|
| NC_019843.3;YP_009047202.1;MERS_1ab;4636 | 0 | 0.05 |
| NC_019843.3;YP_009047202.1;MERS_1ab;4637 | 0 | 0.05 |
| NC_019843.3;YP_009047202.1;MERS_1ab;4638 | 0.05 | 0.025 |
| NC_019843.3;YP_009047202.1;MERS_1ab;4639 | 0.05 | 0.05 |
| NC_019843.3;YP_009047202.1;MERS_1ab;4640 | 0 | 0.025 |
| NC_019843.3;YP_009047202.1;MERS_1ab;4641 | 0 | 0.025 |
| NC_019843.3;YP_009047202.1;MERS_1ab;4642 | 0.1 | 0.05 |
| NC_019843.3;YP_009047202.1;MERS_1ab;4643 | 0.1 | 0.025 |
| NC_019843.3;YP_009047202.1;MERS_1ab;4644 | 0.1 | 0.025 |
| NC_019843.3;YP_009047202.1;MERS_1ab;4645 | 0.15 | 0.025 |
| NC_019843.3;YP_009047202.1;MERS_1ab;4646 | 0.05 | 0 |
| NC_019843.3;YP_009047202.1;MERS_1ab;4647 | 0.05 | 0 |
| NC_019843.3;YP_009047202.1;MERS_1ab;4648 | 0.05 | 0 |
| NC_019843.3;YP_009047202.1;MERS_1ab;4649 | 0.05 | 0.025 |
| NC_019843.3;YP_009047202.1;MERS_1ab;4650 | 0.05 | 0.025 |
| NC_019843.3;YP_009047202.1;MERS_1ab;4651 | 0 | 0 |
| NC_019843.3;YP_009047202.1;MERS_1ab;4652 | 0 | 0 |
| NC_019843.3;YP_009047202.1;MERS_1ab;4653 | 0 | 0 |
| NC_019843.3;YP_009047202.1;MERS_1ab;4654 | 0 | 0 |
| NC_019843.3;YP_009047202.1;MERS_1ab;4655 | 0 | 0 |
| NC_019843.3;YP_009047202.1;MERS_1ab;4656 | 0 | 0 |
| NC_019843.3;YP_009047202.1;MERS_1ab;4657 | 0 | 0 |
| NC_019843.3;YP_009047202.1;MERS_1ab;4658 | 0 | 0.025 |
| NC_019843.3;YP_009047202.1;MERS_1ab;4659 | 0 | 0 |
| NC_019843.3;YP_009047202.1;MERS_1ab;4660 | 0 | 0 |
| NC_019843.3;YP_009047202.1;MERS_1ab;4661 | 0 | 0.05 |
| NC_019843.3;YP_009047202.1;MERS_1ab;4662 | 0 | 0.025 |
| NC_019843.3;YP_009047202.1;MERS_1ab;4663 | 0 | 0 |
| NC_019843.3;YP_009047202.1;MERS_1ab;4664 | 0 | 0 |
| NC_019843.3;YP_009047202.1;MERS_1ab;4665 | 0 | 0.05 |
| NC_019843.3;YP_009047202.1;MERS_1ab;4666 | 0.05 | 0.025 |
| NC_019843.3;YP_009047202.1;MERS_1ab;4667 | 0 | 0.025 |
| NC_019843.3;YP_009047202.1;MERS_1ab;4668 | 0 | 0 |
| NC_019843.3;YP_009047202.1;MERS_1ab;4669 | 0 | 0 |
| NC_019843.3;YP_009047202.1;MERS_1ab;4670 | 0 | 0 |
| NC_019843.3;YP_009047202.1;MERS_1ab;4671 | 0 | 0 |
| NC_019843.3;YP_009047202.1;MERS_1ab;4672 | 0 | 0 |
| NC_019843.3;YP_009047202.1;MERS_1ab;4673 | 0 | 0 |
| NC_019843.3;YP_009047202.1;MERS_1ab;4674 | 0 | 0 |
| NC_019843.3;YP_009047202.1;MERS_1ab;4675 | 0 | 0 |
| NC_019843.3;YP_009047202.1;MERS_1ab;4676 | 0 | 0 |
| NC_019843.3;YP_009047202.1;MERS_1ab;4677 | 0 | 0 |
| NC_019843.3;YP_009047202.1;MERS_1ab;4678 | 0 | 0 |

| | | |
|---|---|---|
| NC_019843.3;YP_009047202.1;MERS_1ab;4679 | 0 | 0 |
| NC_019843.3;YP_009047202.1;MERS_1ab;4680 | 0 | 0 |
| NC_019843.3;YP_009047202.1;MERS_1ab;4681 | 0 | 0 |
| NC_019843.3;YP_009047202.1;MERS_1ab;4682 | 0 | 0 |
| NC_019843.3;YP_009047202.1;MERS_1ab;4683 | 0 | 0 |
| NC_019843.3;YP_009047202.1;MERS_1ab;4684 | 0 | 0 |
| NC_019843.3;YP_009047202.1;MERS_1ab;4685 | 0 | 0 |
| NC_019843.3;YP_009047202.1;MERS_1ab;4686 | 0 | 0 |
| NC_019843.3;YP_009047202.1;MERS_1ab;4687 | 0 | 0 |
| NC_019843.3;YP_009047202.1;MERS_1ab;4688 | 0 | 0 |
| NC_019843.3;YP_009047202.1;MERS_1ab;4689 | 0 | 0 |
| NC_019843.3;YP_009047202.1;MERS_1ab;4690 | 0 | 0 |
| NC_019843.3;YP_009047202.1;MERS_1ab;4691 | 0 | 0 |
| NC_019843.3;YP_009047202.1;MERS_1ab;4692 | 0 | 0 |
| NC_019843.3;YP_009047202.1;MERS_1ab;4693 | 0 | 0 |
| NC_019843.3;YP_009047202.1;MERS_1ab;4694 | 0 | 0 |
| NC_019843.3;YP_009047202.1;MERS_1ab;4695 | 0 | 0 |
| NC_019843.3;YP_009047202.1;MERS_1ab;4696 | 0 | 0 |
| NC_019843.3;YP_009047202.1;MERS_1ab;4697 | 0 | 0 |
| NC_019843.3;YP_009047202.1;MERS_1ab;4698 | 0 | 0 |
| NC_019843.3;YP_009047202.1;MERS_1ab;4699 | 0 | 0 |
| NC_019843.3;YP_009047202.1;MERS_1ab;4700 | 0 | 0 |
| NC_019843.3;YP_009047202.1;MERS_1ab;4701 | 0 | 0 |
| NC_019843.3;YP_009047202.1;MERS_1ab;4702 | 0 | 0 |
| NC_019843.3;YP_009047202.1;MERS_1ab;4703 | 0 | 0 |
| NC_019843.3;YP_009047202.1;MERS_1ab;4704 | 0 | 0 |
| NC_019843.3;YP_009047202.1;MERS_1ab;4705 | 0 | 0 |
| NC_019843.3;YP_009047202.1;MERS_1ab;4706 | 0 | 0 |
| NC_019843.3;YP_009047202.1;MERS_1ab;4707 | 0 | 0 |
| NC_019843.3;YP_009047202.1;MERS_1ab;4708 | 0 | 0 |
| NC_019843.3;YP_009047202.1;MERS_1ab;4709 | 0 | 0 |
| NC_019843.3;YP_009047202.1;MERS_1ab;4710 | 0 | 0 |
| NC_019843.3;YP_009047202.1;MERS_1ab;4711 | 0 | 0 |
| NC_019843.3;YP_009047202.1;MERS_1ab;4712 | 0 | 0 |
| NC_019843.3;YP_009047202.1;MERS_1ab;4713 | 0 | 0 |
| NC_019843.3;YP_009047202.1;MERS_1ab;4714 | 0 | 0 |
| NC_019843.3;YP_009047202.1;MERS_1ab;4715 | 0 | 0 |
| NC_019843.3;YP_009047202.1;MERS_1ab;4716 | 0 | 0 |
| NC_019843.3;YP_009047202.1;MERS_1ab;4717 | 0 | 0 |
| NC_019843.3;YP_009047202.1;MERS_1ab;4718 | 0 | 0 |
| NC_019843.3;YP_009047202.1;MERS_1ab;4719 | 0 | 0 |
| NC_019843.3;YP_009047202.1;MERS_1ab;4720 | 0 | 0 |
| NC_019843.3;YP_009047202.1;MERS_1ab;4721 | 0 | 0 |

| | | |
|---|---|---|
| NC_019843.3;YP_009047202.1;MERS_1ab;4722 | 0 | 0 |
| NC_019843.3;YP_009047202.1;MERS_1ab;4723 | 0 | 0 |
| NC_019843.3;YP_009047202.1;MERS_1ab;4724 | 0 | 0 |
| NC_019843.3;YP_009047202.1;MERS_1ab;4725 | 0.05 | 0 |
| NC_019843.3;YP_009047202.1;MERS_1ab;4726 | 0 | 0 |
| NC_019843.3;YP_009047202.1;MERS_1ab;4727 | 0 | 0 |
| NC_019843.3;YP_009047202.1;MERS_1ab;4728 | 0 | 0 |
| NC_019843.3;YP_009047202.1;MERS_1ab;4729 | 0 | 0 |
| NC_019843.3;YP_009047202.1;MERS_1ab;4730 | 0.05 | 0 |
| NC_019843.3;YP_009047202.1;MERS_1ab;4731 | 0.05 | 0 |
| NC_019843.3;YP_009047202.1;MERS_1ab;4732 | 0 | 0 |
| NC_019843.3;YP_009047202.1;MERS_1ab;4733 | 0.05 | 0 |
| NC_019843.3;YP_009047202.1;MERS_1ab;4734 | 0.1 | 0 |
| NC_019843.3;YP_009047202.1;MERS_1ab;4735 | 0.1 | 0 |
| NC_019843.3;YP_009047202.1;MERS_1ab;4736 | 0.15 | 0 |
| NC_019843.3;YP_009047202.1;MERS_1ab;4737 | 0.1 | 0 |
| NC_019843.3;YP_009047202.1;MERS_1ab;4738 | 0.1 | 0 |
| NC_019843.3;YP_009047202.1;MERS_1ab;4739 | 0.15 | 0 |
| NC_019843.3;YP_009047202.1;MERS_1ab;4740 | 0.15 | 0 |
| NC_019843.3;YP_009047202.1;MERS_1ab;4741 | 0.1 | 0 |
| NC_019843.3;YP_009047202.1;MERS_1ab;4742 | 0.1 | 0 |
| NC_019843.3;YP_009047202.1;MERS_1ab;4743 | 0.1 | 0.025 |
| NC_019843.3;YP_009047202.1;MERS_1ab;4744 | 0.1 | 0.075 |
| NC_019843.3;YP_009047202.1;MERS_1ab;4745 | 0 | 0 |
| NC_019843.3;YP_009047202.1;MERS_1ab;4746 | 0 | 0 |
| NC_019843.3;YP_009047202.1;MERS_1ab;4747 | 0 | 0 |
| NC_019843.3;YP_009047202.1;MERS_1ab;4748 | 0 | 0 |
| NC_019843.3;YP_009047202.1;MERS_1ab;4749 | 0.05 | 0 |
| NC_019843.3;YP_009047202.1;MERS_1ab;4750 | 0.05 | 0 |
| NC_019843.3;YP_009047202.1;MERS_1ab;4751 | 0.05 | 0.025 |
| NC_019843.3;YP_009047202.1;MERS_1ab;4752 | 0.05 | 0.05 |
| NC_019843.3;YP_009047202.1;MERS_1ab;4753 | 0.1 | 0.075 |
| NC_019843.3;YP_009047202.1;MERS_1ab;4754 | 0.1 | 0.05 |
| NC_019843.3;YP_009047202.1;MERS_1ab;4755 | 0.1 | 0.05 |
| NC_019843.3;YP_009047202.1;MERS_1ab;4756 | 0.05 | 0.05 |
| NC_019843.3;YP_009047202.1;MERS_1ab;4757 | 0.05 | 0.05 |
| NC_019843.3;YP_009047202.1;MERS_1ab;4758 | 0.05 | 0.025 |
| NC_019843.3;YP_009047202.1;MERS_1ab;4759 | 0.05 | 0.025 |
| NC_019843.3;YP_009047202.1;MERS_1ab;4760 | 0.05 | 0 |
| NC_019843.3;YP_009047202.1;MERS_1ab;4761 | 0 | 0 |
| NC_019843.3;YP_009047202.1;MERS_1ab;4762 | 0 | 0 |
| NC_019843.3;YP_009047202.1;MERS_1ab;4763 | 0.05 | 0 |
| NC_019843.3;YP_009047202.1;MERS_1ab;4764 | 0 | 0 |

| | | |
|---|---|---|
| NC_019843.3;YP_009047202.1;MERS_1ab;4765 | 0 | 0 |
| NC_019843.3;YP_009047202.1;MERS_1ab;4766 | 0 | 0 |
| NC_019843.3;YP_009047202.1;MERS_1ab;4767 | 0 | 0 |
| NC_019843.3;YP_009047202.1;MERS_1ab;4768 | 0 | 0 |
| NC_019843.3;YP_009047202.1;MERS_1ab;4769 | 0 | 0 |
| NC_019843.3;YP_009047202.1;MERS_1ab;4770 | 0 | 0 |
| NC_019843.3;YP_009047202.1;MERS_1ab;4771 | 0 | 0 |
| NC_019843.3;YP_009047202.1;MERS_1ab;4772 | 0 | 0 |
| NC_019843.3;YP_009047202.1;MERS_1ab;4773 | 0 | 0 |
| NC_019843.3;YP_009047202.1;MERS_1ab;4774 | 0 | 0 |
| NC_019843.3;YP_009047202.1;MERS_1ab;4775 | 0 | 0 |
| NC_019843.3;YP_009047202.1;MERS_1ab;4776 | 0 | 0 |
| NC_019843.3;YP_009047202.1;MERS_1ab;4777 | 0 | 0 |
| NC_019843.3;YP_009047202.1;MERS_1ab;4778 | 0.05 | 0 |
| NC_019843.3;YP_009047202.1;MERS_1ab;4779 | 0 | 0 |
| NC_019843.3;YP_009047202.1;MERS_1ab;4780 | 0 | 0 |
| NC_019843.3;YP_009047202.1;MERS_1ab;4781 | 0.05 | 0 |
| NC_019843.3;YP_009047202.1;MERS_1ab;4782 | 0.05 | 0 |
| NC_019843.3;YP_009047202.1;MERS_1ab;4783 | 0 | 0 |
| NC_019843.3;YP_009047202.1;MERS_1ab;4784 | 0.05 | 0 |
| NC_019843.3;YP_009047202.1;MERS_1ab;4785 | 0.05 | 0 |
| NC_019843.3;YP_009047202.1;MERS_1ab;4786 | 0.2 | 0.025 |
| NC_019843.3;YP_009047202.1;MERS_1ab;4787 | 0.05 | 0 |
| NC_019843.3;YP_009047202.1;MERS_1ab;4788 | 0.05 | 0 |
| NC_019843.3;YP_009047202.1;MERS_1ab;4789 | 0.2 | 0.025 |
| NC_019843.3;YP_009047202.1;MERS_1ab;4790 | 0.15 | 0.025 |
| NC_019843.3;YP_009047202.1;MERS_1ab;4791 | 0 | 0 |
| NC_019843.3;YP_009047202.1;MERS_1ab;4792 | 0 | 0 |
| NC_019843.3;YP_009047202.1;MERS_1ab;4793 | 0 | 0 |
| NC_019843.3;YP_009047202.1;MERS_1ab;4794 | 0 | 0 |
| NC_019843.3;YP_009047202.1;MERS_1ab;4795 | 0.05 | 0 |
| NC_019843.3;YP_009047202.1;MERS_1ab;4796 | 0 | 0 |
| NC_019843.3;YP_009047202.1;MERS_1ab;4797 | 0 | 0 |
| NC_019843.3;YP_009047202.1;MERS_1ab;4798 | 0 | 0 |
| NC_019843.3;YP_009047202.1;MERS_1ab;4799 | 0 | 0 |
| NC_019843.3;YP_009047202.1;MERS_1ab;4800 | 0 | 0 |
| NC_019843.3;YP_009047202.1;MERS_1ab;4801 | 0 | 0 |
| NC_019843.3;YP_009047202.1;MERS_1ab;4802 | 0 | 0 |
| NC_019843.3;YP_009047202.1;MERS_1ab;4803 | 0 | 0 |
| NC_019843.3;YP_009047202.1;MERS_1ab;4804 | 0 | 0 |
| NC_019843.3;YP_009047202.1;MERS_1ab;4805 | 0 | 0 |
| NC_019843.3;YP_009047202.1;MERS_1ab;4806 | 0 | 0 |
| NC_019843.3;YP_009047202.1;MERS_1ab;4807 | 0 | 0 |

| | | |
|---|---|---|
| NC_019843.3;YP_009047202.1;MERS_1ab;4808 | 0 | 0 |
| NC_019843.3;YP_009047202.1;MERS_1ab;4809 | 0 | 0 |
| NC_019843.3;YP_009047202.1;MERS_1ab;4810 | 0 | 0 |
| NC_019843.3;YP_009047202.1;MERS_1ab;4811 | 0 | 0 |
| NC_019843.3;YP_009047202.1;MERS_1ab;4812 | 0.05 | 0 |
| NC_019843.3;YP_009047202.1;MERS_1ab;4813 | 0 | 0 |
| NC_019843.3;YP_009047202.1;MERS_1ab;4814 | 0 | 0 |
| NC_019843.3;YP_009047202.1;MERS_1ab;4815 | 0 | 0 |
| NC_019843.3;YP_009047202.1;MERS_1ab;4816 | 0 | 0 |
| NC_019843.3;YP_009047202.1;MERS_1ab;4817 | 0 | 0 |
| NC_019843.3;YP_009047202.1;MERS_1ab;4818 | 0 | 0 |
| NC_019843.3;YP_009047202.1;MERS_1ab;4819 | 0.05 | 0 |
| NC_019843.3;YP_009047202.1;MERS_1ab;4820 | 0 | 0 |
| NC_019843.3;YP_009047202.1;MERS_1ab;4821 | 0 | 0 |
| NC_019843.3;YP_009047202.1;MERS_1ab;4822 | 0.05 | 0 |
| NC_019843.3;YP_009047202.1;MERS_1ab;4823 | 0 | 0 |
| NC_019843.3;YP_009047202.1;MERS_1ab;4824 | 0.05 | 0 |
| NC_019843.3;YP_009047202.1;MERS_1ab;4825 | 0 | 0 |
| NC_019843.3;YP_009047202.1;MERS_1ab;4826 | 0.1 | 0.05 |
| NC_019843.3;YP_009047202.1;MERS_1ab;4827 | 0.1 | 0.05 |
| NC_019843.3;YP_009047202.1;MERS_1ab;4828 | 0.05 | 0 |
| NC_019843.3;YP_009047202.1;MERS_1ab;4829 | 0.05 | 0 |
| NC_019843.3;YP_009047202.1;MERS_1ab;4830 | 0.1 | 0 |
| NC_019843.3;YP_009047202.1;MERS_1ab;4831 | 0 | 0 |
| NC_019843.3;YP_009047202.1;MERS_1ab;4832 | 0 | 0 |
| NC_019843.3;YP_009047202.1;MERS_1ab;4833 | 0 | 0 |
| NC_019843.3;YP_009047202.1;MERS_1ab;4834 | 0 | 0 |
| NC_019843.3;YP_009047202.1;MERS_1ab;4835 | 0 | 0 |
| NC_019843.3;YP_009047202.1;MERS_1ab;4836 | 0 | 0 |
| NC_019843.3;YP_009047202.1;MERS_1ab;4837 | 0 | 0 |
| NC_019843.3;YP_009047202.1;MERS_1ab;4838 | 0 | 0 |
| NC_019843.3;YP_009047202.1;MERS_1ab;4839 | 0 | 0 |
| NC_019843.3;YP_009047202.1;MERS_1ab;4840 | 0 | 0 |
| NC_019843.3;YP_009047202.1;MERS_1ab;4841 | 0 | 0.025 |
| NC_019843.3;YP_009047202.1;MERS_1ab;4842 | 0.05 | 0 |
| NC_019843.3;YP_009047202.1;MERS_1ab;4843 | 0 | 0 |
| NC_019843.3;YP_009047202.1;MERS_1ab;4844 | 0 | 0 |
| NC_019843.3;YP_009047202.1;MERS_1ab;4845 | 0 | 0 |
| NC_019843.3;YP_009047202.1;MERS_1ab;4846 | 0 | 0 |
| NC_019843.3;YP_009047202.1;MERS_1ab;4847 | 0 | 0 |
| NC_019843.3;YP_009047202.1;MERS_1ab;4848 | 0 | 0 |
| NC_019843.3;YP_009047202.1;MERS_1ab;4849 | 0 | 0 |
| NC_019843.3;YP_009047202.1;MERS_1ab;4850 | 0 | 0 |

| | | |
|---|---|---|
| NC_019843.3;YP_009047202.1;MERS_1ab;4851 | 0 | 0 |
| NC_019843.3;YP_009047202.1;MERS_1ab;4852 | 0 | 0 |
| NC_019843.3;YP_009047202.1;MERS_1ab;4853 | 0 | 0 |
| NC_019843.3;YP_009047202.1;MERS_1ab;4854 | 0 | 0.025 |
| NC_019843.3;YP_009047202.1;MERS_1ab;4855 | 0 | 0 |
| NC_019843.3;YP_009047202.1;MERS_1ab;4856 | 0 | 0.05 |
| NC_019843.3;YP_009047202.1;MERS_1ab;4857 | 0 | 0.025 |
| NC_019843.3;YP_009047202.1;MERS_1ab;4858 | 0 | 0.025 |
| NC_019843.3;YP_009047202.1;MERS_1ab;4859 | 0 | 0 |
| NC_019843.3;YP_009047202.1;MERS_1ab;4860 | 0 | 0 |
| NC_019843.3;YP_009047202.1;MERS_1ab;4861 | 0 | 0.025 |
| NC_019843.3;YP_009047202.1;MERS_1ab;4862 | 0 | 0.025 |
| NC_019843.3;YP_009047202.1;MERS_1ab;4863 | 0 | 0.05 |
| NC_019843.3;YP_009047202.1;MERS_1ab;4864 | 0 | 0.05 |
| NC_019843.3;YP_009047202.1;MERS_1ab;4865 | 0.05 | 0.05 |
| NC_019843.3;YP_009047202.1;MERS_1ab;4866 | 0 | 0.05 |
| NC_019843.3;YP_009047202.1;MERS_1ab;4867 | 0 | 0.025 |
| NC_019843.3;YP_009047202.1;MERS_1ab;4868 | 0.05 | 0.075 |
| NC_019843.3;YP_009047202.1;MERS_1ab;4869 | 0 | 0.05 |
| NC_019843.3;YP_009047202.1;MERS_1ab;4870 | 0 | 0.025 |
| NC_019843.3;YP_009047202.1;MERS_1ab;4871 | 0.05 | 0.025 |
| NC_019843.3;YP_009047202.1;MERS_1ab;4872 | 0 | 0.025 |
| NC_019843.3;YP_009047202.1;MERS_1ab;4873 | 0 | 0 |
| NC_019843.3;YP_009047202.1;MERS_1ab;4874 | 0 | 0 |
| NC_019843.3;YP_009047202.1;MERS_1ab;4875 | 0 | 0 |
| NC_019843.3;YP_009047202.1;MERS_1ab;4876 | 0 | 0 |
| NC_019843.3;YP_009047202.1;MERS_1ab;4877 | 0 | 0.025 |
| NC_019843.3;YP_009047202.1;MERS_1ab;4878 | 0 | 0 |
| NC_019843.3;YP_009047202.1;MERS_1ab;4879 | 0 | 0 |
| NC_019843.3;YP_009047202.1;MERS_1ab;4880 | 0 | 0 |
| NC_019843.3;YP_009047202.1;MERS_1ab;4881 | 0 | 0 |
| NC_019843.3;YP_009047202.1;MERS_1ab;4882 | 0 | 0 |
| NC_019843.3;YP_009047202.1;MERS_1ab;4883 | 0 | 0 |
| NC_019843.3;YP_009047202.1;MERS_1ab;4884 | 0 | 0 |
| NC_019843.3;YP_009047202.1;MERS_1ab;4885 | 0 | 0 |
| NC_019843.3;YP_009047202.1;MERS_1ab;4886 | 0 | 0 |
| NC_019843.3;YP_009047202.1;MERS_1ab;4887 | 0 | 0 |
| NC_019843.3;YP_009047202.1;MERS_1ab;4888 | 0 | 0 |
| NC_019843.3;YP_009047202.1;MERS_1ab;4889 | 0 | 0.025 |
| NC_019843.3;YP_009047202.1;MERS_1ab;4890 | 0 | 0 |
| NC_019843.3;YP_009047202.1;MERS_1ab;4891 | 0 | 0.025 |
| NC_019843.3;YP_009047202.1;MERS_1ab;4892 | 0 | 0 |
| NC_019843.3;YP_009047202.1;MERS_1ab;4893 | 0 | 0 |

| | | |
|---|---|---|
| NC_019843.3;YP_009047202.1;MERS_1ab;4894 | 0 | 0 |
| NC_019843.3;YP_009047202.1;MERS_1ab;4895 | 0.05 | 0.025 |
| NC_019843.3;YP_009047202.1;MERS_1ab;4896 | 0 | 0 |
| NC_019843.3;YP_009047202.1;MERS_1ab;4897 | 0 | 0 |
| NC_019843.3;YP_009047202.1;MERS_1ab;4898 | 0 | 0 |
| NC_019843.3;YP_009047202.1;MERS_1ab;4899 | 0 | 0 |
| NC_019843.3;YP_009047202.1;MERS_1ab;4900 | 0 | 0 |
| NC_019843.3;YP_009047202.1;MERS_1ab;4901 | 0 | 0.025 |
| NC_019843.3;YP_009047202.1;MERS_1ab;4902 | 0.05 | 0.025 |
| NC_019843.3;YP_009047202.1;MERS_1ab;4903 | 0.05 | 0.025 |
| NC_019843.3;YP_009047202.1;MERS_1ab;4904 | 0.05 | 0 |
| NC_019843.3;YP_009047202.1;MERS_1ab;4905 | 0 | 0 |
| NC_019843.3;YP_009047202.1;MERS_1ab;4906 | 0 | 0 |
| NC_019843.3;YP_009047202.1;MERS_1ab;4907 | 0 | 0 |
| NC_019843.3;YP_009047202.1;MERS_1ab;4908 | 0 | 0 |
| NC_019843.3;YP_009047202.1;MERS_1ab;4909 | 0.05 | 0.05 |
| NC_019843.3;YP_009047202.1;MERS_1ab;4910 | 0.1 | 0.15 |
| NC_019843.3;YP_009047202.1;MERS_1ab;4911 | 0 | 0 |
| NC_019843.3;YP_009047202.1;MERS_1ab;4912 | 0 | 0 |
| NC_019843.3;YP_009047202.1;MERS_1ab;4913 | 0 | 0 |
| NC_019843.3;YP_009047202.1;MERS_1ab;4914 | 0 | 0.025 |
| NC_019843.3;YP_009047202.1;MERS_1ab;4915 | 0 | 0.025 |
| NC_019843.3;YP_009047202.1;MERS_1ab;4916 | 0 | 0.025 |
| NC_019843.3;YP_009047202.1;MERS_1ab;4917 | 0 | 0.025 |
| NC_019843.3;YP_009047202.1;MERS_1ab;4918 | 0 | 0.025 |
| NC_019843.3;YP_009047202.1;MERS_1ab;4919 | 0 | 0.025 |
| NC_019843.3;YP_009047202.1;MERS_1ab;4920 | 0 | 0 |
| NC_019843.3;YP_009047202.1;MERS_1ab;4921 | 0 | 0 |
| NC_019843.3;YP_009047202.1;MERS_1ab;4922 | 0 | 0 |
| NC_019843.3;YP_009047202.1;MERS_1ab;4923 | 0 | 0 |
| NC_019843.3;YP_009047202.1;MERS_1ab;4924 | 0 | 0 |
| NC_019843.3;YP_009047202.1;MERS_1ab;4925 | 0 | 0 |
| NC_019843.3;YP_009047202.1;MERS_1ab;4926 | 0 | 0 |
| NC_019843.3;YP_009047202.1;MERS_1ab;4927 | 0 | 0 |
| NC_019843.3;YP_009047202.1;MERS_1ab;4928 | 0 | 0 |
| NC_019843.3;YP_009047202.1;MERS_1ab;4929 | 0 | 0 |
| NC_019843.3;YP_009047202.1;MERS_1ab;4930 | 0 | 0 |
| NC_019843.3;YP_009047202.1;MERS_1ab;4931 | 0 | 0 |
| NC_019843.3;YP_009047202.1;MERS_1ab;4932 | 0 | 0 |
| NC_019843.3;YP_009047202.1;MERS_1ab;4933 | 0 | 0 |
| NC_019843.3;YP_009047202.1;MERS_1ab;4934 | 0 | 0 |
| NC_019843.3;YP_009047202.1;MERS_1ab;4935 | 0 | 0 |
| NC_019843.3;YP_009047202.1;MERS_1ab;4936 | 0 | 0 |

| | | |
|---|---|---|
| NC_019843.3;YP_009047202.1;MERS_1ab;4937 | 0 | 0 |
| NC_019843.3;YP_009047202.1;MERS_1ab;4938 | 0 | 0 |
| NC_019843.3;YP_009047202.1;MERS_1ab;4939 | 0 | 0 |
| NC_019843.3;YP_009047202.1;MERS_1ab;4940 | 0 | 0 |
| NC_019843.3;YP_009047202.1;MERS_1ab;4941 | 0 | 0 |
| NC_019843.3;YP_009047202.1;MERS_1ab;4942 | 0 | 0 |
| NC_019843.3;YP_009047202.1;MERS_1ab;4943 | 0 | 0 |
| NC_019843.3;YP_009047202.1;MERS_1ab;4944 | 0.05 | 0 |
| NC_019843.3;YP_009047202.1;MERS_1ab;4945 | 0 | 0 |
| NC_019843.3;YP_009047202.1;MERS_1ab;4946 | 0 | 0 |
| NC_019843.3;YP_009047202.1;MERS_1ab;4947 | 0 | 0 |
| NC_019843.3;YP_009047202.1;MERS_1ab;4948 | 0 | 0 |
| NC_019843.3;YP_009047202.1;MERS_1ab;4949 | 0.4 | 0.35 |
| NC_019843.3;YP_009047202.1;MERS_1ab;4950 | 0.35 | 0.175 |
| NC_019843.3;YP_009047202.1;MERS_1ab;4951 | 0.25 | 0.2 |
| NC_019843.3;YP_009047202.1;MERS_1ab;4952 | 0.2 | 0.025 |
| NC_019843.3;YP_009047202.1;MERS_1ab;4953 | 0.25 | 0.025 |
| NC_019843.3;YP_009047202.1;MERS_1ab;4954 | 0.25 | 0.025 |
| NC_019843.3;YP_009047202.1;MERS_1ab;4955 | 0.2 | 0.025 |
| NC_019843.3;YP_009047202.1;MERS_1ab;4956 | 0.2 | 0.025 |
| NC_019843.3;YP_009047202.1;MERS_1ab;4957 | 0.1 | 0.025 |
| NC_019843.3;YP_009047202.1;MERS_1ab;4958 | 0.05 | 0 |
| NC_019843.3;YP_009047202.1;MERS_1ab;4959 | 0 | 0 |
| NC_019843.3;YP_009047202.1;MERS_1ab;4960 | 0 | 0 |
| NC_019843.3;YP_009047202.1;MERS_1ab;4961 | 0 | 0 |
| NC_019843.3;YP_009047202.1;MERS_1ab;4962 | 0 | 0 |
| NC_019843.3;YP_009047202.1;MERS_1ab;4963 | 0 | 0 |
| NC_019843.3;YP_009047202.1;MERS_1ab;4964 | 0 | 0 |
| NC_019843.3;YP_009047202.1;MERS_1ab;4965 | 0 | 0 |
| NC_019843.3;YP_009047202.1;MERS_1ab;4966 | 0 | 0 |
| NC_019843.3;YP_009047202.1;MERS_1ab;4967 | 0 | 0 |
| NC_019843.3;YP_009047202.1;MERS_1ab;4968 | 0 | 0 |
| NC_019843.3;YP_009047202.1;MERS_1ab;4969 | 0 | 0 |
| NC_019843.3;YP_009047202.1;MERS_1ab;4970 | 0 | 0 |
| NC_019843.3;YP_009047202.1;MERS_1ab;4971 | 0 | 0 |
| NC_019843.3;YP_009047202.1;MERS_1ab;4972 | 0 | 0 |
| NC_019843.3;YP_009047202.1;MERS_1ab;4973 | 0 | 0.025 |
| NC_019843.3;YP_009047202.1;MERS_1ab;4974 | 0 | 0.025 |
| NC_019843.3;YP_009047202.1;MERS_1ab;4975 | 0.05 | 0.125 |
| NC_019843.3;YP_009047202.1;MERS_1ab;4976 | 0 | 0.025 |
| NC_019843.3;YP_009047202.1;MERS_1ab;4977 | 0 | 0.025 |
| NC_019843.3;YP_009047202.1;MERS_1ab;4978 | 0 | 0.025 |
| NC_019843.3;YP_009047202.1;MERS_1ab;4979 | 0 | 0 |

| | | |
|---|---|---|
| NC_019843.3;YP_009047202.1;MERS_1ab;4980 | 0 | 0 |
| NC_019843.3;YP_009047202.1;MERS_1ab;4981 | 0 | 0.025 |
| NC_019843.3;YP_009047202.1;MERS_1ab;4982 | 0 | 0 |
| NC_019843.3;YP_009047202.1;MERS_1ab;4983 | 0 | 0.025 |
| NC_019843.3;YP_009047202.1;MERS_1ab;4984 | 0 | 0.075 |
| NC_019843.3;YP_009047202.1;MERS_1ab;4985 | 0.05 | 0.05 |
| NC_019843.3;YP_009047202.1;MERS_1ab;4986 | 0 | 0.025 |
| NC_019843.3;YP_009047202.1;MERS_1ab;4987 | 0 | 0.025 |
| NC_019843.3;YP_009047202.1;MERS_1ab;4988 | 0 | 0.025 |
| NC_019843.3;YP_009047202.1;MERS_1ab;4989 | 0 | 0.025 |
| NC_019843.3;YP_009047202.1;MERS_1ab;4990 | 0 | 0.025 |
| NC_019843.3;YP_009047202.1;MERS_1ab;4991 | 0 | 0.025 |
| NC_019843.3;YP_009047202.1;MERS_1ab;4992 | 0 | 0.025 |
| NC_019843.3;YP_009047202.1;MERS_1ab;4993 | 0 | 0.025 |
| NC_019843.3;YP_009047202.1;MERS_1ab;4994 | 0 | 0.025 |
| NC_019843.3;YP_009047202.1;MERS_1ab;4995 | 0 | 0.025 |
| NC_019843.3;YP_009047202.1;MERS_1ab;4996 | 0 | 0 |
| NC_019843.3;YP_009047202.1;MERS_1ab;4997 | 0 | 0 |
| NC_019843.3;YP_009047202.1;MERS_1ab;4998 | 0 | 0 |
| NC_019843.3;YP_009047202.1;MERS_1ab;4999 | 0 | 0 |
| NC_019843.3;YP_009047202.1;MERS_1ab;5000 | 0 | 0 |
| NC_019843.3;YP_009047202.1;MERS_1ab;5001 | 0 | 0 |
| NC_019843.3;YP_009047202.1;MERS_1ab;5002 | 0 | 0 |
| NC_019843.3;YP_009047202.1;MERS_1ab;5003 | 0 | 0 |
| NC_019843.3;YP_009047202.1;MERS_1ab;5004 | 0 | 0 |
| NC_019843.3;YP_009047202.1;MERS_1ab;5005 | 0 | 0 |
| NC_019843.3;YP_009047202.1;MERS_1ab;5006 | 0 | 0 |
| NC_019843.3;YP_009047202.1;MERS_1ab;5007 | 0 | 0 |
| NC_019843.3;YP_009047202.1;MERS_1ab;5008 | 0 | 0 |
| NC_019843.3;YP_009047202.1;MERS_1ab;5009 | 0 | 0 |
| NC_019843.3;YP_009047202.1;MERS_1ab;5010 | 0 | 0 |
| NC_019843.3;YP_009047202.1;MERS_1ab;5011 | 0 | 0 |
| NC_019843.3;YP_009047202.1;MERS_1ab;5012 | 0 | 0 |
| NC_019843.3;YP_009047202.1;MERS_1ab;5013 | 0 | 0 |
| NC_019843.3;YP_009047202.1;MERS_1ab;5014 | 0 | 0 |
| NC_019843.3;YP_009047202.1;MERS_1ab;5015 | 0 | 0 |
| NC_019843.3;YP_009047202.1;MERS_1ab;5016 | 0 | 0 |
| NC_019843.3;YP_009047202.1;MERS_1ab;5017 | 0 | 0 |
| NC_019843.3;YP_009047202.1;MERS_1ab;5018 | 0 | 0 |
| NC_019843.3;YP_009047202.1;MERS_1ab;5019 | 0 | 0 |
| NC_019843.3;YP_009047202.1;MERS_1ab;5020 | 0 | 0 |
| NC_019843.3;YP_009047202.1;MERS_1ab;5021 | 0 | 0 |
| NC_019843.3;YP_009047202.1;MERS_1ab;5022 | 0 | 0 |

| | | |
|---|---|---|
| NC_019843.3;YP_009047202.1;MERS_1ab;5023 | 0 | 0 |
| NC_019843.3;YP_009047202.1;MERS_1ab;5024 | 0.05 | 0 |
| NC_019843.3;YP_009047202.1;MERS_1ab;5025 | 0.05 | 0 |
| NC_019843.3;YP_009047202.1;MERS_1ab;5026 | 0.1 | 0.025 |
| NC_019843.3;YP_009047202.1;MERS_1ab;5027 | 0 | 0 |
| NC_019843.3;YP_009047202.1;MERS_1ab;5028 | 0 | 0 |
| NC_019843.3;YP_009047202.1;MERS_1ab;5029 | 0 | 0.025 |
| NC_019843.3;YP_009047202.1;MERS_1ab;5030 | 0 | 0.025 |
| NC_019843.3;YP_009047202.1;MERS_1ab;5031 | 0 | 0.025 |
| NC_019843.3;YP_009047202.1;MERS_1ab;5032 | 0 | 0.025 |
| NC_019843.3;YP_009047202.1;MERS_1ab;5033 | 0 | 0.025 |
| NC_019843.3;YP_009047202.1;MERS_1ab;5034 | 0 | 0.025 |
| NC_019843.3;YP_009047202.1;MERS_1ab;5035 | 0.05 | 0.025 |
| NC_019843.3;YP_009047202.1;MERS_1ab;5036 | 0 | 0.025 |
| NC_019843.3;YP_009047202.1;MERS_1ab;5037 | 0 | 0.025 |
| NC_019843.3;YP_009047202.1;MERS_1ab;5038 | 0 | 0 |
| NC_019843.3;YP_009047202.1;MERS_1ab;5039 | 0 | 0 |
| NC_019843.3;YP_009047202.1;MERS_1ab;5040 | 0 | 0 |
| NC_019843.3;YP_009047202.1;MERS_1ab;5041 | 0 | 0 |
| NC_019843.3;YP_009047202.1;MERS_1ab;5042 | 0.05 | 0 |
| NC_019843.3;YP_009047202.1;MERS_1ab;5043 | 0 | 0 |
| NC_019843.3;YP_009047202.1;MERS_1ab;5044 | 0 | 0 |
| NC_019843.3;YP_009047202.1;MERS_1ab;5045 | 0 | 0 |
| NC_019843.3;YP_009047202.1;MERS_1ab;5046 | 0 | 0 |
| NC_019843.3;YP_009047202.1;MERS_1ab;5047 | 0 | 0.025 |
| NC_019843.3;YP_009047202.1;MERS_1ab;5048 | 0 | 0.025 |
| NC_019843.3;YP_009047202.1;MERS_1ab;5049 | 0 | 0.025 |
| NC_019843.3;YP_009047202.1;MERS_1ab;5050 | 0.05 | 0.025 |
| NC_019843.3;YP_009047202.1;MERS_1ab;5051 | 0 | 0.05 |
| NC_019843.3;YP_009047202.1;MERS_1ab;5052 | 0 | 0.025 |
| NC_019843.3;YP_009047202.1;MERS_1ab;5053 | 0 | 0.025 |
| NC_019843.3;YP_009047202.1;MERS_1ab;5054 | 0 | 0.05 |
| NC_019843.3;YP_009047202.1;MERS_1ab;5055 | 0.05 | 0.025 |
| NC_019843.3;YP_009047202.1;MERS_1ab;5056 | 0.1 | 0.05 |
| NC_019843.3;YP_009047202.1;MERS_1ab;5057 | 0 | 0 |
| NC_019843.3;YP_009047202.1;MERS_1ab;5058 | 0.05 | 0 |
| NC_019843.3;YP_009047202.1;MERS_1ab;5059 | 0 | 0 |
| NC_019843.3;YP_009047202.1;MERS_1ab;5060 | 0 | 0 |
| NC_019843.3;YP_009047202.1;MERS_1ab;5061 | 0 | 0 |
| NC_019843.3;YP_009047202.1;MERS_1ab;5062 | 0 | 0 |
| NC_019843.3;YP_009047202.1;MERS_1ab;5063 | 0 | 0 |
| NC_019843.3;YP_009047202.1;MERS_1ab;5064 | 0 | 0 |
| NC_019843.3;YP_009047202.1;MERS_1ab;5065 | 0 | 0 |

| | | |
|---|---|---|
| NC_019843.3;YP_009047202.1;MERS_1ab;5066 | 0 | 0.05 |
| NC_019843.3;YP_009047202.1;MERS_1ab;5067 | 0 | 0 |
| NC_019843.3;YP_009047202.1;MERS_1ab;5068 | 0 | 0 |
| NC_019843.3;YP_009047202.1;MERS_1ab;5069 | 0 | 0 |
| NC_019843.3;YP_009047202.1;MERS_1ab;5070 | 0 | 0 |
| NC_019843.3;YP_009047202.1;MERS_1ab;5071 | 0 | 0 |
| NC_019843.3;YP_009047202.1;MERS_1ab;5072 | 0.05 | 0 |
| NC_019843.3;YP_009047202.1;MERS_1ab;5073 | 0.05 | 0 |
| NC_019843.3;YP_009047202.1;MERS_1ab;5074 | 0 | 0 |
| NC_019843.3;YP_009047202.1;MERS_1ab;5075 | 0 | 0 |
| NC_019843.3;YP_009047202.1;MERS_1ab;5076 | 0 | 0.025 |
| NC_019843.3;YP_009047202.1;MERS_1ab;5077 | 0.05 | 0.1 |
| NC_019843.3;YP_009047202.1;MERS_1ab;5078 | 0.05 | 0.1 |
| NC_019843.3;YP_009047202.1;MERS_1ab;5079 | 0.05 | 0.1 |
| NC_019843.3;YP_009047202.1;MERS_1ab;5080 | 0.05 | 0.1 |
| NC_019843.3;YP_009047202.1;MERS_1ab;5081 | 0.1 | 0.1 |
| NC_019843.3;YP_009047202.1;MERS_1ab;5082 | 0.1 | 0.2 |
| NC_019843.3;YP_009047202.1;MERS_1ab;5083 | 0.1 | 0.15 |
| NC_019843.3;YP_009047202.1;MERS_1ab;5084 | 0.05 | 0.125 |
| NC_019843.3;YP_009047202.1;MERS_1ab;5085 | 0.05 | 0.125 |
| NC_019843.3;YP_009047202.1;MERS_1ab;5086 | 0.05 | 0.1 |
| NC_019843.3;YP_009047202.1;MERS_1ab;5087 | 0.05 | 0.075 |
| NC_019843.3;YP_009047202.1;MERS_1ab;5088 | 0.05 | 0.05 |
| NC_019843.3;YP_009047202.1;MERS_1ab;5089 | 0 | 0.025 |
| NC_019843.3;YP_009047202.1;MERS_1ab;5090 | 0 | 0.05 |
| NC_019843.3;YP_009047202.1;MERS_1ab;5091 | 0 | 0.025 |
| NC_019843.3;YP_009047202.1;MERS_1ab;5092 | 0 | 0.025 |
| NC_019843.3;YP_009047202.1;MERS_1ab;5093 | 0 | 0.05 |
| NC_019843.3;YP_009047202.1;MERS_1ab;5094 | 0 | 0.025 |
| NC_019843.3;YP_009047202.1;MERS_1ab;5095 | 0 | 0 |
| NC_019843.3;YP_009047202.1;MERS_1ab;5096 | 0 | 0 |
| NC_019843.3;YP_009047202.1;MERS_1ab;5097 | 0 | 0 |
| NC_019843.3;YP_009047202.1;MERS_1ab;5098 | 0 | 0 |
| NC_019843.3;YP_009047202.1;MERS_1ab;5099 | 0 | 0 |
| NC_019843.3;YP_009047202.1;MERS_1ab;5100 | 0 | 0 |
| NC_019843.3;YP_009047202.1;MERS_1ab;5101 | 0 | 0 |
| NC_019843.3;YP_009047202.1;MERS_1ab;5102 | 0 | 0 |
| NC_019843.3;YP_009047202.1;MERS_1ab;5103 | 0 | 0.025 |
| NC_019843.3;YP_009047202.1;MERS_1ab;5104 | 0 | 0.05 |
| NC_019843.3;YP_009047202.1;MERS_1ab;5105 | 0 | 0.025 |
| NC_019843.3;YP_009047202.1;MERS_1ab;5106 | 0.1 | 0.15 |
| NC_019843.3;YP_009047202.1;MERS_1ab;5107 | 0 | 0.1 |
| NC_019843.3;YP_009047202.1;MERS_1ab;5108 | 0.05 | 0.15 |

| | | |
|---|---|---|
| NC_019843.3;YP_009047202.1;MERS_1ab;5109 | 0.1 | 0.175 |
| NC_019843.3;YP_009047202.1;MERS_1ab;5110 | 0.05 | 0.125 |
| NC_019843.3;YP_009047202.1;MERS_1ab;5111 | 0 | 0.025 |
| NC_019843.3;YP_009047202.1;MERS_1ab;5112 | 0 | 0.025 |
| NC_019843.3;YP_009047202.1;MERS_1ab;5113 | 0 | 0.075 |
| NC_019843.3;YP_009047202.1;MERS_1ab;5114 | 0.05 | 0.025 |
| NC_019843.3;YP_009047202.1;MERS_1ab;5115 | 0 | 0.025 |
| NC_019843.3;YP_009047202.1;MERS_1ab;5116 | 0 | 0.025 |
| NC_019843.3;YP_009047202.1;MERS_1ab;5117 | 0 | 0 |
| NC_019843.3;YP_009047202.1;MERS_1ab;5118 | 0 | 0 |
| NC_019843.3;YP_009047202.1;MERS_1ab;5119 | 0 | 0 |
| NC_019843.3;YP_009047202.1;MERS_1ab;5120 | 0 | 0 |
| NC_019843.3;YP_009047202.1;MERS_1ab;5121 | 0 | 0 |
| NC_019843.3;YP_009047202.1;MERS_1ab;5122 | 0 | 0 |
| NC_019843.3;YP_009047202.1;MERS_1ab;5123 | 0 | 0 |
| NC_019843.3;YP_009047202.1;MERS_1ab;5124 | 0 | 0 |
| NC_019843.3;YP_009047202.1;MERS_1ab;5125 | 0.05 | 0 |
| NC_019843.3;YP_009047202.1;MERS_1ab;5126 | 0 | 0 |
| NC_019843.3;YP_009047202.1;MERS_1ab;5127 | 0 | 0.025 |
| NC_019843.3;YP_009047202.1;MERS_1ab;5128 | 0 | 0 |
| NC_019843.3;YP_009047202.1;MERS_1ab;5129 | 0 | 0 |
| NC_019843.3;YP_009047202.1;MERS_1ab;5130 | 0 | 0 |
| NC_019843.3;YP_009047202.1;MERS_1ab;5131 | 0 | 0 |
| NC_019843.3;YP_009047202.1;MERS_1ab;5132 | 0 | 0 |
| NC_019843.3;YP_009047202.1;MERS_1ab;5133 | 0 | 0 |
| NC_019843.3;YP_009047202.1;MERS_1ab;5134 | 0 | 0.025 |
| NC_019843.3;YP_009047202.1;MERS_1ab;5135 | 0.05 | 0.025 |
| NC_019843.3;YP_009047202.1;MERS_1ab;5136 | 0.05 | 0.025 |
| NC_019843.3;YP_009047202.1;MERS_1ab;5137 | 0.05 | 0 |
| NC_019843.3;YP_009047202.1;MERS_1ab;5138 | 0 | 0.05 |
| NC_019843.3;YP_009047202.1;MERS_1ab;5139 | 0 | 0.05 |
| NC_019843.3;YP_009047202.1;MERS_1ab;5140 | 0 | 0.05 |
| NC_019843.3;YP_009047202.1;MERS_1ab;5141 | 0 | 0 |
| NC_019843.3;YP_009047202.1;MERS_1ab;5142 | 0 | 0 |
| NC_019843.3;YP_009047202.1;MERS_1ab;5143 | 0 | 0 |
| NC_019843.3;YP_009047202.1;MERS_1ab;5144 | 0 | 0 |
| NC_019843.3;YP_009047202.1;MERS_1ab;5145 | 0 | 0 |
| NC_019843.3;YP_009047202.1;MERS_1ab;5146 | 0 | 0 |
| NC_019843.3;YP_009047202.1;MERS_1ab;5147 | 0 | 0.05 |
| NC_019843.3;YP_009047202.1;MERS_1ab;5148 | 0.05 | 0.05 |
| NC_019843.3;YP_009047202.1;MERS_1ab;5149 | 0.05 | 0.05 |
| NC_019843.3;YP_009047202.1;MERS_1ab;5150 | 0.05 | 0.025 |
| NC_019843.3;YP_009047202.1;MERS_1ab;5151 | 0.05 | 0.025 |

| | | |
|---|---|---|
| NC_019843.3;YP_009047202.1;MERS_1ab;5152 | 0.05 | 0 |
| NC_019843.3;YP_009047202.1;MERS_1ab;5153 | 0.05 | 0 |
| NC_019843.3;YP_009047202.1;MERS_1ab;5154 | 0.05 | 0.05 |
| NC_019843.3;YP_009047202.1;MERS_1ab;5155 | 0.1 | 0.075 |
| NC_019843.3;YP_009047202.1;MERS_1ab;5156 | 0.05 | 0 |
| NC_019843.3;YP_009047202.1;MERS_1ab;5157 | 0.05 | 0 |
| NC_019843.3;YP_009047202.1;MERS_1ab;5158 | 0.05 | 0 |
| NC_019843.3;YP_009047202.1;MERS_1ab;5159 | 0.05 | 0 |
| NC_019843.3;YP_009047202.1;MERS_1ab;5160 | 0.05 | 0 |
| NC_019843.3;YP_009047202.1;MERS_1ab;5161 | 0.15 | 0 |
| NC_019843.3;YP_009047202.1;MERS_1ab;5162 | 0.05 | 0 |
| NC_019843.3;YP_009047202.1;MERS_1ab;5163 | 0 | 0 |
| NC_019843.3;YP_009047202.1;MERS_1ab;5164 | 0 | 0 |
| NC_019843.3;YP_009047202.1;MERS_1ab;5165 | 0 | 0.025 |
| NC_019843.3;YP_009047202.1;MERS_1ab;5166 | 0 | 0.025 |
| NC_019843.3;YP_009047202.1;MERS_1ab;5167 | 0 | 0.025 |
| NC_019843.3;YP_009047202.1;MERS_1ab;5168 | 0 | 0.025 |
| NC_019843.3;YP_009047202.1;MERS_1ab;5169 | 0 | 0.025 |
| NC_019843.3;YP_009047202.1;MERS_1ab;5170 | 0 | 0.025 |
| NC_019843.3;YP_009047202.1;MERS_1ab;5171 | 0 | 0 |
| NC_019843.3;YP_009047202.1;MERS_1ab;5172 | 0.05 | 0 |
| NC_019843.3;YP_009047202.1;MERS_1ab;5173 | 0 | 0 |
| NC_019843.3;YP_009047202.1;MERS_1ab;5174 | 0 | 0 |
| NC_019843.3;YP_009047202.1;MERS_1ab;5175 | 0 | 0 |
| NC_019843.3;YP_009047202.1;MERS_1ab;5176 | 0 | 0 |
| NC_019843.3;YP_009047202.1;MERS_1ab;5177 | 0 | 0 |
| NC_019843.3;YP_009047202.1;MERS_1ab;5178 | 0 | 0 |
| NC_019843.3;YP_009047202.1;MERS_1ab;5179 | 0.05 | 0 |
| NC_019843.3;YP_009047202.1;MERS_1ab;5180 | 0 | 0.025 |
| NC_019843.3;YP_009047202.1;MERS_1ab;5181 | 0 | 0 |
| NC_019843.3;YP_009047202.1;MERS_1ab;5182 | 0 | 0 |
| NC_019843.3;YP_009047202.1;MERS_1ab;5183 | 0 | 0 |
| NC_019843.3;YP_009047202.1;MERS_1ab;5184 | 0 | 0 |
| NC_019843.3;YP_009047202.1;MERS_1ab;5185 | 0 | 0 |
| NC_019843.3;YP_009047202.1;MERS_1ab;5186 | 0 | 0 |
| NC_019843.3;YP_009047202.1;MERS_1ab;5187 | 0 | 0.025 |
| NC_019843.3;YP_009047202.1;MERS_1ab;5188 | 0 | 0 |
| NC_019843.3;YP_009047202.1;MERS_1ab;5189 | 0 | 0 |
| NC_019843.3;YP_009047202.1;MERS_1ab;5190 | 0 | 0.025 |
| NC_019843.3;YP_009047202.1;MERS_1ab;5191 | 0 | 0.025 |
| NC_019843.3;YP_009047202.1;MERS_1ab;5192 | 0 | 0.025 |
| NC_019843.3;YP_009047202.1;MERS_1ab;5193 | 0 | 0.025 |
| NC_019843.3;YP_009047202.1;MERS_1ab;5194 | 0 | 0 |

| | | |
|---|---|---|
| NC_019843.3;YP_009047202.1;MERS_1ab;5195 | 0 | 0.025 |
| NC_019843.3;YP_009047202.1;MERS_1ab;5196 | 0 | 0.05 |
| NC_019843.3;YP_009047202.1;MERS_1ab;5197 | 0 | 0.025 |
| NC_019843.3;YP_009047202.1;MERS_1ab;5198 | 0 | 0.025 |
| NC_019843.3;YP_009047202.1;MERS_1ab;5199 | 0.05 | 0.05 |
| NC_019843.3;YP_009047202.1;MERS_1ab;5200 | 0 | 0 |
| NC_019843.3;YP_009047202.1;MERS_1ab;5201 | 0 | 0 |
| NC_019843.3;YP_009047202.1;MERS_1ab;5202 | 0 | 0 |
| NC_019843.3;YP_009047202.1;MERS_1ab;5203 | 0 | 0 |
| NC_019843.3;YP_009047202.1;MERS_1ab;5204 | 0 | 0 |
| NC_019843.3;YP_009047202.1;MERS_1ab;5205 | 0 | 0 |
| NC_019843.3;YP_009047202.1;MERS_1ab;5206 | 0 | 0 |
| NC_019843.3;YP_009047202.1;MERS_1ab;5207 | 0 | 0 |
| NC_019843.3;YP_009047202.1;MERS_1ab;5208 | 0 | 0 |
| NC_019843.3;YP_009047202.1;MERS_1ab;5209 | 0 | 0 |
| NC_019843.3;YP_009047202.1;MERS_1ab;5210 | 0 | 0 |
| NC_019843.3;YP_009047202.1;MERS_1ab;5211 | 0 | 0 |
| NC_019843.3;YP_009047202.1;MERS_1ab;5212 | 0 | 0 |
| NC_019843.3;YP_009047202.1;MERS_1ab;5213 | 0 | 0 |
| NC_019843.3;YP_009047202.1;MERS_1ab;5214 | 0 | 0 |
| NC_019843.3;YP_009047202.1;MERS_1ab;5215 | 0.05 | 0 |
| NC_019843.3;YP_009047202.1;MERS_1ab;5216 | 0 | 0 |
| NC_019843.3;YP_009047202.1;MERS_1ab;5217 | 0 | 0 |
| NC_019843.3;YP_009047202.1;MERS_1ab;5218 | 0 | 0 |
| NC_019843.3;YP_009047202.1;MERS_1ab;5219 | 0 | 0 |
| NC_019843.3;YP_009047202.1;MERS_1ab;5220 | 0 | 0 |
| NC_019843.3;YP_009047202.1;MERS_1ab;5221 | 0 | 0 |
| NC_019843.3;YP_009047202.1;MERS_1ab;5222 | 0 | 0 |
| NC_019843.3;YP_009047202.1;MERS_1ab;5223 | 0 | 0 |
| NC_019843.3;YP_009047202.1;MERS_1ab;5224 | 0 | 0.025 |
| NC_019843.3;YP_009047202.1;MERS_1ab;5225 | 0 | 0 |
| NC_019843.3;YP_009047202.1;MERS_1ab;5226 | 0 | 0 |
| NC_019843.3;YP_009047202.1;MERS_1ab;5227 | 0 | 0 |
| NC_019843.3;YP_009047202.1;MERS_1ab;5228 | 0 | 0 |
| NC_019843.3;YP_009047202.1;MERS_1ab;5229 | 0 | 0 |
| NC_019843.3;YP_009047202.1;MERS_1ab;5230 | 0 | 0 |
| NC_019843.3;YP_009047202.1;MERS_1ab;5231 | 0 | 0 |
| NC_019843.3;YP_009047202.1;MERS_1ab;5232 | 0 | 0 |
| NC_019843.3;YP_009047202.1;MERS_1ab;5233 | 0 | 0 |
| NC_019843.3;YP_009047202.1;MERS_1ab;5234 | 0 | 0 |
| NC_019843.3;YP_009047202.1;MERS_1ab;5235 | 0 | 0 |
| NC_019843.3;YP_009047202.1;MERS_1ab;5236 | 0 | 0 |
| NC_019843.3;YP_009047202.1;MERS_1ab;5237 | 0 | 0 |

| | | |
|---|---|---|
| NC_019843.3;YP_009047202.1;MERS_1ab;5238 | 0 | 0 |
| NC_019843.3;YP_009047202.1;MERS_1ab;5239 | 0 | 0.025 |
| NC_019843.3;YP_009047202.1;MERS_1ab;5240 | 0 | 0.025 |
| NC_019843.3;YP_009047202.1;MERS_1ab;5241 | 0 | 0 |
| NC_019843.3;YP_009047202.1;MERS_1ab;5242 | 0 | 0.025 |
| NC_019843.3;YP_009047202.1;MERS_1ab;5243 | 0 | 0 |
| NC_019843.3;YP_009047202.1;MERS_1ab;5244 | 0 | 0.025 |
| NC_019843.3;YP_009047202.1;MERS_1ab;5245 | 0 | 0 |
| NC_019843.3;YP_009047202.1;MERS_1ab;5246 | 0 | 0 |
| NC_019843.3;YP_009047202.1;MERS_1ab;5247 | 0 | 0 |
| NC_019843.3;YP_009047202.1;MERS_1ab;5248 | 0 | 0 |
| NC_019843.3;YP_009047202.1;MERS_1ab;5249 | 0 | 0 |
| NC_019843.3;YP_009047202.1;MERS_1ab;5250 | 0 | 0 |
| NC_019843.3;YP_009047202.1;MERS_1ab;5251 | 0 | 0 |
| NC_019843.3;YP_009047202.1;MERS_1ab;5252 | 0 | 0 |
| NC_019843.3;YP_009047202.1;MERS_1ab;5253 | 0 | 0 |
| NC_019843.3;YP_009047202.1;MERS_1ab;5254 | 0 | 0 |
| NC_019843.3;YP_009047202.1;MERS_1ab;5255 | 0 | 0 |
| NC_019843.3;YP_009047202.1;MERS_1ab;5256 | 0 | 0 |
| NC_019843.3;YP_009047202.1;MERS_1ab;5257 | 0 | 0 |
| NC_019843.3;YP_009047202.1;MERS_1ab;5258 | 0 | 0 |
| NC_019843.3;YP_009047202.1;MERS_1ab;5259 | 0 | 0 |
| NC_019843.3;YP_009047202.1;MERS_1ab;5260 | 0.05 | 0 |
| NC_019843.3;YP_009047202.1;MERS_1ab;5261 | 0 | 0 |
| NC_019843.3;YP_009047202.1;MERS_1ab;5262 | 0 | 0 |
| NC_019843.3;YP_009047202.1;MERS_1ab;5263 | 0.05 | 0 |
| NC_019843.3;YP_009047202.1;MERS_1ab;5264 | 0 | 0 |
| NC_019843.3;YP_009047202.1;MERS_1ab;5265 | 0.05 | 0.025 |
| NC_019843.3;YP_009047202.1;MERS_1ab;5266 | 0 | 0.075 |
| NC_019843.3;YP_009047202.1;MERS_1ab;5267 | 0 | 0 |
| NC_019843.3;YP_009047202.1;MERS_1ab;5268 | 0 | 0 |
| NC_019843.3;YP_009047202.1;MERS_1ab;5269 | 0 | 0 |
| NC_019843.3;YP_009047202.1;MERS_1ab;5270 | 0 | 0 |
| NC_019843.3;YP_009047202.1;MERS_1ab;5271 | 0 | 0 |
| NC_019843.3;YP_009047202.1;MERS_1ab;5272 | 0 | 0 |
| NC_019843.3;YP_009047202.1;MERS_1ab;5273 | 0 | 0 |
| NC_019843.3;YP_009047202.1;MERS_1ab;5274 | 0 | 0 |
| NC_019843.3;YP_009047202.1;MERS_1ab;5275 | 0 | 0 |
| NC_019843.3;YP_009047202.1;MERS_1ab;5276 | 0 | 0 |
| NC_019843.3;YP_009047202.1;MERS_1ab;5277 | 0.05 | 0 |
| NC_019843.3;YP_009047202.1;MERS_1ab;5278 | 0 | 0 |
| NC_019843.3;YP_009047202.1;MERS_1ab;5279 | 0 | 0 |
| NC_019843.3;YP_009047202.1;MERS_1ab;5280 | 0 | 0 |

| | | |
|---|---|---|
| NC_019843.3;YP_009047202.1;MERS_1ab;5281 | 0 | 0 |
| NC_019843.3;YP_009047202.1;MERS_1ab;5282 | 0 | 0 |
| NC_019843.3;YP_009047202.1;MERS_1ab;5283 | 0 | 0 |
| NC_019843.3;YP_009047202.1;MERS_1ab;5284 | 0 | 0 |
| NC_019843.3;YP_009047202.1;MERS_1ab;5285 | 0.05 | 0 |
| NC_019843.3;YP_009047202.1;MERS_1ab;5286 | 0 | 0 |
| NC_019843.3;YP_009047202.1;MERS_1ab;5287 | 0 | 0.025 |
| NC_019843.3;YP_009047202.1;MERS_1ab;5288 | 0 | 0.025 |
| NC_019843.3;YP_009047202.1;MERS_1ab;5289 | 0.05 | 0 |
| NC_019843.3;YP_009047202.1;MERS_1ab;5290 | 0 | 0 |
| NC_019843.3;YP_009047202.1;MERS_1ab;5291 | 0 | 0 |
| NC_019843.3;YP_009047202.1;MERS_1ab;5292 | 0 | 0 |
| NC_019843.3;YP_009047202.1;MERS_1ab;5293 | 0 | 0 |
| NC_019843.3;YP_009047202.1;MERS_1ab;5294 | 0 | 0 |
| NC_019843.3;YP_009047202.1;MERS_1ab;5295 | 0 | 0 |
| NC_019843.3;YP_009047202.1;MERS_1ab;5296 | 0 | 0 |
| NC_019843.3;YP_009047202.1;MERS_1ab;5297 | 0 | 0 |
| NC_019843.3;YP_009047202.1;MERS_1ab;5298 | 0 | 0 |
| NC_019843.3;YP_009047202.1;MERS_1ab;5299 | 0 | 0 |
| NC_019843.3;YP_009047202.1;MERS_1ab;5300 | 0 | 0 |
| NC_019843.3;YP_009047202.1;MERS_1ab;5301 | 0 | 0 |
| NC_019843.3;YP_009047202.1;MERS_1ab;5302 | 0 | 0 |
| NC_019843.3;YP_009047202.1;MERS_1ab;5303 | 0 | 0 |
| NC_019843.3;YP_009047202.1;MERS_1ab;5304 | 0 | 0 |
| NC_019843.3;YP_009047202.1;MERS_1ab;5305 | 0 | 0 |
| NC_019843.3;YP_009047202.1;MERS_1ab;5306 | 0 | 0 |
| NC_019843.3;YP_009047202.1;MERS_1ab;5307 | 0 | 0 |
| NC_019843.3;YP_009047202.1;MERS_1ab;5308 | 0 | 0 |
| NC_019843.3;YP_009047202.1;MERS_1ab;5309 | 0 | 0 |
| NC_019843.3;YP_009047202.1;MERS_1ab;5310 | 0 | 0 |
| NC_019843.3;YP_009047202.1;MERS_1ab;5311 | 0 | 0 |
| NC_019843.3;YP_009047202.1;MERS_1ab;5312 | 0 | 0 |
| NC_019843.3;YP_009047202.1;MERS_1ab;5313 | 0 | 0 |
| NC_019843.3;YP_009047202.1;MERS_1ab;5314 | 0 | 0 |
| NC_019843.3;YP_009047202.1;MERS_1ab;5315 | 0 | 0 |
| NC_019843.3;YP_009047202.1;MERS_1ab;5316 | 0 | 0 |
| NC_019843.3;YP_009047202.1;MERS_1ab;5317 | 0 | 0 |
| NC_019843.3;YP_009047202.1;MERS_1ab;5318 | 0 | 0 |
| NC_019843.3;YP_009047202.1;MERS_1ab;5319 | 0 | 0 |
| NC_019843.3;YP_009047202.1;MERS_1ab;5320 | 0 | 0 |
| NC_019843.3;YP_009047202.1;MERS_1ab;5321 | 0 | 0 |
| NC_019843.3;YP_009047202.1;MERS_1ab;5322 | 0 | 0 |
| NC_019843.3;YP_009047202.1;MERS_1ab;5323 | 0 | 0 |

| | | |
|---|---|---|
| NC_019843.3;YP_009047202.1;MERS_1ab;5324 | 0 | 0 |
| NC_019843.3;YP_009047202.1;MERS_1ab;5325 | 0 | 0 |
| NC_019843.3;YP_009047202.1;MERS_1ab;5326 | 0 | 0 |
| NC_019843.3;YP_009047202.1;MERS_1ab;5327 | 0 | 0 |
| NC_019843.3;YP_009047202.1;MERS_1ab;5328 | 0 | 0 |
| NC_019843.3;YP_009047202.1;MERS_1ab;5329 | 0 | 0 |
| NC_019843.3;YP_009047202.1;MERS_1ab;5330 | 0 | 0 |
| NC_019843.3;YP_009047202.1;MERS_1ab;5331 | 0 | 0 |
| NC_019843.3;YP_009047202.1;MERS_1ab;5332 | 0 | 0 |
| NC_019843.3;YP_009047202.1;MERS_1ab;5333 | 0 | 0 |
| NC_019843.3;YP_009047202.1;MERS_1ab;5334 | 0 | 0 |
| NC_019843.3;YP_009047202.1;MERS_1ab;5335 | 0 | 0 |
| NC_019843.3;YP_009047202.1;MERS_1ab;5336 | 0 | 0 |
| NC_019843.3;YP_009047202.1;MERS_1ab;5337 | 0 | 0 |
| NC_019843.3;YP_009047202.1;MERS_1ab;5338 | 0 | 0 |
| NC_019843.3;YP_009047202.1;MERS_1ab;5339 | 0 | 0.025 |
| NC_019843.3;YP_009047202.1;MERS_1ab;5340 | 0 | 0 |
| NC_019843.3;YP_009047202.1;MERS_1ab;5341 | 0 | 0 |
| NC_019843.3;YP_009047202.1;MERS_1ab;5342 | 0 | 0 |
| NC_019843.3;YP_009047202.1;MERS_1ab;5343 | 0 | 0 |
| NC_019843.3;YP_009047202.1;MERS_1ab;5344 | 0 | 0 |
| NC_019843.3;YP_009047202.1;MERS_1ab;5345 | 0 | 0 |
| NC_019843.3;YP_009047202.1;MERS_1ab;5346 | 0 | 0 |
| NC_019843.3;YP_009047202.1;MERS_1ab;5347 | 0 | 0 |
| NC_019843.3;YP_009047202.1;MERS_1ab;5348 | 0 | 0 |
| NC_019843.3;YP_009047202.1;MERS_1ab;5349 | 0 | 0 |
| NC_019843.3;YP_009047202.1;MERS_1ab;5350 | 0 | 0 |
| NC_019843.3;YP_009047202.1;MERS_1ab;5351 | 0 | 0 |
| NC_019843.3;YP_009047202.1;MERS_1ab;5352 | 0 | 0 |
| NC_019843.3;YP_009047202.1;MERS_1ab;5353 | 0 | 0.025 |
| NC_019843.3;YP_009047202.1;MERS_1ab;5354 | 0 | 0 |
| NC_019843.3;YP_009047202.1;MERS_1ab;5355 | 0 | 0 |
| NC_019843.3;YP_009047202.1;MERS_1ab;5356 | 0 | 0 |
| NC_019843.3;YP_009047202.1;MERS_1ab;5357 | 0.05 | 0 |
| NC_019843.3;YP_009047202.1;MERS_1ab;5358 | 0 | 0 |
| NC_019843.3;YP_009047202.1;MERS_1ab;5359 | 0 | 0 |
| NC_019843.3;YP_009047202.1;MERS_1ab;5360 | 0 | 0 |
| NC_019843.3;YP_009047202.1;MERS_1ab;5361 | 0 | 0 |
| NC_019843.3;YP_009047202.1;MERS_1ab;5362 | 0 | 0 |
| NC_019843.3;YP_009047202.1;MERS_1ab;5363 | 0 | 0 |
| NC_019843.3;YP_009047202.1;MERS_1ab;5364 | 0 | 0.025 |
| NC_019843.3;YP_009047202.1;MERS_1ab;5365 | 0 | 0.025 |
| NC_019843.3;YP_009047202.1;MERS_1ab;5366 | 0.05 | 0 |

| | | |
|---|---|---|
| NC_019843.3;YP_009047202.1;MERS_1ab;5367 | 0 | 0 |
| NC_019843.3;YP_009047202.1;MERS_1ab;5368 | 0 | 0 |
| NC_019843.3;YP_009047202.1;MERS_1ab;5369 | 0 | 0 |
| NC_019843.3;YP_009047202.1;MERS_1ab;5370 | 0 | 0 |
| NC_019843.3;YP_009047202.1;MERS_1ab;5371 | 0 | 0 |
| NC_019843.3;YP_009047202.1;MERS_1ab;5372 | 0 | 0 |
| NC_019843.3;YP_009047202.1;MERS_1ab;5373 | 0 | 0 |
| NC_019843.3;YP_009047202.1;MERS_1ab;5374 | 0 | 0 |
| NC_019843.3;YP_009047202.1;MERS_1ab;5375 | 0 | 0 |
| NC_019843.3;YP_009047202.1;MERS_1ab;5376 | 0 | 0 |
| NC_019843.3;YP_009047202.1;MERS_1ab;5377 | 0 | 0 |
| NC_019843.3;YP_009047202.1;MERS_1ab;5378 | 0 | 0 |
| NC_019843.3;YP_009047202.1;MERS_1ab;5379 | 0 | 0 |
| NC_019843.3;YP_009047202.1;MERS_1ab;5380 | 0 | 0 |
| NC_019843.3;YP_009047202.1;MERS_1ab;5381 | 0 | 0 |
| NC_019843.3;YP_009047202.1;MERS_1ab;5382 | 0 | 0 |
| NC_019843.3;YP_009047202.1;MERS_1ab;5383 | 0 | 0 |
| NC_019843.3;YP_009047202.1;MERS_1ab;5384 | 0 | 0 |
| NC_019843.3;YP_009047202.1;MERS_1ab;5385 | 0 | 0 |
| NC_019843.3;YP_009047202.1;MERS_1ab;5386 | 0 | 0 |
| NC_019843.3;YP_009047202.1;MERS_1ab;5387 | 0 | 0 |
| NC_019843.3;YP_009047202.1;MERS_1ab;5388 | 0 | 0 |
| NC_019843.3;YP_009047202.1;MERS_1ab;5389 | 0 | 0 |
| NC_019843.3;YP_009047202.1;MERS_1ab;5390 | 0 | 0 |
| NC_019843.3;YP_009047202.1;MERS_1ab;5391 | 0 | 0 |
| NC_019843.3;YP_009047202.1;MERS_1ab;5392 | 0 | 0 |
| NC_019843.3;YP_009047202.1;MERS_1ab;5393 | 0 | 0 |
| NC_019843.3;YP_009047202.1;MERS_1ab;5394 | 0 | 0 |
| NC_019843.3;YP_009047202.1;MERS_1ab;5395 | 0 | 0 |
| NC_019843.3;YP_009047202.1;MERS_1ab;5396 | 0 | 0 |
| NC_019843.3;YP_009047202.1;MERS_1ab;5397 | 0 | 0 |
| NC_019843.3;YP_009047202.1;MERS_1ab;5398 | 0 | 0 |
| NC_019843.3;YP_009047202.1;MERS_1ab;5399 | 0 | 0 |
| NC_019843.3;YP_009047202.1;MERS_1ab;5400 | 0 | 0 |
| NC_019843.3;YP_009047202.1;MERS_1ab;5401 | 0 | 0 |
| NC_019843.3;YP_009047202.1;MERS_1ab;5402 | 0 | 0.025 |
| NC_019843.3;YP_009047202.1;MERS_1ab;5403 | 0 | 0.025 |
| NC_019843.3;YP_009047202.1;MERS_1ab;5404 | 0 | 0.025 |
| NC_019843.3;YP_009047202.1;MERS_1ab;5405 | 0 | 0 |
| NC_019843.3;YP_009047202.1;MERS_1ab;5406 | 0 | 0.025 |
| NC_019843.3;YP_009047202.1;MERS_1ab;5407 | 0 | 0 |
| NC_019843.3;YP_009047202.1;MERS_1ab;5408 | 0 | 0 |
| NC_019843.3;YP_009047202.1;MERS_1ab;5409 | 0 | 0 |

| | | |
|---|---|---|
| NC_019843.3;YP_009047202.1;MERS_1ab;5410 | 0 | 0 |
| NC_019843.3;YP_009047202.1;MERS_1ab;5411 | 0 | 0 |
| NC_019843.3;YP_009047202.1;MERS_1ab;5412 | 0 | 0 |
| NC_019843.3;YP_009047202.1;MERS_1ab;5413 | 0 | 0 |
| NC_019843.3;YP_009047202.1;MERS_1ab;5414 | 0 | 0.025 |
| NC_019843.3;YP_009047202.1;MERS_1ab;5415 | 0 | 0.025 |
| NC_019843.3;YP_009047202.1;MERS_1ab;5416 | 0 | 0.025 |
| NC_019843.3;YP_009047202.1;MERS_1ab;5417 | 0 | 0 |
| NC_019843.3;YP_009047202.1;MERS_1ab;5418 | 0 | 0 |
| NC_019843.3;YP_009047202.1;MERS_1ab;5419 | 0 | 0 |
| NC_019843.3;YP_009047202.1;MERS_1ab;5420 | 0 | 0.025 |
| NC_019843.3;YP_009047202.1;MERS_1ab;5421 | 0 | 0 |
| NC_019843.3;YP_009047202.1;MERS_1ab;5422 | 0 | 0.025 |
| NC_019843.3;YP_009047202.1;MERS_1ab;5423 | 0 | 0.025 |
| NC_019843.3;YP_009047202.1;MERS_1ab;5424 | 0 | 0.025 |
| NC_019843.3;YP_009047202.1;MERS_1ab;5425 | 0 | 0 |
| NC_019843.3;YP_009047202.1;MERS_1ab;5426 | 0 | 0.025 |
| NC_019843.3;YP_009047202.1;MERS_1ab;5427 | 0.05 | 0.025 |
| NC_019843.3;YP_009047202.1;MERS_1ab;5428 | 0.05 | 0 |
| NC_019843.3;YP_009047202.1;MERS_1ab;5429 | 0 | 0.025 |
| NC_019843.3;YP_009047202.1;MERS_1ab;5430 | 0.05 | 0.025 |
| NC_019843.3;YP_009047202.1;MERS_1ab;5431 | 0 | 0 |
| NC_019843.3;YP_009047202.1;MERS_1ab;5432 | 0 | 0.025 |
| NC_019843.3;YP_009047202.1;MERS_1ab;5433 | 0 | 0 |
| NC_019843.3;YP_009047202.1;MERS_1ab;5434 | 0 | 0 |
| NC_019843.3;YP_009047202.1;MERS_1ab;5435 | 0 | 0 |
| NC_019843.3;YP_009047202.1;MERS_1ab;5436 | 0 | 0 |
| NC_019843.3;YP_009047202.1;MERS_1ab;5437 | 0 | 0 |
| NC_019843.3;YP_009047202.1;MERS_1ab;5438 | 0 | 0 |
| NC_019843.3;YP_009047202.1;MERS_1ab;5439 | 0 | 0 |
| NC_019843.3;YP_009047202.1;MERS_1ab;5440 | 0 | 0 |
| NC_019843.3;YP_009047202.1;MERS_1ab;5441 | 0 | 0 |
| NC_019843.3;YP_009047202.1;MERS_1ab;5442 | 0 | 0 |
| NC_019843.3;YP_009047202.1;MERS_1ab;5443 | 0 | 0 |
| NC_019843.3;YP_009047202.1;MERS_1ab;5444 | 0 | 0 |
| NC_019843.3;YP_009047202.1;MERS_1ab;5445 | 0 | 0.025 |
| NC_019843.3;YP_009047202.1;MERS_1ab;5446 | 0 | 0 |
| NC_019843.3;YP_009047202.1;MERS_1ab;5447 | 0 | 0 |
| NC_019843.3;YP_009047202.1;MERS_1ab;5448 | 0 | 0.025 |
| NC_019843.3;YP_009047202.1;MERS_1ab;5449 | 0 | 0 |
| NC_019843.3;YP_009047202.1;MERS_1ab;5450 | 0 | 0 |
| NC_019843.3;YP_009047202.1;MERS_1ab;5451 | 0 | 0 |
| NC_019843.3;YP_009047202.1;MERS_1ab;5452 | 0 | 0 |

| | | |
|---|---|---|
| NC_019843.3;YP_009047202.1;MERS_1ab;5453 | 0 | 0 |
| NC_019843.3;YP_009047202.1;MERS_1ab;5454 | 0 | 0 |
| NC_019843.3;YP_009047202.1;MERS_1ab;5455 | 0 | 0 |
| NC_019843.3;YP_009047202.1;MERS_1ab;5456 | 0 | 0 |
| NC_019843.3;YP_009047202.1;MERS_1ab;5457 | 0 | 0 |
| NC_019843.3;YP_009047202.1;MERS_1ab;5458 | 0 | 0 |
| NC_019843.3;YP_009047202.1;MERS_1ab;5459 | 0 | 0 |
| NC_019843.3;YP_009047202.1;MERS_1ab;5460 | 0 | 0 |
| NC_019843.3;YP_009047202.1;MERS_1ab;5461 | 0 | 0 |
| NC_019843.3;YP_009047202.1;MERS_1ab;5462 | 0 | 0 |
| NC_019843.3;YP_009047202.1;MERS_1ab;5463 | 0 | 0 |
| NC_019843.3;YP_009047202.1;MERS_1ab;5464 | 0 | 0 |
| NC_019843.3;YP_009047202.1;MERS_1ab;5465 | 0.05 | 0 |
| NC_019843.3;YP_009047202.1;MERS_1ab;5466 | 0 | 0 |
| NC_019843.3;YP_009047202.1;MERS_1ab;5467 | 0 | 0.025 |
| NC_019843.3;YP_009047202.1;MERS_1ab;5468 | 0 | 0 |
| NC_019843.3;YP_009047202.1;MERS_1ab;5469 | 0 | 0.025 |
| NC_019843.3;YP_009047202.1;MERS_1ab;5470 | 0.05 | 0.025 |
| NC_019843.3;YP_009047202.1;MERS_1ab;5471 | 0 | 0 |
| NC_019843.3;YP_009047202.1;MERS_1ab;5472 | 0 | 0.025 |
| NC_019843.3;YP_009047202.1;MERS_1ab;5473 | 0 | 0 |
| NC_019843.3;YP_009047202.1;MERS_1ab;5474 | 0 | 0 |
| NC_019843.3;YP_009047202.1;MERS_1ab;5475 | 0.05 | 0.025 |
| NC_019843.3;YP_009047202.1;MERS_1ab;5476 | 0 | 0.025 |
| NC_019843.3;YP_009047202.1;MERS_1ab;5477 | 0 | 0.025 |
| NC_019843.3;YP_009047202.1;MERS_1ab;5478 | 0 | 0 |
| NC_019843.3;YP_009047202.1;MERS_1ab;5479 | 0 | 0 |
| NC_019843.3;YP_009047202.1;MERS_1ab;5480 | 0 | 0 |
| NC_019843.3;YP_009047202.1;MERS_1ab;5481 | 0 | 0 |
| NC_019843.3;YP_009047202.1;MERS_1ab;5482 | 0 | 0 |
| NC_019843.3;YP_009047202.1;MERS_1ab;5483 | 0 | 0 |
| NC_019843.3;YP_009047202.1;MERS_1ab;5484 | 0 | 0 |
| NC_019843.3;YP_009047202.1;MERS_1ab;5485 | 0 | 0 |
| NC_019843.3;YP_009047202.1;MERS_1ab;5486 | 0 | 0 |
| NC_019843.3;YP_009047202.1;MERS_1ab;5487 | 0 | 0 |
| NC_019843.3;YP_009047202.1;MERS_1ab;5488 | 0 | 0 |
| NC_019843.3;YP_009047202.1;MERS_1ab;5489 | 0 | 0 |
| NC_019843.3;YP_009047202.1;MERS_1ab;5490 | 0 | 0 |
| NC_019843.3;YP_009047202.1;MERS_1ab;5491 | 0 | 0.025 |
| NC_019843.3;YP_009047202.1;MERS_1ab;5492 | 0.05 | 0.025 |
| NC_019843.3;YP_009047202.1;MERS_1ab;5493 | 0.1 | 0.025 |
| NC_019843.3;YP_009047202.1;MERS_1ab;5494 | 0.1 | 0 |
| NC_019843.3;YP_009047202.1;MERS_1ab;5495 | 0 | 0 |

| | | |
|---|---|---|
| NC_019843.3;YP_009047202.1;MERS_1ab;5496 | 0 | 0 |
| NC_019843.3;YP_009047202.1;MERS_1ab;5497 | 0.05 | 0 |
| NC_019843.3;YP_009047202.1;MERS_1ab;5498 | 0 | 0 |
| NC_019843.3;YP_009047202.1;MERS_1ab;5499 | 0 | 0 |
| NC_019843.3;YP_009047202.1;MERS_1ab;5500 | 0 | 0 |
| NC_019843.3;YP_009047202.1;MERS_1ab;5501 | 0 | 0 |
| NC_019843.3;YP_009047202.1;MERS_1ab;5502 | 0 | 0 |
| NC_019843.3;YP_009047202.1;MERS_1ab;5503 | 0 | 0 |
| NC_019843.3;YP_009047202.1;MERS_1ab;5504 | 0 | 0.025 |
| NC_019843.3;YP_009047202.1;MERS_1ab;5505 | 0 | 0.025 |
| NC_019843.3;YP_009047202.1;MERS_1ab;5506 | 0 | 0.025 |
| NC_019843.3;YP_009047202.1;MERS_1ab;5507 | 0.1 | 0.025 |
| NC_019843.3;YP_009047202.1;MERS_1ab;5508 | 0.05 | 0.025 |
| NC_019843.3;YP_009047202.1;MERS_1ab;5509 | 0.05 | 0.05 |
| NC_019843.3;YP_009047202.1;MERS_1ab;5510 | 0 | 0.05 |
| NC_019843.3;YP_009047202.1;MERS_1ab;5511 | 0 | 0.025 |
| NC_019843.3;YP_009047202.1;MERS_1ab;5512 | 0.05 | 0 |
| NC_019843.3;YP_009047202.1;MERS_1ab;5513 | 0.05 | 0 |
| NC_019843.3;YP_009047202.1;MERS_1ab;5514 | 0 | 0.025 |
| NC_019843.3;YP_009047202.1;MERS_1ab;5515 | 0 | 0 |
| NC_019843.3;YP_009047202.1;MERS_1ab;5516 | 0 | 0 |
| NC_019843.3;YP_009047202.1;MERS_1ab;5517 | 0 | 0 |
| NC_019843.3;YP_009047202.1;MERS_1ab;5518 | 0 | 0 |
| NC_019843.3;YP_009047202.1;MERS_1ab;5519 | 0.05 | 0 |
| NC_019843.3;YP_009047202.1;MERS_1ab;5520 | 0 | 0 |
| NC_019843.3;YP_009047202.1;MERS_1ab;5521 | 0 | 0 |
| NC_019843.3;YP_009047202.1;MERS_1ab;5522 | 0 | 0 |
| NC_019843.3;YP_009047202.1;MERS_1ab;5523 | 0 | 0 |
| NC_019843.3;YP_009047202.1;MERS_1ab;5524 | 0 | 0 |
| NC_019843.3;YP_009047202.1;MERS_1ab;5525 | 0 | 0 |
| NC_019843.3;YP_009047202.1;MERS_1ab;5526 | 0 | 0 |
| NC_019843.3;YP_009047202.1;MERS_1ab;5527 | 0 | 0 |
| NC_019843.3;YP_009047202.1;MERS_1ab;5528 | 0 | 0 |
| NC_019843.3;YP_009047202.1;MERS_1ab;5529 | 0 | 0.05 |
| NC_019843.3;YP_009047202.1;MERS_1ab;5530 | 0 | 0 |
| NC_019843.3;YP_009047202.1;MERS_1ab;5531 | 0 | 0 |
| NC_019843.3;YP_009047202.1;MERS_1ab;5532 | 0 | 0 |
| NC_019843.3;YP_009047202.1;MERS_1ab;5533 | 0 | 0 |
| NC_019843.3;YP_009047202.1;MERS_1ab;5534 | 0 | 0 |
| NC_019843.3;YP_009047202.1;MERS_1ab;5535 | 0 | 0 |
| NC_019843.3;YP_009047202.1;MERS_1ab;5536 | 0 | 0 |
| NC_019843.3;YP_009047202.1;MERS_1ab;5537 | 0 | 0 |
| NC_019843.3;YP_009047202.1;MERS_1ab;5538 | 0 | 0 |

| | | |
|---|---|---|
| NC_019843.3;YP_009047202.1;MERS_1ab;5539 | 0 | 0 |
| NC_019843.3;YP_009047202.1;MERS_1ab;5540 | 0 | 0 |
| NC_019843.3;YP_009047202.1;MERS_1ab;5541 | 0 | 0 |
| NC_019843.3;YP_009047202.1;MERS_1ab;5542 | 0 | 0 |
| NC_019843.3;YP_009047202.1;MERS_1ab;5543 | 0 | 0.025 |
| NC_019843.3;YP_009047202.1;MERS_1ab;5544 | 0 | 0.025 |
| NC_019843.3;YP_009047202.1;MERS_1ab;5545 | 0 | 0 |
| NC_019843.3;YP_009047202.1;MERS_1ab;5546 | 0.05 | 0.025 |
| NC_019843.3;YP_009047202.1;MERS_1ab;5547 | 0 | 0 |
| NC_019843.3;YP_009047202.1;MERS_1ab;5548 | 0 | 0 |
| NC_019843.3;YP_009047202.1;MERS_1ab;5549 | 0 | 0 |
| NC_019843.3;YP_009047202.1;MERS_1ab;5550 | 0 | 0 |
| NC_019843.3;YP_009047202.1;MERS_1ab;5551 | 0 | 0 |
| NC_019843.3;YP_009047202.1;MERS_1ab;5552 | 0 | 0 |
| NC_019843.3;YP_009047202.1;MERS_1ab;5553 | 0 | 0 |
| NC_019843.3;YP_009047202.1;MERS_1ab;5554 | 0 | 0 |
| NC_019843.3;YP_009047202.1;MERS_1ab;5555 | 0 | 0 |
| NC_019843.3;YP_009047202.1;MERS_1ab;5556 | 0 | 0 |
| NC_019843.3;YP_009047202.1;MERS_1ab;5557 | 0 | 0 |
| NC_019843.3;YP_009047202.1;MERS_1ab;5558 | 0 | 0 |
| NC_019843.3;YP_009047202.1;MERS_1ab;5559 | 0 | 0 |
| NC_019843.3;YP_009047202.1;MERS_1ab;5560 | 0 | 0 |
| NC_019843.3;YP_009047202.1;MERS_1ab;5561 | 0 | 0 |
| NC_019843.3;YP_009047202.1;MERS_1ab;5562 | 0 | 0.025 |
| NC_019843.3;YP_009047202.1;MERS_1ab;5563 | 0 | 0 |
| NC_019843.3;YP_009047202.1;MERS_1ab;5564 | 0 | 0 |
| NC_019843.3;YP_009047202.1;MERS_1ab;5565 | 0 | 0 |
| NC_019843.3;YP_009047202.1;MERS_1ab;5566 | 0 | 0.1 |
| NC_019843.3;YP_009047202.1;MERS_1ab;5567 | 0 | 0.025 |
| NC_019843.3;YP_009047202.1;MERS_1ab;5568 | 0 | 0.025 |
| NC_019843.3;YP_009047202.1;MERS_1ab;5569 | 0 | 0.075 |
| NC_019843.3;YP_009047202.1;MERS_1ab;5570 | 0 | 0.05 |
| NC_019843.3;YP_009047202.1;MERS_1ab;5571 | 0 | 0 |
| NC_019843.3;YP_009047202.1;MERS_1ab;5572 | 0 | 0 |
| NC_019843.3;YP_009047202.1;MERS_1ab;5573 | 0 | 0 |
| NC_019843.3;YP_009047202.1;MERS_1ab;5574 | 0 | 0 |
| NC_019843.3;YP_009047202.1;MERS_1ab;5575 | 0 | 0 |
| NC_019843.3;YP_009047202.1;MERS_1ab;5576 | 0 | 0 |
| NC_019843.3;YP_009047202.1;MERS_1ab;5577 | 0 | 0 |
| NC_019843.3;YP_009047202.1;MERS_1ab;5578 | 0 | 0 |
| NC_019843.3;YP_009047202.1;MERS_1ab;5579 | 0.05 | 0 |
| NC_019843.3;YP_009047202.1;MERS_1ab;5580 | 0.05 | 0 |
| NC_019843.3;YP_009047202.1;MERS_1ab;5581 | 0.05 | 0 |

| | | |
|---|---|---|
| NC_019843.3;YP_009047202.1;MERS_1ab;5582 | 0 | 0 |
| NC_019843.3;YP_009047202.1;MERS_1ab;5583 | 0 | 0 |
| NC_019843.3;YP_009047202.1;MERS_1ab;5584 | 0 | 0 |
| NC_019843.3;YP_009047202.1;MERS_1ab;5585 | 0 | 0 |
| NC_019843.3;YP_009047202.1;MERS_1ab;5586 | 0 | 0 |
| NC_019843.3;YP_009047202.1;MERS_1ab;5587 | 0 | 0 |
| NC_019843.3;YP_009047202.1;MERS_1ab;5588 | 0 | 0 |
| NC_019843.3;YP_009047202.1;MERS_1ab;5589 | 0 | 0 |
| NC_019843.3;YP_009047202.1;MERS_1ab;5590 | 0 | 0 |
| NC_019843.3;YP_009047202.1;MERS_1ab;5591 | 0 | 0 |
| NC_019843.3;YP_009047202.1;MERS_1ab;5592 | 0 | 0 |
| NC_019843.3;YP_009047202.1;MERS_1ab;5593 | 0 | 0 |
| NC_019843.3;YP_009047202.1;MERS_1ab;5594 | 0 | 0 |
| NC_019843.3;YP_009047202.1;MERS_1ab;5595 | 0 | 0 |
| NC_019843.3;YP_009047202.1;MERS_1ab;5596 | 0 | 0 |
| NC_019843.3;YP_009047202.1;MERS_1ab;5597 | 0 | 0 |
| NC_019843.3;YP_009047202.1;MERS_1ab;5598 | 0 | 0 |
| NC_019843.3;YP_009047202.1;MERS_1ab;5599 | 0 | 0 |
| NC_019843.3;YP_009047202.1;MERS_1ab;5600 | 0 | 0 |
| NC_019843.3;YP_009047202.1;MERS_1ab;5601 | 0 | 0 |
| NC_019843.3;YP_009047202.1;MERS_1ab;5602 | 0 | 0 |
| NC_019843.3;YP_009047202.1;MERS_1ab;5603 | 0 | 0 |
| NC_019843.3;YP_009047202.1;MERS_1ab;5604 | 0 | 0 |
| NC_019843.3;YP_009047202.1;MERS_1ab;5605 | 0 | 0 |
| NC_019843.3;YP_009047202.1;MERS_1ab;5606 | 0 | 0 |
| NC_019843.3;YP_009047202.1;MERS_1ab;5607 | 0 | 0 |
| NC_019843.3;YP_009047202.1;MERS_1ab;5608 | 0.05 | 0 |
| NC_019843.3;YP_009047202.1;MERS_1ab;5609 | 0 | 0 |
| NC_019843.3;YP_009047202.1;MERS_1ab;5610 | 0 | 0.025 |
| NC_019843.3;YP_009047202.1;MERS_1ab;5611 | 0 | 0.025 |
| NC_019843.3;YP_009047202.1;MERS_1ab;5612 | 0 | 0.05 |
| NC_019843.3;YP_009047202.1;MERS_1ab;5613 | 0 | 0 |
| NC_019843.3;YP_009047202.1;MERS_1ab;5614 | 0 | 0 |
| NC_019843.3;YP_009047202.1;MERS_1ab;5615 | 0 | 0 |
| NC_019843.3;YP_009047202.1;MERS_1ab;5616 | 0 | 0 |
| NC_019843.3;YP_009047202.1;MERS_1ab;5617 | 0 | 0 |
| NC_019843.3;YP_009047202.1;MERS_1ab;5618 | 0 | 0 |
| NC_019843.3;YP_009047202.1;MERS_1ab;5619 | 0 | 0 |
| NC_019843.3;YP_009047202.1;MERS_1ab;5620 | 0 | 0 |
| NC_019843.3;YP_009047202.1;MERS_1ab;5621 | 0 | 0 |
| NC_019843.3;YP_009047202.1;MERS_1ab;5622 | 0 | 0 |
| NC_019843.3;YP_009047202.1;MERS_1ab;5623 | 0 | 0 |
| NC_019843.3;YP_009047202.1;MERS_1ab;5624 | 0 | 0 |

| | | |
|---|---|---|
| NC_019843.3;YP_009047202.1;MERS_1ab;5625 | 0 | 0 |
| NC_019843.3;YP_009047202.1;MERS_1ab;5626 | 0 | 0 |
| NC_019843.3;YP_009047202.1;MERS_1ab;5627 | 0 | 0 |
| NC_019843.3;YP_009047202.1;MERS_1ab;5628 | 0 | 0 |
| NC_019843.3;YP_009047202.1;MERS_1ab;5629 | 0 | 0 |
| NC_019843.3;YP_009047202.1;MERS_1ab;5630 | 0 | 0 |
| NC_019843.3;YP_009047202.1;MERS_1ab;5631 | 0 | 0 |
| NC_019843.3;YP_009047202.1;MERS_1ab;5632 | 0 | 0 |
| NC_019843.3;YP_009047202.1;MERS_1ab;5633 | 0 | 0 |
| NC_019843.3;YP_009047202.1;MERS_1ab;5634 | 0 | 0 |
| NC_019843.3;YP_009047202.1;MERS_1ab;5635 | 0 | 0 |
| NC_019843.3;YP_009047202.1;MERS_1ab;5636 | 0 | 0 |
| NC_019843.3;YP_009047202.1;MERS_1ab;5637 | 0 | 0 |
| NC_019843.3;YP_009047202.1;MERS_1ab;5638 | 0 | 0 |
| NC_019843.3;YP_009047202.1;MERS_1ab;5639 | 0 | 0 |
| NC_019843.3;YP_009047202.1;MERS_1ab;5640 | 0 | 0 |
| NC_019843.3;YP_009047202.1;MERS_1ab;5641 | 0 | 0 |
| NC_019843.3;YP_009047202.1;MERS_1ab;5642 | 0 | 0 |
| NC_019843.3;YP_009047202.1;MERS_1ab;5643 | 0 | 0 |
| NC_019843.3;YP_009047202.1;MERS_1ab;5644 | 0 | 0 |
| NC_019843.3;YP_009047202.1;MERS_1ab;5645 | 0.05 | 0 |
| NC_019843.3;YP_009047202.1;MERS_1ab;5646 | 0 | 0 |
| NC_019843.3;YP_009047202.1;MERS_1ab;5647 | 0.2 | 0.05 |
| NC_019843.3;YP_009047202.1;MERS_1ab;5648 | 0.3 | 0.225 |
| NC_019843.3;YP_009047202.1;MERS_1ab;5649 | 0.25 | 0.225 |
| NC_019843.3;YP_009047202.1;MERS_1ab;5650 | 0.4 | 0.25 |
| NC_019843.3;YP_009047202.1;MERS_1ab;5651 | 0.4 | 0.25 |
| NC_019843.3;YP_009047202.1;MERS_1ab;5652 | 0.25 | 0.025 |
| NC_019843.3;YP_009047202.1;MERS_1ab;5653 | 0.2 | 0.025 |
| NC_019843.3;YP_009047202.1;MERS_1ab;5654 | 0.2 | 0.075 |
| NC_019843.3;YP_009047202.1;MERS_1ab;5655 | 0.05 | 0.075 |
| NC_019843.3;YP_009047202.1;MERS_1ab;5656 | 0.15 | 0.025 |
| NC_019843.3;YP_009047202.1;MERS_1ab;5657 | 0.2 | 0.125 |
| NC_019843.3;YP_009047202.1;MERS_1ab;5658 | 0.05 | 0.05 |
| NC_019843.3;YP_009047202.1;MERS_1ab;5659 | 0.05 | 0.025 |
| NC_019843.3;YP_009047202.1;MERS_1ab;5660 | 0.05 | 0.05 |
| NC_019843.3;YP_009047202.1;MERS_1ab;5661 | 0.05 | 0.025 |
| NC_019843.3;YP_009047202.1;MERS_1ab;5662 | 0.05 | 0.025 |
| NC_019843.3;YP_009047202.1;MERS_1ab;5663 | 0 | 0.025 |
| NC_019843.3;YP_009047202.1;MERS_1ab;5664 | 0 | 0.025 |
| NC_019843.3;YP_009047202.1;MERS_1ab;5665 | 0 | 0.025 |
| NC_019843.3;YP_009047202.1;MERS_1ab;5666 | 0 | 0 |
| NC_019843.3;YP_009047202.1;MERS_1ab;5667 | 0 | 0 |

| | | |
|---|---|---|
| NC_019843.3;YP_009047202.1;MERS_1ab;5668 | 0 | 0 |
| NC_019843.3;YP_009047202.1;MERS_1ab;5669 | 0.05 | 0 |
| NC_019843.3;YP_009047202.1;MERS_1ab;5670 | 0.05 | 0 |
| NC_019843.3;YP_009047202.1;MERS_1ab;5671 | 0.05 | 0 |
| NC_019843.3;YP_009047202.1;MERS_1ab;5672 | 0.1 | 0 |
| NC_019843.3;YP_009047202.1;MERS_1ab;5673 | 0.05 | 0 |
| NC_019843.3;YP_009047202.1;MERS_1ab;5674 | 0.1 | 0 |
| NC_019843.3;YP_009047202.1;MERS_1ab;5675 | 0.1 | 0 |
| NC_019843.3;YP_009047202.1;MERS_1ab;5676 | 0.1 | 0 |
| NC_019843.3;YP_009047202.1;MERS_1ab;5677 | 0.05 | 0 |
| NC_019843.3;YP_009047202.1;MERS_1ab;5678 | 0 | 0 |
| NC_019843.3;YP_009047202.1;MERS_1ab;5679 | 0 | 0 |
| NC_019843.3;YP_009047202.1;MERS_1ab;5680 | 0 | 0 |
| NC_019843.3;YP_009047202.1;MERS_1ab;5681 | 0 | 0 |
| NC_019843.3;YP_009047202.1;MERS_1ab;5682 | 0.05 | 0.025 |
| NC_019843.3;YP_009047202.1;MERS_1ab;5683 | 0 | 0.025 |
| NC_019843.3;YP_009047202.1;MERS_1ab;5684 | 0 | 0 |
| NC_019843.3;YP_009047202.1;MERS_1ab;5685 | 0 | 0.025 |
| NC_019843.3;YP_009047202.1;MERS_1ab;5686 | 0 | 0 |
| NC_019843.3;YP_009047202.1;MERS_1ab;5687 | 0 | 0 |
| NC_019843.3;YP_009047202.1;MERS_1ab;5688 | 0 | 0 |
| NC_019843.3;YP_009047202.1;MERS_1ab;5689 | 0 | 0 |
| NC_019843.3;YP_009047202.1;MERS_1ab;5690 | 0 | 0 |
| NC_019843.3;YP_009047202.1;MERS_1ab;5691 | 0 | 0 |
| NC_019843.3;YP_009047202.1;MERS_1ab;5692 | 0 | 0 |
| NC_019843.3;YP_009047202.1;MERS_1ab;5693 | 0 | 0 |
| NC_019843.3;YP_009047202.1;MERS_1ab;5694 | 0 | 0 |
| NC_019843.3;YP_009047202.1;MERS_1ab;5695 | 0 | 0 |
| NC_019843.3;YP_009047202.1;MERS_1ab;5696 | 0.05 | 0.05 |
| NC_019843.3;YP_009047202.1;MERS_1ab;5697 | 0 | 0 |
| NC_019843.3;YP_009047202.1;MERS_1ab;5698 | 0 | 0 |
| NC_019843.3;YP_009047202.1;MERS_1ab;5699 | 0.05 | 0 |
| NC_019843.3;YP_009047202.1;MERS_1ab;5700 | 0 | 0 |
| NC_019843.3;YP_009047202.1;MERS_1ab;5701 | 0 | 0 |
| NC_019843.3;YP_009047202.1;MERS_1ab;5702 | 0 | 0 |
| NC_019843.3;YP_009047202.1;MERS_1ab;5703 | 0 | 0.025 |
| NC_019843.3;YP_009047202.1;MERS_1ab;5704 | 0 | 0 |
| NC_019843.3;YP_009047202.1;MERS_1ab;5705 | 0 | 0 |
| NC_019843.3;YP_009047202.1;MERS_1ab;5706 | 0 | 0 |
| NC_019843.3;YP_009047202.1;MERS_1ab;5707 | 0 | 0.025 |
| NC_019843.3;YP_009047202.1;MERS_1ab;5708 | 0 | 0 |
| NC_019843.3;YP_009047202.1;MERS_1ab;5709 | 0 | 0 |
| NC_019843.3;YP_009047202.1;MERS_1ab;5710 | 0 | 0 |

| | | |
|---|---|---|
| NC_019843.3;YP_009047202.1;MERS_1ab;5711 | 0 | 0 |
| NC_019843.3;YP_009047202.1;MERS_1ab;5712 | 0 | 0 |
| NC_019843.3;YP_009047202.1;MERS_1ab;5713 | 0 | 0 |
| NC_019843.3;YP_009047202.1;MERS_1ab;5714 | 0 | 0 |
| NC_019843.3;YP_009047202.1;MERS_1ab;5715 | 0 | 0 |
| NC_019843.3;YP_009047202.1;MERS_1ab;5716 | 0 | 0.025 |
| NC_019843.3;YP_009047202.1;MERS_1ab;5717 | 0 | 0 |
| NC_019843.3;YP_009047202.1;MERS_1ab;5718 | 0.05 | 0 |
| NC_019843.3;YP_009047202.1;MERS_1ab;5719 | 0 | 0 |
| NC_019843.3;YP_009047202.1;MERS_1ab;5720 | 0.05 | 0 |
| NC_019843.3;YP_009047202.1;MERS_1ab;5721 | 0.05 | 0.025 |
| NC_019843.3;YP_009047202.1;MERS_1ab;5722 | 0.05 | 0.025 |
| NC_019843.3;YP_009047202.1;MERS_1ab;5723 | 0 | 0 |
| NC_019843.3;YP_009047202.1;MERS_1ab;5724 | 0 | 0.025 |
| NC_019843.3;YP_009047202.1;MERS_1ab;5725 | 0 | 0 |
| NC_019843.3;YP_009047202.1;MERS_1ab;5726 | 0 | 0 |
| NC_019843.3;YP_009047202.1;MERS_1ab;5727 | 0 | 0 |
| NC_019843.3;YP_009047202.1;MERS_1ab;5728 | 0 | 0 |
| NC_019843.3;YP_009047202.1;MERS_1ab;5729 | 0 | 0 |
| NC_019843.3;YP_009047202.1;MERS_1ab;5730 | 0 | 0 |
| NC_019843.3;YP_009047202.1;MERS_1ab;5731 | 0 | 0 |
| NC_019843.3;YP_009047202.1;MERS_1ab;5732 | 0 | 0 |
| NC_019843.3;YP_009047202.1;MERS_1ab;5733 | 0 | 0 |
| NC_019843.3;YP_009047202.1;MERS_1ab;5734 | 0 | 0.025 |
| NC_019843.3;YP_009047202.1;MERS_1ab;5735 | 0 | 0 |
| NC_019843.3;YP_009047202.1;MERS_1ab;5736 | 0 | 0 |
| NC_019843.3;YP_009047202.1;MERS_1ab;5737 | 0 | 0 |
| NC_019843.3;YP_009047202.1;MERS_1ab;5738 | 0 | 0 |
| NC_019843.3;YP_009047202.1;MERS_1ab;5739 | 0 | 0 |
| NC_019843.3;YP_009047202.1;MERS_1ab;5740 | 0 | 0 |
| NC_019843.3;YP_009047202.1;MERS_1ab;5741 | 0 | 0 |
| NC_019843.3;YP_009047202.1;MERS_1ab;5742 | 0 | 0.025 |
| NC_019843.3;YP_009047202.1;MERS_1ab;5743 | 0 | 0 |
| NC_019843.3;YP_009047202.1;MERS_1ab;5744 | 0 | 0 |
| NC_019843.3;YP_009047202.1;MERS_1ab;5745 | 0 | 0 |
| NC_019843.3;YP_009047202.1;MERS_1ab;5746 | 0 | 0 |
| NC_019843.3;YP_009047202.1;MERS_1ab;5747 | 0 | 0 |
| NC_019843.3;YP_009047202.1;MERS_1ab;5748 | 0.05 | 0 |
| NC_019843.3;YP_009047202.1;MERS_1ab;5749 | 0.05 | 0.025 |
| NC_019843.3;YP_009047202.1;MERS_1ab;5750 | 0.05 | 0.05 |
| NC_019843.3;YP_009047202.1;MERS_1ab;5751 | 0 | 0.025 |
| NC_019843.3;YP_009047202.1;MERS_1ab;5752 | 0 | 0 |
| NC_019843.3;YP_009047202.1;MERS_1ab;5753 | 0 | 0 |

| | | |
|---|---|---|
| NC_019843.3;YP_009047202.1;MERS_1ab;5754 | 0.05 | 0.025 |
| NC_019843.3;YP_009047202.1;MERS_1ab;5755 | 0 | 0 |
| NC_019843.3;YP_009047202.1;MERS_1ab;5756 | 0 | 0 |
| NC_019843.3;YP_009047202.1;MERS_1ab;5757 | 0 | 0 |
| NC_019843.3;YP_009047202.1;MERS_1ab;5758 | 0 | 0 |
| NC_019843.3;YP_009047202.1;MERS_1ab;5759 | 0 | 0 |
| NC_019843.3;YP_009047202.1;MERS_1ab;5760 | 0 | 0 |
| NC_019843.3;YP_009047202.1;MERS_1ab;5761 | 0 | 0.025 |
| NC_019843.3;YP_009047202.1;MERS_1ab;5762 | 0 | 0.025 |
| NC_019843.3;YP_009047202.1;MERS_1ab;5763 | 0 | 0 |
| NC_019843.3;YP_009047202.1;MERS_1ab;5764 | 0 | 0 |
| NC_019843.3;YP_009047202.1;MERS_1ab;5765 | 0 | 0 |
| NC_019843.3;YP_009047202.1;MERS_1ab;5766 | 0 | 0 |
| NC_019843.3;YP_009047202.1;MERS_1ab;5767 | 0.05 | 0 |
| NC_019843.3;YP_009047202.1;MERS_1ab;5768 | 0 | 0 |
| NC_019843.3;YP_009047202.1;MERS_1ab;5769 | 0.05 | 0 |
| NC_019843.3;YP_009047202.1;MERS_1ab;5770 | 0 | 0 |
| NC_019843.3;YP_009047202.1;MERS_1ab;5771 | 0 | 0 |
| NC_019843.3;YP_009047202.1;MERS_1ab;5772 | 0 | 0 |
| NC_019843.3;YP_009047202.1;MERS_1ab;5773 | 0 | 0 |
| NC_019843.3;YP_009047202.1;MERS_1ab;5774 | 0 | 0 |
| NC_019843.3;YP_009047202.1;MERS_1ab;5775 | 0 | 0 |
| NC_019843.3;YP_009047202.1;MERS_1ab;5776 | 0 | 0 |
| NC_019843.3;YP_009047202.1;MERS_1ab;5777 | 0 | 0 |
| NC_019843.3;YP_009047202.1;MERS_1ab;5778 | 0 | 0 |
| NC_019843.3;YP_009047202.1;MERS_1ab;5779 | 0 | 0 |
| NC_019843.3;YP_009047202.1;MERS_1ab;5780 | 0 | 0 |
| NC_019843.3;YP_009047202.1;MERS_1ab;5781 | 0 | 0 |
| NC_019843.3;YP_009047202.1;MERS_1ab;5782 | 0 | 0 |
| NC_019843.3;YP_009047202.1;MERS_1ab;5783 | 0 | 0 |
| NC_019843.3;YP_009047202.1;MERS_1ab;5784 | 0 | 0 |
| NC_019843.3;YP_009047202.1;MERS_1ab;5785 | 0 | 0 |
| NC_019843.3;YP_009047202.1;MERS_1ab;5786 | 0.05 | 0.05 |
| NC_019843.3;YP_009047202.1;MERS_1ab;5787 | 0.05 | 0.05 |
| NC_019843.3;YP_009047202.1;MERS_1ab;5788 | 0 | 0 |
| NC_019843.3;YP_009047202.1;MERS_1ab;5789 | 0 | 0 |
| NC_019843.3;YP_009047202.1;MERS_1ab;5790 | 0 | 0 |
| NC_019843.3;YP_009047202.1;MERS_1ab;5791 | 0 | 0 |
| NC_019843.3;YP_009047202.1;MERS_1ab;5792 | 0.2 | 0.025 |
| NC_019843.3;YP_009047202.1;MERS_1ab;5793 | 0.05 | 0 |
| NC_019843.3;YP_009047202.1;MERS_1ab;5794 | 0 | 0 |
| NC_019843.3;YP_009047202.1;MERS_1ab;5795 | 0 | 0 |
| NC_019843.3;YP_009047202.1;MERS_1ab;5796 | 0 | 0 |

| | | |
|---|---|---|
| NC_019843.3;YP_009047202.1;MERS_1ab;5797 | 0 | 0 |
| NC_019843.3;YP_009047202.1;MERS_1ab;5798 | 0 | 0 |
| NC_019843.3;YP_009047202.1;MERS_1ab;5799 | 0 | 0 |
| NC_019843.3;YP_009047202.1;MERS_1ab;5800 | 0 | 0 |
| NC_019843.3;YP_009047202.1;MERS_1ab;5801 | 0 | 0 |
| NC_019843.3;YP_009047202.1;MERS_1ab;5802 | 0 | 0 |
| NC_019843.3;YP_009047202.1;MERS_1ab;5803 | 0 | 0 |
| NC_019843.3;YP_009047202.1;MERS_1ab;5804 | 0 | 0 |
| NC_019843.3;YP_009047202.1;MERS_1ab;5805 | 0 | 0 |
| NC_019843.3;YP_009047202.1;MERS_1ab;5806 | 0 | 0 |
| NC_019843.3;YP_009047202.1;MERS_1ab;5807 | 0 | 0 |
| NC_019843.3;YP_009047202.1;MERS_1ab;5808 | 0 | 0 |
| NC_019843.3;YP_009047202.1;MERS_1ab;5809 | 0 | 0 |
| NC_019843.3;YP_009047202.1;MERS_1ab;5810 | 0 | 0 |
| NC_019843.3;YP_009047202.1;MERS_1ab;5811 | 0 | 0 |
| NC_019843.3;YP_009047202.1;MERS_1ab;5812 | 0 | 0 |
| NC_019843.3;YP_009047202.1;MERS_1ab;5813 | 0 | 0 |
| NC_019843.3;YP_009047202.1;MERS_1ab;5814 | 0.15 | 0 |
| NC_019843.3;YP_009047202.1;MERS_1ab;5815 | 0 | 0 |
| NC_019843.3;YP_009047202.1;MERS_1ab;5816 | 0 | 0 |
| NC_019843.3;YP_009047202.1;MERS_1ab;5817 | 0.1 | 0 |
| NC_019843.3;YP_009047202.1;MERS_1ab;5818 | 0 | 0 |
| NC_019843.3;YP_009047202.1;MERS_1ab;5819 | 0.05 | 0 |
| NC_019843.3;YP_009047202.1;MERS_1ab;5820 | 0 | 0 |
| NC_019843.3;YP_009047202.1;MERS_1ab;5821 | 0 | 0 |
| NC_019843.3;YP_009047202.1;MERS_1ab;5822 | 0.15 | 0.025 |
| NC_019843.3;YP_009047202.1;MERS_1ab;5823 | 0.05 | 0 |
| NC_019843.3;YP_009047202.1;MERS_1ab;5824 | 0.05 | 0.025 |
| NC_019843.3;YP_009047202.1;MERS_1ab;5825 | 0 | 0.025 |
| NC_019843.3;YP_009047202.1;MERS_1ab;5826 | 0 | 0.025 |
| NC_019843.3;YP_009047202.1;MERS_1ab;5827 | 0 | 0.025 |
| NC_019843.3;YP_009047202.1;MERS_1ab;5828 | 0 | 0.025 |
| NC_019843.3;YP_009047202.1;MERS_1ab;5829 | 0 | 0.025 |
| NC_019843.3;YP_009047202.1;MERS_1ab;5830 | 0 | 0.075 |
| NC_019843.3;YP_009047202.1;MERS_1ab;5831 | 0 | 0 |
| NC_019843.3;YP_009047202.1;MERS_1ab;5832 | 0 | 0 |
| NC_019843.3;YP_009047202.1;MERS_1ab;5833 | 0 | 0 |
| NC_019843.3;YP_009047202.1;MERS_1ab;5834 | 0 | 0.025 |
| NC_019843.3;YP_009047202.1;MERS_1ab;5835 | 0 | 0.025 |
| NC_019843.3;YP_009047202.1;MERS_1ab;5836 | 0 | 0.025 |
| NC_019843.3;YP_009047202.1;MERS_1ab;5837 | 0 | 0.025 |
| NC_019843.3;YP_009047202.1;MERS_1ab;5838 | 0 | 0.025 |
| NC_019843.3;YP_009047202.1;MERS_1ab;5839 | 0 | 0.025 |

| | | |
|---|---|---|
| NC_019843.3;YP_009047202.1;MERS_1ab;5840 | 0 | 0.025 |
| NC_019843.3;YP_009047202.1;MERS_1ab;5841 | 0 | 0.025 |
| NC_019843.3;YP_009047202.1;MERS_1ab;5842 | 0 | 0.025 |
| NC_019843.3;YP_009047202.1;MERS_1ab;5843 | 0 | 0.025 |
| NC_019843.3;YP_009047202.1;MERS_1ab;5844 | 0 | 0 |
| NC_019843.3;YP_009047202.1;MERS_1ab;5845 | 0 | 0 |
| NC_019843.3;YP_009047202.1;MERS_1ab;5846 | 0 | 0 |
| NC_019843.3;YP_009047202.1;MERS_1ab;5847 | 0 | 0 |
| NC_019843.3;YP_009047202.1;MERS_1ab;5848 | 0 | 0 |
| NC_019843.3;YP_009047202.1;MERS_1ab;5849 | 0 | 0 |
| NC_019843.3;YP_009047202.1;MERS_1ab;5850 | 0 | 0 |
| NC_019843.3;YP_009047202.1;MERS_1ab;5851 | 0 | 0 |
| NC_019843.3;YP_009047202.1;MERS_1ab;5852 | 0.05 | 0 |
| NC_019843.3;YP_009047202.1;MERS_1ab;5853 | 0 | 0 |
| NC_019843.3;YP_009047202.1;MERS_1ab;5854 | 0.15 | 0.025 |
| NC_019843.3;YP_009047202.1;MERS_1ab;5855 | 0.1 | 0.025 |
| NC_019843.3;YP_009047202.1;MERS_1ab;5856 | 0.1 | 0.025 |
| NC_019843.3;YP_009047202.1;MERS_1ab;5857 | 0.2 | 0.025 |
| NC_019843.3;YP_009047202.1;MERS_1ab;5858 | 0.2 | 0.05 |
| NC_019843.3;YP_009047202.1;MERS_1ab;5859 | 0.2 | 0.075 |
| NC_019843.3;YP_009047202.1;MERS_1ab;5860 | 0.1 | 0.05 |
| NC_019843.3;YP_009047202.1;MERS_1ab;5861 | 0.1 | 0.025 |
| NC_019843.3;YP_009047202.1;MERS_1ab;5862 | 0.05 | 0 |
| NC_019843.3;YP_009047202.1;MERS_1ab;5863 | 0.05 | 0 |
| NC_019843.3;YP_009047202.1;MERS_1ab;5864 | 0.05 | 0 |
| NC_019843.3;YP_009047202.1;MERS_1ab;5865 | 0.1 | 0 |
| NC_019843.3;YP_009047202.1;MERS_1ab;5866 | 0.05 | 0 |
| NC_019843.3;YP_009047202.1;MERS_1ab;5867 | 0 | 0 |
| NC_019843.3;YP_009047202.1;MERS_1ab;5868 | 0 | 0 |
| NC_019843.3;YP_009047202.1;MERS_1ab;5869 | 0 | 0 |
| NC_019843.3;YP_009047202.1;MERS_1ab;5870 | 0 | 0 |
| NC_019843.3;YP_009047202.1;MERS_1ab;5871 | 0 | 0 |
| NC_019843.3;YP_009047202.1;MERS_1ab;5872 | 0 | 0 |
| NC_019843.3;YP_009047202.1;MERS_1ab;5873 | 0 | 0 |
| NC_019843.3;YP_009047202.1;MERS_1ab;5874 | 0 | 0 |
| NC_019843.3;YP_009047202.1;MERS_1ab;5875 | 0 | 0 |
| NC_019843.3;YP_009047202.1;MERS_1ab;5876 | 0 | 0 |
| NC_019843.3;YP_009047202.1;MERS_1ab;5877 | 0 | 0 |
| NC_019843.3;YP_009047202.1;MERS_1ab;5878 | 0 | 0 |
| NC_019843.3;YP_009047202.1;MERS_1ab;5879 | 0 | 0 |
| NC_019843.3;YP_009047202.1;MERS_1ab;5880 | 0 | 0 |
| NC_019843.3;YP_009047202.1;MERS_1ab;5881 | 0.05 | 0 |
| NC_019843.3;YP_009047202.1;MERS_1ab;5882 | 0.05 | 0 |

| | | |
|---|---|---|
| NC_019843.3;YP_009047202.1;MERS_1ab;5883 | 0 | 0 |
| NC_019843.3;YP_009047202.1;MERS_1ab;5884 | 0 | 0 |
| NC_019843.3;YP_009047202.1;MERS_1ab;5885 | 0 | 0 |
| NC_019843.3;YP_009047202.1;MERS_1ab;5886 | 0 | 0 |
| NC_019843.3;YP_009047202.1;MERS_1ab;5887 | 0 | 0 |
| NC_019843.3;YP_009047202.1;MERS_1ab;5888 | 0 | 0 |
| NC_019843.3;YP_009047202.1;MERS_1ab;5889 | 0 | 0 |
| NC_019843.3;YP_009047202.1;MERS_1ab;5890 | 0 | 0 |
| NC_019843.3;YP_009047202.1;MERS_1ab;5891 | 0 | 0 |
| NC_019843.3;YP_009047202.1;MERS_1ab;5892 | 0 | 0 |
| NC_019843.3;YP_009047202.1;MERS_1ab;5893 | 0 | 0 |
| NC_019843.3;YP_009047202.1;MERS_1ab;5894 | 0 | 0 |
| NC_019843.3;YP_009047202.1;MERS_1ab;5895 | 0 | 0 |
| NC_019843.3;YP_009047202.1;MERS_1ab;5896 | 0 | 0 |
| NC_019843.3;YP_009047202.1;MERS_1ab;5897 | 0 | 0 |
| NC_019843.3;YP_009047202.1;MERS_1ab;5898 | 0 | 0 |
| NC_019843.3;YP_009047202.1;MERS_1ab;5899 | 0 | 0 |
| NC_019843.3;YP_009047202.1;MERS_1ab;5900 | 0 | 0 |
| NC_019843.3;YP_009047202.1;MERS_1ab;5901 | 0 | 0 |
| NC_019843.3;YP_009047202.1;MERS_1ab;5902 | 0 | 0 |
| NC_019843.3;YP_009047202.1;MERS_1ab;5903 | 0 | 0 |
| NC_019843.3;YP_009047202.1;MERS_1ab;5904 | 0 | 0 |
| NC_019843.3;YP_009047202.1;MERS_1ab;5905 | 0 | 0 |
| NC_019843.3;YP_009047202.1;MERS_1ab;5906 | 0 | 0 |
| NC_019843.3;YP_009047202.1;MERS_1ab;5907 | 0 | 0 |
| NC_019843.3;YP_009047202.1;MERS_1ab;5908 | 0 | 0 |
| NC_019843.3;YP_009047202.1;MERS_1ab;5909 | 0 | 0.025 |
| NC_019843.3;YP_009047202.1;MERS_1ab;5910 | 0 | 0 |
| NC_019843.3;YP_009047202.1;MERS_1ab;5911 | 0 | 0.025 |
| NC_019843.3;YP_009047202.1;MERS_1ab;5912 | 0 | 0 |
| NC_019843.3;YP_009047202.1;MERS_1ab;5913 | 0 | 0.025 |
| NC_019843.3;YP_009047202.1;MERS_1ab;5914 | 0 | 0.05 |
| NC_019843.3;YP_009047202.1;MERS_1ab;5915 | 0 | 0 |
| NC_019843.3;YP_009047202.1;MERS_1ab;5916 | 0 | 0.05 |
| NC_019843.3;YP_009047202.1;MERS_1ab;5917 | 0.05 | 0 |
| NC_019843.3;YP_009047202.1;MERS_1ab;5918 | 0 | 0.025 |
| NC_019843.3;YP_009047202.1;MERS_1ab;5919 | 0.05 | 0.025 |
| NC_019843.3;YP_009047202.1;MERS_1ab;5920 | 0 | 0 |
| NC_019843.3;YP_009047202.1;MERS_1ab;5921 | 0 | 0 |
| NC_019843.3;YP_009047202.1;MERS_1ab;5922 | 0 | 0 |
| NC_019843.3;YP_009047202.1;MERS_1ab;5923 | 0 | 0.025 |
| NC_019843.3;YP_009047202.1;MERS_1ab;5924 | 0.05 | 0.025 |
| NC_019843.3;YP_009047202.1;MERS_1ab;5925 | 0.05 | 0.05 |

| | | |
|---|---|---|
| NC_019843.3;YP_009047202.1;MERS_1ab;5926 | 0.05 | 0.05 |
| NC_019843.3;YP_009047202.1;MERS_1ab;5927 | 0.1 | 0.05 |
| NC_019843.3;YP_009047202.1;MERS_1ab;5928 | 0.1 | 0.05 |
| NC_019843.3;YP_009047202.1;MERS_1ab;5929 | 0.15 | 0.05 |
| NC_019843.3;YP_009047202.1;MERS_1ab;5930 | 0.15 | 0.1 |
| NC_019843.3;YP_009047202.1;MERS_1ab;5931 | 0.05 | 0.025 |
| NC_019843.3;YP_009047202.1;MERS_1ab;5932 | 0.05 | 0.025 |
| NC_019843.3;YP_009047202.1;MERS_1ab;5933 | 0 | 0 |
| NC_019843.3;YP_009047202.1;MERS_1ab;5934 | 0.05 | 0 |
| NC_019843.3;YP_009047202.1;MERS_1ab;5935 | 0.05 | 0 |
| NC_019843.3;YP_009047202.1;MERS_1ab;5936 | 0.05 | 0 |
| NC_019843.3;YP_009047202.1;MERS_1ab;5937 | 0 | 0 |
| NC_019843.3;YP_009047202.1;MERS_1ab;5938 | 0 | 0 |
| NC_019843.3;YP_009047202.1;MERS_1ab;5939 | 0 | 0 |
| NC_019843.3;YP_009047202.1;MERS_1ab;5940 | 0 | 0.025 |
| NC_019843.3;YP_009047202.1;MERS_1ab;5941 | 0.05 | 0 |
| NC_019843.3;YP_009047202.1;MERS_1ab;5942 | 0.05 | 0 |
| NC_019843.3;YP_009047202.1;MERS_1ab;5943 | 0 | 0.025 |
| NC_019843.3;YP_009047202.1;MERS_1ab;5944 | 0 | 0 |
| NC_019843.3;YP_009047202.1;MERS_1ab;5945 | 0 | 0 |
| NC_019843.3;YP_009047202.1;MERS_1ab;5946 | 0 | 0 |
| NC_019843.3;YP_009047202.1;MERS_1ab;5947 | 0 | 0 |
| NC_019843.3;YP_009047202.1;MERS_1ab;5948 | 0 | 0 |
| NC_019843.3;YP_009047202.1;MERS_1ab;5949 | 0 | 0 |
| NC_019843.3;YP_009047202.1;MERS_1ab;5950 | 0 | 0 |
| NC_019843.3;YP_009047202.1;MERS_1ab;5951 | 0 | 0 |
| NC_019843.3;YP_009047202.1;MERS_1ab;5952 | 0 | 0 |
| NC_019843.3;YP_009047202.1;MERS_1ab;5953 | 0 | 0 |
| NC_019843.3;YP_009047202.1;MERS_1ab;5954 | 0 | 0 |
| NC_019843.3;YP_009047202.1;MERS_1ab;5955 | 0 | 0 |
| NC_019843.3;YP_009047202.1;MERS_1ab;5956 | 0 | 0 |
| NC_019843.3;YP_009047202.1;MERS_1ab;5957 | 0 | 0 |
| NC_019843.3;YP_009047202.1;MERS_1ab;5958 | 0 | 0 |
| NC_019843.3;YP_009047202.1;MERS_1ab;5959 | 0 | 0.025 |
| NC_019843.3;YP_009047202.1;MERS_1ab;5960 | 0 | 0.025 |
| NC_019843.3;YP_009047202.1;MERS_1ab;5961 | 0.05 | 0.025 |
| NC_019843.3;YP_009047202.1;MERS_1ab;5962 | 0 | 0.025 |
| NC_019843.3;YP_009047202.1;MERS_1ab;5963 | 0 | 0.025 |
| NC_019843.3;YP_009047202.1;MERS_1ab;5964 | 0.05 | 0.025 |
| NC_019843.3;YP_009047202.1;MERS_1ab;5965 | 0 | 0.025 |
| NC_019843.3;YP_009047202.1;MERS_1ab;5966 | 0 | 0 |
| NC_019843.3;YP_009047202.1;MERS_1ab;5967 | 0 | 0 |
| NC_019843.3;YP_009047202.1;MERS_1ab;5968 | 0 | 0 |

| | | |
|---|---|---|
| NC_019843.3;YP_009047202.1;MERS_1ab;5969 | 0 | 0 |
| NC_019843.3;YP_009047202.1;MERS_1ab;5970 | 0 | 0 |
| NC_019843.3;YP_009047202.1;MERS_1ab;5971 | 0 | 0 |
| NC_019843.3;YP_009047202.1;MERS_1ab;5972 | 0 | 0 |
| NC_019843.3;YP_009047202.1;MERS_1ab;5973 | 0 | 0 |
| NC_019843.3;YP_009047202.1;MERS_1ab;5974 | 0 | 0 |
| NC_019843.3;YP_009047202.1;MERS_1ab;5975 | 0 | 0 |
| NC_019843.3;YP_009047202.1;MERS_1ab;5976 | 0 | 0 |
| NC_019843.3;YP_009047202.1;MERS_1ab;5977 | 0 | 0 |
| NC_019843.3;YP_009047202.1;MERS_1ab;5978 | 0 | 0 |
| NC_019843.3;YP_009047202.1;MERS_1ab;5979 | 0 | 0 |
| NC_019843.3;YP_009047202.1;MERS_1ab;5980 | 0 | 0 |
| NC_019843.3;YP_009047202.1;MERS_1ab;5981 | 0 | 0 |
| NC_019843.3;YP_009047202.1;MERS_1ab;5982 | 0 | 0.025 |
| NC_019843.3;YP_009047202.1;MERS_1ab;5983 | 0 | 0.025 |
| NC_019843.3;YP_009047202.1;MERS_1ab;5984 | 0 | 0 |
| NC_019843.3;YP_009047202.1;MERS_1ab;5985 | 0 | 0 |
| NC_019843.3;YP_009047202.1;MERS_1ab;5986 | 0 | 0 |
| NC_019843.3;YP_009047202.1;MERS_1ab;5987 | 0 | 0.025 |
| NC_019843.3;YP_009047202.1;MERS_1ab;5988 | 0 | 0.025 |
| NC_019843.3;YP_009047202.1;MERS_1ab;5989 | 0 | 0.025 |
| NC_019843.3;YP_009047202.1;MERS_1ab;5990 | 0 | 0.025 |
| NC_019843.3;YP_009047202.1;MERS_1ab;5991 | 0 | 0.025 |
| NC_019843.3;YP_009047202.1;MERS_1ab;5992 | 0 | 0.025 |
| NC_019843.3;YP_009047202.1;MERS_1ab;5993 | 0 | 0.05 |
| NC_019843.3;YP_009047202.1;MERS_1ab;5994 | 0 | 0.025 |
| NC_019843.3;YP_009047202.1;MERS_1ab;5995 | 0 | 0 |
| NC_019843.3;YP_009047202.1;MERS_1ab;5996 | 0 | 0 |
| NC_019843.3;YP_009047202.1;MERS_1ab;5997 | 0 | 0 |
| NC_019843.3;YP_009047202.1;MERS_1ab;5998 | 0 | 0 |
| NC_019843.3;YP_009047202.1;MERS_1ab;5999 | 0 | 0 |
| NC_019843.3;YP_009047202.1;MERS_1ab;6000 | 0 | 0.025 |
| NC_019843.3;YP_009047202.1;MERS_1ab;6001 | 0 | 0 |
| NC_019843.3;YP_009047202.1;MERS_1ab;6002 | 0 | 0 |
| NC_019843.3;YP_009047202.1;MERS_1ab;6003 | 0 | 0 |
| NC_019843.3;YP_009047202.1;MERS_1ab;6004 | 0 | 0 |
| NC_019843.3;YP_009047202.1;MERS_1ab;6005 | 0 | 0 |
| NC_019843.3;YP_009047202.1;MERS_1ab;6006 | 0 | 0 |
| NC_019843.3;YP_009047202.1;MERS_1ab;6007 | 0 | 0 |
| NC_019843.3;YP_009047202.1;MERS_1ab;6008 | 0 | 0.025 |
| NC_019843.3;YP_009047202.1;MERS_1ab;6009 | 0 | 0 |
| NC_019843.3;YP_009047202.1;MERS_1ab;6010 | 0 | 0 |
| NC_019843.3;YP_009047202.1;MERS_1ab;6011 | 0 | 0 |

| | | |
|---|---|---|
| NC_019843.3;YP_009047202.1;MERS_1ab;6012 | 0 | 0 |
| NC_019843.3;YP_009047202.1;MERS_1ab;6013 | 0 | 0 |
| NC_019843.3;YP_009047202.1;MERS_1ab;6014 | 0 | 0 |
| NC_019843.3;YP_009047202.1;MERS_1ab;6015 | 0 | 0 |
| NC_019843.3;YP_009047202.1;MERS_1ab;6016 | 0 | 0 |
| NC_019843.3;YP_009047202.1;MERS_1ab;6017 | 0 | 0 |
| NC_019843.3;YP_009047202.1;MERS_1ab;6018 | 0 | 0 |
| NC_019843.3;YP_009047202.1;MERS_1ab;6019 | 0 | 0 |
| NC_019843.3;YP_009047202.1;MERS_1ab;6020 | 0 | 0 |
| NC_019843.3;YP_009047202.1;MERS_1ab;6021 | 0 | 0 |
| NC_019843.3;YP_009047202.1;MERS_1ab;6022 | 0 | 0.025 |
| NC_019843.3;YP_009047202.1;MERS_1ab;6023 | 0 | 0.025 |
| NC_019843.3;YP_009047202.1;MERS_1ab;6024 | 0 | 0.025 |
| NC_019843.3;YP_009047202.1;MERS_1ab;6025 | 0 | 0.025 |
| NC_019843.3;YP_009047202.1;MERS_1ab;6026 | 0 | 0.025 |
| NC_019843.3;YP_009047202.1;MERS_1ab;6027 | 0 | 0.025 |
| NC_019843.3;YP_009047202.1;MERS_1ab;6028 | 0 | 0.025 |
| NC_019843.3;YP_009047202.1;MERS_1ab;6029 | 0 | 0.025 |
| NC_019843.3;YP_009047202.1;MERS_1ab;6030 | 0 | 0.05 |
| NC_019843.3;YP_009047202.1;MERS_1ab;6031 | 0.1 | 0.15 |
| NC_019843.3;YP_009047202.1;MERS_1ab;6032 | 0 | 0 |
| NC_019843.3;YP_009047202.1;MERS_1ab;6033 | 0 | 0 |
| NC_019843.3;YP_009047202.1;MERS_1ab;6034 | 0 | 0 |
| NC_019843.3;YP_009047202.1;MERS_1ab;6035 | 0 | 0 |
| NC_019843.3;YP_009047202.1;MERS_1ab;6036 | 0 | 0 |
| NC_019843.3;YP_009047202.1;MERS_1ab;6037 | 0 | 0 |
| NC_019843.3;YP_009047202.1;MERS_1ab;6038 | 0 | 0 |
| NC_019843.3;YP_009047202.1;MERS_1ab;6039 | 0 | 0 |
| NC_019843.3;YP_009047202.1;MERS_1ab;6040 | 0 | 0 |
| NC_019843.3;YP_009047202.1;MERS_1ab;6041 | 0 | 0 |
| NC_019843.3;YP_009047202.1;MERS_1ab;6042 | 0 | 0 |
| NC_019843.3;YP_009047202.1;MERS_1ab;6043 | 0 | 0 |
| NC_019843.3;YP_009047202.1;MERS_1ab;6044 | 0 | 0.025 |
| NC_019843.3;YP_009047202.1;MERS_1ab;6045 | 0 | 0 |
| NC_019843.3;YP_009047202.1;MERS_1ab;6046 | 0 | 0 |
| NC_019843.3;YP_009047202.1;MERS_1ab;6047 | 0 | 0 |
| NC_019843.3;YP_009047202.1;MERS_1ab;6048 | 0 | 0.025 |
| NC_019843.3;YP_009047202.1;MERS_1ab;6049 | 0 | 0 |
| NC_019843.3;YP_009047202.1;MERS_1ab;6050 | 0 | 0 |
| NC_019843.3;YP_009047202.1;MERS_1ab;6051 | 0.05 | 0 |
| NC_019843.3;YP_009047202.1;MERS_1ab;6052 | 0 | 0 |
| NC_019843.3;YP_009047202.1;MERS_1ab;6053 | 0 | 0 |
| NC_019843.3;YP_009047202.1;MERS_1ab;6054 | 0 | 0 |

| | | |
|---|---|---|
| NC_019843.3;YP_009047202.1;MERS_1ab;6055 | 0 | 0 |
| NC_019843.3;YP_009047202.1;MERS_1ab;6056 | 0 | 0 |
| NC_019843.3;YP_009047202.1;MERS_1ab;6057 | 0 | 0 |
| NC_019843.3;YP_009047202.1;MERS_1ab;6058 | 0 | 0 |
| NC_019843.3;YP_009047202.1;MERS_1ab;6059 | 0 | 0 |
| NC_019843.3;YP_009047202.1;MERS_1ab;6060 | 0 | 0 |
| NC_019843.3;YP_009047202.1;MERS_1ab;6061 | 0 | 0 |
| NC_019843.3;YP_009047202.1;MERS_1ab;6062 | 0 | 0 |
| NC_019843.3;YP_009047202.1;MERS_1ab;6063 | 0 | 0 |
| NC_019843.3;YP_009047202.1;MERS_1ab;6064 | 0 | 0 |
| NC_019843.3;YP_009047202.1;MERS_1ab;6065 | 0 | 0 |
| NC_019843.3;YP_009047202.1;MERS_1ab;6066 | 0 | 0 |
| NC_019843.3;YP_009047202.1;MERS_1ab;6067 | 0 | 0 |
| NC_019843.3;YP_009047202.1;MERS_1ab;6068 | 0 | 0 |
| NC_019843.3;YP_009047202.1;MERS_1ab;6069 | 0.05 | 0 |
| NC_019843.3;YP_009047202.1;MERS_1ab;6070 | 0.05 | 0 |
| NC_019843.3;YP_009047202.1;MERS_1ab;6071 | 0 | 0 |
| NC_019843.3;YP_009047202.1;MERS_1ab;6072 | 0 | 0.025 |
| NC_019843.3;YP_009047202.1;MERS_1ab;6073 | 0 | 0.025 |
| NC_019843.3;YP_009047202.1;MERS_1ab;6074 | 0 | 0 |
| NC_019843.3;YP_009047202.1;MERS_1ab;6075 | 0 | 0 |
| NC_019843.3;YP_009047202.1;MERS_1ab;6076 | 0 | 0 |
| NC_019843.3;YP_009047202.1;MERS_1ab;6077 | 0 | 0 |
| NC_019843.3;YP_009047202.1;MERS_1ab;6078 | 0 | 0 |
| NC_019843.3;YP_009047202.1;MERS_1ab;6079 | 0 | 0 |
| NC_019843.3;YP_009047202.1;MERS_1ab;6080 | 0 | 0 |
| NC_019843.3;YP_009047202.1;MERS_1ab;6081 | 0 | 0 |
| NC_019843.3;YP_009047202.1;MERS_1ab;6082 | 0 | 0 |
| NC_019843.3;YP_009047202.1;MERS_1ab;6083 | 0 | 0 |
| NC_019843.3;YP_009047202.1;MERS_1ab;6084 | 0 | 0 |
| NC_019843.3;YP_009047202.1;MERS_1ab;6085 | 0 | 0 |
| NC_019843.3;YP_009047202.1;MERS_1ab;6086 | 0 | 0 |
| NC_019843.3;YP_009047202.1;MERS_1ab;6087 | 0 | 0 |
| NC_019843.3;YP_009047202.1;MERS_1ab;6088 | 0 | 0 |
| NC_019843.3;YP_009047202.1;MERS_1ab;6089 | 0 | 0 |
| NC_019843.3;YP_009047202.1;MERS_1ab;6090 | 0 | 0 |
| NC_019843.3;YP_009047202.1;MERS_1ab;6091 | 0 | 0 |
| NC_019843.3;YP_009047202.1;MERS_1ab;6092 | 0 | 0 |
| NC_019843.3;YP_009047202.1;MERS_1ab;6093 | 0 | 0 |
| NC_019843.3;YP_009047202.1;MERS_1ab;6094 | 0 | 0 |
| NC_019843.3;YP_009047202.1;MERS_1ab;6095 | 0 | 0 |
| NC_019843.3;YP_009047202.1;MERS_1ab;6096 | 0 | 0 |
| NC_019843.3;YP_009047202.1;MERS_1ab;6097 | 0 | 0 |

| | | |
|---|---|---|
| NC_019843.3;YP_009047202.1;MERS_1ab;6098 | 0 | 0 |
| NC_019843.3;YP_009047202.1;MERS_1ab;6099 | 0.05 | 0.025 |
| NC_019843.3;YP_009047202.1;MERS_1ab;6100 | 0 | 0 |
| NC_019843.3;YP_009047202.1;MERS_1ab;6101 | 0 | 0 |
| NC_019843.3;YP_009047202.1;MERS_1ab;6102 | 0 | 0 |
| NC_019843.3;YP_009047202.1;MERS_1ab;6103 | 0 | 0 |
| NC_019843.3;YP_009047202.1;MERS_1ab;6104 | 0.05 | 0.025 |
| NC_019843.3;YP_009047202.1;MERS_1ab;6105 | 0 | 0 |
| NC_019843.3;YP_009047202.1;MERS_1ab;6106 | 0 | 0 |
| NC_019843.3;YP_009047202.1;MERS_1ab;6107 | 0 | 0 |
| NC_019843.3;YP_009047202.1;MERS_1ab;6108 | 0 | 0 |
| NC_019843.3;YP_009047202.1;MERS_1ab;6109 | 0.05 | 0 |
| NC_019843.3;YP_009047202.1;MERS_1ab;6110 | 0 | 0 |
| NC_019843.3;YP_009047202.1;MERS_1ab;6111 | 0 | 0 |
| NC_019843.3;YP_009047202.1;MERS_1ab;6112 | 0 | 0 |
| NC_019843.3;YP_009047202.1;MERS_1ab;6113 | 0 | 0 |
| NC_019843.3;YP_009047202.1;MERS_1ab;6114 | 0 | 0 |
| NC_019843.3;YP_009047202.1;MERS_1ab;6115 | 0 | 0 |
| NC_019843.3;YP_009047202.1;MERS_1ab;6116 | 0 | 0 |
| NC_019843.3;YP_009047202.1;MERS_1ab;6117 | 0 | 0 |
| NC_019843.3;YP_009047202.1;MERS_1ab;6118 | 0 | 0 |
| NC_019843.3;YP_009047202.1;MERS_1ab;6119 | 0 | 0.025 |
| NC_019843.3;YP_009047202.1;MERS_1ab;6120 | 0 | 0 |
| NC_019843.3;YP_009047202.1;MERS_1ab;6121 | 0 | 0 |
| NC_019843.3;YP_009047202.1;MERS_1ab;6122 | 0 | 0 |
| NC_019843.3;YP_009047202.1;MERS_1ab;6123 | 0 | 0 |
| NC_019843.3;YP_009047202.1;MERS_1ab;6124 | 0 | 0.05 |
| NC_019843.3;YP_009047202.1;MERS_1ab;6125 | 0 | 0 |
| NC_019843.3;YP_009047202.1;MERS_1ab;6126 | 0 | 0 |
| NC_019843.3;YP_009047202.1;MERS_1ab;6127 | 0 | 0 |
| NC_019843.3;YP_009047202.1;MERS_1ab;6128 | 0 | 0 |
| NC_019843.3;YP_009047202.1;MERS_1ab;6129 | 0 | 0 |
| NC_019843.3;YP_009047202.1;MERS_1ab;6130 | 0 | 0 |
| NC_019843.3;YP_009047202.1;MERS_1ab;6131 | 0 | 0 |
| NC_019843.3;YP_009047202.1;MERS_1ab;6132 | 0 | 0 |
| NC_019843.3;YP_009047202.1;MERS_1ab;6133 | 0 | 0 |
| NC_019843.3;YP_009047202.1;MERS_1ab;6134 | 0 | 0 |
| NC_019843.3;YP_009047202.1;MERS_1ab;6135 | 0 | 0 |
| NC_019843.3;YP_009047202.1;MERS_1ab;6136 | 0 | 0 |
| NC_019843.3;YP_009047202.1;MERS_1ab;6137 | 0 | 0 |
| NC_019843.3;YP_009047202.1;MERS_1ab;6138 | 0 | 0 |
| NC_019843.3;YP_009047202.1;MERS_1ab;6139 | 0 | 0 |
| NC_019843.3;YP_009047202.1;MERS_1ab;6140 | 0 | 0 |

| | | |
|---|---|---|
| NC_019843.3;YP_009047202.1;MERS_1ab;6141 | 0 | 0 |
| NC_019843.3;YP_009047202.1;MERS_1ab;6142 | 0 | 0 |
| NC_019843.3;YP_009047202.1;MERS_1ab;6143 | 0 | 0 |
| NC_019843.3;YP_009047202.1;MERS_1ab;6144 | 0 | 0 |
| NC_019843.3;YP_009047202.1;MERS_1ab;6145 | 0 | 0 |
| NC_019843.3;YP_009047202.1;MERS_1ab;6146 | 0 | 0 |
| NC_019843.3;YP_009047202.1;MERS_1ab;6147 | 0 | 0 |
| NC_019843.3;YP_009047202.1;MERS_1ab;6148 | 0 | 0 |
| NC_019843.3;YP_009047202.1;MERS_1ab;6149 | 0 | 0 |
| NC_019843.3;YP_009047202.1;MERS_1ab;6150 | 0 | 0 |
| NC_019843.3;YP_009047202.1;MERS_1ab;6151 | 0 | 0 |
| NC_019843.3;YP_009047202.1;MERS_1ab;6152 | 0 | 0 |
| NC_019843.3;YP_009047202.1;MERS_1ab;6153 | 0 | 0 |
| NC_019843.3;YP_009047202.1;MERS_1ab;6154 | 0 | 0 |
| NC_019843.3;YP_009047202.1;MERS_1ab;6155 | 0 | 0 |
| NC_019843.3;YP_009047202.1;MERS_1ab;6156 | 0 | 0 |
| NC_019843.3;YP_009047202.1;MERS_1ab;6157 | 0 | 0 |
| NC_019843.3;YP_009047202.1;MERS_1ab;6158 | 0 | 0 |
| NC_019843.3;YP_009047202.1;MERS_1ab;6159 | 0 | 0 |
| NC_019843.3;YP_009047202.1;MERS_1ab;6160 | 0 | 0 |
| NC_019843.3;YP_009047202.1;MERS_1ab;6161 | 0 | 0 |
| NC_019843.3;YP_009047202.1;MERS_1ab;6162 | 0 | 0 |
| NC_019843.3;YP_009047202.1;MERS_1ab;6163 | 0.05 | 0.05 |
| NC_019843.3;YP_009047202.1;MERS_1ab;6164 | 0 | 0 |
| NC_019843.3;YP_009047202.1;MERS_1ab;6165 | 0 | 0 |
| NC_019843.3;YP_009047202.1;MERS_1ab;6166 | 0 | 0 |
| NC_019843.3;YP_009047202.1;MERS_1ab;6167 | 0 | 0 |
| NC_019843.3;YP_009047202.1;MERS_1ab;6168 | 0.05 | 0 |
| NC_019843.3;YP_009047202.1;MERS_1ab;6169 | 0.05 | 0 |
| NC_019843.3;YP_009047202.1;MERS_1ab;6170 | 0.05 | 0.05 |
| NC_019843.3;YP_009047202.1;MERS_1ab;6171 | 0 | 0 |
| NC_019843.3;YP_009047202.1;MERS_1ab;6172 | 0 | 0 |
| NC_019843.3;YP_009047202.1;MERS_1ab;6173 | 0 | 0 |
| NC_019843.3;YP_009047202.1;MERS_1ab;6174 | 0 | 0 |
| NC_019843.3;YP_009047202.1;MERS_1ab;6175 | 0 | 0 |
| NC_019843.3;YP_009047202.1;MERS_1ab;6176 | 0 | 0 |
| NC_019843.3;YP_009047202.1;MERS_1ab;6177 | 0 | 0 |
| NC_019843.3;YP_009047202.1;MERS_1ab;6178 | 0 | 0 |
| NC_019843.3;YP_009047202.1;MERS_1ab;6179 | 0 | 0 |
| NC_019843.3;YP_009047202.1;MERS_1ab;6180 | 0 | 0 |
| NC_019843.3;YP_009047202.1;MERS_1ab;6181 | 0 | 0 |
| NC_019843.3;YP_009047202.1;MERS_1ab;6182 | 0 | 0 |
| NC_019843.3;YP_009047202.1;MERS_1ab;6183 | 0 | 0 |

| | | |
|---|---|---|
| NC_019843.3;YP_009047202.1;MERS_1ab;6184 | 0 | 0 |
| NC_019843.3;YP_009047202.1;MERS_1ab;6185 | 0 | 0 |
| NC_019843.3;YP_009047202.1;MERS_1ab;6186 | 0 | 0 |
| NC_019843.3;YP_009047202.1;MERS_1ab;6187 | 0 | 0 |
| NC_019843.3;YP_009047202.1;MERS_1ab;6188 | 0 | 0 |
| NC_019843.3;YP_009047202.1;MERS_1ab;6189 | 0 | 0 |
| NC_019843.3;YP_009047202.1;MERS_1ab;6190 | 0 | 0 |
| NC_019843.3;YP_009047202.1;MERS_1ab;6191 | 0 | 0 |
| NC_019843.3;YP_009047202.1;MERS_1ab;6192 | 0 | 0 |
| NC_019843.3;YP_009047202.1;MERS_1ab;6193 | 0 | 0 |
| NC_019843.3;YP_009047202.1;MERS_1ab;6194 | 0 | 0 |
| NC_019843.3;YP_009047202.1;MERS_1ab;6195 | 0 | 0.025 |
| NC_019843.3;YP_009047202.1;MERS_1ab;6196 | 0.15 | 0.075 |
| NC_019843.3;YP_009047202.1;MERS_1ab;6197 | 0.1 | 0.1 |
| NC_019843.3;YP_009047202.1;MERS_1ab;6198 | 0.1 | 0.1 |
| NC_019843.3;YP_009047202.1;MERS_1ab;6199 | 0.1 | 0.1 |
| NC_019843.3;YP_009047202.1;MERS_1ab;6200 | 0.1 | 0.05 |
| NC_019843.3;YP_009047202.1;MERS_1ab;6201 | 0.05 | 0.025 |
| NC_019843.3;YP_009047202.1;MERS_1ab;6202 | 0.05 | 0 |
| NC_019843.3;YP_009047202.1;MERS_1ab;6203 | 0.05 | 0 |
| NC_019843.3;YP_009047202.1;MERS_1ab;6204 | 0 | 0 |
| NC_019843.3;YP_009047202.1;MERS_1ab;6205 | 0 | 0 |
| NC_019843.3;YP_009047202.1;MERS_1ab;6206 | 0 | 0 |
| NC_019843.3;YP_009047202.1;MERS_1ab;6207 | 0 | 0 |
| NC_019843.3;YP_009047202.1;MERS_1ab;6208 | 0 | 0 |
| NC_019843.3;YP_009047202.1;MERS_1ab;6209 | 0 | 0 |
| NC_019843.3;YP_009047202.1;MERS_1ab;6210 | 0 | 0 |
| NC_019843.3;YP_009047202.1;MERS_1ab;6211 | 0 | 0 |
| NC_019843.3;YP_009047202.1;MERS_1ab;6212 | 0 | 0 |
| NC_019843.3;YP_009047202.1;MERS_1ab;6213 | 0.05 | 0.025 |
| NC_019843.3;YP_009047202.1;MERS_1ab;6214 | 0 | 0 |
| NC_019843.3;YP_009047202.1;MERS_1ab;6215 | 0 | 0 |
| NC_019843.3;YP_009047202.1;MERS_1ab;6216 | 0 | 0 |
| NC_019843.3;YP_009047202.1;MERS_1ab;6217 | 0 | 0 |
| NC_019843.3;YP_009047202.1;MERS_1ab;6218 | 0 | 0 |
| NC_019843.3;YP_009047202.1;MERS_1ab;6219 | 0 | 0 |
| NC_019843.3;YP_009047202.1;MERS_1ab;6220 | 0 | 0 |
| NC_019843.3;YP_009047202.1;MERS_1ab;6221 | 0 | 0 |
| NC_019843.3;YP_009047202.1;MERS_1ab;6222 | 0 | 0 |
| NC_019843.3;YP_009047202.1;MERS_1ab;6223 | 0 | 0 |
| NC_019843.3;YP_009047202.1;MERS_1ab;6224 | 0 | 0 |
| NC_019843.3;YP_009047202.1;MERS_1ab;6225 | 0 | 0 |
| NC_019843.3;YP_009047202.1;MERS_1ab;6226 | 0 | 0 |

| | | |
|---|---|---|
| NC_019843.3;YP_009047202.1;MERS_1ab;6227 | 0 | 0.05 |
| NC_019843.3;YP_009047202.1;MERS_1ab;6228 | 0 | 0.025 |
| NC_019843.3;YP_009047202.1;MERS_1ab;6229 | 0 | 0.025 |
| NC_019843.3;YP_009047202.1;MERS_1ab;6230 | 0.05 | 0 |
| NC_019843.3;YP_009047202.1;MERS_1ab;6231 | 0.05 | 0.025 |
| NC_019843.3;YP_009047202.1;MERS_1ab;6232 | 0.05 | 0 |
| NC_019843.3;YP_009047202.1;MERS_1ab;6233 | 0.1 | 0 |
| NC_019843.3;YP_009047202.1;MERS_1ab;6234 | 0.1 | 0.05 |
| NC_019843.3;YP_009047202.1;MERS_1ab;6235 | 0.15 | 0 |
| NC_019843.3;YP_009047202.1;MERS_1ab;6236 | 0.15 | 0 |
| NC_019843.3;YP_009047202.1;MERS_1ab;6237 | 0.05 | 0.025 |
| NC_019843.3;YP_009047202.1;MERS_1ab;6238 | 0.05 | 0.025 |
| NC_019843.3;YP_009047202.1;MERS_1ab;6239 | 0.05 | 0 |
| NC_019843.3;YP_009047202.1;MERS_1ab;6240 | 0.05 | 0.025 |
| NC_019843.3;YP_009047202.1;MERS_1ab;6241 | 0.05 | 0.025 |
| NC_019843.3;YP_009047202.1;MERS_1ab;6242 | 0.05 | 0.025 |
| NC_019843.3;YP_009047202.1;MERS_1ab;6243 | 0 | 0 |
| NC_019843.3;YP_009047202.1;MERS_1ab;6244 | 0 | 0 |
| NC_019843.3;YP_009047202.1;MERS_1ab;6245 | 0 | 0 |
| NC_019843.3;YP_009047202.1;MERS_1ab;6246 | 0 | 0 |
| NC_019843.3;YP_009047202.1;MERS_1ab;6247 | 0 | 0 |
| NC_019843.3;YP_009047202.1;MERS_1ab;6248 | 0 | 0 |
| NC_019843.3;YP_009047202.1;MERS_1ab;6249 | 0 | 0 |
| NC_019843.3;YP_009047202.1;MERS_1ab;6250 | 0 | 0 |
| NC_019843.3;YP_009047202.1;MERS_1ab;6251 | 0.05 | 0.025 |
| NC_019843.3;YP_009047202.1;MERS_1ab;6252 | 0 | 0 |
| NC_019843.3;YP_009047202.1;MERS_1ab;6253 | 0 | 0 |
| NC_019843.3;YP_009047202.1;MERS_1ab;6254 | 0 | 0 |
| NC_019843.3;YP_009047202.1;MERS_1ab;6255 | 0 | 0 |
| NC_019843.3;YP_009047202.1;MERS_1ab;6256 | 0 | 0 |
| NC_019843.3;YP_009047202.1;MERS_1ab;6257 | 0 | 0 |
| NC_019843.3;YP_009047202.1;MERS_1ab;6258 | 0 | 0 |
| NC_019843.3;YP_009047202.1;MERS_1ab;6259 | 0 | 0 |
| NC_019843.3;YP_009047202.1;MERS_1ab;6260 | 0 | 0 |
| NC_019843.3;YP_009047202.1;MERS_1ab;6261 | 0.05 | 0 |
| NC_019843.3;YP_009047202.1;MERS_1ab;6262 | 0 | 0 |
| NC_019843.3;YP_009047202.1;MERS_1ab;6263 | 0.05 | 0 |
| NC_019843.3;YP_009047202.1;MERS_1ab;6264 | 0 | 0 |
| NC_019843.3;YP_009047202.1;MERS_1ab;6265 | 0 | 0 |
| NC_019843.3;YP_009047202.1;MERS_1ab;6266 | 0 | 0 |
| NC_019843.3;YP_009047202.1;MERS_1ab;6267 | 0 | 0 |
| NC_019843.3;YP_009047202.1;MERS_1ab;6268 | 0 | 0.025 |
| NC_019843.3;YP_009047202.1;MERS_1ab;6269 | 0 | 0.025 |

| | | |
|---|---|---|
| NC_019843.3;YP_009047202.1;MERS_1ab;6270 | 0 | 0.025 |
| NC_019843.3;YP_009047202.1;MERS_1ab;6271 | 0 | 0 |
| NC_019843.3;YP_009047202.1;MERS_1ab;6272 | 0 | 0 |
| NC_019843.3;YP_009047202.1;MERS_1ab;6273 | 0 | 0 |
| NC_019843.3;YP_009047202.1;MERS_1ab;6274 | 0 | 0 |
| NC_019843.3;YP_009047202.1;MERS_1ab;6275 | 0.05 | 0 |
| NC_019843.3;YP_009047202.1;MERS_1ab;6276 | 0 | 0 |
| NC_019843.3;YP_009047202.1;MERS_1ab;6277 | 0 | 0 |
| NC_019843.3;YP_009047202.1;MERS_1ab;6278 | 0 | 0 |
| NC_019843.3;YP_009047202.1;MERS_1ab;6279 | 0 | 0 |
| NC_019843.3;YP_009047202.1;MERS_1ab;6280 | 0 | 0 |
| NC_019843.3;YP_009047202.1;MERS_1ab;6281 | 0.05 | 0 |
| NC_019843.3;YP_009047202.1;MERS_1ab;6282 | 0 | 0 |
| NC_019843.3;YP_009047202.1;MERS_1ab;6283 | 0 | 0 |
| NC_019843.3;YP_009047202.1;MERS_1ab;6284 | 0 | 0 |
| NC_019843.3;YP_009047202.1;MERS_1ab;6285 | 0 | 0 |
| NC_019843.3;YP_009047202.1;MERS_1ab;6286 | 0 | 0 |
| NC_019843.3;YP_009047202.1;MERS_1ab;6287 | 0 | 0 |
| NC_019843.3;YP_009047202.1;MERS_1ab;6288 | 0 | 0 |
| NC_019843.3;YP_009047202.1;MERS_1ab;6289 | 0 | 0 |
| NC_019843.3;YP_009047202.1;MERS_1ab;6290 | 0 | 0 |
| NC_019843.3;YP_009047202.1;MERS_1ab;6291 | 0 | 0 |
| NC_019843.3;YP_009047202.1;MERS_1ab;6292 | 0 | 0 |
| NC_019843.3;YP_009047202.1;MERS_1ab;6293 | 0 | 0 |
| NC_019843.3;YP_009047202.1;MERS_1ab;6294 | 0 | 0 |
| NC_019843.3;YP_009047202.1;MERS_1ab;6295 | 0 | 0 |
| NC_019843.3;YP_009047202.1;MERS_1ab;6296 | 0 | 0 |
| NC_019843.3;YP_009047202.1;MERS_1ab;6297 | 0 | 0 |
| NC_019843.3;YP_009047202.1;MERS_1ab;6298 | 0 | 0 |
| NC_019843.3;YP_009047202.1;MERS_1ab;6299 | 0 | 0 |
| NC_019843.3;YP_009047202.1;MERS_1ab;6300 | 0 | 0 |
| NC_019843.3;YP_009047202.1;MERS_1ab;6301 | 0 | 0 |
| NC_019843.3;YP_009047202.1;MERS_1ab;6302 | 0 | 0 |
| NC_019843.3;YP_009047202.1;MERS_1ab;6303 | 0 | 0 |
| NC_019843.3;YP_009047202.1;MERS_1ab;6304 | 0 | 0 |
| NC_019843.3;YP_009047202.1;MERS_1ab;6305 | 0 | 0 |
| NC_019843.3;YP_009047202.1;MERS_1ab;6306 | 0 | 0 |
| NC_019843.3;YP_009047202.1;MERS_1ab;6307 | 0 | 0 |
| NC_019843.3;YP_009047202.1;MERS_1ab;6308 | 0 | 0.025 |
| NC_019843.3;YP_009047202.1;MERS_1ab;6309 | 0 | 0 |
| NC_019843.3;YP_009047202.1;MERS_1ab;6310 | 0 | 0 |
| NC_019843.3;YP_009047202.1;MERS_1ab;6311 | 0 | 0.025 |
| NC_019843.3;YP_009047202.1;MERS_1ab;6312 | 0 | 0 |

| | | |
|---|---|---|
| NC_019843.3;YP_009047202.1;MERS_1ab;6313 | 0 | 0 |
| NC_019843.3;YP_009047202.1;MERS_1ab;6314 | 0 | 0 |
| NC_019843.3;YP_009047202.1;MERS_1ab;6315 | 0 | 0 |
| NC_019843.3;YP_009047202.1;MERS_1ab;6316 | 0 | 0 |
| NC_019843.3;YP_009047202.1;MERS_1ab;6317 | 0 | 0 |
| NC_019843.3;YP_009047202.1;MERS_1ab;6318 | 0 | 0 |
| NC_019843.3;YP_009047202.1;MERS_1ab;6319 | 0 | 0 |
| NC_019843.3;YP_009047202.1;MERS_1ab;6320 | 0 | 0 |
| NC_019843.3;YP_009047202.1;MERS_1ab;6321 | 0 | 0 |
| NC_019843.3;YP_009047202.1;MERS_1ab;6322 | 0 | 0 |
| NC_019843.3;YP_009047202.1;MERS_1ab;6323 | 0 | 0 |
| NC_019843.3;YP_009047202.1;MERS_1ab;6324 | 0 | 0 |
| NC_019843.3;YP_009047202.1;MERS_1ab;6325 | 0 | 0 |
| NC_019843.3;YP_009047202.1;MERS_1ab;6326 | 0 | 0 |
| NC_019843.3;YP_009047202.1;MERS_1ab;6327 | 0 | 0 |
| NC_019843.3;YP_009047202.1;MERS_1ab;6328 | 0 | 0 |
| NC_019843.3;YP_009047202.1;MERS_1ab;6329 | 0 | 0 |
| NC_019843.3;YP_009047202.1;MERS_1ab;6330 | 0.1 | 0 |
| NC_019843.3;YP_009047202.1;MERS_1ab;6331 | 0 | 0 |
| NC_019843.3;YP_009047202.1;MERS_1ab;6332 | 0.05 | 0 |
| NC_019843.3;YP_009047202.1;MERS_1ab;6333 | 0 | 0 |
| NC_019843.3;YP_009047202.1;MERS_1ab;6334 | 0 | 0.05 |
| NC_019843.3;YP_009047202.1;MERS_1ab;6335 | 0 | 0 |
| NC_019843.3;YP_009047202.1;MERS_1ab;6336 | 0 | 0 |
| NC_019843.3;YP_009047202.1;MERS_1ab;6337 | 0 | 0 |
| NC_019843.3;YP_009047202.1;MERS_1ab;6338 | 0 | 0 |
| NC_019843.3;YP_009047202.1;MERS_1ab;6339 | 0 | 0 |
| NC_019843.3;YP_009047202.1;MERS_1ab;6340 | 0 | 0 |
| NC_019843.3;YP_009047202.1;MERS_1ab;6341 | 0 | 0 |
| NC_019843.3;YP_009047202.1;MERS_1ab;6342 | 0 | 0 |
| NC_019843.3;YP_009047202.1;MERS_1ab;6343 | 0 | 0 |
| NC_019843.3;YP_009047202.1;MERS_1ab;6344 | 0 | 0 |
| NC_019843.3;YP_009047202.1;MERS_1ab;6345 | 0 | 0 |
| NC_019843.3;YP_009047202.1;MERS_1ab;6346 | 0 | 0 |
| NC_019843.3;YP_009047202.1;MERS_1ab;6347 | 0 | 0 |
| NC_019843.3;YP_009047202.1;MERS_1ab;6348 | 0 | 0 |
| NC_019843.3;YP_009047202.1;MERS_1ab;6349 | 0 | 0 |
| NC_019843.3;YP_009047202.1;MERS_1ab;6350 | 0 | 0.05 |
| NC_019843.3;YP_009047202.1;MERS_1ab;6351 | 0 | 0.025 |
| NC_019843.3;YP_009047202.1;MERS_1ab;6352 | 0 | 0.025 |
| NC_019843.3;YP_009047202.1;MERS_1ab;6353 | 0 | 0.025 |
| NC_019843.3;YP_009047202.1;MERS_1ab;6354 | 0 | 0.025 |
| NC_019843.3;YP_009047202.1;MERS_1ab;6355 | 0 | 0.025 |

| | | |
|---|---|---|
| NC_019843.3;YP_009047202.1;MERS_1ab;6356 | 0 | 0 |
| NC_019843.3;YP_009047202.1;MERS_1ab;6357 | 0.05 | 0.025 |
| NC_019843.3;YP_009047202.1;MERS_1ab;6358 | 0 | 0 |
| NC_019843.3;YP_009047202.1;MERS_1ab;6359 | 0 | 0.025 |
| NC_019843.3;YP_009047202.1;MERS_1ab;6360 | 0 | 0.025 |
| NC_019843.3;YP_009047202.1;MERS_1ab;6361 | 0.05 | 0.025 |
| NC_019843.3;YP_009047202.1;MERS_1ab;6362 | 0.05 | 0.025 |
| NC_019843.3;YP_009047202.1;MERS_1ab;6363 | 0 | 0 |
| NC_019843.3;YP_009047202.1;MERS_1ab;6364 | 0 | 0 |
| NC_019843.3;YP_009047202.1;MERS_1ab;6365 | 0 | 0.025 |
| NC_019843.3;YP_009047202.1;MERS_1ab;6366 | 0 | 0 |
| NC_019843.3;YP_009047202.1;MERS_1ab;6367 | 0 | 0 |
| NC_019843.3;YP_009047202.1;MERS_1ab;6368 | 0 | 0 |
| NC_019843.3;YP_009047202.1;MERS_1ab;6369 | 0 | 0 |
| NC_019843.3;YP_009047202.1;MERS_1ab;6370 | 0 | 0 |
| NC_019843.3;YP_009047202.1;MERS_1ab;6371 | 0 | 0 |
| NC_019843.3;YP_009047202.1;MERS_1ab;6372 | 0 | 0 |
| NC_019843.3;YP_009047202.1;MERS_1ab;6373 | 0 | 0 |
| NC_019843.3;YP_009047202.1;MERS_1ab;6374 | 0 | 0 |
| NC_019843.3;YP_009047202.1;MERS_1ab;6375 | 0 | 0 |
| NC_019843.3;YP_009047202.1;MERS_1ab;6376 | 0 | 0 |
| NC_019843.3;YP_009047202.1;MERS_1ab;6377 | 0 | 0 |
| NC_019843.3;YP_009047202.1;MERS_1ab;6378 | 0 | 0 |
| NC_019843.3;YP_009047202.1;MERS_1ab;6379 | 0 | 0 |
| NC_019843.3;YP_009047202.1;MERS_1ab;6380 | 0 | 0 |
| NC_019843.3;YP_009047202.1;MERS_1ab;6381 | 0 | 0.025 |
| NC_019843.3;YP_009047202.1;MERS_1ab;6382 | 0.05 | 0.025 |
| NC_019843.3;YP_009047202.1;MERS_1ab;6383 | 0.05 | 0 |
| NC_019843.3;YP_009047202.1;MERS_1ab;6384 | 0.05 | 0 |
| NC_019843.3;YP_009047202.1;MERS_1ab;6385 | 0.1 | 0 |
| NC_019843.3;YP_009047202.1;MERS_1ab;6386 | 0.1 | 0 |
| NC_019843.3;YP_009047202.1;MERS_1ab;6387 | 0.1 | 0 |
| NC_019843.3;YP_009047202.1;MERS_1ab;6388 | 0.05 | 0 |
| NC_019843.3;YP_009047202.1;MERS_1ab;6389 | 0 | 0.05 |
| NC_019843.3;YP_009047202.1;MERS_1ab;6390 | 0 | 0 |
| NC_019843.3;YP_009047202.1;MERS_1ab;6391 | 0 | 0 |
| NC_019843.3;YP_009047202.1;MERS_1ab;6392 | 0 | 0 |
| NC_019843.3;YP_009047202.1;MERS_1ab;6393 | 0.05 | 0 |
| NC_019843.3;YP_009047202.1;MERS_1ab;6394 | 0 | 0 |
| NC_019843.3;YP_009047202.1;MERS_1ab;6395 | 0 | 0 |
| NC_019843.3;YP_009047202.1;MERS_1ab;6396 | 0 | 0 |
| NC_019843.3;YP_009047202.1;MERS_1ab;6397 | 0 | 0 |
| NC_019843.3;YP_009047202.1;MERS_1ab;6398 | 0 | 0 |

| | | |
|---|---|---|
| NC_019843.3;YP_009047202.1;MERS_1ab;6399 | 0 | 0 |
| NC_019843.3;YP_009047202.1;MERS_1ab;6400 | 0 | 0 |
| NC_019843.3;YP_009047202.1;MERS_1ab;6401 | 0 | 0 |
| NC_019843.3;YP_009047202.1;MERS_1ab;6402 | 0 | 0 |
| NC_019843.3;YP_009047202.1;MERS_1ab;6403 | 0 | 0 |
| NC_019843.3;YP_009047202.1;MERS_1ab;6404 | 0 | 0 |
| NC_019843.3;YP_009047202.1;MERS_1ab;6405 | 0 | 0 |
| NC_019843.3;YP_009047202.1;MERS_1ab;6406 | 0 | 0 |
| NC_019843.3;YP_009047202.1;MERS_1ab;6407 | 0 | 0 |
| NC_019843.3;YP_009047202.1;MERS_1ab;6408 | 0 | 0 |
| NC_019843.3;YP_009047202.1;MERS_1ab;6409 | 0 | 0 |
| NC_019843.3;YP_009047202.1;MERS_1ab;6410 | 0 | 0 |
| NC_019843.3;YP_009047202.1;MERS_1ab;6411 | 0 | 0 |
| NC_019843.3;YP_009047202.1;MERS_1ab;6412 | 0 | 0 |
| NC_019843.3;YP_009047202.1;MERS_1ab;6413 | 0 | 0 |
| NC_019843.3;YP_009047202.1;MERS_1ab;6414 | 0 | 0 |
| NC_019843.3;YP_009047202.1;MERS_1ab;6415 | 0 | 0 |
| NC_019843.3;YP_009047202.1;MERS_1ab;6416 | 0 | 0 |
| NC_019843.3;YP_009047202.1;MERS_1ab;6417 | 0 | 0 |
| NC_019843.3;YP_009047202.1;MERS_1ab;6418 | 0 | 0 |
| NC_019843.3;YP_009047202.1;MERS_1ab;6419 | 0 | 0 |
| NC_019843.3;YP_009047202.1;MERS_1ab;6420 | 0 | 0 |
| NC_019843.3;YP_009047202.1;MERS_1ab;6421 | 0 | 0 |
| NC_019843.3;YP_009047202.1;MERS_1ab;6422 | 0 | 0 |
| NC_019843.3;YP_009047202.1;MERS_1ab;6423 | 0 | 0 |
| NC_019843.3;YP_009047202.1;MERS_1ab;6424 | 0 | 0 |
| NC_019843.3;YP_009047202.1;MERS_1ab;6425 | 0 | 0 |
| NC_019843.3;YP_009047202.1;MERS_1ab;6426 | 0 | 0 |
| NC_019843.3;YP_009047202.1;MERS_1ab;6427 | 0 | 0 |
| NC_019843.3;YP_009047202.1;MERS_1ab;6428 | 0 | 0 |
| NC_019843.3;YP_009047202.1;MERS_1ab;6429 | 0 | 0 |
| NC_019843.3;YP_009047202.1;MERS_1ab;6430 | 0 | 0 |
| NC_019843.3;YP_009047202.1;MERS_1ab;6431 | 0 | 0 |
| NC_019843.3;YP_009047202.1;MERS_1ab;6432 | 0 | 0 |
| NC_019843.3;YP_009047202.1;MERS_1ab;6433 | 0.1 | 0.075 |
| NC_019843.3;YP_009047202.1;MERS_1ab;6434 | 0 | 0 |
| NC_019843.3;YP_009047202.1;MERS_1ab;6435 | 0 | 0 |
| NC_019843.3;YP_009047202.1;MERS_1ab;6436 | 0 | 0 |
| NC_019843.3;YP_009047202.1;MERS_1ab;6437 | 0 | 0 |
| NC_019843.3;YP_009047202.1;MERS_1ab;6438 | 0 | 0 |
| NC_019843.3;YP_009047202.1;MERS_1ab;6439 | 0 | 0 |
| NC_019843.3;YP_009047202.1;MERS_1ab;6440 | 0.05 | 0 |
| NC_019843.3;YP_009047202.1;MERS_1ab;6441 | 0.05 | 0.05 |

| | | |
|---|---|---|
| NC_019843.3;YP_009047202.1;MERS_1ab;6442 | 0.05 | 0.025 |
| NC_019843.3;YP_009047202.1;MERS_1ab;6443 | 0 | 0 |
| NC_019843.3;YP_009047202.1;MERS_1ab;6444 | 0 | 0 |
| NC_019843.3;YP_009047202.1;MERS_1ab;6445 | 0 | 0 |
| NC_019843.3;YP_009047202.1;MERS_1ab;6446 | 0 | 0.025 |
| NC_019843.3;YP_009047202.1;MERS_1ab;6447 | 0.1 | 0.05 |
| NC_019843.3;YP_009047202.1;MERS_1ab;6448 | 0.1 | 0.025 |
| NC_019843.3;YP_009047202.1;MERS_1ab;6449 | 0.15 | 0.05 |
| NC_019843.3;YP_009047202.1;MERS_1ab;6450 | 0.1 | 0.025 |
| NC_019843.3;YP_009047202.1;MERS_1ab;6451 | 0.1 | 0.075 |
| NC_019843.3;YP_009047202.1;MERS_1ab;6452 | 0.1 | 0.05 |
| NC_019843.3;YP_009047202.1;MERS_1ab;6453 | 0.1 | 0.075 |
| NC_019843.3;YP_009047202.1;MERS_1ab;6454 | 0.05 | 0 |
| NC_019843.3;YP_009047202.1;MERS_1ab;6455 | 0 | 0.025 |
| NC_019843.3;YP_009047202.1;MERS_1ab;6456 | 0 | 0.05 |
| NC_019843.3;YP_009047202.1;MERS_1ab;6457 | 0 | 0.05 |
| NC_019843.3;YP_009047202.1;MERS_1ab;6458 | 0 | 0.025 |
| NC_019843.3;YP_009047202.1;MERS_1ab;6459 | 0 | 0 |
| NC_019843.3;YP_009047202.1;MERS_1ab;6460 | 0 | 0 |
| NC_019843.3;YP_009047202.1;MERS_1ab;6461 | 0 | 0 |
| NC_019843.3;YP_009047202.1;MERS_1ab;6462 | 0 | 0 |
| NC_019843.3;YP_009047202.1;MERS_1ab;6463 | 0 | 0 |
| NC_019843.3;YP_009047202.1;MERS_1ab;6464 | 0 | 0.05 |
| NC_019843.3;YP_009047202.1;MERS_1ab;6465 | 0 | 0.05 |
| NC_019843.3;YP_009047202.1;MERS_1ab;6466 | 0 | 0.075 |
| NC_019843.3;YP_009047202.1;MERS_1ab;6467 | 0 | 0.025 |
| NC_019843.3;YP_009047202.1;MERS_1ab;6468 | 0.35 | 0.2 |
| NC_019843.3;YP_009047202.1;MERS_1ab;6469 | 0.05 | 0.05 |
| NC_019843.3;YP_009047202.1;MERS_1ab;6470 | 0 | 0.075 |
| NC_019843.3;YP_009047202.1;MERS_1ab;6471 | 0.1 | 0.025 |
| NC_019843.3;YP_009047202.1;MERS_1ab;6472 | 0.05 | 0.05 |
| NC_019843.3;YP_009047202.1;MERS_1ab;6473 | 0.05 | 0.025 |
| NC_019843.3;YP_009047202.1;MERS_1ab;6474 | 0.1 | 0.025 |
| NC_019843.3;YP_009047202.1;MERS_1ab;6475 | 0 | 0.025 |
| NC_019843.3;YP_009047202.1;MERS_1ab;6476 | 0.1 | 0.025 |
| NC_019843.3;YP_009047202.1;MERS_1ab;6477 | 0 | 0 |
| NC_019843.3;YP_009047202.1;MERS_1ab;6478 | 0 | 0 |
| NC_019843.3;YP_009047202.1;MERS_1ab;6479 | 0 | 0 |
| NC_019843.3;YP_009047202.1;MERS_1ab;6480 | 0 | 0 |
| NC_019843.3;YP_009047202.1;MERS_1ab;6481 | 0 | 0 |
| NC_019843.3;YP_009047202.1;MERS_1ab;6482 | 0 | 0 |
| NC_019843.3;YP_009047202.1;MERS_1ab;6483 | 0 | 0 |
| NC_019843.3;YP_009047202.1;MERS_1ab;6484 | 0 | 0 |

| | | |
|---|---|---|
| NC_019843.3;YP_009047202.1;MERS_1ab;6485 | 0 | 0 |
| NC_019843.3;YP_009047202.1;MERS_1ab;6486 | 0 | 0 |
| NC_019843.3;YP_009047202.1;MERS_1ab;6487 | 0 | 0.025 |
| NC_019843.3;YP_009047202.1;MERS_1ab;6488 | 0 | 0.025 |
| NC_019843.3;YP_009047202.1;MERS_1ab;6489 | 0.05 | 0.05 |
| NC_019843.3;YP_009047202.1;MERS_1ab;6490 | 0.05 | 0.05 |
| NC_019843.3;YP_009047202.1;MERS_1ab;6491 | 0 | 0.025 |
| NC_019843.3;YP_009047202.1;MERS_1ab;6492 | 0 | 0.075 |
| NC_019843.3;YP_009047202.1;MERS_1ab;6493 | 0 | 0.025 |
| NC_019843.3;YP_009047202.1;MERS_1ab;6494 | 0 | 0.025 |
| NC_019843.3;YP_009047202.1;MERS_1ab;6495 | 0 | 0.075 |
| NC_019843.3;YP_009047202.1;MERS_1ab;6496 | 0 | 0.05 |
| NC_019843.3;YP_009047202.1;MERS_1ab;6497 | 0.05 | 0.05 |
| NC_019843.3;YP_009047202.1;MERS_1ab;6498 | 0 | 0.025 |
| NC_019843.3;YP_009047202.1;MERS_1ab;6499 | 0.05 | 0 |
| NC_019843.3;YP_009047202.1;MERS_1ab;6500 | 0 | 0 |
| NC_019843.3;YP_009047202.1;MERS_1ab;6501 | 0 | 0 |
| NC_019843.3;YP_009047202.1;MERS_1ab;6502 | 0 | 0 |
| NC_019843.3;YP_009047202.1;MERS_1ab;6503 | 0 | 0 |
| NC_019843.3;YP_009047202.1;MERS_1ab;6504 | 0 | 0 |
| NC_019843.3;YP_009047202.1;MERS_1ab;6505 | 0 | 0 |
| NC_019843.3;YP_009047202.1;MERS_1ab;6506 | 0 | 0 |
| NC_019843.3;YP_009047202.1;MERS_1ab;6507 | 0 | 0 |
| NC_019843.3;YP_009047202.1;MERS_1ab;6508 | 0 | 0 |
| NC_019843.3;YP_009047202.1;MERS_1ab;6509 | 0 | 0.025 |
| NC_019843.3;YP_009047202.1;MERS_1ab;6510 | 0 | 0 |
| NC_019843.3;YP_009047202.1;MERS_1ab;6511 | 0 | 0 |
| NC_019843.3;YP_009047202.1;MERS_1ab;6512 | 0 | 0 |
| NC_019843.3;YP_009047202.1;MERS_1ab;6513 | 0 | 0 |
| NC_019843.3;YP_009047202.1;MERS_1ab;6514 | 0 | 0 |
| NC_019843.3;YP_009047202.1;MERS_1ab;6515 | 0 | 0 |
| NC_019843.3;YP_009047202.1;MERS_1ab;6516 | 0 | 0 |
| NC_019843.3;YP_009047202.1;MERS_1ab;6517 | 0 | 0 |
| NC_019843.3;YP_009047202.1;MERS_1ab;6518 | 0 | 0 |
| NC_019843.3;YP_009047202.1;MERS_1ab;6519 | 0 | 0 |
| NC_019843.3;YP_009047202.1;MERS_1ab;6520 | 0 | 0 |
| NC_019843.3;YP_009047202.1;MERS_1ab;6521 | 0 | 0 |
| NC_019843.3;YP_009047202.1;MERS_1ab;6522 | 0 | 0 |
| NC_019843.3;YP_009047202.1;MERS_1ab;6523 | 0 | 0 |
| NC_019843.3;YP_009047202.1;MERS_1ab;6524 | 0 | 0 |
| NC_019843.3;YP_009047202.1;MERS_1ab;6525 | 0 | 0 |
| NC_019843.3;YP_009047202.1;MERS_1ab;6526 | 0 | 0 |
| NC_019843.3;YP_009047202.1;MERS_1ab;6527 | 0 | 0 |

| | | |
|---|---|---|
| NC_019843.3;YP_009047202.1;MERS_1ab;6528 | 0.05 | 0 |
| NC_019843.3;YP_009047202.1;MERS_1ab;6529 | 0 | 0.075 |
| NC_019843.3;YP_009047202.1;MERS_1ab;6530 | 0 | 0 |
| NC_019843.3;YP_009047202.1;MERS_1ab;6531 | 0 | 0.05 |
| NC_019843.3;YP_009047202.1;MERS_1ab;6532 | 0 | 0.025 |
| NC_019843.3;YP_009047202.1;MERS_1ab;6533 | 0 | 0.025 |
| NC_019843.3;YP_009047202.1;MERS_1ab;6534 | 0 | 0 |
| NC_019843.3;YP_009047202.1;MERS_1ab;6535 | 0 | 0 |
| NC_019843.3;YP_009047202.1;MERS_1ab;6536 | 0 | 0 |
| NC_019843.3;YP_009047202.1;MERS_1ab;6537 | 0 | 0 |
| NC_019843.3;YP_009047202.1;MERS_1ab;6538 | 0 | 0 |
| NC_019843.3;YP_009047202.1;MERS_1ab;6539 | 0 | 0.05 |
| NC_019843.3;YP_009047202.1;MERS_1ab;6540 | 0 | 0 |
| NC_019843.3;YP_009047202.1;MERS_1ab;6541 | 0 | 0 |
| NC_019843.3;YP_009047202.1;MERS_1ab;6542 | 0 | 0 |
| NC_019843.3;YP_009047202.1;MERS_1ab;6543 | 0 | 0 |
| NC_019843.3;YP_009047202.1;MERS_1ab;6544 | 0 | 0 |
| NC_019843.3;YP_009047202.1;MERS_1ab;6545 | 0 | 0.025 |
| NC_019843.3;YP_009047202.1;MERS_1ab;6546 | 0 | 0 |
| NC_019843.3;YP_009047202.1;MERS_1ab;6547 | 0 | 0 |
| NC_019843.3;YP_009047202.1;MERS_1ab;6548 | 0 | 0 |
| NC_019843.3;YP_009047202.1;MERS_1ab;6549 | 0 | 0 |
| NC_019843.3;YP_009047202.1;MERS_1ab;6550 | 0.05 | 0 |
| NC_019843.3;YP_009047202.1;MERS_1ab;6551 | 0.05 | 0 |
| NC_019843.3;YP_009047202.1;MERS_1ab;6552 | 0 | 0 |
| NC_019843.3;YP_009047202.1;MERS_1ab;6553 | 0 | 0.025 |
| NC_019843.3;YP_009047202.1;MERS_1ab;6554 | 0 | 0 |
| NC_019843.3;YP_009047202.1;MERS_1ab;6555 | 0 | 0 |
| NC_019843.3;YP_009047202.1;MERS_1ab;6556 | 0 | 0 |
| NC_019843.3;YP_009047202.1;MERS_1ab;6557 | 0 | 0 |
| NC_019843.3;YP_009047202.1;MERS_1ab;6558 | 0 | 0 |
| NC_019843.3;YP_009047202.1;MERS_1ab;6559 | 0 | 0 |
| NC_019843.3;YP_009047202.1;MERS_1ab;6560 | 0.05 | 0 |
| NC_019843.3;YP_009047202.1;MERS_1ab;6561 | 0.05 | 0 |
| NC_019843.3;YP_009047202.1;MERS_1ab;6562 | 0.05 | 0 |
| NC_019843.3;YP_009047202.1;MERS_1ab;6563 | 0.05 | 0 |
| NC_019843.3;YP_009047202.1;MERS_1ab;6564 | 0.05 | 0 |
| NC_019843.3;YP_009047202.1;MERS_1ab;6565 | 0.05 | 0 |
| NC_019843.3;YP_009047202.1;MERS_1ab;6566 | 0.05 | 0 |
| NC_019843.3;YP_009047202.1;MERS_1ab;6567 | 0.1 | 0 |
| NC_019843.3;YP_009047202.1;MERS_1ab;6568 | 0 | 0 |
| NC_019843.3;YP_009047202.1;MERS_1ab;6569 | 0 | 0 |
| NC_019843.3;YP_009047202.1;MERS_1ab;6570 | 0 | 0 |

| | | |
|---|---|---|
| NC_019843.3;YP_009047202.1;MERS_1ab;6571 | 0 | 0.025 |
| NC_019843.3;YP_009047202.1;MERS_1ab;6572 | 0 | 0.05 |
| NC_019843.3;YP_009047202.1;MERS_1ab;6573 | 0 | 0.025 |
| NC_019843.3;YP_009047202.1;MERS_1ab;6574 | 0 | 0 |
| NC_019843.3;YP_009047202.1;MERS_1ab;6575 | 0 | 0.025 |
| NC_019843.3;YP_009047202.1;MERS_1ab;6576 | 0 | 0.025 |
| NC_019843.3;YP_009047202.1;MERS_1ab;6577 | 0 | 0.025 |
| NC_019843.3;YP_009047202.1;MERS_1ab;6578 | 0 | 0 |
| NC_019843.3;YP_009047202.1;MERS_1ab;6579 | 0 | 0 |
| NC_019843.3;YP_009047202.1;MERS_1ab;6580 | 0 | 0 |
| NC_019843.3;YP_009047202.1;MERS_1ab;6581 | 0 | 0 |
| NC_019843.3;YP_009047202.1;MERS_1ab;6582 | 0 | 0 |
| NC_019843.3;YP_009047202.1;MERS_1ab;6583 | 0 | 0.025 |
| NC_019843.3;YP_009047202.1;MERS_1ab;6584 | 0 | 0.025 |
| NC_019843.3;YP_009047202.1;MERS_1ab;6585 | 0 | 0.025 |
| NC_019843.3;YP_009047202.1;MERS_1ab;6586 | 0 | 0.025 |
| NC_019843.3;YP_009047202.1;MERS_1ab;6587 | 0 | 0.025 |
| NC_019843.3;YP_009047202.1;MERS_1ab;6588 | 0 | 0.05 |
| NC_019843.3;YP_009047202.1;MERS_1ab;6589 | 0 | 0.05 |
| NC_019843.3;YP_009047202.1;MERS_1ab;6590 | 0.05 | 0.025 |
| NC_019843.3;YP_009047202.1;MERS_1ab;6591 | 0 | 0 |
| NC_019843.3;YP_009047202.1;MERS_1ab;6592 | 0 | 0 |
| NC_019843.3;YP_009047202.1;MERS_1ab;6593 | 0 | 0 |
| NC_019843.3;YP_009047202.1;MERS_1ab;6594 | 0 | 0 |
| NC_019843.3;YP_009047202.1;MERS_1ab;6595 | 0 | 0 |
| NC_019843.3;YP_009047202.1;MERS_1ab;6596 | 0 | 0 |
| NC_019843.3;YP_009047202.1;MERS_1ab;6597 | 0 | 0 |
| NC_019843.3;YP_009047202.1;MERS_1ab;6598 | 0 | 0 |
| NC_019843.3;YP_009047202.1;MERS_1ab;6599 | 0 | 0 |
| NC_019843.3;YP_009047202.1;MERS_1ab;6600 | 0 | 0 |
| NC_019843.3;YP_009047202.1;MERS_1ab;6601 | 0 | 0 |
| NC_019843.3;YP_009047202.1;MERS_1ab;6602 | 0.05 | 0 |
| NC_019843.3;YP_009047202.1;MERS_1ab;6603 | 0 | 0 |
| NC_019843.3;YP_009047202.1;MERS_1ab;6604 | 0 | 0.05 |
| NC_019843.3;YP_009047202.1;MERS_1ab;6605 | 0.15 | 0 |
| NC_019843.3;YP_009047202.1;MERS_1ab;6606 | 0.3 | 0.1 |
| NC_019843.3;YP_009047202.1;MERS_1ab;6607 | 0.35 | 0.1 |
| NC_019843.3;YP_009047202.1;MERS_1ab;6608 | 0.25 | 0.05 |
| NC_019843.3;YP_009047202.1;MERS_1ab;6609 | 0.15 | 0.05 |
| NC_019843.3;YP_009047202.1;MERS_1ab;6610 | 0.1 | 0.05 |
| NC_019843.3;YP_009047202.1;MERS_1ab;6611 | 0.05 | 0 |
| NC_019843.3;YP_009047202.1;MERS_1ab;6612 | 0 | 0.025 |
| NC_019843.3;YP_009047202.1;MERS_1ab;6613 | 0 | 0 |

| | | |
|---|---|---|
| NC_019843.3;YP_009047202.1;MERS_1ab;6614 | 0 | 0 |
| NC_019843.3;YP_009047202.1;MERS_1ab;6615 | 0 | 0 |
| NC_019843.3;YP_009047202.1;MERS_1ab;6616 | 0 | 0 |
| NC_019843.3;YP_009047202.1;MERS_1ab;6617 | 0 | 0 |
| NC_019843.3;YP_009047202.1;MERS_1ab;6618 | 0 | 0 |
| NC_019843.3;YP_009047202.1;MERS_1ab;6619 | 0 | 0 |
| NC_019843.3;YP_009047202.1;MERS_1ab;6620 | 0 | 0 |
| NC_019843.3;YP_009047202.1;MERS_1ab;6621 | 0 | 0 |
| NC_019843.3;YP_009047202.1;MERS_1ab;6622 | 0 | 0 |
| NC_019843.3;YP_009047202.1;MERS_1ab;6623 | 0 | 0 |
| NC_019843.3;YP_009047202.1;MERS_1ab;6624 | 0 | 0 |
| NC_019843.3;YP_009047202.1;MERS_1ab;6625 | 0 | 0 |
| NC_019843.3;YP_009047202.1;MERS_1ab;6626 | 0.05 | 0 |
| NC_019843.3;YP_009047202.1;MERS_1ab;6627 | 0.05 | 0 |
| NC_019843.3;YP_009047202.1;MERS_1ab;6628 | 0 | 0.05 |
| NC_019843.3;YP_009047202.1;MERS_1ab;6629 | 0 | 0.025 |
| NC_019843.3;YP_009047202.1;MERS_1ab;6630 | 0 | 0.025 |
| NC_019843.3;YP_009047202.1;MERS_1ab;6631 | 0 | 0 |
| NC_019843.3;YP_009047202.1;MERS_1ab;6632 | 0 | 0 |
| NC_019843.3;YP_009047202.1;MERS_1ab;6633 | 0 | 0 |
| NC_019843.3;YP_009047202.1;MERS_1ab;6634 | 0 | 0.025 |
| NC_019843.3;YP_009047202.1;MERS_1ab;6635 | 0 | 0.025 |
| NC_019843.3;YP_009047202.1;MERS_1ab;6636 | 0 | 0.025 |
| NC_019843.3;YP_009047202.1;MERS_1ab;6637 | 0 | 0.025 |
| NC_019843.3;YP_009047202.1;MERS_1ab;6638 | 0.05 | 0.025 |
| NC_019843.3;YP_009047202.1;MERS_1ab;6639 | 0 | 0.025 |
| NC_019843.3;YP_009047202.1;MERS_1ab;6640 | 0 | 0 |
| NC_019843.3;YP_009047202.1;MERS_1ab;6641 | 0 | 0 |
| NC_019843.3;YP_009047202.1;MERS_1ab;6642 | 0 | 0.025 |
| NC_019843.3;YP_009047202.1;MERS_1ab;6643 | 0.05 | 0.025 |
| NC_019843.3;YP_009047202.1;MERS_1ab;6644 | 0 | 0.025 |
| NC_019843.3;YP_009047202.1;MERS_1ab;6645 | 0 | 0 |
| NC_019843.3;YP_009047202.1;MERS_1ab;6646 | 0 | 0.025 |
| NC_019843.3;YP_009047202.1;MERS_1ab;6647 | 0 | 0.025 |
| NC_019843.3;YP_009047202.1;MERS_1ab;6648 | 0 | 0 |
| NC_019843.3;YP_009047202.1;MERS_1ab;6649 | 0 | 0 |
| NC_019843.3;YP_009047202.1;MERS_1ab;6650 | 0 | 0 |
| NC_019843.3;YP_009047202.1;MERS_1ab;6651 | 0 | 0 |
| NC_019843.3;YP_009047202.1;MERS_1ab;6652 | 0 | 0 |
| NC_019843.3;YP_009047202.1;MERS_1ab;6653 | 0 | 0 |
| NC_019843.3;YP_009047202.1;MERS_1ab;6654 | 0 | 0 |
| NC_019843.3;YP_009047202.1;MERS_1ab;6655 | 0.05 | 0 |
| NC_019843.3;YP_009047202.1;MERS_1ab;6656 | 0 | 0 |

| | | |
|---|---|---|
| NC_019843.3;YP_009047202.1;MERS_1ab;6657 | 0 | 0 |
| NC_019843.3;YP_009047202.1;MERS_1ab;6658 | 0 | 0 |
| NC_019843.3;YP_009047202.1;MERS_1ab;6659 | 0 | 0 |
| NC_019843.3;YP_009047202.1;MERS_1ab;6660 | 0 | 0 |
| NC_019843.3;YP_009047202.1;MERS_1ab;6661 | 0 | 0 |
| NC_019843.3;YP_009047202.1;MERS_1ab;6662 | 0 | 0 |
| NC_019843.3;YP_009047202.1;MERS_1ab;6663 | 0 | 0 |
| NC_019843.3;YP_009047202.1;MERS_1ab;6664 | 0 | 0 |
| NC_019843.3;YP_009047202.1;MERS_1ab;6665 | 0 | 0 |
| NC_019843.3;YP_009047202.1;MERS_1ab;6666 | 0 | 0 |
| NC_019843.3;YP_009047202.1;MERS_1ab;6667 | 0 | 0 |
| NC_019843.3;YP_009047202.1;MERS_1ab;6668 | 0 | 0 |
| NC_019843.3;YP_009047202.1;MERS_1ab;6669 | 0 | 0 |
| NC_019843.3;YP_009047202.1;MERS_1ab;6670 | 0 | 0 |
| NC_019843.3;YP_009047202.1;MERS_1ab;6671 | 0 | 0 |
| NC_019843.3;YP_009047202.1;MERS_1ab;6672 | 0 | 0 |
| NC_019843.3;YP_009047202.1;MERS_1ab;6673 | 0 | 0 |
| NC_019843.3;YP_009047202.1;MERS_1ab;6674 | 0 | 0 |
| NC_019843.3;YP_009047202.1;MERS_1ab;6675 | 0 | 0 |
| NC_019843.3;YP_009047202.1;MERS_1ab;6676 | 0 | 0 |
| NC_019843.3;YP_009047202.1;MERS_1ab;6677 | 0 | 0 |
| NC_019843.3;YP_009047202.1;MERS_1ab;6678 | 0 | 0 |
| NC_019843.3;YP_009047202.1;MERS_1ab;6679 | 0 | 0 |
| NC_019843.3;YP_009047202.1;MERS_1ab;6680 | 0 | 0 |
| NC_019843.3;YP_009047202.1;MERS_1ab;6681 | 0 | 0 |
| NC_019843.3;YP_009047202.1;MERS_1ab;6682 | 0.05 | 0 |
| NC_019843.3;YP_009047202.1;MERS_1ab;6683 | 0 | 0 |
| NC_019843.3;YP_009047202.1;MERS_1ab;6684 | 0.05 | 0 |
| NC_019843.3;YP_009047202.1;MERS_1ab;6685 | 0 | 0 |
| NC_019843.3;YP_009047202.1;MERS_1ab;6686 | 0 | 0 |
| NC_019843.3;YP_009047202.1;MERS_1ab;6687 | 0 | 0 |
| NC_019843.3;YP_009047202.1;MERS_1ab;6688 | 0 | 0 |
| NC_019843.3;YP_009047202.1;MERS_1ab;6689 | 0 | 0.025 |
| NC_019843.3;YP_009047202.1;MERS_1ab;6690 | 0 | 0 |
| NC_019843.3;YP_009047202.1;MERS_1ab;6691 | 0.05 | 0 |
| NC_019843.3;YP_009047202.1;MERS_1ab;6692 | 0 | 0 |
| NC_019843.3;YP_009047202.1;MERS_1ab;6693 | 0 | 0 |
| NC_019843.3;YP_009047202.1;MERS_1ab;6694 | 0 | 0 |
| NC_019843.3;YP_009047202.1;MERS_1ab;6695 | 0 | 0 |
| NC_019843.3;YP_009047202.1;MERS_1ab;6696 | 0 | 0 |
| NC_019843.3;YP_009047202.1;MERS_1ab;6697 | 0 | 0 |
| NC_019843.3;YP_009047202.1;MERS_1ab;6698 | 0 | 0 |
| NC_019843.3;YP_009047202.1;MERS_1ab;6699 | 0 | 0 |

| | | |
|---|---|---|
| NC_019843.3;YP_009047202.1;MERS_1ab;6700 | 0 | 0 |
| NC_019843.3;YP_009047202.1;MERS_1ab;6701 | 0.05 | 0.025 |
| NC_019843.3;YP_009047202.1;MERS_1ab;6702 | 0 | 0 |
| NC_019843.3;YP_009047202.1;MERS_1ab;6703 | 0 | 0 |
| NC_019843.3;YP_009047202.1;MERS_1ab;6704 | 0.05 | 0 |
| NC_019843.3;YP_009047202.1;MERS_1ab;6705 | 0 | 0 |
| NC_019843.3;YP_009047202.1;MERS_1ab;6706 | 0 | 0 |
| NC_019843.3;YP_009047202.1;MERS_1ab;6707 | 0 | 0 |
| NC_019843.3;YP_009047202.1;MERS_1ab;6708 | 0 | 0 |
| NC_019843.3;YP_009047202.1;MERS_1ab;6709 | 0 | 0 |
| NC_019843.3;YP_009047202.1;MERS_1ab;6710 | 0 | 0 |
| NC_019843.3;YP_009047202.1;MERS_1ab;6711 | 0 | 0 |
| NC_019843.3;YP_009047202.1;MERS_1ab;6712 | 0 | 0 |
| NC_019843.3;YP_009047202.1;MERS_1ab;6713 | 0 | 0 |
| NC_019843.3;YP_009047202.1;MERS_1ab;6714 | 0 | 0 |
| NC_019843.3;YP_009047202.1;MERS_1ab;6715 | 0 | 0 |
| NC_019843.3;YP_009047202.1;MERS_1ab;6716 | 0 | 0 |
| NC_019843.3;YP_009047202.1;MERS_1ab;6717 | 0 | 0 |
| NC_019843.3;YP_009047202.1;MERS_1ab;6718 | 0 | 0 |
| NC_019843.3;YP_009047202.1;MERS_1ab;6719 | 0 | 0 |
| NC_019843.3;YP_009047202.1;MERS_1ab;6720 | 0 | 0 |
| NC_019843.3;YP_009047202.1;MERS_1ab;6721 | 0 | 0 |
| NC_019843.3;YP_009047202.1;MERS_1ab;6722 | 0.05 | 0.025 |
| NC_019843.3;YP_009047202.1;MERS_1ab;6723 | 0.05 | 0 |
| NC_019843.3;YP_009047202.1;MERS_1ab;6724 | 0 | 0 |
| NC_019843.3;YP_009047202.1;MERS_1ab;6725 | 0 | 0 |
| NC_019843.3;YP_009047202.1;MERS_1ab;6726 | 0 | 0 |
| NC_019843.3;YP_009047202.1;MERS_1ab;6727 | 0.05 | 0 |
| NC_019843.3;YP_009047202.1;MERS_1ab;6728 | 0 | 0 |
| NC_019843.3;YP_009047202.1;MERS_1ab;6729 | 0 | 0 |
| NC_019843.3;YP_009047202.1;MERS_1ab;6730 | 0 | 0 |
| NC_019843.3;YP_009047202.1;MERS_1ab;6731 | 0 | 0 |
| NC_019843.3;YP_009047202.1;MERS_1ab;6732 | 0 | 0 |
| NC_019843.3;YP_009047202.1;MERS_1ab;6733 | 0 | 0 |
| NC_019843.3;YP_009047202.1;MERS_1ab;6734 | 0 | 0 |
| NC_019843.3;YP_009047202.1;MERS_1ab;6735 | 0 | 0 |
| NC_019843.3;YP_009047202.1;MERS_1ab;6736 | 0 | 0 |
| NC_019843.3;YP_009047202.1;MERS_1ab;6737 | 0 | 0 |
| NC_019843.3;YP_009047202.1;MERS_1ab;6738 | 0 | 0 |
| NC_019843.3;YP_009047202.1;MERS_1ab;6739 | 0 | 0 |
| NC_019843.3;YP_009047202.1;MERS_1ab;6740 | 0 | 0 |
| NC_019843.3;YP_009047202.1;MERS_1ab;6741 | 0 | 0 |
| NC_019843.3;YP_009047202.1;MERS_1ab;6742 | 0 | 0 |

| | | |
|---|---|---|
| NC_019843.3;YP_009047202.1;MERS_1ab;6743 | 0 | 0 |
| NC_019843.3;YP_009047202.1;MERS_1ab;6744 | 0 | 0 |
| NC_019843.3;YP_009047202.1;MERS_1ab;6745 | 0 | 0 |
| NC_019843.3;YP_009047202.1;MERS_1ab;6746 | 0 | 0 |
| NC_019843.3;YP_009047202.1;MERS_1ab;6747 | 0 | 0 |
| NC_019843.3;YP_009047202.1;MERS_1ab;6748 | 0 | 0 |
| NC_019843.3;YP_009047202.1;MERS_1ab;6749 | 0 | 0 |
| NC_019843.3;YP_009047202.1;MERS_1ab;6750 | 0 | 0 |
| NC_019843.3;YP_009047202.1;MERS_1ab;6751 | 0 | 0 |
| NC_019843.3;YP_009047202.1;MERS_1ab;6752 | 0 | 0 |
| NC_019843.3;YP_009047202.1;MERS_1ab;6753 | 0 | 0.025 |
| NC_019843.3;YP_009047202.1;MERS_1ab;6754 | 0.05 | 0.05 |
| NC_019843.3;YP_009047202.1;MERS_1ab;6755 | 0 | 0.05 |
| NC_019843.3;YP_009047202.1;MERS_1ab;6756 | 0 | 0.025 |
| NC_019843.3;YP_009047202.1;MERS_1ab;6757 | 0 | 0.05 |
| NC_019843.3;YP_009047202.1;MERS_1ab;6758 | 0 | 0.025 |
| NC_019843.3;YP_009047202.1;MERS_1ab;6759 | 0 | 0.025 |
| NC_019843.3;YP_009047202.1;MERS_1ab;6760 | 0 | 0.025 |
| NC_019843.3;YP_009047202.1;MERS_1ab;6761 | 0 | 0.05 |
| NC_019843.3;YP_009047202.1;MERS_1ab;6762 | 0.05 | 0.05 |
| NC_019843.3;YP_009047202.1;MERS_1ab;6763 | 0.05 | 0.05 |
| NC_019843.3;YP_009047202.1;MERS_1ab;6764 | 0 | 0 |
| NC_019843.3;YP_009047202.1;MERS_1ab;6765 | 0 | 0 |
| NC_019843.3;YP_009047202.1;MERS_1ab;6766 | 0 | 0 |
| NC_019843.3;YP_009047202.1;MERS_1ab;6767 | 0 | 0.025 |
| NC_019843.3;YP_009047202.1;MERS_1ab;6768 | 0 | 0.075 |
| NC_019843.3;YP_009047202.1;MERS_1ab;6769 | 0 | 0 |
| NC_019843.3;YP_009047202.1;MERS_1ab;6770 | 0 | 0 |
| NC_019843.3;YP_009047202.1;MERS_1ab;6771 | 0 | 0.025 |
| NC_019843.3;YP_009047202.1;MERS_1ab;6772 | 0.05 | 0.025 |
| NC_019843.3;YP_009047202.1;MERS_1ab;6773 | 0.05 | 0.1 |
| NC_019843.3;YP_009047202.1;MERS_1ab;6774 | 0 | 0.1 |
| NC_019843.3;YP_009047202.1;MERS_1ab;6775 | 0 | 0.025 |
| NC_019843.3;YP_009047202.1;MERS_1ab;6776 | 0 | 0 |
| NC_019843.3;YP_009047202.1;MERS_1ab;6777 | 0 | 0.025 |
| NC_019843.3;YP_009047202.1;MERS_1ab;6778 | 0 | 0 |
| NC_019843.3;YP_009047202.1;MERS_1ab;6779 | 0 | 0 |
| NC_019843.3;YP_009047202.1;MERS_1ab;6780 | 0 | 0 |
| NC_019843.3;YP_009047202.1;MERS_1ab;6781 | 0 | 0 |
| NC_019843.3;YP_009047202.1;MERS_1ab;6782 | 0 | 0 |
| NC_019843.3;YP_009047202.1;MERS_1ab;6783 | 0 | 0.025 |
| NC_019843.3;YP_009047202.1;MERS_1ab;6784 | 0 | 0 |
| NC_019843.3;YP_009047202.1;MERS_1ab;6785 | 0 | 0 |

| | | |
|---|---|---|
| NC_019843.3;YP_009047202.1;MERS_1ab;6786 | 0 | 0.025 |
| NC_019843.3;YP_009047202.1;MERS_1ab;6787 | 0 | 0 |
| NC_019843.3;YP_009047202.1;MERS_1ab;6788 | 0 | 0 |
| NC_019843.3;YP_009047202.1;MERS_1ab;6789 | 0 | 0 |
| NC_019843.3;YP_009047202.1;MERS_1ab;6790 | 0.05 | 0.025 |
| NC_019843.3;YP_009047202.1;MERS_1ab;6791 | 0 | 0.025 |
| NC_019843.3;YP_009047202.1;MERS_1ab;6792 | 0.05 | 0.05 |
| NC_019843.3;YP_009047202.1;MERS_1ab;6793 | 0 | 0 |
| NC_019843.3;YP_009047202.1;MERS_1ab;6794 | 0 | 0 |
| NC_019843.3;YP_009047202.1;MERS_1ab;6795 | 0 | 0 |
| NC_019843.3;YP_009047202.1;MERS_1ab;6796 | 0 | 0 |
| NC_019843.3;YP_009047202.1;MERS_1ab;6797 | 0 | 0.025 |
| NC_019843.3;YP_009047202.1;MERS_1ab;6798 | 0.05 | 0.025 |
| NC_019843.3;YP_009047202.1;MERS_1ab;6799 | 0 | 0 |
| NC_019843.3;YP_009047202.1;MERS_1ab;6800 | 0 | 0 |
| NC_019843.3;YP_009047202.1;MERS_1ab;6801 | 0 | 0 |
| NC_019843.3;YP_009047202.1;MERS_1ab;6802 | 0 | 0 |
| NC_019843.3;YP_009047202.1;MERS_1ab;6803 | 0 | 0 |
| NC_019843.3;YP_009047202.1;MERS_1ab;6804 | 0 | 0 |
| NC_019843.3;YP_009047202.1;MERS_1ab;6805 | 0 | 0 |
| NC_019843.3;YP_009047202.1;MERS_1ab;6806 | 0 | 0 |
| NC_019843.3;YP_009047202.1;MERS_1ab;6807 | 0 | 0 |
| NC_019843.3;YP_009047202.1;MERS_1ab;6808 | 0 | 0 |
| NC_019843.3;YP_009047202.1;MERS_1ab;6809 | 0 | 0 |
| NC_019843.3;YP_009047202.1;MERS_1ab;6810 | 0.05 | 0 |
| NC_019843.3;YP_009047202.1;MERS_1ab;6811 | 0 | 0 |
| NC_019843.3;YP_009047202.1;MERS_1ab;6812 | 0 | 0 |
| NC_019843.3;YP_009047202.1;MERS_1ab;6813 | 0 | 0 |
| NC_019843.3;YP_009047202.1;MERS_1ab;6814 | 0 | 0 |
| NC_019843.3;YP_009047202.1;MERS_1ab;6815 | 0 | 0 |
| NC_019843.3;YP_009047202.1;MERS_1ab;6816 | 0 | 0 |
| NC_019843.3;YP_009047202.1;MERS_1ab;6817 | 0 | 0 |
| NC_019843.3;YP_009047202.1;MERS_1ab;6818 | 0 | 0 |
| NC_019843.3;YP_009047202.1;MERS_1ab;6819 | 0 | 0 |
| NC_019843.3;YP_009047202.1;MERS_1ab;6820 | 0 | 0 |
| NC_019843.3;YP_009047202.1;MERS_1ab;6821 | 0 | 0 |
| NC_019843.3;YP_009047202.1;MERS_1ab;6822 | 0 | 0 |
| NC_019843.3;YP_009047202.1;MERS_1ab;6823 | 0 | 0 |
| NC_019843.3;YP_009047202.1;MERS_1ab;6824 | 0 | 0 |
| NC_019843.3;YP_009047202.1;MERS_1ab;6825 | 0 | 0 |
| NC_019843.3;YP_009047202.1;MERS_1ab;6826 | 0 | 0 |
| NC_019843.3;YP_009047202.1;MERS_1ab;6827 | 0 | 0 |
| NC_019843.3;YP_009047202.1;MERS_1ab;6828 | 0 | 0 |

| | | |
|---|---|---|
| NC_019843.3;YP_009047202.1;MERS_1ab;6829 | 0 | 0 |
| NC_019843.3;YP_009047202.1;MERS_1ab;6830 | 0 | 0 |
| NC_019843.3;YP_009047202.1;MERS_1ab;6831 | 0 | 0 |
| NC_019843.3;YP_009047202.1;MERS_1ab;6832 | 0 | 0 |
| NC_019843.3;YP_009047202.1;MERS_1ab;6833 | 0 | 0 |
| NC_019843.3;YP_009047202.1;MERS_1ab;6834 | 0 | 0 |
| NC_019843.3;YP_009047202.1;MERS_1ab;6835 | 0 | 0 |
| NC_019843.3;YP_009047202.1;MERS_1ab;6836 | 0 | 0 |
| NC_019843.3;YP_009047202.1;MERS_1ab;6837 | 0 | 0 |
| NC_019843.3;YP_009047202.1;MERS_1ab;6838 | 0 | 0 |
| NC_019843.3;YP_009047202.1;MERS_1ab;6839 | 0 | 0.025 |
| NC_019843.3;YP_009047202.1;MERS_1ab;6840 | 0 | 0 |
| NC_019843.3;YP_009047202.1;MERS_1ab;6841 | 0 | 0 |
| NC_019843.3;YP_009047202.1;MERS_1ab;6842 | 0 | 0 |
| NC_019843.3;YP_009047202.1;MERS_1ab;6843 | 0 | 0.025 |
| NC_019843.3;YP_009047202.1;MERS_1ab;6844 | 0 | 0.025 |
| NC_019843.3;YP_009047202.1;MERS_1ab;6845 | 0 | 0.05 |
| NC_019843.3;YP_009047202.1;MERS_1ab;6846 | 0 | 0.025 |
| NC_019843.3;YP_009047202.1;MERS_1ab;6847 | 0 | 0.025 |
| NC_019843.3;YP_009047202.1;MERS_1ab;6848 | 0.05 | 0.025 |
| NC_019843.3;YP_009047202.1;MERS_1ab;6849 | 0 | 0 |
| NC_019843.3;YP_009047202.1;MERS_1ab;6850 | 0 | 0.025 |
| NC_019843.3;YP_009047202.1;MERS_1ab;6851 | 0 | 0.05 |
| NC_019843.3;YP_009047202.1;MERS_1ab;6852 | 0 | 0 |
| NC_019843.3;YP_009047202.1;MERS_1ab;6853 | 0 | 0 |
| NC_019843.3;YP_009047202.1;MERS_1ab;6854 | 0 | 0 |
| NC_019843.3;YP_009047202.1;MERS_1ab;6855 | 0 | 0 |
| NC_019843.3;YP_009047202.1;MERS_1ab;6856 | 0 | 0 |
| NC_019843.3;YP_009047202.1;MERS_1ab;6857 | 0 | 0 |
| NC_019843.3;YP_009047202.1;MERS_1ab;6858 | 0 | 0 |
| NC_019843.3;YP_009047202.1;MERS_1ab;6859 | 0 | 0 |
| NC_019843.3;YP_009047202.1;MERS_1ab;6860 | 0 | 0 |
| NC_019843.3;YP_009047202.1;MERS_1ab;6861 | 0 | 0.05 |
| NC_019843.3;YP_009047202.1;MERS_1ab;6862 | 0 | 0.075 |
| NC_019843.3;YP_009047202.1;MERS_1ab;6863 | 0 | 0.05 |
| NC_019843.3;YP_009047202.1;MERS_1ab;6864 | 0 | 0.05 |
| NC_019843.3;YP_009047202.1;MERS_1ab;6865 | 0 | 0 |
| NC_019843.3;YP_009047202.1;MERS_1ab;6866 | 0.05 | 0.025 |
| NC_019843.3;YP_009047202.1;MERS_1ab;6867 | 0.05 | 0.025 |
| NC_019843.3;YP_009047202.1;MERS_1ab;6868 | 0.05 | 0.025 |
| NC_019843.3;YP_009047202.1;MERS_1ab;6869 | 0.15 | 0.025 |
| NC_019843.3;YP_009047202.1;MERS_1ab;6870 | 0 | 0 |
| NC_019843.3;YP_009047202.1;MERS_1ab;6871 | 0.05 | 0 |

| | | |
|---|---|---|
| NC_019843.3;YP_009047202.1;MERS_1ab;6872 | 0 | 0 |
| NC_019843.3;YP_009047202.1;MERS_1ab;6873 | 0 | 0 |
| NC_019843.3;YP_009047202.1;MERS_1ab;6874 | 0 | 0.025 |
| NC_019843.3;YP_009047202.1;MERS_1ab;6875 | 0 | 0 |
| NC_019843.3;YP_009047202.1;MERS_1ab;6876 | 0 | 0 |
| NC_019843.3;YP_009047202.1;MERS_1ab;6877 | 0.05 | 0 |
| NC_019843.3;YP_009047202.1;MERS_1ab;6878 | 0 | 0 |
| NC_019843.3;YP_009047202.1;MERS_1ab;6879 | 0 | 0 |
| NC_019843.3;YP_009047202.1;MERS_1ab;6880 | 0 | 0 |
| NC_019843.3;YP_009047202.1;MERS_1ab;6881 | 0 | 0 |
| NC_019843.3;YP_009047202.1;MERS_1ab;6882 | 0 | 0 |
| NC_019843.3;YP_009047202.1;MERS_1ab;6883 | 0 | 0 |
| NC_019843.3;YP_009047202.1;MERS_1ab;6884 | 0 | 0 |
| NC_019843.3;YP_009047202.1;MERS_1ab;6885 | 0.05 | 0.025 |
| NC_019843.3;YP_009047202.1;MERS_1ab;6886 | 0 | 0 |
| NC_019843.3;YP_009047202.1;MERS_1ab;6887 | 0 | 0 |
| NC_019843.3;YP_009047202.1;MERS_1ab;6888 | 0 | 0 |
| NC_019843.3;YP_009047202.1;MERS_1ab;6889 | 0 | 0 |
| NC_019843.3;YP_009047202.1;MERS_1ab;6890 | 0 | 0 |
| NC_019843.3;YP_009047202.1;MERS_1ab;6891 | 0 | 0 |
| NC_019843.3;YP_009047202.1;MERS_1ab;6892 | 0 | 0 |
| NC_019843.3;YP_009047202.1;MERS_1ab;6893 | 0 | 0 |
| NC_019843.3;YP_009047202.1;MERS_1ab;6894 | 0 | 0 |
| NC_019843.3;YP_009047202.1;MERS_1ab;6895 | 0 | 0 |
| NC_019843.3;YP_009047202.1;MERS_1ab;6896 | 0 | 0.025 |
| NC_019843.3;YP_009047202.1;MERS_1ab;6897 | 0 | 0.025 |
| NC_019843.3;YP_009047202.1;MERS_1ab;6898 | 0 | 0.025 |
| NC_019843.3;YP_009047202.1;MERS_1ab;6899 | 0.1 | 0 |
| NC_019843.3;YP_009047202.1;MERS_1ab;6900 | 0.05 | 0 |
| NC_019843.3;YP_009047202.1;MERS_1ab;6901 | 0.05 | 0 |
| NC_019843.3;YP_009047202.1;MERS_1ab;6902 | 0.05 | 0 |
| NC_019843.3;YP_009047202.1;MERS_1ab;6903 | 0.3 | 0.2 |
| NC_019843.3;YP_009047202.1;MERS_1ab;6904 | 0.25 | 0.125 |
| NC_019843.3;YP_009047202.1;MERS_1ab;6905 | 0.25 | 0.175 |
| NC_019843.3;YP_009047202.1;MERS_1ab;6906 | 0.3 | 0.175 |
| NC_019843.3;YP_009047202.1;MERS_1ab;6907 | 0.3 | 0.15 |
| NC_019843.3;YP_009047202.1;MERS_1ab;6908 | 0.25 | 0.125 |
| NC_019843.3;YP_009047202.1;MERS_1ab;6909 | 0.2 | 0.1 |
| NC_019843.3;YP_009047202.1;MERS_1ab;6910 | 0.15 | 0.075 |
| NC_019843.3;YP_009047202.1;MERS_1ab;6911 | 0.1 | 0.1 |
| NC_019843.3;YP_009047202.1;MERS_1ab;6912 | 0.05 | 0.05 |
| NC_019843.3;YP_009047202.1;MERS_1ab;6913 | 0 | 0.025 |
| NC_019843.3;YP_009047202.1;MERS_1ab;6914 | 0 | 0 |

| | | |
|---|---|---|
| NC_019843.3;YP_009047202.1;MERS_1ab;6915 | 0 | 0 |
| NC_019843.3;YP_009047202.1;MERS_1ab;6916 | 0 | 0 |
| NC_019843.3;YP_009047202.1;MERS_1ab;6917 | 0 | 0 |
| NC_019843.3;YP_009047202.1;MERS_1ab;6918 | 0 | 0 |
| NC_019843.3;YP_009047202.1;MERS_1ab;6919 | 0 | 0 |
| NC_019843.3;YP_009047202.1;MERS_1ab;6920 | 0 | 0 |
| NC_019843.3;YP_009047202.1;MERS_1ab;6921 | 0 | 0 |
| NC_019843.3;YP_009047202.1;MERS_1ab;6922 | 0 | 0 |
| NC_019843.3;YP_009047202.1;MERS_1ab;6923 | 0 | 0 |
| NC_019843.3;YP_009047202.1;MERS_1ab;6924 | 0 | 0 |
| NC_019843.3;YP_009047202.1;MERS_1ab;6925 | 0 | 0 |
| NC_019843.3;YP_009047202.1;MERS_1ab;6926 | 0 | 0 |
| NC_019843.3;YP_009047202.1;MERS_1ab;6927 | 0 | 0 |
| NC_019843.3;YP_009047202.1;MERS_1ab;6928 | 0.05 | 0.05 |
| NC_019843.3;YP_009047202.1;MERS_1ab;6929 | 0 | 0 |
| NC_019843.3;YP_009047202.1;MERS_1ab;6930 | 0 | 0 |
| NC_019843.3;YP_009047202.1;MERS_1ab;6931 | 0 | 0 |
| NC_019843.3;YP_009047202.1;MERS_1ab;6932 | 0 | 0 |
| NC_019843.3;YP_009047202.1;MERS_1ab;6933 | 0 | 0 |
| NC_019843.3;YP_009047202.1;MERS_1ab;6934 | 0 | 0 |
| NC_019843.3;YP_009047202.1;MERS_1ab;6935 | 0 | 0 |
| NC_019843.3;YP_009047202.1;MERS_1ab;6936 | 0 | 0 |
| NC_019843.3;YP_009047202.1;MERS_1ab;6937 | 0 | 0 |
| NC_019843.3;YP_009047202.1;MERS_1ab;6938 | 0 | 0 |
| NC_019843.3;YP_009047202.1;MERS_1ab;6939 | 0 | 0 |
| NC_019843.3;YP_009047202.1;MERS_1ab;6940 | 0 | 0 |
| NC_019843.3;YP_009047202.1;MERS_1ab;6941 | 0 | 0 |
| NC_019843.3;YP_009047202.1;MERS_1ab;6942 | 0 | 0 |
| NC_019843.3;YP_009047202.1;MERS_1ab;6943 | 0 | 0 |
| NC_019843.3;YP_009047202.1;MERS_1ab;6944 | 0 | 0 |
| NC_019843.3;YP_009047202.1;MERS_1ab;6945 | 0 | 0 |
| NC_019843.3;YP_009047202.1;MERS_1ab;6946 | 0 | 0 |
| NC_019843.3;YP_009047202.1;MERS_1ab;6947 | 0 | 0 |
| NC_019843.3;YP_009047202.1;MERS_1ab;6948 | 0 | 0 |
| NC_019843.3;YP_009047202.1;MERS_1ab;6949 | 0 | 0 |
| NC_019843.3;YP_009047202.1;MERS_1ab;6950 | 0 | 0 |
| NC_019843.3;YP_009047202.1;MERS_1ab;6951 | 0 | 0 |
| NC_019843.3;YP_009047202.1;MERS_1ab;6952 | 0 | 0 |
| NC_019843.3;YP_009047202.1;MERS_1ab;6953 | 0 | 0 |
| NC_019843.3;YP_009047202.1;MERS_1ab;6954 | 0 | 0 |
| NC_019843.3;YP_009047202.1;MERS_1ab;6955 | 0 | 0 |
| NC_019843.3;YP_009047202.1;MERS_1ab;6956 | 0 | 0 |
| NC_019843.3;YP_009047202.1;MERS_1ab;6957 | 0 | 0 |

| | | |
|---|---|---|
| NC_019843.3;YP_009047202.1;MERS_1ab;6958 | 0 | 0 |
| NC_019843.3;YP_009047202.1;MERS_1ab;6959 | 0 | 0 |
| NC_019843.3;YP_009047202.1;MERS_1ab;6960 | 0 | 0 |
| NC_019843.3;YP_009047202.1;MERS_1ab;6961 | 0 | 0 |
| NC_019843.3;YP_009047202.1;MERS_1ab;6962 | 0 | 0 |
| NC_019843.3;YP_009047202.1;MERS_1ab;6963 | 0 | 0 |
| NC_019843.3;YP_009047202.1;MERS_1ab;6964 | 0 | 0.025 |
| NC_019843.3;YP_009047202.1;MERS_1ab;6965 | 0 | 0.025 |
| NC_019843.3;YP_009047202.1;MERS_1ab;6966 | 0 | 0 |
| NC_019843.3;YP_009047202.1;MERS_1ab;6967 | 0 | 0 |
| NC_019843.3;YP_009047202.1;MERS_1ab;6968 | 0 | 0 |
| NC_019843.3;YP_009047202.1;MERS_1ab;6969 | 0 | 0 |
| NC_019843.3;YP_009047202.1;MERS_1ab;6970 | 0.05 | 0.05 |
| NC_019843.3;YP_009047202.1;MERS_1ab;6971 | 0 | 0 |
| NC_019843.3;YP_009047202.1;MERS_1ab;6972 | 0 | 0.025 |
| NC_019843.3;YP_009047202.1;MERS_1ab;6973 | 0 | 0 |
| NC_019843.3;YP_009047202.1;MERS_1ab;6974 | 0 | 0 |
| NC_019843.3;YP_009047202.1;MERS_1ab;6975 | 0 | 0 |
| NC_019843.3;YP_009047202.1;MERS_1ab;6976 | 0 | 0.025 |
| NC_019843.3;YP_009047202.1;MERS_1ab;6977 | 0 | 0.025 |
| NC_019843.3;YP_009047202.1;MERS_1ab;6978 | 0 | 0.025 |
| NC_019843.3;YP_009047202.1;MERS_1ab;6979 | 0 | 0 |
| NC_019843.3;YP_009047202.1;MERS_1ab;6980 | 0 | 0.025 |
| NC_019843.3;YP_009047202.1;MERS_1ab;6981 | 0 | 0.05 |
| NC_019843.3;YP_009047202.1;MERS_1ab;6982 | 0 | 0.05 |
| NC_019843.3;YP_009047202.1;MERS_1ab;6983 | 0.05 | 0.1 |
| NC_019843.3;YP_009047202.1;MERS_1ab;6984 | 0 | 0.05 |
| NC_019843.3;YP_009047202.1;MERS_1ab;6985 | 0 | 0.025 |
| NC_019843.3;YP_009047202.1;MERS_1ab;6986 | 0 | 0.05 |
| NC_019843.3;YP_009047202.1;MERS_1ab;6987 | 0 | 0.025 |
| NC_019843.3;YP_009047202.1;MERS_1ab;6988 | 0 | 0.025 |
| NC_019843.3;YP_009047202.1;MERS_1ab;6989 | 0 | 0.025 |
| NC_019843.3;YP_009047202.1;MERS_1ab;6990 | 0 | 0 |
| NC_019843.3;YP_009047202.1;MERS_1ab;6991 | 0 | 0 |
| NC_019843.3;YP_009047202.1;MERS_1ab;6992 | 0 | 0 |
| NC_019843.3;YP_009047202.1;MERS_1ab;6993 | 0 | 0 |
| NC_019843.3;YP_009047202.1;MERS_1ab;6994 | 0 | 0 |
| NC_019843.3;YP_009047202.1;MERS_1ab;6995 | 0 | 0 |
| NC_019843.3;YP_009047202.1;MERS_1ab;6996 | 0 | 0 |
| NC_019843.3;YP_009047202.1;MERS_1ab;6997 | 0 | 0 |
| NC_019843.3;YP_009047202.1;MERS_1ab;6998 | 0 | 0 |
| NC_019843.3;YP_009047202.1;MERS_1ab;6999 | 0 | 0 |
| NC_019843.3;YP_009047202.1;MERS_1ab;7000 | 0 | 0 |

| | | |
|---|---|---|
| NC_019843.3;YP_009047202.1;MERS_1ab;7001 | 0 | 0 |
| NC_019843.3;YP_009047202.1;MERS_1ab;7002 | 0 | 0 |
| NC_019843.3;YP_009047202.1;MERS_1ab;7003 | 0 | 0 |
| NC_019843.3;YP_009047202.1;MERS_1ab;7004 | 0 | 0 |
| NC_019843.3;YP_009047202.1;MERS_1ab;7005 | 0 | 0 |
| NC_019843.3;YP_009047202.1;MERS_1ab;7006 | 0 | 0 |
| NC_019843.3;YP_009047202.1;MERS_1ab;7007 | 0 | 0 |
| NC_019843.3;YP_009047202.1;MERS_1ab;7008 | 0 | 0 |
| NC_019843.3;YP_009047202.1;MERS_1ab;7009 | 0 | 0 |
| NC_019843.3;YP_009047202.1;MERS_1ab;7010 | 0 | 0 |
| NC_019843.3;YP_009047202.1;MERS_1ab;7011 | 0 | 0 |
| NC_019843.3;YP_009047202.1;MERS_1ab;7012 | 0 | 0.025 |
| NC_019843.3;YP_009047202.1;MERS_1ab;7013 | 0 | 0 |
| NC_019843.3;YP_009047202.1;MERS_1ab;7014 | 0 | 0 |
| NC_019843.3;YP_009047202.1;MERS_1ab;7015 | 0 | 0 |
| NC_019843.3;YP_009047202.1;MERS_1ab;7016 | 0 | 0 |
| NC_019843.3;YP_009047202.1;MERS_1ab;7017 | 0 | 0 |
| NC_019843.3;YP_009047202.1;MERS_1ab;7018 | 0 | 0 |
| NC_019843.3;YP_009047202.1;MERS_1ab;7019 | 0 | 0.025 |
| NC_019843.3;YP_009047202.1;MERS_1ab;7020 | 0 | 0.025 |
| NC_019843.3;YP_009047202.1;MERS_1ab;7021 | 0 | 0.025 |
| NC_019843.3;YP_009047202.1;MERS_1ab;7022 | 0 | 0.025 |
| NC_019843.3;YP_009047202.1;MERS_1ab;7023 | 0 | 0 |
| NC_019843.3;YP_009047202.1;MERS_1ab;7024 | 0 | 0 |
| NC_019843.3;YP_009047202.1;MERS_1ab;7025 | 0 | 0.025 |
| NC_019843.3;YP_009047202.1;MERS_1ab;7026 | 0 | 0 |
| NC_019843.3;YP_009047202.1;MERS_1ab;7027 | 0 | 0 |
| NC_019843.3;YP_009047202.1;MERS_1ab;7028 | 0 | 0 |
| NC_019843.3;YP_009047202.1;MERS_1ab;7029 | 0 | 0 |
| NC_019843.3;YP_009047202.1;MERS_1ab;7030 | 0.05 | 0.025 |
| NC_019843.3;YP_009047202.1;MERS_1ab;7031 | 0.05 | 0 |
| NC_019843.3;YP_009047202.1;MERS_1ab;7032 | 0 | 0 |
| NC_019843.3;YP_009047202.1;MERS_1ab;7033 | 0 | 0 |
| NC_019843.3;YP_009047202.1;MERS_1ab;7034 | 0 | 0 |
| NC_019843.3;YP_009047202.1;MERS_1ab;7035 | 0.05 | 0 |
| NC_019843.3;YP_009047202.1;MERS_1ab;7036 | 0 | 0 |
| NC_019843.3;YP_009047202.1;MERS_1ab;7037 | 0 | 0 |
| NC_019843.3;YP_009047202.1;MERS_1ab;7038 | 0 | 0 |
| NC_019843.3;YP_009047202.1;MERS_1ab;7039 | 0 | 0 |
| NC_019843.3;YP_009047202.1;MERS_1ab;7040 | 0 | 0 |
| NC_019843.3;YP_009047202.1;MERS_1ab;7041 | 0 | 0 |
| NC_019843.3;YP_009047202.1;MERS_1ab;7042 | 0 | 0 |
| NC_019843.3;YP_009047202.1;MERS_1ab;7043 | 0 | 0 |

| | | |
|---|---|---|
| NC_019843.3;YP_009047202.1;MERS_1ab;7044 | 0 | 0 |
| NC_019843.3;YP_009047202.1;MERS_1ab;7045 | 0 | 0 |
| NC_019843.3;YP_009047202.1;MERS_1ab;7046 | 0 | 0 |
| NC_019843.3;YP_009047202.1;MERS_1ab;7047 | 0.05 | 0 |
| NC_019843.3;YP_009047202.1;MERS_1ab;7048 | 0.05 | 0 |
| NC_019843.3;YP_009047202.1;MERS_1ab;7049 | 0.05 | 0.025 |
| NC_019843.3;YP_009047202.1;MERS_1ab;7050 | 0.1 | 0 |
| NC_019843.3;YP_009047202.1;MERS_1ab;7051 | 0.1 | 0 |
| NC_019843.3;YP_009047202.1;MERS_1ab;7052 | 0.1 | 0 |
| NC_019843.3;YP_009047202.1;MERS_1ab;7053 | 0.15 | 0 |
| NC_019843.3;YP_009047202.1;MERS_1ab;7054 | 0.05 | 0 |
| NC_019843.3;YP_009047202.1;MERS_1ab;7055 | 0 | 0 |
| NC_019843.3;YP_009047202.1;MERS_1ab;7056 | 0 | 0 |
| NC_019843.3;YP_009047202.1;MERS_1ab;7057 | 0 | 0 |
| NC_019843.3;YP_009047202.1;MERS_1ab;7058 | 0 | 0 |
| NC_019843.3;YP_009047202.1;MERS_1ab;7059 | 0 | 0 |
| NC_019843.3;YP_009047202.1;MERS_1ab;7060 | 0.05 | 0 |
| NC_019843.3;YP_009047202.1;MERS_1ab;7061 | 0 | 0 |
| NC_019843.3;YP_009047202.1;MERS_1ab;7062 | 0 | 0 |
| NC_019843.3;YP_009047202.1;MERS_1ab;7063 | 0 | 0 |
| NC_019843.3;YP_009047202.1;MERS_1ab;7064 | 0 | 0 |
| NC_019843.3;YP_009047202.1;MERS_1ab;7065 | 0 | 0 |
| NC_019843.3;YP_009047202.1;MERS_1ab;7066 | 0 | 0 |
| NC_019843.3;YP_009047202.1;MERS_1ab;7067 | 0 | 0 |
| NC_019843.3;YP_009047203.1;MERS_1a;1 | 0 | 0 |
| NC_019843.3;YP_009047203.1;MERS_1a;2 | 0 | 0 |
| NC_019843.3;YP_009047203.1;MERS_1a;3 | 0 | 0 |
| NC_019843.3;YP_009047203.1;MERS_1a;4 | 0.05 | 0 |
| NC_019843.3;YP_009047203.1;MERS_1a;5 | 0 | 0 |
| NC_019843.3;YP_009047203.1;MERS_1a;6 | 0 | 0 |
| NC_019843.3;YP_009047203.1;MERS_1a;7 | 0 | 0 |
| NC_019843.3;YP_009047203.1;MERS_1a;8 | 0.05 | 0 |
| NC_019843.3;YP_009047203.1;MERS_1a;9 | 0 | 0.025 |
| NC_019843.3;YP_009047203.1;MERS_1a;10 | 0 | 0.025 |
| NC_019843.3;YP_009047203.1;MERS_1a;11 | 0 | 0.025 |
| NC_019843.3;YP_009047203.1;MERS_1a;12 | 0 | 0.025 |
| NC_019843.3;YP_009047203.1;MERS_1a;13 | 0 | 0.05 |
| NC_019843.3;YP_009047203.1;MERS_1a;14 | 0.05 | 0 |
| NC_019843.3;YP_009047203.1;MERS_1a;15 | 0 | 0 |
| NC_019843.3;YP_009047203.1;MERS_1a;16 | 0.05 | 0 |
| NC_019843.3;YP_009047203.1;MERS_1a;17 | 0 | 0 |
| NC_019843.3;YP_009047203.1;MERS_1a;18 | 0 | 0 |
| NC_019843.3;YP_009047203.1;MERS_1a;19 | 0.1 | 0.025 |

| | | |
|---|---|---|
| NC_019843.3;YP_009047203.1;MERS_1a;20 | 0 | 0.025 |
| NC_019843.3;YP_009047203.1;MERS_1a;21 | 0 | 0.025 |
| NC_019843.3;YP_009047203.1;MERS_1a;22 | 0 | 0.025 |
| NC_019843.3;YP_009047203.1;MERS_1a;23 | 0 | 0.025 |
| NC_019843.3;YP_009047203.1;MERS_1a;24 | 0 | 0 |
| NC_019843.3;YP_009047203.1;MERS_1a;25 | 0 | 0 |
| NC_019843.3;YP_009047203.1;MERS_1a;26 | 0 | 0 |
| NC_019843.3;YP_009047203.1;MERS_1a;27 | 0 | 0 |
| NC_019843.3;YP_009047203.1;MERS_1a;28 | 0 | 0 |
| NC_019843.3;YP_009047203.1;MERS_1a;29 | 0.05 | 0 |
| NC_019843.3;YP_009047203.1;MERS_1a;30 | 0.05 | 0 |
| NC_019843.3;YP_009047203.1;MERS_1a;31 | 0.05 | 0 |
| NC_019843.3;YP_009047203.1;MERS_1a;32 | 0.05 | 0 |
| NC_019843.3;YP_009047203.1;MERS_1a;33 | 0.05 | 0 |
| NC_019843.3;YP_009047203.1;MERS_1a;34 | 0.05 | 0.025 |
| NC_019843.3;YP_009047203.1;MERS_1a;35 | 0.05 | 0 |
| NC_019843.3;YP_009047203.1;MERS_1a;36 | 0.15 | 0.025 |
| NC_019843.3;YP_009047203.1;MERS_1a;37 | 0.05 | 0.05 |
| NC_019843.3;YP_009047203.1;MERS_1a;38 | 0.05 | 0.025 |
| NC_019843.3;YP_009047203.1;MERS_1a;39 | 0.05 | 0.025 |
| NC_019843.3;YP_009047203.1;MERS_1a;40 | 0 | 0.025 |
| NC_019843.3;YP_009047203.1;MERS_1a;41 | 0 | 0.025 |
| NC_019843.3;YP_009047203.1;MERS_1a;42 | 0 | 0.025 |
| NC_019843.3;YP_009047203.1;MERS_1a;43 | 0 | 0.025 |
| NC_019843.3;YP_009047203.1;MERS_1a;44 | 0 | 0.025 |
| NC_019843.3;YP_009047203.1;MERS_1a;45 | 0 | 0.025 |
| NC_019843.3;YP_009047203.1;MERS_1a;46 | 0.05 | 0 |
| NC_019843.3;YP_009047203.1;MERS_1a;47 | 0 | 0.025 |
| NC_019843.3;YP_009047203.1;MERS_1a;48 | 0 | 0.025 |
| NC_019843.3;YP_009047203.1;MERS_1a;49 | 0 | 0 |
| NC_019843.3;YP_009047203.1;MERS_1a;50 | 0 | 0 |
| NC_019843.3;YP_009047203.1;MERS_1a;51 | 0 | 0 |
| NC_019843.3;YP_009047203.1;MERS_1a;52 | 0 | 0 |
| NC_019843.3;YP_009047203.1;MERS_1a;53 | 0 | 0 |
| NC_019843.3;YP_009047203.1;MERS_1a;54 | 0 | 0 |
| NC_019843.3;YP_009047203.1;MERS_1a;55 | 0 | 0 |
| NC_019843.3;YP_009047203.1;MERS_1a;56 | 0 | 0 |
| NC_019843.3;YP_009047203.1;MERS_1a;57 | 0 | 0 |
| NC_019843.3;YP_009047203.1;MERS_1a;58 | 0 | 0 |
| NC_019843.3;YP_009047203.1;MERS_1a;59 | 0 | 0 |
| NC_019843.3;YP_009047203.1;MERS_1a;60 | 0 | 0 |
| NC_019843.3;YP_009047203.1;MERS_1a;61 | 0 | 0 |
| NC_019843.3;YP_009047203.1;MERS_1a;62 | 0 | 0 |

| | | |
|---|---|---|
| NC_019843.3;YP_009047203.1;MERS_1a;63 | 0 | 0 |
| NC_019843.3;YP_009047203.1;MERS_1a;64 | 0 | 0 |
| NC_019843.3;YP_009047203.1;MERS_1a;65 | 0 | 0 |
| NC_019843.3;YP_009047203.1;MERS_1a;66 | 0 | 0 |
| NC_019843.3;YP_009047203.1;MERS_1a;67 | 0 | 0 |
| NC_019843.3;YP_009047203.1;MERS_1a;68 | 0 | 0 |
| NC_019843.3;YP_009047203.1;MERS_1a;69 | 0 | 0 |
| NC_019843.3;YP_009047203.1;MERS_1a;70 | 0 | 0 |
| NC_019843.3;YP_009047203.1;MERS_1a;71 | 0 | 0 |
| NC_019843.3;YP_009047203.1;MERS_1a;72 | 0 | 0 |
| NC_019843.3;YP_009047203.1;MERS_1a;73 | 0 | 0 |
| NC_019843.3;YP_009047203.1;MERS_1a;74 | 0 | 0 |
| NC_019843.3;YP_009047203.1;MERS_1a;75 | 0 | 0 |
| NC_019843.3;YP_009047203.1;MERS_1a;76 | 0 | 0 |
| NC_019843.3;YP_009047203.1;MERS_1a;77 | 0 | 0 |
| NC_019843.3;YP_009047203.1;MERS_1a;78 | 0 | 0 |
| NC_019843.3;YP_009047203.1;MERS_1a;79 | 0 | 0 |
| NC_019843.3;YP_009047203.1;MERS_1a;80 | 0 | 0 |
| NC_019843.3;YP_009047203.1;MERS_1a;81 | 0 | 0 |
| NC_019843.3;YP_009047203.1;MERS_1a;82 | 0 | 0 |
| NC_019843.3;YP_009047203.1;MERS_1a;83 | 0 | 0 |
| NC_019843.3;YP_009047203.1;MERS_1a;84 | 0 | 0 |
| NC_019843.3;YP_009047203.1;MERS_1a;85 | 0 | 0 |
| NC_019843.3;YP_009047203.1;MERS_1a;86 | 0 | 0 |
| NC_019843.3;YP_009047203.1;MERS_1a;87 | 0 | 0 |
| NC_019843.3;YP_009047203.1;MERS_1a;88 | 0 | 0 |
| NC_019843.3;YP_009047203.1;MERS_1a;89 | 0 | 0 |
| NC_019843.3;YP_009047203.1;MERS_1a;90 | 0 | 0 |
| NC_019843.3;YP_009047203.1;MERS_1a;91 | 0 | 0 |
| NC_019843.3;YP_009047203.1;MERS_1a;92 | 0.1 | 0.025 |
| NC_019843.3;YP_009047203.1;MERS_1a;93 | 0 | 0 |
| NC_019843.3;YP_009047203.1;MERS_1a;94 | 0 | 0 |
| NC_019843.3;YP_009047203.1;MERS_1a;95 | 0 | 0 |
| NC_019843.3;YP_009047203.1;MERS_1a;96 | 0 | 0 |
| NC_019843.3;YP_009047203.1;MERS_1a;97 | 0 | 0 |
| NC_019843.3;YP_009047203.1;MERS_1a;98 | 0.05 | 0.075 |
| NC_019843.3;YP_009047203.1;MERS_1a;99 | 0 | 0.025 |
| NC_019843.3;YP_009047203.1;MERS_1a;100 | 0 | 0.025 |
| NC_019843.3;YP_009047203.1;MERS_1a;101 | 0.05 | 0.025 |
| NC_019843.3;YP_009047203.1;MERS_1a;102 | 0.05 | 0 |
| NC_019843.3;YP_009047203.1;MERS_1a;103 | 0.05 | 0.025 |
| NC_019843.3;YP_009047203.1;MERS_1a;104 | 0.05 | 0.05 |
| NC_019843.3;YP_009047203.1;MERS_1a;105 | 0 | 0 |

| | | |
|---|---|---|
| NC_019843.3;YP_009047203.1;MERS_1a;106 | 0 | 0 |
| NC_019843.3;YP_009047203.1;MERS_1a;107 | 0.05 | 0.1 |
| NC_019843.3;YP_009047203.1;MERS_1a;108 | 0 | 0 |
| NC_019843.3;YP_009047203.1;MERS_1a;109 | 0 | 0 |
| NC_019843.3;YP_009047203.1;MERS_1a;110 | 0 | 0 |
| NC_019843.3;YP_009047203.1;MERS_1a;111 | 0 | 0 |
| NC_019843.3;YP_009047203.1;MERS_1a;112 | 0 | 0 |
| NC_019843.3;YP_009047203.1;MERS_1a;113 | 0 | 0 |
| NC_019843.3;YP_009047203.1;MERS_1a;114 | 0 | 0 |
| NC_019843.3;YP_009047203.1;MERS_1a;115 | 0 | 0.025 |
| NC_019843.3;YP_009047203.1;MERS_1a;116 | 0 | 0 |
| NC_019843.3;YP_009047203.1;MERS_1a;117 | 0 | 0 |
| NC_019843.3;YP_009047203.1;MERS_1a;118 | 0 | 0 |
| NC_019843.3;YP_009047203.1;MERS_1a;119 | 0 | 0 |
| NC_019843.3;YP_009047203.1;MERS_1a;120 | 0.05 | 0 |
| NC_019843.3;YP_009047203.1;MERS_1a;121 | 0 | 0 |
| NC_019843.3;YP_009047203.1;MERS_1a;122 | 0 | 0 |
| NC_019843.3;YP_009047203.1;MERS_1a;123 | 0 | 0 |
| NC_019843.3;YP_009047203.1;MERS_1a;124 | 0 | 0 |
| NC_019843.3;YP_009047203.1;MERS_1a;125 | 0 | 0.025 |
| NC_019843.3;YP_009047203.1;MERS_1a;126 | 0.05 | 0 |
| NC_019843.3;YP_009047203.1;MERS_1a;127 | 0.05 | 0 |
| NC_019843.3;YP_009047203.1;MERS_1a;128 | 0.1 | 0 |
| NC_019843.3;YP_009047203.1;MERS_1a;129 | 0 | 0 |
| NC_019843.3;YP_009047203.1;MERS_1a;130 | 0 | 0 |
| NC_019843.3;YP_009047203.1;MERS_1a;131 | 0 | 0 |
| NC_019843.3;YP_009047203.1;MERS_1a;132 | 0 | 0 |
| NC_019843.3;YP_009047203.1;MERS_1a;133 | 0 | 0 |
| NC_019843.3;YP_009047203.1;MERS_1a;134 | 0 | 0 |
| NC_019843.3;YP_009047203.1;MERS_1a;135 | 0 | 0 |
| NC_019843.3;YP_009047203.1;MERS_1a;136 | 0 | 0 |
| NC_019843.3;YP_009047203.1;MERS_1a;137 | 0 | 0 |
| NC_019843.3;YP_009047203.1;MERS_1a;138 | 0 | 0 |
| NC_019843.3;YP_009047203.1;MERS_1a;139 | 0 | 0 |
| NC_019843.3;YP_009047203.1;MERS_1a;140 | 0 | 0 |
| NC_019843.3;YP_009047203.1;MERS_1a;141 | 0 | 0 |
| NC_019843.3;YP_009047203.1;MERS_1a;142 | 0 | 0 |
| NC_019843.3;YP_009047203.1;MERS_1a;143 | 0 | 0 |
| NC_019843.3;YP_009047203.1;MERS_1a;144 | 0 | 0 |
| NC_019843.3;YP_009047203.1;MERS_1a;145 | 0 | 0 |
| NC_019843.3;YP_009047203.1;MERS_1a;146 | 0 | 0 |
| NC_019843.3;YP_009047203.1;MERS_1a;147 | 0 | 0 |
| NC_019843.3;YP_009047203.1;MERS_1a;148 | 0 | 0 |

| | | |
|---|---|---|
| NC_019843.3;YP_009047203.1;MERS_1a;149 | 0 | 0.025 |
| NC_019843.3;YP_009047203.1;MERS_1a;150 | 0 | 0.025 |
| NC_019843.3;YP_009047203.1;MERS_1a;151 | 0 | 0.025 |
| NC_019843.3;YP_009047203.1;MERS_1a;152 | 0 | 0.025 |
| NC_019843.3;YP_009047203.1;MERS_1a;153 | 0 | 0.025 |
| NC_019843.3;YP_009047203.1;MERS_1a;154 | 0 | 0.025 |
| NC_019843.3;YP_009047203.1;MERS_1a;155 | 0 | 0.025 |
| NC_019843.3;YP_009047203.1;MERS_1a;156 | 0 | 0.025 |
| NC_019843.3;YP_009047203.1;MERS_1a;157 | 0 | 0.025 |
| NC_019843.3;YP_009047203.1;MERS_1a;158 | 0 | 0.025 |
| NC_019843.3;YP_009047203.1;MERS_1a;159 | 0 | 0 |
| NC_019843.3;YP_009047203.1;MERS_1a;160 | 0 | 0.05 |
| NC_019843.3;YP_009047203.1;MERS_1a;161 | 0 | 0 |
| NC_019843.3;YP_009047203.1;MERS_1a;162 | 0 | 0 |
| NC_019843.3;YP_009047203.1;MERS_1a;163 | 0.05 | 0.025 |
| NC_019843.3;YP_009047203.1;MERS_1a;164 | 0.1 | 0.025 |
| NC_019843.3;YP_009047203.1;MERS_1a;165 | 0.1 | 0.025 |
| NC_019843.3;YP_009047203.1;MERS_1a;166 | 0.1 | 0.05 |
| NC_019843.3;YP_009047203.1;MERS_1a;167 | 0.1 | 0.075 |
| NC_019843.3;YP_009047203.1;MERS_1a;168 | 0.1 | 0.05 |
| NC_019843.3;YP_009047203.1;MERS_1a;169 | 0.1 | 0.025 |
| NC_019843.3;YP_009047203.1;MERS_1a;170 | 0.1 | 0.05 |
| NC_019843.3;YP_009047203.1;MERS_1a;171 | 0 | 0 |
| NC_019843.3;YP_009047203.1;MERS_1a;172 | 0 | 0 |
| NC_019843.3;YP_009047203.1;MERS_1a;173 | 0 | 0.075 |
| NC_019843.3;YP_009047203.1;MERS_1a;174 | 0 | 0 |
| NC_019843.3;YP_009047203.1;MERS_1a;175 | 0 | 0 |
| NC_019843.3;YP_009047203.1;MERS_1a;176 | 0.05 | 0 |
| NC_019843.3;YP_009047203.1;MERS_1a;177 | 0 | 0 |
| NC_019843.3;YP_009047203.1;MERS_1a;178 | 0 | 0 |
| NC_019843.3;YP_009047203.1;MERS_1a;179 | 0 | 0.05 |
| NC_019843.3;YP_009047203.1;MERS_1a;180 | 0 | 0.025 |
| NC_019843.3;YP_009047203.1;MERS_1a;181 | 0 | 0 |
| NC_019843.3;YP_009047203.1;MERS_1a;182 | 0 | 0 |
| NC_019843.3;YP_009047203.1;MERS_1a;183 | 0 | 0 |
| NC_019843.3;YP_009047203.1;MERS_1a;184 | 0.05 | 0 |
| NC_019843.3;YP_009047203.1;MERS_1a;185 | 0 | 0 |
| NC_019843.3;YP_009047203.1;MERS_1a;186 | 0.1 | 0 |
| NC_019843.3;YP_009047203.1;MERS_1a;187 | 0.1 | 0.025 |
| NC_019843.3;YP_009047203.1;MERS_1a;188 | 0.05 | 0.025 |
| NC_019843.3;YP_009047203.1;MERS_1a;189 | 0 | 0 |
| NC_019843.3;YP_009047203.1;MERS_1a;190 | 0 | 0 |
| NC_019843.3;YP_009047203.1;MERS_1a;191 | 0 | 0 |

| | | |
|---|---|---|
| NC_019843.3;YP_009047203.1;MERS_1a;192 | 0 | 0 |
| NC_019843.3;YP_009047203.1;MERS_1a;193 | 0 | 0 |
| NC_019843.3;YP_009047203.1;MERS_1a;194 | 0 | 0 |
| NC_019843.3;YP_009047203.1;MERS_1a;195 | 0 | 0 |
| NC_019843.3;YP_009047203.1;MERS_1a;196 | 0.05 | 0 |
| NC_019843.3;YP_009047203.1;MERS_1a;197 | 0.1 | 0.025 |
| NC_019843.3;YP_009047203.1;MERS_1a;198 | 0 | 0.025 |
| NC_019843.3;YP_009047203.1;MERS_1a;199 | 0.05 | 0.05 |
| NC_019843.3;YP_009047203.1;MERS_1a;200 | 0 | 0.025 |
| NC_019843.3;YP_009047203.1;MERS_1a;201 | 0 | 0 |
| NC_019843.3;YP_009047203.1;MERS_1a;202 | 0 | 0 |
| NC_019843.3;YP_009047203.1;MERS_1a;203 | 0 | 0 |
| NC_019843.3;YP_009047203.1;MERS_1a;204 | 0 | 0 |
| NC_019843.3;YP_009047203.1;MERS_1a;205 | 0 | 0 |
| NC_019843.3;YP_009047203.1;MERS_1a;206 | 0 | 0 |
| NC_019843.3;YP_009047203.1;MERS_1a;207 | 0 | 0 |
| NC_019843.3;YP_009047203.1;MERS_1a;208 | 0 | 0 |
| NC_019843.3;YP_009047203.1;MERS_1a;209 | 0 | 0 |
| NC_019843.3;YP_009047203.1;MERS_1a;210 | 0 | 0 |
| NC_019843.3;YP_009047203.1;MERS_1a;211 | 0 | 0 |
| NC_019843.3;YP_009047203.1;MERS_1a;212 | 0 | 0 |
| NC_019843.3;YP_009047203.1;MERS_1a;213 | 0 | 0 |
| NC_019843.3;YP_009047203.1;MERS_1a;214 | 0 | 0 |
| NC_019843.3;YP_009047203.1;MERS_1a;215 | 0 | 0 |
| NC_019843.3;YP_009047203.1;MERS_1a;216 | 0 | 0 |
| NC_019843.3;YP_009047203.1;MERS_1a;217 | 0 | 0 |
| NC_019843.3;YP_009047203.1;MERS_1a;218 | 0 | 0 |
| NC_019843.3;YP_009047203.1;MERS_1a;219 | 0.05 | 0 |
| NC_019843.3;YP_009047203.1;MERS_1a;220 | 0 | 0 |
| NC_019843.3;YP_009047203.1;MERS_1a;221 | 0 | 0 |
| NC_019843.3;YP_009047203.1;MERS_1a;222 | 0 | 0 |
| NC_019843.3;YP_009047203.1;MERS_1a;223 | 0 | 0 |
| NC_019843.3;YP_009047203.1;MERS_1a;224 | 0 | 0 |
| NC_019843.3;YP_009047203.1;MERS_1a;225 | 0 | 0 |
| NC_019843.3;YP_009047203.1;MERS_1a;226 | 0 | 0 |
| NC_019843.3;YP_009047203.1;MERS_1a;227 | 0 | 0 |
| NC_019843.3;YP_009047203.1;MERS_1a;228 | 0 | 0 |
| NC_019843.3;YP_009047203.1;MERS_1a;229 | 0 | 0 |
| NC_019843.3;YP_009047203.1;MERS_1a;230 | 0 | 0 |
| NC_019843.3;YP_009047203.1;MERS_1a;231 | 0 | 0 |
| NC_019843.3;YP_009047203.1;MERS_1a;232 | 0.05 | 0 |
| NC_019843.3;YP_009047203.1;MERS_1a;233 | 0 | 0 |
| NC_019843.3;YP_009047203.1;MERS_1a;234 | 0 | 0 |

| | | |
|---|---|---|
| NC_019843.3;YP_009047203.1;MERS_1a;235 | 0 | 0 |
| NC_019843.3;YP_009047203.1;MERS_1a;236 | 0 | 0 |
| NC_019843.3;YP_009047203.1;MERS_1a;237 | 0 | 0 |
| NC_019843.3;YP_009047203.1;MERS_1a;238 | 0 | 0 |
| NC_019843.3;YP_009047203.1;MERS_1a;239 | 0 | 0 |
| NC_019843.3;YP_009047203.1;MERS_1a;240 | 0 | 0 |
| NC_019843.3;YP_009047203.1;MERS_1a;241 | 0 | 0 |
| NC_019843.3;YP_009047203.1;MERS_1a;242 | 0 | 0 |
| NC_019843.3;YP_009047203.1;MERS_1a;243 | 0 | 0 |
| NC_019843.3;YP_009047203.1;MERS_1a;244 | 0 | 0.025 |
| NC_019843.3;YP_009047203.1;MERS_1a;245 | 0 | 0 |
| NC_019843.3;YP_009047203.1;MERS_1a;246 | 0 | 0 |
| NC_019843.3;YP_009047203.1;MERS_1a;247 | 0 | 0.025 |
| NC_019843.3;YP_009047203.1;MERS_1a;248 | 0 | 0 |
| NC_019843.3;YP_009047203.1;MERS_1a;249 | 0 | 0 |
| NC_019843.3;YP_009047203.1;MERS_1a;250 | 0 | 0 |
| NC_019843.3;YP_009047203.1;MERS_1a;251 | 0 | 0.05 |
| NC_019843.3;YP_009047203.1;MERS_1a;252 | 0 | 0.05 |
| NC_019843.3;YP_009047203.1;MERS_1a;253 | 0 | 0.025 |
| NC_019843.3;YP_009047203.1;MERS_1a;254 | 0 | 0 |
| NC_019843.3;YP_009047203.1;MERS_1a;255 | 0 | 0 |
| NC_019843.3;YP_009047203.1;MERS_1a;256 | 0 | 0 |
| NC_019843.3;YP_009047203.1;MERS_1a;257 | 0 | 0 |
| NC_019843.3;YP_009047203.1;MERS_1a;258 | 0 | 0 |
| NC_019843.3;YP_009047203.1;MERS_1a;259 | 0.05 | 0.025 |
| NC_019843.3;YP_009047203.1;MERS_1a;260 | 0.05 | 0.025 |
| NC_019843.3;YP_009047203.1;MERS_1a;261 | 0 | 0 |
| NC_019843.3;YP_009047203.1;MERS_1a;262 | 0 | 0.025 |
| NC_019843.3;YP_009047203.1;MERS_1a;263 | 0 | 0.025 |
| NC_019843.3;YP_009047203.1;MERS_1a;264 | 0 | 0.025 |
| NC_019843.3;YP_009047203.1;MERS_1a;265 | 0 | 0.025 |
| NC_019843.3;YP_009047203.1;MERS_1a;266 | 0 | 0.025 |
| NC_019843.3;YP_009047203.1;MERS_1a;267 | 0.05 | 0.05 |
| NC_019843.3;YP_009047203.1;MERS_1a;268 | 0 | 0 |
| NC_019843.3;YP_009047203.1;MERS_1a;269 | 0 | 0.025 |
| NC_019843.3;YP_009047203.1;MERS_1a;270 | 0 | 0 |
| NC_019843.3;YP_009047203.1;MERS_1a;271 | 0 | 0 |
| NC_019843.3;YP_009047203.1;MERS_1a;272 | 0 | 0 |
| NC_019843.3;YP_009047203.1;MERS_1a;273 | 0 | 0 |
| NC_019843.3;YP_009047203.1;MERS_1a;274 | 0 | 0 |
| NC_019843.3;YP_009047203.1;MERS_1a;275 | 0 | 0 |
| NC_019843.3;YP_009047203.1;MERS_1a;276 | 0 | 0 |
| NC_019843.3;YP_009047203.1;MERS_1a;277 | 0 | 0 |

| | | |
|---|---|---|
| NC_019843.3;YP_009047203.1;MERS_1a;278 | 0 | 0 |
| NC_019843.3;YP_009047203.1;MERS_1a;279 | 0 | 0 |
| NC_019843.3;YP_009047203.1;MERS_1a;280 | 0 | 0 |
| NC_019843.3;YP_009047203.1;MERS_1a;281 | 0 | 0 |
| NC_019843.3;YP_009047203.1;MERS_1a;282 | 0 | 0 |
| NC_019843.3;YP_009047203.1;MERS_1a;283 | 0 | 0 |
| NC_019843.3;YP_009047203.1;MERS_1a;284 | 0 | 0 |
| NC_019843.3;YP_009047203.1;MERS_1a;285 | 0 | 0 |
| NC_019843.3;YP_009047203.1;MERS_1a;286 | 0 | 0 |
| NC_019843.3;YP_009047203.1;MERS_1a;287 | 0 | 0 |
| NC_019843.3;YP_009047203.1;MERS_1a;288 | 0 | 0 |
| NC_019843.3;YP_009047203.1;MERS_1a;289 | 0 | 0 |
| NC_019843.3;YP_009047203.1;MERS_1a;290 | 0 | 0 |
| NC_019843.3;YP_009047203.1;MERS_1a;291 | 0 | 0 |
| NC_019843.3;YP_009047203.1;MERS_1a;292 | 0 | 0 |
| NC_019843.3;YP_009047203.1;MERS_1a;293 | 0 | 0 |
| NC_019843.3;YP_009047203.1;MERS_1a;294 | 0 | 0 |
| NC_019843.3;YP_009047203.1;MERS_1a;295 | 0 | 0 |
| NC_019843.3;YP_009047203.1;MERS_1a;296 | 0 | 0 |
| NC_019843.3;YP_009047203.1;MERS_1a;297 | 0 | 0 |
| NC_019843.3;YP_009047203.1;MERS_1a;298 | 0 | 0 |
| NC_019843.3;YP_009047203.1;MERS_1a;299 | 0 | 0 |
| NC_019843.3;YP_009047203.1;MERS_1a;300 | 0 | 0 |
| NC_019843.3;YP_009047203.1;MERS_1a;301 | 0 | 0 |
| NC_019843.3;YP_009047203.1;MERS_1a;302 | 0 | 0 |
| NC_019843.3;YP_009047203.1;MERS_1a;303 | 0 | 0 |
| NC_019843.3;YP_009047203.1;MERS_1a;304 | 0.05 | 0.05 |
| NC_019843.3;YP_009047203.1;MERS_1a;305 | 0 | 0 |
| NC_019843.3;YP_009047203.1;MERS_1a;306 | 0 | 0 |
| NC_019843.3;YP_009047203.1;MERS_1a;307 | 0 | 0 |
| NC_019843.3;YP_009047203.1;MERS_1a;308 | 0 | 0 |
| NC_019843.3;YP_009047203.1;MERS_1a;309 | 0 | 0 |
| NC_019843.3;YP_009047203.1;MERS_1a;310 | 0 | 0 |
| NC_019843.3;YP_009047203.1;MERS_1a;311 | 0 | 0 |
| NC_019843.3;YP_009047203.1;MERS_1a;312 | 0 | 0 |
| NC_019843.3;YP_009047203.1;MERS_1a;313 | 0 | 0 |
| NC_019843.3;YP_009047203.1;MERS_1a;314 | 0 | 0 |
| NC_019843.3;YP_009047203.1;MERS_1a;315 | 0 | 0 |
| NC_019843.3;YP_009047203.1;MERS_1a;316 | 0 | 0 |
| NC_019843.3;YP_009047203.1;MERS_1a;317 | 0 | 0 |
| NC_019843.3;YP_009047203.1;MERS_1a;318 | 0 | 0 |
| NC_019843.3;YP_009047203.1;MERS_1a;319 | 0 | 0 |
| NC_019843.3;YP_009047203.1;MERS_1a;320 | 0 | 0 |

| | | |
|---|---|---|
| NC_019843.3;YP_009047203.1;MERS_1a;321 | 0 | 0 |
| NC_019843.3;YP_009047203.1;MERS_1a;322 | 0 | 0 |
| NC_019843.3;YP_009047203.1;MERS_1a;323 | 0 | 0 |
| NC_019843.3;YP_009047203.1;MERS_1a;324 | 0 | 0 |
| NC_019843.3;YP_009047203.1;MERS_1a;325 | 0 | 0 |
| NC_019843.3;YP_009047203.1;MERS_1a;326 | 0 | 0 |
| NC_019843.3;YP_009047203.1;MERS_1a;327 | 0 | 0 |
| NC_019843.3;YP_009047203.1;MERS_1a;328 | 0 | 0 |
| NC_019843.3;YP_009047203.1;MERS_1a;329 | 0 | 0 |
| NC_019843.3;YP_009047203.1;MERS_1a;330 | 0 | 0 |
| NC_019843.3;YP_009047203.1;MERS_1a;331 | 0 | 0 |
| NC_019843.3;YP_009047203.1;MERS_1a;332 | 0 | 0 |
| NC_019843.3;YP_009047203.1;MERS_1a;333 | 0 | 0 |
| NC_019843.3;YP_009047203.1;MERS_1a;334 | 0.05 | 0 |
| NC_019843.3;YP_009047203.1;MERS_1a;335 | 0 | 0 |
| NC_019843.3;YP_009047203.1;MERS_1a;336 | 0 | 0.025 |
| NC_019843.3;YP_009047203.1;MERS_1a;337 | 0 | 0.025 |
| NC_019843.3;YP_009047203.1;MERS_1a;338 | 0 | 0.025 |
| NC_019843.3;YP_009047203.1;MERS_1a;339 | 0 | 0 |
| NC_019843.3;YP_009047203.1;MERS_1a;340 | 0.05 | 0.025 |
| NC_019843.3;YP_009047203.1;MERS_1a;341 | 0 | 0.05 |
| NC_019843.3;YP_009047203.1;MERS_1a;342 | 0 | 0 |
| NC_019843.3;YP_009047203.1;MERS_1a;343 | 0 | 0 |
| NC_019843.3;YP_009047203.1;MERS_1a;344 | 0 | 0 |
| NC_019843.3;YP_009047203.1;MERS_1a;345 | 0 | 0 |
| NC_019843.3;YP_009047203.1;MERS_1a;346 | 0 | 0 |
| NC_019843.3;YP_009047203.1;MERS_1a;347 | 0.05 | 0 |
| NC_019843.3;YP_009047203.1;MERS_1a;348 | 0 | 0 |
| NC_019843.3;YP_009047203.1;MERS_1a;349 | 0 | 0 |
| NC_019843.3;YP_009047203.1;MERS_1a;350 | 0 | 0 |
| NC_019843.3;YP_009047203.1;MERS_1a;351 | 0 | 0 |
| NC_019843.3;YP_009047203.1;MERS_1a;352 | 0 | 0.025 |
| NC_019843.3;YP_009047203.1;MERS_1a;353 | 0 | 0.025 |
| NC_019843.3;YP_009047203.1;MERS_1a;354 | 0 | 0.025 |
| NC_019843.3;YP_009047203.1;MERS_1a;355 | 0 | 0.025 |
| NC_019843.3;YP_009047203.1;MERS_1a;356 | 0.05 | 0.05 |
| NC_019843.3;YP_009047203.1;MERS_1a;357 | 0 | 0.05 |
| NC_019843.3;YP_009047203.1;MERS_1a;358 | 0 | 0.05 |
| NC_019843.3;YP_009047203.1;MERS_1a;359 | 0 | 0.025 |
| NC_019843.3;YP_009047203.1;MERS_1a;360 | 0 | 0 |
| NC_019843.3;YP_009047203.1;MERS_1a;361 | 0 | 0.025 |
| NC_019843.3;YP_009047203.1;MERS_1a;362 | 0 | 0 |
| NC_019843.3;YP_009047203.1;MERS_1a;363 | 0 | 0.025 |

| | | |
|---|---|---|
| NC_019843.3;YP_009047203.1;MERS_1a;364 | 0 | 0 |
| NC_019843.3;YP_009047203.1;MERS_1a;365 | 0 | 0 |
| NC_019843.3;YP_009047203.1;MERS_1a;366 | 0 | 0 |
| NC_019843.3;YP_009047203.1;MERS_1a;367 | 0 | 0 |
| NC_019843.3;YP_009047203.1;MERS_1a;368 | 0 | 0 |
| NC_019843.3;YP_009047203.1;MERS_1a;369 | 0 | 0 |
| NC_019843.3;YP_009047203.1;MERS_1a;370 | 0 | 0 |
| NC_019843.3;YP_009047203.1;MERS_1a;371 | 0 | 0 |
| NC_019843.3;YP_009047203.1;MERS_1a;372 | 0 | 0 |
| NC_019843.3;YP_009047203.1;MERS_1a;373 | 0 | 0 |
| NC_019843.3;YP_009047203.1;MERS_1a;374 | 0 | 0 |
| NC_019843.3;YP_009047203.1;MERS_1a;375 | 0 | 0 |
| NC_019843.3;YP_009047203.1;MERS_1a;376 | 0 | 0 |
| NC_019843.3;YP_009047203.1;MERS_1a;377 | 0 | 0 |
| NC_019843.3;YP_009047203.1;MERS_1a;378 | 0 | 0.025 |
| NC_019843.3;YP_009047203.1;MERS_1a;379 | 0 | 0 |
| NC_019843.3;YP_009047203.1;MERS_1a;380 | 0 | 0 |
| NC_019843.3;YP_009047203.1;MERS_1a;381 | 0 | 0.025 |
| NC_019843.3;YP_009047203.1;MERS_1a;382 | 0 | 0.025 |
| NC_019843.3;YP_009047203.1;MERS_1a;383 | 0 | 0.025 |
| NC_019843.3;YP_009047203.1;MERS_1a;384 | 0 | 0.025 |
| NC_019843.3;YP_009047203.1;MERS_1a;385 | 0 | 0 |
| NC_019843.3;YP_009047203.1;MERS_1a;386 | 0 | 0.025 |
| NC_019843.3;YP_009047203.1;MERS_1a;387 | 0 | 0 |
| NC_019843.3;YP_009047203.1;MERS_1a;388 | 0.05 | 0.05 |
| NC_019843.3;YP_009047203.1;MERS_1a;389 | 0 | 0.025 |
| NC_019843.3;YP_009047203.1;MERS_1a;390 | 0 | 0.025 |
| NC_019843.3;YP_009047203.1;MERS_1a;391 | 0 | 0 |
| NC_019843.3;YP_009047203.1;MERS_1a;392 | 0.05 | 0 |
| NC_019843.3;YP_009047203.1;MERS_1a;393 | 0 | 0.025 |
| NC_019843.3;YP_009047203.1;MERS_1a;394 | 0 | 0 |
| NC_019843.3;YP_009047203.1;MERS_1a;395 | 0 | 0 |
| NC_019843.3;YP_009047203.1;MERS_1a;396 | 0 | 0 |
| NC_019843.3;YP_009047203.1;MERS_1a;397 | 0 | 0 |
| NC_019843.3;YP_009047203.1;MERS_1a;398 | 0 | 0 |
| NC_019843.3;YP_009047203.1;MERS_1a;399 | 0 | 0 |
| NC_019843.3;YP_009047203.1;MERS_1a;400 | 0 | 0 |
| NC_019843.3;YP_009047203.1;MERS_1a;401 | 0 | 0 |
| NC_019843.3;YP_009047203.1;MERS_1a;402 | 0 | 0 |
| NC_019843.3;YP_009047203.1;MERS_1a;403 | 0 | 0 |
| NC_019843.3;YP_009047203.1;MERS_1a;404 | 0 | 0 |
| NC_019843.3;YP_009047203.1;MERS_1a;405 | 0.05 | 0 |
| NC_019843.3;YP_009047203.1;MERS_1a;406 | 0.1 | 0.025 |

| | | |
|---|---|---|
| NC_019843.3;YP_009047203.1;MERS_1a;407 | 0.1 | 0.05 |
| NC_019843.3;YP_009047203.1;MERS_1a;408 | 0.05 | 0.025 |
| NC_019843.3;YP_009047203.1;MERS_1a;409 | 0 | 0.05 |
| NC_019843.3;YP_009047203.1;MERS_1a;410 | 0 | 0 |
| NC_019843.3;YP_009047203.1;MERS_1a;411 | 0.05 | 0 |
| NC_019843.3;YP_009047203.1;MERS_1a;412 | 0 | 0 |
| NC_019843.3;YP_009047203.1;MERS_1a;413 | 0 | 0 |
| NC_019843.3;YP_009047203.1;MERS_1a;414 | 0 | 0 |
| NC_019843.3;YP_009047203.1;MERS_1a;415 | 0 | 0 |
| NC_019843.3;YP_009047203.1;MERS_1a;416 | 0 | 0 |
| NC_019843.3;YP_009047203.1;MERS_1a;417 | 0 | 0 |
| NC_019843.3;YP_009047203.1;MERS_1a;418 | 0 | 0 |
| NC_019843.3;YP_009047203.1;MERS_1a;419 | 0 | 0 |
| NC_019843.3;YP_009047203.1;MERS_1a;420 | 0 | 0 |
| NC_019843.3;YP_009047203.1;MERS_1a;421 | 0 | 0 |
| NC_019843.3;YP_009047203.1;MERS_1a;422 | 0 | 0 |
| NC_019843.3;YP_009047203.1;MERS_1a;423 | 0 | 0 |
| NC_019843.3;YP_009047203.1;MERS_1a;424 | 0 | 0 |
| NC_019843.3;YP_009047203.1;MERS_1a;425 | 0 | 0 |
| NC_019843.3;YP_009047203.1;MERS_1a;426 | 0 | 0 |
| NC_019843.3;YP_009047203.1;MERS_1a;427 | 0 | 0 |
| NC_019843.3;YP_009047203.1;MERS_1a;428 | 0 | 0 |
| NC_019843.3;YP_009047203.1;MERS_1a;429 | 0 | 0 |
| NC_019843.3;YP_009047203.1;MERS_1a;430 | 0 | 0 |
| NC_019843.3;YP_009047203.1;MERS_1a;431 | 0 | 0 |
| NC_019843.3;YP_009047203.1;MERS_1a;432 | 0 | 0 |
| NC_019843.3;YP_009047203.1;MERS_1a;433 | 0 | 0 |
| NC_019843.3;YP_009047203.1;MERS_1a;434 | 0 | 0 |
| NC_019843.3;YP_009047203.1;MERS_1a;435 | 0 | 0.025 |
| NC_019843.3;YP_009047203.1;MERS_1a;436 | 0 | 0.025 |
| NC_019843.3;YP_009047203.1;MERS_1a;437 | 0 | 0 |
| NC_019843.3;YP_009047203.1;MERS_1a;438 | 0 | 0 |
| NC_019843.3;YP_009047203.1;MERS_1a;439 | 0 | 0 |
| NC_019843.3;YP_009047203.1;MERS_1a;440 | 0 | 0 |
| NC_019843.3;YP_009047203.1;MERS_1a;441 | 0 | 0 |
| NC_019843.3;YP_009047203.1;MERS_1a;442 | 0 | 0 |
| NC_019843.3;YP_009047203.1;MERS_1a;443 | 0 | 0 |
| NC_019843.3;YP_009047203.1;MERS_1a;444 | 0 | 0 |
| NC_019843.3;YP_009047203.1;MERS_1a;445 | 0 | 0 |
| NC_019843.3;YP_009047203.1;MERS_1a;446 | 0 | 0 |
| NC_019843.3;YP_009047203.1;MERS_1a;447 | 0 | 0 |
| NC_019843.3;YP_009047203.1;MERS_1a;448 | 0 | 0 |
| NC_019843.3;YP_009047203.1;MERS_1a;449 | 0 | 0 |

| | | |
|---|---|---|
| NC_019843.3;YP_009047203.1;MERS_1a;450 | 0 | 0 |
| NC_019843.3;YP_009047203.1;MERS_1a;451 | 0 | 0 |
| NC_019843.3;YP_009047203.1;MERS_1a;452 | 0 | 0 |
| NC_019843.3;YP_009047203.1;MERS_1a;453 | 0 | 0 |
| NC_019843.3;YP_009047203.1;MERS_1a;454 | 0 | 0 |
| NC_019843.3;YP_009047203.1;MERS_1a;455 | 0 | 0 |
| NC_019843.3;YP_009047203.1;MERS_1a;456 | 0 | 0 |
| NC_019843.3;YP_009047203.1;MERS_1a;457 | 0 | 0 |
| NC_019843.3;YP_009047203.1;MERS_1a;458 | 0 | 0 |
| NC_019843.3;YP_009047203.1;MERS_1a;459 | 0 | 0 |
| NC_019843.3;YP_009047203.1;MERS_1a;460 | 0 | 0 |
| NC_019843.3;YP_009047203.1;MERS_1a;461 | 0 | 0 |
| NC_019843.3;YP_009047203.1;MERS_1a;462 | 0 | 0 |
| NC_019843.3;YP_009047203.1;MERS_1a;463 | 0 | 0 |
| NC_019843.3;YP_009047203.1;MERS_1a;464 | 0 | 0 |
| NC_019843.3;YP_009047203.1;MERS_1a;465 | 0 | 0 |
| NC_019843.3;YP_009047203.1;MERS_1a;466 | 0 | 0.025 |
| NC_019843.3;YP_009047203.1;MERS_1a;467 | 0.05 | 0 |
| NC_019843.3;YP_009047203.1;MERS_1a;468 | 0 | 0 |
| NC_019843.3;YP_009047203.1;MERS_1a;469 | 0 | 0 |
| NC_019843.3;YP_009047203.1;MERS_1a;470 | 0 | 0 |
| NC_019843.3;YP_009047203.1;MERS_1a;471 | 0 | 0 |
| NC_019843.3;YP_009047203.1;MERS_1a;472 | 0 | 0.025 |
| NC_019843.3;YP_009047203.1;MERS_1a;473 | 0 | 0 |
| NC_019843.3;YP_009047203.1;MERS_1a;474 | 0 | 0 |
| NC_019843.3;YP_009047203.1;MERS_1a;475 | 0 | 0 |
| NC_019843.3;YP_009047203.1;MERS_1a;476 | 0 | 0 |
| NC_019843.3;YP_009047203.1;MERS_1a;477 | 0 | 0 |
| NC_019843.3;YP_009047203.1;MERS_1a;478 | 0 | 0 |
| NC_019843.3;YP_009047203.1;MERS_1a;479 | 0 | 0 |
| NC_019843.3;YP_009047203.1;MERS_1a;480 | 0 | 0 |
| NC_019843.3;YP_009047203.1;MERS_1a;481 | 0 | 0 |
| NC_019843.3;YP_009047203.1;MERS_1a;482 | 0 | 0 |
| NC_019843.3;YP_009047203.1;MERS_1a;483 | 0 | 0 |
| NC_019843.3;YP_009047203.1;MERS_1a;484 | 0 | 0 |
| NC_019843.3;YP_009047203.1;MERS_1a;485 | 0 | 0 |
| NC_019843.3;YP_009047203.1;MERS_1a;486 | 0 | 0 |
| NC_019843.3;YP_009047203.1;MERS_1a;487 | 0 | 0 |
| NC_019843.3;YP_009047203.1;MERS_1a;488 | 0 | 0 |
| NC_019843.3;YP_009047203.1;MERS_1a;489 | 0.05 | 0 |
| NC_019843.3;YP_009047203.1;MERS_1a;490 | 0 | 0 |
| NC_019843.3;YP_009047203.1;MERS_1a;491 | 0 | 0 |
| NC_019843.3;YP_009047203.1;MERS_1a;492 | 0 | 0 |

| | | |
|---|---|---|
| NC_019843.3;YP_009047203.1;MERS_1a;493 | 0 | 0 |
| NC_019843.3;YP_009047203.1;MERS_1a;494 | 0 | 0 |
| NC_019843.3;YP_009047203.1;MERS_1a;495 | 0 | 0 |
| NC_019843.3;YP_009047203.1;MERS_1a;496 | 0 | 0 |
| NC_019843.3;YP_009047203.1;MERS_1a;497 | 0.05 | 0 |
| NC_019843.3;YP_009047203.1;MERS_1a;498 | 0.05 | 0 |
| NC_019843.3;YP_009047203.1;MERS_1a;499 | 0.05 | 0 |
| NC_019843.3;YP_009047203.1;MERS_1a;500 | 0.05 | 0 |
| NC_019843.3;YP_009047203.1;MERS_1a;501 | 0.05 | 0 |
| NC_019843.3;YP_009047203.1;MERS_1a;502 | 0.05 | 0 |
| NC_019843.3;YP_009047203.1;MERS_1a;503 | 0 | 0 |
| NC_019843.3;YP_009047203.1;MERS_1a;504 | 0 | 0 |
| NC_019843.3;YP_009047203.1;MERS_1a;505 | 0.05 | 0 |
| NC_019843.3;YP_009047203.1;MERS_1a;506 | 0 | 0 |
| NC_019843.3;YP_009047203.1;MERS_1a;507 | 0 | 0 |
| NC_019843.3;YP_009047203.1;MERS_1a;508 | 0 | 0 |
| NC_019843.3;YP_009047203.1;MERS_1a;509 | 0 | 0 |
| NC_019843.3;YP_009047203.1;MERS_1a;510 | 0 | 0 |
| NC_019843.3;YP_009047203.1;MERS_1a;511 | 0 | 0 |
| NC_019843.3;YP_009047203.1;MERS_1a;512 | 0.05 | 0 |
| NC_019843.3;YP_009047203.1;MERS_1a;513 | 0.1 | 0 |
| NC_019843.3;YP_009047203.1;MERS_1a;514 | 0.1 | 0.025 |
| NC_019843.3;YP_009047203.1;MERS_1a;515 | 0.05 | 0.05 |
| NC_019843.3;YP_009047203.1;MERS_1a;516 | 0.1 | 0.05 |
| NC_019843.3;YP_009047203.1;MERS_1a;517 | 0.05 | 0.3 |
| NC_019843.3;YP_009047203.1;MERS_1a;518 | 0.05 | 0 |
| NC_019843.3;YP_009047203.1;MERS_1a;519 | 0.05 | 0 |
| NC_019843.3;YP_009047203.1;MERS_1a;520 | 0 | 0.05 |
| NC_019843.3;YP_009047203.1;MERS_1a;521 | 0.05 | 0.025 |
| NC_019843.3;YP_009047203.1;MERS_1a;522 | 0 | 0.025 |
| NC_019843.3;YP_009047203.1;MERS_1a;523 | 0 | 0 |
| NC_019843.3;YP_009047203.1;MERS_1a;524 | 0 | 0 |
| NC_019843.3;YP_009047203.1;MERS_1a;525 | 0 | 0.025 |
| NC_019843.3;YP_009047203.1;MERS_1a;526 | 0.05 | 0.025 |
| NC_019843.3;YP_009047203.1;MERS_1a;527 | 0 | 0.025 |
| NC_019843.3;YP_009047203.1;MERS_1a;528 | 0 | 0.025 |
| NC_019843.3;YP_009047203.1;MERS_1a;529 | 0.05 | 0.025 |
| NC_019843.3;YP_009047203.1;MERS_1a;530 | 0.05 | 0.025 |
| NC_019843.3;YP_009047203.1;MERS_1a;531 | 0 | 0 |
| NC_019843.3;YP_009047203.1;MERS_1a;532 | 0 | 0 |
| NC_019843.3;YP_009047203.1;MERS_1a;533 | 0 | 0 |
| NC_019843.3;YP_009047203.1;MERS_1a;534 | 0 | 0 |
| NC_019843.3;YP_009047203.1;MERS_1a;535 | 0.05 | 0.025 |

| | | |
|---|---|---|
| NC_019843.3;YP_009047203.1;MERS_1a;536 | 0.05 | 0.025 |
| NC_019843.3;YP_009047203.1;MERS_1a;537 | 0 | 0.025 |
| NC_019843.3;YP_009047203.1;MERS_1a;538 | 0 | 0.025 |
| NC_019843.3;YP_009047203.1;MERS_1a;539 | 0.1 | 0.05 |
| NC_019843.3;YP_009047203.1;MERS_1a;540 | 0 | 0 |
| NC_019843.3;YP_009047203.1;MERS_1a;541 | 0 | 0 |
| NC_019843.3;YP_009047203.1;MERS_1a;542 | 0 | 0 |
| NC_019843.3;YP_009047203.1;MERS_1a;543 | 0 | 0 |
| NC_019843.3;YP_009047203.1;MERS_1a;544 | 0 | 0 |
| NC_019843.3;YP_009047203.1;MERS_1a;545 | 0 | 0 |
| NC_019843.3;YP_009047203.1;MERS_1a;546 | 0 | 0 |
| NC_019843.3;YP_009047203.1;MERS_1a;547 | 0 | 0 |
| NC_019843.3;YP_009047203.1;MERS_1a;548 | 0 | 0 |
| NC_019843.3;YP_009047203.1;MERS_1a;549 | 0 | 0 |
| NC_019843.3;YP_009047203.1;MERS_1a;550 | 0 | 0 |
| NC_019843.3;YP_009047203.1;MERS_1a;551 | 0 | 0 |
| NC_019843.3;YP_009047203.1;MERS_1a;552 | 0 | 0 |
| NC_019843.3;YP_009047203.1;MERS_1a;553 | 0 | 0.025 |
| NC_019843.3;YP_009047203.1;MERS_1a;554 | 0 | 0 |
| NC_019843.3;YP_009047203.1;MERS_1a;555 | 0 | 0.075 |
| NC_019843.3;YP_009047203.1;MERS_1a;556 | 0 | 0.025 |
| NC_019843.3;YP_009047203.1;MERS_1a;557 | 0 | 0 |
| NC_019843.3;YP_009047203.1;MERS_1a;558 | 0 | 0 |
| NC_019843.3;YP_009047203.1;MERS_1a;559 | 0 | 0 |
| NC_019843.3;YP_009047203.1;MERS_1a;560 | 0 | 0 |
| NC_019843.3;YP_009047203.1;MERS_1a;561 | 0 | 0 |
| NC_019843.3;YP_009047203.1;MERS_1a;562 | 0 | 0 |
| NC_019843.3;YP_009047203.1;MERS_1a;563 | 0 | 0 |
| NC_019843.3;YP_009047203.1;MERS_1a;564 | 0.05 | 0 |
| NC_019843.3;YP_009047203.1;MERS_1a;565 | 0 | 0 |
| NC_019843.3;YP_009047203.1;MERS_1a;566 | 0 | 0 |
| NC_019843.3;YP_009047203.1;MERS_1a;567 | 0.05 | 0.025 |
| NC_019843.3;YP_009047203.1;MERS_1a;568 | 0 | 0 |
| NC_019843.3;YP_009047203.1;MERS_1a;569 | 0 | 0 |
| NC_019843.3;YP_009047203.1;MERS_1a;570 | 0 | 0 |
| NC_019843.3;YP_009047203.1;MERS_1a;571 | 0 | 0 |
| NC_019843.3;YP_009047203.1;MERS_1a;572 | 0 | 0.025 |
| NC_019843.3;YP_009047203.1;MERS_1a;573 | 0 | 0.025 |
| NC_019843.3;YP_009047203.1;MERS_1a;574 | 0.05 | 0 |
| NC_019843.3;YP_009047203.1;MERS_1a;575 | 0 | 0 |
| NC_019843.3;YP_009047203.1;MERS_1a;576 | 0 | 0.025 |
| NC_019843.3;YP_009047203.1;MERS_1a;577 | 0 | 0 |
| NC_019843.3;YP_009047203.1;MERS_1a;578 | 0 | 0 |

| | | |
|---|---|---|
| NC_019843.3;YP_009047203.1;MERS_1a;579 | 0 | 0 |
| NC_019843.3;YP_009047203.1;MERS_1a;580 | 0 | 0 |
| NC_019843.3;YP_009047203.1;MERS_1a;581 | 0 | 0 |
| NC_019843.3;YP_009047203.1;MERS_1a;582 | 0 | 0 |
| NC_019843.3;YP_009047203.1;MERS_1a;583 | 0 | 0 |
| NC_019843.3;YP_009047203.1;MERS_1a;584 | 0 | 0 |
| NC_019843.3;YP_009047203.1;MERS_1a;585 | 0 | 0 |
| NC_019843.3;YP_009047203.1;MERS_1a;586 | 0 | 0 |
| NC_019843.3;YP_009047203.1;MERS_1a;587 | 0 | 0 |
| NC_019843.3;YP_009047203.1;MERS_1a;588 | 0 | 0 |
| NC_019843.3;YP_009047203.1;MERS_1a;589 | 0 | 0 |
| NC_019843.3;YP_009047203.1;MERS_1a;590 | 0 | 0 |
| NC_019843.3;YP_009047203.1;MERS_1a;591 | 0 | 0 |
| NC_019843.3;YP_009047203.1;MERS_1a;592 | 0 | 0 |
| NC_019843.3;YP_009047203.1;MERS_1a;593 | 0 | 0 |
| NC_019843.3;YP_009047203.1;MERS_1a;594 | 0.05 | 0 |
| NC_019843.3;YP_009047203.1;MERS_1a;595 | 0.05 | 0 |
| NC_019843.3;YP_009047203.1;MERS_1a;596 | 0.05 | 0 |
| NC_019843.3;YP_009047203.1;MERS_1a;597 | 0.15 | 0.075 |
| NC_019843.3;YP_009047203.1;MERS_1a;598 | 0.05 | 0.025 |
| NC_019843.3;YP_009047203.1;MERS_1a;599 | 0 | 0 |
| NC_019843.3;YP_009047203.1;MERS_1a;600 | 0 | 0 |
| NC_019843.3;YP_009047203.1;MERS_1a;601 | 0 | 0 |
| NC_019843.3;YP_009047203.1;MERS_1a;602 | 0 | 0 |
| NC_019843.3;YP_009047203.1;MERS_1a;603 | 0 | 0 |
| NC_019843.3;YP_009047203.1;MERS_1a;604 | 0 | 0 |
| NC_019843.3;YP_009047203.1;MERS_1a;605 | 0 | 0 |
| NC_019843.3;YP_009047203.1;MERS_1a;606 | 0 | 0 |
| NC_019843.3;YP_009047203.1;MERS_1a;607 | 0 | 0 |
| NC_019843.3;YP_009047203.1;MERS_1a;608 | 0 | 0 |
| NC_019843.3;YP_009047203.1;MERS_1a;609 | 0 | 0 |
| NC_019843.3;YP_009047203.1;MERS_1a;610 | 0 | 0 |
| NC_019843.3;YP_009047203.1;MERS_1a;611 | 0 | 0 |
| NC_019843.3;YP_009047203.1;MERS_1a;612 | 0 | 0 |
| NC_019843.3;YP_009047203.1;MERS_1a;613 | 0 | 0 |
| NC_019843.3;YP_009047203.1;MERS_1a;614 | 0 | 0 |
| NC_019843.3;YP_009047203.1;MERS_1a;615 | 0 | 0 |
| NC_019843.3;YP_009047203.1;MERS_1a;616 | 0 | 0 |
| NC_019843.3;YP_009047203.1;MERS_1a;617 | 0 | 0 |
| NC_019843.3;YP_009047203.1;MERS_1a;618 | 0 | 0 |
| NC_019843.3;YP_009047203.1;MERS_1a;619 | 0 | 0 |
| NC_019843.3;YP_009047203.1;MERS_1a;620 | 0 | 0 |
| NC_019843.3;YP_009047203.1;MERS_1a;621 | 0 | 0 |

| | | |
|---|---|---|
| NC_019843.3;YP_009047203.1;MERS_1a;622 | 0 | 0 |
| NC_019843.3;YP_009047203.1;MERS_1a;623 | 0 | 0 |
| NC_019843.3;YP_009047203.1;MERS_1a;624 | 0 | 0 |
| NC_019843.3;YP_009047203.1;MERS_1a;625 | 0 | 0 |
| NC_019843.3;YP_009047203.1;MERS_1a;626 | 0 | 0 |
| NC_019843.3;YP_009047203.1;MERS_1a;627 | 0 | 0 |
| NC_019843.3;YP_009047203.1;MERS_1a;628 | 0 | 0 |
| NC_019843.3;YP_009047203.1;MERS_1a;629 | 0 | 0 |
| NC_019843.3;YP_009047203.1;MERS_1a;630 | 0 | 0 |
| NC_019843.3;YP_009047203.1;MERS_1a;631 | 0 | 0 |
| NC_019843.3;YP_009047203.1;MERS_1a;632 | 0 | 0 |
| NC_019843.3;YP_009047203.1;MERS_1a;633 | 0 | 0 |
| NC_019843.3;YP_009047203.1;MERS_1a;634 | 0 | 0 |
| NC_019843.3;YP_009047203.1;MERS_1a;635 | 0 | 0 |
| NC_019843.3;YP_009047203.1;MERS_1a;636 | 0 | 0 |
| NC_019843.3;YP_009047203.1;MERS_1a;637 | 0 | 0 |
| NC_019843.3;YP_009047203.1;MERS_1a;638 | 0 | 0 |
| NC_019843.3;YP_009047203.1;MERS_1a;639 | 0 | 0 |
| NC_019843.3;YP_009047203.1;MERS_1a;640 | 0 | 0 |
| NC_019843.3;YP_009047203.1;MERS_1a;641 | 0 | 0 |
| NC_019843.3;YP_009047203.1;MERS_1a;642 | 0 | 0 |
| NC_019843.3;YP_009047203.1;MERS_1a;643 | 0 | 0 |
| NC_019843.3;YP_009047203.1;MERS_1a;644 | 0 | 0 |
| NC_019843.3;YP_009047203.1;MERS_1a;645 | 0 | 0 |
| NC_019843.3;YP_009047203.1;MERS_1a;646 | 0 | 0 |
| NC_019843.3;YP_009047203.1;MERS_1a;647 | 0 | 0 |
| NC_019843.3;YP_009047203.1;MERS_1a;648 | 0 | 0 |
| NC_019843.3;YP_009047203.1;MERS_1a;649 | 0 | 0 |
| NC_019843.3;YP_009047203.1;MERS_1a;650 | 0 | 0 |
| NC_019843.3;YP_009047203.1;MERS_1a;651 | 0 | 0 |
| NC_019843.3;YP_009047203.1;MERS_1a;652 | 0 | 0.075 |
| NC_019843.3;YP_009047203.1;MERS_1a;653 | 0 | 0 |
| NC_019843.3;YP_009047203.1;MERS_1a;654 | 0 | 0 |
| NC_019843.3;YP_009047203.1;MERS_1a;655 | 0 | 0 |
| NC_019843.3;YP_009047203.1;MERS_1a;656 | 0 | 0 |
| NC_019843.3;YP_009047203.1;MERS_1a;657 | 0 | 0 |
| NC_019843.3;YP_009047203.1;MERS_1a;658 | 0 | 0 |
| NC_019843.3;YP_009047203.1;MERS_1a;659 | 0 | 0 |
| NC_019843.3;YP_009047203.1;MERS_1a;660 | 0 | 0 |
| NC_019843.3;YP_009047203.1;MERS_1a;661 | 0 | 0 |
| NC_019843.3;YP_009047203.1;MERS_1a;662 | 0 | 0 |
| NC_019843.3;YP_009047203.1;MERS_1a;663 | 0 | 0 |
| NC_019843.3;YP_009047203.1;MERS_1a;664 | 0 | 0 |

| | | |
|---|---|---|
| NC_019843.3;YP_009047203.1;MERS_1a;665 | 0 | 0 |
| NC_019843.3;YP_009047203.1;MERS_1a;666 | 0 | 0 |
| NC_019843.3;YP_009047203.1;MERS_1a;667 | 0 | 0 |
| NC_019843.3;YP_009047203.1;MERS_1a;668 | 0 | 0 |
| NC_019843.3;YP_009047203.1;MERS_1a;669 | 0 | 0 |
| NC_019843.3;YP_009047203.1;MERS_1a;670 | 0 | 0 |
| NC_019843.3;YP_009047203.1;MERS_1a;671 | 0 | 0 |
| NC_019843.3;YP_009047203.1;MERS_1a;672 | 0 | 0 |
| NC_019843.3;YP_009047203.1;MERS_1a;673 | 0 | 0 |
| NC_019843.3;YP_009047203.1;MERS_1a;674 | 0 | 0 |
| NC_019843.3;YP_009047203.1;MERS_1a;675 | 0 | 0 |
| NC_019843.3;YP_009047203.1;MERS_1a;676 | 0 | 0 |
| NC_019843.3;YP_009047203.1;MERS_1a;677 | 0 | 0 |
| NC_019843.3;YP_009047203.1;MERS_1a;678 | 0 | 0 |
| NC_019843.3;YP_009047203.1;MERS_1a;679 | 0 | 0 |
| NC_019843.3;YP_009047203.1;MERS_1a;680 | 0 | 0 |
| NC_019843.3;YP_009047203.1;MERS_1a;681 | 0 | 0 |
| NC_019843.3;YP_009047203.1;MERS_1a;682 | 0 | 0 |
| NC_019843.3;YP_009047203.1;MERS_1a;683 | 0 | 0 |
| NC_019843.3;YP_009047203.1;MERS_1a;684 | 0 | 0 |
| NC_019843.3;YP_009047203.1;MERS_1a;685 | 0 | 0 |
| NC_019843.3;YP_009047203.1;MERS_1a;686 | 0 | 0 |
| NC_019843.3;YP_009047203.1;MERS_1a;687 | 0 | 0 |
| NC_019843.3;YP_009047203.1;MERS_1a;688 | 0 | 0 |
| NC_019843.3;YP_009047203.1;MERS_1a;689 | 0 | 0 |
| NC_019843.3;YP_009047203.1;MERS_1a;690 | 0 | 0 |
| NC_019843.3;YP_009047203.1;MERS_1a;691 | 0 | 0 |
| NC_019843.3;YP_009047203.1;MERS_1a;692 | 0.05 | 0 |
| NC_019843.3;YP_009047203.1;MERS_1a;693 | 0 | 0 |
| NC_019843.3;YP_009047203.1;MERS_1a;694 | 0 | 0 |
| NC_019843.3;YP_009047203.1;MERS_1a;695 | 0 | 0 |
| NC_019843.3;YP_009047203.1;MERS_1a;696 | 0 | 0 |
| NC_019843.3;YP_009047203.1;MERS_1a;697 | 0 | 0 |
| NC_019843.3;YP_009047203.1;MERS_1a;698 | 0 | 0 |
| NC_019843.3;YP_009047203.1;MERS_1a;699 | 0 | 0 |
| NC_019843.3;YP_009047203.1;MERS_1a;700 | 0 | 0 |
| NC_019843.3;YP_009047203.1;MERS_1a;701 | 0.1 | 0 |
| NC_019843.3;YP_009047203.1;MERS_1a;702 | 0.1 | 0 |
| NC_019843.3;YP_009047203.1;MERS_1a;703 | 0.15 | 0.025 |
| NC_019843.3;YP_009047203.1;MERS_1a;704 | 0 | 0 |
| NC_019843.3;YP_009047203.1;MERS_1a;705 | 0 | 0 |
| NC_019843.3;YP_009047203.1;MERS_1a;706 | 0 | 0 |
| NC_019843.3;YP_009047203.1;MERS_1a;707 | 0 | 0 |

| | | |
|---|---|---|
| NC_019843.3;YP_009047203.1;MERS_1a;708 | 0 | 0 |
| NC_019843.3;YP_009047203.1;MERS_1a;709 | 0 | 0 |
| NC_019843.3;YP_009047203.1;MERS_1a;710 | 0 | 0 |
| NC_019843.3;YP_009047203.1;MERS_1a;711 | 0 | 0 |
| NC_019843.3;YP_009047203.1;MERS_1a;712 | 0 | 0 |
| NC_019843.3;YP_009047203.1;MERS_1a;713 | 0 | 0 |
| NC_019843.3;YP_009047203.1;MERS_1a;714 | 0 | 0 |
| NC_019843.3;YP_009047203.1;MERS_1a;715 | 0 | 0 |
| NC_019843.3;YP_009047203.1;MERS_1a;716 | 0 | 0 |
| NC_019843.3;YP_009047203.1;MERS_1a;717 | 0 | 0 |
| NC_019843.3;YP_009047203.1;MERS_1a;718 | 0 | 0 |
| NC_019843.3;YP_009047203.1;MERS_1a;719 | 0 | 0 |
| NC_019843.3;YP_009047203.1;MERS_1a;720 | 0 | 0 |
| NC_019843.3;YP_009047203.1;MERS_1a;721 | 0 | 0 |
| NC_019843.3;YP_009047203.1;MERS_1a;722 | 0 | 0 |
| NC_019843.3;YP_009047203.1;MERS_1a;723 | 0 | 0 |
| NC_019843.3;YP_009047203.1;MERS_1a;724 | 0 | 0 |
| NC_019843.3;YP_009047203.1;MERS_1a;725 | 0 | 0 |
| NC_019843.3;YP_009047203.1;MERS_1a;726 | 0 | 0 |
| NC_019843.3;YP_009047203.1;MERS_1a;727 | 0 | 0.025 |
| NC_019843.3;YP_009047203.1;MERS_1a;728 | 0 | 0 |
| NC_019843.3;YP_009047203.1;MERS_1a;729 | 0.05 | 0.025 |
| NC_019843.3;YP_009047203.1;MERS_1a;730 | 0 | 0 |
| NC_019843.3;YP_009047203.1;MERS_1a;731 | 0 | 0 |
| NC_019843.3;YP_009047203.1;MERS_1a;732 | 0 | 0 |
| NC_019843.3;YP_009047203.1;MERS_1a;733 | 0 | 0 |
| NC_019843.3;YP_009047203.1;MERS_1a;734 | 0 | 0 |
| NC_019843.3;YP_009047203.1;MERS_1a;735 | 0 | 0 |
| NC_019843.3;YP_009047203.1;MERS_1a;736 | 0 | 0 |
| NC_019843.3;YP_009047203.1;MERS_1a;737 | 0 | 0 |
| NC_019843.3;YP_009047203.1;MERS_1a;738 | 0 | 0 |
| NC_019843.3;YP_009047203.1;MERS_1a;739 | 0 | 0 |
| NC_019843.3;YP_009047203.1;MERS_1a;740 | 0 | 0 |
| NC_019843.3;YP_009047203.1;MERS_1a;741 | 0 | 0 |
| NC_019843.3;YP_009047203.1;MERS_1a;742 | 0 | 0 |
| NC_019843.3;YP_009047203.1;MERS_1a;743 | 0 | 0.025 |
| NC_019843.3;YP_009047203.1;MERS_1a;744 | 0 | 0.025 |
| NC_019843.3;YP_009047203.1;MERS_1a;745 | 0.05 | 0.025 |
| NC_019843.3;YP_009047203.1;MERS_1a;746 | 0.05 | 0.025 |
| NC_019843.3;YP_009047203.1;MERS_1a;747 | 0 | 0 |
| NC_019843.3;YP_009047203.1;MERS_1a;748 | 0 | 0.025 |
| NC_019843.3;YP_009047203.1;MERS_1a;749 | 0.05 | 0 |
| NC_019843.3;YP_009047203.1;MERS_1a;750 | 0 | 0 |

| | | |
|---|---|---|
| NC_019843.3;YP_009047203.1;MERS_1a;751 | 0.1 | 0 |
| NC_019843.3;YP_009047203.1;MERS_1a;752 | 0 | 0 |
| NC_019843.3;YP_009047203.1;MERS_1a;753 | 0 | 0 |
| NC_019843.3;YP_009047203.1;MERS_1a;754 | 0 | 0 |
| NC_019843.3;YP_009047203.1;MERS_1a;755 | 0 | 0 |
| NC_019843.3;YP_009047203.1;MERS_1a;756 | 0 | 0 |
| NC_019843.3;YP_009047203.1;MERS_1a;757 | 0.05 | 0 |
| NC_019843.3;YP_009047203.1;MERS_1a;758 | 0 | 0.025 |
| NC_019843.3;YP_009047203.1;MERS_1a;759 | 0 | 0 |
| NC_019843.3;YP_009047203.1;MERS_1a;760 | 0 | 0 |
| NC_019843.3;YP_009047203.1;MERS_1a;761 | 0 | 0.025 |
| NC_019843.3;YP_009047203.1;MERS_1a;762 | 0 | 0.025 |
| NC_019843.3;YP_009047203.1;MERS_1a;763 | 0 | 0.025 |
| NC_019843.3;YP_009047203.1;MERS_1a;764 | 0 | 0.025 |
| NC_019843.3;YP_009047203.1;MERS_1a;765 | 0 | 0.025 |
| NC_019843.3;YP_009047203.1;MERS_1a;766 | 0.05 | 0.075 |
| NC_019843.3;YP_009047203.1;MERS_1a;767 | 0 | 0.075 |
| NC_019843.3;YP_009047203.1;MERS_1a;768 | 0 | 0 |
| NC_019843.3;YP_009047203.1;MERS_1a;769 | 0 | 0 |
| NC_019843.3;YP_009047203.1;MERS_1a;770 | 0 | 0 |
| NC_019843.3;YP_009047203.1;MERS_1a;771 | 0.05 | 0 |
| NC_019843.3;YP_009047203.1;MERS_1a;772 | 0.05 | 0 |
| NC_019843.3;YP_009047203.1;MERS_1a;773 | 0 | 0 |
| NC_019843.3;YP_009047203.1;MERS_1a;774 | 0 | 0 |
| NC_019843.3;YP_009047203.1;MERS_1a;775 | 0 | 0 |
| NC_019843.3;YP_009047203.1;MERS_1a;776 | 0 | 0.025 |
| NC_019843.3;YP_009047203.1;MERS_1a;777 | 0 | 0.025 |
| NC_019843.3;YP_009047203.1;MERS_1a;778 | 0.05 | 0.05 |
| NC_019843.3;YP_009047203.1;MERS_1a;779 | 0.05 | 0.05 |
| NC_019843.3;YP_009047203.1;MERS_1a;780 | 0.1 | 0.025 |
| NC_019843.3;YP_009047203.1;MERS_1a;781 | 0.1 | 0.025 |
| NC_019843.3;YP_009047203.1;MERS_1a;782 | 0 | 0.05 |
| NC_019843.3;YP_009047203.1;MERS_1a;783 | 0.05 | 0.05 |
| NC_019843.3;YP_009047203.1;MERS_1a;784 | 0.05 | 0.05 |
| NC_019843.3;YP_009047203.1;MERS_1a;785 | 0.05 | 0.05 |
| NC_019843.3;YP_009047203.1;MERS_1a;786 | 0.05 | 0.075 |
| NC_019843.3;YP_009047203.1;MERS_1a;787 | 0.05 | 0.05 |
| NC_019843.3;YP_009047203.1;MERS_1a;788 | 0 | 0.025 |
| NC_019843.3;YP_009047203.1;MERS_1a;789 | 0 | 0.05 |
| NC_019843.3;YP_009047203.1;MERS_1a;790 | 0.05 | 0.075 |
| NC_019843.3;YP_009047203.1;MERS_1a;791 | 0 | 0.025 |
| NC_019843.3;YP_009047203.1;MERS_1a;792 | 0 | 0 |
| NC_019843.3;YP_009047203.1;MERS_1a;793 | 0 | 0 |

| | | |
|---|---|---|
| NC_019843.3;YP_009047203.1;MERS_1a;794 | 0 | 0 |
| NC_019843.3;YP_009047203.1;MERS_1a;795 | 0 | 0 |
| NC_019843.3;YP_009047203.1;MERS_1a;796 | 0 | 0 |
| NC_019843.3;YP_009047203.1;MERS_1a;797 | 0 | 0 |
| NC_019843.3;YP_009047203.1;MERS_1a;798 | 0 | 0 |
| NC_019843.3;YP_009047203.1;MERS_1a;799 | 0 | 0 |
| NC_019843.3;YP_009047203.1;MERS_1a;800 | 0 | 0 |
| NC_019843.3;YP_009047203.1;MERS_1a;801 | 0 | 0 |
| NC_019843.3;YP_009047203.1;MERS_1a;802 | 0 | 0 |
| NC_019843.3;YP_009047203.1;MERS_1a;803 | 0 | 0 |
| NC_019843.3;YP_009047203.1;MERS_1a;804 | 0 | 0 |
| NC_019843.3;YP_009047203.1;MERS_1a;805 | 0 | 0 |
| NC_019843.3;YP_009047203.1;MERS_1a;806 | 0 | 0 |
| NC_019843.3;YP_009047203.1;MERS_1a;807 | 0 | 0 |
| NC_019843.3;YP_009047203.1;MERS_1a;808 | 0 | 0 |
| NC_019843.3;YP_009047203.1;MERS_1a;809 | 0 | 0 |
| NC_019843.3;YP_009047203.1;MERS_1a;810 | 0 | 0 |
| NC_019843.3;YP_009047203.1;MERS_1a;811 | 0 | 0 |
| NC_019843.3;YP_009047203.1;MERS_1a;812 | 0 | 0 |
| NC_019843.3;YP_009047203.1;MERS_1a;813 | 0 | 0 |
| NC_019843.3;YP_009047203.1;MERS_1a;814 | 0 | 0 |
| NC_019843.3;YP_009047203.1;MERS_1a;815 | 0 | 0 |
| NC_019843.3;YP_009047203.1;MERS_1a;816 | 0 | 0 |
| NC_019843.3;YP_009047203.1;MERS_1a;817 | 0.05 | 0 |
| NC_019843.3;YP_009047203.1;MERS_1a;818 | 0.05 | 0 |
| NC_019843.3;YP_009047203.1;MERS_1a;819 | 0 | 0 |
| NC_019843.3;YP_009047203.1;MERS_1a;820 | 0.05 | 0 |
| NC_019843.3;YP_009047203.1;MERS_1a;821 | 0.05 | 0 |
| NC_019843.3;YP_009047203.1;MERS_1a;822 | 0.1 | 0 |
| NC_019843.3;YP_009047203.1;MERS_1a;823 | 0 | 0 |
| NC_019843.3;YP_009047203.1;MERS_1a;824 | 0.05 | 0 |
| NC_019843.3;YP_009047203.1;MERS_1a;825 | 0.05 | 0 |
| NC_019843.3;YP_009047203.1;MERS_1a;826 | 0.1 | 0 |
| NC_019843.3;YP_009047203.1;MERS_1a;827 | 0.05 | 0 |
| NC_019843.3;YP_009047203.1;MERS_1a;828 | 0.05 | 0.025 |
| NC_019843.3;YP_009047203.1;MERS_1a;829 | 0 | 0 |
| NC_019843.3;YP_009047203.1;MERS_1a;830 | 0 | 0 |
| NC_019843.3;YP_009047203.1;MERS_1a;831 | 0 | 0 |
| NC_019843.3;YP_009047203.1;MERS_1a;832 | 0 | 0 |
| NC_019843.3;YP_009047203.1;MERS_1a;833 | 0 | 0 |
| NC_019843.3;YP_009047203.1;MERS_1a;834 | 0 | 0 |
| NC_019843.3;YP_009047203.1;MERS_1a;835 | 0 | 0 |
| NC_019843.3;YP_009047203.1;MERS_1a;836 | 0 | 0 |

| | | |
|---|---|---|
| NC_019843.3;YP_009047203.1;MERS_1a;837 | 0 | 0 |
| NC_019843.3;YP_009047203.1;MERS_1a;838 | 0 | 0 |
| NC_019843.3;YP_009047203.1;MERS_1a;839 | 0 | 0 |
| NC_019843.3;YP_009047203.1;MERS_1a;840 | 0 | 0 |
| NC_019843.3;YP_009047203.1;MERS_1a;841 | 0 | 0 |
| NC_019843.3;YP_009047203.1;MERS_1a;842 | 0 | 0 |
| NC_019843.3;YP_009047203.1;MERS_1a;843 | 0 | 0 |
| NC_019843.3;YP_009047203.1;MERS_1a;844 | 0 | 0 |
| NC_019843.3;YP_009047203.1;MERS_1a;845 | 0 | 0 |
| NC_019843.3;YP_009047203.1;MERS_1a;846 | 0 | 0 |
| NC_019843.3;YP_009047203.1;MERS_1a;847 | 0 | 0 |
| NC_019843.3;YP_009047203.1;MERS_1a;848 | 0 | 0 |
| NC_019843.3;YP_009047203.1;MERS_1a;849 | 0 | 0 |
| NC_019843.3;YP_009047203.1;MERS_1a;850 | 0 | 0.025 |
| NC_019843.3;YP_009047203.1;MERS_1a;851 | 0 | 0.025 |
| NC_019843.3;YP_009047203.1;MERS_1a;852 | 0.05 | 0.025 |
| NC_019843.3;YP_009047203.1;MERS_1a;853 | 0 | 0.05 |
| NC_019843.3;YP_009047203.1;MERS_1a;854 | 0 | 0 |
| NC_019843.3;YP_009047203.1;MERS_1a;855 | 0 | 0.075 |
| NC_019843.3;YP_009047203.1;MERS_1a;856 | 0.05 | 0.025 |
| NC_019843.3;YP_009047203.1;MERS_1a;857 | 0 | 0 |
| NC_019843.3;YP_009047203.1;MERS_1a;858 | 0 | 0.025 |
| NC_019843.3;YP_009047203.1;MERS_1a;859 | 0 | 0.025 |
| NC_019843.3;YP_009047203.1;MERS_1a;860 | 0 | 0.025 |
| NC_019843.3;YP_009047203.1;MERS_1a;861 | 0 | 0.025 |
| NC_019843.3;YP_009047203.1;MERS_1a;862 | 0 | 0 |
| NC_019843.3;YP_009047203.1;MERS_1a;863 | 0 | 0 |
| NC_019843.3;YP_009047203.1;MERS_1a;864 | 0 | 0 |
| NC_019843.3;YP_009047203.1;MERS_1a;865 | 0 | 0 |
| NC_019843.3;YP_009047203.1;MERS_1a;866 | 0 | 0 |
| NC_019843.3;YP_009047203.1;MERS_1a;867 | 0 | 0 |
| NC_019843.3;YP_009047203.1;MERS_1a;868 | 0 | 0 |
| NC_019843.3;YP_009047203.1;MERS_1a;869 | 0 | 0 |
| NC_019843.3;YP_009047203.1;MERS_1a;870 | 0 | 0 |
| NC_019843.3;YP_009047203.1;MERS_1a;871 | 0 | 0 |
| NC_019843.3;YP_009047203.1;MERS_1a;872 | 0 | 0 |
| NC_019843.3;YP_009047203.1;MERS_1a;873 | 0 | 0 |
| NC_019843.3;YP_009047203.1;MERS_1a;874 | 0 | 0 |
| NC_019843.3;YP_009047203.1;MERS_1a;875 | 0 | 0 |
| NC_019843.3;YP_009047203.1;MERS_1a;876 | 0 | 0 |
| NC_019843.3;YP_009047203.1;MERS_1a;877 | 0 | 0 |
| NC_019843.3;YP_009047203.1;MERS_1a;878 | 0 | 0 |
| NC_019843.3;YP_009047203.1;MERS_1a;879 | 0 | 0 |

| | | |
|---|---|---|
| NC_019843.3;YP_009047203.1;MERS_1a;880 | 0 | 0 |
| NC_019843.3;YP_009047203.1;MERS_1a;881 | 0.05 | 0 |
| NC_019843.3;YP_009047203.1;MERS_1a;882 | 0 | 0 |
| NC_019843.3;YP_009047203.1;MERS_1a;883 | 0 | 0 |
| NC_019843.3;YP_009047203.1;MERS_1a;884 | 0 | 0 |
| NC_019843.3;YP_009047203.1;MERS_1a;885 | 0 | 0 |
| NC_019843.3;YP_009047203.1;MERS_1a;886 | 0 | 0 |
| NC_019843.3;YP_009047203.1;MERS_1a;887 | 0 | 0 |
| NC_019843.3;YP_009047203.1;MERS_1a;888 | 0 | 0 |
| NC_019843.3;YP_009047203.1;MERS_1a;889 | 0.05 | 0 |
| NC_019843.3;YP_009047203.1;MERS_1a;890 | 0.05 | 0 |
| NC_019843.3;YP_009047203.1;MERS_1a;891 | 0 | 0 |
| NC_019843.3;YP_009047203.1;MERS_1a;892 | 0 | 0 |
| NC_019843.3;YP_009047203.1;MERS_1a;893 | 0 | 0 |
| NC_019843.3;YP_009047203.1;MERS_1a;894 | 0 | 0 |
| NC_019843.3;YP_009047203.1;MERS_1a;895 | 0.05 | 0.025 |
| NC_019843.3;YP_009047203.1;MERS_1a;896 | 0 | 0.025 |
| NC_019843.3;YP_009047203.1;MERS_1a;897 | 0 | 0.025 |
| NC_019843.3;YP_009047203.1;MERS_1a;898 | 0 | 0 |
| NC_019843.3;YP_009047203.1;MERS_1a;899 | 0 | 0 |
| NC_019843.3;YP_009047203.1;MERS_1a;900 | 0 | 0 |
| NC_019843.3;YP_009047203.1;MERS_1a;901 | 0 | 0 |
| NC_019843.3;YP_009047203.1;MERS_1a;902 | 0 | 0.1 |
| NC_019843.3;YP_009047203.1;MERS_1a;903 | 0 | 0.05 |
| NC_019843.3;YP_009047203.1;MERS_1a;904 | 0 | 0 |
| NC_019843.3;YP_009047203.1;MERS_1a;905 | 0 | 0.025 |
| NC_019843.3;YP_009047203.1;MERS_1a;906 | 0 | 0 |
| NC_019843.3;YP_009047203.1;MERS_1a;907 | 0 | 0 |
| NC_019843.3;YP_009047203.1;MERS_1a;908 | 0 | 0 |
| NC_019843.3;YP_009047203.1;MERS_1a;909 | 0 | 0.025 |
| NC_019843.3;YP_009047203.1;MERS_1a;910 | 0 | 0.025 |
| NC_019843.3;YP_009047203.1;MERS_1a;911 | 0 | 0 |
| NC_019843.3;YP_009047203.1;MERS_1a;912 | 0 | 0 |
| NC_019843.3;YP_009047203.1;MERS_1a;913 | 0 | 0 |
| NC_019843.3;YP_009047203.1;MERS_1a;914 | 0 | 0.025 |
| NC_019843.3;YP_009047203.1;MERS_1a;915 | 0.05 | 0 |
| NC_019843.3;YP_009047203.1;MERS_1a;916 | 0 | 0 |
| NC_019843.3;YP_009047203.1;MERS_1a;917 | 0 | 0 |
| NC_019843.3;YP_009047203.1;MERS_1a;918 | 0 | 0 |
| NC_019843.3;YP_009047203.1;MERS_1a;919 | 0 | 0 |
| NC_019843.3;YP_009047203.1;MERS_1a;920 | 0.05 | 0 |
| NC_019843.3;YP_009047203.1;MERS_1a;921 | 0.05 | 0 |
| NC_019843.3;YP_009047203.1;MERS_1a;922 | 0.05 | 0 |

| | | |
|---|---|---|
| NC_019843.3;YP_009047203.1;MERS_1a;923 | 0.1 | 0 |
| NC_019843.3;YP_009047203.1;MERS_1a;924 | 0.1 | 0.025 |
| NC_019843.3;YP_009047203.1;MERS_1a;925 | 0.15 | 0.05 |
| NC_019843.3;YP_009047203.1;MERS_1a;926 | 0.1 | 0.05 |
| NC_019843.3;YP_009047203.1;MERS_1a;927 | 0.1 | 0.025 |
| NC_019843.3;YP_009047203.1;MERS_1a;928 | 0.05 | 0.025 |
| NC_019843.3;YP_009047203.1;MERS_1a;929 | 0 | 0.025 |
| NC_019843.3;YP_009047203.1;MERS_1a;930 | 0.05 | 0 |
| NC_019843.3;YP_009047203.1;MERS_1a;931 | 0.05 | 0 |
| NC_019843.3;YP_009047203.1;MERS_1a;932 | 0 | 0 |
| NC_019843.3;YP_009047203.1;MERS_1a;933 | 0 | 0 |
| NC_019843.3;YP_009047203.1;MERS_1a;934 | 0 | 0 |
| NC_019843.3;YP_009047203.1;MERS_1a;935 | 0 | 0 |
| NC_019843.3;YP_009047203.1;MERS_1a;936 | 0 | 0 |
| NC_019843.3;YP_009047203.1;MERS_1a;937 | 0 | 0 |
| NC_019843.3;YP_009047203.1;MERS_1a;938 | 0 | 0 |
| NC_019843.3;YP_009047203.1;MERS_1a;939 | 0 | 0 |
| NC_019843.3;YP_009047203.1;MERS_1a;940 | 0 | 0 |
| NC_019843.3;YP_009047203.1;MERS_1a;941 | 0 | 0 |
| NC_019843.3;YP_009047203.1;MERS_1a;942 | 0 | 0 |
| NC_019843.3;YP_009047203.1;MERS_1a;943 | 0 | 0 |
| NC_019843.3;YP_009047203.1;MERS_1a;944 | 0 | 0 |
| NC_019843.3;YP_009047203.1;MERS_1a;945 | 0 | 0 |
| NC_019843.3;YP_009047203.1;MERS_1a;946 | 0 | 0 |
| NC_019843.3;YP_009047203.1;MERS_1a;947 | 0 | 0 |
| NC_019843.3;YP_009047203.1;MERS_1a;948 | 0 | 0 |
| NC_019843.3;YP_009047203.1;MERS_1a;949 | 0 | 0 |
| NC_019843.3;YP_009047203.1;MERS_1a;950 | 0 | 0 |
| NC_019843.3;YP_009047203.1;MERS_1a;951 | 0 | 0 |
| NC_019843.3;YP_009047203.1;MERS_1a;952 | 0 | 0 |
| NC_019843.3;YP_009047203.1;MERS_1a;953 | 0 | 0 |
| NC_019843.3;YP_009047203.1;MERS_1a;954 | 0 | 0 |
| NC_019843.3;YP_009047203.1;MERS_1a;955 | 0 | 0 |
| NC_019843.3;YP_009047203.1;MERS_1a;956 | 0 | 0 |
| NC_019843.3;YP_009047203.1;MERS_1a;957 | 0 | 0 |
| NC_019843.3;YP_009047203.1;MERS_1a;958 | 0 | 0 |
| NC_019843.3;YP_009047203.1;MERS_1a;959 | 0.05 | 0 |
| NC_019843.3;YP_009047203.1;MERS_1a;960 | 0 | 0 |
| NC_019843.3;YP_009047203.1;MERS_1a;961 | 0 | 0 |
| NC_019843.3;YP_009047203.1;MERS_1a;962 | 0 | 0 |
| NC_019843.3;YP_009047203.1;MERS_1a;963 | 0 | 0 |
| NC_019843.3;YP_009047203.1;MERS_1a;964 | 0 | 0 |
| NC_019843.3;YP_009047203.1;MERS_1a;965 | 0 | 0.025 |

| | | |
|---|---|---|
| NC_019843.3;YP_009047203.1;MERS_1a;966 | 0 | 0.025 |
| NC_019843.3;YP_009047203.1;MERS_1a;967 | 0 | 0 |
| NC_019843.3;YP_009047203.1;MERS_1a;968 | 0 | 0 |
| NC_019843.3;YP_009047203.1;MERS_1a;969 | 0 | 0 |
| NC_019843.3;YP_009047203.1;MERS_1a;970 | 0 | 0 |
| NC_019843.3;YP_009047203.1;MERS_1a;971 | 0 | 0 |
| NC_019843.3;YP_009047203.1;MERS_1a;972 | 0 | 0 |
| NC_019843.3;YP_009047203.1;MERS_1a;973 | 0 | 0 |
| NC_019843.3;YP_009047203.1;MERS_1a;974 | 0 | 0 |
| NC_019843.3;YP_009047203.1;MERS_1a;975 | 0 | 0 |
| NC_019843.3;YP_009047203.1;MERS_1a;976 | 0 | 0 |
| NC_019843.3;YP_009047203.1;MERS_1a;977 | 0 | 0 |
| NC_019843.3;YP_009047203.1;MERS_1a;978 | 0 | 0 |
| NC_019843.3;YP_009047203.1;MERS_1a;979 | 0 | 0 |
| NC_019843.3;YP_009047203.1;MERS_1a;980 | 0 | 0 |
| NC_019843.3;YP_009047203.1;MERS_1a;981 | 0 | 0 |
| NC_019843.3;YP_009047203.1;MERS_1a;982 | 0.05 | 0 |
| NC_019843.3;YP_009047203.1;MERS_1a;983 | 0 | 0 |
| NC_019843.3;YP_009047203.1;MERS_1a;984 | 0 | 0 |
| NC_019843.3;YP_009047203.1;MERS_1a;985 | 0 | 0 |
| NC_019843.3;YP_009047203.1;MERS_1a;986 | 0 | 0 |
| NC_019843.3;YP_009047203.1;MERS_1a;987 | 0 | 0 |
| NC_019843.3;YP_009047203.1;MERS_1a;988 | 0 | 0 |
| NC_019843.3;YP_009047203.1;MERS_1a;989 | 0 | 0 |
| NC_019843.3;YP_009047203.1;MERS_1a;990 | 0 | 0 |
| NC_019843.3;YP_009047203.1;MERS_1a;991 | 0 | 0 |
| NC_019843.3;YP_009047203.1;MERS_1a;992 | 0.05 | 0 |
| NC_019843.3;YP_009047203.1;MERS_1a;993 | 0.05 | 0 |
| NC_019843.3;YP_009047203.1;MERS_1a;994 | 0 | 0.025 |
| NC_019843.3;YP_009047203.1;MERS_1a;995 | 0.05 | 0 |
| NC_019843.3;YP_009047203.1;MERS_1a;996 | 0.05 | 0 |
| NC_019843.3;YP_009047203.1;MERS_1a;997 | 0.05 | 0 |
| NC_019843.3;YP_009047203.1;MERS_1a;998 | 0.05 | 0 |
| NC_019843.3;YP_009047203.1;MERS_1a;999 | 0.05 | 0 |
| NC_019843.3;YP_009047203.1;MERS_1a;1000 | 0.1 | 0 |
| NC_019843.3;YP_009047203.1;MERS_1a;1001 | 0.05 | 0.025 |
| NC_019843.3;YP_009047203.1;MERS_1a;1002 | 0.1 | 0.025 |
| NC_019843.3;YP_009047203.1;MERS_1a;1003 | 0.05 | 0.025 |
| NC_019843.3;YP_009047203.1;MERS_1a;1004 | 0.15 | 0.075 |
| NC_019843.3;YP_009047203.1;MERS_1a;1005 | 0.15 | 0.05 |
| NC_019843.3;YP_009047203.1;MERS_1a;1006 | 0.15 | 0.05 |
| NC_019843.3;YP_009047203.1;MERS_1a;1007 | 0.1 | 0.05 |
| NC_019843.3;YP_009047203.1;MERS_1a;1008 | 0.1 | 0.025 |

| | | |
|---|---|---|
| NC_019843.3;YP_009047203.1;MERS_1a;1009 | 0.05 | 0.025 |
| NC_019843.3;YP_009047203.1;MERS_1a;1010 | 0.05 | 0.025 |
| NC_019843.3;YP_009047203.1;MERS_1a;1011 | 0 | 0.025 |
| NC_019843.3;YP_009047203.1;MERS_1a;1012 | 0 | 0.025 |
| NC_019843.3;YP_009047203.1;MERS_1a;1013 | 0 | 0 |
| NC_019843.3;YP_009047203.1;MERS_1a;1014 | 0 | 0 |
| NC_019843.3;YP_009047203.1;MERS_1a;1015 | 0 | 0 |
| NC_019843.3;YP_009047203.1;MERS_1a;1016 | 0 | 0 |
| NC_019843.3;YP_009047203.1;MERS_1a;1017 | 0 | 0 |
| NC_019843.3;YP_009047203.1;MERS_1a;1018 | 0 | 0 |
| NC_019843.3;YP_009047203.1;MERS_1a;1019 | 0.1 | 0.025 |
| NC_019843.3;YP_009047203.1;MERS_1a;1020 | 0 | 0 |
| NC_019843.3;YP_009047203.1;MERS_1a;1021 | 0 | 0 |
| NC_019843.3;YP_009047203.1;MERS_1a;1022 | 0 | 0 |
| NC_019843.3;YP_009047203.1;MERS_1a;1023 | 0 | 0 |
| NC_019843.3;YP_009047203.1;MERS_1a;1024 | 0 | 0 |
| NC_019843.3;YP_009047203.1;MERS_1a;1025 | 0 | 0 |
| NC_019843.3;YP_009047203.1;MERS_1a;1026 | 0 | 0 |
| NC_019843.3;YP_009047203.1;MERS_1a;1027 | 0 | 0 |
| NC_019843.3;YP_009047203.1;MERS_1a;1028 | 0 | 0 |
| NC_019843.3;YP_009047203.1;MERS_1a;1029 | 0 | 0 |
| NC_019843.3;YP_009047203.1;MERS_1a;1030 | 0 | 0 |
| NC_019843.3;YP_009047203.1;MERS_1a;1031 | 0 | 0 |
| NC_019843.3;YP_009047203.1;MERS_1a;1032 | 0 | 0 |
| NC_019843.3;YP_009047203.1;MERS_1a;1033 | 0 | 0 |
| NC_019843.3;YP_009047203.1;MERS_1a;1034 | 0.05 | 0.05 |
| NC_019843.3;YP_009047203.1;MERS_1a;1035 | 0.05 | 0 |
| NC_019843.3;YP_009047203.1;MERS_1a;1036 | 0 | 0.05 |
| NC_019843.3;YP_009047203.1;MERS_1a;1037 | 0 | 0.05 |
| NC_019843.3;YP_009047203.1;MERS_1a;1038 | 0 | 0.05 |
| NC_019843.3;YP_009047203.1;MERS_1a;1039 | 0 | 0.05 |
| NC_019843.3;YP_009047203.1;MERS_1a;1040 | 0.05 | 0.05 |
| NC_019843.3;YP_009047203.1;MERS_1a;1041 | 0.05 | 0.05 |
| NC_019843.3;YP_009047203.1;MERS_1a;1042 | 0.05 | 0.075 |
| NC_019843.3;YP_009047203.1;MERS_1a;1043 | 0.05 | 0.075 |
| NC_019843.3;YP_009047203.1;MERS_1a;1044 | 0.05 | 0.025 |
| NC_019843.3;YP_009047203.1;MERS_1a;1045 | 0 | 0.025 |
| NC_019843.3;YP_009047203.1;MERS_1a;1046 | 0 | 0.025 |
| NC_019843.3;YP_009047203.1;MERS_1a;1047 | 0 | 0.025 |
| NC_019843.3;YP_009047203.1;MERS_1a;1048 | 0 | 0.025 |
| NC_019843.3;YP_009047203.1;MERS_1a;1049 | 0 | 0.025 |
| NC_019843.3;YP_009047203.1;MERS_1a;1050 | 0 | 0.05 |
| NC_019843.3;YP_009047203.1;MERS_1a;1051 | 0.05 | 0.025 |

| | | |
|---|---|---|
| NC_019843.3;YP_009047203.1;MERS_1a;1052 | 0 | 0.025 |
| NC_019843.3;YP_009047203.1;MERS_1a;1053 | 0 | 0.025 |
| NC_019843.3;YP_009047203.1;MERS_1a;1054 | 0 | 0.025 |
| NC_019843.3;YP_009047203.1;MERS_1a;1055 | 0 | 0 |
| NC_019843.3;YP_009047203.1;MERS_1a;1056 | 0.05 | 0 |
| NC_019843.3;YP_009047203.1;MERS_1a;1057 | 0.1 | 0 |
| NC_019843.3;YP_009047203.1;MERS_1a;1058 | 0.05 | 0 |
| NC_019843.3;YP_009047203.1;MERS_1a;1059 | 0.05 | 0.025 |
| NC_019843.3;YP_009047203.1;MERS_1a;1060 | 0.05 | 0 |
| NC_019843.3;YP_009047203.1;MERS_1a;1061 | 0 | 0.025 |
| NC_019843.3;YP_009047203.1;MERS_1a;1062 | 0.05 | 0.075 |
| NC_019843.3;YP_009047203.1;MERS_1a;1063 | 0.1 | 0.05 |
| NC_019843.3;YP_009047203.1;MERS_1a;1064 | 0 | 0.025 |
| NC_019843.3;YP_009047203.1;MERS_1a;1065 | 0 | 0.05 |
| NC_019843.3;YP_009047203.1;MERS_1a;1066 | 0.05 | 0.075 |
| NC_019843.3;YP_009047203.1;MERS_1a;1067 | 0 | 0.05 |
| NC_019843.3;YP_009047203.1;MERS_1a;1068 | 0 | 0 |
| NC_019843.3;YP_009047203.1;MERS_1a;1069 | 0 | 0.025 |
| NC_019843.3;YP_009047203.1;MERS_1a;1070 | 0 | 0.025 |
| NC_019843.3;YP_009047203.1;MERS_1a;1071 | 0 | 0 |
| NC_019843.3;YP_009047203.1;MERS_1a;1072 | 0 | 0 |
| NC_019843.3;YP_009047203.1;MERS_1a;1073 | 0 | 0 |
| NC_019843.3;YP_009047203.1;MERS_1a;1074 | 0 | 0 |
| NC_019843.3;YP_009047203.1;MERS_1a;1075 | 0 | 0 |
| NC_019843.3;YP_009047203.1;MERS_1a;1076 | 0 | 0 |
| NC_019843.3;YP_009047203.1;MERS_1a;1077 | 0 | 0 |
| NC_019843.3;YP_009047203.1;MERS_1a;1078 | 0 | 0 |
| NC_019843.3;YP_009047203.1;MERS_1a;1079 | 0 | 0 |
| NC_019843.3;YP_009047203.1;MERS_1a;1080 | 0 | 0 |
| NC_019843.3;YP_009047203.1;MERS_1a;1081 | 0 | 0.025 |
| NC_019843.3;YP_009047203.1;MERS_1a;1082 | 0.2 | 0.25 |
| NC_019843.3;YP_009047203.1;MERS_1a;1083 | 0.1 | 0.075 |
| NC_019843.3;YP_009047203.1;MERS_1a;1084 | 0.15 | 0.125 |
| NC_019843.3;YP_009047203.1;MERS_1a;1085 | 0.2 | 0.175 |
| NC_019843.3;YP_009047203.1;MERS_1a;1086 | 0.15 | 0.15 |
| NC_019843.3;YP_009047203.1;MERS_1a;1087 | 0.1 | 0.075 |
| NC_019843.3;YP_009047203.1;MERS_1a;1088 | 0.05 | 0.075 |
| NC_019843.3;YP_009047203.1;MERS_1a;1089 | 0.05 | 0.025 |
| NC_019843.3;YP_009047203.1;MERS_1a;1090 | 0.05 | 0.05 |
| NC_019843.3;YP_009047203.1;MERS_1a;1091 | 0.05 | 0 |
| NC_019843.3;YP_009047203.1;MERS_1a;1092 | 0.1 | 0.025 |
| NC_019843.3;YP_009047203.1;MERS_1a;1093 | 0.1 | 0.025 |
| NC_019843.3;YP_009047203.1;MERS_1a;1094 | 0.05 | 0 |

| | | |
|---|---|---|
| NC_019843.3;YP_009047203.1;MERS_1a;1095 | 0 | 0 |
| NC_019843.3;YP_009047203.1;MERS_1a;1096 | 0 | 0 |
| NC_019843.3;YP_009047203.1;MERS_1a;1097 | 0 | 0.025 |
| NC_019843.3;YP_009047203.1;MERS_1a;1098 | 0 | 0.025 |
| NC_019843.3;YP_009047203.1;MERS_1a;1099 | 0.05 | 0.025 |
| NC_019843.3;YP_009047203.1;MERS_1a;1100 | 0 | 0.075 |
| NC_019843.3;YP_009047203.1;MERS_1a;1101 | 0 | 0.05 |
| NC_019843.3;YP_009047203.1;MERS_1a;1102 | 0 | 0.025 |
| NC_019843.3;YP_009047203.1;MERS_1a;1103 | 0 | 0.025 |
| NC_019843.3;YP_009047203.1;MERS_1a;1104 | 0 | 0.125 |
| NC_019843.3;YP_009047203.1;MERS_1a;1105 | 0.15 | 0.15 |
| NC_019843.3;YP_009047203.1;MERS_1a;1106 | 0.05 | 0 |
| NC_019843.3;YP_009047203.1;MERS_1a;1107 | 0 | 0.025 |
| NC_019843.3;YP_009047203.1;MERS_1a;1108 | 0 | 0 |
| NC_019843.3;YP_009047203.1;MERS_1a;1109 | 0 | 0 |
| NC_019843.3;YP_009047203.1;MERS_1a;1110 | 0 | 0 |
| NC_019843.3;YP_009047203.1;MERS_1a;1111 | 0 | 0 |
| NC_019843.3;YP_009047203.1;MERS_1a;1112 | 0 | 0 |
| NC_019843.3;YP_009047203.1;MERS_1a;1113 | 0 | 0 |
| NC_019843.3;YP_009047203.1;MERS_1a;1114 | 0 | 0 |
| NC_019843.3;YP_009047203.1;MERS_1a;1115 | 0 | 0 |
| NC_019843.3;YP_009047203.1;MERS_1a;1116 | 0 | 0 |
| NC_019843.3;YP_009047203.1;MERS_1a;1117 | 0 | 0 |
| NC_019843.3;YP_009047203.1;MERS_1a;1118 | 0 | 0 |
| NC_019843.3;YP_009047203.1;MERS_1a;1119 | 0 | 0 |
| NC_019843.3;YP_009047203.1;MERS_1a;1120 | 0 | 0 |
| NC_019843.3;YP_009047203.1;MERS_1a;1121 | 0 | 0 |
| NC_019843.3;YP_009047203.1;MERS_1a;1122 | 0 | 0 |
| NC_019843.3;YP_009047203.1;MERS_1a;1123 | 0 | 0 |
| NC_019843.3;YP_009047203.1;MERS_1a;1124 | 0 | 0 |
| NC_019843.3;YP_009047203.1;MERS_1a;1125 | 0 | 0 |
| NC_019843.3;YP_009047203.1;MERS_1a;1126 | 0.05 | 0 |
| NC_019843.3;YP_009047203.1;MERS_1a;1127 | 0 | 0 |
| NC_019843.3;YP_009047203.1;MERS_1a;1128 | 0 | 0 |
| NC_019843.3;YP_009047203.1;MERS_1a;1129 | 0 | 0 |
| NC_019843.3;YP_009047203.1;MERS_1a;1130 | 0 | 0 |
| NC_019843.3;YP_009047203.1;MERS_1a;1131 | 0.05 | 0.025 |
| NC_019843.3;YP_009047203.1;MERS_1a;1132 | 0 | 0 |
| NC_019843.3;YP_009047203.1;MERS_1a;1133 | 0 | 0 |
| NC_019843.3;YP_009047203.1;MERS_1a;1134 | 0 | 0 |
| NC_019843.3;YP_009047203.1;MERS_1a;1135 | 0 | 0 |
| NC_019843.3;YP_009047203.1;MERS_1a;1136 | 0 | 0 |
| NC_019843.3;YP_009047203.1;MERS_1a;1137 | 0 | 0 |

| | | |
|---|---|---|
| NC_019843.3;YP_009047203.1;MERS_1a;1138 | 0 | 0 |
| NC_019843.3;YP_009047203.1;MERS_1a;1139 | 0.05 | 0 |
| NC_019843.3;YP_009047203.1;MERS_1a;1140 | 0.05 | 0 |
| NC_019843.3;YP_009047203.1;MERS_1a;1141 | 0 | 0 |
| NC_019843.3;YP_009047203.1;MERS_1a;1142 | 0.4 | 0.3 |
| NC_019843.3;YP_009047203.1;MERS_1a;1143 | 0 | 0.025 |
| NC_019843.3;YP_009047203.1;MERS_1a;1144 | 0 | 0.025 |
| NC_019843.3;YP_009047203.1;MERS_1a;1145 | 0 | 0.025 |
| NC_019843.3;YP_009047203.1;MERS_1a;1146 | 0 | 0.025 |
| NC_019843.3;YP_009047203.1;MERS_1a;1147 | 0 | 0.025 |
| NC_019843.3;YP_009047203.1;MERS_1a;1148 | 0.05 | 0.05 |
| NC_019843.3;YP_009047203.1;MERS_1a;1149 | 0 | 0.025 |
| NC_019843.3;YP_009047203.1;MERS_1a;1150 | 0 | 0 |
| NC_019843.3;YP_009047203.1;MERS_1a;1151 | 0 | 0 |
| NC_019843.3;YP_009047203.1;MERS_1a;1152 | 0 | 0 |
| NC_019843.3;YP_009047203.1;MERS_1a;1153 | 0.05 | 0.05 |
| NC_019843.3;YP_009047203.1;MERS_1a;1154 | 0.05 | 0.025 |
| NC_019843.3;YP_009047203.1;MERS_1a;1155 | 0.05 | 0 |
| NC_019843.3;YP_009047203.1;MERS_1a;1156 | 0.05 | 0 |
| NC_019843.3;YP_009047203.1;MERS_1a;1157 | 0.05 | 0.025 |
| NC_019843.3;YP_009047203.1;MERS_1a;1158 | 0.1 | 0.05 |
| NC_019843.3;YP_009047203.1;MERS_1a;1159 | 0.05 | 0.025 |
| NC_019843.3;YP_009047203.1;MERS_1a;1160 | 0.05 | 0.025 |
| NC_019843.3;YP_009047203.1;MERS_1a;1161 | 0.05 | 0 |
| NC_019843.3;YP_009047203.1;MERS_1a;1162 | 0.05 | 0 |
| NC_019843.3;YP_009047203.1;MERS_1a;1163 | 0 | 0.025 |
| NC_019843.3;YP_009047203.1;MERS_1a;1164 | 0 | 0.025 |
| NC_019843.3;YP_009047203.1;MERS_1a;1165 | 0 | 0.05 |
| NC_019843.3;YP_009047203.1;MERS_1a;1166 | 0 | 0.025 |
| NC_019843.3;YP_009047203.1;MERS_1a;1167 | 0.05 | 0.025 |
| NC_019843.3;YP_009047203.1;MERS_1a;1168 | 0.05 | 0.025 |
| NC_019843.3;YP_009047203.1;MERS_1a;1169 | 0 | 0.025 |
| NC_019843.3;YP_009047203.1;MERS_1a;1170 | 0.05 | 0.05 |
| NC_019843.3;YP_009047203.1;MERS_1a;1171 | 0 | 0.075 |
| NC_019843.3;YP_009047203.1;MERS_1a;1172 | 0.1 | 0.075 |
| NC_019843.3;YP_009047203.1;MERS_1a;1173 | 0 | 0.05 |
| NC_019843.3;YP_009047203.1;MERS_1a;1174 | 0 | 0 |
| NC_019843.3;YP_009047203.1;MERS_1a;1175 | 0 | 0 |
| NC_019843.3;YP_009047203.1;MERS_1a;1176 | 0 | 0 |
| NC_019843.3;YP_009047203.1;MERS_1a;1177 | 0 | 0 |
| NC_019843.3;YP_009047203.1;MERS_1a;1178 | 0 | 0 |
| NC_019843.3;YP_009047203.1;MERS_1a;1179 | 0 | 0 |
| NC_019843.3;YP_009047203.1;MERS_1a;1180 | 0 | 0 |

| | | |
|---|---|---|
| NC_019843.3;YP_009047203.1;MERS_1a;1181 | 0 | 0 |
| NC_019843.3;YP_009047203.1;MERS_1a;1182 | 0 | 0 |
| NC_019843.3;YP_009047203.1;MERS_1a;1183 | 0 | 0 |
| NC_019843.3;YP_009047203.1;MERS_1a;1184 | 0 | 0 |
| NC_019843.3;YP_009047203.1;MERS_1a;1185 | 0 | 0 |
| NC_019843.3;YP_009047203.1;MERS_1a;1186 | 0 | 0 |
| NC_019843.3;YP_009047203.1;MERS_1a;1187 | 0 | 0 |
| NC_019843.3;YP_009047203.1;MERS_1a;1188 | 0 | 0 |
| NC_019843.3;YP_009047203.1;MERS_1a;1189 | 0 | 0 |
| NC_019843.3;YP_009047203.1;MERS_1a;1190 | 0 | 0 |
| NC_019843.3;YP_009047203.1;MERS_1a;1191 | 0 | 0 |
| NC_019843.3;YP_009047203.1;MERS_1a;1192 | 0 | 0 |
| NC_019843.3;YP_009047203.1;MERS_1a;1193 | 0 | 0 |
| NC_019843.3;YP_009047203.1;MERS_1a;1194 | 0 | 0 |
| NC_019843.3;YP_009047203.1;MERS_1a;1195 | 0 | 0 |
| NC_019843.3;YP_009047203.1;MERS_1a;1196 | 0 | 0 |
| NC_019843.3;YP_009047203.1;MERS_1a;1197 | 0 | 0 |
| NC_019843.3;YP_009047203.1;MERS_1a;1198 | 0 | 0.025 |
| NC_019843.3;YP_009047203.1;MERS_1a;1199 | 0 | 0 |
| NC_019843.3;YP_009047203.1;MERS_1a;1200 | 0 | 0.025 |
| NC_019843.3;YP_009047203.1;MERS_1a;1201 | 0 | 0.025 |
| NC_019843.3;YP_009047203.1;MERS_1a;1202 | 0 | 0.05 |
| NC_019843.3;YP_009047203.1;MERS_1a;1203 | 0.05 | 0.025 |
| NC_019843.3;YP_009047203.1;MERS_1a;1204 | 0.05 | 0 |
| NC_019843.3;YP_009047203.1;MERS_1a;1205 | 0.05 | 0.025 |
| NC_019843.3;YP_009047203.1;MERS_1a;1206 | 0.05 | 0 |
| NC_019843.3;YP_009047203.1;MERS_1a;1207 | 0.05 | 0 |
| NC_019843.3;YP_009047203.1;MERS_1a;1208 | 0.05 | 0 |
| NC_019843.3;YP_009047203.1;MERS_1a;1209 | 0.05 | 0.025 |
| NC_019843.3;YP_009047203.1;MERS_1a;1210 | 0.1 | 0.025 |
| NC_019843.3;YP_009047203.1;MERS_1a;1211 | 0 | 0 |
| NC_019843.3;YP_009047203.1;MERS_1a;1212 | 0 | 0 |
| NC_019843.3;YP_009047203.1;MERS_1a;1213 | 0 | 0 |
| NC_019843.3;YP_009047203.1;MERS_1a;1214 | 0 | 0 |
| NC_019843.3;YP_009047203.1;MERS_1a;1215 | 0 | 0 |
| NC_019843.3;YP_009047203.1;MERS_1a;1216 | 0 | 0 |
| NC_019843.3;YP_009047203.1;MERS_1a;1217 | 0 | 0 |
| NC_019843.3;YP_009047203.1;MERS_1a;1218 | 0 | 0 |
| NC_019843.3;YP_009047203.1;MERS_1a;1219 | 0 | 0 |
| NC_019843.3;YP_009047203.1;MERS_1a;1220 | 0 | 0 |
| NC_019843.3;YP_009047203.1;MERS_1a;1221 | 0 | 0 |
| NC_019843.3;YP_009047203.1;MERS_1a;1222 | 0 | 0 |
| NC_019843.3;YP_009047203.1;MERS_1a;1223 | 0 | 0 |

| | | |
|---|---|---|
| NC_019843.3;YP_009047203.1;MERS_1a;1224 | 0 | 0 |
| NC_019843.3;YP_009047203.1;MERS_1a;1225 | 0 | 0 |
| NC_019843.3;YP_009047203.1;MERS_1a;1226 | 0 | 0 |
| NC_019843.3;YP_009047203.1;MERS_1a;1227 | 0 | 0 |
| NC_019843.3;YP_009047203.1;MERS_1a;1228 | 0 | 0 |
| NC_019843.3;YP_009047203.1;MERS_1a;1229 | 0 | 0 |
| NC_019843.3;YP_009047203.1;MERS_1a;1230 | 0 | 0 |
| NC_019843.3;YP_009047203.1;MERS_1a;1231 | 0 | 0 |
| NC_019843.3;YP_009047203.1;MERS_1a;1232 | 0 | 0 |
| NC_019843.3;YP_009047203.1;MERS_1a;1233 | 0 | 0 |
| NC_019843.3;YP_009047203.1;MERS_1a;1234 | 0 | 0 |
| NC_019843.3;YP_009047203.1;MERS_1a;1235 | 0 | 0 |
| NC_019843.3;YP_009047203.1;MERS_1a;1236 | 0 | 0 |
| NC_019843.3;YP_009047203.1;MERS_1a;1237 | 0 | 0 |
| NC_019843.3;YP_009047203.1;MERS_1a;1238 | 0 | 0 |
| NC_019843.3;YP_009047203.1;MERS_1a;1239 | 0 | 0 |
| NC_019843.3;YP_009047203.1;MERS_1a;1240 | 0 | 0 |
| NC_019843.3;YP_009047203.1;MERS_1a;1241 | 0 | 0 |
| NC_019843.3;YP_009047203.1;MERS_1a;1242 | 0 | 0 |
| NC_019843.3;YP_009047203.1;MERS_1a;1243 | 0 | 0 |
| NC_019843.3;YP_009047203.1;MERS_1a;1244 | 0 | 0.025 |
| NC_019843.3;YP_009047203.1;MERS_1a;1245 | 0 | 0 |
| NC_019843.3;YP_009047203.1;MERS_1a;1246 | 0 | 0 |
| NC_019843.3;YP_009047203.1;MERS_1a;1247 | 0 | 0 |
| NC_019843.3;YP_009047203.1;MERS_1a;1248 | 0 | 0 |
| NC_019843.3;YP_009047203.1;MERS_1a;1249 | 0 | 0 |
| NC_019843.3;YP_009047203.1;MERS_1a;1250 | 0 | 0 |
| NC_019843.3;YP_009047203.1;MERS_1a;1251 | 0 | 0 |
| NC_019843.3;YP_009047203.1;MERS_1a;1252 | 0 | 0 |
| NC_019843.3;YP_009047203.1;MERS_1a;1253 | 0 | 0 |
| NC_019843.3;YP_009047203.1;MERS_1a;1254 | 0 | 0 |
| NC_019843.3;YP_009047203.1;MERS_1a;1255 | 0 | 0 |
| NC_019843.3;YP_009047203.1;MERS_1a;1256 | 0 | 0 |
| NC_019843.3;YP_009047203.1;MERS_1a;1257 | 0 | 0 |
| NC_019843.3;YP_009047203.1;MERS_1a;1258 | 0 | 0 |
| NC_019843.3;YP_009047203.1;MERS_1a;1259 | 0 | 0 |
| NC_019843.3;YP_009047203.1;MERS_1a;1260 | 0 | 0.025 |
| NC_019843.3;YP_009047203.1;MERS_1a;1261 | 0 | 0 |
| NC_019843.3;YP_009047203.1;MERS_1a;1262 | 0 | 0 |
| NC_019843.3;YP_009047203.1;MERS_1a;1263 | 0 | 0 |
| NC_019843.3;YP_009047203.1;MERS_1a;1264 | 0 | 0.025 |
| NC_019843.3;YP_009047203.1;MERS_1a;1265 | 0 | 0.025 |
| NC_019843.3;YP_009047203.1;MERS_1a;1266 | 0 | 0 |

| | | |
|---|---|---|
| NC_019843.3;YP_009047203.1;MERS_1a;1267 | 0 | 0 |
| NC_019843.3;YP_009047203.1;MERS_1a;1268 | 0 | 0 |
| NC_019843.3;YP_009047203.1;MERS_1a;1269 | 0 | 0 |
| NC_019843.3;YP_009047203.1;MERS_1a;1270 | 0 | 0 |
| NC_019843.3;YP_009047203.1;MERS_1a;1271 | 0 | 0.025 |
| NC_019843.3;YP_009047203.1;MERS_1a;1272 | 0 | 0.025 |
| NC_019843.3;YP_009047203.1;MERS_1a;1273 | 0 | 0 |
| NC_019843.3;YP_009047203.1;MERS_1a;1274 | 0 | 0 |
| NC_019843.3;YP_009047203.1;MERS_1a;1275 | 0 | 0 |
| NC_019843.3;YP_009047203.1;MERS_1a;1276 | 0 | 0 |
| NC_019843.3;YP_009047203.1;MERS_1a;1277 | 0 | 0 |
| NC_019843.3;YP_009047203.1;MERS_1a;1278 | 0 | 0.025 |
| NC_019843.3;YP_009047203.1;MERS_1a;1279 | 0 | 0.025 |
| NC_019843.3;YP_009047203.1;MERS_1a;1280 | 0 | 0.025 |
| NC_019843.3;YP_009047203.1;MERS_1a;1281 | 0 | 0.025 |
| NC_019843.3;YP_009047203.1;MERS_1a;1282 | 0 | 0.05 |
| NC_019843.3;YP_009047203.1;MERS_1a;1283 | 0 | 0.075 |
| NC_019843.3;YP_009047203.1;MERS_1a;1284 | 0 | 0.025 |
| NC_019843.3;YP_009047203.1;MERS_1a;1285 | 0 | 0.1 |
| NC_019843.3;YP_009047203.1;MERS_1a;1286 | 0 | 0.075 |
| NC_019843.3;YP_009047203.1;MERS_1a;1287 | 0 | 0 |
| NC_019843.3;YP_009047203.1;MERS_1a;1288 | 0 | 0 |
| NC_019843.3;YP_009047203.1;MERS_1a;1289 | 0.05 | 0 |
| NC_019843.3;YP_009047203.1;MERS_1a;1290 | 0 | 0 |
| NC_019843.3;YP_009047203.1;MERS_1a;1291 | 0 | 0 |
| NC_019843.3;YP_009047203.1;MERS_1a;1292 | 0 | 0 |
| NC_019843.3;YP_009047203.1;MERS_1a;1293 | 0 | 0 |
| NC_019843.3;YP_009047203.1;MERS_1a;1294 | 0 | 0 |
| NC_019843.3;YP_009047203.1;MERS_1a;1295 | 0 | 0 |
| NC_019843.3;YP_009047203.1;MERS_1a;1296 | 0 | 0 |
| NC_019843.3;YP_009047203.1;MERS_1a;1297 | 0 | 0 |
| NC_019843.3;YP_009047203.1;MERS_1a;1298 | 0 | 0 |
| NC_019843.3;YP_009047203.1;MERS_1a;1299 | 0 | 0 |
| NC_019843.3;YP_009047203.1;MERS_1a;1300 | 0 | 0 |
| NC_019843.3;YP_009047203.1;MERS_1a;1301 | 0 | 0.05 |
| NC_019843.3;YP_009047203.1;MERS_1a;1302 | 0 | 0.025 |
| NC_019843.3;YP_009047203.1;MERS_1a;1303 | 0 | 0.05 |
| NC_019843.3;YP_009047203.1;MERS_1a;1304 | 0 | 0.025 |
| NC_019843.3;YP_009047203.1;MERS_1a;1305 | 0 | 0.1 |
| NC_019843.3;YP_009047203.1;MERS_1a;1306 | 0.1 | 0.05 |
| NC_019843.3;YP_009047203.1;MERS_1a;1307 | 0 | 0 |
| NC_019843.3;YP_009047203.1;MERS_1a;1308 | 0 | 0 |
| NC_019843.3;YP_009047203.1;MERS_1a;1309 | 0 | 0 |

| | | |
|---|---|---|
| NC_019843.3;YP_009047203.1;MERS_1a;1310 | 0 | 0 |
| NC_019843.3;YP_009047203.1;MERS_1a;1311 | 0 | 0 |
| NC_019843.3;YP_009047203.1;MERS_1a;1312 | 0 | 0 |
| NC_019843.3;YP_009047203.1;MERS_1a;1313 | 0 | 0 |
| NC_019843.3;YP_009047203.1;MERS_1a;1314 | 0 | 0 |
| NC_019843.3;YP_009047203.1;MERS_1a;1315 | 0 | 0 |
| NC_019843.3;YP_009047203.1;MERS_1a;1316 | 0 | 0 |
| NC_019843.3;YP_009047203.1;MERS_1a;1317 | 0 | 0.025 |
| NC_019843.3;YP_009047203.1;MERS_1a;1318 | 0.1 | 0.075 |
| NC_019843.3;YP_009047203.1;MERS_1a;1319 | 0.05 | 0 |
| NC_019843.3;YP_009047203.1;MERS_1a;1320 | 0 | 0 |
| NC_019843.3;YP_009047203.1;MERS_1a;1321 | 0 | 0 |
| NC_019843.3;YP_009047203.1;MERS_1a;1322 | 0 | 0 |
| NC_019843.3;YP_009047203.1;MERS_1a;1323 | 0 | 0 |
| NC_019843.3;YP_009047203.1;MERS_1a;1324 | 0 | 0 |
| NC_019843.3;YP_009047203.1;MERS_1a;1325 | 0 | 0 |
| NC_019843.3;YP_009047203.1;MERS_1a;1326 | 0 | 0 |
| NC_019843.3;YP_009047203.1;MERS_1a;1327 | 0 | 0 |
| NC_019843.3;YP_009047203.1;MERS_1a;1328 | 0 | 0 |
| NC_019843.3;YP_009047203.1;MERS_1a;1329 | 0 | 0 |
| NC_019843.3;YP_009047203.1;MERS_1a;1330 | 0 | 0 |
| NC_019843.3;YP_009047203.1;MERS_1a;1331 | 0 | 0 |
| NC_019843.3;YP_009047203.1;MERS_1a;1332 | 0 | 0 |
| NC_019843.3;YP_009047203.1;MERS_1a;1333 | 0 | 0 |
| NC_019843.3;YP_009047203.1;MERS_1a;1334 | 0 | 0 |
| NC_019843.3;YP_009047203.1;MERS_1a;1335 | 0 | 0 |
| NC_019843.3;YP_009047203.1;MERS_1a;1336 | 0.15 | 0.075 |
| NC_019843.3;YP_009047203.1;MERS_1a;1337 | 0.1 | 0.025 |
| NC_019843.3;YP_009047203.1;MERS_1a;1338 | 0.1 | 0.025 |
| NC_019843.3;YP_009047203.1;MERS_1a;1339 | 0.15 | 0.025 |
| NC_019843.3;YP_009047203.1;MERS_1a;1340 | 0.15 | 0.075 |
| NC_019843.3;YP_009047203.1;MERS_1a;1341 | 0 | 0.05 |
| NC_019843.3;YP_009047203.1;MERS_1a;1342 | 0 | 0.025 |
| NC_019843.3;YP_009047203.1;MERS_1a;1343 | 0.05 | 0.025 |
| NC_019843.3;YP_009047203.1;MERS_1a;1344 | 0.05 | 0.05 |
| NC_019843.3;YP_009047203.1;MERS_1a;1345 | 0.1 | 0.075 |
| NC_019843.3;YP_009047203.1;MERS_1a;1346 | 0.1 | 0 |
| NC_019843.3;YP_009047203.1;MERS_1a;1347 | 0 | 0 |
| NC_019843.3;YP_009047203.1;MERS_1a;1348 | 0 | 0 |
| NC_019843.3;YP_009047203.1;MERS_1a;1349 | 0 | 0.025 |
| NC_019843.3;YP_009047203.1;MERS_1a;1350 | 0 | 0 |
| NC_019843.3;YP_009047203.1;MERS_1a;1351 | 0 | 0 |
| NC_019843.3;YP_009047203.1;MERS_1a;1352 | 0 | 0 |

| | | |
|---|---|---|
| NC_019843.3;YP_009047203.1;MERS_1a;1353 | 0 | 0 |
| NC_019843.3;YP_009047203.1;MERS_1a;1354 | 0 | 0 |
| NC_019843.3;YP_009047203.1;MERS_1a;1355 | 0 | 0 |
| NC_019843.3;YP_009047203.1;MERS_1a;1356 | 0 | 0 |
| NC_019843.3;YP_009047203.1;MERS_1a;1357 | 0 | 0 |
| NC_019843.3;YP_009047203.1;MERS_1a;1358 | 0 | 0 |
| NC_019843.3;YP_009047203.1;MERS_1a;1359 | 0 | 0 |
| NC_019843.3;YP_009047203.1;MERS_1a;1360 | 0 | 0 |
| NC_019843.3;YP_009047203.1;MERS_1a;1361 | 0 | 0 |
| NC_019843.3;YP_009047203.1;MERS_1a;1362 | 0 | 0 |
| NC_019843.3;YP_009047203.1;MERS_1a;1363 | 0 | 0 |
| NC_019843.3;YP_009047203.1;MERS_1a;1364 | 0 | 0 |
| NC_019843.3;YP_009047203.1;MERS_1a;1365 | 0 | 0 |
| NC_019843.3;YP_009047203.1;MERS_1a;1366 | 0 | 0 |
| NC_019843.3;YP_009047203.1;MERS_1a;1367 | 0 | 0 |
| NC_019843.3;YP_009047203.1;MERS_1a;1368 | 0 | 0 |
| NC_019843.3;YP_009047203.1;MERS_1a;1369 | 0 | 0 |
| NC_019843.3;YP_009047203.1;MERS_1a;1370 | 0 | 0 |
| NC_019843.3;YP_009047203.1;MERS_1a;1371 | 0 | 0 |
| NC_019843.3;YP_009047203.1;MERS_1a;1372 | 0 | 0 |
| NC_019843.3;YP_009047203.1;MERS_1a;1373 | 0 | 0 |
| NC_019843.3;YP_009047203.1;MERS_1a;1374 | 0 | 0 |
| NC_019843.3;YP_009047203.1;MERS_1a;1375 | 0 | 0 |
| NC_019843.3;YP_009047203.1;MERS_1a;1376 | 0 | 0.025 |
| NC_019843.3;YP_009047203.1;MERS_1a;1377 | 0 | 0 |
| NC_019843.3;YP_009047203.1;MERS_1a;1378 | 0.05 | 0.025 |
| NC_019843.3;YP_009047203.1;MERS_1a;1379 | 0.05 | 0 |
| NC_019843.3;YP_009047203.1;MERS_1a;1380 | 0.05 | 0 |
| NC_019843.3;YP_009047203.1;MERS_1a;1381 | 0.05 | 0 |
| NC_019843.3;YP_009047203.1;MERS_1a;1382 | 0 | 0 |
| NC_019843.3;YP_009047203.1;MERS_1a;1383 | 0 | 0 |
| NC_019843.3;YP_009047203.1;MERS_1a;1384 | 0 | 0 |
| NC_019843.3;YP_009047203.1;MERS_1a;1385 | 0 | 0 |
| NC_019843.3;YP_009047203.1;MERS_1a;1386 | 0 | 0 |
| NC_019843.3;YP_009047203.1;MERS_1a;1387 | 0 | 0.025 |
| NC_019843.3;YP_009047203.1;MERS_1a;1388 | 0 | 0 |
| NC_019843.3;YP_009047203.1;MERS_1a;1389 | 0 | 0.025 |
| NC_019843.3;YP_009047203.1;MERS_1a;1390 | 0 | 0 |
| NC_019843.3;YP_009047203.1;MERS_1a;1391 | 0 | 0.025 |
| NC_019843.3;YP_009047203.1;MERS_1a;1392 | 0 | 0.025 |
| NC_019843.3;YP_009047203.1;MERS_1a;1393 | 0 | 0.05 |
| NC_019843.3;YP_009047203.1;MERS_1a;1394 | 0.05 | 0.05 |
| NC_019843.3;YP_009047203.1;MERS_1a;1395 | 0.05 | 0.075 |

| | | |
|---|---|---|
| NC_019843.3;YP_009047203.1;MERS_1a;1396 | 0 | 0.025 |
| NC_019843.3;YP_009047203.1;MERS_1a;1397 | 0 | 0.025 |
| NC_019843.3;YP_009047203.1;MERS_1a;1398 | 0.05 | 0.025 |
| NC_019843.3;YP_009047203.1;MERS_1a;1399 | 0 | 0 |
| NC_019843.3;YP_009047203.1;MERS_1a;1400 | 0 | 0 |
| NC_019843.3;YP_009047203.1;MERS_1a;1401 | 0.05 | 0 |
| NC_019843.3;YP_009047203.1;MERS_1a;1402 | 0 | 0 |
| NC_019843.3;YP_009047203.1;MERS_1a;1403 | 0.05 | 0.025 |
| NC_019843.3;YP_009047203.1;MERS_1a;1404 | 0.4 | 0.425 |
| NC_019843.3;YP_009047203.1;MERS_1a;1405 | 0 | 0.025 |
| NC_019843.3;YP_009047203.1;MERS_1a;1406 | 0 | 0.025 |
| NC_019843.3;YP_009047203.1;MERS_1a;1407 | 0 | 0 |
| NC_019843.3;YP_009047203.1;MERS_1a;1408 | 0 | 0 |
| NC_019843.3;YP_009047203.1;MERS_1a;1409 | 0 | 0 |
| NC_019843.3;YP_009047203.1;MERS_1a;1410 | 0 | 0 |
| NC_019843.3;YP_009047203.1;MERS_1a;1411 | 0 | 0 |
| NC_019843.3;YP_009047203.1;MERS_1a;1412 | 0 | 0 |
| NC_019843.3;YP_009047203.1;MERS_1a;1413 | 0 | 0.025 |
| NC_019843.3;YP_009047203.1;MERS_1a;1414 | 0 | 0.025 |
| NC_019843.3;YP_009047203.1;MERS_1a;1415 | 0 | 0 |
| NC_019843.3;YP_009047203.1;MERS_1a;1416 | 0 | 0 |
| NC_019843.3;YP_009047203.1;MERS_1a;1417 | 0 | 0 |
| NC_019843.3;YP_009047203.1;MERS_1a;1418 | 0 | 0 |
| NC_019843.3;YP_009047203.1;MERS_1a;1419 | 0 | 0 |
| NC_019843.3;YP_009047203.1;MERS_1a;1420 | 0 | 0 |
| NC_019843.3;YP_009047203.1;MERS_1a;1421 | 0 | 0 |
| NC_019843.3;YP_009047203.1;MERS_1a;1422 | 0 | 0 |
| NC_019843.3;YP_009047203.1;MERS_1a;1423 | 0 | 0 |
| NC_019843.3;YP_009047203.1;MERS_1a;1424 | 0.05 | 0 |
| NC_019843.3;YP_009047203.1;MERS_1a;1425 | 0 | 0 |
| NC_019843.3;YP_009047203.1;MERS_1a;1426 | 0 | 0 |
| NC_019843.3;YP_009047203.1;MERS_1a;1427 | 0 | 0 |
| NC_019843.3;YP_009047203.1;MERS_1a;1428 | 0 | 0 |
| NC_019843.3;YP_009047203.1;MERS_1a;1429 | 0 | 0 |
| NC_019843.3;YP_009047203.1;MERS_1a;1430 | 0 | 0 |
| NC_019843.3;YP_009047203.1;MERS_1a;1431 | 0 | 0 |
| NC_019843.3;YP_009047203.1;MERS_1a;1432 | 0 | 0 |
| NC_019843.3;YP_009047203.1;MERS_1a;1433 | 0 | 0 |
| NC_019843.3;YP_009047203.1;MERS_1a;1434 | 0 | 0 |
| NC_019843.3;YP_009047203.1;MERS_1a;1435 | 0 | 0 |
| NC_019843.3;YP_009047203.1;MERS_1a;1436 | 0 | 0.025 |
| NC_019843.3;YP_009047203.1;MERS_1a;1437 | 0.05 | 0.05 |
| NC_019843.3;YP_009047203.1;MERS_1a;1438 | 0 | 0.025 |

| | | |
|---|---|---|
| NC_019843.3;YP_009047203.1;MERS_1a;1439 | 0 | 0 |
| NC_019843.3;YP_009047203.1;MERS_1a;1440 | 0 | 0 |
| NC_019843.3;YP_009047203.1;MERS_1a;1441 | 0 | 0 |
| NC_019843.3;YP_009047203.1;MERS_1a;1442 | 0 | 0 |
| NC_019843.3;YP_009047203.1;MERS_1a;1443 | 0 | 0 |
| NC_019843.3;YP_009047203.1;MERS_1a;1444 | 0 | 0 |
| NC_019843.3;YP_009047203.1;MERS_1a;1445 | 0 | 0 |
| NC_019843.3;YP_009047203.1;MERS_1a;1446 | 0 | 0 |
| NC_019843.3;YP_009047203.1;MERS_1a;1447 | 0 | 0 |
| NC_019843.3;YP_009047203.1;MERS_1a;1448 | 0 | 0 |
| NC_019843.3;YP_009047203.1;MERS_1a;1449 | 0 | 0.025 |
| NC_019843.3;YP_009047203.1;MERS_1a;1450 | 0 | 0 |
| NC_019843.3;YP_009047203.1;MERS_1a;1451 | 0 | 0.025 |
| NC_019843.3;YP_009047203.1;MERS_1a;1452 | 0 | 0 |
| NC_019843.3;YP_009047203.1;MERS_1a;1453 | 0 | 0 |
| NC_019843.3;YP_009047203.1;MERS_1a;1454 | 0 | 0.025 |
| NC_019843.3;YP_009047203.1;MERS_1a;1455 | 0 | 0.025 |
| NC_019843.3;YP_009047203.1;MERS_1a;1456 | 0 | 0.025 |
| NC_019843.3;YP_009047203.1;MERS_1a;1457 | 0 | 0.05 |
| NC_019843.3;YP_009047203.1;MERS_1a;1458 | 0 | 0 |
| NC_019843.3;YP_009047203.1;MERS_1a;1459 | 0 | 0 |
| NC_019843.3;YP_009047203.1;MERS_1a;1460 | 0 | 0 |
| NC_019843.3;YP_009047203.1;MERS_1a;1461 | 0 | 0 |
| NC_019843.3;YP_009047203.1;MERS_1a;1462 | 0 | 0 |
| NC_019843.3;YP_009047203.1;MERS_1a;1463 | 0 | 0 |
| NC_019843.3;YP_009047203.1;MERS_1a;1464 | 0 | 0 |
| NC_019843.3;YP_009047203.1;MERS_1a;1465 | 0 | 0 |
| NC_019843.3;YP_009047203.1;MERS_1a;1466 | 0 | 0 |
| NC_019843.3;YP_009047203.1;MERS_1a;1467 | 0 | 0 |
| NC_019843.3;YP_009047203.1;MERS_1a;1468 | 0 | 0 |
| NC_019843.3;YP_009047203.1;MERS_1a;1469 | 0 | 0 |
| NC_019843.3;YP_009047203.1;MERS_1a;1470 | 0 | 0 |
| NC_019843.3;YP_009047203.1;MERS_1a;1471 | 0 | 0 |
| NC_019843.3;YP_009047203.1;MERS_1a;1472 | 0 | 0 |
| NC_019843.3;YP_009047203.1;MERS_1a;1473 | 0.05 | 0 |
| NC_019843.3;YP_009047203.1;MERS_1a;1474 | 0 | 0 |
| NC_019843.3;YP_009047203.1;MERS_1a;1475 | 0 | 0 |
| NC_019843.3;YP_009047203.1;MERS_1a;1476 | 0 | 0 |
| NC_019843.3;YP_009047203.1;MERS_1a;1477 | 0 | 0 |
| NC_019843.3;YP_009047203.1;MERS_1a;1478 | 0 | 0 |
| NC_019843.3;YP_009047203.1;MERS_1a;1479 | 0 | 0 |
| NC_019843.3;YP_009047203.1;MERS_1a;1480 | 0 | 0 |
| NC_019843.3;YP_009047203.1;MERS_1a;1481 | 0 | 0 |

| | | |
|---|---|---|
| NC_019843.3;YP_009047203.1;MERS_1a;1482 | 0 | 0 |
| NC_019843.3;YP_009047203.1;MERS_1a;1483 | 0 | 0 |
| NC_019843.3;YP_009047203.1;MERS_1a;1484 | 0 | 0 |
| NC_019843.3;YP_009047203.1;MERS_1a;1485 | 0 | 0 |
| NC_019843.3;YP_009047203.1;MERS_1a;1486 | 0 | 0 |
| NC_019843.3;YP_009047203.1;MERS_1a;1487 | 0 | 0 |
| NC_019843.3;YP_009047203.1;MERS_1a;1488 | 0 | 0 |
| NC_019843.3;YP_009047203.1;MERS_1a;1489 | 0 | 0.025 |
| NC_019843.3;YP_009047203.1;MERS_1a;1490 | 0.1 | 0.075 |
| NC_019843.3;YP_009047203.1;MERS_1a;1491 | 0 | 0.075 |
| NC_019843.3;YP_009047203.1;MERS_1a;1492 | 0 | 0 |
| NC_019843.3;YP_009047203.1;MERS_1a;1493 | 0 | 0.025 |
| NC_019843.3;YP_009047203.1;MERS_1a;1494 | 0 | 0 |
| NC_019843.3;YP_009047203.1;MERS_1a;1495 | 0 | 0 |
| NC_019843.3;YP_009047203.1;MERS_1a;1496 | 0 | 0 |
| NC_019843.3;YP_009047203.1;MERS_1a;1497 | 0 | 0 |
| NC_019843.3;YP_009047203.1;MERS_1a;1498 | 0 | 0 |
| NC_019843.3;YP_009047203.1;MERS_1a;1499 | 0 | 0 |
| NC_019843.3;YP_009047203.1;MERS_1a;1500 | 0 | 0 |
| NC_019843.3;YP_009047203.1;MERS_1a;1501 | 0 | 0 |
| NC_019843.3;YP_009047203.1;MERS_1a;1502 | 0 | 0 |
| NC_019843.3;YP_009047203.1;MERS_1a;1503 | 0 | 0 |
| NC_019843.3;YP_009047203.1;MERS_1a;1504 | 0 | 0 |
| NC_019843.3;YP_009047203.1;MERS_1a;1505 | 0 | 0 |
| NC_019843.3;YP_009047203.1;MERS_1a;1506 | 0 | 0 |
| NC_019843.3;YP_009047203.1;MERS_1a;1507 | 0 | 0 |
| NC_019843.3;YP_009047203.1;MERS_1a;1508 | 0 | 0 |
| NC_019843.3;YP_009047203.1;MERS_1a;1509 | 0 | 0 |
| NC_019843.3;YP_009047203.1;MERS_1a;1510 | 0 | 0 |
| NC_019843.3;YP_009047203.1;MERS_1a;1511 | 0 | 0 |
| NC_019843.3;YP_009047203.1;MERS_1a;1512 | 0 | 0 |
| NC_019843.3;YP_009047203.1;MERS_1a;1513 | 0 | 0 |
| NC_019843.3;YP_009047203.1;MERS_1a;1514 | 0.05 | 0 |
| NC_019843.3;YP_009047203.1;MERS_1a;1515 | 0.05 | 0 |
| NC_019843.3;YP_009047203.1;MERS_1a;1516 | 0.05 | 0 |
| NC_019843.3;YP_009047203.1;MERS_1a;1517 | 0 | 0 |
| NC_019843.3;YP_009047203.1;MERS_1a;1518 | 0.05 | 0.025 |
| NC_019843.3;YP_009047203.1;MERS_1a;1519 | 0 | 0 |
| NC_019843.3;YP_009047203.1;MERS_1a;1520 | 0.05 | 0 |
| NC_019843.3;YP_009047203.1;MERS_1a;1521 | 0.05 | 0 |
| NC_019843.3;YP_009047203.1;MERS_1a;1522 | 0 | 0 |
| NC_019843.3;YP_009047203.1;MERS_1a;1523 | 0.05 | 0 |
| NC_019843.3;YP_009047203.1;MERS_1a;1524 | 0 | 0 |

| | | |
|---|---|---|
| NC_019843.3;YP_009047203.1;MERS_1a;1525 | 0 | 0 |
| NC_019843.3;YP_009047203.1;MERS_1a;1526 | 0 | 0 |
| NC_019843.3;YP_009047203.1;MERS_1a;1527 | 0 | 0.025 |
| NC_019843.3;YP_009047203.1;MERS_1a;1528 | 0 | 0.025 |
| NC_019843.3;YP_009047203.1;MERS_1a;1529 | 0 | 0.025 |
| NC_019843.3;YP_009047203.1;MERS_1a;1530 | 0.05 | 0 |
| NC_019843.3;YP_009047203.1;MERS_1a;1531 | 0.1 | 0 |
| NC_019843.3;YP_009047203.1;MERS_1a;1532 | 0.05 | 0 |
| NC_019843.3;YP_009047203.1;MERS_1a;1533 | 0.05 | 0 |
| NC_019843.3;YP_009047203.1;MERS_1a;1534 | 0.05 | 0 |
| NC_019843.3;YP_009047203.1;MERS_1a;1535 | 0.15 | 0 |
| NC_019843.3;YP_009047203.1;MERS_1a;1536 | 0.1 | 0 |
| NC_019843.3;YP_009047203.1;MERS_1a;1537 | 0.05 | 0 |
| NC_019843.3;YP_009047203.1;MERS_1a;1538 | 0.05 | 0 |
| NC_019843.3;YP_009047203.1;MERS_1a;1539 | 0.05 | 0 |
| NC_019843.3;YP_009047203.1;MERS_1a;1540 | 0.1 | 0 |
| NC_019843.3;YP_009047203.1;MERS_1a;1541 | 0.1 | 0.05 |
| NC_019843.3;YP_009047203.1;MERS_1a;1542 | 0.15 | 0 |
| NC_019843.3;YP_009047203.1;MERS_1a;1543 | 0.1 | 0 |
| NC_019843.3;YP_009047203.1;MERS_1a;1544 | 0.1 | 0 |
| NC_019843.3;YP_009047203.1;MERS_1a;1545 | 0.1 | 0.05 |
| NC_019843.3;YP_009047203.1;MERS_1a;1546 | 0 | 0.025 |
| NC_019843.3;YP_009047203.1;MERS_1a;1547 | 0 | 0 |
| NC_019843.3;YP_009047203.1;MERS_1a;1548 | 0 | 0 |
| NC_019843.3;YP_009047203.1;MERS_1a;1549 | 0 | 0.025 |
| NC_019843.3;YP_009047203.1;MERS_1a;1550 | 0 | 0 |
| NC_019843.3;YP_009047203.1;MERS_1a;1551 | 0 | 0 |
| NC_019843.3;YP_009047203.1;MERS_1a;1552 | 0 | 0 |
| NC_019843.3;YP_009047203.1;MERS_1a;1553 | 0 | 0 |
| NC_019843.3;YP_009047203.1;MERS_1a;1554 | 0 | 0.025 |
| NC_019843.3;YP_009047203.1;MERS_1a;1555 | 0 | 0 |
| NC_019843.3;YP_009047203.1;MERS_1a;1556 | 0 | 0 |
| NC_019843.3;YP_009047203.1;MERS_1a;1557 | 0 | 0 |
| NC_019843.3;YP_009047203.1;MERS_1a;1558 | 0 | 0 |
| NC_019843.3;YP_009047203.1;MERS_1a;1559 | 0 | 0 |
| NC_019843.3;YP_009047203.1;MERS_1a;1560 | 0 | 0 |
| NC_019843.3;YP_009047203.1;MERS_1a;1561 | 0 | 0 |
| NC_019843.3;YP_009047203.1;MERS_1a;1562 | 0 | 0 |
| NC_019843.3;YP_009047203.1;MERS_1a;1563 | 0 | 0 |
| NC_019843.3;YP_009047203.1;MERS_1a;1564 | 0 | 0 |
| NC_019843.3;YP_009047203.1;MERS_1a;1565 | 0 | 0 |
| NC_019843.3;YP_009047203.1;MERS_1a;1566 | 0 | 0 |
| NC_019843.3;YP_009047203.1;MERS_1a;1567 | 0 | 0.025 |

| | | |
|---|---|---|
| NC_019843.3;YP_009047203.1;MERS_1a;1568 | 0 | 0 |
| NC_019843.3;YP_009047203.1;MERS_1a;1569 | 0 | 0.025 |
| NC_019843.3;YP_009047203.1;MERS_1a;1570 | 0 | 0 |
| NC_019843.3;YP_009047203.1;MERS_1a;1571 | 0 | 0 |
| NC_019843.3;YP_009047203.1;MERS_1a;1572 | 0 | 0 |
| NC_019843.3;YP_009047203.1;MERS_1a;1573 | 0 | 0 |
| NC_019843.3;YP_009047203.1;MERS_1a;1574 | 0 | 0 |
| NC_019843.3;YP_009047203.1;MERS_1a;1575 | 0 | 0 |
| NC_019843.3;YP_009047203.1;MERS_1a;1576 | 0 | 0.075 |
| NC_019843.3;YP_009047203.1;MERS_1a;1577 | 0 | 0.025 |
| NC_019843.3;YP_009047203.1;MERS_1a;1578 | 0 | 0 |
| NC_019843.3;YP_009047203.1;MERS_1a;1579 | 0 | 0.025 |
| NC_019843.3;YP_009047203.1;MERS_1a;1580 | 0 | 0.025 |
| NC_019843.3;YP_009047203.1;MERS_1a;1581 | 0 | 0.05 |
| NC_019843.3;YP_009047203.1;MERS_1a;1582 | 0 | 0.025 |
| NC_019843.3;YP_009047203.1;MERS_1a;1583 | 0 | 0 |
| NC_019843.3;YP_009047203.1;MERS_1a;1584 | 0 | 0.025 |
| NC_019843.3;YP_009047203.1;MERS_1a;1585 | 0 | 0.025 |
| NC_019843.3;YP_009047203.1;MERS_1a;1586 | 0 | 0.025 |
| NC_019843.3;YP_009047203.1;MERS_1a;1587 | 0 | 0 |
| NC_019843.3;YP_009047203.1;MERS_1a;1588 | 0 | 0 |
| NC_019843.3;YP_009047203.1;MERS_1a;1589 | 0 | 0 |
| NC_019843.3;YP_009047203.1;MERS_1a;1590 | 0 | 0 |
| NC_019843.3;YP_009047203.1;MERS_1a;1591 | 0 | 0 |
| NC_019843.3;YP_009047203.1;MERS_1a;1592 | 0 | 0 |
| NC_019843.3;YP_009047203.1;MERS_1a;1593 | 0 | 0 |
| NC_019843.3;YP_009047203.1;MERS_1a;1594 | 0 | 0 |
| NC_019843.3;YP_009047203.1;MERS_1a;1595 | 0 | 0 |
| NC_019843.3;YP_009047203.1;MERS_1a;1596 | 0 | 0.025 |
| NC_019843.3;YP_009047203.1;MERS_1a;1597 | 0 | 0 |
| NC_019843.3;YP_009047203.1;MERS_1a;1598 | 0 | 0 |
| NC_019843.3;YP_009047203.1;MERS_1a;1599 | 0 | 0.15 |
| NC_019843.3;YP_009047203.1;MERS_1a;1600 | 0 | 0 |
| NC_019843.3;YP_009047203.1;MERS_1a;1601 | 0 | 0 |
| NC_019843.3;YP_009047203.1;MERS_1a;1602 | 0 | 0.025 |
| NC_019843.3;YP_009047203.1;MERS_1a;1603 | 0.05 | 0.025 |
| NC_019843.3;YP_009047203.1;MERS_1a;1604 | 0.15 | 0.025 |
| NC_019843.3;YP_009047203.1;MERS_1a;1605 | 0 | 0 |
| NC_019843.3;YP_009047203.1;MERS_1a;1606 | 0 | 0 |
| NC_019843.3;YP_009047203.1;MERS_1a;1607 | 0 | 0 |
| NC_019843.3;YP_009047203.1;MERS_1a;1608 | 0 | 0 |
| NC_019843.3;YP_009047203.1;MERS_1a;1609 | 0 | 0 |
| NC_019843.3;YP_009047203.1;MERS_1a;1610 | 0 | 0 |

| | | |
|---|---|---|
| NC_019843.3;YP_009047203.1;MERS_1a;1611 | 0 | 0 |
| NC_019843.3;YP_009047203.1;MERS_1a;1612 | 0 | 0.025 |
| NC_019843.3;YP_009047203.1;MERS_1a;1613 | 0.05 | 0.025 |
| NC_019843.3;YP_009047203.1;MERS_1a;1614 | 0.05 | 0.025 |
| NC_019843.3;YP_009047203.1;MERS_1a;1615 | 0.05 | 0.025 |
| NC_019843.3;YP_009047203.1;MERS_1a;1616 | 0.1 | 0.025 |
| NC_019843.3;YP_009047203.1;MERS_1a;1617 | 0.1 | 0.025 |
| NC_019843.3;YP_009047203.1;MERS_1a;1618 | 0 | 0.025 |
| NC_019843.3;YP_009047203.1;MERS_1a;1619 | 0 | 0 |
| NC_019843.3;YP_009047203.1;MERS_1a;1620 | 0 | 0 |
| NC_019843.3;YP_009047203.1;MERS_1a;1621 | 0 | 0 |
| NC_019843.3;YP_009047203.1;MERS_1a;1622 | 0 | 0 |
| NC_019843.3;YP_009047203.1;MERS_1a;1623 | 0 | 0 |
| NC_019843.3;YP_009047203.1;MERS_1a;1624 | 0 | 0 |
| NC_019843.3;YP_009047203.1;MERS_1a;1625 | 0 | 0 |
| NC_019843.3;YP_009047203.1;MERS_1a;1626 | 0 | 0 |
| NC_019843.3;YP_009047203.1;MERS_1a;1627 | 0 | 0 |
| NC_019843.3;YP_009047203.1;MERS_1a;1628 | 0 | 0 |
| NC_019843.3;YP_009047203.1;MERS_1a;1629 | 0 | 0 |
| NC_019843.3;YP_009047203.1;MERS_1a;1630 | 0 | 0 |
| NC_019843.3;YP_009047203.1;MERS_1a;1631 | 0 | 0 |
| NC_019843.3;YP_009047203.1;MERS_1a;1632 | 0 | 0.025 |
| NC_019843.3;YP_009047203.1;MERS_1a;1633 | 0.1 | 0 |
| NC_019843.3;YP_009047203.1;MERS_1a;1634 | 0.15 | 0 |
| NC_019843.3;YP_009047203.1;MERS_1a;1635 | 0.15 | 0.05 |
| NC_019843.3;YP_009047203.1;MERS_1a;1636 | 0.1 | 0 |
| NC_019843.3;YP_009047203.1;MERS_1a;1637 | 0.05 | 0 |
| NC_019843.3;YP_009047203.1;MERS_1a;1638 | 0.05 | 0 |
| NC_019843.3;YP_009047203.1;MERS_1a;1639 | 0.1 | 0 |
| NC_019843.3;YP_009047203.1;MERS_1a;1640 | 0.05 | 0 |
| NC_019843.3;YP_009047203.1;MERS_1a;1641 | 0.05 | 0 |
| NC_019843.3;YP_009047203.1;MERS_1a;1642 | 0 | 0 |
| NC_019843.3;YP_009047203.1;MERS_1a;1643 | 0 | 0 |
| NC_019843.3;YP_009047203.1;MERS_1a;1644 | 0 | 0.025 |
| NC_019843.3;YP_009047203.1;MERS_1a;1645 | 0.15 | 0.075 |
| NC_019843.3;YP_009047203.1;MERS_1a;1646 | 0 | 0 |
| NC_019843.3;YP_009047203.1;MERS_1a;1647 | 0 | 0 |
| NC_019843.3;YP_009047203.1;MERS_1a;1648 | 0 | 0 |
| NC_019843.3;YP_009047203.1;MERS_1a;1649 | 0 | 0 |
| NC_019843.3;YP_009047203.1;MERS_1a;1650 | 0 | 0 |
| NC_019843.3;YP_009047203.1;MERS_1a;1651 | 0 | 0 |
| NC_019843.3;YP_009047203.1;MERS_1a;1652 | 0 | 0 |
| NC_019843.3;YP_009047203.1;MERS_1a;1653 | 0 | 0 |

| | | |
|---|---|---|
| NC_019843.3;YP_009047203.1;MERS_1a;1654 | 0 | 0 |
| NC_019843.3;YP_009047203.1;MERS_1a;1655 | 0 | 0 |
| NC_019843.3;YP_009047203.1;MERS_1a;1656 | 0 | 0 |
| NC_019843.3;YP_009047203.1;MERS_1a;1657 | 0 | 0 |
| NC_019843.3;YP_009047203.1;MERS_1a;1658 | 0 | 0 |
| NC_019843.3;YP_009047203.1;MERS_1a;1659 | 0 | 0 |
| NC_019843.3;YP_009047203.1;MERS_1a;1660 | 0 | 0 |
| NC_019843.3;YP_009047203.1;MERS_1a;1661 | 0 | 0 |
| NC_019843.3;YP_009047203.1;MERS_1a;1662 | 0 | 0 |
| NC_019843.3;YP_009047203.1;MERS_1a;1663 | 0 | 0 |
| NC_019843.3;YP_009047203.1;MERS_1a;1664 | 0 | 0 |
| NC_019843.3;YP_009047203.1;MERS_1a;1665 | 0 | 0 |
| NC_019843.3;YP_009047203.1;MERS_1a;1666 | 0 | 0 |
| NC_019843.3;YP_009047203.1;MERS_1a;1667 | 0 | 0 |
| NC_019843.3;YP_009047203.1;MERS_1a;1668 | 0 | 0 |
| NC_019843.3;YP_009047203.1;MERS_1a;1669 | 0 | 0 |
| NC_019843.3;YP_009047203.1;MERS_1a;1670 | 0 | 0 |
| NC_019843.3;YP_009047203.1;MERS_1a;1671 | 0 | 0 |
| NC_019843.3;YP_009047203.1;MERS_1a;1672 | 0 | 0 |
| NC_019843.3;YP_009047203.1;MERS_1a;1673 | 0 | 0 |
| NC_019843.3;YP_009047203.1;MERS_1a;1674 | 0 | 0 |
| NC_019843.3;YP_009047203.1;MERS_1a;1675 | 0 | 0 |
| NC_019843.3;YP_009047203.1;MERS_1a;1676 | 0.15 | 0.125 |
| NC_019843.3;YP_009047203.1;MERS_1a;1677 | 0 | 0 |
| NC_019843.3;YP_009047203.1;MERS_1a;1678 | 0 | 0 |
| NC_019843.3;YP_009047203.1;MERS_1a;1679 | 0 | 0 |
| NC_019843.3;YP_009047203.1;MERS_1a;1680 | 0 | 0 |
| NC_019843.3;YP_009047203.1;MERS_1a;1681 | 0 | 0 |
| NC_019843.3;YP_009047203.1;MERS_1a;1682 | 0 | 0 |
| NC_019843.3;YP_009047203.1;MERS_1a;1683 | 0 | 0 |
| NC_019843.3;YP_009047203.1;MERS_1a;1684 | 0 | 0 |
| NC_019843.3;YP_009047203.1;MERS_1a;1685 | 0 | 0 |
| NC_019843.3;YP_009047203.1;MERS_1a;1686 | 0 | 0 |
| NC_019843.3;YP_009047203.1;MERS_1a;1687 | 0 | 0 |
| NC_019843.3;YP_009047203.1;MERS_1a;1688 | 0 | 0.025 |
| NC_019843.3;YP_009047203.1;MERS_1a;1689 | 0 | 0.025 |
| NC_019843.3;YP_009047203.1;MERS_1a;1690 | 0 | 0.025 |
| NC_019843.3;YP_009047203.1;MERS_1a;1691 | 0 | 0 |
| NC_019843.3;YP_009047203.1;MERS_1a;1692 | 0 | 0.025 |
| NC_019843.3;YP_009047203.1;MERS_1a;1693 | 0 | 0.175 |
| NC_019843.3;YP_009047203.1;MERS_1a;1694 | 0 | 0.225 |
| NC_019843.3;YP_009047203.1;MERS_1a;1695 | 0 | 0.025 |
| NC_019843.3;YP_009047203.1;MERS_1a;1696 | 0 | 0 |

| | | |
|---|---|---|
| NC_019843.3;YP_009047203.1;MERS_1a;1697 | 0 | 0.025 |
| NC_019843.3;YP_009047203.1;MERS_1a;1698 | 0 | 0 |
| NC_019843.3;YP_009047203.1;MERS_1a;1699 | 0 | 0 |
| NC_019843.3;YP_009047203.1;MERS_1a;1700 | 0 | 0 |
| NC_019843.3;YP_009047203.1;MERS_1a;1701 | 0 | 0 |
| NC_019843.3;YP_009047203.1;MERS_1a;1702 | 0 | 0 |
| NC_019843.3;YP_009047203.1;MERS_1a;1703 | 0 | 0 |
| NC_019843.3;YP_009047203.1;MERS_1a;1704 | 0 | 0 |
| NC_019843.3;YP_009047203.1;MERS_1a;1705 | 0 | 0 |
| NC_019843.3;YP_009047203.1;MERS_1a;1706 | 0 | 0 |
| NC_019843.3;YP_009047203.1;MERS_1a;1707 | 0 | 0 |
| NC_019843.3;YP_009047203.1;MERS_1a;1708 | 0 | 0 |
| NC_019843.3;YP_009047203.1;MERS_1a;1709 | 0 | 0 |
| NC_019843.3;YP_009047203.1;MERS_1a;1710 | 0 | 0.025 |
| NC_019843.3;YP_009047203.1;MERS_1a;1711 | 0 | 0 |
| NC_019843.3;YP_009047203.1;MERS_1a;1712 | 0 | 0 |
| NC_019843.3;YP_009047203.1;MERS_1a;1713 | 0 | 0 |
| NC_019843.3;YP_009047203.1;MERS_1a;1714 | 0 | 0 |
| NC_019843.3;YP_009047203.1;MERS_1a;1715 | 0 | 0 |
| NC_019843.3;YP_009047203.1;MERS_1a;1716 | 0 | 0 |
| NC_019843.3;YP_009047203.1;MERS_1a;1717 | 0 | 0 |
| NC_019843.3;YP_009047203.1;MERS_1a;1718 | 0 | 0 |
| NC_019843.3;YP_009047203.1;MERS_1a;1719 | 0 | 0 |
| NC_019843.3;YP_009047203.1;MERS_1a;1720 | 0 | 0 |
| NC_019843.3;YP_009047203.1;MERS_1a;1721 | 0 | 0 |
| NC_019843.3;YP_009047203.1;MERS_1a;1722 | 0 | 0 |
| NC_019843.3;YP_009047203.1;MERS_1a;1723 | 0 | 0 |
| NC_019843.3;YP_009047203.1;MERS_1a;1724 | 0 | 0 |
| NC_019843.3;YP_009047203.1;MERS_1a;1725 | 0 | 0 |
| NC_019843.3;YP_009047203.1;MERS_1a;1726 | 0 | 0 |
| NC_019843.3;YP_009047203.1;MERS_1a;1727 | 0 | 0 |
| NC_019843.3;YP_009047203.1;MERS_1a;1728 | 0 | 0 |
| NC_019843.3;YP_009047203.1;MERS_1a;1729 | 0 | 0 |
| NC_019843.3;YP_009047203.1;MERS_1a;1730 | 0 | 0 |
| NC_019843.3;YP_009047203.1;MERS_1a;1731 | 0 | 0 |
| NC_019843.3;YP_009047203.1;MERS_1a;1732 | 0 | 0 |
| NC_019843.3;YP_009047203.1;MERS_1a;1733 | 0 | 0 |
| NC_019843.3;YP_009047203.1;MERS_1a;1734 | 0 | 0 |
| NC_019843.3;YP_009047203.1;MERS_1a;1735 | 0 | 0 |
| NC_019843.3;YP_009047203.1;MERS_1a;1736 | 0 | 0 |
| NC_019843.3;YP_009047203.1;MERS_1a;1737 | 0 | 0 |
| NC_019843.3;YP_009047203.1;MERS_1a;1738 | 0 | 0 |
| NC_019843.3;YP_009047203.1;MERS_1a;1739 | 0 | 0 |

| | | |
|---|---|---|
| NC_019843.3;YP_009047203.1;MERS_1a;1740 | 0 | 0 |
| NC_019843.3;YP_009047203.1;MERS_1a;1741 | 0 | 0 |
| NC_019843.3;YP_009047203.1;MERS_1a;1742 | 0 | 0 |
| NC_019843.3;YP_009047203.1;MERS_1a;1743 | 0 | 0 |
| NC_019843.3;YP_009047203.1;MERS_1a;1744 | 0 | 0 |
| NC_019843.3;YP_009047203.1;MERS_1a;1745 | 0 | 0 |
| NC_019843.3;YP_009047203.1;MERS_1a;1746 | 0 | 0 |
| NC_019843.3;YP_009047203.1;MERS_1a;1747 | 0 | 0 |
| NC_019843.3;YP_009047203.1;MERS_1a;1748 | 0 | 0 |
| NC_019843.3;YP_009047203.1;MERS_1a;1749 | 0 | 0 |
| NC_019843.3;YP_009047203.1;MERS_1a;1750 | 0 | 0 |
| NC_019843.3;YP_009047203.1;MERS_1a;1751 | 0 | 0 |
| NC_019843.3;YP_009047203.1;MERS_1a;1752 | 0.1 | 0.025 |
| NC_019843.3;YP_009047203.1;MERS_1a;1753 | 0 | 0 |
| NC_019843.3;YP_009047203.1;MERS_1a;1754 | 0 | 0 |
| NC_019843.3;YP_009047203.1;MERS_1a;1755 | 0 | 0 |
| NC_019843.3;YP_009047203.1;MERS_1a;1756 | 0 | 0 |
| NC_019843.3;YP_009047203.1;MERS_1a;1757 | 0 | 0 |
| NC_019843.3;YP_009047203.1;MERS_1a;1758 | 0 | 0 |
| NC_019843.3;YP_009047203.1;MERS_1a;1759 | 0 | 0.025 |
| NC_019843.3;YP_009047203.1;MERS_1a;1760 | 0 | 0.025 |
| NC_019843.3;YP_009047203.1;MERS_1a;1761 | 0 | 0.025 |
| NC_019843.3;YP_009047203.1;MERS_1a;1762 | 0 | 0.025 |
| NC_019843.3;YP_009047203.1;MERS_1a;1763 | 0 | 0.025 |
| NC_019843.3;YP_009047203.1;MERS_1a;1764 | 0 | 0.025 |
| NC_019843.3;YP_009047203.1;MERS_1a;1765 | 0 | 0.025 |
| NC_019843.3;YP_009047203.1;MERS_1a;1766 | 0 | 0.05 |
| NC_019843.3;YP_009047203.1;MERS_1a;1767 | 0 | 0 |
| NC_019843.3;YP_009047203.1;MERS_1a;1768 | 0 | 0 |
| NC_019843.3;YP_009047203.1;MERS_1a;1769 | 0 | 0 |
| NC_019843.3;YP_009047203.1;MERS_1a;1770 | 0 | 0.025 |
| NC_019843.3;YP_009047203.1;MERS_1a;1771 | 0 | 0.025 |
| NC_019843.3;YP_009047203.1;MERS_1a;1772 | 0 | 0.025 |
| NC_019843.3;YP_009047203.1;MERS_1a;1773 | 0.05 | 0.025 |
| NC_019843.3;YP_009047203.1;MERS_1a;1774 | 0.05 | 0.025 |
| NC_019843.3;YP_009047203.1;MERS_1a;1775 | 0 | 0.025 |
| NC_019843.3;YP_009047203.1;MERS_1a;1776 | 0 | 0.025 |
| NC_019843.3;YP_009047203.1;MERS_1a;1777 | 0 | 0.025 |
| NC_019843.3;YP_009047203.1;MERS_1a;1778 | 0 | 0.025 |
| NC_019843.3;YP_009047203.1;MERS_1a;1779 | 0 | 0.025 |
| NC_019843.3;YP_009047203.1;MERS_1a;1780 | 0 | 0 |
| NC_019843.3;YP_009047203.1;MERS_1a;1781 | 0 | 0 |
| NC_019843.3;YP_009047203.1;MERS_1a;1782 | 0 | 0 |

| | | |
|---|---|---|
| NC_019843.3;YP_009047203.1;MERS_1a;1783 | 0 | 0 |
| NC_019843.3;YP_009047203.1;MERS_1a;1784 | 0 | 0 |
| NC_019843.3;YP_009047203.1;MERS_1a;1785 | 0 | 0.025 |
| NC_019843.3;YP_009047203.1;MERS_1a;1786 | 0 | 0.05 |
| NC_019843.3;YP_009047203.1;MERS_1a;1787 | 0.05 | 0.025 |
| NC_019843.3;YP_009047203.1;MERS_1a;1788 | 0 | 0.025 |
| NC_019843.3;YP_009047203.1;MERS_1a;1789 | 0.05 | 0.025 |
| NC_019843.3;YP_009047203.1;MERS_1a;1790 | 0 | 0 |
| NC_019843.3;YP_009047203.1;MERS_1a;1791 | 0 | 0 |
| NC_019843.3;YP_009047203.1;MERS_1a;1792 | 0 | 0 |
| NC_019843.3;YP_009047203.1;MERS_1a;1793 | 0 | 0 |
| NC_019843.3;YP_009047203.1;MERS_1a;1794 | 0 | 0 |
| NC_019843.3;YP_009047203.1;MERS_1a;1795 | 0 | 0 |
| NC_019843.3;YP_009047203.1;MERS_1a;1796 | 0 | 0 |
| NC_019843.3;YP_009047203.1;MERS_1a;1797 | 0 | 0 |
| NC_019843.3;YP_009047203.1;MERS_1a;1798 | 0 | 0 |
| NC_019843.3;YP_009047203.1;MERS_1a;1799 | 0 | 0 |
| NC_019843.3;YP_009047203.1;MERS_1a;1800 | 0 | 0 |
| NC_019843.3;YP_009047203.1;MERS_1a;1801 | 0 | 0 |
| NC_019843.3;YP_009047203.1;MERS_1a;1802 | 0 | 0 |
| NC_019843.3;YP_009047203.1;MERS_1a;1803 | 0.2 | 0.025 |
| NC_019843.3;YP_009047203.1;MERS_1a;1804 | 0 | 0 |
| NC_019843.3;YP_009047203.1;MERS_1a;1805 | 0.05 | 0 |
| NC_019843.3;YP_009047203.1;MERS_1a;1806 | 0.1 | 0 |
| NC_019843.3;YP_009047203.1;MERS_1a;1807 | 0.1 | 0 |
| NC_019843.3;YP_009047203.1;MERS_1a;1808 | 0.05 | 0 |
| NC_019843.3;YP_009047203.1;MERS_1a;1809 | 0.05 | 0 |
| NC_019843.3;YP_009047203.1;MERS_1a;1810 | 0.05 | 0 |
| NC_019843.3;YP_009047203.1;MERS_1a;1811 | 0 | 0 |
| NC_019843.3;YP_009047203.1;MERS_1a;1812 | 0 | 0 |
| NC_019843.3;YP_009047203.1;MERS_1a;1813 | 0 | 0 |
| NC_019843.3;YP_009047203.1;MERS_1a;1814 | 0 | 0 |
| NC_019843.3;YP_009047203.1;MERS_1a;1815 | 0.05 | 0 |
| NC_019843.3;YP_009047203.1;MERS_1a;1816 | 0.1 | 0.025 |
| NC_019843.3;YP_009047203.1;MERS_1a;1817 | 0.15 | 0.05 |
| NC_019843.3;YP_009047203.1;MERS_1a;1818 | 0.2 | 0.025 |
| NC_019843.3;YP_009047203.1;MERS_1a;1819 | 0.15 | 0 |
| NC_019843.3;YP_009047203.1;MERS_1a;1820 | 0.15 | 0.025 |
| NC_019843.3;YP_009047203.1;MERS_1a;1821 | 0.2 | 0 |
| NC_019843.3;YP_009047203.1;MERS_1a;1822 | 0.2 | 0.075 |
| NC_019843.3;YP_009047203.1;MERS_1a;1823 | 0.3 | 0.1 |
| NC_019843.3;YP_009047203.1;MERS_1a;1824 | 0.25 | 0.05 |
| NC_019843.3;YP_009047203.1;MERS_1a;1825 | 0.25 | 0.05 |

| | | |
|---|---|---|
| NC_019843.3;YP_009047203.1;MERS_1a;1826 | 0 | 0.025 |
| NC_019843.3;YP_009047203.1;MERS_1a;1827 | 0 | 0.025 |
| NC_019843.3;YP_009047203.1;MERS_1a;1828 | 0.05 | 0 |
| NC_019843.3;YP_009047203.1;MERS_1a;1829 | 0 | 0 |
| NC_019843.3;YP_009047203.1;MERS_1a;1830 | 0 | 0 |
| NC_019843.3;YP_009047203.1;MERS_1a;1831 | 0 | 0 |
| NC_019843.3;YP_009047203.1;MERS_1a;1832 | 0 | 0.025 |
| NC_019843.3;YP_009047203.1;MERS_1a;1833 | 0 | 0 |
| NC_019843.3;YP_009047203.1;MERS_1a;1834 | 0 | 0 |
| NC_019843.3;YP_009047203.1;MERS_1a;1835 | 0 | 0 |
| NC_019843.3;YP_009047203.1;MERS_1a;1836 | 0 | 0 |
| NC_019843.3;YP_009047203.1;MERS_1a;1837 | 0 | 0 |
| NC_019843.3;YP_009047203.1;MERS_1a;1838 | 0 | 0 |
| NC_019843.3;YP_009047203.1;MERS_1a;1839 | 0 | 0 |
| NC_019843.3;YP_009047203.1;MERS_1a;1840 | 0 | 0 |
| NC_019843.3;YP_009047203.1;MERS_1a;1841 | 0 | 0 |
| NC_019843.3;YP_009047203.1;MERS_1a;1842 | 0 | 0 |
| NC_019843.3;YP_009047203.1;MERS_1a;1843 | 0 | 0 |
| NC_019843.3;YP_009047203.1;MERS_1a;1844 | 0 | 0 |
| NC_019843.3;YP_009047203.1;MERS_1a;1845 | 0 | 0 |
| NC_019843.3;YP_009047203.1;MERS_1a;1846 | 0 | 0 |
| NC_019843.3;YP_009047203.1;MERS_1a;1847 | 0 | 0 |
| NC_019843.3;YP_009047203.1;MERS_1a;1848 | 0 | 0.025 |
| NC_019843.3;YP_009047203.1;MERS_1a;1849 | 0 | 0.025 |
| NC_019843.3;YP_009047203.1;MERS_1a;1850 | 0.1 | 0.025 |
| NC_019843.3;YP_009047203.1;MERS_1a;1851 | 0 | 0.025 |
| NC_019843.3;YP_009047203.1;MERS_1a;1852 | 0 | 0.025 |
| NC_019843.3;YP_009047203.1;MERS_1a;1853 | 0 | 0.025 |
| NC_019843.3;YP_009047203.1;MERS_1a;1854 | 0 | 0.025 |
| NC_019843.3;YP_009047203.1;MERS_1a;1855 | 0 | 0.025 |
| NC_019843.3;YP_009047203.1;MERS_1a;1856 | 0 | 0.025 |
| NC_019843.3;YP_009047203.1;MERS_1a;1857 | 0 | 0 |
| NC_019843.3;YP_009047203.1;MERS_1a;1858 | 0.05 | 0.05 |
| NC_019843.3;YP_009047203.1;MERS_1a;1859 | 0 | 0 |
| NC_019843.3;YP_009047203.1;MERS_1a;1860 | 0 | 0 |
| NC_019843.3;YP_009047203.1;MERS_1a;1861 | 0 | 0 |
| NC_019843.3;YP_009047203.1;MERS_1a;1862 | 0 | 0 |
| NC_019843.3;YP_009047203.1;MERS_1a;1863 | 0.05 | 0.025 |
| NC_019843.3;YP_009047203.1;MERS_1a;1864 | 0 | 0.025 |
| NC_019843.3;YP_009047203.1;MERS_1a;1865 | 0 | 0 |
| NC_019843.3;YP_009047203.1;MERS_1a;1866 | 0 | 0 |
| NC_019843.3;YP_009047203.1;MERS_1a;1867 | 0 | 0.025 |
| NC_019843.3;YP_009047203.1;MERS_1a;1868 | 0.05 | 0.025 |

| | | |
|---|---|---|
| NC_019843.3;YP_009047203.1;MERS_1a;1869 | 0 | 0.025 |
| NC_019843.3;YP_009047203.1;MERS_1a;1870 | 0 | 0.05 |
| NC_019843.3;YP_009047203.1;MERS_1a;1871 | 0.05 | 0.075 |
| NC_019843.3;YP_009047203.1;MERS_1a;1872 | 0 | 0.025 |
| NC_019843.3;YP_009047203.1;MERS_1a;1873 | 0.05 | 0.025 |
| NC_019843.3;YP_009047203.1;MERS_1a;1874 | 0.05 | 0.025 |
| NC_019843.3;YP_009047203.1;MERS_1a;1875 | 0 | 0.025 |
| NC_019843.3;YP_009047203.1;MERS_1a;1876 | 0 | 0.025 |
| NC_019843.3;YP_009047203.1;MERS_1a;1877 | 0 | 0.075 |
| NC_019843.3;YP_009047203.1;MERS_1a;1878 | 0 | 0 |
| NC_019843.3;YP_009047203.1;MERS_1a;1879 | 0 | 0 |
| NC_019843.3;YP_009047203.1;MERS_1a;1880 | 0 | 0 |
| NC_019843.3;YP_009047203.1;MERS_1a;1881 | 0 | 0 |
| NC_019843.3;YP_009047203.1;MERS_1a;1882 | 0 | 0 |
| NC_019843.3;YP_009047203.1;MERS_1a;1883 | 0 | 0.025 |
| NC_019843.3;YP_009047203.1;MERS_1a;1884 | 0 | 0.025 |
| NC_019843.3;YP_009047203.1;MERS_1a;1885 | 0 | 0.025 |
| NC_019843.3;YP_009047203.1;MERS_1a;1886 | 0 | 0.05 |
| NC_019843.3;YP_009047203.1;MERS_1a;1887 | 0 | 0 |
| NC_019843.3;YP_009047203.1;MERS_1a;1888 | 0.05 | 0.025 |
| NC_019843.3;YP_009047203.1;MERS_1a;1889 | 0.05 | 0.05 |
| NC_019843.3;YP_009047203.1;MERS_1a;1890 | 0 | 0.025 |
| NC_019843.3;YP_009047203.1;MERS_1a;1891 | 0 | 0.025 |
| NC_019843.3;YP_009047203.1;MERS_1a;1892 | 0 | 0.025 |
| NC_019843.3;YP_009047203.1;MERS_1a;1893 | 0 | 0.075 |
| NC_019843.3;YP_009047203.1;MERS_1a;1894 | 0 | 0.025 |
| NC_019843.3;YP_009047203.1;MERS_1a;1895 | 0.05 | 0.05 |
| NC_019843.3;YP_009047203.1;MERS_1a;1896 | 0 | 0.025 |
| NC_019843.3;YP_009047203.1;MERS_1a;1897 | 0 | 0.025 |
| NC_019843.3;YP_009047203.1;MERS_1a;1898 | 0 | 0.075 |
| NC_019843.3;YP_009047203.1;MERS_1a;1899 | 0 | 0 |
| NC_019843.3;YP_009047203.1;MERS_1a;1900 | 0 | 0 |
| NC_019843.3;YP_009047203.1;MERS_1a;1901 | 0.05 | 0 |
| NC_019843.3;YP_009047203.1;MERS_1a;1902 | 0 | 0.025 |
| NC_019843.3;YP_009047203.1;MERS_1a;1903 | 0 | 0.05 |
| NC_019843.3;YP_009047203.1;MERS_1a;1904 | 0 | 0.025 |
| NC_019843.3;YP_009047203.1;MERS_1a;1905 | 0.05 | 0 |
| NC_019843.3;YP_009047203.1;MERS_1a;1906 | 0 | 0 |
| NC_019843.3;YP_009047203.1;MERS_1a;1907 | 0 | 0 |
| NC_019843.3;YP_009047203.1;MERS_1a;1908 | 0 | 0 |
| NC_019843.3;YP_009047203.1;MERS_1a;1909 | 0 | 0 |
| NC_019843.3;YP_009047203.1;MERS_1a;1910 | 0 | 0.025 |
| NC_019843.3;YP_009047203.1;MERS_1a;1911 | 0 | 0 |

| | | |
|---|---|---|
| NC_019843.3;YP_009047203.1;MERS_1a;1912 | 0 | 0 |
| NC_019843.3;YP_009047203.1;MERS_1a;1913 | 0 | 0 |
| NC_019843.3;YP_009047203.1;MERS_1a;1914 | 0 | 0 |
| NC_019843.3;YP_009047203.1;MERS_1a;1915 | 0 | 0 |
| NC_019843.3;YP_009047203.1;MERS_1a;1916 | 0 | 0 |
| NC_019843.3;YP_009047203.1;MERS_1a;1917 | 0 | 0 |
| NC_019843.3;YP_009047203.1;MERS_1a;1918 | 0 | 0 |
| NC_019843.3;YP_009047203.1;MERS_1a;1919 | 0 | 0.025 |
| NC_019843.3;YP_009047203.1;MERS_1a;1920 | 0.1 | 0.05 |
| NC_019843.3;YP_009047203.1;MERS_1a;1921 | 0.1 | 0.1 |
| NC_019843.3;YP_009047203.1;MERS_1a;1922 | 0.05 | 0.05 |
| NC_019843.3;YP_009047203.1;MERS_1a;1923 | 0 | 0.025 |
| NC_019843.3;YP_009047203.1;MERS_1a;1924 | 0.1 | 0.025 |
| NC_019843.3;YP_009047203.1;MERS_1a;1925 | 0.1 | 0.05 |
| NC_019843.3;YP_009047203.1;MERS_1a;1926 | 0.15 | 0.05 |
| NC_019843.3;YP_009047203.1;MERS_1a;1927 | 0.15 | 0.075 |
| NC_019843.3;YP_009047203.1;MERS_1a;1928 | 0.1 | 0.1 |
| NC_019843.3;YP_009047203.1;MERS_1a;1929 | 0.05 | 0 |
| NC_019843.3;YP_009047203.1;MERS_1a;1930 | 0.1 | 0 |
| NC_019843.3;YP_009047203.1;MERS_1a;1931 | 0 | 0 |
| NC_019843.3;YP_009047203.1;MERS_1a;1932 | 0.05 | 0 |
| NC_019843.3;YP_009047203.1;MERS_1a;1933 | 0 | 0 |
| NC_019843.3;YP_009047203.1;MERS_1a;1934 | 0 | 0 |
| NC_019843.3;YP_009047203.1;MERS_1a;1935 | 0 | 0.025 |
| NC_019843.3;YP_009047203.1;MERS_1a;1936 | 0 | 0 |
| NC_019843.3;YP_009047203.1;MERS_1a;1937 | 0 | 0 |
| NC_019843.3;YP_009047203.1;MERS_1a;1938 | 0.05 | 0 |
| NC_019843.3;YP_009047203.1;MERS_1a;1939 | 0 | 0 |
| NC_019843.3;YP_009047203.1;MERS_1a;1940 | 0 | 0.025 |
| NC_019843.3;YP_009047203.1;MERS_1a;1941 | 0.05 | 0.025 |
| NC_019843.3;YP_009047203.1;MERS_1a;1942 | 0.1 | 0.075 |
| NC_019843.3;YP_009047203.1;MERS_1a;1943 | 0.1 | 0.15 |
| NC_019843.3;YP_009047203.1;MERS_1a;1944 | 0.3 | 0.175 |
| NC_019843.3;YP_009047203.1;MERS_1a;1945 | 0.1 | 0.1 |
| NC_019843.3;YP_009047203.1;MERS_1a;1946 | 0.05 | 0.05 |
| NC_019843.3;YP_009047203.1;MERS_1a;1947 | 0.2 | 0.075 |
| NC_019843.3;YP_009047203.1;MERS_1a;1948 | 0.15 | 0 |
| NC_019843.3;YP_009047203.1;MERS_1a;1949 | 0.15 | 0.05 |
| NC_019843.3;YP_009047203.1;MERS_1a;1950 | 0 | 0.125 |
| NC_019843.3;YP_009047203.1;MERS_1a;1951 | 0 | 0 |
| NC_019843.3;YP_009047203.1;MERS_1a;1952 | 0 | 0 |
| NC_019843.3;YP_009047203.1;MERS_1a;1953 | 0 | 0 |
| NC_019843.3;YP_009047203.1;MERS_1a;1954 | 0 | 0 |

| | | |
|---|---|---|
| NC_019843.3;YP_009047203.1;MERS_1a;1955 | 0.05 | 0 |
| NC_019843.3;YP_009047203.1;MERS_1a;1956 | 0 | 0 |
| NC_019843.3;YP_009047203.1;MERS_1a;1957 | 0 | 0 |
| NC_019843.3;YP_009047203.1;MERS_1a;1958 | 0 | 0 |
| NC_019843.3;YP_009047203.1;MERS_1a;1959 | 0 | 0 |
| NC_019843.3;YP_009047203.1;MERS_1a;1960 | 0 | 0 |
| NC_019843.3;YP_009047203.1;MERS_1a;1961 | 0 | 0 |
| NC_019843.3;YP_009047203.1;MERS_1a;1962 | 0 | 0 |
| NC_019843.3;YP_009047203.1;MERS_1a;1963 | 0 | 0 |
| NC_019843.3;YP_009047203.1;MERS_1a;1964 | 0.05 | 0 |
| NC_019843.3;YP_009047203.1;MERS_1a;1965 | 0.05 | 0 |
| NC_019843.3;YP_009047203.1;MERS_1a;1966 | 0.05 | 0 |
| NC_019843.3;YP_009047203.1;MERS_1a;1967 | 0.05 | 0 |
| NC_019843.3;YP_009047203.1;MERS_1a;1968 | 0.05 | 0 |
| NC_019843.3;YP_009047203.1;MERS_1a;1969 | 0.05 | 0 |
| NC_019843.3;YP_009047203.1;MERS_1a;1970 | 0.05 | 0 |
| NC_019843.3;YP_009047203.1;MERS_1a;1971 | 0.05 | 0 |
| NC_019843.3;YP_009047203.1;MERS_1a;1972 | 0.05 | 0 |
| NC_019843.3;YP_009047203.1;MERS_1a;1973 | 0.05 | 0.025 |
| NC_019843.3;YP_009047203.1;MERS_1a;1974 | 0 | 0 |
| NC_019843.3;YP_009047203.1;MERS_1a;1975 | 0 | 0 |
| NC_019843.3;YP_009047203.1;MERS_1a;1976 | 0 | 0 |
| NC_019843.3;YP_009047203.1;MERS_1a;1977 | 0 | 0 |
| NC_019843.3;YP_009047203.1;MERS_1a;1978 | 0 | 0.025 |
| NC_019843.3;YP_009047203.1;MERS_1a;1979 | 0.05 | 0 |
| NC_019843.3;YP_009047203.1;MERS_1a;1980 | 0.05 | 0 |
| NC_019843.3;YP_009047203.1;MERS_1a;1981 | 0 | 0 |
| NC_019843.3;YP_009047203.1;MERS_1a;1982 | 0.05 | 0.025 |
| NC_019843.3;YP_009047203.1;MERS_1a;1983 | 0.05 | 0.025 |
| NC_019843.3;YP_009047203.1;MERS_1a;1984 | 0.1 | 0.025 |
| NC_019843.3;YP_009047203.1;MERS_1a;1985 | 0.1 | 0.025 |
| NC_019843.3;YP_009047203.1;MERS_1a;1986 | 0.1 | 0.05 |
| NC_019843.3;YP_009047203.1;MERS_1a;1987 | 0.1 | 0 |
| NC_019843.3;YP_009047203.1;MERS_1a;1988 | 0.05 | 0 |
| NC_019843.3;YP_009047203.1;MERS_1a;1989 | 0 | 0.025 |
| NC_019843.3;YP_009047203.1;MERS_1a;1990 | 0 | 0 |
| NC_019843.3;YP_009047203.1;MERS_1a;1991 | 0 | 0 |
| NC_019843.3;YP_009047203.1;MERS_1a;1992 | 0.05 | 0 |
| NC_019843.3;YP_009047203.1;MERS_1a;1993 | 0 | 0 |
| NC_019843.3;YP_009047203.1;MERS_1a;1994 | 0 | 0 |
| NC_019843.3;YP_009047203.1;MERS_1a;1995 | 0 | 0.025 |
| NC_019843.3;YP_009047203.1;MERS_1a;1996 | 0 | 0 |
| NC_019843.3;YP_009047203.1;MERS_1a;1997 | 0 | 0 |

| | | |
|---|---|---|
| NC_019843.3;YP_009047203.1;MERS_1a;1998 | 0 | 0 |
| NC_019843.3;YP_009047203.1;MERS_1a;1999 | 0 | 0 |
| NC_019843.3;YP_009047203.1;MERS_1a;2000 | 0 | 0 |
| NC_019843.3;YP_009047203.1;MERS_1a;2001 | 0.05 | 0 |
| NC_019843.3;YP_009047203.1;MERS_1a;2002 | 0.05 | 0 |
| NC_019843.3;YP_009047203.1;MERS_1a;2003 | 0.05 | 0 |
| NC_019843.3;YP_009047203.1;MERS_1a;2004 | 0.05 | 0 |
| NC_019843.3;YP_009047203.1;MERS_1a;2005 | 0.05 | 0 |
| NC_019843.3;YP_009047203.1;MERS_1a;2006 | 0.05 | 0.025 |
| NC_019843.3;YP_009047203.1;MERS_1a;2007 | 0.05 | 0.025 |
| NC_019843.3;YP_009047203.1;MERS_1a;2008 | 0.05 | 0.025 |
| NC_019843.3;YP_009047203.1;MERS_1a;2009 | 0 | 0.025 |
| NC_019843.3;YP_009047203.1;MERS_1a;2010 | 0 | 0.025 |
| NC_019843.3;YP_009047203.1;MERS_1a;2011 | 0 | 0 |
| NC_019843.3;YP_009047203.1;MERS_1a;2012 | 0 | 0 |
| NC_019843.3;YP_009047203.1;MERS_1a;2013 | 0.05 | 0 |
| NC_019843.3;YP_009047203.1;MERS_1a;2014 | 0 | 0 |
| NC_019843.3;YP_009047203.1;MERS_1a;2015 | 0.05 | 0 |
| NC_019843.3;YP_009047203.1;MERS_1a;2016 | 0 | 0 |
| NC_019843.3;YP_009047203.1;MERS_1a;2017 | 0 | 0 |
| NC_019843.3;YP_009047203.1;MERS_1a;2018 | 0 | 0 |
| NC_019843.3;YP_009047203.1;MERS_1a;2019 | 0 | 0 |
| NC_019843.3;YP_009047203.1;MERS_1a;2020 | 0 | 0 |
| NC_019843.3;YP_009047203.1;MERS_1a;2021 | 0 | 0 |
| NC_019843.3;YP_009047203.1;MERS_1a;2022 | 0 | 0 |
| NC_019843.3;YP_009047203.1;MERS_1a;2023 | 0 | 0 |
| NC_019843.3;YP_009047203.1;MERS_1a;2024 | 0 | 0.025 |
| NC_019843.3;YP_009047203.1;MERS_1a;2025 | 0 | 0 |
| NC_019843.3;YP_009047203.1;MERS_1a;2026 | 0 | 0 |
| NC_019843.3;YP_009047203.1;MERS_1a;2027 | 0 | 0 |
| NC_019843.3;YP_009047203.1;MERS_1a;2028 | 0 | 0 |
| NC_019843.3;YP_009047203.1;MERS_1a;2029 | 0 | 0 |
| NC_019843.3;YP_009047203.1;MERS_1a;2030 | 0 | 0 |
| NC_019843.3;YP_009047203.1;MERS_1a;2031 | 0 | 0 |
| NC_019843.3;YP_009047203.1;MERS_1a;2032 | 0 | 0 |
| NC_019843.3;YP_009047203.1;MERS_1a;2033 | 0 | 0 |
| NC_019843.3;YP_009047203.1;MERS_1a;2034 | 0 | 0 |
| NC_019843.3;YP_009047203.1;MERS_1a;2035 | 0 | 0 |
| NC_019843.3;YP_009047203.1;MERS_1a;2036 | 0 | 0 |
| NC_019843.3;YP_009047203.1;MERS_1a;2037 | 0 | 0 |
| NC_019843.3;YP_009047203.1;MERS_1a;2038 | 0 | 0 |
| NC_019843.3;YP_009047203.1;MERS_1a;2039 | 0 | 0 |
| NC_019843.3;YP_009047203.1;MERS_1a;2040 | 0 | 0 |

| | | |
|---|---|---|
| NC_019843.3;YP_009047203.1;MERS_1a;2041 | 0 | 0.025 |
| NC_019843.3;YP_009047203.1;MERS_1a;2042 | 0.05 | 0.025 |
| NC_019843.3;YP_009047203.1;MERS_1a;2043 | 0 | 0 |
| NC_019843.3;YP_009047203.1;MERS_1a;2044 | 0 | 0 |
| NC_019843.3;YP_009047203.1;MERS_1a;2045 | 0 | 0 |
| NC_019843.3;YP_009047203.1;MERS_1a;2046 | 0 | 0 |
| NC_019843.3;YP_009047203.1;MERS_1a;2047 | 0 | 0 |
| NC_019843.3;YP_009047203.1;MERS_1a;2048 | 0 | 0 |
| NC_019843.3;YP_009047203.1;MERS_1a;2049 | 0 | 0 |
| NC_019843.3;YP_009047203.1;MERS_1a;2050 | 0 | 0 |
| NC_019843.3;YP_009047203.1;MERS_1a;2051 | 0 | 0 |
| NC_019843.3;YP_009047203.1;MERS_1a;2052 | 0 | 0 |
| NC_019843.3;YP_009047203.1;MERS_1a;2053 | 0 | 0 |
| NC_019843.3;YP_009047203.1;MERS_1a;2054 | 0 | 0 |
| NC_019843.3;YP_009047203.1;MERS_1a;2055 | 0 | 0 |
| NC_019843.3;YP_009047203.1;MERS_1a;2056 | 0 | 0 |
| NC_019843.3;YP_009047203.1;MERS_1a;2057 | 0 | 0 |
| NC_019843.3;YP_009047203.1;MERS_1a;2058 | 0 | 0 |
| NC_019843.3;YP_009047203.1;MERS_1a;2059 | 0 | 0 |
| NC_019843.3;YP_009047203.1;MERS_1a;2060 | 0 | 0 |
| NC_019843.3;YP_009047203.1;MERS_1a;2061 | 0 | 0 |
| NC_019843.3;YP_009047203.1;MERS_1a;2062 | 0 | 0 |
| NC_019843.3;YP_009047203.1;MERS_1a;2063 | 0 | 0 |
| NC_019843.3;YP_009047203.1;MERS_1a;2064 | 0 | 0 |
| NC_019843.3;YP_009047203.1;MERS_1a;2065 | 0 | 0 |
| NC_019843.3;YP_009047203.1;MERS_1a;2066 | 0 | 0 |
| NC_019843.3;YP_009047203.1;MERS_1a;2067 | 0 | 0 |
| NC_019843.3;YP_009047203.1;MERS_1a;2068 | 0 | 0 |
| NC_019843.3;YP_009047203.1;MERS_1a;2069 | 0 | 0 |
| NC_019843.3;YP_009047203.1;MERS_1a;2070 | 0 | 0 |
| NC_019843.3;YP_009047203.1;MERS_1a;2071 | 0 | 0 |
| NC_019843.3;YP_009047203.1;MERS_1a;2072 | 0 | 0 |
| NC_019843.3;YP_009047203.1;MERS_1a;2073 | 0 | 0 |
| NC_019843.3;YP_009047203.1;MERS_1a;2074 | 0 | 0 |
| NC_019843.3;YP_009047203.1;MERS_1a;2075 | 0 | 0 |
| NC_019843.3;YP_009047203.1;MERS_1a;2076 | 0 | 0 |
| NC_019843.3;YP_009047203.1;MERS_1a;2077 | 0 | 0.025 |
| NC_019843.3;YP_009047203.1;MERS_1a;2078 | 0 | 0 |
| NC_019843.3;YP_009047203.1;MERS_1a;2079 | 0 | 0 |
| NC_019843.3;YP_009047203.1;MERS_1a;2080 | 0 | 0 |
| NC_019843.3;YP_009047203.1;MERS_1a;2081 | 0 | 0 |
| NC_019843.3;YP_009047203.1;MERS_1a;2082 | 0 | 0 |
| NC_019843.3;YP_009047203.1;MERS_1a;2083 | 0 | 0 |

| | | |
|---|---|---|
| NC_019843.3;YP_009047203.1;MERS_1a;2084 | 0 | 0 |
| NC_019843.3;YP_009047203.1;MERS_1a;2085 | 0 | 0 |
| NC_019843.3;YP_009047203.1;MERS_1a;2086 | 0 | 0 |
| NC_019843.3;YP_009047203.1;MERS_1a;2087 | 0 | 0 |
| NC_019843.3;YP_009047203.1;MERS_1a;2088 | 0 | 0 |
| NC_019843.3;YP_009047203.1;MERS_1a;2089 | 0 | 0 |
| NC_019843.3;YP_009047203.1;MERS_1a;2090 | 0 | 0 |
| NC_019843.3;YP_009047203.1;MERS_1a;2091 | 0 | 0 |
| NC_019843.3;YP_009047203.1;MERS_1a;2092 | 0 | 0 |
| NC_019843.3;YP_009047203.1;MERS_1a;2093 | 0 | 0 |
| NC_019843.3;YP_009047203.1;MERS_1a;2094 | 0 | 0 |
| NC_019843.3;YP_009047203.1;MERS_1a;2095 | 0 | 0 |
| NC_019843.3;YP_009047203.1;MERS_1a;2096 | 0 | 0 |
| NC_019843.3;YP_009047203.1;MERS_1a;2097 | 0 | 0 |
| NC_019843.3;YP_009047203.1;MERS_1a;2098 | 0 | 0 |
| NC_019843.3;YP_009047203.1;MERS_1a;2099 | 0 | 0 |
| NC_019843.3;YP_009047203.1;MERS_1a;2100 | 0 | 0 |
| NC_019843.3;YP_009047203.1;MERS_1a;2101 | 0 | 0 |
| NC_019843.3;YP_009047203.1;MERS_1a;2102 | 0 | 0 |
| NC_019843.3;YP_009047203.1;MERS_1a;2103 | 0 | 0 |
| NC_019843.3;YP_009047203.1;MERS_1a;2104 | 0 | 0 |
| NC_019843.3;YP_009047203.1;MERS_1a;2105 | 0 | 0 |
| NC_019843.3;YP_009047203.1;MERS_1a;2106 | 0 | 0 |
| NC_019843.3;YP_009047203.1;MERS_1a;2107 | 0 | 0 |
| NC_019843.3;YP_009047203.1;MERS_1a;2108 | 0 | 0 |
| NC_019843.3;YP_009047203.1;MERS_1a;2109 | 0 | 0 |
| NC_019843.3;YP_009047203.1;MERS_1a;2110 | 0 | 0 |
| NC_019843.3;YP_009047203.1;MERS_1a;2111 | 0 | 0 |
| NC_019843.3;YP_009047203.1;MERS_1a;2112 | 0 | 0 |
| NC_019843.3;YP_009047203.1;MERS_1a;2113 | 0 | 0 |
| NC_019843.3;YP_009047203.1;MERS_1a;2114 | 0 | 0 |
| NC_019843.3;YP_009047203.1;MERS_1a;2115 | 0 | 0 |
| NC_019843.3;YP_009047203.1;MERS_1a;2116 | 0 | 0 |
| NC_019843.3;YP_009047203.1;MERS_1a;2117 | 0 | 0 |
| NC_019843.3;YP_009047203.1;MERS_1a;2118 | 0 | 0 |
| NC_019843.3;YP_009047203.1;MERS_1a;2119 | 0.05 | 0.025 |
| NC_019843.3;YP_009047203.1;MERS_1a;2120 | 0.05 | 0.025 |
| NC_019843.3;YP_009047203.1;MERS_1a;2121 | 0.05 | 0 |
| NC_019843.3;YP_009047203.1;MERS_1a;2122 | 0 | 0 |
| NC_019843.3;YP_009047203.1;MERS_1a;2123 | 0 | 0 |
| NC_019843.3;YP_009047203.1;MERS_1a;2124 | 0 | 0 |
| NC_019843.3;YP_009047203.1;MERS_1a;2125 | 0 | 0 |
| NC_019843.3;YP_009047203.1;MERS_1a;2126 | 0 | 0 |

| | | |
|---|---|---|
| NC_019843.3;YP_009047203.1;MERS_1a;2127 | 0 | 0 |
| NC_019843.3;YP_009047203.1;MERS_1a;2128 | 0 | 0 |
| NC_019843.3;YP_009047203.1;MERS_1a;2129 | 0 | 0 |
| NC_019843.3;YP_009047203.1;MERS_1a;2130 | 0 | 0 |
| NC_019843.3;YP_009047203.1;MERS_1a;2131 | 0 | 0 |
| NC_019843.3;YP_009047203.1;MERS_1a;2132 | 0 | 0 |
| NC_019843.3;YP_009047203.1;MERS_1a;2133 | 0 | 0 |
| NC_019843.3;YP_009047203.1;MERS_1a;2134 | 0 | 0 |
| NC_019843.3;YP_009047203.1;MERS_1a;2135 | 0 | 0 |
| NC_019843.3;YP_009047203.1;MERS_1a;2136 | 0 | 0.025 |
| NC_019843.3;YP_009047203.1;MERS_1a;2137 | 0 | 0.025 |
| NC_019843.3;YP_009047203.1;MERS_1a;2138 | 0 | 0.025 |
| NC_019843.3;YP_009047203.1;MERS_1a;2139 | 0 | 0.025 |
| NC_019843.3;YP_009047203.1;MERS_1a;2140 | 0 | 0.025 |
| NC_019843.3;YP_009047203.1;MERS_1a;2141 | 0 | 0.025 |
| NC_019843.3;YP_009047203.1;MERS_1a;2142 | 0 | 0.025 |
| NC_019843.3;YP_009047203.1;MERS_1a;2143 | 0.05 | 0.1 |
| NC_019843.3;YP_009047203.1;MERS_1a;2144 | 0 | 0.075 |
| NC_019843.3;YP_009047203.1;MERS_1a;2145 | 0 | 0.05 |
| NC_019843.3;YP_009047203.1;MERS_1a;2146 | 0 | 0 |
| NC_019843.3;YP_009047203.1;MERS_1a;2147 | 0 | 0 |
| NC_019843.3;YP_009047203.1;MERS_1a;2148 | 0 | 0 |
| NC_019843.3;YP_009047203.1;MERS_1a;2149 | 0 | 0 |
| NC_019843.3;YP_009047203.1;MERS_1a;2150 | 0 | 0 |
| NC_019843.3;YP_009047203.1;MERS_1a;2151 | 0 | 0 |
| NC_019843.3;YP_009047203.1;MERS_1a;2152 | 0 | 0 |
| NC_019843.3;YP_009047203.1;MERS_1a;2153 | 0 | 0 |
| NC_019843.3;YP_009047203.1;MERS_1a;2154 | 0 | 0 |
| NC_019843.3;YP_009047203.1;MERS_1a;2155 | 0 | 0 |
| NC_019843.3;YP_009047203.1;MERS_1a;2156 | 0 | 0 |
| NC_019843.3;YP_009047203.1;MERS_1a;2157 | 0 | 0.05 |
| NC_019843.3;YP_009047203.1;MERS_1a;2158 | 0 | 0 |
| NC_019843.3;YP_009047203.1;MERS_1a;2159 | 0 | 0 |
| NC_019843.3;YP_009047203.1;MERS_1a;2160 | 0 | 0 |
| NC_019843.3;YP_009047203.1;MERS_1a;2161 | 0 | 0 |
| NC_019843.3;YP_009047203.1;MERS_1a;2162 | 0 | 0 |
| NC_019843.3;YP_009047203.1;MERS_1a;2163 | 0 | 0 |
| NC_019843.3;YP_009047203.1;MERS_1a;2164 | 0 | 0 |
| NC_019843.3;YP_009047203.1;MERS_1a;2165 | 0 | 0 |
| NC_019843.3;YP_009047203.1;MERS_1a;2166 | 0 | 0 |
| NC_019843.3;YP_009047203.1;MERS_1a;2167 | 0 | 0 |
| NC_019843.3;YP_009047203.1;MERS_1a;2168 | 0 | 0 |
| NC_019843.3;YP_009047203.1;MERS_1a;2169 | 0 | 0 |

| | | |
|---|---|---|
| NC_019843.3;YP_009047203.1;MERS_1a;2170 | 0 | 0 |
| NC_019843.3;YP_009047203.1;MERS_1a;2171 | 0 | 0 |
| NC_019843.3;YP_009047203.1;MERS_1a;2172 | 0 | 0 |
| NC_019843.3;YP_009047203.1;MERS_1a;2173 | 0 | 0 |
| NC_019843.3;YP_009047203.1;MERS_1a;2174 | 0 | 0 |
| NC_019843.3;YP_009047203.1;MERS_1a;2175 | 0 | 0 |
| NC_019843.3;YP_009047203.1;MERS_1a;2176 | 0 | 0 |
| NC_019843.3;YP_009047203.1;MERS_1a;2177 | 0 | 0 |
| NC_019843.3;YP_009047203.1;MERS_1a;2178 | 0 | 0 |
| NC_019843.3;YP_009047203.1;MERS_1a;2179 | 0 | 0 |
| NC_019843.3;YP_009047203.1;MERS_1a;2180 | 0 | 0 |
| NC_019843.3;YP_009047203.1;MERS_1a;2181 | 0 | 0 |
| NC_019843.3;YP_009047203.1;MERS_1a;2182 | 0 | 0 |
| NC_019843.3;YP_009047203.1;MERS_1a;2183 | 0 | 0 |
| NC_019843.3;YP_009047203.1;MERS_1a;2184 | 0 | 0 |
| NC_019843.3;YP_009047203.1;MERS_1a;2185 | 0 | 0 |
| NC_019843.3;YP_009047203.1;MERS_1a;2186 | 0 | 0 |
| NC_019843.3;YP_009047203.1;MERS_1a;2187 | 0 | 0 |
| NC_019843.3;YP_009047203.1;MERS_1a;2188 | 0 | 0 |
| NC_019843.3;YP_009047203.1;MERS_1a;2189 | 0 | 0 |
| NC_019843.3;YP_009047203.1;MERS_1a;2190 | 0 | 0 |
| NC_019843.3;YP_009047203.1;MERS_1a;2191 | 0 | 0 |
| NC_019843.3;YP_009047203.1;MERS_1a;2192 | 0 | 0 |
| NC_019843.3;YP_009047203.1;MERS_1a;2193 | 0 | 0 |
| NC_019843.3;YP_009047203.1;MERS_1a;2194 | 0 | 0 |
| NC_019843.3;YP_009047203.1;MERS_1a;2195 | 0 | 0 |
| NC_019843.3;YP_009047203.1;MERS_1a;2196 | 0 | 0 |
| NC_019843.3;YP_009047203.1;MERS_1a;2197 | 0 | 0 |
| NC_019843.3;YP_009047203.1;MERS_1a;2198 | 0 | 0 |
| NC_019843.3;YP_009047203.1;MERS_1a;2199 | 0 | 0 |
| NC_019843.3;YP_009047203.1;MERS_1a;2200 | 0 | 0 |
| NC_019843.3;YP_009047203.1;MERS_1a;2201 | 0 | 0 |
| NC_019843.3;YP_009047203.1;MERS_1a;2202 | 0 | 0 |
| NC_019843.3;YP_009047203.1;MERS_1a;2203 | 0 | 0 |
| NC_019843.3;YP_009047203.1;MERS_1a;2204 | 0 | 0 |
| NC_019843.3;YP_009047203.1;MERS_1a;2205 | 0 | 0 |
| NC_019843.3;YP_009047203.1;MERS_1a;2206 | 0 | 0 |
| NC_019843.3;YP_009047203.1;MERS_1a;2207 | 0 | 0 |
| NC_019843.3;YP_009047203.1;MERS_1a;2208 | 0.05 | 0 |
| NC_019843.3;YP_009047203.1;MERS_1a;2209 | 0 | 0 |
| NC_019843.3;YP_009047203.1;MERS_1a;2210 | 0 | 0 |
| NC_019843.3;YP_009047203.1;MERS_1a;2211 | 0 | 0 |
| NC_019843.3;YP_009047203.1;MERS_1a;2212 | 0 | 0 |

| | | |
|---|---|---|
| NC_019843.3;YP_009047203.1;MERS_1a;2213 | 0 | 0 |
| NC_019843.3;YP_009047203.1;MERS_1a;2214 | 0 | 0 |
| NC_019843.3;YP_009047203.1;MERS_1a;2215 | 0 | 0 |
| NC_019843.3;YP_009047203.1;MERS_1a;2216 | 0 | 0 |
| NC_019843.3;YP_009047203.1;MERS_1a;2217 | 0 | 0 |
| NC_019843.3;YP_009047203.1;MERS_1a;2218 | 0 | 0 |
| NC_019843.3;YP_009047203.1;MERS_1a;2219 | 0 | 0 |
| NC_019843.3;YP_009047203.1;MERS_1a;2220 | 0.05 | 0 |
| NC_019843.3;YP_009047203.1;MERS_1a;2221 | 0 | 0 |
| NC_019843.3;YP_009047203.1;MERS_1a;2222 | 0.05 | 0 |
| NC_019843.3;YP_009047203.1;MERS_1a;2223 | 0 | 0 |
| NC_019843.3;YP_009047203.1;MERS_1a;2224 | 0 | 0 |
| NC_019843.3;YP_009047203.1;MERS_1a;2225 | 0 | 0 |
| NC_019843.3;YP_009047203.1;MERS_1a;2226 | 0 | 0 |
| NC_019843.3;YP_009047203.1;MERS_1a;2227 | 0 | 0 |
| NC_019843.3;YP_009047203.1;MERS_1a;2228 | 0.15 | 0.05 |
| NC_019843.3;YP_009047203.1;MERS_1a;2229 | 0.05 | 0.05 |
| NC_019843.3;YP_009047203.1;MERS_1a;2230 | 0.1 | 0.05 |
| NC_019843.3;YP_009047203.1;MERS_1a;2231 | 0.15 | 0.05 |
| NC_019843.3;YP_009047203.1;MERS_1a;2232 | 0.15 | 0.05 |
| NC_019843.3;YP_009047203.1;MERS_1a;2233 | 0.15 | 0.05 |
| NC_019843.3;YP_009047203.1;MERS_1a;2234 | 0.1 | 0.075 |
| NC_019843.3;YP_009047203.1;MERS_1a;2235 | 0.1 | 0.05 |
| NC_019843.3;YP_009047203.1;MERS_1a;2236 | 0 | 0 |
| NC_019843.3;YP_009047203.1;MERS_1a;2237 | 0 | 0 |
| NC_019843.3;YP_009047203.1;MERS_1a;2238 | 0 | 0 |
| NC_019843.3;YP_009047203.1;MERS_1a;2239 | 0.05 | 0 |
| NC_019843.3;YP_009047203.1;MERS_1a;2240 | 0 | 0.025 |
| NC_019843.3;YP_009047203.1;MERS_1a;2241 | 0 | 0.025 |
| NC_019843.3;YP_009047203.1;MERS_1a;2242 | 0 | 0 |
| NC_019843.3;YP_009047203.1;MERS_1a;2243 | 0 | 0 |
| NC_019843.3;YP_009047203.1;MERS_1a;2244 | 0 | 0 |
| NC_019843.3;YP_009047203.1;MERS_1a;2245 | 0 | 0 |
| NC_019843.3;YP_009047203.1;MERS_1a;2246 | 0 | 0 |
| NC_019843.3;YP_009047203.1;MERS_1a;2247 | 0 | 0 |
| NC_019843.3;YP_009047203.1;MERS_1a;2248 | 0 | 0 |
| NC_019843.3;YP_009047203.1;MERS_1a;2249 | 0 | 0 |
| NC_019843.3;YP_009047203.1;MERS_1a;2250 | 0 | 0 |
| NC_019843.3;YP_009047203.1;MERS_1a;2251 | 0 | 0 |
| NC_019843.3;YP_009047203.1;MERS_1a;2252 | 0 | 0 |
| NC_019843.3;YP_009047203.1;MERS_1a;2253 | 0 | 0 |
| NC_019843.3;YP_009047203.1;MERS_1a;2254 | 0 | 0 |
| NC_019843.3;YP_009047203.1;MERS_1a;2255 | 0 | 0 |

| | | |
|---|---|---|
| NC_019843.3;YP_009047203.1;MERS_1a;2256 | 0 | 0 |
| NC_019843.3;YP_009047203.1;MERS_1a;2257 | 0 | 0 |
| NC_019843.3;YP_009047203.1;MERS_1a;2258 | 0 | 0 |
| NC_019843.3;YP_009047203.1;MERS_1a;2259 | 0.15 | 0.075 |
| NC_019843.3;YP_009047203.1;MERS_1a;2260 | 0 | 0.025 |
| NC_019843.3;YP_009047203.1;MERS_1a;2261 | 0 | 0.15 |
| NC_019843.3;YP_009047203.1;MERS_1a;2262 | 0 | 0 |
| NC_019843.3;YP_009047203.1;MERS_1a;2263 | 0 | 0 |
| NC_019843.3;YP_009047203.1;MERS_1a;2264 | 0 | 0 |
| NC_019843.3;YP_009047203.1;MERS_1a;2265 | 0 | 0 |
| NC_019843.3;YP_009047203.1;MERS_1a;2266 | 0 | 0 |
| NC_019843.3;YP_009047203.1;MERS_1a;2267 | 0 | 0 |
| NC_019843.3;YP_009047203.1;MERS_1a;2268 | 0 | 0 |
| NC_019843.3;YP_009047203.1;MERS_1a;2269 | 0 | 0 |
| NC_019843.3;YP_009047203.1;MERS_1a;2270 | 0 | 0 |
| NC_019843.3;YP_009047203.1;MERS_1a;2271 | 0 | 0 |
| NC_019843.3;YP_009047203.1;MERS_1a;2272 | 0 | 0 |
| NC_019843.3;YP_009047203.1;MERS_1a;2273 | 0 | 0 |
| NC_019843.3;YP_009047203.1;MERS_1a;2274 | 0 | 0 |
| NC_019843.3;YP_009047203.1;MERS_1a;2275 | 0 | 0 |
| NC_019843.3;YP_009047203.1;MERS_1a;2276 | 0 | 0 |
| NC_019843.3;YP_009047203.1;MERS_1a;2277 | 0 | 0 |
| NC_019843.3;YP_009047203.1;MERS_1a;2278 | 0 | 0 |
| NC_019843.3;YP_009047203.1;MERS_1a;2279 | 0 | 0 |
| NC_019843.3;YP_009047203.1;MERS_1a;2280 | 0 | 0 |
| NC_019843.3;YP_009047203.1;MERS_1a;2281 | 0 | 0 |
| NC_019843.3;YP_009047203.1;MERS_1a;2282 | 0 | 0 |
| NC_019843.3;YP_009047203.1;MERS_1a;2283 | 0 | 0 |
| NC_019843.3;YP_009047203.1;MERS_1a;2284 | 0 | 0 |
| NC_019843.3;YP_009047203.1;MERS_1a;2285 | 0 | 0 |
| NC_019843.3;YP_009047203.1;MERS_1a;2286 | 0 | 0 |
| NC_019843.3;YP_009047203.1;MERS_1a;2287 | 0 | 0 |
| NC_019843.3;YP_009047203.1;MERS_1a;2288 | 0 | 0 |
| NC_019843.3;YP_009047203.1;MERS_1a;2289 | 0 | 0 |
| NC_019843.3;YP_009047203.1;MERS_1a;2290 | 0 | 0 |
| NC_019843.3;YP_009047203.1;MERS_1a;2291 | 0 | 0 |
| NC_019843.3;YP_009047203.1;MERS_1a;2292 | 0 | 0 |
| NC_019843.3;YP_009047203.1;MERS_1a;2293 | 0 | 0 |
| NC_019843.3;YP_009047203.1;MERS_1a;2294 | 0 | 0 |
| NC_019843.3;YP_009047203.1;MERS_1a;2295 | 0 | 0 |
| NC_019843.3;YP_009047203.1;MERS_1a;2296 | 0 | 0 |
| NC_019843.3;YP_009047203.1;MERS_1a;2297 | 0 | 0 |
| NC_019843.3;YP_009047203.1;MERS_1a;2298 | 0 | 0 |

| | | |
|---|---|---|
| NC_019843.3;YP_009047203.1;MERS_1a;2299 | 0 | 0 |
| NC_019843.3;YP_009047203.1;MERS_1a;2300 | 0 | 0 |
| NC_019843.3;YP_009047203.1;MERS_1a;2301 | 0 | 0 |
| NC_019843.3;YP_009047203.1;MERS_1a;2302 | 0 | 0 |
| NC_019843.3;YP_009047203.1;MERS_1a;2303 | 0 | 0 |
| NC_019843.3;YP_009047203.1;MERS_1a;2304 | 0 | 0 |
| NC_019843.3;YP_009047203.1;MERS_1a;2305 | 0 | 0 |
| NC_019843.3;YP_009047203.1;MERS_1a;2306 | 0 | 0 |
| NC_019843.3;YP_009047203.1;MERS_1a;2307 | 0 | 0 |
| NC_019843.3;YP_009047203.1;MERS_1a;2308 | 0 | 0 |
| NC_019843.3;YP_009047203.1;MERS_1a;2309 | 0 | 0 |
| NC_019843.3;YP_009047203.1;MERS_1a;2310 | 0 | 0 |
| NC_019843.3;YP_009047203.1;MERS_1a;2311 | 0 | 0 |
| NC_019843.3;YP_009047203.1;MERS_1a;2312 | 0 | 0 |
| NC_019843.3;YP_009047203.1;MERS_1a;2313 | 0 | 0 |
| NC_019843.3;YP_009047203.1;MERS_1a;2314 | 0 | 0 |
| NC_019843.3;YP_009047203.1;MERS_1a;2315 | 0 | 0 |
| NC_019843.3;YP_009047203.1;MERS_1a;2316 | 0 | 0 |
| NC_019843.3;YP_009047203.1;MERS_1a;2317 | 0 | 0 |
| NC_019843.3;YP_009047203.1;MERS_1a;2318 | 0 | 0 |
| NC_019843.3;YP_009047203.1;MERS_1a;2319 | 0 | 0 |
| NC_019843.3;YP_009047203.1;MERS_1a;2320 | 0 | 0 |
| NC_019843.3;YP_009047203.1;MERS_1a;2321 | 0 | 0 |
| NC_019843.3;YP_009047203.1;MERS_1a;2322 | 0 | 0 |
| NC_019843.3;YP_009047203.1;MERS_1a;2323 | 0 | 0 |
| NC_019843.3;YP_009047203.1;MERS_1a;2324 | 0 | 0 |
| NC_019843.3;YP_009047203.1;MERS_1a;2325 | 0 | 0 |
| NC_019843.3;YP_009047203.1;MERS_1a;2326 | 0 | 0 |
| NC_019843.3;YP_009047203.1;MERS_1a;2327 | 0 | 0 |
| NC_019843.3;YP_009047203.1;MERS_1a;2328 | 0 | 0 |
| NC_019843.3;YP_009047203.1;MERS_1a;2329 | 0 | 0 |
| NC_019843.3;YP_009047203.1;MERS_1a;2330 | 0 | 0 |
| NC_019843.3;YP_009047203.1;MERS_1a;2331 | 0 | 0 |
| NC_019843.3;YP_009047203.1;MERS_1a;2332 | 0 | 0 |
| NC_019843.3;YP_009047203.1;MERS_1a;2333 | 0.05 | 0 |
| NC_019843.3;YP_009047203.1;MERS_1a;2334 | 0 | 0.025 |
| NC_019843.3;YP_009047203.1;MERS_1a;2335 | 0 | 0 |
| NC_019843.3;YP_009047203.1;MERS_1a;2336 | 0 | 0 |
| NC_019843.3;YP_009047203.1;MERS_1a;2337 | 0 | 0 |
| NC_019843.3;YP_009047203.1;MERS_1a;2338 | 0 | 0 |
| NC_019843.3;YP_009047203.1;MERS_1a;2339 | 0 | 0 |
| NC_019843.3;YP_009047203.1;MERS_1a;2340 | 0 | 0 |
| NC_019843.3;YP_009047203.1;MERS_1a;2341 | 0 | 0 |

| | | |
|---|---|---|
| NC_019843.3;YP_009047203.1;MERS_1a;2342 | 0 | 0 |
| NC_019843.3;YP_009047203.1;MERS_1a;2343 | 0 | 0 |
| NC_019843.3;YP_009047203.1;MERS_1a;2344 | 0 | 0 |
| NC_019843.3;YP_009047203.1;MERS_1a;2345 | 0 | 0 |
| NC_019843.3;YP_009047203.1;MERS_1a;2346 | 0 | 0 |
| NC_019843.3;YP_009047203.1;MERS_1a;2347 | 0 | 0 |
| NC_019843.3;YP_009047203.1;MERS_1a;2348 | 0 | 0 |
| NC_019843.3;YP_009047203.1;MERS_1a;2349 | 0 | 0 |
| NC_019843.3;YP_009047203.1;MERS_1a;2350 | 0 | 0 |
| NC_019843.3;YP_009047203.1;MERS_1a;2351 | 0 | 0 |
| NC_019843.3;YP_009047203.1;MERS_1a;2352 | 0 | 0 |
| NC_019843.3;YP_009047203.1;MERS_1a;2353 | 0 | 0 |
| NC_019843.3;YP_009047203.1;MERS_1a;2354 | 0 | 0 |
| NC_019843.3;YP_009047203.1;MERS_1a;2355 | 0 | 0 |
| NC_019843.3;YP_009047203.1;MERS_1a;2356 | 0 | 0 |
| NC_019843.3;YP_009047203.1;MERS_1a;2357 | 0 | 0 |
| NC_019843.3;YP_009047203.1;MERS_1a;2358 | 0 | 0 |
| NC_019843.3;YP_009047203.1;MERS_1a;2359 | 0 | 0 |
| NC_019843.3;YP_009047203.1;MERS_1a;2360 | 0 | 0 |
| NC_019843.3;YP_009047203.1;MERS_1a;2361 | 0 | 0 |
| NC_019843.3;YP_009047203.1;MERS_1a;2362 | 0 | 0 |
| NC_019843.3;YP_009047203.1;MERS_1a;2363 | 0 | 0 |
| NC_019843.3;YP_009047203.1;MERS_1a;2364 | 0 | 0.025 |
| NC_019843.3;YP_009047203.1;MERS_1a;2365 | 0 | 0 |
| NC_019843.3;YP_009047203.1;MERS_1a;2366 | 0 | 0 |
| NC_019843.3;YP_009047203.1;MERS_1a;2367 | 0 | 0 |
| NC_019843.3;YP_009047203.1;MERS_1a;2368 | 0 | 0 |
| NC_019843.3;YP_009047203.1;MERS_1a;2369 | 0 | 0 |
| NC_019843.3;YP_009047203.1;MERS_1a;2370 | 0 | 0 |
| NC_019843.3;YP_009047203.1;MERS_1a;2371 | 0 | 0 |
| NC_019843.3;YP_009047203.1;MERS_1a;2372 | 0 | 0 |
| NC_019843.3;YP_009047203.1;MERS_1a;2373 | 0 | 0 |
| NC_019843.3;YP_009047203.1;MERS_1a;2374 | 0 | 0 |
| NC_019843.3;YP_009047203.1;MERS_1a;2375 | 0 | 0 |
| NC_019843.3;YP_009047203.1;MERS_1a;2376 | 0 | 0 |
| NC_019843.3;YP_009047203.1;MERS_1a;2377 | 0 | 0 |
| NC_019843.3;YP_009047203.1;MERS_1a;2378 | 0 | 0 |
| NC_019843.3;YP_009047203.1;MERS_1a;2379 | 0 | 0 |
| NC_019843.3;YP_009047203.1;MERS_1a;2380 | 0 | 0 |
| NC_019843.3;YP_009047203.1;MERS_1a;2381 | 0 | 0 |
| NC_019843.3;YP_009047203.1;MERS_1a;2382 | 0 | 0 |
| NC_019843.3;YP_009047203.1;MERS_1a;2383 | 0 | 0 |
| NC_019843.3;YP_009047203.1;MERS_1a;2384 | 0 | 0 |

| | | |
|---|---|---|
| NC_019843.3;YP_009047203.1;MERS_1a;2385 | 0 | 0 |
| NC_019843.3;YP_009047203.1;MERS_1a;2386 | 0 | 0 |
| NC_019843.3;YP_009047203.1;MERS_1a;2387 | 0 | 0 |
| NC_019843.3;YP_009047203.1;MERS_1a;2388 | 0 | 0 |
| NC_019843.3;YP_009047203.1;MERS_1a;2389 | 0 | 0 |
| NC_019843.3;YP_009047203.1;MERS_1a;2390 | 0 | 0 |
| NC_019843.3;YP_009047203.1;MERS_1a;2391 | 0 | 0 |
| NC_019843.3;YP_009047203.1;MERS_1a;2392 | 0 | 0 |
| NC_019843.3;YP_009047203.1;MERS_1a;2393 | 0 | 0 |
| NC_019843.3;YP_009047203.1;MERS_1a;2394 | 0 | 0 |
| NC_019843.3;YP_009047203.1;MERS_1a;2395 | 0 | 0 |
| NC_019843.3;YP_009047203.1;MERS_1a;2396 | 0 | 0 |
| NC_019843.3;YP_009047203.1;MERS_1a;2397 | 0 | 0 |
| NC_019843.3;YP_009047203.1;MERS_1a;2398 | 0 | 0 |
| NC_019843.3;YP_009047203.1;MERS_1a;2399 | 0 | 0 |
| NC_019843.3;YP_009047203.1;MERS_1a;2400 | 0 | 0 |
| NC_019843.3;YP_009047203.1;MERS_1a;2401 | 0 | 0 |
| NC_019843.3;YP_009047203.1;MERS_1a;2402 | 0 | 0 |
| NC_019843.3;YP_009047203.1;MERS_1a;2403 | 0 | 0 |
| NC_019843.3;YP_009047203.1;MERS_1a;2404 | 0 | 0 |
| NC_019843.3;YP_009047203.1;MERS_1a;2405 | 0 | 0 |
| NC_019843.3;YP_009047203.1;MERS_1a;2406 | 0 | 0 |
| NC_019843.3;YP_009047203.1;MERS_1a;2407 | 0 | 0 |
| NC_019843.3;YP_009047203.1;MERS_1a;2408 | 0 | 0 |
| NC_019843.3;YP_009047203.1;MERS_1a;2409 | 0 | 0 |
| NC_019843.3;YP_009047203.1;MERS_1a;2410 | 0 | 0.025 |
| NC_019843.3;YP_009047203.1;MERS_1a;2411 | 0 | 0 |
| NC_019843.3;YP_009047203.1;MERS_1a;2412 | 0 | 0 |
| NC_019843.3;YP_009047203.1;MERS_1a;2413 | 0 | 0 |
| NC_019843.3;YP_009047203.1;MERS_1a;2414 | 0 | 0 |
| NC_019843.3;YP_009047203.1;MERS_1a;2415 | 0 | 0 |
| NC_019843.3;YP_009047203.1;MERS_1a;2416 | 0 | 0 |
| NC_019843.3;YP_009047203.1;MERS_1a;2417 | 0 | 0 |
| NC_019843.3;YP_009047203.1;MERS_1a;2418 | 0 | 0 |
| NC_019843.3;YP_009047203.1;MERS_1a;2419 | 0 | 0 |
| NC_019843.3;YP_009047203.1;MERS_1a;2420 | 0 | 0 |
| NC_019843.3;YP_009047203.1;MERS_1a;2421 | 0 | 0 |
| NC_019843.3;YP_009047203.1;MERS_1a;2422 | 0 | 0 |
| NC_019843.3;YP_009047203.1;MERS_1a;2423 | 0 | 0 |
| NC_019843.3;YP_009047203.1;MERS_1a;2424 | 0 | 0 |
| NC_019843.3;YP_009047203.1;MERS_1a;2425 | 0 | 0 |
| NC_019843.3;YP_009047203.1;MERS_1a;2426 | 0 | 0 |
| NC_019843.3;YP_009047203.1;MERS_1a;2427 | 0 | 0 |

| | | |
|---|---|---|
| NC_019843.3;YP_009047203.1;MERS_1a;2428 | 0 | 0.075 |
| NC_019843.3;YP_009047203.1;MERS_1a;2429 | 0 | 0 |
| NC_019843.3;YP_009047203.1;MERS_1a;2430 | 0 | 0 |
| NC_019843.3;YP_009047203.1;MERS_1a;2431 | 0 | 0 |
| NC_019843.3;YP_009047203.1;MERS_1a;2432 | 0 | 0 |
| NC_019843.3;YP_009047203.1;MERS_1a;2433 | 0 | 0.025 |
| NC_019843.3;YP_009047203.1;MERS_1a;2434 | 0 | 0 |
| NC_019843.3;YP_009047203.1;MERS_1a;2435 | 0 | 0.025 |
| NC_019843.3;YP_009047203.1;MERS_1a;2436 | 0 | 0 |
| NC_019843.3;YP_009047203.1;MERS_1a;2437 | 0.05 | 0 |
| NC_019843.3;YP_009047203.1;MERS_1a;2438 | 0 | 0 |
| NC_019843.3;YP_009047203.1;MERS_1a;2439 | 0.05 | 0 |
| NC_019843.3;YP_009047203.1;MERS_1a;2440 | 0.05 | 0 |
| NC_019843.3;YP_009047203.1;MERS_1a;2441 | 0.05 | 0 |
| NC_019843.3;YP_009047203.1;MERS_1a;2442 | 0.05 | 0 |
| NC_019843.3;YP_009047203.1;MERS_1a;2443 | 0.05 | 0 |
| NC_019843.3;YP_009047203.1;MERS_1a;2444 | 0.05 | 0 |
| NC_019843.3;YP_009047203.1;MERS_1a;2445 | 0.05 | 0 |
| NC_019843.3;YP_009047203.1;MERS_1a;2446 | 0.1 | 0 |
| NC_019843.3;YP_009047203.1;MERS_1a;2447 | 0.05 | 0 |
| NC_019843.3;YP_009047203.1;MERS_1a;2448 | 0 | 0 |
| NC_019843.3;YP_009047203.1;MERS_1a;2449 | 0 | 0.025 |
| NC_019843.3;YP_009047203.1;MERS_1a;2450 | 0 | 0 |
| NC_019843.3;YP_009047203.1;MERS_1a;2451 | 0 | 0 |
| NC_019843.3;YP_009047203.1;MERS_1a;2452 | 0 | 0 |
| NC_019843.3;YP_009047203.1;MERS_1a;2453 | 0 | 0 |
| NC_019843.3;YP_009047203.1;MERS_1a;2454 | 0 | 0 |
| NC_019843.3;YP_009047203.1;MERS_1a;2455 | 0 | 0 |
| NC_019843.3;YP_009047203.1;MERS_1a;2456 | 0 | 0 |
| NC_019843.3;YP_009047203.1;MERS_1a;2457 | 0 | 0 |
| NC_019843.3;YP_009047203.1;MERS_1a;2458 | 0 | 0 |
| NC_019843.3;YP_009047203.1;MERS_1a;2459 | 0 | 0 |
| NC_019843.3;YP_009047203.1;MERS_1a;2460 | 0 | 0 |
| NC_019843.3;YP_009047203.1;MERS_1a;2461 | 0 | 0 |
| NC_019843.3;YP_009047203.1;MERS_1a;2462 | 0 | 0 |
| NC_019843.3;YP_009047203.1;MERS_1a;2463 | 0 | 0.025 |
| NC_019843.3;YP_009047203.1;MERS_1a;2464 | 0 | 0 |
| NC_019843.3;YP_009047203.1;MERS_1a;2465 | 0 | 0 |
| NC_019843.3;YP_009047203.1;MERS_1a;2466 | 0 | 0 |
| NC_019843.3;YP_009047203.1;MERS_1a;2467 | 0 | 0 |
| NC_019843.3;YP_009047203.1;MERS_1a;2468 | 0 | 0 |
| NC_019843.3;YP_009047203.1;MERS_1a;2469 | 0 | 0 |
| NC_019843.3;YP_009047203.1;MERS_1a;2470 | 0 | 0 |

| | | |
|---|---|---|
| NC_019843.3;YP_009047203.1;MERS_1a;2471 | 0 | 0 |
| NC_019843.3;YP_009047203.1;MERS_1a;2472 | 0 | 0 |
| NC_019843.3;YP_009047203.1;MERS_1a;2473 | 0 | 0 |
| NC_019843.3;YP_009047203.1;MERS_1a;2474 | 0 | 0 |
| NC_019843.3;YP_009047203.1;MERS_1a;2475 | 0 | 0 |
| NC_019843.3;YP_009047203.1;MERS_1a;2476 | 0 | 0 |
| NC_019843.3;YP_009047203.1;MERS_1a;2477 | 0 | 0 |
| NC_019843.3;YP_009047203.1;MERS_1a;2478 | 0 | 0 |
| NC_019843.3;YP_009047203.1;MERS_1a;2479 | 0 | 0 |
| NC_019843.3;YP_009047203.1;MERS_1a;2480 | 0 | 0 |
| NC_019843.3;YP_009047203.1;MERS_1a;2481 | 0 | 0 |
| NC_019843.3;YP_009047203.1;MERS_1a;2482 | 0 | 0 |
| NC_019843.3;YP_009047203.1;MERS_1a;2483 | 0.05 | 0 |
| NC_019843.3;YP_009047203.1;MERS_1a;2484 | 0 | 0 |
| NC_019843.3;YP_009047203.1;MERS_1a;2485 | 0 | 0 |
| NC_019843.3;YP_009047203.1;MERS_1a;2486 | 0 | 0 |
| NC_019843.3;YP_009047203.1;MERS_1a;2487 | 0 | 0 |
| NC_019843.3;YP_009047203.1;MERS_1a;2488 | 0 | 0 |
| NC_019843.3;YP_009047203.1;MERS_1a;2489 | 0 | 0 |
| NC_019843.3;YP_009047203.1;MERS_1a;2490 | 0 | 0 |
| NC_019843.3;YP_009047203.1;MERS_1a;2491 | 0 | 0 |
| NC_019843.3;YP_009047203.1;MERS_1a;2492 | 0 | 0 |
| NC_019843.3;YP_009047203.1;MERS_1a;2493 | 0 | 0 |
| NC_019843.3;YP_009047203.1;MERS_1a;2494 | 0 | 0 |
| NC_019843.3;YP_009047203.1;MERS_1a;2495 | 0 | 0 |
| NC_019843.3;YP_009047203.1;MERS_1a;2496 | 0 | 0 |
| NC_019843.3;YP_009047203.1;MERS_1a;2497 | 0 | 0.025 |
| NC_019843.3;YP_009047203.1;MERS_1a;2498 | 0.05 | 0 |
| NC_019843.3;YP_009047203.1;MERS_1a;2499 | 0.05 | 0.025 |
| NC_019843.3;YP_009047203.1;MERS_1a;2500 | 0 | 0 |
| NC_019843.3;YP_009047203.1;MERS_1a;2501 | 0 | 0 |
| NC_019843.3;YP_009047203.1;MERS_1a;2502 | 0 | 0 |
| NC_019843.3;YP_009047203.1;MERS_1a;2503 | 0 | 0 |
| NC_019843.3;YP_009047203.1;MERS_1a;2504 | 0 | 0 |
| NC_019843.3;YP_009047203.1;MERS_1a;2505 | 0 | 0 |
| NC_019843.3;YP_009047203.1;MERS_1a;2506 | 0 | 0 |
| NC_019843.3;YP_009047203.1;MERS_1a;2507 | 0 | 0 |
| NC_019843.3;YP_009047203.1;MERS_1a;2508 | 0 | 0 |
| NC_019843.3;YP_009047203.1;MERS_1a;2509 | 0 | 0 |
| NC_019843.3;YP_009047203.1;MERS_1a;2510 | 0 | 0.025 |
| NC_019843.3;YP_009047203.1;MERS_1a;2511 | 0 | 0.025 |
| NC_019843.3;YP_009047203.1;MERS_1a;2512 | 0 | 0.025 |
| NC_019843.3;YP_009047203.1;MERS_1a;2513 | 0 | 0.025 |

| | | |
|---|---|---|
| NC_019843.3;YP_009047203.1;MERS_1a;2514 | 0 | 0.05 |
| NC_019843.3;YP_009047203.1;MERS_1a;2515 | 0 | 0.05 |
| NC_019843.3;YP_009047203.1;MERS_1a;2516 | 0 | 0 |
| NC_019843.3;YP_009047203.1;MERS_1a;2517 | 0 | 0 |
| NC_019843.3;YP_009047203.1;MERS_1a;2518 | 0 | 0 |
| NC_019843.3;YP_009047203.1;MERS_1a;2519 | 0 | 0 |
| NC_019843.3;YP_009047203.1;MERS_1a;2520 | 0 | 0 |
| NC_019843.3;YP_009047203.1;MERS_1a;2521 | 0 | 0 |
| NC_019843.3;YP_009047203.1;MERS_1a;2522 | 0 | 0 |
| NC_019843.3;YP_009047203.1;MERS_1a;2523 | 0 | 0.025 |
| NC_019843.3;YP_009047203.1;MERS_1a;2524 | 0 | 0 |
| NC_019843.3;YP_009047203.1;MERS_1a;2525 | 0 | 0 |
| NC_019843.3;YP_009047203.1;MERS_1a;2526 | 0 | 0 |
| NC_019843.3;YP_009047203.1;MERS_1a;2527 | 0 | 0 |
| NC_019843.3;YP_009047203.1;MERS_1a;2528 | 0 | 0 |
| NC_019843.3;YP_009047203.1;MERS_1a;2529 | 0 | 0 |
| NC_019843.3;YP_009047203.1;MERS_1a;2530 | 0 | 0 |
| NC_019843.3;YP_009047203.1;MERS_1a;2531 | 0 | 0 |
| NC_019843.3;YP_009047203.1;MERS_1a;2532 | 0 | 0 |
| NC_019843.3;YP_009047203.1;MERS_1a;2533 | 0 | 0 |
| NC_019843.3;YP_009047203.1;MERS_1a;2534 | 0 | 0 |
| NC_019843.3;YP_009047203.1;MERS_1a;2535 | 0 | 0 |
| NC_019843.3;YP_009047203.1;MERS_1a;2536 | 0 | 0 |
| NC_019843.3;YP_009047203.1;MERS_1a;2537 | 0 | 0 |
| NC_019843.3;YP_009047203.1;MERS_1a;2538 | 0 | 0 |
| NC_019843.3;YP_009047203.1;MERS_1a;2539 | 0 | 0 |
| NC_019843.3;YP_009047203.1;MERS_1a;2540 | 0 | 0 |
| NC_019843.3;YP_009047203.1;MERS_1a;2541 | 0 | 0 |
| NC_019843.3;YP_009047203.1;MERS_1a;2542 | 0 | 0 |
| NC_019843.3;YP_009047203.1;MERS_1a;2543 | 0 | 0 |
| NC_019843.3;YP_009047203.1;MERS_1a;2544 | 0.25 | 0.2 |
| NC_019843.3;YP_009047203.1;MERS_1a;2545 | 0 | 0 |
| NC_019843.3;YP_009047203.1;MERS_1a;2546 | 0 | 0 |
| NC_019843.3;YP_009047203.1;MERS_1a;2547 | 0 | 0 |
| NC_019843.3;YP_009047203.1;MERS_1a;2548 | 0 | 0 |
| NC_019843.3;YP_009047203.1;MERS_1a;2549 | 0 | 0 |
| NC_019843.3;YP_009047203.1;MERS_1a;2550 | 0 | 0 |
| NC_019843.3;YP_009047203.1;MERS_1a;2551 | 0 | 0 |
| NC_019843.3;YP_009047203.1;MERS_1a;2552 | 0 | 0 |
| NC_019843.3;YP_009047203.1;MERS_1a;2553 | 0 | 0 |
| NC_019843.3;YP_009047203.1;MERS_1a;2554 | 0 | 0 |
| NC_019843.3;YP_009047203.1;MERS_1a;2555 | 0 | 0 |
| NC_019843.3;YP_009047203.1;MERS_1a;2556 | 0 | 0 |

| | | |
|---|---|---|
| NC_019843.3;YP_009047203.1;MERS_1a;2557 | 0 | 0.05 |
| NC_019843.3;YP_009047203.1;MERS_1a;2558 | 0 | 0.025 |
| NC_019843.3;YP_009047203.1;MERS_1a;2559 | 0 | 0.025 |
| NC_019843.3;YP_009047203.1;MERS_1a;2560 | 0 | 0.025 |
| NC_019843.3;YP_009047203.1;MERS_1a;2561 | 0 | 0.025 |
| NC_019843.3;YP_009047203.1;MERS_1a;2562 | 0 | 0.025 |
| NC_019843.3;YP_009047203.1;MERS_1a;2563 | 0 | 0.05 |
| NC_019843.3;YP_009047203.1;MERS_1a;2564 | 0 | 0 |
| NC_019843.3;YP_009047203.1;MERS_1a;2565 | 0.05 | 0.025 |
| NC_019843.3;YP_009047203.1;MERS_1a;2566 | 0 | 0 |
| NC_019843.3;YP_009047203.1;MERS_1a;2567 | 0 | 0.025 |
| NC_019843.3;YP_009047203.1;MERS_1a;2568 | 0 | 0 |
| NC_019843.3;YP_009047203.1;MERS_1a;2569 | 0 | 0 |
| NC_019843.3;YP_009047203.1;MERS_1a;2570 | 0 | 0 |
| NC_019843.3;YP_009047203.1;MERS_1a;2571 | 0 | 0 |
| NC_019843.3;YP_009047203.1;MERS_1a;2572 | 0 | 0 |
| NC_019843.3;YP_009047203.1;MERS_1a;2573 | 0 | 0 |
| NC_019843.3;YP_009047203.1;MERS_1a;2574 | 0 | 0 |
| NC_019843.3;YP_009047203.1;MERS_1a;2575 | 0 | 0 |
| NC_019843.3;YP_009047203.1;MERS_1a;2576 | 0 | 0 |
| NC_019843.3;YP_009047203.1;MERS_1a;2577 | 0.05 | 0.025 |
| NC_019843.3;YP_009047203.1;MERS_1a;2578 | 0.05 | 0.025 |
| NC_019843.3;YP_009047203.1;MERS_1a;2579 | 0.05 | 0.025 |
| NC_019843.3;YP_009047203.1;MERS_1a;2580 | 0.05 | 0.025 |
| NC_019843.3;YP_009047203.1;MERS_1a;2581 | 0.05 | 0.025 |
| NC_019843.3;YP_009047203.1;MERS_1a;2582 | 0 | 0.075 |
| NC_019843.3;YP_009047203.1;MERS_1a;2583 | 0 | 0.1 |
| NC_019843.3;YP_009047203.1;MERS_1a;2584 | 0 | 0.05 |
| NC_019843.3;YP_009047203.1;MERS_1a;2585 | 0 | 0.025 |
| NC_019843.3;YP_009047203.1;MERS_1a;2586 | 0 | 0.05 |
| NC_019843.3;YP_009047203.1;MERS_1a;2587 | 0 | 0.075 |
| NC_019843.3;YP_009047203.1;MERS_1a;2588 | 0.05 | 0.025 |
| NC_019843.3;YP_009047203.1;MERS_1a;2589 | 0.05 | 0.075 |
| NC_019843.3;YP_009047203.1;MERS_1a;2590 | 0 | 0.025 |
| NC_019843.3;YP_009047203.1;MERS_1a;2591 | 0 | 0 |
| NC_019843.3;YP_009047203.1;MERS_1a;2592 | 0 | 0 |
| NC_019843.3;YP_009047203.1;MERS_1a;2593 | 0 | 0 |
| NC_019843.3;YP_009047203.1;MERS_1a;2594 | 0 | 0 |
| NC_019843.3;YP_009047203.1;MERS_1a;2595 | 0 | 0 |
| NC_019843.3;YP_009047203.1;MERS_1a;2596 | 0 | 0 |
| NC_019843.3;YP_009047203.1;MERS_1a;2597 | 0 | 0 |
| NC_019843.3;YP_009047203.1;MERS_1a;2598 | 0 | 0 |
| NC_019843.3;YP_009047203.1;MERS_1a;2599 | 0 | 0.025 |

| | | |
|---|---|---|
| NC_019843.3;YP_009047203.1;MERS_1a;2600 | 0 | 0.025 |
| NC_019843.3;YP_009047203.1;MERS_1a;2601 | 0 | 0.025 |
| NC_019843.3;YP_009047203.1;MERS_1a;2602 | 0 | 0.05 |
| NC_019843.3;YP_009047203.1;MERS_1a;2603 | 0 | 0.025 |
| NC_019843.3;YP_009047203.1;MERS_1a;2604 | 0 | 0.025 |
| NC_019843.3;YP_009047203.1;MERS_1a;2605 | 0 | 0.025 |
| NC_019843.3;YP_009047203.1;MERS_1a;2606 | 0 | 0.025 |
| NC_019843.3;YP_009047203.1;MERS_1a;2607 | 0 | 0 |
| NC_019843.3;YP_009047203.1;MERS_1a;2608 | 0.05 | 0 |
| NC_019843.3;YP_009047203.1;MERS_1a;2609 | 0 | 0 |
| NC_019843.3;YP_009047203.1;MERS_1a;2610 | 0.05 | 0 |
| NC_019843.3;YP_009047203.1;MERS_1a;2611 | 0.05 | 0 |
| NC_019843.3;YP_009047203.1;MERS_1a;2612 | 0.1 | 0.025 |
| NC_019843.3;YP_009047203.1;MERS_1a;2613 | 0.05 | 0 |
| NC_019843.3;YP_009047203.1;MERS_1a;2614 | 0.05 | 0 |
| NC_019843.3;YP_009047203.1;MERS_1a;2615 | 0.05 | 0.05 |
| NC_019843.3;YP_009047203.1;MERS_1a;2616 | 0.05 | 0 |
| NC_019843.3;YP_009047203.1;MERS_1a;2617 | 0 | 0 |
| NC_019843.3;YP_009047203.1;MERS_1a;2618 | 0 | 0 |
| NC_019843.3;YP_009047203.1;MERS_1a;2619 | 0 | 0 |
| NC_019843.3;YP_009047203.1;MERS_1a;2620 | 0 | 0 |
| NC_019843.3;YP_009047203.1;MERS_1a;2621 | 0 | 0 |
| NC_019843.3;YP_009047203.1;MERS_1a;2622 | 0 | 0 |
| NC_019843.3;YP_009047203.1;MERS_1a;2623 | 0 | 0 |
| NC_019843.3;YP_009047203.1;MERS_1a;2624 | 0 | 0 |
| NC_019843.3;YP_009047203.1;MERS_1a;2625 | 0 | 0 |
| NC_019843.3;YP_009047203.1;MERS_1a;2626 | 0.1 | 0.05 |
| NC_019843.3;YP_009047203.1;MERS_1a;2627 | 0.05 | 0.025 |
| NC_019843.3;YP_009047203.1;MERS_1a;2628 | 0.1 | 0.025 |
| NC_019843.3;YP_009047203.1;MERS_1a;2629 | 0.15 | 0.075 |
| NC_019843.3;YP_009047203.1;MERS_1a;2630 | 0.2 | 0.075 |
| NC_019843.3;YP_009047203.1;MERS_1a;2631 | 0.15 | 0.075 |
| NC_019843.3;YP_009047203.1;MERS_1a;2632 | 0.15 | 0.075 |
| NC_019843.3;YP_009047203.1;MERS_1a;2633 | 0.15 | 0.075 |
| NC_019843.3;YP_009047203.1;MERS_1a;2634 | 0.2 | 0.05 |
| NC_019843.3;YP_009047203.1;MERS_1a;2635 | 0 | 0.175 |
| NC_019843.3;YP_009047203.1;MERS_1a;2636 | 0 | 0.175 |
| NC_019843.3;YP_009047203.1;MERS_1a;2637 | 0.1 | 0.15 |
| NC_019843.3;YP_009047203.1;MERS_1a;2638 | 0.1 | 0.025 |
| NC_019843.3;YP_009047203.1;MERS_1a;2639 | 0.1 | 0.1 |
| NC_019843.3;YP_009047203.1;MERS_1a;2640 | 0 | 0.075 |
| NC_019843.3;YP_009047203.1;MERS_1a;2641 | 0.05 | 0.025 |
| NC_019843.3;YP_009047203.1;MERS_1a;2642 | 0.1 | 0.075 |

| | | |
|---|---|---|
| NC_019843.3;YP_009047203.1;MERS_1a;2643 | 0.45 | 0.175 |
| NC_019843.3;YP_009047203.1;MERS_1a;2644 | 0.15 | 0.125 |
| NC_019843.3;YP_009047203.1;MERS_1a;2645 | 0 | 0.075 |
| NC_019843.3;YP_009047203.1;MERS_1a;2646 | 0 | 0.05 |
| NC_019843.3;YP_009047203.1;MERS_1a;2647 | 0 | 0.025 |
| NC_019843.3;YP_009047203.1;MERS_1a;2648 | 0.05 | 0.05 |
| NC_019843.3;YP_009047203.1;MERS_1a;2649 | 0 | 0 |
| NC_019843.3;YP_009047203.1;MERS_1a;2650 | 0 | 0 |
| NC_019843.3;YP_009047203.1;MERS_1a;2651 | 0 | 0 |
| NC_019843.3;YP_009047203.1;MERS_1a;2652 | 0 | 0 |
| NC_019843.3;YP_009047203.1;MERS_1a;2653 | 0 | 0 |
| NC_019843.3;YP_009047203.1;MERS_1a;2654 | 0 | 0 |
| NC_019843.3;YP_009047203.1;MERS_1a;2655 | 0 | 0 |
| NC_019843.3;YP_009047203.1;MERS_1a;2656 | 0 | 0.025 |
| NC_019843.3;YP_009047203.1;MERS_1a;2657 | 0.05 | 0 |
| NC_019843.3;YP_009047203.1;MERS_1a;2658 | 0 | 0 |
| NC_019843.3;YP_009047203.1;MERS_1a;2659 | 0 | 0 |
| NC_019843.3;YP_009047203.1;MERS_1a;2660 | 0 | 0 |
| NC_019843.3;YP_009047203.1;MERS_1a;2661 | 0 | 0 |
| NC_019843.3;YP_009047203.1;MERS_1a;2662 | 0 | 0.025 |
| NC_019843.3;YP_009047203.1;MERS_1a;2663 | 0 | 0 |
| NC_019843.3;YP_009047203.1;MERS_1a;2664 | 0 | 0.025 |
| NC_019843.3;YP_009047203.1;MERS_1a;2665 | 0 | 0 |
| NC_019843.3;YP_009047203.1;MERS_1a;2666 | 0 | 0 |
| NC_019843.3;YP_009047203.1;MERS_1a;2667 | 0 | 0 |
| NC_019843.3;YP_009047203.1;MERS_1a;2668 | 0 | 0.025 |
| NC_019843.3;YP_009047203.1;MERS_1a;2669 | 0 | 0.025 |
| NC_019843.3;YP_009047203.1;MERS_1a;2670 | 0 | 0 |
| NC_019843.3;YP_009047203.1;MERS_1a;2671 | 0 | 0 |
| NC_019843.3;YP_009047203.1;MERS_1a;2672 | 0 | 0 |
| NC_019843.3;YP_009047203.1;MERS_1a;2673 | 0 | 0 |
| NC_019843.3;YP_009047203.1;MERS_1a;2674 | 0 | 0 |
| NC_019843.3;YP_009047203.1;MERS_1a;2675 | 0 | 0 |
| NC_019843.3;YP_009047203.1;MERS_1a;2676 | 0.05 | 0 |
| NC_019843.3;YP_009047203.1;MERS_1a;2677 | 0 | 0 |
| NC_019843.3;YP_009047203.1;MERS_1a;2678 | 0 | 0 |
| NC_019843.3;YP_009047203.1;MERS_1a;2679 | 0 | 0 |
| NC_019843.3;YP_009047203.1;MERS_1a;2680 | 0 | 0 |
| NC_019843.3;YP_009047203.1;MERS_1a;2681 | 0 | 0 |
| NC_019843.3;YP_009047203.1;MERS_1a;2682 | 0.05 | 0 |
| NC_019843.3;YP_009047203.1;MERS_1a;2683 | 0.05 | 0 |
| NC_019843.3;YP_009047203.1;MERS_1a;2684 | 0.05 | 0 |
| NC_019843.3;YP_009047203.1;MERS_1a;2685 | 0 | 0 |

| | | |
|---|---|---|
| NC_019843.3;YP_009047203.1;MERS_1a;2686 | 0.05 | 0 |
| NC_019843.3;YP_009047203.1;MERS_1a;2687 | 0 | 0 |
| NC_019843.3;YP_009047203.1;MERS_1a;2688 | 0 | 0 |
| NC_019843.3;YP_009047203.1;MERS_1a;2689 | 0 | 0 |
| NC_019843.3;YP_009047203.1;MERS_1a;2690 | 0 | 0 |
| NC_019843.3;YP_009047203.1;MERS_1a;2691 | 0 | 0 |
| NC_019843.3;YP_009047203.1;MERS_1a;2692 | 0 | 0 |
| NC_019843.3;YP_009047203.1;MERS_1a;2693 | 0 | 0 |
| NC_019843.3;YP_009047203.1;MERS_1a;2694 | 0 | 0 |
| NC_019843.3;YP_009047203.1;MERS_1a;2695 | 0 | 0 |
| NC_019843.3;YP_009047203.1;MERS_1a;2696 | 0 | 0 |
| NC_019843.3;YP_009047203.1;MERS_1a;2697 | 0 | 0 |
| NC_019843.3;YP_009047203.1;MERS_1a;2698 | 0 | 0 |
| NC_019843.3;YP_009047203.1;MERS_1a;2699 | 0 | 0 |
| NC_019843.3;YP_009047203.1;MERS_1a;2700 | 0 | 0.025 |
| NC_019843.3;YP_009047203.1;MERS_1a;2701 | 0 | 0 |
| NC_019843.3;YP_009047203.1;MERS_1a;2702 | 0 | 0 |
| NC_019843.3;YP_009047203.1;MERS_1a;2703 | 0 | 0 |
| NC_019843.3;YP_009047203.1;MERS_1a;2704 | 0 | 0 |
| NC_019843.3;YP_009047203.1;MERS_1a;2705 | 0 | 0 |
| NC_019843.3;YP_009047203.1;MERS_1a;2706 | 0 | 0 |
| NC_019843.3;YP_009047203.1;MERS_1a;2707 | 0 | 0 |
| NC_019843.3;YP_009047203.1;MERS_1a;2708 | 0 | 0 |
| NC_019843.3;YP_009047203.1;MERS_1a;2709 | 0 | 0 |
| NC_019843.3;YP_009047203.1;MERS_1a;2710 | 0 | 0 |
| NC_019843.3;YP_009047203.1;MERS_1a;2711 | 0 | 0 |
| NC_019843.3;YP_009047203.1;MERS_1a;2712 | 0 | 0 |
| NC_019843.3;YP_009047203.1;MERS_1a;2713 | 0 | 0 |
| NC_019843.3;YP_009047203.1;MERS_1a;2714 | 0 | 0 |
| NC_019843.3;YP_009047203.1;MERS_1a;2715 | 0.05 | 0 |
| NC_019843.3;YP_009047203.1;MERS_1a;2716 | 0 | 0 |
| NC_019843.3;YP_009047203.1;MERS_1a;2717 | 0 | 0 |
| NC_019843.3;YP_009047203.1;MERS_1a;2718 | 0 | 0 |
| NC_019843.3;YP_009047203.1;MERS_1a;2719 | 0 | 0 |
| NC_019843.3;YP_009047203.1;MERS_1a;2720 | 0 | 0 |
| NC_019843.3;YP_009047203.1;MERS_1a;2721 | 0 | 0 |
| NC_019843.3;YP_009047203.1;MERS_1a;2722 | 0 | 0 |
| NC_019843.3;YP_009047203.1;MERS_1a;2723 | 0 | 0 |
| NC_019843.3;YP_009047203.1;MERS_1a;2724 | 0 | 0 |
| NC_019843.3;YP_009047203.1;MERS_1a;2725 | 0 | 0 |
| NC_019843.3;YP_009047203.1;MERS_1a;2726 | 0 | 0 |
| NC_019843.3;YP_009047203.1;MERS_1a;2727 | 0 | 0 |
| NC_019843.3;YP_009047203.1;MERS_1a;2728 | 0 | 0 |

| | | |
|---|---|---|
| NC_019843.3;YP_009047203.1;MERS_1a;2729 | 0 | 0 |
| NC_019843.3;YP_009047203.1;MERS_1a;2730 | 0 | 0 |
| NC_019843.3;YP_009047203.1;MERS_1a;2731 | 0.05 | 0 |
| NC_019843.3;YP_009047203.1;MERS_1a;2732 | 0 | 0.025 |
| NC_019843.3;YP_009047203.1;MERS_1a;2733 | 0 | 0.025 |
| NC_019843.3;YP_009047203.1;MERS_1a;2734 | 0 | 0.025 |
| NC_019843.3;YP_009047203.1;MERS_1a;2735 | 0 | 0.025 |
| NC_019843.3;YP_009047203.1;MERS_1a;2736 | 0 | 0.025 |
| NC_019843.3;YP_009047203.1;MERS_1a;2737 | 0 | 0 |
| NC_019843.3;YP_009047203.1;MERS_1a;2738 | 0 | 0 |
| NC_019843.3;YP_009047203.1;MERS_1a;2739 | 0 | 0 |
| NC_019843.3;YP_009047203.1;MERS_1a;2740 | 0 | 0 |
| NC_019843.3;YP_009047203.1;MERS_1a;2741 | 0 | 0 |
| NC_019843.3;YP_009047203.1;MERS_1a;2742 | 0 | 0 |
| NC_019843.3;YP_009047203.1;MERS_1a;2743 | 0 | 0 |
| NC_019843.3;YP_009047203.1;MERS_1a;2744 | 0 | 0 |
| NC_019843.3;YP_009047203.1;MERS_1a;2745 | 0 | 0 |
| NC_019843.3;YP_009047203.1;MERS_1a;2746 | 0 | 0 |
| NC_019843.3;YP_009047203.1;MERS_1a;2747 | 0 | 0 |
| NC_019843.3;YP_009047203.1;MERS_1a;2748 | 0 | 0 |
| NC_019843.3;YP_009047203.1;MERS_1a;2749 | 0 | 0 |
| NC_019843.3;YP_009047203.1;MERS_1a;2750 | 0 | 0 |
| NC_019843.3;YP_009047203.1;MERS_1a;2751 | 0 | 0 |
| NC_019843.3;YP_009047203.1;MERS_1a;2752 | 0 | 0 |
| NC_019843.3;YP_009047203.1;MERS_1a;2753 | 0 | 0 |
| NC_019843.3;YP_009047203.1;MERS_1a;2754 | 0 | 0 |
| NC_019843.3;YP_009047203.1;MERS_1a;2755 | 0 | 0 |
| NC_019843.3;YP_009047203.1;MERS_1a;2756 | 0 | 0 |
| NC_019843.3;YP_009047203.1;MERS_1a;2757 | 0 | 0 |
| NC_019843.3;YP_009047203.1;MERS_1a;2758 | 0 | 0 |
| NC_019843.3;YP_009047203.1;MERS_1a;2759 | 0 | 0 |
| NC_019843.3;YP_009047203.1;MERS_1a;2760 | 0 | 0 |
| NC_019843.3;YP_009047203.1;MERS_1a;2761 | 0 | 0 |
| NC_019843.3;YP_009047203.1;MERS_1a;2762 | 0 | 0 |
| NC_019843.3;YP_009047203.1;MERS_1a;2763 | 0 | 0 |
| NC_019843.3;YP_009047203.1;MERS_1a;2764 | 0 | 0 |
| NC_019843.3;YP_009047203.1;MERS_1a;2765 | 0 | 0 |
| NC_019843.3;YP_009047203.1;MERS_1a;2766 | 0 | 0 |
| NC_019843.3;YP_009047203.1;MERS_1a;2767 | 0 | 0 |
| NC_019843.3;YP_009047203.1;MERS_1a;2768 | 0 | 0 |
| NC_019843.3;YP_009047203.1;MERS_1a;2769 | 0 | 0 |
| NC_019843.3;YP_009047203.1;MERS_1a;2770 | 0 | 0 |
| NC_019843.3;YP_009047203.1;MERS_1a;2771 | 0 | 0 |

| | | |
|---|---|---|
| NC_019843.3;YP_009047203.1;MERS_1a;2772 | 0 | 0 |
| NC_019843.3;YP_009047203.1;MERS_1a;2773 | 0 | 0.025 |
| NC_019843.3;YP_009047203.1;MERS_1a;2774 | 0 | 0.075 |
| NC_019843.3;YP_009047203.1;MERS_1a;2775 | 0 | 0.025 |
| NC_019843.3;YP_009047203.1;MERS_1a;2776 | 0 | 0.025 |
| NC_019843.3;YP_009047203.1;MERS_1a;2777 | 0 | 0.05 |
| NC_019843.3;YP_009047203.1;MERS_1a;2778 | 0 | 0.125 |
| NC_019843.3;YP_009047203.1;MERS_1a;2779 | 0 | 0.05 |
| NC_019843.3;YP_009047203.1;MERS_1a;2780 | 0.2 | 0.25 |
| NC_019843.3;YP_009047203.1;MERS_1a;2781 | 0.05 | 0.05 |
| NC_019843.3;YP_009047203.1;MERS_1a;2782 | 0 | 0.025 |
| NC_019843.3;YP_009047203.1;MERS_1a;2783 | 0 | 0.05 |
| NC_019843.3;YP_009047203.1;MERS_1a;2784 | 0 | 0 |
| NC_019843.3;YP_009047203.1;MERS_1a;2785 | 0 | 0 |
| NC_019843.3;YP_009047203.1;MERS_1a;2786 | 0 | 0 |
| NC_019843.3;YP_009047203.1;MERS_1a;2787 | 0 | 0 |
| NC_019843.3;YP_009047203.1;MERS_1a;2788 | 0.05 | 0 |
| NC_019843.3;YP_009047203.1;MERS_1a;2789 | 0.05 | 0.05 |
| NC_019843.3;YP_009047203.1;MERS_1a;2790 | 0 | 0 |
| NC_019843.3;YP_009047203.1;MERS_1a;2791 | 0 | 0 |
| NC_019843.3;YP_009047203.1;MERS_1a;2792 | 0 | 0.025 |
| NC_019843.3;YP_009047203.1;MERS_1a;2793 | 0 | 0.05 |
| NC_019843.3;YP_009047203.1;MERS_1a;2794 | 0.05 | 0 |
| NC_019843.3;YP_009047203.1;MERS_1a;2795 | 0 | 0 |
| NC_019843.3;YP_009047203.1;MERS_1a;2796 | 0 | 0 |
| NC_019843.3;YP_009047203.1;MERS_1a;2797 | 0 | 0.05 |
| NC_019843.3;YP_009047203.1;MERS_1a;2798 | 0 | 0.075 |
| NC_019843.3;YP_009047203.1;MERS_1a;2799 | 0 | 0.05 |
| NC_019843.3;YP_009047203.1;MERS_1a;2800 | 0 | 0 |
| NC_019843.3;YP_009047203.1;MERS_1a;2801 | 0 | 0 |
| NC_019843.3;YP_009047203.1;MERS_1a;2802 | 0 | 0 |
| NC_019843.3;YP_009047203.1;MERS_1a;2803 | 0 | 0.025 |
| NC_019843.3;YP_009047203.1;MERS_1a;2804 | 0 | 0.025 |
| NC_019843.3;YP_009047203.1;MERS_1a;2805 | 0 | 0 |
| NC_019843.3;YP_009047203.1;MERS_1a;2806 | 0 | 0 |
| NC_019843.3;YP_009047203.1;MERS_1a;2807 | 0 | 0 |
| NC_019843.3;YP_009047203.1;MERS_1a;2808 | 0 | 0 |
| NC_019843.3;YP_009047203.1;MERS_1a;2809 | 0 | 0 |
| NC_019843.3;YP_009047203.1;MERS_1a;2810 | 0 | 0 |
| NC_019843.3;YP_009047203.1;MERS_1a;2811 | 0 | 0 |
| NC_019843.3;YP_009047203.1;MERS_1a;2812 | 0 | 0 |
| NC_019843.3;YP_009047203.1;MERS_1a;2813 | 0 | 0 |
| NC_019843.3;YP_009047203.1;MERS_1a;2814 | 0 | 0 |

| | | |
|---|---|---|
| NC_019843.3;YP_009047203.1;MERS_1a;2815 | 0.05 | 0 |
| NC_019843.3;YP_009047203.1;MERS_1a;2816 | 0.05 | 0 |
| NC_019843.3;YP_009047203.1;MERS_1a;2817 | 0.1 | 0 |
| NC_019843.3;YP_009047203.1;MERS_1a;2818 | 0.05 | 0.025 |
| NC_019843.3;YP_009047203.1;MERS_1a;2819 | 0.05 | 0.05 |
| NC_019843.3;YP_009047203.1;MERS_1a;2820 | 0 | 0 |
| NC_019843.3;YP_009047203.1;MERS_1a;2821 | 0.05 | 0 |
| NC_019843.3;YP_009047203.1;MERS_1a;2822 | 0.05 | 0 |
| NC_019843.3;YP_009047203.1;MERS_1a;2823 | 0.05 | 0 |
| NC_019843.3;YP_009047203.1;MERS_1a;2824 | 0 | 0 |
| NC_019843.3;YP_009047203.1;MERS_1a;2825 | 0 | 0.025 |
| NC_019843.3;YP_009047203.1;MERS_1a;2826 | 0 | 0 |
| NC_019843.3;YP_009047203.1;MERS_1a;2827 | 0 | 0 |
| NC_019843.3;YP_009047203.1;MERS_1a;2828 | 0 | 0 |
| NC_019843.3;YP_009047203.1;MERS_1a;2829 | 0 | 0 |
| NC_019843.3;YP_009047203.1;MERS_1a;2830 | 0 | 0 |
| NC_019843.3;YP_009047203.1;MERS_1a;2831 | 0 | 0 |
| NC_019843.3;YP_009047203.1;MERS_1a;2832 | 0 | 0 |
| NC_019843.3;YP_009047203.1;MERS_1a;2833 | 0 | 0 |
| NC_019843.3;YP_009047203.1;MERS_1a;2834 | 0 | 0 |
| NC_019843.3;YP_009047203.1;MERS_1a;2835 | 0 | 0 |
| NC_019843.3;YP_009047203.1;MERS_1a;2836 | 0 | 0 |
| NC_019843.3;YP_009047203.1;MERS_1a;2837 | 0 | 0 |
| NC_019843.3;YP_009047203.1;MERS_1a;2838 | 0 | 0 |
| NC_019843.3;YP_009047203.1;MERS_1a;2839 | 0 | 0 |
| NC_019843.3;YP_009047203.1;MERS_1a;2840 | 0.05 | 0 |
| NC_019843.3;YP_009047203.1;MERS_1a;2841 | 0.05 | 0 |
| NC_019843.3;YP_009047203.1;MERS_1a;2842 | 0.15 | 0.025 |
| NC_019843.3;YP_009047203.1;MERS_1a;2843 | 0 | 0 |
| NC_019843.3;YP_009047203.1;MERS_1a;2844 | 0.05 | 0 |
| NC_019843.3;YP_009047203.1;MERS_1a;2845 | 0 | 0 |
| NC_019843.3;YP_009047203.1;MERS_1a;2846 | 0 | 0 |
| NC_019843.3;YP_009047203.1;MERS_1a;2847 | 0.05 | 0.025 |
| NC_019843.3;YP_009047203.1;MERS_1a;2848 | 0.05 | 0 |
| NC_019843.3;YP_009047203.1;MERS_1a;2849 | 0.1 | 0 |
| NC_019843.3;YP_009047203.1;MERS_1a;2850 | 0 | 0 |
| NC_019843.3;YP_009047203.1;MERS_1a;2851 | 0 | 0 |
| NC_019843.3;YP_009047203.1;MERS_1a;2852 | 0 | 0 |
| NC_019843.3;YP_009047203.1;MERS_1a;2853 | 0 | 0 |
| NC_019843.3;YP_009047203.1;MERS_1a;2854 | 0 | 0 |
| NC_019843.3;YP_009047203.1;MERS_1a;2855 | 0 | 0 |
| NC_019843.3;YP_009047203.1;MERS_1a;2856 | 0 | 0 |
| NC_019843.3;YP_009047203.1;MERS_1a;2857 | 0 | 0 |

| | | |
|---|---|---|
| NC_019843.3;YP_009047203.1;MERS_1a;2858 | 0 | 0 |
| NC_019843.3;YP_009047203.1;MERS_1a;2859 | 0 | 0 |
| NC_019843.3;YP_009047203.1;MERS_1a;2860 | 0 | 0 |
| NC_019843.3;YP_009047203.1;MERS_1a;2861 | 0 | 0 |
| NC_019843.3;YP_009047203.1;MERS_1a;2862 | 0 | 0 |
| NC_019843.3;YP_009047203.1;MERS_1a;2863 | 0 | 0 |
| NC_019843.3;YP_009047203.1;MERS_1a;2864 | 0 | 0 |
| NC_019843.3;YP_009047203.1;MERS_1a;2865 | 0 | 0 |
| NC_019843.3;YP_009047203.1;MERS_1a;2866 | 0 | 0 |
| NC_019843.3;YP_009047203.1;MERS_1a;2867 | 0 | 0 |
| NC_019843.3;YP_009047203.1;MERS_1a;2868 | 0 | 0 |
| NC_019843.3;YP_009047203.1;MERS_1a;2869 | 0 | 0 |
| NC_019843.3;YP_009047203.1;MERS_1a;2870 | 0.05 | 0 |
| NC_019843.3;YP_009047203.1;MERS_1a;2871 | 0 | 0 |
| NC_019843.3;YP_009047203.1;MERS_1a;2872 | 0 | 0.025 |
| NC_019843.3;YP_009047203.1;MERS_1a;2873 | 0 | 0.025 |
| NC_019843.3;YP_009047203.1;MERS_1a;2874 | 0 | 0.025 |
| NC_019843.3;YP_009047203.1;MERS_1a;2875 | 0 | 0 |
| NC_019843.3;YP_009047203.1;MERS_1a;2876 | 0 | 0 |
| NC_019843.3;YP_009047203.1;MERS_1a;2877 | 0 | 0 |
| NC_019843.3;YP_009047203.1;MERS_1a;2878 | 0 | 0 |
| NC_019843.3;YP_009047203.1;MERS_1a;2879 | 0.05 | 0 |
| NC_019843.3;YP_009047203.1;MERS_1a;2880 | 0.05 | 0 |
| NC_019843.3;YP_009047203.1;MERS_1a;2881 | 0 | 0 |
| NC_019843.3;YP_009047203.1;MERS_1a;2882 | 0 | 0 |
| NC_019843.3;YP_009047203.1;MERS_1a;2883 | 0 | 0 |
| NC_019843.3;YP_009047203.1;MERS_1a;2884 | 0.05 | 0 |
| NC_019843.3;YP_009047203.1;MERS_1a;2885 | 0 | 0.05 |
| NC_019843.3;YP_009047203.1;MERS_1a;2886 | 0 | 0.025 |
| NC_019843.3;YP_009047203.1;MERS_1a;2887 | 0 | 0 |
| NC_019843.3;YP_009047203.1;MERS_1a;2888 | 0 | 0 |
| NC_019843.3;YP_009047203.1;MERS_1a;2889 | 0 | 0 |
| NC_019843.3;YP_009047203.1;MERS_1a;2890 | 0 | 0 |
| NC_019843.3;YP_009047203.1;MERS_1a;2891 | 0 | 0 |
| NC_019843.3;YP_009047203.1;MERS_1a;2892 | 0 | 0.025 |
| NC_019843.3;YP_009047203.1;MERS_1a;2893 | 0 | 0 |
| NC_019843.3;YP_009047203.1;MERS_1a;2894 | 0 | 0 |
| NC_019843.3;YP_009047203.1;MERS_1a;2895 | 0 | 0 |
| NC_019843.3;YP_009047203.1;MERS_1a;2896 | 0 | 0 |
| NC_019843.3;YP_009047203.1;MERS_1a;2897 | 0.05 | 0 |
| NC_019843.3;YP_009047203.1;MERS_1a;2898 | 0.05 | 0 |
| NC_019843.3;YP_009047203.1;MERS_1a;2899 | 0.05 | 0 |
| NC_019843.3;YP_009047203.1;MERS_1a;2900 | 0.05 | 0 |

| | | |
|---|---|---|
| NC_019843.3;YP_009047203.1;MERS_1a;2901 | 0.05 | 0 |
| NC_019843.3;YP_009047203.1;MERS_1a;2902 | 0.05 | 0 |
| NC_019843.3;YP_009047203.1;MERS_1a;2903 | 0.05 | 0 |
| NC_019843.3;YP_009047203.1;MERS_1a;2904 | 0.05 | 0 |
| NC_019843.3;YP_009047203.1;MERS_1a;2905 | 0.05 | 0.025 |
| NC_019843.3;YP_009047203.1;MERS_1a;2906 | 0.05 | 0 |
| NC_019843.3;YP_009047203.1;MERS_1a;2907 | 0 | 0 |
| NC_019843.3;YP_009047203.1;MERS_1a;2908 | 0 | 0 |
| NC_019843.3;YP_009047203.1;MERS_1a;2909 | 0 | 0 |
| NC_019843.3;YP_009047203.1;MERS_1a;2910 | 0.05 | 0 |
| NC_019843.3;YP_009047203.1;MERS_1a;2911 | 0 | 0 |
| NC_019843.3;YP_009047203.1;MERS_1a;2912 | 0.05 | 0 |
| NC_019843.3;YP_009047203.1;MERS_1a;2913 | 0 | 0 |
| NC_019843.3;YP_009047203.1;MERS_1a;2914 | 0 | 0 |
| NC_019843.3;YP_009047203.1;MERS_1a;2915 | 0 | 0 |
| NC_019843.3;YP_009047203.1;MERS_1a;2916 | 0 | 0 |
| NC_019843.3;YP_009047203.1;MERS_1a;2917 | 0 | 0 |
| NC_019843.3;YP_009047203.1;MERS_1a;2918 | 0.1 | 0 |
| NC_019843.3;YP_009047203.1;MERS_1a;2919 | 0 | 0 |
| NC_019843.3;YP_009047203.1;MERS_1a;2920 | 0 | 0 |
| NC_019843.3;YP_009047203.1;MERS_1a;2921 | 0 | 0 |
| NC_019843.3;YP_009047203.1;MERS_1a;2922 | 0 | 0 |
| NC_019843.3;YP_009047203.1;MERS_1a;2923 | 0 | 0 |
| NC_019843.3;YP_009047203.1;MERS_1a;2924 | 0.05 | 0 |
| NC_019843.3;YP_009047203.1;MERS_1a;2925 | 0 | 0 |
| NC_019843.3;YP_009047203.1;MERS_1a;2926 | 0 | 0 |
| NC_019843.3;YP_009047203.1;MERS_1a;2927 | 0 | 0.075 |
| NC_019843.3;YP_009047203.1;MERS_1a;2928 | 0 | 0.025 |
| NC_019843.3;YP_009047203.1;MERS_1a;2929 | 0 | 0 |
| NC_019843.3;YP_009047203.1;MERS_1a;2930 | 0.05 | 0.025 |
| NC_019843.3;YP_009047203.1;MERS_1a;2931 | 0 | 0 |
| NC_019843.3;YP_009047203.1;MERS_1a;2932 | 0 | 0 |
| NC_019843.3;YP_009047203.1;MERS_1a;2933 | 0 | 0 |
| NC_019843.3;YP_009047203.1;MERS_1a;2934 | 0 | 0 |
| NC_019843.3;YP_009047203.1;MERS_1a;2935 | 0.05 | 0 |
| NC_019843.3;YP_009047203.1;MERS_1a;2936 | 0.05 | 0 |
| NC_019843.3;YP_009047203.1;MERS_1a;2937 | 0.05 | 0 |
| NC_019843.3;YP_009047203.1;MERS_1a;2938 | 0.05 | 0 |
| NC_019843.3;YP_009047203.1;MERS_1a;2939 | 0 | 0.025 |
| NC_019843.3;YP_009047203.1;MERS_1a;2940 | 0 | 0.025 |
| NC_019843.3;YP_009047203.1;MERS_1a;2941 | 0 | 0 |
| NC_019843.3;YP_009047203.1;MERS_1a;2942 | 0 | 0 |
| NC_019843.3;YP_009047203.1;MERS_1a;2943 | 0 | 0 |

| | | |
|---|---|---|
| NC_019843.3;YP_009047203.1;MERS_1a;2944 | 0 | 0 |
| NC_019843.3;YP_009047203.1;MERS_1a;2945 | 0 | 0 |
| NC_019843.3;YP_009047203.1;MERS_1a;2946 | 0 | 0 |
| NC_019843.3;YP_009047203.1;MERS_1a;2947 | 0 | 0 |
| NC_019843.3;YP_009047203.1;MERS_1a;2948 | 0 | 0 |
| NC_019843.3;YP_009047203.1;MERS_1a;2949 | 0 | 0 |
| NC_019843.3;YP_009047203.1;MERS_1a;2950 | 0 | 0 |
| NC_019843.3;YP_009047203.1;MERS_1a;2951 | 0 | 0 |
| NC_019843.3;YP_009047203.1;MERS_1a;2952 | 0 | 0 |
| NC_019843.3;YP_009047203.1;MERS_1a;2953 | 0 | 0 |
| NC_019843.3;YP_009047203.1;MERS_1a;2954 | 0 | 0 |
| NC_019843.3;YP_009047203.1;MERS_1a;2955 | 0 | 0 |
| NC_019843.3;YP_009047203.1;MERS_1a;2956 | 0 | 0 |
| NC_019843.3;YP_009047203.1;MERS_1a;2957 | 0 | 0 |
| NC_019843.3;YP_009047203.1;MERS_1a;2958 | 0 | 0 |
| NC_019843.3;YP_009047203.1;MERS_1a;2959 | 0 | 0 |
| NC_019843.3;YP_009047203.1;MERS_1a;2960 | 0 | 0 |
| NC_019843.3;YP_009047203.1;MERS_1a;2961 | 0 | 0 |
| NC_019843.3;YP_009047203.1;MERS_1a;2962 | 0 | 0 |
| NC_019843.3;YP_009047203.1;MERS_1a;2963 | 0 | 0 |
| NC_019843.3;YP_009047203.1;MERS_1a;2964 | 0 | 0 |
| NC_019843.3;YP_009047203.1;MERS_1a;2965 | 0 | 0 |
| NC_019843.3;YP_009047203.1;MERS_1a;2966 | 0 | 0 |
| NC_019843.3;YP_009047203.1;MERS_1a;2967 | 0 | 0 |
| NC_019843.3;YP_009047203.1;MERS_1a;2968 | 0 | 0 |
| NC_019843.3;YP_009047203.1;MERS_1a;2969 | 0 | 0 |
| NC_019843.3;YP_009047203.1;MERS_1a;2970 | 0 | 0 |
| NC_019843.3;YP_009047203.1;MERS_1a;2971 | 0 | 0 |
| NC_019843.3;YP_009047203.1;MERS_1a;2972 | 0 | 0 |
| NC_019843.3;YP_009047203.1;MERS_1a;2973 | 0 | 0 |
| NC_019843.3;YP_009047203.1;MERS_1a;2974 | 0 | 0 |
| NC_019843.3;YP_009047203.1;MERS_1a;2975 | 0 | 0 |
| NC_019843.3;YP_009047203.1;MERS_1a;2976 | 0 | 0 |
| NC_019843.3;YP_009047203.1;MERS_1a;2977 | 0 | 0 |
| NC_019843.3;YP_009047203.1;MERS_1a;2978 | 0 | 0.025 |
| NC_019843.3;YP_009047203.1;MERS_1a;2979 | 0 | 0.025 |
| NC_019843.3;YP_009047203.1;MERS_1a;2980 | 0 | 0.025 |
| NC_019843.3;YP_009047203.1;MERS_1a;2981 | 0 | 0.025 |
| NC_019843.3;YP_009047203.1;MERS_1a;2982 | 0 | 0.025 |
| NC_019843.3;YP_009047203.1;MERS_1a;2983 | 0 | 0.025 |
| NC_019843.3;YP_009047203.1;MERS_1a;2984 | 0 | 0.025 |
| NC_019843.3;YP_009047203.1;MERS_1a;2985 | 0 | 0.025 |
| NC_019843.3;YP_009047203.1;MERS_1a;2986 | 0 | 0.025 |

| | | |
|---|---|---|
| NC_019843.3;YP_009047203.1;MERS_1a;2987 | 0 | 0.025 |
| NC_019843.3;YP_009047203.1;MERS_1a;2988 | 0 | 0.025 |
| NC_019843.3;YP_009047203.1;MERS_1a;2989 | 0 | 0.025 |
| NC_019843.3;YP_009047203.1;MERS_1a;2990 | 0 | 0 |
| NC_019843.3;YP_009047203.1;MERS_1a;2991 | 0 | 0 |
| NC_019843.3;YP_009047203.1;MERS_1a;2992 | 0 | 0 |
| NC_019843.3;YP_009047203.1;MERS_1a;2993 | 0 | 0 |
| NC_019843.3;YP_009047203.1;MERS_1a;2994 | 0 | 0 |
| NC_019843.3;YP_009047203.1;MERS_1a;2995 | 0 | 0 |
| NC_019843.3;YP_009047203.1;MERS_1a;2996 | 0 | 0 |
| NC_019843.3;YP_009047203.1;MERS_1a;2997 | 0 | 0 |
| NC_019843.3;YP_009047203.1;MERS_1a;2998 | 0 | 0 |
| NC_019843.3;YP_009047203.1;MERS_1a;2999 | 0 | 0 |
| NC_019843.3;YP_009047203.1;MERS_1a;3000 | 0 | 0 |
| NC_019843.3;YP_009047203.1;MERS_1a;3001 | 0 | 0 |
| NC_019843.3;YP_009047203.1;MERS_1a;3002 | 0 | 0 |
| NC_019843.3;YP_009047203.1;MERS_1a;3003 | 0 | 0 |
| NC_019843.3;YP_009047203.1;MERS_1a;3004 | 0 | 0 |
| NC_019843.3;YP_009047203.1;MERS_1a;3005 | 0 | 0 |
| NC_019843.3;YP_009047203.1;MERS_1a;3006 | 0 | 0 |
| NC_019843.3;YP_009047203.1;MERS_1a;3007 | 0 | 0 |
| NC_019843.3;YP_009047203.1;MERS_1a;3008 | 0 | 0 |
| NC_019843.3;YP_009047203.1;MERS_1a;3009 | 0 | 0 |
| NC_019843.3;YP_009047203.1;MERS_1a;3010 | 0 | 0 |
| NC_019843.3;YP_009047203.1;MERS_1a;3011 | 0 | 0 |
| NC_019843.3;YP_009047203.1;MERS_1a;3012 | 0 | 0 |
| NC_019843.3;YP_009047203.1;MERS_1a;3013 | 0 | 0 |
| NC_019843.3;YP_009047203.1;MERS_1a;3014 | 0 | 0 |
| NC_019843.3;YP_009047203.1;MERS_1a;3015 | 0 | 0 |
| NC_019843.3;YP_009047203.1;MERS_1a;3016 | 0 | 0 |
| NC_019843.3;YP_009047203.1;MERS_1a;3017 | 0 | 0 |
| NC_019843.3;YP_009047203.1;MERS_1a;3018 | 0 | 0 |
| NC_019843.3;YP_009047203.1;MERS_1a;3019 | 0 | 0 |
| NC_019843.3;YP_009047203.1;MERS_1a;3020 | 0 | 0 |
| NC_019843.3;YP_009047203.1;MERS_1a;3021 | 0 | 0 |
| NC_019843.3;YP_009047203.1;MERS_1a;3022 | 0 | 0 |
| NC_019843.3;YP_009047203.1;MERS_1a;3023 | 0 | 0 |
| NC_019843.3;YP_009047203.1;MERS_1a;3024 | 0 | 0 |
| NC_019843.3;YP_009047203.1;MERS_1a;3025 | 0 | 0 |
| NC_019843.3;YP_009047203.1;MERS_1a;3026 | 0 | 0 |
| NC_019843.3;YP_009047203.1;MERS_1a;3027 | 0 | 0 |
| NC_019843.3;YP_009047203.1;MERS_1a;3028 | 0 | 0 |
| NC_019843.3;YP_009047203.1;MERS_1a;3029 | 0 | 0 |

| | | |
|---|---|---|
| NC_019843.3;YP_009047203.1;MERS_1a;3030 | 0 | 0 |
| NC_019843.3;YP_009047203.1;MERS_1a;3031 | 0 | 0 |
| NC_019843.3;YP_009047203.1;MERS_1a;3032 | 0 | 0 |
| NC_019843.3;YP_009047203.1;MERS_1a;3033 | 0 | 0 |
| NC_019843.3;YP_009047203.1;MERS_1a;3034 | 0 | 0 |
| NC_019843.3;YP_009047203.1;MERS_1a;3035 | 0 | 0 |
| NC_019843.3;YP_009047203.1;MERS_1a;3036 | 0 | 0 |
| NC_019843.3;YP_009047203.1;MERS_1a;3037 | 0 | 0 |
| NC_019843.3;YP_009047203.1;MERS_1a;3038 | 0 | 0 |
| NC_019843.3;YP_009047203.1;MERS_1a;3039 | 0 | 0 |
| NC_019843.3;YP_009047203.1;MERS_1a;3040 | 0 | 0 |
| NC_019843.3;YP_009047203.1;MERS_1a;3041 | 0 | 0 |
| NC_019843.3;YP_009047203.1;MERS_1a;3042 | 0 | 0 |
| NC_019843.3;YP_009047203.1;MERS_1a;3043 | 0 | 0 |
| NC_019843.3;YP_009047203.1;MERS_1a;3044 | 0 | 0 |
| NC_019843.3;YP_009047203.1;MERS_1a;3045 | 0 | 0 |
| NC_019843.3;YP_009047203.1;MERS_1a;3046 | 0.05 | 0 |
| NC_019843.3;YP_009047203.1;MERS_1a;3047 | 0.05 | 0 |
| NC_019843.3;YP_009047203.1;MERS_1a;3048 | 0 | 0 |
| NC_019843.3;YP_009047203.1;MERS_1a;3049 | 0 | 0 |
| NC_019843.3;YP_009047203.1;MERS_1a;3050 | 0 | 0 |
| NC_019843.3;YP_009047203.1;MERS_1a;3051 | 0 | 0 |
| NC_019843.3;YP_009047203.1;MERS_1a;3052 | 0 | 0 |
| NC_019843.3;YP_009047203.1;MERS_1a;3053 | 0 | 0 |
| NC_019843.3;YP_009047203.1;MERS_1a;3054 | 0 | 0 |
| NC_019843.3;YP_009047203.1;MERS_1a;3055 | 0 | 0 |
| NC_019843.3;YP_009047203.1;MERS_1a;3056 | 0 | 0 |
| NC_019843.3;YP_009047203.1;MERS_1a;3057 | 0 | 0 |
| NC_019843.3;YP_009047203.1;MERS_1a;3058 | 0 | 0.025 |
| NC_019843.3;YP_009047203.1;MERS_1a;3059 | 0 | 0 |
| NC_019843.3;YP_009047203.1;MERS_1a;3060 | 0 | 0 |
| NC_019843.3;YP_009047203.1;MERS_1a;3061 | 0 | 0 |
| NC_019843.3;YP_009047203.1;MERS_1a;3062 | 0 | 0 |
| NC_019843.3;YP_009047203.1;MERS_1a;3063 | 0 | 0 |
| NC_019843.3;YP_009047203.1;MERS_1a;3064 | 0 | 0 |
| NC_019843.3;YP_009047203.1;MERS_1a;3065 | 0 | 0 |
| NC_019843.3;YP_009047203.1;MERS_1a;3066 | 0 | 0 |
| NC_019843.3;YP_009047203.1;MERS_1a;3067 | 0 | 0 |
| NC_019843.3;YP_009047203.1;MERS_1a;3068 | 0 | 0 |
| NC_019843.3;YP_009047203.1;MERS_1a;3069 | 0 | 0 |
| NC_019843.3;YP_009047203.1;MERS_1a;3070 | 0 | 0 |
| NC_019843.3;YP_009047203.1;MERS_1a;3071 | 0 | 0 |
| NC_019843.3;YP_009047203.1;MERS_1a;3072 | 0 | 0 |

| | | |
|---|---|---|
| NC_019843.3;YP_009047203.1;MERS_1a;3073 | 0 | 0 |
| NC_019843.3;YP_009047203.1;MERS_1a;3074 | 0 | 0 |
| NC_019843.3;YP_009047203.1;MERS_1a;3075 | 0 | 0 |
| NC_019843.3;YP_009047203.1;MERS_1a;3076 | 0 | 0 |
| NC_019843.3;YP_009047203.1;MERS_1a;3077 | 0 | 0 |
| NC_019843.3;YP_009047203.1;MERS_1a;3078 | 0 | 0 |
| NC_019843.3;YP_009047203.1;MERS_1a;3079 | 0 | 0 |
| NC_019843.3;YP_009047203.1;MERS_1a;3080 | 0 | 0 |
| NC_019843.3;YP_009047203.1;MERS_1a;3081 | 0 | 0 |
| NC_019843.3;YP_009047203.1;MERS_1a;3082 | 0 | 0 |
| NC_019843.3;YP_009047203.1;MERS_1a;3083 | 0 | 0 |
| NC_019843.3;YP_009047203.1;MERS_1a;3084 | 0 | 0 |
| NC_019843.3;YP_009047203.1;MERS_1a;3085 | 0 | 0 |
| NC_019843.3;YP_009047203.1;MERS_1a;3086 | 0 | 0 |
| NC_019843.3;YP_009047203.1;MERS_1a;3087 | 0 | 0 |
| NC_019843.3;YP_009047203.1;MERS_1a;3088 | 0 | 0 |
| NC_019843.3;YP_009047203.1;MERS_1a;3089 | 0 | 0 |
| NC_019843.3;YP_009047203.1;MERS_1a;3090 | 0 | 0 |
| NC_019843.3;YP_009047203.1;MERS_1a;3091 | 0 | 0 |
| NC_019843.3;YP_009047203.1;MERS_1a;3092 | 0 | 0 |
| NC_019843.3;YP_009047203.1;MERS_1a;3093 | 0 | 0 |
| NC_019843.3;YP_009047203.1;MERS_1a;3094 | 0 | 0 |
| NC_019843.3;YP_009047203.1;MERS_1a;3095 | 0 | 0 |
| NC_019843.3;YP_009047203.1;MERS_1a;3096 | 0 | 0 |
| NC_019843.3;YP_009047203.1;MERS_1a;3097 | 0 | 0 |
| NC_019843.3;YP_009047203.1;MERS_1a;3098 | 0 | 0 |
| NC_019843.3;YP_009047203.1;MERS_1a;3099 | 0 | 0 |
| NC_019843.3;YP_009047203.1;MERS_1a;3100 | 0 | 0.05 |
| NC_019843.3;YP_009047203.1;MERS_1a;3101 | 0 | 0 |
| NC_019843.3;YP_009047203.1;MERS_1a;3102 | 0 | 0 |
| NC_019843.3;YP_009047203.1;MERS_1a;3103 | 0 | 0 |
| NC_019843.3;YP_009047203.1;MERS_1a;3104 | 0 | 0 |
| NC_019843.3;YP_009047203.1;MERS_1a;3105 | 0 | 0 |
| NC_019843.3;YP_009047203.1;MERS_1a;3106 | 0 | 0 |
| NC_019843.3;YP_009047203.1;MERS_1a;3107 | 0 | 0 |
| NC_019843.3;YP_009047203.1;MERS_1a;3108 | 0 | 0 |
| NC_019843.3;YP_009047203.1;MERS_1a;3109 | 0 | 0 |
| NC_019843.3;YP_009047203.1;MERS_1a;3110 | 0 | 0 |
| NC_019843.3;YP_009047203.1;MERS_1a;3111 | 0 | 0 |
| NC_019843.3;YP_009047203.1;MERS_1a;3112 | 0 | 0 |
| NC_019843.3;YP_009047203.1;MERS_1a;3113 | 0 | 0 |
| NC_019843.3;YP_009047203.1;MERS_1a;3114 | 0 | 0 |
| NC_019843.3;YP_009047203.1;MERS_1a;3115 | 0 | 0 |

| | | |
|---|---|---|
| NC_019843.3;YP_009047203.1;MERS_1a;3116 | 0 | 0 |
| NC_019843.3;YP_009047203.1;MERS_1a;3117 | 0 | 0 |
| NC_019843.3;YP_009047203.1;MERS_1a;3118 | 0 | 0 |
| NC_019843.3;YP_009047203.1;MERS_1a;3119 | 0 | 0 |
| NC_019843.3;YP_009047203.1;MERS_1a;3120 | 0 | 0 |
| NC_019843.3;YP_009047203.1;MERS_1a;3121 | 0 | 0 |
| NC_019843.3;YP_009047203.1;MERS_1a;3122 | 0 | 0 |
| NC_019843.3;YP_009047203.1;MERS_1a;3123 | 0 | 0 |
| NC_019843.3;YP_009047203.1;MERS_1a;3124 | 0 | 0 |
| NC_019843.3;YP_009047203.1;MERS_1a;3125 | 0 | 0 |
| NC_019843.3;YP_009047203.1;MERS_1a;3126 | 0 | 0 |
| NC_019843.3;YP_009047203.1;MERS_1a;3127 | 0 | 0 |
| NC_019843.3;YP_009047203.1;MERS_1a;3128 | 0 | 0 |
| NC_019843.3;YP_009047203.1;MERS_1a;3129 | 0 | 0 |
| NC_019843.3;YP_009047203.1;MERS_1a;3130 | 0 | 0 |
| NC_019843.3;YP_009047203.1;MERS_1a;3131 | 0 | 0 |
| NC_019843.3;YP_009047203.1;MERS_1a;3132 | 0 | 0 |
| NC_019843.3;YP_009047203.1;MERS_1a;3133 | 0 | 0 |
| NC_019843.3;YP_009047203.1;MERS_1a;3134 | 0 | 0 |
| NC_019843.3;YP_009047203.1;MERS_1a;3135 | 0 | 0 |
| NC_019843.3;YP_009047203.1;MERS_1a;3136 | 0 | 0 |
| NC_019843.3;YP_009047203.1;MERS_1a;3137 | 0 | 0 |
| NC_019843.3;YP_009047203.1;MERS_1a;3138 | 0 | 0 |
| NC_019843.3;YP_009047203.1;MERS_1a;3139 | 0 | 0 |
| NC_019843.3;YP_009047203.1;MERS_1a;3140 | 0 | 0.025 |
| NC_019843.3;YP_009047203.1;MERS_1a;3141 | 0 | 0.025 |
| NC_019843.3;YP_009047203.1;MERS_1a;3142 | 0.05 | 0.05 |
| NC_019843.3;YP_009047203.1;MERS_1a;3143 | 0.05 | 0.075 |
| NC_019843.3;YP_009047203.1;MERS_1a;3144 | 0 | 0.05 |
| NC_019843.3;YP_009047203.1;MERS_1a;3145 | 0 | 0.05 |
| NC_019843.3;YP_009047203.1;MERS_1a;3146 | 0 | 0 |
| NC_019843.3;YP_009047203.1;MERS_1a;3147 | 0 | 0 |
| NC_019843.3;YP_009047203.1;MERS_1a;3148 | 0 | 0.05 |
| NC_019843.3;YP_009047203.1;MERS_1a;3149 | 0 | 0.025 |
| NC_019843.3;YP_009047203.1;MERS_1a;3150 | 0.05 | 0.05 |
| NC_019843.3;YP_009047203.1;MERS_1a;3151 | 0 | 0.05 |
| NC_019843.3;YP_009047203.1;MERS_1a;3152 | 0.05 | 0.05 |
| NC_019843.3;YP_009047203.1;MERS_1a;3153 | 0.05 | 0.05 |
| NC_019843.3;YP_009047203.1;MERS_1a;3154 | 0.05 | 0.05 |
| NC_019843.3;YP_009047203.1;MERS_1a;3155 | 0.05 | 0.025 |
| NC_019843.3;YP_009047203.1;MERS_1a;3156 | 0 | 0 |
| NC_019843.3;YP_009047203.1;MERS_1a;3157 | 0 | 0.025 |
| NC_019843.3;YP_009047203.1;MERS_1a;3158 | 0 | 0.025 |

| | | |
|---|---|---|
| NC_019843.3;YP_009047203.1;MERS_1a;3159 | 0 | 0 |
| NC_019843.3;YP_009047203.1;MERS_1a;3160 | 0 | 0.05 |
| NC_019843.3;YP_009047203.1;MERS_1a;3161 | 0 | 0.025 |
| NC_019843.3;YP_009047203.1;MERS_1a;3162 | 0 | 0 |
| NC_019843.3;YP_009047203.1;MERS_1a;3163 | 0.05 | 0.025 |
| NC_019843.3;YP_009047203.1;MERS_1a;3164 | 0 | 0 |
| NC_019843.3;YP_009047203.1;MERS_1a;3165 | 0 | 0 |
| NC_019843.3;YP_009047203.1;MERS_1a;3166 | 0 | 0.025 |
| NC_019843.3;YP_009047203.1;MERS_1a;3167 | 0 | 0.025 |
| NC_019843.3;YP_009047203.1;MERS_1a;3168 | 0 | 0 |
| NC_019843.3;YP_009047203.1;MERS_1a;3169 | 0 | 0 |
| NC_019843.3;YP_009047203.1;MERS_1a;3170 | 0 | 0 |
| NC_019843.3;YP_009047203.1;MERS_1a;3171 | 0.05 | 0 |
| NC_019843.3;YP_009047203.1;MERS_1a;3172 | 0 | 0 |
| NC_019843.3;YP_009047203.1;MERS_1a;3173 | 0.05 | 0 |
| NC_019843.3;YP_009047203.1;MERS_1a;3174 | 0 | 0 |
| NC_019843.3;YP_009047203.1;MERS_1a;3175 | 0 | 0.025 |
| NC_019843.3;YP_009047203.1;MERS_1a;3176 | 0 | 0 |
| NC_019843.3;YP_009047203.1;MERS_1a;3177 | 0 | 0 |
| NC_019843.3;YP_009047203.1;MERS_1a;3178 | 0 | 0.025 |
| NC_019843.3;YP_009047203.1;MERS_1a;3179 | 0 | 0 |
| NC_019843.3;YP_009047203.1;MERS_1a;3180 | 0 | 0 |
| NC_019843.3;YP_009047203.1;MERS_1a;3181 | 0.05 | 0 |
| NC_019843.3;YP_009047203.1;MERS_1a;3182 | 0 | 0 |
| NC_019843.3;YP_009047203.1;MERS_1a;3183 | 0 | 0 |
| NC_019843.3;YP_009047203.1;MERS_1a;3184 | 0 | 0 |
| NC_019843.3;YP_009047203.1;MERS_1a;3185 | 0 | 0 |
| NC_019843.3;YP_009047203.1;MERS_1a;3186 | 0 | 0 |
| NC_019843.3;YP_009047203.1;MERS_1a;3187 | 0 | 0 |
| NC_019843.3;YP_009047203.1;MERS_1a;3188 | 0 | 0 |
| NC_019843.3;YP_009047203.1;MERS_1a;3189 | 0 | 0 |
| NC_019843.3;YP_009047203.1;MERS_1a;3190 | 0 | 0 |
| NC_019843.3;YP_009047203.1;MERS_1a;3191 | 0 | 0 |
| NC_019843.3;YP_009047203.1;MERS_1a;3192 | 0 | 0 |
| NC_019843.3;YP_009047203.1;MERS_1a;3193 | 0 | 0 |
| NC_019843.3;YP_009047203.1;MERS_1a;3194 | 0.05 | 0.025 |
| NC_019843.3;YP_009047203.1;MERS_1a;3195 | 0 | 0.025 |
| NC_019843.3;YP_009047203.1;MERS_1a;3196 | 0 | 0 |
| NC_019843.3;YP_009047203.1;MERS_1a;3197 | 0 | 0 |
| NC_019843.3;YP_009047203.1;MERS_1a;3198 | 0 | 0 |
| NC_019843.3;YP_009047203.1;MERS_1a;3199 | 0.05 | 0.05 |
| NC_019843.3;YP_009047203.1;MERS_1a;3200 | 0.05 | 0.025 |
| NC_019843.3;YP_009047203.1;MERS_1a;3201 | 0 | 0 |

| | | |
|---|---|---|
| NC_019843.3;YP_009047203.1;MERS_1a;3202 | 0 | 0 |
| NC_019843.3;YP_009047203.1;MERS_1a;3203 | 0 | 0 |
| NC_019843.3;YP_009047203.1;MERS_1a;3204 | 0 | 0 |
| NC_019843.3;YP_009047203.1;MERS_1a;3205 | 0 | 0 |
| NC_019843.3;YP_009047203.1;MERS_1a;3206 | 0 | 0 |
| NC_019843.3;YP_009047203.1;MERS_1a;3207 | 0 | 0 |
| NC_019843.3;YP_009047203.1;MERS_1a;3208 | 0 | 0 |
| NC_019843.3;YP_009047203.1;MERS_1a;3209 | 0 | 0 |
| NC_019843.3;YP_009047203.1;MERS_1a;3210 | 0 | 0 |
| NC_019843.3;YP_009047203.1;MERS_1a;3211 | 0 | 0 |
| NC_019843.3;YP_009047203.1;MERS_1a;3212 | 0 | 0 |
| NC_019843.3;YP_009047203.1;MERS_1a;3213 | 0 | 0 |
| NC_019843.3;YP_009047203.1;MERS_1a;3214 | 0.2 | 0.025 |
| NC_019843.3;YP_009047203.1;MERS_1a;3215 | 0.1 | 0 |
| NC_019843.3;YP_009047203.1;MERS_1a;3216 | 0.05 | 0 |
| NC_019843.3;YP_009047203.1;MERS_1a;3217 | 0.05 | 0 |
| NC_019843.3;YP_009047203.1;MERS_1a;3218 | 0.05 | 0 |
| NC_019843.3;YP_009047203.1;MERS_1a;3219 | 0.05 | 0 |
| NC_019843.3;YP_009047203.1;MERS_1a;3220 | 0.1 | 0 |
| NC_019843.3;YP_009047203.1;MERS_1a;3221 | 0.05 | 0 |
| NC_019843.3;YP_009047203.1;MERS_1a;3222 | 0.05 | 0 |
| NC_019843.3;YP_009047203.1;MERS_1a;3223 | 0.05 | 0 |
| NC_019843.3;YP_009047203.1;MERS_1a;3224 | 0.05 | 0 |
| NC_019843.3;YP_009047203.1;MERS_1a;3225 | 0 | 0 |
| NC_019843.3;YP_009047203.1;MERS_1a;3226 | 0 | 0 |
| NC_019843.3;YP_009047203.1;MERS_1a;3227 | 0 | 0.025 |
| NC_019843.3;YP_009047203.1;MERS_1a;3228 | 0 | 0 |
| NC_019843.3;YP_009047203.1;MERS_1a;3229 | 0.05 | 0 |
| NC_019843.3;YP_009047203.1;MERS_1a;3230 | 0 | 0 |
| NC_019843.3;YP_009047203.1;MERS_1a;3231 | 0 | 0 |
| NC_019843.3;YP_009047203.1;MERS_1a;3232 | 0 | 0.025 |
| NC_019843.3;YP_009047203.1;MERS_1a;3233 | 0 | 0 |
| NC_019843.3;YP_009047203.1;MERS_1a;3234 | 0.05 | 0 |
| NC_019843.3;YP_009047203.1;MERS_1a;3235 | 0 | 0 |
| NC_019843.3;YP_009047203.1;MERS_1a;3236 | 0 | 0 |
| NC_019843.3;YP_009047203.1;MERS_1a;3237 | 0 | 0 |
| NC_019843.3;YP_009047203.1;MERS_1a;3238 | 0 | 0 |
| NC_019843.3;YP_009047203.1;MERS_1a;3239 | 0 | 0 |
| NC_019843.3;YP_009047203.1;MERS_1a;3240 | 0 | 0 |
| NC_019843.3;YP_009047203.1;MERS_1a;3241 | 0 | 0 |
| NC_019843.3;YP_009047203.1;MERS_1a;3242 | 0 | 0 |
| NC_019843.3;YP_009047203.1;MERS_1a;3243 | 0 | 0.025 |
| NC_019843.3;YP_009047203.1;MERS_1a;3244 | 0.05 | 0.025 |

| | | |
|---|---|---|
| NC_019843.3;YP_009047203.1;MERS_1a;3245 | 0.05 | 0 |
| NC_019843.3;YP_009047203.1;MERS_1a;3246 | 0.1 | 0 |
| NC_019843.3;YP_009047203.1;MERS_1a;3247 | 0.05 | 0.025 |
| NC_019843.3;YP_009047203.1;MERS_1a;3248 | 0 | 0 |
| NC_019843.3;YP_009047203.1;MERS_1a;3249 | 0.05 | 0 |
| NC_019843.3;YP_009047203.1;MERS_1a;3250 | 0 | 0 |
| NC_019843.3;YP_009047203.1;MERS_1a;3251 | 0 | 0 |
| NC_019843.3;YP_009047203.1;MERS_1a;3252 | 0 | 0 |
| NC_019843.3;YP_009047203.1;MERS_1a;3253 | 0 | 0 |
| NC_019843.3;YP_009047203.1;MERS_1a;3254 | 0 | 0 |
| NC_019843.3;YP_009047203.1;MERS_1a;3255 | 0 | 0 |
| NC_019843.3;YP_009047203.1;MERS_1a;3256 | 0 | 0 |
| NC_019843.3;YP_009047203.1;MERS_1a;3257 | 0 | 0 |
| NC_019843.3;YP_009047203.1;MERS_1a;3258 | 0 | 0.05 |
| NC_019843.3;YP_009047203.1;MERS_1a;3259 | 0 | 0 |
| NC_019843.3;YP_009047203.1;MERS_1a;3260 | 0 | 0 |
| NC_019843.3;YP_009047203.1;MERS_1a;3261 | 0 | 0 |
| NC_019843.3;YP_009047203.1;MERS_1a;3262 | 0 | 0 |
| NC_019843.3;YP_009047203.1;MERS_1a;3263 | 0 | 0 |
| NC_019843.3;YP_009047203.1;MERS_1a;3264 | 0 | 0 |
| NC_019843.3;YP_009047203.1;MERS_1a;3265 | 0 | 0 |
| NC_019843.3;YP_009047203.1;MERS_1a;3266 | 0 | 0 |
| NC_019843.3;YP_009047203.1;MERS_1a;3267 | 0 | 0 |
| NC_019843.3;YP_009047203.1;MERS_1a;3268 | 0 | 0 |
| NC_019843.3;YP_009047203.1;MERS_1a;3269 | 0 | 0 |
| NC_019843.3;YP_009047203.1;MERS_1a;3270 | 0 | 0 |
| NC_019843.3;YP_009047203.1;MERS_1a;3271 | 0 | 0 |
| NC_019843.3;YP_009047203.1;MERS_1a;3272 | 0 | 0 |
| NC_019843.3;YP_009047203.1;MERS_1a;3273 | 0 | 0 |
| NC_019843.3;YP_009047203.1;MERS_1a;3274 | 0 | 0 |
| NC_019843.3;YP_009047203.1;MERS_1a;3275 | 0 | 0 |
| NC_019843.3;YP_009047203.1;MERS_1a;3276 | 0 | 0 |
| NC_019843.3;YP_009047203.1;MERS_1a;3277 | 0 | 0 |
| NC_019843.3;YP_009047203.1;MERS_1a;3278 | 0 | 0 |
| NC_019843.3;YP_009047203.1;MERS_1a;3279 | 0 | 0 |
| NC_019843.3;YP_009047203.1;MERS_1a;3280 | 0 | 0 |
| NC_019843.3;YP_009047203.1;MERS_1a;3281 | 0 | 0 |
| NC_019843.3;YP_009047203.1;MERS_1a;3282 | 0 | 0 |
| NC_019843.3;YP_009047203.1;MERS_1a;3283 | 0.05 | 0 |
| NC_019843.3;YP_009047203.1;MERS_1a;3284 | 0 | 0 |
| NC_019843.3;YP_009047203.1;MERS_1a;3285 | 0 | 0 |
| NC_019843.3;YP_009047203.1;MERS_1a;3286 | 0 | 0 |
| NC_019843.3;YP_009047203.1;MERS_1a;3287 | 0 | 0 |

| | | |
|---|---|---|
| NC_019843.3;YP_009047203.1;MERS_1a;3288 | 0 | 0 |
| NC_019843.3;YP_009047203.1;MERS_1a;3289 | 0 | 0 |
| NC_019843.3;YP_009047203.1;MERS_1a;3290 | 0 | 0 |
| NC_019843.3;YP_009047203.1;MERS_1a;3291 | 0 | 0 |
| NC_019843.3;YP_009047203.1;MERS_1a;3292 | 0 | 0 |
| NC_019843.3;YP_009047203.1;MERS_1a;3293 | 0 | 0 |
| NC_019843.3;YP_009047203.1;MERS_1a;3294 | 0 | 0 |
| NC_019843.3;YP_009047203.1;MERS_1a;3295 | 0 | 0 |
| NC_019843.3;YP_009047203.1;MERS_1a;3296 | 0 | 0 |
| NC_019843.3;YP_009047203.1;MERS_1a;3297 | 0 | 0 |
| NC_019843.3;YP_009047203.1;MERS_1a;3298 | 0 | 0 |
| NC_019843.3;YP_009047203.1;MERS_1a;3299 | 0 | 0 |
| NC_019843.3;YP_009047203.1;MERS_1a;3300 | 0 | 0 |
| NC_019843.3;YP_009047203.1;MERS_1a;3301 | 0 | 0 |
| NC_019843.3;YP_009047203.1;MERS_1a;3302 | 0 | 0 |
| NC_019843.3;YP_009047203.1;MERS_1a;3303 | 0 | 0 |
| NC_019843.3;YP_009047203.1;MERS_1a;3304 | 0 | 0 |
| NC_019843.3;YP_009047203.1;MERS_1a;3305 | 0 | 0 |
| NC_019843.3;YP_009047203.1;MERS_1a;3306 | 0.05 | 0 |
| NC_019843.3;YP_009047203.1;MERS_1a;3307 | 0.15 | 0 |
| NC_019843.3;YP_009047203.1;MERS_1a;3308 | 0.1 | 0.025 |
| NC_019843.3;YP_009047203.1;MERS_1a;3309 | 0.1 | 0.05 |
| NC_019843.3;YP_009047203.1;MERS_1a;3310 | 0.1 | 0.025 |
| NC_019843.3;YP_009047203.1;MERS_1a;3311 | 0.1 | 0.025 |
| NC_019843.3;YP_009047203.1;MERS_1a;3312 | 0.15 | 0.025 |
| NC_019843.3;YP_009047203.1;MERS_1a;3313 | 0.1 | 0.025 |
| NC_019843.3;YP_009047203.1;MERS_1a;3314 | 0.1 | 0.05 |
| NC_019843.3;YP_009047203.1;MERS_1a;3315 | 0.1 | 0.025 |
| NC_019843.3;YP_009047203.1;MERS_1a;3316 | 0.1 | 0.025 |
| NC_019843.3;YP_009047203.1;MERS_1a;3317 | 0.1 | 0.025 |
| NC_019843.3;YP_009047203.1;MERS_1a;3318 | 0.15 | 0.05 |
| NC_019843.3;YP_009047203.1;MERS_1a;3319 | 0 | 0.025 |
| NC_019843.3;YP_009047203.1;MERS_1a;3320 | 0 | 0 |
| NC_019843.3;YP_009047203.1;MERS_1a;3321 | 0 | 0 |
| NC_019843.3;YP_009047203.1;MERS_1a;3322 | 0 | 0 |
| NC_019843.3;YP_009047203.1;MERS_1a;3323 | 0 | 0.025 |
| NC_019843.3;YP_009047203.1;MERS_1a;3324 | 0 | 0.025 |
| NC_019843.3;YP_009047203.1;MERS_1a;3325 | 0 | 0 |
| NC_019843.3;YP_009047203.1;MERS_1a;3326 | 0 | 0 |
| NC_019843.3;YP_009047203.1;MERS_1a;3327 | 0 | 0 |
| NC_019843.3;YP_009047203.1;MERS_1a;3328 | 0 | 0 |
| NC_019843.3;YP_009047203.1;MERS_1a;3329 | 0 | 0 |
| NC_019843.3;YP_009047203.1;MERS_1a;3330 | 0 | 0 |

| | | |
|---|---|---|
| NC_019843.3;YP_009047203.1;MERS_1a;3331 | 0 | 0 |
| NC_019843.3;YP_009047203.1;MERS_1a;3332 | 0 | 0 |
| NC_019843.3;YP_009047203.1;MERS_1a;3333 | 0 | 0 |
| NC_019843.3;YP_009047203.1;MERS_1a;3334 | 0 | 0 |
| NC_019843.3;YP_009047203.1;MERS_1a;3335 | 0 | 0 |
| NC_019843.3;YP_009047203.1;MERS_1a;3336 | 0 | 0 |
| NC_019843.3;YP_009047203.1;MERS_1a;3337 | 0 | 0 |
| NC_019843.3;YP_009047203.1;MERS_1a;3338 | 0 | 0 |
| NC_019843.3;YP_009047203.1;MERS_1a;3339 | 0 | 0.05 |
| NC_019843.3;YP_009047203.1;MERS_1a;3340 | 0 | 0.025 |
| NC_019843.3;YP_009047203.1;MERS_1a;3341 | 0.05 | 0.075 |
| NC_019843.3;YP_009047203.1;MERS_1a;3342 | 0 | 0 |
| NC_019843.3;YP_009047203.1;MERS_1a;3343 | 0 | 0.025 |
| NC_019843.3;YP_009047203.1;MERS_1a;3344 | 0 | 0.025 |
| NC_019843.3;YP_009047203.1;MERS_1a;3345 | 0.05 | 0.025 |
| NC_019843.3;YP_009047203.1;MERS_1a;3346 | 0.1 | 0.075 |
| NC_019843.3;YP_009047203.1;MERS_1a;3347 | 0 | 0 |
| NC_019843.3;YP_009047203.1;MERS_1a;3348 | 0 | 0 |
| NC_019843.3;YP_009047203.1;MERS_1a;3349 | 0 | 0 |
| NC_019843.3;YP_009047203.1;MERS_1a;3350 | 0 | 0 |
| NC_019843.3;YP_009047203.1;MERS_1a;3351 | 0 | 0 |
| NC_019843.3;YP_009047203.1;MERS_1a;3352 | 0 | 0 |
| NC_019843.3;YP_009047203.1;MERS_1a;3353 | 0 | 0 |
| NC_019843.3;YP_009047203.1;MERS_1a;3354 | 0 | 0 |
| NC_019843.3;YP_009047203.1;MERS_1a;3355 | 0 | 0 |
| NC_019843.3;YP_009047203.1;MERS_1a;3356 | 0 | 0 |
| NC_019843.3;YP_009047203.1;MERS_1a;3357 | 0 | 0 |
| NC_019843.3;YP_009047203.1;MERS_1a;3358 | 0 | 0 |
| NC_019843.3;YP_009047203.1;MERS_1a;3359 | 0 | 0 |
| NC_019843.3;YP_009047203.1;MERS_1a;3360 | 0 | 0 |
| NC_019843.3;YP_009047203.1;MERS_1a;3361 | 0 | 0 |
| NC_019843.3;YP_009047203.1;MERS_1a;3362 | 0 | 0 |
| NC_019843.3;YP_009047203.1;MERS_1a;3363 | 0 | 0 |
| NC_019843.3;YP_009047203.1;MERS_1a;3364 | 0 | 0 |
| NC_019843.3;YP_009047203.1;MERS_1a;3365 | 0 | 0 |
| NC_019843.3;YP_009047203.1;MERS_1a;3366 | 0 | 0 |
| NC_019843.3;YP_009047203.1;MERS_1a;3367 | 0 | 0 |
| NC_019843.3;YP_009047203.1;MERS_1a;3368 | 0 | 0 |
| NC_019843.3;YP_009047203.1;MERS_1a;3369 | 0 | 0 |
| NC_019843.3;YP_009047203.1;MERS_1a;3370 | 0 | 0 |
| NC_019843.3;YP_009047203.1;MERS_1a;3371 | 0 | 0 |
| NC_019843.3;YP_009047203.1;MERS_1a;3372 | 0 | 0.025 |
| NC_019843.3;YP_009047203.1;MERS_1a;3373 | 0 | 0.025 |

| | | |
|---|---|---|
| NC_019843.3;YP_009047203.1;MERS_1a;3374 | 0 | 0.05 |
| NC_019843.3;YP_009047203.1;MERS_1a;3375 | 0 | 0 |
| NC_019843.3;YP_009047203.1;MERS_1a;3376 | 0 | 0 |
| NC_019843.3;YP_009047203.1;MERS_1a;3377 | 0 | 0 |
| NC_019843.3;YP_009047203.1;MERS_1a;3378 | 0 | 0 |
| NC_019843.3;YP_009047203.1;MERS_1a;3379 | 0 | 0 |
| NC_019843.3;YP_009047203.1;MERS_1a;3380 | 0 | 0 |
| NC_019843.3;YP_009047203.1;MERS_1a;3381 | 0 | 0 |
| NC_019843.3;YP_009047203.1;MERS_1a;3382 | 0 | 0 |
| NC_019843.3;YP_009047203.1;MERS_1a;3383 | 0 | 0 |
| NC_019843.3;YP_009047203.1;MERS_1a;3384 | 0 | 0 |
| NC_019843.3;YP_009047203.1;MERS_1a;3385 | 0 | 0 |
| NC_019843.3;YP_009047203.1;MERS_1a;3386 | 0 | 0 |
| NC_019843.3;YP_009047203.1;MERS_1a;3387 | 0 | 0.025 |
| NC_019843.3;YP_009047203.1;MERS_1a;3388 | 0 | 0 |
| NC_019843.3;YP_009047203.1;MERS_1a;3389 | 0.05 | 0 |
| NC_019843.3;YP_009047203.1;MERS_1a;3390 | 0.1 | 0 |
| NC_019843.3;YP_009047203.1;MERS_1a;3391 | 0.1 | 0 |
| NC_019843.3;YP_009047203.1;MERS_1a;3392 | 0.05 | 0.025 |
| NC_019843.3;YP_009047203.1;MERS_1a;3393 | 0.1 | 0.025 |
| NC_019843.3;YP_009047203.1;MERS_1a;3394 | 0 | 0 |
| NC_019843.3;YP_009047203.1;MERS_1a;3395 | 0 | 0 |
| NC_019843.3;YP_009047203.1;MERS_1a;3396 | 0 | 0 |
| NC_019843.3;YP_009047203.1;MERS_1a;3397 | 0 | 0 |
| NC_019843.3;YP_009047203.1;MERS_1a;3398 | 0 | 0 |
| NC_019843.3;YP_009047203.1;MERS_1a;3399 | 0 | 0 |
| NC_019843.3;YP_009047203.1;MERS_1a;3400 | 0.05 | 0 |
| NC_019843.3;YP_009047203.1;MERS_1a;3401 | 0 | 0 |
| NC_019843.3;YP_009047203.1;MERS_1a;3402 | 0 | 0 |
| NC_019843.3;YP_009047203.1;MERS_1a;3403 | 0 | 0 |
| NC_019843.3;YP_009047203.1;MERS_1a;3404 | 0 | 0 |
| NC_019843.3;YP_009047203.1;MERS_1a;3405 | 0 | 0 |
| NC_019843.3;YP_009047203.1;MERS_1a;3406 | 0 | 0 |
| NC_019843.3;YP_009047203.1;MERS_1a;3407 | 0 | 0 |
| NC_019843.3;YP_009047203.1;MERS_1a;3408 | 0 | 0 |
| NC_019843.3;YP_009047203.1;MERS_1a;3409 | 0 | 0 |
| NC_019843.3;YP_009047203.1;MERS_1a;3410 | 0 | 0 |
| NC_019843.3;YP_009047203.1;MERS_1a;3411 | 0 | 0 |
| NC_019843.3;YP_009047203.1;MERS_1a;3412 | 0 | 0 |
| NC_019843.3;YP_009047203.1;MERS_1a;3413 | 0.05 | 0 |
| NC_019843.3;YP_009047203.1;MERS_1a;3414 | 0.05 | 0 |
| NC_019843.3;YP_009047203.1;MERS_1a;3415 | 0.05 | 0 |
| NC_019843.3;YP_009047203.1;MERS_1a;3416 | 0.05 | 0 |

| | | |
|---|---|---|
| NC_019843.3;YP_009047203.1;MERS_1a;3417 | 0.05 | 0 |
| NC_019843.3;YP_009047203.1;MERS_1a;3418 | 0.05 | 0.05 |
| NC_019843.3;YP_009047203.1;MERS_1a;3419 | 0.05 | 0 |
| NC_019843.3;YP_009047203.1;MERS_1a;3420 | 0.05 | 0 |
| NC_019843.3;YP_009047203.1;MERS_1a;3421 | 0.05 | 0 |
| NC_019843.3;YP_009047203.1;MERS_1a;3422 | 0 | 0 |
| NC_019843.3;YP_009047203.1;MERS_1a;3423 | 0.05 | 0 |
| NC_019843.3;YP_009047203.1;MERS_1a;3424 | 0 | 0 |
| NC_019843.3;YP_009047203.1;MERS_1a;3425 | 0 | 0 |
| NC_019843.3;YP_009047203.1;MERS_1a;3426 | 0 | 0 |
| NC_019843.3;YP_009047203.1;MERS_1a;3427 | 0 | 0 |
| NC_019843.3;YP_009047203.1;MERS_1a;3428 | 0 | 0 |
| NC_019843.3;YP_009047203.1;MERS_1a;3429 | 0 | 0 |
| NC_019843.3;YP_009047203.1;MERS_1a;3430 | 0 | 0 |
| NC_019843.3;YP_009047203.1;MERS_1a;3431 | 0 | 0 |
| NC_019843.3;YP_009047203.1;MERS_1a;3432 | 0 | 0.025 |
| NC_019843.3;YP_009047203.1;MERS_1a;3433 | 0 | 0 |
| NC_019843.3;YP_009047203.1;MERS_1a;3434 | 0 | 0.025 |
| NC_019843.3;YP_009047203.1;MERS_1a;3435 | 0 | 0 |
| NC_019843.3;YP_009047203.1;MERS_1a;3436 | 0 | 0 |
| NC_019843.3;YP_009047203.1;MERS_1a;3437 | 0 | 0 |
| NC_019843.3;YP_009047203.1;MERS_1a;3438 | 0 | 0 |
| NC_019843.3;YP_009047203.1;MERS_1a;3439 | 0 | 0 |
| NC_019843.3;YP_009047203.1;MERS_1a;3440 | 0 | 0 |
| NC_019843.3;YP_009047203.1;MERS_1a;3441 | 0 | 0 |
| NC_019843.3;YP_009047203.1;MERS_1a;3442 | 0 | 0 |
| NC_019843.3;YP_009047203.1;MERS_1a;3443 | 0 | 0 |
| NC_019843.3;YP_009047203.1;MERS_1a;3444 | 0 | 0 |
| NC_019843.3;YP_009047203.1;MERS_1a;3445 | 0 | 0 |
| NC_019843.3;YP_009047203.1;MERS_1a;3446 | 0 | 0 |
| NC_019843.3;YP_009047203.1;MERS_1a;3447 | 0 | 0 |
| NC_019843.3;YP_009047203.1;MERS_1a;3448 | 0 | 0 |
| NC_019843.3;YP_009047203.1;MERS_1a;3449 | 0 | 0 |
| NC_019843.3;YP_009047203.1;MERS_1a;3450 | 0 | 0 |
| NC_019843.3;YP_009047203.1;MERS_1a;3451 | 0 | 0 |
| NC_019843.3;YP_009047203.1;MERS_1a;3452 | 0 | 0 |
| NC_019843.3;YP_009047203.1;MERS_1a;3453 | 0 | 0 |
| NC_019843.3;YP_009047203.1;MERS_1a;3454 | 0 | 0 |
| NC_019843.3;YP_009047203.1;MERS_1a;3455 | 0 | 0 |
| NC_019843.3;YP_009047203.1;MERS_1a;3456 | 0 | 0 |
| NC_019843.3;YP_009047203.1;MERS_1a;3457 | 0 | 0 |
| NC_019843.3;YP_009047203.1;MERS_1a;3458 | 0 | 0 |
| NC_019843.3;YP_009047203.1;MERS_1a;3459 | 0 | 0 |

| | | |
|---|---|---|
| NC_019843.3;YP_009047203.1;MERS_1a;3460 | 0 | 0.025 |
| NC_019843.3;YP_009047203.1;MERS_1a;3461 | 0 | 0 |
| NC_019843.3;YP_009047203.1;MERS_1a;3462 | 0 | 0 |
| NC_019843.3;YP_009047203.1;MERS_1a;3463 | 0 | 0 |
| NC_019843.3;YP_009047203.1;MERS_1a;3464 | 0 | 0 |
| NC_019843.3;YP_009047203.1;MERS_1a;3465 | 0 | 0 |
| NC_019843.3;YP_009047203.1;MERS_1a;3466 | 0 | 0 |
| NC_019843.3;YP_009047203.1;MERS_1a;3467 | 0 | 0 |
| NC_019843.3;YP_009047203.1;MERS_1a;3468 | 0 | 0 |
| NC_019843.3;YP_009047203.1;MERS_1a;3469 | 0 | 0 |
| NC_019843.3;YP_009047203.1;MERS_1a;3470 | 0 | 0.025 |
| NC_019843.3;YP_009047203.1;MERS_1a;3471 | 0 | 0.025 |
| NC_019843.3;YP_009047203.1;MERS_1a;3472 | 0 | 0 |
| NC_019843.3;YP_009047203.1;MERS_1a;3473 | 0 | 0 |
| NC_019843.3;YP_009047203.1;MERS_1a;3474 | 0 | 0 |
| NC_019843.3;YP_009047203.1;MERS_1a;3475 | 0 | 0 |
| NC_019843.3;YP_009047203.1;MERS_1a;3476 | 0 | 0.025 |
| NC_019843.3;YP_009047203.1;MERS_1a;3477 | 0 | 0.025 |
| NC_019843.3;YP_009047203.1;MERS_1a;3478 | 0 | 0 |
| NC_019843.3;YP_009047203.1;MERS_1a;3479 | 0 | 0.025 |
| NC_019843.3;YP_009047203.1;MERS_1a;3480 | 0 | 0 |
| NC_019843.3;YP_009047203.1;MERS_1a;3481 | 0 | 0 |
| NC_019843.3;YP_009047203.1;MERS_1a;3482 | 0 | 0 |
| NC_019843.3;YP_009047203.1;MERS_1a;3483 | 0 | 0 |
| NC_019843.3;YP_009047203.1;MERS_1a;3484 | 0 | 0.025 |
| NC_019843.3;YP_009047203.1;MERS_1a;3485 | 0 | 0.025 |
| NC_019843.3;YP_009047203.1;MERS_1a;3486 | 0 | 0.05 |
| NC_019843.3;YP_009047203.1;MERS_1a;3487 | 0 | 0.025 |
| NC_019843.3;YP_009047203.1;MERS_1a;3488 | 0 | 0.025 |
| NC_019843.3;YP_009047203.1;MERS_1a;3489 | 0 | 0 |
| NC_019843.3;YP_009047203.1;MERS_1a;3490 | 0 | 0 |
| NC_019843.3;YP_009047203.1;MERS_1a;3491 | 0.05 | 0 |
| NC_019843.3;YP_009047203.1;MERS_1a;3492 | 0 | 0.025 |
| NC_019843.3;YP_009047203.1;MERS_1a;3493 | 0 | 0.025 |
| NC_019843.3;YP_009047203.1;MERS_1a;3494 | 0 | 0 |
| NC_019843.3;YP_009047203.1;MERS_1a;3495 | 0 | 0 |
| NC_019843.3;YP_009047203.1;MERS_1a;3496 | 0 | 0 |
| NC_019843.3;YP_009047203.1;MERS_1a;3497 | 0 | 0 |
| NC_019843.3;YP_009047203.1;MERS_1a;3498 | 0 | 0 |
| NC_019843.3;YP_009047203.1;MERS_1a;3499 | 0 | 0 |
| NC_019843.3;YP_009047203.1;MERS_1a;3500 | 0 | 0 |
| NC_019843.3;YP_009047203.1;MERS_1a;3501 | 0 | 0 |
| NC_019843.3;YP_009047203.1;MERS_1a;3502 | 0 | 0 |

| | | |
|---|---|---|
| NC_019843.3;YP_009047203.1;MERS_1a;3503 | 0 | 0 |
| NC_019843.3;YP_009047203.1;MERS_1a;3504 | 0 | 0 |
| NC_019843.3;YP_009047203.1;MERS_1a;3505 | 0 | 0 |
| NC_019843.3;YP_009047203.1;MERS_1a;3506 | 0 | 0 |
| NC_019843.3;YP_009047203.1;MERS_1a;3507 | 0 | 0.225 |
| NC_019843.3;YP_009047203.1;MERS_1a;3508 | 0 | 0 |
| NC_019843.3;YP_009047203.1;MERS_1a;3509 | 0 | 0 |
| NC_019843.3;YP_009047203.1;MERS_1a;3510 | 0 | 0 |
| NC_019843.3;YP_009047203.1;MERS_1a;3511 | 0 | 0 |
| NC_019843.3;YP_009047203.1;MERS_1a;3512 | 0 | 0 |
| NC_019843.3;YP_009047203.1;MERS_1a;3513 | 0 | 0 |
| NC_019843.3;YP_009047203.1;MERS_1a;3514 | 0 | 0 |
| NC_019843.3;YP_009047203.1;MERS_1a;3515 | 0 | 0 |
| NC_019843.3;YP_009047203.1;MERS_1a;3516 | 0 | 0 |
| NC_019843.3;YP_009047203.1;MERS_1a;3517 | 0 | 0 |
| NC_019843.3;YP_009047203.1;MERS_1a;3518 | 0 | 0 |
| NC_019843.3;YP_009047203.1;MERS_1a;3519 | 0 | 0 |
| NC_019843.3;YP_009047203.1;MERS_1a;3520 | 0 | 0 |
| NC_019843.3;YP_009047203.1;MERS_1a;3521 | 0 | 0 |
| NC_019843.3;YP_009047203.1;MERS_1a;3522 | 0 | 0 |
| NC_019843.3;YP_009047203.1;MERS_1a;3523 | 0 | 0 |
| NC_019843.3;YP_009047203.1;MERS_1a;3524 | 0 | 0 |
| NC_019843.3;YP_009047203.1;MERS_1a;3525 | 0 | 0 |
| NC_019843.3;YP_009047203.1;MERS_1a;3526 | 0 | 0 |
| NC_019843.3;YP_009047203.1;MERS_1a;3527 | 0 | 0 |
| NC_019843.3;YP_009047203.1;MERS_1a;3528 | 0 | 0 |
| NC_019843.3;YP_009047203.1;MERS_1a;3529 | 0 | 0 |
| NC_019843.3;YP_009047203.1;MERS_1a;3530 | 0 | 0 |
| NC_019843.3;YP_009047203.1;MERS_1a;3531 | 0 | 0 |
| NC_019843.3;YP_009047203.1;MERS_1a;3532 | 0 | 0 |
| NC_019843.3;YP_009047203.1;MERS_1a;3533 | 0 | 0 |
| NC_019843.3;YP_009047203.1;MERS_1a;3534 | 0 | 0.025 |
| NC_019843.3;YP_009047203.1;MERS_1a;3535 | 0.1 | 0.025 |
| NC_019843.3;YP_009047203.1;MERS_1a;3536 | 0 | 0 |
| NC_019843.3;YP_009047203.1;MERS_1a;3537 | 0.05 | 0 |
| NC_019843.3;YP_009047203.1;MERS_1a;3538 | 0 | 0.05 |
| NC_019843.3;YP_009047203.1;MERS_1a;3539 | 0 | 0 |
| NC_019843.3;YP_009047203.1;MERS_1a;3540 | 0 | 0 |
| NC_019843.3;YP_009047203.1;MERS_1a;3541 | 0 | 0 |
| NC_019843.3;YP_009047203.1;MERS_1a;3542 | 0 | 0.025 |
| NC_019843.3;YP_009047203.1;MERS_1a;3543 | 0 | 0 |
| NC_019843.3;YP_009047203.1;MERS_1a;3544 | 0 | 0 |
| NC_019843.3;YP_009047203.1;MERS_1a;3545 | 0 | 0 |

| | | |
|---|---|---|
| NC_019843.3;YP_009047203.1;MERS_1a;3546 | 0 | 0 |
| NC_019843.3;YP_009047203.1;MERS_1a;3547 | 0 | 0 |
| NC_019843.3;YP_009047203.1;MERS_1a;3548 | 0 | 0 |
| NC_019843.3;YP_009047203.1;MERS_1a;3549 | 0 | 0 |
| NC_019843.3;YP_009047203.1;MERS_1a;3550 | 0.05 | 0 |
| NC_019843.3;YP_009047203.1;MERS_1a;3551 | 0 | 0 |
| NC_019843.3;YP_009047203.1;MERS_1a;3552 | 0 | 0 |
| NC_019843.3;YP_009047203.1;MERS_1a;3553 | 0 | 0 |
| NC_019843.3;YP_009047203.1;MERS_1a;3554 | 0 | 0 |
| NC_019843.3;YP_009047203.1;MERS_1a;3555 | 0 | 0 |
| NC_019843.3;YP_009047203.1;MERS_1a;3556 | 0 | 0 |
| NC_019843.3;YP_009047203.1;MERS_1a;3557 | 0 | 0 |
| NC_019843.3;YP_009047203.1;MERS_1a;3558 | 0 | 0 |
| NC_019843.3;YP_009047203.1;MERS_1a;3559 | 0 | 0 |
| NC_019843.3;YP_009047203.1;MERS_1a;3560 | 0 | 0 |
| NC_019843.3;YP_009047203.1;MERS_1a;3561 | 0 | 0 |
| NC_019843.3;YP_009047203.1;MERS_1a;3562 | 0 | 0 |
| NC_019843.3;YP_009047203.1;MERS_1a;3563 | 0 | 0 |
| NC_019843.3;YP_009047203.1;MERS_1a;3564 | 0 | 0 |
| NC_019843.3;YP_009047203.1;MERS_1a;3565 | 0 | 0 |
| NC_019843.3;YP_009047203.1;MERS_1a;3566 | 0 | 0 |
| NC_019843.3;YP_009047203.1;MERS_1a;3567 | 0 | 0 |
| NC_019843.3;YP_009047203.1;MERS_1a;3568 | 0 | 0 |
| NC_019843.3;YP_009047203.1;MERS_1a;3569 | 0 | 0 |
| NC_019843.3;YP_009047203.1;MERS_1a;3570 | 0 | 0 |
| NC_019843.3;YP_009047203.1;MERS_1a;3571 | 0 | 0 |
| NC_019843.3;YP_009047203.1;MERS_1a;3572 | 0 | 0 |
| NC_019843.3;YP_009047203.1;MERS_1a;3573 | 0 | 0 |
| NC_019843.3;YP_009047203.1;MERS_1a;3574 | 0 | 0 |
| NC_019843.3;YP_009047203.1;MERS_1a;3575 | 0 | 0 |
| NC_019843.3;YP_009047203.1;MERS_1a;3576 | 0 | 0 |
| NC_019843.3;YP_009047203.1;MERS_1a;3577 | 0 | 0 |
| NC_019843.3;YP_009047203.1;MERS_1a;3578 | 0 | 0 |
| NC_019843.3;YP_009047203.1;MERS_1a;3579 | 0 | 0 |
| NC_019843.3;YP_009047203.1;MERS_1a;3580 | 0 | 0 |
| NC_019843.3;YP_009047203.1;MERS_1a;3581 | 0 | 0 |
| NC_019843.3;YP_009047203.1;MERS_1a;3582 | 0 | 0 |
| NC_019843.3;YP_009047203.1;MERS_1a;3583 | 0 | 0 |
| NC_019843.3;YP_009047203.1;MERS_1a;3584 | 0 | 0 |
| NC_019843.3;YP_009047203.1;MERS_1a;3585 | 0 | 0 |
| NC_019843.3;YP_009047203.1;MERS_1a;3586 | 0 | 0 |
| NC_019843.3;YP_009047203.1;MERS_1a;3587 | 0 | 0 |
| NC_019843.3;YP_009047203.1;MERS_1a;3588 | 0 | 0 |

| | | |
|---|---|---|
| NC_019843.3;YP_009047203.1;MERS_1a;3589 | 0 | 0 |
| NC_019843.3;YP_009047203.1;MERS_1a;3590 | 0 | 0 |
| NC_019843.3;YP_009047203.1;MERS_1a;3591 | 0 | 0 |
| NC_019843.3;YP_009047203.1;MERS_1a;3592 | 0 | 0 |
| NC_019843.3;YP_009047203.1;MERS_1a;3593 | 0 | 0 |
| NC_019843.3;YP_009047203.1;MERS_1a;3594 | 0 | 0 |
| NC_019843.3;YP_009047203.1;MERS_1a;3595 | 0 | 0 |
| NC_019843.3;YP_009047203.1;MERS_1a;3596 | 0 | 0 |
| NC_019843.3;YP_009047203.1;MERS_1a;3597 | 0 | 0 |
| NC_019843.3;YP_009047203.1;MERS_1a;3598 | 0 | 0 |
| NC_019843.3;YP_009047203.1;MERS_1a;3599 | 0 | 0 |
| NC_019843.3;YP_009047203.1;MERS_1a;3600 | 0 | 0 |
| NC_019843.3;YP_009047203.1;MERS_1a;3601 | 0 | 0 |
| NC_019843.3;YP_009047203.1;MERS_1a;3602 | 0 | 0 |
| NC_019843.3;YP_009047203.1;MERS_1a;3603 | 0 | 0 |
| NC_019843.3;YP_009047203.1;MERS_1a;3604 | 0 | 0 |
| NC_019843.3;YP_009047203.1;MERS_1a;3605 | 0 | 0 |
| NC_019843.3;YP_009047203.1;MERS_1a;3606 | 0 | 0 |
| NC_019843.3;YP_009047203.1;MERS_1a;3607 | 0 | 0 |
| NC_019843.3;YP_009047203.1;MERS_1a;3608 | 0 | 0 |
| NC_019843.3;YP_009047203.1;MERS_1a;3609 | 0 | 0 |
| NC_019843.3;YP_009047203.1;MERS_1a;3610 | 0 | 0 |
| NC_019843.3;YP_009047203.1;MERS_1a;3611 | 0 | 0 |
| NC_019843.3;YP_009047203.1;MERS_1a;3612 | 0 | 0 |
| NC_019843.3;YP_009047203.1;MERS_1a;3613 | 0 | 0 |
| NC_019843.3;YP_009047203.1;MERS_1a;3614 | 0 | 0 |
| NC_019843.3;YP_009047203.1;MERS_1a;3615 | 0 | 0 |
| NC_019843.3;YP_009047203.1;MERS_1a;3616 | 0 | 0 |
| NC_019843.3;YP_009047203.1;MERS_1a;3617 | 0 | 0 |
| NC_019843.3;YP_009047203.1;MERS_1a;3618 | 0 | 0 |
| NC_019843.3;YP_009047203.1;MERS_1a;3619 | 0 | 0 |
| NC_019843.3;YP_009047203.1;MERS_1a;3620 | 0 | 0 |
| NC_019843.3;YP_009047203.1;MERS_1a;3621 | 0 | 0 |
| NC_019843.3;YP_009047203.1;MERS_1a;3622 | 0 | 0 |
| NC_019843.3;YP_009047203.1;MERS_1a;3623 | 0 | 0 |
| NC_019843.3;YP_009047203.1;MERS_1a;3624 | 0 | 0 |
| NC_019843.3;YP_009047203.1;MERS_1a;3625 | 0 | 0 |
| NC_019843.3;YP_009047203.1;MERS_1a;3626 | 0 | 0 |
| NC_019843.3;YP_009047203.1;MERS_1a;3627 | 0 | 0 |
| NC_019843.3;YP_009047203.1;MERS_1a;3628 | 0 | 0 |
| NC_019843.3;YP_009047203.1;MERS_1a;3629 | 0 | 0 |
| NC_019843.3;YP_009047203.1;MERS_1a;3630 | 0 | 0 |
| NC_019843.3;YP_009047203.1;MERS_1a;3631 | 0 | 0 |

| | | |
|---|---|---|
| NC_019843.3;YP_009047203.1;MERS_1a;3632 | 0 | 0 |
| NC_019843.3;YP_009047203.1;MERS_1a;3633 | 0 | 0 |
| NC_019843.3;YP_009047203.1;MERS_1a;3634 | 0 | 0.025 |
| NC_019843.3;YP_009047203.1;MERS_1a;3635 | 0 | 0 |
| NC_019843.3;YP_009047203.1;MERS_1a;3636 | 0 | 0 |
| NC_019843.3;YP_009047203.1;MERS_1a;3637 | 0 | 0 |
| NC_019843.3;YP_009047203.1;MERS_1a;3638 | 0 | 0 |
| NC_019843.3;YP_009047203.1;MERS_1a;3639 | 0 | 0 |
| NC_019843.3;YP_009047203.1;MERS_1a;3640 | 0 | 0 |
| NC_019843.3;YP_009047203.1;MERS_1a;3641 | 0 | 0 |
| NC_019843.3;YP_009047203.1;MERS_1a;3642 | 0 | 0 |
| NC_019843.3;YP_009047203.1;MERS_1a;3643 | 0 | 0 |
| NC_019843.3;YP_009047203.1;MERS_1a;3644 | 0 | 0 |
| NC_019843.3;YP_009047203.1;MERS_1a;3645 | 0 | 0 |
| NC_019843.3;YP_009047203.1;MERS_1a;3646 | 0.05 | 0.075 |
| NC_019843.3;YP_009047203.1;MERS_1a;3647 | 0.15 | 0.175 |
| NC_019843.3;YP_009047203.1;MERS_1a;3648 | 0.15 | 0.1 |
| NC_019843.3;YP_009047203.1;MERS_1a;3649 | 0.15 | 0.225 |
| NC_019843.3;YP_009047203.1;MERS_1a;3650 | 0.25 | 0.3 |
| NC_019843.3;YP_009047203.1;MERS_1a;3651 | 0.2 | 0.25 |
| NC_019843.3;YP_009047203.1;MERS_1a;3652 | 0.2 | 0.25 |
| NC_019843.3;YP_009047203.1;MERS_1a;3653 | 0.25 | 0.3 |
| NC_019843.3;YP_009047203.1;MERS_1a;3654 | 0.2 | 0.225 |
| NC_019843.3;YP_009047203.1;MERS_1a;3655 | 0.35 | 0.325 |
| NC_019843.3;YP_009047203.1;MERS_1a;3656 | 0.3 | 0.35 |
| NC_019843.3;YP_009047203.1;MERS_1a;3657 | 0.05 | 0.1 |
| NC_019843.3;YP_009047203.1;MERS_1a;3658 | 0 | 0 |
| NC_019843.3;YP_009047203.1;MERS_1a;3659 | 0 | 0 |
| NC_019843.3;YP_009047203.1;MERS_1a;3660 | 0 | 0 |
| NC_019843.3;YP_009047203.1;MERS_1a;3661 | 0 | 0 |
| NC_019843.3;YP_009047203.1;MERS_1a;3662 | 0 | 0 |
| NC_019843.3;YP_009047203.1;MERS_1a;3663 | 0 | 0 |
| NC_019843.3;YP_009047203.1;MERS_1a;3664 | 0 | 0 |
| NC_019843.3;YP_009047203.1;MERS_1a;3665 | 0 | 0 |
| NC_019843.3;YP_009047203.1;MERS_1a;3666 | 0 | 0 |
| NC_019843.3;YP_009047203.1;MERS_1a;3667 | 0 | 0 |
| NC_019843.3;YP_009047203.1;MERS_1a;3668 | 0 | 0 |
| NC_019843.3;YP_009047203.1;MERS_1a;3669 | 0 | 0 |
| NC_019843.3;YP_009047203.1;MERS_1a;3670 | 0 | 0 |
| NC_019843.3;YP_009047203.1;MERS_1a;3671 | 0 | 0 |
| NC_019843.3;YP_009047203.1;MERS_1a;3672 | 0 | 0 |
| NC_019843.3;YP_009047203.1;MERS_1a;3673 | 0 | 0 |
| NC_019843.3;YP_009047203.1;MERS_1a;3674 | 0 | 0 |

| | | |
|---|---|---|
| NC_019843.3;YP_009047203.1;MERS_1a;3675 | 0 | 0 |
| NC_019843.3;YP_009047203.1;MERS_1a;3676 | 0 | 0 |
| NC_019843.3;YP_009047203.1;MERS_1a;3677 | 0 | 0 |
| NC_019843.3;YP_009047203.1;MERS_1a;3678 | 0 | 0 |
| NC_019843.3;YP_009047203.1;MERS_1a;3679 | 0 | 0 |
| NC_019843.3;YP_009047203.1;MERS_1a;3680 | 0 | 0 |
| NC_019843.3;YP_009047203.1;MERS_1a;3681 | 0 | 0 |
| NC_019843.3;YP_009047203.1;MERS_1a;3682 | 0 | 0 |
| NC_019843.3;YP_009047203.1;MERS_1a;3683 | 0 | 0 |
| NC_019843.3;YP_009047203.1;MERS_1a;3684 | 0 | 0 |
| NC_019843.3;YP_009047203.1;MERS_1a;3685 | 0 | 0 |
| NC_019843.3;YP_009047203.1;MERS_1a;3686 | 0 | 0 |
| NC_019843.3;YP_009047203.1;MERS_1a;3687 | 0 | 0 |
| NC_019843.3;YP_009047203.1;MERS_1a;3688 | 0 | 0 |
| NC_019843.3;YP_009047203.1;MERS_1a;3689 | 0 | 0 |
| NC_019843.3;YP_009047203.1;MERS_1a;3690 | 0 | 0 |
| NC_019843.3;YP_009047203.1;MERS_1a;3691 | 0 | 0 |
| NC_019843.3;YP_009047203.1;MERS_1a;3692 | 0 | 0 |
| NC_019843.3;YP_009047203.1;MERS_1a;3693 | 0 | 0 |
| NC_019843.3;YP_009047203.1;MERS_1a;3694 | 0 | 0 |
| NC_019843.3;YP_009047203.1;MERS_1a;3695 | 0 | 0 |
| NC_019843.3;YP_009047203.1;MERS_1a;3696 | 0 | 0 |
| NC_019843.3;YP_009047203.1;MERS_1a;3697 | 0 | 0 |
| NC_019843.3;YP_009047203.1;MERS_1a;3698 | 0 | 0 |
| NC_019843.3;YP_009047203.1;MERS_1a;3699 | 0.05 | 0.025 |
| NC_019843.3;YP_009047203.1;MERS_1a;3700 | 0.05 | 0.025 |
| NC_019843.3;YP_009047203.1;MERS_1a;3701 | 0.05 | 0.025 |
| NC_019843.3;YP_009047203.1;MERS_1a;3702 | 0.05 | 0.025 |
| NC_019843.3;YP_009047203.1;MERS_1a;3703 | 0.05 | 0.025 |
| NC_019843.3;YP_009047203.1;MERS_1a;3704 | 0.05 | 0.025 |
| NC_019843.3;YP_009047203.1;MERS_1a;3705 | 0.1 | 0 |
| NC_019843.3;YP_009047203.1;MERS_1a;3706 | 0 | 0 |
| NC_019843.3;YP_009047203.1;MERS_1a;3707 | 0 | 0 |
| NC_019843.3;YP_009047203.1;MERS_1a;3708 | 0 | 0 |
| NC_019843.3;YP_009047203.1;MERS_1a;3709 | 0 | 0 |
| NC_019843.3;YP_009047203.1;MERS_1a;3710 | 0 | 0 |
| NC_019843.3;YP_009047203.1;MERS_1a;3711 | 0 | 0 |
| NC_019843.3;YP_009047203.1;MERS_1a;3712 | 0 | 0 |
| NC_019843.3;YP_009047203.1;MERS_1a;3713 | 0 | 0 |
| NC_019843.3;YP_009047203.1;MERS_1a;3714 | 0 | 0 |
| NC_019843.3;YP_009047203.1;MERS_1a;3715 | 0 | 0 |
| NC_019843.3;YP_009047203.1;MERS_1a;3716 | 0 | 0 |
| NC_019843.3;YP_009047203.1;MERS_1a;3717 | 0 | 0 |

| | | |
|---|---|---|
| NC_019843.3;YP_009047203.1;MERS_1a;3718 | 0 | 0 |
| NC_019843.3;YP_009047203.1;MERS_1a;3719 | 0 | 0 |
| NC_019843.3;YP_009047203.1;MERS_1a;3720 | 0 | 0 |
| NC_019843.3;YP_009047203.1;MERS_1a;3721 | 0 | 0 |
| NC_019843.3;YP_009047203.1;MERS_1a;3722 | 0 | 0 |
| NC_019843.3;YP_009047203.1;MERS_1a;3723 | 0 | 0 |
| NC_019843.3;YP_009047203.1;MERS_1a;3724 | 0 | 0 |
| NC_019843.3;YP_009047203.1;MERS_1a;3725 | 0 | 0 |
| NC_019843.3;YP_009047203.1;MERS_1a;3726 | 0 | 0 |
| NC_019843.3;YP_009047203.1;MERS_1a;3727 | 0 | 0 |
| NC_019843.3;YP_009047203.1;MERS_1a;3728 | 0 | 0 |
| NC_019843.3;YP_009047203.1;MERS_1a;3729 | 0 | 0 |
| NC_019843.3;YP_009047203.1;MERS_1a;3730 | 0 | 0 |
| NC_019843.3;YP_009047203.1;MERS_1a;3731 | 0 | 0 |
| NC_019843.3;YP_009047203.1;MERS_1a;3732 | 0 | 0 |
| NC_019843.3;YP_009047203.1;MERS_1a;3733 | 0 | 0 |
| NC_019843.3;YP_009047203.1;MERS_1a;3734 | 0 | 0 |
| NC_019843.3;YP_009047203.1;MERS_1a;3735 | 0 | 0 |
| NC_019843.3;YP_009047203.1;MERS_1a;3736 | 0 | 0 |
| NC_019843.3;YP_009047203.1;MERS_1a;3737 | 0 | 0 |
| NC_019843.3;YP_009047203.1;MERS_1a;3738 | 0 | 0 |
| NC_019843.3;YP_009047203.1;MERS_1a;3739 | 0 | 0 |
| NC_019843.3;YP_009047203.1;MERS_1a;3740 | 0 | 0 |
| NC_019843.3;YP_009047203.1;MERS_1a;3741 | 0 | 0 |
| NC_019843.3;YP_009047203.1;MERS_1a;3742 | 0 | 0 |
| NC_019843.3;YP_009047203.1;MERS_1a;3743 | 0 | 0 |
| NC_019843.3;YP_009047203.1;MERS_1a;3744 | 0 | 0 |
| NC_019843.3;YP_009047203.1;MERS_1a;3745 | 0 | 0 |
| NC_019843.3;YP_009047203.1;MERS_1a;3746 | 0 | 0 |
| NC_019843.3;YP_009047203.1;MERS_1a;3747 | 0 | 0 |
| NC_019843.3;YP_009047203.1;MERS_1a;3748 | 0 | 0 |
| NC_019843.3;YP_009047203.1;MERS_1a;3749 | 0 | 0 |
| NC_019843.3;YP_009047203.1;MERS_1a;3750 | 0 | 0.025 |
| NC_019843.3;YP_009047203.1;MERS_1a;3751 | 0 | 0 |
| NC_019843.3;YP_009047203.1;MERS_1a;3752 | 0 | 0 |
| NC_019843.3;YP_009047203.1;MERS_1a;3753 | 0 | 0 |
| NC_019843.3;YP_009047203.1;MERS_1a;3754 | 0 | 0 |
| NC_019843.3;YP_009047203.1;MERS_1a;3755 | 0 | 0 |
| NC_019843.3;YP_009047203.1;MERS_1a;3756 | 0 | 0 |
| NC_019843.3;YP_009047203.1;MERS_1a;3757 | 0 | 0 |
| NC_019843.3;YP_009047203.1;MERS_1a;3758 | 0 | 0 |
| NC_019843.3;YP_009047203.1;MERS_1a;3759 | 0 | 0 |
| NC_019843.3;YP_009047203.1;MERS_1a;3760 | 0 | 0 |

| | | |
|---|---|---|
| NC_019843.3;YP_009047203.1;MERS_1a;3761 | 0 | 0 |
| NC_019843.3;YP_009047203.1;MERS_1a;3762 | 0 | 0 |
| NC_019843.3;YP_009047203.1;MERS_1a;3763 | 0 | 0 |
| NC_019843.3;YP_009047203.1;MERS_1a;3764 | 0 | 0 |
| NC_019843.3;YP_009047203.1;MERS_1a;3765 | 0 | 0 |
| NC_019843.3;YP_009047203.1;MERS_1a;3766 | 0 | 0 |
| NC_019843.3;YP_009047203.1;MERS_1a;3767 | 0 | 0 |
| NC_019843.3;YP_009047203.1;MERS_1a;3768 | 0 | 0 |
| NC_019843.3;YP_009047203.1;MERS_1a;3769 | 0 | 0 |
| NC_019843.3;YP_009047203.1;MERS_1a;3770 | 0 | 0 |
| NC_019843.3;YP_009047203.1;MERS_1a;3771 | 0 | 0 |
| NC_019843.3;YP_009047203.1;MERS_1a;3772 | 0 | 0 |
| NC_019843.3;YP_009047203.1;MERS_1a;3773 | 0 | 0 |
| NC_019843.3;YP_009047203.1;MERS_1a;3774 | 0 | 0 |
| NC_019843.3;YP_009047203.1;MERS_1a;3775 | 0 | 0 |
| NC_019843.3;YP_009047203.1;MERS_1a;3776 | 0 | 0 |
| NC_019843.3;YP_009047203.1;MERS_1a;3777 | 0 | 0 |
| NC_019843.3;YP_009047203.1;MERS_1a;3778 | 0 | 0 |
| NC_019843.3;YP_009047203.1;MERS_1a;3779 | 0 | 0 |
| NC_019843.3;YP_009047203.1;MERS_1a;3780 | 0 | 0 |
| NC_019843.3;YP_009047203.1;MERS_1a;3781 | 0 | 0 |
| NC_019843.3;YP_009047203.1;MERS_1a;3782 | 0 | 0 |
| NC_019843.3;YP_009047203.1;MERS_1a;3783 | 0 | 0 |
| NC_019843.3;YP_009047203.1;MERS_1a;3784 | 0 | 0 |
| NC_019843.3;YP_009047203.1;MERS_1a;3785 | 0 | 0 |
| NC_019843.3;YP_009047203.1;MERS_1a;3786 | 0 | 0 |
| NC_019843.3;YP_009047203.1;MERS_1a;3787 | 0 | 0 |
| NC_019843.3;YP_009047203.1;MERS_1a;3788 | 0 | 0 |
| NC_019843.3;YP_009047203.1;MERS_1a;3789 | 0 | 0 |
| NC_019843.3;YP_009047203.1;MERS_1a;3790 | 0 | 0 |
| NC_019843.3;YP_009047203.1;MERS_1a;3791 | 0 | 0 |
| NC_019843.3;YP_009047203.1;MERS_1a;3792 | 0 | 0 |
| NC_019843.3;YP_009047203.1;MERS_1a;3793 | 0 | 0 |
| NC_019843.3;YP_009047203.1;MERS_1a;3794 | 0 | 0 |
| NC_019843.3;YP_009047203.1;MERS_1a;3795 | 0 | 0.075 |
| NC_019843.3;YP_009047203.1;MERS_1a;3796 | 0 | 0 |
| NC_019843.3;YP_009047203.1;MERS_1a;3797 | 0 | 0 |
| NC_019843.3;YP_009047203.1;MERS_1a;3798 | 0 | 0 |
| NC_019843.3;YP_009047203.1;MERS_1a;3799 | 0 | 0 |
| NC_019843.3;YP_009047203.1;MERS_1a;3800 | 0 | 0.025 |
| NC_019843.3;YP_009047203.1;MERS_1a;3801 | 0 | 0.025 |
| NC_019843.3;YP_009047203.1;MERS_1a;3802 | 0 | 0 |
| NC_019843.3;YP_009047203.1;MERS_1a;3803 | 0 | 0.025 |

| | | |
|---|---|---|
| NC_019843.3;YP_009047203.1;MERS_1a;3804 | 0 | 0 |
| NC_019843.3;YP_009047203.1;MERS_1a;3805 | 0.05 | 0.025 |
| NC_019843.3;YP_009047203.1;MERS_1a;3806 | 0 | 0 |
| NC_019843.3;YP_009047203.1;MERS_1a;3807 | 0 | 0.025 |
| NC_019843.3;YP_009047203.1;MERS_1a;3808 | 0 | 0 |
| NC_019843.3;YP_009047203.1;MERS_1a;3809 | 0.05 | 0 |
| NC_019843.3;YP_009047203.1;MERS_1a;3810 | 0 | 0 |
| NC_019843.3;YP_009047203.1;MERS_1a;3811 | 0.15 | 0.025 |
| NC_019843.3;YP_009047203.1;MERS_1a;3812 | 0 | 0.025 |
| NC_019843.3;YP_009047203.1;MERS_1a;3813 | 0 | 0 |
| NC_019843.3;YP_009047203.1;MERS_1a;3814 | 0.05 | 0 |
| NC_019843.3;YP_009047203.1;MERS_1a;3815 | 0.05 | 0.025 |
| NC_019843.3;YP_009047203.1;MERS_1a;3816 | 0 | 0 |
| NC_019843.3;YP_009047203.1;MERS_1a;3817 | 0.05 | 0.05 |
| NC_019843.3;YP_009047203.1;MERS_1a;3818 | 0 | 0 |
| NC_019843.3;YP_009047203.1;MERS_1a;3819 | 0 | 0 |
| NC_019843.3;YP_009047203.1;MERS_1a;3820 | 0 | 0 |
| NC_019843.3;YP_009047203.1;MERS_1a;3821 | 0 | 0 |
| NC_019843.3;YP_009047203.1;MERS_1a;3822 | 0 | 0 |
| NC_019843.3;YP_009047203.1;MERS_1a;3823 | 0 | 0 |
| NC_019843.3;YP_009047203.1;MERS_1a;3824 | 0 | 0 |
| NC_019843.3;YP_009047203.1;MERS_1a;3825 | 0 | 0 |
| NC_019843.3;YP_009047203.1;MERS_1a;3826 | 0 | 0 |
| NC_019843.3;YP_009047203.1;MERS_1a;3827 | 0 | 0 |
| NC_019843.3;YP_009047203.1;MERS_1a;3828 | 0 | 0 |
| NC_019843.3;YP_009047203.1;MERS_1a;3829 | 0 | 0 |
| NC_019843.3;YP_009047203.1;MERS_1a;3830 | 0 | 0 |
| NC_019843.3;YP_009047203.1;MERS_1a;3831 | 0 | 0 |
| NC_019843.3;YP_009047203.1;MERS_1a;3832 | 0.05 | 0 |
| NC_019843.3;YP_009047203.1;MERS_1a;3833 | 0.05 | 0 |
| NC_019843.3;YP_009047203.1;MERS_1a;3834 | 0.05 | 0 |
| NC_019843.3;YP_009047203.1;MERS_1a;3835 | 0.05 | 0 |
| NC_019843.3;YP_009047203.1;MERS_1a;3836 | 0.1 | 0 |
| NC_019843.3;YP_009047203.1;MERS_1a;3837 | 0.05 | 0 |
| NC_019843.3;YP_009047203.1;MERS_1a;3838 | 0.05 | 0 |
| NC_019843.3;YP_009047203.1;MERS_1a;3839 | 0.05 | 0 |
| NC_019843.3;YP_009047203.1;MERS_1a;3840 | 0.05 | 0 |
| NC_019843.3;YP_009047203.1;MERS_1a;3841 | 0 | 0 |
| NC_019843.3;YP_009047203.1;MERS_1a;3842 | 0 | 0 |
| NC_019843.3;YP_009047203.1;MERS_1a;3843 | 0 | 0 |
| NC_019843.3;YP_009047203.1;MERS_1a;3844 | 0 | 0 |
| NC_019843.3;YP_009047203.1;MERS_1a;3845 | 0 | 0 |
| NC_019843.3;YP_009047203.1;MERS_1a;3846 | 0 | 0 |

| | | |
|---|---|---|
| NC_019843.3;YP_009047203.1;MERS_1a;3847 | 0 | 0 |
| NC_019843.3;YP_009047203.1;MERS_1a;3848 | 0 | 0 |
| NC_019843.3;YP_009047203.1;MERS_1a;3849 | 0 | 0 |
| NC_019843.3;YP_009047203.1;MERS_1a;3850 | 0 | 0 |
| NC_019843.3;YP_009047203.1;MERS_1a;3851 | 0 | 0 |
| NC_019843.3;YP_009047203.1;MERS_1a;3852 | 0 | 0 |
| NC_019843.3;YP_009047203.1;MERS_1a;3853 | 0 | 0 |
| NC_019843.3;YP_009047203.1;MERS_1a;3854 | 0 | 0 |
| NC_019843.3;YP_009047203.1;MERS_1a;3855 | 0 | 0 |
| NC_019843.3;YP_009047203.1;MERS_1a;3856 | 0 | 0 |
| NC_019843.3;YP_009047203.1;MERS_1a;3857 | 0 | 0 |
| NC_019843.3;YP_009047203.1;MERS_1a;3858 | 0 | 0 |
| NC_019843.3;YP_009047203.1;MERS_1a;3859 | 0 | 0 |
| NC_019843.3;YP_009047203.1;MERS_1a;3860 | 0 | 0 |
| NC_019843.3;YP_009047203.1;MERS_1a;3861 | 0 | 0 |
| NC_019843.3;YP_009047203.1;MERS_1a;3862 | 0 | 0 |
| NC_019843.3;YP_009047203.1;MERS_1a;3863 | 0 | 0 |
| NC_019843.3;YP_009047203.1;MERS_1a;3864 | 0 | 0 |
| NC_019843.3;YP_009047203.1;MERS_1a;3865 | 0 | 0 |
| NC_019843.3;YP_009047203.1;MERS_1a;3866 | 0 | 0 |
| NC_019843.3;YP_009047203.1;MERS_1a;3867 | 0 | 0 |
| NC_019843.3;YP_009047203.1;MERS_1a;3868 | 0 | 0 |
| NC_019843.3;YP_009047203.1;MERS_1a;3869 | 0 | 0 |
| NC_019843.3;YP_009047203.1;MERS_1a;3870 | 0 | 0 |
| NC_019843.3;YP_009047203.1;MERS_1a;3871 | 0 | 0 |
| NC_019843.3;YP_009047203.1;MERS_1a;3872 | 0.05 | 0.025 |
| NC_019843.3;YP_009047203.1;MERS_1a;3873 | 0 | 0 |
| NC_019843.3;YP_009047203.1;MERS_1a;3874 | 0 | 0 |
| NC_019843.3;YP_009047203.1;MERS_1a;3875 | 0 | 0 |
| NC_019843.3;YP_009047203.1;MERS_1a;3876 | 0 | 0 |
| NC_019843.3;YP_009047203.1;MERS_1a;3877 | 0 | 0 |
| NC_019843.3;YP_009047203.1;MERS_1a;3878 | 0 | 0 |
| NC_019843.3;YP_009047203.1;MERS_1a;3879 | 0 | 0 |
| NC_019843.3;YP_009047203.1;MERS_1a;3880 | 0 | 0 |
| NC_019843.3;YP_009047203.1;MERS_1a;3881 | 0 | 0 |
| NC_019843.3;YP_009047203.1;MERS_1a;3882 | 0 | 0 |
| NC_019843.3;YP_009047203.1;MERS_1a;3883 | 0 | 0 |
| NC_019843.3;YP_009047203.1;MERS_1a;3884 | 0 | 0 |
| NC_019843.3;YP_009047203.1;MERS_1a;3885 | 0 | 0 |
| NC_019843.3;YP_009047203.1;MERS_1a;3886 | 0 | 0 |
| NC_019843.3;YP_009047203.1;MERS_1a;3887 | 0 | 0 |
| NC_019843.3;YP_009047203.1;MERS_1a;3888 | 0 | 0 |
| NC_019843.3;YP_009047203.1;MERS_1a;3889 | 0 | 0 |

| | | |
|---|---|---|
| NC_019843.3;YP_009047203.1;MERS_1a;3890 | 0 | 0 |
| NC_019843.3;YP_009047203.1;MERS_1a;3891 | 0 | 0 |
| NC_019843.3;YP_009047203.1;MERS_1a;3892 | 0 | 0 |
| NC_019843.3;YP_009047203.1;MERS_1a;3893 | 0 | 0 |
| NC_019843.3;YP_009047203.1;MERS_1a;3894 | 0 | 0 |
| NC_019843.3;YP_009047203.1;MERS_1a;3895 | 0 | 0 |
| NC_019843.3;YP_009047203.1;MERS_1a;3896 | 0 | 0 |
| NC_019843.3;YP_009047203.1;MERS_1a;3897 | 0 | 0 |
| NC_019843.3;YP_009047203.1;MERS_1a;3898 | 0 | 0 |
| NC_019843.3;YP_009047203.1;MERS_1a;3899 | 0 | 0.025 |
| NC_019843.3;YP_009047203.1;MERS_1a;3900 | 0 | 0.025 |
| NC_019843.3;YP_009047203.1;MERS_1a;3901 | 0 | 0.05 |
| NC_019843.3;YP_009047203.1;MERS_1a;3902 | 0 | 0.05 |
| NC_019843.3;YP_009047203.1;MERS_1a;3903 | 0 | 0.075 |
| NC_019843.3;YP_009047203.1;MERS_1a;3904 | 0 | 0.05 |
| NC_019843.3;YP_009047203.1;MERS_1a;3905 | 0 | 0.075 |
| NC_019843.3;YP_009047203.1;MERS_1a;3906 | 0 | 0.1 |
| NC_019843.3;YP_009047203.1;MERS_1a;3907 | 0 | 0.075 |
| NC_019843.3;YP_009047203.1;MERS_1a;3908 | 0 | 0.05 |
| NC_019843.3;YP_009047203.1;MERS_1a;3909 | 0 | 0.05 |
| NC_019843.3;YP_009047203.1;MERS_1a;3910 | 0 | 0.025 |
| NC_019843.3;YP_009047203.1;MERS_1a;3911 | 0 | 0 |
| NC_019843.3;YP_009047203.1;MERS_1a;3912 | 0 | 0 |
| NC_019843.3;YP_009047203.1;MERS_1a;3913 | 0 | 0 |
| NC_019843.3;YP_009047203.1;MERS_1a;3914 | 0 | 0 |
| NC_019843.3;YP_009047203.1;MERS_1a;3915 | 0 | 0.025 |
| NC_019843.3;YP_009047203.1;MERS_1a;3916 | 0 | 0 |
| NC_019843.3;YP_009047203.1;MERS_1a;3917 | 0.05 | 0 |
| NC_019843.3;YP_009047203.1;MERS_1a;3918 | 0 | 0 |
| NC_019843.3;YP_009047203.1;MERS_1a;3919 | 0 | 0 |
| NC_019843.3;YP_009047203.1;MERS_1a;3920 | 0 | 0 |
| NC_019843.3;YP_009047203.1;MERS_1a;3921 | 0 | 0 |
| NC_019843.3;YP_009047203.1;MERS_1a;3922 | 0 | 0 |
| NC_019843.3;YP_009047203.1;MERS_1a;3923 | 0 | 0 |
| NC_019843.3;YP_009047203.1;MERS_1a;3924 | 0 | 0 |
| NC_019843.3;YP_009047203.1;MERS_1a;3925 | 0 | 0 |
| NC_019843.3;YP_009047203.1;MERS_1a;3926 | 0 | 0 |
| NC_019843.3;YP_009047203.1;MERS_1a;3927 | 0 | 0 |
| NC_019843.3;YP_009047203.1;MERS_1a;3928 | 0 | 0 |
| NC_019843.3;YP_009047203.1;MERS_1a;3929 | 0 | 0 |
| NC_019843.3;YP_009047203.1;MERS_1a;3930 | 0 | 0 |
| NC_019843.3;YP_009047203.1;MERS_1a;3931 | 0 | 0.025 |
| NC_019843.3;YP_009047203.1;MERS_1a;3932 | 0 | 0 |

| | | |
|---|---|---|
| NC_019843.3;YP_009047203.1;MERS_1a;3933 | 0 | 0.025 |
| NC_019843.3;YP_009047203.1;MERS_1a;3934 | 0 | 0 |
| NC_019843.3;YP_009047203.1;MERS_1a;3935 | 0 | 0.05 |
| NC_019843.3;YP_009047203.1;MERS_1a;3936 | 0.05 | 0.05 |
| NC_019843.3;YP_009047203.1;MERS_1a;3937 | 0 | 0.075 |
| NC_019843.3;YP_009047203.1;MERS_1a;3938 | 0.05 | 0.15 |
| NC_019843.3;YP_009047203.1;MERS_1a;3939 | 0.05 | 0.125 |
| NC_019843.3;YP_009047203.1;MERS_1a;3940 | 0.05 | 0.075 |
| NC_019843.3;YP_009047203.1;MERS_1a;3941 | 0.05 | 0 |
| NC_019843.3;YP_009047203.1;MERS_1a;3942 | 0.05 | 0 |
| NC_019843.3;YP_009047203.1;MERS_1a;3943 | 0 | 0 |
| NC_019843.3;YP_009047203.1;MERS_1a;3944 | 0 | 0 |
| NC_019843.3;YP_009047203.1;MERS_1a;3945 | 0 | 0.025 |
| NC_019843.3;YP_009047203.1;MERS_1a;3946 | 0 | 0 |
| NC_019843.3;YP_009047203.1;MERS_1a;3947 | 0 | 0 |
| NC_019843.3;YP_009047203.1;MERS_1a;3948 | 0 | 0 |
| NC_019843.3;YP_009047203.1;MERS_1a;3949 | 0 | 0 |
| NC_019843.3;YP_009047203.1;MERS_1a;3950 | 0 | 0 |
| NC_019843.3;YP_009047203.1;MERS_1a;3951 | 0 | 0 |
| NC_019843.3;YP_009047203.1;MERS_1a;3952 | 0 | 0 |
| NC_019843.3;YP_009047203.1;MERS_1a;3953 | 0 | 0 |
| NC_019843.3;YP_009047203.1;MERS_1a;3954 | 0 | 0 |
| NC_019843.3;YP_009047203.1;MERS_1a;3955 | 0 | 0 |
| NC_019843.3;YP_009047203.1;MERS_1a;3956 | 0 | 0 |
| NC_019843.3;YP_009047203.1;MERS_1a;3957 | 0 | 0 |
| NC_019843.3;YP_009047203.1;MERS_1a;3958 | 0 | 0.025 |
| NC_019843.3;YP_009047203.1;MERS_1a;3959 | 0 | 0 |
| NC_019843.3;YP_009047203.1;MERS_1a;3960 | 0 | 0 |
| NC_019843.3;YP_009047203.1;MERS_1a;3961 | 0 | 0 |
| NC_019843.3;YP_009047203.1;MERS_1a;3962 | 0 | 0 |
| NC_019843.3;YP_009047203.1;MERS_1a;3963 | 0 | 0 |
| NC_019843.3;YP_009047203.1;MERS_1a;3964 | 0 | 0 |
| NC_019843.3;YP_009047203.1;MERS_1a;3965 | 0 | 0 |
| NC_019843.3;YP_009047203.1;MERS_1a;3966 | 0.1 | 0 |
| NC_019843.3;YP_009047203.1;MERS_1a;3967 | 0.05 | 0 |
| NC_019843.3;YP_009047203.1;MERS_1a;3968 | 0.05 | 0 |
| NC_019843.3;YP_009047203.1;MERS_1a;3969 | 0.05 | 0.025 |
| NC_019843.3;YP_009047203.1;MERS_1a;3970 | 0.1 | 0.025 |
| NC_019843.3;YP_009047203.1;MERS_1a;3971 | 0.05 | 0.025 |
| NC_019843.3;YP_009047203.1;MERS_1a;3972 | 0 | 0 |
| NC_019843.3;YP_009047203.1;MERS_1a;3973 | 0 | 0 |
| NC_019843.3;YP_009047203.1;MERS_1a;3974 | 0 | 0.025 |
| NC_019843.3;YP_009047203.1;MERS_1a;3975 | 0 | 0.025 |

| | | |
|---|---|---|
| NC_019843.3;YP_009047203.1;MERS_1a;3976 | 0 | 0.025 |
| NC_019843.3;YP_009047203.1;MERS_1a;3977 | 0 | 0.05 |
| NC_019843.3;YP_009047203.1;MERS_1a;3978 | 0 | 0.025 |
| NC_019843.3;YP_009047203.1;MERS_1a;3979 | 0 | 0 |
| NC_019843.3;YP_009047203.1;MERS_1a;3980 | 0 | 0 |
| NC_019843.3;YP_009047203.1;MERS_1a;3981 | 0.05 | 0 |
| NC_019843.3;YP_009047203.1;MERS_1a;3982 | 0 | 0.025 |
| NC_019843.3;YP_009047203.1;MERS_1a;3983 | 0 | 0 |
| NC_019843.3;YP_009047203.1;MERS_1a;3984 | 0 | 0 |
| NC_019843.3;YP_009047203.1;MERS_1a;3985 | 0 | 0 |
| NC_019843.3;YP_009047203.1;MERS_1a;3986 | 0 | 0.025 |
| NC_019843.3;YP_009047203.1;MERS_1a;3987 | 0 | 0.025 |
| NC_019843.3;YP_009047203.1;MERS_1a;3988 | 0 | 0 |
| NC_019843.3;YP_009047203.1;MERS_1a;3989 | 0 | 0 |
| NC_019843.3;YP_009047203.1;MERS_1a;3990 | 0 | 0 |
| NC_019843.3;YP_009047203.1;MERS_1a;3991 | 0 | 0 |
| NC_019843.3;YP_009047203.1;MERS_1a;3992 | 0.05 | 0.025 |
| NC_019843.3;YP_009047203.1;MERS_1a;3993 | 0.1 | 0.025 |
| NC_019843.3;YP_009047203.1;MERS_1a;3994 | 0.05 | 0.025 |
| NC_019843.3;YP_009047203.1;MERS_1a;3995 | 0.05 | 0.025 |
| NC_019843.3;YP_009047203.1;MERS_1a;3996 | 0.05 | 0.025 |
| NC_019843.3;YP_009047203.1;MERS_1a;3997 | 0 | 0.025 |
| NC_019843.3;YP_009047203.1;MERS_1a;3998 | 0 | 0.025 |
| NC_019843.3;YP_009047203.1;MERS_1a;3999 | 0 | 0.025 |
| NC_019843.3;YP_009047203.1;MERS_1a;4000 | 0 | 0.025 |
| NC_019843.3;YP_009047203.1;MERS_1a;4001 | 0 | 0.025 |
| NC_019843.3;YP_009047203.1;MERS_1a;4002 | 0 | 0 |
| NC_019843.3;YP_009047203.1;MERS_1a;4003 | 0 | 0.025 |
| NC_019843.3;YP_009047203.1;MERS_1a;4004 | 0 | 0 |
| NC_019843.3;YP_009047203.1;MERS_1a;4005 | 0 | 0 |
| NC_019843.3;YP_009047203.1;MERS_1a;4006 | 0 | 0 |
| NC_019843.3;YP_009047203.1;MERS_1a;4007 | 0 | 0 |
| NC_019843.3;YP_009047203.1;MERS_1a;4008 | 0 | 0.025 |
| NC_019843.3;YP_009047203.1;MERS_1a;4009 | 0 | 0 |
| NC_019843.3;YP_009047203.1;MERS_1a;4010 | 0 | 0 |
| NC_019843.3;YP_009047203.1;MERS_1a;4011 | 0 | 0 |
| NC_019843.3;YP_009047203.1;MERS_1a;4012 | 0 | 0 |
| NC_019843.3;YP_009047203.1;MERS_1a;4013 | 0 | 0 |
| NC_019843.3;YP_009047203.1;MERS_1a;4014 | 0 | 0 |
| NC_019843.3;YP_009047203.1;MERS_1a;4015 | 0 | 0 |
| NC_019843.3;YP_009047203.1;MERS_1a;4016 | 0 | 0 |
| NC_019843.3;YP_009047203.1;MERS_1a;4017 | 0 | 0 |
| NC_019843.3;YP_009047203.1;MERS_1a;4018 | 0.2 | 0.125 |

| | | |
|---|---|---|
| NC_019843.3;YP_009047203.1;MERS_1a;4019 | 0.15 | 0.125 |
| NC_019843.3;YP_009047203.1;MERS_1a;4020 | 0.2 | 0.125 |
| NC_019843.3;YP_009047203.1;MERS_1a;4021 | 0.25 | 0.15 |
| NC_019843.3;YP_009047203.1;MERS_1a;4022 | 0.2 | 0.175 |
| NC_019843.3;YP_009047203.1;MERS_1a;4023 | 0.25 | 0.225 |
| NC_019843.3;YP_009047203.1;MERS_1a;4024 | 0.3 | 0.225 |
| NC_019843.3;YP_009047203.1;MERS_1a;4025 | 0.2 | 0.075 |
| NC_019843.3;YP_009047203.1;MERS_1a;4026 | 0.2 | 0.175 |
| NC_019843.3;YP_009047203.1;MERS_1a;4027 | 0.1 | 0.025 |
| NC_019843.3;YP_009047203.1;MERS_1a;4028 | 0.05 | 0 |
| NC_019843.3;YP_009047203.1;MERS_1a;4029 | 0.05 | 0 |
| NC_019843.3;YP_009047203.1;MERS_1a;4030 | 0.05 | 0 |
| NC_019843.3;YP_009047203.1;MERS_1a;4031 | 0.05 | 0 |
| NC_019843.3;YP_009047203.1;MERS_1a;4032 | 0.05 | 0.025 |
| NC_019843.3;YP_009047203.1;MERS_1a;4033 | 0.1 | 0.025 |
| NC_019843.3;YP_009047203.1;MERS_1a;4034 | 0.05 | 0.025 |
| NC_019843.3;YP_009047203.1;MERS_1a;4035 | 0 | 0 |
| NC_019843.3;YP_009047203.1;MERS_1a;4036 | 0.05 | 0 |
| NC_019843.3;YP_009047203.1;MERS_1a;4037 | 0 | 0 |
| NC_019843.3;YP_009047203.1;MERS_1a;4038 | 0 | 0 |
| NC_019843.3;YP_009047203.1;MERS_1a;4039 | 0 | 0 |
| NC_019843.3;YP_009047203.1;MERS_1a;4040 | 0 | 0 |
| NC_019843.3;YP_009047203.1;MERS_1a;4041 | 0 | 0 |
| NC_019843.3;YP_009047203.1;MERS_1a;4042 | 0 | 0 |
| NC_019843.3;YP_009047203.1;MERS_1a;4043 | 0 | 0 |
| NC_019843.3;YP_009047203.1;MERS_1a;4044 | 0 | 0 |
| NC_019843.3;YP_009047203.1;MERS_1a;4045 | 0 | 0 |
| NC_019843.3;YP_009047203.1;MERS_1a;4046 | 0 | 0 |
| NC_019843.3;YP_009047203.1;MERS_1a;4047 | 0 | 0 |
| NC_019843.3;YP_009047203.1;MERS_1a;4048 | 0 | 0 |
| NC_019843.3;YP_009047203.1;MERS_1a;4049 | 0.05 | 0 |
| NC_019843.3;YP_009047203.1;MERS_1a;4050 | 0.05 | 0.05 |
| NC_019843.3;YP_009047203.1;MERS_1a;4051 | 0 | 0 |
| NC_019843.3;YP_009047203.1;MERS_1a;4052 | 0 | 0 |
| NC_019843.3;YP_009047203.1;MERS_1a;4053 | 0.05 | 0 |
| NC_019843.3;YP_009047203.1;MERS_1a;4054 | 0 | 0 |
| NC_019843.3;YP_009047203.1;MERS_1a;4055 | 0 | 0 |
| NC_019843.3;YP_009047203.1;MERS_1a;4056 | 0 | 0 |
| NC_019843.3;YP_009047203.1;MERS_1a;4057 | 0 | 0 |
| NC_019843.3;YP_009047203.1;MERS_1a;4058 | 0 | 0 |
| NC_019843.3;YP_009047203.1;MERS_1a;4059 | 0 | 0 |
| NC_019843.3;YP_009047203.1;MERS_1a;4060 | 0 | 0.025 |
| NC_019843.3;YP_009047203.1;MERS_1a;4061 | 0 | 0 |

| | | |
|---|---|---|
| NC_019843.3;YP_009047203.1;MERS_1a;4062 | 0 | 0 |
| NC_019843.3;YP_009047203.1;MERS_1a;4063 | 0 | 0 |
| NC_019843.3;YP_009047203.1;MERS_1a;4064 | 0 | 0 |
| NC_019843.3;YP_009047203.1;MERS_1a;4065 | 0 | 0 |
| NC_019843.3;YP_009047203.1;MERS_1a;4066 | 0 | 0 |
| NC_019843.3;YP_009047203.1;MERS_1a;4067 | 0 | 0 |
| NC_019843.3;YP_009047203.1;MERS_1a;4068 | 0 | 0 |
| NC_019843.3;YP_009047203.1;MERS_1a;4069 | 0 | 0.025 |
| NC_019843.3;YP_009047203.1;MERS_1a;4070 | 0 | 0.025 |
| NC_019843.3;YP_009047203.1;MERS_1a;4071 | 0 | 0 |
| NC_019843.3;YP_009047203.1;MERS_1a;4072 | 0 | 0 |
| NC_019843.3;YP_009047203.1;MERS_1a;4073 | 0 | 0 |
| NC_019843.3;YP_009047203.1;MERS_1a;4074 | 0 | 0 |
| NC_019843.3;YP_009047203.1;MERS_1a;4075 | 0.05 | 0 |
| NC_019843.3;YP_009047203.1;MERS_1a;4076 | 0 | 0 |
| NC_019843.3;YP_009047203.1;MERS_1a;4077 | 0.05 | 0 |
| NC_019843.3;YP_009047203.1;MERS_1a;4078 | 0.05 | 0 |
| NC_019843.3;YP_009047203.1;MERS_1a;4079 | 0.05 | 0 |
| NC_019843.3;YP_009047203.1;MERS_1a;4080 | 0 | 0 |
| NC_019843.3;YP_009047203.1;MERS_1a;4081 | 0 | 0 |
| NC_019843.3;YP_009047203.1;MERS_1a;4082 | 0 | 0.05 |
| NC_019843.3;YP_009047203.1;MERS_1a;4083 | 0 | 0.025 |
| NC_019843.3;YP_009047203.1;MERS_1a;4084 | 0 | 0 |
| NC_019843.3;YP_009047203.1;MERS_1a;4085 | 0 | 0 |
| NC_019843.3;YP_009047203.1;MERS_1a;4086 | 0 | 0 |
| NC_019843.3;YP_009047203.1;MERS_1a;4087 | 0 | 0 |
| NC_019843.3;YP_009047203.1;MERS_1a;4088 | 0 | 0 |
| NC_019843.3;YP_009047203.1;MERS_1a;4089 | 0 | 0 |
| NC_019843.3;YP_009047203.1;MERS_1a;4090 | 0 | 0 |
| NC_019843.3;YP_009047203.1;MERS_1a;4091 | 0 | 0 |
| NC_019843.3;YP_009047203.1;MERS_1a;4092 | 0 | 0 |
| NC_019843.3;YP_009047203.1;MERS_1a;4093 | 0 | 0 |
| NC_019843.3;YP_009047203.1;MERS_1a;4094 | 0.05 | 0.05 |
| NC_019843.3;YP_009047203.1;MERS_1a;4095 | 0 | 0.025 |
| NC_019843.3;YP_009047203.1;MERS_1a;4096 | 0.1 | 0.075 |
| NC_019843.3;YP_009047203.1;MERS_1a;4097 | 0.05 | 0.075 |
| NC_019843.3;YP_009047203.1;MERS_1a;4098 | 0.05 | 0 |
| NC_019843.3;YP_009047203.1;MERS_1a;4099 | 0 | 0 |
| NC_019843.3;YP_009047203.1;MERS_1a;4100 | 0 | 0 |
| NC_019843.3;YP_009047203.1;MERS_1a;4101 | 0 | 0 |
| NC_019843.3;YP_009047203.1;MERS_1a;4102 | 0 | 0 |
| NC_019843.3;YP_009047203.1;MERS_1a;4103 | 0 | 0 |
| NC_019843.3;YP_009047203.1;MERS_1a;4104 | 0 | 0 |

| | | |
|---|---|---|
| NC_019843.3;YP_009047203.1;MERS_1a;4105 | 0 | 0 |
| NC_019843.3;YP_009047203.1;MERS_1a;4106 | 0 | 0 |
| NC_019843.3;YP_009047203.1;MERS_1a;4107 | 0 | 0 |
| NC_019843.3;YP_009047203.1;MERS_1a;4108 | 0 | 0 |
| NC_019843.3;YP_009047203.1;MERS_1a;4109 | 0 | 0 |
| NC_019843.3;YP_009047203.1;MERS_1a;4110 | 0 | 0 |
| NC_019843.3;YP_009047203.1;MERS_1a;4111 | 0 | 0 |
| NC_019843.3;YP_009047203.1;MERS_1a;4112 | 0 | 0 |
| NC_019843.3;YP_009047203.1;MERS_1a;4113 | 0 | 0 |
| NC_019843.3;YP_009047203.1;MERS_1a;4114 | 0.05 | 0 |
| NC_019843.3;YP_009047203.1;MERS_1a;4115 | 0.05 | 0.05 |
| NC_019843.3;YP_009047203.1;MERS_1a;4116 | 0.05 | 0.05 |
| NC_019843.3;YP_009047203.1;MERS_1a;4117 | 0 | 0 |
| NC_019843.3;YP_009047203.1;MERS_1a;4118 | 0 | 0 |
| NC_019843.3;YP_009047203.1;MERS_1a;4119 | 0 | 0.1 |
| NC_019843.3;YP_009047203.1;MERS_1a;4120 | 0 | 0.025 |
| NC_019843.3;YP_009047203.1;MERS_1a;4121 | 0 | 0 |
| NC_019843.3;YP_009047203.1;MERS_1a;4122 | 0 | 0 |
| NC_019843.3;YP_009047203.1;MERS_1a;4123 | 0 | 0 |
| NC_019843.3;YP_009047203.1;MERS_1a;4124 | 0 | 0 |
| NC_019843.3;YP_009047203.1;MERS_1a;4125 | 0 | 0 |
| NC_019843.3;YP_009047203.1;MERS_1a;4126 | 0.05 | 0.05 |
| NC_019843.3;YP_009047203.1;MERS_1a;4127 | 0 | 0.025 |
| NC_019843.3;YP_009047203.1;MERS_1a;4128 | 0 | 0.025 |
| NC_019843.3;YP_009047203.1;MERS_1a;4129 | 0 | 0.025 |
| NC_019843.3;YP_009047203.1;MERS_1a;4130 | 0 | 0 |
| NC_019843.3;YP_009047203.1;MERS_1a;4131 | 0 | 0 |
| NC_019843.3;YP_009047203.1;MERS_1a;4132 | 0 | 0 |
| NC_019843.3;YP_009047203.1;MERS_1a;4133 | 0 | 0 |
| NC_019843.3;YP_009047203.1;MERS_1a;4134 | 0 | 0 |
| NC_019843.3;YP_009047203.1;MERS_1a;4135 | 0 | 0 |
| NC_019843.3;YP_009047203.1;MERS_1a;4136 | 0 | 0 |
| NC_019843.3;YP_009047203.1;MERS_1a;4137 | 0 | 0 |
| NC_019843.3;YP_009047203.1;MERS_1a;4138 | 0 | 0 |
| NC_019843.3;YP_009047203.1;MERS_1a;4139 | 0 | 0 |
| NC_019843.3;YP_009047203.1;MERS_1a;4140 | 0 | 0 |
| NC_019843.3;YP_009047203.1;MERS_1a;4141 | 0 | 0 |
| NC_019843.3;YP_009047203.1;MERS_1a;4142 | 0 | 0 |
| NC_019843.3;YP_009047203.1;MERS_1a;4143 | 0 | 0 |
| NC_019843.3;YP_009047203.1;MERS_1a;4144 | 0 | 0 |
| NC_019843.3;YP_009047203.1;MERS_1a;4145 | 0 | 0 |
| NC_019843.3;YP_009047203.1;MERS_1a;4146 | 0 | 0.075 |
| NC_019843.3;YP_009047203.1;MERS_1a;4147 | 0 | 0 |

| | | |
|---|---|---|
| NC_019843.3;YP_009047203.1;MERS_1a;4148 | 0 | 0 |
| NC_019843.3;YP_009047203.1;MERS_1a;4149 | 0 | 0 |
| NC_019843.3;YP_009047203.1;MERS_1a;4150 | 0 | 0 |
| NC_019843.3;YP_009047203.1;MERS_1a;4151 | 0 | 0 |
| NC_019843.3;YP_009047203.1;MERS_1a;4152 | 0 | 0 |
| NC_019843.3;YP_009047203.1;MERS_1a;4153 | 0 | 0 |
| NC_019843.3;YP_009047203.1;MERS_1a;4154 | 0 | 0 |
| NC_019843.3;YP_009047203.1;MERS_1a;4155 | 0 | 0 |
| NC_019843.3;YP_009047203.1;MERS_1a;4156 | 0 | 0.025 |
| NC_019843.3;YP_009047203.1;MERS_1a;4157 | 0 | 0 |
| NC_019843.3;YP_009047203.1;MERS_1a;4158 | 0 | 0 |
| NC_019843.3;YP_009047203.1;MERS_1a;4159 | 0.05 | 0 |
| NC_019843.3;YP_009047203.1;MERS_1a;4160 | 0.05 | 0 |
| NC_019843.3;YP_009047203.1;MERS_1a;4161 | 0 | 0 |
| NC_019843.3;YP_009047203.1;MERS_1a;4162 | 0 | 0 |
| NC_019843.3;YP_009047203.1;MERS_1a;4163 | 0.05 | 0.025 |
| NC_019843.3;YP_009047203.1;MERS_1a;4164 | 0 | 0 |
| NC_019843.3;YP_009047203.1;MERS_1a;4165 | 0 | 0 |
| NC_019843.3;YP_009047203.1;MERS_1a;4166 | 0 | 0 |
| NC_019843.3;YP_009047203.1;MERS_1a;4167 | 0 | 0 |
| NC_019843.3;YP_009047203.1;MERS_1a;4168 | 0 | 0 |
| NC_019843.3;YP_009047203.1;MERS_1a;4169 | 0 | 0 |
| NC_019843.3;YP_009047203.1;MERS_1a;4170 | 0 | 0 |
| NC_019843.3;YP_009047203.1;MERS_1a;4171 | 0 | 0.025 |
| NC_019843.3;YP_009047203.1;MERS_1a;4172 | 0.05 | 0 |
| NC_019843.3;YP_009047203.1;MERS_1a;4173 | 0.15 | 0.025 |
| NC_019843.3;YP_009047203.1;MERS_1a;4174 | 0.05 | 0 |
| NC_019843.3;YP_009047203.1;MERS_1a;4175 | 0.1 | 0 |
| NC_019843.3;YP_009047203.1;MERS_1a;4176 | 0.05 | 0 |
| NC_019843.3;YP_009047203.1;MERS_1a;4177 | 0.05 | 0 |
| NC_019843.3;YP_009047203.1;MERS_1a;4178 | 0 | 0 |
| NC_019843.3;YP_009047203.1;MERS_1a;4179 | 0 | 0 |
| NC_019843.3;YP_009047203.1;MERS_1a;4180 | 0.05 | 0 |
| NC_019843.3;YP_009047203.1;MERS_1a;4181 | 0.05 | 0 |
| NC_019843.3;YP_009047203.1;MERS_1a;4182 | 0.05 | 0 |
| NC_019843.3;YP_009047203.1;MERS_1a;4183 | 0.05 | 0 |
| NC_019843.3;YP_009047203.1;MERS_1a;4184 | 0 | 0 |
| NC_019843.3;YP_009047203.1;MERS_1a;4185 | 0 | 0 |
| NC_019843.3;YP_009047203.1;MERS_1a;4186 | 0 | 0 |
| NC_019843.3;YP_009047203.1;MERS_1a;4187 | 0 | 0 |
| NC_019843.3;YP_009047203.1;MERS_1a;4188 | 0 | 0 |
| NC_019843.3;YP_009047203.1;MERS_1a;4189 | 0 | 0 |
| NC_019843.3;YP_009047203.1;MERS_1a;4190 | 0 | 0 |

| | | |
|---|---|---|
| NC_019843.3;YP_009047203.1;MERS_1a;4191 | 0 | 0 |
| NC_019843.3;YP_009047203.1;MERS_1a;4192 | 0 | 0 |
| NC_019843.3;YP_009047203.1;MERS_1a;4193 | 0 | 0 |
| NC_019843.3;YP_009047203.1;MERS_1a;4194 | 0 | 0 |
| NC_019843.3;YP_009047203.1;MERS_1a;4195 | 0 | 0 |
| NC_019843.3;YP_009047203.1;MERS_1a;4196 | 0 | 0 |
| NC_019843.3;YP_009047203.1;MERS_1a;4197 | 0 | 0 |
| NC_019843.3;YP_009047203.1;MERS_1a;4198 | 0 | 0 |
| NC_019843.3;YP_009047203.1;MERS_1a;4199 | 0 | 0 |
| NC_019843.3;YP_009047203.1;MERS_1a;4200 | 0 | 0 |
| NC_019843.3;YP_009047203.1;MERS_1a;4201 | 0 | 0 |
| NC_019843.3;YP_009047203.1;MERS_1a;4202 | 0 | 0 |
| NC_019843.3;YP_009047203.1;MERS_1a;4203 | 0 | 0 |
| NC_019843.3;YP_009047203.1;MERS_1a;4204 | 0 | 0 |
| NC_019843.3;YP_009047203.1;MERS_1a;4205 | 0 | 0 |
| NC_019843.3;YP_009047203.1;MERS_1a;4206 | 0 | 0 |
| NC_019843.3;YP_009047203.1;MERS_1a;4207 | 0 | 0 |
| NC_019843.3;YP_009047203.1;MERS_1a;4208 | 0 | 0 |
| NC_019843.3;YP_009047203.1;MERS_1a;4209 | 0 | 0 |
| NC_019843.3;YP_009047203.1;MERS_1a;4210 | 0 | 0 |
| NC_019843.3;YP_009047203.1;MERS_1a;4211 | 0 | 0 |
| NC_019843.3;YP_009047203.1;MERS_1a;4212 | 0 | 0 |
| NC_019843.3;YP_009047203.1;MERS_1a;4213 | 0 | 0 |
| NC_019843.3;YP_009047203.1;MERS_1a;4214 | 0 | 0 |
| NC_019843.3;YP_009047203.1;MERS_1a;4215 | 0 | 0 |
| NC_019843.3;YP_009047203.1;MERS_1a;4216 | 0 | 0 |
| NC_019843.3;YP_009047203.1;MERS_1a;4217 | 0.1 | 0 |
| NC_019843.3;YP_009047203.1;MERS_1a;4218 | 0 | 0 |
| NC_019843.3;YP_009047203.1;MERS_1a;4219 | 0 | 0 |
| NC_019843.3;YP_009047203.1;MERS_1a;4220 | 0 | 0 |
| NC_019843.3;YP_009047203.1;MERS_1a;4221 | 0 | 0 |
| NC_019843.3;YP_009047203.1;MERS_1a;4222 | 0 | 0 |
| NC_019843.3;YP_009047203.1;MERS_1a;4223 | 0 | 0 |
| NC_019843.3;YP_009047203.1;MERS_1a;4224 | 0 | 0 |
| NC_019843.3;YP_009047203.1;MERS_1a;4225 | 0 | 0 |
| NC_019843.3;YP_009047203.1;MERS_1a;4226 | 0 | 0 |
| NC_019843.3;YP_009047203.1;MERS_1a;4227 | 0 | 0 |
| NC_019843.3;YP_009047203.1;MERS_1a;4228 | 0 | 0 |
| NC_019843.3;YP_009047203.1;MERS_1a;4229 | 0 | 0 |
| NC_019843.3;YP_009047203.1;MERS_1a;4230 | 0 | 0 |
| NC_019843.3;YP_009047203.1;MERS_1a;4231 | 0.1 | 0.025 |
| NC_019843.3;YP_009047203.1;MERS_1a;4232 | 0 | 0 |
| NC_019843.3;YP_009047203.1;MERS_1a;4233 | 0 | 0 |

| | | |
|---|---|---|
| NC_019843.3;YP_009047203.1;MERS_1a;4234 | 0 | 0.025 |
| NC_019843.3;YP_009047203.1;MERS_1a;4235 | 0 | 0.075 |
| NC_019843.3;YP_009047203.1;MERS_1a;4236 | 0.05 | 0.025 |
| NC_019843.3;YP_009047203.1;MERS_1a;4237 | 0 | 0 |
| NC_019843.3;YP_009047203.1;MERS_1a;4238 | 0 | 0 |
| NC_019843.3;YP_009047203.1;MERS_1a;4239 | 0 | 0 |
| NC_019843.3;YP_009047203.1;MERS_1a;4240 | 0 | 0 |
| NC_019843.3;YP_009047203.1;MERS_1a;4241 | 0 | 0 |
| NC_019843.3;YP_009047203.1;MERS_1a;4242 | 0 | 0 |
| NC_019843.3;YP_009047203.1;MERS_1a;4243 | 0 | 0 |
| NC_019843.3;YP_009047203.1;MERS_1a;4244 | 0 | 0 |
| NC_019843.3;YP_009047203.1;MERS_1a;4245 | 0.1 | 0 |
| NC_019843.3;YP_009047203.1;MERS_1a;4246 | 0 | 0.025 |
| NC_019843.3;YP_009047203.1;MERS_1a;4247 | 0 | 0 |
| NC_019843.3;YP_009047203.1;MERS_1a;4248 | 0 | 0 |
| NC_019843.3;YP_009047203.1;MERS_1a;4249 | 0 | 0 |
| NC_019843.3;YP_009047203.1;MERS_1a;4250 | 0 | 0.025 |
| NC_019843.3;YP_009047203.1;MERS_1a;4251 | 0 | 0.025 |
| NC_019843.3;YP_009047203.1;MERS_1a;4252 | 0 | 0.025 |
| NC_019843.3;YP_009047203.1;MERS_1a;4253 | 0 | 0.025 |
| NC_019843.3;YP_009047203.1;MERS_1a;4254 | 0.05 | 0.025 |
| NC_019843.3;YP_009047203.1;MERS_1a;4255 | 0.05 | 0.025 |
| NC_019843.3;YP_009047203.1;MERS_1a;4256 | 0.15 | 0.05 |
| NC_019843.3;YP_009047203.1;MERS_1a;4257 | 0.1 | 0.1 |
| NC_019843.3;YP_009047203.1;MERS_1a;4258 | 0.1 | 0.075 |
| NC_019843.3;YP_009047203.1;MERS_1a;4259 | 0.15 | 0.075 |
| NC_019843.3;YP_009047203.1;MERS_1a;4260 | 0.1 | 0.05 |
| NC_019843.3;YP_009047203.1;MERS_1a;4261 | 0.1 | 0.05 |
| NC_019843.3;YP_009047203.1;MERS_1a;4262 | 0.1 | 0.05 |
| NC_019843.3;YP_009047203.1;MERS_1a;4263 | 0.1 | 0.075 |
| NC_019843.3;YP_009047203.1;MERS_1a;4264 | 0 | 0.025 |
| NC_019843.3;YP_009047203.1;MERS_1a;4265 | 0 | 0.075 |
| NC_019843.3;YP_009047203.1;MERS_1a;4266 | 0 | 0 |
| NC_019843.3;YP_009047203.1;MERS_1a;4267 | 0 | 0.025 |
| NC_019843.3;YP_009047203.1;MERS_1a;4268 | 0 | 0.025 |
| NC_019843.3;YP_009047203.1;MERS_1a;4269 | 0 | 0 |
| NC_019843.3;YP_009047203.1;MERS_1a;4270 | 0 | 0 |
| NC_019843.3;YP_009047203.1;MERS_1a;4271 | 0 | 0 |
| NC_019843.3;YP_009047203.1;MERS_1a;4272 | 0 | 0 |
| NC_019843.3;YP_009047203.1;MERS_1a;4273 | 0.05 | 0 |
| NC_019843.3;YP_009047203.1;MERS_1a;4274 | 0 | 0 |
| NC_019843.3;YP_009047203.1;MERS_1a;4275 | 0 | 0 |
| NC_019843.3;YP_009047203.1;MERS_1a;4276 | 0.05 | 0 |

| | | |
|---|---|---|
| NC_019843.3;YP_009047203.1;MERS_1a;4277 | 0 | 0 |
| NC_019843.3;YP_009047203.1;MERS_1a;4278 | 0.05 | 0 |
| NC_019843.3;YP_009047203.1;MERS_1a;4279 | 0 | 0 |
| NC_019843.3;YP_009047203.1;MERS_1a;4280 | 0 | 0 |
| NC_019843.3;YP_009047203.1;MERS_1a;4281 | 0 | 0 |
| NC_019843.3;YP_009047203.1;MERS_1a;4282 | 0 | 0 |
| NC_019843.3;YP_009047203.1;MERS_1a;4283 | 0.05 | 0 |
| NC_019843.3;YP_009047203.1;MERS_1a;4284 | 0.05 | 0 |
| NC_019843.3;YP_009047203.1;MERS_1a;4285 | 0.05 | 0 |
| NC_019843.3;YP_009047203.1;MERS_1a;4286 | 0 | 0 |
| NC_019843.3;YP_009047203.1;MERS_1a;4287 | 0 | 0 |
| NC_019843.3;YP_009047203.1;MERS_1a;4288 | 0 | 0 |
| NC_019843.3;YP_009047203.1;MERS_1a;4289 | 0 | 0 |
| NC_019843.3;YP_009047203.1;MERS_1a;4290 | 0 | 0 |
| NC_019843.3;YP_009047203.1;MERS_1a;4291 | 0 | 0 |
| NC_019843.3;YP_009047203.1;MERS_1a;4292 | 0 | 0 |
| NC_019843.3;YP_009047203.1;MERS_1a;4293 | 0 | 0 |
| NC_019843.3;YP_009047203.1;MERS_1a;4294 | 0 | 0 |
| NC_019843.3;YP_009047203.1;MERS_1a;4295 | 0 | 0 |
| NC_019843.3;YP_009047203.1;MERS_1a;4296 | 0 | 0 |
| NC_019843.3;YP_009047203.1;MERS_1a;4297 | 0 | 0.075 |
| NC_019843.3;YP_009047203.1;MERS_1a;4298 | 0 | 0 |
| NC_019843.3;YP_009047203.1;MERS_1a;4299 | 0 | 0 |
| NC_019843.3;YP_009047203.1;MERS_1a;4300 | 0 | 0 |
| NC_019843.3;YP_009047203.1;MERS_1a;4301 | 0 | 0 |
| NC_019843.3;YP_009047203.1;MERS_1a;4302 | 0 | 0 |
| NC_019843.3;YP_009047203.1;MERS_1a;4303 | 0 | 0 |
| NC_019843.3;YP_009047203.1;MERS_1a;4304 | 0 | 0 |
| NC_019843.3;YP_009047203.1;MERS_1a;4305 | 0 | 0 |
| NC_019843.3;YP_009047203.1;MERS_1a;4306 | 0 | 0 |
| NC_019843.3;YP_009047203.1;MERS_1a;4307 | 0 | 0 |
| NC_019843.3;YP_009047203.1;MERS_1a;4308 | 0 | 0 |
| NC_019843.3;YP_009047203.1;MERS_1a;4309 | 0 | 0 |
| NC_019843.3;YP_009047203.1;MERS_1a;4310 | 0 | 0 |
| NC_019843.3;YP_009047203.1;MERS_1a;4311 | 0 | 0 |
| NC_019843.3;YP_009047203.1;MERS_1a;4312 | 0 | 0.025 |
| NC_019843.3;YP_009047203.1;MERS_1a;4313 | 0 | 0 |
| NC_019843.3;YP_009047203.1;MERS_1a;4314 | 0 | 0 |
| NC_019843.3;YP_009047203.1;MERS_1a;4315 | 0 | 0 |
| NC_019843.3;YP_009047203.1;MERS_1a;4316 | 0 | 0 |
| NC_019843.3;YP_009047203.1;MERS_1a;4317 | 0 | 0 |
| NC_019843.3;YP_009047203.1;MERS_1a;4318 | 0 | 0 |
| NC_019843.3;YP_009047203.1;MERS_1a;4319 | 0 | 0 |

| | | |
|---|---|---|
| NC_019843.3;YP_009047203.1;MERS_1a;4320 | 0 | 0 |
| NC_019843.3;YP_009047203.1;MERS_1a;4321 | 0 | 0 |
| NC_019843.3;YP_009047203.1;MERS_1a;4322 | 0 | 0 |
| NC_019843.3;YP_009047203.1;MERS_1a;4323 | 0 | 0 |
| NC_019843.3;YP_009047203.1;MERS_1a;4324 | 0 | 0 |
| NC_019843.3;YP_009047203.1;MERS_1a;4325 | 0 | 0 |
| NC_019843.3;YP_009047203.1;MERS_1a;4326 | 0 | 0 |
| NC_019843.3;YP_009047203.1;MERS_1a;4327 | 0 | 0 |
| NC_019843.3;YP_009047203.1;MERS_1a;4328 | 0 | 0 |
| NC_019843.3;YP_009047203.1;MERS_1a;4329 | 0.05 | 0 |
| NC_019843.3;YP_009047203.1;MERS_1a;4330 | 0 | 0.05 |
| NC_019843.3;YP_009047203.1;MERS_1a;4331 | 0.05 | 0 |
| NC_019843.3;YP_009047203.1;MERS_1a;4332 | 0.05 | 0 |
| NC_019843.3;YP_009047203.1;MERS_1a;4333 | 0 | 0.025 |
| NC_019843.3;YP_009047203.1;MERS_1a;4334 | 0 | 0 |
| NC_019843.3;YP_009047203.1;MERS_1a;4335 | 0 | 0 |
| NC_019843.3;YP_009047203.1;MERS_1a;4336 | 0 | 0 |
| NC_019843.3;YP_009047203.1;MERS_1a;4337 | 0 | 0 |
| NC_019843.3;YP_009047203.1;MERS_1a;4338 | 0 | 0 |
| NC_019843.3;YP_009047203.1;MERS_1a;4339 | 0 | 0 |
| NC_019843.3;YP_009047203.1;MERS_1a;4340 | 0 | 0 |
| NC_019843.3;YP_009047203.1;MERS_1a;4341 | 0 | 0 |
| NC_019843.3;YP_009047203.1;MERS_1a;4342 | 0 | 0 |
| NC_019843.3;YP_009047203.1;MERS_1a;4343 | 0.05 | 0.025 |
| NC_019843.3;YP_009047203.1;MERS_1a;4344 | 0 | 0 |
| NC_019843.3;YP_009047203.1;MERS_1a;4345 | 0 | 0 |
| NC_019843.3;YP_009047203.1;MERS_1a;4346 | 0 | 0 |
| NC_019843.3;YP_009047203.1;MERS_1a;4347 | 0 | 0 |
| NC_019843.3;YP_009047203.1;MERS_1a;4348 | 0 | 0 |
| NC_019843.3;YP_009047203.1;MERS_1a;4349 | 0 | 0 |
| NC_019843.3;YP_009047203.1;MERS_1a;4350 | 0 | 0 |
| NC_019843.3;YP_009047203.1;MERS_1a;4351 | 0 | 0 |
| NC_019843.3;YP_009047203.1;MERS_1a;4352 | 0 | 0 |
| NC_019843.3;YP_009047203.1;MERS_1a;4353 | 0 | 0 |
| NC_019843.3;YP_009047203.1;MERS_1a;4354 | 0 | 0 |
| NC_019843.3;YP_009047203.1;MERS_1a;4355 | 0 | 0 |
| NC_019843.3;YP_009047203.1;MERS_1a;4356 | 0 | 0 |
| NC_019843.3;YP_009047203.1;MERS_1a;4357 | 0 | 0 |
| NC_019843.3;YP_009047203.1;MERS_1a;4358 | 0 | 0 |
| NC_019843.3;YP_009047203.1;MERS_1a;4359 | 0.05 | 0.025 |
| NC_019843.3;YP_009047203.1;MERS_1a;4360 | 0 | 0 |
| NC_019843.3;YP_009047203.1;MERS_1a;4361 | 0 | 0 |
| NC_019843.3;YP_009047203.1;MERS_1a;4362 | 0 | 0 |

| | | |
|---|---|---|
| NC_019843.3;YP_009047203.1;MERS_1a;4363 | 0 | 0 |
| NC_019843.3;YP_009047203.1;MERS_1a;4364 | 0 | 0 |
| NC_019843.3;YP_009047203.1;MERS_1a;4365 | 0 | 0 |
| NC_019843.3;YP_009047203.1;MERS_1a;4366 | 0 | 0 |
| NC_019843.3;YP_009047203.1;MERS_1a;4367 | 0 | 0 |
| NC_019843.3;YP_009047203.1;MERS_1a;4368 | 0.1 | 0.05 |
| NC_019843.3;YP_009047203.1;MERS_1a;4369 | 0 | 0 |
| NC_019843.3;YP_009047203.1;MERS_1a;4370 | 0.05 | 0.075 |
| NC_019843.3;YP_009047203.1;MERS_1a;4371 | 0.1 | 0.1 |
| NC_019843.3;YP_009047203.1;MERS_1a;4372 | 0.1 | 0.1 |
| NC_019843.3;YP_009047203.1;MERS_1a;4373 | 0.15 | 0.075 |
| NC_019843.3;YP_009047203.1;MERS_1a;4374 | 0.15 | 0.1 |
| NC_019843.3;YP_009047203.1;MERS_1a;4375 | 0.15 | 0.175 |
| NC_019843.3;YP_009047203.1;MERS_1a;4376 | 0.2 | 0.125 |
| NC_019843.3;YP_009047203.1;MERS_1a;4377 | 0.25 | 0.2 |
| NC_019843.3;YP_009047203.1;MERS_1a;4378 | 0.2 | 0.15 |
| NC_019843.3;YP_009047203.1;MERS_1a;4379 | 0.2 | 0.15 |
| NC_019843.3;YP_009047203.1;MERS_1a;4380 | 0.05 | 0.025 |
| NC_019843.3;YP_009047204.1;MERS_spike;1 | 0 | 0 |
| NC_019843.3;YP_009047204.1;MERS_spike;2 | 0 | 0 |
| NC_019843.3;YP_009047204.1;MERS_spike;3 | 0 | 0 |
| NC_019843.3;YP_009047204.1;MERS_spike;4 | 0 | 0 |
| NC_019843.3;YP_009047204.1;MERS_spike;5 | 0 | 0 |
| NC_019843.3;YP_009047204.1;MERS_spike;6 | 0 | 0 |
| NC_019843.3;YP_009047204.1;MERS_spike;7 | 0 | 0 |
| NC_019843.3;YP_009047204.1;MERS_spike;8 | 0 | 0 |
| NC_019843.3;YP_009047204.1;MERS_spike;9 | 0 | 0 |
| NC_019843.3;YP_009047204.1;MERS_spike;10 | 0 | 0 |
| NC_019843.3;YP_009047204.1;MERS_spike;11 | 0 | 0 |
| NC_019843.3;YP_009047204.1;MERS_spike;12 | 0.05 | 0.025 |
| NC_019843.3;YP_009047204.1;MERS_spike;13 | 0.1 | 0.175 |
| NC_019843.3;YP_009047204.1;MERS_spike;14 | 0 | 0 |
| NC_019843.3;YP_009047204.1;MERS_spike;15 | 0 | 0 |
| NC_019843.3;YP_009047204.1;MERS_spike;16 | 0 | 0 |
| NC_019843.3;YP_009047204.1;MERS_spike;17 | 0 | 0 |
| NC_019843.3;YP_009047204.1;MERS_spike;18 | 0 | 0 |
| NC_019843.3;YP_009047204.1;MERS_spike;19 | 0 | 0.025 |
| NC_019843.3;YP_009047204.1;MERS_spike;20 | 0.05 | 0 |
| NC_019843.3;YP_009047204.1;MERS_spike;21 | 0 | 0 |
| NC_019843.3;YP_009047204.1;MERS_spike;22 | 0 | 0 |
| NC_019843.3;YP_009047204.1;MERS_spike;23 | 0 | 0 |
| NC_019843.3;YP_009047204.1;MERS_spike;24 | 0 | 0 |
| NC_019843.3;YP_009047204.1;MERS_spike;25 | 0 | 0 |

| | | |
|---|---|---|
| NC_019843.3;YP_009047204.1;MERS_spike;26 | 0 | 0 |
| NC_019843.3;YP_009047204.1;MERS_spike;27 | 0 | 0 |
| NC_019843.3;YP_009047204.1;MERS_spike;28 | 0 | 0 |
| NC_019843.3;YP_009047204.1;MERS_spike;29 | 0 | 0 |
| NC_019843.3;YP_009047204.1;MERS_spike;30 | 0 | 0 |
| NC_019843.3;YP_009047204.1;MERS_spike;31 | 0 | 0.025 |
| NC_019843.3;YP_009047204.1;MERS_spike;32 | 0 | 0.025 |
| NC_019843.3;YP_009047204.1;MERS_spike;33 | 0 | 0.05 |
| NC_019843.3;YP_009047204.1;MERS_spike;34 | 0 | 0.025 |
| NC_019843.3;YP_009047204.1;MERS_spike;35 | 0 | 0 |
| NC_019843.3;YP_009047204.1;MERS_spike;36 | 0 | 0 |
| NC_019843.3;YP_009047204.1;MERS_spike;37 | 0.1 | 0 |
| NC_019843.3;YP_009047204.1;MERS_spike;38 | 0.1 | 0 |
| NC_019843.3;YP_009047204.1;MERS_spike;39 | 0.05 | 0 |
| NC_019843.3;YP_009047204.1;MERS_spike;40 | 0.05 | 0 |
| NC_019843.3;YP_009047204.1;MERS_spike;41 | 0.05 | 0 |
| NC_019843.3;YP_009047204.1;MERS_spike;42 | 0.05 | 0.025 |
| NC_019843.3;YP_009047204.1;MERS_spike;43 | 0 | 0.05 |
| NC_019843.3;YP_009047204.1;MERS_spike;44 | 0 | 0.025 |
| NC_019843.3;YP_009047204.1;MERS_spike;45 | 0.05 | 0.025 |
| NC_019843.3;YP_009047204.1;MERS_spike;46 | 0.1 | 0.075 |
| NC_019843.3;YP_009047204.1;MERS_spike;47 | 0.05 | 0 |
| NC_019843.3;YP_009047204.1;MERS_spike;48 | 0 | 0.025 |
| NC_019843.3;YP_009047204.1;MERS_spike;49 | 0 | 0.05 |
| NC_019843.3;YP_009047204.1;MERS_spike;50 | 0 | 0.025 |
| NC_019843.3;YP_009047204.1;MERS_spike;51 | 0.05 | 0.025 |
| NC_019843.3;YP_009047204.1;MERS_spike;52 | 0.05 | 0.025 |
| NC_019843.3;YP_009047204.1;MERS_spike;53 | 0 | 0 |
| NC_019843.3;YP_009047204.1;MERS_spike;54 | 0 | 0 |
| NC_019843.3;YP_009047204.1;MERS_spike;55 | 0 | 0 |
| NC_019843.3;YP_009047204.1;MERS_spike;56 | 0 | 0 |
| NC_019843.3;YP_009047204.1;MERS_spike;57 | 0 | 0 |
| NC_019843.3;YP_009047204.1;MERS_spike;58 | 0 | 0 |
| NC_019843.3;YP_009047204.1;MERS_spike;59 | 0 | 0 |
| NC_019843.3;YP_009047204.1;MERS_spike;60 | 0 | 0 |
| NC_019843.3;YP_009047204.1;MERS_spike;61 | 0 | 0 |
| NC_019843.3;YP_009047204.1;MERS_spike;62 | 0 | 0 |
| NC_019843.3;YP_009047204.1;MERS_spike;63 | 0 | 0 |
| NC_019843.3;YP_009047204.1;MERS_spike;64 | 0 | 0 |
| NC_019843.3;YP_009047204.1;MERS_spike;65 | 0 | 0.025 |
| NC_019843.3;YP_009047204.1;MERS_spike;66 | 0 | 0 |
| NC_019843.3;YP_009047204.1;MERS_spike;67 | 0 | 0.025 |
| NC_019843.3;YP_009047204.1;MERS_spike;68 | 0 | 0 |

| | | |
|---|---|---|
| NC_019843.3;YP_009047204.1;MERS_spike;69 | 0 | 0 |
| NC_019843.3;YP_009047204.1;MERS_spike;70 | 0.05 | 0 |
| NC_019843.3;YP_009047204.1;MERS_spike;71 | 0 | 0 |
| NC_019843.3;YP_009047204.1;MERS_spike;72 | 0 | 0 |
| NC_019843.3;YP_009047204.1;MERS_spike;73 | 0 | 0 |
| NC_019843.3;YP_009047204.1;MERS_spike;74 | 0 | 0 |
| NC_019843.3;YP_009047204.1;MERS_spike;75 | 0 | 0 |
| NC_019843.3;YP_009047204.1;MERS_spike;76 | 0 | 0 |
| NC_019843.3;YP_009047204.1;MERS_spike;77 | 0 | 0 |
| NC_019843.3;YP_009047204.1;MERS_spike;78 | 0 | 0 |
| NC_019843.3;YP_009047204.1;MERS_spike;79 | 0 | 0 |
| NC_019843.3;YP_009047204.1;MERS_spike;80 | 0 | 0.025 |
| NC_019843.3;YP_009047204.1;MERS_spike;81 | 0 | 0 |
| NC_019843.3;YP_009047204.1;MERS_spike;82 | 0 | 0 |
| NC_019843.3;YP_009047204.1;MERS_spike;83 | 0 | 0 |
| NC_019843.3;YP_009047204.1;MERS_spike;84 | 0.05 | 0.025 |
| NC_019843.3;YP_009047204.1;MERS_spike;85 | 0.05 | 0.025 |
| NC_019843.3;YP_009047204.1;MERS_spike;86 | 0.05 | 0.025 |
| NC_019843.3;YP_009047204.1;MERS_spike;87 | 0.05 | 0.025 |
| NC_019843.3;YP_009047204.1;MERS_spike;88 | 0.05 | 0.025 |
| NC_019843.3;YP_009047204.1;MERS_spike;89 | 0 | 0.025 |
| NC_019843.3;YP_009047204.1;MERS_spike;90 | 0 | 0 |
| NC_019843.3;YP_009047204.1;MERS_spike;91 | 0 | 0.025 |
| NC_019843.3;YP_009047204.1;MERS_spike;92 | 0 | 0.025 |
| NC_019843.3;YP_009047204.1;MERS_spike;93 | 0 | 0 |
| NC_019843.3;YP_009047204.1;MERS_spike;94 | 0 | 0 |
| NC_019843.3;YP_009047204.1;MERS_spike;95 | 0 | 0 |
| NC_019843.3;YP_009047204.1;MERS_spike;96 | 0 | 0.025 |
| NC_019843.3;YP_009047204.1;MERS_spike;97 | 0 | 0 |
| NC_019843.3;YP_009047204.1;MERS_spike;98 | 0 | 0 |
| NC_019843.3;YP_009047204.1;MERS_spike;99 | 0.05 | 0 |
| NC_019843.3;YP_009047204.1;MERS_spike;100 | 0 | 0.025 |
| NC_019843.3;YP_009047204.1;MERS_spike;101 | 0.1 | 0.025 |
| NC_019843.3;YP_009047204.1;MERS_spike;102 | 0.1 | 0 |
| NC_019843.3;YP_009047204.1;MERS_spike;103 | 0 | 0 |
| NC_019843.3;YP_009047204.1;MERS_spike;104 | 0 | 0 |
| NC_019843.3;YP_009047204.1;MERS_spike;105 | 0 | 0 |
| NC_019843.3;YP_009047204.1;MERS_spike;106 | 0 | 0.025 |
| NC_019843.3;YP_009047204.1;MERS_spike;107 | 0 | 0 |
| NC_019843.3;YP_009047204.1;MERS_spike;108 | 0.05 | 0 |
| NC_019843.3;YP_009047204.1;MERS_spike;109 | 0.05 | 0 |
| NC_019843.3;YP_009047204.1;MERS_spike;110 | 0 | 0 |
| NC_019843.3;YP_009047204.1;MERS_spike;111 | 0.05 | 0 |

| | | |
|---|---|---|
| NC_019843.3;YP_009047204.1;MERS_spike;112 | 0.05 | 0.025 |
| NC_019843.3;YP_009047204.1;MERS_spike;113 | 0.1 | 0 |
| NC_019843.3;YP_009047204.1;MERS_spike;114 | 0.1 | 0 |
| NC_019843.3;YP_009047204.1;MERS_spike;115 | 0.1 | 0.025 |
| NC_019843.3;YP_009047204.1;MERS_spike;116 | 0.1 | 0 |
| NC_019843.3;YP_009047204.1;MERS_spike;117 | 0.1 | 0.025 |
| NC_019843.3;YP_009047204.1;MERS_spike;118 | 0.05 | 0 |
| NC_019843.3;YP_009047204.1;MERS_spike;119 | 0 | 0 |
| NC_019843.3;YP_009047204.1;MERS_spike;120 | 0 | 0 |
| NC_019843.3;YP_009047204.1;MERS_spike;121 | 0 | 0 |
| NC_019843.3;YP_009047204.1;MERS_spike;122 | 0 | 0 |
| NC_019843.3;YP_009047204.1;MERS_spike;123 | 0.05 | 0 |
| NC_019843.3;YP_009047204.1;MERS_spike;124 | 0 | 0.025 |
| NC_019843.3;YP_009047204.1;MERS_spike;125 | 0 | 0 |
| NC_019843.3;YP_009047204.1;MERS_spike;126 | 0 | 0 |
| NC_019843.3;YP_009047204.1;MERS_spike;127 | 0 | 0 |
| NC_019843.3;YP_009047204.1;MERS_spike;128 | 0 | 0 |
| NC_019843.3;YP_009047204.1;MERS_spike;129 | 0 | 0 |
| NC_019843.3;YP_009047204.1;MERS_spike;130 | 0 | 0 |
| NC_019843.3;YP_009047204.1;MERS_spike;131 | 0 | 0 |
| NC_019843.3;YP_009047204.1;MERS_spike;132 | 0 | 0 |
| NC_019843.3;YP_009047204.1;MERS_spike;133 | 0 | 0 |
| NC_019843.3;YP_009047204.1;MERS_spike;134 | 0 | 0 |
| NC_019843.3;YP_009047204.1;MERS_spike;135 | 0 | 0 |
| NC_019843.3;YP_009047204.1;MERS_spike;136 | 0 | 0 |
| NC_019843.3;YP_009047204.1;MERS_spike;137 | 0 | 0 |
| NC_019843.3;YP_009047204.1;MERS_spike;138 | 0 | 0 |
| NC_019843.3;YP_009047204.1;MERS_spike;139 | 0 | 0 |
| NC_019843.3;YP_009047204.1;MERS_spike;140 | 0 | 0 |
| NC_019843.3;YP_009047204.1;MERS_spike;141 | 0 | 0 |
| NC_019843.3;YP_009047204.1;MERS_spike;142 | 0 | 0 |
| NC_019843.3;YP_009047204.1;MERS_spike;143 | 0 | 0 |
| NC_019843.3;YP_009047204.1;MERS_spike;144 | 0 | 0 |
| NC_019843.3;YP_009047204.1;MERS_spike;145 | 0 | 0.025 |
| NC_019843.3;YP_009047204.1;MERS_spike;146 | 0 | 0.05 |
| NC_019843.3;YP_009047204.1;MERS_spike;147 | 0 | 0.075 |
| NC_019843.3;YP_009047204.1;MERS_spike;148 | 0 | 0.025 |
| NC_019843.3;YP_009047204.1;MERS_spike;149 | 0 | 0.05 |
| NC_019843.3;YP_009047204.1;MERS_spike;150 | 0 | 0.025 |
| NC_019843.3;YP_009047204.1;MERS_spike;151 | 0 | 0 |
| NC_019843.3;YP_009047204.1;MERS_spike;152 | 0 | 0 |
| NC_019843.3;YP_009047204.1;MERS_spike;153 | 0 | 0 |
| NC_019843.3;YP_009047204.1;MERS_spike;154 | 0 | 0 |

| | | |
|---|---|---|
| NC_019843.3;YP_009047204.1;MERS_spike;155 | 0 | 0.025 |
| NC_019843.3;YP_009047204.1;MERS_spike;156 | 0 | 0 |
| NC_019843.3;YP_009047204.1;MERS_spike;157 | 0 | 0 |
| NC_019843.3;YP_009047204.1;MERS_spike;158 | 0 | 0 |
| NC_019843.3;YP_009047204.1;MERS_spike;159 | 0 | 0 |
| NC_019843.3;YP_009047204.1;MERS_spike;160 | 0 | 0 |
| NC_019843.3;YP_009047204.1;MERS_spike;161 | 0 | 0 |
| NC_019843.3;YP_009047204.1;MERS_spike;162 | 0 | 0 |
| NC_019843.3;YP_009047204.1;MERS_spike;163 | 0 | 0 |
| NC_019843.3;YP_009047204.1;MERS_spike;164 | 0 | 0 |
| NC_019843.3;YP_009047204.1;MERS_spike;165 | 0 | 0 |
| NC_019843.3;YP_009047204.1;MERS_spike;166 | 0 | 0 |
| NC_019843.3;YP_009047204.1;MERS_spike;167 | 0 | 0 |
| NC_019843.3;YP_009047204.1;MERS_spike;168 | 0 | 0 |
| NC_019843.3;YP_009047204.1;MERS_spike;169 | 0 | 0 |
| NC_019843.3;YP_009047204.1;MERS_spike;170 | 0 | 0 |
| NC_019843.3;YP_009047204.1;MERS_spike;171 | 0 | 0 |
| NC_019843.3;YP_009047204.1;MERS_spike;172 | 0 | 0 |
| NC_019843.3;YP_009047204.1;MERS_spike;173 | 0 | 0 |
| NC_019843.3;YP_009047204.1;MERS_spike;174 | 0 | 0 |
| NC_019843.3;YP_009047204.1;MERS_spike;175 | 0 | 0 |
| NC_019843.3;YP_009047204.1;MERS_spike;176 | 0 | 0 |
| NC_019843.3;YP_009047204.1;MERS_spike;177 | 0 | 0 |
| NC_019843.3;YP_009047204.1;MERS_spike;178 | 0 | 0 |
| NC_019843.3;YP_009047204.1;MERS_spike;179 | 0 | 0 |
| NC_019843.3;YP_009047204.1;MERS_spike;180 | 0 | 0 |
| NC_019843.3;YP_009047204.1;MERS_spike;181 | 0 | 0 |
| NC_019843.3;YP_009047204.1;MERS_spike;182 | 0 | 0 |
| NC_019843.3;YP_009047204.1;MERS_spike;183 | 0 | 0 |
| NC_019843.3;YP_009047204.1;MERS_spike;184 | 0 | 0.025 |
| NC_019843.3;YP_009047204.1;MERS_spike;185 | 0 | 0 |
| NC_019843.3;YP_009047204.1;MERS_spike;186 | 0 | 0 |
| NC_019843.3;YP_009047204.1;MERS_spike;187 | 0 | 0 |
| NC_019843.3;YP_009047204.1;MERS_spike;188 | 0 | 0 |
| NC_019843.3;YP_009047204.1;MERS_spike;189 | 0.05 | 0 |
| NC_019843.3;YP_009047204.1;MERS_spike;190 | 0.05 | 0 |
| NC_019843.3;YP_009047204.1;MERS_spike;191 | 0.1 | 0 |
| NC_019843.3;YP_009047204.1;MERS_spike;192 | 0.05 | 0 |
| NC_019843.3;YP_009047204.1;MERS_spike;193 | 0.05 | 0 |
| NC_019843.3;YP_009047204.1;MERS_spike;194 | 0.05 | 0 |
| NC_019843.3;YP_009047204.1;MERS_spike;195 | 0.05 | 0 |
| NC_019843.3;YP_009047204.1;MERS_spike;196 | 0.05 | 0.025 |
| NC_019843.3;YP_009047204.1;MERS_spike;197 | 0.1 | 0.025 |

| | | |
|---|---|---|
| NC_019843.3;YP_009047204.1;MERS_spike;198 | 0.05 | 0.025 |
| NC_019843.3;YP_009047204.1;MERS_spike;199 | 0 | 0.025 |
| NC_019843.3;YP_009047204.1;MERS_spike;200 | 0 | 0 |
| NC_019843.3;YP_009047204.1;MERS_spike;201 | 0 | 0 |
| NC_019843.3;YP_009047204.1;MERS_spike;202 | 0 | 0.025 |
| NC_019843.3;YP_009047204.1;MERS_spike;203 | 0 | 0.05 |
| NC_019843.3;YP_009047204.1;MERS_spike;204 | 0 | 0.05 |
| NC_019843.3;YP_009047204.1;MERS_spike;205 | 0.05 | 0.05 |
| NC_019843.3;YP_009047204.1;MERS_spike;206 | 0 | 0.05 |
| NC_019843.3;YP_009047204.1;MERS_spike;207 | 0 | 0.1 |
| NC_019843.3;YP_009047204.1;MERS_spike;208 | 0.05 | 0.025 |
| NC_019843.3;YP_009047204.1;MERS_spike;209 | 0.1 | 0.025 |
| NC_019843.3;YP_009047204.1;MERS_spike;210 | 0.05 | 0.025 |
| NC_019843.3;YP_009047204.1;MERS_spike;211 | 0.25 | 0.35 |
| NC_019843.3;YP_009047204.1;MERS_spike;212 | 0.3 | 0.375 |
| NC_019843.3;YP_009047204.1;MERS_spike;213 | 0.45 | 0.375 |
| NC_019843.3;YP_009047204.1;MERS_spike;214 | 0.2 | 0.15 |
| NC_019843.3;YP_009047204.1;MERS_spike;215 | 0.25 | 0.375 |
| NC_019843.3;YP_009047204.1;MERS_spike;216 | 0.25 | 0.4 |
| NC_019843.3;YP_009047204.1;MERS_spike;217 | 0.1 | 0.125 |
| NC_019843.3;YP_009047204.1;MERS_spike;218 | 0.15 | 0.15 |
| NC_019843.3;YP_009047204.1;MERS_spike;219 | 0.1 | 0.05 |
| NC_019843.3;YP_009047204.1;MERS_spike;220 | 0.1 | 0.05 |
| NC_019843.3;YP_009047204.1;MERS_spike;221 | 0.1 | 0.05 |
| NC_019843.3;YP_009047204.1;MERS_spike;222 | 0.05 | 0.025 |
| NC_019843.3;YP_009047204.1;MERS_spike;223 | 0 | 0 |
| NC_019843.3;YP_009047204.1;MERS_spike;224 | 0 | 0 |
| NC_019843.3;YP_009047204.1;MERS_spike;225 | 0 | 0 |
| NC_019843.3;YP_009047204.1;MERS_spike;226 | 0.05 | 0.025 |
| NC_019843.3;YP_009047204.1;MERS_spike;227 | 0.1 | 0.025 |
| NC_019843.3;YP_009047204.1;MERS_spike;228 | 0 | 0 |
| NC_019843.3;YP_009047204.1;MERS_spike;229 | 0 | 0 |
| NC_019843.3;YP_009047204.1;MERS_spike;230 | 0 | 0 |
| NC_019843.3;YP_009047204.1;MERS_spike;231 | 0 | 0 |
| NC_019843.3;YP_009047204.1;MERS_spike;232 | 0.05 | 0 |
| NC_019843.3;YP_009047204.1;MERS_spike;233 | 0 | 0 |
| NC_019843.3;YP_009047204.1;MERS_spike;234 | 0 | 0.025 |
| NC_019843.3;YP_009047204.1;MERS_spike;235 | 0 | 0 |
| NC_019843.3;YP_009047204.1;MERS_spike;236 | 0 | 0 |
| NC_019843.3;YP_009047204.1;MERS_spike;237 | 0 | 0 |
| NC_019843.3;YP_009047204.1;MERS_spike;238 | 0 | 0 |
| NC_019843.3;YP_009047204.1;MERS_spike;239 | 0 | 0 |
| NC_019843.3;YP_009047204.1;MERS_spike;240 | 0 | 0 |

| | | |
|---|---|---|
| NC_019843.3;YP_009047204.1;MERS_spike;241 | 0 | 0 |
| NC_019843.3;YP_009047204.1;MERS_spike;242 | 0 | 0 |
| NC_019843.3;YP_009047204.1;MERS_spike;243 | 0 | 0 |
| NC_019843.3;YP_009047204.1;MERS_spike;244 | 0 | 0 |
| NC_019843.3;YP_009047204.1;MERS_spike;245 | 0 | 0 |
| NC_019843.3;YP_009047204.1;MERS_spike;246 | 0.05 | 0 |
| NC_019843.3;YP_009047204.1;MERS_spike;247 | 0.15 | 0.05 |
| NC_019843.3;YP_009047204.1;MERS_spike;248 | 0.05 | 0 |
| NC_019843.3;YP_009047204.1;MERS_spike;249 | 0.05 | 0 |
| NC_019843.3;YP_009047204.1;MERS_spike;250 | 0.05 | 0 |
| NC_019843.3;YP_009047204.1;MERS_spike;251 | 0 | 0 |
| NC_019843.3;YP_009047204.1;MERS_spike;252 | 0 | 0 |
| NC_019843.3;YP_009047204.1;MERS_spike;253 | 0 | 0 |
| NC_019843.3;YP_009047204.1;MERS_spike;254 | 0 | 0 |
| NC_019843.3;YP_009047204.1;MERS_spike;255 | 0 | 0 |
| NC_019843.3;YP_009047204.1;MERS_spike;256 | 0 | 0 |
| NC_019843.3;YP_009047204.1;MERS_spike;257 | 0 | 0 |
| NC_019843.3;YP_009047204.1;MERS_spike;258 | 0 | 0 |
| NC_019843.3;YP_009047204.1;MERS_spike;259 | 0 | 0 |
| NC_019843.3;YP_009047204.1;MERS_spike;260 | 0 | 0 |
| NC_019843.3;YP_009047204.1;MERS_spike;261 | 0 | 0 |
| NC_019843.3;YP_009047204.1;MERS_spike;262 | 0 | 0 |
| NC_019843.3;YP_009047204.1;MERS_spike;263 | 0 | 0 |
| NC_019843.3;YP_009047204.1;MERS_spike;264 | 0 | 0 |
| NC_019843.3;YP_009047204.1;MERS_spike;265 | 0.1 | 0 |
| NC_019843.3;YP_009047204.1;MERS_spike;266 | 0.05 | 0 |
| NC_019843.3;YP_009047204.1;MERS_spike;267 | 0.05 | 0 |
| NC_019843.3;YP_009047204.1;MERS_spike;268 | 0.1 | 0 |
| NC_019843.3;YP_009047204.1;MERS_spike;269 | 0.05 | 0 |
| NC_019843.3;YP_009047204.1;MERS_spike;270 | 0.1 | 0 |
| NC_019843.3;YP_009047204.1;MERS_spike;271 | 0.1 | 0.1 |
| NC_019843.3;YP_009047204.1;MERS_spike;272 | 0.05 | 0 |
| NC_019843.3;YP_009047204.1;MERS_spike;273 | 0 | 0 |
| NC_019843.3;YP_009047204.1;MERS_spike;274 | 0 | 0 |
| NC_019843.3;YP_009047204.1;MERS_spike;275 | 0 | 0 |
| NC_019843.3;YP_009047204.1;MERS_spike;276 | 0 | 0.025 |
| NC_019843.3;YP_009047204.1;MERS_spike;277 | 0 | 0.025 |
| NC_019843.3;YP_009047204.1;MERS_spike;278 | 0 | 0 |
| NC_019843.3;YP_009047204.1;MERS_spike;279 | 0 | 0 |
| NC_019843.3;YP_009047204.1;MERS_spike;280 | 0 | 0 |
| NC_019843.3;YP_009047204.1;MERS_spike;281 | 0 | 0 |
| NC_019843.3;YP_009047204.1;MERS_spike;282 | 0 | 0 |
| NC_019843.3;YP_009047204.1;MERS_spike;283 | 0 | 0 |

| | | |
|---|---|---|
| NC_019843.3;YP_009047204.1;MERS_spike;284 | 0 | 0 |
| NC_019843.3;YP_009047204.1;MERS_spike;285 | 0 | 0 |
| NC_019843.3;YP_009047204.1;MERS_spike;286 | 0 | 0 |
| NC_019843.3;YP_009047204.1;MERS_spike;287 | 0 | 0 |
| NC_019843.3;YP_009047204.1;MERS_spike;288 | 0 | 0 |
| NC_019843.3;YP_009047204.1;MERS_spike;289 | 0 | 0 |
| NC_019843.3;YP_009047204.1;MERS_spike;290 | 0 | 0 |
| NC_019843.3;YP_009047204.1;MERS_spike;291 | 0 | 0 |
| NC_019843.3;YP_009047204.1;MERS_spike;292 | 0 | 0 |
| NC_019843.3;YP_009047204.1;MERS_spike;293 | 0 | 0 |
| NC_019843.3;YP_009047204.1;MERS_spike;294 | 0 | 0 |
| NC_019843.3;YP_009047204.1;MERS_spike;295 | 0 | 0 |
| NC_019843.3;YP_009047204.1;MERS_spike;296 | 0 | 0 |
| NC_019843.3;YP_009047204.1;MERS_spike;297 | 0 | 0 |
| NC_019843.3;YP_009047204.1;MERS_spike;298 | 0 | 0 |
| NC_019843.3;YP_009047204.1;MERS_spike;299 | 0 | 0 |
| NC_019843.3;YP_009047204.1;MERS_spike;300 | 0 | 0 |
| NC_019843.3;YP_009047204.1;MERS_spike;301 | 0 | 0 |
| NC_019843.3;YP_009047204.1;MERS_spike;302 | 0 | 0 |
| NC_019843.3;YP_009047204.1;MERS_spike;303 | 0 | 0 |
| NC_019843.3;YP_009047204.1;MERS_spike;304 | 0 | 0 |
| NC_019843.3;YP_009047204.1;MERS_spike;305 | 0 | 0 |
| NC_019843.3;YP_009047204.1;MERS_spike;306 | 0 | 0 |
| NC_019843.3;YP_009047204.1;MERS_spike;307 | 0 | 0 |
| NC_019843.3;YP_009047204.1;MERS_spike;308 | 0 | 0 |
| NC_019843.3;YP_009047204.1;MERS_spike;309 | 0 | 0 |
| NC_019843.3;YP_009047204.1;MERS_spike;310 | 0 | 0 |
| NC_019843.3;YP_009047204.1;MERS_spike;311 | 0 | 0 |
| NC_019843.3;YP_009047204.1;MERS_spike;312 | 0 | 0 |
| NC_019843.3;YP_009047204.1;MERS_spike;313 | 0 | 0 |
| NC_019843.3;YP_009047204.1;MERS_spike;314 | 0 | 0 |
| NC_019843.3;YP_009047204.1;MERS_spike;315 | 0 | 0 |
| NC_019843.3;YP_009047204.1;MERS_spike;316 | 0 | 0 |
| NC_019843.3;YP_009047204.1;MERS_spike;317 | 0 | 0 |
| NC_019843.3;YP_009047204.1;MERS_spike;318 | 0 | 0 |
| NC_019843.3;YP_009047204.1;MERS_spike;319 | 0 | 0 |
| NC_019843.3;YP_009047204.1;MERS_spike;320 | 0 | 0 |
| NC_019843.3;YP_009047204.1;MERS_spike;321 | 0 | 0 |
| NC_019843.3;YP_009047204.1;MERS_spike;322 | 0 | 0 |
| NC_019843.3;YP_009047204.1;MERS_spike;323 | 0 | 0 |
| NC_019843.3;YP_009047204.1;MERS_spike;324 | 0 | 0 |
| NC_019843.3;YP_009047204.1;MERS_spike;325 | 0 | 0 |
| NC_019843.3;YP_009047204.1;MERS_spike;326 | 0 | 0 |

| | | |
|---|---|---|
| NC_019843.3;YP_009047204.1;MERS_spike;327 | 0 | 0 |
| NC_019843.3;YP_009047204.1;MERS_spike;328 | 0 | 0 |
| NC_019843.3;YP_009047204.1;MERS_spike;329 | 0 | 0 |
| NC_019843.3;YP_009047204.1;MERS_spike;330 | 0 | 0 |
| NC_019843.3;YP_009047204.1;MERS_spike;331 | 0 | 0 |
| NC_019843.3;YP_009047204.1;MERS_spike;332 | 0 | 0 |
| NC_019843.3;YP_009047204.1;MERS_spike;333 | 0 | 0.025 |
| NC_019843.3;YP_009047204.1;MERS_spike;334 | 0 | 0.025 |
| NC_019843.3;YP_009047204.1;MERS_spike;335 | 0 | 0.025 |
| NC_019843.3;YP_009047204.1;MERS_spike;336 | 0.05 | 0 |
| NC_019843.3;YP_009047204.1;MERS_spike;337 | 0 | 0 |
| NC_019843.3;YP_009047204.1;MERS_spike;338 | 0 | 0 |
| NC_019843.3;YP_009047204.1;MERS_spike;339 | 0 | 0 |
| NC_019843.3;YP_009047204.1;MERS_spike;340 | 0.05 | 0.05 |
| NC_019843.3;YP_009047204.1;MERS_spike;341 | 0 | 0 |
| NC_019843.3;YP_009047204.1;MERS_spike;342 | 0 | 0 |
| NC_019843.3;YP_009047204.1;MERS_spike;343 | 0 | 0 |
| NC_019843.3;YP_009047204.1;MERS_spike;344 | 0 | 0 |
| NC_019843.3;YP_009047204.1;MERS_spike;345 | 0 | 0 |
| NC_019843.3;YP_009047204.1;MERS_spike;346 | 0 | 0 |
| NC_019843.3;YP_009047204.1;MERS_spike;347 | 0 | 0 |
| NC_019843.3;YP_009047204.1;MERS_spike;348 | 0 | 0 |
| NC_019843.3;YP_009047204.1;MERS_spike;349 | 0 | 0 |
| NC_019843.3;YP_009047204.1;MERS_spike;350 | 0 | 0 |
| NC_019843.3;YP_009047204.1;MERS_spike;351 | 0 | 0 |
| NC_019843.3;YP_009047204.1;MERS_spike;352 | 0 | 0 |
| NC_019843.3;YP_009047204.1;MERS_spike;353 | 0 | 0 |
| NC_019843.3;YP_009047204.1;MERS_spike;354 | 0 | 0.05 |
| NC_019843.3;YP_009047204.1;MERS_spike;355 | 0 | 0.025 |
| NC_019843.3;YP_009047204.1;MERS_spike;356 | 0 | 0.075 |
| NC_019843.3;YP_009047204.1;MERS_spike;357 | 0.05 | 0.075 |
| NC_019843.3;YP_009047204.1;MERS_spike;358 | 0.1 | 0.05 |
| NC_019843.3;YP_009047204.1;MERS_spike;359 | 0.05 | 0.05 |
| NC_019843.3;YP_009047204.1;MERS_spike;360 | 0 | 0.075 |
| NC_019843.3;YP_009047204.1;MERS_spike;361 | 0 | 0.025 |
| NC_019843.3;YP_009047204.1;MERS_spike;362 | 0 | 0.1 |
| NC_019843.3;YP_009047204.1;MERS_spike;363 | 0.05 | 0 |
| NC_019843.3;YP_009047204.1;MERS_spike;364 | 0 | 0.025 |
| NC_019843.3;YP_009047204.1;MERS_spike;365 | 0 | 0 |
| NC_019843.3;YP_009047204.1;MERS_spike;366 | 0 | 0 |
| NC_019843.3;YP_009047204.1;MERS_spike;367 | 0 | 0 |
| NC_019843.3;YP_009047204.1;MERS_spike;368 | 0 | 0 |
| NC_019843.3;YP_009047204.1;MERS_spike;369 | 0 | 0 |

| | | |
|---|---|---|
| NC_019843.3;YP_009047204.1;MERS_spike;370 | 0 | 0 |
| NC_019843.3;YP_009047204.1;MERS_spike;371 | 0 | 0 |
| NC_019843.3;YP_009047204.1;MERS_spike;372 | 0 | 0.025 |
| NC_019843.3;YP_009047204.1;MERS_spike;373 | 0 | 0 |
| NC_019843.3;YP_009047204.1;MERS_spike;374 | 0 | 0 |
| NC_019843.3;YP_009047204.1;MERS_spike;375 | 0 | 0 |
| NC_019843.3;YP_009047204.1;MERS_spike;376 | 0 | 0 |
| NC_019843.3;YP_009047204.1;MERS_spike;377 | 0 | 0 |
| NC_019843.3;YP_009047204.1;MERS_spike;378 | 0 | 0 |
| NC_019843.3;YP_009047204.1;MERS_spike;379 | 0 | 0 |
| NC_019843.3;YP_009047204.1;MERS_spike;380 | 0.1 | 0.05 |
| NC_019843.3;YP_009047204.1;MERS_spike;381 | 0 | 0 |
| NC_019843.3;YP_009047204.1;MERS_spike;382 | 0 | 0 |
| NC_019843.3;YP_009047204.1;MERS_spike;383 | 0 | 0 |
| NC_019843.3;YP_009047204.1;MERS_spike;384 | 0 | 0 |
| NC_019843.3;YP_009047204.1;MERS_spike;385 | 0 | 0 |
| NC_019843.3;YP_009047204.1;MERS_spike;386 | 0 | 0 |
| NC_019843.3;YP_009047204.1;MERS_spike;387 | 0 | 0 |
| NC_019843.3;YP_009047204.1;MERS_spike;388 | 0 | 0 |
| NC_019843.3;YP_009047204.1;MERS_spike;389 | 0 | 0 |
| NC_019843.3;YP_009047204.1;MERS_spike;390 | 0 | 0 |
| NC_019843.3;YP_009047204.1;MERS_spike;391 | 0 | 0 |
| NC_019843.3;YP_009047204.1;MERS_spike;392 | 0 | 0 |
| NC_019843.3;YP_009047204.1;MERS_spike;393 | 0 | 0 |
| NC_019843.3;YP_009047204.1;MERS_spike;394 | 0 | 0 |
| NC_019843.3;YP_009047204.1;MERS_spike;395 | 0 | 0 |
| NC_019843.3;YP_009047204.1;MERS_spike;396 | 0 | 0 |
| NC_019843.3;YP_009047204.1;MERS_spike;397 | 0 | 0 |
| NC_019843.3;YP_009047204.1;MERS_spike;398 | 0 | 0 |
| NC_019843.3;YP_009047204.1;MERS_spike;399 | 0 | 0 |
| NC_019843.3;YP_009047204.1;MERS_spike;400 | 0 | 0.125 |
| NC_019843.3;YP_009047204.1;MERS_spike;401 | 0 | 0 |
| NC_019843.3;YP_009047204.1;MERS_spike;402 | 0 | 0 |
| NC_019843.3;YP_009047204.1;MERS_spike;403 | 0 | 0 |
| NC_019843.3;YP_009047204.1;MERS_spike;404 | 0 | 0 |
| NC_019843.3;YP_009047204.1;MERS_spike;405 | 0 | 0 |
| NC_019843.3;YP_009047204.1;MERS_spike;406 | 0 | 0 |
| NC_019843.3;YP_009047204.1;MERS_spike;407 | 0 | 0 |
| NC_019843.3;YP_009047204.1;MERS_spike;408 | 0 | 0 |
| NC_019843.3;YP_009047204.1;MERS_spike;409 | 0 | 0 |
| NC_019843.3;YP_009047204.1;MERS_spike;410 | 0 | 0 |
| NC_019843.3;YP_009047204.1;MERS_spike;411 | 0 | 0 |
| NC_019843.3;YP_009047204.1;MERS_spike;412 | 0 | 0 |

| | | |
|---|---|---|
| NC_019843.3;YP_009047204.1;MERS_spike;413 | 0 | 0 |
| NC_019843.3;YP_009047204.1;MERS_spike;414 | 0 | 0 |
| NC_019843.3;YP_009047204.1;MERS_spike;415 | 0 | 0 |
| NC_019843.3;YP_009047204.1;MERS_spike;416 | 0 | 0 |
| NC_019843.3;YP_009047204.1;MERS_spike;417 | 0.1 | 0.05 |
| NC_019843.3;YP_009047204.1;MERS_spike;418 | 0 | 0.05 |
| NC_019843.3;YP_009047204.1;MERS_spike;419 | 0.15 | 0.125 |
| NC_019843.3;YP_009047204.1;MERS_spike;420 | 0 | 0 |
| NC_019843.3;YP_009047204.1;MERS_spike;421 | 0 | 0 |
| NC_019843.3;YP_009047204.1;MERS_spike;422 | 0 | 0 |
| NC_019843.3;YP_009047204.1;MERS_spike;423 | 0 | 0 |
| NC_019843.3;YP_009047204.1;MERS_spike;424 | 0 | 0 |
| NC_019843.3;YP_009047204.1;MERS_spike;425 | 0 | 0 |
| NC_019843.3;YP_009047204.1;MERS_spike;426 | 0 | 0.025 |
| NC_019843.3;YP_009047204.1;MERS_spike;427 | 0 | 0 |
| NC_019843.3;YP_009047204.1;MERS_spike;428 | 0 | 0 |
| NC_019843.3;YP_009047204.1;MERS_spike;429 | 0 | 0 |
| NC_019843.3;YP_009047204.1;MERS_spike;430 | 0 | 0 |
| NC_019843.3;YP_009047204.1;MERS_spike;431 | 0 | 0 |
| NC_019843.3;YP_009047204.1;MERS_spike;432 | 0 | 0 |
| NC_019843.3;YP_009047204.1;MERS_spike;433 | 0 | 0 |
| NC_019843.3;YP_009047204.1;MERS_spike;434 | 0 | 0 |
| NC_019843.3;YP_009047204.1;MERS_spike;435 | 0 | 0 |
| NC_019843.3;YP_009047204.1;MERS_spike;436 | 0 | 0 |
| NC_019843.3;YP_009047204.1;MERS_spike;437 | 0 | 0 |
| NC_019843.3;YP_009047204.1;MERS_spike;438 | 0 | 0 |
| NC_019843.3;YP_009047204.1;MERS_spike;439 | 0 | 0 |
| NC_019843.3;YP_009047204.1;MERS_spike;440 | 0 | 0 |
| NC_019843.3;YP_009047204.1;MERS_spike;441 | 0 | 0 |
| NC_019843.3;YP_009047204.1;MERS_spike;442 | 0 | 0 |
| NC_019843.3;YP_009047204.1;MERS_spike;443 | 0 | 0 |
| NC_019843.3;YP_009047204.1;MERS_spike;444 | 0 | 0 |
| NC_019843.3;YP_009047204.1;MERS_spike;445 | 0.05 | 0 |
| NC_019843.3;YP_009047204.1;MERS_spike;446 | 0.05 | 0 |
| NC_019843.3;YP_009047204.1;MERS_spike;447 | 0 | 0 |
| NC_019843.3;YP_009047204.1;MERS_spike;448 | 0 | 0 |
| NC_019843.3;YP_009047204.1;MERS_spike;449 | 0 | 0 |
| NC_019843.3;YP_009047204.1;MERS_spike;450 | 0 | 0 |
| NC_019843.3;YP_009047204.1;MERS_spike;451 | 0 | 0.025 |
| NC_019843.3;YP_009047204.1;MERS_spike;452 | 0 | 0 |
| NC_019843.3;YP_009047204.1;MERS_spike;453 | 0.1 | 0 |
| NC_019843.3;YP_009047204.1;MERS_spike;454 | 0 | 0 |
| NC_019843.3;YP_009047204.1;MERS_spike;455 | 0 | 0 |

| | | |
|---|---|---|
| NC_019843.3;YP_009047204.1;MERS_spike;456 | 0 | 0 |
| NC_019843.3;YP_009047204.1;MERS_spike;457 | 0 | 0.15 |
| NC_019843.3;YP_009047204.1;MERS_spike;458 | 0 | 0 |
| NC_019843.3;YP_009047204.1;MERS_spike;459 | 0 | 0 |
| NC_019843.3;YP_009047204.1;MERS_spike;460 | 0 | 0 |
| NC_019843.3;YP_009047204.1;MERS_spike;461 | 0 | 0 |
| NC_019843.3;YP_009047204.1;MERS_spike;462 | 0.35 | 0.1 |
| NC_019843.3;YP_009047204.1;MERS_spike;463 | 0 | 0 |
| NC_019843.3;YP_009047204.1;MERS_spike;464 | 0 | 0 |
| NC_019843.3;YP_009047204.1;MERS_spike;465 | 0 | 0 |
| NC_019843.3;YP_009047204.1;MERS_spike;466 | 0 | 0 |
| NC_019843.3;YP_009047204.1;MERS_spike;467 | 0 | 0 |
| NC_019843.3;YP_009047204.1;MERS_spike;468 | 0 | 0 |
| NC_019843.3;YP_009047204.1;MERS_spike;469 | 0 | 0 |
| NC_019843.3;YP_009047204.1;MERS_spike;470 | 0 | 0 |
| NC_019843.3;YP_009047204.1;MERS_spike;471 | 0 | 0 |
| NC_019843.3;YP_009047204.1;MERS_spike;472 | 0 | 0 |
| NC_019843.3;YP_009047204.1;MERS_spike;473 | 0 | 0 |
| NC_019843.3;YP_009047204.1;MERS_spike;474 | 0 | 0 |
| NC_019843.3;YP_009047204.1;MERS_spike;475 | 0 | 0 |
| NC_019843.3;YP_009047204.1;MERS_spike;476 | 0 | 0 |
| NC_019843.3;YP_009047204.1;MERS_spike;477 | 0 | 0 |
| NC_019843.3;YP_009047204.1;MERS_spike;478 | 0 | 0 |
| NC_019843.3;YP_009047204.1;MERS_spike;479 | 0.05 | 0 |
| NC_019843.3;YP_009047204.1;MERS_spike;480 | 0.05 | 0 |
| NC_019843.3;YP_009047204.1;MERS_spike;481 | 0.05 | 0 |
| NC_019843.3;YP_009047204.1;MERS_spike;482 | 0.05 | 0 |
| NC_019843.3;YP_009047204.1;MERS_spike;483 | 0.05 | 0 |
| NC_019843.3;YP_009047204.1;MERS_spike;484 | 0 | 0 |
| NC_019843.3;YP_009047204.1;MERS_spike;485 | 0 | 0 |
| NC_019843.3;YP_009047204.1;MERS_spike;486 | 0 | 0 |
| NC_019843.3;YP_009047204.1;MERS_spike;487 | 0 | 0 |
| NC_019843.3;YP_009047204.1;MERS_spike;488 | 0 | 0 |
| NC_019843.3;YP_009047204.1;MERS_spike;489 | 0 | 0 |
| NC_019843.3;YP_009047204.1;MERS_spike;490 | 0 | 0 |
| NC_019843.3;YP_009047204.1;MERS_spike;491 | 0 | 0 |
| NC_019843.3;YP_009047204.1;MERS_spike;492 | 0 | 0 |
| NC_019843.3;YP_009047204.1;MERS_spike;493 | 0 | 0 |
| NC_019843.3;YP_009047204.1;MERS_spike;494 | 0 | 0 |
| NC_019843.3;YP_009047204.1;MERS_spike;495 | 0 | 0.025 |
| NC_019843.3;YP_009047204.1;MERS_spike;496 | 0.05 | 0.025 |
| NC_019843.3;YP_009047204.1;MERS_spike;497 | 0 | 0 |
| NC_019843.3;YP_009047204.1;MERS_spike;498 | 0.05 | 0 |

| | | |
|---|---|---|
| NC_019843.3;YP_009047204.1;MERS_spike;499 | 0.05 | 0 |
| NC_019843.3;YP_009047204.1;MERS_spike;500 | 0.05 | 0.025 |
| NC_019843.3;YP_009047204.1;MERS_spike;501 | 0 | 0.025 |
| NC_019843.3;YP_009047204.1;MERS_spike;502 | 0 | 0.025 |
| NC_019843.3;YP_009047204.1;MERS_spike;503 | 0 | 0.025 |
| NC_019843.3;YP_009047204.1;MERS_spike;504 | 0 | 0.025 |
| NC_019843.3;YP_009047204.1;MERS_spike;505 | 0.05 | 0.025 |
| NC_019843.3;YP_009047204.1;MERS_spike;506 | 0 | 0.025 |
| NC_019843.3;YP_009047204.1;MERS_spike;507 | 0 | 0 |
| NC_019843.3;YP_009047204.1;MERS_spike;508 | 0 | 0 |
| NC_019843.3;YP_009047204.1;MERS_spike;509 | 0 | 0 |
| NC_019843.3;YP_009047204.1;MERS_spike;510 | 0 | 0 |
| NC_019843.3;YP_009047204.1;MERS_spike;511 | 0 | 0.025 |
| NC_019843.3;YP_009047204.1;MERS_spike;512 | 0 | 0 |
| NC_019843.3;YP_009047204.1;MERS_spike;513 | 0 | 0 |
| NC_019843.3;YP_009047204.1;MERS_spike;514 | 0 | 0 |
| NC_019843.3;YP_009047204.1;MERS_spike;515 | 0 | 0 |
| NC_019843.3;YP_009047204.1;MERS_spike;516 | 0 | 0 |
| NC_019843.3;YP_009047204.1;MERS_spike;517 | 0.25 | 0.075 |
| NC_019843.3;YP_009047204.1;MERS_spike;518 | 0 | 0 |
| NC_019843.3;YP_009047204.1;MERS_spike;519 | 0 | 0 |
| NC_019843.3;YP_009047204.1;MERS_spike;520 | 0 | 0 |
| NC_019843.3;YP_009047204.1;MERS_spike;521 | 0 | 0 |
| NC_019843.3;YP_009047204.1;MERS_spike;522 | 0 | 0.025 |
| NC_019843.3;YP_009047204.1;MERS_spike;523 | 0 | 0.025 |
| NC_019843.3;YP_009047204.1;MERS_spike;524 | 0 | 0.025 |
| NC_019843.3;YP_009047204.1;MERS_spike;525 | 0 | 0.025 |
| NC_019843.3;YP_009047204.1;MERS_spike;526 | 0 | 0 |
| NC_019843.3;YP_009047204.1;MERS_spike;527 | 0 | 0.025 |
| NC_019843.3;YP_009047204.1;MERS_spike;528 | 0 | 0.025 |
| NC_019843.3;YP_009047204.1;MERS_spike;529 | 0 | 0.025 |
| NC_019843.3;YP_009047204.1;MERS_spike;530 | 0 | 0.025 |
| NC_019843.3;YP_009047204.1;MERS_spike;531 | 0.05 | 0.05 |
| NC_019843.3;YP_009047204.1;MERS_spike;532 | 0 | 0.025 |
| NC_019843.3;YP_009047204.1;MERS_spike;533 | 0 | 0.025 |
| NC_019843.3;YP_009047204.1;MERS_spike;534 | 0 | 0 |
| NC_019843.3;YP_009047204.1;MERS_spike;535 | 0.1 | 0.025 |
| NC_019843.3;YP_009047204.1;MERS_spike;536 | 0.15 | 0 |
| NC_019843.3;YP_009047204.1;MERS_spike;537 | 0.35 | 0.05 |
| NC_019843.3;YP_009047204.1;MERS_spike;538 | 0.1 | 0.025 |
| NC_019843.3;YP_009047204.1;MERS_spike;539 | 0.05 | 0 |
| NC_019843.3;YP_009047204.1;MERS_spike;540 | 0 | 0 |
| NC_019843.3;YP_009047204.1;MERS_spike;541 | 0 | 0.025 |

| | | |
|---|---|---|
| NC_019843.3;YP_009047204.1;MERS_spike;542 | 0 | 0 |
| NC_019843.3;YP_009047204.1;MERS_spike;543 | 0.05 | 0 |
| NC_019843.3;YP_009047204.1;MERS_spike;544 | 0 | 0 |
| NC_019843.3;YP_009047204.1;MERS_spike;545 | 0 | 0 |
| NC_019843.3;YP_009047204.1;MERS_spike;546 | 0 | 0 |
| NC_019843.3;YP_009047204.1;MERS_spike;547 | 0 | 0.025 |
| NC_019843.3;YP_009047204.1;MERS_spike;548 | 0 | 0 |
| NC_019843.3;YP_009047204.1;MERS_spike;549 | 0.05 | 0 |
| NC_019843.3;YP_009047204.1;MERS_spike;550 | 0 | 0 |
| NC_019843.3;YP_009047204.1;MERS_spike;551 | 0 | 0 |
| NC_019843.3;YP_009047204.1;MERS_spike;552 | 0 | 0 |
| NC_019843.3;YP_009047204.1;MERS_spike;553 | 0 | 0 |
| NC_019843.3;YP_009047204.1;MERS_spike;554 | 0 | 0 |
| NC_019843.3;YP_009047204.1;MERS_spike;555 | 0 | 0 |
| NC_019843.3;YP_009047204.1;MERS_spike;556 | 0 | 0 |
| NC_019843.3;YP_009047204.1;MERS_spike;557 | 0 | 0 |
| NC_019843.3;YP_009047204.1;MERS_spike;558 | 0 | 0 |
| NC_019843.3;YP_009047204.1;MERS_spike;559 | 0 | 0 |
| NC_019843.3;YP_009047204.1;MERS_spike;560 | 0 | 0 |
| NC_019843.3;YP_009047204.1;MERS_spike;561 | 0 | 0 |
| NC_019843.3;YP_009047204.1;MERS_spike;562 | 0 | 0 |
| NC_019843.3;YP_009047204.1;MERS_spike;563 | 0 | 0 |
| NC_019843.3;YP_009047204.1;MERS_spike;564 | 0 | 0 |
| NC_019843.3;YP_009047204.1;MERS_spike;565 | 0 | 0 |
| NC_019843.3;YP_009047204.1;MERS_spike;566 | 0 | 0 |
| NC_019843.3;YP_009047204.1;MERS_spike;567 | 0 | 0.025 |
| NC_019843.3;YP_009047204.1;MERS_spike;568 | 0 | 0 |
| NC_019843.3;YP_009047204.1;MERS_spike;569 | 0 | 0 |
| NC_019843.3;YP_009047204.1;MERS_spike;570 | 0 | 0.025 |
| NC_019843.3;YP_009047204.1;MERS_spike;571 | 0.05 | 0 |
| NC_019843.3;YP_009047204.1;MERS_spike;572 | 0.05 | 0.05 |
| NC_019843.3;YP_009047204.1;MERS_spike;573 | 0.05 | 0 |
| NC_019843.3;YP_009047204.1;MERS_spike;574 | 0.05 | 0.05 |
| NC_019843.3;YP_009047204.1;MERS_spike;575 | 0.05 | 0.05 |
| NC_019843.3;YP_009047204.1;MERS_spike;576 | 0.05 | 0.05 |
| NC_019843.3;YP_009047204.1;MERS_spike;577 | 0.05 | 0.025 |
| NC_019843.3;YP_009047204.1;MERS_spike;578 | 0.1 | 0.025 |
| NC_019843.3;YP_009047204.1;MERS_spike;579 | 0.05 | 0.025 |
| NC_019843.3;YP_009047204.1;MERS_spike;580 | 0.2 | 0.1 |
| NC_019843.3;YP_009047204.1;MERS_spike;581 | 0.1 | 0.125 |
| NC_019843.3;YP_009047204.1;MERS_spike;582 | 0.1 | 0.125 |
| NC_019843.3;YP_009047204.1;MERS_spike;583 | 0.1 | 0.1 |
| NC_019843.3;YP_009047204.1;MERS_spike;584 | 0.1 | 0.1 |

| | | |
|---|---|---|
| NC_019843.3;YP_009047204.1;MERS_spike;585 | 0.1 | 0.1 |
| NC_019843.3;YP_009047204.1;MERS_spike;586 | 0.2 | 0.1 |
| NC_019843.3;YP_009047204.1;MERS_spike;587 | 0.3 | 0.1 |
| NC_019843.3;YP_009047204.1;MERS_spike;588 | 0.1 | 0.1 |
| NC_019843.3;YP_009047204.1;MERS_spike;589 | 0.15 | 0.075 |
| NC_019843.3;YP_009047204.1;MERS_spike;590 | 0.05 | 0.075 |
| NC_019843.3;YP_009047204.1;MERS_spike;591 | 0 | 0 |
| NC_019843.3;YP_009047204.1;MERS_spike;592 | 0.05 | 0 |
| NC_019843.3;YP_009047204.1;MERS_spike;593 | 0 | 0 |
| NC_019843.3;YP_009047204.1;MERS_spike;594 | 0 | 0 |
| NC_019843.3;YP_009047204.1;MERS_spike;595 | 0 | 0 |
| NC_019843.3;YP_009047204.1;MERS_spike;596 | 0 | 0 |
| NC_019843.3;YP_009047204.1;MERS_spike;597 | 0 | 0 |
| NC_019843.3;YP_009047204.1;MERS_spike;598 | 0 | 0 |
| NC_019843.3;YP_009047204.1;MERS_spike;599 | 0 | 0 |
| NC_019843.3;YP_009047204.1;MERS_spike;600 | 0 | 0 |
| NC_019843.3;YP_009047204.1;MERS_spike;601 | 0 | 0 |
| NC_019843.3;YP_009047204.1;MERS_spike;602 | 0 | 0 |
| NC_019843.3;YP_009047204.1;MERS_spike;603 | 0 | 0 |
| NC_019843.3;YP_009047204.1;MERS_spike;604 | 0 | 0 |
| NC_019843.3;YP_009047204.1;MERS_spike;605 | 0 | 0 |
| NC_019843.3;YP_009047204.1;MERS_spike;606 | 0 | 0 |
| NC_019843.3;YP_009047204.1;MERS_spike;607 | 0 | 0 |
| NC_019843.3;YP_009047204.1;MERS_spike;608 | 0 | 0 |
| NC_019843.3;YP_009047204.1;MERS_spike;609 | 0 | 0 |
| NC_019843.3;YP_009047204.1;MERS_spike;610 | 0 | 0.025 |
| NC_019843.3;YP_009047204.1;MERS_spike;611 | 0 | 0 |
| NC_019843.3;YP_009047204.1;MERS_spike;612 | 0 | 0 |
| NC_019843.3;YP_009047204.1;MERS_spike;613 | 0.05 | 0 |
| NC_019843.3;YP_009047204.1;MERS_spike;614 | 0 | 0 |
| NC_019843.3;YP_009047204.1;MERS_spike;615 | 0 | 0 |
| NC_019843.3;YP_009047204.1;MERS_spike;616 | 0 | 0 |
| NC_019843.3;YP_009047204.1;MERS_spike;617 | 0 | 0 |
| NC_019843.3;YP_009047204.1;MERS_spike;618 | 0 | 0 |
| NC_019843.3;YP_009047204.1;MERS_spike;619 | 0 | 0 |
| NC_019843.3;YP_009047204.1;MERS_spike;620 | 0 | 0 |
| NC_019843.3;YP_009047204.1;MERS_spike;621 | 0 | 0 |
| NC_019843.3;YP_009047204.1;MERS_spike;622 | 0 | 0.025 |
| NC_019843.3;YP_009047204.1;MERS_spike;623 | 0 | 0.025 |
| NC_019843.3;YP_009047204.1;MERS_spike;624 | 0 | 0.025 |
| NC_019843.3;YP_009047204.1;MERS_spike;625 | 0 | 0.025 |
| NC_019843.3;YP_009047204.1;MERS_spike;626 | 0 | 0.025 |
| NC_019843.3;YP_009047204.1;MERS_spike;627 | 0 | 0.025 |

| | | |
|---|---|---|
| NC_019843.3;YP_009047204.1;MERS_spike;628 | 0 | 0.025 |
| NC_019843.3;YP_009047204.1;MERS_spike;629 | 0 | 0.025 |
| NC_019843.3;YP_009047204.1;MERS_spike;630 | 0 | 0.025 |
| NC_019843.3;YP_009047204.1;MERS_spike;631 | 0 | 0.025 |
| NC_019843.3;YP_009047204.1;MERS_spike;632 | 0 | 0.025 |
| NC_019843.3;YP_009047204.1;MERS_spike;633 | 0 | 0.05 |
| NC_019843.3;YP_009047204.1;MERS_spike;634 | 0 | 0.05 |
| NC_019843.3;YP_009047204.1;MERS_spike;635 | 0 | 0 |
| NC_019843.3;YP_009047204.1;MERS_spike;636 | 0 | 0 |
| NC_019843.3;YP_009047204.1;MERS_spike;637 | 0 | 0 |
| NC_019843.3;YP_009047204.1;MERS_spike;638 | 0 | 0 |
| NC_019843.3;YP_009047204.1;MERS_spike;639 | 0 | 0 |
| NC_019843.3;YP_009047204.1;MERS_spike;640 | 0 | 0 |
| NC_019843.3;YP_009047204.1;MERS_spike;641 | 0 | 0 |
| NC_019843.3;YP_009047204.1;MERS_spike;642 | 0 | 0 |
| NC_019843.3;YP_009047204.1;MERS_spike;643 | 0 | 0 |
| NC_019843.3;YP_009047204.1;MERS_spike;644 | 0 | 0 |
| NC_019843.3;YP_009047204.1;MERS_spike;645 | 0 | 0 |
| NC_019843.3;YP_009047204.1;MERS_spike;646 | 0 | 0 |
| NC_019843.3;YP_009047204.1;MERS_spike;647 | 0 | 0 |
| NC_019843.3;YP_009047204.1;MERS_spike;648 | 0 | 0 |
| NC_019843.3;YP_009047204.1;MERS_spike;649 | 0 | 0 |
| NC_019843.3;YP_009047204.1;MERS_spike;650 | 0 | 0 |
| NC_019843.3;YP_009047204.1;MERS_spike;651 | 0 | 0 |
| NC_019843.3;YP_009047204.1;MERS_spike;652 | 0 | 0 |
| NC_019843.3;YP_009047204.1;MERS_spike;653 | 0 | 0.05 |
| NC_019843.3;YP_009047204.1;MERS_spike;654 | 0 | 0.1 |
| NC_019843.3;YP_009047204.1;MERS_spike;655 | 0 | 0.05 |
| NC_019843.3;YP_009047204.1;MERS_spike;656 | 0 | 0.1 |
| NC_019843.3;YP_009047204.1;MERS_spike;657 | 0 | 0.1 |
| NC_019843.3;YP_009047204.1;MERS_spike;658 | 0 | 0.075 |
| NC_019843.3;YP_009047204.1;MERS_spike;659 | 0 | 0 |
| NC_019843.3;YP_009047204.1;MERS_spike;660 | 0 | 0.075 |
| NC_019843.3;YP_009047204.1;MERS_spike;661 | 0 | 0.025 |
| NC_019843.3;YP_009047204.1;MERS_spike;662 | 0 | 0 |
| NC_019843.3;YP_009047204.1;MERS_spike;663 | 0 | 0 |
| NC_019843.3;YP_009047204.1;MERS_spike;664 | 0 | 0 |
| NC_019843.3;YP_009047204.1;MERS_spike;665 | 0 | 0 |
| NC_019843.3;YP_009047204.1;MERS_spike;666 | 0 | 0 |
| NC_019843.3;YP_009047204.1;MERS_spike;667 | 0 | 0 |
| NC_019843.3;YP_009047204.1;MERS_spike;668 | 0 | 0 |
| NC_019843.3;YP_009047204.1;MERS_spike;669 | 0 | 0 |
| NC_019843.3;YP_009047204.1;MERS_spike;670 | 0 | 0 |

| | | |
|---|---|---|
| NC_019843.3;YP_009047204.1;MERS_spike;671 | 0 | 0 |
| NC_019843.3;YP_009047204.1;MERS_spike;672 | 0 | 0 |
| NC_019843.3;YP_009047204.1;MERS_spike;673 | 0 | 0 |
| NC_019843.3;YP_009047204.1;MERS_spike;674 | 0 | 0.025 |
| NC_019843.3;YP_009047204.1;MERS_spike;675 | 0 | 0.025 |
| NC_019843.3;YP_009047204.1;MERS_spike;676 | 0 | 0.025 |
| NC_019843.3;YP_009047204.1;MERS_spike;677 | 0 | 0.025 |
| NC_019843.3;YP_009047204.1;MERS_spike;678 | 0 | 0.025 |
| NC_019843.3;YP_009047204.1;MERS_spike;679 | 0 | 0 |
| NC_019843.3;YP_009047204.1;MERS_spike;680 | 0 | 0 |
| NC_019843.3;YP_009047204.1;MERS_spike;681 | 0 | 0 |
| NC_019843.3;YP_009047204.1;MERS_spike;682 | 0 | 0 |
| NC_019843.3;YP_009047204.1;MERS_spike;683 | 0 | 0 |
| NC_019843.3;YP_009047204.1;MERS_spike;684 | 0 | 0 |
| NC_019843.3;YP_009047204.1;MERS_spike;685 | 0 | 0 |
| NC_019843.3;YP_009047204.1;MERS_spike;686 | 0 | 0 |
| NC_019843.3;YP_009047204.1;MERS_spike;687 | 0 | 0 |
| NC_019843.3;YP_009047204.1;MERS_spike;688 | 0.05 | 0 |
| NC_019843.3;YP_009047204.1;MERS_spike;689 | 0.05 | 0.05 |
| NC_019843.3;YP_009047204.1;MERS_spike;690 | 0 | 0.05 |
| NC_019843.3;YP_009047204.1;MERS_spike;691 | 0.05 | 0.025 |
| NC_019843.3;YP_009047204.1;MERS_spike;692 | 0 | 0.025 |
| NC_019843.3;YP_009047204.1;MERS_spike;693 | 0.1 | 0.025 |
| NC_019843.3;YP_009047204.1;MERS_spike;694 | 0.1 | 0.025 |
| NC_019843.3;YP_009047204.1;MERS_spike;695 | 0.3 | 0.025 |
| NC_019843.3;YP_009047204.1;MERS_spike;696 | 0.25 | 0.025 |
| NC_019843.3;YP_009047204.1;MERS_spike;697 | 0.25 | 0.025 |
| NC_019843.3;YP_009047204.1;MERS_spike;698 | 0.15 | 0.1 |
| NC_019843.3;YP_009047204.1;MERS_spike;699 | 0.05 | 0.025 |
| NC_019843.3;YP_009047204.1;MERS_spike;700 | 0 | 0.025 |
| NC_019843.3;YP_009047204.1;MERS_spike;701 | 0 | 0.025 |
| NC_019843.3;YP_009047204.1;MERS_spike;702 | 0 | 0 |
| NC_019843.3;YP_009047204.1;MERS_spike;703 | 0 | 0 |
| NC_019843.3;YP_009047204.1;MERS_spike;704 | 0 | 0 |
| NC_019843.3;YP_009047204.1;MERS_spike;705 | 0 | 0 |
| NC_019843.3;YP_009047204.1;MERS_spike;706 | 0 | 0 |
| NC_019843.3;YP_009047204.1;MERS_spike;707 | 0 | 0 |
| NC_019843.3;YP_009047204.1;MERS_spike;708 | 0 | 0 |
| NC_019843.3;YP_009047204.1;MERS_spike;709 | 0 | 0 |
| NC_019843.3;YP_009047204.1;MERS_spike;710 | 0 | 0 |
| NC_019843.3;YP_009047204.1;MERS_spike;711 | 0 | 0 |
| NC_019843.3;YP_009047204.1;MERS_spike;712 | 0 | 0 |
| NC_019843.3;YP_009047204.1;MERS_spike;713 | 0 | 0 |

| | | |
|---|---|---|
| NC_019843.3;YP_009047204.1;MERS_spike;714 | 0 | 0 |
| NC_019843.3;YP_009047204.1;MERS_spike;715 | 0 | 0 |
| NC_019843.3;YP_009047204.1;MERS_spike;716 | 0 | 0 |
| NC_019843.3;YP_009047204.1;MERS_spike;717 | 0.2 | 0.125 |
| NC_019843.3;YP_009047204.1;MERS_spike;718 | 0 | 0 |
| NC_019843.3;YP_009047204.1;MERS_spike;719 | 0 | 0 |
| NC_019843.3;YP_009047204.1;MERS_spike;720 | 0.05 | 0.05 |
| NC_019843.3;YP_009047204.1;MERS_spike;721 | 0.05 | 0.025 |
| NC_019843.3;YP_009047204.1;MERS_spike;722 | 0 | 0.025 |
| NC_019843.3;YP_009047204.1;MERS_spike;723 | 0 | 0.025 |
| NC_019843.3;YP_009047204.1;MERS_spike;724 | 0.05 | 0.025 |
| NC_019843.3;YP_009047204.1;MERS_spike;725 | 0 | 0.025 |
| NC_019843.3;YP_009047204.1;MERS_spike;726 | 0 | 0.05 |
| NC_019843.3;YP_009047204.1;MERS_spike;727 | 0.05 | 0.05 |
| NC_019843.3;YP_009047204.1;MERS_spike;728 | 0.15 | 0.1 |
| NC_019843.3;YP_009047204.1;MERS_spike;729 | 0.05 | 0 |
| NC_019843.3;YP_009047204.1;MERS_spike;730 | 0 | 0 |
| NC_019843.3;YP_009047204.1;MERS_spike;731 | 0 | 0.05 |
| NC_019843.3;YP_009047204.1;MERS_spike;732 | 0 | 0.025 |
| NC_019843.3;YP_009047204.1;MERS_spike;733 | 0 | 0.075 |
| NC_019843.3;YP_009047204.1;MERS_spike;734 | 0 | 0.05 |
| NC_019843.3;YP_009047204.1;MERS_spike;735 | 0.05 | 0.05 |
| NC_019843.3;YP_009047204.1;MERS_spike;736 | 0.05 | 0.075 |
| NC_019843.3;YP_009047204.1;MERS_spike;737 | 0.05 | 0.075 |
| NC_019843.3;YP_009047204.1;MERS_spike;738 | 0 | 0 |
| NC_019843.3;YP_009047204.1;MERS_spike;739 | 0.1 | 0 |
| NC_019843.3;YP_009047204.1;MERS_spike;740 | 0.05 | 0 |
| NC_019843.3;YP_009047204.1;MERS_spike;741 | 0.05 | 0.025 |
| NC_019843.3;YP_009047204.1;MERS_spike;742 | 0.05 | 0.075 |
| NC_019843.3;YP_009047204.1;MERS_spike;743 | 0.05 | 0 |
| NC_019843.3;YP_009047204.1;MERS_spike;744 | 0.05 | 0.025 |
| NC_019843.3;YP_009047204.1;MERS_spike;745 | 0.05 | 0 |
| NC_019843.3;YP_009047204.1;MERS_spike;746 | 0 | 0.025 |
| NC_019843.3;YP_009047204.1;MERS_spike;747 | 0 | 0 |
| NC_019843.3;YP_009047204.1;MERS_spike;748 | 0 | 0.05 |
| NC_019843.3;YP_009047204.1;MERS_spike;749 | 0 | 0 |
| NC_019843.3;YP_009047204.1;MERS_spike;750 | 0 | 0 |
| NC_019843.3;YP_009047204.1;MERS_spike;751 | 0 | 0 |
| NC_019843.3;YP_009047204.1;MERS_spike;752 | 0 | 0 |
| NC_019843.3;YP_009047204.1;MERS_spike;753 | 0 | 0 |
| NC_019843.3;YP_009047204.1;MERS_spike;754 | 0 | 0 |
| NC_019843.3;YP_009047204.1;MERS_spike;755 | 0 | 0 |
| NC_019843.3;YP_009047204.1;MERS_spike;756 | 0 | 0 |

| | | |
|---|---|---|
| NC_019843.3;YP_009047204.1;MERS_spike;757 | 0 | 0 |
| NC_019843.3;YP_009047204.1;MERS_spike;758 | 0 | 0 |
| NC_019843.3;YP_009047204.1;MERS_spike;759 | 0.05 | 0 |
| NC_019843.3;YP_009047204.1;MERS_spike;760 | 0.05 | 0 |
| NC_019843.3;YP_009047204.1;MERS_spike;761 | 0.1 | 0.05 |
| NC_019843.3;YP_009047204.1;MERS_spike;762 | 0.1 | 0.025 |
| NC_019843.3;YP_009047204.1;MERS_spike;763 | 0.1 | 0.025 |
| NC_019843.3;YP_009047204.1;MERS_spike;764 | 0 | 0.05 |
| NC_019843.3;YP_009047204.1;MERS_spike;765 | 0 | 0.025 |
| NC_019843.3;YP_009047204.1;MERS_spike;766 | 0 | 0.025 |
| NC_019843.3;YP_009047204.1;MERS_spike;767 | 0 | 0 |
| NC_019843.3;YP_009047204.1;MERS_spike;768 | 0 | 0.025 |
| NC_019843.3;YP_009047204.1;MERS_spike;769 | 0 | 0.05 |
| NC_019843.3;YP_009047204.1;MERS_spike;770 | 0.05 | 0.05 |
| NC_019843.3;YP_009047204.1;MERS_spike;771 | 0 | 0.025 |
| NC_019843.3;YP_009047204.1;MERS_spike;772 | 0 | 0 |
| NC_019843.3;YP_009047204.1;MERS_spike;773 | 0 | 0 |
| NC_019843.3;YP_009047204.1;MERS_spike;774 | 0 | 0 |
| NC_019843.3;YP_009047204.1;MERS_spike;775 | 0 | 0 |
| NC_019843.3;YP_009047204.1;MERS_spike;776 | 0 | 0 |
| NC_019843.3;YP_009047204.1;MERS_spike;777 | 0 | 0 |
| NC_019843.3;YP_009047204.1;MERS_spike;778 | 0 | 0 |
| NC_019843.3;YP_009047204.1;MERS_spike;779 | 0 | 0 |
| NC_019843.3;YP_009047204.1;MERS_spike;780 | 0 | 0 |
| NC_019843.3;YP_009047204.1;MERS_spike;781 | 0 | 0 |
| NC_019843.3;YP_009047204.1;MERS_spike;782 | 0 | 0 |
| NC_019843.3;YP_009047204.1;MERS_spike;783 | 0 | 0 |
| NC_019843.3;YP_009047204.1;MERS_spike;784 | 0 | 0 |
| NC_019843.3;YP_009047204.1;MERS_spike;785 | 0 | 0 |
| NC_019843.3;YP_009047204.1;MERS_spike;786 | 0 | 0.075 |
| NC_019843.3;YP_009047204.1;MERS_spike;787 | 0.05 | 0.05 |
| NC_019843.3;YP_009047204.1;MERS_spike;788 | 0 | 0.025 |
| NC_019843.3;YP_009047204.1;MERS_spike;789 | 0 | 0 |
| NC_019843.3;YP_009047204.1;MERS_spike;790 | 0 | 0.025 |
| NC_019843.3;YP_009047204.1;MERS_spike;791 | 0 | 0.025 |
| NC_019843.3;YP_009047204.1;MERS_spike;792 | 0 | 0.05 |
| NC_019843.3;YP_009047204.1;MERS_spike;793 | 0 | 0.025 |
| NC_019843.3;YP_009047204.1;MERS_spike;794 | 0 | 0 |
| NC_019843.3;YP_009047204.1;MERS_spike;795 | 0 | 0.025 |
| NC_019843.3;YP_009047204.1;MERS_spike;796 | 0 | 0 |
| NC_019843.3;YP_009047204.1;MERS_spike;797 | 0 | 0 |
| NC_019843.3;YP_009047204.1;MERS_spike;798 | 0 | 0 |
| NC_019843.3;YP_009047204.1;MERS_spike;799 | 0 | 0.025 |

| | | |
|---|---|---|
| NC_019843.3;YP_009047204.1;MERS_spike;800 | 0 | 0.025 |
| NC_019843.3;YP_009047204.1;MERS_spike;801 | 0 | 0 |
| NC_019843.3;YP_009047204.1;MERS_spike;802 | 0 | 0 |
| NC_019843.3;YP_009047204.1;MERS_spike;803 | 0 | 0.025 |
| NC_019843.3;YP_009047204.1;MERS_spike;804 | 0 | 0 |
| NC_019843.3;YP_009047204.1;MERS_spike;805 | 0 | 0 |
| NC_019843.3;YP_009047204.1;MERS_spike;806 | 0 | 0 |
| NC_019843.3;YP_009047204.1;MERS_spike;807 | 0 | 0 |
| NC_019843.3;YP_009047204.1;MERS_spike;808 | 0 | 0 |
| NC_019843.3;YP_009047204.1;MERS_spike;809 | 0 | 0 |
| NC_019843.3;YP_009047204.1;MERS_spike;810 | 0 | 0 |
| NC_019843.3;YP_009047204.1;MERS_spike;811 | 0 | 0 |
| NC_019843.3;YP_009047204.1;MERS_spike;812 | 0 | 0 |
| NC_019843.3;YP_009047204.1;MERS_spike;813 | 0 | 0.025 |
| NC_019843.3;YP_009047204.1;MERS_spike;814 | 0 | 0.025 |
| NC_019843.3;YP_009047204.1;MERS_spike;815 | 0 | 0 |
| NC_019843.3;YP_009047204.1;MERS_spike;816 | 0 | 0 |
| NC_019843.3;YP_009047204.1;MERS_spike;817 | 0 | 0 |
| NC_019843.3;YP_009047204.1;MERS_spike;818 | 0 | 0 |
| NC_019843.3;YP_009047204.1;MERS_spike;819 | 0 | 0 |
| NC_019843.3;YP_009047204.1;MERS_spike;820 | 0 | 0 |
| NC_019843.3;YP_009047204.1;MERS_spike;821 | 0 | 0 |
| NC_019843.3;YP_009047204.1;MERS_spike;822 | 0 | 0 |
| NC_019843.3;YP_009047204.1;MERS_spike;823 | 0 | 0 |
| NC_019843.3;YP_009047204.1;MERS_spike;824 | 0 | 0 |
| NC_019843.3;YP_009047204.1;MERS_spike;825 | 0 | 0 |
| NC_019843.3;YP_009047204.1;MERS_spike;826 | 0 | 0 |
| NC_019843.3;YP_009047204.1;MERS_spike;827 | 0 | 0 |
| NC_019843.3;YP_009047204.1;MERS_spike;828 | 0 | 0 |
| NC_019843.3;YP_009047204.1;MERS_spike;829 | 0 | 0 |
| NC_019843.3;YP_009047204.1;MERS_spike;830 | 0 | 0 |
| NC_019843.3;YP_009047204.1;MERS_spike;831 | 0.05 | 0 |
| NC_019843.3;YP_009047204.1;MERS_spike;832 | 0 | 0 |
| NC_019843.3;YP_009047204.1;MERS_spike;833 | 0 | 0.025 |
| NC_019843.3;YP_009047204.1;MERS_spike;834 | 0 | 0 |
| NC_019843.3;YP_009047204.1;MERS_spike;835 | 0 | 0 |
| NC_019843.3;YP_009047204.1;MERS_spike;836 | 0 | 0 |
| NC_019843.3;YP_009047204.1;MERS_spike;837 | 0 | 0.025 |
| NC_019843.3;YP_009047204.1;MERS_spike;838 | 0.05 | 0.05 |
| NC_019843.3;YP_009047204.1;MERS_spike;839 | 0 | 0.025 |
| NC_019843.3;YP_009047204.1;MERS_spike;840 | 0 | 0 |
| NC_019843.3;YP_009047204.1;MERS_spike;841 | 0 | 0 |
| NC_019843.3;YP_009047204.1;MERS_spike;842 | 0 | 0 |

| | | |
|---|---|---|
| NC_019843.3;YP_009047204.1;MERS_spike;843 | 0 | 0 |
| NC_019843.3;YP_009047204.1;MERS_spike;844 | 0 | 0 |
| NC_019843.3;YP_009047204.1;MERS_spike;845 | 0 | 0 |
| NC_019843.3;YP_009047204.1;MERS_spike;846 | 0 | 0 |
| NC_019843.3;YP_009047204.1;MERS_spike;847 | 0 | 0 |
| NC_019843.3;YP_009047204.1;MERS_spike;848 | 0 | 0 |
| NC_019843.3;YP_009047204.1;MERS_spike;849 | 0 | 0.025 |
| NC_019843.3;YP_009047204.1;MERS_spike;850 | 0 | 0.025 |
| NC_019843.3;YP_009047204.1;MERS_spike;851 | 0 | 0.075 |
| NC_019843.3;YP_009047204.1;MERS_spike;852 | 0.05 | 0.05 |
| NC_019843.3;YP_009047204.1;MERS_spike;853 | 0 | 0.05 |
| NC_019843.3;YP_009047204.1;MERS_spike;854 | 0 | 0 |
| NC_019843.3;YP_009047204.1;MERS_spike;855 | 0 | 0.05 |
| NC_019843.3;YP_009047204.1;MERS_spike;856 | 0 | 0.025 |
| NC_019843.3;YP_009047204.1;MERS_spike;857 | 0 | 0.025 |
| NC_019843.3;YP_009047204.1;MERS_spike;858 | 0 | 0 |
| NC_019843.3;YP_009047204.1;MERS_spike;859 | 0 | 0.025 |
| NC_019843.3;YP_009047204.1;MERS_spike;860 | 0 | 0 |
| NC_019843.3;YP_009047204.1;MERS_spike;861 | 0 | 0 |
| NC_019843.3;YP_009047204.1;MERS_spike;862 | 0 | 0 |
| NC_019843.3;YP_009047204.1;MERS_spike;863 | 0.05 | 0 |
| NC_019843.3;YP_009047204.1;MERS_spike;864 | 0 | 0 |
| NC_019843.3;YP_009047204.1;MERS_spike;865 | 0 | 0 |
| NC_019843.3;YP_009047204.1;MERS_spike;866 | 0 | 0.025 |
| NC_019843.3;YP_009047204.1;MERS_spike;867 | 0 | 0 |
| NC_019843.3;YP_009047204.1;MERS_spike;868 | 0 | 0 |
| NC_019843.3;YP_009047204.1;MERS_spike;869 | 0 | 0 |
| NC_019843.3;YP_009047204.1;MERS_spike;870 | 0 | 0 |
| NC_019843.3;YP_009047204.1;MERS_spike;871 | 0 | 0 |
| NC_019843.3;YP_009047204.1;MERS_spike;872 | 0 | 0 |
| NC_019843.3;YP_009047204.1;MERS_spike;873 | 0 | 0 |
| NC_019843.3;YP_009047204.1;MERS_spike;874 | 0 | 0 |
| NC_019843.3;YP_009047204.1;MERS_spike;875 | 0.05 | 0 |
| NC_019843.3;YP_009047204.1;MERS_spike;876 | 0 | 0 |
| NC_019843.3;YP_009047204.1;MERS_spike;877 | 0 | 0.05 |
| NC_019843.3;YP_009047204.1;MERS_spike;878 | 0.05 | 0.05 |
| NC_019843.3;YP_009047204.1;MERS_spike;879 | 0.05 | 0.025 |
| NC_019843.3;YP_009047204.1;MERS_spike;880 | 0.2 | 0.575 |
| NC_019843.3;YP_009047204.1;MERS_spike;881 | 0.2 | 0.5 |
| NC_019843.3;YP_009047204.1;MERS_spike;882 | 0.35 | 0.775 |
| NC_019843.3;YP_009047204.1;MERS_spike;883 | 0.2 | 0.75 |
| NC_019843.3;YP_009047204.1;MERS_spike;884 | 0.15 | 0.75 |
| NC_019843.3;YP_009047204.1;MERS_spike;885 | 0 | 0.65 |

| | | |
|---|---|---|
| NC_019843.3;YP_009047204.1;MERS_spike;886 | 0 | 0.65 |
| NC_019843.3;YP_009047204.1;MERS_spike;887 | 0 | 0.575 |
| NC_019843.3;YP_009047204.1;MERS_spike;888 | 0.1 | 0.325 |
| NC_019843.3;YP_009047204.1;MERS_spike;889 | 0.35 | 0.5 |
| NC_019843.3;YP_009047204.1;MERS_spike;890 | 0.1 | 0.275 |
| NC_019843.3;YP_009047204.1;MERS_spike;891 | 0 | 0.175 |
| NC_019843.3;YP_009047204.1;MERS_spike;892 | 0 | 0.025 |
| NC_019843.3;YP_009047204.1;MERS_spike;893 | 0 | 0 |
| NC_019843.3;YP_009047204.1;MERS_spike;894 | 0 | 0 |
| NC_019843.3;YP_009047204.1;MERS_spike;895 | 0 | 0 |
| NC_019843.3;YP_009047204.1;MERS_spike;896 | 0 | 0 |
| NC_019843.3;YP_009047204.1;MERS_spike;897 | 0 | 0 |
| NC_019843.3;YP_009047204.1;MERS_spike;898 | 0 | 0 |
| NC_019843.3;YP_009047204.1;MERS_spike;899 | 0 | 0 |
| NC_019843.3;YP_009047204.1;MERS_spike;900 | 0 | 0 |
| NC_019843.3;YP_009047204.1;MERS_spike;901 | 0 | 0 |
| NC_019843.3;YP_009047204.1;MERS_spike;902 | 0.05 | 0.075 |
| NC_019843.3;YP_009047204.1;MERS_spike;903 | 0 | 0.025 |
| NC_019843.3;YP_009047204.1;MERS_spike;904 | 0 | 0 |
| NC_019843.3;YP_009047204.1;MERS_spike;905 | 0 | 0 |
| NC_019843.3;YP_009047204.1;MERS_spike;906 | 0.05 | 0 |
| NC_019843.3;YP_009047204.1;MERS_spike;907 | 0.05 | 0 |
| NC_019843.3;YP_009047204.1;MERS_spike;908 | 0.1 | 0.075 |
| NC_019843.3;YP_009047204.1;MERS_spike;909 | 0.1 | 0 |
| NC_019843.3;YP_009047204.1;MERS_spike;910 | 0.05 | 0 |
| NC_019843.3;YP_009047204.1;MERS_spike;911 | 0.05 | 0 |
| NC_019843.3;YP_009047204.1;MERS_spike;912 | 0.05 | 0 |
| NC_019843.3;YP_009047204.1;MERS_spike;913 | 0.05 | 0 |
| NC_019843.3;YP_009047204.1;MERS_spike;914 | 0.05 | 0 |
| NC_019843.3;YP_009047204.1;MERS_spike;915 | 0.1 | 0.025 |
| NC_019843.3;YP_009047204.1;MERS_spike;916 | 0.1 | 0.075 |
| NC_019843.3;YP_009047204.1;MERS_spike;917 | 0.05 | 0 |
| NC_019843.3;YP_009047204.1;MERS_spike;918 | 0 | 0 |
| NC_019843.3;YP_009047204.1;MERS_spike;919 | 0 | 0 |
| NC_019843.3;YP_009047204.1;MERS_spike;920 | 0 | 0 |
| NC_019843.3;YP_009047204.1;MERS_spike;921 | 0 | 0 |
| NC_019843.3;YP_009047204.1;MERS_spike;922 | 0 | 0 |
| NC_019843.3;YP_009047204.1;MERS_spike;923 | 0 | 0 |
| NC_019843.3;YP_009047204.1;MERS_spike;924 | 0 | 0 |
| NC_019843.3;YP_009047204.1;MERS_spike;925 | 0 | 0 |
| NC_019843.3;YP_009047204.1;MERS_spike;926 | 0 | 0 |
| NC_019843.3;YP_009047204.1;MERS_spike;927 | 0 | 0.025 |
| NC_019843.3;YP_009047204.1;MERS_spike;928 | 0 | 0.025 |

| | | |
|---|---|---|
| NC_019843.3;YP_009047204.1;MERS_spike;929 | 0 | 0.025 |
| NC_019843.3;YP_009047204.1;MERS_spike;930 | 0 | 0.025 |
| NC_019843.3;YP_009047204.1;MERS_spike;931 | 0.05 | 0.05 |
| NC_019843.3;YP_009047204.1;MERS_spike;932 | 0 | 0.025 |
| NC_019843.3;YP_009047204.1;MERS_spike;933 | 0 | 0 |
| NC_019843.3;YP_009047204.1;MERS_spike;934 | 0 | 0.025 |
| NC_019843.3;YP_009047204.1;MERS_spike;935 | 0 | 0 |
| NC_019843.3;YP_009047204.1;MERS_spike;936 | 0 | 0 |
| NC_019843.3;YP_009047204.1;MERS_spike;937 | 0 | 0 |
| NC_019843.3;YP_009047204.1;MERS_spike;938 | 0 | 0 |
| NC_019843.3;YP_009047204.1;MERS_spike;939 | 0 | 0 |
| NC_019843.3;YP_009047204.1;MERS_spike;940 | 0 | 0 |
| NC_019843.3;YP_009047204.1;MERS_spike;941 | 0 | 0 |
| NC_019843.3;YP_009047204.1;MERS_spike;942 | 0 | 0 |
| NC_019843.3;YP_009047204.1;MERS_spike;943 | 0 | 0 |
| NC_019843.3;YP_009047204.1;MERS_spike;944 | 0 | 0 |
| NC_019843.3;YP_009047204.1;MERS_spike;945 | 0 | 0 |
| NC_019843.3;YP_009047204.1;MERS_spike;946 | 0 | 0 |
| NC_019843.3;YP_009047204.1;MERS_spike;947 | 0 | 0 |
| NC_019843.3;YP_009047204.1;MERS_spike;948 | 0 | 0 |
| NC_019843.3;YP_009047204.1;MERS_spike;949 | 0 | 0 |
| NC_019843.3;YP_009047204.1;MERS_spike;950 | 0 | 0 |
| NC_019843.3;YP_009047204.1;MERS_spike;951 | 0 | 0 |
| NC_019843.3;YP_009047204.1;MERS_spike;952 | 0 | 0 |
| NC_019843.3;YP_009047204.1;MERS_spike;953 | 0 | 0 |
| NC_019843.3;YP_009047204.1;MERS_spike;954 | 0.05 | 0.025 |
| NC_019843.3;YP_009047204.1;MERS_spike;955 | 0 | 0 |
| NC_019843.3;YP_009047204.1;MERS_spike;956 | 0 | 0 |
| NC_019843.3;YP_009047204.1;MERS_spike;957 | 0 | 0 |
| NC_019843.3;YP_009047204.1;MERS_spike;958 | 0 | 0 |
| NC_019843.3;YP_009047204.1;MERS_spike;959 | 0 | 0 |
| NC_019843.3;YP_009047204.1;MERS_spike;960 | 0 | 0 |
| NC_019843.3;YP_009047204.1;MERS_spike;961 | 0 | 0 |
| NC_019843.3;YP_009047204.1;MERS_spike;962 | 0 | 0 |
| NC_019843.3;YP_009047204.1;MERS_spike;963 | 0 | 0 |
| NC_019843.3;YP_009047204.1;MERS_spike;964 | 0 | 0 |
| NC_019843.3;YP_009047204.1;MERS_spike;965 | 0 | 0 |
| NC_019843.3;YP_009047204.1;MERS_spike;966 | 0 | 0 |
| NC_019843.3;YP_009047204.1;MERS_spike;967 | 0 | 0 |
| NC_019843.3;YP_009047204.1;MERS_spike;968 | 0 | 0 |
| NC_019843.3;YP_009047204.1;MERS_spike;969 | 0 | 0 |
| NC_019843.3;YP_009047204.1;MERS_spike;970 | 0 | 0 |
| NC_019843.3;YP_009047204.1;MERS_spike;971 | 0 | 0 |

| | | |
|---|---|---|
| NC_019843.3;YP_009047204.1;MERS_spike;972 | 0 | 0 |
| NC_019843.3;YP_009047204.1;MERS_spike;973 | 0 | 0 |
| NC_019843.3;YP_009047204.1;MERS_spike;974 | 0 | 0 |
| NC_019843.3;YP_009047204.1;MERS_spike;975 | 0 | 0 |
| NC_019843.3;YP_009047204.1;MERS_spike;976 | 0 | 0 |
| NC_019843.3;YP_009047204.1;MERS_spike;977 | 0 | 0 |
| NC_019843.3;YP_009047204.1;MERS_spike;978 | 0 | 0 |
| NC_019843.3;YP_009047204.1;MERS_spike;979 | 0 | 0 |
| NC_019843.3;YP_009047204.1;MERS_spike;980 | 0 | 0 |
| NC_019843.3;YP_009047204.1;MERS_spike;981 | 0 | 0 |
| NC_019843.3;YP_009047204.1;MERS_spike;982 | 0 | 0 |
| NC_019843.3;YP_009047204.1;MERS_spike;983 | 0 | 0.025 |
| NC_019843.3;YP_009047204.1;MERS_spike;984 | 0.1 | 0 |
| NC_019843.3;YP_009047204.1;MERS_spike;985 | 0 | 0.025 |
| NC_019843.3;YP_009047204.1;MERS_spike;986 | 0 | 0.025 |
| NC_019843.3;YP_009047204.1;MERS_spike;987 | 0 | 0.025 |
| NC_019843.3;YP_009047204.1;MERS_spike;988 | 0 | 0.05 |
| NC_019843.3;YP_009047204.1;MERS_spike;989 | 0 | 0.05 |
| NC_019843.3;YP_009047204.1;MERS_spike;990 | 0 | 0.025 |
| NC_019843.3;YP_009047204.1;MERS_spike;991 | 0 | 0.075 |
| NC_019843.3;YP_009047204.1;MERS_spike;992 | 0 | 0.05 |
| NC_019843.3;YP_009047204.1;MERS_spike;993 | 0 | 0.05 |
| NC_019843.3;YP_009047204.1;MERS_spike;994 | 0.05 | 0.025 |
| NC_019843.3;YP_009047204.1;MERS_spike;995 | 0 | 0 |
| NC_019843.3;YP_009047204.1;MERS_spike;996 | 0.05 | 0.05 |
| NC_019843.3;YP_009047204.1;MERS_spike;997 | 0.15 | 0.1 |
| NC_019843.3;YP_009047204.1;MERS_spike;998 | 0.2 | 0.15 |
| NC_019843.3;YP_009047204.1;MERS_spike;999 | 0.05 | 0.1 |
| NC_019843.3;YP_009047204.1;MERS_spike;1000 | 0.05 | 0.1 |
| NC_019843.3;YP_009047204.1;MERS_spike;1001 | 0.05 | 0.15 |
| NC_019843.3;YP_009047204.1;MERS_spike;1002 | 0.15 | 0.15 |
| NC_019843.3;YP_009047204.1;MERS_spike;1003 | 0.15 | 0.15 |
| NC_019843.3;YP_009047204.1;MERS_spike;1004 | 0.15 | 0.15 |
| NC_019843.3;YP_009047204.1;MERS_spike;1005 | 0.1 | 0.05 |
| NC_019843.3;YP_009047204.1;MERS_spike;1006 | 0.05 | 0.1 |
| NC_019843.3;YP_009047204.1;MERS_spike;1007 | 0.05 | 0.05 |
| NC_019843.3;YP_009047204.1;MERS_spike;1008 | 0.05 | 0.075 |
| NC_019843.3;YP_009047204.1;MERS_spike;1009 | 0.1 | 0.15 |
| NC_019843.3;YP_009047204.1;MERS_spike;1010 | 0.15 | 0.1 |
| NC_019843.3;YP_009047204.1;MERS_spike;1011 | 0.1 | 0.1 |
| NC_019843.3;YP_009047204.1;MERS_spike;1012 | 0.05 | 0.1 |
| NC_019843.3;YP_009047204.1;MERS_spike;1013 | 0 | 0 |
| NC_019843.3;YP_009047204.1;MERS_spike;1014 | 0.05 | 0.025 |

| | | |
|---|---|---|
| NC_019843.3;YP_009047204.1;MERS_spike;1015 | 0.1 | 0.05 |
| NC_019843.3;YP_009047204.1;MERS_spike;1016 | 0.05 | 0.075 |
| NC_019843.3;YP_009047204.1;MERS_spike;1017 | 0.05 | 0.05 |
| NC_019843.3;YP_009047204.1;MERS_spike;1018 | 0.1 | 0.1 |
| NC_019843.3;YP_009047204.1;MERS_spike;1019 | 0.15 | 0.175 |
| NC_019843.3;YP_009047204.1;MERS_spike;1020 | 0.1 | 0.05 |
| NC_019843.3;YP_009047204.1;MERS_spike;1021 | 0.1 | 0.025 |
| NC_019843.3;YP_009047204.1;MERS_spike;1022 | 0.1 | 0.125 |
| NC_019843.3;YP_009047204.1;MERS_spike;1023 | 0.05 | 0.025 |
| NC_019843.3;YP_009047204.1;MERS_spike;1024 | 0.05 | 0.025 |
| NC_019843.3;YP_009047204.1;MERS_spike;1025 | 0.2 | 0.05 |
| NC_019843.3;YP_009047204.1;MERS_spike;1026 | 0.1 | 0 |
| NC_019843.3;YP_009047204.1;MERS_spike;1027 | 0.1 | 0 |
| NC_019843.3;YP_009047204.1;MERS_spike;1028 | 0.1 | 0 |
| NC_019843.3;YP_009047204.1;MERS_spike;1029 | 0.05 | 0.05 |
| NC_019843.3;YP_009047204.1;MERS_spike;1030 | 0.05 | 0.05 |
| NC_019843.3;YP_009047204.1;MERS_spike;1031 | 0.05 | 0.05 |
| NC_019843.3;YP_009047204.1;MERS_spike;1032 | 0 | 0 |
| NC_019843.3;YP_009047204.1;MERS_spike;1033 | 0 | 0.025 |
| NC_019843.3;YP_009047204.1;MERS_spike;1034 | 0 | 0.05 |
| NC_019843.3;YP_009047204.1;MERS_spike;1035 | 0.05 | 0.025 |
| NC_019843.3;YP_009047204.1;MERS_spike;1036 | 0 | 0 |
| NC_019843.3;YP_009047204.1;MERS_spike;1037 | 0 | 0 |
| NC_019843.3;YP_009047204.1;MERS_spike;1038 | 0 | 0 |
| NC_019843.3;YP_009047204.1;MERS_spike;1039 | 0 | 0.025 |
| NC_019843.3;YP_009047204.1;MERS_spike;1040 | 0 | 0 |
| NC_019843.3;YP_009047204.1;MERS_spike;1041 | 0 | 0 |
| NC_019843.3;YP_009047204.1;MERS_spike;1042 | 0 | 0 |
| NC_019843.3;YP_009047204.1;MERS_spike;1043 | 0 | 0.025 |
| NC_019843.3;YP_009047204.1;MERS_spike;1044 | 0 | 0.05 |
| NC_019843.3;YP_009047204.1;MERS_spike;1045 | 0.05 | 0.05 |
| NC_019843.3;YP_009047204.1;MERS_spike;1046 | 0 | 0.025 |
| NC_019843.3;YP_009047204.1;MERS_spike;1047 | 0 | 0.025 |
| NC_019843.3;YP_009047204.1;MERS_spike;1048 | 0 | 0.025 |
| NC_019843.3;YP_009047204.1;MERS_spike;1049 | 0 | 0.025 |
| NC_019843.3;YP_009047204.1;MERS_spike;1050 | 0.1 | 0 |
| NC_019843.3;YP_009047204.1;MERS_spike;1051 | 0 | 0.025 |
| NC_019843.3;YP_009047204.1;MERS_spike;1052 | 0 | 0.025 |
| NC_019843.3;YP_009047204.1;MERS_spike;1053 | 0 | 0 |
| NC_019843.3;YP_009047204.1;MERS_spike;1054 | 0 | 0 |
| NC_019843.3;YP_009047204.1;MERS_spike;1055 | 0 | 0 |
| NC_019843.3;YP_009047204.1;MERS_spike;1056 | 0 | 0.025 |
| NC_019843.3;YP_009047204.1;MERS_spike;1057 | 0 | 0 |

| | | |
|---|---|---|
| NC_019843.3;YP_009047204.1;MERS_spike;1058 | 0 | 0 |
| NC_019843.3;YP_009047204.1;MERS_spike;1059 | 0.05 | 0 |
| NC_019843.3;YP_009047204.1;MERS_spike;1060 | 0 | 0.025 |
| NC_019843.3;YP_009047204.1;MERS_spike;1061 | 0 | 0.025 |
| NC_019843.3;YP_009047204.1;MERS_spike;1062 | 0 | 0.025 |
| NC_019843.3;YP_009047204.1;MERS_spike;1063 | 0.05 | 0.025 |
| NC_019843.3;YP_009047204.1;MERS_spike;1064 | 0.05 | 0.025 |
| NC_019843.3;YP_009047204.1;MERS_spike;1065 | 0 | 0.05 |
| NC_019843.3;YP_009047204.1;MERS_spike;1066 | 0 | 0 |
| NC_019843.3;YP_009047204.1;MERS_spike;1067 | 0 | 0 |
| NC_019843.3;YP_009047204.1;MERS_spike;1068 | 0 | 0 |
| NC_019843.3;YP_009047204.1;MERS_spike;1069 | 0 | 0 |
| NC_019843.3;YP_009047204.1;MERS_spike;1070 | 0 | 0 |
| NC_019843.3;YP_009047204.1;MERS_spike;1071 | 0 | 0 |
| NC_019843.3;YP_009047204.1;MERS_spike;1072 | 0 | 0 |
| NC_019843.3;YP_009047204.1;MERS_spike;1073 | 0 | 0 |
| NC_019843.3;YP_009047204.1;MERS_spike;1074 | 0 | 0 |
| NC_019843.3;YP_009047204.1;MERS_spike;1075 | 0 | 0 |
| NC_019843.3;YP_009047204.1;MERS_spike;1076 | 0 | 0.025 |
| NC_019843.3;YP_009047204.1;MERS_spike;1077 | 0 | 0 |
| NC_019843.3;YP_009047204.1;MERS_spike;1078 | 0.05 | 0.025 |
| NC_019843.3;YP_009047204.1;MERS_spike;1079 | 0 | 0 |
| NC_019843.3;YP_009047204.1;MERS_spike;1080 | 0 | 0 |
| NC_019843.3;YP_009047204.1;MERS_spike;1081 | 0 | 0 |
| NC_019843.3;YP_009047204.1;MERS_spike;1082 | 0 | 0 |
| NC_019843.3;YP_009047204.1;MERS_spike;1083 | 0 | 0 |
| NC_019843.3;YP_009047204.1;MERS_spike;1084 | 0 | 0 |
| NC_019843.3;YP_009047204.1;MERS_spike;1085 | 0 | 0 |
| NC_019843.3;YP_009047204.1;MERS_spike;1086 | 0 | 0 |
| NC_019843.3;YP_009047204.1;MERS_spike;1087 | 0 | 0 |
| NC_019843.3;YP_009047204.1;MERS_spike;1088 | 0 | 0 |
| NC_019843.3;YP_009047204.1;MERS_spike;1089 | 0 | 0 |
| NC_019843.3;YP_009047204.1;MERS_spike;1090 | 0 | 0.025 |
| NC_019843.3;YP_009047204.1;MERS_spike;1091 | 0 | 0.025 |
| NC_019843.3;YP_009047204.1;MERS_spike;1092 | 0 | 0.025 |
| NC_019843.3;YP_009047204.1;MERS_spike;1093 | 0 | 0 |
| NC_019843.3;YP_009047204.1;MERS_spike;1094 | 0 | 0 |
| NC_019843.3;YP_009047204.1;MERS_spike;1095 | 0.05 | 0 |
| NC_019843.3;YP_009047204.1;MERS_spike;1096 | 0.1 | 0 |
| NC_019843.3;YP_009047204.1;MERS_spike;1097 | 0 | 0 |
| NC_019843.3;YP_009047204.1;MERS_spike;1098 | 0 | 0 |
| NC_019843.3;YP_009047204.1;MERS_spike;1099 | 0 | 0 |
| NC_019843.3;YP_009047204.1;MERS_spike;1100 | 0 | 0 |

| | | |
|---|---|---|
| NC_019843.3;YP_009047204.1;MERS_spike;1101 | 0 | 0 |
| NC_019843.3;YP_009047204.1;MERS_spike;1102 | 0 | 0 |
| NC_019843.3;YP_009047204.1;MERS_spike;1103 | 0 | 0 |
| NC_019843.3;YP_009047204.1;MERS_spike;1104 | 0 | 0 |
| NC_019843.3;YP_009047204.1;MERS_spike;1105 | 0 | 0 |
| NC_019843.3;YP_009047204.1;MERS_spike;1106 | 0 | 0 |
| NC_019843.3;YP_009047204.1;MERS_spike;1107 | 0 | 0 |
| NC_019843.3;YP_009047204.1;MERS_spike;1108 | 0 | 0 |
| NC_019843.3;YP_009047204.1;MERS_spike;1109 | 0 | 0 |
| NC_019843.3;YP_009047204.1;MERS_spike;1110 | 0 | 0 |
| NC_019843.3;YP_009047204.1;MERS_spike;1111 | 0 | 0 |
| NC_019843.3;YP_009047204.1;MERS_spike;1112 | 0 | 0 |
| NC_019843.3;YP_009047204.1;MERS_spike;1113 | 0 | 0 |
| NC_019843.3;YP_009047204.1;MERS_spike;1114 | 0 | 0 |
| NC_019843.3;YP_009047204.1;MERS_spike;1115 | 0 | 0 |
| NC_019843.3;YP_009047204.1;MERS_spike;1116 | 0 | 0 |
| NC_019843.3;YP_009047204.1;MERS_spike;1117 | 0 | 0 |
| NC_019843.3;YP_009047204.1;MERS_spike;1118 | 0 | 0 |
| NC_019843.3;YP_009047204.1;MERS_spike;1119 | 0 | 0 |
| NC_019843.3;YP_009047204.1;MERS_spike;1120 | 0 | 0 |
| NC_019843.3;YP_009047204.1;MERS_spike;1121 | 0 | 0 |
| NC_019843.3;YP_009047204.1;MERS_spike;1122 | 0 | 0 |
| NC_019843.3;YP_009047204.1;MERS_spike;1123 | 0 | 0 |
| NC_019843.3;YP_009047204.1;MERS_spike;1124 | 0 | 0 |
| NC_019843.3;YP_009047204.1;MERS_spike;1125 | 0 | 0 |
| NC_019843.3;YP_009047204.1;MERS_spike;1126 | 0 | 0 |
| NC_019843.3;YP_009047204.1;MERS_spike;1127 | 0 | 0 |
| NC_019843.3;YP_009047204.1;MERS_spike;1128 | 0 | 0 |
| NC_019843.3;YP_009047204.1;MERS_spike;1129 | 0 | 0 |
| NC_019843.3;YP_009047204.1;MERS_spike;1130 | 0 | 0 |
| NC_019843.3;YP_009047204.1;MERS_spike;1131 | 0 | 0 |
| NC_019843.3;YP_009047204.1;MERS_spike;1132 | 0 | 0 |
| NC_019843.3;YP_009047204.1;MERS_spike;1133 | 0 | 0 |
| NC_019843.3;YP_009047204.1;MERS_spike;1134 | 0 | 0 |
| NC_019843.3;YP_009047204.1;MERS_spike;1135 | 0 | 0 |
| NC_019843.3;YP_009047204.1;MERS_spike;1136 | 0 | 0 |
| NC_019843.3;YP_009047204.1;MERS_spike;1137 | 0 | 0 |
| NC_019843.3;YP_009047204.1;MERS_spike;1138 | 0 | 0 |
| NC_019843.3;YP_009047204.1;MERS_spike;1139 | 0 | 0 |
| NC_019843.3;YP_009047204.1;MERS_spike;1140 | 0 | 0 |
| NC_019843.3;YP_009047204.1;MERS_spike;1141 | 0 | 0 |
| NC_019843.3;YP_009047204.1;MERS_spike;1142 | 0 | 0 |
| NC_019843.3;YP_009047204.1;MERS_spike;1143 | 0 | 0 |

| | | |
|---|---|---|
| NC_019843.3;YP_009047204.1;MERS_spike;1144 | 0 | 0 |
| NC_019843.3;YP_009047204.1;MERS_spike;1145 | 0 | 0 |
| NC_019843.3;YP_009047204.1;MERS_spike;1146 | 0 | 0 |
| NC_019843.3;YP_009047204.1;MERS_spike;1147 | 0 | 0 |
| NC_019843.3;YP_009047204.1;MERS_spike;1148 | 0 | 0 |
| NC_019843.3;YP_009047204.1;MERS_spike;1149 | 0 | 0 |
| NC_019843.3;YP_009047204.1;MERS_spike;1150 | 0 | 0 |
| NC_019843.3;YP_009047204.1;MERS_spike;1151 | 0 | 0.075 |
| NC_019843.3;YP_009047204.1;MERS_spike;1152 | 0 | 0 |
| NC_019843.3;YP_009047204.1;MERS_spike;1153 | 0 | 0 |
| NC_019843.3;YP_009047204.1;MERS_spike;1154 | 0 | 0.05 |
| NC_019843.3;YP_009047204.1;MERS_spike;1155 | 0.1 | 0 |
| NC_019843.3;YP_009047204.1;MERS_spike;1156 | 0 | 0 |
| NC_019843.3;YP_009047204.1;MERS_spike;1157 | 0 | 0 |
| NC_019843.3;YP_009047204.1;MERS_spike;1158 | 0 | 0 |
| NC_019843.3;YP_009047204.1;MERS_spike;1159 | 0 | 0.025 |
| NC_019843.3;YP_009047204.1;MERS_spike;1160 | 0 | 0 |
| NC_019843.3;YP_009047204.1;MERS_spike;1161 | 0 | 0 |
| NC_019843.3;YP_009047204.1;MERS_spike;1162 | 0 | 0 |
| NC_019843.3;YP_009047204.1;MERS_spike;1163 | 0 | 0 |
| NC_019843.3;YP_009047204.1;MERS_spike;1164 | 0 | 0 |
| NC_019843.3;YP_009047204.1;MERS_spike;1165 | 0 | 0.025 |
| NC_019843.3;YP_009047204.1;MERS_spike;1166 | 0 | 0 |
| NC_019843.3;YP_009047204.1;MERS_spike;1167 | 0 | 0 |
| NC_019843.3;YP_009047204.1;MERS_spike;1168 | 0 | 0.025 |
| NC_019843.3;YP_009047204.1;MERS_spike;1169 | 0 | 0.025 |
| NC_019843.3;YP_009047204.1;MERS_spike;1170 | 0 | 0.025 |
| NC_019843.3;YP_009047204.1;MERS_spike;1171 | 0 | 0.025 |
| NC_019843.3;YP_009047204.1;MERS_spike;1172 | 0 | 0.05 |
| NC_019843.3;YP_009047204.1;MERS_spike;1173 | 0 | 0.05 |
| NC_019843.3;YP_009047204.1;MERS_spike;1174 | 0 | 0.05 |
| NC_019843.3;YP_009047204.1;MERS_spike;1175 | 0 | 0.05 |
| NC_019843.3;YP_009047204.1;MERS_spike;1176 | 0 | 0.05 |
| NC_019843.3;YP_009047204.1;MERS_spike;1177 | 0 | 0 |
| NC_019843.3;YP_009047204.1;MERS_spike;1178 | 0 | 0 |
| NC_019843.3;YP_009047204.1;MERS_spike;1179 | 0 | 0.025 |
| NC_019843.3;YP_009047204.1;MERS_spike;1180 | 0 | 0.025 |
| NC_019843.3;YP_009047204.1;MERS_spike;1181 | 0 | 0.025 |
| NC_019843.3;YP_009047204.1;MERS_spike;1182 | 0 | 0.025 |
| NC_019843.3;YP_009047204.1;MERS_spike;1183 | 0 | 0.025 |
| NC_019843.3;YP_009047204.1;MERS_spike;1184 | 0 | 0.025 |
| NC_019843.3;YP_009047204.1;MERS_spike;1185 | 0 | 0.025 |
| NC_019843.3;YP_009047204.1;MERS_spike;1186 | 0 | 0.075 |

| | | |
|---|---|---|
| NC_019843.3;YP_009047204.1;MERS_spike;1187 | 0.05 | 0.025 |
| NC_019843.3;YP_009047204.1;MERS_spike;1188 | 0.1 | 0.025 |
| NC_019843.3;YP_009047204.1;MERS_spike;1189 | 0.05 | 0 |
| NC_019843.3;YP_009047204.1;MERS_spike;1190 | 0.1 | 0 |
| NC_019843.3;YP_009047204.1;MERS_spike;1191 | 0.1 | 0.025 |
| NC_019843.3;YP_009047204.1;MERS_spike;1192 | 0.1 | 0.05 |
| NC_019843.3;YP_009047204.1;MERS_spike;1193 | 0.1 | 0 |
| NC_019843.3;YP_009047204.1;MERS_spike;1194 | 0.1 | 0 |
| NC_019843.3;YP_009047204.1;MERS_spike;1195 | 0.15 | 0.025 |
| NC_019843.3;YP_009047204.1;MERS_spike;1196 | 0 | 0.025 |
| NC_019843.3;YP_009047204.1;MERS_spike;1197 | 0 | 0.025 |
| NC_019843.3;YP_009047204.1;MERS_spike;1198 | 0.05 | 0 |
| NC_019843.3;YP_009047204.1;MERS_spike;1199 | 0.05 | 0 |
| NC_019843.3;YP_009047204.1;MERS_spike;1200 | 0 | 0 |
| NC_019843.3;YP_009047204.1;MERS_spike;1201 | 0 | 0.025 |
| NC_019843.3;YP_009047204.1;MERS_spike;1202 | 0.1 | 0.075 |
| NC_019843.3;YP_009047204.1;MERS_spike;1203 | 0.05 | 0 |
| NC_019843.3;YP_009047204.1;MERS_spike;1204 | 0.2 | 0.1 |
| NC_019843.3;YP_009047204.1;MERS_spike;1205 | 0.2 | 0.175 |
| NC_019843.3;YP_009047204.1;MERS_spike;1206 | 0.15 | 0.2 |
| NC_019843.3;YP_009047204.1;MERS_spike;1207 | 0.15 | 0.1 |
| NC_019843.3;YP_009047204.1;MERS_spike;1208 | 0.25 | 0.125 |
| NC_019843.3;YP_009047204.1;MERS_spike;1209 | 0.2 | 0.125 |
| NC_019843.3;YP_009047204.1;MERS_spike;1210 | 0.25 | 0.175 |
| NC_019843.3;YP_009047204.1;MERS_spike;1211 | 0.35 | 0.325 |
| NC_019843.3;YP_009047204.1;MERS_spike;1212 | 0.05 | 0.025 |
| NC_019843.3;YP_009047204.1;MERS_spike;1213 | 0.05 | 0 |
| NC_019843.3;YP_009047204.1;MERS_spike;1214 | 0 | 0 |
| NC_019843.3;YP_009047204.1;MERS_spike;1215 | 0 | 0.025 |
| NC_019843.3;YP_009047204.1;MERS_spike;1216 | 0 | 0 |
| NC_019843.3;YP_009047204.1;MERS_spike;1217 | 0 | 0 |
| NC_019843.3;YP_009047204.1;MERS_spike;1218 | 0 | 0 |
| NC_019843.3;YP_009047204.1;MERS_spike;1219 | 0 | 0 |
| NC_019843.3;YP_009047204.1;MERS_spike;1220 | 0 | 0 |
| NC_019843.3;YP_009047204.1;MERS_spike;1221 | 0 | 0 |
| NC_019843.3;YP_009047204.1;MERS_spike;1222 | 0 | 0 |
| NC_019843.3;YP_009047204.1;MERS_spike;1223 | 0 | 0.125 |
| NC_019843.3;YP_009047204.1;MERS_spike;1224 | 0 | 0.225 |
| NC_019843.3;YP_009047204.1;MERS_spike;1225 | 0 | 0.4 |
| NC_019843.3;YP_009047204.1;MERS_spike;1226 | 0 | 0.5 |
| NC_019843.3;YP_009047204.1;MERS_spike;1227 | 0 | 0.525 |
| NC_019843.3;YP_009047204.1;MERS_spike;1228 | 0.05 | 0.75 |
| NC_019843.3;YP_009047204.1;MERS_spike;1229 | 0.05 | 0.625 |

| | | |
|---|---|---|
| NC_019843.3;YP_009047204.1;MERS_spike;1230 | 0.05 | 0.625 |
| NC_019843.3;YP_009047204.1;MERS_spike;1231 | 0.05 | 0.475 |
| NC_019843.3;YP_009047204.1;MERS_spike;1232 | 0 | 0.1 |
| NC_019843.3;YP_009047204.1;MERS_spike;1233 | 0.05 | 0.1 |
| NC_019843.3;YP_009047204.1;MERS_spike;1234 | 0 | 0.05 |
| NC_019843.3;YP_009047204.1;MERS_spike;1235 | 0 | 0 |
| NC_019843.3;YP_009047204.1;MERS_spike;1236 | 0.1 | 0.025 |
| NC_019843.3;YP_009047204.1;MERS_spike;1237 | 0 | 0 |
| NC_019843.3;YP_009047204.1;MERS_spike;1238 | 0 | 0 |
| NC_019843.3;YP_009047204.1;MERS_spike;1239 | 0 | 0 |
| NC_019843.3;YP_009047204.1;MERS_spike;1240 | 0 | 0 |
| NC_019843.3;YP_009047204.1;MERS_spike;1241 | 0.05 | 0 |
| NC_019843.3;YP_009047204.1;MERS_spike;1242 | 0 | 0.025 |
| NC_019843.3;YP_009047204.1;MERS_spike;1243 | 0 | 0.025 |
| NC_019843.3;YP_009047204.1;MERS_spike;1244 | 0 | 0.025 |
| NC_019843.3;YP_009047204.1;MERS_spike;1245 | 0 | 0.025 |
| NC_019843.3;YP_009047204.1;MERS_spike;1246 | 0 | 0.025 |
| NC_019843.3;YP_009047204.1;MERS_spike;1247 | 0 | 0 |
| NC_019843.3;YP_009047204.1;MERS_spike;1248 | 0 | 0 |
| NC_019843.3;YP_009047204.1;MERS_spike;1249 | 0 | 0 |
| NC_019843.3;YP_009047204.1;MERS_spike;1250 | 0 | 0 |
| NC_019843.3;YP_009047204.1;MERS_spike;1251 | 0 | 0.025 |
| NC_019843.3;YP_009047204.1;MERS_spike;1252 | 0 | 0 |
| NC_019843.3;YP_009047204.1;MERS_spike;1253 | 0 | 0 |
| NC_019843.3;YP_009047204.1;MERS_spike;1254 | 0 | 0 |
| NC_019843.3;YP_009047204.1;MERS_spike;1255 | 0 | 0 |
| NC_019843.3;YP_009047204.1;MERS_spike;1256 | 0 | 0 |
| NC_019843.3;YP_009047204.1;MERS_spike;1257 | 0 | 0 |
| NC_019843.3;YP_009047204.1;MERS_spike;1258 | 0 | 0 |
| NC_019843.3;YP_009047204.1;MERS_spike;1259 | 0 | 0 |
| NC_019843.3;YP_009047204.1;MERS_spike;1260 | 0 | 0 |
| NC_019843.3;YP_009047204.1;MERS_spike;1261 | 0 | 0 |
| NC_019843.3;YP_009047204.1;MERS_spike;1262 | 0 | 0 |
| NC_019843.3;YP_009047204.1;MERS_spike;1263 | 0 | 0 |
| NC_019843.3;YP_009047204.1;MERS_spike;1264 | 0 | 0.025 |
| NC_019843.3;YP_009047204.1;MERS_spike;1265 | 0 | 0.025 |
| NC_019843.3;YP_009047204.1;MERS_spike;1266 | 0 | 0.075 |
| NC_019843.3;YP_009047204.1;MERS_spike;1267 | 0.15 | 0.125 |
| NC_019843.3;YP_009047204.1;MERS_spike;1268 | 0.05 | 0.15 |
| NC_019843.3;YP_009047204.1;MERS_spike;1269 | 0.15 | 0.175 |
| NC_019843.3;YP_009047204.1;MERS_spike;1270 | 0.15 | 0.175 |
| NC_019843.3;YP_009047204.1;MERS_spike;1271 | 0.1 | 0.125 |
| NC_019843.3;YP_009047204.1;MERS_spike;1272 | 0.1 | 0.125 |

| | | |
|---|---|---|
| NC_019843.3;YP_009047204.1;MERS_spike;1273 | 0.2 | 0.2 |
| NC_019843.3;YP_009047204.1;MERS_spike;1274 | 0.15 | 0.25 |
| NC_019843.3;YP_009047204.1;MERS_spike;1275 | 0.1 | 0.2 |
| NC_019843.3;YP_009047204.1;MERS_spike;1276 | 0.05 | 0.2 |
| NC_019843.3;YP_009047204.1;MERS_spike;1277 | 0 | 0 |
| NC_019843.3;YP_009047204.1;MERS_spike;1278 | 0.05 | 0.05 |
| NC_019843.3;YP_009047204.1;MERS_spike;1279 | 0 | 0.05 |
| NC_019843.3;YP_009047204.1;MERS_spike;1280 | 0 | 0 |
| NC_019843.3;YP_009047204.1;MERS_spike;1281 | 0.05 | 0 |
| NC_019843.3;YP_009047204.1;MERS_spike;1282 | 0 | 0 |
| NC_019843.3;YP_009047204.1;MERS_spike;1283 | 0 | 0 |
| NC_019843.3;YP_009047204.1;MERS_spike;1284 | 0 | 0 |
| NC_019843.3;YP_009047204.1;MERS_spike;1285 | 0 | 0 |
| NC_019843.3;YP_009047204.1;MERS_spike;1286 | 0 | 0 |
| NC_019843.3;YP_009047204.1;MERS_spike;1287 | 0 | 0 |
| NC_019843.3;YP_009047204.1;MERS_spike;1288 | 0 | 0 |
| NC_019843.3;YP_009047204.1;MERS_spike;1289 | 0 | 0 |
| NC_019843.3;YP_009047204.1;MERS_spike;1290 | 0 | 0 |
| NC_019843.3;YP_009047204.1;MERS_spike;1291 | 0 | 0 |
| NC_019843.3;YP_009047204.1;MERS_spike;1292 | 0 | 0 |
| NC_019843.3;YP_009047204.1;MERS_spike;1293 | 0 | 0 |
| NC_019843.3;YP_009047204.1;MERS_spike;1294 | 0 | 0 |
| NC_019843.3;YP_009047204.1;MERS_spike;1295 | 0 | 0 |
| NC_019843.3;YP_009047204.1;MERS_spike;1296 | 0 | 0 |
| NC_019843.3;YP_009047204.1;MERS_spike;1297 | 0 | 0 |
| NC_019843.3;YP_009047204.1;MERS_spike;1298 | 0 | 0 |
| NC_019843.3;YP_009047204.1;MERS_spike;1299 | 0 | 0 |
| NC_019843.3;YP_009047204.1;MERS_spike;1300 | 0 | 0 |
| NC_019843.3;YP_009047204.1;MERS_spike;1301 | 0 | 0 |
| NC_019843.3;YP_009047204.1;MERS_spike;1302 | 0 | 0 |
| NC_019843.3;YP_009047204.1;MERS_spike;1303 | 0 | 0 |
| NC_019843.3;YP_009047204.1;MERS_spike;1304 | 0 | 0 |
| NC_019843.3;YP_009047204.1;MERS_spike;1305 | 0 | 0 |
| NC_019843.3;YP_009047204.1;MERS_spike;1306 | 0 | 0 |
| NC_019843.3;YP_009047204.1;MERS_spike;1307 | 0 | 0 |
| NC_019843.3;YP_009047204.1;MERS_spike;1308 | 0 | 0 |
| NC_019843.3;YP_009047204.1;MERS_spike;1309 | 0 | 0 |
| NC_019843.3;YP_009047204.1;MERS_spike;1310 | 0 | 0 |
| NC_019843.3;YP_009047204.1;MERS_spike;1311 | 0 | 0 |
| NC_019843.3;YP_009047204.1;MERS_spike;1312 | 0 | 0 |
| NC_019843.3;YP_009047204.1;MERS_spike;1313 | 0 | 0 |
| NC_019843.3;YP_009047204.1;MERS_spike;1314 | 0 | 0 |
| NC_019843.3;YP_009047204.1;MERS_spike;1315 | 0 | 0 |

| | | |
|---|---|---|
| NC_019843.3;YP_009047204.1;MERS_spike;1316 | 0 | 0 |
| NC_019843.3;YP_009047204.1;MERS_spike;1317 | 0 | 0 |
| NC_019843.3;YP_009047204.1;MERS_spike;1318 | 0 | 0 |
| NC_019843.3;YP_009047204.1;MERS_spike;1319 | 0 | 0 |
| NC_019843.3;YP_009047204.1;MERS_spike;1320 | 0 | 0 |
| NC_019843.3;YP_009047204.1;MERS_spike;1321 | 0 | 0 |
| NC_019843.3;YP_009047204.1;MERS_spike;1322 | 0 | 0.025 |
| NC_019843.3;YP_009047204.1;MERS_spike;1323 | 0 | 0 |
| NC_019843.3;YP_009047204.1;MERS_spike;1324 | 0 | 0 |
| NC_019843.3;YP_009047204.1;MERS_spike;1325 | 0 | 0 |
| NC_019843.3;YP_009047204.1;MERS_spike;1326 | 0 | 0 |
| NC_019843.3;YP_009047204.1;MERS_spike;1327 | 0 | 0 |
| NC_019843.3;YP_009047204.1;MERS_spike;1328 | 0 | 0 |
| NC_019843.3;YP_009047204.1;MERS_spike;1329 | 0 | 0.025 |
| NC_019843.3;YP_009047204.1;MERS_spike;1330 | 0 | 0.025 |
| NC_019843.3;YP_009047204.1;MERS_spike;1331 | 0 | 0.05 |
| NC_019843.3;YP_009047204.1;MERS_spike;1332 | 0 | 0.05 |
| NC_019843.3;YP_009047204.1;MERS_spike;1333 | 0 | 0.025 |
| NC_019843.3;YP_009047204.1;MERS_spike;1334 | 0 | 0 |
| NC_019843.3;YP_009047204.1;MERS_spike;1335 | 0 | 0 |
| NC_019843.3;YP_009047204.1;MERS_spike;1336 | 0 | 0 |
| NC_019843.3;YP_009047204.1;MERS_spike;1337 | 0 | 0 |
| NC_019843.3;YP_009047204.1;MERS_spike;1338 | 0 | 0 |
| NC_019843.3;YP_009047204.1;MERS_spike;1339 | 0 | 0 |
| NC_019843.3;YP_009047204.1;MERS_spike;1340 | 0 | 0 |
| NC_019843.3;YP_009047204.1;MERS_spike;1341 | 0 | 0 |
| NC_019843.3;YP_009047204.1;MERS_spike;1342 | 0 | 0 |
| NC_019843.3;YP_009047205.1;MERS_ns3;1 | 0 | 0 |
| NC_019843.3;YP_009047205.1;MERS_ns3;2 | 0 | 0 |
| NC_019843.3;YP_009047205.1;MERS_ns3;3 | 0 | 0 |
| NC_019843.3;YP_009047205.1;MERS_ns3;4 | 0 | 0 |
| NC_019843.3;YP_009047205.1;MERS_ns3;5 | 0 | 0 |
| NC_019843.3;YP_009047205.1;MERS_ns3;6 | 0 | 0 |
| NC_019843.3;YP_009047205.1;MERS_ns3;7 | 0 | 0 |
| NC_019843.3;YP_009047205.1;MERS_ns3;8 | 0 | 0 |
| NC_019843.3;YP_009047205.1;MERS_ns3;9 | 0 | 0 |
| NC_019843.3;YP_009047205.1;MERS_ns3;10 | 0 | 0 |
| NC_019843.3;YP_009047205.1;MERS_ns3;11 | 0 | 0 |
| NC_019843.3;YP_009047205.1;MERS_ns3;12 | 0 | 0 |
| NC_019843.3;YP_009047205.1;MERS_ns3;13 | 0 | 0 |
| NC_019843.3;YP_009047205.1;MERS_ns3;14 | 0 | 0 |
| NC_019843.3;YP_009047205.1;MERS_ns3;15 | 0 | 0 |
| NC_019843.3;YP_009047205.1;MERS_ns3;16 | 0 | 0 |

| | | |
|---|---|---|
| NC_019843.3;YP_009047205.1;MERS_ns3;17 | 0.05 | 0.025 |
| NC_019843.3;YP_009047205.1;MERS_ns3;18 | 0.8 | 0.6 |
| NC_019843.3;YP_009047205.1;MERS_ns3;19 | 0.15 | 0.075 |
| NC_019843.3;YP_009047205.1;MERS_ns3;20 | 0.05 | 0 |
| NC_019843.3;YP_009047205.1;MERS_ns3;21 | 0.05 | 0.025 |
| NC_019843.3;YP_009047205.1;MERS_ns3;22 | 0.05 | 0.075 |
| NC_019843.3;YP_009047205.1;MERS_ns3;23 | 0 | 0 |
| NC_019843.3;YP_009047205.1;MERS_ns3;24 | 0 | 0 |
| NC_019843.3;YP_009047205.1;MERS_ns3;25 | 0 | 0 |
| NC_019843.3;YP_009047205.1;MERS_ns3;26 | 0 | 0 |
| NC_019843.3;YP_009047205.1;MERS_ns3;27 | 0 | 0 |
| NC_019843.3;YP_009047205.1;MERS_ns3;28 | 0 | 0 |
| NC_019843.3;YP_009047205.1;MERS_ns3;29 | 0 | 0 |
| NC_019843.3;YP_009047205.1;MERS_ns3;30 | 0 | 0 |
| NC_019843.3;YP_009047205.1;MERS_ns3;31 | 0 | 0 |
| NC_019843.3;YP_009047205.1;MERS_ns3;32 | 0 | 0 |
| NC_019843.3;YP_009047205.1;MERS_ns3;33 | 0 | 0 |
| NC_019843.3;YP_009047205.1;MERS_ns3;34 | 0 | 0 |
| NC_019843.3;YP_009047205.1;MERS_ns3;35 | 0 | 0 |
| NC_019843.3;YP_009047205.1;MERS_ns3;36 | 0.05 | 0.075 |
| NC_019843.3;YP_009047205.1;MERS_ns3;37 | 0.05 | 0.075 |
| NC_019843.3;YP_009047205.1;MERS_ns3;38 | 0.05 | 0.05 |
| NC_019843.3;YP_009047205.1;MERS_ns3;39 | 0.05 | 0.05 |
| NC_019843.3;YP_009047205.1;MERS_ns3;40 | 0.05 | 0.075 |
| NC_019843.3;YP_009047205.1;MERS_ns3;41 | 0.05 | 0.05 |
| NC_019843.3;YP_009047205.1;MERS_ns3;42 | 0.05 | 0.025 |
| NC_019843.3;YP_009047205.1;MERS_ns3;43 | 0.05 | 0.025 |
| NC_019843.3;YP_009047205.1;MERS_ns3;44 | 0.05 | 0.025 |
| NC_019843.3;YP_009047205.1;MERS_ns3;45 | 0.05 | 0 |
| NC_019843.3;YP_009047205.1;MERS_ns3;46 | 0.05 | 0.025 |
| NC_019843.3;YP_009047205.1;MERS_ns3;47 | 0.05 | 0.025 |
| NC_019843.3;YP_009047205.1;MERS_ns3;48 | 0.05 | 0.025 |
| NC_019843.3;YP_009047205.1;MERS_ns3;49 | 0 | 0.025 |
| NC_019843.3;YP_009047205.1;MERS_ns3;50 | 0 | 0 |
| NC_019843.3;YP_009047205.1;MERS_ns3;51 | 0 | 0 |
| NC_019843.3;YP_009047205.1;MERS_ns3;52 | 0.05 | 0 |
| NC_019843.3;YP_009047205.1;MERS_ns3;53 | 0 | 0 |
| NC_019843.3;YP_009047205.1;MERS_ns3;54 | 0.05 | 0 |
| NC_019843.3;YP_009047205.1;MERS_ns3;55 | 0 | 0 |
| NC_019843.3;YP_009047205.1;MERS_ns3;56 | 0 | 0 |
| NC_019843.3;YP_009047205.1;MERS_ns3;57 | 0 | 0.025 |
| NC_019843.3;YP_009047205.1;MERS_ns3;58 | 0 | 0.025 |
| NC_019843.3;YP_009047205.1;MERS_ns3;59 | 0 | 0.05 |

| | | |
|---|---|---|
| NC_019843.3;YP_009047205.1;MERS_ns3;60 | 0 | 0.025 |
| NC_019843.3;YP_009047205.1;MERS_ns3;61 | 0 | 0 |
| NC_019843.3;YP_009047205.1;MERS_ns3;62 | 0 | 0 |
| NC_019843.3;YP_009047205.1;MERS_ns3;63 | 0 | 0 |
| NC_019843.3;YP_009047205.1;MERS_ns3;64 | 0 | 0 |
| NC_019843.3;YP_009047205.1;MERS_ns3;65 | 0 | 0.025 |
| NC_019843.3;YP_009047205.1;MERS_ns3;66 | 0 | 0.125 |
| NC_019843.3;YP_009047205.1;MERS_ns3;67 | 0 | 0.025 |
| NC_019843.3;YP_009047205.1;MERS_ns3;68 | 0 | 0 |
| NC_019843.3;YP_009047205.1;MERS_ns3;69 | 0 | 0 |
| NC_019843.3;YP_009047205.1;MERS_ns3;70 | 0 | 0 |
| NC_019843.3;YP_009047205.1;MERS_ns3;71 | 0 | 0 |
| NC_019843.3;YP_009047205.1;MERS_ns3;72 | 0 | 0 |
| NC_019843.3;YP_009047205.1;MERS_ns3;73 | 0 | 0 |
| NC_019843.3;YP_009047205.1;MERS_ns3;74 | 0 | 0 |
| NC_019843.3;YP_009047205.1;MERS_ns3;75 | 0 | 0 |
| NC_019843.3;YP_009047205.1;MERS_ns3;76 | 0 | 0 |
| NC_019843.3;YP_009047205.1;MERS_ns3;77 | 0 | 0.025 |
| NC_019843.3;YP_009047205.1;MERS_ns3;78 | 0 | 0.15 |
| NC_019843.3;YP_009047205.1;MERS_ns3;79 | 0.05 | 0.05 |
| NC_019843.3;YP_009047205.1;MERS_ns3;80 | 0.05 | 0.025 |
| NC_019843.3;YP_009047205.1;MERS_ns3;81 | 0 | 0 |
| NC_019843.3;YP_009047205.1;MERS_ns3;82 | 0 | 0 |
| NC_019843.3;YP_009047205.1;MERS_ns3;83 | 0 | 0 |
| NC_019843.3;YP_009047205.1;MERS_ns3;84 | 0.05 | 0 |
| NC_019843.3;YP_009047205.1;MERS_ns3;85 | 0.05 | 0.025 |
| NC_019843.3;YP_009047205.1;MERS_ns3;86 | 0.05 | 0.025 |
| NC_019843.3;YP_009047205.1;MERS_ns3;87 | 0.05 | 0.025 |
| NC_019843.3;YP_009047205.1;MERS_ns3;88 | 0.05 | 0.05 |
| NC_019843.3;YP_009047205.1;MERS_ns3;89 | 0.05 | 0.025 |
| NC_019843.3;YP_009047205.1;MERS_ns3;90 | 0.05 | 0.075 |
| NC_019843.3;YP_009047205.1;MERS_ns3;91 | 0 | 0.025 |
| NC_019843.3;YP_009047205.1;MERS_ns3;92 | 0 | 0 |
| NC_019843.3;YP_009047206.1;MERS_ns4a;1 | 0 | 0 |
| NC_019843.3;YP_009047206.1;MERS_ns4a;2 | 0 | 0 |
| NC_019843.3;YP_009047206.1;MERS_ns4a;3 | 0 | 0 |
| NC_019843.3;YP_009047206.1;MERS_ns4a;4 | 0 | 0 |
| NC_019843.3;YP_009047206.1;MERS_ns4a;5 | 0 | 0.025 |
| NC_019843.3;YP_009047206.1;MERS_ns4a;6 | 0 | 0 |
| NC_019843.3;YP_009047206.1;MERS_ns4a;7 | 0 | 0 |
| NC_019843.3;YP_009047206.1;MERS_ns4a;8 | 0 | 0 |
| NC_019843.3;YP_009047206.1;MERS_ns4a;9 | 0 | 0 |
| NC_019843.3;YP_009047206.1;MERS_ns4a;10 | 0 | 0 |

| | | |
|---|---|---|
| NC_019843.3;YP_009047206.1;MERS_ns4a;11 | 0 | 0 |
| NC_019843.3;YP_009047206.1;MERS_ns4a;12 | 0 | 0 |
| NC_019843.3;YP_009047206.1;MERS_ns4a;13 | 0 | 0 |
| NC_019843.3;YP_009047206.1;MERS_ns4a;14 | 0 | 0 |
| NC_019843.3;YP_009047206.1;MERS_ns4a;15 | 0 | 0 |
| NC_019843.3;YP_009047206.1;MERS_ns4a;16 | 0 | 0 |
| NC_019843.3;YP_009047206.1;MERS_ns4a;17 | 0 | 0 |
| NC_019843.3;YP_009047206.1;MERS_ns4a;18 | 0 | 0 |
| NC_019843.3;YP_009047206.1;MERS_ns4a;19 | 0 | 0 |
| NC_019843.3;YP_009047206.1;MERS_ns4a;20 | 0 | 0 |
| NC_019843.3;YP_009047206.1;MERS_ns4a;21 | 0 | 0 |
| NC_019843.3;YP_009047206.1;MERS_ns4a;22 | 0 | 0 |
| NC_019843.3;YP_009047206.1;MERS_ns4a;23 | 0 | 0 |
| NC_019843.3;YP_009047206.1;MERS_ns4a;24 | 0.05 | 0 |
| NC_019843.3;YP_009047206.1;MERS_ns4a;25 | 0 | 0 |
| NC_019843.3;YP_009047206.1;MERS_ns4a;26 | 0 | 0 |
| NC_019843.3;YP_009047206.1;MERS_ns4a;27 | 0 | 0 |
| NC_019843.3;YP_009047206.1;MERS_ns4a;28 | 0.05 | 0 |
| NC_019843.3;YP_009047206.1;MERS_ns4a;29 | 0 | 0 |
| NC_019843.3;YP_009047206.1;MERS_ns4a;30 | 0 | 0 |
| NC_019843.3;YP_009047206.1;MERS_ns4a;31 | 0 | 0 |
| NC_019843.3;YP_009047206.1;MERS_ns4a;32 | 0 | 0 |
| NC_019843.3;YP_009047206.1;MERS_ns4a;33 | 0 | 0 |
| NC_019843.3;YP_009047206.1;MERS_ns4a;34 | 0 | 0 |
| NC_019843.3;YP_009047206.1;MERS_ns4a;35 | 0 | 0 |
| NC_019843.3;YP_009047206.1;MERS_ns4a;36 | 0 | 0 |
| NC_019843.3;YP_009047206.1;MERS_ns4a;37 | 0 | 0 |
| NC_019843.3;YP_009047206.1;MERS_ns4a;38 | 0 | 0 |
| NC_019843.3;YP_009047206.1;MERS_ns4a;39 | 0 | 0 |
| NC_019843.3;YP_009047206.1;MERS_ns4a;40 | 0 | 0 |
| NC_019843.3;YP_009047206.1;MERS_ns4a;41 | 0 | 0 |
| NC_019843.3;YP_009047206.1;MERS_ns4a;42 | 0 | 0 |
| NC_019843.3;YP_009047206.1;MERS_ns4a;43 | 0 | 0 |
| NC_019843.3;YP_009047206.1;MERS_ns4a;44 | 0 | 0.025 |
| NC_019843.3;YP_009047206.1;MERS_ns4a;45 | 0 | 0.025 |
| NC_019843.3;YP_009047206.1;MERS_ns4a;46 | 0 | 0.025 |
| NC_019843.3;YP_009047206.1;MERS_ns4a;47 | 0 | 0.025 |
| NC_019843.3;YP_009047206.1;MERS_ns4a;48 | 0.1 | 0.025 |
| NC_019843.3;YP_009047206.1;MERS_ns4a;49 | 0.05 | 0.025 |
| NC_019843.3;YP_009047206.1;MERS_ns4a;50 | 0.15 | 0.025 |
| NC_019843.3;YP_009047206.1;MERS_ns4a;51 | 0.05 | 0.05 |
| NC_019843.3;YP_009047206.1;MERS_ns4a;52 | 0 | 0.025 |
| NC_019843.3;YP_009047206.1;MERS_ns4a;53 | 0.1 | 0.025 |

| | | |
|---|---|---|
| NC_019843.3;YP_009047206.1;MERS_ns4a;54 | 0.15 | 0.05 |
| NC_019843.3;YP_009047206.1;MERS_ns4a;55 | 0.1 | 0.025 |
| NC_019843.3;YP_009047206.1;MERS_ns4a;56 | 0.15 | 0.025 |
| NC_019843.3;YP_009047206.1;MERS_ns4a;57 | 0 | 0.025 |
| NC_019843.3;YP_009047206.1;MERS_ns4a;58 | 0 | 0 |
| NC_019843.3;YP_009047206.1;MERS_ns4a;59 | 0 | 0 |
| NC_019843.3;YP_009047206.1;MERS_ns4a;60 | 0 | 0 |
| NC_019843.3;YP_009047206.1;MERS_ns4a;61 | 0 | 0.025 |
| NC_019843.3;YP_009047206.1;MERS_ns4a;62 | 0 | 0 |
| NC_019843.3;YP_009047206.1;MERS_ns4a;63 | 0 | 0 |
| NC_019843.3;YP_009047206.1;MERS_ns4a;64 | 0 | 0 |
| NC_019843.3;YP_009047206.1;MERS_ns4a;65 | 0 | 0 |
| NC_019843.3;YP_009047206.1;MERS_ns4a;66 | 0 | 0 |
| NC_019843.3;YP_009047206.1;MERS_ns4a;67 | 0.05 | 0 |
| NC_019843.3;YP_009047206.1;MERS_ns4a;68 | 0 | 0 |
| NC_019843.3;YP_009047206.1;MERS_ns4a;69 | 0.05 | 0 |
| NC_019843.3;YP_009047206.1;MERS_ns4a;70 | 0 | 0 |
| NC_019843.3;YP_009047206.1;MERS_ns4a;71 | 0 | 0 |
| NC_019843.3;YP_009047206.1;MERS_ns4a;72 | 0 | 0 |
| NC_019843.3;YP_009047206.1;MERS_ns4a;73 | 0 | 0 |
| NC_019843.3;YP_009047206.1;MERS_ns4a;74 | 0 | 0 |
| NC_019843.3;YP_009047206.1;MERS_ns4a;75 | 0 | 0 |
| NC_019843.3;YP_009047206.1;MERS_ns4a;76 | 0 | 0 |
| NC_019843.3;YP_009047206.1;MERS_ns4a;77 | 0 | 0 |
| NC_019843.3;YP_009047206.1;MERS_ns4a;78 | 0 | 0 |
| NC_019843.3;YP_009047206.1;MERS_ns4a;79 | 0 | 0 |
| NC_019843.3;YP_009047206.1;MERS_ns4a;80 | 0 | 0 |
| NC_019843.3;YP_009047206.1;MERS_ns4a;81 | 0 | 0 |
| NC_019843.3;YP_009047206.1;MERS_ns4a;82 | 0 | 0 |
| NC_019843.3;YP_009047206.1;MERS_ns4a;83 | 0 | 0 |
| NC_019843.3;YP_009047206.1;MERS_ns4a;84 | 0 | 0 |
| NC_019843.3;YP_009047206.1;MERS_ns4a;85 | 0 | 0 |
| NC_019843.3;YP_009047206.1;MERS_ns4a;86 | 0 | 0 |
| NC_019843.3;YP_009047206.1;MERS_ns4a;87 | 0 | 0 |
| NC_019843.3;YP_009047206.1;MERS_ns4a;88 | 0 | 0 |
| NC_019843.3;YP_009047206.1;MERS_ns4a;89 | 0 | 0 |
| NC_019843.3;YP_009047206.1;MERS_ns4a;90 | 0 | 0 |
| NC_019843.3;YP_009047206.1;MERS_ns4a;91 | 0 | 0 |
| NC_019843.3;YP_009047206.1;MERS_ns4a;92 | 0 | 0 |
| NC_019843.3;YP_009047206.1;MERS_ns4a;93 | 0 | 0 |
| NC_019843.3;YP_009047206.1;MERS_ns4a;94 | 0 | 0 |
| NC_019843.3;YP_009047206.1;MERS_ns4a;95 | 0 | 0 |
| NC_019843.3;YP_009047206.1;MERS_ns4a;96 | 0 | 0.025 |

| | | |
|---|---|---|
| NC_019843.3;YP_009047206.1;MERS_ns4a;97 | 0 | 0 |
| NC_019843.3;YP_009047206.1;MERS_ns4a;98 | 0 | 0 |
| NC_019843.3;YP_009047207.1;MERS_ns4b;1 | 0 | 0 |
| NC_019843.3;YP_009047207.1;MERS_ns4b;2 | 0 | 0 |
| NC_019843.3;YP_009047207.1;MERS_ns4b;3 | 0.05 | 0 |
| NC_019843.3;YP_009047207.1;MERS_ns4b;4 | 0.05 | 0 |
| NC_019843.3;YP_009047207.1;MERS_ns4b;5 | 0 | 0 |
| NC_019843.3;YP_009047207.1;MERS_ns4b;6 | 0 | 0 |
| NC_019843.3;YP_009047207.1;MERS_ns4b;7 | 0 | 0 |
| NC_019843.3;YP_009047207.1;MERS_ns4b;8 | 0 | 0 |
| NC_019843.3;YP_009047207.1;MERS_ns4b;9 | 0 | 0 |
| NC_019843.3;YP_009047207.1;MERS_ns4b;10 | 0 | 0 |
| NC_019843.3;YP_009047207.1;MERS_ns4b;11 | 0 | 0 |
| NC_019843.3;YP_009047207.1;MERS_ns4b;12 | 0 | 0 |
| NC_019843.3;YP_009047207.1;MERS_ns4b;13 | 0 | 0 |
| NC_019843.3;YP_009047207.1;MERS_ns4b;14 | 0 | 0 |
| NC_019843.3;YP_009047207.1;MERS_ns4b;15 | 0 | 0 |
| NC_019843.3;YP_009047207.1;MERS_ns4b;16 | 0 | 0.05 |
| NC_019843.3;YP_009047207.1;MERS_ns4b;17 | 0 | 0 |
| NC_019843.3;YP_009047207.1;MERS_ns4b;18 | 0 | 0 |
| NC_019843.3;YP_009047207.1;MERS_ns4b;19 | 0 | 0 |
| NC_019843.3;YP_009047207.1;MERS_ns4b;20 | 0.05 | 0 |
| NC_019843.3;YP_009047207.1;MERS_ns4b;21 | 0 | 0 |
| NC_019843.3;YP_009047207.1;MERS_ns4b;22 | 0 | 0 |
| NC_019843.3;YP_009047207.1;MERS_ns4b;23 | 0 | 0 |
| NC_019843.3;YP_009047207.1;MERS_ns4b;24 | 0 | 0 |
| NC_019843.3;YP_009047207.1;MERS_ns4b;25 | 0 | 0 |
| NC_019843.3;YP_009047207.1;MERS_ns4b;26 | 0 | 0 |
| NC_019843.3;YP_009047207.1;MERS_ns4b;27 | 0 | 0 |
| NC_019843.3;YP_009047207.1;MERS_ns4b;28 | 0 | 0 |
| NC_019843.3;YP_009047207.1;MERS_ns4b;29 | 0 | 0 |
| NC_019843.3;YP_009047207.1;MERS_ns4b;30 | 0 | 0 |
| NC_019843.3;YP_009047207.1;MERS_ns4b;31 | 0 | 0 |
| NC_019843.3;YP_009047207.1;MERS_ns4b;32 | 0 | 0 |
| NC_019843.3;YP_009047207.1;MERS_ns4b;33 | 0 | 0 |
| NC_019843.3;YP_009047207.1;MERS_ns4b;34 | 0 | 0.025 |
| NC_019843.3;YP_009047207.1;MERS_ns4b;35 | 0 | 0.025 |
| NC_019843.3;YP_009047207.1;MERS_ns4b;36 | 0 | 0.05 |
| NC_019843.3;YP_009047207.1;MERS_ns4b;37 | 0 | 0 |
| NC_019843.3;YP_009047207.1;MERS_ns4b;38 | 0 | 0 |
| NC_019843.3;YP_009047207.1;MERS_ns4b;39 | 0 | 0 |
| NC_019843.3;YP_009047207.1;MERS_ns4b;40 | 0 | 0 |
| NC_019843.3;YP_009047207.1;MERS_ns4b;41 | 0 | 0 |

| | | |
|---|---|---|
| NC_019843.3;YP_009047207.1;MERS_ns4b;42 | 0 | 0 |
| NC_019843.3;YP_009047207.1;MERS_ns4b;43 | 0 | 0 |
| NC_019843.3;YP_009047207.1;MERS_ns4b;44 | 0 | 0 |
| NC_019843.3;YP_009047207.1;MERS_ns4b;45 | 0 | 0 |
| NC_019843.3;YP_009047207.1;MERS_ns4b;46 | 0 | 0 |
| NC_019843.3;YP_009047207.1;MERS_ns4b;47 | 0 | 0 |
| NC_019843.3;YP_009047207.1;MERS_ns4b;48 | 0 | 0 |
| NC_019843.3;YP_009047207.1;MERS_ns4b;49 | 0 | 0 |
| NC_019843.3;YP_009047207.1;MERS_ns4b;50 | 0 | 0 |
| NC_019843.3;YP_009047207.1;MERS_ns4b;51 | 0 | 0 |
| NC_019843.3;YP_009047207.1;MERS_ns4b;52 | 0 | 0 |
| NC_019843.3;YP_009047207.1;MERS_ns4b;53 | 0 | 0 |
| NC_019843.3;YP_009047207.1;MERS_ns4b;54 | 0 | 0 |
| NC_019843.3;YP_009047207.1;MERS_ns4b;55 | 0 | 0 |
| NC_019843.3;YP_009047207.1;MERS_ns4b;56 | 0 | 0 |
| NC_019843.3;YP_009047207.1;MERS_ns4b;57 | 0.05 | 0 |
| NC_019843.3;YP_009047207.1;MERS_ns4b;58 | 0 | 0 |
| NC_019843.3;YP_009047207.1;MERS_ns4b;59 | 0 | 0 |
| NC_019843.3;YP_009047207.1;MERS_ns4b;60 | 0 | 0 |
| NC_019843.3;YP_009047207.1;MERS_ns4b;61 | 0 | 0 |
| NC_019843.3;YP_009047207.1;MERS_ns4b;62 | 0 | 0 |
| NC_019843.3;YP_009047207.1;MERS_ns4b;63 | 0 | 0 |
| NC_019843.3;YP_009047207.1;MERS_ns4b;64 | 0 | 0 |
| NC_019843.3;YP_009047207.1;MERS_ns4b;65 | 0 | 0 |
| NC_019843.3;YP_009047207.1;MERS_ns4b;66 | 0 | 0 |
| NC_019843.3;YP_009047207.1;MERS_ns4b;67 | 0 | 0 |
| NC_019843.3;YP_009047207.1;MERS_ns4b;68 | 0 | 0 |
| NC_019843.3;YP_009047207.1;MERS_ns4b;69 | 0 | 0 |
| NC_019843.3;YP_009047207.1;MERS_ns4b;70 | 0 | 0 |
| NC_019843.3;YP_009047207.1;MERS_ns4b;71 | 0 | 0 |
| NC_019843.3;YP_009047207.1;MERS_ns4b;72 | 0 | 0 |
| NC_019843.3;YP_009047207.1;MERS_ns4b;73 | 0 | 0 |
| NC_019843.3;YP_009047207.1;MERS_ns4b;74 | 0 | 0 |
| NC_019843.3;YP_009047207.1;MERS_ns4b;75 | 0 | 0 |
| NC_019843.3;YP_009047207.1;MERS_ns4b;76 | 0 | 0 |
| NC_019843.3;YP_009047207.1;MERS_ns4b;77 | 0 | 0 |
| NC_019843.3;YP_009047207.1;MERS_ns4b;78 | 0 | 0.025 |
| NC_019843.3;YP_009047207.1;MERS_ns4b;79 | 0 | 0 |
| NC_019843.3;YP_009047207.1;MERS_ns4b;80 | 0 | 0.025 |
| NC_019843.3;YP_009047207.1;MERS_ns4b;81 | 0 | 0.025 |
| NC_019843.3;YP_009047207.1;MERS_ns4b;82 | 0 | 0 |
| NC_019843.3;YP_009047207.1;MERS_ns4b;83 | 0 | 0 |
| NC_019843.3;YP_009047207.1;MERS_ns4b;84 | 0 | 0 |

| | | |
|---|---|---|
| NC_019843.3;YP_009047207.1;MERS_ns4b;85 | 0 | 0 |
| NC_019843.3;YP_009047207.1;MERS_ns4b;86 | 0 | 0 |
| NC_019843.3;YP_009047207.1;MERS_ns4b;87 | 0 | 0 |
| NC_019843.3;YP_009047207.1;MERS_ns4b;88 | 0 | 0 |
| NC_019843.3;YP_009047207.1;MERS_ns4b;89 | 0 | 0 |
| NC_019843.3;YP_009047207.1;MERS_ns4b;90 | 0 | 0 |
| NC_019843.3;YP_009047207.1;MERS_ns4b;91 | 0 | 0 |
| NC_019843.3;YP_009047207.1;MERS_ns4b;92 | 0 | 0 |
| NC_019843.3;YP_009047207.1;MERS_ns4b;93 | 0 | 0 |
| NC_019843.3;YP_009047207.1;MERS_ns4b;94 | 0 | 0 |
| NC_019843.3;YP_009047207.1;MERS_ns4b;95 | 0 | 0 |
| NC_019843.3;YP_009047207.1;MERS_ns4b;96 | 0 | 0 |
| NC_019843.3;YP_009047207.1;MERS_ns4b;97 | 0 | 0 |
| NC_019843.3;YP_009047207.1;MERS_ns4b;98 | 0 | 0.025 |
| NC_019843.3;YP_009047207.1;MERS_ns4b;99 | 0 | 0.05 |
| NC_019843.3;YP_009047207.1;MERS_ns4b;100 | 0 | 0.025 |
| NC_019843.3;YP_009047207.1;MERS_ns4b;101 | 0 | 0.025 |
| NC_019843.3;YP_009047207.1;MERS_ns4b;102 | 0.05 | 0 |
| NC_019843.3;YP_009047207.1;MERS_ns4b;103 | 0 | 0 |
| NC_019843.3;YP_009047207.1;MERS_ns4b;104 | 0 | 0 |
| NC_019843.3;YP_009047207.1;MERS_ns4b;105 | 0 | 0 |
| NC_019843.3;YP_009047207.1;MERS_ns4b;106 | 0 | 0 |
| NC_019843.3;YP_009047207.1;MERS_ns4b;107 | 0 | 0 |
| NC_019843.3;YP_009047207.1;MERS_ns4b;108 | 0 | 0 |
| NC_019843.3;YP_009047207.1;MERS_ns4b;109 | 0 | 0 |
| NC_019843.3;YP_009047207.1;MERS_ns4b;110 | 0 | 0 |
| NC_019843.3;YP_009047207.1;MERS_ns4b;111 | 0 | 0 |
| NC_019843.3;YP_009047207.1;MERS_ns4b;112 | 0 | 0 |
| NC_019843.3;YP_009047207.1;MERS_ns4b;113 | 0 | 0 |
| NC_019843.3;YP_009047207.1;MERS_ns4b;114 | 0 | 0 |
| NC_019843.3;YP_009047207.1;MERS_ns4b;115 | 0 | 0 |
| NC_019843.3;YP_009047207.1;MERS_ns4b;116 | 0 | 0 |
| NC_019843.3;YP_009047207.1;MERS_ns4b;117 | 0 | 0 |
| NC_019843.3;YP_009047207.1;MERS_ns4b;118 | 0 | 0 |
| NC_019843.3;YP_009047207.1;MERS_ns4b;119 | 0 | 0 |
| NC_019843.3;YP_009047207.1;MERS_ns4b;120 | 0 | 0 |
| NC_019843.3;YP_009047207.1;MERS_ns4b;121 | 0 | 0 |
| NC_019843.3;YP_009047207.1;MERS_ns4b;122 | 0 | 0 |
| NC_019843.3;YP_009047207.1;MERS_ns4b;123 | 0 | 0 |
| NC_019843.3;YP_009047207.1;MERS_ns4b;124 | 0 | 0 |
| NC_019843.3;YP_009047207.1;MERS_ns4b;125 | 0 | 0 |
| NC_019843.3;YP_009047207.1;MERS_ns4b;126 | 0 | 0 |
| NC_019843.3;YP_009047207.1;MERS_ns4b;127 | 0 | 0 |

| | | |
|---|---|---|
| NC_019843.3;YP_009047207.1;MERS_ns4b;128 | 0 | 0 |
| NC_019843.3;YP_009047207.1;MERS_ns4b;129 | 0 | 0 |
| NC_019843.3;YP_009047207.1;MERS_ns4b;130 | 0 | 0 |
| NC_019843.3;YP_009047207.1;MERS_ns4b;131 | 0 | 0 |
| NC_019843.3;YP_009047207.1;MERS_ns4b;132 | 0 | 0 |
| NC_019843.3;YP_009047207.1;MERS_ns4b;133 | 0 | 0 |
| NC_019843.3;YP_009047207.1;MERS_ns4b;134 | 0 | 0 |
| NC_019843.3;YP_009047207.1;MERS_ns4b;135 | 0 | 0 |
| NC_019843.3;YP_009047207.1;MERS_ns4b;136 | 0.05 | 0 |
| NC_019843.3;YP_009047207.1;MERS_ns4b;137 | 0 | 0 |
| NC_019843.3;YP_009047207.1;MERS_ns4b;138 | 0 | 0 |
| NC_019843.3;YP_009047207.1;MERS_ns4b;139 | 0 | 0 |
| NC_019843.3;YP_009047207.1;MERS_ns4b;140 | 0 | 0.05 |
| NC_019843.3;YP_009047207.1;MERS_ns4b;141 | 0 | 0 |
| NC_019843.3;YP_009047207.1;MERS_ns4b;142 | 0 | 0 |
| NC_019843.3;YP_009047207.1;MERS_ns4b;143 | 0 | 0 |
| NC_019843.3;YP_009047207.1;MERS_ns4b;144 | 0 | 0 |
| NC_019843.3;YP_009047207.1;MERS_ns4b;145 | 0 | 0 |
| NC_019843.3;YP_009047207.1;MERS_ns4b;146 | 0 | 0 |
| NC_019843.3;YP_009047207.1;MERS_ns4b;147 | 0 | 0 |
| NC_019843.3;YP_009047207.1;MERS_ns4b;148 | 0 | 0 |
| NC_019843.3;YP_009047207.1;MERS_ns4b;149 | 0 | 0 |
| NC_019843.3;YP_009047207.1;MERS_ns4b;150 | 0 | 0 |
| NC_019843.3;YP_009047207.1;MERS_ns4b;151 | 0 | 0 |
| NC_019843.3;YP_009047207.1;MERS_ns4b;152 | 0.05 | 0 |
| NC_019843.3;YP_009047207.1;MERS_ns4b;153 | 0 | 0 |
| NC_019843.3;YP_009047207.1;MERS_ns4b;154 | 0 | 0 |
| NC_019843.3;YP_009047207.1;MERS_ns4b;155 | 0.05 | 0 |
| NC_019843.3;YP_009047207.1;MERS_ns4b;156 | 0 | 0 |
| NC_019843.3;YP_009047207.1;MERS_ns4b;157 | 0 | 0 |
| NC_019843.3;YP_009047207.1;MERS_ns4b;158 | 0 | 0 |
| NC_019843.3;YP_009047207.1;MERS_ns4b;159 | 0 | 0 |
| NC_019843.3;YP_009047207.1;MERS_ns4b;160 | 0 | 0 |
| NC_019843.3;YP_009047207.1;MERS_ns4b;161 | 0 | 0 |
| NC_019843.3;YP_009047207.1;MERS_ns4b;162 | 0 | 0 |
| NC_019843.3;YP_009047207.1;MERS_ns4b;163 | 0 | 0 |
| NC_019843.3;YP_009047207.1;MERS_ns4b;164 | 0 | 0 |
| NC_019843.3;YP_009047207.1;MERS_ns4b;165 | 0 | 0 |
| NC_019843.3;YP_009047207.1;MERS_ns4b;166 | 0 | 0 |
| NC_019843.3;YP_009047207.1;MERS_ns4b;167 | 0 | 0 |
| NC_019843.3;YP_009047207.1;MERS_ns4b;168 | 0 | 0 |
| NC_019843.3;YP_009047207.1;MERS_ns4b;169 | 0 | 0 |
| NC_019843.3;YP_009047207.1;MERS_ns4b;170 | 0 | 0 |

| | | |
|---|---|---|
| NC_019843.3;YP_009047207.1;MERS_ns4b;171 | 0 | 0 |
| NC_019843.3;YP_009047207.1;MERS_ns4b;172 | 0 | 0 |
| NC_019843.3;YP_009047207.1;MERS_ns4b;173 | 0 | 0 |
| NC_019843.3;YP_009047207.1;MERS_ns4b;174 | 0 | 0 |
| NC_019843.3;YP_009047207.1;MERS_ns4b;175 | 0 | 0 |
| NC_019843.3;YP_009047207.1;MERS_ns4b;176 | 0 | 0 |
| NC_019843.3;YP_009047207.1;MERS_ns4b;177 | 0.05 | 0 |
| NC_019843.3;YP_009047207.1;MERS_ns4b;178 | 0.05 | 0 |
| NC_019843.3;YP_009047207.1;MERS_ns4b;179 | 0.05 | 0 |
| NC_019843.3;YP_009047207.1;MERS_ns4b;180 | 0.05 | 0 |
| NC_019843.3;YP_009047207.1;MERS_ns4b;181 | 0.05 | 0 |
| NC_019843.3;YP_009047207.1;MERS_ns4b;182 | 0.05 | 0 |
| NC_019843.3;YP_009047207.1;MERS_ns4b;183 | 0.05 | 0 |
| NC_019843.3;YP_009047207.1;MERS_ns4b;184 | 0.05 | 0 |
| NC_019843.3;YP_009047207.1;MERS_ns4b;185 | 0 | 0 |
| NC_019843.3;YP_009047207.1;MERS_ns4b;186 | 0 | 0 |
| NC_019843.3;YP_009047207.1;MERS_ns4b;187 | 0 | 0 |
| NC_019843.3;YP_009047207.1;MERS_ns4b;188 | 0 | 0 |
| NC_019843.3;YP_009047207.1;MERS_ns4b;189 | 0 | 0 |
| NC_019843.3;YP_009047207.1;MERS_ns4b;190 | 0 | 0 |
| NC_019843.3;YP_009047207.1;MERS_ns4b;191 | 0 | 0 |
| NC_019843.3;YP_009047207.1;MERS_ns4b;192 | 0.05 | 0 |
| NC_019843.3;YP_009047207.1;MERS_ns4b;193 | 0 | 0 |
| NC_019843.3;YP_009047207.1;MERS_ns4b;194 | 0 | 0 |
| NC_019843.3;YP_009047207.1;MERS_ns4b;195 | 0 | 0 |
| NC_019843.3;YP_009047207.1;MERS_ns4b;196 | 0 | 0 |
| NC_019843.3;YP_009047207.1;MERS_ns4b;197 | 0 | 0 |
| NC_019843.3;YP_009047207.1;MERS_ns4b;198 | 0 | 0 |
| NC_019843.3;YP_009047207.1;MERS_ns4b;199 | 0 | 0 |
| NC_019843.3;YP_009047207.1;MERS_ns4b;200 | 0 | 0 |
| NC_019843.3;YP_009047207.1;MERS_ns4b;201 | 0 | 0 |
| NC_019843.3;YP_009047207.1;MERS_ns4b;202 | 0 | 0 |
| NC_019843.3;YP_009047207.1;MERS_ns4b;203 | 0 | 0 |
| NC_019843.3;YP_009047207.1;MERS_ns4b;204 | 0 | 0 |
| NC_019843.3;YP_009047207.1;MERS_ns4b;205 | 0 | 0 |
| NC_019843.3;YP_009047207.1;MERS_ns4b;206 | 0 | 0 |
| NC_019843.3;YP_009047207.1;MERS_ns4b;207 | 0 | 0 |
| NC_019843.3;YP_009047207.1;MERS_ns4b;208 | 0.05 | 0 |
| NC_019843.3;YP_009047207.1;MERS_ns4b;209 | 0 | 0 |
| NC_019843.3;YP_009047207.1;MERS_ns4b;210 | 0 | 0 |
| NC_019843.3;YP_009047207.1;MERS_ns4b;211 | 0 | 0 |
| NC_019843.3;YP_009047207.1;MERS_ns4b;212 | 0 | 0 |
| NC_019843.3;YP_009047207.1;MERS_ns4b;213 | 0 | 0 |

| | | |
|---|---|---|
| NC_019843.3;YP_009047207.1;MERS_ns4b;214 | 0 | 0 |
| NC_019843.3;YP_009047207.1;MERS_ns4b;215 | 0.05 | 0 |
| NC_019843.3;YP_009047207.1;MERS_ns4b;216 | 0.05 | 0.025 |
| NC_019843.3;YP_009047207.1;MERS_ns4b;217 | 0 | 0.025 |
| NC_019843.3;YP_009047207.1;MERS_ns4b;218 | 0.1 | 0.075 |
| NC_019843.3;YP_009047207.1;MERS_ns4b;219 | 0 | 0.075 |
| NC_019843.3;YP_009047207.1;MERS_ns4b;220 | 0 | 0.05 |
| NC_019843.3;YP_009047207.1;MERS_ns4b;221 | 0 | 0.025 |
| NC_019843.3;YP_009047207.1;MERS_ns4b;222 | 0.05 | 0.125 |
| NC_019843.3;YP_009047207.1;MERS_ns4b;223 | 0.2 | 0.1 |
| NC_019843.3;YP_009047207.1;MERS_ns4b;224 | 0.1 | 0.025 |
| NC_019843.3;YP_009047207.1;MERS_ns4b;225 | 0.05 | 0 |
| NC_019843.3;YP_009047207.1;MERS_ns4b;226 | 0.05 | 0.025 |
| NC_019843.3;YP_009047207.1;MERS_ns4b;227 | 0 | 0 |
| NC_019843.3;YP_009047207.1;MERS_ns4b;228 | 0 | 0 |
| NC_019843.3;YP_009047207.1;MERS_ns4b;229 | 0 | 0.05 |
| NC_019843.3;YP_009047207.1;MERS_ns4b;230 | 0 | 0.025 |
| NC_019843.3;YP_009047207.1;MERS_ns4b;231 | 0 | 0 |
| NC_019843.3;YP_009047207.1;MERS_ns4b;232 | 0 | 0 |
| NC_019843.3;YP_009047207.1;MERS_ns4b;233 | 0.05 | 0 |
| NC_019843.3;YP_009047207.1;MERS_ns4b;234 | 0 | 0 |
| NC_019843.3;YP_009047207.1;MERS_ns4b;235 | 0.05 | 0 |
| NC_019843.3;YP_009047208.1;MERS_ns5;1 | 0 | 0.025 |
| NC_019843.3;YP_009047208.1;MERS_ns5;2 | 0 | 0.05 |
| NC_019843.3;YP_009047208.1;MERS_ns5;3 | 0.05 | 0.075 |
| NC_019843.3;YP_009047208.1;MERS_ns5;4 | 0.05 | 0.075 |
| NC_019843.3;YP_009047208.1;MERS_ns5;5 | 0.05 | 0.075 |
| NC_019843.3;YP_009047208.1;MERS_ns5;6 | 0.05 | 0.05 |
| NC_019843.3;YP_009047208.1;MERS_ns5;7 | 0 | 0.05 |
| NC_019843.3;YP_009047208.1;MERS_ns5;8 | 0 | 0.075 |
| NC_019843.3;YP_009047208.1;MERS_ns5;9 | 0.05 | 0 |
| NC_019843.3;YP_009047208.1;MERS_ns5;10 | 0 | 0 |
| NC_019843.3;YP_009047208.1;MERS_ns5;11 | 0.05 | 0 |
| NC_019843.3;YP_009047208.1;MERS_ns5;12 | 0.05 | 0.025 |
| NC_019843.3;YP_009047208.1;MERS_ns5;13 | 0.05 | 0 |
| NC_019843.3;YP_009047208.1;MERS_ns5;14 | 0 | 0.025 |
| NC_019843.3;YP_009047208.1;MERS_ns5;15 | 0 | 0 |
| NC_019843.3;YP_009047208.1;MERS_ns5;16 | 0 | 0 |
| NC_019843.3;YP_009047208.1;MERS_ns5;17 | 0 | 0 |
| NC_019843.3;YP_009047208.1;MERS_ns5;18 | 0 | 0 |
| NC_019843.3;YP_009047208.1;MERS_ns5;19 | 0 | 0 |
| NC_019843.3;YP_009047208.1;MERS_ns5;20 | 0 | 0 |
| NC_019843.3;YP_009047208.1;MERS_ns5;21 | 0 | 0 |

| | | |
|---|---|---|
| NC_019843.3;YP_009047208.1;MERS_ns5;22 | 0 | 0 |
| NC_019843.3;YP_009047208.1;MERS_ns5;23 | 0 | 0 |
| NC_019843.3;YP_009047208.1;MERS_ns5;24 | 0 | 0.05 |
| NC_019843.3;YP_009047208.1;MERS_ns5;25 | 0.05 | 0 |
| NC_019843.3;YP_009047208.1;MERS_ns5;26 | 0 | 0.025 |
| NC_019843.3;YP_009047208.1;MERS_ns5;27 | 0.05 | 0 |
| NC_019843.3;YP_009047208.1;MERS_ns5;28 | 0 | 0 |
| NC_019843.3;YP_009047208.1;MERS_ns5;29 | 0 | 0 |
| NC_019843.3;YP_009047208.1;MERS_ns5;30 | 0 | 0 |
| NC_019843.3;YP_009047208.1;MERS_ns5;31 | 0 | 0 |
| NC_019843.3;YP_009047208.1;MERS_ns5;32 | 0 | 0 |
| NC_019843.3;YP_009047208.1;MERS_ns5;33 | 0 | 0 |
| NC_019843.3;YP_009047208.1;MERS_ns5;34 | 0 | 0 |
| NC_019843.3;YP_009047208.1;MERS_ns5;35 | 0 | 0 |
| NC_019843.3;YP_009047208.1;MERS_ns5;36 | 0 | 0 |
| NC_019843.3;YP_009047208.1;MERS_ns5;37 | 0 | 0 |
| NC_019843.3;YP_009047208.1;MERS_ns5;38 | 0 | 0 |
| NC_019843.3;YP_009047208.1;MERS_ns5;39 | 0 | 0 |
| NC_019843.3;YP_009047208.1;MERS_ns5;40 | 0 | 0 |
| NC_019843.3;YP_009047208.1;MERS_ns5;41 | 0 | 0.025 |
| NC_019843.3;YP_009047208.1;MERS_ns5;42 | 0 | 0 |
| NC_019843.3;YP_009047208.1;MERS_ns5;43 | 0 | 0 |
| NC_019843.3;YP_009047208.1;MERS_ns5;44 | 0 | 0 |
| NC_019843.3;YP_009047208.1;MERS_ns5;45 | 0 | 0 |
| NC_019843.3;YP_009047208.1;MERS_ns5;46 | 0 | 0 |
| NC_019843.3;YP_009047208.1;MERS_ns5;47 | 0 | 0 |
| NC_019843.3;YP_009047208.1;MERS_ns5;48 | 0 | 0 |
| NC_019843.3;YP_009047208.1;MERS_ns5;49 | 0 | 0 |
| NC_019843.3;YP_009047208.1;MERS_ns5;50 | 0 | 0 |
| NC_019843.3;YP_009047208.1;MERS_ns5;51 | 0 | 0 |
| NC_019843.3;YP_009047208.1;MERS_ns5;52 | 0 | 0 |
| NC_019843.3;YP_009047208.1;MERS_ns5;53 | 0 | 0 |
| NC_019843.3;YP_009047208.1;MERS_ns5;54 | 0 | 0 |
| NC_019843.3;YP_009047208.1;MERS_ns5;55 | 0 | 0 |
| NC_019843.3;YP_009047208.1;MERS_ns5;56 | 0 | 0 |
| NC_019843.3;YP_009047208.1;MERS_ns5;57 | 0 | 0 |
| NC_019843.3;YP_009047208.1;MERS_ns5;58 | 0 | 0 |
| NC_019843.3;YP_009047208.1;MERS_ns5;59 | 0 | 0 |
| NC_019843.3;YP_009047208.1;MERS_ns5;60 | 0 | 0 |
| NC_019843.3;YP_009047208.1;MERS_ns5;61 | 0 | 0 |
| NC_019843.3;YP_009047208.1;MERS_ns5;62 | 0 | 0 |
| NC_019843.3;YP_009047208.1;MERS_ns5;63 | 0 | 0 |
| NC_019843.3;YP_009047208.1;MERS_ns5;64 | 0 | 0 |

| | | |
|---|---|---|
| NC_019843.3;YP_009047208.1;MERS_ns5;65 | 0 | 0 |
| NC_019843.3;YP_009047208.1;MERS_ns5;66 | 0 | 0 |
| NC_019843.3;YP_009047208.1;MERS_ns5;67 | 0 | 0 |
| NC_019843.3;YP_009047208.1;MERS_ns5;68 | 0 | 0 |
| NC_019843.3;YP_009047208.1;MERS_ns5;69 | 0 | 0 |
| NC_019843.3;YP_009047208.1;MERS_ns5;70 | 0 | 0 |
| NC_019843.3;YP_009047208.1;MERS_ns5;71 | 0 | 0 |
| NC_019843.3;YP_009047208.1;MERS_ns5;72 | 0 | 0 |
| NC_019843.3;YP_009047208.1;MERS_ns5;73 | 0 | 0 |
| NC_019843.3;YP_009047208.1;MERS_ns5;74 | 0 | 0 |
| NC_019843.3;YP_009047208.1;MERS_ns5;75 | 0 | 0 |
| NC_019843.3;YP_009047208.1;MERS_ns5;76 | 0 | 0 |
| NC_019843.3;YP_009047208.1;MERS_ns5;77 | 0 | 0 |
| NC_019843.3;YP_009047208.1;MERS_ns5;78 | 0 | 0 |
| NC_019843.3;YP_009047208.1;MERS_ns5;79 | 0 | 0 |
| NC_019843.3;YP_009047208.1;MERS_ns5;80 | 0 | 0 |
| NC_019843.3;YP_009047208.1;MERS_ns5;81 | 0 | 0 |
| NC_019843.3;YP_009047208.1;MERS_ns5;82 | 0 | 0 |
| NC_019843.3;YP_009047208.1;MERS_ns5;83 | 0 | 0 |
| NC_019843.3;YP_009047208.1;MERS_ns5;84 | 0 | 0 |
| NC_019843.3;YP_009047208.1;MERS_ns5;85 | 0 | 0 |
| NC_019843.3;YP_009047208.1;MERS_ns5;86 | 0 | 0 |
| NC_019843.3;YP_009047208.1;MERS_ns5;87 | 0 | 0 |
| NC_019843.3;YP_009047208.1;MERS_ns5;88 | 0 | 0 |
| NC_019843.3;YP_009047208.1;MERS_ns5;89 | 0 | 0 |
| NC_019843.3;YP_009047208.1;MERS_ns5;90 | 0 | 0 |
| NC_019843.3;YP_009047208.1;MERS_ns5;91 | 0 | 0 |
| NC_019843.3;YP_009047208.1;MERS_ns5;92 | 0 | 0 |
| NC_019843.3;YP_009047208.1;MERS_ns5;93 | 0 | 0 |
| NC_019843.3;YP_009047208.1;MERS_ns5;94 | 0 | 0 |
| NC_019843.3;YP_009047208.1;MERS_ns5;95 | 0 | 0 |
| NC_019843.3;YP_009047208.1;MERS_ns5;96 | 0 | 0 |
| NC_019843.3;YP_009047208.1;MERS_ns5;97 | 0 | 0 |
| NC_019843.3;YP_009047208.1;MERS_ns5;98 | 0 | 0 |
| NC_019843.3;YP_009047208.1;MERS_ns5;99 | 0 | 0 |
| NC_019843.3;YP_009047208.1;MERS_ns5;100 | 0 | 0 |
| NC_019843.3;YP_009047208.1;MERS_ns5;101 | 0 | 0 |
| NC_019843.3;YP_009047208.1;MERS_ns5;102 | 0 | 0 |
| NC_019843.3;YP_009047208.1;MERS_ns5;103 | 0 | 0 |
| NC_019843.3;YP_009047208.1;MERS_ns5;104 | 0 | 0 |
| NC_019843.3;YP_009047208.1;MERS_ns5;105 | 0 | 0 |
| NC_019843.3;YP_009047208.1;MERS_ns5;106 | 0 | 0 |
| NC_019843.3;YP_009047208.1;MERS_ns5;107 | 0 | 0 |

| | | |
|---|---|---|
| NC_019843.3;YP_009047208.1;MERS_ns5;108 | 0 | 0 |
| NC_019843.3;YP_009047208.1;MERS_ns5;109 | 0 | 0 |
| NC_019843.3;YP_009047208.1;MERS_ns5;110 | 0 | 0 |
| NC_019843.3;YP_009047208.1;MERS_ns5;111 | 0 | 0 |
| NC_019843.3;YP_009047208.1;MERS_ns5;112 | 0 | 0 |
| NC_019843.3;YP_009047208.1;MERS_ns5;113 | 0 | 0 |
| NC_019843.3;YP_009047208.1;MERS_ns5;114 | 0 | 0 |
| NC_019843.3;YP_009047208.1;MERS_ns5;115 | 0 | 0 |
| NC_019843.3;YP_009047208.1;MERS_ns5;116 | 0 | 0 |
| NC_019843.3;YP_009047208.1;MERS_ns5;117 | 0 | 0 |
| NC_019843.3;YP_009047208.1;MERS_ns5;118 | 0 | 0 |
| NC_019843.3;YP_009047208.1;MERS_ns5;119 | 0 | 0 |
| NC_019843.3;YP_009047208.1;MERS_ns5;120 | 0 | 0 |
| NC_019843.3;YP_009047208.1;MERS_ns5;121 | 0 | 0 |
| NC_019843.3;YP_009047208.1;MERS_ns5;122 | 0 | 0 |
| NC_019843.3;YP_009047208.1;MERS_ns5;123 | 0 | 0 |
| NC_019843.3;YP_009047208.1;MERS_ns5;124 | 0 | 0 |
| NC_019843.3;YP_009047208.1;MERS_ns5;125 | 0 | 0 |
| NC_019843.3;YP_009047208.1;MERS_ns5;126 | 0 | 0 |
| NC_019843.3;YP_009047208.1;MERS_ns5;127 | 0 | 0 |
| NC_019843.3;YP_009047208.1;MERS_ns5;128 | 0 | 0 |
| NC_019843.3;YP_009047208.1;MERS_ns5;129 | 0 | 0 |
| NC_019843.3;YP_009047208.1;MERS_ns5;130 | 0 | 0 |
| NC_019843.3;YP_009047208.1;MERS_ns5;131 | 0 | 0 |
| NC_019843.3;YP_009047208.1;MERS_ns5;132 | 0 | 0 |
| NC_019843.3;YP_009047208.1;MERS_ns5;133 | 0 | 0 |
| NC_019843.3;YP_009047208.1;MERS_ns5;134 | 0 | 0 |
| NC_019843.3;YP_009047208.1;MERS_ns5;135 | 0.05 | 0 |
| NC_019843.3;YP_009047208.1;MERS_ns5;136 | 0.05 | 0 |
| NC_019843.3;YP_009047208.1;MERS_ns5;137 | 0.05 | 0 |
| NC_019843.3;YP_009047208.1;MERS_ns5;138 | 0.05 | 0 |
| NC_019843.3;YP_009047208.1;MERS_ns5;139 | 0.05 | 0 |
| NC_019843.3;YP_009047208.1;MERS_ns5;140 | 0.05 | 0 |
| NC_019843.3;YP_009047208.1;MERS_ns5;141 | 0.05 | 0 |
| NC_019843.3;YP_009047208.1;MERS_ns5;142 | 0.05 | 0 |
| NC_019843.3;YP_009047208.1;MERS_ns5;143 | 0.05 | 0 |
| NC_019843.3;YP_009047208.1;MERS_ns5;144 | 0.05 | 0 |
| NC_019843.3;YP_009047208.1;MERS_ns5;145 | 0 | 0 |
| NC_019843.3;YP_009047208.1;MERS_ns5;146 | 0 | 0 |
| NC_019843.3;YP_009047208.1;MERS_ns5;147 | 0 | 0 |
| NC_019843.3;YP_009047208.1;MERS_ns5;148 | 0 | 0 |
| NC_019843.3;YP_009047208.1;MERS_ns5;149 | 0 | 0 |
| NC_019843.3;YP_009047208.1;MERS_ns5;150 | 0 | 0 |

| | | |
|---|---|---|
| NC_019843.3;YP_009047208.1;MERS_ns5;151 | 0 | 0 |
| NC_019843.3;YP_009047208.1;MERS_ns5;152 | 0 | 0 |
| NC_019843.3;YP_009047208.1;MERS_ns5;153 | 0 | 0 |
| NC_019843.3;YP_009047208.1;MERS_ns5;154 | 0 | 0 |
| NC_019843.3;YP_009047208.1;MERS_ns5;155 | 0 | 0 |
| NC_019843.3;YP_009047208.1;MERS_ns5;156 | 0 | 0 |
| NC_019843.3;YP_009047208.1;MERS_ns5;157 | 0 | 0 |
| NC_019843.3;YP_009047208.1;MERS_ns5;158 | 0 | 0 |
| NC_019843.3;YP_009047208.1;MERS_ns5;159 | 0 | 0 |
| NC_019843.3;YP_009047208.1;MERS_ns5;160 | 0 | 0 |
| NC_019843.3;YP_009047208.1;MERS_ns5;161 | 0 | 0 |
| NC_019843.3;YP_009047208.1;MERS_ns5;162 | 0 | 0 |
| NC_019843.3;YP_009047208.1;MERS_ns5;163 | 0 | 0 |
| NC_019843.3;YP_009047208.1;MERS_ns5;164 | 0 | 0 |
| NC_019843.3;YP_009047208.1;MERS_ns5;165 | 0 | 0 |
| NC_019843.3;YP_009047208.1;MERS_ns5;166 | 0 | 0 |
| NC_019843.3;YP_009047208.1;MERS_ns5;167 | 0 | 0 |
| NC_019843.3;YP_009047208.1;MERS_ns5;168 | 0 | 0 |
| NC_019843.3;YP_009047208.1;MERS_ns5;169 | 0 | 0 |
| NC_019843.3;YP_009047208.1;MERS_ns5;170 | 0 | 0 |
| NC_019843.3;YP_009047208.1;MERS_ns5;171 | 0 | 0 |
| NC_019843.3;YP_009047208.1;MERS_ns5;172 | 0 | 0 |
| NC_019843.3;YP_009047208.1;MERS_ns5;173 | 0 | 0 |
| NC_019843.3;YP_009047208.1;MERS_ns5;174 | 0 | 0 |
| NC_019843.3;YP_009047208.1;MERS_ns5;175 | 0 | 0 |
| NC_019843.3;YP_009047208.1;MERS_ns5;176 | 0 | 0 |
| NC_019843.3;YP_009047208.1;MERS_ns5;177 | 0 | 0 |
| NC_019843.3;YP_009047208.1;MERS_ns5;178 | 0 | 0 |
| NC_019843.3;YP_009047208.1;MERS_ns5;179 | 0 | 0 |
| NC_019843.3;YP_009047208.1;MERS_ns5;180 | 0 | 0 |
| NC_019843.3;YP_009047208.1;MERS_ns5;181 | 0 | 0 |
| NC_019843.3;YP_009047208.1;MERS_ns5;182 | 0 | 0 |
| NC_019843.3;YP_009047208.1;MERS_ns5;183 | 0 | 0 |
| NC_019843.3;YP_009047208.1;MERS_ns5;184 | 0 | 0.025 |
| NC_019843.3;YP_009047208.1;MERS_ns5;185 | 0 | 0 |
| NC_019843.3;YP_009047208.1;MERS_ns5;186 | 0 | 0.05 |
| NC_019843.3;YP_009047208.1;MERS_ns5;187 | 0 | 0.05 |
| NC_019843.3;YP_009047208.1;MERS_ns5;188 | 0 | 0.05 |
| NC_019843.3;YP_009047208.1;MERS_ns5;189 | 0.05 | 0 |
| NC_019843.3;YP_009047208.1;MERS_ns5;190 | 0 | 0 |
| NC_019843.3;YP_009047208.1;MERS_ns5;191 | 0 | 0 |
| NC_019843.3;YP_009047208.1;MERS_ns5;192 | 0 | 0 |
| NC_019843.3;YP_009047208.1;MERS_ns5;193 | 0 | 0 |

| | | |
|---|---|---|
| NC_019843.3;YP_009047208.1;MERS_ns5;194 | 0 | 0 |
| NC_019843.3;YP_009047208.1;MERS_ns5;195 | 0.05 | 0.05 |
| NC_019843.3;YP_009047208.1;MERS_ns5;196 | 0 | 0.025 |
| NC_019843.3;YP_009047208.1;MERS_ns5;197 | 0 | 0.025 |
| NC_019843.3;YP_009047208.1;MERS_ns5;198 | 0.05 | 0.025 |
| NC_019843.3;YP_009047208.1;MERS_ns5;199 | 0 | 0 |
| NC_019843.3;YP_009047208.1;MERS_ns5;200 | 0 | 0 |
| NC_019843.3;YP_009047208.1;MERS_ns5;201 | 0 | 0 |
| NC_019843.3;YP_009047208.1;MERS_ns5;202 | 0 | 0 |
| NC_019843.3;YP_009047208.1;MERS_ns5;203 | 0 | 0 |
| NC_019843.3;YP_009047208.1;MERS_ns5;204 | 0 | 0 |
| NC_019843.3;YP_009047208.1;MERS_ns5;205 | 0.05 | 0.025 |
| NC_019843.3;YP_009047208.1;MERS_ns5;206 | 0 | 0 |
| NC_019843.3;YP_009047208.1;MERS_ns5;207 | 0 | 0 |
| NC_019843.3;YP_009047208.1;MERS_ns5;208 | 0 | 0.025 |
| NC_019843.3;YP_009047208.1;MERS_ns5;209 | 0 | 0.025 |
| NC_019843.3;YP_009047208.1;MERS_ns5;210 | 0 | 0 |
| NC_019843.3;YP_009047208.1;MERS_ns5;211 | 0 | 0 |
| NC_019843.3;YP_009047208.1;MERS_ns5;212 | 0 | 0 |
| NC_019843.3;YP_009047208.1;MERS_ns5;213 | 0 | 0 |
| NC_019843.3;YP_009047209.1;MERS_envelope;1 | 0 | 0 |
| NC_019843.3;YP_009047209.1;MERS_envelope;2 | 0 | 0 |
| NC_019843.3;YP_009047209.1;MERS_envelope;3 | 0 | 0 |
| NC_019843.3;YP_009047209.1;MERS_envelope;4 | 0 | 0 |
| NC_019843.3;YP_009047209.1;MERS_envelope;5 | 0 | 0 |
| NC_019843.3;YP_009047209.1;MERS_envelope;6 | 0 | 0 |
| NC_019843.3;YP_009047209.1;MERS_envelope;7 | 0 | 0 |
| NC_019843.3;YP_009047209.1;MERS_envelope;8 | 0 | 0 |
| NC_019843.3;YP_009047209.1;MERS_envelope;9 | 0 | 0 |
| NC_019843.3;YP_009047209.1;MERS_envelope;10 | 0 | 0 |
| NC_019843.3;YP_009047209.1;MERS_envelope;11 | 0 | 0 |
| NC_019843.3;YP_009047209.1;MERS_envelope;12 | 0 | 0 |
| NC_019843.3;YP_009047209.1;MERS_envelope;13 | 0 | 0 |
| NC_019843.3;YP_009047209.1;MERS_envelope;14 | 0 | 0 |
| NC_019843.3;YP_009047209.1;MERS_envelope;15 | 0 | 0 |
| NC_019843.3;YP_009047209.1;MERS_envelope;16 | 0 | 0 |
| NC_019843.3;YP_009047209.1;MERS_envelope;17 | 0 | 0 |
| NC_019843.3;YP_009047209.1;MERS_envelope;18 | 0 | 0 |
| NC_019843.3;YP_009047209.1;MERS_envelope;19 | 0 | 0 |
| NC_019843.3;YP_009047209.1;MERS_envelope;20 | 0 | 0 |
| NC_019843.3;YP_009047209.1;MERS_envelope;21 | 0 | 0 |
| NC_019843.3;YP_009047209.1;MERS_envelope;22 | 0 | 0 |
| NC_019843.3;YP_009047209.1;MERS_envelope;23 | 0 | 0 |

| | | |
|---|---|---|
| NC_019843.3;YP_009047209.1;MERS_envelope;24 | 0 | 0 |
| NC_019843.3;YP_009047209.1;MERS_envelope;25 | 0 | 0 |
| NC_019843.3;YP_009047209.1;MERS_envelope;26 | 0 | 0 |
| NC_019843.3;YP_009047209.1;MERS_envelope;27 | 0 | 0 |
| NC_019843.3;YP_009047209.1;MERS_envelope;28 | 0 | 0 |
| NC_019843.3;YP_009047209.1;MERS_envelope;29 | 0 | 0 |
| NC_019843.3;YP_009047209.1;MERS_envelope;30 | 0 | 0 |
| NC_019843.3;YP_009047209.1;MERS_envelope;31 | 0 | 0 |
| NC_019843.3;YP_009047209.1;MERS_envelope;32 | 0 | 0 |
| NC_019843.3;YP_009047209.1;MERS_envelope;33 | 0 | 0 |
| NC_019843.3;YP_009047209.1;MERS_envelope;34 | 0 | 0 |
| NC_019843.3;YP_009047209.1;MERS_envelope;35 | 0 | 0 |
| NC_019843.3;YP_009047209.1;MERS_envelope;36 | 0 | 0 |
| NC_019843.3;YP_009047209.1;MERS_envelope;37 | 0 | 0 |
| NC_019843.3;YP_009047209.1;MERS_envelope;38 | 0 | 0 |
| NC_019843.3;YP_009047209.1;MERS_envelope;39 | 0 | 0 |
| NC_019843.3;YP_009047209.1;MERS_envelope;40 | 0.05 | 0 |
| NC_019843.3;YP_009047209.1;MERS_envelope;41 | 0 | 0 |
| NC_019843.3;YP_009047209.1;MERS_envelope;42 | 0 | 0 |
| NC_019843.3;YP_009047209.1;MERS_envelope;43 | 0 | 0 |
| NC_019843.3;YP_009047209.1;MERS_envelope;44 | 0 | 0 |
| NC_019843.3;YP_009047209.1;MERS_envelope;45 | 0 | 0 |
| NC_019843.3;YP_009047209.1;MERS_envelope;46 | 0 | 0 |
| NC_019843.3;YP_009047209.1;MERS_envelope;47 | 0 | 0 |
| NC_019843.3;YP_009047209.1;MERS_envelope;48 | 0 | 0 |
| NC_019843.3;YP_009047209.1;MERS_envelope;49 | 0 | 0 |
| NC_019843.3;YP_009047209.1;MERS_envelope;50 | 0 | 0 |
| NC_019843.3;YP_009047209.1;MERS_envelope;51 | 0 | 0 |
| NC_019843.3;YP_009047209.1;MERS_envelope;52 | 0 | 0 |
| NC_019843.3;YP_009047209.1;MERS_envelope;53 | 0 | 0 |
| NC_019843.3;YP_009047209.1;MERS_envelope;54 | 0 | 0 |
| NC_019843.3;YP_009047209.1;MERS_envelope;55 | 0 | 0 |
| NC_019843.3;YP_009047209.1;MERS_envelope;56 | 0 | 0 |
| NC_019843.3;YP_009047209.1;MERS_envelope;57 | 0 | 0.025 |
| NC_019843.3;YP_009047209.1;MERS_envelope;58 | 0 | 0.025 |
| NC_019843.3;YP_009047209.1;MERS_envelope;59 | 0 | 0.025 |
| NC_019843.3;YP_009047209.1;MERS_envelope;60 | 0 | 0.025 |
| NC_019843.3;YP_009047209.1;MERS_envelope;61 | 0 | 0.025 |
| NC_019843.3;YP_009047209.1;MERS_envelope;62 | 0 | 0.025 |
| NC_019843.3;YP_009047209.1;MERS_envelope;63 | 0.05 | 0.025 |
| NC_019843.3;YP_009047209.1;MERS_envelope;64 | 0 | 0 |
| NC_019843.3;YP_009047209.1;MERS_envelope;65 | 0 | 0.025 |
| NC_019843.3;YP_009047209.1;MERS_envelope;66 | 0 | 0 |

| | | |
|---|---|---|
| NC_019843.3;YP_009047209.1;MERS_envelope;67 | 0.1 | 0.075 |
| NC_019843.3;YP_009047209.1;MERS_envelope;68 | 0.1 | 0.075 |
| NC_019843.3;YP_009047209.1;MERS_envelope;69 | 0.1 | 0.05 |
| NC_019843.3;YP_009047209.1;MERS_envelope;70 | 0.05 | 0.05 |
| NC_019843.3;YP_009047209.1;MERS_envelope;71 | 0.1 | 0.025 |
| NC_019843.3;YP_009047210.1;MERS_membrane;1 | 0 | 0 |
| NC_019843.3;YP_009047210.1;MERS_membrane;2 | 0 | 0 |
| NC_019843.3;YP_009047210.1;MERS_membrane;3 | 0 | 0 |
| NC_019843.3;YP_009047210.1;MERS_membrane;4 | 0 | 0 |
| NC_019843.3;YP_009047210.1;MERS_membrane;5 | 0 | 0 |
| NC_019843.3;YP_009047210.1;MERS_membrane;6 | 0 | 0 |
| NC_019843.3;YP_009047210.1;MERS_membrane;7 | 0 | 0 |
| NC_019843.3;YP_009047210.1;MERS_membrane;8 | 0 | 0 |
| NC_019843.3;YP_009047210.1;MERS_membrane;9 | 0 | 0 |
| NC_019843.3;YP_009047210.1;MERS_membrane;10 | 0 | 0.025 |
| NC_019843.3;YP_009047210.1;MERS_membrane;11 | 0 | 0 |
| NC_019843.3;YP_009047210.1;MERS_membrane;12 | 0 | 0 |
| NC_019843.3;YP_009047210.1;MERS_membrane;13 | 0 | 0 |
| NC_019843.3;YP_009047210.1;MERS_membrane;14 | 0 | 0 |
| NC_019843.3;YP_009047210.1;MERS_membrane;15 | 0 | 0 |
| NC_019843.3;YP_009047210.1;MERS_membrane;16 | 0 | 0 |
| NC_019843.3;YP_009047210.1;MERS_membrane;17 | 0 | 0 |
| NC_019843.3;YP_009047210.1;MERS_membrane;18 | 0 | 0 |
| NC_019843.3;YP_009047210.1;MERS_membrane;19 | 0 | 0 |
| NC_019843.3;YP_009047210.1;MERS_membrane;20 | 0 | 0 |
| NC_019843.3;YP_009047210.1;MERS_membrane;21 | 0 | 0 |
| NC_019843.3;YP_009047210.1;MERS_membrane;22 | 0 | 0 |
| NC_019843.3;YP_009047210.1;MERS_membrane;23 | 0 | 0 |
| NC_019843.3;YP_009047210.1;MERS_membrane;24 | 0 | 0 |
| NC_019843.3;YP_009047210.1;MERS_membrane;25 | 0 | 0 |
| NC_019843.3;YP_009047210.1;MERS_membrane;26 | 0 | 0 |
| NC_019843.3;YP_009047210.1;MERS_membrane;27 | 0 | 0 |
| NC_019843.3;YP_009047210.1;MERS_membrane;28 | 0 | 0 |
| NC_019843.3;YP_009047210.1;MERS_membrane;29 | 0 | 0 |
| NC_019843.3;YP_009047210.1;MERS_membrane;30 | 0 | 0 |
| NC_019843.3;YP_009047210.1;MERS_membrane;31 | 0 | 0 |
| NC_019843.3;YP_009047210.1;MERS_membrane;32 | 0 | 0 |
| NC_019843.3;YP_009047210.1;MERS_membrane;33 | 0 | 0 |
| NC_019843.3;YP_009047210.1;MERS_membrane;34 | 0 | 0 |
| NC_019843.3;YP_009047210.1;MERS_membrane;35 | 0 | 0 |
| NC_019843.3;YP_009047210.1;MERS_membrane;36 | 0 | 0 |
| NC_019843.3;YP_009047210.1;MERS_membrane;37 | 0 | 0 |
| NC_019843.3;YP_009047210.1;MERS_membrane;38 | 0 | 0 |

| | | |
|---|---|---|
| NC_019843.3;YP_009047210.1;MERS_membrane;39 | 0 | 0 |
| NC_019843.3;YP_009047210.1;MERS_membrane;40 | 0 | 0 |
| NC_019843.3;YP_009047210.1;MERS_membrane;41 | 0 | 0 |
| NC_019843.3;YP_009047210.1;MERS_membrane;42 | 0 | 0 |
| NC_019843.3;YP_009047210.1;MERS_membrane;43 | 0 | 0 |
| NC_019843.3;YP_009047210.1;MERS_membrane;44 | 0 | 0 |
| NC_019843.3;YP_009047210.1;MERS_membrane;45 | 0 | 0 |
| NC_019843.3;YP_009047210.1;MERS_membrane;46 | 0 | 0 |
| NC_019843.3;YP_009047210.1;MERS_membrane;47 | 0 | 0 |
| NC_019843.3;YP_009047210.1;MERS_membrane;48 | 0 | 0 |
| NC_019843.3;YP_009047210.1;MERS_membrane;49 | 0 | 0 |
| NC_019843.3;YP_009047210.1;MERS_membrane;50 | 0 | 0 |
| NC_019843.3;YP_009047210.1;MERS_membrane;51 | 0 | 0 |
| NC_019843.3;YP_009047210.1;MERS_membrane;52 | 0 | 0 |
| NC_019843.3;YP_009047210.1;MERS_membrane;53 | 0 | 0 |
| NC_019843.3;YP_009047210.1;MERS_membrane;54 | 0 | 0 |
| NC_019843.3;YP_009047210.1;MERS_membrane;55 | 0 | 0 |
| NC_019843.3;YP_009047210.1;MERS_membrane;56 | 0 | 0 |
| NC_019843.3;YP_009047210.1;MERS_membrane;57 | 0 | 0 |
| NC_019843.3;YP_009047210.1;MERS_membrane;58 | 0 | 0 |
| NC_019843.3;YP_009047210.1;MERS_membrane;59 | 0 | 0 |
| NC_019843.3;YP_009047210.1;MERS_membrane;60 | 0 | 0 |
| NC_019843.3;YP_009047210.1;MERS_membrane;61 | 0 | 0 |
| NC_019843.3;YP_009047210.1;MERS_membrane;62 | 0 | 0 |
| NC_019843.3;YP_009047210.1;MERS_membrane;63 | 0 | 0 |
| NC_019843.3;YP_009047210.1;MERS_membrane;64 | 0 | 0 |
| NC_019843.3;YP_009047210.1;MERS_membrane;65 | 0 | 0 |
| NC_019843.3;YP_009047210.1;MERS_membrane;66 | 0 | 0 |
| NC_019843.3;YP_009047210.1;MERS_membrane;67 | 0 | 0 |
| NC_019843.3;YP_009047210.1;MERS_membrane;68 | 0 | 0 |
| NC_019843.3;YP_009047210.1;MERS_membrane;69 | 0 | 0 |
| NC_019843.3;YP_009047210.1;MERS_membrane;70 | 0.05 | 0 |
| NC_019843.3;YP_009047210.1;MERS_membrane;71 | 0 | 0 |
| NC_019843.3;YP_009047210.1;MERS_membrane;72 | 0 | 0 |
| NC_019843.3;YP_009047210.1;MERS_membrane;73 | 0 | 0 |
| NC_019843.3;YP_009047210.1;MERS_membrane;74 | 0 | 0 |
| NC_019843.3;YP_009047210.1;MERS_membrane;75 | 0 | 0 |
| NC_019843.3;YP_009047210.1;MERS_membrane;76 | 0 | 0 |
| NC_019843.3;YP_009047210.1;MERS_membrane;77 | 0 | 0 |
| NC_019843.3;YP_009047210.1;MERS_membrane;78 | 0 | 0 |
| NC_019843.3;YP_009047210.1;MERS_membrane;79 | 0 | 0 |
| NC_019843.3;YP_009047210.1;MERS_membrane;80 | 0 | 0 |
| NC_019843.3;YP_009047210.1;MERS_membrane;81 | 0 | 0 |

| | | |
|---|---|---|
| NC_019843.3;YP_009047210.1;MERS_membrane;82 | 0 | 0 |
| NC_019843.3;YP_009047210.1;MERS_membrane;83 | 0 | 0 |
| NC_019843.3;YP_009047210.1;MERS_membrane;84 | 0 | 0 |
| NC_019843.3;YP_009047210.1;MERS_membrane;85 | 0 | 0 |
| NC_019843.3;YP_009047210.1;MERS_membrane;86 | 0 | 0 |
| NC_019843.3;YP_009047210.1;MERS_membrane;87 | 0 | 0 |
| NC_019843.3;YP_009047210.1;MERS_membrane;88 | 0 | 0 |
| NC_019843.3;YP_009047210.1;MERS_membrane;89 | 0 | 0 |
| NC_019843.3;YP_009047210.1;MERS_membrane;90 | 0 | 0 |
| NC_019843.3;YP_009047210.1;MERS_membrane;91 | 0 | 0 |
| NC_019843.3;YP_009047210.1;MERS_membrane;92 | 0 | 0 |
| NC_019843.3;YP_009047210.1;MERS_membrane;93 | 0 | 0 |
| NC_019843.3;YP_009047210.1;MERS_membrane;94 | 0 | 0 |
| NC_019843.3;YP_009047210.1;MERS_membrane;95 | 0 | 0 |
| NC_019843.3;YP_009047210.1;MERS_membrane;96 | 0 | 0 |
| NC_019843.3;YP_009047210.1;MERS_membrane;97 | 0 | 0 |
| NC_019843.3;YP_009047210.1;MERS_membrane;98 | 0 | 0 |
| NC_019843.3;YP_009047210.1;MERS_membrane;99 | 0 | 0 |
| NC_019843.3;YP_009047210.1;MERS_membrane;100 | 0 | 0 |
| NC_019843.3;YP_009047210.1;MERS_membrane;101 | 0 | 0 |
| NC_019843.3;YP_009047210.1;MERS_membrane;102 | 0 | 0 |
| NC_019843.3;YP_009047210.1;MERS_membrane;103 | 0 | 0 |
| NC_019843.3;YP_009047210.1;MERS_membrane;104 | 0 | 0 |
| NC_019843.3;YP_009047210.1;MERS_membrane;105 | 0 | 0 |
| NC_019843.3;YP_009047210.1;MERS_membrane;106 | 0 | 0 |
| NC_019843.3;YP_009047210.1;MERS_membrane;107 | 0 | 0 |
| NC_019843.3;YP_009047210.1;MERS_membrane;108 | 0 | 0 |
| NC_019843.3;YP_009047210.1;MERS_membrane;109 | 0 | 0 |
| NC_019843.3;YP_009047210.1;MERS_membrane;110 | 0.05 | 0.025 |
| NC_019843.3;YP_009047210.1;MERS_membrane;111 | 0 | 0.025 |
| NC_019843.3;YP_009047210.1;MERS_membrane;112 | 0 | 0.025 |
| NC_019843.3;YP_009047210.1;MERS_membrane;113 | 0 | 0 |
| NC_019843.3;YP_009047210.1;MERS_membrane;114 | 0 | 0 |
| NC_019843.3;YP_009047210.1;MERS_membrane;115 | 0 | 0 |
| NC_019843.3;YP_009047210.1;MERS_membrane;116 | 0 | 0 |
| NC_019843.3;YP_009047210.1;MERS_membrane;117 | 0 | 0 |
| NC_019843.3;YP_009047210.1;MERS_membrane;118 | 0 | 0 |
| NC_019843.3;YP_009047210.1;MERS_membrane;119 | 0 | 0 |
| NC_019843.3;YP_009047210.1;MERS_membrane;120 | 0 | 0 |
| NC_019843.3;YP_009047210.1;MERS_membrane;121 | 0 | 0.05 |
| NC_019843.3;YP_009047210.1;MERS_membrane;122 | 0 | 0.05 |
| NC_019843.3;YP_009047210.1;MERS_membrane;123 | 0 | 0.05 |
| NC_019843.3;YP_009047210.1;MERS_membrane;124 | 0 | 0.05 |

| | | |
|---|---|---|
| NC_019843.3;YP_009047210.1;MERS_membrane;125 | 0 | 0.05 |
| NC_019843.3;YP_009047210.1;MERS_membrane;126 | 0 | 0.05 |
| NC_019843.3;YP_009047210.1;MERS_membrane;127 | 0 | 0 |
| NC_019843.3;YP_009047210.1;MERS_membrane;128 | 0 | 0 |
| NC_019843.3;YP_009047210.1;MERS_membrane;129 | 0 | 0 |
| NC_019843.3;YP_009047210.1;MERS_membrane;130 | 0 | 0 |
| NC_019843.3;YP_009047210.1;MERS_membrane;131 | 0 | 0 |
| NC_019843.3;YP_009047210.1;MERS_membrane;132 | 0.05 | 0 |
| NC_019843.3;YP_009047210.1;MERS_membrane;133 | 0 | 0 |
| NC_019843.3;YP_009047210.1;MERS_membrane;134 | 0.05 | 0 |
| NC_019843.3;YP_009047210.1;MERS_membrane;135 | 0 | 0 |
| NC_019843.3;YP_009047210.1;MERS_membrane;136 | 0 | 0 |
| NC_019843.3;YP_009047210.1;MERS_membrane;137 | 0 | 0 |
| NC_019843.3;YP_009047210.1;MERS_membrane;138 | 0 | 0.025 |
| NC_019843.3;YP_009047210.1;MERS_membrane;139 | 0 | 0 |
| NC_019843.3;YP_009047210.1;MERS_membrane;140 | 0 | 0 |
| NC_019843.3;YP_009047210.1;MERS_membrane;141 | 0 | 0 |
| NC_019843.3;YP_009047210.1;MERS_membrane;142 | 0 | 0 |
| NC_019843.3;YP_009047210.1;MERS_membrane;143 | 0 | 0 |
| NC_019843.3;YP_009047210.1;MERS_membrane;144 | 0 | 0 |
| NC_019843.3;YP_009047210.1;MERS_membrane;145 | 0 | 0 |
| NC_019843.3;YP_009047210.1;MERS_membrane;146 | 0.05 | 0 |
| NC_019843.3;YP_009047210.1;MERS_membrane;147 | 0 | 0 |
| NC_019843.3;YP_009047210.1;MERS_membrane;148 | 0 | 0 |
| NC_019843.3;YP_009047210.1;MERS_membrane;149 | 0 | 0 |
| NC_019843.3;YP_009047210.1;MERS_membrane;150 | 0 | 0 |
| NC_019843.3;YP_009047210.1;MERS_membrane;151 | 0 | 0.025 |
| NC_019843.3;YP_009047210.1;MERS_membrane;152 | 0 | 0.025 |
| NC_019843.3;YP_009047210.1;MERS_membrane;153 | 0 | 0.025 |
| NC_019843.3;YP_009047210.1;MERS_membrane;154 | 0 | 0.025 |
| NC_019843.3;YP_009047210.1;MERS_membrane;155 | 0.05 | 0.025 |
| NC_019843.3;YP_009047210.1;MERS_membrane;156 | 0.1 | 0.075 |
| NC_019843.3;YP_009047210.1;MERS_membrane;157 | 0.05 | 0 |
| NC_019843.3;YP_009047210.1;MERS_membrane;158 | 0.05 | 0 |
| NC_019843.3;YP_009047210.1;MERS_membrane;159 | 0.1 | 0 |
| NC_019843.3;YP_009047210.1;MERS_membrane;160 | 0.1 | 0.025 |
| NC_019843.3;YP_009047210.1;MERS_membrane;161 | 0.1 | 0 |
| NC_019843.3;YP_009047210.1;MERS_membrane;162 | 0.05 | 0 |
| NC_019843.3;YP_009047210.1;MERS_membrane;163 | 0 | 0 |
| NC_019843.3;YP_009047210.1;MERS_membrane;164 | 0 | 0 |
| NC_019843.3;YP_009047210.1;MERS_membrane;165 | 0 | 0 |
| NC_019843.3;YP_009047210.1;MERS_membrane;166 | 0 | 0 |
| NC_019843.3;YP_009047210.1;MERS_membrane;167 | 0 | 0 |

| | | |
|---|---|---|
| NC_019843.3;YP_009047210.1;MERS_membrane;168 | 0 | 0 |
| NC_019843.3;YP_009047210.1;MERS_membrane;169 | 0 | 0 |
| NC_019843.3;YP_009047210.1;MERS_membrane;170 | 0 | 0 |
| NC_019843.3;YP_009047210.1;MERS_membrane;171 | 0 | 0 |
| NC_019843.3;YP_009047210.1;MERS_membrane;172 | 0 | 0 |
| NC_019843.3;YP_009047210.1;MERS_membrane;173 | 0 | 0 |
| NC_019843.3;YP_009047210.1;MERS_membrane;174 | 0 | 0 |
| NC_019843.3;YP_009047210.1;MERS_membrane;175 | 0 | 0 |
| NC_019843.3;YP_009047210.1;MERS_membrane;176 | 0 | 0 |
| NC_019843.3;YP_009047210.1;MERS_membrane;177 | 0 | 0 |
| NC_019843.3;YP_009047210.1;MERS_membrane;178 | 0.05 | 0.025 |
| NC_019843.3;YP_009047210.1;MERS_membrane;179 | 0 | 0.025 |
| NC_019843.3;YP_009047210.1;MERS_membrane;180 | 0 | 0 |
| NC_019843.3;YP_009047210.1;MERS_membrane;181 | 0 | 0 |
| NC_019843.3;YP_009047210.1;MERS_membrane;182 | 0 | 0.025 |
| NC_019843.3;YP_009047210.1;MERS_membrane;183 | 0 | 0 |
| NC_019843.3;YP_009047210.1;MERS_membrane;184 | 0 | 0 |
| NC_019843.3;YP_009047210.1;MERS_membrane;185 | 0 | 0 |
| NC_019843.3;YP_009047210.1;MERS_membrane;186 | 0 | 0 |
| NC_019843.3;YP_009047210.1;MERS_membrane;187 | 0 | 0 |
| NC_019843.3;YP_009047210.1;MERS_membrane;188 | 0 | 0 |
| NC_019843.3;YP_009047210.1;MERS_membrane;189 | 0 | 0 |
| NC_019843.3;YP_009047210.1;MERS_membrane;190 | 0 | 0 |
| NC_019843.3;YP_009047210.1;MERS_membrane;191 | 0 | 0 |
| NC_019843.3;YP_009047210.1;MERS_membrane;192 | 0 | 0 |
| NC_019843.3;YP_009047210.1;MERS_membrane;193 | 0 | 0 |
| NC_019843.3;YP_009047210.1;MERS_membrane;194 | 0 | 0 |
| NC_019843.3;YP_009047210.1;MERS_membrane;195 | 0 | 0 |
| NC_019843.3;YP_009047210.1;MERS_membrane;196 | 0 | 0.025 |
| NC_019843.3;YP_009047210.1;MERS_membrane;197 | 0 | 0.025 |
| NC_019843.3;YP_009047210.1;MERS_membrane;198 | 0.05 | 0 |
| NC_019843.3;YP_009047210.1;MERS_membrane;199 | 0.05 | 0 |
| NC_019843.3;YP_009047210.1;MERS_membrane;200 | 0.05 | 0 |
| NC_019843.3;YP_009047210.1;MERS_membrane;201 | 0 | 0 |
| NC_019843.3;YP_009047210.1;MERS_membrane;202 | 0 | 0 |
| NC_019843.3;YP_009047210.1;MERS_membrane;203 | 0 | 0 |
| NC_019843.3;YP_009047210.1;MERS_membrane;204 | 0 | 0 |
| NC_019843.3;YP_009047210.1;MERS_membrane;205 | 0 | 0 |
| NC_019843.3;YP_009047210.1;MERS_membrane;206 | 0 | 0 |
| NC_019843.3;YP_009047210.1;MERS_membrane;207 | 0 | 0 |
| NC_019843.3;YP_009047210.1;MERS_membrane;208 | 0 | 0.125 |
| NC_019843.3;YP_009047211.1;MERS_nucleoprote;1 | 0.05 | 0 |
| NC_019843.3;YP_009047211.1;MERS_nucleoprote;2 | 0.1 | 0 |

| | | |
|---|---|---|
| NC_019843.3;YP_009047211.1;MERS_nucleoprote;3 | 0.15 | 0.05 |
| NC_019843.3;YP_009047211.1;MERS_nucleoprote;4 | 0.05 | 0.05 |
| NC_019843.3;YP_009047211.1;MERS_nucleoprote;5 | 0.05 | 0.05 |
| NC_019843.3;YP_009047211.1;MERS_nucleoprote;6 | 0.05 | 0.05 |
| NC_019843.3;YP_009047211.1;MERS_nucleoprote;7 | 0.1 | 0.05 |
| NC_019843.3;YP_009047211.1;MERS_nucleoprote;8 | 0.15 | 0.35 |
| NC_019843.3;YP_009047211.1;MERS_nucleoprote;9 | 0.2 | 0.3 |
| NC_019843.3;YP_009047211.1;MERS_nucleoprote;10 | 0.15 | 0.125 |
| NC_019843.3;YP_009047211.1;MERS_nucleoprote;11 | 0.2 | 0.225 |
| NC_019843.3;YP_009047211.1;MERS_nucleoprote;12 | 0.1 | 0.125 |
| NC_019843.3;YP_009047211.1;MERS_nucleoprote;13 | 0.1 | 0.1 |
| NC_019843.3;YP_009047211.1;MERS_nucleoprote;14 | 0.05 | 0.05 |
| NC_019843.3;YP_009047211.1;MERS_nucleoprote;15 | 0.05 | 0.05 |
| NC_019843.3;YP_009047211.1;MERS_nucleoprote;16 | 0.05 | 0.05 |
| NC_019843.3;YP_009047211.1;MERS_nucleoprote;17 | 0.05 | 0.05 |
| NC_019843.3;YP_009047211.1;MERS_nucleoprote;18 | 0.05 | 0.05 |
| NC_019843.3;YP_009047211.1;MERS_nucleoprote;19 | 0.05 | 0.05 |
| NC_019843.3;YP_009047211.1;MERS_nucleoprote;20 | 0.05 | 0.05 |
| NC_019843.3;YP_009047211.1;MERS_nucleoprote;21 | 0.05 | 0.025 |
| NC_019843.3;YP_009047211.1;MERS_nucleoprote;22 | 0 | 0 |
| NC_019843.3;YP_009047211.1;MERS_nucleoprote;23 | 0.05 | 0 |
| NC_019843.3;YP_009047211.1;MERS_nucleoprote;24 | 0 | 0 |
| NC_019843.3;YP_009047211.1;MERS_nucleoprote;25 | 0 | 0 |
| NC_019843.3;YP_009047211.1;MERS_nucleoprote;26 | 0 | 0 |
| NC_019843.3;YP_009047211.1;MERS_nucleoprote;27 | 0 | 0 |
| NC_019843.3;YP_009047211.1;MERS_nucleoprote;28 | 0 | 0 |
| NC_019843.3;YP_009047211.1;MERS_nucleoprote;29 | 0 | 0 |
| NC_019843.3;YP_009047211.1;MERS_nucleoprote;30 | 0 | 0 |
| NC_019843.3;YP_009047211.1;MERS_nucleoprote;31 | 0 | 0 |
| NC_019843.3;YP_009047211.1;MERS_nucleoprote;32 | 0 | 0 |
| NC_019843.3;YP_009047211.1;MERS_nucleoprote;33 | 0 | 0 |
| NC_019843.3;YP_009047211.1;MERS_nucleoprote;34 | 0 | 0 |
| NC_019843.3;YP_009047211.1;MERS_nucleoprote;35 | 0 | 0 |
| NC_019843.3;YP_009047211.1;MERS_nucleoprote;36 | 0.05 | 0 |
| NC_019843.3;YP_009047211.1;MERS_nucleoprote;37 | 0 | 0 |
| NC_019843.3;YP_009047211.1;MERS_nucleoprote;38 | 0 | 0 |
| NC_019843.3;YP_009047211.1;MERS_nucleoprote;39 | 0 | 0 |
| NC_019843.3;YP_009047211.1;MERS_nucleoprote;40 | 0 | 0 |
| NC_019843.3;YP_009047211.1;MERS_nucleoprote;41 | 0 | 0 |
| NC_019843.3;YP_009047211.1;MERS_nucleoprote;42 | 0 | 0 |
| NC_019843.3;YP_009047211.1;MERS_nucleoprote;43 | 0 | 0 |
| NC_019843.3;YP_009047211.1;MERS_nucleoprote;44 | 0 | 0 |
| NC_019843.3;YP_009047211.1;MERS_nucleoprote;45 | 0 | 0 |

| | | |
|---|---|---|
| NC_019843.3;YP_009047211.1;MERS_nucleoprote;46 | 0 | 0 |
| NC_019843.3;YP_009047211.1;MERS_nucleoprote;47 | 0 | 0 |
| NC_019843.3;YP_009047211.1;MERS_nucleoprote;48 | 0 | 0 |
| NC_019843.3;YP_009047211.1;MERS_nucleoprote;49 | 0.05 | 0.05 |
| NC_019843.3;YP_009047211.1;MERS_nucleoprote;50 | 0 | 0 |
| NC_019843.3;YP_009047211.1;MERS_nucleoprote;51 | 0 | 0 |
| NC_019843.3;YP_009047211.1;MERS_nucleoprote;52 | 0 | 0 |
| NC_019843.3;YP_009047211.1;MERS_nucleoprote;53 | 0 | 0 |
| NC_019843.3;YP_009047211.1;MERS_nucleoprote;54 | 0 | 0 |
| NC_019843.3;YP_009047211.1;MERS_nucleoprote;55 | 0.15 | 0.05 |
| NC_019843.3;YP_009047211.1;MERS_nucleoprote;56 | 0 | 0.025 |
| NC_019843.3;YP_009047211.1;MERS_nucleoprote;57 | 0 | 0.05 |
| NC_019843.3;YP_009047211.1;MERS_nucleoprote;58 | 0 | 0.025 |
| NC_019843.3;YP_009047211.1;MERS_nucleoprote;59 | 0 | 0.025 |
| NC_019843.3;YP_009047211.1;MERS_nucleoprote;60 | 0.05 | 0.05 |
| NC_019843.3;YP_009047211.1;MERS_nucleoprote;61 | 0.1 | 0.05 |
| NC_019843.3;YP_009047211.1;MERS_nucleoprote;62 | 0.1 | 0.05 |
| NC_019843.3;YP_009047211.1;MERS_nucleoprote;63 | 0.1 | 0.05 |
| NC_019843.3;YP_009047211.1;MERS_nucleoprote;64 | 0.05 | 0.025 |
| NC_019843.3;YP_009047211.1;MERS_nucleoprote;65 | 0 | 0 |
| NC_019843.3;YP_009047211.1;MERS_nucleoprote;66 | 0 | 0 |
| NC_019843.3;YP_009047211.1;MERS_nucleoprote;67 | 0.05 | 0.025 |
| NC_019843.3;YP_009047211.1;MERS_nucleoprote;68 | 0 | 0 |
| NC_019843.3;YP_009047211.1;MERS_nucleoprote;69 | 0.05 | 0.025 |
| NC_019843.3;YP_009047211.1;MERS_nucleoprote;70 | 0.05 | 0.025 |
| NC_019843.3;YP_009047211.1;MERS_nucleoprote;71 | 0.05 | 0.025 |
| NC_019843.3;YP_009047211.1;MERS_nucleoprote;72 | 0.05 | 0.025 |
| NC_019843.3;YP_009047211.1;MERS_nucleoprote;73 | 0.05 | 0.025 |
| NC_019843.3;YP_009047211.1;MERS_nucleoprote;74 | 0.05 | 0.025 |
| NC_019843.3;YP_009047211.1;MERS_nucleoprote;75 | 0.05 | 0.025 |
| NC_019843.3;YP_009047211.1;MERS_nucleoprote;76 | 0.1 | 0.025 |
| NC_019843.3;YP_009047211.1;MERS_nucleoprote;77 | 0.05 | 0.025 |
| NC_019843.3;YP_009047211.1;MERS_nucleoprote;78 | 0.05 | 0.025 |
| NC_019843.3;YP_009047211.1;MERS_nucleoprote;79 | 0.05 | 0 |
| NC_019843.3;YP_009047211.1;MERS_nucleoprote;80 | 0.05 | 0 |
| NC_019843.3;YP_009047211.1;MERS_nucleoprote;81 | 0.1 | 0 |
| NC_019843.3;YP_009047211.1;MERS_nucleoprote;82 | 0.1 | 0 |
| NC_019843.3;YP_009047211.1;MERS_nucleoprote;83 | 0.05 | 0 |
| NC_019843.3;YP_009047211.1;MERS_nucleoprote;84 | 0 | 0 |
| NC_019843.3;YP_009047211.1;MERS_nucleoprote;85 | 0 | 0 |
| NC_019843.3;YP_009047211.1;MERS_nucleoprote;86 | 0 | 0 |
| NC_019843.3;YP_009047211.1;MERS_nucleoprote;87 | 0.05 | 0 |
| NC_019843.3;YP_009047211.1;MERS_nucleoprote;88 | 0 | 0 |

| | | |
|---|---|---|
| NC_019843.3;YP_009047211.1;MERS_nucleoprote;89 | 0 | 0 |
| NC_019843.3;YP_009047211.1;MERS_nucleoprote;90 | 0 | 0 |
| NC_019843.3;YP_009047211.1;MERS_nucleoprote;91 | 0 | 0 |
| NC_019843.3;YP_009047211.1;MERS_nucleoprote;92 | 0 | 0 |
| NC_019843.3;YP_009047211.1;MERS_nucleoprote;93 | 0 | 0 |
| NC_019843.3;YP_009047211.1;MERS_nucleoprote;94 | 0 | 0 |
| NC_019843.3;YP_009047211.1;MERS_nucleoprote;95 | 0.05 | 0.025 |
| NC_019843.3;YP_009047211.1;MERS_nucleoprote;96 | 0 | 0 |
| NC_019843.3;YP_009047211.1;MERS_nucleoprote;97 | 0 | 0 |
| NC_019843.3;YP_009047211.1;MERS_nucleoprote;98 | 0 | 0 |
| NC_019843.3;YP_009047211.1;MERS_nucleoprote;99 | 0 | 0 |
| NC_019843.3;YP_009047211.1;MERS_nucleoprote;100 | 0 | 0 |
| NC_019843.3;YP_009047211.1;MERS_nucleoprote;101 | 0 | 0 |
| NC_019843.3;YP_009047211.1;MERS_nucleoprote;102 | 0 | 0 |
| NC_019843.3;YP_009047211.1;MERS_nucleoprote;103 | 0 | 0.025 |
| NC_019843.3;YP_009047211.1;MERS_nucleoprote;104 | 0 | 0.025 |
| NC_019843.3;YP_009047211.1;MERS_nucleoprote;105 | 0 | 0.025 |
| NC_019843.3;YP_009047211.1;MERS_nucleoprote;106 | 0 | 0.025 |
| NC_019843.3;YP_009047211.1;MERS_nucleoprote;107 | 0 | 0.025 |
| NC_019843.3;YP_009047211.1;MERS_nucleoprote;108 | 0 | 0 |
| NC_019843.3;YP_009047211.1;MERS_nucleoprote;109 | 0 | 0 |
| NC_019843.3;YP_009047211.1;MERS_nucleoprote;110 | 0 | 0 |
| NC_019843.3;YP_009047211.1;MERS_nucleoprote;111 | 0 | 0 |
| NC_019843.3;YP_009047211.1;MERS_nucleoprote;112 | 0 | 0 |
| NC_019843.3;YP_009047211.1;MERS_nucleoprote;113 | 0 | 0 |
| NC_019843.3;YP_009047211.1;MERS_nucleoprote;114 | 0 | 0.05 |
| NC_019843.3;YP_009047211.1;MERS_nucleoprote;115 | 0 | 0 |
| NC_019843.3;YP_009047211.1;MERS_nucleoprote;116 | 0 | 0 |
| NC_019843.3;YP_009047211.1;MERS_nucleoprote;117 | 0 | 0 |
| NC_019843.3;YP_009047211.1;MERS_nucleoprote;118 | 0 | 0 |
| NC_019843.3;YP_009047211.1;MERS_nucleoprote;119 | 0 | 0 |
| NC_019843.3;YP_009047211.1;MERS_nucleoprote;120 | 0 | 0 |
| NC_019843.3;YP_009047211.1;MERS_nucleoprote;121 | 0 | 0.025 |
| NC_019843.3;YP_009047211.1;MERS_nucleoprote;122 | 0 | 0 |
| NC_019843.3;YP_009047211.1;MERS_nucleoprote;123 | 0 | 0 |
| NC_019843.3;YP_009047211.1;MERS_nucleoprote;124 | 0 | 0 |
| NC_019843.3;YP_009047211.1;MERS_nucleoprote;125 | 0 | 0 |
| NC_019843.3;YP_009047211.1;MERS_nucleoprote;126 | 0 | 0 |
| NC_019843.3;YP_009047211.1;MERS_nucleoprote;127 | 0 | 0 |
| NC_019843.3;YP_009047211.1;MERS_nucleoprote;128 | 0.05 | 0 |
| NC_019843.3;YP_009047211.1;MERS_nucleoprote;129 | 0.05 | 0 |
| NC_019843.3;YP_009047211.1;MERS_nucleoprote;130 | 0.05 | 0 |
| NC_019843.3;YP_009047211.1;MERS_nucleoprote;131 | 0.2 | 0.05 |

| | | |
|---|---|---|
| NC_019843.3;YP_009047211.1;MERS_nucleoprote;132 | 0.1 | 0.05 |
| NC_019843.3;YP_009047211.1;MERS_nucleoprote;133 | 0.05 | 0 |
| NC_019843.3;YP_009047211.1;MERS_nucleoprote;134 | 0.05 | 0 |
| NC_019843.3;YP_009047211.1;MERS_nucleoprote;135 | 0.05 | 0 |
| NC_019843.3;YP_009047211.1;MERS_nucleoprote;136 | 0.05 | 0 |
| NC_019843.3;YP_009047211.1;MERS_nucleoprote;137 | 0.05 | 0.025 |
| NC_019843.3;YP_009047211.1;MERS_nucleoprote;138 | 0 | 0 |
| NC_019843.3;YP_009047211.1;MERS_nucleoprote;139 | 0.05 | 0 |
| NC_019843.3;YP_009047211.1;MERS_nucleoprote;140 | 0 | 0.025 |
| NC_019843.3;YP_009047211.1;MERS_nucleoprote;141 | 0 | 0 |
| NC_019843.3;YP_009047211.1;MERS_nucleoprote;142 | 0 | 0 |
| NC_019843.3;YP_009047211.1;MERS_nucleoprote;143 | 0 | 0 |
| NC_019843.3;YP_009047211.1;MERS_nucleoprote;144 | 0 | 0.025 |
| NC_019843.3;YP_009047211.1;MERS_nucleoprote;145 | 0 | 0 |
| NC_019843.3;YP_009047211.1;MERS_nucleoprote;146 | 0 | 0 |
| NC_019843.3;YP_009047211.1;MERS_nucleoprote;147 | 0 | 0 |
| NC_019843.3;YP_009047211.1;MERS_nucleoprote;148 | 0 | 0 |
| NC_019843.3;YP_009047211.1;MERS_nucleoprote;149 | 0.05 | 0 |
| NC_019843.3;YP_009047211.1;MERS_nucleoprote;150 | 0.05 | 0 |
| NC_019843.3;YP_009047211.1;MERS_nucleoprote;151 | 0 | 0 |
| NC_019843.3;YP_009047211.1;MERS_nucleoprote;152 | 0.05 | 0 |
| NC_019843.3;YP_009047211.1;MERS_nucleoprote;153 | 0 | 0 |
| NC_019843.3;YP_009047211.1;MERS_nucleoprote;154 | 0 | 0 |
| NC_019843.3;YP_009047211.1;MERS_nucleoprote;155 | 0 | 0 |
| NC_019843.3;YP_009047211.1;MERS_nucleoprote;156 | 0.05 | 0 |
| NC_019843.3;YP_009047211.1;MERS_nucleoprote;157 | 0.05 | 0 |
| NC_019843.3;YP_009047211.1;MERS_nucleoprote;158 | 0 | 0.025 |
| NC_019843.3;YP_009047211.1;MERS_nucleoprote;159 | 0 | 0 |
| NC_019843.3;YP_009047211.1;MERS_nucleoprote;160 | 0 | 0.025 |
| NC_019843.3;YP_009047211.1;MERS_nucleoprote;161 | 0 | 0 |
| NC_019843.3;YP_009047211.1;MERS_nucleoprote;162 | 0 | 0.025 |
| NC_019843.3;YP_009047211.1;MERS_nucleoprote;163 | 0.05 | 0.1 |
| NC_019843.3;YP_009047211.1;MERS_nucleoprote;164 | 0.2 | 0.2 |
| NC_019843.3;YP_009047211.1;MERS_nucleoprote;165 | 0.05 | 0.05 |
| NC_019843.3;YP_009047211.1;MERS_nucleoprote;166 | 0.05 | 0.075 |
| NC_019843.3;YP_009047211.1;MERS_nucleoprote;167 | 0.05 | 0.025 |
| NC_019843.3;YP_009047211.1;MERS_nucleoprote;168 | 0.05 | 0.05 |
| NC_019843.3;YP_009047211.1;MERS_nucleoprote;169 | 0.05 | 0.05 |
| NC_019843.3;YP_009047211.1;MERS_nucleoprote;170 | 0.05 | 0.025 |
| NC_019843.3;YP_009047211.1;MERS_nucleoprote;171 | 0.05 | 0.025 |
| NC_019843.3;YP_009047211.1;MERS_nucleoprote;172 | 0.05 | 0.1 |
| NC_019843.3;YP_009047211.1;MERS_nucleoprote;173 | 0.05 | 0.075 |
| NC_019843.3;YP_009047211.1;MERS_nucleoprote;174 | 0 | 0.025 |

| | | |
|---|---|---|
| NC_019843.3;YP_009047211.1;MERS_nucleoprote;175 | 0 | 0.075 |
| NC_019843.3;YP_009047211.1;MERS_nucleoprote;176 | 0 | 0.05 |
| NC_019843.3;YP_009047211.1;MERS_nucleoprote;177 | 0 | 0.05 |
| NC_019843.3;YP_009047211.1;MERS_nucleoprote;178 | 0 | 0.025 |
| NC_019843.3;YP_009047211.1;MERS_nucleoprote;179 | 0 | 0.05 |
| NC_019843.3;YP_009047211.1;MERS_nucleoprote;180 | 0 | 0.05 |
| NC_019843.3;YP_009047211.1;MERS_nucleoprote;181 | 0 | 0.075 |
| NC_019843.3;YP_009047211.1;MERS_nucleoprote;182 | 0 | 0.05 |
| NC_019843.3;YP_009047211.1;MERS_nucleoprote;183 | 0 | 0.025 |
| NC_019843.3;YP_009047211.1;MERS_nucleoprote;184 | 0 | 0.025 |
| NC_019843.3;YP_009047211.1;MERS_nucleoprote;185 | 0 | 0.1 |
| NC_019843.3;YP_009047211.1;MERS_nucleoprote;186 | 0 | 0.05 |
| NC_019843.3;YP_009047211.1;MERS_nucleoprote;187 | 0 | 0.025 |
| NC_019843.3;YP_009047211.1;MERS_nucleoprote;188 | 0 | 0.025 |
| NC_019843.3;YP_009047211.1;MERS_nucleoprote;189 | 0 | 0.05 |
| NC_019843.3;YP_009047211.1;MERS_nucleoprote;190 | 0 | 0.05 |
| NC_019843.3;YP_009047211.1;MERS_nucleoprote;191 | 0 | 0.025 |
| NC_019843.3;YP_009047211.1;MERS_nucleoprote;192 | 0 | 0.075 |
| NC_019843.3;YP_009047211.1;MERS_nucleoprote;193 | 0.1 | 0.15 |
| NC_019843.3;YP_009047211.1;MERS_nucleoprote;194 | 0 | 0.1 |
| NC_019843.3;YP_009047211.1;MERS_nucleoprote;195 | 0 | 0.15 |
| NC_019843.3;YP_009047211.1;MERS_nucleoprote;196 | 0.1 | 0 |
| NC_019843.3;YP_009047211.1;MERS_nucleoprote;197 | 0 | 0.05 |
| NC_019843.3;YP_009047211.1;MERS_nucleoprote;198 | 0 | 0.025 |
| NC_019843.3;YP_009047211.1;MERS_nucleoprote;199 | 0 | 0.025 |
| NC_019843.3;YP_009047211.1;MERS_nucleoprote;200 | 0 | 0 |
| NC_019843.3;YP_009047211.1;MERS_nucleoprote;201 | 0 | 0 |
| NC_019843.3;YP_009047211.1;MERS_nucleoprote;202 | 0 | 0 |
| NC_019843.3;YP_009047211.1;MERS_nucleoprote;203 | 0 | 0.025 |
| NC_019843.3;YP_009047211.1;MERS_nucleoprote;204 | 0 | 0.025 |
| NC_019843.3;YP_009047211.1;MERS_nucleoprote;205 | 0 | 0.05 |
| NC_019843.3;YP_009047211.1;MERS_nucleoprote;206 | 0 | 0.05 |
| NC_019843.3;YP_009047211.1;MERS_nucleoprote;207 | 0 | 0 |
| NC_019843.3;YP_009047211.1;MERS_nucleoprote;208 | 0.05 | 0 |
| NC_019843.3;YP_009047211.1;MERS_nucleoprote;209 | 0.05 | 0 |
| NC_019843.3;YP_009047211.1;MERS_nucleoprote;210 | 0 | 0.025 |
| NC_019843.3;YP_009047211.1;MERS_nucleoprote;211 | 0.05 | 0 |
| NC_019843.3;YP_009047211.1;MERS_nucleoprote;212 | 0 | 0 |
| NC_019843.3;YP_009047211.1;MERS_nucleoprote;213 | 0 | 0.025 |
| NC_019843.3;YP_009047211.1;MERS_nucleoprote;214 | 0.1 | 0.075 |
| NC_019843.3;YP_009047211.1;MERS_nucleoprote;215 | 0 | 0.025 |
| NC_019843.3;YP_009047211.1;MERS_nucleoprote;216 | 0 | 0.025 |
| NC_019843.3;YP_009047211.1;MERS_nucleoprote;217 | 0 | 0 |

| | | |
|---|---|---|
| NC_019843.3;YP_009047211.1;MERS_nucleoprote;218 | 0 | 0 |
| NC_019843.3;YP_009047211.1;MERS_nucleoprote;219 | 0 | 0 |
| NC_019843.3;YP_009047211.1;MERS_nucleoprote;220 | 0 | 0 |
| NC_019843.3;YP_009047211.1;MERS_nucleoprote;221 | 0 | 0 |
| NC_019843.3;YP_009047211.1;MERS_nucleoprote;222 | 0 | 0 |
| NC_019843.3;YP_009047211.1;MERS_nucleoprote;223 | 0 | 0 |
| NC_019843.3;YP_009047211.1;MERS_nucleoprote;224 | 0 | 0 |
| NC_019843.3;YP_009047211.1;MERS_nucleoprote;225 | 0.05 | 0 |
| NC_019843.3;YP_009047211.1;MERS_nucleoprote;226 | 0.05 | 0 |
| NC_019843.3;YP_009047211.1;MERS_nucleoprote;227 | 0.05 | 0 |
| NC_019843.3;YP_009047211.1;MERS_nucleoprote;228 | 0.05 | 0 |
| NC_019843.3;YP_009047211.1;MERS_nucleoprote;229 | 0.05 | 0 |
| NC_019843.3;YP_009047211.1;MERS_nucleoprote;230 | 0.05 | 0 |
| NC_019843.3;YP_009047211.1;MERS_nucleoprote;231 | 0.05 | 0 |
| NC_019843.3;YP_009047211.1;MERS_nucleoprote;232 | 0.05 | 0 |
| NC_019843.3;YP_009047211.1;MERS_nucleoprote;233 | 0 | 0 |
| NC_019843.3;YP_009047211.1;MERS_nucleoprote;234 | 0 | 0 |
| NC_019843.3;YP_009047211.1;MERS_nucleoprote;235 | 0 | 0 |
| NC_019843.3;YP_009047211.1;MERS_nucleoprote;236 | 0 | 0 |
| NC_019843.3;YP_009047211.1;MERS_nucleoprote;237 | 0 | 0 |
| NC_019843.3;YP_009047211.1;MERS_nucleoprote;238 | 0 | 0 |
| NC_019843.3;YP_009047211.1;MERS_nucleoprote;239 | 0 | 0 |
| NC_019843.3;YP_009047211.1;MERS_nucleoprote;240 | 0.05 | 0 |
| NC_019843.3;YP_009047211.1;MERS_nucleoprote;241 | 0 | 0 |
| NC_019843.3;YP_009047211.1;MERS_nucleoprote;242 | 0 | 0 |
| NC_019843.3;YP_009047211.1;MERS_nucleoprote;243 | 0 | 0 |
| NC_019843.3;YP_009047211.1;MERS_nucleoprote;244 | 0 | 0 |
| NC_019843.3;YP_009047211.1;MERS_nucleoprote;245 | 0 | 0 |
| NC_019843.3;YP_009047211.1;MERS_nucleoprote;246 | 0 | 0 |
| NC_019843.3;YP_009047211.1;MERS_nucleoprote;247 | 0 | 0 |
| NC_019843.3;YP_009047211.1;MERS_nucleoprote;248 | 0 | 0 |
| NC_019843.3;YP_009047211.1;MERS_nucleoprote;249 | 0 | 0 |
| NC_019843.3;YP_009047211.1;MERS_nucleoprote;250 | 0 | 0 |
| NC_019843.3;YP_009047211.1;MERS_nucleoprote;251 | 0 | 0 |
| NC_019843.3;YP_009047211.1;MERS_nucleoprote;252 | 0 | 0 |
| NC_019843.3;YP_009047211.1;MERS_nucleoprote;253 | 0 | 0 |
| NC_019843.3;YP_009047211.1;MERS_nucleoprote;254 | 0 | 0 |
| NC_019843.3;YP_009047211.1;MERS_nucleoprote;255 | 0 | 0 |
| NC_019843.3;YP_009047211.1;MERS_nucleoprote;256 | 0 | 0 |
| NC_019843.3;YP_009047211.1;MERS_nucleoprote;257 | 0 | 0 |
| NC_019843.3;YP_009047211.1;MERS_nucleoprote;258 | 0 | 0 |
| NC_019843.3;YP_009047211.1;MERS_nucleoprote;259 | 0 | 0 |
| NC_019843.3;YP_009047211.1;MERS_nucleoprote;260 | 0 | 0 |

| | | |
|---|---|---|
| NC_019843.3;YP_009047211.1;MERS_nucleoprote;261 | 0.05 | 0 |
| NC_019843.3;YP_009047211.1;MERS_nucleoprote;262 | 0.05 | 0 |
| NC_019843.3;YP_009047211.1;MERS_nucleoprote;263 | 0.05 | 0 |
| NC_019843.3;YP_009047211.1;MERS_nucleoprote;264 | 0.05 | 0.025 |
| NC_019843.3;YP_009047211.1;MERS_nucleoprote;265 | 0.1 | 0.025 |
| NC_019843.3;YP_009047211.1;MERS_nucleoprote;266 | 0.15 | 0.075 |
| NC_019843.3;YP_009047211.1;MERS_nucleoprote;267 | 0.1 | 0.05 |
| NC_019843.3;YP_009047211.1;MERS_nucleoprote;268 | 0.15 | 0.075 |
| NC_019843.3;YP_009047211.1;MERS_nucleoprote;269 | 0.1 | 0.1 |
| NC_019843.3;YP_009047211.1;MERS_nucleoprote;270 | 0.1 | 0.125 |
| NC_019843.3;YP_009047211.1;MERS_nucleoprote;271 | 0.05 | 0.025 |
| NC_019843.3;YP_009047211.1;MERS_nucleoprote;272 | 0.15 | 0.05 |
| NC_019843.3;YP_009047211.1;MERS_nucleoprote;273 | 0.05 | 0.1 |
| NC_019843.3;YP_009047211.1;MERS_nucleoprote;274 | 0.15 | 0.15 |
| NC_019843.3;YP_009047211.1;MERS_nucleoprote;275 | 0.05 | 0.1 |
| NC_019843.3;YP_009047211.1;MERS_nucleoprote;276 | 0.05 | 0.125 |
| NC_019843.3;YP_009047211.1;MERS_nucleoprote;277 | 0 | 0.075 |
| NC_019843.3;YP_009047211.1;MERS_nucleoprote;278 | 0.05 | 0.075 |
| NC_019843.3;YP_009047211.1;MERS_nucleoprote;279 | 0 | 0.05 |
| NC_019843.3;YP_009047211.1;MERS_nucleoprote;280 | 0 | 0.05 |
| NC_019843.3;YP_009047211.1;MERS_nucleoprote;281 | 0 | 0.025 |
| NC_019843.3;YP_009047211.1;MERS_nucleoprote;282 | 0.05 | 0.05 |
| NC_019843.3;YP_009047211.1;MERS_nucleoprote;283 | 0.05 | 0.025 |
| NC_019843.3;YP_009047211.1;MERS_nucleoprote;284 | 0.05 | 0 |
| NC_019843.3;YP_009047211.1;MERS_nucleoprote;285 | 0 | 0 |
| NC_019843.3;YP_009047211.1;MERS_nucleoprote;286 | 0 | 0 |
| NC_019843.3;YP_009047211.1;MERS_nucleoprote;287 | 0 | 0.025 |
| NC_019843.3;YP_009047211.1;MERS_nucleoprote;288 | 0 | 0.025 |
| NC_019843.3;YP_009047211.1;MERS_nucleoprote;289 | 0.05 | 0 |
| NC_019843.3;YP_009047211.1;MERS_nucleoprote;290 | 0 | 0.025 |
| NC_019843.3;YP_009047211.1;MERS_nucleoprote;291 | 0.05 | 0 |
| NC_019843.3;YP_009047211.1;MERS_nucleoprote;292 | 0 | 0.05 |
| NC_019843.3;YP_009047211.1;MERS_nucleoprote;293 | 0 | 0 |
| NC_019843.3;YP_009047211.1;MERS_nucleoprote;294 | 0 | 0 |
| NC_019843.3;YP_009047211.1;MERS_nucleoprote;295 | 0 | 0 |
| NC_019843.3;YP_009047211.1;MERS_nucleoprote;296 | 0 | 0 |
| NC_019843.3;YP_009047211.1;MERS_nucleoprote;297 | 0 | 0 |
| NC_019843.3;YP_009047211.1;MERS_nucleoprote;298 | 0 | 0 |
| NC_019843.3;YP_009047211.1;MERS_nucleoprote;299 | 0 | 0 |
| NC_019843.3;YP_009047211.1;MERS_nucleoprote;300 | 0 | 0 |
| NC_019843.3;YP_009047211.1;MERS_nucleoprote;301 | 0 | 0 |
| NC_019843.3;YP_009047211.1;MERS_nucleoprote;302 | 0 | 0 |
| NC_019843.3;YP_009047211.1;MERS_nucleoprote;303 | 0 | 0 |

| | | |
|---|---|---|
| NC_019843.3;YP_009047211.1;MERS_nucleoprote;304 | 0 | 0 |
| NC_019843.3;YP_009047211.1;MERS_nucleoprote;305 | 0 | 0 |
| NC_019843.3;YP_009047211.1;MERS_nucleoprote;306 | 0 | 0 |
| NC_019843.3;YP_009047211.1;MERS_nucleoprote;307 | 0 | 0 |
| NC_019843.3;YP_009047211.1;MERS_nucleoprote;308 | 0 | 0 |
| NC_019843.3;YP_009047211.1;MERS_nucleoprote;309 | 0 | 0.025 |
| NC_019843.3;YP_009047211.1;MERS_nucleoprote;310 | 0 | 0 |
| NC_019843.3;YP_009047211.1;MERS_nucleoprote;311 | 0 | 0 |
| NC_019843.3;YP_009047211.1;MERS_nucleoprote;312 | 0 | 0 |
| NC_019843.3;YP_009047211.1;MERS_nucleoprote;313 | 0 | 0 |
| NC_019843.3;YP_009047211.1;MERS_nucleoprote;314 | 0 | 0 |
| NC_019843.3;YP_009047211.1;MERS_nucleoprote;315 | 0 | 0 |
| NC_019843.3;YP_009047211.1;MERS_nucleoprote;316 | 0 | 0 |
| NC_019843.3;YP_009047211.1;MERS_nucleoprote;317 | 0 | 0 |
| NC_019843.3;YP_009047211.1;MERS_nucleoprote;318 | 0 | 0 |
| NC_019843.3;YP_009047211.1;MERS_nucleoprote;319 | 0 | 0 |
| NC_019843.3;YP_009047211.1;MERS_nucleoprote;320 | 0 | 0 |
| NC_019843.3;YP_009047211.1;MERS_nucleoprote;321 | 0 | 0 |
| NC_019843.3;YP_009047211.1;MERS_nucleoprote;322 | 0 | 0 |
| NC_019843.3;YP_009047211.1;MERS_nucleoprote;323 | 0 | 0 |
| NC_019843.3;YP_009047211.1;MERS_nucleoprote;324 | 0 | 0 |
| NC_019843.3;YP_009047211.1;MERS_nucleoprote;325 | 0 | 0 |
| NC_019843.3;YP_009047211.1;MERS_nucleoprote;326 | 0 | 0 |
| NC_019843.3;YP_009047211.1;MERS_nucleoprote;327 | 0 | 0 |
| NC_019843.3;YP_009047211.1;MERS_nucleoprote;328 | 0 | 0 |
| NC_019843.3;YP_009047211.1;MERS_nucleoprote;329 | 0 | 0 |
| NC_019843.3;YP_009047211.1;MERS_nucleoprote;330 | 0 | 0 |
| NC_019843.3;YP_009047211.1;MERS_nucleoprote;331 | 0 | 0 |
| NC_019843.3;YP_009047211.1;MERS_nucleoprote;332 | 0.05 | 0.025 |
| NC_019843.3;YP_009047211.1;MERS_nucleoprote;333 | 0.05 | 0 |
| NC_019843.3;YP_009047211.1;MERS_nucleoprote;334 | 0.25 | 0.05 |
| NC_019843.3;YP_009047211.1;MERS_nucleoprote;335 | 0.2 | 0.025 |
| NC_019843.3;YP_009047211.1;MERS_nucleoprote;336 | 0.05 | 0 |
| NC_019843.3;YP_009047211.1;MERS_nucleoprote;337 | 0.1 | 0.1 |
| NC_019843.3;YP_009047211.1;MERS_nucleoprote;338 | 0.1 | 0.05 |
| NC_019843.3;YP_009047211.1;MERS_nucleoprote;339 | 0.05 | 0 |
| NC_019843.3;YP_009047211.1;MERS_nucleoprote;340 | 0.15 | 0 |
| NC_019843.3;YP_009047211.1;MERS_nucleoprote;341 | 0 | 0 |
| NC_019843.3;YP_009047211.1;MERS_nucleoprote;342 | 0 | 0 |
| NC_019843.3;YP_009047211.1;MERS_nucleoprote;343 | 0 | 0 |
| NC_019843.3;YP_009047211.1;MERS_nucleoprote;344 | 0.05 | 0.025 |
| NC_019843.3;YP_009047211.1;MERS_nucleoprote;345 | 0 | 0 |
| NC_019843.3;YP_009047211.1;MERS_nucleoprote;346 | 0 | 0.025 |

| | | |
|---|---|---|
| NC_019843.3;YP_009047211.1;MERS_nucleoprote;347 | 0 | 0 |
| NC_019843.3;YP_009047211.1;MERS_nucleoprote;348 | 0.1 | 0.075 |
| NC_019843.3;YP_009047211.1;MERS_nucleoprote;349 | 0.1 | 0.05 |
| NC_019843.3;YP_009047211.1;MERS_nucleoprote;350 | 0.05 | 0.075 |
| NC_019843.3;YP_009047211.1;MERS_nucleoprote;351 | 0.05 | 0.025 |
| NC_019843.3;YP_009047211.1;MERS_nucleoprote;352 | 0.05 | 0.025 |
| NC_019843.3;YP_009047211.1;MERS_nucleoprote;353 | 0.05 | 0.025 |
| NC_019843.3;YP_009047211.1;MERS_nucleoprote;354 | 0 | 0 |
| NC_019843.3;YP_009047211.1;MERS_nucleoprote;355 | 0.05 | 0 |
| NC_019843.3;YP_009047211.1;MERS_nucleoprote;356 | 0.05 | 0 |
| NC_019843.3;YP_009047211.1;MERS_nucleoprote;357 | 0.05 | 0 |
| NC_019843.3;YP_009047211.1;MERS_nucleoprote;358 | 0.05 | 0 |
| NC_019843.3;YP_009047211.1;MERS_nucleoprote;359 | 0 | 0 |
| NC_019843.3;YP_009047211.1;MERS_nucleoprote;360 | 0 | 0 |
| NC_019843.3;YP_009047211.1;MERS_nucleoprote;361 | 0 | 0 |
| NC_019843.3;YP_009047211.1;MERS_nucleoprote;362 | 0 | 0.025 |
| NC_019843.3;YP_009047211.1;MERS_nucleoprote;363 | 0 | 0.05 |
| NC_019843.3;YP_009047211.1;MERS_nucleoprote;364 | 0 | 0.025 |
| NC_019843.3;YP_009047211.1;MERS_nucleoprote;365 | 0.05 | 0 |
| NC_019843.3;YP_009047211.1;MERS_nucleoprote;366 | 0 | 0 |
| NC_019843.3;YP_009047211.1;MERS_nucleoprote;367 | 0 | 0 |
| NC_019843.3;YP_009047211.1;MERS_nucleoprote;368 | 0 | 0 |
| NC_019843.3;YP_009047211.1;MERS_nucleoprote;369 | 0 | 0 |
| NC_019843.3;YP_009047211.1;MERS_nucleoprote;370 | 0.05 | 0.025 |
| NC_019843.3;YP_009047211.1;MERS_nucleoprote;371 | 0 | 0 |
| NC_019843.3;YP_009047211.1;MERS_nucleoprote;372 | 0 | 0 |
| NC_019843.3;YP_009047211.1;MERS_nucleoprote;373 | 0 | 0 |
| NC_019843.3;YP_009047211.1;MERS_nucleoprote;374 | 0 | 0.05 |
| NC_019843.3;YP_009047211.1;MERS_nucleoprote;375 | 0.05 | 0 |
| NC_019843.3;YP_009047211.1;MERS_nucleoprote;376 | 0 | 0 |
| NC_019843.3;YP_009047211.1;MERS_nucleoprote;377 | 0 | 0.025 |
| NC_019843.3;YP_009047211.1;MERS_nucleoprote;378 | 0 | 0 |
| NC_019843.3;YP_009047211.1;MERS_nucleoprote;379 | 0 | 0.025 |
| NC_019843.3;YP_009047211.1;MERS_nucleoprote;380 | 0.05 | 0.075 |
| NC_019843.3;YP_009047211.1;MERS_nucleoprote;381 | 0.1 | 0.15 |
| NC_019843.3;YP_009047211.1;MERS_nucleoprote;382 | 0 | 0.05 |
| NC_019843.3;YP_009047211.1;MERS_nucleoprote;383 | 0 | 0.05 |
| NC_019843.3;YP_009047211.1;MERS_nucleoprote;384 | 0 | 0.025 |
| NC_019843.3;YP_009047211.1;MERS_nucleoprote;385 | 0.05 | 0.05 |
| NC_019843.3;YP_009047211.1;MERS_nucleoprote;386 | 0 | 0.05 |
| NC_019843.3;YP_009047211.1;MERS_nucleoprote;387 | 0 | 0.05 |
| NC_019843.3;YP_009047211.1;MERS_nucleoprote;388 | 0 | 0.05 |
| NC_019843.3;YP_009047211.1;MERS_nucleoprote;389 | 0 | 0.1 |

| | | |
|---|---|---|
| NC_019843.3;YP_009047211.1;MERS_nucleoprote;390 | 0 | 0.05 |
| NC_019843.3;YP_009047211.1;MERS_nucleoprote;391 | 0 | 0.05 |
| NC_019843.3;YP_009047211.1;MERS_nucleoprote;392 | 0 | 0.025 |
| NC_019843.3;YP_009047211.1;MERS_nucleoprote;393 | 0 | 0 |
| NC_019843.3;YP_009047211.1;MERS_nucleoprote;394 | 0 | 0 |
| NC_019843.3;YP_009047211.1;MERS_nucleoprote;395 | 0 | 0 |
| NC_019843.3;YP_009047211.1;MERS_nucleoprote;396 | 0 | 0 |
| NC_019843.3;YP_009047211.1;MERS_nucleoprote;397 | 0 | 0 |
| NC_019843.3;YP_009047211.1;MERS_nucleoprote;398 | 0 | 0 |
| NC_019843.3;YP_009047211.1;MERS_nucleoprote;399 | 0 | 0 |
| NC_019843.3;YP_009047211.1;MERS_nucleoprote;400 | 0 | 0 |
| NC_019843.3;YP_009047211.1;MERS_nucleoprote;401 | 0 | 0 |
| NC_019843.3;YP_009047211.1;MERS_nucleoprote;402 | 0 | 0 |
| NC_019843.3;YP_009047212.1;MERS_orf8b;1 | 0 | 0 |
| NC_019843.3;YP_009047212.1;MERS_orf8b;2 | 0 | 0 |
| NC_019843.3;YP_009047212.1;MERS_orf8b;3 | 0 | 0.025 |
| NC_019843.3;YP_009047212.1;MERS_orf8b;4 | 0 | 0 |
| NC_019843.3;YP_009047212.1;MERS_orf8b;5 | 0 | 0 |
| NC_019843.3;YP_009047212.1;MERS_orf8b;6 | 0 | 0 |
| NC_019843.3;YP_009047212.1;MERS_orf8b;7 | 0 | 0 |
| NC_019843.3;YP_009047212.1;MERS_orf8b;8 | 0 | 0 |
| NC_019843.3;YP_009047212.1;MERS_orf8b;9 | 0 | 0.05 |
| NC_019843.3;YP_009047212.1;MERS_orf8b;10 | 0 | 0.025 |
| NC_019843.3;YP_009047212.1;MERS_orf8b;11 | 0 | 0.025 |
| NC_019843.3;YP_009047212.1;MERS_orf8b;12 | 0 | 0 |
| NC_019843.3;YP_009047212.1;MERS_orf8b;13 | 0 | 0.025 |
| NC_019843.3;YP_009047212.1;MERS_orf8b;14 | 0 | 0 |
| NC_019843.3;YP_009047212.1;MERS_orf8b;15 | 0 | 0.025 |
| NC_019843.3;YP_009047212.1;MERS_orf8b;16 | 0 | 0.025 |
| NC_019843.3;YP_009047212.1;MERS_orf8b;17 | 0 | 0.025 |
| NC_019843.3;YP_009047212.1;MERS_orf8b;18 | 0 | 0.05 |
| NC_019843.3;YP_009047212.1;MERS_orf8b;19 | 0 | 0.025 |
| NC_019843.3;YP_009047212.1;MERS_orf8b;20 | 0 | 0.075 |
| NC_019843.3;YP_009047212.1;MERS_orf8b;21 | 0 | 0.025 |
| NC_019843.3;YP_009047212.1;MERS_orf8b;22 | 0 | 0 |
| NC_019843.3;YP_009047212.1;MERS_orf8b;23 | 0 | 0 |
| NC_019843.3;YP_009047212.1;MERS_orf8b;24 | 0 | 0.025 |
| NC_019843.3;YP_009047212.1;MERS_orf8b;25 | 0.05 | 0 |
| NC_019843.3;YP_009047212.1;MERS_orf8b;26 | 0 | 0 |
| NC_019843.3;YP_009047212.1;MERS_orf8b;27 | 0 | 0 |
| NC_019843.3;YP_009047212.1;MERS_orf8b;28 | 0 | 0 |
| NC_019843.3;YP_009047212.1;MERS_orf8b;29 | 0 | 0 |
| NC_019843.3;YP_009047212.1;MERS_orf8b;30 | 0 | 0 |

| | | |
|---|---|---|
| NC_019843.3;YP_009047212.1;MERS_orf8b;31 | 0 | 0 |
| NC_019843.3;YP_009047212.1;MERS_orf8b;32 | 0 | 0 |
| NC_019843.3;YP_009047212.1;MERS_orf8b;33 | 0 | 0 |
| NC_019843.3;YP_009047212.1;MERS_orf8b;34 | 0 | 0.025 |
| NC_019843.3;YP_009047212.1;MERS_orf8b;35 | 0 | 0 |
| NC_019843.3;YP_009047212.1;MERS_orf8b;36 | 0 | 0 |
| NC_019843.3;YP_009047212.1;MERS_orf8b;37 | 0 | 0 |
| NC_019843.3;YP_009047212.1;MERS_orf8b;38 | 0 | 0 |
| NC_019843.3;YP_009047212.1;MERS_orf8b;39 | 0 | 0 |
| NC_019843.3;YP_009047212.1;MERS_orf8b;40 | 0 | 0 |
| NC_019843.3;YP_009047212.1;MERS_orf8b;41 | 0 | 0 |
| NC_019843.3;YP_009047212.1;MERS_orf8b;42 | 0 | 0 |
| NC_019843.3;YP_009047212.1;MERS_orf8b;43 | 0 | 0 |
| NC_019843.3;YP_009047212.1;MERS_orf8b;44 | 0 | 0 |
| NC_019843.3;YP_009047212.1;MERS_orf8b;45 | 0 | 0 |
| NC_019843.3;YP_009047212.1;MERS_orf8b;46 | 0 | 0 |
| NC_019843.3;YP_009047212.1;MERS_orf8b;47 | 0 | 0 |
| NC_019843.3;YP_009047212.1;MERS_orf8b;48 | 0 | 0 |
| NC_019843.3;YP_009047212.1;MERS_orf8b;49 | 0.05 | 0 |
| NC_019843.3;YP_009047212.1;MERS_orf8b;50 | 0 | 0 |
| NC_019843.3;YP_009047212.1;MERS_orf8b;51 | 0 | 0 |
| NC_019843.3;YP_009047212.1;MERS_orf8b;52 | 0 | 0 |
| NC_019843.3;YP_009047212.1;MERS_orf8b;53 | 0 | 0 |
| NC_019843.3;YP_009047212.1;MERS_orf8b;54 | 0 | 0 |
| NC_019843.3;YP_009047212.1;MERS_orf8b;55 | 0 | 0 |
| NC_019843.3;YP_009047212.1;MERS_orf8b;56 | 0 | 0 |
| NC_019843.3;YP_009047212.1;MERS_orf8b;57 | 0 | 0 |
| NC_019843.3;YP_009047212.1;MERS_orf8b;58 | 0 | 0 |
| NC_019843.3;YP_009047212.1;MERS_orf8b;59 | 0 | 0 |
| NC_019843.3;YP_009047212.1;MERS_orf8b;60 | 0 | 0 |
| NC_019843.3;YP_009047212.1;MERS_orf8b;61 | 0 | 0 |
| NC_019843.3;YP_009047212.1;MERS_orf8b;62 | 0 | 0 |
| NC_019843.3;YP_009047212.1;MERS_orf8b;63 | 0 | 0 |
| NC_019843.3;YP_009047212.1;MERS_orf8b;64 | 0 | 0 |
| NC_019843.3;YP_009047212.1;MERS_orf8b;65 | 0 | 0 |
| NC_019843.3;YP_009047212.1;MERS_orf8b;66 | 0 | 0 |
| NC_019843.3;YP_009047212.1;MERS_orf8b;67 | 0 | 0 |
| NC_019843.3;YP_009047212.1;MERS_orf8b;68 | 0 | 0 |
| NC_019843.3;YP_009047212.1;MERS_orf8b;69 | 0 | 0 |
| NC_019843.3;YP_009047212.1;MERS_orf8b;70 | 0 | 0 |
| NC_019843.3;YP_009047212.1;MERS_orf8b;71 | 0 | 0 |
| NC_019843.3;YP_009047212.1;MERS_orf8b;72 | 0 | 0 |
| NC_019843.3;YP_009047212.1;MERS_orf8b;73 | 0 | 0 |

| | | |
|---|---|---|
| NC_019843.3;YP_009047212.1;MERS_orf8b;74 | 0 | 0 |
| NC_019843.3;YP_009047212.1;MERS_orf8b;75 | 0 | 0 |
| NC_019843.3;YP_009047212.1;MERS_orf8b;76 | 0 | 0 |
| NC_019843.3;YP_009047212.1;MERS_orf8b;77 | 0 | 0 |
| NC_019843.3;YP_009047212.1;MERS_orf8b;78 | 0 | 0 |
| NC_019843.3;YP_009047212.1;MERS_orf8b;79 | 0 | 0 |
| NC_019843.3;YP_009047212.1;MERS_orf8b;80 | 0 | 0 |
| NC_019843.3;YP_009047212.1;MERS_orf8b;81 | 0 | 0 |
| NC_019843.3;YP_009047212.1;MERS_orf8b;82 | 0 | 0 |
| NC_019843.3;YP_009047212.1;MERS_orf8b;83 | 0 | 0 |
| NC_019843.3;YP_009047212.1;MERS_orf8b;84 | 0 | 0 |
| NC_019843.3;YP_009047212.1;MERS_orf8b;85 | 0 | 0 |
| NC_019843.3;YP_009047212.1;MERS_orf8b;86 | 0 | 0 |
| NC_019843.3;YP_009047212.1;MERS_orf8b;87 | 0 | 0 |
| NC_019843.3;YP_009047212.1;MERS_orf8b;88 | 0 | 0 |
| NC_019843.3;YP_009047212.1;MERS_orf8b;89 | 0 | 0 |
| NC_019843.3;YP_009047212.1;MERS_orf8b;90 | 0 | 0 |
| NC_019843.3;YP_009047212.1;MERS_orf8b;91 | 0 | 0 |
| NC_019843.3;YP_009047212.1;MERS_orf8b;92 | 0 | 0 |
| NC_019843.3;YP_009047212.1;MERS_orf8b;93 | 0 | 0 |
| NC_019843.3;YP_009047212.1;MERS_orf8b;94 | 0 | 0 |
| NC_019843.3;YP_009047212.1;MERS_orf8b;95 | 0 | 0 |
| NC_019843.3;YP_009047212.1;MERS_orf8b;96 | 0 | 0 |
| NC_019843.3;YP_009047212.1;MERS_orf8b;97 | 0 | 0.025 |
| NC_019843.3;YP_009047212.1;MERS_orf8b;98 | 0 | 0 |
| NC_019843.3;YP_009047212.1;MERS_orf8b;99 | 0.1 | 0.075 |
| NC_019843.3;YP_009047212.1;MERS_orf8b;100 | 0 | 0 |
| NC_019843.3;YP_009047212.1;MERS_orf8b;101 | 0 | 0 |
| NC_006577.2;YP_173236.1;HCoV-HKU1_orf1ab;1 | 0 | 0 |
| NC_006577.2;YP_173236.1;HCoV-HKU1_orf1ab;2 | 0 | 0 |
| NC_006577.2;YP_173236.1;HCoV-HKU1_orf1ab;3 | 0.05 | 0 |
| NC_006577.2;YP_173236.1;HCoV-HKU1_orf1ab;4 | 0 | 0 |
| NC_006577.2;YP_173236.1;HCoV-HKU1_orf1ab;5 | 0 | 0 |
| NC_006577.2;YP_173236.1;HCoV-HKU1_orf1ab;6 | 0 | 0 |
| NC_006577.2;YP_173236.1;HCoV-HKU1_orf1ab;7 | 0 | 0 |
| NC_006577.2;YP_173236.1;HCoV-HKU1_orf1ab;8 | 0 | 0 |
| NC_006577.2;YP_173236.1;HCoV-HKU1_orf1ab;9 | 0 | 0 |
| NC_006577.2;YP_173236.1;HCoV-HKU1_orf1ab;10 | 0 | 0 |
| NC_006577.2;YP_173236.1;HCoV-HKU1_orf1ab;11 | 0 | 0.025 |
| NC_006577.2;YP_173236.1;HCoV-HKU1_orf1ab;12 | 0 | 0 |
| NC_006577.2;YP_173236.1;HCoV-HKU1_orf1ab;13 | 0 | 0 |
| NC_006577.2;YP_173236.1;HCoV-HKU1_orf1ab;14 | 0 | 0 |
| NC_006577.2;YP_173236.1;HCoV-HKU1_orf1ab;15 | 0 | 0 |

| | | |
|---|---|---|
| NC_006577.2;YP_173236.1;HCoV-HKU1_orf1ab;16 | 0 | 0 |
| NC_006577.2;YP_173236.1;HCoV-HKU1_orf1ab;17 | 0 | 0 |
| NC_006577.2;YP_173236.1;HCoV-HKU1_orf1ab;18 | 0.05 | 0.025 |
| NC_006577.2;YP_173236.1;HCoV-HKU1_orf1ab;19 | 0.05 | 0 |
| NC_006577.2;YP_173236.1;HCoV-HKU1_orf1ab;20 | 0 | 0 |
| NC_006577.2;YP_173236.1;HCoV-HKU1_orf1ab;21 | 0 | 0 |
| NC_006577.2;YP_173236.1;HCoV-HKU1_orf1ab;22 | 0 | 0 |
| NC_006577.2;YP_173236.1;HCoV-HKU1_orf1ab;23 | 0 | 0 |
| NC_006577.2;YP_173236.1;HCoV-HKU1_orf1ab;24 | 0 | 0 |
| NC_006577.2;YP_173236.1;HCoV-HKU1_orf1ab;25 | 0 | 0 |
| NC_006577.2;YP_173236.1;HCoV-HKU1_orf1ab;26 | 0.05 | 0 |
| NC_006577.2;YP_173236.1;HCoV-HKU1_orf1ab;27 | 0.1 | 0 |
| NC_006577.2;YP_173236.1;HCoV-HKU1_orf1ab;28 | 0.05 | 0 |
| NC_006577.2;YP_173236.1;HCoV-HKU1_orf1ab;29 | 0.05 | 0.025 |
| NC_006577.2;YP_173236.1;HCoV-HKU1_orf1ab;30 | 0.05 | 0.025 |
| NC_006577.2;YP_173236.1;HCoV-HKU1_orf1ab;31 | 0.05 | 0 |
| NC_006577.2;YP_173236.1;HCoV-HKU1_orf1ab;32 | 0.05 | 0.025 |
| NC_006577.2;YP_173236.1;HCoV-HKU1_orf1ab;33 | 0.05 | 0.025 |
| NC_006577.2;YP_173236.1;HCoV-HKU1_orf1ab;34 | 0.05 | 0 |
| NC_006577.2;YP_173236.1;HCoV-HKU1_orf1ab;35 | 0.05 | 0 |
| NC_006577.2;YP_173236.1;HCoV-HKU1_orf1ab;36 | 0.05 | 0.025 |
| NC_006577.2;YP_173236.1;HCoV-HKU1_orf1ab;37 | 0 | 0.025 |
| NC_006577.2;YP_173236.1;HCoV-HKU1_orf1ab;38 | 0 | 0 |
| NC_006577.2;YP_173236.1;HCoV-HKU1_orf1ab;39 | 0 | 0 |
| NC_006577.2;YP_173236.1;HCoV-HKU1_orf1ab;40 | 0 | 0 |
| NC_006577.2;YP_173236.1;HCoV-HKU1_orf1ab;41 | 0 | 0 |
| NC_006577.2;YP_173236.1;HCoV-HKU1_orf1ab;42 | 0 | 0 |
| NC_006577.2;YP_173236.1;HCoV-HKU1_orf1ab;43 | 0 | 0 |
| NC_006577.2;YP_173236.1;HCoV-HKU1_orf1ab;44 | 0 | 0 |
| NC_006577.2;YP_173236.1;HCoV-HKU1_orf1ab;45 | 0 | 0 |
| NC_006577.2;YP_173236.1;HCoV-HKU1_orf1ab;46 | 0 | 0 |
| NC_006577.2;YP_173236.1;HCoV-HKU1_orf1ab;47 | 0.05 | 0 |
| NC_006577.2;YP_173236.1;HCoV-HKU1_orf1ab;48 | 0 | 0 |
| NC_006577.2;YP_173236.1;HCoV-HKU1_orf1ab;49 | 0 | 0 |
| NC_006577.2;YP_173236.1;HCoV-HKU1_orf1ab;50 | 0 | 0 |
| NC_006577.2;YP_173236.1;HCoV-HKU1_orf1ab;51 | 0 | 0 |
| NC_006577.2;YP_173236.1;HCoV-HKU1_orf1ab;52 | 0 | 0 |
| NC_006577.2;YP_173236.1;HCoV-HKU1_orf1ab;53 | 0 | 0 |
| NC_006577.2;YP_173236.1;HCoV-HKU1_orf1ab;54 | 0 | 0 |
| NC_006577.2;YP_173236.1;HCoV-HKU1_orf1ab;55 | 0 | 0 |
| NC_006577.2;YP_173236.1;HCoV-HKU1_orf1ab;56 | 0 | 0 |
| NC_006577.2;YP_173236.1;HCoV-HKU1_orf1ab;57 | 0 | 0 |
| NC_006577.2;YP_173236.1;HCoV-HKU1_orf1ab;58 | 0 | 0 |

| | | |
|---|---|---|
| NC_006577.2;YP_173236.1;HCoV-HKU1_orf1ab;59 | 0 | 0 |
| NC_006577.2;YP_173236.1;HCoV-HKU1_orf1ab;60 | 0 | 0 |
| NC_006577.2;YP_173236.1;HCoV-HKU1_orf1ab;61 | 0 | 0 |
| NC_006577.2;YP_173236.1;HCoV-HKU1_orf1ab;62 | 0 | 0 |
| NC_006577.2;YP_173236.1;HCoV-HKU1_orf1ab;63 | 0 | 0 |
| NC_006577.2;YP_173236.1;HCoV-HKU1_orf1ab;64 | 0 | 0 |
| NC_006577.2;YP_173236.1;HCoV-HKU1_orf1ab;65 | 0 | 0 |
| NC_006577.2;YP_173236.1;HCoV-HKU1_orf1ab;66 | 0 | 0 |
| NC_006577.2;YP_173236.1;HCoV-HKU1_orf1ab;67 | 0 | 0 |
| NC_006577.2;YP_173236.1;HCoV-HKU1_orf1ab;68 | 0 | 0 |
| NC_006577.2;YP_173236.1;HCoV-HKU1_orf1ab;69 | 0 | 0 |
| NC_006577.2;YP_173236.1;HCoV-HKU1_orf1ab;70 | 0 | 0 |
| NC_006577.2;YP_173236.1;HCoV-HKU1_orf1ab;71 | 0.05 | 0 |
| NC_006577.2;YP_173236.1;HCoV-HKU1_orf1ab;72 | 0 | 0 |
| NC_006577.2;YP_173236.1;HCoV-HKU1_orf1ab;73 | 0 | 0 |
| NC_006577.2;YP_173236.1;HCoV-HKU1_orf1ab;74 | 0 | 0 |
| NC_006577.2;YP_173236.1;HCoV-HKU1_orf1ab;75 | 0 | 0 |
| NC_006577.2;YP_173236.1;HCoV-HKU1_orf1ab;76 | 0 | 0 |
| NC_006577.2;YP_173236.1;HCoV-HKU1_orf1ab;77 | 0 | 0 |
| NC_006577.2;YP_173236.1;HCoV-HKU1_orf1ab;78 | 0 | 0 |
| NC_006577.2;YP_173236.1;HCoV-HKU1_orf1ab;79 | 0 | 0.025 |
| NC_006577.2;YP_173236.1;HCoV-HKU1_orf1ab;80 | 0 | 0.025 |
| NC_006577.2;YP_173236.1;HCoV-HKU1_orf1ab;81 | 0 | 0.025 |
| NC_006577.2;YP_173236.1;HCoV-HKU1_orf1ab;82 | 0 | 0 |
| NC_006577.2;YP_173236.1;HCoV-HKU1_orf1ab;83 | 0.1 | 0.025 |
| NC_006577.2;YP_173236.1;HCoV-HKU1_orf1ab;84 | 0.1 | 0 |
| NC_006577.2;YP_173236.1;HCoV-HKU1_orf1ab;85 | 0.1 | 0.025 |
| NC_006577.2;YP_173236.1;HCoV-HKU1_orf1ab;86 | 0.1 | 0 |
| NC_006577.2;YP_173236.1;HCoV-HKU1_orf1ab;87 | 0.05 | 0 |
| NC_006577.2;YP_173236.1;HCoV-HKU1_orf1ab;88 | 0 | 0 |
| NC_006577.2;YP_173236.1;HCoV-HKU1_orf1ab;89 | 0.05 | 0 |
| NC_006577.2;YP_173236.1;HCoV-HKU1_orf1ab;90 | 0.05 | 0 |
| NC_006577.2;YP_173236.1;HCoV-HKU1_orf1ab;91 | 0.05 | 0 |
| NC_006577.2;YP_173236.1;HCoV-HKU1_orf1ab;92 | 0 | 0 |
| NC_006577.2;YP_173236.1;HCoV-HKU1_orf1ab;93 | 0.1 | 0 |
| NC_006577.2;YP_173236.1;HCoV-HKU1_orf1ab;94 | 0.1 | 0 |
| NC_006577.2;YP_173236.1;HCoV-HKU1_orf1ab;95 | 0 | 0 |
| NC_006577.2;YP_173236.1;HCoV-HKU1_orf1ab;96 | 0.1 | 0 |
| NC_006577.2;YP_173236.1;HCoV-HKU1_orf1ab;97 | 0.1 | 0 |
| NC_006577.2;YP_173236.1;HCoV-HKU1_orf1ab;98 | 0.1 | 0.025 |
| NC_006577.2;YP_173236.1;HCoV-HKU1_orf1ab;99 | 0 | 0 |
| NC_006577.2;YP_173236.1;HCoV-HKU1_orf1ab;100 | 0 | 0 |
| NC_006577.2;YP_173236.1;HCoV-HKU1_orf1ab;101 | 0 | 0 |

| | | |
|---|---|---|
| NC_006577.2;YP_173236.1;HCoV-HKU1_orf1ab;102 | 0 | 0 |
| NC_006577.2;YP_173236.1;HCoV-HKU1_orf1ab;103 | 0 | 0 |
| NC_006577.2;YP_173236.1;HCoV-HKU1_orf1ab;104 | 0 | 0 |
| NC_006577.2;YP_173236.1;HCoV-HKU1_orf1ab;105 | 0 | 0 |
| NC_006577.2;YP_173236.1;HCoV-HKU1_orf1ab;106 | 0 | 0 |
| NC_006577.2;YP_173236.1;HCoV-HKU1_orf1ab;107 | 0 | 0 |
| NC_006577.2;YP_173236.1;HCoV-HKU1_orf1ab;108 | 0 | 0.025 |
| NC_006577.2;YP_173236.1;HCoV-HKU1_orf1ab;109 | 0 | 0 |
| NC_006577.2;YP_173236.1;HCoV-HKU1_orf1ab;110 | 0 | 0 |
| NC_006577.2;YP_173236.1;HCoV-HKU1_orf1ab;111 | 0 | 0 |
| NC_006577.2;YP_173236.1;HCoV-HKU1_orf1ab;112 | 0 | 0 |
| NC_006577.2;YP_173236.1;HCoV-HKU1_orf1ab;113 | 0 | 0 |
| NC_006577.2;YP_173236.1;HCoV-HKU1_orf1ab;114 | 0 | 0 |
| NC_006577.2;YP_173236.1;HCoV-HKU1_orf1ab;115 | 0 | 0 |
| NC_006577.2;YP_173236.1;HCoV-HKU1_orf1ab;116 | 0 | 0 |
| NC_006577.2;YP_173236.1;HCoV-HKU1_orf1ab;117 | 0 | 0 |
| NC_006577.2;YP_173236.1;HCoV-HKU1_orf1ab;118 | 0 | 0 |
| NC_006577.2;YP_173236.1;HCoV-HKU1_orf1ab;119 | 0 | 0 |
| NC_006577.2;YP_173236.1;HCoV-HKU1_orf1ab;120 | 0 | 0 |
| NC_006577.2;YP_173236.1;HCoV-HKU1_orf1ab;121 | 0 | 0 |
| NC_006577.2;YP_173236.1;HCoV-HKU1_orf1ab;122 | 0 | 0 |
| NC_006577.2;YP_173236.1;HCoV-HKU1_orf1ab;123 | 0 | 0 |
| NC_006577.2;YP_173236.1;HCoV-HKU1_orf1ab;124 | 0 | 0 |
| NC_006577.2;YP_173236.1;HCoV-HKU1_orf1ab;125 | 0 | 0 |
| NC_006577.2;YP_173236.1;HCoV-HKU1_orf1ab;126 | 0 | 0 |
| NC_006577.2;YP_173236.1;HCoV-HKU1_orf1ab;127 | 0 | 0 |
| NC_006577.2;YP_173236.1;HCoV-HKU1_orf1ab;128 | 0 | 0 |
| NC_006577.2;YP_173236.1;HCoV-HKU1_orf1ab;129 | 0 | 0 |
| NC_006577.2;YP_173236.1;HCoV-HKU1_orf1ab;130 | 0 | 0.075 |
| NC_006577.2;YP_173236.1;HCoV-HKU1_orf1ab;131 | 0 | 0 |
| NC_006577.2;YP_173236.1;HCoV-HKU1_orf1ab;132 | 0 | 0 |
| NC_006577.2;YP_173236.1;HCoV-HKU1_orf1ab;133 | 0 | 0 |
| NC_006577.2;YP_173236.1;HCoV-HKU1_orf1ab;134 | 0 | 0 |
| NC_006577.2;YP_173236.1;HCoV-HKU1_orf1ab;135 | 0 | 0 |
| NC_006577.2;YP_173236.1;HCoV-HKU1_orf1ab;136 | 0 | 0.025 |
| NC_006577.2;YP_173236.1;HCoV-HKU1_orf1ab;137 | 0 | 0.05 |
| NC_006577.2;YP_173236.1;HCoV-HKU1_orf1ab;138 | 0 | 0 |
| NC_006577.2;YP_173236.1;HCoV-HKU1_orf1ab;139 | 0 | 0 |
| NC_006577.2;YP_173236.1;HCoV-HKU1_orf1ab;140 | 0 | 0 |
| NC_006577.2;YP_173236.1;HCoV-HKU1_orf1ab;141 | 0 | 0 |
| NC_006577.2;YP_173236.1;HCoV-HKU1_orf1ab;142 | 0 | 0 |
| NC_006577.2;YP_173236.1;HCoV-HKU1_orf1ab;143 | 0 | 0 |
| NC_006577.2;YP_173236.1;HCoV-HKU1_orf1ab;144 | 0 | 0 |

| | | |
|---|---|---|
| NC_006577.2;YP_173236.1;HCoV-HKU1_orf1ab;145 | 0 | 0 |
| NC_006577.2;YP_173236.1;HCoV-HKU1_orf1ab;146 | 0 | 0 |
| NC_006577.2;YP_173236.1;HCoV-HKU1_orf1ab;147 | 0 | 0.025 |
| NC_006577.2;YP_173236.1;HCoV-HKU1_orf1ab;148 | 0 | 0 |
| NC_006577.2;YP_173236.1;HCoV-HKU1_orf1ab;149 | 0 | 0 |
| NC_006577.2;YP_173236.1;HCoV-HKU1_orf1ab;150 | 0 | 0 |
| NC_006577.2;YP_173236.1;HCoV-HKU1_orf1ab;151 | 0 | 0 |
| NC_006577.2;YP_173236.1;HCoV-HKU1_orf1ab;152 | 0 | 0 |
| NC_006577.2;YP_173236.1;HCoV-HKU1_orf1ab;153 | 0 | 0 |
| NC_006577.2;YP_173236.1;HCoV-HKU1_orf1ab;154 | 0 | 0 |
| NC_006577.2;YP_173236.1;HCoV-HKU1_orf1ab;155 | 0 | 0 |
| NC_006577.2;YP_173236.1;HCoV-HKU1_orf1ab;156 | 0 | 0 |
| NC_006577.2;YP_173236.1;HCoV-HKU1_orf1ab;157 | 0 | 0 |
| NC_006577.2;YP_173236.1;HCoV-HKU1_orf1ab;158 | 0 | 0 |
| NC_006577.2;YP_173236.1;HCoV-HKU1_orf1ab;159 | 0 | 0 |
| NC_006577.2;YP_173236.1;HCoV-HKU1_orf1ab;160 | 0 | 0 |
| NC_006577.2;YP_173236.1;HCoV-HKU1_orf1ab;161 | 0 | 0 |
| NC_006577.2;YP_173236.1;HCoV-HKU1_orf1ab;162 | 0 | 0 |
| NC_006577.2;YP_173236.1;HCoV-HKU1_orf1ab;163 | 0 | 0 |
| NC_006577.2;YP_173236.1;HCoV-HKU1_orf1ab;164 | 0 | 0 |
| NC_006577.2;YP_173236.1;HCoV-HKU1_orf1ab;165 | 0 | 0 |
| NC_006577.2;YP_173236.1;HCoV-HKU1_orf1ab;166 | 0 | 0 |
| NC_006577.2;YP_173236.1;HCoV-HKU1_orf1ab;167 | 0 | 0 |
| NC_006577.2;YP_173236.1;HCoV-HKU1_orf1ab;168 | 0 | 0 |
| NC_006577.2;YP_173236.1;HCoV-HKU1_orf1ab;169 | 0 | 0 |
| NC_006577.2;YP_173236.1;HCoV-HKU1_orf1ab;170 | 0 | 0 |
| NC_006577.2;YP_173236.1;HCoV-HKU1_orf1ab;171 | 0 | 0.025 |
| NC_006577.2;YP_173236.1;HCoV-HKU1_orf1ab;172 | 0 | 0 |
| NC_006577.2;YP_173236.1;HCoV-HKU1_orf1ab;173 | 0 | 0 |
| NC_006577.2;YP_173236.1;HCoV-HKU1_orf1ab;174 | 0 | 0 |
| NC_006577.2;YP_173236.1;HCoV-HKU1_orf1ab;175 | 0 | 0.025 |
| NC_006577.2;YP_173236.1;HCoV-HKU1_orf1ab;176 | 0 | 0.05 |
| NC_006577.2;YP_173236.1;HCoV-HKU1_orf1ab;177 | 0 | 0.05 |
| NC_006577.2;YP_173236.1;HCoV-HKU1_orf1ab;178 | 0 | 0.025 |
| NC_006577.2;YP_173236.1;HCoV-HKU1_orf1ab;179 | 0 | 0 |
| NC_006577.2;YP_173236.1;HCoV-HKU1_orf1ab;180 | 0 | 0 |
| NC_006577.2;YP_173236.1;HCoV-HKU1_orf1ab;181 | 0 | 0.025 |
| NC_006577.2;YP_173236.1;HCoV-HKU1_orf1ab;182 | 0.1 | 0.05 |
| NC_006577.2;YP_173236.1;HCoV-HKU1_orf1ab;183 | 0.05 | 0 |
| NC_006577.2;YP_173236.1;HCoV-HKU1_orf1ab;184 | 0 | 0 |
| NC_006577.2;YP_173236.1;HCoV-HKU1_orf1ab;185 | 0 | 0 |
| NC_006577.2;YP_173236.1;HCoV-HKU1_orf1ab;186 | 0 | 0 |
| NC_006577.2;YP_173236.1;HCoV-HKU1_orf1ab;187 | 0 | 0 |

| | | |
|---|---|---|
| NC_006577.2;YP_173236.1;HCoV-HKU1_orf1ab;188 | 0 | 0 |
| NC_006577.2;YP_173236.1;HCoV-HKU1_orf1ab;189 | 0 | 0 |
| NC_006577.2;YP_173236.1;HCoV-HKU1_orf1ab;190 | 0 | 0 |
| NC_006577.2;YP_173236.1;HCoV-HKU1_orf1ab;191 | 0.05 | 0 |
| NC_006577.2;YP_173236.1;HCoV-HKU1_orf1ab;192 | 0 | 0 |
| NC_006577.2;YP_173236.1;HCoV-HKU1_orf1ab;193 | 0.05 | 0 |
| NC_006577.2;YP_173236.1;HCoV-HKU1_orf1ab;194 | 0.05 | 0.05 |
| NC_006577.2;YP_173236.1;HCoV-HKU1_orf1ab;195 | 0 | 0.025 |
| NC_006577.2;YP_173236.1;HCoV-HKU1_orf1ab;196 | 0 | 0.025 |
| NC_006577.2;YP_173236.1;HCoV-HKU1_orf1ab;197 | 0 | 0 |
| NC_006577.2;YP_173236.1;HCoV-HKU1_orf1ab;198 | 0 | 0 |
| NC_006577.2;YP_173236.1;HCoV-HKU1_orf1ab;199 | 0 | 0 |
| NC_006577.2;YP_173236.1;HCoV-HKU1_orf1ab;200 | 0 | 0 |
| NC_006577.2;YP_173236.1;HCoV-HKU1_orf1ab;201 | 0 | 0.025 |
| NC_006577.2;YP_173236.1;HCoV-HKU1_orf1ab;202 | 0 | 0.025 |
| NC_006577.2;YP_173236.1;HCoV-HKU1_orf1ab;203 | 0 | 0.025 |
| NC_006577.2;YP_173236.1;HCoV-HKU1_orf1ab;204 | 0 | 0.025 |
| NC_006577.2;YP_173236.1;HCoV-HKU1_orf1ab;205 | 0 | 0.025 |
| NC_006577.2;YP_173236.1;HCoV-HKU1_orf1ab;206 | 0 | 0 |
| NC_006577.2;YP_173236.1;HCoV-HKU1_orf1ab;207 | 0 | 0.025 |
| NC_006577.2;YP_173236.1;HCoV-HKU1_orf1ab;208 | 0 | 0 |
| NC_006577.2;YP_173236.1;HCoV-HKU1_orf1ab;209 | 0 | 0 |
| NC_006577.2;YP_173236.1;HCoV-HKU1_orf1ab;210 | 0 | 0 |
| NC_006577.2;YP_173236.1;HCoV-HKU1_orf1ab;211 | 0 | 0 |
| NC_006577.2;YP_173236.1;HCoV-HKU1_orf1ab;212 | 0 | 0 |
| NC_006577.2;YP_173236.1;HCoV-HKU1_orf1ab;213 | 0 | 0 |
| NC_006577.2;YP_173236.1;HCoV-HKU1_orf1ab;214 | 0 | 0 |
| NC_006577.2;YP_173236.1;HCoV-HKU1_orf1ab;215 | 0 | 0 |
| NC_006577.2;YP_173236.1;HCoV-HKU1_orf1ab;216 | 0 | 0 |
| NC_006577.2;YP_173236.1;HCoV-HKU1_orf1ab;217 | 0 | 0 |
| NC_006577.2;YP_173236.1;HCoV-HKU1_orf1ab;218 | 0 | 0 |
| NC_006577.2;YP_173236.1;HCoV-HKU1_orf1ab;219 | 0 | 0 |
| NC_006577.2;YP_173236.1;HCoV-HKU1_orf1ab;220 | 0 | 0 |
| NC_006577.2;YP_173236.1;HCoV-HKU1_orf1ab;221 | 0 | 0 |
| NC_006577.2;YP_173236.1;HCoV-HKU1_orf1ab;222 | 0 | 0 |
| NC_006577.2;YP_173236.1;HCoV-HKU1_orf1ab;223 | 0 | 0 |
| NC_006577.2;YP_173236.1;HCoV-HKU1_orf1ab;224 | 0 | 0 |
| NC_006577.2;YP_173236.1;HCoV-HKU1_orf1ab;225 | 0 | 0 |
| NC_006577.2;YP_173236.1;HCoV-HKU1_orf1ab;226 | 0 | 0 |
| NC_006577.2;YP_173236.1;HCoV-HKU1_orf1ab;227 | 0.05 | 0 |
| NC_006577.2;YP_173236.1;HCoV-HKU1_orf1ab;228 | 0 | 0.025 |
| NC_006577.2;YP_173236.1;HCoV-HKU1_orf1ab;229 | 0 | 0 |
| NC_006577.2;YP_173236.1;HCoV-HKU1_orf1ab;230 | 0 | 0.025 |

| | | |
|---|---|---|
| NC_006577.2;YP_173236.1;HCoV-HKU1_orf1ab;231 | 0 | 0.025 |
| NC_006577.2;YP_173236.1;HCoV-HKU1_orf1ab;232 | 0 | 0.025 |
| NC_006577.2;YP_173236.1;HCoV-HKU1_orf1ab;233 | 0.05 | 0.025 |
| NC_006577.2;YP_173236.1;HCoV-HKU1_orf1ab;234 | 0 | 0 |
| NC_006577.2;YP_173236.1;HCoV-HKU1_orf1ab;235 | 0.05 | 0 |
| NC_006577.2;YP_173236.1;HCoV-HKU1_orf1ab;236 | 0.05 | 0 |
| NC_006577.2;YP_173236.1;HCoV-HKU1_orf1ab;237 | 0.05 | 0 |
| NC_006577.2;YP_173236.1;HCoV-HKU1_orf1ab;238 | 0 | 0 |
| NC_006577.2;YP_173236.1;HCoV-HKU1_orf1ab;239 | 0 | 0.05 |
| NC_006577.2;YP_173236.1;HCoV-HKU1_orf1ab;240 | 0 | 0 |
| NC_006577.2;YP_173236.1;HCoV-HKU1_orf1ab;241 | 0 | 0 |
| NC_006577.2;YP_173236.1;HCoV-HKU1_orf1ab;242 | 0 | 0 |
| NC_006577.2;YP_173236.1;HCoV-HKU1_orf1ab;243 | 0 | 0 |
| NC_006577.2;YP_173236.1;HCoV-HKU1_orf1ab;244 | 0 | 0 |
| NC_006577.2;YP_173236.1;HCoV-HKU1_orf1ab;245 | 0 | 0 |
| NC_006577.2;YP_173236.1;HCoV-HKU1_orf1ab;246 | 0 | 0 |
| NC_006577.2;YP_173236.1;HCoV-HKU1_orf1ab;247 | 0 | 0 |
| NC_006577.2;YP_173236.1;HCoV-HKU1_orf1ab;248 | 0 | 0 |
| NC_006577.2;YP_173236.1;HCoV-HKU1_orf1ab;249 | 0 | 0 |
| NC_006577.2;YP_173236.1;HCoV-HKU1_orf1ab;250 | 0 | 0 |
| NC_006577.2;YP_173236.1;HCoV-HKU1_orf1ab;251 | 0 | 0 |
| NC_006577.2;YP_173236.1;HCoV-HKU1_orf1ab;252 | 0.05 | 0 |
| NC_006577.2;YP_173236.1;HCoV-HKU1_orf1ab;253 | 0.05 | 0 |
| NC_006577.2;YP_173236.1;HCoV-HKU1_orf1ab;254 | 0.05 | 0 |
| NC_006577.2;YP_173236.1;HCoV-HKU1_orf1ab;255 | 0.05 | 0 |
| NC_006577.2;YP_173236.1;HCoV-HKU1_orf1ab;256 | 0.05 | 0.025 |
| NC_006577.2;YP_173236.1;HCoV-HKU1_orf1ab;257 | 0 | 0 |
| NC_006577.2;YP_173236.1;HCoV-HKU1_orf1ab;258 | 0 | 0 |
| NC_006577.2;YP_173236.1;HCoV-HKU1_orf1ab;259 | 0 | 0 |
| NC_006577.2;YP_173236.1;HCoV-HKU1_orf1ab;260 | 0 | 0 |
| NC_006577.2;YP_173236.1;HCoV-HKU1_orf1ab;261 | 0 | 0 |
| NC_006577.2;YP_173236.1;HCoV-HKU1_orf1ab;262 | 0 | 0 |
| NC_006577.2;YP_173236.1;HCoV-HKU1_orf1ab;263 | 0 | 0 |
| NC_006577.2;YP_173236.1;HCoV-HKU1_orf1ab;264 | 0 | 0 |
| NC_006577.2;YP_173236.1;HCoV-HKU1_orf1ab;265 | 0 | 0 |
| NC_006577.2;YP_173236.1;HCoV-HKU1_orf1ab;266 | 0 | 0 |
| NC_006577.2;YP_173236.1;HCoV-HKU1_orf1ab;267 | 0 | 0.025 |
| NC_006577.2;YP_173236.1;HCoV-HKU1_orf1ab;268 | 0 | 0.025 |
| NC_006577.2;YP_173236.1;HCoV-HKU1_orf1ab;269 | 0 | 0.05 |
| NC_006577.2;YP_173236.1;HCoV-HKU1_orf1ab;270 | 0 | 0.025 |
| NC_006577.2;YP_173236.1;HCoV-HKU1_orf1ab;271 | 0 | 0.025 |
| NC_006577.2;YP_173236.1;HCoV-HKU1_orf1ab;272 | 0 | 0.025 |
| NC_006577.2;YP_173236.1;HCoV-HKU1_orf1ab;273 | 0 | 0.025 |

| | | |
|---|---|---|
| NC_006577.2;YP_173236.1;HCoV-HKU1_orf1ab;274 | 0 | 0 |
| NC_006577.2;YP_173236.1;HCoV-HKU1_orf1ab;275 | 0 | 0.025 |
| NC_006577.2;YP_173236.1;HCoV-HKU1_orf1ab;276 | 0 | 0.025 |
| NC_006577.2;YP_173236.1;HCoV-HKU1_orf1ab;277 | 0 | 0.025 |
| NC_006577.2;YP_173236.1;HCoV-HKU1_orf1ab;278 | 0 | 0.05 |
| NC_006577.2;YP_173236.1;HCoV-HKU1_orf1ab;279 | 0 | 0 |
| NC_006577.2;YP_173236.1;HCoV-HKU1_orf1ab;280 | 0 | 0 |
| NC_006577.2;YP_173236.1;HCoV-HKU1_orf1ab;281 | 0 | 0 |
| NC_006577.2;YP_173236.1;HCoV-HKU1_orf1ab;282 | 0 | 0 |
| NC_006577.2;YP_173236.1;HCoV-HKU1_orf1ab;283 | 0 | 0 |
| NC_006577.2;YP_173236.1;HCoV-HKU1_orf1ab;284 | 0 | 0 |
| NC_006577.2;YP_173236.1;HCoV-HKU1_orf1ab;285 | 0 | 0 |
| NC_006577.2;YP_173236.1;HCoV-HKU1_orf1ab;286 | 0 | 0 |
| NC_006577.2;YP_173236.1;HCoV-HKU1_orf1ab;287 | 0 | 0 |
| NC_006577.2;YP_173236.1;HCoV-HKU1_orf1ab;288 | 0 | 0 |
| NC_006577.2;YP_173236.1;HCoV-HKU1_orf1ab;289 | 0 | 0 |
| NC_006577.2;YP_173236.1;HCoV-HKU1_orf1ab;290 | 0 | 0 |
| NC_006577.2;YP_173236.1;HCoV-HKU1_orf1ab;291 | 0 | 0.025 |
| NC_006577.2;YP_173236.1;HCoV-HKU1_orf1ab;292 | 0 | 0 |
| NC_006577.2;YP_173236.1;HCoV-HKU1_orf1ab;293 | 0 | 0 |
| NC_006577.2;YP_173236.1;HCoV-HKU1_orf1ab;294 | 0.1 | 0.1 |
| NC_006577.2;YP_173236.1;HCoV-HKU1_orf1ab;295 | 0 | 0 |
| NC_006577.2;YP_173236.1;HCoV-HKU1_orf1ab;296 | 0 | 0 |
| NC_006577.2;YP_173236.1;HCoV-HKU1_orf1ab;297 | 0 | 0 |
| NC_006577.2;YP_173236.1;HCoV-HKU1_orf1ab;298 | 0 | 0 |
| NC_006577.2;YP_173236.1;HCoV-HKU1_orf1ab;299 | 0 | 0 |
| NC_006577.2;YP_173236.1;HCoV-HKU1_orf1ab;300 | 0.05 | 0 |
| NC_006577.2;YP_173236.1;HCoV-HKU1_orf1ab;301 | 0 | 0 |
| NC_006577.2;YP_173236.1;HCoV-HKU1_orf1ab;302 | 0 | 0 |
| NC_006577.2;YP_173236.1;HCoV-HKU1_orf1ab;303 | 0.05 | 0 |
| NC_006577.2;YP_173236.1;HCoV-HKU1_orf1ab;304 | 0 | 0 |
| NC_006577.2;YP_173236.1;HCoV-HKU1_orf1ab;305 | 0 | 0 |
| NC_006577.2;YP_173236.1;HCoV-HKU1_orf1ab;306 | 0 | 0 |
| NC_006577.2;YP_173236.1;HCoV-HKU1_orf1ab;307 | 0 | 0 |
| NC_006577.2;YP_173236.1;HCoV-HKU1_orf1ab;308 | 0 | 0 |
| NC_006577.2;YP_173236.1;HCoV-HKU1_orf1ab;309 | 0 | 0 |
| NC_006577.2;YP_173236.1;HCoV-HKU1_orf1ab;310 | 0 | 0 |
| NC_006577.2;YP_173236.1;HCoV-HKU1_orf1ab;311 | 0 | 0 |
| NC_006577.2;YP_173236.1;HCoV-HKU1_orf1ab;312 | 0 | 0 |
| NC_006577.2;YP_173236.1;HCoV-HKU1_orf1ab;313 | 0 | 0 |
| NC_006577.2;YP_173236.1;HCoV-HKU1_orf1ab;314 | 0 | 0 |
| NC_006577.2;YP_173236.1;HCoV-HKU1_orf1ab;315 | 0 | 0 |
| NC_006577.2;YP_173236.1;HCoV-HKU1_orf1ab;316 | 0 | 0 |

| | | |
|---|---|---|
| NC_006577.2;YP_173236.1;HCoV-HKU1_orf1ab;317 | 0 | 0 |
| NC_006577.2;YP_173236.1;HCoV-HKU1_orf1ab;318 | 0 | 0 |
| NC_006577.2;YP_173236.1;HCoV-HKU1_orf1ab;319 | 0 | 0 |
| NC_006577.2;YP_173236.1;HCoV-HKU1_orf1ab;320 | 0 | 0.025 |
| NC_006577.2;YP_173236.1;HCoV-HKU1_orf1ab;321 | 0 | 0 |
| NC_006577.2;YP_173236.1;HCoV-HKU1_orf1ab;322 | 0 | 0 |
| NC_006577.2;YP_173236.1;HCoV-HKU1_orf1ab;323 | 0 | 0 |
| NC_006577.2;YP_173236.1;HCoV-HKU1_orf1ab;324 | 0 | 0 |
| NC_006577.2;YP_173236.1;HCoV-HKU1_orf1ab;325 | 0 | 0.025 |
| NC_006577.2;YP_173236.1;HCoV-HKU1_orf1ab;326 | 0 | 0 |
| NC_006577.2;YP_173236.1;HCoV-HKU1_orf1ab;327 | 0.05 | 0 |
| NC_006577.2;YP_173236.1;HCoV-HKU1_orf1ab;328 | 0 | 0 |
| NC_006577.2;YP_173236.1;HCoV-HKU1_orf1ab;329 | 0 | 0 |
| NC_006577.2;YP_173236.1;HCoV-HKU1_orf1ab;330 | 0 | 0.025 |
| NC_006577.2;YP_173236.1;HCoV-HKU1_orf1ab;331 | 0 | 0 |
| NC_006577.2;YP_173236.1;HCoV-HKU1_orf1ab;332 | 0 | 0 |
| NC_006577.2;YP_173236.1;HCoV-HKU1_orf1ab;333 | 0 | 0 |
| NC_006577.2;YP_173236.1;HCoV-HKU1_orf1ab;334 | 0 | 0 |
| NC_006577.2;YP_173236.1;HCoV-HKU1_orf1ab;335 | 0 | 0 |
| NC_006577.2;YP_173236.1;HCoV-HKU1_orf1ab;336 | 0 | 0 |
| NC_006577.2;YP_173236.1;HCoV-HKU1_orf1ab;337 | 0 | 0 |
| NC_006577.2;YP_173236.1;HCoV-HKU1_orf1ab;338 | 0 | 0 |
| NC_006577.2;YP_173236.1;HCoV-HKU1_orf1ab;339 | 0 | 0 |
| NC_006577.2;YP_173236.1;HCoV-HKU1_orf1ab;340 | 0 | 0 |
| NC_006577.2;YP_173236.1;HCoV-HKU1_orf1ab;341 | 0 | 0 |
| NC_006577.2;YP_173236.1;HCoV-HKU1_orf1ab;342 | 0 | 0 |
| NC_006577.2;YP_173236.1;HCoV-HKU1_orf1ab;343 | 0 | 0 |
| NC_006577.2;YP_173236.1;HCoV-HKU1_orf1ab;344 | 0 | 0 |
| NC_006577.2;YP_173236.1;HCoV-HKU1_orf1ab;345 | 0 | 0 |
| NC_006577.2;YP_173236.1;HCoV-HKU1_orf1ab;346 | 0 | 0 |
| NC_006577.2;YP_173236.1;HCoV-HKU1_orf1ab;347 | 0 | 0 |
| NC_006577.2;YP_173236.1;HCoV-HKU1_orf1ab;348 | 0 | 0 |
| NC_006577.2;YP_173236.1;HCoV-HKU1_orf1ab;349 | 0 | 0 |
| NC_006577.2;YP_173236.1;HCoV-HKU1_orf1ab;350 | 0 | 0 |
| NC_006577.2;YP_173236.1;HCoV-HKU1_orf1ab;351 | 0 | 0 |
| NC_006577.2;YP_173236.1;HCoV-HKU1_orf1ab;352 | 0 | 0 |
| NC_006577.2;YP_173236.1;HCoV-HKU1_orf1ab;353 | 0 | 0 |
| NC_006577.2;YP_173236.1;HCoV-HKU1_orf1ab;354 | 0 | 0 |
| NC_006577.2;YP_173236.1;HCoV-HKU1_orf1ab;355 | 0 | 0 |
| NC_006577.2;YP_173236.1;HCoV-HKU1_orf1ab;356 | 0.05 | 0.025 |
| NC_006577.2;YP_173236.1;HCoV-HKU1_orf1ab;357 | 0 | 0 |
| NC_006577.2;YP_173236.1;HCoV-HKU1_orf1ab;358 | 0 | 0 |
| NC_006577.2;YP_173236.1;HCoV-HKU1_orf1ab;359 | 0 | 0 |

| | | |
|---|---|---|
| NC_006577.2;YP_173236.1;HCoV-HKU1_orf1ab;360 | 0 | 0 |
| NC_006577.2;YP_173236.1;HCoV-HKU1_orf1ab;361 | 0 | 0.025 |
| NC_006577.2;YP_173236.1;HCoV-HKU1_orf1ab;362 | 0 | 0 |
| NC_006577.2;YP_173236.1;HCoV-HKU1_orf1ab;363 | 0 | 0 |
| NC_006577.2;YP_173236.1;HCoV-HKU1_orf1ab;364 | 0 | 0 |
| NC_006577.2;YP_173236.1;HCoV-HKU1_orf1ab;365 | 0 | 0 |
| NC_006577.2;YP_173236.1;HCoV-HKU1_orf1ab;366 | 0 | 0 |
| NC_006577.2;YP_173236.1;HCoV-HKU1_orf1ab;367 | 0 | 0.025 |
| NC_006577.2;YP_173236.1;HCoV-HKU1_orf1ab;368 | 0 | 0.025 |
| NC_006577.2;YP_173236.1;HCoV-HKU1_orf1ab;369 | 0 | 0.025 |
| NC_006577.2;YP_173236.1;HCoV-HKU1_orf1ab;370 | 0 | 0.025 |
| NC_006577.2;YP_173236.1;HCoV-HKU1_orf1ab;371 | 0 | 0.025 |
| NC_006577.2;YP_173236.1;HCoV-HKU1_orf1ab;372 | 0 | 0.025 |
| NC_006577.2;YP_173236.1;HCoV-HKU1_orf1ab;373 | 0.05 | 0.025 |
| NC_006577.2;YP_173236.1;HCoV-HKU1_orf1ab;374 | 0.05 | 0.025 |
| NC_006577.2;YP_173236.1;HCoV-HKU1_orf1ab;375 | 0 | 0.025 |
| NC_006577.2;YP_173236.1;HCoV-HKU1_orf1ab;376 | 0 | 0 |
| NC_006577.2;YP_173236.1;HCoV-HKU1_orf1ab;377 | 0 | 0 |
| NC_006577.2;YP_173236.1;HCoV-HKU1_orf1ab;378 | 0 | 0 |
| NC_006577.2;YP_173236.1;HCoV-HKU1_orf1ab;379 | 0 | 0 |
| NC_006577.2;YP_173236.1;HCoV-HKU1_orf1ab;380 | 0 | 0 |
| NC_006577.2;YP_173236.1;HCoV-HKU1_orf1ab;381 | 0 | 0.025 |
| NC_006577.2;YP_173236.1;HCoV-HKU1_orf1ab;382 | 0 | 0 |
| NC_006577.2;YP_173236.1;HCoV-HKU1_orf1ab;383 | 0 | 0.05 |
| NC_006577.2;YP_173236.1;HCoV-HKU1_orf1ab;384 | 0 | 0.05 |
| NC_006577.2;YP_173236.1;HCoV-HKU1_orf1ab;385 | 0 | 0.05 |
| NC_006577.2;YP_173236.1;HCoV-HKU1_orf1ab;386 | 0 | 0.05 |
| NC_006577.2;YP_173236.1;HCoV-HKU1_orf1ab;387 | 0 | 0 |
| NC_006577.2;YP_173236.1;HCoV-HKU1_orf1ab;388 | 0 | 0.05 |
| NC_006577.2;YP_173236.1;HCoV-HKU1_orf1ab;389 | 0 | 0.075 |
| NC_006577.2;YP_173236.1;HCoV-HKU1_orf1ab;390 | 0 | 0.05 |
| NC_006577.2;YP_173236.1;HCoV-HKU1_orf1ab;391 | 0 | 0.05 |
| NC_006577.2;YP_173236.1;HCoV-HKU1_orf1ab;392 | 0 | 0.025 |
| NC_006577.2;YP_173236.1;HCoV-HKU1_orf1ab;393 | 0.05 | 0.025 |
| NC_006577.2;YP_173236.1;HCoV-HKU1_orf1ab;394 | 0.05 | 0.025 |
| NC_006577.2;YP_173236.1;HCoV-HKU1_orf1ab;395 | 0.05 | 0.025 |
| NC_006577.2;YP_173236.1;HCoV-HKU1_orf1ab;396 | 0.05 | 0.025 |
| NC_006577.2;YP_173236.1;HCoV-HKU1_orf1ab;397 | 0.05 | 0.025 |
| NC_006577.2;YP_173236.1;HCoV-HKU1_orf1ab;398 | 0.05 | 0.025 |
| NC_006577.2;YP_173236.1;HCoV-HKU1_orf1ab;399 | 0.05 | 0.05 |
| NC_006577.2;YP_173236.1;HCoV-HKU1_orf1ab;400 | 0.05 | 0 |
| NC_006577.2;YP_173236.1;HCoV-HKU1_orf1ab;401 | 0 | 0 |
| NC_006577.2;YP_173236.1;HCoV-HKU1_orf1ab;402 | 0.05 | 0 |

| | | |
|---|---|---|
| NC_006577.2;YP_173236.1;HCoV-HKU1_orf1ab;403 | 0 | 0 |
| NC_006577.2;YP_173236.1;HCoV-HKU1_orf1ab;404 | 0.05 | 0 |
| NC_006577.2;YP_173236.1;HCoV-HKU1_orf1ab;405 | 0 | 0 |
| NC_006577.2;YP_173236.1;HCoV-HKU1_orf1ab;406 | 0 | 0 |
| NC_006577.2;YP_173236.1;HCoV-HKU1_orf1ab;407 | 0 | 0 |
| NC_006577.2;YP_173236.1;HCoV-HKU1_orf1ab;408 | 0 | 0 |
| NC_006577.2;YP_173236.1;HCoV-HKU1_orf1ab;409 | 0.05 | 0.05 |
| NC_006577.2;YP_173236.1;HCoV-HKU1_orf1ab;410 | 0 | 0 |
| NC_006577.2;YP_173236.1;HCoV-HKU1_orf1ab;411 | 0.05 | 0.025 |
| NC_006577.2;YP_173236.1;HCoV-HKU1_orf1ab;412 | 0.05 | 0.05 |
| NC_006577.2;YP_173236.1;HCoV-HKU1_orf1ab;413 | 0.05 | 0.025 |
| NC_006577.2;YP_173236.1;HCoV-HKU1_orf1ab;414 | 0.05 | 0.025 |
| NC_006577.2;YP_173236.1;HCoV-HKU1_orf1ab;415 | 0.05 | 0.025 |
| NC_006577.2;YP_173236.1;HCoV-HKU1_orf1ab;416 | 0.05 | 0.025 |
| NC_006577.2;YP_173236.1;HCoV-HKU1_orf1ab;417 | 0 | 0.025 |
| NC_006577.2;YP_173236.1;HCoV-HKU1_orf1ab;418 | 0 | 0 |
| NC_006577.2;YP_173236.1;HCoV-HKU1_orf1ab;419 | 0 | 0 |
| NC_006577.2;YP_173236.1;HCoV-HKU1_orf1ab;420 | 0 | 0 |
| NC_006577.2;YP_173236.1;HCoV-HKU1_orf1ab;421 | 0 | 0 |
| NC_006577.2;YP_173236.1;HCoV-HKU1_orf1ab;422 | 0 | 0 |
| NC_006577.2;YP_173236.1;HCoV-HKU1_orf1ab;423 | 0.1 | 0 |
| NC_006577.2;YP_173236.1;HCoV-HKU1_orf1ab;424 | 0 | 0 |
| NC_006577.2;YP_173236.1;HCoV-HKU1_orf1ab;425 | 0 | 0 |
| NC_006577.2;YP_173236.1;HCoV-HKU1_orf1ab;426 | 0 | 0 |
| NC_006577.2;YP_173236.1;HCoV-HKU1_orf1ab;427 | 0 | 0 |
| NC_006577.2;YP_173236.1;HCoV-HKU1_orf1ab;428 | 0 | 0 |
| NC_006577.2;YP_173236.1;HCoV-HKU1_orf1ab;429 | 0 | 0 |
| NC_006577.2;YP_173236.1;HCoV-HKU1_orf1ab;430 | 0 | 0 |
| NC_006577.2;YP_173236.1;HCoV-HKU1_orf1ab;431 | 0 | 0 |
| NC_006577.2;YP_173236.1;HCoV-HKU1_orf1ab;432 | 0 | 0 |
| NC_006577.2;YP_173236.1;HCoV-HKU1_orf1ab;433 | 0 | 0 |
| NC_006577.2;YP_173236.1;HCoV-HKU1_orf1ab;434 | 0 | 0 |
| NC_006577.2;YP_173236.1;HCoV-HKU1_orf1ab;435 | 0 | 0 |
| NC_006577.2;YP_173236.1;HCoV-HKU1_orf1ab;436 | 0 | 0 |
| NC_006577.2;YP_173236.1;HCoV-HKU1_orf1ab;437 | 0 | 0 |
| NC_006577.2;YP_173236.1;HCoV-HKU1_orf1ab;438 | 0 | 0 |
| NC_006577.2;YP_173236.1;HCoV-HKU1_orf1ab;439 | 0 | 0 |
| NC_006577.2;YP_173236.1;HCoV-HKU1_orf1ab;440 | 0 | 0 |
| NC_006577.2;YP_173236.1;HCoV-HKU1_orf1ab;441 | 0 | 0 |
| NC_006577.2;YP_173236.1;HCoV-HKU1_orf1ab;442 | 0 | 0 |
| NC_006577.2;YP_173236.1;HCoV-HKU1_orf1ab;443 | 0 | 0 |
| NC_006577.2;YP_173236.1;HCoV-HKU1_orf1ab;444 | 0 | 0 |
| NC_006577.2;YP_173236.1;HCoV-HKU1_orf1ab;445 | 0 | 0 |

| | | |
|---|---|---|
| NC_006577.2;YP_173236.1;HCoV-HKU1_orf1ab;446 | 0 | 0 |
| NC_006577.2;YP_173236.1;HCoV-HKU1_orf1ab;447 | 0 | 0.025 |
| NC_006577.2;YP_173236.1;HCoV-HKU1_orf1ab;448 | 0 | 0 |
| NC_006577.2;YP_173236.1;HCoV-HKU1_orf1ab;449 | 0 | 0 |
| NC_006577.2;YP_173236.1;HCoV-HKU1_orf1ab;450 | 0 | 0.075 |
| NC_006577.2;YP_173236.1;HCoV-HKU1_orf1ab;451 | 0 | 0.025 |
| NC_006577.2;YP_173236.1;HCoV-HKU1_orf1ab;452 | 0 | 0.025 |
| NC_006577.2;YP_173236.1;HCoV-HKU1_orf1ab;453 | 0 | 0 |
| NC_006577.2;YP_173236.1;HCoV-HKU1_orf1ab;454 | 0 | 0.025 |
| NC_006577.2;YP_173236.1;HCoV-HKU1_orf1ab;455 | 0 | 0 |
| NC_006577.2;YP_173236.1;HCoV-HKU1_orf1ab;456 | 0 | 0 |
| NC_006577.2;YP_173236.1;HCoV-HKU1_orf1ab;457 | 0 | 0 |
| NC_006577.2;YP_173236.1;HCoV-HKU1_orf1ab;458 | 0 | 0 |
| NC_006577.2;YP_173236.1;HCoV-HKU1_orf1ab;459 | 0 | 0 |
| NC_006577.2;YP_173236.1;HCoV-HKU1_orf1ab;460 | 0 | 0 |
| NC_006577.2;YP_173236.1;HCoV-HKU1_orf1ab;461 | 0 | 0 |
| NC_006577.2;YP_173236.1;HCoV-HKU1_orf1ab;462 | 0 | 0 |
| NC_006577.2;YP_173236.1;HCoV-HKU1_orf1ab;463 | 0 | 0 |
| NC_006577.2;YP_173236.1;HCoV-HKU1_orf1ab;464 | 0 | 0 |
| NC_006577.2;YP_173236.1;HCoV-HKU1_orf1ab;465 | 0 | 0 |
| NC_006577.2;YP_173236.1;HCoV-HKU1_orf1ab;466 | 0 | 0 |
| NC_006577.2;YP_173236.1;HCoV-HKU1_orf1ab;467 | 0 | 0 |
| NC_006577.2;YP_173236.1;HCoV-HKU1_orf1ab;468 | 0 | 0 |
| NC_006577.2;YP_173236.1;HCoV-HKU1_orf1ab;469 | 0 | 0 |
| NC_006577.2;YP_173236.1;HCoV-HKU1_orf1ab;470 | 0 | 0 |
| NC_006577.2;YP_173236.1;HCoV-HKU1_orf1ab;471 | 0 | 0 |
| NC_006577.2;YP_173236.1;HCoV-HKU1_orf1ab;472 | 0 | 0 |
| NC_006577.2;YP_173236.1;HCoV-HKU1_orf1ab;473 | 0 | 0 |
| NC_006577.2;YP_173236.1;HCoV-HKU1_orf1ab;474 | 0 | 0 |
| NC_006577.2;YP_173236.1;HCoV-HKU1_orf1ab;475 | 0 | 0 |
| NC_006577.2;YP_173236.1;HCoV-HKU1_orf1ab;476 | 0 | 0 |
| NC_006577.2;YP_173236.1;HCoV-HKU1_orf1ab;477 | 0 | 0 |
| NC_006577.2;YP_173236.1;HCoV-HKU1_orf1ab;478 | 0 | 0 |
| NC_006577.2;YP_173236.1;HCoV-HKU1_orf1ab;479 | 0 | 0 |
| NC_006577.2;YP_173236.1;HCoV-HKU1_orf1ab;480 | 0 | 0 |
| NC_006577.2;YP_173236.1;HCoV-HKU1_orf1ab;481 | 0 | 0 |
| NC_006577.2;YP_173236.1;HCoV-HKU1_orf1ab;482 | 0 | 0 |
| NC_006577.2;YP_173236.1;HCoV-HKU1_orf1ab;483 | 0 | 0.025 |
| NC_006577.2;YP_173236.1;HCoV-HKU1_orf1ab;484 | 0 | 0.025 |
| NC_006577.2;YP_173236.1;HCoV-HKU1_orf1ab;485 | 0 | 0 |
| NC_006577.2;YP_173236.1;HCoV-HKU1_orf1ab;486 | 0 | 0 |
| NC_006577.2;YP_173236.1;HCoV-HKU1_orf1ab;487 | 0 | 0 |
| NC_006577.2;YP_173236.1;HCoV-HKU1_orf1ab;488 | 0.05 | 0 |

| | | |
|---|---|---|
| NC_006577.2;YP_173236.1;HCoV-HKU1_orf1ab;489 | 0 | 0 |
| NC_006577.2;YP_173236.1;HCoV-HKU1_orf1ab;490 | 0 | 0 |
| NC_006577.2;YP_173236.1;HCoV-HKU1_orf1ab;491 | 0.05 | 0 |
| NC_006577.2;YP_173236.1;HCoV-HKU1_orf1ab;492 | 0.1 | 0.075 |
| NC_006577.2;YP_173236.1;HCoV-HKU1_orf1ab;493 | 0 | 0 |
| NC_006577.2;YP_173236.1;HCoV-HKU1_orf1ab;494 | 0 | 0 |
| NC_006577.2;YP_173236.1;HCoV-HKU1_orf1ab;495 | 0 | 0 |
| NC_006577.2;YP_173236.1;HCoV-HKU1_orf1ab;496 | 0 | 0 |
| NC_006577.2;YP_173236.1;HCoV-HKU1_orf1ab;497 | 0 | 0 |
| NC_006577.2;YP_173236.1;HCoV-HKU1_orf1ab;498 | 0 | 0 |
| NC_006577.2;YP_173236.1;HCoV-HKU1_orf1ab;499 | 0 | 0 |
| NC_006577.2;YP_173236.1;HCoV-HKU1_orf1ab;500 | 0 | 0 |
| NC_006577.2;YP_173236.1;HCoV-HKU1_orf1ab;501 | 0 | 0 |
| NC_006577.2;YP_173236.1;HCoV-HKU1_orf1ab;502 | 0 | 0 |
| NC_006577.2;YP_173236.1;HCoV-HKU1_orf1ab;503 | 0 | 0 |
| NC_006577.2;YP_173236.1;HCoV-HKU1_orf1ab;504 | 0 | 0 |
| NC_006577.2;YP_173236.1;HCoV-HKU1_orf1ab;505 | 0 | 0 |
| NC_006577.2;YP_173236.1;HCoV-HKU1_orf1ab;506 | 0 | 0 |
| NC_006577.2;YP_173236.1;HCoV-HKU1_orf1ab;507 | 0 | 0 |
| NC_006577.2;YP_173236.1;HCoV-HKU1_orf1ab;508 | 0 | 0 |
| NC_006577.2;YP_173236.1;HCoV-HKU1_orf1ab;509 | 0 | 0 |
| NC_006577.2;YP_173236.1;HCoV-HKU1_orf1ab;510 | 0 | 0 |
| NC_006577.2;YP_173236.1;HCoV-HKU1_orf1ab;511 | 0 | 0 |
| NC_006577.2;YP_173236.1;HCoV-HKU1_orf1ab;512 | 0.05 | 0 |
| NC_006577.2;YP_173236.1;HCoV-HKU1_orf1ab;513 | 0 | 0.05 |
| NC_006577.2;YP_173236.1;HCoV-HKU1_orf1ab;514 | 0 | 0.025 |
| NC_006577.2;YP_173236.1;HCoV-HKU1_orf1ab;515 | 0 | 0 |
| NC_006577.2;YP_173236.1;HCoV-HKU1_orf1ab;516 | 0.05 | 0.025 |
| NC_006577.2;YP_173236.1;HCoV-HKU1_orf1ab;517 | 0.05 | 0 |
| NC_006577.2;YP_173236.1;HCoV-HKU1_orf1ab;518 | 0.05 | 0 |
| NC_006577.2;YP_173236.1;HCoV-HKU1_orf1ab;519 | 0.05 | 0 |
| NC_006577.2;YP_173236.1;HCoV-HKU1_orf1ab;520 | 0.05 | 0 |
| NC_006577.2;YP_173236.1;HCoV-HKU1_orf1ab;521 | 0.05 | 0 |
| NC_006577.2;YP_173236.1;HCoV-HKU1_orf1ab;522 | 0.05 | 0 |
| NC_006577.2;YP_173236.1;HCoV-HKU1_orf1ab;523 | 0.05 | 0 |
| NC_006577.2;YP_173236.1;HCoV-HKU1_orf1ab;524 | 0.05 | 0 |
| NC_006577.2;YP_173236.1;HCoV-HKU1_orf1ab;525 | 0 | 0 |
| NC_006577.2;YP_173236.1;HCoV-HKU1_orf1ab;526 | 0 | 0 |
| NC_006577.2;YP_173236.1;HCoV-HKU1_orf1ab;527 | 0.05 | 0 |
| NC_006577.2;YP_173236.1;HCoV-HKU1_orf1ab;528 | 0 | 0 |
| NC_006577.2;YP_173236.1;HCoV-HKU1_orf1ab;529 | 0 | 0 |
| NC_006577.2;YP_173236.1;HCoV-HKU1_orf1ab;530 | 0 | 0 |
| NC_006577.2;YP_173236.1;HCoV-HKU1_orf1ab;531 | 0 | 0 |

| | | |
|---|---|---|
| NC_006577.2;YP_173236.1;HCoV-HKU1_orf1ab;532 | 0 | 0 |
| NC_006577.2;YP_173236.1;HCoV-HKU1_orf1ab;533 | 0 | 0 |
| NC_006577.2;YP_173236.1;HCoV-HKU1_orf1ab;534 | 0.05 | 0 |
| NC_006577.2;YP_173236.1;HCoV-HKU1_orf1ab;535 | 0 | 0 |
| NC_006577.2;YP_173236.1;HCoV-HKU1_orf1ab;536 | 0 | 0 |
| NC_006577.2;YP_173236.1;HCoV-HKU1_orf1ab;537 | 0 | 0 |
| NC_006577.2;YP_173236.1;HCoV-HKU1_orf1ab;538 | 0 | 0 |
| NC_006577.2;YP_173236.1;HCoV-HKU1_orf1ab;539 | 0 | 0 |
| NC_006577.2;YP_173236.1;HCoV-HKU1_orf1ab;540 | 0 | 0 |
| NC_006577.2;YP_173236.1;HCoV-HKU1_orf1ab;541 | 0 | 0 |
| NC_006577.2;YP_173236.1;HCoV-HKU1_orf1ab;542 | 0 | 0 |
| NC_006577.2;YP_173236.1;HCoV-HKU1_orf1ab;543 | 0 | 0 |
| NC_006577.2;YP_173236.1;HCoV-HKU1_orf1ab;544 | 0 | 0.025 |
| NC_006577.2;YP_173236.1;HCoV-HKU1_orf1ab;545 | 0 | 0 |
| NC_006577.2;YP_173236.1;HCoV-HKU1_orf1ab;546 | 0 | 0 |
| NC_006577.2;YP_173236.1;HCoV-HKU1_orf1ab;547 | 0 | 0 |
| NC_006577.2;YP_173236.1;HCoV-HKU1_orf1ab;548 | 0 | 0 |
| NC_006577.2;YP_173236.1;HCoV-HKU1_orf1ab;549 | 0 | 0 |
| NC_006577.2;YP_173236.1;HCoV-HKU1_orf1ab;550 | 0 | 0 |
| NC_006577.2;YP_173236.1;HCoV-HKU1_orf1ab;551 | 0.05 | 0 |
| NC_006577.2;YP_173236.1;HCoV-HKU1_orf1ab;552 | 0 | 0 |
| NC_006577.2;YP_173236.1;HCoV-HKU1_orf1ab;553 | 0 | 0 |
| NC_006577.2;YP_173236.1;HCoV-HKU1_orf1ab;554 | 0 | 0 |
| NC_006577.2;YP_173236.1;HCoV-HKU1_orf1ab;555 | 0 | 0 |
| NC_006577.2;YP_173236.1;HCoV-HKU1_orf1ab;556 | 0 | 0 |
| NC_006577.2;YP_173236.1;HCoV-HKU1_orf1ab;557 | 0 | 0 |
| NC_006577.2;YP_173236.1;HCoV-HKU1_orf1ab;558 | 0 | 0 |
| NC_006577.2;YP_173236.1;HCoV-HKU1_orf1ab;559 | 0 | 0 |
| NC_006577.2;YP_173236.1;HCoV-HKU1_orf1ab;560 | 0 | 0 |
| NC_006577.2;YP_173236.1;HCoV-HKU1_orf1ab;561 | 0 | 0 |
| NC_006577.2;YP_173236.1;HCoV-HKU1_orf1ab;562 | 0 | 0 |
| NC_006577.2;YP_173236.1;HCoV-HKU1_orf1ab;563 | 0 | 0 |
| NC_006577.2;YP_173236.1;HCoV-HKU1_orf1ab;564 | 0 | 0 |
| NC_006577.2;YP_173236.1;HCoV-HKU1_orf1ab;565 | 0 | 0 |
| NC_006577.2;YP_173236.1;HCoV-HKU1_orf1ab;566 | 0 | 0 |
| NC_006577.2;YP_173236.1;HCoV-HKU1_orf1ab;567 | 0 | 0 |
| NC_006577.2;YP_173236.1;HCoV-HKU1_orf1ab;568 | 0 | 0 |
| NC_006577.2;YP_173236.1;HCoV-HKU1_orf1ab;569 | 0 | 0 |
| NC_006577.2;YP_173236.1;HCoV-HKU1_orf1ab;570 | 0.05 | 0 |
| NC_006577.2;YP_173236.1;HCoV-HKU1_orf1ab;571 | 0 | 0 |
| NC_006577.2;YP_173236.1;HCoV-HKU1_orf1ab;572 | 0 | 0 |
| NC_006577.2;YP_173236.1;HCoV-HKU1_orf1ab;573 | 0 | 0 |
| NC_006577.2;YP_173236.1;HCoV-HKU1_orf1ab;574 | 0 | 0 |

| | | |
|---|---|---|
| NC_006577.2;YP_173236.1;HCoV-HKU1_orf1ab;575 | 0 | 0 |
| NC_006577.2;YP_173236.1;HCoV-HKU1_orf1ab;576 | 0 | 0 |
| NC_006577.2;YP_173236.1;HCoV-HKU1_orf1ab;577 | 0 | 0 |
| NC_006577.2;YP_173236.1;HCoV-HKU1_orf1ab;578 | 0 | 0 |
| NC_006577.2;YP_173236.1;HCoV-HKU1_orf1ab;579 | 0 | 0 |
| NC_006577.2;YP_173236.1;HCoV-HKU1_orf1ab;580 | 0 | 0 |
| NC_006577.2;YP_173236.1;HCoV-HKU1_orf1ab;581 | 0 | 0 |
| NC_006577.2;YP_173236.1;HCoV-HKU1_orf1ab;582 | 0 | 0 |
| NC_006577.2;YP_173236.1;HCoV-HKU1_orf1ab;583 | 0 | 0 |
| NC_006577.2;YP_173236.1;HCoV-HKU1_orf1ab;584 | 0 | 0 |
| NC_006577.2;YP_173236.1;HCoV-HKU1_orf1ab;585 | 0 | 0 |
| NC_006577.2;YP_173236.1;HCoV-HKU1_orf1ab;586 | 0 | 0 |
| NC_006577.2;YP_173236.1;HCoV-HKU1_orf1ab;587 | 0 | 0 |
| NC_006577.2;YP_173236.1;HCoV-HKU1_orf1ab;588 | 0 | 0 |
| NC_006577.2;YP_173236.1;HCoV-HKU1_orf1ab;589 | 0 | 0 |
| NC_006577.2;YP_173236.1;HCoV-HKU1_orf1ab;590 | 0 | 0 |
| NC_006577.2;YP_173236.1;HCoV-HKU1_orf1ab;591 | 0 | 0 |
| NC_006577.2;YP_173236.1;HCoV-HKU1_orf1ab;592 | 0 | 0 |
| NC_006577.2;YP_173236.1;HCoV-HKU1_orf1ab;593 | 0 | 0 |
| NC_006577.2;YP_173236.1;HCoV-HKU1_orf1ab;594 | 0 | 0 |
| NC_006577.2;YP_173236.1;HCoV-HKU1_orf1ab;595 | 0 | 0 |
| NC_006577.2;YP_173236.1;HCoV-HKU1_orf1ab;596 | 0 | 0 |
| NC_006577.2;YP_173236.1;HCoV-HKU1_orf1ab;597 | 0 | 0 |
| NC_006577.2;YP_173236.1;HCoV-HKU1_orf1ab;598 | 0 | 0 |
| NC_006577.2;YP_173236.1;HCoV-HKU1_orf1ab;599 | 0 | 0 |
| NC_006577.2;YP_173236.1;HCoV-HKU1_orf1ab;600 | 0 | 0 |
| NC_006577.2;YP_173236.1;HCoV-HKU1_orf1ab;601 | 0 | 0 |
| NC_006577.2;YP_173236.1;HCoV-HKU1_orf1ab;602 | 0 | 0 |
| NC_006577.2;YP_173236.1;HCoV-HKU1_orf1ab;603 | 0 | 0 |
| NC_006577.2;YP_173236.1;HCoV-HKU1_orf1ab;604 | 0 | 0 |
| NC_006577.2;YP_173236.1;HCoV-HKU1_orf1ab;605 | 0 | 0 |
| NC_006577.2;YP_173236.1;HCoV-HKU1_orf1ab;606 | 0 | 0 |
| NC_006577.2;YP_173236.1;HCoV-HKU1_orf1ab;607 | 0 | 0 |
| NC_006577.2;YP_173236.1;HCoV-HKU1_orf1ab;608 | 0 | 0 |
| NC_006577.2;YP_173236.1;HCoV-HKU1_orf1ab;609 | 0 | 0 |
| NC_006577.2;YP_173236.1;HCoV-HKU1_orf1ab;610 | 0 | 0 |
| NC_006577.2;YP_173236.1;HCoV-HKU1_orf1ab;611 | 0 | 0 |
| NC_006577.2;YP_173236.1;HCoV-HKU1_orf1ab;612 | 0 | 0 |
| NC_006577.2;YP_173236.1;HCoV-HKU1_orf1ab;613 | 0 | 0 |
| NC_006577.2;YP_173236.1;HCoV-HKU1_orf1ab;614 | 0 | 0 |
| NC_006577.2;YP_173236.1;HCoV-HKU1_orf1ab;615 | 0 | 0 |
| NC_006577.2;YP_173236.1;HCoV-HKU1_orf1ab;616 | 0 | 0 |
| NC_006577.2;YP_173236.1;HCoV-HKU1_orf1ab;617 | 0 | 0 |

| | | |
|---|---|---|
| NC_006577.2;YP_173236.1;HCoV-HKU1_orf1ab;618 | 0 | 0 |
| NC_006577.2;YP_173236.1;HCoV-HKU1_orf1ab;619 | 0.05 | 0.025 |
| NC_006577.2;YP_173236.1;HCoV-HKU1_orf1ab;620 | 0 | 0.025 |
| NC_006577.2;YP_173236.1;HCoV-HKU1_orf1ab;621 | 0.1 | 0.05 |
| NC_006577.2;YP_173236.1;HCoV-HKU1_orf1ab;622 | 0.1 | 0.025 |
| NC_006577.2;YP_173236.1;HCoV-HKU1_orf1ab;623 | 0.15 | 0.05 |
| NC_006577.2;YP_173236.1;HCoV-HKU1_orf1ab;624 | 0.15 | 0.025 |
| NC_006577.2;YP_173236.1;HCoV-HKU1_orf1ab;625 | 0.15 | 0.025 |
| NC_006577.2;YP_173236.1;HCoV-HKU1_orf1ab;626 | 0 | 0.025 |
| NC_006577.2;YP_173236.1;HCoV-HKU1_orf1ab;627 | 0 | 0.025 |
| NC_006577.2;YP_173236.1;HCoV-HKU1_orf1ab;628 | 0 | 0.025 |
| NC_006577.2;YP_173236.1;HCoV-HKU1_orf1ab;629 | 0 | 0 |
| NC_006577.2;YP_173236.1;HCoV-HKU1_orf1ab;630 | 0 | 0 |
| NC_006577.2;YP_173236.1;HCoV-HKU1_orf1ab;631 | 0 | 0 |
| NC_006577.2;YP_173236.1;HCoV-HKU1_orf1ab;632 | 0 | 0 |
| NC_006577.2;YP_173236.1;HCoV-HKU1_orf1ab;633 | 0 | 0 |
| NC_006577.2;YP_173236.1;HCoV-HKU1_orf1ab;634 | 0 | 0 |
| NC_006577.2;YP_173236.1;HCoV-HKU1_orf1ab;635 | 0 | 0 |
| NC_006577.2;YP_173236.1;HCoV-HKU1_orf1ab;636 | 0 | 0 |
| NC_006577.2;YP_173236.1;HCoV-HKU1_orf1ab;637 | 0 | 0 |
| NC_006577.2;YP_173236.1;HCoV-HKU1_orf1ab;638 | 0 | 0 |
| NC_006577.2;YP_173236.1;HCoV-HKU1_orf1ab;639 | 0 | 0 |
| NC_006577.2;YP_173236.1;HCoV-HKU1_orf1ab;640 | 0 | 0 |
| NC_006577.2;YP_173236.1;HCoV-HKU1_orf1ab;641 | 0 | 0 |
| NC_006577.2;YP_173236.1;HCoV-HKU1_orf1ab;642 | 0 | 0 |
| NC_006577.2;YP_173236.1;HCoV-HKU1_orf1ab;643 | 0 | 0 |
| NC_006577.2;YP_173236.1;HCoV-HKU1_orf1ab;644 | 0 | 0 |
| NC_006577.2;YP_173236.1;HCoV-HKU1_orf1ab;645 | 0 | 0 |
| NC_006577.2;YP_173236.1;HCoV-HKU1_orf1ab;646 | 0 | 0 |
| NC_006577.2;YP_173236.1;HCoV-HKU1_orf1ab;647 | 0 | 0 |
| NC_006577.2;YP_173236.1;HCoV-HKU1_orf1ab;648 | 0 | 0 |
| NC_006577.2;YP_173236.1;HCoV-HKU1_orf1ab;649 | 0 | 0 |
| NC_006577.2;YP_173236.1;HCoV-HKU1_orf1ab;650 | 0 | 0 |
| NC_006577.2;YP_173236.1;HCoV-HKU1_orf1ab;651 | 0 | 0 |
| NC_006577.2;YP_173236.1;HCoV-HKU1_orf1ab;652 | 0.05 | 0 |
| NC_006577.2;YP_173236.1;HCoV-HKU1_orf1ab;653 | 0.1 | 0 |
| NC_006577.2;YP_173236.1;HCoV-HKU1_orf1ab;654 | 0 | 0 |
| NC_006577.2;YP_173236.1;HCoV-HKU1_orf1ab;655 | 0 | 0 |
| NC_006577.2;YP_173236.1;HCoV-HKU1_orf1ab;656 | 0 | 0 |
| NC_006577.2;YP_173236.1;HCoV-HKU1_orf1ab;657 | 0 | 0 |
| NC_006577.2;YP_173236.1;HCoV-HKU1_orf1ab;658 | 0 | 0 |
| NC_006577.2;YP_173236.1;HCoV-HKU1_orf1ab;659 | 0 | 0 |
| NC_006577.2;YP_173236.1;HCoV-HKU1_orf1ab;660 | 0 | 0 |

| | | |
|---|---|---|
| NC_006577.2;YP_173236.1;HCoV-HKU1_orf1ab;661 | 0 | 0 |
| NC_006577.2;YP_173236.1;HCoV-HKU1_orf1ab;662 | 0 | 0 |
| NC_006577.2;YP_173236.1;HCoV-HKU1_orf1ab;663 | 0 | 0 |
| NC_006577.2;YP_173236.1;HCoV-HKU1_orf1ab;664 | 0 | 0 |
| NC_006577.2;YP_173236.1;HCoV-HKU1_orf1ab;665 | 0 | 0 |
| NC_006577.2;YP_173236.1;HCoV-HKU1_orf1ab;666 | 0 | 0 |
| NC_006577.2;YP_173236.1;HCoV-HKU1_orf1ab;667 | 0 | 0 |
| NC_006577.2;YP_173236.1;HCoV-HKU1_orf1ab;668 | 0 | 0 |
| NC_006577.2;YP_173236.1;HCoV-HKU1_orf1ab;669 | 0 | 0 |
| NC_006577.2;YP_173236.1;HCoV-HKU1_orf1ab;670 | 0 | 0 |
| NC_006577.2;YP_173236.1;HCoV-HKU1_orf1ab;671 | 0 | 0 |
| NC_006577.2;YP_173236.1;HCoV-HKU1_orf1ab;672 | 0 | 0 |
| NC_006577.2;YP_173236.1;HCoV-HKU1_orf1ab;673 | 0 | 0 |
| NC_006577.2;YP_173236.1;HCoV-HKU1_orf1ab;674 | 0 | 0 |
| NC_006577.2;YP_173236.1;HCoV-HKU1_orf1ab;675 | 0 | 0 |
| NC_006577.2;YP_173236.1;HCoV-HKU1_orf1ab;676 | 0 | 0 |
| NC_006577.2;YP_173236.1;HCoV-HKU1_orf1ab;677 | 0 | 0 |
| NC_006577.2;YP_173236.1;HCoV-HKU1_orf1ab;678 | 0 | 0 |
| NC_006577.2;YP_173236.1;HCoV-HKU1_orf1ab;679 | 0 | 0 |
| NC_006577.2;YP_173236.1;HCoV-HKU1_orf1ab;680 | 0 | 0 |
| NC_006577.2;YP_173236.1;HCoV-HKU1_orf1ab;681 | 0 | 0 |
| NC_006577.2;YP_173236.1;HCoV-HKU1_orf1ab;682 | 0 | 0 |
| NC_006577.2;YP_173236.1;HCoV-HKU1_orf1ab;683 | 0 | 0 |
| NC_006577.2;YP_173236.1;HCoV-HKU1_orf1ab;684 | 0 | 0 |
| NC_006577.2;YP_173236.1;HCoV-HKU1_orf1ab;685 | 0 | 0 |
| NC_006577.2;YP_173236.1;HCoV-HKU1_orf1ab;686 | 0 | 0.125 |
| NC_006577.2;YP_173236.1;HCoV-HKU1_orf1ab;687 | 0 | 0 |
| NC_006577.2;YP_173236.1;HCoV-HKU1_orf1ab;688 | 0 | 0.025 |
| NC_006577.2;YP_173236.1;HCoV-HKU1_orf1ab;689 | 0 | 0 |
| NC_006577.2;YP_173236.1;HCoV-HKU1_orf1ab;690 | 0 | 0 |
| NC_006577.2;YP_173236.1;HCoV-HKU1_orf1ab;691 | 0 | 0 |
| NC_006577.2;YP_173236.1;HCoV-HKU1_orf1ab;692 | 0 | 0 |
| NC_006577.2;YP_173236.1;HCoV-HKU1_orf1ab;693 | 0 | 0 |
| NC_006577.2;YP_173236.1;HCoV-HKU1_orf1ab;694 | 0 | 0 |
| NC_006577.2;YP_173236.1;HCoV-HKU1_orf1ab;695 | 0 | 0.025 |
| NC_006577.2;YP_173236.1;HCoV-HKU1_orf1ab;696 | 0 | 0 |
| NC_006577.2;YP_173236.1;HCoV-HKU1_orf1ab;697 | 0 | 0 |
| NC_006577.2;YP_173236.1;HCoV-HKU1_orf1ab;698 | 0 | 0 |
| NC_006577.2;YP_173236.1;HCoV-HKU1_orf1ab;699 | 0 | 0 |
| NC_006577.2;YP_173236.1;HCoV-HKU1_orf1ab;700 | 0 | 0 |
| NC_006577.2;YP_173236.1;HCoV-HKU1_orf1ab;701 | 0 | 0 |
| NC_006577.2;YP_173236.1;HCoV-HKU1_orf1ab;702 | 0 | 0 |
| NC_006577.2;YP_173236.1;HCoV-HKU1_orf1ab;703 | 0 | 0 |

| | | |
|---|---|---|
| NC_006577.2;YP_173236.1;HCoV-HKU1_orf1ab;704 | 0 | 0 |
| NC_006577.2;YP_173236.1;HCoV-HKU1_orf1ab;705 | 0 | 0.025 |
| NC_006577.2;YP_173236.1;HCoV-HKU1_orf1ab;706 | 0 | 0.05 |
| NC_006577.2;YP_173236.1;HCoV-HKU1_orf1ab;707 | 0 | 0.025 |
| NC_006577.2;YP_173236.1;HCoV-HKU1_orf1ab;708 | 0 | 0.025 |
| NC_006577.2;YP_173236.1;HCoV-HKU1_orf1ab;709 | 0 | 0 |
| NC_006577.2;YP_173236.1;HCoV-HKU1_orf1ab;710 | 0 | 0 |
| NC_006577.2;YP_173236.1;HCoV-HKU1_orf1ab;711 | 0 | 0 |
| NC_006577.2;YP_173236.1;HCoV-HKU1_orf1ab;712 | 0 | 0 |
| NC_006577.2;YP_173236.1;HCoV-HKU1_orf1ab;713 | 0 | 0 |
| NC_006577.2;YP_173236.1;HCoV-HKU1_orf1ab;714 | 0 | 0 |
| NC_006577.2;YP_173236.1;HCoV-HKU1_orf1ab;715 | 0 | 0 |
| NC_006577.2;YP_173236.1;HCoV-HKU1_orf1ab;716 | 0 | 0 |
| NC_006577.2;YP_173236.1;HCoV-HKU1_orf1ab;717 | 0 | 0 |
| NC_006577.2;YP_173236.1;HCoV-HKU1_orf1ab;718 | 0 | 0.025 |
| NC_006577.2;YP_173236.1;HCoV-HKU1_orf1ab;719 | 0 | 0 |
| NC_006577.2;YP_173236.1;HCoV-HKU1_orf1ab;720 | 0 | 0 |
| NC_006577.2;YP_173236.1;HCoV-HKU1_orf1ab;721 | 0 | 0 |
| NC_006577.2;YP_173236.1;HCoV-HKU1_orf1ab;722 | 0 | 0 |
| NC_006577.2;YP_173236.1;HCoV-HKU1_orf1ab;723 | 0 | 0 |
| NC_006577.2;YP_173236.1;HCoV-HKU1_orf1ab;724 | 0 | 0 |
| NC_006577.2;YP_173236.1;HCoV-HKU1_orf1ab;725 | 0 | 0.025 |
| NC_006577.2;YP_173236.1;HCoV-HKU1_orf1ab;726 | 0 | 0 |
| NC_006577.2;YP_173236.1;HCoV-HKU1_orf1ab;727 | 0 | 0 |
| NC_006577.2;YP_173236.1;HCoV-HKU1_orf1ab;728 | 0 | 0 |
| NC_006577.2;YP_173236.1;HCoV-HKU1_orf1ab;729 | 0 | 0 |
| NC_006577.2;YP_173236.1;HCoV-HKU1_orf1ab;730 | 0 | 0 |
| NC_006577.2;YP_173236.1;HCoV-HKU1_orf1ab;731 | 0 | 0 |
| NC_006577.2;YP_173236.1;HCoV-HKU1_orf1ab;732 | 0 | 0 |
| NC_006577.2;YP_173236.1;HCoV-HKU1_orf1ab;733 | 0 | 0 |
| NC_006577.2;YP_173236.1;HCoV-HKU1_orf1ab;734 | 0 | 0 |
| NC_006577.2;YP_173236.1;HCoV-HKU1_orf1ab;735 | 0 | 0 |
| NC_006577.2;YP_173236.1;HCoV-HKU1_orf1ab;736 | 0 | 0 |
| NC_006577.2;YP_173236.1;HCoV-HKU1_orf1ab;737 | 0 | 0 |
| NC_006577.2;YP_173236.1;HCoV-HKU1_orf1ab;738 | 0 | 0 |
| NC_006577.2;YP_173236.1;HCoV-HKU1_orf1ab;739 | 0 | 0 |
| NC_006577.2;YP_173236.1;HCoV-HKU1_orf1ab;740 | 0 | 0.025 |
| NC_006577.2;YP_173236.1;HCoV-HKU1_orf1ab;741 | 0 | 0 |
| NC_006577.2;YP_173236.1;HCoV-HKU1_orf1ab;742 | 0 | 0.025 |
| NC_006577.2;YP_173236.1;HCoV-HKU1_orf1ab;743 | 0.05 | 0.05 |
| NC_006577.2;YP_173236.1;HCoV-HKU1_orf1ab;744 | 0 | 0.025 |
| NC_006577.2;YP_173236.1;HCoV-HKU1_orf1ab;745 | 0 | 0 |
| NC_006577.2;YP_173236.1;HCoV-HKU1_orf1ab;746 | 0.1 | 0 |

| | | |
|---|---|---|
| NC_006577.2;YP_173236.1;HCoV-HKU1_orf1ab;747 | 0 | 0 |
| NC_006577.2;YP_173236.1;HCoV-HKU1_orf1ab;748 | 0 | 0 |
| NC_006577.2;YP_173236.1;HCoV-HKU1_orf1ab;749 | 0.05 | 0 |
| NC_006577.2;YP_173236.1;HCoV-HKU1_orf1ab;750 | 0.05 | 0 |
| NC_006577.2;YP_173236.1;HCoV-HKU1_orf1ab;751 | 0 | 0 |
| NC_006577.2;YP_173236.1;HCoV-HKU1_orf1ab;752 | 0 | 0.025 |
| NC_006577.2;YP_173236.1;HCoV-HKU1_orf1ab;753 | 0 | 0.025 |
| NC_006577.2;YP_173236.1;HCoV-HKU1_orf1ab;754 | 0.1 | 0 |
| NC_006577.2;YP_173236.1;HCoV-HKU1_orf1ab;755 | 0 | 0.025 |
| NC_006577.2;YP_173236.1;HCoV-HKU1_orf1ab;756 | 0 | 0.025 |
| NC_006577.2;YP_173236.1;HCoV-HKU1_orf1ab;757 | 0.1 | 0.025 |
| NC_006577.2;YP_173236.1;HCoV-HKU1_orf1ab;758 | 0 | 0 |
| NC_006577.2;YP_173236.1;HCoV-HKU1_orf1ab;759 | 0 | 0 |
| NC_006577.2;YP_173236.1;HCoV-HKU1_orf1ab;760 | 0 | 0 |
| NC_006577.2;YP_173236.1;HCoV-HKU1_orf1ab;761 | 0 | 0 |
| NC_006577.2;YP_173236.1;HCoV-HKU1_orf1ab;762 | 0 | 0 |
| NC_006577.2;YP_173236.1;HCoV-HKU1_orf1ab;763 | 0 | 0 |
| NC_006577.2;YP_173236.1;HCoV-HKU1_orf1ab;764 | 0.05 | 0.025 |
| NC_006577.2;YP_173236.1;HCoV-HKU1_orf1ab;765 | 0 | 0 |
| NC_006577.2;YP_173236.1;HCoV-HKU1_orf1ab;766 | 0 | 0 |
| NC_006577.2;YP_173236.1;HCoV-HKU1_orf1ab;767 | 0 | 0 |
| NC_006577.2;YP_173236.1;HCoV-HKU1_orf1ab;768 | 0.05 | 0 |
| NC_006577.2;YP_173236.1;HCoV-HKU1_orf1ab;769 | 0 | 0 |
| NC_006577.2;YP_173236.1;HCoV-HKU1_orf1ab;770 | 0 | 0.025 |
| NC_006577.2;YP_173236.1;HCoV-HKU1_orf1ab;771 | 0 | 0 |
| NC_006577.2;YP_173236.1;HCoV-HKU1_orf1ab;772 | 0 | 0 |
| NC_006577.2;YP_173236.1;HCoV-HKU1_orf1ab;773 | 0.05 | 0 |
| NC_006577.2;YP_173236.1;HCoV-HKU1_orf1ab;774 | 0.05 | 0 |
| NC_006577.2;YP_173236.1;HCoV-HKU1_orf1ab;775 | 0.05 | 0 |
| NC_006577.2;YP_173236.1;HCoV-HKU1_orf1ab;776 | 0 | 0 |
| NC_006577.2;YP_173236.1;HCoV-HKU1_orf1ab;777 | 0 | 0 |
| NC_006577.2;YP_173236.1;HCoV-HKU1_orf1ab;778 | 0.05 | 0 |
| NC_006577.2;YP_173236.1;HCoV-HKU1_orf1ab;779 | 0 | 0.025 |
| NC_006577.2;YP_173236.1;HCoV-HKU1_orf1ab;780 | 0 | 0.05 |
| NC_006577.2;YP_173236.1;HCoV-HKU1_orf1ab;781 | 0 | 0.05 |
| NC_006577.2;YP_173236.1;HCoV-HKU1_orf1ab;782 | 0.05 | 0.025 |
| NC_006577.2;YP_173236.1;HCoV-HKU1_orf1ab;783 | 0.05 | 0.025 |
| NC_006577.2;YP_173236.1;HCoV-HKU1_orf1ab;784 | 0.05 | 0 |
| NC_006577.2;YP_173236.1;HCoV-HKU1_orf1ab;785 | 0 | 0 |
| NC_006577.2;YP_173236.1;HCoV-HKU1_orf1ab;786 | 0 | 0 |
| NC_006577.2;YP_173236.1;HCoV-HKU1_orf1ab;787 | 0 | 0 |
| NC_006577.2;YP_173236.1;HCoV-HKU1_orf1ab;788 | 0 | 0 |
| NC_006577.2;YP_173236.1;HCoV-HKU1_orf1ab;789 | 0.05 | 0.025 |

| | | |
|---|---|---|
| NC_006577.2;YP_173236.1;HCoV-HKU1_orf1ab;790 | 0 | 0 |
| NC_006577.2;YP_173236.1;HCoV-HKU1_orf1ab;791 | 0 | 0 |
| NC_006577.2;YP_173236.1;HCoV-HKU1_orf1ab;792 | 0 | 0 |
| NC_006577.2;YP_173236.1;HCoV-HKU1_orf1ab;793 | 0 | 0 |
| NC_006577.2;YP_173236.1;HCoV-HKU1_orf1ab;794 | 0 | 0 |
| NC_006577.2;YP_173236.1;HCoV-HKU1_orf1ab;795 | 0 | 0 |
| NC_006577.2;YP_173236.1;HCoV-HKU1_orf1ab;796 | 0 | 0 |
| NC_006577.2;YP_173236.1;HCoV-HKU1_orf1ab;797 | 0 | 0 |
| NC_006577.2;YP_173236.1;HCoV-HKU1_orf1ab;798 | 0 | 0 |
| NC_006577.2;YP_173236.1;HCoV-HKU1_orf1ab;799 | 0.05 | 0.025 |
| NC_006577.2;YP_173236.1;HCoV-HKU1_orf1ab;800 | 0 | 0 |
| NC_006577.2;YP_173236.1;HCoV-HKU1_orf1ab;801 | 0 | 0 |
| NC_006577.2;YP_173236.1;HCoV-HKU1_orf1ab;802 | 0 | 0.05 |
| NC_006577.2;YP_173236.1;HCoV-HKU1_orf1ab;803 | 0 | 0 |
| NC_006577.2;YP_173236.1;HCoV-HKU1_orf1ab;804 | 0 | 0.025 |
| NC_006577.2;YP_173236.1;HCoV-HKU1_orf1ab;805 | 0 | 0.025 |
| NC_006577.2;YP_173236.1;HCoV-HKU1_orf1ab;806 | 0 | 0.1 |
| NC_006577.2;YP_173236.1;HCoV-HKU1_orf1ab;807 | 0 | 0 |
| NC_006577.2;YP_173236.1;HCoV-HKU1_orf1ab;808 | 0.1 | 0.125 |
| NC_006577.2;YP_173236.1;HCoV-HKU1_orf1ab;809 | 0 | 0.025 |
| NC_006577.2;YP_173236.1;HCoV-HKU1_orf1ab;810 | 0 | 0.025 |
| NC_006577.2;YP_173236.1;HCoV-HKU1_orf1ab;811 | 0 | 0 |
| NC_006577.2;YP_173236.1;HCoV-HKU1_orf1ab;812 | 0.1 | 0.075 |
| NC_006577.2;YP_173236.1;HCoV-HKU1_orf1ab;813 | 0 | 0 |
| NC_006577.2;YP_173236.1;HCoV-HKU1_orf1ab;814 | 0.05 | 0.025 |
| NC_006577.2;YP_173236.1;HCoV-HKU1_orf1ab;815 | 0 | 0 |
| NC_006577.2;YP_173236.1;HCoV-HKU1_orf1ab;816 | 0 | 0 |
| NC_006577.2;YP_173236.1;HCoV-HKU1_orf1ab;817 | 0 | 0 |
| NC_006577.2;YP_173236.1;HCoV-HKU1_orf1ab;818 | 0 | 0 |
| NC_006577.2;YP_173236.1;HCoV-HKU1_orf1ab;819 | 0 | 0 |
| NC_006577.2;YP_173236.1;HCoV-HKU1_orf1ab;820 | 0 | 0 |
| NC_006577.2;YP_173236.1;HCoV-HKU1_orf1ab;821 | 0 | 0 |
| NC_006577.2;YP_173236.1;HCoV-HKU1_orf1ab;822 | 0 | 0 |
| NC_006577.2;YP_173236.1;HCoV-HKU1_orf1ab;823 | 0 | 0 |
| NC_006577.2;YP_173236.1;HCoV-HKU1_orf1ab;824 | 0 | 0 |
| NC_006577.2;YP_173236.1;HCoV-HKU1_orf1ab;825 | 0 | 0 |
| NC_006577.2;YP_173236.1;HCoV-HKU1_orf1ab;826 | 0 | 0 |
| NC_006577.2;YP_173236.1;HCoV-HKU1_orf1ab;827 | 0 | 0 |
| NC_006577.2;YP_173236.1;HCoV-HKU1_orf1ab;828 | 0 | 0 |
| NC_006577.2;YP_173236.1;HCoV-HKU1_orf1ab;829 | 0 | 0 |
| NC_006577.2;YP_173236.1;HCoV-HKU1_orf1ab;830 | 0.05 | 0 |
| NC_006577.2;YP_173236.1;HCoV-HKU1_orf1ab;831 | 0 | 0 |
| NC_006577.2;YP_173236.1;HCoV-HKU1_orf1ab;832 | 0 | 0 |

| | | |
|---|---|---|
| NC_006577.2;YP_173236.1;HCoV-HKU1_orf1ab;833 | 0 | 0.025 |
| NC_006577.2;YP_173236.1;HCoV-HKU1_orf1ab;834 | 0 | 0 |
| NC_006577.2;YP_173236.1;HCoV-HKU1_orf1ab;835 | 0 | 0 |
| NC_006577.2;YP_173236.1;HCoV-HKU1_orf1ab;836 | 0 | 0 |
| NC_006577.2;YP_173236.1;HCoV-HKU1_orf1ab;837 | 0.05 | 0 |
| NC_006577.2;YP_173236.1;HCoV-HKU1_orf1ab;838 | 0.05 | 0 |
| NC_006577.2;YP_173236.1;HCoV-HKU1_orf1ab;839 | 0 | 0 |
| NC_006577.2;YP_173236.1;HCoV-HKU1_orf1ab;840 | 0 | 0 |
| NC_006577.2;YP_173236.1;HCoV-HKU1_orf1ab;841 | 0 | 0 |
| NC_006577.2;YP_173236.1;HCoV-HKU1_orf1ab;842 | 0.05 | 0 |
| NC_006577.2;YP_173236.1;HCoV-HKU1_orf1ab;843 | 0 | 0 |
| NC_006577.2;YP_173236.1;HCoV-HKU1_orf1ab;844 | 0 | 0 |
| NC_006577.2;YP_173236.1;HCoV-HKU1_orf1ab;845 | 0 | 0 |
| NC_006577.2;YP_173236.1;HCoV-HKU1_orf1ab;846 | 0 | 0 |
| NC_006577.2;YP_173236.1;HCoV-HKU1_orf1ab;847 | 0 | 0 |
| NC_006577.2;YP_173236.1;HCoV-HKU1_orf1ab;848 | 0 | 0 |
| NC_006577.2;YP_173236.1;HCoV-HKU1_orf1ab;849 | 0 | 0 |
| NC_006577.2;YP_173236.1;HCoV-HKU1_orf1ab;850 | 0 | 0 |
| NC_006577.2;YP_173236.1;HCoV-HKU1_orf1ab;851 | 0 | 0 |
| NC_006577.2;YP_173236.1;HCoV-HKU1_orf1ab;852 | 0 | 0 |
| NC_006577.2;YP_173236.1;HCoV-HKU1_orf1ab;853 | 0 | 0 |
| NC_006577.2;YP_173236.1;HCoV-HKU1_orf1ab;854 | 0 | 0 |
| NC_006577.2;YP_173236.1;HCoV-HKU1_orf1ab;855 | 0 | 0 |
| NC_006577.2;YP_173236.1;HCoV-HKU1_orf1ab;856 | 0 | 0 |
| NC_006577.2;YP_173236.1;HCoV-HKU1_orf1ab;857 | 0 | 0 |
| NC_006577.2;YP_173236.1;HCoV-HKU1_orf1ab;858 | 0 | 0 |
| NC_006577.2;YP_173236.1;HCoV-HKU1_orf1ab;859 | 0 | 0 |
| NC_006577.2;YP_173236.1;HCoV-HKU1_orf1ab;860 | 0 | 0 |
| NC_006577.2;YP_173236.1;HCoV-HKU1_orf1ab;861 | 0 | 0 |
| NC_006577.2;YP_173236.1;HCoV-HKU1_orf1ab;862 | 0 | 0 |
| NC_006577.2;YP_173236.1;HCoV-HKU1_orf1ab;863 | 0 | 0 |
| NC_006577.2;YP_173236.1;HCoV-HKU1_orf1ab;864 | 0.05 | 0.025 |
| NC_006577.2;YP_173236.1;HCoV-HKU1_orf1ab;865 | 0.05 | 0.025 |
| NC_006577.2;YP_173236.1;HCoV-HKU1_orf1ab;866 | 0.05 | 0.05 |
| NC_006577.2;YP_173236.1;HCoV-HKU1_orf1ab;867 | 0 | 0.025 |
| NC_006577.2;YP_173236.1;HCoV-HKU1_orf1ab;868 | 0.1 | 0.15 |
| NC_006577.2;YP_173236.1;HCoV-HKU1_orf1ab;869 | 0.05 | 0 |
| NC_006577.2;YP_173236.1;HCoV-HKU1_orf1ab;870 | 0 | 0 |
| NC_006577.2;YP_173236.1;HCoV-HKU1_orf1ab;871 | 0 | 0 |
| NC_006577.2;YP_173236.1;HCoV-HKU1_orf1ab;872 | 0.05 | 0 |
| NC_006577.2;YP_173236.1;HCoV-HKU1_orf1ab;873 | 0.05 | 0 |
| NC_006577.2;YP_173236.1;HCoV-HKU1_orf1ab;874 | 0.05 | 0 |
| NC_006577.2;YP_173236.1;HCoV-HKU1_orf1ab;875 | 0 | 0 |

| | | |
|---|---|---|
| NC_006577.2;YP_173236.1;HCoV-HKU1_orf1ab;876 | 0 | 0 |
| NC_006577.2;YP_173236.1;HCoV-HKU1_orf1ab;877 | 0 | 0 |
| NC_006577.2;YP_173236.1;HCoV-HKU1_orf1ab;878 | 0 | 0 |
| NC_006577.2;YP_173236.1;HCoV-HKU1_orf1ab;879 | 0 | 0 |
| NC_006577.2;YP_173236.1;HCoV-HKU1_orf1ab;880 | 0 | 0 |
| NC_006577.2;YP_173236.1;HCoV-HKU1_orf1ab;881 | 0 | 0 |
| NC_006577.2;YP_173236.1;HCoV-HKU1_orf1ab;882 | 0 | 0 |
| NC_006577.2;YP_173236.1;HCoV-HKU1_orf1ab;883 | 0 | 0.025 |
| NC_006577.2;YP_173236.1;HCoV-HKU1_orf1ab;884 | 0.05 | 0.025 |
| NC_006577.2;YP_173236.1;HCoV-HKU1_orf1ab;885 | 0 | 0 |
| NC_006577.2;YP_173236.1;HCoV-HKU1_orf1ab;886 | 0.05 | 0 |
| NC_006577.2;YP_173236.1;HCoV-HKU1_orf1ab;887 | 0 | 0 |
| NC_006577.2;YP_173236.1;HCoV-HKU1_orf1ab;888 | 0 | 0 |
| NC_006577.2;YP_173236.1;HCoV-HKU1_orf1ab;889 | 0.05 | 0 |
| NC_006577.2;YP_173236.1;HCoV-HKU1_orf1ab;890 | 0 | 0 |
| NC_006577.2;YP_173236.1;HCoV-HKU1_orf1ab;891 | 0 | 0 |
| NC_006577.2;YP_173236.1;HCoV-HKU1_orf1ab;892 | 0 | 0 |
| NC_006577.2;YP_173236.1;HCoV-HKU1_orf1ab;893 | 0.1 | 0 |
| NC_006577.2;YP_173236.1;HCoV-HKU1_orf1ab;894 | 0.05 | 0 |
| NC_006577.2;YP_173236.1;HCoV-HKU1_orf1ab;895 | 0 | 0 |
| NC_006577.2;YP_173236.1;HCoV-HKU1_orf1ab;896 | 0.05 | 0 |
| NC_006577.2;YP_173236.1;HCoV-HKU1_orf1ab;897 | 0.05 | 0 |
| NC_006577.2;YP_173236.1;HCoV-HKU1_orf1ab;898 | 0.05 | 0 |
| NC_006577.2;YP_173236.1;HCoV-HKU1_orf1ab;899 | 0.05 | 0 |
| NC_006577.2;YP_173236.1;HCoV-HKU1_orf1ab;900 | 0.05 | 0 |
| NC_006577.2;YP_173236.1;HCoV-HKU1_orf1ab;901 | 0 | 0 |
| NC_006577.2;YP_173236.1;HCoV-HKU1_orf1ab;902 | 0 | 0 |
| NC_006577.2;YP_173236.1;HCoV-HKU1_orf1ab;903 | 0 | 0 |
| NC_006577.2;YP_173236.1;HCoV-HKU1_orf1ab;904 | 0 | 0 |
| NC_006577.2;YP_173236.1;HCoV-HKU1_orf1ab;905 | 0 | 0 |
| NC_006577.2;YP_173236.1;HCoV-HKU1_orf1ab;906 | 0 | 0 |
| NC_006577.2;YP_173236.1;HCoV-HKU1_orf1ab;907 | 0 | 0 |
| NC_006577.2;YP_173236.1;HCoV-HKU1_orf1ab;908 | 0 | 0 |
| NC_006577.2;YP_173236.1;HCoV-HKU1_orf1ab;909 | 0 | 0 |
| NC_006577.2;YP_173236.1;HCoV-HKU1_orf1ab;910 | 0 | 0 |
| NC_006577.2;YP_173236.1;HCoV-HKU1_orf1ab;911 | 0 | 0.025 |
| NC_006577.2;YP_173236.1;HCoV-HKU1_orf1ab;912 | 0 | 0 |
| NC_006577.2;YP_173236.1;HCoV-HKU1_orf1ab;913 | 0 | 0.025 |
| NC_006577.2;YP_173236.1;HCoV-HKU1_orf1ab;914 | 0 | 0.025 |
| NC_006577.2;YP_173236.1;HCoV-HKU1_orf1ab;915 | 0 | 0.025 |
| NC_006577.2;YP_173236.1;HCoV-HKU1_orf1ab;916 | 0 | 0 |
| NC_006577.2;YP_173236.1;HCoV-HKU1_orf1ab;917 | 0 | 0 |
| NC_006577.2;YP_173236.1;HCoV-HKU1_orf1ab;918 | 0 | 0 |

| | | |
|---|---|---|
| NC_006577.2;YP_173236.1;HCoV-HKU1_orf1ab;919 | 0.1 | 0.025 |
| NC_006577.2;YP_173236.1;HCoV-HKU1_orf1ab;920 | 0 | 0 |
| NC_006577.2;YP_173236.1;HCoV-HKU1_orf1ab;921 | 0 | 0.05 |
| NC_006577.2;YP_173236.1;HCoV-HKU1_orf1ab;922 | 0 | 0 |
| NC_006577.2;YP_173236.1;HCoV-HKU1_orf1ab;923 | 0.05 | 0 |
| NC_006577.2;YP_173236.1;HCoV-HKU1_orf1ab;924 | 0.05 | 0 |
| NC_006577.2;YP_173236.1;HCoV-HKU1_orf1ab;925 | 0 | 0.025 |
| NC_006577.2;YP_173236.1;HCoV-HKU1_orf1ab;926 | 0 | 0 |
| NC_006577.2;YP_173236.1;HCoV-HKU1_orf1ab;927 | 0 | 0 |
| NC_006577.2;YP_173236.1;HCoV-HKU1_orf1ab;928 | 0 | 0 |
| NC_006577.2;YP_173236.1;HCoV-HKU1_orf1ab;929 | 0 | 0.025 |
| NC_006577.2;YP_173236.1;HCoV-HKU1_orf1ab;930 | 0.1 | 0.025 |
| NC_006577.2;YP_173236.1;HCoV-HKU1_orf1ab;931 | 0.2 | 0.125 |
| NC_006577.2;YP_173236.1;HCoV-HKU1_orf1ab;932 | 0.1 | 0.025 |
| NC_006577.2;YP_173236.1;HCoV-HKU1_orf1ab;933 | 0.1 | 0 |
| NC_006577.2;YP_173236.1;HCoV-HKU1_orf1ab;934 | 0.05 | 0 |
| NC_006577.2;YP_173236.1;HCoV-HKU1_orf1ab;935 | 0.1 | 0 |
| NC_006577.2;YP_173236.1;HCoV-HKU1_orf1ab;936 | 0 | 0 |
| NC_006577.2;YP_173236.1;HCoV-HKU1_orf1ab;937 | 0.05 | 0 |
| NC_006577.2;YP_173236.1;HCoV-HKU1_orf1ab;938 | 0.25 | 0.15 |
| NC_006577.2;YP_173236.1;HCoV-HKU1_orf1ab;939 | 0.05 | 0.075 |
| NC_006577.2;YP_173236.1;HCoV-HKU1_orf1ab;940 | 0.2 | 0.025 |
| NC_006577.2;YP_173236.1;HCoV-HKU1_orf1ab;941 | 0.1 | 0.025 |
| NC_006577.2;YP_173236.1;HCoV-HKU1_orf1ab;942 | 0.15 | 0.1 |
| NC_006577.2;YP_173236.1;HCoV-HKU1_orf1ab;943 | 0.2 | 0.1 |
| NC_006577.2;YP_173236.1;HCoV-HKU1_orf1ab;944 | 0.05 | 0.075 |
| NC_006577.2;YP_173236.1;HCoV-HKU1_orf1ab;945 | 0.05 | 0.05 |
| NC_006577.2;YP_173236.1;HCoV-HKU1_orf1ab;946 | 0.05 | 0.075 |
| NC_006577.2;YP_173236.1;HCoV-HKU1_orf1ab;947 | 0.05 | 0.075 |
| NC_006577.2;YP_173236.1;HCoV-HKU1_orf1ab;948 | 0.15 | 0.075 |
| NC_006577.2;YP_173236.1;HCoV-HKU1_orf1ab;949 | 0.1 | 0.075 |
| NC_006577.2;YP_173236.1;HCoV-HKU1_orf1ab;950 | 0.1 | 0.075 |
| NC_006577.2;YP_173236.1;HCoV-HKU1_orf1ab;951 | 0.1 | 0.05 |
| NC_006577.2;YP_173236.1;HCoV-HKU1_orf1ab;952 | 0.1 | 0.075 |
| NC_006577.2;YP_173236.1;HCoV-HKU1_orf1ab;953 | 0.1 | 0.075 |
| NC_006577.2;YP_173236.1;HCoV-HKU1_orf1ab;954 | 0.1 | 0.075 |
| NC_006577.2;YP_173236.1;HCoV-HKU1_orf1ab;955 | 0.05 | 0.05 |
| NC_006577.2;YP_173236.1;HCoV-HKU1_orf1ab;956 | 0.05 | 0.075 |
| NC_006577.2;YP_173236.1;HCoV-HKU1_orf1ab;957 | 0.05 | 0.075 |
| NC_006577.2;YP_173236.1;HCoV-HKU1_orf1ab;958 | 0.15 | 0.075 |
| NC_006577.2;YP_173236.1;HCoV-HKU1_orf1ab;959 | 0.1 | 0.075 |
| NC_006577.2;YP_173236.1;HCoV-HKU1_orf1ab;960 | 0.1 | 0.075 |
| NC_006577.2;YP_173236.1;HCoV-HKU1_orf1ab;961 | 0.1 | 0.05 |

| | | |
|---|---|---|
| NC_006577.2;YP_173236.1;HCoV-HKU1_orf1ab;962 | 0.1 | 0.075 |
| NC_006577.2;YP_173236.1;HCoV-HKU1_orf1ab;963 | 0.1 | 0.075 |
| NC_006577.2;YP_173236.1;HCoV-HKU1_orf1ab;964 | 0.1 | 0.075 |
| NC_006577.2;YP_173236.1;HCoV-HKU1_orf1ab;965 | 0.05 | 0.05 |
| NC_006577.2;YP_173236.1;HCoV-HKU1_orf1ab;966 | 0.05 | 0.075 |
| NC_006577.2;YP_173236.1;HCoV-HKU1_orf1ab;967 | 0.05 | 0.075 |
| NC_006577.2;YP_173236.1;HCoV-HKU1_orf1ab;968 | 0.15 | 0.075 |
| NC_006577.2;YP_173236.1;HCoV-HKU1_orf1ab;969 | 0.1 | 0.075 |
| NC_006577.2;YP_173236.1;HCoV-HKU1_orf1ab;970 | 0.1 | 0.075 |
| NC_006577.2;YP_173236.1;HCoV-HKU1_orf1ab;971 | 0.1 | 0.05 |
| NC_006577.2;YP_173236.1;HCoV-HKU1_orf1ab;972 | 0.1 | 0.075 |
| NC_006577.2;YP_173236.1;HCoV-HKU1_orf1ab;973 | 0.1 | 0.075 |
| NC_006577.2;YP_173236.1;HCoV-HKU1_orf1ab;974 | 0.1 | 0.075 |
| NC_006577.2;YP_173236.1;HCoV-HKU1_orf1ab;975 | 0.05 | 0.05 |
| NC_006577.2;YP_173236.1;HCoV-HKU1_orf1ab;976 | 0.05 | 0.075 |
| NC_006577.2;YP_173236.1;HCoV-HKU1_orf1ab;977 | 0.05 | 0.075 |
| NC_006577.2;YP_173236.1;HCoV-HKU1_orf1ab;978 | 0.15 | 0.075 |
| NC_006577.2;YP_173236.1;HCoV-HKU1_orf1ab;979 | 0.1 | 0.075 |
| NC_006577.2;YP_173236.1;HCoV-HKU1_orf1ab;980 | 0.1 | 0.075 |
| NC_006577.2;YP_173236.1;HCoV-HKU1_orf1ab;981 | 0.1 | 0.05 |
| NC_006577.2;YP_173236.1;HCoV-HKU1_orf1ab;982 | 0.1 | 0.075 |
| NC_006577.2;YP_173236.1;HCoV-HKU1_orf1ab;983 | 0.1 | 0.075 |
| NC_006577.2;YP_173236.1;HCoV-HKU1_orf1ab;984 | 0.1 | 0.075 |
| NC_006577.2;YP_173236.1;HCoV-HKU1_orf1ab;985 | 0.05 | 0.05 |
| NC_006577.2;YP_173236.1;HCoV-HKU1_orf1ab;986 | 0.05 | 0.075 |
| NC_006577.2;YP_173236.1;HCoV-HKU1_orf1ab;987 | 0.05 | 0.075 |
| NC_006577.2;YP_173236.1;HCoV-HKU1_orf1ab;988 | 0.15 | 0.075 |
| NC_006577.2;YP_173236.1;HCoV-HKU1_orf1ab;989 | 0.1 | 0.075 |
| NC_006577.2;YP_173236.1;HCoV-HKU1_orf1ab;990 | 0.1 | 0.075 |
| NC_006577.2;YP_173236.1;HCoV-HKU1_orf1ab;991 | 0.1 | 0.05 |
| NC_006577.2;YP_173236.1;HCoV-HKU1_orf1ab;992 | 0.1 | 0.075 |
| NC_006577.2;YP_173236.1;HCoV-HKU1_orf1ab;993 | 0.1 | 0.075 |
| NC_006577.2;YP_173236.1;HCoV-HKU1_orf1ab;994 | 0.1 | 0.075 |
| NC_006577.2;YP_173236.1;HCoV-HKU1_orf1ab;995 | 0.05 | 0.05 |
| NC_006577.2;YP_173236.1;HCoV-HKU1_orf1ab;996 | 0.05 | 0.075 |
| NC_006577.2;YP_173236.1;HCoV-HKU1_orf1ab;997 | 0.05 | 0.075 |
| NC_006577.2;YP_173236.1;HCoV-HKU1_orf1ab;998 | 0.15 | 0.075 |
| NC_006577.2;YP_173236.1;HCoV-HKU1_orf1ab;999 | 0.1 | 0.075 |
| NC_006577.2;YP_173236.1;HCoV-HKU1_orf1ab;1000 | 0.1 | 0.075 |
| NC_006577.2;YP_173236.1;HCoV-HKU1_orf1ab;1001 | 0.1 | 0.05 |
| NC_006577.2;YP_173236.1;HCoV-HKU1_orf1ab;1002 | 0.1 | 0.075 |
| NC_006577.2;YP_173236.1;HCoV-HKU1_orf1ab;1003 | 0.1 | 0.075 |
| NC_006577.2;YP_173236.1;HCoV-HKU1_orf1ab;1004 | 0.1 | 0.075 |

| | | |
|---|---|---|
| NC_006577.2;YP_173236.1;HCoV-HKU1_orf1ab;1005 | 0.05 | 0.05 |
| NC_006577.2;YP_173236.1;HCoV-HKU1_orf1ab;1006 | 0.05 | 0.075 |
| NC_006577.2;YP_173236.1;HCoV-HKU1_orf1ab;1007 | 0.05 | 0.075 |
| NC_006577.2;YP_173236.1;HCoV-HKU1_orf1ab;1008 | 0.15 | 0.075 |
| NC_006577.2;YP_173236.1;HCoV-HKU1_orf1ab;1009 | 0.1 | 0.075 |
| NC_006577.2;YP_173236.1;HCoV-HKU1_orf1ab;1010 | 0.1 | 0.075 |
| NC_006577.2;YP_173236.1;HCoV-HKU1_orf1ab;1011 | 0.1 | 0.05 |
| NC_006577.2;YP_173236.1;HCoV-HKU1_orf1ab;1012 | 0.1 | 0.075 |
| NC_006577.2;YP_173236.1;HCoV-HKU1_orf1ab;1013 | 0.1 | 0.075 |
| NC_006577.2;YP_173236.1;HCoV-HKU1_orf1ab;1014 | 0.1 | 0.075 |
| NC_006577.2;YP_173236.1;HCoV-HKU1_orf1ab;1015 | 0.05 | 0.05 |
| NC_006577.2;YP_173236.1;HCoV-HKU1_orf1ab;1016 | 0.05 | 0.075 |
| NC_006577.2;YP_173236.1;HCoV-HKU1_orf1ab;1017 | 0.05 | 0.075 |
| NC_006577.2;YP_173236.1;HCoV-HKU1_orf1ab;1018 | 0.15 | 0.075 |
| NC_006577.2;YP_173236.1;HCoV-HKU1_orf1ab;1019 | 0.1 | 0.075 |
| NC_006577.2;YP_173236.1;HCoV-HKU1_orf1ab;1020 | 0.1 | 0.075 |
| NC_006577.2;YP_173236.1;HCoV-HKU1_orf1ab;1021 | 0.1 | 0.05 |
| NC_006577.2;YP_173236.1;HCoV-HKU1_orf1ab;1022 | 0.1 | 0.075 |
| NC_006577.2;YP_173236.1;HCoV-HKU1_orf1ab;1023 | 0.1 | 0.075 |
| NC_006577.2;YP_173236.1;HCoV-HKU1_orf1ab;1024 | 0.1 | 0.075 |
| NC_006577.2;YP_173236.1;HCoV-HKU1_orf1ab;1025 | 0.05 | 0.05 |
| NC_006577.2;YP_173236.1;HCoV-HKU1_orf1ab;1026 | 0.05 | 0.075 |
| NC_006577.2;YP_173236.1;HCoV-HKU1_orf1ab;1027 | 0.05 | 0.075 |
| NC_006577.2;YP_173236.1;HCoV-HKU1_orf1ab;1028 | 0.15 | 0.075 |
| NC_006577.2;YP_173236.1;HCoV-HKU1_orf1ab;1029 | 0.1 | 0.075 |
| NC_006577.2;YP_173236.1;HCoV-HKU1_orf1ab;1030 | 0.1 | 0.075 |
| NC_006577.2;YP_173236.1;HCoV-HKU1_orf1ab;1031 | 0.1 | 0.05 |
| NC_006577.2;YP_173236.1;HCoV-HKU1_orf1ab;1032 | 0.1 | 0.075 |
| NC_006577.2;YP_173236.1;HCoV-HKU1_orf1ab;1033 | 0.1 | 0.075 |
| NC_006577.2;YP_173236.1;HCoV-HKU1_orf1ab;1034 | 0.1 | 0.075 |
| NC_006577.2;YP_173236.1;HCoV-HKU1_orf1ab;1035 | 0.05 | 0.05 |
| NC_006577.2;YP_173236.1;HCoV-HKU1_orf1ab;1036 | 0.05 | 0.075 |
| NC_006577.2;YP_173236.1;HCoV-HKU1_orf1ab;1037 | 0.05 | 0.075 |
| NC_006577.2;YP_173236.1;HCoV-HKU1_orf1ab;1038 | 0.15 | 0.075 |
| NC_006577.2;YP_173236.1;HCoV-HKU1_orf1ab;1039 | 0.1 | 0.075 |
| NC_006577.2;YP_173236.1;HCoV-HKU1_orf1ab;1040 | 0.1 | 0.075 |
| NC_006577.2;YP_173236.1;HCoV-HKU1_orf1ab;1041 | 0.1 | 0.05 |
| NC_006577.2;YP_173236.1;HCoV-HKU1_orf1ab;1042 | 0.1 | 0.075 |
| NC_006577.2;YP_173236.1;HCoV-HKU1_orf1ab;1043 | 0.1 | 0.075 |
| NC_006577.2;YP_173236.1;HCoV-HKU1_orf1ab;1044 | 0.1 | 0.075 |
| NC_006577.2;YP_173236.1;HCoV-HKU1_orf1ab;1045 | 0.05 | 0.05 |
| NC_006577.2;YP_173236.1;HCoV-HKU1_orf1ab;1046 | 0.05 | 0.075 |
| NC_006577.2;YP_173236.1;HCoV-HKU1_orf1ab;1047 | 0.05 | 0.075 |

| | | |
|---|---|---|
| NC_006577.2;YP_173236.1;HCoV-HKU1_orf1ab;1048 | 0.15 | 0.075 |
| NC_006577.2;YP_173236.1;HCoV-HKU1_orf1ab;1049 | 0.1 | 0.075 |
| NC_006577.2;YP_173236.1;HCoV-HKU1_orf1ab;1050 | 0.1 | 0.075 |
| NC_006577.2;YP_173236.1;HCoV-HKU1_orf1ab;1051 | 0.1 | 0.05 |
| NC_006577.2;YP_173236.1;HCoV-HKU1_orf1ab;1052 | 0.1 | 0.075 |
| NC_006577.2;YP_173236.1;HCoV-HKU1_orf1ab;1053 | 0.1 | 0.075 |
| NC_006577.2;YP_173236.1;HCoV-HKU1_orf1ab;1054 | 0.1 | 0.075 |
| NC_006577.2;YP_173236.1;HCoV-HKU1_orf1ab;1055 | 0.05 | 0.05 |
| NC_006577.2;YP_173236.1;HCoV-HKU1_orf1ab;1056 | 0.05 | 0.075 |
| NC_006577.2;YP_173236.1;HCoV-HKU1_orf1ab;1057 | 0.05 | 0.075 |
| NC_006577.2;YP_173236.1;HCoV-HKU1_orf1ab;1058 | 0.15 | 0.075 |
| NC_006577.2;YP_173236.1;HCoV-HKU1_orf1ab;1059 | 0.1 | 0.075 |
| NC_006577.2;YP_173236.1;HCoV-HKU1_orf1ab;1060 | 0.1 | 0.075 |
| NC_006577.2;YP_173236.1;HCoV-HKU1_orf1ab;1061 | 0.1 | 0.05 |
| NC_006577.2;YP_173236.1;HCoV-HKU1_orf1ab;1062 | 0.1 | 0.075 |
| NC_006577.2;YP_173236.1;HCoV-HKU1_orf1ab;1063 | 0.1 | 0.075 |
| NC_006577.2;YP_173236.1;HCoV-HKU1_orf1ab;1064 | 0.1 | 0.075 |
| NC_006577.2;YP_173236.1;HCoV-HKU1_orf1ab;1065 | 0.05 | 0.05 |
| NC_006577.2;YP_173236.1;HCoV-HKU1_orf1ab;1066 | 0.05 | 0.075 |
| NC_006577.2;YP_173236.1;HCoV-HKU1_orf1ab;1067 | 0.05 | 0.075 |
| NC_006577.2;YP_173236.1;HCoV-HKU1_orf1ab;1068 | 0.15 | 0.075 |
| NC_006577.2;YP_173236.1;HCoV-HKU1_orf1ab;1069 | 0.1 | 0.075 |
| NC_006577.2;YP_173236.1;HCoV-HKU1_orf1ab;1070 | 0.1 | 0.075 |
| NC_006577.2;YP_173236.1;HCoV-HKU1_orf1ab;1071 | 0.1 | 0.05 |
| NC_006577.2;YP_173236.1;HCoV-HKU1_orf1ab;1072 | 0.1 | 0.05 |
| NC_006577.2;YP_173236.1;HCoV-HKU1_orf1ab;1073 | 0.05 | 0.025 |
| NC_006577.2;YP_173236.1;HCoV-HKU1_orf1ab;1074 | 0.05 | 0.075 |
| NC_006577.2;YP_173236.1;HCoV-HKU1_orf1ab;1075 | 0.05 | 0.05 |
| NC_006577.2;YP_173236.1;HCoV-HKU1_orf1ab;1076 | 0.05 | 0.025 |
| NC_006577.2;YP_173236.1;HCoV-HKU1_orf1ab;1077 | 0.05 | 0.025 |
| NC_006577.2;YP_173236.1;HCoV-HKU1_orf1ab;1078 | 0.1 | 0.025 |
| NC_006577.2;YP_173236.1;HCoV-HKU1_orf1ab;1079 | 0.05 | 0.025 |
| NC_006577.2;YP_173236.1;HCoV-HKU1_orf1ab;1080 | 0.05 | 0.025 |
| NC_006577.2;YP_173236.1;HCoV-HKU1_orf1ab;1081 | 0.05 | 0.025 |
| NC_006577.2;YP_173236.1;HCoV-HKU1_orf1ab;1082 | 0.05 | 0.05 |
| NC_006577.2;YP_173236.1;HCoV-HKU1_orf1ab;1083 | 0.05 | 0.025 |
| NC_006577.2;YP_173236.1;HCoV-HKU1_orf1ab;1084 | 0.05 | 0.025 |
| NC_006577.2;YP_173236.1;HCoV-HKU1_orf1ab;1085 | 0.05 | 0.025 |
| NC_006577.2;YP_173236.1;HCoV-HKU1_orf1ab;1086 | 0.05 | 0.025 |
| NC_006577.2;YP_173236.1;HCoV-HKU1_orf1ab;1087 | 0.05 | 0.025 |
| NC_006577.2;YP_173236.1;HCoV-HKU1_orf1ab;1088 | 0.05 | 0.05 |
| NC_006577.2;YP_173236.1;HCoV-HKU1_orf1ab;1089 | 0.05 | 0.025 |
| NC_006577.2;YP_173236.1;HCoV-HKU1_orf1ab;1090 | 0.05 | 0.025 |

| | | |
|---|---|---|
| NC_006577.2;YP_173236.1;HCoV-HKU1_orf1ab;1091 | 0.05 | 0 |
| NC_006577.2;YP_173236.1;HCoV-HKU1_orf1ab;1092 | 0 | 0 |
| NC_006577.2;YP_173236.1;HCoV-HKU1_orf1ab;1093 | 0 | 0.025 |
| NC_006577.2;YP_173236.1;HCoV-HKU1_orf1ab;1094 | 0 | 0.05 |
| NC_006577.2;YP_173236.1;HCoV-HKU1_orf1ab;1095 | 0.05 | 0.05 |
| NC_006577.2;YP_173236.1;HCoV-HKU1_orf1ab;1096 | 0 | 0.025 |
| NC_006577.2;YP_173236.1;HCoV-HKU1_orf1ab;1097 | 0 | 0.025 |
| NC_006577.2;YP_173236.1;HCoV-HKU1_orf1ab;1098 | 0 | 0.025 |
| NC_006577.2;YP_173236.1;HCoV-HKU1_orf1ab;1099 | 0 | 0.025 |
| NC_006577.2;YP_173236.1;HCoV-HKU1_orf1ab;1100 | 0 | 0.025 |
| NC_006577.2;YP_173236.1;HCoV-HKU1_orf1ab;1101 | 0.05 | 0.025 |
| NC_006577.2;YP_173236.1;HCoV-HKU1_orf1ab;1102 | 0.05 | 0 |
| NC_006577.2;YP_173236.1;HCoV-HKU1_orf1ab;1103 | 0.05 | 0 |
| NC_006577.2;YP_173236.1;HCoV-HKU1_orf1ab;1104 | 0.2 | 0 |
| NC_006577.2;YP_173236.1;HCoV-HKU1_orf1ab;1105 | 0.25 | 0.025 |
| NC_006577.2;YP_173236.1;HCoV-HKU1_orf1ab;1106 | 0.2 | 0.025 |
| NC_006577.2;YP_173236.1;HCoV-HKU1_orf1ab;1107 | 0.1 | 0 |
| NC_006577.2;YP_173236.1;HCoV-HKU1_orf1ab;1108 | 0.05 | 0 |
| NC_006577.2;YP_173236.1;HCoV-HKU1_orf1ab;1109 | 0.05 | 0 |
| NC_006577.2;YP_173236.1;HCoV-HKU1_orf1ab;1110 | 0.05 | 0 |
| NC_006577.2;YP_173236.1;HCoV-HKU1_orf1ab;1111 | 0.05 | 0 |
| NC_006577.2;YP_173236.1;HCoV-HKU1_orf1ab;1112 | 0.05 | 0 |
| NC_006577.2;YP_173236.1;HCoV-HKU1_orf1ab;1113 | 0 | 0 |
| NC_006577.2;YP_173236.1;HCoV-HKU1_orf1ab;1114 | 0 | 0 |
| NC_006577.2;YP_173236.1;HCoV-HKU1_orf1ab;1115 | 0 | 0 |
| NC_006577.2;YP_173236.1;HCoV-HKU1_orf1ab;1116 | 0 | 0 |
| NC_006577.2;YP_173236.1;HCoV-HKU1_orf1ab;1117 | 0 | 0 |
| NC_006577.2;YP_173236.1;HCoV-HKU1_orf1ab;1118 | 0 | 0 |
| NC_006577.2;YP_173236.1;HCoV-HKU1_orf1ab;1119 | 0 | 0 |
| NC_006577.2;YP_173236.1;HCoV-HKU1_orf1ab;1120 | 0 | 0 |
| NC_006577.2;YP_173236.1;HCoV-HKU1_orf1ab;1121 | 0 | 0 |
| NC_006577.2;YP_173236.1;HCoV-HKU1_orf1ab;1122 | 0 | 0 |
| NC_006577.2;YP_173236.1;HCoV-HKU1_orf1ab;1123 | 0 | 0.05 |
| NC_006577.2;YP_173236.1;HCoV-HKU1_orf1ab;1124 | 0 | 0 |
| NC_006577.2;YP_173236.1;HCoV-HKU1_orf1ab;1125 | 0 | 0 |
| NC_006577.2;YP_173236.1;HCoV-HKU1_orf1ab;1126 | 0 | 0.025 |
| NC_006577.2;YP_173236.1;HCoV-HKU1_orf1ab;1127 | 0 | 0.025 |
| NC_006577.2;YP_173236.1;HCoV-HKU1_orf1ab;1128 | 0 | 0 |
| NC_006577.2;YP_173236.1;HCoV-HKU1_orf1ab;1129 | 0 | 0 |
| NC_006577.2;YP_173236.1;HCoV-HKU1_orf1ab;1130 | 0 | 0.025 |
| NC_006577.2;YP_173236.1;HCoV-HKU1_orf1ab;1131 | 0 | 0.025 |
| NC_006577.2;YP_173236.1;HCoV-HKU1_orf1ab;1132 | 0.05 | 0.05 |
| NC_006577.2;YP_173236.1;HCoV-HKU1_orf1ab;1133 | 0 | 0.1 |

| | | |
|---|---|---|
| NC_006577.2;YP_173236.1;HCoV-HKU1_orf1ab;1134 | 0.05 | 0.1 |
| NC_006577.2;YP_173236.1;HCoV-HKU1_orf1ab;1135 | 0.05 | 0.125 |
| NC_006577.2;YP_173236.1;HCoV-HKU1_orf1ab;1136 | 0 | 0.075 |
| NC_006577.2;YP_173236.1;HCoV-HKU1_orf1ab;1137 | 0.05 | 0.05 |
| NC_006577.2;YP_173236.1;HCoV-HKU1_orf1ab;1138 | 0 | 0.025 |
| NC_006577.2;YP_173236.1;HCoV-HKU1_orf1ab;1139 | 0 | 0.05 |
| NC_006577.2;YP_173236.1;HCoV-HKU1_orf1ab;1140 | 0.05 | 0.05 |
| NC_006577.2;YP_173236.1;HCoV-HKU1_orf1ab;1141 | 0 | 0.025 |
| NC_006577.2;YP_173236.1;HCoV-HKU1_orf1ab;1142 | 0 | 0 |
| NC_006577.2;YP_173236.1;HCoV-HKU1_orf1ab;1143 | 0 | 0 |
| NC_006577.2;YP_173236.1;HCoV-HKU1_orf1ab;1144 | 0 | 0 |
| NC_006577.2;YP_173236.1;HCoV-HKU1_orf1ab;1145 | 0 | 0 |
| NC_006577.2;YP_173236.1;HCoV-HKU1_orf1ab;1146 | 0 | 0 |
| NC_006577.2;YP_173236.1;HCoV-HKU1_orf1ab;1147 | 0 | 0 |
| NC_006577.2;YP_173236.1;HCoV-HKU1_orf1ab;1148 | 0 | 0 |
| NC_006577.2;YP_173236.1;HCoV-HKU1_orf1ab;1149 | 0 | 0 |
| NC_006577.2;YP_173236.1;HCoV-HKU1_orf1ab;1150 | 0 | 0 |
| NC_006577.2;YP_173236.1;HCoV-HKU1_orf1ab;1151 | 0 | 0 |
| NC_006577.2;YP_173236.1;HCoV-HKU1_orf1ab;1152 | 0 | 0 |
| NC_006577.2;YP_173236.1;HCoV-HKU1_orf1ab;1153 | 0 | 0 |
| NC_006577.2;YP_173236.1;HCoV-HKU1_orf1ab;1154 | 0 | 0 |
| NC_006577.2;YP_173236.1;HCoV-HKU1_orf1ab;1155 | 0 | 0 |
| NC_006577.2;YP_173236.1;HCoV-HKU1_orf1ab;1156 | 0 | 0 |
| NC_006577.2;YP_173236.1;HCoV-HKU1_orf1ab;1157 | 0 | 0 |
| NC_006577.2;YP_173236.1;HCoV-HKU1_orf1ab;1158 | 0 | 0 |
| NC_006577.2;YP_173236.1;HCoV-HKU1_orf1ab;1159 | 0 | 0 |
| NC_006577.2;YP_173236.1;HCoV-HKU1_orf1ab;1160 | 0 | 0 |
| NC_006577.2;YP_173236.1;HCoV-HKU1_orf1ab;1161 | 0 | 0 |
| NC_006577.2;YP_173236.1;HCoV-HKU1_orf1ab;1162 | 0 | 0 |
| NC_006577.2;YP_173236.1;HCoV-HKU1_orf1ab;1163 | 0 | 0 |
| NC_006577.2;YP_173236.1;HCoV-HKU1_orf1ab;1164 | 0 | 0 |
| NC_006577.2;YP_173236.1;HCoV-HKU1_orf1ab;1165 | 0 | 0 |
| NC_006577.2;YP_173236.1;HCoV-HKU1_orf1ab;1166 | 0 | 0 |
| NC_006577.2;YP_173236.1;HCoV-HKU1_orf1ab;1167 | 0 | 0 |
| NC_006577.2;YP_173236.1;HCoV-HKU1_orf1ab;1168 | 0 | 0.05 |
| NC_006577.2;YP_173236.1;HCoV-HKU1_orf1ab;1169 | 0 | 0.025 |
| NC_006577.2;YP_173236.1;HCoV-HKU1_orf1ab;1170 | 0.1 | 0.1 |
| NC_006577.2;YP_173236.1;HCoV-HKU1_orf1ab;1171 | 0 | 0.025 |
| NC_006577.2;YP_173236.1;HCoV-HKU1_orf1ab;1172 | 0 | 0.025 |
| NC_006577.2;YP_173236.1;HCoV-HKU1_orf1ab;1173 | 0 | 0.025 |
| NC_006577.2;YP_173236.1;HCoV-HKU1_orf1ab;1174 | 0 | 0 |
| NC_006577.2;YP_173236.1;HCoV-HKU1_orf1ab;1175 | 0 | 0 |
| NC_006577.2;YP_173236.1;HCoV-HKU1_orf1ab;1176 | 0 | 0 |

| | | |
|---|---|---|
| NC_006577.2;YP_173236.1;HCoV-HKU1_orf1ab;1177 | 0 | 0 |
| NC_006577.2;YP_173236.1;HCoV-HKU1_orf1ab;1178 | 0 | 0 |
| NC_006577.2;YP_173236.1;HCoV-HKU1_orf1ab;1179 | 0 | 0 |
| NC_006577.2;YP_173236.1;HCoV-HKU1_orf1ab;1180 | 0 | 0 |
| NC_006577.2;YP_173236.1;HCoV-HKU1_orf1ab;1181 | 0 | 0 |
| NC_006577.2;YP_173236.1;HCoV-HKU1_orf1ab;1182 | 0.05 | 0 |
| NC_006577.2;YP_173236.1;HCoV-HKU1_orf1ab;1183 | 0 | 0 |
| NC_006577.2;YP_173236.1;HCoV-HKU1_orf1ab;1184 | 0 | 0 |
| NC_006577.2;YP_173236.1;HCoV-HKU1_orf1ab;1185 | 0 | 0 |
| NC_006577.2;YP_173236.1;HCoV-HKU1_orf1ab;1186 | 0 | 0 |
| NC_006577.2;YP_173236.1;HCoV-HKU1_orf1ab;1187 | 0 | 0 |
| NC_006577.2;YP_173236.1;HCoV-HKU1_orf1ab;1188 | 0 | 0 |
| NC_006577.2;YP_173236.1;HCoV-HKU1_orf1ab;1189 | 0 | 0 |
| NC_006577.2;YP_173236.1;HCoV-HKU1_orf1ab;1190 | 0 | 0 |
| NC_006577.2;YP_173236.1;HCoV-HKU1_orf1ab;1191 | 0 | 0 |
| NC_006577.2;YP_173236.1;HCoV-HKU1_orf1ab;1192 | 0 | 0 |
| NC_006577.2;YP_173236.1;HCoV-HKU1_orf1ab;1193 | 0 | 0 |
| NC_006577.2;YP_173236.1;HCoV-HKU1_orf1ab;1194 | 0 | 0 |
| NC_006577.2;YP_173236.1;HCoV-HKU1_orf1ab;1195 | 0 | 0 |
| NC_006577.2;YP_173236.1;HCoV-HKU1_orf1ab;1196 | 0 | 0.025 |
| NC_006577.2;YP_173236.1;HCoV-HKU1_orf1ab;1197 | 0 | 0 |
| NC_006577.2;YP_173236.1;HCoV-HKU1_orf1ab;1198 | 0 | 0 |
| NC_006577.2;YP_173236.1;HCoV-HKU1_orf1ab;1199 | 0 | 0 |
| NC_006577.2;YP_173236.1;HCoV-HKU1_orf1ab;1200 | 0 | 0 |
| NC_006577.2;YP_173236.1;HCoV-HKU1_orf1ab;1201 | 0 | 0 |
| NC_006577.2;YP_173236.1;HCoV-HKU1_orf1ab;1202 | 0 | 0 |
| NC_006577.2;YP_173236.1;HCoV-HKU1_orf1ab;1203 | 0 | 0 |
| NC_006577.2;YP_173236.1;HCoV-HKU1_orf1ab;1204 | 0 | 0 |
| NC_006577.2;YP_173236.1;HCoV-HKU1_orf1ab;1205 | 0 | 0 |
| NC_006577.2;YP_173236.1;HCoV-HKU1_orf1ab;1206 | 0 | 0 |
| NC_006577.2;YP_173236.1;HCoV-HKU1_orf1ab;1207 | 0 | 0 |
| NC_006577.2;YP_173236.1;HCoV-HKU1_orf1ab;1208 | 0 | 0 |
| NC_006577.2;YP_173236.1;HCoV-HKU1_orf1ab;1209 | 0 | 0 |
| NC_006577.2;YP_173236.1;HCoV-HKU1_orf1ab;1210 | 0 | 0 |
| NC_006577.2;YP_173236.1;HCoV-HKU1_orf1ab;1211 | 0 | 0 |
| NC_006577.2;YP_173236.1;HCoV-HKU1_orf1ab;1212 | 0 | 0 |
| NC_006577.2;YP_173236.1;HCoV-HKU1_orf1ab;1213 | 0 | 0 |
| NC_006577.2;YP_173236.1;HCoV-HKU1_orf1ab;1214 | 0 | 0 |
| NC_006577.2;YP_173236.1;HCoV-HKU1_orf1ab;1215 | 0 | 0 |
| NC_006577.2;YP_173236.1;HCoV-HKU1_orf1ab;1216 | 0 | 0 |
| NC_006577.2;YP_173236.1;HCoV-HKU1_orf1ab;1217 | 0 | 0 |
| NC_006577.2;YP_173236.1;HCoV-HKU1_orf1ab;1218 | 0 | 0 |
| NC_006577.2;YP_173236.1;HCoV-HKU1_orf1ab;1219 | 0 | 0 |

| | | |
|---|---|---|
| NC_006577.2;YP_173236.1;HCoV-HKU1_orf1ab;1220 | 0 | 0 |
| NC_006577.2;YP_173236.1;HCoV-HKU1_orf1ab;1221 | 0 | 0 |
| NC_006577.2;YP_173236.1;HCoV-HKU1_orf1ab;1222 | 0 | 0 |
| NC_006577.2;YP_173236.1;HCoV-HKU1_orf1ab;1223 | 0 | 0 |
| NC_006577.2;YP_173236.1;HCoV-HKU1_orf1ab;1224 | 0 | 0 |
| NC_006577.2;YP_173236.1;HCoV-HKU1_orf1ab;1225 | 0 | 0 |
| NC_006577.2;YP_173236.1;HCoV-HKU1_orf1ab;1226 | 0 | 0 |
| NC_006577.2;YP_173236.1;HCoV-HKU1_orf1ab;1227 | 0 | 0.025 |
| NC_006577.2;YP_173236.1;HCoV-HKU1_orf1ab;1228 | 0 | 0 |
| NC_006577.2;YP_173236.1;HCoV-HKU1_orf1ab;1229 | 0 | 0.025 |
| NC_006577.2;YP_173236.1;HCoV-HKU1_orf1ab;1230 | 0 | 0 |
| NC_006577.2;YP_173236.1;HCoV-HKU1_orf1ab;1231 | 0 | 0.025 |
| NC_006577.2;YP_173236.1;HCoV-HKU1_orf1ab;1232 | 0 | 0 |
| NC_006577.2;YP_173236.1;HCoV-HKU1_orf1ab;1233 | 0 | 0 |
| NC_006577.2;YP_173236.1;HCoV-HKU1_orf1ab;1234 | 0.05 | 0 |
| NC_006577.2;YP_173236.1;HCoV-HKU1_orf1ab;1235 | 0 | 0 |
| NC_006577.2;YP_173236.1;HCoV-HKU1_orf1ab;1236 | 0 | 0 |
| NC_006577.2;YP_173236.1;HCoV-HKU1_orf1ab;1237 | 0 | 0.05 |
| NC_006577.2;YP_173236.1;HCoV-HKU1_orf1ab;1238 | 0 | 0.025 |
| NC_006577.2;YP_173236.1;HCoV-HKU1_orf1ab;1239 | 0 | 0.05 |
| NC_006577.2;YP_173236.1;HCoV-HKU1_orf1ab;1240 | 0 | 0.025 |
| NC_006577.2;YP_173236.1;HCoV-HKU1_orf1ab;1241 | 0 | 0 |
| NC_006577.2;YP_173236.1;HCoV-HKU1_orf1ab;1242 | 0 | 0 |
| NC_006577.2;YP_173236.1;HCoV-HKU1_orf1ab;1243 | 0 | 0 |
| NC_006577.2;YP_173236.1;HCoV-HKU1_orf1ab;1244 | 0 | 0 |
| NC_006577.2;YP_173236.1;HCoV-HKU1_orf1ab;1245 | 0 | 0 |
| NC_006577.2;YP_173236.1;HCoV-HKU1_orf1ab;1246 | 0 | 0 |
| NC_006577.2;YP_173236.1;HCoV-HKU1_orf1ab;1247 | 0 | 0 |
| NC_006577.2;YP_173236.1;HCoV-HKU1_orf1ab;1248 | 0 | 0 |
| NC_006577.2;YP_173236.1;HCoV-HKU1_orf1ab;1249 | 0 | 0 |
| NC_006577.2;YP_173236.1;HCoV-HKU1_orf1ab;1250 | 0 | 0 |
| NC_006577.2;YP_173236.1;HCoV-HKU1_orf1ab;1251 | 0 | 0 |
| NC_006577.2;YP_173236.1;HCoV-HKU1_orf1ab;1252 | 0 | 0 |
| NC_006577.2;YP_173236.1;HCoV-HKU1_orf1ab;1253 | 0 | 0 |
| NC_006577.2;YP_173236.1;HCoV-HKU1_orf1ab;1254 | 0 | 0 |
| NC_006577.2;YP_173236.1;HCoV-HKU1_orf1ab;1255 | 0 | 0 |
| NC_006577.2;YP_173236.1;HCoV-HKU1_orf1ab;1256 | 0 | 0 |
| NC_006577.2;YP_173236.1;HCoV-HKU1_orf1ab;1257 | 0 | 0 |
| NC_006577.2;YP_173236.1;HCoV-HKU1_orf1ab;1258 | 0 | 0 |
| NC_006577.2;YP_173236.1;HCoV-HKU1_orf1ab;1259 | 0 | 0 |
| NC_006577.2;YP_173236.1;HCoV-HKU1_orf1ab;1260 | 0.05 | 0.025 |
| NC_006577.2;YP_173236.1;HCoV-HKU1_orf1ab;1261 | 0 | 0 |
| NC_006577.2;YP_173236.1;HCoV-HKU1_orf1ab;1262 | 0 | 0 |

| | | |
|---|---|---|
| NC_006577.2;YP_173236.1;HCoV-HKU1_orf1ab;1263 | 0 | 0 |
| NC_006577.2;YP_173236.1;HCoV-HKU1_orf1ab;1264 | 0 | 0 |
| NC_006577.2;YP_173236.1;HCoV-HKU1_orf1ab;1265 | 0 | 0 |
| NC_006577.2;YP_173236.1;HCoV-HKU1_orf1ab;1266 | 0 | 0 |
| NC_006577.2;YP_173236.1;HCoV-HKU1_orf1ab;1267 | 0 | 0 |
| NC_006577.2;YP_173236.1;HCoV-HKU1_orf1ab;1268 | 0 | 0 |
| NC_006577.2;YP_173236.1;HCoV-HKU1_orf1ab;1269 | 0 | 0 |
| NC_006577.2;YP_173236.1;HCoV-HKU1_orf1ab;1270 | 0 | 0 |
| NC_006577.2;YP_173236.1;HCoV-HKU1_orf1ab;1271 | 0 | 0 |
| NC_006577.2;YP_173236.1;HCoV-HKU1_orf1ab;1272 | 0 | 0 |
| NC_006577.2;YP_173236.1;HCoV-HKU1_orf1ab;1273 | 0 | 0 |
| NC_006577.2;YP_173236.1;HCoV-HKU1_orf1ab;1274 | 0 | 0 |
| NC_006577.2;YP_173236.1;HCoV-HKU1_orf1ab;1275 | 0 | 0 |
| NC_006577.2;YP_173236.1;HCoV-HKU1_orf1ab;1276 | 0 | 0 |
| NC_006577.2;YP_173236.1;HCoV-HKU1_orf1ab;1277 | 0 | 0 |
| NC_006577.2;YP_173236.1;HCoV-HKU1_orf1ab;1278 | 0 | 0 |
| NC_006577.2;YP_173236.1;HCoV-HKU1_orf1ab;1279 | 0 | 0 |
| NC_006577.2;YP_173236.1;HCoV-HKU1_orf1ab;1280 | 0 | 0 |
| NC_006577.2;YP_173236.1;HCoV-HKU1_orf1ab;1281 | 0 | 0 |
| NC_006577.2;YP_173236.1;HCoV-HKU1_orf1ab;1282 | 0 | 0 |
| NC_006577.2;YP_173236.1;HCoV-HKU1_orf1ab;1283 | 0 | 0 |
| NC_006577.2;YP_173236.1;HCoV-HKU1_orf1ab;1284 | 0 | 0 |
| NC_006577.2;YP_173236.1;HCoV-HKU1_orf1ab;1285 | 0 | 0 |
| NC_006577.2;YP_173236.1;HCoV-HKU1_orf1ab;1286 | 0 | 0 |
| NC_006577.2;YP_173236.1;HCoV-HKU1_orf1ab;1287 | 0 | 0 |
| NC_006577.2;YP_173236.1;HCoV-HKU1_orf1ab;1288 | 0 | 0 |
| NC_006577.2;YP_173236.1;HCoV-HKU1_orf1ab;1289 | 0 | 0 |
| NC_006577.2;YP_173236.1;HCoV-HKU1_orf1ab;1290 | 0 | 0 |
| NC_006577.2;YP_173236.1;HCoV-HKU1_orf1ab;1291 | 0 | 0 |
| NC_006577.2;YP_173236.1;HCoV-HKU1_orf1ab;1292 | 0 | 0 |
| NC_006577.2;YP_173236.1;HCoV-HKU1_orf1ab;1293 | 0 | 0 |
| NC_006577.2;YP_173236.1;HCoV-HKU1_orf1ab;1294 | 0 | 0 |
| NC_006577.2;YP_173236.1;HCoV-HKU1_orf1ab;1295 | 0.05 | 0 |
| NC_006577.2;YP_173236.1;HCoV-HKU1_orf1ab;1296 | 0 | 0 |
| NC_006577.2;YP_173236.1;HCoV-HKU1_orf1ab;1297 | 0 | 0 |
| NC_006577.2;YP_173236.1;HCoV-HKU1_orf1ab;1298 | 0 | 0 |
| NC_006577.2;YP_173236.1;HCoV-HKU1_orf1ab;1299 | 0 | 0 |
| NC_006577.2;YP_173236.1;HCoV-HKU1_orf1ab;1300 | 0.05 | 0 |
| NC_006577.2;YP_173236.1;HCoV-HKU1_orf1ab;1301 | 0 | 0 |
| NC_006577.2;YP_173236.1;HCoV-HKU1_orf1ab;1302 | 0 | 0 |
| NC_006577.2;YP_173236.1;HCoV-HKU1_orf1ab;1303 | 0 | 0 |
| NC_006577.2;YP_173236.1;HCoV-HKU1_orf1ab;1304 | 0 | 0 |
| NC_006577.2;YP_173236.1;HCoV-HKU1_orf1ab;1305 | 0 | 0 |

| | | |
|---|---|---|
| NC_006577.2;YP_173236.1;HCoV-HKU1_orf1ab;1306 | 0 | 0 |
| NC_006577.2;YP_173236.1;HCoV-HKU1_orf1ab;1307 | 0 | 0 |
| NC_006577.2;YP_173236.1;HCoV-HKU1_orf1ab;1308 | 0 | 0 |
| NC_006577.2;YP_173236.1;HCoV-HKU1_orf1ab;1309 | 0.05 | 0.025 |
| NC_006577.2;YP_173236.1;HCoV-HKU1_orf1ab;1310 | 0.05 | 0.025 |
| NC_006577.2;YP_173236.1;HCoV-HKU1_orf1ab;1311 | 0.05 | 0.025 |
| NC_006577.2;YP_173236.1;HCoV-HKU1_orf1ab;1312 | 0 | 0.05 |
| NC_006577.2;YP_173236.1;HCoV-HKU1_orf1ab;1313 | 0 | 0.05 |
| NC_006577.2;YP_173236.1;HCoV-HKU1_orf1ab;1314 | 0 | 0.05 |
| NC_006577.2;YP_173236.1;HCoV-HKU1_orf1ab;1315 | 0 | 0 |
| NC_006577.2;YP_173236.1;HCoV-HKU1_orf1ab;1316 | 0 | 0 |
| NC_006577.2;YP_173236.1;HCoV-HKU1_orf1ab;1317 | 0 | 0 |
| NC_006577.2;YP_173236.1;HCoV-HKU1_orf1ab;1318 | 0 | 0 |
| NC_006577.2;YP_173236.1;HCoV-HKU1_orf1ab;1319 | 0.1 | 0.025 |
| NC_006577.2;YP_173236.1;HCoV-HKU1_orf1ab;1320 | 0.1 | 0.05 |
| NC_006577.2;YP_173236.1;HCoV-HKU1_orf1ab;1321 | 0.15 | 0.025 |
| NC_006577.2;YP_173236.1;HCoV-HKU1_orf1ab;1322 | 0.1 | 0.05 |
| NC_006577.2;YP_173236.1;HCoV-HKU1_orf1ab;1323 | 0.05 | 0.025 |
| NC_006577.2;YP_173236.1;HCoV-HKU1_orf1ab;1324 | 0.1 | 0 |
| NC_006577.2;YP_173236.1;HCoV-HKU1_orf1ab;1325 | 0.15 | 0 |
| NC_006577.2;YP_173236.1;HCoV-HKU1_orf1ab;1326 | 0.05 | 0 |
| NC_006577.2;YP_173236.1;HCoV-HKU1_orf1ab;1327 | 0.1 | 0.025 |
| NC_006577.2;YP_173236.1;HCoV-HKU1_orf1ab;1328 | 0.1 | 0 |
| NC_006577.2;YP_173236.1;HCoV-HKU1_orf1ab;1329 | 0.05 | 0 |
| NC_006577.2;YP_173236.1;HCoV-HKU1_orf1ab;1330 | 0 | 0 |
| NC_006577.2;YP_173236.1;HCoV-HKU1_orf1ab;1331 | 0 | 0 |
| NC_006577.2;YP_173236.1;HCoV-HKU1_orf1ab;1332 | 0 | 0 |
| NC_006577.2;YP_173236.1;HCoV-HKU1_orf1ab;1333 | 0.05 | 0 |
| NC_006577.2;YP_173236.1;HCoV-HKU1_orf1ab;1334 | 0 | 0 |
| NC_006577.2;YP_173236.1;HCoV-HKU1_orf1ab;1335 | 0 | 0 |
| NC_006577.2;YP_173236.1;HCoV-HKU1_orf1ab;1336 | 0 | 0 |
| NC_006577.2;YP_173236.1;HCoV-HKU1_orf1ab;1337 | 0 | 0 |
| NC_006577.2;YP_173236.1;HCoV-HKU1_orf1ab;1338 | 0 | 0 |
| NC_006577.2;YP_173236.1;HCoV-HKU1_orf1ab;1339 | 0 | 0 |
| NC_006577.2;YP_173236.1;HCoV-HKU1_orf1ab;1340 | 0 | 0 |
| NC_006577.2;YP_173236.1;HCoV-HKU1_orf1ab;1341 | 0 | 0 |
| NC_006577.2;YP_173236.1;HCoV-HKU1_orf1ab;1342 | 0 | 0 |
| NC_006577.2;YP_173236.1;HCoV-HKU1_orf1ab;1343 | 0 | 0 |
| NC_006577.2;YP_173236.1;HCoV-HKU1_orf1ab;1344 | 0 | 0 |
| NC_006577.2;YP_173236.1;HCoV-HKU1_orf1ab;1345 | 0 | 0 |
| NC_006577.2;YP_173236.1;HCoV-HKU1_orf1ab;1346 | 0.05 | 0.075 |
| NC_006577.2;YP_173236.1;HCoV-HKU1_orf1ab;1347 | 0 | 0 |
| NC_006577.2;YP_173236.1;HCoV-HKU1_orf1ab;1348 | 0 | 0 |

| | | |
|---|---|---|
| NC_006577.2;YP_173236.1;HCoV-HKU1_orf1ab;1349 | 0 | 0 |
| NC_006577.2;YP_173236.1;HCoV-HKU1_orf1ab;1350 | 0 | 0 |
| NC_006577.2;YP_173236.1;HCoV-HKU1_orf1ab;1351 | 0 | 0 |
| NC_006577.2;YP_173236.1;HCoV-HKU1_orf1ab;1352 | 0 | 0 |
| NC_006577.2;YP_173236.1;HCoV-HKU1_orf1ab;1353 | 0 | 0 |
| NC_006577.2;YP_173236.1;HCoV-HKU1_orf1ab;1354 | 0 | 0 |
| NC_006577.2;YP_173236.1;HCoV-HKU1_orf1ab;1355 | 0 | 0 |
| NC_006577.2;YP_173236.1;HCoV-HKU1_orf1ab;1356 | 0 | 0 |
| NC_006577.2;YP_173236.1;HCoV-HKU1_orf1ab;1357 | 0 | 0 |
| NC_006577.2;YP_173236.1;HCoV-HKU1_orf1ab;1358 | 0 | 0 |
| NC_006577.2;YP_173236.1;HCoV-HKU1_orf1ab;1359 | 0 | 0 |
| NC_006577.2;YP_173236.1;HCoV-HKU1_orf1ab;1360 | 0 | 0.025 |
| NC_006577.2;YP_173236.1;HCoV-HKU1_orf1ab;1361 | 0 | 0 |
| NC_006577.2;YP_173236.1;HCoV-HKU1_orf1ab;1362 | 0 | 0 |
| NC_006577.2;YP_173236.1;HCoV-HKU1_orf1ab;1363 | 0 | 0 |
| NC_006577.2;YP_173236.1;HCoV-HKU1_orf1ab;1364 | 0 | 0 |
| NC_006577.2;YP_173236.1;HCoV-HKU1_orf1ab;1365 | 0 | 0 |
| NC_006577.2;YP_173236.1;HCoV-HKU1_orf1ab;1366 | 0 | 0 |
| NC_006577.2;YP_173236.1;HCoV-HKU1_orf1ab;1367 | 0 | 0 |
| NC_006577.2;YP_173236.1;HCoV-HKU1_orf1ab;1368 | 0 | 0 |
| NC_006577.2;YP_173236.1;HCoV-HKU1_orf1ab;1369 | 0 | 0 |
| NC_006577.2;YP_173236.1;HCoV-HKU1_orf1ab;1370 | 0 | 0 |
| NC_006577.2;YP_173236.1;HCoV-HKU1_orf1ab;1371 | 0 | 0 |
| NC_006577.2;YP_173236.1;HCoV-HKU1_orf1ab;1372 | 0 | 0 |
| NC_006577.2;YP_173236.1;HCoV-HKU1_orf1ab;1373 | 0 | 0 |
| NC_006577.2;YP_173236.1;HCoV-HKU1_orf1ab;1374 | 0 | 0 |
| NC_006577.2;YP_173236.1;HCoV-HKU1_orf1ab;1375 | 0 | 0 |
| NC_006577.2;YP_173236.1;HCoV-HKU1_orf1ab;1376 | 0 | 0 |
| NC_006577.2;YP_173236.1;HCoV-HKU1_orf1ab;1377 | 0 | 0 |
| NC_006577.2;YP_173236.1;HCoV-HKU1_orf1ab;1378 | 0 | 0 |
| NC_006577.2;YP_173236.1;HCoV-HKU1_orf1ab;1379 | 0.05 | 0 |
| NC_006577.2;YP_173236.1;HCoV-HKU1_orf1ab;1380 | 0 | 0.05 |
| NC_006577.2;YP_173236.1;HCoV-HKU1_orf1ab;1381 | 0.05 | 0.025 |
| NC_006577.2;YP_173236.1;HCoV-HKU1_orf1ab;1382 | 0 | 0 |
| NC_006577.2;YP_173236.1;HCoV-HKU1_orf1ab;1383 | 0.05 | 0.075 |
| NC_006577.2;YP_173236.1;HCoV-HKU1_orf1ab;1384 | 0.05 | 0 |
| NC_006577.2;YP_173236.1;HCoV-HKU1_orf1ab;1385 | 0.05 | 0 |
| NC_006577.2;YP_173236.1;HCoV-HKU1_orf1ab;1386 | 0 | 0 |
| NC_006577.2;YP_173236.1;HCoV-HKU1_orf1ab;1387 | 0.05 | 0 |
| NC_006577.2;YP_173236.1;HCoV-HKU1_orf1ab;1388 | 0.05 | 0 |
| NC_006577.2;YP_173236.1;HCoV-HKU1_orf1ab;1389 | 0 | 0 |
| NC_006577.2;YP_173236.1;HCoV-HKU1_orf1ab;1390 | 0.05 | 0 |
| NC_006577.2;YP_173236.1;HCoV-HKU1_orf1ab;1391 | 0.05 | 0 |

| | | |
|---|---|---|
| NC_006577.2;YP_173236.1;HCoV-HKU1_orf1ab;1392 | 0 | 0 |
| NC_006577.2;YP_173236.1;HCoV-HKU1_orf1ab;1393 | 0 | 0 |
| NC_006577.2;YP_173236.1;HCoV-HKU1_orf1ab;1394 | 0 | 0 |
| NC_006577.2;YP_173236.1;HCoV-HKU1_orf1ab;1395 | 0 | 0 |
| NC_006577.2;YP_173236.1;HCoV-HKU1_orf1ab;1396 | 0 | 0 |
| NC_006577.2;YP_173236.1;HCoV-HKU1_orf1ab;1397 | 0 | 0.025 |
| NC_006577.2;YP_173236.1;HCoV-HKU1_orf1ab;1398 | 0 | 0 |
| NC_006577.2;YP_173236.1;HCoV-HKU1_orf1ab;1399 | 0 | 0.025 |
| NC_006577.2;YP_173236.1;HCoV-HKU1_orf1ab;1400 | 0 | 0.025 |
| NC_006577.2;YP_173236.1;HCoV-HKU1_orf1ab;1401 | 0 | 0.025 |
| NC_006577.2;YP_173236.1;HCoV-HKU1_orf1ab;1402 | 0 | 0 |
| NC_006577.2;YP_173236.1;HCoV-HKU1_orf1ab;1403 | 0 | 0 |
| NC_006577.2;YP_173236.1;HCoV-HKU1_orf1ab;1404 | 0 | 0 |
| NC_006577.2;YP_173236.1;HCoV-HKU1_orf1ab;1405 | 0 | 0 |
| NC_006577.2;YP_173236.1;HCoV-HKU1_orf1ab;1406 | 0 | 0 |
| NC_006577.2;YP_173236.1;HCoV-HKU1_orf1ab;1407 | 0 | 0 |
| NC_006577.2;YP_173236.1;HCoV-HKU1_orf1ab;1408 | 0 | 0 |
| NC_006577.2;YP_173236.1;HCoV-HKU1_orf1ab;1409 | 0 | 0 |
| NC_006577.2;YP_173236.1;HCoV-HKU1_orf1ab;1410 | 0 | 0 |
| NC_006577.2;YP_173236.1;HCoV-HKU1_orf1ab;1411 | 0 | 0 |
| NC_006577.2;YP_173236.1;HCoV-HKU1_orf1ab;1412 | 0 | 0 |
| NC_006577.2;YP_173236.1;HCoV-HKU1_orf1ab;1413 | 0 | 0 |
| NC_006577.2;YP_173236.1;HCoV-HKU1_orf1ab;1414 | 0 | 0 |
| NC_006577.2;YP_173236.1;HCoV-HKU1_orf1ab;1415 | 0 | 0 |
| NC_006577.2;YP_173236.1;HCoV-HKU1_orf1ab;1416 | 0 | 0.025 |
| NC_006577.2;YP_173236.1;HCoV-HKU1_orf1ab;1417 | 0 | 0 |
| NC_006577.2;YP_173236.1;HCoV-HKU1_orf1ab;1418 | 0 | 0.025 |
| NC_006577.2;YP_173236.1;HCoV-HKU1_orf1ab;1419 | 0.05 | 0 |
| NC_006577.2;YP_173236.1;HCoV-HKU1_orf1ab;1420 | 0 | 0.05 |
| NC_006577.2;YP_173236.1;HCoV-HKU1_orf1ab;1421 | 0 | 0 |
| NC_006577.2;YP_173236.1;HCoV-HKU1_orf1ab;1422 | 0 | 0 |
| NC_006577.2;YP_173236.1;HCoV-HKU1_orf1ab;1423 | 0 | 0 |
| NC_006577.2;YP_173236.1;HCoV-HKU1_orf1ab;1424 | 0 | 0 |
| NC_006577.2;YP_173236.1;HCoV-HKU1_orf1ab;1425 | 0 | 0 |
| NC_006577.2;YP_173236.1;HCoV-HKU1_orf1ab;1426 | 0 | 0 |
| NC_006577.2;YP_173236.1;HCoV-HKU1_orf1ab;1427 | 0 | 0 |
| NC_006577.2;YP_173236.1;HCoV-HKU1_orf1ab;1428 | 0 | 0 |
| NC_006577.2;YP_173236.1;HCoV-HKU1_orf1ab;1429 | 0 | 0 |
| NC_006577.2;YP_173236.1;HCoV-HKU1_orf1ab;1430 | 0 | 0 |
| NC_006577.2;YP_173236.1;HCoV-HKU1_orf1ab;1431 | 0 | 0 |
| NC_006577.2;YP_173236.1;HCoV-HKU1_orf1ab;1432 | 0 | 0 |
| NC_006577.2;YP_173236.1;HCoV-HKU1_orf1ab;1433 | 0 | 0 |
| NC_006577.2;YP_173236.1;HCoV-HKU1_orf1ab;1434 | 0 | 0 |

| | | |
|---|---|---|
| NC_006577.2;YP_173236.1;HCoV-HKU1_orf1ab;1435 | 0 | 0 |
| NC_006577.2;YP_173236.1;HCoV-HKU1_orf1ab;1436 | 0 | 0 |
| NC_006577.2;YP_173236.1;HCoV-HKU1_orf1ab;1437 | 0 | 0.025 |
| NC_006577.2;YP_173236.1;HCoV-HKU1_orf1ab;1438 | 0 | 0.025 |
| NC_006577.2;YP_173236.1;HCoV-HKU1_orf1ab;1439 | 0 | 0.075 |
| NC_006577.2;YP_173236.1;HCoV-HKU1_orf1ab;1440 | 0.05 | 0.075 |
| NC_006577.2;YP_173236.1;HCoV-HKU1_orf1ab;1441 | 0.05 | 0.075 |
| NC_006577.2;YP_173236.1;HCoV-HKU1_orf1ab;1442 | 0.05 | 0.025 |
| NC_006577.2;YP_173236.1;HCoV-HKU1_orf1ab;1443 | 0.05 | 0.025 |
| NC_006577.2;YP_173236.1;HCoV-HKU1_orf1ab;1444 | 0 | 0 |
| NC_006577.2;YP_173236.1;HCoV-HKU1_orf1ab;1445 | 0 | 0 |
| NC_006577.2;YP_173236.1;HCoV-HKU1_orf1ab;1446 | 0 | 0 |
| NC_006577.2;YP_173236.1;HCoV-HKU1_orf1ab;1447 | 0 | 0 |
| NC_006577.2;YP_173236.1;HCoV-HKU1_orf1ab;1448 | 0 | 0 |
| NC_006577.2;YP_173236.1;HCoV-HKU1_orf1ab;1449 | 0 | 0 |
| NC_006577.2;YP_173236.1;HCoV-HKU1_orf1ab;1450 | 0 | 0 |
| NC_006577.2;YP_173236.1;HCoV-HKU1_orf1ab;1451 | 0 | 0 |
| NC_006577.2;YP_173236.1;HCoV-HKU1_orf1ab;1452 | 0 | 0 |
| NC_006577.2;YP_173236.1;HCoV-HKU1_orf1ab;1453 | 0 | 0 |
| NC_006577.2;YP_173236.1;HCoV-HKU1_orf1ab;1454 | 0 | 0 |
| NC_006577.2;YP_173236.1;HCoV-HKU1_orf1ab;1455 | 0 | 0 |
| NC_006577.2;YP_173236.1;HCoV-HKU1_orf1ab;1456 | 0 | 0 |
| NC_006577.2;YP_173236.1;HCoV-HKU1_orf1ab;1457 | 0 | 0 |
| NC_006577.2;YP_173236.1;HCoV-HKU1_orf1ab;1458 | 0.1 | 0 |
| NC_006577.2;YP_173236.1;HCoV-HKU1_orf1ab;1459 | 0 | 0 |
| NC_006577.2;YP_173236.1;HCoV-HKU1_orf1ab;1460 | 0 | 0 |
| NC_006577.2;YP_173236.1;HCoV-HKU1_orf1ab;1461 | 0 | 0 |
| NC_006577.2;YP_173236.1;HCoV-HKU1_orf1ab;1462 | 0 | 0 |
| NC_006577.2;YP_173236.1;HCoV-HKU1_orf1ab;1463 | 0 | 0 |
| NC_006577.2;YP_173236.1;HCoV-HKU1_orf1ab;1464 | 0 | 0 |
| NC_006577.2;YP_173236.1;HCoV-HKU1_orf1ab;1465 | 0 | 0 |
| NC_006577.2;YP_173236.1;HCoV-HKU1_orf1ab;1466 | 0 | 0 |
| NC_006577.2;YP_173236.1;HCoV-HKU1_orf1ab;1467 | 0 | 0 |
| NC_006577.2;YP_173236.1;HCoV-HKU1_orf1ab;1468 | 0 | 0 |
| NC_006577.2;YP_173236.1;HCoV-HKU1_orf1ab;1469 | 0 | 0 |
| NC_006577.2;YP_173236.1;HCoV-HKU1_orf1ab;1470 | 0 | 0 |
| NC_006577.2;YP_173236.1;HCoV-HKU1_orf1ab;1471 | 0 | 0 |
| NC_006577.2;YP_173236.1;HCoV-HKU1_orf1ab;1472 | 0 | 0 |
| NC_006577.2;YP_173236.1;HCoV-HKU1_orf1ab;1473 | 0 | 0 |
| NC_006577.2;YP_173236.1;HCoV-HKU1_orf1ab;1474 | 0 | 0 |
| NC_006577.2;YP_173236.1;HCoV-HKU1_orf1ab;1475 | 0 | 0 |
| NC_006577.2;YP_173236.1;HCoV-HKU1_orf1ab;1476 | 0 | 0.025 |
| NC_006577.2;YP_173236.1;HCoV-HKU1_orf1ab;1477 | 0 | 0.025 |

| | | |
|---|---|---|
| NC_006577.2;YP_173236.1;HCoV-HKU1_orf1ab;1478 | 0 | 0 |
| NC_006577.2;YP_173236.1;HCoV-HKU1_orf1ab;1479 | 0 | 0 |
| NC_006577.2;YP_173236.1;HCoV-HKU1_orf1ab;1480 | 0 | 0 |
| NC_006577.2;YP_173236.1;HCoV-HKU1_orf1ab;1481 | 0 | 0 |
| NC_006577.2;YP_173236.1;HCoV-HKU1_orf1ab;1482 | 0 | 0 |
| NC_006577.2;YP_173236.1;HCoV-HKU1_orf1ab;1483 | 0 | 0 |
| NC_006577.2;YP_173236.1;HCoV-HKU1_orf1ab;1484 | 0.05 | 0 |
| NC_006577.2;YP_173236.1;HCoV-HKU1_orf1ab;1485 | 0 | 0 |
| NC_006577.2;YP_173236.1;HCoV-HKU1_orf1ab;1486 | 0 | 0 |
| NC_006577.2;YP_173236.1;HCoV-HKU1_orf1ab;1487 | 0 | 0 |
| NC_006577.2;YP_173236.1;HCoV-HKU1_orf1ab;1488 | 0 | 0 |
| NC_006577.2;YP_173236.1;HCoV-HKU1_orf1ab;1489 | 0 | 0 |
| NC_006577.2;YP_173236.1;HCoV-HKU1_orf1ab;1490 | 0 | 0 |
| NC_006577.2;YP_173236.1;HCoV-HKU1_orf1ab;1491 | 0 | 0 |
| NC_006577.2;YP_173236.1;HCoV-HKU1_orf1ab;1492 | 0 | 0 |
| NC_006577.2;YP_173236.1;HCoV-HKU1_orf1ab;1493 | 0 | 0 |
| NC_006577.2;YP_173236.1;HCoV-HKU1_orf1ab;1494 | 0.05 | 0 |
| NC_006577.2;YP_173236.1;HCoV-HKU1_orf1ab;1495 | 0.05 | 0.025 |
| NC_006577.2;YP_173236.1;HCoV-HKU1_orf1ab;1496 | 0.05 | 0 |
| NC_006577.2;YP_173236.1;HCoV-HKU1_orf1ab;1497 | 0.05 | 0 |
| NC_006577.2;YP_173236.1;HCoV-HKU1_orf1ab;1498 | 0 | 0 |
| NC_006577.2;YP_173236.1;HCoV-HKU1_orf1ab;1499 | 0 | 0 |
| NC_006577.2;YP_173236.1;HCoV-HKU1_orf1ab;1500 | 0.05 | 0 |
| NC_006577.2;YP_173236.1;HCoV-HKU1_orf1ab;1501 | 0.1 | 0.025 |
| NC_006577.2;YP_173236.1;HCoV-HKU1_orf1ab;1502 | 0.25 | 0.175 |
| NC_006577.2;YP_173236.1;HCoV-HKU1_orf1ab;1503 | 0.6 | 0.5 |
| NC_006577.2;YP_173236.1;HCoV-HKU1_orf1ab;1504 | 0.1 | 0.025 |
| NC_006577.2;YP_173236.1;HCoV-HKU1_orf1ab;1505 | 0 | 0.025 |
| NC_006577.2;YP_173236.1;HCoV-HKU1_orf1ab;1506 | 0 | 0.075 |
| NC_006577.2;YP_173236.1;HCoV-HKU1_orf1ab;1507 | 0 | 0.025 |
| NC_006577.2;YP_173236.1;HCoV-HKU1_orf1ab;1508 | 0 | 0 |
| NC_006577.2;YP_173236.1;HCoV-HKU1_orf1ab;1509 | 0 | 0 |
| NC_006577.2;YP_173236.1;HCoV-HKU1_orf1ab;1510 | 0.05 | 0.025 |
| NC_006577.2;YP_173236.1;HCoV-HKU1_orf1ab;1511 | 0 | 0.025 |
| NC_006577.2;YP_173236.1;HCoV-HKU1_orf1ab;1512 | 0 | 0.05 |
| NC_006577.2;YP_173236.1;HCoV-HKU1_orf1ab;1513 | 0 | 0 |
| NC_006577.2;YP_173236.1;HCoV-HKU1_orf1ab;1514 | 0 | 0 |
| NC_006577.2;YP_173236.1;HCoV-HKU1_orf1ab;1515 | 0 | 0 |
| NC_006577.2;YP_173236.1;HCoV-HKU1_orf1ab;1516 | 0 | 0 |
| NC_006577.2;YP_173236.1;HCoV-HKU1_orf1ab;1517 | 0 | 0 |
| NC_006577.2;YP_173236.1;HCoV-HKU1_orf1ab;1518 | 0 | 0 |
| NC_006577.2;YP_173236.1;HCoV-HKU1_orf1ab;1519 | 0 | 0 |
| NC_006577.2;YP_173236.1;HCoV-HKU1_orf1ab;1520 | 0 | 0 |

| | | |
|---|---|---|
| NC_006577.2;YP_173236.1;HCoV-HKU1_orf1ab;1521 | 0 | 0 |
| NC_006577.2;YP_173236.1;HCoV-HKU1_orf1ab;1522 | 0 | 0 |
| NC_006577.2;YP_173236.1;HCoV-HKU1_orf1ab;1523 | 0 | 0 |
| NC_006577.2;YP_173236.1;HCoV-HKU1_orf1ab;1524 | 0 | 0 |
| NC_006577.2;YP_173236.1;HCoV-HKU1_orf1ab;1525 | 0 | 0 |
| NC_006577.2;YP_173236.1;HCoV-HKU1_orf1ab;1526 | 0 | 0 |
| NC_006577.2;YP_173236.1;HCoV-HKU1_orf1ab;1527 | 0 | 0 |
| NC_006577.2;YP_173236.1;HCoV-HKU1_orf1ab;1528 | 0 | 0 |
| NC_006577.2;YP_173236.1;HCoV-HKU1_orf1ab;1529 | 0 | 0 |
| NC_006577.2;YP_173236.1;HCoV-HKU1_orf1ab;1530 | 0 | 0 |
| NC_006577.2;YP_173236.1;HCoV-HKU1_orf1ab;1531 | 0 | 0.025 |
| NC_006577.2;YP_173236.1;HCoV-HKU1_orf1ab;1532 | 0 | 0 |
| NC_006577.2;YP_173236.1;HCoV-HKU1_orf1ab;1533 | 0.05 | 0.025 |
| NC_006577.2;YP_173236.1;HCoV-HKU1_orf1ab;1534 | 0.05 | 0.05 |
| NC_006577.2;YP_173236.1;HCoV-HKU1_orf1ab;1535 | 0 | 0 |
| NC_006577.2;YP_173236.1;HCoV-HKU1_orf1ab;1536 | 0 | 0 |
| NC_006577.2;YP_173236.1;HCoV-HKU1_orf1ab;1537 | 0 | 0 |
| NC_006577.2;YP_173236.1;HCoV-HKU1_orf1ab;1538 | 0 | 0 |
| NC_006577.2;YP_173236.1;HCoV-HKU1_orf1ab;1539 | 0 | 0 |
| NC_006577.2;YP_173236.1;HCoV-HKU1_orf1ab;1540 | 0 | 0 |
| NC_006577.2;YP_173236.1;HCoV-HKU1_orf1ab;1541 | 0 | 0 |
| NC_006577.2;YP_173236.1;HCoV-HKU1_orf1ab;1542 | 0 | 0 |
| NC_006577.2;YP_173236.1;HCoV-HKU1_orf1ab;1543 | 0 | 0 |
| NC_006577.2;YP_173236.1;HCoV-HKU1_orf1ab;1544 | 0 | 0 |
| NC_006577.2;YP_173236.1;HCoV-HKU1_orf1ab;1545 | 0 | 0 |
| NC_006577.2;YP_173236.1;HCoV-HKU1_orf1ab;1546 | 0 | 0 |
| NC_006577.2;YP_173236.1;HCoV-HKU1_orf1ab;1547 | 0 | 0.025 |
| NC_006577.2;YP_173236.1;HCoV-HKU1_orf1ab;1548 | 0 | 0.05 |
| NC_006577.2;YP_173236.1;HCoV-HKU1_orf1ab;1549 | 0 | 0.05 |
| NC_006577.2;YP_173236.1;HCoV-HKU1_orf1ab;1550 | 0.1 | 0.1 |
| NC_006577.2;YP_173236.1;HCoV-HKU1_orf1ab;1551 | 0.2 | 0.175 |
| NC_006577.2;YP_173236.1;HCoV-HKU1_orf1ab;1552 | 0.1 | 0.1 |
| NC_006577.2;YP_173236.1;HCoV-HKU1_orf1ab;1553 | 0.15 | 0.1 |
| NC_006577.2;YP_173236.1;HCoV-HKU1_orf1ab;1554 | 0 | 0 |
| NC_006577.2;YP_173236.1;HCoV-HKU1_orf1ab;1555 | 0 | 0 |
| NC_006577.2;YP_173236.1;HCoV-HKU1_orf1ab;1556 | 0 | 0 |
| NC_006577.2;YP_173236.1;HCoV-HKU1_orf1ab;1557 | 0 | 0 |
| NC_006577.2;YP_173236.1;HCoV-HKU1_orf1ab;1558 | 0 | 0 |
| NC_006577.2;YP_173236.1;HCoV-HKU1_orf1ab;1559 | 0.05 | 0.025 |
| NC_006577.2;YP_173236.1;HCoV-HKU1_orf1ab;1560 | 0.05 | 0 |
| NC_006577.2;YP_173236.1;HCoV-HKU1_orf1ab;1561 | 0 | 0 |
| NC_006577.2;YP_173236.1;HCoV-HKU1_orf1ab;1562 | 0 | 0 |
| NC_006577.2;YP_173236.1;HCoV-HKU1_orf1ab;1563 | 0 | 0 |

| | | |
|---|---|---|
| NC_006577.2;YP_173236.1;HCoV-HKU1_orf1ab;1564 | 0 | 0 |
| NC_006577.2;YP_173236.1;HCoV-HKU1_orf1ab;1565 | 0 | 0.025 |
| NC_006577.2;YP_173236.1;HCoV-HKU1_orf1ab;1566 | 0.05 | 0 |
| NC_006577.2;YP_173236.1;HCoV-HKU1_orf1ab;1567 | 0 | 0 |
| NC_006577.2;YP_173236.1;HCoV-HKU1_orf1ab;1568 | 0 | 0.075 |
| NC_006577.2;YP_173236.1;HCoV-HKU1_orf1ab;1569 | 0.05 | 0 |
| NC_006577.2;YP_173236.1;HCoV-HKU1_orf1ab;1570 | 0 | 0 |
| NC_006577.2;YP_173236.1;HCoV-HKU1_orf1ab;1571 | 0 | 0 |
| NC_006577.2;YP_173236.1;HCoV-HKU1_orf1ab;1572 | 0 | 0 |
| NC_006577.2;YP_173236.1;HCoV-HKU1_orf1ab;1573 | 0 | 0 |
| NC_006577.2;YP_173236.1;HCoV-HKU1_orf1ab;1574 | 0 | 0.025 |
| NC_006577.2;YP_173236.1;HCoV-HKU1_orf1ab;1575 | 0.05 | 0 |
| NC_006577.2;YP_173236.1;HCoV-HKU1_orf1ab;1576 | 0.05 | 0 |
| NC_006577.2;YP_173236.1;HCoV-HKU1_orf1ab;1577 | 0.05 | 0 |
| NC_006577.2;YP_173236.1;HCoV-HKU1_orf1ab;1578 | 0.05 | 0 |
| NC_006577.2;YP_173236.1;HCoV-HKU1_orf1ab;1579 | 0.15 | 0.05 |
| NC_006577.2;YP_173236.1;HCoV-HKU1_orf1ab;1580 | 0.05 | 0.025 |
| NC_006577.2;YP_173236.1;HCoV-HKU1_orf1ab;1581 | 0.05 | 0 |
| NC_006577.2;YP_173236.1;HCoV-HKU1_orf1ab;1582 | 0 | 0 |
| NC_006577.2;YP_173236.1;HCoV-HKU1_orf1ab;1583 | 0.05 | 0 |
| NC_006577.2;YP_173236.1;HCoV-HKU1_orf1ab;1584 | 0 | 0 |
| NC_006577.2;YP_173236.1;HCoV-HKU1_orf1ab;1585 | 0 | 0 |
| NC_006577.2;YP_173236.1;HCoV-HKU1_orf1ab;1586 | 0 | 0 |
| NC_006577.2;YP_173236.1;HCoV-HKU1_orf1ab;1587 | 0 | 0.025 |
| NC_006577.2;YP_173236.1;HCoV-HKU1_orf1ab;1588 | 0.1 | 0.075 |
| NC_006577.2;YP_173236.1;HCoV-HKU1_orf1ab;1589 | 0.05 | 0.025 |
| NC_006577.2;YP_173236.1;HCoV-HKU1_orf1ab;1590 | 0.1 | 0.05 |
| NC_006577.2;YP_173236.1;HCoV-HKU1_orf1ab;1591 | 0.1 | 0.05 |
| NC_006577.2;YP_173236.1;HCoV-HKU1_orf1ab;1592 | 0.1 | 0.05 |
| NC_006577.2;YP_173236.1;HCoV-HKU1_orf1ab;1593 | 0 | 0.025 |
| NC_006577.2;YP_173236.1;HCoV-HKU1_orf1ab;1594 | 0.05 | 0.05 |
| NC_006577.2;YP_173236.1;HCoV-HKU1_orf1ab;1595 | 0 | 0.05 |
| NC_006577.2;YP_173236.1;HCoV-HKU1_orf1ab;1596 | 0 | 0.05 |
| NC_006577.2;YP_173236.1;HCoV-HKU1_orf1ab;1597 | 0.05 | 0.025 |
| NC_006577.2;YP_173236.1;HCoV-HKU1_orf1ab;1598 | 0 | 0 |
| NC_006577.2;YP_173236.1;HCoV-HKU1_orf1ab;1599 | 0 | 0 |
| NC_006577.2;YP_173236.1;HCoV-HKU1_orf1ab;1600 | 0 | 0 |
| NC_006577.2;YP_173236.1;HCoV-HKU1_orf1ab;1601 | 0 | 0.025 |
| NC_006577.2;YP_173236.1;HCoV-HKU1_orf1ab;1602 | 0 | 0 |
| NC_006577.2;YP_173236.1;HCoV-HKU1_orf1ab;1603 | 0 | 0 |
| NC_006577.2;YP_173236.1;HCoV-HKU1_orf1ab;1604 | 0 | 0 |
| NC_006577.2;YP_173236.1;HCoV-HKU1_orf1ab;1605 | 0 | 0 |
| NC_006577.2;YP_173236.1;HCoV-HKU1_orf1ab;1606 | 0 | 0 |

| | | |
|---|---|---|
| NC_006577.2;YP_173236.1;HCoV-HKU1_orf1ab;1607 | 0 | 0 |
| NC_006577.2;YP_173236.1;HCoV-HKU1_orf1ab;1608 | 0 | 0 |
| NC_006577.2;YP_173236.1;HCoV-HKU1_orf1ab;1609 | 0 | 0 |
| NC_006577.2;YP_173236.1;HCoV-HKU1_orf1ab;1610 | 0 | 0 |
| NC_006577.2;YP_173236.1;HCoV-HKU1_orf1ab;1611 | 0 | 0 |
| NC_006577.2;YP_173236.1;HCoV-HKU1_orf1ab;1612 | 0 | 0 |
| NC_006577.2;YP_173236.1;HCoV-HKU1_orf1ab;1613 | 0 | 0 |
| NC_006577.2;YP_173236.1;HCoV-HKU1_orf1ab;1614 | 0 | 0 |
| NC_006577.2;YP_173236.1;HCoV-HKU1_orf1ab;1615 | 0 | 0 |
| NC_006577.2;YP_173236.1;HCoV-HKU1_orf1ab;1616 | 0 | 0 |
| NC_006577.2;YP_173236.1;HCoV-HKU1_orf1ab;1617 | 0 | 0 |
| NC_006577.2;YP_173236.1;HCoV-HKU1_orf1ab;1618 | 0 | 0 |
| NC_006577.2;YP_173236.1;HCoV-HKU1_orf1ab;1619 | 0 | 0 |
| NC_006577.2;YP_173236.1;HCoV-HKU1_orf1ab;1620 | 0 | 0 |
| NC_006577.2;YP_173236.1;HCoV-HKU1_orf1ab;1621 | 0 | 0 |
| NC_006577.2;YP_173236.1;HCoV-HKU1_orf1ab;1622 | 0 | 0 |
| NC_006577.2;YP_173236.1;HCoV-HKU1_orf1ab;1623 | 0 | 0 |
| NC_006577.2;YP_173236.1;HCoV-HKU1_orf1ab;1624 | 0.05 | 0 |
| NC_006577.2;YP_173236.1;HCoV-HKU1_orf1ab;1625 | 0 | 0 |
| NC_006577.2;YP_173236.1;HCoV-HKU1_orf1ab;1626 | 0 | 0 |
| NC_006577.2;YP_173236.1;HCoV-HKU1_orf1ab;1627 | 0 | 0 |
| NC_006577.2;YP_173236.1;HCoV-HKU1_orf1ab;1628 | 0 | 0 |
| NC_006577.2;YP_173236.1;HCoV-HKU1_orf1ab;1629 | 0 | 0 |
| NC_006577.2;YP_173236.1;HCoV-HKU1_orf1ab;1630 | 0 | 0 |
| NC_006577.2;YP_173236.1;HCoV-HKU1_orf1ab;1631 | 0 | 0 |
| NC_006577.2;YP_173236.1;HCoV-HKU1_orf1ab;1632 | 0 | 0 |
| NC_006577.2;YP_173236.1;HCoV-HKU1_orf1ab;1633 | 0 | 0 |
| NC_006577.2;YP_173236.1;HCoV-HKU1_orf1ab;1634 | 0 | 0 |
| NC_006577.2;YP_173236.1;HCoV-HKU1_orf1ab;1635 | 0 | 0 |
| NC_006577.2;YP_173236.1;HCoV-HKU1_orf1ab;1636 | 0 | 0 |
| NC_006577.2;YP_173236.1;HCoV-HKU1_orf1ab;1637 | 0 | 0 |
| NC_006577.2;YP_173236.1;HCoV-HKU1_orf1ab;1638 | 0.05 | 0 |
| NC_006577.2;YP_173236.1;HCoV-HKU1_orf1ab;1639 | 0 | 0.025 |
| NC_006577.2;YP_173236.1;HCoV-HKU1_orf1ab;1640 | 0 | 0 |
| NC_006577.2;YP_173236.1;HCoV-HKU1_orf1ab;1641 | 0 | 0 |
| NC_006577.2;YP_173236.1;HCoV-HKU1_orf1ab;1642 | 0 | 0 |
| NC_006577.2;YP_173236.1;HCoV-HKU1_orf1ab;1643 | 0 | 0 |
| NC_006577.2;YP_173236.1;HCoV-HKU1_orf1ab;1644 | 0 | 0 |
| NC_006577.2;YP_173236.1;HCoV-HKU1_orf1ab;1645 | 0 | 0 |
| NC_006577.2;YP_173236.1;HCoV-HKU1_orf1ab;1646 | 0 | 0 |
| NC_006577.2;YP_173236.1;HCoV-HKU1_orf1ab;1647 | 0 | 0 |
| NC_006577.2;YP_173236.1;HCoV-HKU1_orf1ab;1648 | 0.05 | 0.025 |
| NC_006577.2;YP_173236.1;HCoV-HKU1_orf1ab;1649 | 0 | 0 |

| | | |
|---|---|---|
| NC_006577.2;YP_173236.1;HCoV-HKU1_orf1ab;1650 | 0.05 | 0.025 |
| NC_006577.2;YP_173236.1;HCoV-HKU1_orf1ab;1651 | 0.05 | 0.05 |
| NC_006577.2;YP_173236.1;HCoV-HKU1_orf1ab;1652 | 0.05 | 0.05 |
| NC_006577.2;YP_173236.1;HCoV-HKU1_orf1ab;1653 | 0 | 0 |
| NC_006577.2;YP_173236.1;HCoV-HKU1_orf1ab;1654 | 0 | 0 |
| NC_006577.2;YP_173236.1;HCoV-HKU1_orf1ab;1655 | 0 | 0 |
| NC_006577.2;YP_173236.1;HCoV-HKU1_orf1ab;1656 | 0 | 0 |
| NC_006577.2;YP_173236.1;HCoV-HKU1_orf1ab;1657 | 0 | 0.025 |
| NC_006577.2;YP_173236.1;HCoV-HKU1_orf1ab;1658 | 0 | 0.025 |
| NC_006577.2;YP_173236.1;HCoV-HKU1_orf1ab;1659 | 0 | 0 |
| NC_006577.2;YP_173236.1;HCoV-HKU1_orf1ab;1660 | 0 | 0.025 |
| NC_006577.2;YP_173236.1;HCoV-HKU1_orf1ab;1661 | 0 | 0.025 |
| NC_006577.2;YP_173236.1;HCoV-HKU1_orf1ab;1662 | 0 | 0.025 |
| NC_006577.2;YP_173236.1;HCoV-HKU1_orf1ab;1663 | 0 | 0.025 |
| NC_006577.2;YP_173236.1;HCoV-HKU1_orf1ab;1664 | 0 | 0.025 |
| NC_006577.2;YP_173236.1;HCoV-HKU1_orf1ab;1665 | 0 | 0.025 |
| NC_006577.2;YP_173236.1;HCoV-HKU1_orf1ab;1666 | 0 | 0.025 |
| NC_006577.2;YP_173236.1;HCoV-HKU1_orf1ab;1667 | 0 | 0 |
| NC_006577.2;YP_173236.1;HCoV-HKU1_orf1ab;1668 | 0 | 0 |
| NC_006577.2;YP_173236.1;HCoV-HKU1_orf1ab;1669 | 0 | 0.025 |
| NC_006577.2;YP_173236.1;HCoV-HKU1_orf1ab;1670 | 0 | 0 |
| NC_006577.2;YP_173236.1;HCoV-HKU1_orf1ab;1671 | 0 | 0 |
| NC_006577.2;YP_173236.1;HCoV-HKU1_orf1ab;1672 | 0 | 0 |
| NC_006577.2;YP_173236.1;HCoV-HKU1_orf1ab;1673 | 0 | 0 |
| NC_006577.2;YP_173236.1;HCoV-HKU1_orf1ab;1674 | 0 | 0 |
| NC_006577.2;YP_173236.1;HCoV-HKU1_orf1ab;1675 | 0 | 0 |
| NC_006577.2;YP_173236.1;HCoV-HKU1_orf1ab;1676 | 0 | 0 |
| NC_006577.2;YP_173236.1;HCoV-HKU1_orf1ab;1677 | 0.05 | 0 |
| NC_006577.2;YP_173236.1;HCoV-HKU1_orf1ab;1678 | 0.05 | 0 |
| NC_006577.2;YP_173236.1;HCoV-HKU1_orf1ab;1679 | 0.05 | 0 |
| NC_006577.2;YP_173236.1;HCoV-HKU1_orf1ab;1680 | 0 | 0 |
| NC_006577.2;YP_173236.1;HCoV-HKU1_orf1ab;1681 | 0 | 0 |
| NC_006577.2;YP_173236.1;HCoV-HKU1_orf1ab;1682 | 0 | 0 |
| NC_006577.2;YP_173236.1;HCoV-HKU1_orf1ab;1683 | 0 | 0 |
| NC_006577.2;YP_173236.1;HCoV-HKU1_orf1ab;1684 | 0 | 0 |
| NC_006577.2;YP_173236.1;HCoV-HKU1_orf1ab;1685 | 0.05 | 0.025 |
| NC_006577.2;YP_173236.1;HCoV-HKU1_orf1ab;1686 | 0.05 | 0 |
| NC_006577.2;YP_173236.1;HCoV-HKU1_orf1ab;1687 | 0 | 0 |
| NC_006577.2;YP_173236.1;HCoV-HKU1_orf1ab;1688 | 0 | 0 |
| NC_006577.2;YP_173236.1;HCoV-HKU1_orf1ab;1689 | 0 | 0 |
| NC_006577.2;YP_173236.1;HCoV-HKU1_orf1ab;1690 | 0 | 0 |
| NC_006577.2;YP_173236.1;HCoV-HKU1_orf1ab;1691 | 0 | 0 |
| NC_006577.2;YP_173236.1;HCoV-HKU1_orf1ab;1692 | 0 | 0 |

| | | |
|---|---|---|
| NC_006577.2;YP_173236.1;HCoV-HKU1_orf1ab;1693 | 0 | 0 |
| NC_006577.2;YP_173236.1;HCoV-HKU1_orf1ab;1694 | 0 | 0 |
| NC_006577.2;YP_173236.1;HCoV-HKU1_orf1ab;1695 | 0 | 0 |
| NC_006577.2;YP_173236.1;HCoV-HKU1_orf1ab;1696 | 0 | 0 |
| NC_006577.2;YP_173236.1;HCoV-HKU1_orf1ab;1697 | 0.1 | 0 |
| NC_006577.2;YP_173236.1;HCoV-HKU1_orf1ab;1698 | 0.05 | 0.025 |
| NC_006577.2;YP_173236.1;HCoV-HKU1_orf1ab;1699 | 0 | 0 |
| NC_006577.2;YP_173236.1;HCoV-HKU1_orf1ab;1700 | 0.05 | 0 |
| NC_006577.2;YP_173236.1;HCoV-HKU1_orf1ab;1701 | 0 | 0 |
| NC_006577.2;YP_173236.1;HCoV-HKU1_orf1ab;1702 | 0.05 | 0 |
| NC_006577.2;YP_173236.1;HCoV-HKU1_orf1ab;1703 | 0.05 | 0 |
| NC_006577.2;YP_173236.1;HCoV-HKU1_orf1ab;1704 | 0 | 0.025 |
| NC_006577.2;YP_173236.1;HCoV-HKU1_orf1ab;1705 | 0 | 0.025 |
| NC_006577.2;YP_173236.1;HCoV-HKU1_orf1ab;1706 | 0 | 0 |
| NC_006577.2;YP_173236.1;HCoV-HKU1_orf1ab;1707 | 0 | 0 |
| NC_006577.2;YP_173236.1;HCoV-HKU1_orf1ab;1708 | 0 | 0 |
| NC_006577.2;YP_173236.1;HCoV-HKU1_orf1ab;1709 | 0.05 | 0.025 |
| NC_006577.2;YP_173236.1;HCoV-HKU1_orf1ab;1710 | 0 | 0 |
| NC_006577.2;YP_173236.1;HCoV-HKU1_orf1ab;1711 | 0 | 0.025 |
| NC_006577.2;YP_173236.1;HCoV-HKU1_orf1ab;1712 | 0 | 0.05 |
| NC_006577.2;YP_173236.1;HCoV-HKU1_orf1ab;1713 | 0 | 0 |
| NC_006577.2;YP_173236.1;HCoV-HKU1_orf1ab;1714 | 0 | 0 |
| NC_006577.2;YP_173236.1;HCoV-HKU1_orf1ab;1715 | 0.1 | 0 |
| NC_006577.2;YP_173236.1;HCoV-HKU1_orf1ab;1716 | 0 | 0 |
| NC_006577.2;YP_173236.1;HCoV-HKU1_orf1ab;1717 | 0 | 0 |
| NC_006577.2;YP_173236.1;HCoV-HKU1_orf1ab;1718 | 0 | 0 |
| NC_006577.2;YP_173236.1;HCoV-HKU1_orf1ab;1719 | 0 | 0 |
| NC_006577.2;YP_173236.1;HCoV-HKU1_orf1ab;1720 | 0 | 0 |
| NC_006577.2;YP_173236.1;HCoV-HKU1_orf1ab;1721 | 0 | 0 |
| NC_006577.2;YP_173236.1;HCoV-HKU1_orf1ab;1722 | 0 | 0 |
| NC_006577.2;YP_173236.1;HCoV-HKU1_orf1ab;1723 | 0 | 0 |
| NC_006577.2;YP_173236.1;HCoV-HKU1_orf1ab;1724 | 0 | 0 |
| NC_006577.2;YP_173236.1;HCoV-HKU1_orf1ab;1725 | 0 | 0 |
| NC_006577.2;YP_173236.1;HCoV-HKU1_orf1ab;1726 | 0 | 0 |
| NC_006577.2;YP_173236.1;HCoV-HKU1_orf1ab;1727 | 0 | 0 |
| NC_006577.2;YP_173236.1;HCoV-HKU1_orf1ab;1728 | 0 | 0 |
| NC_006577.2;YP_173236.1;HCoV-HKU1_orf1ab;1729 | 0 | 0 |
| NC_006577.2;YP_173236.1;HCoV-HKU1_orf1ab;1730 | 0 | 0 |
| NC_006577.2;YP_173236.1;HCoV-HKU1_orf1ab;1731 | 0 | 0 |
| NC_006577.2;YP_173236.1;HCoV-HKU1_orf1ab;1732 | 0 | 0 |
| NC_006577.2;YP_173236.1;HCoV-HKU1_orf1ab;1733 | 0 | 0 |
| NC_006577.2;YP_173236.1;HCoV-HKU1_orf1ab;1734 | 0 | 0 |
| NC_006577.2;YP_173236.1;HCoV-HKU1_orf1ab;1735 | 0 | 0 |

| | | |
|---|---|---|
| NC_006577.2;YP_173236.1;HCoV-HKU1_orf1ab;1736 | 0 | 0 |
| NC_006577.2;YP_173236.1;HCoV-HKU1_orf1ab;1737 | 0 | 0 |
| NC_006577.2;YP_173236.1;HCoV-HKU1_orf1ab;1738 | 0 | 0 |
| NC_006577.2;YP_173236.1;HCoV-HKU1_orf1ab;1739 | 0 | 0 |
| NC_006577.2;YP_173236.1;HCoV-HKU1_orf1ab;1740 | 0 | 0 |
| NC_006577.2;YP_173236.1;HCoV-HKU1_orf1ab;1741 | 0 | 0.025 |
| NC_006577.2;YP_173236.1;HCoV-HKU1_orf1ab;1742 | 0.05 | 0 |
| NC_006577.2;YP_173236.1;HCoV-HKU1_orf1ab;1743 | 0.05 | 0 |
| NC_006577.2;YP_173236.1;HCoV-HKU1_orf1ab;1744 | 0 | 0 |
| NC_006577.2;YP_173236.1;HCoV-HKU1_orf1ab;1745 | 0 | 0 |
| NC_006577.2;YP_173236.1;HCoV-HKU1_orf1ab;1746 | 0 | 0.025 |
| NC_006577.2;YP_173236.1;HCoV-HKU1_orf1ab;1747 | 0.05 | 0 |
| NC_006577.2;YP_173236.1;HCoV-HKU1_orf1ab;1748 | 0.15 | 0.05 |
| NC_006577.2;YP_173236.1;HCoV-HKU1_orf1ab;1749 | 0 | 0 |
| NC_006577.2;YP_173236.1;HCoV-HKU1_orf1ab;1750 | 0 | 0 |
| NC_006577.2;YP_173236.1;HCoV-HKU1_orf1ab;1751 | 0.05 | 0 |
| NC_006577.2;YP_173236.1;HCoV-HKU1_orf1ab;1752 | 0 | 0 |
| NC_006577.2;YP_173236.1;HCoV-HKU1_orf1ab;1753 | 0 | 0 |
| NC_006577.2;YP_173236.1;HCoV-HKU1_orf1ab;1754 | 0 | 0 |
| NC_006577.2;YP_173236.1;HCoV-HKU1_orf1ab;1755 | 0 | 0 |
| NC_006577.2;YP_173236.1;HCoV-HKU1_orf1ab;1756 | 0 | 0 |
| NC_006577.2;YP_173236.1;HCoV-HKU1_orf1ab;1757 | 0 | 0 |
| NC_006577.2;YP_173236.1;HCoV-HKU1_orf1ab;1758 | 0 | 0 |
| NC_006577.2;YP_173236.1;HCoV-HKU1_orf1ab;1759 | 0 | 0 |
| NC_006577.2;YP_173236.1;HCoV-HKU1_orf1ab;1760 | 0 | 0 |
| NC_006577.2;YP_173236.1;HCoV-HKU1_orf1ab;1761 | 0 | 0 |
| NC_006577.2;YP_173236.1;HCoV-HKU1_orf1ab;1762 | 0 | 0 |
| NC_006577.2;YP_173236.1;HCoV-HKU1_orf1ab;1763 | 0 | 0 |
| NC_006577.2;YP_173236.1;HCoV-HKU1_orf1ab;1764 | 0 | 0 |
| NC_006577.2;YP_173236.1;HCoV-HKU1_orf1ab;1765 | 0 | 0 |
| NC_006577.2;YP_173236.1;HCoV-HKU1_orf1ab;1766 | 0 | 0 |
| NC_006577.2;YP_173236.1;HCoV-HKU1_orf1ab;1767 | 0 | 0 |
| NC_006577.2;YP_173236.1;HCoV-HKU1_orf1ab;1768 | 0 | 0 |
| NC_006577.2;YP_173236.1;HCoV-HKU1_orf1ab;1769 | 0 | 0 |
| NC_006577.2;YP_173236.1;HCoV-HKU1_orf1ab;1770 | 0 | 0 |
| NC_006577.2;YP_173236.1;HCoV-HKU1_orf1ab;1771 | 0 | 0 |
| NC_006577.2;YP_173236.1;HCoV-HKU1_orf1ab;1772 | 0.05 | 0 |
| NC_006577.2;YP_173236.1;HCoV-HKU1_orf1ab;1773 | 0 | 0 |
| NC_006577.2;YP_173236.1;HCoV-HKU1_orf1ab;1774 | 0 | 0 |
| NC_006577.2;YP_173236.1;HCoV-HKU1_orf1ab;1775 | 0 | 0 |
| NC_006577.2;YP_173236.1;HCoV-HKU1_orf1ab;1776 | 0 | 0 |
| NC_006577.2;YP_173236.1;HCoV-HKU1_orf1ab;1777 | 0 | 0 |
| NC_006577.2;YP_173236.1;HCoV-HKU1_orf1ab;1778 | 0 | 0 |

| | | |
|---|---|---|
| NC_006577.2;YP_173236.1;HCoV-HKU1_orf1ab;1779 | 0 | 0 |
| NC_006577.2;YP_173236.1;HCoV-HKU1_orf1ab;1780 | 0 | 0 |
| NC_006577.2;YP_173236.1;HCoV-HKU1_orf1ab;1781 | 0 | 0 |
| NC_006577.2;YP_173236.1;HCoV-HKU1_orf1ab;1782 | 0 | 0 |
| NC_006577.2;YP_173236.1;HCoV-HKU1_orf1ab;1783 | 0 | 0 |
| NC_006577.2;YP_173236.1;HCoV-HKU1_orf1ab;1784 | 0 | 0 |
| NC_006577.2;YP_173236.1;HCoV-HKU1_orf1ab;1785 | 0 | 0 |
| NC_006577.2;YP_173236.1;HCoV-HKU1_orf1ab;1786 | 0 | 0 |
| NC_006577.2;YP_173236.1;HCoV-HKU1_orf1ab;1787 | 0 | 0 |
| NC_006577.2;YP_173236.1;HCoV-HKU1_orf1ab;1788 | 0 | 0 |
| NC_006577.2;YP_173236.1;HCoV-HKU1_orf1ab;1789 | 0 | 0 |
| NC_006577.2;YP_173236.1;HCoV-HKU1_orf1ab;1790 | 0 | 0 |
| NC_006577.2;YP_173236.1;HCoV-HKU1_orf1ab;1791 | 0 | 0 |
| NC_006577.2;YP_173236.1;HCoV-HKU1_orf1ab;1792 | 0 | 0 |
| NC_006577.2;YP_173236.1;HCoV-HKU1_orf1ab;1793 | 0 | 0 |
| NC_006577.2;YP_173236.1;HCoV-HKU1_orf1ab;1794 | 0 | 0 |
| NC_006577.2;YP_173236.1;HCoV-HKU1_orf1ab;1795 | 0.15 | 0 |
| NC_006577.2;YP_173236.1;HCoV-HKU1_orf1ab;1796 | 0.1 | 0.025 |
| NC_006577.2;YP_173236.1;HCoV-HKU1_orf1ab;1797 | 0.3 | 0.05 |
| NC_006577.2;YP_173236.1;HCoV-HKU1_orf1ab;1798 | 0.2 | 0.05 |
| NC_006577.2;YP_173236.1;HCoV-HKU1_orf1ab;1799 | 0.2 | 0.025 |
| NC_006577.2;YP_173236.1;HCoV-HKU1_orf1ab;1800 | 0.05 | 0.05 |
| NC_006577.2;YP_173236.1;HCoV-HKU1_orf1ab;1801 | 0 | 0 |
| NC_006577.2;YP_173236.1;HCoV-HKU1_orf1ab;1802 | 0 | 0 |
| NC_006577.2;YP_173236.1;HCoV-HKU1_orf1ab;1803 | 0 | 0 |
| NC_006577.2;YP_173236.1;HCoV-HKU1_orf1ab;1804 | 0 | 0 |
| NC_006577.2;YP_173236.1;HCoV-HKU1_orf1ab;1805 | 0 | 0 |
| NC_006577.2;YP_173236.1;HCoV-HKU1_orf1ab;1806 | 0 | 0 |
| NC_006577.2;YP_173236.1;HCoV-HKU1_orf1ab;1807 | 0 | 0 |
| NC_006577.2;YP_173236.1;HCoV-HKU1_orf1ab;1808 | 0 | 0 |
| NC_006577.2;YP_173236.1;HCoV-HKU1_orf1ab;1809 | 0 | 0 |
| NC_006577.2;YP_173236.1;HCoV-HKU1_orf1ab;1810 | 0 | 0 |
| NC_006577.2;YP_173236.1;HCoV-HKU1_orf1ab;1811 | 0 | 0 |
| NC_006577.2;YP_173236.1;HCoV-HKU1_orf1ab;1812 | 0 | 0 |
| NC_006577.2;YP_173236.1;HCoV-HKU1_orf1ab;1813 | 0 | 0 |
| NC_006577.2;YP_173236.1;HCoV-HKU1_orf1ab;1814 | 0 | 0 |
| NC_006577.2;YP_173236.1;HCoV-HKU1_orf1ab;1815 | 0 | 0 |
| NC_006577.2;YP_173236.1;HCoV-HKU1_orf1ab;1816 | 0 | 0 |
| NC_006577.2;YP_173236.1;HCoV-HKU1_orf1ab;1817 | 0 | 0 |
| NC_006577.2;YP_173236.1;HCoV-HKU1_orf1ab;1818 | 0 | 0 |
| NC_006577.2;YP_173236.1;HCoV-HKU1_orf1ab;1819 | 0 | 0 |
| NC_006577.2;YP_173236.1;HCoV-HKU1_orf1ab;1820 | 0 | 0 |
| NC_006577.2;YP_173236.1;HCoV-HKU1_orf1ab;1821 | 0 | 0 |

| | | |
|---|---|---|
| NC_006577.2;YP_173236.1;HCoV-HKU1_orf1ab;1822 | 0 | 0 |
| NC_006577.2;YP_173236.1;HCoV-HKU1_orf1ab;1823 | 0 | 0 |
| NC_006577.2;YP_173236.1;HCoV-HKU1_orf1ab;1824 | 0 | 0 |
| NC_006577.2;YP_173236.1;HCoV-HKU1_orf1ab;1825 | 0 | 0 |
| NC_006577.2;YP_173236.1;HCoV-HKU1_orf1ab;1826 | 0 | 0 |
| NC_006577.2;YP_173236.1;HCoV-HKU1_orf1ab;1827 | 0 | 0 |
| NC_006577.2;YP_173236.1;HCoV-HKU1_orf1ab;1828 | 0 | 0 |
| NC_006577.2;YP_173236.1;HCoV-HKU1_orf1ab;1829 | 0 | 0 |
| NC_006577.2;YP_173236.1;HCoV-HKU1_orf1ab;1830 | 0.05 | 0 |
| NC_006577.2;YP_173236.1;HCoV-HKU1_orf1ab;1831 | 0.05 | 0 |
| NC_006577.2;YP_173236.1;HCoV-HKU1_orf1ab;1832 | 0.05 | 0 |
| NC_006577.2;YP_173236.1;HCoV-HKU1_orf1ab;1833 | 0.05 | 0 |
| NC_006577.2;YP_173236.1;HCoV-HKU1_orf1ab;1834 | 0.05 | 0 |
| NC_006577.2;YP_173236.1;HCoV-HKU1_orf1ab;1835 | 0.05 | 0 |
| NC_006577.2;YP_173236.1;HCoV-HKU1_orf1ab;1836 | 0.1 | 0 |
| NC_006577.2;YP_173236.1;HCoV-HKU1_orf1ab;1837 | 0.05 | 0 |
| NC_006577.2;YP_173236.1;HCoV-HKU1_orf1ab;1838 | 0.05 | 0 |
| NC_006577.2;YP_173236.1;HCoV-HKU1_orf1ab;1839 | 0.05 | 0 |
| NC_006577.2;YP_173236.1;HCoV-HKU1_orf1ab;1840 | 0.05 | 0 |
| NC_006577.2;YP_173236.1;HCoV-HKU1_orf1ab;1841 | 0 | 0 |
| NC_006577.2;YP_173236.1;HCoV-HKU1_orf1ab;1842 | 0.05 | 0 |
| NC_006577.2;YP_173236.1;HCoV-HKU1_orf1ab;1843 | 0 | 0 |
| NC_006577.2;YP_173236.1;HCoV-HKU1_orf1ab;1844 | 0 | 0 |
| NC_006577.2;YP_173236.1;HCoV-HKU1_orf1ab;1845 | 0 | 0 |
| NC_006577.2;YP_173236.1;HCoV-HKU1_orf1ab;1846 | 0 | 0 |
| NC_006577.2;YP_173236.1;HCoV-HKU1_orf1ab;1847 | 0 | 0 |
| NC_006577.2;YP_173236.1;HCoV-HKU1_orf1ab;1848 | 0 | 0.025 |
| NC_006577.2;YP_173236.1;HCoV-HKU1_orf1ab;1849 | 0 | 0 |
| NC_006577.2;YP_173236.1;HCoV-HKU1_orf1ab;1850 | 0 | 0 |
| NC_006577.2;YP_173236.1;HCoV-HKU1_orf1ab;1851 | 0 | 0.025 |
| NC_006577.2;YP_173236.1;HCoV-HKU1_orf1ab;1852 | 0 | 0 |
| NC_006577.2;YP_173236.1;HCoV-HKU1_orf1ab;1853 | 0 | 0 |
| NC_006577.2;YP_173236.1;HCoV-HKU1_orf1ab;1854 | 0 | 0 |
| NC_006577.2;YP_173236.1;HCoV-HKU1_orf1ab;1855 | 0.05 | 0.025 |
| NC_006577.2;YP_173236.1;HCoV-HKU1_orf1ab;1856 | 0 | 0 |
| NC_006577.2;YP_173236.1;HCoV-HKU1_orf1ab;1857 | 0 | 0.075 |
| NC_006577.2;YP_173236.1;HCoV-HKU1_orf1ab;1858 | 0 | 0.075 |
| NC_006577.2;YP_173236.1;HCoV-HKU1_orf1ab;1859 | 0 | 0 |
| NC_006577.2;YP_173236.1;HCoV-HKU1_orf1ab;1860 | 0 | 0 |
| NC_006577.2;YP_173236.1;HCoV-HKU1_orf1ab;1861 | 0 | 0 |
| NC_006577.2;YP_173236.1;HCoV-HKU1_orf1ab;1862 | 0 | 0 |
| NC_006577.2;YP_173236.1;HCoV-HKU1_orf1ab;1863 | 0 | 0 |
| NC_006577.2;YP_173236.1;HCoV-HKU1_orf1ab;1864 | 0 | 0 |

| | | |
|---|---|---|
| NC_006577.2;YP_173236.1;HCoV-HKU1_orf1ab;1865 | 0 | 0 |
| NC_006577.2;YP_173236.1;HCoV-HKU1_orf1ab;1866 | 0 | 0 |
| NC_006577.2;YP_173236.1;HCoV-HKU1_orf1ab;1867 | 0 | 0 |
| NC_006577.2;YP_173236.1;HCoV-HKU1_orf1ab;1868 | 0 | 0 |
| NC_006577.2;YP_173236.1;HCoV-HKU1_orf1ab;1869 | 0 | 0 |
| NC_006577.2;YP_173236.1;HCoV-HKU1_orf1ab;1870 | 0 | 0 |
| NC_006577.2;YP_173236.1;HCoV-HKU1_orf1ab;1871 | 0 | 0 |
| NC_006577.2;YP_173236.1;HCoV-HKU1_orf1ab;1872 | 0 | 0 |
| NC_006577.2;YP_173236.1;HCoV-HKU1_orf1ab;1873 | 0 | 0 |
| NC_006577.2;YP_173236.1;HCoV-HKU1_orf1ab;1874 | 0 | 0 |
| NC_006577.2;YP_173236.1;HCoV-HKU1_orf1ab;1875 | 0 | 0 |
| NC_006577.2;YP_173236.1;HCoV-HKU1_orf1ab;1876 | 0 | 0 |
| NC_006577.2;YP_173236.1;HCoV-HKU1_orf1ab;1877 | 0 | 0 |
| NC_006577.2;YP_173236.1;HCoV-HKU1_orf1ab;1878 | 0 | 0 |
| NC_006577.2;YP_173236.1;HCoV-HKU1_orf1ab;1879 | 0 | 0 |
| NC_006577.2;YP_173236.1;HCoV-HKU1_orf1ab;1880 | 0 | 0 |
| NC_006577.2;YP_173236.1;HCoV-HKU1_orf1ab;1881 | 0 | 0 |
| NC_006577.2;YP_173236.1;HCoV-HKU1_orf1ab;1882 | 0 | 0 |
| NC_006577.2;YP_173236.1;HCoV-HKU1_orf1ab;1883 | 0 | 0 |
| NC_006577.2;YP_173236.1;HCoV-HKU1_orf1ab;1884 | 0.05 | 0 |
| NC_006577.2;YP_173236.1;HCoV-HKU1_orf1ab;1885 | 0.05 | 0 |
| NC_006577.2;YP_173236.1;HCoV-HKU1_orf1ab;1886 | 0.1 | 0 |
| NC_006577.2;YP_173236.1;HCoV-HKU1_orf1ab;1887 | 0.1 | 0.025 |
| NC_006577.2;YP_173236.1;HCoV-HKU1_orf1ab;1888 | 0.1 | 0 |
| NC_006577.2;YP_173236.1;HCoV-HKU1_orf1ab;1889 | 0.15 | 0 |
| NC_006577.2;YP_173236.1;HCoV-HKU1_orf1ab;1890 | 0.1 | 0 |
| NC_006577.2;YP_173236.1;HCoV-HKU1_orf1ab;1891 | 0.1 | 0 |
| NC_006577.2;YP_173236.1;HCoV-HKU1_orf1ab;1892 | 0.1 | 0.025 |
| NC_006577.2;YP_173236.1;HCoV-HKU1_orf1ab;1893 | 0.1 | 0.05 |
| NC_006577.2;YP_173236.1;HCoV-HKU1_orf1ab;1894 | 0.1 | 0 |
| NC_006577.2;YP_173236.1;HCoV-HKU1_orf1ab;1895 | 0.2 | 0 |
| NC_006577.2;YP_173236.1;HCoV-HKU1_orf1ab;1896 | 0 | 0.025 |
| NC_006577.2;YP_173236.1;HCoV-HKU1_orf1ab;1897 | 0 | 0 |
| NC_006577.2;YP_173236.1;HCoV-HKU1_orf1ab;1898 | 0 | 0 |
| NC_006577.2;YP_173236.1;HCoV-HKU1_orf1ab;1899 | 0 | 0 |
| NC_006577.2;YP_173236.1;HCoV-HKU1_orf1ab;1900 | 0 | 0.025 |
| NC_006577.2;YP_173236.1;HCoV-HKU1_orf1ab;1901 | 0 | 0 |
| NC_006577.2;YP_173236.1;HCoV-HKU1_orf1ab;1902 | 0 | 0 |
| NC_006577.2;YP_173236.1;HCoV-HKU1_orf1ab;1903 | 0 | 0 |
| NC_006577.2;YP_173236.1;HCoV-HKU1_orf1ab;1904 | 0 | 0 |
| NC_006577.2;YP_173236.1;HCoV-HKU1_orf1ab;1905 | 0 | 0 |
| NC_006577.2;YP_173236.1;HCoV-HKU1_orf1ab;1906 | 0 | 0 |
| NC_006577.2;YP_173236.1;HCoV-HKU1_orf1ab;1907 | 0 | 0 |

| | | |
|---|---|---|
| NC_006577.2;YP_173236.1;HCoV-HKU1_orf1ab;1908 | 0 | 0 |
| NC_006577.2;YP_173236.1;HCoV-HKU1_orf1ab;1909 | 0 | 0 |
| NC_006577.2;YP_173236.1;HCoV-HKU1_orf1ab;1910 | 0 | 0 |
| NC_006577.2;YP_173236.1;HCoV-HKU1_orf1ab;1911 | 0 | 0 |
| NC_006577.2;YP_173236.1;HCoV-HKU1_orf1ab;1912 | 0 | 0 |
| NC_006577.2;YP_173236.1;HCoV-HKU1_orf1ab;1913 | 0 | 0 |
| NC_006577.2;YP_173236.1;HCoV-HKU1_orf1ab;1914 | 0 | 0 |
| NC_006577.2;YP_173236.1;HCoV-HKU1_orf1ab;1915 | 0 | 0 |
| NC_006577.2;YP_173236.1;HCoV-HKU1_orf1ab;1916 | 0 | 0 |
| NC_006577.2;YP_173236.1;HCoV-HKU1_orf1ab;1917 | 0 | 0 |
| NC_006577.2;YP_173236.1;HCoV-HKU1_orf1ab;1918 | 0 | 0.025 |
| NC_006577.2;YP_173236.1;HCoV-HKU1_orf1ab;1919 | 0 | 0 |
| NC_006577.2;YP_173236.1;HCoV-HKU1_orf1ab;1920 | 0 | 0 |
| NC_006577.2;YP_173236.1;HCoV-HKU1_orf1ab;1921 | 0 | 0 |
| NC_006577.2;YP_173236.1;HCoV-HKU1_orf1ab;1922 | 0 | 0 |
| NC_006577.2;YP_173236.1;HCoV-HKU1_orf1ab;1923 | 0 | 0.025 |
| NC_006577.2;YP_173236.1;HCoV-HKU1_orf1ab;1924 | 0 | 0 |
| NC_006577.2;YP_173236.1;HCoV-HKU1_orf1ab;1925 | 0 | 0 |
| NC_006577.2;YP_173236.1;HCoV-HKU1_orf1ab;1926 | 0 | 0 |
| NC_006577.2;YP_173236.1;HCoV-HKU1_orf1ab;1927 | 0 | 0 |
| NC_006577.2;YP_173236.1;HCoV-HKU1_orf1ab;1928 | 0 | 0 |
| NC_006577.2;YP_173236.1;HCoV-HKU1_orf1ab;1929 | 0 | 0 |
| NC_006577.2;YP_173236.1;HCoV-HKU1_orf1ab;1930 | 0 | 0 |
| NC_006577.2;YP_173236.1;HCoV-HKU1_orf1ab;1931 | 0 | 0 |
| NC_006577.2;YP_173236.1;HCoV-HKU1_orf1ab;1932 | 0 | 0 |
| NC_006577.2;YP_173236.1;HCoV-HKU1_orf1ab;1933 | 0 | 0 |
| NC_006577.2;YP_173236.1;HCoV-HKU1_orf1ab;1934 | 0 | 0 |
| NC_006577.2;YP_173236.1;HCoV-HKU1_orf1ab;1935 | 0 | 0 |
| NC_006577.2;YP_173236.1;HCoV-HKU1_orf1ab;1936 | 0 | 0 |
| NC_006577.2;YP_173236.1;HCoV-HKU1_orf1ab;1937 | 0 | 0.025 |
| NC_006577.2;YP_173236.1;HCoV-HKU1_orf1ab;1938 | 0 | 0 |
| NC_006577.2;YP_173236.1;HCoV-HKU1_orf1ab;1939 | 0 | 0 |
| NC_006577.2;YP_173236.1;HCoV-HKU1_orf1ab;1940 | 0 | 0 |
| NC_006577.2;YP_173236.1;HCoV-HKU1_orf1ab;1941 | 0 | 0.025 |
| NC_006577.2;YP_173236.1;HCoV-HKU1_orf1ab;1942 | 0 | 0.025 |
| NC_006577.2;YP_173236.1;HCoV-HKU1_orf1ab;1943 | 0 | 0.025 |
| NC_006577.2;YP_173236.1;HCoV-HKU1_orf1ab;1944 | 0 | 0.025 |
| NC_006577.2;YP_173236.1;HCoV-HKU1_orf1ab;1945 | 0 | 0.025 |
| NC_006577.2;YP_173236.1;HCoV-HKU1_orf1ab;1946 | 0 | 0.025 |
| NC_006577.2;YP_173236.1;HCoV-HKU1_orf1ab;1947 | 0 | 0.025 |
| NC_006577.2;YP_173236.1;HCoV-HKU1_orf1ab;1948 | 0 | 0.025 |
| NC_006577.2;YP_173236.1;HCoV-HKU1_orf1ab;1949 | 0 | 0 |
| NC_006577.2;YP_173236.1;HCoV-HKU1_orf1ab;1950 | 0 | 0 |

| | | |
|---|---|---|
| NC_006577.2;YP_173236.1;HCoV-HKU1_orf1ab;1951 | 0 | 0 |
| NC_006577.2;YP_173236.1;HCoV-HKU1_orf1ab;1952 | 0 | 0 |
| NC_006577.2;YP_173236.1;HCoV-HKU1_orf1ab;1953 | 0 | 0.025 |
| NC_006577.2;YP_173236.1;HCoV-HKU1_orf1ab;1954 | 0 | 0.025 |
| NC_006577.2;YP_173236.1;HCoV-HKU1_orf1ab;1955 | 0 | 0 |
| NC_006577.2;YP_173236.1;HCoV-HKU1_orf1ab;1956 | 0 | 0.025 |
| NC_006577.2;YP_173236.1;HCoV-HKU1_orf1ab;1957 | 0 | 0 |
| NC_006577.2;YP_173236.1;HCoV-HKU1_orf1ab;1958 | 0 | 0 |
| NC_006577.2;YP_173236.1;HCoV-HKU1_orf1ab;1959 | 0 | 0 |
| NC_006577.2;YP_173236.1;HCoV-HKU1_orf1ab;1960 | 0 | 0 |
| NC_006577.2;YP_173236.1;HCoV-HKU1_orf1ab;1961 | 0 | 0 |
| NC_006577.2;YP_173236.1;HCoV-HKU1_orf1ab;1962 | 0.05 | 0.025 |
| NC_006577.2;YP_173236.1;HCoV-HKU1_orf1ab;1963 | 0 | 0.025 |
| NC_006577.2;YP_173236.1;HCoV-HKU1_orf1ab;1964 | 0 | 0 |
| NC_006577.2;YP_173236.1;HCoV-HKU1_orf1ab;1965 | 0 | 0 |
| NC_006577.2;YP_173236.1;HCoV-HKU1_orf1ab;1966 | 0 | 0 |
| NC_006577.2;YP_173236.1;HCoV-HKU1_orf1ab;1967 | 0 | 0.05 |
| NC_006577.2;YP_173236.1;HCoV-HKU1_orf1ab;1968 | 0.05 | 0 |
| NC_006577.2;YP_173236.1;HCoV-HKU1_orf1ab;1969 | 0 | 0 |
| NC_006577.2;YP_173236.1;HCoV-HKU1_orf1ab;1970 | 0.1 | 0 |
| NC_006577.2;YP_173236.1;HCoV-HKU1_orf1ab;1971 | 0.05 | 0.075 |
| NC_006577.2;YP_173236.1;HCoV-HKU1_orf1ab;1972 | 0.05 | 0 |
| NC_006577.2;YP_173236.1;HCoV-HKU1_orf1ab;1973 | 0 | 0 |
| NC_006577.2;YP_173236.1;HCoV-HKU1_orf1ab;1974 | 0 | 0.025 |
| NC_006577.2;YP_173236.1;HCoV-HKU1_orf1ab;1975 | 0.1 | 0.025 |
| NC_006577.2;YP_173236.1;HCoV-HKU1_orf1ab;1976 | 0 | 0 |
| NC_006577.2;YP_173236.1;HCoV-HKU1_orf1ab;1977 | 0 | 0 |
| NC_006577.2;YP_173236.1;HCoV-HKU1_orf1ab;1978 | 0 | 0 |
| NC_006577.2;YP_173236.1;HCoV-HKU1_orf1ab;1979 | 0 | 0 |
| NC_006577.2;YP_173236.1;HCoV-HKU1_orf1ab;1980 | 0 | 0.025 |
| NC_006577.2;YP_173236.1;HCoV-HKU1_orf1ab;1981 | 0 | 0 |
| NC_006577.2;YP_173236.1;HCoV-HKU1_orf1ab;1982 | 0.15 | 0.1 |
| NC_006577.2;YP_173236.1;HCoV-HKU1_orf1ab;1983 | 0 | 0.025 |
| NC_006577.2;YP_173236.1;HCoV-HKU1_orf1ab;1984 | 0 | 0.025 |
| NC_006577.2;YP_173236.1;HCoV-HKU1_orf1ab;1985 | 0 | 0 |
| NC_006577.2;YP_173236.1;HCoV-HKU1_orf1ab;1986 | 0 | 0 |
| NC_006577.2;YP_173236.1;HCoV-HKU1_orf1ab;1987 | 0 | 0 |
| NC_006577.2;YP_173236.1;HCoV-HKU1_orf1ab;1988 | 0.05 | 0.025 |
| NC_006577.2;YP_173236.1;HCoV-HKU1_orf1ab;1989 | 0 | 0.025 |
| NC_006577.2;YP_173236.1;HCoV-HKU1_orf1ab;1990 | 0.05 | 0.1 |
| NC_006577.2;YP_173236.1;HCoV-HKU1_orf1ab;1991 | 0 | 0 |
| NC_006577.2;YP_173236.1;HCoV-HKU1_orf1ab;1992 | 0 | 0 |
| NC_006577.2;YP_173236.1;HCoV-HKU1_orf1ab;1993 | 0 | 0.025 |

| | | |
|---|---|---|
| NC_006577.2;YP_173236.1;HCoV-HKU1_orf1ab;1994 | 0 | 0 |
| NC_006577.2;YP_173236.1;HCoV-HKU1_orf1ab;1995 | 0 | 0 |
| NC_006577.2;YP_173236.1;HCoV-HKU1_orf1ab;1996 | 0 | 0 |
| NC_006577.2;YP_173236.1;HCoV-HKU1_orf1ab;1997 | 0 | 0 |
| NC_006577.2;YP_173236.1;HCoV-HKU1_orf1ab;1998 | 0 | 0 |
| NC_006577.2;YP_173236.1;HCoV-HKU1_orf1ab;1999 | 0 | 0 |
| NC_006577.2;YP_173236.1;HCoV-HKU1_orf1ab;2000 | 0.05 | 0 |
| NC_006577.2;YP_173236.1;HCoV-HKU1_orf1ab;2001 | 0.05 | 0 |
| NC_006577.2;YP_173236.1;HCoV-HKU1_orf1ab;2002 | 0.05 | 0 |
| NC_006577.2;YP_173236.1;HCoV-HKU1_orf1ab;2003 | 0.05 | 0 |
| NC_006577.2;YP_173236.1;HCoV-HKU1_orf1ab;2004 | 0.05 | 0 |
| NC_006577.2;YP_173236.1;HCoV-HKU1_orf1ab;2005 | 0.05 | 0 |
| NC_006577.2;YP_173236.1;HCoV-HKU1_orf1ab;2006 | 0.05 | 0 |
| NC_006577.2;YP_173236.1;HCoV-HKU1_orf1ab;2007 | 0.05 | 0 |
| NC_006577.2;YP_173236.1;HCoV-HKU1_orf1ab;2008 | 0 | 0 |
| NC_006577.2;YP_173236.1;HCoV-HKU1_orf1ab;2009 | 0 | 0 |
| NC_006577.2;YP_173236.1;HCoV-HKU1_orf1ab;2010 | 0 | 0 |
| NC_006577.2;YP_173236.1;HCoV-HKU1_orf1ab;2011 | 0 | 0 |
| NC_006577.2;YP_173236.1;HCoV-HKU1_orf1ab;2012 | 0 | 0 |
| NC_006577.2;YP_173236.1;HCoV-HKU1_orf1ab;2013 | 0 | 0 |
| NC_006577.2;YP_173236.1;HCoV-HKU1_orf1ab;2014 | 0 | 0.025 |
| NC_006577.2;YP_173236.1;HCoV-HKU1_orf1ab;2015 | 0 | 0 |
| NC_006577.2;YP_173236.1;HCoV-HKU1_orf1ab;2016 | 0 | 0 |
| NC_006577.2;YP_173236.1;HCoV-HKU1_orf1ab;2017 | 0 | 0 |
| NC_006577.2;YP_173236.1;HCoV-HKU1_orf1ab;2018 | 0.05 | 0 |
| NC_006577.2;YP_173236.1;HCoV-HKU1_orf1ab;2019 | 0 | 0 |
| NC_006577.2;YP_173236.1;HCoV-HKU1_orf1ab;2020 | 0 | 0 |
| NC_006577.2;YP_173236.1;HCoV-HKU1_orf1ab;2021 | 0 | 0 |
| NC_006577.2;YP_173236.1;HCoV-HKU1_orf1ab;2022 | 0 | 0.025 |
| NC_006577.2;YP_173236.1;HCoV-HKU1_orf1ab;2023 | 0 | 0 |
| NC_006577.2;YP_173236.1;HCoV-HKU1_orf1ab;2024 | 0 | 0 |
| NC_006577.2;YP_173236.1;HCoV-HKU1_orf1ab;2025 | 0 | 0 |
| NC_006577.2;YP_173236.1;HCoV-HKU1_orf1ab;2026 | 0 | 0 |
| NC_006577.2;YP_173236.1;HCoV-HKU1_orf1ab;2027 | 0 | 0 |
| NC_006577.2;YP_173236.1;HCoV-HKU1_orf1ab;2028 | 0 | 0 |
| NC_006577.2;YP_173236.1;HCoV-HKU1_orf1ab;2029 | 0 | 0 |
| NC_006577.2;YP_173236.1;HCoV-HKU1_orf1ab;2030 | 0 | 0 |
| NC_006577.2;YP_173236.1;HCoV-HKU1_orf1ab;2031 | 0 | 0 |
| NC_006577.2;YP_173236.1;HCoV-HKU1_orf1ab;2032 | 0 | 0 |
| NC_006577.2;YP_173236.1;HCoV-HKU1_orf1ab;2033 | 0 | 0 |
| NC_006577.2;YP_173236.1;HCoV-HKU1_orf1ab;2034 | 0 | 0 |
| NC_006577.2;YP_173236.1;HCoV-HKU1_orf1ab;2035 | 0 | 0 |
| NC_006577.2;YP_173236.1;HCoV-HKU1_orf1ab;2036 | 0 | 0 |

| | | |
|---|---|---|
| NC_006577.2;YP_173236.1;HCoV-HKU1_orf1ab;2037 | 0 | 0 |
| NC_006577.2;YP_173236.1;HCoV-HKU1_orf1ab;2038 | 0.05 | 0 |
| NC_006577.2;YP_173236.1;HCoV-HKU1_orf1ab;2039 | 0 | 0 |
| NC_006577.2;YP_173236.1;HCoV-HKU1_orf1ab;2040 | 0 | 0.025 |
| NC_006577.2;YP_173236.1;HCoV-HKU1_orf1ab;2041 | 0 | 0.025 |
| NC_006577.2;YP_173236.1;HCoV-HKU1_orf1ab;2042 | 0 | 0.025 |
| NC_006577.2;YP_173236.1;HCoV-HKU1_orf1ab;2043 | 0.05 | 0.025 |
| NC_006577.2;YP_173236.1;HCoV-HKU1_orf1ab;2044 | 0 | 0.05 |
| NC_006577.2;YP_173236.1;HCoV-HKU1_orf1ab;2045 | 0 | 0.1 |
| NC_006577.2;YP_173236.1;HCoV-HKU1_orf1ab;2046 | 0 | 0.05 |
| NC_006577.2;YP_173236.1;HCoV-HKU1_orf1ab;2047 | 0 | 0.025 |
| NC_006577.2;YP_173236.1;HCoV-HKU1_orf1ab;2048 | 0 | 0.025 |
| NC_006577.2;YP_173236.1;HCoV-HKU1_orf1ab;2049 | 0 | 0.025 |
| NC_006577.2;YP_173236.1;HCoV-HKU1_orf1ab;2050 | 0.05 | 0.05 |
| NC_006577.2;YP_173236.1;HCoV-HKU1_orf1ab;2051 | 0 | 0 |
| NC_006577.2;YP_173236.1;HCoV-HKU1_orf1ab;2052 | 0 | 0 |
| NC_006577.2;YP_173236.1;HCoV-HKU1_orf1ab;2053 | 0 | 0 |
| NC_006577.2;YP_173236.1;HCoV-HKU1_orf1ab;2054 | 0 | 0 |
| NC_006577.2;YP_173236.1;HCoV-HKU1_orf1ab;2055 | 0 | 0 |
| NC_006577.2;YP_173236.1;HCoV-HKU1_orf1ab;2056 | 0 | 0 |
| NC_006577.2;YP_173236.1;HCoV-HKU1_orf1ab;2057 | 0 | 0 |
| NC_006577.2;YP_173236.1;HCoV-HKU1_orf1ab;2058 | 0 | 0 |
| NC_006577.2;YP_173236.1;HCoV-HKU1_orf1ab;2059 | 0 | 0 |
| NC_006577.2;YP_173236.1;HCoV-HKU1_orf1ab;2060 | 0 | 0 |
| NC_006577.2;YP_173236.1;HCoV-HKU1_orf1ab;2061 | 0 | 0 |
| NC_006577.2;YP_173236.1;HCoV-HKU1_orf1ab;2062 | 0 | 0 |
| NC_006577.2;YP_173236.1;HCoV-HKU1_orf1ab;2063 | 0 | 0 |
| NC_006577.2;YP_173236.1;HCoV-HKU1_orf1ab;2064 | 0 | 0 |
| NC_006577.2;YP_173236.1;HCoV-HKU1_orf1ab;2065 | 0 | 0 |
| NC_006577.2;YP_173236.1;HCoV-HKU1_orf1ab;2066 | 0 | 0 |
| NC_006577.2;YP_173236.1;HCoV-HKU1_orf1ab;2067 | 0 | 0 |
| NC_006577.2;YP_173236.1;HCoV-HKU1_orf1ab;2068 | 0 | 0 |
| NC_006577.2;YP_173236.1;HCoV-HKU1_orf1ab;2069 | 0 | 0 |
| NC_006577.2;YP_173236.1;HCoV-HKU1_orf1ab;2070 | 0 | 0 |
| NC_006577.2;YP_173236.1;HCoV-HKU1_orf1ab;2071 | 0 | 0 |
| NC_006577.2;YP_173236.1;HCoV-HKU1_orf1ab;2072 | 0 | 0 |
| NC_006577.2;YP_173236.1;HCoV-HKU1_orf1ab;2073 | 0 | 0 |
| NC_006577.2;YP_173236.1;HCoV-HKU1_orf1ab;2074 | 0 | 0 |
| NC_006577.2;YP_173236.1;HCoV-HKU1_orf1ab;2075 | 0 | 0 |
| NC_006577.2;YP_173236.1;HCoV-HKU1_orf1ab;2076 | 0 | 0 |
| NC_006577.2;YP_173236.1;HCoV-HKU1_orf1ab;2077 | 0 | 0 |
| NC_006577.2;YP_173236.1;HCoV-HKU1_orf1ab;2078 | 0 | 0 |
| NC_006577.2;YP_173236.1;HCoV-HKU1_orf1ab;2079 | 0 | 0 |

| | | |
|---|---|---|
| NC_006577.2;YP_173236.1;HCoV-HKU1_orf1ab;2080 | 0 | 0.025 |
| NC_006577.2;YP_173236.1;HCoV-HKU1_orf1ab;2081 | 0 | 0 |
| NC_006577.2;YP_173236.1;HCoV-HKU1_orf1ab;2082 | 0 | 0 |
| NC_006577.2;YP_173236.1;HCoV-HKU1_orf1ab;2083 | 0 | 0.025 |
| NC_006577.2;YP_173236.1;HCoV-HKU1_orf1ab;2084 | 0.05 | 0 |
| NC_006577.2;YP_173236.1;HCoV-HKU1_orf1ab;2085 | 0 | 0 |
| NC_006577.2;YP_173236.1;HCoV-HKU1_orf1ab;2086 | 0.05 | 0 |
| NC_006577.2;YP_173236.1;HCoV-HKU1_orf1ab;2087 | 0 | 0 |
| NC_006577.2;YP_173236.1;HCoV-HKU1_orf1ab;2088 | 0.2 | 0 |
| NC_006577.2;YP_173236.1;HCoV-HKU1_orf1ab;2089 | 0.1 | 0 |
| NC_006577.2;YP_173236.1;HCoV-HKU1_orf1ab;2090 | 0.05 | 0.05 |
| NC_006577.2;YP_173236.1;HCoV-HKU1_orf1ab;2091 | 0.25 | 0.1 |
| NC_006577.2;YP_173236.1;HCoV-HKU1_orf1ab;2092 | 0.15 | 0.1 |
| NC_006577.2;YP_173236.1;HCoV-HKU1_orf1ab;2093 | 0.05 | 0.025 |
| NC_006577.2;YP_173236.1;HCoV-HKU1_orf1ab;2094 | 0.15 | 0.075 |
| NC_006577.2;YP_173236.1;HCoV-HKU1_orf1ab;2095 | 0.1 | 0.1 |
| NC_006577.2;YP_173236.1;HCoV-HKU1_orf1ab;2096 | 0.1 | 0.125 |
| NC_006577.2;YP_173236.1;HCoV-HKU1_orf1ab;2097 | 0.1 | 0.125 |
| NC_006577.2;YP_173236.1;HCoV-HKU1_orf1ab;2098 | 0.15 | 0.15 |
| NC_006577.2;YP_173236.1;HCoV-HKU1_orf1ab;2099 | 0.15 | 0.075 |
| NC_006577.2;YP_173236.1;HCoV-HKU1_orf1ab;2100 | 0.15 | 0.125 |
| NC_006577.2;YP_173236.1;HCoV-HKU1_orf1ab;2101 | 0.15 | 0.1 |
| NC_006577.2;YP_173236.1;HCoV-HKU1_orf1ab;2102 | 0.05 | 0 |
| NC_006577.2;YP_173236.1;HCoV-HKU1_orf1ab;2103 | 0 | 0 |
| NC_006577.2;YP_173236.1;HCoV-HKU1_orf1ab;2104 | 0 | 0 |
| NC_006577.2;YP_173236.1;HCoV-HKU1_orf1ab;2105 | 0.05 | 0.05 |
| NC_006577.2;YP_173236.1;HCoV-HKU1_orf1ab;2106 | 0.05 | 0.05 |
| NC_006577.2;YP_173236.1;HCoV-HKU1_orf1ab;2107 | 0 | 0.025 |
| NC_006577.2;YP_173236.1;HCoV-HKU1_orf1ab;2108 | 0 | 0.025 |
| NC_006577.2;YP_173236.1;HCoV-HKU1_orf1ab;2109 | 0 | 0.05 |
| NC_006577.2;YP_173236.1;HCoV-HKU1_orf1ab;2110 | 0 | 0.025 |
| NC_006577.2;YP_173236.1;HCoV-HKU1_orf1ab;2111 | 0 | 0.025 |
| NC_006577.2;YP_173236.1;HCoV-HKU1_orf1ab;2112 | 0 | 0 |
| NC_006577.2;YP_173236.1;HCoV-HKU1_orf1ab;2113 | 0.05 | 0.025 |
| NC_006577.2;YP_173236.1;HCoV-HKU1_orf1ab;2114 | 0.05 | 0.025 |
| NC_006577.2;YP_173236.1;HCoV-HKU1_orf1ab;2115 | 0 | 0.025 |
| NC_006577.2;YP_173236.1;HCoV-HKU1_orf1ab;2116 | 0 | 0.025 |
| NC_006577.2;YP_173236.1;HCoV-HKU1_orf1ab;2117 | 0 | 0.025 |
| NC_006577.2;YP_173236.1;HCoV-HKU1_orf1ab;2118 | 0 | 0.025 |
| NC_006577.2;YP_173236.1;HCoV-HKU1_orf1ab;2119 | 0 | 0 |
| NC_006577.2;YP_173236.1;HCoV-HKU1_orf1ab;2120 | 0 | 0 |
| NC_006577.2;YP_173236.1;HCoV-HKU1_orf1ab;2121 | 0 | 0 |
| NC_006577.2;YP_173236.1;HCoV-HKU1_orf1ab;2122 | 0 | 0 |

| | | |
|---|---|---|
| NC_006577.2;YP_173236.1;HCoV-HKU1_orf1ab;2123 | 0 | 0 |
| NC_006577.2;YP_173236.1;HCoV-HKU1_orf1ab;2124 | 0 | 0 |
| NC_006577.2;YP_173236.1;HCoV-HKU1_orf1ab;2125 | 0 | 0 |
| NC_006577.2;YP_173236.1;HCoV-HKU1_orf1ab;2126 | 0 | 0 |
| NC_006577.2;YP_173236.1;HCoV-HKU1_orf1ab;2127 | 0 | 0 |
| NC_006577.2;YP_173236.1;HCoV-HKU1_orf1ab;2128 | 0 | 0 |
| NC_006577.2;YP_173236.1;HCoV-HKU1_orf1ab;2129 | 0 | 0 |
| NC_006577.2;YP_173236.1;HCoV-HKU1_orf1ab;2130 | 0 | 0 |
| NC_006577.2;YP_173236.1;HCoV-HKU1_orf1ab;2131 | 0 | 0 |
| NC_006577.2;YP_173236.1;HCoV-HKU1_orf1ab;2132 | 0 | 0 |
| NC_006577.2;YP_173236.1;HCoV-HKU1_orf1ab;2133 | 0 | 0 |
| NC_006577.2;YP_173236.1;HCoV-HKU1_orf1ab;2134 | 0 | 0.025 |
| NC_006577.2;YP_173236.1;HCoV-HKU1_orf1ab;2135 | 0 | 0.05 |
| NC_006577.2;YP_173236.1;HCoV-HKU1_orf1ab;2136 | 0.05 | 0.025 |
| NC_006577.2;YP_173236.1;HCoV-HKU1_orf1ab;2137 | 0.1 | 0.025 |
| NC_006577.2;YP_173236.1;HCoV-HKU1_orf1ab;2138 | 0.1 | 0.025 |
| NC_006577.2;YP_173236.1;HCoV-HKU1_orf1ab;2139 | 0.05 | 0 |
| NC_006577.2;YP_173236.1;HCoV-HKU1_orf1ab;2140 | 0.05 | 0 |
| NC_006577.2;YP_173236.1;HCoV-HKU1_orf1ab;2141 | 0 | 0 |
| NC_006577.2;YP_173236.1;HCoV-HKU1_orf1ab;2142 | 0 | 0 |
| NC_006577.2;YP_173236.1;HCoV-HKU1_orf1ab;2143 | 0 | 0 |
| NC_006577.2;YP_173236.1;HCoV-HKU1_orf1ab;2144 | 0 | 0 |
| NC_006577.2;YP_173236.1;HCoV-HKU1_orf1ab;2145 | 0 | 0 |
| NC_006577.2;YP_173236.1;HCoV-HKU1_orf1ab;2146 | 0 | 0 |
| NC_006577.2;YP_173236.1;HCoV-HKU1_orf1ab;2147 | 0 | 0 |
| NC_006577.2;YP_173236.1;HCoV-HKU1_orf1ab;2148 | 0 | 0 |
| NC_006577.2;YP_173236.1;HCoV-HKU1_orf1ab;2149 | 0 | 0 |
| NC_006577.2;YP_173236.1;HCoV-HKU1_orf1ab;2150 | 0 | 0 |
| NC_006577.2;YP_173236.1;HCoV-HKU1_orf1ab;2151 | 0 | 0 |
| NC_006577.2;YP_173236.1;HCoV-HKU1_orf1ab;2152 | 0 | 0 |
| NC_006577.2;YP_173236.1;HCoV-HKU1_orf1ab;2153 | 0 | 0 |
| NC_006577.2;YP_173236.1;HCoV-HKU1_orf1ab;2154 | 0 | 0 |
| NC_006577.2;YP_173236.1;HCoV-HKU1_orf1ab;2155 | 0 | 0 |
| NC_006577.2;YP_173236.1;HCoV-HKU1_orf1ab;2156 | 0 | 0 |
| NC_006577.2;YP_173236.1;HCoV-HKU1_orf1ab;2157 | 0 | 0 |
| NC_006577.2;YP_173236.1;HCoV-HKU1_orf1ab;2158 | 0 | 0 |
| NC_006577.2;YP_173236.1;HCoV-HKU1_orf1ab;2159 | 0 | 0 |
| NC_006577.2;YP_173236.1;HCoV-HKU1_orf1ab;2160 | 0 | 0 |
| NC_006577.2;YP_173236.1;HCoV-HKU1_orf1ab;2161 | 0 | 0 |
| NC_006577.2;YP_173236.1;HCoV-HKU1_orf1ab;2162 | 0 | 0 |
| NC_006577.2;YP_173236.1;HCoV-HKU1_orf1ab;2163 | 0 | 0 |
| NC_006577.2;YP_173236.1;HCoV-HKU1_orf1ab;2164 | 0 | 0 |
| NC_006577.2;YP_173236.1;HCoV-HKU1_orf1ab;2165 | 0 | 0 |

| | | |
|---|---|---|
| NC_006577.2;YP_173236.1;HCoV-HKU1_orf1ab;2166 | 0 | 0 |
| NC_006577.2;YP_173236.1;HCoV-HKU1_orf1ab;2167 | 0 | 0 |
| NC_006577.2;YP_173236.1;HCoV-HKU1_orf1ab;2168 | 0 | 0 |
| NC_006577.2;YP_173236.1;HCoV-HKU1_orf1ab;2169 | 0 | 0 |
| NC_006577.2;YP_173236.1;HCoV-HKU1_orf1ab;2170 | 0 | 0 |
| NC_006577.2;YP_173236.1;HCoV-HKU1_orf1ab;2171 | 0 | 0 |
| NC_006577.2;YP_173236.1;HCoV-HKU1_orf1ab;2172 | 0 | 0 |
| NC_006577.2;YP_173236.1;HCoV-HKU1_orf1ab;2173 | 0 | 0 |
| NC_006577.2;YP_173236.1;HCoV-HKU1_orf1ab;2174 | 0 | 0.025 |
| NC_006577.2;YP_173236.1;HCoV-HKU1_orf1ab;2175 | 0 | 0 |
| NC_006577.2;YP_173236.1;HCoV-HKU1_orf1ab;2176 | 0 | 0 |
| NC_006577.2;YP_173236.1;HCoV-HKU1_orf1ab;2177 | 0 | 0 |
| NC_006577.2;YP_173236.1;HCoV-HKU1_orf1ab;2178 | 0 | 0 |
| NC_006577.2;YP_173236.1;HCoV-HKU1_orf1ab;2179 | 0 | 0 |
| NC_006577.2;YP_173236.1;HCoV-HKU1_orf1ab;2180 | 0 | 0 |
| NC_006577.2;YP_173236.1;HCoV-HKU1_orf1ab;2181 | 0 | 0 |
| NC_006577.2;YP_173236.1;HCoV-HKU1_orf1ab;2182 | 0 | 0 |
| NC_006577.2;YP_173236.1;HCoV-HKU1_orf1ab;2183 | 0 | 0 |
| NC_006577.2;YP_173236.1;HCoV-HKU1_orf1ab;2184 | 0 | 0 |
| NC_006577.2;YP_173236.1;HCoV-HKU1_orf1ab;2185 | 0 | 0 |
| NC_006577.2;YP_173236.1;HCoV-HKU1_orf1ab;2186 | 0 | 0 |
| NC_006577.2;YP_173236.1;HCoV-HKU1_orf1ab;2187 | 0 | 0 |
| NC_006577.2;YP_173236.1;HCoV-HKU1_orf1ab;2188 | 0 | 0 |
| NC_006577.2;YP_173236.1;HCoV-HKU1_orf1ab;2189 | 0 | 0 |
| NC_006577.2;YP_173236.1;HCoV-HKU1_orf1ab;2190 | 0 | 0 |
| NC_006577.2;YP_173236.1;HCoV-HKU1_orf1ab;2191 | 0 | 0 |
| NC_006577.2;YP_173236.1;HCoV-HKU1_orf1ab;2192 | 0 | 0 |
| NC_006577.2;YP_173236.1;HCoV-HKU1_orf1ab;2193 | 0 | 0 |
| NC_006577.2;YP_173236.1;HCoV-HKU1_orf1ab;2194 | 0 | 0 |
| NC_006577.2;YP_173236.1;HCoV-HKU1_orf1ab;2195 | 0 | 0 |
| NC_006577.2;YP_173236.1;HCoV-HKU1_orf1ab;2196 | 0 | 0 |
| NC_006577.2;YP_173236.1;HCoV-HKU1_orf1ab;2197 | 0 | 0 |
| NC_006577.2;YP_173236.1;HCoV-HKU1_orf1ab;2198 | 0 | 0 |
| NC_006577.2;YP_173236.1;HCoV-HKU1_orf1ab;2199 | 0 | 0 |
| NC_006577.2;YP_173236.1;HCoV-HKU1_orf1ab;2200 | 0 | 0 |
| NC_006577.2;YP_173236.1;HCoV-HKU1_orf1ab;2201 | 0 | 0 |
| NC_006577.2;YP_173236.1;HCoV-HKU1_orf1ab;2202 | 0 | 0 |
| NC_006577.2;YP_173236.1;HCoV-HKU1_orf1ab;2203 | 0 | 0 |
| NC_006577.2;YP_173236.1;HCoV-HKU1_orf1ab;2204 | 0 | 0.025 |
| NC_006577.2;YP_173236.1;HCoV-HKU1_orf1ab;2205 | 0 | 0.025 |
| NC_006577.2;YP_173236.1;HCoV-HKU1_orf1ab;2206 | 0 | 0.025 |
| NC_006577.2;YP_173236.1;HCoV-HKU1_orf1ab;2207 | 0 | 0.025 |
| NC_006577.2;YP_173236.1;HCoV-HKU1_orf1ab;2208 | 0 | 0 |

| | | |
|---|---|---|
| NC_006577.2;YP_173236.1;HCoV-HKU1_orf1ab;2209 | 0 | 0 |
| NC_006577.2;YP_173236.1;HCoV-HKU1_orf1ab;2210 | 0 | 0 |
| NC_006577.2;YP_173236.1;HCoV-HKU1_orf1ab;2211 | 0.05 | 0 |
| NC_006577.2;YP_173236.1;HCoV-HKU1_orf1ab;2212 | 0 | 0 |
| NC_006577.2;YP_173236.1;HCoV-HKU1_orf1ab;2213 | 0 | 0 |
| NC_006577.2;YP_173236.1;HCoV-HKU1_orf1ab;2214 | 0 | 0 |
| NC_006577.2;YP_173236.1;HCoV-HKU1_orf1ab;2215 | 0 | 0 |
| NC_006577.2;YP_173236.1;HCoV-HKU1_orf1ab;2216 | 0 | 0 |
| NC_006577.2;YP_173236.1;HCoV-HKU1_orf1ab;2217 | 0 | 0 |
| NC_006577.2;YP_173236.1;HCoV-HKU1_orf1ab;2218 | 0 | 0 |
| NC_006577.2;YP_173236.1;HCoV-HKU1_orf1ab;2219 | 0 | 0 |
| NC_006577.2;YP_173236.1;HCoV-HKU1_orf1ab;2220 | 0 | 0 |
| NC_006577.2;YP_173236.1;HCoV-HKU1_orf1ab;2221 | 0 | 0 |
| NC_006577.2;YP_173236.1;HCoV-HKU1_orf1ab;2222 | 0 | 0 |
| NC_006577.2;YP_173236.1;HCoV-HKU1_orf1ab;2223 | 0 | 0 |
| NC_006577.2;YP_173236.1;HCoV-HKU1_orf1ab;2224 | 0 | 0 |
| NC_006577.2;YP_173236.1;HCoV-HKU1_orf1ab;2225 | 0 | 0 |
| NC_006577.2;YP_173236.1;HCoV-HKU1_orf1ab;2226 | 0 | 0 |
| NC_006577.2;YP_173236.1;HCoV-HKU1_orf1ab;2227 | 0 | 0 |
| NC_006577.2;YP_173236.1;HCoV-HKU1_orf1ab;2228 | 0 | 0 |
| NC_006577.2;YP_173236.1;HCoV-HKU1_orf1ab;2229 | 0 | 0 |
| NC_006577.2;YP_173236.1;HCoV-HKU1_orf1ab;2230 | 0 | 0 |
| NC_006577.2;YP_173236.1;HCoV-HKU1_orf1ab;2231 | 0 | 0 |
| NC_006577.2;YP_173236.1;HCoV-HKU1_orf1ab;2232 | 0 | 0 |
| NC_006577.2;YP_173236.1;HCoV-HKU1_orf1ab;2233 | 0 | 0 |
| NC_006577.2;YP_173236.1;HCoV-HKU1_orf1ab;2234 | 0 | 0 |
| NC_006577.2;YP_173236.1;HCoV-HKU1_orf1ab;2235 | 0 | 0 |
| NC_006577.2;YP_173236.1;HCoV-HKU1_orf1ab;2236 | 0 | 0 |
| NC_006577.2;YP_173236.1;HCoV-HKU1_orf1ab;2237 | 0 | 0 |
| NC_006577.2;YP_173236.1;HCoV-HKU1_orf1ab;2238 | 0 | 0 |
| NC_006577.2;YP_173236.1;HCoV-HKU1_orf1ab;2239 | 0 | 0 |
| NC_006577.2;YP_173236.1;HCoV-HKU1_orf1ab;2240 | 0 | 0 |
| NC_006577.2;YP_173236.1;HCoV-HKU1_orf1ab;2241 | 0 | 0 |
| NC_006577.2;YP_173236.1;HCoV-HKU1_orf1ab;2242 | 0 | 0 |
| NC_006577.2;YP_173236.1;HCoV-HKU1_orf1ab;2243 | 0 | 0 |
| NC_006577.2;YP_173236.1;HCoV-HKU1_orf1ab;2244 | 0 | 0 |
| NC_006577.2;YP_173236.1;HCoV-HKU1_orf1ab;2245 | 0 | 0 |
| NC_006577.2;YP_173236.1;HCoV-HKU1_orf1ab;2246 | 0 | 0.025 |
| NC_006577.2;YP_173236.1;HCoV-HKU1_orf1ab;2247 | 0 | 0 |
| NC_006577.2;YP_173236.1;HCoV-HKU1_orf1ab;2248 | 0 | 0 |
| NC_006577.2;YP_173236.1;HCoV-HKU1_orf1ab;2249 | 0 | 0 |
| NC_006577.2;YP_173236.1;HCoV-HKU1_orf1ab;2250 | 0 | 0 |
| NC_006577.2;YP_173236.1;HCoV-HKU1_orf1ab;2251 | 0 | 0 |

| | | |
|---|---|---|
| NC_006577.2;YP_173236.1;HCoV-HKU1_orf1ab;2252 | 0 | 0 |
| NC_006577.2;YP_173236.1;HCoV-HKU1_orf1ab;2253 | 0 | 0 |
| NC_006577.2;YP_173236.1;HCoV-HKU1_orf1ab;2254 | 0 | 0 |
| NC_006577.2;YP_173236.1;HCoV-HKU1_orf1ab;2255 | 0 | 0 |
| NC_006577.2;YP_173236.1;HCoV-HKU1_orf1ab;2256 | 0 | 0 |
| NC_006577.2;YP_173236.1;HCoV-HKU1_orf1ab;2257 | 0 | 0 |
| NC_006577.2;YP_173236.1;HCoV-HKU1_orf1ab;2258 | 0 | 0 |
| NC_006577.2;YP_173236.1;HCoV-HKU1_orf1ab;2259 | 0 | 0 |
| NC_006577.2;YP_173236.1;HCoV-HKU1_orf1ab;2260 | 0 | 0 |
| NC_006577.2;YP_173236.1;HCoV-HKU1_orf1ab;2261 | 0 | 0 |
| NC_006577.2;YP_173236.1;HCoV-HKU1_orf1ab;2262 | 0 | 0 |
| NC_006577.2;YP_173236.1;HCoV-HKU1_orf1ab;2263 | 0 | 0 |
| NC_006577.2;YP_173236.1;HCoV-HKU1_orf1ab;2264 | 0 | 0 |
| NC_006577.2;YP_173236.1;HCoV-HKU1_orf1ab;2265 | 0 | 0 |
| NC_006577.2;YP_173236.1;HCoV-HKU1_orf1ab;2266 | 0 | 0 |
| NC_006577.2;YP_173236.1;HCoV-HKU1_orf1ab;2267 | 0 | 0 |
| NC_006577.2;YP_173236.1;HCoV-HKU1_orf1ab;2268 | 0 | 0 |
| NC_006577.2;YP_173236.1;HCoV-HKU1_orf1ab;2269 | 0 | 0 |
| NC_006577.2;YP_173236.1;HCoV-HKU1_orf1ab;2270 | 0 | 0 |
| NC_006577.2;YP_173236.1;HCoV-HKU1_orf1ab;2271 | 0 | 0 |
| NC_006577.2;YP_173236.1;HCoV-HKU1_orf1ab;2272 | 0 | 0 |
| NC_006577.2;YP_173236.1;HCoV-HKU1_orf1ab;2273 | 0 | 0 |
| NC_006577.2;YP_173236.1;HCoV-HKU1_orf1ab;2274 | 0 | 0 |
| NC_006577.2;YP_173236.1;HCoV-HKU1_orf1ab;2275 | 0 | 0 |
| NC_006577.2;YP_173236.1;HCoV-HKU1_orf1ab;2276 | 0 | 0 |
| NC_006577.2;YP_173236.1;HCoV-HKU1_orf1ab;2277 | 0 | 0 |
| NC_006577.2;YP_173236.1;HCoV-HKU1_orf1ab;2278 | 0 | 0 |
| NC_006577.2;YP_173236.1;HCoV-HKU1_orf1ab;2279 | 0 | 0 |
| NC_006577.2;YP_173236.1;HCoV-HKU1_orf1ab;2280 | 0 | 0 |
| NC_006577.2;YP_173236.1;HCoV-HKU1_orf1ab;2281 | 0 | 0 |
| NC_006577.2;YP_173236.1;HCoV-HKU1_orf1ab;2282 | 0 | 0 |
| NC_006577.2;YP_173236.1;HCoV-HKU1_orf1ab;2283 | 0 | 0 |
| NC_006577.2;YP_173236.1;HCoV-HKU1_orf1ab;2284 | 0 | 0 |
| NC_006577.2;YP_173236.1;HCoV-HKU1_orf1ab;2285 | 0 | 0 |
| NC_006577.2;YP_173236.1;HCoV-HKU1_orf1ab;2286 | 0 | 0 |
| NC_006577.2;YP_173236.1;HCoV-HKU1_orf1ab;2287 | 0 | 0 |
| NC_006577.2;YP_173236.1;HCoV-HKU1_orf1ab;2288 | 0 | 0 |
| NC_006577.2;YP_173236.1;HCoV-HKU1_orf1ab;2289 | 0 | 0 |
| NC_006577.2;YP_173236.1;HCoV-HKU1_orf1ab;2290 | 0 | 0 |
| NC_006577.2;YP_173236.1;HCoV-HKU1_orf1ab;2291 | 0 | 0 |
| NC_006577.2;YP_173236.1;HCoV-HKU1_orf1ab;2292 | 0 | 0 |
| NC_006577.2;YP_173236.1;HCoV-HKU1_orf1ab;2293 | 0 | 0 |
| NC_006577.2;YP_173236.1;HCoV-HKU1_orf1ab;2294 | 0 | 0 |

| | | |
|---|---|---|
| NC_006577.2;YP_173236.1;HCoV-HKU1_orf1ab;2295 | 0 | 0 |
| NC_006577.2;YP_173236.1;HCoV-HKU1_orf1ab;2296 | 0 | 0 |
| NC_006577.2;YP_173236.1;HCoV-HKU1_orf1ab;2297 | 0 | 0 |
| NC_006577.2;YP_173236.1;HCoV-HKU1_orf1ab;2298 | 0 | 0 |
| NC_006577.2;YP_173236.1;HCoV-HKU1_orf1ab;2299 | 0 | 0 |
| NC_006577.2;YP_173236.1;HCoV-HKU1_orf1ab;2300 | 0 | 0 |
| NC_006577.2;YP_173236.1;HCoV-HKU1_orf1ab;2301 | 0 | 0 |
| NC_006577.2;YP_173236.1;HCoV-HKU1_orf1ab;2302 | 0 | 0 |
| NC_006577.2;YP_173236.1;HCoV-HKU1_orf1ab;2303 | 0 | 0 |
| NC_006577.2;YP_173236.1;HCoV-HKU1_orf1ab;2304 | 0 | 0 |
| NC_006577.2;YP_173236.1;HCoV-HKU1_orf1ab;2305 | 0 | 0 |
| NC_006577.2;YP_173236.1;HCoV-HKU1_orf1ab;2306 | 0 | 0 |
| NC_006577.2;YP_173236.1;HCoV-HKU1_orf1ab;2307 | 0 | 0 |
| NC_006577.2;YP_173236.1;HCoV-HKU1_orf1ab;2308 | 0 | 0 |
| NC_006577.2;YP_173236.1;HCoV-HKU1_orf1ab;2309 | 0 | 0 |
| NC_006577.2;YP_173236.1;HCoV-HKU1_orf1ab;2310 | 0 | 0 |
| NC_006577.2;YP_173236.1;HCoV-HKU1_orf1ab;2311 | 0 | 0 |
| NC_006577.2;YP_173236.1;HCoV-HKU1_orf1ab;2312 | 0 | 0 |
| NC_006577.2;YP_173236.1;HCoV-HKU1_orf1ab;2313 | 0 | 0 |
| NC_006577.2;YP_173236.1;HCoV-HKU1_orf1ab;2314 | 0 | 0 |
| NC_006577.2;YP_173236.1;HCoV-HKU1_orf1ab;2315 | 0 | 0 |
| NC_006577.2;YP_173236.1;HCoV-HKU1_orf1ab;2316 | 0 | 0 |
| NC_006577.2;YP_173236.1;HCoV-HKU1_orf1ab;2317 | 0 | 0 |
| NC_006577.2;YP_173236.1;HCoV-HKU1_orf1ab;2318 | 0 | 0 |
| NC_006577.2;YP_173236.1;HCoV-HKU1_orf1ab;2319 | 0 | 0 |
| NC_006577.2;YP_173236.1;HCoV-HKU1_orf1ab;2320 | 0 | 0 |
| NC_006577.2;YP_173236.1;HCoV-HKU1_orf1ab;2321 | 0 | 0 |
| NC_006577.2;YP_173236.1;HCoV-HKU1_orf1ab;2322 | 0 | 0 |
| NC_006577.2;YP_173236.1;HCoV-HKU1_orf1ab;2323 | 0 | 0 |
| NC_006577.2;YP_173236.1;HCoV-HKU1_orf1ab;2324 | 0 | 0 |
| NC_006577.2;YP_173236.1;HCoV-HKU1_orf1ab;2325 | 0 | 0 |
| NC_006577.2;YP_173236.1;HCoV-HKU1_orf1ab;2326 | 0 | 0 |
| NC_006577.2;YP_173236.1;HCoV-HKU1_orf1ab;2327 | 0 | 0 |
| NC_006577.2;YP_173236.1;HCoV-HKU1_orf1ab;2328 | 0 | 0 |
| NC_006577.2;YP_173236.1;HCoV-HKU1_orf1ab;2329 | 0 | 0 |
| NC_006577.2;YP_173236.1;HCoV-HKU1_orf1ab;2330 | 0 | 0 |
| NC_006577.2;YP_173236.1;HCoV-HKU1_orf1ab;2331 | 0 | 0 |
| NC_006577.2;YP_173236.1;HCoV-HKU1_orf1ab;2332 | 0 | 0 |
| NC_006577.2;YP_173236.1;HCoV-HKU1_orf1ab;2333 | 0 | 0.025 |
| NC_006577.2;YP_173236.1;HCoV-HKU1_orf1ab;2334 | 0 | 0.025 |
| NC_006577.2;YP_173236.1;HCoV-HKU1_orf1ab;2335 | 0 | 0.025 |
| NC_006577.2;YP_173236.1;HCoV-HKU1_orf1ab;2336 | 0 | 0.025 |
| NC_006577.2;YP_173236.1;HCoV-HKU1_orf1ab;2337 | 0 | 0.025 |

| | | |
|---|---|---|
| NC_006577.2;YP_173236.1;HCoV-HKU1_orf1ab;2338 | 0 | 0 |
| NC_006577.2;YP_173236.1;HCoV-HKU1_orf1ab;2339 | 0 | 0 |
| NC_006577.2;YP_173236.1;HCoV-HKU1_orf1ab;2340 | 0 | 0.025 |
| NC_006577.2;YP_173236.1;HCoV-HKU1_orf1ab;2341 | 0 | 0 |
| NC_006577.2;YP_173236.1;HCoV-HKU1_orf1ab;2342 | 0 | 0 |
| NC_006577.2;YP_173236.1;HCoV-HKU1_orf1ab;2343 | 0 | 0 |
| NC_006577.2;YP_173236.1;HCoV-HKU1_orf1ab;2344 | 0 | 0 |
| NC_006577.2;YP_173236.1;HCoV-HKU1_orf1ab;2345 | 0 | 0 |
| NC_006577.2;YP_173236.1;HCoV-HKU1_orf1ab;2346 | 0 | 0 |
| NC_006577.2;YP_173236.1;HCoV-HKU1_orf1ab;2347 | 0 | 0 |
| NC_006577.2;YP_173236.1;HCoV-HKU1_orf1ab;2348 | 0 | 0 |
| NC_006577.2;YP_173236.1;HCoV-HKU1_orf1ab;2349 | 0 | 0 |
| NC_006577.2;YP_173236.1;HCoV-HKU1_orf1ab;2350 | 0 | 0 |
| NC_006577.2;YP_173236.1;HCoV-HKU1_orf1ab;2351 | 0 | 0 |
| NC_006577.2;YP_173236.1;HCoV-HKU1_orf1ab;2352 | 0 | 0 |
| NC_006577.2;YP_173236.1;HCoV-HKU1_orf1ab;2353 | 0 | 0 |
| NC_006577.2;YP_173236.1;HCoV-HKU1_orf1ab;2354 | 0 | 0 |
| NC_006577.2;YP_173236.1;HCoV-HKU1_orf1ab;2355 | 0 | 0 |
| NC_006577.2;YP_173236.1;HCoV-HKU1_orf1ab;2356 | 0 | 0 |
| NC_006577.2;YP_173236.1;HCoV-HKU1_orf1ab;2357 | 0 | 0 |
| NC_006577.2;YP_173236.1;HCoV-HKU1_orf1ab;2358 | 0 | 0 |
| NC_006577.2;YP_173236.1;HCoV-HKU1_orf1ab;2359 | 0 | 0 |
| NC_006577.2;YP_173236.1;HCoV-HKU1_orf1ab;2360 | 0.05 | 0.025 |
| NC_006577.2;YP_173236.1;HCoV-HKU1_orf1ab;2361 | 0 | 0 |
| NC_006577.2;YP_173236.1;HCoV-HKU1_orf1ab;2362 | 0 | 0 |
| NC_006577.2;YP_173236.1;HCoV-HKU1_orf1ab;2363 | 0 | 0 |
| NC_006577.2;YP_173236.1;HCoV-HKU1_orf1ab;2364 | 0 | 0 |
| NC_006577.2;YP_173236.1;HCoV-HKU1_orf1ab;2365 | 0 | 0 |
| NC_006577.2;YP_173236.1;HCoV-HKU1_orf1ab;2366 | 0 | 0 |
| NC_006577.2;YP_173236.1;HCoV-HKU1_orf1ab;2367 | 0 | 0 |
| NC_006577.2;YP_173236.1;HCoV-HKU1_orf1ab;2368 | 0 | 0 |
| NC_006577.2;YP_173236.1;HCoV-HKU1_orf1ab;2369 | 0 | 0 |
| NC_006577.2;YP_173236.1;HCoV-HKU1_orf1ab;2370 | 0 | 0 |
| NC_006577.2;YP_173236.1;HCoV-HKU1_orf1ab;2371 | 0 | 0 |
| NC_006577.2;YP_173236.1;HCoV-HKU1_orf1ab;2372 | 0 | 0 |
| NC_006577.2;YP_173236.1;HCoV-HKU1_orf1ab;2373 | 0 | 0 |
| NC_006577.2;YP_173236.1;HCoV-HKU1_orf1ab;2374 | 0 | 0.05 |
| NC_006577.2;YP_173236.1;HCoV-HKU1_orf1ab;2375 | 0 | 0 |
| NC_006577.2;YP_173236.1;HCoV-HKU1_orf1ab;2376 | 0 | 0 |
| NC_006577.2;YP_173236.1;HCoV-HKU1_orf1ab;2377 | 0 | 0 |
| NC_006577.2;YP_173236.1;HCoV-HKU1_orf1ab;2378 | 0 | 0 |
| NC_006577.2;YP_173236.1;HCoV-HKU1_orf1ab;2379 | 0 | 0 |
| NC_006577.2;YP_173236.1;HCoV-HKU1_orf1ab;2380 | 0 | 0 |

| | | |
|---|---|---|
| NC_006577.2;YP_173236.1;HCoV-HKU1_orf1ab;2381 | 0 | 0 |
| NC_006577.2;YP_173236.1;HCoV-HKU1_orf1ab;2382 | 0 | 0 |
| NC_006577.2;YP_173236.1;HCoV-HKU1_orf1ab;2383 | 0 | 0 |
| NC_006577.2;YP_173236.1;HCoV-HKU1_orf1ab;2384 | 0 | 0 |
| NC_006577.2;YP_173236.1;HCoV-HKU1_orf1ab;2385 | 0 | 0 |
| NC_006577.2;YP_173236.1;HCoV-HKU1_orf1ab;2386 | 0 | 0 |
| NC_006577.2;YP_173236.1;HCoV-HKU1_orf1ab;2387 | 0 | 0 |
| NC_006577.2;YP_173236.1;HCoV-HKU1_orf1ab;2388 | 0 | 0 |
| NC_006577.2;YP_173236.1;HCoV-HKU1_orf1ab;2389 | 0 | 0 |
| NC_006577.2;YP_173236.1;HCoV-HKU1_orf1ab;2390 | 0 | 0 |
| NC_006577.2;YP_173236.1;HCoV-HKU1_orf1ab;2391 | 0 | 0 |
| NC_006577.2;YP_173236.1;HCoV-HKU1_orf1ab;2392 | 0 | 0 |
| NC_006577.2;YP_173236.1;HCoV-HKU1_orf1ab;2393 | 0 | 0 |
| NC_006577.2;YP_173236.1;HCoV-HKU1_orf1ab;2394 | 0 | 0 |
| NC_006577.2;YP_173236.1;HCoV-HKU1_orf1ab;2395 | 0 | 0 |
| NC_006577.2;YP_173236.1;HCoV-HKU1_orf1ab;2396 | 0 | 0 |
| NC_006577.2;YP_173236.1;HCoV-HKU1_orf1ab;2397 | 0 | 0 |
| NC_006577.2;YP_173236.1;HCoV-HKU1_orf1ab;2398 | 0 | 0 |
| NC_006577.2;YP_173236.1;HCoV-HKU1_orf1ab;2399 | 0 | 0 |
| NC_006577.2;YP_173236.1;HCoV-HKU1_orf1ab;2400 | 0 | 0 |
| NC_006577.2;YP_173236.1;HCoV-HKU1_orf1ab;2401 | 0 | 0 |
| NC_006577.2;YP_173236.1;HCoV-HKU1_orf1ab;2402 | 0 | 0 |
| NC_006577.2;YP_173236.1;HCoV-HKU1_orf1ab;2403 | 0 | 0 |
| NC_006577.2;YP_173236.1;HCoV-HKU1_orf1ab;2404 | 0 | 0 |
| NC_006577.2;YP_173236.1;HCoV-HKU1_orf1ab;2405 | 0 | 0 |
| NC_006577.2;YP_173236.1;HCoV-HKU1_orf1ab;2406 | 0 | 0 |
| NC_006577.2;YP_173236.1;HCoV-HKU1_orf1ab;2407 | 0 | 0 |
| NC_006577.2;YP_173236.1;HCoV-HKU1_orf1ab;2408 | 0 | 0 |
| NC_006577.2;YP_173236.1;HCoV-HKU1_orf1ab;2409 | 0 | 0 |
| NC_006577.2;YP_173236.1;HCoV-HKU1_orf1ab;2410 | 0 | 0 |
| NC_006577.2;YP_173236.1;HCoV-HKU1_orf1ab;2411 | 0 | 0 |
| NC_006577.2;YP_173236.1;HCoV-HKU1_orf1ab;2412 | 0 | 0 |
| NC_006577.2;YP_173236.1;HCoV-HKU1_orf1ab;2413 | 0 | 0 |
| NC_006577.2;YP_173236.1;HCoV-HKU1_orf1ab;2414 | 0 | 0 |
| NC_006577.2;YP_173236.1;HCoV-HKU1_orf1ab;2415 | 0 | 0 |
| NC_006577.2;YP_173236.1;HCoV-HKU1_orf1ab;2416 | 0 | 0 |
| NC_006577.2;YP_173236.1;HCoV-HKU1_orf1ab;2417 | 0 | 0 |
| NC_006577.2;YP_173236.1;HCoV-HKU1_orf1ab;2418 | 0 | 0 |
| NC_006577.2;YP_173236.1;HCoV-HKU1_orf1ab;2419 | 0 | 0 |
| NC_006577.2;YP_173236.1;HCoV-HKU1_orf1ab;2420 | 0 | 0 |
| NC_006577.2;YP_173236.1;HCoV-HKU1_orf1ab;2421 | 0 | 0 |
| NC_006577.2;YP_173236.1;HCoV-HKU1_orf1ab;2422 | 0 | 0 |
| NC_006577.2;YP_173236.1;HCoV-HKU1_orf1ab;2423 | 0 | 0 |

| | | |
|---|---|---|
| NC_006577.2;YP_173236.1;HCoV-HKU1_orf1ab;2424 | 0 | 0 |
| NC_006577.2;YP_173236.1;HCoV-HKU1_orf1ab;2425 | 0 | 0 |
| NC_006577.2;YP_173236.1;HCoV-HKU1_orf1ab;2426 | 0 | 0 |
| NC_006577.2;YP_173236.1;HCoV-HKU1_orf1ab;2427 | 0 | 0 |
| NC_006577.2;YP_173236.1;HCoV-HKU1_orf1ab;2428 | 0 | 0 |
| NC_006577.2;YP_173236.1;HCoV-HKU1_orf1ab;2429 | 0 | 0 |
| NC_006577.2;YP_173236.1;HCoV-HKU1_orf1ab;2430 | 0 | 0 |
| NC_006577.2;YP_173236.1;HCoV-HKU1_orf1ab;2431 | 0 | 0 |
| NC_006577.2;YP_173236.1;HCoV-HKU1_orf1ab;2432 | 0 | 0 |
| NC_006577.2;YP_173236.1;HCoV-HKU1_orf1ab;2433 | 0 | 0 |
| NC_006577.2;YP_173236.1;HCoV-HKU1_orf1ab;2434 | 0 | 0 |
| NC_006577.2;YP_173236.1;HCoV-HKU1_orf1ab;2435 | 0 | 0 |
| NC_006577.2;YP_173236.1;HCoV-HKU1_orf1ab;2436 | 0 | 0 |
| NC_006577.2;YP_173236.1;HCoV-HKU1_orf1ab;2437 | 0 | 0 |
| NC_006577.2;YP_173236.1;HCoV-HKU1_orf1ab;2438 | 0 | 0 |
| NC_006577.2;YP_173236.1;HCoV-HKU1_orf1ab;2439 | 0 | 0 |
| NC_006577.2;YP_173236.1;HCoV-HKU1_orf1ab;2440 | 0 | 0 |
| NC_006577.2;YP_173236.1;HCoV-HKU1_orf1ab;2441 | 0 | 0 |
| NC_006577.2;YP_173236.1;HCoV-HKU1_orf1ab;2442 | 0 | 0 |
| NC_006577.2;YP_173236.1;HCoV-HKU1_orf1ab;2443 | 0 | 0 |
| NC_006577.2;YP_173236.1;HCoV-HKU1_orf1ab;2444 | 0 | 0 |
| NC_006577.2;YP_173236.1;HCoV-HKU1_orf1ab;2445 | 0 | 0 |
| NC_006577.2;YP_173236.1;HCoV-HKU1_orf1ab;2446 | 0 | 0 |
| NC_006577.2;YP_173236.1;HCoV-HKU1_orf1ab;2447 | 0 | 0 |
| NC_006577.2;YP_173236.1;HCoV-HKU1_orf1ab;2448 | 0 | 0 |
| NC_006577.2;YP_173236.1;HCoV-HKU1_orf1ab;2449 | 0 | 0 |
| NC_006577.2;YP_173236.1;HCoV-HKU1_orf1ab;2450 | 0 | 0 |
| NC_006577.2;YP_173236.1;HCoV-HKU1_orf1ab;2451 | 0 | 0 |
| NC_006577.2;YP_173236.1;HCoV-HKU1_orf1ab;2452 | 0 | 0 |
| NC_006577.2;YP_173236.1;HCoV-HKU1_orf1ab;2453 | 0 | 0 |
| NC_006577.2;YP_173236.1;HCoV-HKU1_orf1ab;2454 | 0 | 0 |
| NC_006577.2;YP_173236.1;HCoV-HKU1_orf1ab;2455 | 0 | 0 |
| NC_006577.2;YP_173236.1;HCoV-HKU1_orf1ab;2456 | 0 | 0 |
| NC_006577.2;YP_173236.1;HCoV-HKU1_orf1ab;2457 | 0 | 0 |
| NC_006577.2;YP_173236.1;HCoV-HKU1_orf1ab;2458 | 0 | 0 |
| NC_006577.2;YP_173236.1;HCoV-HKU1_orf1ab;2459 | 0 | 0 |
| NC_006577.2;YP_173236.1;HCoV-HKU1_orf1ab;2460 | 0 | 0 |
| NC_006577.2;YP_173236.1;HCoV-HKU1_orf1ab;2461 | 0 | 0 |
| NC_006577.2;YP_173236.1;HCoV-HKU1_orf1ab;2462 | 0 | 0 |
| NC_006577.2;YP_173236.1;HCoV-HKU1_orf1ab;2463 | 0 | 0 |
| NC_006577.2;YP_173236.1;HCoV-HKU1_orf1ab;2464 | 0 | 0 |
| NC_006577.2;YP_173236.1;HCoV-HKU1_orf1ab;2465 | 0 | 0 |
| NC_006577.2;YP_173236.1;HCoV-HKU1_orf1ab;2466 | 0 | 0 |

| | | |
|---|---|---|
| NC_006577.2;YP_173236.1;HCoV-HKU1_orf1ab;2467 | 0 | 0 |
| NC_006577.2;YP_173236.1;HCoV-HKU1_orf1ab;2468 | 0 | 0 |
| NC_006577.2;YP_173236.1;HCoV-HKU1_orf1ab;2469 | 0 | 0 |
| NC_006577.2;YP_173236.1;HCoV-HKU1_orf1ab;2470 | 0 | 0 |
| NC_006577.2;YP_173236.1;HCoV-HKU1_orf1ab;2471 | 0 | 0 |
| NC_006577.2;YP_173236.1;HCoV-HKU1_orf1ab;2472 | 0 | 0 |
| NC_006577.2;YP_173236.1;HCoV-HKU1_orf1ab;2473 | 0 | 0 |
| NC_006577.2;YP_173236.1;HCoV-HKU1_orf1ab;2474 | 0 | 0 |
| NC_006577.2;YP_173236.1;HCoV-HKU1_orf1ab;2475 | 0 | 0 |
| NC_006577.2;YP_173236.1;HCoV-HKU1_orf1ab;2476 | 0 | 0 |
| NC_006577.2;YP_173236.1;HCoV-HKU1_orf1ab;2477 | 0 | 0 |
| NC_006577.2;YP_173236.1;HCoV-HKU1_orf1ab;2478 | 0 | 0 |
| NC_006577.2;YP_173236.1;HCoV-HKU1_orf1ab;2479 | 0 | 0 |
| NC_006577.2;YP_173236.1;HCoV-HKU1_orf1ab;2480 | 0 | 0 |
| NC_006577.2;YP_173236.1;HCoV-HKU1_orf1ab;2481 | 0 | 0 |
| NC_006577.2;YP_173236.1;HCoV-HKU1_orf1ab;2482 | 0 | 0 |
| NC_006577.2;YP_173236.1;HCoV-HKU1_orf1ab;2483 | 0 | 0 |
| NC_006577.2;YP_173236.1;HCoV-HKU1_orf1ab;2484 | 0 | 0 |
| NC_006577.2;YP_173236.1;HCoV-HKU1_orf1ab;2485 | 0 | 0 |
| NC_006577.2;YP_173236.1;HCoV-HKU1_orf1ab;2486 | 0 | 0 |
| NC_006577.2;YP_173236.1;HCoV-HKU1_orf1ab;2487 | 0 | 0 |
| NC_006577.2;YP_173236.1;HCoV-HKU1_orf1ab;2488 | 0 | 0 |
| NC_006577.2;YP_173236.1;HCoV-HKU1_orf1ab;2489 | 0 | 0 |
| NC_006577.2;YP_173236.1;HCoV-HKU1_orf1ab;2490 | 0 | 0 |
| NC_006577.2;YP_173236.1;HCoV-HKU1_orf1ab;2491 | 0 | 0 |
| NC_006577.2;YP_173236.1;HCoV-HKU1_orf1ab;2492 | 0 | 0 |
| NC_006577.2;YP_173236.1;HCoV-HKU1_orf1ab;2493 | 0 | 0 |
| NC_006577.2;YP_173236.1;HCoV-HKU1_orf1ab;2494 | 0 | 0 |
| NC_006577.2;YP_173236.1;HCoV-HKU1_orf1ab;2495 | 0 | 0 |
| NC_006577.2;YP_173236.1;HCoV-HKU1_orf1ab;2496 | 0 | 0 |
| NC_006577.2;YP_173236.1;HCoV-HKU1_orf1ab;2497 | 0 | 0 |
| NC_006577.2;YP_173236.1;HCoV-HKU1_orf1ab;2498 | 0 | 0 |
| NC_006577.2;YP_173236.1;HCoV-HKU1_orf1ab;2499 | 0 | 0 |
| NC_006577.2;YP_173236.1;HCoV-HKU1_orf1ab;2500 | 0 | 0 |
| NC_006577.2;YP_173236.1;HCoV-HKU1_orf1ab;2501 | 0 | 0 |
| NC_006577.2;YP_173236.1;HCoV-HKU1_orf1ab;2502 | 0.05 | 0 |
| NC_006577.2;YP_173236.1;HCoV-HKU1_orf1ab;2503 | 0 | 0 |
| NC_006577.2;YP_173236.1;HCoV-HKU1_orf1ab;2504 | 0 | 0 |
| NC_006577.2;YP_173236.1;HCoV-HKU1_orf1ab;2505 | 0.1 | 0.125 |
| NC_006577.2;YP_173236.1;HCoV-HKU1_orf1ab;2506 | 0 | 0 |
| NC_006577.2;YP_173236.1;HCoV-HKU1_orf1ab;2507 | 0 | 0 |
| NC_006577.2;YP_173236.1;HCoV-HKU1_orf1ab;2508 | 0 | 0 |
| NC_006577.2;YP_173236.1;HCoV-HKU1_orf1ab;2509 | 0 | 0 |

| | | |
|---|---|---|
| NC_006577.2;YP_173236.1;HCoV-HKU1_orf1ab;2510 | 0 | 0 |
| NC_006577.2;YP_173236.1;HCoV-HKU1_orf1ab;2511 | 0 | 0 |
| NC_006577.2;YP_173236.1;HCoV-HKU1_orf1ab;2512 | 0 | 0 |
| NC_006577.2;YP_173236.1;HCoV-HKU1_orf1ab;2513 | 0 | 0 |
| NC_006577.2;YP_173236.1;HCoV-HKU1_orf1ab;2514 | 0 | 0 |
| NC_006577.2;YP_173236.1;HCoV-HKU1_orf1ab;2515 | 0 | 0 |
| NC_006577.2;YP_173236.1;HCoV-HKU1_orf1ab;2516 | 0 | 0 |
| NC_006577.2;YP_173236.1;HCoV-HKU1_orf1ab;2517 | 0 | 0 |
| NC_006577.2;YP_173236.1;HCoV-HKU1_orf1ab;2518 | 0.05 | 0 |
| NC_006577.2;YP_173236.1;HCoV-HKU1_orf1ab;2519 | 0 | 0 |
| NC_006577.2;YP_173236.1;HCoV-HKU1_orf1ab;2520 | 0 | 0 |
| NC_006577.2;YP_173236.1;HCoV-HKU1_orf1ab;2521 | 0 | 0 |
| NC_006577.2;YP_173236.1;HCoV-HKU1_orf1ab;2522 | 0 | 0 |
| NC_006577.2;YP_173236.1;HCoV-HKU1_orf1ab;2523 | 0 | 0 |
| NC_006577.2;YP_173236.1;HCoV-HKU1_orf1ab;2524 | 0 | 0 |
| NC_006577.2;YP_173236.1;HCoV-HKU1_orf1ab;2525 | 0 | 0 |
| NC_006577.2;YP_173236.1;HCoV-HKU1_orf1ab;2526 | 0 | 0 |
| NC_006577.2;YP_173236.1;HCoV-HKU1_orf1ab;2527 | 0 | 0 |
| NC_006577.2;YP_173236.1;HCoV-HKU1_orf1ab;2528 | 0 | 0 |
| NC_006577.2;YP_173236.1;HCoV-HKU1_orf1ab;2529 | 0 | 0 |
| NC_006577.2;YP_173236.1;HCoV-HKU1_orf1ab;2530 | 0 | 0 |
| NC_006577.2;YP_173236.1;HCoV-HKU1_orf1ab;2531 | 0 | 0 |
| NC_006577.2;YP_173236.1;HCoV-HKU1_orf1ab;2532 | 0 | 0 |
| NC_006577.2;YP_173236.1;HCoV-HKU1_orf1ab;2533 | 0 | 0 |
| NC_006577.2;YP_173236.1;HCoV-HKU1_orf1ab;2534 | 0 | 0 |
| NC_006577.2;YP_173236.1;HCoV-HKU1_orf1ab;2535 | 0 | 0 |
| NC_006577.2;YP_173236.1;HCoV-HKU1_orf1ab;2536 | 0.05 | 0 |
| NC_006577.2;YP_173236.1;HCoV-HKU1_orf1ab;2537 | 0.05 | 0 |
| NC_006577.2;YP_173236.1;HCoV-HKU1_orf1ab;2538 | 0 | 0 |
| NC_006577.2;YP_173236.1;HCoV-HKU1_orf1ab;2539 | 0.05 | 0.025 |
| NC_006577.2;YP_173236.1;HCoV-HKU1_orf1ab;2540 | 0.05 | 0.025 |
| NC_006577.2;YP_173236.1;HCoV-HKU1_orf1ab;2541 | 0.05 | 0.275 |
| NC_006577.2;YP_173236.1;HCoV-HKU1_orf1ab;2542 | 0.05 | 0.025 |
| NC_006577.2;YP_173236.1;HCoV-HKU1_orf1ab;2543 | 0 | 0.025 |
| NC_006577.2;YP_173236.1;HCoV-HKU1_orf1ab;2544 | 0 | 0.025 |
| NC_006577.2;YP_173236.1;HCoV-HKU1_orf1ab;2545 | 0 | 0.05 |
| NC_006577.2;YP_173236.1;HCoV-HKU1_orf1ab;2546 | 0 | 0.025 |
| NC_006577.2;YP_173236.1;HCoV-HKU1_orf1ab;2547 | 0 | 0.025 |
| NC_006577.2;YP_173236.1;HCoV-HKU1_orf1ab;2548 | 0 | 0.025 |
| NC_006577.2;YP_173236.1;HCoV-HKU1_orf1ab;2549 | 0.05 | 0.025 |
| NC_006577.2;YP_173236.1;HCoV-HKU1_orf1ab;2550 | 0 | 0.025 |
| NC_006577.2;YP_173236.1;HCoV-HKU1_orf1ab;2551 | 0.05 | 0 |
| NC_006577.2;YP_173236.1;HCoV-HKU1_orf1ab;2552 | 0 | 0 |

| | | |
|---|---|---|
| NC_006577.2;YP_173236.1;HCoV-HKU1_orf1ab;2553 | 0.05 | 0 |
| NC_006577.2;YP_173236.1;HCoV-HKU1_orf1ab;2554 | 0 | 0.1 |
| NC_006577.2;YP_173236.1;HCoV-HKU1_orf1ab;2555 | 0 | 0 |
| NC_006577.2;YP_173236.1;HCoV-HKU1_orf1ab;2556 | 0 | 0 |
| NC_006577.2;YP_173236.1;HCoV-HKU1_orf1ab;2557 | 0 | 0 |
| NC_006577.2;YP_173236.1;HCoV-HKU1_orf1ab;2558 | 0 | 0 |
| NC_006577.2;YP_173236.1;HCoV-HKU1_orf1ab;2559 | 0 | 0 |
| NC_006577.2;YP_173236.1;HCoV-HKU1_orf1ab;2560 | 0 | 0 |
| NC_006577.2;YP_173236.1;HCoV-HKU1_orf1ab;2561 | 0 | 0 |
| NC_006577.2;YP_173236.1;HCoV-HKU1_orf1ab;2562 | 0 | 0 |
| NC_006577.2;YP_173236.1;HCoV-HKU1_orf1ab;2563 | 0 | 0 |
| NC_006577.2;YP_173236.1;HCoV-HKU1_orf1ab;2564 | 0 | 0 |
| NC_006577.2;YP_173236.1;HCoV-HKU1_orf1ab;2565 | 0 | 0 |
| NC_006577.2;YP_173236.1;HCoV-HKU1_orf1ab;2566 | 0 | 0 |
| NC_006577.2;YP_173236.1;HCoV-HKU1_orf1ab;2567 | 0.05 | 0 |
| NC_006577.2;YP_173236.1;HCoV-HKU1_orf1ab;2568 | 0 | 0 |
| NC_006577.2;YP_173236.1;HCoV-HKU1_orf1ab;2569 | 0 | 0 |
| NC_006577.2;YP_173236.1;HCoV-HKU1_orf1ab;2570 | 0 | 0 |
| NC_006577.2;YP_173236.1;HCoV-HKU1_orf1ab;2571 | 0 | 0.025 |
| NC_006577.2;YP_173236.1;HCoV-HKU1_orf1ab;2572 | 0 | 0 |
| NC_006577.2;YP_173236.1;HCoV-HKU1_orf1ab;2573 | 0 | 0 |
| NC_006577.2;YP_173236.1;HCoV-HKU1_orf1ab;2574 | 0 | 0 |
| NC_006577.2;YP_173236.1;HCoV-HKU1_orf1ab;2575 | 0.05 | 0 |
| NC_006577.2;YP_173236.1;HCoV-HKU1_orf1ab;2576 | 0.05 | 0.025 |
| NC_006577.2;YP_173236.1;HCoV-HKU1_orf1ab;2577 | 0.05 | 0.025 |
| NC_006577.2;YP_173236.1;HCoV-HKU1_orf1ab;2578 | 0.05 | 0.05 |
| NC_006577.2;YP_173236.1;HCoV-HKU1_orf1ab;2579 | 0.05 | 0 |
| NC_006577.2;YP_173236.1;HCoV-HKU1_orf1ab;2580 | 0 | 0.05 |
| NC_006577.2;YP_173236.1;HCoV-HKU1_orf1ab;2581 | 0 | 0.025 |
| NC_006577.2;YP_173236.1;HCoV-HKU1_orf1ab;2582 | 0 | 0.025 |
| NC_006577.2;YP_173236.1;HCoV-HKU1_orf1ab;2583 | 0.05 | 0.025 |
| NC_006577.2;YP_173236.1;HCoV-HKU1_orf1ab;2584 | 0 | 0.025 |
| NC_006577.2;YP_173236.1;HCoV-HKU1_orf1ab;2585 | 0 | 0.025 |
| NC_006577.2;YP_173236.1;HCoV-HKU1_orf1ab;2586 | 0 | 0.025 |
| NC_006577.2;YP_173236.1;HCoV-HKU1_orf1ab;2587 | 0 | 0.025 |
| NC_006577.2;YP_173236.1;HCoV-HKU1_orf1ab;2588 | 0 | 0.05 |
| NC_006577.2;YP_173236.1;HCoV-HKU1_orf1ab;2589 | 0 | 0.025 |
| NC_006577.2;YP_173236.1;HCoV-HKU1_orf1ab;2590 | 0 | 0 |
| NC_006577.2;YP_173236.1;HCoV-HKU1_orf1ab;2591 | 0 | 0 |
| NC_006577.2;YP_173236.1;HCoV-HKU1_orf1ab;2592 | 0 | 0 |
| NC_006577.2;YP_173236.1;HCoV-HKU1_orf1ab;2593 | 0 | 0 |
| NC_006577.2;YP_173236.1;HCoV-HKU1_orf1ab;2594 | 0 | 0 |
| NC_006577.2;YP_173236.1;HCoV-HKU1_orf1ab;2595 | 0 | 0 |

| | | |
|---|---|---|
| NC_006577.2;YP_173236.1;HCoV-HKU1_orf1ab;2596 | 0 | 0 |
| NC_006577.2;YP_173236.1;HCoV-HKU1_orf1ab;2597 | 0 | 0 |
| NC_006577.2;YP_173236.1;HCoV-HKU1_orf1ab;2598 | 0 | 0 |
| NC_006577.2;YP_173236.1;HCoV-HKU1_orf1ab;2599 | 0.1 | 0 |
| NC_006577.2;YP_173236.1;HCoV-HKU1_orf1ab;2600 | 0 | 0 |
| NC_006577.2;YP_173236.1;HCoV-HKU1_orf1ab;2601 | 0.05 | 0 |
| NC_006577.2;YP_173236.1;HCoV-HKU1_orf1ab;2602 | 0 | 0 |
| NC_006577.2;YP_173236.1;HCoV-HKU1_orf1ab;2603 | 0 | 0 |
| NC_006577.2;YP_173236.1;HCoV-HKU1_orf1ab;2604 | 0 | 0 |
| NC_006577.2;YP_173236.1;HCoV-HKU1_orf1ab;2605 | 0.05 | 0 |
| NC_006577.2;YP_173236.1;HCoV-HKU1_orf1ab;2606 | 0 | 0.025 |
| NC_006577.2;YP_173236.1;HCoV-HKU1_orf1ab;2607 | 0 | 0 |
| NC_006577.2;YP_173236.1;HCoV-HKU1_orf1ab;2608 | 0 | 0 |
| NC_006577.2;YP_173236.1;HCoV-HKU1_orf1ab;2609 | 0 | 0 |
| NC_006577.2;YP_173236.1;HCoV-HKU1_orf1ab;2610 | 0 | 0 |
| NC_006577.2;YP_173236.1;HCoV-HKU1_orf1ab;2611 | 0 | 0 |
| NC_006577.2;YP_173236.1;HCoV-HKU1_orf1ab;2612 | 0 | 0.025 |
| NC_006577.2;YP_173236.1;HCoV-HKU1_orf1ab;2613 | 0 | 0.05 |
| NC_006577.2;YP_173236.1;HCoV-HKU1_orf1ab;2614 | 0 | 0.025 |
| NC_006577.2;YP_173236.1;HCoV-HKU1_orf1ab;2615 | 0 | 0.025 |
| NC_006577.2;YP_173236.1;HCoV-HKU1_orf1ab;2616 | 0 | 0.05 |
| NC_006577.2;YP_173236.1;HCoV-HKU1_orf1ab;2617 | 0.05 | 0 |
| NC_006577.2;YP_173236.1;HCoV-HKU1_orf1ab;2618 | 0 | 0 |
| NC_006577.2;YP_173236.1;HCoV-HKU1_orf1ab;2619 | 0 | 0 |
| NC_006577.2;YP_173236.1;HCoV-HKU1_orf1ab;2620 | 0 | 0.025 |
| NC_006577.2;YP_173236.1;HCoV-HKU1_orf1ab;2621 | 0 | 0 |
| NC_006577.2;YP_173236.1;HCoV-HKU1_orf1ab;2622 | 0 | 0 |
| NC_006577.2;YP_173236.1;HCoV-HKU1_orf1ab;2623 | 0 | 0 |
| NC_006577.2;YP_173236.1;HCoV-HKU1_orf1ab;2624 | 0 | 0 |
| NC_006577.2;YP_173236.1;HCoV-HKU1_orf1ab;2625 | 0 | 0 |
| NC_006577.2;YP_173236.1;HCoV-HKU1_orf1ab;2626 | 0 | 0 |
| NC_006577.2;YP_173236.1;HCoV-HKU1_orf1ab;2627 | 0 | 0 |
| NC_006577.2;YP_173236.1;HCoV-HKU1_orf1ab;2628 | 0 | 0 |
| NC_006577.2;YP_173236.1;HCoV-HKU1_orf1ab;2629 | 0 | 0 |
| NC_006577.2;YP_173236.1;HCoV-HKU1_orf1ab;2630 | 0 | 0 |
| NC_006577.2;YP_173236.1;HCoV-HKU1_orf1ab;2631 | 0 | 0 |
| NC_006577.2;YP_173236.1;HCoV-HKU1_orf1ab;2632 | 0 | 0 |
| NC_006577.2;YP_173236.1;HCoV-HKU1_orf1ab;2633 | 0 | 0 |
| NC_006577.2;YP_173236.1;HCoV-HKU1_orf1ab;2634 | 0 | 0 |
| NC_006577.2;YP_173236.1;HCoV-HKU1_orf1ab;2635 | 0 | 0 |
| NC_006577.2;YP_173236.1;HCoV-HKU1_orf1ab;2636 | 0 | 0 |
| NC_006577.2;YP_173236.1;HCoV-HKU1_orf1ab;2637 | 0 | 0 |
| NC_006577.2;YP_173236.1;HCoV-HKU1_orf1ab;2638 | 0 | 0 |

| | | |
|---|---|---|
| NC_006577.2;YP_173236.1;HCoV-HKU1_orf1ab;2639 | 0 | 0 |
| NC_006577.2;YP_173236.1;HCoV-HKU1_orf1ab;2640 | 0 | 0 |
| NC_006577.2;YP_173236.1;HCoV-HKU1_orf1ab;2641 | 0 | 0 |
| NC_006577.2;YP_173236.1;HCoV-HKU1_orf1ab;2642 | 0 | 0 |
| NC_006577.2;YP_173236.1;HCoV-HKU1_orf1ab;2643 | 0 | 0.025 |
| NC_006577.2;YP_173236.1;HCoV-HKU1_orf1ab;2644 | 0.05 | 0.15 |
| NC_006577.2;YP_173236.1;HCoV-HKU1_orf1ab;2645 | 0 | 0 |
| NC_006577.2;YP_173236.1;HCoV-HKU1_orf1ab;2646 | 0 | 0 |
| NC_006577.2;YP_173236.1;HCoV-HKU1_orf1ab;2647 | 0 | 0 |
| NC_006577.2;YP_173236.1;HCoV-HKU1_orf1ab;2648 | 0 | 0 |
| NC_006577.2;YP_173236.1;HCoV-HKU1_orf1ab;2649 | 0 | 0 |
| NC_006577.2;YP_173236.1;HCoV-HKU1_orf1ab;2650 | 0 | 0 |
| NC_006577.2;YP_173236.1;HCoV-HKU1_orf1ab;2651 | 0 | 0 |
| NC_006577.2;YP_173236.1;HCoV-HKU1_orf1ab;2652 | 0 | 0 |
| NC_006577.2;YP_173236.1;HCoV-HKU1_orf1ab;2653 | 0 | 0 |
| NC_006577.2;YP_173236.1;HCoV-HKU1_orf1ab;2654 | 0 | 0 |
| NC_006577.2;YP_173236.1;HCoV-HKU1_orf1ab;2655 | 0 | 0 |
| NC_006577.2;YP_173236.1;HCoV-HKU1_orf1ab;2656 | 0 | 0 |
| NC_006577.2;YP_173236.1;HCoV-HKU1_orf1ab;2657 | 0 | 0 |
| NC_006577.2;YP_173236.1;HCoV-HKU1_orf1ab;2658 | 0 | 0 |
| NC_006577.2;YP_173236.1;HCoV-HKU1_orf1ab;2659 | 0 | 0 |
| NC_006577.2;YP_173236.1;HCoV-HKU1_orf1ab;2660 | 0 | 0 |
| NC_006577.2;YP_173236.1;HCoV-HKU1_orf1ab;2661 | 0 | 0 |
| NC_006577.2;YP_173236.1;HCoV-HKU1_orf1ab;2662 | 0 | 0 |
| NC_006577.2;YP_173236.1;HCoV-HKU1_orf1ab;2663 | 0 | 0 |
| NC_006577.2;YP_173236.1;HCoV-HKU1_orf1ab;2664 | 0 | 0 |
| NC_006577.2;YP_173236.1;HCoV-HKU1_orf1ab;2665 | 0 | 0.025 |
| NC_006577.2;YP_173236.1;HCoV-HKU1_orf1ab;2666 | 0 | 0 |
| NC_006577.2;YP_173236.1;HCoV-HKU1_orf1ab;2667 | 0 | 0.075 |
| NC_006577.2;YP_173236.1;HCoV-HKU1_orf1ab;2668 | 0 | 0 |
| NC_006577.2;YP_173236.1;HCoV-HKU1_orf1ab;2669 | 0 | 0.025 |
| NC_006577.2;YP_173236.1;HCoV-HKU1_orf1ab;2670 | 0.05 | 0 |
| NC_006577.2;YP_173236.1;HCoV-HKU1_orf1ab;2671 | 0 | 0 |
| NC_006577.2;YP_173236.1;HCoV-HKU1_orf1ab;2672 | 0.05 | 0.05 |
| NC_006577.2;YP_173236.1;HCoV-HKU1_orf1ab;2673 | 0.05 | 0.075 |
| NC_006577.2;YP_173236.1;HCoV-HKU1_orf1ab;2674 | 0 | 0 |
| NC_006577.2;YP_173236.1;HCoV-HKU1_orf1ab;2675 | 0 | 0.025 |
| NC_006577.2;YP_173236.1;HCoV-HKU1_orf1ab;2676 | 0 | 0 |
| NC_006577.2;YP_173236.1;HCoV-HKU1_orf1ab;2677 | 0 | 0 |
| NC_006577.2;YP_173236.1;HCoV-HKU1_orf1ab;2678 | 0 | 0 |
| NC_006577.2;YP_173236.1;HCoV-HKU1_orf1ab;2679 | 0 | 0 |
| NC_006577.2;YP_173236.1;HCoV-HKU1_orf1ab;2680 | 0 | 0 |
| NC_006577.2;YP_173236.1;HCoV-HKU1_orf1ab;2681 | 0 | 0 |

| | | |
|---|---|---|
| NC_006577.2;YP_173236.1;HCoV-HKU1_orf1ab;2682 | 0 | 0 |
| NC_006577.2;YP_173236.1;HCoV-HKU1_orf1ab;2683 | 0 | 0 |
| NC_006577.2;YP_173236.1;HCoV-HKU1_orf1ab;2684 | 0 | 0 |
| NC_006577.2;YP_173236.1;HCoV-HKU1_orf1ab;2685 | 0 | 0 |
| NC_006577.2;YP_173236.1;HCoV-HKU1_orf1ab;2686 | 0 | 0.025 |
| NC_006577.2;YP_173236.1;HCoV-HKU1_orf1ab;2687 | 0 | 0 |
| NC_006577.2;YP_173236.1;HCoV-HKU1_orf1ab;2688 | 0 | 0 |
| NC_006577.2;YP_173236.1;HCoV-HKU1_orf1ab;2689 | 0 | 0.025 |
| NC_006577.2;YP_173236.1;HCoV-HKU1_orf1ab;2690 | 0 | 0 |
| NC_006577.2;YP_173236.1;HCoV-HKU1_orf1ab;2691 | 0 | 0 |
| NC_006577.2;YP_173236.1;HCoV-HKU1_orf1ab;2692 | 0 | 0 |
| NC_006577.2;YP_173236.1;HCoV-HKU1_orf1ab;2693 | 0 | 0 |
| NC_006577.2;YP_173236.1;HCoV-HKU1_orf1ab;2694 | 0 | 0 |
| NC_006577.2;YP_173236.1;HCoV-HKU1_orf1ab;2695 | 0 | 0 |
| NC_006577.2;YP_173236.1;HCoV-HKU1_orf1ab;2696 | 0 | 0 |
| NC_006577.2;YP_173236.1;HCoV-HKU1_orf1ab;2697 | 0 | 0 |
| NC_006577.2;YP_173236.1;HCoV-HKU1_orf1ab;2698 | 0 | 0 |
| NC_006577.2;YP_173236.1;HCoV-HKU1_orf1ab;2699 | 0 | 0 |
| NC_006577.2;YP_173236.1;HCoV-HKU1_orf1ab;2700 | 0 | 0 |
| NC_006577.2;YP_173236.1;HCoV-HKU1_orf1ab;2701 | 0 | 0 |
| NC_006577.2;YP_173236.1;HCoV-HKU1_orf1ab;2702 | 0 | 0 |
| NC_006577.2;YP_173236.1;HCoV-HKU1_orf1ab;2703 | 0 | 0 |
| NC_006577.2;YP_173236.1;HCoV-HKU1_orf1ab;2704 | 0 | 0 |
| NC_006577.2;YP_173236.1;HCoV-HKU1_orf1ab;2705 | 0 | 0 |
| NC_006577.2;YP_173236.1;HCoV-HKU1_orf1ab;2706 | 0 | 0 |
| NC_006577.2;YP_173236.1;HCoV-HKU1_orf1ab;2707 | 0 | 0 |
| NC_006577.2;YP_173236.1;HCoV-HKU1_orf1ab;2708 | 0 | 0 |
| NC_006577.2;YP_173236.1;HCoV-HKU1_orf1ab;2709 | 0 | 0 |
| NC_006577.2;YP_173236.1;HCoV-HKU1_orf1ab;2710 | 0 | 0 |
| NC_006577.2;YP_173236.1;HCoV-HKU1_orf1ab;2711 | 0 | 0 |
| NC_006577.2;YP_173236.1;HCoV-HKU1_orf1ab;2712 | 0 | 0 |
| NC_006577.2;YP_173236.1;HCoV-HKU1_orf1ab;2713 | 0 | 0.025 |
| NC_006577.2;YP_173236.1;HCoV-HKU1_orf1ab;2714 | 0 | 0 |
| NC_006577.2;YP_173236.1;HCoV-HKU1_orf1ab;2715 | 0 | 0 |
| NC_006577.2;YP_173236.1;HCoV-HKU1_orf1ab;2716 | 0 | 0 |
| NC_006577.2;YP_173236.1;HCoV-HKU1_orf1ab;2717 | 0 | 0.025 |
| NC_006577.2;YP_173236.1;HCoV-HKU1_orf1ab;2718 | 0 | 0.025 |
| NC_006577.2;YP_173236.1;HCoV-HKU1_orf1ab;2719 | 0 | 0 |
| NC_006577.2;YP_173236.1;HCoV-HKU1_orf1ab;2720 | 0 | 0 |
| NC_006577.2;YP_173236.1;HCoV-HKU1_orf1ab;2721 | 0.05 | 0 |
| NC_006577.2;YP_173236.1;HCoV-HKU1_orf1ab;2722 | 0 | 0 |
| NC_006577.2;YP_173236.1;HCoV-HKU1_orf1ab;2723 | 0 | 0 |
| NC_006577.2;YP_173236.1;HCoV-HKU1_orf1ab;2724 | 0 | 0 |

| | | |
|---|---|---|
| NC_006577.2;YP_173236.1;HCoV-HKU1_orf1ab;2725 | 0 | 0 |
| NC_006577.2;YP_173236.1;HCoV-HKU1_orf1ab;2726 | 0 | 0 |
| NC_006577.2;YP_173236.1;HCoV-HKU1_orf1ab;2727 | 0 | 0 |
| NC_006577.2;YP_173236.1;HCoV-HKU1_orf1ab;2728 | 0 | 0 |
| NC_006577.2;YP_173236.1;HCoV-HKU1_orf1ab;2729 | 0 | 0 |
| NC_006577.2;YP_173236.1;HCoV-HKU1_orf1ab;2730 | 0 | 0 |
| NC_006577.2;YP_173236.1;HCoV-HKU1_orf1ab;2731 | 0 | 0 |
| NC_006577.2;YP_173236.1;HCoV-HKU1_orf1ab;2732 | 0 | 0 |
| NC_006577.2;YP_173236.1;HCoV-HKU1_orf1ab;2733 | 0.05 | 0.075 |
| NC_006577.2;YP_173236.1;HCoV-HKU1_orf1ab;2734 | 0 | 0 |
| NC_006577.2;YP_173236.1;HCoV-HKU1_orf1ab;2735 | 0 | 0.05 |
| NC_006577.2;YP_173236.1;HCoV-HKU1_orf1ab;2736 | 0.05 | 0.05 |
| NC_006577.2;YP_173236.1;HCoV-HKU1_orf1ab;2737 | 0 | 0.025 |
| NC_006577.2;YP_173236.1;HCoV-HKU1_orf1ab;2738 | 0.05 | 0.05 |
| NC_006577.2;YP_173236.1;HCoV-HKU1_orf1ab;2739 | 0.05 | 0.075 |
| NC_006577.2;YP_173236.1;HCoV-HKU1_orf1ab;2740 | 0 | 0 |
| NC_006577.2;YP_173236.1;HCoV-HKU1_orf1ab;2741 | 0 | 0.025 |
| NC_006577.2;YP_173236.1;HCoV-HKU1_orf1ab;2742 | 0.1 | 0.025 |
| NC_006577.2;YP_173236.1;HCoV-HKU1_orf1ab;2743 | 0.1 | 0.025 |
| NC_006577.2;YP_173236.1;HCoV-HKU1_orf1ab;2744 | 0 | 0.025 |
| NC_006577.2;YP_173236.1;HCoV-HKU1_orf1ab;2745 | 0.1 | 0.15 |
| NC_006577.2;YP_173236.1;HCoV-HKU1_orf1ab;2746 | 0.05 | 0.15 |
| NC_006577.2;YP_173236.1;HCoV-HKU1_orf1ab;2747 | 0 | 0.05 |
| NC_006577.2;YP_173236.1;HCoV-HKU1_orf1ab;2748 | 0 | 0.075 |
| NC_006577.2;YP_173236.1;HCoV-HKU1_orf1ab;2749 | 0 | 0.1 |
| NC_006577.2;YP_173236.1;HCoV-HKU1_orf1ab;2750 | 0.05 | 0.05 |
| NC_006577.2;YP_173236.1;HCoV-HKU1_orf1ab;2751 | 0 | 0.025 |
| NC_006577.2;YP_173236.1;HCoV-HKU1_orf1ab;2752 | 0 | 0 |
| NC_006577.2;YP_173236.1;HCoV-HKU1_orf1ab;2753 | 0 | 0 |
| NC_006577.2;YP_173236.1;HCoV-HKU1_orf1ab;2754 | 0 | 0 |
| NC_006577.2;YP_173236.1;HCoV-HKU1_orf1ab;2755 | 0 | 0 |
| NC_006577.2;YP_173236.1;HCoV-HKU1_orf1ab;2756 | 0 | 0 |
| NC_006577.2;YP_173236.1;HCoV-HKU1_orf1ab;2757 | 0 | 0 |
| NC_006577.2;YP_173236.1;HCoV-HKU1_orf1ab;2758 | 0 | 0 |
| NC_006577.2;YP_173236.1;HCoV-HKU1_orf1ab;2759 | 0 | 0 |
| NC_006577.2;YP_173236.1;HCoV-HKU1_orf1ab;2760 | 0 | 0.025 |
| NC_006577.2;YP_173236.1;HCoV-HKU1_orf1ab;2761 | 0.05 | 0.05 |
| NC_006577.2;YP_173236.1;HCoV-HKU1_orf1ab;2762 | 0 | 0.025 |
| NC_006577.2;YP_173236.1;HCoV-HKU1_orf1ab;2763 | 0 | 0 |
| NC_006577.2;YP_173236.1;HCoV-HKU1_orf1ab;2764 | 0 | 0.025 |
| NC_006577.2;YP_173236.1;HCoV-HKU1_orf1ab;2765 | 0 | 0.025 |
| NC_006577.2;YP_173236.1;HCoV-HKU1_orf1ab;2766 | 0 | 0.025 |
| NC_006577.2;YP_173236.1;HCoV-HKU1_orf1ab;2767 | 0 | 0.025 |

| | | |
|---|---|---|
| NC_006577.2;YP_173236.1;HCoV-HKU1_orf1ab;2768 | 0 | 0.025 |
| NC_006577.2;YP_173236.1;HCoV-HKU1_orf1ab;2769 | 0 | 0 |
| NC_006577.2;YP_173236.1;HCoV-HKU1_orf1ab;2770 | 0 | 0 |
| NC_006577.2;YP_173236.1;HCoV-HKU1_orf1ab;2771 | 0 | 0 |
| NC_006577.2;YP_173236.1;HCoV-HKU1_orf1ab;2772 | 0 | 0 |
| NC_006577.2;YP_173236.1;HCoV-HKU1_orf1ab;2773 | 0 | 0 |
| NC_006577.2;YP_173236.1;HCoV-HKU1_orf1ab;2774 | 0 | 0 |
| NC_006577.2;YP_173236.1;HCoV-HKU1_orf1ab;2775 | 0 | 0 |
| NC_006577.2;YP_173236.1;HCoV-HKU1_orf1ab;2776 | 0 | 0 |
| NC_006577.2;YP_173236.1;HCoV-HKU1_orf1ab;2777 | 0 | 0 |
| NC_006577.2;YP_173236.1;HCoV-HKU1_orf1ab;2778 | 0 | 0 |
| NC_006577.2;YP_173236.1;HCoV-HKU1_orf1ab;2779 | 0 | 0 |
| NC_006577.2;YP_173236.1;HCoV-HKU1_orf1ab;2780 | 0 | 0 |
| NC_006577.2;YP_173236.1;HCoV-HKU1_orf1ab;2781 | 0 | 0 |
| NC_006577.2;YP_173236.1;HCoV-HKU1_orf1ab;2782 | 0 | 0 |
| NC_006577.2;YP_173236.1;HCoV-HKU1_orf1ab;2783 | 0 | 0 |
| NC_006577.2;YP_173236.1;HCoV-HKU1_orf1ab;2784 | 0 | 0 |
| NC_006577.2;YP_173236.1;HCoV-HKU1_orf1ab;2785 | 0 | 0 |
| NC_006577.2;YP_173236.1;HCoV-HKU1_orf1ab;2786 | 0 | 0 |
| NC_006577.2;YP_173236.1;HCoV-HKU1_orf1ab;2787 | 0 | 0 |
| NC_006577.2;YP_173236.1;HCoV-HKU1_orf1ab;2788 | 0 | 0 |
| NC_006577.2;YP_173236.1;HCoV-HKU1_orf1ab;2789 | 0 | 0 |
| NC_006577.2;YP_173236.1;HCoV-HKU1_orf1ab;2790 | 0 | 0 |
| NC_006577.2;YP_173236.1;HCoV-HKU1_orf1ab;2791 | 0 | 0 |
| NC_006577.2;YP_173236.1;HCoV-HKU1_orf1ab;2792 | 0 | 0 |
| NC_006577.2;YP_173236.1;HCoV-HKU1_orf1ab;2793 | 0 | 0 |
| NC_006577.2;YP_173236.1;HCoV-HKU1_orf1ab;2794 | 0 | 0.025 |
| NC_006577.2;YP_173236.1;HCoV-HKU1_orf1ab;2795 | 0 | 0 |
| NC_006577.2;YP_173236.1;HCoV-HKU1_orf1ab;2796 | 0 | 0 |
| NC_006577.2;YP_173236.1;HCoV-HKU1_orf1ab;2797 | 0 | 0 |
| NC_006577.2;YP_173236.1;HCoV-HKU1_orf1ab;2798 | 0 | 0 |
| NC_006577.2;YP_173236.1;HCoV-HKU1_orf1ab;2799 | 0 | 0 |
| NC_006577.2;YP_173236.1;HCoV-HKU1_orf1ab;2800 | 0 | 0 |
| NC_006577.2;YP_173236.1;HCoV-HKU1_orf1ab;2801 | 0 | 0 |
| NC_006577.2;YP_173236.1;HCoV-HKU1_orf1ab;2802 | 0 | 0 |
| NC_006577.2;YP_173236.1;HCoV-HKU1_orf1ab;2803 | 0 | 0 |
| NC_006577.2;YP_173236.1;HCoV-HKU1_orf1ab;2804 | 0 | 0 |
| NC_006577.2;YP_173236.1;HCoV-HKU1_orf1ab;2805 | 0 | 0 |
| NC_006577.2;YP_173236.1;HCoV-HKU1_orf1ab;2806 | 0 | 0 |
| NC_006577.2;YP_173236.1;HCoV-HKU1_orf1ab;2807 | 0 | 0 |
| NC_006577.2;YP_173236.1;HCoV-HKU1_orf1ab;2808 | 0 | 0 |
| NC_006577.2;YP_173236.1;HCoV-HKU1_orf1ab;2809 | 0 | 0 |
| NC_006577.2;YP_173236.1;HCoV-HKU1_orf1ab;2810 | 0 | 0 |

| | | |
|---|---|---|
| NC_006577.2;YP_173236.1;HCoV-HKU1_orf1ab;2811 | 0 | 0 |
| NC_006577.2;YP_173236.1;HCoV-HKU1_orf1ab;2812 | 0 | 0 |
| NC_006577.2;YP_173236.1;HCoV-HKU1_orf1ab;2813 | 0.05 | 0 |
| NC_006577.2;YP_173236.1;HCoV-HKU1_orf1ab;2814 | 0.05 | 0 |
| NC_006577.2;YP_173236.1;HCoV-HKU1_orf1ab;2815 | 0.05 | 0.025 |
| NC_006577.2;YP_173236.1;HCoV-HKU1_orf1ab;2816 | 0.05 | 0.025 |
| NC_006577.2;YP_173236.1;HCoV-HKU1_orf1ab;2817 | 0.05 | 0.025 |
| NC_006577.2;YP_173236.1;HCoV-HKU1_orf1ab;2818 | 0.05 | 0.05 |
| NC_006577.2;YP_173236.1;HCoV-HKU1_orf1ab;2819 | 0.05 | 0 |
| NC_006577.2;YP_173236.1;HCoV-HKU1_orf1ab;2820 | 0.05 | 0.025 |
| NC_006577.2;YP_173236.1;HCoV-HKU1_orf1ab;2821 | 0.05 | 0.025 |
| NC_006577.2;YP_173236.1;HCoV-HKU1_orf1ab;2822 | 0 | 0 |
| NC_006577.2;YP_173236.1;HCoV-HKU1_orf1ab;2823 | 0 | 0 |
| NC_006577.2;YP_173236.1;HCoV-HKU1_orf1ab;2824 | 0 | 0 |
| NC_006577.2;YP_173236.1;HCoV-HKU1_orf1ab;2825 | 0 | 0 |
| NC_006577.2;YP_173236.1;HCoV-HKU1_orf1ab;2826 | 0 | 0 |
| NC_006577.2;YP_173236.1;HCoV-HKU1_orf1ab;2827 | 0 | 0 |
| NC_006577.2;YP_173236.1;HCoV-HKU1_orf1ab;2828 | 0 | 0 |
| NC_006577.2;YP_173236.1;HCoV-HKU1_orf1ab;2829 | 0 | 0 |
| NC_006577.2;YP_173236.1;HCoV-HKU1_orf1ab;2830 | 0 | 0 |
| NC_006577.2;YP_173236.1;HCoV-HKU1_orf1ab;2831 | 0 | 0 |
| NC_006577.2;YP_173236.1;HCoV-HKU1_orf1ab;2832 | 0 | 0 |
| NC_006577.2;YP_173236.1;HCoV-HKU1_orf1ab;2833 | 0 | 0 |
| NC_006577.2;YP_173236.1;HCoV-HKU1_orf1ab;2834 | 0 | 0 |
| NC_006577.2;YP_173236.1;HCoV-HKU1_orf1ab;2835 | 0 | 0 |
| NC_006577.2;YP_173236.1;HCoV-HKU1_orf1ab;2836 | 0 | 0 |
| NC_006577.2;YP_173236.1;HCoV-HKU1_orf1ab;2837 | 0 | 0 |
| NC_006577.2;YP_173236.1;HCoV-HKU1_orf1ab;2838 | 0 | 0 |
| NC_006577.2;YP_173236.1;HCoV-HKU1_orf1ab;2839 | 0 | 0 |
| NC_006577.2;YP_173236.1;HCoV-HKU1_orf1ab;2840 | 0 | 0 |
| NC_006577.2;YP_173236.1;HCoV-HKU1_orf1ab;2841 | 0 | 0 |
| NC_006577.2;YP_173236.1;HCoV-HKU1_orf1ab;2842 | 0 | 0 |
| NC_006577.2;YP_173236.1;HCoV-HKU1_orf1ab;2843 | 0 | 0 |
| NC_006577.2;YP_173236.1;HCoV-HKU1_orf1ab;2844 | 0 | 0 |
| NC_006577.2;YP_173236.1;HCoV-HKU1_orf1ab;2845 | 0 | 0 |
| NC_006577.2;YP_173236.1;HCoV-HKU1_orf1ab;2846 | 0 | 0 |
| NC_006577.2;YP_173236.1;HCoV-HKU1_orf1ab;2847 | 0 | 0 |
| NC_006577.2;YP_173236.1;HCoV-HKU1_orf1ab;2848 | 0 | 0 |
| NC_006577.2;YP_173236.1;HCoV-HKU1_orf1ab;2849 | 0 | 0 |
| NC_006577.2;YP_173236.1;HCoV-HKU1_orf1ab;2850 | 0 | 0 |
| NC_006577.2;YP_173236.1;HCoV-HKU1_orf1ab;2851 | 0 | 0 |
| NC_006577.2;YP_173236.1;HCoV-HKU1_orf1ab;2852 | 0 | 0 |
| NC_006577.2;YP_173236.1;HCoV-HKU1_orf1ab;2853 | 0 | 0 |

| | | |
|---|---|---|
| NC_006577.2;YP_173236.1;HCoV-HKU1_orf1ab;2854 | 0 | 0 |
| NC_006577.2;YP_173236.1;HCoV-HKU1_orf1ab;2855 | 0 | 0 |
| NC_006577.2;YP_173236.1;HCoV-HKU1_orf1ab;2856 | 0 | 0 |
| NC_006577.2;YP_173236.1;HCoV-HKU1_orf1ab;2857 | 0 | 0 |
| NC_006577.2;YP_173236.1;HCoV-HKU1_orf1ab;2858 | 0 | 0 |
| NC_006577.2;YP_173236.1;HCoV-HKU1_orf1ab;2859 | 0 | 0 |
| NC_006577.2;YP_173236.1;HCoV-HKU1_orf1ab;2860 | 0 | 0 |
| NC_006577.2;YP_173236.1;HCoV-HKU1_orf1ab;2861 | 0 | 0 |
| NC_006577.2;YP_173236.1;HCoV-HKU1_orf1ab;2862 | 0 | 0 |
| NC_006577.2;YP_173236.1;HCoV-HKU1_orf1ab;2863 | 0 | 0 |
| NC_006577.2;YP_173236.1;HCoV-HKU1_orf1ab;2864 | 0 | 0 |
| NC_006577.2;YP_173236.1;HCoV-HKU1_orf1ab;2865 | 0 | 0 |
| NC_006577.2;YP_173236.1;HCoV-HKU1_orf1ab;2866 | 0 | 0 |
| NC_006577.2;YP_173236.1;HCoV-HKU1_orf1ab;2867 | 0 | 0.025 |
| NC_006577.2;YP_173236.1;HCoV-HKU1_orf1ab;2868 | 0 | 0.025 |
| NC_006577.2;YP_173236.1;HCoV-HKU1_orf1ab;2869 | 0 | 0 |
| NC_006577.2;YP_173236.1;HCoV-HKU1_orf1ab;2870 | 0 | 0.025 |
| NC_006577.2;YP_173236.1;HCoV-HKU1_orf1ab;2871 | 0 | 0.025 |
| NC_006577.2;YP_173236.1;HCoV-HKU1_orf1ab;2872 | 0 | 0.025 |
| NC_006577.2;YP_173236.1;HCoV-HKU1_orf1ab;2873 | 0 | 0.025 |
| NC_006577.2;YP_173236.1;HCoV-HKU1_orf1ab;2874 | 0 | 0.025 |
| NC_006577.2;YP_173236.1;HCoV-HKU1_orf1ab;2875 | 0 | 0.025 |
| NC_006577.2;YP_173236.1;HCoV-HKU1_orf1ab;2876 | 0 | 0 |
| NC_006577.2;YP_173236.1;HCoV-HKU1_orf1ab;2877 | 0 | 0 |
| NC_006577.2;YP_173236.1;HCoV-HKU1_orf1ab;2878 | 0 | 0 |
| NC_006577.2;YP_173236.1;HCoV-HKU1_orf1ab;2879 | 0.05 | 0.025 |
| NC_006577.2;YP_173236.1;HCoV-HKU1_orf1ab;2880 | 0 | 0 |
| NC_006577.2;YP_173236.1;HCoV-HKU1_orf1ab;2881 | 0 | 0 |
| NC_006577.2;YP_173236.1;HCoV-HKU1_orf1ab;2882 | 0 | 0 |
| NC_006577.2;YP_173236.1;HCoV-HKU1_orf1ab;2883 | 0 | 0 |
| NC_006577.2;YP_173236.1;HCoV-HKU1_orf1ab;2884 | 0 | 0 |
| NC_006577.2;YP_173236.1;HCoV-HKU1_orf1ab;2885 | 0 | 0 |
| NC_006577.2;YP_173236.1;HCoV-HKU1_orf1ab;2886 | 0 | 0 |
| NC_006577.2;YP_173236.1;HCoV-HKU1_orf1ab;2887 | 0.3 | 0.4 |
| NC_006577.2;YP_173236.1;HCoV-HKU1_orf1ab;2888 | 0 | 0.025 |
| NC_006577.2;YP_173236.1;HCoV-HKU1_orf1ab;2889 | 0 | 0 |
| NC_006577.2;YP_173236.1;HCoV-HKU1_orf1ab;2890 | 0 | 0 |
| NC_006577.2;YP_173236.1;HCoV-HKU1_orf1ab;2891 | 0 | 0 |
| NC_006577.2;YP_173236.1;HCoV-HKU1_orf1ab;2892 | 0.05 | 0.025 |
| NC_006577.2;YP_173236.1;HCoV-HKU1_orf1ab;2893 | 0.2 | 0.15 |
| NC_006577.2;YP_173236.1;HCoV-HKU1_orf1ab;2894 | 0 | 0.025 |
| NC_006577.2;YP_173236.1;HCoV-HKU1_orf1ab;2895 | 0 | 0 |
| NC_006577.2;YP_173236.1;HCoV-HKU1_orf1ab;2896 | 0 | 0 |

| | | |
|---|---|---|
| NC_006577.2;YP_173236.1;HCoV-HKU1_orf1ab;2897 | 0 | 0 |
| NC_006577.2;YP_173236.1;HCoV-HKU1_orf1ab;2898 | 0 | 0 |
| NC_006577.2;YP_173236.1;HCoV-HKU1_orf1ab;2899 | 0 | 0.025 |
| NC_006577.2;YP_173236.1;HCoV-HKU1_orf1ab;2900 | 0 | 0.025 |
| NC_006577.2;YP_173236.1;HCoV-HKU1_orf1ab;2901 | 0 | 0 |
| NC_006577.2;YP_173236.1;HCoV-HKU1_orf1ab;2902 | 0 | 0 |
| NC_006577.2;YP_173236.1;HCoV-HKU1_orf1ab;2903 | 0 | 0 |
| NC_006577.2;YP_173236.1;HCoV-HKU1_orf1ab;2904 | 0 | 0 |
| NC_006577.2;YP_173236.1;HCoV-HKU1_orf1ab;2905 | 0 | 0 |
| NC_006577.2;YP_173236.1;HCoV-HKU1_orf1ab;2906 | 0 | 0 |
| NC_006577.2;YP_173236.1;HCoV-HKU1_orf1ab;2907 | 0 | 0 |
| NC_006577.2;YP_173236.1;HCoV-HKU1_orf1ab;2908 | 0 | 0 |
| NC_006577.2;YP_173236.1;HCoV-HKU1_orf1ab;2909 | 0 | 0 |
| NC_006577.2;YP_173236.1;HCoV-HKU1_orf1ab;2910 | 0 | 0 |
| NC_006577.2;YP_173236.1;HCoV-HKU1_orf1ab;2911 | 0 | 0.05 |
| NC_006577.2;YP_173236.1;HCoV-HKU1_orf1ab;2912 | 0 | 0 |
| NC_006577.2;YP_173236.1;HCoV-HKU1_orf1ab;2913 | 0 | 0 |
| NC_006577.2;YP_173236.1;HCoV-HKU1_orf1ab;2914 | 0 | 0 |
| NC_006577.2;YP_173236.1;HCoV-HKU1_orf1ab;2915 | 0 | 0.025 |
| NC_006577.2;YP_173236.1;HCoV-HKU1_orf1ab;2916 | 0 | 0 |
| NC_006577.2;YP_173236.1;HCoV-HKU1_orf1ab;2917 | 0 | 0 |
| NC_006577.2;YP_173236.1;HCoV-HKU1_orf1ab;2918 | 0 | 0 |
| NC_006577.2;YP_173236.1;HCoV-HKU1_orf1ab;2919 | 0 | 0 |
| NC_006577.2;YP_173236.1;HCoV-HKU1_orf1ab;2920 | 0 | 0 |
| NC_006577.2;YP_173236.1;HCoV-HKU1_orf1ab;2921 | 0 | 0 |
| NC_006577.2;YP_173236.1;HCoV-HKU1_orf1ab;2922 | 0 | 0 |
| NC_006577.2;YP_173236.1;HCoV-HKU1_orf1ab;2923 | 0 | 0 |
| NC_006577.2;YP_173236.1;HCoV-HKU1_orf1ab;2924 | 0 | 0 |
| NC_006577.2;YP_173236.1;HCoV-HKU1_orf1ab;2925 | 0 | 0 |
| NC_006577.2;YP_173236.1;HCoV-HKU1_orf1ab;2926 | 0 | 0 |
| NC_006577.2;YP_173236.1;HCoV-HKU1_orf1ab;2927 | 0 | 0.025 |
| NC_006577.2;YP_173236.1;HCoV-HKU1_orf1ab;2928 | 0 | 0 |
| NC_006577.2;YP_173236.1;HCoV-HKU1_orf1ab;2929 | 0 | 0 |
| NC_006577.2;YP_173236.1;HCoV-HKU1_orf1ab;2930 | 0 | 0 |
| NC_006577.2;YP_173236.1;HCoV-HKU1_orf1ab;2931 | 0 | 0 |
| NC_006577.2;YP_173236.1;HCoV-HKU1_orf1ab;2932 | 0.05 | 0 |
| NC_006577.2;YP_173236.1;HCoV-HKU1_orf1ab;2933 | 0 | 0 |
| NC_006577.2;YP_173236.1;HCoV-HKU1_orf1ab;2934 | 0 | 0 |
| NC_006577.2;YP_173236.1;HCoV-HKU1_orf1ab;2935 | 0 | 0 |
| NC_006577.2;YP_173236.1;HCoV-HKU1_orf1ab;2936 | 0 | 0 |
| NC_006577.2;YP_173236.1;HCoV-HKU1_orf1ab;2937 | 0 | 0 |
| NC_006577.2;YP_173236.1;HCoV-HKU1_orf1ab;2938 | 0 | 0 |
| NC_006577.2;YP_173236.1;HCoV-HKU1_orf1ab;2939 | 0 | 0 |

| | | |
|---|---|---|
| NC_006577.2;YP_173236.1;HCoV-HKU1_orf1ab;2940 | 0 | 0 |
| NC_006577.2;YP_173236.1;HCoV-HKU1_orf1ab;2941 | 0 | 0 |
| NC_006577.2;YP_173236.1;HCoV-HKU1_orf1ab;2942 | 0 | 0 |
| NC_006577.2;YP_173236.1;HCoV-HKU1_orf1ab;2943 | 0 | 0 |
| NC_006577.2;YP_173236.1;HCoV-HKU1_orf1ab;2944 | 0 | 0 |
| NC_006577.2;YP_173236.1;HCoV-HKU1_orf1ab;2945 | 0 | 0 |
| NC_006577.2;YP_173236.1;HCoV-HKU1_orf1ab;2946 | 0 | 0 |
| NC_006577.2;YP_173236.1;HCoV-HKU1_orf1ab;2947 | 0 | 0 |
| NC_006577.2;YP_173236.1;HCoV-HKU1_orf1ab;2948 | 0 | 0 |
| NC_006577.2;YP_173236.1;HCoV-HKU1_orf1ab;2949 | 0 | 0 |
| NC_006577.2;YP_173236.1;HCoV-HKU1_orf1ab;2950 | 0 | 0 |
| NC_006577.2;YP_173236.1;HCoV-HKU1_orf1ab;2951 | 0 | 0 |
| NC_006577.2;YP_173236.1;HCoV-HKU1_orf1ab;2952 | 0 | 0 |
| NC_006577.2;YP_173236.1;HCoV-HKU1_orf1ab;2953 | 0 | 0 |
| NC_006577.2;YP_173236.1;HCoV-HKU1_orf1ab;2954 | 0 | 0 |
| NC_006577.2;YP_173236.1;HCoV-HKU1_orf1ab;2955 | 0 | 0 |
| NC_006577.2;YP_173236.1;HCoV-HKU1_orf1ab;2956 | 0 | 0 |
| NC_006577.2;YP_173236.1;HCoV-HKU1_orf1ab;2957 | 0 | 0 |
| NC_006577.2;YP_173236.1;HCoV-HKU1_orf1ab;2958 | 0 | 0 |
| NC_006577.2;YP_173236.1;HCoV-HKU1_orf1ab;2959 | 0 | 0 |
| NC_006577.2;YP_173236.1;HCoV-HKU1_orf1ab;2960 | 0 | 0 |
| NC_006577.2;YP_173236.1;HCoV-HKU1_orf1ab;2961 | 0 | 0 |
| NC_006577.2;YP_173236.1;HCoV-HKU1_orf1ab;2962 | 0 | 0 |
| NC_006577.2;YP_173236.1;HCoV-HKU1_orf1ab;2963 | 0 | 0 |
| NC_006577.2;YP_173236.1;HCoV-HKU1_orf1ab;2964 | 0 | 0.025 |
| NC_006577.2;YP_173236.1;HCoV-HKU1_orf1ab;2965 | 0 | 0 |
| NC_006577.2;YP_173236.1;HCoV-HKU1_orf1ab;2966 | 0 | 0 |
| NC_006577.2;YP_173236.1;HCoV-HKU1_orf1ab;2967 | 0 | 0 |
| NC_006577.2;YP_173236.1;HCoV-HKU1_orf1ab;2968 | 0 | 0 |
| NC_006577.2;YP_173236.1;HCoV-HKU1_orf1ab;2969 | 0 | 0 |
| NC_006577.2;YP_173236.1;HCoV-HKU1_orf1ab;2970 | 0 | 0 |
| NC_006577.2;YP_173236.1;HCoV-HKU1_orf1ab;2971 | 0 | 0 |
| NC_006577.2;YP_173236.1;HCoV-HKU1_orf1ab;2972 | 0 | 0 |
| NC_006577.2;YP_173236.1;HCoV-HKU1_orf1ab;2973 | 0 | 0 |
| NC_006577.2;YP_173236.1;HCoV-HKU1_orf1ab;2974 | 0 | 0.025 |
| NC_006577.2;YP_173236.1;HCoV-HKU1_orf1ab;2975 | 0 | 0 |
| NC_006577.2;YP_173236.1;HCoV-HKU1_orf1ab;2976 | 0 | 0 |
| NC_006577.2;YP_173236.1;HCoV-HKU1_orf1ab;2977 | 0 | 0.05 |
| NC_006577.2;YP_173236.1;HCoV-HKU1_orf1ab;2978 | 0 | 0 |
| NC_006577.2;YP_173236.1;HCoV-HKU1_orf1ab;2979 | 0 | 0 |
| NC_006577.2;YP_173236.1;HCoV-HKU1_orf1ab;2980 | 0 | 0 |
| NC_006577.2;YP_173236.1;HCoV-HKU1_orf1ab;2981 | 0 | 0 |
| NC_006577.2;YP_173236.1;HCoV-HKU1_orf1ab;2982 | 0 | 0 |

| | | |
|---|---|---|
| NC_006577.2;YP_173236.1;HCoV-HKU1_orf1ab;2983 | 0 | 0 |
| NC_006577.2;YP_173236.1;HCoV-HKU1_orf1ab;2984 | 0 | 0 |
| NC_006577.2;YP_173236.1;HCoV-HKU1_orf1ab;2985 | 0 | 0 |
| NC_006577.2;YP_173236.1;HCoV-HKU1_orf1ab;2986 | 0 | 0 |
| NC_006577.2;YP_173236.1;HCoV-HKU1_orf1ab;2987 | 0 | 0.025 |
| NC_006577.2;YP_173236.1;HCoV-HKU1_orf1ab;2988 | 0 | 0 |
| NC_006577.2;YP_173236.1;HCoV-HKU1_orf1ab;2989 | 0.05 | 0.025 |
| NC_006577.2;YP_173236.1;HCoV-HKU1_orf1ab;2990 | 0 | 0 |
| NC_006577.2;YP_173236.1;HCoV-HKU1_orf1ab;2991 | 0 | 0 |
| NC_006577.2;YP_173236.1;HCoV-HKU1_orf1ab;2992 | 0 | 0 |
| NC_006577.2;YP_173236.1;HCoV-HKU1_orf1ab;2993 | 0 | 0 |
| NC_006577.2;YP_173236.1;HCoV-HKU1_orf1ab;2994 | 0 | 0.025 |
| NC_006577.2;YP_173236.1;HCoV-HKU1_orf1ab;2995 | 0 | 0.025 |
| NC_006577.2;YP_173236.1;HCoV-HKU1_orf1ab;2996 | 0.05 | 0 |
| NC_006577.2;YP_173236.1;HCoV-HKU1_orf1ab;2997 | 0 | 0 |
| NC_006577.2;YP_173236.1;HCoV-HKU1_orf1ab;2998 | 0 | 0 |
| NC_006577.2;YP_173236.1;HCoV-HKU1_orf1ab;2999 | 0 | 0 |
| NC_006577.2;YP_173236.1;HCoV-HKU1_orf1ab;3000 | 0 | 0 |
| NC_006577.2;YP_173236.1;HCoV-HKU1_orf1ab;3001 | 0.1 | 0.05 |
| NC_006577.2;YP_173236.1;HCoV-HKU1_orf1ab;3002 | 0.1 | 0.05 |
| NC_006577.2;YP_173236.1;HCoV-HKU1_orf1ab;3003 | 0.1 | 0.075 |
| NC_006577.2;YP_173236.1;HCoV-HKU1_orf1ab;3004 | 0.15 | 0.075 |
| NC_006577.2;YP_173236.1;HCoV-HKU1_orf1ab;3005 | 0.1 | 0.075 |
| NC_006577.2;YP_173236.1;HCoV-HKU1_orf1ab;3006 | 0.05 | 0.025 |
| NC_006577.2;YP_173236.1;HCoV-HKU1_orf1ab;3007 | 0 | 0 |
| NC_006577.2;YP_173236.1;HCoV-HKU1_orf1ab;3008 | 0.05 | 0.025 |
| NC_006577.2;YP_173236.1;HCoV-HKU1_orf1ab;3009 | 0.05 | 0 |
| NC_006577.2;YP_173236.1;HCoV-HKU1_orf1ab;3010 | 0.1 | 0.05 |
| NC_006577.2;YP_173236.1;HCoV-HKU1_orf1ab;3011 | 0 | 0 |
| NC_006577.2;YP_173236.1;HCoV-HKU1_orf1ab;3012 | 0 | 0 |
| NC_006577.2;YP_173236.1;HCoV-HKU1_orf1ab;3013 | 0 | 0.025 |
| NC_006577.2;YP_173236.1;HCoV-HKU1_orf1ab;3014 | 0 | 0.025 |
| NC_006577.2;YP_173236.1;HCoV-HKU1_orf1ab;3015 | 0 | 0 |
| NC_006577.2;YP_173236.1;HCoV-HKU1_orf1ab;3016 | 0 | 0 |
| NC_006577.2;YP_173236.1;HCoV-HKU1_orf1ab;3017 | 0 | 0 |
| NC_006577.2;YP_173236.1;HCoV-HKU1_orf1ab;3018 | 0 | 0 |
| NC_006577.2;YP_173236.1;HCoV-HKU1_orf1ab;3019 | 0 | 0 |
| NC_006577.2;YP_173236.1;HCoV-HKU1_orf1ab;3020 | 0 | 0 |
| NC_006577.2;YP_173236.1;HCoV-HKU1_orf1ab;3021 | 0 | 0 |
| NC_006577.2;YP_173236.1;HCoV-HKU1_orf1ab;3022 | 0 | 0 |
| NC_006577.2;YP_173236.1;HCoV-HKU1_orf1ab;3023 | 0 | 0 |
| NC_006577.2;YP_173236.1;HCoV-HKU1_orf1ab;3024 | 0 | 0 |
| NC_006577.2;YP_173236.1;HCoV-HKU1_orf1ab;3025 | 0 | 0 |

| | | |
|---|---|---|
| NC_006577.2;YP_173236.1;HCoV-HKU1_orf1ab;3026 | 0 | 0 |
| NC_006577.2;YP_173236.1;HCoV-HKU1_orf1ab;3027 | 0 | 0 |
| NC_006577.2;YP_173236.1;HCoV-HKU1_orf1ab;3028 | 0 | 0.025 |
| NC_006577.2;YP_173236.1;HCoV-HKU1_orf1ab;3029 | 0.05 | 0.025 |
| NC_006577.2;YP_173236.1;HCoV-HKU1_orf1ab;3030 | 0.05 | 0.05 |
| NC_006577.2;YP_173236.1;HCoV-HKU1_orf1ab;3031 | 0 | 0 |
| NC_006577.2;YP_173236.1;HCoV-HKU1_orf1ab;3032 | 0 | 0 |
| NC_006577.2;YP_173236.1;HCoV-HKU1_orf1ab;3033 | 0 | 0 |
| NC_006577.2;YP_173236.1;HCoV-HKU1_orf1ab;3034 | 0 | 0 |
| NC_006577.2;YP_173236.1;HCoV-HKU1_orf1ab;3035 | 0 | 0 |
| NC_006577.2;YP_173236.1;HCoV-HKU1_orf1ab;3036 | 0 | 0 |
| NC_006577.2;YP_173236.1;HCoV-HKU1_orf1ab;3037 | 0 | 0 |
| NC_006577.2;YP_173236.1;HCoV-HKU1_orf1ab;3038 | 0 | 0 |
| NC_006577.2;YP_173236.1;HCoV-HKU1_orf1ab;3039 | 0 | 0 |
| NC_006577.2;YP_173236.1;HCoV-HKU1_orf1ab;3040 | 0 | 0 |
| NC_006577.2;YP_173236.1;HCoV-HKU1_orf1ab;3041 | 0 | 0 |
| NC_006577.2;YP_173236.1;HCoV-HKU1_orf1ab;3042 | 0 | 0 |
| NC_006577.2;YP_173236.1;HCoV-HKU1_orf1ab;3043 | 0 | 0 |
| NC_006577.2;YP_173236.1;HCoV-HKU1_orf1ab;3044 | 0 | 0 |
| NC_006577.2;YP_173236.1;HCoV-HKU1_orf1ab;3045 | 0 | 0 |
| NC_006577.2;YP_173236.1;HCoV-HKU1_orf1ab;3046 | 0 | 0 |
| NC_006577.2;YP_173236.1;HCoV-HKU1_orf1ab;3047 | 0 | 0 |
| NC_006577.2;YP_173236.1;HCoV-HKU1_orf1ab;3048 | 0 | 0 |
| NC_006577.2;YP_173236.1;HCoV-HKU1_orf1ab;3049 | 0 | 0 |
| NC_006577.2;YP_173236.1;HCoV-HKU1_orf1ab;3050 | 0 | 0 |
| NC_006577.2;YP_173236.1;HCoV-HKU1_orf1ab;3051 | 0 | 0 |
| NC_006577.2;YP_173236.1;HCoV-HKU1_orf1ab;3052 | 0 | 0 |
| NC_006577.2;YP_173236.1;HCoV-HKU1_orf1ab;3053 | 0 | 0 |
| NC_006577.2;YP_173236.1;HCoV-HKU1_orf1ab;3054 | 0 | 0 |
| NC_006577.2;YP_173236.1;HCoV-HKU1_orf1ab;3055 | 0 | 0 |
| NC_006577.2;YP_173236.1;HCoV-HKU1_orf1ab;3056 | 0 | 0 |
| NC_006577.2;YP_173236.1;HCoV-HKU1_orf1ab;3057 | 0 | 0 |
| NC_006577.2;YP_173236.1;HCoV-HKU1_orf1ab;3058 | 0 | 0 |
| NC_006577.2;YP_173236.1;HCoV-HKU1_orf1ab;3059 | 0 | 0 |
| NC_006577.2;YP_173236.1;HCoV-HKU1_orf1ab;3060 | 0 | 0 |
| NC_006577.2;YP_173236.1;HCoV-HKU1_orf1ab;3061 | 0 | 0 |
| NC_006577.2;YP_173236.1;HCoV-HKU1_orf1ab;3062 | 0 | 0 |
| NC_006577.2;YP_173236.1;HCoV-HKU1_orf1ab;3063 | 0 | 0 |
| NC_006577.2;YP_173236.1;HCoV-HKU1_orf1ab;3064 | 0 | 0 |
| NC_006577.2;YP_173236.1;HCoV-HKU1_orf1ab;3065 | 0 | 0 |
| NC_006577.2;YP_173236.1;HCoV-HKU1_orf1ab;3066 | 0 | 0 |
| NC_006577.2;YP_173236.1;HCoV-HKU1_orf1ab;3067 | 0 | 0 |
| NC_006577.2;YP_173236.1;HCoV-HKU1_orf1ab;3068 | 0 | 0 |

| | | |
|---|---|---|
| NC_006577.2;YP_173236.1;HCoV-HKU1_orf1ab;3069 | 0 | 0 |
| NC_006577.2;YP_173236.1;HCoV-HKU1_orf1ab;3070 | 0 | 0 |
| NC_006577.2;YP_173236.1;HCoV-HKU1_orf1ab;3071 | 0 | 0 |
| NC_006577.2;YP_173236.1;HCoV-HKU1_orf1ab;3072 | 0 | 0 |
| NC_006577.2;YP_173236.1;HCoV-HKU1_orf1ab;3073 | 0 | 0 |
| NC_006577.2;YP_173236.1;HCoV-HKU1_orf1ab;3074 | 0 | 0 |
| NC_006577.2;YP_173236.1;HCoV-HKU1_orf1ab;3075 | 0 | 0 |
| NC_006577.2;YP_173236.1;HCoV-HKU1_orf1ab;3076 | 0 | 0 |
| NC_006577.2;YP_173236.1;HCoV-HKU1_orf1ab;3077 | 0 | 0 |
| NC_006577.2;YP_173236.1;HCoV-HKU1_orf1ab;3078 | 0 | 0 |
| NC_006577.2;YP_173236.1;HCoV-HKU1_orf1ab;3079 | 0 | 0 |
| NC_006577.2;YP_173236.1;HCoV-HKU1_orf1ab;3080 | 0 | 0 |
| NC_006577.2;YP_173236.1;HCoV-HKU1_orf1ab;3081 | 0 | 0 |
| NC_006577.2;YP_173236.1;HCoV-HKU1_orf1ab;3082 | 0 | 0 |
| NC_006577.2;YP_173236.1;HCoV-HKU1_orf1ab;3083 | 0 | 0 |
| NC_006577.2;YP_173236.1;HCoV-HKU1_orf1ab;3084 | 0 | 0 |
| NC_006577.2;YP_173236.1;HCoV-HKU1_orf1ab;3085 | 0 | 0 |
| NC_006577.2;YP_173236.1;HCoV-HKU1_orf1ab;3086 | 0 | 0 |
| NC_006577.2;YP_173236.1;HCoV-HKU1_orf1ab;3087 | 0 | 0 |
| NC_006577.2;YP_173236.1;HCoV-HKU1_orf1ab;3088 | 0 | 0 |
| NC_006577.2;YP_173236.1;HCoV-HKU1_orf1ab;3089 | 0 | 0 |
| NC_006577.2;YP_173236.1;HCoV-HKU1_orf1ab;3090 | 0 | 0 |
| NC_006577.2;YP_173236.1;HCoV-HKU1_orf1ab;3091 | 0 | 0 |
| NC_006577.2;YP_173236.1;HCoV-HKU1_orf1ab;3092 | 0 | 0 |
| NC_006577.2;YP_173236.1;HCoV-HKU1_orf1ab;3093 | 0 | 0 |
| NC_006577.2;YP_173236.1;HCoV-HKU1_orf1ab;3094 | 0 | 0 |
| NC_006577.2;YP_173236.1;HCoV-HKU1_orf1ab;3095 | 0 | 0 |
| NC_006577.2;YP_173236.1;HCoV-HKU1_orf1ab;3096 | 0 | 0 |
| NC_006577.2;YP_173236.1;HCoV-HKU1_orf1ab;3097 | 0 | 0 |
| NC_006577.2;YP_173236.1;HCoV-HKU1_orf1ab;3098 | 0 | 0 |
| NC_006577.2;YP_173236.1;HCoV-HKU1_orf1ab;3099 | 0 | 0 |
| NC_006577.2;YP_173236.1;HCoV-HKU1_orf1ab;3100 | 0 | 0 |
| NC_006577.2;YP_173236.1;HCoV-HKU1_orf1ab;3101 | 0 | 0 |
| NC_006577.2;YP_173236.1;HCoV-HKU1_orf1ab;3102 | 0 | 0 |
| NC_006577.2;YP_173236.1;HCoV-HKU1_orf1ab;3103 | 0 | 0 |
| NC_006577.2;YP_173236.1;HCoV-HKU1_orf1ab;3104 | 0 | 0 |
| NC_006577.2;YP_173236.1;HCoV-HKU1_orf1ab;3105 | 0 | 0 |
| NC_006577.2;YP_173236.1;HCoV-HKU1_orf1ab;3106 | 0 | 0 |
| NC_006577.2;YP_173236.1;HCoV-HKU1_orf1ab;3107 | 0 | 0 |
| NC_006577.2;YP_173236.1;HCoV-HKU1_orf1ab;3108 | 0 | 0 |
| NC_006577.2;YP_173236.1;HCoV-HKU1_orf1ab;3109 | 0 | 0 |
| NC_006577.2;YP_173236.1;HCoV-HKU1_orf1ab;3110 | 0 | 0 |
| NC_006577.2;YP_173236.1;HCoV-HKU1_orf1ab;3111 | 0 | 0 |

| | | |
|---|---|---|
| NC_006577.2;YP_173236.1;HCoV-HKU1_orf1ab;3112 | 0 | 0 |
| NC_006577.2;YP_173236.1;HCoV-HKU1_orf1ab;3113 | 0 | 0 |
| NC_006577.2;YP_173236.1;HCoV-HKU1_orf1ab;3114 | 0 | 0 |
| NC_006577.2;YP_173236.1;HCoV-HKU1_orf1ab;3115 | 0 | 0 |
| NC_006577.2;YP_173236.1;HCoV-HKU1_orf1ab;3116 | 0 | 0 |
| NC_006577.2;YP_173236.1;HCoV-HKU1_orf1ab;3117 | 0 | 0 |
| NC_006577.2;YP_173236.1;HCoV-HKU1_orf1ab;3118 | 0 | 0 |
| NC_006577.2;YP_173236.1;HCoV-HKU1_orf1ab;3119 | 0 | 0 |
| NC_006577.2;YP_173236.1;HCoV-HKU1_orf1ab;3120 | 0 | 0 |
| NC_006577.2;YP_173236.1;HCoV-HKU1_orf1ab;3121 | 0 | 0 |
| NC_006577.2;YP_173236.1;HCoV-HKU1_orf1ab;3122 | 0 | 0 |
| NC_006577.2;YP_173236.1;HCoV-HKU1_orf1ab;3123 | 0 | 0 |
| NC_006577.2;YP_173236.1;HCoV-HKU1_orf1ab;3124 | 0 | 0 |
| NC_006577.2;YP_173236.1;HCoV-HKU1_orf1ab;3125 | 0 | 0 |
| NC_006577.2;YP_173236.1;HCoV-HKU1_orf1ab;3126 | 0 | 0 |
| NC_006577.2;YP_173236.1;HCoV-HKU1_orf1ab;3127 | 0 | 0 |
| NC_006577.2;YP_173236.1;HCoV-HKU1_orf1ab;3128 | 0 | 0 |
| NC_006577.2;YP_173236.1;HCoV-HKU1_orf1ab;3129 | 0 | 0 |
| NC_006577.2;YP_173236.1;HCoV-HKU1_orf1ab;3130 | 0 | 0 |
| NC_006577.2;YP_173236.1;HCoV-HKU1_orf1ab;3131 | 0 | 0 |
| NC_006577.2;YP_173236.1;HCoV-HKU1_orf1ab;3132 | 0 | 0 |
| NC_006577.2;YP_173236.1;HCoV-HKU1_orf1ab;3133 | 0 | 0 |
| NC_006577.2;YP_173236.1;HCoV-HKU1_orf1ab;3134 | 0 | 0 |
| NC_006577.2;YP_173236.1;HCoV-HKU1_orf1ab;3135 | 0 | 0 |
| NC_006577.2;YP_173236.1;HCoV-HKU1_orf1ab;3136 | 0 | 0 |
| NC_006577.2;YP_173236.1;HCoV-HKU1_orf1ab;3137 | 0 | 0 |
| NC_006577.2;YP_173236.1;HCoV-HKU1_orf1ab;3138 | 0 | 0 |
| NC_006577.2;YP_173236.1;HCoV-HKU1_orf1ab;3139 | 0 | 0 |
| NC_006577.2;YP_173236.1;HCoV-HKU1_orf1ab;3140 | 0 | 0 |
| NC_006577.2;YP_173236.1;HCoV-HKU1_orf1ab;3141 | 0 | 0 |
| NC_006577.2;YP_173236.1;HCoV-HKU1_orf1ab;3142 | 0 | 0 |
| NC_006577.2;YP_173236.1;HCoV-HKU1_orf1ab;3143 | 0 | 0 |
| NC_006577.2;YP_173236.1;HCoV-HKU1_orf1ab;3144 | 0 | 0 |
| NC_006577.2;YP_173236.1;HCoV-HKU1_orf1ab;3145 | 0 | 0 |
| NC_006577.2;YP_173236.1;HCoV-HKU1_orf1ab;3146 | 0 | 0 |
| NC_006577.2;YP_173236.1;HCoV-HKU1_orf1ab;3147 | 0 | 0 |
| NC_006577.2;YP_173236.1;HCoV-HKU1_orf1ab;3148 | 0 | 0 |
| NC_006577.2;YP_173236.1;HCoV-HKU1_orf1ab;3149 | 0 | 0 |
| NC_006577.2;YP_173236.1;HCoV-HKU1_orf1ab;3150 | 0 | 0 |
| NC_006577.2;YP_173236.1;HCoV-HKU1_orf1ab;3151 | 0 | 0 |
| NC_006577.2;YP_173236.1;HCoV-HKU1_orf1ab;3152 | 0 | 0 |
| NC_006577.2;YP_173236.1;HCoV-HKU1_orf1ab;3153 | 0 | 0 |
| NC_006577.2;YP_173236.1;HCoV-HKU1_orf1ab;3154 | 0 | 0 |

| | | |
|---|---|---|
| NC_006577.2;YP_173236.1;HCoV-HKU1_orf1ab;3155 | 0 | 0 |
| NC_006577.2;YP_173236.1;HCoV-HKU1_orf1ab;3156 | 0 | 0 |
| NC_006577.2;YP_173236.1;HCoV-HKU1_orf1ab;3157 | 0 | 0 |
| NC_006577.2;YP_173236.1;HCoV-HKU1_orf1ab;3158 | 0 | 0 |
| NC_006577.2;YP_173236.1;HCoV-HKU1_orf1ab;3159 | 0 | 0 |
| NC_006577.2;YP_173236.1;HCoV-HKU1_orf1ab;3160 | 0 | 0 |
| NC_006577.2;YP_173236.1;HCoV-HKU1_orf1ab;3161 | 0 | 0 |
| NC_006577.2;YP_173236.1;HCoV-HKU1_orf1ab;3162 | 0 | 0 |
| NC_006577.2;YP_173236.1;HCoV-HKU1_orf1ab;3163 | 0 | 0 |
| NC_006577.2;YP_173236.1;HCoV-HKU1_orf1ab;3164 | 0 | 0 |
| NC_006577.2;YP_173236.1;HCoV-HKU1_orf1ab;3165 | 0 | 0 |
| NC_006577.2;YP_173236.1;HCoV-HKU1_orf1ab;3166 | 0 | 0 |
| NC_006577.2;YP_173236.1;HCoV-HKU1_orf1ab;3167 | 0 | 0 |
| NC_006577.2;YP_173236.1;HCoV-HKU1_orf1ab;3168 | 0 | 0 |
| NC_006577.2;YP_173236.1;HCoV-HKU1_orf1ab;3169 | 0 | 0 |
| NC_006577.2;YP_173236.1;HCoV-HKU1_orf1ab;3170 | 0 | 0 |
| NC_006577.2;YP_173236.1;HCoV-HKU1_orf1ab;3171 | 0 | 0 |
| NC_006577.2;YP_173236.1;HCoV-HKU1_orf1ab;3172 | 0 | 0 |
| NC_006577.2;YP_173236.1;HCoV-HKU1_orf1ab;3173 | 0 | 0 |
| NC_006577.2;YP_173236.1;HCoV-HKU1_orf1ab;3174 | 0 | 0 |
| NC_006577.2;YP_173236.1;HCoV-HKU1_orf1ab;3175 | 0 | 0 |
| NC_006577.2;YP_173236.1;HCoV-HKU1_orf1ab;3176 | 0 | 0 |
| NC_006577.2;YP_173236.1;HCoV-HKU1_orf1ab;3177 | 0 | 0 |
| NC_006577.2;YP_173236.1;HCoV-HKU1_orf1ab;3178 | 0 | 0 |
| NC_006577.2;YP_173236.1;HCoV-HKU1_orf1ab;3179 | 0 | 0 |
| NC_006577.2;YP_173236.1;HCoV-HKU1_orf1ab;3180 | 0 | 0 |
| NC_006577.2;YP_173236.1;HCoV-HKU1_orf1ab;3181 | 0 | 0 |
| NC_006577.2;YP_173236.1;HCoV-HKU1_orf1ab;3182 | 0 | 0 |
| NC_006577.2;YP_173236.1;HCoV-HKU1_orf1ab;3183 | 0 | 0 |
| NC_006577.2;YP_173236.1;HCoV-HKU1_orf1ab;3184 | 0 | 0 |
| NC_006577.2;YP_173236.1;HCoV-HKU1_orf1ab;3185 | 0 | 0 |
| NC_006577.2;YP_173236.1;HCoV-HKU1_orf1ab;3186 | 0 | 0 |
| NC_006577.2;YP_173236.1;HCoV-HKU1_orf1ab;3187 | 0 | 0 |
| NC_006577.2;YP_173236.1;HCoV-HKU1_orf1ab;3188 | 0 | 0 |
| NC_006577.2;YP_173236.1;HCoV-HKU1_orf1ab;3189 | 0 | 0 |
| NC_006577.2;YP_173236.1;HCoV-HKU1_orf1ab;3190 | 0 | 0 |
| NC_006577.2;YP_173236.1;HCoV-HKU1_orf1ab;3191 | 0 | 0 |
| NC_006577.2;YP_173236.1;HCoV-HKU1_orf1ab;3192 | 0 | 0.025 |
| NC_006577.2;YP_173236.1;HCoV-HKU1_orf1ab;3193 | 0 | 0 |
| NC_006577.2;YP_173236.1;HCoV-HKU1_orf1ab;3194 | 0 | 0 |
| NC_006577.2;YP_173236.1;HCoV-HKU1_orf1ab;3195 | 0 | 0 |
| NC_006577.2;YP_173236.1;HCoV-HKU1_orf1ab;3196 | 0 | 0 |
| NC_006577.2;YP_173236.1;HCoV-HKU1_orf1ab;3197 | 0 | 0 |

| | | |
|---|---|---|
| NC_006577.2;YP_173236.1;HCoV-HKU1_orf1ab;3198 | 0 | 0 |
| NC_006577.2;YP_173236.1;HCoV-HKU1_orf1ab;3199 | 0 | 0 |
| NC_006577.2;YP_173236.1;HCoV-HKU1_orf1ab;3200 | 0 | 0 |
| NC_006577.2;YP_173236.1;HCoV-HKU1_orf1ab;3201 | 0 | 0 |
| NC_006577.2;YP_173236.1;HCoV-HKU1_orf1ab;3202 | 0 | 0 |
| NC_006577.2;YP_173236.1;HCoV-HKU1_orf1ab;3203 | 0 | 0 |
| NC_006577.2;YP_173236.1;HCoV-HKU1_orf1ab;3204 | 0 | 0 |
| NC_006577.2;YP_173236.1;HCoV-HKU1_orf1ab;3205 | 0 | 0 |
| NC_006577.2;YP_173236.1;HCoV-HKU1_orf1ab;3206 | 0 | 0 |
| NC_006577.2;YP_173236.1;HCoV-HKU1_orf1ab;3207 | 0 | 0 |
| NC_006577.2;YP_173236.1;HCoV-HKU1_orf1ab;3208 | 0 | 0 |
| NC_006577.2;YP_173236.1;HCoV-HKU1_orf1ab;3209 | 0 | 0 |
| NC_006577.2;YP_173236.1;HCoV-HKU1_orf1ab;3210 | 0 | 0 |
| NC_006577.2;YP_173236.1;HCoV-HKU1_orf1ab;3211 | 0 | 0 |
| NC_006577.2;YP_173236.1;HCoV-HKU1_orf1ab;3212 | 0 | 0 |
| NC_006577.2;YP_173236.1;HCoV-HKU1_orf1ab;3213 | 0 | 0 |
| NC_006577.2;YP_173236.1;HCoV-HKU1_orf1ab;3214 | 0 | 0 |
| NC_006577.2;YP_173236.1;HCoV-HKU1_orf1ab;3215 | 0 | 0 |
| NC_006577.2;YP_173236.1;HCoV-HKU1_orf1ab;3216 | 0 | 0 |
| NC_006577.2;YP_173236.1;HCoV-HKU1_orf1ab;3217 | 0 | 0 |
| NC_006577.2;YP_173236.1;HCoV-HKU1_orf1ab;3218 | 0 | 0 |
| NC_006577.2;YP_173236.1;HCoV-HKU1_orf1ab;3219 | 0 | 0 |
| NC_006577.2;YP_173236.1;HCoV-HKU1_orf1ab;3220 | 0 | 0 |
| NC_006577.2;YP_173236.1;HCoV-HKU1_orf1ab;3221 | 0 | 0 |
| NC_006577.2;YP_173236.1;HCoV-HKU1_orf1ab;3222 | 0 | 0 |
| NC_006577.2;YP_173236.1;HCoV-HKU1_orf1ab;3223 | 0 | 0 |
| NC_006577.2;YP_173236.1;HCoV-HKU1_orf1ab;3224 | 0 | 0 |
| NC_006577.2;YP_173236.1;HCoV-HKU1_orf1ab;3225 | 0 | 0 |
| NC_006577.2;YP_173236.1;HCoV-HKU1_orf1ab;3226 | 0 | 0 |
| NC_006577.2;YP_173236.1;HCoV-HKU1_orf1ab;3227 | 0 | 0 |
| NC_006577.2;YP_173236.1;HCoV-HKU1_orf1ab;3228 | 0 | 0 |
| NC_006577.2;YP_173236.1;HCoV-HKU1_orf1ab;3229 | 0 | 0 |
| NC_006577.2;YP_173236.1;HCoV-HKU1_orf1ab;3230 | 0 | 0 |
| NC_006577.2;YP_173236.1;HCoV-HKU1_orf1ab;3231 | 0.05 | 0.025 |
| NC_006577.2;YP_173236.1;HCoV-HKU1_orf1ab;3232 | 0.05 | 0 |
| NC_006577.2;YP_173236.1;HCoV-HKU1_orf1ab;3233 | 0 | 0.025 |
| NC_006577.2;YP_173236.1;HCoV-HKU1_orf1ab;3234 | 0 | 0.025 |
| NC_006577.2;YP_173236.1;HCoV-HKU1_orf1ab;3235 | 0 | 0 |
| NC_006577.2;YP_173236.1;HCoV-HKU1_orf1ab;3236 | 0 | 0 |
| NC_006577.2;YP_173236.1;HCoV-HKU1_orf1ab;3237 | 0 | 0.025 |
| NC_006577.2;YP_173236.1;HCoV-HKU1_orf1ab;3238 | 0 | 0 |
| NC_006577.2;YP_173236.1;HCoV-HKU1_orf1ab;3239 | 0 | 0 |
| NC_006577.2;YP_173236.1;HCoV-HKU1_orf1ab;3240 | 0 | 0 |

| | | |
|---|---|---|
| NC_006577.2;YP_173236.1;HCoV-HKU1_orf1ab;3241 | 0 | 0 |
| NC_006577.2;YP_173236.1;HCoV-HKU1_orf1ab;3242 | 0 | 0 |
| NC_006577.2;YP_173236.1;HCoV-HKU1_orf1ab;3243 | 0 | 0 |
| NC_006577.2;YP_173236.1;HCoV-HKU1_orf1ab;3244 | 0 | 0 |
| NC_006577.2;YP_173236.1;HCoV-HKU1_orf1ab;3245 | 0 | 0 |
| NC_006577.2;YP_173236.1;HCoV-HKU1_orf1ab;3246 | 0 | 0 |
| NC_006577.2;YP_173236.1;HCoV-HKU1_orf1ab;3247 | 0 | 0 |
| NC_006577.2;YP_173236.1;HCoV-HKU1_orf1ab;3248 | 0 | 0 |
| NC_006577.2;YP_173236.1;HCoV-HKU1_orf1ab;3249 | 0 | 0 |
| NC_006577.2;YP_173236.1;HCoV-HKU1_orf1ab;3250 | 0 | 0 |
| NC_006577.2;YP_173236.1;HCoV-HKU1_orf1ab;3251 | 0 | 0 |
| NC_006577.2;YP_173236.1;HCoV-HKU1_orf1ab;3252 | 0 | 0 |
| NC_006577.2;YP_173236.1;HCoV-HKU1_orf1ab;3253 | 0 | 0 |
| NC_006577.2;YP_173236.1;HCoV-HKU1_orf1ab;3254 | 0 | 0 |
| NC_006577.2;YP_173236.1;HCoV-HKU1_orf1ab;3255 | 0 | 0 |
| NC_006577.2;YP_173236.1;HCoV-HKU1_orf1ab;3256 | 0.1 | 0 |
| NC_006577.2;YP_173236.1;HCoV-HKU1_orf1ab;3257 | 0.05 | 0 |
| NC_006577.2;YP_173236.1;HCoV-HKU1_orf1ab;3258 | 0.15 | 0.025 |
| NC_006577.2;YP_173236.1;HCoV-HKU1_orf1ab;3259 | 0.05 | 0.025 |
| NC_006577.2;YP_173236.1;HCoV-HKU1_orf1ab;3260 | 0.1 | 0.05 |
| NC_006577.2;YP_173236.1;HCoV-HKU1_orf1ab;3261 | 0.15 | 0.025 |
| NC_006577.2;YP_173236.1;HCoV-HKU1_orf1ab;3262 | 0.1 | 0 |
| NC_006577.2;YP_173236.1;HCoV-HKU1_orf1ab;3263 | 0.1 | 0.025 |
| NC_006577.2;YP_173236.1;HCoV-HKU1_orf1ab;3264 | 0.1 | 0.025 |
| NC_006577.2;YP_173236.1;HCoV-HKU1_orf1ab;3265 | 0.1 | 0.025 |
| NC_006577.2;YP_173236.1;HCoV-HKU1_orf1ab;3266 | 0.1 | 0.025 |
| NC_006577.2;YP_173236.1;HCoV-HKU1_orf1ab;3267 | 0 | 0 |
| NC_006577.2;YP_173236.1;HCoV-HKU1_orf1ab;3268 | 0.05 | 0 |
| NC_006577.2;YP_173236.1;HCoV-HKU1_orf1ab;3269 | 0 | 0 |
| NC_006577.2;YP_173236.1;HCoV-HKU1_orf1ab;3270 | 0 | 0 |
| NC_006577.2;YP_173236.1;HCoV-HKU1_orf1ab;3271 | 0 | 0 |
| NC_006577.2;YP_173236.1;HCoV-HKU1_orf1ab;3272 | 0 | 0 |
| NC_006577.2;YP_173236.1;HCoV-HKU1_orf1ab;3273 | 0 | 0 |
| NC_006577.2;YP_173236.1;HCoV-HKU1_orf1ab;3274 | 0 | 0 |
| NC_006577.2;YP_173236.1;HCoV-HKU1_orf1ab;3275 | 0 | 0 |
| NC_006577.2;YP_173236.1;HCoV-HKU1_orf1ab;3276 | 0 | 0 |
| NC_006577.2;YP_173236.1;HCoV-HKU1_orf1ab;3277 | 0 | 0 |
| NC_006577.2;YP_173236.1;HCoV-HKU1_orf1ab;3278 | 0 | 0.025 |
| NC_006577.2;YP_173236.1;HCoV-HKU1_orf1ab;3279 | 0 | 0.025 |
| NC_006577.2;YP_173236.1;HCoV-HKU1_orf1ab;3280 | 0.05 | 0.05 |
| NC_006577.2;YP_173236.1;HCoV-HKU1_orf1ab;3281 | 0 | 0 |
| NC_006577.2;YP_173236.1;HCoV-HKU1_orf1ab;3282 | 0 | 0 |
| NC_006577.2;YP_173236.1;HCoV-HKU1_orf1ab;3283 | 0 | 0 |

| | | |
|---|---|---|
| NC_006577.2;YP_173236.1;HCoV-HKU1_orf1ab;3284 | 0.05 | 0 |
| NC_006577.2;YP_173236.1;HCoV-HKU1_orf1ab;3285 | 0.1 | 0.025 |
| NC_006577.2;YP_173236.1;HCoV-HKU1_orf1ab;3286 | 0 | 0 |
| NC_006577.2;YP_173236.1;HCoV-HKU1_orf1ab;3287 | 0 | 0 |
| NC_006577.2;YP_173236.1;HCoV-HKU1_orf1ab;3288 | 0 | 0 |
| NC_006577.2;YP_173236.1;HCoV-HKU1_orf1ab;3289 | 0 | 0.025 |
| NC_006577.2;YP_173236.1;HCoV-HKU1_orf1ab;3290 | 0.05 | 0.025 |
| NC_006577.2;YP_173236.1;HCoV-HKU1_orf1ab;3291 | 0 | 0 |
| NC_006577.2;YP_173236.1;HCoV-HKU1_orf1ab;3292 | 0 | 0.025 |
| NC_006577.2;YP_173236.1;HCoV-HKU1_orf1ab;3293 | 0 | 0 |
| NC_006577.2;YP_173236.1;HCoV-HKU1_orf1ab;3294 | 0 | 0 |
| NC_006577.2;YP_173236.1;HCoV-HKU1_orf1ab;3295 | 0 | 0 |
| NC_006577.2;YP_173236.1;HCoV-HKU1_orf1ab;3296 | 0.05 | 0 |
| NC_006577.2;YP_173236.1;HCoV-HKU1_orf1ab;3297 | 0.05 | 0 |
| NC_006577.2;YP_173236.1;HCoV-HKU1_orf1ab;3298 | 0 | 0 |
| NC_006577.2;YP_173236.1;HCoV-HKU1_orf1ab;3299 | 0.05 | 0 |
| NC_006577.2;YP_173236.1;HCoV-HKU1_orf1ab;3300 | 0 | 0 |
| NC_006577.2;YP_173236.1;HCoV-HKU1_orf1ab;3301 | 0.1 | 0 |
| NC_006577.2;YP_173236.1;HCoV-HKU1_orf1ab;3302 | 0.05 | 0 |
| NC_006577.2;YP_173236.1;HCoV-HKU1_orf1ab;3303 | 0 | 0 |
| NC_006577.2;YP_173236.1;HCoV-HKU1_orf1ab;3304 | 0 | 0.025 |
| NC_006577.2;YP_173236.1;HCoV-HKU1_orf1ab;3305 | 0 | 0.025 |
| NC_006577.2;YP_173236.1;HCoV-HKU1_orf1ab;3306 | 0 | 0.025 |
| NC_006577.2;YP_173236.1;HCoV-HKU1_orf1ab;3307 | 0 | 0.025 |
| NC_006577.2;YP_173236.1;HCoV-HKU1_orf1ab;3308 | 0 | 0.025 |
| NC_006577.2;YP_173236.1;HCoV-HKU1_orf1ab;3309 | 0.05 | 0.025 |
| NC_006577.2;YP_173236.1;HCoV-HKU1_orf1ab;3310 | 0.05 | 0.025 |
| NC_006577.2;YP_173236.1;HCoV-HKU1_orf1ab;3311 | 0.05 | 0.025 |
| NC_006577.2;YP_173236.1;HCoV-HKU1_orf1ab;3312 | 0.1 | 0.025 |
| NC_006577.2;YP_173236.1;HCoV-HKU1_orf1ab;3313 | 0 | 0.025 |
| NC_006577.2;YP_173236.1;HCoV-HKU1_orf1ab;3314 | 0.05 | 0.025 |
| NC_006577.2;YP_173236.1;HCoV-HKU1_orf1ab;3315 | 0 | 0 |
| NC_006577.2;YP_173236.1;HCoV-HKU1_orf1ab;3316 | 0.05 | 0.025 |
| NC_006577.2;YP_173236.1;HCoV-HKU1_orf1ab;3317 | 0.05 | 0 |
| NC_006577.2;YP_173236.1;HCoV-HKU1_orf1ab;3318 | 0 | 0 |
| NC_006577.2;YP_173236.1;HCoV-HKU1_orf1ab;3319 | 0 | 0 |
| NC_006577.2;YP_173236.1;HCoV-HKU1_orf1ab;3320 | 0 | 0.025 |
| NC_006577.2;YP_173236.1;HCoV-HKU1_orf1ab;3321 | 0.1 | 0.025 |
| NC_006577.2;YP_173236.1;HCoV-HKU1_orf1ab;3322 | 0 | 0.025 |
| NC_006577.2;YP_173236.1;HCoV-HKU1_orf1ab;3323 | 0 | 0 |
| NC_006577.2;YP_173236.1;HCoV-HKU1_orf1ab;3324 | 0 | 0 |
| NC_006577.2;YP_173236.1;HCoV-HKU1_orf1ab;3325 | 0 | 0 |
| NC_006577.2;YP_173236.1;HCoV-HKU1_orf1ab;3326 | 0 | 0 |

| | | |
|---|---|---|
| NC_006577.2;YP_173236.1;HCoV-HKU1_orf1ab;3327 | 0 | 0 |
| NC_006577.2;YP_173236.1;HCoV-HKU1_orf1ab;3328 | 0 | 0 |
| NC_006577.2;YP_173236.1;HCoV-HKU1_orf1ab;3329 | 0 | 0 |
| NC_006577.2;YP_173236.1;HCoV-HKU1_orf1ab;3330 | 0 | 0 |
| NC_006577.2;YP_173236.1;HCoV-HKU1_orf1ab;3331 | 0 | 0 |
| NC_006577.2;YP_173236.1;HCoV-HKU1_orf1ab;3332 | 0.05 | 0 |
| NC_006577.2;YP_173236.1;HCoV-HKU1_orf1ab;3333 | 0.1 | 0 |
| NC_006577.2;YP_173236.1;HCoV-HKU1_orf1ab;3334 | 0.1 | 0 |
| NC_006577.2;YP_173236.1;HCoV-HKU1_orf1ab;3335 | 0.1 | 0 |
| NC_006577.2;YP_173236.1;HCoV-HKU1_orf1ab;3336 | 0.1 | 0 |
| NC_006577.2;YP_173236.1;HCoV-HKU1_orf1ab;3337 | 0.1 | 0 |
| NC_006577.2;YP_173236.1;HCoV-HKU1_orf1ab;3338 | 0.1 | 0 |
| NC_006577.2;YP_173236.1;HCoV-HKU1_orf1ab;3339 | 0.05 | 0 |
| NC_006577.2;YP_173236.1;HCoV-HKU1_orf1ab;3340 | 0 | 0 |
| NC_006577.2;YP_173236.1;HCoV-HKU1_orf1ab;3341 | 0 | 0 |
| NC_006577.2;YP_173236.1;HCoV-HKU1_orf1ab;3342 | 0.05 | 0 |
| NC_006577.2;YP_173236.1;HCoV-HKU1_orf1ab;3343 | 0 | 0 |
| NC_006577.2;YP_173236.1;HCoV-HKU1_orf1ab;3344 | 0 | 0 |
| NC_006577.2;YP_173236.1;HCoV-HKU1_orf1ab;3345 | 0 | 0 |
| NC_006577.2;YP_173236.1;HCoV-HKU1_orf1ab;3346 | 0 | 0 |
| NC_006577.2;YP_173236.1;HCoV-HKU1_orf1ab;3347 | 0 | 0 |
| NC_006577.2;YP_173236.1;HCoV-HKU1_orf1ab;3348 | 0 | 0 |
| NC_006577.2;YP_173236.1;HCoV-HKU1_orf1ab;3349 | 0 | 0 |
| NC_006577.2;YP_173236.1;HCoV-HKU1_orf1ab;3350 | 0 | 0 |
| NC_006577.2;YP_173236.1;HCoV-HKU1_orf1ab;3351 | 0 | 0 |
| NC_006577.2;YP_173236.1;HCoV-HKU1_orf1ab;3352 | 0 | 0 |
| NC_006577.2;YP_173236.1;HCoV-HKU1_orf1ab;3353 | 0 | 0 |
| NC_006577.2;YP_173236.1;HCoV-HKU1_orf1ab;3354 | 0 | 0.025 |
| NC_006577.2;YP_173236.1;HCoV-HKU1_orf1ab;3355 | 0 | 0 |
| NC_006577.2;YP_173236.1;HCoV-HKU1_orf1ab;3356 | 0 | 0 |
| NC_006577.2;YP_173236.1;HCoV-HKU1_orf1ab;3357 | 0 | 0 |
| NC_006577.2;YP_173236.1;HCoV-HKU1_orf1ab;3358 | 0 | 0 |
| NC_006577.2;YP_173236.1;HCoV-HKU1_orf1ab;3359 | 0 | 0 |
| NC_006577.2;YP_173236.1;HCoV-HKU1_orf1ab;3360 | 0 | 0 |
| NC_006577.2;YP_173236.1;HCoV-HKU1_orf1ab;3361 | 0 | 0 |
| NC_006577.2;YP_173236.1;HCoV-HKU1_orf1ab;3362 | 0 | 0 |
| NC_006577.2;YP_173236.1;HCoV-HKU1_orf1ab;3363 | 0 | 0 |
| NC_006577.2;YP_173236.1;HCoV-HKU1_orf1ab;3364 | 0 | 0 |
| NC_006577.2;YP_173236.1;HCoV-HKU1_orf1ab;3365 | 0 | 0 |
| NC_006577.2;YP_173236.1;HCoV-HKU1_orf1ab;3366 | 0 | 0 |
| NC_006577.2;YP_173236.1;HCoV-HKU1_orf1ab;3367 | 0 | 0 |
| NC_006577.2;YP_173236.1;HCoV-HKU1_orf1ab;3368 | 0 | 0.025 |
| NC_006577.2;YP_173236.1;HCoV-HKU1_orf1ab;3369 | 0 | 0 |

| | | |
|---|---|---|
| NC_006577.2;YP_173236.1;HCoV-HKU1_orf1ab;3370 | 0 | 0.025 |
| NC_006577.2;YP_173236.1;HCoV-HKU1_orf1ab;3371 | 0 | 0.025 |
| NC_006577.2;YP_173236.1;HCoV-HKU1_orf1ab;3372 | 0 | 0 |
| NC_006577.2;YP_173236.1;HCoV-HKU1_orf1ab;3373 | 0 | 0.025 |
| NC_006577.2;YP_173236.1;HCoV-HKU1_orf1ab;3374 | 0 | 0 |
| NC_006577.2;YP_173236.1;HCoV-HKU1_orf1ab;3375 | 0.05 | 0 |
| NC_006577.2;YP_173236.1;HCoV-HKU1_orf1ab;3376 | 0 | 0.025 |
| NC_006577.2;YP_173236.1;HCoV-HKU1_orf1ab;3377 | 0.1 | 0.025 |
| NC_006577.2;YP_173236.1;HCoV-HKU1_orf1ab;3378 | 0 | 0 |
| NC_006577.2;YP_173236.1;HCoV-HKU1_orf1ab;3379 | 0 | 0 |
| NC_006577.2;YP_173236.1;HCoV-HKU1_orf1ab;3380 | 0 | 0 |
| NC_006577.2;YP_173236.1;HCoV-HKU1_orf1ab;3381 | 0 | 0 |
| NC_006577.2;YP_173236.1;HCoV-HKU1_orf1ab;3382 | 0 | 0 |
| NC_006577.2;YP_173236.1;HCoV-HKU1_orf1ab;3383 | 0 | 0 |
| NC_006577.2;YP_173236.1;HCoV-HKU1_orf1ab;3384 | 0 | 0 |
| NC_006577.2;YP_173236.1;HCoV-HKU1_orf1ab;3385 | 0 | 0 |
| NC_006577.2;YP_173236.1;HCoV-HKU1_orf1ab;3386 | 0 | 0 |
| NC_006577.2;YP_173236.1;HCoV-HKU1_orf1ab;3387 | 0 | 0 |
| NC_006577.2;YP_173236.1;HCoV-HKU1_orf1ab;3388 | 0 | 0 |
| NC_006577.2;YP_173236.1;HCoV-HKU1_orf1ab;3389 | 0 | 0 |
| NC_006577.2;YP_173236.1;HCoV-HKU1_orf1ab;3390 | 0 | 0 |
| NC_006577.2;YP_173236.1;HCoV-HKU1_orf1ab;3391 | 0 | 0 |
| NC_006577.2;YP_173236.1;HCoV-HKU1_orf1ab;3392 | 0 | 0 |
| NC_006577.2;YP_173236.1;HCoV-HKU1_orf1ab;3393 | 0 | 0 |
| NC_006577.2;YP_173236.1;HCoV-HKU1_orf1ab;3394 | 0 | 0 |
| NC_006577.2;YP_173236.1;HCoV-HKU1_orf1ab;3395 | 0 | 0.025 |
| NC_006577.2;YP_173236.1;HCoV-HKU1_orf1ab;3396 | 0 | 0 |
| NC_006577.2;YP_173236.1;HCoV-HKU1_orf1ab;3397 | 0 | 0 |
| NC_006577.2;YP_173236.1;HCoV-HKU1_orf1ab;3398 | 0 | 0 |
| NC_006577.2;YP_173236.1;HCoV-HKU1_orf1ab;3399 | 0 | 0 |
| NC_006577.2;YP_173236.1;HCoV-HKU1_orf1ab;3400 | 0 | 0 |
| NC_006577.2;YP_173236.1;HCoV-HKU1_orf1ab;3401 | 0 | 0 |
| NC_006577.2;YP_173236.1;HCoV-HKU1_orf1ab;3402 | 0 | 0 |
| NC_006577.2;YP_173236.1;HCoV-HKU1_orf1ab;3403 | 0 | 0 |
| NC_006577.2;YP_173236.1;HCoV-HKU1_orf1ab;3404 | 0 | 0 |
| NC_006577.2;YP_173236.1;HCoV-HKU1_orf1ab;3405 | 0 | 0 |
| NC_006577.2;YP_173236.1;HCoV-HKU1_orf1ab;3406 | 0 | 0 |
| NC_006577.2;YP_173236.1;HCoV-HKU1_orf1ab;3407 | 0 | 0 |
| NC_006577.2;YP_173236.1;HCoV-HKU1_orf1ab;3408 | 0 | 0 |
| NC_006577.2;YP_173236.1;HCoV-HKU1_orf1ab;3409 | 0 | 0 |
| NC_006577.2;YP_173236.1;HCoV-HKU1_orf1ab;3410 | 0 | 0 |
| NC_006577.2;YP_173236.1;HCoV-HKU1_orf1ab;3411 | 0 | 0 |
| NC_006577.2;YP_173236.1;HCoV-HKU1_orf1ab;3412 | 0 | 0 |

| | | |
|---|---|---|
| NC_006577.2;YP_173236.1;HCoV-HKU1_orf1ab;3413 | 0 | 0 |
| NC_006577.2;YP_173236.1;HCoV-HKU1_orf1ab;3414 | 0 | 0 |
| NC_006577.2;YP_173236.1;HCoV-HKU1_orf1ab;3415 | 0 | 0 |
| NC_006577.2;YP_173236.1;HCoV-HKU1_orf1ab;3416 | 0 | 0 |
| NC_006577.2;YP_173236.1;HCoV-HKU1_orf1ab;3417 | 0 | 0 |
| NC_006577.2;YP_173236.1;HCoV-HKU1_orf1ab;3418 | 0 | 0 |
| NC_006577.2;YP_173236.1;HCoV-HKU1_orf1ab;3419 | 0 | 0 |
| NC_006577.2;YP_173236.1;HCoV-HKU1_orf1ab;3420 | 0 | 0 |
| NC_006577.2;YP_173236.1;HCoV-HKU1_orf1ab;3421 | 0 | 0 |
| NC_006577.2;YP_173236.1;HCoV-HKU1_orf1ab;3422 | 0 | 0 |
| NC_006577.2;YP_173236.1;HCoV-HKU1_orf1ab;3423 | 0 | 0 |
| NC_006577.2;YP_173236.1;HCoV-HKU1_orf1ab;3424 | 0 | 0 |
| NC_006577.2;YP_173236.1;HCoV-HKU1_orf1ab;3425 | 0 | 0 |
| NC_006577.2;YP_173236.1;HCoV-HKU1_orf1ab;3426 | 0 | 0 |
| NC_006577.2;YP_173236.1;HCoV-HKU1_orf1ab;3427 | 0 | 0 |
| NC_006577.2;YP_173236.1;HCoV-HKU1_orf1ab;3428 | 0 | 0.025 |
| NC_006577.2;YP_173236.1;HCoV-HKU1_orf1ab;3429 | 0.15 | 0.075 |
| NC_006577.2;YP_173236.1;HCoV-HKU1_orf1ab;3430 | 0.15 | 0.075 |
| NC_006577.2;YP_173236.1;HCoV-HKU1_orf1ab;3431 | 0.05 | 0 |
| NC_006577.2;YP_173236.1;HCoV-HKU1_orf1ab;3432 | 0.1 | 0.075 |
| NC_006577.2;YP_173236.1;HCoV-HKU1_orf1ab;3433 | 0.05 | 0.025 |
| NC_006577.2;YP_173236.1;HCoV-HKU1_orf1ab;3434 | 0 | 0.025 |
| NC_006577.2;YP_173236.1;HCoV-HKU1_orf1ab;3435 | 0 | 0 |
| NC_006577.2;YP_173236.1;HCoV-HKU1_orf1ab;3436 | 0.05 | 0.025 |
| NC_006577.2;YP_173236.1;HCoV-HKU1_orf1ab;3437 | 0 | 0 |
| NC_006577.2;YP_173236.1;HCoV-HKU1_orf1ab;3438 | 0 | 0 |
| NC_006577.2;YP_173236.1;HCoV-HKU1_orf1ab;3439 | 0 | 0 |
| NC_006577.2;YP_173236.1;HCoV-HKU1_orf1ab;3440 | 0 | 0 |
| NC_006577.2;YP_173236.1;HCoV-HKU1_orf1ab;3441 | 0 | 0 |
| NC_006577.2;YP_173236.1;HCoV-HKU1_orf1ab;3442 | 0 | 0 |
| NC_006577.2;YP_173236.1;HCoV-HKU1_orf1ab;3443 | 0 | 0 |
| NC_006577.2;YP_173236.1;HCoV-HKU1_orf1ab;3444 | 0 | 0 |
| NC_006577.2;YP_173236.1;HCoV-HKU1_orf1ab;3445 | 0 | 0 |
| NC_006577.2;YP_173236.1;HCoV-HKU1_orf1ab;3446 | 0 | 0 |
| NC_006577.2;YP_173236.1;HCoV-HKU1_orf1ab;3447 | 0 | 0 |
| NC_006577.2;YP_173236.1;HCoV-HKU1_orf1ab;3448 | 0 | 0 |
| NC_006577.2;YP_173236.1;HCoV-HKU1_orf1ab;3449 | 0 | 0 |
| NC_006577.2;YP_173236.1;HCoV-HKU1_orf1ab;3450 | 0 | 0 |
| NC_006577.2;YP_173236.1;HCoV-HKU1_orf1ab;3451 | 0 | 0 |
| NC_006577.2;YP_173236.1;HCoV-HKU1_orf1ab;3452 | 0 | 0 |
| NC_006577.2;YP_173236.1;HCoV-HKU1_orf1ab;3453 | 0 | 0 |
| NC_006577.2;YP_173236.1;HCoV-HKU1_orf1ab;3454 | 0 | 0 |
| NC_006577.2;YP_173236.1;HCoV-HKU1_orf1ab;3455 | 0 | 0 |

| | | |
|---|---|---|
| NC_006577.2;YP_173236.1;HCoV-HKU1_orf1ab;3456 | 0 | 0 |
| NC_006577.2;YP_173236.1;HCoV-HKU1_orf1ab;3457 | 0 | 0 |
| NC_006577.2;YP_173236.1;HCoV-HKU1_orf1ab;3458 | 0 | 0.025 |
| NC_006577.2;YP_173236.1;HCoV-HKU1_orf1ab;3459 | 0 | 0 |
| NC_006577.2;YP_173236.1;HCoV-HKU1_orf1ab;3460 | 0 | 0 |
| NC_006577.2;YP_173236.1;HCoV-HKU1_orf1ab;3461 | 0 | 0 |
| NC_006577.2;YP_173236.1;HCoV-HKU1_orf1ab;3462 | 0 | 0 |
| NC_006577.2;YP_173236.1;HCoV-HKU1_orf1ab;3463 | 0 | 0 |
| NC_006577.2;YP_173236.1;HCoV-HKU1_orf1ab;3464 | 0 | 0 |
| NC_006577.2;YP_173236.1;HCoV-HKU1_orf1ab;3465 | 0 | 0 |
| NC_006577.2;YP_173236.1;HCoV-HKU1_orf1ab;3466 | 0 | 0 |
| NC_006577.2;YP_173236.1;HCoV-HKU1_orf1ab;3467 | 0 | 0 |
| NC_006577.2;YP_173236.1;HCoV-HKU1_orf1ab;3468 | 0 | 0 |
| NC_006577.2;YP_173236.1;HCoV-HKU1_orf1ab;3469 | 0 | 0 |
| NC_006577.2;YP_173236.1;HCoV-HKU1_orf1ab;3470 | 0 | 0 |
| NC_006577.2;YP_173236.1;HCoV-HKU1_orf1ab;3471 | 0 | 0 |
| NC_006577.2;YP_173236.1;HCoV-HKU1_orf1ab;3472 | 0 | 0 |
| NC_006577.2;YP_173236.1;HCoV-HKU1_orf1ab;3473 | 0 | 0 |
| NC_006577.2;YP_173236.1;HCoV-HKU1_orf1ab;3474 | 0 | 0 |
| NC_006577.2;YP_173236.1;HCoV-HKU1_orf1ab;3475 | 0 | 0 |
| NC_006577.2;YP_173236.1;HCoV-HKU1_orf1ab;3476 | 0 | 0 |
| NC_006577.2;YP_173236.1;HCoV-HKU1_orf1ab;3477 | 0 | 0 |
| NC_006577.2;YP_173236.1;HCoV-HKU1_orf1ab;3478 | 0.1 | 0 |
| NC_006577.2;YP_173236.1;HCoV-HKU1_orf1ab;3479 | 0.1 | 0.025 |
| NC_006577.2;YP_173236.1;HCoV-HKU1_orf1ab;3480 | 0.1 | 0.025 |
| NC_006577.2;YP_173236.1;HCoV-HKU1_orf1ab;3481 | 0.1 | 0.025 |
| NC_006577.2;YP_173236.1;HCoV-HKU1_orf1ab;3482 | 0.1 | 0 |
| NC_006577.2;YP_173236.1;HCoV-HKU1_orf1ab;3483 | 0.1 | 0.025 |
| NC_006577.2;YP_173236.1;HCoV-HKU1_orf1ab;3484 | 0.1 | 0.025 |
| NC_006577.2;YP_173236.1;HCoV-HKU1_orf1ab;3485 | 0.1 | 0.025 |
| NC_006577.2;YP_173236.1;HCoV-HKU1_orf1ab;3486 | 0.1 | 0.025 |
| NC_006577.2;YP_173236.1;HCoV-HKU1_orf1ab;3487 | 0.1 | 0.025 |
| NC_006577.2;YP_173236.1;HCoV-HKU1_orf1ab;3488 | 0 | 0 |
| NC_006577.2;YP_173236.1;HCoV-HKU1_orf1ab;3489 | 0 | 0 |
| NC_006577.2;YP_173236.1;HCoV-HKU1_orf1ab;3490 | 0 | 0 |
| NC_006577.2;YP_173236.1;HCoV-HKU1_orf1ab;3491 | 0 | 0 |
| NC_006577.2;YP_173236.1;HCoV-HKU1_orf1ab;3492 | 0 | 0 |
| NC_006577.2;YP_173236.1;HCoV-HKU1_orf1ab;3493 | 0 | 0 |
| NC_006577.2;YP_173236.1;HCoV-HKU1_orf1ab;3494 | 0 | 0 |
| NC_006577.2;YP_173236.1;HCoV-HKU1_orf1ab;3495 | 0 | 0 |
| NC_006577.2;YP_173236.1;HCoV-HKU1_orf1ab;3496 | 0 | 0 |
| NC_006577.2;YP_173236.1;HCoV-HKU1_orf1ab;3497 | 0 | 0 |
| NC_006577.2;YP_173236.1;HCoV-HKU1_orf1ab;3498 | 0 | 0 |

| | | |
|---|---|---|
| NC_006577.2;YP_173236.1;HCoV-HKU1_orf1ab;3499 | 0 | 0.025 |
| NC_006577.2;YP_173236.1;HCoV-HKU1_orf1ab;3500 | 0.05 | 0.1 |
| NC_006577.2;YP_173236.1;HCoV-HKU1_orf1ab;3501 | 0.05 | 0 |
| NC_006577.2;YP_173236.1;HCoV-HKU1_orf1ab;3502 | 0.05 | 0 |
| NC_006577.2;YP_173236.1;HCoV-HKU1_orf1ab;3503 | 0.05 | 0 |
| NC_006577.2;YP_173236.1;HCoV-HKU1_orf1ab;3504 | 0.05 | 0 |
| NC_006577.2;YP_173236.1;HCoV-HKU1_orf1ab;3505 | 0.05 | 0 |
| NC_006577.2;YP_173236.1;HCoV-HKU1_orf1ab;3506 | 0.05 | 0 |
| NC_006577.2;YP_173236.1;HCoV-HKU1_orf1ab;3507 | 0.05 | 0 |
| NC_006577.2;YP_173236.1;HCoV-HKU1_orf1ab;3508 | 0.05 | 0.025 |
| NC_006577.2;YP_173236.1;HCoV-HKU1_orf1ab;3509 | 0 | 0 |
| NC_006577.2;YP_173236.1;HCoV-HKU1_orf1ab;3510 | 0 | 0 |
| NC_006577.2;YP_173236.1;HCoV-HKU1_orf1ab;3511 | 0 | 0 |
| NC_006577.2;YP_173236.1;HCoV-HKU1_orf1ab;3512 | 0 | 0 |
| NC_006577.2;YP_173236.1;HCoV-HKU1_orf1ab;3513 | 0 | 0 |
| NC_006577.2;YP_173236.1;HCoV-HKU1_orf1ab;3514 | 0 | 0 |
| NC_006577.2;YP_173236.1;HCoV-HKU1_orf1ab;3515 | 0 | 0 |
| NC_006577.2;YP_173236.1;HCoV-HKU1_orf1ab;3516 | 0 | 0.025 |
| NC_006577.2;YP_173236.1;HCoV-HKU1_orf1ab;3517 | 0 | 0.075 |
| NC_006577.2;YP_173236.1;HCoV-HKU1_orf1ab;3518 | 0 | 0 |
| NC_006577.2;YP_173236.1;HCoV-HKU1_orf1ab;3519 | 0 | 0 |
| NC_006577.2;YP_173236.1;HCoV-HKU1_orf1ab;3520 | 0.05 | 0.025 |
| NC_006577.2;YP_173236.1;HCoV-HKU1_orf1ab;3521 | 0 | 0.025 |
| NC_006577.2;YP_173236.1;HCoV-HKU1_orf1ab;3522 | 0 | 0.025 |
| NC_006577.2;YP_173236.1;HCoV-HKU1_orf1ab;3523 | 0 | 0.025 |
| NC_006577.2;YP_173236.1;HCoV-HKU1_orf1ab;3524 | 0 | 0.025 |
| NC_006577.2;YP_173236.1;HCoV-HKU1_orf1ab;3525 | 0 | 0.025 |
| NC_006577.2;YP_173236.1;HCoV-HKU1_orf1ab;3526 | 0 | 0.025 |
| NC_006577.2;YP_173236.1;HCoV-HKU1_orf1ab;3527 | 0 | 0 |
| NC_006577.2;YP_173236.1;HCoV-HKU1_orf1ab;3528 | 0 | 0.025 |
| NC_006577.2;YP_173236.1;HCoV-HKU1_orf1ab;3529 | 0 | 0 |
| NC_006577.2;YP_173236.1;HCoV-HKU1_orf1ab;3530 | 0 | 0 |
| NC_006577.2;YP_173236.1;HCoV-HKU1_orf1ab;3531 | 0 | 0 |
| NC_006577.2;YP_173236.1;HCoV-HKU1_orf1ab;3532 | 0 | 0 |
| NC_006577.2;YP_173236.1;HCoV-HKU1_orf1ab;3533 | 0 | 0.025 |
| NC_006577.2;YP_173236.1;HCoV-HKU1_orf1ab;3534 | 0 | 0 |
| NC_006577.2;YP_173236.1;HCoV-HKU1_orf1ab;3535 | 0 | 0 |
| NC_006577.2;YP_173236.1;HCoV-HKU1_orf1ab;3536 | 0 | 0 |
| NC_006577.2;YP_173236.1;HCoV-HKU1_orf1ab;3537 | 0 | 0 |
| NC_006577.2;YP_173236.1;HCoV-HKU1_orf1ab;3538 | 0 | 0 |
| NC_006577.2;YP_173236.1;HCoV-HKU1_orf1ab;3539 | 0 | 0 |
| NC_006577.2;YP_173236.1;HCoV-HKU1_orf1ab;3540 | 0 | 0 |
| NC_006577.2;YP_173236.1;HCoV-HKU1_orf1ab;3541 | 0 | 0 |

| | | |
|---|---|---|
| NC_006577.2;YP_173236.1;HCoV-HKU1_orf1ab;3542 | 0 | 0 |
| NC_006577.2;YP_173236.1;HCoV-HKU1_orf1ab;3543 | 0 | 0.025 |
| NC_006577.2;YP_173236.1;HCoV-HKU1_orf1ab;3544 | 0 | 0 |
| NC_006577.2;YP_173236.1;HCoV-HKU1_orf1ab;3545 | 0 | 0 |
| NC_006577.2;YP_173236.1;HCoV-HKU1_orf1ab;3546 | 0 | 0 |
| NC_006577.2;YP_173236.1;HCoV-HKU1_orf1ab;3547 | 0 | 0 |
| NC_006577.2;YP_173236.1;HCoV-HKU1_orf1ab;3548 | 0 | 0 |
| NC_006577.2;YP_173236.1;HCoV-HKU1_orf1ab;3549 | 0 | 0 |
| NC_006577.2;YP_173236.1;HCoV-HKU1_orf1ab;3550 | 0 | 0 |
| NC_006577.2;YP_173236.1;HCoV-HKU1_orf1ab;3551 | 0 | 0 |
| NC_006577.2;YP_173236.1;HCoV-HKU1_orf1ab;3552 | 0 | 0 |
| NC_006577.2;YP_173236.1;HCoV-HKU1_orf1ab;3553 | 0 | 0.025 |
| NC_006577.2;YP_173236.1;HCoV-HKU1_orf1ab;3554 | 0 | 0.05 |
| NC_006577.2;YP_173236.1;HCoV-HKU1_orf1ab;3555 | 0 | 0.025 |
| NC_006577.2;YP_173236.1;HCoV-HKU1_orf1ab;3556 | 0 | 0 |
| NC_006577.2;YP_173236.1;HCoV-HKU1_orf1ab;3557 | 0 | 0 |
| NC_006577.2;YP_173236.1;HCoV-HKU1_orf1ab;3558 | 0 | 0 |
| NC_006577.2;YP_173236.1;HCoV-HKU1_orf1ab;3559 | 0 | 0 |
| NC_006577.2;YP_173236.1;HCoV-HKU1_orf1ab;3560 | 0.05 | 0.025 |
| NC_006577.2;YP_173236.1;HCoV-HKU1_orf1ab;3561 | 0 | 0 |
| NC_006577.2;YP_173236.1;HCoV-HKU1_orf1ab;3562 | 0 | 0 |
| NC_006577.2;YP_173236.1;HCoV-HKU1_orf1ab;3563 | 0.1 | 0 |
| NC_006577.2;YP_173236.1;HCoV-HKU1_orf1ab;3564 | 0 | 0 |
| NC_006577.2;YP_173236.1;HCoV-HKU1_orf1ab;3565 | 0 | 0 |
| NC_006577.2;YP_173236.1;HCoV-HKU1_orf1ab;3566 | 0 | 0 |
| NC_006577.2;YP_173236.1;HCoV-HKU1_orf1ab;3567 | 0 | 0 |
| NC_006577.2;YP_173236.1;HCoV-HKU1_orf1ab;3568 | 0 | 0 |
| NC_006577.2;YP_173236.1;HCoV-HKU1_orf1ab;3569 | 0 | 0 |
| NC_006577.2;YP_173236.1;HCoV-HKU1_orf1ab;3570 | 0 | 0 |
| NC_006577.2;YP_173236.1;HCoV-HKU1_orf1ab;3571 | 0.05 | 0 |
| NC_006577.2;YP_173236.1;HCoV-HKU1_orf1ab;3572 | 0.05 | 0 |
| NC_006577.2;YP_173236.1;HCoV-HKU1_orf1ab;3573 | 0.05 | 0 |
| NC_006577.2;YP_173236.1;HCoV-HKU1_orf1ab;3574 | 0 | 0 |
| NC_006577.2;YP_173236.1;HCoV-HKU1_orf1ab;3575 | 0 | 0 |
| NC_006577.2;YP_173236.1;HCoV-HKU1_orf1ab;3576 | 0 | 0 |
| NC_006577.2;YP_173236.1;HCoV-HKU1_orf1ab;3577 | 0 | 0 |
| NC_006577.2;YP_173236.1;HCoV-HKU1_orf1ab;3578 | 0 | 0 |
| NC_006577.2;YP_173236.1;HCoV-HKU1_orf1ab;3579 | 0 | 0 |
| NC_006577.2;YP_173236.1;HCoV-HKU1_orf1ab;3580 | 0 | 0 |
| NC_006577.2;YP_173236.1;HCoV-HKU1_orf1ab;3581 | 0 | 0 |
| NC_006577.2;YP_173236.1;HCoV-HKU1_orf1ab;3582 | 0 | 0.025 |
| NC_006577.2;YP_173236.1;HCoV-HKU1_orf1ab;3583 | 0 | 0.025 |
| NC_006577.2;YP_173236.1;HCoV-HKU1_orf1ab;3584 | 0.05 | 0.05 |

| | | |
|---|---|---|
| NC_006577.2;YP_173236.1;HCoV-HKU1_orf1ab;3585 | 0 | 0.025 |
| NC_006577.2;YP_173236.1;HCoV-HKU1_orf1ab;3586 | 0 | 0 |
| NC_006577.2;YP_173236.1;HCoV-HKU1_orf1ab;3587 | 0 | 0 |
| NC_006577.2;YP_173236.1;HCoV-HKU1_orf1ab;3588 | 0 | 0 |
| NC_006577.2;YP_173236.1;HCoV-HKU1_orf1ab;3589 | 0 | 0 |
| NC_006577.2;YP_173236.1;HCoV-HKU1_orf1ab;3590 | 0 | 0 |
| NC_006577.2;YP_173236.1;HCoV-HKU1_orf1ab;3591 | 0 | 0 |
| NC_006577.2;YP_173236.1;HCoV-HKU1_orf1ab;3592 | 0 | 0 |
| NC_006577.2;YP_173236.1;HCoV-HKU1_orf1ab;3593 | 0 | 0 |
| NC_006577.2;YP_173236.1;HCoV-HKU1_orf1ab;3594 | 0 | 0 |
| NC_006577.2;YP_173236.1;HCoV-HKU1_orf1ab;3595 | 0 | 0 |
| NC_006577.2;YP_173236.1;HCoV-HKU1_orf1ab;3596 | 0 | 0 |
| NC_006577.2;YP_173236.1;HCoV-HKU1_orf1ab;3597 | 0 | 0 |
| NC_006577.2;YP_173236.1;HCoV-HKU1_orf1ab;3598 | 0 | 0 |
| NC_006577.2;YP_173236.1;HCoV-HKU1_orf1ab;3599 | 0 | 0 |
| NC_006577.2;YP_173236.1;HCoV-HKU1_orf1ab;3600 | 0 | 0 |
| NC_006577.2;YP_173236.1;HCoV-HKU1_orf1ab;3601 | 0 | 0 |
| NC_006577.2;YP_173236.1;HCoV-HKU1_orf1ab;3602 | 0 | 0 |
| NC_006577.2;YP_173236.1;HCoV-HKU1_orf1ab;3603 | 0 | 0 |
| NC_006577.2;YP_173236.1;HCoV-HKU1_orf1ab;3604 | 0 | 0 |
| NC_006577.2;YP_173236.1;HCoV-HKU1_orf1ab;3605 | 0 | 0 |
| NC_006577.2;YP_173236.1;HCoV-HKU1_orf1ab;3606 | 0 | 0.05 |
| NC_006577.2;YP_173236.1;HCoV-HKU1_orf1ab;3607 | 0 | 0 |
| NC_006577.2;YP_173236.1;HCoV-HKU1_orf1ab;3608 | 0 | 0 |
| NC_006577.2;YP_173236.1;HCoV-HKU1_orf1ab;3609 | 0 | 0 |
| NC_006577.2;YP_173236.1;HCoV-HKU1_orf1ab;3610 | 0 | 0.025 |
| NC_006577.2;YP_173236.1;HCoV-HKU1_orf1ab;3611 | 0 | 0.025 |
| NC_006577.2;YP_173236.1;HCoV-HKU1_orf1ab;3612 | 0 | 0.025 |
| NC_006577.2;YP_173236.1;HCoV-HKU1_orf1ab;3613 | 0.05 | 0.025 |
| NC_006577.2;YP_173236.1;HCoV-HKU1_orf1ab;3614 | 0.05 | 0.025 |
| NC_006577.2;YP_173236.1;HCoV-HKU1_orf1ab;3615 | 0.05 | 0.1 |
| NC_006577.2;YP_173236.1;HCoV-HKU1_orf1ab;3616 | 0 | 0.05 |
| NC_006577.2;YP_173236.1;HCoV-HKU1_orf1ab;3617 | 0.1 | 0.075 |
| NC_006577.2;YP_173236.1;HCoV-HKU1_orf1ab;3618 | 0 | 0.05 |
| NC_006577.2;YP_173236.1;HCoV-HKU1_orf1ab;3619 | 0.1 | 0.05 |
| NC_006577.2;YP_173236.1;HCoV-HKU1_orf1ab;3620 | 0 | 0.025 |
| NC_006577.2;YP_173236.1;HCoV-HKU1_orf1ab;3621 | 0 | 0.05 |
| NC_006577.2;YP_173236.1;HCoV-HKU1_orf1ab;3622 | 0 | 0.05 |
| NC_006577.2;YP_173236.1;HCoV-HKU1_orf1ab;3623 | 0 | 0.1 |
| NC_006577.2;YP_173236.1;HCoV-HKU1_orf1ab;3624 | 0 | 0 |
| NC_006577.2;YP_173236.1;HCoV-HKU1_orf1ab;3625 | 0 | 0 |
| NC_006577.2;YP_173236.1;HCoV-HKU1_orf1ab;3626 | 0 | 0 |
| NC_006577.2;YP_173236.1;HCoV-HKU1_orf1ab;3627 | 0 | 0 |

| | | |
|---|---|---|
| NC_006577.2;YP_173236.1;HCoV-HKU1_orf1ab;3628 | 0 | 0 |
| NC_006577.2;YP_173236.1;HCoV-HKU1_orf1ab;3629 | 0 | 0 |
| NC_006577.2;YP_173236.1;HCoV-HKU1_orf1ab;3630 | 0 | 0 |
| NC_006577.2;YP_173236.1;HCoV-HKU1_orf1ab;3631 | 0 | 0 |
| NC_006577.2;YP_173236.1;HCoV-HKU1_orf1ab;3632 | 0 | 0 |
| NC_006577.2;YP_173236.1;HCoV-HKU1_orf1ab;3633 | 0.05 | 0 |
| NC_006577.2;YP_173236.1;HCoV-HKU1_orf1ab;3634 | 0.05 | 0 |
| NC_006577.2;YP_173236.1;HCoV-HKU1_orf1ab;3635 | 0 | 0 |
| NC_006577.2;YP_173236.1;HCoV-HKU1_orf1ab;3636 | 0 | 0 |
| NC_006577.2;YP_173236.1;HCoV-HKU1_orf1ab;3637 | 0 | 0 |
| NC_006577.2;YP_173236.1;HCoV-HKU1_orf1ab;3638 | 0 | 0 |
| NC_006577.2;YP_173236.1;HCoV-HKU1_orf1ab;3639 | 0 | 0 |
| NC_006577.2;YP_173236.1;HCoV-HKU1_orf1ab;3640 | 0 | 0 |
| NC_006577.2;YP_173236.1;HCoV-HKU1_orf1ab;3641 | 0 | 0 |
| NC_006577.2;YP_173236.1;HCoV-HKU1_orf1ab;3642 | 0 | 0 |
| NC_006577.2;YP_173236.1;HCoV-HKU1_orf1ab;3643 | 0 | 0 |
| NC_006577.2;YP_173236.1;HCoV-HKU1_orf1ab;3644 | 0 | 0 |
| NC_006577.2;YP_173236.1;HCoV-HKU1_orf1ab;3645 | 0 | 0 |
| NC_006577.2;YP_173236.1;HCoV-HKU1_orf1ab;3646 | 0 | 0 |
| NC_006577.2;YP_173236.1;HCoV-HKU1_orf1ab;3647 | 0 | 0 |
| NC_006577.2;YP_173236.1;HCoV-HKU1_orf1ab;3648 | 0 | 0 |
| NC_006577.2;YP_173236.1;HCoV-HKU1_orf1ab;3649 | 0 | 0 |
| NC_006577.2;YP_173236.1;HCoV-HKU1_orf1ab;3650 | 0 | 0 |
| NC_006577.2;YP_173236.1;HCoV-HKU1_orf1ab;3651 | 0 | 0 |
| NC_006577.2;YP_173236.1;HCoV-HKU1_orf1ab;3652 | 0 | 0 |
| NC_006577.2;YP_173236.1;HCoV-HKU1_orf1ab;3653 | 0 | 0 |
| NC_006577.2;YP_173236.1;HCoV-HKU1_orf1ab;3654 | 0 | 0 |
| NC_006577.2;YP_173236.1;HCoV-HKU1_orf1ab;3655 | 0 | 0 |
| NC_006577.2;YP_173236.1;HCoV-HKU1_orf1ab;3656 | 0 | 0 |
| NC_006577.2;YP_173236.1;HCoV-HKU1_orf1ab;3657 | 0 | 0 |
| NC_006577.2;YP_173236.1;HCoV-HKU1_orf1ab;3658 | 0 | 0 |
| NC_006577.2;YP_173236.1;HCoV-HKU1_orf1ab;3659 | 0 | 0 |
| NC_006577.2;YP_173236.1;HCoV-HKU1_orf1ab;3660 | 0 | 0 |
| NC_006577.2;YP_173236.1;HCoV-HKU1_orf1ab;3661 | 0 | 0 |
| NC_006577.2;YP_173236.1;HCoV-HKU1_orf1ab;3662 | 0 | 0 |
| NC_006577.2;YP_173236.1;HCoV-HKU1_orf1ab;3663 | 0 | 0 |
| NC_006577.2;YP_173236.1;HCoV-HKU1_orf1ab;3664 | 0 | 0 |
| NC_006577.2;YP_173236.1;HCoV-HKU1_orf1ab;3665 | 0 | 0 |
| NC_006577.2;YP_173236.1;HCoV-HKU1_orf1ab;3666 | 0 | 0 |
| NC_006577.2;YP_173236.1;HCoV-HKU1_orf1ab;3667 | 0 | 0 |
| NC_006577.2;YP_173236.1;HCoV-HKU1_orf1ab;3668 | 0 | 0 |
| NC_006577.2;YP_173236.1;HCoV-HKU1_orf1ab;3669 | 0 | 0 |
| NC_006577.2;YP_173236.1;HCoV-HKU1_orf1ab;3670 | 0 | 0 |

| | | |
|---|---|---|
| NC_006577.2;YP_173236.1;HCoV-HKU1_orf1ab;3671 | 0 | 0 |
| NC_006577.2;YP_173236.1;HCoV-HKU1_orf1ab;3672 | 0 | 0 |
| NC_006577.2;YP_173236.1;HCoV-HKU1_orf1ab;3673 | 0 | 0 |
| NC_006577.2;YP_173236.1;HCoV-HKU1_orf1ab;3674 | 0 | 0 |
| NC_006577.2;YP_173236.1;HCoV-HKU1_orf1ab;3675 | 0 | 0 |
| NC_006577.2;YP_173236.1;HCoV-HKU1_orf1ab;3676 | 0 | 0 |
| NC_006577.2;YP_173236.1;HCoV-HKU1_orf1ab;3677 | 0 | 0 |
| NC_006577.2;YP_173236.1;HCoV-HKU1_orf1ab;3678 | 0 | 0 |
| NC_006577.2;YP_173236.1;HCoV-HKU1_orf1ab;3679 | 0 | 0 |
| NC_006577.2;YP_173236.1;HCoV-HKU1_orf1ab;3680 | 0 | 0 |
| NC_006577.2;YP_173236.1;HCoV-HKU1_orf1ab;3681 | 0 | 0 |
| NC_006577.2;YP_173236.1;HCoV-HKU1_orf1ab;3682 | 0 | 0 |
| NC_006577.2;YP_173236.1;HCoV-HKU1_orf1ab;3683 | 0 | 0 |
| NC_006577.2;YP_173236.1;HCoV-HKU1_orf1ab;3684 | 0 | 0 |
| NC_006577.2;YP_173236.1;HCoV-HKU1_orf1ab;3685 | 0 | 0 |
| NC_006577.2;YP_173236.1;HCoV-HKU1_orf1ab;3686 | 0 | 0 |
| NC_006577.2;YP_173236.1;HCoV-HKU1_orf1ab;3687 | 0 | 0 |
| NC_006577.2;YP_173236.1;HCoV-HKU1_orf1ab;3688 | 0 | 0 |
| NC_006577.2;YP_173236.1;HCoV-HKU1_orf1ab;3689 | 0 | 0 |
| NC_006577.2;YP_173236.1;HCoV-HKU1_orf1ab;3690 | 0 | 0 |
| NC_006577.2;YP_173236.1;HCoV-HKU1_orf1ab;3691 | 0 | 0 |
| NC_006577.2;YP_173236.1;HCoV-HKU1_orf1ab;3692 | 0 | 0 |
| NC_006577.2;YP_173236.1;HCoV-HKU1_orf1ab;3693 | 0 | 0 |
| NC_006577.2;YP_173236.1;HCoV-HKU1_orf1ab;3694 | 0 | 0 |
| NC_006577.2;YP_173236.1;HCoV-HKU1_orf1ab;3695 | 0 | 0 |
| NC_006577.2;YP_173236.1;HCoV-HKU1_orf1ab;3696 | 0 | 0 |
| NC_006577.2;YP_173236.1;HCoV-HKU1_orf1ab;3697 | 0 | 0 |
| NC_006577.2;YP_173236.1;HCoV-HKU1_orf1ab;3698 | 0 | 0 |
| NC_006577.2;YP_173236.1;HCoV-HKU1_orf1ab;3699 | 0 | 0 |
| NC_006577.2;YP_173236.1;HCoV-HKU1_orf1ab;3700 | 0 | 0 |
| NC_006577.2;YP_173236.1;HCoV-HKU1_orf1ab;3701 | 0 | 0 |
| NC_006577.2;YP_173236.1;HCoV-HKU1_orf1ab;3702 | 0 | 0 |
| NC_006577.2;YP_173236.1;HCoV-HKU1_orf1ab;3703 | 0 | 0 |
| NC_006577.2;YP_173236.1;HCoV-HKU1_orf1ab;3704 | 0 | 0 |
| NC_006577.2;YP_173236.1;HCoV-HKU1_orf1ab;3705 | 0 | 0 |
| NC_006577.2;YP_173236.1;HCoV-HKU1_orf1ab;3706 | 0 | 0 |
| NC_006577.2;YP_173236.1;HCoV-HKU1_orf1ab;3707 | 0 | 0 |
| NC_006577.2;YP_173236.1;HCoV-HKU1_orf1ab;3708 | 0 | 0 |
| NC_006577.2;YP_173236.1;HCoV-HKU1_orf1ab;3709 | 0 | 0 |
| NC_006577.2;YP_173236.1;HCoV-HKU1_orf1ab;3710 | 0 | 0 |
| NC_006577.2;YP_173236.1;HCoV-HKU1_orf1ab;3711 | 0 | 0 |
| NC_006577.2;YP_173236.1;HCoV-HKU1_orf1ab;3712 | 0 | 0 |
| NC_006577.2;YP_173236.1;HCoV-HKU1_orf1ab;3713 | 0 | 0 |

| | | |
|---|---|---|
| NC_006577.2;YP_173236.1;HCoV-HKU1_orf1ab;3714 | 0 | 0 |
| NC_006577.2;YP_173236.1;HCoV-HKU1_orf1ab;3715 | 0 | 0 |
| NC_006577.2;YP_173236.1;HCoV-HKU1_orf1ab;3716 | 0 | 0 |
| NC_006577.2;YP_173236.1;HCoV-HKU1_orf1ab;3717 | 0 | 0 |
| NC_006577.2;YP_173236.1;HCoV-HKU1_orf1ab;3718 | 0 | 0 |
| NC_006577.2;YP_173236.1;HCoV-HKU1_orf1ab;3719 | 0 | 0 |
| NC_006577.2;YP_173236.1;HCoV-HKU1_orf1ab;3720 | 0 | 0 |
| NC_006577.2;YP_173236.1;HCoV-HKU1_orf1ab;3721 | 0 | 0 |
| NC_006577.2;YP_173236.1;HCoV-HKU1_orf1ab;3722 | 0 | 0 |
| NC_006577.2;YP_173236.1;HCoV-HKU1_orf1ab;3723 | 0 | 0 |
| NC_006577.2;YP_173236.1;HCoV-HKU1_orf1ab;3724 | 0 | 0 |
| NC_006577.2;YP_173236.1;HCoV-HKU1_orf1ab;3725 | 0 | 0 |
| NC_006577.2;YP_173236.1;HCoV-HKU1_orf1ab;3726 | 0 | 0 |
| NC_006577.2;YP_173236.1;HCoV-HKU1_orf1ab;3727 | 0 | 0 |
| NC_006577.2;YP_173236.1;HCoV-HKU1_orf1ab;3728 | 0 | 0 |
| NC_006577.2;YP_173236.1;HCoV-HKU1_orf1ab;3729 | 0 | 0 |
| NC_006577.2;YP_173236.1;HCoV-HKU1_orf1ab;3730 | 0 | 0 |
| NC_006577.2;YP_173236.1;HCoV-HKU1_orf1ab;3731 | 0 | 0 |
| NC_006577.2;YP_173236.1;HCoV-HKU1_orf1ab;3732 | 0 | 0 |
| NC_006577.2;YP_173236.1;HCoV-HKU1_orf1ab;3733 | 0 | 0 |
| NC_006577.2;YP_173236.1;HCoV-HKU1_orf1ab;3734 | 0 | 0 |
| NC_006577.2;YP_173236.1;HCoV-HKU1_orf1ab;3735 | 0 | 0 |
| NC_006577.2;YP_173236.1;HCoV-HKU1_orf1ab;3736 | 0 | 0 |
| NC_006577.2;YP_173236.1;HCoV-HKU1_orf1ab;3737 | 0 | 0 |
| NC_006577.2;YP_173236.1;HCoV-HKU1_orf1ab;3738 | 0 | 0 |
| NC_006577.2;YP_173236.1;HCoV-HKU1_orf1ab;3739 | 0 | 0 |
| NC_006577.2;YP_173236.1;HCoV-HKU1_orf1ab;3740 | 0 | 0 |
| NC_006577.2;YP_173236.1;HCoV-HKU1_orf1ab;3741 | 0 | 0 |
| NC_006577.2;YP_173236.1;HCoV-HKU1_orf1ab;3742 | 0 | 0 |
| NC_006577.2;YP_173236.1;HCoV-HKU1_orf1ab;3743 | 0 | 0 |
| NC_006577.2;YP_173236.1;HCoV-HKU1_orf1ab;3744 | 0 | 0 |
| NC_006577.2;YP_173236.1;HCoV-HKU1_orf1ab;3745 | 0 | 0 |
| NC_006577.2;YP_173236.1;HCoV-HKU1_orf1ab;3746 | 0 | 0 |
| NC_006577.2;YP_173236.1;HCoV-HKU1_orf1ab;3747 | 0 | 0 |
| NC_006577.2;YP_173236.1;HCoV-HKU1_orf1ab;3748 | 0 | 0 |
| NC_006577.2;YP_173236.1;HCoV-HKU1_orf1ab;3749 | 0 | 0 |
| NC_006577.2;YP_173236.1;HCoV-HKU1_orf1ab;3750 | 0 | 0 |
| NC_006577.2;YP_173236.1;HCoV-HKU1_orf1ab;3751 | 0 | 0 |
| NC_006577.2;YP_173236.1;HCoV-HKU1_orf1ab;3752 | 0 | 0 |
| NC_006577.2;YP_173236.1;HCoV-HKU1_orf1ab;3753 | 0 | 0 |
| NC_006577.2;YP_173236.1;HCoV-HKU1_orf1ab;3754 | 0 | 0 |
| NC_006577.2;YP_173236.1;HCoV-HKU1_orf1ab;3755 | 0 | 0 |
| NC_006577.2;YP_173236.1;HCoV-HKU1_orf1ab;3756 | 0 | 0 |

| | | |
|---|---|---|
| NC_006577.2;YP_173236.1;HCoV-HKU1_orf1ab;3757 | 0 | 0 |
| NC_006577.2;YP_173236.1;HCoV-HKU1_orf1ab;3758 | 0 | 0 |
| NC_006577.2;YP_173236.1;HCoV-HKU1_orf1ab;3759 | 0 | 0 |
| NC_006577.2;YP_173236.1;HCoV-HKU1_orf1ab;3760 | 0 | 0 |
| NC_006577.2;YP_173236.1;HCoV-HKU1_orf1ab;3761 | 0 | 0 |
| NC_006577.2;YP_173236.1;HCoV-HKU1_orf1ab;3762 | 0 | 0 |
| NC_006577.2;YP_173236.1;HCoV-HKU1_orf1ab;3763 | 0 | 0 |
| NC_006577.2;YP_173236.1;HCoV-HKU1_orf1ab;3764 | 0 | 0 |
| NC_006577.2;YP_173236.1;HCoV-HKU1_orf1ab;3765 | 0 | 0 |
| NC_006577.2;YP_173236.1;HCoV-HKU1_orf1ab;3766 | 0 | 0 |
| NC_006577.2;YP_173236.1;HCoV-HKU1_orf1ab;3767 | 0 | 0 |
| NC_006577.2;YP_173236.1;HCoV-HKU1_orf1ab;3768 | 0 | 0 |
| NC_006577.2;YP_173236.1;HCoV-HKU1_orf1ab;3769 | 0 | 0 |
| NC_006577.2;YP_173236.1;HCoV-HKU1_orf1ab;3770 | 0 | 0 |
| NC_006577.2;YP_173236.1;HCoV-HKU1_orf1ab;3771 | 0 | 0 |
| NC_006577.2;YP_173236.1;HCoV-HKU1_orf1ab;3772 | 0 | 0 |
| NC_006577.2;YP_173236.1;HCoV-HKU1_orf1ab;3773 | 0 | 0 |
| NC_006577.2;YP_173236.1;HCoV-HKU1_orf1ab;3774 | 0 | 0 |
| NC_006577.2;YP_173236.1;HCoV-HKU1_orf1ab;3775 | 0 | 0 |
| NC_006577.2;YP_173236.1;HCoV-HKU1_orf1ab;3776 | 0 | 0 |
| NC_006577.2;YP_173236.1;HCoV-HKU1_orf1ab;3777 | 0 | 0 |
| NC_006577.2;YP_173236.1;HCoV-HKU1_orf1ab;3778 | 0 | 0 |
| NC_006577.2;YP_173236.1;HCoV-HKU1_orf1ab;3779 | 0 | 0 |
| NC_006577.2;YP_173236.1;HCoV-HKU1_orf1ab;3780 | 0 | 0 |
| NC_006577.2;YP_173236.1;HCoV-HKU1_orf1ab;3781 | 0 | 0 |
| NC_006577.2;YP_173236.1;HCoV-HKU1_orf1ab;3782 | 0 | 0 |
| NC_006577.2;YP_173236.1;HCoV-HKU1_orf1ab;3783 | 0 | 0 |
| NC_006577.2;YP_173236.1;HCoV-HKU1_orf1ab;3784 | 0 | 0 |
| NC_006577.2;YP_173236.1;HCoV-HKU1_orf1ab;3785 | 0 | 0 |
| NC_006577.2;YP_173236.1;HCoV-HKU1_orf1ab;3786 | 0 | 0 |
| NC_006577.2;YP_173236.1;HCoV-HKU1_orf1ab;3787 | 0 | 0 |
| NC_006577.2;YP_173236.1;HCoV-HKU1_orf1ab;3788 | 0 | 0 |
| NC_006577.2;YP_173236.1;HCoV-HKU1_orf1ab;3789 | 0 | 0 |
| NC_006577.2;YP_173236.1;HCoV-HKU1_orf1ab;3790 | 0 | 0 |
| NC_006577.2;YP_173236.1;HCoV-HKU1_orf1ab;3791 | 0 | 0 |
| NC_006577.2;YP_173236.1;HCoV-HKU1_orf1ab;3792 | 0 | 0 |
| NC_006577.2;YP_173236.1;HCoV-HKU1_orf1ab;3793 | 0 | 0 |
| NC_006577.2;YP_173236.1;HCoV-HKU1_orf1ab;3794 | 0 | 0 |
| NC_006577.2;YP_173236.1;HCoV-HKU1_orf1ab;3795 | 0 | 0 |
| NC_006577.2;YP_173236.1;HCoV-HKU1_orf1ab;3796 | 0 | 0 |
| NC_006577.2;YP_173236.1;HCoV-HKU1_orf1ab;3797 | 0 | 0 |
| NC_006577.2;YP_173236.1;HCoV-HKU1_orf1ab;3798 | 0 | 0 |
| NC_006577.2;YP_173236.1;HCoV-HKU1_orf1ab;3799 | 0 | 0 |

| | | |
|---|---|---|
| NC_006577.2;YP_173236.1;HCoV-HKU1_orf1ab;3800 | 0 | 0 |
| NC_006577.2;YP_173236.1;HCoV-HKU1_orf1ab;3801 | 0 | 0 |
| NC_006577.2;YP_173236.1;HCoV-HKU1_orf1ab;3802 | 0 | 0 |
| NC_006577.2;YP_173236.1;HCoV-HKU1_orf1ab;3803 | 0 | 0 |
| NC_006577.2;YP_173236.1;HCoV-HKU1_orf1ab;3804 | 0 | 0 |
| NC_006577.2;YP_173236.1;HCoV-HKU1_orf1ab;3805 | 0 | 0 |
| NC_006577.2;YP_173236.1;HCoV-HKU1_orf1ab;3806 | 0 | 0 |
| NC_006577.2;YP_173236.1;HCoV-HKU1_orf1ab;3807 | 0 | 0 |
| NC_006577.2;YP_173236.1;HCoV-HKU1_orf1ab;3808 | 0 | 0 |
| NC_006577.2;YP_173236.1;HCoV-HKU1_orf1ab;3809 | 0 | 0 |
| NC_006577.2;YP_173236.1;HCoV-HKU1_orf1ab;3810 | 0 | 0 |
| NC_006577.2;YP_173236.1;HCoV-HKU1_orf1ab;3811 | 0 | 0 |
| NC_006577.2;YP_173236.1;HCoV-HKU1_orf1ab;3812 | 0 | 0 |
| NC_006577.2;YP_173236.1;HCoV-HKU1_orf1ab;3813 | 0 | 0 |
| NC_006577.2;YP_173236.1;HCoV-HKU1_orf1ab;3814 | 0 | 0 |
| NC_006577.2;YP_173236.1;HCoV-HKU1_orf1ab;3815 | 0 | 0 |
| NC_006577.2;YP_173236.1;HCoV-HKU1_orf1ab;3816 | 0 | 0 |
| NC_006577.2;YP_173236.1;HCoV-HKU1_orf1ab;3817 | 0 | 0 |
| NC_006577.2;YP_173236.1;HCoV-HKU1_orf1ab;3818 | 0 | 0 |
| NC_006577.2;YP_173236.1;HCoV-HKU1_orf1ab;3819 | 0 | 0 |
| NC_006577.2;YP_173236.1;HCoV-HKU1_orf1ab;3820 | 0 | 0 |
| NC_006577.2;YP_173236.1;HCoV-HKU1_orf1ab;3821 | 0 | 0 |
| NC_006577.2;YP_173236.1;HCoV-HKU1_orf1ab;3822 | 0 | 0 |
| NC_006577.2;YP_173236.1;HCoV-HKU1_orf1ab;3823 | 0 | 0 |
| NC_006577.2;YP_173236.1;HCoV-HKU1_orf1ab;3824 | 0 | 0 |
| NC_006577.2;YP_173236.1;HCoV-HKU1_orf1ab;3825 | 0 | 0 |
| NC_006577.2;YP_173236.1;HCoV-HKU1_orf1ab;3826 | 0 | 0 |
| NC_006577.2;YP_173236.1;HCoV-HKU1_orf1ab;3827 | 0 | 0 |
| NC_006577.2;YP_173236.1;HCoV-HKU1_orf1ab;3828 | 0 | 0 |
| NC_006577.2;YP_173236.1;HCoV-HKU1_orf1ab;3829 | 0 | 0 |
| NC_006577.2;YP_173236.1;HCoV-HKU1_orf1ab;3830 | 0 | 0 |
| NC_006577.2;YP_173236.1;HCoV-HKU1_orf1ab;3831 | 0 | 0 |
| NC_006577.2;YP_173236.1;HCoV-HKU1_orf1ab;3832 | 0 | 0 |
| NC_006577.2;YP_173236.1;HCoV-HKU1_orf1ab;3833 | 0 | 0 |
| NC_006577.2;YP_173236.1;HCoV-HKU1_orf1ab;3834 | 0 | 0 |
| NC_006577.2;YP_173236.1;HCoV-HKU1_orf1ab;3835 | 0 | 0 |
| NC_006577.2;YP_173236.1;HCoV-HKU1_orf1ab;3836 | 0 | 0 |
| NC_006577.2;YP_173236.1;HCoV-HKU1_orf1ab;3837 | 0 | 0 |
| NC_006577.2;YP_173236.1;HCoV-HKU1_orf1ab;3838 | 0 | 0 |
| NC_006577.2;YP_173236.1;HCoV-HKU1_orf1ab;3839 | 0 | 0 |
| NC_006577.2;YP_173236.1;HCoV-HKU1_orf1ab;3840 | 0 | 0 |
| NC_006577.2;YP_173236.1;HCoV-HKU1_orf1ab;3841 | 0 | 0 |
| NC_006577.2;YP_173236.1;HCoV-HKU1_orf1ab;3842 | 0 | 0 |

| | | |
|---|---|---|
| NC_006577.2;YP_173236.1;HCoV-HKU1_orf1ab;3843 | 0 | 0 |
| NC_006577.2;YP_173236.1;HCoV-HKU1_orf1ab;3844 | 0 | 0 |
| NC_006577.2;YP_173236.1;HCoV-HKU1_orf1ab;3845 | 0 | 0 |
| NC_006577.2;YP_173236.1;HCoV-HKU1_orf1ab;3846 | 0 | 0 |
| NC_006577.2;YP_173236.1;HCoV-HKU1_orf1ab;3847 | 0 | 0 |
| NC_006577.2;YP_173236.1;HCoV-HKU1_orf1ab;3848 | 0 | 0 |
| NC_006577.2;YP_173236.1;HCoV-HKU1_orf1ab;3849 | 0 | 0 |
| NC_006577.2;YP_173236.1;HCoV-HKU1_orf1ab;3850 | 0 | 0 |
| NC_006577.2;YP_173236.1;HCoV-HKU1_orf1ab;3851 | 0 | 0 |
| NC_006577.2;YP_173236.1;HCoV-HKU1_orf1ab;3852 | 0 | 0 |
| NC_006577.2;YP_173236.1;HCoV-HKU1_orf1ab;3853 | 0 | 0 |
| NC_006577.2;YP_173236.1;HCoV-HKU1_orf1ab;3854 | 0 | 0 |
| NC_006577.2;YP_173236.1;HCoV-HKU1_orf1ab;3855 | 0 | 0 |
| NC_006577.2;YP_173236.1;HCoV-HKU1_orf1ab;3856 | 0 | 0 |
| NC_006577.2;YP_173236.1;HCoV-HKU1_orf1ab;3857 | 0 | 0 |
| NC_006577.2;YP_173236.1;HCoV-HKU1_orf1ab;3858 | 0 | 0 |
| NC_006577.2;YP_173236.1;HCoV-HKU1_orf1ab;3859 | 0 | 0 |
| NC_006577.2;YP_173236.1;HCoV-HKU1_orf1ab;3860 | 0 | 0 |
| NC_006577.2;YP_173236.1;HCoV-HKU1_orf1ab;3861 | 0 | 0 |
| NC_006577.2;YP_173236.1;HCoV-HKU1_orf1ab;3862 | 0 | 0 |
| NC_006577.2;YP_173236.1;HCoV-HKU1_orf1ab;3863 | 0 | 0 |
| NC_006577.2;YP_173236.1;HCoV-HKU1_orf1ab;3864 | 0 | 0 |
| NC_006577.2;YP_173236.1;HCoV-HKU1_orf1ab;3865 | 0 | 0 |
| NC_006577.2;YP_173236.1;HCoV-HKU1_orf1ab;3866 | 0 | 0 |
| NC_006577.2;YP_173236.1;HCoV-HKU1_orf1ab;3867 | 0 | 0 |
| NC_006577.2;YP_173236.1;HCoV-HKU1_orf1ab;3868 | 0 | 0 |
| NC_006577.2;YP_173236.1;HCoV-HKU1_orf1ab;3869 | 0 | 0 |
| NC_006577.2;YP_173236.1;HCoV-HKU1_orf1ab;3870 | 0 | 0 |
| NC_006577.2;YP_173236.1;HCoV-HKU1_orf1ab;3871 | 0 | 0 |
| NC_006577.2;YP_173236.1;HCoV-HKU1_orf1ab;3872 | 0 | 0 |
| NC_006577.2;YP_173236.1;HCoV-HKU1_orf1ab;3873 | 0 | 0 |
| NC_006577.2;YP_173236.1;HCoV-HKU1_orf1ab;3874 | 0 | 0 |
| NC_006577.2;YP_173236.1;HCoV-HKU1_orf1ab;3875 | 0 | 0 |
| NC_006577.2;YP_173236.1;HCoV-HKU1_orf1ab;3876 | 0 | 0 |
| NC_006577.2;YP_173236.1;HCoV-HKU1_orf1ab;3877 | 0 | 0 |
| NC_006577.2;YP_173236.1;HCoV-HKU1_orf1ab;3878 | 0 | 0 |
| NC_006577.2;YP_173236.1;HCoV-HKU1_orf1ab;3879 | 0 | 0 |
| NC_006577.2;YP_173236.1;HCoV-HKU1_orf1ab;3880 | 0.05 | 0.1 |
| NC_006577.2;YP_173236.1;HCoV-HKU1_orf1ab;3881 | 0 | 0 |
| NC_006577.2;YP_173236.1;HCoV-HKU1_orf1ab;3882 | 0 | 0 |
| NC_006577.2;YP_173236.1;HCoV-HKU1_orf1ab;3883 | 0 | 0 |
| NC_006577.2;YP_173236.1;HCoV-HKU1_orf1ab;3884 | 0.1 | 0.025 |
| NC_006577.2;YP_173236.1;HCoV-HKU1_orf1ab;3885 | 0 | 0 |

| | | |
|---|---|---|
| NC_006577.2;YP_173236.1;HCoV-HKU1_orf1ab;3886 | 0 | 0.025 |
| NC_006577.2;YP_173236.1;HCoV-HKU1_orf1ab;3887 | 0 | 0 |
| NC_006577.2;YP_173236.1;HCoV-HKU1_orf1ab;3888 | 0 | 0 |
| NC_006577.2;YP_173236.1;HCoV-HKU1_orf1ab;3889 | 0 | 0 |
| NC_006577.2;YP_173236.1;HCoV-HKU1_orf1ab;3890 | 0.05 | 0 |
| NC_006577.2;YP_173236.1;HCoV-HKU1_orf1ab;3891 | 0 | 0 |
| NC_006577.2;YP_173236.1;HCoV-HKU1_orf1ab;3892 | 0 | 0 |
| NC_006577.2;YP_173236.1;HCoV-HKU1_orf1ab;3893 | 0 | 0 |
| NC_006577.2;YP_173236.1;HCoV-HKU1_orf1ab;3894 | 0 | 0 |
| NC_006577.2;YP_173236.1;HCoV-HKU1_orf1ab;3895 | 0 | 0 |
| NC_006577.2;YP_173236.1;HCoV-HKU1_orf1ab;3896 | 0 | 0 |
| NC_006577.2;YP_173236.1;HCoV-HKU1_orf1ab;3897 | 0 | 0 |
| NC_006577.2;YP_173236.1;HCoV-HKU1_orf1ab;3898 | 0 | 0 |
| NC_006577.2;YP_173236.1;HCoV-HKU1_orf1ab;3899 | 0 | 0 |
| NC_006577.2;YP_173236.1;HCoV-HKU1_orf1ab;3900 | 0 | 0 |
| NC_006577.2;YP_173236.1;HCoV-HKU1_orf1ab;3901 | 0 | 0 |
| NC_006577.2;YP_173236.1;HCoV-HKU1_orf1ab;3902 | 0 | 0 |
| NC_006577.2;YP_173236.1;HCoV-HKU1_orf1ab;3903 | 0 | 0 |
| NC_006577.2;YP_173236.1;HCoV-HKU1_orf1ab;3904 | 0 | 0 |
| NC_006577.2;YP_173236.1;HCoV-HKU1_orf1ab;3905 | 0 | 0 |
| NC_006577.2;YP_173236.1;HCoV-HKU1_orf1ab;3906 | 0 | 0 |
| NC_006577.2;YP_173236.1;HCoV-HKU1_orf1ab;3907 | 0 | 0 |
| NC_006577.2;YP_173236.1;HCoV-HKU1_orf1ab;3908 | 0 | 0 |
| NC_006577.2;YP_173236.1;HCoV-HKU1_orf1ab;3909 | 0 | 0 |
| NC_006577.2;YP_173236.1;HCoV-HKU1_orf1ab;3910 | 0.05 | 0 |
| NC_006577.2;YP_173236.1;HCoV-HKU1_orf1ab;3911 | 0.4 | 0.225 |
| NC_006577.2;YP_173236.1;HCoV-HKU1_orf1ab;3912 | 0.35 | 0.2 |
| NC_006577.2;YP_173236.1;HCoV-HKU1_orf1ab;3913 | 0.5 | 0.2 |
| NC_006577.2;YP_173236.1;HCoV-HKU1_orf1ab;3914 | 0.4 | 0.2 |
| NC_006577.2;YP_173236.1;HCoV-HKU1_orf1ab;3915 | 0.4 | 0.225 |
| NC_006577.2;YP_173236.1;HCoV-HKU1_orf1ab;3916 | 0.45 | 0.25 |
| NC_006577.2;YP_173236.1;HCoV-HKU1_orf1ab;3917 | 0.35 | 0.225 |
| NC_006577.2;YP_173236.1;HCoV-HKU1_orf1ab;3918 | 0.45 | 0.2 |
| NC_006577.2;YP_173236.1;HCoV-HKU1_orf1ab;3919 | 0.4 | 0.225 |
| NC_006577.2;YP_173236.1;HCoV-HKU1_orf1ab;3920 | 0.05 | 0.025 |
| NC_006577.2;YP_173236.1;HCoV-HKU1_orf1ab;3921 | 0 | 0.025 |
| NC_006577.2;YP_173236.1;HCoV-HKU1_orf1ab;3922 | 0.05 | 0.025 |
| NC_006577.2;YP_173236.1;HCoV-HKU1_orf1ab;3923 | 0 | 0 |
| NC_006577.2;YP_173236.1;HCoV-HKU1_orf1ab;3924 | 0 | 0 |
| NC_006577.2;YP_173236.1;HCoV-HKU1_orf1ab;3925 | 0 | 0 |
| NC_006577.2;YP_173236.1;HCoV-HKU1_orf1ab;3926 | 0.05 | 0.025 |
| NC_006577.2;YP_173236.1;HCoV-HKU1_orf1ab;3927 | 0.15 | 0.025 |
| NC_006577.2;YP_173236.1;HCoV-HKU1_orf1ab;3928 | 0 | 0 |

| | | |
|---|---|---|
| NC_006577.2;YP_173236.1;HCoV-HKU1_orf1ab;3929 | 0 | 0 |
| NC_006577.2;YP_173236.1;HCoV-HKU1_orf1ab;3930 | 0 | 0 |
| NC_006577.2;YP_173236.1;HCoV-HKU1_orf1ab;3931 | 0 | 0 |
| NC_006577.2;YP_173236.1;HCoV-HKU1_orf1ab;3932 | 0 | 0 |
| NC_006577.2;YP_173236.1;HCoV-HKU1_orf1ab;3933 | 0 | 0 |
| NC_006577.2;YP_173236.1;HCoV-HKU1_orf1ab;3934 | 0 | 0 |
| NC_006577.2;YP_173236.1;HCoV-HKU1_orf1ab;3935 | 0 | 0 |
| NC_006577.2;YP_173236.1;HCoV-HKU1_orf1ab;3936 | 0 | 0.05 |
| NC_006577.2;YP_173236.1;HCoV-HKU1_orf1ab;3937 | 0 | 0.075 |
| NC_006577.2;YP_173236.1;HCoV-HKU1_orf1ab;3938 | 0.05 | 0.05 |
| NC_006577.2;YP_173236.1;HCoV-HKU1_orf1ab;3939 | 0.05 | 0.025 |
| NC_006577.2;YP_173236.1;HCoV-HKU1_orf1ab;3940 | 0.05 | 0.025 |
| NC_006577.2;YP_173236.1;HCoV-HKU1_orf1ab;3941 | 0.05 | 0.025 |
| NC_006577.2;YP_173236.1;HCoV-HKU1_orf1ab;3942 | 0.05 | 0.025 |
| NC_006577.2;YP_173236.1;HCoV-HKU1_orf1ab;3943 | 0.05 | 0.025 |
| NC_006577.2;YP_173236.1;HCoV-HKU1_orf1ab;3944 | 0.05 | 0 |
| NC_006577.2;YP_173236.1;HCoV-HKU1_orf1ab;3945 | 0.05 | 0 |
| NC_006577.2;YP_173236.1;HCoV-HKU1_orf1ab;3946 | 0.05 | 0.025 |
| NC_006577.2;YP_173236.1;HCoV-HKU1_orf1ab;3947 | 0.1 | 0 |
| NC_006577.2;YP_173236.1;HCoV-HKU1_orf1ab;3948 | 0 | 0 |
| NC_006577.2;YP_173236.1;HCoV-HKU1_orf1ab;3949 | 0 | 0 |
| NC_006577.2;YP_173236.1;HCoV-HKU1_orf1ab;3950 | 0 | 0 |
| NC_006577.2;YP_173236.1;HCoV-HKU1_orf1ab;3951 | 0 | 0 |
| NC_006577.2;YP_173236.1;HCoV-HKU1_orf1ab;3952 | 0 | 0 |
| NC_006577.2;YP_173236.1;HCoV-HKU1_orf1ab;3953 | 0 | 0 |
| NC_006577.2;YP_173236.1;HCoV-HKU1_orf1ab;3954 | 0 | 0 |
| NC_006577.2;YP_173236.1;HCoV-HKU1_orf1ab;3955 | 0 | 0 |
| NC_006577.2;YP_173236.1;HCoV-HKU1_orf1ab;3956 | 0 | 0 |
| NC_006577.2;YP_173236.1;HCoV-HKU1_orf1ab;3957 | 0.05 | 0 |
| NC_006577.2;YP_173236.1;HCoV-HKU1_orf1ab;3958 | 0 | 0 |
| NC_006577.2;YP_173236.1;HCoV-HKU1_orf1ab;3959 | 0 | 0 |
| NC_006577.2;YP_173236.1;HCoV-HKU1_orf1ab;3960 | 0 | 0 |
| NC_006577.2;YP_173236.1;HCoV-HKU1_orf1ab;3961 | 0 | 0 |
| NC_006577.2;YP_173236.1;HCoV-HKU1_orf1ab;3962 | 0 | 0 |
| NC_006577.2;YP_173236.1;HCoV-HKU1_orf1ab;3963 | 0 | 0.025 |
| NC_006577.2;YP_173236.1;HCoV-HKU1_orf1ab;3964 | 0 | 0 |
| NC_006577.2;YP_173236.1;HCoV-HKU1_orf1ab;3965 | 0 | 0.025 |
| NC_006577.2;YP_173236.1;HCoV-HKU1_orf1ab;3966 | 0 | 0 |
| NC_006577.2;YP_173236.1;HCoV-HKU1_orf1ab;3967 | 0 | 0 |
| NC_006577.2;YP_173236.1;HCoV-HKU1_orf1ab;3968 | 0 | 0 |
| NC_006577.2;YP_173236.1;HCoV-HKU1_orf1ab;3969 | 0 | 0 |
| NC_006577.2;YP_173236.1;HCoV-HKU1_orf1ab;3970 | 0 | 0 |
| NC_006577.2;YP_173236.1;HCoV-HKU1_orf1ab;3971 | 0 | 0 |

| | | |
|---|---|---|
| NC_006577.2;YP_173236.1;HCoV-HKU1_orf1ab;3972 | 0 | 0 |
| NC_006577.2;YP_173236.1;HCoV-HKU1_orf1ab;3973 | 0 | 0 |
| NC_006577.2;YP_173236.1;HCoV-HKU1_orf1ab;3974 | 0 | 0 |
| NC_006577.2;YP_173236.1;HCoV-HKU1_orf1ab;3975 | 0 | 0 |
| NC_006577.2;YP_173236.1;HCoV-HKU1_orf1ab;3976 | 0 | 0 |
| NC_006577.2;YP_173236.1;HCoV-HKU1_orf1ab;3977 | 0 | 0 |
| NC_006577.2;YP_173236.1;HCoV-HKU1_orf1ab;3978 | 0 | 0 |
| NC_006577.2;YP_173236.1;HCoV-HKU1_orf1ab;3979 | 0 | 0 |
| NC_006577.2;YP_173236.1;HCoV-HKU1_orf1ab;3980 | 0 | 0 |
| NC_006577.2;YP_173236.1;HCoV-HKU1_orf1ab;3981 | 0 | 0 |
| NC_006577.2;YP_173236.1;HCoV-HKU1_orf1ab;3982 | 0 | 0 |
| NC_006577.2;YP_173236.1;HCoV-HKU1_orf1ab;3983 | 0 | 0 |
| NC_006577.2;YP_173236.1;HCoV-HKU1_orf1ab;3984 | 0 | 0 |
| NC_006577.2;YP_173236.1;HCoV-HKU1_orf1ab;3985 | 0 | 0 |
| NC_006577.2;YP_173236.1;HCoV-HKU1_orf1ab;3986 | 0 | 0 |
| NC_006577.2;YP_173236.1;HCoV-HKU1_orf1ab;3987 | 0.15 | 0.025 |
| NC_006577.2;YP_173236.1;HCoV-HKU1_orf1ab;3988 | 0.05 | 0 |
| NC_006577.2;YP_173236.1;HCoV-HKU1_orf1ab;3989 | 0 | 0 |
| NC_006577.2;YP_173236.1;HCoV-HKU1_orf1ab;3990 | 0 | 0 |
| NC_006577.2;YP_173236.1;HCoV-HKU1_orf1ab;3991 | 0 | 0 |
| NC_006577.2;YP_173236.1;HCoV-HKU1_orf1ab;3992 | 0 | 0 |
| NC_006577.2;YP_173236.1;HCoV-HKU1_orf1ab;3993 | 0 | 0 |
| NC_006577.2;YP_173236.1;HCoV-HKU1_orf1ab;3994 | 0 | 0 |
| NC_006577.2;YP_173236.1;HCoV-HKU1_orf1ab;3995 | 0 | 0 |
| NC_006577.2;YP_173236.1;HCoV-HKU1_orf1ab;3996 | 0 | 0 |
| NC_006577.2;YP_173236.1;HCoV-HKU1_orf1ab;3997 | 0.05 | 0.025 |
| NC_006577.2;YP_173236.1;HCoV-HKU1_orf1ab;3998 | 0 | 0 |
| NC_006577.2;YP_173236.1;HCoV-HKU1_orf1ab;3999 | 0 | 0 |
| NC_006577.2;YP_173236.1;HCoV-HKU1_orf1ab;4000 | 0 | 0 |
| NC_006577.2;YP_173236.1;HCoV-HKU1_orf1ab;4001 | 0 | 0 |
| NC_006577.2;YP_173236.1;HCoV-HKU1_orf1ab;4002 | 0.1 | 0 |
| NC_006577.2;YP_173236.1;HCoV-HKU1_orf1ab;4003 | 0.05 | 0 |
| NC_006577.2;YP_173236.1;HCoV-HKU1_orf1ab;4004 | 0.05 | 0 |
| NC_006577.2;YP_173236.1;HCoV-HKU1_orf1ab;4005 | 0 | 0 |
| NC_006577.2;YP_173236.1;HCoV-HKU1_orf1ab;4006 | 0 | 0 |
| NC_006577.2;YP_173236.1;HCoV-HKU1_orf1ab;4007 | 0 | 0 |
| NC_006577.2;YP_173236.1;HCoV-HKU1_orf1ab;4008 | 0.05 | 0 |
| NC_006577.2;YP_173236.1;HCoV-HKU1_orf1ab;4009 | 0 | 0 |
| NC_006577.2;YP_173236.1;HCoV-HKU1_orf1ab;4010 | 0 | 0 |
| NC_006577.2;YP_173236.1;HCoV-HKU1_orf1ab;4011 | 0 | 0 |
| NC_006577.2;YP_173236.1;HCoV-HKU1_orf1ab;4012 | 0 | 0.025 |
| NC_006577.2;YP_173236.1;HCoV-HKU1_orf1ab;4013 | 0 | 0 |
| NC_006577.2;YP_173236.1;HCoV-HKU1_orf1ab;4014 | 0 | 0 |

| | | |
|---|---|---|
| NC_006577.2;YP_173236.1;HCoV-HKU1_orf1ab;4015 | 0 | 0 |
| NC_006577.2;YP_173236.1;HCoV-HKU1_orf1ab;4016 | 0 | 0 |
| NC_006577.2;YP_173236.1;HCoV-HKU1_orf1ab;4017 | 0 | 0 |
| NC_006577.2;YP_173236.1;HCoV-HKU1_orf1ab;4018 | 0.05 | 0 |
| NC_006577.2;YP_173236.1;HCoV-HKU1_orf1ab;4019 | 0.05 | 0.025 |
| NC_006577.2;YP_173236.1;HCoV-HKU1_orf1ab;4020 | 0 | 0 |
| NC_006577.2;YP_173236.1;HCoV-HKU1_orf1ab;4021 | 0.05 | 0.025 |
| NC_006577.2;YP_173236.1;HCoV-HKU1_orf1ab;4022 | 0.05 | 0.025 |
| NC_006577.2;YP_173236.1;HCoV-HKU1_orf1ab;4023 | 0.1 | 0.075 |
| NC_006577.2;YP_173236.1;HCoV-HKU1_orf1ab;4024 | 0.1 | 0.075 |
| NC_006577.2;YP_173236.1;HCoV-HKU1_orf1ab;4025 | 0.15 | 0.05 |
| NC_006577.2;YP_173236.1;HCoV-HKU1_orf1ab;4026 | 0.1 | 0.025 |
| NC_006577.2;YP_173236.1;HCoV-HKU1_orf1ab;4027 | 0.05 | 0.05 |
| NC_006577.2;YP_173236.1;HCoV-HKU1_orf1ab;4028 | 0.05 | 0 |
| NC_006577.2;YP_173236.1;HCoV-HKU1_orf1ab;4029 | 0.05 | 0 |
| NC_006577.2;YP_173236.1;HCoV-HKU1_orf1ab;4030 | 0.1 | 0 |
| NC_006577.2;YP_173236.1;HCoV-HKU1_orf1ab;4031 | 0.05 | 0 |
| NC_006577.2;YP_173236.1;HCoV-HKU1_orf1ab;4032 | 0.05 | 0.025 |
| NC_006577.2;YP_173236.1;HCoV-HKU1_orf1ab;4033 | 0 | 0 |
| NC_006577.2;YP_173236.1;HCoV-HKU1_orf1ab;4034 | 0.05 | 0 |
| NC_006577.2;YP_173236.1;HCoV-HKU1_orf1ab;4035 | 0.15 | 0 |
| NC_006577.2;YP_173236.1;HCoV-HKU1_orf1ab;4036 | 0.1 | 0 |
| NC_006577.2;YP_173236.1;HCoV-HKU1_orf1ab;4037 | 0.05 | 0 |
| NC_006577.2;YP_173236.1;HCoV-HKU1_orf1ab;4038 | 0.05 | 0 |
| NC_006577.2;YP_173236.1;HCoV-HKU1_orf1ab;4039 | 0 | 0 |
| NC_006577.2;YP_173236.1;HCoV-HKU1_orf1ab;4040 | 0 | 0 |
| NC_006577.2;YP_173236.1;HCoV-HKU1_orf1ab;4041 | 0 | 0 |
| NC_006577.2;YP_173236.1;HCoV-HKU1_orf1ab;4042 | 0 | 0 |
| NC_006577.2;YP_173236.1;HCoV-HKU1_orf1ab;4043 | 0 | 0 |
| NC_006577.2;YP_173236.1;HCoV-HKU1_orf1ab;4044 | 0 | 0 |
| NC_006577.2;YP_173236.1;HCoV-HKU1_orf1ab;4045 | 0 | 0 |
| NC_006577.2;YP_173236.1;HCoV-HKU1_orf1ab;4046 | 0 | 0 |
| NC_006577.2;YP_173236.1;HCoV-HKU1_orf1ab;4047 | 0 | 0 |
| NC_006577.2;YP_173236.1;HCoV-HKU1_orf1ab;4048 | 0 | 0 |
| NC_006577.2;YP_173236.1;HCoV-HKU1_orf1ab;4049 | 0 | 0 |
| NC_006577.2;YP_173236.1;HCoV-HKU1_orf1ab;4050 | 0 | 0 |
| NC_006577.2;YP_173236.1;HCoV-HKU1_orf1ab;4051 | 0 | 0 |
| NC_006577.2;YP_173236.1;HCoV-HKU1_orf1ab;4052 | 0 | 0 |
| NC_006577.2;YP_173236.1;HCoV-HKU1_orf1ab;4053 | 0 | 0.025 |
| NC_006577.2;YP_173236.1;HCoV-HKU1_orf1ab;4054 | 0.05 | 0.025 |
| NC_006577.2;YP_173236.1;HCoV-HKU1_orf1ab;4055 | 0.05 | 0.05 |
| NC_006577.2;YP_173236.1;HCoV-HKU1_orf1ab;4056 | 0 | 0 |
| NC_006577.2;YP_173236.1;HCoV-HKU1_orf1ab;4057 | 0 | 0 |

| | | |
|---|---|---|
| NC_006577.2;YP_173236.1;HCoV-HKU1_orf1ab;4058 | 0 | 0 |
| NC_006577.2;YP_173236.1;HCoV-HKU1_orf1ab;4059 | 0 | 0 |
| NC_006577.2;YP_173236.1;HCoV-HKU1_orf1ab;4060 | 0 | 0.05 |
| NC_006577.2;YP_173236.1;HCoV-HKU1_orf1ab;4061 | 0 | 0.025 |
| NC_006577.2;YP_173236.1;HCoV-HKU1_orf1ab;4062 | 0 | 0.025 |
| NC_006577.2;YP_173236.1;HCoV-HKU1_orf1ab;4063 | 0 | 0.05 |
| NC_006577.2;YP_173236.1;HCoV-HKU1_orf1ab;4064 | 0 | 0 |
| NC_006577.2;YP_173236.1;HCoV-HKU1_orf1ab;4065 | 0.05 | 0.025 |
| NC_006577.2;YP_173236.1;HCoV-HKU1_orf1ab;4066 | 0.05 | 0 |
| NC_006577.2;YP_173236.1;HCoV-HKU1_orf1ab;4067 | 0 | 0 |
| NC_006577.2;YP_173236.1;HCoV-HKU1_orf1ab;4068 | 0 | 0.025 |
| NC_006577.2;YP_173236.1;HCoV-HKU1_orf1ab;4069 | 0 | 0 |
| NC_006577.2;YP_173236.1;HCoV-HKU1_orf1ab;4070 | 0 | 0.025 |
| NC_006577.2;YP_173236.1;HCoV-HKU1_orf1ab;4071 | 0 | 0.025 |
| NC_006577.2;YP_173236.1;HCoV-HKU1_orf1ab;4072 | 0 | 0.05 |
| NC_006577.2;YP_173236.1;HCoV-HKU1_orf1ab;4073 | 0 | 0 |
| NC_006577.2;YP_173236.1;HCoV-HKU1_orf1ab;4074 | 0.05 | 0.025 |
| NC_006577.2;YP_173236.1;HCoV-HKU1_orf1ab;4075 | 0 | 0.025 |
| NC_006577.2;YP_173236.1;HCoV-HKU1_orf1ab;4076 | 0 | 0.025 |
| NC_006577.2;YP_173236.1;HCoV-HKU1_orf1ab;4077 | 0 | 0.025 |
| NC_006577.2;YP_173236.1;HCoV-HKU1_orf1ab;4078 | 0.05 | 0.025 |
| NC_006577.2;YP_173236.1;HCoV-HKU1_orf1ab;4079 | 0 | 0.025 |
| NC_006577.2;YP_173236.1;HCoV-HKU1_orf1ab;4080 | 0 | 0 |
| NC_006577.2;YP_173236.1;HCoV-HKU1_orf1ab;4081 | 0 | 0 |
| NC_006577.2;YP_173236.1;HCoV-HKU1_orf1ab;4082 | 0 | 0 |
| NC_006577.2;YP_173236.1;HCoV-HKU1_orf1ab;4083 | 0 | 0 |
| NC_006577.2;YP_173236.1;HCoV-HKU1_orf1ab;4084 | 0 | 0 |
| NC_006577.2;YP_173236.1;HCoV-HKU1_orf1ab;4085 | 0 | 0 |
| NC_006577.2;YP_173236.1;HCoV-HKU1_orf1ab;4086 | 0 | 0 |
| NC_006577.2;YP_173236.1;HCoV-HKU1_orf1ab;4087 | 0.05 | 0 |
| NC_006577.2;YP_173236.1;HCoV-HKU1_orf1ab;4088 | 0 | 0 |
| NC_006577.2;YP_173236.1;HCoV-HKU1_orf1ab;4089 | 0 | 0 |
| NC_006577.2;YP_173236.1;HCoV-HKU1_orf1ab;4090 | 0 | 0 |
| NC_006577.2;YP_173236.1;HCoV-HKU1_orf1ab;4091 | 0 | 0 |
| NC_006577.2;YP_173236.1;HCoV-HKU1_orf1ab;4092 | 0 | 0 |
| NC_006577.2;YP_173236.1;HCoV-HKU1_orf1ab;4093 | 0 | 0.025 |
| NC_006577.2;YP_173236.1;HCoV-HKU1_orf1ab;4094 | 0 | 0 |
| NC_006577.2;YP_173236.1;HCoV-HKU1_orf1ab;4095 | 0 | 0 |
| NC_006577.2;YP_173236.1;HCoV-HKU1_orf1ab;4096 | 0 | 0 |
| NC_006577.2;YP_173236.1;HCoV-HKU1_orf1ab;4097 | 0 | 0 |
| NC_006577.2;YP_173236.1;HCoV-HKU1_orf1ab;4098 | 0 | 0 |
| NC_006577.2;YP_173236.1;HCoV-HKU1_orf1ab;4099 | 0 | 0 |
| NC_006577.2;YP_173236.1;HCoV-HKU1_orf1ab;4100 | 0 | 0 |

| | | |
|---|---|---|
| NC_006577.2;YP_173236.1;HCoV-HKU1_orf1ab;4101 | 0 | 0 |
| NC_006577.2;YP_173236.1;HCoV-HKU1_orf1ab;4102 | 0 | 0 |
| NC_006577.2;YP_173236.1;HCoV-HKU1_orf1ab;4103 | 0.2 | 0.175 |
| NC_006577.2;YP_173236.1;HCoV-HKU1_orf1ab;4104 | 0.1 | 0.1 |
| NC_006577.2;YP_173236.1;HCoV-HKU1_orf1ab;4105 | 0.15 | 0.1 |
| NC_006577.2;YP_173236.1;HCoV-HKU1_orf1ab;4106 | 0.2 | 0.1 |
| NC_006577.2;YP_173236.1;HCoV-HKU1_orf1ab;4107 | 0.25 | 0.3 |
| NC_006577.2;YP_173236.1;HCoV-HKU1_orf1ab;4108 | 0.25 | 0.425 |
| NC_006577.2;YP_173236.1;HCoV-HKU1_orf1ab;4109 | 0.3 | 0.3 |
| NC_006577.2;YP_173236.1;HCoV-HKU1_orf1ab;4110 | 0.35 | 0.4 |
| NC_006577.2;YP_173236.1;HCoV-HKU1_orf1ab;4111 | 0.3 | 0.3 |
| NC_006577.2;YP_173236.1;HCoV-HKU1_orf1ab;4112 | 0.1 | 0 |
| NC_006577.2;YP_173236.1;HCoV-HKU1_orf1ab;4113 | 0.1 | 0.075 |
| NC_006577.2;YP_173236.1;HCoV-HKU1_orf1ab;4114 | 0.05 | 0.025 |
| NC_006577.2;YP_173236.1;HCoV-HKU1_orf1ab;4115 | 0.1 | 0.05 |
| NC_006577.2;YP_173236.1;HCoV-HKU1_orf1ab;4116 | 0.15 | 0.05 |
| NC_006577.2;YP_173236.1;HCoV-HKU1_orf1ab;4117 | 0.05 | 0.025 |
| NC_006577.2;YP_173236.1;HCoV-HKU1_orf1ab;4118 | 0.05 | 0.025 |
| NC_006577.2;YP_173236.1;HCoV-HKU1_orf1ab;4119 | 0 | 0.025 |
| NC_006577.2;YP_173236.1;HCoV-HKU1_orf1ab;4120 | 0 | 0.025 |
| NC_006577.2;YP_173236.1;HCoV-HKU1_orf1ab;4121 | 0 | 0 |
| NC_006577.2;YP_173236.1;HCoV-HKU1_orf1ab;4122 | 0.15 | 0.125 |
| NC_006577.2;YP_173236.1;HCoV-HKU1_orf1ab;4123 | 0 | 0 |
| NC_006577.2;YP_173236.1;HCoV-HKU1_orf1ab;4124 | 0.05 | 0.025 |
| NC_006577.2;YP_173236.1;HCoV-HKU1_orf1ab;4125 | 0 | 0 |
| NC_006577.2;YP_173236.1;HCoV-HKU1_orf1ab;4126 | 0.05 | 0.15 |
| NC_006577.2;YP_173236.1;HCoV-HKU1_orf1ab;4127 | 0 | 0.05 |
| NC_006577.2;YP_173236.1;HCoV-HKU1_orf1ab;4128 | 0 | 0.025 |
| NC_006577.2;YP_173236.1;HCoV-HKU1_orf1ab;4129 | 0 | 0.025 |
| NC_006577.2;YP_173236.1;HCoV-HKU1_orf1ab;4130 | 0 | 0 |
| NC_006577.2;YP_173236.1;HCoV-HKU1_orf1ab;4131 | 0 | 0 |
| NC_006577.2;YP_173236.1;HCoV-HKU1_orf1ab;4132 | 0 | 0 |
| NC_006577.2;YP_173236.1;HCoV-HKU1_orf1ab;4133 | 0 | 0 |
| NC_006577.2;YP_173236.1;HCoV-HKU1_orf1ab;4134 | 0 | 0 |
| NC_006577.2;YP_173236.1;HCoV-HKU1_orf1ab;4135 | 0 | 0.025 |
| NC_006577.2;YP_173236.1;HCoV-HKU1_orf1ab;4136 | 0 | 0 |
| NC_006577.2;YP_173236.1;HCoV-HKU1_orf1ab;4137 | 0 | 0 |
| NC_006577.2;YP_173236.1;HCoV-HKU1_orf1ab;4138 | 0.05 | 0 |
| NC_006577.2;YP_173236.1;HCoV-HKU1_orf1ab;4139 | 0 | 0 |
| NC_006577.2;YP_173236.1;HCoV-HKU1_orf1ab;4140 | 0 | 0 |
| NC_006577.2;YP_173236.1;HCoV-HKU1_orf1ab;4141 | 0.05 | 0 |
| NC_006577.2;YP_173236.1;HCoV-HKU1_orf1ab;4142 | 0.1 | 0.025 |
| NC_006577.2;YP_173236.1;HCoV-HKU1_orf1ab;4143 | 0.1 | 0.025 |

| | | |
|---|---|---|
| NC_006577.2;YP_173236.1;HCoV-HKU1_orf1ab;4144 | 0.1 | 0.025 |
| NC_006577.2;YP_173236.1;HCoV-HKU1_orf1ab;4145 | 0.05 | 0.025 |
| NC_006577.2;YP_173236.1;HCoV-HKU1_orf1ab;4146 | 0.05 | 0.025 |
| NC_006577.2;YP_173236.1;HCoV-HKU1_orf1ab;4147 | 0.05 | 0.1 |
| NC_006577.2;YP_173236.1;HCoV-HKU1_orf1ab;4148 | 0.05 | 0 |
| NC_006577.2;YP_173236.1;HCoV-HKU1_orf1ab;4149 | 0 | 0 |
| NC_006577.2;YP_173236.1;HCoV-HKU1_orf1ab;4150 | 0 | 0.025 |
| NC_006577.2;YP_173236.1;HCoV-HKU1_orf1ab;4151 | 0 | 0.025 |
| NC_006577.2;YP_173236.1;HCoV-HKU1_orf1ab;4152 | 0 | 0.025 |
| NC_006577.2;YP_173236.1;HCoV-HKU1_orf1ab;4153 | 0 | 0 |
| NC_006577.2;YP_173236.1;HCoV-HKU1_orf1ab;4154 | 0 | 0 |
| NC_006577.2;YP_173236.1;HCoV-HKU1_orf1ab;4155 | 0 | 0 |
| NC_006577.2;YP_173236.1;HCoV-HKU1_orf1ab;4156 | 0 | 0 |
| NC_006577.2;YP_173236.1;HCoV-HKU1_orf1ab;4157 | 0 | 0 |
| NC_006577.2;YP_173236.1;HCoV-HKU1_orf1ab;4158 | 0 | 0 |
| NC_006577.2;YP_173236.1;HCoV-HKU1_orf1ab;4159 | 0 | 0 |
| NC_006577.2;YP_173236.1;HCoV-HKU1_orf1ab;4160 | 0.05 | 0.05 |
| NC_006577.2;YP_173236.1;HCoV-HKU1_orf1ab;4161 | 0.1 | 0.05 |
| NC_006577.2;YP_173236.1;HCoV-HKU1_orf1ab;4162 | 0.1 | 0.075 |
| NC_006577.2;YP_173236.1;HCoV-HKU1_orf1ab;4163 | 0.1 | 0.075 |
| NC_006577.2;YP_173236.1;HCoV-HKU1_orf1ab;4164 | 0.1 | 0.1 |
| NC_006577.2;YP_173236.1;HCoV-HKU1_orf1ab;4165 | 0.15 | 0.175 |
| NC_006577.2;YP_173236.1;HCoV-HKU1_orf1ab;4166 | 0.15 | 0.15 |
| NC_006577.2;YP_173236.1;HCoV-HKU1_orf1ab;4167 | 0.15 | 0.15 |
| NC_006577.2;YP_173236.1;HCoV-HKU1_orf1ab;4168 | 0.2 | 0.1 |
| NC_006577.2;YP_173236.1;HCoV-HKU1_orf1ab;4169 | 0.1 | 0.075 |
| NC_006577.2;YP_173236.1;HCoV-HKU1_orf1ab;4170 | 0.15 | 0.225 |
| NC_006577.2;YP_173236.1;HCoV-HKU1_orf1ab;4171 | 0.1 | 0.075 |
| NC_006577.2;YP_173236.1;HCoV-HKU1_orf1ab;4172 | 0.1 | 0.075 |
| NC_006577.2;YP_173236.1;HCoV-HKU1_orf1ab;4173 | 0.15 | 0.05 |
| NC_006577.2;YP_173236.1;HCoV-HKU1_orf1ab;4174 | 0.05 | 0.05 |
| NC_006577.2;YP_173236.1;HCoV-HKU1_orf1ab;4175 | 0.1 | 0.025 |
| NC_006577.2;YP_173236.1;HCoV-HKU1_orf1ab;4176 | 0 | 0 |
| NC_006577.2;YP_173236.1;HCoV-HKU1_orf1ab;4177 | 0 | 0 |
| NC_006577.2;YP_173236.1;HCoV-HKU1_orf1ab;4178 | 0 | 0 |
| NC_006577.2;YP_173236.1;HCoV-HKU1_orf1ab;4179 | 0.05 | 0.025 |
| NC_006577.2;YP_173236.1;HCoV-HKU1_orf1ab;4180 | 0 | 0 |
| NC_006577.2;YP_173236.1;HCoV-HKU1_orf1ab;4181 | 0 | 0 |
| NC_006577.2;YP_173236.1;HCoV-HKU1_orf1ab;4182 | 0 | 0 |
| NC_006577.2;YP_173236.1;HCoV-HKU1_orf1ab;4183 | 0 | 0 |
| NC_006577.2;YP_173236.1;HCoV-HKU1_orf1ab;4184 | 0 | 0 |
| NC_006577.2;YP_173236.1;HCoV-HKU1_orf1ab;4185 | 0 | 0 |
| NC_006577.2;YP_173236.1;HCoV-HKU1_orf1ab;4186 | 0 | 0 |

| | | |
|---|---|---|
| NC_006577.2;YP_173236.1;HCoV-HKU1_orf1ab;4187 | 0 | 0 |
| NC_006577.2;YP_173236.1;HCoV-HKU1_orf1ab;4188 | 0 | 0.025 |
| NC_006577.2;YP_173236.1;HCoV-HKU1_orf1ab;4189 | 0.05 | 0 |
| NC_006577.2;YP_173236.1;HCoV-HKU1_orf1ab;4190 | 0 | 0 |
| NC_006577.2;YP_173236.1;HCoV-HKU1_orf1ab;4191 | 0 | 0 |
| NC_006577.2;YP_173236.1;HCoV-HKU1_orf1ab;4192 | 0 | 0 |
| NC_006577.2;YP_173236.1;HCoV-HKU1_orf1ab;4193 | 0 | 0 |
| NC_006577.2;YP_173236.1;HCoV-HKU1_orf1ab;4194 | 0.3 | 0.175 |
| NC_006577.2;YP_173236.1;HCoV-HKU1_orf1ab;4195 | 0 | 0.05 |
| NC_006577.2;YP_173236.1;HCoV-HKU1_orf1ab;4196 | 0 | 0.075 |
| NC_006577.2;YP_173236.1;HCoV-HKU1_orf1ab;4197 | 0.1 | 0.075 |
| NC_006577.2;YP_173236.1;HCoV-HKU1_orf1ab;4198 | 0.15 | 0.075 |
| NC_006577.2;YP_173236.1;HCoV-HKU1_orf1ab;4199 | 0 | 0.025 |
| NC_006577.2;YP_173236.1;HCoV-HKU1_orf1ab;4200 | 0 | 0.025 |
| NC_006577.2;YP_173236.1;HCoV-HKU1_orf1ab;4201 | 0 | 0 |
| NC_006577.2;YP_173236.1;HCoV-HKU1_orf1ab;4202 | 0 | 0 |
| NC_006577.2;YP_173236.1;HCoV-HKU1_orf1ab;4203 | 0 | 0 |
| NC_006577.2;YP_173236.1;HCoV-HKU1_orf1ab;4204 | 0 | 0 |
| NC_006577.2;YP_173236.1;HCoV-HKU1_orf1ab;4205 | 0 | 0 |
| NC_006577.2;YP_173236.1;HCoV-HKU1_orf1ab;4206 | 0 | 0 |
| NC_006577.2;YP_173236.1;HCoV-HKU1_orf1ab;4207 | 0 | 0 |
| NC_006577.2;YP_173236.1;HCoV-HKU1_orf1ab;4208 | 0.05 | 0 |
| NC_006577.2;YP_173236.1;HCoV-HKU1_orf1ab;4209 | 0 | 0 |
| NC_006577.2;YP_173236.1;HCoV-HKU1_orf1ab;4210 | 0 | 0 |
| NC_006577.2;YP_173236.1;HCoV-HKU1_orf1ab;4211 | 0 | 0 |
| NC_006577.2;YP_173236.1;HCoV-HKU1_orf1ab;4212 | 0 | 0 |
| NC_006577.2;YP_173236.1;HCoV-HKU1_orf1ab;4213 | 0 | 0 |
| NC_006577.2;YP_173236.1;HCoV-HKU1_orf1ab;4214 | 0.05 | 0.05 |
| NC_006577.2;YP_173236.1;HCoV-HKU1_orf1ab;4215 | 0 | 0.05 |
| NC_006577.2;YP_173236.1;HCoV-HKU1_orf1ab;4216 | 0 | 0.05 |
| NC_006577.2;YP_173236.1;HCoV-HKU1_orf1ab;4217 | 0 | 0 |
| NC_006577.2;YP_173236.1;HCoV-HKU1_orf1ab;4218 | 0 | 0.025 |
| NC_006577.2;YP_173236.1;HCoV-HKU1_orf1ab;4219 | 0 | 0 |
| NC_006577.2;YP_173236.1;HCoV-HKU1_orf1ab;4220 | 0 | 0 |
| NC_006577.2;YP_173236.1;HCoV-HKU1_orf1ab;4221 | 0.05 | 0.1 |
| NC_006577.2;YP_173236.1;HCoV-HKU1_orf1ab;4222 | 0 | 0 |
| NC_006577.2;YP_173236.1;HCoV-HKU1_orf1ab;4223 | 0 | 0 |
| NC_006577.2;YP_173236.1;HCoV-HKU1_orf1ab;4224 | 0 | 0 |
| NC_006577.2;YP_173236.1;HCoV-HKU1_orf1ab;4225 | 0 | 0.025 |
| NC_006577.2;YP_173236.1;HCoV-HKU1_orf1ab;4226 | 0.05 | 0 |
| NC_006577.2;YP_173236.1;HCoV-HKU1_orf1ab;4227 | 0 | 0 |
| NC_006577.2;YP_173236.1;HCoV-HKU1_orf1ab;4228 | 0 | 0 |
| NC_006577.2;YP_173236.1;HCoV-HKU1_orf1ab;4229 | 0 | 0 |

| | | |
|---|---|---|
| NC_006577.2;YP_173236.1;HCoV-HKU1_orf1ab;4230 | 0 | 0 |
| NC_006577.2;YP_173236.1;HCoV-HKU1_orf1ab;4231 | 0 | 0.025 |
| NC_006577.2;YP_173236.1;HCoV-HKU1_orf1ab;4232 | 0 | 0 |
| NC_006577.2;YP_173236.1;HCoV-HKU1_orf1ab;4233 | 0.05 | 0.025 |
| NC_006577.2;YP_173236.1;HCoV-HKU1_orf1ab;4234 | 0.05 | 0.025 |
| NC_006577.2;YP_173236.1;HCoV-HKU1_orf1ab;4235 | 0 | 0 |
| NC_006577.2;YP_173236.1;HCoV-HKU1_orf1ab;4236 | 0.05 | 0 |
| NC_006577.2;YP_173236.1;HCoV-HKU1_orf1ab;4237 | 0 | 0 |
| NC_006577.2;YP_173236.1;HCoV-HKU1_orf1ab;4238 | 0 | 0 |
| NC_006577.2;YP_173236.1;HCoV-HKU1_orf1ab;4239 | 0 | 0 |
| NC_006577.2;YP_173236.1;HCoV-HKU1_orf1ab;4240 | 0 | 0 |
| NC_006577.2;YP_173236.1;HCoV-HKU1_orf1ab;4241 | 0 | 0 |
| NC_006577.2;YP_173236.1;HCoV-HKU1_orf1ab;4242 | 0 | 0 |
| NC_006577.2;YP_173236.1;HCoV-HKU1_orf1ab;4243 | 0 | 0 |
| NC_006577.2;YP_173236.1;HCoV-HKU1_orf1ab;4244 | 0 | 0 |
| NC_006577.2;YP_173236.1;HCoV-HKU1_orf1ab;4245 | 0 | 0 |
| NC_006577.2;YP_173236.1;HCoV-HKU1_orf1ab;4246 | 0 | 0 |
| NC_006577.2;YP_173236.1;HCoV-HKU1_orf1ab;4247 | 0.05 | 0.05 |
| NC_006577.2;YP_173236.1;HCoV-HKU1_orf1ab;4248 | 0 | 0.025 |
| NC_006577.2;YP_173236.1;HCoV-HKU1_orf1ab;4249 | 0 | 0.05 |
| NC_006577.2;YP_173236.1;HCoV-HKU1_orf1ab;4250 | 0.05 | 0 |
| NC_006577.2;YP_173236.1;HCoV-HKU1_orf1ab;4251 | 0 | 0.025 |
| NC_006577.2;YP_173236.1;HCoV-HKU1_orf1ab;4252 | 0 | 0.025 |
| NC_006577.2;YP_173236.1;HCoV-HKU1_orf1ab;4253 | 0 | 0.05 |
| NC_006577.2;YP_173236.1;HCoV-HKU1_orf1ab;4254 | 0 | 0.025 |
| NC_006577.2;YP_173236.1;HCoV-HKU1_orf1ab;4255 | 0.15 | 0.075 |
| NC_006577.2;YP_173236.1;HCoV-HKU1_orf1ab;4256 | 0.1 | 0.125 |
| NC_006577.2;YP_173236.1;HCoV-HKU1_orf1ab;4257 | 0.15 | 0.15 |
| NC_006577.2;YP_173236.1;HCoV-HKU1_orf1ab;4258 | 0.15 | 0.05 |
| NC_006577.2;YP_173236.1;HCoV-HKU1_orf1ab;4259 | 0.1 | 0.025 |
| NC_006577.2;YP_173236.1;HCoV-HKU1_orf1ab;4260 | 0.15 | 0.05 |
| NC_006577.2;YP_173236.1;HCoV-HKU1_orf1ab;4261 | 0.1 | 0 |
| NC_006577.2;YP_173236.1;HCoV-HKU1_orf1ab;4262 | 0.1 | 0 |
| NC_006577.2;YP_173236.1;HCoV-HKU1_orf1ab;4263 | 0.05 | 0 |
| NC_006577.2;YP_173236.1;HCoV-HKU1_orf1ab;4264 | 0.15 | 0.025 |
| NC_006577.2;YP_173236.1;HCoV-HKU1_orf1ab;4265 | 0.1 | 0.05 |
| NC_006577.2;YP_173236.1;HCoV-HKU1_orf1ab;4266 | 0.2 | 0.025 |
| NC_006577.2;YP_173236.1;HCoV-HKU1_orf1ab;4267 | 0 | 0 |
| NC_006577.2;YP_173236.1;HCoV-HKU1_orf1ab;4268 | 0 | 0 |
| NC_006577.2;YP_173236.1;HCoV-HKU1_orf1ab;4269 | 0 | 0 |
| NC_006577.2;YP_173236.1;HCoV-HKU1_orf1ab;4270 | 0 | 0 |
| NC_006577.2;YP_173236.1;HCoV-HKU1_orf1ab;4271 | 0 | 0 |
| NC_006577.2;YP_173236.1;HCoV-HKU1_orf1ab;4272 | 0 | 0 |

| | | |
|---|---|---|
| NC_006577.2;YP_173236.1;HCoV-HKU1_orf1ab;4273 | 0 | 0 |
| NC_006577.2;YP_173236.1;HCoV-HKU1_orf1ab;4274 | 0 | 0 |
| NC_006577.2;YP_173236.1;HCoV-HKU1_orf1ab;4275 | 0 | 0.025 |
| NC_006577.2;YP_173236.1;HCoV-HKU1_orf1ab;4276 | 0 | 0.025 |
| NC_006577.2;YP_173236.1;HCoV-HKU1_orf1ab;4277 | 0 | 0 |
| NC_006577.2;YP_173236.1;HCoV-HKU1_orf1ab;4278 | 0 | 0 |
| NC_006577.2;YP_173236.1;HCoV-HKU1_orf1ab;4279 | 0 | 0 |
| NC_006577.2;YP_173236.1;HCoV-HKU1_orf1ab;4280 | 0 | 0 |
| NC_006577.2;YP_173236.1;HCoV-HKU1_orf1ab;4281 | 0 | 0 |
| NC_006577.2;YP_173236.1;HCoV-HKU1_orf1ab;4282 | 0 | 0 |
| NC_006577.2;YP_173236.1;HCoV-HKU1_orf1ab;4283 | 0 | 0 |
| NC_006577.2;YP_173236.1;HCoV-HKU1_orf1ab;4284 | 0.05 | 0.025 |
| NC_006577.2;YP_173236.1;HCoV-HKU1_orf1ab;4285 | 0 | 0 |
| NC_006577.2;YP_173236.1;HCoV-HKU1_orf1ab;4286 | 0 | 0 |
| NC_006577.2;YP_173236.1;HCoV-HKU1_orf1ab;4287 | 0 | 0 |
| NC_006577.2;YP_173236.1;HCoV-HKU1_orf1ab;4288 | 0 | 0 |
| NC_006577.2;YP_173236.1;HCoV-HKU1_orf1ab;4289 | 0 | 0 |
| NC_006577.2;YP_173236.1;HCoV-HKU1_orf1ab;4290 | 0 | 0 |
| NC_006577.2;YP_173236.1;HCoV-HKU1_orf1ab;4291 | 0 | 0 |
| NC_006577.2;YP_173236.1;HCoV-HKU1_orf1ab;4292 | 0 | 0 |
| NC_006577.2;YP_173236.1;HCoV-HKU1_orf1ab;4293 | 0 | 0 |
| NC_006577.2;YP_173236.1;HCoV-HKU1_orf1ab;4294 | 0 | 0 |
| NC_006577.2;YP_173236.1;HCoV-HKU1_orf1ab;4295 | 0 | 0 |
| NC_006577.2;YP_173236.1;HCoV-HKU1_orf1ab;4296 | 0 | 0 |
| NC_006577.2;YP_173236.1;HCoV-HKU1_orf1ab;4297 | 0 | 0 |
| NC_006577.2;YP_173236.1;HCoV-HKU1_orf1ab;4298 | 0 | 0 |
| NC_006577.2;YP_173236.1;HCoV-HKU1_orf1ab;4299 | 0 | 0 |
| NC_006577.2;YP_173236.1;HCoV-HKU1_orf1ab;4300 | 0 | 0 |
| NC_006577.2;YP_173236.1;HCoV-HKU1_orf1ab;4301 | 0 | 0 |
| NC_006577.2;YP_173236.1;HCoV-HKU1_orf1ab;4302 | 0 | 0 |
| NC_006577.2;YP_173236.1;HCoV-HKU1_orf1ab;4303 | 0 | 0 |
| NC_006577.2;YP_173236.1;HCoV-HKU1_orf1ab;4304 | 0 | 0 |
| NC_006577.2;YP_173236.1;HCoV-HKU1_orf1ab;4305 | 0 | 0 |
| NC_006577.2;YP_173236.1;HCoV-HKU1_orf1ab;4306 | 0 | 0 |
| NC_006577.2;YP_173236.1;HCoV-HKU1_orf1ab;4307 | 0 | 0 |
| NC_006577.2;YP_173236.1;HCoV-HKU1_orf1ab;4308 | 0 | 0 |
| NC_006577.2;YP_173236.1;HCoV-HKU1_orf1ab;4309 | 0 | 0.025 |
| NC_006577.2;YP_173236.1;HCoV-HKU1_orf1ab;4310 | 0 | 0 |
| NC_006577.2;YP_173236.1;HCoV-HKU1_orf1ab;4311 | 0 | 0 |
| NC_006577.2;YP_173236.1;HCoV-HKU1_orf1ab;4312 | 0 | 0 |
| NC_006577.2;YP_173236.1;HCoV-HKU1_orf1ab;4313 | 0 | 0 |
| NC_006577.2;YP_173236.1;HCoV-HKU1_orf1ab;4314 | 0.05 | 0.075 |
| NC_006577.2;YP_173236.1;HCoV-HKU1_orf1ab;4315 | 0 | 0.025 |

| | | |
|---|---|---|
| NC_006577.2;YP_173236.1;HCoV-HKU1_orf1ab;4316 | 0 | 0.025 |
| NC_006577.2;YP_173236.1;HCoV-HKU1_orf1ab;4317 | 0 | 0.025 |
| NC_006577.2;YP_173236.1;HCoV-HKU1_orf1ab;4318 | 0 | 0.025 |
| NC_006577.2;YP_173236.1;HCoV-HKU1_orf1ab;4319 | 0 | 0 |
| NC_006577.2;YP_173236.1;HCoV-HKU1_orf1ab;4320 | 0 | 0 |
| NC_006577.2;YP_173236.1;HCoV-HKU1_orf1ab;4321 | 0 | 0 |
| NC_006577.2;YP_173236.1;HCoV-HKU1_orf1ab;4322 | 0 | 0 |
| NC_006577.2;YP_173236.1;HCoV-HKU1_orf1ab;4323 | 0 | 0 |
| NC_006577.2;YP_173236.1;HCoV-HKU1_orf1ab;4324 | 0 | 0 |
| NC_006577.2;YP_173236.1;HCoV-HKU1_orf1ab;4325 | 0 | 0 |
| NC_006577.2;YP_173236.1;HCoV-HKU1_orf1ab;4326 | 0 | 0 |
| NC_006577.2;YP_173236.1;HCoV-HKU1_orf1ab;4327 | 0 | 0 |
| NC_006577.2;YP_173236.1;HCoV-HKU1_orf1ab;4328 | 0.05 | 0 |
| NC_006577.2;YP_173236.1;HCoV-HKU1_orf1ab;4329 | 0 | 0 |
| NC_006577.2;YP_173236.1;HCoV-HKU1_orf1ab;4330 | 0 | 0 |
| NC_006577.2;YP_173236.1;HCoV-HKU1_orf1ab;4331 | 0 | 0 |
| NC_006577.2;YP_173236.1;HCoV-HKU1_orf1ab;4332 | 0 | 0 |
| NC_006577.2;YP_173236.1;HCoV-HKU1_orf1ab;4333 | 0 | 0 |
| NC_006577.2;YP_173236.1;HCoV-HKU1_orf1ab;4334 | 0 | 0.025 |
| NC_006577.2;YP_173236.1;HCoV-HKU1_orf1ab;4335 | 0 | 0.025 |
| NC_006577.2;YP_173236.1;HCoV-HKU1_orf1ab;4336 | 0 | 0 |
| NC_006577.2;YP_173236.1;HCoV-HKU1_orf1ab;4337 | 0.05 | 0.05 |
| NC_006577.2;YP_173236.1;HCoV-HKU1_orf1ab;4338 | 0.05 | 0.025 |
| NC_006577.2;YP_173236.1;HCoV-HKU1_orf1ab;4339 | 0.05 | 0.1 |
| NC_006577.2;YP_173236.1;HCoV-HKU1_orf1ab;4340 | 0.1 | 0.15 |
| NC_006577.2;YP_173236.1;HCoV-HKU1_orf1ab;4341 | 0.1 | 0.15 |
| NC_006577.2;YP_173236.1;HCoV-HKU1_orf1ab;4342 | 0.05 | 0.125 |
| NC_006577.2;YP_173236.1;HCoV-HKU1_orf1ab;4343 | 0 | 0.05 |
| NC_006577.2;YP_173236.1;HCoV-HKU1_orf1ab;4344 | 0 | 0.025 |
| NC_006577.2;YP_173236.1;HCoV-HKU1_orf1ab;4345 | 0 | 0.05 |
| NC_006577.2;YP_173236.1;HCoV-HKU1_orf1ab;4346 | 0 | 0.025 |
| NC_006577.2;YP_173236.1;HCoV-HKU1_orf1ab;4347 | 0 | 0 |
| NC_006577.2;YP_173236.1;HCoV-HKU1_orf1ab;4348 | 0 | 0 |
| NC_006577.2;YP_173236.1;HCoV-HKU1_orf1ab;4349 | 0 | 0 |
| NC_006577.2;YP_173236.1;HCoV-HKU1_orf1ab;4350 | 0 | 0 |
| NC_006577.2;YP_173236.1;HCoV-HKU1_orf1ab;4351 | 0 | 0 |
| NC_006577.2;YP_173236.1;HCoV-HKU1_orf1ab;4352 | 0 | 0 |
| NC_006577.2;YP_173236.1;HCoV-HKU1_orf1ab;4353 | 0 | 0 |
| NC_006577.2;YP_173236.1;HCoV-HKU1_orf1ab;4354 | 0 | 0 |
| NC_006577.2;YP_173236.1;HCoV-HKU1_orf1ab;4355 | 0 | 0.025 |
| NC_006577.2;YP_173236.1;HCoV-HKU1_orf1ab;4356 | 0 | 0.025 |
| NC_006577.2;YP_173236.1;HCoV-HKU1_orf1ab;4357 | 0 | 0 |
| NC_006577.2;YP_173236.1;HCoV-HKU1_orf1ab;4358 | 0 | 0 |

| | | |
|---|---|---|
| NC_006577.2;YP_173236.1;HCoV-HKU1_orf1ab;4359 | 0 | 0 |
| NC_006577.2;YP_173236.1;HCoV-HKU1_orf1ab;4360 | 0 | 0 |
| NC_006577.2;YP_173236.1;HCoV-HKU1_orf1ab;4361 | 0 | 0 |
| NC_006577.2;YP_173236.1;HCoV-HKU1_orf1ab;4362 | 0 | 0 |
| NC_006577.2;YP_173236.1;HCoV-HKU1_orf1ab;4363 | 0 | 0 |
| NC_006577.2;YP_173236.1;HCoV-HKU1_orf1ab;4364 | 0 | 0 |
| NC_006577.2;YP_173236.1;HCoV-HKU1_orf1ab;4365 | 0 | 0 |
| NC_006577.2;YP_173236.1;HCoV-HKU1_orf1ab;4366 | 0 | 0 |
| NC_006577.2;YP_173236.1;HCoV-HKU1_orf1ab;4367 | 0 | 0 |
| NC_006577.2;YP_173236.1;HCoV-HKU1_orf1ab;4368 | 0 | 0 |
| NC_006577.2;YP_173236.1;HCoV-HKU1_orf1ab;4369 | 0 | 0 |
| NC_006577.2;YP_173236.1;HCoV-HKU1_orf1ab;4370 | 0 | 0 |
| NC_006577.2;YP_173236.1;HCoV-HKU1_orf1ab;4371 | 0 | 0 |
| NC_006577.2;YP_173236.1;HCoV-HKU1_orf1ab;4372 | 0 | 0 |
| NC_006577.2;YP_173236.1;HCoV-HKU1_orf1ab;4373 | 0 | 0 |
| NC_006577.2;YP_173236.1;HCoV-HKU1_orf1ab;4374 | 0 | 0 |
| NC_006577.2;YP_173236.1;HCoV-HKU1_orf1ab;4375 | 0.05 | 0 |
| NC_006577.2;YP_173236.1;HCoV-HKU1_orf1ab;4376 | 0 | 0 |
| NC_006577.2;YP_173236.1;HCoV-HKU1_orf1ab;4377 | 0 | 0 |
| NC_006577.2;YP_173236.1;HCoV-HKU1_orf1ab;4378 | 0 | 0 |
| NC_006577.2;YP_173236.1;HCoV-HKU1_orf1ab;4379 | 0 | 0 |
| NC_006577.2;YP_173236.1;HCoV-HKU1_orf1ab;4380 | 0.05 | 0 |
| NC_006577.2;YP_173236.1;HCoV-HKU1_orf1ab;4381 | 0 | 0 |
| NC_006577.2;YP_173236.1;HCoV-HKU1_orf1ab;4382 | 0 | 0 |
| NC_006577.2;YP_173236.1;HCoV-HKU1_orf1ab;4383 | 0 | 0 |
| NC_006577.2;YP_173236.1;HCoV-HKU1_orf1ab;4384 | 0 | 0 |
| NC_006577.2;YP_173236.1;HCoV-HKU1_orf1ab;4385 | 0 | 0 |
| NC_006577.2;YP_173236.1;HCoV-HKU1_orf1ab;4386 | 0 | 0 |
| NC_006577.2;YP_173236.1;HCoV-HKU1_orf1ab;4387 | 0 | 0 |
| NC_006577.2;YP_173236.1;HCoV-HKU1_orf1ab;4388 | 0 | 0 |
| NC_006577.2;YP_173236.1;HCoV-HKU1_orf1ab;4389 | 0 | 0 |
| NC_006577.2;YP_173236.1;HCoV-HKU1_orf1ab;4390 | 0 | 0 |
| NC_006577.2;YP_173236.1;HCoV-HKU1_orf1ab;4391 | 0 | 0 |
| NC_006577.2;YP_173236.1;HCoV-HKU1_orf1ab;4392 | 0 | 0 |
| NC_006577.2;YP_173236.1;HCoV-HKU1_orf1ab;4393 | 0 | 0 |
| NC_006577.2;YP_173236.1;HCoV-HKU1_orf1ab;4394 | 0 | 0 |
| NC_006577.2;YP_173236.1;HCoV-HKU1_orf1ab;4395 | 0 | 0 |
| NC_006577.2;YP_173236.1;HCoV-HKU1_orf1ab;4396 | 0 | 0 |
| NC_006577.2;YP_173236.1;HCoV-HKU1_orf1ab;4397 | 0 | 0 |
| NC_006577.2;YP_173236.1;HCoV-HKU1_orf1ab;4398 | 0 | 0 |
| NC_006577.2;YP_173236.1;HCoV-HKU1_orf1ab;4399 | 0 | 0.025 |
| NC_006577.2;YP_173236.1;HCoV-HKU1_orf1ab;4400 | 0 | 0 |
| NC_006577.2;YP_173236.1;HCoV-HKU1_orf1ab;4401 | 0 | 0 |

| | | |
|---|---|---|
| NC_006577.2;YP_173236.1;HCoV-HKU1_orf1ab;4402 | 0 | 0 |
| NC_006577.2;YP_173236.1;HCoV-HKU1_orf1ab;4403 | 0 | 0 |
| NC_006577.2;YP_173236.1;HCoV-HKU1_orf1ab;4404 | 0 | 0 |
| NC_006577.2;YP_173236.1;HCoV-HKU1_orf1ab;4405 | 0 | 0 |
| NC_006577.2;YP_173236.1;HCoV-HKU1_orf1ab;4406 | 0 | 0 |
| NC_006577.2;YP_173236.1;HCoV-HKU1_orf1ab;4407 | 0 | 0 |
| NC_006577.2;YP_173236.1;HCoV-HKU1_orf1ab;4408 | 0 | 0 |
| NC_006577.2;YP_173236.1;HCoV-HKU1_orf1ab;4409 | 0 | 0 |
| NC_006577.2;YP_173236.1;HCoV-HKU1_orf1ab;4410 | 0 | 0 |
| NC_006577.2;YP_173236.1;HCoV-HKU1_orf1ab;4411 | 0 | 0 |
| NC_006577.2;YP_173236.1;HCoV-HKU1_orf1ab;4412 | 0 | 0 |
| NC_006577.2;YP_173236.1;HCoV-HKU1_orf1ab;4413 | 0 | 0.025 |
| NC_006577.2;YP_173236.1;HCoV-HKU1_orf1ab;4414 | 0.05 | 0 |
| NC_006577.2;YP_173236.1;HCoV-HKU1_orf1ab;4415 | 0 | 0 |
| NC_006577.2;YP_173236.1;HCoV-HKU1_orf1ab;4416 | 0 | 0 |
| NC_006577.2;YP_173236.1;HCoV-HKU1_orf1ab;4417 | 0 | 0 |
| NC_006577.2;YP_173236.1;HCoV-HKU1_orf1ab;4418 | 0 | 0 |
| NC_006577.2;YP_173236.1;HCoV-HKU1_orf1ab;4419 | 0 | 0 |
| NC_006577.2;YP_173236.1;HCoV-HKU1_orf1ab;4420 | 0 | 0 |
| NC_006577.2;YP_173236.1;HCoV-HKU1_orf1ab;4421 | 0 | 0 |
| NC_006577.2;YP_173236.1;HCoV-HKU1_orf1ab;4422 | 0 | 0 |
| NC_006577.2;YP_173236.1;HCoV-HKU1_orf1ab;4423 | 0 | 0 |
| NC_006577.2;YP_173236.1;HCoV-HKU1_orf1ab;4424 | 0 | 0 |
| NC_006577.2;YP_173236.1;HCoV-HKU1_orf1ab;4425 | 0 | 0 |
| NC_006577.2;YP_173236.1;HCoV-HKU1_orf1ab;4426 | 0 | 0 |
| NC_006577.2;YP_173236.1;HCoV-HKU1_orf1ab;4427 | 0 | 0 |
| NC_006577.2;YP_173236.1;HCoV-HKU1_orf1ab;4428 | 0 | 0 |
| NC_006577.2;YP_173236.1;HCoV-HKU1_orf1ab;4429 | 0 | 0 |
| NC_006577.2;YP_173236.1;HCoV-HKU1_orf1ab;4430 | 0 | 0 |
| NC_006577.2;YP_173236.1;HCoV-HKU1_orf1ab;4431 | 0 | 0 |
| NC_006577.2;YP_173236.1;HCoV-HKU1_orf1ab;4432 | 0 | 0 |
| NC_006577.2;YP_173236.1;HCoV-HKU1_orf1ab;4433 | 0 | 0 |
| NC_006577.2;YP_173236.1;HCoV-HKU1_orf1ab;4434 | 0 | 0 |
| NC_006577.2;YP_173236.1;HCoV-HKU1_orf1ab;4435 | 0 | 0 |
| NC_006577.2;YP_173236.1;HCoV-HKU1_orf1ab;4436 | 0 | 0 |
| NC_006577.2;YP_173236.1;HCoV-HKU1_orf1ab;4437 | 0 | 0 |
| NC_006577.2;YP_173236.1;HCoV-HKU1_orf1ab;4438 | 0 | 0 |
| NC_006577.2;YP_173236.1;HCoV-HKU1_orf1ab;4439 | 0 | 0 |
| NC_006577.2;YP_173236.1;HCoV-HKU1_orf1ab;4440 | 0.05 | 0.05 |
| NC_006577.2;YP_173236.1;HCoV-HKU1_orf1ab;4441 | 0 | 0 |
| NC_006577.2;YP_173236.1;HCoV-HKU1_orf1ab;4442 | 0.05 | 0 |
| NC_006577.2;YP_173236.1;HCoV-HKU1_orf1ab;4443 | 0 | 0 |
| NC_006577.2;YP_173236.1;HCoV-HKU1_orf1ab;4444 | 0 | 0 |

| | | |
|---|---|---|
| NC_006577.2;YP_173236.1;HCoV-HKU1_orf1ab;4445 | 0 | 0 |
| NC_006577.2;YP_173236.1;HCoV-HKU1_orf1ab;4446 | 0 | 0 |
| NC_006577.2;YP_173236.1;HCoV-HKU1_orf1ab;4447 | 0 | 0.025 |
| NC_006577.2;YP_173236.1;HCoV-HKU1_orf1ab;4448 | 0.05 | 0.025 |
| NC_006577.2;YP_173236.1;HCoV-HKU1_orf1ab;4449 | 0 | 0.025 |
| NC_006577.2;YP_173236.1;HCoV-HKU1_orf1ab;4450 | 0 | 0.025 |
| NC_006577.2;YP_173236.1;HCoV-HKU1_orf1ab;4451 | 0 | 0 |
| NC_006577.2;YP_173236.1;HCoV-HKU1_orf1ab;4452 | 0 | 0.025 |
| NC_006577.2;YP_173236.1;HCoV-HKU1_orf1ab;4453 | 0 | 0 |
| NC_006577.2;YP_173236.1;HCoV-HKU1_orf1ab;4454 | 0 | 0 |
| NC_006577.2;YP_173236.1;HCoV-HKU1_orf1ab;4455 | 0 | 0.025 |
| NC_006577.2;YP_173236.1;HCoV-HKU1_orf1ab;4456 | 0 | 0 |
| NC_006577.2;YP_173236.1;HCoV-HKU1_orf1ab;4457 | 0 | 0 |
| NC_006577.2;YP_173236.1;HCoV-HKU1_orf1ab;4458 | 0 | 0 |
| NC_006577.2;YP_173236.1;HCoV-HKU1_orf1ab;4459 | 0 | 0.025 |
| NC_006577.2;YP_173236.1;HCoV-HKU1_orf1ab;4460 | 0 | 0 |
| NC_006577.2;YP_173236.1;HCoV-HKU1_orf1ab;4461 | 0 | 0 |
| NC_006577.2;YP_173236.1;HCoV-HKU1_orf1ab;4462 | 0 | 0 |
| NC_006577.2;YP_173236.1;HCoV-HKU1_orf1ab;4463 | 0 | 0 |
| NC_006577.2;YP_173236.1;HCoV-HKU1_orf1ab;4464 | 0 | 0 |
| NC_006577.2;YP_173236.1;HCoV-HKU1_orf1ab;4465 | 0 | 0 |
| NC_006577.2;YP_173236.1;HCoV-HKU1_orf1ab;4466 | 0 | 0 |
| NC_006577.2;YP_173236.1;HCoV-HKU1_orf1ab;4467 | 0 | 0 |
| NC_006577.2;YP_173236.1;HCoV-HKU1_orf1ab;4468 | 0 | 0 |
| NC_006577.2;YP_173236.1;HCoV-HKU1_orf1ab;4469 | 0 | 0 |
| NC_006577.2;YP_173236.1;HCoV-HKU1_orf1ab;4470 | 0 | 0 |
| NC_006577.2;YP_173236.1;HCoV-HKU1_orf1ab;4471 | 0 | 0 |
| NC_006577.2;YP_173236.1;HCoV-HKU1_orf1ab;4472 | 0 | 0 |
| NC_006577.2;YP_173236.1;HCoV-HKU1_orf1ab;4473 | 0 | 0 |
| NC_006577.2;YP_173236.1;HCoV-HKU1_orf1ab;4474 | 0 | 0 |
| NC_006577.2;YP_173236.1;HCoV-HKU1_orf1ab;4475 | 0 | 0 |
| NC_006577.2;YP_173236.1;HCoV-HKU1_orf1ab;4476 | 0.2 | 0.075 |
| NC_006577.2;YP_173236.1;HCoV-HKU1_orf1ab;4477 | 0 | 0 |
| NC_006577.2;YP_173236.1;HCoV-HKU1_orf1ab;4478 | 0 | 0 |
| NC_006577.2;YP_173236.1;HCoV-HKU1_orf1ab;4479 | 0 | 0 |
| NC_006577.2;YP_173236.1;HCoV-HKU1_orf1ab;4480 | 0 | 0 |
| NC_006577.2;YP_173236.1;HCoV-HKU1_orf1ab;4481 | 0 | 0 |
| NC_006577.2;YP_173236.1;HCoV-HKU1_orf1ab;4482 | 0 | 0 |
| NC_006577.2;YP_173236.1;HCoV-HKU1_orf1ab;4483 | 0 | 0 |
| NC_006577.2;YP_173236.1;HCoV-HKU1_orf1ab;4484 | 0 | 0 |
| NC_006577.2;YP_173236.1;HCoV-HKU1_orf1ab;4485 | 0 | 0 |
| NC_006577.2;YP_173236.1;HCoV-HKU1_orf1ab;4486 | 0 | 0.025 |
| NC_006577.2;YP_173236.1;HCoV-HKU1_orf1ab;4487 | 0 | 0.025 |

| | | |
|---|---|---|
| NC_006577.2;YP_173236.1;HCoV-HKU1_orf1ab;4488 | 0 | 0 |
| NC_006577.2;YP_173236.1;HCoV-HKU1_orf1ab;4489 | 0 | 0 |
| NC_006577.2;YP_173236.1;HCoV-HKU1_orf1ab;4490 | 0 | 0 |
| NC_006577.2;YP_173236.1;HCoV-HKU1_orf1ab;4491 | 0 | 0 |
| NC_006577.2;YP_173236.1;HCoV-HKU1_orf1ab;4492 | 0 | 0 |
| NC_006577.2;YP_173236.1;HCoV-HKU1_orf1ab;4493 | 0 | 0 |
| NC_006577.2;YP_173236.1;HCoV-HKU1_orf1ab;4494 | 0 | 0.025 |
| NC_006577.2;YP_173236.1;HCoV-HKU1_orf1ab;4495 | 0 | 0 |
| NC_006577.2;YP_173236.1;HCoV-HKU1_orf1ab;4496 | 0 | 0 |
| NC_006577.2;YP_173236.1;HCoV-HKU1_orf1ab;4497 | 0 | 0 |
| NC_006577.2;YP_173236.1;HCoV-HKU1_orf1ab;4498 | 0 | 0 |
| NC_006577.2;YP_173236.1;HCoV-HKU1_orf1ab;4499 | 0 | 0 |
| NC_006577.2;YP_173236.1;HCoV-HKU1_orf1ab;4500 | 0 | 0 |
| NC_006577.2;YP_173236.1;HCoV-HKU1_orf1ab;4501 | 0 | 0 |
| NC_006577.2;YP_173236.1;HCoV-HKU1_orf1ab;4502 | 0 | 0 |
| NC_006577.2;YP_173236.1;HCoV-HKU1_orf1ab;4503 | 0 | 0 |
| NC_006577.2;YP_173236.1;HCoV-HKU1_orf1ab;4504 | 0 | 0.025 |
| NC_006577.2;YP_173236.1;HCoV-HKU1_orf1ab;4505 | 0 | 0.025 |
| NC_006577.2;YP_173236.1;HCoV-HKU1_orf1ab;4506 | 0 | 0.025 |
| NC_006577.2;YP_173236.1;HCoV-HKU1_orf1ab;4507 | 0 | 0.075 |
| NC_006577.2;YP_173236.1;HCoV-HKU1_orf1ab;4508 | 0.15 | 0.075 |
| NC_006577.2;YP_173236.1;HCoV-HKU1_orf1ab;4509 | 0.1 | 0.1 |
| NC_006577.2;YP_173236.1;HCoV-HKU1_orf1ab;4510 | 0.15 | 0.1 |
| NC_006577.2;YP_173236.1;HCoV-HKU1_orf1ab;4511 | 0.2 | 0.125 |
| NC_006577.2;YP_173236.1;HCoV-HKU1_orf1ab;4512 | 0.35 | 0.25 |
| NC_006577.2;YP_173236.1;HCoV-HKU1_orf1ab;4513 | 0.15 | 0.05 |
| NC_006577.2;YP_173236.1;HCoV-HKU1_orf1ab;4514 | 0.05 | 0.05 |
| NC_006577.2;YP_173236.1;HCoV-HKU1_orf1ab;4515 | 0.1 | 0.05 |
| NC_006577.2;YP_173236.1;HCoV-HKU1_orf1ab;4516 | 0 | 0 |
| NC_006577.2;YP_173236.1;HCoV-HKU1_orf1ab;4517 | 0 | 0 |
| NC_006577.2;YP_173236.1;HCoV-HKU1_orf1ab;4518 | 0.15 | 0.075 |
| NC_006577.2;YP_173236.1;HCoV-HKU1_orf1ab;4519 | 0 | 0.025 |
| NC_006577.2;YP_173236.1;HCoV-HKU1_orf1ab;4520 | 0.1 | 0 |
| NC_006577.2;YP_173236.1;HCoV-HKU1_orf1ab;4521 | 0.05 | 0 |
| NC_006577.2;YP_173236.1;HCoV-HKU1_orf1ab;4522 | 0.05 | 0 |
| NC_006577.2;YP_173236.1;HCoV-HKU1_orf1ab;4523 | 0.05 | 0 |
| NC_006577.2;YP_173236.1;HCoV-HKU1_orf1ab;4524 | 0.05 | 0 |
| NC_006577.2;YP_173236.1;HCoV-HKU1_orf1ab;4525 | 0.05 | 0 |
| NC_006577.2;YP_173236.1;HCoV-HKU1_orf1ab;4526 | 0.05 | 0 |
| NC_006577.2;YP_173236.1;HCoV-HKU1_orf1ab;4527 | 0.05 | 0 |
| NC_006577.2;YP_173236.1;HCoV-HKU1_orf1ab;4528 | 0.15 | 0 |
| NC_006577.2;YP_173236.1;HCoV-HKU1_orf1ab;4529 | 0.1 | 0.05 |
| NC_006577.2;YP_173236.1;HCoV-HKU1_orf1ab;4530 | 0.1 | 0.075 |

| | | |
|---|---|---|
| NC_006577.2;YP_173236.1;HCoV-HKU1_orf1ab;4531 | 0.05 | 0.05 |
| NC_006577.2;YP_173236.1;HCoV-HKU1_orf1ab;4532 | 0 | 0 |
| NC_006577.2;YP_173236.1;HCoV-HKU1_orf1ab;4533 | 0.05 | 0 |
| NC_006577.2;YP_173236.1;HCoV-HKU1_orf1ab;4534 | 0 | 0.025 |
| NC_006577.2;YP_173236.1;HCoV-HKU1_orf1ab;4535 | 0 | 0 |
| NC_006577.2;YP_173236.1;HCoV-HKU1_orf1ab;4536 | 0 | 0 |
| NC_006577.2;YP_173236.1;HCoV-HKU1_orf1ab;4537 | 0 | 0 |
| NC_006577.2;YP_173236.1;HCoV-HKU1_orf1ab;4538 | 0 | 0 |
| NC_006577.2;YP_173236.1;HCoV-HKU1_orf1ab;4539 | 0 | 0.025 |
| NC_006577.2;YP_173236.1;HCoV-HKU1_orf1ab;4540 | 0 | 0 |
| NC_006577.2;YP_173236.1;HCoV-HKU1_orf1ab;4541 | 0 | 0 |
| NC_006577.2;YP_173236.1;HCoV-HKU1_orf1ab;4542 | 0 | 0 |
| NC_006577.2;YP_173236.1;HCoV-HKU1_orf1ab;4543 | 0 | 0 |
| NC_006577.2;YP_173236.1;HCoV-HKU1_orf1ab;4544 | 0 | 0 |
| NC_006577.2;YP_173236.1;HCoV-HKU1_orf1ab;4545 | 0 | 0 |
| NC_006577.2;YP_173236.1;HCoV-HKU1_orf1ab;4546 | 0 | 0 |
| NC_006577.2;YP_173236.1;HCoV-HKU1_orf1ab;4547 | 0 | 0 |
| NC_006577.2;YP_173236.1;HCoV-HKU1_orf1ab;4548 | 0 | 0 |
| NC_006577.2;YP_173236.1;HCoV-HKU1_orf1ab;4549 | 0 | 0 |
| NC_006577.2;YP_173236.1;HCoV-HKU1_orf1ab;4550 | 0 | 0 |
| NC_006577.2;YP_173236.1;HCoV-HKU1_orf1ab;4551 | 0 | 0 |
| NC_006577.2;YP_173236.1;HCoV-HKU1_orf1ab;4552 | 0 | 0 |
| NC_006577.2;YP_173236.1;HCoV-HKU1_orf1ab;4553 | 0 | 0 |
| NC_006577.2;YP_173236.1;HCoV-HKU1_orf1ab;4554 | 0 | 0.025 |
| NC_006577.2;YP_173236.1;HCoV-HKU1_orf1ab;4555 | 0 | 0 |
| NC_006577.2;YP_173236.1;HCoV-HKU1_orf1ab;4556 | 0 | 0 |
| NC_006577.2;YP_173236.1;HCoV-HKU1_orf1ab;4557 | 0 | 0 |
| NC_006577.2;YP_173236.1;HCoV-HKU1_orf1ab;4558 | 0 | 0 |
| NC_006577.2;YP_173236.1;HCoV-HKU1_orf1ab;4559 | 0 | 0.05 |
| NC_006577.2;YP_173236.1;HCoV-HKU1_orf1ab;4560 | 0 | 0 |
| NC_006577.2;YP_173236.1;HCoV-HKU1_orf1ab;4561 | 0 | 0 |
| NC_006577.2;YP_173236.1;HCoV-HKU1_orf1ab;4562 | 0 | 0 |
| NC_006577.2;YP_173236.1;HCoV-HKU1_orf1ab;4563 | 0 | 0 |
| NC_006577.2;YP_173236.1;HCoV-HKU1_orf1ab;4564 | 0 | 0 |
| NC_006577.2;YP_173236.1;HCoV-HKU1_orf1ab;4565 | 0 | 0 |
| NC_006577.2;YP_173236.1;HCoV-HKU1_orf1ab;4566 | 0 | 0 |
| NC_006577.2;YP_173236.1;HCoV-HKU1_orf1ab;4567 | 0 | 0 |
| NC_006577.2;YP_173236.1;HCoV-HKU1_orf1ab;4568 | 0 | 0 |
| NC_006577.2;YP_173236.1;HCoV-HKU1_orf1ab;4569 | 0 | 0 |
| NC_006577.2;YP_173236.1;HCoV-HKU1_orf1ab;4570 | 0.05 | 0 |
| NC_006577.2;YP_173236.1;HCoV-HKU1_orf1ab;4571 | 0.05 | 0 |
| NC_006577.2;YP_173236.1;HCoV-HKU1_orf1ab;4572 | 0 | 0 |
| NC_006577.2;YP_173236.1;HCoV-HKU1_orf1ab;4573 | 0 | 0 |

| | | |
|---|---|---|
| NC_006577.2;YP_173236.1;HCoV-HKU1_orf1ab;4574 | 0 | 0 |
| NC_006577.2;YP_173236.1;HCoV-HKU1_orf1ab;4575 | 0 | 0 |
| NC_006577.2;YP_173236.1;HCoV-HKU1_orf1ab;4576 | 0 | 0 |
| NC_006577.2;YP_173236.1;HCoV-HKU1_orf1ab;4577 | 0 | 0 |
| NC_006577.2;YP_173236.1;HCoV-HKU1_orf1ab;4578 | 0 | 0 |
| NC_006577.2;YP_173236.1;HCoV-HKU1_orf1ab;4579 | 0 | 0.025 |
| NC_006577.2;YP_173236.1;HCoV-HKU1_orf1ab;4580 | 0 | 0.025 |
| NC_006577.2;YP_173236.1;HCoV-HKU1_orf1ab;4581 | 0 | 0 |
| NC_006577.2;YP_173236.1;HCoV-HKU1_orf1ab;4582 | 0 | 0 |
| NC_006577.2;YP_173236.1;HCoV-HKU1_orf1ab;4583 | 0 | 0 |
| NC_006577.2;YP_173236.1;HCoV-HKU1_orf1ab;4584 | 0 | 0 |
| NC_006577.2;YP_173236.1;HCoV-HKU1_orf1ab;4585 | 0 | 0 |
| NC_006577.2;YP_173236.1;HCoV-HKU1_orf1ab;4586 | 0 | 0 |
| NC_006577.2;YP_173236.1;HCoV-HKU1_orf1ab;4587 | 0 | 0 |
| NC_006577.2;YP_173236.1;HCoV-HKU1_orf1ab;4588 | 0 | 0 |
| NC_006577.2;YP_173236.1;HCoV-HKU1_orf1ab;4589 | 0 | 0 |
| NC_006577.2;YP_173236.1;HCoV-HKU1_orf1ab;4590 | 0 | 0 |
| NC_006577.2;YP_173236.1;HCoV-HKU1_orf1ab;4591 | 0 | 0 |
| NC_006577.2;YP_173236.1;HCoV-HKU1_orf1ab;4592 | 0 | 0 |
| NC_006577.2;YP_173236.1;HCoV-HKU1_orf1ab;4593 | 0 | 0.05 |
| NC_006577.2;YP_173236.1;HCoV-HKU1_orf1ab;4594 | 0 | 0 |
| NC_006577.2;YP_173236.1;HCoV-HKU1_orf1ab;4595 | 0 | 0.025 |
| NC_006577.2;YP_173236.1;HCoV-HKU1_orf1ab;4596 | 0 | 0 |
| NC_006577.2;YP_173236.1;HCoV-HKU1_orf1ab;4597 | 0 | 0 |
| NC_006577.2;YP_173236.1;HCoV-HKU1_orf1ab;4598 | 0 | 0 |
| NC_006577.2;YP_173236.1;HCoV-HKU1_orf1ab;4599 | 0 | 0.025 |
| NC_006577.2;YP_173236.1;HCoV-HKU1_orf1ab;4600 | 0 | 0.025 |
| NC_006577.2;YP_173236.1;HCoV-HKU1_orf1ab;4601 | 0 | 0.025 |
| NC_006577.2;YP_173236.1;HCoV-HKU1_orf1ab;4602 | 0 | 0.025 |
| NC_006577.2;YP_173236.1;HCoV-HKU1_orf1ab;4603 | 0 | 0 |
| NC_006577.2;YP_173236.1;HCoV-HKU1_orf1ab;4604 | 0 | 0 |
| NC_006577.2;YP_173236.1;HCoV-HKU1_orf1ab;4605 | 0 | 0 |
| NC_006577.2;YP_173236.1;HCoV-HKU1_orf1ab;4606 | 0 | 0 |
| NC_006577.2;YP_173236.1;HCoV-HKU1_orf1ab;4607 | 0 | 0 |
| NC_006577.2;YP_173236.1;HCoV-HKU1_orf1ab;4608 | 0 | 0 |
| NC_006577.2;YP_173236.1;HCoV-HKU1_orf1ab;4609 | 0 | 0 |
| NC_006577.2;YP_173236.1;HCoV-HKU1_orf1ab;4610 | 0.25 | 0.05 |
| NC_006577.2;YP_173236.1;HCoV-HKU1_orf1ab;4611 | 0.05 | 0 |
| NC_006577.2;YP_173236.1;HCoV-HKU1_orf1ab;4612 | 0 | 0 |
| NC_006577.2;YP_173236.1;HCoV-HKU1_orf1ab;4613 | 0 | 0.075 |
| NC_006577.2;YP_173236.1;HCoV-HKU1_orf1ab;4614 | 0.05 | 0.075 |
| NC_006577.2;YP_173236.1;HCoV-HKU1_orf1ab;4615 | 0.1 | 0.125 |
| NC_006577.2;YP_173236.1;HCoV-HKU1_orf1ab;4616 | 0.1 | 0.125 |

| | | |
|---|---|---|
| NC_006577.2;YP_173236.1;HCoV-HKU1_orf1ab;4617 | 0 | 0.025 |
| NC_006577.2;YP_173236.1;HCoV-HKU1_orf1ab;4618 | 0 | 0.05 |
| NC_006577.2;YP_173236.1;HCoV-HKU1_orf1ab;4619 | 0 | 0 |
| NC_006577.2;YP_173236.1;HCoV-HKU1_orf1ab;4620 | 0 | 0 |
| NC_006577.2;YP_173236.1;HCoV-HKU1_orf1ab;4621 | 0 | 0.05 |
| NC_006577.2;YP_173236.1;HCoV-HKU1_orf1ab;4622 | 0 | 0.025 |
| NC_006577.2;YP_173236.1;HCoV-HKU1_orf1ab;4623 | 0 | 0.025 |
| NC_006577.2;YP_173236.1;HCoV-HKU1_orf1ab;4624 | 0 | 0.025 |
| NC_006577.2;YP_173236.1;HCoV-HKU1_orf1ab;4625 | 0 | 0.025 |
| NC_006577.2;YP_173236.1;HCoV-HKU1_orf1ab;4626 | 0 | 0.05 |
| NC_006577.2;YP_173236.1;HCoV-HKU1_orf1ab;4627 | 0 | 0 |
| NC_006577.2;YP_173236.1;HCoV-HKU1_orf1ab;4628 | 0 | 0 |
| NC_006577.2;YP_173236.1;HCoV-HKU1_orf1ab;4629 | 0 | 0 |
| NC_006577.2;YP_173236.1;HCoV-HKU1_orf1ab;4630 | 0 | 0 |
| NC_006577.2;YP_173236.1;HCoV-HKU1_orf1ab;4631 | 0 | 0 |
| NC_006577.2;YP_173236.1;HCoV-HKU1_orf1ab;4632 | 0 | 0 |
| NC_006577.2;YP_173236.1;HCoV-HKU1_orf1ab;4633 | 0 | 0 |
| NC_006577.2;YP_173236.1;HCoV-HKU1_orf1ab;4634 | 0 | 0 |
| NC_006577.2;YP_173236.1;HCoV-HKU1_orf1ab;4635 | 0 | 0 |
| NC_006577.2;YP_173236.1;HCoV-HKU1_orf1ab;4636 | 0 | 0 |
| NC_006577.2;YP_173236.1;HCoV-HKU1_orf1ab;4637 | 0 | 0 |
| NC_006577.2;YP_173236.1;HCoV-HKU1_orf1ab;4638 | 0 | 0 |
| NC_006577.2;YP_173236.1;HCoV-HKU1_orf1ab;4639 | 0 | 0 |
| NC_006577.2;YP_173236.1;HCoV-HKU1_orf1ab;4640 | 0 | 0 |
| NC_006577.2;YP_173236.1;HCoV-HKU1_orf1ab;4641 | 0 | 0 |
| NC_006577.2;YP_173236.1;HCoV-HKU1_orf1ab;4642 | 0 | 0 |
| NC_006577.2;YP_173236.1;HCoV-HKU1_orf1ab;4643 | 0 | 0 |
| NC_006577.2;YP_173236.1;HCoV-HKU1_orf1ab;4644 | 0 | 0 |
| NC_006577.2;YP_173236.1;HCoV-HKU1_orf1ab;4645 | 0 | 0 |
| NC_006577.2;YP_173236.1;HCoV-HKU1_orf1ab;4646 | 0 | 0 |
| NC_006577.2;YP_173236.1;HCoV-HKU1_orf1ab;4647 | 0 | 0 |
| NC_006577.2;YP_173236.1;HCoV-HKU1_orf1ab;4648 | 0 | 0.025 |
| NC_006577.2;YP_173236.1;HCoV-HKU1_orf1ab;4649 | 0 | 0 |
| NC_006577.2;YP_173236.1;HCoV-HKU1_orf1ab;4650 | 0 | 0 |
| NC_006577.2;YP_173236.1;HCoV-HKU1_orf1ab;4651 | 0 | 0 |
| NC_006577.2;YP_173236.1;HCoV-HKU1_orf1ab;4652 | 0 | 0 |
| NC_006577.2;YP_173236.1;HCoV-HKU1_orf1ab;4653 | 0 | 0 |
| NC_006577.2;YP_173236.1;HCoV-HKU1_orf1ab;4654 | 0 | 0 |
| NC_006577.2;YP_173236.1;HCoV-HKU1_orf1ab;4655 | 0 | 0 |
| NC_006577.2;YP_173236.1;HCoV-HKU1_orf1ab;4656 | 0 | 0 |
| NC_006577.2;YP_173236.1;HCoV-HKU1_orf1ab;4657 | 0 | 0 |
| NC_006577.2;YP_173236.1;HCoV-HKU1_orf1ab;4658 | 0 | 0 |
| NC_006577.2;YP_173236.1;HCoV-HKU1_orf1ab;4659 | 0 | 0 |

| | | |
|---|---|---|
| NC_006577.2;YP_173236.1;HCoV-HKU1_orf1ab;4660 | 0 | 0 |
| NC_006577.2;YP_173236.1;HCoV-HKU1_orf1ab;4661 | 0 | 0 |
| NC_006577.2;YP_173236.1;HCoV-HKU1_orf1ab;4662 | 0 | 0 |
| NC_006577.2;YP_173236.1;HCoV-HKU1_orf1ab;4663 | 0 | 0 |
| NC_006577.2;YP_173236.1;HCoV-HKU1_orf1ab;4664 | 0 | 0 |
| NC_006577.2;YP_173236.1;HCoV-HKU1_orf1ab;4665 | 0 | 0 |
| NC_006577.2;YP_173236.1;HCoV-HKU1_orf1ab;4666 | 0 | 0 |
| NC_006577.2;YP_173236.1;HCoV-HKU1_orf1ab;4667 | 0 | 0 |
| NC_006577.2;YP_173236.1;HCoV-HKU1_orf1ab;4668 | 0 | 0 |
| NC_006577.2;YP_173236.1;HCoV-HKU1_orf1ab;4669 | 0 | 0 |
| NC_006577.2;YP_173236.1;HCoV-HKU1_orf1ab;4670 | 0 | 0 |
| NC_006577.2;YP_173236.1;HCoV-HKU1_orf1ab;4671 | 0 | 0 |
| NC_006577.2;YP_173236.1;HCoV-HKU1_orf1ab;4672 | 0 | 0 |
| NC_006577.2;YP_173236.1;HCoV-HKU1_orf1ab;4673 | 0 | 0 |
| NC_006577.2;YP_173236.1;HCoV-HKU1_orf1ab;4674 | 0.1 | 0.05 |
| NC_006577.2;YP_173236.1;HCoV-HKU1_orf1ab;4675 | 0 | 0 |
| NC_006577.2;YP_173236.1;HCoV-HKU1_orf1ab;4676 | 0 | 0 |
| NC_006577.2;YP_173236.1;HCoV-HKU1_orf1ab;4677 | 0 | 0 |
| NC_006577.2;YP_173236.1;HCoV-HKU1_orf1ab;4678 | 0 | 0 |
| NC_006577.2;YP_173236.1;HCoV-HKU1_orf1ab;4679 | 0 | 0.025 |
| NC_006577.2;YP_173236.1;HCoV-HKU1_orf1ab;4680 | 0 | 0 |
| NC_006577.2;YP_173236.1;HCoV-HKU1_orf1ab;4681 | 0.05 | 0 |
| NC_006577.2;YP_173236.1;HCoV-HKU1_orf1ab;4682 | 0 | 0 |
| NC_006577.2;YP_173236.1;HCoV-HKU1_orf1ab;4683 | 0 | 0 |
| NC_006577.2;YP_173236.1;HCoV-HKU1_orf1ab;4684 | 0 | 0.025 |
| NC_006577.2;YP_173236.1;HCoV-HKU1_orf1ab;4685 | 0 | 0.025 |
| NC_006577.2;YP_173236.1;HCoV-HKU1_orf1ab;4686 | 0 | 0 |
| NC_006577.2;YP_173236.1;HCoV-HKU1_orf1ab;4687 | 0 | 0.025 |
| NC_006577.2;YP_173236.1;HCoV-HKU1_orf1ab;4688 | 0.05 | 0.025 |
| NC_006577.2;YP_173236.1;HCoV-HKU1_orf1ab;4689 | 0.05 | 0.05 |
| NC_006577.2;YP_173236.1;HCoV-HKU1_orf1ab;4690 | 0.05 | 0 |
| NC_006577.2;YP_173236.1;HCoV-HKU1_orf1ab;4691 | 0 | 0.025 |
| NC_006577.2;YP_173236.1;HCoV-HKU1_orf1ab;4692 | 0 | 0 |
| NC_006577.2;YP_173236.1;HCoV-HKU1_orf1ab;4693 | 0 | 0 |
| NC_006577.2;YP_173236.1;HCoV-HKU1_orf1ab;4694 | 0 | 0 |
| NC_006577.2;YP_173236.1;HCoV-HKU1_orf1ab;4695 | 0 | 0 |
| NC_006577.2;YP_173236.1;HCoV-HKU1_orf1ab;4696 | 0 | 0 |
| NC_006577.2;YP_173236.1;HCoV-HKU1_orf1ab;4697 | 0 | 0 |
| NC_006577.2;YP_173236.1;HCoV-HKU1_orf1ab;4698 | 0 | 0 |
| NC_006577.2;YP_173236.1;HCoV-HKU1_orf1ab;4699 | 0 | 0 |
| NC_006577.2;YP_173236.1;HCoV-HKU1_orf1ab;4700 | 0 | 0 |
| NC_006577.2;YP_173236.1;HCoV-HKU1_orf1ab;4701 | 0 | 0 |
| NC_006577.2;YP_173236.1;HCoV-HKU1_orf1ab;4702 | 0 | 0 |

| | | |
|---|---|---|
| NC_006577.2;YP_173236.1;HCoV-HKU1_orf1ab;4703 | 0 | 0 |
| NC_006577.2;YP_173236.1;HCoV-HKU1_orf1ab;4704 | 0 | 0 |
| NC_006577.2;YP_173236.1;HCoV-HKU1_orf1ab;4705 | 0 | 0.025 |
| NC_006577.2;YP_173236.1;HCoV-HKU1_orf1ab;4706 | 0 | 0.025 |
| NC_006577.2;YP_173236.1;HCoV-HKU1_orf1ab;4707 | 0 | 0 |
| NC_006577.2;YP_173236.1;HCoV-HKU1_orf1ab;4708 | 0 | 0 |
| NC_006577.2;YP_173236.1;HCoV-HKU1_orf1ab;4709 | 0 | 0 |
| NC_006577.2;YP_173236.1;HCoV-HKU1_orf1ab;4710 | 0.05 | 0.025 |
| NC_006577.2;YP_173236.1;HCoV-HKU1_orf1ab;4711 | 0 | 0 |
| NC_006577.2;YP_173236.1;HCoV-HKU1_orf1ab;4712 | 0 | 0 |
| NC_006577.2;YP_173236.1;HCoV-HKU1_orf1ab;4713 | 0.05 | 0.025 |
| NC_006577.2;YP_173236.1;HCoV-HKU1_orf1ab;4714 | 0 | 0.025 |
| NC_006577.2;YP_173236.1;HCoV-HKU1_orf1ab;4715 | 0 | 0 |
| NC_006577.2;YP_173236.1;HCoV-HKU1_orf1ab;4716 | 0 | 0 |
| NC_006577.2;YP_173236.1;HCoV-HKU1_orf1ab;4717 | 0.05 | 0 |
| NC_006577.2;YP_173236.1;HCoV-HKU1_orf1ab;4718 | 0.05 | 0 |
| NC_006577.2;YP_173236.1;HCoV-HKU1_orf1ab;4719 | 0.05 | 0 |
| NC_006577.2;YP_173236.1;HCoV-HKU1_orf1ab;4720 | 0.05 | 0 |
| NC_006577.2;YP_173236.1;HCoV-HKU1_orf1ab;4721 | 0 | 0 |
| NC_006577.2;YP_173236.1;HCoV-HKU1_orf1ab;4722 | 0.05 | 0 |
| NC_006577.2;YP_173236.1;HCoV-HKU1_orf1ab;4723 | 0.1 | 0.05 |
| NC_006577.2;YP_173236.1;HCoV-HKU1_orf1ab;4724 | 0.1 | 0.075 |
| NC_006577.2;YP_173236.1;HCoV-HKU1_orf1ab;4725 | 0.05 | 0.025 |
| NC_006577.2;YP_173236.1;HCoV-HKU1_orf1ab;4726 | 0 | 0.025 |
| NC_006577.2;YP_173236.1;HCoV-HKU1_orf1ab;4727 | 0 | 0.025 |
| NC_006577.2;YP_173236.1;HCoV-HKU1_orf1ab;4728 | 0 | 0 |
| NC_006577.2;YP_173236.1;HCoV-HKU1_orf1ab;4729 | 0 | 0.025 |
| NC_006577.2;YP_173236.1;HCoV-HKU1_orf1ab;4730 | 0 | 0 |
| NC_006577.2;YP_173236.1;HCoV-HKU1_orf1ab;4731 | 0 | 0 |
| NC_006577.2;YP_173236.1;HCoV-HKU1_orf1ab;4732 | 0 | 0 |
| NC_006577.2;YP_173236.1;HCoV-HKU1_orf1ab;4733 | 0 | 0 |
| NC_006577.2;YP_173236.1;HCoV-HKU1_orf1ab;4734 | 0 | 0 |
| NC_006577.2;YP_173236.1;HCoV-HKU1_orf1ab;4735 | 0 | 0 |
| NC_006577.2;YP_173236.1;HCoV-HKU1_orf1ab;4736 | 0 | 0 |
| NC_006577.2;YP_173236.1;HCoV-HKU1_orf1ab;4737 | 0 | 0 |
| NC_006577.2;YP_173236.1;HCoV-HKU1_orf1ab;4738 | 0 | 0 |
| NC_006577.2;YP_173236.1;HCoV-HKU1_orf1ab;4739 | 0 | 0 |
| NC_006577.2;YP_173236.1;HCoV-HKU1_orf1ab;4740 | 0.1 | 0.025 |
| NC_006577.2;YP_173236.1;HCoV-HKU1_orf1ab;4741 | 0.05 | 0 |
| NC_006577.2;YP_173236.1;HCoV-HKU1_orf1ab;4742 | 0 | 0 |
| NC_006577.2;YP_173236.1;HCoV-HKU1_orf1ab;4743 | 0 | 0.025 |
| NC_006577.2;YP_173236.1;HCoV-HKU1_orf1ab;4744 | 0.05 | 0 |
| NC_006577.2;YP_173236.1;HCoV-HKU1_orf1ab;4745 | 0.05 | 0 |

| | | |
|---|---|---|
| NC_006577.2;YP_173236.1;HCoV-HKU1_orf1ab;4746 | 0 | 0 |
| NC_006577.2;YP_173236.1;HCoV-HKU1_orf1ab;4747 | 0 | 0 |
| NC_006577.2;YP_173236.1;HCoV-HKU1_orf1ab;4748 | 0 | 0 |
| NC_006577.2;YP_173236.1;HCoV-HKU1_orf1ab;4749 | 0 | 0.025 |
| NC_006577.2;YP_173236.1;HCoV-HKU1_orf1ab;4750 | 0 | 0 |
| NC_006577.2;YP_173236.1;HCoV-HKU1_orf1ab;4751 | 0 | 0 |
| NC_006577.2;YP_173236.1;HCoV-HKU1_orf1ab;4752 | 0 | 0 |
| NC_006577.2;YP_173236.1;HCoV-HKU1_orf1ab;4753 | 0 | 0 |
| NC_006577.2;YP_173236.1;HCoV-HKU1_orf1ab;4754 | 0 | 0 |
| NC_006577.2;YP_173236.1;HCoV-HKU1_orf1ab;4755 | 0 | 0 |
| NC_006577.2;YP_173236.1;HCoV-HKU1_orf1ab;4756 | 0 | 0 |
| NC_006577.2;YP_173236.1;HCoV-HKU1_orf1ab;4757 | 0 | 0 |
| NC_006577.2;YP_173236.1;HCoV-HKU1_orf1ab;4758 | 0 | 0 |
| NC_006577.2;YP_173236.1;HCoV-HKU1_orf1ab;4759 | 0 | 0 |
| NC_006577.2;YP_173236.1;HCoV-HKU1_orf1ab;4760 | 0 | 0 |
| NC_006577.2;YP_173236.1;HCoV-HKU1_orf1ab;4761 | 0 | 0 |
| NC_006577.2;YP_173236.1;HCoV-HKU1_orf1ab;4762 | 0 | 0 |
| NC_006577.2;YP_173236.1;HCoV-HKU1_orf1ab;4763 | 0 | 0 |
| NC_006577.2;YP_173236.1;HCoV-HKU1_orf1ab;4764 | 0 | 0 |
| NC_006577.2;YP_173236.1;HCoV-HKU1_orf1ab;4765 | 0 | 0 |
| NC_006577.2;YP_173236.1;HCoV-HKU1_orf1ab;4766 | 0 | 0 |
| NC_006577.2;YP_173236.1;HCoV-HKU1_orf1ab;4767 | 0 | 0 |
| NC_006577.2;YP_173236.1;HCoV-HKU1_orf1ab;4768 | 0 | 0 |
| NC_006577.2;YP_173236.1;HCoV-HKU1_orf1ab;4769 | 0 | 0 |
| NC_006577.2;YP_173236.1;HCoV-HKU1_orf1ab;4770 | 0 | 0 |
| NC_006577.2;YP_173236.1;HCoV-HKU1_orf1ab;4771 | 0 | 0 |
| NC_006577.2;YP_173236.1;HCoV-HKU1_orf1ab;4772 | 0 | 0 |
| NC_006577.2;YP_173236.1;HCoV-HKU1_orf1ab;4773 | 0 | 0 |
| NC_006577.2;YP_173236.1;HCoV-HKU1_orf1ab;4774 | 0 | 0 |
| NC_006577.2;YP_173236.1;HCoV-HKU1_orf1ab;4775 | 0 | 0 |
| NC_006577.2;YP_173236.1;HCoV-HKU1_orf1ab;4776 | 0 | 0 |
| NC_006577.2;YP_173236.1;HCoV-HKU1_orf1ab;4777 | 0 | 0 |
| NC_006577.2;YP_173236.1;HCoV-HKU1_orf1ab;4778 | 0 | 0 |
| NC_006577.2;YP_173236.1;HCoV-HKU1_orf1ab;4779 | 0 | 0 |
| NC_006577.2;YP_173236.1;HCoV-HKU1_orf1ab;4780 | 0 | 0 |
| NC_006577.2;YP_173236.1;HCoV-HKU1_orf1ab;4781 | 0 | 0 |
| NC_006577.2;YP_173236.1;HCoV-HKU1_orf1ab;4782 | 0 | 0 |
| NC_006577.2;YP_173236.1;HCoV-HKU1_orf1ab;4783 | 0 | 0 |
| NC_006577.2;YP_173236.1;HCoV-HKU1_orf1ab;4784 | 0 | 0 |
| NC_006577.2;YP_173236.1;HCoV-HKU1_orf1ab;4785 | 0 | 0 |
| NC_006577.2;YP_173236.1;HCoV-HKU1_orf1ab;4786 | 0 | 0 |
| NC_006577.2;YP_173236.1;HCoV-HKU1_orf1ab;4787 | 0 | 0 |
| NC_006577.2;YP_173236.1;HCoV-HKU1_orf1ab;4788 | 0 | 0 |

| | | |
|---|---|---|
| NC_006577.2;YP_173236.1;HCoV-HKU1_orf1ab;4789 | 0 | 0 |
| NC_006577.2;YP_173236.1;HCoV-HKU1_orf1ab;4790 | 0 | 0 |
| NC_006577.2;YP_173236.1;HCoV-HKU1_orf1ab;4791 | 0 | 0 |
| NC_006577.2;YP_173236.1;HCoV-HKU1_orf1ab;4792 | 0 | 0 |
| NC_006577.2;YP_173236.1;HCoV-HKU1_orf1ab;4793 | 0 | 0 |
| NC_006577.2;YP_173236.1;HCoV-HKU1_orf1ab;4794 | 0 | 0.025 |
| NC_006577.2;YP_173236.1;HCoV-HKU1_orf1ab;4795 | 0 | 0 |
| NC_006577.2;YP_173236.1;HCoV-HKU1_orf1ab;4796 | 0 | 0 |
| NC_006577.2;YP_173236.1;HCoV-HKU1_orf1ab;4797 | 0 | 0.025 |
| NC_006577.2;YP_173236.1;HCoV-HKU1_orf1ab;4798 | 0 | 0 |
| NC_006577.2;YP_173236.1;HCoV-HKU1_orf1ab;4799 | 0 | 0.025 |
| NC_006577.2;YP_173236.1;HCoV-HKU1_orf1ab;4800 | 0.05 | 0.025 |
| NC_006577.2;YP_173236.1;HCoV-HKU1_orf1ab;4801 | 0.05 | 0.025 |
| NC_006577.2;YP_173236.1;HCoV-HKU1_orf1ab;4802 | 0.05 | 0.025 |
| NC_006577.2;YP_173236.1;HCoV-HKU1_orf1ab;4803 | 0 | 0 |
| NC_006577.2;YP_173236.1;HCoV-HKU1_orf1ab;4804 | 0 | 0 |
| NC_006577.2;YP_173236.1;HCoV-HKU1_orf1ab;4805 | 0 | 0 |
| NC_006577.2;YP_173236.1;HCoV-HKU1_orf1ab;4806 | 0 | 0 |
| NC_006577.2;YP_173236.1;HCoV-HKU1_orf1ab;4807 | 0 | 0 |
| NC_006577.2;YP_173236.1;HCoV-HKU1_orf1ab;4808 | 0 | 0 |
| NC_006577.2;YP_173236.1;HCoV-HKU1_orf1ab;4809 | 0 | 0 |
| NC_006577.2;YP_173236.1;HCoV-HKU1_orf1ab;4810 | 0 | 0 |
| NC_006577.2;YP_173236.1;HCoV-HKU1_orf1ab;4811 | 0 | 0 |
| NC_006577.2;YP_173236.1;HCoV-HKU1_orf1ab;4812 | 0 | 0 |
| NC_006577.2;YP_173236.1;HCoV-HKU1_orf1ab;4813 | 0 | 0 |
| NC_006577.2;YP_173236.1;HCoV-HKU1_orf1ab;4814 | 0 | 0 |
| NC_006577.2;YP_173236.1;HCoV-HKU1_orf1ab;4815 | 0 | 0 |
| NC_006577.2;YP_173236.1;HCoV-HKU1_orf1ab;4816 | 0 | 0 |
| NC_006577.2;YP_173236.1;HCoV-HKU1_orf1ab;4817 | 0 | 0 |
| NC_006577.2;YP_173236.1;HCoV-HKU1_orf1ab;4818 | 0 | 0 |
| NC_006577.2;YP_173236.1;HCoV-HKU1_orf1ab;4819 | 0 | 0 |
| NC_006577.2;YP_173236.1;HCoV-HKU1_orf1ab;4820 | 0 | 0.025 |
| NC_006577.2;YP_173236.1;HCoV-HKU1_orf1ab;4821 | 0 | 0.05 |
| NC_006577.2;YP_173236.1;HCoV-HKU1_orf1ab;4822 | 0 | 0 |
| NC_006577.2;YP_173236.1;HCoV-HKU1_orf1ab;4823 | 0 | 0 |
| NC_006577.2;YP_173236.1;HCoV-HKU1_orf1ab;4824 | 0 | 0 |
| NC_006577.2;YP_173236.1;HCoV-HKU1_orf1ab;4825 | 0.05 | 0 |
| NC_006577.2;YP_173236.1;HCoV-HKU1_orf1ab;4826 | 0 | 0 |
| NC_006577.2;YP_173236.1;HCoV-HKU1_orf1ab;4827 | 0 | 0 |
| NC_006577.2;YP_173236.1;HCoV-HKU1_orf1ab;4828 | 0 | 0 |
| NC_006577.2;YP_173236.1;HCoV-HKU1_orf1ab;4829 | 0 | 0 |
| NC_006577.2;YP_173236.1;HCoV-HKU1_orf1ab;4830 | 0 | 0 |
| NC_006577.2;YP_173236.1;HCoV-HKU1_orf1ab;4831 | 0 | 0 |

| | | |
|---|---|---|
| NC_006577.2;YP_173236.1;HCoV-HKU1_orf1ab;4832 | 0 | 0 |
| NC_006577.2;YP_173236.1;HCoV-HKU1_orf1ab;4833 | 0 | 0 |
| NC_006577.2;YP_173236.1;HCoV-HKU1_orf1ab;4834 | 0 | 0 |
| NC_006577.2;YP_173236.1;HCoV-HKU1_orf1ab;4835 | 0 | 0 |
| NC_006577.2;YP_173236.1;HCoV-HKU1_orf1ab;4836 | 0 | 0 |
| NC_006577.2;YP_173236.1;HCoV-HKU1_orf1ab;4837 | 0 | 0 |
| NC_006577.2;YP_173236.1;HCoV-HKU1_orf1ab;4838 | 0 | 0 |
| NC_006577.2;YP_173236.1;HCoV-HKU1_orf1ab;4839 | 0 | 0 |
| NC_006577.2;YP_173236.1;HCoV-HKU1_orf1ab;4840 | 0 | 0 |
| NC_006577.2;YP_173236.1;HCoV-HKU1_orf1ab;4841 | 0 | 0 |
| NC_006577.2;YP_173236.1;HCoV-HKU1_orf1ab;4842 | 0 | 0 |
| NC_006577.2;YP_173236.1;HCoV-HKU1_orf1ab;4843 | 0 | 0 |
| NC_006577.2;YP_173236.1;HCoV-HKU1_orf1ab;4844 | 0 | 0 |
| NC_006577.2;YP_173236.1;HCoV-HKU1_orf1ab;4845 | 0 | 0 |
| NC_006577.2;YP_173236.1;HCoV-HKU1_orf1ab;4846 | 0 | 0 |
| NC_006577.2;YP_173236.1;HCoV-HKU1_orf1ab;4847 | 0 | 0 |
| NC_006577.2;YP_173236.1;HCoV-HKU1_orf1ab;4848 | 0 | 0 |
| NC_006577.2;YP_173236.1;HCoV-HKU1_orf1ab;4849 | 0 | 0 |
| NC_006577.2;YP_173236.1;HCoV-HKU1_orf1ab;4850 | 0 | 0 |
| NC_006577.2;YP_173236.1;HCoV-HKU1_orf1ab;4851 | 0 | 0 |
| NC_006577.2;YP_173236.1;HCoV-HKU1_orf1ab;4852 | 0.05 | 0.025 |
| NC_006577.2;YP_173236.1;HCoV-HKU1_orf1ab;4853 | 0.05 | 0.025 |
| NC_006577.2;YP_173236.1;HCoV-HKU1_orf1ab;4854 | 0.05 | 0.025 |
| NC_006577.2;YP_173236.1;HCoV-HKU1_orf1ab;4855 | 0 | 0.025 |
| NC_006577.2;YP_173236.1;HCoV-HKU1_orf1ab;4856 | 0 | 0.025 |
| NC_006577.2;YP_173236.1;HCoV-HKU1_orf1ab;4857 | 0 | 0.025 |
| NC_006577.2;YP_173236.1;HCoV-HKU1_orf1ab;4858 | 0 | 0.05 |
| NC_006577.2;YP_173236.1;HCoV-HKU1_orf1ab;4859 | 0 | 0.025 |
| NC_006577.2;YP_173236.1;HCoV-HKU1_orf1ab;4860 | 0.05 | 0.025 |
| NC_006577.2;YP_173236.1;HCoV-HKU1_orf1ab;4861 | 0.05 | 0.05 |
| NC_006577.2;YP_173236.1;HCoV-HKU1_orf1ab;4862 | 0.15 | 0.075 |
| NC_006577.2;YP_173236.1;HCoV-HKU1_orf1ab;4863 | 0.15 | 0.05 |
| NC_006577.2;YP_173236.1;HCoV-HKU1_orf1ab;4864 | 0.05 | 0.025 |
| NC_006577.2;YP_173236.1;HCoV-HKU1_orf1ab;4865 | 0.05 | 0.025 |
| NC_006577.2;YP_173236.1;HCoV-HKU1_orf1ab;4866 | 0.05 | 0.025 |
| NC_006577.2;YP_173236.1;HCoV-HKU1_orf1ab;4867 | 0 | 0 |
| NC_006577.2;YP_173236.1;HCoV-HKU1_orf1ab;4868 | 0 | 0 |
| NC_006577.2;YP_173236.1;HCoV-HKU1_orf1ab;4869 | 0 | 0 |
| NC_006577.2;YP_173236.1;HCoV-HKU1_orf1ab;4870 | 0.05 | 0 |
| NC_006577.2;YP_173236.1;HCoV-HKU1_orf1ab;4871 | 0 | 0.025 |
| NC_006577.2;YP_173236.1;HCoV-HKU1_orf1ab;4872 | 0 | 0 |
| NC_006577.2;YP_173236.1;HCoV-HKU1_orf1ab;4873 | 0 | 0 |
| NC_006577.2;YP_173236.1;HCoV-HKU1_orf1ab;4874 | 0.05 | 0 |

| | | |
|---|---|---|
| NC_006577.2;YP_173236.1;HCoV-HKU1_orf1ab;4875 | 0.05 | 0 |
| NC_006577.2;YP_173236.1;HCoV-HKU1_orf1ab;4876 | 0 | 0 |
| NC_006577.2;YP_173236.1;HCoV-HKU1_orf1ab;4877 | 0 | 0.025 |
| NC_006577.2;YP_173236.1;HCoV-HKU1_orf1ab;4878 | 0 | 0 |
| NC_006577.2;YP_173236.1;HCoV-HKU1_orf1ab;4879 | 0 | 0.025 |
| NC_006577.2;YP_173236.1;HCoV-HKU1_orf1ab;4880 | 0 | 0 |
| NC_006577.2;YP_173236.1;HCoV-HKU1_orf1ab;4881 | 0 | 0 |
| NC_006577.2;YP_173236.1;HCoV-HKU1_orf1ab;4882 | 0 | 0 |
| NC_006577.2;YP_173236.1;HCoV-HKU1_orf1ab;4883 | 0 | 0 |
| NC_006577.2;YP_173236.1;HCoV-HKU1_orf1ab;4884 | 0 | 0 |
| NC_006577.2;YP_173236.1;HCoV-HKU1_orf1ab;4885 | 0 | 0 |
| NC_006577.2;YP_173236.1;HCoV-HKU1_orf1ab;4886 | 0 | 0 |
| NC_006577.2;YP_173236.1;HCoV-HKU1_orf1ab;4887 | 0.05 | 0.05 |
| NC_006577.2;YP_173236.1;HCoV-HKU1_orf1ab;4888 | 0 | 0 |
| NC_006577.2;YP_173236.1;HCoV-HKU1_orf1ab;4889 | 0.05 | 0 |
| NC_006577.2;YP_173236.1;HCoV-HKU1_orf1ab;4890 | 0 | 0 |
| NC_006577.2;YP_173236.1;HCoV-HKU1_orf1ab;4891 | 0 | 0 |
| NC_006577.2;YP_173236.1;HCoV-HKU1_orf1ab;4892 | 0 | 0 |
| NC_006577.2;YP_173236.1;HCoV-HKU1_orf1ab;4893 | 0 | 0 |
| NC_006577.2;YP_173236.1;HCoV-HKU1_orf1ab;4894 | 0 | 0.025 |
| NC_006577.2;YP_173236.1;HCoV-HKU1_orf1ab;4895 | 0 | 0.075 |
| NC_006577.2;YP_173236.1;HCoV-HKU1_orf1ab;4896 | 0 | 0.025 |
| NC_006577.2;YP_173236.1;HCoV-HKU1_orf1ab;4897 | 0.15 | 0.025 |
| NC_006577.2;YP_173236.1;HCoV-HKU1_orf1ab;4898 | 0.15 | 0.025 |
| NC_006577.2;YP_173236.1;HCoV-HKU1_orf1ab;4899 | 0.2 | 0.025 |
| NC_006577.2;YP_173236.1;HCoV-HKU1_orf1ab;4900 | 0.2 | 0.025 |
| NC_006577.2;YP_173236.1;HCoV-HKU1_orf1ab;4901 | 0.2 | 0.075 |
| NC_006577.2;YP_173236.1;HCoV-HKU1_orf1ab;4902 | 0.25 | 0.025 |
| NC_006577.2;YP_173236.1;HCoV-HKU1_orf1ab;4903 | 0.25 | 0.05 |
| NC_006577.2;YP_173236.1;HCoV-HKU1_orf1ab;4904 | 0.25 | 0.125 |
| NC_006577.2;YP_173236.1;HCoV-HKU1_orf1ab;4905 | 0.25 | 0.05 |
| NC_006577.2;YP_173236.1;HCoV-HKU1_orf1ab;4906 | 0.05 | 0.025 |
| NC_006577.2;YP_173236.1;HCoV-HKU1_orf1ab;4907 | 0 | 0 |
| NC_006577.2;YP_173236.1;HCoV-HKU1_orf1ab;4908 | 0 | 0 |
| NC_006577.2;YP_173236.1;HCoV-HKU1_orf1ab;4909 | 0 | 0 |
| NC_006577.2;YP_173236.1;HCoV-HKU1_orf1ab;4910 | 0 | 0 |
| NC_006577.2;YP_173236.1;HCoV-HKU1_orf1ab;4911 | 0 | 0 |
| NC_006577.2;YP_173236.1;HCoV-HKU1_orf1ab;4912 | 0 | 0 |
| NC_006577.2;YP_173236.1;HCoV-HKU1_orf1ab;4913 | 0 | 0 |
| NC_006577.2;YP_173236.1;HCoV-HKU1_orf1ab;4914 | 0 | 0 |
| NC_006577.2;YP_173236.1;HCoV-HKU1_orf1ab;4915 | 0 | 0 |
| NC_006577.2;YP_173236.1;HCoV-HKU1_orf1ab;4916 | 0 | 0 |
| NC_006577.2;YP_173236.1;HCoV-HKU1_orf1ab;4917 | 0 | 0 |

| | | |
|---|---|---|
| NC_006577.2;YP_173236.1;HCoV-HKU1_orf1ab;4918 | 0 | 0 |
| NC_006577.2;YP_173236.1;HCoV-HKU1_orf1ab;4919 | 0 | 0 |
| NC_006577.2;YP_173236.1;HCoV-HKU1_orf1ab;4920 | 0 | 0 |
| NC_006577.2;YP_173236.1;HCoV-HKU1_orf1ab;4921 | 0 | 0 |
| NC_006577.2;YP_173236.1;HCoV-HKU1_orf1ab;4922 | 0 | 0 |
| NC_006577.2;YP_173236.1;HCoV-HKU1_orf1ab;4923 | 0 | 0 |
| NC_006577.2;YP_173236.1;HCoV-HKU1_orf1ab;4924 | 0 | 0 |
| NC_006577.2;YP_173236.1;HCoV-HKU1_orf1ab;4925 | 0 | 0 |
| NC_006577.2;YP_173236.1;HCoV-HKU1_orf1ab;4926 | 0 | 0 |
| NC_006577.2;YP_173236.1;HCoV-HKU1_orf1ab;4927 | 0 | 0 |
| NC_006577.2;YP_173236.1;HCoV-HKU1_orf1ab;4928 | 0 | 0 |
| NC_006577.2;YP_173236.1;HCoV-HKU1_orf1ab;4929 | 0 | 0 |
| NC_006577.2;YP_173236.1;HCoV-HKU1_orf1ab;4930 | 0 | 0 |
| NC_006577.2;YP_173236.1;HCoV-HKU1_orf1ab;4931 | 0 | 0.025 |
| NC_006577.2;YP_173236.1;HCoV-HKU1_orf1ab;4932 | 0 | 0.025 |
| NC_006577.2;YP_173236.1;HCoV-HKU1_orf1ab;4933 | 0 | 0.025 |
| NC_006577.2;YP_173236.1;HCoV-HKU1_orf1ab;4934 | 0 | 0 |
| NC_006577.2;YP_173236.1;HCoV-HKU1_orf1ab;4935 | 0 | 0 |
| NC_006577.2;YP_173236.1;HCoV-HKU1_orf1ab;4936 | 0 | 0 |
| NC_006577.2;YP_173236.1;HCoV-HKU1_orf1ab;4937 | 0 | 0 |
| NC_006577.2;YP_173236.1;HCoV-HKU1_orf1ab;4938 | 0 | 0.025 |
| NC_006577.2;YP_173236.1;HCoV-HKU1_orf1ab;4939 | 0 | 0.025 |
| NC_006577.2;YP_173236.1;HCoV-HKU1_orf1ab;4940 | 0.05 | 0.05 |
| NC_006577.2;YP_173236.1;HCoV-HKU1_orf1ab;4941 | 0.05 | 0.025 |
| NC_006577.2;YP_173236.1;HCoV-HKU1_orf1ab;4942 | 0 | 0 |
| NC_006577.2;YP_173236.1;HCoV-HKU1_orf1ab;4943 | 0.1 | 0.075 |
| NC_006577.2;YP_173236.1;HCoV-HKU1_orf1ab;4944 | 0.15 | 0.1 |
| NC_006577.2;YP_173236.1;HCoV-HKU1_orf1ab;4945 | 0.15 | 0.05 |
| NC_006577.2;YP_173236.1;HCoV-HKU1_orf1ab;4946 | 0.25 | 0.05 |
| NC_006577.2;YP_173236.1;HCoV-HKU1_orf1ab;4947 | 0.1 | 0.025 |
| NC_006577.2;YP_173236.1;HCoV-HKU1_orf1ab;4948 | 0.15 | 0.025 |
| NC_006577.2;YP_173236.1;HCoV-HKU1_orf1ab;4949 | 0.1 | 0.025 |
| NC_006577.2;YP_173236.1;HCoV-HKU1_orf1ab;4950 | 0.1 | 0 |
| NC_006577.2;YP_173236.1;HCoV-HKU1_orf1ab;4951 | 0.1 | 0 |
| NC_006577.2;YP_173236.1;HCoV-HKU1_orf1ab;4952 | 0.15 | 0 |
| NC_006577.2;YP_173236.1;HCoV-HKU1_orf1ab;4953 | 0.05 | 0 |
| NC_006577.2;YP_173236.1;HCoV-HKU1_orf1ab;4954 | 0 | 0 |
| NC_006577.2;YP_173236.1;HCoV-HKU1_orf1ab;4955 | 0 | 0 |
| NC_006577.2;YP_173236.1;HCoV-HKU1_orf1ab;4956 | 0 | 0 |
| NC_006577.2;YP_173236.1;HCoV-HKU1_orf1ab;4957 | 0 | 0 |
| NC_006577.2;YP_173236.1;HCoV-HKU1_orf1ab;4958 | 0 | 0 |
| NC_006577.2;YP_173236.1;HCoV-HKU1_orf1ab;4959 | 0 | 0 |
| NC_006577.2;YP_173236.1;HCoV-HKU1_orf1ab;4960 | 0 | 0 |

| | | |
|---|---|---|
| NC_006577.2;YP_173236.1;HCoV-HKU1_orf1ab;4961 | 0 | 0 |
| NC_006577.2;YP_173236.1;HCoV-HKU1_orf1ab;4962 | 0 | 0 |
| NC_006577.2;YP_173236.1;HCoV-HKU1_orf1ab;4963 | 0 | 0 |
| NC_006577.2;YP_173236.1;HCoV-HKU1_orf1ab;4964 | 0 | 0 |
| NC_006577.2;YP_173236.1;HCoV-HKU1_orf1ab;4965 | 0 | 0 |
| NC_006577.2;YP_173236.1;HCoV-HKU1_orf1ab;4966 | 0 | 0 |
| NC_006577.2;YP_173236.1;HCoV-HKU1_orf1ab;4967 | 0 | 0 |
| NC_006577.2;YP_173236.1;HCoV-HKU1_orf1ab;4968 | 0 | 0 |
| NC_006577.2;YP_173236.1;HCoV-HKU1_orf1ab;4969 | 0 | 0 |
| NC_006577.2;YP_173236.1;HCoV-HKU1_orf1ab;4970 | 0.05 | 0.025 |
| NC_006577.2;YP_173236.1;HCoV-HKU1_orf1ab;4971 | 0 | 0.025 |
| NC_006577.2;YP_173236.1;HCoV-HKU1_orf1ab;4972 | 0 | 0.05 |
| NC_006577.2;YP_173236.1;HCoV-HKU1_orf1ab;4973 | 0 | 0.025 |
| NC_006577.2;YP_173236.1;HCoV-HKU1_orf1ab;4974 | 0 | 0 |
| NC_006577.2;YP_173236.1;HCoV-HKU1_orf1ab;4975 | 0 | 0.05 |
| NC_006577.2;YP_173236.1;HCoV-HKU1_orf1ab;4976 | 0 | 0.075 |
| NC_006577.2;YP_173236.1;HCoV-HKU1_orf1ab;4977 | 0 | 0 |
| NC_006577.2;YP_173236.1;HCoV-HKU1_orf1ab;4978 | 0 | 0 |
| NC_006577.2;YP_173236.1;HCoV-HKU1_orf1ab;4979 | 0 | 0 |
| NC_006577.2;YP_173236.1;HCoV-HKU1_orf1ab;4980 | 0 | 0 |
| NC_006577.2;YP_173236.1;HCoV-HKU1_orf1ab;4981 | 0 | 0 |
| NC_006577.2;YP_173236.1;HCoV-HKU1_orf1ab;4982 | 0 | 0 |
| NC_006577.2;YP_173236.1;HCoV-HKU1_orf1ab;4983 | 0 | 0 |
| NC_006577.2;YP_173236.1;HCoV-HKU1_orf1ab;4984 | 0.05 | 0.075 |
| NC_006577.2;YP_173236.1;HCoV-HKU1_orf1ab;4985 | 0.2 | 0.2 |
| NC_006577.2;YP_173236.1;HCoV-HKU1_orf1ab;4986 | 0 | 0 |
| NC_006577.2;YP_173236.1;HCoV-HKU1_orf1ab;4987 | 0 | 0 |
| NC_006577.2;YP_173236.1;HCoV-HKU1_orf1ab;4988 | 0 | 0 |
| NC_006577.2;YP_173236.1;HCoV-HKU1_orf1ab;4989 | 0 | 0.025 |
| NC_006577.2;YP_173236.1;HCoV-HKU1_orf1ab;4990 | 0 | 0.05 |
| NC_006577.2;YP_173236.1;HCoV-HKU1_orf1ab;4991 | 0 | 0.025 |
| NC_006577.2;YP_173236.1;HCoV-HKU1_orf1ab;4992 | 0 | 0.025 |
| NC_006577.2;YP_173236.1;HCoV-HKU1_orf1ab;4993 | 0 | 0.025 |
| NC_006577.2;YP_173236.1;HCoV-HKU1_orf1ab;4994 | 0 | 0.025 |
| NC_006577.2;YP_173236.1;HCoV-HKU1_orf1ab;4995 | 0 | 0 |
| NC_006577.2;YP_173236.1;HCoV-HKU1_orf1ab;4996 | 0 | 0 |
| NC_006577.2;YP_173236.1;HCoV-HKU1_orf1ab;4997 | 0 | 0 |
| NC_006577.2;YP_173236.1;HCoV-HKU1_orf1ab;4998 | 0 | 0 |
| NC_006577.2;YP_173236.1;HCoV-HKU1_orf1ab;4999 | 0 | 0 |
| NC_006577.2;YP_173236.1;HCoV-HKU1_orf1ab;5000 | 0 | 0 |
| NC_006577.2;YP_173236.1;HCoV-HKU1_orf1ab;5001 | 0 | 0 |
| NC_006577.2;YP_173236.1;HCoV-HKU1_orf1ab;5002 | 0 | 0 |
| NC_006577.2;YP_173236.1;HCoV-HKU1_orf1ab;5003 | 0 | 0 |

| | | |
|---|---|---|
| NC_006577.2;YP_173236.1;HCoV-HKU1_orf1ab;5004 | 0 | 0 |
| NC_006577.2;YP_173236.1;HCoV-HKU1_orf1ab;5005 | 0 | 0 |
| NC_006577.2;YP_173236.1;HCoV-HKU1_orf1ab;5006 | 0 | 0 |
| NC_006577.2;YP_173236.1;HCoV-HKU1_orf1ab;5007 | 0 | 0 |
| NC_006577.2;YP_173236.1;HCoV-HKU1_orf1ab;5008 | 0 | 0 |
| NC_006577.2;YP_173236.1;HCoV-HKU1_orf1ab;5009 | 0 | 0 |
| NC_006577.2;YP_173236.1;HCoV-HKU1_orf1ab;5010 | 0 | 0 |
| NC_006577.2;YP_173236.1;HCoV-HKU1_orf1ab;5011 | 0 | 0 |
| NC_006577.2;YP_173236.1;HCoV-HKU1_orf1ab;5012 | 0 | 0 |
| NC_006577.2;YP_173236.1;HCoV-HKU1_orf1ab;5013 | 0 | 0 |
| NC_006577.2;YP_173236.1;HCoV-HKU1_orf1ab;5014 | 0 | 0 |
| NC_006577.2;YP_173236.1;HCoV-HKU1_orf1ab;5015 | 0 | 0 |
| NC_006577.2;YP_173236.1;HCoV-HKU1_orf1ab;5016 | 0 | 0 |
| NC_006577.2;YP_173236.1;HCoV-HKU1_orf1ab;5017 | 0 | 0 |
| NC_006577.2;YP_173236.1;HCoV-HKU1_orf1ab;5018 | 0 | 0 |
| NC_006577.2;YP_173236.1;HCoV-HKU1_orf1ab;5019 | 0 | 0 |
| NC_006577.2;YP_173236.1;HCoV-HKU1_orf1ab;5020 | 0 | 0 |
| NC_006577.2;YP_173236.1;HCoV-HKU1_orf1ab;5021 | 0 | 0 |
| NC_006577.2;YP_173236.1;HCoV-HKU1_orf1ab;5022 | 0 | 0 |
| NC_006577.2;YP_173236.1;HCoV-HKU1_orf1ab;5023 | 0 | 0 |
| NC_006577.2;YP_173236.1;HCoV-HKU1_orf1ab;5024 | 0 | 0 |
| NC_006577.2;YP_173236.1;HCoV-HKU1_orf1ab;5025 | 0 | 0 |
| NC_006577.2;YP_173236.1;HCoV-HKU1_orf1ab;5026 | 0 | 0 |
| NC_006577.2;YP_173236.1;HCoV-HKU1_orf1ab;5027 | 0 | 0 |
| NC_006577.2;YP_173236.1;HCoV-HKU1_orf1ab;5028 | 0 | 0 |
| NC_006577.2;YP_173236.1;HCoV-HKU1_orf1ab;5029 | 0 | 0 |
| NC_006577.2;YP_173236.1;HCoV-HKU1_orf1ab;5030 | 0 | 0 |
| NC_006577.2;YP_173236.1;HCoV-HKU1_orf1ab;5031 | 0 | 0 |
| NC_006577.2;YP_173236.1;HCoV-HKU1_orf1ab;5032 | 0 | 0 |
| NC_006577.2;YP_173236.1;HCoV-HKU1_orf1ab;5033 | 0 | 0 |
| NC_006577.2;YP_173236.1;HCoV-HKU1_orf1ab;5034 | 0 | 0 |
| NC_006577.2;YP_173236.1;HCoV-HKU1_orf1ab;5035 | 0 | 0 |
| NC_006577.2;YP_173236.1;HCoV-HKU1_orf1ab;5036 | 0 | 0 |
| NC_006577.2;YP_173236.1;HCoV-HKU1_orf1ab;5037 | 0 | 0 |
| NC_006577.2;YP_173236.1;HCoV-HKU1_orf1ab;5038 | 0 | 0 |
| NC_006577.2;YP_173236.1;HCoV-HKU1_orf1ab;5039 | 0 | 0 |
| NC_006577.2;YP_173236.1;HCoV-HKU1_orf1ab;5040 | 0 | 0 |
| NC_006577.2;YP_173236.1;HCoV-HKU1_orf1ab;5041 | 0 | 0 |
| NC_006577.2;YP_173236.1;HCoV-HKU1_orf1ab;5042 | 0 | 0 |
| NC_006577.2;YP_173236.1;HCoV-HKU1_orf1ab;5043 | 0 | 0 |
| NC_006577.2;YP_173236.1;HCoV-HKU1_orf1ab;5044 | 0 | 0 |
| NC_006577.2;YP_173236.1;HCoV-HKU1_orf1ab;5045 | 0 | 0 |
| NC_006577.2;YP_173236.1;HCoV-HKU1_orf1ab;5046 | 0.05 | 0.025 |

| | | |
|---|---|---|
| NC_006577.2;YP_173236.1;HCoV-HKU1_orf1ab;5047 | 0.05 | 0 |
| NC_006577.2;YP_173236.1;HCoV-HKU1_orf1ab;5048 | 0.05 | 0 |
| NC_006577.2;YP_173236.1;HCoV-HKU1_orf1ab;5049 | 0 | 0 |
| NC_006577.2;YP_173236.1;HCoV-HKU1_orf1ab;5050 | 0.15 | 0.075 |
| NC_006577.2;YP_173236.1;HCoV-HKU1_orf1ab;5051 | 0 | 0.025 |
| NC_006577.2;YP_173236.1;HCoV-HKU1_orf1ab;5052 | 0 | 0.025 |
| NC_006577.2;YP_173236.1;HCoV-HKU1_orf1ab;5053 | 0 | 0.025 |
| NC_006577.2;YP_173236.1;HCoV-HKU1_orf1ab;5054 | 0.1 | 0.075 |
| NC_006577.2;YP_173236.1;HCoV-HKU1_orf1ab;5055 | 0 | 0.025 |
| NC_006577.2;YP_173236.1;HCoV-HKU1_orf1ab;5056 | 0.05 | 0.025 |
| NC_006577.2;YP_173236.1;HCoV-HKU1_orf1ab;5057 | 0.05 | 0.025 |
| NC_006577.2;YP_173236.1;HCoV-HKU1_orf1ab;5058 | 0.1 | 0.15 |
| NC_006577.2;YP_173236.1;HCoV-HKU1_orf1ab;5059 | 0.1 | 0.175 |
| NC_006577.2;YP_173236.1;HCoV-HKU1_orf1ab;5060 | 0.1 | 0.15 |
| NC_006577.2;YP_173236.1;HCoV-HKU1_orf1ab;5061 | 0 | 0.025 |
| NC_006577.2;YP_173236.1;HCoV-HKU1_orf1ab;5062 | 0 | 0.025 |
| NC_006577.2;YP_173236.1;HCoV-HKU1_orf1ab;5063 | 0 | 0.025 |
| NC_006577.2;YP_173236.1;HCoV-HKU1_orf1ab;5064 | 0 | 0.025 |
| NC_006577.2;YP_173236.1;HCoV-HKU1_orf1ab;5065 | 0 | 0.025 |
| NC_006577.2;YP_173236.1;HCoV-HKU1_orf1ab;5066 | 0 | 0.025 |
| NC_006577.2;YP_173236.1;HCoV-HKU1_orf1ab;5067 | 0 | 0.025 |
| NC_006577.2;YP_173236.1;HCoV-HKU1_orf1ab;5068 | 0 | 0.025 |
| NC_006577.2;YP_173236.1;HCoV-HKU1_orf1ab;5069 | 0 | 0 |
| NC_006577.2;YP_173236.1;HCoV-HKU1_orf1ab;5070 | 0 | 0 |
| NC_006577.2;YP_173236.1;HCoV-HKU1_orf1ab;5071 | 0 | 0 |
| NC_006577.2;YP_173236.1;HCoV-HKU1_orf1ab;5072 | 0 | 0 |
| NC_006577.2;YP_173236.1;HCoV-HKU1_orf1ab;5073 | 0 | 0 |
| NC_006577.2;YP_173236.1;HCoV-HKU1_orf1ab;5074 | 0 | 0 |
| NC_006577.2;YP_173236.1;HCoV-HKU1_orf1ab;5075 | 0 | 0 |
| NC_006577.2;YP_173236.1;HCoV-HKU1_orf1ab;5076 | 0 | 0 |
| NC_006577.2;YP_173236.1;HCoV-HKU1_orf1ab;5077 | 0 | 0 |
| NC_006577.2;YP_173236.1;HCoV-HKU1_orf1ab;5078 | 0 | 0 |
| NC_006577.2;YP_173236.1;HCoV-HKU1_orf1ab;5079 | 0 | 0 |
| NC_006577.2;YP_173236.1;HCoV-HKU1_orf1ab;5080 | 0 | 0 |
| NC_006577.2;YP_173236.1;HCoV-HKU1_orf1ab;5081 | 0 | 0 |
| NC_006577.2;YP_173236.1;HCoV-HKU1_orf1ab;5082 | 0 | 0 |
| NC_006577.2;YP_173236.1;HCoV-HKU1_orf1ab;5083 | 0 | 0 |
| NC_006577.2;YP_173236.1;HCoV-HKU1_orf1ab;5084 | 0 | 0 |
| NC_006577.2;YP_173236.1;HCoV-HKU1_orf1ab;5085 | 0 | 0 |
| NC_006577.2;YP_173236.1;HCoV-HKU1_orf1ab;5086 | 0 | 0 |
| NC_006577.2;YP_173236.1;HCoV-HKU1_orf1ab;5087 | 0 | 0 |
| NC_006577.2;YP_173236.1;HCoV-HKU1_orf1ab;5088 | 0 | 0 |
| NC_006577.2;YP_173236.1;HCoV-HKU1_orf1ab;5089 | 0 | 0 |

| | | |
|---|---|---|
| NC_006577.2;YP_173236.1;HCoV-HKU1_orf1ab;5090 | 0 | 0 |
| NC_006577.2;YP_173236.1;HCoV-HKU1_orf1ab;5091 | 0 | 0 |
| NC_006577.2;YP_173236.1;HCoV-HKU1_orf1ab;5092 | 0 | 0 |
| NC_006577.2;YP_173236.1;HCoV-HKU1_orf1ab;5093 | 0 | 0 |
| NC_006577.2;YP_173236.1;HCoV-HKU1_orf1ab;5094 | 0 | 0 |
| NC_006577.2;YP_173236.1;HCoV-HKU1_orf1ab;5095 | 0 | 0 |
| NC_006577.2;YP_173236.1;HCoV-HKU1_orf1ab;5096 | 0.05 | 0 |
| NC_006577.2;YP_173236.1;HCoV-HKU1_orf1ab;5097 | 0.05 | 0 |
| NC_006577.2;YP_173236.1;HCoV-HKU1_orf1ab;5098 | 0.05 | 0 |
| NC_006577.2;YP_173236.1;HCoV-HKU1_orf1ab;5099 | 0.05 | 0 |
| NC_006577.2;YP_173236.1;HCoV-HKU1_orf1ab;5100 | 0.05 | 0 |
| NC_006577.2;YP_173236.1;HCoV-HKU1_orf1ab;5101 | 0.05 | 0 |
| NC_006577.2;YP_173236.1;HCoV-HKU1_orf1ab;5102 | 0 | 0 |
| NC_006577.2;YP_173236.1;HCoV-HKU1_orf1ab;5103 | 0 | 0 |
| NC_006577.2;YP_173236.1;HCoV-HKU1_orf1ab;5104 | 0 | 0 |
| NC_006577.2;YP_173236.1;HCoV-HKU1_orf1ab;5105 | 0 | 0 |
| NC_006577.2;YP_173236.1;HCoV-HKU1_orf1ab;5106 | 0 | 0 |
| NC_006577.2;YP_173236.1;HCoV-HKU1_orf1ab;5107 | 0 | 0 |
| NC_006577.2;YP_173236.1;HCoV-HKU1_orf1ab;5108 | 0 | 0 |
| NC_006577.2;YP_173236.1;HCoV-HKU1_orf1ab;5109 | 0 | 0 |
| NC_006577.2;YP_173236.1;HCoV-HKU1_orf1ab;5110 | 0 | 0 |
| NC_006577.2;YP_173236.1;HCoV-HKU1_orf1ab;5111 | 0 | 0 |
| NC_006577.2;YP_173236.1;HCoV-HKU1_orf1ab;5112 | 0 | 0 |
| NC_006577.2;YP_173236.1;HCoV-HKU1_orf1ab;5113 | 0 | 0 |
| NC_006577.2;YP_173236.1;HCoV-HKU1_orf1ab;5114 | 0 | 0 |
| NC_006577.2;YP_173236.1;HCoV-HKU1_orf1ab;5115 | 0 | 0 |
| NC_006577.2;YP_173236.1;HCoV-HKU1_orf1ab;5116 | 0 | 0 |
| NC_006577.2;YP_173236.1;HCoV-HKU1_orf1ab;5117 | 0 | 0.025 |
| NC_006577.2;YP_173236.1;HCoV-HKU1_orf1ab;5118 | 0 | 0 |
| NC_006577.2;YP_173236.1;HCoV-HKU1_orf1ab;5119 | 0 | 0 |
| NC_006577.2;YP_173236.1;HCoV-HKU1_orf1ab;5120 | 0 | 0 |
| NC_006577.2;YP_173236.1;HCoV-HKU1_orf1ab;5121 | 0 | 0 |
| NC_006577.2;YP_173236.1;HCoV-HKU1_orf1ab;5122 | 0 | 0.025 |
| NC_006577.2;YP_173236.1;HCoV-HKU1_orf1ab;5123 | 0 | 0.025 |
| NC_006577.2;YP_173236.1;HCoV-HKU1_orf1ab;5124 | 0 | 0.025 |
| NC_006577.2;YP_173236.1;HCoV-HKU1_orf1ab;5125 | 0 | 0.025 |
| NC_006577.2;YP_173236.1;HCoV-HKU1_orf1ab;5126 | 0 | 0.05 |
| NC_006577.2;YP_173236.1;HCoV-HKU1_orf1ab;5127 | 0.05 | 0.075 |
| NC_006577.2;YP_173236.1;HCoV-HKU1_orf1ab;5128 | 0.05 | 0.025 |
| NC_006577.2;YP_173236.1;HCoV-HKU1_orf1ab;5129 | 0.05 | 0.05 |
| NC_006577.2;YP_173236.1;HCoV-HKU1_orf1ab;5130 | 0.05 | 0.025 |
| NC_006577.2;YP_173236.1;HCoV-HKU1_orf1ab;5131 | 0.05 | 0.05 |
| NC_006577.2;YP_173236.1;HCoV-HKU1_orf1ab;5132 | 0.05 | 0 |

| | | |
|---|---|---|
| NC_006577.2;YP_173236.1;HCoV-HKU1_orf1ab;5133 | 0.05 | 0 |
| NC_006577.2;YP_173236.1;HCoV-HKU1_orf1ab;5134 | 0 | 0 |
| NC_006577.2;YP_173236.1;HCoV-HKU1_orf1ab;5135 | 0 | 0 |
| NC_006577.2;YP_173236.1;HCoV-HKU1_orf1ab;5136 | 0 | 0 |
| NC_006577.2;YP_173236.1;HCoV-HKU1_orf1ab;5137 | 0 | 0 |
| NC_006577.2;YP_173236.1;HCoV-HKU1_orf1ab;5138 | 0 | 0 |
| NC_006577.2;YP_173236.1;HCoV-HKU1_orf1ab;5139 | 0 | 0 |
| NC_006577.2;YP_173236.1;HCoV-HKU1_orf1ab;5140 | 0 | 0 |
| NC_006577.2;YP_173236.1;HCoV-HKU1_orf1ab;5141 | 0 | 0 |
| NC_006577.2;YP_173236.1;HCoV-HKU1_orf1ab;5142 | 0 | 0.025 |
| NC_006577.2;YP_173236.1;HCoV-HKU1_orf1ab;5143 | 0 | 0 |
| NC_006577.2;YP_173236.1;HCoV-HKU1_orf1ab;5144 | 0 | 0 |
| NC_006577.2;YP_173236.1;HCoV-HKU1_orf1ab;5145 | 0 | 0 |
| NC_006577.2;YP_173236.1;HCoV-HKU1_orf1ab;5146 | 0 | 0 |
| NC_006577.2;YP_173236.1;HCoV-HKU1_orf1ab;5147 | 0 | 0 |
| NC_006577.2;YP_173236.1;HCoV-HKU1_orf1ab;5148 | 0 | 0 |
| NC_006577.2;YP_173236.1;HCoV-HKU1_orf1ab;5149 | 0 | 0 |
| NC_006577.2;YP_173236.1;HCoV-HKU1_orf1ab;5150 | 0 | 0 |
| NC_006577.2;YP_173236.1;HCoV-HKU1_orf1ab;5151 | 0 | 0 |
| NC_006577.2;YP_173236.1;HCoV-HKU1_orf1ab;5152 | 0 | 0.025 |
| NC_006577.2;YP_173236.1;HCoV-HKU1_orf1ab;5153 | 0 | 0 |
| NC_006577.2;YP_173236.1;HCoV-HKU1_orf1ab;5154 | 0 | 0 |
| NC_006577.2;YP_173236.1;HCoV-HKU1_orf1ab;5155 | 0 | 0 |
| NC_006577.2;YP_173236.1;HCoV-HKU1_orf1ab;5156 | 0 | 0 |
| NC_006577.2;YP_173236.1;HCoV-HKU1_orf1ab;5157 | 0 | 0 |
| NC_006577.2;YP_173236.1;HCoV-HKU1_orf1ab;5158 | 0 | 0 |
| NC_006577.2;YP_173236.1;HCoV-HKU1_orf1ab;5159 | 0 | 0 |
| NC_006577.2;YP_173236.1;HCoV-HKU1_orf1ab;5160 | 0 | 0 |
| NC_006577.2;YP_173236.1;HCoV-HKU1_orf1ab;5161 | 0 | 0 |
| NC_006577.2;YP_173236.1;HCoV-HKU1_orf1ab;5162 | 0 | 0 |
| NC_006577.2;YP_173236.1;HCoV-HKU1_orf1ab;5163 | 0 | 0.025 |
| NC_006577.2;YP_173236.1;HCoV-HKU1_orf1ab;5164 | 0 | 0.025 |
| NC_006577.2;YP_173236.1;HCoV-HKU1_orf1ab;5165 | 0 | 0 |
| NC_006577.2;YP_173236.1;HCoV-HKU1_orf1ab;5166 | 0 | 0 |
| NC_006577.2;YP_173236.1;HCoV-HKU1_orf1ab;5167 | 0 | 0 |
| NC_006577.2;YP_173236.1;HCoV-HKU1_orf1ab;5168 | 0 | 0.025 |
| NC_006577.2;YP_173236.1;HCoV-HKU1_orf1ab;5169 | 0 | 0.05 |
| NC_006577.2;YP_173236.1;HCoV-HKU1_orf1ab;5170 | 0 | 0 |
| NC_006577.2;YP_173236.1;HCoV-HKU1_orf1ab;5171 | 0 | 0 |
| NC_006577.2;YP_173236.1;HCoV-HKU1_orf1ab;5172 | 0 | 0 |
| NC_006577.2;YP_173236.1;HCoV-HKU1_orf1ab;5173 | 0 | 0 |
| NC_006577.2;YP_173236.1;HCoV-HKU1_orf1ab;5174 | 0.05 | 0 |
| NC_006577.2;YP_173236.1;HCoV-HKU1_orf1ab;5175 | 0 | 0 |

| | | |
|---|---|---|
| NC_006577.2;YP_173236.1;HCoV-HKU1_orf1ab;5176 | 0 | 0 |
| NC_006577.2;YP_173236.1;HCoV-HKU1_orf1ab;5177 | 0 | 0 |
| NC_006577.2;YP_173236.1;HCoV-HKU1_orf1ab;5178 | 0 | 0 |
| NC_006577.2;YP_173236.1;HCoV-HKU1_orf1ab;5179 | 0 | 0 |
| NC_006577.2;YP_173236.1;HCoV-HKU1_orf1ab;5180 | 0 | 0.025 |
| NC_006577.2;YP_173236.1;HCoV-HKU1_orf1ab;5181 | 0.05 | 0.025 |
| NC_006577.2;YP_173236.1;HCoV-HKU1_orf1ab;5182 | 0.05 | 0.025 |
| NC_006577.2;YP_173236.1;HCoV-HKU1_orf1ab;5183 | 0 | 0.025 |
| NC_006577.2;YP_173236.1;HCoV-HKU1_orf1ab;5184 | 0 | 0 |
| NC_006577.2;YP_173236.1;HCoV-HKU1_orf1ab;5185 | 0 | 0 |
| NC_006577.2;YP_173236.1;HCoV-HKU1_orf1ab;5186 | 0 | 0 |
| NC_006577.2;YP_173236.1;HCoV-HKU1_orf1ab;5187 | 0 | 0 |
| NC_006577.2;YP_173236.1;HCoV-HKU1_orf1ab;5188 | 0 | 0 |
| NC_006577.2;YP_173236.1;HCoV-HKU1_orf1ab;5189 | 0 | 0 |
| NC_006577.2;YP_173236.1;HCoV-HKU1_orf1ab;5190 | 0 | 0.05 |
| NC_006577.2;YP_173236.1;HCoV-HKU1_orf1ab;5191 | 0 | 0 |
| NC_006577.2;YP_173236.1;HCoV-HKU1_orf1ab;5192 | 0 | 0 |
| NC_006577.2;YP_173236.1;HCoV-HKU1_orf1ab;5193 | 0 | 0.025 |
| NC_006577.2;YP_173236.1;HCoV-HKU1_orf1ab;5194 | 0.05 | 0.05 |
| NC_006577.2;YP_173236.1;HCoV-HKU1_orf1ab;5195 | 0 | 0 |
| NC_006577.2;YP_173236.1;HCoV-HKU1_orf1ab;5196 | 0 | 0 |
| NC_006577.2;YP_173236.1;HCoV-HKU1_orf1ab;5197 | 0 | 0 |
| NC_006577.2;YP_173236.1;HCoV-HKU1_orf1ab;5198 | 0 | 0 |
| NC_006577.2;YP_173236.1;HCoV-HKU1_orf1ab;5199 | 0 | 0 |
| NC_006577.2;YP_173236.1;HCoV-HKU1_orf1ab;5200 | 0 | 0 |
| NC_006577.2;YP_173236.1;HCoV-HKU1_orf1ab;5201 | 0 | 0 |
| NC_006577.2;YP_173236.1;HCoV-HKU1_orf1ab;5202 | 0 | 0 |
| NC_006577.2;YP_173236.1;HCoV-HKU1_orf1ab;5203 | 0 | 0 |
| NC_006577.2;YP_173236.1;HCoV-HKU1_orf1ab;5204 | 0 | 0 |
| NC_006577.2;YP_173236.1;HCoV-HKU1_orf1ab;5205 | 0 | 0 |
| NC_006577.2;YP_173236.1;HCoV-HKU1_orf1ab;5206 | 0 | 0 |
| NC_006577.2;YP_173236.1;HCoV-HKU1_orf1ab;5207 | 0 | 0 |
| NC_006577.2;YP_173236.1;HCoV-HKU1_orf1ab;5208 | 0 | 0 |
| NC_006577.2;YP_173236.1;HCoV-HKU1_orf1ab;5209 | 0 | 0.025 |
| NC_006577.2;YP_173236.1;HCoV-HKU1_orf1ab;5210 | 0 | 0.025 |
| NC_006577.2;YP_173236.1;HCoV-HKU1_orf1ab;5211 | 0 | 0.025 |
| NC_006577.2;YP_173236.1;HCoV-HKU1_orf1ab;5212 | 0 | 0.025 |
| NC_006577.2;YP_173236.1;HCoV-HKU1_orf1ab;5213 | 0 | 0.05 |
| NC_006577.2;YP_173236.1;HCoV-HKU1_orf1ab;5214 | 0 | 0.025 |
| NC_006577.2;YP_173236.1;HCoV-HKU1_orf1ab;5215 | 0 | 0.05 |
| NC_006577.2;YP_173236.1;HCoV-HKU1_orf1ab;5216 | 0.05 | 0.025 |
| NC_006577.2;YP_173236.1;HCoV-HKU1_orf1ab;5217 | 0 | 0.025 |
| NC_006577.2;YP_173236.1;HCoV-HKU1_orf1ab;5218 | 0.05 | 0.025 |

| | | |
|---|---|---|
| NC_006577.2;YP_173236.1;HCoV-HKU1_orf1ab;5219 | 0 | 0 |
| NC_006577.2;YP_173236.1;HCoV-HKU1_orf1ab;5220 | 0 | 0 |
| NC_006577.2;YP_173236.1;HCoV-HKU1_orf1ab;5221 | 0 | 0 |
| NC_006577.2;YP_173236.1;HCoV-HKU1_orf1ab;5222 | 0 | 0 |
| NC_006577.2;YP_173236.1;HCoV-HKU1_orf1ab;5223 | 0 | 0 |
| NC_006577.2;YP_173236.1;HCoV-HKU1_orf1ab;5224 | 0 | 0 |
| NC_006577.2;YP_173236.1;HCoV-HKU1_orf1ab;5225 | 0 | 0 |
| NC_006577.2;YP_173236.1;HCoV-HKU1_orf1ab;5226 | 0 | 0 |
| NC_006577.2;YP_173236.1;HCoV-HKU1_orf1ab;5227 | 0 | 0 |
| NC_006577.2;YP_173236.1;HCoV-HKU1_orf1ab;5228 | 0 | 0 |
| NC_006577.2;YP_173236.1;HCoV-HKU1_orf1ab;5229 | 0 | 0 |
| NC_006577.2;YP_173236.1;HCoV-HKU1_orf1ab;5230 | 0 | 0 |
| NC_006577.2;YP_173236.1;HCoV-HKU1_orf1ab;5231 | 0 | 0 |
| NC_006577.2;YP_173236.1;HCoV-HKU1_orf1ab;5232 | 0 | 0 |
| NC_006577.2;YP_173236.1;HCoV-HKU1_orf1ab;5233 | 0 | 0 |
| NC_006577.2;YP_173236.1;HCoV-HKU1_orf1ab;5234 | 0 | 0 |
| NC_006577.2;YP_173236.1;HCoV-HKU1_orf1ab;5235 | 0 | 0 |
| NC_006577.2;YP_173236.1;HCoV-HKU1_orf1ab;5236 | 0.05 | 0.025 |
| NC_006577.2;YP_173236.1;HCoV-HKU1_orf1ab;5237 | 0 | 0 |
| NC_006577.2;YP_173236.1;HCoV-HKU1_orf1ab;5238 | 0 | 0 |
| NC_006577.2;YP_173236.1;HCoV-HKU1_orf1ab;5239 | 0 | 0 |
| NC_006577.2;YP_173236.1;HCoV-HKU1_orf1ab;5240 | 0 | 0.025 |
| NC_006577.2;YP_173236.1;HCoV-HKU1_orf1ab;5241 | 0 | 0.025 |
| NC_006577.2;YP_173236.1;HCoV-HKU1_orf1ab;5242 | 0 | 0.025 |
| NC_006577.2;YP_173236.1;HCoV-HKU1_orf1ab;5243 | 0.05 | 0.025 |
| NC_006577.2;YP_173236.1;HCoV-HKU1_orf1ab;5244 | 0.05 | 0.025 |
| NC_006577.2;YP_173236.1;HCoV-HKU1_orf1ab;5245 | 0.05 | 0.025 |
| NC_006577.2;YP_173236.1;HCoV-HKU1_orf1ab;5246 | 0.05 | 0 |
| NC_006577.2;YP_173236.1;HCoV-HKU1_orf1ab;5247 | 0.1 | 0.025 |
| NC_006577.2;YP_173236.1;HCoV-HKU1_orf1ab;5248 | 0.05 | 0 |
| NC_006577.2;YP_173236.1;HCoV-HKU1_orf1ab;5249 | 0.05 | 0 |
| NC_006577.2;YP_173236.1;HCoV-HKU1_orf1ab;5250 | 0.05 | 0 |
| NC_006577.2;YP_173236.1;HCoV-HKU1_orf1ab;5251 | 0.05 | 0 |
| NC_006577.2;YP_173236.1;HCoV-HKU1_orf1ab;5252 | 0.05 | 0 |
| NC_006577.2;YP_173236.1;HCoV-HKU1_orf1ab;5253 | 0.05 | 0.025 |
| NC_006577.2;YP_173236.1;HCoV-HKU1_orf1ab;5254 | 0 | 0 |
| NC_006577.2;YP_173236.1;HCoV-HKU1_orf1ab;5255 | 0 | 0.025 |
| NC_006577.2;YP_173236.1;HCoV-HKU1_orf1ab;5256 | 0.05 | 0 |
| NC_006577.2;YP_173236.1;HCoV-HKU1_orf1ab;5257 | 0 | 0 |
| NC_006577.2;YP_173236.1;HCoV-HKU1_orf1ab;5258 | 0 | 0 |
| NC_006577.2;YP_173236.1;HCoV-HKU1_orf1ab;5259 | 0 | 0 |
| NC_006577.2;YP_173236.1;HCoV-HKU1_orf1ab;5260 | 0 | 0 |
| NC_006577.2;YP_173236.1;HCoV-HKU1_orf1ab;5261 | 0 | 0 |

| | | |
|---|---|---|
| NC_006577.2;YP_173236.1;HCoV-HKU1_orf1ab;5262 | 0 | 0 |
| NC_006577.2;YP_173236.1;HCoV-HKU1_orf1ab;5263 | 0 | 0 |
| NC_006577.2;YP_173236.1;HCoV-HKU1_orf1ab;5264 | 0 | 0 |
| NC_006577.2;YP_173236.1;HCoV-HKU1_orf1ab;5265 | 0 | 0 |
| NC_006577.2;YP_173236.1;HCoV-HKU1_orf1ab;5266 | 0 | 0 |
| NC_006577.2;YP_173236.1;HCoV-HKU1_orf1ab;5267 | 0 | 0 |
| NC_006577.2;YP_173236.1;HCoV-HKU1_orf1ab;5268 | 0 | 0 |
| NC_006577.2;YP_173236.1;HCoV-HKU1_orf1ab;5269 | 0 | 0 |
| NC_006577.2;YP_173236.1;HCoV-HKU1_orf1ab;5270 | 0 | 0 |
| NC_006577.2;YP_173236.1;HCoV-HKU1_orf1ab;5271 | 0 | 0 |
| NC_006577.2;YP_173236.1;HCoV-HKU1_orf1ab;5272 | 0 | 0 |
| NC_006577.2;YP_173236.1;HCoV-HKU1_orf1ab;5273 | 0 | 0.075 |
| NC_006577.2;YP_173236.1;HCoV-HKU1_orf1ab;5274 | 0 | 0.075 |
| NC_006577.2;YP_173236.1;HCoV-HKU1_orf1ab;5275 | 0 | 0 |
| NC_006577.2;YP_173236.1;HCoV-HKU1_orf1ab;5276 | 0 | 0 |
| NC_006577.2;YP_173236.1;HCoV-HKU1_orf1ab;5277 | 0 | 0 |
| NC_006577.2;YP_173236.1;HCoV-HKU1_orf1ab;5278 | 0 | 0 |
| NC_006577.2;YP_173236.1;HCoV-HKU1_orf1ab;5279 | 0 | 0 |
| NC_006577.2;YP_173236.1;HCoV-HKU1_orf1ab;5280 | 0 | 0 |
| NC_006577.2;YP_173236.1;HCoV-HKU1_orf1ab;5281 | 0 | 0 |
| NC_006577.2;YP_173236.1;HCoV-HKU1_orf1ab;5282 | 0 | 0 |
| NC_006577.2;YP_173236.1;HCoV-HKU1_orf1ab;5283 | 0 | 0 |
| NC_006577.2;YP_173236.1;HCoV-HKU1_orf1ab;5284 | 0 | 0 |
| NC_006577.2;YP_173236.1;HCoV-HKU1_orf1ab;5285 | 0 | 0 |
| NC_006577.2;YP_173236.1;HCoV-HKU1_orf1ab;5286 | 0 | 0 |
| NC_006577.2;YP_173236.1;HCoV-HKU1_orf1ab;5287 | 0 | 0 |
| NC_006577.2;YP_173236.1;HCoV-HKU1_orf1ab;5288 | 0 | 0 |
| NC_006577.2;YP_173236.1;HCoV-HKU1_orf1ab;5289 | 0 | 0 |
| NC_006577.2;YP_173236.1;HCoV-HKU1_orf1ab;5290 | 0.05 | 0 |
| NC_006577.2;YP_173236.1;HCoV-HKU1_orf1ab;5291 | 0.05 | 0 |
| NC_006577.2;YP_173236.1;HCoV-HKU1_orf1ab;5292 | 0 | 0 |
| NC_006577.2;YP_173236.1;HCoV-HKU1_orf1ab;5293 | 0 | 0 |
| NC_006577.2;YP_173236.1;HCoV-HKU1_orf1ab;5294 | 0 | 0 |
| NC_006577.2;YP_173236.1;HCoV-HKU1_orf1ab;5295 | 0 | 0 |
| NC_006577.2;YP_173236.1;HCoV-HKU1_orf1ab;5296 | 0 | 0 |
| NC_006577.2;YP_173236.1;HCoV-HKU1_orf1ab;5297 | 0 | 0.025 |
| NC_006577.2;YP_173236.1;HCoV-HKU1_orf1ab;5298 | 0 | 0.025 |
| NC_006577.2;YP_173236.1;HCoV-HKU1_orf1ab;5299 | 0 | 0.025 |
| NC_006577.2;YP_173236.1;HCoV-HKU1_orf1ab;5300 | 0 | 0.025 |
| NC_006577.2;YP_173236.1;HCoV-HKU1_orf1ab;5301 | 0 | 0.025 |
| NC_006577.2;YP_173236.1;HCoV-HKU1_orf1ab;5302 | 0 | 0.025 |
| NC_006577.2;YP_173236.1;HCoV-HKU1_orf1ab;5303 | 0 | 0.025 |
| NC_006577.2;YP_173236.1;HCoV-HKU1_orf1ab;5304 | 0 | 0.025 |

| | | |
|---|---|---|
| NC_006577.2;YP_173236.1;HCoV-HKU1_orf1ab;5305 | 0 | 0.025 |
| NC_006577.2;YP_173236.1;HCoV-HKU1_orf1ab;5306 | 0 | 0.025 |
| NC_006577.2;YP_173236.1;HCoV-HKU1_orf1ab;5307 | 0 | 0.025 |
| NC_006577.2;YP_173236.1;HCoV-HKU1_orf1ab;5308 | 0 | 0.025 |
| NC_006577.2;YP_173236.1;HCoV-HKU1_orf1ab;5309 | 0 | 0 |
| NC_006577.2;YP_173236.1;HCoV-HKU1_orf1ab;5310 | 0 | 0 |
| NC_006577.2;YP_173236.1;HCoV-HKU1_orf1ab;5311 | 0 | 0 |
| NC_006577.2;YP_173236.1;HCoV-HKU1_orf1ab;5312 | 0 | 0 |
| NC_006577.2;YP_173236.1;HCoV-HKU1_orf1ab;5313 | 0 | 0 |
| NC_006577.2;YP_173236.1;HCoV-HKU1_orf1ab;5314 | 0 | 0 |
| NC_006577.2;YP_173236.1;HCoV-HKU1_orf1ab;5315 | 0 | 0 |
| NC_006577.2;YP_173236.1;HCoV-HKU1_orf1ab;5316 | 0 | 0 |
| NC_006577.2;YP_173236.1;HCoV-HKU1_orf1ab;5317 | 0 | 0 |
| NC_006577.2;YP_173236.1;HCoV-HKU1_orf1ab;5318 | 0 | 0.025 |
| NC_006577.2;YP_173236.1;HCoV-HKU1_orf1ab;5319 | 0 | 0.025 |
| NC_006577.2;YP_173236.1;HCoV-HKU1_orf1ab;5320 | 0 | 0 |
| NC_006577.2;YP_173236.1;HCoV-HKU1_orf1ab;5321 | 0 | 0 |
| NC_006577.2;YP_173236.1;HCoV-HKU1_orf1ab;5322 | 0 | 0.025 |
| NC_006577.2;YP_173236.1;HCoV-HKU1_orf1ab;5323 | 0.05 | 0 |
| NC_006577.2;YP_173236.1;HCoV-HKU1_orf1ab;5324 | 0 | 0 |
| NC_006577.2;YP_173236.1;HCoV-HKU1_orf1ab;5325 | 0 | 0 |
| NC_006577.2;YP_173236.1;HCoV-HKU1_orf1ab;5326 | 0 | 0 |
| NC_006577.2;YP_173236.1;HCoV-HKU1_orf1ab;5327 | 0 | 0 |
| NC_006577.2;YP_173236.1;HCoV-HKU1_orf1ab;5328 | 0 | 0 |
| NC_006577.2;YP_173236.1;HCoV-HKU1_orf1ab;5329 | 0 | 0 |
| NC_006577.2;YP_173236.1;HCoV-HKU1_orf1ab;5330 | 0 | 0 |
| NC_006577.2;YP_173236.1;HCoV-HKU1_orf1ab;5331 | 0 | 0 |
| NC_006577.2;YP_173236.1;HCoV-HKU1_orf1ab;5332 | 0 | 0 |
| NC_006577.2;YP_173236.1;HCoV-HKU1_orf1ab;5333 | 0 | 0 |
| NC_006577.2;YP_173236.1;HCoV-HKU1_orf1ab;5334 | 0 | 0 |
| NC_006577.2;YP_173236.1;HCoV-HKU1_orf1ab;5335 | 0 | 0 |
| NC_006577.2;YP_173236.1;HCoV-HKU1_orf1ab;5336 | 0 | 0 |
| NC_006577.2;YP_173236.1;HCoV-HKU1_orf1ab;5337 | 0 | 0.025 |
| NC_006577.2;YP_173236.1;HCoV-HKU1_orf1ab;5338 | 0 | 0.075 |
| NC_006577.2;YP_173236.1;HCoV-HKU1_orf1ab;5339 | 0.05 | 0.025 |
| NC_006577.2;YP_173236.1;HCoV-HKU1_orf1ab;5340 | 0.1 | 0.025 |
| NC_006577.2;YP_173236.1;HCoV-HKU1_orf1ab;5341 | 0.05 | 0 |
| NC_006577.2;YP_173236.1;HCoV-HKU1_orf1ab;5342 | 0.05 | 0.025 |
| NC_006577.2;YP_173236.1;HCoV-HKU1_orf1ab;5343 | 0 | 0 |
| NC_006577.2;YP_173236.1;HCoV-HKU1_orf1ab;5344 | 0 | 0 |
| NC_006577.2;YP_173236.1;HCoV-HKU1_orf1ab;5345 | 0 | 0 |
| NC_006577.2;YP_173236.1;HCoV-HKU1_orf1ab;5346 | 0 | 0 |
| NC_006577.2;YP_173236.1;HCoV-HKU1_orf1ab;5347 | 0 | 0 |

| | | |
|---|---|---|
| NC_006577.2;YP_173236.1;HCoV-HKU1_orf1ab;5348 | 0 | 0 |
| NC_006577.2;YP_173236.1;HCoV-HKU1_orf1ab;5349 | 0 | 0 |
| NC_006577.2;YP_173236.1;HCoV-HKU1_orf1ab;5350 | 0 | 0 |
| NC_006577.2;YP_173236.1;HCoV-HKU1_orf1ab;5351 | 0 | 0 |
| NC_006577.2;YP_173236.1;HCoV-HKU1_orf1ab;5352 | 0 | 0 |
| NC_006577.2;YP_173236.1;HCoV-HKU1_orf1ab;5353 | 0 | 0 |
| NC_006577.2;YP_173236.1;HCoV-HKU1_orf1ab;5354 | 0 | 0 |
| NC_006577.2;YP_173236.1;HCoV-HKU1_orf1ab;5355 | 0 | 0 |
| NC_006577.2;YP_173236.1;HCoV-HKU1_orf1ab;5356 | 0.05 | 0.025 |
| NC_006577.2;YP_173236.1;HCoV-HKU1_orf1ab;5357 | 0 | 0.025 |
| NC_006577.2;YP_173236.1;HCoV-HKU1_orf1ab;5358 | 0 | 0 |
| NC_006577.2;YP_173236.1;HCoV-HKU1_orf1ab;5359 | 0 | 0 |
| NC_006577.2;YP_173236.1;HCoV-HKU1_orf1ab;5360 | 0.1 | 0.05 |
| NC_006577.2;YP_173236.1;HCoV-HKU1_orf1ab;5361 | 0.05 | 0.025 |
| NC_006577.2;YP_173236.1;HCoV-HKU1_orf1ab;5362 | 0.05 | 0.075 |
| NC_006577.2;YP_173236.1;HCoV-HKU1_orf1ab;5363 | 0.2 | 0.125 |
| NC_006577.2;YP_173236.1;HCoV-HKU1_orf1ab;5364 | 0.1 | 0.075 |
| NC_006577.2;YP_173236.1;HCoV-HKU1_orf1ab;5365 | 0.1 | 0.025 |
| NC_006577.2;YP_173236.1;HCoV-HKU1_orf1ab;5366 | 0.15 | 0.025 |
| NC_006577.2;YP_173236.1;HCoV-HKU1_orf1ab;5367 | 0.15 | 0.05 |
| NC_006577.2;YP_173236.1;HCoV-HKU1_orf1ab;5368 | 0.1 | 0.05 |
| NC_006577.2;YP_173236.1;HCoV-HKU1_orf1ab;5369 | 0.15 | 0.075 |
| NC_006577.2;YP_173236.1;HCoV-HKU1_orf1ab;5370 | 0.15 | 0.15 |
| NC_006577.2;YP_173236.1;HCoV-HKU1_orf1ab;5371 | 0.05 | 0.025 |
| NC_006577.2;YP_173236.1;HCoV-HKU1_orf1ab;5372 | 0.05 | 0 |
| NC_006577.2;YP_173236.1;HCoV-HKU1_orf1ab;5373 | 0 | 0.025 |
| NC_006577.2;YP_173236.1;HCoV-HKU1_orf1ab;5374 | 0.05 | 0.025 |
| NC_006577.2;YP_173236.1;HCoV-HKU1_orf1ab;5375 | 0 | 0.025 |
| NC_006577.2;YP_173236.1;HCoV-HKU1_orf1ab;5376 | 0 | 0 |
| NC_006577.2;YP_173236.1;HCoV-HKU1_orf1ab;5377 | 0 | 0 |
| NC_006577.2;YP_173236.1;HCoV-HKU1_orf1ab;5378 | 0 | 0 |
| NC_006577.2;YP_173236.1;HCoV-HKU1_orf1ab;5379 | 0 | 0 |
| NC_006577.2;YP_173236.1;HCoV-HKU1_orf1ab;5380 | 0 | 0 |
| NC_006577.2;YP_173236.1;HCoV-HKU1_orf1ab;5381 | 0 | 0 |
| NC_006577.2;YP_173236.1;HCoV-HKU1_orf1ab;5382 | 0 | 0 |
| NC_006577.2;YP_173236.1;HCoV-HKU1_orf1ab;5383 | 0 | 0 |
| NC_006577.2;YP_173236.1;HCoV-HKU1_orf1ab;5384 | 0 | 0 |
| NC_006577.2;YP_173236.1;HCoV-HKU1_orf1ab;5385 | 0 | 0 |
| NC_006577.2;YP_173236.1;HCoV-HKU1_orf1ab;5386 | 0 | 0 |
| NC_006577.2;YP_173236.1;HCoV-HKU1_orf1ab;5387 | 0 | 0 |
| NC_006577.2;YP_173236.1;HCoV-HKU1_orf1ab;5388 | 0 | 0 |
| NC_006577.2;YP_173236.1;HCoV-HKU1_orf1ab;5389 | 0 | 0 |
| NC_006577.2;YP_173236.1;HCoV-HKU1_orf1ab;5390 | 0 | 0 |

| | | |
|---|---|---|
| NC_006577.2;YP_173236.1;HCoV-HKU1_orf1ab;5391 | 0 | 0 |
| NC_006577.2;YP_173236.1;HCoV-HKU1_orf1ab;5392 | 0 | 0 |
| NC_006577.2;YP_173236.1;HCoV-HKU1_orf1ab;5393 | 0 | 0 |
| NC_006577.2;YP_173236.1;HCoV-HKU1_orf1ab;5394 | 0 | 0 |
| NC_006577.2;YP_173236.1;HCoV-HKU1_orf1ab;5395 | 0 | 0 |
| NC_006577.2;YP_173236.1;HCoV-HKU1_orf1ab;5396 | 0 | 0 |
| NC_006577.2;YP_173236.1;HCoV-HKU1_orf1ab;5397 | 0 | 0 |
| NC_006577.2;YP_173236.1;HCoV-HKU1_orf1ab;5398 | 0 | 0 |
| NC_006577.2;YP_173236.1;HCoV-HKU1_orf1ab;5399 | 0 | 0 |
| NC_006577.2;YP_173236.1;HCoV-HKU1_orf1ab;5400 | 0 | 0 |
| NC_006577.2;YP_173236.1;HCoV-HKU1_orf1ab;5401 | 0 | 0 |
| NC_006577.2;YP_173236.1;HCoV-HKU1_orf1ab;5402 | 0 | 0 |
| NC_006577.2;YP_173236.1;HCoV-HKU1_orf1ab;5403 | 0 | 0 |
| NC_006577.2;YP_173236.1;HCoV-HKU1_orf1ab;5404 | 0 | 0 |
| NC_006577.2;YP_173236.1;HCoV-HKU1_orf1ab;5405 | 0 | 0 |
| NC_006577.2;YP_173236.1;HCoV-HKU1_orf1ab;5406 | 0 | 0 |
| NC_006577.2;YP_173236.1;HCoV-HKU1_orf1ab;5407 | 0 | 0 |
| NC_006577.2;YP_173236.1;HCoV-HKU1_orf1ab;5408 | 0 | 0 |
| NC_006577.2;YP_173236.1;HCoV-HKU1_orf1ab;5409 | 0 | 0 |
| NC_006577.2;YP_173236.1;HCoV-HKU1_orf1ab;5410 | 0.05 | 0 |
| NC_006577.2;YP_173236.1;HCoV-HKU1_orf1ab;5411 | 0.05 | 0 |
| NC_006577.2;YP_173236.1;HCoV-HKU1_orf1ab;5412 | 0.05 | 0 |
| NC_006577.2;YP_173236.1;HCoV-HKU1_orf1ab;5413 | 0.05 | 0 |
| NC_006577.2;YP_173236.1;HCoV-HKU1_orf1ab;5414 | 0 | 0.025 |
| NC_006577.2;YP_173236.1;HCoV-HKU1_orf1ab;5415 | 0 | 0 |
| NC_006577.2;YP_173236.1;HCoV-HKU1_orf1ab;5416 | 0 | 0 |
| NC_006577.2;YP_173236.1;HCoV-HKU1_orf1ab;5417 | 0 | 0 |
| NC_006577.2;YP_173236.1;HCoV-HKU1_orf1ab;5418 | 0 | 0 |
| NC_006577.2;YP_173236.1;HCoV-HKU1_orf1ab;5419 | 0 | 0 |
| NC_006577.2;YP_173236.1;HCoV-HKU1_orf1ab;5420 | 0 | 0 |
| NC_006577.2;YP_173236.1;HCoV-HKU1_orf1ab;5421 | 0 | 0 |
| NC_006577.2;YP_173236.1;HCoV-HKU1_orf1ab;5422 | 0 | 0 |
| NC_006577.2;YP_173236.1;HCoV-HKU1_orf1ab;5423 | 0 | 0 |
| NC_006577.2;YP_173236.1;HCoV-HKU1_orf1ab;5424 | 0 | 0 |
| NC_006577.2;YP_173236.1;HCoV-HKU1_orf1ab;5425 | 0 | 0 |
| NC_006577.2;YP_173236.1;HCoV-HKU1_orf1ab;5426 | 0 | 0 |
| NC_006577.2;YP_173236.1;HCoV-HKU1_orf1ab;5427 | 0 | 0 |
| NC_006577.2;YP_173236.1;HCoV-HKU1_orf1ab;5428 | 0 | 0 |
| NC_006577.2;YP_173236.1;HCoV-HKU1_orf1ab;5429 | 0 | 0 |
| NC_006577.2;YP_173236.1;HCoV-HKU1_orf1ab;5430 | 0 | 0 |
| NC_006577.2;YP_173236.1;HCoV-HKU1_orf1ab;5431 | 0 | 0 |
| NC_006577.2;YP_173236.1;HCoV-HKU1_orf1ab;5432 | 0.05 | 0 |
| NC_006577.2;YP_173236.1;HCoV-HKU1_orf1ab;5433 | 0 | 0 |

| | | |
|---|---|---|
| NC_006577.2;YP_173236.1;HCoV-HKU1_orf1ab;5434 | 0.05 | 0 |
| NC_006577.2;YP_173236.1;HCoV-HKU1_orf1ab;5435 | 0 | 0 |
| NC_006577.2;YP_173236.1;HCoV-HKU1_orf1ab;5436 | 0 | 0 |
| NC_006577.2;YP_173236.1;HCoV-HKU1_orf1ab;5437 | 0 | 0 |
| NC_006577.2;YP_173236.1;HCoV-HKU1_orf1ab;5438 | 0 | 0 |
| NC_006577.2;YP_173236.1;HCoV-HKU1_orf1ab;5439 | 0 | 0.025 |
| NC_006577.2;YP_173236.1;HCoV-HKU1_orf1ab;5440 | 0 | 0.025 |
| NC_006577.2;YP_173236.1;HCoV-HKU1_orf1ab;5441 | 0 | 0 |
| NC_006577.2;YP_173236.1;HCoV-HKU1_orf1ab;5442 | 0 | 0 |
| NC_006577.2;YP_173236.1;HCoV-HKU1_orf1ab;5443 | 0 | 0 |
| NC_006577.2;YP_173236.1;HCoV-HKU1_orf1ab;5444 | 0 | 0 |
| NC_006577.2;YP_173236.1;HCoV-HKU1_orf1ab;5445 | 0 | 0 |
| NC_006577.2;YP_173236.1;HCoV-HKU1_orf1ab;5446 | 0 | 0 |
| NC_006577.2;YP_173236.1;HCoV-HKU1_orf1ab;5447 | 0 | 0 |
| NC_006577.2;YP_173236.1;HCoV-HKU1_orf1ab;5448 | 0 | 0 |
| NC_006577.2;YP_173236.1;HCoV-HKU1_orf1ab;5449 | 0 | 0 |
| NC_006577.2;YP_173236.1;HCoV-HKU1_orf1ab;5450 | 0.05 | 0 |
| NC_006577.2;YP_173236.1;HCoV-HKU1_orf1ab;5451 | 0.05 | 0 |
| NC_006577.2;YP_173236.1;HCoV-HKU1_orf1ab;5452 | 0 | 0 |
| NC_006577.2;YP_173236.1;HCoV-HKU1_orf1ab;5453 | 0 | 0 |
| NC_006577.2;YP_173236.1;HCoV-HKU1_orf1ab;5454 | 0 | 0 |
| NC_006577.2;YP_173236.1;HCoV-HKU1_orf1ab;5455 | 0 | 0 |
| NC_006577.2;YP_173236.1;HCoV-HKU1_orf1ab;5456 | 0 | 0 |
| NC_006577.2;YP_173236.1;HCoV-HKU1_orf1ab;5457 | 0 | 0 |
| NC_006577.2;YP_173236.1;HCoV-HKU1_orf1ab;5458 | 0 | 0 |
| NC_006577.2;YP_173236.1;HCoV-HKU1_orf1ab;5459 | 0 | 0 |
| NC_006577.2;YP_173236.1;HCoV-HKU1_orf1ab;5460 | 0 | 0 |
| NC_006577.2;YP_173236.1;HCoV-HKU1_orf1ab;5461 | 0 | 0 |
| NC_006577.2;YP_173236.1;HCoV-HKU1_orf1ab;5462 | 0 | 0 |
| NC_006577.2;YP_173236.1;HCoV-HKU1_orf1ab;5463 | 0 | 0 |
| NC_006577.2;YP_173236.1;HCoV-HKU1_orf1ab;5464 | 0 | 0 |
| NC_006577.2;YP_173236.1;HCoV-HKU1_orf1ab;5465 | 0 | 0 |
| NC_006577.2;YP_173236.1;HCoV-HKU1_orf1ab;5466 | 0 | 0 |
| NC_006577.2;YP_173236.1;HCoV-HKU1_orf1ab;5467 | 0 | 0 |
| NC_006577.2;YP_173236.1;HCoV-HKU1_orf1ab;5468 | 0 | 0 |
| NC_006577.2;YP_173236.1;HCoV-HKU1_orf1ab;5469 | 0 | 0 |
| NC_006577.2;YP_173236.1;HCoV-HKU1_orf1ab;5470 | 0 | 0 |
| NC_006577.2;YP_173236.1;HCoV-HKU1_orf1ab;5471 | 0 | 0 |
| NC_006577.2;YP_173236.1;HCoV-HKU1_orf1ab;5472 | 0 | 0 |
| NC_006577.2;YP_173236.1;HCoV-HKU1_orf1ab;5473 | 0 | 0 |
| NC_006577.2;YP_173236.1;HCoV-HKU1_orf1ab;5474 | 0 | 0 |
| NC_006577.2;YP_173236.1;HCoV-HKU1_orf1ab;5475 | 0 | 0.025 |
| NC_006577.2;YP_173236.1;HCoV-HKU1_orf1ab;5476 | 0 | 0.025 |

| | | |
|---|---|---|
| NC_006577.2;YP_173236.1;HCoV-HKU1_orf1ab;5477 | 0.05 | 0.05 |
| NC_006577.2;YP_173236.1;HCoV-HKU1_orf1ab;5478 | 0.1 | 0.1 |
| NC_006577.2;YP_173236.1;HCoV-HKU1_orf1ab;5479 | 0.25 | 0.15 |
| NC_006577.2;YP_173236.1;HCoV-HKU1_orf1ab;5480 | 0.25 | 0.2 |
| NC_006577.2;YP_173236.1;HCoV-HKU1_orf1ab;5481 | 0.5 | 0.25 |
| NC_006577.2;YP_173236.1;HCoV-HKU1_orf1ab;5482 | 0.3 | 0.1 |
| NC_006577.2;YP_173236.1;HCoV-HKU1_orf1ab;5483 | 0.15 | 0.05 |
| NC_006577.2;YP_173236.1;HCoV-HKU1_orf1ab;5484 | 0.05 | 0.025 |
| NC_006577.2;YP_173236.1;HCoV-HKU1_orf1ab;5485 | 0.05 | 0.025 |
| NC_006577.2;YP_173236.1;HCoV-HKU1_orf1ab;5486 | 0.1 | 0 |
| NC_006577.2;YP_173236.1;HCoV-HKU1_orf1ab;5487 | 0.05 | 0 |
| NC_006577.2;YP_173236.1;HCoV-HKU1_orf1ab;5488 | 0.15 | 0 |
| NC_006577.2;YP_173236.1;HCoV-HKU1_orf1ab;5489 | 0.05 | 0 |
| NC_006577.2;YP_173236.1;HCoV-HKU1_orf1ab;5490 | 0 | 0 |
| NC_006577.2;YP_173236.1;HCoV-HKU1_orf1ab;5491 | 0.05 | 0 |
| NC_006577.2;YP_173236.1;HCoV-HKU1_orf1ab;5492 | 0 | 0 |
| NC_006577.2;YP_173236.1;HCoV-HKU1_orf1ab;5493 | 0 | 0 |
| NC_006577.2;YP_173236.1;HCoV-HKU1_orf1ab;5494 | 0 | 0 |
| NC_006577.2;YP_173236.1;HCoV-HKU1_orf1ab;5495 | 0 | 0 |
| NC_006577.2;YP_173236.1;HCoV-HKU1_orf1ab;5496 | 0 | 0 |
| NC_006577.2;YP_173236.1;HCoV-HKU1_orf1ab;5497 | 0 | 0 |
| NC_006577.2;YP_173236.1;HCoV-HKU1_orf1ab;5498 | 0 | 0 |
| NC_006577.2;YP_173236.1;HCoV-HKU1_orf1ab;5499 | 0 | 0 |
| NC_006577.2;YP_173236.1;HCoV-HKU1_orf1ab;5500 | 0 | 0.025 |
| NC_006577.2;YP_173236.1;HCoV-HKU1_orf1ab;5501 | 0 | 0 |
| NC_006577.2;YP_173236.1;HCoV-HKU1_orf1ab;5502 | 0 | 0 |
| NC_006577.2;YP_173236.1;HCoV-HKU1_orf1ab;5503 | 0 | 0 |
| NC_006577.2;YP_173236.1;HCoV-HKU1_orf1ab;5504 | 0 | 0 |
| NC_006577.2;YP_173236.1;HCoV-HKU1_orf1ab;5505 | 0.05 | 0 |
| NC_006577.2;YP_173236.1;HCoV-HKU1_orf1ab;5506 | 0 | 0 |
| NC_006577.2;YP_173236.1;HCoV-HKU1_orf1ab;5507 | 0 | 0 |
| NC_006577.2;YP_173236.1;HCoV-HKU1_orf1ab;5508 | 0 | 0 |
| NC_006577.2;YP_173236.1;HCoV-HKU1_orf1ab;5509 | 0 | 0 |
| NC_006577.2;YP_173236.1;HCoV-HKU1_orf1ab;5510 | 0 | 0 |
| NC_006577.2;YP_173236.1;HCoV-HKU1_orf1ab;5511 | 0 | 0 |
| NC_006577.2;YP_173236.1;HCoV-HKU1_orf1ab;5512 | 0.05 | 0 |
| NC_006577.2;YP_173236.1;HCoV-HKU1_orf1ab;5513 | 0.05 | 0 |
| NC_006577.2;YP_173236.1;HCoV-HKU1_orf1ab;5514 | 0 | 0 |
| NC_006577.2;YP_173236.1;HCoV-HKU1_orf1ab;5515 | 0 | 0 |
| NC_006577.2;YP_173236.1;HCoV-HKU1_orf1ab;5516 | 0 | 0 |
| NC_006577.2;YP_173236.1;HCoV-HKU1_orf1ab;5517 | 0 | 0 |
| NC_006577.2;YP_173236.1;HCoV-HKU1_orf1ab;5518 | 0 | 0 |
| NC_006577.2;YP_173236.1;HCoV-HKU1_orf1ab;5519 | 0 | 0.025 |

| | | |
|---|---|---|
| NC_006577.2;YP_173236.1;HCoV-HKU1_orf1ab;5520 | 0 | 0.025 |
| NC_006577.2;YP_173236.1;HCoV-HKU1_orf1ab;5521 | 0.05 | 0 |
| NC_006577.2;YP_173236.1;HCoV-HKU1_orf1ab;5522 | 0.05 | 0 |
| NC_006577.2;YP_173236.1;HCoV-HKU1_orf1ab;5523 | 0 | 0 |
| NC_006577.2;YP_173236.1;HCoV-HKU1_orf1ab;5524 | 0 | 0 |
| NC_006577.2;YP_173236.1;HCoV-HKU1_orf1ab;5525 | 0 | 0 |
| NC_006577.2;YP_173236.1;HCoV-HKU1_orf1ab;5526 | 0 | 0 |
| NC_006577.2;YP_173236.1;HCoV-HKU1_orf1ab;5527 | 0 | 0 |
| NC_006577.2;YP_173236.1;HCoV-HKU1_orf1ab;5528 | 0 | 0 |
| NC_006577.2;YP_173236.1;HCoV-HKU1_orf1ab;5529 | 0 | 0 |
| NC_006577.2;YP_173236.1;HCoV-HKU1_orf1ab;5530 | 0 | 0 |
| NC_006577.2;YP_173236.1;HCoV-HKU1_orf1ab;5531 | 0 | 0 |
| NC_006577.2;YP_173236.1;HCoV-HKU1_orf1ab;5532 | 0 | 0 |
| NC_006577.2;YP_173236.1;HCoV-HKU1_orf1ab;5533 | 0 | 0 |
| NC_006577.2;YP_173236.1;HCoV-HKU1_orf1ab;5534 | 0 | 0 |
| NC_006577.2;YP_173236.1;HCoV-HKU1_orf1ab;5535 | 0 | 0.025 |
| NC_006577.2;YP_173236.1;HCoV-HKU1_orf1ab;5536 | 0 | 0 |
| NC_006577.2;YP_173236.1;HCoV-HKU1_orf1ab;5537 | 0 | 0 |
| NC_006577.2;YP_173236.1;HCoV-HKU1_orf1ab;5538 | 0 | 0 |
| NC_006577.2;YP_173236.1;HCoV-HKU1_orf1ab;5539 | 0 | 0 |
| NC_006577.2;YP_173236.1;HCoV-HKU1_orf1ab;5540 | 0 | 0 |
| NC_006577.2;YP_173236.1;HCoV-HKU1_orf1ab;5541 | 0 | 0.025 |
| NC_006577.2;YP_173236.1;HCoV-HKU1_orf1ab;5542 | 0 | 0.025 |
| NC_006577.2;YP_173236.1;HCoV-HKU1_orf1ab;5543 | 0 | 0.025 |
| NC_006577.2;YP_173236.1;HCoV-HKU1_orf1ab;5544 | 0 | 0.025 |
| NC_006577.2;YP_173236.1;HCoV-HKU1_orf1ab;5545 | 0 | 0.025 |
| NC_006577.2;YP_173236.1;HCoV-HKU1_orf1ab;5546 | 0 | 0.025 |
| NC_006577.2;YP_173236.1;HCoV-HKU1_orf1ab;5547 | 0 | 0.025 |
| NC_006577.2;YP_173236.1;HCoV-HKU1_orf1ab;5548 | 0.05 | 0.1 |
| NC_006577.2;YP_173236.1;HCoV-HKU1_orf1ab;5549 | 0.05 | 0.125 |
| NC_006577.2;YP_173236.1;HCoV-HKU1_orf1ab;5550 | 0.1 | 0.15 |
| NC_006577.2;YP_173236.1;HCoV-HKU1_orf1ab;5551 | 0.15 | 0.225 |
| NC_006577.2;YP_173236.1;HCoV-HKU1_orf1ab;5552 | 0.05 | 0.1 |
| NC_006577.2;YP_173236.1;HCoV-HKU1_orf1ab;5553 | 0 | 0.1 |
| NC_006577.2;YP_173236.1;HCoV-HKU1_orf1ab;5554 | 0.05 | 0.1 |
| NC_006577.2;YP_173236.1;HCoV-HKU1_orf1ab;5555 | 0.05 | 0 |
| NC_006577.2;YP_173236.1;HCoV-HKU1_orf1ab;5556 | 0 | 0 |
| NC_006577.2;YP_173236.1;HCoV-HKU1_orf1ab;5557 | 0 | 0 |
| NC_006577.2;YP_173236.1;HCoV-HKU1_orf1ab;5558 | 0 | 0 |
| NC_006577.2;YP_173236.1;HCoV-HKU1_orf1ab;5559 | 0 | 0 |
| NC_006577.2;YP_173236.1;HCoV-HKU1_orf1ab;5560 | 0 | 0 |
| NC_006577.2;YP_173236.1;HCoV-HKU1_orf1ab;5561 | 0 | 0 |
| NC_006577.2;YP_173236.1;HCoV-HKU1_orf1ab;5562 | 0 | 0 |

| | | |
|---|---|---|
| NC_006577.2;YP_173236.1;HCoV-HKU1_orf1ab;5563 | 0 | 0 |
| NC_006577.2;YP_173236.1;HCoV-HKU1_orf1ab;5564 | 0 | 0 |
| NC_006577.2;YP_173236.1;HCoV-HKU1_orf1ab;5565 | 0 | 0 |
| NC_006577.2;YP_173236.1;HCoV-HKU1_orf1ab;5566 | 0 | 0 |
| NC_006577.2;YP_173236.1;HCoV-HKU1_orf1ab;5567 | 0 | 0.05 |
| NC_006577.2;YP_173236.1;HCoV-HKU1_orf1ab;5568 | 0 | 0.05 |
| NC_006577.2;YP_173236.1;HCoV-HKU1_orf1ab;5569 | 0 | 0 |
| NC_006577.2;YP_173236.1;HCoV-HKU1_orf1ab;5570 | 0 | 0 |
| NC_006577.2;YP_173236.1;HCoV-HKU1_orf1ab;5571 | 0 | 0 |
| NC_006577.2;YP_173236.1;HCoV-HKU1_orf1ab;5572 | 0 | 0.075 |
| NC_006577.2;YP_173236.1;HCoV-HKU1_orf1ab;5573 | 0 | 0.075 |
| NC_006577.2;YP_173236.1;HCoV-HKU1_orf1ab;5574 | 0 | 0.075 |
| NC_006577.2;YP_173236.1;HCoV-HKU1_orf1ab;5575 | 0 | 0.05 |
| NC_006577.2;YP_173236.1;HCoV-HKU1_orf1ab;5576 | 0 | 0.075 |
| NC_006577.2;YP_173236.1;HCoV-HKU1_orf1ab;5577 | 0 | 0.1 |
| NC_006577.2;YP_173236.1;HCoV-HKU1_orf1ab;5578 | 0 | 0.1 |
| NC_006577.2;YP_173236.1;HCoV-HKU1_orf1ab;5579 | 0 | 0.075 |
| NC_006577.2;YP_173236.1;HCoV-HKU1_orf1ab;5580 | 0 | 0.1 |
| NC_006577.2;YP_173236.1;HCoV-HKU1_orf1ab;5581 | 0 | 0 |
| NC_006577.2;YP_173236.1;HCoV-HKU1_orf1ab;5582 | 0.05 | 0 |
| NC_006577.2;YP_173236.1;HCoV-HKU1_orf1ab;5583 | 0 | 0 |
| NC_006577.2;YP_173236.1;HCoV-HKU1_orf1ab;5584 | 0 | 0 |
| NC_006577.2;YP_173236.1;HCoV-HKU1_orf1ab;5585 | 0 | 0 |
| NC_006577.2;YP_173236.1;HCoV-HKU1_orf1ab;5586 | 0 | 0 |
| NC_006577.2;YP_173236.1;HCoV-HKU1_orf1ab;5587 | 0 | 0 |
| NC_006577.2;YP_173236.1;HCoV-HKU1_orf1ab;5588 | 0.05 | 0 |
| NC_006577.2;YP_173236.1;HCoV-HKU1_orf1ab;5589 | 0 | 0 |
| NC_006577.2;YP_173236.1;HCoV-HKU1_orf1ab;5590 | 0 | 0 |
| NC_006577.2;YP_173236.1;HCoV-HKU1_orf1ab;5591 | 0 | 0 |
| NC_006577.2;YP_173236.1;HCoV-HKU1_orf1ab;5592 | 0 | 0 |
| NC_006577.2;YP_173236.1;HCoV-HKU1_orf1ab;5593 | 0.05 | 0 |
| NC_006577.2;YP_173236.1;HCoV-HKU1_orf1ab;5594 | 0 | 0 |
| NC_006577.2;YP_173236.1;HCoV-HKU1_orf1ab;5595 | 0 | 0 |
| NC_006577.2;YP_173236.1;HCoV-HKU1_orf1ab;5596 | 0 | 0 |
| NC_006577.2;YP_173236.1;HCoV-HKU1_orf1ab;5597 | 0 | 0 |
| NC_006577.2;YP_173236.1;HCoV-HKU1_orf1ab;5598 | 0 | 0 |
| NC_006577.2;YP_173236.1;HCoV-HKU1_orf1ab;5599 | 0 | 0 |
| NC_006577.2;YP_173236.1;HCoV-HKU1_orf1ab;5600 | 0 | 0 |
| NC_006577.2;YP_173236.1;HCoV-HKU1_orf1ab;5601 | 0 | 0 |
| NC_006577.2;YP_173236.1;HCoV-HKU1_orf1ab;5602 | 0 | 0 |
| NC_006577.2;YP_173236.1;HCoV-HKU1_orf1ab;5603 | 0 | 0 |
| NC_006577.2;YP_173236.1;HCoV-HKU1_orf1ab;5604 | 0 | 0 |
| NC_006577.2;YP_173236.1;HCoV-HKU1_orf1ab;5605 | 0 | 0 |

| | | |
|---|---|---|
| NC_006577.2;YP_173236.1;HCoV-HKU1_orf1ab;5606 | 0 | 0 |
| NC_006577.2;YP_173236.1;HCoV-HKU1_orf1ab;5607 | 0 | 0 |
| NC_006577.2;YP_173236.1;HCoV-HKU1_orf1ab;5608 | 0.05 | 0 |
| NC_006577.2;YP_173236.1;HCoV-HKU1_orf1ab;5609 | 0 | 0 |
| NC_006577.2;YP_173236.1;HCoV-HKU1_orf1ab;5610 | 0 | 0 |
| NC_006577.2;YP_173236.1;HCoV-HKU1_orf1ab;5611 | 0 | 0 |
| NC_006577.2;YP_173236.1;HCoV-HKU1_orf1ab;5612 | 0 | 0.025 |
| NC_006577.2;YP_173236.1;HCoV-HKU1_orf1ab;5613 | 0 | 0 |
| NC_006577.2;YP_173236.1;HCoV-HKU1_orf1ab;5614 | 0 | 0 |
| NC_006577.2;YP_173236.1;HCoV-HKU1_orf1ab;5615 | 0 | 0.025 |
| NC_006577.2;YP_173236.1;HCoV-HKU1_orf1ab;5616 | 0 | 0.275 |
| NC_006577.2;YP_173236.1;HCoV-HKU1_orf1ab;5617 | 0 | 0.05 |
| NC_006577.2;YP_173236.1;HCoV-HKU1_orf1ab;5618 | 0.1 | 0.025 |
| NC_006577.2;YP_173236.1;HCoV-HKU1_orf1ab;5619 | 0.05 | 0.05 |
| NC_006577.2;YP_173236.1;HCoV-HKU1_orf1ab;5620 | 0 | 0.025 |
| NC_006577.2;YP_173236.1;HCoV-HKU1_orf1ab;5621 | 0 | 0.025 |
| NC_006577.2;YP_173236.1;HCoV-HKU1_orf1ab;5622 | 0 | 0.025 |
| NC_006577.2;YP_173236.1;HCoV-HKU1_orf1ab;5623 | 0 | 0.025 |
| NC_006577.2;YP_173236.1;HCoV-HKU1_orf1ab;5624 | 0 | 0 |
| NC_006577.2;YP_173236.1;HCoV-HKU1_orf1ab;5625 | 0 | 0 |
| NC_006577.2;YP_173236.1;HCoV-HKU1_orf1ab;5626 | 0 | 0 |
| NC_006577.2;YP_173236.1;HCoV-HKU1_orf1ab;5627 | 0 | 0 |
| NC_006577.2;YP_173236.1;HCoV-HKU1_orf1ab;5628 | 0.05 | 0 |
| NC_006577.2;YP_173236.1;HCoV-HKU1_orf1ab;5629 | 0 | 0 |
| NC_006577.2;YP_173236.1;HCoV-HKU1_orf1ab;5630 | 0 | 0 |
| NC_006577.2;YP_173236.1;HCoV-HKU1_orf1ab;5631 | 0 | 0 |
| NC_006577.2;YP_173236.1;HCoV-HKU1_orf1ab;5632 | 0 | 0 |
| NC_006577.2;YP_173236.1;HCoV-HKU1_orf1ab;5633 | 0 | 0 |
| NC_006577.2;YP_173236.1;HCoV-HKU1_orf1ab;5634 | 0 | 0 |
| NC_006577.2;YP_173236.1;HCoV-HKU1_orf1ab;5635 | 0 | 0 |
| NC_006577.2;YP_173236.1;HCoV-HKU1_orf1ab;5636 | 0 | 0 |
| NC_006577.2;YP_173236.1;HCoV-HKU1_orf1ab;5637 | 0.05 | 0 |
| NC_006577.2;YP_173236.1;HCoV-HKU1_orf1ab;5638 | 0 | 0 |
| NC_006577.2;YP_173236.1;HCoV-HKU1_orf1ab;5639 | 0 | 0 |
| NC_006577.2;YP_173236.1;HCoV-HKU1_orf1ab;5640 | 0 | 0 |
| NC_006577.2;YP_173236.1;HCoV-HKU1_orf1ab;5641 | 0.05 | 0.025 |
| NC_006577.2;YP_173236.1;HCoV-HKU1_orf1ab;5642 | 0 | 0 |
| NC_006577.2;YP_173236.1;HCoV-HKU1_orf1ab;5643 | 0 | 0 |
| NC_006577.2;YP_173236.1;HCoV-HKU1_orf1ab;5644 | 0 | 0.025 |
| NC_006577.2;YP_173236.1;HCoV-HKU1_orf1ab;5645 | 0 | 0 |
| NC_006577.2;YP_173236.1;HCoV-HKU1_orf1ab;5646 | 0 | 0 |
| NC_006577.2;YP_173236.1;HCoV-HKU1_orf1ab;5647 | 0 | 0 |
| NC_006577.2;YP_173236.1;HCoV-HKU1_orf1ab;5648 | 0 | 0 |

| | | |
|---|---|---|
| NC_006577.2;YP_173236.1;HCoV-HKU1_orf1ab;5649 | 0 | 0 |
| NC_006577.2;YP_173236.1;HCoV-HKU1_orf1ab;5650 | 0 | 0 |
| NC_006577.2;YP_173236.1;HCoV-HKU1_orf1ab;5651 | 0 | 0 |
| NC_006577.2;YP_173236.1;HCoV-HKU1_orf1ab;5652 | 0 | 0 |
| NC_006577.2;YP_173236.1;HCoV-HKU1_orf1ab;5653 | 0 | 0.025 |
| NC_006577.2;YP_173236.1;HCoV-HKU1_orf1ab;5654 | 0 | 0.025 |
| NC_006577.2;YP_173236.1;HCoV-HKU1_orf1ab;5655 | 0 | 0 |
| NC_006577.2;YP_173236.1;HCoV-HKU1_orf1ab;5656 | 0 | 0 |
| NC_006577.2;YP_173236.1;HCoV-HKU1_orf1ab;5657 | 0 | 0 |
| NC_006577.2;YP_173236.1;HCoV-HKU1_orf1ab;5658 | 0 | 0 |
| NC_006577.2;YP_173236.1;HCoV-HKU1_orf1ab;5659 | 0 | 0.025 |
| NC_006577.2;YP_173236.1;HCoV-HKU1_orf1ab;5660 | 0 | 0 |
| NC_006577.2;YP_173236.1;HCoV-HKU1_orf1ab;5661 | 0 | 0.05 |
| NC_006577.2;YP_173236.1;HCoV-HKU1_orf1ab;5662 | 0 | 0.025 |
| NC_006577.2;YP_173236.1;HCoV-HKU1_orf1ab;5663 | 0 | 0.025 |
| NC_006577.2;YP_173236.1;HCoV-HKU1_orf1ab;5664 | 0 | 0 |
| NC_006577.2;YP_173236.1;HCoV-HKU1_orf1ab;5665 | 0 | 0 |
| NC_006577.2;YP_173236.1;HCoV-HKU1_orf1ab;5666 | 0 | 0 |
| NC_006577.2;YP_173236.1;HCoV-HKU1_orf1ab;5667 | 0 | 0 |
| NC_006577.2;YP_173236.1;HCoV-HKU1_orf1ab;5668 | 0 | 0 |
| NC_006577.2;YP_173236.1;HCoV-HKU1_orf1ab;5669 | 0 | 0 |
| NC_006577.2;YP_173236.1;HCoV-HKU1_orf1ab;5670 | 0 | 0 |
| NC_006577.2;YP_173236.1;HCoV-HKU1_orf1ab;5671 | 0 | 0 |
| NC_006577.2;YP_173236.1;HCoV-HKU1_orf1ab;5672 | 0 | 0 |
| NC_006577.2;YP_173236.1;HCoV-HKU1_orf1ab;5673 | 0 | 0 |
| NC_006577.2;YP_173236.1;HCoV-HKU1_orf1ab;5674 | 0 | 0 |
| NC_006577.2;YP_173236.1;HCoV-HKU1_orf1ab;5675 | 0 | 0 |
| NC_006577.2;YP_173236.1;HCoV-HKU1_orf1ab;5676 | 0 | 0 |
| NC_006577.2;YP_173236.1;HCoV-HKU1_orf1ab;5677 | 0 | 0 |
| NC_006577.2;YP_173236.1;HCoV-HKU1_orf1ab;5678 | 0 | 0 |
| NC_006577.2;YP_173236.1;HCoV-HKU1_orf1ab;5679 | 0 | 0 |
| NC_006577.2;YP_173236.1;HCoV-HKU1_orf1ab;5680 | 0 | 0 |
| NC_006577.2;YP_173236.1;HCoV-HKU1_orf1ab;5681 | 0 | 0 |
| NC_006577.2;YP_173236.1;HCoV-HKU1_orf1ab;5682 | 0.05 | 0 |
| NC_006577.2;YP_173236.1;HCoV-HKU1_orf1ab;5683 | 0 | 0 |
| NC_006577.2;YP_173236.1;HCoV-HKU1_orf1ab;5684 | 0 | 0.025 |
| NC_006577.2;YP_173236.1;HCoV-HKU1_orf1ab;5685 | 0 | 0.025 |
| NC_006577.2;YP_173236.1;HCoV-HKU1_orf1ab;5686 | 0 | 0.05 |
| NC_006577.2;YP_173236.1;HCoV-HKU1_orf1ab;5687 | 0 | 0.025 |
| NC_006577.2;YP_173236.1;HCoV-HKU1_orf1ab;5688 | 0 | 0 |
| NC_006577.2;YP_173236.1;HCoV-HKU1_orf1ab;5689 | 0 | 0 |
| NC_006577.2;YP_173236.1;HCoV-HKU1_orf1ab;5690 | 0 | 0 |
| NC_006577.2;YP_173236.1;HCoV-HKU1_orf1ab;5691 | 0 | 0 |

| | | |
|---|---|---|
| NC_006577.2;YP_173236.1;HCoV-HKU1_orf1ab;5692 | 0 | 0 |
| NC_006577.2;YP_173236.1;HCoV-HKU1_orf1ab;5693 | 0 | 0 |
| NC_006577.2;YP_173236.1;HCoV-HKU1_orf1ab;5694 | 0 | 0 |
| NC_006577.2;YP_173236.1;HCoV-HKU1_orf1ab;5695 | 0 | 0 |
| NC_006577.2;YP_173236.1;HCoV-HKU1_orf1ab;5696 | 0 | 0 |
| NC_006577.2;YP_173236.1;HCoV-HKU1_orf1ab;5697 | 0.05 | 0 |
| NC_006577.2;YP_173236.1;HCoV-HKU1_orf1ab;5698 | 0.05 | 0.025 |
| NC_006577.2;YP_173236.1;HCoV-HKU1_orf1ab;5699 | 0 | 0 |
| NC_006577.2;YP_173236.1;HCoV-HKU1_orf1ab;5700 | 0.05 | 0 |
| NC_006577.2;YP_173236.1;HCoV-HKU1_orf1ab;5701 | 0 | 0 |
| NC_006577.2;YP_173236.1;HCoV-HKU1_orf1ab;5702 | 0.05 | 0 |
| NC_006577.2;YP_173236.1;HCoV-HKU1_orf1ab;5703 | 0.05 | 0 |
| NC_006577.2;YP_173236.1;HCoV-HKU1_orf1ab;5704 | 0.05 | 0 |
| NC_006577.2;YP_173236.1;HCoV-HKU1_orf1ab;5705 | 0 | 0 |
| NC_006577.2;YP_173236.1;HCoV-HKU1_orf1ab;5706 | 0 | 0 |
| NC_006577.2;YP_173236.1;HCoV-HKU1_orf1ab;5707 | 0 | 0 |
| NC_006577.2;YP_173236.1;HCoV-HKU1_orf1ab;5708 | 0 | 0 |
| NC_006577.2;YP_173236.1;HCoV-HKU1_orf1ab;5709 | 0 | 0 |
| NC_006577.2;YP_173236.1;HCoV-HKU1_orf1ab;5710 | 0 | 0 |
| NC_006577.2;YP_173236.1;HCoV-HKU1_orf1ab;5711 | 0 | 0 |
| NC_006577.2;YP_173236.1;HCoV-HKU1_orf1ab;5712 | 0 | 0 |
| NC_006577.2;YP_173236.1;HCoV-HKU1_orf1ab;5713 | 0 | 0 |
| NC_006577.2;YP_173236.1;HCoV-HKU1_orf1ab;5714 | 0 | 0 |
| NC_006577.2;YP_173236.1;HCoV-HKU1_orf1ab;5715 | 0 | 0 |
| NC_006577.2;YP_173236.1;HCoV-HKU1_orf1ab;5716 | 0 | 0 |
| NC_006577.2;YP_173236.1;HCoV-HKU1_orf1ab;5717 | 0 | 0 |
| NC_006577.2;YP_173236.1;HCoV-HKU1_orf1ab;5718 | 0 | 0.025 |
| NC_006577.2;YP_173236.1;HCoV-HKU1_orf1ab;5719 | 0.05 | 0.025 |
| NC_006577.2;YP_173236.1;HCoV-HKU1_orf1ab;5720 | 0.05 | 0.075 |
| NC_006577.2;YP_173236.1;HCoV-HKU1_orf1ab;5721 | 0.1 | 0.125 |
| NC_006577.2;YP_173236.1;HCoV-HKU1_orf1ab;5722 | 0.3 | 0.15 |
| NC_006577.2;YP_173236.1;HCoV-HKU1_orf1ab;5723 | 0.3 | 0.15 |
| NC_006577.2;YP_173236.1;HCoV-HKU1_orf1ab;5724 | 0.5 | 0.225 |
| NC_006577.2;YP_173236.1;HCoV-HKU1_orf1ab;5725 | 0.45 | 0.35 |
| NC_006577.2;YP_173236.1;HCoV-HKU1_orf1ab;5726 | 0.25 | 0.15 |
| NC_006577.2;YP_173236.1;HCoV-HKU1_orf1ab;5727 | 0.05 | 0.1 |
| NC_006577.2;YP_173236.1;HCoV-HKU1_orf1ab;5728 | 0.05 | 0.025 |
| NC_006577.2;YP_173236.1;HCoV-HKU1_orf1ab;5729 | 0 | 0 |
| NC_006577.2;YP_173236.1;HCoV-HKU1_orf1ab;5730 | 0 | 0 |
| NC_006577.2;YP_173236.1;HCoV-HKU1_orf1ab;5731 | 0 | 0 |
| NC_006577.2;YP_173236.1;HCoV-HKU1_orf1ab;5732 | 0 | 0 |
| NC_006577.2;YP_173236.1;HCoV-HKU1_orf1ab;5733 | 0 | 0 |
| NC_006577.2;YP_173236.1;HCoV-HKU1_orf1ab;5734 | 0 | 0 |

| | | |
|---|---|---|
| NC_006577.2;YP_173236.1;HCoV-HKU1_orf1ab;5735 | 0 | 0 |
| NC_006577.2;YP_173236.1;HCoV-HKU1_orf1ab;5736 | 0 | 0 |
| NC_006577.2;YP_173236.1;HCoV-HKU1_orf1ab;5737 | 0 | 0 |
| NC_006577.2;YP_173236.1;HCoV-HKU1_orf1ab;5738 | 0 | 0 |
| NC_006577.2;YP_173236.1;HCoV-HKU1_orf1ab;5739 | 0 | 0 |
| NC_006577.2;YP_173236.1;HCoV-HKU1_orf1ab;5740 | 0 | 0 |
| NC_006577.2;YP_173236.1;HCoV-HKU1_orf1ab;5741 | 0 | 0 |
| NC_006577.2;YP_173236.1;HCoV-HKU1_orf1ab;5742 | 0 | 0 |
| NC_006577.2;YP_173236.1;HCoV-HKU1_orf1ab;5743 | 0 | 0 |
| NC_006577.2;YP_173236.1;HCoV-HKU1_orf1ab;5744 | 0 | 0 |
| NC_006577.2;YP_173236.1;HCoV-HKU1_orf1ab;5745 | 0 | 0 |
| NC_006577.2;YP_173236.1;HCoV-HKU1_orf1ab;5746 | 0 | 0 |
| NC_006577.2;YP_173236.1;HCoV-HKU1_orf1ab;5747 | 0 | 0 |
| NC_006577.2;YP_173236.1;HCoV-HKU1_orf1ab;5748 | 0.05 | 0 |
| NC_006577.2;YP_173236.1;HCoV-HKU1_orf1ab;5749 | 0.05 | 0.025 |
| NC_006577.2;YP_173236.1;HCoV-HKU1_orf1ab;5750 | 0.05 | 0.05 |
| NC_006577.2;YP_173236.1;HCoV-HKU1_orf1ab;5751 | 0.05 | 0.05 |
| NC_006577.2;YP_173236.1;HCoV-HKU1_orf1ab;5752 | 0.05 | 0.05 |
| NC_006577.2;YP_173236.1;HCoV-HKU1_orf1ab;5753 | 0.05 | 0 |
| NC_006577.2;YP_173236.1;HCoV-HKU1_orf1ab;5754 | 0.05 | 0 |
| NC_006577.2;YP_173236.1;HCoV-HKU1_orf1ab;5755 | 0.05 | 0.05 |
| NC_006577.2;YP_173236.1;HCoV-HKU1_orf1ab;5756 | 0.05 | 0.025 |
| NC_006577.2;YP_173236.1;HCoV-HKU1_orf1ab;5757 | 0 | 0 |
| NC_006577.2;YP_173236.1;HCoV-HKU1_orf1ab;5758 | 0 | 0 |
| NC_006577.2;YP_173236.1;HCoV-HKU1_orf1ab;5759 | 0.05 | 0 |
| NC_006577.2;YP_173236.1;HCoV-HKU1_orf1ab;5760 | 0 | 0 |
| NC_006577.2;YP_173236.1;HCoV-HKU1_orf1ab;5761 | 0 | 0 |
| NC_006577.2;YP_173236.1;HCoV-HKU1_orf1ab;5762 | 0 | 0 |
| NC_006577.2;YP_173236.1;HCoV-HKU1_orf1ab;5763 | 0 | 0 |
| NC_006577.2;YP_173236.1;HCoV-HKU1_orf1ab;5764 | 0 | 0 |
| NC_006577.2;YP_173236.1;HCoV-HKU1_orf1ab;5765 | 0 | 0 |
| NC_006577.2;YP_173236.1;HCoV-HKU1_orf1ab;5766 | 0 | 0 |
| NC_006577.2;YP_173236.1;HCoV-HKU1_orf1ab;5767 | 0 | 0 |
| NC_006577.2;YP_173236.1;HCoV-HKU1_orf1ab;5768 | 0 | 0 |
| NC_006577.2;YP_173236.1;HCoV-HKU1_orf1ab;5769 | 0 | 0 |
| NC_006577.2;YP_173236.1;HCoV-HKU1_orf1ab;5770 | 0.05 | 0 |
| NC_006577.2;YP_173236.1;HCoV-HKU1_orf1ab;5771 | 0.05 | 0 |
| NC_006577.2;YP_173236.1;HCoV-HKU1_orf1ab;5772 | 0 | 0 |
| NC_006577.2;YP_173236.1;HCoV-HKU1_orf1ab;5773 | 0.05 | 0 |
| NC_006577.2;YP_173236.1;HCoV-HKU1_orf1ab;5774 | 0 | 0 |
| NC_006577.2;YP_173236.1;HCoV-HKU1_orf1ab;5775 | 0 | 0 |
| NC_006577.2;YP_173236.1;HCoV-HKU1_orf1ab;5776 | 0 | 0 |
| NC_006577.2;YP_173236.1;HCoV-HKU1_orf1ab;5777 | 0 | 0 |

| | | |
|---|---|---|
| NC_006577.2;YP_173236.1;HCoV-HKU1_orf1ab;5778 | 0 | 0 |
| NC_006577.2;YP_173236.1;HCoV-HKU1_orf1ab;5779 | 0 | 0 |
| NC_006577.2;YP_173236.1;HCoV-HKU1_orf1ab;5780 | 0 | 0 |
| NC_006577.2;YP_173236.1;HCoV-HKU1_orf1ab;5781 | 0 | 0 |
| NC_006577.2;YP_173236.1;HCoV-HKU1_orf1ab;5782 | 0 | 0.05 |
| NC_006577.2;YP_173236.1;HCoV-HKU1_orf1ab;5783 | 0 | 0 |
| NC_006577.2;YP_173236.1;HCoV-HKU1_orf1ab;5784 | 0 | 0.025 |
| NC_006577.2;YP_173236.1;HCoV-HKU1_orf1ab;5785 | 0 | 0.025 |
| NC_006577.2;YP_173236.1;HCoV-HKU1_orf1ab;5786 | 0 | 0 |
| NC_006577.2;YP_173236.1;HCoV-HKU1_orf1ab;5787 | 0.05 | 0.05 |
| NC_006577.2;YP_173236.1;HCoV-HKU1_orf1ab;5788 | 0 | 0.05 |
| NC_006577.2;YP_173236.1;HCoV-HKU1_orf1ab;5789 | 0 | 0.025 |
| NC_006577.2;YP_173236.1;HCoV-HKU1_orf1ab;5790 | 0 | 0.025 |
| NC_006577.2;YP_173236.1;HCoV-HKU1_orf1ab;5791 | 0 | 0 |
| NC_006577.2;YP_173236.1;HCoV-HKU1_orf1ab;5792 | 0 | 0 |
| NC_006577.2;YP_173236.1;HCoV-HKU1_orf1ab;5793 | 0 | 0 |
| NC_006577.2;YP_173236.1;HCoV-HKU1_orf1ab;5794 | 0 | 0.025 |
| NC_006577.2;YP_173236.1;HCoV-HKU1_orf1ab;5795 | 0 | 0.025 |
| NC_006577.2;YP_173236.1;HCoV-HKU1_orf1ab;5796 | 0.05 | 0.025 |
| NC_006577.2;YP_173236.1;HCoV-HKU1_orf1ab;5797 | 0 | 0.025 |
| NC_006577.2;YP_173236.1;HCoV-HKU1_orf1ab;5798 | 0 | 0 |
| NC_006577.2;YP_173236.1;HCoV-HKU1_orf1ab;5799 | 0 | 0 |
| NC_006577.2;YP_173236.1;HCoV-HKU1_orf1ab;5800 | 0 | 0 |
| NC_006577.2;YP_173236.1;HCoV-HKU1_orf1ab;5801 | 0 | 0 |
| NC_006577.2;YP_173236.1;HCoV-HKU1_orf1ab;5802 | 0 | 0 |
| NC_006577.2;YP_173236.1;HCoV-HKU1_orf1ab;5803 | 0 | 0 |
| NC_006577.2;YP_173236.1;HCoV-HKU1_orf1ab;5804 | 0 | 0 |
| NC_006577.2;YP_173236.1;HCoV-HKU1_orf1ab;5805 | 0 | 0 |
| NC_006577.2;YP_173236.1;HCoV-HKU1_orf1ab;5806 | 0 | 0 |
| NC_006577.2;YP_173236.1;HCoV-HKU1_orf1ab;5807 | 0 | 0 |
| NC_006577.2;YP_173236.1;HCoV-HKU1_orf1ab;5808 | 0 | 0.025 |
| NC_006577.2;YP_173236.1;HCoV-HKU1_orf1ab;5809 | 0 | 0 |
| NC_006577.2;YP_173236.1;HCoV-HKU1_orf1ab;5810 | 0 | 0 |
| NC_006577.2;YP_173236.1;HCoV-HKU1_orf1ab;5811 | 0 | 0 |
| NC_006577.2;YP_173236.1;HCoV-HKU1_orf1ab;5812 | 0 | 0 |
| NC_006577.2;YP_173236.1;HCoV-HKU1_orf1ab;5813 | 0 | 0 |
| NC_006577.2;YP_173236.1;HCoV-HKU1_orf1ab;5814 | 0 | 0 |
| NC_006577.2;YP_173236.1;HCoV-HKU1_orf1ab;5815 | 0 | 0 |
| NC_006577.2;YP_173236.1;HCoV-HKU1_orf1ab;5816 | 0 | 0.025 |
| NC_006577.2;YP_173236.1;HCoV-HKU1_orf1ab;5817 | 0 | 0 |
| NC_006577.2;YP_173236.1;HCoV-HKU1_orf1ab;5818 | 0 | 0 |
| NC_006577.2;YP_173236.1;HCoV-HKU1_orf1ab;5819 | 0 | 0 |
| NC_006577.2;YP_173236.1;HCoV-HKU1_orf1ab;5820 | 0 | 0 |

| | | |
|---|---|---|
| NC_006577.2;YP_173236.1;HCoV-HKU1_orf1ab;5821 | 0 | 0 |
| NC_006577.2;YP_173236.1;HCoV-HKU1_orf1ab;5822 | 0.1 | 0.075 |
| NC_006577.2;YP_173236.1;HCoV-HKU1_orf1ab;5823 | 0.15 | 0.125 |
| NC_006577.2;YP_173236.1;HCoV-HKU1_orf1ab;5824 | 0.15 | 0.125 |
| NC_006577.2;YP_173236.1;HCoV-HKU1_orf1ab;5825 | 0.05 | 0.1 |
| NC_006577.2;YP_173236.1;HCoV-HKU1_orf1ab;5826 | 0 | 0.025 |
| NC_006577.2;YP_173236.1;HCoV-HKU1_orf1ab;5827 | 0 | 0 |
| NC_006577.2;YP_173236.1;HCoV-HKU1_orf1ab;5828 | 0.05 | 0.125 |
| NC_006577.2;YP_173236.1;HCoV-HKU1_orf1ab;5829 | 0.15 | 0.125 |
| NC_006577.2;YP_173236.1;HCoV-HKU1_orf1ab;5830 | 0.05 | 0.125 |
| NC_006577.2;YP_173236.1;HCoV-HKU1_orf1ab;5831 | 0.05 | 0.1 |
| NC_006577.2;YP_173236.1;HCoV-HKU1_orf1ab;5832 | 0.05 | 0.025 |
| NC_006577.2;YP_173236.1;HCoV-HKU1_orf1ab;5833 | 0 | 0.075 |
| NC_006577.2;YP_173236.1;HCoV-HKU1_orf1ab;5834 | 0 | 0 |
| NC_006577.2;YP_173236.1;HCoV-HKU1_orf1ab;5835 | 0 | 0 |
| NC_006577.2;YP_173236.1;HCoV-HKU1_orf1ab;5836 | 0 | 0 |
| NC_006577.2;YP_173236.1;HCoV-HKU1_orf1ab;5837 | 0.1 | 0 |
| NC_006577.2;YP_173236.1;HCoV-HKU1_orf1ab;5838 | 0 | 0 |
| NC_006577.2;YP_173236.1;HCoV-HKU1_orf1ab;5839 | 0 | 0.025 |
| NC_006577.2;YP_173236.1;HCoV-HKU1_orf1ab;5840 | 0 | 0 |
| NC_006577.2;YP_173236.1;HCoV-HKU1_orf1ab;5841 | 0 | 0 |
| NC_006577.2;YP_173236.1;HCoV-HKU1_orf1ab;5842 | 0 | 0 |
| NC_006577.2;YP_173236.1;HCoV-HKU1_orf1ab;5843 | 0 | 0 |
| NC_006577.2;YP_173236.1;HCoV-HKU1_orf1ab;5844 | 0 | 0.025 |
| NC_006577.2;YP_173236.1;HCoV-HKU1_orf1ab;5845 | 0 | 0 |
| NC_006577.2;YP_173236.1;HCoV-HKU1_orf1ab;5846 | 0 | 0 |
| NC_006577.2;YP_173236.1;HCoV-HKU1_orf1ab;5847 | 0 | 0 |
| NC_006577.2;YP_173236.1;HCoV-HKU1_orf1ab;5848 | 0 | 0 |
| NC_006577.2;YP_173236.1;HCoV-HKU1_orf1ab;5849 | 0 | 0 |
| NC_006577.2;YP_173236.1;HCoV-HKU1_orf1ab;5850 | 0 | 0 |
| NC_006577.2;YP_173236.1;HCoV-HKU1_orf1ab;5851 | 0 | 0 |
| NC_006577.2;YP_173236.1;HCoV-HKU1_orf1ab;5852 | 0 | 0.05 |
| NC_006577.2;YP_173236.1;HCoV-HKU1_orf1ab;5853 | 0 | 0.05 |
| NC_006577.2;YP_173236.1;HCoV-HKU1_orf1ab;5854 | 0.05 | 0.05 |
| NC_006577.2;YP_173236.1;HCoV-HKU1_orf1ab;5855 | 0.1 | 0.1 |
| NC_006577.2;YP_173236.1;HCoV-HKU1_orf1ab;5856 | 0.1 | 0.1 |
| NC_006577.2;YP_173236.1;HCoV-HKU1_orf1ab;5857 | 0.3 | 0.15 |
| NC_006577.2;YP_173236.1;HCoV-HKU1_orf1ab;5858 | 0 | 0 |
| NC_006577.2;YP_173236.1;HCoV-HKU1_orf1ab;5859 | 0 | 0 |
| NC_006577.2;YP_173236.1;HCoV-HKU1_orf1ab;5860 | 0 | 0 |
| NC_006577.2;YP_173236.1;HCoV-HKU1_orf1ab;5861 | 0 | 0.05 |
| NC_006577.2;YP_173236.1;HCoV-HKU1_orf1ab;5862 | 0 | 0 |
| NC_006577.2;YP_173236.1;HCoV-HKU1_orf1ab;5863 | 0 | 0 |

| | | |
|---|---|---|
| NC_006577.2;YP_173236.1;HCoV-HKU1_orf1ab;5864 | 0 | 0 |
| NC_006577.2;YP_173236.1;HCoV-HKU1_orf1ab;5865 | 0 | 0 |
| NC_006577.2;YP_173236.1;HCoV-HKU1_orf1ab;5866 | 0 | 0 |
| NC_006577.2;YP_173236.1;HCoV-HKU1_orf1ab;5867 | 0 | 0 |
| NC_006577.2;YP_173236.1;HCoV-HKU1_orf1ab;5868 | 0 | 0.05 |
| NC_006577.2;YP_173236.1;HCoV-HKU1_orf1ab;5869 | 0 | 0 |
| NC_006577.2;YP_173236.1;HCoV-HKU1_orf1ab;5870 | 0 | 0 |
| NC_006577.2;YP_173236.1;HCoV-HKU1_orf1ab;5871 | 0 | 0 |
| NC_006577.2;YP_173236.1;HCoV-HKU1_orf1ab;5872 | 0 | 0 |
| NC_006577.2;YP_173236.1;HCoV-HKU1_orf1ab;5873 | 0 | 0 |
| NC_006577.2;YP_173236.1;HCoV-HKU1_orf1ab;5874 | 0 | 0 |
| NC_006577.2;YP_173236.1;HCoV-HKU1_orf1ab;5875 | 0 | 0 |
| NC_006577.2;YP_173236.1;HCoV-HKU1_orf1ab;5876 | 0 | 0 |
| NC_006577.2;YP_173236.1;HCoV-HKU1_orf1ab;5877 | 0 | 0 |
| NC_006577.2;YP_173236.1;HCoV-HKU1_orf1ab;5878 | 0 | 0.025 |
| NC_006577.2;YP_173236.1;HCoV-HKU1_orf1ab;5879 | 0 | 0 |
| NC_006577.2;YP_173236.1;HCoV-HKU1_orf1ab;5880 | 0 | 0 |
| NC_006577.2;YP_173236.1;HCoV-HKU1_orf1ab;5881 | 0 | 0 |
| NC_006577.2;YP_173236.1;HCoV-HKU1_orf1ab;5882 | 0.05 | 0 |
| NC_006577.2;YP_173236.1;HCoV-HKU1_orf1ab;5883 | 0 | 0 |
| NC_006577.2;YP_173236.1;HCoV-HKU1_orf1ab;5884 | 0 | 0 |
| NC_006577.2;YP_173236.1;HCoV-HKU1_orf1ab;5885 | 0 | 0 |
| NC_006577.2;YP_173236.1;HCoV-HKU1_orf1ab;5886 | 0 | 0 |
| NC_006577.2;YP_173236.1;HCoV-HKU1_orf1ab;5887 | 0 | 0 |
| NC_006577.2;YP_173236.1;HCoV-HKU1_orf1ab;5888 | 0.05 | 0 |
| NC_006577.2;YP_173236.1;HCoV-HKU1_orf1ab;5889 | 0 | 0 |
| NC_006577.2;YP_173236.1;HCoV-HKU1_orf1ab;5890 | 0 | 0 |
| NC_006577.2;YP_173236.1;HCoV-HKU1_orf1ab;5891 | 0.05 | 0.025 |
| NC_006577.2;YP_173236.1;HCoV-HKU1_orf1ab;5892 | 0 | 0.075 |
| NC_006577.2;YP_173236.1;HCoV-HKU1_orf1ab;5893 | 0 | 0.025 |
| NC_006577.2;YP_173236.1;HCoV-HKU1_orf1ab;5894 | 0 | 0.025 |
| NC_006577.2;YP_173236.1;HCoV-HKU1_orf1ab;5895 | 0 | 0.05 |
| NC_006577.2;YP_173236.1;HCoV-HKU1_orf1ab;5896 | 0.05 | 0.05 |
| NC_006577.2;YP_173236.1;HCoV-HKU1_orf1ab;5897 | 0 | 0.05 |
| NC_006577.2;YP_173236.1;HCoV-HKU1_orf1ab;5898 | 0 | 0.025 |
| NC_006577.2;YP_173236.1;HCoV-HKU1_orf1ab;5899 | 0 | 0.05 |
| NC_006577.2;YP_173236.1;HCoV-HKU1_orf1ab;5900 | 0 | 0.05 |
| NC_006577.2;YP_173236.1;HCoV-HKU1_orf1ab;5901 | 0 | 0.075 |
| NC_006577.2;YP_173236.1;HCoV-HKU1_orf1ab;5902 | 0 | 0.025 |
| NC_006577.2;YP_173236.1;HCoV-HKU1_orf1ab;5903 | 0 | 0.05 |
| NC_006577.2;YP_173236.1;HCoV-HKU1_orf1ab;5904 | 0.1 | 0.05 |
| NC_006577.2;YP_173236.1;HCoV-HKU1_orf1ab;5905 | 0 | 0.05 |
| NC_006577.2;YP_173236.1;HCoV-HKU1_orf1ab;5906 | 0 | 0.1 |

| | | |
|---|---|---|
| NC_006577.2;YP_173236.1;HCoV-HKU1_orf1ab;5907 | 0.05 | 0.225 |
| NC_006577.2;YP_173236.1;HCoV-HKU1_orf1ab;5908 | 0 | 0.025 |
| NC_006577.2;YP_173236.1;HCoV-HKU1_orf1ab;5909 | 0 | 0 |
| NC_006577.2;YP_173236.1;HCoV-HKU1_orf1ab;5910 | 0 | 0 |
| NC_006577.2;YP_173236.1;HCoV-HKU1_orf1ab;5911 | 0 | 0 |
| NC_006577.2;YP_173236.1;HCoV-HKU1_orf1ab;5912 | 0.05 | 0 |
| NC_006577.2;YP_173236.1;HCoV-HKU1_orf1ab;5913 | 0.05 | 0 |
| NC_006577.2;YP_173236.1;HCoV-HKU1_orf1ab;5914 | 0 | 0 |
| NC_006577.2;YP_173236.1;HCoV-HKU1_orf1ab;5915 | 0 | 0 |
| NC_006577.2;YP_173236.1;HCoV-HKU1_orf1ab;5916 | 0 | 0 |
| NC_006577.2;YP_173236.1;HCoV-HKU1_orf1ab;5917 | 0 | 0.025 |
| NC_006577.2;YP_173236.1;HCoV-HKU1_orf1ab;5918 | 0 | 0 |
| NC_006577.2;YP_173236.1;HCoV-HKU1_orf1ab;5919 | 0 | 0 |
| NC_006577.2;YP_173236.1;HCoV-HKU1_orf1ab;5920 | 0 | 0 |
| NC_006577.2;YP_173236.1;HCoV-HKU1_orf1ab;5921 | 0 | 0 |
| NC_006577.2;YP_173236.1;HCoV-HKU1_orf1ab;5922 | 0 | 0 |
| NC_006577.2;YP_173236.1;HCoV-HKU1_orf1ab;5923 | 0 | 0 |
| NC_006577.2;YP_173236.1;HCoV-HKU1_orf1ab;5924 | 0.05 | 0.075 |
| NC_006577.2;YP_173236.1;HCoV-HKU1_orf1ab;5925 | 0 | 0 |
| NC_006577.2;YP_173236.1;HCoV-HKU1_orf1ab;5926 | 0 | 0 |
| NC_006577.2;YP_173236.1;HCoV-HKU1_orf1ab;5927 | 0 | 0 |
| NC_006577.2;YP_173236.1;HCoV-HKU1_orf1ab;5928 | 0 | 0.025 |
| NC_006577.2;YP_173236.1;HCoV-HKU1_orf1ab;5929 | 0 | 0.025 |
| NC_006577.2;YP_173236.1;HCoV-HKU1_orf1ab;5930 | 0 | 0 |
| NC_006577.2;YP_173236.1;HCoV-HKU1_orf1ab;5931 | 0.05 | 0 |
| NC_006577.2;YP_173236.1;HCoV-HKU1_orf1ab;5932 | 0.1 | 0 |
| NC_006577.2;YP_173236.1;HCoV-HKU1_orf1ab;5933 | 0 | 0.025 |
| NC_006577.2;YP_173236.1;HCoV-HKU1_orf1ab;5934 | 0 | 0 |
| NC_006577.2;YP_173236.1;HCoV-HKU1_orf1ab;5935 | 0 | 0 |
| NC_006577.2;YP_173236.1;HCoV-HKU1_orf1ab;5936 | 0 | 0 |
| NC_006577.2;YP_173236.1;HCoV-HKU1_orf1ab;5937 | 0 | 0 |
| NC_006577.2;YP_173236.1;HCoV-HKU1_orf1ab;5938 | 0 | 0 |
| NC_006577.2;YP_173236.1;HCoV-HKU1_orf1ab;5939 | 0 | 0 |
| NC_006577.2;YP_173236.1;HCoV-HKU1_orf1ab;5940 | 0 | 0.025 |
| NC_006577.2;YP_173236.1;HCoV-HKU1_orf1ab;5941 | 0 | 0 |
| NC_006577.2;YP_173236.1;HCoV-HKU1_orf1ab;5942 | 0.05 | 0 |
| NC_006577.2;YP_173236.1;HCoV-HKU1_orf1ab;5943 | 0.05 | 0 |
| NC_006577.2;YP_173236.1;HCoV-HKU1_orf1ab;5944 | 0 | 0 |
| NC_006577.2;YP_173236.1;HCoV-HKU1_orf1ab;5945 | 0 | 0 |
| NC_006577.2;YP_173236.1;HCoV-HKU1_orf1ab;5946 | 0 | 0 |
| NC_006577.2;YP_173236.1;HCoV-HKU1_orf1ab;5947 | 0 | 0 |
| NC_006577.2;YP_173236.1;HCoV-HKU1_orf1ab;5948 | 0 | 0 |
| NC_006577.2;YP_173236.1;HCoV-HKU1_orf1ab;5949 | 0 | 0 |

| | | |
|---|---|---|
| NC_006577.2;YP_173236.1;HCoV-HKU1_orf1ab;5950 | 0 | 0 |
| NC_006577.2;YP_173236.1;HCoV-HKU1_orf1ab;5951 | 0 | 0 |
| NC_006577.2;YP_173236.1;HCoV-HKU1_orf1ab;5952 | 0 | 0 |
| NC_006577.2;YP_173236.1;HCoV-HKU1_orf1ab;5953 | 0 | 0 |
| NC_006577.2;YP_173236.1;HCoV-HKU1_orf1ab;5954 | 0 | 0 |
| NC_006577.2;YP_173236.1;HCoV-HKU1_orf1ab;5955 | 0 | 0 |
| NC_006577.2;YP_173236.1;HCoV-HKU1_orf1ab;5956 | 0 | 0 |
| NC_006577.2;YP_173236.1;HCoV-HKU1_orf1ab;5957 | 0 | 0 |
| NC_006577.2;YP_173236.1;HCoV-HKU1_orf1ab;5958 | 0 | 0.025 |
| NC_006577.2;YP_173236.1;HCoV-HKU1_orf1ab;5959 | 0 | 0 |
| NC_006577.2;YP_173236.1;HCoV-HKU1_orf1ab;5960 | 0 | 0.025 |
| NC_006577.2;YP_173236.1;HCoV-HKU1_orf1ab;5961 | 0 | 0 |
| NC_006577.2;YP_173236.1;HCoV-HKU1_orf1ab;5962 | 0 | 0 |
| NC_006577.2;YP_173236.1;HCoV-HKU1_orf1ab;5963 | 0 | 0 |
| NC_006577.2;YP_173236.1;HCoV-HKU1_orf1ab;5964 | 0 | 0 |
| NC_006577.2;YP_173236.1;HCoV-HKU1_orf1ab;5965 | 0 | 0 |
| NC_006577.2;YP_173236.1;HCoV-HKU1_orf1ab;5966 | 0 | 0 |
| NC_006577.2;YP_173236.1;HCoV-HKU1_orf1ab;5967 | 0 | 0 |
| NC_006577.2;YP_173236.1;HCoV-HKU1_orf1ab;5968 | 0 | 0 |
| NC_006577.2;YP_173236.1;HCoV-HKU1_orf1ab;5969 | 0 | 0 |
| NC_006577.2;YP_173236.1;HCoV-HKU1_orf1ab;5970 | 0 | 0 |
| NC_006577.2;YP_173236.1;HCoV-HKU1_orf1ab;5971 | 0 | 0 |
| NC_006577.2;YP_173236.1;HCoV-HKU1_orf1ab;5972 | 0 | 0 |
| NC_006577.2;YP_173236.1;HCoV-HKU1_orf1ab;5973 | 0 | 0 |
| NC_006577.2;YP_173236.1;HCoV-HKU1_orf1ab;5974 | 0 | 0 |
| NC_006577.2;YP_173236.1;HCoV-HKU1_orf1ab;5975 | 0 | 0 |
| NC_006577.2;YP_173236.1;HCoV-HKU1_orf1ab;5976 | 0 | 0 |
| NC_006577.2;YP_173236.1;HCoV-HKU1_orf1ab;5977 | 0.05 | 0 |
| NC_006577.2;YP_173236.1;HCoV-HKU1_orf1ab;5978 | 0.05 | 0 |
| NC_006577.2;YP_173236.1;HCoV-HKU1_orf1ab;5979 | 0 | 0 |
| NC_006577.2;YP_173236.1;HCoV-HKU1_orf1ab;5980 | 0 | 0 |
| NC_006577.2;YP_173236.1;HCoV-HKU1_orf1ab;5981 | 0.05 | 0 |
| NC_006577.2;YP_173236.1;HCoV-HKU1_orf1ab;5982 | 0 | 0 |
| NC_006577.2;YP_173236.1;HCoV-HKU1_orf1ab;5983 | 0.05 | 0 |
| NC_006577.2;YP_173236.1;HCoV-HKU1_orf1ab;5984 | 0 | 0 |
| NC_006577.2;YP_173236.1;HCoV-HKU1_orf1ab;5985 | 0 | 0 |
| NC_006577.2;YP_173236.1;HCoV-HKU1_orf1ab;5986 | 0 | 0 |
| NC_006577.2;YP_173236.1;HCoV-HKU1_orf1ab;5987 | 0 | 0 |
| NC_006577.2;YP_173236.1;HCoV-HKU1_orf1ab;5988 | 0 | 0 |
| NC_006577.2;YP_173236.1;HCoV-HKU1_orf1ab;5989 | 0 | 0 |
| NC_006577.2;YP_173236.1;HCoV-HKU1_orf1ab;5990 | 0 | 0 |
| NC_006577.2;YP_173236.1;HCoV-HKU1_orf1ab;5991 | 0 | 0 |
| NC_006577.2;YP_173236.1;HCoV-HKU1_orf1ab;5992 | 0 | 0.025 |

| | | |
|---|---|---|
| NC_006577.2;YP_173236.1;HCoV-HKU1_orf1ab;5993 | 0 | 0.025 |
| NC_006577.2;YP_173236.1;HCoV-HKU1_orf1ab;5994 | 0 | 0.025 |
| NC_006577.2;YP_173236.1;HCoV-HKU1_orf1ab;5995 | 0 | 0 |
| NC_006577.2;YP_173236.1;HCoV-HKU1_orf1ab;5996 | 0 | 0.025 |
| NC_006577.2;YP_173236.1;HCoV-HKU1_orf1ab;5997 | 0 | 0.025 |
| NC_006577.2;YP_173236.1;HCoV-HKU1_orf1ab;5998 | 0 | 0.025 |
| NC_006577.2;YP_173236.1;HCoV-HKU1_orf1ab;5999 | 0 | 0.025 |
| NC_006577.2;YP_173236.1;HCoV-HKU1_orf1ab;6000 | 0 | 0 |
| NC_006577.2;YP_173236.1;HCoV-HKU1_orf1ab;6001 | 0 | 0 |
| NC_006577.2;YP_173236.1;HCoV-HKU1_orf1ab;6002 | 0 | 0.025 |
| NC_006577.2;YP_173236.1;HCoV-HKU1_orf1ab;6003 | 0 | 0.025 |
| NC_006577.2;YP_173236.1;HCoV-HKU1_orf1ab;6004 | 0 | 0.05 |
| NC_006577.2;YP_173236.1;HCoV-HKU1_orf1ab;6005 | 0 | 0.025 |
| NC_006577.2;YP_173236.1;HCoV-HKU1_orf1ab;6006 | 0.15 | 0.125 |
| NC_006577.2;YP_173236.1;HCoV-HKU1_orf1ab;6007 | 0.15 | 0.05 |
| NC_006577.2;YP_173236.1;HCoV-HKU1_orf1ab;6008 | 0.15 | 0.1 |
| NC_006577.2;YP_173236.1;HCoV-HKU1_orf1ab;6009 | 0.2 | 0.125 |
| NC_006577.2;YP_173236.1;HCoV-HKU1_orf1ab;6010 | 0.2 | 0.15 |
| NC_006577.2;YP_173236.1;HCoV-HKU1_orf1ab;6011 | 0.15 | 0.15 |
| NC_006577.2;YP_173236.1;HCoV-HKU1_orf1ab;6012 | 0.15 | 0.15 |
| NC_006577.2;YP_173236.1;HCoV-HKU1_orf1ab;6013 | 0.2 | 0.075 |
| NC_006577.2;YP_173236.1;HCoV-HKU1_orf1ab;6014 | 0.2 | 0.05 |
| NC_006577.2;YP_173236.1;HCoV-HKU1_orf1ab;6015 | 0.05 | 0 |
| NC_006577.2;YP_173236.1;HCoV-HKU1_orf1ab;6016 | 0 | 0 |
| NC_006577.2;YP_173236.1;HCoV-HKU1_orf1ab;6017 | 0 | 0 |
| NC_006577.2;YP_173236.1;HCoV-HKU1_orf1ab;6018 | 0 | 0 |
| NC_006577.2;YP_173236.1;HCoV-HKU1_orf1ab;6019 | 0 | 0.025 |
| NC_006577.2;YP_173236.1;HCoV-HKU1_orf1ab;6020 | 0 | 0 |
| NC_006577.2;YP_173236.1;HCoV-HKU1_orf1ab;6021 | 0 | 0 |
| NC_006577.2;YP_173236.1;HCoV-HKU1_orf1ab;6022 | 0 | 0 |
| NC_006577.2;YP_173236.1;HCoV-HKU1_orf1ab;6023 | 0 | 0 |
| NC_006577.2;YP_173236.1;HCoV-HKU1_orf1ab;6024 | 0 | 0 |
| NC_006577.2;YP_173236.1;HCoV-HKU1_orf1ab;6025 | 0 | 0 |
| NC_006577.2;YP_173236.1;HCoV-HKU1_orf1ab;6026 | 0 | 0 |
| NC_006577.2;YP_173236.1;HCoV-HKU1_orf1ab;6027 | 0 | 0 |
| NC_006577.2;YP_173236.1;HCoV-HKU1_orf1ab;6028 | 0 | 0 |
| NC_006577.2;YP_173236.1;HCoV-HKU1_orf1ab;6029 | 0 | 0 |
| NC_006577.2;YP_173236.1;HCoV-HKU1_orf1ab;6030 | 0 | 0 |
| NC_006577.2;YP_173236.1;HCoV-HKU1_orf1ab;6031 | 0 | 0 |
| NC_006577.2;YP_173236.1;HCoV-HKU1_orf1ab;6032 | 0 | 0 |
| NC_006577.2;YP_173236.1;HCoV-HKU1_orf1ab;6033 | 0 | 0 |
| NC_006577.2;YP_173236.1;HCoV-HKU1_orf1ab;6034 | 0 | 0 |
| NC_006577.2;YP_173236.1;HCoV-HKU1_orf1ab;6035 | 0 | 0 |

| | | |
|---|---|---|
| NC_006577.2;YP_173236.1;HCoV-HKU1_orf1ab;6036 | 0 | 0 |
| NC_006577.2;YP_173236.1;HCoV-HKU1_orf1ab;6037 | 0 | 0 |
| NC_006577.2;YP_173236.1;HCoV-HKU1_orf1ab;6038 | 0 | 0 |
| NC_006577.2;YP_173236.1;HCoV-HKU1_orf1ab;6039 | 0 | 0 |
| NC_006577.2;YP_173236.1;HCoV-HKU1_orf1ab;6040 | 0 | 0 |
| NC_006577.2;YP_173236.1;HCoV-HKU1_orf1ab;6041 | 0 | 0 |
| NC_006577.2;YP_173236.1;HCoV-HKU1_orf1ab;6042 | 0 | 0 |
| NC_006577.2;YP_173236.1;HCoV-HKU1_orf1ab;6043 | 0 | 0 |
| NC_006577.2;YP_173236.1;HCoV-HKU1_orf1ab;6044 | 0 | 0.025 |
| NC_006577.2;YP_173236.1;HCoV-HKU1_orf1ab;6045 | 0 | 0 |
| NC_006577.2;YP_173236.1;HCoV-HKU1_orf1ab;6046 | 0 | 0 |
| NC_006577.2;YP_173236.1;HCoV-HKU1_orf1ab;6047 | 0 | 0 |
| NC_006577.2;YP_173236.1;HCoV-HKU1_orf1ab;6048 | 0 | 0 |
| NC_006577.2;YP_173236.1;HCoV-HKU1_orf1ab;6049 | 0 | 0.025 |
| NC_006577.2;YP_173236.1;HCoV-HKU1_orf1ab;6050 | 0 | 0.025 |
| NC_006577.2;YP_173236.1;HCoV-HKU1_orf1ab;6051 | 0 | 0.05 |
| NC_006577.2;YP_173236.1;HCoV-HKU1_orf1ab;6052 | 0 | 0.05 |
| NC_006577.2;YP_173236.1;HCoV-HKU1_orf1ab;6053 | 0 | 0.075 |
| NC_006577.2;YP_173236.1;HCoV-HKU1_orf1ab;6054 | 0 | 0.025 |
| NC_006577.2;YP_173236.1;HCoV-HKU1_orf1ab;6055 | 0 | 0.025 |
| NC_006577.2;YP_173236.1;HCoV-HKU1_orf1ab;6056 | 0 | 0.025 |
| NC_006577.2;YP_173236.1;HCoV-HKU1_orf1ab;6057 | 0 | 0.025 |
| NC_006577.2;YP_173236.1;HCoV-HKU1_orf1ab;6058 | 0 | 0.025 |
| NC_006577.2;YP_173236.1;HCoV-HKU1_orf1ab;6059 | 0 | 0.025 |
| NC_006577.2;YP_173236.1;HCoV-HKU1_orf1ab;6060 | 0 | 0.025 |
| NC_006577.2;YP_173236.1;HCoV-HKU1_orf1ab;6061 | 0 | 0.075 |
| NC_006577.2;YP_173236.1;HCoV-HKU1_orf1ab;6062 | 0 | 0.025 |
| NC_006577.2;YP_173236.1;HCoV-HKU1_orf1ab;6063 | 0 | 0 |
| NC_006577.2;YP_173236.1;HCoV-HKU1_orf1ab;6064 | 0.05 | 0 |
| NC_006577.2;YP_173236.1;HCoV-HKU1_orf1ab;6065 | 0.05 | 0 |
| NC_006577.2;YP_173236.1;HCoV-HKU1_orf1ab;6066 | 0 | 0 |
| NC_006577.2;YP_173236.1;HCoV-HKU1_orf1ab;6067 | 0 | 0.025 |
| NC_006577.2;YP_173236.1;HCoV-HKU1_orf1ab;6068 | 0 | 0.025 |
| NC_006577.2;YP_173236.1;HCoV-HKU1_orf1ab;6069 | 0.05 | 0.025 |
| NC_006577.2;YP_173236.1;HCoV-HKU1_orf1ab;6070 | 0 | 0.025 |
| NC_006577.2;YP_173236.1;HCoV-HKU1_orf1ab;6071 | 0 | 0.025 |
| NC_006577.2;YP_173236.1;HCoV-HKU1_orf1ab;6072 | 0 | 0.025 |
| NC_006577.2;YP_173236.1;HCoV-HKU1_orf1ab;6073 | 0 | 0.025 |
| NC_006577.2;YP_173236.1;HCoV-HKU1_orf1ab;6074 | 0 | 0 |
| NC_006577.2;YP_173236.1;HCoV-HKU1_orf1ab;6075 | 0 | 0 |
| NC_006577.2;YP_173236.1;HCoV-HKU1_orf1ab;6076 | 0 | 0.025 |
| NC_006577.2;YP_173236.1;HCoV-HKU1_orf1ab;6077 | 0 | 0.05 |
| NC_006577.2;YP_173236.1;HCoV-HKU1_orf1ab;6078 | 0.05 | 0.075 |

| | | |
|---|---|---|
| NC_006577.2;YP_173236.1;HCoV-HKU1_orf1ab;6079 | 0 | 0.075 |
| NC_006577.2;YP_173236.1;HCoV-HKU1_orf1ab;6080 | 0 | 0.075 |
| NC_006577.2;YP_173236.1;HCoV-HKU1_orf1ab;6081 | 0 | 0.05 |
| NC_006577.2;YP_173236.1;HCoV-HKU1_orf1ab;6082 | 0 | 0.05 |
| NC_006577.2;YP_173236.1;HCoV-HKU1_orf1ab;6083 | 0 | 0.075 |
| NC_006577.2;YP_173236.1;HCoV-HKU1_orf1ab;6084 | 0 | 0.075 |
| NC_006577.2;YP_173236.1;HCoV-HKU1_orf1ab;6085 | 0 | 0.025 |
| NC_006577.2;YP_173236.1;HCoV-HKU1_orf1ab;6086 | 0.05 | 0.05 |
| NC_006577.2;YP_173236.1;HCoV-HKU1_orf1ab;6087 | 0 | 0.025 |
| NC_006577.2;YP_173236.1;HCoV-HKU1_orf1ab;6088 | 0.05 | 0.05 |
| NC_006577.2;YP_173236.1;HCoV-HKU1_orf1ab;6089 | 0 | 0 |
| NC_006577.2;YP_173236.1;HCoV-HKU1_orf1ab;6090 | 0 | 0 |
| NC_006577.2;YP_173236.1;HCoV-HKU1_orf1ab;6091 | 0 | 0 |
| NC_006577.2;YP_173236.1;HCoV-HKU1_orf1ab;6092 | 0 | 0 |
| NC_006577.2;YP_173236.1;HCoV-HKU1_orf1ab;6093 | 0 | 0 |
| NC_006577.2;YP_173236.1;HCoV-HKU1_orf1ab;6094 | 0 | 0 |
| NC_006577.2;YP_173236.1;HCoV-HKU1_orf1ab;6095 | 0 | 0 |
| NC_006577.2;YP_173236.1;HCoV-HKU1_orf1ab;6096 | 0 | 0 |
| NC_006577.2;YP_173236.1;HCoV-HKU1_orf1ab;6097 | 0 | 0 |
| NC_006577.2;YP_173236.1;HCoV-HKU1_orf1ab;6098 | 0 | 0 |
| NC_006577.2;YP_173236.1;HCoV-HKU1_orf1ab;6099 | 0 | 0 |
| NC_006577.2;YP_173236.1;HCoV-HKU1_orf1ab;6100 | 0 | 0 |
| NC_006577.2;YP_173236.1;HCoV-HKU1_orf1ab;6101 | 0 | 0 |
| NC_006577.2;YP_173236.1;HCoV-HKU1_orf1ab;6102 | 0 | 0 |
| NC_006577.2;YP_173236.1;HCoV-HKU1_orf1ab;6103 | 0 | 0 |
| NC_006577.2;YP_173236.1;HCoV-HKU1_orf1ab;6104 | 0 | 0 |
| NC_006577.2;YP_173236.1;HCoV-HKU1_orf1ab;6105 | 0 | 0 |
| NC_006577.2;YP_173236.1;HCoV-HKU1_orf1ab;6106 | 0 | 0 |
| NC_006577.2;YP_173236.1;HCoV-HKU1_orf1ab;6107 | 0 | 0.05 |
| NC_006577.2;YP_173236.1;HCoV-HKU1_orf1ab;6108 | 0 | 0 |
| NC_006577.2;YP_173236.1;HCoV-HKU1_orf1ab;6109 | 0 | 0 |
| NC_006577.2;YP_173236.1;HCoV-HKU1_orf1ab;6110 | 0 | 0 |
| NC_006577.2;YP_173236.1;HCoV-HKU1_orf1ab;6111 | 0 | 0 |
| NC_006577.2;YP_173236.1;HCoV-HKU1_orf1ab;6112 | 0 | 0 |
| NC_006577.2;YP_173236.1;HCoV-HKU1_orf1ab;6113 | 0 | 0.025 |
| NC_006577.2;YP_173236.1;HCoV-HKU1_orf1ab;6114 | 0 | 0 |
| NC_006577.2;YP_173236.1;HCoV-HKU1_orf1ab;6115 | 0 | 0 |
| NC_006577.2;YP_173236.1;HCoV-HKU1_orf1ab;6116 | 0 | 0 |
| NC_006577.2;YP_173236.1;HCoV-HKU1_orf1ab;6117 | 0 | 0 |
| NC_006577.2;YP_173236.1;HCoV-HKU1_orf1ab;6118 | 0 | 0 |
| NC_006577.2;YP_173236.1;HCoV-HKU1_orf1ab;6119 | 0 | 0 |
| NC_006577.2;YP_173236.1;HCoV-HKU1_orf1ab;6120 | 0 | 0 |
| NC_006577.2;YP_173236.1;HCoV-HKU1_orf1ab;6121 | 0 | 0 |

| | | |
|---|---|---|
| NC_006577.2;YP_173236.1;HCoV-HKU1_orf1ab;6122 | 0 | 0 |
| NC_006577.2;YP_173236.1;HCoV-HKU1_orf1ab;6123 | 0.1 | 0 |
| NC_006577.2;YP_173236.1;HCoV-HKU1_orf1ab;6124 | 0.05 | 0 |
| NC_006577.2;YP_173236.1;HCoV-HKU1_orf1ab;6125 | 0 | 0 |
| NC_006577.2;YP_173236.1;HCoV-HKU1_orf1ab;6126 | 0 | 0 |
| NC_006577.2;YP_173236.1;HCoV-HKU1_orf1ab;6127 | 0.05 | 0 |
| NC_006577.2;YP_173236.1;HCoV-HKU1_orf1ab;6128 | 0.05 | 0 |
| NC_006577.2;YP_173236.1;HCoV-HKU1_orf1ab;6129 | 0.05 | 0 |
| NC_006577.2;YP_173236.1;HCoV-HKU1_orf1ab;6130 | 0 | 0 |
| NC_006577.2;YP_173236.1;HCoV-HKU1_orf1ab;6131 | 0 | 0 |
| NC_006577.2;YP_173236.1;HCoV-HKU1_orf1ab;6132 | 0 | 0 |
| NC_006577.2;YP_173236.1;HCoV-HKU1_orf1ab;6133 | 0 | 0 |
| NC_006577.2;YP_173236.1;HCoV-HKU1_orf1ab;6134 | 0 | 0 |
| NC_006577.2;YP_173236.1;HCoV-HKU1_orf1ab;6135 | 0 | 0 |
| NC_006577.2;YP_173236.1;HCoV-HKU1_orf1ab;6136 | 0 | 0 |
| NC_006577.2;YP_173236.1;HCoV-HKU1_orf1ab;6137 | 0 | 0 |
| NC_006577.2;YP_173236.1;HCoV-HKU1_orf1ab;6138 | 0 | 0 |
| NC_006577.2;YP_173236.1;HCoV-HKU1_orf1ab;6139 | 0 | 0 |
| NC_006577.2;YP_173236.1;HCoV-HKU1_orf1ab;6140 | 0 | 0 |
| NC_006577.2;YP_173236.1;HCoV-HKU1_orf1ab;6141 | 0 | 0 |
| NC_006577.2;YP_173236.1;HCoV-HKU1_orf1ab;6142 | 0 | 0 |
| NC_006577.2;YP_173236.1;HCoV-HKU1_orf1ab;6143 | 0 | 0 |
| NC_006577.2;YP_173236.1;HCoV-HKU1_orf1ab;6144 | 0 | 0 |
| NC_006577.2;YP_173236.1;HCoV-HKU1_orf1ab;6145 | 0 | 0 |
| NC_006577.2;YP_173236.1;HCoV-HKU1_orf1ab;6146 | 0 | 0 |
| NC_006577.2;YP_173236.1;HCoV-HKU1_orf1ab;6147 | 0 | 0 |
| NC_006577.2;YP_173236.1;HCoV-HKU1_orf1ab;6148 | 0 | 0 |
| NC_006577.2;YP_173236.1;HCoV-HKU1_orf1ab;6149 | 0 | 0 |
| NC_006577.2;YP_173236.1;HCoV-HKU1_orf1ab;6150 | 0 | 0 |
| NC_006577.2;YP_173236.1;HCoV-HKU1_orf1ab;6151 | 0 | 0 |
| NC_006577.2;YP_173236.1;HCoV-HKU1_orf1ab;6152 | 0 | 0 |
| NC_006577.2;YP_173236.1;HCoV-HKU1_orf1ab;6153 | 0 | 0.025 |
| NC_006577.2;YP_173236.1;HCoV-HKU1_orf1ab;6154 | 0 | 0 |
| NC_006577.2;YP_173236.1;HCoV-HKU1_orf1ab;6155 | 0 | 0 |
| NC_006577.2;YP_173236.1;HCoV-HKU1_orf1ab;6156 | 0 | 0 |
| NC_006577.2;YP_173236.1;HCoV-HKU1_orf1ab;6157 | 0 | 0.025 |
| NC_006577.2;YP_173236.1;HCoV-HKU1_orf1ab;6158 | 0 | 0 |
| NC_006577.2;YP_173236.1;HCoV-HKU1_orf1ab;6159 | 0 | 0 |
| NC_006577.2;YP_173236.1;HCoV-HKU1_orf1ab;6160 | 0 | 0 |
| NC_006577.2;YP_173236.1;HCoV-HKU1_orf1ab;6161 | 0 | 0 |
| NC_006577.2;YP_173236.1;HCoV-HKU1_orf1ab;6162 | 0 | 0 |
| NC_006577.2;YP_173236.1;HCoV-HKU1_orf1ab;6163 | 0 | 0 |
| NC_006577.2;YP_173236.1;HCoV-HKU1_orf1ab;6164 | 0 | 0 |

| | | |
|---|---|---|
| NC_006577.2;YP_173236.1;HCoV-HKU1_orf1ab;6165 | 0 | 0 |
| NC_006577.2;YP_173236.1;HCoV-HKU1_orf1ab;6166 | 0 | 0 |
| NC_006577.2;YP_173236.1;HCoV-HKU1_orf1ab;6167 | 0 | 0 |
| NC_006577.2;YP_173236.1;HCoV-HKU1_orf1ab;6168 | 0 | 0 |
| NC_006577.2;YP_173236.1;HCoV-HKU1_orf1ab;6169 | 0 | 0 |
| NC_006577.2;YP_173236.1;HCoV-HKU1_orf1ab;6170 | 0 | 0 |
| NC_006577.2;YP_173236.1;HCoV-HKU1_orf1ab;6171 | 0 | 0 |
| NC_006577.2;YP_173236.1;HCoV-HKU1_orf1ab;6172 | 0 | 0 |
| NC_006577.2;YP_173236.1;HCoV-HKU1_orf1ab;6173 | 0 | 0 |
| NC_006577.2;YP_173236.1;HCoV-HKU1_orf1ab;6174 | 0 | 0 |
| NC_006577.2;YP_173236.1;HCoV-HKU1_orf1ab;6175 | 0 | 0 |
| NC_006577.2;YP_173236.1;HCoV-HKU1_orf1ab;6176 | 0 | 0 |
| NC_006577.2;YP_173236.1;HCoV-HKU1_orf1ab;6177 | 0 | 0 |
| NC_006577.2;YP_173236.1;HCoV-HKU1_orf1ab;6178 | 0 | 0 |
| NC_006577.2;YP_173236.1;HCoV-HKU1_orf1ab;6179 | 0 | 0 |
| NC_006577.2;YP_173236.1;HCoV-HKU1_orf1ab;6180 | 0 | 0 |
| NC_006577.2;YP_173236.1;HCoV-HKU1_orf1ab;6181 | 0 | 0 |
| NC_006577.2;YP_173236.1;HCoV-HKU1_orf1ab;6182 | 0 | 0 |
| NC_006577.2;YP_173236.1;HCoV-HKU1_orf1ab;6183 | 0 | 0 |
| NC_006577.2;YP_173236.1;HCoV-HKU1_orf1ab;6184 | 0 | 0.025 |
| NC_006577.2;YP_173236.1;HCoV-HKU1_orf1ab;6185 | 0 | 0 |
| NC_006577.2;YP_173236.1;HCoV-HKU1_orf1ab;6186 | 0 | 0 |
| NC_006577.2;YP_173236.1;HCoV-HKU1_orf1ab;6187 | 0 | 0 |
| NC_006577.2;YP_173236.1;HCoV-HKU1_orf1ab;6188 | 0 | 0.025 |
| NC_006577.2;YP_173236.1;HCoV-HKU1_orf1ab;6189 | 0 | 0 |
| NC_006577.2;YP_173236.1;HCoV-HKU1_orf1ab;6190 | 0 | 0 |
| NC_006577.2;YP_173236.1;HCoV-HKU1_orf1ab;6191 | 0 | 0 |
| NC_006577.2;YP_173236.1;HCoV-HKU1_orf1ab;6192 | 0 | 0 |
| NC_006577.2;YP_173236.1;HCoV-HKU1_orf1ab;6193 | 0 | 0 |
| NC_006577.2;YP_173236.1;HCoV-HKU1_orf1ab;6194 | 0 | 0 |
| NC_006577.2;YP_173236.1;HCoV-HKU1_orf1ab;6195 | 0 | 0 |
| NC_006577.2;YP_173236.1;HCoV-HKU1_orf1ab;6196 | 0 | 0 |
| NC_006577.2;YP_173236.1;HCoV-HKU1_orf1ab;6197 | 0.05 | 0 |
| NC_006577.2;YP_173236.1;HCoV-HKU1_orf1ab;6198 | 0.1 | 0 |
| NC_006577.2;YP_173236.1;HCoV-HKU1_orf1ab;6199 | 0 | 0 |
| NC_006577.2;YP_173236.1;HCoV-HKU1_orf1ab;6200 | 0 | 0 |
| NC_006577.2;YP_173236.1;HCoV-HKU1_orf1ab;6201 | 0 | 0 |
| NC_006577.2;YP_173236.1;HCoV-HKU1_orf1ab;6202 | 0 | 0 |
| NC_006577.2;YP_173236.1;HCoV-HKU1_orf1ab;6203 | 0 | 0 |
| NC_006577.2;YP_173236.1;HCoV-HKU1_orf1ab;6204 | 0 | 0 |
| NC_006577.2;YP_173236.1;HCoV-HKU1_orf1ab;6205 | 0 | 0 |
| NC_006577.2;YP_173236.1;HCoV-HKU1_orf1ab;6206 | 0 | 0 |
| NC_006577.2;YP_173236.1;HCoV-HKU1_orf1ab;6207 | 0 | 0 |

| | | |
|---|---|---|
| NC_006577.2;YP_173236.1;HCoV-HKU1_orf1ab;6208 | 0 | 0 |
| NC_006577.2;YP_173236.1;HCoV-HKU1_orf1ab;6209 | 0 | 0 |
| NC_006577.2;YP_173236.1;HCoV-HKU1_orf1ab;6210 | 0 | 0 |
| NC_006577.2;YP_173236.1;HCoV-HKU1_orf1ab;6211 | 0 | 0 |
| NC_006577.2;YP_173236.1;HCoV-HKU1_orf1ab;6212 | 0 | 0 |
| NC_006577.2;YP_173236.1;HCoV-HKU1_orf1ab;6213 | 0 | 0 |
| NC_006577.2;YP_173236.1;HCoV-HKU1_orf1ab;6214 | 0 | 0 |
| NC_006577.2;YP_173236.1;HCoV-HKU1_orf1ab;6215 | 0 | 0 |
| NC_006577.2;YP_173236.1;HCoV-HKU1_orf1ab;6216 | 0 | 0 |
| NC_006577.2;YP_173236.1;HCoV-HKU1_orf1ab;6217 | 0 | 0 |
| NC_006577.2;YP_173236.1;HCoV-HKU1_orf1ab;6218 | 0 | 0 |
| NC_006577.2;YP_173236.1;HCoV-HKU1_orf1ab;6219 | 0 | 0 |
| NC_006577.2;YP_173236.1;HCoV-HKU1_orf1ab;6220 | 0 | 0 |
| NC_006577.2;YP_173236.1;HCoV-HKU1_orf1ab;6221 | 0 | 0 |
| NC_006577.2;YP_173236.1;HCoV-HKU1_orf1ab;6222 | 0 | 0 |
| NC_006577.2;YP_173236.1;HCoV-HKU1_orf1ab;6223 | 0 | 0 |
| NC_006577.2;YP_173236.1;HCoV-HKU1_orf1ab;6224 | 0 | 0 |
| NC_006577.2;YP_173236.1;HCoV-HKU1_orf1ab;6225 | 0 | 0 |
| NC_006577.2;YP_173236.1;HCoV-HKU1_orf1ab;6226 | 0 | 0 |
| NC_006577.2;YP_173236.1;HCoV-HKU1_orf1ab;6227 | 0 | 0 |
| NC_006577.2;YP_173236.1;HCoV-HKU1_orf1ab;6228 | 0 | 0 |
| NC_006577.2;YP_173236.1;HCoV-HKU1_orf1ab;6229 | 0 | 0 |
| NC_006577.2;YP_173236.1;HCoV-HKU1_orf1ab;6230 | 0 | 0 |
| NC_006577.2;YP_173236.1;HCoV-HKU1_orf1ab;6231 | 0 | 0 |
| NC_006577.2;YP_173236.1;HCoV-HKU1_orf1ab;6232 | 0 | 0 |
| NC_006577.2;YP_173236.1;HCoV-HKU1_orf1ab;6233 | 0 | 0 |
| NC_006577.2;YP_173236.1;HCoV-HKU1_orf1ab;6234 | 0 | 0 |
| NC_006577.2;YP_173236.1;HCoV-HKU1_orf1ab;6235 | 0 | 0 |
| NC_006577.2;YP_173236.1;HCoV-HKU1_orf1ab;6236 | 0 | 0 |
| NC_006577.2;YP_173236.1;HCoV-HKU1_orf1ab;6237 | 0 | 0 |
| NC_006577.2;YP_173236.1;HCoV-HKU1_orf1ab;6238 | 0 | 0 |
| NC_006577.2;YP_173236.1;HCoV-HKU1_orf1ab;6239 | 0.05 | 0 |
| NC_006577.2;YP_173236.1;HCoV-HKU1_orf1ab;6240 | 0 | 0 |
| NC_006577.2;YP_173236.1;HCoV-HKU1_orf1ab;6241 | 0 | 0 |
| NC_006577.2;YP_173236.1;HCoV-HKU1_orf1ab;6242 | 0.05 | 0.025 |
| NC_006577.2;YP_173236.1;HCoV-HKU1_orf1ab;6243 | 0 | 0 |
| NC_006577.2;YP_173236.1;HCoV-HKU1_orf1ab;6244 | 0 | 0 |
| NC_006577.2;YP_173236.1;HCoV-HKU1_orf1ab;6245 | 0 | 0 |
| NC_006577.2;YP_173236.1;HCoV-HKU1_orf1ab;6246 | 0 | 0 |
| NC_006577.2;YP_173236.1;HCoV-HKU1_orf1ab;6247 | 0.05 | 0 |
| NC_006577.2;YP_173236.1;HCoV-HKU1_orf1ab;6248 | 0 | 0 |
| NC_006577.2;YP_173236.1;HCoV-HKU1_orf1ab;6249 | 0 | 0 |
| NC_006577.2;YP_173236.1;HCoV-HKU1_orf1ab;6250 | 0 | 0 |

| | | |
|---|---|---|
| NC_006577.2;YP_173236.1;HCoV-HKU1_orf1ab;6251 | 0 | 0 |
| NC_006577.2;YP_173236.1;HCoV-HKU1_orf1ab;6252 | 0 | 0.025 |
| NC_006577.2;YP_173236.1;HCoV-HKU1_orf1ab;6253 | 0 | 0 |
| NC_006577.2;YP_173236.1;HCoV-HKU1_orf1ab;6254 | 0 | 0 |
| NC_006577.2;YP_173236.1;HCoV-HKU1_orf1ab;6255 | 0 | 0 |
| NC_006577.2;YP_173236.1;HCoV-HKU1_orf1ab;6256 | 0 | 0 |
| NC_006577.2;YP_173236.1;HCoV-HKU1_orf1ab;6257 | 0 | 0 |
| NC_006577.2;YP_173236.1;HCoV-HKU1_orf1ab;6258 | 0 | 0 |
| NC_006577.2;YP_173236.1;HCoV-HKU1_orf1ab;6259 | 0 | 0 |
| NC_006577.2;YP_173236.1;HCoV-HKU1_orf1ab;6260 | 0 | 0 |
| NC_006577.2;YP_173236.1;HCoV-HKU1_orf1ab;6261 | 0 | 0 |
| NC_006577.2;YP_173236.1;HCoV-HKU1_orf1ab;6262 | 0 | 0 |
| NC_006577.2;YP_173236.1;HCoV-HKU1_orf1ab;6263 | 0 | 0 |
| NC_006577.2;YP_173236.1;HCoV-HKU1_orf1ab;6264 | 0 | 0 |
| NC_006577.2;YP_173236.1;HCoV-HKU1_orf1ab;6265 | 0 | 0 |
| NC_006577.2;YP_173236.1;HCoV-HKU1_orf1ab;6266 | 0 | 0 |
| NC_006577.2;YP_173236.1;HCoV-HKU1_orf1ab;6267 | 0 | 0 |
| NC_006577.2;YP_173236.1;HCoV-HKU1_orf1ab;6268 | 0 | 0 |
| NC_006577.2;YP_173236.1;HCoV-HKU1_orf1ab;6269 | 0 | 0 |
| NC_006577.2;YP_173236.1;HCoV-HKU1_orf1ab;6270 | 0 | 0 |
| NC_006577.2;YP_173236.1;HCoV-HKU1_orf1ab;6271 | 0 | 0 |
| NC_006577.2;YP_173236.1;HCoV-HKU1_orf1ab;6272 | 0 | 0 |
| NC_006577.2;YP_173236.1;HCoV-HKU1_orf1ab;6273 | 0 | 0 |
| NC_006577.2;YP_173236.1;HCoV-HKU1_orf1ab;6274 | 0.1 | 0.025 |
| NC_006577.2;YP_173236.1;HCoV-HKU1_orf1ab;6275 | 0.15 | 0.05 |
| NC_006577.2;YP_173236.1;HCoV-HKU1_orf1ab;6276 | 0.05 | 0 |
| NC_006577.2;YP_173236.1;HCoV-HKU1_orf1ab;6277 | 0 | 0.025 |
| NC_006577.2;YP_173236.1;HCoV-HKU1_orf1ab;6278 | 0 | 0.025 |
| NC_006577.2;YP_173236.1;HCoV-HKU1_orf1ab;6279 | 0 | 0.05 |
| NC_006577.2;YP_173236.1;HCoV-HKU1_orf1ab;6280 | 0 | 0.05 |
| NC_006577.2;YP_173236.1;HCoV-HKU1_orf1ab;6281 | 0.05 | 0.05 |
| NC_006577.2;YP_173236.1;HCoV-HKU1_orf1ab;6282 | 0 | 0.05 |
| NC_006577.2;YP_173236.1;HCoV-HKU1_orf1ab;6283 | 0 | 0.025 |
| NC_006577.2;YP_173236.1;HCoV-HKU1_orf1ab;6284 | 0.05 | 0 |
| NC_006577.2;YP_173236.1;HCoV-HKU1_orf1ab;6285 | 0 | 0 |
| NC_006577.2;YP_173236.1;HCoV-HKU1_orf1ab;6286 | 0 | 0 |
| NC_006577.2;YP_173236.1;HCoV-HKU1_orf1ab;6287 | 0 | 0 |
| NC_006577.2;YP_173236.1;HCoV-HKU1_orf1ab;6288 | 0 | 0 |
| NC_006577.2;YP_173236.1;HCoV-HKU1_orf1ab;6289 | 0 | 0 |
| NC_006577.2;YP_173236.1;HCoV-HKU1_orf1ab;6290 | 0 | 0 |
| NC_006577.2;YP_173236.1;HCoV-HKU1_orf1ab;6291 | 0 | 0 |
| NC_006577.2;YP_173236.1;HCoV-HKU1_orf1ab;6292 | 0 | 0 |
| NC_006577.2;YP_173236.1;HCoV-HKU1_orf1ab;6293 | 0 | 0 |

| | | |
|---|---|---|
| NC_006577.2;YP_173236.1;HCoV-HKU1_orf1ab;6294 | 0 | 0 |
| NC_006577.2;YP_173236.1;HCoV-HKU1_orf1ab;6295 | 0 | 0 |
| NC_006577.2;YP_173236.1;HCoV-HKU1_orf1ab;6296 | 0 | 0 |
| NC_006577.2;YP_173236.1;HCoV-HKU1_orf1ab;6297 | 0 | 0 |
| NC_006577.2;YP_173236.1;HCoV-HKU1_orf1ab;6298 | 0 | 0 |
| NC_006577.2;YP_173236.1;HCoV-HKU1_orf1ab;6299 | 0 | 0 |
| NC_006577.2;YP_173236.1;HCoV-HKU1_orf1ab;6300 | 0 | 0 |
| NC_006577.2;YP_173236.1;HCoV-HKU1_orf1ab;6301 | 0 | 0 |
| NC_006577.2;YP_173236.1;HCoV-HKU1_orf1ab;6302 | 0 | 0 |
| NC_006577.2;YP_173236.1;HCoV-HKU1_orf1ab;6303 | 0 | 0 |
| NC_006577.2;YP_173236.1;HCoV-HKU1_orf1ab;6304 | 0 | 0 |
| NC_006577.2;YP_173236.1;HCoV-HKU1_orf1ab;6305 | 0 | 0 |
| NC_006577.2;YP_173236.1;HCoV-HKU1_orf1ab;6306 | 0 | 0 |
| NC_006577.2;YP_173236.1;HCoV-HKU1_orf1ab;6307 | 0 | 0 |
| NC_006577.2;YP_173236.1;HCoV-HKU1_orf1ab;6308 | 0 | 0 |
| NC_006577.2;YP_173236.1;HCoV-HKU1_orf1ab;6309 | 0 | 0 |
| NC_006577.2;YP_173236.1;HCoV-HKU1_orf1ab;6310 | 0 | 0 |
| NC_006577.2;YP_173236.1;HCoV-HKU1_orf1ab;6311 | 0 | 0 |
| NC_006577.2;YP_173236.1;HCoV-HKU1_orf1ab;6312 | 0 | 0 |
| NC_006577.2;YP_173236.1;HCoV-HKU1_orf1ab;6313 | 0 | 0 |
| NC_006577.2;YP_173236.1;HCoV-HKU1_orf1ab;6314 | 0 | 0 |
| NC_006577.2;YP_173236.1;HCoV-HKU1_orf1ab;6315 | 0 | 0 |
| NC_006577.2;YP_173236.1;HCoV-HKU1_orf1ab;6316 | 0 | 0 |
| NC_006577.2;YP_173236.1;HCoV-HKU1_orf1ab;6317 | 0 | 0 |
| NC_006577.2;YP_173236.1;HCoV-HKU1_orf1ab;6318 | 0 | 0 |
| NC_006577.2;YP_173236.1;HCoV-HKU1_orf1ab;6319 | 0 | 0 |
| NC_006577.2;YP_173236.1;HCoV-HKU1_orf1ab;6320 | 0 | 0 |
| NC_006577.2;YP_173236.1;HCoV-HKU1_orf1ab;6321 | 0 | 0 |
| NC_006577.2;YP_173236.1;HCoV-HKU1_orf1ab;6322 | 0.05 | 0 |
| NC_006577.2;YP_173236.1;HCoV-HKU1_orf1ab;6323 | 0 | 0.025 |
| NC_006577.2;YP_173236.1;HCoV-HKU1_orf1ab;6324 | 0 | 0 |
| NC_006577.2;YP_173236.1;HCoV-HKU1_orf1ab;6325 | 0 | 0 |
| NC_006577.2;YP_173236.1;HCoV-HKU1_orf1ab;6326 | 0 | 0 |
| NC_006577.2;YP_173236.1;HCoV-HKU1_orf1ab;6327 | 0.05 | 0.025 |
| NC_006577.2;YP_173236.1;HCoV-HKU1_orf1ab;6328 | 0 | 0 |
| NC_006577.2;YP_173236.1;HCoV-HKU1_orf1ab;6329 | 0 | 0.025 |
| NC_006577.2;YP_173236.1;HCoV-HKU1_orf1ab;6330 | 0 | 0.05 |
| NC_006577.2;YP_173236.1;HCoV-HKU1_orf1ab;6331 | 0 | 0 |
| NC_006577.2;YP_173236.1;HCoV-HKU1_orf1ab;6332 | 0 | 0 |
| NC_006577.2;YP_173236.1;HCoV-HKU1_orf1ab;6333 | 0 | 0 |
| NC_006577.2;YP_173236.1;HCoV-HKU1_orf1ab;6334 | 0 | 0.025 |
| NC_006577.2;YP_173236.1;HCoV-HKU1_orf1ab;6335 | 0 | 0 |
| NC_006577.2;YP_173236.1;HCoV-HKU1_orf1ab;6336 | 0.05 | 0 |

| | | |
|---|---|---|
| NC_006577.2;YP_173236.1;HCoV-HKU1_orf1ab;6337 | 0 | 0 |
| NC_006577.2;YP_173236.1;HCoV-HKU1_orf1ab;6338 | 0 | 0 |
| NC_006577.2;YP_173236.1;HCoV-HKU1_orf1ab;6339 | 0 | 0 |
| NC_006577.2;YP_173236.1;HCoV-HKU1_orf1ab;6340 | 0 | 0 |
| NC_006577.2;YP_173236.1;HCoV-HKU1_orf1ab;6341 | 0 | 0 |
| NC_006577.2;YP_173236.1;HCoV-HKU1_orf1ab;6342 | 0 | 0 |
| NC_006577.2;YP_173236.1;HCoV-HKU1_orf1ab;6343 | 0 | 0 |
| NC_006577.2;YP_173236.1;HCoV-HKU1_orf1ab;6344 | 0 | 0 |
| NC_006577.2;YP_173236.1;HCoV-HKU1_orf1ab;6345 | 0 | 0.025 |
| NC_006577.2;YP_173236.1;HCoV-HKU1_orf1ab;6346 | 0.05 | 0.025 |
| NC_006577.2;YP_173236.1;HCoV-HKU1_orf1ab;6347 | 0.05 | 0.05 |
| NC_006577.2;YP_173236.1;HCoV-HKU1_orf1ab;6348 | 0.1 | 0.05 |
| NC_006577.2;YP_173236.1;HCoV-HKU1_orf1ab;6349 | 0.1 | 0.05 |
| NC_006577.2;YP_173236.1;HCoV-HKU1_orf1ab;6350 | 0.15 | 0.05 |
| NC_006577.2;YP_173236.1;HCoV-HKU1_orf1ab;6351 | 0.1 | 0.05 |
| NC_006577.2;YP_173236.1;HCoV-HKU1_orf1ab;6352 | 0.1 | 0.025 |
| NC_006577.2;YP_173236.1;HCoV-HKU1_orf1ab;6353 | 0.1 | 0.025 |
| NC_006577.2;YP_173236.1;HCoV-HKU1_orf1ab;6354 | 0.05 | 0.025 |
| NC_006577.2;YP_173236.1;HCoV-HKU1_orf1ab;6355 | 0.1 | 0 |
| NC_006577.2;YP_173236.1;HCoV-HKU1_orf1ab;6356 | 0 | 0.025 |
| NC_006577.2;YP_173236.1;HCoV-HKU1_orf1ab;6357 | 0 | 0 |
| NC_006577.2;YP_173236.1;HCoV-HKU1_orf1ab;6358 | 0 | 0 |
| NC_006577.2;YP_173236.1;HCoV-HKU1_orf1ab;6359 | 0.05 | 0 |
| NC_006577.2;YP_173236.1;HCoV-HKU1_orf1ab;6360 | 0.05 | 0 |
| NC_006577.2;YP_173236.1;HCoV-HKU1_orf1ab;6361 | 0 | 0.025 |
| NC_006577.2;YP_173236.1;HCoV-HKU1_orf1ab;6362 | 0 | 0 |
| NC_006577.2;YP_173236.1;HCoV-HKU1_orf1ab;6363 | 0.1 | 0.025 |
| NC_006577.2;YP_173236.1;HCoV-HKU1_orf1ab;6364 | 0 | 0 |
| NC_006577.2;YP_173236.1;HCoV-HKU1_orf1ab;6365 | 0 | 0 |
| NC_006577.2;YP_173236.1;HCoV-HKU1_orf1ab;6366 | 0 | 0 |
| NC_006577.2;YP_173236.1;HCoV-HKU1_orf1ab;6367 | 0 | 0 |
| NC_006577.2;YP_173236.1;HCoV-HKU1_orf1ab;6368 | 0 | 0 |
| NC_006577.2;YP_173236.1;HCoV-HKU1_orf1ab;6369 | 0 | 0 |
| NC_006577.2;YP_173236.1;HCoV-HKU1_orf1ab;6370 | 0 | 0 |
| NC_006577.2;YP_173236.1;HCoV-HKU1_orf1ab;6371 | 0 | 0 |
| NC_006577.2;YP_173236.1;HCoV-HKU1_orf1ab;6372 | 0 | 0 |
| NC_006577.2;YP_173236.1;HCoV-HKU1_orf1ab;6373 | 0 | 0 |
| NC_006577.2;YP_173236.1;HCoV-HKU1_orf1ab;6374 | 0 | 0 |
| NC_006577.2;YP_173236.1;HCoV-HKU1_orf1ab;6375 | 0 | 0 |
| NC_006577.2;YP_173236.1;HCoV-HKU1_orf1ab;6376 | 0 | 0 |
| NC_006577.2;YP_173236.1;HCoV-HKU1_orf1ab;6377 | 0 | 0 |
| NC_006577.2;YP_173236.1;HCoV-HKU1_orf1ab;6378 | 0 | 0 |
| NC_006577.2;YP_173236.1;HCoV-HKU1_orf1ab;6379 | 0 | 0 |

| | | |
|---|---|---|
| NC_006577.2;YP_173236.1;HCoV-HKU1_orf1ab;6380 | 0 | 0 |
| NC_006577.2;YP_173236.1;HCoV-HKU1_orf1ab;6381 | 0 | 0 |
| NC_006577.2;YP_173236.1;HCoV-HKU1_orf1ab;6382 | 0 | 0 |
| NC_006577.2;YP_173236.1;HCoV-HKU1_orf1ab;6383 | 0 | 0.025 |
| NC_006577.2;YP_173236.1;HCoV-HKU1_orf1ab;6384 | 0 | 0.025 |
| NC_006577.2;YP_173236.1;HCoV-HKU1_orf1ab;6385 | 0 | 0 |
| NC_006577.2;YP_173236.1;HCoV-HKU1_orf1ab;6386 | 0 | 0 |
| NC_006577.2;YP_173236.1;HCoV-HKU1_orf1ab;6387 | 0 | 0 |
| NC_006577.2;YP_173236.1;HCoV-HKU1_orf1ab;6388 | 0 | 0.025 |
| NC_006577.2;YP_173236.1;HCoV-HKU1_orf1ab;6389 | 0 | 0.05 |
| NC_006577.2;YP_173236.1;HCoV-HKU1_orf1ab;6390 | 0 | 0 |
| NC_006577.2;YP_173236.1;HCoV-HKU1_orf1ab;6391 | 0 | 0 |
| NC_006577.2;YP_173236.1;HCoV-HKU1_orf1ab;6392 | 0 | 0 |
| NC_006577.2;YP_173236.1;HCoV-HKU1_orf1ab;6393 | 0 | 0 |
| NC_006577.2;YP_173236.1;HCoV-HKU1_orf1ab;6394 | 0 | 0 |
| NC_006577.2;YP_173236.1;HCoV-HKU1_orf1ab;6395 | 0 | 0 |
| NC_006577.2;YP_173236.1;HCoV-HKU1_orf1ab;6396 | 0 | 0 |
| NC_006577.2;YP_173236.1;HCoV-HKU1_orf1ab;6397 | 0 | 0 |
| NC_006577.2;YP_173236.1;HCoV-HKU1_orf1ab;6398 | 0 | 0 |
| NC_006577.2;YP_173236.1;HCoV-HKU1_orf1ab;6399 | 0 | 0 |
| NC_006577.2;YP_173236.1;HCoV-HKU1_orf1ab;6400 | 0 | 0 |
| NC_006577.2;YP_173236.1;HCoV-HKU1_orf1ab;6401 | 0 | 0 |
| NC_006577.2;YP_173236.1;HCoV-HKU1_orf1ab;6402 | 0 | 0 |
| NC_006577.2;YP_173236.1;HCoV-HKU1_orf1ab;6403 | 0 | 0 |
| NC_006577.2;YP_173236.1;HCoV-HKU1_orf1ab;6404 | 0 | 0 |
| NC_006577.2;YP_173236.1;HCoV-HKU1_orf1ab;6405 | 0 | 0 |
| NC_006577.2;YP_173236.1;HCoV-HKU1_orf1ab;6406 | 0 | 0 |
| NC_006577.2;YP_173236.1;HCoV-HKU1_orf1ab;6407 | 0 | 0 |
| NC_006577.2;YP_173236.1;HCoV-HKU1_orf1ab;6408 | 0 | 0 |
| NC_006577.2;YP_173236.1;HCoV-HKU1_orf1ab;6409 | 0 | 0 |
| NC_006577.2;YP_173236.1;HCoV-HKU1_orf1ab;6410 | 0.05 | 0.025 |
| NC_006577.2;YP_173236.1;HCoV-HKU1_orf1ab;6411 | 0 | 0.025 |
| NC_006577.2;YP_173236.1;HCoV-HKU1_orf1ab;6412 | 0.05 | 0 |
| NC_006577.2;YP_173236.1;HCoV-HKU1_orf1ab;6413 | 0 | 0 |
| NC_006577.2;YP_173236.1;HCoV-HKU1_orf1ab;6414 | 0 | 0 |
| NC_006577.2;YP_173236.1;HCoV-HKU1_orf1ab;6415 | 0 | 0.025 |
| NC_006577.2;YP_173236.1;HCoV-HKU1_orf1ab;6416 | 0 | 0 |
| NC_006577.2;YP_173236.1;HCoV-HKU1_orf1ab;6417 | 0 | 0 |
| NC_006577.2;YP_173236.1;HCoV-HKU1_orf1ab;6418 | 0 | 0 |
| NC_006577.2;YP_173236.1;HCoV-HKU1_orf1ab;6419 | 0 | 0 |
| NC_006577.2;YP_173236.1;HCoV-HKU1_orf1ab;6420 | 0 | 0 |
| NC_006577.2;YP_173236.1;HCoV-HKU1_orf1ab;6421 | 0 | 0 |
| NC_006577.2;YP_173236.1;HCoV-HKU1_orf1ab;6422 | 0 | 0 |

| | | |
|---|---|---|
| NC_006577.2;YP_173236.1;HCoV-HKU1_orf1ab;6423 | 0 | 0 |
| NC_006577.2;YP_173236.1;HCoV-HKU1_orf1ab;6424 | 0 | 0 |
| NC_006577.2;YP_173236.1;HCoV-HKU1_orf1ab;6425 | 0 | 0 |
| NC_006577.2;YP_173236.1;HCoV-HKU1_orf1ab;6426 | 0 | 0 |
| NC_006577.2;YP_173236.1;HCoV-HKU1_orf1ab;6427 | 0 | 0 |
| NC_006577.2;YP_173236.1;HCoV-HKU1_orf1ab;6428 | 0.05 | 0 |
| NC_006577.2;YP_173236.1;HCoV-HKU1_orf1ab;6429 | 0.05 | 0 |
| NC_006577.2;YP_173236.1;HCoV-HKU1_orf1ab;6430 | 0.05 | 0 |
| NC_006577.2;YP_173236.1;HCoV-HKU1_orf1ab;6431 | 0 | 0 |
| NC_006577.2;YP_173236.1;HCoV-HKU1_orf1ab;6432 | 0 | 0.025 |
| NC_006577.2;YP_173236.1;HCoV-HKU1_orf1ab;6433 | 0 | 0.025 |
| NC_006577.2;YP_173236.1;HCoV-HKU1_orf1ab;6434 | 0 | 0.05 |
| NC_006577.2;YP_173236.1;HCoV-HKU1_orf1ab;6435 | 0.05 | 0.1 |
| NC_006577.2;YP_173236.1;HCoV-HKU1_orf1ab;6436 | 0.05 | 0.075 |
| NC_006577.2;YP_173236.1;HCoV-HKU1_orf1ab;6437 | 0.05 | 0.025 |
| NC_006577.2;YP_173236.1;HCoV-HKU1_orf1ab;6438 | 0.05 | 0.075 |
| NC_006577.2;YP_173236.1;HCoV-HKU1_orf1ab;6439 | 0.2 | 0.075 |
| NC_006577.2;YP_173236.1;HCoV-HKU1_orf1ab;6440 | 0.1 | 0.075 |
| NC_006577.2;YP_173236.1;HCoV-HKU1_orf1ab;6441 | 0.05 | 0.05 |
| NC_006577.2;YP_173236.1;HCoV-HKU1_orf1ab;6442 | 0 | 0 |
| NC_006577.2;YP_173236.1;HCoV-HKU1_orf1ab;6443 | 0 | 0 |
| NC_006577.2;YP_173236.1;HCoV-HKU1_orf1ab;6444 | 0 | 0 |
| NC_006577.2;YP_173236.1;HCoV-HKU1_orf1ab;6445 | 0 | 0 |
| NC_006577.2;YP_173236.1;HCoV-HKU1_orf1ab;6446 | 0 | 0 |
| NC_006577.2;YP_173236.1;HCoV-HKU1_orf1ab;6447 | 0 | 0 |
| NC_006577.2;YP_173236.1;HCoV-HKU1_orf1ab;6448 | 0 | 0 |
| NC_006577.2;YP_173236.1;HCoV-HKU1_orf1ab;6449 | 0 | 0 |
| NC_006577.2;YP_173236.1;HCoV-HKU1_orf1ab;6450 | 0 | 0 |
| NC_006577.2;YP_173236.1;HCoV-HKU1_orf1ab;6451 | 0 | 0 |
| NC_006577.2;YP_173236.1;HCoV-HKU1_orf1ab;6452 | 0 | 0 |
| NC_006577.2;YP_173236.1;HCoV-HKU1_orf1ab;6453 | 0 | 0 |
| NC_006577.2;YP_173236.1;HCoV-HKU1_orf1ab;6454 | 0 | 0 |
| NC_006577.2;YP_173236.1;HCoV-HKU1_orf1ab;6455 | 0 | 0 |
| NC_006577.2;YP_173236.1;HCoV-HKU1_orf1ab;6456 | 0 | 0 |
| NC_006577.2;YP_173236.1;HCoV-HKU1_orf1ab;6457 | 0 | 0 |
| NC_006577.2;YP_173236.1;HCoV-HKU1_orf1ab;6458 | 0 | 0 |
| NC_006577.2;YP_173236.1;HCoV-HKU1_orf1ab;6459 | 0 | 0 |
| NC_006577.2;YP_173236.1;HCoV-HKU1_orf1ab;6460 | 0 | 0 |
| NC_006577.2;YP_173236.1;HCoV-HKU1_orf1ab;6461 | 0 | 0 |
| NC_006577.2;YP_173236.1;HCoV-HKU1_orf1ab;6462 | 0 | 0 |
| NC_006577.2;YP_173236.1;HCoV-HKU1_orf1ab;6463 | 0 | 0 |
| NC_006577.2;YP_173236.1;HCoV-HKU1_orf1ab;6464 | 0 | 0 |
| NC_006577.2;YP_173236.1;HCoV-HKU1_orf1ab;6465 | 0 | 0.025 |

| | | |
|---|---|---|
| NC_006577.2;YP_173236.1;HCoV-HKU1_orf1ab;6466 | 0 | 0.025 |
| NC_006577.2;YP_173236.1;HCoV-HKU1_orf1ab;6467 | 0 | 0.025 |
| NC_006577.2;YP_173236.1;HCoV-HKU1_orf1ab;6468 | 0 | 0 |
| NC_006577.2;YP_173236.1;HCoV-HKU1_orf1ab;6469 | 0 | 0 |
| NC_006577.2;YP_173236.1;HCoV-HKU1_orf1ab;6470 | 0 | 0.025 |
| NC_006577.2;YP_173236.1;HCoV-HKU1_orf1ab;6471 | 0 | 0.025 |
| NC_006577.2;YP_173236.1;HCoV-HKU1_orf1ab;6472 | 0 | 0 |
| NC_006577.2;YP_173236.1;HCoV-HKU1_orf1ab;6473 | 0 | 0 |
| NC_006577.2;YP_173236.1;HCoV-HKU1_orf1ab;6474 | 0 | 0.025 |
| NC_006577.2;YP_173236.1;HCoV-HKU1_orf1ab;6475 | 0 | 0 |
| NC_006577.2;YP_173236.1;HCoV-HKU1_orf1ab;6476 | 0 | 0 |
| NC_006577.2;YP_173236.1;HCoV-HKU1_orf1ab;6477 | 0 | 0 |
| NC_006577.2;YP_173236.1;HCoV-HKU1_orf1ab;6478 | 0 | 0 |
| NC_006577.2;YP_173236.1;HCoV-HKU1_orf1ab;6479 | 0 | 0 |
| NC_006577.2;YP_173236.1;HCoV-HKU1_orf1ab;6480 | 0 | 0 |
| NC_006577.2;YP_173236.1;HCoV-HKU1_orf1ab;6481 | 0 | 0 |
| NC_006577.2;YP_173236.1;HCoV-HKU1_orf1ab;6482 | 0 | 0 |
| NC_006577.2;YP_173236.1;HCoV-HKU1_orf1ab;6483 | 0 | 0 |
| NC_006577.2;YP_173236.1;HCoV-HKU1_orf1ab;6484 | 0 | 0 |
| NC_006577.2;YP_173236.1;HCoV-HKU1_orf1ab;6485 | 0 | 0 |
| NC_006577.2;YP_173236.1;HCoV-HKU1_orf1ab;6486 | 0 | 0 |
| NC_006577.2;YP_173236.1;HCoV-HKU1_orf1ab;6487 | 0 | 0 |
| NC_006577.2;YP_173236.1;HCoV-HKU1_orf1ab;6488 | 0 | 0 |
| NC_006577.2;YP_173236.1;HCoV-HKU1_orf1ab;6489 | 0 | 0 |
| NC_006577.2;YP_173236.1;HCoV-HKU1_orf1ab;6490 | 0 | 0 |
| NC_006577.2;YP_173236.1;HCoV-HKU1_orf1ab;6491 | 0 | 0 |
| NC_006577.2;YP_173236.1;HCoV-HKU1_orf1ab;6492 | 0 | 0 |
| NC_006577.2;YP_173236.1;HCoV-HKU1_orf1ab;6493 | 0 | 0 |
| NC_006577.2;YP_173236.1;HCoV-HKU1_orf1ab;6494 | 0 | 0 |
| NC_006577.2;YP_173236.1;HCoV-HKU1_orf1ab;6495 | 0 | 0 |
| NC_006577.2;YP_173236.1;HCoV-HKU1_orf1ab;6496 | 0 | 0 |
| NC_006577.2;YP_173236.1;HCoV-HKU1_orf1ab;6497 | 0 | 0 |
| NC_006577.2;YP_173236.1;HCoV-HKU1_orf1ab;6498 | 0 | 0 |
| NC_006577.2;YP_173236.1;HCoV-HKU1_orf1ab;6499 | 0 | 0 |
| NC_006577.2;YP_173236.1;HCoV-HKU1_orf1ab;6500 | 0 | 0 |
| NC_006577.2;YP_173236.1;HCoV-HKU1_orf1ab;6501 | 0.05 | 0 |
| NC_006577.2;YP_173236.1;HCoV-HKU1_orf1ab;6502 | 0.05 | 0 |
| NC_006577.2;YP_173236.1;HCoV-HKU1_orf1ab;6503 | 0.05 | 0 |
| NC_006577.2;YP_173236.1;HCoV-HKU1_orf1ab;6504 | 0 | 0 |
| NC_006577.2;YP_173236.1;HCoV-HKU1_orf1ab;6505 | 0 | 0 |
| NC_006577.2;YP_173236.1;HCoV-HKU1_orf1ab;6506 | 0 | 0.025 |
| NC_006577.2;YP_173236.1;HCoV-HKU1_orf1ab;6507 | 0 | 0.025 |
| NC_006577.2;YP_173236.1;HCoV-HKU1_orf1ab;6508 | 0 | 0 |

| | | |
|---|---|---|
| NC_006577.2;YP_173236.1;HCoV-HKU1_orf1ab;6509 | 0 | 0.025 |
| NC_006577.2;YP_173236.1;HCoV-HKU1_orf1ab;6510 | 0 | 0 |
| NC_006577.2;YP_173236.1;HCoV-HKU1_orf1ab;6511 | 0 | 0 |
| NC_006577.2;YP_173236.1;HCoV-HKU1_orf1ab;6512 | 0 | 0 |
| NC_006577.2;YP_173236.1;HCoV-HKU1_orf1ab;6513 | 0 | 0 |
| NC_006577.2;YP_173236.1;HCoV-HKU1_orf1ab;6514 | 0 | 0 |
| NC_006577.2;YP_173236.1;HCoV-HKU1_orf1ab;6515 | 0 | 0 |
| NC_006577.2;YP_173236.1;HCoV-HKU1_orf1ab;6516 | 0 | 0.025 |
| NC_006577.2;YP_173236.1;HCoV-HKU1_orf1ab;6517 | 0.05 | 0.025 |
| NC_006577.2;YP_173236.1;HCoV-HKU1_orf1ab;6518 | 0 | 0.025 |
| NC_006577.2;YP_173236.1;HCoV-HKU1_orf1ab;6519 | 0 | 0.025 |
| NC_006577.2;YP_173236.1;HCoV-HKU1_orf1ab;6520 | 0 | 0.025 |
| NC_006577.2;YP_173236.1;HCoV-HKU1_orf1ab;6521 | 0 | 0.025 |
| NC_006577.2;YP_173236.1;HCoV-HKU1_orf1ab;6522 | 0 | 0 |
| NC_006577.2;YP_173236.1;HCoV-HKU1_orf1ab;6523 | 0 | 0.025 |
| NC_006577.2;YP_173236.1;HCoV-HKU1_orf1ab;6524 | 0 | 0.025 |
| NC_006577.2;YP_173236.1;HCoV-HKU1_orf1ab;6525 | 0 | 0 |
| NC_006577.2;YP_173236.1;HCoV-HKU1_orf1ab;6526 | 0 | 0.05 |
| NC_006577.2;YP_173236.1;HCoV-HKU1_orf1ab;6527 | 0 | 0 |
| NC_006577.2;YP_173236.1;HCoV-HKU1_orf1ab;6528 | 0 | 0 |
| NC_006577.2;YP_173236.1;HCoV-HKU1_orf1ab;6529 | 0 | 0 |
| NC_006577.2;YP_173236.1;HCoV-HKU1_orf1ab;6530 | 0 | 0 |
| NC_006577.2;YP_173236.1;HCoV-HKU1_orf1ab;6531 | 0 | 0 |
| NC_006577.2;YP_173236.1;HCoV-HKU1_orf1ab;6532 | 0 | 0 |
| NC_006577.2;YP_173236.1;HCoV-HKU1_orf1ab;6533 | 0 | 0 |
| NC_006577.2;YP_173236.1;HCoV-HKU1_orf1ab;6534 | 0 | 0.025 |
| NC_006577.2;YP_173236.1;HCoV-HKU1_orf1ab;6535 | 0 | 0.025 |
| NC_006577.2;YP_173236.1;HCoV-HKU1_orf1ab;6536 | 0 | 0 |
| NC_006577.2;YP_173236.1;HCoV-HKU1_orf1ab;6537 | 0 | 0 |
| NC_006577.2;YP_173236.1;HCoV-HKU1_orf1ab;6538 | 0 | 0 |
| NC_006577.2;YP_173236.1;HCoV-HKU1_orf1ab;6539 | 0 | 0 |
| NC_006577.2;YP_173236.1;HCoV-HKU1_orf1ab;6540 | 0 | 0 |
| NC_006577.2;YP_173236.1;HCoV-HKU1_orf1ab;6541 | 0.05 | 0 |
| NC_006577.2;YP_173236.1;HCoV-HKU1_orf1ab;6542 | 0.05 | 0.05 |
| NC_006577.2;YP_173236.1;HCoV-HKU1_orf1ab;6543 | 0 | 0 |
| NC_006577.2;YP_173236.1;HCoV-HKU1_orf1ab;6544 | 0.05 | 0 |
| NC_006577.2;YP_173236.1;HCoV-HKU1_orf1ab;6545 | 0.05 | 0.05 |
| NC_006577.2;YP_173236.1;HCoV-HKU1_orf1ab;6546 | 0.05 | 0 |
| NC_006577.2;YP_173236.1;HCoV-HKU1_orf1ab;6547 | 0.2 | 0.1 |
| NC_006577.2;YP_173236.1;HCoV-HKU1_orf1ab;6548 | 0.1 | 0 |
| NC_006577.2;YP_173236.1;HCoV-HKU1_orf1ab;6549 | 0 | 0 |
| NC_006577.2;YP_173236.1;HCoV-HKU1_orf1ab;6550 | 0.05 | 0.05 |
| NC_006577.2;YP_173236.1;HCoV-HKU1_orf1ab;6551 | 0.05 | 0.1 |

| | | |
|---|---|---|
| NC_006577.2;YP_173236.1;HCoV-HKU1_orf1ab;6552 | 0 | 0 |
| NC_006577.2;YP_173236.1;HCoV-HKU1_orf1ab;6553 | 0 | 0.025 |
| NC_006577.2;YP_173236.1;HCoV-HKU1_orf1ab;6554 | 0 | 0 |
| NC_006577.2;YP_173236.1;HCoV-HKU1_orf1ab;6555 | 0 | 0 |
| NC_006577.2;YP_173236.1;HCoV-HKU1_orf1ab;6556 | 0 | 0 |
| NC_006577.2;YP_173236.1;HCoV-HKU1_orf1ab;6557 | 0 | 0 |
| NC_006577.2;YP_173236.1;HCoV-HKU1_orf1ab;6558 | 0 | 0 |
| NC_006577.2;YP_173236.1;HCoV-HKU1_orf1ab;6559 | 0 | 0 |
| NC_006577.2;YP_173236.1;HCoV-HKU1_orf1ab;6560 | 0 | 0 |
| NC_006577.2;YP_173236.1;HCoV-HKU1_orf1ab;6561 | 0 | 0 |
| NC_006577.2;YP_173236.1;HCoV-HKU1_orf1ab;6562 | 0 | 0 |
| NC_006577.2;YP_173236.1;HCoV-HKU1_orf1ab;6563 | 0 | 0 |
| NC_006577.2;YP_173236.1;HCoV-HKU1_orf1ab;6564 | 0 | 0 |
| NC_006577.2;YP_173236.1;HCoV-HKU1_orf1ab;6565 | 0 | 0 |
| NC_006577.2;YP_173236.1;HCoV-HKU1_orf1ab;6566 | 0 | 0 |
| NC_006577.2;YP_173236.1;HCoV-HKU1_orf1ab;6567 | 0 | 0 |
| NC_006577.2;YP_173236.1;HCoV-HKU1_orf1ab;6568 | 0 | 0 |
| NC_006577.2;YP_173236.1;HCoV-HKU1_orf1ab;6569 | 0 | 0.025 |
| NC_006577.2;YP_173236.1;HCoV-HKU1_orf1ab;6570 | 0.05 | 0.025 |
| NC_006577.2;YP_173236.1;HCoV-HKU1_orf1ab;6571 | 0 | 0.025 |
| NC_006577.2;YP_173236.1;HCoV-HKU1_orf1ab;6572 | 0.05 | 0 |
| NC_006577.2;YP_173236.1;HCoV-HKU1_orf1ab;6573 | 0 | 0 |
| NC_006577.2;YP_173236.1;HCoV-HKU1_orf1ab;6574 | 0 | 0 |
| NC_006577.2;YP_173236.1;HCoV-HKU1_orf1ab;6575 | 0 | 0 |
| NC_006577.2;YP_173236.1;HCoV-HKU1_orf1ab;6576 | 0 | 0 |
| NC_006577.2;YP_173236.1;HCoV-HKU1_orf1ab;6577 | 0 | 0 |
| NC_006577.2;YP_173236.1;HCoV-HKU1_orf1ab;6578 | 0 | 0 |
| NC_006577.2;YP_173236.1;HCoV-HKU1_orf1ab;6579 | 0.05 | 0 |
| NC_006577.2;YP_173236.1;HCoV-HKU1_orf1ab;6580 | 0 | 0 |
| NC_006577.2;YP_173236.1;HCoV-HKU1_orf1ab;6581 | 0 | 0 |
| NC_006577.2;YP_173236.1;HCoV-HKU1_orf1ab;6582 | 0 | 0 |
| NC_006577.2;YP_173236.1;HCoV-HKU1_orf1ab;6583 | 0 | 0 |
| NC_006577.2;YP_173236.1;HCoV-HKU1_orf1ab;6584 | 0 | 0 |
| NC_006577.2;YP_173236.1;HCoV-HKU1_orf1ab;6585 | 0 | 0 |
| NC_006577.2;YP_173236.1;HCoV-HKU1_orf1ab;6586 | 0 | 0 |
| NC_006577.2;YP_173236.1;HCoV-HKU1_orf1ab;6587 | 0 | 0 |
| NC_006577.2;YP_173236.1;HCoV-HKU1_orf1ab;6588 | 0 | 0 |
| NC_006577.2;YP_173236.1;HCoV-HKU1_orf1ab;6589 | 0 | 0 |
| NC_006577.2;YP_173236.1;HCoV-HKU1_orf1ab;6590 | 0 | 0 |
| NC_006577.2;YP_173236.1;HCoV-HKU1_orf1ab;6591 | 0 | 0 |
| NC_006577.2;YP_173236.1;HCoV-HKU1_orf1ab;6592 | 0 | 0 |
| NC_006577.2;YP_173236.1;HCoV-HKU1_orf1ab;6593 | 0 | 0 |
| NC_006577.2;YP_173236.1;HCoV-HKU1_orf1ab;6594 | 0 | 0 |

| | | |
|---|---|---|
| NC_006577.2;YP_173236.1;HCoV-HKU1_orf1ab;6595 | 0 | 0 |
| NC_006577.2;YP_173236.1;HCoV-HKU1_orf1ab;6596 | 0 | 0 |
| NC_006577.2;YP_173236.1;HCoV-HKU1_orf1ab;6597 | 0 | 0 |
| NC_006577.2;YP_173236.1;HCoV-HKU1_orf1ab;6598 | 0 | 0 |
| NC_006577.2;YP_173236.1;HCoV-HKU1_orf1ab;6599 | 0 | 0 |
| NC_006577.2;YP_173236.1;HCoV-HKU1_orf1ab;6600 | 0 | 0 |
| NC_006577.2;YP_173236.1;HCoV-HKU1_orf1ab;6601 | 0 | 0 |
| NC_006577.2;YP_173236.1;HCoV-HKU1_orf1ab;6602 | 0 | 0 |
| NC_006577.2;YP_173236.1;HCoV-HKU1_orf1ab;6603 | 0 | 0 |
| NC_006577.2;YP_173236.1;HCoV-HKU1_orf1ab;6604 | 0 | 0 |
| NC_006577.2;YP_173236.1;HCoV-HKU1_orf1ab;6605 | 0.05 | 0 |
| NC_006577.2;YP_173236.1;HCoV-HKU1_orf1ab;6606 | 0.05 | 0.025 |
| NC_006577.2;YP_173236.1;HCoV-HKU1_orf1ab;6607 | 0.05 | 0 |
| NC_006577.2;YP_173236.1;HCoV-HKU1_orf1ab;6608 | 0.15 | 0.025 |
| NC_006577.2;YP_173236.1;HCoV-HKU1_orf1ab;6609 | 0 | 0 |
| NC_006577.2;YP_173236.1;HCoV-HKU1_orf1ab;6610 | 0 | 0 |
| NC_006577.2;YP_173236.1;HCoV-HKU1_orf1ab;6611 | 0 | 0 |
| NC_006577.2;YP_173236.1;HCoV-HKU1_orf1ab;6612 | 0 | 0 |
| NC_006577.2;YP_173236.1;HCoV-HKU1_orf1ab;6613 | 0 | 0 |
| NC_006577.2;YP_173236.1;HCoV-HKU1_orf1ab;6614 | 0 | 0 |
| NC_006577.2;YP_173236.1;HCoV-HKU1_orf1ab;6615 | 0 | 0 |
| NC_006577.2;YP_173236.1;HCoV-HKU1_orf1ab;6616 | 0 | 0 |
| NC_006577.2;YP_173236.1;HCoV-HKU1_orf1ab;6617 | 0 | 0 |
| NC_006577.2;YP_173236.1;HCoV-HKU1_orf1ab;6618 | 0 | 0 |
| NC_006577.2;YP_173236.1;HCoV-HKU1_orf1ab;6619 | 0 | 0 |
| NC_006577.2;YP_173236.1;HCoV-HKU1_orf1ab;6620 | 0 | 0 |
| NC_006577.2;YP_173236.1;HCoV-HKU1_orf1ab;6621 | 0 | 0 |
| NC_006577.2;YP_173236.1;HCoV-HKU1_orf1ab;6622 | 0 | 0.025 |
| NC_006577.2;YP_173236.1;HCoV-HKU1_orf1ab;6623 | 0 | 0 |
| NC_006577.2;YP_173236.1;HCoV-HKU1_orf1ab;6624 | 0 | 0 |
| NC_006577.2;YP_173236.1;HCoV-HKU1_orf1ab;6625 | 0.05 | 0 |
| NC_006577.2;YP_173236.1;HCoV-HKU1_orf1ab;6626 | 0 | 0 |
| NC_006577.2;YP_173236.1;HCoV-HKU1_orf1ab;6627 | 0 | 0.025 |
| NC_006577.2;YP_173236.1;HCoV-HKU1_orf1ab;6628 | 0 | 0.05 |
| NC_006577.2;YP_173236.1;HCoV-HKU1_orf1ab;6629 | 0 | 0 |
| NC_006577.2;YP_173236.1;HCoV-HKU1_orf1ab;6630 | 0 | 0 |
| NC_006577.2;YP_173236.1;HCoV-HKU1_orf1ab;6631 | 0 | 0 |
| NC_006577.2;YP_173236.1;HCoV-HKU1_orf1ab;6632 | 0 | 0 |
| NC_006577.2;YP_173236.1;HCoV-HKU1_orf1ab;6633 | 0 | 0 |
| NC_006577.2;YP_173236.1;HCoV-HKU1_orf1ab;6634 | 0 | 0 |
| NC_006577.2;YP_173236.1;HCoV-HKU1_orf1ab;6635 | 0 | 0.05 |
| NC_006577.2;YP_173236.1;HCoV-HKU1_orf1ab;6636 | 0 | 0.025 |
| NC_006577.2;YP_173236.1;HCoV-HKU1_orf1ab;6637 | 0 | 0.025 |

| | | |
|---|---|---|
| NC_006577.2;YP_173236.1;HCoV-HKU1_orf1ab;6638 | 0 | 0.025 |
| NC_006577.2;YP_173236.1;HCoV-HKU1_orf1ab;6639 | 0 | 0.025 |
| NC_006577.2;YP_173236.1;HCoV-HKU1_orf1ab;6640 | 0 | 0 |
| NC_006577.2;YP_173236.1;HCoV-HKU1_orf1ab;6641 | 0 | 0 |
| NC_006577.2;YP_173236.1;HCoV-HKU1_orf1ab;6642 | 0 | 0 |
| NC_006577.2;YP_173236.1;HCoV-HKU1_orf1ab;6643 | 0 | 0 |
| NC_006577.2;YP_173236.1;HCoV-HKU1_orf1ab;6644 | 0 | 0 |
| NC_006577.2;YP_173236.1;HCoV-HKU1_orf1ab;6645 | 0 | 0 |
| NC_006577.2;YP_173236.1;HCoV-HKU1_orf1ab;6646 | 0 | 0.025 |
| NC_006577.2;YP_173236.1;HCoV-HKU1_orf1ab;6647 | 0 | 0.025 |
| NC_006577.2;YP_173236.1;HCoV-HKU1_orf1ab;6648 | 0 | 0.025 |
| NC_006577.2;YP_173236.1;HCoV-HKU1_orf1ab;6649 | 0 | 0.025 |
| NC_006577.2;YP_173236.1;HCoV-HKU1_orf1ab;6650 | 0 | 0.025 |
| NC_006577.2;YP_173236.1;HCoV-HKU1_orf1ab;6651 | 0 | 0 |
| NC_006577.2;YP_173236.1;HCoV-HKU1_orf1ab;6652 | 0 | 0 |
| NC_006577.2;YP_173236.1;HCoV-HKU1_orf1ab;6653 | 0 | 0 |
| NC_006577.2;YP_173236.1;HCoV-HKU1_orf1ab;6654 | 0 | 0.025 |
| NC_006577.2;YP_173236.1;HCoV-HKU1_orf1ab;6655 | 0 | 0.025 |
| NC_006577.2;YP_173236.1;HCoV-HKU1_orf1ab;6656 | 0 | 0.05 |
| NC_006577.2;YP_173236.1;HCoV-HKU1_orf1ab;6657 | 0 | 0.075 |
| NC_006577.2;YP_173236.1;HCoV-HKU1_orf1ab;6658 | 0 | 0.05 |
| NC_006577.2;YP_173236.1;HCoV-HKU1_orf1ab;6659 | 0.05 | 0.1 |
| NC_006577.2;YP_173236.1;HCoV-HKU1_orf1ab;6660 | 0.05 | 0.15 |
| NC_006577.2;YP_173236.1;HCoV-HKU1_orf1ab;6661 | 0 | 0.125 |
| NC_006577.2;YP_173236.1;HCoV-HKU1_orf1ab;6662 | 0.05 | 0.15 |
| NC_006577.2;YP_173236.1;HCoV-HKU1_orf1ab;6663 | 0 | 0.125 |
| NC_006577.2;YP_173236.1;HCoV-HKU1_orf1ab;6664 | 0.05 | 0.125 |
| NC_006577.2;YP_173236.1;HCoV-HKU1_orf1ab;6665 | 0 | 0.05 |
| NC_006577.2;YP_173236.1;HCoV-HKU1_orf1ab;6666 | 0 | 0.025 |
| NC_006577.2;YP_173236.1;HCoV-HKU1_orf1ab;6667 | 0 | 0 |
| NC_006577.2;YP_173236.1;HCoV-HKU1_orf1ab;6668 | 0 | 0 |
| NC_006577.2;YP_173236.1;HCoV-HKU1_orf1ab;6669 | 0 | 0 |
| NC_006577.2;YP_173236.1;HCoV-HKU1_orf1ab;6670 | 0 | 0 |
| NC_006577.2;YP_173236.1;HCoV-HKU1_orf1ab;6671 | 0 | 0.025 |
| NC_006577.2;YP_173236.1;HCoV-HKU1_orf1ab;6672 | 0 | 0.05 |
| NC_006577.2;YP_173236.1;HCoV-HKU1_orf1ab;6673 | 0 | 0 |
| NC_006577.2;YP_173236.1;HCoV-HKU1_orf1ab;6674 | 0 | 0.025 |
| NC_006577.2;YP_173236.1;HCoV-HKU1_orf1ab;6675 | 0 | 0 |
| NC_006577.2;YP_173236.1;HCoV-HKU1_orf1ab;6676 | 0 | 0.025 |
| NC_006577.2;YP_173236.1;HCoV-HKU1_orf1ab;6677 | 0 | 0.025 |
| NC_006577.2;YP_173236.1;HCoV-HKU1_orf1ab;6678 | 0 | 0.05 |
| NC_006577.2;YP_173236.1;HCoV-HKU1_orf1ab;6679 | 0 | 0.05 |
| NC_006577.2;YP_173236.1;HCoV-HKU1_orf1ab;6680 | 0 | 0.025 |

| | | |
|---|---|---|
| NC_006577.2;YP_173236.1;HCoV-HKU1_orf1ab;6681 | 0 | 0.025 |
| NC_006577.2;YP_173236.1;HCoV-HKU1_orf1ab;6682 | 0 | 0 |
| NC_006577.2;YP_173236.1;HCoV-HKU1_orf1ab;6683 | 0 | 0 |
| NC_006577.2;YP_173236.1;HCoV-HKU1_orf1ab;6684 | 0 | 0 |
| NC_006577.2;YP_173236.1;HCoV-HKU1_orf1ab;6685 | 0 | 0 |
| NC_006577.2;YP_173236.1;HCoV-HKU1_orf1ab;6686 | 0 | 0 |
| NC_006577.2;YP_173236.1;HCoV-HKU1_orf1ab;6687 | 0 | 0 |
| NC_006577.2;YP_173236.1;HCoV-HKU1_orf1ab;6688 | 0 | 0 |
| NC_006577.2;YP_173236.1;HCoV-HKU1_orf1ab;6689 | 0 | 0 |
| NC_006577.2;YP_173236.1;HCoV-HKU1_orf1ab;6690 | 0 | 0 |
| NC_006577.2;YP_173236.1;HCoV-HKU1_orf1ab;6691 | 0 | 0 |
| NC_006577.2;YP_173236.1;HCoV-HKU1_orf1ab;6692 | 0 | 0 |
| NC_006577.2;YP_173236.1;HCoV-HKU1_orf1ab;6693 | 0 | 0 |
| NC_006577.2;YP_173236.1;HCoV-HKU1_orf1ab;6694 | 0.05 | 0 |
| NC_006577.2;YP_173236.1;HCoV-HKU1_orf1ab;6695 | 0 | 0.025 |
| NC_006577.2;YP_173236.1;HCoV-HKU1_orf1ab;6696 | 0 | 0.025 |
| NC_006577.2;YP_173236.1;HCoV-HKU1_orf1ab;6697 | 0.05 | 0.025 |
| NC_006577.2;YP_173236.1;HCoV-HKU1_orf1ab;6698 | 0.1 | 0.025 |
| NC_006577.2;YP_173236.1;HCoV-HKU1_orf1ab;6699 | 0.05 | 0.05 |
| NC_006577.2;YP_173236.1;HCoV-HKU1_orf1ab;6700 | 0 | 0.025 |
| NC_006577.2;YP_173236.1;HCoV-HKU1_orf1ab;6701 | 0.1 | 0.025 |
| NC_006577.2;YP_173236.1;HCoV-HKU1_orf1ab;6702 | 0.05 | 0.025 |
| NC_006577.2;YP_173236.1;HCoV-HKU1_orf1ab;6703 | 0.1 | 0.025 |
| NC_006577.2;YP_173236.1;HCoV-HKU1_orf1ab;6704 | 0.1 | 0.025 |
| NC_006577.2;YP_173236.1;HCoV-HKU1_orf1ab;6705 | 0.15 | 0.025 |
| NC_006577.2;YP_173236.1;HCoV-HKU1_orf1ab;6706 | 0 | 0.075 |
| NC_006577.2;YP_173236.1;HCoV-HKU1_orf1ab;6707 | 0.05 | 0.025 |
| NC_006577.2;YP_173236.1;HCoV-HKU1_orf1ab;6708 | 0.1 | 0.025 |
| NC_006577.2;YP_173236.1;HCoV-HKU1_orf1ab;6709 | 0.05 | 0 |
| NC_006577.2;YP_173236.1;HCoV-HKU1_orf1ab;6710 | 0.05 | 0 |
| NC_006577.2;YP_173236.1;HCoV-HKU1_orf1ab;6711 | 0.1 | 0 |
| NC_006577.2;YP_173236.1;HCoV-HKU1_orf1ab;6712 | 0.15 | 0.025 |
| NC_006577.2;YP_173236.1;HCoV-HKU1_orf1ab;6713 | 0.1 | 0.05 |
| NC_006577.2;YP_173236.1;HCoV-HKU1_orf1ab;6714 | 0.1 | 0.025 |
| NC_006577.2;YP_173236.1;HCoV-HKU1_orf1ab;6715 | 0.05 | 0.025 |
| NC_006577.2;YP_173236.1;HCoV-HKU1_orf1ab;6716 | 0 | 0.025 |
| NC_006577.2;YP_173236.1;HCoV-HKU1_orf1ab;6717 | 0 | 0 |
| NC_006577.2;YP_173236.1;HCoV-HKU1_orf1ab;6718 | 0 | 0 |
| NC_006577.2;YP_173236.1;HCoV-HKU1_orf1ab;6719 | 0 | 0 |
| NC_006577.2;YP_173236.1;HCoV-HKU1_orf1ab;6720 | 0 | 0 |
| NC_006577.2;YP_173236.1;HCoV-HKU1_orf1ab;6721 | 0 | 0 |
| NC_006577.2;YP_173236.1;HCoV-HKU1_orf1ab;6722 | 0 | 0 |
| NC_006577.2;YP_173236.1;HCoV-HKU1_orf1ab;6723 | 0 | 0 |

| | | |
|---|---|---|
| NC_006577.2;YP_173236.1;HCoV-HKU1_orf1ab;6724 | 0 | 0 |
| NC_006577.2;YP_173236.1;HCoV-HKU1_orf1ab;6725 | 0 | 0 |
| NC_006577.2;YP_173236.1;HCoV-HKU1_orf1ab;6726 | 0 | 0 |
| NC_006577.2;YP_173236.1;HCoV-HKU1_orf1ab;6727 | 0 | 0 |
| NC_006577.2;YP_173236.1;HCoV-HKU1_orf1ab;6728 | 0 | 0 |
| NC_006577.2;YP_173236.1;HCoV-HKU1_orf1ab;6729 | 0 | 0 |
| NC_006577.2;YP_173236.1;HCoV-HKU1_orf1ab;6730 | 0 | 0 |
| NC_006577.2;YP_173236.1;HCoV-HKU1_orf1ab;6731 | 0 | 0 |
| NC_006577.2;YP_173236.1;HCoV-HKU1_orf1ab;6732 | 0 | 0 |
| NC_006577.2;YP_173236.1;HCoV-HKU1_orf1ab;6733 | 0 | 0 |
| NC_006577.2;YP_173236.1;HCoV-HKU1_orf1ab;6734 | 0 | 0.025 |
| NC_006577.2;YP_173236.1;HCoV-HKU1_orf1ab;6735 | 0.05 | 0 |
| NC_006577.2;YP_173236.1;HCoV-HKU1_orf1ab;6736 | 0 | 0 |
| NC_006577.2;YP_173236.1;HCoV-HKU1_orf1ab;6737 | 0 | 0 |
| NC_006577.2;YP_173236.1;HCoV-HKU1_orf1ab;6738 | 0 | 0 |
| NC_006577.2;YP_173236.1;HCoV-HKU1_orf1ab;6739 | 0 | 0 |
| NC_006577.2;YP_173236.1;HCoV-HKU1_orf1ab;6740 | 0 | 0 |
| NC_006577.2;YP_173236.1;HCoV-HKU1_orf1ab;6741 | 0 | 0 |
| NC_006577.2;YP_173236.1;HCoV-HKU1_orf1ab;6742 | 0 | 0 |
| NC_006577.2;YP_173236.1;HCoV-HKU1_orf1ab;6743 | 0 | 0 |
| NC_006577.2;YP_173236.1;HCoV-HKU1_orf1ab;6744 | 0 | 0 |
| NC_006577.2;YP_173236.1;HCoV-HKU1_orf1ab;6745 | 0.05 | 0.05 |
| NC_006577.2;YP_173236.1;HCoV-HKU1_orf1ab;6746 | 0 | 0.05 |
| NC_006577.2;YP_173236.1;HCoV-HKU1_orf1ab;6747 | 0.05 | 0.025 |
| NC_006577.2;YP_173236.1;HCoV-HKU1_orf1ab;6748 | 0 | 0 |
| NC_006577.2;YP_173236.1;HCoV-HKU1_orf1ab;6749 | 0 | 0 |
| NC_006577.2;YP_173236.1;HCoV-HKU1_orf1ab;6750 | 0 | 0 |
| NC_006577.2;YP_173236.1;HCoV-HKU1_orf1ab;6751 | 0 | 0 |
| NC_006577.2;YP_173236.1;HCoV-HKU1_orf1ab;6752 | 0.05 | 0 |
| NC_006577.2;YP_173236.1;HCoV-HKU1_orf1ab;6753 | 0 | 0 |
| NC_006577.2;YP_173236.1;HCoV-HKU1_orf1ab;6754 | 0 | 0 |
| NC_006577.2;YP_173236.1;HCoV-HKU1_orf1ab;6755 | 0 | 0 |
| NC_006577.2;YP_173236.1;HCoV-HKU1_orf1ab;6756 | 0 | 0 |
| NC_006577.2;YP_173236.1;HCoV-HKU1_orf1ab;6757 | 0 | 0 |
| NC_006577.2;YP_173236.1;HCoV-HKU1_orf1ab;6758 | 0 | 0 |
| NC_006577.2;YP_173236.1;HCoV-HKU1_orf1ab;6759 | 0 | 0 |
| NC_006577.2;YP_173236.1;HCoV-HKU1_orf1ab;6760 | 0 | 0 |
| NC_006577.2;YP_173236.1;HCoV-HKU1_orf1ab;6761 | 0 | 0 |
| NC_006577.2;YP_173236.1;HCoV-HKU1_orf1ab;6762 | 0 | 0 |
| NC_006577.2;YP_173236.1;HCoV-HKU1_orf1ab;6763 | 0.1 | 0.15 |
| NC_006577.2;YP_173236.1;HCoV-HKU1_orf1ab;6764 | 0 | 0 |
| NC_006577.2;YP_173236.1;HCoV-HKU1_orf1ab;6765 | 0 | 0.025 |
| NC_006577.2;YP_173236.1;HCoV-HKU1_orf1ab;6766 | 0 | 0 |

| | | |
|---|---|---|
| NC_006577.2;YP_173236.1;HCoV-HKU1_orf1ab;6767 | 0 | 0 |
| NC_006577.2;YP_173236.1;HCoV-HKU1_orf1ab;6768 | 0 | 0 |
| NC_006577.2;YP_173236.1;HCoV-HKU1_orf1ab;6769 | 0 | 0 |
| NC_006577.2;YP_173236.1;HCoV-HKU1_orf1ab;6770 | 0 | 0 |
| NC_006577.2;YP_173236.1;HCoV-HKU1_orf1ab;6771 | 0 | 0 |
| NC_006577.2;YP_173236.1;HCoV-HKU1_orf1ab;6772 | 0 | 0 |
| NC_006577.2;YP_173236.1;HCoV-HKU1_orf1ab;6773 | 0 | 0 |
| NC_006577.2;YP_173236.1;HCoV-HKU1_orf1ab;6774 | 0 | 0 |
| NC_006577.2;YP_173236.1;HCoV-HKU1_orf1ab;6775 | 0 | 0 |
| NC_006577.2;YP_173236.1;HCoV-HKU1_orf1ab;6776 | 0 | 0 |
| NC_006577.2;YP_173236.1;HCoV-HKU1_orf1ab;6777 | 0 | 0 |
| NC_006577.2;YP_173236.1;HCoV-HKU1_orf1ab;6778 | 0 | 0 |
| NC_006577.2;YP_173236.1;HCoV-HKU1_orf1ab;6779 | 0 | 0 |
| NC_006577.2;YP_173236.1;HCoV-HKU1_orf1ab;6780 | 0 | 0 |
| NC_006577.2;YP_173236.1;HCoV-HKU1_orf1ab;6781 | 0 | 0 |
| NC_006577.2;YP_173236.1;HCoV-HKU1_orf1ab;6782 | 0 | 0 |
| NC_006577.2;YP_173236.1;HCoV-HKU1_orf1ab;6783 | 0 | 0 |
| NC_006577.2;YP_173236.1;HCoV-HKU1_orf1ab;6784 | 0 | 0 |
| NC_006577.2;YP_173236.1;HCoV-HKU1_orf1ab;6785 | 0 | 0 |
| NC_006577.2;YP_173236.1;HCoV-HKU1_orf1ab;6786 | 0 | 0 |
| NC_006577.2;YP_173236.1;HCoV-HKU1_orf1ab;6787 | 0 | 0 |
| NC_006577.2;YP_173236.1;HCoV-HKU1_orf1ab;6788 | 0 | 0 |
| NC_006577.2;YP_173236.1;HCoV-HKU1_orf1ab;6789 | 0 | 0 |
| NC_006577.2;YP_173236.1;HCoV-HKU1_orf1ab;6790 | 0 | 0 |
| NC_006577.2;YP_173236.1;HCoV-HKU1_orf1ab;6791 | 0 | 0 |
| NC_006577.2;YP_173236.1;HCoV-HKU1_orf1ab;6792 | 0 | 0 |
| NC_006577.2;YP_173236.1;HCoV-HKU1_orf1ab;6793 | 0 | 0.1 |
| NC_006577.2;YP_173236.1;HCoV-HKU1_orf1ab;6794 | 0 | 0.025 |
| NC_006577.2;YP_173236.1;HCoV-HKU1_orf1ab;6795 | 0 | 0 |
| NC_006577.2;YP_173236.1;HCoV-HKU1_orf1ab;6796 | 0.05 | 0.05 |
| NC_006577.2;YP_173236.1;HCoV-HKU1_orf1ab;6797 | 0.05 | 0.025 |
| NC_006577.2;YP_173236.1;HCoV-HKU1_orf1ab;6798 | 0.05 | 0 |
| NC_006577.2;YP_173236.1;HCoV-HKU1_orf1ab;6799 | 0 | 0.025 |
| NC_006577.2;YP_173236.1;HCoV-HKU1_orf1ab;6800 | 0 | 0.025 |
| NC_006577.2;YP_173236.1;HCoV-HKU1_orf1ab;6801 | 0 | 0 |
| NC_006577.2;YP_173236.1;HCoV-HKU1_orf1ab;6802 | 0 | 0 |
| NC_006577.2;YP_173236.1;HCoV-HKU1_orf1ab;6803 | 0 | 0 |
| NC_006577.2;YP_173236.1;HCoV-HKU1_orf1ab;6804 | 0 | 0 |
| NC_006577.2;YP_173236.1;HCoV-HKU1_orf1ab;6805 | 0 | 0 |
| NC_006577.2;YP_173236.1;HCoV-HKU1_orf1ab;6806 | 0 | 0.025 |
| NC_006577.2;YP_173236.1;HCoV-HKU1_orf1ab;6807 | 0 | 0 |
| NC_006577.2;YP_173236.1;HCoV-HKU1_orf1ab;6808 | 0 | 0 |
| NC_006577.2;YP_173236.1;HCoV-HKU1_orf1ab;6809 | 0 | 0 |

| | | |
|---|---|---|
| NC_006577.2;YP_173236.1;HCoV-HKU1_orf1ab;6810 | 0 | 0 |
| NC_006577.2;YP_173236.1;HCoV-HKU1_orf1ab;6811 | 0 | 0 |
| NC_006577.2;YP_173236.1;HCoV-HKU1_orf1ab;6812 | 0 | 0 |
| NC_006577.2;YP_173236.1;HCoV-HKU1_orf1ab;6813 | 0.05 | 0 |
| NC_006577.2;YP_173236.1;HCoV-HKU1_orf1ab;6814 | 0.05 | 0 |
| NC_006577.2;YP_173236.1;HCoV-HKU1_orf1ab;6815 | 0.05 | 0 |
| NC_006577.2;YP_173236.1;HCoV-HKU1_orf1ab;6816 | 0 | 0 |
| NC_006577.2;YP_173236.1;HCoV-HKU1_orf1ab;6817 | 0 | 0 |
| NC_006577.2;YP_173236.1;HCoV-HKU1_orf1ab;6818 | 0 | 0 |
| NC_006577.2;YP_173236.1;HCoV-HKU1_orf1ab;6819 | 0 | 0 |
| NC_006577.2;YP_173236.1;HCoV-HKU1_orf1ab;6820 | 0 | 0 |
| NC_006577.2;YP_173236.1;HCoV-HKU1_orf1ab;6821 | 0 | 0 |
| NC_006577.2;YP_173236.1;HCoV-HKU1_orf1ab;6822 | 0 | 0 |
| NC_006577.2;YP_173236.1;HCoV-HKU1_orf1ab;6823 | 0 | 0 |
| NC_006577.2;YP_173236.1;HCoV-HKU1_orf1ab;6824 | 0 | 0 |
| NC_006577.2;YP_173236.1;HCoV-HKU1_orf1ab;6825 | 0 | 0 |
| NC_006577.2;YP_173236.1;HCoV-HKU1_orf1ab;6826 | 0 | 0 |
| NC_006577.2;YP_173236.1;HCoV-HKU1_orf1ab;6827 | 0 | 0 |
| NC_006577.2;YP_173236.1;HCoV-HKU1_orf1ab;6828 | 0 | 0 |
| NC_006577.2;YP_173236.1;HCoV-HKU1_orf1ab;6829 | 0 | 0 |
| NC_006577.2;YP_173236.1;HCoV-HKU1_orf1ab;6830 | 0 | 0 |
| NC_006577.2;YP_173236.1;HCoV-HKU1_orf1ab;6831 | 0 | 0 |
| NC_006577.2;YP_173236.1;HCoV-HKU1_orf1ab;6832 | 0 | 0 |
| NC_006577.2;YP_173236.1;HCoV-HKU1_orf1ab;6833 | 0 | 0 |
| NC_006577.2;YP_173236.1;HCoV-HKU1_orf1ab;6834 | 0 | 0 |
| NC_006577.2;YP_173236.1;HCoV-HKU1_orf1ab;6835 | 0 | 0 |
| NC_006577.2;YP_173236.1;HCoV-HKU1_orf1ab;6836 | 0 | 0 |
| NC_006577.2;YP_173236.1;HCoV-HKU1_orf1ab;6837 | 0 | 0 |
| NC_006577.2;YP_173236.1;HCoV-HKU1_orf1ab;6838 | 0 | 0 |
| NC_006577.2;YP_173236.1;HCoV-HKU1_orf1ab;6839 | 0 | 0 |
| NC_006577.2;YP_173236.1;HCoV-HKU1_orf1ab;6840 | 0 | 0 |
| NC_006577.2;YP_173236.1;HCoV-HKU1_orf1ab;6841 | 0 | 0 |
| NC_006577.2;YP_173236.1;HCoV-HKU1_orf1ab;6842 | 0 | 0 |
| NC_006577.2;YP_173236.1;HCoV-HKU1_orf1ab;6843 | 0 | 0 |
| NC_006577.2;YP_173236.1;HCoV-HKU1_orf1ab;6844 | 0 | 0 |
| NC_006577.2;YP_173236.1;HCoV-HKU1_orf1ab;6845 | 0 | 0.025 |
| NC_006577.2;YP_173236.1;HCoV-HKU1_orf1ab;6846 | 0 | 0 |
| NC_006577.2;YP_173236.1;HCoV-HKU1_orf1ab;6847 | 0 | 0 |
| NC_006577.2;YP_173236.1;HCoV-HKU1_orf1ab;6848 | 0 | 0 |
| NC_006577.2;YP_173236.1;HCoV-HKU1_orf1ab;6849 | 0 | 0 |
| NC_006577.2;YP_173236.1;HCoV-HKU1_orf1ab;6850 | 0.05 | 0 |
| NC_006577.2;YP_173236.1;HCoV-HKU1_orf1ab;6851 | 0.05 | 0.025 |
| NC_006577.2;YP_173236.1;HCoV-HKU1_orf1ab;6852 | 0.05 | 0.05 |

| | | |
|---|---|---|
| NC_006577.2;YP_173236.1;HCoV-HKU1_orf1ab;6853 | 0 | 0.025 |
| NC_006577.2;YP_173236.1;HCoV-HKU1_orf1ab;6854 | 0.05 | 0.025 |
| NC_006577.2;YP_173236.1;HCoV-HKU1_orf1ab;6855 | 0 | 0 |
| NC_006577.2;YP_173236.1;HCoV-HKU1_orf1ab;6856 | 0 | 0 |
| NC_006577.2;YP_173236.1;HCoV-HKU1_orf1ab;6857 | 0 | 0 |
| NC_006577.2;YP_173236.1;HCoV-HKU1_orf1ab;6858 | 0 | 0 |
| NC_006577.2;YP_173236.1;HCoV-HKU1_orf1ab;6859 | 0.1 | 0 |
| NC_006577.2;YP_173236.1;HCoV-HKU1_orf1ab;6860 | 0.05 | 0 |
| NC_006577.2;YP_173236.1;HCoV-HKU1_orf1ab;6861 | 0.1 | 0.025 |
| NC_006577.2;YP_173236.1;HCoV-HKU1_orf1ab;6862 | 0.1 | 0 |
| NC_006577.2;YP_173236.1;HCoV-HKU1_orf1ab;6863 | 0 | 0 |
| NC_006577.2;YP_173236.1;HCoV-HKU1_orf1ab;6864 | 0 | 0 |
| NC_006577.2;YP_173236.1;HCoV-HKU1_orf1ab;6865 | 0 | 0 |
| NC_006577.2;YP_173236.1;HCoV-HKU1_orf1ab;6866 | 0 | 0 |
| NC_006577.2;YP_173236.1;HCoV-HKU1_orf1ab;6867 | 0 | 0 |
| NC_006577.2;YP_173236.1;HCoV-HKU1_orf1ab;6868 | 0 | 0 |
| NC_006577.2;YP_173236.1;HCoV-HKU1_orf1ab;6869 | 0 | 0 |
| NC_006577.2;YP_173236.1;HCoV-HKU1_orf1ab;6870 | 0 | 0 |
| NC_006577.2;YP_173236.1;HCoV-HKU1_orf1ab;6871 | 0 | 0 |
| NC_006577.2;YP_173236.1;HCoV-HKU1_orf1ab;6872 | 0 | 0 |
| NC_006577.2;YP_173236.1;HCoV-HKU1_orf1ab;6873 | 0 | 0 |
| NC_006577.2;YP_173236.1;HCoV-HKU1_orf1ab;6874 | 0 | 0 |
| NC_006577.2;YP_173236.1;HCoV-HKU1_orf1ab;6875 | 0 | 0 |
| NC_006577.2;YP_173236.1;HCoV-HKU1_orf1ab;6876 | 0 | 0 |
| NC_006577.2;YP_173236.1;HCoV-HKU1_orf1ab;6877 | 0 | 0 |
| NC_006577.2;YP_173236.1;HCoV-HKU1_orf1ab;6878 | 0 | 0 |
| NC_006577.2;YP_173236.1;HCoV-HKU1_orf1ab;6879 | 0 | 0 |
| NC_006577.2;YP_173236.1;HCoV-HKU1_orf1ab;6880 | 0.05 | 0 |
| NC_006577.2;YP_173236.1;HCoV-HKU1_orf1ab;6881 | 0.05 | 0 |
| NC_006577.2;YP_173236.1;HCoV-HKU1_orf1ab;6882 | 0 | 0 |
| NC_006577.2;YP_173236.1;HCoV-HKU1_orf1ab;6883 | 0 | 0.025 |
| NC_006577.2;YP_173236.1;HCoV-HKU1_orf1ab;6884 | 0 | 0 |
| NC_006577.2;YP_173236.1;HCoV-HKU1_orf1ab;6885 | 0 | 0 |
| NC_006577.2;YP_173236.1;HCoV-HKU1_orf1ab;6886 | 0 | 0 |
| NC_006577.2;YP_173236.1;HCoV-HKU1_orf1ab;6887 | 0 | 0 |
| NC_006577.2;YP_173236.1;HCoV-HKU1_orf1ab;6888 | 0 | 0 |
| NC_006577.2;YP_173236.1;HCoV-HKU1_orf1ab;6889 | 0 | 0 |
| NC_006577.2;YP_173236.1;HCoV-HKU1_orf1ab;6890 | 0 | 0 |
| NC_006577.2;YP_173236.1;HCoV-HKU1_orf1ab;6891 | 0 | 0 |
| NC_006577.2;YP_173236.1;HCoV-HKU1_orf1ab;6892 | 0 | 0 |
| NC_006577.2;YP_173236.1;HCoV-HKU1_orf1ab;6893 | 0 | 0 |
| NC_006577.2;YP_173236.1;HCoV-HKU1_orf1ab;6894 | 0 | 0 |
| NC_006577.2;YP_173236.1;HCoV-HKU1_orf1ab;6895 | 0 | 0 |

| | | |
|---|---|---|
| NC_006577.2;YP_173236.1;HCoV-HKU1_orf1ab;6896 | 0 | 0 |
| NC_006577.2;YP_173236.1;HCoV-HKU1_orf1ab;6897 | 0 | 0 |
| NC_006577.2;YP_173236.1;HCoV-HKU1_orf1ab;6898 | 0 | 0 |
| NC_006577.2;YP_173236.1;HCoV-HKU1_orf1ab;6899 | 0 | 0 |
| NC_006577.2;YP_173236.1;HCoV-HKU1_orf1ab;6900 | 0 | 0 |
| NC_006577.2;YP_173236.1;HCoV-HKU1_orf1ab;6901 | 0 | 0 |
| NC_006577.2;YP_173236.1;HCoV-HKU1_orf1ab;6902 | 0 | 0 |
| NC_006577.2;YP_173236.1;HCoV-HKU1_orf1ab;6903 | 0 | 0 |
| NC_006577.2;YP_173236.1;HCoV-HKU1_orf1ab;6904 | 0 | 0 |
| NC_006577.2;YP_173236.1;HCoV-HKU1_orf1ab;6905 | 0 | 0.025 |
| NC_006577.2;YP_173236.1;HCoV-HKU1_orf1ab;6906 | 0.05 | 0 |
| NC_006577.2;YP_173236.1;HCoV-HKU1_orf1ab;6907 | 0 | 0 |
| NC_006577.2;YP_173236.1;HCoV-HKU1_orf1ab;6908 | 0 | 0 |
| NC_006577.2;YP_173236.1;HCoV-HKU1_orf1ab;6909 | 0 | 0 |
| NC_006577.2;YP_173236.1;HCoV-HKU1_orf1ab;6910 | 0 | 0 |
| NC_006577.2;YP_173236.1;HCoV-HKU1_orf1ab;6911 | 0 | 0 |
| NC_006577.2;YP_173236.1;HCoV-HKU1_orf1ab;6912 | 0 | 0 |
| NC_006577.2;YP_173236.1;HCoV-HKU1_orf1ab;6913 | 0.05 | 0.05 |
| NC_006577.2;YP_173236.1;HCoV-HKU1_orf1ab;6914 | 0 | 0 |
| NC_006577.2;YP_173236.1;HCoV-HKU1_orf1ab;6915 | 0 | 0 |
| NC_006577.2;YP_173236.1;HCoV-HKU1_orf1ab;6916 | 0 | 0 |
| NC_006577.2;YP_173236.1;HCoV-HKU1_orf1ab;6917 | 0 | 0 |
| NC_006577.2;YP_173236.1;HCoV-HKU1_orf1ab;6918 | 0 | 0 |
| NC_006577.2;YP_173236.1;HCoV-HKU1_orf1ab;6919 | 0 | 0 |
| NC_006577.2;YP_173236.1;HCoV-HKU1_orf1ab;6920 | 0 | 0 |
| NC_006577.2;YP_173236.1;HCoV-HKU1_orf1ab;6921 | 0 | 0 |
| NC_006577.2;YP_173236.1;HCoV-HKU1_orf1ab;6922 | 0 | 0 |
| NC_006577.2;YP_173236.1;HCoV-HKU1_orf1ab;6923 | 0 | 0 |
| NC_006577.2;YP_173236.1;HCoV-HKU1_orf1ab;6924 | 0 | 0 |
| NC_006577.2;YP_173236.1;HCoV-HKU1_orf1ab;6925 | 0 | 0 |
| NC_006577.2;YP_173236.1;HCoV-HKU1_orf1ab;6926 | 0.05 | 0 |
| NC_006577.2;YP_173236.1;HCoV-HKU1_orf1ab;6927 | 0 | 0 |
| NC_006577.2;YP_173236.1;HCoV-HKU1_orf1ab;6928 | 0 | 0 |
| NC_006577.2;YP_173236.1;HCoV-HKU1_orf1ab;6929 | 0 | 0 |
| NC_006577.2;YP_173236.1;HCoV-HKU1_orf1ab;6930 | 0.05 | 0 |
| NC_006577.2;YP_173236.1;HCoV-HKU1_orf1ab;6931 | 0.05 | 0 |
| NC_006577.2;YP_173236.1;HCoV-HKU1_orf1ab;6932 | 0.05 | 0 |
| NC_006577.2;YP_173236.1;HCoV-HKU1_orf1ab;6933 | 0.05 | 0 |
| NC_006577.2;YP_173236.1;HCoV-HKU1_orf1ab;6934 | 0 | 0 |
| NC_006577.2;YP_173236.1;HCoV-HKU1_orf1ab;6935 | 0 | 0 |
| NC_006577.2;YP_173236.1;HCoV-HKU1_orf1ab;6936 | 0 | 0 |
| NC_006577.2;YP_173236.1;HCoV-HKU1_orf1ab;6937 | 0 | 0 |
| NC_006577.2;YP_173236.1;HCoV-HKU1_orf1ab;6938 | 0 | 0 |

| | | |
|---|---|---|
| NC_006577.2;YP_173236.1;HCoV-HKU1_orf1ab;6939 | 0 | 0 |
| NC_006577.2;YP_173236.1;HCoV-HKU1_orf1ab;6940 | 0 | 0 |
| NC_006577.2;YP_173236.1;HCoV-HKU1_orf1ab;6941 | 0 | 0 |
| NC_006577.2;YP_173236.1;HCoV-HKU1_orf1ab;6942 | 0 | 0 |
| NC_006577.2;YP_173236.1;HCoV-HKU1_orf1ab;6943 | 0 | 0 |
| NC_006577.2;YP_173236.1;HCoV-HKU1_orf1ab;6944 | 0 | 0 |
| NC_006577.2;YP_173236.1;HCoV-HKU1_orf1ab;6945 | 0 | 0 |
| NC_006577.2;YP_173236.1;HCoV-HKU1_orf1ab;6946 | 0 | 0.025 |
| NC_006577.2;YP_173236.1;HCoV-HKU1_orf1ab;6947 | 0 | 0.025 |
| NC_006577.2;YP_173236.1;HCoV-HKU1_orf1ab;6948 | 0 | 0.025 |
| NC_006577.2;YP_173236.1;HCoV-HKU1_orf1ab;6949 | 0 | 0.025 |
| NC_006577.2;YP_173236.1;HCoV-HKU1_orf1ab;6950 | 0 | 0.025 |
| NC_006577.2;YP_173236.1;HCoV-HKU1_orf1ab;6951 | 0 | 0 |
| NC_006577.2;YP_173236.1;HCoV-HKU1_orf1ab;6952 | 0 | 0 |
| NC_006577.2;YP_173236.1;HCoV-HKU1_orf1ab;6953 | 0 | 0 |
| NC_006577.2;YP_173236.1;HCoV-HKU1_orf1ab;6954 | 0 | 0 |
| NC_006577.2;YP_173236.1;HCoV-HKU1_orf1ab;6955 | 0 | 0 |
| NC_006577.2;YP_173236.1;HCoV-HKU1_orf1ab;6956 | 0.05 | 0 |
| NC_006577.2;YP_173236.1;HCoV-HKU1_orf1ab;6957 | 0 | 0 |
| NC_006577.2;YP_173236.1;HCoV-HKU1_orf1ab;6958 | 0 | 0 |
| NC_006577.2;YP_173236.1;HCoV-HKU1_orf1ab;6959 | 0 | 0 |
| NC_006577.2;YP_173236.1;HCoV-HKU1_orf1ab;6960 | 0 | 0 |
| NC_006577.2;YP_173236.1;HCoV-HKU1_orf1ab;6961 | 0 | 0 |
| NC_006577.2;YP_173236.1;HCoV-HKU1_orf1ab;6962 | 0 | 0 |
| NC_006577.2;YP_173236.1;HCoV-HKU1_orf1ab;6963 | 0 | 0 |
| NC_006577.2;YP_173236.1;HCoV-HKU1_orf1ab;6964 | 0 | 0 |
| NC_006577.2;YP_173236.1;HCoV-HKU1_orf1ab;6965 | 0 | 0 |
| NC_006577.2;YP_173236.1;HCoV-HKU1_orf1ab;6966 | 0 | 0 |
| NC_006577.2;YP_173236.1;HCoV-HKU1_orf1ab;6967 | 0 | 0 |
| NC_006577.2;YP_173236.1;HCoV-HKU1_orf1ab;6968 | 0 | 0 |
| NC_006577.2;YP_173236.1;HCoV-HKU1_orf1ab;6969 | 0 | 0 |
| NC_006577.2;YP_173236.1;HCoV-HKU1_orf1ab;6970 | 0 | 0 |
| NC_006577.2;YP_173236.1;HCoV-HKU1_orf1ab;6971 | 0 | 0 |
| NC_006577.2;YP_173236.1;HCoV-HKU1_orf1ab;6972 | 0 | 0 |
| NC_006577.2;YP_173236.1;HCoV-HKU1_orf1ab;6973 | 0 | 0 |
| NC_006577.2;YP_173236.1;HCoV-HKU1_orf1ab;6974 | 0 | 0 |
| NC_006577.2;YP_173236.1;HCoV-HKU1_orf1ab;6975 | 0 | 0 |
| NC_006577.2;YP_173236.1;HCoV-HKU1_orf1ab;6976 | 0 | 0 |
| NC_006577.2;YP_173236.1;HCoV-HKU1_orf1ab;6977 | 0 | 0 |
| NC_006577.2;YP_173236.1;HCoV-HKU1_orf1ab;6978 | 0 | 0 |
| NC_006577.2;YP_173236.1;HCoV-HKU1_orf1ab;6979 | 0 | 0 |
| NC_006577.2;YP_173236.1;HCoV-HKU1_orf1ab;6980 | 0 | 0 |
| NC_006577.2;YP_173236.1;HCoV-HKU1_orf1ab;6981 | 0 | 0 |

| | | |
|---|---|---|
| NC_006577.2;YP_173236.1;HCoV-HKU1_orf1ab;6982 | 0 | 0 |
| NC_006577.2;YP_173236.1;HCoV-HKU1_orf1ab;6983 | 0 | 0 |
| NC_006577.2;YP_173236.1;HCoV-HKU1_orf1ab;6984 | 0 | 0 |
| NC_006577.2;YP_173236.1;HCoV-HKU1_orf1ab;6985 | 0 | 0 |
| NC_006577.2;YP_173236.1;HCoV-HKU1_orf1ab;6986 | 0 | 0 |
| NC_006577.2;YP_173236.1;HCoV-HKU1_orf1ab;6987 | 0 | 0 |
| NC_006577.2;YP_173236.1;HCoV-HKU1_orf1ab;6988 | 0 | 0 |
| NC_006577.2;YP_173236.1;HCoV-HKU1_orf1ab;6989 | 0 | 0 |
| NC_006577.2;YP_173236.1;HCoV-HKU1_orf1ab;6990 | 0 | 0 |
| NC_006577.2;YP_173236.1;HCoV-HKU1_orf1ab;6991 | 0 | 0 |
| NC_006577.2;YP_173236.1;HCoV-HKU1_orf1ab;6992 | 0 | 0 |
| NC_006577.2;YP_173236.1;HCoV-HKU1_orf1ab;6993 | 0 | 0 |
| NC_006577.2;YP_173236.1;HCoV-HKU1_orf1ab;6994 | 0 | 0 |
| NC_006577.2;YP_173236.1;HCoV-HKU1_orf1ab;6995 | 0 | 0 |
| NC_006577.2;YP_173236.1;HCoV-HKU1_orf1ab;6996 | 0 | 0 |
| NC_006577.2;YP_173236.1;HCoV-HKU1_orf1ab;6997 | 0 | 0 |
| NC_006577.2;YP_173236.1;HCoV-HKU1_orf1ab;6998 | 0 | 0 |
| NC_006577.2;YP_173236.1;HCoV-HKU1_orf1ab;6999 | 0 | 0 |
| NC_006577.2;YP_173236.1;HCoV-HKU1_orf1ab;7000 | 0 | 0 |
| NC_006577.2;YP_173236.1;HCoV-HKU1_orf1ab;7001 | 0 | 0 |
| NC_006577.2;YP_173236.1;HCoV-HKU1_orf1ab;7002 | 0 | 0 |
| NC_006577.2;YP_173236.1;HCoV-HKU1_orf1ab;7003 | 0 | 0 |
| NC_006577.2;YP_173236.1;HCoV-HKU1_orf1ab;7004 | 0 | 0 |
| NC_006577.2;YP_173236.1;HCoV-HKU1_orf1ab;7005 | 0 | 0 |
| NC_006577.2;YP_173236.1;HCoV-HKU1_orf1ab;7006 | 0 | 0 |
| NC_006577.2;YP_173236.1;HCoV-HKU1_orf1ab;7007 | 0 | 0 |
| NC_006577.2;YP_173236.1;HCoV-HKU1_orf1ab;7008 | 0 | 0 |
| NC_006577.2;YP_173236.1;HCoV-HKU1_orf1ab;7009 | 0 | 0 |
| NC_006577.2;YP_173236.1;HCoV-HKU1_orf1ab;7010 | 0.05 | 0 |
| NC_006577.2;YP_173236.1;HCoV-HKU1_orf1ab;7011 | 0.1 | 0.025 |
| NC_006577.2;YP_173236.1;HCoV-HKU1_orf1ab;7012 | 0.05 | 0.025 |
| NC_006577.2;YP_173236.1;HCoV-HKU1_orf1ab;7013 | 0.05 | 0.05 |
| NC_006577.2;YP_173236.1;HCoV-HKU1_orf1ab;7014 | 0.1 | 0.125 |
| NC_006577.2;YP_173236.1;HCoV-HKU1_orf1ab;7015 | 0.1 | 0.075 |
| NC_006577.2;YP_173236.1;HCoV-HKU1_orf1ab;7016 | 0.05 | 0.05 |
| NC_006577.2;YP_173236.1;HCoV-HKU1_orf1ab;7017 | 0.2 | 0.1 |
| NC_006577.2;YP_173236.1;HCoV-HKU1_orf1ab;7018 | 0.15 | 0 |
| NC_006577.2;YP_173236.1;HCoV-HKU1_orf1ab;7019 | 0.15 | 0.025 |
| NC_006577.2;YP_173236.1;HCoV-HKU1_orf1ab;7020 | 0.1 | 0.025 |
| NC_006577.2;YP_173236.1;HCoV-HKU1_orf1ab;7021 | 0.05 | 0 |
| NC_006577.2;YP_173236.1;HCoV-HKU1_orf1ab;7022 | 0 | 0 |
| NC_006577.2;YP_173236.1;HCoV-HKU1_orf1ab;7023 | 0 | 0 |
| NC_006577.2;YP_173236.1;HCoV-HKU1_orf1ab;7024 | 0 | 0 |

| | | |
|---|---|---|
| NC_006577.2;YP_173236.1;HCoV-HKU1_orf1ab;7025 | 0 | 0 |
| NC_006577.2;YP_173236.1;HCoV-HKU1_orf1ab;7026 | 0 | 0 |
| NC_006577.2;YP_173236.1;HCoV-HKU1_orf1ab;7027 | 0 | 0 |
| NC_006577.2;YP_173236.1;HCoV-HKU1_orf1ab;7028 | 0 | 0 |
| NC_006577.2;YP_173236.1;HCoV-HKU1_orf1ab;7029 | 0 | 0 |
| NC_006577.2;YP_173236.1;HCoV-HKU1_orf1ab;7030 | 0 | 0 |
| NC_006577.2;YP_173236.1;HCoV-HKU1_orf1ab;7031 | 0 | 0 |
| NC_006577.2;YP_173236.1;HCoV-HKU1_orf1ab;7032 | 0 | 0 |
| NC_006577.2;YP_173236.1;HCoV-HKU1_orf1ab;7033 | 0 | 0 |
| NC_006577.2;YP_173236.1;HCoV-HKU1_orf1ab;7034 | 0 | 0 |
| NC_006577.2;YP_173236.1;HCoV-HKU1_orf1ab;7035 | 0 | 0 |
| NC_006577.2;YP_173236.1;HCoV-HKU1_orf1ab;7036 | 0.05 | 0 |
| NC_006577.2;YP_173236.1;HCoV-HKU1_orf1ab;7037 | 0 | 0 |
| NC_006577.2;YP_173236.1;HCoV-HKU1_orf1ab;7038 | 0 | 0 |
| NC_006577.2;YP_173236.1;HCoV-HKU1_orf1ab;7039 | 0 | 0 |
| NC_006577.2;YP_173236.1;HCoV-HKU1_orf1ab;7040 | 0 | 0 |
| NC_006577.2;YP_173236.1;HCoV-HKU1_orf1ab;7041 | 0 | 0 |
| NC_006577.2;YP_173236.1;HCoV-HKU1_orf1ab;7042 | 0 | 0 |
| NC_006577.2;YP_173236.1;HCoV-HKU1_orf1ab;7043 | 0 | 0 |
| NC_006577.2;YP_173236.1;HCoV-HKU1_orf1ab;7044 | 0 | 0 |
| NC_006577.2;YP_173236.1;HCoV-HKU1_orf1ab;7045 | 0 | 0 |
| NC_006577.2;YP_173236.1;HCoV-HKU1_orf1ab;7046 | 0 | 0 |
| NC_006577.2;YP_173236.1;HCoV-HKU1_orf1ab;7047 | 0 | 0 |
| NC_006577.2;YP_173236.1;HCoV-HKU1_orf1ab;7048 | 0 | 0 |
| NC_006577.2;YP_173236.1;HCoV-HKU1_orf1ab;7049 | 0 | 0 |
| NC_006577.2;YP_173236.1;HCoV-HKU1_orf1ab;7050 | 0 | 0 |
| NC_006577.2;YP_173236.1;HCoV-HKU1_orf1ab;7051 | 0 | 0.025 |
| NC_006577.2;YP_173236.1;HCoV-HKU1_orf1ab;7052 | 0 | 0.075 |
| NC_006577.2;YP_173236.1;HCoV-HKU1_orf1ab;7053 | 0 | 0.025 |
| NC_006577.2;YP_173236.1;HCoV-HKU1_orf1ab;7054 | 0 | 0.025 |
| NC_006577.2;YP_173236.1;HCoV-HKU1_orf1ab;7055 | 0 | 0.075 |
| NC_006577.2;YP_173236.1;HCoV-HKU1_orf1ab;7056 | 0.05 | 0.025 |
| NC_006577.2;YP_173236.1;HCoV-HKU1_orf1ab;7057 | 0 | 0.025 |
| NC_006577.2;YP_173236.1;HCoV-HKU1_orf1ab;7058 | 0 | 0 |
| NC_006577.2;YP_173236.1;HCoV-HKU1_orf1ab;7059 | 0 | 0 |
| NC_006577.2;YP_173236.1;HCoV-HKU1_orf1ab;7060 | 0 | 0 |
| NC_006577.2;YP_173236.1;HCoV-HKU1_orf1ab;7061 | 0 | 0 |
| NC_006577.2;YP_173236.1;HCoV-HKU1_orf1ab;7062 | 0 | 0 |
| NC_006577.2;YP_173236.1;HCoV-HKU1_orf1ab;7063 | 0 | 0 |
| NC_006577.2;YP_173236.1;HCoV-HKU1_orf1ab;7064 | 0 | 0 |
| NC_006577.2;YP_173236.1;HCoV-HKU1_orf1ab;7065 | 0 | 0 |
| NC_006577.2;YP_173236.1;HCoV-HKU1_orf1ab;7066 | 0 | 0 |
| NC_006577.2;YP_173236.1;HCoV-HKU1_orf1ab;7067 | 0 | 0 |

| | | |
|---|---|---|
| NC_006577.2;YP_173236.1;HCoV-HKU1_orf1ab;7068 | 0 | 0 |
| NC_006577.2;YP_173236.1;HCoV-HKU1_orf1ab;7069 | 0 | 0 |
| NC_006577.2;YP_173236.1;HCoV-HKU1_orf1ab;7070 | 0 | 0 |
| NC_006577.2;YP_173236.1;HCoV-HKU1_orf1ab;7071 | 0 | 0 |
| NC_006577.2;YP_173236.1;HCoV-HKU1_orf1ab;7072 | 0 | 0.025 |
| NC_006577.2;YP_173236.1;HCoV-HKU1_orf1ab;7073 | 0 | 0.025 |
| NC_006577.2;YP_173236.1;HCoV-HKU1_orf1ab;7074 | 0 | 0 |
| NC_006577.2;YP_173236.1;HCoV-HKU1_orf1ab;7075 | 0 | 0 |
| NC_006577.2;YP_173236.1;HCoV-HKU1_orf1ab;7076 | 0 | 0 |
| NC_006577.2;YP_173236.1;HCoV-HKU1_orf1ab;7077 | 0 | 0 |
| NC_006577.2;YP_173236.1;HCoV-HKU1_orf1ab;7078 | 0.1 | 0 |
| NC_006577.2;YP_173236.1;HCoV-HKU1_orf1ab;7079 | 0 | 0 |
| NC_006577.2;YP_173236.1;HCoV-HKU1_orf1ab;7080 | 0 | 0 |
| NC_006577.2;YP_173236.1;HCoV-HKU1_orf1ab;7081 | 0 | 0 |
| NC_006577.2;YP_173236.1;HCoV-HKU1_orf1ab;7082 | 0 | 0 |
| NC_006577.2;YP_173236.1;HCoV-HKU1_orf1ab;7083 | 0 | 0 |
| NC_006577.2;YP_173236.1;HCoV-HKU1_orf1ab;7084 | 0 | 0 |
| NC_006577.2;YP_173236.1;HCoV-HKU1_orf1ab;7085 | 0 | 0 |
| NC_006577.2;YP_173236.1;HCoV-HKU1_orf1ab;7086 | 0 | 0 |
| NC_006577.2;YP_173236.1;HCoV-HKU1_orf1ab;7087 | 0 | 0 |
| NC_006577.2;YP_173236.1;HCoV-HKU1_orf1ab;7088 | 0 | 0 |
| NC_006577.2;YP_173236.1;HCoV-HKU1_orf1ab;7089 | 0 | 0 |
| NC_006577.2;YP_173236.1;HCoV-HKU1_orf1ab;7090 | 0 | 0.025 |
| NC_006577.2;YP_173236.1;HCoV-HKU1_orf1ab;7091 | 0 | 0 |
| NC_006577.2;YP_173236.1;HCoV-HKU1_orf1ab;7092 | 0.05 | 0 |
| NC_006577.2;YP_173236.1;HCoV-HKU1_orf1ab;7093 | 0 | 0 |
| NC_006577.2;YP_173236.1;HCoV-HKU1_orf1ab;7094 | 0.05 | 0 |
| NC_006577.2;YP_173236.1;HCoV-HKU1_orf1ab;7095 | 0.05 | 0 |
| NC_006577.2;YP_173236.1;HCoV-HKU1_orf1ab;7096 | 0.15 | 0 |
| NC_006577.2;YP_173236.1;HCoV-HKU1_orf1ab;7097 | 0.2 | 0.075 |
| NC_006577.2;YP_173236.1;HCoV-HKU1_orf1ab;7098 | 0.05 | 0 |
| NC_006577.2;YP_173236.1;HCoV-HKU1_orf1ab;7099 | 0 | 0 |
| NC_006577.2;YP_173236.1;HCoV-HKU1_orf1ab;7100 | 0.15 | 0.175 |
| NC_006577.2;YP_173236.1;HCoV-HKU1_orf1ab;7101 | 0.25 | 0.275 |
| NC_006577.2;YP_173236.1;HCoV-HKU1_orf1ab;7102 | 0 | 0 |
| NC_006577.2;YP_173236.1;HCoV-HKU1_orf1ab;7103 | 0 | 0 |
| NC_006577.2;YP_173236.1;HCoV-HKU1_orf1ab;7104 | 0 | 0 |
| NC_006577.2;YP_173236.1;HCoV-HKU1_orf1ab;7105 | 0 | 0 |
| NC_006577.2;YP_173236.1;HCoV-HKU1_orf1ab;7106 | 0 | 0 |
| NC_006577.2;YP_173236.1;HCoV-HKU1_orf1ab;7107 | 0 | 0 |
| NC_006577.2;YP_173236.1;HCoV-HKU1_orf1ab;7108 | 0 | 0 |
| NC_006577.2;YP_173236.1;HCoV-HKU1_orf1ab;7109 | 0 | 0 |
| NC_006577.2;YP_173236.1;HCoV-HKU1_orf1ab;7110 | 0 | 0 |

| | | |
|---|---|---|
| NC_006577.2;YP_173236.1;HCoV-HKU1_orf1ab;7111 | 0 | 0 |
| NC_006577.2;YP_173236.1;HCoV-HKU1_orf1ab;7112 | 0 | 0 |
| NC_006577.2;YP_173236.1;HCoV-HKU1_orf1ab;7113 | 0 | 0 |
| NC_006577.2;YP_173236.1;HCoV-HKU1_orf1ab;7114 | 0 | 0 |
| NC_006577.2;YP_173236.1;HCoV-HKU1_orf1ab;7115 | 0 | 0 |
| NC_006577.2;YP_173236.1;HCoV-HKU1_orf1ab;7116 | 0 | 0 |
| NC_006577.2;YP_173236.1;HCoV-HKU1_orf1ab;7117 | 0.05 | 0.05 |
| NC_006577.2;YP_173236.1;HCoV-HKU1_orf1ab;7118 | 0.05 | 0.025 |
| NC_006577.2;YP_173236.1;HCoV-HKU1_orf1ab;7119 | 0.05 | 0.025 |
| NC_006577.2;YP_173236.1;HCoV-HKU1_orf1ab;7120 | 0 | 0.025 |
| NC_006577.2;YP_173236.1;HCoV-HKU1_orf1ab;7121 | 0 | 0.025 |
| NC_006577.2;YP_173236.1;HCoV-HKU1_orf1ab;7122 | 0 | 0.025 |
| NC_006577.2;YP_173236.1;HCoV-HKU1_orf1ab;7123 | 0 | 0 |
| NC_006577.2;YP_173236.1;HCoV-HKU1_orf1ab;7124 | 0 | 0 |
| NC_006577.2;YP_173236.1;HCoV-HKU1_orf1ab;7125 | 0 | 0 |
| NC_006577.2;YP_173236.1;HCoV-HKU1_orf1ab;7126 | 0 | 0 |
| NC_006577.2;YP_173236.1;HCoV-HKU1_orf1ab;7127 | 0 | 0 |
| NC_006577.2;YP_173236.1;HCoV-HKU1_orf1ab;7128 | 0 | 0 |
| NC_006577.2;YP_173236.1;HCoV-HKU1_orf1ab;7129 | 0 | 0.025 |
| NC_006577.2;YP_173236.1;HCoV-HKU1_orf1ab;7130 | 0 | 0 |
| NC_006577.2;YP_173236.1;HCoV-HKU1_orf1ab;7131 | 0 | 0 |
| NC_006577.2;YP_173236.1;HCoV-HKU1_orf1ab;7132 | 0 | 0 |
| NC_006577.2;YP_173236.1;HCoV-HKU1_orf1ab;7133 | 0 | 0.025 |
| NC_006577.2;YP_173236.1;HCoV-HKU1_orf1ab;7134 | 0 | 0.025 |
| NC_006577.2;YP_173236.1;HCoV-HKU1_orf1ab;7135 | 0 | 0.025 |
| NC_006577.2;YP_173236.1;HCoV-HKU1_orf1ab;7136 | 0 | 0.025 |
| NC_006577.2;YP_173236.1;HCoV-HKU1_orf1ab;7137 | 0 | 0 |
| NC_006577.2;YP_173236.1;HCoV-HKU1_orf1ab;7138 | 0 | 0 |
| NC_006577.2;YP_173236.1;HCoV-HKU1_orf1ab;7139 | 0 | 0.025 |
| NC_006577.2;YP_173236.1;HCoV-HKU1_orf1ab;7140 | 0 | 0.05 |
| NC_006577.2;YP_173236.1;HCoV-HKU1_orf1ab;7141 | 0 | 0.075 |
| NC_006577.2;YP_173236.1;HCoV-HKU1_orf1ab;7142 | 0 | 0 |
| NC_006577.2;YP_173236.1;HCoV-HKU1_orf1ab;7143 | 0 | 0 |
| NC_006577.2;YP_173236.1;HCoV-HKU1_orf1ab;7144 | 0 | 0 |
| NC_006577.2;YP_173236.1;HCoV-HKU1_orf1ab;7145 | 0 | 0 |
| NC_006577.2;YP_173236.1;HCoV-HKU1_orf1ab;7146 | 0 | 0 |
| NC_006577.2;YP_173236.1;HCoV-HKU1_orf1ab;7147 | 0 | 0 |
| NC_006577.2;YP_173236.1;HCoV-HKU1_orf1ab;7148 | 0 | 0.025 |
| NC_006577.2;YP_173236.1;HCoV-HKU1_orf1ab;7149 | 0 | 0 |
| NC_006577.2;YP_173236.1;HCoV-HKU1_orf1ab;7150 | 0 | 0 |
| NC_006577.2;YP_173236.1;HCoV-HKU1_orf1ab;7151 | 0 | 0 |
| NC_006577.2;YP_173236.1;HCoV-HKU1_orf1ab;7152 | 0 | 0 |
| NC_006577.2;YP_173236.1;HCoV-HKU1_orf1ab;7153 | 0 | 0 |

| | | |
|---|---|---|
| NC_006577.2;YP_173236.1;HCoV-HKU1_orf1ab;7154 | 0 | 0 |
| NC_006577.2;YP_173236.1;HCoV-HKU1_orf1ab;7155 | 0 | 0 |
| NC_006577.2;YP_173236.1;HCoV-HKU1_orf1ab;7156 | 0.05 | 0.075 |
| NC_006577.2;YP_173236.1;HCoV-HKU1_orf1ab;7157 | 0.1 | 0.1 |
| NC_006577.2;YP_173236.1;HCoV-HKU1_orf1ab;7158 | 0.1 | 0.075 |
| NC_006577.2;YP_173236.1;HCoV-HKU1_orf1ab;7159 | 0.1 | 0.075 |
| NC_006577.2;YP_173236.1;HCoV-HKU1_orf1ab;7160 | 0.15 | 0.1 |
| NC_006577.2;YP_173236.1;HCoV-HKU1_orf1ab;7161 | 0 | 0 |
| NC_006577.2;YP_173236.1;HCoV-HKU1_orf1ab;7162 | 0 | 0 |
| NC_006577.2;YP_173236.1;HCoV-HKU1_orf1ab;7163 | 0 | 0 |
| NC_006577.2;YP_173236.1;HCoV-HKU1_orf1ab;7164 | 0 | 0 |
| NC_006577.2;YP_173236.1;HCoV-HKU1_orf1ab;7165 | 0 | 0 |
| NC_006577.2;YP_173236.1;HCoV-HKU1_orf1ab;7166 | 0 | 0 |
| NC_006577.2;YP_173236.1;HCoV-HKU1_orf1ab;7167 | 0 | 0 |
| NC_006577.2;YP_173236.1;HCoV-HKU1_orf1ab;7168 | 0 | 0 |
| NC_006577.2;YP_173236.1;HCoV-HKU1_orf1ab;7169 | 0 | 0 |
| NC_006577.2;YP_173236.1;HCoV-HKU1_orf1ab;7170 | 0 | 0 |
| NC_006577.2;YP_173236.1;HCoV-HKU1_orf1ab;7171 | 0 | 0 |
| NC_006577.2;YP_173237.1;HCoV-HKU1_HA-E-glycop;1 | 0 | 0 |
| NC_006577.2;YP_173237.1;HCoV-HKU1_HA-E-glycop;2 | 0 | 0 |
| NC_006577.2;YP_173237.1;HCoV-HKU1_HA-E-glycop;3 | 0 | 0 |
| NC_006577.2;YP_173237.1;HCoV-HKU1_HA-E-glycop;4 | 0 | 0 |
| NC_006577.2;YP_173237.1;HCoV-HKU1_HA-E-glycop;5 | 0 | 0 |
| NC_006577.2;YP_173237.1;HCoV-HKU1_HA-E-glycop;6 | 0 | 0 |
| NC_006577.2;YP_173237.1;HCoV-HKU1_HA-E-glycop;7 | 0 | 0 |
| NC_006577.2;YP_173237.1;HCoV-HKU1_HA-E-glycop;8 | 0 | 0 |
| NC_006577.2;YP_173237.1;HCoV-HKU1_HA-E-glycop;9 | 0 | 0 |
| NC_006577.2;YP_173237.1;HCoV-HKU1_HA-E-glycop;10 | 0 | 0 |
| NC_006577.2;YP_173237.1;HCoV-HKU1_HA-E-glycop;11 | 0 | 0 |
| NC_006577.2;YP_173237.1;HCoV-HKU1_HA-E-glycop;12 | 0 | 0 |
| NC_006577.2;YP_173237.1;HCoV-HKU1_HA-E-glycop;13 | 0.05 | 0 |
| NC_006577.2;YP_173237.1;HCoV-HKU1_HA-E-glycop;14 | 0 | 0 |
| NC_006577.2;YP_173237.1;HCoV-HKU1_HA-E-glycop;15 | 0 | 0 |
| NC_006577.2;YP_173237.1;HCoV-HKU1_HA-E-glycop;16 | 0 | 0 |
| NC_006577.2;YP_173237.1;HCoV-HKU1_HA-E-glycop;17 | 0 | 0 |
| NC_006577.2;YP_173237.1;HCoV-HKU1_HA-E-glycop;18 | 0 | 0 |
| NC_006577.2;YP_173237.1;HCoV-HKU1_HA-E-glycop;19 | 0 | 0 |
| NC_006577.2;YP_173237.1;HCoV-HKU1_HA-E-glycop;20 | 0 | 0 |
| NC_006577.2;YP_173237.1;HCoV-HKU1_HA-E-glycop;21 | 0 | 0 |
| NC_006577.2;YP_173237.1;HCoV-HKU1_HA-E-glycop;22 | 0 | 0 |
| NC_006577.2;YP_173237.1;HCoV-HKU1_HA-E-glycop;23 | 0 | 0 |
| NC_006577.2;YP_173237.1;HCoV-HKU1_HA-E-glycop;24 | 0 | 0 |
| NC_006577.2;YP_173237.1;HCoV-HKU1_HA-E-glycop;25 | 0 | 0 |

| | | |
|---|---|---|
| NC_006577.2;YP_173237.1;HCoV-HKU1_HA-E-glycop;26 | 0 | 0 |
| NC_006577.2;YP_173237.1;HCoV-HKU1_HA-E-glycop;27 | 0 | 0 |
| NC_006577.2;YP_173237.1;HCoV-HKU1_HA-E-glycop;28 | 0 | 0 |
| NC_006577.2;YP_173237.1;HCoV-HKU1_HA-E-glycop;29 | 0.05 | 0.025 |
| NC_006577.2;YP_173237.1;HCoV-HKU1_HA-E-glycop;30 | 0.1 | 0.025 |
| NC_006577.2;YP_173237.1;HCoV-HKU1_HA-E-glycop;31 | 0.1 | 0.025 |
| NC_006577.2;YP_173237.1;HCoV-HKU1_HA-E-glycop;32 | 0 | 0 |
| NC_006577.2;YP_173237.1;HCoV-HKU1_HA-E-glycop;33 | 0 | 0.05 |
| NC_006577.2;YP_173237.1;HCoV-HKU1_HA-E-glycop;34 | 0.05 | 0 |
| NC_006577.2;YP_173237.1;HCoV-HKU1_HA-E-glycop;35 | 0 | 0.05 |
| NC_006577.2;YP_173237.1;HCoV-HKU1_HA-E-glycop;36 | 0.05 | 0.05 |
| NC_006577.2;YP_173237.1;HCoV-HKU1_HA-E-glycop;37 | 0.1 | 0.05 |
| NC_006577.2;YP_173237.1;HCoV-HKU1_HA-E-glycop;38 | 0 | 0.05 |
| NC_006577.2;YP_173237.1;HCoV-HKU1_HA-E-glycop;39 | 0 | 0 |
| NC_006577.2;YP_173237.1;HCoV-HKU1_HA-E-glycop;40 | 0 | 0 |
| NC_006577.2;YP_173237.1;HCoV-HKU1_HA-E-glycop;41 | 0.05 | 0.1 |
| NC_006577.2;YP_173237.1;HCoV-HKU1_HA-E-glycop;42 | 0 | 0.025 |
| NC_006577.2;YP_173237.1;HCoV-HKU1_HA-E-glycop;43 | 0 | 0 |
| NC_006577.2;YP_173237.1;HCoV-HKU1_HA-E-glycop;44 | 0 | 0 |
| NC_006577.2;YP_173237.1;HCoV-HKU1_HA-E-glycop;45 | 0 | 0 |
| NC_006577.2;YP_173237.1;HCoV-HKU1_HA-E-glycop;46 | 0.05 | 0 |
| NC_006577.2;YP_173237.1;HCoV-HKU1_HA-E-glycop;47 | 0.05 | 0 |
| NC_006577.2;YP_173237.1;HCoV-HKU1_HA-E-glycop;48 | 0 | 0.025 |
| NC_006577.2;YP_173237.1;HCoV-HKU1_HA-E-glycop;49 | 0 | 0.025 |
| NC_006577.2;YP_173237.1;HCoV-HKU1_HA-E-glycop;50 | 0 | 0.05 |
| NC_006577.2;YP_173237.1;HCoV-HKU1_HA-E-glycop;51 | 0 | 0.025 |
| NC_006577.2;YP_173237.1;HCoV-HKU1_HA-E-glycop;52 | 0 | 0.025 |
| NC_006577.2;YP_173237.1;HCoV-HKU1_HA-E-glycop;53 | 0 | 0 |
| NC_006577.2;YP_173237.1;HCoV-HKU1_HA-E-glycop;54 | 0 | 0 |
| NC_006577.2;YP_173237.1;HCoV-HKU1_HA-E-glycop;55 | 0 | 0.025 |
| NC_006577.2;YP_173237.1;HCoV-HKU1_HA-E-glycop;56 | 0 | 0.025 |
| NC_006577.2;YP_173237.1;HCoV-HKU1_HA-E-glycop;57 | 0 | 0.025 |
| NC_006577.2;YP_173237.1;HCoV-HKU1_HA-E-glycop;58 | 0 | 0.025 |
| NC_006577.2;YP_173237.1;HCoV-HKU1_HA-E-glycop;59 | 0 | 0 |
| NC_006577.2;YP_173237.1;HCoV-HKU1_HA-E-glycop;60 | 0 | 0 |
| NC_006577.2;YP_173237.1;HCoV-HKU1_HA-E-glycop;61 | 0 | 0 |
| NC_006577.2;YP_173237.1;HCoV-HKU1_HA-E-glycop;62 | 0 | 0 |
| NC_006577.2;YP_173237.1;HCoV-HKU1_HA-E-glycop;63 | 0 | 0 |
| NC_006577.2;YP_173237.1;HCoV-HKU1_HA-E-glycop;64 | 0 | 0 |
| NC_006577.2;YP_173237.1;HCoV-HKU1_HA-E-glycop;65 | 0 | 0 |
| NC_006577.2;YP_173237.1;HCoV-HKU1_HA-E-glycop;66 | 0 | 0 |
| NC_006577.2;YP_173237.1;HCoV-HKU1_HA-E-glycop;67 | 0 | 0 |
| NC_006577.2;YP_173237.1;HCoV-HKU1_HA-E-glycop;68 | 0 | 0 |

| | | |
|---|---|---|
| NC_006577.2;YP_173237.1;HCoV-HKU1_HA-E-glycop;69 | 0 | 0 |
| NC_006577.2;YP_173237.1;HCoV-HKU1_HA-E-glycop;70 | 0 | 0 |
| NC_006577.2;YP_173237.1;HCoV-HKU1_HA-E-glycop;71 | 0 | 0 |
| NC_006577.2;YP_173237.1;HCoV-HKU1_HA-E-glycop;72 | 0 | 0 |
| NC_006577.2;YP_173237.1;HCoV-HKU1_HA-E-glycop;73 | 0 | 0 |
| NC_006577.2;YP_173237.1;HCoV-HKU1_HA-E-glycop;74 | 0 | 0 |
| NC_006577.2;YP_173237.1;HCoV-HKU1_HA-E-glycop;75 | 0 | 0 |
| NC_006577.2;YP_173237.1;HCoV-HKU1_HA-E-glycop;76 | 0 | 0 |
| NC_006577.2;YP_173237.1;HCoV-HKU1_HA-E-glycop;77 | 0 | 0 |
| NC_006577.2;YP_173237.1;HCoV-HKU1_HA-E-glycop;78 | 0 | 0 |
| NC_006577.2;YP_173237.1;HCoV-HKU1_HA-E-glycop;79 | 0 | 0 |
| NC_006577.2;YP_173237.1;HCoV-HKU1_HA-E-glycop;80 | 0 | 0 |
| NC_006577.2;YP_173237.1;HCoV-HKU1_HA-E-glycop;81 | 0 | 0 |
| NC_006577.2;YP_173237.1;HCoV-HKU1_HA-E-glycop;82 | 0 | 0 |
| NC_006577.2;YP_173237.1;HCoV-HKU1_HA-E-glycop;83 | 0 | 0 |
| NC_006577.2;YP_173237.1;HCoV-HKU1_HA-E-glycop;84 | 0 | 0 |
| NC_006577.2;YP_173237.1;HCoV-HKU1_HA-E-glycop;85 | 0 | 0 |
| NC_006577.2;YP_173237.1;HCoV-HKU1_HA-E-glycop;86 | 0 | 0 |
| NC_006577.2;YP_173237.1;HCoV-HKU1_HA-E-glycop;87 | 0 | 0.025 |
| NC_006577.2;YP_173237.1;HCoV-HKU1_HA-E-glycop;88 | 0 | 0 |
| NC_006577.2;YP_173237.1;HCoV-HKU1_HA-E-glycop;89 | 0 | 0 |
| NC_006577.2;YP_173237.1;HCoV-HKU1_HA-E-glycop;90 | 0 | 0 |
| NC_006577.2;YP_173237.1;HCoV-HKU1_HA-E-glycop;91 | 0 | 0 |
| NC_006577.2;YP_173237.1;HCoV-HKU1_HA-E-glycop;92 | 0 | 0 |
| NC_006577.2;YP_173237.1;HCoV-HKU1_HA-E-glycop;93 | 0 | 0 |
| NC_006577.2;YP_173237.1;HCoV-HKU1_HA-E-glycop;94 | 0 | 0 |
| NC_006577.2;YP_173237.1;HCoV-HKU1_HA-E-glycop;95 | 0 | 0 |
| NC_006577.2;YP_173237.1;HCoV-HKU1_HA-E-glycop;96 | 0 | 0.025 |
| NC_006577.2;YP_173237.1;HCoV-HKU1_HA-E-glycop;97 | 0 | 0 |
| NC_006577.2;YP_173237.1;HCoV-HKU1_HA-E-glycop;98 | 0 | 0 |
| NC_006577.2;YP_173237.1;HCoV-HKU1_HA-E-glycop;99 | 0 | 0 |
| NC_006577.2;YP_173237.1;HCoV-HKU1_HA-E-glycop;100 | 0 | 0.025 |
| NC_006577.2;YP_173237.1;HCoV-HKU1_HA-E-glycop;101 | 0 | 0.025 |
| NC_006577.2;YP_173237.1;HCoV-HKU1_HA-E-glycop;102 | 0 | 0 |
| NC_006577.2;YP_173237.1;HCoV-HKU1_HA-E-glycop;103 | 0 | 0 |
| NC_006577.2;YP_173237.1;HCoV-HKU1_HA-E-glycop;104 | 0 | 0 |
| NC_006577.2;YP_173237.1;HCoV-HKU1_HA-E-glycop;105 | 0.05 | 0 |
| NC_006577.2;YP_173237.1;HCoV-HKU1_HA-E-glycop;106 | 0.05 | 0 |
| NC_006577.2;YP_173237.1;HCoV-HKU1_HA-E-glycop;107 | 0.05 | 0 |
| NC_006577.2;YP_173237.1;HCoV-HKU1_HA-E-glycop;108 | 0.05 | 0 |
| NC_006577.2;YP_173237.1;HCoV-HKU1_HA-E-glycop;109 | 0 | 0 |
| NC_006577.2;YP_173237.1;HCoV-HKU1_HA-E-glycop;110 | 0 | 0 |
| NC_006577.2;YP_173237.1;HCoV-HKU1_HA-E-glycop;111 | 0 | 0 |

| | | |
|---|---|---|
| NC_006577.2;YP_173237.1;HCoV-HKU1_HA-E-glycop;112 | 0 | 0 |
| NC_006577.2;YP_173237.1;HCoV-HKU1_HA-E-glycop;113 | 0 | 0 |
| NC_006577.2;YP_173237.1;HCoV-HKU1_HA-E-glycop;114 | 0 | 0 |
| NC_006577.2;YP_173237.1;HCoV-HKU1_HA-E-glycop;115 | 0 | 0 |
| NC_006577.2;YP_173237.1;HCoV-HKU1_HA-E-glycop;116 | 0 | 0 |
| NC_006577.2;YP_173237.1;HCoV-HKU1_HA-E-glycop;117 | 0 | 0 |
| NC_006577.2;YP_173237.1;HCoV-HKU1_HA-E-glycop;118 | 0 | 0 |
| NC_006577.2;YP_173237.1;HCoV-HKU1_HA-E-glycop;119 | 0 | 0 |
| NC_006577.2;YP_173237.1;HCoV-HKU1_HA-E-glycop;120 | 0 | 0 |
| NC_006577.2;YP_173237.1;HCoV-HKU1_HA-E-glycop;121 | 0 | 0 |
| NC_006577.2;YP_173237.1;HCoV-HKU1_HA-E-glycop;122 | 0 | 0 |
| NC_006577.2;YP_173237.1;HCoV-HKU1_HA-E-glycop;123 | 0 | 0 |
| NC_006577.2;YP_173237.1;HCoV-HKU1_HA-E-glycop;124 | 0 | 0 |
| NC_006577.2;YP_173237.1;HCoV-HKU1_HA-E-glycop;125 | 0 | 0 |
| NC_006577.2;YP_173237.1;HCoV-HKU1_HA-E-glycop;126 | 0 | 0 |
| NC_006577.2;YP_173237.1;HCoV-HKU1_HA-E-glycop;127 | 0 | 0 |
| NC_006577.2;YP_173237.1;HCoV-HKU1_HA-E-glycop;128 | 0 | 0 |
| NC_006577.2;YP_173237.1;HCoV-HKU1_HA-E-glycop;129 | 0 | 0 |
| NC_006577.2;YP_173237.1;HCoV-HKU1_HA-E-glycop;130 | 0 | 0 |
| NC_006577.2;YP_173237.1;HCoV-HKU1_HA-E-glycop;131 | 0 | 0 |
| NC_006577.2;YP_173237.1;HCoV-HKU1_HA-E-glycop;132 | 0 | 0 |
| NC_006577.2;YP_173237.1;HCoV-HKU1_HA-E-glycop;133 | 0 | 0 |
| NC_006577.2;YP_173237.1;HCoV-HKU1_HA-E-glycop;134 | 0 | 0 |
| NC_006577.2;YP_173237.1;HCoV-HKU1_HA-E-glycop;135 | 0 | 0 |
| NC_006577.2;YP_173237.1;HCoV-HKU1_HA-E-glycop;136 | 0 | 0 |
| NC_006577.2;YP_173237.1;HCoV-HKU1_HA-E-glycop;137 | 0 | 0 |
| NC_006577.2;YP_173237.1;HCoV-HKU1_HA-E-glycop;138 | 0 | 0 |
| NC_006577.2;YP_173237.1;HCoV-HKU1_HA-E-glycop;139 | 0 | 0 |
| NC_006577.2;YP_173237.1;HCoV-HKU1_HA-E-glycop;140 | 0 | 0 |
| NC_006577.2;YP_173237.1;HCoV-HKU1_HA-E-glycop;141 | 0 | 0.025 |
| NC_006577.2;YP_173237.1;HCoV-HKU1_HA-E-glycop;142 | 0.05 | 0.025 |
| NC_006577.2;YP_173237.1;HCoV-HKU1_HA-E-glycop;143 | 0 | 0.025 |
| NC_006577.2;YP_173237.1;HCoV-HKU1_HA-E-glycop;144 | 0 | 0.025 |
| NC_006577.2;YP_173237.1;HCoV-HKU1_HA-E-glycop;145 | 0 | 0 |
| NC_006577.2;YP_173237.1;HCoV-HKU1_HA-E-glycop;146 | 0 | 0 |
| NC_006577.2;YP_173237.1;HCoV-HKU1_HA-E-glycop;147 | 0 | 0.025 |
| NC_006577.2;YP_173237.1;HCoV-HKU1_HA-E-glycop;148 | 0 | 0 |
| NC_006577.2;YP_173237.1;HCoV-HKU1_HA-E-glycop;149 | 0 | 0.05 |
| NC_006577.2;YP_173237.1;HCoV-HKU1_HA-E-glycop;150 | 0 | 0.025 |
| NC_006577.2;YP_173237.1;HCoV-HKU1_HA-E-glycop;151 | 0.05 | 0.025 |
| NC_006577.2;YP_173237.1;HCoV-HKU1_HA-E-glycop;152 | 0 | 0 |
| NC_006577.2;YP_173237.1;HCoV-HKU1_HA-E-glycop;153 | 0.05 | 0 |
| NC_006577.2;YP_173237.1;HCoV-HKU1_HA-E-glycop;154 | 0 | 0 |

| | | |
|---|---|---|
| NC_006577.2;YP_173237.1;HCoV-HKU1_HA-E-glycop;155 | 0 | 0.025 |
| NC_006577.2;YP_173237.1;HCoV-HKU1_HA-E-glycop;156 | 0 | 0.025 |
| NC_006577.2;YP_173237.1;HCoV-HKU1_HA-E-glycop;157 | 0 | 0.025 |
| NC_006577.2;YP_173237.1;HCoV-HKU1_HA-E-glycop;158 | 0 | 0.025 |
| NC_006577.2;YP_173237.1;HCoV-HKU1_HA-E-glycop;159 | 0 | 0 |
| NC_006577.2;YP_173237.1;HCoV-HKU1_HA-E-glycop;160 | 0.05 | 0.025 |
| NC_006577.2;YP_173237.1;HCoV-HKU1_HA-E-glycop;161 | 0 | 0 |
| NC_006577.2;YP_173237.1;HCoV-HKU1_HA-E-glycop;162 | 0 | 0 |
| NC_006577.2;YP_173237.1;HCoV-HKU1_HA-E-glycop;163 | 0 | 0 |
| NC_006577.2;YP_173237.1;HCoV-HKU1_HA-E-glycop;164 | 0 | 0 |
| NC_006577.2;YP_173237.1;HCoV-HKU1_HA-E-glycop;165 | 0 | 0 |
| NC_006577.2;YP_173237.1;HCoV-HKU1_HA-E-glycop;166 | 0 | 0 |
| NC_006577.2;YP_173237.1;HCoV-HKU1_HA-E-glycop;167 | 0 | 0 |
| NC_006577.2;YP_173237.1;HCoV-HKU1_HA-E-glycop;168 | 0 | 0 |
| NC_006577.2;YP_173237.1;HCoV-HKU1_HA-E-glycop;169 | 0 | 0 |
| NC_006577.2;YP_173237.1;HCoV-HKU1_HA-E-glycop;170 | 0 | 0 |
| NC_006577.2;YP_173237.1;HCoV-HKU1_HA-E-glycop;171 | 0 | 0 |
| NC_006577.2;YP_173237.1;HCoV-HKU1_HA-E-glycop;172 | 0 | 0 |
| NC_006577.2;YP_173237.1;HCoV-HKU1_HA-E-glycop;173 | 0 | 0 |
| NC_006577.2;YP_173237.1;HCoV-HKU1_HA-E-glycop;174 | 0 | 0 |
| NC_006577.2;YP_173237.1;HCoV-HKU1_HA-E-glycop;175 | 0 | 0 |
| NC_006577.2;YP_173237.1;HCoV-HKU1_HA-E-glycop;176 | 0 | 0 |
| NC_006577.2;YP_173237.1;HCoV-HKU1_HA-E-glycop;177 | 0 | 0 |
| NC_006577.2;YP_173237.1;HCoV-HKU1_HA-E-glycop;178 | 0 | 0 |
| NC_006577.2;YP_173237.1;HCoV-HKU1_HA-E-glycop;179 | 0 | 0 |
| NC_006577.2;YP_173237.1;HCoV-HKU1_HA-E-glycop;180 | 0 | 0 |
| NC_006577.2;YP_173237.1;HCoV-HKU1_HA-E-glycop;181 | 0 | 0 |
| NC_006577.2;YP_173237.1;HCoV-HKU1_HA-E-glycop;182 | 0 | 0 |
| NC_006577.2;YP_173237.1;HCoV-HKU1_HA-E-glycop;183 | 0 | 0 |
| NC_006577.2;YP_173237.1;HCoV-HKU1_HA-E-glycop;184 | 0 | 0 |
| NC_006577.2;YP_173237.1;HCoV-HKU1_HA-E-glycop;185 | 0 | 0 |
| NC_006577.2;YP_173237.1;HCoV-HKU1_HA-E-glycop;186 | 0 | 0 |
| NC_006577.2;YP_173237.1;HCoV-HKU1_HA-E-glycop;187 | 0 | 0 |
| NC_006577.2;YP_173237.1;HCoV-HKU1_HA-E-glycop;188 | 0 | 0 |
| NC_006577.2;YP_173237.1;HCoV-HKU1_HA-E-glycop;189 | 0 | 0 |
| NC_006577.2;YP_173237.1;HCoV-HKU1_HA-E-glycop;190 | 0 | 0.025 |
| NC_006577.2;YP_173237.1;HCoV-HKU1_HA-E-glycop;191 | 0.05 | 0.025 |
| NC_006577.2;YP_173237.1;HCoV-HKU1_HA-E-glycop;192 | 0 | 0.025 |
| NC_006577.2;YP_173237.1;HCoV-HKU1_HA-E-glycop;193 | 0 | 0.025 |
| NC_006577.2;YP_173237.1;HCoV-HKU1_HA-E-glycop;194 | 0 | 0 |
| NC_006577.2;YP_173237.1;HCoV-HKU1_HA-E-glycop;195 | 0 | 0 |
| NC_006577.2;YP_173237.1;HCoV-HKU1_HA-E-glycop;196 | 0 | 0 |
| NC_006577.2;YP_173237.1;HCoV-HKU1_HA-E-glycop;197 | 0 | 0 |

| | | |
|---|---|---|
| NC_006577.2;YP_173237.1;HCoV-HKU1_HA-E-glycop;198 | 0 | 0 |
| NC_006577.2;YP_173237.1;HCoV-HKU1_HA-E-glycop;199 | 0 | 0 |
| NC_006577.2;YP_173237.1;HCoV-HKU1_HA-E-glycop;200 | 0 | 0 |
| NC_006577.2;YP_173237.1;HCoV-HKU1_HA-E-glycop;201 | 0 | 0 |
| NC_006577.2;YP_173237.1;HCoV-HKU1_HA-E-glycop;202 | 0 | 0 |
| NC_006577.2;YP_173237.1;HCoV-HKU1_HA-E-glycop;203 | 0.05 | 0 |
| NC_006577.2;YP_173237.1;HCoV-HKU1_HA-E-glycop;204 | 0.05 | 0 |
| NC_006577.2;YP_173237.1;HCoV-HKU1_HA-E-glycop;205 | 0 | 0 |
| NC_006577.2;YP_173237.1;HCoV-HKU1_HA-E-glycop;206 | 0 | 0 |
| NC_006577.2;YP_173237.1;HCoV-HKU1_HA-E-glycop;207 | 0 | 0 |
| NC_006577.2;YP_173237.1;HCoV-HKU1_HA-E-glycop;208 | 0 | 0 |
| NC_006577.2;YP_173237.1;HCoV-HKU1_HA-E-glycop;209 | 0 | 0 |
| NC_006577.2;YP_173237.1;HCoV-HKU1_HA-E-glycop;210 | 0 | 0.025 |
| NC_006577.2;YP_173237.1;HCoV-HKU1_HA-E-glycop;211 | 0 | 0 |
| NC_006577.2;YP_173237.1;HCoV-HKU1_HA-E-glycop;212 | 0 | 0 |
| NC_006577.2;YP_173237.1;HCoV-HKU1_HA-E-glycop;213 | 0 | 0 |
| NC_006577.2;YP_173237.1;HCoV-HKU1_HA-E-glycop;214 | 0 | 0 |
| NC_006577.2;YP_173237.1;HCoV-HKU1_HA-E-glycop;215 | 0 | 0 |
| NC_006577.2;YP_173237.1;HCoV-HKU1_HA-E-glycop;216 | 0 | 0 |
| NC_006577.2;YP_173237.1;HCoV-HKU1_HA-E-glycop;217 | 0 | 0 |
| NC_006577.2;YP_173237.1;HCoV-HKU1_HA-E-glycop;218 | 0 | 0 |
| NC_006577.2;YP_173237.1;HCoV-HKU1_HA-E-glycop;219 | 0 | 0 |
| NC_006577.2;YP_173237.1;HCoV-HKU1_HA-E-glycop;220 | 0 | 0 |
| NC_006577.2;YP_173237.1;HCoV-HKU1_HA-E-glycop;221 | 0.05 | 0 |
| NC_006577.2;YP_173237.1;HCoV-HKU1_HA-E-glycop;222 | 0 | 0 |
| NC_006577.2;YP_173237.1;HCoV-HKU1_HA-E-glycop;223 | 0 | 0 |
| NC_006577.2;YP_173237.1;HCoV-HKU1_HA-E-glycop;224 | 0 | 0 |
| NC_006577.2;YP_173237.1;HCoV-HKU1_HA-E-glycop;225 | 0 | 0 |
| NC_006577.2;YP_173237.1;HCoV-HKU1_HA-E-glycop;226 | 0 | 0 |
| NC_006577.2;YP_173237.1;HCoV-HKU1_HA-E-glycop;227 | 0.05 | 0 |
| NC_006577.2;YP_173237.1;HCoV-HKU1_HA-E-glycop;228 | 0 | 0 |
| NC_006577.2;YP_173237.1;HCoV-HKU1_HA-E-glycop;229 | 0 | 0 |
| NC_006577.2;YP_173237.1;HCoV-HKU1_HA-E-glycop;230 | 0 | 0 |
| NC_006577.2;YP_173237.1;HCoV-HKU1_HA-E-glycop;231 | 0 | 0 |
| NC_006577.2;YP_173237.1;HCoV-HKU1_HA-E-glycop;232 | 0 | 0 |
| NC_006577.2;YP_173237.1;HCoV-HKU1_HA-E-glycop;233 | 0 | 0 |
| NC_006577.2;YP_173237.1;HCoV-HKU1_HA-E-glycop;234 | 0 | 0 |
| NC_006577.2;YP_173237.1;HCoV-HKU1_HA-E-glycop;235 | 0 | 0 |
| NC_006577.2;YP_173237.1;HCoV-HKU1_HA-E-glycop;236 | 0 | 0 |
| NC_006577.2;YP_173237.1;HCoV-HKU1_HA-E-glycop;237 | 0 | 0 |
| NC_006577.2;YP_173237.1;HCoV-HKU1_HA-E-glycop;238 | 0 | 0 |
| NC_006577.2;YP_173237.1;HCoV-HKU1_HA-E-glycop;239 | 0 | 0 |
| NC_006577.2;YP_173237.1;HCoV-HKU1_HA-E-glycop;240 | 0 | 0.025 |

| | | |
|---|---|---|
| NC_006577.2;YP_173237.1;HCoV-HKU1_HA-E-glycop;241 | 0 | 0.025 |
| NC_006577.2;YP_173237.1;HCoV-HKU1_HA-E-glycop;242 | 0 | 0.025 |
| NC_006577.2;YP_173237.1;HCoV-HKU1_HA-E-glycop;243 | 0 | 0.025 |
| NC_006577.2;YP_173237.1;HCoV-HKU1_HA-E-glycop;244 | 0 | 0.025 |
| NC_006577.2;YP_173237.1;HCoV-HKU1_HA-E-glycop;245 | 0 | 0.025 |
| NC_006577.2;YP_173237.1;HCoV-HKU1_HA-E-glycop;246 | 0 | 0.025 |
| NC_006577.2;YP_173237.1;HCoV-HKU1_HA-E-glycop;247 | 0 | 0 |
| NC_006577.2;YP_173237.1;HCoV-HKU1_HA-E-glycop;248 | 0 | 0 |
| NC_006577.2;YP_173237.1;HCoV-HKU1_HA-E-glycop;249 | 0 | 0 |
| NC_006577.2;YP_173237.1;HCoV-HKU1_HA-E-glycop;250 | 0.05 | 0 |
| NC_006577.2;YP_173237.1;HCoV-HKU1_HA-E-glycop;251 | 0 | 0 |
| NC_006577.2;YP_173237.1;HCoV-HKU1_HA-E-glycop;252 | 0 | 0 |
| NC_006577.2;YP_173237.1;HCoV-HKU1_HA-E-glycop;253 | 0 | 0 |
| NC_006577.2;YP_173237.1;HCoV-HKU1_HA-E-glycop;254 | 0.05 | 0.025 |
| NC_006577.2;YP_173237.1;HCoV-HKU1_HA-E-glycop;255 | 0.05 | 0 |
| NC_006577.2;YP_173237.1;HCoV-HKU1_HA-E-glycop;256 | 0 | 0 |
| NC_006577.2;YP_173237.1;HCoV-HKU1_HA-E-glycop;257 | 0.1 | 0 |
| NC_006577.2;YP_173237.1;HCoV-HKU1_HA-E-glycop;258 | 0 | 0 |
| NC_006577.2;YP_173237.1;HCoV-HKU1_HA-E-glycop;259 | 0 | 0.075 |
| NC_006577.2;YP_173237.1;HCoV-HKU1_HA-E-glycop;260 | 0 | 0 |
| NC_006577.2;YP_173237.1;HCoV-HKU1_HA-E-glycop;261 | 0.05 | 0.075 |
| NC_006577.2;YP_173237.1;HCoV-HKU1_HA-E-glycop;262 | 0 | 0.025 |
| NC_006577.2;YP_173237.1;HCoV-HKU1_HA-E-glycop;263 | 0 | 0 |
| NC_006577.2;YP_173237.1;HCoV-HKU1_HA-E-glycop;264 | 0 | 0 |
| NC_006577.2;YP_173237.1;HCoV-HKU1_HA-E-glycop;265 | 0 | 0 |
| NC_006577.2;YP_173237.1;HCoV-HKU1_HA-E-glycop;266 | 0 | 0 |
| NC_006577.2;YP_173237.1;HCoV-HKU1_HA-E-glycop;267 | 0 | 0 |
| NC_006577.2;YP_173237.1;HCoV-HKU1_HA-E-glycop;268 | 0 | 0 |
| NC_006577.2;YP_173237.1;HCoV-HKU1_HA-E-glycop;269 | 0 | 0.025 |
| NC_006577.2;YP_173237.1;HCoV-HKU1_HA-E-glycop;270 | 0 | 0 |
| NC_006577.2;YP_173237.1;HCoV-HKU1_HA-E-glycop;271 | 0 | 0 |
| NC_006577.2;YP_173237.1;HCoV-HKU1_HA-E-glycop;272 | 0 | 0 |
| NC_006577.2;YP_173237.1;HCoV-HKU1_HA-E-glycop;273 | 0 | 0 |
| NC_006577.2;YP_173237.1;HCoV-HKU1_HA-E-glycop;274 | 0 | 0 |
| NC_006577.2;YP_173237.1;HCoV-HKU1_HA-E-glycop;275 | 0 | 0.025 |
| NC_006577.2;YP_173237.1;HCoV-HKU1_HA-E-glycop;276 | 0 | 0.025 |
| NC_006577.2;YP_173237.1;HCoV-HKU1_HA-E-glycop;277 | 0 | 0.025 |
| NC_006577.2;YP_173237.1;HCoV-HKU1_HA-E-glycop;278 | 0 | 0.025 |
| NC_006577.2;YP_173237.1;HCoV-HKU1_HA-E-glycop;279 | 0 | 0.025 |
| NC_006577.2;YP_173237.1;HCoV-HKU1_HA-E-glycop;280 | 0 | 0.025 |
| NC_006577.2;YP_173237.1;HCoV-HKU1_HA-E-glycop;281 | 0.05 | 0.025 |
| NC_006577.2;YP_173237.1;HCoV-HKU1_HA-E-glycop;282 | 0.05 | 0.05 |
| NC_006577.2;YP_173237.1;HCoV-HKU1_HA-E-glycop;283 | 0.2 | 0.075 |

| | | |
|---|---|---|
| NC_006577.2;YP_173237.1;HCoV-HKU1_HA-E-glycop;284 | 0.15 | 0.025 |
| NC_006577.2;YP_173237.1;HCoV-HKU1_HA-E-glycop;285 | 0.1 | 0.025 |
| NC_006577.2;YP_173237.1;HCoV-HKU1_HA-E-glycop;286 | 0.1 | 0.075 |
| NC_006577.2;YP_173237.1;HCoV-HKU1_HA-E-glycop;287 | 0.05 | 0.025 |
| NC_006577.2;YP_173237.1;HCoV-HKU1_HA-E-glycop;288 | 0 | 0.075 |
| NC_006577.2;YP_173237.1;HCoV-HKU1_HA-E-glycop;289 | 0 | 0.025 |
| NC_006577.2;YP_173237.1;HCoV-HKU1_HA-E-glycop;290 | 0 | 0.025 |
| NC_006577.2;YP_173237.1;HCoV-HKU1_HA-E-glycop;291 | 0.05 | 0.025 |
| NC_006577.2;YP_173237.1;HCoV-HKU1_HA-E-glycop;292 | 0 | 0 |
| NC_006577.2;YP_173237.1;HCoV-HKU1_HA-E-glycop;293 | 0 | 0 |
| NC_006577.2;YP_173237.1;HCoV-HKU1_HA-E-glycop;294 | 0 | 0 |
| NC_006577.2;YP_173237.1;HCoV-HKU1_HA-E-glycop;295 | 0 | 0 |
| NC_006577.2;YP_173237.1;HCoV-HKU1_HA-E-glycop;296 | 0 | 0 |
| NC_006577.2;YP_173237.1;HCoV-HKU1_HA-E-glycop;297 | 0 | 0 |
| NC_006577.2;YP_173237.1;HCoV-HKU1_HA-E-glycop;298 | 0 | 0 |
| NC_006577.2;YP_173237.1;HCoV-HKU1_HA-E-glycop;299 | 0 | 0 |
| NC_006577.2;YP_173237.1;HCoV-HKU1_HA-E-glycop;300 | 0 | 0 |
| NC_006577.2;YP_173237.1;HCoV-HKU1_HA-E-glycop;301 | 0 | 0 |
| NC_006577.2;YP_173237.1;HCoV-HKU1_HA-E-glycop;302 | 0 | 0 |
| NC_006577.2;YP_173237.1;HCoV-HKU1_HA-E-glycop;303 | 0 | 0 |
| NC_006577.2;YP_173237.1;HCoV-HKU1_HA-E-glycop;304 | 0 | 0 |
| NC_006577.2;YP_173237.1;HCoV-HKU1_HA-E-glycop;305 | 0 | 0 |
| NC_006577.2;YP_173237.1;HCoV-HKU1_HA-E-glycop;306 | 0 | 0 |
| NC_006577.2;YP_173237.1;HCoV-HKU1_HA-E-glycop;307 | 0 | 0 |
| NC_006577.2;YP_173237.1;HCoV-HKU1_HA-E-glycop;308 | 0 | 0 |
| NC_006577.2;YP_173237.1;HCoV-HKU1_HA-E-glycop;309 | 0 | 0 |
| NC_006577.2;YP_173237.1;HCoV-HKU1_HA-E-glycop;310 | 0 | 0 |
| NC_006577.2;YP_173237.1;HCoV-HKU1_HA-E-glycop;311 | 0 | 0 |
| NC_006577.2;YP_173237.1;HCoV-HKU1_HA-E-glycop;312 | 0 | 0 |
| NC_006577.2;YP_173237.1;HCoV-HKU1_HA-E-glycop;313 | 0 | 0 |
| NC_006577.2;YP_173237.1;HCoV-HKU1_HA-E-glycop;314 | 0 | 0 |
| NC_006577.2;YP_173237.1;HCoV-HKU1_HA-E-glycop;315 | 0 | 0 |
| NC_006577.2;YP_173237.1;HCoV-HKU1_HA-E-glycop;316 | 0 | 0 |
| NC_006577.2;YP_173237.1;HCoV-HKU1_HA-E-glycop;317 | 0 | 0 |
| NC_006577.2;YP_173237.1;HCoV-HKU1_HA-E-glycop;318 | 0 | 0 |
| NC_006577.2;YP_173237.1;HCoV-HKU1_HA-E-glycop;319 | 0 | 0 |
| NC_006577.2;YP_173237.1;HCoV-HKU1_HA-E-glycop;320 | 0 | 0 |
| NC_006577.2;YP_173237.1;HCoV-HKU1_HA-E-glycop;321 | 0 | 0 |
| NC_006577.2;YP_173237.1;HCoV-HKU1_HA-E-glycop;322 | 0 | 0 |
| NC_006577.2;YP_173237.1;HCoV-HKU1_HA-E-glycop;323 | 0 | 0 |
| NC_006577.2;YP_173237.1;HCoV-HKU1_HA-E-glycop;324 | 0 | 0 |
| NC_006577.2;YP_173237.1;HCoV-HKU1_HA-E-glycop;325 | 0 | 0 |
| NC_006577.2;YP_173237.1;HCoV-HKU1_HA-E-glycop;326 | 0 | 0 |

| | | |
|---|---|---|
| NC_006577.2;YP_173237.1;HCoV-HKU1_HA-E-glycop;327 | 0 | 0 |
| NC_006577.2;YP_173237.1;HCoV-HKU1_HA-E-glycop;328 | 0 | 0 |
| NC_006577.2;YP_173237.1;HCoV-HKU1_HA-E-glycop;329 | 0.05 | 0 |
| NC_006577.2;YP_173237.1;HCoV-HKU1_HA-E-glycop;330 | 0 | 0 |
| NC_006577.2;YP_173237.1;HCoV-HKU1_HA-E-glycop;331 | 0 | 0 |
| NC_006577.2;YP_173237.1;HCoV-HKU1_HA-E-glycop;332 | 0 | 0 |
| NC_006577.2;YP_173237.1;HCoV-HKU1_HA-E-glycop;333 | 0 | 0 |
| NC_006577.2;YP_173237.1;HCoV-HKU1_HA-E-glycop;334 | 0 | 0 |
| NC_006577.2;YP_173237.1;HCoV-HKU1_HA-E-glycop;335 | 0 | 0 |
| NC_006577.2;YP_173237.1;HCoV-HKU1_HA-E-glycop;336 | 0 | 0 |
| NC_006577.2;YP_173237.1;HCoV-HKU1_HA-E-glycop;337 | 0 | 0 |
| NC_006577.2;YP_173237.1;HCoV-HKU1_HA-E-glycop;338 | 0 | 0 |
| NC_006577.2;YP_173237.1;HCoV-HKU1_HA-E-glycop;339 | 0 | 0 |
| NC_006577.2;YP_173237.1;HCoV-HKU1_HA-E-glycop;340 | 0 | 0 |
| NC_006577.2;YP_173237.1;HCoV-HKU1_HA-E-glycop;341 | 0 | 0 |
| NC_006577.2;YP_173237.1;HCoV-HKU1_HA-E-glycop;342 | 0 | 0 |
| NC_006577.2;YP_173237.1;HCoV-HKU1_HA-E-glycop;343 | 0 | 0 |
| NC_006577.2;YP_173237.1;HCoV-HKU1_HA-E-glycop;344 | 0 | 0 |
| NC_006577.2;YP_173237.1;HCoV-HKU1_HA-E-glycop;345 | 0 | 0 |
| NC_006577.2;YP_173237.1;HCoV-HKU1_HA-E-glycop;346 | 0 | 0 |
| NC_006577.2;YP_173237.1;HCoV-HKU1_HA-E-glycop;347 | 0 | 0 |
| NC_006577.2;YP_173237.1;HCoV-HKU1_HA-E-glycop;348 | 0 | 0 |
| NC_006577.2;YP_173237.1;HCoV-HKU1_HA-E-glycop;349 | 0 | 0 |
| NC_006577.2;YP_173237.1;HCoV-HKU1_HA-E-glycop;350 | 0 | 0 |
| NC_006577.2;YP_173237.1;HCoV-HKU1_HA-E-glycop;351 | 0 | 0 |
| NC_006577.2;YP_173237.1;HCoV-HKU1_HA-E-glycop;352 | 0 | 0 |
| NC_006577.2;YP_173237.1;HCoV-HKU1_HA-E-glycop;353 | 0 | 0 |
| NC_006577.2;YP_173237.1;HCoV-HKU1_HA-E-glycop;354 | 0 | 0 |
| NC_006577.2;YP_173237.1;HCoV-HKU1_HA-E-glycop;355 | 0 | 0 |
| NC_006577.2;YP_173237.1;HCoV-HKU1_HA-E-glycop;356 | 0 | 0 |
| NC_006577.2;YP_173237.1;HCoV-HKU1_HA-E-glycop;357 | 0 | 0 |
| NC_006577.2;YP_173237.1;HCoV-HKU1_HA-E-glycop;358 | 0 | 0 |
| NC_006577.2;YP_173237.1;HCoV-HKU1_HA-E-glycop;359 | 0 | 0 |
| NC_006577.2;YP_173237.1;HCoV-HKU1_HA-E-glycop;360 | 0 | 0 |
| NC_006577.2;YP_173237.1;HCoV-HKU1_HA-E-glycop;361 | 0 | 0 |
| NC_006577.2;YP_173237.1;HCoV-HKU1_HA-E-glycop;362 | 0 | 0 |
| NC_006577.2;YP_173237.1;HCoV-HKU1_HA-E-glycop;363 | 0 | 0 |
| NC_006577.2;YP_173237.1;HCoV-HKU1_HA-E-glycop;364 | 0 | 0 |
| NC_006577.2;YP_173237.1;HCoV-HKU1_HA-E-glycop;365 | 0 | 0 |
| NC_006577.2;YP_173237.1;HCoV-HKU1_HA-E-glycop;366 | 0 | 0 |
| NC_006577.2;YP_173237.1;HCoV-HKU1_HA-E-glycop;367 | 0 | 0 |
| NC_006577.2;YP_173237.1;HCoV-HKU1_HA-E-glycop;368 | 0 | 0 |
| NC_006577.2;YP_173237.1;HCoV-HKU1_HA-E-glycop;369 | 0 | 0 |

| | | |
|---|---|---|
| NC_006577.2;YP_173237.1;HCoV-HKU1_HA-E-glycop;370 | 0 | 0 |
| NC_006577.2;YP_173237.1;HCoV-HKU1_HA-E-glycop;371 | 0 | 0 |
| NC_006577.2;YP_173237.1;HCoV-HKU1_HA-E-glycop;372 | 0 | 0 |
| NC_006577.2;YP_173237.1;HCoV-HKU1_HA-E-glycop;373 | 0 | 0 |
| NC_006577.2;YP_173237.1;HCoV-HKU1_HA-E-glycop;374 | 0 | 0 |
| NC_006577.2;YP_173237.1;HCoV-HKU1_HA-E-glycop;375 | 0 | 0 |
| NC_006577.2;YP_173238.1;HCoV-HKU1_spike;1 | 0 | 0 |
| NC_006577.2;YP_173238.1;HCoV-HKU1_spike;2 | 0 | 0 |
| NC_006577.2;YP_173238.1;HCoV-HKU1_spike;3 | 0 | 0 |
| NC_006577.2;YP_173238.1;HCoV-HKU1_spike;4 | 0 | 0 |
| NC_006577.2;YP_173238.1;HCoV-HKU1_spike;5 | 0 | 0 |
| NC_006577.2;YP_173238.1;HCoV-HKU1_spike;6 | 0 | 0 |
| NC_006577.2;YP_173238.1;HCoV-HKU1_spike;7 | 0 | 0 |
| NC_006577.2;YP_173238.1;HCoV-HKU1_spike;8 | 0 | 0 |
| NC_006577.2;YP_173238.1;HCoV-HKU1_spike;9 | 0 | 0 |
| NC_006577.2;YP_173238.1;HCoV-HKU1_spike;10 | 0 | 0 |
| NC_006577.2;YP_173238.1;HCoV-HKU1_spike;11 | 0 | 0 |
| NC_006577.2;YP_173238.1;HCoV-HKU1_spike;12 | 0 | 0.075 |
| NC_006577.2;YP_173238.1;HCoV-HKU1_spike;13 | 0 | 0.075 |
| NC_006577.2;YP_173238.1;HCoV-HKU1_spike;14 | 0.05 | 0.175 |
| NC_006577.2;YP_173238.1;HCoV-HKU1_spike;15 | 0.05 | 0.175 |
| NC_006577.2;YP_173238.1;HCoV-HKU1_spike;16 | 0.3 | 0.25 |
| NC_006577.2;YP_173238.1;HCoV-HKU1_spike;17 | 0.05 | 0.25 |
| NC_006577.2;YP_173238.1;HCoV-HKU1_spike;18 | 0 | 0.1 |
| NC_006577.2;YP_173238.1;HCoV-HKU1_spike;19 | 0.05 | 0.025 |
| NC_006577.2;YP_173238.1;HCoV-HKU1_spike;20 | 0.1 | 0.05 |
| NC_006577.2;YP_173238.1;HCoV-HKU1_spike;21 | 0.1 | 0.2 |
| NC_006577.2;YP_173238.1;HCoV-HKU1_spike;22 | 0.05 | 0.075 |
| NC_006577.2;YP_173238.1;HCoV-HKU1_spike;23 | 0.05 | 0.225 |
| NC_006577.2;YP_173238.1;HCoV-HKU1_spike;24 | 0.1 | 0 |
| NC_006577.2;YP_173238.1;HCoV-HKU1_spike;25 | 0 | 0 |
| NC_006577.2;YP_173238.1;HCoV-HKU1_spike;26 | 0 | 0 |
| NC_006577.2;YP_173238.1;HCoV-HKU1_spike;27 | 0 | 0 |
| NC_006577.2;YP_173238.1;HCoV-HKU1_spike;28 | 0 | 0 |
| NC_006577.2;YP_173238.1;HCoV-HKU1_spike;29 | 0 | 0 |
| NC_006577.2;YP_173238.1;HCoV-HKU1_spike;30 | 0 | 0 |
| NC_006577.2;YP_173238.1;HCoV-HKU1_spike;31 | 0 | 0 |
| NC_006577.2;YP_173238.1;HCoV-HKU1_spike;32 | 0 | 0 |
| NC_006577.2;YP_173238.1;HCoV-HKU1_spike;33 | 0 | 0 |
| NC_006577.2;YP_173238.1;HCoV-HKU1_spike;34 | 0 | 0 |
| NC_006577.2;YP_173238.1;HCoV-HKU1_spike;35 | 0 | 0 |
| NC_006577.2;YP_173238.1;HCoV-HKU1_spike;36 | 0 | 0 |
| NC_006577.2;YP_173238.1;HCoV-HKU1_spike;37 | 0 | 0 |

| | | |
|---|---|---|
| NC_006577.2;YP_173238.1;HCoV-HKU1_spike;38 | 0 | 0 |
| NC_006577.2;YP_173238.1;HCoV-HKU1_spike;39 | 0 | 0 |
| NC_006577.2;YP_173238.1;HCoV-HKU1_spike;40 | 0 | 0 |
| NC_006577.2;YP_173238.1;HCoV-HKU1_spike;41 | 0 | 0 |
| NC_006577.2;YP_173238.1;HCoV-HKU1_spike;42 | 0 | 0 |
| NC_006577.2;YP_173238.1;HCoV-HKU1_spike;43 | 0 | 0 |
| NC_006577.2;YP_173238.1;HCoV-HKU1_spike;44 | 0 | 0 |
| NC_006577.2;YP_173238.1;HCoV-HKU1_spike;45 | 0 | 0 |
| NC_006577.2;YP_173238.1;HCoV-HKU1_spike;46 | 0 | 0 |
| NC_006577.2;YP_173238.1;HCoV-HKU1_spike;47 | 0 | 0 |
| NC_006577.2;YP_173238.1;HCoV-HKU1_spike;48 | 0 | 0 |
| NC_006577.2;YP_173238.1;HCoV-HKU1_spike;49 | 0 | 0 |
| NC_006577.2;YP_173238.1;HCoV-HKU1_spike;50 | 0 | 0 |
| NC_006577.2;YP_173238.1;HCoV-HKU1_spike;51 | 0 | 0 |
| NC_006577.2;YP_173238.1;HCoV-HKU1_spike;52 | 0 | 0 |
| NC_006577.2;YP_173238.1;HCoV-HKU1_spike;53 | 0 | 0 |
| NC_006577.2;YP_173238.1;HCoV-HKU1_spike;54 | 0 | 0 |
| NC_006577.2;YP_173238.1;HCoV-HKU1_spike;55 | 0 | 0 |
| NC_006577.2;YP_173238.1;HCoV-HKU1_spike;56 | 0 | 0 |
| NC_006577.2;YP_173238.1;HCoV-HKU1_spike;57 | 0 | 0 |
| NC_006577.2;YP_173238.1;HCoV-HKU1_spike;58 | 0 | 0 |
| NC_006577.2;YP_173238.1;HCoV-HKU1_spike;59 | 0 | 0 |
| NC_006577.2;YP_173238.1;HCoV-HKU1_spike;60 | 0 | 0 |
| NC_006577.2;YP_173238.1;HCoV-HKU1_spike;61 | 0 | 0 |
| NC_006577.2;YP_173238.1;HCoV-HKU1_spike;62 | 0 | 0 |
| NC_006577.2;YP_173238.1;HCoV-HKU1_spike;63 | 0 | 0 |
| NC_006577.2;YP_173238.1;HCoV-HKU1_spike;64 | 0.05 | 0 |
| NC_006577.2;YP_173238.1;HCoV-HKU1_spike;65 | 0 | 0 |
| NC_006577.2;YP_173238.1;HCoV-HKU1_spike;66 | 0 | 0 |
| NC_006577.2;YP_173238.1;HCoV-HKU1_spike;67 | 0 | 0 |
| NC_006577.2;YP_173238.1;HCoV-HKU1_spike;68 | 0 | 0 |
| NC_006577.2;YP_173238.1;HCoV-HKU1_spike;69 | 0 | 0 |
| NC_006577.2;YP_173238.1;HCoV-HKU1_spike;70 | 0 | 0 |
| NC_006577.2;YP_173238.1;HCoV-HKU1_spike;71 | 0 | 0.025 |
| NC_006577.2;YP_173238.1;HCoV-HKU1_spike;72 | 0 | 0 |
| NC_006577.2;YP_173238.1;HCoV-HKU1_spike;73 | 0 | 0 |
| NC_006577.2;YP_173238.1;HCoV-HKU1_spike;74 | 0 | 0 |
| NC_006577.2;YP_173238.1;HCoV-HKU1_spike;75 | 0 | 0 |
| NC_006577.2;YP_173238.1;HCoV-HKU1_spike;76 | 0 | 0 |
| NC_006577.2;YP_173238.1;HCoV-HKU1_spike;77 | 0 | 0 |
| NC_006577.2;YP_173238.1;HCoV-HKU1_spike;78 | 0 | 0 |
| NC_006577.2;YP_173238.1;HCoV-HKU1_spike;79 | 0 | 0 |
| NC_006577.2;YP_173238.1;HCoV-HKU1_spike;80 | 0 | 0 |

| | | |
|---|---|---|
| NC_006577.2;YP_173238.1;HCoV-HKU1_spike;81 | 0 | 0 |
| NC_006577.2;YP_173238.1;HCoV-HKU1_spike;82 | 0 | 0 |
| NC_006577.2;YP_173238.1;HCoV-HKU1_spike;83 | 0 | 0 |
| NC_006577.2;YP_173238.1;HCoV-HKU1_spike;84 | 0 | 0 |
| NC_006577.2;YP_173238.1;HCoV-HKU1_spike;85 | 0 | 0 |
| NC_006577.2;YP_173238.1;HCoV-HKU1_spike;86 | 0.05 | 0 |
| NC_006577.2;YP_173238.1;HCoV-HKU1_spike;87 | 0 | 0 |
| NC_006577.2;YP_173238.1;HCoV-HKU1_spike;88 | 0 | 0 |
| NC_006577.2;YP_173238.1;HCoV-HKU1_spike;89 | 0 | 0 |
| NC_006577.2;YP_173238.1;HCoV-HKU1_spike;90 | 0 | 0 |
| NC_006577.2;YP_173238.1;HCoV-HKU1_spike;91 | 0 | 0 |
| NC_006577.2;YP_173238.1;HCoV-HKU1_spike;92 | 0 | 0 |
| NC_006577.2;YP_173238.1;HCoV-HKU1_spike;93 | 0 | 0 |
| NC_006577.2;YP_173238.1;HCoV-HKU1_spike;94 | 0 | 0 |
| NC_006577.2;YP_173238.1;HCoV-HKU1_spike;95 | 0 | 0 |
| NC_006577.2;YP_173238.1;HCoV-HKU1_spike;96 | 0 | 0.025 |
| NC_006577.2;YP_173238.1;HCoV-HKU1_spike;97 | 0 | 0 |
| NC_006577.2;YP_173238.1;HCoV-HKU1_spike;98 | 0 | 0 |
| NC_006577.2;YP_173238.1;HCoV-HKU1_spike;99 | 0 | 0 |
| NC_006577.2;YP_173238.1;HCoV-HKU1_spike;100 | 0 | 0 |
| NC_006577.2;YP_173238.1;HCoV-HKU1_spike;101 | 0 | 0 |
| NC_006577.2;YP_173238.1;HCoV-HKU1_spike;102 | 0 | 0 |
| NC_006577.2;YP_173238.1;HCoV-HKU1_spike;103 | 0 | 0 |
| NC_006577.2;YP_173238.1;HCoV-HKU1_spike;104 | 0 | 0 |
| NC_006577.2;YP_173238.1;HCoV-HKU1_spike;105 | 0.05 | 0.025 |
| NC_006577.2;YP_173238.1;HCoV-HKU1_spike;106 | 0 | 0.025 |
| NC_006577.2;YP_173238.1;HCoV-HKU1_spike;107 | 0 | 0.025 |
| NC_006577.2;YP_173238.1;HCoV-HKU1_spike;108 | 0 | 0 |
| NC_006577.2;YP_173238.1;HCoV-HKU1_spike;109 | 0 | 0 |
| NC_006577.2;YP_173238.1;HCoV-HKU1_spike;110 | 0 | 0 |
| NC_006577.2;YP_173238.1;HCoV-HKU1_spike;111 | 0 | 0 |
| NC_006577.2;YP_173238.1;HCoV-HKU1_spike;112 | 0 | 0 |
| NC_006577.2;YP_173238.1;HCoV-HKU1_spike;113 | 0 | 0.025 |
| NC_006577.2;YP_173238.1;HCoV-HKU1_spike;114 | 0 | 0 |
| NC_006577.2;YP_173238.1;HCoV-HKU1_spike;115 | 0 | 0 |
| NC_006577.2;YP_173238.1;HCoV-HKU1_spike;116 | 0 | 0 |
| NC_006577.2;YP_173238.1;HCoV-HKU1_spike;117 | 0 | 0 |
| NC_006577.2;YP_173238.1;HCoV-HKU1_spike;118 | 0 | 0 |
| NC_006577.2;YP_173238.1;HCoV-HKU1_spike;119 | 0 | 0 |
| NC_006577.2;YP_173238.1;HCoV-HKU1_spike;120 | 0 | 0 |
| NC_006577.2;YP_173238.1;HCoV-HKU1_spike;121 | 0 | 0 |
| NC_006577.2;YP_173238.1;HCoV-HKU1_spike;122 | 0 | 0 |
| NC_006577.2;YP_173238.1;HCoV-HKU1_spike;123 | 0 | 0 |

| | | |
|---|---|---|
| NC_006577.2;YP_173238.1;HCoV-HKU1_spike;124 | 0 | 0 |
| NC_006577.2;YP_173238.1;HCoV-HKU1_spike;125 | 0 | 0 |
| NC_006577.2;YP_173238.1;HCoV-HKU1_spike;126 | 0 | 0 |
| NC_006577.2;YP_173238.1;HCoV-HKU1_spike;127 | 0 | 0 |
| NC_006577.2;YP_173238.1;HCoV-HKU1_spike;128 | 0 | 0 |
| NC_006577.2;YP_173238.1;HCoV-HKU1_spike;129 | 0 | 0 |
| NC_006577.2;YP_173238.1;HCoV-HKU1_spike;130 | 0 | 0 |
| NC_006577.2;YP_173238.1;HCoV-HKU1_spike;131 | 0 | 0 |
| NC_006577.2;YP_173238.1;HCoV-HKU1_spike;132 | 0 | 0 |
| NC_006577.2;YP_173238.1;HCoV-HKU1_spike;133 | 0 | 0 |
| NC_006577.2;YP_173238.1;HCoV-HKU1_spike;134 | 0 | 0 |
| NC_006577.2;YP_173238.1;HCoV-HKU1_spike;135 | 0 | 0 |
| NC_006577.2;YP_173238.1;HCoV-HKU1_spike;136 | 0 | 0 |
| NC_006577.2;YP_173238.1;HCoV-HKU1_spike;137 | 0 | 0 |
| NC_006577.2;YP_173238.1;HCoV-HKU1_spike;138 | 0 | 0 |
| NC_006577.2;YP_173238.1;HCoV-HKU1_spike;139 | 0 | 0 |
| NC_006577.2;YP_173238.1;HCoV-HKU1_spike;140 | 0 | 0 |
| NC_006577.2;YP_173238.1;HCoV-HKU1_spike;141 | 0 | 0 |
| NC_006577.2;YP_173238.1;HCoV-HKU1_spike;142 | 0 | 0 |
| NC_006577.2;YP_173238.1;HCoV-HKU1_spike;143 | 0 | 0 |
| NC_006577.2;YP_173238.1;HCoV-HKU1_spike;144 | 0 | 0 |
| NC_006577.2;YP_173238.1;HCoV-HKU1_spike;145 | 0 | 0 |
| NC_006577.2;YP_173238.1;HCoV-HKU1_spike;146 | 0 | 0.05 |
| NC_006577.2;YP_173238.1;HCoV-HKU1_spike;147 | 0 | 0 |
| NC_006577.2;YP_173238.1;HCoV-HKU1_spike;148 | 0 | 0 |
| NC_006577.2;YP_173238.1;HCoV-HKU1_spike;149 | 0 | 0 |
| NC_006577.2;YP_173238.1;HCoV-HKU1_spike;150 | 0 | 0 |
| NC_006577.2;YP_173238.1;HCoV-HKU1_spike;151 | 0 | 0 |
| NC_006577.2;YP_173238.1;HCoV-HKU1_spike;152 | 0 | 0 |
| NC_006577.2;YP_173238.1;HCoV-HKU1_spike;153 | 0 | 0 |
| NC_006577.2;YP_173238.1;HCoV-HKU1_spike;154 | 0 | 0 |
| NC_006577.2;YP_173238.1;HCoV-HKU1_spike;155 | 0 | 0 |
| NC_006577.2;YP_173238.1;HCoV-HKU1_spike;156 | 0 | 0 |
| NC_006577.2;YP_173238.1;HCoV-HKU1_spike;157 | 0 | 0 |
| NC_006577.2;YP_173238.1;HCoV-HKU1_spike;158 | 0 | 0 |
| NC_006577.2;YP_173238.1;HCoV-HKU1_spike;159 | 0 | 0 |
| NC_006577.2;YP_173238.1;HCoV-HKU1_spike;160 | 0 | 0 |
| NC_006577.2;YP_173238.1;HCoV-HKU1_spike;161 | 0 | 0 |
| NC_006577.2;YP_173238.1;HCoV-HKU1_spike;162 | 0 | 0 |
| NC_006577.2;YP_173238.1;HCoV-HKU1_spike;163 | 0 | 0.025 |
| NC_006577.2;YP_173238.1;HCoV-HKU1_spike;164 | 0 | 0.05 |
| NC_006577.2;YP_173238.1;HCoV-HKU1_spike;165 | 0.05 | 0.05 |
| NC_006577.2;YP_173238.1;HCoV-HKU1_spike;166 | 0.05 | 0.05 |

| | | |
|---|---|---|
| NC_006577.2;YP_173238.1;HCoV-HKU1_spike;167 | 0.15 | 0.25 |
| NC_006577.2;YP_173238.1;HCoV-HKU1_spike;168 | 0.1 | 0.1 |
| NC_006577.2;YP_173238.1;HCoV-HKU1_spike;169 | 0.1 | 0.125 |
| NC_006577.2;YP_173238.1;HCoV-HKU1_spike;170 | 0 | 0.05 |
| NC_006577.2;YP_173238.1;HCoV-HKU1_spike;171 | 0.05 | 0.025 |
| NC_006577.2;YP_173238.1;HCoV-HKU1_spike;172 | 0 | 0 |
| NC_006577.2;YP_173238.1;HCoV-HKU1_spike;173 | 0 | 0 |
| NC_006577.2;YP_173238.1;HCoV-HKU1_spike;174 | 0 | 0 |
| NC_006577.2;YP_173238.1;HCoV-HKU1_spike;175 | 0 | 0 |
| NC_006577.2;YP_173238.1;HCoV-HKU1_spike;176 | 0 | 0 |
| NC_006577.2;YP_173238.1;HCoV-HKU1_spike;177 | 0.05 | 0.025 |
| NC_006577.2;YP_173238.1;HCoV-HKU1_spike;178 | 0.05 | 0 |
| NC_006577.2;YP_173238.1;HCoV-HKU1_spike;179 | 0.05 | 0 |
| NC_006577.2;YP_173238.1;HCoV-HKU1_spike;180 | 0.05 | 0 |
| NC_006577.2;YP_173238.1;HCoV-HKU1_spike;181 | 0.05 | 0 |
| NC_006577.2;YP_173238.1;HCoV-HKU1_spike;182 | 0.05 | 0 |
| NC_006577.2;YP_173238.1;HCoV-HKU1_spike;183 | 0.05 | 0 |
| NC_006577.2;YP_173238.1;HCoV-HKU1_spike;184 | 0 | 0 |
| NC_006577.2;YP_173238.1;HCoV-HKU1_spike;185 | 0 | 0 |
| NC_006577.2;YP_173238.1;HCoV-HKU1_spike;186 | 0.05 | 0 |
| NC_006577.2;YP_173238.1;HCoV-HKU1_spike;187 | 0 | 0 |
| NC_006577.2;YP_173238.1;HCoV-HKU1_spike;188 | 0 | 0 |
| NC_006577.2;YP_173238.1;HCoV-HKU1_spike;189 | 0 | 0 |
| NC_006577.2;YP_173238.1;HCoV-HKU1_spike;190 | 0 | 0 |
| NC_006577.2;YP_173238.1;HCoV-HKU1_spike;191 | 0 | 0 |
| NC_006577.2;YP_173238.1;HCoV-HKU1_spike;192 | 0 | 0 |
| NC_006577.2;YP_173238.1;HCoV-HKU1_spike;193 | 0 | 0 |
| NC_006577.2;YP_173238.1;HCoV-HKU1_spike;194 | 0 | 0 |
| NC_006577.2;YP_173238.1;HCoV-HKU1_spike;195 | 0 | 0 |
| NC_006577.2;YP_173238.1;HCoV-HKU1_spike;196 | 0 | 0 |
| NC_006577.2;YP_173238.1;HCoV-HKU1_spike;197 | 0 | 0 |
| NC_006577.2;YP_173238.1;HCoV-HKU1_spike;198 | 0 | 0 |
| NC_006577.2;YP_173238.1;HCoV-HKU1_spike;199 | 0 | 0 |
| NC_006577.2;YP_173238.1;HCoV-HKU1_spike;200 | 0 | 0 |
| NC_006577.2;YP_173238.1;HCoV-HKU1_spike;201 | 0 | 0 |
| NC_006577.2;YP_173238.1;HCoV-HKU1_spike;202 | 0 | 0 |
| NC_006577.2;YP_173238.1;HCoV-HKU1_spike;203 | 0 | 0 |
| NC_006577.2;YP_173238.1;HCoV-HKU1_spike;204 | 0 | 0 |
| NC_006577.2;YP_173238.1;HCoV-HKU1_spike;205 | 0 | 0 |
| NC_006577.2;YP_173238.1;HCoV-HKU1_spike;206 | 0 | 0 |
| NC_006577.2;YP_173238.1;HCoV-HKU1_spike;207 | 0 | 0 |
| NC_006577.2;YP_173238.1;HCoV-HKU1_spike;208 | 0 | 0 |
| NC_006577.2;YP_173238.1;HCoV-HKU1_spike;209 | 0 | 0 |

| | | |
|---|---|---|
| NC_006577.2;YP_173238.1;HCoV-HKU1_spike;210 | 0 | 0 |
| NC_006577.2;YP_173238.1;HCoV-HKU1_spike;211 | 0 | 0 |
| NC_006577.2;YP_173238.1;HCoV-HKU1_spike;212 | 0 | 0 |
| NC_006577.2;YP_173238.1;HCoV-HKU1_spike;213 | 0 | 0 |
| NC_006577.2;YP_173238.1;HCoV-HKU1_spike;214 | 0 | 0 |
| NC_006577.2;YP_173238.1;HCoV-HKU1_spike;215 | 0 | 0 |
| NC_006577.2;YP_173238.1;HCoV-HKU1_spike;216 | 0 | 0 |
| NC_006577.2;YP_173238.1;HCoV-HKU1_spike;217 | 0 | 0 |
| NC_006577.2;YP_173238.1;HCoV-HKU1_spike;218 | 0 | 0 |
| NC_006577.2;YP_173238.1;HCoV-HKU1_spike;219 | 0 | 0 |
| NC_006577.2;YP_173238.1;HCoV-HKU1_spike;220 | 0 | 0 |
| NC_006577.2;YP_173238.1;HCoV-HKU1_spike;221 | 0 | 0 |
| NC_006577.2;YP_173238.1;HCoV-HKU1_spike;222 | 0 | 0 |
| NC_006577.2;YP_173238.1;HCoV-HKU1_spike;223 | 0 | 0 |
| NC_006577.2;YP_173238.1;HCoV-HKU1_spike;224 | 0 | 0 |
| NC_006577.2;YP_173238.1;HCoV-HKU1_spike;225 | 0 | 0 |
| NC_006577.2;YP_173238.1;HCoV-HKU1_spike;226 | 0 | 0 |
| NC_006577.2;YP_173238.1;HCoV-HKU1_spike;227 | 0 | 0 |
| NC_006577.2;YP_173238.1;HCoV-HKU1_spike;228 | 0 | 0 |
| NC_006577.2;YP_173238.1;HCoV-HKU1_spike;229 | 0 | 0 |
| NC_006577.2;YP_173238.1;HCoV-HKU1_spike;230 | 0 | 0 |
| NC_006577.2;YP_173238.1;HCoV-HKU1_spike;231 | 0 | 0 |
| NC_006577.2;YP_173238.1;HCoV-HKU1_spike;232 | 0 | 0 |
| NC_006577.2;YP_173238.1;HCoV-HKU1_spike;233 | 0 | 0 |
| NC_006577.2;YP_173238.1;HCoV-HKU1_spike;234 | 0 | 0 |
| NC_006577.2;YP_173238.1;HCoV-HKU1_spike;235 | 0 | 0 |
| NC_006577.2;YP_173238.1;HCoV-HKU1_spike;236 | 0.1 | 0.125 |
| NC_006577.2;YP_173238.1;HCoV-HKU1_spike;237 | 0.1 | 0.125 |
| NC_006577.2;YP_173238.1;HCoV-HKU1_spike;238 | 0 | 0.125 |
| NC_006577.2;YP_173238.1;HCoV-HKU1_spike;239 | 0 | 0.125 |
| NC_006577.2;YP_173238.1;HCoV-HKU1_spike;240 | 0 | 0.15 |
| NC_006577.2;YP_173238.1;HCoV-HKU1_spike;241 | 0.05 | 0.175 |
| NC_006577.2;YP_173238.1;HCoV-HKU1_spike;242 | 0 | 0.075 |
| NC_006577.2;YP_173238.1;HCoV-HKU1_spike;243 | 0 | 0.075 |
| NC_006577.2;YP_173238.1;HCoV-HKU1_spike;244 | 0.15 | 0.05 |
| NC_006577.2;YP_173238.1;HCoV-HKU1_spike;245 | 0.05 | 0.05 |
| NC_006577.2;YP_173238.1;HCoV-HKU1_spike;246 | 0 | 0.025 |
| NC_006577.2;YP_173238.1;HCoV-HKU1_spike;247 | 0 | 0.025 |
| NC_006577.2;YP_173238.1;HCoV-HKU1_spike;248 | 0 | 0 |
| NC_006577.2;YP_173238.1;HCoV-HKU1_spike;249 | 0 | 0 |
| NC_006577.2;YP_173238.1;HCoV-HKU1_spike;250 | 0 | 0 |
| NC_006577.2;YP_173238.1;HCoV-HKU1_spike;251 | 0 | 0.025 |
| NC_006577.2;YP_173238.1;HCoV-HKU1_spike;252 | 0 | 0 |

| | | |
|---|---|---|
| NC_006577.2;YP_173238.1;HCoV-HKU1_spike;253 | 0 | 0 |
| NC_006577.2;YP_173238.1;HCoV-HKU1_spike;254 | 0 | 0 |
| NC_006577.2;YP_173238.1;HCoV-HKU1_spike;255 | 0 | 0 |
| NC_006577.2;YP_173238.1;HCoV-HKU1_spike;256 | 0 | 0 |
| NC_006577.2;YP_173238.1;HCoV-HKU1_spike;257 | 0 | 0 |
| NC_006577.2;YP_173238.1;HCoV-HKU1_spike;258 | 0 | 0 |
| NC_006577.2;YP_173238.1;HCoV-HKU1_spike;259 | 0 | 0 |
| NC_006577.2;YP_173238.1;HCoV-HKU1_spike;260 | 0 | 0 |
| NC_006577.2;YP_173238.1;HCoV-HKU1_spike;261 | 0 | 0 |
| NC_006577.2;YP_173238.1;HCoV-HKU1_spike;262 | 0 | 0 |
| NC_006577.2;YP_173238.1;HCoV-HKU1_spike;263 | 0 | 0.025 |
| NC_006577.2;YP_173238.1;HCoV-HKU1_spike;264 | 0 | 0 |
| NC_006577.2;YP_173238.1;HCoV-HKU1_spike;265 | 0 | 0.025 |
| NC_006577.2;YP_173238.1;HCoV-HKU1_spike;266 | 0.1 | 0.05 |
| NC_006577.2;YP_173238.1;HCoV-HKU1_spike;267 | 0.05 | 0.05 |
| NC_006577.2;YP_173238.1;HCoV-HKU1_spike;268 | 0.1 | 0.05 |
| NC_006577.2;YP_173238.1;HCoV-HKU1_spike;269 | 0.1 | 0.05 |
| NC_006577.2;YP_173238.1;HCoV-HKU1_spike;270 | 0 | 0 |
| NC_006577.2;YP_173238.1;HCoV-HKU1_spike;271 | 0 | 0 |
| NC_006577.2;YP_173238.1;HCoV-HKU1_spike;272 | 0 | 0 |
| NC_006577.2;YP_173238.1;HCoV-HKU1_spike;273 | 0 | 0 |
| NC_006577.2;YP_173238.1;HCoV-HKU1_spike;274 | 0 | 0 |
| NC_006577.2;YP_173238.1;HCoV-HKU1_spike;275 | 0 | 0 |
| NC_006577.2;YP_173238.1;HCoV-HKU1_spike;276 | 0.05 | 0 |
| NC_006577.2;YP_173238.1;HCoV-HKU1_spike;277 | 0 | 0 |
| NC_006577.2;YP_173238.1;HCoV-HKU1_spike;278 | 0.05 | 0.025 |
| NC_006577.2;YP_173238.1;HCoV-HKU1_spike;279 | 0.05 | 0 |
| NC_006577.2;YP_173238.1;HCoV-HKU1_spike;280 | 0 | 0 |
| NC_006577.2;YP_173238.1;HCoV-HKU1_spike;281 | 0 | 0 |
| NC_006577.2;YP_173238.1;HCoV-HKU1_spike;282 | 0 | 0 |
| NC_006577.2;YP_173238.1;HCoV-HKU1_spike;283 | 0 | 0 |
| NC_006577.2;YP_173238.1;HCoV-HKU1_spike;284 | 0 | 0 |
| NC_006577.2;YP_173238.1;HCoV-HKU1_spike;285 | 0 | 0 |
| NC_006577.2;YP_173238.1;HCoV-HKU1_spike;286 | 0 | 0 |
| NC_006577.2;YP_173238.1;HCoV-HKU1_spike;287 | 0.1 | 0 |
| NC_006577.2;YP_173238.1;HCoV-HKU1_spike;288 | 0 | 0 |
| NC_006577.2;YP_173238.1;HCoV-HKU1_spike;289 | 0.05 | 0 |
| NC_006577.2;YP_173238.1;HCoV-HKU1_spike;290 | 0.05 | 0.025 |
| NC_006577.2;YP_173238.1;HCoV-HKU1_spike;291 | 0.05 | 0.025 |
| NC_006577.2;YP_173238.1;HCoV-HKU1_spike;292 | 0.05 | 0.025 |
| NC_006577.2;YP_173238.1;HCoV-HKU1_spike;293 | 0 | 0.025 |
| NC_006577.2;YP_173238.1;HCoV-HKU1_spike;294 | 0.05 | 0.025 |
| NC_006577.2;YP_173238.1;HCoV-HKU1_spike;295 | 0 | 0.025 |

| | | |
|---|---|---|
| NC_006577.2;YP_173238.1;HCoV-HKU1_spike;296 | 0 | 0 |
| NC_006577.2;YP_173238.1;HCoV-HKU1_spike;297 | 0 | 0 |
| NC_006577.2;YP_173238.1;HCoV-HKU1_spike;298 | 0 | 0 |
| NC_006577.2;YP_173238.1;HCoV-HKU1_spike;299 | 0 | 0 |
| NC_006577.2;YP_173238.1;HCoV-HKU1_spike;300 | 0.1 | 0.025 |
| NC_006577.2;YP_173238.1;HCoV-HKU1_spike;301 | 0 | 0 |
| NC_006577.2;YP_173238.1;HCoV-HKU1_spike;302 | 0 | 0 |
| NC_006577.2;YP_173238.1;HCoV-HKU1_spike;303 | 0 | 0 |
| NC_006577.2;YP_173238.1;HCoV-HKU1_spike;304 | 0 | 0 |
| NC_006577.2;YP_173238.1;HCoV-HKU1_spike;305 | 0 | 0 |
| NC_006577.2;YP_173238.1;HCoV-HKU1_spike;306 | 0 | 0 |
| NC_006577.2;YP_173238.1;HCoV-HKU1_spike;307 | 0 | 0 |
| NC_006577.2;YP_173238.1;HCoV-HKU1_spike;308 | 0 | 0.025 |
| NC_006577.2;YP_173238.1;HCoV-HKU1_spike;309 | 0 | 0 |
| NC_006577.2;YP_173238.1;HCoV-HKU1_spike;310 | 0 | 0.025 |
| NC_006577.2;YP_173238.1;HCoV-HKU1_spike;311 | 0 | 0.025 |
| NC_006577.2;YP_173238.1;HCoV-HKU1_spike;312 | 0 | 0.025 |
| NC_006577.2;YP_173238.1;HCoV-HKU1_spike;313 | 0 | 0 |
| NC_006577.2;YP_173238.1;HCoV-HKU1_spike;314 | 0 | 0 |
| NC_006577.2;YP_173238.1;HCoV-HKU1_spike;315 | 0 | 0 |
| NC_006577.2;YP_173238.1;HCoV-HKU1_spike;316 | 0 | 0 |
| NC_006577.2;YP_173238.1;HCoV-HKU1_spike;317 | 0 | 0 |
| NC_006577.2;YP_173238.1;HCoV-HKU1_spike;318 | 0 | 0 |
| NC_006577.2;YP_173238.1;HCoV-HKU1_spike;319 | 0 | 0 |
| NC_006577.2;YP_173238.1;HCoV-HKU1_spike;320 | 0 | 0 |
| NC_006577.2;YP_173238.1;HCoV-HKU1_spike;321 | 0 | 0 |
| NC_006577.2;YP_173238.1;HCoV-HKU1_spike;322 | 0.1 | 0.025 |
| NC_006577.2;YP_173238.1;HCoV-HKU1_spike;323 | 0.1 | 0 |
| NC_006577.2;YP_173238.1;HCoV-HKU1_spike;324 | 0.1 | 0.025 |
| NC_006577.2;YP_173238.1;HCoV-HKU1_spike;325 | 0.1 | 0 |
| NC_006577.2;YP_173238.1;HCoV-HKU1_spike;326 | 0.1 | 0.025 |
| NC_006577.2;YP_173238.1;HCoV-HKU1_spike;327 | 0.15 | 0.025 |
| NC_006577.2;YP_173238.1;HCoV-HKU1_spike;328 | 0.1 | 0.025 |
| NC_006577.2;YP_173238.1;HCoV-HKU1_spike;329 | 0.05 | 0.025 |
| NC_006577.2;YP_173238.1;HCoV-HKU1_spike;330 | 0.15 | 0.1 |
| NC_006577.2;YP_173238.1;HCoV-HKU1_spike;331 | 0.05 | 0.025 |
| NC_006577.2;YP_173238.1;HCoV-HKU1_spike;332 | 0.05 | 0 |
| NC_006577.2;YP_173238.1;HCoV-HKU1_spike;333 | 0.2 | 0 |
| NC_006577.2;YP_173238.1;HCoV-HKU1_spike;334 | 0.05 | 0 |
| NC_006577.2;YP_173238.1;HCoV-HKU1_spike;335 | 0 | 0 |
| NC_006577.2;YP_173238.1;HCoV-HKU1_spike;336 | 0 | 0 |
| NC_006577.2;YP_173238.1;HCoV-HKU1_spike;337 | 0 | 0 |
| NC_006577.2;YP_173238.1;HCoV-HKU1_spike;338 | 0.05 | 0 |

| | | |
|---|---|---|
| NC_006577.2;YP_173238.1;HCoV-HKU1_spike;339 | 0 | 0 |
| NC_006577.2;YP_173238.1;HCoV-HKU1_spike;340 | 0 | 0 |
| NC_006577.2;YP_173238.1;HCoV-HKU1_spike;341 | 0 | 0 |
| NC_006577.2;YP_173238.1;HCoV-HKU1_spike;342 | 0 | 0 |
| NC_006577.2;YP_173238.1;HCoV-HKU1_spike;343 | 0 | 0 |
| NC_006577.2;YP_173238.1;HCoV-HKU1_spike;344 | 0 | 0 |
| NC_006577.2;YP_173238.1;HCoV-HKU1_spike;345 | 0.05 | 0 |
| NC_006577.2;YP_173238.1;HCoV-HKU1_spike;346 | 0 | 0 |
| NC_006577.2;YP_173238.1;HCoV-HKU1_spike;347 | 0 | 0 |
| NC_006577.2;YP_173238.1;HCoV-HKU1_spike;348 | 0 | 0 |
| NC_006577.2;YP_173238.1;HCoV-HKU1_spike;349 | 0 | 0 |
| NC_006577.2;YP_173238.1;HCoV-HKU1_spike;350 | 0 | 0 |
| NC_006577.2;YP_173238.1;HCoV-HKU1_spike;351 | 0 | 0 |
| NC_006577.2;YP_173238.1;HCoV-HKU1_spike;352 | 0 | 0 |
| NC_006577.2;YP_173238.1;HCoV-HKU1_spike;353 | 0 | 0 |
| NC_006577.2;YP_173238.1;HCoV-HKU1_spike;354 | 0 | 0 |
| NC_006577.2;YP_173238.1;HCoV-HKU1_spike;355 | 0 | 0 |
| NC_006577.2;YP_173238.1;HCoV-HKU1_spike;356 | 0 | 0 |
| NC_006577.2;YP_173238.1;HCoV-HKU1_spike;357 | 0 | 0 |
| NC_006577.2;YP_173238.1;HCoV-HKU1_spike;358 | 0 | 0 |
| NC_006577.2;YP_173238.1;HCoV-HKU1_spike;359 | 0 | 0 |
| NC_006577.2;YP_173238.1;HCoV-HKU1_spike;360 | 0 | 0 |
| NC_006577.2;YP_173238.1;HCoV-HKU1_spike;361 | 0 | 0 |
| NC_006577.2;YP_173238.1;HCoV-HKU1_spike;362 | 0 | 0 |
| NC_006577.2;YP_173238.1;HCoV-HKU1_spike;363 | 0 | 0 |
| NC_006577.2;YP_173238.1;HCoV-HKU1_spike;364 | 0 | 0 |
| NC_006577.2;YP_173238.1;HCoV-HKU1_spike;365 | 0.15 | 0.05 |
| NC_006577.2;YP_173238.1;HCoV-HKU1_spike;366 | 0.2 | 0.075 |
| NC_006577.2;YP_173238.1;HCoV-HKU1_spike;367 | 0.1 | 0.025 |
| NC_006577.2;YP_173238.1;HCoV-HKU1_spike;368 | 0.1 | 0.025 |
| NC_006577.2;YP_173238.1;HCoV-HKU1_spike;369 | 0.05 | 0 |
| NC_006577.2;YP_173238.1;HCoV-HKU1_spike;370 | 0.05 | 0 |
| NC_006577.2;YP_173238.1;HCoV-HKU1_spike;371 | 0 | 0 |
| NC_006577.2;YP_173238.1;HCoV-HKU1_spike;372 | 0 | 0 |
| NC_006577.2;YP_173238.1;HCoV-HKU1_spike;373 | 0 | 0 |
| NC_006577.2;YP_173238.1;HCoV-HKU1_spike;374 | 0 | 0 |
| NC_006577.2;YP_173238.1;HCoV-HKU1_spike;375 | 0 | 0.025 |
| NC_006577.2;YP_173238.1;HCoV-HKU1_spike;376 | 0 | 0.025 |
| NC_006577.2;YP_173238.1;HCoV-HKU1_spike;377 | 0 | 0 |
| NC_006577.2;YP_173238.1;HCoV-HKU1_spike;378 | 0 | 0 |
| NC_006577.2;YP_173238.1;HCoV-HKU1_spike;379 | 0 | 0 |
| NC_006577.2;YP_173238.1;HCoV-HKU1_spike;380 | 0 | 0 |
| NC_006577.2;YP_173238.1;HCoV-HKU1_spike;381 | 0 | 0 |

| | | |
|---|---|---|
| NC_006577.2;YP_173238.1;HCoV-HKU1_spike;382 | 0 | 0 |
| NC_006577.2;YP_173238.1;HCoV-HKU1_spike;383 | 0 | 0 |
| NC_006577.2;YP_173238.1;HCoV-HKU1_spike;384 | 0 | 0 |
| NC_006577.2;YP_173238.1;HCoV-HKU1_spike;385 | 0 | 0 |
| NC_006577.2;YP_173238.1;HCoV-HKU1_spike;386 | 0.05 | 0.025 |
| NC_006577.2;YP_173238.1;HCoV-HKU1_spike;387 | 0.05 | 0 |
| NC_006577.2;YP_173238.1;HCoV-HKU1_spike;388 | 0 | 0 |
| NC_006577.2;YP_173238.1;HCoV-HKU1_spike;389 | 0 | 0 |
| NC_006577.2;YP_173238.1;HCoV-HKU1_spike;390 | 0 | 0 |
| NC_006577.2;YP_173238.1;HCoV-HKU1_spike;391 | 0 | 0 |
| NC_006577.2;YP_173238.1;HCoV-HKU1_spike;392 | 0 | 0 |
| NC_006577.2;YP_173238.1;HCoV-HKU1_spike;393 | 0 | 0 |
| NC_006577.2;YP_173238.1;HCoV-HKU1_spike;394 | 0 | 0 |
| NC_006577.2;YP_173238.1;HCoV-HKU1_spike;395 | 0 | 0 |
| NC_006577.2;YP_173238.1;HCoV-HKU1_spike;396 | 0 | 0 |
| NC_006577.2;YP_173238.1;HCoV-HKU1_spike;397 | 0 | 0 |
| NC_006577.2;YP_173238.1;HCoV-HKU1_spike;398 | 0 | 0.025 |
| NC_006577.2;YP_173238.1;HCoV-HKU1_spike;399 | 0 | 0.075 |
| NC_006577.2;YP_173238.1;HCoV-HKU1_spike;400 | 0 | 0.025 |
| NC_006577.2;YP_173238.1;HCoV-HKU1_spike;401 | 0 | 0.025 |
| NC_006577.2;YP_173238.1;HCoV-HKU1_spike;402 | 0 | 0.1 |
| NC_006577.2;YP_173238.1;HCoV-HKU1_spike;403 | 0 | 0.05 |
| NC_006577.2;YP_173238.1;HCoV-HKU1_spike;404 | 0.1 | 0.075 |
| NC_006577.2;YP_173238.1;HCoV-HKU1_spike;405 | 0.05 | 0.05 |
| NC_006577.2;YP_173238.1;HCoV-HKU1_spike;406 | 0.1 | 0.15 |
| NC_006577.2;YP_173238.1;HCoV-HKU1_spike;407 | 0.1 | 0.2 |
| NC_006577.2;YP_173238.1;HCoV-HKU1_spike;408 | 0.1 | 0.075 |
| NC_006577.2;YP_173238.1;HCoV-HKU1_spike;409 | 0.2 | 0.1 |
| NC_006577.2;YP_173238.1;HCoV-HKU1_spike;410 | 0.1 | 0.075 |
| NC_006577.2;YP_173238.1;HCoV-HKU1_spike;411 | 0.05 | 0.05 |
| NC_006577.2;YP_173238.1;HCoV-HKU1_spike;412 | 0 | 0.025 |
| NC_006577.2;YP_173238.1;HCoV-HKU1_spike;413 | 0 | 0.025 |
| NC_006577.2;YP_173238.1;HCoV-HKU1_spike;414 | 0 | 0.025 |
| NC_006577.2;YP_173238.1;HCoV-HKU1_spike;415 | 0.05 | 0.05 |
| NC_006577.2;YP_173238.1;HCoV-HKU1_spike;416 | 0.05 | 0 |
| NC_006577.2;YP_173238.1;HCoV-HKU1_spike;417 | 0 | 0 |
| NC_006577.2;YP_173238.1;HCoV-HKU1_spike;418 | 0 | 0 |
| NC_006577.2;YP_173238.1;HCoV-HKU1_spike;419 | 0 | 0 |
| NC_006577.2;YP_173238.1;HCoV-HKU1_spike;420 | 0 | 0 |
| NC_006577.2;YP_173238.1;HCoV-HKU1_spike;421 | 0 | 0 |
| NC_006577.2;YP_173238.1;HCoV-HKU1_spike;422 | 0.05 | 0 |
| NC_006577.2;YP_173238.1;HCoV-HKU1_spike;423 | 0.1 | 0 |
| NC_006577.2;YP_173238.1;HCoV-HKU1_spike;424 | 0.05 | 0 |

| | | |
|---|---|---|
| NC_006577.2;YP_173238.1;HCoV-HKU1_spike;425 | 0.1 | 0 |
| NC_006577.2;YP_173238.1;HCoV-HKU1_spike;426 | 0.1 | 0.025 |
| NC_006577.2;YP_173238.1;HCoV-HKU1_spike;427 | 0.1 | 0.025 |
| NC_006577.2;YP_173238.1;HCoV-HKU1_spike;428 | 0.1 | 0.025 |
| NC_006577.2;YP_173238.1;HCoV-HKU1_spike;429 | 0.1 | 0.025 |
| NC_006577.2;YP_173238.1;HCoV-HKU1_spike;430 | 0.1 | 0.05 |
| NC_006577.2;YP_173238.1;HCoV-HKU1_spike;431 | 0.1 | 0.05 |
| NC_006577.2;YP_173238.1;HCoV-HKU1_spike;432 | 0.1 | 0 |
| NC_006577.2;YP_173238.1;HCoV-HKU1_spike;433 | 0.05 | 0 |
| NC_006577.2;YP_173238.1;HCoV-HKU1_spike;434 | 0 | 0 |
| NC_006577.2;YP_173238.1;HCoV-HKU1_spike;435 | 0 | 0 |
| NC_006577.2;YP_173238.1;HCoV-HKU1_spike;436 | 0 | 0 |
| NC_006577.2;YP_173238.1;HCoV-HKU1_spike;437 | 0 | 0 |
| NC_006577.2;YP_173238.1;HCoV-HKU1_spike;438 | 0 | 0 |
| NC_006577.2;YP_173238.1;HCoV-HKU1_spike;439 | 0 | 0 |
| NC_006577.2;YP_173238.1;HCoV-HKU1_spike;440 | 0 | 0 |
| NC_006577.2;YP_173238.1;HCoV-HKU1_spike;441 | 0 | 0 |
| NC_006577.2;YP_173238.1;HCoV-HKU1_spike;442 | 0 | 0 |
| NC_006577.2;YP_173238.1;HCoV-HKU1_spike;443 | 0 | 0 |
| NC_006577.2;YP_173238.1;HCoV-HKU1_spike;444 | 0 | 0 |
| NC_006577.2;YP_173238.1;HCoV-HKU1_spike;445 | 0 | 0 |
| NC_006577.2;YP_173238.1;HCoV-HKU1_spike;446 | 0 | 0 |
| NC_006577.2;YP_173238.1;HCoV-HKU1_spike;447 | 0 | 0 |
| NC_006577.2;YP_173238.1;HCoV-HKU1_spike;448 | 0 | 0 |
| NC_006577.2;YP_173238.1;HCoV-HKU1_spike;449 | 0 | 0 |
| NC_006577.2;YP_173238.1;HCoV-HKU1_spike;450 | 0 | 0 |
| NC_006577.2;YP_173238.1;HCoV-HKU1_spike;451 | 0 | 0 |
| NC_006577.2;YP_173238.1;HCoV-HKU1_spike;452 | 0 | 0 |
| NC_006577.2;YP_173238.1;HCoV-HKU1_spike;453 | 0 | 0 |
| NC_006577.2;YP_173238.1;HCoV-HKU1_spike;454 | 0 | 0 |
| NC_006577.2;YP_173238.1;HCoV-HKU1_spike;455 | 0 | 0 |
| NC_006577.2;YP_173238.1;HCoV-HKU1_spike;456 | 0 | 0 |
| NC_006577.2;YP_173238.1;HCoV-HKU1_spike;457 | 0 | 0 |
| NC_006577.2;YP_173238.1;HCoV-HKU1_spike;458 | 0 | 0 |
| NC_006577.2;YP_173238.1;HCoV-HKU1_spike;459 | 0 | 0 |
| NC_006577.2;YP_173238.1;HCoV-HKU1_spike;460 | 0 | 0 |
| NC_006577.2;YP_173238.1;HCoV-HKU1_spike;461 | 0 | 0 |
| NC_006577.2;YP_173238.1;HCoV-HKU1_spike;462 | 0 | 0 |
| NC_006577.2;YP_173238.1;HCoV-HKU1_spike;463 | 0 | 0 |
| NC_006577.2;YP_173238.1;HCoV-HKU1_spike;464 | 0 | 0 |
| NC_006577.2;YP_173238.1;HCoV-HKU1_spike;465 | 0 | 0.025 |
| NC_006577.2;YP_173238.1;HCoV-HKU1_spike;466 | 0 | 0 |
| NC_006577.2;YP_173238.1;HCoV-HKU1_spike;467 | 0 | 0.025 |

| | | |
|---|---|---|
| NC_006577.2;YP_173238.1;HCoV-HKU1_spike;468 | 0.05 | 0 |
| NC_006577.2;YP_173238.1;HCoV-HKU1_spike;469 | 0 | 0 |
| NC_006577.2;YP_173238.1;HCoV-HKU1_spike;470 | 0 | 0 |
| NC_006577.2;YP_173238.1;HCoV-HKU1_spike;471 | 0.05 | 0 |
| NC_006577.2;YP_173238.1;HCoV-HKU1_spike;472 | 0.05 | 0 |
| NC_006577.2;YP_173238.1;HCoV-HKU1_spike;473 | 0.05 | 0 |
| NC_006577.2;YP_173238.1;HCoV-HKU1_spike;474 | 0 | 0 |
| NC_006577.2;YP_173238.1;HCoV-HKU1_spike;475 | 0 | 0 |
| NC_006577.2;YP_173238.1;HCoV-HKU1_spike;476 | 0 | 0 |
| NC_006577.2;YP_173238.1;HCoV-HKU1_spike;477 | 0 | 0 |
| NC_006577.2;YP_173238.1;HCoV-HKU1_spike;478 | 0 | 0 |
| NC_006577.2;YP_173238.1;HCoV-HKU1_spike;479 | 0.2 | 0.1 |
| NC_006577.2;YP_173238.1;HCoV-HKU1_spike;480 | 0.05 | 0.05 |
| NC_006577.2;YP_173238.1;HCoV-HKU1_spike;481 | 0.05 | 0 |
| NC_006577.2;YP_173238.1;HCoV-HKU1_spike;482 | 0 | 0 |
| NC_006577.2;YP_173238.1;HCoV-HKU1_spike;483 | 0 | 0 |
| NC_006577.2;YP_173238.1;HCoV-HKU1_spike;484 | 0 | 0 |
| NC_006577.2;YP_173238.1;HCoV-HKU1_spike;485 | 0 | 0.075 |
| NC_006577.2;YP_173238.1;HCoV-HKU1_spike;486 | 0 | 0.05 |
| NC_006577.2;YP_173238.1;HCoV-HKU1_spike;487 | 0 | 0.05 |
| NC_006577.2;YP_173238.1;HCoV-HKU1_spike;488 | 0 | 0.05 |
| NC_006577.2;YP_173238.1;HCoV-HKU1_spike;489 | 0 | 0.05 |
| NC_006577.2;YP_173238.1;HCoV-HKU1_spike;490 | 0 | 0.05 |
| NC_006577.2;YP_173238.1;HCoV-HKU1_spike;491 | 0.05 | 0.05 |
| NC_006577.2;YP_173238.1;HCoV-HKU1_spike;492 | 0 | 0.075 |
| NC_006577.2;YP_173238.1;HCoV-HKU1_spike;493 | 0 | 0.05 |
| NC_006577.2;YP_173238.1;HCoV-HKU1_spike;494 | 0 | 0.025 |
| NC_006577.2;YP_173238.1;HCoV-HKU1_spike;495 | 0 | 0.05 |
| NC_006577.2;YP_173238.1;HCoV-HKU1_spike;496 | 0 | 0.025 |
| NC_006577.2;YP_173238.1;HCoV-HKU1_spike;497 | 0.05 | 0.05 |
| NC_006577.2;YP_173238.1;HCoV-HKU1_spike;498 | 0 | 0 |
| NC_006577.2;YP_173238.1;HCoV-HKU1_spike;499 | 0 | 0 |
| NC_006577.2;YP_173238.1;HCoV-HKU1_spike;500 | 0 | 0 |
| NC_006577.2;YP_173238.1;HCoV-HKU1_spike;501 | 0 | 0 |
| NC_006577.2;YP_173238.1;HCoV-HKU1_spike;502 | 0 | 0 |
| NC_006577.2;YP_173238.1;HCoV-HKU1_spike;503 | 0 | 0 |
| NC_006577.2;YP_173238.1;HCoV-HKU1_spike;504 | 0 | 0 |
| NC_006577.2;YP_173238.1;HCoV-HKU1_spike;505 | 0 | 0.075 |
| NC_006577.2;YP_173238.1;HCoV-HKU1_spike;506 | 0.05 | 0 |
| NC_006577.2;YP_173238.1;HCoV-HKU1_spike;507 | 0 | 0 |
| NC_006577.2;YP_173238.1;HCoV-HKU1_spike;508 | 0 | 0 |
| NC_006577.2;YP_173238.1;HCoV-HKU1_spike;509 | 0 | 0 |
| NC_006577.2;YP_173238.1;HCoV-HKU1_spike;510 | 0 | 0 |

| | | |
|---|---|---|
| NC_006577.2;YP_173238.1;HCoV-HKU1_spike;511 | 0 | 0 |
| NC_006577.2;YP_173238.1;HCoV-HKU1_spike;512 | 0 | 0 |
| NC_006577.2;YP_173238.1;HCoV-HKU1_spike;513 | 0 | 0 |
| NC_006577.2;YP_173238.1;HCoV-HKU1_spike;514 | 0 | 0 |
| NC_006577.2;YP_173238.1;HCoV-HKU1_spike;515 | 0.05 | 0 |
| NC_006577.2;YP_173238.1;HCoV-HKU1_spike;516 | 0 | 0 |
| NC_006577.2;YP_173238.1;HCoV-HKU1_spike;517 | 0 | 0 |
| NC_006577.2;YP_173238.1;HCoV-HKU1_spike;518 | 0 | 0 |
| NC_006577.2;YP_173238.1;HCoV-HKU1_spike;519 | 0 | 0 |
| NC_006577.2;YP_173238.1;HCoV-HKU1_spike;520 | 0 | 0.025 |
| NC_006577.2;YP_173238.1;HCoV-HKU1_spike;521 | 0 | 0 |
| NC_006577.2;YP_173238.1;HCoV-HKU1_spike;522 | 0 | 0 |
| NC_006577.2;YP_173238.1;HCoV-HKU1_spike;523 | 0 | 0.025 |
| NC_006577.2;YP_173238.1;HCoV-HKU1_spike;524 | 0 | 0 |
| NC_006577.2;YP_173238.1;HCoV-HKU1_spike;525 | 0 | 0 |
| NC_006577.2;YP_173238.1;HCoV-HKU1_spike;526 | 0 | 0 |
| NC_006577.2;YP_173238.1;HCoV-HKU1_spike;527 | 0 | 0 |
| NC_006577.2;YP_173238.1;HCoV-HKU1_spike;528 | 0 | 0 |
| NC_006577.2;YP_173238.1;HCoV-HKU1_spike;529 | 0.05 | 0.025 |
| NC_006577.2;YP_173238.1;HCoV-HKU1_spike;530 | 0 | 0 |
| NC_006577.2;YP_173238.1;HCoV-HKU1_spike;531 | 0 | 0 |
| NC_006577.2;YP_173238.1;HCoV-HKU1_spike;532 | 0 | 0 |
| NC_006577.2;YP_173238.1;HCoV-HKU1_spike;533 | 0 | 0 |
| NC_006577.2;YP_173238.1;HCoV-HKU1_spike;534 | 0 | 0 |
| NC_006577.2;YP_173238.1;HCoV-HKU1_spike;535 | 0 | 0 |
| NC_006577.2;YP_173238.1;HCoV-HKU1_spike;536 | 0 | 0 |
| NC_006577.2;YP_173238.1;HCoV-HKU1_spike;537 | 0 | 0 |
| NC_006577.2;YP_173238.1;HCoV-HKU1_spike;538 | 0 | 0 |
| NC_006577.2;YP_173238.1;HCoV-HKU1_spike;539 | 0 | 0 |
| NC_006577.2;YP_173238.1;HCoV-HKU1_spike;540 | 0 | 0 |
| NC_006577.2;YP_173238.1;HCoV-HKU1_spike;541 | 0 | 0 |
| NC_006577.2;YP_173238.1;HCoV-HKU1_spike;542 | 0 | 0 |
| NC_006577.2;YP_173238.1;HCoV-HKU1_spike;543 | 0 | 0 |
| NC_006577.2;YP_173238.1;HCoV-HKU1_spike;544 | 0 | 0 |
| NC_006577.2;YP_173238.1;HCoV-HKU1_spike;545 | 0 | 0 |
| NC_006577.2;YP_173238.1;HCoV-HKU1_spike;546 | 0 | 0 |
| NC_006577.2;YP_173238.1;HCoV-HKU1_spike;547 | 0 | 0 |
| NC_006577.2;YP_173238.1;HCoV-HKU1_spike;548 | 0.15 | 0.125 |
| NC_006577.2;YP_173238.1;HCoV-HKU1_spike;549 | 0.05 | 0 |
| NC_006577.2;YP_173238.1;HCoV-HKU1_spike;550 | 0 | 0 |
| NC_006577.2;YP_173238.1;HCoV-HKU1_spike;551 | 0 | 0 |
| NC_006577.2;YP_173238.1;HCoV-HKU1_spike;552 | 0 | 0 |
| NC_006577.2;YP_173238.1;HCoV-HKU1_spike;553 | 0 | 0 |

| | | |
|---|---|---|
| NC_006577.2;YP_173238.1;HCoV-HKU1_spike;554 | 0 | 0 |
| NC_006577.2;YP_173238.1;HCoV-HKU1_spike;555 | 0 | 0 |
| NC_006577.2;YP_173238.1;HCoV-HKU1_spike;556 | 0 | 0 |
| NC_006577.2;YP_173238.1;HCoV-HKU1_spike;557 | 0 | 0.025 |
| NC_006577.2;YP_173238.1;HCoV-HKU1_spike;558 | 0 | 0 |
| NC_006577.2;YP_173238.1;HCoV-HKU1_spike;559 | 0 | 0 |
| NC_006577.2;YP_173238.1;HCoV-HKU1_spike;560 | 0 | 0 |
| NC_006577.2;YP_173238.1;HCoV-HKU1_spike;561 | 0 | 0 |
| NC_006577.2;YP_173238.1;HCoV-HKU1_spike;562 | 0 | 0 |
| NC_006577.2;YP_173238.1;HCoV-HKU1_spike;563 | 0 | 0 |
| NC_006577.2;YP_173238.1;HCoV-HKU1_spike;564 | 0 | 0 |
| NC_006577.2;YP_173238.1;HCoV-HKU1_spike;565 | 0 | 0 |
| NC_006577.2;YP_173238.1;HCoV-HKU1_spike;566 | 0 | 0 |
| NC_006577.2;YP_173238.1;HCoV-HKU1_spike;567 | 0 | 0 |
| NC_006577.2;YP_173238.1;HCoV-HKU1_spike;568 | 0 | 0 |
| NC_006577.2;YP_173238.1;HCoV-HKU1_spike;569 | 0 | 0 |
| NC_006577.2;YP_173238.1;HCoV-HKU1_spike;570 | 0 | 0 |
| NC_006577.2;YP_173238.1;HCoV-HKU1_spike;571 | 0 | 0 |
| NC_006577.2;YP_173238.1;HCoV-HKU1_spike;572 | 0 | 0 |
| NC_006577.2;YP_173238.1;HCoV-HKU1_spike;573 | 0 | 0 |
| NC_006577.2;YP_173238.1;HCoV-HKU1_spike;574 | 0 | 0 |
| NC_006577.2;YP_173238.1;HCoV-HKU1_spike;575 | 0 | 0 |
| NC_006577.2;YP_173238.1;HCoV-HKU1_spike;576 | 0 | 0 |
| NC_006577.2;YP_173238.1;HCoV-HKU1_spike;577 | 0 | 0 |
| NC_006577.2;YP_173238.1;HCoV-HKU1_spike;578 | 0 | 0 |
| NC_006577.2;YP_173238.1;HCoV-HKU1_spike;579 | 0 | 0 |
| NC_006577.2;YP_173238.1;HCoV-HKU1_spike;580 | 0 | 0 |
| NC_006577.2;YP_173238.1;HCoV-HKU1_spike;581 | 0 | 0 |
| NC_006577.2;YP_173238.1;HCoV-HKU1_spike;582 | 0 | 0 |
| NC_006577.2;YP_173238.1;HCoV-HKU1_spike;583 | 0.05 | 0 |
| NC_006577.2;YP_173238.1;HCoV-HKU1_spike;584 | 0 | 0 |
| NC_006577.2;YP_173238.1;HCoV-HKU1_spike;585 | 0 | 0 |
| NC_006577.2;YP_173238.1;HCoV-HKU1_spike;586 | 0 | 0 |
| NC_006577.2;YP_173238.1;HCoV-HKU1_spike;587 | 0.05 | 0 |
| NC_006577.2;YP_173238.1;HCoV-HKU1_spike;588 | 0 | 0 |
| NC_006577.2;YP_173238.1;HCoV-HKU1_spike;589 | 0 | 0 |
| NC_006577.2;YP_173238.1;HCoV-HKU1_spike;590 | 0 | 0 |
| NC_006577.2;YP_173238.1;HCoV-HKU1_spike;591 | 0 | 0 |
| NC_006577.2;YP_173238.1;HCoV-HKU1_spike;592 | 0 | 0 |
| NC_006577.2;YP_173238.1;HCoV-HKU1_spike;593 | 0.05 | 0 |
| NC_006577.2;YP_173238.1;HCoV-HKU1_spike;594 | 0 | 0 |
| NC_006577.2;YP_173238.1;HCoV-HKU1_spike;595 | 0.1 | 0 |
| NC_006577.2;YP_173238.1;HCoV-HKU1_spike;596 | 0 | 0 |

| | | |
|---|---|---|
| NC_006577.2;YP_173238.1;HCoV-HKU1_spike;597 | 0.05 | 0 |
| NC_006577.2;YP_173238.1;HCoV-HKU1_spike;598 | 0 | 0.025 |
| NC_006577.2;YP_173238.1;HCoV-HKU1_spike;599 | 0 | 0 |
| NC_006577.2;YP_173238.1;HCoV-HKU1_spike;600 | 0.05 | 0 |
| NC_006577.2;YP_173238.1;HCoV-HKU1_spike;601 | 0.05 | 0.025 |
| NC_006577.2;YP_173238.1;HCoV-HKU1_spike;602 | 0.05 | 0.025 |
| NC_006577.2;YP_173238.1;HCoV-HKU1_spike;603 | 0 | 0.025 |
| NC_006577.2;YP_173238.1;HCoV-HKU1_spike;604 | 0.05 | 0.025 |
| NC_006577.2;YP_173238.1;HCoV-HKU1_spike;605 | 0 | 0.025 |
| NC_006577.2;YP_173238.1;HCoV-HKU1_spike;606 | 0.05 | 0.1 |
| NC_006577.2;YP_173238.1;HCoV-HKU1_spike;607 | 0 | 0 |
| NC_006577.2;YP_173238.1;HCoV-HKU1_spike;608 | 0 | 0 |
| NC_006577.2;YP_173238.1;HCoV-HKU1_spike;609 | 0 | 0 |
| NC_006577.2;YP_173238.1;HCoV-HKU1_spike;610 | 0 | 0 |
| NC_006577.2;YP_173238.1;HCoV-HKU1_spike;611 | 0 | 0 |
| NC_006577.2;YP_173238.1;HCoV-HKU1_spike;612 | 0 | 0 |
| NC_006577.2;YP_173238.1;HCoV-HKU1_spike;613 | 0 | 0 |
| NC_006577.2;YP_173238.1;HCoV-HKU1_spike;614 | 0 | 0 |
| NC_006577.2;YP_173238.1;HCoV-HKU1_spike;615 | 0 | 0 |
| NC_006577.2;YP_173238.1;HCoV-HKU1_spike;616 | 0 | 0 |
| NC_006577.2;YP_173238.1;HCoV-HKU1_spike;617 | 0 | 0 |
| NC_006577.2;YP_173238.1;HCoV-HKU1_spike;618 | 0 | 0 |
| NC_006577.2;YP_173238.1;HCoV-HKU1_spike;619 | 0 | 0 |
| NC_006577.2;YP_173238.1;HCoV-HKU1_spike;620 | 0 | 0 |
| NC_006577.2;YP_173238.1;HCoV-HKU1_spike;621 | 0 | 0 |
| NC_006577.2;YP_173238.1;HCoV-HKU1_spike;622 | 0 | 0 |
| NC_006577.2;YP_173238.1;HCoV-HKU1_spike;623 | 0 | 0 |
| NC_006577.2;YP_173238.1;HCoV-HKU1_spike;624 | 0 | 0 |
| NC_006577.2;YP_173238.1;HCoV-HKU1_spike;625 | 0 | 0 |
| NC_006577.2;YP_173238.1;HCoV-HKU1_spike;626 | 0 | 0 |
| NC_006577.2;YP_173238.1;HCoV-HKU1_spike;627 | 0 | 0 |
| NC_006577.2;YP_173238.1;HCoV-HKU1_spike;628 | 0 | 0 |
| NC_006577.2;YP_173238.1;HCoV-HKU1_spike;629 | 0 | 0 |
| NC_006577.2;YP_173238.1;HCoV-HKU1_spike;630 | 0 | 0 |
| NC_006577.2;YP_173238.1;HCoV-HKU1_spike;631 | 0 | 0 |
| NC_006577.2;YP_173238.1;HCoV-HKU1_spike;632 | 0 | 0 |
| NC_006577.2;YP_173238.1;HCoV-HKU1_spike;633 | 0 | 0 |
| NC_006577.2;YP_173238.1;HCoV-HKU1_spike;634 | 0 | 0 |
| NC_006577.2;YP_173238.1;HCoV-HKU1_spike;635 | 0 | 0 |
| NC_006577.2;YP_173238.1;HCoV-HKU1_spike;636 | 0 | 0 |
| NC_006577.2;YP_173238.1;HCoV-HKU1_spike;637 | 0 | 0 |
| NC_006577.2;YP_173238.1;HCoV-HKU1_spike;638 | 0 | 0 |
| NC_006577.2;YP_173238.1;HCoV-HKU1_spike;639 | 0 | 0 |

| | | |
|---|---|---|
| NC_006577.2;YP_173238.1;HCoV-HKU1_spike;640 | 0 | 0 |
| NC_006577.2;YP_173238.1;HCoV-HKU1_spike;641 | 0 | 0.025 |
| NC_006577.2;YP_173238.1;HCoV-HKU1_spike;642 | 0.1 | 0.1 |
| NC_006577.2;YP_173238.1;HCoV-HKU1_spike;643 | 0.15 | 0.175 |
| NC_006577.2;YP_173238.1;HCoV-HKU1_spike;644 | 0.15 | 0.175 |
| NC_006577.2;YP_173238.1;HCoV-HKU1_spike;645 | 0 | 0.1 |
| NC_006577.2;YP_173238.1;HCoV-HKU1_spike;646 | 0 | 0.125 |
| NC_006577.2;YP_173238.1;HCoV-HKU1_spike;647 | 0.15 | 0.175 |
| NC_006577.2;YP_173238.1;HCoV-HKU1_spike;648 | 0 | 0.15 |
| NC_006577.2;YP_173238.1;HCoV-HKU1_spike;649 | 0.05 | 0.125 |
| NC_006577.2;YP_173238.1;HCoV-HKU1_spike;650 | 0.1 | 0.15 |
| NC_006577.2;YP_173238.1;HCoV-HKU1_spike;651 | 0 | 0.125 |
| NC_006577.2;YP_173238.1;HCoV-HKU1_spike;652 | 0 | 0.025 |
| NC_006577.2;YP_173238.1;HCoV-HKU1_spike;653 | 0 | 0.025 |
| NC_006577.2;YP_173238.1;HCoV-HKU1_spike;654 | 0 | 0 |
| NC_006577.2;YP_173238.1;HCoV-HKU1_spike;655 | 0.05 | 0.025 |
| NC_006577.2;YP_173238.1;HCoV-HKU1_spike;656 | 0.05 | 0 |
| NC_006577.2;YP_173238.1;HCoV-HKU1_spike;657 | 0.05 | 0 |
| NC_006577.2;YP_173238.1;HCoV-HKU1_spike;658 | 0.05 | 0.025 |
| NC_006577.2;YP_173238.1;HCoV-HKU1_spike;659 | 0.15 | 0.1 |
| NC_006577.2;YP_173238.1;HCoV-HKU1_spike;660 | 0 | 0 |
| NC_006577.2;YP_173238.1;HCoV-HKU1_spike;661 | 0.05 | 0 |
| NC_006577.2;YP_173238.1;HCoV-HKU1_spike;662 | 0.05 | 0 |
| NC_006577.2;YP_173238.1;HCoV-HKU1_spike;663 | 0 | 0 |
| NC_006577.2;YP_173238.1;HCoV-HKU1_spike;664 | 0.05 | 0 |
| NC_006577.2;YP_173238.1;HCoV-HKU1_spike;665 | 0.05 | 0 |
| NC_006577.2;YP_173238.1;HCoV-HKU1_spike;666 | 0 | 0 |
| NC_006577.2;YP_173238.1;HCoV-HKU1_spike;667 | 0.05 | 0 |
| NC_006577.2;YP_173238.1;HCoV-HKU1_spike;668 | 0 | 0 |
| NC_006577.2;YP_173238.1;HCoV-HKU1_spike;669 | 0 | 0 |
| NC_006577.2;YP_173238.1;HCoV-HKU1_spike;670 | 0 | 0 |
| NC_006577.2;YP_173238.1;HCoV-HKU1_spike;671 | 0 | 0 |
| NC_006577.2;YP_173238.1;HCoV-HKU1_spike;672 | 0 | 0 |
| NC_006577.2;YP_173238.1;HCoV-HKU1_spike;673 | 0 | 0 |
| NC_006577.2;YP_173238.1;HCoV-HKU1_spike;674 | 0 | 0 |
| NC_006577.2;YP_173238.1;HCoV-HKU1_spike;675 | 0 | 0 |
| NC_006577.2;YP_173238.1;HCoV-HKU1_spike;676 | 0 | 0 |
| NC_006577.2;YP_173238.1;HCoV-HKU1_spike;677 | 0 | 0 |
| NC_006577.2;YP_173238.1;HCoV-HKU1_spike;678 | 0 | 0 |
| NC_006577.2;YP_173238.1;HCoV-HKU1_spike;679 | 0 | 0 |
| NC_006577.2;YP_173238.1;HCoV-HKU1_spike;680 | 0 | 0 |
| NC_006577.2;YP_173238.1;HCoV-HKU1_spike;681 | 0 | 0 |
| NC_006577.2;YP_173238.1;HCoV-HKU1_spike;682 | 0 | 0 |

| | | |
|---|---|---|
| NC_006577.2;YP_173238.1;HCoV-HKU1_spike;683 | 0 | 0 |
| NC_006577.2;YP_173238.1;HCoV-HKU1_spike;684 | 0 | 0 |
| NC_006577.2;YP_173238.1;HCoV-HKU1_spike;685 | 0 | 0 |
| NC_006577.2;YP_173238.1;HCoV-HKU1_spike;686 | 0 | 0 |
| NC_006577.2;YP_173238.1;HCoV-HKU1_spike;687 | 0 | 0 |
| NC_006577.2;YP_173238.1;HCoV-HKU1_spike;688 | 0 | 0 |
| NC_006577.2;YP_173238.1;HCoV-HKU1_spike;689 | 0 | 0 |
| NC_006577.2;YP_173238.1;HCoV-HKU1_spike;690 | 0 | 0 |
| NC_006577.2;YP_173238.1;HCoV-HKU1_spike;691 | 0 | 0 |
| NC_006577.2;YP_173238.1;HCoV-HKU1_spike;692 | 0 | 0 |
| NC_006577.2;YP_173238.1;HCoV-HKU1_spike;693 | 0 | 0 |
| NC_006577.2;YP_173238.1;HCoV-HKU1_spike;694 | 0 | 0 |
| NC_006577.2;YP_173238.1;HCoV-HKU1_spike;695 | 0 | 0 |
| NC_006577.2;YP_173238.1;HCoV-HKU1_spike;696 | 0 | 0 |
| NC_006577.2;YP_173238.1;HCoV-HKU1_spike;697 | 0 | 0 |
| NC_006577.2;YP_173238.1;HCoV-HKU1_spike;698 | 0 | 0 |
| NC_006577.2;YP_173238.1;HCoV-HKU1_spike;699 | 0 | 0 |
| NC_006577.2;YP_173238.1;HCoV-HKU1_spike;700 | 0 | 0.025 |
| NC_006577.2;YP_173238.1;HCoV-HKU1_spike;701 | 0 | 0.05 |
| NC_006577.2;YP_173238.1;HCoV-HKU1_spike;702 | 0 | 0.05 |
| NC_006577.2;YP_173238.1;HCoV-HKU1_spike;703 | 0 | 0 |
| NC_006577.2;YP_173238.1;HCoV-HKU1_spike;704 | 0 | 0.025 |
| NC_006577.2;YP_173238.1;HCoV-HKU1_spike;705 | 0 | 0 |
| NC_006577.2;YP_173238.1;HCoV-HKU1_spike;706 | 0 | 0 |
| NC_006577.2;YP_173238.1;HCoV-HKU1_spike;707 | 0 | 0 |
| NC_006577.2;YP_173238.1;HCoV-HKU1_spike;708 | 0 | 0 |
| NC_006577.2;YP_173238.1;HCoV-HKU1_spike;709 | 0 | 0 |
| NC_006577.2;YP_173238.1;HCoV-HKU1_spike;710 | 0 | 0 |
| NC_006577.2;YP_173238.1;HCoV-HKU1_spike;711 | 0 | 0 |
| NC_006577.2;YP_173238.1;HCoV-HKU1_spike;712 | 0 | 0 |
| NC_006577.2;YP_173238.1;HCoV-HKU1_spike;713 | 0.05 | 0 |
| NC_006577.2;YP_173238.1;HCoV-HKU1_spike;714 | 0.05 | 0 |
| NC_006577.2;YP_173238.1;HCoV-HKU1_spike;715 | 0.05 | 0 |
| NC_006577.2;YP_173238.1;HCoV-HKU1_spike;716 | 0.05 | 0 |
| NC_006577.2;YP_173238.1;HCoV-HKU1_spike;717 | 0.05 | 0 |
| NC_006577.2;YP_173238.1;HCoV-HKU1_spike;718 | 0.05 | 0 |
| NC_006577.2;YP_173238.1;HCoV-HKU1_spike;719 | 0.05 | 0 |
| NC_006577.2;YP_173238.1;HCoV-HKU1_spike;720 | 0.05 | 0 |
| NC_006577.2;YP_173238.1;HCoV-HKU1_spike;721 | 0.05 | 0 |
| NC_006577.2;YP_173238.1;HCoV-HKU1_spike;722 | 0.05 | 0 |
| NC_006577.2;YP_173238.1;HCoV-HKU1_spike;723 | 0.05 | 0 |
| NC_006577.2;YP_173238.1;HCoV-HKU1_spike;724 | 0 | 0 |
| NC_006577.2;YP_173238.1;HCoV-HKU1_spike;725 | 0 | 0 |

| | | |
|---|---|---|
| NC_006577.2;YP_173238.1;HCoV-HKU1_spike;726 | 0 | 0 |
| NC_006577.2;YP_173238.1;HCoV-HKU1_spike;727 | 0 | 0 |
| NC_006577.2;YP_173238.1;HCoV-HKU1_spike;728 | 0 | 0 |
| NC_006577.2;YP_173238.1;HCoV-HKU1_spike;729 | 0 | 0 |
| NC_006577.2;YP_173238.1;HCoV-HKU1_spike;730 | 0 | 0 |
| NC_006577.2;YP_173238.1;HCoV-HKU1_spike;731 | 0 | 0.05 |
| NC_006577.2;YP_173238.1;HCoV-HKU1_spike;732 | 0 | 0 |
| NC_006577.2;YP_173238.1;HCoV-HKU1_spike;733 | 0 | 0 |
| NC_006577.2;YP_173238.1;HCoV-HKU1_spike;734 | 0 | 0 |
| NC_006577.2;YP_173238.1;HCoV-HKU1_spike;735 | 0 | 0 |
| NC_006577.2;YP_173238.1;HCoV-HKU1_spike;736 | 0 | 0.025 |
| NC_006577.2;YP_173238.1;HCoV-HKU1_spike;737 | 0 | 0.05 |
| NC_006577.2;YP_173238.1;HCoV-HKU1_spike;738 | 0 | 0.05 |
| NC_006577.2;YP_173238.1;HCoV-HKU1_spike;739 | 0 | 0.05 |
| NC_006577.2;YP_173238.1;HCoV-HKU1_spike;740 | 0.05 | 0.025 |
| NC_006577.2;YP_173238.1;HCoV-HKU1_spike;741 | 0 | 0 |
| NC_006577.2;YP_173238.1;HCoV-HKU1_spike;742 | 0 | 0 |
| NC_006577.2;YP_173238.1;HCoV-HKU1_spike;743 | 0 | 0 |
| NC_006577.2;YP_173238.1;HCoV-HKU1_spike;744 | 0 | 0 |
| NC_006577.2;YP_173238.1;HCoV-HKU1_spike;745 | 0 | 0.05 |
| NC_006577.2;YP_173238.1;HCoV-HKU1_spike;746 | 0 | 0.125 |
| NC_006577.2;YP_173238.1;HCoV-HKU1_spike;747 | 0 | 0.075 |
| NC_006577.2;YP_173238.1;HCoV-HKU1_spike;748 | 0 | 0.075 |
| NC_006577.2;YP_173238.1;HCoV-HKU1_spike;749 | 0 | 0.075 |
| NC_006577.2;YP_173238.1;HCoV-HKU1_spike;750 | 0.1 | 0.125 |
| NC_006577.2;YP_173238.1;HCoV-HKU1_spike;751 | 0 | 0.1 |
| NC_006577.2;YP_173238.1;HCoV-HKU1_spike;752 | 0 | 0.1 |
| NC_006577.2;YP_173238.1;HCoV-HKU1_spike;753 | 0 | 0.075 |
| NC_006577.2;YP_173238.1;HCoV-HKU1_spike;754 | 0 | 0.05 |
| NC_006577.2;YP_173238.1;HCoV-HKU1_spike;755 | 0 | 0.025 |
| NC_006577.2;YP_173238.1;HCoV-HKU1_spike;756 | 0 | 0 |
| NC_006577.2;YP_173238.1;HCoV-HKU1_spike;757 | 0 | 0.025 |
| NC_006577.2;YP_173238.1;HCoV-HKU1_spike;758 | 0.15 | 0.075 |
| NC_006577.2;YP_173238.1;HCoV-HKU1_spike;759 | 0.05 | 0.05 |
| NC_006577.2;YP_173238.1;HCoV-HKU1_spike;760 | 0.2 | 0.075 |
| NC_006577.2;YP_173238.1;HCoV-HKU1_spike;761 | 0.2 | 0.075 |
| NC_006577.2;YP_173238.1;HCoV-HKU1_spike;762 | 0.25 | 0.15 |
| NC_006577.2;YP_173238.1;HCoV-HKU1_spike;763 | 0.05 | 0.05 |
| NC_006577.2;YP_173238.1;HCoV-HKU1_spike;764 | 0.05 | 0.05 |
| NC_006577.2;YP_173238.1;HCoV-HKU1_spike;765 | 0.05 | 0.05 |
| NC_006577.2;YP_173238.1;HCoV-HKU1_spike;766 | 0.15 | 0.05 |
| NC_006577.2;YP_173238.1;HCoV-HKU1_spike;767 | 0.2 | 0.175 |
| NC_006577.2;YP_173238.1;HCoV-HKU1_spike;768 | 0.15 | 0.05 |

| | | |
|---|---|---|
| NC_006577.2;YP_173238.1;HCoV-HKU1_spike;769 | 0.05 | 0.15 |
| NC_006577.2;YP_173238.1;HCoV-HKU1_spike;770 | 0.2 | 0.2 |
| NC_006577.2;YP_173238.1;HCoV-HKU1_spike;771 | 0.05 | 0.05 |
| NC_006577.2;YP_173238.1;HCoV-HKU1_spike;772 | 0.05 | 0.025 |
| NC_006577.2;YP_173238.1;HCoV-HKU1_spike;773 | 0.05 | 0.025 |
| NC_006577.2;YP_173238.1;HCoV-HKU1_spike;774 | 0 | 0.05 |
| NC_006577.2;YP_173238.1;HCoV-HKU1_spike;775 | 0.05 | 0.075 |
| NC_006577.2;YP_173238.1;HCoV-HKU1_spike;776 | 0 | 0.075 |
| NC_006577.2;YP_173238.1;HCoV-HKU1_spike;777 | 0 | 0.05 |
| NC_006577.2;YP_173238.1;HCoV-HKU1_spike;778 | 0.05 | 0.025 |
| NC_006577.2;YP_173238.1;HCoV-HKU1_spike;779 | 0 | 0 |
| NC_006577.2;YP_173238.1;HCoV-HKU1_spike;780 | 0 | 0 |
| NC_006577.2;YP_173238.1;HCoV-HKU1_spike;781 | 0 | 0.075 |
| NC_006577.2;YP_173238.1;HCoV-HKU1_spike;782 | 0.05 | 0.05 |
| NC_006577.2;YP_173238.1;HCoV-HKU1_spike;783 | 0 | 0 |
| NC_006577.2;YP_173238.1;HCoV-HKU1_spike;784 | 0 | 0 |
| NC_006577.2;YP_173238.1;HCoV-HKU1_spike;785 | 0 | 0 |
| NC_006577.2;YP_173238.1;HCoV-HKU1_spike;786 | 0 | 0 |
| NC_006577.2;YP_173238.1;HCoV-HKU1_spike;787 | 0 | 0 |
| NC_006577.2;YP_173238.1;HCoV-HKU1_spike;788 | 0 | 0 |
| NC_006577.2;YP_173238.1;HCoV-HKU1_spike;789 | 0 | 0 |
| NC_006577.2;YP_173238.1;HCoV-HKU1_spike;790 | 0 | 0 |
| NC_006577.2;YP_173238.1;HCoV-HKU1_spike;791 | 0 | 0 |
| NC_006577.2;YP_173238.1;HCoV-HKU1_spike;792 | 0 | 0.025 |
| NC_006577.2;YP_173238.1;HCoV-HKU1_spike;793 | 0 | 0.025 |
| NC_006577.2;YP_173238.1;HCoV-HKU1_spike;794 | 0 | 0.025 |
| NC_006577.2;YP_173238.1;HCoV-HKU1_spike;795 | 0 | 0.025 |
| NC_006577.2;YP_173238.1;HCoV-HKU1_spike;796 | 0 | 0.025 |
| NC_006577.2;YP_173238.1;HCoV-HKU1_spike;797 | 0 | 0.025 |
| NC_006577.2;YP_173238.1;HCoV-HKU1_spike;798 | 0.05 | 0.15 |
| NC_006577.2;YP_173238.1;HCoV-HKU1_spike;799 | 0.05 | 0.1 |
| NC_006577.2;YP_173238.1;HCoV-HKU1_spike;800 | 0.05 | 0.15 |
| NC_006577.2;YP_173238.1;HCoV-HKU1_spike;801 | 0.1 | 0.1 |
| NC_006577.2;YP_173238.1;HCoV-HKU1_spike;802 | 0.15 | 0.1 |
| NC_006577.2;YP_173238.1;HCoV-HKU1_spike;803 | 0.05 | 0.05 |
| NC_006577.2;YP_173238.1;HCoV-HKU1_spike;804 | 0.1 | 0.05 |
| NC_006577.2;YP_173238.1;HCoV-HKU1_spike;805 | 0.05 | 0.025 |
| NC_006577.2;YP_173238.1;HCoV-HKU1_spike;806 | 0 | 0 |
| NC_006577.2;YP_173238.1;HCoV-HKU1_spike;807 | 0 | 0.05 |
| NC_006577.2;YP_173238.1;HCoV-HKU1_spike;808 | 0 | 0 |
| NC_006577.2;YP_173238.1;HCoV-HKU1_spike;809 | 0 | 0 |
| NC_006577.2;YP_173238.1;HCoV-HKU1_spike;810 | 0 | 0 |
| NC_006577.2;YP_173238.1;HCoV-HKU1_spike;811 | 0 | 0 |

| | | |
|---|---|---|
| NC_006577.2;YP_173238.1;HCoV-HKU1_spike;812 | 0 | 0 |
| NC_006577.2;YP_173238.1;HCoV-HKU1_spike;813 | 0.05 | 0 |
| NC_006577.2;YP_173238.1;HCoV-HKU1_spike;814 | 0 | 0 |
| NC_006577.2;YP_173238.1;HCoV-HKU1_spike;815 | 0 | 0 |
| NC_006577.2;YP_173238.1;HCoV-HKU1_spike;816 | 0 | 0 |
| NC_006577.2;YP_173238.1;HCoV-HKU1_spike;817 | 0 | 0 |
| NC_006577.2;YP_173238.1;HCoV-HKU1_spike;818 | 0 | 0 |
| NC_006577.2;YP_173238.1;HCoV-HKU1_spike;819 | 0 | 0 |
| NC_006577.2;YP_173238.1;HCoV-HKU1_spike;820 | 0.1 | 0 |
| NC_006577.2;YP_173238.1;HCoV-HKU1_spike;821 | 0 | 0 |
| NC_006577.2;YP_173238.1;HCoV-HKU1_spike;822 | 0 | 0 |
| NC_006577.2;YP_173238.1;HCoV-HKU1_spike;823 | 0 | 0 |
| NC_006577.2;YP_173238.1;HCoV-HKU1_spike;824 | 0 | 0 |
| NC_006577.2;YP_173238.1;HCoV-HKU1_spike;825 | 0 | 0 |
| NC_006577.2;YP_173238.1;HCoV-HKU1_spike;826 | 0 | 0 |
| NC_006577.2;YP_173238.1;HCoV-HKU1_spike;827 | 0 | 0 |
| NC_006577.2;YP_173238.1;HCoV-HKU1_spike;828 | 0 | 0 |
| NC_006577.2;YP_173238.1;HCoV-HKU1_spike;829 | 0 | 0 |
| NC_006577.2;YP_173238.1;HCoV-HKU1_spike;830 | 0 | 0 |
| NC_006577.2;YP_173238.1;HCoV-HKU1_spike;831 | 0.05 | 0 |
| NC_006577.2;YP_173238.1;HCoV-HKU1_spike;832 | 0 | 0 |
| NC_006577.2;YP_173238.1;HCoV-HKU1_spike;833 | 0.05 | 0.025 |
| NC_006577.2;YP_173238.1;HCoV-HKU1_spike;834 | 0.1 | 0.025 |
| NC_006577.2;YP_173238.1;HCoV-HKU1_spike;835 | 0.1 | 0.025 |
| NC_006577.2;YP_173238.1;HCoV-HKU1_spike;836 | 0.15 | 0.075 |
| NC_006577.2;YP_173238.1;HCoV-HKU1_spike;837 | 0.15 | 0.025 |
| NC_006577.2;YP_173238.1;HCoV-HKU1_spike;838 | 0.1 | 0.025 |
| NC_006577.2;YP_173238.1;HCoV-HKU1_spike;839 | 0.1 | 0.05 |
| NC_006577.2;YP_173238.1;HCoV-HKU1_spike;840 | 0.05 | 0.025 |
| NC_006577.2;YP_173238.1;HCoV-HKU1_spike;841 | 0.05 | 0.025 |
| NC_006577.2;YP_173238.1;HCoV-HKU1_spike;842 | 0 | 0 |
| NC_006577.2;YP_173238.1;HCoV-HKU1_spike;843 | 0.1 | 0.05 |
| NC_006577.2;YP_173238.1;HCoV-HKU1_spike;844 | 0.05 | 0.05 |
| NC_006577.2;YP_173238.1;HCoV-HKU1_spike;845 | 0 | 0.025 |
| NC_006577.2;YP_173238.1;HCoV-HKU1_spike;846 | 0.05 | 0.025 |
| NC_006577.2;YP_173238.1;HCoV-HKU1_spike;847 | 0.05 | 0 |
| NC_006577.2;YP_173238.1;HCoV-HKU1_spike;848 | 0.15 | 0.1 |
| NC_006577.2;YP_173238.1;HCoV-HKU1_spike;849 | 0.1 | 0.025 |
| NC_006577.2;YP_173238.1;HCoV-HKU1_spike;850 | 0.05 | 0.05 |
| NC_006577.2;YP_173238.1;HCoV-HKU1_spike;851 | 0.05 | 0.05 |
| NC_006577.2;YP_173238.1;HCoV-HKU1_spike;852 | 0.15 | 0.125 |
| NC_006577.2;YP_173238.1;HCoV-HKU1_spike;853 | 0.05 | 0.05 |
| NC_006577.2;YP_173238.1;HCoV-HKU1_spike;854 | 0.05 | 0.05 |

| | | |
|---|---|---|
| NC_006577.2;YP_173238.1;HCoV-HKU1_spike;855 | 0.05 | 0.05 |
| NC_006577.2;YP_173238.1;HCoV-HKU1_spike;856 | 0.05 | 0.05 |
| NC_006577.2;YP_173238.1;HCoV-HKU1_spike;857 | 0 | 0.05 |
| NC_006577.2;YP_173238.1;HCoV-HKU1_spike;858 | 0 | 0.075 |
| NC_006577.2;YP_173238.1;HCoV-HKU1_spike;859 | 0.05 | 0.075 |
| NC_006577.2;YP_173238.1;HCoV-HKU1_spike;860 | 0.05 | 0.05 |
| NC_006577.2;YP_173238.1;HCoV-HKU1_spike;861 | 0.05 | 0.05 |
| NC_006577.2;YP_173238.1;HCoV-HKU1_spike;862 | 0.1 | 0.05 |
| NC_006577.2;YP_173238.1;HCoV-HKU1_spike;863 | 0.05 | 0.075 |
| NC_006577.2;YP_173238.1;HCoV-HKU1_spike;864 | 0.05 | 0.075 |
| NC_006577.2;YP_173238.1;HCoV-HKU1_spike;865 | 0.05 | 0.05 |
| NC_006577.2;YP_173238.1;HCoV-HKU1_spike;866 | 0.05 | 0.075 |
| NC_006577.2;YP_173238.1;HCoV-HKU1_spike;867 | 0.05 | 0.05 |
| NC_006577.2;YP_173238.1;HCoV-HKU1_spike;868 | 0.05 | 0.025 |
| NC_006577.2;YP_173238.1;HCoV-HKU1_spike;869 | 0 | 0.05 |
| NC_006577.2;YP_173238.1;HCoV-HKU1_spike;870 | 0 | 0.025 |
| NC_006577.2;YP_173238.1;HCoV-HKU1_spike;871 | 0.1 | 0.05 |
| NC_006577.2;YP_173238.1;HCoV-HKU1_spike;872 | 0.15 | 0.25 |
| NC_006577.2;YP_173238.1;HCoV-HKU1_spike;873 | 0.1 | 0.05 |
| NC_006577.2;YP_173238.1;HCoV-HKU1_spike;874 | 0.1 | 0.1 |
| NC_006577.2;YP_173238.1;HCoV-HKU1_spike;875 | 0.1 | 0.075 |
| NC_006577.2;YP_173238.1;HCoV-HKU1_spike;876 | 0.15 | 0.025 |
| NC_006577.2;YP_173238.1;HCoV-HKU1_spike;877 | 0.05 | 0.05 |
| NC_006577.2;YP_173238.1;HCoV-HKU1_spike;878 | 0 | 0.025 |
| NC_006577.2;YP_173238.1;HCoV-HKU1_spike;879 | 0 | 0 |
| NC_006577.2;YP_173238.1;HCoV-HKU1_spike;880 | 0 | 0 |
| NC_006577.2;YP_173238.1;HCoV-HKU1_spike;881 | 0 | 0 |
| NC_006577.2;YP_173238.1;HCoV-HKU1_spike;882 | 0.05 | 0.025 |
| NC_006577.2;YP_173238.1;HCoV-HKU1_spike;883 | 0 | 0 |
| NC_006577.2;YP_173238.1;HCoV-HKU1_spike;884 | 0 | 0 |
| NC_006577.2;YP_173238.1;HCoV-HKU1_spike;885 | 0 | 0 |
| NC_006577.2;YP_173238.1;HCoV-HKU1_spike;886 | 0 | 0 |
| NC_006577.2;YP_173238.1;HCoV-HKU1_spike;887 | 0 | 0 |
| NC_006577.2;YP_173238.1;HCoV-HKU1_spike;888 | 0 | 0 |
| NC_006577.2;YP_173238.1;HCoV-HKU1_spike;889 | 0 | 0 |
| NC_006577.2;YP_173238.1;HCoV-HKU1_spike;890 | 0 | 0 |
| NC_006577.2;YP_173238.1;HCoV-HKU1_spike;891 | 0 | 0 |
| NC_006577.2;YP_173238.1;HCoV-HKU1_spike;892 | 0 | 0 |
| NC_006577.2;YP_173238.1;HCoV-HKU1_spike;893 | 0 | 0 |
| NC_006577.2;YP_173238.1;HCoV-HKU1_spike;894 | 0 | 0.05 |
| NC_006577.2;YP_173238.1;HCoV-HKU1_spike;895 | 0 | 0.05 |
| NC_006577.2;YP_173238.1;HCoV-HKU1_spike;896 | 0 | 0 |
| NC_006577.2;YP_173238.1;HCoV-HKU1_spike;897 | 0 | 0.25 |

| | | |
|---|---|---|
| NC_006577.2;YP_173238.1;HCoV-HKU1_spike;898 | 0.05 | 0.3 |
| NC_006577.2;YP_173238.1;HCoV-HKU1_spike;899 | 0.15 | 0.6 |
| NC_006577.2;YP_173238.1;HCoV-HKU1_spike;900 | 0.05 | 0.625 |
| NC_006577.2;YP_173238.1;HCoV-HKU1_spike;901 | 0.05 | 0.625 |
| NC_006577.2;YP_173238.1;HCoV-HKU1_spike;902 | 0.1 | 0.575 |
| NC_006577.2;YP_173238.1;HCoV-HKU1_spike;903 | 0 | 0.525 |
| NC_006577.2;YP_173238.1;HCoV-HKU1_spike;904 | 0 | 0.55 |
| NC_006577.2;YP_173238.1;HCoV-HKU1_spike;905 | 0 | 0.325 |
| NC_006577.2;YP_173238.1;HCoV-HKU1_spike;906 | 0 | 0.35 |
| NC_006577.2;YP_173238.1;HCoV-HKU1_spike;907 | 0 | 0.275 |
| NC_006577.2;YP_173238.1;HCoV-HKU1_spike;908 | 0 | 0.125 |
| NC_006577.2;YP_173238.1;HCoV-HKU1_spike;909 | 0 | 0 |
| NC_006577.2;YP_173238.1;HCoV-HKU1_spike;910 | 0 | 0 |
| NC_006577.2;YP_173238.1;HCoV-HKU1_spike;911 | 0 | 0 |
| NC_006577.2;YP_173238.1;HCoV-HKU1_spike;912 | 0 | 0.025 |
| NC_006577.2;YP_173238.1;HCoV-HKU1_spike;913 | 0 | 0.025 |
| NC_006577.2;YP_173238.1;HCoV-HKU1_spike;914 | 0.05 | 0.025 |
| NC_006577.2;YP_173238.1;HCoV-HKU1_spike;915 | 0.05 | 0.025 |
| NC_006577.2;YP_173238.1;HCoV-HKU1_spike;916 | 0.05 | 0.075 |
| NC_006577.2;YP_173238.1;HCoV-HKU1_spike;917 | 0 | 0.025 |
| NC_006577.2;YP_173238.1;HCoV-HKU1_spike;918 | 0.05 | 0.05 |
| NC_006577.2;YP_173238.1;HCoV-HKU1_spike;919 | 0 | 0.075 |
| NC_006577.2;YP_173238.1;HCoV-HKU1_spike;920 | 0 | 0.075 |
| NC_006577.2;YP_173238.1;HCoV-HKU1_spike;921 | 0 | 0 |
| NC_006577.2;YP_173238.1;HCoV-HKU1_spike;922 | 0 | 0.025 |
| NC_006577.2;YP_173238.1;HCoV-HKU1_spike;923 | 0 | 0 |
| NC_006577.2;YP_173238.1;HCoV-HKU1_spike;924 | 0 | 0 |
| NC_006577.2;YP_173238.1;HCoV-HKU1_spike;925 | 0 | 0 |
| NC_006577.2;YP_173238.1;HCoV-HKU1_spike;926 | 0 | 0 |
| NC_006577.2;YP_173238.1;HCoV-HKU1_spike;927 | 0 | 0 |
| NC_006577.2;YP_173238.1;HCoV-HKU1_spike;928 | 0 | 0 |
| NC_006577.2;YP_173238.1;HCoV-HKU1_spike;929 | 0 | 0 |
| NC_006577.2;YP_173238.1;HCoV-HKU1_spike;930 | 0 | 0 |
| NC_006577.2;YP_173238.1;HCoV-HKU1_spike;931 | 0.05 | 0 |
| NC_006577.2;YP_173238.1;HCoV-HKU1_spike;932 | 0 | 0 |
| NC_006577.2;YP_173238.1;HCoV-HKU1_spike;933 | 0 | 0.025 |
| NC_006577.2;YP_173238.1;HCoV-HKU1_spike;934 | 0 | 0 |
| NC_006577.2;YP_173238.1;HCoV-HKU1_spike;935 | 0 | 0 |
| NC_006577.2;YP_173238.1;HCoV-HKU1_spike;936 | 0 | 0 |
| NC_006577.2;YP_173238.1;HCoV-HKU1_spike;937 | 0 | 0 |
| NC_006577.2;YP_173238.1;HCoV-HKU1_spike;938 | 0 | 0 |
| NC_006577.2;YP_173238.1;HCoV-HKU1_spike;939 | 0 | 0 |
| NC_006577.2;YP_173238.1;HCoV-HKU1_spike;940 | 0 | 0 |

| | | |
|---|---|---|
| NC_006577.2;YP_173238.1;HCoV-HKU1_spike;941 | 0 | 0 |
| NC_006577.2;YP_173238.1;HCoV-HKU1_spike;942 | 0 | 0 |
| NC_006577.2;YP_173238.1;HCoV-HKU1_spike;943 | 0 | 0 |
| NC_006577.2;YP_173238.1;HCoV-HKU1_spike;944 | 0 | 0 |
| NC_006577.2;YP_173238.1;HCoV-HKU1_spike;945 | 0 | 0 |
| NC_006577.2;YP_173238.1;HCoV-HKU1_spike;946 | 0 | 0 |
| NC_006577.2;YP_173238.1;HCoV-HKU1_spike;947 | 0 | 0 |
| NC_006577.2;YP_173238.1;HCoV-HKU1_spike;948 | 0 | 0 |
| NC_006577.2;YP_173238.1;HCoV-HKU1_spike;949 | 0 | 0 |
| NC_006577.2;YP_173238.1;HCoV-HKU1_spike;950 | 0 | 0 |
| NC_006577.2;YP_173238.1;HCoV-HKU1_spike;951 | 0.05 | 0.025 |
| NC_006577.2;YP_173238.1;HCoV-HKU1_spike;952 | 0 | 0 |
| NC_006577.2;YP_173238.1;HCoV-HKU1_spike;953 | 0 | 0 |
| NC_006577.2;YP_173238.1;HCoV-HKU1_spike;954 | 0.05 | 0.05 |
| NC_006577.2;YP_173238.1;HCoV-HKU1_spike;955 | 0.05 | 0.025 |
| NC_006577.2;YP_173238.1;HCoV-HKU1_spike;956 | 0 | 0 |
| NC_006577.2;YP_173238.1;HCoV-HKU1_spike;957 | 0 | 0.025 |
| NC_006577.2;YP_173238.1;HCoV-HKU1_spike;958 | 0 | 0 |
| NC_006577.2;YP_173238.1;HCoV-HKU1_spike;959 | 0 | 0 |
| NC_006577.2;YP_173238.1;HCoV-HKU1_spike;960 | 0 | 0 |
| NC_006577.2;YP_173238.1;HCoV-HKU1_spike;961 | 0 | 0 |
| NC_006577.2;YP_173238.1;HCoV-HKU1_spike;962 | 0 | 0 |
| NC_006577.2;YP_173238.1;HCoV-HKU1_spike;963 | 0.05 | 0.025 |
| NC_006577.2;YP_173238.1;HCoV-HKU1_spike;964 | 0.05 | 0.025 |
| NC_006577.2;YP_173238.1;HCoV-HKU1_spike;965 | 0.05 | 0.025 |
| NC_006577.2;YP_173238.1;HCoV-HKU1_spike;966 | 0.05 | 0.025 |
| NC_006577.2;YP_173238.1;HCoV-HKU1_spike;967 | 0 | 0 |
| NC_006577.2;YP_173238.1;HCoV-HKU1_spike;968 | 0 | 0 |
| NC_006577.2;YP_173238.1;HCoV-HKU1_spike;969 | 0 | 0 |
| NC_006577.2;YP_173238.1;HCoV-HKU1_spike;970 | 0 | 0 |
| NC_006577.2;YP_173238.1;HCoV-HKU1_spike;971 | 0 | 0 |
| NC_006577.2;YP_173238.1;HCoV-HKU1_spike;972 | 0 | 0 |
| NC_006577.2;YP_173238.1;HCoV-HKU1_spike;973 | 0 | 0 |
| NC_006577.2;YP_173238.1;HCoV-HKU1_spike;974 | 0 | 0 |
| NC_006577.2;YP_173238.1;HCoV-HKU1_spike;975 | 0 | 0 |
| NC_006577.2;YP_173238.1;HCoV-HKU1_spike;976 | 0 | 0 |
| NC_006577.2;YP_173238.1;HCoV-HKU1_spike;977 | 0 | 0 |
| NC_006577.2;YP_173238.1;HCoV-HKU1_spike;978 | 0 | 0 |
| NC_006577.2;YP_173238.1;HCoV-HKU1_spike;979 | 0 | 0 |
| NC_006577.2;YP_173238.1;HCoV-HKU1_spike;980 | 0 | 0 |
| NC_006577.2;YP_173238.1;HCoV-HKU1_spike;981 | 0 | 0 |
| NC_006577.2;YP_173238.1;HCoV-HKU1_spike;982 | 0 | 0 |
| NC_006577.2;YP_173238.1;HCoV-HKU1_spike;983 | 0 | 0 |

| | | |
|---|---|---|
| NC_006577.2;YP_173238.1;HCoV-HKU1_spike;984 | 0.05 | 0.025 |
| NC_006577.2;YP_173238.1;HCoV-HKU1_spike;985 | 0 | 0.025 |
| NC_006577.2;YP_173238.1;HCoV-HKU1_spike;986 | 0 | 0.025 |
| NC_006577.2;YP_173238.1;HCoV-HKU1_spike;987 | 0 | 0.025 |
| NC_006577.2;YP_173238.1;HCoV-HKU1_spike;988 | 0 | 0.05 |
| NC_006577.2;YP_173238.1;HCoV-HKU1_spike;989 | 0 | 0.05 |
| NC_006577.2;YP_173238.1;HCoV-HKU1_spike;990 | 0 | 0.075 |
| NC_006577.2;YP_173238.1;HCoV-HKU1_spike;991 | 0 | 0.025 |
| NC_006577.2;YP_173238.1;HCoV-HKU1_spike;992 | 0 | 0.025 |
| NC_006577.2;YP_173238.1;HCoV-HKU1_spike;993 | 0 | 0.025 |
| NC_006577.2;YP_173238.1;HCoV-HKU1_spike;994 | 0 | 0.025 |
| NC_006577.2;YP_173238.1;HCoV-HKU1_spike;995 | 0 | 0.025 |
| NC_006577.2;YP_173238.1;HCoV-HKU1_spike;996 | 0 | 0.025 |
| NC_006577.2;YP_173238.1;HCoV-HKU1_spike;997 | 0 | 0 |
| NC_006577.2;YP_173238.1;HCoV-HKU1_spike;998 | 0 | 0 |
| NC_006577.2;YP_173238.1;HCoV-HKU1_spike;999 | 0.1 | 0 |
| NC_006577.2;YP_173238.1;HCoV-HKU1_spike;1000 | 0 | 0 |
| NC_006577.2;YP_173238.1;HCoV-HKU1_spike;1001 | 0 | 0.025 |
| NC_006577.2;YP_173238.1;HCoV-HKU1_spike;1002 | 0 | 0 |
| NC_006577.2;YP_173238.1;HCoV-HKU1_spike;1003 | 0 | 0.025 |
| NC_006577.2;YP_173238.1;HCoV-HKU1_spike;1004 | 0 | 0 |
| NC_006577.2;YP_173238.1;HCoV-HKU1_spike;1005 | 0 | 0 |
| NC_006577.2;YP_173238.1;HCoV-HKU1_spike;1006 | 0 | 0 |
| NC_006577.2;YP_173238.1;HCoV-HKU1_spike;1007 | 0 | 0 |
| NC_006577.2;YP_173238.1;HCoV-HKU1_spike;1008 | 0 | 0.025 |
| NC_006577.2;YP_173238.1;HCoV-HKU1_spike;1009 | 0 | 0.025 |
| NC_006577.2;YP_173238.1;HCoV-HKU1_spike;1010 | 0.05 | 0.025 |
| NC_006577.2;YP_173238.1;HCoV-HKU1_spike;1011 | 0 | 0 |
| NC_006577.2;YP_173238.1;HCoV-HKU1_spike;1012 | 0 | 0 |
| NC_006577.2;YP_173238.1;HCoV-HKU1_spike;1013 | 0 | 0 |
| NC_006577.2;YP_173238.1;HCoV-HKU1_spike;1014 | 0.05 | 0.025 |
| NC_006577.2;YP_173238.1;HCoV-HKU1_spike;1015 | 0 | 0 |
| NC_006577.2;YP_173238.1;HCoV-HKU1_spike;1016 | 0 | 0 |
| NC_006577.2;YP_173238.1;HCoV-HKU1_spike;1017 | 0 | 0 |
| NC_006577.2;YP_173238.1;HCoV-HKU1_spike;1018 | 0 | 0 |
| NC_006577.2;YP_173238.1;HCoV-HKU1_spike;1019 | 0 | 0 |
| NC_006577.2;YP_173238.1;HCoV-HKU1_spike;1020 | 0 | 0 |
| NC_006577.2;YP_173238.1;HCoV-HKU1_spike;1021 | 0 | 0 |
| NC_006577.2;YP_173238.1;HCoV-HKU1_spike;1022 | 0 | 0 |
| NC_006577.2;YP_173238.1;HCoV-HKU1_spike;1023 | 0 | 0 |
| NC_006577.2;YP_173238.1;HCoV-HKU1_spike;1024 | 0 | 0 |
| NC_006577.2;YP_173238.1;HCoV-HKU1_spike;1025 | 0.05 | 0 |
| NC_006577.2;YP_173238.1;HCoV-HKU1_spike;1026 | 0 | 0.025 |

| | | |
|---|---|---|
| NC_006577.2;YP_173238.1;HCoV-HKU1_spike;1027 | 0 | 0.025 |
| NC_006577.2;YP_173238.1;HCoV-HKU1_spike;1028 | 0 | 0 |
| NC_006577.2;YP_173238.1;HCoV-HKU1_spike;1029 | 0 | 0 |
| NC_006577.2;YP_173238.1;HCoV-HKU1_spike;1030 | 0 | 0 |
| NC_006577.2;YP_173238.1;HCoV-HKU1_spike;1031 | 0.05 | 0.075 |
| NC_006577.2;YP_173238.1;HCoV-HKU1_spike;1032 | 0 | 0 |
| NC_006577.2;YP_173238.1;HCoV-HKU1_spike;1033 | 0.05 | 0.05 |
| NC_006577.2;YP_173238.1;HCoV-HKU1_spike;1034 | 0.05 | 0.025 |
| NC_006577.2;YP_173238.1;HCoV-HKU1_spike;1035 | 0.15 | 0.05 |
| NC_006577.2;YP_173238.1;HCoV-HKU1_spike;1036 | 0.1 | 0.1 |
| NC_006577.2;YP_173238.1;HCoV-HKU1_spike;1037 | 0 | 0.025 |
| NC_006577.2;YP_173238.1;HCoV-HKU1_spike;1038 | 0 | 0 |
| NC_006577.2;YP_173238.1;HCoV-HKU1_spike;1039 | 0 | 0 |
| NC_006577.2;YP_173238.1;HCoV-HKU1_spike;1040 | 0 | 0 |
| NC_006577.2;YP_173238.1;HCoV-HKU1_spike;1041 | 0 | 0 |
| NC_006577.2;YP_173238.1;HCoV-HKU1_spike;1042 | 0 | 0 |
| NC_006577.2;YP_173238.1;HCoV-HKU1_spike;1043 | 0 | 0 |
| NC_006577.2;YP_173238.1;HCoV-HKU1_spike;1044 | 0 | 0 |
| NC_006577.2;YP_173238.1;HCoV-HKU1_spike;1045 | 0 | 0 |
| NC_006577.2;YP_173238.1;HCoV-HKU1_spike;1046 | 0 | 0 |
| NC_006577.2;YP_173238.1;HCoV-HKU1_spike;1047 | 0 | 0 |
| NC_006577.2;YP_173238.1;HCoV-HKU1_spike;1048 | 0 | 0 |
| NC_006577.2;YP_173238.1;HCoV-HKU1_spike;1049 | 0 | 0 |
| NC_006577.2;YP_173238.1;HCoV-HKU1_spike;1050 | 0 | 0 |
| NC_006577.2;YP_173238.1;HCoV-HKU1_spike;1051 | 0 | 0 |
| NC_006577.2;YP_173238.1;HCoV-HKU1_spike;1052 | 0 | 0 |
| NC_006577.2;YP_173238.1;HCoV-HKU1_spike;1053 | 0 | 0 |
| NC_006577.2;YP_173238.1;HCoV-HKU1_spike;1054 | 0 | 0 |
| NC_006577.2;YP_173238.1;HCoV-HKU1_spike;1055 | 0 | 0 |
| NC_006577.2;YP_173238.1;HCoV-HKU1_spike;1056 | 0 | 0 |
| NC_006577.2;YP_173238.1;HCoV-HKU1_spike;1057 | 0.05 | 0 |
| NC_006577.2;YP_173238.1;HCoV-HKU1_spike;1058 | 0 | 0 |
| NC_006577.2;YP_173238.1;HCoV-HKU1_spike;1059 | 0 | 0.05 |
| NC_006577.2;YP_173238.1;HCoV-HKU1_spike;1060 | 0 | 0.025 |
| NC_006577.2;YP_173238.1;HCoV-HKU1_spike;1061 | 0 | 0 |
| NC_006577.2;YP_173238.1;HCoV-HKU1_spike;1062 | 0 | 0 |
| NC_006577.2;YP_173238.1;HCoV-HKU1_spike;1063 | 0 | 0 |
| NC_006577.2;YP_173238.1;HCoV-HKU1_spike;1064 | 0 | 0 |
| NC_006577.2;YP_173238.1;HCoV-HKU1_spike;1065 | 0.1 | 0 |
| NC_006577.2;YP_173238.1;HCoV-HKU1_spike;1066 | 0.05 | 0 |
| NC_006577.2;YP_173238.1;HCoV-HKU1_spike;1067 | 0 | 0.05 |
| NC_006577.2;YP_173238.1;HCoV-HKU1_spike;1068 | 0 | 0.025 |
| NC_006577.2;YP_173238.1;HCoV-HKU1_spike;1069 | 0 | 0.05 |

| | | |
|---|---|---|
| NC_006577.2;YP_173238.1;HCoV-HKU1_spike;1070 | 0 | 0.05 |
| NC_006577.2;YP_173238.1;HCoV-HKU1_spike;1071 | 0.05 | 0.025 |
| NC_006577.2;YP_173238.1;HCoV-HKU1_spike;1072 | 0 | 0.025 |
| NC_006577.2;YP_173238.1;HCoV-HKU1_spike;1073 | 0.05 | 0.025 |
| NC_006577.2;YP_173238.1;HCoV-HKU1_spike;1074 | 0 | 0.025 |
| NC_006577.2;YP_173238.1;HCoV-HKU1_spike;1075 | 0 | 0 |
| NC_006577.2;YP_173238.1;HCoV-HKU1_spike;1076 | 0 | 0 |
| NC_006577.2;YP_173238.1;HCoV-HKU1_spike;1077 | 0 | 0 |
| NC_006577.2;YP_173238.1;HCoV-HKU1_spike;1078 | 0 | 0 |
| NC_006577.2;YP_173238.1;HCoV-HKU1_spike;1079 | 0 | 0 |
| NC_006577.2;YP_173238.1;HCoV-HKU1_spike;1080 | 0 | 0 |
| NC_006577.2;YP_173238.1;HCoV-HKU1_spike;1081 | 0 | 0 |
| NC_006577.2;YP_173238.1;HCoV-HKU1_spike;1082 | 0 | 0 |
| NC_006577.2;YP_173238.1;HCoV-HKU1_spike;1083 | 0 | 0 |
| NC_006577.2;YP_173238.1;HCoV-HKU1_spike;1084 | 0 | 0 |
| NC_006577.2;YP_173238.1;HCoV-HKU1_spike;1085 | 0 | 0 |
| NC_006577.2;YP_173238.1;HCoV-HKU1_spike;1086 | 0 | 0.025 |
| NC_006577.2;YP_173238.1;HCoV-HKU1_spike;1087 | 0 | 0.025 |
| NC_006577.2;YP_173238.1;HCoV-HKU1_spike;1088 | 0 | 0.025 |
| NC_006577.2;YP_173238.1;HCoV-HKU1_spike;1089 | 0 | 0 |
| NC_006577.2;YP_173238.1;HCoV-HKU1_spike;1090 | 0 | 0 |
| NC_006577.2;YP_173238.1;HCoV-HKU1_spike;1091 | 0 | 0.025 |
| NC_006577.2;YP_173238.1;HCoV-HKU1_spike;1092 | 0 | 0 |
| NC_006577.2;YP_173238.1;HCoV-HKU1_spike;1093 | 0.05 | 0 |
| NC_006577.2;YP_173238.1;HCoV-HKU1_spike;1094 | 0 | 0 |
| NC_006577.2;YP_173238.1;HCoV-HKU1_spike;1095 | 0 | 0 |
| NC_006577.2;YP_173238.1;HCoV-HKU1_spike;1096 | 0 | 0 |
| NC_006577.2;YP_173238.1;HCoV-HKU1_spike;1097 | 0 | 0 |
| NC_006577.2;YP_173238.1;HCoV-HKU1_spike;1098 | 0.05 | 0.025 |
| NC_006577.2;YP_173238.1;HCoV-HKU1_spike;1099 | 0 | 0 |
| NC_006577.2;YP_173238.1;HCoV-HKU1_spike;1100 | 0 | 0 |
| NC_006577.2;YP_173238.1;HCoV-HKU1_spike;1101 | 0.05 | 0 |
| NC_006577.2;YP_173238.1;HCoV-HKU1_spike;1102 | 0.05 | 0 |
| NC_006577.2;YP_173238.1;HCoV-HKU1_spike;1103 | 0.05 | 0.025 |
| NC_006577.2;YP_173238.1;HCoV-HKU1_spike;1104 | 0.05 | 0 |
| NC_006577.2;YP_173238.1;HCoV-HKU1_spike;1105 | 0 | 0.025 |
| NC_006577.2;YP_173238.1;HCoV-HKU1_spike;1106 | 0 | 0.025 |
| NC_006577.2;YP_173238.1;HCoV-HKU1_spike;1107 | 0 | 0.025 |
| NC_006577.2;YP_173238.1;HCoV-HKU1_spike;1108 | 0 | 0 |
| NC_006577.2;YP_173238.1;HCoV-HKU1_spike;1109 | 0 | 0 |
| NC_006577.2;YP_173238.1;HCoV-HKU1_spike;1110 | 0 | 0 |
| NC_006577.2;YP_173238.1;HCoV-HKU1_spike;1111 | 0 | 0 |
| NC_006577.2;YP_173238.1;HCoV-HKU1_spike;1112 | 0 | 0 |

| | | |
|---|---|---|
| NC_006577.2;YP_173238.1;HCoV-HKU1_spike;1113 | 0 | 0.025 |
| NC_006577.2;YP_173238.1;HCoV-HKU1_spike;1114 | 0 | 0 |
| NC_006577.2;YP_173238.1;HCoV-HKU1_spike;1115 | 0 | 0 |
| NC_006577.2;YP_173238.1;HCoV-HKU1_spike;1116 | 0 | 0 |
| NC_006577.2;YP_173238.1;HCoV-HKU1_spike;1117 | 0 | 0.025 |
| NC_006577.2;YP_173238.1;HCoV-HKU1_spike;1118 | 0 | 0 |
| NC_006577.2;YP_173238.1;HCoV-HKU1_spike;1119 | 0 | 0.025 |
| NC_006577.2;YP_173238.1;HCoV-HKU1_spike;1120 | 0 | 0 |
| NC_006577.2;YP_173238.1;HCoV-HKU1_spike;1121 | 0 | 0 |
| NC_006577.2;YP_173238.1;HCoV-HKU1_spike;1122 | 0 | 0 |
| NC_006577.2;YP_173238.1;HCoV-HKU1_spike;1123 | 0 | 0 |
| NC_006577.2;YP_173238.1;HCoV-HKU1_spike;1124 | 0 | 0 |
| NC_006577.2;YP_173238.1;HCoV-HKU1_spike;1125 | 0 | 0 |
| NC_006577.2;YP_173238.1;HCoV-HKU1_spike;1126 | 0 | 0 |
| NC_006577.2;YP_173238.1;HCoV-HKU1_spike;1127 | 0 | 0 |
| NC_006577.2;YP_173238.1;HCoV-HKU1_spike;1128 | 0 | 0 |
| NC_006577.2;YP_173238.1;HCoV-HKU1_spike;1129 | 0 | 0 |
| NC_006577.2;YP_173238.1;HCoV-HKU1_spike;1130 | 0 | 0 |
| NC_006577.2;YP_173238.1;HCoV-HKU1_spike;1131 | 0 | 0 |
| NC_006577.2;YP_173238.1;HCoV-HKU1_spike;1132 | 0 | 0 |
| NC_006577.2;YP_173238.1;HCoV-HKU1_spike;1133 | 0 | 0 |
| NC_006577.2;YP_173238.1;HCoV-HKU1_spike;1134 | 0 | 0 |
| NC_006577.2;YP_173238.1;HCoV-HKU1_spike;1135 | 0 | 0 |
| NC_006577.2;YP_173238.1;HCoV-HKU1_spike;1136 | 0 | 0 |
| NC_006577.2;YP_173238.1;HCoV-HKU1_spike;1137 | 0 | 0 |
| NC_006577.2;YP_173238.1;HCoV-HKU1_spike;1138 | 0 | 0 |
| NC_006577.2;YP_173238.1;HCoV-HKU1_spike;1139 | 0 | 0 |
| NC_006577.2;YP_173238.1;HCoV-HKU1_spike;1140 | 0 | 0 |
| NC_006577.2;YP_173238.1;HCoV-HKU1_spike;1141 | 0 | 0 |
| NC_006577.2;YP_173238.1;HCoV-HKU1_spike;1142 | 0 | 0 |
| NC_006577.2;YP_173238.1;HCoV-HKU1_spike;1143 | 0 | 0 |
| NC_006577.2;YP_173238.1;HCoV-HKU1_spike;1144 | 0 | 0 |
| NC_006577.2;YP_173238.1;HCoV-HKU1_spike;1145 | 0 | 0 |
| NC_006577.2;YP_173238.1;HCoV-HKU1_spike;1146 | 0 | 0 |
| NC_006577.2;YP_173238.1;HCoV-HKU1_spike;1147 | 0 | 0 |
| NC_006577.2;YP_173238.1;HCoV-HKU1_spike;1148 | 0 | 0 |
| NC_006577.2;YP_173238.1;HCoV-HKU1_spike;1149 | 0 | 0 |
| NC_006577.2;YP_173238.1;HCoV-HKU1_spike;1150 | 0 | 0 |
| NC_006577.2;YP_173238.1;HCoV-HKU1_spike;1151 | 0 | 0 |
| NC_006577.2;YP_173238.1;HCoV-HKU1_spike;1152 | 0 | 0 |
| NC_006577.2;YP_173238.1;HCoV-HKU1_spike;1153 | 0 | 0 |
| NC_006577.2;YP_173238.1;HCoV-HKU1_spike;1154 | 0 | 0 |
| NC_006577.2;YP_173238.1;HCoV-HKU1_spike;1155 | 0 | 0 |

| | | |
|---|---|---|
| NC_006577.2;YP_173238.1;HCoV-HKU1_spike;1156 | 0 | 0 |
| NC_006577.2;YP_173238.1;HCoV-HKU1_spike;1157 | 0 | 0 |
| NC_006577.2;YP_173238.1;HCoV-HKU1_spike;1158 | 0 | 0 |
| NC_006577.2;YP_173238.1;HCoV-HKU1_spike;1159 | 0 | 0 |
| NC_006577.2;YP_173238.1;HCoV-HKU1_spike;1160 | 0 | 0.025 |
| NC_006577.2;YP_173238.1;HCoV-HKU1_spike;1161 | 0 | 0 |
| NC_006577.2;YP_173238.1;HCoV-HKU1_spike;1162 | 0 | 0 |
| NC_006577.2;YP_173238.1;HCoV-HKU1_spike;1163 | 0 | 0 |
| NC_006577.2;YP_173238.1;HCoV-HKU1_spike;1164 | 0 | 0 |
| NC_006577.2;YP_173238.1;HCoV-HKU1_spike;1165 | 0 | 0 |
| NC_006577.2;YP_173238.1;HCoV-HKU1_spike;1166 | 0 | 0 |
| NC_006577.2;YP_173238.1;HCoV-HKU1_spike;1167 | 0.05 | 0 |
| NC_006577.2;YP_173238.1;HCoV-HKU1_spike;1168 | 0 | 0 |
| NC_006577.2;YP_173238.1;HCoV-HKU1_spike;1169 | 0 | 0 |
| NC_006577.2;YP_173238.1;HCoV-HKU1_spike;1170 | 0 | 0.025 |
| NC_006577.2;YP_173238.1;HCoV-HKU1_spike;1171 | 0.05 | 0 |
| NC_006577.2;YP_173238.1;HCoV-HKU1_spike;1172 | 0 | 0 |
| NC_006577.2;YP_173238.1;HCoV-HKU1_spike;1173 | 0 | 0 |
| NC_006577.2;YP_173238.1;HCoV-HKU1_spike;1174 | 0 | 0 |
| NC_006577.2;YP_173238.1;HCoV-HKU1_spike;1175 | 0 | 0 |
| NC_006577.2;YP_173238.1;HCoV-HKU1_spike;1176 | 0 | 0 |
| NC_006577.2;YP_173238.1;HCoV-HKU1_spike;1177 | 0 | 0 |
| NC_006577.2;YP_173238.1;HCoV-HKU1_spike;1178 | 0 | 0 |
| NC_006577.2;YP_173238.1;HCoV-HKU1_spike;1179 | 0 | 0 |
| NC_006577.2;YP_173238.1;HCoV-HKU1_spike;1180 | 0 | 0 |
| NC_006577.2;YP_173238.1;HCoV-HKU1_spike;1181 | 0 | 0 |
| NC_006577.2;YP_173238.1;HCoV-HKU1_spike;1182 | 0 | 0 |
| NC_006577.2;YP_173238.1;HCoV-HKU1_spike;1183 | 0 | 0 |
| NC_006577.2;YP_173238.1;HCoV-HKU1_spike;1184 | 0 | 0 |
| NC_006577.2;YP_173238.1;HCoV-HKU1_spike;1185 | 0 | 0 |
| NC_006577.2;YP_173238.1;HCoV-HKU1_spike;1186 | 0 | 0 |
| NC_006577.2;YP_173238.1;HCoV-HKU1_spike;1187 | 0 | 0 |
| NC_006577.2;YP_173238.1;HCoV-HKU1_spike;1188 | 0 | 0 |
| NC_006577.2;YP_173238.1;HCoV-HKU1_spike;1189 | 0 | 0.025 |
| NC_006577.2;YP_173238.1;HCoV-HKU1_spike;1190 | 0 | 0.025 |
| NC_006577.2;YP_173238.1;HCoV-HKU1_spike;1191 | 0 | 0 |
| NC_006577.2;YP_173238.1;HCoV-HKU1_spike;1192 | 0 | 0.025 |
| NC_006577.2;YP_173238.1;HCoV-HKU1_spike;1193 | 0 | 0.025 |
| NC_006577.2;YP_173238.1;HCoV-HKU1_spike;1194 | 0 | 0 |
| NC_006577.2;YP_173238.1;HCoV-HKU1_spike;1195 | 0.05 | 0 |
| NC_006577.2;YP_173238.1;HCoV-HKU1_spike;1196 | 0.05 | 0.05 |
| NC_006577.2;YP_173238.1;HCoV-HKU1_spike;1197 | 0.05 | 0 |
| NC_006577.2;YP_173238.1;HCoV-HKU1_spike;1198 | 0.05 | 0.025 |

| | | |
|---|---|---|
| NC_006577.2;YP_173238.1;HCoV-HKU1_spike;1199 | 0.1 | 0 |
| NC_006577.2;YP_173238.1;HCoV-HKU1_spike;1200 | 0.05 | 0 |
| NC_006577.2;YP_173238.1;HCoV-HKU1_spike;1201 | 0.05 | 0 |
| NC_006577.2;YP_173238.1;HCoV-HKU1_spike;1202 | 0 | 0 |
| NC_006577.2;YP_173238.1;HCoV-HKU1_spike;1203 | 0 | 0 |
| NC_006577.2;YP_173238.1;HCoV-HKU1_spike;1204 | 0 | 0 |
| NC_006577.2;YP_173238.1;HCoV-HKU1_spike;1205 | 0 | 0 |
| NC_006577.2;YP_173238.1;HCoV-HKU1_spike;1206 | 0 | 0 |
| NC_006577.2;YP_173238.1;HCoV-HKU1_spike;1207 | 0 | 0 |
| NC_006577.2;YP_173238.1;HCoV-HKU1_spike;1208 | 0 | 0 |
| NC_006577.2;YP_173238.1;HCoV-HKU1_spike;1209 | 0 | 0 |
| NC_006577.2;YP_173238.1;HCoV-HKU1_spike;1210 | 0 | 0 |
| NC_006577.2;YP_173238.1;HCoV-HKU1_spike;1211 | 0 | 0 |
| NC_006577.2;YP_173238.1;HCoV-HKU1_spike;1212 | 0 | 0 |
| NC_006577.2;YP_173238.1;HCoV-HKU1_spike;1213 | 0 | 0 |
| NC_006577.2;YP_173238.1;HCoV-HKU1_spike;1214 | 0 | 0 |
| NC_006577.2;YP_173238.1;HCoV-HKU1_spike;1215 | 0 | 0 |
| NC_006577.2;YP_173238.1;HCoV-HKU1_spike;1216 | 0 | 0 |
| NC_006577.2;YP_173238.1;HCoV-HKU1_spike;1217 | 0 | 0.025 |
| NC_006577.2;YP_173238.1;HCoV-HKU1_spike;1218 | 0 | 0.025 |
| NC_006577.2;YP_173238.1;HCoV-HKU1_spike;1219 | 0 | 0.025 |
| NC_006577.2;YP_173238.1;HCoV-HKU1_spike;1220 | 0 | 0.025 |
| NC_006577.2;YP_173238.1;HCoV-HKU1_spike;1221 | 0.05 | 0.05 |
| NC_006577.2;YP_173238.1;HCoV-HKU1_spike;1222 | 0 | 0.075 |
| NC_006577.2;YP_173238.1;HCoV-HKU1_spike;1223 | 0 | 0.05 |
| NC_006577.2;YP_173238.1;HCoV-HKU1_spike;1224 | 0 | 0.05 |
| NC_006577.2;YP_173238.1;HCoV-HKU1_spike;1225 | 0 | 0.025 |
| NC_006577.2;YP_173238.1;HCoV-HKU1_spike;1226 | 0 | 0.025 |
| NC_006577.2;YP_173238.1;HCoV-HKU1_spike;1227 | 0 | 0.125 |
| NC_006577.2;YP_173238.1;HCoV-HKU1_spike;1228 | 0.05 | 0.275 |
| NC_006577.2;YP_173238.1;HCoV-HKU1_spike;1229 | 0.1 | 0.275 |
| NC_006577.2;YP_173238.1;HCoV-HKU1_spike;1230 | 0.05 | 0.325 |
| NC_006577.2;YP_173238.1;HCoV-HKU1_spike;1231 | 0.1 | 0.35 |
| NC_006577.2;YP_173238.1;HCoV-HKU1_spike;1232 | 0.05 | 0.35 |
| NC_006577.2;YP_173238.1;HCoV-HKU1_spike;1233 | 0.05 | 0.275 |
| NC_006577.2;YP_173238.1;HCoV-HKU1_spike;1234 | 0.05 | 0.175 |
| NC_006577.2;YP_173238.1;HCoV-HKU1_spike;1235 | 0.05 | 0 |
| NC_006577.2;YP_173238.1;HCoV-HKU1_spike;1236 | 0.05 | 0 |
| NC_006577.2;YP_173238.1;HCoV-HKU1_spike;1237 | 0 | 0 |
| NC_006577.2;YP_173238.1;HCoV-HKU1_spike;1238 | 0 | 0 |
| NC_006577.2;YP_173238.1;HCoV-HKU1_spike;1239 | 0 | 0 |
| NC_006577.2;YP_173238.1;HCoV-HKU1_spike;1240 | 0.1 | 0 |
| NC_006577.2;YP_173238.1;HCoV-HKU1_spike;1241 | 0.05 | 0 |

| | | |
|---|---|---|
| NC_006577.2;YP_173238.1;HCoV-HKU1_spike;1242 | 0.05 | 0 |
| NC_006577.2;YP_173238.1;HCoV-HKU1_spike;1243 | 0.05 | 0 |
| NC_006577.2;YP_173238.1;HCoV-HKU1_spike;1244 | 0.05 | 0 |
| NC_006577.2;YP_173238.1;HCoV-HKU1_spike;1245 | 0.1 | 0 |
| NC_006577.2;YP_173238.1;HCoV-HKU1_spike;1246 | 0.1 | 0 |
| NC_006577.2;YP_173238.1;HCoV-HKU1_spike;1247 | 0.1 | 0 |
| NC_006577.2;YP_173238.1;HCoV-HKU1_spike;1248 | 0.05 | 0 |
| NC_006577.2;YP_173238.1;HCoV-HKU1_spike;1249 | 0.05 | 0 |
| NC_006577.2;YP_173238.1;HCoV-HKU1_spike;1250 | 0.05 | 0 |
| NC_006577.2;YP_173238.1;HCoV-HKU1_spike;1251 | 0 | 0 |
| NC_006577.2;YP_173238.1;HCoV-HKU1_spike;1252 | 0 | 0 |
| NC_006577.2;YP_173238.1;HCoV-HKU1_spike;1253 | 0 | 0 |
| NC_006577.2;YP_173238.1;HCoV-HKU1_spike;1254 | 0 | 0 |
| NC_006577.2;YP_173238.1;HCoV-HKU1_spike;1255 | 0 | 0 |
| NC_006577.2;YP_173238.1;HCoV-HKU1_spike;1256 | 0 | 0 |
| NC_006577.2;YP_173238.1;HCoV-HKU1_spike;1257 | 0 | 0 |
| NC_006577.2;YP_173238.1;HCoV-HKU1_spike;1258 | 0 | 0 |
| NC_006577.2;YP_173238.1;HCoV-HKU1_spike;1259 | 0 | 0 |
| NC_006577.2;YP_173238.1;HCoV-HKU1_spike;1260 | 0 | 0 |
| NC_006577.2;YP_173238.1;HCoV-HKU1_spike;1261 | 0 | 0.05 |
| NC_006577.2;YP_173238.1;HCoV-HKU1_spike;1262 | 0 | 0 |
| NC_006577.2;YP_173238.1;HCoV-HKU1_spike;1263 | 0 | 0.025 |
| NC_006577.2;YP_173238.1;HCoV-HKU1_spike;1264 | 0 | 0 |
| NC_006577.2;YP_173238.1;HCoV-HKU1_spike;1265 | 0 | 0 |
| NC_006577.2;YP_173238.1;HCoV-HKU1_spike;1266 | 0 | 0 |
| NC_006577.2;YP_173238.1;HCoV-HKU1_spike;1267 | 0 | 0 |
| NC_006577.2;YP_173238.1;HCoV-HKU1_spike;1268 | 0 | 0 |
| NC_006577.2;YP_173238.1;HCoV-HKU1_spike;1269 | 0.05 | 0 |
| NC_006577.2;YP_173238.1;HCoV-HKU1_spike;1270 | 0 | 0 |
| NC_006577.2;YP_173238.1;HCoV-HKU1_spike;1271 | 0.15 | 0 |
| NC_006577.2;YP_173238.1;HCoV-HKU1_spike;1272 | 0.05 | 0.05 |
| NC_006577.2;YP_173238.1;HCoV-HKU1_spike;1273 | 0.05 | 0 |
| NC_006577.2;YP_173238.1;HCoV-HKU1_spike;1274 | 0.3 | 0.05 |
| NC_006577.2;YP_173238.1;HCoV-HKU1_spike;1275 | 0.15 | 0.05 |
| NC_006577.2;YP_173238.1;HCoV-HKU1_spike;1276 | 0.15 | 0.025 |
| NC_006577.2;YP_173238.1;HCoV-HKU1_spike;1277 | 0.1 | 0.075 |
| NC_006577.2;YP_173238.1;HCoV-HKU1_spike;1278 | 0.15 | 0 |
| NC_006577.2;YP_173238.1;HCoV-HKU1_spike;1279 | 0.25 | 0.1 |
| NC_006577.2;YP_173238.1;HCoV-HKU1_spike;1280 | 0.05 | 0.05 |
| NC_006577.2;YP_173238.1;HCoV-HKU1_spike;1281 | 0 | 0 |
| NC_006577.2;YP_173238.1;HCoV-HKU1_spike;1282 | 0 | 0 |
| NC_006577.2;YP_173238.1;HCoV-HKU1_spike;1283 | 0 | 0 |
| NC_006577.2;YP_173238.1;HCoV-HKU1_spike;1284 | 0 | 0 |

| | | |
|---|---|---|
| NC_006577.2;YP_173238.1;HCoV-HKU1_spike;1285 | 0 | 0 |
| NC_006577.2;YP_173238.1;HCoV-HKU1_spike;1286 | 0 | 0 |
| NC_006577.2;YP_173238.1;HCoV-HKU1_spike;1287 | 0.05 | 0 |
| NC_006577.2;YP_173238.1;HCoV-HKU1_spike;1288 | 0 | 0 |
| NC_006577.2;YP_173238.1;HCoV-HKU1_spike;1289 | 0 | 0 |
| NC_006577.2;YP_173238.1;HCoV-HKU1_spike;1290 | 0 | 0 |
| NC_006577.2;YP_173238.1;HCoV-HKU1_spike;1291 | 0 | 0 |
| NC_006577.2;YP_173238.1;HCoV-HKU1_spike;1292 | 0 | 0 |
| NC_006577.2;YP_173238.1;HCoV-HKU1_spike;1293 | 0 | 0.025 |
| NC_006577.2;YP_173238.1;HCoV-HKU1_spike;1294 | 0 | 0 |
| NC_006577.2;YP_173238.1;HCoV-HKU1_spike;1295 | 0 | 0 |
| NC_006577.2;YP_173238.1;HCoV-HKU1_spike;1296 | 0 | 0 |
| NC_006577.2;YP_173238.1;HCoV-HKU1_spike;1297 | 0 | 0 |
| NC_006577.2;YP_173238.1;HCoV-HKU1_spike;1298 | 0 | 0 |
| NC_006577.2;YP_173238.1;HCoV-HKU1_spike;1299 | 0 | 0 |
| NC_006577.2;YP_173238.1;HCoV-HKU1_spike;1300 | 0 | 0 |
| NC_006577.2;YP_173238.1;HCoV-HKU1_spike;1301 | 0 | 0 |
| NC_006577.2;YP_173238.1;HCoV-HKU1_spike;1302 | 0 | 0 |
| NC_006577.2;YP_173238.1;HCoV-HKU1_spike;1303 | 0 | 0 |
| NC_006577.2;YP_173238.1;HCoV-HKU1_spike;1304 | 0 | 0 |
| NC_006577.2;YP_173238.1;HCoV-HKU1_spike;1305 | 0 | 0 |
| NC_006577.2;YP_173238.1;HCoV-HKU1_spike;1306 | 0 | 0 |
| NC_006577.2;YP_173238.1;HCoV-HKU1_spike;1307 | 0 | 0 |
| NC_006577.2;YP_173238.1;HCoV-HKU1_spike;1308 | 0 | 0 |
| NC_006577.2;YP_173238.1;HCoV-HKU1_spike;1309 | 0 | 0 |
| NC_006577.2;YP_173238.1;HCoV-HKU1_spike;1310 | 0 | 0 |
| NC_006577.2;YP_173238.1;HCoV-HKU1_spike;1311 | 0 | 0 |
| NC_006577.2;YP_173238.1;HCoV-HKU1_spike;1312 | 0 | 0 |
| NC_006577.2;YP_173238.1;HCoV-HKU1_spike;1313 | 0 | 0 |
| NC_006577.2;YP_173238.1;HCoV-HKU1_spike;1314 | 0 | 0 |
| NC_006577.2;YP_173238.1;HCoV-HKU1_spike;1315 | 0 | 0 |
| NC_006577.2;YP_173238.1;HCoV-HKU1_spike;1316 | 0 | 0 |
| NC_006577.2;YP_173238.1;HCoV-HKU1_spike;1317 | 0 | 0 |
| NC_006577.2;YP_173238.1;HCoV-HKU1_spike;1318 | 0 | 0 |
| NC_006577.2;YP_173238.1;HCoV-HKU1_spike;1319 | 0 | 0 |
| NC_006577.2;YP_173238.1;HCoV-HKU1_spike;1320 | 0 | 0 |
| NC_006577.2;YP_173238.1;HCoV-HKU1_spike;1321 | 0 | 0 |
| NC_006577.2;YP_173238.1;HCoV-HKU1_spike;1322 | 0 | 0 |
| NC_006577.2;YP_173238.1;HCoV-HKU1_spike;1323 | 0 | 0 |
| NC_006577.2;YP_173238.1;HCoV-HKU1_spike;1324 | 0 | 0 |
| NC_006577.2;YP_173238.1;HCoV-HKU1_spike;1325 | 0 | 0 |
| NC_006577.2;YP_173238.1;HCoV-HKU1_spike;1326 | 0 | 0 |
| NC_006577.2;YP_173238.1;HCoV-HKU1_spike;1327 | 0 | 0 |

| | | |
|---|---|---|
| NC_006577.2;YP_173238.1;HCoV-HKU1_spike;1328 | 0 | 0 |
| NC_006577.2;YP_173238.1;HCoV-HKU1_spike;1329 | 0 | 0 |
| NC_006577.2;YP_173238.1;HCoV-HKU1_spike;1330 | 0 | 0.025 |
| NC_006577.2;YP_173238.1;HCoV-HKU1_spike;1331 | 0 | 0.025 |
| NC_006577.2;YP_173238.1;HCoV-HKU1_spike;1332 | 0 | 0.025 |
| NC_006577.2;YP_173238.1;HCoV-HKU1_spike;1333 | 0 | 0.025 |
| NC_006577.2;YP_173238.1;HCoV-HKU1_spike;1334 | 0 | 0.025 |
| NC_006577.2;YP_173238.1;HCoV-HKU1_spike;1335 | 0 | 0.025 |
| NC_006577.2;YP_173238.1;HCoV-HKU1_spike;1336 | 0 | 0.025 |
| NC_006577.2;YP_173238.1;HCoV-HKU1_spike;1337 | 0 | 0.025 |
| NC_006577.2;YP_173238.1;HCoV-HKU1_spike;1338 | 0 | 0.025 |
| NC_006577.2;YP_173238.1;HCoV-HKU1_spike;1339 | 0 | 0.025 |
| NC_006577.2;YP_173238.1;HCoV-HKU1_spike;1340 | 0 | 0.05 |
| NC_006577.2;YP_173238.1;HCoV-HKU1_spike;1341 | 0.15 | 0.05 |
| NC_006577.2;YP_173238.1;HCoV-HKU1_spike;1342 | 0.15 | 0.025 |
| NC_006577.2;YP_173238.1;HCoV-HKU1_spike;1343 | 0.15 | 0.025 |
| NC_006577.2;YP_173238.1;HCoV-HKU1_spike;1344 | 0.15 | 0.025 |
| NC_006577.2;YP_173238.1;HCoV-HKU1_spike;1345 | 0.15 | 0.025 |
| NC_006577.2;YP_173239.1;HCoV-HKU1_non-structu;1 | 0 | 0 |
| NC_006577.2;YP_173239.1;HCoV-HKU1_non-structu;2 | 0 | 0 |
| NC_006577.2;YP_173239.1;HCoV-HKU1_non-structu;3 | 0 | 0 |
| NC_006577.2;YP_173239.1;HCoV-HKU1_non-structu;4 | 0 | 0 |
| NC_006577.2;YP_173239.1;HCoV-HKU1_non-structu;5 | 0 | 0 |
| NC_006577.2;YP_173239.1;HCoV-HKU1_non-structu;6 | 0 | 0 |
| NC_006577.2;YP_173239.1;HCoV-HKU1_non-structu;7 | 0 | 0 |
| NC_006577.2;YP_173239.1;HCoV-HKU1_non-structu;8 | 0 | 0 |
| NC_006577.2;YP_173239.1;HCoV-HKU1_non-structu;9 | 0 | 0 |
| NC_006577.2;YP_173239.1;HCoV-HKU1_non-structu;10 | 0 | 0 |
| NC_006577.2;YP_173239.1;HCoV-HKU1_non-structu;11 | 0 | 0 |
| NC_006577.2;YP_173239.1;HCoV-HKU1_non-structu;12 | 0.05 | 0 |
| NC_006577.2;YP_173239.1;HCoV-HKU1_non-structu;13 | 0.05 | 0 |
| NC_006577.2;YP_173239.1;HCoV-HKU1_non-structu;14 | 0.05 | 0 |
| NC_006577.2;YP_173239.1;HCoV-HKU1_non-structu;15 | 0.1 | 0 |
| NC_006577.2;YP_173239.1;HCoV-HKU1_non-structu;16 | 0.05 | 0 |
| NC_006577.2;YP_173239.1;HCoV-HKU1_non-structu;17 | 0.05 | 0.025 |
| NC_006577.2;YP_173239.1;HCoV-HKU1_non-structu;18 | 0.15 | 0.1 |
| NC_006577.2;YP_173239.1;HCoV-HKU1_non-structu;19 | 0.05 | 0 |
| NC_006577.2;YP_173239.1;HCoV-HKU1_non-structu;20 | 0.15 | 0 |
| NC_006577.2;YP_173239.1;HCoV-HKU1_non-structu;21 | 0 | 0.025 |
| NC_006577.2;YP_173239.1;HCoV-HKU1_non-structu;22 | 0 | 0 |
| NC_006577.2;YP_173239.1;HCoV-HKU1_non-structu;23 | 0 | 0 |
| NC_006577.2;YP_173239.1;HCoV-HKU1_non-structu;24 | 0 | 0 |
| NC_006577.2;YP_173239.1;HCoV-HKU1_non-structu;25 | 0 | 0 |

| | | |
|---|---|---|
| NC_006577.2;YP_173239.1;HCoV-HKU1_non-structu;26 | 0 | 0 |
| NC_006577.2;YP_173239.1;HCoV-HKU1_non-structu;27 | 0 | 0 |
| NC_006577.2;YP_173239.1;HCoV-HKU1_non-structu;28 | 0 | 0 |
| NC_006577.2;YP_173239.1;HCoV-HKU1_non-structu;29 | 0 | 0 |
| NC_006577.2;YP_173239.1;HCoV-HKU1_non-structu;30 | 0 | 0 |
| NC_006577.2;YP_173239.1;HCoV-HKU1_non-structu;31 | 0 | 0 |
| NC_006577.2;YP_173239.1;HCoV-HKU1_non-structu;32 | 0 | 0 |
| NC_006577.2;YP_173239.1;HCoV-HKU1_non-structu;33 | 0 | 0 |
| NC_006577.2;YP_173239.1;HCoV-HKU1_non-structu;34 | 0 | 0 |
| NC_006577.2;YP_173239.1;HCoV-HKU1_non-structu;35 | 0 | 0 |
| NC_006577.2;YP_173239.1;HCoV-HKU1_non-structu;36 | 0 | 0 |
| NC_006577.2;YP_173239.1;HCoV-HKU1_non-structu;37 | 0 | 0 |
| NC_006577.2;YP_173239.1;HCoV-HKU1_non-structu;38 | 0 | 0 |
| NC_006577.2;YP_173239.1;HCoV-HKU1_non-structu;39 | 0 | 0 |
| NC_006577.2;YP_173239.1;HCoV-HKU1_non-structu;40 | 0 | 0 |
| NC_006577.2;YP_173239.1;HCoV-HKU1_non-structu;41 | 0 | 0 |
| NC_006577.2;YP_173239.1;HCoV-HKU1_non-structu;42 | 0 | 0 |
| NC_006577.2;YP_173239.1;HCoV-HKU1_non-structu;43 | 0 | 0 |
| NC_006577.2;YP_173239.1;HCoV-HKU1_non-structu;44 | 0 | 0 |
| NC_006577.2;YP_173239.1;HCoV-HKU1_non-structu;45 | 0 | 0 |
| NC_006577.2;YP_173239.1;HCoV-HKU1_non-structu;46 | 0 | 0 |
| NC_006577.2;YP_173239.1;HCoV-HKU1_non-structu;47 | 0 | 0 |
| NC_006577.2;YP_173239.1;HCoV-HKU1_non-structu;48 | 0 | 0 |
| NC_006577.2;YP_173239.1;HCoV-HKU1_non-structu;49 | 0 | 0 |
| NC_006577.2;YP_173239.1;HCoV-HKU1_non-structu;50 | 0 | 0 |
| NC_006577.2;YP_173239.1;HCoV-HKU1_non-structu;51 | 0 | 0 |
| NC_006577.2;YP_173239.1;HCoV-HKU1_non-structu;52 | 0 | 0 |
| NC_006577.2;YP_173239.1;HCoV-HKU1_non-structu;53 | 0 | 0 |
| NC_006577.2;YP_173239.1;HCoV-HKU1_non-structu;54 | 0 | 0 |
| NC_006577.2;YP_173239.1;HCoV-HKU1_non-structu;55 | 0 | 0 |
| NC_006577.2;YP_173239.1;HCoV-HKU1_non-structu;56 | 0 | 0 |
| NC_006577.2;YP_173239.1;HCoV-HKU1_non-structu;57 | 0 | 0 |
| NC_006577.2;YP_173239.1;HCoV-HKU1_non-structu;58 | 0 | 0 |
| NC_006577.2;YP_173239.1;HCoV-HKU1_non-structu;59 | 0 | 0 |
| NC_006577.2;YP_173239.1;HCoV-HKU1_non-structu;60 | 0.05 | 0 |
| NC_006577.2;YP_173239.1;HCoV-HKU1_non-structu;61 | 0 | 0 |
| NC_006577.2;YP_173239.1;HCoV-HKU1_non-structu;62 | 0 | 0 |
| NC_006577.2;YP_173239.1;HCoV-HKU1_non-structu;63 | 0 | 0 |
| NC_006577.2;YP_173239.1;HCoV-HKU1_non-structu;64 | 0 | 0 |
| NC_006577.2;YP_173239.1;HCoV-HKU1_non-structu;65 | 0 | 0 |
| NC_006577.2;YP_173239.1;HCoV-HKU1_non-structu;66 | 0 | 0 |
| NC_006577.2;YP_173239.1;HCoV-HKU1_non-structu;67 | 0 | 0 |
| NC_006577.2;YP_173239.1;HCoV-HKU1_non-structu;68 | 0 | 0 |

| | | |
|---|---|---|
| NC_006577.2;YP_173239.1;HCoV-HKU1_non-structu;69 | 0 | 0 |
| NC_006577.2;YP_173239.1;HCoV-HKU1_non-structu;70 | 0 | 0 |
| NC_006577.2;YP_173239.1;HCoV-HKU1_non-structu;71 | 0 | 0 |
| NC_006577.2;YP_173239.1;HCoV-HKU1_non-structu;72 | 0 | 0 |
| NC_006577.2;YP_173239.1;HCoV-HKU1_non-structu;73 | 0 | 0.025 |
| NC_006577.2;YP_173239.1;HCoV-HKU1_non-structu;74 | 0.05 | 0.025 |
| NC_006577.2;YP_173239.1;HCoV-HKU1_non-structu;75 | 0 | 0.025 |
| NC_006577.2;YP_173239.1;HCoV-HKU1_non-structu;76 | 0 | 0 |
| NC_006577.2;YP_173239.1;HCoV-HKU1_non-structu;77 | 0.05 | 0 |
| NC_006577.2;YP_173239.1;HCoV-HKU1_non-structu;78 | 0 | 0 |
| NC_006577.2;YP_173239.1;HCoV-HKU1_non-structu;79 | 0 | 0 |
| NC_006577.2;YP_173239.1;HCoV-HKU1_non-structu;80 | 0.05 | 0 |
| NC_006577.2;YP_173239.1;HCoV-HKU1_non-structu;81 | 0 | 0 |
| NC_006577.2;YP_173239.1;HCoV-HKU1_non-structu;82 | 0 | 0 |
| NC_006577.2;YP_173239.1;HCoV-HKU1_non-structu;83 | 0 | 0 |
| NC_006577.2;YP_173239.1;HCoV-HKU1_non-structu;84 | 0 | 0 |
| NC_006577.2;YP_173239.1;HCoV-HKU1_non-structu;85 | 0 | 0 |
| NC_006577.2;YP_173239.1;HCoV-HKU1_non-structu;86 | 0 | 0 |
| NC_006577.2;YP_173239.1;HCoV-HKU1_non-structu;87 | 0 | 0 |
| NC_006577.2;YP_173239.1;HCoV-HKU1_non-structu;88 | 0 | 0 |
| NC_006577.2;YP_173239.1;HCoV-HKU1_non-structu;89 | 0 | 0 |
| NC_006577.2;YP_173239.1;HCoV-HKU1_non-structu;90 | 0 | 0 |
| NC_006577.2;YP_173239.1;HCoV-HKU1_non-structu;91 | 0 | 0 |
| NC_006577.2;YP_173239.1;HCoV-HKU1_non-structu;92 | 0 | 0 |
| NC_006577.2;YP_173239.1;HCoV-HKU1_non-structu;93 | 0 | 0 |
| NC_006577.2;YP_173239.1;HCoV-HKU1_non-structu;94 | 0 | 0 |
| NC_006577.2;YP_173239.1;HCoV-HKU1_non-structu;95 | 0 | 0 |
| NC_006577.2;YP_173239.1;HCoV-HKU1_non-structu;96 | 0 | 0 |
| NC_006577.2;YP_173239.1;HCoV-HKU1_non-structu;97 | 0 | 0 |
| NC_006577.2;YP_173239.1;HCoV-HKU1_non-structu;98 | 0 | 0 |
| NC_006577.2;YP_173240.1;HCoV-HKU1_small-membr;1 | 0 | 0 |
| NC_006577.2;YP_173240.1;HCoV-HKU1_small-membr;2 | 0 | 0 |
| NC_006577.2;YP_173240.1;HCoV-HKU1_small-membr;3 | 0 | 0 |
| NC_006577.2;YP_173240.1;HCoV-HKU1_small-membr;4 | 0 | 0 |
| NC_006577.2;YP_173240.1;HCoV-HKU1_small-membr;5 | 0 | 0 |
| NC_006577.2;YP_173240.1;HCoV-HKU1_small-membr;6 | 0 | 0 |
| NC_006577.2;YP_173240.1;HCoV-HKU1_small-membr;7 | 0 | 0 |
| NC_006577.2;YP_173240.1;HCoV-HKU1_small-membr;8 | 0 | 0 |
| NC_006577.2;YP_173240.1;HCoV-HKU1_small-membr;9 | 0 | 0 |
| NC_006577.2;YP_173240.1;HCoV-HKU1_small-membr;10 | 0 | 0 |
| NC_006577.2;YP_173240.1;HCoV-HKU1_small-membr;11 | 0 | 0 |
| NC_006577.2;YP_173240.1;HCoV-HKU1_small-membr;12 | 0 | 0 |
| NC_006577.2;YP_173240.1;HCoV-HKU1_small-membr;13 | 0 | 0 |

| | | |
|---|---|---|
| NC_006577.2;YP_173240.1;HCoV-HKU1_small-membr;14 | 0 | 0 |
| NC_006577.2;YP_173240.1;HCoV-HKU1_small-membr;15 | 0 | 0 |
| NC_006577.2;YP_173240.1;HCoV-HKU1_small-membr;16 | 0 | 0 |
| NC_006577.2;YP_173240.1;HCoV-HKU1_small-membr;17 | 0 | 0 |
| NC_006577.2;YP_173240.1;HCoV-HKU1_small-membr;18 | 0 | 0 |
| NC_006577.2;YP_173240.1;HCoV-HKU1_small-membr;19 | 0 | 0 |
| NC_006577.2;YP_173240.1;HCoV-HKU1_small-membr;20 | 0 | 0 |
| NC_006577.2;YP_173240.1;HCoV-HKU1_small-membr;21 | 0 | 0 |
| NC_006577.2;YP_173240.1;HCoV-HKU1_small-membr;22 | 0 | 0 |
| NC_006577.2;YP_173240.1;HCoV-HKU1_small-membr;23 | 0 | 0 |
| NC_006577.2;YP_173240.1;HCoV-HKU1_small-membr;24 | 0 | 0 |
| NC_006577.2;YP_173240.1;HCoV-HKU1_small-membr;25 | 0 | 0 |
| NC_006577.2;YP_173240.1;HCoV-HKU1_small-membr;26 | 0 | 0 |
| NC_006577.2;YP_173240.1;HCoV-HKU1_small-membr;27 | 0 | 0 |
| NC_006577.2;YP_173240.1;HCoV-HKU1_small-membr;28 | 0 | 0 |
| NC_006577.2;YP_173240.1;HCoV-HKU1_small-membr;29 | 0 | 0 |
| NC_006577.2;YP_173240.1;HCoV-HKU1_small-membr;30 | 0 | 0 |
| NC_006577.2;YP_173240.1;HCoV-HKU1_small-membr;31 | 0 | 0 |
| NC_006577.2;YP_173240.1;HCoV-HKU1_small-membr;32 | 0 | 0 |
| NC_006577.2;YP_173240.1;HCoV-HKU1_small-membr;33 | 0 | 0 |
| NC_006577.2;YP_173240.1;HCoV-HKU1_small-membr;34 | 0 | 0 |
| NC_006577.2;YP_173240.1;HCoV-HKU1_small-membr;35 | 0 | 0 |
| NC_006577.2;YP_173240.1;HCoV-HKU1_small-membr;36 | 0 | 0 |
| NC_006577.2;YP_173240.1;HCoV-HKU1_small-membr;37 | 0 | 0 |
| NC_006577.2;YP_173240.1;HCoV-HKU1_small-membr;38 | 0 | 0 |
| NC_006577.2;YP_173240.1;HCoV-HKU1_small-membr;39 | 0 | 0 |
| NC_006577.2;YP_173240.1;HCoV-HKU1_small-membr;40 | 0 | 0 |
| NC_006577.2;YP_173240.1;HCoV-HKU1_small-membr;41 | 0 | 0 |
| NC_006577.2;YP_173240.1;HCoV-HKU1_small-membr;42 | 0 | 0 |
| NC_006577.2;YP_173240.1;HCoV-HKU1_small-membr;43 | 0 | 0 |
| NC_006577.2;YP_173240.1;HCoV-HKU1_small-membr;44 | 0 | 0 |
| NC_006577.2;YP_173240.1;HCoV-HKU1_small-membr;45 | 0 | 0 |
| NC_006577.2;YP_173240.1;HCoV-HKU1_small-membr;46 | 0 | 0 |
| NC_006577.2;YP_173240.1;HCoV-HKU1_small-membr;47 | 0 | 0 |
| NC_006577.2;YP_173240.1;HCoV-HKU1_small-membr;48 | 0 | 0 |
| NC_006577.2;YP_173240.1;HCoV-HKU1_small-membr;49 | 0 | 0 |
| NC_006577.2;YP_173240.1;HCoV-HKU1_small-membr;50 | 0 | 0 |
| NC_006577.2;YP_173240.1;HCoV-HKU1_small-membr;51 | 0 | 0 |
| NC_006577.2;YP_173240.1;HCoV-HKU1_small-membr;52 | 0 | 0 |
| NC_006577.2;YP_173240.1;HCoV-HKU1_small-membr;53 | 0 | 0 |
| NC_006577.2;YP_173240.1;HCoV-HKU1_small-membr;54 | 0 | 0 |
| NC_006577.2;YP_173240.1;HCoV-HKU1_small-membr;55 | 0 | 0 |
| NC_006577.2;YP_173240.1;HCoV-HKU1_small-membr;56 | 0 | 0 |

| | | |
|---|---|---|
| NC_006577.2;YP_173240.1;HCoV-HKU1_small-membr;57 | 0.05 | 0 |
| NC_006577.2;YP_173240.1;HCoV-HKU1_small-membr;58 | 0 | 0 |
| NC_006577.2;YP_173240.1;HCoV-HKU1_small-membr;59 | 0 | 0 |
| NC_006577.2;YP_173240.1;HCoV-HKU1_small-membr;60 | 0.05 | 0.075 |
| NC_006577.2;YP_173240.1;HCoV-HKU1_small-membr;61 | 0 | 0 |
| NC_006577.2;YP_173240.1;HCoV-HKU1_small-membr;62 | 0 | 0 |
| NC_006577.2;YP_173240.1;HCoV-HKU1_small-membr;63 | 0 | 0 |
| NC_006577.2;YP_173240.1;HCoV-HKU1_small-membr;64 | 0 | 0 |
| NC_006577.2;YP_173240.1;HCoV-HKU1_small-membr;65 | 0 | 0.025 |
| NC_006577.2;YP_173240.1;HCoV-HKU1_small-membr;66 | 0.05 | 0 |
| NC_006577.2;YP_173240.1;HCoV-HKU1_small-membr;67 | 0.05 | 0 |
| NC_006577.2;YP_173240.1;HCoV-HKU1_small-membr;68 | 0.05 | 0 |
| NC_006577.2;YP_173240.1;HCoV-HKU1_small-membr;69 | 0.05 | 0 |
| NC_006577.2;YP_173240.1;HCoV-HKU1_small-membr;70 | 0.05 | 0 |
| NC_006577.2;YP_173240.1;HCoV-HKU1_small-membr;71 | 0.05 | 0 |
| NC_006577.2;YP_173241.1;HCoV-HKU1_membrane;1 | 1 | 0.8 |
| NC_006577.2;YP_173241.1;HCoV-HKU1_membrane;2 | 0.95 | 0.775 |
| NC_006577.2;YP_173241.1;HCoV-HKU1_membrane;3 | 0.85 | 0.625 |
| NC_006577.2;YP_173241.1;HCoV-HKU1_membrane;4 | 0.05 | 0.05 |
| NC_006577.2;YP_173241.1;HCoV-HKU1_membrane;5 | 0.05 | 0.075 |
| NC_006577.2;YP_173241.1;HCoV-HKU1_membrane;6 | 0.1 | 0.075 |
| NC_006577.2;YP_173241.1;HCoV-HKU1_membrane;7 | 0.1 | 0.05 |
| NC_006577.2;YP_173241.1;HCoV-HKU1_membrane;8 | 0.05 | 0.075 |
| NC_006577.2;YP_173241.1;HCoV-HKU1_membrane;9 | 0.05 | 0.05 |
| NC_006577.2;YP_173241.1;HCoV-HKU1_membrane;10 | 0 | 0.05 |
| NC_006577.2;YP_173241.1;HCoV-HKU1_membrane;11 | 0 | 0 |
| NC_006577.2;YP_173241.1;HCoV-HKU1_membrane;12 | 0 | 0 |
| NC_006577.2;YP_173241.1;HCoV-HKU1_membrane;13 | 0 | 0 |
| NC_006577.2;YP_173241.1;HCoV-HKU1_membrane;14 | 0 | 0 |
| NC_006577.2;YP_173241.1;HCoV-HKU1_membrane;15 | 0 | 0 |
| NC_006577.2;YP_173241.1;HCoV-HKU1_membrane;16 | 0 | 0 |
| NC_006577.2;YP_173241.1;HCoV-HKU1_membrane;17 | 0 | 0 |
| NC_006577.2;YP_173241.1;HCoV-HKU1_membrane;18 | 0 | 0 |
| NC_006577.2;YP_173241.1;HCoV-HKU1_membrane;19 | 0 | 0 |
| NC_006577.2;YP_173241.1;HCoV-HKU1_membrane;20 | 0 | 0 |
| NC_006577.2;YP_173241.1;HCoV-HKU1_membrane;21 | 0 | 0 |
| NC_006577.2;YP_173241.1;HCoV-HKU1_membrane;22 | 0 | 0 |
| NC_006577.2;YP_173241.1;HCoV-HKU1_membrane;23 | 0 | 0 |
| NC_006577.2;YP_173241.1;HCoV-HKU1_membrane;24 | 0 | 0 |
| NC_006577.2;YP_173241.1;HCoV-HKU1_membrane;25 | 0 | 0 |
| NC_006577.2;YP_173241.1;HCoV-HKU1_membrane;26 | 0 | 0 |
| NC_006577.2;YP_173241.1;HCoV-HKU1_membrane;27 | 0 | 0 |
| NC_006577.2;YP_173241.1;HCoV-HKU1_membrane;28 | 0 | 0 |

| | | |
|---|---|---|
| NC_006577.2;YP_173241.1;HCoV-HKU1_membrane;29 | 0 | 0 |
| NC_006577.2;YP_173241.1;HCoV-HKU1_membrane;30 | 0 | 0 |
| NC_006577.2;YP_173241.1;HCoV-HKU1_membrane;31 | 0 | 0 |
| NC_006577.2;YP_173241.1;HCoV-HKU1_membrane;32 | 0 | 0 |
| NC_006577.2;YP_173241.1;HCoV-HKU1_membrane;33 | 0 | 0 |
| NC_006577.2;YP_173241.1;HCoV-HKU1_membrane;34 | 0 | 0 |
| NC_006577.2;YP_173241.1;HCoV-HKU1_membrane;35 | 0 | 0 |
| NC_006577.2;YP_173241.1;HCoV-HKU1_membrane;36 | 0 | 0 |
| NC_006577.2;YP_173241.1;HCoV-HKU1_membrane;37 | 0 | 0 |
| NC_006577.2;YP_173241.1;HCoV-HKU1_membrane;38 | 0 | 0 |
| NC_006577.2;YP_173241.1;HCoV-HKU1_membrane;39 | 0 | 0 |
| NC_006577.2;YP_173241.1;HCoV-HKU1_membrane;40 | 0 | 0 |
| NC_006577.2;YP_173241.1;HCoV-HKU1_membrane;41 | 0 | 0 |
| NC_006577.2;YP_173241.1;HCoV-HKU1_membrane;42 | 0 | 0 |
| NC_006577.2;YP_173241.1;HCoV-HKU1_membrane;43 | 0 | 0 |
| NC_006577.2;YP_173241.1;HCoV-HKU1_membrane;44 | 0 | 0 |
| NC_006577.2;YP_173241.1;HCoV-HKU1_membrane;45 | 0 | 0 |
| NC_006577.2;YP_173241.1;HCoV-HKU1_membrane;46 | 0 | 0 |
| NC_006577.2;YP_173241.1;HCoV-HKU1_membrane;47 | 0 | 0 |
| NC_006577.2;YP_173241.1;HCoV-HKU1_membrane;48 | 0 | 0 |
| NC_006577.2;YP_173241.1;HCoV-HKU1_membrane;49 | 0 | 0 |
| NC_006577.2;YP_173241.1;HCoV-HKU1_membrane;50 | 0 | 0 |
| NC_006577.2;YP_173241.1;HCoV-HKU1_membrane;51 | 0 | 0 |
| NC_006577.2;YP_173241.1;HCoV-HKU1_membrane;52 | 0 | 0 |
| NC_006577.2;YP_173241.1;HCoV-HKU1_membrane;53 | 0 | 0 |
| NC_006577.2;YP_173241.1;HCoV-HKU1_membrane;54 | 0 | 0 |
| NC_006577.2;YP_173241.1;HCoV-HKU1_membrane;55 | 0 | 0 |
| NC_006577.2;YP_173241.1;HCoV-HKU1_membrane;56 | 0 | 0 |
| NC_006577.2;YP_173241.1;HCoV-HKU1_membrane;57 | 0 | 0 |
| NC_006577.2;YP_173241.1;HCoV-HKU1_membrane;58 | 0 | 0 |
| NC_006577.2;YP_173241.1;HCoV-HKU1_membrane;59 | 0 | 0 |
| NC_006577.2;YP_173241.1;HCoV-HKU1_membrane;60 | 0 | 0 |
| NC_006577.2;YP_173241.1;HCoV-HKU1_membrane;61 | 0 | 0 |
| NC_006577.2;YP_173241.1;HCoV-HKU1_membrane;62 | 0 | 0 |
| NC_006577.2;YP_173241.1;HCoV-HKU1_membrane;63 | 0 | 0 |
| NC_006577.2;YP_173241.1;HCoV-HKU1_membrane;64 | 0 | 0 |
| NC_006577.2;YP_173241.1;HCoV-HKU1_membrane;65 | 0 | 0 |
| NC_006577.2;YP_173241.1;HCoV-HKU1_membrane;66 | 0 | 0.025 |
| NC_006577.2;YP_173241.1;HCoV-HKU1_membrane;67 | 0 | 0.025 |
| NC_006577.2;YP_173241.1;HCoV-HKU1_membrane;68 | 0 | 0.025 |
| NC_006577.2;YP_173241.1;HCoV-HKU1_membrane;69 | 0 | 0 |
| NC_006577.2;YP_173241.1;HCoV-HKU1_membrane;70 | 0 | 0 |
| NC_006577.2;YP_173241.1;HCoV-HKU1_membrane;71 | 0 | 0 |

| | | |
|---|---|---|
| NC_006577.2;YP_173241.1;HCoV-HKU1_membrane;72 | 0 | 0 |
| NC_006577.2;YP_173241.1;HCoV-HKU1_membrane;73 | 0 | 0 |
| NC_006577.2;YP_173241.1;HCoV-HKU1_membrane;74 | 0 | 0 |
| NC_006577.2;YP_173241.1;HCoV-HKU1_membrane;75 | 0 | 0 |
| NC_006577.2;YP_173241.1;HCoV-HKU1_membrane;76 | 0 | 0 |
| NC_006577.2;YP_173241.1;HCoV-HKU1_membrane;77 | 0 | 0 |
| NC_006577.2;YP_173241.1;HCoV-HKU1_membrane;78 | 0 | 0 |
| NC_006577.2;YP_173241.1;HCoV-HKU1_membrane;79 | 0 | 0 |
| NC_006577.2;YP_173241.1;HCoV-HKU1_membrane;80 | 0 | 0 |
| NC_006577.2;YP_173241.1;HCoV-HKU1_membrane;81 | 0 | 0 |
| NC_006577.2;YP_173241.1;HCoV-HKU1_membrane;82 | 0 | 0 |
| NC_006577.2;YP_173241.1;HCoV-HKU1_membrane;83 | 0 | 0 |
| NC_006577.2;YP_173241.1;HCoV-HKU1_membrane;84 | 0 | 0 |
| NC_006577.2;YP_173241.1;HCoV-HKU1_membrane;85 | 0 | 0 |
| NC_006577.2;YP_173241.1;HCoV-HKU1_membrane;86 | 0 | 0 |
| NC_006577.2;YP_173241.1;HCoV-HKU1_membrane;87 | 0 | 0 |
| NC_006577.2;YP_173241.1;HCoV-HKU1_membrane;88 | 0 | 0 |
| NC_006577.2;YP_173241.1;HCoV-HKU1_membrane;89 | 0 | 0 |
| NC_006577.2;YP_173241.1;HCoV-HKU1_membrane;90 | 0 | 0 |
| NC_006577.2;YP_173241.1;HCoV-HKU1_membrane;91 | 0 | 0 |
| NC_006577.2;YP_173241.1;HCoV-HKU1_membrane;92 | 0 | 0 |
| NC_006577.2;YP_173241.1;HCoV-HKU1_membrane;93 | 0 | 0 |
| NC_006577.2;YP_173241.1;HCoV-HKU1_membrane;94 | 0 | 0 |
| NC_006577.2;YP_173241.1;HCoV-HKU1_membrane;95 | 0 | 0 |
| NC_006577.2;YP_173241.1;HCoV-HKU1_membrane;96 | 0 | 0 |
| NC_006577.2;YP_173241.1;HCoV-HKU1_membrane;97 | 0 | 0 |
| NC_006577.2;YP_173241.1;HCoV-HKU1_membrane;98 | 0 | 0 |
| NC_006577.2;YP_173241.1;HCoV-HKU1_membrane;99 | 0 | 0 |
| NC_006577.2;YP_173241.1;HCoV-HKU1_membrane;100 | 0 | 0 |
| NC_006577.2;YP_173241.1;HCoV-HKU1_membrane;101 | 0 | 0 |
| NC_006577.2;YP_173241.1;HCoV-HKU1_membrane;102 | 0 | 0 |
| NC_006577.2;YP_173241.1;HCoV-HKU1_membrane;103 | 0 | 0 |
| NC_006577.2;YP_173241.1;HCoV-HKU1_membrane;104 | 0 | 0 |
| NC_006577.2;YP_173241.1;HCoV-HKU1_membrane;105 | 0 | 0 |
| NC_006577.2;YP_173241.1;HCoV-HKU1_membrane;106 | 0 | 0 |
| NC_006577.2;YP_173241.1;HCoV-HKU1_membrane;107 | 0 | 0 |
| NC_006577.2;YP_173241.1;HCoV-HKU1_membrane;108 | 0 | 0 |
| NC_006577.2;YP_173241.1;HCoV-HKU1_membrane;109 | 0 | 0 |
| NC_006577.2;YP_173241.1;HCoV-HKU1_membrane;110 | 0 | 0 |
| NC_006577.2;YP_173241.1;HCoV-HKU1_membrane;111 | 0.1 | 0.025 |
| NC_006577.2;YP_173241.1;HCoV-HKU1_membrane;112 | 0.05 | 0 |
| NC_006577.2;YP_173241.1;HCoV-HKU1_membrane;113 | 0.05 | 0 |
| NC_006577.2;YP_173241.1;HCoV-HKU1_membrane;114 | 0.05 | 0 |

| | | |
|---|---|---|
| NC_006577.2;YP_173241.1;HCoV-HKU1_membrane;115 | 0.05 | 0.025 |
| NC_006577.2;YP_173241.1;HCoV-HKU1_membrane;116 | 0 | 0 |
| NC_006577.2;YP_173241.1;HCoV-HKU1_membrane;117 | 0 | 0 |
| NC_006577.2;YP_173241.1;HCoV-HKU1_membrane;118 | 0 | 0 |
| NC_006577.2;YP_173241.1;HCoV-HKU1_membrane;119 | 0 | 0 |
| NC_006577.2;YP_173241.1;HCoV-HKU1_membrane;120 | 0 | 0 |
| NC_006577.2;YP_173241.1;HCoV-HKU1_membrane;121 | 0 | 0 |
| NC_006577.2;YP_173241.1;HCoV-HKU1_membrane;122 | 0 | 0.025 |
| NC_006577.2;YP_173241.1;HCoV-HKU1_membrane;123 | 0.05 | 0.05 |
| NC_006577.2;YP_173241.1;HCoV-HKU1_membrane;124 | 0 | 0.025 |
| NC_006577.2;YP_173241.1;HCoV-HKU1_membrane;125 | 0 | 0.025 |
| NC_006577.2;YP_173241.1;HCoV-HKU1_membrane;126 | 0 | 0.05 |
| NC_006577.2;YP_173241.1;HCoV-HKU1_membrane;127 | 0 | 0.025 |
| NC_006577.2;YP_173241.1;HCoV-HKU1_membrane;128 | 0 | 0.025 |
| NC_006577.2;YP_173241.1;HCoV-HKU1_membrane;129 | 0 | 0.025 |
| NC_006577.2;YP_173241.1;HCoV-HKU1_membrane;130 | 0 | 0.025 |
| NC_006577.2;YP_173241.1;HCoV-HKU1_membrane;131 | 0 | 0.025 |
| NC_006577.2;YP_173241.1;HCoV-HKU1_membrane;132 | 0 | 0.025 |
| NC_006577.2;YP_173241.1;HCoV-HKU1_membrane;133 | 0 | 0 |
| NC_006577.2;YP_173241.1;HCoV-HKU1_membrane;134 | 0 | 0 |
| NC_006577.2;YP_173241.1;HCoV-HKU1_membrane;135 | 0 | 0 |
| NC_006577.2;YP_173241.1;HCoV-HKU1_membrane;136 | 0 | 0 |
| NC_006577.2;YP_173241.1;HCoV-HKU1_membrane;137 | 0 | 0 |
| NC_006577.2;YP_173241.1;HCoV-HKU1_membrane;138 | 0 | 0 |
| NC_006577.2;YP_173241.1;HCoV-HKU1_membrane;139 | 0 | 0 |
| NC_006577.2;YP_173241.1;HCoV-HKU1_membrane;140 | 0 | 0 |
| NC_006577.2;YP_173241.1;HCoV-HKU1_membrane;141 | 0 | 0 |
| NC_006577.2;YP_173241.1;HCoV-HKU1_membrane;142 | 0 | 0 |
| NC_006577.2;YP_173241.1;HCoV-HKU1_membrane;143 | 0 | 0.025 |
| NC_006577.2;YP_173241.1;HCoV-HKU1_membrane;144 | 0 | 0.05 |
| NC_006577.2;YP_173241.1;HCoV-HKU1_membrane;145 | 0 | 0.05 |
| NC_006577.2;YP_173241.1;HCoV-HKU1_membrane;146 | 0 | 0.05 |
| NC_006577.2;YP_173241.1;HCoV-HKU1_membrane;147 | 0 | 0.05 |
| NC_006577.2;YP_173241.1;HCoV-HKU1_membrane;148 | 0 | 0.05 |
| NC_006577.2;YP_173241.1;HCoV-HKU1_membrane;149 | 0 | 0.05 |
| NC_006577.2;YP_173241.1;HCoV-HKU1_membrane;150 | 0 | 0.05 |
| NC_006577.2;YP_173241.1;HCoV-HKU1_membrane;151 | 0 | 0.05 |
| NC_006577.2;YP_173241.1;HCoV-HKU1_membrane;152 | 0 | 0.1 |
| NC_006577.2;YP_173241.1;HCoV-HKU1_membrane;153 | 0.05 | 0.1 |
| NC_006577.2;YP_173241.1;HCoV-HKU1_membrane;154 | 0 | 0.05 |
| NC_006577.2;YP_173241.1;HCoV-HKU1_membrane;155 | 0 | 0.025 |
| NC_006577.2;YP_173241.1;HCoV-HKU1_membrane;156 | 0 | 0 |
| NC_006577.2;YP_173241.1;HCoV-HKU1_membrane;157 | 0 | 0 |

| | | |
|---|---|---|
| NC_006577.2;YP_173241.1;HCoV-HKU1_membrane;158 | 0 | 0 |
| NC_006577.2;YP_173241.1;HCoV-HKU1_membrane;159 | 0 | 0 |
| NC_006577.2;YP_173241.1;HCoV-HKU1_membrane;160 | 0 | 0 |
| NC_006577.2;YP_173241.1;HCoV-HKU1_membrane;161 | 0 | 0.025 |
| NC_006577.2;YP_173241.1;HCoV-HKU1_membrane;162 | 0 | 0 |
| NC_006577.2;YP_173241.1;HCoV-HKU1_membrane;163 | 0 | 0 |
| NC_006577.2;YP_173241.1;HCoV-HKU1_membrane;164 | 0 | 0 |
| NC_006577.2;YP_173241.1;HCoV-HKU1_membrane;165 | 0 | 0 |
| NC_006577.2;YP_173241.1;HCoV-HKU1_membrane;166 | 0 | 0 |
| NC_006577.2;YP_173241.1;HCoV-HKU1_membrane;167 | 0 | 0 |
| NC_006577.2;YP_173241.1;HCoV-HKU1_membrane;168 | 0 | 0 |
| NC_006577.2;YP_173241.1;HCoV-HKU1_membrane;169 | 0 | 0 |
| NC_006577.2;YP_173241.1;HCoV-HKU1_membrane;170 | 0 | 0 |
| NC_006577.2;YP_173241.1;HCoV-HKU1_membrane;171 | 0 | 0 |
| NC_006577.2;YP_173241.1;HCoV-HKU1_membrane;172 | 0 | 0 |
| NC_006577.2;YP_173241.1;HCoV-HKU1_membrane;173 | 0 | 0 |
| NC_006577.2;YP_173241.1;HCoV-HKU1_membrane;174 | 0 | 0 |
| NC_006577.2;YP_173241.1;HCoV-HKU1_membrane;175 | 0 | 0 |
| NC_006577.2;YP_173241.1;HCoV-HKU1_membrane;176 | 0 | 0 |
| NC_006577.2;YP_173241.1;HCoV-HKU1_membrane;177 | 0 | 0 |
| NC_006577.2;YP_173241.1;HCoV-HKU1_membrane;178 | 0 | 0 |
| NC_006577.2;YP_173241.1;HCoV-HKU1_membrane;179 | 0.05 | 0.025 |
| NC_006577.2;YP_173241.1;HCoV-HKU1_membrane;180 | 0 | 0 |
| NC_006577.2;YP_173241.1;HCoV-HKU1_membrane;181 | 0 | 0.025 |
| NC_006577.2;YP_173241.1;HCoV-HKU1_membrane;182 | 0.05 | 0.075 |
| NC_006577.2;YP_173241.1;HCoV-HKU1_membrane;183 | 0 | 0 |
| NC_006577.2;YP_173241.1;HCoV-HKU1_membrane;184 | 0 | 0 |
| NC_006577.2;YP_173241.1;HCoV-HKU1_membrane;185 | 0 | 0 |
| NC_006577.2;YP_173241.1;HCoV-HKU1_membrane;186 | 0 | 0 |
| NC_006577.2;YP_173241.1;HCoV-HKU1_membrane;187 | 0 | 0 |
| NC_006577.2;YP_173241.1;HCoV-HKU1_membrane;188 | 0 | 0 |
| NC_006577.2;YP_173241.1;HCoV-HKU1_membrane;189 | 0 | 0 |
| NC_006577.2;YP_173241.1;HCoV-HKU1_membrane;190 | 0 | 0 |
| NC_006577.2;YP_173241.1;HCoV-HKU1_membrane;191 | 0 | 0 |
| NC_006577.2;YP_173241.1;HCoV-HKU1_membrane;192 | 0 | 0 |
| NC_006577.2;YP_173241.1;HCoV-HKU1_membrane;193 | 0 | 0 |
| NC_006577.2;YP_173241.1;HCoV-HKU1_membrane;194 | 0 | 0 |
| NC_006577.2;YP_173241.1;HCoV-HKU1_membrane;195 | 0 | 0 |
| NC_006577.2;YP_173241.1;HCoV-HKU1_membrane;196 | 0 | 0 |
| NC_006577.2;YP_173241.1;HCoV-HKU1_membrane;197 | 0 | 0 |
| NC_006577.2;YP_173241.1;HCoV-HKU1_membrane;198 | 0 | 0.025 |
| NC_006577.2;YP_173241.1;HCoV-HKU1_membrane;199 | 0 | 0.025 |
| NC_006577.2;YP_173241.1;HCoV-HKU1_membrane;200 | 0 | 0.025 |

| | | |
|---|---|---|
| NC_006577.2;YP_173241.1;HCoV-HKU1_membrane;201 | 0 | 0.025 |
| NC_006577.2;YP_173241.1;HCoV-HKU1_membrane;202 | 0.05 | 0.075 |
| NC_006577.2;YP_173241.1;HCoV-HKU1_membrane;203 | 0 | 0.05 |
| NC_006577.2;YP_173241.1;HCoV-HKU1_membrane;204 | 0 | 0.05 |
| NC_006577.2;YP_173241.1;HCoV-HKU1_membrane;205 | 0 | 0 |
| NC_006577.2;YP_173241.1;HCoV-HKU1_membrane;206 | 0 | 0.025 |
| NC_006577.2;YP_173241.1;HCoV-HKU1_membrane;207 | 0 | 0 |
| NC_006577.2;YP_173241.1;HCoV-HKU1_membrane;208 | 0 | 0 |
| NC_006577.2;YP_173241.1;HCoV-HKU1_membrane;209 | 0 | 0 |
| NC_006577.2;YP_173241.1;HCoV-HKU1_membrane;210 | 0 | 0 |
| NC_006577.2;YP_173241.1;HCoV-HKU1_membrane;211 | 0 | 0 |
| NC_006577.2;YP_173241.1;HCoV-HKU1_membrane;212 | 0 | 0.025 |
| NC_006577.2;YP_173242.1;HCoV-HKU1_nucleocapsi;1 | 0 | 0.025 |
| NC_006577.2;YP_173242.1;HCoV-HKU1_nucleocapsi;2 | 0.05 | 0.05 |
| NC_006577.2;YP_173242.1;HCoV-HKU1_nucleocapsi;3 | 0 | 0.075 |
| NC_006577.2;YP_173242.1;HCoV-HKU1_nucleocapsi;4 | 0.05 | 0.05 |
| NC_006577.2;YP_173242.1;HCoV-HKU1_nucleocapsi;5 | 0 | 0.05 |
| NC_006577.2;YP_173242.1;HCoV-HKU1_nucleocapsi;6 | 0 | 0.075 |
| NC_006577.2;YP_173242.1;HCoV-HKU1_nucleocapsi;7 | 0 | 0.05 |
| NC_006577.2;YP_173242.1;HCoV-HKU1_nucleocapsi;8 | 0 | 0.05 |
| NC_006577.2;YP_173242.1;HCoV-HKU1_nucleocapsi;9 | 0 | 0 |
| NC_006577.2;YP_173242.1;HCoV-HKU1_nucleocapsi;10 | 0 | 0 |
| NC_006577.2;YP_173242.1;HCoV-HKU1_nucleocapsi;11 | 0 | 0 |
| NC_006577.2;YP_173242.1;HCoV-HKU1_nucleocapsi;12 | 0 | 0 |
| NC_006577.2;YP_173242.1;HCoV-HKU1_nucleocapsi;13 | 0 | 0 |
| NC_006577.2;YP_173242.1;HCoV-HKU1_nucleocapsi;14 | 0 | 0 |
| NC_006577.2;YP_173242.1;HCoV-HKU1_nucleocapsi;15 | 0 | 0 |
| NC_006577.2;YP_173242.1;HCoV-HKU1_nucleocapsi;16 | 0.05 | 0 |
| NC_006577.2;YP_173242.1;HCoV-HKU1_nucleocapsi;17 | 0.05 | 0 |
| NC_006577.2;YP_173242.1;HCoV-HKU1_nucleocapsi;18 | 0.05 | 0.025 |
| NC_006577.2;YP_173242.1;HCoV-HKU1_nucleocapsi;19 | 0.05 | 0.025 |
| NC_006577.2;YP_173242.1;HCoV-HKU1_nucleocapsi;20 | 0.05 | 0.025 |
| NC_006577.2;YP_173242.1;HCoV-HKU1_nucleocapsi;21 | 0.05 | 0.025 |
| NC_006577.2;YP_173242.1;HCoV-HKU1_nucleocapsi;22 | 0.05 | 0.05 |
| NC_006577.2;YP_173242.1;HCoV-HKU1_nucleocapsi;23 | 0.05 | 0.075 |
| NC_006577.2;YP_173242.1;HCoV-HKU1_nucleocapsi;24 | 0 | 0.05 |
| NC_006577.2;YP_173242.1;HCoV-HKU1_nucleocapsi;25 | 0.05 | 0.05 |
| NC_006577.2;YP_173242.1;HCoV-HKU1_nucleocapsi;26 | 0.15 | 0.05 |
| NC_006577.2;YP_173242.1;HCoV-HKU1_nucleocapsi;27 | 0 | 0.025 |
| NC_006577.2;YP_173242.1;HCoV-HKU1_nucleocapsi;28 | 0 | 0 |
| NC_006577.2;YP_173242.1;HCoV-HKU1_nucleocapsi;29 | 0 | 0 |
| NC_006577.2;YP_173242.1;HCoV-HKU1_nucleocapsi;30 | 0 | 0 |
| NC_006577.2;YP_173242.1;HCoV-HKU1_nucleocapsi;31 | 0 | 0 |

| | | |
|---|---|---|
| NC_006577.2;YP_173242.1;HCoV-HKU1_nucleocapsi;32 | 0 | 0 |
| NC_006577.2;YP_173242.1;HCoV-HKU1_nucleocapsi;33 | 0 | 0 |
| NC_006577.2;YP_173242.1;HCoV-HKU1_nucleocapsi;34 | 0 | 0 |
| NC_006577.2;YP_173242.1;HCoV-HKU1_nucleocapsi;35 | 0 | 0 |
| NC_006577.2;YP_173242.1;HCoV-HKU1_nucleocapsi;36 | 0 | 0 |
| NC_006577.2;YP_173242.1;HCoV-HKU1_nucleocapsi;37 | 0 | 0 |
| NC_006577.2;YP_173242.1;HCoV-HKU1_nucleocapsi;38 | 0 | 0 |
| NC_006577.2;YP_173242.1;HCoV-HKU1_nucleocapsi;39 | 0 | 0 |
| NC_006577.2;YP_173242.1;HCoV-HKU1_nucleocapsi;40 | 0 | 0 |
| NC_006577.2;YP_173242.1;HCoV-HKU1_nucleocapsi;41 | 0 | 0 |
| NC_006577.2;YP_173242.1;HCoV-HKU1_nucleocapsi;42 | 0 | 0 |
| NC_006577.2;YP_173242.1;HCoV-HKU1_nucleocapsi;43 | 0 | 0 |
| NC_006577.2;YP_173242.1;HCoV-HKU1_nucleocapsi;44 | 0 | 0.025 |
| NC_006577.2;YP_173242.1;HCoV-HKU1_nucleocapsi;45 | 0 | 0 |
| NC_006577.2;YP_173242.1;HCoV-HKU1_nucleocapsi;46 | 0 | 0 |
| NC_006577.2;YP_173242.1;HCoV-HKU1_nucleocapsi;47 | 0 | 0 |
| NC_006577.2;YP_173242.1;HCoV-HKU1_nucleocapsi;48 | 0 | 0.025 |
| NC_006577.2;YP_173242.1;HCoV-HKU1_nucleocapsi;49 | 0 | 0 |
| NC_006577.2;YP_173242.1;HCoV-HKU1_nucleocapsi;50 | 0 | 0 |
| NC_006577.2;YP_173242.1;HCoV-HKU1_nucleocapsi;51 | 0 | 0 |
| NC_006577.2;YP_173242.1;HCoV-HKU1_nucleocapsi;52 | 0 | 0 |
| NC_006577.2;YP_173242.1;HCoV-HKU1_nucleocapsi;53 | 0 | 0 |
| NC_006577.2;YP_173242.1;HCoV-HKU1_nucleocapsi;54 | 0 | 0 |
| NC_006577.2;YP_173242.1;HCoV-HKU1_nucleocapsi;55 | 0 | 0 |
| NC_006577.2;YP_173242.1;HCoV-HKU1_nucleocapsi;56 | 0 | 0 |
| NC_006577.2;YP_173242.1;HCoV-HKU1_nucleocapsi;57 | 0 | 0 |
| NC_006577.2;YP_173242.1;HCoV-HKU1_nucleocapsi;58 | 0 | 0 |
| NC_006577.2;YP_173242.1;HCoV-HKU1_nucleocapsi;59 | 0 | 0 |
| NC_006577.2;YP_173242.1;HCoV-HKU1_nucleocapsi;60 | 0 | 0 |
| NC_006577.2;YP_173242.1;HCoV-HKU1_nucleocapsi;61 | 0 | 0 |
| NC_006577.2;YP_173242.1;HCoV-HKU1_nucleocapsi;62 | 0 | 0 |
| NC_006577.2;YP_173242.1;HCoV-HKU1_nucleocapsi;63 | 0 | 0 |
| NC_006577.2;YP_173242.1;HCoV-HKU1_nucleocapsi;64 | 0 | 0 |
| NC_006577.2;YP_173242.1;HCoV-HKU1_nucleocapsi;65 | 0 | 0 |
| NC_006577.2;YP_173242.1;HCoV-HKU1_nucleocapsi;66 | 0 | 0 |
| NC_006577.2;YP_173242.1;HCoV-HKU1_nucleocapsi;67 | 0.05 | 0.075 |
| NC_006577.2;YP_173242.1;HCoV-HKU1_nucleocapsi;68 | 0 | 0 |
| NC_006577.2;YP_173242.1;HCoV-HKU1_nucleocapsi;69 | 0 | 0 |
| NC_006577.2;YP_173242.1;HCoV-HKU1_nucleocapsi;70 | 0 | 0 |
| NC_006577.2;YP_173242.1;HCoV-HKU1_nucleocapsi;71 | 0.1 | 0.05 |
| NC_006577.2;YP_173242.1;HCoV-HKU1_nucleocapsi;72 | 0 | 0 |
| NC_006577.2;YP_173242.1;HCoV-HKU1_nucleocapsi;73 | 0 | 0.075 |
| NC_006577.2;YP_173242.1;HCoV-HKU1_nucleocapsi;74 | 0 | 0.025 |

| | | |
|---|---|---|
| NC_006577.2;YP_173242.1;HCoV-HKU1_nucleocapsi;75 | 0 | 0 |
| NC_006577.2;YP_173242.1;HCoV-HKU1_nucleocapsi;76 | 0 | 0 |
| NC_006577.2;YP_173242.1;HCoV-HKU1_nucleocapsi;77 | 0.05 | 0.025 |
| NC_006577.2;YP_173242.1;HCoV-HKU1_nucleocapsi;78 | 0 | 0 |
| NC_006577.2;YP_173242.1;HCoV-HKU1_nucleocapsi;79 | 0 | 0.025 |
| NC_006577.2;YP_173242.1;HCoV-HKU1_nucleocapsi;80 | 0 | 0 |
| NC_006577.2;YP_173242.1;HCoV-HKU1_nucleocapsi;81 | 0 | 0 |
| NC_006577.2;YP_173242.1;HCoV-HKU1_nucleocapsi;82 | 0 | 0 |
| NC_006577.2;YP_173242.1;HCoV-HKU1_nucleocapsi;83 | 0 | 0.025 |
| NC_006577.2;YP_173242.1;HCoV-HKU1_nucleocapsi;84 | 0 | 0.05 |
| NC_006577.2;YP_173242.1;HCoV-HKU1_nucleocapsi;85 | 0 | 0.05 |
| NC_006577.2;YP_173242.1;HCoV-HKU1_nucleocapsi;86 | 0 | 0.025 |
| NC_006577.2;YP_173242.1;HCoV-HKU1_nucleocapsi;87 | 0 | 0 |
| NC_006577.2;YP_173242.1;HCoV-HKU1_nucleocapsi;88 | 0 | 0 |
| NC_006577.2;YP_173242.1;HCoV-HKU1_nucleocapsi;89 | 0 | 0.025 |
| NC_006577.2;YP_173242.1;HCoV-HKU1_nucleocapsi;90 | 0 | 0 |
| NC_006577.2;YP_173242.1;HCoV-HKU1_nucleocapsi;91 | 0 | 0 |
| NC_006577.2;YP_173242.1;HCoV-HKU1_nucleocapsi;92 | 0 | 0 |
| NC_006577.2;YP_173242.1;HCoV-HKU1_nucleocapsi;93 | 0 | 0 |
| NC_006577.2;YP_173242.1;HCoV-HKU1_nucleocapsi;94 | 0 | 0 |
| NC_006577.2;YP_173242.1;HCoV-HKU1_nucleocapsi;95 | 0 | 0 |
| NC_006577.2;YP_173242.1;HCoV-HKU1_nucleocapsi;96 | 0 | 0 |
| NC_006577.2;YP_173242.1;HCoV-HKU1_nucleocapsi;97 | 0 | 0 |
| NC_006577.2;YP_173242.1;HCoV-HKU1_nucleocapsi;98 | 0 | 0 |
| NC_006577.2;YP_173242.1;HCoV-HKU1_nucleocapsi;99 | 0 | 0 |
| NC_006577.2;YP_173242.1;HCoV-HKU1_nucleocapsi;100 | 0 | 0.025 |
| NC_006577.2;YP_173242.1;HCoV-HKU1_nucleocapsi;101 | 0 | 0 |
| NC_006577.2;YP_173242.1;HCoV-HKU1_nucleocapsi;102 | 0 | 0 |
| NC_006577.2;YP_173242.1;HCoV-HKU1_nucleocapsi;103 | 0 | 0 |
| NC_006577.2;YP_173242.1;HCoV-HKU1_nucleocapsi;104 | 0 | 0 |
| NC_006577.2;YP_173242.1;HCoV-HKU1_nucleocapsi;105 | 0 | 0.025 |
| NC_006577.2;YP_173242.1;HCoV-HKU1_nucleocapsi;106 | 0 | 0 |
| NC_006577.2;YP_173242.1;HCoV-HKU1_nucleocapsi;107 | 0 | 0.025 |
| NC_006577.2;YP_173242.1;HCoV-HKU1_nucleocapsi;108 | 0 | 0.025 |
| NC_006577.2;YP_173242.1;HCoV-HKU1_nucleocapsi;109 | 0 | 0 |
| NC_006577.2;YP_173242.1;HCoV-HKU1_nucleocapsi;110 | 0 | 0 |
| NC_006577.2;YP_173242.1;HCoV-HKU1_nucleocapsi;111 | 0.05 | 0 |
| NC_006577.2;YP_173242.1;HCoV-HKU1_nucleocapsi;112 | 0 | 0 |
| NC_006577.2;YP_173242.1;HCoV-HKU1_nucleocapsi;113 | 0 | 0 |
| NC_006577.2;YP_173242.1;HCoV-HKU1_nucleocapsi;114 | 0 | 0 |
| NC_006577.2;YP_173242.1;HCoV-HKU1_nucleocapsi;115 | 0 | 0 |
| NC_006577.2;YP_173242.1;HCoV-HKU1_nucleocapsi;116 | 0 | 0 |
| NC_006577.2;YP_173242.1;HCoV-HKU1_nucleocapsi;117 | 0 | 0 |

| | | |
|---|---|---|
| NC_006577.2;YP_173242.1;HCoV-HKU1_nucleocapsi;118 | 0 | 0 |
| NC_006577.2;YP_173242.1;HCoV-HKU1_nucleocapsi;119 | 0 | 0 |
| NC_006577.2;YP_173242.1;HCoV-HKU1_nucleocapsi;120 | 0 | 0 |
| NC_006577.2;YP_173242.1;HCoV-HKU1_nucleocapsi;121 | 0 | 0 |
| NC_006577.2;YP_173242.1;HCoV-HKU1_nucleocapsi;122 | 0 | 0 |
| NC_006577.2;YP_173242.1;HCoV-HKU1_nucleocapsi;123 | 0 | 0 |
| NC_006577.2;YP_173242.1;HCoV-HKU1_nucleocapsi;124 | 0 | 0 |
| NC_006577.2;YP_173242.1;HCoV-HKU1_nucleocapsi;125 | 0 | 0.025 |
| NC_006577.2;YP_173242.1;HCoV-HKU1_nucleocapsi;126 | 0 | 0 |
| NC_006577.2;YP_173242.1;HCoV-HKU1_nucleocapsi;127 | 0 | 0 |
| NC_006577.2;YP_173242.1;HCoV-HKU1_nucleocapsi;128 | 0.1 | 0.025 |
| NC_006577.2;YP_173242.1;HCoV-HKU1_nucleocapsi;129 | 0.05 | 0 |
| NC_006577.2;YP_173242.1;HCoV-HKU1_nucleocapsi;130 | 0 | 0 |
| NC_006577.2;YP_173242.1;HCoV-HKU1_nucleocapsi;131 | 0 | 0 |
| NC_006577.2;YP_173242.1;HCoV-HKU1_nucleocapsi;132 | 0 | 0 |
| NC_006577.2;YP_173242.1;HCoV-HKU1_nucleocapsi;133 | 0 | 0 |
| NC_006577.2;YP_173242.1;HCoV-HKU1_nucleocapsi;134 | 0 | 0 |
| NC_006577.2;YP_173242.1;HCoV-HKU1_nucleocapsi;135 | 0 | 0 |
| NC_006577.2;YP_173242.1;HCoV-HKU1_nucleocapsi;136 | 0 | 0 |
| NC_006577.2;YP_173242.1;HCoV-HKU1_nucleocapsi;137 | 0 | 0 |
| NC_006577.2;YP_173242.1;HCoV-HKU1_nucleocapsi;138 | 0 | 0 |
| NC_006577.2;YP_173242.1;HCoV-HKU1_nucleocapsi;139 | 0.05 | 0 |
| NC_006577.2;YP_173242.1;HCoV-HKU1_nucleocapsi;140 | 0.05 | 0.025 |
| NC_006577.2;YP_173242.1;HCoV-HKU1_nucleocapsi;141 | 0.05 | 0 |
| NC_006577.2;YP_173242.1;HCoV-HKU1_nucleocapsi;142 | 0.05 | 0.025 |
| NC_006577.2;YP_173242.1;HCoV-HKU1_nucleocapsi;143 | 0.05 | 0.05 |
| NC_006577.2;YP_173242.1;HCoV-HKU1_nucleocapsi;144 | 0.05 | 0.025 |
| NC_006577.2;YP_173242.1;HCoV-HKU1_nucleocapsi;145 | 0.05 | 0.025 |
| NC_006577.2;YP_173242.1;HCoV-HKU1_nucleocapsi;146 | 0.05 | 0.05 |
| NC_006577.2;YP_173242.1;HCoV-HKU1_nucleocapsi;147 | 0.05 | 0.05 |
| NC_006577.2;YP_173242.1;HCoV-HKU1_nucleocapsi;148 | 0.05 | 0.025 |
| NC_006577.2;YP_173242.1;HCoV-HKU1_nucleocapsi;149 | 0.05 | 0.025 |
| NC_006577.2;YP_173242.1;HCoV-HKU1_nucleocapsi;150 | 0.05 | 0.05 |
| NC_006577.2;YP_173242.1;HCoV-HKU1_nucleocapsi;151 | 0.05 | 0.05 |
| NC_006577.2;YP_173242.1;HCoV-HKU1_nucleocapsi;152 | 0 | 0.025 |
| NC_006577.2;YP_173242.1;HCoV-HKU1_nucleocapsi;153 | 0 | 0.025 |
| NC_006577.2;YP_173242.1;HCoV-HKU1_nucleocapsi;154 | 0 | 0.025 |
| NC_006577.2;YP_173242.1;HCoV-HKU1_nucleocapsi;155 | 0.05 | 0.05 |
| NC_006577.2;YP_173242.1;HCoV-HKU1_nucleocapsi;156 | 0.05 | 0.025 |
| NC_006577.2;YP_173242.1;HCoV-HKU1_nucleocapsi;157 | 0 | 0 |
| NC_006577.2;YP_173242.1;HCoV-HKU1_nucleocapsi;158 | 0.05 | 0.025 |
| NC_006577.2;YP_173242.1;HCoV-HKU1_nucleocapsi;159 | 0.05 | 0 |
| NC_006577.2;YP_173242.1;HCoV-HKU1_nucleocapsi;160 | 0.05 | 0 |

| | | |
|---|---|---|
| NC_006577.2;YP_173242.1;HCoV-HKU1_nucleocapsi;161 | 0.05 | 0 |
| NC_006577.2;YP_173242.1;HCoV-HKU1_nucleocapsi;162 | 0.05 | 0.025 |
| NC_006577.2;YP_173242.1;HCoV-HKU1_nucleocapsi;163 | 0.05 | 0 |
| NC_006577.2;YP_173242.1;HCoV-HKU1_nucleocapsi;164 | 0.1 | 0 |
| NC_006577.2;YP_173242.1;HCoV-HKU1_nucleocapsi;165 | 0 | 0 |
| NC_006577.2;YP_173242.1;HCoV-HKU1_nucleocapsi;166 | 0 | 0 |
| NC_006577.2;YP_173242.1;HCoV-HKU1_nucleocapsi;167 | 0 | 0 |
| NC_006577.2;YP_173242.1;HCoV-HKU1_nucleocapsi;168 | 0 | 0 |
| NC_006577.2;YP_173242.1;HCoV-HKU1_nucleocapsi;169 | 0.05 | 0.025 |
| NC_006577.2;YP_173242.1;HCoV-HKU1_nucleocapsi;170 | 0 | 0 |
| NC_006577.2;YP_173242.1;HCoV-HKU1_nucleocapsi;171 | 0 | 0 |
| NC_006577.2;YP_173242.1;HCoV-HKU1_nucleocapsi;172 | 0 | 0 |
| NC_006577.2;YP_173242.1;HCoV-HKU1_nucleocapsi;173 | 0 | 0 |
| NC_006577.2;YP_173242.1;HCoV-HKU1_nucleocapsi;174 | 0 | 0 |
| NC_006577.2;YP_173242.1;HCoV-HKU1_nucleocapsi;175 | 0 | 0 |
| NC_006577.2;YP_173242.1;HCoV-HKU1_nucleocapsi;176 | 0 | 0 |
| NC_006577.2;YP_173242.1;HCoV-HKU1_nucleocapsi;177 | 0 | 0 |
| NC_006577.2;YP_173242.1;HCoV-HKU1_nucleocapsi;178 | 0 | 0 |
| NC_006577.2;YP_173242.1;HCoV-HKU1_nucleocapsi;179 | 0 | 0 |
| NC_006577.2;YP_173242.1;HCoV-HKU1_nucleocapsi;180 | 0.1 | 0 |
| NC_006577.2;YP_173242.1;HCoV-HKU1_nucleocapsi;181 | 0 | 0 |
| NC_006577.2;YP_173242.1;HCoV-HKU1_nucleocapsi;182 | 0 | 0 |
| NC_006577.2;YP_173242.1;HCoV-HKU1_nucleocapsi;183 | 0 | 0 |
| NC_006577.2;YP_173242.1;HCoV-HKU1_nucleocapsi;184 | 0 | 0 |
| NC_006577.2;YP_173242.1;HCoV-HKU1_nucleocapsi;185 | 0 | 0 |
| NC_006577.2;YP_173242.1;HCoV-HKU1_nucleocapsi;186 | 0.05 | 0 |
| NC_006577.2;YP_173242.1;HCoV-HKU1_nucleocapsi;187 | 0.05 | 0.025 |
| NC_006577.2;YP_173242.1;HCoV-HKU1_nucleocapsi;188 | 0 | 0.05 |
| NC_006577.2;YP_173242.1;HCoV-HKU1_nucleocapsi;189 | 0 | 0.025 |
| NC_006577.2;YP_173242.1;HCoV-HKU1_nucleocapsi;190 | 0 | 0.025 |
| NC_006577.2;YP_173242.1;HCoV-HKU1_nucleocapsi;191 | 0 | 0 |
| NC_006577.2;YP_173242.1;HCoV-HKU1_nucleocapsi;192 | 0 | 0 |
| NC_006577.2;YP_173242.1;HCoV-HKU1_nucleocapsi;193 | 0 | 0.05 |
| NC_006577.2;YP_173242.1;HCoV-HKU1_nucleocapsi;194 | 0.1 | 0.025 |
| NC_006577.2;YP_173242.1;HCoV-HKU1_nucleocapsi;195 | 0 | 0.05 |
| NC_006577.2;YP_173242.1;HCoV-HKU1_nucleocapsi;196 | 0 | 0 |
| NC_006577.2;YP_173242.1;HCoV-HKU1_nucleocapsi;197 | 0 | 0 |
| NC_006577.2;YP_173242.1;HCoV-HKU1_nucleocapsi;198 | 0 | 0 |
| NC_006577.2;YP_173242.1;HCoV-HKU1_nucleocapsi;199 | 0 | 0 |
| NC_006577.2;YP_173242.1;HCoV-HKU1_nucleocapsi;200 | 0 | 0 |
| NC_006577.2;YP_173242.1;HCoV-HKU1_nucleocapsi;201 | 0 | 0 |
| NC_006577.2;YP_173242.1;HCoV-HKU1_nucleocapsi;202 | 0 | 0 |
| NC_006577.2;YP_173242.1;HCoV-HKU1_nucleocapsi;203 | 0 | 0 |

| | | |
|---|---|---|
| NC_006577.2;YP_173242.1;HCoV-HKU1_nucleocapsi;204 | 0 | 0.025 |
| NC_006577.2;YP_173242.1;HCoV-HKU1_nucleocapsi;205 | 0.05 | 0.025 |
| NC_006577.2;YP_173242.1;HCoV-HKU1_nucleocapsi;206 | 0 | 0 |
| NC_006577.2;YP_173242.1;HCoV-HKU1_nucleocapsi;207 | 0 | 0.025 |
| NC_006577.2;YP_173242.1;HCoV-HKU1_nucleocapsi;208 | 0 | 0.025 |
| NC_006577.2;YP_173242.1;HCoV-HKU1_nucleocapsi;209 | 0 | 0.025 |
| NC_006577.2;YP_173242.1;HCoV-HKU1_nucleocapsi;210 | 0 | 0.05 |
| NC_006577.2;YP_173242.1;HCoV-HKU1_nucleocapsi;211 | 0 | 0.025 |
| NC_006577.2;YP_173242.1;HCoV-HKU1_nucleocapsi;212 | 0.05 | 0.05 |
| NC_006577.2;YP_173242.1;HCoV-HKU1_nucleocapsi;213 | 0.05 | 0.025 |
| NC_006577.2;YP_173242.1;HCoV-HKU1_nucleocapsi;214 | 0 | 0 |
| NC_006577.2;YP_173242.1;HCoV-HKU1_nucleocapsi;215 | 0 | 0.1 |
| NC_006577.2;YP_173242.1;HCoV-HKU1_nucleocapsi;216 | 0.05 | 0 |
| NC_006577.2;YP_173242.1;HCoV-HKU1_nucleocapsi;217 | 0 | 0 |
| NC_006577.2;YP_173242.1;HCoV-HKU1_nucleocapsi;218 | 0 | 0 |
| NC_006577.2;YP_173242.1;HCoV-HKU1_nucleocapsi;219 | 0 | 0 |
| NC_006577.2;YP_173242.1;HCoV-HKU1_nucleocapsi;220 | 0 | 0 |
| NC_006577.2;YP_173242.1;HCoV-HKU1_nucleocapsi;221 | 0 | 0 |
| NC_006577.2;YP_173242.1;HCoV-HKU1_nucleocapsi;222 | 0 | 0 |
| NC_006577.2;YP_173242.1;HCoV-HKU1_nucleocapsi;223 | 0 | 0 |
| NC_006577.2;YP_173242.1;HCoV-HKU1_nucleocapsi;224 | 0 | 0 |
| NC_006577.2;YP_173242.1;HCoV-HKU1_nucleocapsi;225 | 0 | 0 |
| NC_006577.2;YP_173242.1;HCoV-HKU1_nucleocapsi;226 | 0 | 0 |
| NC_006577.2;YP_173242.1;HCoV-HKU1_nucleocapsi;227 | 0 | 0 |
| NC_006577.2;YP_173242.1;HCoV-HKU1_nucleocapsi;228 | 0 | 0 |
| NC_006577.2;YP_173242.1;HCoV-HKU1_nucleocapsi;229 | 0 | 0 |
| NC_006577.2;YP_173242.1;HCoV-HKU1_nucleocapsi;230 | 0 | 0 |
| NC_006577.2;YP_173242.1;HCoV-HKU1_nucleocapsi;231 | 0 | 0 |
| NC_006577.2;YP_173242.1;HCoV-HKU1_nucleocapsi;232 | 0.05 | 0.1 |
| NC_006577.2;YP_173242.1;HCoV-HKU1_nucleocapsi;233 | 0.05 | 0.075 |
| NC_006577.2;YP_173242.1;HCoV-HKU1_nucleocapsi;234 | 0.05 | 0.075 |
| NC_006577.2;YP_173242.1;HCoV-HKU1_nucleocapsi;235 | 0.05 | 0.05 |
| NC_006577.2;YP_173242.1;HCoV-HKU1_nucleocapsi;236 | 0.05 | 0.075 |
| NC_006577.2;YP_173242.1;HCoV-HKU1_nucleocapsi;237 | 0.05 | 0.075 |
| NC_006577.2;YP_173242.1;HCoV-HKU1_nucleocapsi;238 | 0.05 | 0.075 |
| NC_006577.2;YP_173242.1;HCoV-HKU1_nucleocapsi;239 | 0.05 | 0 |
| NC_006577.2;YP_173242.1;HCoV-HKU1_nucleocapsi;240 | 0.05 | 0 |
| NC_006577.2;YP_173242.1;HCoV-HKU1_nucleocapsi;241 | 0.05 | 0 |
| NC_006577.2;YP_173242.1;HCoV-HKU1_nucleocapsi;242 | 0.05 | 0 |
| NC_006577.2;YP_173242.1;HCoV-HKU1_nucleocapsi;243 | 0.05 | 0 |
| NC_006577.2;YP_173242.1;HCoV-HKU1_nucleocapsi;244 | 0.05 | 0 |
| NC_006577.2;YP_173242.1;HCoV-HKU1_nucleocapsi;245 | 0.05 | 0 |
| NC_006577.2;YP_173242.1;HCoV-HKU1_nucleocapsi;246 | 0.1 | 0 |

| | | |
|---|---|---|
| NC_006577.2;YP_173242.1;HCoV-HKU1_nucleocapsi;247 | 0.05 | 0 |
| NC_006577.2;YP_173242.1;HCoV-HKU1_nucleocapsi;248 | 0.05 | 0 |
| NC_006577.2;YP_173242.1;HCoV-HKU1_nucleocapsi;249 | 0 | 0 |
| NC_006577.2;YP_173242.1;HCoV-HKU1_nucleocapsi;250 | 0.05 | 0 |
| NC_006577.2;YP_173242.1;HCoV-HKU1_nucleocapsi;251 | 0.05 | 0 |
| NC_006577.2;YP_173242.1;HCoV-HKU1_nucleocapsi;252 | 0.05 | 0 |
| NC_006577.2;YP_173242.1;HCoV-HKU1_nucleocapsi;253 | 0.05 | 0 |
| NC_006577.2;YP_173242.1;HCoV-HKU1_nucleocapsi;254 | 0.05 | 0 |
| NC_006577.2;YP_173242.1;HCoV-HKU1_nucleocapsi;255 | 0.05 | 0.025 |
| NC_006577.2;YP_173242.1;HCoV-HKU1_nucleocapsi;256 | 0 | 0 |
| NC_006577.2;YP_173242.1;HCoV-HKU1_nucleocapsi;257 | 0 | 0 |
| NC_006577.2;YP_173242.1;HCoV-HKU1_nucleocapsi;258 | 0 | 0 |
| NC_006577.2;YP_173242.1;HCoV-HKU1_nucleocapsi;259 | 0 | 0 |
| NC_006577.2;YP_173242.1;HCoV-HKU1_nucleocapsi;260 | 0 | 0 |
| NC_006577.2;YP_173242.1;HCoV-HKU1_nucleocapsi;261 | 0 | 0 |
| NC_006577.2;YP_173242.1;HCoV-HKU1_nucleocapsi;262 | 0 | 0 |
| NC_006577.2;YP_173242.1;HCoV-HKU1_nucleocapsi;263 | 0 | 0 |
| NC_006577.2;YP_173242.1;HCoV-HKU1_nucleocapsi;264 | 0 | 0 |
| NC_006577.2;YP_173242.1;HCoV-HKU1_nucleocapsi;265 | 0 | 0 |
| NC_006577.2;YP_173242.1;HCoV-HKU1_nucleocapsi;266 | 0 | 0 |
| NC_006577.2;YP_173242.1;HCoV-HKU1_nucleocapsi;267 | 0 | 0 |
| NC_006577.2;YP_173242.1;HCoV-HKU1_nucleocapsi;268 | 0.05 | 0 |
| NC_006577.2;YP_173242.1;HCoV-HKU1_nucleocapsi;269 | 0 | 0 |
| NC_006577.2;YP_173242.1;HCoV-HKU1_nucleocapsi;270 | 0 | 0.025 |
| NC_006577.2;YP_173242.1;HCoV-HKU1_nucleocapsi;271 | 0 | 0 |
| NC_006577.2;YP_173242.1;HCoV-HKU1_nucleocapsi;272 | 0 | 0.025 |
| NC_006577.2;YP_173242.1;HCoV-HKU1_nucleocapsi;273 | 0 | 0.025 |
| NC_006577.2;YP_173242.1;HCoV-HKU1_nucleocapsi;274 | 0 | 0.025 |
| NC_006577.2;YP_173242.1;HCoV-HKU1_nucleocapsi;275 | 0 | 0.025 |
| NC_006577.2;YP_173242.1;HCoV-HKU1_nucleocapsi;276 | 0 | 0 |
| NC_006577.2;YP_173242.1;HCoV-HKU1_nucleocapsi;277 | 0 | 0 |
| NC_006577.2;YP_173242.1;HCoV-HKU1_nucleocapsi;278 | 0 | 0 |
| NC_006577.2;YP_173242.1;HCoV-HKU1_nucleocapsi;279 | 0 | 0 |
| NC_006577.2;YP_173242.1;HCoV-HKU1_nucleocapsi;280 | 0.05 | 0.025 |
| NC_006577.2;YP_173242.1;HCoV-HKU1_nucleocapsi;281 | 0.05 | 0.025 |
| NC_006577.2;YP_173242.1;HCoV-HKU1_nucleocapsi;282 | 0.05 | 0.025 |
| NC_006577.2;YP_173242.1;HCoV-HKU1_nucleocapsi;283 | 0.05 | 0.025 |
| NC_006577.2;YP_173242.1;HCoV-HKU1_nucleocapsi;284 | 0.05 | 0.025 |
| NC_006577.2;YP_173242.1;HCoV-HKU1_nucleocapsi;285 | 0.05 | 0 |
| NC_006577.2;YP_173242.1;HCoV-HKU1_nucleocapsi;286 | 0.05 | 0 |
| NC_006577.2;YP_173242.1;HCoV-HKU1_nucleocapsi;287 | 0 | 0 |
| NC_006577.2;YP_173242.1;HCoV-HKU1_nucleocapsi;288 | 0 | 0.025 |
| NC_006577.2;YP_173242.1;HCoV-HKU1_nucleocapsi;289 | 0 | 0 |

| | | |
|---|---|---|
| NC_006577.2;YP_173242.1;HCoV-HKU1_nucleocapsi;290 | 0 | 0.025 |
| NC_006577.2;YP_173242.1;HCoV-HKU1_nucleocapsi;291 | 0 | 0 |
| NC_006577.2;YP_173242.1;HCoV-HKU1_nucleocapsi;292 | 0 | 0 |
| NC_006577.2;YP_173242.1;HCoV-HKU1_nucleocapsi;293 | 0 | 0 |
| NC_006577.2;YP_173242.1;HCoV-HKU1_nucleocapsi;294 | 0 | 0 |
| NC_006577.2;YP_173242.1;HCoV-HKU1_nucleocapsi;295 | 0 | 0 |
| NC_006577.2;YP_173242.1;HCoV-HKU1_nucleocapsi;296 | 0 | 0.05 |
| NC_006577.2;YP_173242.1;HCoV-HKU1_nucleocapsi;297 | 0 | 0.025 |
| NC_006577.2;YP_173242.1;HCoV-HKU1_nucleocapsi;298 | 0 | 0 |
| NC_006577.2;YP_173242.1;HCoV-HKU1_nucleocapsi;299 | 0 | 0.025 |
| NC_006577.2;YP_173242.1;HCoV-HKU1_nucleocapsi;300 | 0 | 0 |
| NC_006577.2;YP_173242.1;HCoV-HKU1_nucleocapsi;301 | 0.05 | 0.025 |
| NC_006577.2;YP_173242.1;HCoV-HKU1_nucleocapsi;302 | 0.05 | 0 |
| NC_006577.2;YP_173242.1;HCoV-HKU1_nucleocapsi;303 | 0.05 | 0 |
| NC_006577.2;YP_173242.1;HCoV-HKU1_nucleocapsi;304 | 0.05 | 0 |
| NC_006577.2;YP_173242.1;HCoV-HKU1_nucleocapsi;305 | 0.1 | 0 |
| NC_006577.2;YP_173242.1;HCoV-HKU1_nucleocapsi;306 | 0 | 0 |
| NC_006577.2;YP_173242.1;HCoV-HKU1_nucleocapsi;307 | 0 | 0 |
| NC_006577.2;YP_173242.1;HCoV-HKU1_nucleocapsi;308 | 0 | 0 |
| NC_006577.2;YP_173242.1;HCoV-HKU1_nucleocapsi;309 | 0.05 | 0 |
| NC_006577.2;YP_173242.1;HCoV-HKU1_nucleocapsi;310 | 0 | 0 |
| NC_006577.2;YP_173242.1;HCoV-HKU1_nucleocapsi;311 | 0 | 0 |
| NC_006577.2;YP_173242.1;HCoV-HKU1_nucleocapsi;312 | 0 | 0.025 |
| NC_006577.2;YP_173242.1;HCoV-HKU1_nucleocapsi;313 | 0 | 0.025 |
| NC_006577.2;YP_173242.1;HCoV-HKU1_nucleocapsi;314 | 0 | 0.025 |
| NC_006577.2;YP_173242.1;HCoV-HKU1_nucleocapsi;315 | 0 | 0.025 |
| NC_006577.2;YP_173242.1;HCoV-HKU1_nucleocapsi;316 | 0 | 0.025 |
| NC_006577.2;YP_173242.1;HCoV-HKU1_nucleocapsi;317 | 0 | 0.025 |
| NC_006577.2;YP_173242.1;HCoV-HKU1_nucleocapsi;318 | 0 | 0.025 |
| NC_006577.2;YP_173242.1;HCoV-HKU1_nucleocapsi;319 | 0 | 0.025 |
| NC_006577.2;YP_173242.1;HCoV-HKU1_nucleocapsi;320 | 0 | 0.025 |
| NC_006577.2;YP_173242.1;HCoV-HKU1_nucleocapsi;321 | 0 | 0 |
| NC_006577.2;YP_173242.1;HCoV-HKU1_nucleocapsi;322 | 0 | 0 |
| NC_006577.2;YP_173242.1;HCoV-HKU1_nucleocapsi;323 | 0 | 0 |
| NC_006577.2;YP_173242.1;HCoV-HKU1_nucleocapsi;324 | 0 | 0 |
| NC_006577.2;YP_173242.1;HCoV-HKU1_nucleocapsi;325 | 0 | 0 |
| NC_006577.2;YP_173242.1;HCoV-HKU1_nucleocapsi;326 | 0.05 | 0 |
| NC_006577.2;YP_173242.1;HCoV-HKU1_nucleocapsi;327 | 0 | 0 |
| NC_006577.2;YP_173242.1;HCoV-HKU1_nucleocapsi;328 | 0 | 0 |
| NC_006577.2;YP_173242.1;HCoV-HKU1_nucleocapsi;329 | 0 | 0 |
| NC_006577.2;YP_173242.1;HCoV-HKU1_nucleocapsi;330 | 0 | 0 |
| NC_006577.2;YP_173242.1;HCoV-HKU1_nucleocapsi;331 | 0.1 | 0.05 |
| NC_006577.2;YP_173242.1;HCoV-HKU1_nucleocapsi;332 | 0 | 0 |

| | | |
|---|---|---|
| NC_006577.2;YP_173242.1;HCoV-HKU1_nucleocapsi;333 | 0 | 0 |
| NC_006577.2;YP_173242.1;HCoV-HKU1_nucleocapsi;334 | 0 | 0 |
| NC_006577.2;YP_173242.1;HCoV-HKU1_nucleocapsi;335 | 0 | 0 |
| NC_006577.2;YP_173242.1;HCoV-HKU1_nucleocapsi;336 | 0.05 | 0 |
| NC_006577.2;YP_173242.1;HCoV-HKU1_nucleocapsi;337 | 0.05 | 0 |
| NC_006577.2;YP_173242.1;HCoV-HKU1_nucleocapsi;338 | 0 | 0.025 |
| NC_006577.2;YP_173242.1;HCoV-HKU1_nucleocapsi;339 | 0 | 0 |
| NC_006577.2;YP_173242.1;HCoV-HKU1_nucleocapsi;340 | 0 | 0 |
| NC_006577.2;YP_173242.1;HCoV-HKU1_nucleocapsi;341 | 0 | 0 |
| NC_006577.2;YP_173242.1;HCoV-HKU1_nucleocapsi;342 | 0 | 0 |
| NC_006577.2;YP_173242.1;HCoV-HKU1_nucleocapsi;343 | 0 | 0 |
| NC_006577.2;YP_173242.1;HCoV-HKU1_nucleocapsi;344 | 0 | 0 |
| NC_006577.2;YP_173242.1;HCoV-HKU1_nucleocapsi;345 | 0 | 0 |
| NC_006577.2;YP_173242.1;HCoV-HKU1_nucleocapsi;346 | 0 | 0 |
| NC_006577.2;YP_173242.1;HCoV-HKU1_nucleocapsi;347 | 0 | 0 |
| NC_006577.2;YP_173242.1;HCoV-HKU1_nucleocapsi;348 | 0 | 0 |
| NC_006577.2;YP_173242.1;HCoV-HKU1_nucleocapsi;349 | 0 | 0 |
| NC_006577.2;YP_173242.1;HCoV-HKU1_nucleocapsi;350 | 0.1 | 0.3 |
| NC_006577.2;YP_173242.1;HCoV-HKU1_nucleocapsi;351 | 0.1 | 0.025 |
| NC_006577.2;YP_173242.1;HCoV-HKU1_nucleocapsi;352 | 0.05 | 0 |
| NC_006577.2;YP_173242.1;HCoV-HKU1_nucleocapsi;353 | 0 | 0 |
| NC_006577.2;YP_173242.1;HCoV-HKU1_nucleocapsi;354 | 0 | 0 |
| NC_006577.2;YP_173242.1;HCoV-HKU1_nucleocapsi;355 | 0 | 0 |
| NC_006577.2;YP_173242.1;HCoV-HKU1_nucleocapsi;356 | 0 | 0 |
| NC_006577.2;YP_173242.1;HCoV-HKU1_nucleocapsi;357 | 0 | 0 |
| NC_006577.2;YP_173242.1;HCoV-HKU1_nucleocapsi;358 | 0 | 0 |
| NC_006577.2;YP_173242.1;HCoV-HKU1_nucleocapsi;359 | 0 | 0 |
| NC_006577.2;YP_173242.1;HCoV-HKU1_nucleocapsi;360 | 0.05 | 0 |
| NC_006577.2;YP_173242.1;HCoV-HKU1_nucleocapsi;361 | 0 | 0.025 |
| NC_006577.2;YP_173242.1;HCoV-HKU1_nucleocapsi;362 | 0 | 0 |
| NC_006577.2;YP_173242.1;HCoV-HKU1_nucleocapsi;363 | 0 | 0 |
| NC_006577.2;YP_173242.1;HCoV-HKU1_nucleocapsi;364 | 0 | 0 |
| NC_006577.2;YP_173242.1;HCoV-HKU1_nucleocapsi;365 | 0 | 0 |
| NC_006577.2;YP_173242.1;HCoV-HKU1_nucleocapsi;366 | 0.05 | 0.075 |
| NC_006577.2;YP_173242.1;HCoV-HKU1_nucleocapsi;367 | 0 | 0.05 |
| NC_006577.2;YP_173242.1;HCoV-HKU1_nucleocapsi;368 | 0.1 | 0.1 |
| NC_006577.2;YP_173242.1;HCoV-HKU1_nucleocapsi;369 | 0.2 | 0.225 |
| NC_006577.2;YP_173242.1;HCoV-HKU1_nucleocapsi;370 | 0.1 | 0.25 |
| NC_006577.2;YP_173242.1;HCoV-HKU1_nucleocapsi;371 | 0.05 | 0.15 |
| NC_006577.2;YP_173242.1;HCoV-HKU1_nucleocapsi;372 | 0.05 | 0.15 |
| NC_006577.2;YP_173242.1;HCoV-HKU1_nucleocapsi;373 | 0 | 0.175 |
| NC_006577.2;YP_173242.1;HCoV-HKU1_nucleocapsi;374 | 0.1 | 0.175 |
| NC_006577.2;YP_173242.1;HCoV-HKU1_nucleocapsi;375 | 0.05 | 0.05 |

| | | |
|---|---|---|
| NC_006577.2;YP_173242.1;HCoV-HKU1_nucleocapsi;376 | 0.05 | 0.05 |
| NC_006577.2;YP_173242.1;HCoV-HKU1_nucleocapsi;377 | 0 | 0.025 |
| NC_006577.2;YP_173242.1;HCoV-HKU1_nucleocapsi;378 | 0 | 0.025 |
| NC_006577.2;YP_173242.1;HCoV-HKU1_nucleocapsi;379 | 0 | 0 |
| NC_006577.2;YP_173242.1;HCoV-HKU1_nucleocapsi;380 | 0 | 0 |
| NC_006577.2;YP_173242.1;HCoV-HKU1_nucleocapsi;381 | 0 | 0 |
| NC_006577.2;YP_173242.1;HCoV-HKU1_nucleocapsi;382 | 0 | 0.025 |
| NC_006577.2;YP_173242.1;HCoV-HKU1_nucleocapsi;383 | 0 | 0.025 |
| NC_006577.2;YP_173242.1;HCoV-HKU1_nucleocapsi;384 | 0 | 0 |
| NC_006577.2;YP_173242.1;HCoV-HKU1_nucleocapsi;385 | 0 | 0 |
| NC_006577.2;YP_173242.1;HCoV-HKU1_nucleocapsi;386 | 0 | 0.15 |
| NC_006577.2;YP_173242.1;HCoV-HKU1_nucleocapsi;387 | 0 | 0 |
| NC_006577.2;YP_173242.1;HCoV-HKU1_nucleocapsi;388 | 0 | 0 |
| NC_006577.2;YP_173242.1;HCoV-HKU1_nucleocapsi;389 | 0 | 0 |
| NC_006577.2;YP_173242.1;HCoV-HKU1_nucleocapsi;390 | 0 | 0.025 |
| NC_006577.2;YP_173242.1;HCoV-HKU1_nucleocapsi;391 | 0.05 | 0.025 |
| NC_006577.2;YP_173242.1;HCoV-HKU1_nucleocapsi;392 | 0 | 0.025 |
| NC_006577.2;YP_173242.1;HCoV-HKU1_nucleocapsi;393 | 0 | 0.075 |
| NC_006577.2;YP_173242.1;HCoV-HKU1_nucleocapsi;394 | 0.1 | 0.125 |
| NC_006577.2;YP_173242.1;HCoV-HKU1_nucleocapsi;395 | 0.3 | 0.175 |
| NC_006577.2;YP_173242.1;HCoV-HKU1_nucleocapsi;396 | 0.3 | 0.275 |
| NC_006577.2;YP_173242.1;HCoV-HKU1_nucleocapsi;397 | 0.35 | 0.275 |
| NC_006577.2;YP_173242.1;HCoV-HKU1_nucleocapsi;398 | 0.35 | 0.25 |
| NC_006577.2;YP_173242.1;HCoV-HKU1_nucleocapsi;399 | 0.35 | 0.3 |
| NC_006577.2;YP_173242.1;HCoV-HKU1_nucleocapsi;400 | 0.25 | 0.2 |
| NC_006577.2;YP_173242.1;HCoV-HKU1_nucleocapsi;401 | 0.25 | 0.175 |
| NC_006577.2;YP_173242.1;HCoV-HKU1_nucleocapsi;402 | 0.25 | 0.2 |
| NC_006577.2;YP_173242.1;HCoV-HKU1_nucleocapsi;403 | 0.2 | 0.075 |
| NC_006577.2;YP_173242.1;HCoV-HKU1_nucleocapsi;404 | 0.1 | 0.025 |
| NC_006577.2;YP_173242.1;HCoV-HKU1_nucleocapsi;405 | 0 | 0.025 |
| NC_006577.2;YP_173242.1;HCoV-HKU1_nucleocapsi;406 | 0.1 | 0 |
| NC_006577.2;YP_173242.1;HCoV-HKU1_nucleocapsi;407 | 0 | 0.05 |
| NC_006577.2;YP_173242.1;HCoV-HKU1_nucleocapsi;408 | 0 | 0.05 |
| NC_006577.2;YP_173242.1;HCoV-HKU1_nucleocapsi;409 | 0.05 | 0 |
| NC_006577.2;YP_173242.1;HCoV-HKU1_nucleocapsi;410 | 0 | 0.025 |
| NC_006577.2;YP_173242.1;HCoV-HKU1_nucleocapsi;411 | 0 | 0 |
| NC_006577.2;YP_173242.1;HCoV-HKU1_nucleocapsi;412 | 0 | 0.05 |
| NC_006577.2;YP_173242.1;HCoV-HKU1_nucleocapsi;413 | 0 | 0 |
| NC_006577.2;YP_173242.1;HCoV-HKU1_nucleocapsi;414 | 0 | 0 |
| NC_006577.2;YP_173242.1;HCoV-HKU1_nucleocapsi;415 | 0 | 0 |
| NC_006577.2;YP_173242.1;HCoV-HKU1_nucleocapsi;416 | 0 | 0 |
| NC_006577.2;YP_173242.1;HCoV-HKU1_nucleocapsi;417 | 0.1 | 0.125 |
| NC_006577.2;YP_173242.1;HCoV-HKU1_nucleocapsi;418 | 0 | 0 |

| | | |
|---|---|---|
| NC_006577.2;YP_173242.1;HCoV-HKU1_nucleocapsi;419 | 0 | 0.025 |
| NC_006577.2;YP_173242.1;HCoV-HKU1_nucleocapsi;420 | 0 | 0.1 |
| NC_006577.2;YP_173242.1;HCoV-HKU1_nucleocapsi;421 | 0 | 0.025 |
| NC_006577.2;YP_173242.1;HCoV-HKU1_nucleocapsi;422 | 0 | 0.025 |
| NC_006577.2;YP_173242.1;HCoV-HKU1_nucleocapsi;423 | 0 | 0.025 |
| NC_006577.2;YP_173242.1;HCoV-HKU1_nucleocapsi;424 | 0 | 0.05 |
| NC_006577.2;YP_173242.1;HCoV-HKU1_nucleocapsi;425 | 0 | 0.025 |
| NC_006577.2;YP_173242.1;HCoV-HKU1_nucleocapsi;426 | 0.05 | 0.05 |
| NC_006577.2;YP_173242.1;HCoV-HKU1_nucleocapsi;427 | 0.05 | 0.025 |
| NC_006577.2;YP_173242.1;HCoV-HKU1_nucleocapsi;428 | 0.1 | 0.025 |
| NC_006577.2;YP_173242.1;HCoV-HKU1_nucleocapsi;429 | 0.05 | 0.025 |
| NC_006577.2;YP_173242.1;HCoV-HKU1_nucleocapsi;430 | 0.05 | 0.025 |
| NC_006577.2;YP_173243.1;HCoV-HKU1_nucleocap2;1 | 0 | 0 |
| NC_006577.2;YP_173243.1;HCoV-HKU1_nucleocap2;2 | 0 | 0 |
| NC_006577.2;YP_173243.1;HCoV-HKU1_nucleocap2;3 | 0 | 0 |
| NC_006577.2;YP_173243.1;HCoV-HKU1_nucleocap2;4 | 0 | 0 |
| NC_006577.2;YP_173243.1;HCoV-HKU1_nucleocap2;5 | 0 | 0 |
| NC_006577.2;YP_173243.1;HCoV-HKU1_nucleocap2;6 | 0 | 0 |
| NC_006577.2;YP_173243.1;HCoV-HKU1_nucleocap2;7 | 0 | 0 |
| NC_006577.2;YP_173243.1;HCoV-HKU1_nucleocap2;8 | 0 | 0.025 |
| NC_006577.2;YP_173243.1;HCoV-HKU1_nucleocap2;9 | 0 | 0 |
| NC_006577.2;YP_173243.1;HCoV-HKU1_nucleocap2;10 | 0.2 | 0.025 |
| NC_006577.2;YP_173243.1;HCoV-HKU1_nucleocap2;11 | 0.05 | 0 |
| NC_006577.2;YP_173243.1;HCoV-HKU1_nucleocap2;12 | 0 | 0 |
| NC_006577.2;YP_173243.1;HCoV-HKU1_nucleocap2;13 | 0.1 | 0.075 |
| NC_006577.2;YP_173243.1;HCoV-HKU1_nucleocap2;14 | 0 | 0 |
| NC_006577.2;YP_173243.1;HCoV-HKU1_nucleocap2;15 | 0 | 0 |
| NC_006577.2;YP_173243.1;HCoV-HKU1_nucleocap2;16 | 0 | 0 |
| NC_006577.2;YP_173243.1;HCoV-HKU1_nucleocap2;17 | 0 | 0 |
| NC_006577.2;YP_173243.1;HCoV-HKU1_nucleocap2;18 | 0 | 0.025 |
| NC_006577.2;YP_173243.1;HCoV-HKU1_nucleocap2;19 | 0 | 0.075 |
| NC_006577.2;YP_173243.1;HCoV-HKU1_nucleocap2;20 | 0 | 0.075 |
| NC_006577.2;YP_173243.1;HCoV-HKU1_nucleocap2;21 | 0.1 | 0.075 |
| NC_006577.2;YP_173243.1;HCoV-HKU1_nucleocap2;22 | 0.1 | 0.125 |
| NC_006577.2;YP_173243.1;HCoV-HKU1_nucleocap2;23 | 0.05 | 0.1 |
| NC_006577.2;YP_173243.1;HCoV-HKU1_nucleocap2;24 | 0.05 | 0.1 |
| NC_006577.2;YP_173243.1;HCoV-HKU1_nucleocap2;25 | 0.05 | 0.15 |
| NC_006577.2;YP_173243.1;HCoV-HKU1_nucleocap2;26 | 0.05 | 0.125 |
| NC_006577.2;YP_173243.1;HCoV-HKU1_nucleocap2;27 | 0.15 | 0.275 |
| NC_006577.2;YP_173243.1;HCoV-HKU1_nucleocap2;28 | 0.15 | 0.25 |
| NC_006577.2;YP_173243.1;HCoV-HKU1_nucleocap2;29 | 0.05 | 0.25 |
| NC_006577.2;YP_173243.1;HCoV-HKU1_nucleocap2;30 | 0 | 0 |
| NC_006577.2;YP_173243.1;HCoV-HKU1_nucleocap2;31 | 0 | 0 |

| | | |
|---|---|---|
| NC_006577.2;YP_173243.1;HCoV-HKU1_nucleocap2;32 | 0 | 0 |
| NC_006577.2;YP_173243.1;HCoV-HKU1_nucleocap2;33 | 0 | 0 |
| NC_006577.2;YP_173243.1;HCoV-HKU1_nucleocap2;34 | 0 | 0 |
| NC_006577.2;YP_173243.1;HCoV-HKU1_nucleocap2;35 | 0 | 0 |
| NC_006577.2;YP_173243.1;HCoV-HKU1_nucleocap2;36 | 0 | 0 |
| NC_006577.2;YP_173243.1;HCoV-HKU1_nucleocap2;37 | 0 | 0.025 |
| NC_006577.2;YP_173243.1;HCoV-HKU1_nucleocap2;38 | 0 | 0 |
| NC_006577.2;YP_173243.1;HCoV-HKU1_nucleocap2;39 | 0 | 0 |
| NC_006577.2;YP_173243.1;HCoV-HKU1_nucleocap2;40 | 0 | 0 |
| NC_006577.2;YP_173243.1;HCoV-HKU1_nucleocap2;41 | 0 | 0 |
| NC_006577.2;YP_173243.1;HCoV-HKU1_nucleocap2;42 | 0 | 0 |
| NC_006577.2;YP_173243.1;HCoV-HKU1_nucleocap2;43 | 0 | 0 |
| NC_006577.2;YP_173243.1;HCoV-HKU1_nucleocap2;44 | 0 | 0 |
| NC_006577.2;YP_173243.1;HCoV-HKU1_nucleocap2;45 | 0 | 0 |
| NC_006577.2;YP_173243.1;HCoV-HKU1_nucleocap2;46 | 0.05 | 0 |
| NC_006577.2;YP_173243.1;HCoV-HKU1_nucleocap2;47 | 0 | 0 |
| NC_006577.2;YP_173243.1;HCoV-HKU1_nucleocap2;48 | 0 | 0.025 |
| NC_006577.2;YP_173243.1;HCoV-HKU1_nucleocap2;49 | 0.05 | 0 |
| NC_006577.2;YP_173243.1;HCoV-HKU1_nucleocap2;50 | 0 | 0 |
| NC_006577.2;YP_173243.1;HCoV-HKU1_nucleocap2;51 | 0 | 0 |
| NC_006577.2;YP_173243.1;HCoV-HKU1_nucleocap2;52 | 0 | 0 |
| NC_006577.2;YP_173243.1;HCoV-HKU1_nucleocap2;53 | 0 | 0.025 |
| NC_006577.2;YP_173243.1;HCoV-HKU1_nucleocap2;54 | 0 | 0.05 |
| NC_006577.2;YP_173243.1;HCoV-HKU1_nucleocap2;55 | 0 | 0.025 |
| NC_006577.2;YP_173243.1;HCoV-HKU1_nucleocap2;56 | 0 | 0 |
| NC_006577.2;YP_173243.1;HCoV-HKU1_nucleocap2;57 | 0.05 | 0.025 |
| NC_006577.2;YP_173243.1;HCoV-HKU1_nucleocap2;58 | 0.05 | 0.05 |
| NC_006577.2;YP_173243.1;HCoV-HKU1_nucleocap2;59 | 0 | 0.025 |
| NC_006577.2;YP_173243.1;HCoV-HKU1_nucleocap2;60 | 0 | 0 |
| NC_006577.2;YP_173243.1;HCoV-HKU1_nucleocap2;61 | 0 | 0 |
| NC_006577.2;YP_173243.1;HCoV-HKU1_nucleocap2;62 | 0.05 | 0.025 |
| NC_006577.2;YP_173243.1;HCoV-HKU1_nucleocap2;63 | 0.05 | 0.025 |
| NC_006577.2;YP_173243.1;HCoV-HKU1_nucleocap2;64 | 0.15 | 0.075 |
| NC_006577.2;YP_173243.1;HCoV-HKU1_nucleocap2;65 | 0 | 0 |
| NC_006577.2;YP_173243.1;HCoV-HKU1_nucleocap2;66 | 0 | 0.025 |
| NC_006577.2;YP_173243.1;HCoV-HKU1_nucleocap2;67 | 0.05 | 0.075 |
| NC_006577.2;YP_173243.1;HCoV-HKU1_nucleocap2;68 | 0 | 0 |
| NC_006577.2;YP_173243.1;HCoV-HKU1_nucleocap2;69 | 0 | 0 |
| NC_006577.2;YP_173243.1;HCoV-HKU1_nucleocap2;70 | 0 | 0 |
| NC_006577.2;YP_173243.1;HCoV-HKU1_nucleocap2;71 | 0 | 0 |
| NC_006577.2;YP_173243.1;HCoV-HKU1_nucleocap2;72 | 0 | 0 |
| NC_006577.2;YP_173243.1;HCoV-HKU1_nucleocap2;73 | 0.05 | 0 |
| NC_006577.2;YP_173243.1;HCoV-HKU1_nucleocap2;74 | 0 | 0.05 |

| | | |
|---|---|---|
| NC_006577.2;YP_173243.1;HCoV-HKU1_nucleocap2;75 | 0 | 0 |
| NC_006577.2;YP_173243.1;HCoV-HKU1_nucleocap2;76 | 0.05 | 0.025 |
| NC_006577.2;YP_173243.1;HCoV-HKU1_nucleocap2;77 | 0.05 | 0.025 |
| NC_006577.2;YP_173243.1;HCoV-HKU1_nucleocap2;78 | 0.05 | 0 |
| NC_006577.2;YP_173243.1;HCoV-HKU1_nucleocap2;79 | 0.05 | 0.025 |
| NC_006577.2;YP_173243.1;HCoV-HKU1_nucleocap2;80 | 0 | 0.1 |
| NC_006577.2;YP_173243.1;HCoV-HKU1_nucleocap2;81 | 0 | 0.125 |
| NC_006577.2;YP_173243.1;HCoV-HKU1_nucleocap2;82 | 0 | 0.075 |
| NC_006577.2;YP_173243.1;HCoV-HKU1_nucleocap2;83 | 0 | 0.075 |
| NC_006577.2;YP_173243.1;HCoV-HKU1_nucleocap2;84 | 0 | 0.025 |
| NC_006577.2;YP_173243.1;HCoV-HKU1_nucleocap2;85 | 0 | 0.05 |
| NC_006577.2;YP_173243.1;HCoV-HKU1_nucleocap2;86 | 0 | 0 |
| NC_006577.2;YP_173243.1;HCoV-HKU1_nucleocap2;87 | 0 | 0.025 |
| NC_006577.2;YP_173243.1;HCoV-HKU1_nucleocap2;88 | 0 | 0 |
| NC_006577.2;YP_173243.1;HCoV-HKU1_nucleocap2;89 | 0.05 | 0.025 |
| NC_006577.2;YP_173243.1;HCoV-HKU1_nucleocap2;90 | 0 | 0 |
| NC_006577.2;YP_173243.1;HCoV-HKU1_nucleocap2;91 | 0 | 0 |
| NC_006577.2;YP_173243.1;HCoV-HKU1_nucleocap2;92 | 0 | 0.025 |
| NC_006577.2;YP_173243.1;HCoV-HKU1_nucleocap2;93 | 0 | 0 |
| NC_006577.2;YP_173243.1;HCoV-HKU1_nucleocap2;94 | 0 | 0 |
| NC_006577.2;YP_173243.1;HCoV-HKU1_nucleocap2;95 | 0 | 0.025 |
| NC_006577.2;YP_173243.1;HCoV-HKU1_nucleocap2;96 | 0 | 0 |
| NC_006577.2;YP_173243.1;HCoV-HKU1_nucleocap2;97 | 0 | 0.025 |
| NC_006577.2;YP_173243.1;HCoV-HKU1_nucleocap2;98 | 0 | 0.025 |
| NC_006577.2;YP_173243.1;HCoV-HKU1_nucleocap2;99 | 0 | 0 |
| NC_006577.2;YP_173243.1;HCoV-HKU1_nucleocap2;100 | 0 | 0 |
| NC_006577.2;YP_173243.1;HCoV-HKU1_nucleocap2;101 | 0 | 0.025 |
| NC_006577.2;YP_173243.1;HCoV-HKU1_nucleocap2;102 | 0 | 0 |
| NC_006577.2;YP_173243.1;HCoV-HKU1_nucleocap2;103 | 0.05 | 0.025 |
| NC_006577.2;YP_173243.1;HCoV-HKU1_nucleocap2;104 | 0 | 0 |
| NC_006577.2;YP_173243.1;HCoV-HKU1_nucleocap2;105 | 0 | 0 |
| NC_006577.2;YP_173243.1;HCoV-HKU1_nucleocap2;106 | 0 | 0 |
| NC_006577.2;YP_173243.1;HCoV-HKU1_nucleocap2;107 | 0.05 | 0 |
| NC_006577.2;YP_173243.1;HCoV-HKU1_nucleocap2;108 | 0 | 0 |
| NC_006577.2;YP_173243.1;HCoV-HKU1_nucleocap2;109 | 0 | 0 |
| NC_006577.2;YP_173243.1;HCoV-HKU1_nucleocap2;110 | 0 | 0 |
| NC_006577.2;YP_173243.1;HCoV-HKU1_nucleocap2;111 | 0 | 0 |
| NC_006577.2;YP_173243.1;HCoV-HKU1_nucleocap2;112 | 0.1 | 0.05 |
| NC_006577.2;YP_173243.1;HCoV-HKU1_nucleocap2;113 | 0 | 0 |
| NC_006577.2;YP_173243.1;HCoV-HKU1_nucleocap2;114 | 0 | 0 |
| NC_006577.2;YP_173243.1;HCoV-HKU1_nucleocap2;115 | 0 | 0 |
| NC_006577.2;YP_173243.1;HCoV-HKU1_nucleocap2;116 | 0 | 0 |
| NC_006577.2;YP_173243.1;HCoV-HKU1_nucleocap2;117 | 0 | 0 |

| | | |
|---|---|---|
| NC_006577.2;YP_173243.1;HCoV-HKU1_nucleocap2;118 | 0 | 0 |
| NC_006577.2;YP_173243.1;HCoV-HKU1_nucleocap2;119 | 0 | 0 |
| NC_006577.2;YP_173243.1;HCoV-HKU1_nucleocap2;120 | 0 | 0 |
| NC_006577.2;YP_173243.1;HCoV-HKU1_nucleocap2;121 | 0 | 0 |
| NC_006577.2;YP_173243.1;HCoV-HKU1_nucleocap2;122 | 0 | 0 |
| NC_006577.2;YP_173243.1;HCoV-HKU1_nucleocap2;123 | 0 | 0 |
| NC_006577.2;YP_173243.1;HCoV-HKU1_nucleocap2;124 | 0 | 0 |
| NC_006577.2;YP_173243.1;HCoV-HKU1_nucleocap2;125 | 0 | 0 |
| NC_006577.2;YP_173243.1;HCoV-HKU1_nucleocap2;126 | 0 | 0.025 |
| NC_006577.2;YP_173243.1;HCoV-HKU1_nucleocap2;127 | 0 | 0.05 |
| NC_006577.2;YP_173243.1;HCoV-HKU1_nucleocap2;128 | 0 | 0 |
| NC_006577.2;YP_173243.1;HCoV-HKU1_nucleocap2;129 | 0 | 0.05 |
| NC_006577.2;YP_173243.1;HCoV-HKU1_nucleocap2;130 | 0 | 0 |
| NC_006577.2;YP_173243.1;HCoV-HKU1_nucleocap2;131 | 0 | 0 |
| NC_006577.2;YP_173243.1;HCoV-HKU1_nucleocap2;132 | 0 | 0 |
| NC_006577.2;YP_173243.1;HCoV-HKU1_nucleocap2;133 | 0 | 0 |
| NC_006577.2;YP_173243.1;HCoV-HKU1_nucleocap2;134 | 0 | 0 |
| NC_006577.2;YP_173243.1;HCoV-HKU1_nucleocap2;135 | 0 | 0 |
| NC_006577.2;YP_173243.1;HCoV-HKU1_nucleocap2;136 | 0 | 0 |
| NC_006577.2;YP_173243.1;HCoV-HKU1_nucleocap2;137 | 0 | 0 |
| NC_006577.2;YP_173243.1;HCoV-HKU1_nucleocap2;138 | 0 | 0 |
| NC_006577.2;YP_173243.1;HCoV-HKU1_nucleocap2;139 | 0 | 0 |
| NC_006577.2;YP_173243.1;HCoV-HKU1_nucleocap2;140 | 0 | 0 |
| NC_006577.2;YP_173243.1;HCoV-HKU1_nucleocap2;141 | 0 | 0 |
| NC_006577.2;YP_173243.1;HCoV-HKU1_nucleocap2;142 | 0 | 0 |
| NC_006577.2;YP_173243.1;HCoV-HKU1_nucleocap2;143 | 0 | 0 |
| NC_006577.2;YP_173243.1;HCoV-HKU1_nucleocap2;144 | 0 | 0 |
| NC_006577.2;YP_173243.1;HCoV-HKU1_nucleocap2;145 | 0 | 0 |
| NC_006577.2;YP_173243.1;HCoV-HKU1_nucleocap2;146 | 0 | 0 |
| NC_006577.2;YP_173243.1;HCoV-HKU1_nucleocap2;147 | 0 | 0 |
| NC_006577.2;YP_173243.1;HCoV-HKU1_nucleocap2;148 | 0 | 0 |
| NC_006577.2;YP_173243.1;HCoV-HKU1_nucleocap2;149 | 0 | 0 |
| NC_006577.2;YP_173243.1;HCoV-HKU1_nucleocap2;150 | 0 | 0 |
| NC_006577.2;YP_173243.1;HCoV-HKU1_nucleocap2;151 | 0 | 0 |
| NC_006577.2;YP_173243.1;HCoV-HKU1_nucleocap2;152 | 0 | 0 |
| NC_006577.2;YP_173243.1;HCoV-HKU1_nucleocap2;153 | 0 | 0 |
| NC_006577.2;YP_173243.1;HCoV-HKU1_nucleocap2;154 | 0 | 0 |
| NC_006577.2;YP_173243.1;HCoV-HKU1_nucleocap2;155 | 0 | 0 |
| NC_006577.2;YP_173243.1;HCoV-HKU1_nucleocap2;156 | 0 | 0 |
| NC_006577.2;YP_173243.1;HCoV-HKU1_nucleocap2;157 | 0 | 0 |
| NC_006577.2;YP_173243.1;HCoV-HKU1_nucleocap2;158 | 0 | 0 |
| NC_006577.2;YP_173243.1;HCoV-HKU1_nucleocap2;159 | 0 | 0 |
| NC_006577.2;YP_173243.1;HCoV-HKU1_nucleocap2;160 | 0 | 0 |

| | | |
|---|---|---|
| NC_006577.2;YP_173243.1;HCoV-HKU1_nucleocap2;161 | 0 | 0 |
| NC_006577.2;YP_173243.1;HCoV-HKU1_nucleocap2;162 | 0 | 0 |
| NC_006577.2;YP_173243.1;HCoV-HKU1_nucleocap2;163 | 0 | 0 |
| NC_006577.2;YP_173243.1;HCoV-HKU1_nucleocap2;164 | 0 | 0 |
| NC_006577.2;YP_173243.1;HCoV-HKU1_nucleocap2;165 | 0 | 0 |
| NC_006577.2;YP_173243.1;HCoV-HKU1_nucleocap2;166 | 0 | 0 |
| NC_006577.2;YP_173243.1;HCoV-HKU1_nucleocap2;167 | 0 | 0 |
| NC_006577.2;YP_173243.1;HCoV-HKU1_nucleocap2;168 | 0 | 0 |
| NC_006577.2;YP_173243.1;HCoV-HKU1_nucleocap2;169 | 0 | 0 |
| NC_006577.2;YP_173243.1;HCoV-HKU1_nucleocap2;170 | 0 | 0 |
| NC_006577.2;YP_173243.1;HCoV-HKU1_nucleocap2;171 | 0 | 0 |
| NC_006577.2;YP_173243.1;HCoV-HKU1_nucleocap2;172 | 0 | 0 |
| NC_006577.2;YP_173243.1;HCoV-HKU1_nucleocap2;173 | 0 | 0 |
| NC_006577.2;YP_173243.1;HCoV-HKU1_nucleocap2;174 | 0 | 0 |
| NC_006577.2;YP_173243.1;HCoV-HKU1_nucleocap2;175 | 0 | 0 |
| NC_006577.2;YP_173243.1;HCoV-HKU1_nucleocap2;176 | 0 | 0 |
| NC_006577.2;YP_173243.1;HCoV-HKU1_nucleocap2;177 | 0 | 0 |
| NC_006577.2;YP_173243.1;HCoV-HKU1_nucleocap2;178 | 0 | 0 |
| NC_006577.2;YP_173243.1;HCoV-HKU1_nucleocap2;179 | 0 | 0 |
| NC_006577.2;YP_173243.1;HCoV-HKU1_nucleocap2;180 | 0 | 0 |
| NC_006577.2;YP_173243.1;HCoV-HKU1_nucleocap2;181 | 0 | 0 |
| NC_006577.2;YP_173243.1;HCoV-HKU1_nucleocap2;182 | 0 | 0 |
| NC_006577.2;YP_173243.1;HCoV-HKU1_nucleocap2;183 | 0 | 0 |
| NC_006577.2;YP_173243.1;HCoV-HKU1_nucleocap2;184 | 0 | 0 |
| NC_006577.2;YP_173243.1;HCoV-HKU1_nucleocap2;185 | 0 | 0 |
| NC_006577.2;YP_173243.1;HCoV-HKU1_nucleocap2;186 | 0 | 0 |
| NC_006577.2;YP_173243.1;HCoV-HKU1_nucleocap2;187 | 0 | 0 |
| NC_006577.2;YP_173243.1;HCoV-HKU1_nucleocap2;188 | 0.05 | 0.025 |
| NC_006577.2;YP_173243.1;HCoV-HKU1_nucleocap2;189 | 0.15 | 0.125 |
| NC_006577.2;YP_173243.1;HCoV-HKU1_nucleocap2;190 | 0 | 0 |
| NC_006577.2;YP_173243.1;HCoV-HKU1_nucleocap2;191 | 0 | 0 |
| NC_006577.2;YP_173243.1;HCoV-HKU1_nucleocap2;192 | 0 | 0 |
| NC_006577.2;YP_173243.1;HCoV-HKU1_nucleocap2;193 | 0 | 0 |
| NC_006577.2;YP_173243.1;HCoV-HKU1_nucleocap2;194 | 0 | 0 |
| NC_006213.1;YP_009555238.1;HCoV-OC43_orf1ab;1 | 0 | 0.05 |
| NC_006213.1;YP_009555238.1;HCoV-OC43_orf1ab;2 | 0 | 0.05 |
| NC_006213.1;YP_009555238.1;HCoV-OC43_orf1ab;3 | 0 | 0.05 |
| NC_006213.1;YP_009555238.1;HCoV-OC43_orf1ab;4 | 0 | 0.025 |
| NC_006213.1;YP_009555238.1;HCoV-OC43_orf1ab;5 | 0 | 0 |
| NC_006213.1;YP_009555238.1;HCoV-OC43_orf1ab;6 | 0.05 | 0 |
| NC_006213.1;YP_009555238.1;HCoV-OC43_orf1ab;7 | 0 | 0 |
| NC_006213.1;YP_009555238.1;HCoV-OC43_orf1ab;8 | 0 | 0.025 |
| NC_006213.1;YP_009555238.1;HCoV-OC43_orf1ab;9 | 0 | 0.025 |

| | | |
|---|---|---|
| NC_006213.1;YP_009555238.1;HCoV-OC43_orf1ab;10 | 0 | 0.025 |
| NC_006213.1;YP_009555238.1;HCoV-OC43_orf1ab;11 | 0 | 0 |
| NC_006213.1;YP_009555238.1;HCoV-OC43_orf1ab;12 | 0 | 0.025 |
| NC_006213.1;YP_009555238.1;HCoV-OC43_orf1ab;13 | 0 | 0.025 |
| NC_006213.1;YP_009555238.1;HCoV-OC43_orf1ab;14 | 0.05 | 0 |
| NC_006213.1;YP_009555238.1;HCoV-OC43_orf1ab;15 | 0.05 | 0 |
| NC_006213.1;YP_009555238.1;HCoV-OC43_orf1ab;16 | 0 | 0 |
| NC_006213.1;YP_009555238.1;HCoV-OC43_orf1ab;17 | 0 | 0 |
| NC_006213.1;YP_009555238.1;HCoV-OC43_orf1ab;18 | 0 | 0.025 |
| NC_006213.1;YP_009555238.1;HCoV-OC43_orf1ab;19 | 0.05 | 0.05 |
| NC_006213.1;YP_009555238.1;HCoV-OC43_orf1ab;20 | 0.05 | 0 |
| NC_006213.1;YP_009555238.1;HCoV-OC43_orf1ab;21 | 0.05 | 0 |
| NC_006213.1;YP_009555238.1;HCoV-OC43_orf1ab;22 | 0.35 | 0.1 |
| NC_006213.1;YP_009555238.1;HCoV-OC43_orf1ab;23 | 0.05 | 0 |
| NC_006213.1;YP_009555238.1;HCoV-OC43_orf1ab;24 | 0 | 0 |
| NC_006213.1;YP_009555238.1;HCoV-OC43_orf1ab;25 | 0.05 | 0 |
| NC_006213.1;YP_009555238.1;HCoV-OC43_orf1ab;26 | 0.05 | 0 |
| NC_006213.1;YP_009555238.1;HCoV-OC43_orf1ab;27 | 0 | 0 |
| NC_006213.1;YP_009555238.1;HCoV-OC43_orf1ab;28 | 0 | 0 |
| NC_006213.1;YP_009555238.1;HCoV-OC43_orf1ab;29 | 0 | 0.025 |
| NC_006213.1;YP_009555238.1;HCoV-OC43_orf1ab;30 | 0 | 0.025 |
| NC_006213.1;YP_009555238.1;HCoV-OC43_orf1ab;31 | 0.05 | 0 |
| NC_006213.1;YP_009555238.1;HCoV-OC43_orf1ab;32 | 0 | 0.025 |
| NC_006213.1;YP_009555238.1;HCoV-OC43_orf1ab;33 | 0 | 0 |
| NC_006213.1;YP_009555238.1;HCoV-OC43_orf1ab;34 | 0 | 0 |
| NC_006213.1;YP_009555238.1;HCoV-OC43_orf1ab;35 | 0 | 0 |
| NC_006213.1;YP_009555238.1;HCoV-OC43_orf1ab;36 | 0 | 0 |
| NC_006213.1;YP_009555238.1;HCoV-OC43_orf1ab;37 | 0 | 0 |
| NC_006213.1;YP_009555238.1;HCoV-OC43_orf1ab;38 | 0 | 0 |
| NC_006213.1;YP_009555238.1;HCoV-OC43_orf1ab;39 | 0 | 0 |
| NC_006213.1;YP_009555238.1;HCoV-OC43_orf1ab;40 | 0 | 0 |
| NC_006213.1;YP_009555238.1;HCoV-OC43_orf1ab;41 | 0 | 0 |
| NC_006213.1;YP_009555238.1;HCoV-OC43_orf1ab;42 | 0 | 0 |
| NC_006213.1;YP_009555238.1;HCoV-OC43_orf1ab;43 | 0.05 | 0 |
| NC_006213.1;YP_009555238.1;HCoV-OC43_orf1ab;44 | 0.05 | 0 |
| NC_006213.1;YP_009555238.1;HCoV-OC43_orf1ab;45 | 0 | 0.025 |
| NC_006213.1;YP_009555238.1;HCoV-OC43_orf1ab;46 | 0.05 | 0 |
| NC_006213.1;YP_009555238.1;HCoV-OC43_orf1ab;47 | 0.2 | 0.15 |
| NC_006213.1;YP_009555238.1;HCoV-OC43_orf1ab;48 | 0.1 | 0.025 |
| NC_006213.1;YP_009555238.1;HCoV-OC43_orf1ab;49 | 0 | 0 |
| NC_006213.1;YP_009555238.1;HCoV-OC43_orf1ab;50 | 0 | 0 |
| NC_006213.1;YP_009555238.1;HCoV-OC43_orf1ab;51 | 0 | 0.025 |
| NC_006213.1;YP_009555238.1;HCoV-OC43_orf1ab;52 | 0 | 0 |

| | | |
|---|---|---|
| NC_006213.1;YP_009555238.1;HCoV-OC43_orf1ab;53 | 0 | 0 |
| NC_006213.1;YP_009555238.1;HCoV-OC43_orf1ab;54 | 0 | 0 |
| NC_006213.1;YP_009555238.1;HCoV-OC43_orf1ab;55 | 0 | 0 |
| NC_006213.1;YP_009555238.1;HCoV-OC43_orf1ab;56 | 0 | 0 |
| NC_006213.1;YP_009555238.1;HCoV-OC43_orf1ab;57 | 0 | 0 |
| NC_006213.1;YP_009555238.1;HCoV-OC43_orf1ab;58 | 0 | 0 |
| NC_006213.1;YP_009555238.1;HCoV-OC43_orf1ab;59 | 0 | 0 |
| NC_006213.1;YP_009555238.1;HCoV-OC43_orf1ab;60 | 0 | 0 |
| NC_006213.1;YP_009555238.1;HCoV-OC43_orf1ab;61 | 0 | 0 |
| NC_006213.1;YP_009555238.1;HCoV-OC43_orf1ab;62 | 0 | 0 |
| NC_006213.1;YP_009555238.1;HCoV-OC43_orf1ab;63 | 0 | 0 |
| NC_006213.1;YP_009555238.1;HCoV-OC43_orf1ab;64 | 0 | 0 |
| NC_006213.1;YP_009555238.1;HCoV-OC43_orf1ab;65 | 0 | 0 |
| NC_006213.1;YP_009555238.1;HCoV-OC43_orf1ab;66 | 0 | 0 |
| NC_006213.1;YP_009555238.1;HCoV-OC43_orf1ab;67 | 0 | 0 |
| NC_006213.1;YP_009555238.1;HCoV-OC43_orf1ab;68 | 0 | 0 |
| NC_006213.1;YP_009555238.1;HCoV-OC43_orf1ab;69 | 0 | 0 |
| NC_006213.1;YP_009555238.1;HCoV-OC43_orf1ab;70 | 0 | 0 |
| NC_006213.1;YP_009555238.1;HCoV-OC43_orf1ab;71 | 0 | 0 |
| NC_006213.1;YP_009555238.1;HCoV-OC43_orf1ab;72 | 0 | 0 |
| NC_006213.1;YP_009555238.1;HCoV-OC43_orf1ab;73 | 0.05 | 0 |
| NC_006213.1;YP_009555238.1;HCoV-OC43_orf1ab;74 | 0.05 | 0 |
| NC_006213.1;YP_009555238.1;HCoV-OC43_orf1ab;75 | 0.05 | 0 |
| NC_006213.1;YP_009555238.1;HCoV-OC43_orf1ab;76 | 0.05 | 0 |
| NC_006213.1;YP_009555238.1;HCoV-OC43_orf1ab;77 | 0.05 | 0 |
| NC_006213.1;YP_009555238.1;HCoV-OC43_orf1ab;78 | 0.05 | 0 |
| NC_006213.1;YP_009555238.1;HCoV-OC43_orf1ab;79 | 0.1 | 0 |
| NC_006213.1;YP_009555238.1;HCoV-OC43_orf1ab;80 | 0.05 | 0 |
| NC_006213.1;YP_009555238.1;HCoV-OC43_orf1ab;81 | 0.05 | 0 |
| NC_006213.1;YP_009555238.1;HCoV-OC43_orf1ab;82 | 0.05 | 0.025 |
| NC_006213.1;YP_009555238.1;HCoV-OC43_orf1ab;83 | 0.05 | 0.025 |
| NC_006213.1;YP_009555238.1;HCoV-OC43_orf1ab;84 | 0.05 | 0.025 |
| NC_006213.1;YP_009555238.1;HCoV-OC43_orf1ab;85 | 0.1 | 0.1 |
| NC_006213.1;YP_009555238.1;HCoV-OC43_orf1ab;86 | 0.1 | 0.025 |
| NC_006213.1;YP_009555238.1;HCoV-OC43_orf1ab;87 | 0.05 | 0.025 |
| NC_006213.1;YP_009555238.1;HCoV-OC43_orf1ab;88 | 0.05 | 0.05 |
| NC_006213.1;YP_009555238.1;HCoV-OC43_orf1ab;89 | 0 | 0.025 |
| NC_006213.1;YP_009555238.1;HCoV-OC43_orf1ab;90 | 0 | 0 |
| NC_006213.1;YP_009555238.1;HCoV-OC43_orf1ab;91 | 0 | 0 |
| NC_006213.1;YP_009555238.1;HCoV-OC43_orf1ab;92 | 0.05 | 0 |
| NC_006213.1;YP_009555238.1;HCoV-OC43_orf1ab;93 | 0 | 0 |
| NC_006213.1;YP_009555238.1;HCoV-OC43_orf1ab;94 | 0 | 0 |
| NC_006213.1;YP_009555238.1;HCoV-OC43_orf1ab;95 | 0 | 0 |

| | | |
|---|---|---|
| NC_006213.1;YP_009555238.1;HCoV-OC43_orf1ab;96 | 0 | 0 |
| NC_006213.1;YP_009555238.1;HCoV-OC43_orf1ab;97 | 0 | 0 |
| NC_006213.1;YP_009555238.1;HCoV-OC43_orf1ab;98 | 0 | 0 |
| NC_006213.1;YP_009555238.1;HCoV-OC43_orf1ab;99 | 0 | 0 |
| NC_006213.1;YP_009555238.1;HCoV-OC43_orf1ab;100 | 0 | 0 |
| NC_006213.1;YP_009555238.1;HCoV-OC43_orf1ab;101 | 0 | 0 |
| NC_006213.1;YP_009555238.1;HCoV-OC43_orf1ab;102 | 0 | 0 |
| NC_006213.1;YP_009555238.1;HCoV-OC43_orf1ab;103 | 0 | 0 |
| NC_006213.1;YP_009555238.1;HCoV-OC43_orf1ab;104 | 0 | 0 |
| NC_006213.1;YP_009555238.1;HCoV-OC43_orf1ab;105 | 0 | 0 |
| NC_006213.1;YP_009555238.1;HCoV-OC43_orf1ab;106 | 0 | 0 |
| NC_006213.1;YP_009555238.1;HCoV-OC43_orf1ab;107 | 0 | 0 |
| NC_006213.1;YP_009555238.1;HCoV-OC43_orf1ab;108 | 0 | 0.025 |
| NC_006213.1;YP_009555238.1;HCoV-OC43_orf1ab;109 | 0 | 0.025 |
| NC_006213.1;YP_009555238.1;HCoV-OC43_orf1ab;110 | 0 | 0 |
| NC_006213.1;YP_009555238.1;HCoV-OC43_orf1ab;111 | 0 | 0 |
| NC_006213.1;YP_009555238.1;HCoV-OC43_orf1ab;112 | 0 | 0 |
| NC_006213.1;YP_009555238.1;HCoV-OC43_orf1ab;113 | 0 | 0 |
| NC_006213.1;YP_009555238.1;HCoV-OC43_orf1ab;114 | 0 | 0 |
| NC_006213.1;YP_009555238.1;HCoV-OC43_orf1ab;115 | 0 | 0 |
| NC_006213.1;YP_009555238.1;HCoV-OC43_orf1ab;116 | 0 | 0 |
| NC_006213.1;YP_009555238.1;HCoV-OC43_orf1ab;117 | 0 | 0 |
| NC_006213.1;YP_009555238.1;HCoV-OC43_orf1ab;118 | 0 | 0 |
| NC_006213.1;YP_009555238.1;HCoV-OC43_orf1ab;119 | 0 | 0 |
| NC_006213.1;YP_009555238.1;HCoV-OC43_orf1ab;120 | 0 | 0 |
| NC_006213.1;YP_009555238.1;HCoV-OC43_orf1ab;121 | 0 | 0 |
| NC_006213.1;YP_009555238.1;HCoV-OC43_orf1ab;122 | 0 | 0 |
| NC_006213.1;YP_009555238.1;HCoV-OC43_orf1ab;123 | 0 | 0 |
| NC_006213.1;YP_009555238.1;HCoV-OC43_orf1ab;124 | 0 | 0 |
| NC_006213.1;YP_009555238.1;HCoV-OC43_orf1ab;125 | 0 | 0 |
| NC_006213.1;YP_009555238.1;HCoV-OC43_orf1ab;126 | 0 | 0 |
| NC_006213.1;YP_009555238.1;HCoV-OC43_orf1ab;127 | 0 | 0 |
| NC_006213.1;YP_009555238.1;HCoV-OC43_orf1ab;128 | 0 | 0 |
| NC_006213.1;YP_009555238.1;HCoV-OC43_orf1ab;129 | 0 | 0 |
| NC_006213.1;YP_009555238.1;HCoV-OC43_orf1ab;130 | 0 | 0 |
| NC_006213.1;YP_009555238.1;HCoV-OC43_orf1ab;131 | 0 | 0 |
| NC_006213.1;YP_009555238.1;HCoV-OC43_orf1ab;132 | 0 | 0 |
| NC_006213.1;YP_009555238.1;HCoV-OC43_orf1ab;133 | 0 | 0 |
| NC_006213.1;YP_009555238.1;HCoV-OC43_orf1ab;134 | 0 | 0 |
| NC_006213.1;YP_009555238.1;HCoV-OC43_orf1ab;135 | 0 | 0 |
| NC_006213.1;YP_009555238.1;HCoV-OC43_orf1ab;136 | 0 | 0 |
| NC_006213.1;YP_009555238.1;HCoV-OC43_orf1ab;137 | 0 | 0 |
| NC_006213.1;YP_009555238.1;HCoV-OC43_orf1ab;138 | 0 | 0 |

| | | |
|---|---|---|
| NC_006213.1;YP_009555238.1;HCoV-OC43_orf1ab;139 | 0 | 0 |
| NC_006213.1;YP_009555238.1;HCoV-OC43_orf1ab;140 | 0 | 0 |
| NC_006213.1;YP_009555238.1;HCoV-OC43_orf1ab;141 | 0 | 0 |
| NC_006213.1;YP_009555238.1;HCoV-OC43_orf1ab;142 | 0 | 0 |
| NC_006213.1;YP_009555238.1;HCoV-OC43_orf1ab;143 | 0 | 0 |
| NC_006213.1;YP_009555238.1;HCoV-OC43_orf1ab;144 | 0 | 0 |
| NC_006213.1;YP_009555238.1;HCoV-OC43_orf1ab;145 | 0 | 0 |
| NC_006213.1;YP_009555238.1;HCoV-OC43_orf1ab;146 | 0 | 0 |
| NC_006213.1;YP_009555238.1;HCoV-OC43_orf1ab;147 | 0 | 0 |
| NC_006213.1;YP_009555238.1;HCoV-OC43_orf1ab;148 | 0 | 0 |
| NC_006213.1;YP_009555238.1;HCoV-OC43_orf1ab;149 | 0 | 0 |
| NC_006213.1;YP_009555238.1;HCoV-OC43_orf1ab;150 | 0 | 0 |
| NC_006213.1;YP_009555238.1;HCoV-OC43_orf1ab;151 | 0 | 0 |
| NC_006213.1;YP_009555238.1;HCoV-OC43_orf1ab;152 | 0 | 0 |
| NC_006213.1;YP_009555238.1;HCoV-OC43_orf1ab;153 | 0 | 0 |
| NC_006213.1;YP_009555238.1;HCoV-OC43_orf1ab;154 | 0 | 0 |
| NC_006213.1;YP_009555238.1;HCoV-OC43_orf1ab;155 | 0 | 0 |
| NC_006213.1;YP_009555238.1;HCoV-OC43_orf1ab;156 | 0 | 0 |
| NC_006213.1;YP_009555238.1;HCoV-OC43_orf1ab;157 | 0 | 0 |
| NC_006213.1;YP_009555238.1;HCoV-OC43_orf1ab;158 | 0 | 0 |
| NC_006213.1;YP_009555238.1;HCoV-OC43_orf1ab;159 | 0 | 0 |
| NC_006213.1;YP_009555238.1;HCoV-OC43_orf1ab;160 | 0 | 0 |
| NC_006213.1;YP_009555238.1;HCoV-OC43_orf1ab;161 | 0 | 0 |
| NC_006213.1;YP_009555238.1;HCoV-OC43_orf1ab;162 | 0 | 0 |
| NC_006213.1;YP_009555238.1;HCoV-OC43_orf1ab;163 | 0 | 0 |
| NC_006213.1;YP_009555238.1;HCoV-OC43_orf1ab;164 | 0 | 0 |
| NC_006213.1;YP_009555238.1;HCoV-OC43_orf1ab;165 | 0 | 0 |
| NC_006213.1;YP_009555238.1;HCoV-OC43_orf1ab;166 | 0 | 0 |
| NC_006213.1;YP_009555238.1;HCoV-OC43_orf1ab;167 | 0 | 0 |
| NC_006213.1;YP_009555238.1;HCoV-OC43_orf1ab;168 | 0 | 0 |
| NC_006213.1;YP_009555238.1;HCoV-OC43_orf1ab;169 | 0 | 0 |
| NC_006213.1;YP_009555238.1;HCoV-OC43_orf1ab;170 | 0 | 0 |
| NC_006213.1;YP_009555238.1;HCoV-OC43_orf1ab;171 | 0 | 0 |
| NC_006213.1;YP_009555238.1;HCoV-OC43_orf1ab;172 | 0 | 0 |
| NC_006213.1;YP_009555238.1;HCoV-OC43_orf1ab;173 | 0 | 0 |
| NC_006213.1;YP_009555238.1;HCoV-OC43_orf1ab;174 | 0 | 0 |
| NC_006213.1;YP_009555238.1;HCoV-OC43_orf1ab;175 | 0 | 0 |
| NC_006213.1;YP_009555238.1;HCoV-OC43_orf1ab;176 | 0 | 0 |
| NC_006213.1;YP_009555238.1;HCoV-OC43_orf1ab;177 | 0 | 0 |
| NC_006213.1;YP_009555238.1;HCoV-OC43_orf1ab;178 | 0 | 0 |
| NC_006213.1;YP_009555238.1;HCoV-OC43_orf1ab;179 | 0 | 0 |
| NC_006213.1;YP_009555238.1;HCoV-OC43_orf1ab;180 | 0 | 0 |
| NC_006213.1;YP_009555238.1;HCoV-OC43_orf1ab;181 | 0 | 0 |

| | | |
|---|---|---|
| NC_006213.1;YP_009555238.1;HCoV-OC43_orf1ab;182 | 0 | 0 |
| NC_006213.1;YP_009555238.1;HCoV-OC43_orf1ab;183 | 0 | 0 |
| NC_006213.1;YP_009555238.1;HCoV-OC43_orf1ab;184 | 0 | 0 |
| NC_006213.1;YP_009555238.1;HCoV-OC43_orf1ab;185 | 0 | 0 |
| NC_006213.1;YP_009555238.1;HCoV-OC43_orf1ab;186 | 0 | 0 |
| NC_006213.1;YP_009555238.1;HCoV-OC43_orf1ab;187 | 0 | 0 |
| NC_006213.1;YP_009555238.1;HCoV-OC43_orf1ab;188 | 0 | 0 |
| NC_006213.1;YP_009555238.1;HCoV-OC43_orf1ab;189 | 0.05 | 0 |
| NC_006213.1;YP_009555238.1;HCoV-OC43_orf1ab;190 | 0 | 0.025 |
| NC_006213.1;YP_009555238.1;HCoV-OC43_orf1ab;191 | 0.05 | 0 |
| NC_006213.1;YP_009555238.1;HCoV-OC43_orf1ab;192 | 0 | 0 |
| NC_006213.1;YP_009555238.1;HCoV-OC43_orf1ab;193 | 0 | 0.025 |
| NC_006213.1;YP_009555238.1;HCoV-OC43_orf1ab;194 | 0.05 | 0.025 |
| NC_006213.1;YP_009555238.1;HCoV-OC43_orf1ab;195 | 0 | 0.025 |
| NC_006213.1;YP_009555238.1;HCoV-OC43_orf1ab;196 | 0 | 0.025 |
| NC_006213.1;YP_009555238.1;HCoV-OC43_orf1ab;197 | 0 | 0.025 |
| NC_006213.1;YP_009555238.1;HCoV-OC43_orf1ab;198 | 0 | 0.025 |
| NC_006213.1;YP_009555238.1;HCoV-OC43_orf1ab;199 | 0 | 0 |
| NC_006213.1;YP_009555238.1;HCoV-OC43_orf1ab;200 | 0 | 0 |
| NC_006213.1;YP_009555238.1;HCoV-OC43_orf1ab;201 | 0 | 0 |
| NC_006213.1;YP_009555238.1;HCoV-OC43_orf1ab;202 | 0.05 | 0 |
| NC_006213.1;YP_009555238.1;HCoV-OC43_orf1ab;203 | 0 | 0 |
| NC_006213.1;YP_009555238.1;HCoV-OC43_orf1ab;204 | 0.05 | 0 |
| NC_006213.1;YP_009555238.1;HCoV-OC43_orf1ab;205 | 0 | 0 |
| NC_006213.1;YP_009555238.1;HCoV-OC43_orf1ab;206 | 0.05 | 0 |
| NC_006213.1;YP_009555238.1;HCoV-OC43_orf1ab;207 | 0.05 | 0 |
| NC_006213.1;YP_009555238.1;HCoV-OC43_orf1ab;208 | 0 | 0 |
| NC_006213.1;YP_009555238.1;HCoV-OC43_orf1ab;209 | 0 | 0 |
| NC_006213.1;YP_009555238.1;HCoV-OC43_orf1ab;210 | 0.1 | 0.025 |
| NC_006213.1;YP_009555238.1;HCoV-OC43_orf1ab;211 | 0 | 0 |
| NC_006213.1;YP_009555238.1;HCoV-OC43_orf1ab;212 | 0 | 0.025 |
| NC_006213.1;YP_009555238.1;HCoV-OC43_orf1ab;213 | 0 | 0.025 |
| NC_006213.1;YP_009555238.1;HCoV-OC43_orf1ab;214 | 0 | 0.025 |
| NC_006213.1;YP_009555238.1;HCoV-OC43_orf1ab;215 | 0 | 0 |
| NC_006213.1;YP_009555238.1;HCoV-OC43_orf1ab;216 | 0 | 0.025 |
| NC_006213.1;YP_009555238.1;HCoV-OC43_orf1ab;217 | 0 | 0 |
| NC_006213.1;YP_009555238.1;HCoV-OC43_orf1ab;218 | 0.05 | 0.025 |
| NC_006213.1;YP_009555238.1;HCoV-OC43_orf1ab;219 | 0 | 0 |
| NC_006213.1;YP_009555238.1;HCoV-OC43_orf1ab;220 | 0 | 0.025 |
| NC_006213.1;YP_009555238.1;HCoV-OC43_orf1ab;221 | 0 | 0 |
| NC_006213.1;YP_009555238.1;HCoV-OC43_orf1ab;222 | 0 | 0 |
| NC_006213.1;YP_009555238.1;HCoV-OC43_orf1ab;223 | 0 | 0 |
| NC_006213.1;YP_009555238.1;HCoV-OC43_orf1ab;224 | 0 | 0 |

| | | |
|---|---|---|
| NC_006213.1;YP_009555238.1;HCoV-OC43_orf1ab;225 | 0 | 0 |
| NC_006213.1;YP_009555238.1;HCoV-OC43_orf1ab;226 | 0 | 0 |
| NC_006213.1;YP_009555238.1;HCoV-OC43_orf1ab;227 | 0 | 0 |
| NC_006213.1;YP_009555238.1;HCoV-OC43_orf1ab;228 | 0 | 0 |
| NC_006213.1;YP_009555238.1;HCoV-OC43_orf1ab;229 | 0 | 0 |
| NC_006213.1;YP_009555238.1;HCoV-OC43_orf1ab;230 | 0 | 0 |
| NC_006213.1;YP_009555238.1;HCoV-OC43_orf1ab;231 | 0 | 0 |
| NC_006213.1;YP_009555238.1;HCoV-OC43_orf1ab;232 | 0 | 0 |
| NC_006213.1;YP_009555238.1;HCoV-OC43_orf1ab;233 | 0 | 0 |
| NC_006213.1;YP_009555238.1;HCoV-OC43_orf1ab;234 | 0 | 0 |
| NC_006213.1;YP_009555238.1;HCoV-OC43_orf1ab;235 | 0 | 0 |
| NC_006213.1;YP_009555238.1;HCoV-OC43_orf1ab;236 | 0 | 0 |
| NC_006213.1;YP_009555238.1;HCoV-OC43_orf1ab;237 | 0 | 0 |
| NC_006213.1;YP_009555238.1;HCoV-OC43_orf1ab;238 | 0 | 0 |
| NC_006213.1;YP_009555238.1;HCoV-OC43_orf1ab;239 | 0 | 0 |
| NC_006213.1;YP_009555238.1;HCoV-OC43_orf1ab;240 | 0 | 0 |
| NC_006213.1;YP_009555238.1;HCoV-OC43_orf1ab;241 | 0 | 0 |
| NC_006213.1;YP_009555238.1;HCoV-OC43_orf1ab;242 | 0 | 0 |
| NC_006213.1;YP_009555238.1;HCoV-OC43_orf1ab;243 | 0 | 0 |
| NC_006213.1;YP_009555238.1;HCoV-OC43_orf1ab;244 | 0 | 0 |
| NC_006213.1;YP_009555238.1;HCoV-OC43_orf1ab;245 | 0 | 0 |
| NC_006213.1;YP_009555238.1;HCoV-OC43_orf1ab;246 | 0 | 0 |
| NC_006213.1;YP_009555238.1;HCoV-OC43_orf1ab;247 | 0 | 0 |
| NC_006213.1;YP_009555238.1;HCoV-OC43_orf1ab;248 | 0 | 0 |
| NC_006213.1;YP_009555238.1;HCoV-OC43_orf1ab;249 | 0 | 0 |
| NC_006213.1;YP_009555238.1;HCoV-OC43_orf1ab;250 | 0 | 0 |
| NC_006213.1;YP_009555238.1;HCoV-OC43_orf1ab;251 | 0 | 0 |
| NC_006213.1;YP_009555238.1;HCoV-OC43_orf1ab;252 | 0 | 0.025 |
| NC_006213.1;YP_009555238.1;HCoV-OC43_orf1ab;253 | 0 | 0 |
| NC_006213.1;YP_009555238.1;HCoV-OC43_orf1ab;254 | 0 | 0 |
| NC_006213.1;YP_009555238.1;HCoV-OC43_orf1ab;255 | 0 | 0 |
| NC_006213.1;YP_009555238.1;HCoV-OC43_orf1ab;256 | 0 | 0 |
| NC_006213.1;YP_009555238.1;HCoV-OC43_orf1ab;257 | 0 | 0 |
| NC_006213.1;YP_009555238.1;HCoV-OC43_orf1ab;258 | 0 | 0 |
| NC_006213.1;YP_009555238.1;HCoV-OC43_orf1ab;259 | 0.05 | 0.025 |
| NC_006213.1;YP_009555238.1;HCoV-OC43_orf1ab;260 | 0.05 | 0.025 |
| NC_006213.1;YP_009555238.1;HCoV-OC43_orf1ab;261 | 0.05 | 0.075 |
| NC_006213.1;YP_009555238.1;HCoV-OC43_orf1ab;262 | 0.05 | 0.1 |
| NC_006213.1;YP_009555238.1;HCoV-OC43_orf1ab;263 | 0.05 | 0.1 |
| NC_006213.1;YP_009555238.1;HCoV-OC43_orf1ab;264 | 0.1 | 0.075 |
| NC_006213.1;YP_009555238.1;HCoV-OC43_orf1ab;265 | 0.05 | 0.075 |
| NC_006213.1;YP_009555238.1;HCoV-OC43_orf1ab;266 | 0.05 | 0.075 |
| NC_006213.1;YP_009555238.1;HCoV-OC43_orf1ab;267 | 0.1 | 0.075 |

| | | |
|---|---|---|
| NC_006213.1;YP_009555238.1;HCoV-OC43_orf1ab;268 | 0.05 | 0.05 |
| NC_006213.1;YP_009555238.1;HCoV-OC43_orf1ab;269 | 0.05 | 0.05 |
| NC_006213.1;YP_009555238.1;HCoV-OC43_orf1ab;270 | 0.1 | 0 |
| NC_006213.1;YP_009555238.1;HCoV-OC43_orf1ab;271 | 0 | 0.025 |
| NC_006213.1;YP_009555238.1;HCoV-OC43_orf1ab;272 | 0 | 0 |
| NC_006213.1;YP_009555238.1;HCoV-OC43_orf1ab;273 | 0 | 0 |
| NC_006213.1;YP_009555238.1;HCoV-OC43_orf1ab;274 | 0 | 0.05 |
| NC_006213.1;YP_009555238.1;HCoV-OC43_orf1ab;275 | 0 | 0.025 |
| NC_006213.1;YP_009555238.1;HCoV-OC43_orf1ab;276 | 0 | 0 |
| NC_006213.1;YP_009555238.1;HCoV-OC43_orf1ab;277 | 0 | 0 |
| NC_006213.1;YP_009555238.1;HCoV-OC43_orf1ab;278 | 0 | 0 |
| NC_006213.1;YP_009555238.1;HCoV-OC43_orf1ab;279 | 0 | 0 |
| NC_006213.1;YP_009555238.1;HCoV-OC43_orf1ab;280 | 0 | 0 |
| NC_006213.1;YP_009555238.1;HCoV-OC43_orf1ab;281 | 0 | 0 |
| NC_006213.1;YP_009555238.1;HCoV-OC43_orf1ab;282 | 0 | 0 |
| NC_006213.1;YP_009555238.1;HCoV-OC43_orf1ab;283 | 0 | 0 |
| NC_006213.1;YP_009555238.1;HCoV-OC43_orf1ab;284 | 0 | 0 |
| NC_006213.1;YP_009555238.1;HCoV-OC43_orf1ab;285 | 0 | 0 |
| NC_006213.1;YP_009555238.1;HCoV-OC43_orf1ab;286 | 0 | 0 |
| NC_006213.1;YP_009555238.1;HCoV-OC43_orf1ab;287 | 0 | 0 |
| NC_006213.1;YP_009555238.1;HCoV-OC43_orf1ab;288 | 0 | 0 |
| NC_006213.1;YP_009555238.1;HCoV-OC43_orf1ab;289 | 0 | 0 |
| NC_006213.1;YP_009555238.1;HCoV-OC43_orf1ab;290 | 0 | 0 |
| NC_006213.1;YP_009555238.1;HCoV-OC43_orf1ab;291 | 0.1 | 0.1 |
| NC_006213.1;YP_009555238.1;HCoV-OC43_orf1ab;292 | 0.1 | 0.05 |
| NC_006213.1;YP_009555238.1;HCoV-OC43_orf1ab;293 | 0.15 | 0.075 |
| NC_006213.1;YP_009555238.1;HCoV-OC43_orf1ab;294 | 0.1 | 0.05 |
| NC_006213.1;YP_009555238.1;HCoV-OC43_orf1ab;295 | 0.1 | 0.05 |
| NC_006213.1;YP_009555238.1;HCoV-OC43_orf1ab;296 | 0.15 | 0.05 |
| NC_006213.1;YP_009555238.1;HCoV-OC43_orf1ab;297 | 0.15 | 0.05 |
| NC_006213.1;YP_009555238.1;HCoV-OC43_orf1ab;298 | 0.15 | 0.05 |
| NC_006213.1;YP_009555238.1;HCoV-OC43_orf1ab;299 | 0.15 | 0.05 |
| NC_006213.1;YP_009555238.1;HCoV-OC43_orf1ab;300 | 0.15 | 0.125 |
| NC_006213.1;YP_009555238.1;HCoV-OC43_orf1ab;301 | 0.15 | 0.15 |
| NC_006213.1;YP_009555238.1;HCoV-OC43_orf1ab;302 | 0.15 | 0.1 |
| NC_006213.1;YP_009555238.1;HCoV-OC43_orf1ab;303 | 0 | 0 |
| NC_006213.1;YP_009555238.1;HCoV-OC43_orf1ab;304 | 0 | 0 |
| NC_006213.1;YP_009555238.1;HCoV-OC43_orf1ab;305 | 0 | 0 |
| NC_006213.1;YP_009555238.1;HCoV-OC43_orf1ab;306 | 0 | 0 |
| NC_006213.1;YP_009555238.1;HCoV-OC43_orf1ab;307 | 0 | 0 |
| NC_006213.1;YP_009555238.1;HCoV-OC43_orf1ab;308 | 0 | 0 |
| NC_006213.1;YP_009555238.1;HCoV-OC43_orf1ab;309 | 0 | 0 |
| NC_006213.1;YP_009555238.1;HCoV-OC43_orf1ab;310 | 0 | 0 |

| | | |
|---|---|---|
| NC_006213.1;YP_009555238.1;HCoV-OC43_orf1ab;311 | 0 | 0 |
| NC_006213.1;YP_009555238.1;HCoV-OC43_orf1ab;312 | 0 | 0 |
| NC_006213.1;YP_009555238.1;HCoV-OC43_orf1ab;313 | 0 | 0 |
| NC_006213.1;YP_009555238.1;HCoV-OC43_orf1ab;314 | 0 | 0 |
| NC_006213.1;YP_009555238.1;HCoV-OC43_orf1ab;315 | 0 | 0 |
| NC_006213.1;YP_009555238.1;HCoV-OC43_orf1ab;316 | 0 | 0 |
| NC_006213.1;YP_009555238.1;HCoV-OC43_orf1ab;317 | 0.05 | 0 |
| NC_006213.1;YP_009555238.1;HCoV-OC43_orf1ab;318 | 0 | 0 |
| NC_006213.1;YP_009555238.1;HCoV-OC43_orf1ab;319 | 0 | 0 |
| NC_006213.1;YP_009555238.1;HCoV-OC43_orf1ab;320 | 0 | 0.025 |
| NC_006213.1;YP_009555238.1;HCoV-OC43_orf1ab;321 | 0 | 0 |
| NC_006213.1;YP_009555238.1;HCoV-OC43_orf1ab;322 | 0 | 0 |
| NC_006213.1;YP_009555238.1;HCoV-OC43_orf1ab;323 | 0 | 0 |
| NC_006213.1;YP_009555238.1;HCoV-OC43_orf1ab;324 | 0.05 | 0.075 |
| NC_006213.1;YP_009555238.1;HCoV-OC43_orf1ab;325 | 0 | 0 |
| NC_006213.1;YP_009555238.1;HCoV-OC43_orf1ab;326 | 0 | 0 |
| NC_006213.1;YP_009555238.1;HCoV-OC43_orf1ab;327 | 0 | 0 |
| NC_006213.1;YP_009555238.1;HCoV-OC43_orf1ab;328 | 0 | 0 |
| NC_006213.1;YP_009555238.1;HCoV-OC43_orf1ab;329 | 0 | 0 |
| NC_006213.1;YP_009555238.1;HCoV-OC43_orf1ab;330 | 0 | 0 |
| NC_006213.1;YP_009555238.1;HCoV-OC43_orf1ab;331 | 0 | 0 |
| NC_006213.1;YP_009555238.1;HCoV-OC43_orf1ab;332 | 0 | 0.025 |
| NC_006213.1;YP_009555238.1;HCoV-OC43_orf1ab;333 | 0 | 0 |
| NC_006213.1;YP_009555238.1;HCoV-OC43_orf1ab;334 | 0 | 0 |
| NC_006213.1;YP_009555238.1;HCoV-OC43_orf1ab;335 | 0 | 0 |
| NC_006213.1;YP_009555238.1;HCoV-OC43_orf1ab;336 | 0 | 0 |
| NC_006213.1;YP_009555238.1;HCoV-OC43_orf1ab;337 | 0 | 0 |
| NC_006213.1;YP_009555238.1;HCoV-OC43_orf1ab;338 | 0 | 0 |
| NC_006213.1;YP_009555238.1;HCoV-OC43_orf1ab;339 | 0 | 0 |
| NC_006213.1;YP_009555238.1;HCoV-OC43_orf1ab;340 | 0 | 0 |
| NC_006213.1;YP_009555238.1;HCoV-OC43_orf1ab;341 | 0 | 0 |
| NC_006213.1;YP_009555238.1;HCoV-OC43_orf1ab;342 | 0 | 0.025 |
| NC_006213.1;YP_009555238.1;HCoV-OC43_orf1ab;343 | 0 | 0 |
| NC_006213.1;YP_009555238.1;HCoV-OC43_orf1ab;344 | 0 | 0 |
| NC_006213.1;YP_009555238.1;HCoV-OC43_orf1ab;345 | 0 | 0 |
| NC_006213.1;YP_009555238.1;HCoV-OC43_orf1ab;346 | 0 | 0 |
| NC_006213.1;YP_009555238.1;HCoV-OC43_orf1ab;347 | 0 | 0 |
| NC_006213.1;YP_009555238.1;HCoV-OC43_orf1ab;348 | 0 | 0 |
| NC_006213.1;YP_009555238.1;HCoV-OC43_orf1ab;349 | 0 | 0 |
| NC_006213.1;YP_009555238.1;HCoV-OC43_orf1ab;350 | 0 | 0 |
| NC_006213.1;YP_009555238.1;HCoV-OC43_orf1ab;351 | 0 | 0 |
| NC_006213.1;YP_009555238.1;HCoV-OC43_orf1ab;352 | 0 | 0 |
| NC_006213.1;YP_009555238.1;HCoV-OC43_orf1ab;353 | 0 | 0 |

| | | |
|---|---|---|
| NC_006213.1;YP_009555238.1;HCoV-OC43_orf1ab;354 | 0 | 0 |
| NC_006213.1;YP_009555238.1;HCoV-OC43_orf1ab;355 | 0.05 | 0 |
| NC_006213.1;YP_009555238.1;HCoV-OC43_orf1ab;356 | 0 | 0 |
| NC_006213.1;YP_009555238.1;HCoV-OC43_orf1ab;357 | 0 | 0 |
| NC_006213.1;YP_009555238.1;HCoV-OC43_orf1ab;358 | 0 | 0 |
| NC_006213.1;YP_009555238.1;HCoV-OC43_orf1ab;359 | 0 | 0 |
| NC_006213.1;YP_009555238.1;HCoV-OC43_orf1ab;360 | 0 | 0 |
| NC_006213.1;YP_009555238.1;HCoV-OC43_orf1ab;361 | 0 | 0 |
| NC_006213.1;YP_009555238.1;HCoV-OC43_orf1ab;362 | 0 | 0.025 |
| NC_006213.1;YP_009555238.1;HCoV-OC43_orf1ab;363 | 0 | 0 |
| NC_006213.1;YP_009555238.1;HCoV-OC43_orf1ab;364 | 0 | 0 |
| NC_006213.1;YP_009555238.1;HCoV-OC43_orf1ab;365 | 0 | 0 |
| NC_006213.1;YP_009555238.1;HCoV-OC43_orf1ab;366 | 0 | 0 |
| NC_006213.1;YP_009555238.1;HCoV-OC43_orf1ab;367 | 0 | 0 |
| NC_006213.1;YP_009555238.1;HCoV-OC43_orf1ab;368 | 0 | 0 |
| NC_006213.1;YP_009555238.1;HCoV-OC43_orf1ab;369 | 0 | 0 |
| NC_006213.1;YP_009555238.1;HCoV-OC43_orf1ab;370 | 0 | 0 |
| NC_006213.1;YP_009555238.1;HCoV-OC43_orf1ab;371 | 0 | 0 |
| NC_006213.1;YP_009555238.1;HCoV-OC43_orf1ab;372 | 0 | 0 |
| NC_006213.1;YP_009555238.1;HCoV-OC43_orf1ab;373 | 0 | 0 |
| NC_006213.1;YP_009555238.1;HCoV-OC43_orf1ab;374 | 0 | 0 |
| NC_006213.1;YP_009555238.1;HCoV-OC43_orf1ab;375 | 0 | 0.025 |
| NC_006213.1;YP_009555238.1;HCoV-OC43_orf1ab;376 | 0 | 0 |
| NC_006213.1;YP_009555238.1;HCoV-OC43_orf1ab;377 | 0 | 0 |
| NC_006213.1;YP_009555238.1;HCoV-OC43_orf1ab;378 | 0.05 | 0 |
| NC_006213.1;YP_009555238.1;HCoV-OC43_orf1ab;379 | 0 | 0 |
| NC_006213.1;YP_009555238.1;HCoV-OC43_orf1ab;380 | 0 | 0 |
| NC_006213.1;YP_009555238.1;HCoV-OC43_orf1ab;381 | 0 | 0 |
| NC_006213.1;YP_009555238.1;HCoV-OC43_orf1ab;382 | 0 | 0 |
| NC_006213.1;YP_009555238.1;HCoV-OC43_orf1ab;383 | 0 | 0.025 |
| NC_006213.1;YP_009555238.1;HCoV-OC43_orf1ab;384 | 0 | 0.025 |
| NC_006213.1;YP_009555238.1;HCoV-OC43_orf1ab;385 | 0 | 0 |
| NC_006213.1;YP_009555238.1;HCoV-OC43_orf1ab;386 | 0 | 0 |
| NC_006213.1;YP_009555238.1;HCoV-OC43_orf1ab;387 | 0 | 0.025 |
| NC_006213.1;YP_009555238.1;HCoV-OC43_orf1ab;388 | 0 | 0.025 |
| NC_006213.1;YP_009555238.1;HCoV-OC43_orf1ab;389 | 0 | 0 |
| NC_006213.1;YP_009555238.1;HCoV-OC43_orf1ab;390 | 0 | 0 |
| NC_006213.1;YP_009555238.1;HCoV-OC43_orf1ab;391 | 0 | 0 |
| NC_006213.1;YP_009555238.1;HCoV-OC43_orf1ab;392 | 0 | 0 |
| NC_006213.1;YP_009555238.1;HCoV-OC43_orf1ab;393 | 0 | 0 |
| NC_006213.1;YP_009555238.1;HCoV-OC43_orf1ab;394 | 0 | 0 |
| NC_006213.1;YP_009555238.1;HCoV-OC43_orf1ab;395 | 0 | 0 |
| NC_006213.1;YP_009555238.1;HCoV-OC43_orf1ab;396 | 0 | 0 |

| | | |
|---|---|---|
| NC_006213.1;YP_009555238.1;HCoV-OC43_orf1ab;397 | 0 | 0 |
| NC_006213.1;YP_009555238.1;HCoV-OC43_orf1ab;398 | 0.05 | 0 |
| NC_006213.1;YP_009555238.1;HCoV-OC43_orf1ab;399 | 0.1 | 0.025 |
| NC_006213.1;YP_009555238.1;HCoV-OC43_orf1ab;400 | 0.15 | 0.075 |
| NC_006213.1;YP_009555238.1;HCoV-OC43_orf1ab;401 | 0 | 0 |
| NC_006213.1;YP_009555238.1;HCoV-OC43_orf1ab;402 | 0 | 0 |
| NC_006213.1;YP_009555238.1;HCoV-OC43_orf1ab;403 | 0 | 0 |
| NC_006213.1;YP_009555238.1;HCoV-OC43_orf1ab;404 | 0 | 0 |
| NC_006213.1;YP_009555238.1;HCoV-OC43_orf1ab;405 | 0 | 0 |
| NC_006213.1;YP_009555238.1;HCoV-OC43_orf1ab;406 | 0 | 0 |
| NC_006213.1;YP_009555238.1;HCoV-OC43_orf1ab;407 | 0 | 0 |
| NC_006213.1;YP_009555238.1;HCoV-OC43_orf1ab;408 | 0 | 0 |
| NC_006213.1;YP_009555238.1;HCoV-OC43_orf1ab;409 | 0.05 | 0 |
| NC_006213.1;YP_009555238.1;HCoV-OC43_orf1ab;410 | 0.05 | 0 |
| NC_006213.1;YP_009555238.1;HCoV-OC43_orf1ab;411 | 0.05 | 0 |
| NC_006213.1;YP_009555238.1;HCoV-OC43_orf1ab;412 | 0.05 | 0 |
| NC_006213.1;YP_009555238.1;HCoV-OC43_orf1ab;413 | 0.05 | 0 |
| NC_006213.1;YP_009555238.1;HCoV-OC43_orf1ab;414 | 0.05 | 0 |
| NC_006213.1;YP_009555238.1;HCoV-OC43_orf1ab;415 | 0.05 | 0 |
| NC_006213.1;YP_009555238.1;HCoV-OC43_orf1ab;416 | 0.05 | 0 |
| NC_006213.1;YP_009555238.1;HCoV-OC43_orf1ab;417 | 0.05 | 0 |
| NC_006213.1;YP_009555238.1;HCoV-OC43_orf1ab;418 | 0.05 | 0 |
| NC_006213.1;YP_009555238.1;HCoV-OC43_orf1ab;419 | 0.05 | 0 |
| NC_006213.1;YP_009555238.1;HCoV-OC43_orf1ab;420 | 0 | 0 |
| NC_006213.1;YP_009555238.1;HCoV-OC43_orf1ab;421 | 0.05 | 0 |
| NC_006213.1;YP_009555238.1;HCoV-OC43_orf1ab;422 | 0 | 0 |
| NC_006213.1;YP_009555238.1;HCoV-OC43_orf1ab;423 | 0 | 0 |
| NC_006213.1;YP_009555238.1;HCoV-OC43_orf1ab;424 | 0 | 0 |
| NC_006213.1;YP_009555238.1;HCoV-OC43_orf1ab;425 | 0 | 0 |
| NC_006213.1;YP_009555238.1;HCoV-OC43_orf1ab;426 | 0 | 0 |
| NC_006213.1;YP_009555238.1;HCoV-OC43_orf1ab;427 | 0.1 | 0 |
| NC_006213.1;YP_009555238.1;HCoV-OC43_orf1ab;428 | 0.05 | 0 |
| NC_006213.1;YP_009555238.1;HCoV-OC43_orf1ab;429 | 0.05 | 0 |
| NC_006213.1;YP_009555238.1;HCoV-OC43_orf1ab;430 | 0.1 | 0 |
| NC_006213.1;YP_009555238.1;HCoV-OC43_orf1ab;431 | 0.05 | 0 |
| NC_006213.1;YP_009555238.1;HCoV-OC43_orf1ab;432 | 0.05 | 0 |
| NC_006213.1;YP_009555238.1;HCoV-OC43_orf1ab;433 | 0.05 | 0 |
| NC_006213.1;YP_009555238.1;HCoV-OC43_orf1ab;434 | 0.15 | 0 |
| NC_006213.1;YP_009555238.1;HCoV-OC43_orf1ab;435 | 0.05 | 0 |
| NC_006213.1;YP_009555238.1;HCoV-OC43_orf1ab;436 | 0.05 | 0 |
| NC_006213.1;YP_009555238.1;HCoV-OC43_orf1ab;437 | 0.05 | 0 |
| NC_006213.1;YP_009555238.1;HCoV-OC43_orf1ab;438 | 0 | 0 |
| NC_006213.1;YP_009555238.1;HCoV-OC43_orf1ab;439 | 0 | 0 |

| | | |
|---|---|---|
| NC_006213.1;YP_009555238.1;HCoV-OC43_orf1ab;440 | 0 | 0 |
| NC_006213.1;YP_009555238.1;HCoV-OC43_orf1ab;441 | 0 | 0 |
| NC_006213.1;YP_009555238.1;HCoV-OC43_orf1ab;442 | 0 | 0 |
| NC_006213.1;YP_009555238.1;HCoV-OC43_orf1ab;443 | 0 | 0 |
| NC_006213.1;YP_009555238.1;HCoV-OC43_orf1ab;444 | 0 | 0 |
| NC_006213.1;YP_009555238.1;HCoV-OC43_orf1ab;445 | 0 | 0 |
| NC_006213.1;YP_009555238.1;HCoV-OC43_orf1ab;446 | 0 | 0 |
| NC_006213.1;YP_009555238.1;HCoV-OC43_orf1ab;447 | 0 | 0 |
| NC_006213.1;YP_009555238.1;HCoV-OC43_orf1ab;448 | 0 | 0 |
| NC_006213.1;YP_009555238.1;HCoV-OC43_orf1ab;449 | 0 | 0 |
| NC_006213.1;YP_009555238.1;HCoV-OC43_orf1ab;450 | 0 | 0 |
| NC_006213.1;YP_009555238.1;HCoV-OC43_orf1ab;451 | 0 | 0 |
| NC_006213.1;YP_009555238.1;HCoV-OC43_orf1ab;452 | 0 | 0 |
| NC_006213.1;YP_009555238.1;HCoV-OC43_orf1ab;453 | 0 | 0 |
| NC_006213.1;YP_009555238.1;HCoV-OC43_orf1ab;454 | 0 | 0 |
| NC_006213.1;YP_009555238.1;HCoV-OC43_orf1ab;455 | 0 | 0 |
| NC_006213.1;YP_009555238.1;HCoV-OC43_orf1ab;456 | 0 | 0 |
| NC_006213.1;YP_009555238.1;HCoV-OC43_orf1ab;457 | 0 | 0 |
| NC_006213.1;YP_009555238.1;HCoV-OC43_orf1ab;458 | 0 | 0 |
| NC_006213.1;YP_009555238.1;HCoV-OC43_orf1ab;459 | 0 | 0 |
| NC_006213.1;YP_009555238.1;HCoV-OC43_orf1ab;460 | 0 | 0 |
| NC_006213.1;YP_009555238.1;HCoV-OC43_orf1ab;461 | 0 | 0 |
| NC_006213.1;YP_009555238.1;HCoV-OC43_orf1ab;462 | 0 | 0 |
| NC_006213.1;YP_009555238.1;HCoV-OC43_orf1ab;463 | 0 | 0 |
| NC_006213.1;YP_009555238.1;HCoV-OC43_orf1ab;464 | 0 | 0 |
| NC_006213.1;YP_009555238.1;HCoV-OC43_orf1ab;465 | 0 | 0 |
| NC_006213.1;YP_009555238.1;HCoV-OC43_orf1ab;466 | 0 | 0 |
| NC_006213.1;YP_009555238.1;HCoV-OC43_orf1ab;467 | 0 | 0 |
| NC_006213.1;YP_009555238.1;HCoV-OC43_orf1ab;468 | 0 | 0 |
| NC_006213.1;YP_009555238.1;HCoV-OC43_orf1ab;469 | 0 | 0 |
| NC_006213.1;YP_009555238.1;HCoV-OC43_orf1ab;470 | 0 | 0 |
| NC_006213.1;YP_009555238.1;HCoV-OC43_orf1ab;471 | 0 | 0 |
| NC_006213.1;YP_009555238.1;HCoV-OC43_orf1ab;472 | 0 | 0 |
| NC_006213.1;YP_009555238.1;HCoV-OC43_orf1ab;473 | 0 | 0 |
| NC_006213.1;YP_009555238.1;HCoV-OC43_orf1ab;474 | 0 | 0 |
| NC_006213.1;YP_009555238.1;HCoV-OC43_orf1ab;475 | 0 | 0 |
| NC_006213.1;YP_009555238.1;HCoV-OC43_orf1ab;476 | 0 | 0 |
| NC_006213.1;YP_009555238.1;HCoV-OC43_orf1ab;477 | 0 | 0 |
| NC_006213.1;YP_009555238.1;HCoV-OC43_orf1ab;478 | 0 | 0 |
| NC_006213.1;YP_009555238.1;HCoV-OC43_orf1ab;479 | 0 | 0 |
| NC_006213.1;YP_009555238.1;HCoV-OC43_orf1ab;480 | 0 | 0 |
| NC_006213.1;YP_009555238.1;HCoV-OC43_orf1ab;481 | 0 | 0.025 |
| NC_006213.1;YP_009555238.1;HCoV-OC43_orf1ab;482 | 0 | 0 |

| | | |
|---|---|---|
| NC_006213.1;YP_009555238.1;HCoV-OC43_orf1ab;483 | 0 | 0 |
| NC_006213.1;YP_009555238.1;HCoV-OC43_orf1ab;484 | 0 | 0 |
| NC_006213.1;YP_009555238.1;HCoV-OC43_orf1ab;485 | 0 | 0 |
| NC_006213.1;YP_009555238.1;HCoV-OC43_orf1ab;486 | 0 | 0 |
| NC_006213.1;YP_009555238.1;HCoV-OC43_orf1ab;487 | 0 | 0 |
| NC_006213.1;YP_009555238.1;HCoV-OC43_orf1ab;488 | 0 | 0 |
| NC_006213.1;YP_009555238.1;HCoV-OC43_orf1ab;489 | 0 | 0 |
| NC_006213.1;YP_009555238.1;HCoV-OC43_orf1ab;490 | 0 | 0 |
| NC_006213.1;YP_009555238.1;HCoV-OC43_orf1ab;491 | 0 | 0 |
| NC_006213.1;YP_009555238.1;HCoV-OC43_orf1ab;492 | 0 | 0 |
| NC_006213.1;YP_009555238.1;HCoV-OC43_orf1ab;493 | 0 | 0 |
| NC_006213.1;YP_009555238.1;HCoV-OC43_orf1ab;494 | 0 | 0 |
| NC_006213.1;YP_009555238.1;HCoV-OC43_orf1ab;495 | 0 | 0 |
| NC_006213.1;YP_009555238.1;HCoV-OC43_orf1ab;496 | 0 | 0 |
| NC_006213.1;YP_009555238.1;HCoV-OC43_orf1ab;497 | 0.05 | 0.05 |
| NC_006213.1;YP_009555238.1;HCoV-OC43_orf1ab;498 | 0 | 0 |
| NC_006213.1;YP_009555238.1;HCoV-OC43_orf1ab;499 | 0 | 0 |
| NC_006213.1;YP_009555238.1;HCoV-OC43_orf1ab;500 | 0 | 0 |
| NC_006213.1;YP_009555238.1;HCoV-OC43_orf1ab;501 | 0 | 0 |
| NC_006213.1;YP_009555238.1;HCoV-OC43_orf1ab;502 | 0 | 0 |
| NC_006213.1;YP_009555238.1;HCoV-OC43_orf1ab;503 | 0 | 0 |
| NC_006213.1;YP_009555238.1;HCoV-OC43_orf1ab;504 | 0 | 0 |
| NC_006213.1;YP_009555238.1;HCoV-OC43_orf1ab;505 | 0 | 0 |
| NC_006213.1;YP_009555238.1;HCoV-OC43_orf1ab;506 | 0 | 0 |
| NC_006213.1;YP_009555238.1;HCoV-OC43_orf1ab;507 | 0 | 0 |
| NC_006213.1;YP_009555238.1;HCoV-OC43_orf1ab;508 | 0 | 0 |
| NC_006213.1;YP_009555238.1;HCoV-OC43_orf1ab;509 | 0 | 0 |
| NC_006213.1;YP_009555238.1;HCoV-OC43_orf1ab;510 | 0 | 0 |
| NC_006213.1;YP_009555238.1;HCoV-OC43_orf1ab;511 | 0 | 0 |
| NC_006213.1;YP_009555238.1;HCoV-OC43_orf1ab;512 | 0 | 0 |
| NC_006213.1;YP_009555238.1;HCoV-OC43_orf1ab;513 | 0 | 0 |
| NC_006213.1;YP_009555238.1;HCoV-OC43_orf1ab;514 | 0 | 0.025 |
| NC_006213.1;YP_009555238.1;HCoV-OC43_orf1ab;515 | 0 | 0 |
| NC_006213.1;YP_009555238.1;HCoV-OC43_orf1ab;516 | 0 | 0 |
| NC_006213.1;YP_009555238.1;HCoV-OC43_orf1ab;517 | 0 | 0 |
| NC_006213.1;YP_009555238.1;HCoV-OC43_orf1ab;518 | 0.05 | 0 |
| NC_006213.1;YP_009555238.1;HCoV-OC43_orf1ab;519 | 0.05 | 0 |
| NC_006213.1;YP_009555238.1;HCoV-OC43_orf1ab;520 | 0.05 | 0 |
| NC_006213.1;YP_009555238.1;HCoV-OC43_orf1ab;521 | 0.05 | 0 |
| NC_006213.1;YP_009555238.1;HCoV-OC43_orf1ab;522 | 0.05 | 0 |
| NC_006213.1;YP_009555238.1;HCoV-OC43_orf1ab;523 | 0.05 | 0.025 |
| NC_006213.1;YP_009555238.1;HCoV-OC43_orf1ab;524 | 0 | 0.025 |
| NC_006213.1;YP_009555238.1;HCoV-OC43_orf1ab;525 | 0 | 0.025 |

| | | |
|---|---|---|
| NC_006213.1;YP_009555238.1;HCoV-OC43_orf1ab;526 | 0 | 0 |
| NC_006213.1;YP_009555238.1;HCoV-OC43_orf1ab;527 | 0 | 0.025 |
| NC_006213.1;YP_009555238.1;HCoV-OC43_orf1ab;528 | 0 | 0 |
| NC_006213.1;YP_009555238.1;HCoV-OC43_orf1ab;529 | 0 | 0 |
| NC_006213.1;YP_009555238.1;HCoV-OC43_orf1ab;530 | 0 | 0 |
| NC_006213.1;YP_009555238.1;HCoV-OC43_orf1ab;531 | 0 | 0 |
| NC_006213.1;YP_009555238.1;HCoV-OC43_orf1ab;532 | 0 | 0 |
| NC_006213.1;YP_009555238.1;HCoV-OC43_orf1ab;533 | 0 | 0 |
| NC_006213.1;YP_009555238.1;HCoV-OC43_orf1ab;534 | 0.05 | 0.025 |
| NC_006213.1;YP_009555238.1;HCoV-OC43_orf1ab;535 | 0 | 0 |
| NC_006213.1;YP_009555238.1;HCoV-OC43_orf1ab;536 | 0 | 0 |
| NC_006213.1;YP_009555238.1;HCoV-OC43_orf1ab;537 | 0 | 0 |
| NC_006213.1;YP_009555238.1;HCoV-OC43_orf1ab;538 | 0 | 0 |
| NC_006213.1;YP_009555238.1;HCoV-OC43_orf1ab;539 | 0 | 0 |
| NC_006213.1;YP_009555238.1;HCoV-OC43_orf1ab;540 | 0 | 0 |
| NC_006213.1;YP_009555238.1;HCoV-OC43_orf1ab;541 | 0 | 0 |
| NC_006213.1;YP_009555238.1;HCoV-OC43_orf1ab;542 | 0 | 0 |
| NC_006213.1;YP_009555238.1;HCoV-OC43_orf1ab;543 | 0 | 0.025 |
| NC_006213.1;YP_009555238.1;HCoV-OC43_orf1ab;544 | 0 | 0.05 |
| NC_006213.1;YP_009555238.1;HCoV-OC43_orf1ab;545 | 0 | 0.05 |
| NC_006213.1;YP_009555238.1;HCoV-OC43_orf1ab;546 | 0.1 | 0.05 |
| NC_006213.1;YP_009555238.1;HCoV-OC43_orf1ab;547 | 0.1 | 0.125 |
| NC_006213.1;YP_009555238.1;HCoV-OC43_orf1ab;548 | 0.05 | 0.025 |
| NC_006213.1;YP_009555238.1;HCoV-OC43_orf1ab;549 | 0 | 0 |
| NC_006213.1;YP_009555238.1;HCoV-OC43_orf1ab;550 | 0 | 0.025 |
| NC_006213.1;YP_009555238.1;HCoV-OC43_orf1ab;551 | 0 | 0.025 |
| NC_006213.1;YP_009555238.1;HCoV-OC43_orf1ab;552 | 0 | 0 |
| NC_006213.1;YP_009555238.1;HCoV-OC43_orf1ab;553 | 0.05 | 0.025 |
| NC_006213.1;YP_009555238.1;HCoV-OC43_orf1ab;554 | 0.05 | 0.025 |
| NC_006213.1;YP_009555238.1;HCoV-OC43_orf1ab;555 | 0.05 | 0.025 |
| NC_006213.1;YP_009555238.1;HCoV-OC43_orf1ab;556 | 0 | 0 |
| NC_006213.1;YP_009555238.1;HCoV-OC43_orf1ab;557 | 0 | 0 |
| NC_006213.1;YP_009555238.1;HCoV-OC43_orf1ab;558 | 0 | 0 |
| NC_006213.1;YP_009555238.1;HCoV-OC43_orf1ab;559 | 0 | 0 |
| NC_006213.1;YP_009555238.1;HCoV-OC43_orf1ab;560 | 0 | 0 |
| NC_006213.1;YP_009555238.1;HCoV-OC43_orf1ab;561 | 0 | 0 |
| NC_006213.1;YP_009555238.1;HCoV-OC43_orf1ab;562 | 0 | 0 |
| NC_006213.1;YP_009555238.1;HCoV-OC43_orf1ab;563 | 0 | 0 |
| NC_006213.1;YP_009555238.1;HCoV-OC43_orf1ab;564 | 0 | 0 |
| NC_006213.1;YP_009555238.1;HCoV-OC43_orf1ab;565 | 0 | 0 |
| NC_006213.1;YP_009555238.1;HCoV-OC43_orf1ab;566 | 0 | 0 |
| NC_006213.1;YP_009555238.1;HCoV-OC43_orf1ab;567 | 0 | 0 |
| NC_006213.1;YP_009555238.1;HCoV-OC43_orf1ab;568 | 0 | 0 |

| | | |
|---|---|---|
| NC_006213.1;YP_009555238.1;HCoV-OC43_orf1ab;569 | 0 | 0 |
| NC_006213.1;YP_009555238.1;HCoV-OC43_orf1ab;570 | 0 | 0 |
| NC_006213.1;YP_009555238.1;HCoV-OC43_orf1ab;571 | 0 | 0 |
| NC_006213.1;YP_009555238.1;HCoV-OC43_orf1ab;572 | 0 | 0 |
| NC_006213.1;YP_009555238.1;HCoV-OC43_orf1ab;573 | 0 | 0 |
| NC_006213.1;YP_009555238.1;HCoV-OC43_orf1ab;574 | 0 | 0 |
| NC_006213.1;YP_009555238.1;HCoV-OC43_orf1ab;575 | 0 | 0 |
| NC_006213.1;YP_009555238.1;HCoV-OC43_orf1ab;576 | 0 | 0 |
| NC_006213.1;YP_009555238.1;HCoV-OC43_orf1ab;577 | 0 | 0 |
| NC_006213.1;YP_009555238.1;HCoV-OC43_orf1ab;578 | 0 | 0 |
| NC_006213.1;YP_009555238.1;HCoV-OC43_orf1ab;579 | 0 | 0 |
| NC_006213.1;YP_009555238.1;HCoV-OC43_orf1ab;580 | 0 | 0 |
| NC_006213.1;YP_009555238.1;HCoV-OC43_orf1ab;581 | 0 | 0 |
| NC_006213.1;YP_009555238.1;HCoV-OC43_orf1ab;582 | 0 | 0 |
| NC_006213.1;YP_009555238.1;HCoV-OC43_orf1ab;583 | 0 | 0 |
| NC_006213.1;YP_009555238.1;HCoV-OC43_orf1ab;584 | 0 | 0 |
| NC_006213.1;YP_009555238.1;HCoV-OC43_orf1ab;585 | 0 | 0 |
| NC_006213.1;YP_009555238.1;HCoV-OC43_orf1ab;586 | 0 | 0 |
| NC_006213.1;YP_009555238.1;HCoV-OC43_orf1ab;587 | 0 | 0 |
| NC_006213.1;YP_009555238.1;HCoV-OC43_orf1ab;588 | 0 | 0 |
| NC_006213.1;YP_009555238.1;HCoV-OC43_orf1ab;589 | 0 | 0 |
| NC_006213.1;YP_009555238.1;HCoV-OC43_orf1ab;590 | 0 | 0 |
| NC_006213.1;YP_009555238.1;HCoV-OC43_orf1ab;591 | 0 | 0 |
| NC_006213.1;YP_009555238.1;HCoV-OC43_orf1ab;592 | 0.05 | 0 |
| NC_006213.1;YP_009555238.1;HCoV-OC43_orf1ab;593 | 0.1 | 0 |
| NC_006213.1;YP_009555238.1;HCoV-OC43_orf1ab;594 | 0.05 | 0.025 |
| NC_006213.1;YP_009555238.1;HCoV-OC43_orf1ab;595 | 0 | 0 |
| NC_006213.1;YP_009555238.1;HCoV-OC43_orf1ab;596 | 0.05 | 0 |
| NC_006213.1;YP_009555238.1;HCoV-OC43_orf1ab;597 | 0 | 0 |
| NC_006213.1;YP_009555238.1;HCoV-OC43_orf1ab;598 | 0 | 0 |
| NC_006213.1;YP_009555238.1;HCoV-OC43_orf1ab;599 | 0 | 0 |
| NC_006213.1;YP_009555238.1;HCoV-OC43_orf1ab;600 | 0.05 | 0 |
| NC_006213.1;YP_009555238.1;HCoV-OC43_orf1ab;601 | 0.05 | 0 |
| NC_006213.1;YP_009555238.1;HCoV-OC43_orf1ab;602 | 0.05 | 0 |
| NC_006213.1;YP_009555238.1;HCoV-OC43_orf1ab;603 | 0.05 | 0 |
| NC_006213.1;YP_009555238.1;HCoV-OC43_orf1ab;604 | 0.05 | 0 |
| NC_006213.1;YP_009555238.1;HCoV-OC43_orf1ab;605 | 0 | 0 |
| NC_006213.1;YP_009555238.1;HCoV-OC43_orf1ab;606 | 0 | 0 |
| NC_006213.1;YP_009555238.1;HCoV-OC43_orf1ab;607 | 0.05 | 0 |
| NC_006213.1;YP_009555238.1;HCoV-OC43_orf1ab;608 | 0.05 | 0 |
| NC_006213.1;YP_009555238.1;HCoV-OC43_orf1ab;609 | 0 | 0 |
| NC_006213.1;YP_009555238.1;HCoV-OC43_orf1ab;610 | 0 | 0 |
| NC_006213.1;YP_009555238.1;HCoV-OC43_orf1ab;611 | 0 | 0 |

| | | |
|---|---|---|
| NC_006213.1;YP_009555238.1;HCoV-OC43_orf1ab;612 | 0 | 0 |
| NC_006213.1;YP_009555238.1;HCoV-OC43_orf1ab;613 | 0 | 0 |
| NC_006213.1;YP_009555238.1;HCoV-OC43_orf1ab;614 | 0 | 0 |
| NC_006213.1;YP_009555238.1;HCoV-OC43_orf1ab;615 | 0.1 | 0.075 |
| NC_006213.1;YP_009555238.1;HCoV-OC43_orf1ab;616 | 0 | 0 |
| NC_006213.1;YP_009555238.1;HCoV-OC43_orf1ab;617 | 0 | 0 |
| NC_006213.1;YP_009555238.1;HCoV-OC43_orf1ab;618 | 0 | 0 |
| NC_006213.1;YP_009555238.1;HCoV-OC43_orf1ab;619 | 0 | 0 |
| NC_006213.1;YP_009555238.1;HCoV-OC43_orf1ab;620 | 0.05 | 0 |
| NC_006213.1;YP_009555238.1;HCoV-OC43_orf1ab;621 | 0 | 0 |
| NC_006213.1;YP_009555238.1;HCoV-OC43_orf1ab;622 | 0 | 0.025 |
| NC_006213.1;YP_009555238.1;HCoV-OC43_orf1ab;623 | 0 | 0 |
| NC_006213.1;YP_009555238.1;HCoV-OC43_orf1ab;624 | 0 | 0 |
| NC_006213.1;YP_009555238.1;HCoV-OC43_orf1ab;625 | 0 | 0 |
| NC_006213.1;YP_009555238.1;HCoV-OC43_orf1ab;626 | 0 | 0 |
| NC_006213.1;YP_009555238.1;HCoV-OC43_orf1ab;627 | 0 | 0.025 |
| NC_006213.1;YP_009555238.1;HCoV-OC43_orf1ab;628 | 0.05 | 0.025 |
| NC_006213.1;YP_009555238.1;HCoV-OC43_orf1ab;629 | 0.05 | 0.025 |
| NC_006213.1;YP_009555238.1;HCoV-OC43_orf1ab;630 | 0 | 0 |
| NC_006213.1;YP_009555238.1;HCoV-OC43_orf1ab;631 | 0 | 0 |
| NC_006213.1;YP_009555238.1;HCoV-OC43_orf1ab;632 | 0 | 0 |
| NC_006213.1;YP_009555238.1;HCoV-OC43_orf1ab;633 | 0 | 0 |
| NC_006213.1;YP_009555238.1;HCoV-OC43_orf1ab;634 | 0 | 0 |
| NC_006213.1;YP_009555238.1;HCoV-OC43_orf1ab;635 | 0 | 0.025 |
| NC_006213.1;YP_009555238.1;HCoV-OC43_orf1ab;636 | 0 | 0.025 |
| NC_006213.1;YP_009555238.1;HCoV-OC43_orf1ab;637 | 0 | 0 |
| NC_006213.1;YP_009555238.1;HCoV-OC43_orf1ab;638 | 0 | 0 |
| NC_006213.1;YP_009555238.1;HCoV-OC43_orf1ab;639 | 0 | 0 |
| NC_006213.1;YP_009555238.1;HCoV-OC43_orf1ab;640 | 0 | 0 |
| NC_006213.1;YP_009555238.1;HCoV-OC43_orf1ab;641 | 0 | 0 |
| NC_006213.1;YP_009555238.1;HCoV-OC43_orf1ab;642 | 0 | 0 |
| NC_006213.1;YP_009555238.1;HCoV-OC43_orf1ab;643 | 0 | 0 |
| NC_006213.1;YP_009555238.1;HCoV-OC43_orf1ab;644 | 0 | 0 |
| NC_006213.1;YP_009555238.1;HCoV-OC43_orf1ab;645 | 0 | 0 |
| NC_006213.1;YP_009555238.1;HCoV-OC43_orf1ab;646 | 0 | 0 |
| NC_006213.1;YP_009555238.1;HCoV-OC43_orf1ab;647 | 0 | 0 |
| NC_006213.1;YP_009555238.1;HCoV-OC43_orf1ab;648 | 0 | 0 |
| NC_006213.1;YP_009555238.1;HCoV-OC43_orf1ab;649 | 0 | 0 |
| NC_006213.1;YP_009555238.1;HCoV-OC43_orf1ab;650 | 0 | 0 |
| NC_006213.1;YP_009555238.1;HCoV-OC43_orf1ab;651 | 0 | 0 |
| NC_006213.1;YP_009555238.1;HCoV-OC43_orf1ab;652 | 0 | 0 |
| NC_006213.1;YP_009555238.1;HCoV-OC43_orf1ab;653 | 0 | 0 |
| NC_006213.1;YP_009555238.1;HCoV-OC43_orf1ab;654 | 0 | 0 |

| | | |
|---|---|---|
| NC_006213.1;YP_009555238.1;HCoV-OC43_orf1ab;655 | 0 | 0 |
| NC_006213.1;YP_009555238.1;HCoV-OC43_orf1ab;656 | 0 | 0 |
| NC_006213.1;YP_009555238.1;HCoV-OC43_orf1ab;657 | 0 | 0 |
| NC_006213.1;YP_009555238.1;HCoV-OC43_orf1ab;658 | 0 | 0 |
| NC_006213.1;YP_009555238.1;HCoV-OC43_orf1ab;659 | 0 | 0 |
| NC_006213.1;YP_009555238.1;HCoV-OC43_orf1ab;660 | 0 | 0 |
| NC_006213.1;YP_009555238.1;HCoV-OC43_orf1ab;661 | 0 | 0 |
| NC_006213.1;YP_009555238.1;HCoV-OC43_orf1ab;662 | 0 | 0 |
| NC_006213.1;YP_009555238.1;HCoV-OC43_orf1ab;663 | 0 | 0 |
| NC_006213.1;YP_009555238.1;HCoV-OC43_orf1ab;664 | 0 | 0 |
| NC_006213.1;YP_009555238.1;HCoV-OC43_orf1ab;665 | 0 | 0 |
| NC_006213.1;YP_009555238.1;HCoV-OC43_orf1ab;666 | 0 | 0 |
| NC_006213.1;YP_009555238.1;HCoV-OC43_orf1ab;667 | 0 | 0 |
| NC_006213.1;YP_009555238.1;HCoV-OC43_orf1ab;668 | 0 | 0 |
| NC_006213.1;YP_009555238.1;HCoV-OC43_orf1ab;669 | 0 | 0 |
| NC_006213.1;YP_009555238.1;HCoV-OC43_orf1ab;670 | 0 | 0 |
| NC_006213.1;YP_009555238.1;HCoV-OC43_orf1ab;671 | 0 | 0 |
| NC_006213.1;YP_009555238.1;HCoV-OC43_orf1ab;672 | 0 | 0 |
| NC_006213.1;YP_009555238.1;HCoV-OC43_orf1ab;673 | 0 | 0 |
| NC_006213.1;YP_009555238.1;HCoV-OC43_orf1ab;674 | 0 | 0 |
| NC_006213.1;YP_009555238.1;HCoV-OC43_orf1ab;675 | 0 | 0 |
| NC_006213.1;YP_009555238.1;HCoV-OC43_orf1ab;676 | 0 | 0 |
| NC_006213.1;YP_009555238.1;HCoV-OC43_orf1ab;677 | 0 | 0 |
| NC_006213.1;YP_009555238.1;HCoV-OC43_orf1ab;678 | 0 | 0 |
| NC_006213.1;YP_009555238.1;HCoV-OC43_orf1ab;679 | 0 | 0 |
| NC_006213.1;YP_009555238.1;HCoV-OC43_orf1ab;680 | 0 | 0 |
| NC_006213.1;YP_009555238.1;HCoV-OC43_orf1ab;681 | 0 | 0 |
| NC_006213.1;YP_009555238.1;HCoV-OC43_orf1ab;682 | 0 | 0 |
| NC_006213.1;YP_009555238.1;HCoV-OC43_orf1ab;683 | 0 | 0 |
| NC_006213.1;YP_009555238.1;HCoV-OC43_orf1ab;684 | 0 | 0 |
| NC_006213.1;YP_009555238.1;HCoV-OC43_orf1ab;685 | 0 | 0 |
| NC_006213.1;YP_009555238.1;HCoV-OC43_orf1ab;686 | 0 | 0 |
| NC_006213.1;YP_009555238.1;HCoV-OC43_orf1ab;687 | 0 | 0.025 |
| NC_006213.1;YP_009555238.1;HCoV-OC43_orf1ab;688 | 0 | 0.025 |
| NC_006213.1;YP_009555238.1;HCoV-OC43_orf1ab;689 | 0.05 | 0 |
| NC_006213.1;YP_009555238.1;HCoV-OC43_orf1ab;690 | 0 | 0 |
| NC_006213.1;YP_009555238.1;HCoV-OC43_orf1ab;691 | 0 | 0 |
| NC_006213.1;YP_009555238.1;HCoV-OC43_orf1ab;692 | 0 | 0 |
| NC_006213.1;YP_009555238.1;HCoV-OC43_orf1ab;693 | 0.05 | 0 |
| NC_006213.1;YP_009555238.1;HCoV-OC43_orf1ab;694 | 0.05 | 0 |
| NC_006213.1;YP_009555238.1;HCoV-OC43_orf1ab;695 | 0 | 0 |
| NC_006213.1;YP_009555238.1;HCoV-OC43_orf1ab;696 | 0.05 | 0 |
| NC_006213.1;YP_009555238.1;HCoV-OC43_orf1ab;697 | 0 | 0 |

| | | |
|---|---|---|
| NC_006213.1;YP_009555238.1;HCoV-OC43_orf1ab;698 | 0 | 0 |
| NC_006213.1;YP_009555238.1;HCoV-OC43_orf1ab;699 | 0 | 0 |
| NC_006213.1;YP_009555238.1;HCoV-OC43_orf1ab;700 | 0.05 | 0 |
| NC_006213.1;YP_009555238.1;HCoV-OC43_orf1ab;701 | 0 | 0 |
| NC_006213.1;YP_009555238.1;HCoV-OC43_orf1ab;702 | 0 | 0 |
| NC_006213.1;YP_009555238.1;HCoV-OC43_orf1ab;703 | 0 | 0 |
| NC_006213.1;YP_009555238.1;HCoV-OC43_orf1ab;704 | 0 | 0 |
| NC_006213.1;YP_009555238.1;HCoV-OC43_orf1ab;705 | 0 | 0 |
| NC_006213.1;YP_009555238.1;HCoV-OC43_orf1ab;706 | 0 | 0 |
| NC_006213.1;YP_009555238.1;HCoV-OC43_orf1ab;707 | 0 | 0 |
| NC_006213.1;YP_009555238.1;HCoV-OC43_orf1ab;708 | 0 | 0 |
| NC_006213.1;YP_009555238.1;HCoV-OC43_orf1ab;709 | 0 | 0 |
| NC_006213.1;YP_009555238.1;HCoV-OC43_orf1ab;710 | 0 | 0 |
| NC_006213.1;YP_009555238.1;HCoV-OC43_orf1ab;711 | 0 | 0 |
| NC_006213.1;YP_009555238.1;HCoV-OC43_orf1ab;712 | 0 | 0 |
| NC_006213.1;YP_009555238.1;HCoV-OC43_orf1ab;713 | 0 | 0 |
| NC_006213.1;YP_009555238.1;HCoV-OC43_orf1ab;714 | 0 | 0 |
| NC_006213.1;YP_009555238.1;HCoV-OC43_orf1ab;715 | 0 | 0 |
| NC_006213.1;YP_009555238.1;HCoV-OC43_orf1ab;716 | 0 | 0 |
| NC_006213.1;YP_009555238.1;HCoV-OC43_orf1ab;717 | 0 | 0 |
| NC_006213.1;YP_009555238.1;HCoV-OC43_orf1ab;718 | 0 | 0 |
| NC_006213.1;YP_009555238.1;HCoV-OC43_orf1ab;719 | 0 | 0 |
| NC_006213.1;YP_009555238.1;HCoV-OC43_orf1ab;720 | 0 | 0 |
| NC_006213.1;YP_009555238.1;HCoV-OC43_orf1ab;721 | 0 | 0 |
| NC_006213.1;YP_009555238.1;HCoV-OC43_orf1ab;722 | 0 | 0 |
| NC_006213.1;YP_009555238.1;HCoV-OC43_orf1ab;723 | 0 | 0 |
| NC_006213.1;YP_009555238.1;HCoV-OC43_orf1ab;724 | 0 | 0 |
| NC_006213.1;YP_009555238.1;HCoV-OC43_orf1ab;725 | 0 | 0 |
| NC_006213.1;YP_009555238.1;HCoV-OC43_orf1ab;726 | 0 | 0 |
| NC_006213.1;YP_009555238.1;HCoV-OC43_orf1ab;727 | 0 | 0 |
| NC_006213.1;YP_009555238.1;HCoV-OC43_orf1ab;728 | 0 | 0 |
| NC_006213.1;YP_009555238.1;HCoV-OC43_orf1ab;729 | 0 | 0 |
| NC_006213.1;YP_009555238.1;HCoV-OC43_orf1ab;730 | 0 | 0 |
| NC_006213.1;YP_009555238.1;HCoV-OC43_orf1ab;731 | 0 | 0 |
| NC_006213.1;YP_009555238.1;HCoV-OC43_orf1ab;732 | 0 | 0 |
| NC_006213.1;YP_009555238.1;HCoV-OC43_orf1ab;733 | 0 | 0 |
| NC_006213.1;YP_009555238.1;HCoV-OC43_orf1ab;734 | 0 | 0 |
| NC_006213.1;YP_009555238.1;HCoV-OC43_orf1ab;735 | 0 | 0 |
| NC_006213.1;YP_009555238.1;HCoV-OC43_orf1ab;736 | 0 | 0 |
| NC_006213.1;YP_009555238.1;HCoV-OC43_orf1ab;737 | 0 | 0 |
| NC_006213.1;YP_009555238.1;HCoV-OC43_orf1ab;738 | 0 | 0 |
| NC_006213.1;YP_009555238.1;HCoV-OC43_orf1ab;739 | 0 | 0 |
| NC_006213.1;YP_009555238.1;HCoV-OC43_orf1ab;740 | 0 | 0 |

| | | |
|---|---|---|
| NC_006213.1;YP_009555238.1;HCoV-OC43_orf1ab;741 | 0 | 0 |
| NC_006213.1;YP_009555238.1;HCoV-OC43_orf1ab;742 | 0 | 0 |
| NC_006213.1;YP_009555238.1;HCoV-OC43_orf1ab;743 | 0.05 | 0 |
| NC_006213.1;YP_009555238.1;HCoV-OC43_orf1ab;744 | 0.1 | 0 |
| NC_006213.1;YP_009555238.1;HCoV-OC43_orf1ab;745 | 0.1 | 0 |
| NC_006213.1;YP_009555238.1;HCoV-OC43_orf1ab;746 | 0.05 | 0 |
| NC_006213.1;YP_009555238.1;HCoV-OC43_orf1ab;747 | 0.1 | 0 |
| NC_006213.1;YP_009555238.1;HCoV-OC43_orf1ab;748 | 0 | 0.025 |
| NC_006213.1;YP_009555238.1;HCoV-OC43_orf1ab;749 | 0.05 | 0.05 |
| NC_006213.1;YP_009555238.1;HCoV-OC43_orf1ab;750 | 0.05 | 0.025 |
| NC_006213.1;YP_009555238.1;HCoV-OC43_orf1ab;751 | 0.05 | 0.025 |
| NC_006213.1;YP_009555238.1;HCoV-OC43_orf1ab;752 | 0.1 | 0.075 |
| NC_006213.1;YP_009555238.1;HCoV-OC43_orf1ab;753 | 0.05 | 0.075 |
| NC_006213.1;YP_009555238.1;HCoV-OC43_orf1ab;754 | 0.05 | 0.025 |
| NC_006213.1;YP_009555238.1;HCoV-OC43_orf1ab;755 | 0.05 | 0.05 |
| NC_006213.1;YP_009555238.1;HCoV-OC43_orf1ab;756 | 0 | 0 |
| NC_006213.1;YP_009555238.1;HCoV-OC43_orf1ab;757 | 0 | 0 |
| NC_006213.1;YP_009555238.1;HCoV-OC43_orf1ab;758 | 0 | 0 |
| NC_006213.1;YP_009555238.1;HCoV-OC43_orf1ab;759 | 0 | 0 |
| NC_006213.1;YP_009555238.1;HCoV-OC43_orf1ab;760 | 0 | 0 |
| NC_006213.1;YP_009555238.1;HCoV-OC43_orf1ab;761 | 0 | 0 |
| NC_006213.1;YP_009555238.1;HCoV-OC43_orf1ab;762 | 0 | 0 |
| NC_006213.1;YP_009555238.1;HCoV-OC43_orf1ab;763 | 0 | 0 |
| NC_006213.1;YP_009555238.1;HCoV-OC43_orf1ab;764 | 0 | 0 |
| NC_006213.1;YP_009555238.1;HCoV-OC43_orf1ab;765 | 0 | 0 |
| NC_006213.1;YP_009555238.1;HCoV-OC43_orf1ab;766 | 0 | 0 |
| NC_006213.1;YP_009555238.1;HCoV-OC43_orf1ab;767 | 0 | 0 |
| NC_006213.1;YP_009555238.1;HCoV-OC43_orf1ab;768 | 0 | 0 |
| NC_006213.1;YP_009555238.1;HCoV-OC43_orf1ab;769 | 0 | 0 |
| NC_006213.1;YP_009555238.1;HCoV-OC43_orf1ab;770 | 0 | 0.025 |
| NC_006213.1;YP_009555238.1;HCoV-OC43_orf1ab;771 | 0 | 0.05 |
| NC_006213.1;YP_009555238.1;HCoV-OC43_orf1ab;772 | 0.05 | 0.05 |
| NC_006213.1;YP_009555238.1;HCoV-OC43_orf1ab;773 | 0.05 | 0.075 |
| NC_006213.1;YP_009555238.1;HCoV-OC43_orf1ab;774 | 0.1 | 0.05 |
| NC_006213.1;YP_009555238.1;HCoV-OC43_orf1ab;775 | 0.05 | 0.125 |
| NC_006213.1;YP_009555238.1;HCoV-OC43_orf1ab;776 | 0.05 | 0.05 |
| NC_006213.1;YP_009555238.1;HCoV-OC43_orf1ab;777 | 0.05 | 0.05 |
| NC_006213.1;YP_009555238.1;HCoV-OC43_orf1ab;778 | 0.05 | 0.05 |
| NC_006213.1;YP_009555238.1;HCoV-OC43_orf1ab;779 | 0.05 | 0.05 |
| NC_006213.1;YP_009555238.1;HCoV-OC43_orf1ab;780 | 0.05 | 0.05 |
| NC_006213.1;YP_009555238.1;HCoV-OC43_orf1ab;781 | 0 | 0 |
| NC_006213.1;YP_009555238.1;HCoV-OC43_orf1ab;782 | 0 | 0 |
| NC_006213.1;YP_009555238.1;HCoV-OC43_orf1ab;783 | 0 | 0.025 |

| | | |
|---|---|---|
| NC_006213.1;YP_009555238.1;HCoV-OC43_orf1ab;784 | 0 | 0.025 |
| NC_006213.1;YP_009555238.1;HCoV-OC43_orf1ab;785 | 0 | 0.025 |
| NC_006213.1;YP_009555238.1;HCoV-OC43_orf1ab;786 | 0 | 0.025 |
| NC_006213.1;YP_009555238.1;HCoV-OC43_orf1ab;787 | 0 | 0.025 |
| NC_006213.1;YP_009555238.1;HCoV-OC43_orf1ab;788 | 0 | 0.025 |
| NC_006213.1;YP_009555238.1;HCoV-OC43_orf1ab;789 | 0 | 0.025 |
| NC_006213.1;YP_009555238.1;HCoV-OC43_orf1ab;790 | 0 | 0.025 |
| NC_006213.1;YP_009555238.1;HCoV-OC43_orf1ab;791 | 0 | 0.025 |
| NC_006213.1;YP_009555238.1;HCoV-OC43_orf1ab;792 | 0 | 0.025 |
| NC_006213.1;YP_009555238.1;HCoV-OC43_orf1ab;793 | 0 | 0 |
| NC_006213.1;YP_009555238.1;HCoV-OC43_orf1ab;794 | 0 | 0 |
| NC_006213.1;YP_009555238.1;HCoV-OC43_orf1ab;795 | 0 | 0 |
| NC_006213.1;YP_009555238.1;HCoV-OC43_orf1ab;796 | 0 | 0 |
| NC_006213.1;YP_009555238.1;HCoV-OC43_orf1ab;797 | 0.05 | 0.025 |
| NC_006213.1;YP_009555238.1;HCoV-OC43_orf1ab;798 | 0.05 | 0.025 |
| NC_006213.1;YP_009555238.1;HCoV-OC43_orf1ab;799 | 0.05 | 0.05 |
| NC_006213.1;YP_009555238.1;HCoV-OC43_orf1ab;800 | 0.05 | 0.025 |
| NC_006213.1;YP_009555238.1;HCoV-OC43_orf1ab;801 | 0.05 | 0.025 |
| NC_006213.1;YP_009555238.1;HCoV-OC43_orf1ab;802 | 0 | 0.025 |
| NC_006213.1;YP_009555238.1;HCoV-OC43_orf1ab;803 | 0 | 0 |
| NC_006213.1;YP_009555238.1;HCoV-OC43_orf1ab;804 | 0 | 0 |
| NC_006213.1;YP_009555238.1;HCoV-OC43_orf1ab;805 | 0 | 0 |
| NC_006213.1;YP_009555238.1;HCoV-OC43_orf1ab;806 | 0 | 0 |
| NC_006213.1;YP_009555238.1;HCoV-OC43_orf1ab;807 | 0 | 0 |
| NC_006213.1;YP_009555238.1;HCoV-OC43_orf1ab;808 | 0 | 0 |
| NC_006213.1;YP_009555238.1;HCoV-OC43_orf1ab;809 | 0 | 0 |
| NC_006213.1;YP_009555238.1;HCoV-OC43_orf1ab;810 | 0 | 0 |
| NC_006213.1;YP_009555238.1;HCoV-OC43_orf1ab;811 | 0 | 0 |
| NC_006213.1;YP_009555238.1;HCoV-OC43_orf1ab;812 | 0 | 0 |
| NC_006213.1;YP_009555238.1;HCoV-OC43_orf1ab;813 | 0 | 0.025 |
| NC_006213.1;YP_009555238.1;HCoV-OC43_orf1ab;814 | 0 | 0.05 |
| NC_006213.1;YP_009555238.1;HCoV-OC43_orf1ab;815 | 0.05 | 0.025 |
| NC_006213.1;YP_009555238.1;HCoV-OC43_orf1ab;816 | 0.05 | 0.05 |
| NC_006213.1;YP_009555238.1;HCoV-OC43_orf1ab;817 | 0 | 0.075 |
| NC_006213.1;YP_009555238.1;HCoV-OC43_orf1ab;818 | 0.15 | 0.125 |
| NC_006213.1;YP_009555238.1;HCoV-OC43_orf1ab;819 | 0 | 0.1 |
| NC_006213.1;YP_009555238.1;HCoV-OC43_orf1ab;820 | 0 | 0.05 |
| NC_006213.1;YP_009555238.1;HCoV-OC43_orf1ab;821 | 0.05 | 0.1 |
| NC_006213.1;YP_009555238.1;HCoV-OC43_orf1ab;822 | 0.05 | 0.125 |
| NC_006213.1;YP_009555238.1;HCoV-OC43_orf1ab;823 | 0 | 0.05 |
| NC_006213.1;YP_009555238.1;HCoV-OC43_orf1ab;824 | 0 | 0.05 |
| NC_006213.1;YP_009555238.1;HCoV-OC43_orf1ab;825 | 0 | 0.05 |
| NC_006213.1;YP_009555238.1;HCoV-OC43_orf1ab;826 | 0 | 0 |

| | | |
|---|---|---|
| NC_006213.1;YP_009555238.1;HCoV-OC43_orf1ab;827 | 0 | 0 |
| NC_006213.1;YP_009555238.1;HCoV-OC43_orf1ab;828 | 0 | 0 |
| NC_006213.1;YP_009555238.1;HCoV-OC43_orf1ab;829 | 0 | 0 |
| NC_006213.1;YP_009555238.1;HCoV-OC43_orf1ab;830 | 0 | 0 |
| NC_006213.1;YP_009555238.1;HCoV-OC43_orf1ab;831 | 0 | 0 |
| NC_006213.1;YP_009555238.1;HCoV-OC43_orf1ab;832 | 0 | 0.025 |
| NC_006213.1;YP_009555238.1;HCoV-OC43_orf1ab;833 | 0 | 0 |
| NC_006213.1;YP_009555238.1;HCoV-OC43_orf1ab;834 | 0 | 0.05 |
| NC_006213.1;YP_009555238.1;HCoV-OC43_orf1ab;835 | 0 | 0 |
| NC_006213.1;YP_009555238.1;HCoV-OC43_orf1ab;836 | 0 | 0 |
| NC_006213.1;YP_009555238.1;HCoV-OC43_orf1ab;837 | 0 | 0 |
| NC_006213.1;YP_009555238.1;HCoV-OC43_orf1ab;838 | 0 | 0 |
| NC_006213.1;YP_009555238.1;HCoV-OC43_orf1ab;839 | 0 | 0 |
| NC_006213.1;YP_009555238.1;HCoV-OC43_orf1ab;840 | 0 | 0 |
| NC_006213.1;YP_009555238.1;HCoV-OC43_orf1ab;841 | 0.05 | 0.05 |
| NC_006213.1;YP_009555238.1;HCoV-OC43_orf1ab;842 | 0 | 0 |
| NC_006213.1;YP_009555238.1;HCoV-OC43_orf1ab;843 | 0 | 0 |
| NC_006213.1;YP_009555238.1;HCoV-OC43_orf1ab;844 | 0 | 0 |
| NC_006213.1;YP_009555238.1;HCoV-OC43_orf1ab;845 | 0 | 0 |
| NC_006213.1;YP_009555238.1;HCoV-OC43_orf1ab;846 | 0 | 0 |
| NC_006213.1;YP_009555238.1;HCoV-OC43_orf1ab;847 | 0 | 0 |
| NC_006213.1;YP_009555238.1;HCoV-OC43_orf1ab;848 | 0.1 | 0 |
| NC_006213.1;YP_009555238.1;HCoV-OC43_orf1ab;849 | 0 | 0 |
| NC_006213.1;YP_009555238.1;HCoV-OC43_orf1ab;850 | 0 | 0.025 |
| NC_006213.1;YP_009555238.1;HCoV-OC43_orf1ab;851 | 0 | 0 |
| NC_006213.1;YP_009555238.1;HCoV-OC43_orf1ab;852 | 0 | 0 |
| NC_006213.1;YP_009555238.1;HCoV-OC43_orf1ab;853 | 0 | 0 |
| NC_006213.1;YP_009555238.1;HCoV-OC43_orf1ab;854 | 0 | 0 |
| NC_006213.1;YP_009555238.1;HCoV-OC43_orf1ab;855 | 0 | 0 |
| NC_006213.1;YP_009555238.1;HCoV-OC43_orf1ab;856 | 0 | 0 |
| NC_006213.1;YP_009555238.1;HCoV-OC43_orf1ab;857 | 0 | 0 |
| NC_006213.1;YP_009555238.1;HCoV-OC43_orf1ab;858 | 0 | 0 |
| NC_006213.1;YP_009555238.1;HCoV-OC43_orf1ab;859 | 0 | 0 |
| NC_006213.1;YP_009555238.1;HCoV-OC43_orf1ab;860 | 0 | 0 |
| NC_006213.1;YP_009555238.1;HCoV-OC43_orf1ab;861 | 0 | 0.025 |
| NC_006213.1;YP_009555238.1;HCoV-OC43_orf1ab;862 | 0 | 0 |
| NC_006213.1;YP_009555238.1;HCoV-OC43_orf1ab;863 | 0 | 0 |
| NC_006213.1;YP_009555238.1;HCoV-OC43_orf1ab;864 | 0 | 0 |
| NC_006213.1;YP_009555238.1;HCoV-OC43_orf1ab;865 | 0 | 0 |
| NC_006213.1;YP_009555238.1;HCoV-OC43_orf1ab;866 | 0 | 0 |
| NC_006213.1;YP_009555238.1;HCoV-OC43_orf1ab;867 | 0 | 0 |
| NC_006213.1;YP_009555238.1;HCoV-OC43_orf1ab;868 | 0 | 0.025 |
| NC_006213.1;YP_009555238.1;HCoV-OC43_orf1ab;869 | 0 | 0.025 |

| | | |
|---|---|---|
| NC_006213.1;YP_009555238.1;HCoV-OC43_orf1ab;870 | 0 | 0.025 |
| NC_006213.1;YP_009555238.1;HCoV-OC43_orf1ab;871 | 0 | 0 |
| NC_006213.1;YP_009555238.1;HCoV-OC43_orf1ab;872 | 0 | 0 |
| NC_006213.1;YP_009555238.1;HCoV-OC43_orf1ab;873 | 0 | 0.025 |
| NC_006213.1;YP_009555238.1;HCoV-OC43_orf1ab;874 | 0 | 0 |
| NC_006213.1;YP_009555238.1;HCoV-OC43_orf1ab;875 | 0 | 0.05 |
| NC_006213.1;YP_009555238.1;HCoV-OC43_orf1ab;876 | 0 | 0.025 |
| NC_006213.1;YP_009555238.1;HCoV-OC43_orf1ab;877 | 0.05 | 0.05 |
| NC_006213.1;YP_009555238.1;HCoV-OC43_orf1ab;878 | 0.05 | 0.05 |
| NC_006213.1;YP_009555238.1;HCoV-OC43_orf1ab;879 | 0 | 0.025 |
| NC_006213.1;YP_009555238.1;HCoV-OC43_orf1ab;880 | 0 | 0.025 |
| NC_006213.1;YP_009555238.1;HCoV-OC43_orf1ab;881 | 0 | 0.025 |
| NC_006213.1;YP_009555238.1;HCoV-OC43_orf1ab;882 | 0 | 0.025 |
| NC_006213.1;YP_009555238.1;HCoV-OC43_orf1ab;883 | 0 | 0.025 |
| NC_006213.1;YP_009555238.1;HCoV-OC43_orf1ab;884 | 0 | 0.025 |
| NC_006213.1;YP_009555238.1;HCoV-OC43_orf1ab;885 | 0 | 0.025 |
| NC_006213.1;YP_009555238.1;HCoV-OC43_orf1ab;886 | 0 | 0 |
| NC_006213.1;YP_009555238.1;HCoV-OC43_orf1ab;887 | 0 | 0 |
| NC_006213.1;YP_009555238.1;HCoV-OC43_orf1ab;888 | 0 | 0 |
| NC_006213.1;YP_009555238.1;HCoV-OC43_orf1ab;889 | 0 | 0 |
| NC_006213.1;YP_009555238.1;HCoV-OC43_orf1ab;890 | 0 | 0 |
| NC_006213.1;YP_009555238.1;HCoV-OC43_orf1ab;891 | 0 | 0 |
| NC_006213.1;YP_009555238.1;HCoV-OC43_orf1ab;892 | 0 | 0 |
| NC_006213.1;YP_009555238.1;HCoV-OC43_orf1ab;893 | 0 | 0.025 |
| NC_006213.1;YP_009555238.1;HCoV-OC43_orf1ab;894 | 0 | 0 |
| NC_006213.1;YP_009555238.1;HCoV-OC43_orf1ab;895 | 0 | 0 |
| NC_006213.1;YP_009555238.1;HCoV-OC43_orf1ab;896 | 0 | 0 |
| NC_006213.1;YP_009555238.1;HCoV-OC43_orf1ab;897 | 0 | 0 |
| NC_006213.1;YP_009555238.1;HCoV-OC43_orf1ab;898 | 0 | 0 |
| NC_006213.1;YP_009555238.1;HCoV-OC43_orf1ab;899 | 0 | 0 |
| NC_006213.1;YP_009555238.1;HCoV-OC43_orf1ab;900 | 0 | 0 |
| NC_006213.1;YP_009555238.1;HCoV-OC43_orf1ab;901 | 0 | 0 |
| NC_006213.1;YP_009555238.1;HCoV-OC43_orf1ab;902 | 0 | 0 |
| NC_006213.1;YP_009555238.1;HCoV-OC43_orf1ab;903 | 0 | 0 |
| NC_006213.1;YP_009555238.1;HCoV-OC43_orf1ab;904 | 0 | 0 |
| NC_006213.1;YP_009555238.1;HCoV-OC43_orf1ab;905 | 0 | 0 |
| NC_006213.1;YP_009555238.1;HCoV-OC43_orf1ab;906 | 0 | 0 |
| NC_006213.1;YP_009555238.1;HCoV-OC43_orf1ab;907 | 0 | 0 |
| NC_006213.1;YP_009555238.1;HCoV-OC43_orf1ab;908 | 0 | 0 |
| NC_006213.1;YP_009555238.1;HCoV-OC43_orf1ab;909 | 0 | 0 |
| NC_006213.1;YP_009555238.1;HCoV-OC43_orf1ab;910 | 0 | 0 |
| NC_006213.1;YP_009555238.1;HCoV-OC43_orf1ab;911 | 0.05 | 0 |
| NC_006213.1;YP_009555238.1;HCoV-OC43_orf1ab;912 | 0 | 0 |

| | | |
|---|---|---|
| NC_006213.1;YP_009555238.1;HCoV-OC43_orf1ab;913 | 0 | 0 |
| NC_006213.1;YP_009555238.1;HCoV-OC43_orf1ab;914 | 0 | 0 |
| NC_006213.1;YP_009555238.1;HCoV-OC43_orf1ab;915 | 0 | 0 |
| NC_006213.1;YP_009555238.1;HCoV-OC43_orf1ab;916 | 0 | 0 |
| NC_006213.1;YP_009555238.1;HCoV-OC43_orf1ab;917 | 0 | 0 |
| NC_006213.1;YP_009555238.1;HCoV-OC43_orf1ab;918 | 0 | 0.025 |
| NC_006213.1;YP_009555238.1;HCoV-OC43_orf1ab;919 | 0 | 0 |
| NC_006213.1;YP_009555238.1;HCoV-OC43_orf1ab;920 | 0 | 0 |
| NC_006213.1;YP_009555238.1;HCoV-OC43_orf1ab;921 | 0 | 0 |
| NC_006213.1;YP_009555238.1;HCoV-OC43_orf1ab;922 | 0 | 0 |
| NC_006213.1;YP_009555238.1;HCoV-OC43_orf1ab;923 | 0 | 0 |
| NC_006213.1;YP_009555238.1;HCoV-OC43_orf1ab;924 | 0 | 0.025 |
| NC_006213.1;YP_009555238.1;HCoV-OC43_orf1ab;925 | 0 | 0 |
| NC_006213.1;YP_009555238.1;HCoV-OC43_orf1ab;926 | 0.05 | 0 |
| NC_006213.1;YP_009555238.1;HCoV-OC43_orf1ab;927 | 0.05 | 0 |
| NC_006213.1;YP_009555238.1;HCoV-OC43_orf1ab;928 | 0.05 | 0 |
| NC_006213.1;YP_009555238.1;HCoV-OC43_orf1ab;929 | 0.05 | 0 |
| NC_006213.1;YP_009555238.1;HCoV-OC43_orf1ab;930 | 0 | 0 |
| NC_006213.1;YP_009555238.1;HCoV-OC43_orf1ab;931 | 0 | 0 |
| NC_006213.1;YP_009555238.1;HCoV-OC43_orf1ab;932 | 0 | 0 |
| NC_006213.1;YP_009555238.1;HCoV-OC43_orf1ab;933 | 0 | 0 |
| NC_006213.1;YP_009555238.1;HCoV-OC43_orf1ab;934 | 0 | 0 |
| NC_006213.1;YP_009555238.1;HCoV-OC43_orf1ab;935 | 0.05 | 0 |
| NC_006213.1;YP_009555238.1;HCoV-OC43_orf1ab;936 | 0.05 | 0 |
| NC_006213.1;YP_009555238.1;HCoV-OC43_orf1ab;937 | 0.05 | 0 |
| NC_006213.1;YP_009555238.1;HCoV-OC43_orf1ab;938 | 0 | 0 |
| NC_006213.1;YP_009555238.1;HCoV-OC43_orf1ab;939 | 0.05 | 0 |
| NC_006213.1;YP_009555238.1;HCoV-OC43_orf1ab;940 | 0.05 | 0 |
| NC_006213.1;YP_009555238.1;HCoV-OC43_orf1ab;941 | 0.1 | 0.05 |
| NC_006213.1;YP_009555238.1;HCoV-OC43_orf1ab;942 | 0.1 | 0.025 |
| NC_006213.1;YP_009555238.1;HCoV-OC43_orf1ab;943 | 0 | 0.025 |
| NC_006213.1;YP_009555238.1;HCoV-OC43_orf1ab;944 | 0.05 | 0.025 |
| NC_006213.1;YP_009555238.1;HCoV-OC43_orf1ab;945 | 0 | 0.025 |
| NC_006213.1;YP_009555238.1;HCoV-OC43_orf1ab;946 | 0 | 0 |
| NC_006213.1;YP_009555238.1;HCoV-OC43_orf1ab;947 | 0 | 0 |
| NC_006213.1;YP_009555238.1;HCoV-OC43_orf1ab;948 | 0 | 0 |
| NC_006213.1;YP_009555238.1;HCoV-OC43_orf1ab;949 | 0 | 0 |
| NC_006213.1;YP_009555238.1;HCoV-OC43_orf1ab;950 | 0 | 0 |
| NC_006213.1;YP_009555238.1;HCoV-OC43_orf1ab;951 | 0 | 0 |
| NC_006213.1;YP_009555238.1;HCoV-OC43_orf1ab;952 | 0 | 0 |
| NC_006213.1;YP_009555238.1;HCoV-OC43_orf1ab;953 | 0 | 0 |
| NC_006213.1;YP_009555238.1;HCoV-OC43_orf1ab;954 | 0 | 0 |
| NC_006213.1;YP_009555238.1;HCoV-OC43_orf1ab;955 | 0 | 0 |

| | | |
|---|---|---|
| NC_006213.1;YP_009555238.1;HCoV-OC43_orf1ab;956 | 0 | 0.025 |
| NC_006213.1;YP_009555238.1;HCoV-OC43_orf1ab;957 | 0 | 0.025 |
| NC_006213.1;YP_009555238.1;HCoV-OC43_orf1ab;958 | 0 | 0.025 |
| NC_006213.1;YP_009555238.1;HCoV-OC43_orf1ab;959 | 0 | 0 |
| NC_006213.1;YP_009555238.1;HCoV-OC43_orf1ab;960 | 0 | 0 |
| NC_006213.1;YP_009555238.1;HCoV-OC43_orf1ab;961 | 0 | 0 |
| NC_006213.1;YP_009555238.1;HCoV-OC43_orf1ab;962 | 0 | 0 |
| NC_006213.1;YP_009555238.1;HCoV-OC43_orf1ab;963 | 0 | 0 |
| NC_006213.1;YP_009555238.1;HCoV-OC43_orf1ab;964 | 0 | 0 |
| NC_006213.1;YP_009555238.1;HCoV-OC43_orf1ab;965 | 0.1 | 0 |
| NC_006213.1;YP_009555238.1;HCoV-OC43_orf1ab;966 | 0 | 0 |
| NC_006213.1;YP_009555238.1;HCoV-OC43_orf1ab;967 | 0 | 0 |
| NC_006213.1;YP_009555238.1;HCoV-OC43_orf1ab;968 | 0 | 0 |
| NC_006213.1;YP_009555238.1;HCoV-OC43_orf1ab;969 | 0 | 0 |
| NC_006213.1;YP_009555238.1;HCoV-OC43_orf1ab;970 | 0 | 0 |
| NC_006213.1;YP_009555238.1;HCoV-OC43_orf1ab;971 | 0 | 0 |
| NC_006213.1;YP_009555238.1;HCoV-OC43_orf1ab;972 | 0 | 0 |
| NC_006213.1;YP_009555238.1;HCoV-OC43_orf1ab;973 | 0 | 0 |
| NC_006213.1;YP_009555238.1;HCoV-OC43_orf1ab;974 | 0 | 0 |
| NC_006213.1;YP_009555238.1;HCoV-OC43_orf1ab;975 | 0 | 0.025 |
| NC_006213.1;YP_009555238.1;HCoV-OC43_orf1ab;976 | 0 | 0 |
| NC_006213.1;YP_009555238.1;HCoV-OC43_orf1ab;977 | 0.15 | 0 |
| NC_006213.1;YP_009555238.1;HCoV-OC43_orf1ab;978 | 0.05 | 0 |
| NC_006213.1;YP_009555238.1;HCoV-OC43_orf1ab;979 | 0 | 0 |
| NC_006213.1;YP_009555238.1;HCoV-OC43_orf1ab;980 | 0 | 0 |
| NC_006213.1;YP_009555238.1;HCoV-OC43_orf1ab;981 | 0 | 0.025 |
| NC_006213.1;YP_009555238.1;HCoV-OC43_orf1ab;982 | 0 | 0 |
| NC_006213.1;YP_009555238.1;HCoV-OC43_orf1ab;983 | 0 | 0.025 |
| NC_006213.1;YP_009555238.1;HCoV-OC43_orf1ab;984 | 0 | 0 |
| NC_006213.1;YP_009555238.1;HCoV-OC43_orf1ab;985 | 0 | 0 |
| NC_006213.1;YP_009555238.1;HCoV-OC43_orf1ab;986 | 0 | 0 |
| NC_006213.1;YP_009555238.1;HCoV-OC43_orf1ab;987 | 0 | 0 |
| NC_006213.1;YP_009555238.1;HCoV-OC43_orf1ab;988 | 0 | 0 |
| NC_006213.1;YP_009555238.1;HCoV-OC43_orf1ab;989 | 0 | 0 |
| NC_006213.1;YP_009555238.1;HCoV-OC43_orf1ab;990 | 0 | 0 |
| NC_006213.1;YP_009555238.1;HCoV-OC43_orf1ab;991 | 0 | 0 |
| NC_006213.1;YP_009555238.1;HCoV-OC43_orf1ab;992 | 0 | 0 |
| NC_006213.1;YP_009555238.1;HCoV-OC43_orf1ab;993 | 0 | 0 |
| NC_006213.1;YP_009555238.1;HCoV-OC43_orf1ab;994 | 0 | 0 |
| NC_006213.1;YP_009555238.1;HCoV-OC43_orf1ab;995 | 0 | 0 |
| NC_006213.1;YP_009555238.1;HCoV-OC43_orf1ab;996 | 0 | 0 |
| NC_006213.1;YP_009555238.1;HCoV-OC43_orf1ab;997 | 0 | 0 |
| NC_006213.1;YP_009555238.1;HCoV-OC43_orf1ab;998 | 0 | 0 |

| | | |
|---|---|---|
| NC_006213.1;YP_009555238.1;HCoV-OC43_orf1ab;999 | 0 | 0 |
| NC_006213.1;YP_009555238.1;HCoV-OC43_orf1ab;1000 | 0 | 0 |
| NC_006213.1;YP_009555238.1;HCoV-OC43_orf1ab;1001 | 0 | 0 |
| NC_006213.1;YP_009555238.1;HCoV-OC43_orf1ab;1002 | 0.05 | 0 |
| NC_006213.1;YP_009555238.1;HCoV-OC43_orf1ab;1003 | 0.05 | 0 |
| NC_006213.1;YP_009555238.1;HCoV-OC43_orf1ab;1004 | 0.05 | 0 |
| NC_006213.1;YP_009555238.1;HCoV-OC43_orf1ab;1005 | 0.05 | 0 |
| NC_006213.1;YP_009555238.1;HCoV-OC43_orf1ab;1006 | 0.05 | 0 |
| NC_006213.1;YP_009555238.1;HCoV-OC43_orf1ab;1007 | 0.05 | 0.05 |
| NC_006213.1;YP_009555238.1;HCoV-OC43_orf1ab;1008 | 0.05 | 0.025 |
| NC_006213.1;YP_009555238.1;HCoV-OC43_orf1ab;1009 | 0.1 | 0 |
| NC_006213.1;YP_009555238.1;HCoV-OC43_orf1ab;1010 | 0 | 0 |
| NC_006213.1;YP_009555238.1;HCoV-OC43_orf1ab;1011 | 0 | 0 |
| NC_006213.1;YP_009555238.1;HCoV-OC43_orf1ab;1012 | 0 | 0 |
| NC_006213.1;YP_009555238.1;HCoV-OC43_orf1ab;1013 | 0 | 0 |
| NC_006213.1;YP_009555238.1;HCoV-OC43_orf1ab;1014 | 0 | 0 |
| NC_006213.1;YP_009555238.1;HCoV-OC43_orf1ab;1015 | 0 | 0 |
| NC_006213.1;YP_009555238.1;HCoV-OC43_orf1ab;1016 | 0 | 0 |
| NC_006213.1;YP_009555238.1;HCoV-OC43_orf1ab;1017 | 0 | 0 |
| NC_006213.1;YP_009555238.1;HCoV-OC43_orf1ab;1018 | 0 | 0 |
| NC_006213.1;YP_009555238.1;HCoV-OC43_orf1ab;1019 | 0 | 0 |
| NC_006213.1;YP_009555238.1;HCoV-OC43_orf1ab;1020 | 0 | 0 |
| NC_006213.1;YP_009555238.1;HCoV-OC43_orf1ab;1021 | 0 | 0 |
| NC_006213.1;YP_009555238.1;HCoV-OC43_orf1ab;1022 | 0 | 0 |
| NC_006213.1;YP_009555238.1;HCoV-OC43_orf1ab;1023 | 0 | 0 |
| NC_006213.1;YP_009555238.1;HCoV-OC43_orf1ab;1024 | 0 | 0 |
| NC_006213.1;YP_009555238.1;HCoV-OC43_orf1ab;1025 | 0 | 0.025 |
| NC_006213.1;YP_009555238.1;HCoV-OC43_orf1ab;1026 | 0 | 0 |
| NC_006213.1;YP_009555238.1;HCoV-OC43_orf1ab;1027 | 0.05 | 0 |
| NC_006213.1;YP_009555238.1;HCoV-OC43_orf1ab;1028 | 0.1 | 0 |
| NC_006213.1;YP_009555238.1;HCoV-OC43_orf1ab;1029 | 0.1 | 0 |
| NC_006213.1;YP_009555238.1;HCoV-OC43_orf1ab;1030 | 0.1 | 0 |
| NC_006213.1;YP_009555238.1;HCoV-OC43_orf1ab;1031 | 0.1 | 0 |
| NC_006213.1;YP_009555238.1;HCoV-OC43_orf1ab;1032 | 0.1 | 0.025 |
| NC_006213.1;YP_009555238.1;HCoV-OC43_orf1ab;1033 | 0.1 | 0.05 |
| NC_006213.1;YP_009555238.1;HCoV-OC43_orf1ab;1034 | 0.1 | 0 |
| NC_006213.1;YP_009555238.1;HCoV-OC43_orf1ab;1035 | 0.05 | 0 |
| NC_006213.1;YP_009555238.1;HCoV-OC43_orf1ab;1036 | 0.05 | 0.05 |
| NC_006213.1;YP_009555238.1;HCoV-OC43_orf1ab;1037 | 0 | 0 |
| NC_006213.1;YP_009555238.1;HCoV-OC43_orf1ab;1038 | 0 | 0 |
| NC_006213.1;YP_009555238.1;HCoV-OC43_orf1ab;1039 | 0 | 0 |
| NC_006213.1;YP_009555238.1;HCoV-OC43_orf1ab;1040 | 0 | 0 |
| NC_006213.1;YP_009555238.1;HCoV-OC43_orf1ab;1041 | 0 | 0 |

| | | |
|---|---|---|
| NC_006213.1;YP_009555238.1;HCoV-OC43_orf1ab;1042 | 0 | 0 |
| NC_006213.1;YP_009555238.1;HCoV-OC43_orf1ab;1043 | 0 | 0 |
| NC_006213.1;YP_009555238.1;HCoV-OC43_orf1ab;1044 | 0 | 0 |
| NC_006213.1;YP_009555238.1;HCoV-OC43_orf1ab;1045 | 0 | 0 |
| NC_006213.1;YP_009555238.1;HCoV-OC43_orf1ab;1046 | 0 | 0 |
| NC_006213.1;YP_009555238.1;HCoV-OC43_orf1ab;1047 | 0 | 0 |
| NC_006213.1;YP_009555238.1;HCoV-OC43_orf1ab;1048 | 0 | 0 |
| NC_006213.1;YP_009555238.1;HCoV-OC43_orf1ab;1049 | 0 | 0 |
| NC_006213.1;YP_009555238.1;HCoV-OC43_orf1ab;1050 | 0 | 0 |
| NC_006213.1;YP_009555238.1;HCoV-OC43_orf1ab;1051 | 0 | 0 |
| NC_006213.1;YP_009555238.1;HCoV-OC43_orf1ab;1052 | 0 | 0 |
| NC_006213.1;YP_009555238.1;HCoV-OC43_orf1ab;1053 | 0 | 0 |
| NC_006213.1;YP_009555238.1;HCoV-OC43_orf1ab;1054 | 0 | 0 |
| NC_006213.1;YP_009555238.1;HCoV-OC43_orf1ab;1055 | 0 | 0 |
| NC_006213.1;YP_009555238.1;HCoV-OC43_orf1ab;1056 | 0 | 0 |
| NC_006213.1;YP_009555238.1;HCoV-OC43_orf1ab;1057 | 0 | 0 |
| NC_006213.1;YP_009555238.1;HCoV-OC43_orf1ab;1058 | 0 | 0 |
| NC_006213.1;YP_009555238.1;HCoV-OC43_orf1ab;1059 | 0 | 0 |
| NC_006213.1;YP_009555238.1;HCoV-OC43_orf1ab;1060 | 0 | 0 |
| NC_006213.1;YP_009555238.1;HCoV-OC43_orf1ab;1061 | 0 | 0 |
| NC_006213.1;YP_009555238.1;HCoV-OC43_orf1ab;1062 | 0 | 0 |
| NC_006213.1;YP_009555238.1;HCoV-OC43_orf1ab;1063 | 0 | 0 |
| NC_006213.1;YP_009555238.1;HCoV-OC43_orf1ab;1064 | 0 | 0 |
| NC_006213.1;YP_009555238.1;HCoV-OC43_orf1ab;1065 | 0 | 0 |
| NC_006213.1;YP_009555238.1;HCoV-OC43_orf1ab;1066 | 0 | 0 |
| NC_006213.1;YP_009555238.1;HCoV-OC43_orf1ab;1067 | 0 | 0 |
| NC_006213.1;YP_009555238.1;HCoV-OC43_orf1ab;1068 | 0 | 0 |
| NC_006213.1;YP_009555238.1;HCoV-OC43_orf1ab;1069 | 0 | 0 |
| NC_006213.1;YP_009555238.1;HCoV-OC43_orf1ab;1070 | 0 | 0 |
| NC_006213.1;YP_009555238.1;HCoV-OC43_orf1ab;1071 | 0 | 0 |
| NC_006213.1;YP_009555238.1;HCoV-OC43_orf1ab;1072 | 0 | 0 |
| NC_006213.1;YP_009555238.1;HCoV-OC43_orf1ab;1073 | 0 | 0 |
| NC_006213.1;YP_009555238.1;HCoV-OC43_orf1ab;1074 | 0 | 0 |
| NC_006213.1;YP_009555238.1;HCoV-OC43_orf1ab;1075 | 0 | 0 |
| NC_006213.1;YP_009555238.1;HCoV-OC43_orf1ab;1076 | 0 | 0 |
| NC_006213.1;YP_009555238.1;HCoV-OC43_orf1ab;1077 | 0 | 0 |
| NC_006213.1;YP_009555238.1;HCoV-OC43_orf1ab;1078 | 0 | 0 |
| NC_006213.1;YP_009555238.1;HCoV-OC43_orf1ab;1079 | 0 | 0 |
| NC_006213.1;YP_009555238.1;HCoV-OC43_orf1ab;1080 | 0 | 0 |
| NC_006213.1;YP_009555238.1;HCoV-OC43_orf1ab;1081 | 0 | 0 |
| NC_006213.1;YP_009555238.1;HCoV-OC43_orf1ab;1082 | 0 | 0 |
| NC_006213.1;YP_009555238.1;HCoV-OC43_orf1ab;1083 | 0 | 0.025 |
| NC_006213.1;YP_009555238.1;HCoV-OC43_orf1ab;1084 | 0.05 | 0.05 |

| | | |
|---|---|---|
| NC_006213.1;YP_009555238.1;HCoV-OC43_orf1ab;1085 | 0.05 | 0.05 |
| NC_006213.1;YP_009555238.1;HCoV-OC43_orf1ab;1086 | 0 | 0.05 |
| NC_006213.1;YP_009555238.1;HCoV-OC43_orf1ab;1087 | 0 | 0 |
| NC_006213.1;YP_009555238.1;HCoV-OC43_orf1ab;1088 | 0 | 0 |
| NC_006213.1;YP_009555238.1;HCoV-OC43_orf1ab;1089 | 0 | 0 |
| NC_006213.1;YP_009555238.1;HCoV-OC43_orf1ab;1090 | 0 | 0 |
| NC_006213.1;YP_009555238.1;HCoV-OC43_orf1ab;1091 | 0 | 0 |
| NC_006213.1;YP_009555238.1;HCoV-OC43_orf1ab;1092 | 0 | 0 |
| NC_006213.1;YP_009555238.1;HCoV-OC43_orf1ab;1093 | 0.05 | 0 |
| NC_006213.1;YP_009555238.1;HCoV-OC43_orf1ab;1094 | 0 | 0 |
| NC_006213.1;YP_009555238.1;HCoV-OC43_orf1ab;1095 | 0 | 0 |
| NC_006213.1;YP_009555238.1;HCoV-OC43_orf1ab;1096 | 0 | 0 |
| NC_006213.1;YP_009555238.1;HCoV-OC43_orf1ab;1097 | 0 | 0 |
| NC_006213.1;YP_009555238.1;HCoV-OC43_orf1ab;1098 | 0 | 0 |
| NC_006213.1;YP_009555238.1;HCoV-OC43_orf1ab;1099 | 0 | 0 |
| NC_006213.1;YP_009555238.1;HCoV-OC43_orf1ab;1100 | 0 | 0 |
| NC_006213.1;YP_009555238.1;HCoV-OC43_orf1ab;1101 | 0 | 0 |
| NC_006213.1;YP_009555238.1;HCoV-OC43_orf1ab;1102 | 0 | 0 |
| NC_006213.1;YP_009555238.1;HCoV-OC43_orf1ab;1103 | 0 | 0 |
| NC_006213.1;YP_009555238.1;HCoV-OC43_orf1ab;1104 | 0 | 0 |
| NC_006213.1;YP_009555238.1;HCoV-OC43_orf1ab;1105 | 0 | 0 |
| NC_006213.1;YP_009555238.1;HCoV-OC43_orf1ab;1106 | 0 | 0 |
| NC_006213.1;YP_009555238.1;HCoV-OC43_orf1ab;1107 | 0 | 0.025 |
| NC_006213.1;YP_009555238.1;HCoV-OC43_orf1ab;1108 | 0.05 | 0 |
| NC_006213.1;YP_009555238.1;HCoV-OC43_orf1ab;1109 | 0 | 0.025 |
| NC_006213.1;YP_009555238.1;HCoV-OC43_orf1ab;1110 | 0 | 0 |
| NC_006213.1;YP_009555238.1;HCoV-OC43_orf1ab;1111 | 0.1 | 0.025 |
| NC_006213.1;YP_009555238.1;HCoV-OC43_orf1ab;1112 | 0 | 0 |
| NC_006213.1;YP_009555238.1;HCoV-OC43_orf1ab;1113 | 0 | 0 |
| NC_006213.1;YP_009555238.1;HCoV-OC43_orf1ab;1114 | 0 | 0 |
| NC_006213.1;YP_009555238.1;HCoV-OC43_orf1ab;1115 | 0 | 0 |
| NC_006213.1;YP_009555238.1;HCoV-OC43_orf1ab;1116 | 0 | 0 |
| NC_006213.1;YP_009555238.1;HCoV-OC43_orf1ab;1117 | 0 | 0.05 |
| NC_006213.1;YP_009555238.1;HCoV-OC43_orf1ab;1118 | 0 | 0 |
| NC_006213.1;YP_009555238.1;HCoV-OC43_orf1ab;1119 | 0 | 0 |
| NC_006213.1;YP_009555238.1;HCoV-OC43_orf1ab;1120 | 0 | 0 |
| NC_006213.1;YP_009555238.1;HCoV-OC43_orf1ab;1121 | 0 | 0 |
| NC_006213.1;YP_009555238.1;HCoV-OC43_orf1ab;1122 | 0 | 0 |
| NC_006213.1;YP_009555238.1;HCoV-OC43_orf1ab;1123 | 0 | 0 |
| NC_006213.1;YP_009555238.1;HCoV-OC43_orf1ab;1124 | 0 | 0 |
| NC_006213.1;YP_009555238.1;HCoV-OC43_orf1ab;1125 | 0 | 0 |
| NC_006213.1;YP_009555238.1;HCoV-OC43_orf1ab;1126 | 0 | 0 |
| NC_006213.1;YP_009555238.1;HCoV-OC43_orf1ab;1127 | 0 | 0 |

| | | |
|---|---|---|
| NC_006213.1;YP_009555238.1;HCoV-OC43_orf1ab;1128 | 0 | 0 |
| NC_006213.1;YP_009555238.1;HCoV-OC43_orf1ab;1129 | 0 | 0 |
| NC_006213.1;YP_009555238.1;HCoV-OC43_orf1ab;1130 | 0 | 0 |
| NC_006213.1;YP_009555238.1;HCoV-OC43_orf1ab;1131 | 0 | 0 |
| NC_006213.1;YP_009555238.1;HCoV-OC43_orf1ab;1132 | 0 | 0 |
| NC_006213.1;YP_009555238.1;HCoV-OC43_orf1ab;1133 | 0 | 0 |
| NC_006213.1;YP_009555238.1;HCoV-OC43_orf1ab;1134 | 0 | 0 |
| NC_006213.1;YP_009555238.1;HCoV-OC43_orf1ab;1135 | 0 | 0 |
| NC_006213.1;YP_009555238.1;HCoV-OC43_orf1ab;1136 | 0 | 0 |
| NC_006213.1;YP_009555238.1;HCoV-OC43_orf1ab;1137 | 0 | 0 |
| NC_006213.1;YP_009555238.1;HCoV-OC43_orf1ab;1138 | 0 | 0 |
| NC_006213.1;YP_009555238.1;HCoV-OC43_orf1ab;1139 | 0 | 0 |
| NC_006213.1;YP_009555238.1;HCoV-OC43_orf1ab;1140 | 0 | 0 |
| NC_006213.1;YP_009555238.1;HCoV-OC43_orf1ab;1141 | 0 | 0 |
| NC_006213.1;YP_009555238.1;HCoV-OC43_orf1ab;1142 | 0 | 0 |
| NC_006213.1;YP_009555238.1;HCoV-OC43_orf1ab;1143 | 0 | 0 |
| NC_006213.1;YP_009555238.1;HCoV-OC43_orf1ab;1144 | 0 | 0 |
| NC_006213.1;YP_009555238.1;HCoV-OC43_orf1ab;1145 | 0 | 0 |
| NC_006213.1;YP_009555238.1;HCoV-OC43_orf1ab;1146 | 0 | 0 |
| NC_006213.1;YP_009555238.1;HCoV-OC43_orf1ab;1147 | 0 | 0 |
| NC_006213.1;YP_009555238.1;HCoV-OC43_orf1ab;1148 | 0 | 0 |
| NC_006213.1;YP_009555238.1;HCoV-OC43_orf1ab;1149 | 0 | 0 |
| NC_006213.1;YP_009555238.1;HCoV-OC43_orf1ab;1150 | 0 | 0 |
| NC_006213.1;YP_009555238.1;HCoV-OC43_orf1ab;1151 | 0 | 0 |
| NC_006213.1;YP_009555238.1;HCoV-OC43_orf1ab;1152 | 0.05 | 0 |
| NC_006213.1;YP_009555238.1;HCoV-OC43_orf1ab;1153 | 0.05 | 0 |
| NC_006213.1;YP_009555238.1;HCoV-OC43_orf1ab;1154 | 0 | 0 |
| NC_006213.1;YP_009555238.1;HCoV-OC43_orf1ab;1155 | 0 | 0 |
| NC_006213.1;YP_009555238.1;HCoV-OC43_orf1ab;1156 | 0 | 0 |
| NC_006213.1;YP_009555238.1;HCoV-OC43_orf1ab;1157 | 0 | 0 |
| NC_006213.1;YP_009555238.1;HCoV-OC43_orf1ab;1158 | 0 | 0 |
| NC_006213.1;YP_009555238.1;HCoV-OC43_orf1ab;1159 | 0 | 0 |
| NC_006213.1;YP_009555238.1;HCoV-OC43_orf1ab;1160 | 0 | 0 |
| NC_006213.1;YP_009555238.1;HCoV-OC43_orf1ab;1161 | 0 | 0 |
| NC_006213.1;YP_009555238.1;HCoV-OC43_orf1ab;1162 | 0 | 0 |
| NC_006213.1;YP_009555238.1;HCoV-OC43_orf1ab;1163 | 0 | 0 |
| NC_006213.1;YP_009555238.1;HCoV-OC43_orf1ab;1164 | 0 | 0 |
| NC_006213.1;YP_009555238.1;HCoV-OC43_orf1ab;1165 | 0 | 0 |
| NC_006213.1;YP_009555238.1;HCoV-OC43_orf1ab;1166 | 0 | 0 |
| NC_006213.1;YP_009555238.1;HCoV-OC43_orf1ab;1167 | 0 | 0 |
| NC_006213.1;YP_009555238.1;HCoV-OC43_orf1ab;1168 | 0 | 0 |
| NC_006213.1;YP_009555238.1;HCoV-OC43_orf1ab;1169 | 0 | 0 |
| NC_006213.1;YP_009555238.1;HCoV-OC43_orf1ab;1170 | 0 | 0 |

| | | |
|---|---|---|
| NC_006213.1;YP_009555238.1;HCoV-OC43_orf1ab;1171 | 0 | 0 |
| NC_006213.1;YP_009555238.1;HCoV-OC43_orf1ab;1172 | 0 | 0.025 |
| NC_006213.1;YP_009555238.1;HCoV-OC43_orf1ab;1173 | 0 | 0 |
| NC_006213.1;YP_009555238.1;HCoV-OC43_orf1ab;1174 | 0 | 0 |
| NC_006213.1;YP_009555238.1;HCoV-OC43_orf1ab;1175 | 0 | 0 |
| NC_006213.1;YP_009555238.1;HCoV-OC43_orf1ab;1176 | 0 | 0 |
| NC_006213.1;YP_009555238.1;HCoV-OC43_orf1ab;1177 | 0 | 0 |
| NC_006213.1;YP_009555238.1;HCoV-OC43_orf1ab;1178 | 0 | 0 |
| NC_006213.1;YP_009555238.1;HCoV-OC43_orf1ab;1179 | 0 | 0 |
| NC_006213.1;YP_009555238.1;HCoV-OC43_orf1ab;1180 | 0 | 0 |
| NC_006213.1;YP_009555238.1;HCoV-OC43_orf1ab;1181 | 0 | 0 |
| NC_006213.1;YP_009555238.1;HCoV-OC43_orf1ab;1182 | 0 | 0.025 |
| NC_006213.1;YP_009555238.1;HCoV-OC43_orf1ab;1183 | 0 | 0 |
| NC_006213.1;YP_009555238.1;HCoV-OC43_orf1ab;1184 | 0 | 0 |
| NC_006213.1;YP_009555238.1;HCoV-OC43_orf1ab;1185 | 0 | 0 |
| NC_006213.1;YP_009555238.1;HCoV-OC43_orf1ab;1186 | 0 | 0 |
| NC_006213.1;YP_009555238.1;HCoV-OC43_orf1ab;1187 | 0.05 | 0 |
| NC_006213.1;YP_009555238.1;HCoV-OC43_orf1ab;1188 | 0 | 0 |
| NC_006213.1;YP_009555238.1;HCoV-OC43_orf1ab;1189 | 0 | 0 |
| NC_006213.1;YP_009555238.1;HCoV-OC43_orf1ab;1190 | 0 | 0 |
| NC_006213.1;YP_009555238.1;HCoV-OC43_orf1ab;1191 | 0 | 0 |
| NC_006213.1;YP_009555238.1;HCoV-OC43_orf1ab;1192 | 0 | 0 |
| NC_006213.1;YP_009555238.1;HCoV-OC43_orf1ab;1193 | 0 | 0 |
| NC_006213.1;YP_009555238.1;HCoV-OC43_orf1ab;1194 | 0 | 0 |
| NC_006213.1;YP_009555238.1;HCoV-OC43_orf1ab;1195 | 0 | 0 |
| NC_006213.1;YP_009555238.1;HCoV-OC43_orf1ab;1196 | 0 | 0 |
| NC_006213.1;YP_009555238.1;HCoV-OC43_orf1ab;1197 | 0 | 0 |
| NC_006213.1;YP_009555238.1;HCoV-OC43_orf1ab;1198 | 0 | 0 |
| NC_006213.1;YP_009555238.1;HCoV-OC43_orf1ab;1199 | 0 | 0 |
| NC_006213.1;YP_009555238.1;HCoV-OC43_orf1ab;1200 | 0 | 0 |
| NC_006213.1;YP_009555238.1;HCoV-OC43_orf1ab;1201 | 0 | 0 |
| NC_006213.1;YP_009555238.1;HCoV-OC43_orf1ab;1202 | 0 | 0 |
| NC_006213.1;YP_009555238.1;HCoV-OC43_orf1ab;1203 | 0 | 0 |
| NC_006213.1;YP_009555238.1;HCoV-OC43_orf1ab;1204 | 0 | 0 |
| NC_006213.1;YP_009555238.1;HCoV-OC43_orf1ab;1205 | 0 | 0 |
| NC_006213.1;YP_009555238.1;HCoV-OC43_orf1ab;1206 | 0 | 0 |
| NC_006213.1;YP_009555238.1;HCoV-OC43_orf1ab;1207 | 0 | 0 |
| NC_006213.1;YP_009555238.1;HCoV-OC43_orf1ab;1208 | 0 | 0 |
| NC_006213.1;YP_009555238.1;HCoV-OC43_orf1ab;1209 | 0 | 0.05 |
| NC_006213.1;YP_009555238.1;HCoV-OC43_orf1ab;1210 | 0 | 0 |
| NC_006213.1;YP_009555238.1;HCoV-OC43_orf1ab;1211 | 0 | 0.025 |
| NC_006213.1;YP_009555238.1;HCoV-OC43_orf1ab;1212 | 0 | 0 |
| NC_006213.1;YP_009555238.1;HCoV-OC43_orf1ab;1213 | 0 | 0.025 |

| | | |
|---|---|---|
| NC_006213.1;YP_009555238.1;HCoV-OC43_orf1ab;1214 | 0.05 | 0.05 |
| NC_006213.1;YP_009555238.1;HCoV-OC43_orf1ab;1215 | 0 | 0 |
| NC_006213.1;YP_009555238.1;HCoV-OC43_orf1ab;1216 | 0 | 0 |
| NC_006213.1;YP_009555238.1;HCoV-OC43_orf1ab;1217 | 0 | 0 |
| NC_006213.1;YP_009555238.1;HCoV-OC43_orf1ab;1218 | 0 | 0.05 |
| NC_006213.1;YP_009555238.1;HCoV-OC43_orf1ab;1219 | 0 | 0.025 |
| NC_006213.1;YP_009555238.1;HCoV-OC43_orf1ab;1220 | 0 | 0 |
| NC_006213.1;YP_009555238.1;HCoV-OC43_orf1ab;1221 | 0 | 0 |
| NC_006213.1;YP_009555238.1;HCoV-OC43_orf1ab;1222 | 0 | 0 |
| NC_006213.1;YP_009555238.1;HCoV-OC43_orf1ab;1223 | 0 | 0 |
| NC_006213.1;YP_009555238.1;HCoV-OC43_orf1ab;1224 | 0 | 0 |
| NC_006213.1;YP_009555238.1;HCoV-OC43_orf1ab;1225 | 0.05 | 0.025 |
| NC_006213.1;YP_009555238.1;HCoV-OC43_orf1ab;1226 | 0.05 | 0 |
| NC_006213.1;YP_009555238.1;HCoV-OC43_orf1ab;1227 | 0 | 0 |
| NC_006213.1;YP_009555238.1;HCoV-OC43_orf1ab;1228 | 0 | 0 |
| NC_006213.1;YP_009555238.1;HCoV-OC43_orf1ab;1229 | 0 | 0 |
| NC_006213.1;YP_009555238.1;HCoV-OC43_orf1ab;1230 | 0 | 0 |
| NC_006213.1;YP_009555238.1;HCoV-OC43_orf1ab;1231 | 0 | 0 |
| NC_006213.1;YP_009555238.1;HCoV-OC43_orf1ab;1232 | 0.05 | 0.025 |
| NC_006213.1;YP_009555238.1;HCoV-OC43_orf1ab;1233 | 0.05 | 0.025 |
| NC_006213.1;YP_009555238.1;HCoV-OC43_orf1ab;1234 | 0.05 | 0 |
| NC_006213.1;YP_009555238.1;HCoV-OC43_orf1ab;1235 | 0.05 | 0 |
| NC_006213.1;YP_009555238.1;HCoV-OC43_orf1ab;1236 | 0.05 | 0 |
| NC_006213.1;YP_009555238.1;HCoV-OC43_orf1ab;1237 | 0 | 0 |
| NC_006213.1;YP_009555238.1;HCoV-OC43_orf1ab;1238 | 0 | 0 |
| NC_006213.1;YP_009555238.1;HCoV-OC43_orf1ab;1239 | 0 | 0 |
| NC_006213.1;YP_009555238.1;HCoV-OC43_orf1ab;1240 | 0.05 | 0 |
| NC_006213.1;YP_009555238.1;HCoV-OC43_orf1ab;1241 | 0.15 | 0 |
| NC_006213.1;YP_009555238.1;HCoV-OC43_orf1ab;1242 | 0.1 | 0.05 |
| NC_006213.1;YP_009555238.1;HCoV-OC43_orf1ab;1243 | 0.05 | 0 |
| NC_006213.1;YP_009555238.1;HCoV-OC43_orf1ab;1244 | 0.05 | 0 |
| NC_006213.1;YP_009555238.1;HCoV-OC43_orf1ab;1245 | 0.05 | 0.025 |
| NC_006213.1;YP_009555238.1;HCoV-OC43_orf1ab;1246 | 0.1 | 0.025 |
| NC_006213.1;YP_009555238.1;HCoV-OC43_orf1ab;1247 | 0 | 0 |
| NC_006213.1;YP_009555238.1;HCoV-OC43_orf1ab;1248 | 0 | 0 |
| NC_006213.1;YP_009555238.1;HCoV-OC43_orf1ab;1249 | 0 | 0 |
| NC_006213.1;YP_009555238.1;HCoV-OC43_orf1ab;1250 | 0 | 0 |
| NC_006213.1;YP_009555238.1;HCoV-OC43_orf1ab;1251 | 0 | 0 |
| NC_006213.1;YP_009555238.1;HCoV-OC43_orf1ab;1252 | 0 | 0 |
| NC_006213.1;YP_009555238.1;HCoV-OC43_orf1ab;1253 | 0 | 0 |
| NC_006213.1;YP_009555238.1;HCoV-OC43_orf1ab;1254 | 0 | 0 |
| NC_006213.1;YP_009555238.1;HCoV-OC43_orf1ab;1255 | 0 | 0 |
| NC_006213.1;YP_009555238.1;HCoV-OC43_orf1ab;1256 | 0 | 0 |

| | | |
|---|---|---|
| NC_006213.1;YP_009555238.1;HCoV-OC43_orf1ab;1257 | 0 | 0 |
| NC_006213.1;YP_009555238.1;HCoV-OC43_orf1ab;1258 | 0 | 0 |
| NC_006213.1;YP_009555238.1;HCoV-OC43_orf1ab;1259 | 0 | 0 |
| NC_006213.1;YP_009555238.1;HCoV-OC43_orf1ab;1260 | 0 | 0 |
| NC_006213.1;YP_009555238.1;HCoV-OC43_orf1ab;1261 | 0 | 0 |
| NC_006213.1;YP_009555238.1;HCoV-OC43_orf1ab;1262 | 0 | 0 |
| NC_006213.1;YP_009555238.1;HCoV-OC43_orf1ab;1263 | 0 | 0.025 |
| NC_006213.1;YP_009555238.1;HCoV-OC43_orf1ab;1264 | 0 | 0 |
| NC_006213.1;YP_009555238.1;HCoV-OC43_orf1ab;1265 | 0 | 0 |
| NC_006213.1;YP_009555238.1;HCoV-OC43_orf1ab;1266 | 0 | 0 |
| NC_006213.1;YP_009555238.1;HCoV-OC43_orf1ab;1267 | 0 | 0 |
| NC_006213.1;YP_009555238.1;HCoV-OC43_orf1ab;1268 | 0 | 0 |
| NC_006213.1;YP_009555238.1;HCoV-OC43_orf1ab;1269 | 0 | 0 |
| NC_006213.1;YP_009555238.1;HCoV-OC43_orf1ab;1270 | 0 | 0 |
| NC_006213.1;YP_009555238.1;HCoV-OC43_orf1ab;1271 | 0 | 0 |
| NC_006213.1;YP_009555238.1;HCoV-OC43_orf1ab;1272 | 0 | 0 |
| NC_006213.1;YP_009555238.1;HCoV-OC43_orf1ab;1273 | 0 | 0 |
| NC_006213.1;YP_009555238.1;HCoV-OC43_orf1ab;1274 | 0 | 0 |
| NC_006213.1;YP_009555238.1;HCoV-OC43_orf1ab;1275 | 0 | 0 |
| NC_006213.1;YP_009555238.1;HCoV-OC43_orf1ab;1276 | 0 | 0 |
| NC_006213.1;YP_009555238.1;HCoV-OC43_orf1ab;1277 | 0 | 0 |
| NC_006213.1;YP_009555238.1;HCoV-OC43_orf1ab;1278 | 0 | 0 |
| NC_006213.1;YP_009555238.1;HCoV-OC43_orf1ab;1279 | 0 | 0 |
| NC_006213.1;YP_009555238.1;HCoV-OC43_orf1ab;1280 | 0 | 0 |
| NC_006213.1;YP_009555238.1;HCoV-OC43_orf1ab;1281 | 0 | 0 |
| NC_006213.1;YP_009555238.1;HCoV-OC43_orf1ab;1282 | 0 | 0 |
| NC_006213.1;YP_009555238.1;HCoV-OC43_orf1ab;1283 | 0.05 | 0 |
| NC_006213.1;YP_009555238.1;HCoV-OC43_orf1ab;1284 | 0 | 0 |
| NC_006213.1;YP_009555238.1;HCoV-OC43_orf1ab;1285 | 0 | 0 |
| NC_006213.1;YP_009555238.1;HCoV-OC43_orf1ab;1286 | 0 | 0 |
| NC_006213.1;YP_009555238.1;HCoV-OC43_orf1ab;1287 | 0 | 0 |
| NC_006213.1;YP_009555238.1;HCoV-OC43_orf1ab;1288 | 0 | 0 |
| NC_006213.1;YP_009555238.1;HCoV-OC43_orf1ab;1289 | 0 | 0 |
| NC_006213.1;YP_009555238.1;HCoV-OC43_orf1ab;1290 | 0.05 | 0 |
| NC_006213.1;YP_009555238.1;HCoV-OC43_orf1ab;1291 | 0 | 0 |
| NC_006213.1;YP_009555238.1;HCoV-OC43_orf1ab;1292 | 0 | 0.025 |
| NC_006213.1;YP_009555238.1;HCoV-OC43_orf1ab;1293 | 0.15 | 0.15 |
| NC_006213.1;YP_009555238.1;HCoV-OC43_orf1ab;1294 | 0.15 | 0.05 |
| NC_006213.1;YP_009555238.1;HCoV-OC43_orf1ab;1295 | 0 | 0 |
| NC_006213.1;YP_009555238.1;HCoV-OC43_orf1ab;1296 | 0.05 | 0.05 |
| NC_006213.1;YP_009555238.1;HCoV-OC43_orf1ab;1297 | 0.05 | 0 |
| NC_006213.1;YP_009555238.1;HCoV-OC43_orf1ab;1298 | 0.05 | 0 |
| NC_006213.1;YP_009555238.1;HCoV-OC43_orf1ab;1299 | 0.05 | 0 |

| | | |
|---|---|---|
| NC_006213.1;YP_009555238.1;HCoV-OC43_orf1ab;1300 | 0.05 | 0 |
| NC_006213.1;YP_009555238.1;HCoV-OC43_orf1ab;1301 | 0.05 | 0 |
| NC_006213.1;YP_009555238.1;HCoV-OC43_orf1ab;1302 | 0.05 | 0 |
| NC_006213.1;YP_009555238.1;HCoV-OC43_orf1ab;1303 | 0.1 | 0 |
| NC_006213.1;YP_009555238.1;HCoV-OC43_orf1ab;1304 | 0.05 | 0 |
| NC_006213.1;YP_009555238.1;HCoV-OC43_orf1ab;1305 | 0 | 0 |
| NC_006213.1;YP_009555238.1;HCoV-OC43_orf1ab;1306 | 0 | 0 |
| NC_006213.1;YP_009555238.1;HCoV-OC43_orf1ab;1307 | 0 | 0 |
| NC_006213.1;YP_009555238.1;HCoV-OC43_orf1ab;1308 | 0 | 0 |
| NC_006213.1;YP_009555238.1;HCoV-OC43_orf1ab;1309 | 0 | 0 |
| NC_006213.1;YP_009555238.1;HCoV-OC43_orf1ab;1310 | 0 | 0 |
| NC_006213.1;YP_009555238.1;HCoV-OC43_orf1ab;1311 | 0 | 0 |
| NC_006213.1;YP_009555238.1;HCoV-OC43_orf1ab;1312 | 0 | 0 |
| NC_006213.1;YP_009555238.1;HCoV-OC43_orf1ab;1313 | 0 | 0 |
| NC_006213.1;YP_009555238.1;HCoV-OC43_orf1ab;1314 | 0 | 0 |
| NC_006213.1;YP_009555238.1;HCoV-OC43_orf1ab;1315 | 0 | 0 |
| NC_006213.1;YP_009555238.1;HCoV-OC43_orf1ab;1316 | 0 | 0 |
| NC_006213.1;YP_009555238.1;HCoV-OC43_orf1ab;1317 | 0 | 0 |
| NC_006213.1;YP_009555238.1;HCoV-OC43_orf1ab;1318 | 0 | 0 |
| NC_006213.1;YP_009555238.1;HCoV-OC43_orf1ab;1319 | 0 | 0 |
| NC_006213.1;YP_009555238.1;HCoV-OC43_orf1ab;1320 | 0 | 0 |
| NC_006213.1;YP_009555238.1;HCoV-OC43_orf1ab;1321 | 0 | 0 |
| NC_006213.1;YP_009555238.1;HCoV-OC43_orf1ab;1322 | 0 | 0 |
| NC_006213.1;YP_009555238.1;HCoV-OC43_orf1ab;1323 | 0 | 0 |
| NC_006213.1;YP_009555238.1;HCoV-OC43_orf1ab;1324 | 0 | 0 |
| NC_006213.1;YP_009555238.1;HCoV-OC43_orf1ab;1325 | 0 | 0.025 |
| NC_006213.1;YP_009555238.1;HCoV-OC43_orf1ab;1326 | 0 | 0.025 |
| NC_006213.1;YP_009555238.1;HCoV-OC43_orf1ab;1327 | 0 | 0.025 |
| NC_006213.1;YP_009555238.1;HCoV-OC43_orf1ab;1328 | 0 | 0 |
| NC_006213.1;YP_009555238.1;HCoV-OC43_orf1ab;1329 | 0 | 0.025 |
| NC_006213.1;YP_009555238.1;HCoV-OC43_orf1ab;1330 | 0 | 0 |
| NC_006213.1;YP_009555238.1;HCoV-OC43_orf1ab;1331 | 0 | 0 |
| NC_006213.1;YP_009555238.1;HCoV-OC43_orf1ab;1332 | 0.05 | 0 |
| NC_006213.1;YP_009555238.1;HCoV-OC43_orf1ab;1333 | 0 | 0.05 |
| NC_006213.1;YP_009555238.1;HCoV-OC43_orf1ab;1334 | 0 | 0 |
| NC_006213.1;YP_009555238.1;HCoV-OC43_orf1ab;1335 | 0 | 0 |
| NC_006213.1;YP_009555238.1;HCoV-OC43_orf1ab;1336 | 0 | 0 |
| NC_006213.1;YP_009555238.1;HCoV-OC43_orf1ab;1337 | 0 | 0 |
| NC_006213.1;YP_009555238.1;HCoV-OC43_orf1ab;1338 | 0 | 0 |
| NC_006213.1;YP_009555238.1;HCoV-OC43_orf1ab;1339 | 0 | 0 |
| NC_006213.1;YP_009555238.1;HCoV-OC43_orf1ab;1340 | 0 | 0 |
| NC_006213.1;YP_009555238.1;HCoV-OC43_orf1ab;1341 | 0 | 0 |
| NC_006213.1;YP_009555238.1;HCoV-OC43_orf1ab;1342 | 0 | 0 |

| | | |
|---|---|---|
| NC_006213.1;YP_009555238.1;HCoV-OC43_orf1ab;1343 | 0 | 0 |
| NC_006213.1;YP_009555238.1;HCoV-OC43_orf1ab;1344 | 0 | 0 |
| NC_006213.1;YP_009555238.1;HCoV-OC43_orf1ab;1345 | 0 | 0 |
| NC_006213.1;YP_009555238.1;HCoV-OC43_orf1ab;1346 | 0 | 0 |
| NC_006213.1;YP_009555238.1;HCoV-OC43_orf1ab;1347 | 0 | 0 |
| NC_006213.1;YP_009555238.1;HCoV-OC43_orf1ab;1348 | 0 | 0 |
| NC_006213.1;YP_009555238.1;HCoV-OC43_orf1ab;1349 | 0 | 0.025 |
| NC_006213.1;YP_009555238.1;HCoV-OC43_orf1ab;1350 | 0 | 0.025 |
| NC_006213.1;YP_009555238.1;HCoV-OC43_orf1ab;1351 | 0 | 0 |
| NC_006213.1;YP_009555238.1;HCoV-OC43_orf1ab;1352 | 0 | 0.025 |
| NC_006213.1;YP_009555238.1;HCoV-OC43_orf1ab;1353 | 0 | 0.025 |
| NC_006213.1;YP_009555238.1;HCoV-OC43_orf1ab;1354 | 0 | 0.025 |
| NC_006213.1;YP_009555238.1;HCoV-OC43_orf1ab;1355 | 0 | 0.025 |
| NC_006213.1;YP_009555238.1;HCoV-OC43_orf1ab;1356 | 0 | 0.05 |
| NC_006213.1;YP_009555238.1;HCoV-OC43_orf1ab;1357 | 0 | 0.025 |
| NC_006213.1;YP_009555238.1;HCoV-OC43_orf1ab;1358 | 0 | 0 |
| NC_006213.1;YP_009555238.1;HCoV-OC43_orf1ab;1359 | 0 | 0 |
| NC_006213.1;YP_009555238.1;HCoV-OC43_orf1ab;1360 | 0 | 0 |
| NC_006213.1;YP_009555238.1;HCoV-OC43_orf1ab;1361 | 0.05 | 0 |
| NC_006213.1;YP_009555238.1;HCoV-OC43_orf1ab;1362 | 0 | 0 |
| NC_006213.1;YP_009555238.1;HCoV-OC43_orf1ab;1363 | 0 | 0 |
| NC_006213.1;YP_009555238.1;HCoV-OC43_orf1ab;1364 | 0 | 0 |
| NC_006213.1;YP_009555238.1;HCoV-OC43_orf1ab;1365 | 0 | 0 |
| NC_006213.1;YP_009555238.1;HCoV-OC43_orf1ab;1366 | 0 | 0 |
| NC_006213.1;YP_009555238.1;HCoV-OC43_orf1ab;1367 | 0 | 0 |
| NC_006213.1;YP_009555238.1;HCoV-OC43_orf1ab;1368 | 0 | 0 |
| NC_006213.1;YP_009555238.1;HCoV-OC43_orf1ab;1369 | 0 | 0 |
| NC_006213.1;YP_009555238.1;HCoV-OC43_orf1ab;1370 | 0 | 0 |
| NC_006213.1;YP_009555238.1;HCoV-OC43_orf1ab;1371 | 0 | 0 |
| NC_006213.1;YP_009555238.1;HCoV-OC43_orf1ab;1372 | 0.05 | 0.05 |
| NC_006213.1;YP_009555238.1;HCoV-OC43_orf1ab;1373 | 0.1 | 0 |
| NC_006213.1;YP_009555238.1;HCoV-OC43_orf1ab;1374 | 0.05 | 0 |
| NC_006213.1;YP_009555238.1;HCoV-OC43_orf1ab;1375 | 0 | 0 |
| NC_006213.1;YP_009555238.1;HCoV-OC43_orf1ab;1376 | 0 | 0 |
| NC_006213.1;YP_009555238.1;HCoV-OC43_orf1ab;1377 | 0 | 0 |
| NC_006213.1;YP_009555238.1;HCoV-OC43_orf1ab;1378 | 0 | 0 |
| NC_006213.1;YP_009555238.1;HCoV-OC43_orf1ab;1379 | 0 | 0 |
| NC_006213.1;YP_009555238.1;HCoV-OC43_orf1ab;1380 | 0 | 0 |
| NC_006213.1;YP_009555238.1;HCoV-OC43_orf1ab;1381 | 0 | 0 |
| NC_006213.1;YP_009555238.1;HCoV-OC43_orf1ab;1382 | 0 | 0 |
| NC_006213.1;YP_009555238.1;HCoV-OC43_orf1ab;1383 | 0 | 0 |
| NC_006213.1;YP_009555238.1;HCoV-OC43_orf1ab;1384 | 0 | 0 |
| NC_006213.1;YP_009555238.1;HCoV-OC43_orf1ab;1385 | 0 | 0 |

| | | |
|---|---|---|
| NC_006213.1;YP_009555238.1;HCoV-OC43_orf1ab;1386 | 0 | 0 |
| NC_006213.1;YP_009555238.1;HCoV-OC43_orf1ab;1387 | 0 | 0 |
| NC_006213.1;YP_009555238.1;HCoV-OC43_orf1ab;1388 | 0 | 0 |
| NC_006213.1;YP_009555238.1;HCoV-OC43_orf1ab;1389 | 0 | 0.025 |
| NC_006213.1;YP_009555238.1;HCoV-OC43_orf1ab;1390 | 0 | 0.025 |
| NC_006213.1;YP_009555238.1;HCoV-OC43_orf1ab;1391 | 0 | 0 |
| NC_006213.1;YP_009555238.1;HCoV-OC43_orf1ab;1392 | 0 | 0 |
| NC_006213.1;YP_009555238.1;HCoV-OC43_orf1ab;1393 | 0 | 0 |
| NC_006213.1;YP_009555238.1;HCoV-OC43_orf1ab;1394 | 0 | 0 |
| NC_006213.1;YP_009555238.1;HCoV-OC43_orf1ab;1395 | 0 | 0 |
| NC_006213.1;YP_009555238.1;HCoV-OC43_orf1ab;1396 | 0 | 0 |
| NC_006213.1;YP_009555238.1;HCoV-OC43_orf1ab;1397 | 0.05 | 0 |
| NC_006213.1;YP_009555238.1;HCoV-OC43_orf1ab;1398 | 0 | 0.025 |
| NC_006213.1;YP_009555238.1;HCoV-OC43_orf1ab;1399 | 0 | 0.025 |
| NC_006213.1;YP_009555238.1;HCoV-OC43_orf1ab;1400 | 0 | 0.025 |
| NC_006213.1;YP_009555238.1;HCoV-OC43_orf1ab;1401 | 0 | 0.025 |
| NC_006213.1;YP_009555238.1;HCoV-OC43_orf1ab;1402 | 0 | 0 |
| NC_006213.1;YP_009555238.1;HCoV-OC43_orf1ab;1403 | 0 | 0 |
| NC_006213.1;YP_009555238.1;HCoV-OC43_orf1ab;1404 | 0 | 0 |
| NC_006213.1;YP_009555238.1;HCoV-OC43_orf1ab;1405 | 0 | 0 |
| NC_006213.1;YP_009555238.1;HCoV-OC43_orf1ab;1406 | 0 | 0.025 |
| NC_006213.1;YP_009555238.1;HCoV-OC43_orf1ab;1407 | 0 | 0.025 |
| NC_006213.1;YP_009555238.1;HCoV-OC43_orf1ab;1408 | 0 | 0.025 |
| NC_006213.1;YP_009555238.1;HCoV-OC43_orf1ab;1409 | 0.05 | 0.05 |
| NC_006213.1;YP_009555238.1;HCoV-OC43_orf1ab;1410 | 0.05 | 0.05 |
| NC_006213.1;YP_009555238.1;HCoV-OC43_orf1ab;1411 | 0.05 | 0.075 |
| NC_006213.1;YP_009555238.1;HCoV-OC43_orf1ab;1412 | 0 | 0.025 |
| NC_006213.1;YP_009555238.1;HCoV-OC43_orf1ab;1413 | 0 | 0.025 |
| NC_006213.1;YP_009555238.1;HCoV-OC43_orf1ab;1414 | 0 | 0.05 |
| NC_006213.1;YP_009555238.1;HCoV-OC43_orf1ab;1415 | 0.1 | 0.175 |
| NC_006213.1;YP_009555238.1;HCoV-OC43_orf1ab;1416 | 0.15 | 0.125 |
| NC_006213.1;YP_009555238.1;HCoV-OC43_orf1ab;1417 | 0.05 | 0.05 |
| NC_006213.1;YP_009555238.1;HCoV-OC43_orf1ab;1418 | 0 | 0 |
| NC_006213.1;YP_009555238.1;HCoV-OC43_orf1ab;1419 | 0 | 0 |
| NC_006213.1;YP_009555238.1;HCoV-OC43_orf1ab;1420 | 0 | 0 |
| NC_006213.1;YP_009555238.1;HCoV-OC43_orf1ab;1421 | 0 | 0.05 |
| NC_006213.1;YP_009555238.1;HCoV-OC43_orf1ab;1422 | 0.15 | 0.075 |
| NC_006213.1;YP_009555238.1;HCoV-OC43_orf1ab;1423 | 0.15 | 0.1 |
| NC_006213.1;YP_009555238.1;HCoV-OC43_orf1ab;1424 | 0.1 | 0.1 |
| NC_006213.1;YP_009555238.1;HCoV-OC43_orf1ab;1425 | 0.05 | 0.025 |
| NC_006213.1;YP_009555238.1;HCoV-OC43_orf1ab;1426 | 0 | 0 |
| NC_006213.1;YP_009555238.1;HCoV-OC43_orf1ab;1427 | 0 | 0 |
| NC_006213.1;YP_009555238.1;HCoV-OC43_orf1ab;1428 | 0.05 | 0.025 |

| | | |
|---|---|---|
| NC_006213.1;YP_009555238.1;HCoV-OC43_orf1ab;1429 | 0 | 0 |
| NC_006213.1;YP_009555238.1;HCoV-OC43_orf1ab;1430 | 0 | 0 |
| NC_006213.1;YP_009555238.1;HCoV-OC43_orf1ab;1431 | 0 | 0 |
| NC_006213.1;YP_009555238.1;HCoV-OC43_orf1ab;1432 | 0 | 0 |
| NC_006213.1;YP_009555238.1;HCoV-OC43_orf1ab;1433 | 0 | 0 |
| NC_006213.1;YP_009555238.1;HCoV-OC43_orf1ab;1434 | 0 | 0 |
| NC_006213.1;YP_009555238.1;HCoV-OC43_orf1ab;1435 | 0 | 0 |
| NC_006213.1;YP_009555238.1;HCoV-OC43_orf1ab;1436 | 0 | 0.025 |
| NC_006213.1;YP_009555238.1;HCoV-OC43_orf1ab;1437 | 0 | 0 |
| NC_006213.1;YP_009555238.1;HCoV-OC43_orf1ab;1438 | 0 | 0 |
| NC_006213.1;YP_009555238.1;HCoV-OC43_orf1ab;1439 | 0 | 0 |
| NC_006213.1;YP_009555238.1;HCoV-OC43_orf1ab;1440 | 0 | 0 |
| NC_006213.1;YP_009555238.1;HCoV-OC43_orf1ab;1441 | 0 | 0 |
| NC_006213.1;YP_009555238.1;HCoV-OC43_orf1ab;1442 | 0 | 0 |
| NC_006213.1;YP_009555238.1;HCoV-OC43_orf1ab;1443 | 0 | 0 |
| NC_006213.1;YP_009555238.1;HCoV-OC43_orf1ab;1444 | 0 | 0 |
| NC_006213.1;YP_009555238.1;HCoV-OC43_orf1ab;1445 | 0 | 0 |
| NC_006213.1;YP_009555238.1;HCoV-OC43_orf1ab;1446 | 0.05 | 0 |
| NC_006213.1;YP_009555238.1;HCoV-OC43_orf1ab;1447 | 0 | 0 |
| NC_006213.1;YP_009555238.1;HCoV-OC43_orf1ab;1448 | 0 | 0 |
| NC_006213.1;YP_009555238.1;HCoV-OC43_orf1ab;1449 | 0.05 | 0 |
| NC_006213.1;YP_009555238.1;HCoV-OC43_orf1ab;1450 | 0 | 0 |
| NC_006213.1;YP_009555238.1;HCoV-OC43_orf1ab;1451 | 0 | 0 |
| NC_006213.1;YP_009555238.1;HCoV-OC43_orf1ab;1452 | 0 | 0.025 |
| NC_006213.1;YP_009555238.1;HCoV-OC43_orf1ab;1453 | 0 | 0 |
| NC_006213.1;YP_009555238.1;HCoV-OC43_orf1ab;1454 | 0.05 | 0 |
| NC_006213.1;YP_009555238.1;HCoV-OC43_orf1ab;1455 | 0 | 0 |
| NC_006213.1;YP_009555238.1;HCoV-OC43_orf1ab;1456 | 0 | 0 |
| NC_006213.1;YP_009555238.1;HCoV-OC43_orf1ab;1457 | 0 | 0.025 |
| NC_006213.1;YP_009555238.1;HCoV-OC43_orf1ab;1458 | 0 | 0.025 |
| NC_006213.1;YP_009555238.1;HCoV-OC43_orf1ab;1459 | 0 | 0.025 |
| NC_006213.1;YP_009555238.1;HCoV-OC43_orf1ab;1460 | 0 | 0 |
| NC_006213.1;YP_009555238.1;HCoV-OC43_orf1ab;1461 | 0 | 0 |
| NC_006213.1;YP_009555238.1;HCoV-OC43_orf1ab;1462 | 0 | 0 |
| NC_006213.1;YP_009555238.1;HCoV-OC43_orf1ab;1463 | 0 | 0 |
| NC_006213.1;YP_009555238.1;HCoV-OC43_orf1ab;1464 | 0 | 0 |
| NC_006213.1;YP_009555238.1;HCoV-OC43_orf1ab;1465 | 0 | 0 |
| NC_006213.1;YP_009555238.1;HCoV-OC43_orf1ab;1466 | 0 | 0 |
| NC_006213.1;YP_009555238.1;HCoV-OC43_orf1ab;1467 | 0 | 0 |
| NC_006213.1;YP_009555238.1;HCoV-OC43_orf1ab;1468 | 0 | 0 |
| NC_006213.1;YP_009555238.1;HCoV-OC43_orf1ab;1469 | 0 | 0 |
| NC_006213.1;YP_009555238.1;HCoV-OC43_orf1ab;1470 | 0 | 0 |
| NC_006213.1;YP_009555238.1;HCoV-OC43_orf1ab;1471 | 0 | 0.025 |

| | | |
|---|---|---|
| NC_006213.1;YP_009555238.1;HCoV-OC43_orf1ab;1472 | 0 | 0 |
| NC_006213.1;YP_009555238.1;HCoV-OC43_orf1ab;1473 | 0 | 0 |
| NC_006213.1;YP_009555238.1;HCoV-OC43_orf1ab;1474 | 0 | 0 |
| NC_006213.1;YP_009555238.1;HCoV-OC43_orf1ab;1475 | 0 | 0.025 |
| NC_006213.1;YP_009555238.1;HCoV-OC43_orf1ab;1476 | 0.15 | 0 |
| NC_006213.1;YP_009555238.1;HCoV-OC43_orf1ab;1477 | 0 | 0 |
| NC_006213.1;YP_009555238.1;HCoV-OC43_orf1ab;1478 | 0 | 0 |
| NC_006213.1;YP_009555238.1;HCoV-OC43_orf1ab;1479 | 0 | 0 |
| NC_006213.1;YP_009555238.1;HCoV-OC43_orf1ab;1480 | 0 | 0.025 |
| NC_006213.1;YP_009555238.1;HCoV-OC43_orf1ab;1481 | 0 | 0 |
| NC_006213.1;YP_009555238.1;HCoV-OC43_orf1ab;1482 | 0.05 | 0 |
| NC_006213.1;YP_009555238.1;HCoV-OC43_orf1ab;1483 | 0 | 0 |
| NC_006213.1;YP_009555238.1;HCoV-OC43_orf1ab;1484 | 0 | 0 |
| NC_006213.1;YP_009555238.1;HCoV-OC43_orf1ab;1485 | 0 | 0 |
| NC_006213.1;YP_009555238.1;HCoV-OC43_orf1ab;1486 | 0 | 0 |
| NC_006213.1;YP_009555238.1;HCoV-OC43_orf1ab;1487 | 0 | 0 |
| NC_006213.1;YP_009555238.1;HCoV-OC43_orf1ab;1488 | 0 | 0 |
| NC_006213.1;YP_009555238.1;HCoV-OC43_orf1ab;1489 | 0 | 0 |
| NC_006213.1;YP_009555238.1;HCoV-OC43_orf1ab;1490 | 0 | 0 |
| NC_006213.1;YP_009555238.1;HCoV-OC43_orf1ab;1491 | 0 | 0.025 |
| NC_006213.1;YP_009555238.1;HCoV-OC43_orf1ab;1492 | 0 | 0 |
| NC_006213.1;YP_009555238.1;HCoV-OC43_orf1ab;1493 | 0 | 0 |
| NC_006213.1;YP_009555238.1;HCoV-OC43_orf1ab;1494 | 0 | 0 |
| NC_006213.1;YP_009555238.1;HCoV-OC43_orf1ab;1495 | 0 | 0 |
| NC_006213.1;YP_009555238.1;HCoV-OC43_orf1ab;1496 | 0.05 | 0.025 |
| NC_006213.1;YP_009555238.1;HCoV-OC43_orf1ab;1497 | 0 | 0 |
| NC_006213.1;YP_009555238.1;HCoV-OC43_orf1ab;1498 | 0.05 | 0.05 |
| NC_006213.1;YP_009555238.1;HCoV-OC43_orf1ab;1499 | 0.1 | 0.05 |
| NC_006213.1;YP_009555238.1;HCoV-OC43_orf1ab;1500 | 0 | 0 |
| NC_006213.1;YP_009555238.1;HCoV-OC43_orf1ab;1501 | 0 | 0.025 |
| NC_006213.1;YP_009555238.1;HCoV-OC43_orf1ab;1502 | 0 | 0 |
| NC_006213.1;YP_009555238.1;HCoV-OC43_orf1ab;1503 | 0 | 0 |
| NC_006213.1;YP_009555238.1;HCoV-OC43_orf1ab;1504 | 0.05 | 0 |
| NC_006213.1;YP_009555238.1;HCoV-OC43_orf1ab;1505 | 0.05 | 0 |
| NC_006213.1;YP_009555238.1;HCoV-OC43_orf1ab;1506 | 0 | 0 |
| NC_006213.1;YP_009555238.1;HCoV-OC43_orf1ab;1507 | 0 | 0 |
| NC_006213.1;YP_009555238.1;HCoV-OC43_orf1ab;1508 | 0 | 0 |
| NC_006213.1;YP_009555238.1;HCoV-OC43_orf1ab;1509 | 0 | 0 |
| NC_006213.1;YP_009555238.1;HCoV-OC43_orf1ab;1510 | 0 | 0 |
| NC_006213.1;YP_009555238.1;HCoV-OC43_orf1ab;1511 | 0 | 0 |
| NC_006213.1;YP_009555238.1;HCoV-OC43_orf1ab;1512 | 0 | 0 |
| NC_006213.1;YP_009555238.1;HCoV-OC43_orf1ab;1513 | 0 | 0 |
| NC_006213.1;YP_009555238.1;HCoV-OC43_orf1ab;1514 | 0 | 0 |

| | | |
|---|---|---|
| NC_006213.1;YP_009555238.1;HCoV-OC43_orf1ab;1515 | 0 | 0 |
| NC_006213.1;YP_009555238.1;HCoV-OC43_orf1ab;1516 | 0 | 0 |
| NC_006213.1;YP_009555238.1;HCoV-OC43_orf1ab;1517 | 0 | 0 |
| NC_006213.1;YP_009555238.1;HCoV-OC43_orf1ab;1518 | 0 | 0 |
| NC_006213.1;YP_009555238.1;HCoV-OC43_orf1ab;1519 | 0 | 0 |
| NC_006213.1;YP_009555238.1;HCoV-OC43_orf1ab;1520 | 0 | 0 |
| NC_006213.1;YP_009555238.1;HCoV-OC43_orf1ab;1521 | 0 | 0 |
| NC_006213.1;YP_009555238.1;HCoV-OC43_orf1ab;1522 | 0 | 0 |
| NC_006213.1;YP_009555238.1;HCoV-OC43_orf1ab;1523 | 0 | 0 |
| NC_006213.1;YP_009555238.1;HCoV-OC43_orf1ab;1524 | 0 | 0 |
| NC_006213.1;YP_009555238.1;HCoV-OC43_orf1ab;1525 | 0 | 0 |
| NC_006213.1;YP_009555238.1;HCoV-OC43_orf1ab;1526 | 0 | 0 |
| NC_006213.1;YP_009555238.1;HCoV-OC43_orf1ab;1527 | 0 | 0 |
| NC_006213.1;YP_009555238.1;HCoV-OC43_orf1ab;1528 | 0 | 0 |
| NC_006213.1;YP_009555238.1;HCoV-OC43_orf1ab;1529 | 0 | 0 |
| NC_006213.1;YP_009555238.1;HCoV-OC43_orf1ab;1530 | 0 | 0 |
| NC_006213.1;YP_009555238.1;HCoV-OC43_orf1ab;1531 | 0.05 | 0.05 |
| NC_006213.1;YP_009555238.1;HCoV-OC43_orf1ab;1532 | 0 | 0 |
| NC_006213.1;YP_009555238.1;HCoV-OC43_orf1ab;1533 | 0 | 0 |
| NC_006213.1;YP_009555238.1;HCoV-OC43_orf1ab;1534 | 0 | 0 |
| NC_006213.1;YP_009555238.1;HCoV-OC43_orf1ab;1535 | 0 | 0 |
| NC_006213.1;YP_009555238.1;HCoV-OC43_orf1ab;1536 | 0.05 | 0.075 |
| NC_006213.1;YP_009555238.1;HCoV-OC43_orf1ab;1537 | 0.1 | 0.075 |
| NC_006213.1;YP_009555238.1;HCoV-OC43_orf1ab;1538 | 0.15 | 0.125 |
| NC_006213.1;YP_009555238.1;HCoV-OC43_orf1ab;1539 | 0.05 | 0.075 |
| NC_006213.1;YP_009555238.1;HCoV-OC43_orf1ab;1540 | 0.1 | 0.075 |
| NC_006213.1;YP_009555238.1;HCoV-OC43_orf1ab;1541 | 0.15 | 0.1 |
| NC_006213.1;YP_009555238.1;HCoV-OC43_orf1ab;1542 | 0.15 | 0.1 |
| NC_006213.1;YP_009555238.1;HCoV-OC43_orf1ab;1543 | 0.15 | 0.1 |
| NC_006213.1;YP_009555238.1;HCoV-OC43_orf1ab;1544 | 0.15 | 0.1 |
| NC_006213.1;YP_009555238.1;HCoV-OC43_orf1ab;1545 | 0.1 | 0.05 |
| NC_006213.1;YP_009555238.1;HCoV-OC43_orf1ab;1546 | 0 | 0 |
| NC_006213.1;YP_009555238.1;HCoV-OC43_orf1ab;1547 | 0 | 0 |
| NC_006213.1;YP_009555238.1;HCoV-OC43_orf1ab;1548 | 0 | 0.025 |
| NC_006213.1;YP_009555238.1;HCoV-OC43_orf1ab;1549 | 0 | 0 |
| NC_006213.1;YP_009555238.1;HCoV-OC43_orf1ab;1550 | 0 | 0 |
| NC_006213.1;YP_009555238.1;HCoV-OC43_orf1ab;1551 | 0.05 | 0 |
| NC_006213.1;YP_009555238.1;HCoV-OC43_orf1ab;1552 | 0.05 | 0 |
| NC_006213.1;YP_009555238.1;HCoV-OC43_orf1ab;1553 | 0.05 | 0 |
| NC_006213.1;YP_009555238.1;HCoV-OC43_orf1ab;1554 | 0 | 0 |
| NC_006213.1;YP_009555238.1;HCoV-OC43_orf1ab;1555 | 0.05 | 0.05 |
| NC_006213.1;YP_009555238.1;HCoV-OC43_orf1ab;1556 | 0 | 0 |
| NC_006213.1;YP_009555238.1;HCoV-OC43_orf1ab;1557 | 0 | 0 |

| | | |
|---|---|---|
| NC_006213.1;YP_009555238.1;HCoV-OC43_orf1ab;1558 | 0.05 | 0 |
| NC_006213.1;YP_009555238.1;HCoV-OC43_orf1ab;1559 | 0 | 0 |
| NC_006213.1;YP_009555238.1;HCoV-OC43_orf1ab;1560 | 0 | 0 |
| NC_006213.1;YP_009555238.1;HCoV-OC43_orf1ab;1561 | 0 | 0.025 |
| NC_006213.1;YP_009555238.1;HCoV-OC43_orf1ab;1562 | 0 | 0.025 |
| NC_006213.1;YP_009555238.1;HCoV-OC43_orf1ab;1563 | 0.05 | 0.025 |
| NC_006213.1;YP_009555238.1;HCoV-OC43_orf1ab;1564 | 0 | 0 |
| NC_006213.1;YP_009555238.1;HCoV-OC43_orf1ab;1565 | 0 | 0 |
| NC_006213.1;YP_009555238.1;HCoV-OC43_orf1ab;1566 | 0 | 0 |
| NC_006213.1;YP_009555238.1;HCoV-OC43_orf1ab;1567 | 0 | 0 |
| NC_006213.1;YP_009555238.1;HCoV-OC43_orf1ab;1568 | 0.05 | 0 |
| NC_006213.1;YP_009555238.1;HCoV-OC43_orf1ab;1569 | 0.15 | 0.1 |
| NC_006213.1;YP_009555238.1;HCoV-OC43_orf1ab;1570 | 0.2 | 0.25 |
| NC_006213.1;YP_009555238.1;HCoV-OC43_orf1ab;1571 | 0.2 | 0.175 |
| NC_006213.1;YP_009555238.1;HCoV-OC43_orf1ab;1572 | 0.1 | 0.025 |
| NC_006213.1;YP_009555238.1;HCoV-OC43_orf1ab;1573 | 0.05 | 0.025 |
| NC_006213.1;YP_009555238.1;HCoV-OC43_orf1ab;1574 | 0.05 | 0 |
| NC_006213.1;YP_009555238.1;HCoV-OC43_orf1ab;1575 | 0 | 0 |
| NC_006213.1;YP_009555238.1;HCoV-OC43_orf1ab;1576 | 0 | 0 |
| NC_006213.1;YP_009555238.1;HCoV-OC43_orf1ab;1577 | 0 | 0 |
| NC_006213.1;YP_009555238.1;HCoV-OC43_orf1ab;1578 | 0 | 0 |
| NC_006213.1;YP_009555238.1;HCoV-OC43_orf1ab;1579 | 0 | 0 |
| NC_006213.1;YP_009555238.1;HCoV-OC43_orf1ab;1580 | 0 | 0 |
| NC_006213.1;YP_009555238.1;HCoV-OC43_orf1ab;1581 | 0 | 0 |
| NC_006213.1;YP_009555238.1;HCoV-OC43_orf1ab;1582 | 0 | 0 |
| NC_006213.1;YP_009555238.1;HCoV-OC43_orf1ab;1583 | 0 | 0 |
| NC_006213.1;YP_009555238.1;HCoV-OC43_orf1ab;1584 | 0 | 0 |
| NC_006213.1;YP_009555238.1;HCoV-OC43_orf1ab;1585 | 0 | 0 |
| NC_006213.1;YP_009555238.1;HCoV-OC43_orf1ab;1586 | 0 | 0 |
| NC_006213.1;YP_009555238.1;HCoV-OC43_orf1ab;1587 | 0 | 0 |
| NC_006213.1;YP_009555238.1;HCoV-OC43_orf1ab;1588 | 0 | 0.025 |
| NC_006213.1;YP_009555238.1;HCoV-OC43_orf1ab;1589 | 0 | 0 |
| NC_006213.1;YP_009555238.1;HCoV-OC43_orf1ab;1590 | 0 | 0.025 |
| NC_006213.1;YP_009555238.1;HCoV-OC43_orf1ab;1591 | 0 | 0.025 |
| NC_006213.1;YP_009555238.1;HCoV-OC43_orf1ab;1592 | 0 | 0.025 |
| NC_006213.1;YP_009555238.1;HCoV-OC43_orf1ab;1593 | 0 | 0.025 |
| NC_006213.1;YP_009555238.1;HCoV-OC43_orf1ab;1594 | 0 | 0.025 |
| NC_006213.1;YP_009555238.1;HCoV-OC43_orf1ab;1595 | 0 | 0.025 |
| NC_006213.1;YP_009555238.1;HCoV-OC43_orf1ab;1596 | 0 | 0.025 |
| NC_006213.1;YP_009555238.1;HCoV-OC43_orf1ab;1597 | 0 | 0.025 |
| NC_006213.1;YP_009555238.1;HCoV-OC43_orf1ab;1598 | 0 | 0.025 |
| NC_006213.1;YP_009555238.1;HCoV-OC43_orf1ab;1599 | 0 | 0.025 |
| NC_006213.1;YP_009555238.1;HCoV-OC43_orf1ab;1600 | 0.05 | 0.05 |

| | | |
|---|---|---|
| NC_006213.1;YP_009555238.1;HCoV-OC43_orf1ab;1601 | 0.05 | 0 |
| NC_006213.1;YP_009555238.1;HCoV-OC43_orf1ab;1602 | 0.05 | 0.05 |
| NC_006213.1;YP_009555238.1;HCoV-OC43_orf1ab;1603 | 0 | 0 |
| NC_006213.1;YP_009555238.1;HCoV-OC43_orf1ab;1604 | 0 | 0 |
| NC_006213.1;YP_009555238.1;HCoV-OC43_orf1ab;1605 | 0 | 0.025 |
| NC_006213.1;YP_009555238.1;HCoV-OC43_orf1ab;1606 | 0 | 0.025 |
| NC_006213.1;YP_009555238.1;HCoV-OC43_orf1ab;1607 | 0 | 0.025 |
| NC_006213.1;YP_009555238.1;HCoV-OC43_orf1ab;1608 | 0 | 0.025 |
| NC_006213.1;YP_009555238.1;HCoV-OC43_orf1ab;1609 | 0 | 0 |
| NC_006213.1;YP_009555238.1;HCoV-OC43_orf1ab;1610 | 0 | 0 |
| NC_006213.1;YP_009555238.1;HCoV-OC43_orf1ab;1611 | 0.05 | 0.025 |
| NC_006213.1;YP_009555238.1;HCoV-OC43_orf1ab;1612 | 0.05 | 0.1 |
| NC_006213.1;YP_009555238.1;HCoV-OC43_orf1ab;1613 | 0 | 0.025 |
| NC_006213.1;YP_009555238.1;HCoV-OC43_orf1ab;1614 | 0 | 0.025 |
| NC_006213.1;YP_009555238.1;HCoV-OC43_orf1ab;1615 | 0 | 0 |
| NC_006213.1;YP_009555238.1;HCoV-OC43_orf1ab;1616 | 0.05 | 0 |
| NC_006213.1;YP_009555238.1;HCoV-OC43_orf1ab;1617 | 0 | 0.025 |
| NC_006213.1;YP_009555238.1;HCoV-OC43_orf1ab;1618 | 0 | 0 |
| NC_006213.1;YP_009555238.1;HCoV-OC43_orf1ab;1619 | 0 | 0.025 |
| NC_006213.1;YP_009555238.1;HCoV-OC43_orf1ab;1620 | 0 | 0 |
| NC_006213.1;YP_009555238.1;HCoV-OC43_orf1ab;1621 | 0 | 0 |
| NC_006213.1;YP_009555238.1;HCoV-OC43_orf1ab;1622 | 0.05 | 0 |
| NC_006213.1;YP_009555238.1;HCoV-OC43_orf1ab;1623 | 0.1 | 0.025 |
| NC_006213.1;YP_009555238.1;HCoV-OC43_orf1ab;1624 | 0.05 | 0.05 |
| NC_006213.1;YP_009555238.1;HCoV-OC43_orf1ab;1625 | 0 | 0 |
| NC_006213.1;YP_009555238.1;HCoV-OC43_orf1ab;1626 | 0 | 0.025 |
| NC_006213.1;YP_009555238.1;HCoV-OC43_orf1ab;1627 | 0 | 0.025 |
| NC_006213.1;YP_009555238.1;HCoV-OC43_orf1ab;1628 | 0.1 | 0.075 |
| NC_006213.1;YP_009555238.1;HCoV-OC43_orf1ab;1629 | 0 | 0.05 |
| NC_006213.1;YP_009555238.1;HCoV-OC43_orf1ab;1630 | 0.1 | 0.075 |
| NC_006213.1;YP_009555238.1;HCoV-OC43_orf1ab;1631 | 0.05 | 0.1 |
| NC_006213.1;YP_009555238.1;HCoV-OC43_orf1ab;1632 | 0.05 | 0.1 |
| NC_006213.1;YP_009555238.1;HCoV-OC43_orf1ab;1633 | 0 | 0.075 |
| NC_006213.1;YP_009555238.1;HCoV-OC43_orf1ab;1634 | 0.05 | 0.05 |
| NC_006213.1;YP_009555238.1;HCoV-OC43_orf1ab;1635 | 0 | 0 |
| NC_006213.1;YP_009555238.1;HCoV-OC43_orf1ab;1636 | 0 | 0 |
| NC_006213.1;YP_009555238.1;HCoV-OC43_orf1ab;1637 | 0.05 | 0.025 |
| NC_006213.1;YP_009555238.1;HCoV-OC43_orf1ab;1638 | 0 | 0 |
| NC_006213.1;YP_009555238.1;HCoV-OC43_orf1ab;1639 | 0 | 0 |
| NC_006213.1;YP_009555238.1;HCoV-OC43_orf1ab;1640 | 0 | 0 |
| NC_006213.1;YP_009555238.1;HCoV-OC43_orf1ab;1641 | 0 | 0 |
| NC_006213.1;YP_009555238.1;HCoV-OC43_orf1ab;1642 | 0 | 0 |
| NC_006213.1;YP_009555238.1;HCoV-OC43_orf1ab;1643 | 0 | 0 |

| | | |
|---|---|---|
| NC_006213.1;YP_009555238.1;HCoV-OC43_orf1ab;1644 | 0 | 0 |
| NC_006213.1;YP_009555238.1;HCoV-OC43_orf1ab;1645 | 0.05 | 0 |
| NC_006213.1;YP_009555238.1;HCoV-OC43_orf1ab;1646 | 0 | 0 |
| NC_006213.1;YP_009555238.1;HCoV-OC43_orf1ab;1647 | 0 | 0 |
| NC_006213.1;YP_009555238.1;HCoV-OC43_orf1ab;1648 | 0 | 0 |
| NC_006213.1;YP_009555238.1;HCoV-OC43_orf1ab;1649 | 0 | 0 |
| NC_006213.1;YP_009555238.1;HCoV-OC43_orf1ab;1650 | 0 | 0 |
| NC_006213.1;YP_009555238.1;HCoV-OC43_orf1ab;1651 | 0 | 0 |
| NC_006213.1;YP_009555238.1;HCoV-OC43_orf1ab;1652 | 0 | 0 |
| NC_006213.1;YP_009555238.1;HCoV-OC43_orf1ab;1653 | 0 | 0 |
| NC_006213.1;YP_009555238.1;HCoV-OC43_orf1ab;1654 | 0 | 0 |
| NC_006213.1;YP_009555238.1;HCoV-OC43_orf1ab;1655 | 0.05 | 0.05 |
| NC_006213.1;YP_009555238.1;HCoV-OC43_orf1ab;1656 | 0.05 | 0.025 |
| NC_006213.1;YP_009555238.1;HCoV-OC43_orf1ab;1657 | 0.1 | 0 |
| NC_006213.1;YP_009555238.1;HCoV-OC43_orf1ab;1658 | 0.05 | 0 |
| NC_006213.1;YP_009555238.1;HCoV-OC43_orf1ab;1659 | 0.1 | 0.025 |
| NC_006213.1;YP_009555238.1;HCoV-OC43_orf1ab;1660 | 0.05 | 0 |
| NC_006213.1;YP_009555238.1;HCoV-OC43_orf1ab;1661 | 0.05 | 0 |
| NC_006213.1;YP_009555238.1;HCoV-OC43_orf1ab;1662 | 0.1 | 0.05 |
| NC_006213.1;YP_009555238.1;HCoV-OC43_orf1ab;1663 | 0.05 | 0.025 |
| NC_006213.1;YP_009555238.1;HCoV-OC43_orf1ab;1664 | 0.05 | 0.075 |
| NC_006213.1;YP_009555238.1;HCoV-OC43_orf1ab;1665 | 0.05 | 0.075 |
| NC_006213.1;YP_009555238.1;HCoV-OC43_orf1ab;1666 | 0.05 | 0.05 |
| NC_006213.1;YP_009555238.1;HCoV-OC43_orf1ab;1667 | 0 | 0 |
| NC_006213.1;YP_009555238.1;HCoV-OC43_orf1ab;1668 | 0 | 0 |
| NC_006213.1;YP_009555238.1;HCoV-OC43_orf1ab;1669 | 0 | 0 |
| NC_006213.1;YP_009555238.1;HCoV-OC43_orf1ab;1670 | 0 | 0 |
| NC_006213.1;YP_009555238.1;HCoV-OC43_orf1ab;1671 | 0 | 0 |
| NC_006213.1;YP_009555238.1;HCoV-OC43_orf1ab;1672 | 0 | 0 |
| NC_006213.1;YP_009555238.1;HCoV-OC43_orf1ab;1673 | 0 | 0 |
| NC_006213.1;YP_009555238.1;HCoV-OC43_orf1ab;1674 | 0 | 0 |
| NC_006213.1;YP_009555238.1;HCoV-OC43_orf1ab;1675 | 0 | 0 |
| NC_006213.1;YP_009555238.1;HCoV-OC43_orf1ab;1676 | 0 | 0 |
| NC_006213.1;YP_009555238.1;HCoV-OC43_orf1ab;1677 | 0 | 0 |
| NC_006213.1;YP_009555238.1;HCoV-OC43_orf1ab;1678 | 0 | 0 |
| NC_006213.1;YP_009555238.1;HCoV-OC43_orf1ab;1679 | 0 | 0 |
| NC_006213.1;YP_009555238.1;HCoV-OC43_orf1ab;1680 | 0 | 0 |
| NC_006213.1;YP_009555238.1;HCoV-OC43_orf1ab;1681 | 0 | 0 |
| NC_006213.1;YP_009555238.1;HCoV-OC43_orf1ab;1682 | 0 | 0 |
| NC_006213.1;YP_009555238.1;HCoV-OC43_orf1ab;1683 | 0 | 0 |
| NC_006213.1;YP_009555238.1;HCoV-OC43_orf1ab;1684 | 0 | 0 |
| NC_006213.1;YP_009555238.1;HCoV-OC43_orf1ab;1685 | 0 | 0 |
| NC_006213.1;YP_009555238.1;HCoV-OC43_orf1ab;1686 | 0 | 0 |

| | | |
|---|---|---|
| NC_006213.1;YP_009555238.1;HCoV-OC43_orf1ab;1687 | 0 | 0 |
| NC_006213.1;YP_009555238.1;HCoV-OC43_orf1ab;1688 | 0 | 0 |
| NC_006213.1;YP_009555238.1;HCoV-OC43_orf1ab;1689 | 0 | 0 |
| NC_006213.1;YP_009555238.1;HCoV-OC43_orf1ab;1690 | 0 | 0 |
| NC_006213.1;YP_009555238.1;HCoV-OC43_orf1ab;1691 | 0 | 0 |
| NC_006213.1;YP_009555238.1;HCoV-OC43_orf1ab;1692 | 0 | 0 |
| NC_006213.1;YP_009555238.1;HCoV-OC43_orf1ab;1693 | 0 | 0 |
| NC_006213.1;YP_009555238.1;HCoV-OC43_orf1ab;1694 | 0 | 0 |
| NC_006213.1;YP_009555238.1;HCoV-OC43_orf1ab;1695 | 0 | 0 |
| NC_006213.1;YP_009555238.1;HCoV-OC43_orf1ab;1696 | 0 | 0 |
| NC_006213.1;YP_009555238.1;HCoV-OC43_orf1ab;1697 | 0 | 0 |
| NC_006213.1;YP_009555238.1;HCoV-OC43_orf1ab;1698 | 0 | 0 |
| NC_006213.1;YP_009555238.1;HCoV-OC43_orf1ab;1699 | 0 | 0 |
| NC_006213.1;YP_009555238.1;HCoV-OC43_orf1ab;1700 | 0 | 0 |
| NC_006213.1;YP_009555238.1;HCoV-OC43_orf1ab;1701 | 0 | 0 |
| NC_006213.1;YP_009555238.1;HCoV-OC43_orf1ab;1702 | 0 | 0 |
| NC_006213.1;YP_009555238.1;HCoV-OC43_orf1ab;1703 | 0 | 0 |
| NC_006213.1;YP_009555238.1;HCoV-OC43_orf1ab;1704 | 0 | 0 |
| NC_006213.1;YP_009555238.1;HCoV-OC43_orf1ab;1705 | 0 | 0 |
| NC_006213.1;YP_009555238.1;HCoV-OC43_orf1ab;1706 | 0.05 | 0 |
| NC_006213.1;YP_009555238.1;HCoV-OC43_orf1ab;1707 | 0.05 | 0 |
| NC_006213.1;YP_009555238.1;HCoV-OC43_orf1ab;1708 | 0.1 | 0 |
| NC_006213.1;YP_009555238.1;HCoV-OC43_orf1ab;1709 | 0.15 | 0.025 |
| NC_006213.1;YP_009555238.1;HCoV-OC43_orf1ab;1710 | 0.2 | 0.025 |
| NC_006213.1;YP_009555238.1;HCoV-OC43_orf1ab;1711 | 0.1 | 0.05 |
| NC_006213.1;YP_009555238.1;HCoV-OC43_orf1ab;1712 | 0.25 | 0.05 |
| NC_006213.1;YP_009555238.1;HCoV-OC43_orf1ab;1713 | 0.15 | 0.05 |
| NC_006213.1;YP_009555238.1;HCoV-OC43_orf1ab;1714 | 0.1 | 0.1 |
| NC_006213.1;YP_009555238.1;HCoV-OC43_orf1ab;1715 | 0.15 | 0 |
| NC_006213.1;YP_009555238.1;HCoV-OC43_orf1ab;1716 | 0.05 | 0.025 |
| NC_006213.1;YP_009555238.1;HCoV-OC43_orf1ab;1717 | 0 | 0 |
| NC_006213.1;YP_009555238.1;HCoV-OC43_orf1ab;1718 | 0 | 0 |
| NC_006213.1;YP_009555238.1;HCoV-OC43_orf1ab;1719 | 0 | 0 |
| NC_006213.1;YP_009555238.1;HCoV-OC43_orf1ab;1720 | 0.05 | 0 |
| NC_006213.1;YP_009555238.1;HCoV-OC43_orf1ab;1721 | 0 | 0 |
| NC_006213.1;YP_009555238.1;HCoV-OC43_orf1ab;1722 | 0.05 | 0.025 |
| NC_006213.1;YP_009555238.1;HCoV-OC43_orf1ab;1723 | 0 | 0 |
| NC_006213.1;YP_009555238.1;HCoV-OC43_orf1ab;1724 | 0 | 0 |
| NC_006213.1;YP_009555238.1;HCoV-OC43_orf1ab;1725 | 0 | 0 |
| NC_006213.1;YP_009555238.1;HCoV-OC43_orf1ab;1726 | 0 | 0 |
| NC_006213.1;YP_009555238.1;HCoV-OC43_orf1ab;1727 | 0 | 0 |
| NC_006213.1;YP_009555238.1;HCoV-OC43_orf1ab;1728 | 0 | 0 |
| NC_006213.1;YP_009555238.1;HCoV-OC43_orf1ab;1729 | 0 | 0 |

| | | |
|---|---|---|
| NC_006213.1;YP_009555238.1;HCoV-OC43_orf1ab;1730 | 0 | 0 |
| NC_006213.1;YP_009555238.1;HCoV-OC43_orf1ab;1731 | 0 | 0 |
| NC_006213.1;YP_009555238.1;HCoV-OC43_orf1ab;1732 | 0 | 0 |
| NC_006213.1;YP_009555238.1;HCoV-OC43_orf1ab;1733 | 0 | 0 |
| NC_006213.1;YP_009555238.1;HCoV-OC43_orf1ab;1734 | 0 | 0 |
| NC_006213.1;YP_009555238.1;HCoV-OC43_orf1ab;1735 | 0 | 0 |
| NC_006213.1;YP_009555238.1;HCoV-OC43_orf1ab;1736 | 0 | 0 |
| NC_006213.1;YP_009555238.1;HCoV-OC43_orf1ab;1737 | 0 | 0 |
| NC_006213.1;YP_009555238.1;HCoV-OC43_orf1ab;1738 | 0 | 0 |
| NC_006213.1;YP_009555238.1;HCoV-OC43_orf1ab;1739 | 0 | 0.025 |
| NC_006213.1;YP_009555238.1;HCoV-OC43_orf1ab;1740 | 0 | 0 |
| NC_006213.1;YP_009555238.1;HCoV-OC43_orf1ab;1741 | 0 | 0 |
| NC_006213.1;YP_009555238.1;HCoV-OC43_orf1ab;1742 | 0 | 0 |
| NC_006213.1;YP_009555238.1;HCoV-OC43_orf1ab;1743 | 0 | 0.025 |
| NC_006213.1;YP_009555238.1;HCoV-OC43_orf1ab;1744 | 0 | 0.025 |
| NC_006213.1;YP_009555238.1;HCoV-OC43_orf1ab;1745 | 0 | 0.025 |
| NC_006213.1;YP_009555238.1;HCoV-OC43_orf1ab;1746 | 0 | 0.025 |
| NC_006213.1;YP_009555238.1;HCoV-OC43_orf1ab;1747 | 0 | 0.025 |
| NC_006213.1;YP_009555238.1;HCoV-OC43_orf1ab;1748 | 0 | 0.075 |
| NC_006213.1;YP_009555238.1;HCoV-OC43_orf1ab;1749 | 0 | 0.05 |
| NC_006213.1;YP_009555238.1;HCoV-OC43_orf1ab;1750 | 0 | 0.075 |
| NC_006213.1;YP_009555238.1;HCoV-OC43_orf1ab;1751 | 0 | 0 |
| NC_006213.1;YP_009555238.1;HCoV-OC43_orf1ab;1752 | 0 | 0.05 |
| NC_006213.1;YP_009555238.1;HCoV-OC43_orf1ab;1753 | 0 | 0 |
| NC_006213.1;YP_009555238.1;HCoV-OC43_orf1ab;1754 | 0 | 0 |
| NC_006213.1;YP_009555238.1;HCoV-OC43_orf1ab;1755 | 0 | 0 |
| NC_006213.1;YP_009555238.1;HCoV-OC43_orf1ab;1756 | 0.05 | 0 |
| NC_006213.1;YP_009555238.1;HCoV-OC43_orf1ab;1757 | 0 | 0 |
| NC_006213.1;YP_009555238.1;HCoV-OC43_orf1ab;1758 | 0 | 0 |
| NC_006213.1;YP_009555238.1;HCoV-OC43_orf1ab;1759 | 0 | 0 |
| NC_006213.1;YP_009555238.1;HCoV-OC43_orf1ab;1760 | 0.05 | 0 |
| NC_006213.1;YP_009555238.1;HCoV-OC43_orf1ab;1761 | 0 | 0 |
| NC_006213.1;YP_009555238.1;HCoV-OC43_orf1ab;1762 | 0 | 0 |
| NC_006213.1;YP_009555238.1;HCoV-OC43_orf1ab;1763 | 0 | 0 |
| NC_006213.1;YP_009555238.1;HCoV-OC43_orf1ab;1764 | 0 | 0 |
| NC_006213.1;YP_009555238.1;HCoV-OC43_orf1ab;1765 | 0 | 0 |
| NC_006213.1;YP_009555238.1;HCoV-OC43_orf1ab;1766 | 0 | 0 |
| NC_006213.1;YP_009555238.1;HCoV-OC43_orf1ab;1767 | 0 | 0 |
| NC_006213.1;YP_009555238.1;HCoV-OC43_orf1ab;1768 | 0 | 0 |
| NC_006213.1;YP_009555238.1;HCoV-OC43_orf1ab;1769 | 0 | 0 |
| NC_006213.1;YP_009555238.1;HCoV-OC43_orf1ab;1770 | 0 | 0 |
| NC_006213.1;YP_009555238.1;HCoV-OC43_orf1ab;1771 | 0 | 0 |
| NC_006213.1;YP_009555238.1;HCoV-OC43_orf1ab;1772 | 0 | 0 |

| | | |
|---|---|---|
| NC_006213.1;YP_009555238.1;HCoV-OC43_orf1ab;1773 | 0 | 0.025 |
| NC_006213.1;YP_009555238.1;HCoV-OC43_orf1ab;1774 | 0 | 0.025 |
| NC_006213.1;YP_009555238.1;HCoV-OC43_orf1ab;1775 | 0.05 | 0.025 |
| NC_006213.1;YP_009555238.1;HCoV-OC43_orf1ab;1776 | 0.05 | 0 |
| NC_006213.1;YP_009555238.1;HCoV-OC43_orf1ab;1777 | 0 | 0 |
| NC_006213.1;YP_009555238.1;HCoV-OC43_orf1ab;1778 | 0 | 0 |
| NC_006213.1;YP_009555238.1;HCoV-OC43_orf1ab;1779 | 0 | 0 |
| NC_006213.1;YP_009555238.1;HCoV-OC43_orf1ab;1780 | 0 | 0 |
| NC_006213.1;YP_009555238.1;HCoV-OC43_orf1ab;1781 | 0 | 0 |
| NC_006213.1;YP_009555238.1;HCoV-OC43_orf1ab;1782 | 0 | 0 |
| NC_006213.1;YP_009555238.1;HCoV-OC43_orf1ab;1783 | 0 | 0 |
| NC_006213.1;YP_009555238.1;HCoV-OC43_orf1ab;1784 | 0 | 0 |
| NC_006213.1;YP_009555238.1;HCoV-OC43_orf1ab;1785 | 0 | 0 |
| NC_006213.1;YP_009555238.1;HCoV-OC43_orf1ab;1786 | 0 | 0 |
| NC_006213.1;YP_009555238.1;HCoV-OC43_orf1ab;1787 | 0 | 0 |
| NC_006213.1;YP_009555238.1;HCoV-OC43_orf1ab;1788 | 0 | 0 |
| NC_006213.1;YP_009555238.1;HCoV-OC43_orf1ab;1789 | 0 | 0 |
| NC_006213.1;YP_009555238.1;HCoV-OC43_orf1ab;1790 | 0 | 0 |
| NC_006213.1;YP_009555238.1;HCoV-OC43_orf1ab;1791 | 0 | 0 |
| NC_006213.1;YP_009555238.1;HCoV-OC43_orf1ab;1792 | 0 | 0 |
| NC_006213.1;YP_009555238.1;HCoV-OC43_orf1ab;1793 | 0 | 0 |
| NC_006213.1;YP_009555238.1;HCoV-OC43_orf1ab;1794 | 0 | 0 |
| NC_006213.1;YP_009555238.1;HCoV-OC43_orf1ab;1795 | 0 | 0 |
| NC_006213.1;YP_009555238.1;HCoV-OC43_orf1ab;1796 | 0 | 0 |
| NC_006213.1;YP_009555238.1;HCoV-OC43_orf1ab;1797 | 0 | 0 |
| NC_006213.1;YP_009555238.1;HCoV-OC43_orf1ab;1798 | 0 | 0 |
| NC_006213.1;YP_009555238.1;HCoV-OC43_orf1ab;1799 | 0 | 0 |
| NC_006213.1;YP_009555238.1;HCoV-OC43_orf1ab;1800 | 0 | 0 |
| NC_006213.1;YP_009555238.1;HCoV-OC43_orf1ab;1801 | 0 | 0 |
| NC_006213.1;YP_009555238.1;HCoV-OC43_orf1ab;1802 | 0 | 0 |
| NC_006213.1;YP_009555238.1;HCoV-OC43_orf1ab;1803 | 0.05 | 0 |
| NC_006213.1;YP_009555238.1;HCoV-OC43_orf1ab;1804 | 0 | 0 |
| NC_006213.1;YP_009555238.1;HCoV-OC43_orf1ab;1805 | 0 | 0 |
| NC_006213.1;YP_009555238.1;HCoV-OC43_orf1ab;1806 | 0 | 0 |
| NC_006213.1;YP_009555238.1;HCoV-OC43_orf1ab;1807 | 0 | 0 |
| NC_006213.1;YP_009555238.1;HCoV-OC43_orf1ab;1808 | 0 | 0 |
| NC_006213.1;YP_009555238.1;HCoV-OC43_orf1ab;1809 | 0 | 0 |
| NC_006213.1;YP_009555238.1;HCoV-OC43_orf1ab;1810 | 0 | 0.025 |
| NC_006213.1;YP_009555238.1;HCoV-OC43_orf1ab;1811 | 0.05 | 0.025 |
| NC_006213.1;YP_009555238.1;HCoV-OC43_orf1ab;1812 | 0.05 | 0.05 |
| NC_006213.1;YP_009555238.1;HCoV-OC43_orf1ab;1813 | 0 | 0.05 |
| NC_006213.1;YP_009555238.1;HCoV-OC43_orf1ab;1814 | 0 | 0.075 |
| NC_006213.1;YP_009555238.1;HCoV-OC43_orf1ab;1815 | 0 | 0.025 |

| | | |
|---|---|---|
| NC_006213.1;YP_009555238.1;HCoV-OC43_orf1ab;1816 | 0 | 0 |
| NC_006213.1;YP_009555238.1;HCoV-OC43_orf1ab;1817 | 0 | 0 |
| NC_006213.1;YP_009555238.1;HCoV-OC43_orf1ab;1818 | 0 | 0.025 |
| NC_006213.1;YP_009555238.1;HCoV-OC43_orf1ab;1819 | 0 | 0 |
| NC_006213.1;YP_009555238.1;HCoV-OC43_orf1ab;1820 | 0 | 0 |
| NC_006213.1;YP_009555238.1;HCoV-OC43_orf1ab;1821 | 0 | 0 |
| NC_006213.1;YP_009555238.1;HCoV-OC43_orf1ab;1822 | 0 | 0 |
| NC_006213.1;YP_009555238.1;HCoV-OC43_orf1ab;1823 | 0 | 0 |
| NC_006213.1;YP_009555238.1;HCoV-OC43_orf1ab;1824 | 0 | 0.05 |
| NC_006213.1;YP_009555238.1;HCoV-OC43_orf1ab;1825 | 0 | 0 |
| NC_006213.1;YP_009555238.1;HCoV-OC43_orf1ab;1826 | 0 | 0 |
| NC_006213.1;YP_009555238.1;HCoV-OC43_orf1ab;1827 | 0 | 0 |
| NC_006213.1;YP_009555238.1;HCoV-OC43_orf1ab;1828 | 0 | 0 |
| NC_006213.1;YP_009555238.1;HCoV-OC43_orf1ab;1829 | 0 | 0 |
| NC_006213.1;YP_009555238.1;HCoV-OC43_orf1ab;1830 | 0 | 0.025 |
| NC_006213.1;YP_009555238.1;HCoV-OC43_orf1ab;1831 | 0 | 0 |
| NC_006213.1;YP_009555238.1;HCoV-OC43_orf1ab;1832 | 0.05 | 0.025 |
| NC_006213.1;YP_009555238.1;HCoV-OC43_orf1ab;1833 | 0.05 | 0 |
| NC_006213.1;YP_009555238.1;HCoV-OC43_orf1ab;1834 | 0.1 | 0.05 |
| NC_006213.1;YP_009555238.1;HCoV-OC43_orf1ab;1835 | 0.05 | 0 |
| NC_006213.1;YP_009555238.1;HCoV-OC43_orf1ab;1836 | 0.1 | 0.025 |
| NC_006213.1;YP_009555238.1;HCoV-OC43_orf1ab;1837 | 0.1 | 0.025 |
| NC_006213.1;YP_009555238.1;HCoV-OC43_orf1ab;1838 | 0.05 | 0 |
| NC_006213.1;YP_009555238.1;HCoV-OC43_orf1ab;1839 | 0.05 | 0 |
| NC_006213.1;YP_009555238.1;HCoV-OC43_orf1ab;1840 | 0.3 | 0.125 |
| NC_006213.1;YP_009555238.1;HCoV-OC43_orf1ab;1841 | 0.05 | 0 |
| NC_006213.1;YP_009555238.1;HCoV-OC43_orf1ab;1842 | 0 | 0 |
| NC_006213.1;YP_009555238.1;HCoV-OC43_orf1ab;1843 | 0 | 0.05 |
| NC_006213.1;YP_009555238.1;HCoV-OC43_orf1ab;1844 | 0 | 0.025 |
| NC_006213.1;YP_009555238.1;HCoV-OC43_orf1ab;1845 | 0 | 0 |
| NC_006213.1;YP_009555238.1;HCoV-OC43_orf1ab;1846 | 0 | 0 |
| NC_006213.1;YP_009555238.1;HCoV-OC43_orf1ab;1847 | 0 | 0 |
| NC_006213.1;YP_009555238.1;HCoV-OC43_orf1ab;1848 | 0 | 0 |
| NC_006213.1;YP_009555238.1;HCoV-OC43_orf1ab;1849 | 0 | 0 |
| NC_006213.1;YP_009555238.1;HCoV-OC43_orf1ab;1850 | 0.05 | 0 |
| NC_006213.1;YP_009555238.1;HCoV-OC43_orf1ab;1851 | 0 | 0 |
| NC_006213.1;YP_009555238.1;HCoV-OC43_orf1ab;1852 | 0 | 0 |
| NC_006213.1;YP_009555238.1;HCoV-OC43_orf1ab;1853 | 0 | 0 |
| NC_006213.1;YP_009555238.1;HCoV-OC43_orf1ab;1854 | 0 | 0 |
| NC_006213.1;YP_009555238.1;HCoV-OC43_orf1ab;1855 | 0 | 0 |
| NC_006213.1;YP_009555238.1;HCoV-OC43_orf1ab;1856 | 0 | 0 |
| NC_006213.1;YP_009555238.1;HCoV-OC43_orf1ab;1857 | 0 | 0 |
| NC_006213.1;YP_009555238.1;HCoV-OC43_orf1ab;1858 | 0 | 0 |

| | | |
|---|---|---|
| NC_006213.1;YP_009555238.1;HCoV-OC43_orf1ab;1859 | 0 | 0 |
| NC_006213.1;YP_009555238.1;HCoV-OC43_orf1ab;1860 | 0.05 | 0 |
| NC_006213.1;YP_009555238.1;HCoV-OC43_orf1ab;1861 | 0 | 0 |
| NC_006213.1;YP_009555238.1;HCoV-OC43_orf1ab;1862 | 0 | 0 |
| NC_006213.1;YP_009555238.1;HCoV-OC43_orf1ab;1863 | 0 | 0 |
| NC_006213.1;YP_009555238.1;HCoV-OC43_orf1ab;1864 | 0 | 0 |
| NC_006213.1;YP_009555238.1;HCoV-OC43_orf1ab;1865 | 0 | 0 |
| NC_006213.1;YP_009555238.1;HCoV-OC43_orf1ab;1866 | 0 | 0 |
| NC_006213.1;YP_009555238.1;HCoV-OC43_orf1ab;1867 | 0 | 0.025 |
| NC_006213.1;YP_009555238.1;HCoV-OC43_orf1ab;1868 | 0 | 0 |
| NC_006213.1;YP_009555238.1;HCoV-OC43_orf1ab;1869 | 0 | 0 |
| NC_006213.1;YP_009555238.1;HCoV-OC43_orf1ab;1870 | 0.1 | 0 |
| NC_006213.1;YP_009555238.1;HCoV-OC43_orf1ab;1871 | 0.05 | 0.025 |
| NC_006213.1;YP_009555238.1;HCoV-OC43_orf1ab;1872 | 0 | 0 |
| NC_006213.1;YP_009555238.1;HCoV-OC43_orf1ab;1873 | 0.05 | 0.025 |
| NC_006213.1;YP_009555238.1;HCoV-OC43_orf1ab;1874 | 0.05 | 0.075 |
| NC_006213.1;YP_009555238.1;HCoV-OC43_orf1ab;1875 | 0.1 | 0.15 |
| NC_006213.1;YP_009555238.1;HCoV-OC43_orf1ab;1876 | 0.1 | 0.075 |
| NC_006213.1;YP_009555238.1;HCoV-OC43_orf1ab;1877 | 0.05 | 0.025 |
| NC_006213.1;YP_009555238.1;HCoV-OC43_orf1ab;1878 | 0 | 0.025 |
| NC_006213.1;YP_009555238.1;HCoV-OC43_orf1ab;1879 | 0.1 | 0 |
| NC_006213.1;YP_009555238.1;HCoV-OC43_orf1ab;1880 | 0 | 0 |
| NC_006213.1;YP_009555238.1;HCoV-OC43_orf1ab;1881 | 0 | 0 |
| NC_006213.1;YP_009555238.1;HCoV-OC43_orf1ab;1882 | 0 | 0 |
| NC_006213.1;YP_009555238.1;HCoV-OC43_orf1ab;1883 | 0 | 0 |
| NC_006213.1;YP_009555238.1;HCoV-OC43_orf1ab;1884 | 0 | 0 |
| NC_006213.1;YP_009555238.1;HCoV-OC43_orf1ab;1885 | 0 | 0 |
| NC_006213.1;YP_009555238.1;HCoV-OC43_orf1ab;1886 | 0 | 0 |
| NC_006213.1;YP_009555238.1;HCoV-OC43_orf1ab;1887 | 0.1 | 0.075 |
| NC_006213.1;YP_009555238.1;HCoV-OC43_orf1ab;1888 | 0 | 0 |
| NC_006213.1;YP_009555238.1;HCoV-OC43_orf1ab;1889 | 0 | 0 |
| NC_006213.1;YP_009555238.1;HCoV-OC43_orf1ab;1890 | 0 | 0 |
| NC_006213.1;YP_009555238.1;HCoV-OC43_orf1ab;1891 | 0 | 0 |
| NC_006213.1;YP_009555238.1;HCoV-OC43_orf1ab;1892 | 0 | 0 |
| NC_006213.1;YP_009555238.1;HCoV-OC43_orf1ab;1893 | 0 | 0 |
| NC_006213.1;YP_009555238.1;HCoV-OC43_orf1ab;1894 | 0 | 0 |
| NC_006213.1;YP_009555238.1;HCoV-OC43_orf1ab;1895 | 0 | 0 |
| NC_006213.1;YP_009555238.1;HCoV-OC43_orf1ab;1896 | 0 | 0 |
| NC_006213.1;YP_009555238.1;HCoV-OC43_orf1ab;1897 | 0 | 0 |
| NC_006213.1;YP_009555238.1;HCoV-OC43_orf1ab;1898 | 0 | 0 |
| NC_006213.1;YP_009555238.1;HCoV-OC43_orf1ab;1899 | 0 | 0 |
| NC_006213.1;YP_009555238.1;HCoV-OC43_orf1ab;1900 | 0 | 0 |
| NC_006213.1;YP_009555238.1;HCoV-OC43_orf1ab;1901 | 0 | 0 |

| | | |
|---|---|---|
| NC_006213.1;YP_009555238.1;HCoV-OC43_orf1ab;1902 | 0 | 0 |
| NC_006213.1;YP_009555238.1;HCoV-OC43_orf1ab;1903 | 0 | 0 |
| NC_006213.1;YP_009555238.1;HCoV-OC43_orf1ab;1904 | 0.05 | 0 |
| NC_006213.1;YP_009555238.1;HCoV-OC43_orf1ab;1905 | 0 | 0.025 |
| NC_006213.1;YP_009555238.1;HCoV-OC43_orf1ab;1906 | 0 | 0.025 |
| NC_006213.1;YP_009555238.1;HCoV-OC43_orf1ab;1907 | 0 | 0.05 |
| NC_006213.1;YP_009555238.1;HCoV-OC43_orf1ab;1908 | 0 | 0.05 |
| NC_006213.1;YP_009555238.1;HCoV-OC43_orf1ab;1909 | 0.05 | 0.05 |
| NC_006213.1;YP_009555238.1;HCoV-OC43_orf1ab;1910 | 0.05 | 0.025 |
| NC_006213.1;YP_009555238.1;HCoV-OC43_orf1ab;1911 | 0 | 0 |
| NC_006213.1;YP_009555238.1;HCoV-OC43_orf1ab;1912 | 0.05 | 0 |
| NC_006213.1;YP_009555238.1;HCoV-OC43_orf1ab;1913 | 0.05 | 0 |
| NC_006213.1;YP_009555238.1;HCoV-OC43_orf1ab;1914 | 0 | 0 |
| NC_006213.1;YP_009555238.1;HCoV-OC43_orf1ab;1915 | 0 | 0 |
| NC_006213.1;YP_009555238.1;HCoV-OC43_orf1ab;1916 | 0 | 0 |
| NC_006213.1;YP_009555238.1;HCoV-OC43_orf1ab;1917 | 0 | 0 |
| NC_006213.1;YP_009555238.1;HCoV-OC43_orf1ab;1918 | 0 | 0 |
| NC_006213.1;YP_009555238.1;HCoV-OC43_orf1ab;1919 | 0 | 0 |
| NC_006213.1;YP_009555238.1;HCoV-OC43_orf1ab;1920 | 0 | 0 |
| NC_006213.1;YP_009555238.1;HCoV-OC43_orf1ab;1921 | 0.1 | 0 |
| NC_006213.1;YP_009555238.1;HCoV-OC43_orf1ab;1922 | 0 | 0 |
| NC_006213.1;YP_009555238.1;HCoV-OC43_orf1ab;1923 | 0 | 0 |
| NC_006213.1;YP_009555238.1;HCoV-OC43_orf1ab;1924 | 0 | 0 |
| NC_006213.1;YP_009555238.1;HCoV-OC43_orf1ab;1925 | 0 | 0 |
| NC_006213.1;YP_009555238.1;HCoV-OC43_orf1ab;1926 | 0.05 | 0.05 |
| NC_006213.1;YP_009555238.1;HCoV-OC43_orf1ab;1927 | 0.05 | 0.025 |
| NC_006213.1;YP_009555238.1;HCoV-OC43_orf1ab;1928 | 0.05 | 0.025 |
| NC_006213.1;YP_009555238.1;HCoV-OC43_orf1ab;1929 | 0.1 | 0.075 |
| NC_006213.1;YP_009555238.1;HCoV-OC43_orf1ab;1930 | 0.1 | 0.075 |
| NC_006213.1;YP_009555238.1;HCoV-OC43_orf1ab;1931 | 0.1 | 0.125 |
| NC_006213.1;YP_009555238.1;HCoV-OC43_orf1ab;1932 | 0.05 | 0.1 |
| NC_006213.1;YP_009555238.1;HCoV-OC43_orf1ab;1933 | 0.05 | 0.1 |
| NC_006213.1;YP_009555238.1;HCoV-OC43_orf1ab;1934 | 0 | 0.075 |
| NC_006213.1;YP_009555238.1;HCoV-OC43_orf1ab;1935 | 0.05 | 0.05 |
| NC_006213.1;YP_009555238.1;HCoV-OC43_orf1ab;1936 | 0.05 | 0.1 |
| NC_006213.1;YP_009555238.1;HCoV-OC43_orf1ab;1937 | 0.05 | 0.075 |
| NC_006213.1;YP_009555238.1;HCoV-OC43_orf1ab;1938 | 0 | 0.025 |
| NC_006213.1;YP_009555238.1;HCoV-OC43_orf1ab;1939 | 0 | 0.05 |
| NC_006213.1;YP_009555238.1;HCoV-OC43_orf1ab;1940 | 0 | 0 |
| NC_006213.1;YP_009555238.1;HCoV-OC43_orf1ab;1941 | 0.05 | 0 |
| NC_006213.1;YP_009555238.1;HCoV-OC43_orf1ab;1942 | 0 | 0 |
| NC_006213.1;YP_009555238.1;HCoV-OC43_orf1ab;1943 | 0.1 | 0 |
| NC_006213.1;YP_009555238.1;HCoV-OC43_orf1ab;1944 | 0.05 | 0 |

| | | |
|---|---|---|
| NC_006213.1;YP_009555238.1;HCoV-OC43_orf1ab;1945 | 0.05 | 0 |
| NC_006213.1;YP_009555238.1;HCoV-OC43_orf1ab;1946 | 0 | 0 |
| NC_006213.1;YP_009555238.1;HCoV-OC43_orf1ab;1947 | 0 | 0 |
| NC_006213.1;YP_009555238.1;HCoV-OC43_orf1ab;1948 | 0 | 0 |
| NC_006213.1;YP_009555238.1;HCoV-OC43_orf1ab;1949 | 0 | 0 |
| NC_006213.1;YP_009555238.1;HCoV-OC43_orf1ab;1950 | 0 | 0 |
| NC_006213.1;YP_009555238.1;HCoV-OC43_orf1ab;1951 | 0 | 0 |
| NC_006213.1;YP_009555238.1;HCoV-OC43_orf1ab;1952 | 0 | 0 |
| NC_006213.1;YP_009555238.1;HCoV-OC43_orf1ab;1953 | 0 | 0 |
| NC_006213.1;YP_009555238.1;HCoV-OC43_orf1ab;1954 | 0 | 0 |
| NC_006213.1;YP_009555238.1;HCoV-OC43_orf1ab;1955 | 0 | 0 |
| NC_006213.1;YP_009555238.1;HCoV-OC43_orf1ab;1956 | 0 | 0.025 |
| NC_006213.1;YP_009555238.1;HCoV-OC43_orf1ab;1957 | 0.05 | 0.025 |
| NC_006213.1;YP_009555238.1;HCoV-OC43_orf1ab;1958 | 0 | 0.025 |
| NC_006213.1;YP_009555238.1;HCoV-OC43_orf1ab;1959 | 0 | 0.1 |
| NC_006213.1;YP_009555238.1;HCoV-OC43_orf1ab;1960 | 0 | 0.05 |
| NC_006213.1;YP_009555238.1;HCoV-OC43_orf1ab;1961 | 0 | 0.025 |
| NC_006213.1;YP_009555238.1;HCoV-OC43_orf1ab;1962 | 0 | 0.05 |
| NC_006213.1;YP_009555238.1;HCoV-OC43_orf1ab;1963 | 0 | 0 |
| NC_006213.1;YP_009555238.1;HCoV-OC43_orf1ab;1964 | 0 | 0 |
| NC_006213.1;YP_009555238.1;HCoV-OC43_orf1ab;1965 | 0 | 0 |
| NC_006213.1;YP_009555238.1;HCoV-OC43_orf1ab;1966 | 0 | 0 |
| NC_006213.1;YP_009555238.1;HCoV-OC43_orf1ab;1967 | 0 | 0 |
| NC_006213.1;YP_009555238.1;HCoV-OC43_orf1ab;1968 | 0 | 0 |
| NC_006213.1;YP_009555238.1;HCoV-OC43_orf1ab;1969 | 0 | 0 |
| NC_006213.1;YP_009555238.1;HCoV-OC43_orf1ab;1970 | 0 | 0 |
| NC_006213.1;YP_009555238.1;HCoV-OC43_orf1ab;1971 | 0 | 0 |
| NC_006213.1;YP_009555238.1;HCoV-OC43_orf1ab;1972 | 0 | 0 |
| NC_006213.1;YP_009555238.1;HCoV-OC43_orf1ab;1973 | 0 | 0 |
| NC_006213.1;YP_009555238.1;HCoV-OC43_orf1ab;1974 | 0 | 0 |
| NC_006213.1;YP_009555238.1;HCoV-OC43_orf1ab;1975 | 0 | 0 |
| NC_006213.1;YP_009555238.1;HCoV-OC43_orf1ab;1976 | 0 | 0 |
| NC_006213.1;YP_009555238.1;HCoV-OC43_orf1ab;1977 | 0 | 0 |
| NC_006213.1;YP_009555238.1;HCoV-OC43_orf1ab;1978 | 0 | 0 |
| NC_006213.1;YP_009555238.1;HCoV-OC43_orf1ab;1979 | 0 | 0 |
| NC_006213.1;YP_009555238.1;HCoV-OC43_orf1ab;1980 | 0 | 0 |
| NC_006213.1;YP_009555238.1;HCoV-OC43_orf1ab;1981 | 0 | 0 |
| NC_006213.1;YP_009555238.1;HCoV-OC43_orf1ab;1982 | 0 | 0 |
| NC_006213.1;YP_009555238.1;HCoV-OC43_orf1ab;1983 | 0 | 0 |
| NC_006213.1;YP_009555238.1;HCoV-OC43_orf1ab;1984 | 0 | 0 |
| NC_006213.1;YP_009555238.1;HCoV-OC43_orf1ab;1985 | 0 | 0 |
| NC_006213.1;YP_009555238.1;HCoV-OC43_orf1ab;1986 | 0 | 0 |
| NC_006213.1;YP_009555238.1;HCoV-OC43_orf1ab;1987 | 0 | 0 |

| | | |
|---|---|---|
| NC_006213.1;YP_009555238.1;HCoV-OC43_orf1ab;1988 | 0 | 0 |
| NC_006213.1;YP_009555238.1;HCoV-OC43_orf1ab;1989 | 0 | 0 |
| NC_006213.1;YP_009555238.1;HCoV-OC43_orf1ab;1990 | 0 | 0.05 |
| NC_006213.1;YP_009555238.1;HCoV-OC43_orf1ab;1991 | 0 | 0 |
| NC_006213.1;YP_009555238.1;HCoV-OC43_orf1ab;1992 | 0 | 0 |
| NC_006213.1;YP_009555238.1;HCoV-OC43_orf1ab;1993 | 0.05 | 0 |
| NC_006213.1;YP_009555238.1;HCoV-OC43_orf1ab;1994 | 0 | 0.025 |
| NC_006213.1;YP_009555238.1;HCoV-OC43_orf1ab;1995 | 0.05 | 0.025 |
| NC_006213.1;YP_009555238.1;HCoV-OC43_orf1ab;1996 | 0.05 | 0 |
| NC_006213.1;YP_009555238.1;HCoV-OC43_orf1ab;1997 | 0.1 | 0.025 |
| NC_006213.1;YP_009555238.1;HCoV-OC43_orf1ab;1998 | 0.1 | 0.05 |
| NC_006213.1;YP_009555238.1;HCoV-OC43_orf1ab;1999 | 0.1 | 0.025 |
| NC_006213.1;YP_009555238.1;HCoV-OC43_orf1ab;2000 | 0.05 | 0.05 |
| NC_006213.1;YP_009555238.1;HCoV-OC43_orf1ab;2001 | 0.05 | 0 |
| NC_006213.1;YP_009555238.1;HCoV-OC43_orf1ab;2002 | 0.05 | 0 |
| NC_006213.1;YP_009555238.1;HCoV-OC43_orf1ab;2003 | 0.05 | 0 |
| NC_006213.1;YP_009555238.1;HCoV-OC43_orf1ab;2004 | 0.1 | 0 |
| NC_006213.1;YP_009555238.1;HCoV-OC43_orf1ab;2005 | 0 | 0 |
| NC_006213.1;YP_009555238.1;HCoV-OC43_orf1ab;2006 | 0 | 0 |
| NC_006213.1;YP_009555238.1;HCoV-OC43_orf1ab;2007 | 0 | 0 |
| NC_006213.1;YP_009555238.1;HCoV-OC43_orf1ab;2008 | 0.05 | 0.025 |
| NC_006213.1;YP_009555238.1;HCoV-OC43_orf1ab;2009 | 0 | 0 |
| NC_006213.1;YP_009555238.1;HCoV-OC43_orf1ab;2010 | 0 | 0 |
| NC_006213.1;YP_009555238.1;HCoV-OC43_orf1ab;2011 | 0 | 0 |
| NC_006213.1;YP_009555238.1;HCoV-OC43_orf1ab;2012 | 0 | 0 |
| NC_006213.1;YP_009555238.1;HCoV-OC43_orf1ab;2013 | 0 | 0 |
| NC_006213.1;YP_009555238.1;HCoV-OC43_orf1ab;2014 | 0 | 0 |
| NC_006213.1;YP_009555238.1;HCoV-OC43_orf1ab;2015 | 0 | 0 |
| NC_006213.1;YP_009555238.1;HCoV-OC43_orf1ab;2016 | 0 | 0 |
| NC_006213.1;YP_009555238.1;HCoV-OC43_orf1ab;2017 | 0.05 | 0 |
| NC_006213.1;YP_009555238.1;HCoV-OC43_orf1ab;2018 | 0 | 0 |
| NC_006213.1;YP_009555238.1;HCoV-OC43_orf1ab;2019 | 0 | 0 |
| NC_006213.1;YP_009555238.1;HCoV-OC43_orf1ab;2020 | 0 | 0 |
| NC_006213.1;YP_009555238.1;HCoV-OC43_orf1ab;2021 | 0 | 0.025 |
| NC_006213.1;YP_009555238.1;HCoV-OC43_orf1ab;2022 | 0 | 0.025 |
| NC_006213.1;YP_009555238.1;HCoV-OC43_orf1ab;2023 | 0 | 0.025 |
| NC_006213.1;YP_009555238.1;HCoV-OC43_orf1ab;2024 | 0 | 0.025 |
| NC_006213.1;YP_009555238.1;HCoV-OC43_orf1ab;2025 | 0 | 0.025 |
| NC_006213.1;YP_009555238.1;HCoV-OC43_orf1ab;2026 | 0 | 0.025 |
| NC_006213.1;YP_009555238.1;HCoV-OC43_orf1ab;2027 | 0 | 0.025 |
| NC_006213.1;YP_009555238.1;HCoV-OC43_orf1ab;2028 | 0 | 0.025 |
| NC_006213.1;YP_009555238.1;HCoV-OC43_orf1ab;2029 | 0 | 0.025 |
| NC_006213.1;YP_009555238.1;HCoV-OC43_orf1ab;2030 | 0 | 0 |

| | | |
|---|---|---|
| NC_006213.1;YP_009555238.1;HCoV-OC43_orf1ab;2031 | 0 | 0 |
| NC_006213.1;YP_009555238.1;HCoV-OC43_orf1ab;2032 | 0 | 0.025 |
| NC_006213.1;YP_009555238.1;HCoV-OC43_orf1ab;2033 | 0 | 0 |
| NC_006213.1;YP_009555238.1;HCoV-OC43_orf1ab;2034 | 0 | 0 |
| NC_006213.1;YP_009555238.1;HCoV-OC43_orf1ab;2035 | 0 | 0 |
| NC_006213.1;YP_009555238.1;HCoV-OC43_orf1ab;2036 | 0 | 0 |
| NC_006213.1;YP_009555238.1;HCoV-OC43_orf1ab;2037 | 0 | 0 |
| NC_006213.1;YP_009555238.1;HCoV-OC43_orf1ab;2038 | 0 | 0 |
| NC_006213.1;YP_009555238.1;HCoV-OC43_orf1ab;2039 | 0 | 0 |
| NC_006213.1;YP_009555238.1;HCoV-OC43_orf1ab;2040 | 0.05 | 0 |
| NC_006213.1;YP_009555238.1;HCoV-OC43_orf1ab;2041 | 0 | 0 |
| NC_006213.1;YP_009555238.1;HCoV-OC43_orf1ab;2042 | 0.05 | 0 |
| NC_006213.1;YP_009555238.1;HCoV-OC43_orf1ab;2043 | 0 | 0 |
| NC_006213.1;YP_009555238.1;HCoV-OC43_orf1ab;2044 | 0 | 0 |
| NC_006213.1;YP_009555238.1;HCoV-OC43_orf1ab;2045 | 0.05 | 0 |
| NC_006213.1;YP_009555238.1;HCoV-OC43_orf1ab;2046 | 0 | 0.025 |
| NC_006213.1;YP_009555238.1;HCoV-OC43_orf1ab;2047 | 0 | 0 |
| NC_006213.1;YP_009555238.1;HCoV-OC43_orf1ab;2048 | 0 | 0 |
| NC_006213.1;YP_009555238.1;HCoV-OC43_orf1ab;2049 | 0 | 0 |
| NC_006213.1;YP_009555238.1;HCoV-OC43_orf1ab;2050 | 0 | 0 |
| NC_006213.1;YP_009555238.1;HCoV-OC43_orf1ab;2051 | 0 | 0 |
| NC_006213.1;YP_009555238.1;HCoV-OC43_orf1ab;2052 | 0 | 0.025 |
| NC_006213.1;YP_009555238.1;HCoV-OC43_orf1ab;2053 | 0 | 0.025 |
| NC_006213.1;YP_009555238.1;HCoV-OC43_orf1ab;2054 | 0 | 0.025 |
| NC_006213.1;YP_009555238.1;HCoV-OC43_orf1ab;2055 | 0.05 | 0.075 |
| NC_006213.1;YP_009555238.1;HCoV-OC43_orf1ab;2056 | 0 | 0.025 |
| NC_006213.1;YP_009555238.1;HCoV-OC43_orf1ab;2057 | 0 | 0.025 |
| NC_006213.1;YP_009555238.1;HCoV-OC43_orf1ab;2058 | 0 | 0.025 |
| NC_006213.1;YP_009555238.1;HCoV-OC43_orf1ab;2059 | 0 | 0 |
| NC_006213.1;YP_009555238.1;HCoV-OC43_orf1ab;2060 | 0 | 0 |
| NC_006213.1;YP_009555238.1;HCoV-OC43_orf1ab;2061 | 0 | 0 |
| NC_006213.1;YP_009555238.1;HCoV-OC43_orf1ab;2062 | 0 | 0 |
| NC_006213.1;YP_009555238.1;HCoV-OC43_orf1ab;2063 | 0 | 0 |
| NC_006213.1;YP_009555238.1;HCoV-OC43_orf1ab;2064 | 0 | 0 |
| NC_006213.1;YP_009555238.1;HCoV-OC43_orf1ab;2065 | 0 | 0 |
| NC_006213.1;YP_009555238.1;HCoV-OC43_orf1ab;2066 | 0 | 0 |
| NC_006213.1;YP_009555238.1;HCoV-OC43_orf1ab;2067 | 0 | 0 |
| NC_006213.1;YP_009555238.1;HCoV-OC43_orf1ab;2068 | 0 | 0 |
| NC_006213.1;YP_009555238.1;HCoV-OC43_orf1ab;2069 | 0 | 0 |
| NC_006213.1;YP_009555238.1;HCoV-OC43_orf1ab;2070 | 0 | 0 |
| NC_006213.1;YP_009555238.1;HCoV-OC43_orf1ab;2071 | 0 | 0 |
| NC_006213.1;YP_009555238.1;HCoV-OC43_orf1ab;2072 | 0 | 0 |
| NC_006213.1;YP_009555238.1;HCoV-OC43_orf1ab;2073 | 0 | 0 |

| | | |
|---|---|---|
| NC_006213.1;YP_009555238.1;HCoV-OC43_orf1ab;2074 | 0 | 0 |
| NC_006213.1;YP_009555238.1;HCoV-OC43_orf1ab;2075 | 0 | 0 |
| NC_006213.1;YP_009555238.1;HCoV-OC43_orf1ab;2076 | 0 | 0 |
| NC_006213.1;YP_009555238.1;HCoV-OC43_orf1ab;2077 | 0 | 0 |
| NC_006213.1;YP_009555238.1;HCoV-OC43_orf1ab;2078 | 0 | 0 |
| NC_006213.1;YP_009555238.1;HCoV-OC43_orf1ab;2079 | 0 | 0 |
| NC_006213.1;YP_009555238.1;HCoV-OC43_orf1ab;2080 | 0 | 0 |
| NC_006213.1;YP_009555238.1;HCoV-OC43_orf1ab;2081 | 0 | 0 |
| NC_006213.1;YP_009555238.1;HCoV-OC43_orf1ab;2082 | 0 | 0 |
| NC_006213.1;YP_009555238.1;HCoV-OC43_orf1ab;2083 | 0 | 0 |
| NC_006213.1;YP_009555238.1;HCoV-OC43_orf1ab;2084 | 0 | 0 |
| NC_006213.1;YP_009555238.1;HCoV-OC43_orf1ab;2085 | 0 | 0.05 |
| NC_006213.1;YP_009555238.1;HCoV-OC43_orf1ab;2086 | 0 | 0.05 |
| NC_006213.1;YP_009555238.1;HCoV-OC43_orf1ab;2087 | 0 | 0 |
| NC_006213.1;YP_009555238.1;HCoV-OC43_orf1ab;2088 | 0 | 0 |
| NC_006213.1;YP_009555238.1;HCoV-OC43_orf1ab;2089 | 0 | 0.025 |
| NC_006213.1;YP_009555238.1;HCoV-OC43_orf1ab;2090 | 0 | 0.025 |
| NC_006213.1;YP_009555238.1;HCoV-OC43_orf1ab;2091 | 0 | 0 |
| NC_006213.1;YP_009555238.1;HCoV-OC43_orf1ab;2092 | 0 | 0 |
| NC_006213.1;YP_009555238.1;HCoV-OC43_orf1ab;2093 | 0 | 0 |
| NC_006213.1;YP_009555238.1;HCoV-OC43_orf1ab;2094 | 0 | 0 |
| NC_006213.1;YP_009555238.1;HCoV-OC43_orf1ab;2095 | 0 | 0 |
| NC_006213.1;YP_009555238.1;HCoV-OC43_orf1ab;2096 | 0 | 0 |
| NC_006213.1;YP_009555238.1;HCoV-OC43_orf1ab;2097 | 0 | 0 |
| NC_006213.1;YP_009555238.1;HCoV-OC43_orf1ab;2098 | 0 | 0 |
| NC_006213.1;YP_009555238.1;HCoV-OC43_orf1ab;2099 | 0 | 0 |
| NC_006213.1;YP_009555238.1;HCoV-OC43_orf1ab;2100 | 0 | 0 |
| NC_006213.1;YP_009555238.1;HCoV-OC43_orf1ab;2101 | 0 | 0 |
| NC_006213.1;YP_009555238.1;HCoV-OC43_orf1ab;2102 | 0 | 0 |
| NC_006213.1;YP_009555238.1;HCoV-OC43_orf1ab;2103 | 0 | 0 |
| NC_006213.1;YP_009555238.1;HCoV-OC43_orf1ab;2104 | 0 | 0 |
| NC_006213.1;YP_009555238.1;HCoV-OC43_orf1ab;2105 | 0 | 0 |
| NC_006213.1;YP_009555238.1;HCoV-OC43_orf1ab;2106 | 0 | 0 |
| NC_006213.1;YP_009555238.1;HCoV-OC43_orf1ab;2107 | 0 | 0 |
| NC_006213.1;YP_009555238.1;HCoV-OC43_orf1ab;2108 | 0 | 0 |
| NC_006213.1;YP_009555238.1;HCoV-OC43_orf1ab;2109 | 0 | 0 |
| NC_006213.1;YP_009555238.1;HCoV-OC43_orf1ab;2110 | 0 | 0 |
| NC_006213.1;YP_009555238.1;HCoV-OC43_orf1ab;2111 | 0.05 | 0 |
| NC_006213.1;YP_009555238.1;HCoV-OC43_orf1ab;2112 | 0.05 | 0.025 |
| NC_006213.1;YP_009555238.1;HCoV-OC43_orf1ab;2113 | 0.05 | 0.025 |
| NC_006213.1;YP_009555238.1;HCoV-OC43_orf1ab;2114 | 0.15 | 0.025 |
| NC_006213.1;YP_009555238.1;HCoV-OC43_orf1ab;2115 | 0 | 0 |
| NC_006213.1;YP_009555238.1;HCoV-OC43_orf1ab;2116 | 0 | 0.025 |

| | | |
|---|---|---|
| NC_006213.1;YP_009555238.1;HCoV-OC43_orf1ab;2117 | 0 | 0 |
| NC_006213.1;YP_009555238.1;HCoV-OC43_orf1ab;2118 | 0 | 0 |
| NC_006213.1;YP_009555238.1;HCoV-OC43_orf1ab;2119 | 0 | 0 |
| NC_006213.1;YP_009555238.1;HCoV-OC43_orf1ab;2120 | 0 | 0 |
| NC_006213.1;YP_009555238.1;HCoV-OC43_orf1ab;2121 | 0 | 0 |
| NC_006213.1;YP_009555238.1;HCoV-OC43_orf1ab;2122 | 0 | 0 |
| NC_006213.1;YP_009555238.1;HCoV-OC43_orf1ab;2123 | 0 | 0 |
| NC_006213.1;YP_009555238.1;HCoV-OC43_orf1ab;2124 | 0 | 0 |
| NC_006213.1;YP_009555238.1;HCoV-OC43_orf1ab;2125 | 0 | 0 |
| NC_006213.1;YP_009555238.1;HCoV-OC43_orf1ab;2126 | 0 | 0 |
| NC_006213.1;YP_009555238.1;HCoV-OC43_orf1ab;2127 | 0 | 0 |
| NC_006213.1;YP_009555238.1;HCoV-OC43_orf1ab;2128 | 0 | 0 |
| NC_006213.1;YP_009555238.1;HCoV-OC43_orf1ab;2129 | 0 | 0 |
| NC_006213.1;YP_009555238.1;HCoV-OC43_orf1ab;2130 | 0 | 0 |
| NC_006213.1;YP_009555238.1;HCoV-OC43_orf1ab;2131 | 0 | 0 |
| NC_006213.1;YP_009555238.1;HCoV-OC43_orf1ab;2132 | 0 | 0 |
| NC_006213.1;YP_009555238.1;HCoV-OC43_orf1ab;2133 | 0 | 0 |
| NC_006213.1;YP_009555238.1;HCoV-OC43_orf1ab;2134 | 0 | 0 |
| NC_006213.1;YP_009555238.1;HCoV-OC43_orf1ab;2135 | 0 | 0 |
| NC_006213.1;YP_009555238.1;HCoV-OC43_orf1ab;2136 | 0 | 0 |
| NC_006213.1;YP_009555238.1;HCoV-OC43_orf1ab;2137 | 0 | 0 |
| NC_006213.1;YP_009555238.1;HCoV-OC43_orf1ab;2138 | 0 | 0 |
| NC_006213.1;YP_009555238.1;HCoV-OC43_orf1ab;2139 | 0 | 0 |
| NC_006213.1;YP_009555238.1;HCoV-OC43_orf1ab;2140 | 0 | 0 |
| NC_006213.1;YP_009555238.1;HCoV-OC43_orf1ab;2141 | 0 | 0 |
| NC_006213.1;YP_009555238.1;HCoV-OC43_orf1ab;2142 | 0 | 0 |
| NC_006213.1;YP_009555238.1;HCoV-OC43_orf1ab;2143 | 0 | 0 |
| NC_006213.1;YP_009555238.1;HCoV-OC43_orf1ab;2144 | 0 | 0 |
| NC_006213.1;YP_009555238.1;HCoV-OC43_orf1ab;2145 | 0 | 0 |
| NC_006213.1;YP_009555238.1;HCoV-OC43_orf1ab;2146 | 0 | 0 |
| NC_006213.1;YP_009555238.1;HCoV-OC43_orf1ab;2147 | 0 | 0 |
| NC_006213.1;YP_009555238.1;HCoV-OC43_orf1ab;2148 | 0 | 0 |
| NC_006213.1;YP_009555238.1;HCoV-OC43_orf1ab;2149 | 0 | 0 |
| NC_006213.1;YP_009555238.1;HCoV-OC43_orf1ab;2150 | 0 | 0 |
| NC_006213.1;YP_009555238.1;HCoV-OC43_orf1ab;2151 | 0 | 0 |
| NC_006213.1;YP_009555238.1;HCoV-OC43_orf1ab;2152 | 0 | 0 |
| NC_006213.1;YP_009555238.1;HCoV-OC43_orf1ab;2153 | 0 | 0.025 |
| NC_006213.1;YP_009555238.1;HCoV-OC43_orf1ab;2154 | 0 | 0.025 |
| NC_006213.1;YP_009555238.1;HCoV-OC43_orf1ab;2155 | 0 | 0 |
| NC_006213.1;YP_009555238.1;HCoV-OC43_orf1ab;2156 | 0 | 0.05 |
| NC_006213.1;YP_009555238.1;HCoV-OC43_orf1ab;2157 | 0.05 | 0.05 |
| NC_006213.1;YP_009555238.1;HCoV-OC43_orf1ab;2158 | 0 | 0 |
| NC_006213.1;YP_009555238.1;HCoV-OC43_orf1ab;2159 | 0 | 0 |

| | | |
|---|---|---|
| NC_006213.1;YP_009555238.1;HCoV-OC43_orf1ab;2160 | 0 | 0 |
| NC_006213.1;YP_009555238.1;HCoV-OC43_orf1ab;2161 | 0 | 0 |
| NC_006213.1;YP_009555238.1;HCoV-OC43_orf1ab;2162 | 0 | 0.025 |
| NC_006213.1;YP_009555238.1;HCoV-OC43_orf1ab;2163 | 0 | 0 |
| NC_006213.1;YP_009555238.1;HCoV-OC43_orf1ab;2164 | 0 | 0 |
| NC_006213.1;YP_009555238.1;HCoV-OC43_orf1ab;2165 | 0 | 0 |
| NC_006213.1;YP_009555238.1;HCoV-OC43_orf1ab;2166 | 0 | 0 |
| NC_006213.1;YP_009555238.1;HCoV-OC43_orf1ab;2167 | 0 | 0 |
| NC_006213.1;YP_009555238.1;HCoV-OC43_orf1ab;2168 | 0 | 0 |
| NC_006213.1;YP_009555238.1;HCoV-OC43_orf1ab;2169 | 0 | 0 |
| NC_006213.1;YP_009555238.1;HCoV-OC43_orf1ab;2170 | 0 | 0 |
| NC_006213.1;YP_009555238.1;HCoV-OC43_orf1ab;2171 | 0 | 0 |
| NC_006213.1;YP_009555238.1;HCoV-OC43_orf1ab;2172 | 0 | 0 |
| NC_006213.1;YP_009555238.1;HCoV-OC43_orf1ab;2173 | 0 | 0 |
| NC_006213.1;YP_009555238.1;HCoV-OC43_orf1ab;2174 | 0 | 0 |
| NC_006213.1;YP_009555238.1;HCoV-OC43_orf1ab;2175 | 0 | 0 |
| NC_006213.1;YP_009555238.1;HCoV-OC43_orf1ab;2176 | 0 | 0 |
| NC_006213.1;YP_009555238.1;HCoV-OC43_orf1ab;2177 | 0 | 0 |
| NC_006213.1;YP_009555238.1;HCoV-OC43_orf1ab;2178 | 0 | 0 |
| NC_006213.1;YP_009555238.1;HCoV-OC43_orf1ab;2179 | 0 | 0 |
| NC_006213.1;YP_009555238.1;HCoV-OC43_orf1ab;2180 | 0 | 0 |
| NC_006213.1;YP_009555238.1;HCoV-OC43_orf1ab;2181 | 0 | 0 |
| NC_006213.1;YP_009555238.1;HCoV-OC43_orf1ab;2182 | 0 | 0 |
| NC_006213.1;YP_009555238.1;HCoV-OC43_orf1ab;2183 | 0 | 0 |
| NC_006213.1;YP_009555238.1;HCoV-OC43_orf1ab;2184 | 0 | 0 |
| NC_006213.1;YP_009555238.1;HCoV-OC43_orf1ab;2185 | 0 | 0 |
| NC_006213.1;YP_009555238.1;HCoV-OC43_orf1ab;2186 | 0 | 0 |
| NC_006213.1;YP_009555238.1;HCoV-OC43_orf1ab;2187 | 0 | 0 |
| NC_006213.1;YP_009555238.1;HCoV-OC43_orf1ab;2188 | 0 | 0 |
| NC_006213.1;YP_009555238.1;HCoV-OC43_orf1ab;2189 | 0 | 0 |
| NC_006213.1;YP_009555238.1;HCoV-OC43_orf1ab;2190 | 0 | 0 |
| NC_006213.1;YP_009555238.1;HCoV-OC43_orf1ab;2191 | 0 | 0 |
| NC_006213.1;YP_009555238.1;HCoV-OC43_orf1ab;2192 | 0 | 0 |
| NC_006213.1;YP_009555238.1;HCoV-OC43_orf1ab;2193 | 0 | 0 |
| NC_006213.1;YP_009555238.1;HCoV-OC43_orf1ab;2194 | 0 | 0 |
| NC_006213.1;YP_009555238.1;HCoV-OC43_orf1ab;2195 | 0 | 0 |
| NC_006213.1;YP_009555238.1;HCoV-OC43_orf1ab;2196 | 0 | 0 |
| NC_006213.1;YP_009555238.1;HCoV-OC43_orf1ab;2197 | 0 | 0 |
| NC_006213.1;YP_009555238.1;HCoV-OC43_orf1ab;2198 | 0 | 0 |
| NC_006213.1;YP_009555238.1;HCoV-OC43_orf1ab;2199 | 0 | 0 |
| NC_006213.1;YP_009555238.1;HCoV-OC43_orf1ab;2200 | 0 | 0 |
| NC_006213.1;YP_009555238.1;HCoV-OC43_orf1ab;2201 | 0 | 0 |
| NC_006213.1;YP_009555238.1;HCoV-OC43_orf1ab;2202 | 0 | 0 |

| | | |
|---|---|---|
| NC_006213.1;YP_009555238.1;HCoV-OC43_orf1ab;2203 | 0 | 0 |
| NC_006213.1;YP_009555238.1;HCoV-OC43_orf1ab;2204 | 0 | 0 |
| NC_006213.1;YP_009555238.1;HCoV-OC43_orf1ab;2205 | 0 | 0 |
| NC_006213.1;YP_009555238.1;HCoV-OC43_orf1ab;2206 | 0 | 0 |
| NC_006213.1;YP_009555238.1;HCoV-OC43_orf1ab;2207 | 0 | 0 |
| NC_006213.1;YP_009555238.1;HCoV-OC43_orf1ab;2208 | 0 | 0 |
| NC_006213.1;YP_009555238.1;HCoV-OC43_orf1ab;2209 | 0 | 0 |
| NC_006213.1;YP_009555238.1;HCoV-OC43_orf1ab;2210 | 0 | 0 |
| NC_006213.1;YP_009555238.1;HCoV-OC43_orf1ab;2211 | 0 | 0 |
| NC_006213.1;YP_009555238.1;HCoV-OC43_orf1ab;2212 | 0 | 0 |
| NC_006213.1;YP_009555238.1;HCoV-OC43_orf1ab;2213 | 0 | 0 |
| NC_006213.1;YP_009555238.1;HCoV-OC43_orf1ab;2214 | 0.05 | 0 |
| NC_006213.1;YP_009555238.1;HCoV-OC43_orf1ab;2215 | 0.05 | 0 |
| NC_006213.1;YP_009555238.1;HCoV-OC43_orf1ab;2216 | 0.05 | 0 |
| NC_006213.1;YP_009555238.1;HCoV-OC43_orf1ab;2217 | 0.05 | 0 |
| NC_006213.1;YP_009555238.1;HCoV-OC43_orf1ab;2218 | 0.05 | 0 |
| NC_006213.1;YP_009555238.1;HCoV-OC43_orf1ab;2219 | 0.05 | 0 |
| NC_006213.1;YP_009555238.1;HCoV-OC43_orf1ab;2220 | 0.05 | 0 |
| NC_006213.1;YP_009555238.1;HCoV-OC43_orf1ab;2221 | 0.05 | 0 |
| NC_006213.1;YP_009555238.1;HCoV-OC43_orf1ab;2222 | 0.05 | 0 |
| NC_006213.1;YP_009555238.1;HCoV-OC43_orf1ab;2223 | 0.05 | 0 |
| NC_006213.1;YP_009555238.1;HCoV-OC43_orf1ab;2224 | 0 | 0 |
| NC_006213.1;YP_009555238.1;HCoV-OC43_orf1ab;2225 | 0 | 0 |
| NC_006213.1;YP_009555238.1;HCoV-OC43_orf1ab;2226 | 0 | 0 |
| NC_006213.1;YP_009555238.1;HCoV-OC43_orf1ab;2227 | 0 | 0 |
| NC_006213.1;YP_009555238.1;HCoV-OC43_orf1ab;2228 | 0 | 0 |
| NC_006213.1;YP_009555238.1;HCoV-OC43_orf1ab;2229 | 0 | 0 |
| NC_006213.1;YP_009555238.1;HCoV-OC43_orf1ab;2230 | 0 | 0 |
| NC_006213.1;YP_009555238.1;HCoV-OC43_orf1ab;2231 | 0 | 0 |
| NC_006213.1;YP_009555238.1;HCoV-OC43_orf1ab;2232 | 0 | 0 |
| NC_006213.1;YP_009555238.1;HCoV-OC43_orf1ab;2233 | 0 | 0 |
| NC_006213.1;YP_009555238.1;HCoV-OC43_orf1ab;2234 | 0 | 0 |
| NC_006213.1;YP_009555238.1;HCoV-OC43_orf1ab;2235 | 0 | 0 |
| NC_006213.1;YP_009555238.1;HCoV-OC43_orf1ab;2236 | 0 | 0 |
| NC_006213.1;YP_009555238.1;HCoV-OC43_orf1ab;2237 | 0 | 0 |
| NC_006213.1;YP_009555238.1;HCoV-OC43_orf1ab;2238 | 0 | 0 |
| NC_006213.1;YP_009555238.1;HCoV-OC43_orf1ab;2239 | 0 | 0 |
| NC_006213.1;YP_009555238.1;HCoV-OC43_orf1ab;2240 | 0 | 0 |
| NC_006213.1;YP_009555238.1;HCoV-OC43_orf1ab;2241 | 0 | 0 |
| NC_006213.1;YP_009555238.1;HCoV-OC43_orf1ab;2242 | 0 | 0 |
| NC_006213.1;YP_009555238.1;HCoV-OC43_orf1ab;2243 | 0 | 0 |
| NC_006213.1;YP_009555238.1;HCoV-OC43_orf1ab;2244 | 0 | 0 |
| NC_006213.1;YP_009555238.1;HCoV-OC43_orf1ab;2245 | 0 | 0 |

| | | |
|---|---|---|
| NC_006213.1;YP_009555238.1;HCoV-OC43_orf1ab;2246 | 0 | 0 |
| NC_006213.1;YP_009555238.1;HCoV-OC43_orf1ab;2247 | 0 | 0 |
| NC_006213.1;YP_009555238.1;HCoV-OC43_orf1ab;2248 | 0 | 0 |
| NC_006213.1;YP_009555238.1;HCoV-OC43_orf1ab;2249 | 0 | 0 |
| NC_006213.1;YP_009555238.1;HCoV-OC43_orf1ab;2250 | 0 | 0 |
| NC_006213.1;YP_009555238.1;HCoV-OC43_orf1ab;2251 | 0 | 0 |
| NC_006213.1;YP_009555238.1;HCoV-OC43_orf1ab;2252 | 0 | 0 |
| NC_006213.1;YP_009555238.1;HCoV-OC43_orf1ab;2253 | 0.05 | 0 |
| NC_006213.1;YP_009555238.1;HCoV-OC43_orf1ab;2254 | 0.05 | 0.075 |
| NC_006213.1;YP_009555238.1;HCoV-OC43_orf1ab;2255 | 0.1 | 0.05 |
| NC_006213.1;YP_009555238.1;HCoV-OC43_orf1ab;2256 | 0.15 | 0.075 |
| NC_006213.1;YP_009555238.1;HCoV-OC43_orf1ab;2257 | 0.1 | 0.075 |
| NC_006213.1;YP_009555238.1;HCoV-OC43_orf1ab;2258 | 0.05 | 0.025 |
| NC_006213.1;YP_009555238.1;HCoV-OC43_orf1ab;2259 | 0 | 0 |
| NC_006213.1;YP_009555238.1;HCoV-OC43_orf1ab;2260 | 0 | 0 |
| NC_006213.1;YP_009555238.1;HCoV-OC43_orf1ab;2261 | 0 | 0 |
| NC_006213.1;YP_009555238.1;HCoV-OC43_orf1ab;2262 | 0 | 0 |
| NC_006213.1;YP_009555238.1;HCoV-OC43_orf1ab;2263 | 0 | 0 |
| NC_006213.1;YP_009555238.1;HCoV-OC43_orf1ab;2264 | 0 | 0 |
| NC_006213.1;YP_009555238.1;HCoV-OC43_orf1ab;2265 | 0 | 0 |
| NC_006213.1;YP_009555238.1;HCoV-OC43_orf1ab;2266 | 0 | 0 |
| NC_006213.1;YP_009555238.1;HCoV-OC43_orf1ab;2267 | 0 | 0 |
| NC_006213.1;YP_009555238.1;HCoV-OC43_orf1ab;2268 | 0 | 0 |
| NC_006213.1;YP_009555238.1;HCoV-OC43_orf1ab;2269 | 0 | 0 |
| NC_006213.1;YP_009555238.1;HCoV-OC43_orf1ab;2270 | 0 | 0 |
| NC_006213.1;YP_009555238.1;HCoV-OC43_orf1ab;2271 | 0.05 | 0 |
| NC_006213.1;YP_009555238.1;HCoV-OC43_orf1ab;2272 | 0.05 | 0 |
| NC_006213.1;YP_009555238.1;HCoV-OC43_orf1ab;2273 | 0 | 0 |
| NC_006213.1;YP_009555238.1;HCoV-OC43_orf1ab;2274 | 0.05 | 0 |
| NC_006213.1;YP_009555238.1;HCoV-OC43_orf1ab;2275 | 0.05 | 0 |
| NC_006213.1;YP_009555238.1;HCoV-OC43_orf1ab;2276 | 0.05 | 0.025 |
| NC_006213.1;YP_009555238.1;HCoV-OC43_orf1ab;2277 | 0.05 | 0 |
| NC_006213.1;YP_009555238.1;HCoV-OC43_orf1ab;2278 | 0 | 0.025 |
| NC_006213.1;YP_009555238.1;HCoV-OC43_orf1ab;2279 | 0 | 0 |
| NC_006213.1;YP_009555238.1;HCoV-OC43_orf1ab;2280 | 0 | 0 |
| NC_006213.1;YP_009555238.1;HCoV-OC43_orf1ab;2281 | 0 | 0 |
| NC_006213.1;YP_009555238.1;HCoV-OC43_orf1ab;2282 | 0 | 0 |
| NC_006213.1;YP_009555238.1;HCoV-OC43_orf1ab;2283 | 0 | 0 |
| NC_006213.1;YP_009555238.1;HCoV-OC43_orf1ab;2284 | 0 | 0 |
| NC_006213.1;YP_009555238.1;HCoV-OC43_orf1ab;2285 | 0 | 0 |
| NC_006213.1;YP_009555238.1;HCoV-OC43_orf1ab;2286 | 0 | 0 |
| NC_006213.1;YP_009555238.1;HCoV-OC43_orf1ab;2287 | 0.15 | 0 |
| NC_006213.1;YP_009555238.1;HCoV-OC43_orf1ab;2288 | 0.05 | 0.025 |

| | | |
|---|---|---|
| NC_006213.1;YP_009555238.1;HCoV-OC43_orf1ab;2289 | 0.05 | 0.05 |
| NC_006213.1;YP_009555238.1;HCoV-OC43_orf1ab;2290 | 0.05 | 0.025 |
| NC_006213.1;YP_009555238.1;HCoV-OC43_orf1ab;2291 | 0 | 0 |
| NC_006213.1;YP_009555238.1;HCoV-OC43_orf1ab;2292 | 0 | 0 |
| NC_006213.1;YP_009555238.1;HCoV-OC43_orf1ab;2293 | 0 | 0 |
| NC_006213.1;YP_009555238.1;HCoV-OC43_orf1ab;2294 | 0 | 0 |
| NC_006213.1;YP_009555238.1;HCoV-OC43_orf1ab;2295 | 0 | 0 |
| NC_006213.1;YP_009555238.1;HCoV-OC43_orf1ab;2296 | 0 | 0 |
| NC_006213.1;YP_009555238.1;HCoV-OC43_orf1ab;2297 | 0 | 0 |
| NC_006213.1;YP_009555238.1;HCoV-OC43_orf1ab;2298 | 0 | 0 |
| NC_006213.1;YP_009555238.1;HCoV-OC43_orf1ab;2299 | 0 | 0 |
| NC_006213.1;YP_009555238.1;HCoV-OC43_orf1ab;2300 | 0 | 0 |
| NC_006213.1;YP_009555238.1;HCoV-OC43_orf1ab;2301 | 0 | 0 |
| NC_006213.1;YP_009555238.1;HCoV-OC43_orf1ab;2302 | 0 | 0 |
| NC_006213.1;YP_009555238.1;HCoV-OC43_orf1ab;2303 | 0 | 0 |
| NC_006213.1;YP_009555238.1;HCoV-OC43_orf1ab;2304 | 0 | 0 |
| NC_006213.1;YP_009555238.1;HCoV-OC43_orf1ab;2305 | 0 | 0 |
| NC_006213.1;YP_009555238.1;HCoV-OC43_orf1ab;2306 | 0 | 0 |
| NC_006213.1;YP_009555238.1;HCoV-OC43_orf1ab;2307 | 0 | 0 |
| NC_006213.1;YP_009555238.1;HCoV-OC43_orf1ab;2308 | 0 | 0 |
| NC_006213.1;YP_009555238.1;HCoV-OC43_orf1ab;2309 | 0 | 0 |
| NC_006213.1;YP_009555238.1;HCoV-OC43_orf1ab;2310 | 0 | 0 |
| NC_006213.1;YP_009555238.1;HCoV-OC43_orf1ab;2311 | 0 | 0 |
| NC_006213.1;YP_009555238.1;HCoV-OC43_orf1ab;2312 | 0 | 0 |
| NC_006213.1;YP_009555238.1;HCoV-OC43_orf1ab;2313 | 0 | 0 |
| NC_006213.1;YP_009555238.1;HCoV-OC43_orf1ab;2314 | 0 | 0 |
| NC_006213.1;YP_009555238.1;HCoV-OC43_orf1ab;2315 | 0 | 0 |
| NC_006213.1;YP_009555238.1;HCoV-OC43_orf1ab;2316 | 0 | 0 |
| NC_006213.1;YP_009555238.1;HCoV-OC43_orf1ab;2317 | 0 | 0 |
| NC_006213.1;YP_009555238.1;HCoV-OC43_orf1ab;2318 | 0 | 0 |
| NC_006213.1;YP_009555238.1;HCoV-OC43_orf1ab;2319 | 0 | 0 |
| NC_006213.1;YP_009555238.1;HCoV-OC43_orf1ab;2320 | 0 | 0 |
| NC_006213.1;YP_009555238.1;HCoV-OC43_orf1ab;2321 | 0 | 0 |
| NC_006213.1;YP_009555238.1;HCoV-OC43_orf1ab;2322 | 0 | 0 |
| NC_006213.1;YP_009555238.1;HCoV-OC43_orf1ab;2323 | 0 | 0 |
| NC_006213.1;YP_009555238.1;HCoV-OC43_orf1ab;2324 | 0 | 0 |
| NC_006213.1;YP_009555238.1;HCoV-OC43_orf1ab;2325 | 0 | 0 |
| NC_006213.1;YP_009555238.1;HCoV-OC43_orf1ab;2326 | 0 | 0 |
| NC_006213.1;YP_009555238.1;HCoV-OC43_orf1ab;2327 | 0 | 0 |
| NC_006213.1;YP_009555238.1;HCoV-OC43_orf1ab;2328 | 0 | 0 |
| NC_006213.1;YP_009555238.1;HCoV-OC43_orf1ab;2329 | 0 | 0 |
| NC_006213.1;YP_009555238.1;HCoV-OC43_orf1ab;2330 | 0 | 0 |
| NC_006213.1;YP_009555238.1;HCoV-OC43_orf1ab;2331 | 0 | 0 |

| | | |
|---|---|---|
| NC_006213.1;YP_009555238.1;HCoV-OC43_orf1ab;2332 | 0 | 0 |
| NC_006213.1;YP_009555238.1;HCoV-OC43_orf1ab;2333 | 0 | 0 |
| NC_006213.1;YP_009555238.1;HCoV-OC43_orf1ab;2334 | 0 | 0 |
| NC_006213.1;YP_009555238.1;HCoV-OC43_orf1ab;2335 | 0 | 0 |
| NC_006213.1;YP_009555238.1;HCoV-OC43_orf1ab;2336 | 0 | 0 |
| NC_006213.1;YP_009555238.1;HCoV-OC43_orf1ab;2337 | 0 | 0 |
| NC_006213.1;YP_009555238.1;HCoV-OC43_orf1ab;2338 | 0 | 0 |
| NC_006213.1;YP_009555238.1;HCoV-OC43_orf1ab;2339 | 0 | 0 |
| NC_006213.1;YP_009555238.1;HCoV-OC43_orf1ab;2340 | 0 | 0 |
| NC_006213.1;YP_009555238.1;HCoV-OC43_orf1ab;2341 | 0 | 0 |
| NC_006213.1;YP_009555238.1;HCoV-OC43_orf1ab;2342 | 0 | 0 |
| NC_006213.1;YP_009555238.1;HCoV-OC43_orf1ab;2343 | 0 | 0 |
| NC_006213.1;YP_009555238.1;HCoV-OC43_orf1ab;2344 | 0 | 0 |
| NC_006213.1;YP_009555238.1;HCoV-OC43_orf1ab;2345 | 0 | 0 |
| NC_006213.1;YP_009555238.1;HCoV-OC43_orf1ab;2346 | 0 | 0 |
| NC_006213.1;YP_009555238.1;HCoV-OC43_orf1ab;2347 | 0 | 0 |
| NC_006213.1;YP_009555238.1;HCoV-OC43_orf1ab;2348 | 0 | 0 |
| NC_006213.1;YP_009555238.1;HCoV-OC43_orf1ab;2349 | 0 | 0 |
| NC_006213.1;YP_009555238.1;HCoV-OC43_orf1ab;2350 | 0 | 0 |
| NC_006213.1;YP_009555238.1;HCoV-OC43_orf1ab;2351 | 0 | 0 |
| NC_006213.1;YP_009555238.1;HCoV-OC43_orf1ab;2352 | 0 | 0 |
| NC_006213.1;YP_009555238.1;HCoV-OC43_orf1ab;2353 | 0 | 0 |
| NC_006213.1;YP_009555238.1;HCoV-OC43_orf1ab;2354 | 0 | 0 |
| NC_006213.1;YP_009555238.1;HCoV-OC43_orf1ab;2355 | 0 | 0 |
| NC_006213.1;YP_009555238.1;HCoV-OC43_orf1ab;2356 | 0.2 | 0.025 |
| NC_006213.1;YP_009555238.1;HCoV-OC43_orf1ab;2357 | 0 | 0 |
| NC_006213.1;YP_009555238.1;HCoV-OC43_orf1ab;2358 | 0 | 0 |
| NC_006213.1;YP_009555238.1;HCoV-OC43_orf1ab;2359 | 0 | 0 |
| NC_006213.1;YP_009555238.1;HCoV-OC43_orf1ab;2360 | 0 | 0 |
| NC_006213.1;YP_009555238.1;HCoV-OC43_orf1ab;2361 | 0 | 0 |
| NC_006213.1;YP_009555238.1;HCoV-OC43_orf1ab;2362 | 0 | 0 |
| NC_006213.1;YP_009555238.1;HCoV-OC43_orf1ab;2363 | 0 | 0 |
| NC_006213.1;YP_009555238.1;HCoV-OC43_orf1ab;2364 | 0 | 0 |
| NC_006213.1;YP_009555238.1;HCoV-OC43_orf1ab;2365 | 0 | 0 |
| NC_006213.1;YP_009555238.1;HCoV-OC43_orf1ab;2366 | 0 | 0 |
| NC_006213.1;YP_009555238.1;HCoV-OC43_orf1ab;2367 | 0 | 0 |
| NC_006213.1;YP_009555238.1;HCoV-OC43_orf1ab;2368 | 0 | 0 |
| NC_006213.1;YP_009555238.1;HCoV-OC43_orf1ab;2369 | 0 | 0 |
| NC_006213.1;YP_009555238.1;HCoV-OC43_orf1ab;2370 | 0 | 0 |
| NC_006213.1;YP_009555238.1;HCoV-OC43_orf1ab;2371 | 0 | 0 |
| NC_006213.1;YP_009555238.1;HCoV-OC43_orf1ab;2372 | 0 | 0 |
| NC_006213.1;YP_009555238.1;HCoV-OC43_orf1ab;2373 | 0 | 0 |
| NC_006213.1;YP_009555238.1;HCoV-OC43_orf1ab;2374 | 0 | 0 |

| | | |
|---|---|---|
| NC_006213.1;YP_009555238.1;HCoV-OC43_orf1ab;2375 | 0 | 0 |
| NC_006213.1;YP_009555238.1;HCoV-OC43_orf1ab;2376 | 0 | 0 |
| NC_006213.1;YP_009555238.1;HCoV-OC43_orf1ab;2377 | 0 | 0 |
| NC_006213.1;YP_009555238.1;HCoV-OC43_orf1ab;2378 | 0 | 0 |
| NC_006213.1;YP_009555238.1;HCoV-OC43_orf1ab;2379 | 0 | 0 |
| NC_006213.1;YP_009555238.1;HCoV-OC43_orf1ab;2380 | 0 | 0 |
| NC_006213.1;YP_009555238.1;HCoV-OC43_orf1ab;2381 | 0 | 0 |
| NC_006213.1;YP_009555238.1;HCoV-OC43_orf1ab;2382 | 0 | 0 |
| NC_006213.1;YP_009555238.1;HCoV-OC43_orf1ab;2383 | 0 | 0 |
| NC_006213.1;YP_009555238.1;HCoV-OC43_orf1ab;2384 | 0 | 0 |
| NC_006213.1;YP_009555238.1;HCoV-OC43_orf1ab;2385 | 0 | 0 |
| NC_006213.1;YP_009555238.1;HCoV-OC43_orf1ab;2386 | 0 | 0 |
| NC_006213.1;YP_009555238.1;HCoV-OC43_orf1ab;2387 | 0 | 0 |
| NC_006213.1;YP_009555238.1;HCoV-OC43_orf1ab;2388 | 0 | 0 |
| NC_006213.1;YP_009555238.1;HCoV-OC43_orf1ab;2389 | 0 | 0 |
| NC_006213.1;YP_009555238.1;HCoV-OC43_orf1ab;2390 | 0 | 0 |
| NC_006213.1;YP_009555238.1;HCoV-OC43_orf1ab;2391 | 0 | 0 |
| NC_006213.1;YP_009555238.1;HCoV-OC43_orf1ab;2392 | 0 | 0 |
| NC_006213.1;YP_009555238.1;HCoV-OC43_orf1ab;2393 | 0 | 0 |
| NC_006213.1;YP_009555238.1;HCoV-OC43_orf1ab;2394 | 0 | 0 |
| NC_006213.1;YP_009555238.1;HCoV-OC43_orf1ab;2395 | 0 | 0 |
| NC_006213.1;YP_009555238.1;HCoV-OC43_orf1ab;2396 | 0 | 0 |
| NC_006213.1;YP_009555238.1;HCoV-OC43_orf1ab;2397 | 0 | 0 |
| NC_006213.1;YP_009555238.1;HCoV-OC43_orf1ab;2398 | 0 | 0 |
| NC_006213.1;YP_009555238.1;HCoV-OC43_orf1ab;2399 | 0 | 0 |
| NC_006213.1;YP_009555238.1;HCoV-OC43_orf1ab;2400 | 0 | 0 |
| NC_006213.1;YP_009555238.1;HCoV-OC43_orf1ab;2401 | 0 | 0 |
| NC_006213.1;YP_009555238.1;HCoV-OC43_orf1ab;2402 | 0 | 0.025 |
| NC_006213.1;YP_009555238.1;HCoV-OC43_orf1ab;2403 | 0 | 0 |
| NC_006213.1;YP_009555238.1;HCoV-OC43_orf1ab;2404 | 0 | 0 |
| NC_006213.1;YP_009555238.1;HCoV-OC43_orf1ab;2405 | 0 | 0 |
| NC_006213.1;YP_009555238.1;HCoV-OC43_orf1ab;2406 | 0 | 0 |
| NC_006213.1;YP_009555238.1;HCoV-OC43_orf1ab;2407 | 0 | 0 |
| NC_006213.1;YP_009555238.1;HCoV-OC43_orf1ab;2408 | 0 | 0 |
| NC_006213.1;YP_009555238.1;HCoV-OC43_orf1ab;2409 | 0 | 0 |
| NC_006213.1;YP_009555238.1;HCoV-OC43_orf1ab;2410 | 0 | 0 |
| NC_006213.1;YP_009555238.1;HCoV-OC43_orf1ab;2411 | 0 | 0 |
| NC_006213.1;YP_009555238.1;HCoV-OC43_orf1ab;2412 | 0 | 0 |
| NC_006213.1;YP_009555238.1;HCoV-OC43_orf1ab;2413 | 0 | 0 |
| NC_006213.1;YP_009555238.1;HCoV-OC43_orf1ab;2414 | 0.05 | 0 |
| NC_006213.1;YP_009555238.1;HCoV-OC43_orf1ab;2415 | 0 | 0.025 |
| NC_006213.1;YP_009555238.1;HCoV-OC43_orf1ab;2416 | 0 | 0.05 |
| NC_006213.1;YP_009555238.1;HCoV-OC43_orf1ab;2417 | 0 | 0.05 |

| | | |
|---|---|---|
| NC_006213.1;YP_009555238.1;HCoV-OC43_orf1ab;2418 | 0 | 0 |
| NC_006213.1;YP_009555238.1;HCoV-OC43_orf1ab;2419 | 0 | 0 |
| NC_006213.1;YP_009555238.1;HCoV-OC43_orf1ab;2420 | 0 | 0 |
| NC_006213.1;YP_009555238.1;HCoV-OC43_orf1ab;2421 | 0 | 0 |
| NC_006213.1;YP_009555238.1;HCoV-OC43_orf1ab;2422 | 0 | 0 |
| NC_006213.1;YP_009555238.1;HCoV-OC43_orf1ab;2423 | 0 | 0 |
| NC_006213.1;YP_009555238.1;HCoV-OC43_orf1ab;2424 | 0 | 0 |
| NC_006213.1;YP_009555238.1;HCoV-OC43_orf1ab;2425 | 0 | 0 |
| NC_006213.1;YP_009555238.1;HCoV-OC43_orf1ab;2426 | 0 | 0 |
| NC_006213.1;YP_009555238.1;HCoV-OC43_orf1ab;2427 | 0 | 0 |
| NC_006213.1;YP_009555238.1;HCoV-OC43_orf1ab;2428 | 0 | 0 |
| NC_006213.1;YP_009555238.1;HCoV-OC43_orf1ab;2429 | 0 | 0 |
| NC_006213.1;YP_009555238.1;HCoV-OC43_orf1ab;2430 | 0.05 | 0 |
| NC_006213.1;YP_009555238.1;HCoV-OC43_orf1ab;2431 | 0 | 0 |
| NC_006213.1;YP_009555238.1;HCoV-OC43_orf1ab;2432 | 0 | 0 |
| NC_006213.1;YP_009555238.1;HCoV-OC43_orf1ab;2433 | 0 | 0 |
| NC_006213.1;YP_009555238.1;HCoV-OC43_orf1ab;2434 | 0 | 0 |
| NC_006213.1;YP_009555238.1;HCoV-OC43_orf1ab;2435 | 0 | 0 |
| NC_006213.1;YP_009555238.1;HCoV-OC43_orf1ab;2436 | 0 | 0 |
| NC_006213.1;YP_009555238.1;HCoV-OC43_orf1ab;2437 | 0.05 | 0 |
| NC_006213.1;YP_009555238.1;HCoV-OC43_orf1ab;2438 | 0 | 0 |
| NC_006213.1;YP_009555238.1;HCoV-OC43_orf1ab;2439 | 0 | 0.025 |
| NC_006213.1;YP_009555238.1;HCoV-OC43_orf1ab;2440 | 0 | 0 |
| NC_006213.1;YP_009555238.1;HCoV-OC43_orf1ab;2441 | 0 | 0 |
| NC_006213.1;YP_009555238.1;HCoV-OC43_orf1ab;2442 | 0 | 0 |
| NC_006213.1;YP_009555238.1;HCoV-OC43_orf1ab;2443 | 0 | 0 |
| NC_006213.1;YP_009555238.1;HCoV-OC43_orf1ab;2444 | 0 | 0 |
| NC_006213.1;YP_009555238.1;HCoV-OC43_orf1ab;2445 | 0 | 0 |
| NC_006213.1;YP_009555238.1;HCoV-OC43_orf1ab;2446 | 0 | 0 |
| NC_006213.1;YP_009555238.1;HCoV-OC43_orf1ab;2447 | 0 | 0 |
| NC_006213.1;YP_009555238.1;HCoV-OC43_orf1ab;2448 | 0 | 0 |
| NC_006213.1;YP_009555238.1;HCoV-OC43_orf1ab;2449 | 0 | 0 |
| NC_006213.1;YP_009555238.1;HCoV-OC43_orf1ab;2450 | 0 | 0 |
| NC_006213.1;YP_009555238.1;HCoV-OC43_orf1ab;2451 | 0.05 | 0.05 |
| NC_006213.1;YP_009555238.1;HCoV-OC43_orf1ab;2452 | 0.05 | 0.025 |
| NC_006213.1;YP_009555238.1;HCoV-OC43_orf1ab;2453 | 0.05 | 0.275 |
| NC_006213.1;YP_009555238.1;HCoV-OC43_orf1ab;2454 | 0.05 | 0.05 |
| NC_006213.1;YP_009555238.1;HCoV-OC43_orf1ab;2455 | 0.05 | 0.05 |
| NC_006213.1;YP_009555238.1;HCoV-OC43_orf1ab;2456 | 0 | 0.05 |
| NC_006213.1;YP_009555238.1;HCoV-OC43_orf1ab;2457 | 0 | 0.025 |
| NC_006213.1;YP_009555238.1;HCoV-OC43_orf1ab;2458 | 0 | 0.05 |
| NC_006213.1;YP_009555238.1;HCoV-OC43_orf1ab;2459 | 0 | 0.05 |
| NC_006213.1;YP_009555238.1;HCoV-OC43_orf1ab;2460 | 0 | 0.025 |

| | | |
|---|---|---|
| NC_006213.1;YP_009555238.1;HCoV-OC43_orf1ab;2461 | 0 | 0.025 |
| NC_006213.1;YP_009555238.1;HCoV-OC43_orf1ab;2462 | 0 | 0.025 |
| NC_006213.1;YP_009555238.1;HCoV-OC43_orf1ab;2463 | 0 | 0 |
| NC_006213.1;YP_009555238.1;HCoV-OC43_orf1ab;2464 | 0 | 0 |
| NC_006213.1;YP_009555238.1;HCoV-OC43_orf1ab;2465 | 0 | 0 |
| NC_006213.1;YP_009555238.1;HCoV-OC43_orf1ab;2466 | 0 | 0.075 |
| NC_006213.1;YP_009555238.1;HCoV-OC43_orf1ab;2467 | 0 | 0 |
| NC_006213.1;YP_009555238.1;HCoV-OC43_orf1ab;2468 | 0 | 0.025 |
| NC_006213.1;YP_009555238.1;HCoV-OC43_orf1ab;2469 | 0 | 0 |
| NC_006213.1;YP_009555238.1;HCoV-OC43_orf1ab;2470 | 0 | 0 |
| NC_006213.1;YP_009555238.1;HCoV-OC43_orf1ab;2471 | 0 | 0 |
| NC_006213.1;YP_009555238.1;HCoV-OC43_orf1ab;2472 | 0.05 | 0.025 |
| NC_006213.1;YP_009555238.1;HCoV-OC43_orf1ab;2473 | 0.05 | 0 |
| NC_006213.1;YP_009555238.1;HCoV-OC43_orf1ab;2474 | 0.1 | 0 |
| NC_006213.1;YP_009555238.1;HCoV-OC43_orf1ab;2475 | 0 | 0 |
| NC_006213.1;YP_009555238.1;HCoV-OC43_orf1ab;2476 | 0 | 0 |
| NC_006213.1;YP_009555238.1;HCoV-OC43_orf1ab;2477 | 0 | 0 |
| NC_006213.1;YP_009555238.1;HCoV-OC43_orf1ab;2478 | 0 | 0 |
| NC_006213.1;YP_009555238.1;HCoV-OC43_orf1ab;2479 | 0.05 | 0 |
| NC_006213.1;YP_009555238.1;HCoV-OC43_orf1ab;2480 | 0 | 0 |
| NC_006213.1;YP_009555238.1;HCoV-OC43_orf1ab;2481 | 0 | 0 |
| NC_006213.1;YP_009555238.1;HCoV-OC43_orf1ab;2482 | 0 | 0 |
| NC_006213.1;YP_009555238.1;HCoV-OC43_orf1ab;2483 | 0 | 0.025 |
| NC_006213.1;YP_009555238.1;HCoV-OC43_orf1ab;2484 | 0 | 0 |
| NC_006213.1;YP_009555238.1;HCoV-OC43_orf1ab;2485 | 0 | 0 |
| NC_006213.1;YP_009555238.1;HCoV-OC43_orf1ab;2486 | 0 | 0 |
| NC_006213.1;YP_009555238.1;HCoV-OC43_orf1ab;2487 | 0 | 0 |
| NC_006213.1;YP_009555238.1;HCoV-OC43_orf1ab;2488 | 0 | 0.025 |
| NC_006213.1;YP_009555238.1;HCoV-OC43_orf1ab;2489 | 0 | 0.025 |
| NC_006213.1;YP_009555238.1;HCoV-OC43_orf1ab;2490 | 0.05 | 0.025 |
| NC_006213.1;YP_009555238.1;HCoV-OC43_orf1ab;2491 | 0.1 | 0.025 |
| NC_006213.1;YP_009555238.1;HCoV-OC43_orf1ab;2492 | 0.1 | 0.05 |
| NC_006213.1;YP_009555238.1;HCoV-OC43_orf1ab;2493 | 0.05 | 0.05 |
| NC_006213.1;YP_009555238.1;HCoV-OC43_orf1ab;2494 | 0.15 | 0.05 |
| NC_006213.1;YP_009555238.1;HCoV-OC43_orf1ab;2495 | 0.15 | 0 |
| NC_006213.1;YP_009555238.1;HCoV-OC43_orf1ab;2496 | 0 | 0 |
| NC_006213.1;YP_009555238.1;HCoV-OC43_orf1ab;2497 | 0 | 0 |
| NC_006213.1;YP_009555238.1;HCoV-OC43_orf1ab;2498 | 0 | 0 |
| NC_006213.1;YP_009555238.1;HCoV-OC43_orf1ab;2499 | 0 | 0.075 |
| NC_006213.1;YP_009555238.1;HCoV-OC43_orf1ab;2500 | 0 | 0 |
| NC_006213.1;YP_009555238.1;HCoV-OC43_orf1ab;2501 | 0 | 0 |
| NC_006213.1;YP_009555238.1;HCoV-OC43_orf1ab;2502 | 0 | 0.05 |
| NC_006213.1;YP_009555238.1;HCoV-OC43_orf1ab;2503 | 0 | 0 |

| | | |
|---|---|---|
| NC_006213.1;YP_009555238.1;HCoV-OC43_orf1ab;2504 | 0 | 0 |
| NC_006213.1;YP_009555238.1;HCoV-OC43_orf1ab;2505 | 0 | 0 |
| NC_006213.1;YP_009555238.1;HCoV-OC43_orf1ab;2506 | 0 | 0 |
| NC_006213.1;YP_009555238.1;HCoV-OC43_orf1ab;2507 | 0 | 0 |
| NC_006213.1;YP_009555238.1;HCoV-OC43_orf1ab;2508 | 0 | 0 |
| NC_006213.1;YP_009555238.1;HCoV-OC43_orf1ab;2509 | 0.15 | 0.075 |
| NC_006213.1;YP_009555238.1;HCoV-OC43_orf1ab;2510 | 0.15 | 0.025 |
| NC_006213.1;YP_009555238.1;HCoV-OC43_orf1ab;2511 | 0 | 0.025 |
| NC_006213.1;YP_009555238.1;HCoV-OC43_orf1ab;2512 | 0.05 | 0 |
| NC_006213.1;YP_009555238.1;HCoV-OC43_orf1ab;2513 | 0 | 0 |
| NC_006213.1;YP_009555238.1;HCoV-OC43_orf1ab;2514 | 0 | 0 |
| NC_006213.1;YP_009555238.1;HCoV-OC43_orf1ab;2515 | 0 | 0 |
| NC_006213.1;YP_009555238.1;HCoV-OC43_orf1ab;2516 | 0 | 0 |
| NC_006213.1;YP_009555238.1;HCoV-OC43_orf1ab;2517 | 0.05 | 0 |
| NC_006213.1;YP_009555238.1;HCoV-OC43_orf1ab;2518 | 0 | 0 |
| NC_006213.1;YP_009555238.1;HCoV-OC43_orf1ab;2519 | 0 | 0 |
| NC_006213.1;YP_009555238.1;HCoV-OC43_orf1ab;2520 | 0 | 0 |
| NC_006213.1;YP_009555238.1;HCoV-OC43_orf1ab;2521 | 0 | 0 |
| NC_006213.1;YP_009555238.1;HCoV-OC43_orf1ab;2522 | 0.05 | 0 |
| NC_006213.1;YP_009555238.1;HCoV-OC43_orf1ab;2523 | 0 | 0 |
| NC_006213.1;YP_009555238.1;HCoV-OC43_orf1ab;2524 | 0.05 | 0 |
| NC_006213.1;YP_009555238.1;HCoV-OC43_orf1ab;2525 | 0.05 | 0 |
| NC_006213.1;YP_009555238.1;HCoV-OC43_orf1ab;2526 | 0.05 | 0 |
| NC_006213.1;YP_009555238.1;HCoV-OC43_orf1ab;2527 | 0.1 | 0 |
| NC_006213.1;YP_009555238.1;HCoV-OC43_orf1ab;2528 | 0.1 | 0.025 |
| NC_006213.1;YP_009555238.1;HCoV-OC43_orf1ab;2529 | 0.1 | 0 |
| NC_006213.1;YP_009555238.1;HCoV-OC43_orf1ab;2530 | 0.05 | 0.05 |
| NC_006213.1;YP_009555238.1;HCoV-OC43_orf1ab;2531 | 0.05 | 0 |
| NC_006213.1;YP_009555238.1;HCoV-OC43_orf1ab;2532 | 0 | 0 |
| NC_006213.1;YP_009555238.1;HCoV-OC43_orf1ab;2533 | 0 | 0 |
| NC_006213.1;YP_009555238.1;HCoV-OC43_orf1ab;2534 | 0.05 | 0.025 |
| NC_006213.1;YP_009555238.1;HCoV-OC43_orf1ab;2535 | 0 | 0 |
| NC_006213.1;YP_009555238.1;HCoV-OC43_orf1ab;2536 | 0 | 0 |
| NC_006213.1;YP_009555238.1;HCoV-OC43_orf1ab;2537 | 0 | 0 |
| NC_006213.1;YP_009555238.1;HCoV-OC43_orf1ab;2538 | 0 | 0 |
| NC_006213.1;YP_009555238.1;HCoV-OC43_orf1ab;2539 | 0 | 0 |
| NC_006213.1;YP_009555238.1;HCoV-OC43_orf1ab;2540 | 0 | 0 |
| NC_006213.1;YP_009555238.1;HCoV-OC43_orf1ab;2541 | 0 | 0 |
| NC_006213.1;YP_009555238.1;HCoV-OC43_orf1ab;2542 | 0 | 0 |
| NC_006213.1;YP_009555238.1;HCoV-OC43_orf1ab;2543 | 0 | 0 |
| NC_006213.1;YP_009555238.1;HCoV-OC43_orf1ab;2544 | 0 | 0 |
| NC_006213.1;YP_009555238.1;HCoV-OC43_orf1ab;2545 | 0 | 0 |
| NC_006213.1;YP_009555238.1;HCoV-OC43_orf1ab;2546 | 0 | 0 |

| | | |
|---|---|---|
| NC_006213.1;YP_009555238.1;HCoV-OC43_orf1ab;2547 | 0 | 0 |
| NC_006213.1;YP_009555238.1;HCoV-OC43_orf1ab;2548 | 0 | 0 |
| NC_006213.1;YP_009555238.1;HCoV-OC43_orf1ab;2549 | 0 | 0 |
| NC_006213.1;YP_009555238.1;HCoV-OC43_orf1ab;2550 | 0 | 0 |
| NC_006213.1;YP_009555238.1;HCoV-OC43_orf1ab;2551 | 0 | 0 |
| NC_006213.1;YP_009555238.1;HCoV-OC43_orf1ab;2552 | 0 | 0 |
| NC_006213.1;YP_009555238.1;HCoV-OC43_orf1ab;2553 | 0 | 0 |
| NC_006213.1;YP_009555238.1;HCoV-OC43_orf1ab;2554 | 0 | 0.025 |
| NC_006213.1;YP_009555238.1;HCoV-OC43_orf1ab;2555 | 0.05 | 0.025 |
| NC_006213.1;YP_009555238.1;HCoV-OC43_orf1ab;2556 | 0.05 | 0.025 |
| NC_006213.1;YP_009555238.1;HCoV-OC43_orf1ab;2557 | 0.05 | 0 |
| NC_006213.1;YP_009555238.1;HCoV-OC43_orf1ab;2558 | 0.05 | 0 |
| NC_006213.1;YP_009555238.1;HCoV-OC43_orf1ab;2559 | 0.05 | 0.025 |
| NC_006213.1;YP_009555238.1;HCoV-OC43_orf1ab;2560 | 0 | 0 |
| NC_006213.1;YP_009555238.1;HCoV-OC43_orf1ab;2561 | 0 | 0 |
| NC_006213.1;YP_009555238.1;HCoV-OC43_orf1ab;2562 | 0 | 0 |
| NC_006213.1;YP_009555238.1;HCoV-OC43_orf1ab;2563 | 0 | 0 |
| NC_006213.1;YP_009555238.1;HCoV-OC43_orf1ab;2564 | 0 | 0 |
| NC_006213.1;YP_009555238.1;HCoV-OC43_orf1ab;2565 | 0 | 0 |
| NC_006213.1;YP_009555238.1;HCoV-OC43_orf1ab;2566 | 0 | 0 |
| NC_006213.1;YP_009555238.1;HCoV-OC43_orf1ab;2567 | 0 | 0 |
| NC_006213.1;YP_009555238.1;HCoV-OC43_orf1ab;2568 | 0 | 0 |
| NC_006213.1;YP_009555238.1;HCoV-OC43_orf1ab;2569 | 0 | 0 |
| NC_006213.1;YP_009555238.1;HCoV-OC43_orf1ab;2570 | 0 | 0 |
| NC_006213.1;YP_009555238.1;HCoV-OC43_orf1ab;2571 | 0 | 0 |
| NC_006213.1;YP_009555238.1;HCoV-OC43_orf1ab;2572 | 0 | 0 |
| NC_006213.1;YP_009555238.1;HCoV-OC43_orf1ab;2573 | 0 | 0 |
| NC_006213.1;YP_009555238.1;HCoV-OC43_orf1ab;2574 | 0 | 0 |
| NC_006213.1;YP_009555238.1;HCoV-OC43_orf1ab;2575 | 0 | 0 |
| NC_006213.1;YP_009555238.1;HCoV-OC43_orf1ab;2576 | 0 | 0.025 |
| NC_006213.1;YP_009555238.1;HCoV-OC43_orf1ab;2577 | 0 | 0.025 |
| NC_006213.1;YP_009555238.1;HCoV-OC43_orf1ab;2578 | 0 | 0 |
| NC_006213.1;YP_009555238.1;HCoV-OC43_orf1ab;2579 | 0 | 0 |
| NC_006213.1;YP_009555238.1;HCoV-OC43_orf1ab;2580 | 0 | 0 |
| NC_006213.1;YP_009555238.1;HCoV-OC43_orf1ab;2581 | 0.1 | 0.075 |
| NC_006213.1;YP_009555238.1;HCoV-OC43_orf1ab;2582 | 0.1 | 0.075 |
| NC_006213.1;YP_009555238.1;HCoV-OC43_orf1ab;2583 | 0.1 | 0.075 |
| NC_006213.1;YP_009555238.1;HCoV-OC43_orf1ab;2584 | 0.1 | 0.025 |
| NC_006213.1;YP_009555238.1;HCoV-OC43_orf1ab;2585 | 0.1 | 0.05 |
| NC_006213.1;YP_009555238.1;HCoV-OC43_orf1ab;2586 | 0.15 | 0.05 |
| NC_006213.1;YP_009555238.1;HCoV-OC43_orf1ab;2587 | 0.15 | 0.1 |
| NC_006213.1;YP_009555238.1;HCoV-OC43_orf1ab;2588 | 0.1 | 0.025 |
| NC_006213.1;YP_009555238.1;HCoV-OC43_orf1ab;2589 | 0 | 0 |

| | | |
|---|---|---|
| NC_006213.1;YP_009555238.1;HCoV-OC43_orf1ab;2590 | 0 | 0 |
| NC_006213.1;YP_009555238.1;HCoV-OC43_orf1ab;2591 | 0 | 0 |
| NC_006213.1;YP_009555238.1;HCoV-OC43_orf1ab;2592 | 0 | 0 |
| NC_006213.1;YP_009555238.1;HCoV-OC43_orf1ab;2593 | 0 | 0 |
| NC_006213.1;YP_009555238.1;HCoV-OC43_orf1ab;2594 | 0 | 0.025 |
| NC_006213.1;YP_009555238.1;HCoV-OC43_orf1ab;2595 | 0 | 0.025 |
| NC_006213.1;YP_009555238.1;HCoV-OC43_orf1ab;2596 | 0 | 0.025 |
| NC_006213.1;YP_009555238.1;HCoV-OC43_orf1ab;2597 | 0 | 0.025 |
| NC_006213.1;YP_009555238.1;HCoV-OC43_orf1ab;2598 | 0 | 0 |
| NC_006213.1;YP_009555238.1;HCoV-OC43_orf1ab;2599 | 0 | 0 |
| NC_006213.1;YP_009555238.1;HCoV-OC43_orf1ab;2600 | 0 | 0 |
| NC_006213.1;YP_009555238.1;HCoV-OC43_orf1ab;2601 | 0 | 0 |
| NC_006213.1;YP_009555238.1;HCoV-OC43_orf1ab;2602 | 0.05 | 0 |
| NC_006213.1;YP_009555238.1;HCoV-OC43_orf1ab;2603 | 0.05 | 0 |
| NC_006213.1;YP_009555238.1;HCoV-OC43_orf1ab;2604 | 0 | 0 |
| NC_006213.1;YP_009555238.1;HCoV-OC43_orf1ab;2605 | 0 | 0 |
| NC_006213.1;YP_009555238.1;HCoV-OC43_orf1ab;2606 | 0 | 0 |
| NC_006213.1;YP_009555238.1;HCoV-OC43_orf1ab;2607 | 0 | 0 |
| NC_006213.1;YP_009555238.1;HCoV-OC43_orf1ab;2608 | 0 | 0 |
| NC_006213.1;YP_009555238.1;HCoV-OC43_orf1ab;2609 | 0 | 0 |
| NC_006213.1;YP_009555238.1;HCoV-OC43_orf1ab;2610 | 0 | 0 |
| NC_006213.1;YP_009555238.1;HCoV-OC43_orf1ab;2611 | 0 | 0 |
| NC_006213.1;YP_009555238.1;HCoV-OC43_orf1ab;2612 | 0 | 0 |
| NC_006213.1;YP_009555238.1;HCoV-OC43_orf1ab;2613 | 0 | 0 |
| NC_006213.1;YP_009555238.1;HCoV-OC43_orf1ab;2614 | 0 | 0 |
| NC_006213.1;YP_009555238.1;HCoV-OC43_orf1ab;2615 | 0 | 0 |
| NC_006213.1;YP_009555238.1;HCoV-OC43_orf1ab;2616 | 0 | 0 |
| NC_006213.1;YP_009555238.1;HCoV-OC43_orf1ab;2617 | 0 | 0 |
| NC_006213.1;YP_009555238.1;HCoV-OC43_orf1ab;2618 | 0 | 0 |
| NC_006213.1;YP_009555238.1;HCoV-OC43_orf1ab;2619 | 0 | 0 |
| NC_006213.1;YP_009555238.1;HCoV-OC43_orf1ab;2620 | 0 | 0 |
| NC_006213.1;YP_009555238.1;HCoV-OC43_orf1ab;2621 | 0 | 0 |
| NC_006213.1;YP_009555238.1;HCoV-OC43_orf1ab;2622 | 0 | 0 |
| NC_006213.1;YP_009555238.1;HCoV-OC43_orf1ab;2623 | 0 | 0 |
| NC_006213.1;YP_009555238.1;HCoV-OC43_orf1ab;2624 | 0 | 0 |
| NC_006213.1;YP_009555238.1;HCoV-OC43_orf1ab;2625 | 0 | 0 |
| NC_006213.1;YP_009555238.1;HCoV-OC43_orf1ab;2626 | 0 | 0 |
| NC_006213.1;YP_009555238.1;HCoV-OC43_orf1ab;2627 | 0 | 0 |
| NC_006213.1;YP_009555238.1;HCoV-OC43_orf1ab;2628 | 0 | 0 |
| NC_006213.1;YP_009555238.1;HCoV-OC43_orf1ab;2629 | 0 | 0.025 |
| NC_006213.1;YP_009555238.1;HCoV-OC43_orf1ab;2630 | 0 | 0 |
| NC_006213.1;YP_009555238.1;HCoV-OC43_orf1ab;2631 | 0 | 0 |
| NC_006213.1;YP_009555238.1;HCoV-OC43_orf1ab;2632 | 0 | 0 |

| | | |
|---|---|---|
| NC_006213.1;YP_009555238.1;HCoV-OC43_orf1ab;2633 | 0 | 0 |
| NC_006213.1;YP_009555238.1;HCoV-OC43_orf1ab;2634 | 0 | 0 |
| NC_006213.1;YP_009555238.1;HCoV-OC43_orf1ab;2635 | 0 | 0 |
| NC_006213.1;YP_009555238.1;HCoV-OC43_orf1ab;2636 | 0 | 0 |
| NC_006213.1;YP_009555238.1;HCoV-OC43_orf1ab;2637 | 0 | 0 |
| NC_006213.1;YP_009555238.1;HCoV-OC43_orf1ab;2638 | 0 | 0 |
| NC_006213.1;YP_009555238.1;HCoV-OC43_orf1ab;2639 | 0 | 0 |
| NC_006213.1;YP_009555238.1;HCoV-OC43_orf1ab;2640 | 0 | 0 |
| NC_006213.1;YP_009555238.1;HCoV-OC43_orf1ab;2641 | 0.05 | 0 |
| NC_006213.1;YP_009555238.1;HCoV-OC43_orf1ab;2642 | 0 | 0 |
| NC_006213.1;YP_009555238.1;HCoV-OC43_orf1ab;2643 | 0 | 0 |
| NC_006213.1;YP_009555238.1;HCoV-OC43_orf1ab;2644 | 0 | 0 |
| NC_006213.1;YP_009555238.1;HCoV-OC43_orf1ab;2645 | 0 | 0 |
| NC_006213.1;YP_009555238.1;HCoV-OC43_orf1ab;2646 | 0 | 0 |
| NC_006213.1;YP_009555238.1;HCoV-OC43_orf1ab;2647 | 0 | 0.025 |
| NC_006213.1;YP_009555238.1;HCoV-OC43_orf1ab;2648 | 0 | 0 |
| NC_006213.1;YP_009555238.1;HCoV-OC43_orf1ab;2649 | 0 | 0.05 |
| NC_006213.1;YP_009555238.1;HCoV-OC43_orf1ab;2650 | 0 | 0 |
| NC_006213.1;YP_009555238.1;HCoV-OC43_orf1ab;2651 | 0 | 0 |
| NC_006213.1;YP_009555238.1;HCoV-OC43_orf1ab;2652 | 0.1 | 0.125 |
| NC_006213.1;YP_009555238.1;HCoV-OC43_orf1ab;2653 | 0.05 | 0 |
| NC_006213.1;YP_009555238.1;HCoV-OC43_orf1ab;2654 | 0.05 | 0.025 |
| NC_006213.1;YP_009555238.1;HCoV-OC43_orf1ab;2655 | 0.05 | 0.05 |
| NC_006213.1;YP_009555238.1;HCoV-OC43_orf1ab;2656 | 0 | 0.05 |
| NC_006213.1;YP_009555238.1;HCoV-OC43_orf1ab;2657 | 0.05 | 0.15 |
| NC_006213.1;YP_009555238.1;HCoV-OC43_orf1ab;2658 | 0.05 | 0.15 |
| NC_006213.1;YP_009555238.1;HCoV-OC43_orf1ab;2659 | 0 | 0.05 |
| NC_006213.1;YP_009555238.1;HCoV-OC43_orf1ab;2660 | 0 | 0.075 |
| NC_006213.1;YP_009555238.1;HCoV-OC43_orf1ab;2661 | 0 | 0.1 |
| NC_006213.1;YP_009555238.1;HCoV-OC43_orf1ab;2662 | 0.05 | 0.05 |
| NC_006213.1;YP_009555238.1;HCoV-OC43_orf1ab;2663 | 0 | 0.025 |
| NC_006213.1;YP_009555238.1;HCoV-OC43_orf1ab;2664 | 0 | 0 |
| NC_006213.1;YP_009555238.1;HCoV-OC43_orf1ab;2665 | 0 | 0 |
| NC_006213.1;YP_009555238.1;HCoV-OC43_orf1ab;2666 | 0 | 0 |
| NC_006213.1;YP_009555238.1;HCoV-OC43_orf1ab;2667 | 0 | 0 |
| NC_006213.1;YP_009555238.1;HCoV-OC43_orf1ab;2668 | 0 | 0.05 |
| NC_006213.1;YP_009555238.1;HCoV-OC43_orf1ab;2669 | 0 | 0.025 |
| NC_006213.1;YP_009555238.1;HCoV-OC43_orf1ab;2670 | 0 | 0 |
| NC_006213.1;YP_009555238.1;HCoV-OC43_orf1ab;2671 | 0 | 0.025 |
| NC_006213.1;YP_009555238.1;HCoV-OC43_orf1ab;2672 | 0 | 0.05 |
| NC_006213.1;YP_009555238.1;HCoV-OC43_orf1ab;2673 | 0.05 | 0.025 |
| NC_006213.1;YP_009555238.1;HCoV-OC43_orf1ab;2674 | 0 | 0.05 |
| NC_006213.1;YP_009555238.1;HCoV-OC43_orf1ab;2675 | 0 | 0 |

| | | |
|---|---|---|
| NC_006213.1;YP_009555238.1;HCoV-OC43_orf1ab;2676 | 0 | 0 |
| NC_006213.1;YP_009555238.1;HCoV-OC43_orf1ab;2677 | 0 | 0 |
| NC_006213.1;YP_009555238.1;HCoV-OC43_orf1ab;2678 | 0.05 | 0 |
| NC_006213.1;YP_009555238.1;HCoV-OC43_orf1ab;2679 | 0 | 0.025 |
| NC_006213.1;YP_009555238.1;HCoV-OC43_orf1ab;2680 | 0 | 0 |
| NC_006213.1;YP_009555238.1;HCoV-OC43_orf1ab;2681 | 0 | 0 |
| NC_006213.1;YP_009555238.1;HCoV-OC43_orf1ab;2682 | 0 | 0 |
| NC_006213.1;YP_009555238.1;HCoV-OC43_orf1ab;2683 | 0 | 0 |
| NC_006213.1;YP_009555238.1;HCoV-OC43_orf1ab;2684 | 0 | 0 |
| NC_006213.1;YP_009555238.1;HCoV-OC43_orf1ab;2685 | 0 | 0 |
| NC_006213.1;YP_009555238.1;HCoV-OC43_orf1ab;2686 | 0 | 0 |
| NC_006213.1;YP_009555238.1;HCoV-OC43_orf1ab;2687 | 0 | 0 |
| NC_006213.1;YP_009555238.1;HCoV-OC43_orf1ab;2688 | 0 | 0 |
| NC_006213.1;YP_009555238.1;HCoV-OC43_orf1ab;2689 | 0 | 0 |
| NC_006213.1;YP_009555238.1;HCoV-OC43_orf1ab;2690 | 0 | 0 |
| NC_006213.1;YP_009555238.1;HCoV-OC43_orf1ab;2691 | 0 | 0 |
| NC_006213.1;YP_009555238.1;HCoV-OC43_orf1ab;2692 | 0 | 0 |
| NC_006213.1;YP_009555238.1;HCoV-OC43_orf1ab;2693 | 0 | 0 |
| NC_006213.1;YP_009555238.1;HCoV-OC43_orf1ab;2694 | 0 | 0 |
| NC_006213.1;YP_009555238.1;HCoV-OC43_orf1ab;2695 | 0.05 | 0 |
| NC_006213.1;YP_009555238.1;HCoV-OC43_orf1ab;2696 | 0 | 0 |
| NC_006213.1;YP_009555238.1;HCoV-OC43_orf1ab;2697 | 0 | 0 |
| NC_006213.1;YP_009555238.1;HCoV-OC43_orf1ab;2698 | 0 | 0 |
| NC_006213.1;YP_009555238.1;HCoV-OC43_orf1ab;2699 | 0.05 | 0 |
| NC_006213.1;YP_009555238.1;HCoV-OC43_orf1ab;2700 | 0.05 | 0 |
| NC_006213.1;YP_009555238.1;HCoV-OC43_orf1ab;2701 | 0.1 | 0.075 |
| NC_006213.1;YP_009555238.1;HCoV-OC43_orf1ab;2702 | 0.2 | 0.075 |
| NC_006213.1;YP_009555238.1;HCoV-OC43_orf1ab;2703 | 0.1 | 0 |
| NC_006213.1;YP_009555238.1;HCoV-OC43_orf1ab;2704 | 0.05 | 0 |
| NC_006213.1;YP_009555238.1;HCoV-OC43_orf1ab;2705 | 0 | 0 |
| NC_006213.1;YP_009555238.1;HCoV-OC43_orf1ab;2706 | 0 | 0 |
| NC_006213.1;YP_009555238.1;HCoV-OC43_orf1ab;2707 | 0 | 0 |
| NC_006213.1;YP_009555238.1;HCoV-OC43_orf1ab;2708 | 0 | 0 |
| NC_006213.1;YP_009555238.1;HCoV-OC43_orf1ab;2709 | 0 | 0 |
| NC_006213.1;YP_009555238.1;HCoV-OC43_orf1ab;2710 | 0 | 0 |
| NC_006213.1;YP_009555238.1;HCoV-OC43_orf1ab;2711 | 0 | 0 |
| NC_006213.1;YP_009555238.1;HCoV-OC43_orf1ab;2712 | 0 | 0 |
| NC_006213.1;YP_009555238.1;HCoV-OC43_orf1ab;2713 | 0 | 0 |
| NC_006213.1;YP_009555238.1;HCoV-OC43_orf1ab;2714 | 0 | 0 |
| NC_006213.1;YP_009555238.1;HCoV-OC43_orf1ab;2715 | 0 | 0 |
| NC_006213.1;YP_009555238.1;HCoV-OC43_orf1ab;2716 | 0 | 0 |
| NC_006213.1;YP_009555238.1;HCoV-OC43_orf1ab;2717 | 0 | 0 |
| NC_006213.1;YP_009555238.1;HCoV-OC43_orf1ab;2718 | 0 | 0 |

| | | |
|---|---|---|
| NC_006213.1;YP_009555238.1;HCoV-OC43_orf1ab;2719 | 0 | 0 |
| NC_006213.1;YP_009555238.1;HCoV-OC43_orf1ab;2720 | 0 | 0 |
| NC_006213.1;YP_009555238.1;HCoV-OC43_orf1ab;2721 | 0.05 | 0 |
| NC_006213.1;YP_009555238.1;HCoV-OC43_orf1ab;2722 | 0.05 | 0 |
| NC_006213.1;YP_009555238.1;HCoV-OC43_orf1ab;2723 | 0 | 0 |
| NC_006213.1;YP_009555238.1;HCoV-OC43_orf1ab;2724 | 0 | 0 |
| NC_006213.1;YP_009555238.1;HCoV-OC43_orf1ab;2725 | 0 | 0 |
| NC_006213.1;YP_009555238.1;HCoV-OC43_orf1ab;2726 | 0 | 0 |
| NC_006213.1;YP_009555238.1;HCoV-OC43_orf1ab;2727 | 0 | 0 |
| NC_006213.1;YP_009555238.1;HCoV-OC43_orf1ab;2728 | 0 | 0 |
| NC_006213.1;YP_009555238.1;HCoV-OC43_orf1ab;2729 | 0 | 0.025 |
| NC_006213.1;YP_009555238.1;HCoV-OC43_orf1ab;2730 | 0 | 0 |
| NC_006213.1;YP_009555238.1;HCoV-OC43_orf1ab;2731 | 0 | 0 |
| NC_006213.1;YP_009555238.1;HCoV-OC43_orf1ab;2732 | 0 | 0.05 |
| NC_006213.1;YP_009555238.1;HCoV-OC43_orf1ab;2733 | 0 | 0 |
| NC_006213.1;YP_009555238.1;HCoV-OC43_orf1ab;2734 | 0 | 0 |
| NC_006213.1;YP_009555238.1;HCoV-OC43_orf1ab;2735 | 0 | 0 |
| NC_006213.1;YP_009555238.1;HCoV-OC43_orf1ab;2736 | 0 | 0 |
| NC_006213.1;YP_009555238.1;HCoV-OC43_orf1ab;2737 | 0 | 0 |
| NC_006213.1;YP_009555238.1;HCoV-OC43_orf1ab;2738 | 0 | 0 |
| NC_006213.1;YP_009555238.1;HCoV-OC43_orf1ab;2739 | 0 | 0 |
| NC_006213.1;YP_009555238.1;HCoV-OC43_orf1ab;2740 | 0 | 0 |
| NC_006213.1;YP_009555238.1;HCoV-OC43_orf1ab;2741 | 0 | 0 |
| NC_006213.1;YP_009555238.1;HCoV-OC43_orf1ab;2742 | 0 | 0 |
| NC_006213.1;YP_009555238.1;HCoV-OC43_orf1ab;2743 | 0 | 0 |
| NC_006213.1;YP_009555238.1;HCoV-OC43_orf1ab;2744 | 0 | 0 |
| NC_006213.1;YP_009555238.1;HCoV-OC43_orf1ab;2745 | 0 | 0 |
| NC_006213.1;YP_009555238.1;HCoV-OC43_orf1ab;2746 | 0 | 0 |
| NC_006213.1;YP_009555238.1;HCoV-OC43_orf1ab;2747 | 0 | 0 |
| NC_006213.1;YP_009555238.1;HCoV-OC43_orf1ab;2748 | 0 | 0 |
| NC_006213.1;YP_009555238.1;HCoV-OC43_orf1ab;2749 | 0 | 0 |
| NC_006213.1;YP_009555238.1;HCoV-OC43_orf1ab;2750 | 0 | 0 |
| NC_006213.1;YP_009555238.1;HCoV-OC43_orf1ab;2751 | 0 | 0 |
| NC_006213.1;YP_009555238.1;HCoV-OC43_orf1ab;2752 | 0 | 0 |
| NC_006213.1;YP_009555238.1;HCoV-OC43_orf1ab;2753 | 0 | 0 |
| NC_006213.1;YP_009555238.1;HCoV-OC43_orf1ab;2754 | 0 | 0 |
| NC_006213.1;YP_009555238.1;HCoV-OC43_orf1ab;2755 | 0 | 0 |
| NC_006213.1;YP_009555238.1;HCoV-OC43_orf1ab;2756 | 0 | 0 |
| NC_006213.1;YP_009555238.1;HCoV-OC43_orf1ab;2757 | 0 | 0 |
| NC_006213.1;YP_009555238.1;HCoV-OC43_orf1ab;2758 | 0 | 0 |
| NC_006213.1;YP_009555238.1;HCoV-OC43_orf1ab;2759 | 0 | 0 |
| NC_006213.1;YP_009555238.1;HCoV-OC43_orf1ab;2760 | 0 | 0 |
| NC_006213.1;YP_009555238.1;HCoV-OC43_orf1ab;2761 | 0 | 0 |

| | | |
|---|---|---|
| NC_006213.1;YP_009555238.1;HCoV-OC43_orf1ab;2762 | 0 | 0 |
| NC_006213.1;YP_009555238.1;HCoV-OC43_orf1ab;2763 | 0 | 0 |
| NC_006213.1;YP_009555238.1;HCoV-OC43_orf1ab;2764 | 0 | 0 |
| NC_006213.1;YP_009555238.1;HCoV-OC43_orf1ab;2765 | 0 | 0 |
| NC_006213.1;YP_009555238.1;HCoV-OC43_orf1ab;2766 | 0 | 0 |
| NC_006213.1;YP_009555238.1;HCoV-OC43_orf1ab;2767 | 0 | 0 |
| NC_006213.1;YP_009555238.1;HCoV-OC43_orf1ab;2768 | 0 | 0 |
| NC_006213.1;YP_009555238.1;HCoV-OC43_orf1ab;2769 | 0 | 0 |
| NC_006213.1;YP_009555238.1;HCoV-OC43_orf1ab;2770 | 0 | 0 |
| NC_006213.1;YP_009555238.1;HCoV-OC43_orf1ab;2771 | 0 | 0 |
| NC_006213.1;YP_009555238.1;HCoV-OC43_orf1ab;2772 | 0.1 | 0 |
| NC_006213.1;YP_009555238.1;HCoV-OC43_orf1ab;2773 | 0.1 | 0 |
| NC_006213.1;YP_009555238.1;HCoV-OC43_orf1ab;2774 | 0.1 | 0 |
| NC_006213.1;YP_009555238.1;HCoV-OC43_orf1ab;2775 | 0.1 | 0 |
| NC_006213.1;YP_009555238.1;HCoV-OC43_orf1ab;2776 | 0.1 | 0.025 |
| NC_006213.1;YP_009555238.1;HCoV-OC43_orf1ab;2777 | 0.1 | 0 |
| NC_006213.1;YP_009555238.1;HCoV-OC43_orf1ab;2778 | 0.05 | 0 |
| NC_006213.1;YP_009555238.1;HCoV-OC43_orf1ab;2779 | 0.05 | 0 |
| NC_006213.1;YP_009555238.1;HCoV-OC43_orf1ab;2780 | 0.05 | 0 |
| NC_006213.1;YP_009555238.1;HCoV-OC43_orf1ab;2781 | 0.05 | 0 |
| NC_006213.1;YP_009555238.1;HCoV-OC43_orf1ab;2782 | 0.05 | 0 |
| NC_006213.1;YP_009555238.1;HCoV-OC43_orf1ab;2783 | 0 | 0 |
| NC_006213.1;YP_009555238.1;HCoV-OC43_orf1ab;2784 | 0 | 0 |
| NC_006213.1;YP_009555238.1;HCoV-OC43_orf1ab;2785 | 0 | 0 |
| NC_006213.1;YP_009555238.1;HCoV-OC43_orf1ab;2786 | 0 | 0 |
| NC_006213.1;YP_009555238.1;HCoV-OC43_orf1ab;2787 | 0 | 0 |
| NC_006213.1;YP_009555238.1;HCoV-OC43_orf1ab;2788 | 0.05 | 0.025 |
| NC_006213.1;YP_009555238.1;HCoV-OC43_orf1ab;2789 | 0 | 0 |
| NC_006213.1;YP_009555238.1;HCoV-OC43_orf1ab;2790 | 0 | 0 |
| NC_006213.1;YP_009555238.1;HCoV-OC43_orf1ab;2791 | 0 | 0 |
| NC_006213.1;YP_009555238.1;HCoV-OC43_orf1ab;2792 | 0 | 0 |
| NC_006213.1;YP_009555238.1;HCoV-OC43_orf1ab;2793 | 0 | 0 |
| NC_006213.1;YP_009555238.1;HCoV-OC43_orf1ab;2794 | 0 | 0 |
| NC_006213.1;YP_009555238.1;HCoV-OC43_orf1ab;2795 | 0 | 0 |
| NC_006213.1;YP_009555238.1;HCoV-OC43_orf1ab;2796 | 0 | 0.025 |
| NC_006213.1;YP_009555238.1;HCoV-OC43_orf1ab;2797 | 0 | 0.025 |
| NC_006213.1;YP_009555238.1;HCoV-OC43_orf1ab;2798 | 0 | 0.025 |
| NC_006213.1;YP_009555238.1;HCoV-OC43_orf1ab;2799 | 0.05 | 0.05 |
| NC_006213.1;YP_009555238.1;HCoV-OC43_orf1ab;2800 | 0 | 0.025 |
| NC_006213.1;YP_009555238.1;HCoV-OC43_orf1ab;2801 | 0 | 0.025 |
| NC_006213.1;YP_009555238.1;HCoV-OC43_orf1ab;2802 | 0 | 0.025 |
| NC_006213.1;YP_009555238.1;HCoV-OC43_orf1ab;2803 | 0 | 0.025 |
| NC_006213.1;YP_009555238.1;HCoV-OC43_orf1ab;2804 | 0.1 | 0.15 |

| | | |
|---|---|---|
| NC_006213.1;YP_009555238.1;HCoV-OC43_orf1ab;2805 | 0.05 | 0.05 |
| NC_006213.1;YP_009555238.1;HCoV-OC43_orf1ab;2806 | 0 | 0 |
| NC_006213.1;YP_009555238.1;HCoV-OC43_orf1ab;2807 | 0 | 0 |
| NC_006213.1;YP_009555238.1;HCoV-OC43_orf1ab;2808 | 0 | 0 |
| NC_006213.1;YP_009555238.1;HCoV-OC43_orf1ab;2809 | 0 | 0 |
| NC_006213.1;YP_009555238.1;HCoV-OC43_orf1ab;2810 | 0 | 0.025 |
| NC_006213.1;YP_009555238.1;HCoV-OC43_orf1ab;2811 | 0 | 0.025 |
| NC_006213.1;YP_009555238.1;HCoV-OC43_orf1ab;2812 | 0.05 | 0.025 |
| NC_006213.1;YP_009555238.1;HCoV-OC43_orf1ab;2813 | 0 | 0 |
| NC_006213.1;YP_009555238.1;HCoV-OC43_orf1ab;2814 | 0 | 0 |
| NC_006213.1;YP_009555238.1;HCoV-OC43_orf1ab;2815 | 0 | 0 |
| NC_006213.1;YP_009555238.1;HCoV-OC43_orf1ab;2816 | 0 | 0 |
| NC_006213.1;YP_009555238.1;HCoV-OC43_orf1ab;2817 | 0 | 0 |
| NC_006213.1;YP_009555238.1;HCoV-OC43_orf1ab;2818 | 0 | 0 |
| NC_006213.1;YP_009555238.1;HCoV-OC43_orf1ab;2819 | 0 | 0 |
| NC_006213.1;YP_009555238.1;HCoV-OC43_orf1ab;2820 | 0 | 0 |
| NC_006213.1;YP_009555238.1;HCoV-OC43_orf1ab;2821 | 0 | 0 |
| NC_006213.1;YP_009555238.1;HCoV-OC43_orf1ab;2822 | 0 | 0 |
| NC_006213.1;YP_009555238.1;HCoV-OC43_orf1ab;2823 | 0 | 0.1 |
| NC_006213.1;YP_009555238.1;HCoV-OC43_orf1ab;2824 | 0 | 0 |
| NC_006213.1;YP_009555238.1;HCoV-OC43_orf1ab;2825 | 0 | 0 |
| NC_006213.1;YP_009555238.1;HCoV-OC43_orf1ab;2826 | 0 | 0 |
| NC_006213.1;YP_009555238.1;HCoV-OC43_orf1ab;2827 | 0 | 0 |
| NC_006213.1;YP_009555238.1;HCoV-OC43_orf1ab;2828 | 0 | 0 |
| NC_006213.1;YP_009555238.1;HCoV-OC43_orf1ab;2829 | 0 | 0 |
| NC_006213.1;YP_009555238.1;HCoV-OC43_orf1ab;2830 | 0 | 0 |
| NC_006213.1;YP_009555238.1;HCoV-OC43_orf1ab;2831 | 0 | 0 |
| NC_006213.1;YP_009555238.1;HCoV-OC43_orf1ab;2832 | 0 | 0 |
| NC_006213.1;YP_009555238.1;HCoV-OC43_orf1ab;2833 | 0 | 0 |
| NC_006213.1;YP_009555238.1;HCoV-OC43_orf1ab;2834 | 0 | 0 |
| NC_006213.1;YP_009555238.1;HCoV-OC43_orf1ab;2835 | 0 | 0 |
| NC_006213.1;YP_009555238.1;HCoV-OC43_orf1ab;2836 | 0 | 0 |
| NC_006213.1;YP_009555238.1;HCoV-OC43_orf1ab;2837 | 0 | 0 |
| NC_006213.1;YP_009555238.1;HCoV-OC43_orf1ab;2838 | 0 | 0 |
| NC_006213.1;YP_009555238.1;HCoV-OC43_orf1ab;2839 | 0 | 0 |
| NC_006213.1;YP_009555238.1;HCoV-OC43_orf1ab;2840 | 0 | 0.025 |
| NC_006213.1;YP_009555238.1;HCoV-OC43_orf1ab;2841 | 0 | 0 |
| NC_006213.1;YP_009555238.1;HCoV-OC43_orf1ab;2842 | 0 | 0.025 |
| NC_006213.1;YP_009555238.1;HCoV-OC43_orf1ab;2843 | 0.05 | 0.05 |
| NC_006213.1;YP_009555238.1;HCoV-OC43_orf1ab;2844 | 0 | 0.075 |
| NC_006213.1;YP_009555238.1;HCoV-OC43_orf1ab;2845 | 0.05 | 0.075 |
| NC_006213.1;YP_009555238.1;HCoV-OC43_orf1ab;2846 | 0 | 0.025 |
| NC_006213.1;YP_009555238.1;HCoV-OC43_orf1ab;2847 | 0 | 0 |

| | | |
|---|---|---|
| NC_006213.1;YP_009555238.1;HCoV-OC43_orf1ab;2848 | 0 | 0 |
| NC_006213.1;YP_009555238.1;HCoV-OC43_orf1ab;2849 | 0 | 0 |
| NC_006213.1;YP_009555238.1;HCoV-OC43_orf1ab;2850 | 0 | 0 |
| NC_006213.1;YP_009555238.1;HCoV-OC43_orf1ab;2851 | 0 | 0 |
| NC_006213.1;YP_009555238.1;HCoV-OC43_orf1ab;2852 | 0 | 0 |
| NC_006213.1;YP_009555238.1;HCoV-OC43_orf1ab;2853 | 0 | 0 |
| NC_006213.1;YP_009555238.1;HCoV-OC43_orf1ab;2854 | 0 | 0 |
| NC_006213.1;YP_009555238.1;HCoV-OC43_orf1ab;2855 | 0 | 0 |
| NC_006213.1;YP_009555238.1;HCoV-OC43_orf1ab;2856 | 0 | 0 |
| NC_006213.1;YP_009555238.1;HCoV-OC43_orf1ab;2857 | 0 | 0 |
| NC_006213.1;YP_009555238.1;HCoV-OC43_orf1ab;2858 | 0 | 0 |
| NC_006213.1;YP_009555238.1;HCoV-OC43_orf1ab;2859 | 0 | 0 |
| NC_006213.1;YP_009555238.1;HCoV-OC43_orf1ab;2860 | 0 | 0 |
| NC_006213.1;YP_009555238.1;HCoV-OC43_orf1ab;2861 | 0 | 0 |
| NC_006213.1;YP_009555238.1;HCoV-OC43_orf1ab;2862 | 0 | 0 |
| NC_006213.1;YP_009555238.1;HCoV-OC43_orf1ab;2863 | 0 | 0 |
| NC_006213.1;YP_009555238.1;HCoV-OC43_orf1ab;2864 | 0 | 0 |
| NC_006213.1;YP_009555238.1;HCoV-OC43_orf1ab;2865 | 0 | 0 |
| NC_006213.1;YP_009555238.1;HCoV-OC43_orf1ab;2866 | 0 | 0 |
| NC_006213.1;YP_009555238.1;HCoV-OC43_orf1ab;2867 | 0 | 0 |
| NC_006213.1;YP_009555238.1;HCoV-OC43_orf1ab;2868 | 0 | 0 |
| NC_006213.1;YP_009555238.1;HCoV-OC43_orf1ab;2869 | 0 | 0 |
| NC_006213.1;YP_009555238.1;HCoV-OC43_orf1ab;2870 | 0 | 0 |
| NC_006213.1;YP_009555238.1;HCoV-OC43_orf1ab;2871 | 0 | 0 |
| NC_006213.1;YP_009555238.1;HCoV-OC43_orf1ab;2872 | 0 | 0 |
| NC_006213.1;YP_009555238.1;HCoV-OC43_orf1ab;2873 | 0 | 0 |
| NC_006213.1;YP_009555238.1;HCoV-OC43_orf1ab;2874 | 0 | 0 |
| NC_006213.1;YP_009555238.1;HCoV-OC43_orf1ab;2875 | 0 | 0 |
| NC_006213.1;YP_009555238.1;HCoV-OC43_orf1ab;2876 | 0 | 0 |
| NC_006213.1;YP_009555238.1;HCoV-OC43_orf1ab;2877 | 0 | 0 |
| NC_006213.1;YP_009555238.1;HCoV-OC43_orf1ab;2878 | 0 | 0 |
| NC_006213.1;YP_009555238.1;HCoV-OC43_orf1ab;2879 | 0 | 0 |
| NC_006213.1;YP_009555238.1;HCoV-OC43_orf1ab;2880 | 0 | 0 |
| NC_006213.1;YP_009555238.1;HCoV-OC43_orf1ab;2881 | 0 | 0 |
| NC_006213.1;YP_009555238.1;HCoV-OC43_orf1ab;2882 | 0 | 0 |
| NC_006213.1;YP_009555238.1;HCoV-OC43_orf1ab;2883 | 0 | 0 |
| NC_006213.1;YP_009555238.1;HCoV-OC43_orf1ab;2884 | 0 | 0 |
| NC_006213.1;YP_009555238.1;HCoV-OC43_orf1ab;2885 | 0 | 0 |
| NC_006213.1;YP_009555238.1;HCoV-OC43_orf1ab;2886 | 0 | 0.025 |
| NC_006213.1;YP_009555238.1;HCoV-OC43_orf1ab;2887 | 0 | 0 |
| NC_006213.1;YP_009555238.1;HCoV-OC43_orf1ab;2888 | 0 | 0 |
| NC_006213.1;YP_009555238.1;HCoV-OC43_orf1ab;2889 | 0 | 0.025 |
| NC_006213.1;YP_009555238.1;HCoV-OC43_orf1ab;2890 | 0 | 0.025 |

| | | |
|---|---|---|
| NC_006213.1;YP_009555238.1;HCoV-OC43_orf1ab;2891 | 0 | 0.025 |
| NC_006213.1;YP_009555238.1;HCoV-OC43_orf1ab;2892 | 0 | 0 |
| NC_006213.1;YP_009555238.1;HCoV-OC43_orf1ab;2893 | 0 | 0 |
| NC_006213.1;YP_009555238.1;HCoV-OC43_orf1ab;2894 | 0 | 0 |
| NC_006213.1;YP_009555238.1;HCoV-OC43_orf1ab;2895 | 0 | 0 |
| NC_006213.1;YP_009555238.1;HCoV-OC43_orf1ab;2896 | 0 | 0 |
| NC_006213.1;YP_009555238.1;HCoV-OC43_orf1ab;2897 | 0 | 0 |
| NC_006213.1;YP_009555238.1;HCoV-OC43_orf1ab;2898 | 0 | 0 |
| NC_006213.1;YP_009555238.1;HCoV-OC43_orf1ab;2899 | 0 | 0 |
| NC_006213.1;YP_009555238.1;HCoV-OC43_orf1ab;2900 | 0.05 | 0.025 |
| NC_006213.1;YP_009555238.1;HCoV-OC43_orf1ab;2901 | 0 | 0 |
| NC_006213.1;YP_009555238.1;HCoV-OC43_orf1ab;2902 | 0 | 0 |
| NC_006213.1;YP_009555238.1;HCoV-OC43_orf1ab;2903 | 0 | 0 |
| NC_006213.1;YP_009555238.1;HCoV-OC43_orf1ab;2904 | 0 | 0 |
| NC_006213.1;YP_009555238.1;HCoV-OC43_orf1ab;2905 | 0 | 0.025 |
| NC_006213.1;YP_009555238.1;HCoV-OC43_orf1ab;2906 | 0 | 0 |
| NC_006213.1;YP_009555238.1;HCoV-OC43_orf1ab;2907 | 0 | 0 |
| NC_006213.1;YP_009555238.1;HCoV-OC43_orf1ab;2908 | 0 | 0 |
| NC_006213.1;YP_009555238.1;HCoV-OC43_orf1ab;2909 | 0 | 0 |
| NC_006213.1;YP_009555238.1;HCoV-OC43_orf1ab;2910 | 0 | 0 |
| NC_006213.1;YP_009555238.1;HCoV-OC43_orf1ab;2911 | 0 | 0 |
| NC_006213.1;YP_009555238.1;HCoV-OC43_orf1ab;2912 | 0 | 0 |
| NC_006213.1;YP_009555238.1;HCoV-OC43_orf1ab;2913 | 0 | 0 |
| NC_006213.1;YP_009555238.1;HCoV-OC43_orf1ab;2914 | 0 | 0 |
| NC_006213.1;YP_009555238.1;HCoV-OC43_orf1ab;2915 | 0 | 0 |
| NC_006213.1;YP_009555238.1;HCoV-OC43_orf1ab;2916 | 0 | 0 |
| NC_006213.1;YP_009555238.1;HCoV-OC43_orf1ab;2917 | 0 | 0 |
| NC_006213.1;YP_009555238.1;HCoV-OC43_orf1ab;2918 | 0 | 0 |
| NC_006213.1;YP_009555238.1;HCoV-OC43_orf1ab;2919 | 0 | 0 |
| NC_006213.1;YP_009555238.1;HCoV-OC43_orf1ab;2920 | 0.05 | 0.025 |
| NC_006213.1;YP_009555238.1;HCoV-OC43_orf1ab;2921 | 0 | 0 |
| NC_006213.1;YP_009555238.1;HCoV-OC43_orf1ab;2922 | 0 | 0 |
| NC_006213.1;YP_009555238.1;HCoV-OC43_orf1ab;2923 | 0 | 0 |
| NC_006213.1;YP_009555238.1;HCoV-OC43_orf1ab;2924 | 0 | 0 |
| NC_006213.1;YP_009555238.1;HCoV-OC43_orf1ab;2925 | 0 | 0 |
| NC_006213.1;YP_009555238.1;HCoV-OC43_orf1ab;2926 | 0 | 0 |
| NC_006213.1;YP_009555238.1;HCoV-OC43_orf1ab;2927 | 0 | 0 |
| NC_006213.1;YP_009555238.1;HCoV-OC43_orf1ab;2928 | 0 | 0 |
| NC_006213.1;YP_009555238.1;HCoV-OC43_orf1ab;2929 | 0 | 0 |
| NC_006213.1;YP_009555238.1;HCoV-OC43_orf1ab;2930 | 0 | 0 |
| NC_006213.1;YP_009555238.1;HCoV-OC43_orf1ab;2931 | 0 | 0 |
| NC_006213.1;YP_009555238.1;HCoV-OC43_orf1ab;2932 | 0 | 0 |
| NC_006213.1;YP_009555238.1;HCoV-OC43_orf1ab;2933 | 0 | 0 |

| | | |
|---|---|---|
| NC_006213.1;YP_009555238.1;HCoV-OC43_orf1ab;2934 | 0 | 0 |
| NC_006213.1;YP_009555238.1;HCoV-OC43_orf1ab;2935 | 0 | 0 |
| NC_006213.1;YP_009555238.1;HCoV-OC43_orf1ab;2936 | 0 | 0 |
| NC_006213.1;YP_009555238.1;HCoV-OC43_orf1ab;2937 | 0 | 0 |
| NC_006213.1;YP_009555238.1;HCoV-OC43_orf1ab;2938 | 0 | 0 |
| NC_006213.1;YP_009555238.1;HCoV-OC43_orf1ab;2939 | 0 | 0 |
| NC_006213.1;YP_009555238.1;HCoV-OC43_orf1ab;2940 | 0 | 0 |
| NC_006213.1;YP_009555238.1;HCoV-OC43_orf1ab;2941 | 0 | 0 |
| NC_006213.1;YP_009555238.1;HCoV-OC43_orf1ab;2942 | 0 | 0.025 |
| NC_006213.1;YP_009555238.1;HCoV-OC43_orf1ab;2943 | 0 | 0 |
| NC_006213.1;YP_009555238.1;HCoV-OC43_orf1ab;2944 | 0 | 0 |
| NC_006213.1;YP_009555238.1;HCoV-OC43_orf1ab;2945 | 0 | 0.025 |
| NC_006213.1;YP_009555238.1;HCoV-OC43_orf1ab;2946 | 0 | 0 |
| NC_006213.1;YP_009555238.1;HCoV-OC43_orf1ab;2947 | 0 | 0 |
| NC_006213.1;YP_009555238.1;HCoV-OC43_orf1ab;2948 | 0 | 0 |
| NC_006213.1;YP_009555238.1;HCoV-OC43_orf1ab;2949 | 0 | 0 |
| NC_006213.1;YP_009555238.1;HCoV-OC43_orf1ab;2950 | 0 | 0 |
| NC_006213.1;YP_009555238.1;HCoV-OC43_orf1ab;2951 | 0 | 0 |
| NC_006213.1;YP_009555238.1;HCoV-OC43_orf1ab;2952 | 0 | 0 |
| NC_006213.1;YP_009555238.1;HCoV-OC43_orf1ab;2953 | 0 | 0 |
| NC_006213.1;YP_009555238.1;HCoV-OC43_orf1ab;2954 | 0 | 0 |
| NC_006213.1;YP_009555238.1;HCoV-OC43_orf1ab;2955 | 0 | 0 |
| NC_006213.1;YP_009555238.1;HCoV-OC43_orf1ab;2956 | 0 | 0 |
| NC_006213.1;YP_009555238.1;HCoV-OC43_orf1ab;2957 | 0 | 0 |
| NC_006213.1;YP_009555238.1;HCoV-OC43_orf1ab;2958 | 0 | 0 |
| NC_006213.1;YP_009555238.1;HCoV-OC43_orf1ab;2959 | 0 | 0 |
| NC_006213.1;YP_009555238.1;HCoV-OC43_orf1ab;2960 | 0 | 0 |
| NC_006213.1;YP_009555238.1;HCoV-OC43_orf1ab;2961 | 0 | 0 |
| NC_006213.1;YP_009555238.1;HCoV-OC43_orf1ab;2962 | 0 | 0 |
| NC_006213.1;YP_009555238.1;HCoV-OC43_orf1ab;2963 | 0 | 0 |
| NC_006213.1;YP_009555238.1;HCoV-OC43_orf1ab;2964 | 0 | 0 |
| NC_006213.1;YP_009555238.1;HCoV-OC43_orf1ab;2965 | 0 | 0 |
| NC_006213.1;YP_009555238.1;HCoV-OC43_orf1ab;2966 | 0 | 0 |
| NC_006213.1;YP_009555238.1;HCoV-OC43_orf1ab;2967 | 0 | 0 |
| NC_006213.1;YP_009555238.1;HCoV-OC43_orf1ab;2968 | 0 | 0 |
| NC_006213.1;YP_009555238.1;HCoV-OC43_orf1ab;2969 | 0 | 0 |
| NC_006213.1;YP_009555238.1;HCoV-OC43_orf1ab;2970 | 0 | 0 |
| NC_006213.1;YP_009555238.1;HCoV-OC43_orf1ab;2971 | 0 | 0.025 |
| NC_006213.1;YP_009555238.1;HCoV-OC43_orf1ab;2972 | 0 | 0 |
| NC_006213.1;YP_009555238.1;HCoV-OC43_orf1ab;2973 | 0.05 | 0 |
| NC_006213.1;YP_009555238.1;HCoV-OC43_orf1ab;2974 | 0 | 0 |
| NC_006213.1;YP_009555238.1;HCoV-OC43_orf1ab;2975 | 0 | 0 |
| NC_006213.1;YP_009555238.1;HCoV-OC43_orf1ab;2976 | 0 | 0.025 |

| | | |
|---|---|---|
| NC_006213.1;YP_009555238.1;HCoV-OC43_orf1ab;2977 | 0 | 0.025 |
| NC_006213.1;YP_009555238.1;HCoV-OC43_orf1ab;2978 | 0 | 0 |
| NC_006213.1;YP_009555238.1;HCoV-OC43_orf1ab;2979 | 0 | 0 |
| NC_006213.1;YP_009555238.1;HCoV-OC43_orf1ab;2980 | 0 | 0 |
| NC_006213.1;YP_009555238.1;HCoV-OC43_orf1ab;2981 | 0 | 0 |
| NC_006213.1;YP_009555238.1;HCoV-OC43_orf1ab;2982 | 0 | 0 |
| NC_006213.1;YP_009555238.1;HCoV-OC43_orf1ab;2983 | 0 | 0 |
| NC_006213.1;YP_009555238.1;HCoV-OC43_orf1ab;2984 | 0 | 0 |
| NC_006213.1;YP_009555238.1;HCoV-OC43_orf1ab;2985 | 0 | 0 |
| NC_006213.1;YP_009555238.1;HCoV-OC43_orf1ab;2986 | 0 | 0 |
| NC_006213.1;YP_009555238.1;HCoV-OC43_orf1ab;2987 | 0 | 0 |
| NC_006213.1;YP_009555238.1;HCoV-OC43_orf1ab;2988 | 0 | 0 |
| NC_006213.1;YP_009555238.1;HCoV-OC43_orf1ab;2989 | 0 | 0 |
| NC_006213.1;YP_009555238.1;HCoV-OC43_orf1ab;2990 | 0 | 0 |
| NC_006213.1;YP_009555238.1;HCoV-OC43_orf1ab;2991 | 0 | 0 |
| NC_006213.1;YP_009555238.1;HCoV-OC43_orf1ab;2992 | 0 | 0 |
| NC_006213.1;YP_009555238.1;HCoV-OC43_orf1ab;2993 | 0 | 0 |
| NC_006213.1;YP_009555238.1;HCoV-OC43_orf1ab;2994 | 0 | 0 |
| NC_006213.1;YP_009555238.1;HCoV-OC43_orf1ab;2995 | 0 | 0 |
| NC_006213.1;YP_009555238.1;HCoV-OC43_orf1ab;2996 | 0 | 0 |
| NC_006213.1;YP_009555238.1;HCoV-OC43_orf1ab;2997 | 0 | 0 |
| NC_006213.1;YP_009555238.1;HCoV-OC43_orf1ab;2998 | 0 | 0 |
| NC_006213.1;YP_009555238.1;HCoV-OC43_orf1ab;2999 | 0 | 0 |
| NC_006213.1;YP_009555238.1;HCoV-OC43_orf1ab;3000 | 0 | 0 |
| NC_006213.1;YP_009555238.1;HCoV-OC43_orf1ab;3001 | 0 | 0 |
| NC_006213.1;YP_009555238.1;HCoV-OC43_orf1ab;3002 | 0 | 0 |
| NC_006213.1;YP_009555238.1;HCoV-OC43_orf1ab;3003 | 0 | 0 |
| NC_006213.1;YP_009555238.1;HCoV-OC43_orf1ab;3004 | 0 | 0 |
| NC_006213.1;YP_009555238.1;HCoV-OC43_orf1ab;3005 | 0 | 0 |
| NC_006213.1;YP_009555238.1;HCoV-OC43_orf1ab;3006 | 0 | 0 |
| NC_006213.1;YP_009555238.1;HCoV-OC43_orf1ab;3007 | 0 | 0 |
| NC_006213.1;YP_009555238.1;HCoV-OC43_orf1ab;3008 | 0 | 0 |
| NC_006213.1;YP_009555238.1;HCoV-OC43_orf1ab;3009 | 0 | 0 |
| NC_006213.1;YP_009555238.1;HCoV-OC43_orf1ab;3010 | 0 | 0 |
| NC_006213.1;YP_009555238.1;HCoV-OC43_orf1ab;3011 | 0 | 0 |
| NC_006213.1;YP_009555238.1;HCoV-OC43_orf1ab;3012 | 0 | 0 |
| NC_006213.1;YP_009555238.1;HCoV-OC43_orf1ab;3013 | 0 | 0 |
| NC_006213.1;YP_009555238.1;HCoV-OC43_orf1ab;3014 | 0 | 0 |
| NC_006213.1;YP_009555238.1;HCoV-OC43_orf1ab;3015 | 0 | 0 |
| NC_006213.1;YP_009555238.1;HCoV-OC43_orf1ab;3016 | 0 | 0 |
| NC_006213.1;YP_009555238.1;HCoV-OC43_orf1ab;3017 | 0 | 0 |
| NC_006213.1;YP_009555238.1;HCoV-OC43_orf1ab;3018 | 0 | 0 |
| NC_006213.1;YP_009555238.1;HCoV-OC43_orf1ab;3019 | 0 | 0 |

| | | |
|---|---|---|
| NC_006213.1;YP_009555238.1;HCoV-OC43_orf1ab;3020 | 0 | 0.025 |
| NC_006213.1;YP_009555238.1;HCoV-OC43_orf1ab;3021 | 0 | 0 |
| NC_006213.1;YP_009555238.1;HCoV-OC43_orf1ab;3022 | 0 | 0 |
| NC_006213.1;YP_009555238.1;HCoV-OC43_orf1ab;3023 | 0 | 0 |
| NC_006213.1;YP_009555238.1;HCoV-OC43_orf1ab;3024 | 0 | 0 |
| NC_006213.1;YP_009555238.1;HCoV-OC43_orf1ab;3025 | 0 | 0 |
| NC_006213.1;YP_009555238.1;HCoV-OC43_orf1ab;3026 | 0 | 0 |
| NC_006213.1;YP_009555238.1;HCoV-OC43_orf1ab;3027 | 0 | 0 |
| NC_006213.1;YP_009555238.1;HCoV-OC43_orf1ab;3028 | 0 | 0 |
| NC_006213.1;YP_009555238.1;HCoV-OC43_orf1ab;3029 | 0 | 0 |
| NC_006213.1;YP_009555238.1;HCoV-OC43_orf1ab;3030 | 0 | 0 |
| NC_006213.1;YP_009555238.1;HCoV-OC43_orf1ab;3031 | 0 | 0 |
| NC_006213.1;YP_009555238.1;HCoV-OC43_orf1ab;3032 | 0 | 0 |
| NC_006213.1;YP_009555238.1;HCoV-OC43_orf1ab;3033 | 0 | 0 |
| NC_006213.1;YP_009555238.1;HCoV-OC43_orf1ab;3034 | 0 | 0 |
| NC_006213.1;YP_009555238.1;HCoV-OC43_orf1ab;3035 | 0 | 0 |
| NC_006213.1;YP_009555238.1;HCoV-OC43_orf1ab;3036 | 0 | 0 |
| NC_006213.1;YP_009555238.1;HCoV-OC43_orf1ab;3037 | 0 | 0 |
| NC_006213.1;YP_009555238.1;HCoV-OC43_orf1ab;3038 | 0 | 0 |
| NC_006213.1;YP_009555238.1;HCoV-OC43_orf1ab;3039 | 0 | 0 |
| NC_006213.1;YP_009555238.1;HCoV-OC43_orf1ab;3040 | 0 | 0 |
| NC_006213.1;YP_009555238.1;HCoV-OC43_orf1ab;3041 | 0 | 0 |
| NC_006213.1;YP_009555238.1;HCoV-OC43_orf1ab;3042 | 0 | 0 |
| NC_006213.1;YP_009555238.1;HCoV-OC43_orf1ab;3043 | 0 | 0 |
| NC_006213.1;YP_009555238.1;HCoV-OC43_orf1ab;3044 | 0 | 0 |
| NC_006213.1;YP_009555238.1;HCoV-OC43_orf1ab;3045 | 0 | 0 |
| NC_006213.1;YP_009555238.1;HCoV-OC43_orf1ab;3046 | 0 | 0 |
| NC_006213.1;YP_009555238.1;HCoV-OC43_orf1ab;3047 | 0 | 0 |
| NC_006213.1;YP_009555238.1;HCoV-OC43_orf1ab;3048 | 0 | 0 |
| NC_006213.1;YP_009555238.1;HCoV-OC43_orf1ab;3049 | 0 | 0 |
| NC_006213.1;YP_009555238.1;HCoV-OC43_orf1ab;3050 | 0 | 0 |
| NC_006213.1;YP_009555238.1;HCoV-OC43_orf1ab;3051 | 0 | 0 |
| NC_006213.1;YP_009555238.1;HCoV-OC43_orf1ab;3052 | 0 | 0 |
| NC_006213.1;YP_009555238.1;HCoV-OC43_orf1ab;3053 | 0 | 0 |
| NC_006213.1;YP_009555238.1;HCoV-OC43_orf1ab;3054 | 0 | 0 |
| NC_006213.1;YP_009555238.1;HCoV-OC43_orf1ab;3055 | 0 | 0 |
| NC_006213.1;YP_009555238.1;HCoV-OC43_orf1ab;3056 | 0 | 0 |
| NC_006213.1;YP_009555238.1;HCoV-OC43_orf1ab;3057 | 0 | 0 |
| NC_006213.1;YP_009555238.1;HCoV-OC43_orf1ab;3058 | 0 | 0 |
| NC_006213.1;YP_009555238.1;HCoV-OC43_orf1ab;3059 | 0 | 0 |
| NC_006213.1;YP_009555238.1;HCoV-OC43_orf1ab;3060 | 0 | 0 |
| NC_006213.1;YP_009555238.1;HCoV-OC43_orf1ab;3061 | 0 | 0 |
| NC_006213.1;YP_009555238.1;HCoV-OC43_orf1ab;3062 | 0 | 0 |

| | | |
|---|---|---|
| NC_006213.1;YP_009555238.1;HCoV-OC43_orf1ab;3063 | 0 | 0 |
| NC_006213.1;YP_009555238.1;HCoV-OC43_orf1ab;3064 | 0 | 0 |
| NC_006213.1;YP_009555238.1;HCoV-OC43_orf1ab;3065 | 0 | 0 |
| NC_006213.1;YP_009555238.1;HCoV-OC43_orf1ab;3066 | 0 | 0 |
| NC_006213.1;YP_009555238.1;HCoV-OC43_orf1ab;3067 | 0 | 0 |
| NC_006213.1;YP_009555238.1;HCoV-OC43_orf1ab;3068 | 0 | 0 |
| NC_006213.1;YP_009555238.1;HCoV-OC43_orf1ab;3069 | 0 | 0 |
| NC_006213.1;YP_009555238.1;HCoV-OC43_orf1ab;3070 | 0 | 0 |
| NC_006213.1;YP_009555238.1;HCoV-OC43_orf1ab;3071 | 0 | 0 |
| NC_006213.1;YP_009555238.1;HCoV-OC43_orf1ab;3072 | 0 | 0 |
| NC_006213.1;YP_009555238.1;HCoV-OC43_orf1ab;3073 | 0 | 0 |
| NC_006213.1;YP_009555238.1;HCoV-OC43_orf1ab;3074 | 0 | 0 |
| NC_006213.1;YP_009555238.1;HCoV-OC43_orf1ab;3075 | 0 | 0 |
| NC_006213.1;YP_009555238.1;HCoV-OC43_orf1ab;3076 | 0 | 0 |
| NC_006213.1;YP_009555238.1;HCoV-OC43_orf1ab;3077 | 0 | 0 |
| NC_006213.1;YP_009555238.1;HCoV-OC43_orf1ab;3078 | 0 | 0 |
| NC_006213.1;YP_009555238.1;HCoV-OC43_orf1ab;3079 | 0 | 0 |
| NC_006213.1;YP_009555238.1;HCoV-OC43_orf1ab;3080 | 0 | 0 |
| NC_006213.1;YP_009555238.1;HCoV-OC43_orf1ab;3081 | 0 | 0 |
| NC_006213.1;YP_009555238.1;HCoV-OC43_orf1ab;3082 | 0 | 0 |
| NC_006213.1;YP_009555238.1;HCoV-OC43_orf1ab;3083 | 0 | 0 |
| NC_006213.1;YP_009555238.1;HCoV-OC43_orf1ab;3084 | 0 | 0 |
| NC_006213.1;YP_009555238.1;HCoV-OC43_orf1ab;3085 | 0 | 0 |
| NC_006213.1;YP_009555238.1;HCoV-OC43_orf1ab;3086 | 0 | 0 |
| NC_006213.1;YP_009555238.1;HCoV-OC43_orf1ab;3087 | 0 | 0 |
| NC_006213.1;YP_009555238.1;HCoV-OC43_orf1ab;3088 | 0 | 0 |
| NC_006213.1;YP_009555238.1;HCoV-OC43_orf1ab;3089 | 0 | 0 |
| NC_006213.1;YP_009555238.1;HCoV-OC43_orf1ab;3090 | 0 | 0 |
| NC_006213.1;YP_009555238.1;HCoV-OC43_orf1ab;3091 | 0 | 0 |
| NC_006213.1;YP_009555238.1;HCoV-OC43_orf1ab;3092 | 0 | 0 |
| NC_006213.1;YP_009555238.1;HCoV-OC43_orf1ab;3093 | 0 | 0 |
| NC_006213.1;YP_009555238.1;HCoV-OC43_orf1ab;3094 | 0 | 0 |
| NC_006213.1;YP_009555238.1;HCoV-OC43_orf1ab;3095 | 0 | 0 |
| NC_006213.1;YP_009555238.1;HCoV-OC43_orf1ab;3096 | 0 | 0 |
| NC_006213.1;YP_009555238.1;HCoV-OC43_orf1ab;3097 | 0 | 0 |
| NC_006213.1;YP_009555238.1;HCoV-OC43_orf1ab;3098 | 0 | 0 |
| NC_006213.1;YP_009555238.1;HCoV-OC43_orf1ab;3099 | 0 | 0 |
| NC_006213.1;YP_009555238.1;HCoV-OC43_orf1ab;3100 | 0 | 0 |
| NC_006213.1;YP_009555238.1;HCoV-OC43_orf1ab;3101 | 0 | 0 |
| NC_006213.1;YP_009555238.1;HCoV-OC43_orf1ab;3102 | 0 | 0 |
| NC_006213.1;YP_009555238.1;HCoV-OC43_orf1ab;3103 | 0 | 0 |
| NC_006213.1;YP_009555238.1;HCoV-OC43_orf1ab;3104 | 0 | 0.025 |
| NC_006213.1;YP_009555238.1;HCoV-OC43_orf1ab;3105 | 0 | 0 |

| | | |
|---|---|---|
| NC_006213.1;YP_009555238.1;HCoV-OC43_orf1ab;3106 | 0 | 0 |
| NC_006213.1;YP_009555238.1;HCoV-OC43_orf1ab;3107 | 0 | 0 |
| NC_006213.1;YP_009555238.1;HCoV-OC43_orf1ab;3108 | 0 | 0 |
| NC_006213.1;YP_009555238.1;HCoV-OC43_orf1ab;3109 | 0 | 0 |
| NC_006213.1;YP_009555238.1;HCoV-OC43_orf1ab;3110 | 0 | 0 |
| NC_006213.1;YP_009555238.1;HCoV-OC43_orf1ab;3111 | 0 | 0 |
| NC_006213.1;YP_009555238.1;HCoV-OC43_orf1ab;3112 | 0 | 0 |
| NC_006213.1;YP_009555238.1;HCoV-OC43_orf1ab;3113 | 0 | 0 |
| NC_006213.1;YP_009555238.1;HCoV-OC43_orf1ab;3114 | 0 | 0 |
| NC_006213.1;YP_009555238.1;HCoV-OC43_orf1ab;3115 | 0 | 0 |
| NC_006213.1;YP_009555238.1;HCoV-OC43_orf1ab;3116 | 0 | 0 |
| NC_006213.1;YP_009555238.1;HCoV-OC43_orf1ab;3117 | 0 | 0 |
| NC_006213.1;YP_009555238.1;HCoV-OC43_orf1ab;3118 | 0 | 0 |
| NC_006213.1;YP_009555238.1;HCoV-OC43_orf1ab;3119 | 0 | 0 |
| NC_006213.1;YP_009555238.1;HCoV-OC43_orf1ab;3120 | 0 | 0 |
| NC_006213.1;YP_009555238.1;HCoV-OC43_orf1ab;3121 | 0 | 0 |
| NC_006213.1;YP_009555238.1;HCoV-OC43_orf1ab;3122 | 0 | 0 |
| NC_006213.1;YP_009555238.1;HCoV-OC43_orf1ab;3123 | 0 | 0 |
| NC_006213.1;YP_009555238.1;HCoV-OC43_orf1ab;3124 | 0 | 0 |
| NC_006213.1;YP_009555238.1;HCoV-OC43_orf1ab;3125 | 0 | 0 |
| NC_006213.1;YP_009555238.1;HCoV-OC43_orf1ab;3126 | 0 | 0 |
| NC_006213.1;YP_009555238.1;HCoV-OC43_orf1ab;3127 | 0 | 0 |
| NC_006213.1;YP_009555238.1;HCoV-OC43_orf1ab;3128 | 0 | 0 |
| NC_006213.1;YP_009555238.1;HCoV-OC43_orf1ab;3129 | 0 | 0 |
| NC_006213.1;YP_009555238.1;HCoV-OC43_orf1ab;3130 | 0 | 0 |
| NC_006213.1;YP_009555238.1;HCoV-OC43_orf1ab;3131 | 0 | 0 |
| NC_006213.1;YP_009555238.1;HCoV-OC43_orf1ab;3132 | 0 | 0 |
| NC_006213.1;YP_009555238.1;HCoV-OC43_orf1ab;3133 | 0 | 0 |
| NC_006213.1;YP_009555238.1;HCoV-OC43_orf1ab;3134 | 0 | 0 |
| NC_006213.1;YP_009555238.1;HCoV-OC43_orf1ab;3135 | 0 | 0 |
| NC_006213.1;YP_009555238.1;HCoV-OC43_orf1ab;3136 | 0 | 0 |
| NC_006213.1;YP_009555238.1;HCoV-OC43_orf1ab;3137 | 0 | 0 |
| NC_006213.1;YP_009555238.1;HCoV-OC43_orf1ab;3138 | 0 | 0 |
| NC_006213.1;YP_009555238.1;HCoV-OC43_orf1ab;3139 | 0 | 0 |
| NC_006213.1;YP_009555238.1;HCoV-OC43_orf1ab;3140 | 0 | 0 |
| NC_006213.1;YP_009555238.1;HCoV-OC43_orf1ab;3141 | 0 | 0 |
| NC_006213.1;YP_009555238.1;HCoV-OC43_orf1ab;3142 | 0 | 0 |
| NC_006213.1;YP_009555238.1;HCoV-OC43_orf1ab;3143 | 0 | 0 |
| NC_006213.1;YP_009555238.1;HCoV-OC43_orf1ab;3144 | 0 | 0 |
| NC_006213.1;YP_009555238.1;HCoV-OC43_orf1ab;3145 | 0.05 | 0 |
| NC_006213.1;YP_009555238.1;HCoV-OC43_orf1ab;3146 | 0 | 0 |
| NC_006213.1;YP_009555238.1;HCoV-OC43_orf1ab;3147 | 0 | 0 |
| NC_006213.1;YP_009555238.1;HCoV-OC43_orf1ab;3148 | 0 | 0 |

| | | |
|---|---|---|
| NC_006213.1;YP_009555238.1;HCoV-OC43_orf1ab;3149 | 0 | 0 |
| NC_006213.1;YP_009555238.1;HCoV-OC43_orf1ab;3150 | 0 | 0 |
| NC_006213.1;YP_009555238.1;HCoV-OC43_orf1ab;3151 | 0 | 0 |
| NC_006213.1;YP_009555238.1;HCoV-OC43_orf1ab;3152 | 0 | 0 |
| NC_006213.1;YP_009555238.1;HCoV-OC43_orf1ab;3153 | 0 | 0 |
| NC_006213.1;YP_009555238.1;HCoV-OC43_orf1ab;3154 | 0 | 0 |
| NC_006213.1;YP_009555238.1;HCoV-OC43_orf1ab;3155 | 0 | 0 |
| NC_006213.1;YP_009555238.1;HCoV-OC43_orf1ab;3156 | 0 | 0 |
| NC_006213.1;YP_009555238.1;HCoV-OC43_orf1ab;3157 | 0 | 0 |
| NC_006213.1;YP_009555238.1;HCoV-OC43_orf1ab;3158 | 0 | 0 |
| NC_006213.1;YP_009555238.1;HCoV-OC43_orf1ab;3159 | 0 | 0 |
| NC_006213.1;YP_009555238.1;HCoV-OC43_orf1ab;3160 | 0 | 0 |
| NC_006213.1;YP_009555238.1;HCoV-OC43_orf1ab;3161 | 0 | 0 |
| NC_006213.1;YP_009555238.1;HCoV-OC43_orf1ab;3162 | 0 | 0 |
| NC_006213.1;YP_009555238.1;HCoV-OC43_orf1ab;3163 | 0 | 0 |
| NC_006213.1;YP_009555238.1;HCoV-OC43_orf1ab;3164 | 0 | 0 |
| NC_006213.1;YP_009555238.1;HCoV-OC43_orf1ab;3165 | 0 | 0 |
| NC_006213.1;YP_009555238.1;HCoV-OC43_orf1ab;3166 | 0 | 0 |
| NC_006213.1;YP_009555238.1;HCoV-OC43_orf1ab;3167 | 0 | 0 |
| NC_006213.1;YP_009555238.1;HCoV-OC43_orf1ab;3168 | 0 | 0 |
| NC_006213.1;YP_009555238.1;HCoV-OC43_orf1ab;3169 | 0 | 0 |
| NC_006213.1;YP_009555238.1;HCoV-OC43_orf1ab;3170 | 0 | 0 |
| NC_006213.1;YP_009555238.1;HCoV-OC43_orf1ab;3171 | 0 | 0 |
| NC_006213.1;YP_009555238.1;HCoV-OC43_orf1ab;3172 | 0.05 | 0.025 |
| NC_006213.1;YP_009555238.1;HCoV-OC43_orf1ab;3173 | 0 | 0 |
| NC_006213.1;YP_009555238.1;HCoV-OC43_orf1ab;3174 | 0 | 0 |
| NC_006213.1;YP_009555238.1;HCoV-OC43_orf1ab;3175 | 0 | 0 |
| NC_006213.1;YP_009555238.1;HCoV-OC43_orf1ab;3176 | 0 | 0 |
| NC_006213.1;YP_009555238.1;HCoV-OC43_orf1ab;3177 | 0 | 0 |
| NC_006213.1;YP_009555238.1;HCoV-OC43_orf1ab;3178 | 0 | 0 |
| NC_006213.1;YP_009555238.1;HCoV-OC43_orf1ab;3179 | 0 | 0 |
| NC_006213.1;YP_009555238.1;HCoV-OC43_orf1ab;3180 | 0 | 0 |
| NC_006213.1;YP_009555238.1;HCoV-OC43_orf1ab;3181 | 0 | 0 |
| NC_006213.1;YP_009555238.1;HCoV-OC43_orf1ab;3182 | 0 | 0 |
| NC_006213.1;YP_009555238.1;HCoV-OC43_orf1ab;3183 | 0 | 0 |
| NC_006213.1;YP_009555238.1;HCoV-OC43_orf1ab;3184 | 0 | 0 |
| NC_006213.1;YP_009555238.1;HCoV-OC43_orf1ab;3185 | 0 | 0 |
| NC_006213.1;YP_009555238.1;HCoV-OC43_orf1ab;3186 | 0 | 0 |
| NC_006213.1;YP_009555238.1;HCoV-OC43_orf1ab;3187 | 0 | 0 |
| NC_006213.1;YP_009555238.1;HCoV-OC43_orf1ab;3188 | 0 | 0 |
| NC_006213.1;YP_009555238.1;HCoV-OC43_orf1ab;3189 | 0 | 0 |
| NC_006213.1;YP_009555238.1;HCoV-OC43_orf1ab;3190 | 0 | 0.025 |
| NC_006213.1;YP_009555238.1;HCoV-OC43_orf1ab;3191 | 0 | 0.025 |

| | | |
|---|---|---|
| NC_006213.1;YP_009555238.1;HCoV-OC43_orf1ab;3192 | 0.05 | 0.05 |
| NC_006213.1;YP_009555238.1;HCoV-OC43_orf1ab;3193 | 0 | 0 |
| NC_006213.1;YP_009555238.1;HCoV-OC43_orf1ab;3194 | 0 | 0 |
| NC_006213.1;YP_009555238.1;HCoV-OC43_orf1ab;3195 | 0 | 0 |
| NC_006213.1;YP_009555238.1;HCoV-OC43_orf1ab;3196 | 0.05 | 0 |
| NC_006213.1;YP_009555238.1;HCoV-OC43_orf1ab;3197 | 0.1 | 0.025 |
| NC_006213.1;YP_009555238.1;HCoV-OC43_orf1ab;3198 | 0 | 0 |
| NC_006213.1;YP_009555238.1;HCoV-OC43_orf1ab;3199 | 0 | 0 |
| NC_006213.1;YP_009555238.1;HCoV-OC43_orf1ab;3200 | 0 | 0 |
| NC_006213.1;YP_009555238.1;HCoV-OC43_orf1ab;3201 | 0 | 0.025 |
| NC_006213.1;YP_009555238.1;HCoV-OC43_orf1ab;3202 | 0.05 | 0.025 |
| NC_006213.1;YP_009555238.1;HCoV-OC43_orf1ab;3203 | 0 | 0 |
| NC_006213.1;YP_009555238.1;HCoV-OC43_orf1ab;3204 | 0 | 0.025 |
| NC_006213.1;YP_009555238.1;HCoV-OC43_orf1ab;3205 | 0 | 0 |
| NC_006213.1;YP_009555238.1;HCoV-OC43_orf1ab;3206 | 0 | 0 |
| NC_006213.1;YP_009555238.1;HCoV-OC43_orf1ab;3207 | 0 | 0 |
| NC_006213.1;YP_009555238.1;HCoV-OC43_orf1ab;3208 | 0 | 0.025 |
| NC_006213.1;YP_009555238.1;HCoV-OC43_orf1ab;3209 | 0.05 | 0 |
| NC_006213.1;YP_009555238.1;HCoV-OC43_orf1ab;3210 | 0 | 0 |
| NC_006213.1;YP_009555238.1;HCoV-OC43_orf1ab;3211 | 0.05 | 0 |
| NC_006213.1;YP_009555238.1;HCoV-OC43_orf1ab;3212 | 0.05 | 0 |
| NC_006213.1;YP_009555238.1;HCoV-OC43_orf1ab;3213 | 0.1 | 0 |
| NC_006213.1;YP_009555238.1;HCoV-OC43_orf1ab;3214 | 0 | 0 |
| NC_006213.1;YP_009555238.1;HCoV-OC43_orf1ab;3215 | 0 | 0 |
| NC_006213.1;YP_009555238.1;HCoV-OC43_orf1ab;3216 | 0 | 0.05 |
| NC_006213.1;YP_009555238.1;HCoV-OC43_orf1ab;3217 | 0 | 0.05 |
| NC_006213.1;YP_009555238.1;HCoV-OC43_orf1ab;3218 | 0 | 0.025 |
| NC_006213.1;YP_009555238.1;HCoV-OC43_orf1ab;3219 | 0 | 0.025 |
| NC_006213.1;YP_009555238.1;HCoV-OC43_orf1ab;3220 | 0.1 | 0.025 |
| NC_006213.1;YP_009555238.1;HCoV-OC43_orf1ab;3221 | 0.25 | 0.05 |
| NC_006213.1;YP_009555238.1;HCoV-OC43_orf1ab;3222 | 0.05 | 0.1 |
| NC_006213.1;YP_009555238.1;HCoV-OC43_orf1ab;3223 | 0 | 0.075 |
| NC_006213.1;YP_009555238.1;HCoV-OC43_orf1ab;3224 | 0.05 | 0 |
| NC_006213.1;YP_009555238.1;HCoV-OC43_orf1ab;3225 | 0 | 0.05 |
| NC_006213.1;YP_009555238.1;HCoV-OC43_orf1ab;3226 | 0.05 | 0 |
| NC_006213.1;YP_009555238.1;HCoV-OC43_orf1ab;3227 | 0 | 0.025 |
| NC_006213.1;YP_009555238.1;HCoV-OC43_orf1ab;3228 | 0.05 | 0.025 |
| NC_006213.1;YP_009555238.1;HCoV-OC43_orf1ab;3229 | 0.05 | 0 |
| NC_006213.1;YP_009555238.1;HCoV-OC43_orf1ab;3230 | 0 | 0 |
| NC_006213.1;YP_009555238.1;HCoV-OC43_orf1ab;3231 | 0 | 0 |
| NC_006213.1;YP_009555238.1;HCoV-OC43_orf1ab;3232 | 0 | 0.025 |
| NC_006213.1;YP_009555238.1;HCoV-OC43_orf1ab;3233 | 0.1 | 0.025 |
| NC_006213.1;YP_009555238.1;HCoV-OC43_orf1ab;3234 | 0 | 0.025 |

| | | |
|---|---|---|
| NC_006213.1;YP_009555238.1;HCoV-OC43_orf1ab;3235 | 0 | 0 |
| NC_006213.1;YP_009555238.1;HCoV-OC43_orf1ab;3236 | 0 | 0 |
| NC_006213.1;YP_009555238.1;HCoV-OC43_orf1ab;3237 | 0 | 0 |
| NC_006213.1;YP_009555238.1;HCoV-OC43_orf1ab;3238 | 0 | 0 |
| NC_006213.1;YP_009555238.1;HCoV-OC43_orf1ab;3239 | 0 | 0 |
| NC_006213.1;YP_009555238.1;HCoV-OC43_orf1ab;3240 | 0 | 0 |
| NC_006213.1;YP_009555238.1;HCoV-OC43_orf1ab;3241 | 0 | 0.025 |
| NC_006213.1;YP_009555238.1;HCoV-OC43_orf1ab;3242 | 0 | 0 |
| NC_006213.1;YP_009555238.1;HCoV-OC43_orf1ab;3243 | 0 | 0 |
| NC_006213.1;YP_009555238.1;HCoV-OC43_orf1ab;3244 | 0 | 0 |
| NC_006213.1;YP_009555238.1;HCoV-OC43_orf1ab;3245 | 0 | 0 |
| NC_006213.1;YP_009555238.1;HCoV-OC43_orf1ab;3246 | 0 | 0 |
| NC_006213.1;YP_009555238.1;HCoV-OC43_orf1ab;3247 | 0 | 0 |
| NC_006213.1;YP_009555238.1;HCoV-OC43_orf1ab;3248 | 0.05 | 0 |
| NC_006213.1;YP_009555238.1;HCoV-OC43_orf1ab;3249 | 0 | 0 |
| NC_006213.1;YP_009555238.1;HCoV-OC43_orf1ab;3250 | 0 | 0 |
| NC_006213.1;YP_009555238.1;HCoV-OC43_orf1ab;3251 | 0 | 0 |
| NC_006213.1;YP_009555238.1;HCoV-OC43_orf1ab;3252 | 0 | 0 |
| NC_006213.1;YP_009555238.1;HCoV-OC43_orf1ab;3253 | 0 | 0 |
| NC_006213.1;YP_009555238.1;HCoV-OC43_orf1ab;3254 | 0 | 0 |
| NC_006213.1;YP_009555238.1;HCoV-OC43_orf1ab;3255 | 0 | 0 |
| NC_006213.1;YP_009555238.1;HCoV-OC43_orf1ab;3256 | 0 | 0 |
| NC_006213.1;YP_009555238.1;HCoV-OC43_orf1ab;3257 | 0 | 0 |
| NC_006213.1;YP_009555238.1;HCoV-OC43_orf1ab;3258 | 0 | 0 |
| NC_006213.1;YP_009555238.1;HCoV-OC43_orf1ab;3259 | 0.1 | 0 |
| NC_006213.1;YP_009555238.1;HCoV-OC43_orf1ab;3260 | 0.05 | 0 |
| NC_006213.1;YP_009555238.1;HCoV-OC43_orf1ab;3261 | 0.05 | 0 |
| NC_006213.1;YP_009555238.1;HCoV-OC43_orf1ab;3262 | 0.05 | 0 |
| NC_006213.1;YP_009555238.1;HCoV-OC43_orf1ab;3263 | 0.05 | 0 |
| NC_006213.1;YP_009555238.1;HCoV-OC43_orf1ab;3264 | 0.05 | 0 |
| NC_006213.1;YP_009555238.1;HCoV-OC43_orf1ab;3265 | 0.05 | 0 |
| NC_006213.1;YP_009555238.1;HCoV-OC43_orf1ab;3266 | 0.1 | 0.025 |
| NC_006213.1;YP_009555238.1;HCoV-OC43_orf1ab;3267 | 0.1 | 0.025 |
| NC_006213.1;YP_009555238.1;HCoV-OC43_orf1ab;3268 | 0.05 | 0.025 |
| NC_006213.1;YP_009555238.1;HCoV-OC43_orf1ab;3269 | 0.1 | 0 |
| NC_006213.1;YP_009555238.1;HCoV-OC43_orf1ab;3270 | 0.05 | 0 |
| NC_006213.1;YP_009555238.1;HCoV-OC43_orf1ab;3271 | 0 | 0 |
| NC_006213.1;YP_009555238.1;HCoV-OC43_orf1ab;3272 | 0 | 0 |
| NC_006213.1;YP_009555238.1;HCoV-OC43_orf1ab;3273 | 0 | 0 |
| NC_006213.1;YP_009555238.1;HCoV-OC43_orf1ab;3274 | 0 | 0 |
| NC_006213.1;YP_009555238.1;HCoV-OC43_orf1ab;3275 | 0 | 0 |
| NC_006213.1;YP_009555238.1;HCoV-OC43_orf1ab;3276 | 0 | 0 |
| NC_006213.1;YP_009555238.1;HCoV-OC43_orf1ab;3277 | 0 | 0 |

| | | |
|---|---|---|
| NC_006213.1;YP_009555238.1;HCoV-OC43_orf1ab;3278 | 0 | 0 |
| NC_006213.1;YP_009555238.1;HCoV-OC43_orf1ab;3279 | 0 | 0 |
| NC_006213.1;YP_009555238.1;HCoV-OC43_orf1ab;3280 | 0 | 0.025 |
| NC_006213.1;YP_009555238.1;HCoV-OC43_orf1ab;3281 | 0 | 0 |
| NC_006213.1;YP_009555238.1;HCoV-OC43_orf1ab;3282 | 0 | 0 |
| NC_006213.1;YP_009555238.1;HCoV-OC43_orf1ab;3283 | 0 | 0 |
| NC_006213.1;YP_009555238.1;HCoV-OC43_orf1ab;3284 | 0 | 0 |
| NC_006213.1;YP_009555238.1;HCoV-OC43_orf1ab;3285 | 0 | 0 |
| NC_006213.1;YP_009555238.1;HCoV-OC43_orf1ab;3286 | 0 | 0 |
| NC_006213.1;YP_009555238.1;HCoV-OC43_orf1ab;3287 | 0 | 0 |
| NC_006213.1;YP_009555238.1;HCoV-OC43_orf1ab;3288 | 0.05 | 0.025 |
| NC_006213.1;YP_009555238.1;HCoV-OC43_orf1ab;3289 | 0.05 | 0.025 |
| NC_006213.1;YP_009555238.1;HCoV-OC43_orf1ab;3290 | 0 | 0.025 |
| NC_006213.1;YP_009555238.1;HCoV-OC43_orf1ab;3291 | 0 | 0.025 |
| NC_006213.1;YP_009555238.1;HCoV-OC43_orf1ab;3292 | 0.05 | 0 |
| NC_006213.1;YP_009555238.1;HCoV-OC43_orf1ab;3293 | 0 | 0 |
| NC_006213.1;YP_009555238.1;HCoV-OC43_orf1ab;3294 | 0 | 0.025 |
| NC_006213.1;YP_009555238.1;HCoV-OC43_orf1ab;3295 | 0 | 0 |
| NC_006213.1;YP_009555238.1;HCoV-OC43_orf1ab;3296 | 0 | 0 |
| NC_006213.1;YP_009555238.1;HCoV-OC43_orf1ab;3297 | 0.05 | 0 |
| NC_006213.1;YP_009555238.1;HCoV-OC43_orf1ab;3298 | 0.05 | 0.025 |
| NC_006213.1;YP_009555238.1;HCoV-OC43_orf1ab;3299 | 0.05 | 0 |
| NC_006213.1;YP_009555238.1;HCoV-OC43_orf1ab;3300 | 0 | 0 |
| NC_006213.1;YP_009555238.1;HCoV-OC43_orf1ab;3301 | 0 | 0 |
| NC_006213.1;YP_009555238.1;HCoV-OC43_orf1ab;3302 | 0 | 0 |
| NC_006213.1;YP_009555238.1;HCoV-OC43_orf1ab;3303 | 0 | 0 |
| NC_006213.1;YP_009555238.1;HCoV-OC43_orf1ab;3304 | 0 | 0 |
| NC_006213.1;YP_009555238.1;HCoV-OC43_orf1ab;3305 | 0 | 0 |
| NC_006213.1;YP_009555238.1;HCoV-OC43_orf1ab;3306 | 0 | 0 |
| NC_006213.1;YP_009555238.1;HCoV-OC43_orf1ab;3307 | 0 | 0 |
| NC_006213.1;YP_009555238.1;HCoV-OC43_orf1ab;3308 | 0 | 0 |
| NC_006213.1;YP_009555238.1;HCoV-OC43_orf1ab;3309 | 0 | 0 |
| NC_006213.1;YP_009555238.1;HCoV-OC43_orf1ab;3310 | 0 | 0 |
| NC_006213.1;YP_009555238.1;HCoV-OC43_orf1ab;3311 | 0 | 0 |
| NC_006213.1;YP_009555238.1;HCoV-OC43_orf1ab;3312 | 0 | 0 |
| NC_006213.1;YP_009555238.1;HCoV-OC43_orf1ab;3313 | 0 | 0 |
| NC_006213.1;YP_009555238.1;HCoV-OC43_orf1ab;3314 | 0 | 0 |
| NC_006213.1;YP_009555238.1;HCoV-OC43_orf1ab;3315 | 0 | 0 |
| NC_006213.1;YP_009555238.1;HCoV-OC43_orf1ab;3316 | 0 | 0 |
| NC_006213.1;YP_009555238.1;HCoV-OC43_orf1ab;3317 | 0 | 0 |
| NC_006213.1;YP_009555238.1;HCoV-OC43_orf1ab;3318 | 0 | 0 |
| NC_006213.1;YP_009555238.1;HCoV-OC43_orf1ab;3319 | 0 | 0 |
| NC_006213.1;YP_009555238.1;HCoV-OC43_orf1ab;3320 | 0 | 0 |

| | | |
|---|---|---|
| NC_006213.1;YP_009555238.1;HCoV-OC43_orf1ab;3321 | 0 | 0 |
| NC_006213.1;YP_009555238.1;HCoV-OC43_orf1ab;3322 | 0 | 0 |
| NC_006213.1;YP_009555238.1;HCoV-OC43_orf1ab;3323 | 0 | 0 |
| NC_006213.1;YP_009555238.1;HCoV-OC43_orf1ab;3324 | 0 | 0 |
| NC_006213.1;YP_009555238.1;HCoV-OC43_orf1ab;3325 | 0 | 0 |
| NC_006213.1;YP_009555238.1;HCoV-OC43_orf1ab;3326 | 0 | 0 |
| NC_006213.1;YP_009555238.1;HCoV-OC43_orf1ab;3327 | 0 | 0 |
| NC_006213.1;YP_009555238.1;HCoV-OC43_orf1ab;3328 | 0 | 0 |
| NC_006213.1;YP_009555238.1;HCoV-OC43_orf1ab;3329 | 0 | 0 |
| NC_006213.1;YP_009555238.1;HCoV-OC43_orf1ab;3330 | 0 | 0 |
| NC_006213.1;YP_009555238.1;HCoV-OC43_orf1ab;3331 | 0 | 0 |
| NC_006213.1;YP_009555238.1;HCoV-OC43_orf1ab;3332 | 0 | 0 |
| NC_006213.1;YP_009555238.1;HCoV-OC43_orf1ab;3333 | 0 | 0 |
| NC_006213.1;YP_009555238.1;HCoV-OC43_orf1ab;3334 | 0 | 0 |
| NC_006213.1;YP_009555238.1;HCoV-OC43_orf1ab;3335 | 0 | 0 |
| NC_006213.1;YP_009555238.1;HCoV-OC43_orf1ab;3336 | 0 | 0 |
| NC_006213.1;YP_009555238.1;HCoV-OC43_orf1ab;3337 | 0 | 0 |
| NC_006213.1;YP_009555238.1;HCoV-OC43_orf1ab;3338 | 0 | 0 |
| NC_006213.1;YP_009555238.1;HCoV-OC43_orf1ab;3339 | 0 | 0 |
| NC_006213.1;YP_009555238.1;HCoV-OC43_orf1ab;3340 | 0 | 0 |
| NC_006213.1;YP_009555238.1;HCoV-OC43_orf1ab;3341 | 0 | 0 |
| NC_006213.1;YP_009555238.1;HCoV-OC43_orf1ab;3342 | 0 | 0 |
| NC_006213.1;YP_009555238.1;HCoV-OC43_orf1ab;3343 | 0 | 0 |
| NC_006213.1;YP_009555238.1;HCoV-OC43_orf1ab;3344 | 0 | 0 |
| NC_006213.1;YP_009555238.1;HCoV-OC43_orf1ab;3345 | 0 | 0 |
| NC_006213.1;YP_009555238.1;HCoV-OC43_orf1ab;3346 | 0 | 0 |
| NC_006213.1;YP_009555238.1;HCoV-OC43_orf1ab;3347 | 0 | 0 |
| NC_006213.1;YP_009555238.1;HCoV-OC43_orf1ab;3348 | 0.05 | 0.025 |
| NC_006213.1;YP_009555238.1;HCoV-OC43_orf1ab;3349 | 0 | 0 |
| NC_006213.1;YP_009555238.1;HCoV-OC43_orf1ab;3350 | 0 | 0 |
| NC_006213.1;YP_009555238.1;HCoV-OC43_orf1ab;3351 | 0 | 0 |
| NC_006213.1;YP_009555238.1;HCoV-OC43_orf1ab;3352 | 0 | 0 |
| NC_006213.1;YP_009555238.1;HCoV-OC43_orf1ab;3353 | 0 | 0 |
| NC_006213.1;YP_009555238.1;HCoV-OC43_orf1ab;3354 | 0 | 0 |
| NC_006213.1;YP_009555238.1;HCoV-OC43_orf1ab;3355 | 0 | 0 |
| NC_006213.1;YP_009555238.1;HCoV-OC43_orf1ab;3356 | 0 | 0 |
| NC_006213.1;YP_009555238.1;HCoV-OC43_orf1ab;3357 | 0 | 0 |
| NC_006213.1;YP_009555238.1;HCoV-OC43_orf1ab;3358 | 0 | 0 |
| NC_006213.1;YP_009555238.1;HCoV-OC43_orf1ab;3359 | 0 | 0 |
| NC_006213.1;YP_009555238.1;HCoV-OC43_orf1ab;3360 | 0 | 0 |
| NC_006213.1;YP_009555238.1;HCoV-OC43_orf1ab;3361 | 0 | 0 |
| NC_006213.1;YP_009555238.1;HCoV-OC43_orf1ab;3362 | 0.05 | 0 |
| NC_006213.1;YP_009555238.1;HCoV-OC43_orf1ab;3363 | 0.05 | 0 |

| | | |
|---|---|---|
| NC_006213.1;YP_009555238.1;HCoV-OC43_orf1ab;3364 | 0 | 0 |
| NC_006213.1;YP_009555238.1;HCoV-OC43_orf1ab;3365 | 0 | 0 |
| NC_006213.1;YP_009555238.1;HCoV-OC43_orf1ab;3366 | 0 | 0 |
| NC_006213.1;YP_009555238.1;HCoV-OC43_orf1ab;3367 | 0 | 0 |
| NC_006213.1;YP_009555238.1;HCoV-OC43_orf1ab;3368 | 0 | 0 |
| NC_006213.1;YP_009555238.1;HCoV-OC43_orf1ab;3369 | 0 | 0 |
| NC_006213.1;YP_009555238.1;HCoV-OC43_orf1ab;3370 | 0 | 0.025 |
| NC_006213.1;YP_009555238.1;HCoV-OC43_orf1ab;3371 | 0 | 0 |
| NC_006213.1;YP_009555238.1;HCoV-OC43_orf1ab;3372 | 0 | 0 |
| NC_006213.1;YP_009555238.1;HCoV-OC43_orf1ab;3373 | 0 | 0 |
| NC_006213.1;YP_009555238.1;HCoV-OC43_orf1ab;3374 | 0 | 0 |
| NC_006213.1;YP_009555238.1;HCoV-OC43_orf1ab;3375 | 0 | 0 |
| NC_006213.1;YP_009555238.1;HCoV-OC43_orf1ab;3376 | 0 | 0 |
| NC_006213.1;YP_009555238.1;HCoV-OC43_orf1ab;3377 | 0 | 0 |
| NC_006213.1;YP_009555238.1;HCoV-OC43_orf1ab;3378 | 0 | 0 |
| NC_006213.1;YP_009555238.1;HCoV-OC43_orf1ab;3379 | 0 | 0 |
| NC_006213.1;YP_009555238.1;HCoV-OC43_orf1ab;3380 | 0 | 0 |
| NC_006213.1;YP_009555238.1;HCoV-OC43_orf1ab;3381 | 0 | 0 |
| NC_006213.1;YP_009555238.1;HCoV-OC43_orf1ab;3382 | 0 | 0 |
| NC_006213.1;YP_009555238.1;HCoV-OC43_orf1ab;3383 | 0 | 0 |
| NC_006213.1;YP_009555238.1;HCoV-OC43_orf1ab;3384 | 0 | 0 |
| NC_006213.1;YP_009555238.1;HCoV-OC43_orf1ab;3385 | 0 | 0 |
| NC_006213.1;YP_009555238.1;HCoV-OC43_orf1ab;3386 | 0 | 0 |
| NC_006213.1;YP_009555238.1;HCoV-OC43_orf1ab;3387 | 0 | 0 |
| NC_006213.1;YP_009555238.1;HCoV-OC43_orf1ab;3388 | 0 | 0 |
| NC_006213.1;YP_009555238.1;HCoV-OC43_orf1ab;3389 | 0 | 0 |
| NC_006213.1;YP_009555238.1;HCoV-OC43_orf1ab;3390 | 0 | 0 |
| NC_006213.1;YP_009555238.1;HCoV-OC43_orf1ab;3391 | 0.1 | 0.025 |
| NC_006213.1;YP_009555238.1;HCoV-OC43_orf1ab;3392 | 0.05 | 0 |
| NC_006213.1;YP_009555238.1;HCoV-OC43_orf1ab;3393 | 0.05 | 0 |
| NC_006213.1;YP_009555238.1;HCoV-OC43_orf1ab;3394 | 0 | 0 |
| NC_006213.1;YP_009555238.1;HCoV-OC43_orf1ab;3395 | 0.1 | 0 |
| NC_006213.1;YP_009555238.1;HCoV-OC43_orf1ab;3396 | 0 | 0 |
| NC_006213.1;YP_009555238.1;HCoV-OC43_orf1ab;3397 | 0.1 | 0 |
| NC_006213.1;YP_009555238.1;HCoV-OC43_orf1ab;3398 | 0.1 | 0 |
| NC_006213.1;YP_009555238.1;HCoV-OC43_orf1ab;3399 | 0.05 | 0 |
| NC_006213.1;YP_009555238.1;HCoV-OC43_orf1ab;3400 | 0.05 | 0 |
| NC_006213.1;YP_009555238.1;HCoV-OC43_orf1ab;3401 | 0 | 0 |
| NC_006213.1;YP_009555238.1;HCoV-OC43_orf1ab;3402 | 0 | 0 |
| NC_006213.1;YP_009555238.1;HCoV-OC43_orf1ab;3403 | 0 | 0 |
| NC_006213.1;YP_009555238.1;HCoV-OC43_orf1ab;3404 | 0 | 0 |
| NC_006213.1;YP_009555238.1;HCoV-OC43_orf1ab;3405 | 0 | 0 |
| NC_006213.1;YP_009555238.1;HCoV-OC43_orf1ab;3406 | 0 | 0 |

| | | |
|---|---|---|
| NC_006213.1;YP_009555238.1;HCoV-OC43_orf1ab;3407 | 0 | 0 |
| NC_006213.1;YP_009555238.1;HCoV-OC43_orf1ab;3408 | 0 | 0 |
| NC_006213.1;YP_009555238.1;HCoV-OC43_orf1ab;3409 | 0 | 0 |
| NC_006213.1;YP_009555238.1;HCoV-OC43_orf1ab;3410 | 0 | 0 |
| NC_006213.1;YP_009555238.1;HCoV-OC43_orf1ab;3411 | 0 | 0 |
| NC_006213.1;YP_009555238.1;HCoV-OC43_orf1ab;3412 | 0 | 0.025 |
| NC_006213.1;YP_009555238.1;HCoV-OC43_orf1ab;3413 | 0 | 0 |
| NC_006213.1;YP_009555238.1;HCoV-OC43_orf1ab;3414 | 0 | 0 |
| NC_006213.1;YP_009555238.1;HCoV-OC43_orf1ab;3415 | 0 | 0 |
| NC_006213.1;YP_009555238.1;HCoV-OC43_orf1ab;3416 | 0 | 0 |
| NC_006213.1;YP_009555238.1;HCoV-OC43_orf1ab;3417 | 0.05 | 0 |
| NC_006213.1;YP_009555238.1;HCoV-OC43_orf1ab;3418 | 0 | 0 |
| NC_006213.1;YP_009555238.1;HCoV-OC43_orf1ab;3419 | 0.1 | 0.025 |
| NC_006213.1;YP_009555238.1;HCoV-OC43_orf1ab;3420 | 0.05 | 0.1 |
| NC_006213.1;YP_009555238.1;HCoV-OC43_orf1ab;3421 | 0.05 | 0 |
| NC_006213.1;YP_009555238.1;HCoV-OC43_orf1ab;3422 | 0.05 | 0 |
| NC_006213.1;YP_009555238.1;HCoV-OC43_orf1ab;3423 | 0 | 0 |
| NC_006213.1;YP_009555238.1;HCoV-OC43_orf1ab;3424 | 0 | 0 |
| NC_006213.1;YP_009555238.1;HCoV-OC43_orf1ab;3425 | 0 | 0 |
| NC_006213.1;YP_009555238.1;HCoV-OC43_orf1ab;3426 | 0 | 0 |
| NC_006213.1;YP_009555238.1;HCoV-OC43_orf1ab;3427 | 0 | 0 |
| NC_006213.1;YP_009555238.1;HCoV-OC43_orf1ab;3428 | 0 | 0 |
| NC_006213.1;YP_009555238.1;HCoV-OC43_orf1ab;3429 | 0 | 0.05 |
| NC_006213.1;YP_009555238.1;HCoV-OC43_orf1ab;3430 | 0 | 0 |
| NC_006213.1;YP_009555238.1;HCoV-OC43_orf1ab;3431 | 0 | 0 |
| NC_006213.1;YP_009555238.1;HCoV-OC43_orf1ab;3432 | 0 | 0 |
| NC_006213.1;YP_009555238.1;HCoV-OC43_orf1ab;3433 | 0 | 0 |
| NC_006213.1;YP_009555238.1;HCoV-OC43_orf1ab;3434 | 0 | 0 |
| NC_006213.1;YP_009555238.1;HCoV-OC43_orf1ab;3435 | 0 | 0 |
| NC_006213.1;YP_009555238.1;HCoV-OC43_orf1ab;3436 | 0 | 0 |
| NC_006213.1;YP_009555238.1;HCoV-OC43_orf1ab;3437 | 0 | 0 |
| NC_006213.1;YP_009555238.1;HCoV-OC43_orf1ab;3438 | 0 | 0 |
| NC_006213.1;YP_009555238.1;HCoV-OC43_orf1ab;3439 | 0 | 0 |
| NC_006213.1;YP_009555238.1;HCoV-OC43_orf1ab;3440 | 0 | 0 |
| NC_006213.1;YP_009555238.1;HCoV-OC43_orf1ab;3441 | 0 | 0 |
| NC_006213.1;YP_009555238.1;HCoV-OC43_orf1ab;3442 | 0 | 0 |
| NC_006213.1;YP_009555238.1;HCoV-OC43_orf1ab;3443 | 0 | 0 |
| NC_006213.1;YP_009555238.1;HCoV-OC43_orf1ab;3444 | 0 | 0 |
| NC_006213.1;YP_009555238.1;HCoV-OC43_orf1ab;3445 | 0 | 0 |
| NC_006213.1;YP_009555238.1;HCoV-OC43_orf1ab;3446 | 0 | 0 |
| NC_006213.1;YP_009555238.1;HCoV-OC43_orf1ab;3447 | 0 | 0 |
| NC_006213.1;YP_009555238.1;HCoV-OC43_orf1ab;3448 | 0 | 0 |
| NC_006213.1;YP_009555238.1;HCoV-OC43_orf1ab;3449 | 0 | 0 |

| | | |
|---|---|---|
| NC_006213.1;YP_009555238.1;HCoV-OC43_orf1ab;3450 | 0 | 0 |
| NC_006213.1;YP_009555238.1;HCoV-OC43_orf1ab;3451 | 0 | 0 |
| NC_006213.1;YP_009555238.1;HCoV-OC43_orf1ab;3452 | 0 | 0 |
| NC_006213.1;YP_009555238.1;HCoV-OC43_orf1ab;3453 | 0 | 0 |
| NC_006213.1;YP_009555238.1;HCoV-OC43_orf1ab;3454 | 0 | 0 |
| NC_006213.1;YP_009555238.1;HCoV-OC43_orf1ab;3455 | 0 | 0.025 |
| NC_006213.1;YP_009555238.1;HCoV-OC43_orf1ab;3456 | 0.05 | 0 |
| NC_006213.1;YP_009555238.1;HCoV-OC43_orf1ab;3457 | 0 | 0 |
| NC_006213.1;YP_009555238.1;HCoV-OC43_orf1ab;3458 | 0 | 0 |
| NC_006213.1;YP_009555238.1;HCoV-OC43_orf1ab;3459 | 0 | 0 |
| NC_006213.1;YP_009555238.1;HCoV-OC43_orf1ab;3460 | 0 | 0 |
| NC_006213.1;YP_009555238.1;HCoV-OC43_orf1ab;3461 | 0 | 0 |
| NC_006213.1;YP_009555238.1;HCoV-OC43_orf1ab;3462 | 0 | 0 |
| NC_006213.1;YP_009555238.1;HCoV-OC43_orf1ab;3463 | 0 | 0 |
| NC_006213.1;YP_009555238.1;HCoV-OC43_orf1ab;3464 | 0 | 0 |
| NC_006213.1;YP_009555238.1;HCoV-OC43_orf1ab;3465 | 0 | 0.05 |
| NC_006213.1;YP_009555238.1;HCoV-OC43_orf1ab;3466 | 0 | 0 |
| NC_006213.1;YP_009555238.1;HCoV-OC43_orf1ab;3467 | 0.05 | 0 |
| NC_006213.1;YP_009555238.1;HCoV-OC43_orf1ab;3468 | 0 | 0 |
| NC_006213.1;YP_009555238.1;HCoV-OC43_orf1ab;3469 | 0 | 0 |
| NC_006213.1;YP_009555238.1;HCoV-OC43_orf1ab;3470 | 0 | 0 |
| NC_006213.1;YP_009555238.1;HCoV-OC43_orf1ab;3471 | 0 | 0.15 |
| NC_006213.1;YP_009555238.1;HCoV-OC43_orf1ab;3472 | 0.1 | 0.05 |
| NC_006213.1;YP_009555238.1;HCoV-OC43_orf1ab;3473 | 0 | 0 |
| NC_006213.1;YP_009555238.1;HCoV-OC43_orf1ab;3474 | 0 | 0 |
| NC_006213.1;YP_009555238.1;HCoV-OC43_orf1ab;3475 | 0 | 0 |
| NC_006213.1;YP_009555238.1;HCoV-OC43_orf1ab;3476 | 0 | 0 |
| NC_006213.1;YP_009555238.1;HCoV-OC43_orf1ab;3477 | 0 | 0 |
| NC_006213.1;YP_009555238.1;HCoV-OC43_orf1ab;3478 | 0 | 0 |
| NC_006213.1;YP_009555238.1;HCoV-OC43_orf1ab;3479 | 0 | 0 |
| NC_006213.1;YP_009555238.1;HCoV-OC43_orf1ab;3480 | 0.05 | 0 |
| NC_006213.1;YP_009555238.1;HCoV-OC43_orf1ab;3481 | 0 | 0 |
| NC_006213.1;YP_009555238.1;HCoV-OC43_orf1ab;3482 | 0 | 0 |
| NC_006213.1;YP_009555238.1;HCoV-OC43_orf1ab;3483 | 0 | 0 |
| NC_006213.1;YP_009555238.1;HCoV-OC43_orf1ab;3484 | 0.05 | 0 |
| NC_006213.1;YP_009555238.1;HCoV-OC43_orf1ab;3485 | 0.05 | 0 |
| NC_006213.1;YP_009555238.1;HCoV-OC43_orf1ab;3486 | 0 | 0 |
| NC_006213.1;YP_009555238.1;HCoV-OC43_orf1ab;3487 | 0 | 0 |
| NC_006213.1;YP_009555238.1;HCoV-OC43_orf1ab;3488 | 0 | 0 |
| NC_006213.1;YP_009555238.1;HCoV-OC43_orf1ab;3489 | 0 | 0 |
| NC_006213.1;YP_009555238.1;HCoV-OC43_orf1ab;3490 | 0.05 | 0 |
| NC_006213.1;YP_009555238.1;HCoV-OC43_orf1ab;3491 | 0 | 0 |
| NC_006213.1;YP_009555238.1;HCoV-OC43_orf1ab;3492 | 0 | 0 |

| | | |
|---|---|---|
| NC_006213.1;YP_009555238.1;HCoV-OC43_orf1ab;3493 | 0 | 0 |
| NC_006213.1;YP_009555238.1;HCoV-OC43_orf1ab;3494 | 0 | 0 |
| NC_006213.1;YP_009555238.1;HCoV-OC43_orf1ab;3495 | 0 | 0 |
| NC_006213.1;YP_009555238.1;HCoV-OC43_orf1ab;3496 | 0.05 | 0.025 |
| NC_006213.1;YP_009555238.1;HCoV-OC43_orf1ab;3497 | 0 | 0 |
| NC_006213.1;YP_009555238.1;HCoV-OC43_orf1ab;3498 | 0 | 0 |
| NC_006213.1;YP_009555238.1;HCoV-OC43_orf1ab;3499 | 0 | 0 |
| NC_006213.1;YP_009555238.1;HCoV-OC43_orf1ab;3500 | 0 | 0 |
| NC_006213.1;YP_009555238.1;HCoV-OC43_orf1ab;3501 | 0 | 0 |
| NC_006213.1;YP_009555238.1;HCoV-OC43_orf1ab;3502 | 0 | 0 |
| NC_006213.1;YP_009555238.1;HCoV-OC43_orf1ab;3503 | 0 | 0.025 |
| NC_006213.1;YP_009555238.1;HCoV-OC43_orf1ab;3504 | 0 | 0 |
| NC_006213.1;YP_009555238.1;HCoV-OC43_orf1ab;3505 | 0 | 0 |
| NC_006213.1;YP_009555238.1;HCoV-OC43_orf1ab;3506 | 0 | 0 |
| NC_006213.1;YP_009555238.1;HCoV-OC43_orf1ab;3507 | 0 | 0 |
| NC_006213.1;YP_009555238.1;HCoV-OC43_orf1ab;3508 | 0 | 0 |
| NC_006213.1;YP_009555238.1;HCoV-OC43_orf1ab;3509 | 0 | 0 |
| NC_006213.1;YP_009555238.1;HCoV-OC43_orf1ab;3510 | 0 | 0 |
| NC_006213.1;YP_009555238.1;HCoV-OC43_orf1ab;3511 | 0 | 0 |
| NC_006213.1;YP_009555238.1;HCoV-OC43_orf1ab;3512 | 0.05 | 0 |
| NC_006213.1;YP_009555238.1;HCoV-OC43_orf1ab;3513 | 0 | 0.05 |
| NC_006213.1;YP_009555238.1;HCoV-OC43_orf1ab;3514 | 0 | 0 |
| NC_006213.1;YP_009555238.1;HCoV-OC43_orf1ab;3515 | 0 | 0 |
| NC_006213.1;YP_009555238.1;HCoV-OC43_orf1ab;3516 | 0 | 0.025 |
| NC_006213.1;YP_009555238.1;HCoV-OC43_orf1ab;3517 | 0 | 0 |
| NC_006213.1;YP_009555238.1;HCoV-OC43_orf1ab;3518 | 0.05 | 0.05 |
| NC_006213.1;YP_009555238.1;HCoV-OC43_orf1ab;3519 | 0 | 0 |
| NC_006213.1;YP_009555238.1;HCoV-OC43_orf1ab;3520 | 0 | 0 |
| NC_006213.1;YP_009555238.1;HCoV-OC43_orf1ab;3521 | 0 | 0.025 |
| NC_006213.1;YP_009555238.1;HCoV-OC43_orf1ab;3522 | 0 | 0.025 |
| NC_006213.1;YP_009555238.1;HCoV-OC43_orf1ab;3523 | 0 | 0.025 |
| NC_006213.1;YP_009555238.1;HCoV-OC43_orf1ab;3524 | 0 | 0.025 |
| NC_006213.1;YP_009555238.1;HCoV-OC43_orf1ab;3525 | 0 | 0.025 |
| NC_006213.1;YP_009555238.1;HCoV-OC43_orf1ab;3526 | 0.05 | 0.025 |
| NC_006213.1;YP_009555238.1;HCoV-OC43_orf1ab;3527 | 0.05 | 0.075 |
| NC_006213.1;YP_009555238.1;HCoV-OC43_orf1ab;3528 | 0 | 0 |
| NC_006213.1;YP_009555238.1;HCoV-OC43_orf1ab;3529 | 0.05 | 0.025 |
| NC_006213.1;YP_009555238.1;HCoV-OC43_orf1ab;3530 | 0 | 0.05 |
| NC_006213.1;YP_009555238.1;HCoV-OC43_orf1ab;3531 | 0.1 | 0.05 |
| NC_006213.1;YP_009555238.1;HCoV-OC43_orf1ab;3532 | 0 | 0 |
| NC_006213.1;YP_009555238.1;HCoV-OC43_orf1ab;3533 | 0 | 0.025 |
| NC_006213.1;YP_009555238.1;HCoV-OC43_orf1ab;3534 | 0 | 0.075 |
| NC_006213.1;YP_009555238.1;HCoV-OC43_orf1ab;3535 | 0 | 0.025 |

| | | |
|---|---|---|
| NC_006213.1;YP_009555238.1;HCoV-OC43_orf1ab;3536 | 0 | 0 |
| NC_006213.1;YP_009555238.1;HCoV-OC43_orf1ab;3537 | 0 | 0 |
| NC_006213.1;YP_009555238.1;HCoV-OC43_orf1ab;3538 | 0 | 0 |
| NC_006213.1;YP_009555238.1;HCoV-OC43_orf1ab;3539 | 0 | 0 |
| NC_006213.1;YP_009555238.1;HCoV-OC43_orf1ab;3540 | 0 | 0 |
| NC_006213.1;YP_009555238.1;HCoV-OC43_orf1ab;3541 | 0 | 0 |
| NC_006213.1;YP_009555238.1;HCoV-OC43_orf1ab;3542 | 0 | 0 |
| NC_006213.1;YP_009555238.1;HCoV-OC43_orf1ab;3543 | 0 | 0 |
| NC_006213.1;YP_009555238.1;HCoV-OC43_orf1ab;3544 | 0 | 0 |
| NC_006213.1;YP_009555238.1;HCoV-OC43_orf1ab;3545 | 0 | 0 |
| NC_006213.1;YP_009555238.1;HCoV-OC43_orf1ab;3546 | 0 | 0 |
| NC_006213.1;YP_009555238.1;HCoV-OC43_orf1ab;3547 | 0 | 0 |
| NC_006213.1;YP_009555238.1;HCoV-OC43_orf1ab;3548 | 0 | 0 |
| NC_006213.1;YP_009555238.1;HCoV-OC43_orf1ab;3549 | 0 | 0 |
| NC_006213.1;YP_009555238.1;HCoV-OC43_orf1ab;3550 | 0 | 0 |
| NC_006213.1;YP_009555238.1;HCoV-OC43_orf1ab;3551 | 0 | 0 |
| NC_006213.1;YP_009555238.1;HCoV-OC43_orf1ab;3552 | 0 | 0 |
| NC_006213.1;YP_009555238.1;HCoV-OC43_orf1ab;3553 | 0 | 0 |
| NC_006213.1;YP_009555238.1;HCoV-OC43_orf1ab;3554 | 0 | 0 |
| NC_006213.1;YP_009555238.1;HCoV-OC43_orf1ab;3555 | 0 | 0 |
| NC_006213.1;YP_009555238.1;HCoV-OC43_orf1ab;3556 | 0 | 0 |
| NC_006213.1;YP_009555238.1;HCoV-OC43_orf1ab;3557 | 0 | 0 |
| NC_006213.1;YP_009555238.1;HCoV-OC43_orf1ab;3558 | 0 | 0 |
| NC_006213.1;YP_009555238.1;HCoV-OC43_orf1ab;3559 | 0 | 0 |
| NC_006213.1;YP_009555238.1;HCoV-OC43_orf1ab;3560 | 0 | 0 |
| NC_006213.1;YP_009555238.1;HCoV-OC43_orf1ab;3561 | 0 | 0 |
| NC_006213.1;YP_009555238.1;HCoV-OC43_orf1ab;3562 | 0 | 0 |
| NC_006213.1;YP_009555238.1;HCoV-OC43_orf1ab;3563 | 0 | 0 |
| NC_006213.1;YP_009555238.1;HCoV-OC43_orf1ab;3564 | 0 | 0 |
| NC_006213.1;YP_009555238.1;HCoV-OC43_orf1ab;3565 | 0 | 0 |
| NC_006213.1;YP_009555238.1;HCoV-OC43_orf1ab;3566 | 0 | 0 |
| NC_006213.1;YP_009555238.1;HCoV-OC43_orf1ab;3567 | 0 | 0 |
| NC_006213.1;YP_009555238.1;HCoV-OC43_orf1ab;3568 | 0 | 0 |
| NC_006213.1;YP_009555238.1;HCoV-OC43_orf1ab;3569 | 0 | 0 |
| NC_006213.1;YP_009555238.1;HCoV-OC43_orf1ab;3570 | 0 | 0 |
| NC_006213.1;YP_009555238.1;HCoV-OC43_orf1ab;3571 | 0 | 0 |
| NC_006213.1;YP_009555238.1;HCoV-OC43_orf1ab;3572 | 0 | 0 |
| NC_006213.1;YP_009555238.1;HCoV-OC43_orf1ab;3573 | 0 | 0 |
| NC_006213.1;YP_009555238.1;HCoV-OC43_orf1ab;3574 | 0 | 0 |
| NC_006213.1;YP_009555238.1;HCoV-OC43_orf1ab;3575 | 0 | 0 |
| NC_006213.1;YP_009555238.1;HCoV-OC43_orf1ab;3576 | 0 | 0 |
| NC_006213.1;YP_009555238.1;HCoV-OC43_orf1ab;3577 | 0 | 0 |
| NC_006213.1;YP_009555238.1;HCoV-OC43_orf1ab;3578 | 0 | 0 |

| | | |
|---|---|---|
| NC_006213.1;YP_009555238.1;HCoV-OC43_orf1ab;3579 | 0 | 0 |
| NC_006213.1;YP_009555238.1;HCoV-OC43_orf1ab;3580 | 0 | 0 |
| NC_006213.1;YP_009555238.1;HCoV-OC43_orf1ab;3581 | 0 | 0 |
| NC_006213.1;YP_009555238.1;HCoV-OC43_orf1ab;3582 | 0 | 0 |
| NC_006213.1;YP_009555238.1;HCoV-OC43_orf1ab;3583 | 0 | 0 |
| NC_006213.1;YP_009555238.1;HCoV-OC43_orf1ab;3584 | 0 | 0 |
| NC_006213.1;YP_009555238.1;HCoV-OC43_orf1ab;3585 | 0 | 0 |
| NC_006213.1;YP_009555238.1;HCoV-OC43_orf1ab;3586 | 0 | 0 |
| NC_006213.1;YP_009555238.1;HCoV-OC43_orf1ab;3587 | 0 | 0 |
| NC_006213.1;YP_009555238.1;HCoV-OC43_orf1ab;3588 | 0 | 0 |
| NC_006213.1;YP_009555238.1;HCoV-OC43_orf1ab;3589 | 0 | 0 |
| NC_006213.1;YP_009555238.1;HCoV-OC43_orf1ab;3590 | 0 | 0 |
| NC_006213.1;YP_009555238.1;HCoV-OC43_orf1ab;3591 | 0 | 0 |
| NC_006213.1;YP_009555238.1;HCoV-OC43_orf1ab;3592 | 0 | 0 |
| NC_006213.1;YP_009555238.1;HCoV-OC43_orf1ab;3593 | 0 | 0 |
| NC_006213.1;YP_009555238.1;HCoV-OC43_orf1ab;3594 | 0 | 0 |
| NC_006213.1;YP_009555238.1;HCoV-OC43_orf1ab;3595 | 0 | 0 |
| NC_006213.1;YP_009555238.1;HCoV-OC43_orf1ab;3596 | 0 | 0 |
| NC_006213.1;YP_009555238.1;HCoV-OC43_orf1ab;3597 | 0 | 0 |
| NC_006213.1;YP_009555238.1;HCoV-OC43_orf1ab;3598 | 0 | 0 |
| NC_006213.1;YP_009555238.1;HCoV-OC43_orf1ab;3599 | 0 | 0 |
| NC_006213.1;YP_009555238.1;HCoV-OC43_orf1ab;3600 | 0 | 0 |
| NC_006213.1;YP_009555238.1;HCoV-OC43_orf1ab;3601 | 0 | 0 |
| NC_006213.1;YP_009555238.1;HCoV-OC43_orf1ab;3602 | 0 | 0 |
| NC_006213.1;YP_009555238.1;HCoV-OC43_orf1ab;3603 | 0 | 0 |
| NC_006213.1;YP_009555238.1;HCoV-OC43_orf1ab;3604 | 0 | 0 |
| NC_006213.1;YP_009555238.1;HCoV-OC43_orf1ab;3605 | 0 | 0 |
| NC_006213.1;YP_009555238.1;HCoV-OC43_orf1ab;3606 | 0 | 0 |
| NC_006213.1;YP_009555238.1;HCoV-OC43_orf1ab;3607 | 0 | 0 |
| NC_006213.1;YP_009555238.1;HCoV-OC43_orf1ab;3608 | 0 | 0 |
| NC_006213.1;YP_009555238.1;HCoV-OC43_orf1ab;3609 | 0 | 0 |
| NC_006213.1;YP_009555238.1;HCoV-OC43_orf1ab;3610 | 0 | 0 |
| NC_006213.1;YP_009555238.1;HCoV-OC43_orf1ab;3611 | 0 | 0 |
| NC_006213.1;YP_009555238.1;HCoV-OC43_orf1ab;3612 | 0 | 0 |
| NC_006213.1;YP_009555238.1;HCoV-OC43_orf1ab;3613 | 0 | 0 |
| NC_006213.1;YP_009555238.1;HCoV-OC43_orf1ab;3614 | 0 | 0 |
| NC_006213.1;YP_009555238.1;HCoV-OC43_orf1ab;3615 | 0 | 0 |
| NC_006213.1;YP_009555238.1;HCoV-OC43_orf1ab;3616 | 0 | 0 |
| NC_006213.1;YP_009555238.1;HCoV-OC43_orf1ab;3617 | 0 | 0 |
| NC_006213.1;YP_009555238.1;HCoV-OC43_orf1ab;3618 | 0 | 0 |
| NC_006213.1;YP_009555238.1;HCoV-OC43_orf1ab;3619 | 0 | 0 |
| NC_006213.1;YP_009555238.1;HCoV-OC43_orf1ab;3620 | 0 | 0 |
| NC_006213.1;YP_009555238.1;HCoV-OC43_orf1ab;3621 | 0 | 0 |

| | | |
|---|---|---|
| NC_006213.1;YP_009555238.1;HCoV-OC43_orf1ab;3622 | 0 | 0 |
| NC_006213.1;YP_009555238.1;HCoV-OC43_orf1ab;3623 | 0 | 0 |
| NC_006213.1;YP_009555238.1;HCoV-OC43_orf1ab;3624 | 0 | 0 |
| NC_006213.1;YP_009555238.1;HCoV-OC43_orf1ab;3625 | 0 | 0 |
| NC_006213.1;YP_009555238.1;HCoV-OC43_orf1ab;3626 | 0 | 0 |
| NC_006213.1;YP_009555238.1;HCoV-OC43_orf1ab;3627 | 0 | 0 |
| NC_006213.1;YP_009555238.1;HCoV-OC43_orf1ab;3628 | 0 | 0 |
| NC_006213.1;YP_009555238.1;HCoV-OC43_orf1ab;3629 | 0 | 0 |
| NC_006213.1;YP_009555238.1;HCoV-OC43_orf1ab;3630 | 0 | 0 |
| NC_006213.1;YP_009555238.1;HCoV-OC43_orf1ab;3631 | 0 | 0 |
| NC_006213.1;YP_009555238.1;HCoV-OC43_orf1ab;3632 | 0 | 0 |
| NC_006213.1;YP_009555238.1;HCoV-OC43_orf1ab;3633 | 0 | 0 |
| NC_006213.1;YP_009555238.1;HCoV-OC43_orf1ab;3634 | 0 | 0 |
| NC_006213.1;YP_009555238.1;HCoV-OC43_orf1ab;3635 | 0 | 0 |
| NC_006213.1;YP_009555238.1;HCoV-OC43_orf1ab;3636 | 0 | 0 |
| NC_006213.1;YP_009555238.1;HCoV-OC43_orf1ab;3637 | 0 | 0 |
| NC_006213.1;YP_009555238.1;HCoV-OC43_orf1ab;3638 | 0 | 0 |
| NC_006213.1;YP_009555238.1;HCoV-OC43_orf1ab;3639 | 0 | 0 |
| NC_006213.1;YP_009555238.1;HCoV-OC43_orf1ab;3640 | 0 | 0 |
| NC_006213.1;YP_009555238.1;HCoV-OC43_orf1ab;3641 | 0 | 0 |
| NC_006213.1;YP_009555238.1;HCoV-OC43_orf1ab;3642 | 0 | 0 |
| NC_006213.1;YP_009555238.1;HCoV-OC43_orf1ab;3643 | 0 | 0 |
| NC_006213.1;YP_009555238.1;HCoV-OC43_orf1ab;3644 | 0 | 0 |
| NC_006213.1;YP_009555238.1;HCoV-OC43_orf1ab;3645 | 0 | 0 |
| NC_006213.1;YP_009555238.1;HCoV-OC43_orf1ab;3646 | 0 | 0 |
| NC_006213.1;YP_009555238.1;HCoV-OC43_orf1ab;3647 | 0.05 | 0.025 |
| NC_006213.1;YP_009555238.1;HCoV-OC43_orf1ab;3648 | 0 | 0 |
| NC_006213.1;YP_009555238.1;HCoV-OC43_orf1ab;3649 | 0 | 0 |
| NC_006213.1;YP_009555238.1;HCoV-OC43_orf1ab;3650 | 0 | 0 |
| NC_006213.1;YP_009555238.1;HCoV-OC43_orf1ab;3651 | 0 | 0 |
| NC_006213.1;YP_009555238.1;HCoV-OC43_orf1ab;3652 | 0 | 0 |
| NC_006213.1;YP_009555238.1;HCoV-OC43_orf1ab;3653 | 0 | 0 |
| NC_006213.1;YP_009555238.1;HCoV-OC43_orf1ab;3654 | 0 | 0 |
| NC_006213.1;YP_009555238.1;HCoV-OC43_orf1ab;3655 | 0 | 0 |
| NC_006213.1;YP_009555238.1;HCoV-OC43_orf1ab;3656 | 0 | 0 |
| NC_006213.1;YP_009555238.1;HCoV-OC43_orf1ab;3657 | 0 | 0 |
| NC_006213.1;YP_009555238.1;HCoV-OC43_orf1ab;3658 | 0 | 0 |
| NC_006213.1;YP_009555238.1;HCoV-OC43_orf1ab;3659 | 0 | 0 |
| NC_006213.1;YP_009555238.1;HCoV-OC43_orf1ab;3660 | 0 | 0 |
| NC_006213.1;YP_009555238.1;HCoV-OC43_orf1ab;3661 | 0 | 0 |
| NC_006213.1;YP_009555238.1;HCoV-OC43_orf1ab;3662 | 0 | 0 |
| NC_006213.1;YP_009555238.1;HCoV-OC43_orf1ab;3663 | 0 | 0 |
| NC_006213.1;YP_009555238.1;HCoV-OC43_orf1ab;3664 | 0 | 0 |

| | | |
|---|---|---|
| NC_006213.1;YP_009555238.1;HCoV-OC43_orf1ab;3665 | 0 | 0 |
| NC_006213.1;YP_009555238.1;HCoV-OC43_orf1ab;3666 | 0 | 0 |
| NC_006213.1;YP_009555238.1;HCoV-OC43_orf1ab;3667 | 0 | 0 |
| NC_006213.1;YP_009555238.1;HCoV-OC43_orf1ab;3668 | 0 | 0 |
| NC_006213.1;YP_009555238.1;HCoV-OC43_orf1ab;3669 | 0 | 0 |
| NC_006213.1;YP_009555238.1;HCoV-OC43_orf1ab;3670 | 0 | 0 |
| NC_006213.1;YP_009555238.1;HCoV-OC43_orf1ab;3671 | 0 | 0 |
| NC_006213.1;YP_009555238.1;HCoV-OC43_orf1ab;3672 | 0 | 0 |
| NC_006213.1;YP_009555238.1;HCoV-OC43_orf1ab;3673 | 0 | 0 |
| NC_006213.1;YP_009555238.1;HCoV-OC43_orf1ab;3674 | 0 | 0 |
| NC_006213.1;YP_009555238.1;HCoV-OC43_orf1ab;3675 | 0 | 0 |
| NC_006213.1;YP_009555238.1;HCoV-OC43_orf1ab;3676 | 0 | 0 |
| NC_006213.1;YP_009555238.1;HCoV-OC43_orf1ab;3677 | 0 | 0 |
| NC_006213.1;YP_009555238.1;HCoV-OC43_orf1ab;3678 | 0 | 0 |
| NC_006213.1;YP_009555238.1;HCoV-OC43_orf1ab;3679 | 0 | 0 |
| NC_006213.1;YP_009555238.1;HCoV-OC43_orf1ab;3680 | 0 | 0 |
| NC_006213.1;YP_009555238.1;HCoV-OC43_orf1ab;3681 | 0 | 0 |
| NC_006213.1;YP_009555238.1;HCoV-OC43_orf1ab;3682 | 0 | 0 |
| NC_006213.1;YP_009555238.1;HCoV-OC43_orf1ab;3683 | 0 | 0 |
| NC_006213.1;YP_009555238.1;HCoV-OC43_orf1ab;3684 | 0 | 0 |
| NC_006213.1;YP_009555238.1;HCoV-OC43_orf1ab;3685 | 0 | 0 |
| NC_006213.1;YP_009555238.1;HCoV-OC43_orf1ab;3686 | 0 | 0 |
| NC_006213.1;YP_009555238.1;HCoV-OC43_orf1ab;3687 | 0 | 0 |
| NC_006213.1;YP_009555238.1;HCoV-OC43_orf1ab;3688 | 0 | 0 |
| NC_006213.1;YP_009555238.1;HCoV-OC43_orf1ab;3689 | 0 | 0 |
| NC_006213.1;YP_009555238.1;HCoV-OC43_orf1ab;3690 | 0 | 0 |
| NC_006213.1;YP_009555238.1;HCoV-OC43_orf1ab;3691 | 0 | 0 |
| NC_006213.1;YP_009555238.1;HCoV-OC43_orf1ab;3692 | 0 | 0 |
| NC_006213.1;YP_009555238.1;HCoV-OC43_orf1ab;3693 | 0 | 0 |
| NC_006213.1;YP_009555238.1;HCoV-OC43_orf1ab;3694 | 0 | 0 |
| NC_006213.1;YP_009555238.1;HCoV-OC43_orf1ab;3695 | 0 | 0 |
| NC_006213.1;YP_009555238.1;HCoV-OC43_orf1ab;3696 | 0 | 0 |
| NC_006213.1;YP_009555238.1;HCoV-OC43_orf1ab;3697 | 0 | 0 |
| NC_006213.1;YP_009555238.1;HCoV-OC43_orf1ab;3698 | 0 | 0 |
| NC_006213.1;YP_009555238.1;HCoV-OC43_orf1ab;3699 | 0 | 0 |
| NC_006213.1;YP_009555238.1;HCoV-OC43_orf1ab;3700 | 0 | 0 |
| NC_006213.1;YP_009555238.1;HCoV-OC43_orf1ab;3701 | 0 | 0.025 |
| NC_006213.1;YP_009555238.1;HCoV-OC43_orf1ab;3702 | 0.05 | 0 |
| NC_006213.1;YP_009555238.1;HCoV-OC43_orf1ab;3703 | 0 | 0 |
| NC_006213.1;YP_009555238.1;HCoV-OC43_orf1ab;3704 | 0.05 | 0 |
| NC_006213.1;YP_009555238.1;HCoV-OC43_orf1ab;3705 | 0 | 0 |
| NC_006213.1;YP_009555238.1;HCoV-OC43_orf1ab;3706 | 0 | 0 |
| NC_006213.1;YP_009555238.1;HCoV-OC43_orf1ab;3707 | 0 | 0 |

| | | |
|---|---|---|
| NC_006213.1;YP_009555238.1;HCoV-OC43_orf1ab;3708 | 0 | 0 |
| NC_006213.1;YP_009555238.1;HCoV-OC43_orf1ab;3709 | 0 | 0 |
| NC_006213.1;YP_009555238.1;HCoV-OC43_orf1ab;3710 | 0 | 0 |
| NC_006213.1;YP_009555238.1;HCoV-OC43_orf1ab;3711 | 0 | 0 |
| NC_006213.1;YP_009555238.1;HCoV-OC43_orf1ab;3712 | 0 | 0 |
| NC_006213.1;YP_009555238.1;HCoV-OC43_orf1ab;3713 | 0 | 0 |
| NC_006213.1;YP_009555238.1;HCoV-OC43_orf1ab;3714 | 0 | 0 |
| NC_006213.1;YP_009555238.1;HCoV-OC43_orf1ab;3715 | 0 | 0 |
| NC_006213.1;YP_009555238.1;HCoV-OC43_orf1ab;3716 | 0 | 0 |
| NC_006213.1;YP_009555238.1;HCoV-OC43_orf1ab;3717 | 0 | 0 |
| NC_006213.1;YP_009555238.1;HCoV-OC43_orf1ab;3718 | 0 | 0 |
| NC_006213.1;YP_009555238.1;HCoV-OC43_orf1ab;3719 | 0 | 0 |
| NC_006213.1;YP_009555238.1;HCoV-OC43_orf1ab;3720 | 0 | 0 |
| NC_006213.1;YP_009555238.1;HCoV-OC43_orf1ab;3721 | 0 | 0 |
| NC_006213.1;YP_009555238.1;HCoV-OC43_orf1ab;3722 | 0 | 0 |
| NC_006213.1;YP_009555238.1;HCoV-OC43_orf1ab;3723 | 0 | 0 |
| NC_006213.1;YP_009555238.1;HCoV-OC43_orf1ab;3724 | 0 | 0 |
| NC_006213.1;YP_009555238.1;HCoV-OC43_orf1ab;3725 | 0 | 0 |
| NC_006213.1;YP_009555238.1;HCoV-OC43_orf1ab;3726 | 0 | 0 |
| NC_006213.1;YP_009555238.1;HCoV-OC43_orf1ab;3727 | 0 | 0 |
| NC_006213.1;YP_009555238.1;HCoV-OC43_orf1ab;3728 | 0 | 0 |
| NC_006213.1;YP_009555238.1;HCoV-OC43_orf1ab;3729 | 0 | 0 |
| NC_006213.1;YP_009555238.1;HCoV-OC43_orf1ab;3730 | 0 | 0 |
| NC_006213.1;YP_009555238.1;HCoV-OC43_orf1ab;3731 | 0 | 0 |
| NC_006213.1;YP_009555238.1;HCoV-OC43_orf1ab;3732 | 0 | 0 |
| NC_006213.1;YP_009555238.1;HCoV-OC43_orf1ab;3733 | 0 | 0 |
| NC_006213.1;YP_009555238.1;HCoV-OC43_orf1ab;3734 | 0 | 0 |
| NC_006213.1;YP_009555238.1;HCoV-OC43_orf1ab;3735 | 0 | 0 |
| NC_006213.1;YP_009555238.1;HCoV-OC43_orf1ab;3736 | 0 | 0 |
| NC_006213.1;YP_009555238.1;HCoV-OC43_orf1ab;3737 | 0 | 0 |
| NC_006213.1;YP_009555238.1;HCoV-OC43_orf1ab;3738 | 0 | 0 |
| NC_006213.1;YP_009555238.1;HCoV-OC43_orf1ab;3739 | 0 | 0 |
| NC_006213.1;YP_009555238.1;HCoV-OC43_orf1ab;3740 | 0 | 0 |
| NC_006213.1;YP_009555238.1;HCoV-OC43_orf1ab;3741 | 0 | 0 |
| NC_006213.1;YP_009555238.1;HCoV-OC43_orf1ab;3742 | 0 | 0 |
| NC_006213.1;YP_009555238.1;HCoV-OC43_orf1ab;3743 | 0 | 0 |
| NC_006213.1;YP_009555238.1;HCoV-OC43_orf1ab;3744 | 0 | 0 |
| NC_006213.1;YP_009555238.1;HCoV-OC43_orf1ab;3745 | 0 | 0 |
| NC_006213.1;YP_009555238.1;HCoV-OC43_orf1ab;3746 | 0 | 0 |
| NC_006213.1;YP_009555238.1;HCoV-OC43_orf1ab;3747 | 0 | 0 |
| NC_006213.1;YP_009555238.1;HCoV-OC43_orf1ab;3748 | 0 | 0 |
| NC_006213.1;YP_009555238.1;HCoV-OC43_orf1ab;3749 | 0 | 0 |
| NC_006213.1;YP_009555238.1;HCoV-OC43_orf1ab;3750 | 0 | 0 |

| | | |
|---|---|---|
| NC_006213.1;YP_009555238.1;HCoV-OC43_orf1ab;3751 | 0 | 0 |
| NC_006213.1;YP_009555238.1;HCoV-OC43_orf1ab;3752 | 0 | 0 |
| NC_006213.1;YP_009555238.1;HCoV-OC43_orf1ab;3753 | 0 | 0 |
| NC_006213.1;YP_009555238.1;HCoV-OC43_orf1ab;3754 | 0 | 0 |
| NC_006213.1;YP_009555238.1;HCoV-OC43_orf1ab;3755 | 0 | 0 |
| NC_006213.1;YP_009555238.1;HCoV-OC43_orf1ab;3756 | 0 | 0 |
| NC_006213.1;YP_009555238.1;HCoV-OC43_orf1ab;3757 | 0 | 0 |
| NC_006213.1;YP_009555238.1;HCoV-OC43_orf1ab;3758 | 0 | 0 |
| NC_006213.1;YP_009555238.1;HCoV-OC43_orf1ab;3759 | 0 | 0 |
| NC_006213.1;YP_009555238.1;HCoV-OC43_orf1ab;3760 | 0 | 0 |
| NC_006213.1;YP_009555238.1;HCoV-OC43_orf1ab;3761 | 0 | 0 |
| NC_006213.1;YP_009555238.1;HCoV-OC43_orf1ab;3762 | 0 | 0 |
| NC_006213.1;YP_009555238.1;HCoV-OC43_orf1ab;3763 | 0 | 0 |
| NC_006213.1;YP_009555238.1;HCoV-OC43_orf1ab;3764 | 0 | 0 |
| NC_006213.1;YP_009555238.1;HCoV-OC43_orf1ab;3765 | 0 | 0 |
| NC_006213.1;YP_009555238.1;HCoV-OC43_orf1ab;3766 | 0 | 0 |
| NC_006213.1;YP_009555238.1;HCoV-OC43_orf1ab;3767 | 0 | 0 |
| NC_006213.1;YP_009555238.1;HCoV-OC43_orf1ab;3768 | 0 | 0 |
| NC_006213.1;YP_009555238.1;HCoV-OC43_orf1ab;3769 | 0 | 0 |
| NC_006213.1;YP_009555238.1;HCoV-OC43_orf1ab;3770 | 0 | 0 |
| NC_006213.1;YP_009555238.1;HCoV-OC43_orf1ab;3771 | 0 | 0 |
| NC_006213.1;YP_009555238.1;HCoV-OC43_orf1ab;3772 | 0 | 0 |
| NC_006213.1;YP_009555238.1;HCoV-OC43_orf1ab;3773 | 0 | 0 |
| NC_006213.1;YP_009555238.1;HCoV-OC43_orf1ab;3774 | 0 | 0 |
| NC_006213.1;YP_009555238.1;HCoV-OC43_orf1ab;3775 | 0 | 0 |
| NC_006213.1;YP_009555238.1;HCoV-OC43_orf1ab;3776 | 0 | 0 |
| NC_006213.1;YP_009555238.1;HCoV-OC43_orf1ab;3777 | 0 | 0 |
| NC_006213.1;YP_009555238.1;HCoV-OC43_orf1ab;3778 | 0 | 0 |
| NC_006213.1;YP_009555238.1;HCoV-OC43_orf1ab;3779 | 0 | 0 |
| NC_006213.1;YP_009555238.1;HCoV-OC43_orf1ab;3780 | 0 | 0 |
| NC_006213.1;YP_009555238.1;HCoV-OC43_orf1ab;3781 | 0 | 0 |
| NC_006213.1;YP_009555238.1;HCoV-OC43_orf1ab;3782 | 0 | 0 |
| NC_006213.1;YP_009555238.1;HCoV-OC43_orf1ab;3783 | 0 | 0 |
| NC_006213.1;YP_009555238.1;HCoV-OC43_orf1ab;3784 | 0 | 0 |
| NC_006213.1;YP_009555238.1;HCoV-OC43_orf1ab;3785 | 0 | 0 |
| NC_006213.1;YP_009555238.1;HCoV-OC43_orf1ab;3786 | 0 | 0 |
| NC_006213.1;YP_009555238.1;HCoV-OC43_orf1ab;3787 | 0 | 0 |
| NC_006213.1;YP_009555238.1;HCoV-OC43_orf1ab;3788 | 0 | 0 |
| NC_006213.1;YP_009555238.1;HCoV-OC43_orf1ab;3789 | 0 | 0 |
| NC_006213.1;YP_009555238.1;HCoV-OC43_orf1ab;3790 | 0 | 0 |
| NC_006213.1;YP_009555238.1;HCoV-OC43_orf1ab;3791 | 0 | 0 |
| NC_006213.1;YP_009555238.1;HCoV-OC43_orf1ab;3792 | 0.05 | 0.1 |
| NC_006213.1;YP_009555238.1;HCoV-OC43_orf1ab;3793 | 0 | 0 |

| | | |
|---|---|---|
| NC_006213.1;YP_009555238.1;HCoV-OC43_orf1ab;3794 | 0 | 0 |
| NC_006213.1;YP_009555238.1;HCoV-OC43_orf1ab;3795 | 0 | 0 |
| NC_006213.1;YP_009555238.1;HCoV-OC43_orf1ab;3796 | 0.1 | 0.025 |
| NC_006213.1;YP_009555238.1;HCoV-OC43_orf1ab;3797 | 0 | 0 |
| NC_006213.1;YP_009555238.1;HCoV-OC43_orf1ab;3798 | 0 | 0.05 |
| NC_006213.1;YP_009555238.1;HCoV-OC43_orf1ab;3799 | 0 | 0.025 |
| NC_006213.1;YP_009555238.1;HCoV-OC43_orf1ab;3800 | 0 | 0 |
| NC_006213.1;YP_009555238.1;HCoV-OC43_orf1ab;3801 | 0 | 0 |
| NC_006213.1;YP_009555238.1;HCoV-OC43_orf1ab;3802 | 0.05 | 0 |
| NC_006213.1;YP_009555238.1;HCoV-OC43_orf1ab;3803 | 0 | 0 |
| NC_006213.1;YP_009555238.1;HCoV-OC43_orf1ab;3804 | 0 | 0 |
| NC_006213.1;YP_009555238.1;HCoV-OC43_orf1ab;3805 | 0 | 0.025 |
| NC_006213.1;YP_009555238.1;HCoV-OC43_orf1ab;3806 | 0 | 0 |
| NC_006213.1;YP_009555238.1;HCoV-OC43_orf1ab;3807 | 0 | 0 |
| NC_006213.1;YP_009555238.1;HCoV-OC43_orf1ab;3808 | 0 | 0.025 |
| NC_006213.1;YP_009555238.1;HCoV-OC43_orf1ab;3809 | 0 | 0 |
| NC_006213.1;YP_009555238.1;HCoV-OC43_orf1ab;3810 | 0 | 0 |
| NC_006213.1;YP_009555238.1;HCoV-OC43_orf1ab;3811 | 0 | 0 |
| NC_006213.1;YP_009555238.1;HCoV-OC43_orf1ab;3812 | 0 | 0 |
| NC_006213.1;YP_009555238.1;HCoV-OC43_orf1ab;3813 | 0 | 0 |
| NC_006213.1;YP_009555238.1;HCoV-OC43_orf1ab;3814 | 0 | 0 |
| NC_006213.1;YP_009555238.1;HCoV-OC43_orf1ab;3815 | 0 | 0 |
| NC_006213.1;YP_009555238.1;HCoV-OC43_orf1ab;3816 | 0 | 0 |
| NC_006213.1;YP_009555238.1;HCoV-OC43_orf1ab;3817 | 0 | 0 |
| NC_006213.1;YP_009555238.1;HCoV-OC43_orf1ab;3818 | 0 | 0 |
| NC_006213.1;YP_009555238.1;HCoV-OC43_orf1ab;3819 | 0 | 0.025 |
| NC_006213.1;YP_009555238.1;HCoV-OC43_orf1ab;3820 | 0 | 0 |
| NC_006213.1;YP_009555238.1;HCoV-OC43_orf1ab;3821 | 0 | 0 |
| NC_006213.1;YP_009555238.1;HCoV-OC43_orf1ab;3822 | 0 | 0 |
| NC_006213.1;YP_009555238.1;HCoV-OC43_orf1ab;3823 | 0 | 0.025 |
| NC_006213.1;YP_009555238.1;HCoV-OC43_orf1ab;3824 | 0 | 0.025 |
| NC_006213.1;YP_009555238.1;HCoV-OC43_orf1ab;3825 | 0 | 0 |
| NC_006213.1;YP_009555238.1;HCoV-OC43_orf1ab;3826 | 0.05 | 0 |
| NC_006213.1;YP_009555238.1;HCoV-OC43_orf1ab;3827 | 0.05 | 0 |
| NC_006213.1;YP_009555238.1;HCoV-OC43_orf1ab;3828 | 0.05 | 0.025 |
| NC_006213.1;YP_009555238.1;HCoV-OC43_orf1ab;3829 | 0.1 | 0.025 |
| NC_006213.1;YP_009555238.1;HCoV-OC43_orf1ab;3830 | 0.05 | 0.025 |
| NC_006213.1;YP_009555238.1;HCoV-OC43_orf1ab;3831 | 0.1 | 0.05 |
| NC_006213.1;YP_009555238.1;HCoV-OC43_orf1ab;3832 | 0 | 0 |
| NC_006213.1;YP_009555238.1;HCoV-OC43_orf1ab;3833 | 0 | 0 |
| NC_006213.1;YP_009555238.1;HCoV-OC43_orf1ab;3834 | 0 | 0 |
| NC_006213.1;YP_009555238.1;HCoV-OC43_orf1ab;3835 | 0 | 0 |
| NC_006213.1;YP_009555238.1;HCoV-OC43_orf1ab;3836 | 0 | 0 |

| | | |
|---|---|---|
| NC_006213.1;YP_009555238.1;HCoV-OC43_orf1ab;3837 | 0 | 0 |
| NC_006213.1;YP_009555238.1;HCoV-OC43_orf1ab;3838 | 0.05 | 0.025 |
| NC_006213.1;YP_009555238.1;HCoV-OC43_orf1ab;3839 | 0.15 | 0.025 |
| NC_006213.1;YP_009555238.1;HCoV-OC43_orf1ab;3840 | 0 | 0 |
| NC_006213.1;YP_009555238.1;HCoV-OC43_orf1ab;3841 | 0 | 0 |
| NC_006213.1;YP_009555238.1;HCoV-OC43_orf1ab;3842 | 0 | 0 |
| NC_006213.1;YP_009555238.1;HCoV-OC43_orf1ab;3843 | 0 | 0 |
| NC_006213.1;YP_009555238.1;HCoV-OC43_orf1ab;3844 | 0 | 0 |
| NC_006213.1;YP_009555238.1;HCoV-OC43_orf1ab;3845 | 0 | 0 |
| NC_006213.1;YP_009555238.1;HCoV-OC43_orf1ab;3846 | 0 | 0 |
| NC_006213.1;YP_009555238.1;HCoV-OC43_orf1ab;3847 | 0 | 0 |
| NC_006213.1;YP_009555238.1;HCoV-OC43_orf1ab;3848 | 0 | 0.05 |
| NC_006213.1;YP_009555238.1;HCoV-OC43_orf1ab;3849 | 0 | 0.075 |
| NC_006213.1;YP_009555238.1;HCoV-OC43_orf1ab;3850 | 0.05 | 0.05 |
| NC_006213.1;YP_009555238.1;HCoV-OC43_orf1ab;3851 | 0.05 | 0.05 |
| NC_006213.1;YP_009555238.1;HCoV-OC43_orf1ab;3852 | 0.05 | 0.025 |
| NC_006213.1;YP_009555238.1;HCoV-OC43_orf1ab;3853 | 0.05 | 0 |
| NC_006213.1;YP_009555238.1;HCoV-OC43_orf1ab;3854 | 0.05 | 0 |
| NC_006213.1;YP_009555238.1;HCoV-OC43_orf1ab;3855 | 0.05 | 0 |
| NC_006213.1;YP_009555238.1;HCoV-OC43_orf1ab;3856 | 0 | 0 |
| NC_006213.1;YP_009555238.1;HCoV-OC43_orf1ab;3857 | 0 | 0 |
| NC_006213.1;YP_009555238.1;HCoV-OC43_orf1ab;3858 | 0.05 | 0.025 |
| NC_006213.1;YP_009555238.1;HCoV-OC43_orf1ab;3859 | 0.15 | 0 |
| NC_006213.1;YP_009555238.1;HCoV-OC43_orf1ab;3860 | 0 | 0 |
| NC_006213.1;YP_009555238.1;HCoV-OC43_orf1ab;3861 | 0 | 0 |
| NC_006213.1;YP_009555238.1;HCoV-OC43_orf1ab;3862 | 0 | 0 |
| NC_006213.1;YP_009555238.1;HCoV-OC43_orf1ab;3863 | 0 | 0 |
| NC_006213.1;YP_009555238.1;HCoV-OC43_orf1ab;3864 | 0 | 0 |
| NC_006213.1;YP_009555238.1;HCoV-OC43_orf1ab;3865 | 0 | 0 |
| NC_006213.1;YP_009555238.1;HCoV-OC43_orf1ab;3866 | 0 | 0 |
| NC_006213.1;YP_009555238.1;HCoV-OC43_orf1ab;3867 | 0 | 0 |
| NC_006213.1;YP_009555238.1;HCoV-OC43_orf1ab;3868 | 0 | 0 |
| NC_006213.1;YP_009555238.1;HCoV-OC43_orf1ab;3869 | 0.05 | 0 |
| NC_006213.1;YP_009555238.1;HCoV-OC43_orf1ab;3870 | 0 | 0 |
| NC_006213.1;YP_009555238.1;HCoV-OC43_orf1ab;3871 | 0 | 0 |
| NC_006213.1;YP_009555238.1;HCoV-OC43_orf1ab;3872 | 0 | 0 |
| NC_006213.1;YP_009555238.1;HCoV-OC43_orf1ab;3873 | 0 | 0 |
| NC_006213.1;YP_009555238.1;HCoV-OC43_orf1ab;3874 | 0 | 0 |
| NC_006213.1;YP_009555238.1;HCoV-OC43_orf1ab;3875 | 0 | 0 |
| NC_006213.1;YP_009555238.1;HCoV-OC43_orf1ab;3876 | 0 | 0 |
| NC_006213.1;YP_009555238.1;HCoV-OC43_orf1ab;3877 | 0 | 0 |
| NC_006213.1;YP_009555238.1;HCoV-OC43_orf1ab;3878 | 0 | 0 |
| NC_006213.1;YP_009555238.1;HCoV-OC43_orf1ab;3879 | 0 | 0 |

| | | |
|---|---|---|
| NC_006213.1;YP_009555238.1;HCoV-OC43_orf1ab;3880 | 0 | 0 |
| NC_006213.1;YP_009555238.1;HCoV-OC43_orf1ab;3881 | 0 | 0 |
| NC_006213.1;YP_009555238.1;HCoV-OC43_orf1ab;3882 | 0 | 0 |
| NC_006213.1;YP_009555238.1;HCoV-OC43_orf1ab;3883 | 0 | 0 |
| NC_006213.1;YP_009555238.1;HCoV-OC43_orf1ab;3884 | 0 | 0 |
| NC_006213.1;YP_009555238.1;HCoV-OC43_orf1ab;3885 | 0 | 0 |
| NC_006213.1;YP_009555238.1;HCoV-OC43_orf1ab;3886 | 0 | 0 |
| NC_006213.1;YP_009555238.1;HCoV-OC43_orf1ab;3887 | 0 | 0 |
| NC_006213.1;YP_009555238.1;HCoV-OC43_orf1ab;3888 | 0 | 0 |
| NC_006213.1;YP_009555238.1;HCoV-OC43_orf1ab;3889 | 0 | 0 |
| NC_006213.1;YP_009555238.1;HCoV-OC43_orf1ab;3890 | 0 | 0 |
| NC_006213.1;YP_009555238.1;HCoV-OC43_orf1ab;3891 | 0 | 0 |
| NC_006213.1;YP_009555238.1;HCoV-OC43_orf1ab;3892 | 0 | 0 |
| NC_006213.1;YP_009555238.1;HCoV-OC43_orf1ab;3893 | 0 | 0 |
| NC_006213.1;YP_009555238.1;HCoV-OC43_orf1ab;3894 | 0 | 0 |
| NC_006213.1;YP_009555238.1;HCoV-OC43_orf1ab;3895 | 0 | 0 |
| NC_006213.1;YP_009555238.1;HCoV-OC43_orf1ab;3896 | 0 | 0 |
| NC_006213.1;YP_009555238.1;HCoV-OC43_orf1ab;3897 | 0 | 0 |
| NC_006213.1;YP_009555238.1;HCoV-OC43_orf1ab;3898 | 0 | 0 |
| NC_006213.1;YP_009555238.1;HCoV-OC43_orf1ab;3899 | 0 | 0 |
| NC_006213.1;YP_009555238.1;HCoV-OC43_orf1ab;3900 | 0 | 0 |
| NC_006213.1;YP_009555238.1;HCoV-OC43_orf1ab;3901 | 0 | 0 |
| NC_006213.1;YP_009555238.1;HCoV-OC43_orf1ab;3902 | 0 | 0 |
| NC_006213.1;YP_009555238.1;HCoV-OC43_orf1ab;3903 | 0 | 0 |
| NC_006213.1;YP_009555238.1;HCoV-OC43_orf1ab;3904 | 0 | 0 |
| NC_006213.1;YP_009555238.1;HCoV-OC43_orf1ab;3905 | 0 | 0 |
| NC_006213.1;YP_009555238.1;HCoV-OC43_orf1ab;3906 | 0 | 0 |
| NC_006213.1;YP_009555238.1;HCoV-OC43_orf1ab;3907 | 0 | 0 |
| NC_006213.1;YP_009555238.1;HCoV-OC43_orf1ab;3908 | 0 | 0 |
| NC_006213.1;YP_009555238.1;HCoV-OC43_orf1ab;3909 | 0.05 | 0 |
| NC_006213.1;YP_009555238.1;HCoV-OC43_orf1ab;3910 | 0.05 | 0 |
| NC_006213.1;YP_009555238.1;HCoV-OC43_orf1ab;3911 | 0 | 0 |
| NC_006213.1;YP_009555238.1;HCoV-OC43_orf1ab;3912 | 0 | 0 |
| NC_006213.1;YP_009555238.1;HCoV-OC43_orf1ab;3913 | 0 | 0 |
| NC_006213.1;YP_009555238.1;HCoV-OC43_orf1ab;3914 | 0 | 0 |
| NC_006213.1;YP_009555238.1;HCoV-OC43_orf1ab;3915 | 0 | 0 |
| NC_006213.1;YP_009555238.1;HCoV-OC43_orf1ab;3916 | 0 | 0 |
| NC_006213.1;YP_009555238.1;HCoV-OC43_orf1ab;3917 | 0 | 0 |
| NC_006213.1;YP_009555238.1;HCoV-OC43_orf1ab;3918 | 0 | 0 |
| NC_006213.1;YP_009555238.1;HCoV-OC43_orf1ab;3919 | 0 | 0.025 |
| NC_006213.1;YP_009555238.1;HCoV-OC43_orf1ab;3920 | 0 | 0 |
| NC_006213.1;YP_009555238.1;HCoV-OC43_orf1ab;3921 | 0 | 0 |
| NC_006213.1;YP_009555238.1;HCoV-OC43_orf1ab;3922 | 0 | 0 |

| | | |
|---|---|---|
| NC_006213.1;YP_009555238.1;HCoV-OC43_orf1ab;3923 | 0 | 0 |
| NC_006213.1;YP_009555238.1;HCoV-OC43_orf1ab;3924 | 0 | 0.025 |
| NC_006213.1;YP_009555238.1;HCoV-OC43_orf1ab;3925 | 0 | 0 |
| NC_006213.1;YP_009555238.1;HCoV-OC43_orf1ab;3926 | 0 | 0 |
| NC_006213.1;YP_009555238.1;HCoV-OC43_orf1ab;3927 | 0 | 0 |
| NC_006213.1;YP_009555238.1;HCoV-OC43_orf1ab;3928 | 0 | 0 |
| NC_006213.1;YP_009555238.1;HCoV-OC43_orf1ab;3929 | 0 | 0 |
| NC_006213.1;YP_009555238.1;HCoV-OC43_orf1ab;3930 | 0.05 | 0 |
| NC_006213.1;YP_009555238.1;HCoV-OC43_orf1ab;3931 | 0.05 | 0.025 |
| NC_006213.1;YP_009555238.1;HCoV-OC43_orf1ab;3932 | 0 | 0 |
| NC_006213.1;YP_009555238.1;HCoV-OC43_orf1ab;3933 | 0.05 | 0 |
| NC_006213.1;YP_009555238.1;HCoV-OC43_orf1ab;3934 | 0.05 | 0 |
| NC_006213.1;YP_009555238.1;HCoV-OC43_orf1ab;3935 | 0.05 | 0.05 |
| NC_006213.1;YP_009555238.1;HCoV-OC43_orf1ab;3936 | 0.05 | 0.025 |
| NC_006213.1;YP_009555238.1;HCoV-OC43_orf1ab;3937 | 0.05 | 0 |
| NC_006213.1;YP_009555238.1;HCoV-OC43_orf1ab;3938 | 0.05 | 0 |
| NC_006213.1;YP_009555238.1;HCoV-OC43_orf1ab;3939 | 0.1 | 0 |
| NC_006213.1;YP_009555238.1;HCoV-OC43_orf1ab;3940 | 0.1 | 0.025 |
| NC_006213.1;YP_009555238.1;HCoV-OC43_orf1ab;3941 | 0.05 | 0 |
| NC_006213.1;YP_009555238.1;HCoV-OC43_orf1ab;3942 | 0 | 0 |
| NC_006213.1;YP_009555238.1;HCoV-OC43_orf1ab;3943 | 0 | 0 |
| NC_006213.1;YP_009555238.1;HCoV-OC43_orf1ab;3944 | 0 | 0.025 |
| NC_006213.1;YP_009555238.1;HCoV-OC43_orf1ab;3945 | 0 | 0.025 |
| NC_006213.1;YP_009555238.1;HCoV-OC43_orf1ab;3946 | 0.05 | 0.05 |
| NC_006213.1;YP_009555238.1;HCoV-OC43_orf1ab;3947 | 0 | 0 |
| NC_006213.1;YP_009555238.1;HCoV-OC43_orf1ab;3948 | 0 | 0 |
| NC_006213.1;YP_009555238.1;HCoV-OC43_orf1ab;3949 | 0 | 0.05 |
| NC_006213.1;YP_009555238.1;HCoV-OC43_orf1ab;3950 | 0 | 0 |
| NC_006213.1;YP_009555238.1;HCoV-OC43_orf1ab;3951 | 0 | 0 |
| NC_006213.1;YP_009555238.1;HCoV-OC43_orf1ab;3952 | 0 | 0 |
| NC_006213.1;YP_009555238.1;HCoV-OC43_orf1ab;3953 | 0 | 0 |
| NC_006213.1;YP_009555238.1;HCoV-OC43_orf1ab;3954 | 0 | 0 |
| NC_006213.1;YP_009555238.1;HCoV-OC43_orf1ab;3955 | 0 | 0 |
| NC_006213.1;YP_009555238.1;HCoV-OC43_orf1ab;3956 | 0 | 0.025 |
| NC_006213.1;YP_009555238.1;HCoV-OC43_orf1ab;3957 | 0 | 0 |
| NC_006213.1;YP_009555238.1;HCoV-OC43_orf1ab;3958 | 0.05 | 0 |
| NC_006213.1;YP_009555238.1;HCoV-OC43_orf1ab;3959 | 0 | 0 |
| NC_006213.1;YP_009555238.1;HCoV-OC43_orf1ab;3960 | 0 | 0 |
| NC_006213.1;YP_009555238.1;HCoV-OC43_orf1ab;3961 | 0 | 0 |
| NC_006213.1;YP_009555238.1;HCoV-OC43_orf1ab;3962 | 0 | 0 |
| NC_006213.1;YP_009555238.1;HCoV-OC43_orf1ab;3963 | 0 | 0.025 |
| NC_006213.1;YP_009555238.1;HCoV-OC43_orf1ab;3964 | 0 | 0.05 |
| NC_006213.1;YP_009555238.1;HCoV-OC43_orf1ab;3965 | 0.05 | 0.1 |

| | | |
|---|---|---|
| NC_006213.1;YP_009555238.1;HCoV-OC43_orf1ab;3966 | 0 | 0.025 |
| NC_006213.1;YP_009555238.1;HCoV-OC43_orf1ab;3967 | 0.15 | 0.1 |
| NC_006213.1;YP_009555238.1;HCoV-OC43_orf1ab;3968 | 0.05 | 0.025 |
| NC_006213.1;YP_009555238.1;HCoV-OC43_orf1ab;3969 | 0 | 0 |
| NC_006213.1;YP_009555238.1;HCoV-OC43_orf1ab;3970 | 0 | 0 |
| NC_006213.1;YP_009555238.1;HCoV-OC43_orf1ab;3971 | 0 | 0 |
| NC_006213.1;YP_009555238.1;HCoV-OC43_orf1ab;3972 | 0 | 0.025 |
| NC_006213.1;YP_009555238.1;HCoV-OC43_orf1ab;3973 | 0 | 0 |
| NC_006213.1;YP_009555238.1;HCoV-OC43_orf1ab;3974 | 0 | 0 |
| NC_006213.1;YP_009555238.1;HCoV-OC43_orf1ab;3975 | 0 | 0.05 |
| NC_006213.1;YP_009555238.1;HCoV-OC43_orf1ab;3976 | 0 | 0 |
| NC_006213.1;YP_009555238.1;HCoV-OC43_orf1ab;3977 | 0.05 | 0.025 |
| NC_006213.1;YP_009555238.1;HCoV-OC43_orf1ab;3978 | 0 | 0 |
| NC_006213.1;YP_009555238.1;HCoV-OC43_orf1ab;3979 | 0 | 0 |
| NC_006213.1;YP_009555238.1;HCoV-OC43_orf1ab;3980 | 0 | 0 |
| NC_006213.1;YP_009555238.1;HCoV-OC43_orf1ab;3981 | 0 | 0 |
| NC_006213.1;YP_009555238.1;HCoV-OC43_orf1ab;3982 | 0 | 0.05 |
| NC_006213.1;YP_009555238.1;HCoV-OC43_orf1ab;3983 | 0 | 0.05 |
| NC_006213.1;YP_009555238.1;HCoV-OC43_orf1ab;3984 | 0 | 0.05 |
| NC_006213.1;YP_009555238.1;HCoV-OC43_orf1ab;3985 | 0 | 0 |
| NC_006213.1;YP_009555238.1;HCoV-OC43_orf1ab;3986 | 0.05 | 0.025 |
| NC_006213.1;YP_009555238.1;HCoV-OC43_orf1ab;3987 | 0 | 0.025 |
| NC_006213.1;YP_009555238.1;HCoV-OC43_orf1ab;3988 | 0 | 0.025 |
| NC_006213.1;YP_009555238.1;HCoV-OC43_orf1ab;3989 | 0 | 0.025 |
| NC_006213.1;YP_009555238.1;HCoV-OC43_orf1ab;3990 | 0.05 | 0.025 |
| NC_006213.1;YP_009555238.1;HCoV-OC43_orf1ab;3991 | 0 | 0.025 |
| NC_006213.1;YP_009555238.1;HCoV-OC43_orf1ab;3992 | 0 | 0 |
| NC_006213.1;YP_009555238.1;HCoV-OC43_orf1ab;3993 | 0 | 0 |
| NC_006213.1;YP_009555238.1;HCoV-OC43_orf1ab;3994 | 0 | 0 |
| NC_006213.1;YP_009555238.1;HCoV-OC43_orf1ab;3995 | 0 | 0 |
| NC_006213.1;YP_009555238.1;HCoV-OC43_orf1ab;3996 | 0 | 0 |
| NC_006213.1;YP_009555238.1;HCoV-OC43_orf1ab;3997 | 0 | 0 |
| NC_006213.1;YP_009555238.1;HCoV-OC43_orf1ab;3998 | 0 | 0 |
| NC_006213.1;YP_009555238.1;HCoV-OC43_orf1ab;3999 | 0.05 | 0 |
| NC_006213.1;YP_009555238.1;HCoV-OC43_orf1ab;4000 | 0 | 0 |
| NC_006213.1;YP_009555238.1;HCoV-OC43_orf1ab;4001 | 0 | 0 |
| NC_006213.1;YP_009555238.1;HCoV-OC43_orf1ab;4002 | 0 | 0 |
| NC_006213.1;YP_009555238.1;HCoV-OC43_orf1ab;4003 | 0 | 0 |
| NC_006213.1;YP_009555238.1;HCoV-OC43_orf1ab;4004 | 0 | 0 |
| NC_006213.1;YP_009555238.1;HCoV-OC43_orf1ab;4005 | 0 | 0.025 |
| NC_006213.1;YP_009555238.1;HCoV-OC43_orf1ab;4006 | 0 | 0 |
| NC_006213.1;YP_009555238.1;HCoV-OC43_orf1ab;4007 | 0 | 0 |
| NC_006213.1;YP_009555238.1;HCoV-OC43_orf1ab;4008 | 0 | 0 |

| | | |
|---|---|---|
| NC_006213.1;YP_009555238.1;HCoV-OC43_orf1ab;4009 | 0 | 0 |
| NC_006213.1;YP_009555238.1;HCoV-OC43_orf1ab;4010 | 0 | 0 |
| NC_006213.1;YP_009555238.1;HCoV-OC43_orf1ab;4011 | 0 | 0 |
| NC_006213.1;YP_009555238.1;HCoV-OC43_orf1ab;4012 | 0 | 0 |
| NC_006213.1;YP_009555238.1;HCoV-OC43_orf1ab;4013 | 0 | 0 |
| NC_006213.1;YP_009555238.1;HCoV-OC43_orf1ab;4014 | 0 | 0 |
| NC_006213.1;YP_009555238.1;HCoV-OC43_orf1ab;4015 | 0.2 | 0.175 |
| NC_006213.1;YP_009555238.1;HCoV-OC43_orf1ab;4016 | 0.1 | 0.1 |
| NC_006213.1;YP_009555238.1;HCoV-OC43_orf1ab;4017 | 0.15 | 0.1 |
| NC_006213.1;YP_009555238.1;HCoV-OC43_orf1ab;4018 | 0.2 | 0.1 |
| NC_006213.1;YP_009555238.1;HCoV-OC43_orf1ab;4019 | 0.25 | 0.3 |
| NC_006213.1;YP_009555238.1;HCoV-OC43_orf1ab;4020 | 0.25 | 0.425 |
| NC_006213.1;YP_009555238.1;HCoV-OC43_orf1ab;4021 | 0.3 | 0.3 |
| NC_006213.1;YP_009555238.1;HCoV-OC43_orf1ab;4022 | 0.35 | 0.4 |
| NC_006213.1;YP_009555238.1;HCoV-OC43_orf1ab;4023 | 0.3 | 0.3 |
| NC_006213.1;YP_009555238.1;HCoV-OC43_orf1ab;4024 | 0.1 | 0 |
| NC_006213.1;YP_009555238.1;HCoV-OC43_orf1ab;4025 | 0.1 | 0.075 |
| NC_006213.1;YP_009555238.1;HCoV-OC43_orf1ab;4026 | 0.05 | 0.025 |
| NC_006213.1;YP_009555238.1;HCoV-OC43_orf1ab;4027 | 0.1 | 0.05 |
| NC_006213.1;YP_009555238.1;HCoV-OC43_orf1ab;4028 | 0.15 | 0.05 |
| NC_006213.1;YP_009555238.1;HCoV-OC43_orf1ab;4029 | 0.05 | 0.025 |
| NC_006213.1;YP_009555238.1;HCoV-OC43_orf1ab;4030 | 0.05 | 0.025 |
| NC_006213.1;YP_009555238.1;HCoV-OC43_orf1ab;4031 | 0.05 | 0.025 |
| NC_006213.1;YP_009555238.1;HCoV-OC43_orf1ab;4032 | 0 | 0.025 |
| NC_006213.1;YP_009555238.1;HCoV-OC43_orf1ab;4033 | 0 | 0.025 |
| NC_006213.1;YP_009555238.1;HCoV-OC43_orf1ab;4034 | 0.15 | 0.1 |
| NC_006213.1;YP_009555238.1;HCoV-OC43_orf1ab;4035 | 0 | 0.025 |
| NC_006213.1;YP_009555238.1;HCoV-OC43_orf1ab;4036 | 0.05 | 0.025 |
| NC_006213.1;YP_009555238.1;HCoV-OC43_orf1ab;4037 | 0 | 0 |
| NC_006213.1;YP_009555238.1;HCoV-OC43_orf1ab;4038 | 0.1 | 0.125 |
| NC_006213.1;YP_009555238.1;HCoV-OC43_orf1ab;4039 | 0 | 0 |
| NC_006213.1;YP_009555238.1;HCoV-OC43_orf1ab;4040 | 0.1 | 0 |
| NC_006213.1;YP_009555238.1;HCoV-OC43_orf1ab;4041 | 0.05 | 0 |
| NC_006213.1;YP_009555238.1;HCoV-OC43_orf1ab;4042 | 0.05 | 0.1 |
| NC_006213.1;YP_009555238.1;HCoV-OC43_orf1ab;4043 | 0 | 0 |
| NC_006213.1;YP_009555238.1;HCoV-OC43_orf1ab;4044 | 0.05 | 0.025 |
| NC_006213.1;YP_009555238.1;HCoV-OC43_orf1ab;4045 | 0.05 | 0.05 |
| NC_006213.1;YP_009555238.1;HCoV-OC43_orf1ab;4046 | 0.1 | 0.075 |
| NC_006213.1;YP_009555238.1;HCoV-OC43_orf1ab;4047 | 0.1 | 0.1 |
| NC_006213.1;YP_009555238.1;HCoV-OC43_orf1ab;4048 | 0.2 | 0.15 |
| NC_006213.1;YP_009555238.1;HCoV-OC43_orf1ab;4049 | 0.15 | 0.125 |
| NC_006213.1;YP_009555238.1;HCoV-OC43_orf1ab;4050 | 0.25 | 0.3 |
| NC_006213.1;YP_009555238.1;HCoV-OC43_orf1ab;4051 | 0.15 | 0.075 |

| | | |
|---|---|---|
| NC_006213.1;YP_009555238.1;HCoV-OC43_orf1ab;4052 | 0.05 | 0.025 |
| NC_006213.1;YP_009555238.1;HCoV-OC43_orf1ab;4053 | 0.05 | 0.05 |
| NC_006213.1;YP_009555238.1;HCoV-OC43_orf1ab;4054 | 0.05 | 0.05 |
| NC_006213.1;YP_009555238.1;HCoV-OC43_orf1ab;4055 | 0.05 | 0.025 |
| NC_006213.1;YP_009555238.1;HCoV-OC43_orf1ab;4056 | 0.15 | 0.05 |
| NC_006213.1;YP_009555238.1;HCoV-OC43_orf1ab;4057 | 0.05 | 0.025 |
| NC_006213.1;YP_009555238.1;HCoV-OC43_orf1ab;4058 | 0.05 | 0.025 |
| NC_006213.1;YP_009555238.1;HCoV-OC43_orf1ab;4059 | 0.05 | 0.025 |
| NC_006213.1;YP_009555238.1;HCoV-OC43_orf1ab;4060 | 0 | 0 |
| NC_006213.1;YP_009555238.1;HCoV-OC43_orf1ab;4061 | 0 | 0 |
| NC_006213.1;YP_009555238.1;HCoV-OC43_orf1ab;4062 | 0 | 0.025 |
| NC_006213.1;YP_009555238.1;HCoV-OC43_orf1ab;4063 | 0 | 0.025 |
| NC_006213.1;YP_009555238.1;HCoV-OC43_orf1ab;4064 | 0 | 0.025 |
| NC_006213.1;YP_009555238.1;HCoV-OC43_orf1ab;4065 | 0 | 0 |
| NC_006213.1;YP_009555238.1;HCoV-OC43_orf1ab;4066 | 0 | 0 |
| NC_006213.1;YP_009555238.1;HCoV-OC43_orf1ab;4067 | 0 | 0 |
| NC_006213.1;YP_009555238.1;HCoV-OC43_orf1ab;4068 | 0 | 0 |
| NC_006213.1;YP_009555238.1;HCoV-OC43_orf1ab;4069 | 0 | 0 |
| NC_006213.1;YP_009555238.1;HCoV-OC43_orf1ab;4070 | 0 | 0 |
| NC_006213.1;YP_009555238.1;HCoV-OC43_orf1ab;4071 | 0 | 0 |
| NC_006213.1;YP_009555238.1;HCoV-OC43_orf1ab;4072 | 0 | 0.05 |
| NC_006213.1;YP_009555238.1;HCoV-OC43_orf1ab;4073 | 0 | 0.05 |
| NC_006213.1;YP_009555238.1;HCoV-OC43_orf1ab;4074 | 0.1 | 0.05 |
| NC_006213.1;YP_009555238.1;HCoV-OC43_orf1ab;4075 | 0.15 | 0.05 |
| NC_006213.1;YP_009555238.1;HCoV-OC43_orf1ab;4076 | 0.15 | 0.1 |
| NC_006213.1;YP_009555238.1;HCoV-OC43_orf1ab;4077 | 0.15 | 0.075 |
| NC_006213.1;YP_009555238.1;HCoV-OC43_orf1ab;4078 | 0.15 | 0.1 |
| NC_006213.1;YP_009555238.1;HCoV-OC43_orf1ab;4079 | 0.05 | 0.15 |
| NC_006213.1;YP_009555238.1;HCoV-OC43_orf1ab;4080 | 0.1 | 0.15 |
| NC_006213.1;YP_009555238.1;HCoV-OC43_orf1ab;4081 | 0 | 0.075 |
| NC_006213.1;YP_009555238.1;HCoV-OC43_orf1ab;4082 | 0 | 0.025 |
| NC_006213.1;YP_009555238.1;HCoV-OC43_orf1ab;4083 | 0.05 | 0.05 |
| NC_006213.1;YP_009555238.1;HCoV-OC43_orf1ab;4084 | 0 | 0.05 |
| NC_006213.1;YP_009555238.1;HCoV-OC43_orf1ab;4085 | 0.1 | 0.1 |
| NC_006213.1;YP_009555238.1;HCoV-OC43_orf1ab;4086 | 0.1 | 0 |
| NC_006213.1;YP_009555238.1;HCoV-OC43_orf1ab;4087 | 0.1 | 0.025 |
| NC_006213.1;YP_009555238.1;HCoV-OC43_orf1ab;4088 | 0.05 | 0.025 |
| NC_006213.1;YP_009555238.1;HCoV-OC43_orf1ab;4089 | 0.1 | 0.05 |
| NC_006213.1;YP_009555238.1;HCoV-OC43_orf1ab;4090 | 0.05 | 0.05 |
| NC_006213.1;YP_009555238.1;HCoV-OC43_orf1ab;4091 | 0.05 | 0.025 |
| NC_006213.1;YP_009555238.1;HCoV-OC43_orf1ab;4092 | 0.05 | 0.05 |
| NC_006213.1;YP_009555238.1;HCoV-OC43_orf1ab;4093 | 0.05 | 0.025 |
| NC_006213.1;YP_009555238.1;HCoV-OC43_orf1ab;4094 | 0 | 0 |

| | | |
|---|---|---|
| NC_006213.1;YP_009555238.1;HCoV-OC43_orf1ab;4095 | 0 | 0 |
| NC_006213.1;YP_009555238.1;HCoV-OC43_orf1ab;4096 | 0 | 0 |
| NC_006213.1;YP_009555238.1;HCoV-OC43_orf1ab;4097 | 0 | 0 |
| NC_006213.1;YP_009555238.1;HCoV-OC43_orf1ab;4098 | 0 | 0 |
| NC_006213.1;YP_009555238.1;HCoV-OC43_orf1ab;4099 | 0 | 0 |
| NC_006213.1;YP_009555238.1;HCoV-OC43_orf1ab;4100 | 0 | 0 |
| NC_006213.1;YP_009555238.1;HCoV-OC43_orf1ab;4101 | 0 | 0 |
| NC_006213.1;YP_009555238.1;HCoV-OC43_orf1ab;4102 | 0 | 0 |
| NC_006213.1;YP_009555238.1;HCoV-OC43_orf1ab;4103 | 0 | 0 |
| NC_006213.1;YP_009555238.1;HCoV-OC43_orf1ab;4104 | 0 | 0 |
| NC_006213.1;YP_009555238.1;HCoV-OC43_orf1ab;4105 | 0 | 0 |
| NC_006213.1;YP_009555238.1;HCoV-OC43_orf1ab;4106 | 0 | 0 |
| NC_006213.1;YP_009555238.1;HCoV-OC43_orf1ab;4107 | 0 | 0 |
| NC_006213.1;YP_009555238.1;HCoV-OC43_orf1ab;4108 | 0 | 0 |
| NC_006213.1;YP_009555238.1;HCoV-OC43_orf1ab;4109 | 0.1 | 0.025 |
| NC_006213.1;YP_009555238.1;HCoV-OC43_orf1ab;4110 | 0.1 | 0 |
| NC_006213.1;YP_009555238.1;HCoV-OC43_orf1ab;4111 | 0 | 0 |
| NC_006213.1;YP_009555238.1;HCoV-OC43_orf1ab;4112 | 0 | 0 |
| NC_006213.1;YP_009555238.1;HCoV-OC43_orf1ab;4113 | 0 | 0 |
| NC_006213.1;YP_009555238.1;HCoV-OC43_orf1ab;4114 | 0 | 0 |
| NC_006213.1;YP_009555238.1;HCoV-OC43_orf1ab;4115 | 0 | 0 |
| NC_006213.1;YP_009555238.1;HCoV-OC43_orf1ab;4116 | 0 | 0 |
| NC_006213.1;YP_009555238.1;HCoV-OC43_orf1ab;4117 | 0 | 0 |
| NC_006213.1;YP_009555238.1;HCoV-OC43_orf1ab;4118 | 0.05 | 0 |
| NC_006213.1;YP_009555238.1;HCoV-OC43_orf1ab;4119 | 0.1 | 0 |
| NC_006213.1;YP_009555238.1;HCoV-OC43_orf1ab;4120 | 0.1 | 0 |
| NC_006213.1;YP_009555238.1;HCoV-OC43_orf1ab;4121 | 0 | 0 |
| NC_006213.1;YP_009555238.1;HCoV-OC43_orf1ab;4122 | 0 | 0 |
| NC_006213.1;YP_009555238.1;HCoV-OC43_orf1ab;4123 | 0 | 0.025 |
| NC_006213.1;YP_009555238.1;HCoV-OC43_orf1ab;4124 | 0 | 0 |
| NC_006213.1;YP_009555238.1;HCoV-OC43_orf1ab;4125 | 0 | 0 |
| NC_006213.1;YP_009555238.1;HCoV-OC43_orf1ab;4126 | 0 | 0 |
| NC_006213.1;YP_009555238.1;HCoV-OC43_orf1ab;4127 | 0 | 0 |
| NC_006213.1;YP_009555238.1;HCoV-OC43_orf1ab;4128 | 0 | 0.025 |
| NC_006213.1;YP_009555238.1;HCoV-OC43_orf1ab;4129 | 0 | 0.025 |
| NC_006213.1;YP_009555238.1;HCoV-OC43_orf1ab;4130 | 0 | 0 |
| NC_006213.1;YP_009555238.1;HCoV-OC43_orf1ab;4131 | 0 | 0 |
| NC_006213.1;YP_009555238.1;HCoV-OC43_orf1ab;4132 | 0 | 0.025 |
| NC_006213.1;YP_009555238.1;HCoV-OC43_orf1ab;4133 | 0.05 | 0.05 |
| NC_006213.1;YP_009555238.1;HCoV-OC43_orf1ab;4134 | 0 | 0 |
| NC_006213.1;YP_009555238.1;HCoV-OC43_orf1ab;4135 | 0 | 0 |
| NC_006213.1;YP_009555238.1;HCoV-OC43_orf1ab;4136 | 0 | 0 |
| NC_006213.1;YP_009555238.1;HCoV-OC43_orf1ab;4137 | 0 | 0 |

| | | |
|---|---|---|
| NC_006213.1;YP_009555238.1;HCoV-OC43_orf1ab;4138 | 0 | 0 |
| NC_006213.1;YP_009555238.1;HCoV-OC43_orf1ab;4139 | 0 | 0 |
| NC_006213.1;YP_009555238.1;HCoV-OC43_orf1ab;4140 | 0 | 0.025 |
| NC_006213.1;YP_009555238.1;HCoV-OC43_orf1ab;4141 | 0.05 | 0 |
| NC_006213.1;YP_009555238.1;HCoV-OC43_orf1ab;4142 | 0.35 | 0.325 |
| NC_006213.1;YP_009555238.1;HCoV-OC43_orf1ab;4143 | 0.4 | 0.3 |
| NC_006213.1;YP_009555238.1;HCoV-OC43_orf1ab;4144 | 0.2 | 0.25 |
| NC_006213.1;YP_009555238.1;HCoV-OC43_orf1ab;4145 | 0.2 | 0.25 |
| NC_006213.1;YP_009555238.1;HCoV-OC43_orf1ab;4146 | 0.2 | 0.2 |
| NC_006213.1;YP_009555238.1;HCoV-OC43_orf1ab;4147 | 0.05 | 0.05 |
| NC_006213.1;YP_009555238.1;HCoV-OC43_orf1ab;4148 | 0.1 | 0.125 |
| NC_006213.1;YP_009555238.1;HCoV-OC43_orf1ab;4149 | 0.05 | 0.025 |
| NC_006213.1;YP_009555238.1;HCoV-OC43_orf1ab;4150 | 0.05 | 0 |
| NC_006213.1;YP_009555238.1;HCoV-OC43_orf1ab;4151 | 0.05 | 0.025 |
| NC_006213.1;YP_009555238.1;HCoV-OC43_orf1ab;4152 | 0.05 | 0.025 |
| NC_006213.1;YP_009555238.1;HCoV-OC43_orf1ab;4153 | 0 | 0 |
| NC_006213.1;YP_009555238.1;HCoV-OC43_orf1ab;4154 | 0 | 0 |
| NC_006213.1;YP_009555238.1;HCoV-OC43_orf1ab;4155 | 0 | 0 |
| NC_006213.1;YP_009555238.1;HCoV-OC43_orf1ab;4156 | 0 | 0 |
| NC_006213.1;YP_009555238.1;HCoV-OC43_orf1ab;4157 | 0 | 0 |
| NC_006213.1;YP_009555238.1;HCoV-OC43_orf1ab;4158 | 0 | 0 |
| NC_006213.1;YP_009555238.1;HCoV-OC43_orf1ab;4159 | 0.05 | 0.025 |
| NC_006213.1;YP_009555238.1;HCoV-OC43_orf1ab;4160 | 0 | 0 |
| NC_006213.1;YP_009555238.1;HCoV-OC43_orf1ab;4161 | 0 | 0 |
| NC_006213.1;YP_009555238.1;HCoV-OC43_orf1ab;4162 | 0 | 0 |
| NC_006213.1;YP_009555238.1;HCoV-OC43_orf1ab;4163 | 0 | 0 |
| NC_006213.1;YP_009555238.1;HCoV-OC43_orf1ab;4164 | 0 | 0 |
| NC_006213.1;YP_009555238.1;HCoV-OC43_orf1ab;4165 | 0 | 0.05 |
| NC_006213.1;YP_009555238.1;HCoV-OC43_orf1ab;4166 | 0 | 0.05 |
| NC_006213.1;YP_009555238.1;HCoV-OC43_orf1ab;4167 | 0.15 | 0.075 |
| NC_006213.1;YP_009555238.1;HCoV-OC43_orf1ab;4168 | 0.3 | 0.225 |
| NC_006213.1;YP_009555238.1;HCoV-OC43_orf1ab;4169 | 0.25 | 0.275 |
| NC_006213.1;YP_009555238.1;HCoV-OC43_orf1ab;4170 | 0.25 | 0.125 |
| NC_006213.1;YP_009555238.1;HCoV-OC43_orf1ab;4171 | 0.15 | 0.05 |
| NC_006213.1;YP_009555238.1;HCoV-OC43_orf1ab;4172 | 0.2 | 0.1 |
| NC_006213.1;YP_009555238.1;HCoV-OC43_orf1ab;4173 | 0.05 | 0 |
| NC_006213.1;YP_009555238.1;HCoV-OC43_orf1ab;4174 | 0.05 | 0 |
| NC_006213.1;YP_009555238.1;HCoV-OC43_orf1ab;4175 | 0.15 | 0.025 |
| NC_006213.1;YP_009555238.1;HCoV-OC43_orf1ab;4176 | 0.2 | 0 |
| NC_006213.1;YP_009555238.1;HCoV-OC43_orf1ab;4177 | 0.15 | 0.025 |
| NC_006213.1;YP_009555238.1;HCoV-OC43_orf1ab;4178 | 0.25 | 0.075 |
| NC_006213.1;YP_009555238.1;HCoV-OC43_orf1ab;4179 | 0 | 0 |
| NC_006213.1;YP_009555238.1;HCoV-OC43_orf1ab;4180 | 0 | 0 |

| | | |
|---|---|---|
| NC_006213.1;YP_009555238.1;HCoV-OC43_orf1ab;4181 | 0 | 0 |
| NC_006213.1;YP_009555238.1;HCoV-OC43_orf1ab;4182 | 0 | 0 |
| NC_006213.1;YP_009555238.1;HCoV-OC43_orf1ab;4183 | 0 | 0 |
| NC_006213.1;YP_009555238.1;HCoV-OC43_orf1ab;4184 | 0 | 0 |
| NC_006213.1;YP_009555238.1;HCoV-OC43_orf1ab;4185 | 0.05 | 0 |
| NC_006213.1;YP_009555238.1;HCoV-OC43_orf1ab;4186 | 0.05 | 0 |
| NC_006213.1;YP_009555238.1;HCoV-OC43_orf1ab;4187 | 0.05 | 0 |
| NC_006213.1;YP_009555238.1;HCoV-OC43_orf1ab;4188 | 0.05 | 0 |
| NC_006213.1;YP_009555238.1;HCoV-OC43_orf1ab;4189 | 0.05 | 0.05 |
| NC_006213.1;YP_009555238.1;HCoV-OC43_orf1ab;4190 | 0.05 | 0.025 |
| NC_006213.1;YP_009555238.1;HCoV-OC43_orf1ab;4191 | 0 | 0.025 |
| NC_006213.1;YP_009555238.1;HCoV-OC43_orf1ab;4192 | 0 | 0 |
| NC_006213.1;YP_009555238.1;HCoV-OC43_orf1ab;4193 | 0.05 | 0 |
| NC_006213.1;YP_009555238.1;HCoV-OC43_orf1ab;4194 | 0 | 0 |
| NC_006213.1;YP_009555238.1;HCoV-OC43_orf1ab;4195 | 0 | 0.025 |
| NC_006213.1;YP_009555238.1;HCoV-OC43_orf1ab;4196 | 0 | 0.025 |
| NC_006213.1;YP_009555238.1;HCoV-OC43_orf1ab;4197 | 0 | 0 |
| NC_006213.1;YP_009555238.1;HCoV-OC43_orf1ab;4198 | 0 | 0 |
| NC_006213.1;YP_009555238.1;HCoV-OC43_orf1ab;4199 | 0 | 0 |
| NC_006213.1;YP_009555238.1;HCoV-OC43_orf1ab;4200 | 0 | 0 |
| NC_006213.1;YP_009555238.1;HCoV-OC43_orf1ab;4201 | 0 | 0 |
| NC_006213.1;YP_009555238.1;HCoV-OC43_orf1ab;4202 | 0 | 0 |
| NC_006213.1;YP_009555238.1;HCoV-OC43_orf1ab;4203 | 0 | 0 |
| NC_006213.1;YP_009555238.1;HCoV-OC43_orf1ab;4204 | 0 | 0 |
| NC_006213.1;YP_009555238.1;HCoV-OC43_orf1ab;4205 | 0 | 0 |
| NC_006213.1;YP_009555238.1;HCoV-OC43_orf1ab;4206 | 0 | 0 |
| NC_006213.1;YP_009555238.1;HCoV-OC43_orf1ab;4207 | 0 | 0 |
| NC_006213.1;YP_009555238.1;HCoV-OC43_orf1ab;4208 | 0 | 0 |
| NC_006213.1;YP_009555238.1;HCoV-OC43_orf1ab;4209 | 0 | 0 |
| NC_006213.1;YP_009555238.1;HCoV-OC43_orf1ab;4210 | 0 | 0 |
| NC_006213.1;YP_009555238.1;HCoV-OC43_orf1ab;4211 | 0 | 0 |
| NC_006213.1;YP_009555238.1;HCoV-OC43_orf1ab;4212 | 0 | 0 |
| NC_006213.1;YP_009555238.1;HCoV-OC43_orf1ab;4213 | 0 | 0 |
| NC_006213.1;YP_009555238.1;HCoV-OC43_orf1ab;4214 | 0 | 0 |
| NC_006213.1;YP_009555238.1;HCoV-OC43_orf1ab;4215 | 0 | 0 |
| NC_006213.1;YP_009555238.1;HCoV-OC43_orf1ab;4216 | 0 | 0 |
| NC_006213.1;YP_009555238.1;HCoV-OC43_orf1ab;4217 | 0 | 0 |
| NC_006213.1;YP_009555238.1;HCoV-OC43_orf1ab;4218 | 0 | 0 |
| NC_006213.1;YP_009555238.1;HCoV-OC43_orf1ab;4219 | 0 | 0 |
| NC_006213.1;YP_009555238.1;HCoV-OC43_orf1ab;4220 | 0 | 0 |
| NC_006213.1;YP_009555238.1;HCoV-OC43_orf1ab;4221 | 0 | 0 |
| NC_006213.1;YP_009555238.1;HCoV-OC43_orf1ab;4222 | 0 | 0 |
| NC_006213.1;YP_009555238.1;HCoV-OC43_orf1ab;4223 | 0.05 | 0 |

| | | |
|---|---|---|
| NC_006213.1;YP_009555238.1;HCoV-OC43_orf1ab;4224 | 0.05 | 0 |
| NC_006213.1;YP_009555238.1;HCoV-OC43_orf1ab;4225 | 0.1 | 0 |
| NC_006213.1;YP_009555238.1;HCoV-OC43_orf1ab;4226 | 0.1 | 0.025 |
| NC_006213.1;YP_009555238.1;HCoV-OC43_orf1ab;4227 | 0.05 | 0 |
| NC_006213.1;YP_009555238.1;HCoV-OC43_orf1ab;4228 | 0.05 | 0 |
| NC_006213.1;YP_009555238.1;HCoV-OC43_orf1ab;4229 | 0.05 | 0.025 |
| NC_006213.1;YP_009555238.1;HCoV-OC43_orf1ab;4230 | 0 | 0.05 |
| NC_006213.1;YP_009555238.1;HCoV-OC43_orf1ab;4231 | 0.1 | 0.025 |
| NC_006213.1;YP_009555238.1;HCoV-OC43_orf1ab;4232 | 0 | 0 |
| NC_006213.1;YP_009555238.1;HCoV-OC43_orf1ab;4233 | 0 | 0 |
| NC_006213.1;YP_009555238.1;HCoV-OC43_orf1ab;4234 | 0 | 0 |
| NC_006213.1;YP_009555238.1;HCoV-OC43_orf1ab;4235 | 0 | 0 |
| NC_006213.1;YP_009555238.1;HCoV-OC43_orf1ab;4236 | 0 | 0 |
| NC_006213.1;YP_009555238.1;HCoV-OC43_orf1ab;4237 | 0 | 0 |
| NC_006213.1;YP_009555238.1;HCoV-OC43_orf1ab;4238 | 0 | 0 |
| NC_006213.1;YP_009555238.1;HCoV-OC43_orf1ab;4239 | 0 | 0 |
| NC_006213.1;YP_009555238.1;HCoV-OC43_orf1ab;4240 | 0.05 | 0 |
| NC_006213.1;YP_009555238.1;HCoV-OC43_orf1ab;4241 | 0 | 0.025 |
| NC_006213.1;YP_009555238.1;HCoV-OC43_orf1ab;4242 | 0 | 0 |
| NC_006213.1;YP_009555238.1;HCoV-OC43_orf1ab;4243 | 0 | 0 |
| NC_006213.1;YP_009555238.1;HCoV-OC43_orf1ab;4244 | 0 | 0 |
| NC_006213.1;YP_009555238.1;HCoV-OC43_orf1ab;4245 | 0 | 0 |
| NC_006213.1;YP_009555238.1;HCoV-OC43_orf1ab;4246 | 0 | 0.025 |
| NC_006213.1;YP_009555238.1;HCoV-OC43_orf1ab;4247 | 0 | 0.025 |
| NC_006213.1;YP_009555238.1;HCoV-OC43_orf1ab;4248 | 0 | 0 |
| NC_006213.1;YP_009555238.1;HCoV-OC43_orf1ab;4249 | 0 | 0 |
| NC_006213.1;YP_009555238.1;HCoV-OC43_orf1ab;4250 | 0.05 | 0.025 |
| NC_006213.1;YP_009555238.1;HCoV-OC43_orf1ab;4251 | 0.05 | 0.025 |
| NC_006213.1;YP_009555238.1;HCoV-OC43_orf1ab;4252 | 0.1 | 0.025 |
| NC_006213.1;YP_009555238.1;HCoV-OC43_orf1ab;4253 | 0.1 | 0.05 |
| NC_006213.1;YP_009555238.1;HCoV-OC43_orf1ab;4254 | 0 | 0.025 |
| NC_006213.1;YP_009555238.1;HCoV-OC43_orf1ab;4255 | 0 | 0 |
| NC_006213.1;YP_009555238.1;HCoV-OC43_orf1ab;4256 | 0 | 0 |
| NC_006213.1;YP_009555238.1;HCoV-OC43_orf1ab;4257 | 0 | 0.025 |
| NC_006213.1;YP_009555238.1;HCoV-OC43_orf1ab;4258 | 0 | 0.025 |
| NC_006213.1;YP_009555238.1;HCoV-OC43_orf1ab;4259 | 0 | 0.025 |
| NC_006213.1;YP_009555238.1;HCoV-OC43_orf1ab;4260 | 0 | 0 |
| NC_006213.1;YP_009555238.1;HCoV-OC43_orf1ab;4261 | 0.05 | 0 |
| NC_006213.1;YP_009555238.1;HCoV-OC43_orf1ab;4262 | 0 | 0 |
| NC_006213.1;YP_009555238.1;HCoV-OC43_orf1ab;4263 | 0 | 0 |
| NC_006213.1;YP_009555238.1;HCoV-OC43_orf1ab;4264 | 0 | 0 |
| NC_006213.1;YP_009555238.1;HCoV-OC43_orf1ab;4265 | 0 | 0 |
| NC_006213.1;YP_009555238.1;HCoV-OC43_orf1ab;4266 | 0 | 0 |

| | | |
|---|---|---|
| NC_006213.1;YP_009555238.1;HCoV-OC43_orf1ab;4267 | 0 | 0 |
| NC_006213.1;YP_009555238.1;HCoV-OC43_orf1ab;4268 | 0 | 0 |
| NC_006213.1;YP_009555238.1;HCoV-OC43_orf1ab;4269 | 0 | 0 |
| NC_006213.1;YP_009555238.1;HCoV-OC43_orf1ab;4270 | 0 | 0 |
| NC_006213.1;YP_009555238.1;HCoV-OC43_orf1ab;4271 | 0 | 0 |
| NC_006213.1;YP_009555238.1;HCoV-OC43_orf1ab;4272 | 0 | 0 |
| NC_006213.1;YP_009555238.1;HCoV-OC43_orf1ab;4273 | 0 | 0 |
| NC_006213.1;YP_009555238.1;HCoV-OC43_orf1ab;4274 | 0 | 0 |
| NC_006213.1;YP_009555238.1;HCoV-OC43_orf1ab;4275 | 0 | 0 |
| NC_006213.1;YP_009555238.1;HCoV-OC43_orf1ab;4276 | 0 | 0 |
| NC_006213.1;YP_009555238.1;HCoV-OC43_orf1ab;4277 | 0 | 0 |
| NC_006213.1;YP_009555238.1;HCoV-OC43_orf1ab;4278 | 0 | 0 |
| NC_006213.1;YP_009555238.1;HCoV-OC43_orf1ab;4279 | 0 | 0 |
| NC_006213.1;YP_009555238.1;HCoV-OC43_orf1ab;4280 | 0 | 0 |
| NC_006213.1;YP_009555238.1;HCoV-OC43_orf1ab;4281 | 0 | 0 |
| NC_006213.1;YP_009555238.1;HCoV-OC43_orf1ab;4282 | 0 | 0.025 |
| NC_006213.1;YP_009555238.1;HCoV-OC43_orf1ab;4283 | 0 | 0 |
| NC_006213.1;YP_009555238.1;HCoV-OC43_orf1ab;4284 | 0 | 0 |
| NC_006213.1;YP_009555238.1;HCoV-OC43_orf1ab;4285 | 0.05 | 0 |
| NC_006213.1;YP_009555238.1;HCoV-OC43_orf1ab;4286 | 0 | 0 |
| NC_006213.1;YP_009555238.1;HCoV-OC43_orf1ab;4287 | 0.05 | 0 |
| NC_006213.1;YP_009555238.1;HCoV-OC43_orf1ab;4288 | 0 | 0 |
| NC_006213.1;YP_009555238.1;HCoV-OC43_orf1ab;4289 | 0 | 0 |
| NC_006213.1;YP_009555238.1;HCoV-OC43_orf1ab;4290 | 0 | 0 |
| NC_006213.1;YP_009555238.1;HCoV-OC43_orf1ab;4291 | 0 | 0 |
| NC_006213.1;YP_009555238.1;HCoV-OC43_orf1ab;4292 | 0 | 0 |
| NC_006213.1;YP_009555238.1;HCoV-OC43_orf1ab;4293 | 0 | 0 |
| NC_006213.1;YP_009555238.1;HCoV-OC43_orf1ab;4294 | 0 | 0 |
| NC_006213.1;YP_009555238.1;HCoV-OC43_orf1ab;4295 | 0 | 0 |
| NC_006213.1;YP_009555238.1;HCoV-OC43_orf1ab;4296 | 0 | 0 |
| NC_006213.1;YP_009555238.1;HCoV-OC43_orf1ab;4297 | 0 | 0 |
| NC_006213.1;YP_009555238.1;HCoV-OC43_orf1ab;4298 | 0 | 0 |
| NC_006213.1;YP_009555238.1;HCoV-OC43_orf1ab;4299 | 0 | 0 |
| NC_006213.1;YP_009555238.1;HCoV-OC43_orf1ab;4300 | 0 | 0 |
| NC_006213.1;YP_009555238.1;HCoV-OC43_orf1ab;4301 | 0 | 0 |
| NC_006213.1;YP_009555238.1;HCoV-OC43_orf1ab;4302 | 0 | 0 |
| NC_006213.1;YP_009555238.1;HCoV-OC43_orf1ab;4303 | 0 | 0 |
| NC_006213.1;YP_009555238.1;HCoV-OC43_orf1ab;4304 | 0 | 0 |
| NC_006213.1;YP_009555238.1;HCoV-OC43_orf1ab;4305 | 0 | 0 |
| NC_006213.1;YP_009555238.1;HCoV-OC43_orf1ab;4306 | 0 | 0 |
| NC_006213.1;YP_009555238.1;HCoV-OC43_orf1ab;4307 | 0 | 0 |
| NC_006213.1;YP_009555238.1;HCoV-OC43_orf1ab;4308 | 0 | 0 |
| NC_006213.1;YP_009555238.1;HCoV-OC43_orf1ab;4309 | 0 | 0 |

| | | |
|---|---|---|
| NC_006213.1;YP_009555238.1;HCoV-OC43_orf1ab;4310 | 0 | 0.05 |
| NC_006213.1;YP_009555238.1;HCoV-OC43_orf1ab;4311 | 0 | 0 |
| NC_006213.1;YP_009555238.1;HCoV-OC43_orf1ab;4312 | 0 | 0 |
| NC_006213.1;YP_009555238.1;HCoV-OC43_orf1ab;4313 | 0 | 0 |
| NC_006213.1;YP_009555238.1;HCoV-OC43_orf1ab;4314 | 0 | 0 |
| NC_006213.1;YP_009555238.1;HCoV-OC43_orf1ab;4315 | 0 | 0 |
| NC_006213.1;YP_009555238.1;HCoV-OC43_orf1ab;4316 | 0 | 0 |
| NC_006213.1;YP_009555238.1;HCoV-OC43_orf1ab;4317 | 0 | 0 |
| NC_006213.1;YP_009555238.1;HCoV-OC43_orf1ab;4318 | 0 | 0 |
| NC_006213.1;YP_009555238.1;HCoV-OC43_orf1ab;4319 | 0 | 0 |
| NC_006213.1;YP_009555238.1;HCoV-OC43_orf1ab;4320 | 0 | 0 |
| NC_006213.1;YP_009555238.1;HCoV-OC43_orf1ab;4321 | 0 | 0 |
| NC_006213.1;YP_009555238.1;HCoV-OC43_orf1ab;4322 | 0 | 0 |
| NC_006213.1;YP_009555238.1;HCoV-OC43_orf1ab;4323 | 0 | 0 |
| NC_006213.1;YP_009555238.1;HCoV-OC43_orf1ab;4324 | 0 | 0 |
| NC_006213.1;YP_009555238.1;HCoV-OC43_orf1ab;4325 | 0 | 0.025 |
| NC_006213.1;YP_009555238.1;HCoV-OC43_orf1ab;4326 | 0.05 | 0 |
| NC_006213.1;YP_009555238.1;HCoV-OC43_orf1ab;4327 | 0.05 | 0 |
| NC_006213.1;YP_009555238.1;HCoV-OC43_orf1ab;4328 | 0 | 0 |
| NC_006213.1;YP_009555238.1;HCoV-OC43_orf1ab;4329 | 0 | 0 |
| NC_006213.1;YP_009555238.1;HCoV-OC43_orf1ab;4330 | 0 | 0.025 |
| NC_006213.1;YP_009555238.1;HCoV-OC43_orf1ab;4331 | 0 | 0 |
| NC_006213.1;YP_009555238.1;HCoV-OC43_orf1ab;4332 | 0 | 0 |
| NC_006213.1;YP_009555238.1;HCoV-OC43_orf1ab;4333 | 0 | 0 |
| NC_006213.1;YP_009555238.1;HCoV-OC43_orf1ab;4334 | 0 | 0 |
| NC_006213.1;YP_009555238.1;HCoV-OC43_orf1ab;4335 | 0 | 0 |
| NC_006213.1;YP_009555238.1;HCoV-OC43_orf1ab;4336 | 0 | 0 |
| NC_006213.1;YP_009555238.1;HCoV-OC43_orf1ab;4337 | 0 | 0 |
| NC_006213.1;YP_009555238.1;HCoV-OC43_orf1ab;4338 | 0 | 0 |
| NC_006213.1;YP_009555238.1;HCoV-OC43_orf1ab;4339 | 0 | 0 |
| NC_006213.1;YP_009555238.1;HCoV-OC43_orf1ab;4340 | 0 | 0 |
| NC_006213.1;YP_009555238.1;HCoV-OC43_orf1ab;4341 | 0 | 0 |
| NC_006213.1;YP_009555238.1;HCoV-OC43_orf1ab;4342 | 0 | 0 |
| NC_006213.1;YP_009555238.1;HCoV-OC43_orf1ab;4343 | 0 | 0 |
| NC_006213.1;YP_009555238.1;HCoV-OC43_orf1ab;4344 | 0 | 0 |
| NC_006213.1;YP_009555238.1;HCoV-OC43_orf1ab;4345 | 0 | 0 |
| NC_006213.1;YP_009555238.1;HCoV-OC43_orf1ab;4346 | 0 | 0 |
| NC_006213.1;YP_009555238.1;HCoV-OC43_orf1ab;4347 | 0 | 0 |
| NC_006213.1;YP_009555238.1;HCoV-OC43_orf1ab;4348 | 0 | 0 |
| NC_006213.1;YP_009555238.1;HCoV-OC43_orf1ab;4349 | 0 | 0 |
| NC_006213.1;YP_009555238.1;HCoV-OC43_orf1ab;4350 | 0 | 0 |
| NC_006213.1;YP_009555238.1;HCoV-OC43_orf1ab;4351 | 0 | 0 |
| NC_006213.1;YP_009555238.1;HCoV-OC43_orf1ab;4352 | 0 | 0 |

| | | |
|---|---|---|
| NC_006213.1;YP_009555238.1;HCoV-OC43_orf1ab;4353 | 0 | 0 |
| NC_006213.1;YP_009555238.1;HCoV-OC43_orf1ab;4354 | 0 | 0 |
| NC_006213.1;YP_009555238.1;HCoV-OC43_orf1ab;4355 | 0 | 0 |
| NC_006213.1;YP_009555238.1;HCoV-OC43_orf1ab;4356 | 0.05 | 0.025 |
| NC_006213.1;YP_009555238.1;HCoV-OC43_orf1ab;4357 | 0.05 | 0 |
| NC_006213.1;YP_009555238.1;HCoV-OC43_orf1ab;4358 | 0.05 | 0.05 |
| NC_006213.1;YP_009555238.1;HCoV-OC43_orf1ab;4359 | 0.05 | 0.025 |
| NC_006213.1;YP_009555238.1;HCoV-OC43_orf1ab;4360 | 0.05 | 0.025 |
| NC_006213.1;YP_009555238.1;HCoV-OC43_orf1ab;4361 | 0.1 | 0.025 |
| NC_006213.1;YP_009555238.1;HCoV-OC43_orf1ab;4362 | 0.1 | 0.025 |
| NC_006213.1;YP_009555238.1;HCoV-OC43_orf1ab;4363 | 0.15 | 0 |
| NC_006213.1;YP_009555238.1;HCoV-OC43_orf1ab;4364 | 0.2 | 0 |
| NC_006213.1;YP_009555238.1;HCoV-OC43_orf1ab;4365 | 0.2 | 0 |
| NC_006213.1;YP_009555238.1;HCoV-OC43_orf1ab;4366 | 0.1 | 0 |
| NC_006213.1;YP_009555238.1;HCoV-OC43_orf1ab;4367 | 0.1 | 0 |
| NC_006213.1;YP_009555238.1;HCoV-OC43_orf1ab;4368 | 0.05 | 0 |
| NC_006213.1;YP_009555238.1;HCoV-OC43_orf1ab;4369 | 0.1 | 0 |
| NC_006213.1;YP_009555238.1;HCoV-OC43_orf1ab;4370 | 0.05 | 0.025 |
| NC_006213.1;YP_009555238.1;HCoV-OC43_orf1ab;4371 | 0 | 0.025 |
| NC_006213.1;YP_009555238.1;HCoV-OC43_orf1ab;4372 | 0 | 0 |
| NC_006213.1;YP_009555238.1;HCoV-OC43_orf1ab;4373 | 0 | 0 |
| NC_006213.1;YP_009555238.1;HCoV-OC43_orf1ab;4374 | 0 | 0 |
| NC_006213.1;YP_009555238.1;HCoV-OC43_orf1ab;4375 | 0 | 0.025 |
| NC_006213.1;YP_009555238.1;HCoV-OC43_orf1ab;4376 | 0 | 0 |
| NC_006213.1;YP_009555238.1;HCoV-OC43_orf1ab;4377 | 0 | 0 |
| NC_006213.1;YP_009555238.1;HCoV-OC43_orf1ab;4378 | 0 | 0 |
| NC_006213.1;YP_009555238.1;HCoV-OC43_orf1ab;4379 | 0 | 0 |
| NC_006213.1;YP_009555238.1;HCoV-OC43_orf1ab;4380 | 0 | 0 |
| NC_006213.1;YP_009555238.1;HCoV-OC43_orf1ab;4381 | 0 | 0 |
| NC_006213.1;YP_009555238.1;HCoV-OC43_orf1ab;4382 | 0 | 0 |
| NC_006213.1;YP_009555238.1;HCoV-OC43_orf1ab;4383 | 0 | 0 |
| NC_006213.1;YP_009555238.1;HCoV-OC43_orf1ab;4384 | 0 | 0 |
| NC_006213.1;YP_009555238.1;HCoV-OC43_orf1ab;4385 | 0 | 0 |
| NC_006213.1;YP_009555238.1;HCoV-OC43_orf1ab;4386 | 0 | 0 |
| NC_006213.1;YP_009555238.1;HCoV-OC43_orf1ab;4387 | 0 | 0 |
| NC_006213.1;YP_009555238.1;HCoV-OC43_orf1ab;4388 | 0.2 | 0.075 |
| NC_006213.1;YP_009555238.1;HCoV-OC43_orf1ab;4389 | 0 | 0 |
| NC_006213.1;YP_009555238.1;HCoV-OC43_orf1ab;4390 | 0 | 0 |
| NC_006213.1;YP_009555238.1;HCoV-OC43_orf1ab;4391 | 0 | 0 |
| NC_006213.1;YP_009555238.1;HCoV-OC43_orf1ab;4392 | 0 | 0 |
| NC_006213.1;YP_009555238.1;HCoV-OC43_orf1ab;4393 | 0 | 0 |
| NC_006213.1;YP_009555238.1;HCoV-OC43_orf1ab;4394 | 0 | 0 |
| NC_006213.1;YP_009555238.1;HCoV-OC43_orf1ab;4395 | 0 | 0 |

| | | |
|---|---|---|
| NC_006213.1;YP_009555238.1;HCoV-OC43_orf1ab;4396 | 0.05 | 0 |
| NC_006213.1;YP_009555238.1;HCoV-OC43_orf1ab;4397 | 0.05 | 0.025 |
| NC_006213.1;YP_009555238.1;HCoV-OC43_orf1ab;4398 | 0 | 0 |
| NC_006213.1;YP_009555238.1;HCoV-OC43_orf1ab;4399 | 0 | 0 |
| NC_006213.1;YP_009555238.1;HCoV-OC43_orf1ab;4400 | 0 | 0 |
| NC_006213.1;YP_009555238.1;HCoV-OC43_orf1ab;4401 | 0 | 0.025 |
| NC_006213.1;YP_009555238.1;HCoV-OC43_orf1ab;4402 | 0 | 0 |
| NC_006213.1;YP_009555238.1;HCoV-OC43_orf1ab;4403 | 0 | 0 |
| NC_006213.1;YP_009555238.1;HCoV-OC43_orf1ab;4404 | 0 | 0 |
| NC_006213.1;YP_009555238.1;HCoV-OC43_orf1ab;4405 | 0 | 0 |
| NC_006213.1;YP_009555238.1;HCoV-OC43_orf1ab;4406 | 0 | 0 |
| NC_006213.1;YP_009555238.1;HCoV-OC43_orf1ab;4407 | 0 | 0 |
| NC_006213.1;YP_009555238.1;HCoV-OC43_orf1ab;4408 | 0 | 0 |
| NC_006213.1;YP_009555238.1;HCoV-OC43_orf1ab;4409 | 0 | 0 |
| NC_006213.1;YP_009555238.1;HCoV-OC43_orf1ab;4410 | 0 | 0 |
| NC_006213.1;YP_009555238.1;HCoV-OC43_orf1ab;4411 | 0 | 0 |
| NC_006213.1;YP_009555238.1;HCoV-OC43_orf1ab;4412 | 0 | 0 |
| NC_006213.1;YP_009555238.1;HCoV-OC43_orf1ab;4413 | 0 | 0 |
| NC_006213.1;YP_009555238.1;HCoV-OC43_orf1ab;4414 | 0 | 0 |
| NC_006213.1;YP_009555238.1;HCoV-OC43_orf1ab;4415 | 0 | 0 |
| NC_006213.1;YP_009555238.1;HCoV-OC43_orf1ab;4416 | 0 | 0.025 |
| NC_006213.1;YP_009555238.1;HCoV-OC43_orf1ab;4417 | 0 | 0.025 |
| NC_006213.1;YP_009555238.1;HCoV-OC43_orf1ab;4418 | 0 | 0.025 |
| NC_006213.1;YP_009555238.1;HCoV-OC43_orf1ab;4419 | 0 | 0.05 |
| NC_006213.1;YP_009555238.1;HCoV-OC43_orf1ab;4420 | 0.05 | 0.05 |
| NC_006213.1;YP_009555238.1;HCoV-OC43_orf1ab;4421 | 0.05 | 0.075 |
| NC_006213.1;YP_009555238.1;HCoV-OC43_orf1ab;4422 | 0.05 | 0.05 |
| NC_006213.1;YP_009555238.1;HCoV-OC43_orf1ab;4423 | 0.15 | 0.05 |
| NC_006213.1;YP_009555238.1;HCoV-OC43_orf1ab;4424 | 0.05 | 0.1 |
| NC_006213.1;YP_009555238.1;HCoV-OC43_orf1ab;4425 | 0.05 | 0.075 |
| NC_006213.1;YP_009555238.1;HCoV-OC43_orf1ab;4426 | 0 | 0.05 |
| NC_006213.1;YP_009555238.1;HCoV-OC43_orf1ab;4427 | 0 | 0.05 |
| NC_006213.1;YP_009555238.1;HCoV-OC43_orf1ab;4428 | 0 | 0.025 |
| NC_006213.1;YP_009555238.1;HCoV-OC43_orf1ab;4429 | 0 | 0 |
| NC_006213.1;YP_009555238.1;HCoV-OC43_orf1ab;4430 | 0 | 0 |
| NC_006213.1;YP_009555238.1;HCoV-OC43_orf1ab;4431 | 0 | 0 |
| NC_006213.1;YP_009555238.1;HCoV-OC43_orf1ab;4432 | 0 | 0 |
| NC_006213.1;YP_009555238.1;HCoV-OC43_orf1ab;4433 | 0 | 0 |
| NC_006213.1;YP_009555238.1;HCoV-OC43_orf1ab;4434 | 0 | 0 |
| NC_006213.1;YP_009555238.1;HCoV-OC43_orf1ab;4435 | 0 | 0 |
| NC_006213.1;YP_009555238.1;HCoV-OC43_orf1ab;4436 | 0 | 0 |
| NC_006213.1;YP_009555238.1;HCoV-OC43_orf1ab;4437 | 0 | 0 |
| NC_006213.1;YP_009555238.1;HCoV-OC43_orf1ab;4438 | 0 | 0 |

| | | |
|---|---|---|
| NC_006213.1;YP_009555238.1;HCoV-OC43_orf1ab;4439 | 0 | 0 |
| NC_006213.1;YP_009555238.1;HCoV-OC43_orf1ab;4440 | 0 | 0 |
| NC_006213.1;YP_009555238.1;HCoV-OC43_orf1ab;4441 | 0 | 0.025 |
| NC_006213.1;YP_009555238.1;HCoV-OC43_orf1ab;4442 | 0.1 | 0.1 |
| NC_006213.1;YP_009555238.1;HCoV-OC43_orf1ab;4443 | 0 | 0 |
| NC_006213.1;YP_009555238.1;HCoV-OC43_orf1ab;4444 | 0 | 0 |
| NC_006213.1;YP_009555238.1;HCoV-OC43_orf1ab;4445 | 0 | 0 |
| NC_006213.1;YP_009555238.1;HCoV-OC43_orf1ab;4446 | 0 | 0 |
| NC_006213.1;YP_009555238.1;HCoV-OC43_orf1ab;4447 | 0 | 0 |
| NC_006213.1;YP_009555238.1;HCoV-OC43_orf1ab;4448 | 0 | 0 |
| NC_006213.1;YP_009555238.1;HCoV-OC43_orf1ab;4449 | 0 | 0 |
| NC_006213.1;YP_009555238.1;HCoV-OC43_orf1ab;4450 | 0 | 0 |
| NC_006213.1;YP_009555238.1;HCoV-OC43_orf1ab;4451 | 0 | 0 |
| NC_006213.1;YP_009555238.1;HCoV-OC43_orf1ab;4452 | 0 | 0 |
| NC_006213.1;YP_009555238.1;HCoV-OC43_orf1ab;4453 | 0 | 0 |
| NC_006213.1;YP_009555238.1;HCoV-OC43_orf1ab;4454 | 0 | 0 |
| NC_006213.1;YP_009555238.1;HCoV-OC43_orf1ab;4455 | 0 | 0 |
| NC_006213.1;YP_009555238.1;HCoV-OC43_orf1ab;4456 | 0 | 0 |
| NC_006213.1;YP_009555238.1;HCoV-OC43_orf1ab;4457 | 0.05 | 0 |
| NC_006213.1;YP_009555238.1;HCoV-OC43_orf1ab;4458 | 0 | 0 |
| NC_006213.1;YP_009555238.1;HCoV-OC43_orf1ab;4459 | 0 | 0 |
| NC_006213.1;YP_009555238.1;HCoV-OC43_orf1ab;4460 | 0 | 0 |
| NC_006213.1;YP_009555238.1;HCoV-OC43_orf1ab;4461 | 0 | 0 |
| NC_006213.1;YP_009555238.1;HCoV-OC43_orf1ab;4462 | 0 | 0 |
| NC_006213.1;YP_009555238.1;HCoV-OC43_orf1ab;4463 | 0 | 0 |
| NC_006213.1;YP_009555238.1;HCoV-OC43_orf1ab;4464 | 0 | 0 |
| NC_006213.1;YP_009555238.1;HCoV-OC43_orf1ab;4465 | 0 | 0 |
| NC_006213.1;YP_009555238.1;HCoV-OC43_orf1ab;4466 | 0 | 0.025 |
| NC_006213.1;YP_009555238.1;HCoV-OC43_orf1ab;4467 | 0 | 0 |
| NC_006213.1;YP_009555238.1;HCoV-OC43_orf1ab;4468 | 0 | 0 |
| NC_006213.1;YP_009555238.1;HCoV-OC43_orf1ab;4469 | 0 | 0 |
| NC_006213.1;YP_009555238.1;HCoV-OC43_orf1ab;4470 | 0 | 0 |
| NC_006213.1;YP_009555238.1;HCoV-OC43_orf1ab;4471 | 0.05 | 0.05 |
| NC_006213.1;YP_009555238.1;HCoV-OC43_orf1ab;4472 | 0 | 0 |
| NC_006213.1;YP_009555238.1;HCoV-OC43_orf1ab;4473 | 0 | 0 |
| NC_006213.1;YP_009555238.1;HCoV-OC43_orf1ab;4474 | 0 | 0 |
| NC_006213.1;YP_009555238.1;HCoV-OC43_orf1ab;4475 | 0 | 0 |
| NC_006213.1;YP_009555238.1;HCoV-OC43_orf1ab;4476 | 0 | 0.05 |
| NC_006213.1;YP_009555238.1;HCoV-OC43_orf1ab;4477 | 0 | 0.025 |
| NC_006213.1;YP_009555238.1;HCoV-OC43_orf1ab;4478 | 0 | 0.025 |
| NC_006213.1;YP_009555238.1;HCoV-OC43_orf1ab;4479 | 0 | 0.025 |
| NC_006213.1;YP_009555238.1;HCoV-OC43_orf1ab;4480 | 0 | 0.025 |
| NC_006213.1;YP_009555238.1;HCoV-OC43_orf1ab;4481 | 0 | 0 |

| | | |
|---|---|---|
| NC_006213.1;YP_009555238.1;HCoV-OC43_orf1ab;4482 | 0 | 0 |
| NC_006213.1;YP_009555238.1;HCoV-OC43_orf1ab;4483 | 0 | 0 |
| NC_006213.1;YP_009555238.1;HCoV-OC43_orf1ab;4484 | 0 | 0 |
| NC_006213.1;YP_009555238.1;HCoV-OC43_orf1ab;4485 | 0 | 0 |
| NC_006213.1;YP_009555238.1;HCoV-OC43_orf1ab;4486 | 0 | 0 |
| NC_006213.1;YP_009555238.1;HCoV-OC43_orf1ab;4487 | 0 | 0 |
| NC_006213.1;YP_009555238.1;HCoV-OC43_orf1ab;4488 | 0 | 0 |
| NC_006213.1;YP_009555238.1;HCoV-OC43_orf1ab;4489 | 0 | 0 |
| NC_006213.1;YP_009555238.1;HCoV-OC43_orf1ab;4490 | 0 | 0 |
| NC_006213.1;YP_009555238.1;HCoV-OC43_orf1ab;4491 | 0 | 0.025 |
| NC_006213.1;YP_009555238.1;HCoV-OC43_orf1ab;4492 | 0 | 0.025 |
| NC_006213.1;YP_009555238.1;HCoV-OC43_orf1ab;4493 | 0 | 0 |
| NC_006213.1;YP_009555238.1;HCoV-OC43_orf1ab;4494 | 0 | 0 |
| NC_006213.1;YP_009555238.1;HCoV-OC43_orf1ab;4495 | 0 | 0 |
| NC_006213.1;YP_009555238.1;HCoV-OC43_orf1ab;4496 | 0 | 0 |
| NC_006213.1;YP_009555238.1;HCoV-OC43_orf1ab;4497 | 0 | 0 |
| NC_006213.1;YP_009555238.1;HCoV-OC43_orf1ab;4498 | 0 | 0 |
| NC_006213.1;YP_009555238.1;HCoV-OC43_orf1ab;4499 | 0 | 0 |
| NC_006213.1;YP_009555238.1;HCoV-OC43_orf1ab;4500 | 0 | 0 |
| NC_006213.1;YP_009555238.1;HCoV-OC43_orf1ab;4501 | 0 | 0 |
| NC_006213.1;YP_009555238.1;HCoV-OC43_orf1ab;4502 | 0 | 0 |
| NC_006213.1;YP_009555238.1;HCoV-OC43_orf1ab;4503 | 0 | 0 |
| NC_006213.1;YP_009555238.1;HCoV-OC43_orf1ab;4504 | 0 | 0 |
| NC_006213.1;YP_009555238.1;HCoV-OC43_orf1ab;4505 | 0 | 0.025 |
| NC_006213.1;YP_009555238.1;HCoV-OC43_orf1ab;4506 | 0 | 0 |
| NC_006213.1;YP_009555238.1;HCoV-OC43_orf1ab;4507 | 0 | 0 |
| NC_006213.1;YP_009555238.1;HCoV-OC43_orf1ab;4508 | 0 | 0 |
| NC_006213.1;YP_009555238.1;HCoV-OC43_orf1ab;4509 | 0 | 0 |
| NC_006213.1;YP_009555238.1;HCoV-OC43_orf1ab;4510 | 0 | 0 |
| NC_006213.1;YP_009555238.1;HCoV-OC43_orf1ab;4511 | 0 | 0 |
| NC_006213.1;YP_009555238.1;HCoV-OC43_orf1ab;4512 | 0 | 0 |
| NC_006213.1;YP_009555238.1;HCoV-OC43_orf1ab;4513 | 0 | 0 |
| NC_006213.1;YP_009555238.1;HCoV-OC43_orf1ab;4514 | 0 | 0 |
| NC_006213.1;YP_009555238.1;HCoV-OC43_orf1ab;4515 | 0 | 0 |
| NC_006213.1;YP_009555238.1;HCoV-OC43_orf1ab;4516 | 0 | 0 |
| NC_006213.1;YP_009555238.1;HCoV-OC43_orf1ab;4517 | 0 | 0 |
| NC_006213.1;YP_009555238.1;HCoV-OC43_orf1ab;4518 | 0 | 0 |
| NC_006213.1;YP_009555238.1;HCoV-OC43_orf1ab;4519 | 0 | 0 |
| NC_006213.1;YP_009555238.1;HCoV-OC43_orf1ab;4520 | 0 | 0 |
| NC_006213.1;YP_009555238.1;HCoV-OC43_orf1ab;4521 | 0 | 0 |
| NC_006213.1;YP_009555238.1;HCoV-OC43_orf1ab;4522 | 0.2 | 0.025 |
| NC_006213.1;YP_009555238.1;HCoV-OC43_orf1ab;4523 | 0.05 | 0 |
| NC_006213.1;YP_009555238.1;HCoV-OC43_orf1ab;4524 | 0 | 0 |

| | | |
|---|---|---|
| NC_006213.1;YP_009555238.1;HCoV-OC43_orf1ab;4525 | 0 | 0.075 |
| NC_006213.1;YP_009555238.1;HCoV-OC43_orf1ab;4526 | 0.05 | 0.075 |
| NC_006213.1;YP_009555238.1;HCoV-OC43_orf1ab;4527 | 0.05 | 0.15 |
| NC_006213.1;YP_009555238.1;HCoV-OC43_orf1ab;4528 | 0.1 | 0.2 |
| NC_006213.1;YP_009555238.1;HCoV-OC43_orf1ab;4529 | 0 | 0.025 |
| NC_006213.1;YP_009555238.1;HCoV-OC43_orf1ab;4530 | 0.05 | 0.05 |
| NC_006213.1;YP_009555238.1;HCoV-OC43_orf1ab;4531 | 0.05 | 0.025 |
| NC_006213.1;YP_009555238.1;HCoV-OC43_orf1ab;4532 | 0 | 0.025 |
| NC_006213.1;YP_009555238.1;HCoV-OC43_orf1ab;4533 | 0.05 | 0.025 |
| NC_006213.1;YP_009555238.1;HCoV-OC43_orf1ab;4534 | 0 | 0 |
| NC_006213.1;YP_009555238.1;HCoV-OC43_orf1ab;4535 | 0 | 0.025 |
| NC_006213.1;YP_009555238.1;HCoV-OC43_orf1ab;4536 | 0 | 0.025 |
| NC_006213.1;YP_009555238.1;HCoV-OC43_orf1ab;4537 | 0 | 0.025 |
| NC_006213.1;YP_009555238.1;HCoV-OC43_orf1ab;4538 | 0 | 0.025 |
| NC_006213.1;YP_009555238.1;HCoV-OC43_orf1ab;4539 | 0 | 0 |
| NC_006213.1;YP_009555238.1;HCoV-OC43_orf1ab;4540 | 0 | 0 |
| NC_006213.1;YP_009555238.1;HCoV-OC43_orf1ab;4541 | 0 | 0 |
| NC_006213.1;YP_009555238.1;HCoV-OC43_orf1ab;4542 | 0 | 0 |
| NC_006213.1;YP_009555238.1;HCoV-OC43_orf1ab;4543 | 0 | 0 |
| NC_006213.1;YP_009555238.1;HCoV-OC43_orf1ab;4544 | 0 | 0 |
| NC_006213.1;YP_009555238.1;HCoV-OC43_orf1ab;4545 | 0 | 0.025 |
| NC_006213.1;YP_009555238.1;HCoV-OC43_orf1ab;4546 | 0 | 0.025 |
| NC_006213.1;YP_009555238.1;HCoV-OC43_orf1ab;4547 | 0 | 0 |
| NC_006213.1;YP_009555238.1;HCoV-OC43_orf1ab;4548 | 0 | 0 |
| NC_006213.1;YP_009555238.1;HCoV-OC43_orf1ab;4549 | 0 | 0 |
| NC_006213.1;YP_009555238.1;HCoV-OC43_orf1ab;4550 | 0 | 0 |
| NC_006213.1;YP_009555238.1;HCoV-OC43_orf1ab;4551 | 0 | 0.025 |
| NC_006213.1;YP_009555238.1;HCoV-OC43_orf1ab;4552 | 0 | 0.025 |
| NC_006213.1;YP_009555238.1;HCoV-OC43_orf1ab;4553 | 0 | 0 |
| NC_006213.1;YP_009555238.1;HCoV-OC43_orf1ab;4554 | 0 | 0 |
| NC_006213.1;YP_009555238.1;HCoV-OC43_orf1ab;4555 | 0 | 0 |
| NC_006213.1;YP_009555238.1;HCoV-OC43_orf1ab;4556 | 0 | 0 |
| NC_006213.1;YP_009555238.1;HCoV-OC43_orf1ab;4557 | 0 | 0 |
| NC_006213.1;YP_009555238.1;HCoV-OC43_orf1ab;4558 | 0 | 0 |
| NC_006213.1;YP_009555238.1;HCoV-OC43_orf1ab;4559 | 0 | 0 |
| NC_006213.1;YP_009555238.1;HCoV-OC43_orf1ab;4560 | 0 | 0 |
| NC_006213.1;YP_009555238.1;HCoV-OC43_orf1ab;4561 | 0 | 0 |
| NC_006213.1;YP_009555238.1;HCoV-OC43_orf1ab;4562 | 0 | 0 |
| NC_006213.1;YP_009555238.1;HCoV-OC43_orf1ab;4563 | 0 | 0 |
| NC_006213.1;YP_009555238.1;HCoV-OC43_orf1ab;4564 | 0 | 0 |
| NC_006213.1;YP_009555238.1;HCoV-OC43_orf1ab;4565 | 0 | 0 |
| NC_006213.1;YP_009555238.1;HCoV-OC43_orf1ab;4566 | 0 | 0 |
| NC_006213.1;YP_009555238.1;HCoV-OC43_orf1ab;4567 | 0 | 0 |

| | | |
|---|---|---|
| NC_006213.1;YP_009555238.1;HCoV-OC43_orf1ab;4568 | 0.05 | 0.025 |
| NC_006213.1;YP_009555238.1;HCoV-OC43_orf1ab;4569 | 0 | 0.025 |
| NC_006213.1;YP_009555238.1;HCoV-OC43_orf1ab;4570 | 0 | 0.025 |
| NC_006213.1;YP_009555238.1;HCoV-OC43_orf1ab;4571 | 0 | 0 |
| NC_006213.1;YP_009555238.1;HCoV-OC43_orf1ab;4572 | 0.15 | 0.05 |
| NC_006213.1;YP_009555238.1;HCoV-OC43_orf1ab;4573 | 0 | 0.025 |
| NC_006213.1;YP_009555238.1;HCoV-OC43_orf1ab;4574 | 0.05 | 0.025 |
| NC_006213.1;YP_009555238.1;HCoV-OC43_orf1ab;4575 | 0 | 0.025 |
| NC_006213.1;YP_009555238.1;HCoV-OC43_orf1ab;4576 | 0 | 0.025 |
| NC_006213.1;YP_009555238.1;HCoV-OC43_orf1ab;4577 | 0 | 0.025 |
| NC_006213.1;YP_009555238.1;HCoV-OC43_orf1ab;4578 | 0 | 0 |
| NC_006213.1;YP_009555238.1;HCoV-OC43_orf1ab;4579 | 0 | 0.025 |
| NC_006213.1;YP_009555238.1;HCoV-OC43_orf1ab;4580 | 0 | 0 |
| NC_006213.1;YP_009555238.1;HCoV-OC43_orf1ab;4581 | 0 | 0 |
| NC_006213.1;YP_009555238.1;HCoV-OC43_orf1ab;4582 | 0 | 0 |
| NC_006213.1;YP_009555238.1;HCoV-OC43_orf1ab;4583 | 0 | 0 |
| NC_006213.1;YP_009555238.1;HCoV-OC43_orf1ab;4584 | 0 | 0 |
| NC_006213.1;YP_009555238.1;HCoV-OC43_orf1ab;4585 | 0 | 0 |
| NC_006213.1;YP_009555238.1;HCoV-OC43_orf1ab;4586 | 0 | 0.025 |
| NC_006213.1;YP_009555238.1;HCoV-OC43_orf1ab;4587 | 0 | 0 |
| NC_006213.1;YP_009555238.1;HCoV-OC43_orf1ab;4588 | 0 | 0 |
| NC_006213.1;YP_009555238.1;HCoV-OC43_orf1ab;4589 | 0 | 0 |
| NC_006213.1;YP_009555238.1;HCoV-OC43_orf1ab;4590 | 0 | 0 |
| NC_006213.1;YP_009555238.1;HCoV-OC43_orf1ab;4591 | 0 | 0 |
| NC_006213.1;YP_009555238.1;HCoV-OC43_orf1ab;4592 | 0 | 0 |
| NC_006213.1;YP_009555238.1;HCoV-OC43_orf1ab;4593 | 0 | 0 |
| NC_006213.1;YP_009555238.1;HCoV-OC43_orf1ab;4594 | 0 | 0 |
| NC_006213.1;YP_009555238.1;HCoV-OC43_orf1ab;4595 | 0 | 0 |
| NC_006213.1;YP_009555238.1;HCoV-OC43_orf1ab;4596 | 0 | 0 |
| NC_006213.1;YP_009555238.1;HCoV-OC43_orf1ab;4597 | 0 | 0 |
| NC_006213.1;YP_009555238.1;HCoV-OC43_orf1ab;4598 | 0 | 0 |
| NC_006213.1;YP_009555238.1;HCoV-OC43_orf1ab;4599 | 0 | 0 |
| NC_006213.1;YP_009555238.1;HCoV-OC43_orf1ab;4600 | 0 | 0.025 |
| NC_006213.1;YP_009555238.1;HCoV-OC43_orf1ab;4601 | 0.05 | 0 |
| NC_006213.1;YP_009555238.1;HCoV-OC43_orf1ab;4602 | 0 | 0 |
| NC_006213.1;YP_009555238.1;HCoV-OC43_orf1ab;4603 | 0 | 0.025 |
| NC_006213.1;YP_009555238.1;HCoV-OC43_orf1ab;4604 | 0 | 0 |
| NC_006213.1;YP_009555238.1;HCoV-OC43_orf1ab;4605 | 0 | 0 |
| NC_006213.1;YP_009555238.1;HCoV-OC43_orf1ab;4606 | 0 | 0 |
| NC_006213.1;YP_009555238.1;HCoV-OC43_orf1ab;4607 | 0 | 0 |
| NC_006213.1;YP_009555238.1;HCoV-OC43_orf1ab;4608 | 0 | 0 |
| NC_006213.1;YP_009555238.1;HCoV-OC43_orf1ab;4609 | 0 | 0 |
| NC_006213.1;YP_009555238.1;HCoV-OC43_orf1ab;4610 | 0 | 0 |

| | | |
|---|---|---|
| NC_006213.1;YP_009555238.1;HCoV-OC43_orf1ab;4611 | 0 | 0 |
| NC_006213.1;YP_009555238.1;HCoV-OC43_orf1ab;4612 | 0 | 0 |
| NC_006213.1;YP_009555238.1;HCoV-OC43_orf1ab;4613 | 0 | 0 |
| NC_006213.1;YP_009555238.1;HCoV-OC43_orf1ab;4614 | 0 | 0 |
| NC_006213.1;YP_009555238.1;HCoV-OC43_orf1ab;4615 | 0 | 0 |
| NC_006213.1;YP_009555238.1;HCoV-OC43_orf1ab;4616 | 0 | 0 |
| NC_006213.1;YP_009555238.1;HCoV-OC43_orf1ab;4617 | 0 | 0.025 |
| NC_006213.1;YP_009555238.1;HCoV-OC43_orf1ab;4618 | 0 | 0.025 |
| NC_006213.1;YP_009555238.1;HCoV-OC43_orf1ab;4619 | 0 | 0 |
| NC_006213.1;YP_009555238.1;HCoV-OC43_orf1ab;4620 | 0 | 0 |
| NC_006213.1;YP_009555238.1;HCoV-OC43_orf1ab;4621 | 0 | 0 |
| NC_006213.1;YP_009555238.1;HCoV-OC43_orf1ab;4622 | 0 | 0.025 |
| NC_006213.1;YP_009555238.1;HCoV-OC43_orf1ab;4623 | 0 | 0.025 |
| NC_006213.1;YP_009555238.1;HCoV-OC43_orf1ab;4624 | 0 | 0 |
| NC_006213.1;YP_009555238.1;HCoV-OC43_orf1ab;4625 | 0 | 0 |
| NC_006213.1;YP_009555238.1;HCoV-OC43_orf1ab;4626 | 0 | 0 |
| NC_006213.1;YP_009555238.1;HCoV-OC43_orf1ab;4627 | 0 | 0 |
| NC_006213.1;YP_009555238.1;HCoV-OC43_orf1ab;4628 | 0 | 0 |
| NC_006213.1;YP_009555238.1;HCoV-OC43_orf1ab;4629 | 0.05 | 0 |
| NC_006213.1;YP_009555238.1;HCoV-OC43_orf1ab;4630 | 0.05 | 0 |
| NC_006213.1;YP_009555238.1;HCoV-OC43_orf1ab;4631 | 0.05 | 0 |
| NC_006213.1;YP_009555238.1;HCoV-OC43_orf1ab;4632 | 0.05 | 0 |
| NC_006213.1;YP_009555238.1;HCoV-OC43_orf1ab;4633 | 0 | 0 |
| NC_006213.1;YP_009555238.1;HCoV-OC43_orf1ab;4634 | 0.05 | 0 |
| NC_006213.1;YP_009555238.1;HCoV-OC43_orf1ab;4635 | 0.1 | 0.05 |
| NC_006213.1;YP_009555238.1;HCoV-OC43_orf1ab;4636 | 0.1 | 0.075 |
| NC_006213.1;YP_009555238.1;HCoV-OC43_orf1ab;4637 | 0.05 | 0.025 |
| NC_006213.1;YP_009555238.1;HCoV-OC43_orf1ab;4638 | 0 | 0.025 |
| NC_006213.1;YP_009555238.1;HCoV-OC43_orf1ab;4639 | 0 | 0.025 |
| NC_006213.1;YP_009555238.1;HCoV-OC43_orf1ab;4640 | 0 | 0.025 |
| NC_006213.1;YP_009555238.1;HCoV-OC43_orf1ab;4641 | 0 | 0.025 |
| NC_006213.1;YP_009555238.1;HCoV-OC43_orf1ab;4642 | 0 | 0.025 |
| NC_006213.1;YP_009555238.1;HCoV-OC43_orf1ab;4643 | 0 | 0 |
| NC_006213.1;YP_009555238.1;HCoV-OC43_orf1ab;4644 | 0 | 0 |
| NC_006213.1;YP_009555238.1;HCoV-OC43_orf1ab;4645 | 0 | 0 |
| NC_006213.1;YP_009555238.1;HCoV-OC43_orf1ab;4646 | 0 | 0 |
| NC_006213.1;YP_009555238.1;HCoV-OC43_orf1ab;4647 | 0 | 0 |
| NC_006213.1;YP_009555238.1;HCoV-OC43_orf1ab;4648 | 0 | 0.025 |
| NC_006213.1;YP_009555238.1;HCoV-OC43_orf1ab;4649 | 0 | 0.025 |
| NC_006213.1;YP_009555238.1;HCoV-OC43_orf1ab;4650 | 0.1 | 0.025 |
| NC_006213.1;YP_009555238.1;HCoV-OC43_orf1ab;4651 | 0.05 | 0.025 |
| NC_006213.1;YP_009555238.1;HCoV-OC43_orf1ab;4652 | 0 | 0.025 |
| NC_006213.1;YP_009555238.1;HCoV-OC43_orf1ab;4653 | 0 | 0.025 |

| | | |
|---|---|---|
| NC_006213.1;YP_009555238.1;HCoV-OC43_orf1ab;4654 | 0 | 0.025 |
| NC_006213.1;YP_009555238.1;HCoV-OC43_orf1ab;4655 | 0 | 0.05 |
| NC_006213.1;YP_009555238.1;HCoV-OC43_orf1ab;4656 | 0 | 0.05 |
| NC_006213.1;YP_009555238.1;HCoV-OC43_orf1ab;4657 | 0 | 0.05 |
| NC_006213.1;YP_009555238.1;HCoV-OC43_orf1ab;4658 | 0 | 0.05 |
| NC_006213.1;YP_009555238.1;HCoV-OC43_orf1ab;4659 | 0 | 0 |
| NC_006213.1;YP_009555238.1;HCoV-OC43_orf1ab;4660 | 0 | 0 |
| NC_006213.1;YP_009555238.1;HCoV-OC43_orf1ab;4661 | 0 | 0.025 |
| NC_006213.1;YP_009555238.1;HCoV-OC43_orf1ab;4662 | 0 | 0 |
| NC_006213.1;YP_009555238.1;HCoV-OC43_orf1ab;4663 | 0 | 0 |
| NC_006213.1;YP_009555238.1;HCoV-OC43_orf1ab;4664 | 0 | 0 |
| NC_006213.1;YP_009555238.1;HCoV-OC43_orf1ab;4665 | 0 | 0 |
| NC_006213.1;YP_009555238.1;HCoV-OC43_orf1ab;4666 | 0 | 0 |
| NC_006213.1;YP_009555238.1;HCoV-OC43_orf1ab;4667 | 0 | 0 |
| NC_006213.1;YP_009555238.1;HCoV-OC43_orf1ab;4668 | 0 | 0 |
| NC_006213.1;YP_009555238.1;HCoV-OC43_orf1ab;4669 | 0 | 0 |
| NC_006213.1;YP_009555238.1;HCoV-OC43_orf1ab;4670 | 0 | 0 |
| NC_006213.1;YP_009555238.1;HCoV-OC43_orf1ab;4671 | 0 | 0 |
| NC_006213.1;YP_009555238.1;HCoV-OC43_orf1ab;4672 | 0 | 0 |
| NC_006213.1;YP_009555238.1;HCoV-OC43_orf1ab;4673 | 0 | 0 |
| NC_006213.1;YP_009555238.1;HCoV-OC43_orf1ab;4674 | 0 | 0 |
| NC_006213.1;YP_009555238.1;HCoV-OC43_orf1ab;4675 | 0 | 0 |
| NC_006213.1;YP_009555238.1;HCoV-OC43_orf1ab;4676 | 0 | 0 |
| NC_006213.1;YP_009555238.1;HCoV-OC43_orf1ab;4677 | 0 | 0 |
| NC_006213.1;YP_009555238.1;HCoV-OC43_orf1ab;4678 | 0 | 0 |
| NC_006213.1;YP_009555238.1;HCoV-OC43_orf1ab;4679 | 0 | 0 |
| NC_006213.1;YP_009555238.1;HCoV-OC43_orf1ab;4680 | 0 | 0 |
| NC_006213.1;YP_009555238.1;HCoV-OC43_orf1ab;4681 | 0 | 0 |
| NC_006213.1;YP_009555238.1;HCoV-OC43_orf1ab;4682 | 0 | 0 |
| NC_006213.1;YP_009555238.1;HCoV-OC43_orf1ab;4683 | 0 | 0 |
| NC_006213.1;YP_009555238.1;HCoV-OC43_orf1ab;4684 | 0 | 0 |
| NC_006213.1;YP_009555238.1;HCoV-OC43_orf1ab;4685 | 0 | 0 |
| NC_006213.1;YP_009555238.1;HCoV-OC43_orf1ab;4686 | 0 | 0 |
| NC_006213.1;YP_009555238.1;HCoV-OC43_orf1ab;4687 | 0 | 0 |
| NC_006213.1;YP_009555238.1;HCoV-OC43_orf1ab;4688 | 0 | 0 |
| NC_006213.1;YP_009555238.1;HCoV-OC43_orf1ab;4689 | 0 | 0 |
| NC_006213.1;YP_009555238.1;HCoV-OC43_orf1ab;4690 | 0 | 0 |
| NC_006213.1;YP_009555238.1;HCoV-OC43_orf1ab;4691 | 0 | 0 |
| NC_006213.1;YP_009555238.1;HCoV-OC43_orf1ab;4692 | 0 | 0 |
| NC_006213.1;YP_009555238.1;HCoV-OC43_orf1ab;4693 | 0 | 0 |
| NC_006213.1;YP_009555238.1;HCoV-OC43_orf1ab;4694 | 0 | 0 |
| NC_006213.1;YP_009555238.1;HCoV-OC43_orf1ab;4695 | 0 | 0 |
| NC_006213.1;YP_009555238.1;HCoV-OC43_orf1ab;4696 | 0 | 0 |

| | | |
|---|---|---|
| NC_006213.1;YP_009555238.1;HCoV-OC43_orf1ab;4697 | 0 | 0 |
| NC_006213.1;YP_009555238.1;HCoV-OC43_orf1ab;4698 | 0 | 0 |
| NC_006213.1;YP_009555238.1;HCoV-OC43_orf1ab;4699 | 0 | 0 |
| NC_006213.1;YP_009555238.1;HCoV-OC43_orf1ab;4700 | 0 | 0 |
| NC_006213.1;YP_009555238.1;HCoV-OC43_orf1ab;4701 | 0 | 0 |
| NC_006213.1;YP_009555238.1;HCoV-OC43_orf1ab;4702 | 0 | 0 |
| NC_006213.1;YP_009555238.1;HCoV-OC43_orf1ab;4703 | 0 | 0 |
| NC_006213.1;YP_009555238.1;HCoV-OC43_orf1ab;4704 | 0 | 0 |
| NC_006213.1;YP_009555238.1;HCoV-OC43_orf1ab;4705 | 0 | 0 |
| NC_006213.1;YP_009555238.1;HCoV-OC43_orf1ab;4706 | 0 | 0 |
| NC_006213.1;YP_009555238.1;HCoV-OC43_orf1ab;4707 | 0 | 0 |
| NC_006213.1;YP_009555238.1;HCoV-OC43_orf1ab;4708 | 0 | 0 |
| NC_006213.1;YP_009555238.1;HCoV-OC43_orf1ab;4709 | 0 | 0 |
| NC_006213.1;YP_009555238.1;HCoV-OC43_orf1ab;4710 | 0 | 0 |
| NC_006213.1;YP_009555238.1;HCoV-OC43_orf1ab;4711 | 0 | 0 |
| NC_006213.1;YP_009555238.1;HCoV-OC43_orf1ab;4712 | 0.05 | 0 |
| NC_006213.1;YP_009555238.1;HCoV-OC43_orf1ab;4713 | 0 | 0 |
| NC_006213.1;YP_009555238.1;HCoV-OC43_orf1ab;4714 | 0 | 0.025 |
| NC_006213.1;YP_009555238.1;HCoV-OC43_orf1ab;4715 | 0 | 0 |
| NC_006213.1;YP_009555238.1;HCoV-OC43_orf1ab;4716 | 0 | 0 |
| NC_006213.1;YP_009555238.1;HCoV-OC43_orf1ab;4717 | 0 | 0 |
| NC_006213.1;YP_009555238.1;HCoV-OC43_orf1ab;4718 | 0 | 0 |
| NC_006213.1;YP_009555238.1;HCoV-OC43_orf1ab;4719 | 0 | 0 |
| NC_006213.1;YP_009555238.1;HCoV-OC43_orf1ab;4720 | 0 | 0 |
| NC_006213.1;YP_009555238.1;HCoV-OC43_orf1ab;4721 | 0 | 0 |
| NC_006213.1;YP_009555238.1;HCoV-OC43_orf1ab;4722 | 0 | 0 |
| NC_006213.1;YP_009555238.1;HCoV-OC43_orf1ab;4723 | 0 | 0 |
| NC_006213.1;YP_009555238.1;HCoV-OC43_orf1ab;4724 | 0 | 0 |
| NC_006213.1;YP_009555238.1;HCoV-OC43_orf1ab;4725 | 0 | 0 |
| NC_006213.1;YP_009555238.1;HCoV-OC43_orf1ab;4726 | 0 | 0 |
| NC_006213.1;YP_009555238.1;HCoV-OC43_orf1ab;4727 | 0 | 0 |
| NC_006213.1;YP_009555238.1;HCoV-OC43_orf1ab;4728 | 0 | 0 |
| NC_006213.1;YP_009555238.1;HCoV-OC43_orf1ab;4729 | 0 | 0 |
| NC_006213.1;YP_009555238.1;HCoV-OC43_orf1ab;4730 | 0 | 0 |
| NC_006213.1;YP_009555238.1;HCoV-OC43_orf1ab;4731 | 0 | 0 |
| NC_006213.1;YP_009555238.1;HCoV-OC43_orf1ab;4732 | 0 | 0.025 |
| NC_006213.1;YP_009555238.1;HCoV-OC43_orf1ab;4733 | 0 | 0.025 |
| NC_006213.1;YP_009555238.1;HCoV-OC43_orf1ab;4734 | 0 | 0 |
| NC_006213.1;YP_009555238.1;HCoV-OC43_orf1ab;4735 | 0 | 0 |
| NC_006213.1;YP_009555238.1;HCoV-OC43_orf1ab;4736 | 0 | 0 |
| NC_006213.1;YP_009555238.1;HCoV-OC43_orf1ab;4737 | 0 | 0 |
| NC_006213.1;YP_009555238.1;HCoV-OC43_orf1ab;4738 | 0 | 0 |
| NC_006213.1;YP_009555238.1;HCoV-OC43_orf1ab;4739 | 0 | 0 |

| | | |
|---|---|---|
| NC_006213.1;YP_009555238.1;HCoV-OC43_orf1ab;4740 | 0 | 0.025 |
| NC_006213.1;YP_009555238.1;HCoV-OC43_orf1ab;4741 | 0 | 0 |
| NC_006213.1;YP_009555238.1;HCoV-OC43_orf1ab;4742 | 0 | 0 |
| NC_006213.1;YP_009555238.1;HCoV-OC43_orf1ab;4743 | 0 | 0 |
| NC_006213.1;YP_009555238.1;HCoV-OC43_orf1ab;4744 | 0 | 0 |
| NC_006213.1;YP_009555238.1;HCoV-OC43_orf1ab;4745 | 0 | 0 |
| NC_006213.1;YP_009555238.1;HCoV-OC43_orf1ab;4746 | 0 | 0 |
| NC_006213.1;YP_009555238.1;HCoV-OC43_orf1ab;4747 | 0 | 0 |
| NC_006213.1;YP_009555238.1;HCoV-OC43_orf1ab;4748 | 0 | 0 |
| NC_006213.1;YP_009555238.1;HCoV-OC43_orf1ab;4749 | 0 | 0 |
| NC_006213.1;YP_009555238.1;HCoV-OC43_orf1ab;4750 | 0.05 | 0 |
| NC_006213.1;YP_009555238.1;HCoV-OC43_orf1ab;4751 | 0 | 0 |
| NC_006213.1;YP_009555238.1;HCoV-OC43_orf1ab;4752 | 0 | 0 |
| NC_006213.1;YP_009555238.1;HCoV-OC43_orf1ab;4753 | 0 | 0 |
| NC_006213.1;YP_009555238.1;HCoV-OC43_orf1ab;4754 | 0 | 0 |
| NC_006213.1;YP_009555238.1;HCoV-OC43_orf1ab;4755 | 0 | 0 |
| NC_006213.1;YP_009555238.1;HCoV-OC43_orf1ab;4756 | 0 | 0 |
| NC_006213.1;YP_009555238.1;HCoV-OC43_orf1ab;4757 | 0 | 0 |
| NC_006213.1;YP_009555238.1;HCoV-OC43_orf1ab;4758 | 0 | 0 |
| NC_006213.1;YP_009555238.1;HCoV-OC43_orf1ab;4759 | 0 | 0.025 |
| NC_006213.1;YP_009555238.1;HCoV-OC43_orf1ab;4760 | 0 | 0.025 |
| NC_006213.1;YP_009555238.1;HCoV-OC43_orf1ab;4761 | 0 | 0.025 |
| NC_006213.1;YP_009555238.1;HCoV-OC43_orf1ab;4762 | 0 | 0.075 |
| NC_006213.1;YP_009555238.1;HCoV-OC43_orf1ab;4763 | 0 | 0.05 |
| NC_006213.1;YP_009555238.1;HCoV-OC43_orf1ab;4764 | 0.05 | 0.05 |
| NC_006213.1;YP_009555238.1;HCoV-OC43_orf1ab;4765 | 0.05 | 0.05 |
| NC_006213.1;YP_009555238.1;HCoV-OC43_orf1ab;4766 | 0.05 | 0.025 |
| NC_006213.1;YP_009555238.1;HCoV-OC43_orf1ab;4767 | 0 | 0.025 |
| NC_006213.1;YP_009555238.1;HCoV-OC43_orf1ab;4768 | 0 | 0.025 |
| NC_006213.1;YP_009555238.1;HCoV-OC43_orf1ab;4769 | 0 | 0.025 |
| NC_006213.1;YP_009555238.1;HCoV-OC43_orf1ab;4770 | 0 | 0.075 |
| NC_006213.1;YP_009555238.1;HCoV-OC43_orf1ab;4771 | 0 | 0.025 |
| NC_006213.1;YP_009555238.1;HCoV-OC43_orf1ab;4772 | 0.05 | 0 |
| NC_006213.1;YP_009555238.1;HCoV-OC43_orf1ab;4773 | 0.15 | 0.025 |
| NC_006213.1;YP_009555238.1;HCoV-OC43_orf1ab;4774 | 0 | 0.025 |
| NC_006213.1;YP_009555238.1;HCoV-OC43_orf1ab;4775 | 0 | 0.025 |
| NC_006213.1;YP_009555238.1;HCoV-OC43_orf1ab;4776 | 0.2 | 0.05 |
| NC_006213.1;YP_009555238.1;HCoV-OC43_orf1ab;4777 | 0.15 | 0.05 |
| NC_006213.1;YP_009555238.1;HCoV-OC43_orf1ab;4778 | 0.1 | 0 |
| NC_006213.1;YP_009555238.1;HCoV-OC43_orf1ab;4779 | 0.05 | 0 |
| NC_006213.1;YP_009555238.1;HCoV-OC43_orf1ab;4780 | 0.05 | 0 |
| NC_006213.1;YP_009555238.1;HCoV-OC43_orf1ab;4781 | 0.05 | 0 |
| NC_006213.1;YP_009555238.1;HCoV-OC43_orf1ab;4782 | 0.1 | 0 |

| | | |
|---|---|---|
| NC_006213.1;YP_009555238.1;HCoV-OC43_orf1ab;4783 | 0.1 | 0 |
| NC_006213.1;YP_009555238.1;HCoV-OC43_orf1ab;4784 | 0.05 | 0 |
| NC_006213.1;YP_009555238.1;HCoV-OC43_orf1ab;4785 | 0.05 | 0 |
| NC_006213.1;YP_009555238.1;HCoV-OC43_orf1ab;4786 | 0.05 | 0 |
| NC_006213.1;YP_009555238.1;HCoV-OC43_orf1ab;4787 | 0.1 | 0 |
| NC_006213.1;YP_009555238.1;HCoV-OC43_orf1ab;4788 | 0.05 | 0 |
| NC_006213.1;YP_009555238.1;HCoV-OC43_orf1ab;4789 | 0 | 0 |
| NC_006213.1;YP_009555238.1;HCoV-OC43_orf1ab;4790 | 0 | 0 |
| NC_006213.1;YP_009555238.1;HCoV-OC43_orf1ab;4791 | 0 | 0.025 |
| NC_006213.1;YP_009555238.1;HCoV-OC43_orf1ab;4792 | 0.05 | 0 |
| NC_006213.1;YP_009555238.1;HCoV-OC43_orf1ab;4793 | 0 | 0 |
| NC_006213.1;YP_009555238.1;HCoV-OC43_orf1ab;4794 | 0 | 0 |
| NC_006213.1;YP_009555238.1;HCoV-OC43_orf1ab;4795 | 0 | 0 |
| NC_006213.1;YP_009555238.1;HCoV-OC43_orf1ab;4796 | 0 | 0 |
| NC_006213.1;YP_009555238.1;HCoV-OC43_orf1ab;4797 | 0 | 0 |
| NC_006213.1;YP_009555238.1;HCoV-OC43_orf1ab;4798 | 0 | 0.025 |
| NC_006213.1;YP_009555238.1;HCoV-OC43_orf1ab;4799 | 0.05 | 0.05 |
| NC_006213.1;YP_009555238.1;HCoV-OC43_orf1ab;4800 | 0 | 0 |
| NC_006213.1;YP_009555238.1;HCoV-OC43_orf1ab;4801 | 0.05 | 0 |
| NC_006213.1;YP_009555238.1;HCoV-OC43_orf1ab;4802 | 0 | 0 |
| NC_006213.1;YP_009555238.1;HCoV-OC43_orf1ab;4803 | 0 | 0 |
| NC_006213.1;YP_009555238.1;HCoV-OC43_orf1ab;4804 | 0 | 0 |
| NC_006213.1;YP_009555238.1;HCoV-OC43_orf1ab;4805 | 0 | 0 |
| NC_006213.1;YP_009555238.1;HCoV-OC43_orf1ab;4806 | 0 | 0.025 |
| NC_006213.1;YP_009555238.1;HCoV-OC43_orf1ab;4807 | 0 | 0.075 |
| NC_006213.1;YP_009555238.1;HCoV-OC43_orf1ab;4808 | 0 | 0.025 |
| NC_006213.1;YP_009555238.1;HCoV-OC43_orf1ab;4809 | 0.15 | 0.025 |
| NC_006213.1;YP_009555238.1;HCoV-OC43_orf1ab;4810 | 0.15 | 0.025 |
| NC_006213.1;YP_009555238.1;HCoV-OC43_orf1ab;4811 | 0.2 | 0.025 |
| NC_006213.1;YP_009555238.1;HCoV-OC43_orf1ab;4812 | 0.2 | 0.025 |
| NC_006213.1;YP_009555238.1;HCoV-OC43_orf1ab;4813 | 0.2 | 0.075 |
| NC_006213.1;YP_009555238.1;HCoV-OC43_orf1ab;4814 | 0.25 | 0.025 |
| NC_006213.1;YP_009555238.1;HCoV-OC43_orf1ab;4815 | 0.25 | 0.05 |
| NC_006213.1;YP_009555238.1;HCoV-OC43_orf1ab;4816 | 0.25 | 0.125 |
| NC_006213.1;YP_009555238.1;HCoV-OC43_orf1ab;4817 | 0.25 | 0.05 |
| NC_006213.1;YP_009555238.1;HCoV-OC43_orf1ab;4818 | 0.05 | 0.025 |
| NC_006213.1;YP_009555238.1;HCoV-OC43_orf1ab;4819 | 0 | 0 |
| NC_006213.1;YP_009555238.1;HCoV-OC43_orf1ab;4820 | 0 | 0 |
| NC_006213.1;YP_009555238.1;HCoV-OC43_orf1ab;4821 | 0 | 0 |
| NC_006213.1;YP_009555238.1;HCoV-OC43_orf1ab;4822 | 0 | 0 |
| NC_006213.1;YP_009555238.1;HCoV-OC43_orf1ab;4823 | 0 | 0 |
| NC_006213.1;YP_009555238.1;HCoV-OC43_orf1ab;4824 | 0 | 0 |
| NC_006213.1;YP_009555238.1;HCoV-OC43_orf1ab;4825 | 0 | 0 |

| | | |
|---|---|---|
| NC_006213.1;YP_009555238.1;HCoV-OC43_orf1ab;4826 | 0 | 0 |
| NC_006213.1;YP_009555238.1;HCoV-OC43_orf1ab;4827 | 0 | 0 |
| NC_006213.1;YP_009555238.1;HCoV-OC43_orf1ab;4828 | 0 | 0 |
| NC_006213.1;YP_009555238.1;HCoV-OC43_orf1ab;4829 | 0 | 0 |
| NC_006213.1;YP_009555238.1;HCoV-OC43_orf1ab;4830 | 0 | 0 |
| NC_006213.1;YP_009555238.1;HCoV-OC43_orf1ab;4831 | 0 | 0 |
| NC_006213.1;YP_009555238.1;HCoV-OC43_orf1ab;4832 | 0 | 0 |
| NC_006213.1;YP_009555238.1;HCoV-OC43_orf1ab;4833 | 0 | 0 |
| NC_006213.1;YP_009555238.1;HCoV-OC43_orf1ab;4834 | 0 | 0 |
| NC_006213.1;YP_009555238.1;HCoV-OC43_orf1ab;4835 | 0 | 0 |
| NC_006213.1;YP_009555238.1;HCoV-OC43_orf1ab;4836 | 0 | 0 |
| NC_006213.1;YP_009555238.1;HCoV-OC43_orf1ab;4837 | 0 | 0 |
| NC_006213.1;YP_009555238.1;HCoV-OC43_orf1ab;4838 | 0 | 0 |
| NC_006213.1;YP_009555238.1;HCoV-OC43_orf1ab;4839 | 0 | 0 |
| NC_006213.1;YP_009555238.1;HCoV-OC43_orf1ab;4840 | 0 | 0 |
| NC_006213.1;YP_009555238.1;HCoV-OC43_orf1ab;4841 | 0 | 0 |
| NC_006213.1;YP_009555238.1;HCoV-OC43_orf1ab;4842 | 0 | 0 |
| NC_006213.1;YP_009555238.1;HCoV-OC43_orf1ab;4843 | 0 | 0.025 |
| NC_006213.1;YP_009555238.1;HCoV-OC43_orf1ab;4844 | 0 | 0.025 |
| NC_006213.1;YP_009555238.1;HCoV-OC43_orf1ab;4845 | 0 | 0.025 |
| NC_006213.1;YP_009555238.1;HCoV-OC43_orf1ab;4846 | 0 | 0 |
| NC_006213.1;YP_009555238.1;HCoV-OC43_orf1ab;4847 | 0 | 0 |
| NC_006213.1;YP_009555238.1;HCoV-OC43_orf1ab;4848 | 0 | 0 |
| NC_006213.1;YP_009555238.1;HCoV-OC43_orf1ab;4849 | 0 | 0 |
| NC_006213.1;YP_009555238.1;HCoV-OC43_orf1ab;4850 | 0 | 0.025 |
| NC_006213.1;YP_009555238.1;HCoV-OC43_orf1ab;4851 | 0 | 0.025 |
| NC_006213.1;YP_009555238.1;HCoV-OC43_orf1ab;4852 | 0.05 | 0.05 |
| NC_006213.1;YP_009555238.1;HCoV-OC43_orf1ab;4853 | 0.05 | 0.025 |
| NC_006213.1;YP_009555238.1;HCoV-OC43_orf1ab;4854 | 0 | 0 |
| NC_006213.1;YP_009555238.1;HCoV-OC43_orf1ab;4855 | 0.1 | 0.075 |
| NC_006213.1;YP_009555238.1;HCoV-OC43_orf1ab;4856 | 0.15 | 0.1 |
| NC_006213.1;YP_009555238.1;HCoV-OC43_orf1ab;4857 | 0.15 | 0.05 |
| NC_006213.1;YP_009555238.1;HCoV-OC43_orf1ab;4858 | 0.25 | 0.05 |
| NC_006213.1;YP_009555238.1;HCoV-OC43_orf1ab;4859 | 0.1 | 0.025 |
| NC_006213.1;YP_009555238.1;HCoV-OC43_orf1ab;4860 | 0.15 | 0.025 |
| NC_006213.1;YP_009555238.1;HCoV-OC43_orf1ab;4861 | 0.1 | 0.025 |
| NC_006213.1;YP_009555238.1;HCoV-OC43_orf1ab;4862 | 0.1 | 0 |
| NC_006213.1;YP_009555238.1;HCoV-OC43_orf1ab;4863 | 0.1 | 0 |
| NC_006213.1;YP_009555238.1;HCoV-OC43_orf1ab;4864 | 0.15 | 0 |
| NC_006213.1;YP_009555238.1;HCoV-OC43_orf1ab;4865 | 0.05 | 0 |
| NC_006213.1;YP_009555238.1;HCoV-OC43_orf1ab;4866 | 0 | 0 |
| NC_006213.1;YP_009555238.1;HCoV-OC43_orf1ab;4867 | 0 | 0 |
| NC_006213.1;YP_009555238.1;HCoV-OC43_orf1ab;4868 | 0 | 0 |

| | | |
|---|---|---|
| NC_006213.1;YP_009555238.1;HCoV-OC43_orf1ab;4869 | 0 | 0 |
| NC_006213.1;YP_009555238.1;HCoV-OC43_orf1ab;4870 | 0 | 0 |
| NC_006213.1;YP_009555238.1;HCoV-OC43_orf1ab;4871 | 0 | 0 |
| NC_006213.1;YP_009555238.1;HCoV-OC43_orf1ab;4872 | 0 | 0 |
| NC_006213.1;YP_009555238.1;HCoV-OC43_orf1ab;4873 | 0 | 0 |
| NC_006213.1;YP_009555238.1;HCoV-OC43_orf1ab;4874 | 0 | 0 |
| NC_006213.1;YP_009555238.1;HCoV-OC43_orf1ab;4875 | 0 | 0 |
| NC_006213.1;YP_009555238.1;HCoV-OC43_orf1ab;4876 | 0 | 0.025 |
| NC_006213.1;YP_009555238.1;HCoV-OC43_orf1ab;4877 | 0 | 0.025 |
| NC_006213.1;YP_009555238.1;HCoV-OC43_orf1ab;4878 | 0 | 0 |
| NC_006213.1;YP_009555238.1;HCoV-OC43_orf1ab;4879 | 0 | 0 |
| NC_006213.1;YP_009555238.1;HCoV-OC43_orf1ab;4880 | 0 | 0 |
| NC_006213.1;YP_009555238.1;HCoV-OC43_orf1ab;4881 | 0 | 0 |
| NC_006213.1;YP_009555238.1;HCoV-OC43_orf1ab;4882 | 0 | 0.025 |
| NC_006213.1;YP_009555238.1;HCoV-OC43_orf1ab;4883 | 0 | 0.05 |
| NC_006213.1;YP_009555238.1;HCoV-OC43_orf1ab;4884 | 0 | 0.05 |
| NC_006213.1;YP_009555238.1;HCoV-OC43_orf1ab;4885 | 0 | 0.075 |
| NC_006213.1;YP_009555238.1;HCoV-OC43_orf1ab;4886 | 0 | 0.075 |
| NC_006213.1;YP_009555238.1;HCoV-OC43_orf1ab;4887 | 0 | 0.025 |
| NC_006213.1;YP_009555238.1;HCoV-OC43_orf1ab;4888 | 0 | 0.075 |
| NC_006213.1;YP_009555238.1;HCoV-OC43_orf1ab;4889 | 0 | 0 |
| NC_006213.1;YP_009555238.1;HCoV-OC43_orf1ab;4890 | 0 | 0 |
| NC_006213.1;YP_009555238.1;HCoV-OC43_orf1ab;4891 | 0 | 0 |
| NC_006213.1;YP_009555238.1;HCoV-OC43_orf1ab;4892 | 0 | 0 |
| NC_006213.1;YP_009555238.1;HCoV-OC43_orf1ab;4893 | 0 | 0 |
| NC_006213.1;YP_009555238.1;HCoV-OC43_orf1ab;4894 | 0 | 0 |
| NC_006213.1;YP_009555238.1;HCoV-OC43_orf1ab;4895 | 0 | 0 |
| NC_006213.1;YP_009555238.1;HCoV-OC43_orf1ab;4896 | 0.05 | 0.075 |
| NC_006213.1;YP_009555238.1;HCoV-OC43_orf1ab;4897 | 0.2 | 0.2 |
| NC_006213.1;YP_009555238.1;HCoV-OC43_orf1ab;4898 | 0 | 0 |
| NC_006213.1;YP_009555238.1;HCoV-OC43_orf1ab;4899 | 0 | 0 |
| NC_006213.1;YP_009555238.1;HCoV-OC43_orf1ab;4900 | 0 | 0 |
| NC_006213.1;YP_009555238.1;HCoV-OC43_orf1ab;4901 | 0 | 0.025 |
| NC_006213.1;YP_009555238.1;HCoV-OC43_orf1ab;4902 | 0 | 0.05 |
| NC_006213.1;YP_009555238.1;HCoV-OC43_orf1ab;4903 | 0 | 0.025 |
| NC_006213.1;YP_009555238.1;HCoV-OC43_orf1ab;4904 | 0 | 0.025 |
| NC_006213.1;YP_009555238.1;HCoV-OC43_orf1ab;4905 | 0 | 0.025 |
| NC_006213.1;YP_009555238.1;HCoV-OC43_orf1ab;4906 | 0 | 0.025 |
| NC_006213.1;YP_009555238.1;HCoV-OC43_orf1ab;4907 | 0 | 0 |
| NC_006213.1;YP_009555238.1;HCoV-OC43_orf1ab;4908 | 0 | 0 |
| NC_006213.1;YP_009555238.1;HCoV-OC43_orf1ab;4909 | 0 | 0 |
| NC_006213.1;YP_009555238.1;HCoV-OC43_orf1ab;4910 | 0 | 0 |
| NC_006213.1;YP_009555238.1;HCoV-OC43_orf1ab;4911 | 0 | 0 |

| | | |
|---|---|---|
| NC_006213.1;YP_009555238.1;HCoV-OC43_orf1ab;4912 | 0 | 0 |
| NC_006213.1;YP_009555238.1;HCoV-OC43_orf1ab;4913 | 0 | 0 |
| NC_006213.1;YP_009555238.1;HCoV-OC43_orf1ab;4914 | 0 | 0 |
| NC_006213.1;YP_009555238.1;HCoV-OC43_orf1ab;4915 | 0 | 0 |
| NC_006213.1;YP_009555238.1;HCoV-OC43_orf1ab;4916 | 0 | 0 |
| NC_006213.1;YP_009555238.1;HCoV-OC43_orf1ab;4917 | 0 | 0 |
| NC_006213.1;YP_009555238.1;HCoV-OC43_orf1ab;4918 | 0 | 0 |
| NC_006213.1;YP_009555238.1;HCoV-OC43_orf1ab;4919 | 0 | 0 |
| NC_006213.1;YP_009555238.1;HCoV-OC43_orf1ab;4920 | 0 | 0 |
| NC_006213.1;YP_009555238.1;HCoV-OC43_orf1ab;4921 | 0 | 0 |
| NC_006213.1;YP_009555238.1;HCoV-OC43_orf1ab;4922 | 0 | 0 |
| NC_006213.1;YP_009555238.1;HCoV-OC43_orf1ab;4923 | 0 | 0 |
| NC_006213.1;YP_009555238.1;HCoV-OC43_orf1ab;4924 | 0 | 0 |
| NC_006213.1;YP_009555238.1;HCoV-OC43_orf1ab;4925 | 0 | 0 |
| NC_006213.1;YP_009555238.1;HCoV-OC43_orf1ab;4926 | 0 | 0 |
| NC_006213.1;YP_009555238.1;HCoV-OC43_orf1ab;4927 | 0 | 0 |
| NC_006213.1;YP_009555238.1;HCoV-OC43_orf1ab;4928 | 0 | 0 |
| NC_006213.1;YP_009555238.1;HCoV-OC43_orf1ab;4929 | 0 | 0 |
| NC_006213.1;YP_009555238.1;HCoV-OC43_orf1ab;4930 | 0 | 0 |
| NC_006213.1;YP_009555238.1;HCoV-OC43_orf1ab;4931 | 0 | 0 |
| NC_006213.1;YP_009555238.1;HCoV-OC43_orf1ab;4932 | 0 | 0 |
| NC_006213.1;YP_009555238.1;HCoV-OC43_orf1ab;4933 | 0 | 0 |
| NC_006213.1;YP_009555238.1;HCoV-OC43_orf1ab;4934 | 0 | 0 |
| NC_006213.1;YP_009555238.1;HCoV-OC43_orf1ab;4935 | 0 | 0 |
| NC_006213.1;YP_009555238.1;HCoV-OC43_orf1ab;4936 | 0 | 0 |
| NC_006213.1;YP_009555238.1;HCoV-OC43_orf1ab;4937 | 0 | 0 |
| NC_006213.1;YP_009555238.1;HCoV-OC43_orf1ab;4938 | 0 | 0 |
| NC_006213.1;YP_009555238.1;HCoV-OC43_orf1ab;4939 | 0 | 0 |
| NC_006213.1;YP_009555238.1;HCoV-OC43_orf1ab;4940 | 0 | 0 |
| NC_006213.1;YP_009555238.1;HCoV-OC43_orf1ab;4941 | 0 | 0 |
| NC_006213.1;YP_009555238.1;HCoV-OC43_orf1ab;4942 | 0 | 0 |
| NC_006213.1;YP_009555238.1;HCoV-OC43_orf1ab;4943 | 0 | 0 |
| NC_006213.1;YP_009555238.1;HCoV-OC43_orf1ab;4944 | 0 | 0 |
| NC_006213.1;YP_009555238.1;HCoV-OC43_orf1ab;4945 | 0 | 0 |
| NC_006213.1;YP_009555238.1;HCoV-OC43_orf1ab;4946 | 0 | 0 |
| NC_006213.1;YP_009555238.1;HCoV-OC43_orf1ab;4947 | 0 | 0 |
| NC_006213.1;YP_009555238.1;HCoV-OC43_orf1ab;4948 | 0 | 0 |
| NC_006213.1;YP_009555238.1;HCoV-OC43_orf1ab;4949 | 0 | 0 |
| NC_006213.1;YP_009555238.1;HCoV-OC43_orf1ab;4950 | 0 | 0 |
| NC_006213.1;YP_009555238.1;HCoV-OC43_orf1ab;4951 | 0 | 0 |
| NC_006213.1;YP_009555238.1;HCoV-OC43_orf1ab;4952 | 0 | 0 |
| NC_006213.1;YP_009555238.1;HCoV-OC43_orf1ab;4953 | 0 | 0 |
| NC_006213.1;YP_009555238.1;HCoV-OC43_orf1ab;4954 | 0 | 0 |

| | | |
|---|---|---|
| NC_006213.1;YP_009555238.1;HCoV-OC43_orf1ab;4955 | 0 | 0.025 |
| NC_006213.1;YP_009555238.1;HCoV-OC43_orf1ab;4956 | 0 | 0 |
| NC_006213.1;YP_009555238.1;HCoV-OC43_orf1ab;4957 | 0 | 0 |
| NC_006213.1;YP_009555238.1;HCoV-OC43_orf1ab;4958 | 0 | 0 |
| NC_006213.1;YP_009555238.1;HCoV-OC43_orf1ab;4959 | 0 | 0 |
| NC_006213.1;YP_009555238.1;HCoV-OC43_orf1ab;4960 | 0.05 | 0.025 |
| NC_006213.1;YP_009555238.1;HCoV-OC43_orf1ab;4961 | 0.05 | 0 |
| NC_006213.1;YP_009555238.1;HCoV-OC43_orf1ab;4962 | 0.2 | 0.175 |
| NC_006213.1;YP_009555238.1;HCoV-OC43_orf1ab;4963 | 0.1 | 0.1 |
| NC_006213.1;YP_009555238.1;HCoV-OC43_orf1ab;4964 | 0.05 | 0.075 |
| NC_006213.1;YP_009555238.1;HCoV-OC43_orf1ab;4965 | 0.05 | 0.075 |
| NC_006213.1;YP_009555238.1;HCoV-OC43_orf1ab;4966 | 0.05 | 0.025 |
| NC_006213.1;YP_009555238.1;HCoV-OC43_orf1ab;4967 | 0 | 0.025 |
| NC_006213.1;YP_009555238.1;HCoV-OC43_orf1ab;4968 | 0.2 | 0.1 |
| NC_006213.1;YP_009555238.1;HCoV-OC43_orf1ab;4969 | 0.1 | 0.075 |
| NC_006213.1;YP_009555238.1;HCoV-OC43_orf1ab;4970 | 0.1 | 0.15 |
| NC_006213.1;YP_009555238.1;HCoV-OC43_orf1ab;4971 | 0.1 | 0.175 |
| NC_006213.1;YP_009555238.1;HCoV-OC43_orf1ab;4972 | 0.1 | 0.15 |
| NC_006213.1;YP_009555238.1;HCoV-OC43_orf1ab;4973 | 0 | 0.025 |
| NC_006213.1;YP_009555238.1;HCoV-OC43_orf1ab;4974 | 0 | 0.025 |
| NC_006213.1;YP_009555238.1;HCoV-OC43_orf1ab;4975 | 0 | 0.025 |
| NC_006213.1;YP_009555238.1;HCoV-OC43_orf1ab;4976 | 0 | 0.025 |
| NC_006213.1;YP_009555238.1;HCoV-OC43_orf1ab;4977 | 0 | 0.025 |
| NC_006213.1;YP_009555238.1;HCoV-OC43_orf1ab;4978 | 0 | 0.025 |
| NC_006213.1;YP_009555238.1;HCoV-OC43_orf1ab;4979 | 0 | 0.025 |
| NC_006213.1;YP_009555238.1;HCoV-OC43_orf1ab;4980 | 0 | 0.025 |
| NC_006213.1;YP_009555238.1;HCoV-OC43_orf1ab;4981 | 0 | 0.025 |
| NC_006213.1;YP_009555238.1;HCoV-OC43_orf1ab;4982 | 0 | 0 |
| NC_006213.1;YP_009555238.1;HCoV-OC43_orf1ab;4983 | 0 | 0 |
| NC_006213.1;YP_009555238.1;HCoV-OC43_orf1ab;4984 | 0 | 0 |
| NC_006213.1;YP_009555238.1;HCoV-OC43_orf1ab;4985 | 0 | 0 |
| NC_006213.1;YP_009555238.1;HCoV-OC43_orf1ab;4986 | 0 | 0 |
| NC_006213.1;YP_009555238.1;HCoV-OC43_orf1ab;4987 | 0 | 0 |
| NC_006213.1;YP_009555238.1;HCoV-OC43_orf1ab;4988 | 0 | 0 |
| NC_006213.1;YP_009555238.1;HCoV-OC43_orf1ab;4989 | 0 | 0 |
| NC_006213.1;YP_009555238.1;HCoV-OC43_orf1ab;4990 | 0 | 0 |
| NC_006213.1;YP_009555238.1;HCoV-OC43_orf1ab;4991 | 0 | 0 |
| NC_006213.1;YP_009555238.1;HCoV-OC43_orf1ab;4992 | 0 | 0 |
| NC_006213.1;YP_009555238.1;HCoV-OC43_orf1ab;4993 | 0 | 0 |
| NC_006213.1;YP_009555238.1;HCoV-OC43_orf1ab;4994 | 0 | 0 |
| NC_006213.1;YP_009555238.1;HCoV-OC43_orf1ab;4995 | 0 | 0 |
| NC_006213.1;YP_009555238.1;HCoV-OC43_orf1ab;4996 | 0 | 0 |
| NC_006213.1;YP_009555238.1;HCoV-OC43_orf1ab;4997 | 0 | 0 |

| | | |
|---|---|---|
| NC_006213.1;YP_009555238.1;HCoV-OC43_orf1ab;4998 | 0 | 0 |
| NC_006213.1;YP_009555238.1;HCoV-OC43_orf1ab;4999 | 0 | 0 |
| NC_006213.1;YP_009555238.1;HCoV-OC43_orf1ab;5000 | 0 | 0 |
| NC_006213.1;YP_009555238.1;HCoV-OC43_orf1ab;5001 | 0 | 0 |
| NC_006213.1;YP_009555238.1;HCoV-OC43_orf1ab;5002 | 0 | 0 |
| NC_006213.1;YP_009555238.1;HCoV-OC43_orf1ab;5003 | 0 | 0 |
| NC_006213.1;YP_009555238.1;HCoV-OC43_orf1ab;5004 | 0 | 0 |
| NC_006213.1;YP_009555238.1;HCoV-OC43_orf1ab;5005 | 0 | 0 |
| NC_006213.1;YP_009555238.1;HCoV-OC43_orf1ab;5006 | 0 | 0 |
| NC_006213.1;YP_009555238.1;HCoV-OC43_orf1ab;5007 | 0.05 | 0 |
| NC_006213.1;YP_009555238.1;HCoV-OC43_orf1ab;5008 | 0.1 | 0 |
| NC_006213.1;YP_009555238.1;HCoV-OC43_orf1ab;5009 | 0.05 | 0 |
| NC_006213.1;YP_009555238.1;HCoV-OC43_orf1ab;5010 | 0.05 | 0 |
| NC_006213.1;YP_009555238.1;HCoV-OC43_orf1ab;5011 | 0.05 | 0 |
| NC_006213.1;YP_009555238.1;HCoV-OC43_orf1ab;5012 | 0.05 | 0 |
| NC_006213.1;YP_009555238.1;HCoV-OC43_orf1ab;5013 | 0.05 | 0 |
| NC_006213.1;YP_009555238.1;HCoV-OC43_orf1ab;5014 | 0 | 0 |
| NC_006213.1;YP_009555238.1;HCoV-OC43_orf1ab;5015 | 0 | 0 |
| NC_006213.1;YP_009555238.1;HCoV-OC43_orf1ab;5016 | 0 | 0 |
| NC_006213.1;YP_009555238.1;HCoV-OC43_orf1ab;5017 | 0 | 0 |
| NC_006213.1;YP_009555238.1;HCoV-OC43_orf1ab;5018 | 0 | 0 |
| NC_006213.1;YP_009555238.1;HCoV-OC43_orf1ab;5019 | 0 | 0 |
| NC_006213.1;YP_009555238.1;HCoV-OC43_orf1ab;5020 | 0 | 0 |
| NC_006213.1;YP_009555238.1;HCoV-OC43_orf1ab;5021 | 0 | 0 |
| NC_006213.1;YP_009555238.1;HCoV-OC43_orf1ab;5022 | 0 | 0 |
| NC_006213.1;YP_009555238.1;HCoV-OC43_orf1ab;5023 | 0 | 0 |
| NC_006213.1;YP_009555238.1;HCoV-OC43_orf1ab;5024 | 0 | 0 |
| NC_006213.1;YP_009555238.1;HCoV-OC43_orf1ab;5025 | 0 | 0 |
| NC_006213.1;YP_009555238.1;HCoV-OC43_orf1ab;5026 | 0 | 0 |
| NC_006213.1;YP_009555238.1;HCoV-OC43_orf1ab;5027 | 0 | 0 |
| NC_006213.1;YP_009555238.1;HCoV-OC43_orf1ab;5028 | 0 | 0 |
| NC_006213.1;YP_009555238.1;HCoV-OC43_orf1ab;5029 | 0 | 0.025 |
| NC_006213.1;YP_009555238.1;HCoV-OC43_orf1ab;5030 | 0 | 0 |
| NC_006213.1;YP_009555238.1;HCoV-OC43_orf1ab;5031 | 0 | 0 |
| NC_006213.1;YP_009555238.1;HCoV-OC43_orf1ab;5032 | 0 | 0 |
| NC_006213.1;YP_009555238.1;HCoV-OC43_orf1ab;5033 | 0 | 0 |
| NC_006213.1;YP_009555238.1;HCoV-OC43_orf1ab;5034 | 0 | 0.025 |
| NC_006213.1;YP_009555238.1;HCoV-OC43_orf1ab;5035 | 0 | 0.025 |
| NC_006213.1;YP_009555238.1;HCoV-OC43_orf1ab;5036 | 0 | 0.025 |
| NC_006213.1;YP_009555238.1;HCoV-OC43_orf1ab;5037 | 0 | 0.025 |
| NC_006213.1;YP_009555238.1;HCoV-OC43_orf1ab;5038 | 0 | 0.05 |
| NC_006213.1;YP_009555238.1;HCoV-OC43_orf1ab;5039 | 0.05 | 0.075 |
| NC_006213.1;YP_009555238.1;HCoV-OC43_orf1ab;5040 | 0.05 | 0.025 |

| | | |
|---|---|---|
| NC_006213.1;YP_009555238.1;HCoV-OC43_orf1ab;5041 | 0.05 | 0.05 |
| NC_006213.1;YP_009555238.1;HCoV-OC43_orf1ab;5042 | 0.05 | 0.025 |
| NC_006213.1;YP_009555238.1;HCoV-OC43_orf1ab;5043 | 0.05 | 0.05 |
| NC_006213.1;YP_009555238.1;HCoV-OC43_orf1ab;5044 | 0.05 | 0 |
| NC_006213.1;YP_009555238.1;HCoV-OC43_orf1ab;5045 | 0.05 | 0 |
| NC_006213.1;YP_009555238.1;HCoV-OC43_orf1ab;5046 | 0 | 0 |
| NC_006213.1;YP_009555238.1;HCoV-OC43_orf1ab;5047 | 0 | 0 |
| NC_006213.1;YP_009555238.1;HCoV-OC43_orf1ab;5048 | 0 | 0 |
| NC_006213.1;YP_009555238.1;HCoV-OC43_orf1ab;5049 | 0 | 0 |
| NC_006213.1;YP_009555238.1;HCoV-OC43_orf1ab;5050 | 0 | 0 |
| NC_006213.1;YP_009555238.1;HCoV-OC43_orf1ab;5051 | 0 | 0 |
| NC_006213.1;YP_009555238.1;HCoV-OC43_orf1ab;5052 | 0 | 0 |
| NC_006213.1;YP_009555238.1;HCoV-OC43_orf1ab;5053 | 0 | 0.025 |
| NC_006213.1;YP_009555238.1;HCoV-OC43_orf1ab;5054 | 0 | 0.05 |
| NC_006213.1;YP_009555238.1;HCoV-OC43_orf1ab;5055 | 0 | 0.025 |
| NC_006213.1;YP_009555238.1;HCoV-OC43_orf1ab;5056 | 0 | 0.025 |
| NC_006213.1;YP_009555238.1;HCoV-OC43_orf1ab;5057 | 0 | 0.025 |
| NC_006213.1;YP_009555238.1;HCoV-OC43_orf1ab;5058 | 0 | 0.025 |
| NC_006213.1;YP_009555238.1;HCoV-OC43_orf1ab;5059 | 0 | 0.025 |
| NC_006213.1;YP_009555238.1;HCoV-OC43_orf1ab;5060 | 0 | 0.025 |
| NC_006213.1;YP_009555238.1;HCoV-OC43_orf1ab;5061 | 0 | 0.025 |
| NC_006213.1;YP_009555238.1;HCoV-OC43_orf1ab;5062 | 0 | 0.025 |
| NC_006213.1;YP_009555238.1;HCoV-OC43_orf1ab;5063 | 0 | 0 |
| NC_006213.1;YP_009555238.1;HCoV-OC43_orf1ab;5064 | 0.05 | 0 |
| NC_006213.1;YP_009555238.1;HCoV-OC43_orf1ab;5065 | 0 | 0.025 |
| NC_006213.1;YP_009555238.1;HCoV-OC43_orf1ab;5066 | 0 | 0 |
| NC_006213.1;YP_009555238.1;HCoV-OC43_orf1ab;5067 | 0 | 0 |
| NC_006213.1;YP_009555238.1;HCoV-OC43_orf1ab;5068 | 0 | 0.025 |
| NC_006213.1;YP_009555238.1;HCoV-OC43_orf1ab;5069 | 0 | 0.025 |
| NC_006213.1;YP_009555238.1;HCoV-OC43_orf1ab;5070 | 0 | 0 |
| NC_006213.1;YP_009555238.1;HCoV-OC43_orf1ab;5071 | 0.05 | 0 |
| NC_006213.1;YP_009555238.1;HCoV-OC43_orf1ab;5072 | 0 | 0 |
| NC_006213.1;YP_009555238.1;HCoV-OC43_orf1ab;5073 | 0.05 | 0 |
| NC_006213.1;YP_009555238.1;HCoV-OC43_orf1ab;5074 | 0 | 0 |
| NC_006213.1;YP_009555238.1;HCoV-OC43_orf1ab;5075 | 0 | 0 |
| NC_006213.1;YP_009555238.1;HCoV-OC43_orf1ab;5076 | 0 | 0 |
| NC_006213.1;YP_009555238.1;HCoV-OC43_orf1ab;5077 | 0 | 0 |
| NC_006213.1;YP_009555238.1;HCoV-OC43_orf1ab;5078 | 0 | 0 |
| NC_006213.1;YP_009555238.1;HCoV-OC43_orf1ab;5079 | 0 | 0 |
| NC_006213.1;YP_009555238.1;HCoV-OC43_orf1ab;5080 | 0 | 0 |
| NC_006213.1;YP_009555238.1;HCoV-OC43_orf1ab;5081 | 0 | 0 |
| NC_006213.1;YP_009555238.1;HCoV-OC43_orf1ab;5082 | 0 | 0 |
| NC_006213.1;YP_009555238.1;HCoV-OC43_orf1ab;5083 | 0 | 0 |

| | | |
|---|---|---|
| NC_006213.1;YP_009555238.1;HCoV-OC43_orf1ab;5084 | 0 | 0 |
| NC_006213.1;YP_009555238.1;HCoV-OC43_orf1ab;5085 | 0 | 0 |
| NC_006213.1;YP_009555238.1;HCoV-OC43_orf1ab;5086 | 0 | 0 |
| NC_006213.1;YP_009555238.1;HCoV-OC43_orf1ab;5087 | 0 | 0 |
| NC_006213.1;YP_009555238.1;HCoV-OC43_orf1ab;5088 | 0 | 0.025 |
| NC_006213.1;YP_009555238.1;HCoV-OC43_orf1ab;5089 | 0 | 0.025 |
| NC_006213.1;YP_009555238.1;HCoV-OC43_orf1ab;5090 | 0 | 0.05 |
| NC_006213.1;YP_009555238.1;HCoV-OC43_orf1ab;5091 | 0 | 0.025 |
| NC_006213.1;YP_009555238.1;HCoV-OC43_orf1ab;5092 | 0 | 0 |
| NC_006213.1;YP_009555238.1;HCoV-OC43_orf1ab;5093 | 0 | 0 |
| NC_006213.1;YP_009555238.1;HCoV-OC43_orf1ab;5094 | 0 | 0 |
| NC_006213.1;YP_009555238.1;HCoV-OC43_orf1ab;5095 | 0 | 0 |
| NC_006213.1;YP_009555238.1;HCoV-OC43_orf1ab;5096 | 0 | 0 |
| NC_006213.1;YP_009555238.1;HCoV-OC43_orf1ab;5097 | 0 | 0 |
| NC_006213.1;YP_009555238.1;HCoV-OC43_orf1ab;5098 | 0 | 0 |
| NC_006213.1;YP_009555238.1;HCoV-OC43_orf1ab;5099 | 0 | 0 |
| NC_006213.1;YP_009555238.1;HCoV-OC43_orf1ab;5100 | 0 | 0.025 |
| NC_006213.1;YP_009555238.1;HCoV-OC43_orf1ab;5101 | 0 | 0.025 |
| NC_006213.1;YP_009555238.1;HCoV-OC43_orf1ab;5102 | 0 | 0 |
| NC_006213.1;YP_009555238.1;HCoV-OC43_orf1ab;5103 | 0 | 0.025 |
| NC_006213.1;YP_009555238.1;HCoV-OC43_orf1ab;5104 | 0.05 | 0.025 |
| NC_006213.1;YP_009555238.1;HCoV-OC43_orf1ab;5105 | 0 | 0 |
| NC_006213.1;YP_009555238.1;HCoV-OC43_orf1ab;5106 | 0.05 | 0.025 |
| NC_006213.1;YP_009555238.1;HCoV-OC43_orf1ab;5107 | 0 | 0.025 |
| NC_006213.1;YP_009555238.1;HCoV-OC43_orf1ab;5108 | 0 | 0.025 |
| NC_006213.1;YP_009555238.1;HCoV-OC43_orf1ab;5109 | 0 | 0 |
| NC_006213.1;YP_009555238.1;HCoV-OC43_orf1ab;5110 | 0 | 0 |
| NC_006213.1;YP_009555238.1;HCoV-OC43_orf1ab;5111 | 0 | 0 |
| NC_006213.1;YP_009555238.1;HCoV-OC43_orf1ab;5112 | 0.05 | 0 |
| NC_006213.1;YP_009555238.1;HCoV-OC43_orf1ab;5113 | 0 | 0 |
| NC_006213.1;YP_009555238.1;HCoV-OC43_orf1ab;5114 | 0 | 0.025 |
| NC_006213.1;YP_009555238.1;HCoV-OC43_orf1ab;5115 | 0 | 0 |
| NC_006213.1;YP_009555238.1;HCoV-OC43_orf1ab;5116 | 0 | 0 |
| NC_006213.1;YP_009555238.1;HCoV-OC43_orf1ab;5117 | 0 | 0 |
| NC_006213.1;YP_009555238.1;HCoV-OC43_orf1ab;5118 | 0 | 0 |
| NC_006213.1;YP_009555238.1;HCoV-OC43_orf1ab;5119 | 0 | 0 |
| NC_006213.1;YP_009555238.1;HCoV-OC43_orf1ab;5120 | 0 | 0 |
| NC_006213.1;YP_009555238.1;HCoV-OC43_orf1ab;5121 | 0 | 0.025 |
| NC_006213.1;YP_009555238.1;HCoV-OC43_orf1ab;5122 | 0 | 0.025 |
| NC_006213.1;YP_009555238.1;HCoV-OC43_orf1ab;5123 | 0 | 0.025 |
| NC_006213.1;YP_009555238.1;HCoV-OC43_orf1ab;5124 | 0 | 0.025 |
| NC_006213.1;YP_009555238.1;HCoV-OC43_orf1ab;5125 | 0 | 0.05 |
| NC_006213.1;YP_009555238.1;HCoV-OC43_orf1ab;5126 | 0 | 0.025 |

| | | |
|---|---|---|
| NC_006213.1;YP_009555238.1;HCoV-OC43_orf1ab;5127 | 0 | 0.05 |
| NC_006213.1;YP_009555238.1;HCoV-OC43_orf1ab;5128 | 0.05 | 0.025 |
| NC_006213.1;YP_009555238.1;HCoV-OC43_orf1ab;5129 | 0 | 0.025 |
| NC_006213.1;YP_009555238.1;HCoV-OC43_orf1ab;5130 | 0.05 | 0.025 |
| NC_006213.1;YP_009555238.1;HCoV-OC43_orf1ab;5131 | 0 | 0.025 |
| NC_006213.1;YP_009555238.1;HCoV-OC43_orf1ab;5132 | 0 | 0 |
| NC_006213.1;YP_009555238.1;HCoV-OC43_orf1ab;5133 | 0 | 0 |
| NC_006213.1;YP_009555238.1;HCoV-OC43_orf1ab;5134 | 0 | 0 |
| NC_006213.1;YP_009555238.1;HCoV-OC43_orf1ab;5135 | 0 | 0 |
| NC_006213.1;YP_009555238.1;HCoV-OC43_orf1ab;5136 | 0 | 0 |
| NC_006213.1;YP_009555238.1;HCoV-OC43_orf1ab;5137 | 0 | 0 |
| NC_006213.1;YP_009555238.1;HCoV-OC43_orf1ab;5138 | 0 | 0 |
| NC_006213.1;YP_009555238.1;HCoV-OC43_orf1ab;5139 | 0 | 0 |
| NC_006213.1;YP_009555238.1;HCoV-OC43_orf1ab;5140 | 0 | 0 |
| NC_006213.1;YP_009555238.1;HCoV-OC43_orf1ab;5141 | 0 | 0 |
| NC_006213.1;YP_009555238.1;HCoV-OC43_orf1ab;5142 | 0 | 0 |
| NC_006213.1;YP_009555238.1;HCoV-OC43_orf1ab;5143 | 0 | 0 |
| NC_006213.1;YP_009555238.1;HCoV-OC43_orf1ab;5144 | 0 | 0 |
| NC_006213.1;YP_009555238.1;HCoV-OC43_orf1ab;5145 | 0 | 0 |
| NC_006213.1;YP_009555238.1;HCoV-OC43_orf1ab;5146 | 0 | 0 |
| NC_006213.1;YP_009555238.1;HCoV-OC43_orf1ab;5147 | 0 | 0 |
| NC_006213.1;YP_009555238.1;HCoV-OC43_orf1ab;5148 | 0.05 | 0.025 |
| NC_006213.1;YP_009555238.1;HCoV-OC43_orf1ab;5149 | 0 | 0 |
| NC_006213.1;YP_009555238.1;HCoV-OC43_orf1ab;5150 | 0 | 0 |
| NC_006213.1;YP_009555238.1;HCoV-OC43_orf1ab;5151 | 0 | 0 |
| NC_006213.1;YP_009555238.1;HCoV-OC43_orf1ab;5152 | 0 | 0.025 |
| NC_006213.1;YP_009555238.1;HCoV-OC43_orf1ab;5153 | 0 | 0 |
| NC_006213.1;YP_009555238.1;HCoV-OC43_orf1ab;5154 | 0 | 0.025 |
| NC_006213.1;YP_009555238.1;HCoV-OC43_orf1ab;5155 | 0 | 0.025 |
| NC_006213.1;YP_009555238.1;HCoV-OC43_orf1ab;5156 | 0 | 0.025 |
| NC_006213.1;YP_009555238.1;HCoV-OC43_orf1ab;5157 | 0 | 0.025 |
| NC_006213.1;YP_009555238.1;HCoV-OC43_orf1ab;5158 | 0.1 | 0 |
| NC_006213.1;YP_009555238.1;HCoV-OC43_orf1ab;5159 | 0.05 | 0.025 |
| NC_006213.1;YP_009555238.1;HCoV-OC43_orf1ab;5160 | 0 | 0 |
| NC_006213.1;YP_009555238.1;HCoV-OC43_orf1ab;5161 | 0 | 0 |
| NC_006213.1;YP_009555238.1;HCoV-OC43_orf1ab;5162 | 0 | 0 |
| NC_006213.1;YP_009555238.1;HCoV-OC43_orf1ab;5163 | 0 | 0 |
| NC_006213.1;YP_009555238.1;HCoV-OC43_orf1ab;5164 | 0 | 0 |
| NC_006213.1;YP_009555238.1;HCoV-OC43_orf1ab;5165 | 0 | 0 |
| NC_006213.1;YP_009555238.1;HCoV-OC43_orf1ab;5166 | 0 | 0 |
| NC_006213.1;YP_009555238.1;HCoV-OC43_orf1ab;5167 | 0 | 0 |
| NC_006213.1;YP_009555238.1;HCoV-OC43_orf1ab;5168 | 0 | 0 |
| NC_006213.1;YP_009555238.1;HCoV-OC43_orf1ab;5169 | 0 | 0 |

| | | |
|---|---|---|
| NC_006213.1;YP_009555238.1;HCoV-OC43_orf1ab;5170 | 0 | 0 |
| NC_006213.1;YP_009555238.1;HCoV-OC43_orf1ab;5171 | 0 | 0 |
| NC_006213.1;YP_009555238.1;HCoV-OC43_orf1ab;5172 | 0 | 0 |
| NC_006213.1;YP_009555238.1;HCoV-OC43_orf1ab;5173 | 0 | 0 |
| NC_006213.1;YP_009555238.1;HCoV-OC43_orf1ab;5174 | 0 | 0 |
| NC_006213.1;YP_009555238.1;HCoV-OC43_orf1ab;5175 | 0 | 0 |
| NC_006213.1;YP_009555238.1;HCoV-OC43_orf1ab;5176 | 0 | 0.025 |
| NC_006213.1;YP_009555238.1;HCoV-OC43_orf1ab;5177 | 0 | 0 |
| NC_006213.1;YP_009555238.1;HCoV-OC43_orf1ab;5178 | 0.05 | 0.025 |
| NC_006213.1;YP_009555238.1;HCoV-OC43_orf1ab;5179 | 0 | 0 |
| NC_006213.1;YP_009555238.1;HCoV-OC43_orf1ab;5180 | 0 | 0 |
| NC_006213.1;YP_009555238.1;HCoV-OC43_orf1ab;5181 | 0 | 0 |
| NC_006213.1;YP_009555238.1;HCoV-OC43_orf1ab;5182 | 0.1 | 0.025 |
| NC_006213.1;YP_009555238.1;HCoV-OC43_orf1ab;5183 | 0.1 | 0.05 |
| NC_006213.1;YP_009555238.1;HCoV-OC43_orf1ab;5184 | 0.15 | 0.05 |
| NC_006213.1;YP_009555238.1;HCoV-OC43_orf1ab;5185 | 0.15 | 0.05 |
| NC_006213.1;YP_009555238.1;HCoV-OC43_orf1ab;5186 | 0.15 | 0.05 |
| NC_006213.1;YP_009555238.1;HCoV-OC43_orf1ab;5187 | 0 | 0 |
| NC_006213.1;YP_009555238.1;HCoV-OC43_orf1ab;5188 | 0 | 0 |
| NC_006213.1;YP_009555238.1;HCoV-OC43_orf1ab;5189 | 0 | 0 |
| NC_006213.1;YP_009555238.1;HCoV-OC43_orf1ab;5190 | 0 | 0 |
| NC_006213.1;YP_009555238.1;HCoV-OC43_orf1ab;5191 | 0 | 0 |
| NC_006213.1;YP_009555238.1;HCoV-OC43_orf1ab;5192 | 0 | 0 |
| NC_006213.1;YP_009555238.1;HCoV-OC43_orf1ab;5193 | 0 | 0 |
| NC_006213.1;YP_009555238.1;HCoV-OC43_orf1ab;5194 | 0 | 0 |
| NC_006213.1;YP_009555238.1;HCoV-OC43_orf1ab;5195 | 0 | 0 |
| NC_006213.1;YP_009555238.1;HCoV-OC43_orf1ab;5196 | 0 | 0 |
| NC_006213.1;YP_009555238.1;HCoV-OC43_orf1ab;5197 | 0 | 0 |
| NC_006213.1;YP_009555238.1;HCoV-OC43_orf1ab;5198 | 0 | 0 |
| NC_006213.1;YP_009555238.1;HCoV-OC43_orf1ab;5199 | 0 | 0 |
| NC_006213.1;YP_009555238.1;HCoV-OC43_orf1ab;5200 | 0 | 0 |
| NC_006213.1;YP_009555238.1;HCoV-OC43_orf1ab;5201 | 0 | 0 |
| NC_006213.1;YP_009555238.1;HCoV-OC43_orf1ab;5202 | 0.05 | 0 |
| NC_006213.1;YP_009555238.1;HCoV-OC43_orf1ab;5203 | 0.05 | 0 |
| NC_006213.1;YP_009555238.1;HCoV-OC43_orf1ab;5204 | 0 | 0 |
| NC_006213.1;YP_009555238.1;HCoV-OC43_orf1ab;5205 | 0 | 0 |
| NC_006213.1;YP_009555238.1;HCoV-OC43_orf1ab;5206 | 0 | 0 |
| NC_006213.1;YP_009555238.1;HCoV-OC43_orf1ab;5207 | 0 | 0 |
| NC_006213.1;YP_009555238.1;HCoV-OC43_orf1ab;5208 | 0 | 0 |
| NC_006213.1;YP_009555238.1;HCoV-OC43_orf1ab;5209 | 0 | 0.025 |
| NC_006213.1;YP_009555238.1;HCoV-OC43_orf1ab;5210 | 0 | 0.025 |
| NC_006213.1;YP_009555238.1;HCoV-OC43_orf1ab;5211 | 0 | 0.025 |
| NC_006213.1;YP_009555238.1;HCoV-OC43_orf1ab;5212 | 0 | 0.025 |

| | | |
|---|---|---|
| NC_006213.1;YP_009555238.1;HCoV-OC43_orf1ab;5213 | 0 | 0.025 |
| NC_006213.1;YP_009555238.1;HCoV-OC43_orf1ab;5214 | 0 | 0.025 |
| NC_006213.1;YP_009555238.1;HCoV-OC43_orf1ab;5215 | 0 | 0.025 |
| NC_006213.1;YP_009555238.1;HCoV-OC43_orf1ab;5216 | 0 | 0.025 |
| NC_006213.1;YP_009555238.1;HCoV-OC43_orf1ab;5217 | 0 | 0.025 |
| NC_006213.1;YP_009555238.1;HCoV-OC43_orf1ab;5218 | 0 | 0.025 |
| NC_006213.1;YP_009555238.1;HCoV-OC43_orf1ab;5219 | 0 | 0.025 |
| NC_006213.1;YP_009555238.1;HCoV-OC43_orf1ab;5220 | 0 | 0.025 |
| NC_006213.1;YP_009555238.1;HCoV-OC43_orf1ab;5221 | 0 | 0 |
| NC_006213.1;YP_009555238.1;HCoV-OC43_orf1ab;5222 | 0 | 0 |
| NC_006213.1;YP_009555238.1;HCoV-OC43_orf1ab;5223 | 0 | 0 |
| NC_006213.1;YP_009555238.1;HCoV-OC43_orf1ab;5224 | 0 | 0 |
| NC_006213.1;YP_009555238.1;HCoV-OC43_orf1ab;5225 | 0 | 0 |
| NC_006213.1;YP_009555238.1;HCoV-OC43_orf1ab;5226 | 0 | 0 |
| NC_006213.1;YP_009555238.1;HCoV-OC43_orf1ab;5227 | 0 | 0 |
| NC_006213.1;YP_009555238.1;HCoV-OC43_orf1ab;5228 | 0 | 0 |
| NC_006213.1;YP_009555238.1;HCoV-OC43_orf1ab;5229 | 0 | 0 |
| NC_006213.1;YP_009555238.1;HCoV-OC43_orf1ab;5230 | 0 | 0.025 |
| NC_006213.1;YP_009555238.1;HCoV-OC43_orf1ab;5231 | 0.05 | 0.025 |
| NC_006213.1;YP_009555238.1;HCoV-OC43_orf1ab;5232 | 0 | 0.05 |
| NC_006213.1;YP_009555238.1;HCoV-OC43_orf1ab;5233 | 0 | 0.025 |
| NC_006213.1;YP_009555238.1;HCoV-OC43_orf1ab;5234 | 0 | 0.025 |
| NC_006213.1;YP_009555238.1;HCoV-OC43_orf1ab;5235 | 0 | 0 |
| NC_006213.1;YP_009555238.1;HCoV-OC43_orf1ab;5236 | 0 | 0 |
| NC_006213.1;YP_009555238.1;HCoV-OC43_orf1ab;5237 | 0 | 0 |
| NC_006213.1;YP_009555238.1;HCoV-OC43_orf1ab;5238 | 0 | 0 |
| NC_006213.1;YP_009555238.1;HCoV-OC43_orf1ab;5239 | 0 | 0 |
| NC_006213.1;YP_009555238.1;HCoV-OC43_orf1ab;5240 | 0 | 0 |
| NC_006213.1;YP_009555238.1;HCoV-OC43_orf1ab;5241 | 0 | 0 |
| NC_006213.1;YP_009555238.1;HCoV-OC43_orf1ab;5242 | 0 | 0 |
| NC_006213.1;YP_009555238.1;HCoV-OC43_orf1ab;5243 | 0 | 0 |
| NC_006213.1;YP_009555238.1;HCoV-OC43_orf1ab;5244 | 0 | 0 |
| NC_006213.1;YP_009555238.1;HCoV-OC43_orf1ab;5245 | 0 | 0 |
| NC_006213.1;YP_009555238.1;HCoV-OC43_orf1ab;5246 | 0 | 0 |
| NC_006213.1;YP_009555238.1;HCoV-OC43_orf1ab;5247 | 0 | 0 |
| NC_006213.1;YP_009555238.1;HCoV-OC43_orf1ab;5248 | 0 | 0 |
| NC_006213.1;YP_009555238.1;HCoV-OC43_orf1ab;5249 | 0.05 | 0.025 |
| NC_006213.1;YP_009555238.1;HCoV-OC43_orf1ab;5250 | 0.05 | 0 |
| NC_006213.1;YP_009555238.1;HCoV-OC43_orf1ab;5251 | 0.15 | 0.175 |
| NC_006213.1;YP_009555238.1;HCoV-OC43_orf1ab;5252 | 0.15 | 0.2 |
| NC_006213.1;YP_009555238.1;HCoV-OC43_orf1ab;5253 | 0.1 | 0.1 |
| NC_006213.1;YP_009555238.1;HCoV-OC43_orf1ab;5254 | 0.15 | 0.05 |
| NC_006213.1;YP_009555238.1;HCoV-OC43_orf1ab;5255 | 0.05 | 0 |

| | | |
|---|---|---|
| NC_006213.1;YP_009555238.1;HCoV-OC43_orf1ab;5256 | 0 | 0 |
| NC_006213.1;YP_009555238.1;HCoV-OC43_orf1ab;5257 | 0 | 0 |
| NC_006213.1;YP_009555238.1;HCoV-OC43_orf1ab;5258 | 0 | 0 |
| NC_006213.1;YP_009555238.1;HCoV-OC43_orf1ab;5259 | 0 | 0 |
| NC_006213.1;YP_009555238.1;HCoV-OC43_orf1ab;5260 | 0 | 0 |
| NC_006213.1;YP_009555238.1;HCoV-OC43_orf1ab;5261 | 0 | 0 |
| NC_006213.1;YP_009555238.1;HCoV-OC43_orf1ab;5262 | 0 | 0 |
| NC_006213.1;YP_009555238.1;HCoV-OC43_orf1ab;5263 | 0 | 0 |
| NC_006213.1;YP_009555238.1;HCoV-OC43_orf1ab;5264 | 0 | 0 |
| NC_006213.1;YP_009555238.1;HCoV-OC43_orf1ab;5265 | 0 | 0 |
| NC_006213.1;YP_009555238.1;HCoV-OC43_orf1ab;5266 | 0 | 0 |
| NC_006213.1;YP_009555238.1;HCoV-OC43_orf1ab;5267 | 0.05 | 0 |
| NC_006213.1;YP_009555238.1;HCoV-OC43_orf1ab;5268 | 0 | 0 |
| NC_006213.1;YP_009555238.1;HCoV-OC43_orf1ab;5269 | 0 | 0 |
| NC_006213.1;YP_009555238.1;HCoV-OC43_orf1ab;5270 | 0.05 | 0 |
| NC_006213.1;YP_009555238.1;HCoV-OC43_orf1ab;5271 | 0.05 | 0 |
| NC_006213.1;YP_009555238.1;HCoV-OC43_orf1ab;5272 | 0.05 | 0 |
| NC_006213.1;YP_009555238.1;HCoV-OC43_orf1ab;5273 | 0.05 | 0 |
| NC_006213.1;YP_009555238.1;HCoV-OC43_orf1ab;5274 | 0.05 | 0 |
| NC_006213.1;YP_009555238.1;HCoV-OC43_orf1ab;5275 | 0.05 | 0.025 |
| NC_006213.1;YP_009555238.1;HCoV-OC43_orf1ab;5276 | 0.05 | 0.05 |
| NC_006213.1;YP_009555238.1;HCoV-OC43_orf1ab;5277 | 0 | 0 |
| NC_006213.1;YP_009555238.1;HCoV-OC43_orf1ab;5278 | 0 | 0 |
| NC_006213.1;YP_009555238.1;HCoV-OC43_orf1ab;5279 | 0 | 0 |
| NC_006213.1;YP_009555238.1;HCoV-OC43_orf1ab;5280 | 0 | 0 |
| NC_006213.1;YP_009555238.1;HCoV-OC43_orf1ab;5281 | 0 | 0 |
| NC_006213.1;YP_009555238.1;HCoV-OC43_orf1ab;5282 | 0 | 0 |
| NC_006213.1;YP_009555238.1;HCoV-OC43_orf1ab;5283 | 0 | 0 |
| NC_006213.1;YP_009555238.1;HCoV-OC43_orf1ab;5284 | 0 | 0 |
| NC_006213.1;YP_009555238.1;HCoV-OC43_orf1ab;5285 | 0 | 0 |
| NC_006213.1;YP_009555238.1;HCoV-OC43_orf1ab;5286 | 0 | 0 |
| NC_006213.1;YP_009555238.1;HCoV-OC43_orf1ab;5287 | 0 | 0 |
| NC_006213.1;YP_009555238.1;HCoV-OC43_orf1ab;5288 | 0 | 0 |
| NC_006213.1;YP_009555238.1;HCoV-OC43_orf1ab;5289 | 0 | 0 |
| NC_006213.1;YP_009555238.1;HCoV-OC43_orf1ab;5290 | 0 | 0 |
| NC_006213.1;YP_009555238.1;HCoV-OC43_orf1ab;5291 | 0 | 0 |
| NC_006213.1;YP_009555238.1;HCoV-OC43_orf1ab;5292 | 0 | 0 |
| NC_006213.1;YP_009555238.1;HCoV-OC43_orf1ab;5293 | 0 | 0 |
| NC_006213.1;YP_009555238.1;HCoV-OC43_orf1ab;5294 | 0 | 0 |
| NC_006213.1;YP_009555238.1;HCoV-OC43_orf1ab;5295 | 0 | 0 |
| NC_006213.1;YP_009555238.1;HCoV-OC43_orf1ab;5296 | 0 | 0 |
| NC_006213.1;YP_009555238.1;HCoV-OC43_orf1ab;5297 | 0 | 0 |
| NC_006213.1;YP_009555238.1;HCoV-OC43_orf1ab;5298 | 0 | 0 |

| | | |
|---|---|---|
| NC_006213.1;YP_009555238.1;HCoV-OC43_orf1ab;5299 | 0 | 0 |
| NC_006213.1;YP_009555238.1;HCoV-OC43_orf1ab;5300 | 0 | 0 |
| NC_006213.1;YP_009555238.1;HCoV-OC43_orf1ab;5301 | 0 | 0 |
| NC_006213.1;YP_009555238.1;HCoV-OC43_orf1ab;5302 | 0 | 0 |
| NC_006213.1;YP_009555238.1;HCoV-OC43_orf1ab;5303 | 0 | 0 |
| NC_006213.1;YP_009555238.1;HCoV-OC43_orf1ab;5304 | 0 | 0 |
| NC_006213.1;YP_009555238.1;HCoV-OC43_orf1ab;5305 | 0 | 0 |
| NC_006213.1;YP_009555238.1;HCoV-OC43_orf1ab;5306 | 0 | 0 |
| NC_006213.1;YP_009555238.1;HCoV-OC43_orf1ab;5307 | 0 | 0 |
| NC_006213.1;YP_009555238.1;HCoV-OC43_orf1ab;5308 | 0 | 0 |
| NC_006213.1;YP_009555238.1;HCoV-OC43_orf1ab;5309 | 0 | 0 |
| NC_006213.1;YP_009555238.1;HCoV-OC43_orf1ab;5310 | 0 | 0 |
| NC_006213.1;YP_009555238.1;HCoV-OC43_orf1ab;5311 | 0 | 0 |
| NC_006213.1;YP_009555238.1;HCoV-OC43_orf1ab;5312 | 0 | 0 |
| NC_006213.1;YP_009555238.1;HCoV-OC43_orf1ab;5313 | 0 | 0 |
| NC_006213.1;YP_009555238.1;HCoV-OC43_orf1ab;5314 | 0 | 0 |
| NC_006213.1;YP_009555238.1;HCoV-OC43_orf1ab;5315 | 0 | 0 |
| NC_006213.1;YP_009555238.1;HCoV-OC43_orf1ab;5316 | 0 | 0 |
| NC_006213.1;YP_009555238.1;HCoV-OC43_orf1ab;5317 | 0 | 0 |
| NC_006213.1;YP_009555238.1;HCoV-OC43_orf1ab;5318 | 0 | 0 |
| NC_006213.1;YP_009555238.1;HCoV-OC43_orf1ab;5319 | 0 | 0 |
| NC_006213.1;YP_009555238.1;HCoV-OC43_orf1ab;5320 | 0 | 0 |
| NC_006213.1;YP_009555238.1;HCoV-OC43_orf1ab;5321 | 0 | 0 |
| NC_006213.1;YP_009555238.1;HCoV-OC43_orf1ab;5322 | 0 | 0 |
| NC_006213.1;YP_009555238.1;HCoV-OC43_orf1ab;5323 | 0 | 0 |
| NC_006213.1;YP_009555238.1;HCoV-OC43_orf1ab;5324 | 0 | 0 |
| NC_006213.1;YP_009555238.1;HCoV-OC43_orf1ab;5325 | 0 | 0 |
| NC_006213.1;YP_009555238.1;HCoV-OC43_orf1ab;5326 | 0 | 0.025 |
| NC_006213.1;YP_009555238.1;HCoV-OC43_orf1ab;5327 | 0 | 0 |
| NC_006213.1;YP_009555238.1;HCoV-OC43_orf1ab;5328 | 0 | 0 |
| NC_006213.1;YP_009555238.1;HCoV-OC43_orf1ab;5329 | 0 | 0 |
| NC_006213.1;YP_009555238.1;HCoV-OC43_orf1ab;5330 | 0 | 0 |
| NC_006213.1;YP_009555238.1;HCoV-OC43_orf1ab;5331 | 0 | 0 |
| NC_006213.1;YP_009555238.1;HCoV-OC43_orf1ab;5332 | 0 | 0 |
| NC_006213.1;YP_009555238.1;HCoV-OC43_orf1ab;5333 | 0 | 0 |
| NC_006213.1;YP_009555238.1;HCoV-OC43_orf1ab;5334 | 0 | 0 |
| NC_006213.1;YP_009555238.1;HCoV-OC43_orf1ab;5335 | 0 | 0 |
| NC_006213.1;YP_009555238.1;HCoV-OC43_orf1ab;5336 | 0 | 0 |
| NC_006213.1;YP_009555238.1;HCoV-OC43_orf1ab;5337 | 0 | 0 |
| NC_006213.1;YP_009555238.1;HCoV-OC43_orf1ab;5338 | 0 | 0 |
| NC_006213.1;YP_009555238.1;HCoV-OC43_orf1ab;5339 | 0 | 0 |
| NC_006213.1;YP_009555238.1;HCoV-OC43_orf1ab;5340 | 0 | 0 |
| NC_006213.1;YP_009555238.1;HCoV-OC43_orf1ab;5341 | 0.05 | 0 |

| | | |
|---|---|---|
| NC_006213.1;YP_009555238.1;HCoV-OC43_orf1ab;5342 | 0 | 0.025 |
| NC_006213.1;YP_009555238.1;HCoV-OC43_orf1ab;5343 | 0 | 0.05 |
| NC_006213.1;YP_009555238.1;HCoV-OC43_orf1ab;5344 | 0.05 | 0.075 |
| NC_006213.1;YP_009555238.1;HCoV-OC43_orf1ab;5345 | 0 | 0.1 |
| NC_006213.1;YP_009555238.1;HCoV-OC43_orf1ab;5346 | 0.05 | 0.05 |
| NC_006213.1;YP_009555238.1;HCoV-OC43_orf1ab;5347 | 0 | 0.025 |
| NC_006213.1;YP_009555238.1;HCoV-OC43_orf1ab;5348 | 0 | 0.025 |
| NC_006213.1;YP_009555238.1;HCoV-OC43_orf1ab;5349 | 0 | 0.025 |
| NC_006213.1;YP_009555238.1;HCoV-OC43_orf1ab;5350 | 0.05 | 0.025 |
| NC_006213.1;YP_009555238.1;HCoV-OC43_orf1ab;5351 | 0.05 | 0.05 |
| NC_006213.1;YP_009555238.1;HCoV-OC43_orf1ab;5352 | 0 | 0.025 |
| NC_006213.1;YP_009555238.1;HCoV-OC43_orf1ab;5353 | 0 | 0.025 |
| NC_006213.1;YP_009555238.1;HCoV-OC43_orf1ab;5354 | 0 | 0 |
| NC_006213.1;YP_009555238.1;HCoV-OC43_orf1ab;5355 | 0 | 0 |
| NC_006213.1;YP_009555238.1;HCoV-OC43_orf1ab;5356 | 0 | 0 |
| NC_006213.1;YP_009555238.1;HCoV-OC43_orf1ab;5357 | 0 | 0 |
| NC_006213.1;YP_009555238.1;HCoV-OC43_orf1ab;5358 | 0 | 0 |
| NC_006213.1;YP_009555238.1;HCoV-OC43_orf1ab;5359 | 0 | 0.025 |
| NC_006213.1;YP_009555238.1;HCoV-OC43_orf1ab;5360 | 0 | 0 |
| NC_006213.1;YP_009555238.1;HCoV-OC43_orf1ab;5361 | 0 | 0 |
| NC_006213.1;YP_009555238.1;HCoV-OC43_orf1ab;5362 | 0 | 0 |
| NC_006213.1;YP_009555238.1;HCoV-OC43_orf1ab;5363 | 0 | 0.025 |
| NC_006213.1;YP_009555238.1;HCoV-OC43_orf1ab;5364 | 0.05 | 0.025 |
| NC_006213.1;YP_009555238.1;HCoV-OC43_orf1ab;5365 | 0.05 | 0 |
| NC_006213.1;YP_009555238.1;HCoV-OC43_orf1ab;5366 | 0 | 0 |
| NC_006213.1;YP_009555238.1;HCoV-OC43_orf1ab;5367 | 0 | 0 |
| NC_006213.1;YP_009555238.1;HCoV-OC43_orf1ab;5368 | 0 | 0 |
| NC_006213.1;YP_009555238.1;HCoV-OC43_orf1ab;5369 | 0 | 0 |
| NC_006213.1;YP_009555238.1;HCoV-OC43_orf1ab;5370 | 0 | 0 |
| NC_006213.1;YP_009555238.1;HCoV-OC43_orf1ab;5371 | 0 | 0 |
| NC_006213.1;YP_009555238.1;HCoV-OC43_orf1ab;5372 | 0 | 0 |
| NC_006213.1;YP_009555238.1;HCoV-OC43_orf1ab;5373 | 0 | 0 |
| NC_006213.1;YP_009555238.1;HCoV-OC43_orf1ab;5374 | 0 | 0 |
| NC_006213.1;YP_009555238.1;HCoV-OC43_orf1ab;5375 | 0 | 0 |
| NC_006213.1;YP_009555238.1;HCoV-OC43_orf1ab;5376 | 0 | 0 |
| NC_006213.1;YP_009555238.1;HCoV-OC43_orf1ab;5377 | 0 | 0 |
| NC_006213.1;YP_009555238.1;HCoV-OC43_orf1ab;5378 | 0 | 0 |
| NC_006213.1;YP_009555238.1;HCoV-OC43_orf1ab;5379 | 0 | 0 |
| NC_006213.1;YP_009555238.1;HCoV-OC43_orf1ab;5380 | 0 | 0 |
| NC_006213.1;YP_009555238.1;HCoV-OC43_orf1ab;5381 | 0 | 0 |
| NC_006213.1;YP_009555238.1;HCoV-OC43_orf1ab;5382 | 0 | 0 |
| NC_006213.1;YP_009555238.1;HCoV-OC43_orf1ab;5383 | 0 | 0 |
| NC_006213.1;YP_009555238.1;HCoV-OC43_orf1ab;5384 | 0 | 0 |

| | | |
|---|---|---|
| NC_006213.1;YP_009555238.1;HCoV-OC43_orf1ab;5385 | 0 | 0 |
| NC_006213.1;YP_009555238.1;HCoV-OC43_orf1ab;5386 | 0 | 0 |
| NC_006213.1;YP_009555238.1;HCoV-OC43_orf1ab;5387 | 0 | 0.025 |
| NC_006213.1;YP_009555238.1;HCoV-OC43_orf1ab;5388 | 0 | 0.025 |
| NC_006213.1;YP_009555238.1;HCoV-OC43_orf1ab;5389 | 0.05 | 0.05 |
| NC_006213.1;YP_009555238.1;HCoV-OC43_orf1ab;5390 | 0 | 0.125 |
| NC_006213.1;YP_009555238.1;HCoV-OC43_orf1ab;5391 | 0 | 0.025 |
| NC_006213.1;YP_009555238.1;HCoV-OC43_orf1ab;5392 | 0.05 | 0.1 |
| NC_006213.1;YP_009555238.1;HCoV-OC43_orf1ab;5393 | 0.1 | 0.05 |
| NC_006213.1;YP_009555238.1;HCoV-OC43_orf1ab;5394 | 0 | 0 |
| NC_006213.1;YP_009555238.1;HCoV-OC43_orf1ab;5395 | 0.05 | 0.05 |
| NC_006213.1;YP_009555238.1;HCoV-OC43_orf1ab;5396 | 0 | 0 |
| NC_006213.1;YP_009555238.1;HCoV-OC43_orf1ab;5397 | 0 | 0.025 |
| NC_006213.1;YP_009555238.1;HCoV-OC43_orf1ab;5398 | 0 | 0.025 |
| NC_006213.1;YP_009555238.1;HCoV-OC43_orf1ab;5399 | 0 | 0 |
| NC_006213.1;YP_009555238.1;HCoV-OC43_orf1ab;5400 | 0 | 0 |
| NC_006213.1;YP_009555238.1;HCoV-OC43_orf1ab;5401 | 0 | 0 |
| NC_006213.1;YP_009555238.1;HCoV-OC43_orf1ab;5402 | 0 | 0 |
| NC_006213.1;YP_009555238.1;HCoV-OC43_orf1ab;5403 | 0 | 0 |
| NC_006213.1;YP_009555238.1;HCoV-OC43_orf1ab;5404 | 0 | 0 |
| NC_006213.1;YP_009555238.1;HCoV-OC43_orf1ab;5405 | 0 | 0 |
| NC_006213.1;YP_009555238.1;HCoV-OC43_orf1ab;5406 | 0 | 0 |
| NC_006213.1;YP_009555238.1;HCoV-OC43_orf1ab;5407 | 0 | 0 |
| NC_006213.1;YP_009555238.1;HCoV-OC43_orf1ab;5408 | 0 | 0 |
| NC_006213.1;YP_009555238.1;HCoV-OC43_orf1ab;5409 | 0 | 0 |
| NC_006213.1;YP_009555238.1;HCoV-OC43_orf1ab;5410 | 0 | 0 |
| NC_006213.1;YP_009555238.1;HCoV-OC43_orf1ab;5411 | 0 | 0 |
| NC_006213.1;YP_009555238.1;HCoV-OC43_orf1ab;5412 | 0 | 0 |
| NC_006213.1;YP_009555238.1;HCoV-OC43_orf1ab;5413 | 0 | 0 |
| NC_006213.1;YP_009555238.1;HCoV-OC43_orf1ab;5414 | 0.05 | 0 |
| NC_006213.1;YP_009555238.1;HCoV-OC43_orf1ab;5415 | 0.05 | 0 |
| NC_006213.1;YP_009555238.1;HCoV-OC43_orf1ab;5416 | 0 | 0 |
| NC_006213.1;YP_009555238.1;HCoV-OC43_orf1ab;5417 | 0.05 | 0.025 |
| NC_006213.1;YP_009555238.1;HCoV-OC43_orf1ab;5418 | 0 | 0 |
| NC_006213.1;YP_009555238.1;HCoV-OC43_orf1ab;5419 | 0 | 0 |
| NC_006213.1;YP_009555238.1;HCoV-OC43_orf1ab;5420 | 0 | 0 |
| NC_006213.1;YP_009555238.1;HCoV-OC43_orf1ab;5421 | 0 | 0 |
| NC_006213.1;YP_009555238.1;HCoV-OC43_orf1ab;5422 | 0 | 0 |
| NC_006213.1;YP_009555238.1;HCoV-OC43_orf1ab;5423 | 0.05 | 0 |
| NC_006213.1;YP_009555238.1;HCoV-OC43_orf1ab;5424 | 0 | 0 |
| NC_006213.1;YP_009555238.1;HCoV-OC43_orf1ab;5425 | 0.05 | 0 |
| NC_006213.1;YP_009555238.1;HCoV-OC43_orf1ab;5426 | 0 | 0 |
| NC_006213.1;YP_009555238.1;HCoV-OC43_orf1ab;5427 | 0 | 0 |

| | | |
|---|---|---|
| NC_006213.1;YP_009555238.1;HCoV-OC43_orf1ab;5428 | 0 | 0 |
| NC_006213.1;YP_009555238.1;HCoV-OC43_orf1ab;5429 | 0 | 0 |
| NC_006213.1;YP_009555238.1;HCoV-OC43_orf1ab;5430 | 0 | 0 |
| NC_006213.1;YP_009555238.1;HCoV-OC43_orf1ab;5431 | 0 | 0.025 |
| NC_006213.1;YP_009555238.1;HCoV-OC43_orf1ab;5432 | 0 | 0.025 |
| NC_006213.1;YP_009555238.1;HCoV-OC43_orf1ab;5433 | 0.05 | 0 |
| NC_006213.1;YP_009555238.1;HCoV-OC43_orf1ab;5434 | 0.05 | 0 |
| NC_006213.1;YP_009555238.1;HCoV-OC43_orf1ab;5435 | 0 | 0 |
| NC_006213.1;YP_009555238.1;HCoV-OC43_orf1ab;5436 | 0 | 0 |
| NC_006213.1;YP_009555238.1;HCoV-OC43_orf1ab;5437 | 0 | 0 |
| NC_006213.1;YP_009555238.1;HCoV-OC43_orf1ab;5438 | 0 | 0 |
| NC_006213.1;YP_009555238.1;HCoV-OC43_orf1ab;5439 | 0 | 0 |
| NC_006213.1;YP_009555238.1;HCoV-OC43_orf1ab;5440 | 0 | 0 |
| NC_006213.1;YP_009555238.1;HCoV-OC43_orf1ab;5441 | 0 | 0 |
| NC_006213.1;YP_009555238.1;HCoV-OC43_orf1ab;5442 | 0 | 0 |
| NC_006213.1;YP_009555238.1;HCoV-OC43_orf1ab;5443 | 0 | 0 |
| NC_006213.1;YP_009555238.1;HCoV-OC43_orf1ab;5444 | 0 | 0 |
| NC_006213.1;YP_009555238.1;HCoV-OC43_orf1ab;5445 | 0 | 0 |
| NC_006213.1;YP_009555238.1;HCoV-OC43_orf1ab;5446 | 0 | 0 |
| NC_006213.1;YP_009555238.1;HCoV-OC43_orf1ab;5447 | 0 | 0.025 |
| NC_006213.1;YP_009555238.1;HCoV-OC43_orf1ab;5448 | 0 | 0 |
| NC_006213.1;YP_009555238.1;HCoV-OC43_orf1ab;5449 | 0 | 0 |
| NC_006213.1;YP_009555238.1;HCoV-OC43_orf1ab;5450 | 0 | 0 |
| NC_006213.1;YP_009555238.1;HCoV-OC43_orf1ab;5451 | 0 | 0 |
| NC_006213.1;YP_009555238.1;HCoV-OC43_orf1ab;5452 | 0 | 0 |
| NC_006213.1;YP_009555238.1;HCoV-OC43_orf1ab;5453 | 0 | 0.025 |
| NC_006213.1;YP_009555238.1;HCoV-OC43_orf1ab;5454 | 0 | 0.025 |
| NC_006213.1;YP_009555238.1;HCoV-OC43_orf1ab;5455 | 0 | 0 |
| NC_006213.1;YP_009555238.1;HCoV-OC43_orf1ab;5456 | 0.05 | 0.05 |
| NC_006213.1;YP_009555238.1;HCoV-OC43_orf1ab;5457 | 0.05 | 0.05 |
| NC_006213.1;YP_009555238.1;HCoV-OC43_orf1ab;5458 | 0 | 0.025 |
| NC_006213.1;YP_009555238.1;HCoV-OC43_orf1ab;5459 | 0 | 0.025 |
| NC_006213.1;YP_009555238.1;HCoV-OC43_orf1ab;5460 | 0 | 0.1 |
| NC_006213.1;YP_009555238.1;HCoV-OC43_orf1ab;5461 | 0.05 | 0.125 |
| NC_006213.1;YP_009555238.1;HCoV-OC43_orf1ab;5462 | 0.05 | 0.1 |
| NC_006213.1;YP_009555238.1;HCoV-OC43_orf1ab;5463 | 0.05 | 0.2 |
| NC_006213.1;YP_009555238.1;HCoV-OC43_orf1ab;5464 | 0 | 0.1 |
| NC_006213.1;YP_009555238.1;HCoV-OC43_orf1ab;5465 | 0.1 | 0.1 |
| NC_006213.1;YP_009555238.1;HCoV-OC43_orf1ab;5466 | 0.05 | 0.05 |
| NC_006213.1;YP_009555238.1;HCoV-OC43_orf1ab;5467 | 0.05 | 0 |
| NC_006213.1;YP_009555238.1;HCoV-OC43_orf1ab;5468 | 0 | 0 |
| NC_006213.1;YP_009555238.1;HCoV-OC43_orf1ab;5469 | 0 | 0 |
| NC_006213.1;YP_009555238.1;HCoV-OC43_orf1ab;5470 | 0 | 0 |

| | | |
|---|---|---|
| NC_006213.1;YP_009555238.1;HCoV-OC43_orf1ab;5471 | 0 | 0 |
| NC_006213.1;YP_009555238.1;HCoV-OC43_orf1ab;5472 | 0 | 0 |
| NC_006213.1;YP_009555238.1;HCoV-OC43_orf1ab;5473 | 0 | 0 |
| NC_006213.1;YP_009555238.1;HCoV-OC43_orf1ab;5474 | 0 | 0 |
| NC_006213.1;YP_009555238.1;HCoV-OC43_orf1ab;5475 | 0 | 0 |
| NC_006213.1;YP_009555238.1;HCoV-OC43_orf1ab;5476 | 0 | 0 |
| NC_006213.1;YP_009555238.1;HCoV-OC43_orf1ab;5477 | 0 | 0 |
| NC_006213.1;YP_009555238.1;HCoV-OC43_orf1ab;5478 | 0 | 0 |
| NC_006213.1;YP_009555238.1;HCoV-OC43_orf1ab;5479 | 0 | 0.05 |
| NC_006213.1;YP_009555238.1;HCoV-OC43_orf1ab;5480 | 0 | 0.05 |
| NC_006213.1;YP_009555238.1;HCoV-OC43_orf1ab;5481 | 0 | 0 |
| NC_006213.1;YP_009555238.1;HCoV-OC43_orf1ab;5482 | 0 | 0 |
| NC_006213.1;YP_009555238.1;HCoV-OC43_orf1ab;5483 | 0 | 0 |
| NC_006213.1;YP_009555238.1;HCoV-OC43_orf1ab;5484 | 0 | 0.1 |
| NC_006213.1;YP_009555238.1;HCoV-OC43_orf1ab;5485 | 0 | 0.075 |
| NC_006213.1;YP_009555238.1;HCoV-OC43_orf1ab;5486 | 0 | 0.1 |
| NC_006213.1;YP_009555238.1;HCoV-OC43_orf1ab;5487 | 0 | 0.05 |
| NC_006213.1;YP_009555238.1;HCoV-OC43_orf1ab;5488 | 0 | 0.075 |
| NC_006213.1;YP_009555238.1;HCoV-OC43_orf1ab;5489 | 0 | 0.1 |
| NC_006213.1;YP_009555238.1;HCoV-OC43_orf1ab;5490 | 0 | 0.1 |
| NC_006213.1;YP_009555238.1;HCoV-OC43_orf1ab;5491 | 0 | 0.075 |
| NC_006213.1;YP_009555238.1;HCoV-OC43_orf1ab;5492 | 0 | 0.1 |
| NC_006213.1;YP_009555238.1;HCoV-OC43_orf1ab;5493 | 0 | 0 |
| NC_006213.1;YP_009555238.1;HCoV-OC43_orf1ab;5494 | 0.05 | 0 |
| NC_006213.1;YP_009555238.1;HCoV-OC43_orf1ab;5495 | 0 | 0 |
| NC_006213.1;YP_009555238.1;HCoV-OC43_orf1ab;5496 | 0 | 0 |
| NC_006213.1;YP_009555238.1;HCoV-OC43_orf1ab;5497 | 0 | 0 |
| NC_006213.1;YP_009555238.1;HCoV-OC43_orf1ab;5498 | 0 | 0 |
| NC_006213.1;YP_009555238.1;HCoV-OC43_orf1ab;5499 | 0 | 0 |
| NC_006213.1;YP_009555238.1;HCoV-OC43_orf1ab;5500 | 0.05 | 0 |
| NC_006213.1;YP_009555238.1;HCoV-OC43_orf1ab;5501 | 0 | 0 |
| NC_006213.1;YP_009555238.1;HCoV-OC43_orf1ab;5502 | 0 | 0 |
| NC_006213.1;YP_009555238.1;HCoV-OC43_orf1ab;5503 | 0 | 0 |
| NC_006213.1;YP_009555238.1;HCoV-OC43_orf1ab;5504 | 0 | 0 |
| NC_006213.1;YP_009555238.1;HCoV-OC43_orf1ab;5505 | 0 | 0 |
| NC_006213.1;YP_009555238.1;HCoV-OC43_orf1ab;5506 | 0 | 0 |
| NC_006213.1;YP_009555238.1;HCoV-OC43_orf1ab;5507 | 0 | 0 |
| NC_006213.1;YP_009555238.1;HCoV-OC43_orf1ab;5508 | 0 | 0 |
| NC_006213.1;YP_009555238.1;HCoV-OC43_orf1ab;5509 | 0 | 0 |
| NC_006213.1;YP_009555238.1;HCoV-OC43_orf1ab;5510 | 0 | 0 |
| NC_006213.1;YP_009555238.1;HCoV-OC43_orf1ab;5511 | 0 | 0 |
| NC_006213.1;YP_009555238.1;HCoV-OC43_orf1ab;5512 | 0 | 0 |
| NC_006213.1;YP_009555238.1;HCoV-OC43_orf1ab;5513 | 0 | 0 |

| | | |
|---|---|---|
| NC_006213.1;YP_009555238.1;HCoV-OC43_orf1ab;5514 | 0 | 0 |
| NC_006213.1;YP_009555238.1;HCoV-OC43_orf1ab;5515 | 0 | 0 |
| NC_006213.1;YP_009555238.1;HCoV-OC43_orf1ab;5516 | 0 | 0 |
| NC_006213.1;YP_009555238.1;HCoV-OC43_orf1ab;5517 | 0 | 0 |
| NC_006213.1;YP_009555238.1;HCoV-OC43_orf1ab;5518 | 0 | 0 |
| NC_006213.1;YP_009555238.1;HCoV-OC43_orf1ab;5519 | 0 | 0 |
| NC_006213.1;YP_009555238.1;HCoV-OC43_orf1ab;5520 | 0.05 | 0 |
| NC_006213.1;YP_009555238.1;HCoV-OC43_orf1ab;5521 | 0 | 0 |
| NC_006213.1;YP_009555238.1;HCoV-OC43_orf1ab;5522 | 0 | 0 |
| NC_006213.1;YP_009555238.1;HCoV-OC43_orf1ab;5523 | 0 | 0 |
| NC_006213.1;YP_009555238.1;HCoV-OC43_orf1ab;5524 | 0 | 0.025 |
| NC_006213.1;YP_009555238.1;HCoV-OC43_orf1ab;5525 | 0 | 0 |
| NC_006213.1;YP_009555238.1;HCoV-OC43_orf1ab;5526 | 0 | 0 |
| NC_006213.1;YP_009555238.1;HCoV-OC43_orf1ab;5527 | 0 | 0 |
| NC_006213.1;YP_009555238.1;HCoV-OC43_orf1ab;5528 | 0 | 0.025 |
| NC_006213.1;YP_009555238.1;HCoV-OC43_orf1ab;5529 | 0.05 | 0.025 |
| NC_006213.1;YP_009555238.1;HCoV-OC43_orf1ab;5530 | 0.05 | 0.025 |
| NC_006213.1;YP_009555238.1;HCoV-OC43_orf1ab;5531 | 0.05 | 0.05 |
| NC_006213.1;YP_009555238.1;HCoV-OC43_orf1ab;5532 | 0 | 0.025 |
| NC_006213.1;YP_009555238.1;HCoV-OC43_orf1ab;5533 | 0 | 0.025 |
| NC_006213.1;YP_009555238.1;HCoV-OC43_orf1ab;5534 | 0 | 0.025 |
| NC_006213.1;YP_009555238.1;HCoV-OC43_orf1ab;5535 | 0.05 | 0.025 |
| NC_006213.1;YP_009555238.1;HCoV-OC43_orf1ab;5536 | 0.05 | 0 |
| NC_006213.1;YP_009555238.1;HCoV-OC43_orf1ab;5537 | 0.05 | 0 |
| NC_006213.1;YP_009555238.1;HCoV-OC43_orf1ab;5538 | 0.05 | 0 |
| NC_006213.1;YP_009555238.1;HCoV-OC43_orf1ab;5539 | 0 | 0 |
| NC_006213.1;YP_009555238.1;HCoV-OC43_orf1ab;5540 | 0 | 0 |
| NC_006213.1;YP_009555238.1;HCoV-OC43_orf1ab;5541 | 0 | 0 |
| NC_006213.1;YP_009555238.1;HCoV-OC43_orf1ab;5542 | 0 | 0 |
| NC_006213.1;YP_009555238.1;HCoV-OC43_orf1ab;5543 | 0 | 0 |
| NC_006213.1;YP_009555238.1;HCoV-OC43_orf1ab;5544 | 0 | 0 |
| NC_006213.1;YP_009555238.1;HCoV-OC43_orf1ab;5545 | 0 | 0 |
| NC_006213.1;YP_009555238.1;HCoV-OC43_orf1ab;5546 | 0 | 0 |
| NC_006213.1;YP_009555238.1;HCoV-OC43_orf1ab;5547 | 0 | 0 |
| NC_006213.1;YP_009555238.1;HCoV-OC43_orf1ab;5548 | 0 | 0 |
| NC_006213.1;YP_009555238.1;HCoV-OC43_orf1ab;5549 | 0 | 0.025 |
| NC_006213.1;YP_009555238.1;HCoV-OC43_orf1ab;5550 | 0 | 0.025 |
| NC_006213.1;YP_009555238.1;HCoV-OC43_orf1ab;5551 | 0 | 0.05 |
| NC_006213.1;YP_009555238.1;HCoV-OC43_orf1ab;5552 | 0 | 0.025 |
| NC_006213.1;YP_009555238.1;HCoV-OC43_orf1ab;5553 | 0 | 0.075 |
| NC_006213.1;YP_009555238.1;HCoV-OC43_orf1ab;5554 | 0 | 0.025 |
| NC_006213.1;YP_009555238.1;HCoV-OC43_orf1ab;5555 | 0 | 0.1 |
| NC_006213.1;YP_009555238.1;HCoV-OC43_orf1ab;5556 | 0 | 0 |

| | | |
|---|---|---|
| NC_006213.1;YP_009555238.1;HCoV-OC43_orf1ab;5557 | 0 | 0 |
| NC_006213.1;YP_009555238.1;HCoV-OC43_orf1ab;5558 | 0 | 0 |
| NC_006213.1;YP_009555238.1;HCoV-OC43_orf1ab;5559 | 0 | 0 |
| NC_006213.1;YP_009555238.1;HCoV-OC43_orf1ab;5560 | 0 | 0 |
| NC_006213.1;YP_009555238.1;HCoV-OC43_orf1ab;5561 | 0 | 0 |
| NC_006213.1;YP_009555238.1;HCoV-OC43_orf1ab;5562 | 0 | 0 |
| NC_006213.1;YP_009555238.1;HCoV-OC43_orf1ab;5563 | 0 | 0 |
| NC_006213.1;YP_009555238.1;HCoV-OC43_orf1ab;5564 | 0 | 0 |
| NC_006213.1;YP_009555238.1;HCoV-OC43_orf1ab;5565 | 0 | 0.025 |
| NC_006213.1;YP_009555238.1;HCoV-OC43_orf1ab;5566 | 0 | 0.025 |
| NC_006213.1;YP_009555238.1;HCoV-OC43_orf1ab;5567 | 0 | 0 |
| NC_006213.1;YP_009555238.1;HCoV-OC43_orf1ab;5568 | 0 | 0 |
| NC_006213.1;YP_009555238.1;HCoV-OC43_orf1ab;5569 | 0 | 0 |
| NC_006213.1;YP_009555238.1;HCoV-OC43_orf1ab;5570 | 0 | 0 |
| NC_006213.1;YP_009555238.1;HCoV-OC43_orf1ab;5571 | 0 | 0.025 |
| NC_006213.1;YP_009555238.1;HCoV-OC43_orf1ab;5572 | 0 | 0 |
| NC_006213.1;YP_009555238.1;HCoV-OC43_orf1ab;5573 | 0 | 0.05 |
| NC_006213.1;YP_009555238.1;HCoV-OC43_orf1ab;5574 | 0 | 0.025 |
| NC_006213.1;YP_009555238.1;HCoV-OC43_orf1ab;5575 | 0 | 0.025 |
| NC_006213.1;YP_009555238.1;HCoV-OC43_orf1ab;5576 | 0 | 0 |
| NC_006213.1;YP_009555238.1;HCoV-OC43_orf1ab;5577 | 0 | 0 |
| NC_006213.1;YP_009555238.1;HCoV-OC43_orf1ab;5578 | 0 | 0 |
| NC_006213.1;YP_009555238.1;HCoV-OC43_orf1ab;5579 | 0 | 0 |
| NC_006213.1;YP_009555238.1;HCoV-OC43_orf1ab;5580 | 0 | 0 |
| NC_006213.1;YP_009555238.1;HCoV-OC43_orf1ab;5581 | 0 | 0 |
| NC_006213.1;YP_009555238.1;HCoV-OC43_orf1ab;5582 | 0 | 0 |
| NC_006213.1;YP_009555238.1;HCoV-OC43_orf1ab;5583 | 0 | 0 |
| NC_006213.1;YP_009555238.1;HCoV-OC43_orf1ab;5584 | 0 | 0 |
| NC_006213.1;YP_009555238.1;HCoV-OC43_orf1ab;5585 | 0 | 0 |
| NC_006213.1;YP_009555238.1;HCoV-OC43_orf1ab;5586 | 0 | 0 |
| NC_006213.1;YP_009555238.1;HCoV-OC43_orf1ab;5587 | 0 | 0 |
| NC_006213.1;YP_009555238.1;HCoV-OC43_orf1ab;5588 | 0 | 0 |
| NC_006213.1;YP_009555238.1;HCoV-OC43_orf1ab;5589 | 0 | 0 |
| NC_006213.1;YP_009555238.1;HCoV-OC43_orf1ab;5590 | 0 | 0 |
| NC_006213.1;YP_009555238.1;HCoV-OC43_orf1ab;5591 | 0 | 0 |
| NC_006213.1;YP_009555238.1;HCoV-OC43_orf1ab;5592 | 0 | 0 |
| NC_006213.1;YP_009555238.1;HCoV-OC43_orf1ab;5593 | 0 | 0 |
| NC_006213.1;YP_009555238.1;HCoV-OC43_orf1ab;5594 | 0.05 | 0 |
| NC_006213.1;YP_009555238.1;HCoV-OC43_orf1ab;5595 | 0 | 0 |
| NC_006213.1;YP_009555238.1;HCoV-OC43_orf1ab;5596 | 0 | 0.025 |
| NC_006213.1;YP_009555238.1;HCoV-OC43_orf1ab;5597 | 0 | 0.025 |
| NC_006213.1;YP_009555238.1;HCoV-OC43_orf1ab;5598 | 0 | 0.05 |
| NC_006213.1;YP_009555238.1;HCoV-OC43_orf1ab;5599 | 0 | 0.025 |

| | | |
|---|---|---|
| NC_006213.1;YP_009555238.1;HCoV-OC43_orf1ab;5600 | 0 | 0 |
| NC_006213.1;YP_009555238.1;HCoV-OC43_orf1ab;5601 | 0 | 0 |
| NC_006213.1;YP_009555238.1;HCoV-OC43_orf1ab;5602 | 0 | 0 |
| NC_006213.1;YP_009555238.1;HCoV-OC43_orf1ab;5603 | 0 | 0 |
| NC_006213.1;YP_009555238.1;HCoV-OC43_orf1ab;5604 | 0 | 0 |
| NC_006213.1;YP_009555238.1;HCoV-OC43_orf1ab;5605 | 0 | 0 |
| NC_006213.1;YP_009555238.1;HCoV-OC43_orf1ab;5606 | 0 | 0 |
| NC_006213.1;YP_009555238.1;HCoV-OC43_orf1ab;5607 | 0 | 0 |
| NC_006213.1;YP_009555238.1;HCoV-OC43_orf1ab;5608 | 0 | 0 |
| NC_006213.1;YP_009555238.1;HCoV-OC43_orf1ab;5609 | 0.05 | 0 |
| NC_006213.1;YP_009555238.1;HCoV-OC43_orf1ab;5610 | 0.05 | 0.025 |
| NC_006213.1;YP_009555238.1;HCoV-OC43_orf1ab;5611 | 0 | 0 |
| NC_006213.1;YP_009555238.1;HCoV-OC43_orf1ab;5612 | 0.05 | 0 |
| NC_006213.1;YP_009555238.1;HCoV-OC43_orf1ab;5613 | 0 | 0 |
| NC_006213.1;YP_009555238.1;HCoV-OC43_orf1ab;5614 | 0.05 | 0 |
| NC_006213.1;YP_009555238.1;HCoV-OC43_orf1ab;5615 | 0 | 0 |
| NC_006213.1;YP_009555238.1;HCoV-OC43_orf1ab;5616 | 0.05 | 0 |
| NC_006213.1;YP_009555238.1;HCoV-OC43_orf1ab;5617 | 0.05 | 0.025 |
| NC_006213.1;YP_009555238.1;HCoV-OC43_orf1ab;5618 | 0 | 0 |
| NC_006213.1;YP_009555238.1;HCoV-OC43_orf1ab;5619 | 0 | 0 |
| NC_006213.1;YP_009555238.1;HCoV-OC43_orf1ab;5620 | 0 | 0 |
| NC_006213.1;YP_009555238.1;HCoV-OC43_orf1ab;5621 | 0 | 0 |
| NC_006213.1;YP_009555238.1;HCoV-OC43_orf1ab;5622 | 0 | 0 |
| NC_006213.1;YP_009555238.1;HCoV-OC43_orf1ab;5623 | 0 | 0 |
| NC_006213.1;YP_009555238.1;HCoV-OC43_orf1ab;5624 | 0 | 0 |
| NC_006213.1;YP_009555238.1;HCoV-OC43_orf1ab;5625 | 0 | 0 |
| NC_006213.1;YP_009555238.1;HCoV-OC43_orf1ab;5626 | 0 | 0 |
| NC_006213.1;YP_009555238.1;HCoV-OC43_orf1ab;5627 | 0 | 0 |
| NC_006213.1;YP_009555238.1;HCoV-OC43_orf1ab;5628 | 0 | 0 |
| NC_006213.1;YP_009555238.1;HCoV-OC43_orf1ab;5629 | 0 | 0 |
| NC_006213.1;YP_009555238.1;HCoV-OC43_orf1ab;5630 | 0 | 0 |
| NC_006213.1;YP_009555238.1;HCoV-OC43_orf1ab;5631 | 0.1 | 0.05 |
| NC_006213.1;YP_009555238.1;HCoV-OC43_orf1ab;5632 | 0.05 | 0.025 |
| NC_006213.1;YP_009555238.1;HCoV-OC43_orf1ab;5633 | 0.15 | 0.125 |
| NC_006213.1;YP_009555238.1;HCoV-OC43_orf1ab;5634 | 0.35 | 0.1 |
| NC_006213.1;YP_009555238.1;HCoV-OC43_orf1ab;5635 | 0.1 | 0.1 |
| NC_006213.1;YP_009555238.1;HCoV-OC43_orf1ab;5636 | 0.4 | 0.175 |
| NC_006213.1;YP_009555238.1;HCoV-OC43_orf1ab;5637 | 0.5 | 0.275 |
| NC_006213.1;YP_009555238.1;HCoV-OC43_orf1ab;5638 | 0.2 | 0.1 |
| NC_006213.1;YP_009555238.1;HCoV-OC43_orf1ab;5639 | 0.05 | 0.05 |
| NC_006213.1;YP_009555238.1;HCoV-OC43_orf1ab;5640 | 0 | 0.025 |
| NC_006213.1;YP_009555238.1;HCoV-OC43_orf1ab;5641 | 0 | 0 |
| NC_006213.1;YP_009555238.1;HCoV-OC43_orf1ab;5642 | 0 | 0 |

| | | |
|---|---|---|
| NC_006213.1;YP_009555238.1;HCoV-OC43_orf1ab;5643 | 0 | 0 |
| NC_006213.1;YP_009555238.1;HCoV-OC43_orf1ab;5644 | 0 | 0 |
| NC_006213.1;YP_009555238.1;HCoV-OC43_orf1ab;5645 | 0 | 0 |
| NC_006213.1;YP_009555238.1;HCoV-OC43_orf1ab;5646 | 0 | 0 |
| NC_006213.1;YP_009555238.1;HCoV-OC43_orf1ab;5647 | 0 | 0 |
| NC_006213.1;YP_009555238.1;HCoV-OC43_orf1ab;5648 | 0.05 | 0 |
| NC_006213.1;YP_009555238.1;HCoV-OC43_orf1ab;5649 | 0.05 | 0 |
| NC_006213.1;YP_009555238.1;HCoV-OC43_orf1ab;5650 | 0 | 0 |
| NC_006213.1;YP_009555238.1;HCoV-OC43_orf1ab;5651 | 0 | 0 |
| NC_006213.1;YP_009555238.1;HCoV-OC43_orf1ab;5652 | 0 | 0.05 |
| NC_006213.1;YP_009555238.1;HCoV-OC43_orf1ab;5653 | 0 | 0 |
| NC_006213.1;YP_009555238.1;HCoV-OC43_orf1ab;5654 | 0 | 0 |
| NC_006213.1;YP_009555238.1;HCoV-OC43_orf1ab;5655 | 0 | 0 |
| NC_006213.1;YP_009555238.1;HCoV-OC43_orf1ab;5656 | 0 | 0 |
| NC_006213.1;YP_009555238.1;HCoV-OC43_orf1ab;5657 | 0 | 0 |
| NC_006213.1;YP_009555238.1;HCoV-OC43_orf1ab;5658 | 0.05 | 0 |
| NC_006213.1;YP_009555238.1;HCoV-OC43_orf1ab;5659 | 0 | 0 |
| NC_006213.1;YP_009555238.1;HCoV-OC43_orf1ab;5660 | 0.05 | 0 |
| NC_006213.1;YP_009555238.1;HCoV-OC43_orf1ab;5661 | 0.1 | 0 |
| NC_006213.1;YP_009555238.1;HCoV-OC43_orf1ab;5662 | 0.05 | 0.025 |
| NC_006213.1;YP_009555238.1;HCoV-OC43_orf1ab;5663 | 0.05 | 0.05 |
| NC_006213.1;YP_009555238.1;HCoV-OC43_orf1ab;5664 | 0.05 | 0.05 |
| NC_006213.1;YP_009555238.1;HCoV-OC43_orf1ab;5665 | 0.05 | 0 |
| NC_006213.1;YP_009555238.1;HCoV-OC43_orf1ab;5666 | 0.05 | 0 |
| NC_006213.1;YP_009555238.1;HCoV-OC43_orf1ab;5667 | 0.05 | 0.05 |
| NC_006213.1;YP_009555238.1;HCoV-OC43_orf1ab;5668 | 0.05 | 0.025 |
| NC_006213.1;YP_009555238.1;HCoV-OC43_orf1ab;5669 | 0 | 0 |
| NC_006213.1;YP_009555238.1;HCoV-OC43_orf1ab;5670 | 0 | 0 |
| NC_006213.1;YP_009555238.1;HCoV-OC43_orf1ab;5671 | 0.05 | 0 |
| NC_006213.1;YP_009555238.1;HCoV-OC43_orf1ab;5672 | 0 | 0 |
| NC_006213.1;YP_009555238.1;HCoV-OC43_orf1ab;5673 | 0 | 0 |
| NC_006213.1;YP_009555238.1;HCoV-OC43_orf1ab;5674 | 0 | 0 |
| NC_006213.1;YP_009555238.1;HCoV-OC43_orf1ab;5675 | 0 | 0 |
| NC_006213.1;YP_009555238.1;HCoV-OC43_orf1ab;5676 | 0 | 0 |
| NC_006213.1;YP_009555238.1;HCoV-OC43_orf1ab;5677 | 0 | 0 |
| NC_006213.1;YP_009555238.1;HCoV-OC43_orf1ab;5678 | 0 | 0 |
| NC_006213.1;YP_009555238.1;HCoV-OC43_orf1ab;5679 | 0 | 0 |
| NC_006213.1;YP_009555238.1;HCoV-OC43_orf1ab;5680 | 0 | 0 |
| NC_006213.1;YP_009555238.1;HCoV-OC43_orf1ab;5681 | 0 | 0 |
| NC_006213.1;YP_009555238.1;HCoV-OC43_orf1ab;5682 | 0 | 0.025 |
| NC_006213.1;YP_009555238.1;HCoV-OC43_orf1ab;5683 | 0 | 0.025 |
| NC_006213.1;YP_009555238.1;HCoV-OC43_orf1ab;5684 | 0 | 0.025 |
| NC_006213.1;YP_009555238.1;HCoV-OC43_orf1ab;5685 | 0 | 0.025 |

| | | |
|---|---|---|
| NC_006213.1;YP_009555238.1;HCoV-OC43_orf1ab;5686 | 0 | 0.025 |
| NC_006213.1;YP_009555238.1;HCoV-OC43_orf1ab;5687 | 0 | 0 |
| NC_006213.1;YP_009555238.1;HCoV-OC43_orf1ab;5688 | 0 | 0.025 |
| NC_006213.1;YP_009555238.1;HCoV-OC43_orf1ab;5689 | 0 | 0 |
| NC_006213.1;YP_009555238.1;HCoV-OC43_orf1ab;5690 | 0 | 0 |
| NC_006213.1;YP_009555238.1;HCoV-OC43_orf1ab;5691 | 0 | 0 |
| NC_006213.1;YP_009555238.1;HCoV-OC43_orf1ab;5692 | 0 | 0 |
| NC_006213.1;YP_009555238.1;HCoV-OC43_orf1ab;5693 | 0 | 0 |
| NC_006213.1;YP_009555238.1;HCoV-OC43_orf1ab;5694 | 0 | 0.05 |
| NC_006213.1;YP_009555238.1;HCoV-OC43_orf1ab;5695 | 0 | 0 |
| NC_006213.1;YP_009555238.1;HCoV-OC43_orf1ab;5696 | 0 | 0.025 |
| NC_006213.1;YP_009555238.1;HCoV-OC43_orf1ab;5697 | 0 | 0.025 |
| NC_006213.1;YP_009555238.1;HCoV-OC43_orf1ab;5698 | 0 | 0 |
| NC_006213.1;YP_009555238.1;HCoV-OC43_orf1ab;5699 | 0.05 | 0.025 |
| NC_006213.1;YP_009555238.1;HCoV-OC43_orf1ab;5700 | 0.05 | 0.075 |
| NC_006213.1;YP_009555238.1;HCoV-OC43_orf1ab;5701 | 0 | 0.025 |
| NC_006213.1;YP_009555238.1;HCoV-OC43_orf1ab;5702 | 0 | 0.05 |
| NC_006213.1;YP_009555238.1;HCoV-OC43_orf1ab;5703 | 0 | 0 |
| NC_006213.1;YP_009555238.1;HCoV-OC43_orf1ab;5704 | 0 | 0 |
| NC_006213.1;YP_009555238.1;HCoV-OC43_orf1ab;5705 | 0 | 0 |
| NC_006213.1;YP_009555238.1;HCoV-OC43_orf1ab;5706 | 0 | 0 |
| NC_006213.1;YP_009555238.1;HCoV-OC43_orf1ab;5707 | 0.05 | 0 |
| NC_006213.1;YP_009555238.1;HCoV-OC43_orf1ab;5708 | 0 | 0 |
| NC_006213.1;YP_009555238.1;HCoV-OC43_orf1ab;5709 | 0 | 0 |
| NC_006213.1;YP_009555238.1;HCoV-OC43_orf1ab;5710 | 0 | 0 |
| NC_006213.1;YP_009555238.1;HCoV-OC43_orf1ab;5711 | 0 | 0 |
| NC_006213.1;YP_009555238.1;HCoV-OC43_orf1ab;5712 | 0 | 0 |
| NC_006213.1;YP_009555238.1;HCoV-OC43_orf1ab;5713 | 0 | 0 |
| NC_006213.1;YP_009555238.1;HCoV-OC43_orf1ab;5714 | 0 | 0 |
| NC_006213.1;YP_009555238.1;HCoV-OC43_orf1ab;5715 | 0 | 0 |
| NC_006213.1;YP_009555238.1;HCoV-OC43_orf1ab;5716 | 0.05 | 0 |
| NC_006213.1;YP_009555238.1;HCoV-OC43_orf1ab;5717 | 0 | 0 |
| NC_006213.1;YP_009555238.1;HCoV-OC43_orf1ab;5718 | 0 | 0 |
| NC_006213.1;YP_009555238.1;HCoV-OC43_orf1ab;5719 | 0 | 0 |
| NC_006213.1;YP_009555238.1;HCoV-OC43_orf1ab;5720 | 0 | 0 |
| NC_006213.1;YP_009555238.1;HCoV-OC43_orf1ab;5721 | 0 | 0 |
| NC_006213.1;YP_009555238.1;HCoV-OC43_orf1ab;5722 | 0 | 0 |
| NC_006213.1;YP_009555238.1;HCoV-OC43_orf1ab;5723 | 0 | 0 |
| NC_006213.1;YP_009555238.1;HCoV-OC43_orf1ab;5724 | 0 | 0 |
| NC_006213.1;YP_009555238.1;HCoV-OC43_orf1ab;5725 | 0 | 0 |
| NC_006213.1;YP_009555238.1;HCoV-OC43_orf1ab;5726 | 0 | 0 |
| NC_006213.1;YP_009555238.1;HCoV-OC43_orf1ab;5727 | 0 | 0 |
| NC_006213.1;YP_009555238.1;HCoV-OC43_orf1ab;5728 | 0 | 0.025 |

| | | |
|---|---|---|
| NC_006213.1;YP_009555238.1;HCoV-OC43_orf1ab;5729 | 0 | 0 |
| NC_006213.1;YP_009555238.1;HCoV-OC43_orf1ab;5730 | 0 | 0 |
| NC_006213.1;YP_009555238.1;HCoV-OC43_orf1ab;5731 | 0 | 0 |
| NC_006213.1;YP_009555238.1;HCoV-OC43_orf1ab;5732 | 0 | 0 |
| NC_006213.1;YP_009555238.1;HCoV-OC43_orf1ab;5733 | 0 | 0 |
| NC_006213.1;YP_009555238.1;HCoV-OC43_orf1ab;5734 | 0.05 | 0.05 |
| NC_006213.1;YP_009555238.1;HCoV-OC43_orf1ab;5735 | 0.1 | 0.075 |
| NC_006213.1;YP_009555238.1;HCoV-OC43_orf1ab;5736 | 0.1 | 0.1 |
| NC_006213.1;YP_009555238.1;HCoV-OC43_orf1ab;5737 | 0.15 | 0.1 |
| NC_006213.1;YP_009555238.1;HCoV-OC43_orf1ab;5738 | 0.1 | 0.05 |
| NC_006213.1;YP_009555238.1;HCoV-OC43_orf1ab;5739 | 0.1 | 0.025 |
| NC_006213.1;YP_009555238.1;HCoV-OC43_orf1ab;5740 | 0.1 | 0.025 |
| NC_006213.1;YP_009555238.1;HCoV-OC43_orf1ab;5741 | 0.1 | 0.025 |
| NC_006213.1;YP_009555238.1;HCoV-OC43_orf1ab;5742 | 0.1 | 0.05 |
| NC_006213.1;YP_009555238.1;HCoV-OC43_orf1ab;5743 | 0.1 | 0.05 |
| NC_006213.1;YP_009555238.1;HCoV-OC43_orf1ab;5744 | 0.1 | 0.075 |
| NC_006213.1;YP_009555238.1;HCoV-OC43_orf1ab;5745 | 0.15 | 0.125 |
| NC_006213.1;YP_009555238.1;HCoV-OC43_orf1ab;5746 | 0 | 0.025 |
| NC_006213.1;YP_009555238.1;HCoV-OC43_orf1ab;5747 | 0 | 0.025 |
| NC_006213.1;YP_009555238.1;HCoV-OC43_orf1ab;5748 | 0 | 0.025 |
| NC_006213.1;YP_009555238.1;HCoV-OC43_orf1ab;5749 | 0 | 0 |
| NC_006213.1;YP_009555238.1;HCoV-OC43_orf1ab;5750 | 0 | 0 |
| NC_006213.1;YP_009555238.1;HCoV-OC43_orf1ab;5751 | 0 | 0 |
| NC_006213.1;YP_009555238.1;HCoV-OC43_orf1ab;5752 | 0 | 0 |
| NC_006213.1;YP_009555238.1;HCoV-OC43_orf1ab;5753 | 0 | 0 |
| NC_006213.1;YP_009555238.1;HCoV-OC43_orf1ab;5754 | 0 | 0 |
| NC_006213.1;YP_009555238.1;HCoV-OC43_orf1ab;5755 | 0 | 0 |
| NC_006213.1;YP_009555238.1;HCoV-OC43_orf1ab;5756 | 0 | 0.025 |
| NC_006213.1;YP_009555238.1;HCoV-OC43_orf1ab;5757 | 0 | 0 |
| NC_006213.1;YP_009555238.1;HCoV-OC43_orf1ab;5758 | 0 | 0 |
| NC_006213.1;YP_009555238.1;HCoV-OC43_orf1ab;5759 | 0 | 0 |
| NC_006213.1;YP_009555238.1;HCoV-OC43_orf1ab;5760 | 0 | 0 |
| NC_006213.1;YP_009555238.1;HCoV-OC43_orf1ab;5761 | 0 | 0 |
| NC_006213.1;YP_009555238.1;HCoV-OC43_orf1ab;5762 | 0 | 0 |
| NC_006213.1;YP_009555238.1;HCoV-OC43_orf1ab;5763 | 0 | 0 |
| NC_006213.1;YP_009555238.1;HCoV-OC43_orf1ab;5764 | 0 | 0 |
| NC_006213.1;YP_009555238.1;HCoV-OC43_orf1ab;5765 | 0 | 0 |
| NC_006213.1;YP_009555238.1;HCoV-OC43_orf1ab;5766 | 0 | 0 |
| NC_006213.1;YP_009555238.1;HCoV-OC43_orf1ab;5767 | 0 | 0 |
| NC_006213.1;YP_009555238.1;HCoV-OC43_orf1ab;5768 | 0 | 0 |
| NC_006213.1;YP_009555238.1;HCoV-OC43_orf1ab;5769 | 0 | 0 |
| NC_006213.1;YP_009555238.1;HCoV-OC43_orf1ab;5770 | 0 | 0.025 |
| NC_006213.1;YP_009555238.1;HCoV-OC43_orf1ab;5771 | 0 | 0.025 |

| | | |
|---|---|---|
| NC_006213.1;YP_009555238.1;HCoV-OC43_orf1ab;5772 | 0 | 0 |
| NC_006213.1;YP_009555238.1;HCoV-OC43_orf1ab;5773 | 0 | 0.025 |
| NC_006213.1;YP_009555238.1;HCoV-OC43_orf1ab;5774 | 0 | 0 |
| NC_006213.1;YP_009555238.1;HCoV-OC43_orf1ab;5775 | 0 | 0 |
| NC_006213.1;YP_009555238.1;HCoV-OC43_orf1ab;5776 | 0 | 0 |
| NC_006213.1;YP_009555238.1;HCoV-OC43_orf1ab;5777 | 0 | 0 |
| NC_006213.1;YP_009555238.1;HCoV-OC43_orf1ab;5778 | 0 | 0 |
| NC_006213.1;YP_009555238.1;HCoV-OC43_orf1ab;5779 | 0.05 | 0 |
| NC_006213.1;YP_009555238.1;HCoV-OC43_orf1ab;5780 | 0.05 | 0.05 |
| NC_006213.1;YP_009555238.1;HCoV-OC43_orf1ab;5781 | 0 | 0 |
| NC_006213.1;YP_009555238.1;HCoV-OC43_orf1ab;5782 | 0 | 0 |
| NC_006213.1;YP_009555238.1;HCoV-OC43_orf1ab;5783 | 0 | 0 |
| NC_006213.1;YP_009555238.1;HCoV-OC43_orf1ab;5784 | 0 | 0.025 |
| NC_006213.1;YP_009555238.1;HCoV-OC43_orf1ab;5785 | 0 | 0 |
| NC_006213.1;YP_009555238.1;HCoV-OC43_orf1ab;5786 | 0 | 0 |
| NC_006213.1;YP_009555238.1;HCoV-OC43_orf1ab;5787 | 0 | 0 |
| NC_006213.1;YP_009555238.1;HCoV-OC43_orf1ab;5788 | 0 | 0 |
| NC_006213.1;YP_009555238.1;HCoV-OC43_orf1ab;5789 | 0.05 | 0 |
| NC_006213.1;YP_009555238.1;HCoV-OC43_orf1ab;5790 | 0.05 | 0 |
| NC_006213.1;YP_009555238.1;HCoV-OC43_orf1ab;5791 | 0 | 0.025 |
| NC_006213.1;YP_009555238.1;HCoV-OC43_orf1ab;5792 | 0 | 0.025 |
| NC_006213.1;YP_009555238.1;HCoV-OC43_orf1ab;5793 | 0 | 0 |
| NC_006213.1;YP_009555238.1;HCoV-OC43_orf1ab;5794 | 0.05 | 0 |
| NC_006213.1;YP_009555238.1;HCoV-OC43_orf1ab;5795 | 0 | 0 |
| NC_006213.1;YP_009555238.1;HCoV-OC43_orf1ab;5796 | 0 | 0 |
| NC_006213.1;YP_009555238.1;HCoV-OC43_orf1ab;5797 | 0 | 0.025 |
| NC_006213.1;YP_009555238.1;HCoV-OC43_orf1ab;5798 | 0 | 0 |
| NC_006213.1;YP_009555238.1;HCoV-OC43_orf1ab;5799 | 0 | 0 |
| NC_006213.1;YP_009555238.1;HCoV-OC43_orf1ab;5800 | 0 | 0 |
| NC_006213.1;YP_009555238.1;HCoV-OC43_orf1ab;5801 | 0 | 0 |
| NC_006213.1;YP_009555238.1;HCoV-OC43_orf1ab;5802 | 0 | 0 |
| NC_006213.1;YP_009555238.1;HCoV-OC43_orf1ab;5803 | 0.1 | 0.025 |
| NC_006213.1;YP_009555238.1;HCoV-OC43_orf1ab;5804 | 0 | 0.05 |
| NC_006213.1;YP_009555238.1;HCoV-OC43_orf1ab;5805 | 0 | 0.05 |
| NC_006213.1;YP_009555238.1;HCoV-OC43_orf1ab;5806 | 0.05 | 0.05 |
| NC_006213.1;YP_009555238.1;HCoV-OC43_orf1ab;5807 | 0 | 0.05 |
| NC_006213.1;YP_009555238.1;HCoV-OC43_orf1ab;5808 | 0.05 | 0.05 |
| NC_006213.1;YP_009555238.1;HCoV-OC43_orf1ab;5809 | 0.05 | 0.075 |
| NC_006213.1;YP_009555238.1;HCoV-OC43_orf1ab;5810 | 0 | 0.1 |
| NC_006213.1;YP_009555238.1;HCoV-OC43_orf1ab;5811 | 0 | 0.1 |
| NC_006213.1;YP_009555238.1;HCoV-OC43_orf1ab;5812 | 0 | 0.1 |
| NC_006213.1;YP_009555238.1;HCoV-OC43_orf1ab;5813 | 0 | 0.1 |
| NC_006213.1;YP_009555238.1;HCoV-OC43_orf1ab;5814 | 0 | 0.075 |

| | | |
|---|---|---|
| NC_006213.1;YP_009555238.1;HCoV-OC43_orf1ab;5815 | 0 | 0.05 |
| NC_006213.1;YP_009555238.1;HCoV-OC43_orf1ab;5816 | 0 | 0.125 |
| NC_006213.1;YP_009555238.1;HCoV-OC43_orf1ab;5817 | 0 | 0.125 |
| NC_006213.1;YP_009555238.1;HCoV-OC43_orf1ab;5818 | 0.05 | 0.1 |
| NC_006213.1;YP_009555238.1;HCoV-OC43_orf1ab;5819 | 0.15 | 0.175 |
| NC_006213.1;YP_009555238.1;HCoV-OC43_orf1ab;5820 | 0 | 0.05 |
| NC_006213.1;YP_009555238.1;HCoV-OC43_orf1ab;5821 | 0 | 0.025 |
| NC_006213.1;YP_009555238.1;HCoV-OC43_orf1ab;5822 | 0 | 0 |
| NC_006213.1;YP_009555238.1;HCoV-OC43_orf1ab;5823 | 0.05 | 0.025 |
| NC_006213.1;YP_009555238.1;HCoV-OC43_orf1ab;5824 | 0.05 | 0 |
| NC_006213.1;YP_009555238.1;HCoV-OC43_orf1ab;5825 | 0.05 | 0 |
| NC_006213.1;YP_009555238.1;HCoV-OC43_orf1ab;5826 | 0 | 0 |
| NC_006213.1;YP_009555238.1;HCoV-OC43_orf1ab;5827 | 0 | 0 |
| NC_006213.1;YP_009555238.1;HCoV-OC43_orf1ab;5828 | 0 | 0 |
| NC_006213.1;YP_009555238.1;HCoV-OC43_orf1ab;5829 | 0 | 0.025 |
| NC_006213.1;YP_009555238.1;HCoV-OC43_orf1ab;5830 | 0 | 0 |
| NC_006213.1;YP_009555238.1;HCoV-OC43_orf1ab;5831 | 0 | 0 |
| NC_006213.1;YP_009555238.1;HCoV-OC43_orf1ab;5832 | 0 | 0 |
| NC_006213.1;YP_009555238.1;HCoV-OC43_orf1ab;5833 | 0 | 0 |
| NC_006213.1;YP_009555238.1;HCoV-OC43_orf1ab;5834 | 0 | 0 |
| NC_006213.1;YP_009555238.1;HCoV-OC43_orf1ab;5835 | 0 | 0 |
| NC_006213.1;YP_009555238.1;HCoV-OC43_orf1ab;5836 | 0.05 | 0.075 |
| NC_006213.1;YP_009555238.1;HCoV-OC43_orf1ab;5837 | 0 | 0 |
| NC_006213.1;YP_009555238.1;HCoV-OC43_orf1ab;5838 | 0 | 0 |
| NC_006213.1;YP_009555238.1;HCoV-OC43_orf1ab;5839 | 0 | 0 |
| NC_006213.1;YP_009555238.1;HCoV-OC43_orf1ab;5840 | 0 | 0.025 |
| NC_006213.1;YP_009555238.1;HCoV-OC43_orf1ab;5841 | 0 | 0.025 |
| NC_006213.1;YP_009555238.1;HCoV-OC43_orf1ab;5842 | 0 | 0 |
| NC_006213.1;YP_009555238.1;HCoV-OC43_orf1ab;5843 | 0.05 | 0 |
| NC_006213.1;YP_009555238.1;HCoV-OC43_orf1ab;5844 | 0.1 | 0 |
| NC_006213.1;YP_009555238.1;HCoV-OC43_orf1ab;5845 | 0 | 0.025 |
| NC_006213.1;YP_009555238.1;HCoV-OC43_orf1ab;5846 | 0 | 0 |
| NC_006213.1;YP_009555238.1;HCoV-OC43_orf1ab;5847 | 0 | 0 |
| NC_006213.1;YP_009555238.1;HCoV-OC43_orf1ab;5848 | 0 | 0 |
| NC_006213.1;YP_009555238.1;HCoV-OC43_orf1ab;5849 | 0 | 0 |
| NC_006213.1;YP_009555238.1;HCoV-OC43_orf1ab;5850 | 0 | 0 |
| NC_006213.1;YP_009555238.1;HCoV-OC43_orf1ab;5851 | 0 | 0 |
| NC_006213.1;YP_009555238.1;HCoV-OC43_orf1ab;5852 | 0 | 0.025 |
| NC_006213.1;YP_009555238.1;HCoV-OC43_orf1ab;5853 | 0 | 0 |
| NC_006213.1;YP_009555238.1;HCoV-OC43_orf1ab;5854 | 0.05 | 0 |
| NC_006213.1;YP_009555238.1;HCoV-OC43_orf1ab;5855 | 0.05 | 0 |
| NC_006213.1;YP_009555238.1;HCoV-OC43_orf1ab;5856 | 0.05 | 0 |
| NC_006213.1;YP_009555238.1;HCoV-OC43_orf1ab;5857 | 0.05 | 0 |

| | | |
|---|---|---|
| NC_006213.1;YP_009555238.1;HCoV-OC43_orf1ab;5858 | 0 | 0 |
| NC_006213.1;YP_009555238.1;HCoV-OC43_orf1ab;5859 | 0 | 0 |
| NC_006213.1;YP_009555238.1;HCoV-OC43_orf1ab;5860 | 0 | 0 |
| NC_006213.1;YP_009555238.1;HCoV-OC43_orf1ab;5861 | 0 | 0 |
| NC_006213.1;YP_009555238.1;HCoV-OC43_orf1ab;5862 | 0 | 0 |
| NC_006213.1;YP_009555238.1;HCoV-OC43_orf1ab;5863 | 0 | 0 |
| NC_006213.1;YP_009555238.1;HCoV-OC43_orf1ab;5864 | 0 | 0 |
| NC_006213.1;YP_009555238.1;HCoV-OC43_orf1ab;5865 | 0 | 0 |
| NC_006213.1;YP_009555238.1;HCoV-OC43_orf1ab;5866 | 0 | 0 |
| NC_006213.1;YP_009555238.1;HCoV-OC43_orf1ab;5867 | 0 | 0 |
| NC_006213.1;YP_009555238.1;HCoV-OC43_orf1ab;5868 | 0 | 0 |
| NC_006213.1;YP_009555238.1;HCoV-OC43_orf1ab;5869 | 0 | 0 |
| NC_006213.1;YP_009555238.1;HCoV-OC43_orf1ab;5870 | 0 | 0 |
| NC_006213.1;YP_009555238.1;HCoV-OC43_orf1ab;5871 | 0 | 0 |
| NC_006213.1;YP_009555238.1;HCoV-OC43_orf1ab;5872 | 0 | 0 |
| NC_006213.1;YP_009555238.1;HCoV-OC43_orf1ab;5873 | 0 | 0 |
| NC_006213.1;YP_009555238.1;HCoV-OC43_orf1ab;5874 | 0 | 0 |
| NC_006213.1;YP_009555238.1;HCoV-OC43_orf1ab;5875 | 0 | 0 |
| NC_006213.1;YP_009555238.1;HCoV-OC43_orf1ab;5876 | 0 | 0 |
| NC_006213.1;YP_009555238.1;HCoV-OC43_orf1ab;5877 | 0 | 0 |
| NC_006213.1;YP_009555238.1;HCoV-OC43_orf1ab;5878 | 0 | 0 |
| NC_006213.1;YP_009555238.1;HCoV-OC43_orf1ab;5879 | 0 | 0 |
| NC_006213.1;YP_009555238.1;HCoV-OC43_orf1ab;5880 | 0 | 0 |
| NC_006213.1;YP_009555238.1;HCoV-OC43_orf1ab;5881 | 0 | 0 |
| NC_006213.1;YP_009555238.1;HCoV-OC43_orf1ab;5882 | 0 | 0 |
| NC_006213.1;YP_009555238.1;HCoV-OC43_orf1ab;5883 | 0.05 | 0 |
| NC_006213.1;YP_009555238.1;HCoV-OC43_orf1ab;5884 | 0.05 | 0 |
| NC_006213.1;YP_009555238.1;HCoV-OC43_orf1ab;5885 | 0.1 | 0 |
| NC_006213.1;YP_009555238.1;HCoV-OC43_orf1ab;5886 | 0.05 | 0 |
| NC_006213.1;YP_009555238.1;HCoV-OC43_orf1ab;5887 | 0.1 | 0 |
| NC_006213.1;YP_009555238.1;HCoV-OC43_orf1ab;5888 | 0.05 | 0 |
| NC_006213.1;YP_009555238.1;HCoV-OC43_orf1ab;5889 | 0.05 | 0 |
| NC_006213.1;YP_009555238.1;HCoV-OC43_orf1ab;5890 | 0.1 | 0 |
| NC_006213.1;YP_009555238.1;HCoV-OC43_orf1ab;5891 | 0.05 | 0 |
| NC_006213.1;YP_009555238.1;HCoV-OC43_orf1ab;5892 | 0 | 0 |
| NC_006213.1;YP_009555238.1;HCoV-OC43_orf1ab;5893 | 0 | 0 |
| NC_006213.1;YP_009555238.1;HCoV-OC43_orf1ab;5894 | 0 | 0 |
| NC_006213.1;YP_009555238.1;HCoV-OC43_orf1ab;5895 | 0 | 0 |
| NC_006213.1;YP_009555238.1;HCoV-OC43_orf1ab;5896 | 0 | 0.05 |
| NC_006213.1;YP_009555238.1;HCoV-OC43_orf1ab;5897 | 0 | 0 |
| NC_006213.1;YP_009555238.1;HCoV-OC43_orf1ab;5898 | 0 | 0.025 |
| NC_006213.1;YP_009555238.1;HCoV-OC43_orf1ab;5899 | 0 | 0 |
| NC_006213.1;YP_009555238.1;HCoV-OC43_orf1ab;5900 | 0 | 0.025 |

| | | |
|---|---|---|
| NC_006213.1;YP_009555238.1;HCoV-OC43_orf1ab;5901 | 0 | 0.025 |
| NC_006213.1;YP_009555238.1;HCoV-OC43_orf1ab;5902 | 0 | 0 |
| NC_006213.1;YP_009555238.1;HCoV-OC43_orf1ab;5903 | 0 | 0 |
| NC_006213.1;YP_009555238.1;HCoV-OC43_orf1ab;5904 | 0.05 | 0.025 |
| NC_006213.1;YP_009555238.1;HCoV-OC43_orf1ab;5905 | 0 | 0.025 |
| NC_006213.1;YP_009555238.1;HCoV-OC43_orf1ab;5906 | 0 | 0.025 |
| NC_006213.1;YP_009555238.1;HCoV-OC43_orf1ab;5907 | 0 | 0.025 |
| NC_006213.1;YP_009555238.1;HCoV-OC43_orf1ab;5908 | 0 | 0.025 |
| NC_006213.1;YP_009555238.1;HCoV-OC43_orf1ab;5909 | 0.05 | 0.025 |
| NC_006213.1;YP_009555238.1;HCoV-OC43_orf1ab;5910 | 0 | 0.025 |
| NC_006213.1;YP_009555238.1;HCoV-OC43_orf1ab;5911 | 0 | 0.025 |
| NC_006213.1;YP_009555238.1;HCoV-OC43_orf1ab;5912 | 0 | 0.025 |
| NC_006213.1;YP_009555238.1;HCoV-OC43_orf1ab;5913 | 0 | 0.025 |
| NC_006213.1;YP_009555238.1;HCoV-OC43_orf1ab;5914 | 0 | 0.025 |
| NC_006213.1;YP_009555238.1;HCoV-OC43_orf1ab;5915 | 0 | 0.025 |
| NC_006213.1;YP_009555238.1;HCoV-OC43_orf1ab;5916 | 0 | 0.05 |
| NC_006213.1;YP_009555238.1;HCoV-OC43_orf1ab;5917 | 0 | 0.025 |
| NC_006213.1;YP_009555238.1;HCoV-OC43_orf1ab;5918 | 0.05 | 0.025 |
| NC_006213.1;YP_009555238.1;HCoV-OC43_orf1ab;5919 | 0.1 | 0.025 |
| NC_006213.1;YP_009555238.1;HCoV-OC43_orf1ab;5920 | 0.2 | 0.025 |
| NC_006213.1;YP_009555238.1;HCoV-OC43_orf1ab;5921 | 0.2 | 0.025 |
| NC_006213.1;YP_009555238.1;HCoV-OC43_orf1ab;5922 | 0.15 | 0.05 |
| NC_006213.1;YP_009555238.1;HCoV-OC43_orf1ab;5923 | 0.15 | 0.05 |
| NC_006213.1;YP_009555238.1;HCoV-OC43_orf1ab;5924 | 0.15 | 0.025 |
| NC_006213.1;YP_009555238.1;HCoV-OC43_orf1ab;5925 | 0.05 | 0 |
| NC_006213.1;YP_009555238.1;HCoV-OC43_orf1ab;5926 | 0 | 0 |
| NC_006213.1;YP_009555238.1;HCoV-OC43_orf1ab;5927 | 0 | 0 |
| NC_006213.1;YP_009555238.1;HCoV-OC43_orf1ab;5928 | 0 | 0 |
| NC_006213.1;YP_009555238.1;HCoV-OC43_orf1ab;5929 | 0 | 0 |
| NC_006213.1;YP_009555238.1;HCoV-OC43_orf1ab;5930 | 0 | 0.025 |
| NC_006213.1;YP_009555238.1;HCoV-OC43_orf1ab;5931 | 0 | 0.025 |
| NC_006213.1;YP_009555238.1;HCoV-OC43_orf1ab;5932 | 0 | 0.025 |
| NC_006213.1;YP_009555238.1;HCoV-OC43_orf1ab;5933 | 0.05 | 0.025 |
| NC_006213.1;YP_009555238.1;HCoV-OC43_orf1ab;5934 | 0 | 0.025 |
| NC_006213.1;YP_009555238.1;HCoV-OC43_orf1ab;5935 | 0 | 0.025 |
| NC_006213.1;YP_009555238.1;HCoV-OC43_orf1ab;5936 | 0 | 0.025 |
| NC_006213.1;YP_009555238.1;HCoV-OC43_orf1ab;5937 | 0 | 0 |
| NC_006213.1;YP_009555238.1;HCoV-OC43_orf1ab;5938 | 0 | 0 |
| NC_006213.1;YP_009555238.1;HCoV-OC43_orf1ab;5939 | 0 | 0 |
| NC_006213.1;YP_009555238.1;HCoV-OC43_orf1ab;5940 | 0 | 0 |
| NC_006213.1;YP_009555238.1;HCoV-OC43_orf1ab;5941 | 0 | 0 |
| NC_006213.1;YP_009555238.1;HCoV-OC43_orf1ab;5942 | 0.05 | 0.05 |
| NC_006213.1;YP_009555238.1;HCoV-OC43_orf1ab;5943 | 0 | 0 |

| | | |
|---|---|---|
| NC_006213.1;YP_009555238.1;HCoV-OC43_orf1ab;5944 | 0 | 0 |
| NC_006213.1;YP_009555238.1;HCoV-OC43_orf1ab;5945 | 0 | 0 |
| NC_006213.1;YP_009555238.1;HCoV-OC43_orf1ab;5946 | 0 | 0 |
| NC_006213.1;YP_009555238.1;HCoV-OC43_orf1ab;5947 | 0 | 0 |
| NC_006213.1;YP_009555238.1;HCoV-OC43_orf1ab;5948 | 0 | 0 |
| NC_006213.1;YP_009555238.1;HCoV-OC43_orf1ab;5949 | 0 | 0 |
| NC_006213.1;YP_009555238.1;HCoV-OC43_orf1ab;5950 | 0 | 0 |
| NC_006213.1;YP_009555238.1;HCoV-OC43_orf1ab;5951 | 0 | 0 |
| NC_006213.1;YP_009555238.1;HCoV-OC43_orf1ab;5952 | 0 | 0 |
| NC_006213.1;YP_009555238.1;HCoV-OC43_orf1ab;5953 | 0 | 0 |
| NC_006213.1;YP_009555238.1;HCoV-OC43_orf1ab;5954 | 0 | 0 |
| NC_006213.1;YP_009555238.1;HCoV-OC43_orf1ab;5955 | 0 | 0 |
| NC_006213.1;YP_009555238.1;HCoV-OC43_orf1ab;5956 | 0 | 0 |
| NC_006213.1;YP_009555238.1;HCoV-OC43_orf1ab;5957 | 0 | 0 |
| NC_006213.1;YP_009555238.1;HCoV-OC43_orf1ab;5958 | 0 | 0 |
| NC_006213.1;YP_009555238.1;HCoV-OC43_orf1ab;5959 | 0 | 0 |
| NC_006213.1;YP_009555238.1;HCoV-OC43_orf1ab;5960 | 0 | 0 |
| NC_006213.1;YP_009555238.1;HCoV-OC43_orf1ab;5961 | 0 | 0 |
| NC_006213.1;YP_009555238.1;HCoV-OC43_orf1ab;5962 | 0 | 0 |
| NC_006213.1;YP_009555238.1;HCoV-OC43_orf1ab;5963 | 0 | 0 |
| NC_006213.1;YP_009555238.1;HCoV-OC43_orf1ab;5964 | 0 | 0 |
| NC_006213.1;YP_009555238.1;HCoV-OC43_orf1ab;5965 | 0 | 0 |
| NC_006213.1;YP_009555238.1;HCoV-OC43_orf1ab;5966 | 0 | 0.025 |
| NC_006213.1;YP_009555238.1;HCoV-OC43_orf1ab;5967 | 0 | 0 |
| NC_006213.1;YP_009555238.1;HCoV-OC43_orf1ab;5968 | 0 | 0 |
| NC_006213.1;YP_009555238.1;HCoV-OC43_orf1ab;5969 | 0 | 0 |
| NC_006213.1;YP_009555238.1;HCoV-OC43_orf1ab;5970 | 0 | 0 |
| NC_006213.1;YP_009555238.1;HCoV-OC43_orf1ab;5971 | 0 | 0 |
| NC_006213.1;YP_009555238.1;HCoV-OC43_orf1ab;5972 | 0 | 0 |
| NC_006213.1;YP_009555238.1;HCoV-OC43_orf1ab;5973 | 0 | 0 |
| NC_006213.1;YP_009555238.1;HCoV-OC43_orf1ab;5974 | 0 | 0 |
| NC_006213.1;YP_009555238.1;HCoV-OC43_orf1ab;5975 | 0 | 0 |
| NC_006213.1;YP_009555238.1;HCoV-OC43_orf1ab;5976 | 0 | 0 |
| NC_006213.1;YP_009555238.1;HCoV-OC43_orf1ab;5977 | 0 | 0 |
| NC_006213.1;YP_009555238.1;HCoV-OC43_orf1ab;5978 | 0 | 0.025 |
| NC_006213.1;YP_009555238.1;HCoV-OC43_orf1ab;5979 | 0 | 0.025 |
| NC_006213.1;YP_009555238.1;HCoV-OC43_orf1ab;5980 | 0 | 0.025 |
| NC_006213.1;YP_009555238.1;HCoV-OC43_orf1ab;5981 | 0 | 0.025 |
| NC_006213.1;YP_009555238.1;HCoV-OC43_orf1ab;5982 | 0 | 0.025 |
| NC_006213.1;YP_009555238.1;HCoV-OC43_orf1ab;5983 | 0 | 0.05 |
| NC_006213.1;YP_009555238.1;HCoV-OC43_orf1ab;5984 | 0 | 0.025 |
| NC_006213.1;YP_009555238.1;HCoV-OC43_orf1ab;5985 | 0 | 0.025 |
| NC_006213.1;YP_009555238.1;HCoV-OC43_orf1ab;5986 | 0 | 0 |

| | | |
|---|---|---|
| NC_006213.1;YP_009555238.1;HCoV-OC43_orf1ab;5987 | 0 | 0 |
| NC_006213.1;YP_009555238.1;HCoV-OC43_orf1ab;5988 | 0 | 0.1 |
| NC_006213.1;YP_009555238.1;HCoV-OC43_orf1ab;5989 | 0 | 0.075 |
| NC_006213.1;YP_009555238.1;HCoV-OC43_orf1ab;5990 | 0.05 | 0.075 |
| NC_006213.1;YP_009555238.1;HCoV-OC43_orf1ab;5991 | 0.05 | 0.125 |
| NC_006213.1;YP_009555238.1;HCoV-OC43_orf1ab;5992 | 0.05 | 0.15 |
| NC_006213.1;YP_009555238.1;HCoV-OC43_orf1ab;5993 | 0.05 | 0.1 |
| NC_006213.1;YP_009555238.1;HCoV-OC43_orf1ab;5994 | 0.05 | 0.1 |
| NC_006213.1;YP_009555238.1;HCoV-OC43_orf1ab;5995 | 0.05 | 0.05 |
| NC_006213.1;YP_009555238.1;HCoV-OC43_orf1ab;5996 | 0.05 | 0.075 |
| NC_006213.1;YP_009555238.1;HCoV-OC43_orf1ab;5997 | 0.05 | 0.1 |
| NC_006213.1;YP_009555238.1;HCoV-OC43_orf1ab;5998 | 0.05 | 0.025 |
| NC_006213.1;YP_009555238.1;HCoV-OC43_orf1ab;5999 | 0.05 | 0.075 |
| NC_006213.1;YP_009555238.1;HCoV-OC43_orf1ab;6000 | 0.05 | 0.125 |
| NC_006213.1;YP_009555238.1;HCoV-OC43_orf1ab;6001 | 0 | 0 |
| NC_006213.1;YP_009555238.1;HCoV-OC43_orf1ab;6002 | 0 | 0 |
| NC_006213.1;YP_009555238.1;HCoV-OC43_orf1ab;6003 | 0 | 0 |
| NC_006213.1;YP_009555238.1;HCoV-OC43_orf1ab;6004 | 0 | 0 |
| NC_006213.1;YP_009555238.1;HCoV-OC43_orf1ab;6005 | 0 | 0 |
| NC_006213.1;YP_009555238.1;HCoV-OC43_orf1ab;6006 | 0 | 0 |
| NC_006213.1;YP_009555238.1;HCoV-OC43_orf1ab;6007 | 0 | 0 |
| NC_006213.1;YP_009555238.1;HCoV-OC43_orf1ab;6008 | 0 | 0 |
| NC_006213.1;YP_009555238.1;HCoV-OC43_orf1ab;6009 | 0 | 0 |
| NC_006213.1;YP_009555238.1;HCoV-OC43_orf1ab;6010 | 0 | 0 |
| NC_006213.1;YP_009555238.1;HCoV-OC43_orf1ab;6011 | 0 | 0 |
| NC_006213.1;YP_009555238.1;HCoV-OC43_orf1ab;6012 | 0 | 0 |
| NC_006213.1;YP_009555238.1;HCoV-OC43_orf1ab;6013 | 0 | 0 |
| NC_006213.1;YP_009555238.1;HCoV-OC43_orf1ab;6014 | 0 | 0 |
| NC_006213.1;YP_009555238.1;HCoV-OC43_orf1ab;6015 | 0 | 0 |
| NC_006213.1;YP_009555238.1;HCoV-OC43_orf1ab;6016 | 0 | 0 |
| NC_006213.1;YP_009555238.1;HCoV-OC43_orf1ab;6017 | 0 | 0 |
| NC_006213.1;YP_009555238.1;HCoV-OC43_orf1ab;6018 | 0 | 0 |
| NC_006213.1;YP_009555238.1;HCoV-OC43_orf1ab;6019 | 0 | 0.025 |
| NC_006213.1;YP_009555238.1;HCoV-OC43_orf1ab;6020 | 0 | 0 |
| NC_006213.1;YP_009555238.1;HCoV-OC43_orf1ab;6021 | 0 | 0 |
| NC_006213.1;YP_009555238.1;HCoV-OC43_orf1ab;6022 | 0 | 0 |
| NC_006213.1;YP_009555238.1;HCoV-OC43_orf1ab;6023 | 0 | 0 |
| NC_006213.1;YP_009555238.1;HCoV-OC43_orf1ab;6024 | 0 | 0 |
| NC_006213.1;YP_009555238.1;HCoV-OC43_orf1ab;6025 | 0 | 0 |
| NC_006213.1;YP_009555238.1;HCoV-OC43_orf1ab;6026 | 0 | 0 |
| NC_006213.1;YP_009555238.1;HCoV-OC43_orf1ab;6027 | 0 | 0 |
| NC_006213.1;YP_009555238.1;HCoV-OC43_orf1ab;6028 | 0 | 0 |
| NC_006213.1;YP_009555238.1;HCoV-OC43_orf1ab;6029 | 0 | 0 |

| | | |
|---|---|---|
| NC_006213.1;YP_009555238.1;HCoV-OC43_orf1ab;6030 | 0 | 0 |
| NC_006213.1;YP_009555238.1;HCoV-OC43_orf1ab;6031 | 0 | 0 |
| NC_006213.1;YP_009555238.1;HCoV-OC43_orf1ab;6032 | 0 | 0 |
| NC_006213.1;YP_009555238.1;HCoV-OC43_orf1ab;6033 | 0 | 0 |
| NC_006213.1;YP_009555238.1;HCoV-OC43_orf1ab;6034 | 0 | 0 |
| NC_006213.1;YP_009555238.1;HCoV-OC43_orf1ab;6035 | 0 | 0 |
| NC_006213.1;YP_009555238.1;HCoV-OC43_orf1ab;6036 | 0 | 0 |
| NC_006213.1;YP_009555238.1;HCoV-OC43_orf1ab;6037 | 0 | 0 |
| NC_006213.1;YP_009555238.1;HCoV-OC43_orf1ab;6038 | 0 | 0 |
| NC_006213.1;YP_009555238.1;HCoV-OC43_orf1ab;6039 | 0 | 0 |
| NC_006213.1;YP_009555238.1;HCoV-OC43_orf1ab;6040 | 0 | 0 |
| NC_006213.1;YP_009555238.1;HCoV-OC43_orf1ab;6041 | 0 | 0 |
| NC_006213.1;YP_009555238.1;HCoV-OC43_orf1ab;6042 | 0 | 0 |
| NC_006213.1;YP_009555238.1;HCoV-OC43_orf1ab;6043 | 0 | 0 |
| NC_006213.1;YP_009555238.1;HCoV-OC43_orf1ab;6044 | 0 | 0 |
| NC_006213.1;YP_009555238.1;HCoV-OC43_orf1ab;6045 | 0 | 0 |
| NC_006213.1;YP_009555238.1;HCoV-OC43_orf1ab;6046 | 0 | 0 |
| NC_006213.1;YP_009555238.1;HCoV-OC43_orf1ab;6047 | 0 | 0 |
| NC_006213.1;YP_009555238.1;HCoV-OC43_orf1ab;6048 | 0 | 0 |
| NC_006213.1;YP_009555238.1;HCoV-OC43_orf1ab;6049 | 0 | 0 |
| NC_006213.1;YP_009555238.1;HCoV-OC43_orf1ab;6050 | 0 | 0 |
| NC_006213.1;YP_009555238.1;HCoV-OC43_orf1ab;6051 | 0 | 0 |
| NC_006213.1;YP_009555238.1;HCoV-OC43_orf1ab;6052 | 0 | 0 |
| NC_006213.1;YP_009555238.1;HCoV-OC43_orf1ab;6053 | 0 | 0 |
| NC_006213.1;YP_009555238.1;HCoV-OC43_orf1ab;6054 | 0 | 0 |
| NC_006213.1;YP_009555238.1;HCoV-OC43_orf1ab;6055 | 0 | 0 |
| NC_006213.1;YP_009555238.1;HCoV-OC43_orf1ab;6056 | 0 | 0 |
| NC_006213.1;YP_009555238.1;HCoV-OC43_orf1ab;6057 | 0 | 0 |
| NC_006213.1;YP_009555238.1;HCoV-OC43_orf1ab;6058 | 0 | 0 |
| NC_006213.1;YP_009555238.1;HCoV-OC43_orf1ab;6059 | 0 | 0 |
| NC_006213.1;YP_009555238.1;HCoV-OC43_orf1ab;6060 | 0 | 0 |
| NC_006213.1;YP_009555238.1;HCoV-OC43_orf1ab;6061 | 0 | 0 |
| NC_006213.1;YP_009555238.1;HCoV-OC43_orf1ab;6062 | 0 | 0 |
| NC_006213.1;YP_009555238.1;HCoV-OC43_orf1ab;6063 | 0 | 0 |
| NC_006213.1;YP_009555238.1;HCoV-OC43_orf1ab;6064 | 0 | 0 |
| NC_006213.1;YP_009555238.1;HCoV-OC43_orf1ab;6065 | 0 | 0 |
| NC_006213.1;YP_009555238.1;HCoV-OC43_orf1ab;6066 | 0 | 0 |
| NC_006213.1;YP_009555238.1;HCoV-OC43_orf1ab;6067 | 0 | 0 |
| NC_006213.1;YP_009555238.1;HCoV-OC43_orf1ab;6068 | 0 | 0 |
| NC_006213.1;YP_009555238.1;HCoV-OC43_orf1ab;6069 | 0 | 0 |
| NC_006213.1;YP_009555238.1;HCoV-OC43_orf1ab;6070 | 0 | 0 |
| NC_006213.1;YP_009555238.1;HCoV-OC43_orf1ab;6071 | 0 | 0 |
| NC_006213.1;YP_009555238.1;HCoV-OC43_orf1ab;6072 | 0 | 0 |

| | | |
|---|---|---|
| NC_006213.1;YP_009555238.1;HCoV-OC43_orf1ab;6073 | 0 | 0 |
| NC_006213.1;YP_009555238.1;HCoV-OC43_orf1ab;6074 | 0 | 0 |
| NC_006213.1;YP_009555238.1;HCoV-OC43_orf1ab;6075 | 0 | 0 |
| NC_006213.1;YP_009555238.1;HCoV-OC43_orf1ab;6076 | 0 | 0 |
| NC_006213.1;YP_009555238.1;HCoV-OC43_orf1ab;6077 | 0 | 0 |
| NC_006213.1;YP_009555238.1;HCoV-OC43_orf1ab;6078 | 0 | 0 |
| NC_006213.1;YP_009555238.1;HCoV-OC43_orf1ab;6079 | 0 | 0 |
| NC_006213.1;YP_009555238.1;HCoV-OC43_orf1ab;6080 | 0 | 0 |
| NC_006213.1;YP_009555238.1;HCoV-OC43_orf1ab;6081 | 0 | 0 |
| NC_006213.1;YP_009555238.1;HCoV-OC43_orf1ab;6082 | 0 | 0 |
| NC_006213.1;YP_009555238.1;HCoV-OC43_orf1ab;6083 | 0 | 0 |
| NC_006213.1;YP_009555238.1;HCoV-OC43_orf1ab;6084 | 0 | 0 |
| NC_006213.1;YP_009555238.1;HCoV-OC43_orf1ab;6085 | 0 | 0 |
| NC_006213.1;YP_009555238.1;HCoV-OC43_orf1ab;6086 | 0 | 0 |
| NC_006213.1;YP_009555238.1;HCoV-OC43_orf1ab;6087 | 0 | 0 |
| NC_006213.1;YP_009555238.1;HCoV-OC43_orf1ab;6088 | 0 | 0 |
| NC_006213.1;YP_009555238.1;HCoV-OC43_orf1ab;6089 | 0 | 0 |
| NC_006213.1;YP_009555238.1;HCoV-OC43_orf1ab;6090 | 0 | 0 |
| NC_006213.1;YP_009555238.1;HCoV-OC43_orf1ab;6091 | 0 | 0 |
| NC_006213.1;YP_009555238.1;HCoV-OC43_orf1ab;6092 | 0 | 0 |
| NC_006213.1;YP_009555238.1;HCoV-OC43_orf1ab;6093 | 0 | 0 |
| NC_006213.1;YP_009555238.1;HCoV-OC43_orf1ab;6094 | 0 | 0 |
| NC_006213.1;YP_009555238.1;HCoV-OC43_orf1ab;6095 | 0 | 0 |
| NC_006213.1;YP_009555238.1;HCoV-OC43_orf1ab;6096 | 0 | 0.025 |
| NC_006213.1;YP_009555238.1;HCoV-OC43_orf1ab;6097 | 0 | 0 |
| NC_006213.1;YP_009555238.1;HCoV-OC43_orf1ab;6098 | 0 | 0 |
| NC_006213.1;YP_009555238.1;HCoV-OC43_orf1ab;6099 | 0 | 0 |
| NC_006213.1;YP_009555238.1;HCoV-OC43_orf1ab;6100 | 0 | 0 |
| NC_006213.1;YP_009555238.1;HCoV-OC43_orf1ab;6101 | 0 | 0.025 |
| NC_006213.1;YP_009555238.1;HCoV-OC43_orf1ab;6102 | 0 | 0.025 |
| NC_006213.1;YP_009555238.1;HCoV-OC43_orf1ab;6103 | 0 | 0.025 |
| NC_006213.1;YP_009555238.1;HCoV-OC43_orf1ab;6104 | 0 | 0 |
| NC_006213.1;YP_009555238.1;HCoV-OC43_orf1ab;6105 | 0 | 0 |
| NC_006213.1;YP_009555238.1;HCoV-OC43_orf1ab;6106 | 0 | 0 |
| NC_006213.1;YP_009555238.1;HCoV-OC43_orf1ab;6107 | 0 | 0 |
| NC_006213.1;YP_009555238.1;HCoV-OC43_orf1ab;6108 | 0 | 0 |
| NC_006213.1;YP_009555238.1;HCoV-OC43_orf1ab;6109 | 0 | 0 |
| NC_006213.1;YP_009555238.1;HCoV-OC43_orf1ab;6110 | 0 | 0 |
| NC_006213.1;YP_009555238.1;HCoV-OC43_orf1ab;6111 | 0 | 0 |
| NC_006213.1;YP_009555238.1;HCoV-OC43_orf1ab;6112 | 0 | 0 |
| NC_006213.1;YP_009555238.1;HCoV-OC43_orf1ab;6113 | 0 | 0 |
| NC_006213.1;YP_009555238.1;HCoV-OC43_orf1ab;6114 | 0 | 0 |
| NC_006213.1;YP_009555238.1;HCoV-OC43_orf1ab;6115 | 0 | 0 |

| | | |
|---|---|---|
| NC_006213.1;YP_009555238.1;HCoV-OC43_orf1ab;6116 | 0 | 0 |
| NC_006213.1;YP_009555238.1;HCoV-OC43_orf1ab;6117 | 0 | 0 |
| NC_006213.1;YP_009555238.1;HCoV-OC43_orf1ab;6118 | 0 | 0 |
| NC_006213.1;YP_009555238.1;HCoV-OC43_orf1ab;6119 | 0 | 0 |
| NC_006213.1;YP_009555238.1;HCoV-OC43_orf1ab;6120 | 0 | 0 |
| NC_006213.1;YP_009555238.1;HCoV-OC43_orf1ab;6121 | 0 | 0 |
| NC_006213.1;YP_009555238.1;HCoV-OC43_orf1ab;6122 | 0 | 0 |
| NC_006213.1;YP_009555238.1;HCoV-OC43_orf1ab;6123 | 0 | 0 |
| NC_006213.1;YP_009555238.1;HCoV-OC43_orf1ab;6124 | 0 | 0 |
| NC_006213.1;YP_009555238.1;HCoV-OC43_orf1ab;6125 | 0 | 0 |
| NC_006213.1;YP_009555238.1;HCoV-OC43_orf1ab;6126 | 0 | 0 |
| NC_006213.1;YP_009555238.1;HCoV-OC43_orf1ab;6127 | 0 | 0 |
| NC_006213.1;YP_009555238.1;HCoV-OC43_orf1ab;6128 | 0 | 0 |
| NC_006213.1;YP_009555238.1;HCoV-OC43_orf1ab;6129 | 0 | 0 |
| NC_006213.1;YP_009555238.1;HCoV-OC43_orf1ab;6130 | 0 | 0 |
| NC_006213.1;YP_009555238.1;HCoV-OC43_orf1ab;6131 | 0 | 0 |
| NC_006213.1;YP_009555238.1;HCoV-OC43_orf1ab;6132 | 0 | 0 |
| NC_006213.1;YP_009555238.1;HCoV-OC43_orf1ab;6133 | 0 | 0 |
| NC_006213.1;YP_009555238.1;HCoV-OC43_orf1ab;6134 | 0 | 0 |
| NC_006213.1;YP_009555238.1;HCoV-OC43_orf1ab;6135 | 0 | 0 |
| NC_006213.1;YP_009555238.1;HCoV-OC43_orf1ab;6136 | 0 | 0 |
| NC_006213.1;YP_009555238.1;HCoV-OC43_orf1ab;6137 | 0 | 0 |
| NC_006213.1;YP_009555238.1;HCoV-OC43_orf1ab;6138 | 0 | 0 |
| NC_006213.1;YP_009555238.1;HCoV-OC43_orf1ab;6139 | 0 | 0 |
| NC_006213.1;YP_009555238.1;HCoV-OC43_orf1ab;6140 | 0 | 0 |
| NC_006213.1;YP_009555238.1;HCoV-OC43_orf1ab;6141 | 0 | 0 |
| NC_006213.1;YP_009555238.1;HCoV-OC43_orf1ab;6142 | 0 | 0 |
| NC_006213.1;YP_009555238.1;HCoV-OC43_orf1ab;6143 | 0 | 0 |
| NC_006213.1;YP_009555238.1;HCoV-OC43_orf1ab;6144 | 0 | 0 |
| NC_006213.1;YP_009555238.1;HCoV-OC43_orf1ab;6145 | 0 | 0 |
| NC_006213.1;YP_009555238.1;HCoV-OC43_orf1ab;6146 | 0 | 0 |
| NC_006213.1;YP_009555238.1;HCoV-OC43_orf1ab;6147 | 0 | 0 |
| NC_006213.1;YP_009555238.1;HCoV-OC43_orf1ab;6148 | 0 | 0 |
| NC_006213.1;YP_009555238.1;HCoV-OC43_orf1ab;6149 | 0 | 0 |
| NC_006213.1;YP_009555238.1;HCoV-OC43_orf1ab;6150 | 0 | 0 |
| NC_006213.1;YP_009555238.1;HCoV-OC43_orf1ab;6151 | 0 | 0.025 |
| NC_006213.1;YP_009555238.1;HCoV-OC43_orf1ab;6152 | 0 | 0 |
| NC_006213.1;YP_009555238.1;HCoV-OC43_orf1ab;6153 | 0 | 0 |
| NC_006213.1;YP_009555238.1;HCoV-OC43_orf1ab;6154 | 0.05 | 0.025 |
| NC_006213.1;YP_009555238.1;HCoV-OC43_orf1ab;6155 | 0 | 0 |
| NC_006213.1;YP_009555238.1;HCoV-OC43_orf1ab;6156 | 0 | 0 |
| NC_006213.1;YP_009555238.1;HCoV-OC43_orf1ab;6157 | 0 | 0 |
| NC_006213.1;YP_009555238.1;HCoV-OC43_orf1ab;6158 | 0 | 0 |

| | | |
|---|---|---|
| NC_006213.1;YP_009555238.1;HCoV-OC43_orf1ab;6159 | 0 | 0 |
| NC_006213.1;YP_009555238.1;HCoV-OC43_orf1ab;6160 | 0 | 0 |
| NC_006213.1;YP_009555238.1;HCoV-OC43_orf1ab;6161 | 0 | 0 |
| NC_006213.1;YP_009555238.1;HCoV-OC43_orf1ab;6162 | 0 | 0 |
| NC_006213.1;YP_009555238.1;HCoV-OC43_orf1ab;6163 | 0 | 0 |
| NC_006213.1;YP_009555238.1;HCoV-OC43_orf1ab;6164 | 0 | 0.025 |
| NC_006213.1;YP_009555238.1;HCoV-OC43_orf1ab;6165 | 0 | 0 |
| NC_006213.1;YP_009555238.1;HCoV-OC43_orf1ab;6166 | 0 | 0 |
| NC_006213.1;YP_009555238.1;HCoV-OC43_orf1ab;6167 | 0 | 0 |
| NC_006213.1;YP_009555238.1;HCoV-OC43_orf1ab;6168 | 0 | 0 |
| NC_006213.1;YP_009555238.1;HCoV-OC43_orf1ab;6169 | 0 | 0 |
| NC_006213.1;YP_009555238.1;HCoV-OC43_orf1ab;6170 | 0 | 0 |
| NC_006213.1;YP_009555238.1;HCoV-OC43_orf1ab;6171 | 0 | 0 |
| NC_006213.1;YP_009555238.1;HCoV-OC43_orf1ab;6172 | 0 | 0 |
| NC_006213.1;YP_009555238.1;HCoV-OC43_orf1ab;6173 | 0 | 0 |
| NC_006213.1;YP_009555238.1;HCoV-OC43_orf1ab;6174 | 0 | 0 |
| NC_006213.1;YP_009555238.1;HCoV-OC43_orf1ab;6175 | 0 | 0 |
| NC_006213.1;YP_009555238.1;HCoV-OC43_orf1ab;6176 | 0 | 0 |
| NC_006213.1;YP_009555238.1;HCoV-OC43_orf1ab;6177 | 0 | 0 |
| NC_006213.1;YP_009555238.1;HCoV-OC43_orf1ab;6178 | 0 | 0 |
| NC_006213.1;YP_009555238.1;HCoV-OC43_orf1ab;6179 | 0 | 0 |
| NC_006213.1;YP_009555238.1;HCoV-OC43_orf1ab;6180 | 0 | 0 |
| NC_006213.1;YP_009555238.1;HCoV-OC43_orf1ab;6181 | 0 | 0 |
| NC_006213.1;YP_009555238.1;HCoV-OC43_orf1ab;6182 | 0 | 0 |
| NC_006213.1;YP_009555238.1;HCoV-OC43_orf1ab;6183 | 0 | 0 |
| NC_006213.1;YP_009555238.1;HCoV-OC43_orf1ab;6184 | 0 | 0 |
| NC_006213.1;YP_009555238.1;HCoV-OC43_orf1ab;6185 | 0 | 0 |
| NC_006213.1;YP_009555238.1;HCoV-OC43_orf1ab;6186 | 0.05 | 0.025 |
| NC_006213.1;YP_009555238.1;HCoV-OC43_orf1ab;6187 | 0.05 | 0 |
| NC_006213.1;YP_009555238.1;HCoV-OC43_orf1ab;6188 | 0.05 | 0.025 |
| NC_006213.1;YP_009555238.1;HCoV-OC43_orf1ab;6189 | 0 | 0.05 |
| NC_006213.1;YP_009555238.1;HCoV-OC43_orf1ab;6190 | 0 | 0.025 |
| NC_006213.1;YP_009555238.1;HCoV-OC43_orf1ab;6191 | 0 | 0.025 |
| NC_006213.1;YP_009555238.1;HCoV-OC43_orf1ab;6192 | 0.05 | 0.025 |
| NC_006213.1;YP_009555238.1;HCoV-OC43_orf1ab;6193 | 0 | 0.025 |
| NC_006213.1;YP_009555238.1;HCoV-OC43_orf1ab;6194 | 0 | 0.025 |
| NC_006213.1;YP_009555238.1;HCoV-OC43_orf1ab;6195 | 0 | 0 |
| NC_006213.1;YP_009555238.1;HCoV-OC43_orf1ab;6196 | 0.05 | 0 |
| NC_006213.1;YP_009555238.1;HCoV-OC43_orf1ab;6197 | 0 | 0 |
| NC_006213.1;YP_009555238.1;HCoV-OC43_orf1ab;6198 | 0 | 0 |
| NC_006213.1;YP_009555238.1;HCoV-OC43_orf1ab;6199 | 0 | 0 |
| NC_006213.1;YP_009555238.1;HCoV-OC43_orf1ab;6200 | 0 | 0 |
| NC_006213.1;YP_009555238.1;HCoV-OC43_orf1ab;6201 | 0 | 0 |

| | | |
|---|---|---|
| NC_006213.1;YP_009555238.1;HCoV-OC43_orf1ab;6202 | 0 | 0 |
| NC_006213.1;YP_009555238.1;HCoV-OC43_orf1ab;6203 | 0 | 0 |
| NC_006213.1;YP_009555238.1;HCoV-OC43_orf1ab;6204 | 0 | 0 |
| NC_006213.1;YP_009555238.1;HCoV-OC43_orf1ab;6205 | 0 | 0 |
| NC_006213.1;YP_009555238.1;HCoV-OC43_orf1ab;6206 | 0 | 0 |
| NC_006213.1;YP_009555238.1;HCoV-OC43_orf1ab;6207 | 0 | 0 |
| NC_006213.1;YP_009555238.1;HCoV-OC43_orf1ab;6208 | 0 | 0 |
| NC_006213.1;YP_009555238.1;HCoV-OC43_orf1ab;6209 | 0 | 0 |
| NC_006213.1;YP_009555238.1;HCoV-OC43_orf1ab;6210 | 0 | 0 |
| NC_006213.1;YP_009555238.1;HCoV-OC43_orf1ab;6211 | 0 | 0 |
| NC_006213.1;YP_009555238.1;HCoV-OC43_orf1ab;6212 | 0 | 0 |
| NC_006213.1;YP_009555238.1;HCoV-OC43_orf1ab;6213 | 0 | 0 |
| NC_006213.1;YP_009555238.1;HCoV-OC43_orf1ab;6214 | 0 | 0 |
| NC_006213.1;YP_009555238.1;HCoV-OC43_orf1ab;6215 | 0 | 0 |
| NC_006213.1;YP_009555238.1;HCoV-OC43_orf1ab;6216 | 0 | 0 |
| NC_006213.1;YP_009555238.1;HCoV-OC43_orf1ab;6217 | 0 | 0 |
| NC_006213.1;YP_009555238.1;HCoV-OC43_orf1ab;6218 | 0 | 0 |
| NC_006213.1;YP_009555238.1;HCoV-OC43_orf1ab;6219 | 0 | 0 |
| NC_006213.1;YP_009555238.1;HCoV-OC43_orf1ab;6220 | 0 | 0 |
| NC_006213.1;YP_009555238.1;HCoV-OC43_orf1ab;6221 | 0 | 0 |
| NC_006213.1;YP_009555238.1;HCoV-OC43_orf1ab;6222 | 0 | 0 |
| NC_006213.1;YP_009555238.1;HCoV-OC43_orf1ab;6223 | 0 | 0 |
| NC_006213.1;YP_009555238.1;HCoV-OC43_orf1ab;6224 | 0 | 0 |
| NC_006213.1;YP_009555238.1;HCoV-OC43_orf1ab;6225 | 0 | 0 |
| NC_006213.1;YP_009555238.1;HCoV-OC43_orf1ab;6226 | 0 | 0 |
| NC_006213.1;YP_009555238.1;HCoV-OC43_orf1ab;6227 | 0 | 0 |
| NC_006213.1;YP_009555238.1;HCoV-OC43_orf1ab;6228 | 0 | 0 |
| NC_006213.1;YP_009555238.1;HCoV-OC43_orf1ab;6229 | 0 | 0 |
| NC_006213.1;YP_009555238.1;HCoV-OC43_orf1ab;6230 | 0 | 0 |
| NC_006213.1;YP_009555238.1;HCoV-OC43_orf1ab;6231 | 0 | 0 |
| NC_006213.1;YP_009555238.1;HCoV-OC43_orf1ab;6232 | 0 | 0 |
| NC_006213.1;YP_009555238.1;HCoV-OC43_orf1ab;6233 | 0 | 0 |
| NC_006213.1;YP_009555238.1;HCoV-OC43_orf1ab;6234 | 0 | 0 |
| NC_006213.1;YP_009555238.1;HCoV-OC43_orf1ab;6235 | 0 | 0.025 |
| NC_006213.1;YP_009555238.1;HCoV-OC43_orf1ab;6236 | 0 | 0 |
| NC_006213.1;YP_009555238.1;HCoV-OC43_orf1ab;6237 | 0 | 0.025 |
| NC_006213.1;YP_009555238.1;HCoV-OC43_orf1ab;6238 | 0.05 | 0.025 |
| NC_006213.1;YP_009555238.1;HCoV-OC43_orf1ab;6239 | 0 | 0.025 |
| NC_006213.1;YP_009555238.1;HCoV-OC43_orf1ab;6240 | 0.1 | 0.075 |
| NC_006213.1;YP_009555238.1;HCoV-OC43_orf1ab;6241 | 0.3 | 0.075 |
| NC_006213.1;YP_009555238.1;HCoV-OC43_orf1ab;6242 | 0.1 | 0 |
| NC_006213.1;YP_009555238.1;HCoV-OC43_orf1ab;6243 | 0.05 | 0 |
| NC_006213.1;YP_009555238.1;HCoV-OC43_orf1ab;6244 | 0.05 | 0 |

| | | |
|---|---|---|
| NC_006213.1;YP_009555238.1;HCoV-OC43_orf1ab;6245 | 0 | 0 |
| NC_006213.1;YP_009555238.1;HCoV-OC43_orf1ab;6246 | 0.05 | 0 |
| NC_006213.1;YP_009555238.1;HCoV-OC43_orf1ab;6247 | 0 | 0 |
| NC_006213.1;YP_009555238.1;HCoV-OC43_orf1ab;6248 | 0 | 0 |
| NC_006213.1;YP_009555238.1;HCoV-OC43_orf1ab;6249 | 0 | 0 |
| NC_006213.1;YP_009555238.1;HCoV-OC43_orf1ab;6250 | 0 | 0 |
| NC_006213.1;YP_009555238.1;HCoV-OC43_orf1ab;6251 | 0 | 0 |
| NC_006213.1;YP_009555238.1;HCoV-OC43_orf1ab;6252 | 0 | 0 |
| NC_006213.1;YP_009555238.1;HCoV-OC43_orf1ab;6253 | 0 | 0 |
| NC_006213.1;YP_009555238.1;HCoV-OC43_orf1ab;6254 | 0 | 0 |
| NC_006213.1;YP_009555238.1;HCoV-OC43_orf1ab;6255 | 0 | 0 |
| NC_006213.1;YP_009555238.1;HCoV-OC43_orf1ab;6256 | 0 | 0 |
| NC_006213.1;YP_009555238.1;HCoV-OC43_orf1ab;6257 | 0 | 0 |
| NC_006213.1;YP_009555238.1;HCoV-OC43_orf1ab;6258 | 0 | 0.025 |
| NC_006213.1;YP_009555238.1;HCoV-OC43_orf1ab;6259 | 0.1 | 0 |
| NC_006213.1;YP_009555238.1;HCoV-OC43_orf1ab;6260 | 0.1 | 0.025 |
| NC_006213.1;YP_009555238.1;HCoV-OC43_orf1ab;6261 | 0.1 | 0.025 |
| NC_006213.1;YP_009555238.1;HCoV-OC43_orf1ab;6262 | 0.05 | 0 |
| NC_006213.1;YP_009555238.1;HCoV-OC43_orf1ab;6263 | 0.05 | 0.025 |
| NC_006213.1;YP_009555238.1;HCoV-OC43_orf1ab;6264 | 0.05 | 0 |
| NC_006213.1;YP_009555238.1;HCoV-OC43_orf1ab;6265 | 0.05 | 0 |
| NC_006213.1;YP_009555238.1;HCoV-OC43_orf1ab;6266 | 0.05 | 0 |
| NC_006213.1;YP_009555238.1;HCoV-OC43_orf1ab;6267 | 0.05 | 0 |
| NC_006213.1;YP_009555238.1;HCoV-OC43_orf1ab;6268 | 0.05 | 0.025 |
| NC_006213.1;YP_009555238.1;HCoV-OC43_orf1ab;6269 | 0 | 0 |
| NC_006213.1;YP_009555238.1;HCoV-OC43_orf1ab;6270 | 0.05 | 0 |
| NC_006213.1;YP_009555238.1;HCoV-OC43_orf1ab;6271 | 0.05 | 0 |
| NC_006213.1;YP_009555238.1;HCoV-OC43_orf1ab;6272 | 0.05 | 0 |
| NC_006213.1;YP_009555238.1;HCoV-OC43_orf1ab;6273 | 0 | 0 |
| NC_006213.1;YP_009555238.1;HCoV-OC43_orf1ab;6274 | 0 | 0 |
| NC_006213.1;YP_009555238.1;HCoV-OC43_orf1ab;6275 | 0.05 | 0 |
| NC_006213.1;YP_009555238.1;HCoV-OC43_orf1ab;6276 | 0.05 | 0 |
| NC_006213.1;YP_009555238.1;HCoV-OC43_orf1ab;6277 | 0 | 0 |
| NC_006213.1;YP_009555238.1;HCoV-OC43_orf1ab;6278 | 0 | 0 |
| NC_006213.1;YP_009555238.1;HCoV-OC43_orf1ab;6279 | 0 | 0 |
| NC_006213.1;YP_009555238.1;HCoV-OC43_orf1ab;6280 | 0 | 0 |
| NC_006213.1;YP_009555238.1;HCoV-OC43_orf1ab;6281 | 0 | 0 |
| NC_006213.1;YP_009555238.1;HCoV-OC43_orf1ab;6282 | 0 | 0 |
| NC_006213.1;YP_009555238.1;HCoV-OC43_orf1ab;6283 | 0 | 0 |
| NC_006213.1;YP_009555238.1;HCoV-OC43_orf1ab;6284 | 0 | 0 |
| NC_006213.1;YP_009555238.1;HCoV-OC43_orf1ab;6285 | 0 | 0 |
| NC_006213.1;YP_009555238.1;HCoV-OC43_orf1ab;6286 | 0 | 0 |
| NC_006213.1;YP_009555238.1;HCoV-OC43_orf1ab;6287 | 0 | 0 |

| | | |
|---|---|---|
| NC_006213.1;YP_009555238.1;HCoV-OC43_orf1ab;6288 | 0 | 0 |
| NC_006213.1;YP_009555238.1;HCoV-OC43_orf1ab;6289 | 0 | 0 |
| NC_006213.1;YP_009555238.1;HCoV-OC43_orf1ab;6290 | 0 | 0 |
| NC_006213.1;YP_009555238.1;HCoV-OC43_orf1ab;6291 | 0.05 | 0 |
| NC_006213.1;YP_009555238.1;HCoV-OC43_orf1ab;6292 | 0.05 | 0 |
| NC_006213.1;YP_009555238.1;HCoV-OC43_orf1ab;6293 | 0.1 | 0.025 |
| NC_006213.1;YP_009555238.1;HCoV-OC43_orf1ab;6294 | 0.05 | 0.025 |
| NC_006213.1;YP_009555238.1;HCoV-OC43_orf1ab;6295 | 0.1 | 0.05 |
| NC_006213.1;YP_009555238.1;HCoV-OC43_orf1ab;6296 | 0.2 | 0.1 |
| NC_006213.1;YP_009555238.1;HCoV-OC43_orf1ab;6297 | 0.15 | 0.075 |
| NC_006213.1;YP_009555238.1;HCoV-OC43_orf1ab;6298 | 0.2 | 0.075 |
| NC_006213.1;YP_009555238.1;HCoV-OC43_orf1ab;6299 | 0.05 | 0.05 |
| NC_006213.1;YP_009555238.1;HCoV-OC43_orf1ab;6300 | 0.05 | 0 |
| NC_006213.1;YP_009555238.1;HCoV-OC43_orf1ab;6301 | 0.05 | 0 |
| NC_006213.1;YP_009555238.1;HCoV-OC43_orf1ab;6302 | 0 | 0 |
| NC_006213.1;YP_009555238.1;HCoV-OC43_orf1ab;6303 | 0 | 0 |
| NC_006213.1;YP_009555238.1;HCoV-OC43_orf1ab;6304 | 0 | 0 |
| NC_006213.1;YP_009555238.1;HCoV-OC43_orf1ab;6305 | 0 | 0 |
| NC_006213.1;YP_009555238.1;HCoV-OC43_orf1ab;6306 | 0 | 0 |
| NC_006213.1;YP_009555238.1;HCoV-OC43_orf1ab;6307 | 0 | 0 |
| NC_006213.1;YP_009555238.1;HCoV-OC43_orf1ab;6308 | 0 | 0 |
| NC_006213.1;YP_009555238.1;HCoV-OC43_orf1ab;6309 | 0 | 0 |
| NC_006213.1;YP_009555238.1;HCoV-OC43_orf1ab;6310 | 0 | 0 |
| NC_006213.1;YP_009555238.1;HCoV-OC43_orf1ab;6311 | 0 | 0 |
| NC_006213.1;YP_009555238.1;HCoV-OC43_orf1ab;6312 | 0 | 0 |
| NC_006213.1;YP_009555238.1;HCoV-OC43_orf1ab;6313 | 0 | 0 |
| NC_006213.1;YP_009555238.1;HCoV-OC43_orf1ab;6314 | 0 | 0 |
| NC_006213.1;YP_009555238.1;HCoV-OC43_orf1ab;6315 | 0 | 0 |
| NC_006213.1;YP_009555238.1;HCoV-OC43_orf1ab;6316 | 0 | 0 |
| NC_006213.1;YP_009555238.1;HCoV-OC43_orf1ab;6317 | 0 | 0 |
| NC_006213.1;YP_009555238.1;HCoV-OC43_orf1ab;6318 | 0 | 0 |
| NC_006213.1;YP_009555238.1;HCoV-OC43_orf1ab;6319 | 0 | 0 |
| NC_006213.1;YP_009555238.1;HCoV-OC43_orf1ab;6320 | 0 | 0 |
| NC_006213.1;YP_009555238.1;HCoV-OC43_orf1ab;6321 | 0 | 0 |
| NC_006213.1;YP_009555238.1;HCoV-OC43_orf1ab;6322 | 0 | 0 |
| NC_006213.1;YP_009555238.1;HCoV-OC43_orf1ab;6323 | 0 | 0.025 |
| NC_006213.1;YP_009555238.1;HCoV-OC43_orf1ab;6324 | 0 | 0 |
| NC_006213.1;YP_009555238.1;HCoV-OC43_orf1ab;6325 | 0 | 0 |
| NC_006213.1;YP_009555238.1;HCoV-OC43_orf1ab;6326 | 0 | 0 |
| NC_006213.1;YP_009555238.1;HCoV-OC43_orf1ab;6327 | 0 | 0 |
| NC_006213.1;YP_009555238.1;HCoV-OC43_orf1ab;6328 | 0 | 0 |
| NC_006213.1;YP_009555238.1;HCoV-OC43_orf1ab;6329 | 0 | 0 |
| NC_006213.1;YP_009555238.1;HCoV-OC43_orf1ab;6330 | 0 | 0 |

| | | |
|---|---|---|
| NC_006213.1;YP_009555238.1;HCoV-OC43_orf1ab;6331 | 0 | 0 |
| NC_006213.1;YP_009555238.1;HCoV-OC43_orf1ab;6332 | 0 | 0 |
| NC_006213.1;YP_009555238.1;HCoV-OC43_orf1ab;6333 | 0 | 0 |
| NC_006213.1;YP_009555238.1;HCoV-OC43_orf1ab;6334 | 0 | 0 |
| NC_006213.1;YP_009555238.1;HCoV-OC43_orf1ab;6335 | 0 | 0 |
| NC_006213.1;YP_009555238.1;HCoV-OC43_orf1ab;6336 | 0 | 0 |
| NC_006213.1;YP_009555238.1;HCoV-OC43_orf1ab;6337 | 0 | 0 |
| NC_006213.1;YP_009555238.1;HCoV-OC43_orf1ab;6338 | 0 | 0 |
| NC_006213.1;YP_009555238.1;HCoV-OC43_orf1ab;6339 | 0 | 0 |
| NC_006213.1;YP_009555238.1;HCoV-OC43_orf1ab;6340 | 0 | 0 |
| NC_006213.1;YP_009555238.1;HCoV-OC43_orf1ab;6341 | 0 | 0 |
| NC_006213.1;YP_009555238.1;HCoV-OC43_orf1ab;6342 | 0 | 0 |
| NC_006213.1;YP_009555238.1;HCoV-OC43_orf1ab;6343 | 0 | 0 |
| NC_006213.1;YP_009555238.1;HCoV-OC43_orf1ab;6344 | 0 | 0 |
| NC_006213.1;YP_009555238.1;HCoV-OC43_orf1ab;6345 | 0 | 0 |
| NC_006213.1;YP_009555238.1;HCoV-OC43_orf1ab;6346 | 0 | 0 |
| NC_006213.1;YP_009555238.1;HCoV-OC43_orf1ab;6347 | 0 | 0 |
| NC_006213.1;YP_009555238.1;HCoV-OC43_orf1ab;6348 | 0 | 0.075 |
| NC_006213.1;YP_009555238.1;HCoV-OC43_orf1ab;6349 | 0 | 0.075 |
| NC_006213.1;YP_009555238.1;HCoV-OC43_orf1ab;6350 | 0 | 0.075 |
| NC_006213.1;YP_009555238.1;HCoV-OC43_orf1ab;6351 | 0 | 0.05 |
| NC_006213.1;YP_009555238.1;HCoV-OC43_orf1ab;6352 | 0 | 0.05 |
| NC_006213.1;YP_009555238.1;HCoV-OC43_orf1ab;6353 | 0 | 0.05 |
| NC_006213.1;YP_009555238.1;HCoV-OC43_orf1ab;6354 | 0 | 0.05 |
| NC_006213.1;YP_009555238.1;HCoV-OC43_orf1ab;6355 | 0 | 0 |
| NC_006213.1;YP_009555238.1;HCoV-OC43_orf1ab;6356 | 0 | 0 |
| NC_006213.1;YP_009555238.1;HCoV-OC43_orf1ab;6357 | 0 | 0 |
| NC_006213.1;YP_009555238.1;HCoV-OC43_orf1ab;6358 | 0 | 0 |
| NC_006213.1;YP_009555238.1;HCoV-OC43_orf1ab;6359 | 0 | 0 |
| NC_006213.1;YP_009555238.1;HCoV-OC43_orf1ab;6360 | 0 | 0 |
| NC_006213.1;YP_009555238.1;HCoV-OC43_orf1ab;6361 | 0 | 0 |
| NC_006213.1;YP_009555238.1;HCoV-OC43_orf1ab;6362 | 0 | 0 |
| NC_006213.1;YP_009555238.1;HCoV-OC43_orf1ab;6363 | 0 | 0 |
| NC_006213.1;YP_009555238.1;HCoV-OC43_orf1ab;6364 | 0 | 0 |
| NC_006213.1;YP_009555238.1;HCoV-OC43_orf1ab;6365 | 0 | 0 |
| NC_006213.1;YP_009555238.1;HCoV-OC43_orf1ab;6366 | 0 | 0 |
| NC_006213.1;YP_009555238.1;HCoV-OC43_orf1ab;6367 | 0 | 0 |
| NC_006213.1;YP_009555238.1;HCoV-OC43_orf1ab;6368 | 0 | 0 |
| NC_006213.1;YP_009555238.1;HCoV-OC43_orf1ab;6369 | 0 | 0 |
| NC_006213.1;YP_009555238.1;HCoV-OC43_orf1ab;6370 | 0 | 0 |
| NC_006213.1;YP_009555238.1;HCoV-OC43_orf1ab;6371 | 0 | 0 |
| NC_006213.1;YP_009555238.1;HCoV-OC43_orf1ab;6372 | 0 | 0 |
| NC_006213.1;YP_009555238.1;HCoV-OC43_orf1ab;6373 | 0 | 0 |

| | | |
|---|---|---|
| NC_006213.1;YP_009555238.1;HCoV-OC43_orf1ab;6374 | 0 | 0.025 |
| NC_006213.1;YP_009555238.1;HCoV-OC43_orf1ab;6375 | 0 | 0.025 |
| NC_006213.1;YP_009555238.1;HCoV-OC43_orf1ab;6376 | 0 | 0.025 |
| NC_006213.1;YP_009555238.1;HCoV-OC43_orf1ab;6377 | 0 | 0.025 |
| NC_006213.1;YP_009555238.1;HCoV-OC43_orf1ab;6378 | 0 | 0.1 |
| NC_006213.1;YP_009555238.1;HCoV-OC43_orf1ab;6379 | 0 | 0.025 |
| NC_006213.1;YP_009555238.1;HCoV-OC43_orf1ab;6380 | 0 | 0 |
| NC_006213.1;YP_009555238.1;HCoV-OC43_orf1ab;6381 | 0 | 0.05 |
| NC_006213.1;YP_009555238.1;HCoV-OC43_orf1ab;6382 | 0.15 | 0.025 |
| NC_006213.1;YP_009555238.1;HCoV-OC43_orf1ab;6383 | 0.1 | 0.125 |
| NC_006213.1;YP_009555238.1;HCoV-OC43_orf1ab;6384 | 0.15 | 0.025 |
| NC_006213.1;YP_009555238.1;HCoV-OC43_orf1ab;6385 | 0.1 | 0.025 |
| NC_006213.1;YP_009555238.1;HCoV-OC43_orf1ab;6386 | 0.05 | 0.025 |
| NC_006213.1;YP_009555238.1;HCoV-OC43_orf1ab;6387 | 0 | 0.05 |
| NC_006213.1;YP_009555238.1;HCoV-OC43_orf1ab;6388 | 0 | 0 |
| NC_006213.1;YP_009555238.1;HCoV-OC43_orf1ab;6389 | 0 | 0 |
| NC_006213.1;YP_009555238.1;HCoV-OC43_orf1ab;6390 | 0 | 0 |
| NC_006213.1;YP_009555238.1;HCoV-OC43_orf1ab;6391 | 0 | 0 |
| NC_006213.1;YP_009555238.1;HCoV-OC43_orf1ab;6392 | 0 | 0 |
| NC_006213.1;YP_009555238.1;HCoV-OC43_orf1ab;6393 | 0 | 0 |
| NC_006213.1;YP_009555238.1;HCoV-OC43_orf1ab;6394 | 0 | 0 |
| NC_006213.1;YP_009555238.1;HCoV-OC43_orf1ab;6395 | 0 | 0 |
| NC_006213.1;YP_009555238.1;HCoV-OC43_orf1ab;6396 | 0 | 0 |
| NC_006213.1;YP_009555238.1;HCoV-OC43_orf1ab;6397 | 0 | 0 |
| NC_006213.1;YP_009555238.1;HCoV-OC43_orf1ab;6398 | 0 | 0 |
| NC_006213.1;YP_009555238.1;HCoV-OC43_orf1ab;6399 | 0 | 0 |
| NC_006213.1;YP_009555238.1;HCoV-OC43_orf1ab;6400 | 0 | 0 |
| NC_006213.1;YP_009555238.1;HCoV-OC43_orf1ab;6401 | 0 | 0 |
| NC_006213.1;YP_009555238.1;HCoV-OC43_orf1ab;6402 | 0 | 0 |
| NC_006213.1;YP_009555238.1;HCoV-OC43_orf1ab;6403 | 0 | 0 |
| NC_006213.1;YP_009555238.1;HCoV-OC43_orf1ab;6404 | 0 | 0 |
| NC_006213.1;YP_009555238.1;HCoV-OC43_orf1ab;6405 | 0 | 0 |
| NC_006213.1;YP_009555238.1;HCoV-OC43_orf1ab;6406 | 0 | 0 |
| NC_006213.1;YP_009555238.1;HCoV-OC43_orf1ab;6407 | 0 | 0 |
| NC_006213.1;YP_009555238.1;HCoV-OC43_orf1ab;6408 | 0 | 0 |
| NC_006213.1;YP_009555238.1;HCoV-OC43_orf1ab;6409 | 0 | 0 |
| NC_006213.1;YP_009555238.1;HCoV-OC43_orf1ab;6410 | 0 | 0 |
| NC_006213.1;YP_009555238.1;HCoV-OC43_orf1ab;6411 | 0 | 0 |
| NC_006213.1;YP_009555238.1;HCoV-OC43_orf1ab;6412 | 0 | 0 |
| NC_006213.1;YP_009555238.1;HCoV-OC43_orf1ab;6413 | 0 | 0 |
| NC_006213.1;YP_009555238.1;HCoV-OC43_orf1ab;6414 | 0 | 0 |
| NC_006213.1;YP_009555238.1;HCoV-OC43_orf1ab;6415 | 0 | 0 |
| NC_006213.1;YP_009555238.1;HCoV-OC43_orf1ab;6416 | 0 | 0 |

| | | |
|---|---|---|
| NC_006213.1;YP_009555238.1;HCoV-OC43_orf1ab;6417 | 0 | 0 |
| NC_006213.1;YP_009555238.1;HCoV-OC43_orf1ab;6418 | 0 | 0 |
| NC_006213.1;YP_009555238.1;HCoV-OC43_orf1ab;6419 | 0 | 0 |
| NC_006213.1;YP_009555238.1;HCoV-OC43_orf1ab;6420 | 0 | 0 |
| NC_006213.1;YP_009555238.1;HCoV-OC43_orf1ab;6421 | 0 | 0 |
| NC_006213.1;YP_009555238.1;HCoV-OC43_orf1ab;6422 | 0 | 0 |
| NC_006213.1;YP_009555238.1;HCoV-OC43_orf1ab;6423 | 0 | 0 |
| NC_006213.1;YP_009555238.1;HCoV-OC43_orf1ab;6424 | 0 | 0 |
| NC_006213.1;YP_009555238.1;HCoV-OC43_orf1ab;6425 | 0 | 0 |
| NC_006213.1;YP_009555238.1;HCoV-OC43_orf1ab;6426 | 0 | 0 |
| NC_006213.1;YP_009555238.1;HCoV-OC43_orf1ab;6427 | 0.05 | 0.025 |
| NC_006213.1;YP_009555238.1;HCoV-OC43_orf1ab;6428 | 0 | 0.025 |
| NC_006213.1;YP_009555238.1;HCoV-OC43_orf1ab;6429 | 0 | 0.05 |
| NC_006213.1;YP_009555238.1;HCoV-OC43_orf1ab;6430 | 0.05 | 0.05 |
| NC_006213.1;YP_009555238.1;HCoV-OC43_orf1ab;6431 | 0.05 | 0.05 |
| NC_006213.1;YP_009555238.1;HCoV-OC43_orf1ab;6432 | 0.1 | 0.05 |
| NC_006213.1;YP_009555238.1;HCoV-OC43_orf1ab;6433 | 0 | 0.025 |
| NC_006213.1;YP_009555238.1;HCoV-OC43_orf1ab;6434 | 0 | 0.05 |
| NC_006213.1;YP_009555238.1;HCoV-OC43_orf1ab;6435 | 0 | 0.05 |
| NC_006213.1;YP_009555238.1;HCoV-OC43_orf1ab;6436 | 0.05 | 0.05 |
| NC_006213.1;YP_009555238.1;HCoV-OC43_orf1ab;6437 | 0.05 | 0.05 |
| NC_006213.1;YP_009555238.1;HCoV-OC43_orf1ab;6438 | 0.05 | 0.025 |
| NC_006213.1;YP_009555238.1;HCoV-OC43_orf1ab;6439 | 0.05 | 0.025 |
| NC_006213.1;YP_009555238.1;HCoV-OC43_orf1ab;6440 | 0.05 | 0.025 |
| NC_006213.1;YP_009555238.1;HCoV-OC43_orf1ab;6441 | 0.05 | 0.025 |
| NC_006213.1;YP_009555238.1;HCoV-OC43_orf1ab;6442 | 0.1 | 0.075 |
| NC_006213.1;YP_009555238.1;HCoV-OC43_orf1ab;6443 | 0.15 | 0.05 |
| NC_006213.1;YP_009555238.1;HCoV-OC43_orf1ab;6444 | 0.1 | 0.075 |
| NC_006213.1;YP_009555238.1;HCoV-OC43_orf1ab;6445 | 0 | 0.025 |
| NC_006213.1;YP_009555238.1;HCoV-OC43_orf1ab;6446 | 0.05 | 0.075 |
| NC_006213.1;YP_009555238.1;HCoV-OC43_orf1ab;6447 | 0.1 | 0.05 |
| NC_006213.1;YP_009555238.1;HCoV-OC43_orf1ab;6448 | 0 | 0.025 |
| NC_006213.1;YP_009555238.1;HCoV-OC43_orf1ab;6449 | 0 | 0 |
| NC_006213.1;YP_009555238.1;HCoV-OC43_orf1ab;6450 | 0 | 0 |
| NC_006213.1;YP_009555238.1;HCoV-OC43_orf1ab;6451 | 0 | 0.025 |
| NC_006213.1;YP_009555238.1;HCoV-OC43_orf1ab;6452 | 0 | 0 |
| NC_006213.1;YP_009555238.1;HCoV-OC43_orf1ab;6453 | 0 | 0 |
| NC_006213.1;YP_009555238.1;HCoV-OC43_orf1ab;6454 | 0 | 0 |
| NC_006213.1;YP_009555238.1;HCoV-OC43_orf1ab;6455 | 0 | 0 |
| NC_006213.1;YP_009555238.1;HCoV-OC43_orf1ab;6456 | 0 | 0.025 |
| NC_006213.1;YP_009555238.1;HCoV-OC43_orf1ab;6457 | 0 | 0.025 |
| NC_006213.1;YP_009555238.1;HCoV-OC43_orf1ab;6458 | 0 | 0.025 |
| NC_006213.1;YP_009555238.1;HCoV-OC43_orf1ab;6459 | 0 | 0.025 |

| | | |
|---|---|---|
| NC_006213.1;YP_009555238.1;HCoV-OC43_orf1ab;6460 | 0 | 0 |
| NC_006213.1;YP_009555238.1;HCoV-OC43_orf1ab;6461 | 0 | 0.025 |
| NC_006213.1;YP_009555238.1;HCoV-OC43_orf1ab;6462 | 0.1 | 0.025 |
| NC_006213.1;YP_009555238.1;HCoV-OC43_orf1ab;6463 | 0.05 | 0.075 |
| NC_006213.1;YP_009555238.1;HCoV-OC43_orf1ab;6464 | 0.1 | 0.05 |
| NC_006213.1;YP_009555238.1;HCoV-OC43_orf1ab;6465 | 0 | 0 |
| NC_006213.1;YP_009555238.1;HCoV-OC43_orf1ab;6466 | 0 | 0 |
| NC_006213.1;YP_009555238.1;HCoV-OC43_orf1ab;6467 | 0 | 0 |
| NC_006213.1;YP_009555238.1;HCoV-OC43_orf1ab;6468 | 0 | 0 |
| NC_006213.1;YP_009555238.1;HCoV-OC43_orf1ab;6469 | 0 | 0 |
| NC_006213.1;YP_009555238.1;HCoV-OC43_orf1ab;6470 | 0 | 0 |
| NC_006213.1;YP_009555238.1;HCoV-OC43_orf1ab;6471 | 0 | 0 |
| NC_006213.1;YP_009555238.1;HCoV-OC43_orf1ab;6472 | 0.05 | 0 |
| NC_006213.1;YP_009555238.1;HCoV-OC43_orf1ab;6473 | 0 | 0 |
| NC_006213.1;YP_009555238.1;HCoV-OC43_orf1ab;6474 | 0 | 0 |
| NC_006213.1;YP_009555238.1;HCoV-OC43_orf1ab;6475 | 0 | 0 |
| NC_006213.1;YP_009555238.1;HCoV-OC43_orf1ab;6476 | 0 | 0 |
| NC_006213.1;YP_009555238.1;HCoV-OC43_orf1ab;6477 | 0 | 0 |
| NC_006213.1;YP_009555238.1;HCoV-OC43_orf1ab;6478 | 0 | 0 |
| NC_006213.1;YP_009555238.1;HCoV-OC43_orf1ab;6479 | 0 | 0 |
| NC_006213.1;YP_009555238.1;HCoV-OC43_orf1ab;6480 | 0 | 0 |
| NC_006213.1;YP_009555238.1;HCoV-OC43_orf1ab;6481 | 0 | 0.075 |
| NC_006213.1;YP_009555238.1;HCoV-OC43_orf1ab;6482 | 0 | 0.05 |
| NC_006213.1;YP_009555238.1;HCoV-OC43_orf1ab;6483 | 0 | 0.05 |
| NC_006213.1;YP_009555238.1;HCoV-OC43_orf1ab;6484 | 0 | 0.025 |
| NC_006213.1;YP_009555238.1;HCoV-OC43_orf1ab;6485 | 0 | 0 |
| NC_006213.1;YP_009555238.1;HCoV-OC43_orf1ab;6486 | 0 | 0.025 |
| NC_006213.1;YP_009555238.1;HCoV-OC43_orf1ab;6487 | 0 | 0 |
| NC_006213.1;YP_009555238.1;HCoV-OC43_orf1ab;6488 | 0 | 0 |
| NC_006213.1;YP_009555238.1;HCoV-OC43_orf1ab;6489 | 0 | 0 |
| NC_006213.1;YP_009555238.1;HCoV-OC43_orf1ab;6490 | 0 | 0 |
| NC_006213.1;YP_009555238.1;HCoV-OC43_orf1ab;6491 | 0 | 0 |
| NC_006213.1;YP_009555238.1;HCoV-OC43_orf1ab;6492 | 0 | 0 |
| NC_006213.1;YP_009555238.1;HCoV-OC43_orf1ab;6493 | 0 | 0 |
| NC_006213.1;YP_009555238.1;HCoV-OC43_orf1ab;6494 | 0 | 0 |
| NC_006213.1;YP_009555238.1;HCoV-OC43_orf1ab;6495 | 0 | 0 |
| NC_006213.1;YP_009555238.1;HCoV-OC43_orf1ab;6496 | 0 | 0 |
| NC_006213.1;YP_009555238.1;HCoV-OC43_orf1ab;6497 | 0 | 0 |
| NC_006213.1;YP_009555238.1;HCoV-OC43_orf1ab;6498 | 0 | 0 |
| NC_006213.1;YP_009555238.1;HCoV-OC43_orf1ab;6499 | 0 | 0 |
| NC_006213.1;YP_009555238.1;HCoV-OC43_orf1ab;6500 | 0 | 0 |
| NC_006213.1;YP_009555238.1;HCoV-OC43_orf1ab;6501 | 0 | 0 |
| NC_006213.1;YP_009555238.1;HCoV-OC43_orf1ab;6502 | 0 | 0 |

| | | |
|---|---|---|
| NC_006213.1;YP_009555238.1;HCoV-OC43_orf1ab;6503 | 0 | 0 |
| NC_006213.1;YP_009555238.1;HCoV-OC43_orf1ab;6504 | 0 | 0 |
| NC_006213.1;YP_009555238.1;HCoV-OC43_orf1ab;6505 | 0 | 0 |
| NC_006213.1;YP_009555238.1;HCoV-OC43_orf1ab;6506 | 0 | 0 |
| NC_006213.1;YP_009555238.1;HCoV-OC43_orf1ab;6507 | 0 | 0 |
| NC_006213.1;YP_009555238.1;HCoV-OC43_orf1ab;6508 | 0 | 0 |
| NC_006213.1;YP_009555238.1;HCoV-OC43_orf1ab;6509 | 0 | 0 |
| NC_006213.1;YP_009555238.1;HCoV-OC43_orf1ab;6510 | 0 | 0 |
| NC_006213.1;YP_009555238.1;HCoV-OC43_orf1ab;6511 | 0 | 0 |
| NC_006213.1;YP_009555238.1;HCoV-OC43_orf1ab;6512 | 0 | 0 |
| NC_006213.1;YP_009555238.1;HCoV-OC43_orf1ab;6513 | 0 | 0 |
| NC_006213.1;YP_009555238.1;HCoV-OC43_orf1ab;6514 | 0 | 0 |
| NC_006213.1;YP_009555238.1;HCoV-OC43_orf1ab;6515 | 0 | 0 |
| NC_006213.1;YP_009555238.1;HCoV-OC43_orf1ab;6516 | 0.05 | 0 |
| NC_006213.1;YP_009555238.1;HCoV-OC43_orf1ab;6517 | 0.05 | 0 |
| NC_006213.1;YP_009555238.1;HCoV-OC43_orf1ab;6518 | 0.05 | 0 |
| NC_006213.1;YP_009555238.1;HCoV-OC43_orf1ab;6519 | 0.05 | 0 |
| NC_006213.1;YP_009555238.1;HCoV-OC43_orf1ab;6520 | 0.05 | 0.025 |
| NC_006213.1;YP_009555238.1;HCoV-OC43_orf1ab;6521 | 0.05 | 0 |
| NC_006213.1;YP_009555238.1;HCoV-OC43_orf1ab;6522 | 0 | 0 |
| NC_006213.1;YP_009555238.1;HCoV-OC43_orf1ab;6523 | 0 | 0 |
| NC_006213.1;YP_009555238.1;HCoV-OC43_orf1ab;6524 | 0 | 0 |
| NC_006213.1;YP_009555238.1;HCoV-OC43_orf1ab;6525 | 0 | 0 |
| NC_006213.1;YP_009555238.1;HCoV-OC43_orf1ab;6526 | 0 | 0 |
| NC_006213.1;YP_009555238.1;HCoV-OC43_orf1ab;6527 | 0 | 0.025 |
| NC_006213.1;YP_009555238.1;HCoV-OC43_orf1ab;6528 | 0 | 0.025 |
| NC_006213.1;YP_009555238.1;HCoV-OC43_orf1ab;6529 | 0 | 0.025 |
| NC_006213.1;YP_009555238.1;HCoV-OC43_orf1ab;6530 | 0 | 0 |
| NC_006213.1;YP_009555238.1;HCoV-OC43_orf1ab;6531 | 0 | 0.025 |
| NC_006213.1;YP_009555238.1;HCoV-OC43_orf1ab;6532 | 0 | 0 |
| NC_006213.1;YP_009555238.1;HCoV-OC43_orf1ab;6533 | 0 | 0 |
| NC_006213.1;YP_009555238.1;HCoV-OC43_orf1ab;6534 | 0 | 0 |
| NC_006213.1;YP_009555238.1;HCoV-OC43_orf1ab;6535 | 0 | 0.025 |
| NC_006213.1;YP_009555238.1;HCoV-OC43_orf1ab;6536 | 0 | 0.025 |
| NC_006213.1;YP_009555238.1;HCoV-OC43_orf1ab;6537 | 0 | 0.025 |
| NC_006213.1;YP_009555238.1;HCoV-OC43_orf1ab;6538 | 0 | 0.025 |
| NC_006213.1;YP_009555238.1;HCoV-OC43_orf1ab;6539 | 0 | 0.025 |
| NC_006213.1;YP_009555238.1;HCoV-OC43_orf1ab;6540 | 0.05 | 0.025 |
| NC_006213.1;YP_009555238.1;HCoV-OC43_orf1ab;6541 | 0 | 0.025 |
| NC_006213.1;YP_009555238.1;HCoV-OC43_orf1ab;6542 | 0 | 0.025 |
| NC_006213.1;YP_009555238.1;HCoV-OC43_orf1ab;6543 | 0 | 0.025 |
| NC_006213.1;YP_009555238.1;HCoV-OC43_orf1ab;6544 | 0 | 0 |
| NC_006213.1;YP_009555238.1;HCoV-OC43_orf1ab;6545 | 0 | 0 |

| | | |
|---|---|---|
| NC_006213.1;YP_009555238.1;HCoV-OC43_orf1ab;6546 | 0 | 0 |
| NC_006213.1;YP_009555238.1;HCoV-OC43_orf1ab;6547 | 0 | 0.025 |
| NC_006213.1;YP_009555238.1;HCoV-OC43_orf1ab;6548 | 0 | 0 |
| NC_006213.1;YP_009555238.1;HCoV-OC43_orf1ab;6549 | 0 | 0 |
| NC_006213.1;YP_009555238.1;HCoV-OC43_orf1ab;6550 | 0 | 0 |
| NC_006213.1;YP_009555238.1;HCoV-OC43_orf1ab;6551 | 0 | 0 |
| NC_006213.1;YP_009555238.1;HCoV-OC43_orf1ab;6552 | 0.05 | 0.05 |
| NC_006213.1;YP_009555238.1;HCoV-OC43_orf1ab;6553 | 0.05 | 0 |
| NC_006213.1;YP_009555238.1;HCoV-OC43_orf1ab;6554 | 0 | 0 |
| NC_006213.1;YP_009555238.1;HCoV-OC43_orf1ab;6555 | 0 | 0 |
| NC_006213.1;YP_009555238.1;HCoV-OC43_orf1ab;6556 | 0 | 0 |
| NC_006213.1;YP_009555238.1;HCoV-OC43_orf1ab;6557 | 0 | 0 |
| NC_006213.1;YP_009555238.1;HCoV-OC43_orf1ab;6558 | 0 | 0 |
| NC_006213.1;YP_009555238.1;HCoV-OC43_orf1ab;6559 | 0 | 0 |
| NC_006213.1;YP_009555238.1;HCoV-OC43_orf1ab;6560 | 0 | 0 |
| NC_006213.1;YP_009555238.1;HCoV-OC43_orf1ab;6561 | 0 | 0 |
| NC_006213.1;YP_009555238.1;HCoV-OC43_orf1ab;6562 | 0 | 0 |
| NC_006213.1;YP_009555238.1;HCoV-OC43_orf1ab;6563 | 0 | 0.075 |
| NC_006213.1;YP_009555238.1;HCoV-OC43_orf1ab;6564 | 0 | 0 |
| NC_006213.1;YP_009555238.1;HCoV-OC43_orf1ab;6565 | 0 | 0.05 |
| NC_006213.1;YP_009555238.1;HCoV-OC43_orf1ab;6566 | 0 | 0.05 |
| NC_006213.1;YP_009555238.1;HCoV-OC43_orf1ab;6567 | 0 | 0.025 |
| NC_006213.1;YP_009555238.1;HCoV-OC43_orf1ab;6568 | 0 | 0.05 |
| NC_006213.1;YP_009555238.1;HCoV-OC43_orf1ab;6569 | 0.05 | 0.075 |
| NC_006213.1;YP_009555238.1;HCoV-OC43_orf1ab;6570 | 0 | 0.05 |
| NC_006213.1;YP_009555238.1;HCoV-OC43_orf1ab;6571 | 0.05 | 0.125 |
| NC_006213.1;YP_009555238.1;HCoV-OC43_orf1ab;6572 | 0 | 0.05 |
| NC_006213.1;YP_009555238.1;HCoV-OC43_orf1ab;6573 | 0 | 0.05 |
| NC_006213.1;YP_009555238.1;HCoV-OC43_orf1ab;6574 | 0 | 0.025 |
| NC_006213.1;YP_009555238.1;HCoV-OC43_orf1ab;6575 | 0 | 0 |
| NC_006213.1;YP_009555238.1;HCoV-OC43_orf1ab;6576 | 0 | 0.025 |
| NC_006213.1;YP_009555238.1;HCoV-OC43_orf1ab;6577 | 0 | 0 |
| NC_006213.1;YP_009555238.1;HCoV-OC43_orf1ab;6578 | 0 | 0 |
| NC_006213.1;YP_009555238.1;HCoV-OC43_orf1ab;6579 | 0 | 0 |
| NC_006213.1;YP_009555238.1;HCoV-OC43_orf1ab;6580 | 0 | 0 |
| NC_006213.1;YP_009555238.1;HCoV-OC43_orf1ab;6581 | 0 | 0 |
| NC_006213.1;YP_009555238.1;HCoV-OC43_orf1ab;6582 | 0 | 0 |
| NC_006213.1;YP_009555238.1;HCoV-OC43_orf1ab;6583 | 0 | 0 |
| NC_006213.1;YP_009555238.1;HCoV-OC43_orf1ab;6584 | 0 | 0 |
| NC_006213.1;YP_009555238.1;HCoV-OC43_orf1ab;6585 | 0 | 0 |
| NC_006213.1;YP_009555238.1;HCoV-OC43_orf1ab;6586 | 0 | 0 |
| NC_006213.1;YP_009555238.1;HCoV-OC43_orf1ab;6587 | 0 | 0 |
| NC_006213.1;YP_009555238.1;HCoV-OC43_orf1ab;6588 | 0 | 0.025 |

| | | |
|---|---|---|
| NC_006213.1;YP_009555238.1;HCoV-OC43_orf1ab;6589 | 0 | 0.025 |
| NC_006213.1;YP_009555238.1;HCoV-OC43_orf1ab;6590 | 0 | 0.025 |
| NC_006213.1;YP_009555238.1;HCoV-OC43_orf1ab;6591 | 0 | 0.05 |
| NC_006213.1;YP_009555238.1;HCoV-OC43_orf1ab;6592 | 0 | 0.05 |
| NC_006213.1;YP_009555238.1;HCoV-OC43_orf1ab;6593 | 0 | 0.025 |
| NC_006213.1;YP_009555238.1;HCoV-OC43_orf1ab;6594 | 0 | 0.025 |
| NC_006213.1;YP_009555238.1;HCoV-OC43_orf1ab;6595 | 0 | 0.025 |
| NC_006213.1;YP_009555238.1;HCoV-OC43_orf1ab;6596 | 0 | 0.025 |
| NC_006213.1;YP_009555238.1;HCoV-OC43_orf1ab;6597 | 0 | 0 |
| NC_006213.1;YP_009555238.1;HCoV-OC43_orf1ab;6598 | 0 | 0 |
| NC_006213.1;YP_009555238.1;HCoV-OC43_orf1ab;6599 | 0.1 | 0.025 |
| NC_006213.1;YP_009555238.1;HCoV-OC43_orf1ab;6600 | 0.05 | 0.025 |
| NC_006213.1;YP_009555238.1;HCoV-OC43_orf1ab;6601 | 0.1 | 0.025 |
| NC_006213.1;YP_009555238.1;HCoV-OC43_orf1ab;6602 | 0.15 | 0.05 |
| NC_006213.1;YP_009555238.1;HCoV-OC43_orf1ab;6603 | 0.15 | 0.025 |
| NC_006213.1;YP_009555238.1;HCoV-OC43_orf1ab;6604 | 0.2 | 0.025 |
| NC_006213.1;YP_009555238.1;HCoV-OC43_orf1ab;6605 | 0.2 | 0.05 |
| NC_006213.1;YP_009555238.1;HCoV-OC43_orf1ab;6606 | 0.15 | 0.075 |
| NC_006213.1;YP_009555238.1;HCoV-OC43_orf1ab;6607 | 0.15 | 0.075 |
| NC_006213.1;YP_009555238.1;HCoV-OC43_orf1ab;6608 | 0.05 | 0 |
| NC_006213.1;YP_009555238.1;HCoV-OC43_orf1ab;6609 | 0.05 | 0 |
| NC_006213.1;YP_009555238.1;HCoV-OC43_orf1ab;6610 | 0 | 0.025 |
| NC_006213.1;YP_009555238.1;HCoV-OC43_orf1ab;6611 | 0.05 | 0.025 |
| NC_006213.1;YP_009555238.1;HCoV-OC43_orf1ab;6612 | 0.15 | 0.05 |
| NC_006213.1;YP_009555238.1;HCoV-OC43_orf1ab;6613 | 0.05 | 0.05 |
| NC_006213.1;YP_009555238.1;HCoV-OC43_orf1ab;6614 | 0 | 0.1 |
| NC_006213.1;YP_009555238.1;HCoV-OC43_orf1ab;6615 | 0.05 | 0.075 |
| NC_006213.1;YP_009555238.1;HCoV-OC43_orf1ab;6616 | 0.05 | 0.075 |
| NC_006213.1;YP_009555238.1;HCoV-OC43_orf1ab;6617 | 0.05 | 0.1 |
| NC_006213.1;YP_009555238.1;HCoV-OC43_orf1ab;6618 | 0 | 0.1 |
| NC_006213.1;YP_009555238.1;HCoV-OC43_orf1ab;6619 | 0 | 0.1 |
| NC_006213.1;YP_009555238.1;HCoV-OC43_orf1ab;6620 | 0.05 | 0.075 |
| NC_006213.1;YP_009555238.1;HCoV-OC43_orf1ab;6621 | 0.1 | 0.05 |
| NC_006213.1;YP_009555238.1;HCoV-OC43_orf1ab;6622 | 0.1 | 0.025 |
| NC_006213.1;YP_009555238.1;HCoV-OC43_orf1ab;6623 | 0.1 | 0.05 |
| NC_006213.1;YP_009555238.1;HCoV-OC43_orf1ab;6624 | 0.1 | 0.025 |
| NC_006213.1;YP_009555238.1;HCoV-OC43_orf1ab;6625 | 0.1 | 0 |
| NC_006213.1;YP_009555238.1;HCoV-OC43_orf1ab;6626 | 0.1 | 0 |
| NC_006213.1;YP_009555238.1;HCoV-OC43_orf1ab;6627 | 0.1 | 0.025 |
| NC_006213.1;YP_009555238.1;HCoV-OC43_orf1ab;6628 | 0 | 0 |
| NC_006213.1;YP_009555238.1;HCoV-OC43_orf1ab;6629 | 0 | 0 |
| NC_006213.1;YP_009555238.1;HCoV-OC43_orf1ab;6630 | 0 | 0 |
| NC_006213.1;YP_009555238.1;HCoV-OC43_orf1ab;6631 | 0 | 0 |

| | | |
|---|---|---|
| NC_006213.1;YP_009555238.1;HCoV-OC43_orf1ab;6632 | 0 | 0 |
| NC_006213.1;YP_009555238.1;HCoV-OC43_orf1ab;6633 | 0 | 0 |
| NC_006213.1;YP_009555238.1;HCoV-OC43_orf1ab;6634 | 0 | 0 |
| NC_006213.1;YP_009555238.1;HCoV-OC43_orf1ab;6635 | 0 | 0 |
| NC_006213.1;YP_009555238.1;HCoV-OC43_orf1ab;6636 | 0 | 0 |
| NC_006213.1;YP_009555238.1;HCoV-OC43_orf1ab;6637 | 0 | 0 |
| NC_006213.1;YP_009555238.1;HCoV-OC43_orf1ab;6638 | 0 | 0 |
| NC_006213.1;YP_009555238.1;HCoV-OC43_orf1ab;6639 | 0 | 0 |
| NC_006213.1;YP_009555238.1;HCoV-OC43_orf1ab;6640 | 0 | 0 |
| NC_006213.1;YP_009555238.1;HCoV-OC43_orf1ab;6641 | 0 | 0 |
| NC_006213.1;YP_009555238.1;HCoV-OC43_orf1ab;6642 | 0 | 0 |
| NC_006213.1;YP_009555238.1;HCoV-OC43_orf1ab;6643 | 0 | 0 |
| NC_006213.1;YP_009555238.1;HCoV-OC43_orf1ab;6644 | 0 | 0 |
| NC_006213.1;YP_009555238.1;HCoV-OC43_orf1ab;6645 | 0 | 0 |
| NC_006213.1;YP_009555238.1;HCoV-OC43_orf1ab;6646 | 0 | 0 |
| NC_006213.1;YP_009555238.1;HCoV-OC43_orf1ab;6647 | 0 | 0 |
| NC_006213.1;YP_009555238.1;HCoV-OC43_orf1ab;6648 | 0 | 0 |
| NC_006213.1;YP_009555238.1;HCoV-OC43_orf1ab;6649 | 0 | 0 |
| NC_006213.1;YP_009555238.1;HCoV-OC43_orf1ab;6650 | 0 | 0 |
| NC_006213.1;YP_009555238.1;HCoV-OC43_orf1ab;6651 | 0 | 0 |
| NC_006213.1;YP_009555238.1;HCoV-OC43_orf1ab;6652 | 0.05 | 0.025 |
| NC_006213.1;YP_009555238.1;HCoV-OC43_orf1ab;6653 | 0.05 | 0.025 |
| NC_006213.1;YP_009555238.1;HCoV-OC43_orf1ab;6654 | 0.1 | 0.025 |
| NC_006213.1;YP_009555238.1;HCoV-OC43_orf1ab;6655 | 0.1 | 0.025 |
| NC_006213.1;YP_009555238.1;HCoV-OC43_orf1ab;6656 | 0.05 | 0.025 |
| NC_006213.1;YP_009555238.1;HCoV-OC43_orf1ab;6657 | 0.05 | 0 |
| NC_006213.1;YP_009555238.1;HCoV-OC43_orf1ab;6658 | 0.1 | 0 |
| NC_006213.1;YP_009555238.1;HCoV-OC43_orf1ab;6659 | 0.1 | 0.05 |
| NC_006213.1;YP_009555238.1;HCoV-OC43_orf1ab;6660 | 0.1 | 0.025 |
| NC_006213.1;YP_009555238.1;HCoV-OC43_orf1ab;6661 | 0.1 | 0 |
| NC_006213.1;YP_009555238.1;HCoV-OC43_orf1ab;6662 | 0 | 0 |
| NC_006213.1;YP_009555238.1;HCoV-OC43_orf1ab;6663 | 0 | 0 |
| NC_006213.1;YP_009555238.1;HCoV-OC43_orf1ab;6664 | 0.05 | 0 |
| NC_006213.1;YP_009555238.1;HCoV-OC43_orf1ab;6665 | 0.1 | 0 |
| NC_006213.1;YP_009555238.1;HCoV-OC43_orf1ab;6666 | 0.05 | 0.025 |
| NC_006213.1;YP_009555238.1;HCoV-OC43_orf1ab;6667 | 0.05 | 0 |
| NC_006213.1;YP_009555238.1;HCoV-OC43_orf1ab;6668 | 0.15 | 0.025 |
| NC_006213.1;YP_009555238.1;HCoV-OC43_orf1ab;6669 | 0.1 | 0 |
| NC_006213.1;YP_009555238.1;HCoV-OC43_orf1ab;6670 | 0 | 0 |
| NC_006213.1;YP_009555238.1;HCoV-OC43_orf1ab;6671 | 0.05 | 0 |
| NC_006213.1;YP_009555238.1;HCoV-OC43_orf1ab;6672 | 0 | 0 |
| NC_006213.1;YP_009555238.1;HCoV-OC43_orf1ab;6673 | 0 | 0 |
| NC_006213.1;YP_009555238.1;HCoV-OC43_orf1ab;6674 | 0.05 | 0 |

| | | |
|---|---|---|
| NC_006213.1;YP_009555238.1;HCoV-OC43_orf1ab;6675 | 0 | 0 |
| NC_006213.1;YP_009555238.1;HCoV-OC43_orf1ab;6676 | 0.1 | 0.15 |
| NC_006213.1;YP_009555238.1;HCoV-OC43_orf1ab;6677 | 0 | 0 |
| NC_006213.1;YP_009555238.1;HCoV-OC43_orf1ab;6678 | 0 | 0.025 |
| NC_006213.1;YP_009555238.1;HCoV-OC43_orf1ab;6679 | 0 | 0.025 |
| NC_006213.1;YP_009555238.1;HCoV-OC43_orf1ab;6680 | 0 | 0 |
| NC_006213.1;YP_009555238.1;HCoV-OC43_orf1ab;6681 | 0 | 0 |
| NC_006213.1;YP_009555238.1;HCoV-OC43_orf1ab;6682 | 0 | 0 |
| NC_006213.1;YP_009555238.1;HCoV-OC43_orf1ab;6683 | 0 | 0 |
| NC_006213.1;YP_009555238.1;HCoV-OC43_orf1ab;6684 | 0 | 0 |
| NC_006213.1;YP_009555238.1;HCoV-OC43_orf1ab;6685 | 0 | 0 |
| NC_006213.1;YP_009555238.1;HCoV-OC43_orf1ab;6686 | 0 | 0 |
| NC_006213.1;YP_009555238.1;HCoV-OC43_orf1ab;6687 | 0 | 0 |
| NC_006213.1;YP_009555238.1;HCoV-OC43_orf1ab;6688 | 0 | 0 |
| NC_006213.1;YP_009555238.1;HCoV-OC43_orf1ab;6689 | 0 | 0 |
| NC_006213.1;YP_009555238.1;HCoV-OC43_orf1ab;6690 | 0 | 0 |
| NC_006213.1;YP_009555238.1;HCoV-OC43_orf1ab;6691 | 0 | 0 |
| NC_006213.1;YP_009555238.1;HCoV-OC43_orf1ab;6692 | 0 | 0 |
| NC_006213.1;YP_009555238.1;HCoV-OC43_orf1ab;6693 | 0 | 0 |
| NC_006213.1;YP_009555238.1;HCoV-OC43_orf1ab;6694 | 0 | 0 |
| NC_006213.1;YP_009555238.1;HCoV-OC43_orf1ab;6695 | 0 | 0 |
| NC_006213.1;YP_009555238.1;HCoV-OC43_orf1ab;6696 | 0 | 0 |
| NC_006213.1;YP_009555238.1;HCoV-OC43_orf1ab;6697 | 0 | 0.025 |
| NC_006213.1;YP_009555238.1;HCoV-OC43_orf1ab;6698 | 0 | 0 |
| NC_006213.1;YP_009555238.1;HCoV-OC43_orf1ab;6699 | 0 | 0 |
| NC_006213.1;YP_009555238.1;HCoV-OC43_orf1ab;6700 | 0 | 0 |
| NC_006213.1;YP_009555238.1;HCoV-OC43_orf1ab;6701 | 0 | 0.025 |
| NC_006213.1;YP_009555238.1;HCoV-OC43_orf1ab;6702 | 0 | 0.025 |
| NC_006213.1;YP_009555238.1;HCoV-OC43_orf1ab;6703 | 0 | 0.025 |
| NC_006213.1;YP_009555238.1;HCoV-OC43_orf1ab;6704 | 0 | 0.025 |
| NC_006213.1;YP_009555238.1;HCoV-OC43_orf1ab;6705 | 0 | 0.05 |
| NC_006213.1;YP_009555238.1;HCoV-OC43_orf1ab;6706 | 0 | 0.025 |
| NC_006213.1;YP_009555238.1;HCoV-OC43_orf1ab;6707 | 0 | 0.025 |
| NC_006213.1;YP_009555238.1;HCoV-OC43_orf1ab;6708 | 0 | 0.05 |
| NC_006213.1;YP_009555238.1;HCoV-OC43_orf1ab;6709 | 0 | 0.025 |
| NC_006213.1;YP_009555238.1;HCoV-OC43_orf1ab;6710 | 0 | 0 |
| NC_006213.1;YP_009555238.1;HCoV-OC43_orf1ab;6711 | 0 | 0 |
| NC_006213.1;YP_009555238.1;HCoV-OC43_orf1ab;6712 | 0 | 0 |
| NC_006213.1;YP_009555238.1;HCoV-OC43_orf1ab;6713 | 0 | 0 |
| NC_006213.1;YP_009555238.1;HCoV-OC43_orf1ab;6714 | 0 | 0 |
| NC_006213.1;YP_009555238.1;HCoV-OC43_orf1ab;6715 | 0 | 0 |
| NC_006213.1;YP_009555238.1;HCoV-OC43_orf1ab;6716 | 0 | 0 |
| NC_006213.1;YP_009555238.1;HCoV-OC43_orf1ab;6717 | 0 | 0 |

| | | |
|---|---|---|
| NC_006213.1;YP_009555238.1;HCoV-OC43_orf1ab;6718 | 0 | 0 |
| NC_006213.1;YP_009555238.1;HCoV-OC43_orf1ab;6719 | 0 | 0 |
| NC_006213.1;YP_009555238.1;HCoV-OC43_orf1ab;6720 | 0 | 0 |
| NC_006213.1;YP_009555238.1;HCoV-OC43_orf1ab;6721 | 0 | 0 |
| NC_006213.1;YP_009555238.1;HCoV-OC43_orf1ab;6722 | 0.05 | 0 |
| NC_006213.1;YP_009555238.1;HCoV-OC43_orf1ab;6723 | 0 | 0 |
| NC_006213.1;YP_009555238.1;HCoV-OC43_orf1ab;6724 | 0 | 0.025 |
| NC_006213.1;YP_009555238.1;HCoV-OC43_orf1ab;6725 | 0 | 0 |
| NC_006213.1;YP_009555238.1;HCoV-OC43_orf1ab;6726 | 0 | 0 |
| NC_006213.1;YP_009555238.1;HCoV-OC43_orf1ab;6727 | 0 | 0 |
| NC_006213.1;YP_009555238.1;HCoV-OC43_orf1ab;6728 | 0 | 0 |
| NC_006213.1;YP_009555238.1;HCoV-OC43_orf1ab;6729 | 0 | 0 |
| NC_006213.1;YP_009555238.1;HCoV-OC43_orf1ab;6730 | 0 | 0 |
| NC_006213.1;YP_009555238.1;HCoV-OC43_orf1ab;6731 | 0 | 0 |
| NC_006213.1;YP_009555238.1;HCoV-OC43_orf1ab;6732 | 0 | 0 |
| NC_006213.1;YP_009555238.1;HCoV-OC43_orf1ab;6733 | 0 | 0 |
| NC_006213.1;YP_009555238.1;HCoV-OC43_orf1ab;6734 | 0 | 0 |
| NC_006213.1;YP_009555238.1;HCoV-OC43_orf1ab;6735 | 0 | 0 |
| NC_006213.1;YP_009555238.1;HCoV-OC43_orf1ab;6736 | 0 | 0 |
| NC_006213.1;YP_009555238.1;HCoV-OC43_orf1ab;6737 | 0 | 0 |
| NC_006213.1;YP_009555238.1;HCoV-OC43_orf1ab;6738 | 0 | 0 |
| NC_006213.1;YP_009555238.1;HCoV-OC43_orf1ab;6739 | 0 | 0 |
| NC_006213.1;YP_009555238.1;HCoV-OC43_orf1ab;6740 | 0 | 0 |
| NC_006213.1;YP_009555238.1;HCoV-OC43_orf1ab;6741 | 0 | 0 |
| NC_006213.1;YP_009555238.1;HCoV-OC43_orf1ab;6742 | 0 | 0 |
| NC_006213.1;YP_009555238.1;HCoV-OC43_orf1ab;6743 | 0 | 0.025 |
| NC_006213.1;YP_009555238.1;HCoV-OC43_orf1ab;6744 | 0 | 0 |
| NC_006213.1;YP_009555238.1;HCoV-OC43_orf1ab;6745 | 0 | 0 |
| NC_006213.1;YP_009555238.1;HCoV-OC43_orf1ab;6746 | 0 | 0 |
| NC_006213.1;YP_009555238.1;HCoV-OC43_orf1ab;6747 | 0 | 0.075 |
| NC_006213.1;YP_009555238.1;HCoV-OC43_orf1ab;6748 | 0 | 0 |
| NC_006213.1;YP_009555238.1;HCoV-OC43_orf1ab;6749 | 0 | 0 |
| NC_006213.1;YP_009555238.1;HCoV-OC43_orf1ab;6750 | 0 | 0 |
| NC_006213.1;YP_009555238.1;HCoV-OC43_orf1ab;6751 | 0 | 0 |
| NC_006213.1;YP_009555238.1;HCoV-OC43_orf1ab;6752 | 0 | 0 |
| NC_006213.1;YP_009555238.1;HCoV-OC43_orf1ab;6753 | 0 | 0 |
| NC_006213.1;YP_009555238.1;HCoV-OC43_orf1ab;6754 | 0 | 0 |
| NC_006213.1;YP_009555238.1;HCoV-OC43_orf1ab;6755 | 0 | 0 |
| NC_006213.1;YP_009555238.1;HCoV-OC43_orf1ab;6756 | 0 | 0 |
| NC_006213.1;YP_009555238.1;HCoV-OC43_orf1ab;6757 | 0 | 0 |
| NC_006213.1;YP_009555238.1;HCoV-OC43_orf1ab;6758 | 0 | 0 |
| NC_006213.1;YP_009555238.1;HCoV-OC43_orf1ab;6759 | 0 | 0 |
| NC_006213.1;YP_009555238.1;HCoV-OC43_orf1ab;6760 | 0 | 0 |

| | | |
|---|---|---|
| NC_006213.1;YP_009555238.1;HCoV-OC43_orf1ab;6761 | 0 | 0 |
| NC_006213.1;YP_009555238.1;HCoV-OC43_orf1ab;6762 | 0 | 0 |
| NC_006213.1;YP_009555238.1;HCoV-OC43_orf1ab;6763 | 0 | 0 |
| NC_006213.1;YP_009555238.1;HCoV-OC43_orf1ab;6764 | 0 | 0 |
| NC_006213.1;YP_009555238.1;HCoV-OC43_orf1ab;6765 | 0 | 0.025 |
| NC_006213.1;YP_009555238.1;HCoV-OC43_orf1ab;6766 | 0.05 | 0.025 |
| NC_006213.1;YP_009555238.1;HCoV-OC43_orf1ab;6767 | 0.05 | 0.025 |
| NC_006213.1;YP_009555238.1;HCoV-OC43_orf1ab;6768 | 0 | 0.025 |
| NC_006213.1;YP_009555238.1;HCoV-OC43_orf1ab;6769 | 0 | 0 |
| NC_006213.1;YP_009555238.1;HCoV-OC43_orf1ab;6770 | 0 | 0 |
| NC_006213.1;YP_009555238.1;HCoV-OC43_orf1ab;6771 | 0 | 0 |
| NC_006213.1;YP_009555238.1;HCoV-OC43_orf1ab;6772 | 0.05 | 0 |
| NC_006213.1;YP_009555238.1;HCoV-OC43_orf1ab;6773 | 0.05 | 0 |
| NC_006213.1;YP_009555238.1;HCoV-OC43_orf1ab;6774 | 0.1 | 0 |
| NC_006213.1;YP_009555238.1;HCoV-OC43_orf1ab;6775 | 0 | 0 |
| NC_006213.1;YP_009555238.1;HCoV-OC43_orf1ab;6776 | 0 | 0 |
| NC_006213.1;YP_009555238.1;HCoV-OC43_orf1ab;6777 | 0 | 0 |
| NC_006213.1;YP_009555238.1;HCoV-OC43_orf1ab;6778 | 0 | 0 |
| NC_006213.1;YP_009555238.1;HCoV-OC43_orf1ab;6779 | 0 | 0 |
| NC_006213.1;YP_009555238.1;HCoV-OC43_orf1ab;6780 | 0 | 0 |
| NC_006213.1;YP_009555238.1;HCoV-OC43_orf1ab;6781 | 0 | 0 |
| NC_006213.1;YP_009555238.1;HCoV-OC43_orf1ab;6782 | 0 | 0 |
| NC_006213.1;YP_009555238.1;HCoV-OC43_orf1ab;6783 | 0.05 | 0.025 |
| NC_006213.1;YP_009555238.1;HCoV-OC43_orf1ab;6784 | 0.05 | 0.025 |
| NC_006213.1;YP_009555238.1;HCoV-OC43_orf1ab;6785 | 0 | 0 |
| NC_006213.1;YP_009555238.1;HCoV-OC43_orf1ab;6786 | 0 | 0.025 |
| NC_006213.1;YP_009555238.1;HCoV-OC43_orf1ab;6787 | 0 | 0 |
| NC_006213.1;YP_009555238.1;HCoV-OC43_orf1ab;6788 | 0 | 0.025 |
| NC_006213.1;YP_009555238.1;HCoV-OC43_orf1ab;6789 | 0 | 0.075 |
| NC_006213.1;YP_009555238.1;HCoV-OC43_orf1ab;6790 | 0 | 0.025 |
| NC_006213.1;YP_009555238.1;HCoV-OC43_orf1ab;6791 | 0 | 0.025 |
| NC_006213.1;YP_009555238.1;HCoV-OC43_orf1ab;6792 | 0.05 | 0.025 |
| NC_006213.1;YP_009555238.1;HCoV-OC43_orf1ab;6793 | 0.05 | 0.05 |
| NC_006213.1;YP_009555238.1;HCoV-OC43_orf1ab;6794 | 0 | 0.05 |
| NC_006213.1;YP_009555238.1;HCoV-OC43_orf1ab;6795 | 0 | 0.025 |
| NC_006213.1;YP_009555238.1;HCoV-OC43_orf1ab;6796 | 0 | 0.025 |
| NC_006213.1;YP_009555238.1;HCoV-OC43_orf1ab;6797 | 0 | 0 |
| NC_006213.1;YP_009555238.1;HCoV-OC43_orf1ab;6798 | 0 | 0 |
| NC_006213.1;YP_009555238.1;HCoV-OC43_orf1ab;6799 | 0 | 0 |
| NC_006213.1;YP_009555238.1;HCoV-OC43_orf1ab;6800 | 0 | 0 |
| NC_006213.1;YP_009555238.1;HCoV-OC43_orf1ab;6801 | 0 | 0 |
| NC_006213.1;YP_009555238.1;HCoV-OC43_orf1ab;6802 | 0 | 0 |
| NC_006213.1;YP_009555238.1;HCoV-OC43_orf1ab;6803 | 0 | 0 |

| | | |
|---|---|---|
| NC_006213.1;YP_009555238.1;HCoV-OC43_orf1ab;6804 | 0 | 0 |
| NC_006213.1;YP_009555238.1;HCoV-OC43_orf1ab;6805 | 0 | 0 |
| NC_006213.1;YP_009555238.1;HCoV-OC43_orf1ab;6806 | 0 | 0 |
| NC_006213.1;YP_009555238.1;HCoV-OC43_orf1ab;6807 | 0 | 0 |
| NC_006213.1;YP_009555238.1;HCoV-OC43_orf1ab;6808 | 0 | 0 |
| NC_006213.1;YP_009555238.1;HCoV-OC43_orf1ab;6809 | 0 | 0 |
| NC_006213.1;YP_009555238.1;HCoV-OC43_orf1ab;6810 | 0.05 | 0.025 |
| NC_006213.1;YP_009555238.1;HCoV-OC43_orf1ab;6811 | 0.05 | 0.025 |
| NC_006213.1;YP_009555238.1;HCoV-OC43_orf1ab;6812 | 0 | 0 |
| NC_006213.1;YP_009555238.1;HCoV-OC43_orf1ab;6813 | 0 | 0 |
| NC_006213.1;YP_009555238.1;HCoV-OC43_orf1ab;6814 | 0 | 0 |
| NC_006213.1;YP_009555238.1;HCoV-OC43_orf1ab;6815 | 0 | 0 |
| NC_006213.1;YP_009555238.1;HCoV-OC43_orf1ab;6816 | 0 | 0 |
| NC_006213.1;YP_009555238.1;HCoV-OC43_orf1ab;6817 | 0 | 0 |
| NC_006213.1;YP_009555238.1;HCoV-OC43_orf1ab;6818 | 0 | 0 |
| NC_006213.1;YP_009555238.1;HCoV-OC43_orf1ab;6819 | 0.05 | 0 |
| NC_006213.1;YP_009555238.1;HCoV-OC43_orf1ab;6820 | 0 | 0 |
| NC_006213.1;YP_009555238.1;HCoV-OC43_orf1ab;6821 | 0 | 0 |
| NC_006213.1;YP_009555238.1;HCoV-OC43_orf1ab;6822 | 0 | 0 |
| NC_006213.1;YP_009555238.1;HCoV-OC43_orf1ab;6823 | 0 | 0 |
| NC_006213.1;YP_009555238.1;HCoV-OC43_orf1ab;6824 | 0 | 0 |
| NC_006213.1;YP_009555238.1;HCoV-OC43_orf1ab;6825 | 0 | 0 |
| NC_006213.1;YP_009555238.1;HCoV-OC43_orf1ab;6826 | 0.05 | 0.025 |
| NC_006213.1;YP_009555238.1;HCoV-OC43_orf1ab;6827 | 0 | 0 |
| NC_006213.1;YP_009555238.1;HCoV-OC43_orf1ab;6828 | 0 | 0 |
| NC_006213.1;YP_009555238.1;HCoV-OC43_orf1ab;6829 | 0 | 0 |
| NC_006213.1;YP_009555238.1;HCoV-OC43_orf1ab;6830 | 0 | 0 |
| NC_006213.1;YP_009555238.1;HCoV-OC43_orf1ab;6831 | 0 | 0 |
| NC_006213.1;YP_009555238.1;HCoV-OC43_orf1ab;6832 | 0 | 0 |
| NC_006213.1;YP_009555238.1;HCoV-OC43_orf1ab;6833 | 0 | 0 |
| NC_006213.1;YP_009555238.1;HCoV-OC43_orf1ab;6834 | 0 | 0 |
| NC_006213.1;YP_009555238.1;HCoV-OC43_orf1ab;6835 | 0 | 0 |
| NC_006213.1;YP_009555238.1;HCoV-OC43_orf1ab;6836 | 0 | 0 |
| NC_006213.1;YP_009555238.1;HCoV-OC43_orf1ab;6837 | 0 | 0 |
| NC_006213.1;YP_009555238.1;HCoV-OC43_orf1ab;6838 | 0 | 0 |
| NC_006213.1;YP_009555238.1;HCoV-OC43_orf1ab;6839 | 0.05 | 0 |
| NC_006213.1;YP_009555238.1;HCoV-OC43_orf1ab;6840 | 0 | 0 |
| NC_006213.1;YP_009555238.1;HCoV-OC43_orf1ab;6841 | 0 | 0 |
| NC_006213.1;YP_009555238.1;HCoV-OC43_orf1ab;6842 | 0 | 0 |
| NC_006213.1;YP_009555238.1;HCoV-OC43_orf1ab;6843 | 0.05 | 0 |
| NC_006213.1;YP_009555238.1;HCoV-OC43_orf1ab;6844 | 0.05 | 0 |
| NC_006213.1;YP_009555238.1;HCoV-OC43_orf1ab;6845 | 0.05 | 0 |
| NC_006213.1;YP_009555238.1;HCoV-OC43_orf1ab;6846 | 0.05 | 0 |

| | | |
|---|---|---|
| NC_006213.1;YP_009555238.1;HCoV-OC43_orf1ab;6847 | 0 | 0 |
| NC_006213.1;YP_009555238.1;HCoV-OC43_orf1ab;6848 | 0 | 0 |
| NC_006213.1;YP_009555238.1;HCoV-OC43_orf1ab;6849 | 0 | 0 |
| NC_006213.1;YP_009555238.1;HCoV-OC43_orf1ab;6850 | 0 | 0 |
| NC_006213.1;YP_009555238.1;HCoV-OC43_orf1ab;6851 | 0 | 0 |
| NC_006213.1;YP_009555238.1;HCoV-OC43_orf1ab;6852 | 0 | 0 |
| NC_006213.1;YP_009555238.1;HCoV-OC43_orf1ab;6853 | 0 | 0 |
| NC_006213.1;YP_009555238.1;HCoV-OC43_orf1ab;6854 | 0 | 0 |
| NC_006213.1;YP_009555238.1;HCoV-OC43_orf1ab;6855 | 0 | 0 |
| NC_006213.1;YP_009555238.1;HCoV-OC43_orf1ab;6856 | 0 | 0 |
| NC_006213.1;YP_009555238.1;HCoV-OC43_orf1ab;6857 | 0 | 0 |
| NC_006213.1;YP_009555238.1;HCoV-OC43_orf1ab;6858 | 0.05 | 0 |
| NC_006213.1;YP_009555238.1;HCoV-OC43_orf1ab;6859 | 0 | 0.025 |
| NC_006213.1;YP_009555238.1;HCoV-OC43_orf1ab;6860 | 0.05 | 0.05 |
| NC_006213.1;YP_009555238.1;HCoV-OC43_orf1ab;6861 | 0 | 0 |
| NC_006213.1;YP_009555238.1;HCoV-OC43_orf1ab;6862 | 0 | 0 |
| NC_006213.1;YP_009555238.1;HCoV-OC43_orf1ab;6863 | 0 | 0 |
| NC_006213.1;YP_009555238.1;HCoV-OC43_orf1ab;6864 | 0 | 0 |
| NC_006213.1;YP_009555238.1;HCoV-OC43_orf1ab;6865 | 0 | 0 |
| NC_006213.1;YP_009555238.1;HCoV-OC43_orf1ab;6866 | 0 | 0 |
| NC_006213.1;YP_009555238.1;HCoV-OC43_orf1ab;6867 | 0 | 0.025 |
| NC_006213.1;YP_009555238.1;HCoV-OC43_orf1ab;6868 | 0.05 | 0 |
| NC_006213.1;YP_009555238.1;HCoV-OC43_orf1ab;6869 | 0 | 0.025 |
| NC_006213.1;YP_009555238.1;HCoV-OC43_orf1ab;6870 | 0 | 0 |
| NC_006213.1;YP_009555238.1;HCoV-OC43_orf1ab;6871 | 0 | 0 |
| NC_006213.1;YP_009555238.1;HCoV-OC43_orf1ab;6872 | 0 | 0 |
| NC_006213.1;YP_009555238.1;HCoV-OC43_orf1ab;6873 | 0 | 0 |
| NC_006213.1;YP_009555238.1;HCoV-OC43_orf1ab;6874 | 0 | 0 |
| NC_006213.1;YP_009555238.1;HCoV-OC43_orf1ab;6875 | 0 | 0 |
| NC_006213.1;YP_009555238.1;HCoV-OC43_orf1ab;6876 | 0 | 0 |
| NC_006213.1;YP_009555238.1;HCoV-OC43_orf1ab;6877 | 0 | 0 |
| NC_006213.1;YP_009555238.1;HCoV-OC43_orf1ab;6878 | 0 | 0 |
| NC_006213.1;YP_009555238.1;HCoV-OC43_orf1ab;6879 | 0 | 0 |
| NC_006213.1;YP_009555238.1;HCoV-OC43_orf1ab;6880 | 0 | 0 |
| NC_006213.1;YP_009555238.1;HCoV-OC43_orf1ab;6881 | 0 | 0 |
| NC_006213.1;YP_009555238.1;HCoV-OC43_orf1ab;6882 | 0 | 0 |
| NC_006213.1;YP_009555238.1;HCoV-OC43_orf1ab;6883 | 0 | 0.025 |
| NC_006213.1;YP_009555238.1;HCoV-OC43_orf1ab;6884 | 0 | 0 |
| NC_006213.1;YP_009555238.1;HCoV-OC43_orf1ab;6885 | 0 | 0 |
| NC_006213.1;YP_009555238.1;HCoV-OC43_orf1ab;6886 | 0 | 0.025 |
| NC_006213.1;YP_009555238.1;HCoV-OC43_orf1ab;6887 | 0 | 0.05 |
| NC_006213.1;YP_009555238.1;HCoV-OC43_orf1ab;6888 | 0 | 0 |
| NC_006213.1;YP_009555238.1;HCoV-OC43_orf1ab;6889 | 0 | 0 |

| | | |
|---|---|---|
| NC_006213.1;YP_009555238.1;HCoV-OC43_orf1ab;6890 | 0 | 0.025 |
| NC_006213.1;YP_009555238.1;HCoV-OC43_orf1ab;6891 | 0 | 0.025 |
| NC_006213.1;YP_009555238.1;HCoV-OC43_orf1ab;6892 | 0 | 0.025 |
| NC_006213.1;YP_009555238.1;HCoV-OC43_orf1ab;6893 | 0 | 0 |
| NC_006213.1;YP_009555238.1;HCoV-OC43_orf1ab;6894 | 0 | 0 |
| NC_006213.1;YP_009555238.1;HCoV-OC43_orf1ab;6895 | 0 | 0 |
| NC_006213.1;YP_009555238.1;HCoV-OC43_orf1ab;6896 | 0 | 0 |
| NC_006213.1;YP_009555238.1;HCoV-OC43_orf1ab;6897 | 0 | 0 |
| NC_006213.1;YP_009555238.1;HCoV-OC43_orf1ab;6898 | 0 | 0 |
| NC_006213.1;YP_009555238.1;HCoV-OC43_orf1ab;6899 | 0 | 0 |
| NC_006213.1;YP_009555238.1;HCoV-OC43_orf1ab;6900 | 0 | 0 |
| NC_006213.1;YP_009555238.1;HCoV-OC43_orf1ab;6901 | 0 | 0.025 |
| NC_006213.1;YP_009555238.1;HCoV-OC43_orf1ab;6902 | 0 | 0 |
| NC_006213.1;YP_009555238.1;HCoV-OC43_orf1ab;6903 | 0 | 0 |
| NC_006213.1;YP_009555238.1;HCoV-OC43_orf1ab;6904 | 0 | 0 |
| NC_006213.1;YP_009555238.1;HCoV-OC43_orf1ab;6905 | 0 | 0 |
| NC_006213.1;YP_009555238.1;HCoV-OC43_orf1ab;6906 | 0 | 0 |
| NC_006213.1;YP_009555238.1;HCoV-OC43_orf1ab;6907 | 0 | 0 |
| NC_006213.1;YP_009555238.1;HCoV-OC43_orf1ab;6908 | 0 | 0 |
| NC_006213.1;YP_009555238.1;HCoV-OC43_orf1ab;6909 | 0 | 0 |
| NC_006213.1;YP_009555238.1;HCoV-OC43_orf1ab;6910 | 0 | 0 |
| NC_006213.1;YP_009555238.1;HCoV-OC43_orf1ab;6911 | 0 | 0 |
| NC_006213.1;YP_009555238.1;HCoV-OC43_orf1ab;6912 | 0 | 0 |
| NC_006213.1;YP_009555238.1;HCoV-OC43_orf1ab;6913 | 0 | 0 |
| NC_006213.1;YP_009555238.1;HCoV-OC43_orf1ab;6914 | 0 | 0 |
| NC_006213.1;YP_009555238.1;HCoV-OC43_orf1ab;6915 | 0 | 0 |
| NC_006213.1;YP_009555238.1;HCoV-OC43_orf1ab;6916 | 0 | 0 |
| NC_006213.1;YP_009555238.1;HCoV-OC43_orf1ab;6917 | 0 | 0 |
| NC_006213.1;YP_009555238.1;HCoV-OC43_orf1ab;6918 | 0 | 0 |
| NC_006213.1;YP_009555238.1;HCoV-OC43_orf1ab;6919 | 0 | 0.025 |
| NC_006213.1;YP_009555238.1;HCoV-OC43_orf1ab;6920 | 0 | 0 |
| NC_006213.1;YP_009555238.1;HCoV-OC43_orf1ab;6921 | 0 | 0 |
| NC_006213.1;YP_009555238.1;HCoV-OC43_orf1ab;6922 | 0 | 0.025 |
| NC_006213.1;YP_009555238.1;HCoV-OC43_orf1ab;6923 | 0 | 0 |
| NC_006213.1;YP_009555238.1;HCoV-OC43_orf1ab;6924 | 0.05 | 0 |
| NC_006213.1;YP_009555238.1;HCoV-OC43_orf1ab;6925 | 0.05 | 0 |
| NC_006213.1;YP_009555238.1;HCoV-OC43_orf1ab;6926 | 0.05 | 0.025 |
| NC_006213.1;YP_009555238.1;HCoV-OC43_orf1ab;6927 | 0.15 | 0.025 |
| NC_006213.1;YP_009555238.1;HCoV-OC43_orf1ab;6928 | 0.15 | 0.025 |
| NC_006213.1;YP_009555238.1;HCoV-OC43_orf1ab;6929 | 0.2 | 0 |
| NC_006213.1;YP_009555238.1;HCoV-OC43_orf1ab;6930 | 0.1 | 0.025 |
| NC_006213.1;YP_009555238.1;HCoV-OC43_orf1ab;6931 | 0.1 | 0 |
| NC_006213.1;YP_009555238.1;HCoV-OC43_orf1ab;6932 | 0.15 | 0.025 |

| | | |
|---|---|---|
| NC_006213.1;YP_009555238.1;HCoV-OC43_orf1ab;6933 | 0.1 | 0.025 |
| NC_006213.1;YP_009555238.1;HCoV-OC43_orf1ab;6934 | 0.05 | 0 |
| NC_006213.1;YP_009555238.1;HCoV-OC43_orf1ab;6935 | 0.05 | 0 |
| NC_006213.1;YP_009555238.1;HCoV-OC43_orf1ab;6936 | 0 | 0 |
| NC_006213.1;YP_009555238.1;HCoV-OC43_orf1ab;6937 | 0 | 0 |
| NC_006213.1;YP_009555238.1;HCoV-OC43_orf1ab;6938 | 0 | 0 |
| NC_006213.1;YP_009555238.1;HCoV-OC43_orf1ab;6939 | 0 | 0 |
| NC_006213.1;YP_009555238.1;HCoV-OC43_orf1ab;6940 | 0 | 0 |
| NC_006213.1;YP_009555238.1;HCoV-OC43_orf1ab;6941 | 0 | 0 |
| NC_006213.1;YP_009555238.1;HCoV-OC43_orf1ab;6942 | 0 | 0 |
| NC_006213.1;YP_009555238.1;HCoV-OC43_orf1ab;6943 | 0 | 0 |
| NC_006213.1;YP_009555238.1;HCoV-OC43_orf1ab;6944 | 0 | 0 |
| NC_006213.1;YP_009555238.1;HCoV-OC43_orf1ab;6945 | 0 | 0 |
| NC_006213.1;YP_009555238.1;HCoV-OC43_orf1ab;6946 | 0 | 0 |
| NC_006213.1;YP_009555238.1;HCoV-OC43_orf1ab;6947 | 0 | 0 |
| NC_006213.1;YP_009555238.1;HCoV-OC43_orf1ab;6948 | 0 | 0 |
| NC_006213.1;YP_009555238.1;HCoV-OC43_orf1ab;6949 | 0.05 | 0 |
| NC_006213.1;YP_009555238.1;HCoV-OC43_orf1ab;6950 | 0 | 0 |
| NC_006213.1;YP_009555238.1;HCoV-OC43_orf1ab;6951 | 0 | 0 |
| NC_006213.1;YP_009555238.1;HCoV-OC43_orf1ab;6952 | 0 | 0 |
| NC_006213.1;YP_009555238.1;HCoV-OC43_orf1ab;6953 | 0 | 0 |
| NC_006213.1;YP_009555238.1;HCoV-OC43_orf1ab;6954 | 0 | 0 |
| NC_006213.1;YP_009555238.1;HCoV-OC43_orf1ab;6955 | 0 | 0 |
| NC_006213.1;YP_009555238.1;HCoV-OC43_orf1ab;6956 | 0 | 0 |
| NC_006213.1;YP_009555238.1;HCoV-OC43_orf1ab;6957 | 0 | 0 |
| NC_006213.1;YP_009555238.1;HCoV-OC43_orf1ab;6958 | 0 | 0 |
| NC_006213.1;YP_009555238.1;HCoV-OC43_orf1ab;6959 | 0 | 0 |
| NC_006213.1;YP_009555238.1;HCoV-OC43_orf1ab;6960 | 0 | 0 |
| NC_006213.1;YP_009555238.1;HCoV-OC43_orf1ab;6961 | 0 | 0 |
| NC_006213.1;YP_009555238.1;HCoV-OC43_orf1ab;6962 | 0 | 0 |
| NC_006213.1;YP_009555238.1;HCoV-OC43_orf1ab;6963 | 0 | 0 |
| NC_006213.1;YP_009555238.1;HCoV-OC43_orf1ab;6964 | 0 | 0 |
| NC_006213.1;YP_009555238.1;HCoV-OC43_orf1ab;6965 | 0 | 0.025 |
| NC_006213.1;YP_009555238.1;HCoV-OC43_orf1ab;6966 | 0 | 0 |
| NC_006213.1;YP_009555238.1;HCoV-OC43_orf1ab;6967 | 0 | 0 |
| NC_006213.1;YP_009555238.1;HCoV-OC43_orf1ab;6968 | 0 | 0 |
| NC_006213.1;YP_009555238.1;HCoV-OC43_orf1ab;6969 | 0 | 0 |
| NC_006213.1;YP_009555238.1;HCoV-OC43_orf1ab;6970 | 0 | 0 |
| NC_006213.1;YP_009555238.1;HCoV-OC43_orf1ab;6971 | 0 | 0 |
| NC_006213.1;YP_009555238.1;HCoV-OC43_orf1ab;6972 | 0 | 0 |
| NC_006213.1;YP_009555238.1;HCoV-OC43_orf1ab;6973 | 0 | 0 |
| NC_006213.1;YP_009555238.1;HCoV-OC43_orf1ab;6974 | 0 | 0 |
| NC_006213.1;YP_009555238.1;HCoV-OC43_orf1ab;6975 | 0 | 0 |

| | | |
|---|---|---|
| NC_006213.1;YP_009555238.1;HCoV-OC43_orf1ab;6976 | 0 | 0 |
| NC_006213.1;YP_009555238.1;HCoV-OC43_orf1ab;6977 | 0 | 0 |
| NC_006213.1;YP_009555238.1;HCoV-OC43_orf1ab;6978 | 0 | 0 |
| NC_006213.1;YP_009555238.1;HCoV-OC43_orf1ab;6979 | 0 | 0 |
| NC_006213.1;YP_009555238.1;HCoV-OC43_orf1ab;6980 | 0 | 0 |
| NC_006213.1;YP_009555238.1;HCoV-OC43_orf1ab;6981 | 0 | 0 |
| NC_006213.1;YP_009555238.1;HCoV-OC43_orf1ab;6982 | 0 | 0 |
| NC_006213.1;YP_009555238.1;HCoV-OC43_orf1ab;6983 | 0 | 0 |
| NC_006213.1;YP_009555238.1;HCoV-OC43_orf1ab;6984 | 0 | 0 |
| NC_006213.1;YP_009555238.1;HCoV-OC43_orf1ab;6985 | 0 | 0.025 |
| NC_006213.1;YP_009555238.1;HCoV-OC43_orf1ab;6986 | 0 | 0.025 |
| NC_006213.1;YP_009555238.1;HCoV-OC43_orf1ab;6987 | 0 | 0 |
| NC_006213.1;YP_009555238.1;HCoV-OC43_orf1ab;6988 | 0 | 0 |
| NC_006213.1;YP_009555238.1;HCoV-OC43_orf1ab;6989 | 0 | 0 |
| NC_006213.1;YP_009555238.1;HCoV-OC43_orf1ab;6990 | 0 | 0 |
| NC_006213.1;YP_009555238.1;HCoV-OC43_orf1ab;6991 | 0.05 | 0.025 |
| NC_006213.1;YP_009555238.1;HCoV-OC43_orf1ab;6992 | 0 | 0 |
| NC_006213.1;YP_009555238.1;HCoV-OC43_orf1ab;6993 | 0 | 0 |
| NC_006213.1;YP_009555238.1;HCoV-OC43_orf1ab;6994 | 0 | 0 |
| NC_006213.1;YP_009555238.1;HCoV-OC43_orf1ab;6995 | 0 | 0 |
| NC_006213.1;YP_009555238.1;HCoV-OC43_orf1ab;6996 | 0 | 0 |
| NC_006213.1;YP_009555238.1;HCoV-OC43_orf1ab;6997 | 0 | 0 |
| NC_006213.1;YP_009555238.1;HCoV-OC43_orf1ab;6998 | 0 | 0 |
| NC_006213.1;YP_009555238.1;HCoV-OC43_orf1ab;6999 | 0 | 0 |
| NC_006213.1;YP_009555238.1;HCoV-OC43_orf1ab;7000 | 0 | 0 |
| NC_006213.1;YP_009555238.1;HCoV-OC43_orf1ab;7001 | 0 | 0 |
| NC_006213.1;YP_009555238.1;HCoV-OC43_orf1ab;7002 | 0 | 0.025 |
| NC_006213.1;YP_009555238.1;HCoV-OC43_orf1ab;7003 | 0 | 0.025 |
| NC_006213.1;YP_009555238.1;HCoV-OC43_orf1ab;7004 | 0 | 0 |
| NC_006213.1;YP_009555238.1;HCoV-OC43_orf1ab;7005 | 0 | 0 |
| NC_006213.1;YP_009555238.1;HCoV-OC43_orf1ab;7006 | 0 | 0 |
| NC_006213.1;YP_009555238.1;HCoV-OC43_orf1ab;7007 | 0.05 | 0 |
| NC_006213.1;YP_009555238.1;HCoV-OC43_orf1ab;7008 | 0.05 | 0 |
| NC_006213.1;YP_009555238.1;HCoV-OC43_orf1ab;7009 | 0.05 | 0.025 |
| NC_006213.1;YP_009555238.1;HCoV-OC43_orf1ab;7010 | 0.05 | 0.025 |
| NC_006213.1;YP_009555238.1;HCoV-OC43_orf1ab;7011 | 0.05 | 0.025 |
| NC_006213.1;YP_009555238.1;HCoV-OC43_orf1ab;7012 | 0 | 0 |
| NC_006213.1;YP_009555238.1;HCoV-OC43_orf1ab;7013 | 0.25 | 0.3 |
| NC_006213.1;YP_009555238.1;HCoV-OC43_orf1ab;7014 | 0.25 | 0.275 |
| NC_006213.1;YP_009555238.1;HCoV-OC43_orf1ab;7015 | 0 | 0 |
| NC_006213.1;YP_009555238.1;HCoV-OC43_orf1ab;7016 | 0 | 0 |
| NC_006213.1;YP_009555238.1;HCoV-OC43_orf1ab;7017 | 0 | 0 |
| NC_006213.1;YP_009555238.1;HCoV-OC43_orf1ab;7018 | 0 | 0 |

| | | |
|---|---|---|
| NC_006213.1;YP_009555238.1;HCoV-OC43_orf1ab;7019 | 0 | 0 |
| NC_006213.1;YP_009555238.1;HCoV-OC43_orf1ab;7020 | 0 | 0 |
| NC_006213.1;YP_009555238.1;HCoV-OC43_orf1ab;7021 | 0 | 0 |
| NC_006213.1;YP_009555238.1;HCoV-OC43_orf1ab;7022 | 0 | 0 |
| NC_006213.1;YP_009555238.1;HCoV-OC43_orf1ab;7023 | 0 | 0 |
| NC_006213.1;YP_009555238.1;HCoV-OC43_orf1ab;7024 | 0 | 0 |
| NC_006213.1;YP_009555238.1;HCoV-OC43_orf1ab;7025 | 0 | 0 |
| NC_006213.1;YP_009555238.1;HCoV-OC43_orf1ab;7026 | 0 | 0 |
| NC_006213.1;YP_009555238.1;HCoV-OC43_orf1ab;7027 | 0 | 0 |
| NC_006213.1;YP_009555238.1;HCoV-OC43_orf1ab;7028 | 0 | 0 |
| NC_006213.1;YP_009555238.1;HCoV-OC43_orf1ab;7029 | 0 | 0 |
| NC_006213.1;YP_009555238.1;HCoV-OC43_orf1ab;7030 | 0.05 | 0.05 |
| NC_006213.1;YP_009555238.1;HCoV-OC43_orf1ab;7031 | 0.05 | 0.05 |
| NC_006213.1;YP_009555238.1;HCoV-OC43_orf1ab;7032 | 0.05 | 0.05 |
| NC_006213.1;YP_009555238.1;HCoV-OC43_orf1ab;7033 | 0.05 | 0.05 |
| NC_006213.1;YP_009555238.1;HCoV-OC43_orf1ab;7034 | 0.05 | 0.05 |
| NC_006213.1;YP_009555238.1;HCoV-OC43_orf1ab;7035 | 0.05 | 0.05 |
| NC_006213.1;YP_009555238.1;HCoV-OC43_orf1ab;7036 | 0.05 | 0.05 |
| NC_006213.1;YP_009555238.1;HCoV-OC43_orf1ab;7037 | 0.05 | 0.1 |
| NC_006213.1;YP_009555238.1;HCoV-OC43_orf1ab;7038 | 0 | 0.025 |
| NC_006213.1;YP_009555238.1;HCoV-OC43_orf1ab;7039 | 0 | 0 |
| NC_006213.1;YP_009555238.1;HCoV-OC43_orf1ab;7040 | 0 | 0 |
| NC_006213.1;YP_009555238.1;HCoV-OC43_orf1ab;7041 | 0 | 0.025 |
| NC_006213.1;YP_009555238.1;HCoV-OC43_orf1ab;7042 | 0 | 0.025 |
| NC_006213.1;YP_009555238.1;HCoV-OC43_orf1ab;7043 | 0 | 0.025 |
| NC_006213.1;YP_009555238.1;HCoV-OC43_orf1ab;7044 | 0 | 0.025 |
| NC_006213.1;YP_009555238.1;HCoV-OC43_orf1ab;7045 | 0 | 0.025 |
| NC_006213.1;YP_009555238.1;HCoV-OC43_orf1ab;7046 | 0 | 0.025 |
| NC_006213.1;YP_009555238.1;HCoV-OC43_orf1ab;7047 | 0 | 0.025 |
| NC_006213.1;YP_009555238.1;HCoV-OC43_orf1ab;7048 | 0 | 0.025 |
| NC_006213.1;YP_009555238.1;HCoV-OC43_orf1ab;7049 | 0 | 0.025 |
| NC_006213.1;YP_009555238.1;HCoV-OC43_orf1ab;7050 | 0 | 0 |
| NC_006213.1;YP_009555238.1;HCoV-OC43_orf1ab;7051 | 0 | 0.025 |
| NC_006213.1;YP_009555238.1;HCoV-OC43_orf1ab;7052 | 0 | 0.05 |
| NC_006213.1;YP_009555238.1;HCoV-OC43_orf1ab;7053 | 0 | 0.025 |
| NC_006213.1;YP_009555238.1;HCoV-OC43_orf1ab;7054 | 0.05 | 0.075 |
| NC_006213.1;YP_009555238.1;HCoV-OC43_orf1ab;7055 | 0.05 | 0.025 |
| NC_006213.1;YP_009555238.1;HCoV-OC43_orf1ab;7056 | 0 | 0.025 |
| NC_006213.1;YP_009555238.1;HCoV-OC43_orf1ab;7057 | 0 | 0 |
| NC_006213.1;YP_009555238.1;HCoV-OC43_orf1ab;7058 | 0 | 0 |
| NC_006213.1;YP_009555238.1;HCoV-OC43_orf1ab;7059 | 0 | 0.025 |
| NC_006213.1;YP_009555238.1;HCoV-OC43_orf1ab;7060 | 0 | 0 |
| NC_006213.1;YP_009555238.1;HCoV-OC43_orf1ab;7061 | 0.05 | 0 |

| | | |
|---|---|---|
| NC_006213.1;YP_009555238.1;HCoV-OC43_orf1ab;7062 | 0.05 | 0.025 |
| NC_006213.1;YP_009555238.1;HCoV-OC43_orf1ab;7063 | 0 | 0 |
| NC_006213.1;YP_009555238.1;HCoV-OC43_orf1ab;7064 | 0 | 0 |
| NC_006213.1;YP_009555238.1;HCoV-OC43_orf1ab;7065 | 0.05 | 0 |
| NC_006213.1;YP_009555238.1;HCoV-OC43_orf1ab;7066 | 0.05 | 0 |
| NC_006213.1;YP_009555238.1;HCoV-OC43_orf1ab;7067 | 0.05 | 0 |
| NC_006213.1;YP_009555238.1;HCoV-OC43_orf1ab;7068 | 0 | 0 |
| NC_006213.1;YP_009555238.1;HCoV-OC43_orf1ab;7069 | 0.2 | 0 |
| NC_006213.1;YP_009555238.1;HCoV-OC43_orf1ab;7070 | 0.2 | 0.05 |
| NC_006213.1;YP_009555238.1;HCoV-OC43_orf1ab;7071 | 0.25 | 0.05 |
| NC_006213.1;YP_009555238.1;HCoV-OC43_orf1ab;7072 | 0.25 | 0.05 |
| NC_006213.1;YP_009555238.1;HCoV-OC43_orf1ab;7073 | 0.2 | 0.075 |
| NC_006213.1;YP_009555238.1;HCoV-OC43_orf1ab;7074 | 0.2 | 0.075 |
| NC_006213.1;YP_009555238.1;HCoV-OC43_orf1ab;7075 | 0.15 | 0.025 |
| NC_006213.1;YP_009555238.1;HCoV-OC43_orf1ab;7076 | 0.05 | 0 |
| NC_006213.1;YP_009555238.1;HCoV-OC43_orf1ab;7077 | 0 | 0 |
| NC_006213.1;YP_009555238.1;HCoV-OC43_orf1ab;7078 | 0 | 0 |
| NC_006213.1;YP_009555238.1;HCoV-OC43_orf1ab;7079 | 0 | 0 |
| NC_006213.1;YP_009555238.1;HCoV-OC43_orf1ab;7080 | 0 | 0 |
| NC_006213.1;YP_009555238.1;HCoV-OC43_orf1ab;7081 | 0 | 0 |
| NC_006213.1;YP_009555238.1;HCoV-OC43_orf1ab;7082 | 0 | 0 |
| NC_006213.1;YP_009555238.1;HCoV-OC43_orf1ab;7083 | 0 | 0 |
| NC_006213.1;YP_009555238.1;HCoV-OC43_orf1ab;7084 | 0 | 0 |
| NC_006213.1;YP_009555239.1;HCoV-OC43_ns2;1 | 0 | 0.05 |
| NC_006213.1;YP_009555239.1;HCoV-OC43_ns2;2 | 0 | 0.075 |
| NC_006213.1;YP_009555239.1;HCoV-OC43_ns2;3 | 0.05 | 0.05 |
| NC_006213.1;YP_009555239.1;HCoV-OC43_ns2;4 | 0.05 | 0.025 |
| NC_006213.1;YP_009555239.1;HCoV-OC43_ns2;5 | 0 | 0.05 |
| NC_006213.1;YP_009555239.1;HCoV-OC43_ns2;6 | 0 | 0.025 |
| NC_006213.1;YP_009555239.1;HCoV-OC43_ns2;7 | 0 | 0.025 |
| NC_006213.1;YP_009555239.1;HCoV-OC43_ns2;8 | 0 | 0 |
| NC_006213.1;YP_009555239.1;HCoV-OC43_ns2;9 | 0 | 0 |
| NC_006213.1;YP_009555239.1;HCoV-OC43_ns2;10 | 0 | 0 |
| NC_006213.1;YP_009555239.1;HCoV-OC43_ns2;11 | 0 | 0 |
| NC_006213.1;YP_009555239.1;HCoV-OC43_ns2;12 | 0 | 0 |
| NC_006213.1;YP_009555239.1;HCoV-OC43_ns2;13 | 0 | 0 |
| NC_006213.1;YP_009555239.1;HCoV-OC43_ns2;14 | 0 | 0 |
| NC_006213.1;YP_009555239.1;HCoV-OC43_ns2;15 | 0 | 0 |
| NC_006213.1;YP_009555239.1;HCoV-OC43_ns2;16 | 0 | 0 |
| NC_006213.1;YP_009555239.1;HCoV-OC43_ns2;17 | 0 | 0 |
| NC_006213.1;YP_009555239.1;HCoV-OC43_ns2;18 | 0 | 0 |
| NC_006213.1;YP_009555239.1;HCoV-OC43_ns2;19 | 0 | 0 |
| NC_006213.1;YP_009555239.1;HCoV-OC43_ns2;20 | 0 | 0 |

| | | |
|---|---|---|
| NC_006213.1;YP_009555239.1;HCoV-OC43_ns2;21 | 0 | 0 |
| NC_006213.1;YP_009555239.1;HCoV-OC43_ns2;22 | 0 | 0 |
| NC_006213.1;YP_009555239.1;HCoV-OC43_ns2;23 | 0.05 | 0 |
| NC_006213.1;YP_009555239.1;HCoV-OC43_ns2;24 | 0 | 0 |
| NC_006213.1;YP_009555239.1;HCoV-OC43_ns2;25 | 0 | 0 |
| NC_006213.1;YP_009555239.1;HCoV-OC43_ns2;26 | 0 | 0 |
| NC_006213.1;YP_009555239.1;HCoV-OC43_ns2;27 | 0 | 0 |
| NC_006213.1;YP_009555239.1;HCoV-OC43_ns2;28 | 0 | 0 |
| NC_006213.1;YP_009555239.1;HCoV-OC43_ns2;29 | 0 | 0.025 |
| NC_006213.1;YP_009555239.1;HCoV-OC43_ns2;30 | 0 | 0 |
| NC_006213.1;YP_009555239.1;HCoV-OC43_ns2;31 | 0 | 0.025 |
| NC_006213.1;YP_009555239.1;HCoV-OC43_ns2;32 | 0 | 0 |
| NC_006213.1;YP_009555239.1;HCoV-OC43_ns2;33 | 0 | 0 |
| NC_006213.1;YP_009555239.1;HCoV-OC43_ns2;34 | 0 | 0 |
| NC_006213.1;YP_009555239.1;HCoV-OC43_ns2;35 | 0 | 0 |
| NC_006213.1;YP_009555239.1;HCoV-OC43_ns2;36 | 0 | 0 |
| NC_006213.1;YP_009555239.1;HCoV-OC43_ns2;37 | 0 | 0 |
| NC_006213.1;YP_009555239.1;HCoV-OC43_ns2;38 | 0 | 0 |
| NC_006213.1;YP_009555239.1;HCoV-OC43_ns2;39 | 0 | 0 |
| NC_006213.1;YP_009555239.1;HCoV-OC43_ns2;40 | 0 | 0 |
| NC_006213.1;YP_009555239.1;HCoV-OC43_ns2;41 | 0 | 0 |
| NC_006213.1;YP_009555239.1;HCoV-OC43_ns2;42 | 0 | 0 |
| NC_006213.1;YP_009555239.1;HCoV-OC43_ns2;43 | 0 | 0 |
| NC_006213.1;YP_009555239.1;HCoV-OC43_ns2;44 | 0 | 0 |
| NC_006213.1;YP_009555239.1;HCoV-OC43_ns2;45 | 0 | 0 |
| NC_006213.1;YP_009555239.1;HCoV-OC43_ns2;46 | 0 | 0 |
| NC_006213.1;YP_009555239.1;HCoV-OC43_ns2;47 | 0 | 0 |
| NC_006213.1;YP_009555239.1;HCoV-OC43_ns2;48 | 0 | 0 |
| NC_006213.1;YP_009555239.1;HCoV-OC43_ns2;49 | 0 | 0.025 |
| NC_006213.1;YP_009555239.1;HCoV-OC43_ns2;50 | 0 | 0.025 |
| NC_006213.1;YP_009555239.1;HCoV-OC43_ns2;51 | 0 | 0.025 |
| NC_006213.1;YP_009555239.1;HCoV-OC43_ns2;52 | 0 | 0 |
| NC_006213.1;YP_009555239.1;HCoV-OC43_ns2;53 | 0 | 0 |
| NC_006213.1;YP_009555239.1;HCoV-OC43_ns2;54 | 0.05 | 0 |
| NC_006213.1;YP_009555239.1;HCoV-OC43_ns2;55 | 0 | 0 |
| NC_006213.1;YP_009555239.1;HCoV-OC43_ns2;56 | 0 | 0 |
| NC_006213.1;YP_009555239.1;HCoV-OC43_ns2;57 | 0 | 0 |
| NC_006213.1;YP_009555239.1;HCoV-OC43_ns2;58 | 0.05 | 0 |
| NC_006213.1;YP_009555239.1;HCoV-OC43_ns2;59 | 0.05 | 0 |
| NC_006213.1;YP_009555239.1;HCoV-OC43_ns2;60 | 0.05 | 0 |
| NC_006213.1;YP_009555239.1;HCoV-OC43_ns2;61 | 0.05 | 0 |
| NC_006213.1;YP_009555239.1;HCoV-OC43_ns2;62 | 0.1 | 0 |
| NC_006213.1;YP_009555239.1;HCoV-OC43_ns2;63 | 0.15 | 0 |

| | | |
|---|---|---|
| NC_006213.1;YP_009555239.1;HCoV-OC43_ns2;64 | 0.1 | 0.025 |
| NC_006213.1;YP_009555239.1;HCoV-OC43_ns2;65 | 0 | 0 |
| NC_006213.1;YP_009555239.1;HCoV-OC43_ns2;66 | 0.05 | 0.05 |
| NC_006213.1;YP_009555239.1;HCoV-OC43_ns2;67 | 0.05 | 0 |
| NC_006213.1;YP_009555239.1;HCoV-OC43_ns2;68 | 0.05 | 0 |
| NC_006213.1;YP_009555239.1;HCoV-OC43_ns2;69 | 0.1 | 0.05 |
| NC_006213.1;YP_009555239.1;HCoV-OC43_ns2;70 | 0.05 | 0 |
| NC_006213.1;YP_009555239.1;HCoV-OC43_ns2;71 | 0 | 0 |
| NC_006213.1;YP_009555239.1;HCoV-OC43_ns2;72 | 0 | 0 |
| NC_006213.1;YP_009555239.1;HCoV-OC43_ns2;73 | 0 | 0 |
| NC_006213.1;YP_009555239.1;HCoV-OC43_ns2;74 | 0 | 0 |
| NC_006213.1;YP_009555239.1;HCoV-OC43_ns2;75 | 0 | 0 |
| NC_006213.1;YP_009555239.1;HCoV-OC43_ns2;76 | 0 | 0 |
| NC_006213.1;YP_009555239.1;HCoV-OC43_ns2;77 | 0 | 0 |
| NC_006213.1;YP_009555239.1;HCoV-OC43_ns2;78 | 0 | 0 |
| NC_006213.1;YP_009555239.1;HCoV-OC43_ns2;79 | 0 | 0 |
| NC_006213.1;YP_009555239.1;HCoV-OC43_ns2;80 | 0 | 0 |
| NC_006213.1;YP_009555239.1;HCoV-OC43_ns2;81 | 0 | 0 |
| NC_006213.1;YP_009555239.1;HCoV-OC43_ns2;82 | 0 | 0 |
| NC_006213.1;YP_009555239.1;HCoV-OC43_ns2;83 | 0 | 0 |
| NC_006213.1;YP_009555239.1;HCoV-OC43_ns2;84 | 0 | 0 |
| NC_006213.1;YP_009555239.1;HCoV-OC43_ns2;85 | 0 | 0 |
| NC_006213.1;YP_009555239.1;HCoV-OC43_ns2;86 | 0 | 0 |
| NC_006213.1;YP_009555239.1;HCoV-OC43_ns2;87 | 0 | 0 |
| NC_006213.1;YP_009555239.1;HCoV-OC43_ns2;88 | 0 | 0 |
| NC_006213.1;YP_009555239.1;HCoV-OC43_ns2;89 | 0 | 0.025 |
| NC_006213.1;YP_009555239.1;HCoV-OC43_ns2;90 | 0 | 0.025 |
| NC_006213.1;YP_009555239.1;HCoV-OC43_ns2;91 | 0 | 0.025 |
| NC_006213.1;YP_009555239.1;HCoV-OC43_ns2;92 | 0 | 0.025 |
| NC_006213.1;YP_009555239.1;HCoV-OC43_ns2;93 | 0 | 0.025 |
| NC_006213.1;YP_009555239.1;HCoV-OC43_ns2;94 | 0 | 0 |
| NC_006213.1;YP_009555239.1;HCoV-OC43_ns2;95 | 0 | 0.025 |
| NC_006213.1;YP_009555239.1;HCoV-OC43_ns2;96 | 0 | 0.025 |
| NC_006213.1;YP_009555239.1;HCoV-OC43_ns2;97 | 0 | 0.025 |
| NC_006213.1;YP_009555239.1;HCoV-OC43_ns2;98 | 0 | 0 |
| NC_006213.1;YP_009555239.1;HCoV-OC43_ns2;99 | 0 | 0 |
| NC_006213.1;YP_009555239.1;HCoV-OC43_ns2;100 | 0 | 0.05 |
| NC_006213.1;YP_009555239.1;HCoV-OC43_ns2;101 | 0.45 | 0.275 |
| NC_006213.1;YP_009555239.1;HCoV-OC43_ns2;102 | 0 | 0.075 |
| NC_006213.1;YP_009555239.1;HCoV-OC43_ns2;103 | 0 | 0.05 |
| NC_006213.1;YP_009555239.1;HCoV-OC43_ns2;104 | 0 | 0.05 |
| NC_006213.1;YP_009555239.1;HCoV-OC43_ns2;105 | 0.05 | 0.1 |
| NC_006213.1;YP_009555239.1;HCoV-OC43_ns2;106 | 0 | 0.025 |

| | | |
|---|---|---|
| NC_006213.1;YP_009555239.1;HCoV-OC43_ns2;107 | 0.1 | 0.025 |
| NC_006213.1;YP_009555239.1;HCoV-OC43_ns2;108 | 0.25 | 0 |
| NC_006213.1;YP_009555239.1;HCoV-OC43_ns2;109 | 0 | 0 |
| NC_006213.1;YP_009555239.1;HCoV-OC43_ns2;110 | 0 | 0 |
| NC_006213.1;YP_009555239.1;HCoV-OC43_ns2;111 | 0 | 0 |
| NC_006213.1;YP_009555239.1;HCoV-OC43_ns2;112 | 0.05 | 0 |
| NC_006213.1;YP_009555239.1;HCoV-OC43_ns2;113 | 0.05 | 0.025 |
| NC_006213.1;YP_009555239.1;HCoV-OC43_ns2;114 | 0.05 | 0 |
| NC_006213.1;YP_009555239.1;HCoV-OC43_ns2;115 | 0 | 0 |
| NC_006213.1;YP_009555239.1;HCoV-OC43_ns2;116 | 0 | 0.025 |
| NC_006213.1;YP_009555239.1;HCoV-OC43_ns2;117 | 0 | 0 |
| NC_006213.1;YP_009555239.1;HCoV-OC43_ns2;118 | 0 | 0 |
| NC_006213.1;YP_009555239.1;HCoV-OC43_ns2;119 | 0.05 | 0.05 |
| NC_006213.1;YP_009555239.1;HCoV-OC43_ns2;120 | 0.05 | 0 |
| NC_006213.1;YP_009555239.1;HCoV-OC43_ns2;121 | 0 | 0 |
| NC_006213.1;YP_009555239.1;HCoV-OC43_ns2;122 | 0 | 0 |
| NC_006213.1;YP_009555239.1;HCoV-OC43_ns2;123 | 0 | 0 |
| NC_006213.1;YP_009555239.1;HCoV-OC43_ns2;124 | 0 | 0 |
| NC_006213.1;YP_009555239.1;HCoV-OC43_ns2;125 | 0 | 0 |
| NC_006213.1;YP_009555239.1;HCoV-OC43_ns2;126 | 0 | 0 |
| NC_006213.1;YP_009555239.1;HCoV-OC43_ns2;127 | 0 | 0 |
| NC_006213.1;YP_009555239.1;HCoV-OC43_ns2;128 | 0 | 0 |
| NC_006213.1;YP_009555239.1;HCoV-OC43_ns2;129 | 0 | 0 |
| NC_006213.1;YP_009555239.1;HCoV-OC43_ns2;130 | 0 | 0.15 |
| NC_006213.1;YP_009555239.1;HCoV-OC43_ns2;131 | 0 | 0 |
| NC_006213.1;YP_009555239.1;HCoV-OC43_ns2;132 | 0 | 0 |
| NC_006213.1;YP_009555239.1;HCoV-OC43_ns2;133 | 0 | 0 |
| NC_006213.1;YP_009555239.1;HCoV-OC43_ns2;134 | 0 | 0 |
| NC_006213.1;YP_009555239.1;HCoV-OC43_ns2;135 | 0 | 0.025 |
| NC_006213.1;YP_009555239.1;HCoV-OC43_ns2;136 | 0 | 0 |
| NC_006213.1;YP_009555239.1;HCoV-OC43_ns2;137 | 0 | 0 |
| NC_006213.1;YP_009555239.1;HCoV-OC43_ns2;138 | 0 | 0 |
| NC_006213.1;YP_009555239.1;HCoV-OC43_ns2;139 | 0 | 0 |
| NC_006213.1;YP_009555239.1;HCoV-OC43_ns2;140 | 0.05 | 0 |
| NC_006213.1;YP_009555239.1;HCoV-OC43_ns2;141 | 0 | 0.025 |
| NC_006213.1;YP_009555239.1;HCoV-OC43_ns2;142 | 0.15 | 0.025 |
| NC_006213.1;YP_009555239.1;HCoV-OC43_ns2;143 | 0.1 | 0.025 |
| NC_006213.1;YP_009555239.1;HCoV-OC43_ns2;144 | 0.1 | 0 |
| NC_006213.1;YP_009555239.1;HCoV-OC43_ns2;145 | 0 | 0 |
| NC_006213.1;YP_009555239.1;HCoV-OC43_ns2;146 | 0 | 0.025 |
| NC_006213.1;YP_009555239.1;HCoV-OC43_ns2;147 | 0 | 0 |
| NC_006213.1;YP_009555239.1;HCoV-OC43_ns2;148 | 0 | 0 |
| NC_006213.1;YP_009555239.1;HCoV-OC43_ns2;149 | 0 | 0 |

| | | |
|---|---|---|
| NC_006213.1;YP_009555239.1;HCoV-OC43_ns2;150 | 0 | 0 |
| NC_006213.1;YP_009555239.1;HCoV-OC43_ns2;151 | 0.05 | 0 |
| NC_006213.1;YP_009555239.1;HCoV-OC43_ns2;152 | 0 | 0 |
| NC_006213.1;YP_009555239.1;HCoV-OC43_ns2;153 | 0 | 0 |
| NC_006213.1;YP_009555239.1;HCoV-OC43_ns2;154 | 0.05 | 0 |
| NC_006213.1;YP_009555239.1;HCoV-OC43_ns2;155 | 0.05 | 0 |
| NC_006213.1;YP_009555239.1;HCoV-OC43_ns2;156 | 0 | 0 |
| NC_006213.1;YP_009555239.1;HCoV-OC43_ns2;157 | 0.1 | 0 |
| NC_006213.1;YP_009555239.1;HCoV-OC43_ns2;158 | 0.05 | 0 |
| NC_006213.1;YP_009555239.1;HCoV-OC43_ns2;159 | 0.05 | 0 |
| NC_006213.1;YP_009555239.1;HCoV-OC43_ns2;160 | 0.05 | 0 |
| NC_006213.1;YP_009555239.1;HCoV-OC43_ns2;161 | 0.1 | 0 |
| NC_006213.1;YP_009555239.1;HCoV-OC43_ns2;162 | 0.05 | 0 |
| NC_006213.1;YP_009555239.1;HCoV-OC43_ns2;163 | 0.05 | 0.025 |
| NC_006213.1;YP_009555239.1;HCoV-OC43_ns2;164 | 0.05 | 0.05 |
| NC_006213.1;YP_009555239.1;HCoV-OC43_ns2;165 | 0.05 | 0.025 |
| NC_006213.1;YP_009555239.1;HCoV-OC43_ns2;166 | 0 | 0.025 |
| NC_006213.1;YP_009555239.1;HCoV-OC43_ns2;167 | 0 | 0.025 |
| NC_006213.1;YP_009555239.1;HCoV-OC43_ns2;168 | 0.05 | 0.025 |
| NC_006213.1;YP_009555239.1;HCoV-OC43_ns2;169 | 0.05 | 0.05 |
| NC_006213.1;YP_009555239.1;HCoV-OC43_ns2;170 | 0.1 | 0.05 |
| NC_006213.1;YP_009555239.1;HCoV-OC43_ns2;171 | 0.05 | 0.05 |
| NC_006213.1;YP_009555239.1;HCoV-OC43_ns2;172 | 0.1 | 0.05 |
| NC_006213.1;YP_009555239.1;HCoV-OC43_ns2;173 | 0.05 | 0 |
| NC_006213.1;YP_009555239.1;HCoV-OC43_ns2;174 | 0 | 0 |
| NC_006213.1;YP_009555239.1;HCoV-OC43_ns2;175 | 0 | 0 |
| NC_006213.1;YP_009555239.1;HCoV-OC43_ns2;176 | 0 | 0 |
| NC_006213.1;YP_009555239.1;HCoV-OC43_ns2;177 | 0.05 | 0.025 |
| NC_006213.1;YP_009555239.1;HCoV-OC43_ns2;178 | 0 | 0 |
| NC_006213.1;YP_009555239.1;HCoV-OC43_ns2;179 | 0 | 0 |
| NC_006213.1;YP_009555239.1;HCoV-OC43_ns2;180 | 0 | 0 |
| NC_006213.1;YP_009555239.1;HCoV-OC43_ns2;181 | 0 | 0 |
| NC_006213.1;YP_009555239.1;HCoV-OC43_ns2;182 | 0 | 0 |
| NC_006213.1;YP_009555239.1;HCoV-OC43_ns2;183 | 0 | 0 |
| NC_006213.1;YP_009555239.1;HCoV-OC43_ns2;184 | 0 | 0 |
| NC_006213.1;YP_009555239.1;HCoV-OC43_ns2;185 | 0 | 0 |
| NC_006213.1;YP_009555239.1;HCoV-OC43_ns2;186 | 0 | 0 |
| NC_006213.1;YP_009555239.1;HCoV-OC43_ns2;187 | 0.05 | 0.025 |
| NC_006213.1;YP_009555239.1;HCoV-OC43_ns2;188 | 0 | 0.025 |
| NC_006213.1;YP_009555239.1;HCoV-OC43_ns2;189 | 0.05 | 0.05 |
| NC_006213.1;YP_009555239.1;HCoV-OC43_ns2;190 | 0.05 | 0.05 |
| NC_006213.1;YP_009555239.1;HCoV-OC43_ns2;191 | 0.05 | 0.075 |
| NC_006213.1;YP_009555239.1;HCoV-OC43_ns2;192 | 0.15 | 0.125 |

| | | |
|---|---|---|
| NC_006213.1;YP_009555239.1;HCoV-OC43_ns2;193 | 0.15 | 0.075 |
| NC_006213.1;YP_009555239.1;HCoV-OC43_ns2;194 | 0.05 | 0.025 |
| NC_006213.1;YP_009555239.1;HCoV-OC43_ns2;195 | 0 | 0 |
| NC_006213.1;YP_009555239.1;HCoV-OC43_ns2;196 | 0 | 0 |
| NC_006213.1;YP_009555239.1;HCoV-OC43_ns2;197 | 0 | 0 |
| NC_006213.1;YP_009555239.1;HCoV-OC43_ns2;198 | 0 | 0 |
| NC_006213.1;YP_009555239.1;HCoV-OC43_ns2;199 | 0 | 0 |
| NC_006213.1;YP_009555239.1;HCoV-OC43_ns2;200 | 0 | 0 |
| NC_006213.1;YP_009555239.1;HCoV-OC43_ns2;201 | 0 | 0 |
| NC_006213.1;YP_009555239.1;HCoV-OC43_ns2;202 | 0 | 0 |
| NC_006213.1;YP_009555239.1;HCoV-OC43_ns2;203 | 0 | 0 |
| NC_006213.1;YP_009555239.1;HCoV-OC43_ns2;204 | 0 | 0 |
| NC_006213.1;YP_009555239.1;HCoV-OC43_ns2;205 | 0 | 0 |
| NC_006213.1;YP_009555239.1;HCoV-OC43_ns2;206 | 0.05 | 0 |
| NC_006213.1;YP_009555239.1;HCoV-OC43_ns2;207 | 0.05 | 0 |
| NC_006213.1;YP_009555239.1;HCoV-OC43_ns2;208 | 0.05 | 0 |
| NC_006213.1;YP_009555239.1;HCoV-OC43_ns2;209 | 0.05 | 0 |
| NC_006213.1;YP_009555239.1;HCoV-OC43_ns2;210 | 0.05 | 0 |
| NC_006213.1;YP_009555239.1;HCoV-OC43_ns2;211 | 0.05 | 0 |
| NC_006213.1;YP_009555239.1;HCoV-OC43_ns2;212 | 0 | 0 |
| NC_006213.1;YP_009555239.1;HCoV-OC43_ns2;213 | 0 | 0 |
| NC_006213.1;YP_009555239.1;HCoV-OC43_ns2;214 | 0 | 0 |
| NC_006213.1;YP_009555239.1;HCoV-OC43_ns2;215 | 0 | 0 |
| NC_006213.1;YP_009555239.1;HCoV-OC43_ns2;216 | 0 | 0 |
| NC_006213.1;YP_009555239.1;HCoV-OC43_ns2;217 | 0 | 0 |
| NC_006213.1;YP_009555239.1;HCoV-OC43_ns2;218 | 0 | 0 |
| NC_006213.1;YP_009555239.1;HCoV-OC43_ns2;219 | 0 | 0 |
| NC_006213.1;YP_009555239.1;HCoV-OC43_ns2;220 | 0 | 0 |
| NC_006213.1;YP_009555239.1;HCoV-OC43_ns2;221 | 0 | 0 |
| NC_006213.1;YP_009555239.1;HCoV-OC43_ns2;222 | 0 | 0 |
| NC_006213.1;YP_009555239.1;HCoV-OC43_ns2;223 | 0 | 0 |
| NC_006213.1;YP_009555239.1;HCoV-OC43_ns2;224 | 0 | 0 |
| NC_006213.1;YP_009555239.1;HCoV-OC43_ns2;225 | 0 | 0 |
| NC_006213.1;YP_009555239.1;HCoV-OC43_ns2;226 | 0 | 0 |
| NC_006213.1;YP_009555239.1;HCoV-OC43_ns2;227 | 0 | 0.025 |
| NC_006213.1;YP_009555239.1;HCoV-OC43_ns2;228 | 0 | 0 |
| NC_006213.1;YP_009555239.1;HCoV-OC43_ns2;229 | 0 | 0 |
| NC_006213.1;YP_009555239.1;HCoV-OC43_ns2;230 | 0 | 0 |
| NC_006213.1;YP_009555239.1;HCoV-OC43_ns2;231 | 0 | 0 |
| NC_006213.1;YP_009555239.1;HCoV-OC43_ns2;232 | 0 | 0 |
| NC_006213.1;YP_009555239.1;HCoV-OC43_ns2;233 | 0 | 0 |
| NC_006213.1;YP_009555239.1;HCoV-OC43_ns2;234 | 0 | 0 |
| NC_006213.1;YP_009555239.1;HCoV-OC43_ns2;235 | 0 | 0.025 |

| | | |
|---|---|---|
| NC_006213.1;YP_009555239.1;HCoV-OC43_ns2;236 | 0 | 0 |
| NC_006213.1;YP_009555239.1;HCoV-OC43_ns2;237 | 0 | 0 |
| NC_006213.1;YP_009555239.1;HCoV-OC43_ns2;238 | 0 | 0 |
| NC_006213.1;YP_009555239.1;HCoV-OC43_ns2;239 | 0 | 0 |
| NC_006213.1;YP_009555239.1;HCoV-OC43_ns2;240 | 0 | 0 |
| NC_006213.1;YP_009555239.1;HCoV-OC43_ns2;241 | 0 | 0 |
| NC_006213.1;YP_009555239.1;HCoV-OC43_ns2;242 | 0 | 0 |
| NC_006213.1;YP_009555239.1;HCoV-OC43_ns2;243 | 0 | 0 |
| NC_006213.1;YP_009555239.1;HCoV-OC43_ns2;244 | 0 | 0 |
| NC_006213.1;YP_009555239.1;HCoV-OC43_ns2;245 | 0 | 0 |
| NC_006213.1;YP_009555239.1;HCoV-OC43_ns2;246 | 0 | 0 |
| NC_006213.1;YP_009555239.1;HCoV-OC43_ns2;247 | 0 | 0 |
| NC_006213.1;YP_009555239.1;HCoV-OC43_ns2;248 | 0 | 0 |
| NC_006213.1;YP_009555239.1;HCoV-OC43_ns2;249 | 0 | 0 |
| NC_006213.1;YP_009555239.1;HCoV-OC43_ns2;250 | 0 | 0 |
| NC_006213.1;YP_009555239.1;HCoV-OC43_ns2;251 | 0 | 0 |
| NC_006213.1;YP_009555239.1;HCoV-OC43_ns2;252 | 0 | 0 |
| NC_006213.1;YP_009555239.1;HCoV-OC43_ns2;253 | 0 | 0 |
| NC_006213.1;YP_009555239.1;HCoV-OC43_ns2;254 | 0 | 0 |
| NC_006213.1;YP_009555239.1;HCoV-OC43_ns2;255 | 0 | 0 |
| NC_006213.1;YP_009555239.1;HCoV-OC43_ns2;256 | 0 | 0 |
| NC_006213.1;YP_009555239.1;HCoV-OC43_ns2;257 | 0 | 0 |
| NC_006213.1;YP_009555239.1;HCoV-OC43_ns2;258 | 0 | 0 |
| NC_006213.1;YP_009555239.1;HCoV-OC43_ns2;259 | 0 | 0 |
| NC_006213.1;YP_009555239.1;HCoV-OC43_ns2;260 | 0 | 0 |
| NC_006213.1;YP_009555239.1;HCoV-OC43_ns2;261 | 0 | 0 |
| NC_006213.1;YP_009555239.1;HCoV-OC43_ns2;262 | 0 | 0 |
| NC_006213.1;YP_009555239.1;HCoV-OC43_ns2;263 | 0 | 0 |
| NC_006213.1;YP_009555239.1;HCoV-OC43_ns2;264 | 0 | 0 |
| NC_006213.1;YP_009555239.1;HCoV-OC43_ns2;265 | 0 | 0 |
| NC_006213.1;YP_009555239.1;HCoV-OC43_ns2;266 | 0 | 0 |
| NC_006213.1;YP_009555239.1;HCoV-OC43_ns2;267 | 0 | 0 |
| NC_006213.1;YP_009555240.1;HCoV-OC43_HA-E;1 | 0 | 0 |
| NC_006213.1;YP_009555240.1;HCoV-OC43_HA-E;2 | 0 | 0 |
| NC_006213.1;YP_009555240.1;HCoV-OC43_HA-E;3 | 0 | 0 |
| NC_006213.1;YP_009555240.1;HCoV-OC43_HA-E;4 | 0 | 0 |
| NC_006213.1;YP_009555240.1;HCoV-OC43_HA-E;5 | 0 | 0 |
| NC_006213.1;YP_009555240.1;HCoV-OC43_HA-E;6 | 0 | 0 |
| NC_006213.1;YP_009555240.1;HCoV-OC43_HA-E;7 | 0 | 0 |
| NC_006213.1;YP_009555240.1;HCoV-OC43_HA-E;8 | 0 | 0 |
| NC_006213.1;YP_009555240.1;HCoV-OC43_HA-E;9 | 0 | 0 |
| NC_006213.1;YP_009555240.1;HCoV-OC43_HA-E;10 | 0 | 0 |
| NC_006213.1;YP_009555240.1;HCoV-OC43_HA-E;11 | 0 | 0.05 |

| | | |
|---|---|---|
| NC_006213.1;YP_009555240.1;HCoV-OC43_HA-E;12 | 0 | 0 |
| NC_006213.1;YP_009555240.1;HCoV-OC43_HA-E;13 | 0.1 | 0.025 |
| NC_006213.1;YP_009555240.1;HCoV-OC43_HA-E;14 | 0 | 0 |
| NC_006213.1;YP_009555240.1;HCoV-OC43_HA-E;15 | 0 | 0 |
| NC_006213.1;YP_009555240.1;HCoV-OC43_HA-E;16 | 0.05 | 0 |
| NC_006213.1;YP_009555240.1;HCoV-OC43_HA-E;17 | 0.1 | 0 |
| NC_006213.1;YP_009555240.1;HCoV-OC43_HA-E;18 | 0.05 | 0 |
| NC_006213.1;YP_009555240.1;HCoV-OC43_HA-E;19 | 0.05 | 0 |
| NC_006213.1;YP_009555240.1;HCoV-OC43_HA-E;20 | 0.05 | 0 |
| NC_006213.1;YP_009555240.1;HCoV-OC43_HA-E;21 | 0 | 0 |
| NC_006213.1;YP_009555240.1;HCoV-OC43_HA-E;22 | 0 | 0 |
| NC_006213.1;YP_009555240.1;HCoV-OC43_HA-E;23 | 0 | 0 |
| NC_006213.1;YP_009555240.1;HCoV-OC43_HA-E;24 | 0.05 | 0 |
| NC_006213.1;YP_009555240.1;HCoV-OC43_HA-E;25 | 0 | 0.025 |
| NC_006213.1;YP_009555240.1;HCoV-OC43_HA-E;26 | 0 | 0 |
| NC_006213.1;YP_009555240.1;HCoV-OC43_HA-E;27 | 0 | 0 |
| NC_006213.1;YP_009555240.1;HCoV-OC43_HA-E;28 | 0 | 0.025 |
| NC_006213.1;YP_009555240.1;HCoV-OC43_HA-E;29 | 0 | 0 |
| NC_006213.1;YP_009555240.1;HCoV-OC43_HA-E;30 | 0 | 0.025 |
| NC_006213.1;YP_009555240.1;HCoV-OC43_HA-E;31 | 0 | 0 |
| NC_006213.1;YP_009555240.1;HCoV-OC43_HA-E;32 | 0 | 0 |
| NC_006213.1;YP_009555240.1;HCoV-OC43_HA-E;33 | 0 | 0 |
| NC_006213.1;YP_009555240.1;HCoV-OC43_HA-E;34 | 0 | 0 |
| NC_006213.1;YP_009555240.1;HCoV-OC43_HA-E;35 | 0 | 0.025 |
| NC_006213.1;YP_009555240.1;HCoV-OC43_HA-E;36 | 0.05 | 0 |
| NC_006213.1;YP_009555240.1;HCoV-OC43_HA-E;37 | 0 | 0 |
| NC_006213.1;YP_009555240.1;HCoV-OC43_HA-E;38 | 0 | 0 |
| NC_006213.1;YP_009555240.1;HCoV-OC43_HA-E;39 | 0.05 | 0 |
| NC_006213.1;YP_009555240.1;HCoV-OC43_HA-E;40 | 0 | 0 |
| NC_006213.1;YP_009555240.1;HCoV-OC43_HA-E;41 | 0 | 0 |
| NC_006213.1;YP_009555240.1;HCoV-OC43_HA-E;42 | 0 | 0 |
| NC_006213.1;YP_009555240.1;HCoV-OC43_HA-E;43 | 0 | 0 |
| NC_006213.1;YP_009555240.1;HCoV-OC43_HA-E;44 | 0 | 0 |
| NC_006213.1;YP_009555240.1;HCoV-OC43_HA-E;45 | 0 | 0 |
| NC_006213.1;YP_009555240.1;HCoV-OC43_HA-E;46 | 0.05 | 0 |
| NC_006213.1;YP_009555240.1;HCoV-OC43_HA-E;47 | 0.05 | 0 |
| NC_006213.1;YP_009555240.1;HCoV-OC43_HA-E;48 | 0.1 | 0.025 |
| NC_006213.1;YP_009555240.1;HCoV-OC43_HA-E;49 | 0 | 0.025 |
| NC_006213.1;YP_009555240.1;HCoV-OC43_HA-E;50 | 0 | 0 |
| NC_006213.1;YP_009555240.1;HCoV-OC43_HA-E;51 | 0.05 | 0 |
| NC_006213.1;YP_009555240.1;HCoV-OC43_HA-E;52 | 0 | 0 |
| NC_006213.1;YP_009555240.1;HCoV-OC43_HA-E;53 | 0 | 0 |
| NC_006213.1;YP_009555240.1;HCoV-OC43_HA-E;54 | 0.05 | 0 |

| | | |
|---|---|---|
| NC_006213.1;YP_009555240.1;HCoV-OC43_HA-E;55 | 0.05 | 0 |
| NC_006213.1;YP_009555240.1;HCoV-OC43_HA-E;56 | 0 | 0 |
| NC_006213.1;YP_009555240.1;HCoV-OC43_HA-E;57 | 0 | 0 |
| NC_006213.1;YP_009555240.1;HCoV-OC43_HA-E;58 | 0 | 0.025 |
| NC_006213.1;YP_009555240.1;HCoV-OC43_HA-E;59 | 0 | 0 |
| NC_006213.1;YP_009555240.1;HCoV-OC43_HA-E;60 | 0 | 0 |
| NC_006213.1;YP_009555240.1;HCoV-OC43_HA-E;61 | 0 | 0 |
| NC_006213.1;YP_009555240.1;HCoV-OC43_HA-E;62 | 0 | 0 |
| NC_006213.1;YP_009555240.1;HCoV-OC43_HA-E;63 | 0.05 | 0 |
| NC_006213.1;YP_009555240.1;HCoV-OC43_HA-E;64 | 0.05 | 0.05 |
| NC_006213.1;YP_009555240.1;HCoV-OC43_HA-E;65 | 0 | 0 |
| NC_006213.1;YP_009555240.1;HCoV-OC43_HA-E;66 | 0 | 0 |
| NC_006213.1;YP_009555240.1;HCoV-OC43_HA-E;67 | 0 | 0 |
| NC_006213.1;YP_009555240.1;HCoV-OC43_HA-E;68 | 0 | 0 |
| NC_006213.1;YP_009555240.1;HCoV-OC43_HA-E;69 | 0 | 0 |
| NC_006213.1;YP_009555240.1;HCoV-OC43_HA-E;70 | 0 | 0 |
| NC_006213.1;YP_009555240.1;HCoV-OC43_HA-E;71 | 0 | 0 |
| NC_006213.1;YP_009555240.1;HCoV-OC43_HA-E;72 | 0 | 0 |
| NC_006213.1;YP_009555240.1;HCoV-OC43_HA-E;73 | 0 | 0 |
| NC_006213.1;YP_009555240.1;HCoV-OC43_HA-E;74 | 0 | 0 |
| NC_006213.1;YP_009555240.1;HCoV-OC43_HA-E;75 | 0 | 0 |
| NC_006213.1;YP_009555240.1;HCoV-OC43_HA-E;76 | 0 | 0 |
| NC_006213.1;YP_009555240.1;HCoV-OC43_HA-E;77 | 0 | 0 |
| NC_006213.1;YP_009555240.1;HCoV-OC43_HA-E;78 | 0 | 0 |
| NC_006213.1;YP_009555240.1;HCoV-OC43_HA-E;79 | 0 | 0 |
| NC_006213.1;YP_009555240.1;HCoV-OC43_HA-E;80 | 0.05 | 0.025 |
| NC_006213.1;YP_009555240.1;HCoV-OC43_HA-E;81 | 0.05 | 0 |
| NC_006213.1;YP_009555240.1;HCoV-OC43_HA-E;82 | 0 | 0 |
| NC_006213.1;YP_009555240.1;HCoV-OC43_HA-E;83 | 0 | 0 |
| NC_006213.1;YP_009555240.1;HCoV-OC43_HA-E;84 | 0 | 0 |
| NC_006213.1;YP_009555240.1;HCoV-OC43_HA-E;85 | 0 | 0 |
| NC_006213.1;YP_009555240.1;HCoV-OC43_HA-E;86 | 0 | 0 |
| NC_006213.1;YP_009555240.1;HCoV-OC43_HA-E;87 | 0 | 0 |
| NC_006213.1;YP_009555240.1;HCoV-OC43_HA-E;88 | 0 | 0 |
| NC_006213.1;YP_009555240.1;HCoV-OC43_HA-E;89 | 0 | 0 |
| NC_006213.1;YP_009555240.1;HCoV-OC43_HA-E;90 | 0 | 0 |
| NC_006213.1;YP_009555240.1;HCoV-OC43_HA-E;91 | 0 | 0 |
| NC_006213.1;YP_009555240.1;HCoV-OC43_HA-E;92 | 0 | 0 |
| NC_006213.1;YP_009555240.1;HCoV-OC43_HA-E;93 | 0 | 0 |
| NC_006213.1;YP_009555240.1;HCoV-OC43_HA-E;94 | 0 | 0 |
| NC_006213.1;YP_009555240.1;HCoV-OC43_HA-E;95 | 0 | 0 |
| NC_006213.1;YP_009555240.1;HCoV-OC43_HA-E;96 | 0 | 0 |
| NC_006213.1;YP_009555240.1;HCoV-OC43_HA-E;97 | 0 | 0 |

| | | |
|---|---|---|
| NC_006213.1;YP_009555240.1;HCoV-OC43_HA-E;98 | 0 | 0 |
| NC_006213.1;YP_009555240.1;HCoV-OC43_HA-E;99 | 0 | 0 |
| NC_006213.1;YP_009555240.1;HCoV-OC43_HA-E;100 | 0 | 0 |
| NC_006213.1;YP_009555240.1;HCoV-OC43_HA-E;101 | 0 | 0 |
| NC_006213.1;YP_009555240.1;HCoV-OC43_HA-E;102 | 0 | 0.025 |
| NC_006213.1;YP_009555240.1;HCoV-OC43_HA-E;103 | 0 | 0 |
| NC_006213.1;YP_009555240.1;HCoV-OC43_HA-E;104 | 0.05 | 0 |
| NC_006213.1;YP_009555240.1;HCoV-OC43_HA-E;105 | 0.05 | 0.025 |
| NC_006213.1;YP_009555240.1;HCoV-OC43_HA-E;106 | 0.1 | 0.025 |
| NC_006213.1;YP_009555240.1;HCoV-OC43_HA-E;107 | 0.05 | 0 |
| NC_006213.1;YP_009555240.1;HCoV-OC43_HA-E;108 | 0.05 | 0.025 |
| NC_006213.1;YP_009555240.1;HCoV-OC43_HA-E;109 | 0.05 | 0 |
| NC_006213.1;YP_009555240.1;HCoV-OC43_HA-E;110 | 0.05 | 0 |
| NC_006213.1;YP_009555240.1;HCoV-OC43_HA-E;111 | 0.1 | 0 |
| NC_006213.1;YP_009555240.1;HCoV-OC43_HA-E;112 | 0.25 | 0.025 |
| NC_006213.1;YP_009555240.1;HCoV-OC43_HA-E;113 | 0.45 | 0.2 |
| NC_006213.1;YP_009555240.1;HCoV-OC43_HA-E;114 | 0.2 | 0.125 |
| NC_006213.1;YP_009555240.1;HCoV-OC43_HA-E;115 | 0.05 | 0 |
| NC_006213.1;YP_009555240.1;HCoV-OC43_HA-E;116 | 0.2 | 0.125 |
| NC_006213.1;YP_009555240.1;HCoV-OC43_HA-E;117 | 0.25 | 0.125 |
| NC_006213.1;YP_009555240.1;HCoV-OC43_HA-E;118 | 0.1 | 0.025 |
| NC_006213.1;YP_009555240.1;HCoV-OC43_HA-E;119 | 0 | 0 |
| NC_006213.1;YP_009555240.1;HCoV-OC43_HA-E;120 | 0 | 0 |
| NC_006213.1;YP_009555240.1;HCoV-OC43_HA-E;121 | 0 | 0 |
| NC_006213.1;YP_009555240.1;HCoV-OC43_HA-E;122 | 0 | 0 |
| NC_006213.1;YP_009555240.1;HCoV-OC43_HA-E;123 | 0 | 0 |
| NC_006213.1;YP_009555240.1;HCoV-OC43_HA-E;124 | 0 | 0.025 |
| NC_006213.1;YP_009555240.1;HCoV-OC43_HA-E;125 | 0 | 0 |
| NC_006213.1;YP_009555240.1;HCoV-OC43_HA-E;126 | 0 | 0 |
| NC_006213.1;YP_009555240.1;HCoV-OC43_HA-E;127 | 0 | 0 |
| NC_006213.1;YP_009555240.1;HCoV-OC43_HA-E;128 | 0 | 0 |
| NC_006213.1;YP_009555240.1;HCoV-OC43_HA-E;129 | 0 | 0 |
| NC_006213.1;YP_009555240.1;HCoV-OC43_HA-E;130 | 0 | 0 |
| NC_006213.1;YP_009555240.1;HCoV-OC43_HA-E;131 | 0 | 0 |
| NC_006213.1;YP_009555240.1;HCoV-OC43_HA-E;132 | 0 | 0 |
| NC_006213.1;YP_009555240.1;HCoV-OC43_HA-E;133 | 0 | 0 |
| NC_006213.1;YP_009555240.1;HCoV-OC43_HA-E;134 | 0 | 0 |
| NC_006213.1;YP_009555240.1;HCoV-OC43_HA-E;135 | 0 | 0 |
| NC_006213.1;YP_009555240.1;HCoV-OC43_HA-E;136 | 0 | 0 |
| NC_006213.1;YP_009555240.1;HCoV-OC43_HA-E;137 | 0 | 0 |
| NC_006213.1;YP_009555240.1;HCoV-OC43_HA-E;138 | 0 | 0 |
| NC_006213.1;YP_009555240.1;HCoV-OC43_HA-E;139 | 0 | 0 |
| NC_006213.1;YP_009555240.1;HCoV-OC43_HA-E;140 | 0 | 0 |

| | | |
|---|---|---|
| NC_006213.1;YP_009555240.1;HCoV-OC43_HA-E;141 | 0 | 0 |
| NC_006213.1;YP_009555240.1;HCoV-OC43_HA-E;142 | 0 | 0.025 |
| NC_006213.1;YP_009555240.1;HCoV-OC43_HA-E;143 | 0 | 0.025 |
| NC_006213.1;YP_009555240.1;HCoV-OC43_HA-E;144 | 0 | 0 |
| NC_006213.1;YP_009555240.1;HCoV-OC43_HA-E;145 | 0 | 0.025 |
| NC_006213.1;YP_009555240.1;HCoV-OC43_HA-E;146 | 0 | 0 |
| NC_006213.1;YP_009555240.1;HCoV-OC43_HA-E;147 | 0 | 0 |
| NC_006213.1;YP_009555240.1;HCoV-OC43_HA-E;148 | 0 | 0 |
| NC_006213.1;YP_009555240.1;HCoV-OC43_HA-E;149 | 0.05 | 0 |
| NC_006213.1;YP_009555240.1;HCoV-OC43_HA-E;150 | 0 | 0 |
| NC_006213.1;YP_009555240.1;HCoV-OC43_HA-E;151 | 0 | 0 |
| NC_006213.1;YP_009555240.1;HCoV-OC43_HA-E;152 | 0 | 0 |
| NC_006213.1;YP_009555240.1;HCoV-OC43_HA-E;153 | 0 | 0 |
| NC_006213.1;YP_009555240.1;HCoV-OC43_HA-E;154 | 0 | 0 |
| NC_006213.1;YP_009555240.1;HCoV-OC43_HA-E;155 | 0 | 0 |
| NC_006213.1;YP_009555240.1;HCoV-OC43_HA-E;156 | 0.05 | 0 |
| NC_006213.1;YP_009555240.1;HCoV-OC43_HA-E;157 | 0.1 | 0 |
| NC_006213.1;YP_009555240.1;HCoV-OC43_HA-E;158 | 0.1 | 0 |
| NC_006213.1;YP_009555240.1;HCoV-OC43_HA-E;159 | 0 | 0 |
| NC_006213.1;YP_009555240.1;HCoV-OC43_HA-E;160 | 0 | 0 |
| NC_006213.1;YP_009555240.1;HCoV-OC43_HA-E;161 | 0 | 0 |
| NC_006213.1;YP_009555240.1;HCoV-OC43_HA-E;162 | 0.15 | 0 |
| NC_006213.1;YP_009555240.1;HCoV-OC43_HA-E;163 | 0.25 | 0.1 |
| NC_006213.1;YP_009555240.1;HCoV-OC43_HA-E;164 | 0.05 | 0.025 |
| NC_006213.1;YP_009555240.1;HCoV-OC43_HA-E;165 | 0 | 0 |
| NC_006213.1;YP_009555240.1;HCoV-OC43_HA-E;166 | 0 | 0 |
| NC_006213.1;YP_009555240.1;HCoV-OC43_HA-E;167 | 0.05 | 0 |
| NC_006213.1;YP_009555240.1;HCoV-OC43_HA-E;168 | 0.05 | 0 |
| NC_006213.1;YP_009555240.1;HCoV-OC43_HA-E;169 | 0.1 | 0 |
| NC_006213.1;YP_009555240.1;HCoV-OC43_HA-E;170 | 0.05 | 0 |
| NC_006213.1;YP_009555240.1;HCoV-OC43_HA-E;171 | 0 | 0 |
| NC_006213.1;YP_009555240.1;HCoV-OC43_HA-E;172 | 0 | 0 |
| NC_006213.1;YP_009555240.1;HCoV-OC43_HA-E;173 | 0 | 0 |
| NC_006213.1;YP_009555240.1;HCoV-OC43_HA-E;174 | 0 | 0 |
| NC_006213.1;YP_009555240.1;HCoV-OC43_HA-E;175 | 0.1 | 0 |
| NC_006213.1;YP_009555240.1;HCoV-OC43_HA-E;176 | 0 | 0 |
| NC_006213.1;YP_009555240.1;HCoV-OC43_HA-E;177 | 0 | 0 |
| NC_006213.1;YP_009555240.1;HCoV-OC43_HA-E;178 | 0.05 | 0 |
| NC_006213.1;YP_009555240.1;HCoV-OC43_HA-E;179 | 0 | 0 |
| NC_006213.1;YP_009555240.1;HCoV-OC43_HA-E;180 | 0 | 0 |
| NC_006213.1;YP_009555240.1;HCoV-OC43_HA-E;181 | 0 | 0 |
| NC_006213.1;YP_009555240.1;HCoV-OC43_HA-E;182 | 0 | 0 |
| NC_006213.1;YP_009555240.1;HCoV-OC43_HA-E;183 | 0 | 0 |

| | | |
|---|---|---|
| NC_006213.1;YP_009555240.1;HCoV-OC43_HA-E;184 | 0 | 0 |
| NC_006213.1;YP_009555240.1;HCoV-OC43_HA-E;185 | 0 | 0 |
| NC_006213.1;YP_009555240.1;HCoV-OC43_HA-E;186 | 0 | 0 |
| NC_006213.1;YP_009555240.1;HCoV-OC43_HA-E;187 | 0.05 | 0 |
| NC_006213.1;YP_009555240.1;HCoV-OC43_HA-E;188 | 0 | 0 |
| NC_006213.1;YP_009555240.1;HCoV-OC43_HA-E;189 | 0 | 0 |
| NC_006213.1;YP_009555240.1;HCoV-OC43_HA-E;190 | 0 | 0 |
| NC_006213.1;YP_009555240.1;HCoV-OC43_HA-E;191 | 0 | 0 |
| NC_006213.1;YP_009555240.1;HCoV-OC43_HA-E;192 | 0 | 0 |
| NC_006213.1;YP_009555240.1;HCoV-OC43_HA-E;193 | 0 | 0 |
| NC_006213.1;YP_009555240.1;HCoV-OC43_HA-E;194 | 0 | 0 |
| NC_006213.1;YP_009555240.1;HCoV-OC43_HA-E;195 | 0 | 0 |
| NC_006213.1;YP_009555240.1;HCoV-OC43_HA-E;196 | 0 | 0 |
| NC_006213.1;YP_009555240.1;HCoV-OC43_HA-E;197 | 0 | 0 |
| NC_006213.1;YP_009555240.1;HCoV-OC43_HA-E;198 | 0 | 0 |
| NC_006213.1;YP_009555240.1;HCoV-OC43_HA-E;199 | 0 | 0 |
| NC_006213.1;YP_009555240.1;HCoV-OC43_HA-E;200 | 0 | 0 |
| NC_006213.1;YP_009555240.1;HCoV-OC43_HA-E;201 | 0 | 0.025 |
| NC_006213.1;YP_009555240.1;HCoV-OC43_HA-E;202 | 0 | 0.025 |
| NC_006213.1;YP_009555240.1;HCoV-OC43_HA-E;203 | 0 | 0.025 |
| NC_006213.1;YP_009555240.1;HCoV-OC43_HA-E;204 | 0.05 | 0.125 |
| NC_006213.1;YP_009555240.1;HCoV-OC43_HA-E;205 | 0.2 | 0.25 |
| NC_006213.1;YP_009555240.1;HCoV-OC43_HA-E;206 | 0.3 | 0.275 |
| NC_006213.1;YP_009555240.1;HCoV-OC43_HA-E;207 | 0.2 | 0.275 |
| NC_006213.1;YP_009555240.1;HCoV-OC43_HA-E;208 | 0.3 | 0.25 |
| NC_006213.1;YP_009555240.1;HCoV-OC43_HA-E;209 | 0.35 | 0.275 |
| NC_006213.1;YP_009555240.1;HCoV-OC43_HA-E;210 | 0.2 | 0.175 |
| NC_006213.1;YP_009555240.1;HCoV-OC43_HA-E;211 | 0 | 0.05 |
| NC_006213.1;YP_009555240.1;HCoV-OC43_HA-E;212 | 0.1 | 0.025 |
| NC_006213.1;YP_009555240.1;HCoV-OC43_HA-E;213 | 0.15 | 0.025 |
| NC_006213.1;YP_009555240.1;HCoV-OC43_HA-E;214 | 0.05 | 0.075 |
| NC_006213.1;YP_009555240.1;HCoV-OC43_HA-E;215 | 0.15 | 0.1 |
| NC_006213.1;YP_009555240.1;HCoV-OC43_HA-E;216 | 0.15 | 0.05 |
| NC_006213.1;YP_009555240.1;HCoV-OC43_HA-E;217 | 0.1 | 0.15 |
| NC_006213.1;YP_009555240.1;HCoV-OC43_HA-E;218 | 0 | 0.075 |
| NC_006213.1;YP_009555240.1;HCoV-OC43_HA-E;219 | 0.1 | 0.025 |
| NC_006213.1;YP_009555240.1;HCoV-OC43_HA-E;220 | 0 | 0 |
| NC_006213.1;YP_009555240.1;HCoV-OC43_HA-E;221 | 0 | 0.025 |
| NC_006213.1;YP_009555240.1;HCoV-OC43_HA-E;222 | 0 | 0.025 |
| NC_006213.1;YP_009555240.1;HCoV-OC43_HA-E;223 | 0 | 0 |
| NC_006213.1;YP_009555240.1;HCoV-OC43_HA-E;224 | 0 | 0 |
| NC_006213.1;YP_009555240.1;HCoV-OC43_HA-E;225 | 0 | 0.025 |
| NC_006213.1;YP_009555240.1;HCoV-OC43_HA-E;226 | 0 | 0 |

| | | |
|---|---|---|
| NC_006213.1;YP_009555240.1;HCoV-OC43_HA-E;227 | 0.05 | 0 |
| NC_006213.1;YP_009555240.1;HCoV-OC43_HA-E;228 | 0.05 | 0 |
| NC_006213.1;YP_009555240.1;HCoV-OC43_HA-E;229 | 0.05 | 0 |
| NC_006213.1;YP_009555240.1;HCoV-OC43_HA-E;230 | 0.05 | 0 |
| NC_006213.1;YP_009555240.1;HCoV-OC43_HA-E;231 | 0 | 0 |
| NC_006213.1;YP_009555240.1;HCoV-OC43_HA-E;232 | 0 | 0 |
| NC_006213.1;YP_009555240.1;HCoV-OC43_HA-E;233 | 0.05 | 0.025 |
| NC_006213.1;YP_009555240.1;HCoV-OC43_HA-E;234 | 0 | 0 |
| NC_006213.1;YP_009555240.1;HCoV-OC43_HA-E;235 | 0 | 0 |
| NC_006213.1;YP_009555240.1;HCoV-OC43_HA-E;236 | 0 | 0 |
| NC_006213.1;YP_009555240.1;HCoV-OC43_HA-E;237 | 0 | 0 |
| NC_006213.1;YP_009555240.1;HCoV-OC43_HA-E;238 | 0 | 0 |
| NC_006213.1;YP_009555240.1;HCoV-OC43_HA-E;239 | 0 | 0 |
| NC_006213.1;YP_009555240.1;HCoV-OC43_HA-E;240 | 0 | 0 |
| NC_006213.1;YP_009555240.1;HCoV-OC43_HA-E;241 | 0 | 0 |
| NC_006213.1;YP_009555240.1;HCoV-OC43_HA-E;242 | 0 | 0 |
| NC_006213.1;YP_009555240.1;HCoV-OC43_HA-E;243 | 0.05 | 0 |
| NC_006213.1;YP_009555240.1;HCoV-OC43_HA-E;244 | 0 | 0 |
| NC_006213.1;YP_009555240.1;HCoV-OC43_HA-E;245 | 0 | 0 |
| NC_006213.1;YP_009555240.1;HCoV-OC43_HA-E;246 | 0 | 0 |
| NC_006213.1;YP_009555240.1;HCoV-OC43_HA-E;247 | 0 | 0 |
| NC_006213.1;YP_009555240.1;HCoV-OC43_HA-E;248 | 0 | 0 |
| NC_006213.1;YP_009555240.1;HCoV-OC43_HA-E;249 | 0 | 0 |
| NC_006213.1;YP_009555240.1;HCoV-OC43_HA-E;250 | 0 | 0 |
| NC_006213.1;YP_009555240.1;HCoV-OC43_HA-E;251 | 0 | 0 |
| NC_006213.1;YP_009555240.1;HCoV-OC43_HA-E;252 | 0 | 0 |
| NC_006213.1;YP_009555240.1;HCoV-OC43_HA-E;253 | 0 | 0 |
| NC_006213.1;YP_009555240.1;HCoV-OC43_HA-E;254 | 0 | 0 |
| NC_006213.1;YP_009555240.1;HCoV-OC43_HA-E;255 | 0 | 0 |
| NC_006213.1;YP_009555240.1;HCoV-OC43_HA-E;256 | 0 | 0 |
| NC_006213.1;YP_009555240.1;HCoV-OC43_HA-E;257 | 0 | 0 |
| NC_006213.1;YP_009555240.1;HCoV-OC43_HA-E;258 | 0 | 0 |
| NC_006213.1;YP_009555240.1;HCoV-OC43_HA-E;259 | 0 | 0 |
| NC_006213.1;YP_009555240.1;HCoV-OC43_HA-E;260 | 0 | 0 |
| NC_006213.1;YP_009555240.1;HCoV-OC43_HA-E;261 | 0 | 0 |
| NC_006213.1;YP_009555240.1;HCoV-OC43_HA-E;262 | 0 | 0 |
| NC_006213.1;YP_009555240.1;HCoV-OC43_HA-E;263 | 0 | 0 |
| NC_006213.1;YP_009555240.1;HCoV-OC43_HA-E;264 | 0 | 0 |
| NC_006213.1;YP_009555240.1;HCoV-OC43_HA-E;265 | 0 | 0 |
| NC_006213.1;YP_009555240.1;HCoV-OC43_HA-E;266 | 0 | 0 |
| NC_006213.1;YP_009555240.1;HCoV-OC43_HA-E;267 | 0 | 0 |
| NC_006213.1;YP_009555240.1;HCoV-OC43_HA-E;268 | 0 | 0 |
| NC_006213.1;YP_009555240.1;HCoV-OC43_HA-E;269 | 0.05 | 0 |

| | | |
|---|---|---|
| NC_006213.1;YP_009555240.1;HCoV-OC43_HA-E;270 | 0.05 | 0 |
| NC_006213.1;YP_009555240.1;HCoV-OC43_HA-E;271 | 0.05 | 0 |
| NC_006213.1;YP_009555240.1;HCoV-OC43_HA-E;272 | 0 | 0 |
| NC_006213.1;YP_009555240.1;HCoV-OC43_HA-E;273 | 0 | 0 |
| NC_006213.1;YP_009555240.1;HCoV-OC43_HA-E;274 | 0 | 0 |
| NC_006213.1;YP_009555240.1;HCoV-OC43_HA-E;275 | 0 | 0 |
| NC_006213.1;YP_009555240.1;HCoV-OC43_HA-E;276 | 0 | 0 |
| NC_006213.1;YP_009555240.1;HCoV-OC43_HA-E;277 | 0 | 0 |
| NC_006213.1;YP_009555240.1;HCoV-OC43_HA-E;278 | 0 | 0 |
| NC_006213.1;YP_009555240.1;HCoV-OC43_HA-E;279 | 0 | 0 |
| NC_006213.1;YP_009555240.1;HCoV-OC43_HA-E;280 | 0 | 0 |
| NC_006213.1;YP_009555240.1;HCoV-OC43_HA-E;281 | 0 | 0 |
| NC_006213.1;YP_009555240.1;HCoV-OC43_HA-E;282 | 0 | 0.05 |
| NC_006213.1;YP_009555240.1;HCoV-OC43_HA-E;283 | 0 | 0 |
| NC_006213.1;YP_009555240.1;HCoV-OC43_HA-E;284 | 0 | 0.025 |
| NC_006213.1;YP_009555240.1;HCoV-OC43_HA-E;285 | 0.05 | 0 |
| NC_006213.1;YP_009555240.1;HCoV-OC43_HA-E;286 | 0.05 | 0 |
| NC_006213.1;YP_009555240.1;HCoV-OC43_HA-E;287 | 0.05 | 0.025 |
| NC_006213.1;YP_009555240.1;HCoV-OC43_HA-E;288 | 0 | 0 |
| NC_006213.1;YP_009555240.1;HCoV-OC43_HA-E;289 | 0 | 0.075 |
| NC_006213.1;YP_009555240.1;HCoV-OC43_HA-E;290 | 0.15 | 0.05 |
| NC_006213.1;YP_009555240.1;HCoV-OC43_HA-E;291 | 0.05 | 0.075 |
| NC_006213.1;YP_009555240.1;HCoV-OC43_HA-E;292 | 0.05 | 0 |
| NC_006213.1;YP_009555240.1;HCoV-OC43_HA-E;293 | 0 | 0 |
| NC_006213.1;YP_009555240.1;HCoV-OC43_HA-E;294 | 0 | 0.025 |
| NC_006213.1;YP_009555240.1;HCoV-OC43_HA-E;295 | 0 | 0 |
| NC_006213.1;YP_009555240.1;HCoV-OC43_HA-E;296 | 0 | 0.05 |
| NC_006213.1;YP_009555240.1;HCoV-OC43_HA-E;297 | 0.05 | 0.025 |
| NC_006213.1;YP_009555240.1;HCoV-OC43_HA-E;298 | 0 | 0 |
| NC_006213.1;YP_009555240.1;HCoV-OC43_HA-E;299 | 0 | 0 |
| NC_006213.1;YP_009555240.1;HCoV-OC43_HA-E;300 | 0 | 0 |
| NC_006213.1;YP_009555240.1;HCoV-OC43_HA-E;301 | 0 | 0 |
| NC_006213.1;YP_009555240.1;HCoV-OC43_HA-E;302 | 0 | 0 |
| NC_006213.1;YP_009555240.1;HCoV-OC43_HA-E;303 | 0 | 0 |
| NC_006213.1;YP_009555240.1;HCoV-OC43_HA-E;304 | 0 | 0 |
| NC_006213.1;YP_009555240.1;HCoV-OC43_HA-E;305 | 0.1 | 0.025 |
| NC_006213.1;YP_009555240.1;HCoV-OC43_HA-E;306 | 0.1 | 0.025 |
| NC_006213.1;YP_009555240.1;HCoV-OC43_HA-E;307 | 0.1 | 0 |
| NC_006213.1;YP_009555240.1;HCoV-OC43_HA-E;308 | 0.1 | 0 |
| NC_006213.1;YP_009555240.1;HCoV-OC43_HA-E;309 | 0.1 | 0 |
| NC_006213.1;YP_009555240.1;HCoV-OC43_HA-E;310 | 0 | 0 |
| NC_006213.1;YP_009555240.1;HCoV-OC43_HA-E;311 | 0.1 | 0.025 |
| NC_006213.1;YP_009555240.1;HCoV-OC43_HA-E;312 | 0.1 | 0 |

| | | |
|---|---|---|
| NC_006213.1;YP_009555240.1;HCoV-OC43_HA-E;313 | 0.2 | 0.025 |
| NC_006213.1;YP_009555240.1;HCoV-OC43_HA-E;314 | 0.15 | 0.075 |
| NC_006213.1;YP_009555240.1;HCoV-OC43_HA-E;315 | 0.15 | 0.075 |
| NC_006213.1;YP_009555240.1;HCoV-OC43_HA-E;316 | 0.05 | 0.05 |
| NC_006213.1;YP_009555240.1;HCoV-OC43_HA-E;317 | 0 | 0.025 |
| NC_006213.1;YP_009555240.1;HCoV-OC43_HA-E;318 | 0 | 0.025 |
| NC_006213.1;YP_009555240.1;HCoV-OC43_HA-E;319 | 0 | 0.025 |
| NC_006213.1;YP_009555240.1;HCoV-OC43_HA-E;320 | 0 | 0.025 |
| NC_006213.1;YP_009555240.1;HCoV-OC43_HA-E;321 | 0 | 0 |
| NC_006213.1;YP_009555240.1;HCoV-OC43_HA-E;322 | 0 | 0.025 |
| NC_006213.1;YP_009555240.1;HCoV-OC43_HA-E;323 | 0.05 | 0 |
| NC_006213.1;YP_009555240.1;HCoV-OC43_HA-E;324 | 0 | 0 |
| NC_006213.1;YP_009555240.1;HCoV-OC43_HA-E;325 | 0 | 0 |
| NC_006213.1;YP_009555240.1;HCoV-OC43_HA-E;326 | 0 | 0 |
| NC_006213.1;YP_009555240.1;HCoV-OC43_HA-E;327 | 0 | 0 |
| NC_006213.1;YP_009555240.1;HCoV-OC43_HA-E;328 | 0 | 0 |
| NC_006213.1;YP_009555240.1;HCoV-OC43_HA-E;329 | 0 | 0 |
| NC_006213.1;YP_009555240.1;HCoV-OC43_HA-E;330 | 0 | 0 |
| NC_006213.1;YP_009555240.1;HCoV-OC43_HA-E;331 | 0 | 0 |
| NC_006213.1;YP_009555240.1;HCoV-OC43_HA-E;332 | 0 | 0 |
| NC_006213.1;YP_009555240.1;HCoV-OC43_HA-E;333 | 0 | 0 |
| NC_006213.1;YP_009555240.1;HCoV-OC43_HA-E;334 | 0 | 0 |
| NC_006213.1;YP_009555240.1;HCoV-OC43_HA-E;335 | 0 | 0 |
| NC_006213.1;YP_009555240.1;HCoV-OC43_HA-E;336 | 0 | 0 |
| NC_006213.1;YP_009555240.1;HCoV-OC43_HA-E;337 | 0 | 0 |
| NC_006213.1;YP_009555240.1;HCoV-OC43_HA-E;338 | 0 | 0 |
| NC_006213.1;YP_009555240.1;HCoV-OC43_HA-E;339 | 0 | 0 |
| NC_006213.1;YP_009555240.1;HCoV-OC43_HA-E;340 | 0 | 0 |
| NC_006213.1;YP_009555240.1;HCoV-OC43_HA-E;341 | 0 | 0 |
| NC_006213.1;YP_009555240.1;HCoV-OC43_HA-E;342 | 0 | 0 |
| NC_006213.1;YP_009555240.1;HCoV-OC43_HA-E;343 | 0 | 0 |
| NC_006213.1;YP_009555240.1;HCoV-OC43_HA-E;344 | 0 | 0 |
| NC_006213.1;YP_009555240.1;HCoV-OC43_HA-E;345 | 0 | 0 |
| NC_006213.1;YP_009555240.1;HCoV-OC43_HA-E;346 | 0 | 0 |
| NC_006213.1;YP_009555240.1;HCoV-OC43_HA-E;347 | 0 | 0 |
| NC_006213.1;YP_009555240.1;HCoV-OC43_HA-E;348 | 0 | 0 |
| NC_006213.1;YP_009555240.1;HCoV-OC43_HA-E;349 | 0 | 0.025 |
| NC_006213.1;YP_009555240.1;HCoV-OC43_HA-E;350 | 0 | 0 |
| NC_006213.1;YP_009555240.1;HCoV-OC43_HA-E;351 | 0 | 0 |
| NC_006213.1;YP_009555240.1;HCoV-OC43_HA-E;352 | 0 | 0 |
| NC_006213.1;YP_009555240.1;HCoV-OC43_HA-E;353 | 0 | 0 |
| NC_006213.1;YP_009555240.1;HCoV-OC43_HA-E;354 | 0 | 0 |
| NC_006213.1;YP_009555240.1;HCoV-OC43_HA-E;355 | 0 | 0 |

| | | |
|---|---|---|
| NC_006213.1;YP_009555240.1;HCoV-OC43_HA-E;356 | 0 | 0 |
| NC_006213.1;YP_009555240.1;HCoV-OC43_HA-E;357 | 0 | 0.025 |
| NC_006213.1;YP_009555240.1;HCoV-OC43_HA-E;358 | 0 | 0 |
| NC_006213.1;YP_009555240.1;HCoV-OC43_HA-E;359 | 0 | 0 |
| NC_006213.1;YP_009555240.1;HCoV-OC43_HA-E;360 | 0.05 | 0.025 |
| NC_006213.1;YP_009555240.1;HCoV-OC43_HA-E;361 | 0 | 0 |
| NC_006213.1;YP_009555240.1;HCoV-OC43_HA-E;362 | 0.15 | 0 |
| NC_006213.1;YP_009555240.1;HCoV-OC43_HA-E;363 | 0.1 | 0.025 |
| NC_006213.1;YP_009555240.1;HCoV-OC43_HA-E;364 | 0.15 | 0.05 |
| NC_006213.1;YP_009555240.1;HCoV-OC43_HA-E;365 | 0.15 | 0.025 |
| NC_006213.1;YP_009555240.1;HCoV-OC43_HA-E;366 | 0.1 | 0.025 |
| NC_006213.1;YP_009555240.1;HCoV-OC43_HA-E;367 | 0.1 | 0.025 |
| NC_006213.1;YP_009555240.1;HCoV-OC43_HA-E;368 | 0.15 | 0.025 |
| NC_006213.1;YP_009555240.1;HCoV-OC43_HA-E;369 | 0.45 | 0.175 |
| NC_006213.1;YP_009555240.1;HCoV-OC43_HA-E;370 | 0.2 | 0.05 |
| NC_006213.1;YP_009555240.1;HCoV-OC43_HA-E;371 | 0.1 | 0.025 |
| NC_006213.1;YP_009555240.1;HCoV-OC43_HA-E;372 | 0 | 0.025 |
| NC_006213.1;YP_009555240.1;HCoV-OC43_HA-E;373 | 0 | 0 |
| NC_006213.1;YP_009555240.1;HCoV-OC43_HA-E;374 | 0 | 0 |
| NC_006213.1;YP_009555240.1;HCoV-OC43_HA-E;375 | 0 | 0.025 |
| NC_006213.1;YP_009555240.1;HCoV-OC43_HA-E;376 | 0 | 0 |
| NC_006213.1;YP_009555240.1;HCoV-OC43_HA-E;377 | 0 | 0 |
| NC_006213.1;YP_009555240.1;HCoV-OC43_HA-E;378 | 0 | 0 |
| NC_006213.1;YP_009555240.1;HCoV-OC43_HA-E;379 | 0 | 0 |
| NC_006213.1;YP_009555240.1;HCoV-OC43_HA-E;380 | 0 | 0 |
| NC_006213.1;YP_009555240.1;HCoV-OC43_HA-E;381 | 0 | 0 |
| NC_006213.1;YP_009555240.1;HCoV-OC43_HA-E;382 | 0 | 0 |
| NC_006213.1;YP_009555240.1;HCoV-OC43_HA-E;383 | 0 | 0 |
| NC_006213.1;YP_009555240.1;HCoV-OC43_HA-E;384 | 0 | 0 |
| NC_006213.1;YP_009555240.1;HCoV-OC43_HA-E;385 | 0 | 0 |
| NC_006213.1;YP_009555240.1;HCoV-OC43_HA-E;386 | 0 | 0 |
| NC_006213.1;YP_009555240.1;HCoV-OC43_HA-E;387 | 0 | 0 |
| NC_006213.1;YP_009555240.1;HCoV-OC43_HA-E;388 | 0 | 0 |
| NC_006213.1;YP_009555240.1;HCoV-OC43_HA-E;389 | 0 | 0 |
| NC_006213.1;YP_009555240.1;HCoV-OC43_HA-E;390 | 0 | 0 |
| NC_006213.1;YP_009555240.1;HCoV-OC43_HA-E;391 | 0 | 0 |
| NC_006213.1;YP_009555240.1;HCoV-OC43_HA-E;392 | 0 | 0 |
| NC_006213.1;YP_009555240.1;HCoV-OC43_HA-E;393 | 0 | 0 |
| NC_006213.1;YP_009555240.1;HCoV-OC43_HA-E;394 | 0 | 0 |
| NC_006213.1;YP_009555240.1;HCoV-OC43_HA-E;395 | 0 | 0 |
| NC_006213.1;YP_009555240.1;HCoV-OC43_HA-E;396 | 0 | 0 |
| NC_006213.1;YP_009555240.1;HCoV-OC43_HA-E;397 | 0 | 0 |
| NC_006213.1;YP_009555240.1;HCoV-OC43_HA-E;398 | 0 | 0 |

| | | |
|---|---|---|
| NC_006213.1;YP_009555240.1;HCoV-OC43_HA-E;399 | 0 | 0 |
| NC_006213.1;YP_009555240.1;HCoV-OC43_HA-E;400 | 0 | 0 |
| NC_006213.1;YP_009555240.1;HCoV-OC43_HA-E;401 | 0 | 0 |
| NC_006213.1;YP_009555240.1;HCoV-OC43_HA-E;402 | 0 | 0 |
| NC_006213.1;YP_009555240.1;HCoV-OC43_HA-E;403 | 0 | 0 |
| NC_006213.1;YP_009555240.1;HCoV-OC43_HA-E;404 | 0 | 0 |
| NC_006213.1;YP_009555240.1;HCoV-OC43_HA-E;405 | 0 | 0 |
| NC_006213.1;YP_009555240.1;HCoV-OC43_HA-E;406 | 0 | 0 |
| NC_006213.1;YP_009555240.1;HCoV-OC43_HA-E;407 | 0 | 0 |
| NC_006213.1;YP_009555240.1;HCoV-OC43_HA-E;408 | 0 | 0 |
| NC_006213.1;YP_009555240.1;HCoV-OC43_HA-E;409 | 0 | 0 |
| NC_006213.1;YP_009555240.1;HCoV-OC43_HA-E;410 | 0 | 0 |
| NC_006213.1;YP_009555240.1;HCoV-OC43_HA-E;411 | 0 | 0 |
| NC_006213.1;YP_009555240.1;HCoV-OC43_HA-E;412 | 0 | 0.025 |
| NC_006213.1;YP_009555240.1;HCoV-OC43_HA-E;413 | 0 | 0.025 |
| NC_006213.1;YP_009555241.1;HCoV-OC43_spike;1 | 0 | 0 |
| NC_006213.1;YP_009555241.1;HCoV-OC43_spike;2 | 0 | 0 |
| NC_006213.1;YP_009555241.1;HCoV-OC43_spike;3 | 0 | 0 |
| NC_006213.1;YP_009555241.1;HCoV-OC43_spike;4 | 0 | 0 |
| NC_006213.1;YP_009555241.1;HCoV-OC43_spike;5 | 0 | 0 |
| NC_006213.1;YP_009555241.1;HCoV-OC43_spike;6 | 0 | 0 |
| NC_006213.1;YP_009555241.1;HCoV-OC43_spike;7 | 0 | 0 |
| NC_006213.1;YP_009555241.1;HCoV-OC43_spike;8 | 0 | 0 |
| NC_006213.1;YP_009555241.1;HCoV-OC43_spike;9 | 0 | 0.025 |
| NC_006213.1;YP_009555241.1;HCoV-OC43_spike;10 | 0.05 | 0 |
| NC_006213.1;YP_009555241.1;HCoV-OC43_spike;11 | 0.05 | 0 |
| NC_006213.1;YP_009555241.1;HCoV-OC43_spike;12 | 0.05 | 0.025 |
| NC_006213.1;YP_009555241.1;HCoV-OC43_spike;13 | 0.05 | 0.075 |
| NC_006213.1;YP_009555241.1;HCoV-OC43_spike;14 | 0.05 | 0.025 |
| NC_006213.1;YP_009555241.1;HCoV-OC43_spike;15 | 0.05 | 0.05 |
| NC_006213.1;YP_009555241.1;HCoV-OC43_spike;16 | 0.05 | 0.075 |
| NC_006213.1;YP_009555241.1;HCoV-OC43_spike;17 | 0.05 | 0.05 |
| NC_006213.1;YP_009555241.1;HCoV-OC43_spike;18 | 0.05 | 0.075 |
| NC_006213.1;YP_009555241.1;HCoV-OC43_spike;19 | 0.05 | 0.075 |
| NC_006213.1;YP_009555241.1;HCoV-OC43_spike;20 | 0.05 | 0.05 |
| NC_006213.1;YP_009555241.1;HCoV-OC43_spike;21 | 0 | 0.025 |
| NC_006213.1;YP_009555241.1;HCoV-OC43_spike;22 | 0 | 0.05 |
| NC_006213.1;YP_009555241.1;HCoV-OC43_spike;23 | 0.05 | 0.05 |
| NC_006213.1;YP_009555241.1;HCoV-OC43_spike;24 | 0 | 0 |
| NC_006213.1;YP_009555241.1;HCoV-OC43_spike;25 | 0 | 0 |
| NC_006213.1;YP_009555241.1;HCoV-OC43_spike;26 | 0 | 0 |
| NC_006213.1;YP_009555241.1;HCoV-OC43_spike;27 | 0 | 0 |
| NC_006213.1;YP_009555241.1;HCoV-OC43_spike;28 | 0 | 0 |

| | | |
|---|---|---|
| NC_006213.1;YP_009555241.1;HCoV-OC43_spike;29 | 0 | 0 |
| NC_006213.1;YP_009555241.1;HCoV-OC43_spike;30 | 0.1 | 0.025 |
| NC_006213.1;YP_009555241.1;HCoV-OC43_spike;31 | 0.05 | 0.025 |
| NC_006213.1;YP_009555241.1;HCoV-OC43_spike;32 | 0.05 | 0.025 |
| NC_006213.1;YP_009555241.1;HCoV-OC43_spike;33 | 0.05 | 0 |
| NC_006213.1;YP_009555241.1;HCoV-OC43_spike;34 | 0.05 | 0 |
| NC_006213.1;YP_009555241.1;HCoV-OC43_spike;35 | 0 | 0 |
| NC_006213.1;YP_009555241.1;HCoV-OC43_spike;36 | 0.05 | 0 |
| NC_006213.1;YP_009555241.1;HCoV-OC43_spike;37 | 0.1 | 0 |
| NC_006213.1;YP_009555241.1;HCoV-OC43_spike;38 | 0 | 0 |
| NC_006213.1;YP_009555241.1;HCoV-OC43_spike;39 | 0 | 0 |
| NC_006213.1;YP_009555241.1;HCoV-OC43_spike;40 | 0 | 0 |
| NC_006213.1;YP_009555241.1;HCoV-OC43_spike;41 | 0 | 0.025 |
| NC_006213.1;YP_009555241.1;HCoV-OC43_spike;42 | 0 | 0 |
| NC_006213.1;YP_009555241.1;HCoV-OC43_spike;43 | 0 | 0 |
| NC_006213.1;YP_009555241.1;HCoV-OC43_spike;44 | 0 | 0 |
| NC_006213.1;YP_009555241.1;HCoV-OC43_spike;45 | 0 | 0 |
| NC_006213.1;YP_009555241.1;HCoV-OC43_spike;46 | 0 | 0 |
| NC_006213.1;YP_009555241.1;HCoV-OC43_spike;47 | 0 | 0 |
| NC_006213.1;YP_009555241.1;HCoV-OC43_spike;48 | 0 | 0 |
| NC_006213.1;YP_009555241.1;HCoV-OC43_spike;49 | 0 | 0 |
| NC_006213.1;YP_009555241.1;HCoV-OC43_spike;50 | 0 | 0 |
| NC_006213.1;YP_009555241.1;HCoV-OC43_spike;51 | 0 | 0 |
| NC_006213.1;YP_009555241.1;HCoV-OC43_spike;52 | 0 | 0 |
| NC_006213.1;YP_009555241.1;HCoV-OC43_spike;53 | 0 | 0 |
| NC_006213.1;YP_009555241.1;HCoV-OC43_spike;54 | 0 | 0 |
| NC_006213.1;YP_009555241.1;HCoV-OC43_spike;55 | 0 | 0 |
| NC_006213.1;YP_009555241.1;HCoV-OC43_spike;56 | 0 | 0 |
| NC_006213.1;YP_009555241.1;HCoV-OC43_spike;57 | 0 | 0 |
| NC_006213.1;YP_009555241.1;HCoV-OC43_spike;58 | 0.05 | 0.025 |
| NC_006213.1;YP_009555241.1;HCoV-OC43_spike;59 | 0 | 0 |
| NC_006213.1;YP_009555241.1;HCoV-OC43_spike;60 | 0 | 0 |
| NC_006213.1;YP_009555241.1;HCoV-OC43_spike;61 | 0 | 0 |
| NC_006213.1;YP_009555241.1;HCoV-OC43_spike;62 | 0 | 0 |
| NC_006213.1;YP_009555241.1;HCoV-OC43_spike;63 | 0 | 0 |
| NC_006213.1;YP_009555241.1;HCoV-OC43_spike;64 | 0.05 | 0 |
| NC_006213.1;YP_009555241.1;HCoV-OC43_spike;65 | 0 | 0 |
| NC_006213.1;YP_009555241.1;HCoV-OC43_spike;66 | 0 | 0 |
| NC_006213.1;YP_009555241.1;HCoV-OC43_spike;67 | 0 | 0 |
| NC_006213.1;YP_009555241.1;HCoV-OC43_spike;68 | 0 | 0 |
| NC_006213.1;YP_009555241.1;HCoV-OC43_spike;69 | 0 | 0 |
| NC_006213.1;YP_009555241.1;HCoV-OC43_spike;70 | 0 | 0 |
| NC_006213.1;YP_009555241.1;HCoV-OC43_spike;71 | 0 | 0 |

| | | |
|---|---|---|
| NC_006213.1;YP_009555241.1;HCoV-OC43_spike;72 | 0 | 0 |
| NC_006213.1;YP_009555241.1;HCoV-OC43_spike;73 | 0 | 0 |
| NC_006213.1;YP_009555241.1;HCoV-OC43_spike;74 | 0 | 0 |
| NC_006213.1;YP_009555241.1;HCoV-OC43_spike;75 | 0 | 0 |
| NC_006213.1;YP_009555241.1;HCoV-OC43_spike;76 | 0 | 0 |
| NC_006213.1;YP_009555241.1;HCoV-OC43_spike;77 | 0 | 0 |
| NC_006213.1;YP_009555241.1;HCoV-OC43_spike;78 | 0 | 0 |
| NC_006213.1;YP_009555241.1;HCoV-OC43_spike;79 | 0 | 0 |
| NC_006213.1;YP_009555241.1;HCoV-OC43_spike;80 | 0 | 0 |
| NC_006213.1;YP_009555241.1;HCoV-OC43_spike;81 | 0 | 0 |
| NC_006213.1;YP_009555241.1;HCoV-OC43_spike;82 | 0 | 0 |
| NC_006213.1;YP_009555241.1;HCoV-OC43_spike;83 | 0 | 0 |
| NC_006213.1;YP_009555241.1;HCoV-OC43_spike;84 | 0 | 0 |
| NC_006213.1;YP_009555241.1;HCoV-OC43_spike;85 | 0 | 0 |
| NC_006213.1;YP_009555241.1;HCoV-OC43_spike;86 | 0 | 0 |
| NC_006213.1;YP_009555241.1;HCoV-OC43_spike;87 | 0 | 0 |
| NC_006213.1;YP_009555241.1;HCoV-OC43_spike;88 | 0 | 0 |
| NC_006213.1;YP_009555241.1;HCoV-OC43_spike;89 | 0 | 0 |
| NC_006213.1;YP_009555241.1;HCoV-OC43_spike;90 | 0 | 0 |
| NC_006213.1;YP_009555241.1;HCoV-OC43_spike;91 | 0 | 0 |
| NC_006213.1;YP_009555241.1;HCoV-OC43_spike;92 | 0 | 0 |
| NC_006213.1;YP_009555241.1;HCoV-OC43_spike;93 | 0 | 0 |
| NC_006213.1;YP_009555241.1;HCoV-OC43_spike;94 | 0 | 0 |
| NC_006213.1;YP_009555241.1;HCoV-OC43_spike;95 | 0 | 0.025 |
| NC_006213.1;YP_009555241.1;HCoV-OC43_spike;96 | 0 | 0.025 |
| NC_006213.1;YP_009555241.1;HCoV-OC43_spike;97 | 0.05 | 0.025 |
| NC_006213.1;YP_009555241.1;HCoV-OC43_spike;98 | 0.05 | 0.025 |
| NC_006213.1;YP_009555241.1;HCoV-OC43_spike;99 | 0 | 0.025 |
| NC_006213.1;YP_009555241.1;HCoV-OC43_spike;100 | 0 | 0 |
| NC_006213.1;YP_009555241.1;HCoV-OC43_spike;101 | 0 | 0 |
| NC_006213.1;YP_009555241.1;HCoV-OC43_spike;102 | 0 | 0 |
| NC_006213.1;YP_009555241.1;HCoV-OC43_spike;103 | 0 | 0 |
| NC_006213.1;YP_009555241.1;HCoV-OC43_spike;104 | 0 | 0 |
| NC_006213.1;YP_009555241.1;HCoV-OC43_spike;105 | 0 | 0.025 |
| NC_006213.1;YP_009555241.1;HCoV-OC43_spike;106 | 0 | 0 |
| NC_006213.1;YP_009555241.1;HCoV-OC43_spike;107 | 0.05 | 0 |
| NC_006213.1;YP_009555241.1;HCoV-OC43_spike;108 | 0.05 | 0 |
| NC_006213.1;YP_009555241.1;HCoV-OC43_spike;109 | 0.05 | 0.025 |
| NC_006213.1;YP_009555241.1;HCoV-OC43_spike;110 | 0.05 | 0 |
| NC_006213.1;YP_009555241.1;HCoV-OC43_spike;111 | 0.05 | 0 |
| NC_006213.1;YP_009555241.1;HCoV-OC43_spike;112 | 0.05 | 0 |
| NC_006213.1;YP_009555241.1;HCoV-OC43_spike;113 | 0.05 | 0.025 |
| NC_006213.1;YP_009555241.1;HCoV-OC43_spike;114 | 0 | 0 |

| | | |
|---|---|---|
| NC_006213.1;YP_009555241.1;HCoV-OC43_spike;115 | 0 | 0 |
| NC_006213.1;YP_009555241.1;HCoV-OC43_spike;116 | 0 | 0 |
| NC_006213.1;YP_009555241.1;HCoV-OC43_spike;117 | 0 | 0 |
| NC_006213.1;YP_009555241.1;HCoV-OC43_spike;118 | 0 | 0 |
| NC_006213.1;YP_009555241.1;HCoV-OC43_spike;119 | 0 | 0 |
| NC_006213.1;YP_009555241.1;HCoV-OC43_spike;120 | 0 | 0 |
| NC_006213.1;YP_009555241.1;HCoV-OC43_spike;121 | 0 | 0 |
| NC_006213.1;YP_009555241.1;HCoV-OC43_spike;122 | 0 | 0 |
| NC_006213.1;YP_009555241.1;HCoV-OC43_spike;123 | 0 | 0 |
| NC_006213.1;YP_009555241.1;HCoV-OC43_spike;124 | 0 | 0 |
| NC_006213.1;YP_009555241.1;HCoV-OC43_spike;125 | 0 | 0 |
| NC_006213.1;YP_009555241.1;HCoV-OC43_spike;126 | 0 | 0 |
| NC_006213.1;YP_009555241.1;HCoV-OC43_spike;127 | 0 | 0 |
| NC_006213.1;YP_009555241.1;HCoV-OC43_spike;128 | 0 | 0 |
| NC_006213.1;YP_009555241.1;HCoV-OC43_spike;129 | 0 | 0 |
| NC_006213.1;YP_009555241.1;HCoV-OC43_spike;130 | 0 | 0 |
| NC_006213.1;YP_009555241.1;HCoV-OC43_spike;131 | 0 | 0 |
| NC_006213.1;YP_009555241.1;HCoV-OC43_spike;132 | 0.05 | 0 |
| NC_006213.1;YP_009555241.1;HCoV-OC43_spike;133 | 0.05 | 0 |
| NC_006213.1;YP_009555241.1;HCoV-OC43_spike;134 | 0 | 0 |
| NC_006213.1;YP_009555241.1;HCoV-OC43_spike;135 | 0 | 0 |
| NC_006213.1;YP_009555241.1;HCoV-OC43_spike;136 | 0 | 0 |
| NC_006213.1;YP_009555241.1;HCoV-OC43_spike;137 | 0.05 | 0.025 |
| NC_006213.1;YP_009555241.1;HCoV-OC43_spike;138 | 0.05 | 0 |
| NC_006213.1;YP_009555241.1;HCoV-OC43_spike;139 | 0.05 | 0.025 |
| NC_006213.1;YP_009555241.1;HCoV-OC43_spike;140 | 0.1 | 0.05 |
| NC_006213.1;YP_009555241.1;HCoV-OC43_spike;141 | 0.1 | 0.025 |
| NC_006213.1;YP_009555241.1;HCoV-OC43_spike;142 | 0.05 | 0.05 |
| NC_006213.1;YP_009555241.1;HCoV-OC43_spike;143 | 0.05 | 0.025 |
| NC_006213.1;YP_009555241.1;HCoV-OC43_spike;144 | 0 | 0 |
| NC_006213.1;YP_009555241.1;HCoV-OC43_spike;145 | 0 | 0.025 |
| NC_006213.1;YP_009555241.1;HCoV-OC43_spike;146 | 0 | 0.025 |
| NC_006213.1;YP_009555241.1;HCoV-OC43_spike;147 | 0 | 0.025 |
| NC_006213.1;YP_009555241.1;HCoV-OC43_spike;148 | 0 | 0.05 |
| NC_006213.1;YP_009555241.1;HCoV-OC43_spike;149 | 0 | 0 |
| NC_006213.1;YP_009555241.1;HCoV-OC43_spike;150 | 0 | 0 |
| NC_006213.1;YP_009555241.1;HCoV-OC43_spike;151 | 0 | 0 |
| NC_006213.1;YP_009555241.1;HCoV-OC43_spike;152 | 0 | 0 |
| NC_006213.1;YP_009555241.1;HCoV-OC43_spike;153 | 0 | 0 |
| NC_006213.1;YP_009555241.1;HCoV-OC43_spike;154 | 0 | 0 |
| NC_006213.1;YP_009555241.1;HCoV-OC43_spike;155 | 0 | 0 |
| NC_006213.1;YP_009555241.1;HCoV-OC43_spike;156 | 0 | 0 |
| NC_006213.1;YP_009555241.1;HCoV-OC43_spike;157 | 0.05 | 0.025 |

| | | |
|---|---|---|
| NC_006213.1;YP_009555241.1;HCoV-OC43_spike;158 | 0 | 0 |
| NC_006213.1;YP_009555241.1;HCoV-OC43_spike;159 | 0 | 0 |
| NC_006213.1;YP_009555241.1;HCoV-OC43_spike;160 | 0 | 0 |
| NC_006213.1;YP_009555241.1;HCoV-OC43_spike;161 | 0 | 0 |
| NC_006213.1;YP_009555241.1;HCoV-OC43_spike;162 | 0 | 0 |
| NC_006213.1;YP_009555241.1;HCoV-OC43_spike;163 | 0 | 0 |
| NC_006213.1;YP_009555241.1;HCoV-OC43_spike;164 | 0 | 0 |
| NC_006213.1;YP_009555241.1;HCoV-OC43_spike;165 | 0 | 0 |
| NC_006213.1;YP_009555241.1;HCoV-OC43_spike;166 | 0 | 0 |
| NC_006213.1;YP_009555241.1;HCoV-OC43_spike;167 | 0 | 0 |
| NC_006213.1;YP_009555241.1;HCoV-OC43_spike;168 | 0 | 0 |
| NC_006213.1;YP_009555241.1;HCoV-OC43_spike;169 | 0 | 0 |
| NC_006213.1;YP_009555241.1;HCoV-OC43_spike;170 | 0.05 | 0 |
| NC_006213.1;YP_009555241.1;HCoV-OC43_spike;171 | 0.05 | 0 |
| NC_006213.1;YP_009555241.1;HCoV-OC43_spike;172 | 0 | 0 |
| NC_006213.1;YP_009555241.1;HCoV-OC43_spike;173 | 0.05 | 0 |
| NC_006213.1;YP_009555241.1;HCoV-OC43_spike;174 | 0.05 | 0 |
| NC_006213.1;YP_009555241.1;HCoV-OC43_spike;175 | 0 | 0 |
| NC_006213.1;YP_009555241.1;HCoV-OC43_spike;176 | 0.05 | 0 |
| NC_006213.1;YP_009555241.1;HCoV-OC43_spike;177 | 0.05 | 0 |
| NC_006213.1;YP_009555241.1;HCoV-OC43_spike;178 | 0.05 | 0 |
| NC_006213.1;YP_009555241.1;HCoV-OC43_spike;179 | 0 | 0 |
| NC_006213.1;YP_009555241.1;HCoV-OC43_spike;180 | 0 | 0 |
| NC_006213.1;YP_009555241.1;HCoV-OC43_spike;181 | 0 | 0 |
| NC_006213.1;YP_009555241.1;HCoV-OC43_spike;182 | 0 | 0 |
| NC_006213.1;YP_009555241.1;HCoV-OC43_spike;183 | 0 | 0 |
| NC_006213.1;YP_009555241.1;HCoV-OC43_spike;184 | 0 | 0 |
| NC_006213.1;YP_009555241.1;HCoV-OC43_spike;185 | 0 | 0 |
| NC_006213.1;YP_009555241.1;HCoV-OC43_spike;186 | 0 | 0 |
| NC_006213.1;YP_009555241.1;HCoV-OC43_spike;187 | 0 | 0 |
| NC_006213.1;YP_009555241.1;HCoV-OC43_spike;188 | 0 | 0 |
| NC_006213.1;YP_009555241.1;HCoV-OC43_spike;189 | 0 | 0 |
| NC_006213.1;YP_009555241.1;HCoV-OC43_spike;190 | 0 | 0 |
| NC_006213.1;YP_009555241.1;HCoV-OC43_spike;191 | 0 | 0 |
| NC_006213.1;YP_009555241.1;HCoV-OC43_spike;192 | 0 | 0 |
| NC_006213.1;YP_009555241.1;HCoV-OC43_spike;193 | 0 | 0 |
| NC_006213.1;YP_009555241.1;HCoV-OC43_spike;194 | 0 | 0 |
| NC_006213.1;YP_009555241.1;HCoV-OC43_spike;195 | 0 | 0 |
| NC_006213.1;YP_009555241.1;HCoV-OC43_spike;196 | 0 | 0 |
| NC_006213.1;YP_009555241.1;HCoV-OC43_spike;197 | 0 | 0 |
| NC_006213.1;YP_009555241.1;HCoV-OC43_spike;198 | 0 | 0 |
| NC_006213.1;YP_009555241.1;HCoV-OC43_spike;199 | 0 | 0 |
| NC_006213.1;YP_009555241.1;HCoV-OC43_spike;200 | 0.05 | 0.05 |

| | | |
|---|---|---|
| NC_006213.1;YP_009555241.1;HCoV-OC43_spike;201 | 0 | 0 |
| NC_006213.1;YP_009555241.1;HCoV-OC43_spike;202 | 0 | 0 |
| NC_006213.1;YP_009555241.1;HCoV-OC43_spike;203 | 0 | 0 |
| NC_006213.1;YP_009555241.1;HCoV-OC43_spike;204 | 0 | 0 |
| NC_006213.1;YP_009555241.1;HCoV-OC43_spike;205 | 0.05 | 0 |
| NC_006213.1;YP_009555241.1;HCoV-OC43_spike;206 | 0.05 | 0 |
| NC_006213.1;YP_009555241.1;HCoV-OC43_spike;207 | 0.05 | 0 |
| NC_006213.1;YP_009555241.1;HCoV-OC43_spike;208 | 0 | 0 |
| NC_006213.1;YP_009555241.1;HCoV-OC43_spike;209 | 0 | 0 |
| NC_006213.1;YP_009555241.1;HCoV-OC43_spike;210 | 0 | 0 |
| NC_006213.1;YP_009555241.1;HCoV-OC43_spike;211 | 0 | 0 |
| NC_006213.1;YP_009555241.1;HCoV-OC43_spike;212 | 0 | 0 |
| NC_006213.1;YP_009555241.1;HCoV-OC43_spike;213 | 0 | 0 |
| NC_006213.1;YP_009555241.1;HCoV-OC43_spike;214 | 0 | 0 |
| NC_006213.1;YP_009555241.1;HCoV-OC43_spike;215 | 0 | 0 |
| NC_006213.1;YP_009555241.1;HCoV-OC43_spike;216 | 0 | 0 |
| NC_006213.1;YP_009555241.1;HCoV-OC43_spike;217 | 0 | 0 |
| NC_006213.1;YP_009555241.1;HCoV-OC43_spike;218 | 0 | 0 |
| NC_006213.1;YP_009555241.1;HCoV-OC43_spike;219 | 0 | 0 |
| NC_006213.1;YP_009555241.1;HCoV-OC43_spike;220 | 0 | 0 |
| NC_006213.1;YP_009555241.1;HCoV-OC43_spike;221 | 0 | 0 |
| NC_006213.1;YP_009555241.1;HCoV-OC43_spike;222 | 0 | 0 |
| NC_006213.1;YP_009555241.1;HCoV-OC43_spike;223 | 0 | 0 |
| NC_006213.1;YP_009555241.1;HCoV-OC43_spike;224 | 0 | 0 |
| NC_006213.1;YP_009555241.1;HCoV-OC43_spike;225 | 0 | 0 |
| NC_006213.1;YP_009555241.1;HCoV-OC43_spike;226 | 0 | 0 |
| NC_006213.1;YP_009555241.1;HCoV-OC43_spike;227 | 0 | 0 |
| NC_006213.1;YP_009555241.1;HCoV-OC43_spike;228 | 0 | 0 |
| NC_006213.1;YP_009555241.1;HCoV-OC43_spike;229 | 0 | 0 |
| NC_006213.1;YP_009555241.1;HCoV-OC43_spike;230 | 0 | 0 |
| NC_006213.1;YP_009555241.1;HCoV-OC43_spike;231 | 0 | 0 |
| NC_006213.1;YP_009555241.1;HCoV-OC43_spike;232 | 0 | 0 |
| NC_006213.1;YP_009555241.1;HCoV-OC43_spike;233 | 0 | 0 |
| NC_006213.1;YP_009555241.1;HCoV-OC43_spike;234 | 0 | 0 |
| NC_006213.1;YP_009555241.1;HCoV-OC43_spike;235 | 0 | 0 |
| NC_006213.1;YP_009555241.1;HCoV-OC43_spike;236 | 0 | 0 |
| NC_006213.1;YP_009555241.1;HCoV-OC43_spike;237 | 0 | 0 |
| NC_006213.1;YP_009555241.1;HCoV-OC43_spike;238 | 0 | 0 |
| NC_006213.1;YP_009555241.1;HCoV-OC43_spike;239 | 0 | 0 |
| NC_006213.1;YP_009555241.1;HCoV-OC43_spike;240 | 0 | 0 |
| NC_006213.1;YP_009555241.1;HCoV-OC43_spike;241 | 0 | 0 |
| NC_006213.1;YP_009555241.1;HCoV-OC43_spike;242 | 0 | 0 |
| NC_006213.1;YP_009555241.1;HCoV-OC43_spike;243 | 0 | 0 |

| | | |
|---|---|---|
| NC_006213.1;YP_009555241.1;HCoV-OC43_spike;244 | 0 | 0 |
| NC_006213.1;YP_009555241.1;HCoV-OC43_spike;245 | 0 | 0 |
| NC_006213.1;YP_009555241.1;HCoV-OC43_spike;246 | 0 | 0 |
| NC_006213.1;YP_009555241.1;HCoV-OC43_spike;247 | 0 | 0 |
| NC_006213.1;YP_009555241.1;HCoV-OC43_spike;248 | 0 | 0 |
| NC_006213.1;YP_009555241.1;HCoV-OC43_spike;249 | 0 | 0 |
| NC_006213.1;YP_009555241.1;HCoV-OC43_spike;250 | 0 | 0 |
| NC_006213.1;YP_009555241.1;HCoV-OC43_spike;251 | 0 | 0 |
| NC_006213.1;YP_009555241.1;HCoV-OC43_spike;252 | 0 | 0 |
| NC_006213.1;YP_009555241.1;HCoV-OC43_spike;253 | 0 | 0 |
| NC_006213.1;YP_009555241.1;HCoV-OC43_spike;254 | 0 | 0 |
| NC_006213.1;YP_009555241.1;HCoV-OC43_spike;255 | 0 | 0 |
| NC_006213.1;YP_009555241.1;HCoV-OC43_spike;256 | 0 | 0 |
| NC_006213.1;YP_009555241.1;HCoV-OC43_spike;257 | 0 | 0 |
| NC_006213.1;YP_009555241.1;HCoV-OC43_spike;258 | 0 | 0 |
| NC_006213.1;YP_009555241.1;HCoV-OC43_spike;259 | 0 | 0 |
| NC_006213.1;YP_009555241.1;HCoV-OC43_spike;260 | 0 | 0 |
| NC_006213.1;YP_009555241.1;HCoV-OC43_spike;261 | 0 | 0 |
| NC_006213.1;YP_009555241.1;HCoV-OC43_spike;262 | 0 | 0 |
| NC_006213.1;YP_009555241.1;HCoV-OC43_spike;263 | 0 | 0 |
| NC_006213.1;YP_009555241.1;HCoV-OC43_spike;264 | 0 | 0 |
| NC_006213.1;YP_009555241.1;HCoV-OC43_spike;265 | 0 | 0 |
| NC_006213.1;YP_009555241.1;HCoV-OC43_spike;266 | 0 | 0 |
| NC_006213.1;YP_009555241.1;HCoV-OC43_spike;267 | 0 | 0.025 |
| NC_006213.1;YP_009555241.1;HCoV-OC43_spike;268 | 0 | 0 |
| NC_006213.1;YP_009555241.1;HCoV-OC43_spike;269 | 0 | 0 |
| NC_006213.1;YP_009555241.1;HCoV-OC43_spike;270 | 0 | 0 |
| NC_006213.1;YP_009555241.1;HCoV-OC43_spike;271 | 0 | 0 |
| NC_006213.1;YP_009555241.1;HCoV-OC43_spike;272 | 0 | 0.025 |
| NC_006213.1;YP_009555241.1;HCoV-OC43_spike;273 | 0.05 | 0.025 |
| NC_006213.1;YP_009555241.1;HCoV-OC43_spike;274 | 0.05 | 0.025 |
| NC_006213.1;YP_009555241.1;HCoV-OC43_spike;275 | 0.05 | 0.025 |
| NC_006213.1;YP_009555241.1;HCoV-OC43_spike;276 | 0.05 | 0.05 |
| NC_006213.1;YP_009555241.1;HCoV-OC43_spike;277 | 0.05 | 0.05 |
| NC_006213.1;YP_009555241.1;HCoV-OC43_spike;278 | 0 | 0.025 |
| NC_006213.1;YP_009555241.1;HCoV-OC43_spike;279 | 0 | 0 |
| NC_006213.1;YP_009555241.1;HCoV-OC43_spike;280 | 0 | 0 |
| NC_006213.1;YP_009555241.1;HCoV-OC43_spike;281 | 0 | 0 |
| NC_006213.1;YP_009555241.1;HCoV-OC43_spike;282 | 0 | 0 |
| NC_006213.1;YP_009555241.1;HCoV-OC43_spike;283 | 0 | 0.025 |
| NC_006213.1;YP_009555241.1;HCoV-OC43_spike;284 | 0 | 0.025 |
| NC_006213.1;YP_009555241.1;HCoV-OC43_spike;285 | 0 | 0.025 |
| NC_006213.1;YP_009555241.1;HCoV-OC43_spike;286 | 0.05 | 0.025 |

| | | |
|---|---|---|
| NC_006213.1;YP_009555241.1;HCoV-OC43_spike;287 | 0 | 0 |
| NC_006213.1;YP_009555241.1;HCoV-OC43_spike;288 | 0 | 0 |
| NC_006213.1;YP_009555241.1;HCoV-OC43_spike;289 | 0 | 0 |
| NC_006213.1;YP_009555241.1;HCoV-OC43_spike;290 | 0 | 0 |
| NC_006213.1;YP_009555241.1;HCoV-OC43_spike;291 | 0 | 0.05 |
| NC_006213.1;YP_009555241.1;HCoV-OC43_spike;292 | 0.05 | 0 |
| NC_006213.1;YP_009555241.1;HCoV-OC43_spike;293 | 0.05 | 0 |
| NC_006213.1;YP_009555241.1;HCoV-OC43_spike;294 | 0 | 0 |
| NC_006213.1;YP_009555241.1;HCoV-OC43_spike;295 | 0 | 0 |
| NC_006213.1;YP_009555241.1;HCoV-OC43_spike;296 | 0 | 0.025 |
| NC_006213.1;YP_009555241.1;HCoV-OC43_spike;297 | 0 | 0 |
| NC_006213.1;YP_009555241.1;HCoV-OC43_spike;298 | 0 | 0 |
| NC_006213.1;YP_009555241.1;HCoV-OC43_spike;299 | 0 | 0.025 |
| NC_006213.1;YP_009555241.1;HCoV-OC43_spike;300 | 0 | 0 |
| NC_006213.1;YP_009555241.1;HCoV-OC43_spike;301 | 0.05 | 0 |
| NC_006213.1;YP_009555241.1;HCoV-OC43_spike;302 | 0 | 0 |
| NC_006213.1;YP_009555241.1;HCoV-OC43_spike;303 | 0 | 0 |
| NC_006213.1;YP_009555241.1;HCoV-OC43_spike;304 | 0 | 0 |
| NC_006213.1;YP_009555241.1;HCoV-OC43_spike;305 | 0 | 0 |
| NC_006213.1;YP_009555241.1;HCoV-OC43_spike;306 | 0.1 | 0 |
| NC_006213.1;YP_009555241.1;HCoV-OC43_spike;307 | 0.05 | 0 |
| NC_006213.1;YP_009555241.1;HCoV-OC43_spike;308 | 0.05 | 0 |
| NC_006213.1;YP_009555241.1;HCoV-OC43_spike;309 | 0.05 | 0 |
| NC_006213.1;YP_009555241.1;HCoV-OC43_spike;310 | 0.05 | 0 |
| NC_006213.1;YP_009555241.1;HCoV-OC43_spike;311 | 0.05 | 0 |
| NC_006213.1;YP_009555241.1;HCoV-OC43_spike;312 | 0 | 0 |
| NC_006213.1;YP_009555241.1;HCoV-OC43_spike;313 | 0 | 0 |
| NC_006213.1;YP_009555241.1;HCoV-OC43_spike;314 | 0 | 0 |
| NC_006213.1;YP_009555241.1;HCoV-OC43_spike;315 | 0 | 0 |
| NC_006213.1;YP_009555241.1;HCoV-OC43_spike;316 | 0 | 0 |
| NC_006213.1;YP_009555241.1;HCoV-OC43_spike;317 | 0 | 0 |
| NC_006213.1;YP_009555241.1;HCoV-OC43_spike;318 | 0 | 0 |
| NC_006213.1;YP_009555241.1;HCoV-OC43_spike;319 | 0 | 0 |
| NC_006213.1;YP_009555241.1;HCoV-OC43_spike;320 | 0.05 | 0 |
| NC_006213.1;YP_009555241.1;HCoV-OC43_spike;321 | 0 | 0 |
| NC_006213.1;YP_009555241.1;HCoV-OC43_spike;322 | 0 | 0 |
| NC_006213.1;YP_009555241.1;HCoV-OC43_spike;323 | 0 | 0.05 |
| NC_006213.1;YP_009555241.1;HCoV-OC43_spike;324 | 0 | 0.05 |
| NC_006213.1;YP_009555241.1;HCoV-OC43_spike;325 | 0 | 0.025 |
| NC_006213.1;YP_009555241.1;HCoV-OC43_spike;326 | 0 | 0 |
| NC_006213.1;YP_009555241.1;HCoV-OC43_spike;327 | 0 | 0.025 |
| NC_006213.1;YP_009555241.1;HCoV-OC43_spike;328 | 0 | 0.075 |
| NC_006213.1;YP_009555241.1;HCoV-OC43_spike;329 | 0.15 | 0 |

| | | |
|---|---|---|
| NC_006213.1;YP_009555241.1;HCoV-OC43_spike;330 | 0.1 | 0 |
| NC_006213.1;YP_009555241.1;HCoV-OC43_spike;331 | 0 | 0 |
| NC_006213.1;YP_009555241.1;HCoV-OC43_spike;332 | 0.2 | 0.025 |
| NC_006213.1;YP_009555241.1;HCoV-OC43_spike;333 | 0.1 | 0 |
| NC_006213.1;YP_009555241.1;HCoV-OC43_spike;334 | 0.15 | 0 |
| NC_006213.1;YP_009555241.1;HCoV-OC43_spike;335 | 0.15 | 0.05 |
| NC_006213.1;YP_009555241.1;HCoV-OC43_spike;336 | 0.25 | 0.1 |
| NC_006213.1;YP_009555241.1;HCoV-OC43_spike;337 | 0.05 | 0 |
| NC_006213.1;YP_009555241.1;HCoV-OC43_spike;338 | 0.05 | 0.05 |
| NC_006213.1;YP_009555241.1;HCoV-OC43_spike;339 | 0 | 0.05 |
| NC_006213.1;YP_009555241.1;HCoV-OC43_spike;340 | 0.05 | 0.025 |
| NC_006213.1;YP_009555241.1;HCoV-OC43_spike;341 | 0.05 | 0 |
| NC_006213.1;YP_009555241.1;HCoV-OC43_spike;342 | 0.1 | 0 |
| NC_006213.1;YP_009555241.1;HCoV-OC43_spike;343 | 0.05 | 0 |
| NC_006213.1;YP_009555241.1;HCoV-OC43_spike;344 | 0 | 0 |
| NC_006213.1;YP_009555241.1;HCoV-OC43_spike;345 | 0.05 | 0 |
| NC_006213.1;YP_009555241.1;HCoV-OC43_spike;346 | 0.1 | 0 |
| NC_006213.1;YP_009555241.1;HCoV-OC43_spike;347 | 0 | 0 |
| NC_006213.1;YP_009555241.1;HCoV-OC43_spike;348 | 0.05 | 0 |
| NC_006213.1;YP_009555241.1;HCoV-OC43_spike;349 | 0.1 | 0 |
| NC_006213.1;YP_009555241.1;HCoV-OC43_spike;350 | 0 | 0 |
| NC_006213.1;YP_009555241.1;HCoV-OC43_spike;351 | 0 | 0 |
| NC_006213.1;YP_009555241.1;HCoV-OC43_spike;352 | 0 | 0 |
| NC_006213.1;YP_009555241.1;HCoV-OC43_spike;353 | 0.05 | 0 |
| NC_006213.1;YP_009555241.1;HCoV-OC43_spike;354 | 0 | 0 |
| NC_006213.1;YP_009555241.1;HCoV-OC43_spike;355 | 0 | 0 |
| NC_006213.1;YP_009555241.1;HCoV-OC43_spike;356 | 0 | 0 |
| NC_006213.1;YP_009555241.1;HCoV-OC43_spike;357 | 0 | 0.025 |
| NC_006213.1;YP_009555241.1;HCoV-OC43_spike;358 | 0 | 0 |
| NC_006213.1;YP_009555241.1;HCoV-OC43_spike;359 | 0 | 0 |
| NC_006213.1;YP_009555241.1;HCoV-OC43_spike;360 | 0.05 | 0 |
| NC_006213.1;YP_009555241.1;HCoV-OC43_spike;361 | 0 | 0 |
| NC_006213.1;YP_009555241.1;HCoV-OC43_spike;362 | 0 | 0 |
| NC_006213.1;YP_009555241.1;HCoV-OC43_spike;363 | 0 | 0 |
| NC_006213.1;YP_009555241.1;HCoV-OC43_spike;364 | 0 | 0 |
| NC_006213.1;YP_009555241.1;HCoV-OC43_spike;365 | 0 | 0 |
| NC_006213.1;YP_009555241.1;HCoV-OC43_spike;366 | 0 | 0 |
| NC_006213.1;YP_009555241.1;HCoV-OC43_spike;367 | 0 | 0 |
| NC_006213.1;YP_009555241.1;HCoV-OC43_spike;368 | 0 | 0 |
| NC_006213.1;YP_009555241.1;HCoV-OC43_spike;369 | 0.05 | 0.025 |
| NC_006213.1;YP_009555241.1;HCoV-OC43_spike;370 | 0 | 0 |
| NC_006213.1;YP_009555241.1;HCoV-OC43_spike;371 | 0 | 0.025 |
| NC_006213.1;YP_009555241.1;HCoV-OC43_spike;372 | 0 | 0.05 |

| | | |
|---|---|---|
| NC_006213.1;YP_009555241.1;HCoV-OC43_spike;373 | 0.05 | 0.125 |
| NC_006213.1;YP_009555241.1;HCoV-OC43_spike;374 | 0.15 | 0.075 |
| NC_006213.1;YP_009555241.1;HCoV-OC43_spike;375 | 0.05 | 0.05 |
| NC_006213.1;YP_009555241.1;HCoV-OC43_spike;376 | 0.05 | 0.05 |
| NC_006213.1;YP_009555241.1;HCoV-OC43_spike;377 | 0.05 | 0.05 |
| NC_006213.1;YP_009555241.1;HCoV-OC43_spike;378 | 0 | 0.025 |
| NC_006213.1;YP_009555241.1;HCoV-OC43_spike;379 | 0 | 0 |
| NC_006213.1;YP_009555241.1;HCoV-OC43_spike;380 | 0 | 0 |
| NC_006213.1;YP_009555241.1;HCoV-OC43_spike;381 | 0 | 0 |
| NC_006213.1;YP_009555241.1;HCoV-OC43_spike;382 | 0 | 0 |
| NC_006213.1;YP_009555241.1;HCoV-OC43_spike;383 | 0 | 0 |
| NC_006213.1;YP_009555241.1;HCoV-OC43_spike;384 | 0 | 0.025 |
| NC_006213.1;YP_009555241.1;HCoV-OC43_spike;385 | 0 | 0 |
| NC_006213.1;YP_009555241.1;HCoV-OC43_spike;386 | 0 | 0 |
| NC_006213.1;YP_009555241.1;HCoV-OC43_spike;387 | 0 | 0 |
| NC_006213.1;YP_009555241.1;HCoV-OC43_spike;388 | 0 | 0 |
| NC_006213.1;YP_009555241.1;HCoV-OC43_spike;389 | 0 | 0 |
| NC_006213.1;YP_009555241.1;HCoV-OC43_spike;390 | 0 | 0 |
| NC_006213.1;YP_009555241.1;HCoV-OC43_spike;391 | 0 | 0 |
| NC_006213.1;YP_009555241.1;HCoV-OC43_spike;392 | 0 | 0.025 |
| NC_006213.1;YP_009555241.1;HCoV-OC43_spike;393 | 0 | 0 |
| NC_006213.1;YP_009555241.1;HCoV-OC43_spike;394 | 0.05 | 0.325 |
| NC_006213.1;YP_009555241.1;HCoV-OC43_spike;395 | 0 | 0.025 |
| NC_006213.1;YP_009555241.1;HCoV-OC43_spike;396 | 0 | 0.025 |
| NC_006213.1;YP_009555241.1;HCoV-OC43_spike;397 | 0 | 0.025 |
| NC_006213.1;YP_009555241.1;HCoV-OC43_spike;398 | 0 | 0.025 |
| NC_006213.1;YP_009555241.1;HCoV-OC43_spike;399 | 0 | 0.075 |
| NC_006213.1;YP_009555241.1;HCoV-OC43_spike;400 | 0.05 | 0.075 |
| NC_006213.1;YP_009555241.1;HCoV-OC43_spike;401 | 0 | 0.075 |
| NC_006213.1;YP_009555241.1;HCoV-OC43_spike;402 | 0 | 0 |
| NC_006213.1;YP_009555241.1;HCoV-OC43_spike;403 | 0 | 0.05 |
| NC_006213.1;YP_009555241.1;HCoV-OC43_spike;404 | 0 | 0.025 |
| NC_006213.1;YP_009555241.1;HCoV-OC43_spike;405 | 0 | 0 |
| NC_006213.1;YP_009555241.1;HCoV-OC43_spike;406 | 0 | 0.05 |
| NC_006213.1;YP_009555241.1;HCoV-OC43_spike;407 | 0 | 0 |
| NC_006213.1;YP_009555241.1;HCoV-OC43_spike;408 | 0 | 0 |
| NC_006213.1;YP_009555241.1;HCoV-OC43_spike;409 | 0 | 0 |
| NC_006213.1;YP_009555241.1;HCoV-OC43_spike;410 | 0 | 0 |
| NC_006213.1;YP_009555241.1;HCoV-OC43_spike;411 | 0 | 0 |
| NC_006213.1;YP_009555241.1;HCoV-OC43_spike;412 | 0 | 0 |
| NC_006213.1;YP_009555241.1;HCoV-OC43_spike;413 | 0 | 0 |
| NC_006213.1;YP_009555241.1;HCoV-OC43_spike;414 | 0 | 0 |
| NC_006213.1;YP_009555241.1;HCoV-OC43_spike;415 | 0 | 0 |

| | | |
|---|---|---|
| NC_006213.1;YP_009555241.1;HCoV-OC43_spike;416 | 0 | 0 |
| NC_006213.1;YP_009555241.1;HCoV-OC43_spike;417 | 0 | 0.025 |
| NC_006213.1;YP_009555241.1;HCoV-OC43_spike;418 | 0 | 0 |
| NC_006213.1;YP_009555241.1;HCoV-OC43_spike;419 | 0 | 0 |
| NC_006213.1;YP_009555241.1;HCoV-OC43_spike;420 | 0 | 0 |
| NC_006213.1;YP_009555241.1;HCoV-OC43_spike;421 | 0 | 0 |
| NC_006213.1;YP_009555241.1;HCoV-OC43_spike;422 | 0 | 0 |
| NC_006213.1;YP_009555241.1;HCoV-OC43_spike;423 | 0 | 0 |
| NC_006213.1;YP_009555241.1;HCoV-OC43_spike;424 | 0 | 0 |
| NC_006213.1;YP_009555241.1;HCoV-OC43_spike;425 | 0.1 | 0.025 |
| NC_006213.1;YP_009555241.1;HCoV-OC43_spike;426 | 0.05 | 0 |
| NC_006213.1;YP_009555241.1;HCoV-OC43_spike;427 | 0 | 0 |
| NC_006213.1;YP_009555241.1;HCoV-OC43_spike;428 | 0 | 0.05 |
| NC_006213.1;YP_009555241.1;HCoV-OC43_spike;429 | 0 | 0 |
| NC_006213.1;YP_009555241.1;HCoV-OC43_spike;430 | 0 | 0 |
| NC_006213.1;YP_009555241.1;HCoV-OC43_spike;431 | 0 | 0 |
| NC_006213.1;YP_009555241.1;HCoV-OC43_spike;432 | 0 | 0 |
| NC_006213.1;YP_009555241.1;HCoV-OC43_spike;433 | 0 | 0 |
| NC_006213.1;YP_009555241.1;HCoV-OC43_spike;434 | 0 | 0 |
| NC_006213.1;YP_009555241.1;HCoV-OC43_spike;435 | 0 | 0 |
| NC_006213.1;YP_009555241.1;HCoV-OC43_spike;436 | 0 | 0 |
| NC_006213.1;YP_009555241.1;HCoV-OC43_spike;437 | 0 | 0 |
| NC_006213.1;YP_009555241.1;HCoV-OC43_spike;438 | 0 | 0 |
| NC_006213.1;YP_009555241.1;HCoV-OC43_spike;439 | 0 | 0 |
| NC_006213.1;YP_009555241.1;HCoV-OC43_spike;440 | 0 | 0 |
| NC_006213.1;YP_009555241.1;HCoV-OC43_spike;441 | 0 | 0 |
| NC_006213.1;YP_009555241.1;HCoV-OC43_spike;442 | 0 | 0 |
| NC_006213.1;YP_009555241.1;HCoV-OC43_spike;443 | 0 | 0.025 |
| NC_006213.1;YP_009555241.1;HCoV-OC43_spike;444 | 0 | 0 |
| NC_006213.1;YP_009555241.1;HCoV-OC43_spike;445 | 0 | 0 |
| NC_006213.1;YP_009555241.1;HCoV-OC43_spike;446 | 0 | 0.05 |
| NC_006213.1;YP_009555241.1;HCoV-OC43_spike;447 | 0.05 | 0.075 |
| NC_006213.1;YP_009555241.1;HCoV-OC43_spike;448 | 0 | 0.075 |
| NC_006213.1;YP_009555241.1;HCoV-OC43_spike;449 | 0.05 | 0.05 |
| NC_006213.1;YP_009555241.1;HCoV-OC43_spike;450 | 0.1 | 0.025 |
| NC_006213.1;YP_009555241.1;HCoV-OC43_spike;451 | 0.1 | 0.05 |
| NC_006213.1;YP_009555241.1;HCoV-OC43_spike;452 | 0.1 | 0.025 |
| NC_006213.1;YP_009555241.1;HCoV-OC43_spike;453 | 0.1 | 0.05 |
| NC_006213.1;YP_009555241.1;HCoV-OC43_spike;454 | 0.1 | 0.025 |
| NC_006213.1;YP_009555241.1;HCoV-OC43_spike;455 | 0.15 | 0.05 |
| NC_006213.1;YP_009555241.1;HCoV-OC43_spike;456 | 0.15 | 0 |
| NC_006213.1;YP_009555241.1;HCoV-OC43_spike;457 | 0.2 | 0.05 |
| NC_006213.1;YP_009555241.1;HCoV-OC43_spike;458 | 0.15 | 0 |

| | | |
|---|---|---|
| NC_006213.1;YP_009555241.1;HCoV-OC43_spike;459 | 0.15 | 0.05 |
| NC_006213.1;YP_009555241.1;HCoV-OC43_spike;460 | 0 | 0 |
| NC_006213.1;YP_009555241.1;HCoV-OC43_spike;461 | 0 | 0 |
| NC_006213.1;YP_009555241.1;HCoV-OC43_spike;462 | 0 | 0 |
| NC_006213.1;YP_009555241.1;HCoV-OC43_spike;463 | 0 | 0 |
| NC_006213.1;YP_009555241.1;HCoV-OC43_spike;464 | 0 | 0 |
| NC_006213.1;YP_009555241.1;HCoV-OC43_spike;465 | 0.05 | 0 |
| NC_006213.1;YP_009555241.1;HCoV-OC43_spike;466 | 0 | 0 |
| NC_006213.1;YP_009555241.1;HCoV-OC43_spike;467 | 0 | 0 |
| NC_006213.1;YP_009555241.1;HCoV-OC43_spike;468 | 0 | 0 |
| NC_006213.1;YP_009555241.1;HCoV-OC43_spike;469 | 0 | 0 |
| NC_006213.1;YP_009555241.1;HCoV-OC43_spike;470 | 0.05 | 0 |
| NC_006213.1;YP_009555241.1;HCoV-OC43_spike;471 | 0.05 | 0 |
| NC_006213.1;YP_009555241.1;HCoV-OC43_spike;472 | 0.05 | 0.025 |
| NC_006213.1;YP_009555241.1;HCoV-OC43_spike;473 | 0.05 | 0 |
| NC_006213.1;YP_009555241.1;HCoV-OC43_spike;474 | 0.05 | 0 |
| NC_006213.1;YP_009555241.1;HCoV-OC43_spike;475 | 0.05 | 0 |
| NC_006213.1;YP_009555241.1;HCoV-OC43_spike;476 | 0.05 | 0 |
| NC_006213.1;YP_009555241.1;HCoV-OC43_spike;477 | 0.05 | 0 |
| NC_006213.1;YP_009555241.1;HCoV-OC43_spike;478 | 0.05 | 0 |
| NC_006213.1;YP_009555241.1;HCoV-OC43_spike;479 | 0.05 | 0 |
| NC_006213.1;YP_009555241.1;HCoV-OC43_spike;480 | 0 | 0 |
| NC_006213.1;YP_009555241.1;HCoV-OC43_spike;481 | 0 | 0 |
| NC_006213.1;YP_009555241.1;HCoV-OC43_spike;482 | 0 | 0 |
| NC_006213.1;YP_009555241.1;HCoV-OC43_spike;483 | 0 | 0 |
| NC_006213.1;YP_009555241.1;HCoV-OC43_spike;484 | 0 | 0 |
| NC_006213.1;YP_009555241.1;HCoV-OC43_spike;485 | 0 | 0.025 |
| NC_006213.1;YP_009555241.1;HCoV-OC43_spike;486 | 0 | 0 |
| NC_006213.1;YP_009555241.1;HCoV-OC43_spike;487 | 0 | 0.025 |
| NC_006213.1;YP_009555241.1;HCoV-OC43_spike;488 | 0 | 0 |
| NC_006213.1;YP_009555241.1;HCoV-OC43_spike;489 | 0 | 0 |
| NC_006213.1;YP_009555241.1;HCoV-OC43_spike;490 | 0 | 0 |
| NC_006213.1;YP_009555241.1;HCoV-OC43_spike;491 | 0.05 | 0.025 |
| NC_006213.1;YP_009555241.1;HCoV-OC43_spike;492 | 0 | 0 |
| NC_006213.1;YP_009555241.1;HCoV-OC43_spike;493 | 0.05 | 0 |
| NC_006213.1;YP_009555241.1;HCoV-OC43_spike;494 | 0 | 0 |
| NC_006213.1;YP_009555241.1;HCoV-OC43_spike;495 | 0.05 | 0 |
| NC_006213.1;YP_009555241.1;HCoV-OC43_spike;496 | 0.05 | 0 |
| NC_006213.1;YP_009555241.1;HCoV-OC43_spike;497 | 0.05 | 0.025 |
| NC_006213.1;YP_009555241.1;HCoV-OC43_spike;498 | 0.05 | 0 |
| NC_006213.1;YP_009555241.1;HCoV-OC43_spike;499 | 0.05 | 0 |
| NC_006213.1;YP_009555241.1;HCoV-OC43_spike;500 | 0.05 | 0 |
| NC_006213.1;YP_009555241.1;HCoV-OC43_spike;501 | 0.05 | 0 |

| | | |
|---|---|---|
| NC_006213.1;YP_009555241.1;HCoV-OC43_spike;502 | 0.1 | 0 |
| NC_006213.1;YP_009555241.1;HCoV-OC43_spike;503 | 0.05 | 0 |
| NC_006213.1;YP_009555241.1;HCoV-OC43_spike;504 | 0.05 | 0 |
| NC_006213.1;YP_009555241.1;HCoV-OC43_spike;505 | 0.05 | 0 |
| NC_006213.1;YP_009555241.1;HCoV-OC43_spike;506 | 0.05 | 0 |
| NC_006213.1;YP_009555241.1;HCoV-OC43_spike;507 | 0 | 0 |
| NC_006213.1;YP_009555241.1;HCoV-OC43_spike;508 | 0 | 0 |
| NC_006213.1;YP_009555241.1;HCoV-OC43_spike;509 | 0 | 0.025 |
| NC_006213.1;YP_009555241.1;HCoV-OC43_spike;510 | 0 | 0 |
| NC_006213.1;YP_009555241.1;HCoV-OC43_spike;511 | 0 | 0 |
| NC_006213.1;YP_009555241.1;HCoV-OC43_spike;512 | 0 | 0 |
| NC_006213.1;YP_009555241.1;HCoV-OC43_spike;513 | 0 | 0 |
| NC_006213.1;YP_009555241.1;HCoV-OC43_spike;514 | 0 | 0 |
| NC_006213.1;YP_009555241.1;HCoV-OC43_spike;515 | 0 | 0.025 |
| NC_006213.1;YP_009555241.1;HCoV-OC43_spike;516 | 0 | 0.025 |
| NC_006213.1;YP_009555241.1;HCoV-OC43_spike;517 | 0 | 0 |
| NC_006213.1;YP_009555241.1;HCoV-OC43_spike;518 | 0 | 0 |
| NC_006213.1;YP_009555241.1;HCoV-OC43_spike;519 | 0 | 0 |
| NC_006213.1;YP_009555241.1;HCoV-OC43_spike;520 | 0 | 0 |
| NC_006213.1;YP_009555241.1;HCoV-OC43_spike;521 | 0 | 0 |
| NC_006213.1;YP_009555241.1;HCoV-OC43_spike;522 | 0 | 0 |
| NC_006213.1;YP_009555241.1;HCoV-OC43_spike;523 | 0 | 0 |
| NC_006213.1;YP_009555241.1;HCoV-OC43_spike;524 | 0 | 0 |
| NC_006213.1;YP_009555241.1;HCoV-OC43_spike;525 | 0 | 0 |
| NC_006213.1;YP_009555241.1;HCoV-OC43_spike;526 | 0 | 0 |
| NC_006213.1;YP_009555241.1;HCoV-OC43_spike;527 | 0 | 0 |
| NC_006213.1;YP_009555241.1;HCoV-OC43_spike;528 | 0 | 0 |
| NC_006213.1;YP_009555241.1;HCoV-OC43_spike;529 | 0 | 0 |
| NC_006213.1;YP_009555241.1;HCoV-OC43_spike;530 | 0 | 0 |
| NC_006213.1;YP_009555241.1;HCoV-OC43_spike;531 | 0 | 0 |
| NC_006213.1;YP_009555241.1;HCoV-OC43_spike;532 | 0 | 0 |
| NC_006213.1;YP_009555241.1;HCoV-OC43_spike;533 | 0 | 0 |
| NC_006213.1;YP_009555241.1;HCoV-OC43_spike;534 | 0 | 0 |
| NC_006213.1;YP_009555241.1;HCoV-OC43_spike;535 | 0.05 | 0 |
| NC_006213.1;YP_009555241.1;HCoV-OC43_spike;536 | 0 | 0 |
| NC_006213.1;YP_009555241.1;HCoV-OC43_spike;537 | 0 | 0 |
| NC_006213.1;YP_009555241.1;HCoV-OC43_spike;538 | 0 | 0 |
| NC_006213.1;YP_009555241.1;HCoV-OC43_spike;539 | 0 | 0 |
| NC_006213.1;YP_009555241.1;HCoV-OC43_spike;540 | 0 | 0 |
| NC_006213.1;YP_009555241.1;HCoV-OC43_spike;541 | 0 | 0 |
| NC_006213.1;YP_009555241.1;HCoV-OC43_spike;542 | 0 | 0 |
| NC_006213.1;YP_009555241.1;HCoV-OC43_spike;543 | 0 | 0 |
| NC_006213.1;YP_009555241.1;HCoV-OC43_spike;544 | 0 | 0 |

| | | |
|---|---|---|
| NC_006213.1;YP_009555241.1;HCoV-OC43_spike;545 | 0 | 0 |
| NC_006213.1;YP_009555241.1;HCoV-OC43_spike;546 | 0 | 0 |
| NC_006213.1;YP_009555241.1;HCoV-OC43_spike;547 | 0 | 0 |
| NC_006213.1;YP_009555241.1;HCoV-OC43_spike;548 | 0 | 0.025 |
| NC_006213.1;YP_009555241.1;HCoV-OC43_spike;549 | 0 | 0 |
| NC_006213.1;YP_009555241.1;HCoV-OC43_spike;550 | 0 | 0 |
| NC_006213.1;YP_009555241.1;HCoV-OC43_spike;551 | 0 | 0 |
| NC_006213.1;YP_009555241.1;HCoV-OC43_spike;552 | 0 | 0.025 |
| NC_006213.1;YP_009555241.1;HCoV-OC43_spike;553 | 0 | 0 |
| NC_006213.1;YP_009555241.1;HCoV-OC43_spike;554 | 0 | 0 |
| NC_006213.1;YP_009555241.1;HCoV-OC43_spike;555 | 0.05 | 0 |
| NC_006213.1;YP_009555241.1;HCoV-OC43_spike;556 | 0 | 0.025 |
| NC_006213.1;YP_009555241.1;HCoV-OC43_spike;557 | 0 | 0 |
| NC_006213.1;YP_009555241.1;HCoV-OC43_spike;558 | 0 | 0 |
| NC_006213.1;YP_009555241.1;HCoV-OC43_spike;559 | 0 | 0 |
| NC_006213.1;YP_009555241.1;HCoV-OC43_spike;560 | 0 | 0 |
| NC_006213.1;YP_009555241.1;HCoV-OC43_spike;561 | 0 | 0 |
| NC_006213.1;YP_009555241.1;HCoV-OC43_spike;562 | 0 | 0 |
| NC_006213.1;YP_009555241.1;HCoV-OC43_spike;563 | 0 | 0 |
| NC_006213.1;YP_009555241.1;HCoV-OC43_spike;564 | 0 | 0 |
| NC_006213.1;YP_009555241.1;HCoV-OC43_spike;565 | 0 | 0 |
| NC_006213.1;YP_009555241.1;HCoV-OC43_spike;566 | 0.05 | 0 |
| NC_006213.1;YP_009555241.1;HCoV-OC43_spike;567 | 0 | 0 |
| NC_006213.1;YP_009555241.1;HCoV-OC43_spike;568 | 0 | 0 |
| NC_006213.1;YP_009555241.1;HCoV-OC43_spike;569 | 0 | 0 |
| NC_006213.1;YP_009555241.1;HCoV-OC43_spike;570 | 0 | 0 |
| NC_006213.1;YP_009555241.1;HCoV-OC43_spike;571 | 0 | 0 |
| NC_006213.1;YP_009555241.1;HCoV-OC43_spike;572 | 0 | 0 |
| NC_006213.1;YP_009555241.1;HCoV-OC43_spike;573 | 0 | 0 |
| NC_006213.1;YP_009555241.1;HCoV-OC43_spike;574 | 0 | 0 |
| NC_006213.1;YP_009555241.1;HCoV-OC43_spike;575 | 0 | 0 |
| NC_006213.1;YP_009555241.1;HCoV-OC43_spike;576 | 0 | 0 |
| NC_006213.1;YP_009555241.1;HCoV-OC43_spike;577 | 0 | 0.025 |
| NC_006213.1;YP_009555241.1;HCoV-OC43_spike;578 | 0 | 0.025 |
| NC_006213.1;YP_009555241.1;HCoV-OC43_spike;579 | 0 | 0.025 |
| NC_006213.1;YP_009555241.1;HCoV-OC43_spike;580 | 0 | 0 |
| NC_006213.1;YP_009555241.1;HCoV-OC43_spike;581 | 0 | 0 |
| NC_006213.1;YP_009555241.1;HCoV-OC43_spike;582 | 0 | 0 |
| NC_006213.1;YP_009555241.1;HCoV-OC43_spike;583 | 0 | 0 |
| NC_006213.1;YP_009555241.1;HCoV-OC43_spike;584 | 0 | 0.025 |
| NC_006213.1;YP_009555241.1;HCoV-OC43_spike;585 | 0 | 0 |
| NC_006213.1;YP_009555241.1;HCoV-OC43_spike;586 | 0 | 0 |
| NC_006213.1;YP_009555241.1;HCoV-OC43_spike;587 | 0.05 | 0 |

| | | |
|---|---|---|
| NC_006213.1;YP_009555241.1;HCoV-OC43_spike;588 | 0 | 0 |
| NC_006213.1;YP_009555241.1;HCoV-OC43_spike;589 | 0 | 0 |
| NC_006213.1;YP_009555241.1;HCoV-OC43_spike;590 | 0 | 0 |
| NC_006213.1;YP_009555241.1;HCoV-OC43_spike;591 | 0.05 | 0.05 |
| NC_006213.1;YP_009555241.1;HCoV-OC43_spike;592 | 0.05 | 0 |
| NC_006213.1;YP_009555241.1;HCoV-OC43_spike;593 | 0 | 0 |
| NC_006213.1;YP_009555241.1;HCoV-OC43_spike;594 | 0 | 0 |
| NC_006213.1;YP_009555241.1;HCoV-OC43_spike;595 | 0.1 | 0 |
| NC_006213.1;YP_009555241.1;HCoV-OC43_spike;596 | 0.05 | 0 |
| NC_006213.1;YP_009555241.1;HCoV-OC43_spike;597 | 0.05 | 0 |
| NC_006213.1;YP_009555241.1;HCoV-OC43_spike;598 | 0.05 | 0 |
| NC_006213.1;YP_009555241.1;HCoV-OC43_spike;599 | 0.05 | 0 |
| NC_006213.1;YP_009555241.1;HCoV-OC43_spike;600 | 0.05 | 0 |
| NC_006213.1;YP_009555241.1;HCoV-OC43_spike;601 | 0.05 | 0 |
| NC_006213.1;YP_009555241.1;HCoV-OC43_spike;602 | 0.05 | 0 |
| NC_006213.1;YP_009555241.1;HCoV-OC43_spike;603 | 0 | 0 |
| NC_006213.1;YP_009555241.1;HCoV-OC43_spike;604 | 0 | 0 |
| NC_006213.1;YP_009555241.1;HCoV-OC43_spike;605 | 0 | 0 |
| NC_006213.1;YP_009555241.1;HCoV-OC43_spike;606 | 0 | 0 |
| NC_006213.1;YP_009555241.1;HCoV-OC43_spike;607 | 0 | 0 |
| NC_006213.1;YP_009555241.1;HCoV-OC43_spike;608 | 0 | 0 |
| NC_006213.1;YP_009555241.1;HCoV-OC43_spike;609 | 0 | 0 |
| NC_006213.1;YP_009555241.1;HCoV-OC43_spike;610 | 0 | 0 |
| NC_006213.1;YP_009555241.1;HCoV-OC43_spike;611 | 0 | 0 |
| NC_006213.1;YP_009555241.1;HCoV-OC43_spike;612 | 0 | 0 |
| NC_006213.1;YP_009555241.1;HCoV-OC43_spike;613 | 0 | 0 |
| NC_006213.1;YP_009555241.1;HCoV-OC43_spike;614 | 0 | 0 |
| NC_006213.1;YP_009555241.1;HCoV-OC43_spike;615 | 0 | 0 |
| NC_006213.1;YP_009555241.1;HCoV-OC43_spike;616 | 0 | 0 |
| NC_006213.1;YP_009555241.1;HCoV-OC43_spike;617 | 0 | 0 |
| NC_006213.1;YP_009555241.1;HCoV-OC43_spike;618 | 0 | 0 |
| NC_006213.1;YP_009555241.1;HCoV-OC43_spike;619 | 0 | 0 |
| NC_006213.1;YP_009555241.1;HCoV-OC43_spike;620 | 0 | 0 |
| NC_006213.1;YP_009555241.1;HCoV-OC43_spike;621 | 0 | 0 |
| NC_006213.1;YP_009555241.1;HCoV-OC43_spike;622 | 0 | 0 |
| NC_006213.1;YP_009555241.1;HCoV-OC43_spike;623 | 0 | 0 |
| NC_006213.1;YP_009555241.1;HCoV-OC43_spike;624 | 0 | 0 |
| NC_006213.1;YP_009555241.1;HCoV-OC43_spike;625 | 0 | 0 |
| NC_006213.1;YP_009555241.1;HCoV-OC43_spike;626 | 0 | 0 |
| NC_006213.1;YP_009555241.1;HCoV-OC43_spike;627 | 0 | 0 |
| NC_006213.1;YP_009555241.1;HCoV-OC43_spike;628 | 0 | 0 |
| NC_006213.1;YP_009555241.1;HCoV-OC43_spike;629 | 0 | 0 |
| NC_006213.1;YP_009555241.1;HCoV-OC43_spike;630 | 0 | 0 |

| | | |
|---|---|---|
| NC_006213.1;YP_009555241.1;HCoV-OC43_spike;631 | 0 | 0 |
| NC_006213.1;YP_009555241.1;HCoV-OC43_spike;632 | 0 | 0 |
| NC_006213.1;YP_009555241.1;HCoV-OC43_spike;633 | 0 | 0 |
| NC_006213.1;YP_009555241.1;HCoV-OC43_spike;634 | 0 | 0 |
| NC_006213.1;YP_009555241.1;HCoV-OC43_spike;635 | 0 | 0 |
| NC_006213.1;YP_009555241.1;HCoV-OC43_spike;636 | 0 | 0.05 |
| NC_006213.1;YP_009555241.1;HCoV-OC43_spike;637 | 0 | 0 |
| NC_006213.1;YP_009555241.1;HCoV-OC43_spike;638 | 0 | 0 |
| NC_006213.1;YP_009555241.1;HCoV-OC43_spike;639 | 0 | 0 |
| NC_006213.1;YP_009555241.1;HCoV-OC43_spike;640 | 0 | 0.025 |
| NC_006213.1;YP_009555241.1;HCoV-OC43_spike;641 | 0 | 0.025 |
| NC_006213.1;YP_009555241.1;HCoV-OC43_spike;642 | 0.1 | 0.05 |
| NC_006213.1;YP_009555241.1;HCoV-OC43_spike;643 | 0 | 0.1 |
| NC_006213.1;YP_009555241.1;HCoV-OC43_spike;644 | 0 | 0.075 |
| NC_006213.1;YP_009555241.1;HCoV-OC43_spike;645 | 0 | 0.025 |
| NC_006213.1;YP_009555241.1;HCoV-OC43_spike;646 | 0 | 0 |
| NC_006213.1;YP_009555241.1;HCoV-OC43_spike;647 | 0 | 0.05 |
| NC_006213.1;YP_009555241.1;HCoV-OC43_spike;648 | 0 | 0 |
| NC_006213.1;YP_009555241.1;HCoV-OC43_spike;649 | 0 | 0.025 |
| NC_006213.1;YP_009555241.1;HCoV-OC43_spike;650 | 0.05 | 0.075 |
| NC_006213.1;YP_009555241.1;HCoV-OC43_spike;651 | 0 | 0 |
| NC_006213.1;YP_009555241.1;HCoV-OC43_spike;652 | 0.05 | 0.025 |
| NC_006213.1;YP_009555241.1;HCoV-OC43_spike;653 | 0 | 0 |
| NC_006213.1;YP_009555241.1;HCoV-OC43_spike;654 | 0 | 0 |
| NC_006213.1;YP_009555241.1;HCoV-OC43_spike;655 | 0 | 0 |
| NC_006213.1;YP_009555241.1;HCoV-OC43_spike;656 | 0 | 0 |
| NC_006213.1;YP_009555241.1;HCoV-OC43_spike;657 | 0 | 0 |
| NC_006213.1;YP_009555241.1;HCoV-OC43_spike;658 | 0 | 0 |
| NC_006213.1;YP_009555241.1;HCoV-OC43_spike;659 | 0 | 0.05 |
| NC_006213.1;YP_009555241.1;HCoV-OC43_spike;660 | 0 | 0 |
| NC_006213.1;YP_009555241.1;HCoV-OC43_spike;661 | 0 | 0 |
| NC_006213.1;YP_009555241.1;HCoV-OC43_spike;662 | 0 | 0 |
| NC_006213.1;YP_009555241.1;HCoV-OC43_spike;663 | 0 | 0 |
| NC_006213.1;YP_009555241.1;HCoV-OC43_spike;664 | 0 | 0 |
| NC_006213.1;YP_009555241.1;HCoV-OC43_spike;665 | 0 | 0 |
| NC_006213.1;YP_009555241.1;HCoV-OC43_spike;666 | 0 | 0 |
| NC_006213.1;YP_009555241.1;HCoV-OC43_spike;667 | 0.05 | 0 |
| NC_006213.1;YP_009555241.1;HCoV-OC43_spike;668 | 0 | 0 |
| NC_006213.1;YP_009555241.1;HCoV-OC43_spike;669 | 0 | 0 |
| NC_006213.1;YP_009555241.1;HCoV-OC43_spike;670 | 0 | 0 |
| NC_006213.1;YP_009555241.1;HCoV-OC43_spike;671 | 0 | 0 |
| NC_006213.1;YP_009555241.1;HCoV-OC43_spike;672 | 0 | 0 |
| NC_006213.1;YP_009555241.1;HCoV-OC43_spike;673 | 0 | 0.025 |

| | | |
|---|---|---|
| NC_006213.1;YP_009555241.1;HCoV-OC43_spike;674 | 0 | 0.05 |
| NC_006213.1;YP_009555241.1;HCoV-OC43_spike;675 | 0 | 0.05 |
| NC_006213.1;YP_009555241.1;HCoV-OC43_spike;676 | 0 | 0.05 |
| NC_006213.1;YP_009555241.1;HCoV-OC43_spike;677 | 0 | 0.025 |
| NC_006213.1;YP_009555241.1;HCoV-OC43_spike;678 | 0 | 0.025 |
| NC_006213.1;YP_009555241.1;HCoV-OC43_spike;679 | 0 | 0.025 |
| NC_006213.1;YP_009555241.1;HCoV-OC43_spike;680 | 0 | 0.025 |
| NC_006213.1;YP_009555241.1;HCoV-OC43_spike;681 | 0 | 0.025 |
| NC_006213.1;YP_009555241.1;HCoV-OC43_spike;682 | 0 | 0.025 |
| NC_006213.1;YP_009555241.1;HCoV-OC43_spike;683 | 0 | 0.025 |
| NC_006213.1;YP_009555241.1;HCoV-OC43_spike;684 | 0 | 0.025 |
| NC_006213.1;YP_009555241.1;HCoV-OC43_spike;685 | 0 | 0.025 |
| NC_006213.1;YP_009555241.1;HCoV-OC43_spike;686 | 0 | 0 |
| NC_006213.1;YP_009555241.1;HCoV-OC43_spike;687 | 0 | 0 |
| NC_006213.1;YP_009555241.1;HCoV-OC43_spike;688 | 0 | 0 |
| NC_006213.1;YP_009555241.1;HCoV-OC43_spike;689 | 0 | 0 |
| NC_006213.1;YP_009555241.1;HCoV-OC43_spike;690 | 0 | 0 |
| NC_006213.1;YP_009555241.1;HCoV-OC43_spike;691 | 0 | 0 |
| NC_006213.1;YP_009555241.1;HCoV-OC43_spike;692 | 0 | 0 |
| NC_006213.1;YP_009555241.1;HCoV-OC43_spike;693 | 0 | 0 |
| NC_006213.1;YP_009555241.1;HCoV-OC43_spike;694 | 0 | 0 |
| NC_006213.1;YP_009555241.1;HCoV-OC43_spike;695 | 0 | 0 |
| NC_006213.1;YP_009555241.1;HCoV-OC43_spike;696 | 0 | 0 |
| NC_006213.1;YP_009555241.1;HCoV-OC43_spike;697 | 0 | 0 |
| NC_006213.1;YP_009555241.1;HCoV-OC43_spike;698 | 0 | 0 |
| NC_006213.1;YP_009555241.1;HCoV-OC43_spike;699 | 0 | 0 |
| NC_006213.1;YP_009555241.1;HCoV-OC43_spike;700 | 0 | 0 |
| NC_006213.1;YP_009555241.1;HCoV-OC43_spike;701 | 0 | 0 |
| NC_006213.1;YP_009555241.1;HCoV-OC43_spike;702 | 0 | 0 |
| NC_006213.1;YP_009555241.1;HCoV-OC43_spike;703 | 0.05 | 0.025 |
| NC_006213.1;YP_009555241.1;HCoV-OC43_spike;704 | 0.05 | 0.05 |
| NC_006213.1;YP_009555241.1;HCoV-OC43_spike;705 | 0 | 0 |
| NC_006213.1;YP_009555241.1;HCoV-OC43_spike;706 | 0 | 0 |
| NC_006213.1;YP_009555241.1;HCoV-OC43_spike;707 | 0 | 0 |
| NC_006213.1;YP_009555241.1;HCoV-OC43_spike;708 | 0 | 0 |
| NC_006213.1;YP_009555241.1;HCoV-OC43_spike;709 | 0 | 0 |
| NC_006213.1;YP_009555241.1;HCoV-OC43_spike;710 | 0 | 0 |
| NC_006213.1;YP_009555241.1;HCoV-OC43_spike;711 | 0 | 0 |
| NC_006213.1;YP_009555241.1;HCoV-OC43_spike;712 | 0 | 0 |
| NC_006213.1;YP_009555241.1;HCoV-OC43_spike;713 | 0 | 0 |
| NC_006213.1;YP_009555241.1;HCoV-OC43_spike;714 | 0.05 | 0 |
| NC_006213.1;YP_009555241.1;HCoV-OC43_spike;715 | 0 | 0 |
| NC_006213.1;YP_009555241.1;HCoV-OC43_spike;716 | 0 | 0 |

| | | |
|---|---|---|
| NC_006213.1;YP_009555241.1;HCoV-OC43_spike;717 | 0.05 | 0 |
| NC_006213.1;YP_009555241.1;HCoV-OC43_spike;718 | 0.05 | 0 |
| NC_006213.1;YP_009555241.1;HCoV-OC43_spike;719 | 0 | 0 |
| NC_006213.1;YP_009555241.1;HCoV-OC43_spike;720 | 0 | 0 |
| NC_006213.1;YP_009555241.1;HCoV-OC43_spike;721 | 0 | 0 |
| NC_006213.1;YP_009555241.1;HCoV-OC43_spike;722 | 0 | 0 |
| NC_006213.1;YP_009555241.1;HCoV-OC43_spike;723 | 0 | 0 |
| NC_006213.1;YP_009555241.1;HCoV-OC43_spike;724 | 0 | 0 |
| NC_006213.1;YP_009555241.1;HCoV-OC43_spike;725 | 0 | 0 |
| NC_006213.1;YP_009555241.1;HCoV-OC43_spike;726 | 0 | 0 |
| NC_006213.1;YP_009555241.1;HCoV-OC43_spike;727 | 0 | 0 |
| NC_006213.1;YP_009555241.1;HCoV-OC43_spike;728 | 0 | 0 |
| NC_006213.1;YP_009555241.1;HCoV-OC43_spike;729 | 0 | 0 |
| NC_006213.1;YP_009555241.1;HCoV-OC43_spike;730 | 0 | 0 |
| NC_006213.1;YP_009555241.1;HCoV-OC43_spike;731 | 0 | 0 |
| NC_006213.1;YP_009555241.1;HCoV-OC43_spike;732 | 0 | 0.025 |
| NC_006213.1;YP_009555241.1;HCoV-OC43_spike;733 | 0 | 0 |
| NC_006213.1;YP_009555241.1;HCoV-OC43_spike;734 | 0 | 0 |
| NC_006213.1;YP_009555241.1;HCoV-OC43_spike;735 | 0 | 0 |
| NC_006213.1;YP_009555241.1;HCoV-OC43_spike;736 | 0 | 0 |
| NC_006213.1;YP_009555241.1;HCoV-OC43_spike;737 | 0 | 0 |
| NC_006213.1;YP_009555241.1;HCoV-OC43_spike;738 | 0 | 0 |
| NC_006213.1;YP_009555241.1;HCoV-OC43_spike;739 | 0 | 0 |
| NC_006213.1;YP_009555241.1;HCoV-OC43_spike;740 | 0 | 0 |
| NC_006213.1;YP_009555241.1;HCoV-OC43_spike;741 | 0 | 0 |
| NC_006213.1;YP_009555241.1;HCoV-OC43_spike;742 | 0 | 0 |
| NC_006213.1;YP_009555241.1;HCoV-OC43_spike;743 | 0 | 0 |
| NC_006213.1;YP_009555241.1;HCoV-OC43_spike;744 | 0 | 0 |
| NC_006213.1;YP_009555241.1;HCoV-OC43_spike;745 | 0 | 0.025 |
| NC_006213.1;YP_009555241.1;HCoV-OC43_spike;746 | 0 | 0 |
| NC_006213.1;YP_009555241.1;HCoV-OC43_spike;747 | 0.05 | 0 |
| NC_006213.1;YP_009555241.1;HCoV-OC43_spike;748 | 0.1 | 0 |
| NC_006213.1;YP_009555241.1;HCoV-OC43_spike;749 | 0.1 | 0 |
| NC_006213.1;YP_009555241.1;HCoV-OC43_spike;750 | 0 | 0 |
| NC_006213.1;YP_009555241.1;HCoV-OC43_spike;751 | 0 | 0 |
| NC_006213.1;YP_009555241.1;HCoV-OC43_spike;752 | 0 | 0 |
| NC_006213.1;YP_009555241.1;HCoV-OC43_spike;753 | 0 | 0.025 |
| NC_006213.1;YP_009555241.1;HCoV-OC43_spike;754 | 0 | 0.025 |
| NC_006213.1;YP_009555241.1;HCoV-OC43_spike;755 | 0.2 | 0.125 |
| NC_006213.1;YP_009555241.1;HCoV-OC43_spike;756 | 0.55 | 0.3 |
| NC_006213.1;YP_009555241.1;HCoV-OC43_spike;757 | 0.4 | 0.175 |
| NC_006213.1;YP_009555241.1;HCoV-OC43_spike;758 | 0.5 | 0.275 |
| NC_006213.1;YP_009555241.1;HCoV-OC43_spike;759 | 0.45 | 0.2 |

| | | |
|---|---|---|
| NC_006213.1;YP_009555241.1;HCoV-OC43_spike;760 | 0.5 | 0.35 |
| NC_006213.1;YP_009555241.1;HCoV-OC43_spike;761 | 0.7 | 0.425 |
| NC_006213.1;YP_009555241.1;HCoV-OC43_spike;762 | 0.75 | 0.35 |
| NC_006213.1;YP_009555241.1;HCoV-OC43_spike;763 | 0.8 | 0.4 |
| NC_006213.1;YP_009555241.1;HCoV-OC43_spike;764 | 0.85 | 0.575 |
| NC_006213.1;YP_009555241.1;HCoV-OC43_spike;765 | 0.65 | 0.55 |
| NC_006213.1;YP_009555241.1;HCoV-OC43_spike;766 | 0.6 | 0.475 |
| NC_006213.1;YP_009555241.1;HCoV-OC43_spike;767 | 0.35 | 0.175 |
| NC_006213.1;YP_009555241.1;HCoV-OC43_spike;768 | 0.15 | 0.125 |
| NC_006213.1;YP_009555241.1;HCoV-OC43_spike;769 | 0.2 | 0.125 |
| NC_006213.1;YP_009555241.1;HCoV-OC43_spike;770 | 0.3 | 0.15 |
| NC_006213.1;YP_009555241.1;HCoV-OC43_spike;771 | 0.3 | 0.2 |
| NC_006213.1;YP_009555241.1;HCoV-OC43_spike;772 | 0.25 | 0.075 |
| NC_006213.1;YP_009555241.1;HCoV-OC43_spike;773 | 0.2 | 0.125 |
| NC_006213.1;YP_009555241.1;HCoV-OC43_spike;774 | 0.05 | 0.15 |
| NC_006213.1;YP_009555241.1;HCoV-OC43_spike;775 | 0 | 0.05 |
| NC_006213.1;YP_009555241.1;HCoV-OC43_spike;776 | 0.25 | 0.125 |
| NC_006213.1;YP_009555241.1;HCoV-OC43_spike;777 | 0 | 0 |
| NC_006213.1;YP_009555241.1;HCoV-OC43_spike;778 | 0 | 0 |
| NC_006213.1;YP_009555241.1;HCoV-OC43_spike;779 | 0 | 0 |
| NC_006213.1;YP_009555241.1;HCoV-OC43_spike;780 | 0 | 0 |
| NC_006213.1;YP_009555241.1;HCoV-OC43_spike;781 | 0 | 0 |
| NC_006213.1;YP_009555241.1;HCoV-OC43_spike;782 | 0 | 0 |
| NC_006213.1;YP_009555241.1;HCoV-OC43_spike;783 | 0 | 0 |
| NC_006213.1;YP_009555241.1;HCoV-OC43_spike;784 | 0 | 0 |
| NC_006213.1;YP_009555241.1;HCoV-OC43_spike;785 | 0 | 0 |
| NC_006213.1;YP_009555241.1;HCoV-OC43_spike;786 | 0 | 0.025 |
| NC_006213.1;YP_009555241.1;HCoV-OC43_spike;787 | 0 | 0 |
| NC_006213.1;YP_009555241.1;HCoV-OC43_spike;788 | 0 | 0 |
| NC_006213.1;YP_009555241.1;HCoV-OC43_spike;789 | 0 | 0 |
| NC_006213.1;YP_009555241.1;HCoV-OC43_spike;790 | 0 | 0 |
| NC_006213.1;YP_009555241.1;HCoV-OC43_spike;791 | 0 | 0 |
| NC_006213.1;YP_009555241.1;HCoV-OC43_spike;792 | 0 | 0 |
| NC_006213.1;YP_009555241.1;HCoV-OC43_spike;793 | 0 | 0 |
| NC_006213.1;YP_009555241.1;HCoV-OC43_spike;794 | 0 | 0 |
| NC_006213.1;YP_009555241.1;HCoV-OC43_spike;795 | 0 | 0 |
| NC_006213.1;YP_009555241.1;HCoV-OC43_spike;796 | 0 | 0 |
| NC_006213.1;YP_009555241.1;HCoV-OC43_spike;797 | 0.05 | 0 |
| NC_006213.1;YP_009555241.1;HCoV-OC43_spike;798 | 0.05 | 0.025 |
| NC_006213.1;YP_009555241.1;HCoV-OC43_spike;799 | 0.05 | 0.025 |
| NC_006213.1;YP_009555241.1;HCoV-OC43_spike;800 | 0 | 0.025 |
| NC_006213.1;YP_009555241.1;HCoV-OC43_spike;801 | 0 | 0 |
| NC_006213.1;YP_009555241.1;HCoV-OC43_spike;802 | 0 | 0 |

| | | |
|---|---|---|
| NC_006213.1;YP_009555241.1;HCoV-OC43_spike;803 | 0.05 | 0.025 |
| NC_006213.1;YP_009555241.1;HCoV-OC43_spike;804 | 0 | 0 |
| NC_006213.1;YP_009555241.1;HCoV-OC43_spike;805 | 0 | 0 |
| NC_006213.1;YP_009555241.1;HCoV-OC43_spike;806 | 0 | 0 |
| NC_006213.1;YP_009555241.1;HCoV-OC43_spike;807 | 0 | 0 |
| NC_006213.1;YP_009555241.1;HCoV-OC43_spike;808 | 0 | 0 |
| NC_006213.1;YP_009555241.1;HCoV-OC43_spike;809 | 0 | 0 |
| NC_006213.1;YP_009555241.1;HCoV-OC43_spike;810 | 0 | 0 |
| NC_006213.1;YP_009555241.1;HCoV-OC43_spike;811 | 0.05 | 0 |
| NC_006213.1;YP_009555241.1;HCoV-OC43_spike;812 | 0 | 0 |
| NC_006213.1;YP_009555241.1;HCoV-OC43_spike;813 | 0 | 0 |
| NC_006213.1;YP_009555241.1;HCoV-OC43_spike;814 | 0 | 0 |
| NC_006213.1;YP_009555241.1;HCoV-OC43_spike;815 | 0 | 0 |
| NC_006213.1;YP_009555241.1;HCoV-OC43_spike;816 | 0 | 0 |
| NC_006213.1;YP_009555241.1;HCoV-OC43_spike;817 | 0 | 0 |
| NC_006213.1;YP_009555241.1;HCoV-OC43_spike;818 | 0 | 0 |
| NC_006213.1;YP_009555241.1;HCoV-OC43_spike;819 | 0 | 0 |
| NC_006213.1;YP_009555241.1;HCoV-OC43_spike;820 | 0 | 0 |
| NC_006213.1;YP_009555241.1;HCoV-OC43_spike;821 | 0 | 0.025 |
| NC_006213.1;YP_009555241.1;HCoV-OC43_spike;822 | 0 | 0 |
| NC_006213.1;YP_009555241.1;HCoV-OC43_spike;823 | 0 | 0 |
| NC_006213.1;YP_009555241.1;HCoV-OC43_spike;824 | 0 | 0 |
| NC_006213.1;YP_009555241.1;HCoV-OC43_spike;825 | 0 | 0 |
| NC_006213.1;YP_009555241.1;HCoV-OC43_spike;826 | 0 | 0 |
| NC_006213.1;YP_009555241.1;HCoV-OC43_spike;827 | 0 | 0.025 |
| NC_006213.1;YP_009555241.1;HCoV-OC43_spike;828 | 0 | 0.025 |
| NC_006213.1;YP_009555241.1;HCoV-OC43_spike;829 | 0 | 0 |
| NC_006213.1;YP_009555241.1;HCoV-OC43_spike;830 | 0 | 0 |
| NC_006213.1;YP_009555241.1;HCoV-OC43_spike;831 | 0 | 0 |
| NC_006213.1;YP_009555241.1;HCoV-OC43_spike;832 | 0 | 0 |
| NC_006213.1;YP_009555241.1;HCoV-OC43_spike;833 | 0 | 0 |
| NC_006213.1;YP_009555241.1;HCoV-OC43_spike;834 | 0 | 0 |
| NC_006213.1;YP_009555241.1;HCoV-OC43_spike;835 | 0 | 0 |
| NC_006213.1;YP_009555241.1;HCoV-OC43_spike;836 | 0 | 0.025 |
| NC_006213.1;YP_009555241.1;HCoV-OC43_spike;837 | 0.05 | 0 |
| NC_006213.1;YP_009555241.1;HCoV-OC43_spike;838 | 0.05 | 0 |
| NC_006213.1;YP_009555241.1;HCoV-OC43_spike;839 | 0 | 0.025 |
| NC_006213.1;YP_009555241.1;HCoV-OC43_spike;840 | 0 | 0.025 |
| NC_006213.1;YP_009555241.1;HCoV-OC43_spike;841 | 0 | 0 |
| NC_006213.1;YP_009555241.1;HCoV-OC43_spike;842 | 0 | 0.05 |
| NC_006213.1;YP_009555241.1;HCoV-OC43_spike;843 | 0 | 0 |
| NC_006213.1;YP_009555241.1;HCoV-OC43_spike;844 | 0 | 0.025 |
| NC_006213.1;YP_009555241.1;HCoV-OC43_spike;845 | 0 | 0.05 |

| | | |
|---|---|---|
| NC_006213.1;YP_009555241.1;HCoV-OC43_spike;846 | 0.05 | 0.075 |
| NC_006213.1;YP_009555241.1;HCoV-OC43_spike;847 | 0.05 | 0.05 |
| NC_006213.1;YP_009555241.1;HCoV-OC43_spike;848 | 0 | 0.05 |
| NC_006213.1;YP_009555241.1;HCoV-OC43_spike;849 | 0 | 0.025 |
| NC_006213.1;YP_009555241.1;HCoV-OC43_spike;850 | 0 | 0.025 |
| NC_006213.1;YP_009555241.1;HCoV-OC43_spike;851 | 0 | 0 |
| NC_006213.1;YP_009555241.1;HCoV-OC43_spike;852 | 0.05 | 0 |
| NC_006213.1;YP_009555241.1;HCoV-OC43_spike;853 | 0 | 0 |
| NC_006213.1;YP_009555241.1;HCoV-OC43_spike;854 | 0 | 0 |
| NC_006213.1;YP_009555241.1;HCoV-OC43_spike;855 | 0 | 0 |
| NC_006213.1;YP_009555241.1;HCoV-OC43_spike;856 | 0 | 0 |
| NC_006213.1;YP_009555241.1;HCoV-OC43_spike;857 | 0 | 0 |
| NC_006213.1;YP_009555241.1;HCoV-OC43_spike;858 | 0 | 0 |
| NC_006213.1;YP_009555241.1;HCoV-OC43_spike;859 | 0 | 0 |
| NC_006213.1;YP_009555241.1;HCoV-OC43_spike;860 | 0 | 0 |
| NC_006213.1;YP_009555241.1;HCoV-OC43_spike;861 | 0 | 0 |
| NC_006213.1;YP_009555241.1;HCoV-OC43_spike;862 | 0 | 0 |
| NC_006213.1;YP_009555241.1;HCoV-OC43_spike;863 | 0 | 0 |
| NC_006213.1;YP_009555241.1;HCoV-OC43_spike;864 | 0 | 0 |
| NC_006213.1;YP_009555241.1;HCoV-OC43_spike;865 | 0.05 | 0.025 |
| NC_006213.1;YP_009555241.1;HCoV-OC43_spike;866 | 0 | 0.075 |
| NC_006213.1;YP_009555241.1;HCoV-OC43_spike;867 | 0.1 | 0.025 |
| NC_006213.1;YP_009555241.1;HCoV-OC43_spike;868 | 0.1 | 0.025 |
| NC_006213.1;YP_009555241.1;HCoV-OC43_spike;869 | 0.15 | 0.025 |
| NC_006213.1;YP_009555241.1;HCoV-OC43_spike;870 | 0.2 | 0.05 |
| NC_006213.1;YP_009555241.1;HCoV-OC43_spike;871 | 0.2 | 0.075 |
| NC_006213.1;YP_009555241.1;HCoV-OC43_spike;872 | 0.3 | 0.05 |
| NC_006213.1;YP_009555241.1;HCoV-OC43_spike;873 | 0.3 | 0.075 |
| NC_006213.1;YP_009555241.1;HCoV-OC43_spike;874 | 0.2 | 0.125 |
| NC_006213.1;YP_009555241.1;HCoV-OC43_spike;875 | 0.05 | 0.1 |
| NC_006213.1;YP_009555241.1;HCoV-OC43_spike;876 | 0.15 | 0.075 |
| NC_006213.1;YP_009555241.1;HCoV-OC43_spike;877 | 0.05 | 0.05 |
| NC_006213.1;YP_009555241.1;HCoV-OC43_spike;878 | 0 | 0.025 |
| NC_006213.1;YP_009555241.1;HCoV-OC43_spike;879 | 0.05 | 0.05 |
| NC_006213.1;YP_009555241.1;HCoV-OC43_spike;880 | 0.1 | 0.2 |
| NC_006213.1;YP_009555241.1;HCoV-OC43_spike;881 | 0.1 | 0.1 |
| NC_006213.1;YP_009555241.1;HCoV-OC43_spike;882 | 0 | 0 |
| NC_006213.1;YP_009555241.1;HCoV-OC43_spike;883 | 0 | 0 |
| NC_006213.1;YP_009555241.1;HCoV-OC43_spike;884 | 0 | 0 |
| NC_006213.1;YP_009555241.1;HCoV-OC43_spike;885 | 0 | 0 |
| NC_006213.1;YP_009555241.1;HCoV-OC43_spike;886 | 0.05 | 0 |
| NC_006213.1;YP_009555241.1;HCoV-OC43_spike;887 | 0 | 0 |
| NC_006213.1;YP_009555241.1;HCoV-OC43_spike;888 | 0 | 0 |

| | | |
|---|---|---|
| NC_006213.1;YP_009555241.1;HCoV-OC43_spike;889 | 0 | 0 |
| NC_006213.1;YP_009555241.1;HCoV-OC43_spike;890 | 0 | 0 |
| NC_006213.1;YP_009555241.1;HCoV-OC43_spike;891 | 0 | 0 |
| NC_006213.1;YP_009555241.1;HCoV-OC43_spike;892 | 0 | 0 |
| NC_006213.1;YP_009555241.1;HCoV-OC43_spike;893 | 0 | 0.025 |
| NC_006213.1;YP_009555241.1;HCoV-OC43_spike;894 | 0 | 0.025 |
| NC_006213.1;YP_009555241.1;HCoV-OC43_spike;895 | 0 | 0.05 |
| NC_006213.1;YP_009555241.1;HCoV-OC43_spike;896 | 0.1 | 0.575 |
| NC_006213.1;YP_009555241.1;HCoV-OC43_spike;897 | 0.2 | 0.525 |
| NC_006213.1;YP_009555241.1;HCoV-OC43_spike;898 | 0.15 | 0.8 |
| NC_006213.1;YP_009555241.1;HCoV-OC43_spike;899 | 0.1 | 0.725 |
| NC_006213.1;YP_009555241.1;HCoV-OC43_spike;900 | 0.05 | 0.7 |
| NC_006213.1;YP_009555241.1;HCoV-OC43_spike;901 | 0.1 | 0.725 |
| NC_006213.1;YP_009555241.1;HCoV-OC43_spike;902 | 0 | 0.675 |
| NC_006213.1;YP_009555241.1;HCoV-OC43_spike;903 | 0.05 | 0.675 |
| NC_006213.1;YP_009555241.1;HCoV-OC43_spike;904 | 0.1 | 0.4 |
| NC_006213.1;YP_009555241.1;HCoV-OC43_spike;905 | 0.45 | 0.55 |
| NC_006213.1;YP_009555241.1;HCoV-OC43_spike;906 | 0 | 0.275 |
| NC_006213.1;YP_009555241.1;HCoV-OC43_spike;907 | 0 | 0.125 |
| NC_006213.1;YP_009555241.1;HCoV-OC43_spike;908 | 0 | 0 |
| NC_006213.1;YP_009555241.1;HCoV-OC43_spike;909 | 0 | 0 |
| NC_006213.1;YP_009555241.1;HCoV-OC43_spike;910 | 0 | 0 |
| NC_006213.1;YP_009555241.1;HCoV-OC43_spike;911 | 0 | 0.025 |
| NC_006213.1;YP_009555241.1;HCoV-OC43_spike;912 | 0 | 0.025 |
| NC_006213.1;YP_009555241.1;HCoV-OC43_spike;913 | 0.05 | 0.025 |
| NC_006213.1;YP_009555241.1;HCoV-OC43_spike;914 | 0.05 | 0.025 |
| NC_006213.1;YP_009555241.1;HCoV-OC43_spike;915 | 0.05 | 0.075 |
| NC_006213.1;YP_009555241.1;HCoV-OC43_spike;916 | 0 | 0.025 |
| NC_006213.1;YP_009555241.1;HCoV-OC43_spike;917 | 0.05 | 0.025 |
| NC_006213.1;YP_009555241.1;HCoV-OC43_spike;918 | 0 | 0.1 |
| NC_006213.1;YP_009555241.1;HCoV-OC43_spike;919 | 0 | 0.05 |
| NC_006213.1;YP_009555241.1;HCoV-OC43_spike;920 | 0 | 0 |
| NC_006213.1;YP_009555241.1;HCoV-OC43_spike;921 | 0 | 0.025 |
| NC_006213.1;YP_009555241.1;HCoV-OC43_spike;922 | 0 | 0 |
| NC_006213.1;YP_009555241.1;HCoV-OC43_spike;923 | 0 | 0 |
| NC_006213.1;YP_009555241.1;HCoV-OC43_spike;924 | 0 | 0 |
| NC_006213.1;YP_009555241.1;HCoV-OC43_spike;925 | 0 | 0 |
| NC_006213.1;YP_009555241.1;HCoV-OC43_spike;926 | 0 | 0 |
| NC_006213.1;YP_009555241.1;HCoV-OC43_spike;927 | 0 | 0 |
| NC_006213.1;YP_009555241.1;HCoV-OC43_spike;928 | 0 | 0 |
| NC_006213.1;YP_009555241.1;HCoV-OC43_spike;929 | 0 | 0 |
| NC_006213.1;YP_009555241.1;HCoV-OC43_spike;930 | 0.05 | 0 |
| NC_006213.1;YP_009555241.1;HCoV-OC43_spike;931 | 0 | 0 |

| | | |
|---|---|---|
| NC_006213.1;YP_009555241.1;HCoV-OC43_spike;932 | 0 | 0 |
| NC_006213.1;YP_009555241.1;HCoV-OC43_spike;933 | 0.05 | 0 |
| NC_006213.1;YP_009555241.1;HCoV-OC43_spike;934 | 0.05 | 0 |
| NC_006213.1;YP_009555241.1;HCoV-OC43_spike;935 | 0.05 | 0 |
| NC_006213.1;YP_009555241.1;HCoV-OC43_spike;936 | 0.05 | 0 |
| NC_006213.1;YP_009555241.1;HCoV-OC43_spike;937 | 0.05 | 0 |
| NC_006213.1;YP_009555241.1;HCoV-OC43_spike;938 | 0 | 0 |
| NC_006213.1;YP_009555241.1;HCoV-OC43_spike;939 | 0.05 | 0 |
| NC_006213.1;YP_009555241.1;HCoV-OC43_spike;940 | 0.05 | 0 |
| NC_006213.1;YP_009555241.1;HCoV-OC43_spike;941 | 0.1 | 0 |
| NC_006213.1;YP_009555241.1;HCoV-OC43_spike;942 | 0.05 | 0 |
| NC_006213.1;YP_009555241.1;HCoV-OC43_spike;943 | 0.05 | 0 |
| NC_006213.1;YP_009555241.1;HCoV-OC43_spike;944 | 0 | 0.025 |
| NC_006213.1;YP_009555241.1;HCoV-OC43_spike;945 | 0.05 | 0.025 |
| NC_006213.1;YP_009555241.1;HCoV-OC43_spike;946 | 0.05 | 0.025 |
| NC_006213.1;YP_009555241.1;HCoV-OC43_spike;947 | 0.05 | 0.05 |
| NC_006213.1;YP_009555241.1;HCoV-OC43_spike;948 | 0.05 | 0.025 |
| NC_006213.1;YP_009555241.1;HCoV-OC43_spike;949 | 0 | 0.025 |
| NC_006213.1;YP_009555241.1;HCoV-OC43_spike;950 | 0.05 | 0.075 |
| NC_006213.1;YP_009555241.1;HCoV-OC43_spike;951 | 0 | 0.025 |
| NC_006213.1;YP_009555241.1;HCoV-OC43_spike;952 | 0 | 0 |
| NC_006213.1;YP_009555241.1;HCoV-OC43_spike;953 | 0 | 0 |
| NC_006213.1;YP_009555241.1;HCoV-OC43_spike;954 | 0.2 | 0.075 |
| NC_006213.1;YP_009555241.1;HCoV-OC43_spike;955 | 0.2 | 0.15 |
| NC_006213.1;YP_009555241.1;HCoV-OC43_spike;956 | 0 | 0 |
| NC_006213.1;YP_009555241.1;HCoV-OC43_spike;957 | 0 | 0 |
| NC_006213.1;YP_009555241.1;HCoV-OC43_spike;958 | 0.05 | 0.025 |
| NC_006213.1;YP_009555241.1;HCoV-OC43_spike;959 | 0 | 0 |
| NC_006213.1;YP_009555241.1;HCoV-OC43_spike;960 | 0 | 0.025 |
| NC_006213.1;YP_009555241.1;HCoV-OC43_spike;961 | 0 | 0 |
| NC_006213.1;YP_009555241.1;HCoV-OC43_spike;962 | 0.05 | 0.025 |
| NC_006213.1;YP_009555241.1;HCoV-OC43_spike;963 | 0.1 | 0.025 |
| NC_006213.1;YP_009555241.1;HCoV-OC43_spike;964 | 0 | 0 |
| NC_006213.1;YP_009555241.1;HCoV-OC43_spike;965 | 0.05 | 0.025 |
| NC_006213.1;YP_009555241.1;HCoV-OC43_spike;966 | 0.1 | 0.025 |
| NC_006213.1;YP_009555241.1;HCoV-OC43_spike;967 | 0.05 | 0 |
| NC_006213.1;YP_009555241.1;HCoV-OC43_spike;968 | 0 | 0 |
| NC_006213.1;YP_009555241.1;HCoV-OC43_spike;969 | 0 | 0 |
| NC_006213.1;YP_009555241.1;HCoV-OC43_spike;970 | 0 | 0 |
| NC_006213.1;YP_009555241.1;HCoV-OC43_spike;971 | 0 | 0 |
| NC_006213.1;YP_009555241.1;HCoV-OC43_spike;972 | 0 | 0.025 |
| NC_006213.1;YP_009555241.1;HCoV-OC43_spike;973 | 0 | 0 |
| NC_006213.1;YP_009555241.1;HCoV-OC43_spike;974 | 0 | 0 |

| | | |
|---|---|---|
| NC_006213.1;YP_009555241.1;HCoV-OC43_spike;975 | 0 | 0 |
| NC_006213.1;YP_009555241.1;HCoV-OC43_spike;976 | 0 | 0 |
| NC_006213.1;YP_009555241.1;HCoV-OC43_spike;977 | 0 | 0 |
| NC_006213.1;YP_009555241.1;HCoV-OC43_spike;978 | 0 | 0 |
| NC_006213.1;YP_009555241.1;HCoV-OC43_spike;979 | 0 | 0 |
| NC_006213.1;YP_009555241.1;HCoV-OC43_spike;980 | 0 | 0 |
| NC_006213.1;YP_009555241.1;HCoV-OC43_spike;981 | 0 | 0 |
| NC_006213.1;YP_009555241.1;HCoV-OC43_spike;982 | 0 | 0 |
| NC_006213.1;YP_009555241.1;HCoV-OC43_spike;983 | 0 | 0.025 |
| NC_006213.1;YP_009555241.1;HCoV-OC43_spike;984 | 0 | 0.025 |
| NC_006213.1;YP_009555241.1;HCoV-OC43_spike;985 | 0 | 0.025 |
| NC_006213.1;YP_009555241.1;HCoV-OC43_spike;986 | 0 | 0.025 |
| NC_006213.1;YP_009555241.1;HCoV-OC43_spike;987 | 0 | 0.025 |
| NC_006213.1;YP_009555241.1;HCoV-OC43_spike;988 | 0 | 0.05 |
| NC_006213.1;YP_009555241.1;HCoV-OC43_spike;989 | 0 | 0 |
| NC_006213.1;YP_009555241.1;HCoV-OC43_spike;990 | 0 | 0 |
| NC_006213.1;YP_009555241.1;HCoV-OC43_spike;991 | 0 | 0 |
| NC_006213.1;YP_009555241.1;HCoV-OC43_spike;992 | 0 | 0.025 |
| NC_006213.1;YP_009555241.1;HCoV-OC43_spike;993 | 0 | 0.025 |
| NC_006213.1;YP_009555241.1;HCoV-OC43_spike;994 | 0 | 0 |
| NC_006213.1;YP_009555241.1;HCoV-OC43_spike;995 | 0 | 0.025 |
| NC_006213.1;YP_009555241.1;HCoV-OC43_spike;996 | 0 | 0 |
| NC_006213.1;YP_009555241.1;HCoV-OC43_spike;997 | 0.05 | 0.05 |
| NC_006213.1;YP_009555241.1;HCoV-OC43_spike;998 | 0.05 | 0.075 |
| NC_006213.1;YP_009555241.1;HCoV-OC43_spike;999 | 0.05 | 0 |
| NC_006213.1;YP_009555241.1;HCoV-OC43_spike;1000 | 0 | 0 |
| NC_006213.1;YP_009555241.1;HCoV-OC43_spike;1001 | 0 | 0 |
| NC_006213.1;YP_009555241.1;HCoV-OC43_spike;1002 | 0 | 0 |
| NC_006213.1;YP_009555241.1;HCoV-OC43_spike;1003 | 0 | 0 |
| NC_006213.1;YP_009555241.1;HCoV-OC43_spike;1004 | 0 | 0 |
| NC_006213.1;YP_009555241.1;HCoV-OC43_spike;1005 | 0.05 | 0 |
| NC_006213.1;YP_009555241.1;HCoV-OC43_spike;1006 | 0 | 0.025 |
| NC_006213.1;YP_009555241.1;HCoV-OC43_spike;1007 | 0 | 0 |
| NC_006213.1;YP_009555241.1;HCoV-OC43_spike;1008 | 0 | 0 |
| NC_006213.1;YP_009555241.1;HCoV-OC43_spike;1009 | 0 | 0 |
| NC_006213.1;YP_009555241.1;HCoV-OC43_spike;1010 | 0 | 0 |
| NC_006213.1;YP_009555241.1;HCoV-OC43_spike;1011 | 0.05 | 0.025 |
| NC_006213.1;YP_009555241.1;HCoV-OC43_spike;1012 | 0.05 | 0.025 |
| NC_006213.1;YP_009555241.1;HCoV-OC43_spike;1013 | 0.05 | 0.025 |
| NC_006213.1;YP_009555241.1;HCoV-OC43_spike;1014 | 0.1 | 0.025 |
| NC_006213.1;YP_009555241.1;HCoV-OC43_spike;1015 | 0.1 | 0.025 |
| NC_006213.1;YP_009555241.1;HCoV-OC43_spike;1016 | 0.1 | 0.025 |
| NC_006213.1;YP_009555241.1;HCoV-OC43_spike;1017 | 0.05 | 0.025 |

| | | |
|---|---|---|
| NC_006213.1;YP_009555241.1;HCoV-OC43_spike;1018 | 0.05 | 0.025 |
| NC_006213.1;YP_009555241.1;HCoV-OC43_spike;1019 | 0.05 | 0.025 |
| NC_006213.1;YP_009555241.1;HCoV-OC43_spike;1020 | 0 | 0 |
| NC_006213.1;YP_009555241.1;HCoV-OC43_spike;1021 | 0 | 0 |
| NC_006213.1;YP_009555241.1;HCoV-OC43_spike;1022 | 0 | 0.025 |
| NC_006213.1;YP_009555241.1;HCoV-OC43_spike;1023 | 0 | 0 |
| NC_006213.1;YP_009555241.1;HCoV-OC43_spike;1024 | 0.05 | 0.025 |
| NC_006213.1;YP_009555241.1;HCoV-OC43_spike;1025 | 0 | 0.025 |
| NC_006213.1;YP_009555241.1;HCoV-OC43_spike;1026 | 0 | 0.025 |
| NC_006213.1;YP_009555241.1;HCoV-OC43_spike;1027 | 0 | 0.025 |
| NC_006213.1;YP_009555241.1;HCoV-OC43_spike;1028 | 0 | 0.025 |
| NC_006213.1;YP_009555241.1;HCoV-OC43_spike;1029 | 0 | 0.025 |
| NC_006213.1;YP_009555241.1;HCoV-OC43_spike;1030 | 0.05 | 0.025 |
| NC_006213.1;YP_009555241.1;HCoV-OC43_spike;1031 | 0.05 | 0.025 |
| NC_006213.1;YP_009555241.1;HCoV-OC43_spike;1032 | 0.1 | 0 |
| NC_006213.1;YP_009555241.1;HCoV-OC43_spike;1033 | 0.05 | 0 |
| NC_006213.1;YP_009555241.1;HCoV-OC43_spike;1034 | 0.1 | 0.05 |
| NC_006213.1;YP_009555241.1;HCoV-OC43_spike;1035 | 0.05 | 0.025 |
| NC_006213.1;YP_009555241.1;HCoV-OC43_spike;1036 | 0 | 0 |
| NC_006213.1;YP_009555241.1;HCoV-OC43_spike;1037 | 0 | 0 |
| NC_006213.1;YP_009555241.1;HCoV-OC43_spike;1038 | 0 | 0 |
| NC_006213.1;YP_009555241.1;HCoV-OC43_spike;1039 | 0 | 0 |
| NC_006213.1;YP_009555241.1;HCoV-OC43_spike;1040 | 0.05 | 0 |
| NC_006213.1;YP_009555241.1;HCoV-OC43_spike;1041 | 0 | 0.025 |
| NC_006213.1;YP_009555241.1;HCoV-OC43_spike;1042 | 0 | 0 |
| NC_006213.1;YP_009555241.1;HCoV-OC43_spike;1043 | 0 | 0 |
| NC_006213.1;YP_009555241.1;HCoV-OC43_spike;1044 | 0 | 0 |
| NC_006213.1;YP_009555241.1;HCoV-OC43_spike;1045 | 0 | 0 |
| NC_006213.1;YP_009555241.1;HCoV-OC43_spike;1046 | 0.05 | 0 |
| NC_006213.1;YP_009555241.1;HCoV-OC43_spike;1047 | 0.1 | 0 |
| NC_006213.1;YP_009555241.1;HCoV-OC43_spike;1048 | 0 | 0 |
| NC_006213.1;YP_009555241.1;HCoV-OC43_spike;1049 | 0 | 0 |
| NC_006213.1;YP_009555241.1;HCoV-OC43_spike;1050 | 0.05 | 0 |
| NC_006213.1;YP_009555241.1;HCoV-OC43_spike;1051 | 0.05 | 0 |
| NC_006213.1;YP_009555241.1;HCoV-OC43_spike;1052 | 0 | 0 |
| NC_006213.1;YP_009555241.1;HCoV-OC43_spike;1053 | 0 | 0 |
| NC_006213.1;YP_009555241.1;HCoV-OC43_spike;1054 | 0 | 0 |
| NC_006213.1;YP_009555241.1;HCoV-OC43_spike;1055 | 0 | 0 |
| NC_006213.1;YP_009555241.1;HCoV-OC43_spike;1056 | 0 | 0 |
| NC_006213.1;YP_009555241.1;HCoV-OC43_spike;1057 | 0.05 | 0 |
| NC_006213.1;YP_009555241.1;HCoV-OC43_spike;1058 | 0 | 0.05 |
| NC_006213.1;YP_009555241.1;HCoV-OC43_spike;1059 | 0 | 0.025 |
| NC_006213.1;YP_009555241.1;HCoV-OC43_spike;1060 | 0 | 0 |

| | | |
|---|---|---|
| NC_006213.1;YP_009555241.1;HCoV-OC43_spike;1061 | 0 | 0 |
| NC_006213.1;YP_009555241.1;HCoV-OC43_spike;1062 | 0 | 0 |
| NC_006213.1;YP_009555241.1;HCoV-OC43_spike;1063 | 0 | 0 |
| NC_006213.1;YP_009555241.1;HCoV-OC43_spike;1064 | 0.05 | 0 |
| NC_006213.1;YP_009555241.1;HCoV-OC43_spike;1065 | 0 | 0.025 |
| NC_006213.1;YP_009555241.1;HCoV-OC43_spike;1066 | 0.1 | 0.075 |
| NC_006213.1;YP_009555241.1;HCoV-OC43_spike;1067 | 0 | 0 |
| NC_006213.1;YP_009555241.1;HCoV-OC43_spike;1068 | 0 | 0.025 |
| NC_006213.1;YP_009555241.1;HCoV-OC43_spike;1069 | 0 | 0.025 |
| NC_006213.1;YP_009555241.1;HCoV-OC43_spike;1070 | 0.05 | 0 |
| NC_006213.1;YP_009555241.1;HCoV-OC43_spike;1071 | 0 | 0 |
| NC_006213.1;YP_009555241.1;HCoV-OC43_spike;1072 | 0.1 | 0.025 |
| NC_006213.1;YP_009555241.1;HCoV-OC43_spike;1073 | 0.05 | 0.05 |
| NC_006213.1;YP_009555241.1;HCoV-OC43_spike;1074 | 0.05 | 0.025 |
| NC_006213.1;YP_009555241.1;HCoV-OC43_spike;1075 | 0 | 0.05 |
| NC_006213.1;YP_009555241.1;HCoV-OC43_spike;1076 | 0 | 0 |
| NC_006213.1;YP_009555241.1;HCoV-OC43_spike;1077 | 0 | 0 |
| NC_006213.1;YP_009555241.1;HCoV-OC43_spike;1078 | 0 | 0 |
| NC_006213.1;YP_009555241.1;HCoV-OC43_spike;1079 | 0 | 0 |
| NC_006213.1;YP_009555241.1;HCoV-OC43_spike;1080 | 0 | 0 |
| NC_006213.1;YP_009555241.1;HCoV-OC43_spike;1081 | 0 | 0 |
| NC_006213.1;YP_009555241.1;HCoV-OC43_spike;1082 | 0 | 0 |
| NC_006213.1;YP_009555241.1;HCoV-OC43_spike;1083 | 0 | 0 |
| NC_006213.1;YP_009555241.1;HCoV-OC43_spike;1084 | 0 | 0 |
| NC_006213.1;YP_009555241.1;HCoV-OC43_spike;1085 | 0 | 0.025 |
| NC_006213.1;YP_009555241.1;HCoV-OC43_spike;1086 | 0.05 | 0.025 |
| NC_006213.1;YP_009555241.1;HCoV-OC43_spike;1087 | 0.05 | 0.025 |
| NC_006213.1;YP_009555241.1;HCoV-OC43_spike;1088 | 0.05 | 0.025 |
| NC_006213.1;YP_009555241.1;HCoV-OC43_spike;1089 | 0.05 | 0.025 |
| NC_006213.1;YP_009555241.1;HCoV-OC43_spike;1090 | 0.05 | 0.025 |
| NC_006213.1;YP_009555241.1;HCoV-OC43_spike;1091 | 0.05 | 0 |
| NC_006213.1;YP_009555241.1;HCoV-OC43_spike;1092 | 0.05 | 0 |
| NC_006213.1;YP_009555241.1;HCoV-OC43_spike;1093 | 0 | 0 |
| NC_006213.1;YP_009555241.1;HCoV-OC43_spike;1094 | 0 | 0 |
| NC_006213.1;YP_009555241.1;HCoV-OC43_spike;1095 | 0 | 0 |
| NC_006213.1;YP_009555241.1;HCoV-OC43_spike;1096 | 0 | 0 |
| NC_006213.1;YP_009555241.1;HCoV-OC43_spike;1097 | 0 | 0.025 |
| NC_006213.1;YP_009555241.1;HCoV-OC43_spike;1098 | 0 | 0.025 |
| NC_006213.1;YP_009555241.1;HCoV-OC43_spike;1099 | 0 | 0 |
| NC_006213.1;YP_009555241.1;HCoV-OC43_spike;1100 | 0 | 0 |
| NC_006213.1;YP_009555241.1;HCoV-OC43_spike;1101 | 0 | 0 |
| NC_006213.1;YP_009555241.1;HCoV-OC43_spike;1102 | 0 | 0 |
| NC_006213.1;YP_009555241.1;HCoV-OC43_spike;1103 | 0 | 0 |

| | | |
|---|---|---|
| NC_006213.1;YP_009555241.1;HCoV-OC43_spike;1104 | 0 | 0 |
| NC_006213.1;YP_009555241.1;HCoV-OC43_spike;1105 | 0 | 0.025 |
| NC_006213.1;YP_009555241.1;HCoV-OC43_spike;1106 | 0 | 0 |
| NC_006213.1;YP_009555241.1;HCoV-OC43_spike;1107 | 0 | 0 |
| NC_006213.1;YP_009555241.1;HCoV-OC43_spike;1108 | 0 | 0 |
| NC_006213.1;YP_009555241.1;HCoV-OC43_spike;1109 | 0 | 0 |
| NC_006213.1;YP_009555241.1;HCoV-OC43_spike;1110 | 0.05 | 0.025 |
| NC_006213.1;YP_009555241.1;HCoV-OC43_spike;1111 | 0.05 | 0 |
| NC_006213.1;YP_009555241.1;HCoV-OC43_spike;1112 | 0 | 0 |
| NC_006213.1;YP_009555241.1;HCoV-OC43_spike;1113 | 0 | 0 |
| NC_006213.1;YP_009555241.1;HCoV-OC43_spike;1114 | 0 | 0 |
| NC_006213.1;YP_009555241.1;HCoV-OC43_spike;1115 | 0 | 0 |
| NC_006213.1;YP_009555241.1;HCoV-OC43_spike;1116 | 0 | 0 |
| NC_006213.1;YP_009555241.1;HCoV-OC43_spike;1117 | 0 | 0 |
| NC_006213.1;YP_009555241.1;HCoV-OC43_spike;1118 | 0 | 0.025 |
| NC_006213.1;YP_009555241.1;HCoV-OC43_spike;1119 | 0 | 0.1 |
| NC_006213.1;YP_009555241.1;HCoV-OC43_spike;1120 | 0 | 0 |
| NC_006213.1;YP_009555241.1;HCoV-OC43_spike;1121 | 0 | 0 |
| NC_006213.1;YP_009555241.1;HCoV-OC43_spike;1122 | 0 | 0.025 |
| NC_006213.1;YP_009555241.1;HCoV-OC43_spike;1123 | 0 | 0 |
| NC_006213.1;YP_009555241.1;HCoV-OC43_spike;1124 | 0 | 0 |
| NC_006213.1;YP_009555241.1;HCoV-OC43_spike;1125 | 0 | 0 |
| NC_006213.1;YP_009555241.1;HCoV-OC43_spike;1126 | 0 | 0 |
| NC_006213.1;YP_009555241.1;HCoV-OC43_spike;1127 | 0 | 0 |
| NC_006213.1;YP_009555241.1;HCoV-OC43_spike;1128 | 0 | 0 |
| NC_006213.1;YP_009555241.1;HCoV-OC43_spike;1129 | 0 | 0 |
| NC_006213.1;YP_009555241.1;HCoV-OC43_spike;1130 | 0 | 0 |
| NC_006213.1;YP_009555241.1;HCoV-OC43_spike;1131 | 0 | 0 |
| NC_006213.1;YP_009555241.1;HCoV-OC43_spike;1132 | 0 | 0 |
| NC_006213.1;YP_009555241.1;HCoV-OC43_spike;1133 | 0 | 0 |
| NC_006213.1;YP_009555241.1;HCoV-OC43_spike;1134 | 0 | 0 |
| NC_006213.1;YP_009555241.1;HCoV-OC43_spike;1135 | 0 | 0 |
| NC_006213.1;YP_009555241.1;HCoV-OC43_spike;1136 | 0 | 0 |
| NC_006213.1;YP_009555241.1;HCoV-OC43_spike;1137 | 0 | 0 |
| NC_006213.1;YP_009555241.1;HCoV-OC43_spike;1138 | 0 | 0 |
| NC_006213.1;YP_009555241.1;HCoV-OC43_spike;1139 | 0 | 0 |
| NC_006213.1;YP_009555241.1;HCoV-OC43_spike;1140 | 0 | 0 |
| NC_006213.1;YP_009555241.1;HCoV-OC43_spike;1141 | 0 | 0 |
| NC_006213.1;YP_009555241.1;HCoV-OC43_spike;1142 | 0 | 0 |
| NC_006213.1;YP_009555241.1;HCoV-OC43_spike;1143 | 0 | 0 |
| NC_006213.1;YP_009555241.1;HCoV-OC43_spike;1144 | 0 | 0 |
| NC_006213.1;YP_009555241.1;HCoV-OC43_spike;1145 | 0 | 0 |
| NC_006213.1;YP_009555241.1;HCoV-OC43_spike;1146 | 0 | 0 |

| | | |
|---|---|---|
| NC_006213.1;YP_009555241.1;HCoV-OC43_spike;1147 | 0 | 0 |
| NC_006213.1;YP_009555241.1;HCoV-OC43_spike;1148 | 0 | 0 |
| NC_006213.1;YP_009555241.1;HCoV-OC43_spike;1149 | 0 | 0 |
| NC_006213.1;YP_009555241.1;HCoV-OC43_spike;1150 | 0 | 0 |
| NC_006213.1;YP_009555241.1;HCoV-OC43_spike;1151 | 0 | 0 |
| NC_006213.1;YP_009555241.1;HCoV-OC43_spike;1152 | 0 | 0 |
| NC_006213.1;YP_009555241.1;HCoV-OC43_spike;1153 | 0 | 0 |
| NC_006213.1;YP_009555241.1;HCoV-OC43_spike;1154 | 0 | 0 |
| NC_006213.1;YP_009555241.1;HCoV-OC43_spike;1155 | 0 | 0 |
| NC_006213.1;YP_009555241.1;HCoV-OC43_spike;1156 | 0 | 0 |
| NC_006213.1;YP_009555241.1;HCoV-OC43_spike;1157 | 0 | 0 |
| NC_006213.1;YP_009555241.1;HCoV-OC43_spike;1158 | 0 | 0 |
| NC_006213.1;YP_009555241.1;HCoV-OC43_spike;1159 | 0 | 0.025 |
| NC_006213.1;YP_009555241.1;HCoV-OC43_spike;1160 | 0.05 | 0 |
| NC_006213.1;YP_009555241.1;HCoV-OC43_spike;1161 | 0 | 0 |
| NC_006213.1;YP_009555241.1;HCoV-OC43_spike;1162 | 0 | 0 |
| NC_006213.1;YP_009555241.1;HCoV-OC43_spike;1163 | 0 | 0 |
| NC_006213.1;YP_009555241.1;HCoV-OC43_spike;1164 | 0 | 0 |
| NC_006213.1;YP_009555241.1;HCoV-OC43_spike;1165 | 0 | 0 |
| NC_006213.1;YP_009555241.1;HCoV-OC43_spike;1166 | 0.05 | 0 |
| NC_006213.1;YP_009555241.1;HCoV-OC43_spike;1167 | 0 | 0 |
| NC_006213.1;YP_009555241.1;HCoV-OC43_spike;1168 | 0 | 0 |
| NC_006213.1;YP_009555241.1;HCoV-OC43_spike;1169 | 0 | 0.025 |
| NC_006213.1;YP_009555241.1;HCoV-OC43_spike;1170 | 0 | 0.025 |
| NC_006213.1;YP_009555241.1;HCoV-OC43_spike;1171 | 0 | 0.025 |
| NC_006213.1;YP_009555241.1;HCoV-OC43_spike;1172 | 0 | 0 |
| NC_006213.1;YP_009555241.1;HCoV-OC43_spike;1173 | 0.05 | 0.025 |
| NC_006213.1;YP_009555241.1;HCoV-OC43_spike;1174 | 0 | 0.025 |
| NC_006213.1;YP_009555241.1;HCoV-OC43_spike;1175 | 0 | 0.025 |
| NC_006213.1;YP_009555241.1;HCoV-OC43_spike;1176 | 0 | 0.025 |
| NC_006213.1;YP_009555241.1;HCoV-OC43_spike;1177 | 0 | 0 |
| NC_006213.1;YP_009555241.1;HCoV-OC43_spike;1178 | 0 | 0.025 |
| NC_006213.1;YP_009555241.1;HCoV-OC43_spike;1179 | 0.05 | 0.025 |
| NC_006213.1;YP_009555241.1;HCoV-OC43_spike;1180 | 0.05 | 0.05 |
| NC_006213.1;YP_009555241.1;HCoV-OC43_spike;1181 | 0 | 0.025 |
| NC_006213.1;YP_009555241.1;HCoV-OC43_spike;1182 | 0 | 0 |
| NC_006213.1;YP_009555241.1;HCoV-OC43_spike;1183 | 0 | 0 |
| NC_006213.1;YP_009555241.1;HCoV-OC43_spike;1184 | 0 | 0 |
| NC_006213.1;YP_009555241.1;HCoV-OC43_spike;1185 | 0 | 0 |
| NC_006213.1;YP_009555241.1;HCoV-OC43_spike;1186 | 0 | 0 |
| NC_006213.1;YP_009555241.1;HCoV-OC43_spike;1187 | 0 | 0 |
| NC_006213.1;YP_009555241.1;HCoV-OC43_spike;1188 | 0 | 0 |
| NC_006213.1;YP_009555241.1;HCoV-OC43_spike;1189 | 0 | 0 |

| | | |
|---|---|---|
| NC_006213.1;YP_009555241.1;HCoV-OC43_spike;1190 | 0.05 | 0 |
| NC_006213.1;YP_009555241.1;HCoV-OC43_spike;1191 | 0.05 | 0.025 |
| NC_006213.1;YP_009555241.1;HCoV-OC43_spike;1192 | 0 | 0 |
| NC_006213.1;YP_009555241.1;HCoV-OC43_spike;1193 | 0.05 | 0 |
| NC_006213.1;YP_009555241.1;HCoV-OC43_spike;1194 | 0 | 0 |
| NC_006213.1;YP_009555241.1;HCoV-OC43_spike;1195 | 0 | 0.025 |
| NC_006213.1;YP_009555241.1;HCoV-OC43_spike;1196 | 0 | 0 |
| NC_006213.1;YP_009555241.1;HCoV-OC43_spike;1197 | 0 | 0 |
| NC_006213.1;YP_009555241.1;HCoV-OC43_spike;1198 | 0.05 | 0 |
| NC_006213.1;YP_009555241.1;HCoV-OC43_spike;1199 | 0 | 0.025 |
| NC_006213.1;YP_009555241.1;HCoV-OC43_spike;1200 | 0 | 0 |
| NC_006213.1;YP_009555241.1;HCoV-OC43_spike;1201 | 0 | 0 |
| NC_006213.1;YP_009555241.1;HCoV-OC43_spike;1202 | 0 | 0 |
| NC_006213.1;YP_009555241.1;HCoV-OC43_spike;1203 | 0 | 0 |
| NC_006213.1;YP_009555241.1;HCoV-OC43_spike;1204 | 0 | 0 |
| NC_006213.1;YP_009555241.1;HCoV-OC43_spike;1205 | 0 | 0.025 |
| NC_006213.1;YP_009555241.1;HCoV-OC43_spike;1206 | 0 | 0 |
| NC_006213.1;YP_009555241.1;HCoV-OC43_spike;1207 | 0 | 0 |
| NC_006213.1;YP_009555241.1;HCoV-OC43_spike;1208 | 0 | 0 |
| NC_006213.1;YP_009555241.1;HCoV-OC43_spike;1209 | 0 | 0 |
| NC_006213.1;YP_009555241.1;HCoV-OC43_spike;1210 | 0 | 0 |
| NC_006213.1;YP_009555241.1;HCoV-OC43_spike;1211 | 0 | 0 |
| NC_006213.1;YP_009555241.1;HCoV-OC43_spike;1212 | 0 | 0 |
| NC_006213.1;YP_009555241.1;HCoV-OC43_spike;1213 | 0 | 0 |
| NC_006213.1;YP_009555241.1;HCoV-OC43_spike;1214 | 0 | 0 |
| NC_006213.1;YP_009555241.1;HCoV-OC43_spike;1215 | 0 | 0 |
| NC_006213.1;YP_009555241.1;HCoV-OC43_spike;1216 | 0 | 0.025 |
| NC_006213.1;YP_009555241.1;HCoV-OC43_spike;1217 | 0.05 | 0.025 |
| NC_006213.1;YP_009555241.1;HCoV-OC43_spike;1218 | 0.05 | 0.05 |
| NC_006213.1;YP_009555241.1;HCoV-OC43_spike;1219 | 0 | 0.025 |
| NC_006213.1;YP_009555241.1;HCoV-OC43_spike;1220 | 0 | 0.05 |
| NC_006213.1;YP_009555241.1;HCoV-OC43_spike;1221 | 0.05 | 0.05 |
| NC_006213.1;YP_009555241.1;HCoV-OC43_spike;1222 | 0 | 0.1 |
| NC_006213.1;YP_009555241.1;HCoV-OC43_spike;1223 | 0 | 0.1 |
| NC_006213.1;YP_009555241.1;HCoV-OC43_spike;1224 | 0 | 0.025 |
| NC_006213.1;YP_009555241.1;HCoV-OC43_spike;1225 | 0 | 0.175 |
| NC_006213.1;YP_009555241.1;HCoV-OC43_spike;1226 | 0 | 0.4 |
| NC_006213.1;YP_009555241.1;HCoV-OC43_spike;1227 | 0 | 0.55 |
| NC_006213.1;YP_009555241.1;HCoV-OC43_spike;1228 | 0.05 | 0.55 |
| NC_006213.1;YP_009555241.1;HCoV-OC43_spike;1229 | 0.05 | 0.65 |
| NC_006213.1;YP_009555241.1;HCoV-OC43_spike;1230 | 0.05 | 0.675 |
| NC_006213.1;YP_009555241.1;HCoV-OC43_spike;1231 | 0 | 0.675 |
| NC_006213.1;YP_009555241.1;HCoV-OC43_spike;1232 | 0.05 | 0.7 |

| | | |
|---|---|---|
| NC_006213.1;YP_009555241.1;HCoV-OC43_spike;1233 | 0.1 | 0.725 |
| NC_006213.1;YP_009555241.1;HCoV-OC43_spike;1234 | 0.15 | 0.4 |
| NC_006213.1;YP_009555241.1;HCoV-OC43_spike;1235 | 0.05 | 0.1 |
| NC_006213.1;YP_009555241.1;HCoV-OC43_spike;1236 | 0 | 0.05 |
| NC_006213.1;YP_009555241.1;HCoV-OC43_spike;1237 | 0 | 0 |
| NC_006213.1;YP_009555241.1;HCoV-OC43_spike;1238 | 0.05 | 0 |
| NC_006213.1;YP_009555241.1;HCoV-OC43_spike;1239 | 0.1 | 0 |
| NC_006213.1;YP_009555241.1;HCoV-OC43_spike;1240 | 0 | 0.025 |
| NC_006213.1;YP_009555241.1;HCoV-OC43_spike;1241 | 0 | 0 |
| NC_006213.1;YP_009555241.1;HCoV-OC43_spike;1242 | 0 | 0 |
| NC_006213.1;YP_009555241.1;HCoV-OC43_spike;1243 | 0 | 0 |
| NC_006213.1;YP_009555241.1;HCoV-OC43_spike;1244 | 0 | 0 |
| NC_006213.1;YP_009555241.1;HCoV-OC43_spike;1245 | 0 | 0 |
| NC_006213.1;YP_009555241.1;HCoV-OC43_spike;1246 | 0 | 0 |
| NC_006213.1;YP_009555241.1;HCoV-OC43_spike;1247 | 0 | 0 |
| NC_006213.1;YP_009555241.1;HCoV-OC43_spike;1248 | 0 | 0 |
| NC_006213.1;YP_009555241.1;HCoV-OC43_spike;1249 | 0 | 0 |
| NC_006213.1;YP_009555241.1;HCoV-OC43_spike;1250 | 0 | 0 |
| NC_006213.1;YP_009555241.1;HCoV-OC43_spike;1251 | 0 | 0 |
| NC_006213.1;YP_009555241.1;HCoV-OC43_spike;1252 | 0 | 0 |
| NC_006213.1;YP_009555241.1;HCoV-OC43_spike;1253 | 0 | 0 |
| NC_006213.1;YP_009555241.1;HCoV-OC43_spike;1254 | 0 | 0.025 |
| NC_006213.1;YP_009555241.1;HCoV-OC43_spike;1255 | 0 | 0 |
| NC_006213.1;YP_009555241.1;HCoV-OC43_spike;1256 | 0 | 0 |
| NC_006213.1;YP_009555241.1;HCoV-OC43_spike;1257 | 0 | 0 |
| NC_006213.1;YP_009555241.1;HCoV-OC43_spike;1258 | 0 | 0.025 |
| NC_006213.1;YP_009555241.1;HCoV-OC43_spike;1259 | 0 | 0.05 |
| NC_006213.1;YP_009555241.1;HCoV-OC43_spike;1260 | 0 | 0 |
| NC_006213.1;YP_009555241.1;HCoV-OC43_spike;1261 | 0 | 0 |
| NC_006213.1;YP_009555241.1;HCoV-OC43_spike;1262 | 0 | 0 |
| NC_006213.1;YP_009555241.1;HCoV-OC43_spike;1263 | 0 | 0 |
| NC_006213.1;YP_009555241.1;HCoV-OC43_spike;1264 | 0 | 0 |
| NC_006213.1;YP_009555241.1;HCoV-OC43_spike;1265 | 0 | 0 |
| NC_006213.1;YP_009555241.1;HCoV-OC43_spike;1266 | 0 | 0 |
| NC_006213.1;YP_009555241.1;HCoV-OC43_spike;1267 | 0 | 0 |
| NC_006213.1;YP_009555241.1;HCoV-OC43_spike;1268 | 0 | 0 |
| NC_006213.1;YP_009555241.1;HCoV-OC43_spike;1269 | 0 | 0 |
| NC_006213.1;YP_009555241.1;HCoV-OC43_spike;1270 | 0 | 0 |
| NC_006213.1;YP_009555241.1;HCoV-OC43_spike;1271 | 0 | 0 |
| NC_006213.1;YP_009555241.1;HCoV-OC43_spike;1272 | 0.05 | 0 |
| NC_006213.1;YP_009555241.1;HCoV-OC43_spike;1273 | 0.05 | 0 |
| NC_006213.1;YP_009555241.1;HCoV-OC43_spike;1274 | 0.05 | 0 |
| NC_006213.1;YP_009555241.1;HCoV-OC43_spike;1275 | 0.05 | 0 |

| | | |
|---|---|---|
| NC_006213.1;YP_009555241.1;HCoV-OC43_spike;1276 | 0.05 | 0.025 |
| NC_006213.1;YP_009555241.1;HCoV-OC43_spike;1277 | 0 | 0.05 |
| NC_006213.1;YP_009555241.1;HCoV-OC43_spike;1278 | 0 | 0 |
| NC_006213.1;YP_009555241.1;HCoV-OC43_spike;1279 | 0 | 0 |
| NC_006213.1;YP_009555241.1;HCoV-OC43_spike;1280 | 0 | 0 |
| NC_006213.1;YP_009555241.1;HCoV-OC43_spike;1281 | 0 | 0 |
| NC_006213.1;YP_009555241.1;HCoV-OC43_spike;1282 | 0 | 0 |
| NC_006213.1;YP_009555241.1;HCoV-OC43_spike;1283 | 0 | 0 |
| NC_006213.1;YP_009555241.1;HCoV-OC43_spike;1284 | 0.05 | 0 |
| NC_006213.1;YP_009555241.1;HCoV-OC43_spike;1285 | 0 | 0 |
| NC_006213.1;YP_009555241.1;HCoV-OC43_spike;1286 | 0 | 0 |
| NC_006213.1;YP_009555241.1;HCoV-OC43_spike;1287 | 0 | 0 |
| NC_006213.1;YP_009555241.1;HCoV-OC43_spike;1288 | 0 | 0 |
| NC_006213.1;YP_009555241.1;HCoV-OC43_spike;1289 | 0 | 0 |
| NC_006213.1;YP_009555241.1;HCoV-OC43_spike;1290 | 0 | 0 |
| NC_006213.1;YP_009555241.1;HCoV-OC43_spike;1291 | 0 | 0 |
| NC_006213.1;YP_009555241.1;HCoV-OC43_spike;1292 | 0 | 0 |
| NC_006213.1;YP_009555241.1;HCoV-OC43_spike;1293 | 0 | 0 |
| NC_006213.1;YP_009555241.1;HCoV-OC43_spike;1294 | 0 | 0 |
| NC_006213.1;YP_009555241.1;HCoV-OC43_spike;1295 | 0 | 0 |
| NC_006213.1;YP_009555241.1;HCoV-OC43_spike;1296 | 0 | 0 |
| NC_006213.1;YP_009555241.1;HCoV-OC43_spike;1297 | 0 | 0 |
| NC_006213.1;YP_009555241.1;HCoV-OC43_spike;1298 | 0 | 0 |
| NC_006213.1;YP_009555241.1;HCoV-OC43_spike;1299 | 0 | 0 |
| NC_006213.1;YP_009555241.1;HCoV-OC43_spike;1300 | 0 | 0 |
| NC_006213.1;YP_009555241.1;HCoV-OC43_spike;1301 | 0 | 0 |
| NC_006213.1;YP_009555241.1;HCoV-OC43_spike;1302 | 0 | 0 |
| NC_006213.1;YP_009555241.1;HCoV-OC43_spike;1303 | 0 | 0 |
| NC_006213.1;YP_009555241.1;HCoV-OC43_spike;1304 | 0 | 0 |
| NC_006213.1;YP_009555241.1;HCoV-OC43_spike;1305 | 0 | 0 |
| NC_006213.1;YP_009555241.1;HCoV-OC43_spike;1306 | 0 | 0 |
| NC_006213.1;YP_009555241.1;HCoV-OC43_spike;1307 | 0 | 0 |
| NC_006213.1;YP_009555241.1;HCoV-OC43_spike;1308 | 0 | 0 |
| NC_006213.1;YP_009555241.1;HCoV-OC43_spike;1309 | 0 | 0 |
| NC_006213.1;YP_009555241.1;HCoV-OC43_spike;1310 | 0 | 0 |
| NC_006213.1;YP_009555241.1;HCoV-OC43_spike;1311 | 0 | 0 |
| NC_006213.1;YP_009555241.1;HCoV-OC43_spike;1312 | 0 | 0 |
| NC_006213.1;YP_009555241.1;HCoV-OC43_spike;1313 | 0 | 0 |
| NC_006213.1;YP_009555241.1;HCoV-OC43_spike;1314 | 0 | 0 |
| NC_006213.1;YP_009555241.1;HCoV-OC43_spike;1315 | 0 | 0 |
| NC_006213.1;YP_009555241.1;HCoV-OC43_spike;1316 | 0 | 0 |
| NC_006213.1;YP_009555241.1;HCoV-OC43_spike;1317 | 0 | 0 |
| NC_006213.1;YP_009555241.1;HCoV-OC43_spike;1318 | 0 | 0 |

| | | |
|---|---|---|
| NC_006213.1;YP_009555241.1;HCoV-OC43_spike;1319 | 0 | 0 |
| NC_006213.1;YP_009555241.1;HCoV-OC43_spike;1320 | 0 | 0 |
| NC_006213.1;YP_009555241.1;HCoV-OC43_spike;1321 | 0 | 0 |
| NC_006213.1;YP_009555241.1;HCoV-OC43_spike;1322 | 0 | 0 |
| NC_006213.1;YP_009555241.1;HCoV-OC43_spike;1323 | 0 | 0 |
| NC_006213.1;YP_009555241.1;HCoV-OC43_spike;1324 | 0 | 0 |
| NC_006213.1;YP_009555241.1;HCoV-OC43_spike;1325 | 0 | 0 |
| NC_006213.1;YP_009555241.1;HCoV-OC43_spike;1326 | 0 | 0 |
| NC_006213.1;YP_009555241.1;HCoV-OC43_spike;1327 | 0 | 0 |
| NC_006213.1;YP_009555241.1;HCoV-OC43_spike;1328 | 0 | 0 |
| NC_006213.1;YP_009555241.1;HCoV-OC43_spike;1329 | 0.05 | 0.025 |
| NC_006213.1;YP_009555241.1;HCoV-OC43_spike;1330 | 0.05 | 0.025 |
| NC_006213.1;YP_009555241.1;HCoV-OC43_spike;1331 | 0.05 | 0.025 |
| NC_006213.1;YP_009555241.1;HCoV-OC43_spike;1332 | 0.05 | 0.025 |
| NC_006213.1;YP_009555241.1;HCoV-OC43_spike;1333 | 0.3 | 0.2 |
| NC_006213.1;YP_009555241.1;HCoV-OC43_spike;1334 | 0.15 | 0.125 |
| NC_006213.1;YP_009555241.1;HCoV-OC43_spike;1335 | 0.15 | 0.1 |
| NC_006213.1;YP_009555241.1;HCoV-OC43_spike;1336 | 0.05 | 0.025 |
| NC_006213.1;YP_009555241.1;HCoV-OC43_spike;1337 | 0.05 | 0.075 |
| NC_006213.1;YP_009555241.1;HCoV-OC43_spike;1338 | 0.2 | 0.075 |
| NC_006213.1;YP_009555241.1;HCoV-OC43_spike;1339 | 0.2 | 0.05 |
| NC_006213.1;YP_009555241.1;HCoV-OC43_spike;1340 | 0.15 | 0.05 |
| NC_006213.1;YP_009555241.1;HCoV-OC43_spike;1341 | 0.15 | 0.05 |
| NC_006213.1;YP_009555241.1;HCoV-OC43_spike;1342 | 0.15 | 0.05 |
| NC_006213.1;YP_009555242.1;HCoV-OC43_ns12.9;1 | 0 | 0 |
| NC_006213.1;YP_009555242.1;HCoV-OC43_ns12.9;2 | 0 | 0 |
| NC_006213.1;YP_009555242.1;HCoV-OC43_ns12.9;3 | 0 | 0 |
| NC_006213.1;YP_009555242.1;HCoV-OC43_ns12.9;4 | 0 | 0 |
| NC_006213.1;YP_009555242.1;HCoV-OC43_ns12.9;5 | 0 | 0 |
| NC_006213.1;YP_009555242.1;HCoV-OC43_ns12.9;6 | 0.05 | 0 |
| NC_006213.1;YP_009555242.1;HCoV-OC43_ns12.9;7 | 0.05 | 0 |
| NC_006213.1;YP_009555242.1;HCoV-OC43_ns12.9;8 | 0.1 | 0 |
| NC_006213.1;YP_009555242.1;HCoV-OC43_ns12.9;9 | 0.15 | 0 |
| NC_006213.1;YP_009555242.1;HCoV-OC43_ns12.9;10 | 0.15 | 0.025 |
| NC_006213.1;YP_009555242.1;HCoV-OC43_ns12.9;11 | 0.15 | 0 |
| NC_006213.1;YP_009555242.1;HCoV-OC43_ns12.9;12 | 0.1 | 0 |
| NC_006213.1;YP_009555242.1;HCoV-OC43_ns12.9;13 | 0.15 | 0.025 |
| NC_006213.1;YP_009555242.1;HCoV-OC43_ns12.9;14 | 0.15 | 0.025 |
| NC_006213.1;YP_009555242.1;HCoV-OC43_ns12.9;15 | 0.1 | 0.05 |
| NC_006213.1;YP_009555242.1;HCoV-OC43_ns12.9;16 | 0.1 | 0.05 |
| NC_006213.1;YP_009555242.1;HCoV-OC43_ns12.9;17 | 0.1 | 0 |
| NC_006213.1;YP_009555242.1;HCoV-OC43_ns12.9;18 | 0.05 | 0 |
| NC_006213.1;YP_009555242.1;HCoV-OC43_ns12.9;19 | 0.05 | 0 |

| | | |
|---|---|---|
| NC_006213.1;YP_009555242.1;HCoV-OC43_ns12.9;20 | 0.1 | 0 |
| NC_006213.1;YP_009555242.1;HCoV-OC43_ns12.9;21 | 0.05 | 0 |
| NC_006213.1;YP_009555242.1;HCoV-OC43_ns12.9;22 | 0 | 0 |
| NC_006213.1;YP_009555242.1;HCoV-OC43_ns12.9;23 | 0.05 | 0 |
| NC_006213.1;YP_009555242.1;HCoV-OC43_ns12.9;24 | 0.05 | 0 |
| NC_006213.1;YP_009555242.1;HCoV-OC43_ns12.9;25 | 0 | 0 |
| NC_006213.1;YP_009555242.1;HCoV-OC43_ns12.9;26 | 0 | 0 |
| NC_006213.1;YP_009555242.1;HCoV-OC43_ns12.9;27 | 0 | 0 |
| NC_006213.1;YP_009555242.1;HCoV-OC43_ns12.9;28 | 0 | 0.025 |
| NC_006213.1;YP_009555242.1;HCoV-OC43_ns12.9;29 | 0.05 | 0 |
| NC_006213.1;YP_009555242.1;HCoV-OC43_ns12.9;30 | 0 | 0 |
| NC_006213.1;YP_009555242.1;HCoV-OC43_ns12.9;31 | 0 | 0 |
| NC_006213.1;YP_009555242.1;HCoV-OC43_ns12.9;32 | 0 | 0 |
| NC_006213.1;YP_009555242.1;HCoV-OC43_ns12.9;33 | 0 | 0 |
| NC_006213.1;YP_009555242.1;HCoV-OC43_ns12.9;34 | 0 | 0 |
| NC_006213.1;YP_009555242.1;HCoV-OC43_ns12.9;35 | 0 | 0 |
| NC_006213.1;YP_009555242.1;HCoV-OC43_ns12.9;36 | 0 | 0 |
| NC_006213.1;YP_009555242.1;HCoV-OC43_ns12.9;37 | 0 | 0 |
| NC_006213.1;YP_009555242.1;HCoV-OC43_ns12.9;38 | 0 | 0 |
| NC_006213.1;YP_009555242.1;HCoV-OC43_ns12.9;39 | 0 | 0 |
| NC_006213.1;YP_009555242.1;HCoV-OC43_ns12.9;40 | 0 | 0 |
| NC_006213.1;YP_009555242.1;HCoV-OC43_ns12.9;41 | 0 | 0 |
| NC_006213.1;YP_009555242.1;HCoV-OC43_ns12.9;42 | 0 | 0 |
| NC_006213.1;YP_009555242.1;HCoV-OC43_ns12.9;43 | 0 | 0 |
| NC_006213.1;YP_009555242.1;HCoV-OC43_ns12.9;44 | 0 | 0 |
| NC_006213.1;YP_009555242.1;HCoV-OC43_ns12.9;45 | 0 | 0 |
| NC_006213.1;YP_009555242.1;HCoV-OC43_ns12.9;46 | 0 | 0 |
| NC_006213.1;YP_009555242.1;HCoV-OC43_ns12.9;47 | 0 | 0 |
| NC_006213.1;YP_009555242.1;HCoV-OC43_ns12.9;48 | 0 | 0 |
| NC_006213.1;YP_009555242.1;HCoV-OC43_ns12.9;49 | 0 | 0 |
| NC_006213.1;YP_009555242.1;HCoV-OC43_ns12.9;50 | 0 | 0 |
| NC_006213.1;YP_009555242.1;HCoV-OC43_ns12.9;51 | 0 | 0 |
| NC_006213.1;YP_009555242.1;HCoV-OC43_ns12.9;52 | 0 | 0 |
| NC_006213.1;YP_009555242.1;HCoV-OC43_ns12.9;53 | 0 | 0 |
| NC_006213.1;YP_009555242.1;HCoV-OC43_ns12.9;54 | 0 | 0 |
| NC_006213.1;YP_009555242.1;HCoV-OC43_ns12.9;55 | 0 | 0 |
| NC_006213.1;YP_009555242.1;HCoV-OC43_ns12.9;56 | 0 | 0 |
| NC_006213.1;YP_009555242.1;HCoV-OC43_ns12.9;57 | 0 | 0 |
| NC_006213.1;YP_009555242.1;HCoV-OC43_ns12.9;58 | 0 | 0 |
| NC_006213.1;YP_009555242.1;HCoV-OC43_ns12.9;59 | 0 | 0 |
| NC_006213.1;YP_009555242.1;HCoV-OC43_ns12.9;60 | 0 | 0 |
| NC_006213.1;YP_009555242.1;HCoV-OC43_ns12.9;61 | 0 | 0 |
| NC_006213.1;YP_009555242.1;HCoV-OC43_ns12.9;62 | 0 | 0 |

| | | |
|---|---|---|
| NC_006213.1;YP_009555242.1;HCoV-OC43_ns12.9;63 | 0 | 0 |
| NC_006213.1;YP_009555242.1;HCoV-OC43_ns12.9;64 | 0 | 0 |
| NC_006213.1;YP_009555242.1;HCoV-OC43_ns12.9;65 | 0 | 0 |
| NC_006213.1;YP_009555242.1;HCoV-OC43_ns12.9;66 | 0 | 0 |
| NC_006213.1;YP_009555242.1;HCoV-OC43_ns12.9;67 | 0 | 0 |
| NC_006213.1;YP_009555242.1;HCoV-OC43_ns12.9;68 | 0 | 0 |
| NC_006213.1;YP_009555242.1;HCoV-OC43_ns12.9;69 | 0 | 0 |
| NC_006213.1;YP_009555242.1;HCoV-OC43_ns12.9;70 | 0 | 0 |
| NC_006213.1;YP_009555242.1;HCoV-OC43_ns12.9;71 | 0 | 0 |
| NC_006213.1;YP_009555242.1;HCoV-OC43_ns12.9;72 | 0 | 0 |
| NC_006213.1;YP_009555242.1;HCoV-OC43_ns12.9;73 | 0 | 0 |
| NC_006213.1;YP_009555242.1;HCoV-OC43_ns12.9;74 | 0.1 | 0 |
| NC_006213.1;YP_009555242.1;HCoV-OC43_ns12.9;75 | 0.1 | 0.05 |
| NC_006213.1;YP_009555242.1;HCoV-OC43_ns12.9;76 | 0.05 | 0.05 |
| NC_006213.1;YP_009555242.1;HCoV-OC43_ns12.9;77 | 0.05 | 0.05 |
| NC_006213.1;YP_009555242.1;HCoV-OC43_ns12.9;78 | 0.05 | 0 |
| NC_006213.1;YP_009555242.1;HCoV-OC43_ns12.9;79 | 0.05 | 0 |
| NC_006213.1;YP_009555242.1;HCoV-OC43_ns12.9;80 | 0.05 | 0.025 |
| NC_006213.1;YP_009555242.1;HCoV-OC43_ns12.9;81 | 0.05 | 0 |
| NC_006213.1;YP_009555242.1;HCoV-OC43_ns12.9;82 | 0.05 | 0 |
| NC_006213.1;YP_009555242.1;HCoV-OC43_ns12.9;83 | 0.05 | 0 |
| NC_006213.1;YP_009555242.1;HCoV-OC43_ns12.9;84 | 0 | 0 |
| NC_006213.1;YP_009555242.1;HCoV-OC43_ns12.9;85 | 0 | 0 |
| NC_006213.1;YP_009555242.1;HCoV-OC43_ns12.9;86 | 0 | 0 |
| NC_006213.1;YP_009555242.1;HCoV-OC43_ns12.9;87 | 0 | 0 |
| NC_006213.1;YP_009555242.1;HCoV-OC43_ns12.9;88 | 0 | 0 |
| NC_006213.1;YP_009555242.1;HCoV-OC43_ns12.9;89 | 0 | 0 |
| NC_006213.1;YP_009555242.1;HCoV-OC43_ns12.9;90 | 0 | 0 |
| NC_006213.1;YP_009555242.1;HCoV-OC43_ns12.9;91 | 0 | 0 |
| NC_006213.1;YP_009555242.1;HCoV-OC43_ns12.9;92 | 0 | 0 |
| NC_006213.1;YP_009555242.1;HCoV-OC43_ns12.9;93 | 0 | 0 |
| NC_006213.1;YP_009555242.1;HCoV-OC43_ns12.9;94 | 0 | 0 |
| NC_006213.1;YP_009555242.1;HCoV-OC43_ns12.9;95 | 0 | 0 |
| NC_006213.1;YP_009555242.1;HCoV-OC43_ns12.9;96 | 0 | 0 |
| NC_006213.1;YP_009555242.1;HCoV-OC43_ns12.9;97 | 0 | 0 |
| NC_006213.1;YP_009555242.1;HCoV-OC43_ns12.9;98 | 0 | 0 |
| NC_006213.1;YP_009555243.1;HCoV-OC43_envelope;1 | 0 | 0 |
| NC_006213.1;YP_009555243.1;HCoV-OC43_envelope;2 | 0 | 0 |
| NC_006213.1;YP_009555243.1;HCoV-OC43_envelope;3 | 0 | 0 |
| NC_006213.1;YP_009555243.1;HCoV-OC43_envelope;4 | 0 | 0 |
| NC_006213.1;YP_009555243.1;HCoV-OC43_envelope;5 | 0 | 0 |
| NC_006213.1;YP_009555243.1;HCoV-OC43_envelope;6 | 0 | 0 |
| NC_006213.1;YP_009555243.1;HCoV-OC43_envelope;7 | 0 | 0 |

| | | |
|---|---|---|
| NC_006213.1;YP_009555243.1;HCoV-OC43_envelope;8 | 0 | 0 |
| NC_006213.1;YP_009555243.1;HCoV-OC43_envelope;9 | 0 | 0 |
| NC_006213.1;YP_009555243.1;HCoV-OC43_envelope;10 | 0 | 0 |
| NC_006213.1;YP_009555243.1;HCoV-OC43_envelope;11 | 0 | 0 |
| NC_006213.1;YP_009555243.1;HCoV-OC43_envelope;12 | 0 | 0 |
| NC_006213.1;YP_009555243.1;HCoV-OC43_envelope;13 | 0 | 0 |
| NC_006213.1;YP_009555243.1;HCoV-OC43_envelope;14 | 0 | 0 |
| NC_006213.1;YP_009555243.1;HCoV-OC43_envelope;15 | 0 | 0 |
| NC_006213.1;YP_009555243.1;HCoV-OC43_envelope;16 | 0 | 0 |
| NC_006213.1;YP_009555243.1;HCoV-OC43_envelope;17 | 0 | 0 |
| NC_006213.1;YP_009555243.1;HCoV-OC43_envelope;18 | 0 | 0 |
| NC_006213.1;YP_009555243.1;HCoV-OC43_envelope;19 | 0 | 0 |
| NC_006213.1;YP_009555243.1;HCoV-OC43_envelope;20 | 0 | 0 |
| NC_006213.1;YP_009555243.1;HCoV-OC43_envelope;21 | 0 | 0 |
| NC_006213.1;YP_009555243.1;HCoV-OC43_envelope;22 | 0 | 0 |
| NC_006213.1;YP_009555243.1;HCoV-OC43_envelope;23 | 0 | 0 |
| NC_006213.1;YP_009555243.1;HCoV-OC43_envelope;24 | 0 | 0 |
| NC_006213.1;YP_009555243.1;HCoV-OC43_envelope;25 | 0 | 0 |
| NC_006213.1;YP_009555243.1;HCoV-OC43_envelope;26 | 0 | 0 |
| NC_006213.1;YP_009555243.1;HCoV-OC43_envelope;27 | 0 | 0 |
| NC_006213.1;YP_009555243.1;HCoV-OC43_envelope;28 | 0 | 0 |
| NC_006213.1;YP_009555243.1;HCoV-OC43_envelope;29 | 0 | 0 |
| NC_006213.1;YP_009555243.1;HCoV-OC43_envelope;30 | 0 | 0 |
| NC_006213.1;YP_009555243.1;HCoV-OC43_envelope;31 | 0 | 0 |
| NC_006213.1;YP_009555243.1;HCoV-OC43_envelope;32 | 0 | 0 |
| NC_006213.1;YP_009555243.1;HCoV-OC43_envelope;33 | 0 | 0 |
| NC_006213.1;YP_009555243.1;HCoV-OC43_envelope;34 | 0 | 0 |
| NC_006213.1;YP_009555243.1;HCoV-OC43_envelope;35 | 0 | 0 |
| NC_006213.1;YP_009555243.1;HCoV-OC43_envelope;36 | 0 | 0 |
| NC_006213.1;YP_009555243.1;HCoV-OC43_envelope;37 | 0 | 0 |
| NC_006213.1;YP_009555243.1;HCoV-OC43_envelope;38 | 0 | 0 |
| NC_006213.1;YP_009555243.1;HCoV-OC43_envelope;39 | 0 | 0 |
| NC_006213.1;YP_009555243.1;HCoV-OC43_envelope;40 | 0 | 0 |
| NC_006213.1;YP_009555243.1;HCoV-OC43_envelope;41 | 0 | 0 |
| NC_006213.1;YP_009555243.1;HCoV-OC43_envelope;42 | 0 | 0 |
| NC_006213.1;YP_009555243.1;HCoV-OC43_envelope;43 | 0 | 0 |
| NC_006213.1;YP_009555243.1;HCoV-OC43_envelope;44 | 0 | 0 |
| NC_006213.1;YP_009555243.1;HCoV-OC43_envelope;45 | 0 | 0 |
| NC_006213.1;YP_009555243.1;HCoV-OC43_envelope;46 | 0 | 0 |
| NC_006213.1;YP_009555243.1;HCoV-OC43_envelope;47 | 0 | 0 |
| NC_006213.1;YP_009555243.1;HCoV-OC43_envelope;48 | 0 | 0 |
| NC_006213.1;YP_009555243.1;HCoV-OC43_envelope;49 | 0 | 0 |
| NC_006213.1;YP_009555243.1;HCoV-OC43_envelope;50 | 0 | 0 |

| | | |
|---|---|---|
| NC_006213.1;YP_009555243.1;HCoV-OC43_envelope;51 | 0 | 0 |
| NC_006213.1;YP_009555243.1;HCoV-OC43_envelope;52 | 0 | 0 |
| NC_006213.1;YP_009555243.1;HCoV-OC43_envelope;53 | 0 | 0 |
| NC_006213.1;YP_009555243.1;HCoV-OC43_envelope;54 | 0 | 0 |
| NC_006213.1;YP_009555243.1;HCoV-OC43_envelope;55 | 0 | 0 |
| NC_006213.1;YP_009555243.1;HCoV-OC43_envelope;56 | 0 | 0 |
| NC_006213.1;YP_009555243.1;HCoV-OC43_envelope;57 | 0 | 0 |
| NC_006213.1;YP_009555243.1;HCoV-OC43_envelope;58 | 0 | 0 |
| NC_006213.1;YP_009555243.1;HCoV-OC43_envelope;59 | 0 | 0 |
| NC_006213.1;YP_009555243.1;HCoV-OC43_envelope;60 | 0 | 0 |
| NC_006213.1;YP_009555243.1;HCoV-OC43_envelope;61 | 0 | 0.025 |
| NC_006213.1;YP_009555243.1;HCoV-OC43_envelope;62 | 0.05 | 0.025 |
| NC_006213.1;YP_009555243.1;HCoV-OC43_envelope;63 | 0.05 | 0.025 |
| NC_006213.1;YP_009555243.1;HCoV-OC43_envelope;64 | 0 | 0.025 |
| NC_006213.1;YP_009555243.1;HCoV-OC43_envelope;65 | 0.1 | 0.05 |
| NC_006213.1;YP_009555243.1;HCoV-OC43_envelope;66 | 0.1 | 0.05 |
| NC_006213.1;YP_009555243.1;HCoV-OC43_envelope;67 | 0.1 | 0.075 |
| NC_006213.1;YP_009555243.1;HCoV-OC43_envelope;68 | 0.05 | 0 |
| NC_006213.1;YP_009555243.1;HCoV-OC43_envelope;69 | 0.05 | 0.025 |
| NC_006213.1;YP_009555243.1;HCoV-OC43_envelope;70 | 0.1 | 0 |
| NC_006213.1;YP_009555243.1;HCoV-OC43_envelope;71 | 0 | 0 |
| NC_006213.1;YP_009555243.1;HCoV-OC43_envelope;72 | 0 | 0 |
| NC_006213.1;YP_009555243.1;HCoV-OC43_envelope;73 | 0 | 0 |
| NC_006213.1;YP_009555244.1;HCoV-OC43_membrane;1 | 0.05 | 0 |
| NC_006213.1;YP_009555244.1;HCoV-OC43_membrane;2 | 0.05 | 0 |
| NC_006213.1;YP_009555244.1;HCoV-OC43_membrane;3 | 0.05 | 0 |
| NC_006213.1;YP_009555244.1;HCoV-OC43_membrane;4 | 0.05 | 0.05 |
| NC_006213.1;YP_009555244.1;HCoV-OC43_membrane;5 | 0.05 | 0.05 |
| NC_006213.1;YP_009555244.1;HCoV-OC43_membrane;6 | 0.05 | 0.05 |
| NC_006213.1;YP_009555244.1;HCoV-OC43_membrane;7 | 0.05 | 0.075 |
| NC_006213.1;YP_009555244.1;HCoV-OC43_membrane;8 | 0 | 0.1 |
| NC_006213.1;YP_009555244.1;HCoV-OC43_membrane;9 | 0.1 | 0.075 |
| NC_006213.1;YP_009555244.1;HCoV-OC43_membrane;10 | 0.1 | 0.125 |
| NC_006213.1;YP_009555244.1;HCoV-OC43_membrane;11 | 0.15 | 0.125 |
| NC_006213.1;YP_009555244.1;HCoV-OC43_membrane;12 | 0.15 | 0.225 |
| NC_006213.1;YP_009555244.1;HCoV-OC43_membrane;13 | 0.15 | 0.55 |
| NC_006213.1;YP_009555244.1;HCoV-OC43_membrane;14 | 0 | 0.05 |
| NC_006213.1;YP_009555244.1;HCoV-OC43_membrane;15 | 0 | 0 |
| NC_006213.1;YP_009555244.1;HCoV-OC43_membrane;16 | 0 | 0 |
| NC_006213.1;YP_009555244.1;HCoV-OC43_membrane;17 | 0 | 0 |
| NC_006213.1;YP_009555244.1;HCoV-OC43_membrane;18 | 0 | 0 |
| NC_006213.1;YP_009555244.1;HCoV-OC43_membrane;19 | 0 | 0 |
| NC_006213.1;YP_009555244.1;HCoV-OC43_membrane;20 | 0 | 0 |

| | | |
|---|---|---|
| NC_006213.1;YP_009555244.1;HCoV-OC43_membrane;21 | 0 | 0 |
| NC_006213.1;YP_009555244.1;HCoV-OC43_membrane;22 | 0.05 | 0.025 |
| NC_006213.1;YP_009555244.1;HCoV-OC43_membrane;23 | 0 | 0 |
| NC_006213.1;YP_009555244.1;HCoV-OC43_membrane;24 | 0 | 0 |
| NC_006213.1;YP_009555244.1;HCoV-OC43_membrane;25 | 0 | 0 |
| NC_006213.1;YP_009555244.1;HCoV-OC43_membrane;26 | 0 | 0 |
| NC_006213.1;YP_009555244.1;HCoV-OC43_membrane;27 | 0 | 0 |
| NC_006213.1;YP_009555244.1;HCoV-OC43_membrane;28 | 0 | 0 |
| NC_006213.1;YP_009555244.1;HCoV-OC43_membrane;29 | 0 | 0 |
| NC_006213.1;YP_009555244.1;HCoV-OC43_membrane;30 | 0 | 0 |
| NC_006213.1;YP_009555244.1;HCoV-OC43_membrane;31 | 0 | 0 |
| NC_006213.1;YP_009555244.1;HCoV-OC43_membrane;32 | 0 | 0 |
| NC_006213.1;YP_009555244.1;HCoV-OC43_membrane;33 | 0 | 0 |
| NC_006213.1;YP_009555244.1;HCoV-OC43_membrane;34 | 0 | 0 |
| NC_006213.1;YP_009555244.1;HCoV-OC43_membrane;35 | 0 | 0 |
| NC_006213.1;YP_009555244.1;HCoV-OC43_membrane;36 | 0 | 0 |
| NC_006213.1;YP_009555244.1;HCoV-OC43_membrane;37 | 0 | 0 |
| NC_006213.1;YP_009555244.1;HCoV-OC43_membrane;38 | 0 | 0 |
| NC_006213.1;YP_009555244.1;HCoV-OC43_membrane;39 | 0 | 0 |
| NC_006213.1;YP_009555244.1;HCoV-OC43_membrane;40 | 0 | 0 |
| NC_006213.1;YP_009555244.1;HCoV-OC43_membrane;41 | 0 | 0 |
| NC_006213.1;YP_009555244.1;HCoV-OC43_membrane;42 | 0 | 0 |
| NC_006213.1;YP_009555244.1;HCoV-OC43_membrane;43 | 0 | 0 |
| NC_006213.1;YP_009555244.1;HCoV-OC43_membrane;44 | 0 | 0 |
| NC_006213.1;YP_009555244.1;HCoV-OC43_membrane;45 | 0 | 0 |
| NC_006213.1;YP_009555244.1;HCoV-OC43_membrane;46 | 0 | 0 |
| NC_006213.1;YP_009555244.1;HCoV-OC43_membrane;47 | 0 | 0 |
| NC_006213.1;YP_009555244.1;HCoV-OC43_membrane;48 | 0 | 0 |
| NC_006213.1;YP_009555244.1;HCoV-OC43_membrane;49 | 0 | 0 |
| NC_006213.1;YP_009555244.1;HCoV-OC43_membrane;50 | 0 | 0 |
| NC_006213.1;YP_009555244.1;HCoV-OC43_membrane;51 | 0 | 0 |
| NC_006213.1;YP_009555244.1;HCoV-OC43_membrane;52 | 0 | 0 |
| NC_006213.1;YP_009555244.1;HCoV-OC43_membrane;53 | 0 | 0 |
| NC_006213.1;YP_009555244.1;HCoV-OC43_membrane;54 | 0 | 0 |
| NC_006213.1;YP_009555244.1;HCoV-OC43_membrane;55 | 0 | 0 |
| NC_006213.1;YP_009555244.1;HCoV-OC43_membrane;56 | 0 | 0 |
| NC_006213.1;YP_009555244.1;HCoV-OC43_membrane;57 | 0 | 0 |
| NC_006213.1;YP_009555244.1;HCoV-OC43_membrane;58 | 0 | 0 |
| NC_006213.1;YP_009555244.1;HCoV-OC43_membrane;59 | 0 | 0 |
| NC_006213.1;YP_009555244.1;HCoV-OC43_membrane;60 | 0 | 0 |
| NC_006213.1;YP_009555244.1;HCoV-OC43_membrane;61 | 0 | 0 |
| NC_006213.1;YP_009555244.1;HCoV-OC43_membrane;62 | 0 | 0 |
| NC_006213.1;YP_009555244.1;HCoV-OC43_membrane;63 | 0 | 0 |

| | | |
|---|---|---|
| NC_006213.1;YP_009555244.1;HCoV-OC43_membrane;64 | 0 | 0 |
| NC_006213.1;YP_009555244.1;HCoV-OC43_membrane;65 | 0 | 0 |
| NC_006213.1;YP_009555244.1;HCoV-OC43_membrane;66 | 0 | 0 |
| NC_006213.1;YP_009555244.1;HCoV-OC43_membrane;67 | 0 | 0 |
| NC_006213.1;YP_009555244.1;HCoV-OC43_membrane;68 | 0 | 0 |
| NC_006213.1;YP_009555244.1;HCoV-OC43_membrane;69 | 0 | 0 |
| NC_006213.1;YP_009555244.1;HCoV-OC43_membrane;70 | 0 | 0 |
| NC_006213.1;YP_009555244.1;HCoV-OC43_membrane;71 | 0 | 0 |
| NC_006213.1;YP_009555244.1;HCoV-OC43_membrane;72 | 0 | 0 |
| NC_006213.1;YP_009555244.1;HCoV-OC43_membrane;73 | 0 | 0 |
| NC_006213.1;YP_009555244.1;HCoV-OC43_membrane;74 | 0 | 0 |
| NC_006213.1;YP_009555244.1;HCoV-OC43_membrane;75 | 0 | 0 |
| NC_006213.1;YP_009555244.1;HCoV-OC43_membrane;76 | 0 | 0 |
| NC_006213.1;YP_009555244.1;HCoV-OC43_membrane;77 | 0 | 0 |
| NC_006213.1;YP_009555244.1;HCoV-OC43_membrane;78 | 0 | 0 |
| NC_006213.1;YP_009555244.1;HCoV-OC43_membrane;79 | 0 | 0 |
| NC_006213.1;YP_009555244.1;HCoV-OC43_membrane;80 | 0 | 0 |
| NC_006213.1;YP_009555244.1;HCoV-OC43_membrane;81 | 0 | 0 |
| NC_006213.1;YP_009555244.1;HCoV-OC43_membrane;82 | 0 | 0 |
| NC_006213.1;YP_009555244.1;HCoV-OC43_membrane;83 | 0 | 0 |
| NC_006213.1;YP_009555244.1;HCoV-OC43_membrane;84 | 0 | 0 |
| NC_006213.1;YP_009555244.1;HCoV-OC43_membrane;85 | 0 | 0 |
| NC_006213.1;YP_009555244.1;HCoV-OC43_membrane;86 | 0 | 0 |
| NC_006213.1;YP_009555244.1;HCoV-OC43_membrane;87 | 0 | 0 |
| NC_006213.1;YP_009555244.1;HCoV-OC43_membrane;88 | 0 | 0 |
| NC_006213.1;YP_009555244.1;HCoV-OC43_membrane;89 | 0 | 0 |
| NC_006213.1;YP_009555244.1;HCoV-OC43_membrane;90 | 0 | 0 |
| NC_006213.1;YP_009555244.1;HCoV-OC43_membrane;91 | 0 | 0 |
| NC_006213.1;YP_009555244.1;HCoV-OC43_membrane;92 | 0 | 0 |
| NC_006213.1;YP_009555244.1;HCoV-OC43_membrane;93 | 0 | 0 |
| NC_006213.1;YP_009555244.1;HCoV-OC43_membrane;94 | 0 | 0 |
| NC_006213.1;YP_009555244.1;HCoV-OC43_membrane;95 | 0 | 0 |
| NC_006213.1;YP_009555244.1;HCoV-OC43_membrane;96 | 0 | 0 |
| NC_006213.1;YP_009555244.1;HCoV-OC43_membrane;97 | 0 | 0 |
| NC_006213.1;YP_009555244.1;HCoV-OC43_membrane;98 | 0 | 0 |
| NC_006213.1;YP_009555244.1;HCoV-OC43_membrane;99 | 0 | 0 |
| NC_006213.1;YP_009555244.1;HCoV-OC43_membrane;100 | 0 | 0 |
| NC_006213.1;YP_009555244.1;HCoV-OC43_membrane;101 | 0 | 0 |
| NC_006213.1;YP_009555244.1;HCoV-OC43_membrane;102 | 0 | 0 |
| NC_006213.1;YP_009555244.1;HCoV-OC43_membrane;103 | 0 | 0 |
| NC_006213.1;YP_009555244.1;HCoV-OC43_membrane;104 | 0 | 0 |
| NC_006213.1;YP_009555244.1;HCoV-OC43_membrane;105 | 0 | 0 |
| NC_006213.1;YP_009555244.1;HCoV-OC43_membrane;106 | 0 | 0 |

| | | |
|---|---|---|
| NC_006213.1;YP_009555244.1;HCoV-OC43_membrane;107 | 0 | 0 |
| NC_006213.1;YP_009555244.1;HCoV-OC43_membrane;108 | 0 | 0 |
| NC_006213.1;YP_009555244.1;HCoV-OC43_membrane;109 | 0 | 0 |
| NC_006213.1;YP_009555244.1;HCoV-OC43_membrane;110 | 0 | 0 |
| NC_006213.1;YP_009555244.1;HCoV-OC43_membrane;111 | 0 | 0 |
| NC_006213.1;YP_009555244.1;HCoV-OC43_membrane;112 | 0 | 0 |
| NC_006213.1;YP_009555244.1;HCoV-OC43_membrane;113 | 0 | 0 |
| NC_006213.1;YP_009555244.1;HCoV-OC43_membrane;114 | 0 | 0.025 |
| NC_006213.1;YP_009555244.1;HCoV-OC43_membrane;115 | 0.1 | 0.025 |
| NC_006213.1;YP_009555244.1;HCoV-OC43_membrane;116 | 0.05 | 0 |
| NC_006213.1;YP_009555244.1;HCoV-OC43_membrane;117 | 0.05 | 0 |
| NC_006213.1;YP_009555244.1;HCoV-OC43_membrane;118 | 0.05 | 0 |
| NC_006213.1;YP_009555244.1;HCoV-OC43_membrane;119 | 0.05 | 0.025 |
| NC_006213.1;YP_009555244.1;HCoV-OC43_membrane;120 | 0.05 | 0.025 |
| NC_006213.1;YP_009555244.1;HCoV-OC43_membrane;121 | 0 | 0 |
| NC_006213.1;YP_009555244.1;HCoV-OC43_membrane;122 | 0 | 0.025 |
| NC_006213.1;YP_009555244.1;HCoV-OC43_membrane;123 | 0 | 0 |
| NC_006213.1;YP_009555244.1;HCoV-OC43_membrane;124 | 0 | 0 |
| NC_006213.1;YP_009555244.1;HCoV-OC43_membrane;125 | 0 | 0.025 |
| NC_006213.1;YP_009555244.1;HCoV-OC43_membrane;126 | 0 | 0.075 |
| NC_006213.1;YP_009555244.1;HCoV-OC43_membrane;127 | 0 | 0.075 |
| NC_006213.1;YP_009555244.1;HCoV-OC43_membrane;128 | 0 | 0.025 |
| NC_006213.1;YP_009555244.1;HCoV-OC43_membrane;129 | 0 | 0.05 |
| NC_006213.1;YP_009555244.1;HCoV-OC43_membrane;130 | 0 | 0.05 |
| NC_006213.1;YP_009555244.1;HCoV-OC43_membrane;131 | 0 | 0.05 |
| NC_006213.1;YP_009555244.1;HCoV-OC43_membrane;132 | 0 | 0.025 |
| NC_006213.1;YP_009555244.1;HCoV-OC43_membrane;133 | 0 | 0.025 |
| NC_006213.1;YP_009555244.1;HCoV-OC43_membrane;134 | 0 | 0.025 |
| NC_006213.1;YP_009555244.1;HCoV-OC43_membrane;135 | 0 | 0.025 |
| NC_006213.1;YP_009555244.1;HCoV-OC43_membrane;136 | 0 | 0.025 |
| NC_006213.1;YP_009555244.1;HCoV-OC43_membrane;137 | 0 | 0 |
| NC_006213.1;YP_009555244.1;HCoV-OC43_membrane;138 | 0 | 0 |
| NC_006213.1;YP_009555244.1;HCoV-OC43_membrane;139 | 0 | 0 |
| NC_006213.1;YP_009555244.1;HCoV-OC43_membrane;140 | 0 | 0 |
| NC_006213.1;YP_009555244.1;HCoV-OC43_membrane;141 | 0 | 0 |
| NC_006213.1;YP_009555244.1;HCoV-OC43_membrane;142 | 0 | 0 |
| NC_006213.1;YP_009555244.1;HCoV-OC43_membrane;143 | 0 | 0 |
| NC_006213.1;YP_009555244.1;HCoV-OC43_membrane;144 | 0 | 0 |
| NC_006213.1;YP_009555244.1;HCoV-OC43_membrane;145 | 0 | 0 |
| NC_006213.1;YP_009555244.1;HCoV-OC43_membrane;146 | 0 | 0 |
| NC_006213.1;YP_009555244.1;HCoV-OC43_membrane;147 | 0 | 0 |
| NC_006213.1;YP_009555244.1;HCoV-OC43_membrane;148 | 0 | 0 |
| NC_006213.1;YP_009555244.1;HCoV-OC43_membrane;149 | 0 | 0.025 |

| | | |
|---|---|---|
| NC_006213.1;YP_009555244.1;HCoV-OC43_membrane;150 | 0 | 0.025 |
| NC_006213.1;YP_009555244.1;HCoV-OC43_membrane;151 | 0 | 0.025 |
| NC_006213.1;YP_009555244.1;HCoV-OC43_membrane;152 | 0 | 0.025 |
| NC_006213.1;YP_009555244.1;HCoV-OC43_membrane;153 | 0 | 0.025 |
| NC_006213.1;YP_009555244.1;HCoV-OC43_membrane;154 | 0 | 0.05 |
| NC_006213.1;YP_009555244.1;HCoV-OC43_membrane;155 | 0 | 0.05 |
| NC_006213.1;YP_009555244.1;HCoV-OC43_membrane;156 | 0 | 0.1 |
| NC_006213.1;YP_009555244.1;HCoV-OC43_membrane;157 | 0 | 0.05 |
| NC_006213.1;YP_009555244.1;HCoV-OC43_membrane;158 | 0 | 0.025 |
| NC_006213.1;YP_009555244.1;HCoV-OC43_membrane;159 | 0 | 0.075 |
| NC_006213.1;YP_009555244.1;HCoV-OC43_membrane;160 | 0 | 0.05 |
| NC_006213.1;YP_009555244.1;HCoV-OC43_membrane;161 | 0 | 0.05 |
| NC_006213.1;YP_009555244.1;HCoV-OC43_membrane;162 | 0 | 0.125 |
| NC_006213.1;YP_009555244.1;HCoV-OC43_membrane;163 | 0 | 0.025 |
| NC_006213.1;YP_009555244.1;HCoV-OC43_membrane;164 | 0 | 0 |
| NC_006213.1;YP_009555244.1;HCoV-OC43_membrane;165 | 0 | 0 |
| NC_006213.1;YP_009555244.1;HCoV-OC43_membrane;166 | 0 | 0 |
| NC_006213.1;YP_009555244.1;HCoV-OC43_membrane;167 | 0 | 0 |
| NC_006213.1;YP_009555244.1;HCoV-OC43_membrane;168 | 0 | 0 |
| NC_006213.1;YP_009555244.1;HCoV-OC43_membrane;169 | 0 | 0 |
| NC_006213.1;YP_009555244.1;HCoV-OC43_membrane;170 | 0 | 0 |
| NC_006213.1;YP_009555244.1;HCoV-OC43_membrane;171 | 0 | 0 |
| NC_006213.1;YP_009555244.1;HCoV-OC43_membrane;172 | 0 | 0 |
| NC_006213.1;YP_009555244.1;HCoV-OC43_membrane;173 | 0 | 0 |
| NC_006213.1;YP_009555244.1;HCoV-OC43_membrane;174 | 0 | 0 |
| NC_006213.1;YP_009555244.1;HCoV-OC43_membrane;175 | 0 | 0 |
| NC_006213.1;YP_009555244.1;HCoV-OC43_membrane;176 | 0 | 0 |
| NC_006213.1;YP_009555244.1;HCoV-OC43_membrane;177 | 0 | 0 |
| NC_006213.1;YP_009555244.1;HCoV-OC43_membrane;178 | 0 | 0 |
| NC_006213.1;YP_009555244.1;HCoV-OC43_membrane;179 | 0 | 0 |
| NC_006213.1;YP_009555244.1;HCoV-OC43_membrane;180 | 0 | 0 |
| NC_006213.1;YP_009555244.1;HCoV-OC43_membrane;181 | 0 | 0 |
| NC_006213.1;YP_009555244.1;HCoV-OC43_membrane;182 | 0 | 0 |
| NC_006213.1;YP_009555244.1;HCoV-OC43_membrane;183 | 0 | 0 |
| NC_006213.1;YP_009555244.1;HCoV-OC43_membrane;184 | 0 | 0 |
| NC_006213.1;YP_009555244.1;HCoV-OC43_membrane;185 | 0 | 0 |
| NC_006213.1;YP_009555244.1;HCoV-OC43_membrane;186 | 0 | 0 |
| NC_006213.1;YP_009555244.1;HCoV-OC43_membrane;187 | 0 | 0 |
| NC_006213.1;YP_009555244.1;HCoV-OC43_membrane;188 | 0.05 | 0 |
| NC_006213.1;YP_009555244.1;HCoV-OC43_membrane;189 | 0 | 0 |
| NC_006213.1;YP_009555244.1;HCoV-OC43_membrane;190 | 0 | 0 |
| NC_006213.1;YP_009555244.1;HCoV-OC43_membrane;191 | 0 | 0 |
| NC_006213.1;YP_009555244.1;HCoV-OC43_membrane;192 | 0 | 0 |

| | | |
|---|---|---|
| NC_006213.1;YP_009555244.1;HCoV-OC43_membrane;193 | 0 | 0 |
| NC_006213.1;YP_009555244.1;HCoV-OC43_membrane;194 | 0 | 0 |
| NC_006213.1;YP_009555244.1;HCoV-OC43_membrane;195 | 0 | 0 |
| NC_006213.1;YP_009555244.1;HCoV-OC43_membrane;196 | 0 | 0 |
| NC_006213.1;YP_009555244.1;HCoV-OC43_membrane;197 | 0 | 0 |
| NC_006213.1;YP_009555244.1;HCoV-OC43_membrane;198 | 0 | 0 |
| NC_006213.1;YP_009555244.1;HCoV-OC43_membrane;199 | 0 | 0 |
| NC_006213.1;YP_009555244.1;HCoV-OC43_membrane;200 | 0 | 0 |
| NC_006213.1;YP_009555244.1;HCoV-OC43_membrane;201 | 0 | 0.025 |
| NC_006213.1;YP_009555244.1;HCoV-OC43_membrane;202 | 0.05 | 0.025 |
| NC_006213.1;YP_009555244.1;HCoV-OC43_membrane;203 | 0 | 0.025 |
| NC_006213.1;YP_009555244.1;HCoV-OC43_membrane;204 | 0 | 0.025 |
| NC_006213.1;YP_009555244.1;HCoV-OC43_membrane;205 | 0 | 0.025 |
| NC_006213.1;YP_009555244.1;HCoV-OC43_membrane;206 | 0 | 0.025 |
| NC_006213.1;YP_009555244.1;HCoV-OC43_membrane;207 | 0.05 | 0.05 |
| NC_006213.1;YP_009555244.1;HCoV-OC43_membrane;208 | 0 | 0.075 |
| NC_006213.1;YP_009555244.1;HCoV-OC43_membrane;209 | 0 | 0.075 |
| NC_006213.1;YP_009555244.1;HCoV-OC43_membrane;210 | 0.1 | 0.15 |
| NC_006213.1;YP_009555244.1;HCoV-OC43_membrane;211 | 0.15 | 0.175 |
| NC_006213.1;YP_009555244.1;HCoV-OC43_membrane;212 | 0.15 | 0.175 |
| NC_006213.1;YP_009555244.1;HCoV-OC43_membrane;213 | 0.2 | 0.175 |
| NC_006213.1;YP_009555244.1;HCoV-OC43_membrane;214 | 0.15 | 0.175 |
| NC_006213.1;YP_009555244.1;HCoV-OC43_membrane;215 | 0.1 | 0.15 |
| NC_006213.1;YP_009555244.1;HCoV-OC43_membrane;216 | 0.15 | 0.175 |
| NC_006213.1;YP_009555244.1;HCoV-OC43_membrane;217 | 0.2 | 0.175 |
| NC_006213.1;YP_009555244.1;HCoV-OC43_membrane;218 | 0 | 0 |
| NC_006213.1;YP_009555244.1;HCoV-OC43_membrane;219 | 0 | 0 |
| NC_006213.1;YP_009555245.1;HCoV-OC43_nucleoca;1 | 0.05 | 0.125 |
| NC_006213.1;YP_009555245.1;HCoV-OC43_nucleoca;2 | 0.05 | 0.15 |
| NC_006213.1;YP_009555245.1;HCoV-OC43_nucleoca;3 | 0.05 | 0.1 |
| NC_006213.1;YP_009555245.1;HCoV-OC43_nucleoca;4 | 0.05 | 0.075 |
| NC_006213.1;YP_009555245.1;HCoV-OC43_nucleoca;5 | 0.05 | 0.075 |
| NC_006213.1;YP_009555245.1;HCoV-OC43_nucleoca;6 | 0.05 | 0.075 |
| NC_006213.1;YP_009555245.1;HCoV-OC43_nucleoca;7 | 0.05 | 0.025 |
| NC_006213.1;YP_009555245.1;HCoV-OC43_nucleoca;8 | 0 | 0.075 |
| NC_006213.1;YP_009555245.1;HCoV-OC43_nucleoca;9 | 0 | 0.05 |
| NC_006213.1;YP_009555245.1;HCoV-OC43_nucleoca;10 | 0 | 0.025 |
| NC_006213.1;YP_009555245.1;HCoV-OC43_nucleoca;11 | 0 | 0.125 |
| NC_006213.1;YP_009555245.1;HCoV-OC43_nucleoca;12 | 0 | 0.025 |
| NC_006213.1;YP_009555245.1;HCoV-OC43_nucleoca;13 | 0.05 | 0.1 |
| NC_006213.1;YP_009555245.1;HCoV-OC43_nucleoca;14 | 0 | 0.025 |
| NC_006213.1;YP_009555245.1;HCoV-OC43_nucleoca;15 | 0 | 0.025 |
| NC_006213.1;YP_009555245.1;HCoV-OC43_nucleoca;16 | 0 | 0.025 |

| | | |
|---|---|---|
| NC_006213.1;YP_009555245.1;HCoV-OC43_nucleoca;17 | 0 | 0 |
| NC_006213.1;YP_009555245.1;HCoV-OC43_nucleoca;18 | 0 | 0.1 |
| NC_006213.1;YP_009555245.1;HCoV-OC43_nucleoca;19 | 0.1 | 0.025 |
| NC_006213.1;YP_009555245.1;HCoV-OC43_nucleoca;20 | 0.1 | 0.025 |
| NC_006213.1;YP_009555245.1;HCoV-OC43_nucleoca;21 | 0.05 | 0 |
| NC_006213.1;YP_009555245.1;HCoV-OC43_nucleoca;22 | 0.35 | 0.075 |
| NC_006213.1;YP_009555245.1;HCoV-OC43_nucleoca;23 | 0.15 | 0.025 |
| NC_006213.1;YP_009555245.1;HCoV-OC43_nucleoca;24 | 0.3 | 0.025 |
| NC_006213.1;YP_009555245.1;HCoV-OC43_nucleoca;25 | 0.3 | 0.025 |
| NC_006213.1;YP_009555245.1;HCoV-OC43_nucleoca;26 | 0 | 0 |
| NC_006213.1;YP_009555245.1;HCoV-OC43_nucleoca;27 | 0 | 0 |
| NC_006213.1;YP_009555245.1;HCoV-OC43_nucleoca;28 | 0 | 0 |
| NC_006213.1;YP_009555245.1;HCoV-OC43_nucleoca;29 | 0 | 0 |
| NC_006213.1;YP_009555245.1;HCoV-OC43_nucleoca;30 | 0 | 0 |
| NC_006213.1;YP_009555245.1;HCoV-OC43_nucleoca;31 | 0 | 0 |
| NC_006213.1;YP_009555245.1;HCoV-OC43_nucleoca;32 | 0 | 0 |
| NC_006213.1;YP_009555245.1;HCoV-OC43_nucleoca;33 | 0 | 0 |
| NC_006213.1;YP_009555245.1;HCoV-OC43_nucleoca;34 | 0 | 0 |
| NC_006213.1;YP_009555245.1;HCoV-OC43_nucleoca;35 | 0.05 | 0 |
| NC_006213.1;YP_009555245.1;HCoV-OC43_nucleoca;36 | 0.05 | 0 |
| NC_006213.1;YP_009555245.1;HCoV-OC43_nucleoca;37 | 0.05 | 0.025 |
| NC_006213.1;YP_009555245.1;HCoV-OC43_nucleoca;38 | 0.05 | 0.025 |
| NC_006213.1;YP_009555245.1;HCoV-OC43_nucleoca;39 | 0.05 | 0 |
| NC_006213.1;YP_009555245.1;HCoV-OC43_nucleoca;40 | 0.05 | 0 |
| NC_006213.1;YP_009555245.1;HCoV-OC43_nucleoca;41 | 0.05 | 0 |
| NC_006213.1;YP_009555245.1;HCoV-OC43_nucleoca;42 | 0.05 | 0 |
| NC_006213.1;YP_009555245.1;HCoV-OC43_nucleoca;43 | 0.15 | 0.025 |
| NC_006213.1;YP_009555245.1;HCoV-OC43_nucleoca;44 | 0.15 | 0 |
| NC_006213.1;YP_009555245.1;HCoV-OC43_nucleoca;45 | 0.1 | 0 |
| NC_006213.1;YP_009555245.1;HCoV-OC43_nucleoca;46 | 0.15 | 0.025 |
| NC_006213.1;YP_009555245.1;HCoV-OC43_nucleoca;47 | 0 | 0.025 |
| NC_006213.1;YP_009555245.1;HCoV-OC43_nucleoca;48 | 0 | 0.025 |
| NC_006213.1;YP_009555245.1;HCoV-OC43_nucleoca;49 | 0 | 0.075 |
| NC_006213.1;YP_009555245.1;HCoV-OC43_nucleoca;50 | 0 | 0 |
| NC_006213.1;YP_009555245.1;HCoV-OC43_nucleoca;51 | 0 | 0 |
| NC_006213.1;YP_009555245.1;HCoV-OC43_nucleoca;52 | 0 | 0 |
| NC_006213.1;YP_009555245.1;HCoV-OC43_nucleoca;53 | 0 | 0 |
| NC_006213.1;YP_009555245.1;HCoV-OC43_nucleoca;54 | 0 | 0 |
| NC_006213.1;YP_009555245.1;HCoV-OC43_nucleoca;55 | 0 | 0 |
| NC_006213.1;YP_009555245.1;HCoV-OC43_nucleoca;56 | 0 | 0 |
| NC_006213.1;YP_009555245.1;HCoV-OC43_nucleoca;57 | 0 | 0 |
| NC_006213.1;YP_009555245.1;HCoV-OC43_nucleoca;58 | 0 | 0 |
| NC_006213.1;YP_009555245.1;HCoV-OC43_nucleoca;59 | 0 | 0 |

| | | |
|---|---|---|
| NC_006213.1;YP_009555245.1;HCoV-OC43_nucleoca;60 | 0 | 0 |
| NC_006213.1;YP_009555245.1;HCoV-OC43_nucleoca;61 | 0 | 0 |
| NC_006213.1;YP_009555245.1;HCoV-OC43_nucleoca;62 | 0 | 0 |
| NC_006213.1;YP_009555245.1;HCoV-OC43_nucleoca;63 | 0 | 0 |
| NC_006213.1;YP_009555245.1;HCoV-OC43_nucleoca;64 | 0 | 0 |
| NC_006213.1;YP_009555245.1;HCoV-OC43_nucleoca;65 | 0 | 0 |
| NC_006213.1;YP_009555245.1;HCoV-OC43_nucleoca;66 | 0 | 0 |
| NC_006213.1;YP_009555245.1;HCoV-OC43_nucleoca;67 | 0 | 0.025 |
| NC_006213.1;YP_009555245.1;HCoV-OC43_nucleoca;68 | 0.1 | 0.05 |
| NC_006213.1;YP_009555245.1;HCoV-OC43_nucleoca;69 | 0 | 0.05 |
| NC_006213.1;YP_009555245.1;HCoV-OC43_nucleoca;70 | 0 | 0.05 |
| NC_006213.1;YP_009555245.1;HCoV-OC43_nucleoca;71 | 0 | 0.025 |
| NC_006213.1;YP_009555245.1;HCoV-OC43_nucleoca;72 | 0.1 | 0.1 |
| NC_006213.1;YP_009555245.1;HCoV-OC43_nucleoca;73 | 0 | 0.025 |
| NC_006213.1;YP_009555245.1;HCoV-OC43_nucleoca;74 | 0 | 0.05 |
| NC_006213.1;YP_009555245.1;HCoV-OC43_nucleoca;75 | 0.1 | 0.05 |
| NC_006213.1;YP_009555245.1;HCoV-OC43_nucleoca;76 | 0.05 | 0.025 |
| NC_006213.1;YP_009555245.1;HCoV-OC43_nucleoca;77 | 0.1 | 0.025 |
| NC_006213.1;YP_009555245.1;HCoV-OC43_nucleoca;78 | 0.15 | 0.025 |
| NC_006213.1;YP_009555245.1;HCoV-OC43_nucleoca;79 | 0.1 | 0.025 |
| NC_006213.1;YP_009555245.1;HCoV-OC43_nucleoca;80 | 0.05 | 0 |
| NC_006213.1;YP_009555245.1;HCoV-OC43_nucleoca;81 | 0.05 | 0 |
| NC_006213.1;YP_009555245.1;HCoV-OC43_nucleoca;82 | 0.05 | 0 |
| NC_006213.1;YP_009555245.1;HCoV-OC43_nucleoca;83 | 0.1 | 0 |
| NC_006213.1;YP_009555245.1;HCoV-OC43_nucleoca;84 | 0.1 | 0 |
| NC_006213.1;YP_009555245.1;HCoV-OC43_nucleoca;85 | 0.15 | 0 |
| NC_006213.1;YP_009555245.1;HCoV-OC43_nucleoca;86 | 0.15 | 0 |
| NC_006213.1;YP_009555245.1;HCoV-OC43_nucleoca;87 | 0.05 | 0 |
| NC_006213.1;YP_009555245.1;HCoV-OC43_nucleoca;88 | 0 | 0 |
| NC_006213.1;YP_009555245.1;HCoV-OC43_nucleoca;89 | 0 | 0 |
| NC_006213.1;YP_009555245.1;HCoV-OC43_nucleoca;90 | 0 | 0 |
| NC_006213.1;YP_009555245.1;HCoV-OC43_nucleoca;91 | 0 | 0 |
| NC_006213.1;YP_009555245.1;HCoV-OC43_nucleoca;92 | 0 | 0 |
| NC_006213.1;YP_009555245.1;HCoV-OC43_nucleoca;93 | 0 | 0 |
| NC_006213.1;YP_009555245.1;HCoV-OC43_nucleoca;94 | 0 | 0 |
| NC_006213.1;YP_009555245.1;HCoV-OC43_nucleoca;95 | 0 | 0 |
| NC_006213.1;YP_009555245.1;HCoV-OC43_nucleoca;96 | 0 | 0 |
| NC_006213.1;YP_009555245.1;HCoV-OC43_nucleoca;97 | 0 | 0 |
| NC_006213.1;YP_009555245.1;HCoV-OC43_nucleoca;98 | 0 | 0 |
| NC_006213.1;YP_009555245.1;HCoV-OC43_nucleoca;99 | 0 | 0 |
| NC_006213.1;YP_009555245.1;HCoV-OC43_nucleoca;100 | 0 | 0 |
| NC_006213.1;YP_009555245.1;HCoV-OC43_nucleoca;101 | 0 | 0 |
| NC_006213.1;YP_009555245.1;HCoV-OC43_nucleoca;102 | 0 | 0 |

| | | |
|---|---|---|
| NC_006213.1;YP_009555245.1;HCoV-OC43_nucleoca;103 | 0 | 0 |
| NC_006213.1;YP_009555245.1;HCoV-OC43_nucleoca;104 | 0 | 0 |
| NC_006213.1;YP_009555245.1;HCoV-OC43_nucleoca;105 | 0 | 0.025 |
| NC_006213.1;YP_009555245.1;HCoV-OC43_nucleoca;106 | 0 | 0 |
| NC_006213.1;YP_009555245.1;HCoV-OC43_nucleoca;107 | 0 | 0.025 |
| NC_006213.1;YP_009555245.1;HCoV-OC43_nucleoca;108 | 0 | 0 |
| NC_006213.1;YP_009555245.1;HCoV-OC43_nucleoca;109 | 0 | 0.025 |
| NC_006213.1;YP_009555245.1;HCoV-OC43_nucleoca;110 | 0 | 0.025 |
| NC_006213.1;YP_009555245.1;HCoV-OC43_nucleoca;111 | 0 | 0 |
| NC_006213.1;YP_009555245.1;HCoV-OC43_nucleoca;112 | 0 | 0 |
| NC_006213.1;YP_009555245.1;HCoV-OC43_nucleoca;113 | 0 | 0 |
| NC_006213.1;YP_009555245.1;HCoV-OC43_nucleoca;114 | 0 | 0 |
| NC_006213.1;YP_009555245.1;HCoV-OC43_nucleoca;115 | 0 | 0 |
| NC_006213.1;YP_009555245.1;HCoV-OC43_nucleoca;116 | 0 | 0 |
| NC_006213.1;YP_009555245.1;HCoV-OC43_nucleoca;117 | 0 | 0 |
| NC_006213.1;YP_009555245.1;HCoV-OC43_nucleoca;118 | 0 | 0 |
| NC_006213.1;YP_009555245.1;HCoV-OC43_nucleoca;119 | 0 | 0 |
| NC_006213.1;YP_009555245.1;HCoV-OC43_nucleoca;120 | 0 | 0 |
| NC_006213.1;YP_009555245.1;HCoV-OC43_nucleoca;121 | 0 | 0 |
| NC_006213.1;YP_009555245.1;HCoV-OC43_nucleoca;122 | 0 | 0 |
| NC_006213.1;YP_009555245.1;HCoV-OC43_nucleoca;123 | 0 | 0 |
| NC_006213.1;YP_009555245.1;HCoV-OC43_nucleoca;124 | 0 | 0 |
| NC_006213.1;YP_009555245.1;HCoV-OC43_nucleoca;125 | 0.05 | 0 |
| NC_006213.1;YP_009555245.1;HCoV-OC43_nucleoca;126 | 0.05 | 0 |
| NC_006213.1;YP_009555245.1;HCoV-OC43_nucleoca;127 | 0.1 | 0 |
| NC_006213.1;YP_009555245.1;HCoV-OC43_nucleoca;128 | 0.05 | 0.025 |
| NC_006213.1;YP_009555245.1;HCoV-OC43_nucleoca;129 | 0 | 0 |
| NC_006213.1;YP_009555245.1;HCoV-OC43_nucleoca;130 | 0 | 0 |
| NC_006213.1;YP_009555245.1;HCoV-OC43_nucleoca;131 | 0 | 0 |
| NC_006213.1;YP_009555245.1;HCoV-OC43_nucleoca;132 | 0 | 0 |
| NC_006213.1;YP_009555245.1;HCoV-OC43_nucleoca;133 | 0 | 0 |
| NC_006213.1;YP_009555245.1;HCoV-OC43_nucleoca;134 | 0 | 0 |
| NC_006213.1;YP_009555245.1;HCoV-OC43_nucleoca;135 | 0 | 0 |
| NC_006213.1;YP_009555245.1;HCoV-OC43_nucleoca;136 | 0 | 0 |
| NC_006213.1;YP_009555245.1;HCoV-OC43_nucleoca;137 | 0 | 0 |
| NC_006213.1;YP_009555245.1;HCoV-OC43_nucleoca;138 | 0 | 0 |
| NC_006213.1;YP_009555245.1;HCoV-OC43_nucleoca;139 | 0.05 | 0 |
| NC_006213.1;YP_009555245.1;HCoV-OC43_nucleoca;140 | 0 | 0 |
| NC_006213.1;YP_009555245.1;HCoV-OC43_nucleoca;141 | 0.05 | 0 |
| NC_006213.1;YP_009555245.1;HCoV-OC43_nucleoca;142 | 0 | 0 |
| NC_006213.1;YP_009555245.1;HCoV-OC43_nucleoca;143 | 0 | 0 |
| NC_006213.1;YP_009555245.1;HCoV-OC43_nucleoca;144 | 0 | 0.025 |
| NC_006213.1;YP_009555245.1;HCoV-OC43_nucleoca;145 | 0 | 0 |

| | | |
|---|---|---|
| NC_006213.1;YP_009555245.1;HCoV-OC43_nucleoca;146 | 0 | 0 |
| NC_006213.1;YP_009555245.1;HCoV-OC43_nucleoca;147 | 0 | 0.05 |
| NC_006213.1;YP_009555245.1;HCoV-OC43_nucleoca;148 | 0 | 0.025 |
| NC_006213.1;YP_009555245.1;HCoV-OC43_nucleoca;149 | 0.05 | 0.05 |
| NC_006213.1;YP_009555245.1;HCoV-OC43_nucleoca;150 | 0 | 0.05 |
| NC_006213.1;YP_009555245.1;HCoV-OC43_nucleoca;151 | 0.05 | 0 |
| NC_006213.1;YP_009555245.1;HCoV-OC43_nucleoca;152 | 0 | 0 |
| NC_006213.1;YP_009555245.1;HCoV-OC43_nucleoca;153 | 0 | 0 |
| NC_006213.1;YP_009555245.1;HCoV-OC43_nucleoca;154 | 0 | 0 |
| NC_006213.1;YP_009555245.1;HCoV-OC43_nucleoca;155 | 0 | 0 |
| NC_006213.1;YP_009555245.1;HCoV-OC43_nucleoca;156 | 0 | 0 |
| NC_006213.1;YP_009555245.1;HCoV-OC43_nucleoca;157 | 0.05 | 0 |
| NC_006213.1;YP_009555245.1;HCoV-OC43_nucleoca;158 | 0 | 0 |
| NC_006213.1;YP_009555245.1;HCoV-OC43_nucleoca;159 | 0 | 0.025 |
| NC_006213.1;YP_009555245.1;HCoV-OC43_nucleoca;160 | 0.05 | 0 |
| NC_006213.1;YP_009555245.1;HCoV-OC43_nucleoca;161 | 0 | 0 |
| NC_006213.1;YP_009555245.1;HCoV-OC43_nucleoca;162 | 0 | 0 |
| NC_006213.1;YP_009555245.1;HCoV-OC43_nucleoca;163 | 0 | 0 |
| NC_006213.1;YP_009555245.1;HCoV-OC43_nucleoca;164 | 0 | 0.05 |
| NC_006213.1;YP_009555245.1;HCoV-OC43_nucleoca;165 | 0.05 | 0.05 |
| NC_006213.1;YP_009555245.1;HCoV-OC43_nucleoca;166 | 0 | 0.025 |
| NC_006213.1;YP_009555245.1;HCoV-OC43_nucleoca;167 | 0 | 0.025 |
| NC_006213.1;YP_009555245.1;HCoV-OC43_nucleoca;168 | 0 | 0.025 |
| NC_006213.1;YP_009555245.1;HCoV-OC43_nucleoca;169 | 0 | 0.025 |
| NC_006213.1;YP_009555245.1;HCoV-OC43_nucleoca;170 | 0.05 | 0.05 |
| NC_006213.1;YP_009555245.1;HCoV-OC43_nucleoca;171 | 0 | 0 |
| NC_006213.1;YP_009555245.1;HCoV-OC43_nucleoca;172 | 0 | 0 |
| NC_006213.1;YP_009555245.1;HCoV-OC43_nucleoca;173 | 0 | 0 |
| NC_006213.1;YP_009555245.1;HCoV-OC43_nucleoca;174 | 0 | 0 |
| NC_006213.1;YP_009555245.1;HCoV-OC43_nucleoca;175 | 0 | 0 |
| NC_006213.1;YP_009555245.1;HCoV-OC43_nucleoca;176 | 0 | 0 |
| NC_006213.1;YP_009555245.1;HCoV-OC43_nucleoca;177 | 0 | 0 |
| NC_006213.1;YP_009555245.1;HCoV-OC43_nucleoca;178 | 0 | 0 |
| NC_006213.1;YP_009555245.1;HCoV-OC43_nucleoca;179 | 0 | 0 |
| NC_006213.1;YP_009555245.1;HCoV-OC43_nucleoca;180 | 0 | 0 |
| NC_006213.1;YP_009555245.1;HCoV-OC43_nucleoca;181 | 0.05 | 0 |
| NC_006213.1;YP_009555245.1;HCoV-OC43_nucleoca;182 | 0 | 0 |
| NC_006213.1;YP_009555245.1;HCoV-OC43_nucleoca;183 | 0 | 0.025 |
| NC_006213.1;YP_009555245.1;HCoV-OC43_nucleoca;184 | 0 | 0 |
| NC_006213.1;YP_009555245.1;HCoV-OC43_nucleoca;185 | 0 | 0 |
| NC_006213.1;YP_009555245.1;HCoV-OC43_nucleoca;186 | 0 | 0 |
| NC_006213.1;YP_009555245.1;HCoV-OC43_nucleoca;187 | 0.05 | 0 |
| NC_006213.1;YP_009555245.1;HCoV-OC43_nucleoca;188 | 0 | 0.025 |

| | | |
|---|---|---|
| NC_006213.1;YP_009555245.1;HCoV-OC43_nucleoca;189 | 0 | 0 |
| NC_006213.1;YP_009555245.1;HCoV-OC43_nucleoca;190 | 0 | 0 |
| NC_006213.1;YP_009555245.1;HCoV-OC43_nucleoca;191 | 0 | 0 |
| NC_006213.1;YP_009555245.1;HCoV-OC43_nucleoca;192 | 0 | 0 |
| NC_006213.1;YP_009555245.1;HCoV-OC43_nucleoca;193 | 0 | 0.025 |
| NC_006213.1;YP_009555245.1;HCoV-OC43_nucleoca;194 | 0 | 0.025 |
| NC_006213.1;YP_009555245.1;HCoV-OC43_nucleoca;195 | 0.05 | 0.025 |
| NC_006213.1;YP_009555245.1;HCoV-OC43_nucleoca;196 | 0 | 0.025 |
| NC_006213.1;YP_009555245.1;HCoV-OC43_nucleoca;197 | 0 | 0.025 |
| NC_006213.1;YP_009555245.1;HCoV-OC43_nucleoca;198 | 0 | 0.025 |
| NC_006213.1;YP_009555245.1;HCoV-OC43_nucleoca;199 | 0 | 0.025 |
| NC_006213.1;YP_009555245.1;HCoV-OC43_nucleoca;200 | 0.05 | 0.025 |
| NC_006213.1;YP_009555245.1;HCoV-OC43_nucleoca;201 | 0 | 0 |
| NC_006213.1;YP_009555245.1;HCoV-OC43_nucleoca;202 | 0 | 0 |
| NC_006213.1;YP_009555245.1;HCoV-OC43_nucleoca;203 | 0.05 | 0.025 |
| NC_006213.1;YP_009555245.1;HCoV-OC43_nucleoca;204 | 0.05 | 0.025 |
| NC_006213.1;YP_009555245.1;HCoV-OC43_nucleoca;205 | 0.05 | 0.025 |
| NC_006213.1;YP_009555245.1;HCoV-OC43_nucleoca;206 | 0.05 | 0.075 |
| NC_006213.1;YP_009555245.1;HCoV-OC43_nucleoca;207 | 0 | 0.05 |
| NC_006213.1;YP_009555245.1;HCoV-OC43_nucleoca;208 | 0.05 | 0 |
| NC_006213.1;YP_009555245.1;HCoV-OC43_nucleoca;209 | 0.05 | 0.025 |
| NC_006213.1;YP_009555245.1;HCoV-OC43_nucleoca;210 | 0.1 | 0.05 |
| NC_006213.1;YP_009555245.1;HCoV-OC43_nucleoca;211 | 0 | 0.05 |
| NC_006213.1;YP_009555245.1;HCoV-OC43_nucleoca;212 | 0.05 | 0.05 |
| NC_006213.1;YP_009555245.1;HCoV-OC43_nucleoca;213 | 0.1 | 0.025 |
| NC_006213.1;YP_009555245.1;HCoV-OC43_nucleoca;214 | 0 | 0 |
| NC_006213.1;YP_009555245.1;HCoV-OC43_nucleoca;215 | 0 | 0.05 |
| NC_006213.1;YP_009555245.1;HCoV-OC43_nucleoca;216 | 0.1 | 0.05 |
| NC_006213.1;YP_009555245.1;HCoV-OC43_nucleoca;217 | 0.1 | 0.075 |
| NC_006213.1;YP_009555245.1;HCoV-OC43_nucleoca;218 | 0.05 | 0.075 |
| NC_006213.1;YP_009555245.1;HCoV-OC43_nucleoca;219 | 0 | 0 |
| NC_006213.1;YP_009555245.1;HCoV-OC43_nucleoca;220 | 0 | 0 |
| NC_006213.1;YP_009555245.1;HCoV-OC43_nucleoca;221 | 0 | 0 |
| NC_006213.1;YP_009555245.1;HCoV-OC43_nucleoca;222 | 0 | 0.025 |
| NC_006213.1;YP_009555245.1;HCoV-OC43_nucleoca;223 | 0 | 0 |
| NC_006213.1;YP_009555245.1;HCoV-OC43_nucleoca;224 | 0 | 0 |
| NC_006213.1;YP_009555245.1;HCoV-OC43_nucleoca;225 | 0 | 0 |
| NC_006213.1;YP_009555245.1;HCoV-OC43_nucleoca;226 | 0 | 0 |
| NC_006213.1;YP_009555245.1;HCoV-OC43_nucleoca;227 | 0 | 0 |
| NC_006213.1;YP_009555245.1;HCoV-OC43_nucleoca;228 | 0 | 0 |
| NC_006213.1;YP_009555245.1;HCoV-OC43_nucleoca;229 | 0 | 0 |
| NC_006213.1;YP_009555245.1;HCoV-OC43_nucleoca;230 | 0 | 0 |
| NC_006213.1;YP_009555245.1;HCoV-OC43_nucleoca;231 | 0 | 0 |

| | | |
|---|---|---|
| NC_006213.1;YP_009555245.1;HCoV-OC43_nucleoca;232 | 0 | 0 |
| NC_006213.1;YP_009555245.1;HCoV-OC43_nucleoca;233 | 0 | 0 |
| NC_006213.1;YP_009555245.1;HCoV-OC43_nucleoca;234 | 0 | 0 |
| NC_006213.1;YP_009555245.1;HCoV-OC43_nucleoca;235 | 0 | 0 |
| NC_006213.1;YP_009555245.1;HCoV-OC43_nucleoca;236 | 0 | 0.025 |
| NC_006213.1;YP_009555245.1;HCoV-OC43_nucleoca;237 | 0.05 | 0.025 |
| NC_006213.1;YP_009555245.1;HCoV-OC43_nucleoca;238 | 0.1 | 0.025 |
| NC_006213.1;YP_009555245.1;HCoV-OC43_nucleoca;239 | 0.1 | 0.025 |
| NC_006213.1;YP_009555245.1;HCoV-OC43_nucleoca;240 | 0.1 | 0.025 |
| NC_006213.1;YP_009555245.1;HCoV-OC43_nucleoca;241 | 0.05 | 0.025 |
| NC_006213.1;YP_009555245.1;HCoV-OC43_nucleoca;242 | 0 | 0 |
| NC_006213.1;YP_009555245.1;HCoV-OC43_nucleoca;243 | 0.05 | 0 |
| NC_006213.1;YP_009555245.1;HCoV-OC43_nucleoca;244 | 0.05 | 0 |
| NC_006213.1;YP_009555245.1;HCoV-OC43_nucleoca;245 | 0 | 0 |
| NC_006213.1;YP_009555245.1;HCoV-OC43_nucleoca;246 | 0.05 | 0.025 |
| NC_006213.1;YP_009555245.1;HCoV-OC43_nucleoca;247 | 0.35 | 0.175 |
| NC_006213.1;YP_009555245.1;HCoV-OC43_nucleoca;248 | 0.05 | 0 |
| NC_006213.1;YP_009555245.1;HCoV-OC43_nucleoca;249 | 0 | 0 |
| NC_006213.1;YP_009555245.1;HCoV-OC43_nucleoca;250 | 0 | 0 |
| NC_006213.1;YP_009555245.1;HCoV-OC43_nucleoca;251 | 0 | 0 |
| NC_006213.1;YP_009555245.1;HCoV-OC43_nucleoca;252 | 0 | 0 |
| NC_006213.1;YP_009555245.1;HCoV-OC43_nucleoca;253 | 0.05 | 0 |
| NC_006213.1;YP_009555245.1;HCoV-OC43_nucleoca;254 | 0.05 | 0 |
| NC_006213.1;YP_009555245.1;HCoV-OC43_nucleoca;255 | 0.05 | 0 |
| NC_006213.1;YP_009555245.1;HCoV-OC43_nucleoca;256 | 0.05 | 0 |
| NC_006213.1;YP_009555245.1;HCoV-OC43_nucleoca;257 | 0.05 | 0 |
| NC_006213.1;YP_009555245.1;HCoV-OC43_nucleoca;258 | 0.05 | 0 |
| NC_006213.1;YP_009555245.1;HCoV-OC43_nucleoca;259 | 0.05 | 0 |
| NC_006213.1;YP_009555245.1;HCoV-OC43_nucleoca;260 | 0.05 | 0 |
| NC_006213.1;YP_009555245.1;HCoV-OC43_nucleoca;261 | 0.05 | 0.025 |
| NC_006213.1;YP_009555245.1;HCoV-OC43_nucleoca;262 | 0.05 | 0 |
| NC_006213.1;YP_009555245.1;HCoV-OC43_nucleoca;263 | 0.05 | 0 |
| NC_006213.1;YP_009555245.1;HCoV-OC43_nucleoca;264 | 0.05 | 0 |
| NC_006213.1;YP_009555245.1;HCoV-OC43_nucleoca;265 | 0 | 0 |
| NC_006213.1;YP_009555245.1;HCoV-OC43_nucleoca;266 | 0 | 0 |
| NC_006213.1;YP_009555245.1;HCoV-OC43_nucleoca;267 | 0 | 0 |
| NC_006213.1;YP_009555245.1;HCoV-OC43_nucleoca;268 | 0 | 0.025 |
| NC_006213.1;YP_009555245.1;HCoV-OC43_nucleoca;269 | 0 | 0 |
| NC_006213.1;YP_009555245.1;HCoV-OC43_nucleoca;270 | 0.05 | 0 |
| NC_006213.1;YP_009555245.1;HCoV-OC43_nucleoca;271 | 0 | 0 |
| NC_006213.1;YP_009555245.1;HCoV-OC43_nucleoca;272 | 0 | 0 |
| NC_006213.1;YP_009555245.1;HCoV-OC43_nucleoca;273 | 0 | 0 |
| NC_006213.1;YP_009555245.1;HCoV-OC43_nucleoca;274 | 0 | 0 |

| | | |
|---|---|---|
| NC_006213.1;YP_009555245.1;HCoV-OC43_nucleoca;275 | 0.05 | 0.025 |
| NC_006213.1;YP_009555245.1;HCoV-OC43_nucleoca;276 | 0 | 0 |
| NC_006213.1;YP_009555245.1;HCoV-OC43_nucleoca;277 | 0 | 0 |
| NC_006213.1;YP_009555245.1;HCoV-OC43_nucleoca;278 | 0 | 0.025 |
| NC_006213.1;YP_009555245.1;HCoV-OC43_nucleoca;279 | 0 | 0.025 |
| NC_006213.1;YP_009555245.1;HCoV-OC43_nucleoca;280 | 0 | 0.025 |
| NC_006213.1;YP_009555245.1;HCoV-OC43_nucleoca;281 | 0 | 0.025 |
| NC_006213.1;YP_009555245.1;HCoV-OC43_nucleoca;282 | 0.05 | 0.05 |
| NC_006213.1;YP_009555245.1;HCoV-OC43_nucleoca;283 | 0.35 | 0.125 |
| NC_006213.1;YP_009555245.1;HCoV-OC43_nucleoca;284 | 0.2 | 0.05 |
| NC_006213.1;YP_009555245.1;HCoV-OC43_nucleoca;285 | 0.15 | 0.05 |
| NC_006213.1;YP_009555245.1;HCoV-OC43_nucleoca;286 | 0.1 | 0.075 |
| NC_006213.1;YP_009555245.1;HCoV-OC43_nucleoca;287 | 0 | 0.05 |
| NC_006213.1;YP_009555245.1;HCoV-OC43_nucleoca;288 | 0 | 0.05 |
| NC_006213.1;YP_009555245.1;HCoV-OC43_nucleoca;289 | 0.05 | 0.025 |
| NC_006213.1;YP_009555245.1;HCoV-OC43_nucleoca;290 | 0 | 0 |
| NC_006213.1;YP_009555245.1;HCoV-OC43_nucleoca;291 | 0 | 0 |
| NC_006213.1;YP_009555245.1;HCoV-OC43_nucleoca;292 | 0 | 0 |
| NC_006213.1;YP_009555245.1;HCoV-OC43_nucleoca;293 | 0 | 0 |
| NC_006213.1;YP_009555245.1;HCoV-OC43_nucleoca;294 | 0 | 0 |
| NC_006213.1;YP_009555245.1;HCoV-OC43_nucleoca;295 | 0 | 0 |
| NC_006213.1;YP_009555245.1;HCoV-OC43_nucleoca;296 | 0 | 0 |
| NC_006213.1;YP_009555245.1;HCoV-OC43_nucleoca;297 | 0 | 0.025 |
| NC_006213.1;YP_009555245.1;HCoV-OC43_nucleoca;298 | 0 | 0 |
| NC_006213.1;YP_009555245.1;HCoV-OC43_nucleoca;299 | 0 | 0 |
| NC_006213.1;YP_009555245.1;HCoV-OC43_nucleoca;300 | 0 | 0 |
| NC_006213.1;YP_009555245.1;HCoV-OC43_nucleoca;301 | 0 | 0 |
| NC_006213.1;YP_009555245.1;HCoV-OC43_nucleoca;302 | 0 | 0 |
| NC_006213.1;YP_009555245.1;HCoV-OC43_nucleoca;303 | 0.05 | 0 |
| NC_006213.1;YP_009555245.1;HCoV-OC43_nucleoca;304 | 0.05 | 0 |
| NC_006213.1;YP_009555245.1;HCoV-OC43_nucleoca;305 | 0.05 | 0 |
| NC_006213.1;YP_009555245.1;HCoV-OC43_nucleoca;306 | 0.05 | 0.075 |
| NC_006213.1;YP_009555245.1;HCoV-OC43_nucleoca;307 | 0.1 | 0 |
| NC_006213.1;YP_009555245.1;HCoV-OC43_nucleoca;308 | 0.05 | 0 |
| NC_006213.1;YP_009555245.1;HCoV-OC43_nucleoca;309 | 0 | 0 |
| NC_006213.1;YP_009555245.1;HCoV-OC43_nucleoca;310 | 0.05 | 0 |
| NC_006213.1;YP_009555245.1;HCoV-OC43_nucleoca;311 | 0.05 | 0 |
| NC_006213.1;YP_009555245.1;HCoV-OC43_nucleoca;312 | 0 | 0 |
| NC_006213.1;YP_009555245.1;HCoV-OC43_nucleoca;313 | 0 | 0 |
| NC_006213.1;YP_009555245.1;HCoV-OC43_nucleoca;314 | 0 | 0 |
| NC_006213.1;YP_009555245.1;HCoV-OC43_nucleoca;315 | 0 | 0 |
| NC_006213.1;YP_009555245.1;HCoV-OC43_nucleoca;316 | 0 | 0 |
| NC_006213.1;YP_009555245.1;HCoV-OC43_nucleoca;317 | 0 | 0 |

| | | |
|---|---|---|
| NC_006213.1;YP_009555245.1;HCoV-OC43_nucleoca;318 | 0 | 0 |
| NC_006213.1;YP_009555245.1;HCoV-OC43_nucleoca;319 | 0 | 0 |
| NC_006213.1;YP_009555245.1;HCoV-OC43_nucleoca;320 | 0 | 0 |
| NC_006213.1;YP_009555245.1;HCoV-OC43_nucleoca;321 | 0 | 0 |
| NC_006213.1;YP_009555245.1;HCoV-OC43_nucleoca;322 | 0 | 0 |
| NC_006213.1;YP_009555245.1;HCoV-OC43_nucleoca;323 | 0 | 0 |
| NC_006213.1;YP_009555245.1;HCoV-OC43_nucleoca;324 | 0 | 0 |
| NC_006213.1;YP_009555245.1;HCoV-OC43_nucleoca;325 | 0 | 0.05 |
| NC_006213.1;YP_009555245.1;HCoV-OC43_nucleoca;326 | 0 | 0 |
| NC_006213.1;YP_009555245.1;HCoV-OC43_nucleoca;327 | 0 | 0.025 |
| NC_006213.1;YP_009555245.1;HCoV-OC43_nucleoca;328 | 0.05 | 0 |
| NC_006213.1;YP_009555245.1;HCoV-OC43_nucleoca;329 | 0.05 | 0 |
| NC_006213.1;YP_009555245.1;HCoV-OC43_nucleoca;330 | 0.05 | 0.05 |
| NC_006213.1;YP_009555245.1;HCoV-OC43_nucleoca;331 | 0.1 | 0.1 |
| NC_006213.1;YP_009555245.1;HCoV-OC43_nucleoca;332 | 0.2 | 0.125 |
| NC_006213.1;YP_009555245.1;HCoV-OC43_nucleoca;333 | 0.15 | 0.15 |
| NC_006213.1;YP_009555245.1;HCoV-OC43_nucleoca;334 | 0.2 | 0.15 |
| NC_006213.1;YP_009555245.1;HCoV-OC43_nucleoca;335 | 0.2 | 0.1 |
| NC_006213.1;YP_009555245.1;HCoV-OC43_nucleoca;336 | 0.2 | 0.1 |
| NC_006213.1;YP_009555245.1;HCoV-OC43_nucleoca;337 | 0.15 | 0.025 |
| NC_006213.1;YP_009555245.1;HCoV-OC43_nucleoca;338 | 0.15 | 0.025 |
| NC_006213.1;YP_009555245.1;HCoV-OC43_nucleoca;339 | 0.1 | 0.025 |
| NC_006213.1;YP_009555245.1;HCoV-OC43_nucleoca;340 | 0.1 | 0.025 |
| NC_006213.1;YP_009555245.1;HCoV-OC43_nucleoca;341 | 0.1 | 0.05 |
| NC_006213.1;YP_009555245.1;HCoV-OC43_nucleoca;342 | 0 | 0.025 |
| NC_006213.1;YP_009555245.1;HCoV-OC43_nucleoca;343 | 0 | 0.05 |
| NC_006213.1;YP_009555245.1;HCoV-OC43_nucleoca;344 | 0 | 0 |
| NC_006213.1;YP_009555245.1;HCoV-OC43_nucleoca;345 | 0 | 0 |
| NC_006213.1;YP_009555245.1;HCoV-OC43_nucleoca;346 | 0 | 0 |
| NC_006213.1;YP_009555245.1;HCoV-OC43_nucleoca;347 | 0 | 0 |
| NC_006213.1;YP_009555245.1;HCoV-OC43_nucleoca;348 | 0 | 0 |
| NC_006213.1;YP_009555245.1;HCoV-OC43_nucleoca;349 | 0 | 0 |
| NC_006213.1;YP_009555245.1;HCoV-OC43_nucleoca;350 | 0.05 | 0 |
| NC_006213.1;YP_009555245.1;HCoV-OC43_nucleoca;351 | 0 | 0 |
| NC_006213.1;YP_009555245.1;HCoV-OC43_nucleoca;352 | 0.05 | 0 |
| NC_006213.1;YP_009555245.1;HCoV-OC43_nucleoca;353 | 0.05 | 0 |
| NC_006213.1;YP_009555245.1;HCoV-OC43_nucleoca;354 | 0 | 0 |
| NC_006213.1;YP_009555245.1;HCoV-OC43_nucleoca;355 | 0.05 | 0 |
| NC_006213.1;YP_009555245.1;HCoV-OC43_nucleoca;356 | 0.05 | 0 |
| NC_006213.1;YP_009555245.1;HCoV-OC43_nucleoca;357 | 0.05 | 0 |
| NC_006213.1;YP_009555245.1;HCoV-OC43_nucleoca;358 | 0.05 | 0 |
| NC_006213.1;YP_009555245.1;HCoV-OC43_nucleoca;359 | 0 | 0 |
| NC_006213.1;YP_009555245.1;HCoV-OC43_nucleoca;360 | 0 | 0 |

| | | |
|---|---|---|
| NC_006213.1;YP_009555245.1;HCoV-OC43_nucleoca;361 | 0 | 0 |
| NC_006213.1;YP_009555245.1;HCoV-OC43_nucleoca;362 | 0 | 0 |
| NC_006213.1;YP_009555245.1;HCoV-OC43_nucleoca;363 | 0.05 | 0.05 |
| NC_006213.1;YP_009555245.1;HCoV-OC43_nucleoca;364 | 0 | 0.025 |
| NC_006213.1;YP_009555245.1;HCoV-OC43_nucleoca;365 | 0.05 | 0.075 |
| NC_006213.1;YP_009555245.1;HCoV-OC43_nucleoca;366 | 0.05 | 0.15 |
| NC_006213.1;YP_009555245.1;HCoV-OC43_nucleoca;367 | 0.05 | 0.125 |
| NC_006213.1;YP_009555245.1;HCoV-OC43_nucleoca;368 | 0.1 | 0.175 |
| NC_006213.1;YP_009555245.1;HCoV-OC43_nucleoca;369 | 0.2 | 0.225 |
| NC_006213.1;YP_009555245.1;HCoV-OC43_nucleoca;370 | 0.2 | 0.175 |
| NC_006213.1;YP_009555245.1;HCoV-OC43_nucleoca;371 | 0.25 | 0.225 |
| NC_006213.1;YP_009555245.1;HCoV-OC43_nucleoca;372 | 0.05 | 0.2 |
| NC_006213.1;YP_009555245.1;HCoV-OC43_nucleoca;373 | 0.05 | 0.2 |
| NC_006213.1;YP_009555245.1;HCoV-OC43_nucleoca;374 | 0.05 | 0.1 |
| NC_006213.1;YP_009555245.1;HCoV-OC43_nucleoca;375 | 0 | 0 |
| NC_006213.1;YP_009555245.1;HCoV-OC43_nucleoca;376 | 0 | 0 |
| NC_006213.1;YP_009555245.1;HCoV-OC43_nucleoca;377 | 0 | 0 |
| NC_006213.1;YP_009555245.1;HCoV-OC43_nucleoca;378 | 0 | 0 |
| NC_006213.1;YP_009555245.1;HCoV-OC43_nucleoca;379 | 0 | 0 |
| NC_006213.1;YP_009555245.1;HCoV-OC43_nucleoca;380 | 0 | 0 |
| NC_006213.1;YP_009555245.1;HCoV-OC43_nucleoca;381 | 0 | 0 |
| NC_006213.1;YP_009555245.1;HCoV-OC43_nucleoca;382 | 0 | 0 |
| NC_006213.1;YP_009555245.1;HCoV-OC43_nucleoca;383 | 0 | 0 |
| NC_006213.1;YP_009555245.1;HCoV-OC43_nucleoca;384 | 0 | 0.025 |
| NC_006213.1;YP_009555245.1;HCoV-OC43_nucleoca;385 | 0 | 0 |
| NC_006213.1;YP_009555245.1;HCoV-OC43_nucleoca;386 | 0 | 0 |
| NC_006213.1;YP_009555245.1;HCoV-OC43_nucleoca;387 | 0 | 0 |
| NC_006213.1;YP_009555245.1;HCoV-OC43_nucleoca;388 | 0 | 0.025 |
| NC_006213.1;YP_009555245.1;HCoV-OC43_nucleoca;389 | 0 | 0 |
| NC_006213.1;YP_009555245.1;HCoV-OC43_nucleoca;390 | 0 | 0 |
| NC_006213.1;YP_009555245.1;HCoV-OC43_nucleoca;391 | 0 | 0 |
| NC_006213.1;YP_009555245.1;HCoV-OC43_nucleoca;392 | 0 | 0.025 |
| NC_006213.1;YP_009555245.1;HCoV-OC43_nucleoca;393 | 0.05 | 0.025 |
| NC_006213.1;YP_009555245.1;HCoV-OC43_nucleoca;394 | 0 | 0.025 |
| NC_006213.1;YP_009555245.1;HCoV-OC43_nucleoca;395 | 0 | 0.025 |
| NC_006213.1;YP_009555245.1;HCoV-OC43_nucleoca;396 | 0.1 | 0.025 |
| NC_006213.1;YP_009555245.1;HCoV-OC43_nucleoca;397 | 0.05 | 0.025 |
| NC_006213.1;YP_009555245.1;HCoV-OC43_nucleoca;398 | 0 | 0.025 |
| NC_006213.1;YP_009555245.1;HCoV-OC43_nucleoca;399 | 0.05 | 0 |
| NC_006213.1;YP_009555245.1;HCoV-OC43_nucleoca;400 | 0.1 | 0 |
| NC_006213.1;YP_009555245.1;HCoV-OC43_nucleoca;401 | 0.1 | 0 |
| NC_006213.1;YP_009555245.1;HCoV-OC43_nucleoca;402 | 0.1 | 0.025 |
| NC_006213.1;YP_009555245.1;HCoV-OC43_nucleoca;403 | 0.1 | 0 |

| | | |
|---|---|---|
| NC_006213.1;YP_009555245.1;HCoV-OC43_nucleoca;404 | 0.1 | 0.025 |
| NC_006213.1;YP_009555245.1;HCoV-OC43_nucleoca;405 | 0.05 | 0 |
| NC_006213.1;YP_009555245.1;HCoV-OC43_nucleoca;406 | 0 | 0 |
| NC_006213.1;YP_009555245.1;HCoV-OC43_nucleoca;407 | 0 | 0.025 |
| NC_006213.1;YP_009555245.1;HCoV-OC43_nucleoca;408 | 0 | 0 |
| NC_006213.1;YP_009555245.1;HCoV-OC43_nucleoca;409 | 0 | 0 |
| NC_006213.1;YP_009555245.1;HCoV-OC43_nucleoca;410 | 0.05 | 0 |
| NC_006213.1;YP_009555245.1;HCoV-OC43_nucleoca;411 | 0.05 | 0 |
| NC_006213.1;YP_009555245.1;HCoV-OC43_nucleoca;412 | 0.1 | 0 |
| NC_006213.1;YP_009555245.1;HCoV-OC43_nucleoca;413 | 0.05 | 0 |
| NC_006213.1;YP_009555245.1;HCoV-OC43_nucleoca;414 | 0.05 | 0.05 |
| NC_006213.1;YP_009555245.1;HCoV-OC43_nucleoca;415 | 0.05 | 0.025 |
| NC_006213.1;YP_009555245.1;HCoV-OC43_nucleoca;416 | 0.05 | 0 |
| NC_006213.1;YP_009555245.1;HCoV-OC43_nucleoca;417 | 0 | 0.025 |
| NC_006213.1;YP_009555245.1;HCoV-OC43_nucleoca;418 | 0 | 0 |
| NC_006213.1;YP_009555245.1;HCoV-OC43_nucleoca;419 | 0 | 0 |
| NC_006213.1;YP_009555245.1;HCoV-OC43_nucleoca;420 | 0.15 | 0.025 |
| NC_006213.1;YP_009555245.1;HCoV-OC43_nucleoca;421 | 0.05 | 0 |
| NC_006213.1;YP_009555245.1;HCoV-OC43_nucleoca;422 | 0.05 | 0 |
| NC_006213.1;YP_009555245.1;HCoV-OC43_nucleoca;423 | 0.1 | 0.025 |
| NC_006213.1;YP_009555245.1;HCoV-OC43_nucleoca;424 | 0.05 | 0.05 |
| NC_006213.1;YP_009555245.1;HCoV-OC43_nucleoca;425 | 0.05 | 0.05 |
| NC_006213.1;YP_009555245.1;HCoV-OC43_nucleoca;426 | 0.1 | 0.325 |
| NC_006213.1;YP_009555245.1;HCoV-OC43_nucleoca;427 | 0.05 | 0.025 |
| NC_006213.1;YP_009555245.1;HCoV-OC43_nucleoca;428 | 0 | 0 |
| NC_006213.1;YP_009555245.1;HCoV-OC43_nucleoca;429 | 0.05 | 0 |
| NC_006213.1;YP_009555245.1;HCoV-OC43_nucleoca;430 | 0 | 0 |
| NC_006213.1;YP_009555245.1;HCoV-OC43_nucleoca;431 | 0 | 0 |
| NC_006213.1;YP_009555245.1;HCoV-OC43_nucleoca;432 | 0 | 0.025 |
| NC_006213.1;YP_009555245.1;HCoV-OC43_nucleoca;433 | 0.05 | 0 |
| NC_006213.1;YP_009555245.1;HCoV-OC43_nucleoca;434 | 0 | 0 |
| NC_006213.1;YP_009555245.1;HCoV-OC43_nucleoca;435 | 0 | 0 |
| NC_006213.1;YP_009555245.1;HCoV-OC43_nucleoca;436 | 0 | 0 |
| NC_006213.1;YP_009555245.1;HCoV-OC43_nucleoca;437 | 0 | 0 |
| NC_005831.2;YP_003766.2;HCoV-NL63_1ab;1 | 0.05 | 0 |
| NC_005831.2;YP_003766.2;HCoV-NL63_1ab;2 | 0 | 0 |
| NC_005831.2;YP_003766.2;HCoV-NL63_1ab;3 | 0 | 0 |
| NC_005831.2;YP_003766.2;HCoV-NL63_1ab;4 | 0 | 0 |
| NC_005831.2;YP_003766.2;HCoV-NL63_1ab;5 | 0 | 0 |
| NC_005831.2;YP_003766.2;HCoV-NL63_1ab;6 | 0 | 0 |
| NC_005831.2;YP_003766.2;HCoV-NL63_1ab;7 | 0.05 | 0 |
| NC_005831.2;YP_003766.2;HCoV-NL63_1ab;8 | 0.05 | 0 |
| NC_005831.2;YP_003766.2;HCoV-NL63_1ab;9 | 0 | 0 |

| | | |
|---|---|---|
| NC_005831.2;YP_003766.2;HCoV-NL63_1ab;10 | 0 | 0.025 |
| NC_005831.2;YP_003766.2;HCoV-NL63_1ab;11 | 0 | 0 |
| NC_005831.2;YP_003766.2;HCoV-NL63_1ab;12 | 0.05 | 0 |
| NC_005831.2;YP_003766.2;HCoV-NL63_1ab;13 | 0 | 0.025 |
| NC_005831.2;YP_003766.2;HCoV-NL63_1ab;14 | 0 | 0.025 |
| NC_005831.2;YP_003766.2;HCoV-NL63_1ab;15 | 0 | 0 |
| NC_005831.2;YP_003766.2;HCoV-NL63_1ab;16 | 0 | 0 |
| NC_005831.2;YP_003766.2;HCoV-NL63_1ab;17 | 0 | 0 |
| NC_005831.2;YP_003766.2;HCoV-NL63_1ab;18 | 0 | 0 |
| NC_005831.2;YP_003766.2;HCoV-NL63_1ab;19 | 0 | 0 |
| NC_005831.2;YP_003766.2;HCoV-NL63_1ab;20 | 0.05 | 0.025 |
| NC_005831.2;YP_003766.2;HCoV-NL63_1ab;21 | 0.05 | 0.025 |
| NC_005831.2;YP_003766.2;HCoV-NL63_1ab;22 | 0 | 0 |
| NC_005831.2;YP_003766.2;HCoV-NL63_1ab;23 | 0 | 0 |
| NC_005831.2;YP_003766.2;HCoV-NL63_1ab;24 | 0 | 0 |
| NC_005831.2;YP_003766.2;HCoV-NL63_1ab;25 | 0 | 0 |
| NC_005831.2;YP_003766.2;HCoV-NL63_1ab;26 | 0 | 0 |
| NC_005831.2;YP_003766.2;HCoV-NL63_1ab;27 | 0 | 0 |
| NC_005831.2;YP_003766.2;HCoV-NL63_1ab;28 | 0 | 0 |
| NC_005831.2;YP_003766.2;HCoV-NL63_1ab;29 | 0 | 0 |
| NC_005831.2;YP_003766.2;HCoV-NL63_1ab;30 | 0 | 0 |
| NC_005831.2;YP_003766.2;HCoV-NL63_1ab;31 | 0 | 0 |
| NC_005831.2;YP_003766.2;HCoV-NL63_1ab;32 | 0 | 0 |
| NC_005831.2;YP_003766.2;HCoV-NL63_1ab;33 | 0 | 0 |
| NC_005831.2;YP_003766.2;HCoV-NL63_1ab;34 | 0 | 0 |
| NC_005831.2;YP_003766.2;HCoV-NL63_1ab;35 | 0 | 0 |
| NC_005831.2;YP_003766.2;HCoV-NL63_1ab;36 | 0 | 0 |
| NC_005831.2;YP_003766.2;HCoV-NL63_1ab;37 | 0 | 0 |
| NC_005831.2;YP_003766.2;HCoV-NL63_1ab;38 | 0 | 0 |
| NC_005831.2;YP_003766.2;HCoV-NL63_1ab;39 | 0 | 0 |
| NC_005831.2;YP_003766.2;HCoV-NL63_1ab;40 | 0 | 0 |
| NC_005831.2;YP_003766.2;HCoV-NL63_1ab;41 | 0 | 0 |
| NC_005831.2;YP_003766.2;HCoV-NL63_1ab;42 | 0 | 0 |
| NC_005831.2;YP_003766.2;HCoV-NL63_1ab;43 | 0 | 0 |
| NC_005831.2;YP_003766.2;HCoV-NL63_1ab;44 | 0 | 0 |
| NC_005831.2;YP_003766.2;HCoV-NL63_1ab;45 | 0 | 0.025 |
| NC_005831.2;YP_003766.2;HCoV-NL63_1ab;46 | 0 | 0.05 |
| NC_005831.2;YP_003766.2;HCoV-NL63_1ab;47 | 0 | 0.05 |
| NC_005831.2;YP_003766.2;HCoV-NL63_1ab;48 | 0 | 0.125 |
| NC_005831.2;YP_003766.2;HCoV-NL63_1ab;49 | 0 | 0.025 |
| NC_005831.2;YP_003766.2;HCoV-NL63_1ab;50 | 0 | 0 |
| NC_005831.2;YP_003766.2;HCoV-NL63_1ab;51 | 0 | 0 |
| NC_005831.2;YP_003766.2;HCoV-NL63_1ab;52 | 0 | 0 |

| | | |
|---|---|---|
| NC_005831.2;YP_003766.2;HCoV-NL63_1ab;53 | 0 | 0.025 |
| NC_005831.2;YP_003766.2;HCoV-NL63_1ab;54 | 0 | 0 |
| NC_005831.2;YP_003766.2;HCoV-NL63_1ab;55 | 0.1 | 0.05 |
| NC_005831.2;YP_003766.2;HCoV-NL63_1ab;56 | 0 | 0 |
| NC_005831.2;YP_003766.2;HCoV-NL63_1ab;57 | 0 | 0 |
| NC_005831.2;YP_003766.2;HCoV-NL63_1ab;58 | 0 | 0 |
| NC_005831.2;YP_003766.2;HCoV-NL63_1ab;59 | 0 | 0 |
| NC_005831.2;YP_003766.2;HCoV-NL63_1ab;60 | 0 | 0 |
| NC_005831.2;YP_003766.2;HCoV-NL63_1ab;61 | 0 | 0 |
| NC_005831.2;YP_003766.2;HCoV-NL63_1ab;62 | 0 | 0 |
| NC_005831.2;YP_003766.2;HCoV-NL63_1ab;63 | 0 | 0 |
| NC_005831.2;YP_003766.2;HCoV-NL63_1ab;64 | 0 | 0 |
| NC_005831.2;YP_003766.2;HCoV-NL63_1ab;65 | 0 | 0.025 |
| NC_005831.2;YP_003766.2;HCoV-NL63_1ab;66 | 0 | 0 |
| NC_005831.2;YP_003766.2;HCoV-NL63_1ab;67 | 0 | 0 |
| NC_005831.2;YP_003766.2;HCoV-NL63_1ab;68 | 0 | 0 |
| NC_005831.2;YP_003766.2;HCoV-NL63_1ab;69 | 0 | 0 |
| NC_005831.2;YP_003766.2;HCoV-NL63_1ab;70 | 0 | 0 |
| NC_005831.2;YP_003766.2;HCoV-NL63_1ab;71 | 0 | 0 |
| NC_005831.2;YP_003766.2;HCoV-NL63_1ab;72 | 0 | 0 |
| NC_005831.2;YP_003766.2;HCoV-NL63_1ab;73 | 0 | 0 |
| NC_005831.2;YP_003766.2;HCoV-NL63_1ab;74 | 0 | 0 |
| NC_005831.2;YP_003766.2;HCoV-NL63_1ab;75 | 0 | 0 |
| NC_005831.2;YP_003766.2;HCoV-NL63_1ab;76 | 0 | 0 |
| NC_005831.2;YP_003766.2;HCoV-NL63_1ab;77 | 0 | 0 |
| NC_005831.2;YP_003766.2;HCoV-NL63_1ab;78 | 0 | 0 |
| NC_005831.2;YP_003766.2;HCoV-NL63_1ab;79 | 0 | 0 |
| NC_005831.2;YP_003766.2;HCoV-NL63_1ab;80 | 0 | 0 |
| NC_005831.2;YP_003766.2;HCoV-NL63_1ab;81 | 0 | 0 |
| NC_005831.2;YP_003766.2;HCoV-NL63_1ab;82 | 0 | 0 |
| NC_005831.2;YP_003766.2;HCoV-NL63_1ab;83 | 0 | 0 |
| NC_005831.2;YP_003766.2;HCoV-NL63_1ab;84 | 0 | 0 |
| NC_005831.2;YP_003766.2;HCoV-NL63_1ab;85 | 0 | 0 |
| NC_005831.2;YP_003766.2;HCoV-NL63_1ab;86 | 0 | 0 |
| NC_005831.2;YP_003766.2;HCoV-NL63_1ab;87 | 0 | 0 |
| NC_005831.2;YP_003766.2;HCoV-NL63_1ab;88 | 0 | 0 |
| NC_005831.2;YP_003766.2;HCoV-NL63_1ab;89 | 0.05 | 0 |
| NC_005831.2;YP_003766.2;HCoV-NL63_1ab;90 | 0 | 0 |
| NC_005831.2;YP_003766.2;HCoV-NL63_1ab;91 | 0 | 0.025 |
| NC_005831.2;YP_003766.2;HCoV-NL63_1ab;92 | 0 | 0 |
| NC_005831.2;YP_003766.2;HCoV-NL63_1ab;93 | 0 | 0 |
| NC_005831.2;YP_003766.2;HCoV-NL63_1ab;94 | 0 | 0 |
| NC_005831.2;YP_003766.2;HCoV-NL63_1ab;95 | 0 | 0 |

| | | |
|---|---|---|
| NC_005831.2;YP_003766.2;HCoV-NL63_1ab;96 | 0 | 0 |
| NC_005831.2;YP_003766.2;HCoV-NL63_1ab;97 | 0 | 0 |
| NC_005831.2;YP_003766.2;HCoV-NL63_1ab;98 | 0 | 0 |
| NC_005831.2;YP_003766.2;HCoV-NL63_1ab;99 | 0 | 0 |
| NC_005831.2;YP_003766.2;HCoV-NL63_1ab;100 | 0 | 0 |
| NC_005831.2;YP_003766.2;HCoV-NL63_1ab;101 | 0 | 0 |
| NC_005831.2;YP_003766.2;HCoV-NL63_1ab;102 | 0 | 0 |
| NC_005831.2;YP_003766.2;HCoV-NL63_1ab;103 | 0 | 0 |
| NC_005831.2;YP_003766.2;HCoV-NL63_1ab;104 | 0 | 0 |
| NC_005831.2;YP_003766.2;HCoV-NL63_1ab;105 | 0 | 0 |
| NC_005831.2;YP_003766.2;HCoV-NL63_1ab;106 | 0 | 0 |
| NC_005831.2;YP_003766.2;HCoV-NL63_1ab;107 | 0 | 0 |
| NC_005831.2;YP_003766.2;HCoV-NL63_1ab;108 | 0 | 0 |
| NC_005831.2;YP_003766.2;HCoV-NL63_1ab;109 | 0 | 0 |
| NC_005831.2;YP_003766.2;HCoV-NL63_1ab;110 | 0 | 0.025 |
| NC_005831.2;YP_003766.2;HCoV-NL63_1ab;111 | 0 | 0 |
| NC_005831.2;YP_003766.2;HCoV-NL63_1ab;112 | 0 | 0.025 |
| NC_005831.2;YP_003766.2;HCoV-NL63_1ab;113 | 0.05 | 0.025 |
| NC_005831.2;YP_003766.2;HCoV-NL63_1ab;114 | 0 | 0 |
| NC_005831.2;YP_003766.2;HCoV-NL63_1ab;115 | 0 | 0.025 |
| NC_005831.2;YP_003766.2;HCoV-NL63_1ab;116 | 0 | 0 |
| NC_005831.2;YP_003766.2;HCoV-NL63_1ab;117 | 0 | 0 |
| NC_005831.2;YP_003766.2;HCoV-NL63_1ab;118 | 0 | 0 |
| NC_005831.2;YP_003766.2;HCoV-NL63_1ab;119 | 0 | 0 |
| NC_005831.2;YP_003766.2;HCoV-NL63_1ab;120 | 0 | 0 |
| NC_005831.2;YP_003766.2;HCoV-NL63_1ab;121 | 0 | 0 |
| NC_005831.2;YP_003766.2;HCoV-NL63_1ab;122 | 0.05 | 0 |
| NC_005831.2;YP_003766.2;HCoV-NL63_1ab;123 | 0 | 0 |
| NC_005831.2;YP_003766.2;HCoV-NL63_1ab;124 | 0 | 0 |
| NC_005831.2;YP_003766.2;HCoV-NL63_1ab;125 | 0.05 | 0 |
| NC_005831.2;YP_003766.2;HCoV-NL63_1ab;126 | 0.05 | 0.025 |
| NC_005831.2;YP_003766.2;HCoV-NL63_1ab;127 | 0.05 | 0 |
| NC_005831.2;YP_003766.2;HCoV-NL63_1ab;128 | 0.05 | 0 |
| NC_005831.2;YP_003766.2;HCoV-NL63_1ab;129 | 0 | 0 |
| NC_005831.2;YP_003766.2;HCoV-NL63_1ab;130 | 0.1 | 0.125 |
| NC_005831.2;YP_003766.2;HCoV-NL63_1ab;131 | 0 | 0.025 |
| NC_005831.2;YP_003766.2;HCoV-NL63_1ab;132 | 0 | 0.075 |
| NC_005831.2;YP_003766.2;HCoV-NL63_1ab;133 | 0 | 0 |
| NC_005831.2;YP_003766.2;HCoV-NL63_1ab;134 | 0 | 0 |
| NC_005831.2;YP_003766.2;HCoV-NL63_1ab;135 | 0 | 0.025 |
| NC_005831.2;YP_003766.2;HCoV-NL63_1ab;136 | 0 | 0 |
| NC_005831.2;YP_003766.2;HCoV-NL63_1ab;137 | 0 | 0 |
| NC_005831.2;YP_003766.2;HCoV-NL63_1ab;138 | 0 | 0 |

| | | |
|---|---|---|
| NC_005831.2;YP_003766.2;HCoV-NL63_1ab;139 | 0 | 0 |
| NC_005831.2;YP_003766.2;HCoV-NL63_1ab;140 | 0 | 0 |
| NC_005831.2;YP_003766.2;HCoV-NL63_1ab;141 | 0 | 0 |
| NC_005831.2;YP_003766.2;HCoV-NL63_1ab;142 | 0 | 0 |
| NC_005831.2;YP_003766.2;HCoV-NL63_1ab;143 | 0 | 0 |
| NC_005831.2;YP_003766.2;HCoV-NL63_1ab;144 | 0 | 0 |
| NC_005831.2;YP_003766.2;HCoV-NL63_1ab;145 | 0 | 0 |
| NC_005831.2;YP_003766.2;HCoV-NL63_1ab;146 | 0 | 0 |
| NC_005831.2;YP_003766.2;HCoV-NL63_1ab;147 | 0 | 0 |
| NC_005831.2;YP_003766.2;HCoV-NL63_1ab;148 | 0 | 0 |
| NC_005831.2;YP_003766.2;HCoV-NL63_1ab;149 | 0 | 0 |
| NC_005831.2;YP_003766.2;HCoV-NL63_1ab;150 | 0.05 | 0.025 |
| NC_005831.2;YP_003766.2;HCoV-NL63_1ab;151 | 0 | 0 |
| NC_005831.2;YP_003766.2;HCoV-NL63_1ab;152 | 0 | 0 |
| NC_005831.2;YP_003766.2;HCoV-NL63_1ab;153 | 0 | 0 |
| NC_005831.2;YP_003766.2;HCoV-NL63_1ab;154 | 0.05 | 0 |
| NC_005831.2;YP_003766.2;HCoV-NL63_1ab;155 | 0.05 | 0.075 |
| NC_005831.2;YP_003766.2;HCoV-NL63_1ab;156 | 0 | 0.075 |
| NC_005831.2;YP_003766.2;HCoV-NL63_1ab;157 | 0 | 0.05 |
| NC_005831.2;YP_003766.2;HCoV-NL63_1ab;158 | 0.05 | 0.05 |
| NC_005831.2;YP_003766.2;HCoV-NL63_1ab;159 | 0 | 0.025 |
| NC_005831.2;YP_003766.2;HCoV-NL63_1ab;160 | 0 | 0.025 |
| NC_005831.2;YP_003766.2;HCoV-NL63_1ab;161 | 0 | 0.025 |
| NC_005831.2;YP_003766.2;HCoV-NL63_1ab;162 | 0 | 0 |
| NC_005831.2;YP_003766.2;HCoV-NL63_1ab;163 | 0 | 0 |
| NC_005831.2;YP_003766.2;HCoV-NL63_1ab;164 | 0 | 0 |
| NC_005831.2;YP_003766.2;HCoV-NL63_1ab;165 | 0 | 0 |
| NC_005831.2;YP_003766.2;HCoV-NL63_1ab;166 | 0 | 0 |
| NC_005831.2;YP_003766.2;HCoV-NL63_1ab;167 | 0 | 0 |
| NC_005831.2;YP_003766.2;HCoV-NL63_1ab;168 | 0 | 0 |
| NC_005831.2;YP_003766.2;HCoV-NL63_1ab;169 | 0 | 0 |
| NC_005831.2;YP_003766.2;HCoV-NL63_1ab;170 | 0 | 0 |
| NC_005831.2;YP_003766.2;HCoV-NL63_1ab;171 | 0 | 0.025 |
| NC_005831.2;YP_003766.2;HCoV-NL63_1ab;172 | 0 | 0 |
| NC_005831.2;YP_003766.2;HCoV-NL63_1ab;173 | 0 | 0 |
| NC_005831.2;YP_003766.2;HCoV-NL63_1ab;174 | 0 | 0.025 |
| NC_005831.2;YP_003766.2;HCoV-NL63_1ab;175 | 0 | 0.025 |
| NC_005831.2;YP_003766.2;HCoV-NL63_1ab;176 | 0 | 0 |
| NC_005831.2;YP_003766.2;HCoV-NL63_1ab;177 | 0 | 0 |
| NC_005831.2;YP_003766.2;HCoV-NL63_1ab;178 | 0 | 0 |
| NC_005831.2;YP_003766.2;HCoV-NL63_1ab;179 | 0 | 0 |
| NC_005831.2;YP_003766.2;HCoV-NL63_1ab;180 | 0 | 0 |
| NC_005831.2;YP_003766.2;HCoV-NL63_1ab;181 | 0 | 0 |

| | | |
|---|---|---|
| NC_005831.2;YP_003766.2;HCoV-NL63_1ab;182 | 0 | 0 |
| NC_005831.2;YP_003766.2;HCoV-NL63_1ab;183 | 0 | 0 |
| NC_005831.2;YP_003766.2;HCoV-NL63_1ab;184 | 0 | 0 |
| NC_005831.2;YP_003766.2;HCoV-NL63_1ab;185 | 0 | 0 |
| NC_005831.2;YP_003766.2;HCoV-NL63_1ab;186 | 0 | 0 |
| NC_005831.2;YP_003766.2;HCoV-NL63_1ab;187 | 0 | 0 |
| NC_005831.2;YP_003766.2;HCoV-NL63_1ab;188 | 0 | 0.025 |
| NC_005831.2;YP_003766.2;HCoV-NL63_1ab;189 | 0 | 0 |
| NC_005831.2;YP_003766.2;HCoV-NL63_1ab;190 | 0 | 0 |
| NC_005831.2;YP_003766.2;HCoV-NL63_1ab;191 | 0 | 0 |
| NC_005831.2;YP_003766.2;HCoV-NL63_1ab;192 | 0 | 0 |
| NC_005831.2;YP_003766.2;HCoV-NL63_1ab;193 | 0 | 0 |
| NC_005831.2;YP_003766.2;HCoV-NL63_1ab;194 | 0 | 0.025 |
| NC_005831.2;YP_003766.2;HCoV-NL63_1ab;195 | 0 | 0 |
| NC_005831.2;YP_003766.2;HCoV-NL63_1ab;196 | 0 | 0 |
| NC_005831.2;YP_003766.2;HCoV-NL63_1ab;197 | 0 | 0 |
| NC_005831.2;YP_003766.2;HCoV-NL63_1ab;198 | 0 | 0 |
| NC_005831.2;YP_003766.2;HCoV-NL63_1ab;199 | 0 | 0 |
| NC_005831.2;YP_003766.2;HCoV-NL63_1ab;200 | 0 | 0 |
| NC_005831.2;YP_003766.2;HCoV-NL63_1ab;201 | 0 | 0 |
| NC_005831.2;YP_003766.2;HCoV-NL63_1ab;202 | 0 | 0 |
| NC_005831.2;YP_003766.2;HCoV-NL63_1ab;203 | 0 | 0 |
| NC_005831.2;YP_003766.2;HCoV-NL63_1ab;204 | 0 | 0 |
| NC_005831.2;YP_003766.2;HCoV-NL63_1ab;205 | 0 | 0 |
| NC_005831.2;YP_003766.2;HCoV-NL63_1ab;206 | 0 | 0 |
| NC_005831.2;YP_003766.2;HCoV-NL63_1ab;207 | 0 | 0 |
| NC_005831.2;YP_003766.2;HCoV-NL63_1ab;208 | 0 | 0 |
| NC_005831.2;YP_003766.2;HCoV-NL63_1ab;209 | 0 | 0 |
| NC_005831.2;YP_003766.2;HCoV-NL63_1ab;210 | 0 | 0 |
| NC_005831.2;YP_003766.2;HCoV-NL63_1ab;211 | 0 | 0 |
| NC_005831.2;YP_003766.2;HCoV-NL63_1ab;212 | 0 | 0 |
| NC_005831.2;YP_003766.2;HCoV-NL63_1ab;213 | 0 | 0 |
| NC_005831.2;YP_003766.2;HCoV-NL63_1ab;214 | 0 | 0 |
| NC_005831.2;YP_003766.2;HCoV-NL63_1ab;215 | 0 | 0 |
| NC_005831.2;YP_003766.2;HCoV-NL63_1ab;216 | 0 | 0 |
| NC_005831.2;YP_003766.2;HCoV-NL63_1ab;217 | 0 | 0 |
| NC_005831.2;YP_003766.2;HCoV-NL63_1ab;218 | 0 | 0 |
| NC_005831.2;YP_003766.2;HCoV-NL63_1ab;219 | 0 | 0 |
| NC_005831.2;YP_003766.2;HCoV-NL63_1ab;220 | 0 | 0 |
| NC_005831.2;YP_003766.2;HCoV-NL63_1ab;221 | 0 | 0 |
| NC_005831.2;YP_003766.2;HCoV-NL63_1ab;222 | 0 | 0 |
| NC_005831.2;YP_003766.2;HCoV-NL63_1ab;223 | 0 | 0 |
| NC_005831.2;YP_003766.2;HCoV-NL63_1ab;224 | 0 | 0 |

| | | |
|---|---|---|
| NC_005831.2;YP_003766.2;HCoV-NL63_1ab;225 | 0 | 0 |
| NC_005831.2;YP_003766.2;HCoV-NL63_1ab;226 | 0 | 0 |
| NC_005831.2;YP_003766.2;HCoV-NL63_1ab;227 | 0 | 0 |
| NC_005831.2;YP_003766.2;HCoV-NL63_1ab;228 | 0 | 0 |
| NC_005831.2;YP_003766.2;HCoV-NL63_1ab;229 | 0 | 0 |
| NC_005831.2;YP_003766.2;HCoV-NL63_1ab;230 | 0 | 0 |
| NC_005831.2;YP_003766.2;HCoV-NL63_1ab;231 | 0 | 0 |
| NC_005831.2;YP_003766.2;HCoV-NL63_1ab;232 | 0 | 0 |
| NC_005831.2;YP_003766.2;HCoV-NL63_1ab;233 | 0 | 0 |
| NC_005831.2;YP_003766.2;HCoV-NL63_1ab;234 | 0 | 0 |
| NC_005831.2;YP_003766.2;HCoV-NL63_1ab;235 | 0 | 0 |
| NC_005831.2;YP_003766.2;HCoV-NL63_1ab;236 | 0 | 0 |
| NC_005831.2;YP_003766.2;HCoV-NL63_1ab;237 | 0 | 0 |
| NC_005831.2;YP_003766.2;HCoV-NL63_1ab;238 | 0.05 | 0 |
| NC_005831.2;YP_003766.2;HCoV-NL63_1ab;239 | 0.05 | 0 |
| NC_005831.2;YP_003766.2;HCoV-NL63_1ab;240 | 0.05 | 0 |
| NC_005831.2;YP_003766.2;HCoV-NL63_1ab;241 | 0.05 | 0 |
| NC_005831.2;YP_003766.2;HCoV-NL63_1ab;242 | 0 | 0 |
| NC_005831.2;YP_003766.2;HCoV-NL63_1ab;243 | 0.05 | 0 |
| NC_005831.2;YP_003766.2;HCoV-NL63_1ab;244 | 0.05 | 0.025 |
| NC_005831.2;YP_003766.2;HCoV-NL63_1ab;245 | 0 | 0 |
| NC_005831.2;YP_003766.2;HCoV-NL63_1ab;246 | 0 | 0 |
| NC_005831.2;YP_003766.2;HCoV-NL63_1ab;247 | 0 | 0 |
| NC_005831.2;YP_003766.2;HCoV-NL63_1ab;248 | 0 | 0 |
| NC_005831.2;YP_003766.2;HCoV-NL63_1ab;249 | 0 | 0 |
| NC_005831.2;YP_003766.2;HCoV-NL63_1ab;250 | 0 | 0 |
| NC_005831.2;YP_003766.2;HCoV-NL63_1ab;251 | 0 | 0 |
| NC_005831.2;YP_003766.2;HCoV-NL63_1ab;252 | 0 | 0 |
| NC_005831.2;YP_003766.2;HCoV-NL63_1ab;253 | 0 | 0 |
| NC_005831.2;YP_003766.2;HCoV-NL63_1ab;254 | 0 | 0.025 |
| NC_005831.2;YP_003766.2;HCoV-NL63_1ab;255 | 0.05 | 0.05 |
| NC_005831.2;YP_003766.2;HCoV-NL63_1ab;256 | 0 | 0 |
| NC_005831.2;YP_003766.2;HCoV-NL63_1ab;257 | 0 | 0 |
| NC_005831.2;YP_003766.2;HCoV-NL63_1ab;258 | 0 | 0 |
| NC_005831.2;YP_003766.2;HCoV-NL63_1ab;259 | 0 | 0 |
| NC_005831.2;YP_003766.2;HCoV-NL63_1ab;260 | 0 | 0 |
| NC_005831.2;YP_003766.2;HCoV-NL63_1ab;261 | 0 | 0 |
| NC_005831.2;YP_003766.2;HCoV-NL63_1ab;262 | 0 | 0 |
| NC_005831.2;YP_003766.2;HCoV-NL63_1ab;263 | 0 | 0 |
| NC_005831.2;YP_003766.2;HCoV-NL63_1ab;264 | 0 | 0 |
| NC_005831.2;YP_003766.2;HCoV-NL63_1ab;265 | 0 | 0 |
| NC_005831.2;YP_003766.2;HCoV-NL63_1ab;266 | 0 | 0.025 |
| NC_005831.2;YP_003766.2;HCoV-NL63_1ab;267 | 0.05 | 0.025 |

| | | |
|---|---|---|
| NC_005831.2;YP_003766.2;HCoV-NL63_1ab;268 | 0.05 | 0 |
| NC_005831.2;YP_003766.2;HCoV-NL63_1ab;269 | 0 | 0 |
| NC_005831.2;YP_003766.2;HCoV-NL63_1ab;270 | 0.1 | 0.025 |
| NC_005831.2;YP_003766.2;HCoV-NL63_1ab;271 | 0.05 | 0 |
| NC_005831.2;YP_003766.2;HCoV-NL63_1ab;272 | 0.05 | 0 |
| NC_005831.2;YP_003766.2;HCoV-NL63_1ab;273 | 0.05 | 0 |
| NC_005831.2;YP_003766.2;HCoV-NL63_1ab;274 | 0.05 | 0 |
| NC_005831.2;YP_003766.2;HCoV-NL63_1ab;275 | 0 | 0 |
| NC_005831.2;YP_003766.2;HCoV-NL63_1ab;276 | 0 | 0 |
| NC_005831.2;YP_003766.2;HCoV-NL63_1ab;277 | 0 | 0 |
| NC_005831.2;YP_003766.2;HCoV-NL63_1ab;278 | 0 | 0 |
| NC_005831.2;YP_003766.2;HCoV-NL63_1ab;279 | 0 | 0 |
| NC_005831.2;YP_003766.2;HCoV-NL63_1ab;280 | 0 | 0 |
| NC_005831.2;YP_003766.2;HCoV-NL63_1ab;281 | 0.05 | 0 |
| NC_005831.2;YP_003766.2;HCoV-NL63_1ab;282 | 0 | 0 |
| NC_005831.2;YP_003766.2;HCoV-NL63_1ab;283 | 0 | 0 |
| NC_005831.2;YP_003766.2;HCoV-NL63_1ab;284 | 0 | 0 |
| NC_005831.2;YP_003766.2;HCoV-NL63_1ab;285 | 0 | 0.025 |
| NC_005831.2;YP_003766.2;HCoV-NL63_1ab;286 | 0 | 0 |
| NC_005831.2;YP_003766.2;HCoV-NL63_1ab;287 | 0 | 0 |
| NC_005831.2;YP_003766.2;HCoV-NL63_1ab;288 | 0 | 0 |
| NC_005831.2;YP_003766.2;HCoV-NL63_1ab;289 | 0 | 0 |
| NC_005831.2;YP_003766.2;HCoV-NL63_1ab;290 | 0 | 0 |
| NC_005831.2;YP_003766.2;HCoV-NL63_1ab;291 | 0 | 0 |
| NC_005831.2;YP_003766.2;HCoV-NL63_1ab;292 | 0 | 0 |
| NC_005831.2;YP_003766.2;HCoV-NL63_1ab;293 | 0 | 0 |
| NC_005831.2;YP_003766.2;HCoV-NL63_1ab;294 | 0.05 | 0 |
| NC_005831.2;YP_003766.2;HCoV-NL63_1ab;295 | 0 | 0 |
| NC_005831.2;YP_003766.2;HCoV-NL63_1ab;296 | 0 | 0 |
| NC_005831.2;YP_003766.2;HCoV-NL63_1ab;297 | 0 | 0 |
| NC_005831.2;YP_003766.2;HCoV-NL63_1ab;298 | 0 | 0 |
| NC_005831.2;YP_003766.2;HCoV-NL63_1ab;299 | 0 | 0 |
| NC_005831.2;YP_003766.2;HCoV-NL63_1ab;300 | 0 | 0 |
| NC_005831.2;YP_003766.2;HCoV-NL63_1ab;301 | 0 | 0 |
| NC_005831.2;YP_003766.2;HCoV-NL63_1ab;302 | 0 | 0 |
| NC_005831.2;YP_003766.2;HCoV-NL63_1ab;303 | 0 | 0 |
| NC_005831.2;YP_003766.2;HCoV-NL63_1ab;304 | 0 | 0 |
| NC_005831.2;YP_003766.2;HCoV-NL63_1ab;305 | 0 | 0 |
| NC_005831.2;YP_003766.2;HCoV-NL63_1ab;306 | 0 | 0 |
| NC_005831.2;YP_003766.2;HCoV-NL63_1ab;307 | 0 | 0 |
| NC_005831.2;YP_003766.2;HCoV-NL63_1ab;308 | 0 | 0 |
| NC_005831.2;YP_003766.2;HCoV-NL63_1ab;309 | 0 | 0 |
| NC_005831.2;YP_003766.2;HCoV-NL63_1ab;310 | 0 | 0 |

| | | |
|---|---|---|
| NC_005831.2;YP_003766.2;HCoV-NL63_1ab;311 | 0 | 0 |
| NC_005831.2;YP_003766.2;HCoV-NL63_1ab;312 | 0 | 0 |
| NC_005831.2;YP_003766.2;HCoV-NL63_1ab;313 | 0 | 0.025 |
| NC_005831.2;YP_003766.2;HCoV-NL63_1ab;314 | 0 | 0 |
| NC_005831.2;YP_003766.2;HCoV-NL63_1ab;315 | 0.05 | 0 |
| NC_005831.2;YP_003766.2;HCoV-NL63_1ab;316 | 0 | 0 |
| NC_005831.2;YP_003766.2;HCoV-NL63_1ab;317 | 0 | 0 |
| NC_005831.2;YP_003766.2;HCoV-NL63_1ab;318 | 0 | 0 |
| NC_005831.2;YP_003766.2;HCoV-NL63_1ab;319 | 0 | 0 |
| NC_005831.2;YP_003766.2;HCoV-NL63_1ab;320 | 0 | 0 |
| NC_005831.2;YP_003766.2;HCoV-NL63_1ab;321 | 0 | 0 |
| NC_005831.2;YP_003766.2;HCoV-NL63_1ab;322 | 0 | 0 |
| NC_005831.2;YP_003766.2;HCoV-NL63_1ab;323 | 0 | 0 |
| NC_005831.2;YP_003766.2;HCoV-NL63_1ab;324 | 0 | 0 |
| NC_005831.2;YP_003766.2;HCoV-NL63_1ab;325 | 0 | 0 |
| NC_005831.2;YP_003766.2;HCoV-NL63_1ab;326 | 0 | 0 |
| NC_005831.2;YP_003766.2;HCoV-NL63_1ab;327 | 0 | 0.05 |
| NC_005831.2;YP_003766.2;HCoV-NL63_1ab;328 | 0.05 | 0.025 |
| NC_005831.2;YP_003766.2;HCoV-NL63_1ab;329 | 0 | 0.075 |
| NC_005831.2;YP_003766.2;HCoV-NL63_1ab;330 | 0 | 0.025 |
| NC_005831.2;YP_003766.2;HCoV-NL63_1ab;331 | 0 | 0 |
| NC_005831.2;YP_003766.2;HCoV-NL63_1ab;332 | 0 | 0.025 |
| NC_005831.2;YP_003766.2;HCoV-NL63_1ab;333 | 0.05 | 0.025 |
| NC_005831.2;YP_003766.2;HCoV-NL63_1ab;334 | 0 | 0 |
| NC_005831.2;YP_003766.2;HCoV-NL63_1ab;335 | 0 | 0 |
| NC_005831.2;YP_003766.2;HCoV-NL63_1ab;336 | 0 | 0 |
| NC_005831.2;YP_003766.2;HCoV-NL63_1ab;337 | 0 | 0 |
| NC_005831.2;YP_003766.2;HCoV-NL63_1ab;338 | 0.05 | 0 |
| NC_005831.2;YP_003766.2;HCoV-NL63_1ab;339 | 0 | 0 |
| NC_005831.2;YP_003766.2;HCoV-NL63_1ab;340 | 0 | 0.025 |
| NC_005831.2;YP_003766.2;HCoV-NL63_1ab;341 | 0.05 | 0.025 |
| NC_005831.2;YP_003766.2;HCoV-NL63_1ab;342 | 0 | 0 |
| NC_005831.2;YP_003766.2;HCoV-NL63_1ab;343 | 0 | 0 |
| NC_005831.2;YP_003766.2;HCoV-NL63_1ab;344 | 0 | 0 |
| NC_005831.2;YP_003766.2;HCoV-NL63_1ab;345 | 0 | 0 |
| NC_005831.2;YP_003766.2;HCoV-NL63_1ab;346 | 0 | 0 |
| NC_005831.2;YP_003766.2;HCoV-NL63_1ab;347 | 0 | 0 |
| NC_005831.2;YP_003766.2;HCoV-NL63_1ab;348 | 0 | 0 |
| NC_005831.2;YP_003766.2;HCoV-NL63_1ab;349 | 0 | 0 |
| NC_005831.2;YP_003766.2;HCoV-NL63_1ab;350 | 0 | 0 |
| NC_005831.2;YP_003766.2;HCoV-NL63_1ab;351 | 0.1 | 0.075 |
| NC_005831.2;YP_003766.2;HCoV-NL63_1ab;352 | 0 | 0.025 |
| NC_005831.2;YP_003766.2;HCoV-NL63_1ab;353 | 0 | 0 |

| | | |
|---|---|---|
| NC_005831.2;YP_003766.2;HCoV-NL63_1ab;354 | 0 | 0 |
| NC_005831.2;YP_003766.2;HCoV-NL63_1ab;355 | 0.05 | 0.025 |
| NC_005831.2;YP_003766.2;HCoV-NL63_1ab;356 | 0.25 | 0.175 |
| NC_005831.2;YP_003766.2;HCoV-NL63_1ab;357 | 0.25 | 0.25 |
| NC_005831.2;YP_003766.2;HCoV-NL63_1ab;358 | 0.4 | 0.225 |
| NC_005831.2;YP_003766.2;HCoV-NL63_1ab;359 | 0.4 | 0.3 |
| NC_005831.2;YP_003766.2;HCoV-NL63_1ab;360 | 0.3 | 0.25 |
| NC_005831.2;YP_003766.2;HCoV-NL63_1ab;361 | 0.25 | 0.2 |
| NC_005831.2;YP_003766.2;HCoV-NL63_1ab;362 | 0.2 | 0.2 |
| NC_005831.2;YP_003766.2;HCoV-NL63_1ab;363 | 0.2 | 0.175 |
| NC_005831.2;YP_003766.2;HCoV-NL63_1ab;364 | 0.2 | 0.225 |
| NC_005831.2;YP_003766.2;HCoV-NL63_1ab;365 | 0.2 | 0.175 |
| NC_005831.2;YP_003766.2;HCoV-NL63_1ab;366 | 0.2 | 0.175 |
| NC_005831.2;YP_003766.2;HCoV-NL63_1ab;367 | 0.2 | 0.2 |
| NC_005831.2;YP_003766.2;HCoV-NL63_1ab;368 | 0.05 | 0 |
| NC_005831.2;YP_003766.2;HCoV-NL63_1ab;369 | 0 | 0 |
| NC_005831.2;YP_003766.2;HCoV-NL63_1ab;370 | 0 | 0 |
| NC_005831.2;YP_003766.2;HCoV-NL63_1ab;371 | 0 | 0 |
| NC_005831.2;YP_003766.2;HCoV-NL63_1ab;372 | 0 | 0 |
| NC_005831.2;YP_003766.2;HCoV-NL63_1ab;373 | 0 | 0 |
| NC_005831.2;YP_003766.2;HCoV-NL63_1ab;374 | 0 | 0 |
| NC_005831.2;YP_003766.2;HCoV-NL63_1ab;375 | 0 | 0 |
| NC_005831.2;YP_003766.2;HCoV-NL63_1ab;376 | 0 | 0.05 |
| NC_005831.2;YP_003766.2;HCoV-NL63_1ab;377 | 0 | 0 |
| NC_005831.2;YP_003766.2;HCoV-NL63_1ab;378 | 0 | 0 |
| NC_005831.2;YP_003766.2;HCoV-NL63_1ab;379 | 0 | 0 |
| NC_005831.2;YP_003766.2;HCoV-NL63_1ab;380 | 0 | 0 |
| NC_005831.2;YP_003766.2;HCoV-NL63_1ab;381 | 0 | 0.05 |
| NC_005831.2;YP_003766.2;HCoV-NL63_1ab;382 | 0 | 0.025 |
| NC_005831.2;YP_003766.2;HCoV-NL63_1ab;383 | 0 | 0.05 |
| NC_005831.2;YP_003766.2;HCoV-NL63_1ab;384 | 0 | 0.025 |
| NC_005831.2;YP_003766.2;HCoV-NL63_1ab;385 | 0.05 | 0 |
| NC_005831.2;YP_003766.2;HCoV-NL63_1ab;386 | 0.05 | 0.05 |
| NC_005831.2;YP_003766.2;HCoV-NL63_1ab;387 | 0 | 0 |
| NC_005831.2;YP_003766.2;HCoV-NL63_1ab;388 | 0 | 0 |
| NC_005831.2;YP_003766.2;HCoV-NL63_1ab;389 | 0 | 0 |
| NC_005831.2;YP_003766.2;HCoV-NL63_1ab;390 | 0 | 0 |
| NC_005831.2;YP_003766.2;HCoV-NL63_1ab;391 | 0.05 | 0.025 |
| NC_005831.2;YP_003766.2;HCoV-NL63_1ab;392 | 0 | 0 |
| NC_005831.2;YP_003766.2;HCoV-NL63_1ab;393 | 0.05 | 0 |
| NC_005831.2;YP_003766.2;HCoV-NL63_1ab;394 | 0 | 0.025 |
| NC_005831.2;YP_003766.2;HCoV-NL63_1ab;395 | 0 | 0 |
| NC_005831.2;YP_003766.2;HCoV-NL63_1ab;396 | 0 | 0 |

| | | |
|---|---|---|
| NC_005831.2;YP_003766.2;HCoV-NL63_1ab;397 | 0 | 0 |
| NC_005831.2;YP_003766.2;HCoV-NL63_1ab;398 | 0.05 | 0 |
| NC_005831.2;YP_003766.2;HCoV-NL63_1ab;399 | 0.05 | 0 |
| NC_005831.2;YP_003766.2;HCoV-NL63_1ab;400 | 0.05 | 0 |
| NC_005831.2;YP_003766.2;HCoV-NL63_1ab;401 | 0.05 | 0 |
| NC_005831.2;YP_003766.2;HCoV-NL63_1ab;402 | 0.1 | 0 |
| NC_005831.2;YP_003766.2;HCoV-NL63_1ab;403 | 0.05 | 0 |
| NC_005831.2;YP_003766.2;HCoV-NL63_1ab;404 | 0.05 | 0 |
| NC_005831.2;YP_003766.2;HCoV-NL63_1ab;405 | 0.05 | 0.025 |
| NC_005831.2;YP_003766.2;HCoV-NL63_1ab;406 | 0 | 0 |
| NC_005831.2;YP_003766.2;HCoV-NL63_1ab;407 | 0 | 0 |
| NC_005831.2;YP_003766.2;HCoV-NL63_1ab;408 | 0 | 0 |
| NC_005831.2;YP_003766.2;HCoV-NL63_1ab;409 | 0 | 0.025 |
| NC_005831.2;YP_003766.2;HCoV-NL63_1ab;410 | 0 | 0 |
| NC_005831.2;YP_003766.2;HCoV-NL63_1ab;411 | 0 | 0 |
| NC_005831.2;YP_003766.2;HCoV-NL63_1ab;412 | 0.05 | 0.025 |
| NC_005831.2;YP_003766.2;HCoV-NL63_1ab;413 | 0 | 0 |
| NC_005831.2;YP_003766.2;HCoV-NL63_1ab;414 | 0 | 0 |
| NC_005831.2;YP_003766.2;HCoV-NL63_1ab;415 | 0 | 0 |
| NC_005831.2;YP_003766.2;HCoV-NL63_1ab;416 | 0 | 0.025 |
| NC_005831.2;YP_003766.2;HCoV-NL63_1ab;417 | 0 | 0 |
| NC_005831.2;YP_003766.2;HCoV-NL63_1ab;418 | 0 | 0 |
| NC_005831.2;YP_003766.2;HCoV-NL63_1ab;419 | 0 | 0 |
| NC_005831.2;YP_003766.2;HCoV-NL63_1ab;420 | 0 | 0 |
| NC_005831.2;YP_003766.2;HCoV-NL63_1ab;421 | 0 | 0 |
| NC_005831.2;YP_003766.2;HCoV-NL63_1ab;422 | 0 | 0 |
| NC_005831.2;YP_003766.2;HCoV-NL63_1ab;423 | 0 | 0 |
| NC_005831.2;YP_003766.2;HCoV-NL63_1ab;424 | 0 | 0 |
| NC_005831.2;YP_003766.2;HCoV-NL63_1ab;425 | 0 | 0 |
| NC_005831.2;YP_003766.2;HCoV-NL63_1ab;426 | 0 | 0 |
| NC_005831.2;YP_003766.2;HCoV-NL63_1ab;427 | 0 | 0 |
| NC_005831.2;YP_003766.2;HCoV-NL63_1ab;428 | 0 | 0 |
| NC_005831.2;YP_003766.2;HCoV-NL63_1ab;429 | 0 | 0 |
| NC_005831.2;YP_003766.2;HCoV-NL63_1ab;430 | 0 | 0 |
| NC_005831.2;YP_003766.2;HCoV-NL63_1ab;431 | 0 | 0 |
| NC_005831.2;YP_003766.2;HCoV-NL63_1ab;432 | 0 | 0 |
| NC_005831.2;YP_003766.2;HCoV-NL63_1ab;433 | 0 | 0 |
| NC_005831.2;YP_003766.2;HCoV-NL63_1ab;434 | 0 | 0 |
| NC_005831.2;YP_003766.2;HCoV-NL63_1ab;435 | 0 | 0 |
| NC_005831.2;YP_003766.2;HCoV-NL63_1ab;436 | 0 | 0 |
| NC_005831.2;YP_003766.2;HCoV-NL63_1ab;437 | 0 | 0 |
| NC_005831.2;YP_003766.2;HCoV-NL63_1ab;438 | 0 | 0 |
| NC_005831.2;YP_003766.2;HCoV-NL63_1ab;439 | 0 | 0 |

| | | |
|---|---|---|
| NC_005831.2;YP_003766.2;HCoV-NL63_1ab;440 | 0 | 0 |
| NC_005831.2;YP_003766.2;HCoV-NL63_1ab;441 | 0 | 0 |
| NC_005831.2;YP_003766.2;HCoV-NL63_1ab;442 | 0 | 0 |
| NC_005831.2;YP_003766.2;HCoV-NL63_1ab;443 | 0 | 0 |
| NC_005831.2;YP_003766.2;HCoV-NL63_1ab;444 | 0 | 0 |
| NC_005831.2;YP_003766.2;HCoV-NL63_1ab;445 | 0 | 0.025 |
| NC_005831.2;YP_003766.2;HCoV-NL63_1ab;446 | 0 | 0 |
| NC_005831.2;YP_003766.2;HCoV-NL63_1ab;447 | 0.05 | 0 |
| NC_005831.2;YP_003766.2;HCoV-NL63_1ab;448 | 0 | 0.025 |
| NC_005831.2;YP_003766.2;HCoV-NL63_1ab;449 | 0 | 0 |
| NC_005831.2;YP_003766.2;HCoV-NL63_1ab;450 | 0 | 0 |
| NC_005831.2;YP_003766.2;HCoV-NL63_1ab;451 | 0 | 0 |
| NC_005831.2;YP_003766.2;HCoV-NL63_1ab;452 | 0 | 0 |
| NC_005831.2;YP_003766.2;HCoV-NL63_1ab;453 | 0 | 0 |
| NC_005831.2;YP_003766.2;HCoV-NL63_1ab;454 | 0 | 0 |
| NC_005831.2;YP_003766.2;HCoV-NL63_1ab;455 | 0 | 0 |
| NC_005831.2;YP_003766.2;HCoV-NL63_1ab;456 | 0 | 0 |
| NC_005831.2;YP_003766.2;HCoV-NL63_1ab;457 | 0 | 0 |
| NC_005831.2;YP_003766.2;HCoV-NL63_1ab;458 | 0 | 0 |
| NC_005831.2;YP_003766.2;HCoV-NL63_1ab;459 | 0 | 0 |
| NC_005831.2;YP_003766.2;HCoV-NL63_1ab;460 | 0 | 0 |
| NC_005831.2;YP_003766.2;HCoV-NL63_1ab;461 | 0 | 0 |
| NC_005831.2;YP_003766.2;HCoV-NL63_1ab;462 | 0 | 0 |
| NC_005831.2;YP_003766.2;HCoV-NL63_1ab;463 | 0 | 0 |
| NC_005831.2;YP_003766.2;HCoV-NL63_1ab;464 | 0 | 0 |
| NC_005831.2;YP_003766.2;HCoV-NL63_1ab;465 | 0 | 0 |
| NC_005831.2;YP_003766.2;HCoV-NL63_1ab;466 | 0 | 0 |
| NC_005831.2;YP_003766.2;HCoV-NL63_1ab;467 | 0 | 0 |
| NC_005831.2;YP_003766.2;HCoV-NL63_1ab;468 | 0 | 0.025 |
| NC_005831.2;YP_003766.2;HCoV-NL63_1ab;469 | 0 | 0 |
| NC_005831.2;YP_003766.2;HCoV-NL63_1ab;470 | 0 | 0.025 |
| NC_005831.2;YP_003766.2;HCoV-NL63_1ab;471 | 0.1 | 0.1 |
| NC_005831.2;YP_003766.2;HCoV-NL63_1ab;472 | 0.1 | 0.075 |
| NC_005831.2;YP_003766.2;HCoV-NL63_1ab;473 | 0.05 | 0.05 |
| NC_005831.2;YP_003766.2;HCoV-NL63_1ab;474 | 0.05 | 0 |
| NC_005831.2;YP_003766.2;HCoV-NL63_1ab;475 | 0.05 | 0.025 |
| NC_005831.2;YP_003766.2;HCoV-NL63_1ab;476 | 0.1 | 0.05 |
| NC_005831.2;YP_003766.2;HCoV-NL63_1ab;477 | 0.1 | 0.125 |
| NC_005831.2;YP_003766.2;HCoV-NL63_1ab;478 | 0.05 | 0.05 |
| NC_005831.2;YP_003766.2;HCoV-NL63_1ab;479 | 0.05 | 0.025 |
| NC_005831.2;YP_003766.2;HCoV-NL63_1ab;480 | 0.05 | 0.025 |
| NC_005831.2;YP_003766.2;HCoV-NL63_1ab;481 | 0.15 | 0.075 |
| NC_005831.2;YP_003766.2;HCoV-NL63_1ab;482 | 0.2 | 0.075 |

| | | |
|---|---|---|
| NC_005831.2;YP_003766.2;HCoV-NL63_1ab;483 | 0.1 | 0 |
| NC_005831.2;YP_003766.2;HCoV-NL63_1ab;484 | 0.05 | 0 |
| NC_005831.2;YP_003766.2;HCoV-NL63_1ab;485 | 0.05 | 0 |
| NC_005831.2;YP_003766.2;HCoV-NL63_1ab;486 | 0 | 0 |
| NC_005831.2;YP_003766.2;HCoV-NL63_1ab;487 | 0 | 0.025 |
| NC_005831.2;YP_003766.2;HCoV-NL63_1ab;488 | 0 | 0 |
| NC_005831.2;YP_003766.2;HCoV-NL63_1ab;489 | 0 | 0 |
| NC_005831.2;YP_003766.2;HCoV-NL63_1ab;490 | 0 | 0 |
| NC_005831.2;YP_003766.2;HCoV-NL63_1ab;491 | 0 | 0 |
| NC_005831.2;YP_003766.2;HCoV-NL63_1ab;492 | 0 | 0 |
| NC_005831.2;YP_003766.2;HCoV-NL63_1ab;493 | 0 | 0.025 |
| NC_005831.2;YP_003766.2;HCoV-NL63_1ab;494 | 0 | 0 |
| NC_005831.2;YP_003766.2;HCoV-NL63_1ab;495 | 0 | 0 |
| NC_005831.2;YP_003766.2;HCoV-NL63_1ab;496 | 0 | 0 |
| NC_005831.2;YP_003766.2;HCoV-NL63_1ab;497 | 0 | 0 |
| NC_005831.2;YP_003766.2;HCoV-NL63_1ab;498 | 0 | 0 |
| NC_005831.2;YP_003766.2;HCoV-NL63_1ab;499 | 0 | 0 |
| NC_005831.2;YP_003766.2;HCoV-NL63_1ab;500 | 0 | 0 |
| NC_005831.2;YP_003766.2;HCoV-NL63_1ab;501 | 0 | 0 |
| NC_005831.2;YP_003766.2;HCoV-NL63_1ab;502 | 0 | 0 |
| NC_005831.2;YP_003766.2;HCoV-NL63_1ab;503 | 0 | 0 |
| NC_005831.2;YP_003766.2;HCoV-NL63_1ab;504 | 0 | 0 |
| NC_005831.2;YP_003766.2;HCoV-NL63_1ab;505 | 0 | 0 |
| NC_005831.2;YP_003766.2;HCoV-NL63_1ab;506 | 0 | 0 |
| NC_005831.2;YP_003766.2;HCoV-NL63_1ab;507 | 0 | 0 |
| NC_005831.2;YP_003766.2;HCoV-NL63_1ab;508 | 0 | 0 |
| NC_005831.2;YP_003766.2;HCoV-NL63_1ab;509 | 0 | 0.025 |
| NC_005831.2;YP_003766.2;HCoV-NL63_1ab;510 | 0 | 0 |
| NC_005831.2;YP_003766.2;HCoV-NL63_1ab;511 | 0 | 0 |
| NC_005831.2;YP_003766.2;HCoV-NL63_1ab;512 | 0 | 0 |
| NC_005831.2;YP_003766.2;HCoV-NL63_1ab;513 | 0 | 0 |
| NC_005831.2;YP_003766.2;HCoV-NL63_1ab;514 | 0 | 0 |
| NC_005831.2;YP_003766.2;HCoV-NL63_1ab;515 | 0 | 0 |
| NC_005831.2;YP_003766.2;HCoV-NL63_1ab;516 | 0.05 | 0.025 |
| NC_005831.2;YP_003766.2;HCoV-NL63_1ab;517 | 0.05 | 0.025 |
| NC_005831.2;YP_003766.2;HCoV-NL63_1ab;518 | 0.05 | 0.025 |
| NC_005831.2;YP_003766.2;HCoV-NL63_1ab;519 | 0.05 | 0.025 |
| NC_005831.2;YP_003766.2;HCoV-NL63_1ab;520 | 0.05 | 0.025 |
| NC_005831.2;YP_003766.2;HCoV-NL63_1ab;521 | 0 | 0.025 |
| NC_005831.2;YP_003766.2;HCoV-NL63_1ab;522 | 0.05 | 0.025 |
| NC_005831.2;YP_003766.2;HCoV-NL63_1ab;523 | 0 | 0 |
| NC_005831.2;YP_003766.2;HCoV-NL63_1ab;524 | 0 | 0 |
| NC_005831.2;YP_003766.2;HCoV-NL63_1ab;525 | 0 | 0 |

| | | |
|---|---|---|
| NC_005831.2;YP_003766.2;HCoV-NL63_1ab;526 | 0.05 | 0 |
| NC_005831.2;YP_003766.2;HCoV-NL63_1ab;527 | 0 | 0 |
| NC_005831.2;YP_003766.2;HCoV-NL63_1ab;528 | 0 | 0 |
| NC_005831.2;YP_003766.2;HCoV-NL63_1ab;529 | 0 | 0 |
| NC_005831.2;YP_003766.2;HCoV-NL63_1ab;530 | 0 | 0 |
| NC_005831.2;YP_003766.2;HCoV-NL63_1ab;531 | 0 | 0 |
| NC_005831.2;YP_003766.2;HCoV-NL63_1ab;532 | 0 | 0 |
| NC_005831.2;YP_003766.2;HCoV-NL63_1ab;533 | 0 | 0 |
| NC_005831.2;YP_003766.2;HCoV-NL63_1ab;534 | 0 | 0.025 |
| NC_005831.2;YP_003766.2;HCoV-NL63_1ab;535 | 0 | 0 |
| NC_005831.2;YP_003766.2;HCoV-NL63_1ab;536 | 0 | 0 |
| NC_005831.2;YP_003766.2;HCoV-NL63_1ab;537 | 0 | 0 |
| NC_005831.2;YP_003766.2;HCoV-NL63_1ab;538 | 0 | 0 |
| NC_005831.2;YP_003766.2;HCoV-NL63_1ab;539 | 0 | 0 |
| NC_005831.2;YP_003766.2;HCoV-NL63_1ab;540 | 0 | 0 |
| NC_005831.2;YP_003766.2;HCoV-NL63_1ab;541 | 0 | 0 |
| NC_005831.2;YP_003766.2;HCoV-NL63_1ab;542 | 0 | 0 |
| NC_005831.2;YP_003766.2;HCoV-NL63_1ab;543 | 0 | 0.025 |
| NC_005831.2;YP_003766.2;HCoV-NL63_1ab;544 | 0 | 0 |
| NC_005831.2;YP_003766.2;HCoV-NL63_1ab;545 | 0 | 0 |
| NC_005831.2;YP_003766.2;HCoV-NL63_1ab;546 | 0 | 0 |
| NC_005831.2;YP_003766.2;HCoV-NL63_1ab;547 | 0 | 0 |
| NC_005831.2;YP_003766.2;HCoV-NL63_1ab;548 | 0 | 0.025 |
| NC_005831.2;YP_003766.2;HCoV-NL63_1ab;549 | 0 | 0 |
| NC_005831.2;YP_003766.2;HCoV-NL63_1ab;550 | 0 | 0 |
| NC_005831.2;YP_003766.2;HCoV-NL63_1ab;551 | 0 | 0 |
| NC_005831.2;YP_003766.2;HCoV-NL63_1ab;552 | 0 | 0 |
| NC_005831.2;YP_003766.2;HCoV-NL63_1ab;553 | 0 | 0 |
| NC_005831.2;YP_003766.2;HCoV-NL63_1ab;554 | 0 | 0 |
| NC_005831.2;YP_003766.2;HCoV-NL63_1ab;555 | 0 | 0 |
| NC_005831.2;YP_003766.2;HCoV-NL63_1ab;556 | 0 | 0 |
| NC_005831.2;YP_003766.2;HCoV-NL63_1ab;557 | 0 | 0 |
| NC_005831.2;YP_003766.2;HCoV-NL63_1ab;558 | 0 | 0 |
| NC_005831.2;YP_003766.2;HCoV-NL63_1ab;559 | 0 | 0 |
| NC_005831.2;YP_003766.2;HCoV-NL63_1ab;560 | 0 | 0 |
| NC_005831.2;YP_003766.2;HCoV-NL63_1ab;561 | 0 | 0 |
| NC_005831.2;YP_003766.2;HCoV-NL63_1ab;562 | 0 | 0 |
| NC_005831.2;YP_003766.2;HCoV-NL63_1ab;563 | 0 | 0 |
| NC_005831.2;YP_003766.2;HCoV-NL63_1ab;564 | 0 | 0 |
| NC_005831.2;YP_003766.2;HCoV-NL63_1ab;565 | 0 | 0 |
| NC_005831.2;YP_003766.2;HCoV-NL63_1ab;566 | 0 | 0 |
| NC_005831.2;YP_003766.2;HCoV-NL63_1ab;567 | 0 | 0 |
| NC_005831.2;YP_003766.2;HCoV-NL63_1ab;568 | 0 | 0 |

| | | |
|---|---|---|
| NC_005831.2;YP_003766.2;HCoV-NL63_1ab;569 | 0 | 0 |
| NC_005831.2;YP_003766.2;HCoV-NL63_1ab;570 | 0 | 0.025 |
| NC_005831.2;YP_003766.2;HCoV-NL63_1ab;571 | 0 | 0 |
| NC_005831.2;YP_003766.2;HCoV-NL63_1ab;572 | 0 | 0 |
| NC_005831.2;YP_003766.2;HCoV-NL63_1ab;573 | 0 | 0 |
| NC_005831.2;YP_003766.2;HCoV-NL63_1ab;574 | 0 | 0 |
| NC_005831.2;YP_003766.2;HCoV-NL63_1ab;575 | 0 | 0 |
| NC_005831.2;YP_003766.2;HCoV-NL63_1ab;576 | 0 | 0 |
| NC_005831.2;YP_003766.2;HCoV-NL63_1ab;577 | 0 | 0.025 |
| NC_005831.2;YP_003766.2;HCoV-NL63_1ab;578 | 0.05 | 0 |
| NC_005831.2;YP_003766.2;HCoV-NL63_1ab;579 | 0 | 0 |
| NC_005831.2;YP_003766.2;HCoV-NL63_1ab;580 | 0 | 0 |
| NC_005831.2;YP_003766.2;HCoV-NL63_1ab;581 | 0 | 0.05 |
| NC_005831.2;YP_003766.2;HCoV-NL63_1ab;582 | 0 | 0 |
| NC_005831.2;YP_003766.2;HCoV-NL63_1ab;583 | 0 | 0 |
| NC_005831.2;YP_003766.2;HCoV-NL63_1ab;584 | 0 | 0 |
| NC_005831.2;YP_003766.2;HCoV-NL63_1ab;585 | 0 | 0 |
| NC_005831.2;YP_003766.2;HCoV-NL63_1ab;586 | 0 | 0 |
| NC_005831.2;YP_003766.2;HCoV-NL63_1ab;587 | 0 | 0 |
| NC_005831.2;YP_003766.2;HCoV-NL63_1ab;588 | 0 | 0 |
| NC_005831.2;YP_003766.2;HCoV-NL63_1ab;589 | 0 | 0.025 |
| NC_005831.2;YP_003766.2;HCoV-NL63_1ab;590 | 0 | 0.025 |
| NC_005831.2;YP_003766.2;HCoV-NL63_1ab;591 | 0 | 0 |
| NC_005831.2;YP_003766.2;HCoV-NL63_1ab;592 | 0 | 0 |
| NC_005831.2;YP_003766.2;HCoV-NL63_1ab;593 | 0 | 0 |
| NC_005831.2;YP_003766.2;HCoV-NL63_1ab;594 | 0 | 0.025 |
| NC_005831.2;YP_003766.2;HCoV-NL63_1ab;595 | 0 | 0.025 |
| NC_005831.2;YP_003766.2;HCoV-NL63_1ab;596 | 0 | 0.05 |
| NC_005831.2;YP_003766.2;HCoV-NL63_1ab;597 | 0 | 0.05 |
| NC_005831.2;YP_003766.2;HCoV-NL63_1ab;598 | 0 | 0.025 |
| NC_005831.2;YP_003766.2;HCoV-NL63_1ab;599 | 0 | 0.05 |
| NC_005831.2;YP_003766.2;HCoV-NL63_1ab;600 | 0 | 0 |
| NC_005831.2;YP_003766.2;HCoV-NL63_1ab;601 | 0 | 0 |
| NC_005831.2;YP_003766.2;HCoV-NL63_1ab;602 | 0 | 0.025 |
| NC_005831.2;YP_003766.2;HCoV-NL63_1ab;603 | 0 | 0 |
| NC_005831.2;YP_003766.2;HCoV-NL63_1ab;604 | 0 | 0 |
| NC_005831.2;YP_003766.2;HCoV-NL63_1ab;605 | 0 | 0 |
| NC_005831.2;YP_003766.2;HCoV-NL63_1ab;606 | 0 | 0 |
| NC_005831.2;YP_003766.2;HCoV-NL63_1ab;607 | 0 | 0 |
| NC_005831.2;YP_003766.2;HCoV-NL63_1ab;608 | 0 | 0 |
| NC_005831.2;YP_003766.2;HCoV-NL63_1ab;609 | 0 | 0 |
| NC_005831.2;YP_003766.2;HCoV-NL63_1ab;610 | 0 | 0 |
| NC_005831.2;YP_003766.2;HCoV-NL63_1ab;611 | 0 | 0 |

| | | |
|---|---|---|
| NC_005831.2;YP_003766.2;HCoV-NL63_1ab;612 | 0 | 0 |
| NC_005831.2;YP_003766.2;HCoV-NL63_1ab;613 | 0 | 0 |
| NC_005831.2;YP_003766.2;HCoV-NL63_1ab;614 | 0 | 0 |
| NC_005831.2;YP_003766.2;HCoV-NL63_1ab;615 | 0 | 0 |
| NC_005831.2;YP_003766.2;HCoV-NL63_1ab;616 | 0 | 0 |
| NC_005831.2;YP_003766.2;HCoV-NL63_1ab;617 | 0 | 0 |
| NC_005831.2;YP_003766.2;HCoV-NL63_1ab;618 | 0 | 0 |
| NC_005831.2;YP_003766.2;HCoV-NL63_1ab;619 | 0 | 0.025 |
| NC_005831.2;YP_003766.2;HCoV-NL63_1ab;620 | 0 | 0 |
| NC_005831.2;YP_003766.2;HCoV-NL63_1ab;621 | 0 | 0 |
| NC_005831.2;YP_003766.2;HCoV-NL63_1ab;622 | 0 | 0 |
| NC_005831.2;YP_003766.2;HCoV-NL63_1ab;623 | 0 | 0 |
| NC_005831.2;YP_003766.2;HCoV-NL63_1ab;624 | 0 | 0 |
| NC_005831.2;YP_003766.2;HCoV-NL63_1ab;625 | 0 | 0 |
| NC_005831.2;YP_003766.2;HCoV-NL63_1ab;626 | 0 | 0 |
| NC_005831.2;YP_003766.2;HCoV-NL63_1ab;627 | 0 | 0 |
| NC_005831.2;YP_003766.2;HCoV-NL63_1ab;628 | 0 | 0 |
| NC_005831.2;YP_003766.2;HCoV-NL63_1ab;629 | 0 | 0 |
| NC_005831.2;YP_003766.2;HCoV-NL63_1ab;630 | 0 | 0 |
| NC_005831.2;YP_003766.2;HCoV-NL63_1ab;631 | 0 | 0 |
| NC_005831.2;YP_003766.2;HCoV-NL63_1ab;632 | 0 | 0 |
| NC_005831.2;YP_003766.2;HCoV-NL63_1ab;633 | 0 | 0 |
| NC_005831.2;YP_003766.2;HCoV-NL63_1ab;634 | 0 | 0 |
| NC_005831.2;YP_003766.2;HCoV-NL63_1ab;635 | 0 | 0 |
| NC_005831.2;YP_003766.2;HCoV-NL63_1ab;636 | 0 | 0 |
| NC_005831.2;YP_003766.2;HCoV-NL63_1ab;637 | 0 | 0 |
| NC_005831.2;YP_003766.2;HCoV-NL63_1ab;638 | 0 | 0 |
| NC_005831.2;YP_003766.2;HCoV-NL63_1ab;639 | 0 | 0 |
| NC_005831.2;YP_003766.2;HCoV-NL63_1ab;640 | 0 | 0 |
| NC_005831.2;YP_003766.2;HCoV-NL63_1ab;641 | 0.05 | 0 |
| NC_005831.2;YP_003766.2;HCoV-NL63_1ab;642 | 0.05 | 0 |
| NC_005831.2;YP_003766.2;HCoV-NL63_1ab;643 | 0.05 | 0 |
| NC_005831.2;YP_003766.2;HCoV-NL63_1ab;644 | 0.05 | 0 |
| NC_005831.2;YP_003766.2;HCoV-NL63_1ab;645 | 0 | 0 |
| NC_005831.2;YP_003766.2;HCoV-NL63_1ab;646 | 0 | 0 |
| NC_005831.2;YP_003766.2;HCoV-NL63_1ab;647 | 0 | 0 |
| NC_005831.2;YP_003766.2;HCoV-NL63_1ab;648 | 0 | 0 |
| NC_005831.2;YP_003766.2;HCoV-NL63_1ab;649 | 0 | 0 |
| NC_005831.2;YP_003766.2;HCoV-NL63_1ab;650 | 0 | 0 |
| NC_005831.2;YP_003766.2;HCoV-NL63_1ab;651 | 0 | 0 |
| NC_005831.2;YP_003766.2;HCoV-NL63_1ab;652 | 0 | 0 |
| NC_005831.2;YP_003766.2;HCoV-NL63_1ab;653 | 0 | 0 |
| NC_005831.2;YP_003766.2;HCoV-NL63_1ab;654 | 0 | 0 |

| | | |
|---|---|---|
| NC_005831.2;YP_003766.2;HCoV-NL63_1ab;655 | 0 | 0 |
| NC_005831.2;YP_003766.2;HCoV-NL63_1ab;656 | 0 | 0 |
| NC_005831.2;YP_003766.2;HCoV-NL63_1ab;657 | 0 | 0 |
| NC_005831.2;YP_003766.2;HCoV-NL63_1ab;658 | 0 | 0 |
| NC_005831.2;YP_003766.2;HCoV-NL63_1ab;659 | 0 | 0 |
| NC_005831.2;YP_003766.2;HCoV-NL63_1ab;660 | 0 | 0 |
| NC_005831.2;YP_003766.2;HCoV-NL63_1ab;661 | 0 | 0 |
| NC_005831.2;YP_003766.2;HCoV-NL63_1ab;662 | 0 | 0 |
| NC_005831.2;YP_003766.2;HCoV-NL63_1ab;663 | 0 | 0 |
| NC_005831.2;YP_003766.2;HCoV-NL63_1ab;664 | 0 | 0 |
| NC_005831.2;YP_003766.2;HCoV-NL63_1ab;665 | 0 | 0 |
| NC_005831.2;YP_003766.2;HCoV-NL63_1ab;666 | 0 | 0 |
| NC_005831.2;YP_003766.2;HCoV-NL63_1ab;667 | 0 | 0 |
| NC_005831.2;YP_003766.2;HCoV-NL63_1ab;668 | 0 | 0 |
| NC_005831.2;YP_003766.2;HCoV-NL63_1ab;669 | 0 | 0 |
| NC_005831.2;YP_003766.2;HCoV-NL63_1ab;670 | 0 | 0 |
| NC_005831.2;YP_003766.2;HCoV-NL63_1ab;671 | 0 | 0 |
| NC_005831.2;YP_003766.2;HCoV-NL63_1ab;672 | 0 | 0 |
| NC_005831.2;YP_003766.2;HCoV-NL63_1ab;673 | 0 | 0 |
| NC_005831.2;YP_003766.2;HCoV-NL63_1ab;674 | 0 | 0 |
| NC_005831.2;YP_003766.2;HCoV-NL63_1ab;675 | 0 | 0.025 |
| NC_005831.2;YP_003766.2;HCoV-NL63_1ab;676 | 0 | 0 |
| NC_005831.2;YP_003766.2;HCoV-NL63_1ab;677 | 0 | 0 |
| NC_005831.2;YP_003766.2;HCoV-NL63_1ab;678 | 0 | 0 |
| NC_005831.2;YP_003766.2;HCoV-NL63_1ab;679 | 0 | 0 |
| NC_005831.2;YP_003766.2;HCoV-NL63_1ab;680 | 0.05 | 0 |
| NC_005831.2;YP_003766.2;HCoV-NL63_1ab;681 | 0.05 | 0.025 |
| NC_005831.2;YP_003766.2;HCoV-NL63_1ab;682 | 0 | 0.05 |
| NC_005831.2;YP_003766.2;HCoV-NL63_1ab;683 | 0 | 0.025 |
| NC_005831.2;YP_003766.2;HCoV-NL63_1ab;684 | 0 | 0 |
| NC_005831.2;YP_003766.2;HCoV-NL63_1ab;685 | 0 | 0 |
| NC_005831.2;YP_003766.2;HCoV-NL63_1ab;686 | 0.1 | 0 |
| NC_005831.2;YP_003766.2;HCoV-NL63_1ab;687 | 0.05 | 0.05 |
| NC_005831.2;YP_003766.2;HCoV-NL63_1ab;688 | 0 | 0.025 |
| NC_005831.2;YP_003766.2;HCoV-NL63_1ab;689 | 0.05 | 0 |
| NC_005831.2;YP_003766.2;HCoV-NL63_1ab;690 | 0 | 0 |
| NC_005831.2;YP_003766.2;HCoV-NL63_1ab;691 | 0 | 0 |
| NC_005831.2;YP_003766.2;HCoV-NL63_1ab;692 | 0 | 0 |
| NC_005831.2;YP_003766.2;HCoV-NL63_1ab;693 | 0 | 0 |
| NC_005831.2;YP_003766.2;HCoV-NL63_1ab;694 | 0 | 0 |
| NC_005831.2;YP_003766.2;HCoV-NL63_1ab;695 | 0.05 | 0.025 |
| NC_005831.2;YP_003766.2;HCoV-NL63_1ab;696 | 0 | 0.025 |
| NC_005831.2;YP_003766.2;HCoV-NL63_1ab;697 | 0 | 0.025 |

| | | |
|---|---|---|
| NC_005831.2;YP_003766.2;HCoV-NL63_1ab;698 | 0 | 0.025 |
| NC_005831.2;YP_003766.2;HCoV-NL63_1ab;699 | 0 | 0.025 |
| NC_005831.2;YP_003766.2;HCoV-NL63_1ab;700 | 0 | 0.025 |
| NC_005831.2;YP_003766.2;HCoV-NL63_1ab;701 | 0 | 0.025 |
| NC_005831.2;YP_003766.2;HCoV-NL63_1ab;702 | 0.1 | 0.05 |
| NC_005831.2;YP_003766.2;HCoV-NL63_1ab;703 | 0 | 0.025 |
| NC_005831.2;YP_003766.2;HCoV-NL63_1ab;704 | 0 | 0.025 |
| NC_005831.2;YP_003766.2;HCoV-NL63_1ab;705 | 0 | 0 |
| NC_005831.2;YP_003766.2;HCoV-NL63_1ab;706 | 0 | 0 |
| NC_005831.2;YP_003766.2;HCoV-NL63_1ab;707 | 0 | 0 |
| NC_005831.2;YP_003766.2;HCoV-NL63_1ab;708 | 0.05 | 0 |
| NC_005831.2;YP_003766.2;HCoV-NL63_1ab;709 | 0 | 0 |
| NC_005831.2;YP_003766.2;HCoV-NL63_1ab;710 | 0 | 0 |
| NC_005831.2;YP_003766.2;HCoV-NL63_1ab;711 | 0 | 0 |
| NC_005831.2;YP_003766.2;HCoV-NL63_1ab;712 | 0 | 0 |
| NC_005831.2;YP_003766.2;HCoV-NL63_1ab;713 | 0 | 0 |
| NC_005831.2;YP_003766.2;HCoV-NL63_1ab;714 | 0 | 0 |
| NC_005831.2;YP_003766.2;HCoV-NL63_1ab;715 | 0 | 0 |
| NC_005831.2;YP_003766.2;HCoV-NL63_1ab;716 | 0 | 0 |
| NC_005831.2;YP_003766.2;HCoV-NL63_1ab;717 | 0 | 0 |
| NC_005831.2;YP_003766.2;HCoV-NL63_1ab;718 | 0 | 0 |
| NC_005831.2;YP_003766.2;HCoV-NL63_1ab;719 | 0 | 0.025 |
| NC_005831.2;YP_003766.2;HCoV-NL63_1ab;720 | 0.05 | 0 |
| NC_005831.2;YP_003766.2;HCoV-NL63_1ab;721 | 0 | 0 |
| NC_005831.2;YP_003766.2;HCoV-NL63_1ab;722 | 0 | 0 |
| NC_005831.2;YP_003766.2;HCoV-NL63_1ab;723 | 0 | 0 |
| NC_005831.2;YP_003766.2;HCoV-NL63_1ab;724 | 0 | 0 |
| NC_005831.2;YP_003766.2;HCoV-NL63_1ab;725 | 0 | 0 |
| NC_005831.2;YP_003766.2;HCoV-NL63_1ab;726 | 0 | 0 |
| NC_005831.2;YP_003766.2;HCoV-NL63_1ab;727 | 0 | 0 |
| NC_005831.2;YP_003766.2;HCoV-NL63_1ab;728 | 0 | 0 |
| NC_005831.2;YP_003766.2;HCoV-NL63_1ab;729 | 0 | 0 |
| NC_005831.2;YP_003766.2;HCoV-NL63_1ab;730 | 0 | 0 |
| NC_005831.2;YP_003766.2;HCoV-NL63_1ab;731 | 0 | 0 |
| NC_005831.2;YP_003766.2;HCoV-NL63_1ab;732 | 0 | 0 |
| NC_005831.2;YP_003766.2;HCoV-NL63_1ab;733 | 0 | 0 |
| NC_005831.2;YP_003766.2;HCoV-NL63_1ab;734 | 0 | 0 |
| NC_005831.2;YP_003766.2;HCoV-NL63_1ab;735 | 0 | 0 |
| NC_005831.2;YP_003766.2;HCoV-NL63_1ab;736 | 0 | 0 |
| NC_005831.2;YP_003766.2;HCoV-NL63_1ab;737 | 0 | 0 |
| NC_005831.2;YP_003766.2;HCoV-NL63_1ab;738 | 0 | 0 |
| NC_005831.2;YP_003766.2;HCoV-NL63_1ab;739 | 0 | 0 |
| NC_005831.2;YP_003766.2;HCoV-NL63_1ab;740 | 0 | 0 |

| | | |
|---|---|---|
| NC_005831.2;YP_003766.2;HCoV-NL63_1ab;741 | 0 | 0 |
| NC_005831.2;YP_003766.2;HCoV-NL63_1ab;742 | 0 | 0 |
| NC_005831.2;YP_003766.2;HCoV-NL63_1ab;743 | 0 | 0 |
| NC_005831.2;YP_003766.2;HCoV-NL63_1ab;744 | 0 | 0 |
| NC_005831.2;YP_003766.2;HCoV-NL63_1ab;745 | 0 | 0 |
| NC_005831.2;YP_003766.2;HCoV-NL63_1ab;746 | 0 | 0 |
| NC_005831.2;YP_003766.2;HCoV-NL63_1ab;747 | 0 | 0 |
| NC_005831.2;YP_003766.2;HCoV-NL63_1ab;748 | 0 | 0 |
| NC_005831.2;YP_003766.2;HCoV-NL63_1ab;749 | 0 | 0 |
| NC_005831.2;YP_003766.2;HCoV-NL63_1ab;750 | 0 | 0 |
| NC_005831.2;YP_003766.2;HCoV-NL63_1ab;751 | 0 | 0 |
| NC_005831.2;YP_003766.2;HCoV-NL63_1ab;752 | 0 | 0 |
| NC_005831.2;YP_003766.2;HCoV-NL63_1ab;753 | 0 | 0 |
| NC_005831.2;YP_003766.2;HCoV-NL63_1ab;754 | 0 | 0 |
| NC_005831.2;YP_003766.2;HCoV-NL63_1ab;755 | 0 | 0 |
| NC_005831.2;YP_003766.2;HCoV-NL63_1ab;756 | 0 | 0 |
| NC_005831.2;YP_003766.2;HCoV-NL63_1ab;757 | 0 | 0 |
| NC_005831.2;YP_003766.2;HCoV-NL63_1ab;758 | 0 | 0 |
| NC_005831.2;YP_003766.2;HCoV-NL63_1ab;759 | 0 | 0 |
| NC_005831.2;YP_003766.2;HCoV-NL63_1ab;760 | 0 | 0 |
| NC_005831.2;YP_003766.2;HCoV-NL63_1ab;761 | 0 | 0 |
| NC_005831.2;YP_003766.2;HCoV-NL63_1ab;762 | 0 | 0 |
| NC_005831.2;YP_003766.2;HCoV-NL63_1ab;763 | 0 | 0 |
| NC_005831.2;YP_003766.2;HCoV-NL63_1ab;764 | 0 | 0 |
| NC_005831.2;YP_003766.2;HCoV-NL63_1ab;765 | 0 | 0 |
| NC_005831.2;YP_003766.2;HCoV-NL63_1ab;766 | 0 | 0 |
| NC_005831.2;YP_003766.2;HCoV-NL63_1ab;767 | 0 | 0 |
| NC_005831.2;YP_003766.2;HCoV-NL63_1ab;768 | 0 | 0 |
| NC_005831.2;YP_003766.2;HCoV-NL63_1ab;769 | 0 | 0 |
| NC_005831.2;YP_003766.2;HCoV-NL63_1ab;770 | 0 | 0 |
| NC_005831.2;YP_003766.2;HCoV-NL63_1ab;771 | 0 | 0 |
| NC_005831.2;YP_003766.2;HCoV-NL63_1ab;772 | 0 | 0 |
| NC_005831.2;YP_003766.2;HCoV-NL63_1ab;773 | 0 | 0 |
| NC_005831.2;YP_003766.2;HCoV-NL63_1ab;774 | 0 | 0 |
| NC_005831.2;YP_003766.2;HCoV-NL63_1ab;775 | 0 | 0 |
| NC_005831.2;YP_003766.2;HCoV-NL63_1ab;776 | 0 | 0 |
| NC_005831.2;YP_003766.2;HCoV-NL63_1ab;777 | 0 | 0 |
| NC_005831.2;YP_003766.2;HCoV-NL63_1ab;778 | 0 | 0 |
| NC_005831.2;YP_003766.2;HCoV-NL63_1ab;779 | 0 | 0 |
| NC_005831.2;YP_003766.2;HCoV-NL63_1ab;780 | 0 | 0 |
| NC_005831.2;YP_003766.2;HCoV-NL63_1ab;781 | 0 | 0 |
| NC_005831.2;YP_003766.2;HCoV-NL63_1ab;782 | 0 | 0 |
| NC_005831.2;YP_003766.2;HCoV-NL63_1ab;783 | 0 | 0 |

| | | |
|---|---|---|
| NC_005831.2;YP_003766.2;HCoV-NL63_1ab;784 | 0 | 0 |
| NC_005831.2;YP_003766.2;HCoV-NL63_1ab;785 | 0 | 0 |
| NC_005831.2;YP_003766.2;HCoV-NL63_1ab;786 | 0 | 0 |
| NC_005831.2;YP_003766.2;HCoV-NL63_1ab;787 | 0 | 0 |
| NC_005831.2;YP_003766.2;HCoV-NL63_1ab;788 | 0 | 0 |
| NC_005831.2;YP_003766.2;HCoV-NL63_1ab;789 | 0 | 0 |
| NC_005831.2;YP_003766.2;HCoV-NL63_1ab;790 | 0 | 0 |
| NC_005831.2;YP_003766.2;HCoV-NL63_1ab;791 | 0 | 0 |
| NC_005831.2;YP_003766.2;HCoV-NL63_1ab;792 | 0 | 0 |
| NC_005831.2;YP_003766.2;HCoV-NL63_1ab;793 | 0 | 0 |
| NC_005831.2;YP_003766.2;HCoV-NL63_1ab;794 | 0 | 0 |
| NC_005831.2;YP_003766.2;HCoV-NL63_1ab;795 | 0 | 0 |
| NC_005831.2;YP_003766.2;HCoV-NL63_1ab;796 | 0 | 0 |
| NC_005831.2;YP_003766.2;HCoV-NL63_1ab;797 | 0 | 0 |
| NC_005831.2;YP_003766.2;HCoV-NL63_1ab;798 | 0 | 0 |
| NC_005831.2;YP_003766.2;HCoV-NL63_1ab;799 | 0 | 0 |
| NC_005831.2;YP_003766.2;HCoV-NL63_1ab;800 | 0 | 0 |
| NC_005831.2;YP_003766.2;HCoV-NL63_1ab;801 | 0 | 0 |
| NC_005831.2;YP_003766.2;HCoV-NL63_1ab;802 | 0 | 0 |
| NC_005831.2;YP_003766.2;HCoV-NL63_1ab;803 | 0 | 0 |
| NC_005831.2;YP_003766.2;HCoV-NL63_1ab;804 | 0 | 0 |
| NC_005831.2;YP_003766.2;HCoV-NL63_1ab;805 | 0 | 0 |
| NC_005831.2;YP_003766.2;HCoV-NL63_1ab;806 | 0 | 0 |
| NC_005831.2;YP_003766.2;HCoV-NL63_1ab;807 | 0 | 0 |
| NC_005831.2;YP_003766.2;HCoV-NL63_1ab;808 | 0 | 0.05 |
| NC_005831.2;YP_003766.2;HCoV-NL63_1ab;809 | 0 | 0 |
| NC_005831.2;YP_003766.2;HCoV-NL63_1ab;810 | 0 | 0 |
| NC_005831.2;YP_003766.2;HCoV-NL63_1ab;811 | 0 | 0 |
| NC_005831.2;YP_003766.2;HCoV-NL63_1ab;812 | 0 | 0 |
| NC_005831.2;YP_003766.2;HCoV-NL63_1ab;813 | 0 | 0 |
| NC_005831.2;YP_003766.2;HCoV-NL63_1ab;814 | 0 | 0 |
| NC_005831.2;YP_003766.2;HCoV-NL63_1ab;815 | 0 | 0 |
| NC_005831.2;YP_003766.2;HCoV-NL63_1ab;816 | 0 | 0 |
| NC_005831.2;YP_003766.2;HCoV-NL63_1ab;817 | 0 | 0 |
| NC_005831.2;YP_003766.2;HCoV-NL63_1ab;818 | 0 | 0 |
| NC_005831.2;YP_003766.2;HCoV-NL63_1ab;819 | 0 | 0 |
| NC_005831.2;YP_003766.2;HCoV-NL63_1ab;820 | 0 | 0 |
| NC_005831.2;YP_003766.2;HCoV-NL63_1ab;821 | 0 | 0 |
| NC_005831.2;YP_003766.2;HCoV-NL63_1ab;822 | 0 | 0 |
| NC_005831.2;YP_003766.2;HCoV-NL63_1ab;823 | 0 | 0 |
| NC_005831.2;YP_003766.2;HCoV-NL63_1ab;824 | 0 | 0 |
| NC_005831.2;YP_003766.2;HCoV-NL63_1ab;825 | 0 | 0 |
| NC_005831.2;YP_003766.2;HCoV-NL63_1ab;826 | 0 | 0 |

| | | |
|---|---|---|
| NC_005831.2;YP_003766.2;HCoV-NL63_1ab;827 | 0 | 0 |
| NC_005831.2;YP_003766.2;HCoV-NL63_1ab;828 | 0 | 0 |
| NC_005831.2;YP_003766.2;HCoV-NL63_1ab;829 | 0.05 | 0 |
| NC_005831.2;YP_003766.2;HCoV-NL63_1ab;830 | 0 | 0 |
| NC_005831.2;YP_003766.2;HCoV-NL63_1ab;831 | 0 | 0 |
| NC_005831.2;YP_003766.2;HCoV-NL63_1ab;832 | 0 | 0 |
| NC_005831.2;YP_003766.2;HCoV-NL63_1ab;833 | 0 | 0 |
| NC_005831.2;YP_003766.2;HCoV-NL63_1ab;834 | 0 | 0 |
| NC_005831.2;YP_003766.2;HCoV-NL63_1ab;835 | 0.05 | 0 |
| NC_005831.2;YP_003766.2;HCoV-NL63_1ab;836 | 0 | 0 |
| NC_005831.2;YP_003766.2;HCoV-NL63_1ab;837 | 0 | 0.05 |
| NC_005831.2;YP_003766.2;HCoV-NL63_1ab;838 | 0 | 0 |
| NC_005831.2;YP_003766.2;HCoV-NL63_1ab;839 | 0 | 0 |
| NC_005831.2;YP_003766.2;HCoV-NL63_1ab;840 | 0 | 0 |
| NC_005831.2;YP_003766.2;HCoV-NL63_1ab;841 | 0 | 0 |
| NC_005831.2;YP_003766.2;HCoV-NL63_1ab;842 | 0 | 0 |
| NC_005831.2;YP_003766.2;HCoV-NL63_1ab;843 | 0 | 0 |
| NC_005831.2;YP_003766.2;HCoV-NL63_1ab;844 | 0 | 0 |
| NC_005831.2;YP_003766.2;HCoV-NL63_1ab;845 | 0 | 0 |
| NC_005831.2;YP_003766.2;HCoV-NL63_1ab;846 | 0 | 0.025 |
| NC_005831.2;YP_003766.2;HCoV-NL63_1ab;847 | 0 | 0 |
| NC_005831.2;YP_003766.2;HCoV-NL63_1ab;848 | 0 | 0 |
| NC_005831.2;YP_003766.2;HCoV-NL63_1ab;849 | 0 | 0 |
| NC_005831.2;YP_003766.2;HCoV-NL63_1ab;850 | 0 | 0 |
| NC_005831.2;YP_003766.2;HCoV-NL63_1ab;851 | 0 | 0 |
| NC_005831.2;YP_003766.2;HCoV-NL63_1ab;852 | 0 | 0 |
| NC_005831.2;YP_003766.2;HCoV-NL63_1ab;853 | 0 | 0 |
| NC_005831.2;YP_003766.2;HCoV-NL63_1ab;854 | 0 | 0 |
| NC_005831.2;YP_003766.2;HCoV-NL63_1ab;855 | 0 | 0 |
| NC_005831.2;YP_003766.2;HCoV-NL63_1ab;856 | 0 | 0 |
| NC_005831.2;YP_003766.2;HCoV-NL63_1ab;857 | 0 | 0 |
| NC_005831.2;YP_003766.2;HCoV-NL63_1ab;858 | 0 | 0 |
| NC_005831.2;YP_003766.2;HCoV-NL63_1ab;859 | 0 | 0 |
| NC_005831.2;YP_003766.2;HCoV-NL63_1ab;860 | 0 | 0 |
| NC_005831.2;YP_003766.2;HCoV-NL63_1ab;861 | 0 | 0 |
| NC_005831.2;YP_003766.2;HCoV-NL63_1ab;862 | 0 | 0 |
| NC_005831.2;YP_003766.2;HCoV-NL63_1ab;863 | 0 | 0 |
| NC_005831.2;YP_003766.2;HCoV-NL63_1ab;864 | 0 | 0 |
| NC_005831.2;YP_003766.2;HCoV-NL63_1ab;865 | 0 | 0 |
| NC_005831.2;YP_003766.2;HCoV-NL63_1ab;866 | 0 | 0 |
| NC_005831.2;YP_003766.2;HCoV-NL63_1ab;867 | 0 | 0 |
| NC_005831.2;YP_003766.2;HCoV-NL63_1ab;868 | 0 | 0 |
| NC_005831.2;YP_003766.2;HCoV-NL63_1ab;869 | 0 | 0 |

| | | |
|---|---|---|
| NC_005831.2;YP_003766.2;HCoV-NL63_1ab;870 | 0 | 0 |
| NC_005831.2;YP_003766.2;HCoV-NL63_1ab;871 | 0 | 0 |
| NC_005831.2;YP_003766.2;HCoV-NL63_1ab;872 | 0 | 0 |
| NC_005831.2;YP_003766.2;HCoV-NL63_1ab;873 | 0 | 0 |
| NC_005831.2;YP_003766.2;HCoV-NL63_1ab;874 | 0 | 0.025 |
| NC_005831.2;YP_003766.2;HCoV-NL63_1ab;875 | 0.05 | 0.05 |
| NC_005831.2;YP_003766.2;HCoV-NL63_1ab;876 | 0 | 0.025 |
| NC_005831.2;YP_003766.2;HCoV-NL63_1ab;877 | 0.1 | 0.05 |
| NC_005831.2;YP_003766.2;HCoV-NL63_1ab;878 | 0 | 0 |
| NC_005831.2;YP_003766.2;HCoV-NL63_1ab;879 | 0 | 0 |
| NC_005831.2;YP_003766.2;HCoV-NL63_1ab;880 | 0 | 0 |
| NC_005831.2;YP_003766.2;HCoV-NL63_1ab;881 | 0 | 0 |
| NC_005831.2;YP_003766.2;HCoV-NL63_1ab;882 | 0 | 0 |
| NC_005831.2;YP_003766.2;HCoV-NL63_1ab;883 | 0 | 0 |
| NC_005831.2;YP_003766.2;HCoV-NL63_1ab;884 | 0.05 | 0 |
| NC_005831.2;YP_003766.2;HCoV-NL63_1ab;885 | 0 | 0 |
| NC_005831.2;YP_003766.2;HCoV-NL63_1ab;886 | 0 | 0 |
| NC_005831.2;YP_003766.2;HCoV-NL63_1ab;887 | 0 | 0 |
| NC_005831.2;YP_003766.2;HCoV-NL63_1ab;888 | 0 | 0 |
| NC_005831.2;YP_003766.2;HCoV-NL63_1ab;889 | 0 | 0 |
| NC_005831.2;YP_003766.2;HCoV-NL63_1ab;890 | 0 | 0.025 |
| NC_005831.2;YP_003766.2;HCoV-NL63_1ab;891 | 0 | 0.025 |
| NC_005831.2;YP_003766.2;HCoV-NL63_1ab;892 | 0 | 0.025 |
| NC_005831.2;YP_003766.2;HCoV-NL63_1ab;893 | 0 | 0.025 |
| NC_005831.2;YP_003766.2;HCoV-NL63_1ab;894 | 0 | 0.025 |
| NC_005831.2;YP_003766.2;HCoV-NL63_1ab;895 | 0 | 0.025 |
| NC_005831.2;YP_003766.2;HCoV-NL63_1ab;896 | 0 | 0 |
| NC_005831.2;YP_003766.2;HCoV-NL63_1ab;897 | 0 | 0 |
| NC_005831.2;YP_003766.2;HCoV-NL63_1ab;898 | 0 | 0 |
| NC_005831.2;YP_003766.2;HCoV-NL63_1ab;899 | 0 | 0 |
| NC_005831.2;YP_003766.2;HCoV-NL63_1ab;900 | 0 | 0 |
| NC_005831.2;YP_003766.2;HCoV-NL63_1ab;901 | 0.05 | 0 |
| NC_005831.2;YP_003766.2;HCoV-NL63_1ab;902 | 0 | 0 |
| NC_005831.2;YP_003766.2;HCoV-NL63_1ab;903 | 0 | 0 |
| NC_005831.2;YP_003766.2;HCoV-NL63_1ab;904 | 0 | 0 |
| NC_005831.2;YP_003766.2;HCoV-NL63_1ab;905 | 0 | 0 |
| NC_005831.2;YP_003766.2;HCoV-NL63_1ab;906 | 0 | 0 |
| NC_005831.2;YP_003766.2;HCoV-NL63_1ab;907 | 0 | 0 |
| NC_005831.2;YP_003766.2;HCoV-NL63_1ab;908 | 0 | 0 |
| NC_005831.2;YP_003766.2;HCoV-NL63_1ab;909 | 0 | 0 |
| NC_005831.2;YP_003766.2;HCoV-NL63_1ab;910 | 0 | 0 |
| NC_005831.2;YP_003766.2;HCoV-NL63_1ab;911 | 0 | 0 |
| NC_005831.2;YP_003766.2;HCoV-NL63_1ab;912 | 0 | 0 |

| | | |
|---|---|---|
| NC_005831.2;YP_003766.2;HCoV-NL63_1ab;913 | 0 | 0 |
| NC_005831.2;YP_003766.2;HCoV-NL63_1ab;914 | 0 | 0 |
| NC_005831.2;YP_003766.2;HCoV-NL63_1ab;915 | 0 | 0 |
| NC_005831.2;YP_003766.2;HCoV-NL63_1ab;916 | 0 | 0 |
| NC_005831.2;YP_003766.2;HCoV-NL63_1ab;917 | 0 | 0.025 |
| NC_005831.2;YP_003766.2;HCoV-NL63_1ab;918 | 0 | 0 |
| NC_005831.2;YP_003766.2;HCoV-NL63_1ab;919 | 0 | 0 |
| NC_005831.2;YP_003766.2;HCoV-NL63_1ab;920 | 0 | 0 |
| NC_005831.2;YP_003766.2;HCoV-NL63_1ab;921 | 0 | 0 |
| NC_005831.2;YP_003766.2;HCoV-NL63_1ab;922 | 0 | 0 |
| NC_005831.2;YP_003766.2;HCoV-NL63_1ab;923 | 0 | 0 |
| NC_005831.2;YP_003766.2;HCoV-NL63_1ab;924 | 0 | 0.025 |
| NC_005831.2;YP_003766.2;HCoV-NL63_1ab;925 | 0 | 0 |
| NC_005831.2;YP_003766.2;HCoV-NL63_1ab;926 | 0 | 0 |
| NC_005831.2;YP_003766.2;HCoV-NL63_1ab;927 | 0 | 0 |
| NC_005831.2;YP_003766.2;HCoV-NL63_1ab;928 | 0 | 0 |
| NC_005831.2;YP_003766.2;HCoV-NL63_1ab;929 | 0 | 0 |
| NC_005831.2;YP_003766.2;HCoV-NL63_1ab;930 | 0 | 0 |
| NC_005831.2;YP_003766.2;HCoV-NL63_1ab;931 | 0 | 0 |
| NC_005831.2;YP_003766.2;HCoV-NL63_1ab;932 | 0 | 0 |
| NC_005831.2;YP_003766.2;HCoV-NL63_1ab;933 | 0.05 | 0 |
| NC_005831.2;YP_003766.2;HCoV-NL63_1ab;934 | 0.1 | 0 |
| NC_005831.2;YP_003766.2;HCoV-NL63_1ab;935 | 0 | 0 |
| NC_005831.2;YP_003766.2;HCoV-NL63_1ab;936 | 0 | 0 |
| NC_005831.2;YP_003766.2;HCoV-NL63_1ab;937 | 0 | 0 |
| NC_005831.2;YP_003766.2;HCoV-NL63_1ab;938 | 0 | 0.025 |
| NC_005831.2;YP_003766.2;HCoV-NL63_1ab;939 | 0 | 0.025 |
| NC_005831.2;YP_003766.2;HCoV-NL63_1ab;940 | 0 | 0.05 |
| NC_005831.2;YP_003766.2;HCoV-NL63_1ab;941 | 0 | 0.025 |
| NC_005831.2;YP_003766.2;HCoV-NL63_1ab;942 | 0 | 0.025 |
| NC_005831.2;YP_003766.2;HCoV-NL63_1ab;943 | 0 | 0 |
| NC_005831.2;YP_003766.2;HCoV-NL63_1ab;944 | 0 | 0 |
| NC_005831.2;YP_003766.2;HCoV-NL63_1ab;945 | 0 | 0 |
| NC_005831.2;YP_003766.2;HCoV-NL63_1ab;946 | 0 | 0 |
| NC_005831.2;YP_003766.2;HCoV-NL63_1ab;947 | 0 | 0 |
| NC_005831.2;YP_003766.2;HCoV-NL63_1ab;948 | 0 | 0 |
| NC_005831.2;YP_003766.2;HCoV-NL63_1ab;949 | 0.05 | 0 |
| NC_005831.2;YP_003766.2;HCoV-NL63_1ab;950 | 0 | 0 |
| NC_005831.2;YP_003766.2;HCoV-NL63_1ab;951 | 0 | 0 |
| NC_005831.2;YP_003766.2;HCoV-NL63_1ab;952 | 0 | 0 |
| NC_005831.2;YP_003766.2;HCoV-NL63_1ab;953 | 0 | 0 |
| NC_005831.2;YP_003766.2;HCoV-NL63_1ab;954 | 0 | 0 |
| NC_005831.2;YP_003766.2;HCoV-NL63_1ab;955 | 0 | 0 |

| | | |
|---|---|---|
| NC_005831.2;YP_003766.2;HCoV-NL63_1ab;956 | 0 | 0 |
| NC_005831.2;YP_003766.2;HCoV-NL63_1ab;957 | 0 | 0 |
| NC_005831.2;YP_003766.2;HCoV-NL63_1ab;958 | 0 | 0 |
| NC_005831.2;YP_003766.2;HCoV-NL63_1ab;959 | 0 | 0 |
| NC_005831.2;YP_003766.2;HCoV-NL63_1ab;960 | 0 | 0 |
| NC_005831.2;YP_003766.2;HCoV-NL63_1ab;961 | 0 | 0 |
| NC_005831.2;YP_003766.2;HCoV-NL63_1ab;962 | 0 | 0 |
| NC_005831.2;YP_003766.2;HCoV-NL63_1ab;963 | 0 | 0 |
| NC_005831.2;YP_003766.2;HCoV-NL63_1ab;964 | 0 | 0 |
| NC_005831.2;YP_003766.2;HCoV-NL63_1ab;965 | 0 | 0.05 |
| NC_005831.2;YP_003766.2;HCoV-NL63_1ab;966 | 0 | 0.025 |
| NC_005831.2;YP_003766.2;HCoV-NL63_1ab;967 | 0 | 0.025 |
| NC_005831.2;YP_003766.2;HCoV-NL63_1ab;968 | 0.05 | 0.05 |
| NC_005831.2;YP_003766.2;HCoV-NL63_1ab;969 | 0.05 | 0.05 |
| NC_005831.2;YP_003766.2;HCoV-NL63_1ab;970 | 0.05 | 0.025 |
| NC_005831.2;YP_003766.2;HCoV-NL63_1ab;971 | 0 | 0.025 |
| NC_005831.2;YP_003766.2;HCoV-NL63_1ab;972 | 0 | 0.025 |
| NC_005831.2;YP_003766.2;HCoV-NL63_1ab;973 | 0 | 0.025 |
| NC_005831.2;YP_003766.2;HCoV-NL63_1ab;974 | 0 | 0 |
| NC_005831.2;YP_003766.2;HCoV-NL63_1ab;975 | 0 | 0 |
| NC_005831.2;YP_003766.2;HCoV-NL63_1ab;976 | 0.05 | 0 |
| NC_005831.2;YP_003766.2;HCoV-NL63_1ab;977 | 0 | 0 |
| NC_005831.2;YP_003766.2;HCoV-NL63_1ab;978 | 0 | 0 |
| NC_005831.2;YP_003766.2;HCoV-NL63_1ab;979 | 0 | 0 |
| NC_005831.2;YP_003766.2;HCoV-NL63_1ab;980 | 0 | 0 |
| NC_005831.2;YP_003766.2;HCoV-NL63_1ab;981 | 0 | 0 |
| NC_005831.2;YP_003766.2;HCoV-NL63_1ab;982 | 0.05 | 0.025 |
| NC_005831.2;YP_003766.2;HCoV-NL63_1ab;983 | 0.1 | 0.05 |
| NC_005831.2;YP_003766.2;HCoV-NL63_1ab;984 | 0.05 | 0.05 |
| NC_005831.2;YP_003766.2;HCoV-NL63_1ab;985 | 0.05 | 0.05 |
| NC_005831.2;YP_003766.2;HCoV-NL63_1ab;986 | 0.05 | 0.1 |
| NC_005831.2;YP_003766.2;HCoV-NL63_1ab;987 | 0.15 | 0.125 |
| NC_005831.2;YP_003766.2;HCoV-NL63_1ab;988 | 0.1 | 0.125 |
| NC_005831.2;YP_003766.2;HCoV-NL63_1ab;989 | 0.05 | 0.05 |
| NC_005831.2;YP_003766.2;HCoV-NL63_1ab;990 | 0 | 0.025 |
| NC_005831.2;YP_003766.2;HCoV-NL63_1ab;991 | 0 | 0.025 |
| NC_005831.2;YP_003766.2;HCoV-NL63_1ab;992 | 0 | 0.025 |
| NC_005831.2;YP_003766.2;HCoV-NL63_1ab;993 | 0 | 0.025 |
| NC_005831.2;YP_003766.2;HCoV-NL63_1ab;994 | 0 | 0.025 |
| NC_005831.2;YP_003766.2;HCoV-NL63_1ab;995 | 0 | 0.025 |
| NC_005831.2;YP_003766.2;HCoV-NL63_1ab;996 | 0 | 0.025 |
| NC_005831.2;YP_003766.2;HCoV-NL63_1ab;997 | 0 | 0 |
| NC_005831.2;YP_003766.2;HCoV-NL63_1ab;998 | 0 | 0 |

| | | |
|---|---|---|
| NC_005831.2;YP_003766.2;HCoV-NL63_1ab;999 | 0 | 0 |
| NC_005831.2;YP_003766.2;HCoV-NL63_1ab;1000 | 0 | 0 |
| NC_005831.2;YP_003766.2;HCoV-NL63_1ab;1001 | 0 | 0 |
| NC_005831.2;YP_003766.2;HCoV-NL63_1ab;1002 | 0 | 0 |
| NC_005831.2;YP_003766.2;HCoV-NL63_1ab;1003 | 0 | 0 |
| NC_005831.2;YP_003766.2;HCoV-NL63_1ab;1004 | 0 | 0 |
| NC_005831.2;YP_003766.2;HCoV-NL63_1ab;1005 | 0 | 0 |
| NC_005831.2;YP_003766.2;HCoV-NL63_1ab;1006 | 0 | 0 |
| NC_005831.2;YP_003766.2;HCoV-NL63_1ab;1007 | 0 | 0 |
| NC_005831.2;YP_003766.2;HCoV-NL63_1ab;1008 | 0 | 0 |
| NC_005831.2;YP_003766.2;HCoV-NL63_1ab;1009 | 0 | 0 |
| NC_005831.2;YP_003766.2;HCoV-NL63_1ab;1010 | 0 | 0 |
| NC_005831.2;YP_003766.2;HCoV-NL63_1ab;1011 | 0 | 0 |
| NC_005831.2;YP_003766.2;HCoV-NL63_1ab;1012 | 0 | 0 |
| NC_005831.2;YP_003766.2;HCoV-NL63_1ab;1013 | 0 | 0 |
| NC_005831.2;YP_003766.2;HCoV-NL63_1ab;1014 | 0 | 0 |
| NC_005831.2;YP_003766.2;HCoV-NL63_1ab;1015 | 0.05 | 0 |
| NC_005831.2;YP_003766.2;HCoV-NL63_1ab;1016 | 0.05 | 0.025 |
| NC_005831.2;YP_003766.2;HCoV-NL63_1ab;1017 | 0.05 | 0 |
| NC_005831.2;YP_003766.2;HCoV-NL63_1ab;1018 | 0.05 | 0.025 |
| NC_005831.2;YP_003766.2;HCoV-NL63_1ab;1019 | 0.05 | 0.025 |
| NC_005831.2;YP_003766.2;HCoV-NL63_1ab;1020 | 0.05 | 0.05 |
| NC_005831.2;YP_003766.2;HCoV-NL63_1ab;1021 | 0.05 | 0.025 |
| NC_005831.2;YP_003766.2;HCoV-NL63_1ab;1022 | 0.05 | 0 |
| NC_005831.2;YP_003766.2;HCoV-NL63_1ab;1023 | 0.05 | 0 |
| NC_005831.2;YP_003766.2;HCoV-NL63_1ab;1024 | 0.05 | 0 |
| NC_005831.2;YP_003766.2;HCoV-NL63_1ab;1025 | 0.05 | 0 |
| NC_005831.2;YP_003766.2;HCoV-NL63_1ab;1026 | 0 | 0 |
| NC_005831.2;YP_003766.2;HCoV-NL63_1ab;1027 | 0 | 0 |
| NC_005831.2;YP_003766.2;HCoV-NL63_1ab;1028 | 0 | 0 |
| NC_005831.2;YP_003766.2;HCoV-NL63_1ab;1029 | 0 | 0 |
| NC_005831.2;YP_003766.2;HCoV-NL63_1ab;1030 | 0 | 0 |
| NC_005831.2;YP_003766.2;HCoV-NL63_1ab;1031 | 0 | 0 |
| NC_005831.2;YP_003766.2;HCoV-NL63_1ab;1032 | 0 | 0 |
| NC_005831.2;YP_003766.2;HCoV-NL63_1ab;1033 | 0 | 0 |
| NC_005831.2;YP_003766.2;HCoV-NL63_1ab;1034 | 0 | 0 |
| NC_005831.2;YP_003766.2;HCoV-NL63_1ab;1035 | 0 | 0 |
| NC_005831.2;YP_003766.2;HCoV-NL63_1ab;1036 | 0 | 0 |
| NC_005831.2;YP_003766.2;HCoV-NL63_1ab;1037 | 0 | 0 |
| NC_005831.2;YP_003766.2;HCoV-NL63_1ab;1038 | 0 | 0.05 |
| NC_005831.2;YP_003766.2;HCoV-NL63_1ab;1039 | 0 | 0.025 |
| NC_005831.2;YP_003766.2;HCoV-NL63_1ab;1040 | 0 | 0 |
| NC_005831.2;YP_003766.2;HCoV-NL63_1ab;1041 | 0 | 0 |

| | | |
|---|---|---|
| NC_005831.2;YP_003766.2;HCoV-NL63_1ab;1042 | 0.05 | 0 |
| NC_005831.2;YP_003766.2;HCoV-NL63_1ab;1043 | 0.1 | 0.025 |
| NC_005831.2;YP_003766.2;HCoV-NL63_1ab;1044 | 0.05 | 0.025 |
| NC_005831.2;YP_003766.2;HCoV-NL63_1ab;1045 | 0.1 | 0.025 |
| NC_005831.2;YP_003766.2;HCoV-NL63_1ab;1046 | 0.2 | 0.05 |
| NC_005831.2;YP_003766.2;HCoV-NL63_1ab;1047 | 0.05 | 0.025 |
| NC_005831.2;YP_003766.2;HCoV-NL63_1ab;1048 | 0 | 0 |
| NC_005831.2;YP_003766.2;HCoV-NL63_1ab;1049 | 0 | 0 |
| NC_005831.2;YP_003766.2;HCoV-NL63_1ab;1050 | 0 | 0 |
| NC_005831.2;YP_003766.2;HCoV-NL63_1ab;1051 | 0.05 | 0 |
| NC_005831.2;YP_003766.2;HCoV-NL63_1ab;1052 | 0 | 0 |
| NC_005831.2;YP_003766.2;HCoV-NL63_1ab;1053 | 0 | 0.025 |
| NC_005831.2;YP_003766.2;HCoV-NL63_1ab;1054 | 0.05 | 0.025 |
| NC_005831.2;YP_003766.2;HCoV-NL63_1ab;1055 | 0 | 0 |
| NC_005831.2;YP_003766.2;HCoV-NL63_1ab;1056 | 0 | 0 |
| NC_005831.2;YP_003766.2;HCoV-NL63_1ab;1057 | 0 | 0 |
| NC_005831.2;YP_003766.2;HCoV-NL63_1ab;1058 | 0 | 0 |
| NC_005831.2;YP_003766.2;HCoV-NL63_1ab;1059 | 0 | 0 |
| NC_005831.2;YP_003766.2;HCoV-NL63_1ab;1060 | 0 | 0 |
| NC_005831.2;YP_003766.2;HCoV-NL63_1ab;1061 | 0 | 0 |
| NC_005831.2;YP_003766.2;HCoV-NL63_1ab;1062 | 0 | 0 |
| NC_005831.2;YP_003766.2;HCoV-NL63_1ab;1063 | 0 | 0 |
| NC_005831.2;YP_003766.2;HCoV-NL63_1ab;1064 | 0 | 0 |
| NC_005831.2;YP_003766.2;HCoV-NL63_1ab;1065 | 0 | 0 |
| NC_005831.2;YP_003766.2;HCoV-NL63_1ab;1066 | 0 | 0 |
| NC_005831.2;YP_003766.2;HCoV-NL63_1ab;1067 | 0 | 0 |
| NC_005831.2;YP_003766.2;HCoV-NL63_1ab;1068 | 0 | 0 |
| NC_005831.2;YP_003766.2;HCoV-NL63_1ab;1069 | 0 | 0 |
| NC_005831.2;YP_003766.2;HCoV-NL63_1ab;1070 | 0 | 0 |
| NC_005831.2;YP_003766.2;HCoV-NL63_1ab;1071 | 0 | 0 |
| NC_005831.2;YP_003766.2;HCoV-NL63_1ab;1072 | 0 | 0 |
| NC_005831.2;YP_003766.2;HCoV-NL63_1ab;1073 | 0 | 0 |
| NC_005831.2;YP_003766.2;HCoV-NL63_1ab;1074 | 0 | 0 |
| NC_005831.2;YP_003766.2;HCoV-NL63_1ab;1075 | 0 | 0 |
| NC_005831.2;YP_003766.2;HCoV-NL63_1ab;1076 | 0 | 0 |
| NC_005831.2;YP_003766.2;HCoV-NL63_1ab;1077 | 0 | 0 |
| NC_005831.2;YP_003766.2;HCoV-NL63_1ab;1078 | 0.05 | 0 |
| NC_005831.2;YP_003766.2;HCoV-NL63_1ab;1079 | 0 | 0.025 |
| NC_005831.2;YP_003766.2;HCoV-NL63_1ab;1080 | 0.05 | 0 |
| NC_005831.2;YP_003766.2;HCoV-NL63_1ab;1081 | 0 | 0.025 |
| NC_005831.2;YP_003766.2;HCoV-NL63_1ab;1082 | 0 | 0 |
| NC_005831.2;YP_003766.2;HCoV-NL63_1ab;1083 | 0 | 0 |
| NC_005831.2;YP_003766.2;HCoV-NL63_1ab;1084 | 0 | 0 |

| | | |
|---|---|---|
| NC_005831.2;YP_003766.2;HCoV-NL63_1ab;1085 | 0 | 0.025 |
| NC_005831.2;YP_003766.2;HCoV-NL63_1ab;1086 | 0 | 0.025 |
| NC_005831.2;YP_003766.2;HCoV-NL63_1ab;1087 | 0 | 0.025 |
| NC_005831.2;YP_003766.2;HCoV-NL63_1ab;1088 | 0 | 0 |
| NC_005831.2;YP_003766.2;HCoV-NL63_1ab;1089 | 0 | 0.025 |
| NC_005831.2;YP_003766.2;HCoV-NL63_1ab;1090 | 0 | 0 |
| NC_005831.2;YP_003766.2;HCoV-NL63_1ab;1091 | 0 | 0 |
| NC_005831.2;YP_003766.2;HCoV-NL63_1ab;1092 | 0 | 0 |
| NC_005831.2;YP_003766.2;HCoV-NL63_1ab;1093 | 0 | 0 |
| NC_005831.2;YP_003766.2;HCoV-NL63_1ab;1094 | 0 | 0 |
| NC_005831.2;YP_003766.2;HCoV-NL63_1ab;1095 | 0 | 0 |
| NC_005831.2;YP_003766.2;HCoV-NL63_1ab;1096 | 0 | 0 |
| NC_005831.2;YP_003766.2;HCoV-NL63_1ab;1097 | 0 | 0 |
| NC_005831.2;YP_003766.2;HCoV-NL63_1ab;1098 | 0 | 0 |
| NC_005831.2;YP_003766.2;HCoV-NL63_1ab;1099 | 0 | 0 |
| NC_005831.2;YP_003766.2;HCoV-NL63_1ab;1100 | 0 | 0 |
| NC_005831.2;YP_003766.2;HCoV-NL63_1ab;1101 | 0 | 0 |
| NC_005831.2;YP_003766.2;HCoV-NL63_1ab;1102 | 0 | 0 |
| NC_005831.2;YP_003766.2;HCoV-NL63_1ab;1103 | 0 | 0 |
| NC_005831.2;YP_003766.2;HCoV-NL63_1ab;1104 | 0 | 0 |
| NC_005831.2;YP_003766.2;HCoV-NL63_1ab;1105 | 0 | 0 |
| NC_005831.2;YP_003766.2;HCoV-NL63_1ab;1106 | 0 | 0 |
| NC_005831.2;YP_003766.2;HCoV-NL63_1ab;1107 | 0 | 0 |
| NC_005831.2;YP_003766.2;HCoV-NL63_1ab;1108 | 0 | 0 |
| NC_005831.2;YP_003766.2;HCoV-NL63_1ab;1109 | 0.05 | 0 |
| NC_005831.2;YP_003766.2;HCoV-NL63_1ab;1110 | 0 | 0.025 |
| NC_005831.2;YP_003766.2;HCoV-NL63_1ab;1111 | 0 | 0 |
| NC_005831.2;YP_003766.2;HCoV-NL63_1ab;1112 | 0 | 0 |
| NC_005831.2;YP_003766.2;HCoV-NL63_1ab;1113 | 0 | 0 |
| NC_005831.2;YP_003766.2;HCoV-NL63_1ab;1114 | 0 | 0 |
| NC_005831.2;YP_003766.2;HCoV-NL63_1ab;1115 | 0 | 0 |
| NC_005831.2;YP_003766.2;HCoV-NL63_1ab;1116 | 0 | 0 |
| NC_005831.2;YP_003766.2;HCoV-NL63_1ab;1117 | 0 | 0.025 |
| NC_005831.2;YP_003766.2;HCoV-NL63_1ab;1118 | 0 | 0 |
| NC_005831.2;YP_003766.2;HCoV-NL63_1ab;1119 | 0 | 0 |
| NC_005831.2;YP_003766.2;HCoV-NL63_1ab;1120 | 0 | 0 |
| NC_005831.2;YP_003766.2;HCoV-NL63_1ab;1121 | 0 | 0 |
| NC_005831.2;YP_003766.2;HCoV-NL63_1ab;1122 | 0 | 0 |
| NC_005831.2;YP_003766.2;HCoV-NL63_1ab;1123 | 0 | 0 |
| NC_005831.2;YP_003766.2;HCoV-NL63_1ab;1124 | 0 | 0.025 |
| NC_005831.2;YP_003766.2;HCoV-NL63_1ab;1125 | 0 | 0 |
| NC_005831.2;YP_003766.2;HCoV-NL63_1ab;1126 | 0 | 0 |
| NC_005831.2;YP_003766.2;HCoV-NL63_1ab;1127 | 0 | 0.025 |

| | | |
|---|---|---|
| NC_005831.2;YP_003766.2;HCoV-NL63_1ab;1128 | 0.05 | 0 |
| NC_005831.2;YP_003766.2;HCoV-NL63_1ab;1129 | 0 | 0.025 |
| NC_005831.2;YP_003766.2;HCoV-NL63_1ab;1130 | 0.05 | 0 |
| NC_005831.2;YP_003766.2;HCoV-NL63_1ab;1131 | 0.05 | 0.025 |
| NC_005831.2;YP_003766.2;HCoV-NL63_1ab;1132 | 0 | 0.025 |
| NC_005831.2;YP_003766.2;HCoV-NL63_1ab;1133 | 0 | 0.05 |
| NC_005831.2;YP_003766.2;HCoV-NL63_1ab;1134 | 0 | 0 |
| NC_005831.2;YP_003766.2;HCoV-NL63_1ab;1135 | 0.05 | 0 |
| NC_005831.2;YP_003766.2;HCoV-NL63_1ab;1136 | 0 | 0 |
| NC_005831.2;YP_003766.2;HCoV-NL63_1ab;1137 | 0.05 | 0 |
| NC_005831.2;YP_003766.2;HCoV-NL63_1ab;1138 | 0 | 0 |
| NC_005831.2;YP_003766.2;HCoV-NL63_1ab;1139 | 0 | 0 |
| NC_005831.2;YP_003766.2;HCoV-NL63_1ab;1140 | 0 | 0 |
| NC_005831.2;YP_003766.2;HCoV-NL63_1ab;1141 | 0 | 0 |
| NC_005831.2;YP_003766.2;HCoV-NL63_1ab;1142 | 0 | 0 |
| NC_005831.2;YP_003766.2;HCoV-NL63_1ab;1143 | 0 | 0 |
| NC_005831.2;YP_003766.2;HCoV-NL63_1ab;1144 | 0 | 0 |
| NC_005831.2;YP_003766.2;HCoV-NL63_1ab;1145 | 0 | 0 |
| NC_005831.2;YP_003766.2;HCoV-NL63_1ab;1146 | 0.05 | 0 |
| NC_005831.2;YP_003766.2;HCoV-NL63_1ab;1147 | 0 | 0 |
| NC_005831.2;YP_003766.2;HCoV-NL63_1ab;1148 | 0.05 | 0 |
| NC_005831.2;YP_003766.2;HCoV-NL63_1ab;1149 | 0.05 | 0 |
| NC_005831.2;YP_003766.2;HCoV-NL63_1ab;1150 | 0.05 | 0 |
| NC_005831.2;YP_003766.2;HCoV-NL63_1ab;1151 | 0.05 | 0.025 |
| NC_005831.2;YP_003766.2;HCoV-NL63_1ab;1152 | 0 | 0 |
| NC_005831.2;YP_003766.2;HCoV-NL63_1ab;1153 | 0 | 0 |
| NC_005831.2;YP_003766.2;HCoV-NL63_1ab;1154 | 0 | 0 |
| NC_005831.2;YP_003766.2;HCoV-NL63_1ab;1155 | 0 | 0 |
| NC_005831.2;YP_003766.2;HCoV-NL63_1ab;1156 | 0 | 0 |
| NC_005831.2;YP_003766.2;HCoV-NL63_1ab;1157 | 0 | 0 |
| NC_005831.2;YP_003766.2;HCoV-NL63_1ab;1158 | 0 | 0 |
| NC_005831.2;YP_003766.2;HCoV-NL63_1ab;1159 | 0 | 0 |
| NC_005831.2;YP_003766.2;HCoV-NL63_1ab;1160 | 0 | 0 |
| NC_005831.2;YP_003766.2;HCoV-NL63_1ab;1161 | 0 | 0 |
| NC_005831.2;YP_003766.2;HCoV-NL63_1ab;1162 | 0 | 0 |
| NC_005831.2;YP_003766.2;HCoV-NL63_1ab;1163 | 0 | 0 |
| NC_005831.2;YP_003766.2;HCoV-NL63_1ab;1164 | 0 | 0 |
| NC_005831.2;YP_003766.2;HCoV-NL63_1ab;1165 | 0 | 0 |
| NC_005831.2;YP_003766.2;HCoV-NL63_1ab;1166 | 0 | 0 |
| NC_005831.2;YP_003766.2;HCoV-NL63_1ab;1167 | 0 | 0 |
| NC_005831.2;YP_003766.2;HCoV-NL63_1ab;1168 | 0 | 0 |
| NC_005831.2;YP_003766.2;HCoV-NL63_1ab;1169 | 0 | 0 |
| NC_005831.2;YP_003766.2;HCoV-NL63_1ab;1170 | 0 | 0 |

| | | |
|---|---|---|
| NC_005831.2;YP_003766.2;HCoV-NL63_1ab;1171 | 0 | 0 |
| NC_005831.2;YP_003766.2;HCoV-NL63_1ab;1172 | 0 | 0.025 |
| NC_005831.2;YP_003766.2;HCoV-NL63_1ab;1173 | 0 | 0 |
| NC_005831.2;YP_003766.2;HCoV-NL63_1ab;1174 | 0 | 0.025 |
| NC_005831.2;YP_003766.2;HCoV-NL63_1ab;1175 | 0 | 0.025 |
| NC_005831.2;YP_003766.2;HCoV-NL63_1ab;1176 | 0 | 0.05 |
| NC_005831.2;YP_003766.2;HCoV-NL63_1ab;1177 | 0 | 0.025 |
| NC_005831.2;YP_003766.2;HCoV-NL63_1ab;1178 | 0 | 0 |
| NC_005831.2;YP_003766.2;HCoV-NL63_1ab;1179 | 0 | 0 |
| NC_005831.2;YP_003766.2;HCoV-NL63_1ab;1180 | 0 | 0.025 |
| NC_005831.2;YP_003766.2;HCoV-NL63_1ab;1181 | 0 | 0 |
| NC_005831.2;YP_003766.2;HCoV-NL63_1ab;1182 | 0 | 0.125 |
| NC_005831.2;YP_003766.2;HCoV-NL63_1ab;1183 | 0 | 0.075 |
| NC_005831.2;YP_003766.2;HCoV-NL63_1ab;1184 | 0 | 0.075 |
| NC_005831.2;YP_003766.2;HCoV-NL63_1ab;1185 | 0.05 | 0.075 |
| NC_005831.2;YP_003766.2;HCoV-NL63_1ab;1186 | 0.05 | 0.075 |
| NC_005831.2;YP_003766.2;HCoV-NL63_1ab;1187 | 0.05 | 0.1 |
| NC_005831.2;YP_003766.2;HCoV-NL63_1ab;1188 | 0 | 0.05 |
| NC_005831.2;YP_003766.2;HCoV-NL63_1ab;1189 | 0 | 0.025 |
| NC_005831.2;YP_003766.2;HCoV-NL63_1ab;1190 | 0 | 0 |
| NC_005831.2;YP_003766.2;HCoV-NL63_1ab;1191 | 0 | 0 |
| NC_005831.2;YP_003766.2;HCoV-NL63_1ab;1192 | 0 | 0 |
| NC_005831.2;YP_003766.2;HCoV-NL63_1ab;1193 | 0 | 0 |
| NC_005831.2;YP_003766.2;HCoV-NL63_1ab;1194 | 0 | 0 |
| NC_005831.2;YP_003766.2;HCoV-NL63_1ab;1195 | 0 | 0.025 |
| NC_005831.2;YP_003766.2;HCoV-NL63_1ab;1196 | 0 | 0 |
| NC_005831.2;YP_003766.2;HCoV-NL63_1ab;1197 | 0 | 0 |
| NC_005831.2;YP_003766.2;HCoV-NL63_1ab;1198 | 0 | 0 |
| NC_005831.2;YP_003766.2;HCoV-NL63_1ab;1199 | 0 | 0 |
| NC_005831.2;YP_003766.2;HCoV-NL63_1ab;1200 | 0 | 0 |
| NC_005831.2;YP_003766.2;HCoV-NL63_1ab;1201 | 0 | 0 |
| NC_005831.2;YP_003766.2;HCoV-NL63_1ab;1202 | 0 | 0.025 |
| NC_005831.2;YP_003766.2;HCoV-NL63_1ab;1203 | 0 | 0 |
| NC_005831.2;YP_003766.2;HCoV-NL63_1ab;1204 | 0 | 0 |
| NC_005831.2;YP_003766.2;HCoV-NL63_1ab;1205 | 0 | 0 |
| NC_005831.2;YP_003766.2;HCoV-NL63_1ab;1206 | 0.05 | 0 |
| NC_005831.2;YP_003766.2;HCoV-NL63_1ab;1207 | 0 | 0 |
| NC_005831.2;YP_003766.2;HCoV-NL63_1ab;1208 | 0 | 0 |
| NC_005831.2;YP_003766.2;HCoV-NL63_1ab;1209 | 0 | 0 |
| NC_005831.2;YP_003766.2;HCoV-NL63_1ab;1210 | 0.05 | 0.025 |
| NC_005831.2;YP_003766.2;HCoV-NL63_1ab;1211 | 0.1 | 0 |
| NC_005831.2;YP_003766.2;HCoV-NL63_1ab;1212 | 0.1 | 0.025 |
| NC_005831.2;YP_003766.2;HCoV-NL63_1ab;1213 | 0.1 | 0.025 |

| | | |
|---|---|---|
| NC_005831.2;YP_003766.2;HCoV-NL63_1ab;1214 | 0.1 | 0.05 |
| NC_005831.2;YP_003766.2;HCoV-NL63_1ab;1215 | 0.05 | 0 |
| NC_005831.2;YP_003766.2;HCoV-NL63_1ab;1216 | 0.05 | 0 |
| NC_005831.2;YP_003766.2;HCoV-NL63_1ab;1217 | 0 | 0 |
| NC_005831.2;YP_003766.2;HCoV-NL63_1ab;1218 | 0.05 | 0 |
| NC_005831.2;YP_003766.2;HCoV-NL63_1ab;1219 | 0.05 | 0 |
| NC_005831.2;YP_003766.2;HCoV-NL63_1ab;1220 | 0.05 | 0.025 |
| NC_005831.2;YP_003766.2;HCoV-NL63_1ab;1221 | 0 | 0.075 |
| NC_005831.2;YP_003766.2;HCoV-NL63_1ab;1222 | 0.05 | 0 |
| NC_005831.2;YP_003766.2;HCoV-NL63_1ab;1223 | 0.25 | 0.3 |
| NC_005831.2;YP_003766.2;HCoV-NL63_1ab;1224 | 0.3 | 0.25 |
| NC_005831.2;YP_003766.2;HCoV-NL63_1ab;1225 | 0.45 | 0.35 |
| NC_005831.2;YP_003766.2;HCoV-NL63_1ab;1226 | 0.2 | 0.225 |
| NC_005831.2;YP_003766.2;HCoV-NL63_1ab;1227 | 0.1 | 0.1 |
| NC_005831.2;YP_003766.2;HCoV-NL63_1ab;1228 | 0.25 | 0.1 |
| NC_005831.2;YP_003766.2;HCoV-NL63_1ab;1229 | 0.15 | 0.075 |
| NC_005831.2;YP_003766.2;HCoV-NL63_1ab;1230 | 0.05 | 0.05 |
| NC_005831.2;YP_003766.2;HCoV-NL63_1ab;1231 | 0.15 | 0.05 |
| NC_005831.2;YP_003766.2;HCoV-NL63_1ab;1232 | 0.05 | 0 |
| NC_005831.2;YP_003766.2;HCoV-NL63_1ab;1233 | 0 | 0.025 |
| NC_005831.2;YP_003766.2;HCoV-NL63_1ab;1234 | 0 | 0 |
| NC_005831.2;YP_003766.2;HCoV-NL63_1ab;1235 | 0 | 0 |
| NC_005831.2;YP_003766.2;HCoV-NL63_1ab;1236 | 0 | 0 |
| NC_005831.2;YP_003766.2;HCoV-NL63_1ab;1237 | 0 | 0.025 |
| NC_005831.2;YP_003766.2;HCoV-NL63_1ab;1238 | 0 | 0 |
| NC_005831.2;YP_003766.2;HCoV-NL63_1ab;1239 | 0 | 0 |
| NC_005831.2;YP_003766.2;HCoV-NL63_1ab;1240 | 0 | 0 |
| NC_005831.2;YP_003766.2;HCoV-NL63_1ab;1241 | 0 | 0 |
| NC_005831.2;YP_003766.2;HCoV-NL63_1ab;1242 | 0 | 0 |
| NC_005831.2;YP_003766.2;HCoV-NL63_1ab;1243 | 0 | 0 |
| NC_005831.2;YP_003766.2;HCoV-NL63_1ab;1244 | 0.05 | 0.025 |
| NC_005831.2;YP_003766.2;HCoV-NL63_1ab;1245 | 0.05 | 0.05 |
| NC_005831.2;YP_003766.2;HCoV-NL63_1ab;1246 | 0.05 | 0.05 |
| NC_005831.2;YP_003766.2;HCoV-NL63_1ab;1247 | 0.05 | 0.075 |
| NC_005831.2;YP_003766.2;HCoV-NL63_1ab;1248 | 0.05 | 0.025 |
| NC_005831.2;YP_003766.2;HCoV-NL63_1ab;1249 | 0.05 | 0.025 |
| NC_005831.2;YP_003766.2;HCoV-NL63_1ab;1250 | 0.05 | 0.05 |
| NC_005831.2;YP_003766.2;HCoV-NL63_1ab;1251 | 0.05 | 0.075 |
| NC_005831.2;YP_003766.2;HCoV-NL63_1ab;1252 | 0 | 0.05 |
| NC_005831.2;YP_003766.2;HCoV-NL63_1ab;1253 | 0 | 0 |
| NC_005831.2;YP_003766.2;HCoV-NL63_1ab;1254 | 0 | 0 |
| NC_005831.2;YP_003766.2;HCoV-NL63_1ab;1255 | 0 | 0 |
| NC_005831.2;YP_003766.2;HCoV-NL63_1ab;1256 | 0 | 0 |

| | | |
|---|---|---|
| NC_005831.2;YP_003766.2;HCoV-NL63_1ab;1257 | 0 | 0 |
| NC_005831.2;YP_003766.2;HCoV-NL63_1ab;1258 | 0 | 0 |
| NC_005831.2;YP_003766.2;HCoV-NL63_1ab;1259 | 0 | 0 |
| NC_005831.2;YP_003766.2;HCoV-NL63_1ab;1260 | 0 | 0 |
| NC_005831.2;YP_003766.2;HCoV-NL63_1ab;1261 | 0 | 0 |
| NC_005831.2;YP_003766.2;HCoV-NL63_1ab;1262 | 0 | 0 |
| NC_005831.2;YP_003766.2;HCoV-NL63_1ab;1263 | 0 | 0 |
| NC_005831.2;YP_003766.2;HCoV-NL63_1ab;1264 | 0 | 0 |
| NC_005831.2;YP_003766.2;HCoV-NL63_1ab;1265 | 0 | 0 |
| NC_005831.2;YP_003766.2;HCoV-NL63_1ab;1266 | 0 | 0 |
| NC_005831.2;YP_003766.2;HCoV-NL63_1ab;1267 | 0 | 0 |
| NC_005831.2;YP_003766.2;HCoV-NL63_1ab;1268 | 0 | 0 |
| NC_005831.2;YP_003766.2;HCoV-NL63_1ab;1269 | 0 | 0 |
| NC_005831.2;YP_003766.2;HCoV-NL63_1ab;1270 | 0 | 0 |
| NC_005831.2;YP_003766.2;HCoV-NL63_1ab;1271 | 0 | 0 |
| NC_005831.2;YP_003766.2;HCoV-NL63_1ab;1272 | 0 | 0 |
| NC_005831.2;YP_003766.2;HCoV-NL63_1ab;1273 | 0 | 0 |
| NC_005831.2;YP_003766.2;HCoV-NL63_1ab;1274 | 0 | 0 |
| NC_005831.2;YP_003766.2;HCoV-NL63_1ab;1275 | 0 | 0 |
| NC_005831.2;YP_003766.2;HCoV-NL63_1ab;1276 | 0 | 0 |
| NC_005831.2;YP_003766.2;HCoV-NL63_1ab;1277 | 0 | 0 |
| NC_005831.2;YP_003766.2;HCoV-NL63_1ab;1278 | 0 | 0 |
| NC_005831.2;YP_003766.2;HCoV-NL63_1ab;1279 | 0 | 0 |
| NC_005831.2;YP_003766.2;HCoV-NL63_1ab;1280 | 0 | 0 |
| NC_005831.2;YP_003766.2;HCoV-NL63_1ab;1281 | 0 | 0 |
| NC_005831.2;YP_003766.2;HCoV-NL63_1ab;1282 | 0 | 0 |
| NC_005831.2;YP_003766.2;HCoV-NL63_1ab;1283 | 0.05 | 0.1 |
| NC_005831.2;YP_003766.2;HCoV-NL63_1ab;1284 | 0 | 0.025 |
| NC_005831.2;YP_003766.2;HCoV-NL63_1ab;1285 | 0.05 | 0.075 |
| NC_005831.2;YP_003766.2;HCoV-NL63_1ab;1286 | 0 | 0 |
| NC_005831.2;YP_003766.2;HCoV-NL63_1ab;1287 | 0 | 0 |
| NC_005831.2;YP_003766.2;HCoV-NL63_1ab;1288 | 0 | 0 |
| NC_005831.2;YP_003766.2;HCoV-NL63_1ab;1289 | 0 | 0 |
| NC_005831.2;YP_003766.2;HCoV-NL63_1ab;1290 | 0 | 0 |
| NC_005831.2;YP_003766.2;HCoV-NL63_1ab;1291 | 0 | 0 |
| NC_005831.2;YP_003766.2;HCoV-NL63_1ab;1292 | 0 | 0 |
| NC_005831.2;YP_003766.2;HCoV-NL63_1ab;1293 | 0 | 0 |
| NC_005831.2;YP_003766.2;HCoV-NL63_1ab;1294 | 0 | 0 |
| NC_005831.2;YP_003766.2;HCoV-NL63_1ab;1295 | 0 | 0 |
| NC_005831.2;YP_003766.2;HCoV-NL63_1ab;1296 | 0.05 | 0 |
| NC_005831.2;YP_003766.2;HCoV-NL63_1ab;1297 | 0 | 0 |
| NC_005831.2;YP_003766.2;HCoV-NL63_1ab;1298 | 0 | 0.025 |
| NC_005831.2;YP_003766.2;HCoV-NL63_1ab;1299 | 0 | 0 |

| | | |
|---|---|---|
| NC_005831.2;YP_003766.2;HCoV-NL63_1ab;1300 | 0 | 0.05 |
| NC_005831.2;YP_003766.2;HCoV-NL63_1ab;1301 | 0.05 | 0 |
| NC_005831.2;YP_003766.2;HCoV-NL63_1ab;1302 | 0 | 0.025 |
| NC_005831.2;YP_003766.2;HCoV-NL63_1ab;1303 | 0 | 0.025 |
| NC_005831.2;YP_003766.2;HCoV-NL63_1ab;1304 | 0 | 0 |
| NC_005831.2;YP_003766.2;HCoV-NL63_1ab;1305 | 0.05 | 0 |
| NC_005831.2;YP_003766.2;HCoV-NL63_1ab;1306 | 0.1 | 0 |
| NC_005831.2;YP_003766.2;HCoV-NL63_1ab;1307 | 0.1 | 0.05 |
| NC_005831.2;YP_003766.2;HCoV-NL63_1ab;1308 | 0.15 | 0.05 |
| NC_005831.2;YP_003766.2;HCoV-NL63_1ab;1309 | 0.2 | 0.15 |
| NC_005831.2;YP_003766.2;HCoV-NL63_1ab;1310 | 0.15 | 0.125 |
| NC_005831.2;YP_003766.2;HCoV-NL63_1ab;1311 | 0.15 | 0.125 |
| NC_005831.2;YP_003766.2;HCoV-NL63_1ab;1312 | 0.15 | 0.15 |
| NC_005831.2;YP_003766.2;HCoV-NL63_1ab;1313 | 0.1 | 0.125 |
| NC_005831.2;YP_003766.2;HCoV-NL63_1ab;1314 | 0.1 | 0.175 |
| NC_005831.2;YP_003766.2;HCoV-NL63_1ab;1315 | 0.05 | 0.1 |
| NC_005831.2;YP_003766.2;HCoV-NL63_1ab;1316 | 0.05 | 0.1 |
| NC_005831.2;YP_003766.2;HCoV-NL63_1ab;1317 | 0 | 0.05 |
| NC_005831.2;YP_003766.2;HCoV-NL63_1ab;1318 | 0.05 | 0.05 |
| NC_005831.2;YP_003766.2;HCoV-NL63_1ab;1319 | 0 | 0.1 |
| NC_005831.2;YP_003766.2;HCoV-NL63_1ab;1320 | 0.05 | 0.025 |
| NC_005831.2;YP_003766.2;HCoV-NL63_1ab;1321 | 0.05 | 0.025 |
| NC_005831.2;YP_003766.2;HCoV-NL63_1ab;1322 | 0 | 0.05 |
| NC_005831.2;YP_003766.2;HCoV-NL63_1ab;1323 | 0 | 0.05 |
| NC_005831.2;YP_003766.2;HCoV-NL63_1ab;1324 | 0 | 0.05 |
| NC_005831.2;YP_003766.2;HCoV-NL63_1ab;1325 | 0.05 | 0.05 |
| NC_005831.2;YP_003766.2;HCoV-NL63_1ab;1326 | 0 | 0.025 |
| NC_005831.2;YP_003766.2;HCoV-NL63_1ab;1327 | 0 | 0 |
| NC_005831.2;YP_003766.2;HCoV-NL63_1ab;1328 | 0.05 | 0.05 |
| NC_005831.2;YP_003766.2;HCoV-NL63_1ab;1329 | 0 | 0 |
| NC_005831.2;YP_003766.2;HCoV-NL63_1ab;1330 | 0.05 | 0 |
| NC_005831.2;YP_003766.2;HCoV-NL63_1ab;1331 | 0.05 | 0 |
| NC_005831.2;YP_003766.2;HCoV-NL63_1ab;1332 | 0.05 | 0 |
| NC_005831.2;YP_003766.2;HCoV-NL63_1ab;1333 | 0.05 | 0 |
| NC_005831.2;YP_003766.2;HCoV-NL63_1ab;1334 | 0.05 | 0 |
| NC_005831.2;YP_003766.2;HCoV-NL63_1ab;1335 | 0 | 0 |
| NC_005831.2;YP_003766.2;HCoV-NL63_1ab;1336 | 0.1 | 0 |
| NC_005831.2;YP_003766.2;HCoV-NL63_1ab;1337 | 0.05 | 0 |
| NC_005831.2;YP_003766.2;HCoV-NL63_1ab;1338 | 0 | 0 |
| NC_005831.2;YP_003766.2;HCoV-NL63_1ab;1339 | 0.05 | 0 |
| NC_005831.2;YP_003766.2;HCoV-NL63_1ab;1340 | 0 | 0 |
| NC_005831.2;YP_003766.2;HCoV-NL63_1ab;1341 | 0 | 0 |
| NC_005831.2;YP_003766.2;HCoV-NL63_1ab;1342 | 0.05 | 0 |

| | | |
|---|---|---|
| NC_005831.2;YP_003766.2;HCoV-NL63_1ab;1343 | 0.05 | 0 |
| NC_005831.2;YP_003766.2;HCoV-NL63_1ab;1344 | 0 | 0 |
| NC_005831.2;YP_003766.2;HCoV-NL63_1ab;1345 | 0 | 0 |
| NC_005831.2;YP_003766.2;HCoV-NL63_1ab;1346 | 0 | 0.025 |
| NC_005831.2;YP_003766.2;HCoV-NL63_1ab;1347 | 0 | 0.025 |
| NC_005831.2;YP_003766.2;HCoV-NL63_1ab;1348 | 0 | 0 |
| NC_005831.2;YP_003766.2;HCoV-NL63_1ab;1349 | 0 | 0.025 |
| NC_005831.2;YP_003766.2;HCoV-NL63_1ab;1350 | 0 | 0.025 |
| NC_005831.2;YP_003766.2;HCoV-NL63_1ab;1351 | 0 | 0.025 |
| NC_005831.2;YP_003766.2;HCoV-NL63_1ab;1352 | 0 | 0.025 |
| NC_005831.2;YP_003766.2;HCoV-NL63_1ab;1353 | 0 | 0 |
| NC_005831.2;YP_003766.2;HCoV-NL63_1ab;1354 | 0 | 0.025 |
| NC_005831.2;YP_003766.2;HCoV-NL63_1ab;1355 | 0 | 0 |
| NC_005831.2;YP_003766.2;HCoV-NL63_1ab;1356 | 0 | 0 |
| NC_005831.2;YP_003766.2;HCoV-NL63_1ab;1357 | 0.1 | 0.05 |
| NC_005831.2;YP_003766.2;HCoV-NL63_1ab;1358 | 0.1 | 0.025 |
| NC_005831.2;YP_003766.2;HCoV-NL63_1ab;1359 | 0.1 | 0.025 |
| NC_005831.2;YP_003766.2;HCoV-NL63_1ab;1360 | 0 | 0 |
| NC_005831.2;YP_003766.2;HCoV-NL63_1ab;1361 | 0 | 0 |
| NC_005831.2;YP_003766.2;HCoV-NL63_1ab;1362 | 0 | 0 |
| NC_005831.2;YP_003766.2;HCoV-NL63_1ab;1363 | 0 | 0 |
| NC_005831.2;YP_003766.2;HCoV-NL63_1ab;1364 | 0 | 0.025 |
| NC_005831.2;YP_003766.2;HCoV-NL63_1ab;1365 | 0 | 0 |
| NC_005831.2;YP_003766.2;HCoV-NL63_1ab;1366 | 0 | 0 |
| NC_005831.2;YP_003766.2;HCoV-NL63_1ab;1367 | 0 | 0 |
| NC_005831.2;YP_003766.2;HCoV-NL63_1ab;1368 | 0 | 0 |
| NC_005831.2;YP_003766.2;HCoV-NL63_1ab;1369 | 0 | 0 |
| NC_005831.2;YP_003766.2;HCoV-NL63_1ab;1370 | 0 | 0 |
| NC_005831.2;YP_003766.2;HCoV-NL63_1ab;1371 | 0 | 0 |
| NC_005831.2;YP_003766.2;HCoV-NL63_1ab;1372 | 0 | 0 |
| NC_005831.2;YP_003766.2;HCoV-NL63_1ab;1373 | 0 | 0 |
| NC_005831.2;YP_003766.2;HCoV-NL63_1ab;1374 | 0 | 0 |
| NC_005831.2;YP_003766.2;HCoV-NL63_1ab;1375 | 0 | 0 |
| NC_005831.2;YP_003766.2;HCoV-NL63_1ab;1376 | 0 | 0 |
| NC_005831.2;YP_003766.2;HCoV-NL63_1ab;1377 | 0 | 0 |
| NC_005831.2;YP_003766.2;HCoV-NL63_1ab;1378 | 0 | 0 |
| NC_005831.2;YP_003766.2;HCoV-NL63_1ab;1379 | 0 | 0 |
| NC_005831.2;YP_003766.2;HCoV-NL63_1ab;1380 | 0 | 0 |
| NC_005831.2;YP_003766.2;HCoV-NL63_1ab;1381 | 0 | 0 |
| NC_005831.2;YP_003766.2;HCoV-NL63_1ab;1382 | 0 | 0 |
| NC_005831.2;YP_003766.2;HCoV-NL63_1ab;1383 | 0 | 0 |
| NC_005831.2;YP_003766.2;HCoV-NL63_1ab;1384 | 0 | 0 |
| NC_005831.2;YP_003766.2;HCoV-NL63_1ab;1385 | 0 | 0 |

| | | |
|---|---|---|
| NC_005831.2;YP_003766.2;HCoV-NL63_1ab;1386 | 0 | 0 |
| NC_005831.2;YP_003766.2;HCoV-NL63_1ab;1387 | 0 | 0 |
| NC_005831.2;YP_003766.2;HCoV-NL63_1ab;1388 | 0 | 0 |
| NC_005831.2;YP_003766.2;HCoV-NL63_1ab;1389 | 0 | 0 |
| NC_005831.2;YP_003766.2;HCoV-NL63_1ab;1390 | 0 | 0 |
| NC_005831.2;YP_003766.2;HCoV-NL63_1ab;1391 | 0 | 0 |
| NC_005831.2;YP_003766.2;HCoV-NL63_1ab;1392 | 0 | 0 |
| NC_005831.2;YP_003766.2;HCoV-NL63_1ab;1393 | 0 | 0 |
| NC_005831.2;YP_003766.2;HCoV-NL63_1ab;1394 | 0 | 0 |
| NC_005831.2;YP_003766.2;HCoV-NL63_1ab;1395 | 0 | 0 |
| NC_005831.2;YP_003766.2;HCoV-NL63_1ab;1396 | 0 | 0 |
| NC_005831.2;YP_003766.2;HCoV-NL63_1ab;1397 | 0 | 0 |
| NC_005831.2;YP_003766.2;HCoV-NL63_1ab;1398 | 0 | 0.025 |
| NC_005831.2;YP_003766.2;HCoV-NL63_1ab;1399 | 0 | 0 |
| NC_005831.2;YP_003766.2;HCoV-NL63_1ab;1400 | 0 | 0.025 |
| NC_005831.2;YP_003766.2;HCoV-NL63_1ab;1401 | 0 | 0 |
| NC_005831.2;YP_003766.2;HCoV-NL63_1ab;1402 | 0 | 0 |
| NC_005831.2;YP_003766.2;HCoV-NL63_1ab;1403 | 0.05 | 0.025 |
| NC_005831.2;YP_003766.2;HCoV-NL63_1ab;1404 | 0.05 | 0.125 |
| NC_005831.2;YP_003766.2;HCoV-NL63_1ab;1405 | 0 | 0.025 |
| NC_005831.2;YP_003766.2;HCoV-NL63_1ab;1406 | 0 | 0.075 |
| NC_005831.2;YP_003766.2;HCoV-NL63_1ab;1407 | 0 | 0.025 |
| NC_005831.2;YP_003766.2;HCoV-NL63_1ab;1408 | 0 | 0 |
| NC_005831.2;YP_003766.2;HCoV-NL63_1ab;1409 | 0 | 0 |
| NC_005831.2;YP_003766.2;HCoV-NL63_1ab;1410 | 0 | 0 |
| NC_005831.2;YP_003766.2;HCoV-NL63_1ab;1411 | 0 | 0 |
| NC_005831.2;YP_003766.2;HCoV-NL63_1ab;1412 | 0 | 0 |
| NC_005831.2;YP_003766.2;HCoV-NL63_1ab;1413 | 0 | 0 |
| NC_005831.2;YP_003766.2;HCoV-NL63_1ab;1414 | 0 | 0 |
| NC_005831.2;YP_003766.2;HCoV-NL63_1ab;1415 | 0 | 0 |
| NC_005831.2;YP_003766.2;HCoV-NL63_1ab;1416 | 0.05 | 0 |
| NC_005831.2;YP_003766.2;HCoV-NL63_1ab;1417 | 0 | 0 |
| NC_005831.2;YP_003766.2;HCoV-NL63_1ab;1418 | 0 | 0 |
| NC_005831.2;YP_003766.2;HCoV-NL63_1ab;1419 | 0 | 0 |
| NC_005831.2;YP_003766.2;HCoV-NL63_1ab;1420 | 0 | 0 |
| NC_005831.2;YP_003766.2;HCoV-NL63_1ab;1421 | 0 | 0 |
| NC_005831.2;YP_003766.2;HCoV-NL63_1ab;1422 | 0 | 0 |
| NC_005831.2;YP_003766.2;HCoV-NL63_1ab;1423 | 0 | 0 |
| NC_005831.2;YP_003766.2;HCoV-NL63_1ab;1424 | 0 | 0 |
| NC_005831.2;YP_003766.2;HCoV-NL63_1ab;1425 | 0 | 0.025 |
| NC_005831.2;YP_003766.2;HCoV-NL63_1ab;1426 | 0 | 0 |
| NC_005831.2;YP_003766.2;HCoV-NL63_1ab;1427 | 0 | 0.025 |
| NC_005831.2;YP_003766.2;HCoV-NL63_1ab;1428 | 0 | 0 |

| | | |
|---|---|---|
| NC_005831.2;YP_003766.2;HCoV-NL63_1ab;1429 | 0 | 0 |
| NC_005831.2;YP_003766.2;HCoV-NL63_1ab;1430 | 0 | 0 |
| NC_005831.2;YP_003766.2;HCoV-NL63_1ab;1431 | 0 | 0 |
| NC_005831.2;YP_003766.2;HCoV-NL63_1ab;1432 | 0 | 0 |
| NC_005831.2;YP_003766.2;HCoV-NL63_1ab;1433 | 0 | 0 |
| NC_005831.2;YP_003766.2;HCoV-NL63_1ab;1434 | 0 | 0 |
| NC_005831.2;YP_003766.2;HCoV-NL63_1ab;1435 | 0 | 0 |
| NC_005831.2;YP_003766.2;HCoV-NL63_1ab;1436 | 0 | 0 |
| NC_005831.2;YP_003766.2;HCoV-NL63_1ab;1437 | 0 | 0 |
| NC_005831.2;YP_003766.2;HCoV-NL63_1ab;1438 | 0 | 0 |
| NC_005831.2;YP_003766.2;HCoV-NL63_1ab;1439 | 0.1 | 0 |
| NC_005831.2;YP_003766.2;HCoV-NL63_1ab;1440 | 0.05 | 0 |
| NC_005831.2;YP_003766.2;HCoV-NL63_1ab;1441 | 0.05 | 0 |
| NC_005831.2;YP_003766.2;HCoV-NL63_1ab;1442 | 0.05 | 0 |
| NC_005831.2;YP_003766.2;HCoV-NL63_1ab;1443 | 0 | 0 |
| NC_005831.2;YP_003766.2;HCoV-NL63_1ab;1444 | 0 | 0 |
| NC_005831.2;YP_003766.2;HCoV-NL63_1ab;1445 | 0 | 0 |
| NC_005831.2;YP_003766.2;HCoV-NL63_1ab;1446 | 0 | 0 |
| NC_005831.2;YP_003766.2;HCoV-NL63_1ab;1447 | 0 | 0 |
| NC_005831.2;YP_003766.2;HCoV-NL63_1ab;1448 | 0 | 0 |
| NC_005831.2;YP_003766.2;HCoV-NL63_1ab;1449 | 0.05 | 0 |
| NC_005831.2;YP_003766.2;HCoV-NL63_1ab;1450 | 0 | 0 |
| NC_005831.2;YP_003766.2;HCoV-NL63_1ab;1451 | 0 | 0 |
| NC_005831.2;YP_003766.2;HCoV-NL63_1ab;1452 | 0 | 0 |
| NC_005831.2;YP_003766.2;HCoV-NL63_1ab;1453 | 0 | 0 |
| NC_005831.2;YP_003766.2;HCoV-NL63_1ab;1454 | 0 | 0 |
| NC_005831.2;YP_003766.2;HCoV-NL63_1ab;1455 | 0 | 0 |
| NC_005831.2;YP_003766.2;HCoV-NL63_1ab;1456 | 0 | 0 |
| NC_005831.2;YP_003766.2;HCoV-NL63_1ab;1457 | 0.05 | 0 |
| NC_005831.2;YP_003766.2;HCoV-NL63_1ab;1458 | 0.1 | 0.025 |
| NC_005831.2;YP_003766.2;HCoV-NL63_1ab;1459 | 0.2 | 0.075 |
| NC_005831.2;YP_003766.2;HCoV-NL63_1ab;1460 | 0.2 | 0.075 |
| NC_005831.2;YP_003766.2;HCoV-NL63_1ab;1461 | 0.15 | 0.075 |
| NC_005831.2;YP_003766.2;HCoV-NL63_1ab;1462 | 0.15 | 0.075 |
| NC_005831.2;YP_003766.2;HCoV-NL63_1ab;1463 | 0.2 | 0.075 |
| NC_005831.2;YP_003766.2;HCoV-NL63_1ab;1464 | 0.25 | 0.075 |
| NC_005831.2;YP_003766.2;HCoV-NL63_1ab;1465 | 0.25 | 0.075 |
| NC_005831.2;YP_003766.2;HCoV-NL63_1ab;1466 | 0.25 | 0.025 |
| NC_005831.2;YP_003766.2;HCoV-NL63_1ab;1467 | 0.15 | 0.025 |
| NC_005831.2;YP_003766.2;HCoV-NL63_1ab;1468 | 0.05 | 0.05 |
| NC_005831.2;YP_003766.2;HCoV-NL63_1ab;1469 | 0.05 | 0.05 |
| NC_005831.2;YP_003766.2;HCoV-NL63_1ab;1470 | 0.15 | 0.1 |
| NC_005831.2;YP_003766.2;HCoV-NL63_1ab;1471 | 0.1 | 0.1 |

| | | |
|---|---|---|
| NC_005831.2;YP_003766.2;HCoV-NL63_1ab;1472 | 0.1 | 0.075 |
| NC_005831.2;YP_003766.2;HCoV-NL63_1ab;1473 | 0.1 | 0.025 |
| NC_005831.2;YP_003766.2;HCoV-NL63_1ab;1474 | 0.05 | 0.025 |
| NC_005831.2;YP_003766.2;HCoV-NL63_1ab;1475 | 0 | 0.05 |
| NC_005831.2;YP_003766.2;HCoV-NL63_1ab;1476 | 0 | 0.025 |
| NC_005831.2;YP_003766.2;HCoV-NL63_1ab;1477 | 0 | 0 |
| NC_005831.2;YP_003766.2;HCoV-NL63_1ab;1478 | 0.05 | 0 |
| NC_005831.2;YP_003766.2;HCoV-NL63_1ab;1479 | 0 | 0.025 |
| NC_005831.2;YP_003766.2;HCoV-NL63_1ab;1480 | 0 | 0 |
| NC_005831.2;YP_003766.2;HCoV-NL63_1ab;1481 | 0 | 0 |
| NC_005831.2;YP_003766.2;HCoV-NL63_1ab;1482 | 0 | 0.025 |
| NC_005831.2;YP_003766.2;HCoV-NL63_1ab;1483 | 0 | 0.025 |
| NC_005831.2;YP_003766.2;HCoV-NL63_1ab;1484 | 0 | 0.05 |
| NC_005831.2;YP_003766.2;HCoV-NL63_1ab;1485 | 0 | 0 |
| NC_005831.2;YP_003766.2;HCoV-NL63_1ab;1486 | 0 | 0 |
| NC_005831.2;YP_003766.2;HCoV-NL63_1ab;1487 | 0 | 0 |
| NC_005831.2;YP_003766.2;HCoV-NL63_1ab;1488 | 0 | 0 |
| NC_005831.2;YP_003766.2;HCoV-NL63_1ab;1489 | 0 | 0 |
| NC_005831.2;YP_003766.2;HCoV-NL63_1ab;1490 | 0 | 0 |
| NC_005831.2;YP_003766.2;HCoV-NL63_1ab;1491 | 0 | 0 |
| NC_005831.2;YP_003766.2;HCoV-NL63_1ab;1492 | 0 | 0 |
| NC_005831.2;YP_003766.2;HCoV-NL63_1ab;1493 | 0 | 0 |
| NC_005831.2;YP_003766.2;HCoV-NL63_1ab;1494 | 0 | 0 |
| NC_005831.2;YP_003766.2;HCoV-NL63_1ab;1495 | 0 | 0 |
| NC_005831.2;YP_003766.2;HCoV-NL63_1ab;1496 | 0 | 0 |
| NC_005831.2;YP_003766.2;HCoV-NL63_1ab;1497 | 0 | 0 |
| NC_005831.2;YP_003766.2;HCoV-NL63_1ab;1498 | 0 | 0 |
| NC_005831.2;YP_003766.2;HCoV-NL63_1ab;1499 | 0 | 0 |
| NC_005831.2;YP_003766.2;HCoV-NL63_1ab;1500 | 0 | 0 |
| NC_005831.2;YP_003766.2;HCoV-NL63_1ab;1501 | 0 | 0 |
| NC_005831.2;YP_003766.2;HCoV-NL63_1ab;1502 | 0 | 0.025 |
| NC_005831.2;YP_003766.2;HCoV-NL63_1ab;1503 | 0 | 0 |
| NC_005831.2;YP_003766.2;HCoV-NL63_1ab;1504 | 0 | 0 |
| NC_005831.2;YP_003766.2;HCoV-NL63_1ab;1505 | 0 | 0 |
| NC_005831.2;YP_003766.2;HCoV-NL63_1ab;1506 | 0 | 0 |
| NC_005831.2;YP_003766.2;HCoV-NL63_1ab;1507 | 0 | 0 |
| NC_005831.2;YP_003766.2;HCoV-NL63_1ab;1508 | 0 | 0 |
| NC_005831.2;YP_003766.2;HCoV-NL63_1ab;1509 | 0 | 0.025 |
| NC_005831.2;YP_003766.2;HCoV-NL63_1ab;1510 | 0.05 | 0.05 |
| NC_005831.2;YP_003766.2;HCoV-NL63_1ab;1511 | 0.05 | 0.1 |
| NC_005831.2;YP_003766.2;HCoV-NL63_1ab;1512 | 0.05 | 0.175 |
| NC_005831.2;YP_003766.2;HCoV-NL63_1ab;1513 | 0.05 | 0.15 |
| NC_005831.2;YP_003766.2;HCoV-NL63_1ab;1514 | 0.1 | 0.1 |

| | | |
|---|---|---|
| NC_005831.2;YP_003766.2;HCoV-NL63_1ab;1515 | 0.15 | 0.125 |
| NC_005831.2;YP_003766.2;HCoV-NL63_1ab;1516 | 0.1 | 0.1 |
| NC_005831.2;YP_003766.2;HCoV-NL63_1ab;1517 | 0.35 | 0.2 |
| NC_005831.2;YP_003766.2;HCoV-NL63_1ab;1518 | 0 | 0 |
| NC_005831.2;YP_003766.2;HCoV-NL63_1ab;1519 | 0 | 0 |
| NC_005831.2;YP_003766.2;HCoV-NL63_1ab;1520 | 0 | 0 |
| NC_005831.2;YP_003766.2;HCoV-NL63_1ab;1521 | 0 | 0 |
| NC_005831.2;YP_003766.2;HCoV-NL63_1ab;1522 | 0 | 0 |
| NC_005831.2;YP_003766.2;HCoV-NL63_1ab;1523 | 0 | 0 |
| NC_005831.2;YP_003766.2;HCoV-NL63_1ab;1524 | 0 | 0 |
| NC_005831.2;YP_003766.2;HCoV-NL63_1ab;1525 | 0 | 0 |
| NC_005831.2;YP_003766.2;HCoV-NL63_1ab;1526 | 0 | 0 |
| NC_005831.2;YP_003766.2;HCoV-NL63_1ab;1527 | 0 | 0 |
| NC_005831.2;YP_003766.2;HCoV-NL63_1ab;1528 | 0 | 0 |
| NC_005831.2;YP_003766.2;HCoV-NL63_1ab;1529 | 0 | 0 |
| NC_005831.2;YP_003766.2;HCoV-NL63_1ab;1530 | 0 | 0 |
| NC_005831.2;YP_003766.2;HCoV-NL63_1ab;1531 | 0 | 0 |
| NC_005831.2;YP_003766.2;HCoV-NL63_1ab;1532 | 0 | 0 |
| NC_005831.2;YP_003766.2;HCoV-NL63_1ab;1533 | 0 | 0 |
| NC_005831.2;YP_003766.2;HCoV-NL63_1ab;1534 | 0 | 0 |
| NC_005831.2;YP_003766.2;HCoV-NL63_1ab;1535 | 0 | 0 |
| NC_005831.2;YP_003766.2;HCoV-NL63_1ab;1536 | 0 | 0 |
| NC_005831.2;YP_003766.2;HCoV-NL63_1ab;1537 | 0 | 0.025 |
| NC_005831.2;YP_003766.2;HCoV-NL63_1ab;1538 | 0.05 | 0.025 |
| NC_005831.2;YP_003766.2;HCoV-NL63_1ab;1539 | 0.05 | 0.05 |
| NC_005831.2;YP_003766.2;HCoV-NL63_1ab;1540 | 0 | 0.025 |
| NC_005831.2;YP_003766.2;HCoV-NL63_1ab;1541 | 0 | 0.025 |
| NC_005831.2;YP_003766.2;HCoV-NL63_1ab;1542 | 0 | 0.025 |
| NC_005831.2;YP_003766.2;HCoV-NL63_1ab;1543 | 0 | 0.025 |
| NC_005831.2;YP_003766.2;HCoV-NL63_1ab;1544 | 0 | 0.025 |
| NC_005831.2;YP_003766.2;HCoV-NL63_1ab;1545 | 0 | 0 |
| NC_005831.2;YP_003766.2;HCoV-NL63_1ab;1546 | 0 | 0 |
| NC_005831.2;YP_003766.2;HCoV-NL63_1ab;1547 | 0 | 0 |
| NC_005831.2;YP_003766.2;HCoV-NL63_1ab;1548 | 0 | 0 |
| NC_005831.2;YP_003766.2;HCoV-NL63_1ab;1549 | 0 | 0 |
| NC_005831.2;YP_003766.2;HCoV-NL63_1ab;1550 | 0 | 0 |
| NC_005831.2;YP_003766.2;HCoV-NL63_1ab;1551 | 0 | 0 |
| NC_005831.2;YP_003766.2;HCoV-NL63_1ab;1552 | 0 | 0 |
| NC_005831.2;YP_003766.2;HCoV-NL63_1ab;1553 | 0 | 0 |
| NC_005831.2;YP_003766.2;HCoV-NL63_1ab;1554 | 0 | 0 |
| NC_005831.2;YP_003766.2;HCoV-NL63_1ab;1555 | 0 | 0 |
| NC_005831.2;YP_003766.2;HCoV-NL63_1ab;1556 | 0 | 0 |
| NC_005831.2;YP_003766.2;HCoV-NL63_1ab;1557 | 0 | 0 |

| | | |
|---|---|---|
| NC_005831.2;YP_003766.2;HCoV-NL63_1ab;1558 | 0 | 0 |
| NC_005831.2;YP_003766.2;HCoV-NL63_1ab;1559 | 0 | 0 |
| NC_005831.2;YP_003766.2;HCoV-NL63_1ab;1560 | 0 | 0 |
| NC_005831.2;YP_003766.2;HCoV-NL63_1ab;1561 | 0 | 0 |
| NC_005831.2;YP_003766.2;HCoV-NL63_1ab;1562 | 0 | 0 |
| NC_005831.2;YP_003766.2;HCoV-NL63_1ab;1563 | 0.05 | 0.025 |
| NC_005831.2;YP_003766.2;HCoV-NL63_1ab;1564 | 0.05 | 0 |
| NC_005831.2;YP_003766.2;HCoV-NL63_1ab;1565 | 0.1 | 0 |
| NC_005831.2;YP_003766.2;HCoV-NL63_1ab;1566 | 0.05 | 0 |
| NC_005831.2;YP_003766.2;HCoV-NL63_1ab;1567 | 0.05 | 0 |
| NC_005831.2;YP_003766.2;HCoV-NL63_1ab;1568 | 0.05 | 0 |
| NC_005831.2;YP_003766.2;HCoV-NL63_1ab;1569 | 0 | 0.025 |
| NC_005831.2;YP_003766.2;HCoV-NL63_1ab;1570 | 0.05 | 0.025 |
| NC_005831.2;YP_003766.2;HCoV-NL63_1ab;1571 | 0 | 0 |
| NC_005831.2;YP_003766.2;HCoV-NL63_1ab;1572 | 0 | 0 |
| NC_005831.2;YP_003766.2;HCoV-NL63_1ab;1573 | 0 | 0 |
| NC_005831.2;YP_003766.2;HCoV-NL63_1ab;1574 | 0 | 0 |
| NC_005831.2;YP_003766.2;HCoV-NL63_1ab;1575 | 0 | 0.025 |
| NC_005831.2;YP_003766.2;HCoV-NL63_1ab;1576 | 0 | 0.025 |
| NC_005831.2;YP_003766.2;HCoV-NL63_1ab;1577 | 0.05 | 0.025 |
| NC_005831.2;YP_003766.2;HCoV-NL63_1ab;1578 | 0 | 0 |
| NC_005831.2;YP_003766.2;HCoV-NL63_1ab;1579 | 0.05 | 0 |
| NC_005831.2;YP_003766.2;HCoV-NL63_1ab;1580 | 0 | 0 |
| NC_005831.2;YP_003766.2;HCoV-NL63_1ab;1581 | 0 | 0 |
| NC_005831.2;YP_003766.2;HCoV-NL63_1ab;1582 | 0 | 0 |
| NC_005831.2;YP_003766.2;HCoV-NL63_1ab;1583 | 0 | 0 |
| NC_005831.2;YP_003766.2;HCoV-NL63_1ab;1584 | 0 | 0 |
| NC_005831.2;YP_003766.2;HCoV-NL63_1ab;1585 | 0 | 0 |
| NC_005831.2;YP_003766.2;HCoV-NL63_1ab;1586 | 0 | 0 |
| NC_005831.2;YP_003766.2;HCoV-NL63_1ab;1587 | 0 | 0 |
| NC_005831.2;YP_003766.2;HCoV-NL63_1ab;1588 | 0 | 0 |
| NC_005831.2;YP_003766.2;HCoV-NL63_1ab;1589 | 0 | 0 |
| NC_005831.2;YP_003766.2;HCoV-NL63_1ab;1590 | 0 | 0 |
| NC_005831.2;YP_003766.2;HCoV-NL63_1ab;1591 | 0 | 0 |
| NC_005831.2;YP_003766.2;HCoV-NL63_1ab;1592 | 0.05 | 0 |
| NC_005831.2;YP_003766.2;HCoV-NL63_1ab;1593 | 0.05 | 0 |
| NC_005831.2;YP_003766.2;HCoV-NL63_1ab;1594 | 0.05 | 0 |
| NC_005831.2;YP_003766.2;HCoV-NL63_1ab;1595 | 0.05 | 0 |
| NC_005831.2;YP_003766.2;HCoV-NL63_1ab;1596 | 0.05 | 0 |
| NC_005831.2;YP_003766.2;HCoV-NL63_1ab;1597 | 0.1 | 0 |
| NC_005831.2;YP_003766.2;HCoV-NL63_1ab;1598 | 0.1 | 0.075 |
| NC_005831.2;YP_003766.2;HCoV-NL63_1ab;1599 | 0.05 | 0.05 |
| NC_005831.2;YP_003766.2;HCoV-NL63_1ab;1600 | 0.05 | 0.025 |

| | | |
|---|---|---|
| NC_005831.2;YP_003766.2;HCoV-NL63_1ab;1601 | 0.05 | 0 |
| NC_005831.2;YP_003766.2;HCoV-NL63_1ab;1602 | 0.05 | 0 |
| NC_005831.2;YP_003766.2;HCoV-NL63_1ab;1603 | 0 | 0 |
| NC_005831.2;YP_003766.2;HCoV-NL63_1ab;1604 | 0 | 0 |
| NC_005831.2;YP_003766.2;HCoV-NL63_1ab;1605 | 0 | 0 |
| NC_005831.2;YP_003766.2;HCoV-NL63_1ab;1606 | 0 | 0 |
| NC_005831.2;YP_003766.2;HCoV-NL63_1ab;1607 | 0 | 0 |
| NC_005831.2;YP_003766.2;HCoV-NL63_1ab;1608 | 0 | 0 |
| NC_005831.2;YP_003766.2;HCoV-NL63_1ab;1609 | 0 | 0 |
| NC_005831.2;YP_003766.2;HCoV-NL63_1ab;1610 | 0 | 0 |
| NC_005831.2;YP_003766.2;HCoV-NL63_1ab;1611 | 0 | 0 |
| NC_005831.2;YP_003766.2;HCoV-NL63_1ab;1612 | 0 | 0.05 |
| NC_005831.2;YP_003766.2;HCoV-NL63_1ab;1613 | 0.1 | 0.1 |
| NC_005831.2;YP_003766.2;HCoV-NL63_1ab;1614 | 0.05 | 0.075 |
| NC_005831.2;YP_003766.2;HCoV-NL63_1ab;1615 | 0.05 | 0.025 |
| NC_005831.2;YP_003766.2;HCoV-NL63_1ab;1616 | 0.1 | 0.075 |
| NC_005831.2;YP_003766.2;HCoV-NL63_1ab;1617 | 0 | 0.025 |
| NC_005831.2;YP_003766.2;HCoV-NL63_1ab;1618 | 0 | 0.025 |
| NC_005831.2;YP_003766.2;HCoV-NL63_1ab;1619 | 0 | 0.025 |
| NC_005831.2;YP_003766.2;HCoV-NL63_1ab;1620 | 0 | 0.025 |
| NC_005831.2;YP_003766.2;HCoV-NL63_1ab;1621 | 0 | 0.025 |
| NC_005831.2;YP_003766.2;HCoV-NL63_1ab;1622 | 0 | 0.025 |
| NC_005831.2;YP_003766.2;HCoV-NL63_1ab;1623 | 0 | 0.025 |
| NC_005831.2;YP_003766.2;HCoV-NL63_1ab;1624 | 0 | 0 |
| NC_005831.2;YP_003766.2;HCoV-NL63_1ab;1625 | 0.05 | 0 |
| NC_005831.2;YP_003766.2;HCoV-NL63_1ab;1626 | 0 | 0 |
| NC_005831.2;YP_003766.2;HCoV-NL63_1ab;1627 | 0 | 0 |
| NC_005831.2;YP_003766.2;HCoV-NL63_1ab;1628 | 0 | 0 |
| NC_005831.2;YP_003766.2;HCoV-NL63_1ab;1629 | 0 | 0 |
| NC_005831.2;YP_003766.2;HCoV-NL63_1ab;1630 | 0 | 0 |
| NC_005831.2;YP_003766.2;HCoV-NL63_1ab;1631 | 0 | 0 |
| NC_005831.2;YP_003766.2;HCoV-NL63_1ab;1632 | 0 | 0 |
| NC_005831.2;YP_003766.2;HCoV-NL63_1ab;1633 | 0.05 | 0 |
| NC_005831.2;YP_003766.2;HCoV-NL63_1ab;1634 | 0 | 0 |
| NC_005831.2;YP_003766.2;HCoV-NL63_1ab;1635 | 0 | 0 |
| NC_005831.2;YP_003766.2;HCoV-NL63_1ab;1636 | 0 | 0 |
| NC_005831.2;YP_003766.2;HCoV-NL63_1ab;1637 | 0 | 0 |
| NC_005831.2;YP_003766.2;HCoV-NL63_1ab;1638 | 0 | 0 |
| NC_005831.2;YP_003766.2;HCoV-NL63_1ab;1639 | 0 | 0 |
| NC_005831.2;YP_003766.2;HCoV-NL63_1ab;1640 | 0 | 0 |
| NC_005831.2;YP_003766.2;HCoV-NL63_1ab;1641 | 0 | 0 |
| NC_005831.2;YP_003766.2;HCoV-NL63_1ab;1642 | 0 | 0 |
| NC_005831.2;YP_003766.2;HCoV-NL63_1ab;1643 | 0 | 0 |

| | | |
|---|---|---|
| NC_005831.2;YP_003766.2;HCoV-NL63_1ab;1644 | 0 | 0 |
| NC_005831.2;YP_003766.2;HCoV-NL63_1ab;1645 | 0 | 0 |
| NC_005831.2;YP_003766.2;HCoV-NL63_1ab;1646 | 0 | 0 |
| NC_005831.2;YP_003766.2;HCoV-NL63_1ab;1647 | 0 | 0 |
| NC_005831.2;YP_003766.2;HCoV-NL63_1ab;1648 | 0 | 0.025 |
| NC_005831.2;YP_003766.2;HCoV-NL63_1ab;1649 | 0 | 0 |
| NC_005831.2;YP_003766.2;HCoV-NL63_1ab;1650 | 0 | 0.05 |
| NC_005831.2;YP_003766.2;HCoV-NL63_1ab;1651 | 0.1 | 0.075 |
| NC_005831.2;YP_003766.2;HCoV-NL63_1ab;1652 | 0.3 | 0.2 |
| NC_005831.2;YP_003766.2;HCoV-NL63_1ab;1653 | 0 | 0.025 |
| NC_005831.2;YP_003766.2;HCoV-NL63_1ab;1654 | 0 | 0.05 |
| NC_005831.2;YP_003766.2;HCoV-NL63_1ab;1655 | 0 | 0 |
| NC_005831.2;YP_003766.2;HCoV-NL63_1ab;1656 | 0 | 0 |
| NC_005831.2;YP_003766.2;HCoV-NL63_1ab;1657 | 0 | 0 |
| NC_005831.2;YP_003766.2;HCoV-NL63_1ab;1658 | 0 | 0 |
| NC_005831.2;YP_003766.2;HCoV-NL63_1ab;1659 | 0 | 0 |
| NC_005831.2;YP_003766.2;HCoV-NL63_1ab;1660 | 0 | 0 |
| NC_005831.2;YP_003766.2;HCoV-NL63_1ab;1661 | 0 | 0 |
| NC_005831.2;YP_003766.2;HCoV-NL63_1ab;1662 | 0 | 0 |
| NC_005831.2;YP_003766.2;HCoV-NL63_1ab;1663 | 0 | 0 |
| NC_005831.2;YP_003766.2;HCoV-NL63_1ab;1664 | 0 | 0 |
| NC_005831.2;YP_003766.2;HCoV-NL63_1ab;1665 | 0 | 0 |
| NC_005831.2;YP_003766.2;HCoV-NL63_1ab;1666 | 0 | 0 |
| NC_005831.2;YP_003766.2;HCoV-NL63_1ab;1667 | 0 | 0 |
| NC_005831.2;YP_003766.2;HCoV-NL63_1ab;1668 | 0 | 0 |
| NC_005831.2;YP_003766.2;HCoV-NL63_1ab;1669 | 0 | 0 |
| NC_005831.2;YP_003766.2;HCoV-NL63_1ab;1670 | 0 | 0 |
| NC_005831.2;YP_003766.2;HCoV-NL63_1ab;1671 | 0 | 0 |
| NC_005831.2;YP_003766.2;HCoV-NL63_1ab;1672 | 0 | 0 |
| NC_005831.2;YP_003766.2;HCoV-NL63_1ab;1673 | 0 | 0 |
| NC_005831.2;YP_003766.2;HCoV-NL63_1ab;1674 | 0 | 0 |
| NC_005831.2;YP_003766.2;HCoV-NL63_1ab;1675 | 0 | 0 |
| NC_005831.2;YP_003766.2;HCoV-NL63_1ab;1676 | 0 | 0 |
| NC_005831.2;YP_003766.2;HCoV-NL63_1ab;1677 | 0 | 0 |
| NC_005831.2;YP_003766.2;HCoV-NL63_1ab;1678 | 0 | 0 |
| NC_005831.2;YP_003766.2;HCoV-NL63_1ab;1679 | 0 | 0 |
| NC_005831.2;YP_003766.2;HCoV-NL63_1ab;1680 | 0 | 0 |
| NC_005831.2;YP_003766.2;HCoV-NL63_1ab;1681 | 0 | 0 |
| NC_005831.2;YP_003766.2;HCoV-NL63_1ab;1682 | 0 | 0 |
| NC_005831.2;YP_003766.2;HCoV-NL63_1ab;1683 | 0 | 0 |
| NC_005831.2;YP_003766.2;HCoV-NL63_1ab;1684 | 0 | 0 |
| NC_005831.2;YP_003766.2;HCoV-NL63_1ab;1685 | 0 | 0 |
| NC_005831.2;YP_003766.2;HCoV-NL63_1ab;1686 | 0 | 0 |

| | | |
|---|---|---|
| NC_005831.2;YP_003766.2;HCoV-NL63_1ab;1687 | 0 | 0 |
| NC_005831.2;YP_003766.2;HCoV-NL63_1ab;1688 | 0 | 0 |
| NC_005831.2;YP_003766.2;HCoV-NL63_1ab;1689 | 0 | 0 |
| NC_005831.2;YP_003766.2;HCoV-NL63_1ab;1690 | 0 | 0 |
| NC_005831.2;YP_003766.2;HCoV-NL63_1ab;1691 | 0 | 0 |
| NC_005831.2;YP_003766.2;HCoV-NL63_1ab;1692 | 0 | 0 |
| NC_005831.2;YP_003766.2;HCoV-NL63_1ab;1693 | 0 | 0 |
| NC_005831.2;YP_003766.2;HCoV-NL63_1ab;1694 | 0 | 0 |
| NC_005831.2;YP_003766.2;HCoV-NL63_1ab;1695 | 0.05 | 0.025 |
| NC_005831.2;YP_003766.2;HCoV-NL63_1ab;1696 | 0 | 0 |
| NC_005831.2;YP_003766.2;HCoV-NL63_1ab;1697 | 0 | 0.025 |
| NC_005831.2;YP_003766.2;HCoV-NL63_1ab;1698 | 0 | 0 |
| NC_005831.2;YP_003766.2;HCoV-NL63_1ab;1699 | 0 | 0 |
| NC_005831.2;YP_003766.2;HCoV-NL63_1ab;1700 | 0 | 0 |
| NC_005831.2;YP_003766.2;HCoV-NL63_1ab;1701 | 0 | 0 |
| NC_005831.2;YP_003766.2;HCoV-NL63_1ab;1702 | 0 | 0 |
| NC_005831.2;YP_003766.2;HCoV-NL63_1ab;1703 | 0 | 0 |
| NC_005831.2;YP_003766.2;HCoV-NL63_1ab;1704 | 0 | 0 |
| NC_005831.2;YP_003766.2;HCoV-NL63_1ab;1705 | 0 | 0 |
| NC_005831.2;YP_003766.2;HCoV-NL63_1ab;1706 | 0 | 0 |
| NC_005831.2;YP_003766.2;HCoV-NL63_1ab;1707 | 0 | 0 |
| NC_005831.2;YP_003766.2;HCoV-NL63_1ab;1708 | 0 | 0 |
| NC_005831.2;YP_003766.2;HCoV-NL63_1ab;1709 | 0 | 0 |
| NC_005831.2;YP_003766.2;HCoV-NL63_1ab;1710 | 0 | 0 |
| NC_005831.2;YP_003766.2;HCoV-NL63_1ab;1711 | 0 | 0 |
| NC_005831.2;YP_003766.2;HCoV-NL63_1ab;1712 | 0 | 0 |
| NC_005831.2;YP_003766.2;HCoV-NL63_1ab;1713 | 0 | 0 |
| NC_005831.2;YP_003766.2;HCoV-NL63_1ab;1714 | 0 | 0 |
| NC_005831.2;YP_003766.2;HCoV-NL63_1ab;1715 | 0 | 0 |
| NC_005831.2;YP_003766.2;HCoV-NL63_1ab;1716 | 0 | 0 |
| NC_005831.2;YP_003766.2;HCoV-NL63_1ab;1717 | 0 | 0 |
| NC_005831.2;YP_003766.2;HCoV-NL63_1ab;1718 | 0 | 0.025 |
| NC_005831.2;YP_003766.2;HCoV-NL63_1ab;1719 | 0 | 0.025 |
| NC_005831.2;YP_003766.2;HCoV-NL63_1ab;1720 | 0 | 0.025 |
| NC_005831.2;YP_003766.2;HCoV-NL63_1ab;1721 | 0 | 0 |
| NC_005831.2;YP_003766.2;HCoV-NL63_1ab;1722 | 0 | 0 |
| NC_005831.2;YP_003766.2;HCoV-NL63_1ab;1723 | 0 | 0 |
| NC_005831.2;YP_003766.2;HCoV-NL63_1ab;1724 | 0 | 0 |
| NC_005831.2;YP_003766.2;HCoV-NL63_1ab;1725 | 0 | 0 |
| NC_005831.2;YP_003766.2;HCoV-NL63_1ab;1726 | 0 | 0.025 |
| NC_005831.2;YP_003766.2;HCoV-NL63_1ab;1727 | 0.05 | 0.025 |
| NC_005831.2;YP_003766.2;HCoV-NL63_1ab;1728 | 0 | 0.05 |
| NC_005831.2;YP_003766.2;HCoV-NL63_1ab;1729 | 0 | 0.075 |

| | | |
|---|---|---|
| NC_005831.2;YP_003766.2;HCoV-NL63_1ab;1730 | 0 | 0.075 |
| NC_005831.2;YP_003766.2;HCoV-NL63_1ab;1731 | 0 | 0.025 |
| NC_005831.2;YP_003766.2;HCoV-NL63_1ab;1732 | 0 | 0.025 |
| NC_005831.2;YP_003766.2;HCoV-NL63_1ab;1733 | 0 | 0.025 |
| NC_005831.2;YP_003766.2;HCoV-NL63_1ab;1734 | 0 | 0 |
| NC_005831.2;YP_003766.2;HCoV-NL63_1ab;1735 | 0 | 0 |
| NC_005831.2;YP_003766.2;HCoV-NL63_1ab;1736 | 0 | 0 |
| NC_005831.2;YP_003766.2;HCoV-NL63_1ab;1737 | 0 | 0 |
| NC_005831.2;YP_003766.2;HCoV-NL63_1ab;1738 | 0 | 0 |
| NC_005831.2;YP_003766.2;HCoV-NL63_1ab;1739 | 0.05 | 0 |
| NC_005831.2;YP_003766.2;HCoV-NL63_1ab;1740 | 0 | 0 |
| NC_005831.2;YP_003766.2;HCoV-NL63_1ab;1741 | 0 | 0 |
| NC_005831.2;YP_003766.2;HCoV-NL63_1ab;1742 | 0 | 0 |
| NC_005831.2;YP_003766.2;HCoV-NL63_1ab;1743 | 0 | 0 |
| NC_005831.2;YP_003766.2;HCoV-NL63_1ab;1744 | 0 | 0 |
| NC_005831.2;YP_003766.2;HCoV-NL63_1ab;1745 | 0 | 0 |
| NC_005831.2;YP_003766.2;HCoV-NL63_1ab;1746 | 0 | 0 |
| NC_005831.2;YP_003766.2;HCoV-NL63_1ab;1747 | 0 | 0 |
| NC_005831.2;YP_003766.2;HCoV-NL63_1ab;1748 | 0 | 0.025 |
| NC_005831.2;YP_003766.2;HCoV-NL63_1ab;1749 | 0 | 0.05 |
| NC_005831.2;YP_003766.2;HCoV-NL63_1ab;1750 | 0 | 0 |
| NC_005831.2;YP_003766.2;HCoV-NL63_1ab;1751 | 0 | 0 |
| NC_005831.2;YP_003766.2;HCoV-NL63_1ab;1752 | 0 | 0 |
| NC_005831.2;YP_003766.2;HCoV-NL63_1ab;1753 | 0 | 0.025 |
| NC_005831.2;YP_003766.2;HCoV-NL63_1ab;1754 | 0 | 0 |
| NC_005831.2;YP_003766.2;HCoV-NL63_1ab;1755 | 0 | 0 |
| NC_005831.2;YP_003766.2;HCoV-NL63_1ab;1756 | 0 | 0.025 |
| NC_005831.2;YP_003766.2;HCoV-NL63_1ab;1757 | 0 | 0.025 |
| NC_005831.2;YP_003766.2;HCoV-NL63_1ab;1758 | 0 | 0 |
| NC_005831.2;YP_003766.2;HCoV-NL63_1ab;1759 | 0 | 0 |
| NC_005831.2;YP_003766.2;HCoV-NL63_1ab;1760 | 0 | 0 |
| NC_005831.2;YP_003766.2;HCoV-NL63_1ab;1761 | 0 | 0 |
| NC_005831.2;YP_003766.2;HCoV-NL63_1ab;1762 | 0 | 0 |
| NC_005831.2;YP_003766.2;HCoV-NL63_1ab;1763 | 0 | 0 |
| NC_005831.2;YP_003766.2;HCoV-NL63_1ab;1764 | 0 | 0 |
| NC_005831.2;YP_003766.2;HCoV-NL63_1ab;1765 | 0.05 | 0 |
| NC_005831.2;YP_003766.2;HCoV-NL63_1ab;1766 | 0 | 0 |
| NC_005831.2;YP_003766.2;HCoV-NL63_1ab;1767 | 0 | 0 |
| NC_005831.2;YP_003766.2;HCoV-NL63_1ab;1768 | 0 | 0 |
| NC_005831.2;YP_003766.2;HCoV-NL63_1ab;1769 | 0 | 0.025 |
| NC_005831.2;YP_003766.2;HCoV-NL63_1ab;1770 | 0 | 0 |
| NC_005831.2;YP_003766.2;HCoV-NL63_1ab;1771 | 0 | 0 |
| NC_005831.2;YP_003766.2;HCoV-NL63_1ab;1772 | 0 | 0 |

| | | |
|---|---|---|
| NC_005831.2;YP_003766.2;HCoV-NL63_1ab;1773 | 0 | 0 |
| NC_005831.2;YP_003766.2;HCoV-NL63_1ab;1774 | 0 | 0 |
| NC_005831.2;YP_003766.2;HCoV-NL63_1ab;1775 | 0 | 0 |
| NC_005831.2;YP_003766.2;HCoV-NL63_1ab;1776 | 0 | 0 |
| NC_005831.2;YP_003766.2;HCoV-NL63_1ab;1777 | 0 | 0 |
| NC_005831.2;YP_003766.2;HCoV-NL63_1ab;1778 | 0 | 0 |
| NC_005831.2;YP_003766.2;HCoV-NL63_1ab;1779 | 0 | 0 |
| NC_005831.2;YP_003766.2;HCoV-NL63_1ab;1780 | 0 | 0 |
| NC_005831.2;YP_003766.2;HCoV-NL63_1ab;1781 | 0 | 0 |
| NC_005831.2;YP_003766.2;HCoV-NL63_1ab;1782 | 0 | 0 |
| NC_005831.2;YP_003766.2;HCoV-NL63_1ab;1783 | 0 | 0 |
| NC_005831.2;YP_003766.2;HCoV-NL63_1ab;1784 | 0 | 0 |
| NC_005831.2;YP_003766.2;HCoV-NL63_1ab;1785 | 0 | 0 |
| NC_005831.2;YP_003766.2;HCoV-NL63_1ab;1786 | 0 | 0 |
| NC_005831.2;YP_003766.2;HCoV-NL63_1ab;1787 | 0 | 0 |
| NC_005831.2;YP_003766.2;HCoV-NL63_1ab;1788 | 0 | 0 |
| NC_005831.2;YP_003766.2;HCoV-NL63_1ab;1789 | 0 | 0 |
| NC_005831.2;YP_003766.2;HCoV-NL63_1ab;1790 | 0 | 0 |
| NC_005831.2;YP_003766.2;HCoV-NL63_1ab;1791 | 0 | 0 |
| NC_005831.2;YP_003766.2;HCoV-NL63_1ab;1792 | 0 | 0 |
| NC_005831.2;YP_003766.2;HCoV-NL63_1ab;1793 | 0 | 0 |
| NC_005831.2;YP_003766.2;HCoV-NL63_1ab;1794 | 0 | 0 |
| NC_005831.2;YP_003766.2;HCoV-NL63_1ab;1795 | 0 | 0 |
| NC_005831.2;YP_003766.2;HCoV-NL63_1ab;1796 | 0 | 0 |
| NC_005831.2;YP_003766.2;HCoV-NL63_1ab;1797 | 0 | 0 |
| NC_005831.2;YP_003766.2;HCoV-NL63_1ab;1798 | 0 | 0 |
| NC_005831.2;YP_003766.2;HCoV-NL63_1ab;1799 | 0 | 0 |
| NC_005831.2;YP_003766.2;HCoV-NL63_1ab;1800 | 0 | 0 |
| NC_005831.2;YP_003766.2;HCoV-NL63_1ab;1801 | 0 | 0 |
| NC_005831.2;YP_003766.2;HCoV-NL63_1ab;1802 | 0 | 0 |
| NC_005831.2;YP_003766.2;HCoV-NL63_1ab;1803 | 0 | 0.025 |
| NC_005831.2;YP_003766.2;HCoV-NL63_1ab;1804 | 0 | 0.05 |
| NC_005831.2;YP_003766.2;HCoV-NL63_1ab;1805 | 0 | 0 |
| NC_005831.2;YP_003766.2;HCoV-NL63_1ab;1806 | 0.05 | 0 |
| NC_005831.2;YP_003766.2;HCoV-NL63_1ab;1807 | 0 | 0 |
| NC_005831.2;YP_003766.2;HCoV-NL63_1ab;1808 | 0 | 0 |
| NC_005831.2;YP_003766.2;HCoV-NL63_1ab;1809 | 0 | 0 |
| NC_005831.2;YP_003766.2;HCoV-NL63_1ab;1810 | 0.05 | 0 |
| NC_005831.2;YP_003766.2;HCoV-NL63_1ab;1811 | 0 | 0 |
| NC_005831.2;YP_003766.2;HCoV-NL63_1ab;1812 | 0 | 0 |
| NC_005831.2;YP_003766.2;HCoV-NL63_1ab;1813 | 0 | 0 |
| NC_005831.2;YP_003766.2;HCoV-NL63_1ab;1814 | 0 | 0 |
| NC_005831.2;YP_003766.2;HCoV-NL63_1ab;1815 | 0 | 0 |

| | | |
|---|---|---|
| NC_005831.2;YP_003766.2;HCoV-NL63_1ab;1816 | 0 | 0.025 |
| NC_005831.2;YP_003766.2;HCoV-NL63_1ab;1817 | 0 | 0 |
| NC_005831.2;YP_003766.2;HCoV-NL63_1ab;1818 | 0 | 0 |
| NC_005831.2;YP_003766.2;HCoV-NL63_1ab;1819 | 0 | 0.025 |
| NC_005831.2;YP_003766.2;HCoV-NL63_1ab;1820 | 0 | 0 |
| NC_005831.2;YP_003766.2;HCoV-NL63_1ab;1821 | 0 | 0.05 |
| NC_005831.2;YP_003766.2;HCoV-NL63_1ab;1822 | 0 | 0 |
| NC_005831.2;YP_003766.2;HCoV-NL63_1ab;1823 | 0 | 0 |
| NC_005831.2;YP_003766.2;HCoV-NL63_1ab;1824 | 0 | 0.025 |
| NC_005831.2;YP_003766.2;HCoV-NL63_1ab;1825 | 0 | 0 |
| NC_005831.2;YP_003766.2;HCoV-NL63_1ab;1826 | 0 | 0 |
| NC_005831.2;YP_003766.2;HCoV-NL63_1ab;1827 | 0 | 0.025 |
| NC_005831.2;YP_003766.2;HCoV-NL63_1ab;1828 | 0 | 0 |
| NC_005831.2;YP_003766.2;HCoV-NL63_1ab;1829 | 0 | 0 |
| NC_005831.2;YP_003766.2;HCoV-NL63_1ab;1830 | 0 | 0.025 |
| NC_005831.2;YP_003766.2;HCoV-NL63_1ab;1831 | 0 | 0 |
| NC_005831.2;YP_003766.2;HCoV-NL63_1ab;1832 | 0 | 0 |
| NC_005831.2;YP_003766.2;HCoV-NL63_1ab;1833 | 0 | 0 |
| NC_005831.2;YP_003766.2;HCoV-NL63_1ab;1834 | 0 | 0 |
| NC_005831.2;YP_003766.2;HCoV-NL63_1ab;1835 | 0 | 0 |
| NC_005831.2;YP_003766.2;HCoV-NL63_1ab;1836 | 0 | 0 |
| NC_005831.2;YP_003766.2;HCoV-NL63_1ab;1837 | 0.05 | 0.025 |
| NC_005831.2;YP_003766.2;HCoV-NL63_1ab;1838 | 0.05 | 0 |
| NC_005831.2;YP_003766.2;HCoV-NL63_1ab;1839 | 0.1 | 0.025 |
| NC_005831.2;YP_003766.2;HCoV-NL63_1ab;1840 | 0 | 0 |
| NC_005831.2;YP_003766.2;HCoV-NL63_1ab;1841 | 0 | 0 |
| NC_005831.2;YP_003766.2;HCoV-NL63_1ab;1842 | 0.05 | 0 |
| NC_005831.2;YP_003766.2;HCoV-NL63_1ab;1843 | 0 | 0 |
| NC_005831.2;YP_003766.2;HCoV-NL63_1ab;1844 | 0 | 0 |
| NC_005831.2;YP_003766.2;HCoV-NL63_1ab;1845 | 0.05 | 0 |
| NC_005831.2;YP_003766.2;HCoV-NL63_1ab;1846 | 0 | 0 |
| NC_005831.2;YP_003766.2;HCoV-NL63_1ab;1847 | 0 | 0 |
| NC_005831.2;YP_003766.2;HCoV-NL63_1ab;1848 | 0 | 0 |
| NC_005831.2;YP_003766.2;HCoV-NL63_1ab;1849 | 0 | 0 |
| NC_005831.2;YP_003766.2;HCoV-NL63_1ab;1850 | 0 | 0 |
| NC_005831.2;YP_003766.2;HCoV-NL63_1ab;1851 | 0 | 0 |
| NC_005831.2;YP_003766.2;HCoV-NL63_1ab;1852 | 0 | 0 |
| NC_005831.2;YP_003766.2;HCoV-NL63_1ab;1853 | 0 | 0 |
| NC_005831.2;YP_003766.2;HCoV-NL63_1ab;1854 | 0 | 0 |
| NC_005831.2;YP_003766.2;HCoV-NL63_1ab;1855 | 0 | 0 |
| NC_005831.2;YP_003766.2;HCoV-NL63_1ab;1856 | 0 | 0 |
| NC_005831.2;YP_003766.2;HCoV-NL63_1ab;1857 | 0 | 0 |
| NC_005831.2;YP_003766.2;HCoV-NL63_1ab;1858 | 0 | 0 |

| | | |
|---|---|---|
| NC_005831.2;YP_003766.2;HCoV-NL63_1ab;1859 | 0 | 0 |
| NC_005831.2;YP_003766.2;HCoV-NL63_1ab;1860 | 0 | 0 |
| NC_005831.2;YP_003766.2;HCoV-NL63_1ab;1861 | 0 | 0 |
| NC_005831.2;YP_003766.2;HCoV-NL63_1ab;1862 | 0 | 0 |
| NC_005831.2;YP_003766.2;HCoV-NL63_1ab;1863 | 0 | 0 |
| NC_005831.2;YP_003766.2;HCoV-NL63_1ab;1864 | 0 | 0 |
| NC_005831.2;YP_003766.2;HCoV-NL63_1ab;1865 | 0 | 0 |
| NC_005831.2;YP_003766.2;HCoV-NL63_1ab;1866 | 0 | 0 |
| NC_005831.2;YP_003766.2;HCoV-NL63_1ab;1867 | 0 | 0 |
| NC_005831.2;YP_003766.2;HCoV-NL63_1ab;1868 | 0 | 0 |
| NC_005831.2;YP_003766.2;HCoV-NL63_1ab;1869 | 0 | 0 |
| NC_005831.2;YP_003766.2;HCoV-NL63_1ab;1870 | 0 | 0 |
| NC_005831.2;YP_003766.2;HCoV-NL63_1ab;1871 | 0 | 0 |
| NC_005831.2;YP_003766.2;HCoV-NL63_1ab;1872 | 0 | 0 |
| NC_005831.2;YP_003766.2;HCoV-NL63_1ab;1873 | 0 | 0 |
| NC_005831.2;YP_003766.2;HCoV-NL63_1ab;1874 | 0 | 0 |
| NC_005831.2;YP_003766.2;HCoV-NL63_1ab;1875 | 0 | 0 |
| NC_005831.2;YP_003766.2;HCoV-NL63_1ab;1876 | 0 | 0 |
| NC_005831.2;YP_003766.2;HCoV-NL63_1ab;1877 | 0 | 0 |
| NC_005831.2;YP_003766.2;HCoV-NL63_1ab;1878 | 0 | 0.025 |
| NC_005831.2;YP_003766.2;HCoV-NL63_1ab;1879 | 0 | 0.025 |
| NC_005831.2;YP_003766.2;HCoV-NL63_1ab;1880 | 0 | 0.025 |
| NC_005831.2;YP_003766.2;HCoV-NL63_1ab;1881 | 0 | 0.025 |
| NC_005831.2;YP_003766.2;HCoV-NL63_1ab;1882 | 0 | 0.025 |
| NC_005831.2;YP_003766.2;HCoV-NL63_1ab;1883 | 0 | 0 |
| NC_005831.2;YP_003766.2;HCoV-NL63_1ab;1884 | 0 | 0 |
| NC_005831.2;YP_003766.2;HCoV-NL63_1ab;1885 | 0 | 0.025 |
| NC_005831.2;YP_003766.2;HCoV-NL63_1ab;1886 | 0 | 0 |
| NC_005831.2;YP_003766.2;HCoV-NL63_1ab;1887 | 0 | 0 |
| NC_005831.2;YP_003766.2;HCoV-NL63_1ab;1888 | 0 | 0.025 |
| NC_005831.2;YP_003766.2;HCoV-NL63_1ab;1889 | 0 | 0.025 |
| NC_005831.2;YP_003766.2;HCoV-NL63_1ab;1890 | 0.1 | 0.025 |
| NC_005831.2;YP_003766.2;HCoV-NL63_1ab;1891 | 0 | 0 |
| NC_005831.2;YP_003766.2;HCoV-NL63_1ab;1892 | 0 | 0 |
| NC_005831.2;YP_003766.2;HCoV-NL63_1ab;1893 | 0 | 0 |
| NC_005831.2;YP_003766.2;HCoV-NL63_1ab;1894 | 0 | 0 |
| NC_005831.2;YP_003766.2;HCoV-NL63_1ab;1895 | 0.05 | 0 |
| NC_005831.2;YP_003766.2;HCoV-NL63_1ab;1896 | 0 | 0 |
| NC_005831.2;YP_003766.2;HCoV-NL63_1ab;1897 | 0 | 0 |
| NC_005831.2;YP_003766.2;HCoV-NL63_1ab;1898 | 0 | 0 |
| NC_005831.2;YP_003766.2;HCoV-NL63_1ab;1899 | 0 | 0 |
| NC_005831.2;YP_003766.2;HCoV-NL63_1ab;1900 | 0 | 0 |
| NC_005831.2;YP_003766.2;HCoV-NL63_1ab;1901 | 0 | 0 |

| | | |
|---|---|---|
| NC_005831.2;YP_003766.2;HCoV-NL63_1ab;1902 | 0 | 0 |
| NC_005831.2;YP_003766.2;HCoV-NL63_1ab;1903 | 0 | 0 |
| NC_005831.2;YP_003766.2;HCoV-NL63_1ab;1904 | 0 | 0 |
| NC_005831.2;YP_003766.2;HCoV-NL63_1ab;1905 | 0 | 0 |
| NC_005831.2;YP_003766.2;HCoV-NL63_1ab;1906 | 0 | 0 |
| NC_005831.2;YP_003766.2;HCoV-NL63_1ab;1907 | 0 | 0 |
| NC_005831.2;YP_003766.2;HCoV-NL63_1ab;1908 | 0 | 0 |
| NC_005831.2;YP_003766.2;HCoV-NL63_1ab;1909 | 0 | 0 |
| NC_005831.2;YP_003766.2;HCoV-NL63_1ab;1910 | 0 | 0 |
| NC_005831.2;YP_003766.2;HCoV-NL63_1ab;1911 | 0 | 0 |
| NC_005831.2;YP_003766.2;HCoV-NL63_1ab;1912 | 0 | 0 |
| NC_005831.2;YP_003766.2;HCoV-NL63_1ab;1913 | 0 | 0 |
| NC_005831.2;YP_003766.2;HCoV-NL63_1ab;1914 | 0 | 0 |
| NC_005831.2;YP_003766.2;HCoV-NL63_1ab;1915 | 0 | 0 |
| NC_005831.2;YP_003766.2;HCoV-NL63_1ab;1916 | 0 | 0 |
| NC_005831.2;YP_003766.2;HCoV-NL63_1ab;1917 | 0 | 0 |
| NC_005831.2;YP_003766.2;HCoV-NL63_1ab;1918 | 0 | 0 |
| NC_005831.2;YP_003766.2;HCoV-NL63_1ab;1919 | 0 | 0 |
| NC_005831.2;YP_003766.2;HCoV-NL63_1ab;1920 | 0 | 0 |
| NC_005831.2;YP_003766.2;HCoV-NL63_1ab;1921 | 0 | 0 |
| NC_005831.2;YP_003766.2;HCoV-NL63_1ab;1922 | 0 | 0 |
| NC_005831.2;YP_003766.2;HCoV-NL63_1ab;1923 | 0 | 0 |
| NC_005831.2;YP_003766.2;HCoV-NL63_1ab;1924 | 0 | 0 |
| NC_005831.2;YP_003766.2;HCoV-NL63_1ab;1925 | 0 | 0 |
| NC_005831.2;YP_003766.2;HCoV-NL63_1ab;1926 | 0 | 0 |
| NC_005831.2;YP_003766.2;HCoV-NL63_1ab;1927 | 0 | 0 |
| NC_005831.2;YP_003766.2;HCoV-NL63_1ab;1928 | 0 | 0 |
| NC_005831.2;YP_003766.2;HCoV-NL63_1ab;1929 | 0 | 0 |
| NC_005831.2;YP_003766.2;HCoV-NL63_1ab;1930 | 0 | 0 |
| NC_005831.2;YP_003766.2;HCoV-NL63_1ab;1931 | 0 | 0 |
| NC_005831.2;YP_003766.2;HCoV-NL63_1ab;1932 | 0 | 0 |
| NC_005831.2;YP_003766.2;HCoV-NL63_1ab;1933 | 0 | 0 |
| NC_005831.2;YP_003766.2;HCoV-NL63_1ab;1934 | 0 | 0 |
| NC_005831.2;YP_003766.2;HCoV-NL63_1ab;1935 | 0 | 0 |
| NC_005831.2;YP_003766.2;HCoV-NL63_1ab;1936 | 0 | 0 |
| NC_005831.2;YP_003766.2;HCoV-NL63_1ab;1937 | 0 | 0 |
| NC_005831.2;YP_003766.2;HCoV-NL63_1ab;1938 | 0 | 0 |
| NC_005831.2;YP_003766.2;HCoV-NL63_1ab;1939 | 0 | 0 |
| NC_005831.2;YP_003766.2;HCoV-NL63_1ab;1940 | 0 | 0 |
| NC_005831.2;YP_003766.2;HCoV-NL63_1ab;1941 | 0 | 0 |
| NC_005831.2;YP_003766.2;HCoV-NL63_1ab;1942 | 0 | 0 |
| NC_005831.2;YP_003766.2;HCoV-NL63_1ab;1943 | 0 | 0 |
| NC_005831.2;YP_003766.2;HCoV-NL63_1ab;1944 | 0 | 0 |

| | | |
|---|---|---|
| NC_005831.2;YP_003766.2;HCoV-NL63_1ab;1945 | 0 | 0 |
| NC_005831.2;YP_003766.2;HCoV-NL63_1ab;1946 | 0 | 0 |
| NC_005831.2;YP_003766.2;HCoV-NL63_1ab;1947 | 0 | 0 |
| NC_005831.2;YP_003766.2;HCoV-NL63_1ab;1948 | 0 | 0 |
| NC_005831.2;YP_003766.2;HCoV-NL63_1ab;1949 | 0 | 0 |
| NC_005831.2;YP_003766.2;HCoV-NL63_1ab;1950 | 0 | 0 |
| NC_005831.2;YP_003766.2;HCoV-NL63_1ab;1951 | 0 | 0 |
| NC_005831.2;YP_003766.2;HCoV-NL63_1ab;1952 | 0 | 0 |
| NC_005831.2;YP_003766.2;HCoV-NL63_1ab;1953 | 0 | 0 |
| NC_005831.2;YP_003766.2;HCoV-NL63_1ab;1954 | 0 | 0 |
| NC_005831.2;YP_003766.2;HCoV-NL63_1ab;1955 | 0 | 0 |
| NC_005831.2;YP_003766.2;HCoV-NL63_1ab;1956 | 0 | 0 |
| NC_005831.2;YP_003766.2;HCoV-NL63_1ab;1957 | 0 | 0 |
| NC_005831.2;YP_003766.2;HCoV-NL63_1ab;1958 | 0 | 0 |
| NC_005831.2;YP_003766.2;HCoV-NL63_1ab;1959 | 0 | 0 |
| NC_005831.2;YP_003766.2;HCoV-NL63_1ab;1960 | 0 | 0 |
| NC_005831.2;YP_003766.2;HCoV-NL63_1ab;1961 | 0 | 0 |
| NC_005831.2;YP_003766.2;HCoV-NL63_1ab;1962 | 0 | 0 |
| NC_005831.2;YP_003766.2;HCoV-NL63_1ab;1963 | 0 | 0 |
| NC_005831.2;YP_003766.2;HCoV-NL63_1ab;1964 | 0 | 0 |
| NC_005831.2;YP_003766.2;HCoV-NL63_1ab;1965 | 0 | 0 |
| NC_005831.2;YP_003766.2;HCoV-NL63_1ab;1966 | 0 | 0 |
| NC_005831.2;YP_003766.2;HCoV-NL63_1ab;1967 | 0 | 0 |
| NC_005831.2;YP_003766.2;HCoV-NL63_1ab;1968 | 0 | 0 |
| NC_005831.2;YP_003766.2;HCoV-NL63_1ab;1969 | 0 | 0 |
| NC_005831.2;YP_003766.2;HCoV-NL63_1ab;1970 | 0 | 0 |
| NC_005831.2;YP_003766.2;HCoV-NL63_1ab;1971 | 0 | 0 |
| NC_005831.2;YP_003766.2;HCoV-NL63_1ab;1972 | 0 | 0 |
| NC_005831.2;YP_003766.2;HCoV-NL63_1ab;1973 | 0 | 0 |
| NC_005831.2;YP_003766.2;HCoV-NL63_1ab;1974 | 0 | 0 |
| NC_005831.2;YP_003766.2;HCoV-NL63_1ab;1975 | 0 | 0 |
| NC_005831.2;YP_003766.2;HCoV-NL63_1ab;1976 | 0 | 0 |
| NC_005831.2;YP_003766.2;HCoV-NL63_1ab;1977 | 0 | 0 |
| NC_005831.2;YP_003766.2;HCoV-NL63_1ab;1978 | 0 | 0 |
| NC_005831.2;YP_003766.2;HCoV-NL63_1ab;1979 | 0 | 0 |
| NC_005831.2;YP_003766.2;HCoV-NL63_1ab;1980 | 0 | 0 |
| NC_005831.2;YP_003766.2;HCoV-NL63_1ab;1981 | 0 | 0 |
| NC_005831.2;YP_003766.2;HCoV-NL63_1ab;1982 | 0 | 0 |
| NC_005831.2;YP_003766.2;HCoV-NL63_1ab;1983 | 0 | 0 |
| NC_005831.2;YP_003766.2;HCoV-NL63_1ab;1984 | 0 | 0 |
| NC_005831.2;YP_003766.2;HCoV-NL63_1ab;1985 | 0 | 0 |
| NC_005831.2;YP_003766.2;HCoV-NL63_1ab;1986 | 0 | 0 |
| NC_005831.2;YP_003766.2;HCoV-NL63_1ab;1987 | 0 | 0 |

| | | |
|---|---|---|
| NC_005831.2;YP_003766.2;HCoV-NL63_1ab;1988 | 0 | 0 |
| NC_005831.2;YP_003766.2;HCoV-NL63_1ab;1989 | 0 | 0 |
| NC_005831.2;YP_003766.2;HCoV-NL63_1ab;1990 | 0 | 0 |
| NC_005831.2;YP_003766.2;HCoV-NL63_1ab;1991 | 0 | 0 |
| NC_005831.2;YP_003766.2;HCoV-NL63_1ab;1992 | 0.05 | 0 |
| NC_005831.2;YP_003766.2;HCoV-NL63_1ab;1993 | 0.05 | 0 |
| NC_005831.2;YP_003766.2;HCoV-NL63_1ab;1994 | 0 | 0 |
| NC_005831.2;YP_003766.2;HCoV-NL63_1ab;1995 | 0 | 0 |
| NC_005831.2;YP_003766.2;HCoV-NL63_1ab;1996 | 0 | 0 |
| NC_005831.2;YP_003766.2;HCoV-NL63_1ab;1997 | 0 | 0 |
| NC_005831.2;YP_003766.2;HCoV-NL63_1ab;1998 | 0 | 0.05 |
| NC_005831.2;YP_003766.2;HCoV-NL63_1ab;1999 | 0 | 0.025 |
| NC_005831.2;YP_003766.2;HCoV-NL63_1ab;2000 | 0 | 0.025 |
| NC_005831.2;YP_003766.2;HCoV-NL63_1ab;2001 | 0 | 0.025 |
| NC_005831.2;YP_003766.2;HCoV-NL63_1ab;2002 | 0 | 0.025 |
| NC_005831.2;YP_003766.2;HCoV-NL63_1ab;2003 | 0.1 | 0.025 |
| NC_005831.2;YP_003766.2;HCoV-NL63_1ab;2004 | 0.05 | 0.075 |
| NC_005831.2;YP_003766.2;HCoV-NL63_1ab;2005 | 0.05 | 0.05 |
| NC_005831.2;YP_003766.2;HCoV-NL63_1ab;2006 | 0.05 | 0.05 |
| NC_005831.2;YP_003766.2;HCoV-NL63_1ab;2007 | 0.05 | 0.025 |
| NC_005831.2;YP_003766.2;HCoV-NL63_1ab;2008 | 0 | 0.025 |
| NC_005831.2;YP_003766.2;HCoV-NL63_1ab;2009 | 0 | 0.025 |
| NC_005831.2;YP_003766.2;HCoV-NL63_1ab;2010 | 0 | 0.025 |
| NC_005831.2;YP_003766.2;HCoV-NL63_1ab;2011 | 0 | 0 |
| NC_005831.2;YP_003766.2;HCoV-NL63_1ab;2012 | 0 | 0 |
| NC_005831.2;YP_003766.2;HCoV-NL63_1ab;2013 | 0 | 0 |
| NC_005831.2;YP_003766.2;HCoV-NL63_1ab;2014 | 0 | 0 |
| NC_005831.2;YP_003766.2;HCoV-NL63_1ab;2015 | 0 | 0 |
| NC_005831.2;YP_003766.2;HCoV-NL63_1ab;2016 | 0 | 0 |
| NC_005831.2;YP_003766.2;HCoV-NL63_1ab;2017 | 0 | 0 |
| NC_005831.2;YP_003766.2;HCoV-NL63_1ab;2018 | 0 | 0 |
| NC_005831.2;YP_003766.2;HCoV-NL63_1ab;2019 | 0 | 0 |
| NC_005831.2;YP_003766.2;HCoV-NL63_1ab;2020 | 0 | 0 |
| NC_005831.2;YP_003766.2;HCoV-NL63_1ab;2021 | 0 | 0 |
| NC_005831.2;YP_003766.2;HCoV-NL63_1ab;2022 | 0 | 0 |
| NC_005831.2;YP_003766.2;HCoV-NL63_1ab;2023 | 0 | 0 |
| NC_005831.2;YP_003766.2;HCoV-NL63_1ab;2024 | 0 | 0 |
| NC_005831.2;YP_003766.2;HCoV-NL63_1ab;2025 | 0 | 0 |
| NC_005831.2;YP_003766.2;HCoV-NL63_1ab;2026 | 0 | 0 |
| NC_005831.2;YP_003766.2;HCoV-NL63_1ab;2027 | 0 | 0 |
| NC_005831.2;YP_003766.2;HCoV-NL63_1ab;2028 | 0 | 0 |
| NC_005831.2;YP_003766.2;HCoV-NL63_1ab;2029 | 0.05 | 0 |
| NC_005831.2;YP_003766.2;HCoV-NL63_1ab;2030 | 0 | 0 |

| | | |
|---|---|---|
| NC_005831.2;YP_003766.2;HCoV-NL63_1ab;2031 | 0 | 0 |
| NC_005831.2;YP_003766.2;HCoV-NL63_1ab;2032 | 0 | 0 |
| NC_005831.2;YP_003766.2;HCoV-NL63_1ab;2033 | 0 | 0 |
| NC_005831.2;YP_003766.2;HCoV-NL63_1ab;2034 | 0 | 0 |
| NC_005831.2;YP_003766.2;HCoV-NL63_1ab;2035 | 0 | 0 |
| NC_005831.2;YP_003766.2;HCoV-NL63_1ab;2036 | 0 | 0 |
| NC_005831.2;YP_003766.2;HCoV-NL63_1ab;2037 | 0 | 0 |
| NC_005831.2;YP_003766.2;HCoV-NL63_1ab;2038 | 0 | 0 |
| NC_005831.2;YP_003766.2;HCoV-NL63_1ab;2039 | 0 | 0 |
| NC_005831.2;YP_003766.2;HCoV-NL63_1ab;2040 | 0 | 0 |
| NC_005831.2;YP_003766.2;HCoV-NL63_1ab;2041 | 0 | 0 |
| NC_005831.2;YP_003766.2;HCoV-NL63_1ab;2042 | 0 | 0 |
| NC_005831.2;YP_003766.2;HCoV-NL63_1ab;2043 | 0 | 0 |
| NC_005831.2;YP_003766.2;HCoV-NL63_1ab;2044 | 0 | 0 |
| NC_005831.2;YP_003766.2;HCoV-NL63_1ab;2045 | 0 | 0 |
| NC_005831.2;YP_003766.2;HCoV-NL63_1ab;2046 | 0 | 0 |
| NC_005831.2;YP_003766.2;HCoV-NL63_1ab;2047 | 0 | 0 |
| NC_005831.2;YP_003766.2;HCoV-NL63_1ab;2048 | 0 | 0 |
| NC_005831.2;YP_003766.2;HCoV-NL63_1ab;2049 | 0 | 0 |
| NC_005831.2;YP_003766.2;HCoV-NL63_1ab;2050 | 0 | 0 |
| NC_005831.2;YP_003766.2;HCoV-NL63_1ab;2051 | 0 | 0 |
| NC_005831.2;YP_003766.2;HCoV-NL63_1ab;2052 | 0 | 0 |
| NC_005831.2;YP_003766.2;HCoV-NL63_1ab;2053 | 0 | 0 |
| NC_005831.2;YP_003766.2;HCoV-NL63_1ab;2054 | 0 | 0 |
| NC_005831.2;YP_003766.2;HCoV-NL63_1ab;2055 | 0 | 0 |
| NC_005831.2;YP_003766.2;HCoV-NL63_1ab;2056 | 0 | 0 |
| NC_005831.2;YP_003766.2;HCoV-NL63_1ab;2057 | 0 | 0 |
| NC_005831.2;YP_003766.2;HCoV-NL63_1ab;2058 | 0 | 0 |
| NC_005831.2;YP_003766.2;HCoV-NL63_1ab;2059 | 0 | 0 |
| NC_005831.2;YP_003766.2;HCoV-NL63_1ab;2060 | 0 | 0 |
| NC_005831.2;YP_003766.2;HCoV-NL63_1ab;2061 | 0 | 0 |
| NC_005831.2;YP_003766.2;HCoV-NL63_1ab;2062 | 0 | 0 |
| NC_005831.2;YP_003766.2;HCoV-NL63_1ab;2063 | 0 | 0 |
| NC_005831.2;YP_003766.2;HCoV-NL63_1ab;2064 | 0 | 0 |
| NC_005831.2;YP_003766.2;HCoV-NL63_1ab;2065 | 0 | 0 |
| NC_005831.2;YP_003766.2;HCoV-NL63_1ab;2066 | 0 | 0 |
| NC_005831.2;YP_003766.2;HCoV-NL63_1ab;2067 | 0 | 0 |
| NC_005831.2;YP_003766.2;HCoV-NL63_1ab;2068 | 0 | 0 |
| NC_005831.2;YP_003766.2;HCoV-NL63_1ab;2069 | 0 | 0 |
| NC_005831.2;YP_003766.2;HCoV-NL63_1ab;2070 | 0 | 0 |
| NC_005831.2;YP_003766.2;HCoV-NL63_1ab;2071 | 0 | 0 |
| NC_005831.2;YP_003766.2;HCoV-NL63_1ab;2072 | 0 | 0 |
| NC_005831.2;YP_003766.2;HCoV-NL63_1ab;2073 | 0 | 0 |

| | | |
|---|---|---|
| NC_005831.2;YP_003766.2;HCoV-NL63_1ab;2074 | 0 | 0 |
| NC_005831.2;YP_003766.2;HCoV-NL63_1ab;2075 | 0 | 0 |
| NC_005831.2;YP_003766.2;HCoV-NL63_1ab;2076 | 0 | 0 |
| NC_005831.2;YP_003766.2;HCoV-NL63_1ab;2077 | 0 | 0 |
| NC_005831.2;YP_003766.2;HCoV-NL63_1ab;2078 | 0 | 0 |
| NC_005831.2;YP_003766.2;HCoV-NL63_1ab;2079 | 0 | 0 |
| NC_005831.2;YP_003766.2;HCoV-NL63_1ab;2080 | 0 | 0 |
| NC_005831.2;YP_003766.2;HCoV-NL63_1ab;2081 | 0 | 0 |
| NC_005831.2;YP_003766.2;HCoV-NL63_1ab;2082 | 0 | 0 |
| NC_005831.2;YP_003766.2;HCoV-NL63_1ab;2083 | 0 | 0 |
| NC_005831.2;YP_003766.2;HCoV-NL63_1ab;2084 | 0 | 0 |
| NC_005831.2;YP_003766.2;HCoV-NL63_1ab;2085 | 0 | 0 |
| NC_005831.2;YP_003766.2;HCoV-NL63_1ab;2086 | 0 | 0 |
| NC_005831.2;YP_003766.2;HCoV-NL63_1ab;2087 | 0 | 0 |
| NC_005831.2;YP_003766.2;HCoV-NL63_1ab;2088 | 0 | 0 |
| NC_005831.2;YP_003766.2;HCoV-NL63_1ab;2089 | 0 | 0 |
| NC_005831.2;YP_003766.2;HCoV-NL63_1ab;2090 | 0 | 0 |
| NC_005831.2;YP_003766.2;HCoV-NL63_1ab;2091 | 0 | 0 |
| NC_005831.2;YP_003766.2;HCoV-NL63_1ab;2092 | 0 | 0 |
| NC_005831.2;YP_003766.2;HCoV-NL63_1ab;2093 | 0 | 0 |
| NC_005831.2;YP_003766.2;HCoV-NL63_1ab;2094 | 0 | 0 |
| NC_005831.2;YP_003766.2;HCoV-NL63_1ab;2095 | 0 | 0 |
| NC_005831.2;YP_003766.2;HCoV-NL63_1ab;2096 | 0 | 0 |
| NC_005831.2;YP_003766.2;HCoV-NL63_1ab;2097 | 0 | 0 |
| NC_005831.2;YP_003766.2;HCoV-NL63_1ab;2098 | 0 | 0 |
| NC_005831.2;YP_003766.2;HCoV-NL63_1ab;2099 | 0 | 0 |
| NC_005831.2;YP_003766.2;HCoV-NL63_1ab;2100 | 0 | 0 |
| NC_005831.2;YP_003766.2;HCoV-NL63_1ab;2101 | 0 | 0 |
| NC_005831.2;YP_003766.2;HCoV-NL63_1ab;2102 | 0 | 0 |
| NC_005831.2;YP_003766.2;HCoV-NL63_1ab;2103 | 0 | 0 |
| NC_005831.2;YP_003766.2;HCoV-NL63_1ab;2104 | 0 | 0 |
| NC_005831.2;YP_003766.2;HCoV-NL63_1ab;2105 | 0 | 0 |
| NC_005831.2;YP_003766.2;HCoV-NL63_1ab;2106 | 0 | 0 |
| NC_005831.2;YP_003766.2;HCoV-NL63_1ab;2107 | 0 | 0 |
| NC_005831.2;YP_003766.2;HCoV-NL63_1ab;2108 | 0 | 0 |
| NC_005831.2;YP_003766.2;HCoV-NL63_1ab;2109 | 0 | 0 |
| NC_005831.2;YP_003766.2;HCoV-NL63_1ab;2110 | 0 | 0 |
| NC_005831.2;YP_003766.2;HCoV-NL63_1ab;2111 | 0 | 0 |
| NC_005831.2;YP_003766.2;HCoV-NL63_1ab;2112 | 0 | 0 |
| NC_005831.2;YP_003766.2;HCoV-NL63_1ab;2113 | 0 | 0 |
| NC_005831.2;YP_003766.2;HCoV-NL63_1ab;2114 | 0 | 0 |
| NC_005831.2;YP_003766.2;HCoV-NL63_1ab;2115 | 0 | 0 |
| NC_005831.2;YP_003766.2;HCoV-NL63_1ab;2116 | 0 | 0 |

| | | |
|---|---|---|
| NC_005831.2;YP_003766.2;HCoV-NL63_1ab;2117 | 0 | 0 |
| NC_005831.2;YP_003766.2;HCoV-NL63_1ab;2118 | 0 | 0 |
| NC_005831.2;YP_003766.2;HCoV-NL63_1ab;2119 | 0 | 0 |
| NC_005831.2;YP_003766.2;HCoV-NL63_1ab;2120 | 0 | 0 |
| NC_005831.2;YP_003766.2;HCoV-NL63_1ab;2121 | 0 | 0 |
| NC_005831.2;YP_003766.2;HCoV-NL63_1ab;2122 | 0 | 0 |
| NC_005831.2;YP_003766.2;HCoV-NL63_1ab;2123 | 0 | 0.025 |
| NC_005831.2;YP_003766.2;HCoV-NL63_1ab;2124 | 0 | 0 |
| NC_005831.2;YP_003766.2;HCoV-NL63_1ab;2125 | 0 | 0 |
| NC_005831.2;YP_003766.2;HCoV-NL63_1ab;2126 | 0 | 0 |
| NC_005831.2;YP_003766.2;HCoV-NL63_1ab;2127 | 0 | 0 |
| NC_005831.2;YP_003766.2;HCoV-NL63_1ab;2128 | 0 | 0 |
| NC_005831.2;YP_003766.2;HCoV-NL63_1ab;2129 | 0.05 | 0 |
| NC_005831.2;YP_003766.2;HCoV-NL63_1ab;2130 | 0 | 0 |
| NC_005831.2;YP_003766.2;HCoV-NL63_1ab;2131 | 0 | 0 |
| NC_005831.2;YP_003766.2;HCoV-NL63_1ab;2132 | 0 | 0 |
| NC_005831.2;YP_003766.2;HCoV-NL63_1ab;2133 | 0 | 0 |
| NC_005831.2;YP_003766.2;HCoV-NL63_1ab;2134 | 0.05 | 0 |
| NC_005831.2;YP_003766.2;HCoV-NL63_1ab;2135 | 0 | 0 |
| NC_005831.2;YP_003766.2;HCoV-NL63_1ab;2136 | 0 | 0 |
| NC_005831.2;YP_003766.2;HCoV-NL63_1ab;2137 | 0 | 0 |
| NC_005831.2;YP_003766.2;HCoV-NL63_1ab;2138 | 0 | 0 |
| NC_005831.2;YP_003766.2;HCoV-NL63_1ab;2139 | 0 | 0 |
| NC_005831.2;YP_003766.2;HCoV-NL63_1ab;2140 | 0 | 0 |
| NC_005831.2;YP_003766.2;HCoV-NL63_1ab;2141 | 0 | 0 |
| NC_005831.2;YP_003766.2;HCoV-NL63_1ab;2142 | 0 | 0 |
| NC_005831.2;YP_003766.2;HCoV-NL63_1ab;2143 | 0 | 0.025 |
| NC_005831.2;YP_003766.2;HCoV-NL63_1ab;2144 | 0 | 0 |
| NC_005831.2;YP_003766.2;HCoV-NL63_1ab;2145 | 0 | 0 |
| NC_005831.2;YP_003766.2;HCoV-NL63_1ab;2146 | 0 | 0 |
| NC_005831.2;YP_003766.2;HCoV-NL63_1ab;2147 | 0 | 0 |
| NC_005831.2;YP_003766.2;HCoV-NL63_1ab;2148 | 0 | 0 |
| NC_005831.2;YP_003766.2;HCoV-NL63_1ab;2149 | 0 | 0 |
| NC_005831.2;YP_003766.2;HCoV-NL63_1ab;2150 | 0 | 0 |
| NC_005831.2;YP_003766.2;HCoV-NL63_1ab;2151 | 0 | 0 |
| NC_005831.2;YP_003766.2;HCoV-NL63_1ab;2152 | 0 | 0 |
| NC_005831.2;YP_003766.2;HCoV-NL63_1ab;2153 | 0.05 | 0 |
| NC_005831.2;YP_003766.2;HCoV-NL63_1ab;2154 | 0.05 | 0.025 |
| NC_005831.2;YP_003766.2;HCoV-NL63_1ab;2155 | 0 | 0 |
| NC_005831.2;YP_003766.2;HCoV-NL63_1ab;2156 | 0 | 0 |
| NC_005831.2;YP_003766.2;HCoV-NL63_1ab;2157 | 0 | 0 |
| NC_005831.2;YP_003766.2;HCoV-NL63_1ab;2158 | 0 | 0 |
| NC_005831.2;YP_003766.2;HCoV-NL63_1ab;2159 | 0.05 | 0 |

| | | |
|---|---|---|
| NC_005831.2;YP_003766.2;HCoV-NL63_1ab;2160 | 0 | 0 |
| NC_005831.2;YP_003766.2;HCoV-NL63_1ab;2161 | 0 | 0 |
| NC_005831.2;YP_003766.2;HCoV-NL63_1ab;2162 | 0 | 0 |
| NC_005831.2;YP_003766.2;HCoV-NL63_1ab;2163 | 0 | 0 |
| NC_005831.2;YP_003766.2;HCoV-NL63_1ab;2164 | 0 | 0 |
| NC_005831.2;YP_003766.2;HCoV-NL63_1ab;2165 | 0 | 0 |
| NC_005831.2;YP_003766.2;HCoV-NL63_1ab;2166 | 0 | 0 |
| NC_005831.2;YP_003766.2;HCoV-NL63_1ab;2167 | 0 | 0 |
| NC_005831.2;YP_003766.2;HCoV-NL63_1ab;2168 | 0 | 0 |
| NC_005831.2;YP_003766.2;HCoV-NL63_1ab;2169 | 0 | 0 |
| NC_005831.2;YP_003766.2;HCoV-NL63_1ab;2170 | 0 | 0 |
| NC_005831.2;YP_003766.2;HCoV-NL63_1ab;2171 | 0 | 0 |
| NC_005831.2;YP_003766.2;HCoV-NL63_1ab;2172 | 0 | 0 |
| NC_005831.2;YP_003766.2;HCoV-NL63_1ab;2173 | 0 | 0 |
| NC_005831.2;YP_003766.2;HCoV-NL63_1ab;2174 | 0 | 0 |
| NC_005831.2;YP_003766.2;HCoV-NL63_1ab;2175 | 0 | 0 |
| NC_005831.2;YP_003766.2;HCoV-NL63_1ab;2176 | 0 | 0 |
| NC_005831.2;YP_003766.2;HCoV-NL63_1ab;2177 | 0 | 0 |
| NC_005831.2;YP_003766.2;HCoV-NL63_1ab;2178 | 0 | 0 |
| NC_005831.2;YP_003766.2;HCoV-NL63_1ab;2179 | 0.05 | 0.025 |
| NC_005831.2;YP_003766.2;HCoV-NL63_1ab;2180 | 0.05 | 0.025 |
| NC_005831.2;YP_003766.2;HCoV-NL63_1ab;2181 | 0.05 | 0.025 |
| NC_005831.2;YP_003766.2;HCoV-NL63_1ab;2182 | 0.05 | 0.025 |
| NC_005831.2;YP_003766.2;HCoV-NL63_1ab;2183 | 0.15 | 0.075 |
| NC_005831.2;YP_003766.2;HCoV-NL63_1ab;2184 | 0.05 | 0.05 |
| NC_005831.2;YP_003766.2;HCoV-NL63_1ab;2185 | 0.05 | 0.025 |
| NC_005831.2;YP_003766.2;HCoV-NL63_1ab;2186 | 0 | 0.025 |
| NC_005831.2;YP_003766.2;HCoV-NL63_1ab;2187 | 0 | 0.025 |
| NC_005831.2;YP_003766.2;HCoV-NL63_1ab;2188 | 0 | 0 |
| NC_005831.2;YP_003766.2;HCoV-NL63_1ab;2189 | 0 | 0 |
| NC_005831.2;YP_003766.2;HCoV-NL63_1ab;2190 | 0 | 0 |
| NC_005831.2;YP_003766.2;HCoV-NL63_1ab;2191 | 0 | 0 |
| NC_005831.2;YP_003766.2;HCoV-NL63_1ab;2192 | 0 | 0 |
| NC_005831.2;YP_003766.2;HCoV-NL63_1ab;2193 | 0 | 0 |
| NC_005831.2;YP_003766.2;HCoV-NL63_1ab;2194 | 0 | 0 |
| NC_005831.2;YP_003766.2;HCoV-NL63_1ab;2195 | 0 | 0 |
| NC_005831.2;YP_003766.2;HCoV-NL63_1ab;2196 | 0 | 0 |
| NC_005831.2;YP_003766.2;HCoV-NL63_1ab;2197 | 0 | 0 |
| NC_005831.2;YP_003766.2;HCoV-NL63_1ab;2198 | 0.1 | 0 |
| NC_005831.2;YP_003766.2;HCoV-NL63_1ab;2199 | 0.1 | 0 |
| NC_005831.2;YP_003766.2;HCoV-NL63_1ab;2200 | 0.1 | 0.025 |
| NC_005831.2;YP_003766.2;HCoV-NL63_1ab;2201 | 0.1 | 0.025 |
| NC_005831.2;YP_003766.2;HCoV-NL63_1ab;2202 | 0.1 | 0 |

| | | |
|---|---|---|
| NC_005831.2;YP_003766.2;HCoV-NL63_1ab;2203 | 0.05 | 0 |
| NC_005831.2;YP_003766.2;HCoV-NL63_1ab;2204 | 0.1 | 0 |
| NC_005831.2;YP_003766.2;HCoV-NL63_1ab;2205 | 0.1 | 0 |
| NC_005831.2;YP_003766.2;HCoV-NL63_1ab;2206 | 0.1 | 0 |
| NC_005831.2;YP_003766.2;HCoV-NL63_1ab;2207 | 0.05 | 0 |
| NC_005831.2;YP_003766.2;HCoV-NL63_1ab;2208 | 0.1 | 0 |
| NC_005831.2;YP_003766.2;HCoV-NL63_1ab;2209 | 0.1 | 0 |
| NC_005831.2;YP_003766.2;HCoV-NL63_1ab;2210 | 0 | 0 |
| NC_005831.2;YP_003766.2;HCoV-NL63_1ab;2211 | 0 | 0 |
| NC_005831.2;YP_003766.2;HCoV-NL63_1ab;2212 | 0 | 0 |
| NC_005831.2;YP_003766.2;HCoV-NL63_1ab;2213 | 0 | 0 |
| NC_005831.2;YP_003766.2;HCoV-NL63_1ab;2214 | 0 | 0.025 |
| NC_005831.2;YP_003766.2;HCoV-NL63_1ab;2215 | 0 | 0 |
| NC_005831.2;YP_003766.2;HCoV-NL63_1ab;2216 | 0 | 0 |
| NC_005831.2;YP_003766.2;HCoV-NL63_1ab;2217 | 0 | 0 |
| NC_005831.2;YP_003766.2;HCoV-NL63_1ab;2218 | 0 | 0 |
| NC_005831.2;YP_003766.2;HCoV-NL63_1ab;2219 | 0 | 0.025 |
| NC_005831.2;YP_003766.2;HCoV-NL63_1ab;2220 | 0 | 0 |
| NC_005831.2;YP_003766.2;HCoV-NL63_1ab;2221 | 0 | 0 |
| NC_005831.2;YP_003766.2;HCoV-NL63_1ab;2222 | 0 | 0 |
| NC_005831.2;YP_003766.2;HCoV-NL63_1ab;2223 | 0 | 0.025 |
| NC_005831.2;YP_003766.2;HCoV-NL63_1ab;2224 | 0 | 0 |
| NC_005831.2;YP_003766.2;HCoV-NL63_1ab;2225 | 0 | 0 |
| NC_005831.2;YP_003766.2;HCoV-NL63_1ab;2226 | 0 | 0 |
| NC_005831.2;YP_003766.2;HCoV-NL63_1ab;2227 | 0 | 0 |
| NC_005831.2;YP_003766.2;HCoV-NL63_1ab;2228 | 0 | 0 |
| NC_005831.2;YP_003766.2;HCoV-NL63_1ab;2229 | 0 | 0 |
| NC_005831.2;YP_003766.2;HCoV-NL63_1ab;2230 | 0 | 0 |
| NC_005831.2;YP_003766.2;HCoV-NL63_1ab;2231 | 0 | 0.025 |
| NC_005831.2;YP_003766.2;HCoV-NL63_1ab;2232 | 0.05 | 0 |
| NC_005831.2;YP_003766.2;HCoV-NL63_1ab;2233 | 0.05 | 0 |
| NC_005831.2;YP_003766.2;HCoV-NL63_1ab;2234 | 0.05 | 0 |
| NC_005831.2;YP_003766.2;HCoV-NL63_1ab;2235 | 0.05 | 0 |
| NC_005831.2;YP_003766.2;HCoV-NL63_1ab;2236 | 0 | 0 |
| NC_005831.2;YP_003766.2;HCoV-NL63_1ab;2237 | 0.05 | 0.025 |
| NC_005831.2;YP_003766.2;HCoV-NL63_1ab;2238 | 0.1 | 0 |
| NC_005831.2;YP_003766.2;HCoV-NL63_1ab;2239 | 0.1 | 0 |
| NC_005831.2;YP_003766.2;HCoV-NL63_1ab;2240 | 0 | 0.025 |
| NC_005831.2;YP_003766.2;HCoV-NL63_1ab;2241 | 0 | 0 |
| NC_005831.2;YP_003766.2;HCoV-NL63_1ab;2242 | 0 | 0.025 |
| NC_005831.2;YP_003766.2;HCoV-NL63_1ab;2243 | 0 | 0 |
| NC_005831.2;YP_003766.2;HCoV-NL63_1ab;2244 | 0.05 | 0 |
| NC_005831.2;YP_003766.2;HCoV-NL63_1ab;2245 | 0.05 | 0.1 |

| | | |
|---|---|---|
| NC_005831.2;YP_003766.2;HCoV-NL63_1ab;2246 | 0 | 0 |
| NC_005831.2;YP_003766.2;HCoV-NL63_1ab;2247 | 0 | 0 |
| NC_005831.2;YP_003766.2;HCoV-NL63_1ab;2248 | 0 | 0 |
| NC_005831.2;YP_003766.2;HCoV-NL63_1ab;2249 | 0 | 0 |
| NC_005831.2;YP_003766.2;HCoV-NL63_1ab;2250 | 0 | 0 |
| NC_005831.2;YP_003766.2;HCoV-NL63_1ab;2251 | 0 | 0 |
| NC_005831.2;YP_003766.2;HCoV-NL63_1ab;2252 | 0 | 0 |
| NC_005831.2;YP_003766.2;HCoV-NL63_1ab;2253 | 0 | 0 |
| NC_005831.2;YP_003766.2;HCoV-NL63_1ab;2254 | 0 | 0 |
| NC_005831.2;YP_003766.2;HCoV-NL63_1ab;2255 | 0 | 0 |
| NC_005831.2;YP_003766.2;HCoV-NL63_1ab;2256 | 0 | 0 |
| NC_005831.2;YP_003766.2;HCoV-NL63_1ab;2257 | 0.05 | 0 |
| NC_005831.2;YP_003766.2;HCoV-NL63_1ab;2258 | 0.05 | 0 |
| NC_005831.2;YP_003766.2;HCoV-NL63_1ab;2259 | 0.05 | 0 |
| NC_005831.2;YP_003766.2;HCoV-NL63_1ab;2260 | 0.05 | 0 |
| NC_005831.2;YP_003766.2;HCoV-NL63_1ab;2261 | 0.05 | 0 |
| NC_005831.2;YP_003766.2;HCoV-NL63_1ab;2262 | 0.05 | 0 |
| NC_005831.2;YP_003766.2;HCoV-NL63_1ab;2263 | 0.05 | 0 |
| NC_005831.2;YP_003766.2;HCoV-NL63_1ab;2264 | 0.05 | 0.025 |
| NC_005831.2;YP_003766.2;HCoV-NL63_1ab;2265 | 0.05 | 0.025 |
| NC_005831.2;YP_003766.2;HCoV-NL63_1ab;2266 | 0.1 | 0.025 |
| NC_005831.2;YP_003766.2;HCoV-NL63_1ab;2267 | 0 | 0.025 |
| NC_005831.2;YP_003766.2;HCoV-NL63_1ab;2268 | 0 | 0.025 |
| NC_005831.2;YP_003766.2;HCoV-NL63_1ab;2269 | 0 | 0 |
| NC_005831.2;YP_003766.2;HCoV-NL63_1ab;2270 | 0 | 0 |
| NC_005831.2;YP_003766.2;HCoV-NL63_1ab;2271 | 0 | 0 |
| NC_005831.2;YP_003766.2;HCoV-NL63_1ab;2272 | 0 | 0 |
| NC_005831.2;YP_003766.2;HCoV-NL63_1ab;2273 | 0 | 0 |
| NC_005831.2;YP_003766.2;HCoV-NL63_1ab;2274 | 0 | 0 |
| NC_005831.2;YP_003766.2;HCoV-NL63_1ab;2275 | 0 | 0 |
| NC_005831.2;YP_003766.2;HCoV-NL63_1ab;2276 | 0 | 0 |
| NC_005831.2;YP_003766.2;HCoV-NL63_1ab;2277 | 0 | 0 |
| NC_005831.2;YP_003766.2;HCoV-NL63_1ab;2278 | 0 | 0 |
| NC_005831.2;YP_003766.2;HCoV-NL63_1ab;2279 | 0 | 0 |
| NC_005831.2;YP_003766.2;HCoV-NL63_1ab;2280 | 0 | 0 |
| NC_005831.2;YP_003766.2;HCoV-NL63_1ab;2281 | 0 | 0 |
| NC_005831.2;YP_003766.2;HCoV-NL63_1ab;2282 | 0 | 0 |
| NC_005831.2;YP_003766.2;HCoV-NL63_1ab;2283 | 0 | 0 |
| NC_005831.2;YP_003766.2;HCoV-NL63_1ab;2284 | 0 | 0 |
| NC_005831.2;YP_003766.2;HCoV-NL63_1ab;2285 | 0 | 0 |
| NC_005831.2;YP_003766.2;HCoV-NL63_1ab;2286 | 0 | 0 |
| NC_005831.2;YP_003766.2;HCoV-NL63_1ab;2287 | 0 | 0 |
| NC_005831.2;YP_003766.2;HCoV-NL63_1ab;2288 | 0 | 0 |

| | | |
|---|---|---|
| NC_005831.2;YP_003766.2;HCoV-NL63_1ab;2289 | 0 | 0 |
| NC_005831.2;YP_003766.2;HCoV-NL63_1ab;2290 | 0 | 0 |
| NC_005831.2;YP_003766.2;HCoV-NL63_1ab;2291 | 0 | 0 |
| NC_005831.2;YP_003766.2;HCoV-NL63_1ab;2292 | 0.05 | 0 |
| NC_005831.2;YP_003766.2;HCoV-NL63_1ab;2293 | 0 | 0 |
| NC_005831.2;YP_003766.2;HCoV-NL63_1ab;2294 | 0 | 0 |
| NC_005831.2;YP_003766.2;HCoV-NL63_1ab;2295 | 0 | 0 |
| NC_005831.2;YP_003766.2;HCoV-NL63_1ab;2296 | 0 | 0 |
| NC_005831.2;YP_003766.2;HCoV-NL63_1ab;2297 | 0 | 0 |
| NC_005831.2;YP_003766.2;HCoV-NL63_1ab;2298 | 0 | 0 |
| NC_005831.2;YP_003766.2;HCoV-NL63_1ab;2299 | 0 | 0 |
| NC_005831.2;YP_003766.2;HCoV-NL63_1ab;2300 | 0 | 0 |
| NC_005831.2;YP_003766.2;HCoV-NL63_1ab;2301 | 0 | 0 |
| NC_005831.2;YP_003766.2;HCoV-NL63_1ab;2302 | 0 | 0 |
| NC_005831.2;YP_003766.2;HCoV-NL63_1ab;2303 | 0 | 0 |
| NC_005831.2;YP_003766.2;HCoV-NL63_1ab;2304 | 0 | 0 |
| NC_005831.2;YP_003766.2;HCoV-NL63_1ab;2305 | 0 | 0 |
| NC_005831.2;YP_003766.2;HCoV-NL63_1ab;2306 | 0 | 0 |
| NC_005831.2;YP_003766.2;HCoV-NL63_1ab;2307 | 0 | 0 |
| NC_005831.2;YP_003766.2;HCoV-NL63_1ab;2308 | 0 | 0 |
| NC_005831.2;YP_003766.2;HCoV-NL63_1ab;2309 | 0 | 0 |
| NC_005831.2;YP_003766.2;HCoV-NL63_1ab;2310 | 0 | 0 |
| NC_005831.2;YP_003766.2;HCoV-NL63_1ab;2311 | 0 | 0 |
| NC_005831.2;YP_003766.2;HCoV-NL63_1ab;2312 | 0 | 0 |
| NC_005831.2;YP_003766.2;HCoV-NL63_1ab;2313 | 0 | 0.025 |
| NC_005831.2;YP_003766.2;HCoV-NL63_1ab;2314 | 0 | 0 |
| NC_005831.2;YP_003766.2;HCoV-NL63_1ab;2315 | 0 | 0 |
| NC_005831.2;YP_003766.2;HCoV-NL63_1ab;2316 | 0 | 0.025 |
| NC_005831.2;YP_003766.2;HCoV-NL63_1ab;2317 | 0 | 0 |
| NC_005831.2;YP_003766.2;HCoV-NL63_1ab;2318 | 0 | 0 |
| NC_005831.2;YP_003766.2;HCoV-NL63_1ab;2319 | 0 | 0.025 |
| NC_005831.2;YP_003766.2;HCoV-NL63_1ab;2320 | 0.05 | 0.025 |
| NC_005831.2;YP_003766.2;HCoV-NL63_1ab;2321 | 0 | 0.025 |
| NC_005831.2;YP_003766.2;HCoV-NL63_1ab;2322 | 0 | 0.025 |
| NC_005831.2;YP_003766.2;HCoV-NL63_1ab;2323 | 0 | 0.025 |
| NC_005831.2;YP_003766.2;HCoV-NL63_1ab;2324 | 0 | 0 |
| NC_005831.2;YP_003766.2;HCoV-NL63_1ab;2325 | 0 | 0 |
| NC_005831.2;YP_003766.2;HCoV-NL63_1ab;2326 | 0 | 0.025 |
| NC_005831.2;YP_003766.2;HCoV-NL63_1ab;2327 | 0 | 0 |
| NC_005831.2;YP_003766.2;HCoV-NL63_1ab;2328 | 0 | 0 |
| NC_005831.2;YP_003766.2;HCoV-NL63_1ab;2329 | 0 | 0 |
| NC_005831.2;YP_003766.2;HCoV-NL63_1ab;2330 | 0 | 0 |
| NC_005831.2;YP_003766.2;HCoV-NL63_1ab;2331 | 0 | 0 |

| | | |
|---|---|---|
| NC_005831.2;YP_003766.2;HCoV-NL63_1ab;2332 | 0 | 0 |
| NC_005831.2;YP_003766.2;HCoV-NL63_1ab;2333 | 0 | 0 |
| NC_005831.2;YP_003766.2;HCoV-NL63_1ab;2334 | 0 | 0 |
| NC_005831.2;YP_003766.2;HCoV-NL63_1ab;2335 | 0 | 0 |
| NC_005831.2;YP_003766.2;HCoV-NL63_1ab;2336 | 0 | 0 |
| NC_005831.2;YP_003766.2;HCoV-NL63_1ab;2337 | 0 | 0 |
| NC_005831.2;YP_003766.2;HCoV-NL63_1ab;2338 | 0 | 0 |
| NC_005831.2;YP_003766.2;HCoV-NL63_1ab;2339 | 0 | 0 |
| NC_005831.2;YP_003766.2;HCoV-NL63_1ab;2340 | 0 | 0 |
| NC_005831.2;YP_003766.2;HCoV-NL63_1ab;2341 | 0 | 0.025 |
| NC_005831.2;YP_003766.2;HCoV-NL63_1ab;2342 | 0 | 0.025 |
| NC_005831.2;YP_003766.2;HCoV-NL63_1ab;2343 | 0 | 0.025 |
| NC_005831.2;YP_003766.2;HCoV-NL63_1ab;2344 | 0 | 0.025 |
| NC_005831.2;YP_003766.2;HCoV-NL63_1ab;2345 | 0 | 0.025 |
| NC_005831.2;YP_003766.2;HCoV-NL63_1ab;2346 | 0 | 0 |
| NC_005831.2;YP_003766.2;HCoV-NL63_1ab;2347 | 0 | 0 |
| NC_005831.2;YP_003766.2;HCoV-NL63_1ab;2348 | 0 | 0 |
| NC_005831.2;YP_003766.2;HCoV-NL63_1ab;2349 | 0 | 0 |
| NC_005831.2;YP_003766.2;HCoV-NL63_1ab;2350 | 0 | 0 |
| NC_005831.2;YP_003766.2;HCoV-NL63_1ab;2351 | 0 | 0.025 |
| NC_005831.2;YP_003766.2;HCoV-NL63_1ab;2352 | 0 | 0 |
| NC_005831.2;YP_003766.2;HCoV-NL63_1ab;2353 | 0 | 0 |
| NC_005831.2;YP_003766.2;HCoV-NL63_1ab;2354 | 0 | 0 |
| NC_005831.2;YP_003766.2;HCoV-NL63_1ab;2355 | 0 | 0 |
| NC_005831.2;YP_003766.2;HCoV-NL63_1ab;2356 | 0 | 0 |
| NC_005831.2;YP_003766.2;HCoV-NL63_1ab;2357 | 0 | 0 |
| NC_005831.2;YP_003766.2;HCoV-NL63_1ab;2358 | 0 | 0 |
| NC_005831.2;YP_003766.2;HCoV-NL63_1ab;2359 | 0 | 0 |
| NC_005831.2;YP_003766.2;HCoV-NL63_1ab;2360 | 0 | 0 |
| NC_005831.2;YP_003766.2;HCoV-NL63_1ab;2361 | 0 | 0 |
| NC_005831.2;YP_003766.2;HCoV-NL63_1ab;2362 | 0 | 0 |
| NC_005831.2;YP_003766.2;HCoV-NL63_1ab;2363 | 0 | 0 |
| NC_005831.2;YP_003766.2;HCoV-NL63_1ab;2364 | 0.05 | 0 |
| NC_005831.2;YP_003766.2;HCoV-NL63_1ab;2365 | 0 | 0 |
| NC_005831.2;YP_003766.2;HCoV-NL63_1ab;2366 | 0 | 0 |
| NC_005831.2;YP_003766.2;HCoV-NL63_1ab;2367 | 0 | 0.025 |
| NC_005831.2;YP_003766.2;HCoV-NL63_1ab;2368 | 0 | 0 |
| NC_005831.2;YP_003766.2;HCoV-NL63_1ab;2369 | 0 | 0.025 |
| NC_005831.2;YP_003766.2;HCoV-NL63_1ab;2370 | 0 | 0 |
| NC_005831.2;YP_003766.2;HCoV-NL63_1ab;2371 | 0 | 0 |
| NC_005831.2;YP_003766.2;HCoV-NL63_1ab;2372 | 0 | 0 |
| NC_005831.2;YP_003766.2;HCoV-NL63_1ab;2373 | 0 | 0 |
| NC_005831.2;YP_003766.2;HCoV-NL63_1ab;2374 | 0 | 0 |

| | | |
|---|---|---|
| NC_005831.2;YP_003766.2;HCoV-NL63_1ab;2375 | 0 | 0 |
| NC_005831.2;YP_003766.2;HCoV-NL63_1ab;2376 | 0 | 0 |
| NC_005831.2;YP_003766.2;HCoV-NL63_1ab;2377 | 0 | 0 |
| NC_005831.2;YP_003766.2;HCoV-NL63_1ab;2378 | 0.05 | 0.025 |
| NC_005831.2;YP_003766.2;HCoV-NL63_1ab;2379 | 0 | 0.025 |
| NC_005831.2;YP_003766.2;HCoV-NL63_1ab;2380 | 0 | 0 |
| NC_005831.2;YP_003766.2;HCoV-NL63_1ab;2381 | 0.05 | 0 |
| NC_005831.2;YP_003766.2;HCoV-NL63_1ab;2382 | 0 | 0.025 |
| NC_005831.2;YP_003766.2;HCoV-NL63_1ab;2383 | 0 | 0.025 |
| NC_005831.2;YP_003766.2;HCoV-NL63_1ab;2384 | 0 | 0 |
| NC_005831.2;YP_003766.2;HCoV-NL63_1ab;2385 | 0 | 0 |
| NC_005831.2;YP_003766.2;HCoV-NL63_1ab;2386 | 0 | 0 |
| NC_005831.2;YP_003766.2;HCoV-NL63_1ab;2387 | 0 | 0 |
| NC_005831.2;YP_003766.2;HCoV-NL63_1ab;2388 | 0 | 0 |
| NC_005831.2;YP_003766.2;HCoV-NL63_1ab;2389 | 0 | 0 |
| NC_005831.2;YP_003766.2;HCoV-NL63_1ab;2390 | 0 | 0.025 |
| NC_005831.2;YP_003766.2;HCoV-NL63_1ab;2391 | 0 | 0 |
| NC_005831.2;YP_003766.2;HCoV-NL63_1ab;2392 | 0 | 0 |
| NC_005831.2;YP_003766.2;HCoV-NL63_1ab;2393 | 0 | 0 |
| NC_005831.2;YP_003766.2;HCoV-NL63_1ab;2394 | 0 | 0 |
| NC_005831.2;YP_003766.2;HCoV-NL63_1ab;2395 | 0 | 0 |
| NC_005831.2;YP_003766.2;HCoV-NL63_1ab;2396 | 0 | 0 |
| NC_005831.2;YP_003766.2;HCoV-NL63_1ab;2397 | 0 | 0 |
| NC_005831.2;YP_003766.2;HCoV-NL63_1ab;2398 | 0 | 0 |
| NC_005831.2;YP_003766.2;HCoV-NL63_1ab;2399 | 0 | 0 |
| NC_005831.2;YP_003766.2;HCoV-NL63_1ab;2400 | 0 | 0.025 |
| NC_005831.2;YP_003766.2;HCoV-NL63_1ab;2401 | 0 | 0 |
| NC_005831.2;YP_003766.2;HCoV-NL63_1ab;2402 | 0 | 0.05 |
| NC_005831.2;YP_003766.2;HCoV-NL63_1ab;2403 | 0 | 0.025 |
| NC_005831.2;YP_003766.2;HCoV-NL63_1ab;2404 | 0 | 0 |
| NC_005831.2;YP_003766.2;HCoV-NL63_1ab;2405 | 0 | 0.025 |
| NC_005831.2;YP_003766.2;HCoV-NL63_1ab;2406 | 0 | 0.025 |
| NC_005831.2;YP_003766.2;HCoV-NL63_1ab;2407 | 0.1 | 0.1 |
| NC_005831.2;YP_003766.2;HCoV-NL63_1ab;2408 | 0.15 | 0.1 |
| NC_005831.2;YP_003766.2;HCoV-NL63_1ab;2409 | 0 | 0.1 |
| NC_005831.2;YP_003766.2;HCoV-NL63_1ab;2410 | 0 | 0.075 |
| NC_005831.2;YP_003766.2;HCoV-NL63_1ab;2411 | 0 | 0.075 |
| NC_005831.2;YP_003766.2;HCoV-NL63_1ab;2412 | 0.15 | 0.25 |
| NC_005831.2;YP_003766.2;HCoV-NL63_1ab;2413 | 0 | 0.05 |
| NC_005831.2;YP_003766.2;HCoV-NL63_1ab;2414 | 0 | 0 |
| NC_005831.2;YP_003766.2;HCoV-NL63_1ab;2415 | 0 | 0 |
| NC_005831.2;YP_003766.2;HCoV-NL63_1ab;2416 | 0 | 0 |
| NC_005831.2;YP_003766.2;HCoV-NL63_1ab;2417 | 0 | 0 |

| | | |
|---|---|---|
| NC_005831.2;YP_003766.2;HCoV-NL63_1ab;2418 | 0 | 0 |
| NC_005831.2;YP_003766.2;HCoV-NL63_1ab;2419 | 0 | 0 |
| NC_005831.2;YP_003766.2;HCoV-NL63_1ab;2420 | 0 | 0.025 |
| NC_005831.2;YP_003766.2;HCoV-NL63_1ab;2421 | 0 | 0 |
| NC_005831.2;YP_003766.2;HCoV-NL63_1ab;2422 | 0 | 0 |
| NC_005831.2;YP_003766.2;HCoV-NL63_1ab;2423 | 0 | 0 |
| NC_005831.2;YP_003766.2;HCoV-NL63_1ab;2424 | 0 | 0 |
| NC_005831.2;YP_003766.2;HCoV-NL63_1ab;2425 | 0 | 0 |
| NC_005831.2;YP_003766.2;HCoV-NL63_1ab;2426 | 0 | 0 |
| NC_005831.2;YP_003766.2;HCoV-NL63_1ab;2427 | 0 | 0 |
| NC_005831.2;YP_003766.2;HCoV-NL63_1ab;2428 | 0 | 0 |
| NC_005831.2;YP_003766.2;HCoV-NL63_1ab;2429 | 0 | 0 |
| NC_005831.2;YP_003766.2;HCoV-NL63_1ab;2430 | 0 | 0 |
| NC_005831.2;YP_003766.2;HCoV-NL63_1ab;2431 | 0 | 0 |
| NC_005831.2;YP_003766.2;HCoV-NL63_1ab;2432 | 0 | 0 |
| NC_005831.2;YP_003766.2;HCoV-NL63_1ab;2433 | 0 | 0 |
| NC_005831.2;YP_003766.2;HCoV-NL63_1ab;2434 | 0 | 0 |
| NC_005831.2;YP_003766.2;HCoV-NL63_1ab;2435 | 0 | 0 |
| NC_005831.2;YP_003766.2;HCoV-NL63_1ab;2436 | 0 | 0 |
| NC_005831.2;YP_003766.2;HCoV-NL63_1ab;2437 | 0 | 0.025 |
| NC_005831.2;YP_003766.2;HCoV-NL63_1ab;2438 | 0 | 0 |
| NC_005831.2;YP_003766.2;HCoV-NL63_1ab;2439 | 0 | 0.025 |
| NC_005831.2;YP_003766.2;HCoV-NL63_1ab;2440 | 0 | 0.025 |
| NC_005831.2;YP_003766.2;HCoV-NL63_1ab;2441 | 0 | 0.025 |
| NC_005831.2;YP_003766.2;HCoV-NL63_1ab;2442 | 0 | 0 |
| NC_005831.2;YP_003766.2;HCoV-NL63_1ab;2443 | 0.05 | 0 |
| NC_005831.2;YP_003766.2;HCoV-NL63_1ab;2444 | 0 | 0 |
| NC_005831.2;YP_003766.2;HCoV-NL63_1ab;2445 | 0 | 0 |
| NC_005831.2;YP_003766.2;HCoV-NL63_1ab;2446 | 0 | 0 |
| NC_005831.2;YP_003766.2;HCoV-NL63_1ab;2447 | 0 | 0 |
| NC_005831.2;YP_003766.2;HCoV-NL63_1ab;2448 | 0 | 0 |
| NC_005831.2;YP_003766.2;HCoV-NL63_1ab;2449 | 0 | 0 |
| NC_005831.2;YP_003766.2;HCoV-NL63_1ab;2450 | 0 | 0.025 |
| NC_005831.2;YP_003766.2;HCoV-NL63_1ab;2451 | 0.05 | 0 |
| NC_005831.2;YP_003766.2;HCoV-NL63_1ab;2452 | 0.05 | 0 |
| NC_005831.2;YP_003766.2;HCoV-NL63_1ab;2453 | 0.05 | 0 |
| NC_005831.2;YP_003766.2;HCoV-NL63_1ab;2454 | 0.05 | 0 |
| NC_005831.2;YP_003766.2;HCoV-NL63_1ab;2455 | 0.05 | 0 |
| NC_005831.2;YP_003766.2;HCoV-NL63_1ab;2456 | 0.05 | 0 |
| NC_005831.2;YP_003766.2;HCoV-NL63_1ab;2457 | 0.05 | 0 |
| NC_005831.2;YP_003766.2;HCoV-NL63_1ab;2458 | 0.05 | 0 |
| NC_005831.2;YP_003766.2;HCoV-NL63_1ab;2459 | 0.05 | 0 |
| NC_005831.2;YP_003766.2;HCoV-NL63_1ab;2460 | 0 | 0 |

| | | |
|---|---|---|
| NC_005831.2;YP_003766.2;HCoV-NL63_1ab;2461 | 0 | 0 |
| NC_005831.2;YP_003766.2;HCoV-NL63_1ab;2462 | 0 | 0 |
| NC_005831.2;YP_003766.2;HCoV-NL63_1ab;2463 | 0 | 0 |
| NC_005831.2;YP_003766.2;HCoV-NL63_1ab;2464 | 0 | 0 |
| NC_005831.2;YP_003766.2;HCoV-NL63_1ab;2465 | 0 | 0 |
| NC_005831.2;YP_003766.2;HCoV-NL63_1ab;2466 | 0 | 0 |
| NC_005831.2;YP_003766.2;HCoV-NL63_1ab;2467 | 0 | 0 |
| NC_005831.2;YP_003766.2;HCoV-NL63_1ab;2468 | 0 | 0 |
| NC_005831.2;YP_003766.2;HCoV-NL63_1ab;2469 | 0 | 0 |
| NC_005831.2;YP_003766.2;HCoV-NL63_1ab;2470 | 0 | 0 |
| NC_005831.2;YP_003766.2;HCoV-NL63_1ab;2471 | 0 | 0 |
| NC_005831.2;YP_003766.2;HCoV-NL63_1ab;2472 | 0 | 0 |
| NC_005831.2;YP_003766.2;HCoV-NL63_1ab;2473 | 0 | 0 |
| NC_005831.2;YP_003766.2;HCoV-NL63_1ab;2474 | 0 | 0 |
| NC_005831.2;YP_003766.2;HCoV-NL63_1ab;2475 | 0 | 0 |
| NC_005831.2;YP_003766.2;HCoV-NL63_1ab;2476 | 0 | 0 |
| NC_005831.2;YP_003766.2;HCoV-NL63_1ab;2477 | 0 | 0 |
| NC_005831.2;YP_003766.2;HCoV-NL63_1ab;2478 | 0 | 0 |
| NC_005831.2;YP_003766.2;HCoV-NL63_1ab;2479 | 0 | 0 |
| NC_005831.2;YP_003766.2;HCoV-NL63_1ab;2480 | 0 | 0 |
| NC_005831.2;YP_003766.2;HCoV-NL63_1ab;2481 | 0 | 0 |
| NC_005831.2;YP_003766.2;HCoV-NL63_1ab;2482 | 0.05 | 0 |
| NC_005831.2;YP_003766.2;HCoV-NL63_1ab;2483 | 0.05 | 0 |
| NC_005831.2;YP_003766.2;HCoV-NL63_1ab;2484 | 0.05 | 0 |
| NC_005831.2;YP_003766.2;HCoV-NL63_1ab;2485 | 0 | 0 |
| NC_005831.2;YP_003766.2;HCoV-NL63_1ab;2486 | 0 | 0.025 |
| NC_005831.2;YP_003766.2;HCoV-NL63_1ab;2487 | 0 | 0 |
| NC_005831.2;YP_003766.2;HCoV-NL63_1ab;2488 | 0 | 0 |
| NC_005831.2;YP_003766.2;HCoV-NL63_1ab;2489 | 0 | 0 |
| NC_005831.2;YP_003766.2;HCoV-NL63_1ab;2490 | 0 | 0 |
| NC_005831.2;YP_003766.2;HCoV-NL63_1ab;2491 | 0 | 0 |
| NC_005831.2;YP_003766.2;HCoV-NL63_1ab;2492 | 0 | 0 |
| NC_005831.2;YP_003766.2;HCoV-NL63_1ab;2493 | 0 | 0 |
| NC_005831.2;YP_003766.2;HCoV-NL63_1ab;2494 | 0 | 0 |
| NC_005831.2;YP_003766.2;HCoV-NL63_1ab;2495 | 0.05 | 0 |
| NC_005831.2;YP_003766.2;HCoV-NL63_1ab;2496 | 0.05 | 0 |
| NC_005831.2;YP_003766.2;HCoV-NL63_1ab;2497 | 0.05 | 0 |
| NC_005831.2;YP_003766.2;HCoV-NL63_1ab;2498 | 0 | 0 |
| NC_005831.2;YP_003766.2;HCoV-NL63_1ab;2499 | 0 | 0 |
| NC_005831.2;YP_003766.2;HCoV-NL63_1ab;2500 | 0 | 0 |
| NC_005831.2;YP_003766.2;HCoV-NL63_1ab;2501 | 0.05 | 0 |
| NC_005831.2;YP_003766.2;HCoV-NL63_1ab;2502 | 0.05 | 0 |
| NC_005831.2;YP_003766.2;HCoV-NL63_1ab;2503 | 0 | 0 |

| | | |
|---|---|---|
| NC_005831.2;YP_003766.2;HCoV-NL63_1ab;2504 | 0 | 0 |
| NC_005831.2;YP_003766.2;HCoV-NL63_1ab;2505 | 0 | 0 |
| NC_005831.2;YP_003766.2;HCoV-NL63_1ab;2506 | 0 | 0 |
| NC_005831.2;YP_003766.2;HCoV-NL63_1ab;2507 | 0 | 0 |
| NC_005831.2;YP_003766.2;HCoV-NL63_1ab;2508 | 0 | 0 |
| NC_005831.2;YP_003766.2;HCoV-NL63_1ab;2509 | 0 | 0 |
| NC_005831.2;YP_003766.2;HCoV-NL63_1ab;2510 | 0 | 0 |
| NC_005831.2;YP_003766.2;HCoV-NL63_1ab;2511 | 0 | 0 |
| NC_005831.2;YP_003766.2;HCoV-NL63_1ab;2512 | 0 | 0 |
| NC_005831.2;YP_003766.2;HCoV-NL63_1ab;2513 | 0.05 | 0 |
| NC_005831.2;YP_003766.2;HCoV-NL63_1ab;2514 | 0.05 | 0 |
| NC_005831.2;YP_003766.2;HCoV-NL63_1ab;2515 | 0 | 0 |
| NC_005831.2;YP_003766.2;HCoV-NL63_1ab;2516 | 0 | 0 |
| NC_005831.2;YP_003766.2;HCoV-NL63_1ab;2517 | 0 | 0 |
| NC_005831.2;YP_003766.2;HCoV-NL63_1ab;2518 | 0 | 0 |
| NC_005831.2;YP_003766.2;HCoV-NL63_1ab;2519 | 0 | 0 |
| NC_005831.2;YP_003766.2;HCoV-NL63_1ab;2520 | 0 | 0 |
| NC_005831.2;YP_003766.2;HCoV-NL63_1ab;2521 | 0 | 0 |
| NC_005831.2;YP_003766.2;HCoV-NL63_1ab;2522 | 0 | 0 |
| NC_005831.2;YP_003766.2;HCoV-NL63_1ab;2523 | 0 | 0 |
| NC_005831.2;YP_003766.2;HCoV-NL63_1ab;2524 | 0 | 0 |
| NC_005831.2;YP_003766.2;HCoV-NL63_1ab;2525 | 0 | 0 |
| NC_005831.2;YP_003766.2;HCoV-NL63_1ab;2526 | 0 | 0 |
| NC_005831.2;YP_003766.2;HCoV-NL63_1ab;2527 | 0 | 0 |
| NC_005831.2;YP_003766.2;HCoV-NL63_1ab;2528 | 0 | 0.025 |
| NC_005831.2;YP_003766.2;HCoV-NL63_1ab;2529 | 0 | 0 |
| NC_005831.2;YP_003766.2;HCoV-NL63_1ab;2530 | 0 | 0 |
| NC_005831.2;YP_003766.2;HCoV-NL63_1ab;2531 | 0 | 0.025 |
| NC_005831.2;YP_003766.2;HCoV-NL63_1ab;2532 | 0.05 | 0 |
| NC_005831.2;YP_003766.2;HCoV-NL63_1ab;2533 | 0.05 | 0 |
| NC_005831.2;YP_003766.2;HCoV-NL63_1ab;2534 | 0 | 0 |
| NC_005831.2;YP_003766.2;HCoV-NL63_1ab;2535 | 0 | 0 |
| NC_005831.2;YP_003766.2;HCoV-NL63_1ab;2536 | 0 | 0 |
| NC_005831.2;YP_003766.2;HCoV-NL63_1ab;2537 | 0 | 0.05 |
| NC_005831.2;YP_003766.2;HCoV-NL63_1ab;2538 | 0 | 0.025 |
| NC_005831.2;YP_003766.2;HCoV-NL63_1ab;2539 | 0 | 0 |
| NC_005831.2;YP_003766.2;HCoV-NL63_1ab;2540 | 0 | 0 |
| NC_005831.2;YP_003766.2;HCoV-NL63_1ab;2541 | 0 | 0 |
| NC_005831.2;YP_003766.2;HCoV-NL63_1ab;2542 | 0 | 0 |
| NC_005831.2;YP_003766.2;HCoV-NL63_1ab;2543 | 0 | 0 |
| NC_005831.2;YP_003766.2;HCoV-NL63_1ab;2544 | 0 | 0 |
| NC_005831.2;YP_003766.2;HCoV-NL63_1ab;2545 | 0 | 0 |
| NC_005831.2;YP_003766.2;HCoV-NL63_1ab;2546 | 0 | 0 |

| | | |
|---|---|---|
| NC_005831.2;YP_003766.2;HCoV-NL63_1ab;2547 | 0 | 0 |
| NC_005831.2;YP_003766.2;HCoV-NL63_1ab;2548 | 0 | 0.025 |
| NC_005831.2;YP_003766.2;HCoV-NL63_1ab;2549 | 0 | 0.05 |
| NC_005831.2;YP_003766.2;HCoV-NL63_1ab;2550 | 0.05 | 0.175 |
| NC_005831.2;YP_003766.2;HCoV-NL63_1ab;2551 | 0.05 | 0.075 |
| NC_005831.2;YP_003766.2;HCoV-NL63_1ab;2552 | 0 | 0 |
| NC_005831.2;YP_003766.2;HCoV-NL63_1ab;2553 | 0.05 | 0 |
| NC_005831.2;YP_003766.2;HCoV-NL63_1ab;2554 | 0 | 0 |
| NC_005831.2;YP_003766.2;HCoV-NL63_1ab;2555 | 0 | 0 |
| NC_005831.2;YP_003766.2;HCoV-NL63_1ab;2556 | 0 | 0 |
| NC_005831.2;YP_003766.2;HCoV-NL63_1ab;2557 | 0 | 0 |
| NC_005831.2;YP_003766.2;HCoV-NL63_1ab;2558 | 0 | 0 |
| NC_005831.2;YP_003766.2;HCoV-NL63_1ab;2559 | 0 | 0 |
| NC_005831.2;YP_003766.2;HCoV-NL63_1ab;2560 | 0 | 0 |
| NC_005831.2;YP_003766.2;HCoV-NL63_1ab;2561 | 0 | 0 |
| NC_005831.2;YP_003766.2;HCoV-NL63_1ab;2562 | 0 | 0 |
| NC_005831.2;YP_003766.2;HCoV-NL63_1ab;2563 | 0 | 0 |
| NC_005831.2;YP_003766.2;HCoV-NL63_1ab;2564 | 0 | 0 |
| NC_005831.2;YP_003766.2;HCoV-NL63_1ab;2565 | 0 | 0 |
| NC_005831.2;YP_003766.2;HCoV-NL63_1ab;2566 | 0 | 0 |
| NC_005831.2;YP_003766.2;HCoV-NL63_1ab;2567 | 0 | 0 |
| NC_005831.2;YP_003766.2;HCoV-NL63_1ab;2568 | 0 | 0 |
| NC_005831.2;YP_003766.2;HCoV-NL63_1ab;2569 | 0 | 0 |
| NC_005831.2;YP_003766.2;HCoV-NL63_1ab;2570 | 0 | 0 |
| NC_005831.2;YP_003766.2;HCoV-NL63_1ab;2571 | 0 | 0 |
| NC_005831.2;YP_003766.2;HCoV-NL63_1ab;2572 | 0 | 0 |
| NC_005831.2;YP_003766.2;HCoV-NL63_1ab;2573 | 0 | 0 |
| NC_005831.2;YP_003766.2;HCoV-NL63_1ab;2574 | 0 | 0 |
| NC_005831.2;YP_003766.2;HCoV-NL63_1ab;2575 | 0 | 0 |
| NC_005831.2;YP_003766.2;HCoV-NL63_1ab;2576 | 0 | 0 |
| NC_005831.2;YP_003766.2;HCoV-NL63_1ab;2577 | 0 | 0 |
| NC_005831.2;YP_003766.2;HCoV-NL63_1ab;2578 | 0 | 0 |
| NC_005831.2;YP_003766.2;HCoV-NL63_1ab;2579 | 0 | 0 |
| NC_005831.2;YP_003766.2;HCoV-NL63_1ab;2580 | 0.05 | 0 |
| NC_005831.2;YP_003766.2;HCoV-NL63_1ab;2581 | 0 | 0 |
| NC_005831.2;YP_003766.2;HCoV-NL63_1ab;2582 | 0.05 | 0.025 |
| NC_005831.2;YP_003766.2;HCoV-NL63_1ab;2583 | 0.05 | 0.025 |
| NC_005831.2;YP_003766.2;HCoV-NL63_1ab;2584 | 0.05 | 0.05 |
| NC_005831.2;YP_003766.2;HCoV-NL63_1ab;2585 | 0.05 | 0 |
| NC_005831.2;YP_003766.2;HCoV-NL63_1ab;2586 | 0.05 | 0.025 |
| NC_005831.2;YP_003766.2;HCoV-NL63_1ab;2587 | 0.05 | 0.05 |
| NC_005831.2;YP_003766.2;HCoV-NL63_1ab;2588 | 0.05 | 0.05 |
| NC_005831.2;YP_003766.2;HCoV-NL63_1ab;2589 | 0.05 | 0.05 |

| | | |
|---|---|---|
| NC_005831.2;YP_003766.2;HCoV-NL63_1ab;2590 | 0 | 0.05 |
| NC_005831.2;YP_003766.2;HCoV-NL63_1ab;2591 | 0 | 0.025 |
| NC_005831.2;YP_003766.2;HCoV-NL63_1ab;2592 | 0 | 0 |
| NC_005831.2;YP_003766.2;HCoV-NL63_1ab;2593 | 0 | 0.025 |
| NC_005831.2;YP_003766.2;HCoV-NL63_1ab;2594 | 0 | 0.025 |
| NC_005831.2;YP_003766.2;HCoV-NL63_1ab;2595 | 0 | 0 |
| NC_005831.2;YP_003766.2;HCoV-NL63_1ab;2596 | 0 | 0.025 |
| NC_005831.2;YP_003766.2;HCoV-NL63_1ab;2597 | 0.1 | 0 |
| NC_005831.2;YP_003766.2;HCoV-NL63_1ab;2598 | 0 | 0 |
| NC_005831.2;YP_003766.2;HCoV-NL63_1ab;2599 | 0 | 0.025 |
| NC_005831.2;YP_003766.2;HCoV-NL63_1ab;2600 | 0 | 0 |
| NC_005831.2;YP_003766.2;HCoV-NL63_1ab;2601 | 0 | 0 |
| NC_005831.2;YP_003766.2;HCoV-NL63_1ab;2602 | 0 | 0 |
| NC_005831.2;YP_003766.2;HCoV-NL63_1ab;2603 | 0 | 0 |
| NC_005831.2;YP_003766.2;HCoV-NL63_1ab;2604 | 0 | 0 |
| NC_005831.2;YP_003766.2;HCoV-NL63_1ab;2605 | 0 | 0.025 |
| NC_005831.2;YP_003766.2;HCoV-NL63_1ab;2606 | 0.05 | 0.025 |
| NC_005831.2;YP_003766.2;HCoV-NL63_1ab;2607 | 0.05 | 0 |
| NC_005831.2;YP_003766.2;HCoV-NL63_1ab;2608 | 0 | 0 |
| NC_005831.2;YP_003766.2;HCoV-NL63_1ab;2609 | 0 | 0 |
| NC_005831.2;YP_003766.2;HCoV-NL63_1ab;2610 | 0.05 | 0 |
| NC_005831.2;YP_003766.2;HCoV-NL63_1ab;2611 | 0 | 0.025 |
| NC_005831.2;YP_003766.2;HCoV-NL63_1ab;2612 | 0 | 0.025 |
| NC_005831.2;YP_003766.2;HCoV-NL63_1ab;2613 | 0 | 0.025 |
| NC_005831.2;YP_003766.2;HCoV-NL63_1ab;2614 | 0 | 0.025 |
| NC_005831.2;YP_003766.2;HCoV-NL63_1ab;2615 | 0 | 0.025 |
| NC_005831.2;YP_003766.2;HCoV-NL63_1ab;2616 | 0 | 0.025 |
| NC_005831.2;YP_003766.2;HCoV-NL63_1ab;2617 | 0 | 0.025 |
| NC_005831.2;YP_003766.2;HCoV-NL63_1ab;2618 | 0 | 0.025 |
| NC_005831.2;YP_003766.2;HCoV-NL63_1ab;2619 | 0 | 0.05 |
| NC_005831.2;YP_003766.2;HCoV-NL63_1ab;2620 | 0 | 0 |
| NC_005831.2;YP_003766.2;HCoV-NL63_1ab;2621 | 0 | 0.025 |
| NC_005831.2;YP_003766.2;HCoV-NL63_1ab;2622 | 0.25 | 0.075 |
| NC_005831.2;YP_003766.2;HCoV-NL63_1ab;2623 | 0.15 | 0.05 |
| NC_005831.2;YP_003766.2;HCoV-NL63_1ab;2624 | 0.1 | 0.1 |
| NC_005831.2;YP_003766.2;HCoV-NL63_1ab;2625 | 0 | 0 |
| NC_005831.2;YP_003766.2;HCoV-NL63_1ab;2626 | 0.1 | 0.075 |
| NC_005831.2;YP_003766.2;HCoV-NL63_1ab;2627 | 0.15 | 0.1 |
| NC_005831.2;YP_003766.2;HCoV-NL63_1ab;2628 | 0.05 | 0 |
| NC_005831.2;YP_003766.2;HCoV-NL63_1ab;2629 | 0.05 | 0 |
| NC_005831.2;YP_003766.2;HCoV-NL63_1ab;2630 | 0.1 | 0.025 |
| NC_005831.2;YP_003766.2;HCoV-NL63_1ab;2631 | 0.05 | 0.05 |
| NC_005831.2;YP_003766.2;HCoV-NL63_1ab;2632 | 0 | 0 |

| | | |
|---|---|---|
| NC_005831.2;YP_003766.2;HCoV-NL63_1ab;2633 | 0 | 0 |
| NC_005831.2;YP_003766.2;HCoV-NL63_1ab;2634 | 0 | 0 |
| NC_005831.2;YP_003766.2;HCoV-NL63_1ab;2635 | 0 | 0.05 |
| NC_005831.2;YP_003766.2;HCoV-NL63_1ab;2636 | 0 | 0.075 |
| NC_005831.2;YP_003766.2;HCoV-NL63_1ab;2637 | 0 | 0 |
| NC_005831.2;YP_003766.2;HCoV-NL63_1ab;2638 | 0.05 | 0.075 |
| NC_005831.2;YP_003766.2;HCoV-NL63_1ab;2639 | 0.1 | 0.125 |
| NC_005831.2;YP_003766.2;HCoV-NL63_1ab;2640 | 0 | 0 |
| NC_005831.2;YP_003766.2;HCoV-NL63_1ab;2641 | 0 | 0 |
| NC_005831.2;YP_003766.2;HCoV-NL63_1ab;2642 | 0 | 0 |
| NC_005831.2;YP_003766.2;HCoV-NL63_1ab;2643 | 0 | 0 |
| NC_005831.2;YP_003766.2;HCoV-NL63_1ab;2644 | 0 | 0 |
| NC_005831.2;YP_003766.2;HCoV-NL63_1ab;2645 | 0 | 0 |
| NC_005831.2;YP_003766.2;HCoV-NL63_1ab;2646 | 0 | 0 |
| NC_005831.2;YP_003766.2;HCoV-NL63_1ab;2647 | 0 | 0 |
| NC_005831.2;YP_003766.2;HCoV-NL63_1ab;2648 | 0 | 0 |
| NC_005831.2;YP_003766.2;HCoV-NL63_1ab;2649 | 0 | 0 |
| NC_005831.2;YP_003766.2;HCoV-NL63_1ab;2650 | 0 | 0 |
| NC_005831.2;YP_003766.2;HCoV-NL63_1ab;2651 | 0 | 0 |
| NC_005831.2;YP_003766.2;HCoV-NL63_1ab;2652 | 0 | 0 |
| NC_005831.2;YP_003766.2;HCoV-NL63_1ab;2653 | 0 | 0 |
| NC_005831.2;YP_003766.2;HCoV-NL63_1ab;2654 | 0 | 0 |
| NC_005831.2;YP_003766.2;HCoV-NL63_1ab;2655 | 0 | 0 |
| NC_005831.2;YP_003766.2;HCoV-NL63_1ab;2656 | 0 | 0 |
| NC_005831.2;YP_003766.2;HCoV-NL63_1ab;2657 | 0 | 0 |
| NC_005831.2;YP_003766.2;HCoV-NL63_1ab;2658 | 0 | 0 |
| NC_005831.2;YP_003766.2;HCoV-NL63_1ab;2659 | 0 | 0 |
| NC_005831.2;YP_003766.2;HCoV-NL63_1ab;2660 | 0 | 0 |
| NC_005831.2;YP_003766.2;HCoV-NL63_1ab;2661 | 0 | 0 |
| NC_005831.2;YP_003766.2;HCoV-NL63_1ab;2662 | 0 | 0 |
| NC_005831.2;YP_003766.2;HCoV-NL63_1ab;2663 | 0 | 0 |
| NC_005831.2;YP_003766.2;HCoV-NL63_1ab;2664 | 0 | 0 |
| NC_005831.2;YP_003766.2;HCoV-NL63_1ab;2665 | 0 | 0 |
| NC_005831.2;YP_003766.2;HCoV-NL63_1ab;2666 | 0 | 0 |
| NC_005831.2;YP_003766.2;HCoV-NL63_1ab;2667 | 0 | 0 |
| NC_005831.2;YP_003766.2;HCoV-NL63_1ab;2668 | 0 | 0 |
| NC_005831.2;YP_003766.2;HCoV-NL63_1ab;2669 | 0 | 0 |
| NC_005831.2;YP_003766.2;HCoV-NL63_1ab;2670 | 0 | 0 |
| NC_005831.2;YP_003766.2;HCoV-NL63_1ab;2671 | 0 | 0 |
| NC_005831.2;YP_003766.2;HCoV-NL63_1ab;2672 | 0 | 0 |
| NC_005831.2;YP_003766.2;HCoV-NL63_1ab;2673 | 0 | 0 |
| NC_005831.2;YP_003766.2;HCoV-NL63_1ab;2674 | 0 | 0 |
| NC_005831.2;YP_003766.2;HCoV-NL63_1ab;2675 | 0 | 0.025 |

| | | |
|---|---|---|
| NC_005831.2;YP_003766.2;HCoV-NL63_1ab;2676 | 0 | 0 |
| NC_005831.2;YP_003766.2;HCoV-NL63_1ab;2677 | 0 | 0 |
| NC_005831.2;YP_003766.2;HCoV-NL63_1ab;2678 | 0 | 0 |
| NC_005831.2;YP_003766.2;HCoV-NL63_1ab;2679 | 0 | 0.05 |
| NC_005831.2;YP_003766.2;HCoV-NL63_1ab;2680 | 0 | 0 |
| NC_005831.2;YP_003766.2;HCoV-NL63_1ab;2681 | 0 | 0 |
| NC_005831.2;YP_003766.2;HCoV-NL63_1ab;2682 | 0 | 0 |
| NC_005831.2;YP_003766.2;HCoV-NL63_1ab;2683 | 0.05 | 0 |
| NC_005831.2;YP_003766.2;HCoV-NL63_1ab;2684 | 0 | 0 |
| NC_005831.2;YP_003766.2;HCoV-NL63_1ab;2685 | 0 | 0 |
| NC_005831.2;YP_003766.2;HCoV-NL63_1ab;2686 | 0 | 0 |
| NC_005831.2;YP_003766.2;HCoV-NL63_1ab;2687 | 0 | 0 |
| NC_005831.2;YP_003766.2;HCoV-NL63_1ab;2688 | 0 | 0.025 |
| NC_005831.2;YP_003766.2;HCoV-NL63_1ab;2689 | 0 | 0 |
| NC_005831.2;YP_003766.2;HCoV-NL63_1ab;2690 | 0 | 0 |
| NC_005831.2;YP_003766.2;HCoV-NL63_1ab;2691 | 0 | 0 |
| NC_005831.2;YP_003766.2;HCoV-NL63_1ab;2692 | 0 | 0 |
| NC_005831.2;YP_003766.2;HCoV-NL63_1ab;2693 | 0 | 0 |
| NC_005831.2;YP_003766.2;HCoV-NL63_1ab;2694 | 0 | 0 |
| NC_005831.2;YP_003766.2;HCoV-NL63_1ab;2695 | 0 | 0 |
| NC_005831.2;YP_003766.2;HCoV-NL63_1ab;2696 | 0 | 0 |
| NC_005831.2;YP_003766.2;HCoV-NL63_1ab;2697 | 0 | 0 |
| NC_005831.2;YP_003766.2;HCoV-NL63_1ab;2698 | 0 | 0 |
| NC_005831.2;YP_003766.2;HCoV-NL63_1ab;2699 | 0.05 | 0 |
| NC_005831.2;YP_003766.2;HCoV-NL63_1ab;2700 | 0 | 0 |
| NC_005831.2;YP_003766.2;HCoV-NL63_1ab;2701 | 0 | 0 |
| NC_005831.2;YP_003766.2;HCoV-NL63_1ab;2702 | 0 | 0 |
| NC_005831.2;YP_003766.2;HCoV-NL63_1ab;2703 | 0 | 0 |
| NC_005831.2;YP_003766.2;HCoV-NL63_1ab;2704 | 0 | 0.05 |
| NC_005831.2;YP_003766.2;HCoV-NL63_1ab;2705 | 0 | 0 |
| NC_005831.2;YP_003766.2;HCoV-NL63_1ab;2706 | 0 | 0 |
| NC_005831.2;YP_003766.2;HCoV-NL63_1ab;2707 | 0 | 0 |
| NC_005831.2;YP_003766.2;HCoV-NL63_1ab;2708 | 0 | 0 |
| NC_005831.2;YP_003766.2;HCoV-NL63_1ab;2709 | 0 | 0 |
| NC_005831.2;YP_003766.2;HCoV-NL63_1ab;2710 | 0 | 0 |
| NC_005831.2;YP_003766.2;HCoV-NL63_1ab;2711 | 0 | 0 |
| NC_005831.2;YP_003766.2;HCoV-NL63_1ab;2712 | 0 | 0 |
| NC_005831.2;YP_003766.2;HCoV-NL63_1ab;2713 | 0 | 0 |
| NC_005831.2;YP_003766.2;HCoV-NL63_1ab;2714 | 0 | 0 |
| NC_005831.2;YP_003766.2;HCoV-NL63_1ab;2715 | 0 | 0 |
| NC_005831.2;YP_003766.2;HCoV-NL63_1ab;2716 | 0 | 0 |
| NC_005831.2;YP_003766.2;HCoV-NL63_1ab;2717 | 0 | 0 |
| NC_005831.2;YP_003766.2;HCoV-NL63_1ab;2718 | 0 | 0 |

| | | |
|---|---|---|
| NC_005831.2;YP_003766.2;HCoV-NL63_1ab;2719 | 0 | 0 |
| NC_005831.2;YP_003766.2;HCoV-NL63_1ab;2720 | 0 | 0 |
| NC_005831.2;YP_003766.2;HCoV-NL63_1ab;2721 | 0 | 0 |
| NC_005831.2;YP_003766.2;HCoV-NL63_1ab;2722 | 0 | 0 |
| NC_005831.2;YP_003766.2;HCoV-NL63_1ab;2723 | 0 | 0 |
| NC_005831.2;YP_003766.2;HCoV-NL63_1ab;2724 | 0 | 0 |
| NC_005831.2;YP_003766.2;HCoV-NL63_1ab;2725 | 0 | 0 |
| NC_005831.2;YP_003766.2;HCoV-NL63_1ab;2726 | 0 | 0 |
| NC_005831.2;YP_003766.2;HCoV-NL63_1ab;2727 | 0 | 0 |
| NC_005831.2;YP_003766.2;HCoV-NL63_1ab;2728 | 0 | 0 |
| NC_005831.2;YP_003766.2;HCoV-NL63_1ab;2729 | 0 | 0 |
| NC_005831.2;YP_003766.2;HCoV-NL63_1ab;2730 | 0 | 0 |
| NC_005831.2;YP_003766.2;HCoV-NL63_1ab;2731 | 0 | 0 |
| NC_005831.2;YP_003766.2;HCoV-NL63_1ab;2732 | 0 | 0 |
| NC_005831.2;YP_003766.2;HCoV-NL63_1ab;2733 | 0 | 0 |
| NC_005831.2;YP_003766.2;HCoV-NL63_1ab;2734 | 0 | 0 |
| NC_005831.2;YP_003766.2;HCoV-NL63_1ab;2735 | 0 | 0 |
| NC_005831.2;YP_003766.2;HCoV-NL63_1ab;2736 | 0 | 0 |
| NC_005831.2;YP_003766.2;HCoV-NL63_1ab;2737 | 0 | 0 |
| NC_005831.2;YP_003766.2;HCoV-NL63_1ab;2738 | 0 | 0 |
| NC_005831.2;YP_003766.2;HCoV-NL63_1ab;2739 | 0 | 0 |
| NC_005831.2;YP_003766.2;HCoV-NL63_1ab;2740 | 0 | 0 |
| NC_005831.2;YP_003766.2;HCoV-NL63_1ab;2741 | 0 | 0 |
| NC_005831.2;YP_003766.2;HCoV-NL63_1ab;2742 | 0 | 0 |
| NC_005831.2;YP_003766.2;HCoV-NL63_1ab;2743 | 0 | 0 |
| NC_005831.2;YP_003766.2;HCoV-NL63_1ab;2744 | 0 | 0 |
| NC_005831.2;YP_003766.2;HCoV-NL63_1ab;2745 | 0 | 0 |
| NC_005831.2;YP_003766.2;HCoV-NL63_1ab;2746 | 0 | 0 |
| NC_005831.2;YP_003766.2;HCoV-NL63_1ab;2747 | 0 | 0 |
| NC_005831.2;YP_003766.2;HCoV-NL63_1ab;2748 | 0 | 0 |
| NC_005831.2;YP_003766.2;HCoV-NL63_1ab;2749 | 0 | 0 |
| NC_005831.2;YP_003766.2;HCoV-NL63_1ab;2750 | 0 | 0 |
| NC_005831.2;YP_003766.2;HCoV-NL63_1ab;2751 | 0 | 0 |
| NC_005831.2;YP_003766.2;HCoV-NL63_1ab;2752 | 0 | 0 |
| NC_005831.2;YP_003766.2;HCoV-NL63_1ab;2753 | 0 | 0 |
| NC_005831.2;YP_003766.2;HCoV-NL63_1ab;2754 | 0 | 0 |
| NC_005831.2;YP_003766.2;HCoV-NL63_1ab;2755 | 0 | 0 |
| NC_005831.2;YP_003766.2;HCoV-NL63_1ab;2756 | 0 | 0 |
| NC_005831.2;YP_003766.2;HCoV-NL63_1ab;2757 | 0 | 0 |
| NC_005831.2;YP_003766.2;HCoV-NL63_1ab;2758 | 0 | 0 |
| NC_005831.2;YP_003766.2;HCoV-NL63_1ab;2759 | 0 | 0 |
| NC_005831.2;YP_003766.2;HCoV-NL63_1ab;2760 | 0 | 0 |
| NC_005831.2;YP_003766.2;HCoV-NL63_1ab;2761 | 0 | 0 |

| | | |
|---|---|---|
| NC_005831.2;YP_003766.2;HCoV-NL63_1ab;2762 | 0 | 0 |
| NC_005831.2;YP_003766.2;HCoV-NL63_1ab;2763 | 0 | 0 |
| NC_005831.2;YP_003766.2;HCoV-NL63_1ab;2764 | 0 | 0 |
| NC_005831.2;YP_003766.2;HCoV-NL63_1ab;2765 | 0 | 0 |
| NC_005831.2;YP_003766.2;HCoV-NL63_1ab;2766 | 0 | 0 |
| NC_005831.2;YP_003766.2;HCoV-NL63_1ab;2767 | 0 | 0 |
| NC_005831.2;YP_003766.2;HCoV-NL63_1ab;2768 | 0 | 0 |
| NC_005831.2;YP_003766.2;HCoV-NL63_1ab;2769 | 0 | 0 |
| NC_005831.2;YP_003766.2;HCoV-NL63_1ab;2770 | 0 | 0 |
| NC_005831.2;YP_003766.2;HCoV-NL63_1ab;2771 | 0 | 0 |
| NC_005831.2;YP_003766.2;HCoV-NL63_1ab;2772 | 0 | 0 |
| NC_005831.2;YP_003766.2;HCoV-NL63_1ab;2773 | 0 | 0 |
| NC_005831.2;YP_003766.2;HCoV-NL63_1ab;2774 | 0 | 0 |
| NC_005831.2;YP_003766.2;HCoV-NL63_1ab;2775 | 0 | 0 |
| NC_005831.2;YP_003766.2;HCoV-NL63_1ab;2776 | 0 | 0 |
| NC_005831.2;YP_003766.2;HCoV-NL63_1ab;2777 | 0 | 0 |
| NC_005831.2;YP_003766.2;HCoV-NL63_1ab;2778 | 0 | 0 |
| NC_005831.2;YP_003766.2;HCoV-NL63_1ab;2779 | 0 | 0 |
| NC_005831.2;YP_003766.2;HCoV-NL63_1ab;2780 | 0 | 0 |
| NC_005831.2;YP_003766.2;HCoV-NL63_1ab;2781 | 0 | 0 |
| NC_005831.2;YP_003766.2;HCoV-NL63_1ab;2782 | 0 | 0 |
| NC_005831.2;YP_003766.2;HCoV-NL63_1ab;2783 | 0 | 0 |
| NC_005831.2;YP_003766.2;HCoV-NL63_1ab;2784 | 0 | 0 |
| NC_005831.2;YP_003766.2;HCoV-NL63_1ab;2785 | 0 | 0 |
| NC_005831.2;YP_003766.2;HCoV-NL63_1ab;2786 | 0 | 0 |
| NC_005831.2;YP_003766.2;HCoV-NL63_1ab;2787 | 0 | 0 |
| NC_005831.2;YP_003766.2;HCoV-NL63_1ab;2788 | 0 | 0 |
| NC_005831.2;YP_003766.2;HCoV-NL63_1ab;2789 | 0 | 0 |
| NC_005831.2;YP_003766.2;HCoV-NL63_1ab;2790 | 0 | 0 |
| NC_005831.2;YP_003766.2;HCoV-NL63_1ab;2791 | 0 | 0 |
| NC_005831.2;YP_003766.2;HCoV-NL63_1ab;2792 | 0 | 0 |
| NC_005831.2;YP_003766.2;HCoV-NL63_1ab;2793 | 0 | 0 |
| NC_005831.2;YP_003766.2;HCoV-NL63_1ab;2794 | 0 | 0 |
| NC_005831.2;YP_003766.2;HCoV-NL63_1ab;2795 | 0 | 0 |
| NC_005831.2;YP_003766.2;HCoV-NL63_1ab;2796 | 0 | 0 |
| NC_005831.2;YP_003766.2;HCoV-NL63_1ab;2797 | 0 | 0 |
| NC_005831.2;YP_003766.2;HCoV-NL63_1ab;2798 | 0 | 0 |
| NC_005831.2;YP_003766.2;HCoV-NL63_1ab;2799 | 0 | 0 |
| NC_005831.2;YP_003766.2;HCoV-NL63_1ab;2800 | 0 | 0 |
| NC_005831.2;YP_003766.2;HCoV-NL63_1ab;2801 | 0 | 0 |
| NC_005831.2;YP_003766.2;HCoV-NL63_1ab;2802 | 0 | 0 |
| NC_005831.2;YP_003766.2;HCoV-NL63_1ab;2803 | 0 | 0 |
| NC_005831.2;YP_003766.2;HCoV-NL63_1ab;2804 | 0 | 0 |

| | | |
|---|---|---|
| NC_005831.2;YP_003766.2;HCoV-NL63_1ab;2805 | 0 | 0 |
| NC_005831.2;YP_003766.2;HCoV-NL63_1ab;2806 | 0 | 0 |
| NC_005831.2;YP_003766.2;HCoV-NL63_1ab;2807 | 0 | 0 |
| NC_005831.2;YP_003766.2;HCoV-NL63_1ab;2808 | 0 | 0 |
| NC_005831.2;YP_003766.2;HCoV-NL63_1ab;2809 | 0 | 0 |
| NC_005831.2;YP_003766.2;HCoV-NL63_1ab;2810 | 0 | 0 |
| NC_005831.2;YP_003766.2;HCoV-NL63_1ab;2811 | 0 | 0 |
| NC_005831.2;YP_003766.2;HCoV-NL63_1ab;2812 | 0 | 0 |
| NC_005831.2;YP_003766.2;HCoV-NL63_1ab;2813 | 0 | 0 |
| NC_005831.2;YP_003766.2;HCoV-NL63_1ab;2814 | 0 | 0 |
| NC_005831.2;YP_003766.2;HCoV-NL63_1ab;2815 | 0 | 0 |
| NC_005831.2;YP_003766.2;HCoV-NL63_1ab;2816 | 0 | 0 |
| NC_005831.2;YP_003766.2;HCoV-NL63_1ab;2817 | 0 | 0 |
| NC_005831.2;YP_003766.2;HCoV-NL63_1ab;2818 | 0 | 0 |
| NC_005831.2;YP_003766.2;HCoV-NL63_1ab;2819 | 0 | 0 |
| NC_005831.2;YP_003766.2;HCoV-NL63_1ab;2820 | 0 | 0 |
| NC_005831.2;YP_003766.2;HCoV-NL63_1ab;2821 | 0 | 0 |
| NC_005831.2;YP_003766.2;HCoV-NL63_1ab;2822 | 0 | 0 |
| NC_005831.2;YP_003766.2;HCoV-NL63_1ab;2823 | 0 | 0 |
| NC_005831.2;YP_003766.2;HCoV-NL63_1ab;2824 | 0.05 | 0 |
| NC_005831.2;YP_003766.2;HCoV-NL63_1ab;2825 | 0 | 0 |
| NC_005831.2;YP_003766.2;HCoV-NL63_1ab;2826 | 0 | 0 |
| NC_005831.2;YP_003766.2;HCoV-NL63_1ab;2827 | 0 | 0 |
| NC_005831.2;YP_003766.2;HCoV-NL63_1ab;2828 | 0 | 0 |
| NC_005831.2;YP_003766.2;HCoV-NL63_1ab;2829 | 0 | 0.025 |
| NC_005831.2;YP_003766.2;HCoV-NL63_1ab;2830 | 0 | 0 |
| NC_005831.2;YP_003766.2;HCoV-NL63_1ab;2831 | 0 | 0 |
| NC_005831.2;YP_003766.2;HCoV-NL63_1ab;2832 | 0 | 0 |
| NC_005831.2;YP_003766.2;HCoV-NL63_1ab;2833 | 0 | 0 |
| NC_005831.2;YP_003766.2;HCoV-NL63_1ab;2834 | 0 | 0.025 |
| NC_005831.2;YP_003766.2;HCoV-NL63_1ab;2835 | 0 | 0 |
| NC_005831.2;YP_003766.2;HCoV-NL63_1ab;2836 | 0.1 | 0.05 |
| NC_005831.2;YP_003766.2;HCoV-NL63_1ab;2837 | 0.05 | 0.025 |
| NC_005831.2;YP_003766.2;HCoV-NL63_1ab;2838 | 0.1 | 0.05 |
| NC_005831.2;YP_003766.2;HCoV-NL63_1ab;2839 | 0.15 | 0.05 |
| NC_005831.2;YP_003766.2;HCoV-NL63_1ab;2840 | 0.15 | 0.025 |
| NC_005831.2;YP_003766.2;HCoV-NL63_1ab;2841 | 0.1 | 0 |
| NC_005831.2;YP_003766.2;HCoV-NL63_1ab;2842 | 0 | 0 |
| NC_005831.2;YP_003766.2;HCoV-NL63_1ab;2843 | 0.05 | 0.025 |
| NC_005831.2;YP_003766.2;HCoV-NL63_1ab;2844 | 0.05 | 0.025 |
| NC_005831.2;YP_003766.2;HCoV-NL63_1ab;2845 | 0 | 0 |
| NC_005831.2;YP_003766.2;HCoV-NL63_1ab;2846 | 0 | 0 |
| NC_005831.2;YP_003766.2;HCoV-NL63_1ab;2847 | 0 | 0 |

| | | |
|---|---|---|
| NC_005831.2;YP_003766.2;HCoV-NL63_1ab;2848 | 0 | 0 |
| NC_005831.2;YP_003766.2;HCoV-NL63_1ab;2849 | 0 | 0 |
| NC_005831.2;YP_003766.2;HCoV-NL63_1ab;2850 | 0 | 0 |
| NC_005831.2;YP_003766.2;HCoV-NL63_1ab;2851 | 0 | 0 |
| NC_005831.2;YP_003766.2;HCoV-NL63_1ab;2852 | 0 | 0 |
| NC_005831.2;YP_003766.2;HCoV-NL63_1ab;2853 | 0 | 0 |
| NC_005831.2;YP_003766.2;HCoV-NL63_1ab;2854 | 0 | 0 |
| NC_005831.2;YP_003766.2;HCoV-NL63_1ab;2855 | 0 | 0 |
| NC_005831.2;YP_003766.2;HCoV-NL63_1ab;2856 | 0 | 0 |
| NC_005831.2;YP_003766.2;HCoV-NL63_1ab;2857 | 0 | 0 |
| NC_005831.2;YP_003766.2;HCoV-NL63_1ab;2858 | 0 | 0 |
| NC_005831.2;YP_003766.2;HCoV-NL63_1ab;2859 | 0 | 0 |
| NC_005831.2;YP_003766.2;HCoV-NL63_1ab;2860 | 0 | 0 |
| NC_005831.2;YP_003766.2;HCoV-NL63_1ab;2861 | 0 | 0 |
| NC_005831.2;YP_003766.2;HCoV-NL63_1ab;2862 | 0 | 0 |
| NC_005831.2;YP_003766.2;HCoV-NL63_1ab;2863 | 0 | 0 |
| NC_005831.2;YP_003766.2;HCoV-NL63_1ab;2864 | 0.05 | 0 |
| NC_005831.2;YP_003766.2;HCoV-NL63_1ab;2865 | 0.05 | 0 |
| NC_005831.2;YP_003766.2;HCoV-NL63_1ab;2866 | 0.05 | 0 |
| NC_005831.2;YP_003766.2;HCoV-NL63_1ab;2867 | 0 | 0 |
| NC_005831.2;YP_003766.2;HCoV-NL63_1ab;2868 | 0.05 | 0 |
| NC_005831.2;YP_003766.2;HCoV-NL63_1ab;2869 | 0 | 0 |
| NC_005831.2;YP_003766.2;HCoV-NL63_1ab;2870 | 0.05 | 0 |
| NC_005831.2;YP_003766.2;HCoV-NL63_1ab;2871 | 0.05 | 0 |
| NC_005831.2;YP_003766.2;HCoV-NL63_1ab;2872 | 0 | 0 |
| NC_005831.2;YP_003766.2;HCoV-NL63_1ab;2873 | 0 | 0.025 |
| NC_005831.2;YP_003766.2;HCoV-NL63_1ab;2874 | 0 | 0.025 |
| NC_005831.2;YP_003766.2;HCoV-NL63_1ab;2875 | 0 | 0 |
| NC_005831.2;YP_003766.2;HCoV-NL63_1ab;2876 | 0 | 0 |
| NC_005831.2;YP_003766.2;HCoV-NL63_1ab;2877 | 0 | 0 |
| NC_005831.2;YP_003766.2;HCoV-NL63_1ab;2878 | 0 | 0 |
| NC_005831.2;YP_003766.2;HCoV-NL63_1ab;2879 | 0 | 0 |
| NC_005831.2;YP_003766.2;HCoV-NL63_1ab;2880 | 0 | 0 |
| NC_005831.2;YP_003766.2;HCoV-NL63_1ab;2881 | 0.05 | 0.025 |
| NC_005831.2;YP_003766.2;HCoV-NL63_1ab;2882 | 0 | 0 |
| NC_005831.2;YP_003766.2;HCoV-NL63_1ab;2883 | 0 | 0 |
| NC_005831.2;YP_003766.2;HCoV-NL63_1ab;2884 | 0 | 0 |
| NC_005831.2;YP_003766.2;HCoV-NL63_1ab;2885 | 0 | 0.075 |
| NC_005831.2;YP_003766.2;HCoV-NL63_1ab;2886 | 0 | 0.025 |
| NC_005831.2;YP_003766.2;HCoV-NL63_1ab;2887 | 0.05 | 0 |
| NC_005831.2;YP_003766.2;HCoV-NL63_1ab;2888 | 0 | 0 |
| NC_005831.2;YP_003766.2;HCoV-NL63_1ab;2889 | 0 | 0.025 |
| NC_005831.2;YP_003766.2;HCoV-NL63_1ab;2890 | 0 | 0 |

| | | |
|---|---|---|
| NC_005831.2;YP_003766.2;HCoV-NL63_1ab;2891 | 0.05 | 0 |
| NC_005831.2;YP_003766.2;HCoV-NL63_1ab;2892 | 0.05 | 0 |
| NC_005831.2;YP_003766.2;HCoV-NL63_1ab;2893 | 0 | 0 |
| NC_005831.2;YP_003766.2;HCoV-NL63_1ab;2894 | 0 | 0 |
| NC_005831.2;YP_003766.2;HCoV-NL63_1ab;2895 | 0 | 0 |
| NC_005831.2;YP_003766.2;HCoV-NL63_1ab;2896 | 0 | 0 |
| NC_005831.2;YP_003766.2;HCoV-NL63_1ab;2897 | 0 | 0.025 |
| NC_005831.2;YP_003766.2;HCoV-NL63_1ab;2898 | 0.05 | 0 |
| NC_005831.2;YP_003766.2;HCoV-NL63_1ab;2899 | 0 | 0.025 |
| NC_005831.2;YP_003766.2;HCoV-NL63_1ab;2900 | 0 | 0 |
| NC_005831.2;YP_003766.2;HCoV-NL63_1ab;2901 | 0 | 0 |
| NC_005831.2;YP_003766.2;HCoV-NL63_1ab;2902 | 0 | 0 |
| NC_005831.2;YP_003766.2;HCoV-NL63_1ab;2903 | 0 | 0 |
| NC_005831.2;YP_003766.2;HCoV-NL63_1ab;2904 | 0 | 0 |
| NC_005831.2;YP_003766.2;HCoV-NL63_1ab;2905 | 0 | 0 |
| NC_005831.2;YP_003766.2;HCoV-NL63_1ab;2906 | 0 | 0 |
| NC_005831.2;YP_003766.2;HCoV-NL63_1ab;2907 | 0 | 0 |
| NC_005831.2;YP_003766.2;HCoV-NL63_1ab;2908 | 0 | 0 |
| NC_005831.2;YP_003766.2;HCoV-NL63_1ab;2909 | 0 | 0 |
| NC_005831.2;YP_003766.2;HCoV-NL63_1ab;2910 | 0 | 0 |
| NC_005831.2;YP_003766.2;HCoV-NL63_1ab;2911 | 0 | 0 |
| NC_005831.2;YP_003766.2;HCoV-NL63_1ab;2912 | 0 | 0.025 |
| NC_005831.2;YP_003766.2;HCoV-NL63_1ab;2913 | 0 | 0.025 |
| NC_005831.2;YP_003766.2;HCoV-NL63_1ab;2914 | 0 | 0 |
| NC_005831.2;YP_003766.2;HCoV-NL63_1ab;2915 | 0 | 0.05 |
| NC_005831.2;YP_003766.2;HCoV-NL63_1ab;2916 | 0 | 0.025 |
| NC_005831.2;YP_003766.2;HCoV-NL63_1ab;2917 | 0 | 0 |
| NC_005831.2;YP_003766.2;HCoV-NL63_1ab;2918 | 0 | 0 |
| NC_005831.2;YP_003766.2;HCoV-NL63_1ab;2919 | 0 | 0 |
| NC_005831.2;YP_003766.2;HCoV-NL63_1ab;2920 | 0.1 | 0 |
| NC_005831.2;YP_003766.2;HCoV-NL63_1ab;2921 | 0.1 | 0 |
| NC_005831.2;YP_003766.2;HCoV-NL63_1ab;2922 | 0.05 | 0.05 |
| NC_005831.2;YP_003766.2;HCoV-NL63_1ab;2923 | 0 | 0.025 |
| NC_005831.2;YP_003766.2;HCoV-NL63_1ab;2924 | 0 | 0.025 |
| NC_005831.2;YP_003766.2;HCoV-NL63_1ab;2925 | 0 | 0.025 |
| NC_005831.2;YP_003766.2;HCoV-NL63_1ab;2926 | 0.05 | 0.05 |
| NC_005831.2;YP_003766.2;HCoV-NL63_1ab;2927 | 0 | 0 |
| NC_005831.2;YP_003766.2;HCoV-NL63_1ab;2928 | 0 | 0 |
| NC_005831.2;YP_003766.2;HCoV-NL63_1ab;2929 | 0 | 0 |
| NC_005831.2;YP_003766.2;HCoV-NL63_1ab;2930 | 0.05 | 0 |
| NC_005831.2;YP_003766.2;HCoV-NL63_1ab;2931 | 0 | 0 |
| NC_005831.2;YP_003766.2;HCoV-NL63_1ab;2932 | 0 | 0 |
| NC_005831.2;YP_003766.2;HCoV-NL63_1ab;2933 | 0 | 0 |

| | | |
|---|---|---|
| NC_005831.2;YP_003766.2;HCoV-NL63_1ab;2934 | 0 | 0 |
| NC_005831.2;YP_003766.2;HCoV-NL63_1ab;2935 | 0 | 0 |
| NC_005831.2;YP_003766.2;HCoV-NL63_1ab;2936 | 0 | 0 |
| NC_005831.2;YP_003766.2;HCoV-NL63_1ab;2937 | 0 | 0 |
| NC_005831.2;YP_003766.2;HCoV-NL63_1ab;2938 | 0 | 0 |
| NC_005831.2;YP_003766.2;HCoV-NL63_1ab;2939 | 0 | 0 |
| NC_005831.2;YP_003766.2;HCoV-NL63_1ab;2940 | 0 | 0 |
| NC_005831.2;YP_003766.2;HCoV-NL63_1ab;2941 | 0 | 0 |
| NC_005831.2;YP_003766.2;HCoV-NL63_1ab;2942 | 0 | 0 |
| NC_005831.2;YP_003766.2;HCoV-NL63_1ab;2943 | 0 | 0.025 |
| NC_005831.2;YP_003766.2;HCoV-NL63_1ab;2944 | 0 | 0 |
| NC_005831.2;YP_003766.2;HCoV-NL63_1ab;2945 | 0 | 0 |
| NC_005831.2;YP_003766.2;HCoV-NL63_1ab;2946 | 0 | 0 |
| NC_005831.2;YP_003766.2;HCoV-NL63_1ab;2947 | 0 | 0 |
| NC_005831.2;YP_003766.2;HCoV-NL63_1ab;2948 | 0 | 0 |
| NC_005831.2;YP_003766.2;HCoV-NL63_1ab;2949 | 0 | 0 |
| NC_005831.2;YP_003766.2;HCoV-NL63_1ab;2950 | 0 | 0 |
| NC_005831.2;YP_003766.2;HCoV-NL63_1ab;2951 | 0 | 0 |
| NC_005831.2;YP_003766.2;HCoV-NL63_1ab;2952 | 0 | 0 |
| NC_005831.2;YP_003766.2;HCoV-NL63_1ab;2953 | 0.05 | 0 |
| NC_005831.2;YP_003766.2;HCoV-NL63_1ab;2954 | 0 | 0 |
| NC_005831.2;YP_003766.2;HCoV-NL63_1ab;2955 | 0 | 0 |
| NC_005831.2;YP_003766.2;HCoV-NL63_1ab;2956 | 0 | 0 |
| NC_005831.2;YP_003766.2;HCoV-NL63_1ab;2957 | 0 | 0 |
| NC_005831.2;YP_003766.2;HCoV-NL63_1ab;2958 | 0 | 0 |
| NC_005831.2;YP_003766.2;HCoV-NL63_1ab;2959 | 0 | 0 |
| NC_005831.2;YP_003766.2;HCoV-NL63_1ab;2960 | 0 | 0 |
| NC_005831.2;YP_003766.2;HCoV-NL63_1ab;2961 | 0 | 0.025 |
| NC_005831.2;YP_003766.2;HCoV-NL63_1ab;2962 | 0 | 0 |
| NC_005831.2;YP_003766.2;HCoV-NL63_1ab;2963 | 0 | 0.025 |
| NC_005831.2;YP_003766.2;HCoV-NL63_1ab;2964 | 0 | 0 |
| NC_005831.2;YP_003766.2;HCoV-NL63_1ab;2965 | 0 | 0 |
| NC_005831.2;YP_003766.2;HCoV-NL63_1ab;2966 | 0 | 0 |
| NC_005831.2;YP_003766.2;HCoV-NL63_1ab;2967 | 0 | 0 |
| NC_005831.2;YP_003766.2;HCoV-NL63_1ab;2968 | 0 | 0 |
| NC_005831.2;YP_003766.2;HCoV-NL63_1ab;2969 | 0 | 0 |
| NC_005831.2;YP_003766.2;HCoV-NL63_1ab;2970 | 0 | 0.025 |
| NC_005831.2;YP_003766.2;HCoV-NL63_1ab;2971 | 0 | 0 |
| NC_005831.2;YP_003766.2;HCoV-NL63_1ab;2972 | 0 | 0 |
| NC_005831.2;YP_003766.2;HCoV-NL63_1ab;2973 | 0 | 0 |
| NC_005831.2;YP_003766.2;HCoV-NL63_1ab;2974 | 0 | 0 |
| NC_005831.2;YP_003766.2;HCoV-NL63_1ab;2975 | 0 | 0 |
| NC_005831.2;YP_003766.2;HCoV-NL63_1ab;2976 | 0 | 0 |

| | | |
|---|---|---|
| NC_005831.2;YP_003766.2;HCoV-NL63_1ab;2977 | 0 | 0 |
| NC_005831.2;YP_003766.2;HCoV-NL63_1ab;2978 | 0 | 0 |
| NC_005831.2;YP_003766.2;HCoV-NL63_1ab;2979 | 0 | 0 |
| NC_005831.2;YP_003766.2;HCoV-NL63_1ab;2980 | 0 | 0 |
| NC_005831.2;YP_003766.2;HCoV-NL63_1ab;2981 | 0 | 0 |
| NC_005831.2;YP_003766.2;HCoV-NL63_1ab;2982 | 0 | 0 |
| NC_005831.2;YP_003766.2;HCoV-NL63_1ab;2983 | 0 | 0 |
| NC_005831.2;YP_003766.2;HCoV-NL63_1ab;2984 | 0 | 0.025 |
| NC_005831.2;YP_003766.2;HCoV-NL63_1ab;2985 | 0 | 0 |
| NC_005831.2;YP_003766.2;HCoV-NL63_1ab;2986 | 0 | 0 |
| NC_005831.2;YP_003766.2;HCoV-NL63_1ab;2987 | 0 | 0 |
| NC_005831.2;YP_003766.2;HCoV-NL63_1ab;2988 | 0 | 0.025 |
| NC_005831.2;YP_003766.2;HCoV-NL63_1ab;2989 | 0 | 0 |
| NC_005831.2;YP_003766.2;HCoV-NL63_1ab;2990 | 0 | 0 |
| NC_005831.2;YP_003766.2;HCoV-NL63_1ab;2991 | 0 | 0 |
| NC_005831.2;YP_003766.2;HCoV-NL63_1ab;2992 | 0 | 0 |
| NC_005831.2;YP_003766.2;HCoV-NL63_1ab;2993 | 0 | 0 |
| NC_005831.2;YP_003766.2;HCoV-NL63_1ab;2994 | 0 | 0 |
| NC_005831.2;YP_003766.2;HCoV-NL63_1ab;2995 | 0 | 0.025 |
| NC_005831.2;YP_003766.2;HCoV-NL63_1ab;2996 | 0 | 0 |
| NC_005831.2;YP_003766.2;HCoV-NL63_1ab;2997 | 0 | 0 |
| NC_005831.2;YP_003766.2;HCoV-NL63_1ab;2998 | 0 | 0 |
| NC_005831.2;YP_003766.2;HCoV-NL63_1ab;2999 | 0 | 0 |
| NC_005831.2;YP_003766.2;HCoV-NL63_1ab;3000 | 0 | 0 |
| NC_005831.2;YP_003766.2;HCoV-NL63_1ab;3001 | 0 | 0 |
| NC_005831.2;YP_003766.2;HCoV-NL63_1ab;3002 | 0 | 0 |
| NC_005831.2;YP_003766.2;HCoV-NL63_1ab;3003 | 0 | 0 |
| NC_005831.2;YP_003766.2;HCoV-NL63_1ab;3004 | 0 | 0 |
| NC_005831.2;YP_003766.2;HCoV-NL63_1ab;3005 | 0 | 0 |
| NC_005831.2;YP_003766.2;HCoV-NL63_1ab;3006 | 0 | 0 |
| NC_005831.2;YP_003766.2;HCoV-NL63_1ab;3007 | 0 | 0 |
| NC_005831.2;YP_003766.2;HCoV-NL63_1ab;3008 | 0 | 0 |
| NC_005831.2;YP_003766.2;HCoV-NL63_1ab;3009 | 0 | 0 |
| NC_005831.2;YP_003766.2;HCoV-NL63_1ab;3010 | 0 | 0 |
| NC_005831.2;YP_003766.2;HCoV-NL63_1ab;3011 | 0 | 0 |
| NC_005831.2;YP_003766.2;HCoV-NL63_1ab;3012 | 0 | 0 |
| NC_005831.2;YP_003766.2;HCoV-NL63_1ab;3013 | 0 | 0 |
| NC_005831.2;YP_003766.2;HCoV-NL63_1ab;3014 | 0 | 0 |
| NC_005831.2;YP_003766.2;HCoV-NL63_1ab;3015 | 0 | 0 |
| NC_005831.2;YP_003766.2;HCoV-NL63_1ab;3016 | 0 | 0 |
| NC_005831.2;YP_003766.2;HCoV-NL63_1ab;3017 | 0 | 0 |
| NC_005831.2;YP_003766.2;HCoV-NL63_1ab;3018 | 0 | 0 |
| NC_005831.2;YP_003766.2;HCoV-NL63_1ab;3019 | 0 | 0 |

| | | |
|---|---|---|
| NC_005831.2;YP_003766.2;HCoV-NL63_1ab;3020 | 0 | 0 |
| NC_005831.2;YP_003766.2;HCoV-NL63_1ab;3021 | 0 | 0 |
| NC_005831.2;YP_003766.2;HCoV-NL63_1ab;3022 | 0 | 0 |
| NC_005831.2;YP_003766.2;HCoV-NL63_1ab;3023 | 0 | 0 |
| NC_005831.2;YP_003766.2;HCoV-NL63_1ab;3024 | 0 | 0 |
| NC_005831.2;YP_003766.2;HCoV-NL63_1ab;3025 | 0 | 0 |
| NC_005831.2;YP_003766.2;HCoV-NL63_1ab;3026 | 0 | 0 |
| NC_005831.2;YP_003766.2;HCoV-NL63_1ab;3027 | 0 | 0 |
| NC_005831.2;YP_003766.2;HCoV-NL63_1ab;3028 | 0.05 | 0 |
| NC_005831.2;YP_003766.2;HCoV-NL63_1ab;3029 | 0.15 | 0 |
| NC_005831.2;YP_003766.2;HCoV-NL63_1ab;3030 | 0 | 0 |
| NC_005831.2;YP_003766.2;HCoV-NL63_1ab;3031 | 0 | 0 |
| NC_005831.2;YP_003766.2;HCoV-NL63_1ab;3032 | 0 | 0 |
| NC_005831.2;YP_003766.2;HCoV-NL63_1ab;3033 | 0 | 0.025 |
| NC_005831.2;YP_003766.2;HCoV-NL63_1ab;3034 | 0 | 0.025 |
| NC_005831.2;YP_003766.2;HCoV-NL63_1ab;3035 | 0 | 0.025 |
| NC_005831.2;YP_003766.2;HCoV-NL63_1ab;3036 | 0 | 0.05 |
| NC_005831.2;YP_003766.2;HCoV-NL63_1ab;3037 | 0 | 0.025 |
| NC_005831.2;YP_003766.2;HCoV-NL63_1ab;3038 | 0 | 0 |
| NC_005831.2;YP_003766.2;HCoV-NL63_1ab;3039 | 0 | 0.025 |
| NC_005831.2;YP_003766.2;HCoV-NL63_1ab;3040 | 0 | 0.025 |
| NC_005831.2;YP_003766.2;HCoV-NL63_1ab;3041 | 0 | 0.025 |
| NC_005831.2;YP_003766.2;HCoV-NL63_1ab;3042 | 0 | 0.025 |
| NC_005831.2;YP_003766.2;HCoV-NL63_1ab;3043 | 0 | 0.025 |
| NC_005831.2;YP_003766.2;HCoV-NL63_1ab;3044 | 0 | 0.025 |
| NC_005831.2;YP_003766.2;HCoV-NL63_1ab;3045 | 0 | 0.05 |
| NC_005831.2;YP_003766.2;HCoV-NL63_1ab;3046 | 0 | 0 |
| NC_005831.2;YP_003766.2;HCoV-NL63_1ab;3047 | 0 | 0 |
| NC_005831.2;YP_003766.2;HCoV-NL63_1ab;3048 | 0.1 | 0.05 |
| NC_005831.2;YP_003766.2;HCoV-NL63_1ab;3049 | 0 | 0 |
| NC_005831.2;YP_003766.2;HCoV-NL63_1ab;3050 | 0 | 0 |
| NC_005831.2;YP_003766.2;HCoV-NL63_1ab;3051 | 0 | 0 |
| NC_005831.2;YP_003766.2;HCoV-NL63_1ab;3052 | 0 | 0 |
| NC_005831.2;YP_003766.2;HCoV-NL63_1ab;3053 | 0 | 0 |
| NC_005831.2;YP_003766.2;HCoV-NL63_1ab;3054 | 0 | 0.025 |
| NC_005831.2;YP_003766.2;HCoV-NL63_1ab;3055 | 0 | 0.025 |
| NC_005831.2;YP_003766.2;HCoV-NL63_1ab;3056 | 0 | 0 |
| NC_005831.2;YP_003766.2;HCoV-NL63_1ab;3057 | 0 | 0 |
| NC_005831.2;YP_003766.2;HCoV-NL63_1ab;3058 | 0 | 0.025 |
| NC_005831.2;YP_003766.2;HCoV-NL63_1ab;3059 | 0 | 0 |
| NC_005831.2;YP_003766.2;HCoV-NL63_1ab;3060 | 0 | 0 |
| NC_005831.2;YP_003766.2;HCoV-NL63_1ab;3061 | 0 | 0.025 |
| NC_005831.2;YP_003766.2;HCoV-NL63_1ab;3062 | 0 | 0 |

| | | |
|---|---|---|
| NC_005831.2;YP_003766.2;HCoV-NL63_1ab;3063 | 0 | 0 |
| NC_005831.2;YP_003766.2;HCoV-NL63_1ab;3064 | 0 | 0 |
| NC_005831.2;YP_003766.2;HCoV-NL63_1ab;3065 | 0 | 0.025 |
| NC_005831.2;YP_003766.2;HCoV-NL63_1ab;3066 | 0 | 0 |
| NC_005831.2;YP_003766.2;HCoV-NL63_1ab;3067 | 0 | 0 |
| NC_005831.2;YP_003766.2;HCoV-NL63_1ab;3068 | 0 | 0.025 |
| NC_005831.2;YP_003766.2;HCoV-NL63_1ab;3069 | 0 | 0 |
| NC_005831.2;YP_003766.2;HCoV-NL63_1ab;3070 | 0 | 0 |
| NC_005831.2;YP_003766.2;HCoV-NL63_1ab;3071 | 0 | 0 |
| NC_005831.2;YP_003766.2;HCoV-NL63_1ab;3072 | 0 | 0 |
| NC_005831.2;YP_003766.2;HCoV-NL63_1ab;3073 | 0 | 0 |
| NC_005831.2;YP_003766.2;HCoV-NL63_1ab;3074 | 0 | 0 |
| NC_005831.2;YP_003766.2;HCoV-NL63_1ab;3075 | 0 | 0.025 |
| NC_005831.2;YP_003766.2;HCoV-NL63_1ab;3076 | 0 | 0 |
| NC_005831.2;YP_003766.2;HCoV-NL63_1ab;3077 | 0 | 0 |
| NC_005831.2;YP_003766.2;HCoV-NL63_1ab;3078 | 0 | 0.025 |
| NC_005831.2;YP_003766.2;HCoV-NL63_1ab;3079 | 0 | 0.025 |
| NC_005831.2;YP_003766.2;HCoV-NL63_1ab;3080 | 0 | 0.025 |
| NC_005831.2;YP_003766.2;HCoV-NL63_1ab;3081 | 0 | 0.05 |
| NC_005831.2;YP_003766.2;HCoV-NL63_1ab;3082 | 0 | 0.025 |
| NC_005831.2;YP_003766.2;HCoV-NL63_1ab;3083 | 0.05 | 0.025 |
| NC_005831.2;YP_003766.2;HCoV-NL63_1ab;3084 | 0 | 0.025 |
| NC_005831.2;YP_003766.2;HCoV-NL63_1ab;3085 | 0 | 0.025 |
| NC_005831.2;YP_003766.2;HCoV-NL63_1ab;3086 | 0 | 0 |
| NC_005831.2;YP_003766.2;HCoV-NL63_1ab;3087 | 0 | 0.025 |
| NC_005831.2;YP_003766.2;HCoV-NL63_1ab;3088 | 0 | 0.025 |
| NC_005831.2;YP_003766.2;HCoV-NL63_1ab;3089 | 0 | 0 |
| NC_005831.2;YP_003766.2;HCoV-NL63_1ab;3090 | 0 | 0.025 |
| NC_005831.2;YP_003766.2;HCoV-NL63_1ab;3091 | 0 | 0 |
| NC_005831.2;YP_003766.2;HCoV-NL63_1ab;3092 | 0 | 0 |
| NC_005831.2;YP_003766.2;HCoV-NL63_1ab;3093 | 0 | 0 |
| NC_005831.2;YP_003766.2;HCoV-NL63_1ab;3094 | 0 | 0 |
| NC_005831.2;YP_003766.2;HCoV-NL63_1ab;3095 | 0 | 0 |
| NC_005831.2;YP_003766.2;HCoV-NL63_1ab;3096 | 0 | 0 |
| NC_005831.2;YP_003766.2;HCoV-NL63_1ab;3097 | 0.05 | 0 |
| NC_005831.2;YP_003766.2;HCoV-NL63_1ab;3098 | 0.05 | 0 |
| NC_005831.2;YP_003766.2;HCoV-NL63_1ab;3099 | 0.05 | 0 |
| NC_005831.2;YP_003766.2;HCoV-NL63_1ab;3100 | 0.05 | 0 |
| NC_005831.2;YP_003766.2;HCoV-NL63_1ab;3101 | 0.05 | 0 |
| NC_005831.2;YP_003766.2;HCoV-NL63_1ab;3102 | 0.05 | 0 |
| NC_005831.2;YP_003766.2;HCoV-NL63_1ab;3103 | 0 | 0 |
| NC_005831.2;YP_003766.2;HCoV-NL63_1ab;3104 | 0 | 0 |
| NC_005831.2;YP_003766.2;HCoV-NL63_1ab;3105 | 0 | 0 |

| | | |
|---|---|---|
| NC_005831.2;YP_003766.2;HCoV-NL63_1ab;3106 | 0 | 0 |
| NC_005831.2;YP_003766.2;HCoV-NL63_1ab;3107 | 0 | 0 |
| NC_005831.2;YP_003766.2;HCoV-NL63_1ab;3108 | 0 | 0 |
| NC_005831.2;YP_003766.2;HCoV-NL63_1ab;3109 | 0 | 0 |
| NC_005831.2;YP_003766.2;HCoV-NL63_1ab;3110 | 0 | 0 |
| NC_005831.2;YP_003766.2;HCoV-NL63_1ab;3111 | 0 | 0 |
| NC_005831.2;YP_003766.2;HCoV-NL63_1ab;3112 | 0.05 | 0 |
| NC_005831.2;YP_003766.2;HCoV-NL63_1ab;3113 | 0 | 0.025 |
| NC_005831.2;YP_003766.2;HCoV-NL63_1ab;3114 | 0.1 | 0.05 |
| NC_005831.2;YP_003766.2;HCoV-NL63_1ab;3115 | 0.1 | 0.15 |
| NC_005831.2;YP_003766.2;HCoV-NL63_1ab;3116 | 0.1 | 0.175 |
| NC_005831.2;YP_003766.2;HCoV-NL63_1ab;3117 | 0.1 | 0.225 |
| NC_005831.2;YP_003766.2;HCoV-NL63_1ab;3118 | 0.1 | 0.325 |
| NC_005831.2;YP_003766.2;HCoV-NL63_1ab;3119 | 0.15 | 0.225 |
| NC_005831.2;YP_003766.2;HCoV-NL63_1ab;3120 | 0.1 | 0.2 |
| NC_005831.2;YP_003766.2;HCoV-NL63_1ab;3121 | 0.1 | 0.2 |
| NC_005831.2;YP_003766.2;HCoV-NL63_1ab;3122 | 0.1 | 0.2 |
| NC_005831.2;YP_003766.2;HCoV-NL63_1ab;3123 | 0.1 | 0.2 |
| NC_005831.2;YP_003766.2;HCoV-NL63_1ab;3124 | 0.1 | 0.15 |
| NC_005831.2;YP_003766.2;HCoV-NL63_1ab;3125 | 0 | 0.025 |
| NC_005831.2;YP_003766.2;HCoV-NL63_1ab;3126 | 0.05 | 0.05 |
| NC_005831.2;YP_003766.2;HCoV-NL63_1ab;3127 | 0 | 0.025 |
| NC_005831.2;YP_003766.2;HCoV-NL63_1ab;3128 | 0 | 0 |
| NC_005831.2;YP_003766.2;HCoV-NL63_1ab;3129 | 0 | 0.025 |
| NC_005831.2;YP_003766.2;HCoV-NL63_1ab;3130 | 0.2 | 0.075 |
| NC_005831.2;YP_003766.2;HCoV-NL63_1ab;3131 | 0.05 | 0.025 |
| NC_005831.2;YP_003766.2;HCoV-NL63_1ab;3132 | 0 | 0.05 |
| NC_005831.2;YP_003766.2;HCoV-NL63_1ab;3133 | 0 | 0 |
| NC_005831.2;YP_003766.2;HCoV-NL63_1ab;3134 | 0 | 0 |
| NC_005831.2;YP_003766.2;HCoV-NL63_1ab;3135 | 0 | 0 |
| NC_005831.2;YP_003766.2;HCoV-NL63_1ab;3136 | 0 | 0 |
| NC_005831.2;YP_003766.2;HCoV-NL63_1ab;3137 | 0 | 0 |
| NC_005831.2;YP_003766.2;HCoV-NL63_1ab;3138 | 0 | 0 |
| NC_005831.2;YP_003766.2;HCoV-NL63_1ab;3139 | 0 | 0 |
| NC_005831.2;YP_003766.2;HCoV-NL63_1ab;3140 | 0 | 0 |
| NC_005831.2;YP_003766.2;HCoV-NL63_1ab;3141 | 0 | 0 |
| NC_005831.2;YP_003766.2;HCoV-NL63_1ab;3142 | 0 | 0 |
| NC_005831.2;YP_003766.2;HCoV-NL63_1ab;3143 | 0 | 0 |
| NC_005831.2;YP_003766.2;HCoV-NL63_1ab;3144 | 0 | 0 |
| NC_005831.2;YP_003766.2;HCoV-NL63_1ab;3145 | 0 | 0 |
| NC_005831.2;YP_003766.2;HCoV-NL63_1ab;3146 | 0 | 0 |
| NC_005831.2;YP_003766.2;HCoV-NL63_1ab;3147 | 0 | 0 |
| NC_005831.2;YP_003766.2;HCoV-NL63_1ab;3148 | 0 | 0 |

| | | |
|---|---|---|
| NC_005831.2;YP_003766.2;HCoV-NL63_1ab;3149 | 0 | 0 |
| NC_005831.2;YP_003766.2;HCoV-NL63_1ab;3150 | 0 | 0 |
| NC_005831.2;YP_003766.2;HCoV-NL63_1ab;3151 | 0 | 0 |
| NC_005831.2;YP_003766.2;HCoV-NL63_1ab;3152 | 0 | 0 |
| NC_005831.2;YP_003766.2;HCoV-NL63_1ab;3153 | 0 | 0 |
| NC_005831.2;YP_003766.2;HCoV-NL63_1ab;3154 | 0 | 0 |
| NC_005831.2;YP_003766.2;HCoV-NL63_1ab;3155 | 0 | 0 |
| NC_005831.2;YP_003766.2;HCoV-NL63_1ab;3156 | 0 | 0 |
| NC_005831.2;YP_003766.2;HCoV-NL63_1ab;3157 | 0 | 0 |
| NC_005831.2;YP_003766.2;HCoV-NL63_1ab;3158 | 0 | 0 |
| NC_005831.2;YP_003766.2;HCoV-NL63_1ab;3159 | 0 | 0.075 |
| NC_005831.2;YP_003766.2;HCoV-NL63_1ab;3160 | 0.05 | 0.075 |
| NC_005831.2;YP_003766.2;HCoV-NL63_1ab;3161 | 0.1 | 0.075 |
| NC_005831.2;YP_003766.2;HCoV-NL63_1ab;3162 | 0.1 | 0.125 |
| NC_005831.2;YP_003766.2;HCoV-NL63_1ab;3163 | 0.1 | 0.175 |
| NC_005831.2;YP_003766.2;HCoV-NL63_1ab;3164 | 0.1 | 0.15 |
| NC_005831.2;YP_003766.2;HCoV-NL63_1ab;3165 | 0.15 | 0.15 |
| NC_005831.2;YP_003766.2;HCoV-NL63_1ab;3166 | 0.1 | 0 |
| NC_005831.2;YP_003766.2;HCoV-NL63_1ab;3167 | 0.05 | 0 |
| NC_005831.2;YP_003766.2;HCoV-NL63_1ab;3168 | 0.05 | 0.075 |
| NC_005831.2;YP_003766.2;HCoV-NL63_1ab;3169 | 0 | 0 |
| NC_005831.2;YP_003766.2;HCoV-NL63_1ab;3170 | 0 | 0 |
| NC_005831.2;YP_003766.2;HCoV-NL63_1ab;3171 | 0.05 | 0.075 |
| NC_005831.2;YP_003766.2;HCoV-NL63_1ab;3172 | 0.05 | 0 |
| NC_005831.2;YP_003766.2;HCoV-NL63_1ab;3173 | 0.05 | 0.075 |
| NC_005831.2;YP_003766.2;HCoV-NL63_1ab;3174 | 0.05 | 0 |
| NC_005831.2;YP_003766.2;HCoV-NL63_1ab;3175 | 0 | 0 |
| NC_005831.2;YP_003766.2;HCoV-NL63_1ab;3176 | 0 | 0 |
| NC_005831.2;YP_003766.2;HCoV-NL63_1ab;3177 | 0 | 0 |
| NC_005831.2;YP_003766.2;HCoV-NL63_1ab;3178 | 0 | 0 |
| NC_005831.2;YP_003766.2;HCoV-NL63_1ab;3179 | 0 | 0 |
| NC_005831.2;YP_003766.2;HCoV-NL63_1ab;3180 | 0 | 0.025 |
| NC_005831.2;YP_003766.2;HCoV-NL63_1ab;3181 | 0 | 0.025 |
| NC_005831.2;YP_003766.2;HCoV-NL63_1ab;3182 | 0 | 0 |
| NC_005831.2;YP_003766.2;HCoV-NL63_1ab;3183 | 0 | 0.025 |
| NC_005831.2;YP_003766.2;HCoV-NL63_1ab;3184 | 0 | 0.025 |
| NC_005831.2;YP_003766.2;HCoV-NL63_1ab;3185 | 0 | 0.025 |
| NC_005831.2;YP_003766.2;HCoV-NL63_1ab;3186 | 0 | 0.025 |
| NC_005831.2;YP_003766.2;HCoV-NL63_1ab;3187 | 0 | 0.025 |
| NC_005831.2;YP_003766.2;HCoV-NL63_1ab;3188 | 0 | 0 |
| NC_005831.2;YP_003766.2;HCoV-NL63_1ab;3189 | 0 | 0 |
| NC_005831.2;YP_003766.2;HCoV-NL63_1ab;3190 | 0 | 0 |
| NC_005831.2;YP_003766.2;HCoV-NL63_1ab;3191 | 0 | 0 |

| | | |
|---|---|---|
| NC_005831.2;YP_003766.2;HCoV-NL63_1ab;3192 | 0 | 0 |
| NC_005831.2;YP_003766.2;HCoV-NL63_1ab;3193 | 0 | 0 |
| NC_005831.2;YP_003766.2;HCoV-NL63_1ab;3194 | 0 | 0 |
| NC_005831.2;YP_003766.2;HCoV-NL63_1ab;3195 | 0 | 0 |
| NC_005831.2;YP_003766.2;HCoV-NL63_1ab;3196 | 0 | 0.225 |
| NC_005831.2;YP_003766.2;HCoV-NL63_1ab;3197 | 0 | 0 |
| NC_005831.2;YP_003766.2;HCoV-NL63_1ab;3198 | 0 | 0 |
| NC_005831.2;YP_003766.2;HCoV-NL63_1ab;3199 | 0.05 | 0 |
| NC_005831.2;YP_003766.2;HCoV-NL63_1ab;3200 | 0 | 0 |
| NC_005831.2;YP_003766.2;HCoV-NL63_1ab;3201 | 0 | 0 |
| NC_005831.2;YP_003766.2;HCoV-NL63_1ab;3202 | 0 | 0 |
| NC_005831.2;YP_003766.2;HCoV-NL63_1ab;3203 | 0 | 0.025 |
| NC_005831.2;YP_003766.2;HCoV-NL63_1ab;3204 | 0.05 | 0 |
| NC_005831.2;YP_003766.2;HCoV-NL63_1ab;3205 | 0.05 | 0 |
| NC_005831.2;YP_003766.2;HCoV-NL63_1ab;3206 | 0.05 | 0 |
| NC_005831.2;YP_003766.2;HCoV-NL63_1ab;3207 | 0.05 | 0 |
| NC_005831.2;YP_003766.2;HCoV-NL63_1ab;3208 | 0 | 0 |
| NC_005831.2;YP_003766.2;HCoV-NL63_1ab;3209 | 0 | 0 |
| NC_005831.2;YP_003766.2;HCoV-NL63_1ab;3210 | 0.05 | 0 |
| NC_005831.2;YP_003766.2;HCoV-NL63_1ab;3211 | 0.05 | 0 |
| NC_005831.2;YP_003766.2;HCoV-NL63_1ab;3212 | 0.05 | 0 |
| NC_005831.2;YP_003766.2;HCoV-NL63_1ab;3213 | 0.05 | 0 |
| NC_005831.2;YP_003766.2;HCoV-NL63_1ab;3214 | 0.05 | 0 |
| NC_005831.2;YP_003766.2;HCoV-NL63_1ab;3215 | 0.05 | 0 |
| NC_005831.2;YP_003766.2;HCoV-NL63_1ab;3216 | 0.05 | 0 |
| NC_005831.2;YP_003766.2;HCoV-NL63_1ab;3217 | 0 | 0 |
| NC_005831.2;YP_003766.2;HCoV-NL63_1ab;3218 | 0 | 0 |
| NC_005831.2;YP_003766.2;HCoV-NL63_1ab;3219 | 0 | 0.025 |
| NC_005831.2;YP_003766.2;HCoV-NL63_1ab;3220 | 0.05 | 0.025 |
| NC_005831.2;YP_003766.2;HCoV-NL63_1ab;3221 | 0.1 | 0.025 |
| NC_005831.2;YP_003766.2;HCoV-NL63_1ab;3222 | 0.1 | 0.025 |
| NC_005831.2;YP_003766.2;HCoV-NL63_1ab;3223 | 0 | 0.025 |
| NC_005831.2;YP_003766.2;HCoV-NL63_1ab;3224 | 0.05 | 0.025 |
| NC_005831.2;YP_003766.2;HCoV-NL63_1ab;3225 | 0 | 0 |
| NC_005831.2;YP_003766.2;HCoV-NL63_1ab;3226 | 0 | 0 |
| NC_005831.2;YP_003766.2;HCoV-NL63_1ab;3227 | 0 | 0.05 |
| NC_005831.2;YP_003766.2;HCoV-NL63_1ab;3228 | 0 | 0 |
| NC_005831.2;YP_003766.2;HCoV-NL63_1ab;3229 | 0.05 | 0 |
| NC_005831.2;YP_003766.2;HCoV-NL63_1ab;3230 | 0.05 | 0.15 |
| NC_005831.2;YP_003766.2;HCoV-NL63_1ab;3231 | 0 | 0 |
| NC_005831.2;YP_003766.2;HCoV-NL63_1ab;3232 | 0 | 0 |
| NC_005831.2;YP_003766.2;HCoV-NL63_1ab;3233 | 0 | 0 |
| NC_005831.2;YP_003766.2;HCoV-NL63_1ab;3234 | 0 | 0 |

| | | |
|---|---|---|
| NC_005831.2;YP_003766.2;HCoV-NL63_1ab;3235 | 0 | 0 |
| NC_005831.2;YP_003766.2;HCoV-NL63_1ab;3236 | 0 | 0 |
| NC_005831.2;YP_003766.2;HCoV-NL63_1ab;3237 | 0 | 0 |
| NC_005831.2;YP_003766.2;HCoV-NL63_1ab;3238 | 0 | 0 |
| NC_005831.2;YP_003766.2;HCoV-NL63_1ab;3239 | 0 | 0 |
| NC_005831.2;YP_003766.2;HCoV-NL63_1ab;3240 | 0 | 0 |
| NC_005831.2;YP_003766.2;HCoV-NL63_1ab;3241 | 0 | 0 |
| NC_005831.2;YP_003766.2;HCoV-NL63_1ab;3242 | 0 | 0 |
| NC_005831.2;YP_003766.2;HCoV-NL63_1ab;3243 | 0 | 0 |
| NC_005831.2;YP_003766.2;HCoV-NL63_1ab;3244 | 0 | 0 |
| NC_005831.2;YP_003766.2;HCoV-NL63_1ab;3245 | 0 | 0 |
| NC_005831.2;YP_003766.2;HCoV-NL63_1ab;3246 | 0 | 0 |
| NC_005831.2;YP_003766.2;HCoV-NL63_1ab;3247 | 0 | 0 |
| NC_005831.2;YP_003766.2;HCoV-NL63_1ab;3248 | 0 | 0 |
| NC_005831.2;YP_003766.2;HCoV-NL63_1ab;3249 | 0 | 0 |
| NC_005831.2;YP_003766.2;HCoV-NL63_1ab;3250 | 0 | 0 |
| NC_005831.2;YP_003766.2;HCoV-NL63_1ab;3251 | 0 | 0 |
| NC_005831.2;YP_003766.2;HCoV-NL63_1ab;3252 | 0 | 0 |
| NC_005831.2;YP_003766.2;HCoV-NL63_1ab;3253 | 0 | 0 |
| NC_005831.2;YP_003766.2;HCoV-NL63_1ab;3254 | 0 | 0 |
| NC_005831.2;YP_003766.2;HCoV-NL63_1ab;3255 | 0 | 0 |
| NC_005831.2;YP_003766.2;HCoV-NL63_1ab;3256 | 0 | 0 |
| NC_005831.2;YP_003766.2;HCoV-NL63_1ab;3257 | 0 | 0 |
| NC_005831.2;YP_003766.2;HCoV-NL63_1ab;3258 | 0 | 0 |
| NC_005831.2;YP_003766.2;HCoV-NL63_1ab;3259 | 0 | 0 |
| NC_005831.2;YP_003766.2;HCoV-NL63_1ab;3260 | 0 | 0 |
| NC_005831.2;YP_003766.2;HCoV-NL63_1ab;3261 | 0 | 0 |
| NC_005831.2;YP_003766.2;HCoV-NL63_1ab;3262 | 0 | 0 |
| NC_005831.2;YP_003766.2;HCoV-NL63_1ab;3263 | 0 | 0 |
| NC_005831.2;YP_003766.2;HCoV-NL63_1ab;3264 | 0 | 0 |
| NC_005831.2;YP_003766.2;HCoV-NL63_1ab;3265 | 0 | 0 |
| NC_005831.2;YP_003766.2;HCoV-NL63_1ab;3266 | 0 | 0 |
| NC_005831.2;YP_003766.2;HCoV-NL63_1ab;3267 | 0 | 0 |
| NC_005831.2;YP_003766.2;HCoV-NL63_1ab;3268 | 0 | 0 |
| NC_005831.2;YP_003766.2;HCoV-NL63_1ab;3269 | 0 | 0 |
| NC_005831.2;YP_003766.2;HCoV-NL63_1ab;3270 | 0 | 0 |
| NC_005831.2;YP_003766.2;HCoV-NL63_1ab;3271 | 0 | 0 |
| NC_005831.2;YP_003766.2;HCoV-NL63_1ab;3272 | 0 | 0 |
| NC_005831.2;YP_003766.2;HCoV-NL63_1ab;3273 | 0 | 0 |
| NC_005831.2;YP_003766.2;HCoV-NL63_1ab;3274 | 0 | 0 |
| NC_005831.2;YP_003766.2;HCoV-NL63_1ab;3275 | 0 | 0 |
| NC_005831.2;YP_003766.2;HCoV-NL63_1ab;3276 | 0 | 0 |
| NC_005831.2;YP_003766.2;HCoV-NL63_1ab;3277 | 0 | 0 |

| | | |
|---|---|---|
| NC_005831.2;YP_003766.2;HCoV-NL63_1ab;3278 | 0 | 0 |
| NC_005831.2;YP_003766.2;HCoV-NL63_1ab;3279 | 0 | 0 |
| NC_005831.2;YP_003766.2;HCoV-NL63_1ab;3280 | 0 | 0 |
| NC_005831.2;YP_003766.2;HCoV-NL63_1ab;3281 | 0 | 0 |
| NC_005831.2;YP_003766.2;HCoV-NL63_1ab;3282 | 0 | 0 |
| NC_005831.2;YP_003766.2;HCoV-NL63_1ab;3283 | 0 | 0 |
| NC_005831.2;YP_003766.2;HCoV-NL63_1ab;3284 | 0 | 0 |
| NC_005831.2;YP_003766.2;HCoV-NL63_1ab;3285 | 0 | 0 |
| NC_005831.2;YP_003766.2;HCoV-NL63_1ab;3286 | 0 | 0 |
| NC_005831.2;YP_003766.2;HCoV-NL63_1ab;3287 | 0 | 0 |
| NC_005831.2;YP_003766.2;HCoV-NL63_1ab;3288 | 0 | 0 |
| NC_005831.2;YP_003766.2;HCoV-NL63_1ab;3289 | 0 | 0 |
| NC_005831.2;YP_003766.2;HCoV-NL63_1ab;3290 | 0 | 0 |
| NC_005831.2;YP_003766.2;HCoV-NL63_1ab;3291 | 0 | 0 |
| NC_005831.2;YP_003766.2;HCoV-NL63_1ab;3292 | 0 | 0 |
| NC_005831.2;YP_003766.2;HCoV-NL63_1ab;3293 | 0 | 0 |
| NC_005831.2;YP_003766.2;HCoV-NL63_1ab;3294 | 0 | 0 |
| NC_005831.2;YP_003766.2;HCoV-NL63_1ab;3295 | 0 | 0 |
| NC_005831.2;YP_003766.2;HCoV-NL63_1ab;3296 | 0 | 0 |
| NC_005831.2;YP_003766.2;HCoV-NL63_1ab;3297 | 0 | 0 |
| NC_005831.2;YP_003766.2;HCoV-NL63_1ab;3298 | 0 | 0 |
| NC_005831.2;YP_003766.2;HCoV-NL63_1ab;3299 | 0 | 0 |
| NC_005831.2;YP_003766.2;HCoV-NL63_1ab;3300 | 0 | 0 |
| NC_005831.2;YP_003766.2;HCoV-NL63_1ab;3301 | 0 | 0 |
| NC_005831.2;YP_003766.2;HCoV-NL63_1ab;3302 | 0 | 0 |
| NC_005831.2;YP_003766.2;HCoV-NL63_1ab;3303 | 0 | 0 |
| NC_005831.2;YP_003766.2;HCoV-NL63_1ab;3304 | 0 | 0 |
| NC_005831.2;YP_003766.2;HCoV-NL63_1ab;3305 | 0 | 0 |
| NC_005831.2;YP_003766.2;HCoV-NL63_1ab;3306 | 0 | 0 |
| NC_005831.2;YP_003766.2;HCoV-NL63_1ab;3307 | 0 | 0 |
| NC_005831.2;YP_003766.2;HCoV-NL63_1ab;3308 | 0 | 0 |
| NC_005831.2;YP_003766.2;HCoV-NL63_1ab;3309 | 0 | 0 |
| NC_005831.2;YP_003766.2;HCoV-NL63_1ab;3310 | 0 | 0 |
| NC_005831.2;YP_003766.2;HCoV-NL63_1ab;3311 | 0 | 0 |
| NC_005831.2;YP_003766.2;HCoV-NL63_1ab;3312 | 0 | 0 |
| NC_005831.2;YP_003766.2;HCoV-NL63_1ab;3313 | 0 | 0 |
| NC_005831.2;YP_003766.2;HCoV-NL63_1ab;3314 | 0 | 0 |
| NC_005831.2;YP_003766.2;HCoV-NL63_1ab;3315 | 0 | 0 |
| NC_005831.2;YP_003766.2;HCoV-NL63_1ab;3316 | 0 | 0 |
| NC_005831.2;YP_003766.2;HCoV-NL63_1ab;3317 | 0 | 0 |
| NC_005831.2;YP_003766.2;HCoV-NL63_1ab;3318 | 0 | 0 |
| NC_005831.2;YP_003766.2;HCoV-NL63_1ab;3319 | 0 | 0 |
| NC_005831.2;YP_003766.2;HCoV-NL63_1ab;3320 | 0 | 0 |

| | | |
|---|---|---|
| NC_005831.2;YP_003766.2;HCoV-NL63_1ab;3321 | 0 | 0 |
| NC_005831.2;YP_003766.2;HCoV-NL63_1ab;3322 | 0 | 0 |
| NC_005831.2;YP_003766.2;HCoV-NL63_1ab;3323 | 0 | 0 |
| NC_005831.2;YP_003766.2;HCoV-NL63_1ab;3324 | 0 | 0 |
| NC_005831.2;YP_003766.2;HCoV-NL63_1ab;3325 | 0 | 0 |
| NC_005831.2;YP_003766.2;HCoV-NL63_1ab;3326 | 0 | 0 |
| NC_005831.2;YP_003766.2;HCoV-NL63_1ab;3327 | 0 | 0 |
| NC_005831.2;YP_003766.2;HCoV-NL63_1ab;3328 | 0 | 0 |
| NC_005831.2;YP_003766.2;HCoV-NL63_1ab;3329 | 0 | 0 |
| NC_005831.2;YP_003766.2;HCoV-NL63_1ab;3330 | 0.05 | 0 |
| NC_005831.2;YP_003766.2;HCoV-NL63_1ab;3331 | 0 | 0 |
| NC_005831.2;YP_003766.2;HCoV-NL63_1ab;3332 | 0 | 0 |
| NC_005831.2;YP_003766.2;HCoV-NL63_1ab;3333 | 0 | 0 |
| NC_005831.2;YP_003766.2;HCoV-NL63_1ab;3334 | 0 | 0 |
| NC_005831.2;YP_003766.2;HCoV-NL63_1ab;3335 | 0 | 0 |
| NC_005831.2;YP_003766.2;HCoV-NL63_1ab;3336 | 0 | 0.025 |
| NC_005831.2;YP_003766.2;HCoV-NL63_1ab;3337 | 0 | 0.025 |
| NC_005831.2;YP_003766.2;HCoV-NL63_1ab;3338 | 0 | 0 |
| NC_005831.2;YP_003766.2;HCoV-NL63_1ab;3339 | 0 | 0 |
| NC_005831.2;YP_003766.2;HCoV-NL63_1ab;3340 | 0 | 0 |
| NC_005831.2;YP_003766.2;HCoV-NL63_1ab;3341 | 0 | 0 |
| NC_005831.2;YP_003766.2;HCoV-NL63_1ab;3342 | 0 | 0 |
| NC_005831.2;YP_003766.2;HCoV-NL63_1ab;3343 | 0 | 0 |
| NC_005831.2;YP_003766.2;HCoV-NL63_1ab;3344 | 0 | 0 |
| NC_005831.2;YP_003766.2;HCoV-NL63_1ab;3345 | 0 | 0 |
| NC_005831.2;YP_003766.2;HCoV-NL63_1ab;3346 | 0 | 0 |
| NC_005831.2;YP_003766.2;HCoV-NL63_1ab;3347 | 0 | 0 |
| NC_005831.2;YP_003766.2;HCoV-NL63_1ab;3348 | 0 | 0 |
| NC_005831.2;YP_003766.2;HCoV-NL63_1ab;3349 | 0 | 0 |
| NC_005831.2;YP_003766.2;HCoV-NL63_1ab;3350 | 0 | 0 |
| NC_005831.2;YP_003766.2;HCoV-NL63_1ab;3351 | 0 | 0 |
| NC_005831.2;YP_003766.2;HCoV-NL63_1ab;3352 | 0 | 0 |
| NC_005831.2;YP_003766.2;HCoV-NL63_1ab;3353 | 0 | 0 |
| NC_005831.2;YP_003766.2;HCoV-NL63_1ab;3354 | 0 | 0 |
| NC_005831.2;YP_003766.2;HCoV-NL63_1ab;3355 | 0 | 0 |
| NC_005831.2;YP_003766.2;HCoV-NL63_1ab;3356 | 0 | 0 |
| NC_005831.2;YP_003766.2;HCoV-NL63_1ab;3357 | 0 | 0 |
| NC_005831.2;YP_003766.2;HCoV-NL63_1ab;3358 | 0 | 0 |
| NC_005831.2;YP_003766.2;HCoV-NL63_1ab;3359 | 0 | 0 |
| NC_005831.2;YP_003766.2;HCoV-NL63_1ab;3360 | 0 | 0 |
| NC_005831.2;YP_003766.2;HCoV-NL63_1ab;3361 | 0 | 0 |
| NC_005831.2;YP_003766.2;HCoV-NL63_1ab;3362 | 0 | 0 |
| NC_005831.2;YP_003766.2;HCoV-NL63_1ab;3363 | 0 | 0 |

| | | |
|---|---|---|
| NC_005831.2;YP_003766.2;HCoV-NL63_1ab;3364 | 0 | 0 |
| NC_005831.2;YP_003766.2;HCoV-NL63_1ab;3365 | 0 | 0 |
| NC_005831.2;YP_003766.2;HCoV-NL63_1ab;3366 | 0 | 0 |
| NC_005831.2;YP_003766.2;HCoV-NL63_1ab;3367 | 0 | 0 |
| NC_005831.2;YP_003766.2;HCoV-NL63_1ab;3368 | 0 | 0 |
| NC_005831.2;YP_003766.2;HCoV-NL63_1ab;3369 | 0 | 0 |
| NC_005831.2;YP_003766.2;HCoV-NL63_1ab;3370 | 0 | 0 |
| NC_005831.2;YP_003766.2;HCoV-NL63_1ab;3371 | 0 | 0 |
| NC_005831.2;YP_003766.2;HCoV-NL63_1ab;3372 | 0 | 0 |
| NC_005831.2;YP_003766.2;HCoV-NL63_1ab;3373 | 0 | 0 |
| NC_005831.2;YP_003766.2;HCoV-NL63_1ab;3374 | 0 | 0 |
| NC_005831.2;YP_003766.2;HCoV-NL63_1ab;3375 | 0 | 0 |
| NC_005831.2;YP_003766.2;HCoV-NL63_1ab;3376 | 0 | 0 |
| NC_005831.2;YP_003766.2;HCoV-NL63_1ab;3377 | 0 | 0 |
| NC_005831.2;YP_003766.2;HCoV-NL63_1ab;3378 | 0 | 0 |
| NC_005831.2;YP_003766.2;HCoV-NL63_1ab;3379 | 0 | 0 |
| NC_005831.2;YP_003766.2;HCoV-NL63_1ab;3380 | 0 | 0 |
| NC_005831.2;YP_003766.2;HCoV-NL63_1ab;3381 | 0 | 0 |
| NC_005831.2;YP_003766.2;HCoV-NL63_1ab;3382 | 0 | 0 |
| NC_005831.2;YP_003766.2;HCoV-NL63_1ab;3383 | 0 | 0 |
| NC_005831.2;YP_003766.2;HCoV-NL63_1ab;3384 | 0 | 0 |
| NC_005831.2;YP_003766.2;HCoV-NL63_1ab;3385 | 0 | 0 |
| NC_005831.2;YP_003766.2;HCoV-NL63_1ab;3386 | 0 | 0 |
| NC_005831.2;YP_003766.2;HCoV-NL63_1ab;3387 | 0 | 0 |
| NC_005831.2;YP_003766.2;HCoV-NL63_1ab;3388 | 0 | 0 |
| NC_005831.2;YP_003766.2;HCoV-NL63_1ab;3389 | 0 | 0 |
| NC_005831.2;YP_003766.2;HCoV-NL63_1ab;3390 | 0 | 0 |
| NC_005831.2;YP_003766.2;HCoV-NL63_1ab;3391 | 0 | 0 |
| NC_005831.2;YP_003766.2;HCoV-NL63_1ab;3392 | 0 | 0 |
| NC_005831.2;YP_003766.2;HCoV-NL63_1ab;3393 | 0 | 0 |
| NC_005831.2;YP_003766.2;HCoV-NL63_1ab;3394 | 0 | 0 |
| NC_005831.2;YP_003766.2;HCoV-NL63_1ab;3395 | 0 | 0 |
| NC_005831.2;YP_003766.2;HCoV-NL63_1ab;3396 | 0 | 0 |
| NC_005831.2;YP_003766.2;HCoV-NL63_1ab;3397 | 0 | 0 |
| NC_005831.2;YP_003766.2;HCoV-NL63_1ab;3398 | 0 | 0 |
| NC_005831.2;YP_003766.2;HCoV-NL63_1ab;3399 | 0 | 0 |
| NC_005831.2;YP_003766.2;HCoV-NL63_1ab;3400 | 0 | 0 |
| NC_005831.2;YP_003766.2;HCoV-NL63_1ab;3401 | 0 | 0 |
| NC_005831.2;YP_003766.2;HCoV-NL63_1ab;3402 | 0 | 0 |
| NC_005831.2;YP_003766.2;HCoV-NL63_1ab;3403 | 0 | 0 |
| NC_005831.2;YP_003766.2;HCoV-NL63_1ab;3404 | 0 | 0 |
| NC_005831.2;YP_003766.2;HCoV-NL63_1ab;3405 | 0 | 0 |
| NC_005831.2;YP_003766.2;HCoV-NL63_1ab;3406 | 0 | 0 |

| | | |
|---|---|---|
| NC_005831.2;YP_003766.2;HCoV-NL63_1ab;3407 | 0 | 0 |
| NC_005831.2;YP_003766.2;HCoV-NL63_1ab;3408 | 0 | 0 |
| NC_005831.2;YP_003766.2;HCoV-NL63_1ab;3409 | 0 | 0 |
| NC_005831.2;YP_003766.2;HCoV-NL63_1ab;3410 | 0 | 0 |
| NC_005831.2;YP_003766.2;HCoV-NL63_1ab;3411 | 0 | 0 |
| NC_005831.2;YP_003766.2;HCoV-NL63_1ab;3412 | 0 | 0 |
| NC_005831.2;YP_003766.2;HCoV-NL63_1ab;3413 | 0 | 0 |
| NC_005831.2;YP_003766.2;HCoV-NL63_1ab;3414 | 0 | 0 |
| NC_005831.2;YP_003766.2;HCoV-NL63_1ab;3415 | 0 | 0 |
| NC_005831.2;YP_003766.2;HCoV-NL63_1ab;3416 | 0 | 0 |
| NC_005831.2;YP_003766.2;HCoV-NL63_1ab;3417 | 0 | 0 |
| NC_005831.2;YP_003766.2;HCoV-NL63_1ab;3418 | 0 | 0 |
| NC_005831.2;YP_003766.2;HCoV-NL63_1ab;3419 | 0 | 0 |
| NC_005831.2;YP_003766.2;HCoV-NL63_1ab;3420 | 0 | 0 |
| NC_005831.2;YP_003766.2;HCoV-NL63_1ab;3421 | 0 | 0 |
| NC_005831.2;YP_003766.2;HCoV-NL63_1ab;3422 | 0 | 0 |
| NC_005831.2;YP_003766.2;HCoV-NL63_1ab;3423 | 0 | 0 |
| NC_005831.2;YP_003766.2;HCoV-NL63_1ab;3424 | 0 | 0.025 |
| NC_005831.2;YP_003766.2;HCoV-NL63_1ab;3425 | 0 | 0.025 |
| NC_005831.2;YP_003766.2;HCoV-NL63_1ab;3426 | 0.1 | 0 |
| NC_005831.2;YP_003766.2;HCoV-NL63_1ab;3427 | 0.05 | 0 |
| NC_005831.2;YP_003766.2;HCoV-NL63_1ab;3428 | 0 | 0 |
| NC_005831.2;YP_003766.2;HCoV-NL63_1ab;3429 | 0 | 0 |
| NC_005831.2;YP_003766.2;HCoV-NL63_1ab;3430 | 0 | 0 |
| NC_005831.2;YP_003766.2;HCoV-NL63_1ab;3431 | 0 | 0 |
| NC_005831.2;YP_003766.2;HCoV-NL63_1ab;3432 | 0 | 0 |
| NC_005831.2;YP_003766.2;HCoV-NL63_1ab;3433 | 0 | 0 |
| NC_005831.2;YP_003766.2;HCoV-NL63_1ab;3434 | 0 | 0 |
| NC_005831.2;YP_003766.2;HCoV-NL63_1ab;3435 | 0 | 0 |
| NC_005831.2;YP_003766.2;HCoV-NL63_1ab;3436 | 0 | 0 |
| NC_005831.2;YP_003766.2;HCoV-NL63_1ab;3437 | 0 | 0 |
| NC_005831.2;YP_003766.2;HCoV-NL63_1ab;3438 | 0 | 0 |
| NC_005831.2;YP_003766.2;HCoV-NL63_1ab;3439 | 0 | 0 |
| NC_005831.2;YP_003766.2;HCoV-NL63_1ab;3440 | 0 | 0 |
| NC_005831.2;YP_003766.2;HCoV-NL63_1ab;3441 | 0 | 0 |
| NC_005831.2;YP_003766.2;HCoV-NL63_1ab;3442 | 0 | 0 |
| NC_005831.2;YP_003766.2;HCoV-NL63_1ab;3443 | 0 | 0 |
| NC_005831.2;YP_003766.2;HCoV-NL63_1ab;3444 | 0 | 0 |
| NC_005831.2;YP_003766.2;HCoV-NL63_1ab;3445 | 0 | 0 |
| NC_005831.2;YP_003766.2;HCoV-NL63_1ab;3446 | 0 | 0 |
| NC_005831.2;YP_003766.2;HCoV-NL63_1ab;3447 | 0 | 0 |
| NC_005831.2;YP_003766.2;HCoV-NL63_1ab;3448 | 0 | 0 |
| NC_005831.2;YP_003766.2;HCoV-NL63_1ab;3449 | 0 | 0 |

| | | |
|---|---|---|
| NC_005831.2;YP_003766.2;HCoV-NL63_1ab;3450 | 0 | 0 |
| NC_005831.2;YP_003766.2;HCoV-NL63_1ab;3451 | 0 | 0 |
| NC_005831.2;YP_003766.2;HCoV-NL63_1ab;3452 | 0 | 0 |
| NC_005831.2;YP_003766.2;HCoV-NL63_1ab;3453 | 0 | 0 |
| NC_005831.2;YP_003766.2;HCoV-NL63_1ab;3454 | 0 | 0 |
| NC_005831.2;YP_003766.2;HCoV-NL63_1ab;3455 | 0 | 0 |
| NC_005831.2;YP_003766.2;HCoV-NL63_1ab;3456 | 0 | 0 |
| NC_005831.2;YP_003766.2;HCoV-NL63_1ab;3457 | 0 | 0 |
| NC_005831.2;YP_003766.2;HCoV-NL63_1ab;3458 | 0 | 0 |
| NC_005831.2;YP_003766.2;HCoV-NL63_1ab;3459 | 0 | 0 |
| NC_005831.2;YP_003766.2;HCoV-NL63_1ab;3460 | 0 | 0 |
| NC_005831.2;YP_003766.2;HCoV-NL63_1ab;3461 | 0 | 0 |
| NC_005831.2;YP_003766.2;HCoV-NL63_1ab;3462 | 0 | 0 |
| NC_005831.2;YP_003766.2;HCoV-NL63_1ab;3463 | 0 | 0 |
| NC_005831.2;YP_003766.2;HCoV-NL63_1ab;3464 | 0 | 0 |
| NC_005831.2;YP_003766.2;HCoV-NL63_1ab;3465 | 0.05 | 0.025 |
| NC_005831.2;YP_003766.2;HCoV-NL63_1ab;3466 | 0 | 0.025 |
| NC_005831.2;YP_003766.2;HCoV-NL63_1ab;3467 | 0 | 0 |
| NC_005831.2;YP_003766.2;HCoV-NL63_1ab;3468 | 0 | 0 |
| NC_005831.2;YP_003766.2;HCoV-NL63_1ab;3469 | 0 | 0.025 |
| NC_005831.2;YP_003766.2;HCoV-NL63_1ab;3470 | 0 | 0.025 |
| NC_005831.2;YP_003766.2;HCoV-NL63_1ab;3471 | 0 | 0.125 |
| NC_005831.2;YP_003766.2;HCoV-NL63_1ab;3472 | 0 | 0.05 |
| NC_005831.2;YP_003766.2;HCoV-NL63_1ab;3473 | 0 | 0 |
| NC_005831.2;YP_003766.2;HCoV-NL63_1ab;3474 | 0.05 | 0 |
| NC_005831.2;YP_003766.2;HCoV-NL63_1ab;3475 | 0.1 | 0 |
| NC_005831.2;YP_003766.2;HCoV-NL63_1ab;3476 | 0.1 | 0 |
| NC_005831.2;YP_003766.2;HCoV-NL63_1ab;3477 | 0 | 0.025 |
| NC_005831.2;YP_003766.2;HCoV-NL63_1ab;3478 | 0.05 | 0.025 |
| NC_005831.2;YP_003766.2;HCoV-NL63_1ab;3479 | 0.05 | 0.025 |
| NC_005831.2;YP_003766.2;HCoV-NL63_1ab;3480 | 0.05 | 0.05 |
| NC_005831.2;YP_003766.2;HCoV-NL63_1ab;3481 | 0.15 | 0.05 |
| NC_005831.2;YP_003766.2;HCoV-NL63_1ab;3482 | 0 | 0 |
| NC_005831.2;YP_003766.2;HCoV-NL63_1ab;3483 | 0.05 | 0.05 |
| NC_005831.2;YP_003766.2;HCoV-NL63_1ab;3484 | 0 | 0 |
| NC_005831.2;YP_003766.2;HCoV-NL63_1ab;3485 | 0 | 0.075 |
| NC_005831.2;YP_003766.2;HCoV-NL63_1ab;3486 | 0 | 0 |
| NC_005831.2;YP_003766.2;HCoV-NL63_1ab;3487 | 0 | 0 |
| NC_005831.2;YP_003766.2;HCoV-NL63_1ab;3488 | 0 | 0 |
| NC_005831.2;YP_003766.2;HCoV-NL63_1ab;3489 | 0 | 0 |
| NC_005831.2;YP_003766.2;HCoV-NL63_1ab;3490 | 0 | 0 |
| NC_005831.2;YP_003766.2;HCoV-NL63_1ab;3491 | 0 | 0 |
| NC_005831.2;YP_003766.2;HCoV-NL63_1ab;3492 | 0 | 0 |

| | | |
|---|---|---|
| NC_005831.2;YP_003766.2;HCoV-NL63_1ab;3493 | 0 | 0 |
| NC_005831.2;YP_003766.2;HCoV-NL63_1ab;3494 | 0 | 0 |
| NC_005831.2;YP_003766.2;HCoV-NL63_1ab;3495 | 0 | 0 |
| NC_005831.2;YP_003766.2;HCoV-NL63_1ab;3496 | 0 | 0 |
| NC_005831.2;YP_003766.2;HCoV-NL63_1ab;3497 | 0 | 0 |
| NC_005831.2;YP_003766.2;HCoV-NL63_1ab;3498 | 0 | 0 |
| NC_005831.2;YP_003766.2;HCoV-NL63_1ab;3499 | 0 | 0 |
| NC_005831.2;YP_003766.2;HCoV-NL63_1ab;3500 | 0 | 0 |
| NC_005831.2;YP_003766.2;HCoV-NL63_1ab;3501 | 0 | 0 |
| NC_005831.2;YP_003766.2;HCoV-NL63_1ab;3502 | 0 | 0 |
| NC_005831.2;YP_003766.2;HCoV-NL63_1ab;3503 | 0 | 0 |
| NC_005831.2;YP_003766.2;HCoV-NL63_1ab;3504 | 0 | 0 |
| NC_005831.2;YP_003766.2;HCoV-NL63_1ab;3505 | 0 | 0 |
| NC_005831.2;YP_003766.2;HCoV-NL63_1ab;3506 | 0 | 0 |
| NC_005831.2;YP_003766.2;HCoV-NL63_1ab;3507 | 0.05 | 0.025 |
| NC_005831.2;YP_003766.2;HCoV-NL63_1ab;3508 | 0.05 | 0.05 |
| NC_005831.2;YP_003766.2;HCoV-NL63_1ab;3509 | 0.05 | 0.025 |
| NC_005831.2;YP_003766.2;HCoV-NL63_1ab;3510 | 0.05 | 0.025 |
| NC_005831.2;YP_003766.2;HCoV-NL63_1ab;3511 | 0.05 | 0.1 |
| NC_005831.2;YP_003766.2;HCoV-NL63_1ab;3512 | 0.1 | 0.075 |
| NC_005831.2;YP_003766.2;HCoV-NL63_1ab;3513 | 0.05 | 0.025 |
| NC_005831.2;YP_003766.2;HCoV-NL63_1ab;3514 | 0.05 | 0.025 |
| NC_005831.2;YP_003766.2;HCoV-NL63_1ab;3515 | 0.05 | 0.025 |
| NC_005831.2;YP_003766.2;HCoV-NL63_1ab;3516 | 0.05 | 0.025 |
| NC_005831.2;YP_003766.2;HCoV-NL63_1ab;3517 | 0.05 | 0.025 |
| NC_005831.2;YP_003766.2;HCoV-NL63_1ab;3518 | 0 | 0 |
| NC_005831.2;YP_003766.2;HCoV-NL63_1ab;3519 | 0 | 0 |
| NC_005831.2;YP_003766.2;HCoV-NL63_1ab;3520 | 0 | 0 |
| NC_005831.2;YP_003766.2;HCoV-NL63_1ab;3521 | 0 | 0 |
| NC_005831.2;YP_003766.2;HCoV-NL63_1ab;3522 | 0 | 0 |
| NC_005831.2;YP_003766.2;HCoV-NL63_1ab;3523 | 0 | 0 |
| NC_005831.2;YP_003766.2;HCoV-NL63_1ab;3524 | 0 | 0.025 |
| NC_005831.2;YP_003766.2;HCoV-NL63_1ab;3525 | 0 | 0 |
| NC_005831.2;YP_003766.2;HCoV-NL63_1ab;3526 | 0 | 0 |
| NC_005831.2;YP_003766.2;HCoV-NL63_1ab;3527 | 0 | 0 |
| NC_005831.2;YP_003766.2;HCoV-NL63_1ab;3528 | 0 | 0 |
| NC_005831.2;YP_003766.2;HCoV-NL63_1ab;3529 | 0 | 0 |
| NC_005831.2;YP_003766.2;HCoV-NL63_1ab;3530 | 0.1 | 0.025 |
| NC_005831.2;YP_003766.2;HCoV-NL63_1ab;3531 | 0 | 0 |
| NC_005831.2;YP_003766.2;HCoV-NL63_1ab;3532 | 0 | 0 |
| NC_005831.2;YP_003766.2;HCoV-NL63_1ab;3533 | 0 | 0 |
| NC_005831.2;YP_003766.2;HCoV-NL63_1ab;3534 | 0 | 0.05 |
| NC_005831.2;YP_003766.2;HCoV-NL63_1ab;3535 | 0 | 0.05 |

| | | |
|---|---|---|
| NC_005831.2;YP_003766.2;HCoV-NL63_1ab;3536 | 0 | 0.05 |
| NC_005831.2;YP_003766.2;HCoV-NL63_1ab;3537 | 0 | 0.025 |
| NC_005831.2;YP_003766.2;HCoV-NL63_1ab;3538 | 0 | 0 |
| NC_005831.2;YP_003766.2;HCoV-NL63_1ab;3539 | 0 | 0 |
| NC_005831.2;YP_003766.2;HCoV-NL63_1ab;3540 | 0 | 0 |
| NC_005831.2;YP_003766.2;HCoV-NL63_1ab;3541 | 0 | 0 |
| NC_005831.2;YP_003766.2;HCoV-NL63_1ab;3542 | 0 | 0 |
| NC_005831.2;YP_003766.2;HCoV-NL63_1ab;3543 | 0 | 0 |
| NC_005831.2;YP_003766.2;HCoV-NL63_1ab;3544 | 0.15 | 0.075 |
| NC_005831.2;YP_003766.2;HCoV-NL63_1ab;3545 | 0.15 | 0.1 |
| NC_005831.2;YP_003766.2;HCoV-NL63_1ab;3546 | 0.05 | 0.1 |
| NC_005831.2;YP_003766.2;HCoV-NL63_1ab;3547 | 0.05 | 0.075 |
| NC_005831.2;YP_003766.2;HCoV-NL63_1ab;3548 | 0.05 | 0.025 |
| NC_005831.2;YP_003766.2;HCoV-NL63_1ab;3549 | 0 | 0.025 |
| NC_005831.2;YP_003766.2;HCoV-NL63_1ab;3550 | 0 | 0.05 |
| NC_005831.2;YP_003766.2;HCoV-NL63_1ab;3551 | 0 | 0.075 |
| NC_005831.2;YP_003766.2;HCoV-NL63_1ab;3552 | 0 | 0.075 |
| NC_005831.2;YP_003766.2;HCoV-NL63_1ab;3553 | 0.05 | 0.1 |
| NC_005831.2;YP_003766.2;HCoV-NL63_1ab;3554 | 0.1 | 0.1 |
| NC_005831.2;YP_003766.2;HCoV-NL63_1ab;3555 | 0 | 0.05 |
| NC_005831.2;YP_003766.2;HCoV-NL63_1ab;3556 | 0 | 0 |
| NC_005831.2;YP_003766.2;HCoV-NL63_1ab;3557 | 0 | 0 |
| NC_005831.2;YP_003766.2;HCoV-NL63_1ab;3558 | 0 | 0.025 |
| NC_005831.2;YP_003766.2;HCoV-NL63_1ab;3559 | 0 | 0 |
| NC_005831.2;YP_003766.2;HCoV-NL63_1ab;3560 | 0 | 0 |
| NC_005831.2;YP_003766.2;HCoV-NL63_1ab;3561 | 0 | 0 |
| NC_005831.2;YP_003766.2;HCoV-NL63_1ab;3562 | 0 | 0 |
| NC_005831.2;YP_003766.2;HCoV-NL63_1ab;3563 | 0 | 0 |
| NC_005831.2;YP_003766.2;HCoV-NL63_1ab;3564 | 0 | 0 |
| NC_005831.2;YP_003766.2;HCoV-NL63_1ab;3565 | 0 | 0 |
| NC_005831.2;YP_003766.2;HCoV-NL63_1ab;3566 | 0 | 0 |
| NC_005831.2;YP_003766.2;HCoV-NL63_1ab;3567 | 0 | 0 |
| NC_005831.2;YP_003766.2;HCoV-NL63_1ab;3568 | 0 | 0 |
| NC_005831.2;YP_003766.2;HCoV-NL63_1ab;3569 | 0 | 0 |
| NC_005831.2;YP_003766.2;HCoV-NL63_1ab;3570 | 0 | 0 |
| NC_005831.2;YP_003766.2;HCoV-NL63_1ab;3571 | 0 | 0 |
| NC_005831.2;YP_003766.2;HCoV-NL63_1ab;3572 | 0 | 0 |
| NC_005831.2;YP_003766.2;HCoV-NL63_1ab;3573 | 0 | 0 |
| NC_005831.2;YP_003766.2;HCoV-NL63_1ab;3574 | 0 | 0 |
| NC_005831.2;YP_003766.2;HCoV-NL63_1ab;3575 | 0 | 0 |
| NC_005831.2;YP_003766.2;HCoV-NL63_1ab;3576 | 0 | 0 |
| NC_005831.2;YP_003766.2;HCoV-NL63_1ab;3577 | 0 | 0 |
| NC_005831.2;YP_003766.2;HCoV-NL63_1ab;3578 | 0.05 | 0 |

| | | |
|---|---|---|
| NC_005831.2;YP_003766.2;HCoV-NL63_1ab;3579 | 0.05 | 0 |
| NC_005831.2;YP_003766.2;HCoV-NL63_1ab;3580 | 0.05 | 0 |
| NC_005831.2;YP_003766.2;HCoV-NL63_1ab;3581 | 0.1 | 0 |
| NC_005831.2;YP_003766.2;HCoV-NL63_1ab;3582 | 0.05 | 0 |
| NC_005831.2;YP_003766.2;HCoV-NL63_1ab;3583 | 0.05 | 0 |
| NC_005831.2;YP_003766.2;HCoV-NL63_1ab;3584 | 0.05 | 0 |
| NC_005831.2;YP_003766.2;HCoV-NL63_1ab;3585 | 0 | 0 |
| NC_005831.2;YP_003766.2;HCoV-NL63_1ab;3586 | 0 | 0 |
| NC_005831.2;YP_003766.2;HCoV-NL63_1ab;3587 | 0 | 0 |
| NC_005831.2;YP_003766.2;HCoV-NL63_1ab;3588 | 0.05 | 0.05 |
| NC_005831.2;YP_003766.2;HCoV-NL63_1ab;3589 | 0.05 | 0 |
| NC_005831.2;YP_003766.2;HCoV-NL63_1ab;3590 | 0 | 0 |
| NC_005831.2;YP_003766.2;HCoV-NL63_1ab;3591 | 0 | 0.1 |
| NC_005831.2;YP_003766.2;HCoV-NL63_1ab;3592 | 0.1 | 0.025 |
| NC_005831.2;YP_003766.2;HCoV-NL63_1ab;3593 | 0 | 0 |
| NC_005831.2;YP_003766.2;HCoV-NL63_1ab;3594 | 0.05 | 0.025 |
| NC_005831.2;YP_003766.2;HCoV-NL63_1ab;3595 | 0 | 0 |
| NC_005831.2;YP_003766.2;HCoV-NL63_1ab;3596 | 0 | 0 |
| NC_005831.2;YP_003766.2;HCoV-NL63_1ab;3597 | 0 | 0 |
| NC_005831.2;YP_003766.2;HCoV-NL63_1ab;3598 | 0 | 0.025 |
| NC_005831.2;YP_003766.2;HCoV-NL63_1ab;3599 | 0 | 0 |
| NC_005831.2;YP_003766.2;HCoV-NL63_1ab;3600 | 0 | 0 |
| NC_005831.2;YP_003766.2;HCoV-NL63_1ab;3601 | 0 | 0 |
| NC_005831.2;YP_003766.2;HCoV-NL63_1ab;3602 | 0 | 0 |
| NC_005831.2;YP_003766.2;HCoV-NL63_1ab;3603 | 0 | 0 |
| NC_005831.2;YP_003766.2;HCoV-NL63_1ab;3604 | 0 | 0 |
| NC_005831.2;YP_003766.2;HCoV-NL63_1ab;3605 | 0 | 0.025 |
| NC_005831.2;YP_003766.2;HCoV-NL63_1ab;3606 | 0 | 0 |
| NC_005831.2;YP_003766.2;HCoV-NL63_1ab;3607 | 0 | 0 |
| NC_005831.2;YP_003766.2;HCoV-NL63_1ab;3608 | 0 | 0 |
| NC_005831.2;YP_003766.2;HCoV-NL63_1ab;3609 | 0 | 0.025 |
| NC_005831.2;YP_003766.2;HCoV-NL63_1ab;3610 | 0 | 0 |
| NC_005831.2;YP_003766.2;HCoV-NL63_1ab;3611 | 0 | 0 |
| NC_005831.2;YP_003766.2;HCoV-NL63_1ab;3612 | 0.05 | 0.025 |
| NC_005831.2;YP_003766.2;HCoV-NL63_1ab;3613 | 0.05 | 0 |
| NC_005831.2;YP_003766.2;HCoV-NL63_1ab;3614 | 0.05 | 0.05 |
| NC_005831.2;YP_003766.2;HCoV-NL63_1ab;3615 | 0.15 | 0.05 |
| NC_005831.2;YP_003766.2;HCoV-NL63_1ab;3616 | 0.1 | 0.05 |
| NC_005831.2;YP_003766.2;HCoV-NL63_1ab;3617 | 0.1 | 0.075 |
| NC_005831.2;YP_003766.2;HCoV-NL63_1ab;3618 | 0.05 | 0.075 |
| NC_005831.2;YP_003766.2;HCoV-NL63_1ab;3619 | 0 | 0.05 |
| NC_005831.2;YP_003766.2;HCoV-NL63_1ab;3620 | 0 | 0 |
| NC_005831.2;YP_003766.2;HCoV-NL63_1ab;3621 | 0 | 0.025 |

| | | |
|---|---|---|
| NC_005831.2;YP_003766.2;HCoV-NL63_1ab;3622 | 0 | 0.025 |
| NC_005831.2;YP_003766.2;HCoV-NL63_1ab;3623 | 0 | 0.025 |
| NC_005831.2;YP_003766.2;HCoV-NL63_1ab;3624 | 0 | 0.025 |
| NC_005831.2;YP_003766.2;HCoV-NL63_1ab;3625 | 0 | 0.025 |
| NC_005831.2;YP_003766.2;HCoV-NL63_1ab;3626 | 0 | 0.025 |
| NC_005831.2;YP_003766.2;HCoV-NL63_1ab;3627 | 0 | 0 |
| NC_005831.2;YP_003766.2;HCoV-NL63_1ab;3628 | 0 | 0 |
| NC_005831.2;YP_003766.2;HCoV-NL63_1ab;3629 | 0 | 0 |
| NC_005831.2;YP_003766.2;HCoV-NL63_1ab;3630 | 0 | 0 |
| NC_005831.2;YP_003766.2;HCoV-NL63_1ab;3631 | 0 | 0 |
| NC_005831.2;YP_003766.2;HCoV-NL63_1ab;3632 | 0 | 0 |
| NC_005831.2;YP_003766.2;HCoV-NL63_1ab;3633 | 0.05 | 0 |
| NC_005831.2;YP_003766.2;HCoV-NL63_1ab;3634 | 0.05 | 0 |
| NC_005831.2;YP_003766.2;HCoV-NL63_1ab;3635 | 0 | 0 |
| NC_005831.2;YP_003766.2;HCoV-NL63_1ab;3636 | 0 | 0 |
| NC_005831.2;YP_003766.2;HCoV-NL63_1ab;3637 | 0 | 0 |
| NC_005831.2;YP_003766.2;HCoV-NL63_1ab;3638 | 0 | 0.025 |
| NC_005831.2;YP_003766.2;HCoV-NL63_1ab;3639 | 0 | 0.025 |
| NC_005831.2;YP_003766.2;HCoV-NL63_1ab;3640 | 0 | 0.025 |
| NC_005831.2;YP_003766.2;HCoV-NL63_1ab;3641 | 0 | 0.05 |
| NC_005831.2;YP_003766.2;HCoV-NL63_1ab;3642 | 0 | 0.05 |
| NC_005831.2;YP_003766.2;HCoV-NL63_1ab;3643 | 0 | 0.075 |
| NC_005831.2;YP_003766.2;HCoV-NL63_1ab;3644 | 0 | 0.025 |
| NC_005831.2;YP_003766.2;HCoV-NL63_1ab;3645 | 0.05 | 0 |
| NC_005831.2;YP_003766.2;HCoV-NL63_1ab;3646 | 0 | 0 |
| NC_005831.2;YP_003766.2;HCoV-NL63_1ab;3647 | 0 | 0 |
| NC_005831.2;YP_003766.2;HCoV-NL63_1ab;3648 | 0 | 0 |
| NC_005831.2;YP_003766.2;HCoV-NL63_1ab;3649 | 0 | 0 |
| NC_005831.2;YP_003766.2;HCoV-NL63_1ab;3650 | 0 | 0 |
| NC_005831.2;YP_003766.2;HCoV-NL63_1ab;3651 | 0 | 0 |
| NC_005831.2;YP_003766.2;HCoV-NL63_1ab;3652 | 0 | 0 |
| NC_005831.2;YP_003766.2;HCoV-NL63_1ab;3653 | 0 | 0 |
| NC_005831.2;YP_003766.2;HCoV-NL63_1ab;3654 | 0 | 0 |
| NC_005831.2;YP_003766.2;HCoV-NL63_1ab;3655 | 0.05 | 0 |
| NC_005831.2;YP_003766.2;HCoV-NL63_1ab;3656 | 0 | 0.025 |
| NC_005831.2;YP_003766.2;HCoV-NL63_1ab;3657 | 0 | 0 |
| NC_005831.2;YP_003766.2;HCoV-NL63_1ab;3658 | 0 | 0 |
| NC_005831.2;YP_003766.2;HCoV-NL63_1ab;3659 | 0 | 0 |
| NC_005831.2;YP_003766.2;HCoV-NL63_1ab;3660 | 0 | 0 |
| NC_005831.2;YP_003766.2;HCoV-NL63_1ab;3661 | 0 | 0 |
| NC_005831.2;YP_003766.2;HCoV-NL63_1ab;3662 | 0 | 0 |
| NC_005831.2;YP_003766.2;HCoV-NL63_1ab;3663 | 0 | 0 |
| NC_005831.2;YP_003766.2;HCoV-NL63_1ab;3664 | 0 | 0 |

| | | |
|---|---|---|
| NC_005831.2;YP_003766.2;HCoV-NL63_1ab;3665 | 0 | 0 |
| NC_005831.2;YP_003766.2;HCoV-NL63_1ab;3666 | 0 | 0 |
| NC_005831.2;YP_003766.2;HCoV-NL63_1ab;3667 | 0 | 0.025 |
| NC_005831.2;YP_003766.2;HCoV-NL63_1ab;3668 | 0 | 0.05 |
| NC_005831.2;YP_003766.2;HCoV-NL63_1ab;3669 | 0 | 0 |
| NC_005831.2;YP_003766.2;HCoV-NL63_1ab;3670 | 0 | 0 |
| NC_005831.2;YP_003766.2;HCoV-NL63_1ab;3671 | 0 | 0 |
| NC_005831.2;YP_003766.2;HCoV-NL63_1ab;3672 | 0 | 0.025 |
| NC_005831.2;YP_003766.2;HCoV-NL63_1ab;3673 | 0 | 0.025 |
| NC_005831.2;YP_003766.2;HCoV-NL63_1ab;3674 | 0 | 0.025 |
| NC_005831.2;YP_003766.2;HCoV-NL63_1ab;3675 | 0 | 0 |
| NC_005831.2;YP_003766.2;HCoV-NL63_1ab;3676 | 0 | 0 |
| NC_005831.2;YP_003766.2;HCoV-NL63_1ab;3677 | 0 | 0.025 |
| NC_005831.2;YP_003766.2;HCoV-NL63_1ab;3678 | 0 | 0 |
| NC_005831.2;YP_003766.2;HCoV-NL63_1ab;3679 | 0.05 | 0 |
| NC_005831.2;YP_003766.2;HCoV-NL63_1ab;3680 | 0 | 0 |
| NC_005831.2;YP_003766.2;HCoV-NL63_1ab;3681 | 0 | 0 |
| NC_005831.2;YP_003766.2;HCoV-NL63_1ab;3682 | 0 | 0 |
| NC_005831.2;YP_003766.2;HCoV-NL63_1ab;3683 | 0 | 0 |
| NC_005831.2;YP_003766.2;HCoV-NL63_1ab;3684 | 0 | 0 |
| NC_005831.2;YP_003766.2;HCoV-NL63_1ab;3685 | 0 | 0 |
| NC_005831.2;YP_003766.2;HCoV-NL63_1ab;3686 | 0 | 0 |
| NC_005831.2;YP_003766.2;HCoV-NL63_1ab;3687 | 0 | 0 |
| NC_005831.2;YP_003766.2;HCoV-NL63_1ab;3688 | 0 | 0 |
| NC_005831.2;YP_003766.2;HCoV-NL63_1ab;3689 | 0 | 0 |
| NC_005831.2;YP_003766.2;HCoV-NL63_1ab;3690 | 0 | 0 |
| NC_005831.2;YP_003766.2;HCoV-NL63_1ab;3691 | 0 | 0 |
| NC_005831.2;YP_003766.2;HCoV-NL63_1ab;3692 | 0 | 0 |
| NC_005831.2;YP_003766.2;HCoV-NL63_1ab;3693 | 0 | 0 |
| NC_005831.2;YP_003766.2;HCoV-NL63_1ab;3694 | 0 | 0 |
| NC_005831.2;YP_003766.2;HCoV-NL63_1ab;3695 | 0 | 0 |
| NC_005831.2;YP_003766.2;HCoV-NL63_1ab;3696 | 0 | 0 |
| NC_005831.2;YP_003766.2;HCoV-NL63_1ab;3697 | 0 | 0 |
| NC_005831.2;YP_003766.2;HCoV-NL63_1ab;3698 | 0 | 0 |
| NC_005831.2;YP_003766.2;HCoV-NL63_1ab;3699 | 0 | 0.025 |
| NC_005831.2;YP_003766.2;HCoV-NL63_1ab;3700 | 0 | 0 |
| NC_005831.2;YP_003766.2;HCoV-NL63_1ab;3701 | 0 | 0 |
| NC_005831.2;YP_003766.2;HCoV-NL63_1ab;3702 | 0 | 0 |
| NC_005831.2;YP_003766.2;HCoV-NL63_1ab;3703 | 0 | 0 |
| NC_005831.2;YP_003766.2;HCoV-NL63_1ab;3704 | 0.1 | 0.05 |
| NC_005831.2;YP_003766.2;HCoV-NL63_1ab;3705 | 0 | 0.05 |
| NC_005831.2;YP_003766.2;HCoV-NL63_1ab;3706 | 0 | 0.025 |
| NC_005831.2;YP_003766.2;HCoV-NL63_1ab;3707 | 0 | 0.075 |

| | | |
|---|---|---|
| NC_005831.2;YP_003766.2;HCoV-NL63_1ab;3708 | 0 | 0 |
| NC_005831.2;YP_003766.2;HCoV-NL63_1ab;3709 | 0 | 0 |
| NC_005831.2;YP_003766.2;HCoV-NL63_1ab;3710 | 0 | 0 |
| NC_005831.2;YP_003766.2;HCoV-NL63_1ab;3711 | 0 | 0 |
| NC_005831.2;YP_003766.2;HCoV-NL63_1ab;3712 | 0 | 0.025 |
| NC_005831.2;YP_003766.2;HCoV-NL63_1ab;3713 | 0 | 0.025 |
| NC_005831.2;YP_003766.2;HCoV-NL63_1ab;3714 | 0 | 0.05 |
| NC_005831.2;YP_003766.2;HCoV-NL63_1ab;3715 | 0.2 | 0.025 |
| NC_005831.2;YP_003766.2;HCoV-NL63_1ab;3716 | 0.05 | 0.05 |
| NC_005831.2;YP_003766.2;HCoV-NL63_1ab;3717 | 0.1 | 0.025 |
| NC_005831.2;YP_003766.2;HCoV-NL63_1ab;3718 | 0.1 | 0.025 |
| NC_005831.2;YP_003766.2;HCoV-NL63_1ab;3719 | 0 | 0.025 |
| NC_005831.2;YP_003766.2;HCoV-NL63_1ab;3720 | 0 | 0 |
| NC_005831.2;YP_003766.2;HCoV-NL63_1ab;3721 | 0 | 0 |
| NC_005831.2;YP_003766.2;HCoV-NL63_1ab;3722 | 0 | 0.05 |
| NC_005831.2;YP_003766.2;HCoV-NL63_1ab;3723 | 0 | 0.05 |
| NC_005831.2;YP_003766.2;HCoV-NL63_1ab;3724 | 0 | 0.05 |
| NC_005831.2;YP_003766.2;HCoV-NL63_1ab;3725 | 0 | 0.05 |
| NC_005831.2;YP_003766.2;HCoV-NL63_1ab;3726 | 0 | 0.025 |
| NC_005831.2;YP_003766.2;HCoV-NL63_1ab;3727 | 0 | 0.025 |
| NC_005831.2;YP_003766.2;HCoV-NL63_1ab;3728 | 0 | 0.05 |
| NC_005831.2;YP_003766.2;HCoV-NL63_1ab;3729 | 0 | 0.025 |
| NC_005831.2;YP_003766.2;HCoV-NL63_1ab;3730 | 0 | 0.025 |
| NC_005831.2;YP_003766.2;HCoV-NL63_1ab;3731 | 0 | 0 |
| NC_005831.2;YP_003766.2;HCoV-NL63_1ab;3732 | 0 | 0 |
| NC_005831.2;YP_003766.2;HCoV-NL63_1ab;3733 | 0 | 0 |
| NC_005831.2;YP_003766.2;HCoV-NL63_1ab;3734 | 0 | 0 |
| NC_005831.2;YP_003766.2;HCoV-NL63_1ab;3735 | 0 | 0 |
| NC_005831.2;YP_003766.2;HCoV-NL63_1ab;3736 | 0 | 0 |
| NC_005831.2;YP_003766.2;HCoV-NL63_1ab;3737 | 0 | 0 |
| NC_005831.2;YP_003766.2;HCoV-NL63_1ab;3738 | 0 | 0 |
| NC_005831.2;YP_003766.2;HCoV-NL63_1ab;3739 | 0 | 0 |
| NC_005831.2;YP_003766.2;HCoV-NL63_1ab;3740 | 0 | 0 |
| NC_005831.2;YP_003766.2;HCoV-NL63_1ab;3741 | 0 | 0.025 |
| NC_005831.2;YP_003766.2;HCoV-NL63_1ab;3742 | 0 | 0 |
| NC_005831.2;YP_003766.2;HCoV-NL63_1ab;3743 | 0 | 0 |
| NC_005831.2;YP_003766.2;HCoV-NL63_1ab;3744 | 0 | 0 |
| NC_005831.2;YP_003766.2;HCoV-NL63_1ab;3745 | 0 | 0 |
| NC_005831.2;YP_003766.2;HCoV-NL63_1ab;3746 | 0 | 0 |
| NC_005831.2;YP_003766.2;HCoV-NL63_1ab;3747 | 0 | 0 |
| NC_005831.2;YP_003766.2;HCoV-NL63_1ab;3748 | 0 | 0 |
| NC_005831.2;YP_003766.2;HCoV-NL63_1ab;3749 | 0 | 0 |
| NC_005831.2;YP_003766.2;HCoV-NL63_1ab;3750 | 0 | 0 |

| | | |
|---|---|---|
| NC_005831.2;YP_003766.2;HCoV-NL63_1ab;3751 | 0 | 0 |
| NC_005831.2;YP_003766.2;HCoV-NL63_1ab;3752 | 0 | 0.025 |
| NC_005831.2;YP_003766.2;HCoV-NL63_1ab;3753 | 0 | 0.1 |
| NC_005831.2;YP_003766.2;HCoV-NL63_1ab;3754 | 0 | 0.1 |
| NC_005831.2;YP_003766.2;HCoV-NL63_1ab;3755 | 0 | 0.1 |
| NC_005831.2;YP_003766.2;HCoV-NL63_1ab;3756 | 0.05 | 0.1 |
| NC_005831.2;YP_003766.2;HCoV-NL63_1ab;3757 | 0 | 0.175 |
| NC_005831.2;YP_003766.2;HCoV-NL63_1ab;3758 | 0.05 | 0.1 |
| NC_005831.2;YP_003766.2;HCoV-NL63_1ab;3759 | 0.15 | 0.025 |
| NC_005831.2;YP_003766.2;HCoV-NL63_1ab;3760 | 0.05 | 0.025 |
| NC_005831.2;YP_003766.2;HCoV-NL63_1ab;3761 | 0 | 0 |
| NC_005831.2;YP_003766.2;HCoV-NL63_1ab;3762 | 0 | 0 |
| NC_005831.2;YP_003766.2;HCoV-NL63_1ab;3763 | 0 | 0 |
| NC_005831.2;YP_003766.2;HCoV-NL63_1ab;3764 | 0 | 0 |
| NC_005831.2;YP_003766.2;HCoV-NL63_1ab;3765 | 0 | 0.025 |
| NC_005831.2;YP_003766.2;HCoV-NL63_1ab;3766 | 0 | 0 |
| NC_005831.2;YP_003766.2;HCoV-NL63_1ab;3767 | 0 | 0 |
| NC_005831.2;YP_003766.2;HCoV-NL63_1ab;3768 | 0.05 | 0 |
| NC_005831.2;YP_003766.2;HCoV-NL63_1ab;3769 | 0 | 0 |
| NC_005831.2;YP_003766.2;HCoV-NL63_1ab;3770 | 0.05 | 0.025 |
| NC_005831.2;YP_003766.2;HCoV-NL63_1ab;3771 | 0 | 0 |
| NC_005831.2;YP_003766.2;HCoV-NL63_1ab;3772 | 0 | 0 |
| NC_005831.2;YP_003766.2;HCoV-NL63_1ab;3773 | 0 | 0 |
| NC_005831.2;YP_003766.2;HCoV-NL63_1ab;3774 | 0 | 0 |
| NC_005831.2;YP_003766.2;HCoV-NL63_1ab;3775 | 0 | 0 |
| NC_005831.2;YP_003766.2;HCoV-NL63_1ab;3776 | 0 | 0.025 |
| NC_005831.2;YP_003766.2;HCoV-NL63_1ab;3777 | 0 | 0 |
| NC_005831.2;YP_003766.2;HCoV-NL63_1ab;3778 | 0 | 0 |
| NC_005831.2;YP_003766.2;HCoV-NL63_1ab;3779 | 0 | 0 |
| NC_005831.2;YP_003766.2;HCoV-NL63_1ab;3780 | 0 | 0 |
| NC_005831.2;YP_003766.2;HCoV-NL63_1ab;3781 | 0 | 0 |
| NC_005831.2;YP_003766.2;HCoV-NL63_1ab;3782 | 0 | 0 |
| NC_005831.2;YP_003766.2;HCoV-NL63_1ab;3783 | 0 | 0 |
| NC_005831.2;YP_003766.2;HCoV-NL63_1ab;3784 | 0 | 0 |
| NC_005831.2;YP_003766.2;HCoV-NL63_1ab;3785 | 0 | 0 |
| NC_005831.2;YP_003766.2;HCoV-NL63_1ab;3786 | 0 | 0 |
| NC_005831.2;YP_003766.2;HCoV-NL63_1ab;3787 | 0 | 0 |
| NC_005831.2;YP_003766.2;HCoV-NL63_1ab;3788 | 0 | 0 |
| NC_005831.2;YP_003766.2;HCoV-NL63_1ab;3789 | 0 | 0 |
| NC_005831.2;YP_003766.2;HCoV-NL63_1ab;3790 | 0 | 0.025 |
| NC_005831.2;YP_003766.2;HCoV-NL63_1ab;3791 | 0 | 0 |
| NC_005831.2;YP_003766.2;HCoV-NL63_1ab;3792 | 0 | 0 |
| NC_005831.2;YP_003766.2;HCoV-NL63_1ab;3793 | 0.05 | 0 |

| | | |
|---|---|---|
| NC_005831.2;YP_003766.2;HCoV-NL63_1ab;3794 | 0 | 0 |
| NC_005831.2;YP_003766.2;HCoV-NL63_1ab;3795 | 0 | 0 |
| NC_005831.2;YP_003766.2;HCoV-NL63_1ab;3796 | 0 | 0 |
| NC_005831.2;YP_003766.2;HCoV-NL63_1ab;3797 | 0 | 0 |
| NC_005831.2;YP_003766.2;HCoV-NL63_1ab;3798 | 0 | 0 |
| NC_005831.2;YP_003766.2;HCoV-NL63_1ab;3799 | 0 | 0 |
| NC_005831.2;YP_003766.2;HCoV-NL63_1ab;3800 | 0 | 0 |
| NC_005831.2;YP_003766.2;HCoV-NL63_1ab;3801 | 0 | 0 |
| NC_005831.2;YP_003766.2;HCoV-NL63_1ab;3802 | 0.05 | 0 |
| NC_005831.2;YP_003766.2;HCoV-NL63_1ab;3803 | 0 | 0 |
| NC_005831.2;YP_003766.2;HCoV-NL63_1ab;3804 | 0 | 0 |
| NC_005831.2;YP_003766.2;HCoV-NL63_1ab;3805 | 0 | 0 |
| NC_005831.2;YP_003766.2;HCoV-NL63_1ab;3806 | 0 | 0 |
| NC_005831.2;YP_003766.2;HCoV-NL63_1ab;3807 | 0 | 0 |
| NC_005831.2;YP_003766.2;HCoV-NL63_1ab;3808 | 0 | 0 |
| NC_005831.2;YP_003766.2;HCoV-NL63_1ab;3809 | 0 | 0 |
| NC_005831.2;YP_003766.2;HCoV-NL63_1ab;3810 | 0 | 0 |
| NC_005831.2;YP_003766.2;HCoV-NL63_1ab;3811 | 0 | 0 |
| NC_005831.2;YP_003766.2;HCoV-NL63_1ab;3812 | 0 | 0 |
| NC_005831.2;YP_003766.2;HCoV-NL63_1ab;3813 | 0 | 0 |
| NC_005831.2;YP_003766.2;HCoV-NL63_1ab;3814 | 0 | 0 |
| NC_005831.2;YP_003766.2;HCoV-NL63_1ab;3815 | 0 | 0 |
| NC_005831.2;YP_003766.2;HCoV-NL63_1ab;3816 | 0 | 0 |
| NC_005831.2;YP_003766.2;HCoV-NL63_1ab;3817 | 0 | 0 |
| NC_005831.2;YP_003766.2;HCoV-NL63_1ab;3818 | 0 | 0 |
| NC_005831.2;YP_003766.2;HCoV-NL63_1ab;3819 | 0.1 | 0.1 |
| NC_005831.2;YP_003766.2;HCoV-NL63_1ab;3820 | 0.2 | 0.3 |
| NC_005831.2;YP_003766.2;HCoV-NL63_1ab;3821 | 0.05 | 0 |
| NC_005831.2;YP_003766.2;HCoV-NL63_1ab;3822 | 0.05 | 0 |
| NC_005831.2;YP_003766.2;HCoV-NL63_1ab;3823 | 0.1 | 0.025 |
| NC_005831.2;YP_003766.2;HCoV-NL63_1ab;3824 | 0.05 | 0 |
| NC_005831.2;YP_003766.2;HCoV-NL63_1ab;3825 | 0.05 | 0 |
| NC_005831.2;YP_003766.2;HCoV-NL63_1ab;3826 | 0.1 | 0 |
| NC_005831.2;YP_003766.2;HCoV-NL63_1ab;3827 | 0 | 0 |
| NC_005831.2;YP_003766.2;HCoV-NL63_1ab;3828 | 0 | 0 |
| NC_005831.2;YP_003766.2;HCoV-NL63_1ab;3829 | 0 | 0 |
| NC_005831.2;YP_003766.2;HCoV-NL63_1ab;3830 | 0 | 0 |
| NC_005831.2;YP_003766.2;HCoV-NL63_1ab;3831 | 0 | 0 |
| NC_005831.2;YP_003766.2;HCoV-NL63_1ab;3832 | 0.05 | 0.025 |
| NC_005831.2;YP_003766.2;HCoV-NL63_1ab;3833 | 0 | 0 |
| NC_005831.2;YP_003766.2;HCoV-NL63_1ab;3834 | 0 | 0 |
| NC_005831.2;YP_003766.2;HCoV-NL63_1ab;3835 | 0 | 0 |
| NC_005831.2;YP_003766.2;HCoV-NL63_1ab;3836 | 0 | 0 |

| | | |
|---|---|---|
| NC_005831.2;YP_003766.2;HCoV-NL63_1ab;3837 | 0 | 0 |
| NC_005831.2;YP_003766.2;HCoV-NL63_1ab;3838 | 0 | 0 |
| NC_005831.2;YP_003766.2;HCoV-NL63_1ab;3839 | 0 | 0 |
| NC_005831.2;YP_003766.2;HCoV-NL63_1ab;3840 | 0.05 | 0.05 |
| NC_005831.2;YP_003766.2;HCoV-NL63_1ab;3841 | 0.15 | 0 |
| NC_005831.2;YP_003766.2;HCoV-NL63_1ab;3842 | 0.1 | 0.05 |
| NC_005831.2;YP_003766.2;HCoV-NL63_1ab;3843 | 0.1 | 0 |
| NC_005831.2;YP_003766.2;HCoV-NL63_1ab;3844 | 0 | 0 |
| NC_005831.2;YP_003766.2;HCoV-NL63_1ab;3845 | 0.05 | 0 |
| NC_005831.2;YP_003766.2;HCoV-NL63_1ab;3846 | 0 | 0 |
| NC_005831.2;YP_003766.2;HCoV-NL63_1ab;3847 | 0 | 0 |
| NC_005831.2;YP_003766.2;HCoV-NL63_1ab;3848 | 0 | 0 |
| NC_005831.2;YP_003766.2;HCoV-NL63_1ab;3849 | 0 | 0 |
| NC_005831.2;YP_003766.2;HCoV-NL63_1ab;3850 | 0.1 | 0 |
| NC_005831.2;YP_003766.2;HCoV-NL63_1ab;3851 | 0 | 0 |
| NC_005831.2;YP_003766.2;HCoV-NL63_1ab;3852 | 0.05 | 0 |
| NC_005831.2;YP_003766.2;HCoV-NL63_1ab;3853 | 0 | 0 |
| NC_005831.2;YP_003766.2;HCoV-NL63_1ab;3854 | 0 | 0 |
| NC_005831.2;YP_003766.2;HCoV-NL63_1ab;3855 | 0 | 0 |
| NC_005831.2;YP_003766.2;HCoV-NL63_1ab;3856 | 0 | 0.025 |
| NC_005831.2;YP_003766.2;HCoV-NL63_1ab;3857 | 0 | 0 |
| NC_005831.2;YP_003766.2;HCoV-NL63_1ab;3858 | 0 | 0 |
| NC_005831.2;YP_003766.2;HCoV-NL63_1ab;3859 | 0 | 0 |
| NC_005831.2;YP_003766.2;HCoV-NL63_1ab;3860 | 0 | 0 |
| NC_005831.2;YP_003766.2;HCoV-NL63_1ab;3861 | 0 | 0 |
| NC_005831.2;YP_003766.2;HCoV-NL63_1ab;3862 | 0 | 0 |
| NC_005831.2;YP_003766.2;HCoV-NL63_1ab;3863 | 0 | 0 |
| NC_005831.2;YP_003766.2;HCoV-NL63_1ab;3864 | 0 | 0 |
| NC_005831.2;YP_003766.2;HCoV-NL63_1ab;3865 | 0.05 | 0 |
| NC_005831.2;YP_003766.2;HCoV-NL63_1ab;3866 | 0 | 0 |
| NC_005831.2;YP_003766.2;HCoV-NL63_1ab;3867 | 0.05 | 0 |
| NC_005831.2;YP_003766.2;HCoV-NL63_1ab;3868 | 0 | 0 |
| NC_005831.2;YP_003766.2;HCoV-NL63_1ab;3869 | 0 | 0 |
| NC_005831.2;YP_003766.2;HCoV-NL63_1ab;3870 | 0 | 0 |
| NC_005831.2;YP_003766.2;HCoV-NL63_1ab;3871 | 0 | 0 |
| NC_005831.2;YP_003766.2;HCoV-NL63_1ab;3872 | 0 | 0 |
| NC_005831.2;YP_003766.2;HCoV-NL63_1ab;3873 | 0 | 0 |
| NC_005831.2;YP_003766.2;HCoV-NL63_1ab;3874 | 0 | 0 |
| NC_005831.2;YP_003766.2;HCoV-NL63_1ab;3875 | 0 | 0 |
| NC_005831.2;YP_003766.2;HCoV-NL63_1ab;3876 | 0 | 0 |
| NC_005831.2;YP_003766.2;HCoV-NL63_1ab;3877 | 0 | 0 |
| NC_005831.2;YP_003766.2;HCoV-NL63_1ab;3878 | 0 | 0 |
| NC_005831.2;YP_003766.2;HCoV-NL63_1ab;3879 | 0 | 0 |

| | | |
|---|---|---|
| NC_005831.2;YP_003766.2;HCoV-NL63_1ab;3880 | 0 | 0 |
| NC_005831.2;YP_003766.2;HCoV-NL63_1ab;3881 | 0 | 0 |
| NC_005831.2;YP_003766.2;HCoV-NL63_1ab;3882 | 0 | 0 |
| NC_005831.2;YP_003766.2;HCoV-NL63_1ab;3883 | 0 | 0 |
| NC_005831.2;YP_003766.2;HCoV-NL63_1ab;3884 | 0 | 0 |
| NC_005831.2;YP_003766.2;HCoV-NL63_1ab;3885 | 0 | 0 |
| NC_005831.2;YP_003766.2;HCoV-NL63_1ab;3886 | 0 | 0 |
| NC_005831.2;YP_003766.2;HCoV-NL63_1ab;3887 | 0 | 0 |
| NC_005831.2;YP_003766.2;HCoV-NL63_1ab;3888 | 0 | 0 |
| NC_005831.2;YP_003766.2;HCoV-NL63_1ab;3889 | 0 | 0 |
| NC_005831.2;YP_003766.2;HCoV-NL63_1ab;3890 | 0 | 0 |
| NC_005831.2;YP_003766.2;HCoV-NL63_1ab;3891 | 0 | 0 |
| NC_005831.2;YP_003766.2;HCoV-NL63_1ab;3892 | 0 | 0 |
| NC_005831.2;YP_003766.2;HCoV-NL63_1ab;3893 | 0 | 0 |
| NC_005831.2;YP_003766.2;HCoV-NL63_1ab;3894 | 0 | 0 |
| NC_005831.2;YP_003766.2;HCoV-NL63_1ab;3895 | 0 | 0 |
| NC_005831.2;YP_003766.2;HCoV-NL63_1ab;3896 | 0 | 0 |
| NC_005831.2;YP_003766.2;HCoV-NL63_1ab;3897 | 0 | 0 |
| NC_005831.2;YP_003766.2;HCoV-NL63_1ab;3898 | 0 | 0 |
| NC_005831.2;YP_003766.2;HCoV-NL63_1ab;3899 | 0.05 | 0 |
| NC_005831.2;YP_003766.2;HCoV-NL63_1ab;3900 | 0.05 | 0.025 |
| NC_005831.2;YP_003766.2;HCoV-NL63_1ab;3901 | 0.05 | 0.025 |
| NC_005831.2;YP_003766.2;HCoV-NL63_1ab;3902 | 0.05 | 0.025 |
| NC_005831.2;YP_003766.2;HCoV-NL63_1ab;3903 | 0.05 | 0.025 |
| NC_005831.2;YP_003766.2;HCoV-NL63_1ab;3904 | 0 | 0.025 |
| NC_005831.2;YP_003766.2;HCoV-NL63_1ab;3905 | 0 | 0 |
| NC_005831.2;YP_003766.2;HCoV-NL63_1ab;3906 | 0 | 0 |
| NC_005831.2;YP_003766.2;HCoV-NL63_1ab;3907 | 0 | 0 |
| NC_005831.2;YP_003766.2;HCoV-NL63_1ab;3908 | 0 | 0 |
| NC_005831.2;YP_003766.2;HCoV-NL63_1ab;3909 | 0 | 0 |
| NC_005831.2;YP_003766.2;HCoV-NL63_1ab;3910 | 0 | 0 |
| NC_005831.2;YP_003766.2;HCoV-NL63_1ab;3911 | 0 | 0 |
| NC_005831.2;YP_003766.2;HCoV-NL63_1ab;3912 | 0 | 0 |
| NC_005831.2;YP_003766.2;HCoV-NL63_1ab;3913 | 0 | 0 |
| NC_005831.2;YP_003766.2;HCoV-NL63_1ab;3914 | 0 | 0 |
| NC_005831.2;YP_003766.2;HCoV-NL63_1ab;3915 | 0 | 0 |
| NC_005831.2;YP_003766.2;HCoV-NL63_1ab;3916 | 0 | 0 |
| NC_005831.2;YP_003766.2;HCoV-NL63_1ab;3917 | 0 | 0 |
| NC_005831.2;YP_003766.2;HCoV-NL63_1ab;3918 | 0 | 0 |
| NC_005831.2;YP_003766.2;HCoV-NL63_1ab;3919 | 0 | 0 |
| NC_005831.2;YP_003766.2;HCoV-NL63_1ab;3920 | 0.05 | 0.025 |
| NC_005831.2;YP_003766.2;HCoV-NL63_1ab;3921 | 0 | 0 |
| NC_005831.2;YP_003766.2;HCoV-NL63_1ab;3922 | 0 | 0 |

| | | |
|---|---|---|
| NC_005831.2;YP_003766.2;HCoV-NL63_1ab;3923 | 0 | 0.025 |
| NC_005831.2;YP_003766.2;HCoV-NL63_1ab;3924 | 0 | 0 |
| NC_005831.2;YP_003766.2;HCoV-NL63_1ab;3925 | 0 | 0 |
| NC_005831.2;YP_003766.2;HCoV-NL63_1ab;3926 | 0 | 0 |
| NC_005831.2;YP_003766.2;HCoV-NL63_1ab;3927 | 0 | 0.025 |
| NC_005831.2;YP_003766.2;HCoV-NL63_1ab;3928 | 0 | 0.025 |
| NC_005831.2;YP_003766.2;HCoV-NL63_1ab;3929 | 0 | 0.025 |
| NC_005831.2;YP_003766.2;HCoV-NL63_1ab;3930 | 0 | 0 |
| NC_005831.2;YP_003766.2;HCoV-NL63_1ab;3931 | 0 | 0 |
| NC_005831.2;YP_003766.2;HCoV-NL63_1ab;3932 | 0 | 0.025 |
| NC_005831.2;YP_003766.2;HCoV-NL63_1ab;3933 | 0 | 0.025 |
| NC_005831.2;YP_003766.2;HCoV-NL63_1ab;3934 | 0 | 0 |
| NC_005831.2;YP_003766.2;HCoV-NL63_1ab;3935 | 0 | 0 |
| NC_005831.2;YP_003766.2;HCoV-NL63_1ab;3936 | 0 | 0 |
| NC_005831.2;YP_003766.2;HCoV-NL63_1ab;3937 | 0 | 0 |
| NC_005831.2;YP_003766.2;HCoV-NL63_1ab;3938 | 0 | 0 |
| NC_005831.2;YP_003766.2;HCoV-NL63_1ab;3939 | 0 | 0 |
| NC_005831.2;YP_003766.2;HCoV-NL63_1ab;3940 | 0 | 0 |
| NC_005831.2;YP_003766.2;HCoV-NL63_1ab;3941 | 0 | 0 |
| NC_005831.2;YP_003766.2;HCoV-NL63_1ab;3942 | 0 | 0 |
| NC_005831.2;YP_003766.2;HCoV-NL63_1ab;3943 | 0 | 0 |
| NC_005831.2;YP_003766.2;HCoV-NL63_1ab;3944 | 0 | 0 |
| NC_005831.2;YP_003766.2;HCoV-NL63_1ab;3945 | 0.05 | 0 |
| NC_005831.2;YP_003766.2;HCoV-NL63_1ab;3946 | 0.05 | 0.1 |
| NC_005831.2;YP_003766.2;HCoV-NL63_1ab;3947 | 0.05 | 0 |
| NC_005831.2;YP_003766.2;HCoV-NL63_1ab;3948 | 0 | 0 |
| NC_005831.2;YP_003766.2;HCoV-NL63_1ab;3949 | 0 | 0 |
| NC_005831.2;YP_003766.2;HCoV-NL63_1ab;3950 | 0.05 | 0 |
| NC_005831.2;YP_003766.2;HCoV-NL63_1ab;3951 | 0 | 0 |
| NC_005831.2;YP_003766.2;HCoV-NL63_1ab;3952 | 0 | 0 |
| NC_005831.2;YP_003766.2;HCoV-NL63_1ab;3953 | 0 | 0 |
| NC_005831.2;YP_003766.2;HCoV-NL63_1ab;3954 | 0 | 0 |
| NC_005831.2;YP_003766.2;HCoV-NL63_1ab;3955 | 0 | 0 |
| NC_005831.2;YP_003766.2;HCoV-NL63_1ab;3956 | 0 | 0.025 |
| NC_005831.2;YP_003766.2;HCoV-NL63_1ab;3957 | 0 | 0 |
| NC_005831.2;YP_003766.2;HCoV-NL63_1ab;3958 | 0 | 0 |
| NC_005831.2;YP_003766.2;HCoV-NL63_1ab;3959 | 0 | 0.025 |
| NC_005831.2;YP_003766.2;HCoV-NL63_1ab;3960 | 0 | 0 |
| NC_005831.2;YP_003766.2;HCoV-NL63_1ab;3961 | 0 | 0 |
| NC_005831.2;YP_003766.2;HCoV-NL63_1ab;3962 | 0.05 | 0 |
| NC_005831.2;YP_003766.2;HCoV-NL63_1ab;3963 | 0.1 | 0.05 |
| NC_005831.2;YP_003766.2;HCoV-NL63_1ab;3964 | 0 | 0 |
| NC_005831.2;YP_003766.2;HCoV-NL63_1ab;3965 | 0 | 0.05 |

| | | |
|---|---|---|
| NC_005831.2;YP_003766.2;HCoV-NL63_1ab;3966 | 0 | 0 |
| NC_005831.2;YP_003766.2;HCoV-NL63_1ab;3967 | 0 | 0.025 |
| NC_005831.2;YP_003766.2;HCoV-NL63_1ab;3968 | 0 | 0 |
| NC_005831.2;YP_003766.2;HCoV-NL63_1ab;3969 | 0 | 0.025 |
| NC_005831.2;YP_003766.2;HCoV-NL63_1ab;3970 | 0 | 0 |
| NC_005831.2;YP_003766.2;HCoV-NL63_1ab;3971 | 0 | 0 |
| NC_005831.2;YP_003766.2;HCoV-NL63_1ab;3972 | 0 | 0 |
| NC_005831.2;YP_003766.2;HCoV-NL63_1ab;3973 | 0 | 0 |
| NC_005831.2;YP_003766.2;HCoV-NL63_1ab;3974 | 0 | 0 |
| NC_005831.2;YP_003766.2;HCoV-NL63_1ab;3975 | 0 | 0 |
| NC_005831.2;YP_003766.2;HCoV-NL63_1ab;3976 | 0 | 0 |
| NC_005831.2;YP_003766.2;HCoV-NL63_1ab;3977 | 0 | 0 |
| NC_005831.2;YP_003766.2;HCoV-NL63_1ab;3978 | 0 | 0 |
| NC_005831.2;YP_003766.2;HCoV-NL63_1ab;3979 | 0 | 0 |
| NC_005831.2;YP_003766.2;HCoV-NL63_1ab;3980 | 0 | 0 |
| NC_005831.2;YP_003766.2;HCoV-NL63_1ab;3981 | 0 | 0 |
| NC_005831.2;YP_003766.2;HCoV-NL63_1ab;3982 | 0 | 0 |
| NC_005831.2;YP_003766.2;HCoV-NL63_1ab;3983 | 0 | 0 |
| NC_005831.2;YP_003766.2;HCoV-NL63_1ab;3984 | 0 | 0 |
| NC_005831.2;YP_003766.2;HCoV-NL63_1ab;3985 | 0 | 0 |
| NC_005831.2;YP_003766.2;HCoV-NL63_1ab;3986 | 0 | 0 |
| NC_005831.2;YP_003766.2;HCoV-NL63_1ab;3987 | 0 | 0 |
| NC_005831.2;YP_003766.2;HCoV-NL63_1ab;3988 | 0 | 0.025 |
| NC_005831.2;YP_003766.2;HCoV-NL63_1ab;3989 | 0 | 0 |
| NC_005831.2;YP_003766.2;HCoV-NL63_1ab;3990 | 0 | 0 |
| NC_005831.2;YP_003766.2;HCoV-NL63_1ab;3991 | 0 | 0 |
| NC_005831.2;YP_003766.2;HCoV-NL63_1ab;3992 | 0 | 0.025 |
| NC_005831.2;YP_003766.2;HCoV-NL63_1ab;3993 | 0 | 0 |
| NC_005831.2;YP_003766.2;HCoV-NL63_1ab;3994 | 0 | 0 |
| NC_005831.2;YP_003766.2;HCoV-NL63_1ab;3995 | 0 | 0 |
| NC_005831.2;YP_003766.2;HCoV-NL63_1ab;3996 | 0 | 0 |
| NC_005831.2;YP_003766.2;HCoV-NL63_1ab;3997 | 0 | 0 |
| NC_005831.2;YP_003766.2;HCoV-NL63_1ab;3998 | 0 | 0.025 |
| NC_005831.2;YP_003766.2;HCoV-NL63_1ab;3999 | 0 | 0.05 |
| NC_005831.2;YP_003766.2;HCoV-NL63_1ab;4000 | 0 | 0 |
| NC_005831.2;YP_003766.2;HCoV-NL63_1ab;4001 | 0 | 0 |
| NC_005831.2;YP_003766.2;HCoV-NL63_1ab;4002 | 0 | 0 |
| NC_005831.2;YP_003766.2;HCoV-NL63_1ab;4003 | 0 | 0 |
| NC_005831.2;YP_003766.2;HCoV-NL63_1ab;4004 | 0 | 0 |
| NC_005831.2;YP_003766.2;HCoV-NL63_1ab;4005 | 0 | 0 |
| NC_005831.2;YP_003766.2;HCoV-NL63_1ab;4006 | 0 | 0 |
| NC_005831.2;YP_003766.2;HCoV-NL63_1ab;4007 | 0 | 0 |
| NC_005831.2;YP_003766.2;HCoV-NL63_1ab;4008 | 0 | 0 |

| | | |
|---|---|---|
| NC_005831.2;YP_003766.2;HCoV-NL63_1ab;4009 | 0 | 0 |
| NC_005831.2;YP_003766.2;HCoV-NL63_1ab;4010 | 0 | 0 |
| NC_005831.2;YP_003766.2;HCoV-NL63_1ab;4011 | 0 | 0 |
| NC_005831.2;YP_003766.2;HCoV-NL63_1ab;4012 | 0 | 0 |
| NC_005831.2;YP_003766.2;HCoV-NL63_1ab;4013 | 0 | 0.025 |
| NC_005831.2;YP_003766.2;HCoV-NL63_1ab;4014 | 0 | 0 |
| NC_005831.2;YP_003766.2;HCoV-NL63_1ab;4015 | 0 | 0 |
| NC_005831.2;YP_003766.2;HCoV-NL63_1ab;4016 | 0 | 0 |
| NC_005831.2;YP_003766.2;HCoV-NL63_1ab;4017 | 0 | 0 |
| NC_005831.2;YP_003766.2;HCoV-NL63_1ab;4018 | 0 | 0 |
| NC_005831.2;YP_003766.2;HCoV-NL63_1ab;4019 | 0 | 0 |
| NC_005831.2;YP_003766.2;HCoV-NL63_1ab;4020 | 0 | 0 |
| NC_005831.2;YP_003766.2;HCoV-NL63_1ab;4021 | 0 | 0 |
| NC_005831.2;YP_003766.2;HCoV-NL63_1ab;4022 | 0 | 0 |
| NC_005831.2;YP_003766.2;HCoV-NL63_1ab;4023 | 0 | 0 |
| NC_005831.2;YP_003766.2;HCoV-NL63_1ab;4024 | 0 | 0 |
| NC_005831.2;YP_003766.2;HCoV-NL63_1ab;4025 | 0 | 0 |
| NC_005831.2;YP_003766.2;HCoV-NL63_1ab;4026 | 0 | 0 |
| NC_005831.2;YP_003766.2;HCoV-NL63_1ab;4027 | 0 | 0 |
| NC_005831.2;YP_003766.2;HCoV-NL63_1ab;4028 | 0 | 0 |
| NC_005831.2;YP_003766.2;HCoV-NL63_1ab;4029 | 0 | 0 |
| NC_005831.2;YP_003766.2;HCoV-NL63_1ab;4030 | 0 | 0 |
| NC_005831.2;YP_003766.2;HCoV-NL63_1ab;4031 | 0.05 | 0.025 |
| NC_005831.2;YP_003766.2;HCoV-NL63_1ab;4032 | 0 | 0 |
| NC_005831.2;YP_003766.2;HCoV-NL63_1ab;4033 | 0.05 | 0.075 |
| NC_005831.2;YP_003766.2;HCoV-NL63_1ab;4034 | 0.05 | 0.05 |
| NC_005831.2;YP_003766.2;HCoV-NL63_1ab;4035 | 0 | 0 |
| NC_005831.2;YP_003766.2;HCoV-NL63_1ab;4036 | 0 | 0 |
| NC_005831.2;YP_003766.2;HCoV-NL63_1ab;4037 | 0 | 0 |
| NC_005831.2;YP_003766.2;HCoV-NL63_1ab;4038 | 0 | 0 |
| NC_005831.2;YP_003766.2;HCoV-NL63_1ab;4039 | 0 | 0 |
| NC_005831.2;YP_003766.2;HCoV-NL63_1ab;4040 | 0 | 0 |
| NC_005831.2;YP_003766.2;HCoV-NL63_1ab;4041 | 0 | 0 |
| NC_005831.2;YP_003766.2;HCoV-NL63_1ab;4042 | 0 | 0 |
| NC_005831.2;YP_003766.2;HCoV-NL63_1ab;4043 | 0 | 0 |
| NC_005831.2;YP_003766.2;HCoV-NL63_1ab;4044 | 0.05 | 0.025 |
| NC_005831.2;YP_003766.2;HCoV-NL63_1ab;4045 | 0 | 0 |
| NC_005831.2;YP_003766.2;HCoV-NL63_1ab;4046 | 0 | 0 |
| NC_005831.2;YP_003766.2;HCoV-NL63_1ab;4047 | 0 | 0 |
| NC_005831.2;YP_003766.2;HCoV-NL63_1ab;4048 | 0.05 | 0 |
| NC_005831.2;YP_003766.2;HCoV-NL63_1ab;4049 | 0.05 | 0 |
| NC_005831.2;YP_003766.2;HCoV-NL63_1ab;4050 | 0 | 0.025 |
| NC_005831.2;YP_003766.2;HCoV-NL63_1ab;4051 | 0 | 0 |

| | | |
|---|---|---|
| NC_005831.2;YP_003766.2;HCoV-NL63_1ab;4052 | 0 | 0 |
| NC_005831.2;YP_003766.2;HCoV-NL63_1ab;4053 | 0.05 | 0 |
| NC_005831.2;YP_003766.2;HCoV-NL63_1ab;4054 | 0 | 0 |
| NC_005831.2;YP_003766.2;HCoV-NL63_1ab;4055 | 0 | 0 |
| NC_005831.2;YP_003766.2;HCoV-NL63_1ab;4056 | 0 | 0 |
| NC_005831.2;YP_003766.2;HCoV-NL63_1ab;4057 | 0 | 0.05 |
| NC_005831.2;YP_003766.2;HCoV-NL63_1ab;4058 | 0 | 0 |
| NC_005831.2;YP_003766.2;HCoV-NL63_1ab;4059 | 0 | 0.025 |
| NC_005831.2;YP_003766.2;HCoV-NL63_1ab;4060 | 0 | 0.025 |
| NC_005831.2;YP_003766.2;HCoV-NL63_1ab;4061 | 0 | 0 |
| NC_005831.2;YP_003766.2;HCoV-NL63_1ab;4062 | 0 | 0 |
| NC_005831.2;YP_003766.2;HCoV-NL63_1ab;4063 | 0 | 0 |
| NC_005831.2;YP_003766.2;HCoV-NL63_1ab;4064 | 0 | 0 |
| NC_005831.2;YP_003766.2;HCoV-NL63_1ab;4065 | 0 | 0.025 |
| NC_005831.2;YP_003766.2;HCoV-NL63_1ab;4066 | 0 | 0 |
| NC_005831.2;YP_003766.2;HCoV-NL63_1ab;4067 | 0 | 0 |
| NC_005831.2;YP_003766.2;HCoV-NL63_1ab;4068 | 0 | 0 |
| NC_005831.2;YP_003766.2;HCoV-NL63_1ab;4069 | 0 | 0 |
| NC_005831.2;YP_003766.2;HCoV-NL63_1ab;4070 | 0 | 0 |
| NC_005831.2;YP_003766.2;HCoV-NL63_1ab;4071 | 0 | 0 |
| NC_005831.2;YP_003766.2;HCoV-NL63_1ab;4072 | 0 | 0 |
| NC_005831.2;YP_003766.2;HCoV-NL63_1ab;4073 | 0 | 0 |
| NC_005831.2;YP_003766.2;HCoV-NL63_1ab;4074 | 0 | 0 |
| NC_005831.2;YP_003766.2;HCoV-NL63_1ab;4075 | 0 | 0 |
| NC_005831.2;YP_003766.2;HCoV-NL63_1ab;4076 | 0 | 0 |
| NC_005831.2;YP_003766.2;HCoV-NL63_1ab;4077 | 0.05 | 0 |
| NC_005831.2;YP_003766.2;HCoV-NL63_1ab;4078 | 0.05 | 0 |
| NC_005831.2;YP_003766.2;HCoV-NL63_1ab;4079 | 0.05 | 0 |
| NC_005831.2;YP_003766.2;HCoV-NL63_1ab;4080 | 0.05 | 0 |
| NC_005831.2;YP_003766.2;HCoV-NL63_1ab;4081 | 0.05 | 0 |
| NC_005831.2;YP_003766.2;HCoV-NL63_1ab;4082 | 0 | 0 |
| NC_005831.2;YP_003766.2;HCoV-NL63_1ab;4083 | 0 | 0 |
| NC_005831.2;YP_003766.2;HCoV-NL63_1ab;4084 | 0.05 | 0 |
| NC_005831.2;YP_003766.2;HCoV-NL63_1ab;4085 | 0.05 | 0 |
| NC_005831.2;YP_003766.2;HCoV-NL63_1ab;4086 | 0 | 0 |
| NC_005831.2;YP_003766.2;HCoV-NL63_1ab;4087 | 0 | 0.025 |
| NC_005831.2;YP_003766.2;HCoV-NL63_1ab;4088 | 0 | 0 |
| NC_005831.2;YP_003766.2;HCoV-NL63_1ab;4089 | 0 | 0 |
| NC_005831.2;YP_003766.2;HCoV-NL63_1ab;4090 | 0 | 0 |
| NC_005831.2;YP_003766.2;HCoV-NL63_1ab;4091 | 0 | 0.025 |
| NC_005831.2;YP_003766.2;HCoV-NL63_1ab;4092 | 0 | 0 |
| NC_005831.2;YP_003766.2;HCoV-NL63_1ab;4093 | 0 | 0 |
| NC_005831.2;YP_003766.2;HCoV-NL63_1ab;4094 | 0 | 0 |

| | | |
|---|---|---|
| NC_005831.2;YP_003766.2;HCoV-NL63_1ab;4095 | 0 | 0 |
| NC_005831.2;YP_003766.2;HCoV-NL63_1ab;4096 | 0 | 0 |
| NC_005831.2;YP_003766.2;HCoV-NL63_1ab;4097 | 0 | 0 |
| NC_005831.2;YP_003766.2;HCoV-NL63_1ab;4098 | 0 | 0 |
| NC_005831.2;YP_003766.2;HCoV-NL63_1ab;4099 | 0 | 0 |
| NC_005831.2;YP_003766.2;HCoV-NL63_1ab;4100 | 0 | 0 |
| NC_005831.2;YP_003766.2;HCoV-NL63_1ab;4101 | 0 | 0 |
| NC_005831.2;YP_003766.2;HCoV-NL63_1ab;4102 | 0 | 0 |
| NC_005831.2;YP_003766.2;HCoV-NL63_1ab;4103 | 0 | 0 |
| NC_005831.2;YP_003766.2;HCoV-NL63_1ab;4104 | 0 | 0 |
| NC_005831.2;YP_003766.2;HCoV-NL63_1ab;4105 | 0 | 0 |
| NC_005831.2;YP_003766.2;HCoV-NL63_1ab;4106 | 0 | 0 |
| NC_005831.2;YP_003766.2;HCoV-NL63_1ab;4107 | 0 | 0 |
| NC_005831.2;YP_003766.2;HCoV-NL63_1ab;4108 | 0 | 0 |
| NC_005831.2;YP_003766.2;HCoV-NL63_1ab;4109 | 0 | 0 |
| NC_005831.2;YP_003766.2;HCoV-NL63_1ab;4110 | 0 | 0.025 |
| NC_005831.2;YP_003766.2;HCoV-NL63_1ab;4111 | 0 | 0 |
| NC_005831.2;YP_003766.2;HCoV-NL63_1ab;4112 | 0 | 0 |
| NC_005831.2;YP_003766.2;HCoV-NL63_1ab;4113 | 0 | 0 |
| NC_005831.2;YP_003766.2;HCoV-NL63_1ab;4114 | 0 | 0.025 |
| NC_005831.2;YP_003766.2;HCoV-NL63_1ab;4115 | 0 | 0 |
| NC_005831.2;YP_003766.2;HCoV-NL63_1ab;4116 | 0 | 0 |
| NC_005831.2;YP_003766.2;HCoV-NL63_1ab;4117 | 0 | 0 |
| NC_005831.2;YP_003766.2;HCoV-NL63_1ab;4118 | 0 | 0 |
| NC_005831.2;YP_003766.2;HCoV-NL63_1ab;4119 | 0 | 0 |
| NC_005831.2;YP_003766.2;HCoV-NL63_1ab;4120 | 0 | 0 |
| NC_005831.2;YP_003766.2;HCoV-NL63_1ab;4121 | 0 | 0 |
| NC_005831.2;YP_003766.2;HCoV-NL63_1ab;4122 | 0 | 0 |
| NC_005831.2;YP_003766.2;HCoV-NL63_1ab;4123 | 0 | 0 |
| NC_005831.2;YP_003766.2;HCoV-NL63_1ab;4124 | 0 | 0 |
| NC_005831.2;YP_003766.2;HCoV-NL63_1ab;4125 | 0 | 0 |
| NC_005831.2;YP_003766.2;HCoV-NL63_1ab;4126 | 0 | 0 |
| NC_005831.2;YP_003766.2;HCoV-NL63_1ab;4127 | 0 | 0 |
| NC_005831.2;YP_003766.2;HCoV-NL63_1ab;4128 | 0 | 0 |
| NC_005831.2;YP_003766.2;HCoV-NL63_1ab;4129 | 0 | 0 |
| NC_005831.2;YP_003766.2;HCoV-NL63_1ab;4130 | 0 | 0 |
| NC_005831.2;YP_003766.2;HCoV-NL63_1ab;4131 | 0 | 0 |
| NC_005831.2;YP_003766.2;HCoV-NL63_1ab;4132 | 0 | 0 |
| NC_005831.2;YP_003766.2;HCoV-NL63_1ab;4133 | 0 | 0 |
| NC_005831.2;YP_003766.2;HCoV-NL63_1ab;4134 | 0 | 0.025 |
| NC_005831.2;YP_003766.2;HCoV-NL63_1ab;4135 | 0 | 0 |
| NC_005831.2;YP_003766.2;HCoV-NL63_1ab;4136 | 0 | 0 |
| NC_005831.2;YP_003766.2;HCoV-NL63_1ab;4137 | 0 | 0 |

| | | |
|---|---|---|
| NC_005831.2;YP_003766.2;HCoV-NL63_1ab;4138 | 0 | 0 |
| NC_005831.2;YP_003766.2;HCoV-NL63_1ab;4139 | 0 | 0 |
| NC_005831.2;YP_003766.2;HCoV-NL63_1ab;4140 | 0 | 0 |
| NC_005831.2;YP_003766.2;HCoV-NL63_1ab;4141 | 0 | 0 |
| NC_005831.2;YP_003766.2;HCoV-NL63_1ab;4142 | 0 | 0 |
| NC_005831.2;YP_003766.2;HCoV-NL63_1ab;4143 | 0 | 0 |
| NC_005831.2;YP_003766.2;HCoV-NL63_1ab;4144 | 0 | 0 |
| NC_005831.2;YP_003766.2;HCoV-NL63_1ab;4145 | 0 | 0 |
| NC_005831.2;YP_003766.2;HCoV-NL63_1ab;4146 | 0 | 0 |
| NC_005831.2;YP_003766.2;HCoV-NL63_1ab;4147 | 0.05 | 0 |
| NC_005831.2;YP_003766.2;HCoV-NL63_1ab;4148 | 0.05 | 0.05 |
| NC_005831.2;YP_003766.2;HCoV-NL63_1ab;4149 | 0.05 | 0.025 |
| NC_005831.2;YP_003766.2;HCoV-NL63_1ab;4150 | 0.05 | 0 |
| NC_005831.2;YP_003766.2;HCoV-NL63_1ab;4151 | 0 | 0 |
| NC_005831.2;YP_003766.2;HCoV-NL63_1ab;4152 | 0 | 0 |
| NC_005831.2;YP_003766.2;HCoV-NL63_1ab;4153 | 0 | 0 |
| NC_005831.2;YP_003766.2;HCoV-NL63_1ab;4154 | 0 | 0 |
| NC_005831.2;YP_003766.2;HCoV-NL63_1ab;4155 | 0 | 0 |
| NC_005831.2;YP_003766.2;HCoV-NL63_1ab;4156 | 0 | 0 |
| NC_005831.2;YP_003766.2;HCoV-NL63_1ab;4157 | 0 | 0 |
| NC_005831.2;YP_003766.2;HCoV-NL63_1ab;4158 | 0.05 | 0 |
| NC_005831.2;YP_003766.2;HCoV-NL63_1ab;4159 | 0.05 | 0.375 |
| NC_005831.2;YP_003766.2;HCoV-NL63_1ab;4160 | 0.05 | 0 |
| NC_005831.2;YP_003766.2;HCoV-NL63_1ab;4161 | 0.05 | 0 |
| NC_005831.2;YP_003766.2;HCoV-NL63_1ab;4162 | 0.05 | 0 |
| NC_005831.2;YP_003766.2;HCoV-NL63_1ab;4163 | 0.05 | 0 |
| NC_005831.2;YP_003766.2;HCoV-NL63_1ab;4164 | 0.05 | 0 |
| NC_005831.2;YP_003766.2;HCoV-NL63_1ab;4165 | 0.1 | 0.05 |
| NC_005831.2;YP_003766.2;HCoV-NL63_1ab;4166 | 0 | 0 |
| NC_005831.2;YP_003766.2;HCoV-NL63_1ab;4167 | 0 | 0 |
| NC_005831.2;YP_003766.2;HCoV-NL63_1ab;4168 | 0.05 | 0.025 |
| NC_005831.2;YP_003766.2;HCoV-NL63_1ab;4169 | 0.15 | 0.2 |
| NC_005831.2;YP_003766.2;HCoV-NL63_1ab;4170 | 0.05 | 0.075 |
| NC_005831.2;YP_003766.2;HCoV-NL63_1ab;4171 | 0.05 | 0.05 |
| NC_005831.2;YP_003766.2;HCoV-NL63_1ab;4172 | 0.05 | 0.025 |
| NC_005831.2;YP_003766.2;HCoV-NL63_1ab;4173 | 0 | 0 |
| NC_005831.2;YP_003766.2;HCoV-NL63_1ab;4174 | 0 | 0 |
| NC_005831.2;YP_003766.2;HCoV-NL63_1ab;4175 | 0 | 0 |
| NC_005831.2;YP_003766.2;HCoV-NL63_1ab;4176 | 0 | 0 |
| NC_005831.2;YP_003766.2;HCoV-NL63_1ab;4177 | 0 | 0 |
| NC_005831.2;YP_003766.2;HCoV-NL63_1ab;4178 | 0 | 0 |
| NC_005831.2;YP_003766.2;HCoV-NL63_1ab;4179 | 0 | 0 |
| NC_005831.2;YP_003766.2;HCoV-NL63_1ab;4180 | 0 | 0 |

| | | |
|---|---|---|
| NC_005831.2;YP_003766.2;HCoV-NL63_1ab;4181 | 0 | 0 |
| NC_005831.2;YP_003766.2;HCoV-NL63_1ab;4182 | 0 | 0 |
| NC_005831.2;YP_003766.2;HCoV-NL63_1ab;4183 | 0 | 0 |
| NC_005831.2;YP_003766.2;HCoV-NL63_1ab;4184 | 0 | 0.025 |
| NC_005831.2;YP_003766.2;HCoV-NL63_1ab;4185 | 0 | 0.025 |
| NC_005831.2;YP_003766.2;HCoV-NL63_1ab;4186 | 0 | 0.025 |
| NC_005831.2;YP_003766.2;HCoV-NL63_1ab;4187 | 0 | 0.025 |
| NC_005831.2;YP_003766.2;HCoV-NL63_1ab;4188 | 0 | 0 |
| NC_005831.2;YP_003766.2;HCoV-NL63_1ab;4189 | 0 | 0.025 |
| NC_005831.2;YP_003766.2;HCoV-NL63_1ab;4190 | 0 | 0.025 |
| NC_005831.2;YP_003766.2;HCoV-NL63_1ab;4191 | 0 | 0.025 |
| NC_005831.2;YP_003766.2;HCoV-NL63_1ab;4192 | 0 | 0.05 |
| NC_005831.2;YP_003766.2;HCoV-NL63_1ab;4193 | 0 | 0.025 |
| NC_005831.2;YP_003766.2;HCoV-NL63_1ab;4194 | 0 | 0.025 |
| NC_005831.2;YP_003766.2;HCoV-NL63_1ab;4195 | 0.05 | 0.025 |
| NC_005831.2;YP_003766.2;HCoV-NL63_1ab;4196 | 0.05 | 0.025 |
| NC_005831.2;YP_003766.2;HCoV-NL63_1ab;4197 | 0 | 0.05 |
| NC_005831.2;YP_003766.2;HCoV-NL63_1ab;4198 | 0 | 0.05 |
| NC_005831.2;YP_003766.2;HCoV-NL63_1ab;4199 | 0 | 0.025 |
| NC_005831.2;YP_003766.2;HCoV-NL63_1ab;4200 | 0 | 0.025 |
| NC_005831.2;YP_003766.2;HCoV-NL63_1ab;4201 | 0.05 | 0 |
| NC_005831.2;YP_003766.2;HCoV-NL63_1ab;4202 | 0 | 0 |
| NC_005831.2;YP_003766.2;HCoV-NL63_1ab;4203 | 0.05 | 0 |
| NC_005831.2;YP_003766.2;HCoV-NL63_1ab;4204 | 0 | 0 |
| NC_005831.2;YP_003766.2;HCoV-NL63_1ab;4205 | 0 | 0 |
| NC_005831.2;YP_003766.2;HCoV-NL63_1ab;4206 | 0 | 0 |
| NC_005831.2;YP_003766.2;HCoV-NL63_1ab;4207 | 0 | 0 |
| NC_005831.2;YP_003766.2;HCoV-NL63_1ab;4208 | 0.05 | 0 |
| NC_005831.2;YP_003766.2;HCoV-NL63_1ab;4209 | 0 | 0 |
| NC_005831.2;YP_003766.2;HCoV-NL63_1ab;4210 | 0 | 0 |
| NC_005831.2;YP_003766.2;HCoV-NL63_1ab;4211 | 0 | 0 |
| NC_005831.2;YP_003766.2;HCoV-NL63_1ab;4212 | 0 | 0 |
| NC_005831.2;YP_003766.2;HCoV-NL63_1ab;4213 | 0 | 0 |
| NC_005831.2;YP_003766.2;HCoV-NL63_1ab;4214 | 0 | 0 |
| NC_005831.2;YP_003766.2;HCoV-NL63_1ab;4215 | 0 | 0 |
| NC_005831.2;YP_003766.2;HCoV-NL63_1ab;4216 | 0 | 0 |
| NC_005831.2;YP_003766.2;HCoV-NL63_1ab;4217 | 0 | 0 |
| NC_005831.2;YP_003766.2;HCoV-NL63_1ab;4218 | 0 | 0 |
| NC_005831.2;YP_003766.2;HCoV-NL63_1ab;4219 | 0 | 0 |
| NC_005831.2;YP_003766.2;HCoV-NL63_1ab;4220 | 0 | 0.025 |
| NC_005831.2;YP_003766.2;HCoV-NL63_1ab;4221 | 0 | 0 |
| NC_005831.2;YP_003766.2;HCoV-NL63_1ab;4222 | 0 | 0 |
| NC_005831.2;YP_003766.2;HCoV-NL63_1ab;4223 | 0 | 0 |

| | | |
|---|---|---|
| NC_005831.2;YP_003766.2;HCoV-NL63_1ab;4224 | 0 | 0 |
| NC_005831.2;YP_003766.2;HCoV-NL63_1ab;4225 | 0 | 0 |
| NC_005831.2;YP_003766.2;HCoV-NL63_1ab;4226 | 0 | 0 |
| NC_005831.2;YP_003766.2;HCoV-NL63_1ab;4227 | 0 | 0 |
| NC_005831.2;YP_003766.2;HCoV-NL63_1ab;4228 | 0 | 0 |
| NC_005831.2;YP_003766.2;HCoV-NL63_1ab;4229 | 0 | 0 |
| NC_005831.2;YP_003766.2;HCoV-NL63_1ab;4230 | 0 | 0 |
| NC_005831.2;YP_003766.2;HCoV-NL63_1ab;4231 | 0 | 0 |
| NC_005831.2;YP_003766.2;HCoV-NL63_1ab;4232 | 0 | 0 |
| NC_005831.2;YP_003766.2;HCoV-NL63_1ab;4233 | 0 | 0 |
| NC_005831.2;YP_003766.2;HCoV-NL63_1ab;4234 | 0 | 0 |
| NC_005831.2;YP_003766.2;HCoV-NL63_1ab;4235 | 0 | 0 |
| NC_005831.2;YP_003766.2;HCoV-NL63_1ab;4236 | 0.05 | 0 |
| NC_005831.2;YP_003766.2;HCoV-NL63_1ab;4237 | 0 | 0 |
| NC_005831.2;YP_003766.2;HCoV-NL63_1ab;4238 | 0 | 0 |
| NC_005831.2;YP_003766.2;HCoV-NL63_1ab;4239 | 0 | 0 |
| NC_005831.2;YP_003766.2;HCoV-NL63_1ab;4240 | 0.1 | 0 |
| NC_005831.2;YP_003766.2;HCoV-NL63_1ab;4241 | 0 | 0 |
| NC_005831.2;YP_003766.2;HCoV-NL63_1ab;4242 | 0 | 0 |
| NC_005831.2;YP_003766.2;HCoV-NL63_1ab;4243 | 0 | 0 |
| NC_005831.2;YP_003766.2;HCoV-NL63_1ab;4244 | 0 | 0 |
| NC_005831.2;YP_003766.2;HCoV-NL63_1ab;4245 | 0 | 0 |
| NC_005831.2;YP_003766.2;HCoV-NL63_1ab;4246 | 0 | 0 |
| NC_005831.2;YP_003766.2;HCoV-NL63_1ab;4247 | 0 | 0 |
| NC_005831.2;YP_003766.2;HCoV-NL63_1ab;4248 | 0 | 0 |
| NC_005831.2;YP_003766.2;HCoV-NL63_1ab;4249 | 0 | 0 |
| NC_005831.2;YP_003766.2;HCoV-NL63_1ab;4250 | 0 | 0 |
| NC_005831.2;YP_003766.2;HCoV-NL63_1ab;4251 | 0 | 0 |
| NC_005831.2;YP_003766.2;HCoV-NL63_1ab;4252 | 0 | 0 |
| NC_005831.2;YP_003766.2;HCoV-NL63_1ab;4253 | 0 | 0 |
| NC_005831.2;YP_003766.2;HCoV-NL63_1ab;4254 | 0.05 | 0.075 |
| NC_005831.2;YP_003766.2;HCoV-NL63_1ab;4255 | 0 | 0 |
| NC_005831.2;YP_003766.2;HCoV-NL63_1ab;4256 | 0 | 0 |
| NC_005831.2;YP_003766.2;HCoV-NL63_1ab;4257 | 0 | 0 |
| NC_005831.2;YP_003766.2;HCoV-NL63_1ab;4258 | 0 | 0 |
| NC_005831.2;YP_003766.2;HCoV-NL63_1ab;4259 | 0 | 0 |
| NC_005831.2;YP_003766.2;HCoV-NL63_1ab;4260 | 0 | 0 |
| NC_005831.2;YP_003766.2;HCoV-NL63_1ab;4261 | 0 | 0 |
| NC_005831.2;YP_003766.2;HCoV-NL63_1ab;4262 | 0 | 0 |
| NC_005831.2;YP_003766.2;HCoV-NL63_1ab;4263 | 0 | 0 |
| NC_005831.2;YP_003766.2;HCoV-NL63_1ab;4264 | 0 | 0 |
| NC_005831.2;YP_003766.2;HCoV-NL63_1ab;4265 | 0 | 0 |
| NC_005831.2;YP_003766.2;HCoV-NL63_1ab;4266 | 0 | 0 |

| | | |
|---|---|---|
| NC_005831.2;YP_003766.2;HCoV-NL63_1ab;4267 | 0 | 0 |
| NC_005831.2;YP_003766.2;HCoV-NL63_1ab;4268 | 0 | 0 |
| NC_005831.2;YP_003766.2;HCoV-NL63_1ab;4269 | 0 | 0 |
| NC_005831.2;YP_003766.2;HCoV-NL63_1ab;4270 | 0 | 0 |
| NC_005831.2;YP_003766.2;HCoV-NL63_1ab;4271 | 0 | 0 |
| NC_005831.2;YP_003766.2;HCoV-NL63_1ab;4272 | 0 | 0 |
| NC_005831.2;YP_003766.2;HCoV-NL63_1ab;4273 | 0 | 0 |
| NC_005831.2;YP_003766.2;HCoV-NL63_1ab;4274 | 0 | 0 |
| NC_005831.2;YP_003766.2;HCoV-NL63_1ab;4275 | 0 | 0 |
| NC_005831.2;YP_003766.2;HCoV-NL63_1ab;4276 | 0 | 0 |
| NC_005831.2;YP_003766.2;HCoV-NL63_1ab;4277 | 0 | 0 |
| NC_005831.2;YP_003766.2;HCoV-NL63_1ab;4278 | 0 | 0 |
| NC_005831.2;YP_003766.2;HCoV-NL63_1ab;4279 | 0 | 0 |
| NC_005831.2;YP_003766.2;HCoV-NL63_1ab;4280 | 0 | 0 |
| NC_005831.2;YP_003766.2;HCoV-NL63_1ab;4281 | 0 | 0 |
| NC_005831.2;YP_003766.2;HCoV-NL63_1ab;4282 | 0 | 0 |
| NC_005831.2;YP_003766.2;HCoV-NL63_1ab;4283 | 0 | 0.025 |
| NC_005831.2;YP_003766.2;HCoV-NL63_1ab;4284 | 0 | 0 |
| NC_005831.2;YP_003766.2;HCoV-NL63_1ab;4285 | 0 | 0 |
| NC_005831.2;YP_003766.2;HCoV-NL63_1ab;4286 | 0 | 0 |
| NC_005831.2;YP_003766.2;HCoV-NL63_1ab;4287 | 0.05 | 0.025 |
| NC_005831.2;YP_003766.2;HCoV-NL63_1ab;4288 | 0.15 | 0.05 |
| NC_005831.2;YP_003766.2;HCoV-NL63_1ab;4289 | 0.15 | 0.125 |
| NC_005831.2;YP_003766.2;HCoV-NL63_1ab;4290 | 0.1 | 0.125 |
| NC_005831.2;YP_003766.2;HCoV-NL63_1ab;4291 | 0.1 | 0.05 |
| NC_005831.2;YP_003766.2;HCoV-NL63_1ab;4292 | 0.1 | 0.05 |
| NC_005831.2;YP_003766.2;HCoV-NL63_1ab;4293 | 0.1 | 0.075 |
| NC_005831.2;YP_003766.2;HCoV-NL63_1ab;4294 | 0.1 | 0.1 |
| NC_005831.2;YP_003766.2;HCoV-NL63_1ab;4295 | 0.1 | 0.1 |
| NC_005831.2;YP_003766.2;HCoV-NL63_1ab;4296 | 0 | 0.025 |
| NC_005831.2;YP_003766.2;HCoV-NL63_1ab;4297 | 0 | 0.025 |
| NC_005831.2;YP_003766.2;HCoV-NL63_1ab;4298 | 0 | 0 |
| NC_005831.2;YP_003766.2;HCoV-NL63_1ab;4299 | 0 | 0 |
| NC_005831.2;YP_003766.2;HCoV-NL63_1ab;4300 | 0 | 0 |
| NC_005831.2;YP_003766.2;HCoV-NL63_1ab;4301 | 0 | 0 |
| NC_005831.2;YP_003766.2;HCoV-NL63_1ab;4302 | 0 | 0 |
| NC_005831.2;YP_003766.2;HCoV-NL63_1ab;4303 | 0 | 0.025 |
| NC_005831.2;YP_003766.2;HCoV-NL63_1ab;4304 | 0 | 0.025 |
| NC_005831.2;YP_003766.2;HCoV-NL63_1ab;4305 | 0 | 0 |
| NC_005831.2;YP_003766.2;HCoV-NL63_1ab;4306 | 0 | 0 |
| NC_005831.2;YP_003766.2;HCoV-NL63_1ab;4307 | 0.15 | 0.025 |
| NC_005831.2;YP_003766.2;HCoV-NL63_1ab;4308 | 0.1 | 0.025 |
| NC_005831.2;YP_003766.2;HCoV-NL63_1ab;4309 | 0.1 | 0 |

| | | |
|---|---|---|
| NC_005831.2;YP_003766.2;HCoV-NL63_1ab;4310 | 0.05 | 0.025 |
| NC_005831.2;YP_003766.2;HCoV-NL63_1ab;4311 | 0 | 0 |
| NC_005831.2;YP_003766.2;HCoV-NL63_1ab;4312 | 0 | 0 |
| NC_005831.2;YP_003766.2;HCoV-NL63_1ab;4313 | 0 | 0 |
| NC_005831.2;YP_003766.2;HCoV-NL63_1ab;4314 | 0 | 0 |
| NC_005831.2;YP_003766.2;HCoV-NL63_1ab;4315 | 0 | 0 |
| NC_005831.2;YP_003766.2;HCoV-NL63_1ab;4316 | 0 | 0 |
| NC_005831.2;YP_003766.2;HCoV-NL63_1ab;4317 | 0 | 0 |
| NC_005831.2;YP_003766.2;HCoV-NL63_1ab;4318 | 0 | 0 |
| NC_005831.2;YP_003766.2;HCoV-NL63_1ab;4319 | 0 | 0 |
| NC_005831.2;YP_003766.2;HCoV-NL63_1ab;4320 | 0 | 0 |
| NC_005831.2;YP_003766.2;HCoV-NL63_1ab;4321 | 0 | 0 |
| NC_005831.2;YP_003766.2;HCoV-NL63_1ab;4322 | 0 | 0 |
| NC_005831.2;YP_003766.2;HCoV-NL63_1ab;4323 | 0 | 0 |
| NC_005831.2;YP_003766.2;HCoV-NL63_1ab;4324 | 0 | 0 |
| NC_005831.2;YP_003766.2;HCoV-NL63_1ab;4325 | 0 | 0 |
| NC_005831.2;YP_003766.2;HCoV-NL63_1ab;4326 | 0 | 0 |
| NC_005831.2;YP_003766.2;HCoV-NL63_1ab;4327 | 0 | 0.025 |
| NC_005831.2;YP_003766.2;HCoV-NL63_1ab;4328 | 0 | 0.025 |
| NC_005831.2;YP_003766.2;HCoV-NL63_1ab;4329 | 0 | 0.025 |
| NC_005831.2;YP_003766.2;HCoV-NL63_1ab;4330 | 0 | 0.025 |
| NC_005831.2;YP_003766.2;HCoV-NL63_1ab;4331 | 0 | 0 |
| NC_005831.2;YP_003766.2;HCoV-NL63_1ab;4332 | 0 | 0 |
| NC_005831.2;YP_003766.2;HCoV-NL63_1ab;4333 | 0.05 | 0 |
| NC_005831.2;YP_003766.2;HCoV-NL63_1ab;4334 | 0.05 | 0 |
| NC_005831.2;YP_003766.2;HCoV-NL63_1ab;4335 | 0.05 | 0 |
| NC_005831.2;YP_003766.2;HCoV-NL63_1ab;4336 | 0.05 | 0 |
| NC_005831.2;YP_003766.2;HCoV-NL63_1ab;4337 | 0.05 | 0 |
| NC_005831.2;YP_003766.2;HCoV-NL63_1ab;4338 | 0.05 | 0 |
| NC_005831.2;YP_003766.2;HCoV-NL63_1ab;4339 | 0.05 | 0 |
| NC_005831.2;YP_003766.2;HCoV-NL63_1ab;4340 | 0 | 0 |
| NC_005831.2;YP_003766.2;HCoV-NL63_1ab;4341 | 0 | 0 |
| NC_005831.2;YP_003766.2;HCoV-NL63_1ab;4342 | 0 | 0 |
| NC_005831.2;YP_003766.2;HCoV-NL63_1ab;4343 | 0 | 0 |
| NC_005831.2;YP_003766.2;HCoV-NL63_1ab;4344 | 0 | 0 |
| NC_005831.2;YP_003766.2;HCoV-NL63_1ab;4345 | 0 | 0 |
| NC_005831.2;YP_003766.2;HCoV-NL63_1ab;4346 | 0 | 0 |
| NC_005831.2;YP_003766.2;HCoV-NL63_1ab;4347 | 0 | 0 |
| NC_005831.2;YP_003766.2;HCoV-NL63_1ab;4348 | 0 | 0 |
| NC_005831.2;YP_003766.2;HCoV-NL63_1ab;4349 | 0 | 0 |
| NC_005831.2;YP_003766.2;HCoV-NL63_1ab;4350 | 0 | 0 |
| NC_005831.2;YP_003766.2;HCoV-NL63_1ab;4351 | 0 | 0 |
| NC_005831.2;YP_003766.2;HCoV-NL63_1ab;4352 | 0 | 0 |

| | | |
|---|---|---|
| NC_005831.2;YP_003766.2;HCoV-NL63_1ab;4353 | 0 | 0 |
| NC_005831.2;YP_003766.2;HCoV-NL63_1ab;4354 | 0 | 0 |
| NC_005831.2;YP_003766.2;HCoV-NL63_1ab;4355 | 0 | 0 |
| NC_005831.2;YP_003766.2;HCoV-NL63_1ab;4356 | 0 | 0 |
| NC_005831.2;YP_003766.2;HCoV-NL63_1ab;4357 | 0 | 0 |
| NC_005831.2;YP_003766.2;HCoV-NL63_1ab;4358 | 0 | 0 |
| NC_005831.2;YP_003766.2;HCoV-NL63_1ab;4359 | 0 | 0 |
| NC_005831.2;YP_003766.2;HCoV-NL63_1ab;4360 | 0 | 0 |
| NC_005831.2;YP_003766.2;HCoV-NL63_1ab;4361 | 0 | 0 |
| NC_005831.2;YP_003766.2;HCoV-NL63_1ab;4362 | 0 | 0 |
| NC_005831.2;YP_003766.2;HCoV-NL63_1ab;4363 | 0 | 0 |
| NC_005831.2;YP_003766.2;HCoV-NL63_1ab;4364 | 0 | 0 |
| NC_005831.2;YP_003766.2;HCoV-NL63_1ab;4365 | 0 | 0 |
| NC_005831.2;YP_003766.2;HCoV-NL63_1ab;4366 | 0 | 0 |
| NC_005831.2;YP_003766.2;HCoV-NL63_1ab;4367 | 0 | 0 |
| NC_005831.2;YP_003766.2;HCoV-NL63_1ab;4368 | 0 | 0 |
| NC_005831.2;YP_003766.2;HCoV-NL63_1ab;4369 | 0 | 0 |
| NC_005831.2;YP_003766.2;HCoV-NL63_1ab;4370 | 0 | 0 |
| NC_005831.2;YP_003766.2;HCoV-NL63_1ab;4371 | 0 | 0 |
| NC_005831.2;YP_003766.2;HCoV-NL63_1ab;4372 | 0 | 0 |
| NC_005831.2;YP_003766.2;HCoV-NL63_1ab;4373 | 0 | 0 |
| NC_005831.2;YP_003766.2;HCoV-NL63_1ab;4374 | 0 | 0 |
| NC_005831.2;YP_003766.2;HCoV-NL63_1ab;4375 | 0 | 0 |
| NC_005831.2;YP_003766.2;HCoV-NL63_1ab;4376 | 0 | 0 |
| NC_005831.2;YP_003766.2;HCoV-NL63_1ab;4377 | 0 | 0 |
| NC_005831.2;YP_003766.2;HCoV-NL63_1ab;4378 | 0 | 0 |
| NC_005831.2;YP_003766.2;HCoV-NL63_1ab;4379 | 0 | 0 |
| NC_005831.2;YP_003766.2;HCoV-NL63_1ab;4380 | 0 | 0 |
| NC_005831.2;YP_003766.2;HCoV-NL63_1ab;4381 | 0 | 0 |
| NC_005831.2;YP_003766.2;HCoV-NL63_1ab;4382 | 0 | 0 |
| NC_005831.2;YP_003766.2;HCoV-NL63_1ab;4383 | 0 | 0 |
| NC_005831.2;YP_003766.2;HCoV-NL63_1ab;4384 | 0 | 0 |
| NC_005831.2;YP_003766.2;HCoV-NL63_1ab;4385 | 0 | 0 |
| NC_005831.2;YP_003766.2;HCoV-NL63_1ab;4386 | 0 | 0 |
| NC_005831.2;YP_003766.2;HCoV-NL63_1ab;4387 | 0 | 0 |
| NC_005831.2;YP_003766.2;HCoV-NL63_1ab;4388 | 0 | 0 |
| NC_005831.2;YP_003766.2;HCoV-NL63_1ab;4389 | 0 | 0 |
| NC_005831.2;YP_003766.2;HCoV-NL63_1ab;4390 | 0 | 0 |
| NC_005831.2;YP_003766.2;HCoV-NL63_1ab;4391 | 0 | 0 |
| NC_005831.2;YP_003766.2;HCoV-NL63_1ab;4392 | 0 | 0 |
| NC_005831.2;YP_003766.2;HCoV-NL63_1ab;4393 | 0 | 0 |
| NC_005831.2;YP_003766.2;HCoV-NL63_1ab;4394 | 0 | 0 |
| NC_005831.2;YP_003766.2;HCoV-NL63_1ab;4395 | 0 | 0 |

| | | |
|---|---|---|
| NC_005831.2;YP_003766.2;HCoV-NL63_1ab;4396 | 0 | 0 |
| NC_005831.2;YP_003766.2;HCoV-NL63_1ab;4397 | 0 | 0 |
| NC_005831.2;YP_003766.2;HCoV-NL63_1ab;4398 | 0 | 0 |
| NC_005831.2;YP_003766.2;HCoV-NL63_1ab;4399 | 0 | 0 |
| NC_005831.2;YP_003766.2;HCoV-NL63_1ab;4400 | 0 | 0 |
| NC_005831.2;YP_003766.2;HCoV-NL63_1ab;4401 | 0 | 0 |
| NC_005831.2;YP_003766.2;HCoV-NL63_1ab;4402 | 0 | 0 |
| NC_005831.2;YP_003766.2;HCoV-NL63_1ab;4403 | 0 | 0 |
| NC_005831.2;YP_003766.2;HCoV-NL63_1ab;4404 | 0 | 0.175 |
| NC_005831.2;YP_003766.2;HCoV-NL63_1ab;4405 | 0 | 0 |
| NC_005831.2;YP_003766.2;HCoV-NL63_1ab;4406 | 0 | 0 |
| NC_005831.2;YP_003766.2;HCoV-NL63_1ab;4407 | 0 | 0 |
| NC_005831.2;YP_003766.2;HCoV-NL63_1ab;4408 | 0 | 0 |
| NC_005831.2;YP_003766.2;HCoV-NL63_1ab;4409 | 0 | 0 |
| NC_005831.2;YP_003766.2;HCoV-NL63_1ab;4410 | 0 | 0 |
| NC_005831.2;YP_003766.2;HCoV-NL63_1ab;4411 | 0 | 0 |
| NC_005831.2;YP_003766.2;HCoV-NL63_1ab;4412 | 0 | 0 |
| NC_005831.2;YP_003766.2;HCoV-NL63_1ab;4413 | 0 | 0 |
| NC_005831.2;YP_003766.2;HCoV-NL63_1ab;4414 | 0 | 0 |
| NC_005831.2;YP_003766.2;HCoV-NL63_1ab;4415 | 0 | 0 |
| NC_005831.2;YP_003766.2;HCoV-NL63_1ab;4416 | 0 | 0 |
| NC_005831.2;YP_003766.2;HCoV-NL63_1ab;4417 | 0 | 0.025 |
| NC_005831.2;YP_003766.2;HCoV-NL63_1ab;4418 | 0 | 0 |
| NC_005831.2;YP_003766.2;HCoV-NL63_1ab;4419 | 0 | 0 |
| NC_005831.2;YP_003766.2;HCoV-NL63_1ab;4420 | 0 | 0 |
| NC_005831.2;YP_003766.2;HCoV-NL63_1ab;4421 | 0 | 0 |
| NC_005831.2;YP_003766.2;HCoV-NL63_1ab;4422 | 0 | 0 |
| NC_005831.2;YP_003766.2;HCoV-NL63_1ab;4423 | 0.05 | 0 |
| NC_005831.2;YP_003766.2;HCoV-NL63_1ab;4424 | 0 | 0 |
| NC_005831.2;YP_003766.2;HCoV-NL63_1ab;4425 | 0 | 0 |
| NC_005831.2;YP_003766.2;HCoV-NL63_1ab;4426 | 0.1 | 0.05 |
| NC_005831.2;YP_003766.2;HCoV-NL63_1ab;4427 | 0 | 0 |
| NC_005831.2;YP_003766.2;HCoV-NL63_1ab;4428 | 0 | 0 |
| NC_005831.2;YP_003766.2;HCoV-NL63_1ab;4429 | 0 | 0 |
| NC_005831.2;YP_003766.2;HCoV-NL63_1ab;4430 | 0 | 0 |
| NC_005831.2;YP_003766.2;HCoV-NL63_1ab;4431 | 0 | 0 |
| NC_005831.2;YP_003766.2;HCoV-NL63_1ab;4432 | 0 | 0 |
| NC_005831.2;YP_003766.2;HCoV-NL63_1ab;4433 | 0 | 0 |
| NC_005831.2;YP_003766.2;HCoV-NL63_1ab;4434 | 0 | 0 |
| NC_005831.2;YP_003766.2;HCoV-NL63_1ab;4435 | 0 | 0 |
| NC_005831.2;YP_003766.2;HCoV-NL63_1ab;4436 | 0 | 0 |
| NC_005831.2;YP_003766.2;HCoV-NL63_1ab;4437 | 0 | 0 |
| NC_005831.2;YP_003766.2;HCoV-NL63_1ab;4438 | 0.1 | 0 |

| | | |
|---|---|---|
| NC_005831.2;YP_003766.2;HCoV-NL63_1ab;4439 | 0 | 0 |
| NC_005831.2;YP_003766.2;HCoV-NL63_1ab;4440 | 0 | 0 |
| NC_005831.2;YP_003766.2;HCoV-NL63_1ab;4441 | 0 | 0 |
| NC_005831.2;YP_003766.2;HCoV-NL63_1ab;4442 | 0.05 | 0.05 |
| NC_005831.2;YP_003766.2;HCoV-NL63_1ab;4443 | 0.15 | 0.05 |
| NC_005831.2;YP_003766.2;HCoV-NL63_1ab;4444 | 0.2 | 0.075 |
| NC_005831.2;YP_003766.2;HCoV-NL63_1ab;4445 | 0.15 | 0.05 |
| NC_005831.2;YP_003766.2;HCoV-NL63_1ab;4446 | 0.15 | 0.075 |
| NC_005831.2;YP_003766.2;HCoV-NL63_1ab;4447 | 0.2 | 0.075 |
| NC_005831.2;YP_003766.2;HCoV-NL63_1ab;4448 | 0.15 | 0.05 |
| NC_005831.2;YP_003766.2;HCoV-NL63_1ab;4449 | 0.05 | 0.025 |
| NC_005831.2;YP_003766.2;HCoV-NL63_1ab;4450 | 0.05 | 0 |
| NC_005831.2;YP_003766.2;HCoV-NL63_1ab;4451 | 0 | 0 |
| NC_005831.2;YP_003766.2;HCoV-NL63_1ab;4452 | 0 | 0 |
| NC_005831.2;YP_003766.2;HCoV-NL63_1ab;4453 | 0.05 | 0 |
| NC_005831.2;YP_003766.2;HCoV-NL63_1ab;4454 | 0 | 0 |
| NC_005831.2;YP_003766.2;HCoV-NL63_1ab;4455 | 0 | 0.025 |
| NC_005831.2;YP_003766.2;HCoV-NL63_1ab;4456 | 0 | 0 |
| NC_005831.2;YP_003766.2;HCoV-NL63_1ab;4457 | 0 | 0 |
| NC_005831.2;YP_003766.2;HCoV-NL63_1ab;4458 | 0 | 0 |
| NC_005831.2;YP_003766.2;HCoV-NL63_1ab;4459 | 0 | 0 |
| NC_005831.2;YP_003766.2;HCoV-NL63_1ab;4460 | 0 | 0 |
| NC_005831.2;YP_003766.2;HCoV-NL63_1ab;4461 | 0.05 | 0 |
| NC_005831.2;YP_003766.2;HCoV-NL63_1ab;4462 | 0 | 0 |
| NC_005831.2;YP_003766.2;HCoV-NL63_1ab;4463 | 0 | 0 |
| NC_005831.2;YP_003766.2;HCoV-NL63_1ab;4464 | 0 | 0 |
| NC_005831.2;YP_003766.2;HCoV-NL63_1ab;4465 | 0.1 | 0 |
| NC_005831.2;YP_003766.2;HCoV-NL63_1ab;4466 | 0 | 0 |
| NC_005831.2;YP_003766.2;HCoV-NL63_1ab;4467 | 0 | 0 |
| NC_005831.2;YP_003766.2;HCoV-NL63_1ab;4468 | 0 | 0 |
| NC_005831.2;YP_003766.2;HCoV-NL63_1ab;4469 | 0 | 0 |
| NC_005831.2;YP_003766.2;HCoV-NL63_1ab;4470 | 0 | 0 |
| NC_005831.2;YP_003766.2;HCoV-NL63_1ab;4471 | 0 | 0 |
| NC_005831.2;YP_003766.2;HCoV-NL63_1ab;4472 | 0 | 0 |
| NC_005831.2;YP_003766.2;HCoV-NL63_1ab;4473 | 0 | 0 |
| NC_005831.2;YP_003766.2;HCoV-NL63_1ab;4474 | 0 | 0 |
| NC_005831.2;YP_003766.2;HCoV-NL63_1ab;4475 | 0 | 0 |
| NC_005831.2;YP_003766.2;HCoV-NL63_1ab;4476 | 0 | 0 |
| NC_005831.2;YP_003766.2;HCoV-NL63_1ab;4477 | 0 | 0 |
| NC_005831.2;YP_003766.2;HCoV-NL63_1ab;4478 | 0 | 0 |
| NC_005831.2;YP_003766.2;HCoV-NL63_1ab;4479 | 0 | 0 |
| NC_005831.2;YP_003766.2;HCoV-NL63_1ab;4480 | 0 | 0 |
| NC_005831.2;YP_003766.2;HCoV-NL63_1ab;4481 | 0 | 0 |

| | | |
|---|---|---|
| NC_005831.2;YP_003766.2;HCoV-NL63_1ab;4482 | 0.15 | 0.05 |
| NC_005831.2;YP_003766.2;HCoV-NL63_1ab;4483 | 0.15 | 0.1 |
| NC_005831.2;YP_003766.2;HCoV-NL63_1ab;4484 | 0.15 | 0.125 |
| NC_005831.2;YP_003766.2;HCoV-NL63_1ab;4485 | 0.2 | 0.1 |
| NC_005831.2;YP_003766.2;HCoV-NL63_1ab;4486 | 0.15 | 0.025 |
| NC_005831.2;YP_003766.2;HCoV-NL63_1ab;4487 | 0.15 | 0.025 |
| NC_005831.2;YP_003766.2;HCoV-NL63_1ab;4488 | 0.15 | 0.1 |
| NC_005831.2;YP_003766.2;HCoV-NL63_1ab;4489 | 0.15 | 0.05 |
| NC_005831.2;YP_003766.2;HCoV-NL63_1ab;4490 | 0.05 | 0 |
| NC_005831.2;YP_003766.2;HCoV-NL63_1ab;4491 | 0 | 0 |
| NC_005831.2;YP_003766.2;HCoV-NL63_1ab;4492 | 0 | 0 |
| NC_005831.2;YP_003766.2;HCoV-NL63_1ab;4493 | 0 | 0 |
| NC_005831.2;YP_003766.2;HCoV-NL63_1ab;4494 | 0 | 0 |
| NC_005831.2;YP_003766.2;HCoV-NL63_1ab;4495 | 0 | 0 |
| NC_005831.2;YP_003766.2;HCoV-NL63_1ab;4496 | 0 | 0 |
| NC_005831.2;YP_003766.2;HCoV-NL63_1ab;4497 | 0 | 0 |
| NC_005831.2;YP_003766.2;HCoV-NL63_1ab;4498 | 0 | 0 |
| NC_005831.2;YP_003766.2;HCoV-NL63_1ab;4499 | 0 | 0 |
| NC_005831.2;YP_003766.2;HCoV-NL63_1ab;4500 | 0 | 0 |
| NC_005831.2;YP_003766.2;HCoV-NL63_1ab;4501 | 0 | 0 |
| NC_005831.2;YP_003766.2;HCoV-NL63_1ab;4502 | 0 | 0 |
| NC_005831.2;YP_003766.2;HCoV-NL63_1ab;4503 | 0 | 0 |
| NC_005831.2;YP_003766.2;HCoV-NL63_1ab;4504 | 0 | 0 |
| NC_005831.2;YP_003766.2;HCoV-NL63_1ab;4505 | 0 | 0 |
| NC_005831.2;YP_003766.2;HCoV-NL63_1ab;4506 | 0 | 0 |
| NC_005831.2;YP_003766.2;HCoV-NL63_1ab;4507 | 0 | 0 |
| NC_005831.2;YP_003766.2;HCoV-NL63_1ab;4508 | 0 | 0 |
| NC_005831.2;YP_003766.2;HCoV-NL63_1ab;4509 | 0 | 0 |
| NC_005831.2;YP_003766.2;HCoV-NL63_1ab;4510 | 0 | 0 |
| NC_005831.2;YP_003766.2;HCoV-NL63_1ab;4511 | 0 | 0 |
| NC_005831.2;YP_003766.2;HCoV-NL63_1ab;4512 | 0 | 0 |
| NC_005831.2;YP_003766.2;HCoV-NL63_1ab;4513 | 0 | 0 |
| NC_005831.2;YP_003766.2;HCoV-NL63_1ab;4514 | 0 | 0 |
| NC_005831.2;YP_003766.2;HCoV-NL63_1ab;4515 | 0 | 0 |
| NC_005831.2;YP_003766.2;HCoV-NL63_1ab;4516 | 0 | 0 |
| NC_005831.2;YP_003766.2;HCoV-NL63_1ab;4517 | 0 | 0 |
| NC_005831.2;YP_003766.2;HCoV-NL63_1ab;4518 | 0 | 0 |
| NC_005831.2;YP_003766.2;HCoV-NL63_1ab;4519 | 0 | 0 |
| NC_005831.2;YP_003766.2;HCoV-NL63_1ab;4520 | 0 | 0 |
| NC_005831.2;YP_003766.2;HCoV-NL63_1ab;4521 | 0.05 | 0.025 |
| NC_005831.2;YP_003766.2;HCoV-NL63_1ab;4522 | 0.05 | 0 |
| NC_005831.2;YP_003766.2;HCoV-NL63_1ab;4523 | 0.05 | 0.075 |
| NC_005831.2;YP_003766.2;HCoV-NL63_1ab;4524 | 0.05 | 0.05 |

| | | |
|---|---|---|
| NC_005831.2;YP_003766.2;HCoV-NL63_1ab;4525 | 0.1 | 0.075 |
| NC_005831.2;YP_003766.2;HCoV-NL63_1ab;4526 | 0.1 | 0.05 |
| NC_005831.2;YP_003766.2;HCoV-NL63_1ab;4527 | 0.1 | 0.025 |
| NC_005831.2;YP_003766.2;HCoV-NL63_1ab;4528 | 0.1 | 0.075 |
| NC_005831.2;YP_003766.2;HCoV-NL63_1ab;4529 | 0.2 | 0.1 |
| NC_005831.2;YP_003766.2;HCoV-NL63_1ab;4530 | 0.25 | 0.15 |
| NC_005831.2;YP_003766.2;HCoV-NL63_1ab;4531 | 0.1 | 0 |
| NC_005831.2;YP_003766.2;HCoV-NL63_1ab;4532 | 0.1 | 0 |
| NC_005831.2;YP_003766.2;HCoV-NL63_1ab;4533 | 0.1 | 0.025 |
| NC_005831.2;YP_003766.2;HCoV-NL63_1ab;4534 | 0.1 | 0 |
| NC_005831.2;YP_003766.2;HCoV-NL63_1ab;4535 | 0.1 | 0 |
| NC_005831.2;YP_003766.2;HCoV-NL63_1ab;4536 | 0.15 | 0.025 |
| NC_005831.2;YP_003766.2;HCoV-NL63_1ab;4537 | 0.1 | 0 |
| NC_005831.2;YP_003766.2;HCoV-NL63_1ab;4538 | 0.1 | 0 |
| NC_005831.2;YP_003766.2;HCoV-NL63_1ab;4539 | 0 | 0 |
| NC_005831.2;YP_003766.2;HCoV-NL63_1ab;4540 | 0 | 0 |
| NC_005831.2;YP_003766.2;HCoV-NL63_1ab;4541 | 0 | 0 |
| NC_005831.2;YP_003766.2;HCoV-NL63_1ab;4542 | 0 | 0 |
| NC_005831.2;YP_003766.2;HCoV-NL63_1ab;4543 | 0 | 0 |
| NC_005831.2;YP_003766.2;HCoV-NL63_1ab;4544 | 0 | 0.025 |
| NC_005831.2;YP_003766.2;HCoV-NL63_1ab;4545 | 0 | 0 |
| NC_005831.2;YP_003766.2;HCoV-NL63_1ab;4546 | 0 | 0 |
| NC_005831.2;YP_003766.2;HCoV-NL63_1ab;4547 | 0 | 0 |
| NC_005831.2;YP_003766.2;HCoV-NL63_1ab;4548 | 0 | 0 |
| NC_005831.2;YP_003766.2;HCoV-NL63_1ab;4549 | 0 | 0 |
| NC_005831.2;YP_003766.2;HCoV-NL63_1ab;4550 | 0 | 0 |
| NC_005831.2;YP_003766.2;HCoV-NL63_1ab;4551 | 0 | 0 |
| NC_005831.2;YP_003766.2;HCoV-NL63_1ab;4552 | 0 | 0 |
| NC_005831.2;YP_003766.2;HCoV-NL63_1ab;4553 | 0 | 0 |
| NC_005831.2;YP_003766.2;HCoV-NL63_1ab;4554 | 0 | 0 |
| NC_005831.2;YP_003766.2;HCoV-NL63_1ab;4555 | 0 | 0 |
| NC_005831.2;YP_003766.2;HCoV-NL63_1ab;4556 | 0 | 0 |
| NC_005831.2;YP_003766.2;HCoV-NL63_1ab;4557 | 0 | 0.025 |
| NC_005831.2;YP_003766.2;HCoV-NL63_1ab;4558 | 0 | 0 |
| NC_005831.2;YP_003766.2;HCoV-NL63_1ab;4559 | 0 | 0 |
| NC_005831.2;YP_003766.2;HCoV-NL63_1ab;4560 | 0 | 0 |
| NC_005831.2;YP_003766.2;HCoV-NL63_1ab;4561 | 0 | 0 |
| NC_005831.2;YP_003766.2;HCoV-NL63_1ab;4562 | 0 | 0 |
| NC_005831.2;YP_003766.2;HCoV-NL63_1ab;4563 | 0 | 0 |
| NC_005831.2;YP_003766.2;HCoV-NL63_1ab;4564 | 0 | 0 |
| NC_005831.2;YP_003766.2;HCoV-NL63_1ab;4565 | 0 | 0 |
| NC_005831.2;YP_003766.2;HCoV-NL63_1ab;4566 | 0 | 0 |
| NC_005831.2;YP_003766.2;HCoV-NL63_1ab;4567 | 0 | 0 |

| | | |
|---|---|---|
| NC_005831.2;YP_003766.2;HCoV-NL63_1ab;4568 | 0 | 0 |
| NC_005831.2;YP_003766.2;HCoV-NL63_1ab;4569 | 0.15 | 0.075 |
| NC_005831.2;YP_003766.2;HCoV-NL63_1ab;4570 | 0.2 | 0.2 |
| NC_005831.2;YP_003766.2;HCoV-NL63_1ab;4571 | 0 | 0 |
| NC_005831.2;YP_003766.2;HCoV-NL63_1ab;4572 | 0 | 0 |
| NC_005831.2;YP_003766.2;HCoV-NL63_1ab;4573 | 0 | 0 |
| NC_005831.2;YP_003766.2;HCoV-NL63_1ab;4574 | 0 | 0 |
| NC_005831.2;YP_003766.2;HCoV-NL63_1ab;4575 | 0.05 | 0.05 |
| NC_005831.2;YP_003766.2;HCoV-NL63_1ab;4576 | 0.05 | 0.025 |
| NC_005831.2;YP_003766.2;HCoV-NL63_1ab;4577 | 0.05 | 0.025 |
| NC_005831.2;YP_003766.2;HCoV-NL63_1ab;4578 | 0 | 0 |
| NC_005831.2;YP_003766.2;HCoV-NL63_1ab;4579 | 0 | 0 |
| NC_005831.2;YP_003766.2;HCoV-NL63_1ab;4580 | 0 | 0 |
| NC_005831.2;YP_003766.2;HCoV-NL63_1ab;4581 | 0 | 0 |
| NC_005831.2;YP_003766.2;HCoV-NL63_1ab;4582 | 0 | 0 |
| NC_005831.2;YP_003766.2;HCoV-NL63_1ab;4583 | 0 | 0 |
| NC_005831.2;YP_003766.2;HCoV-NL63_1ab;4584 | 0 | 0.025 |
| NC_005831.2;YP_003766.2;HCoV-NL63_1ab;4585 | 0 | 0 |
| NC_005831.2;YP_003766.2;HCoV-NL63_1ab;4586 | 0.1 | 0 |
| NC_005831.2;YP_003766.2;HCoV-NL63_1ab;4587 | 0 | 0 |
| NC_005831.2;YP_003766.2;HCoV-NL63_1ab;4588 | 0 | 0 |
| NC_005831.2;YP_003766.2;HCoV-NL63_1ab;4589 | 0 | 0 |
| NC_005831.2;YP_003766.2;HCoV-NL63_1ab;4590 | 0 | 0 |
| NC_005831.2;YP_003766.2;HCoV-NL63_1ab;4591 | 0 | 0 |
| NC_005831.2;YP_003766.2;HCoV-NL63_1ab;4592 | 0 | 0 |
| NC_005831.2;YP_003766.2;HCoV-NL63_1ab;4593 | 0 | 0 |
| NC_005831.2;YP_003766.2;HCoV-NL63_1ab;4594 | 0 | 0 |
| NC_005831.2;YP_003766.2;HCoV-NL63_1ab;4595 | 0 | 0 |
| NC_005831.2;YP_003766.2;HCoV-NL63_1ab;4596 | 0 | 0 |
| NC_005831.2;YP_003766.2;HCoV-NL63_1ab;4597 | 0 | 0 |
| NC_005831.2;YP_003766.2;HCoV-NL63_1ab;4598 | 0 | 0 |
| NC_005831.2;YP_003766.2;HCoV-NL63_1ab;4599 | 0 | 0 |
| NC_005831.2;YP_003766.2;HCoV-NL63_1ab;4600 | 0 | 0 |
| NC_005831.2;YP_003766.2;HCoV-NL63_1ab;4601 | 0 | 0 |
| NC_005831.2;YP_003766.2;HCoV-NL63_1ab;4602 | 0 | 0 |
| NC_005831.2;YP_003766.2;HCoV-NL63_1ab;4603 | 0 | 0 |
| NC_005831.2;YP_003766.2;HCoV-NL63_1ab;4604 | 0 | 0 |
| NC_005831.2;YP_003766.2;HCoV-NL63_1ab;4605 | 0 | 0 |
| NC_005831.2;YP_003766.2;HCoV-NL63_1ab;4606 | 0 | 0 |
| NC_005831.2;YP_003766.2;HCoV-NL63_1ab;4607 | 0 | 0 |
| NC_005831.2;YP_003766.2;HCoV-NL63_1ab;4608 | 0 | 0 |
| NC_005831.2;YP_003766.2;HCoV-NL63_1ab;4609 | 0.05 | 0 |
| NC_005831.2;YP_003766.2;HCoV-NL63_1ab;4610 | 0.05 | 0 |

| | | |
|---|---|---|
| NC_005831.2;YP_003766.2;HCoV-NL63_1ab;4611 | 0.05 | 0 |
| NC_005831.2;YP_003766.2;HCoV-NL63_1ab;4612 | 0.05 | 0 |
| NC_005831.2;YP_003766.2;HCoV-NL63_1ab;4613 | 0.05 | 0 |
| NC_005831.2;YP_003766.2;HCoV-NL63_1ab;4614 | 0.05 | 0.025 |
| NC_005831.2;YP_003766.2;HCoV-NL63_1ab;4615 | 0.05 | 0 |
| NC_005831.2;YP_003766.2;HCoV-NL63_1ab;4616 | 0.05 | 0 |
| NC_005831.2;YP_003766.2;HCoV-NL63_1ab;4617 | 0.05 | 0 |
| NC_005831.2;YP_003766.2;HCoV-NL63_1ab;4618 | 0 | 0 |
| NC_005831.2;YP_003766.2;HCoV-NL63_1ab;4619 | 0 | 0 |
| NC_005831.2;YP_003766.2;HCoV-NL63_1ab;4620 | 0 | 0 |
| NC_005831.2;YP_003766.2;HCoV-NL63_1ab;4621 | 0 | 0 |
| NC_005831.2;YP_003766.2;HCoV-NL63_1ab;4622 | 0 | 0 |
| NC_005831.2;YP_003766.2;HCoV-NL63_1ab;4623 | 0 | 0 |
| NC_005831.2;YP_003766.2;HCoV-NL63_1ab;4624 | 0 | 0 |
| NC_005831.2;YP_003766.2;HCoV-NL63_1ab;4625 | 0 | 0 |
| NC_005831.2;YP_003766.2;HCoV-NL63_1ab;4626 | 0 | 0 |
| NC_005831.2;YP_003766.2;HCoV-NL63_1ab;4627 | 0 | 0 |
| NC_005831.2;YP_003766.2;HCoV-NL63_1ab;4628 | 0 | 0 |
| NC_005831.2;YP_003766.2;HCoV-NL63_1ab;4629 | 0 | 0 |
| NC_005831.2;YP_003766.2;HCoV-NL63_1ab;4630 | 0 | 0 |
| NC_005831.2;YP_003766.2;HCoV-NL63_1ab;4631 | 0 | 0 |
| NC_005831.2;YP_003766.2;HCoV-NL63_1ab;4632 | 0.05 | 0 |
| NC_005831.2;YP_003766.2;HCoV-NL63_1ab;4633 | 0.05 | 0 |
| NC_005831.2;YP_003766.2;HCoV-NL63_1ab;4634 | 0 | 0 |
| NC_005831.2;YP_003766.2;HCoV-NL63_1ab;4635 | 0.05 | 0 |
| NC_005831.2;YP_003766.2;HCoV-NL63_1ab;4636 | 0 | 0 |
| NC_005831.2;YP_003766.2;HCoV-NL63_1ab;4637 | 0 | 0 |
| NC_005831.2;YP_003766.2;HCoV-NL63_1ab;4638 | 0 | 0 |
| NC_005831.2;YP_003766.2;HCoV-NL63_1ab;4639 | 0 | 0 |
| NC_005831.2;YP_003766.2;HCoV-NL63_1ab;4640 | 0 | 0 |
| NC_005831.2;YP_003766.2;HCoV-NL63_1ab;4641 | 0.05 | 0 |
| NC_005831.2;YP_003766.2;HCoV-NL63_1ab;4642 | 0 | 0 |
| NC_005831.2;YP_003766.2;HCoV-NL63_1ab;4643 | 0 | 0.025 |
| NC_005831.2;YP_003766.2;HCoV-NL63_1ab;4644 | 0 | 0.025 |
| NC_005831.2;YP_003766.2;HCoV-NL63_1ab;4645 | 0.05 | 0.025 |
| NC_005831.2;YP_003766.2;HCoV-NL63_1ab;4646 | 0 | 0.05 |
| NC_005831.2;YP_003766.2;HCoV-NL63_1ab;4647 | 0 | 0.025 |
| NC_005831.2;YP_003766.2;HCoV-NL63_1ab;4648 | 0 | 0.025 |
| NC_005831.2;YP_003766.2;HCoV-NL63_1ab;4649 | 0 | 0.05 |
| NC_005831.2;YP_003766.2;HCoV-NL63_1ab;4650 | 0 | 0.025 |
| NC_005831.2;YP_003766.2;HCoV-NL63_1ab;4651 | 0 | 0.025 |
| NC_005831.2;YP_003766.2;HCoV-NL63_1ab;4652 | 0 | 0.025 |
| NC_005831.2;YP_003766.2;HCoV-NL63_1ab;4653 | 0 | 0.025 |

| | | |
|---|---|---|
| NC_005831.2;YP_003766.2;HCoV-NL63_1ab;4654 | 0 | 0.025 |
| NC_005831.2;YP_003766.2;HCoV-NL63_1ab;4655 | 0 | 0.025 |
| NC_005831.2;YP_003766.2;HCoV-NL63_1ab;4656 | 0 | 0 |
| NC_005831.2;YP_003766.2;HCoV-NL63_1ab;4657 | 0 | 0 |
| NC_005831.2;YP_003766.2;HCoV-NL63_1ab;4658 | 0 | 0 |
| NC_005831.2;YP_003766.2;HCoV-NL63_1ab;4659 | 0 | 0 |
| NC_005831.2;YP_003766.2;HCoV-NL63_1ab;4660 | 0 | 0 |
| NC_005831.2;YP_003766.2;HCoV-NL63_1ab;4661 | 0 | 0 |
| NC_005831.2;YP_003766.2;HCoV-NL63_1ab;4662 | 0 | 0 |
| NC_005831.2;YP_003766.2;HCoV-NL63_1ab;4663 | 0 | 0 |
| NC_005831.2;YP_003766.2;HCoV-NL63_1ab;4664 | 0 | 0 |
| NC_005831.2;YP_003766.2;HCoV-NL63_1ab;4665 | 0 | 0 |
| NC_005831.2;YP_003766.2;HCoV-NL63_1ab;4666 | 0 | 0 |
| NC_005831.2;YP_003766.2;HCoV-NL63_1ab;4667 | 0 | 0 |
| NC_005831.2;YP_003766.2;HCoV-NL63_1ab;4668 | 0 | 0 |
| NC_005831.2;YP_003766.2;HCoV-NL63_1ab;4669 | 0 | 0 |
| NC_005831.2;YP_003766.2;HCoV-NL63_1ab;4670 | 0 | 0 |
| NC_005831.2;YP_003766.2;HCoV-NL63_1ab;4671 | 0 | 0 |
| NC_005831.2;YP_003766.2;HCoV-NL63_1ab;4672 | 0 | 0 |
| NC_005831.2;YP_003766.2;HCoV-NL63_1ab;4673 | 0 | 0 |
| NC_005831.2;YP_003766.2;HCoV-NL63_1ab;4674 | 0 | 0 |
| NC_005831.2;YP_003766.2;HCoV-NL63_1ab;4675 | 0 | 0 |
| NC_005831.2;YP_003766.2;HCoV-NL63_1ab;4676 | 0 | 0 |
| NC_005831.2;YP_003766.2;HCoV-NL63_1ab;4677 | 0 | 0 |
| NC_005831.2;YP_003766.2;HCoV-NL63_1ab;4678 | 0 | 0 |
| NC_005831.2;YP_003766.2;HCoV-NL63_1ab;4679 | 0 | 0 |
| NC_005831.2;YP_003766.2;HCoV-NL63_1ab;4680 | 0 | 0 |
| NC_005831.2;YP_003766.2;HCoV-NL63_1ab;4681 | 0 | 0 |
| NC_005831.2;YP_003766.2;HCoV-NL63_1ab;4682 | 0 | 0 |
| NC_005831.2;YP_003766.2;HCoV-NL63_1ab;4683 | 0 | 0 |
| NC_005831.2;YP_003766.2;HCoV-NL63_1ab;4684 | 0 | 0 |
| NC_005831.2;YP_003766.2;HCoV-NL63_1ab;4685 | 0 | 0 |
| NC_005831.2;YP_003766.2;HCoV-NL63_1ab;4686 | 0 | 0 |
| NC_005831.2;YP_003766.2;HCoV-NL63_1ab;4687 | 0 | 0 |
| NC_005831.2;YP_003766.2;HCoV-NL63_1ab;4688 | 0 | 0 |
| NC_005831.2;YP_003766.2;HCoV-NL63_1ab;4689 | 0 | 0 |
| NC_005831.2;YP_003766.2;HCoV-NL63_1ab;4690 | 0 | 0 |
| NC_005831.2;YP_003766.2;HCoV-NL63_1ab;4691 | 0 | 0 |
| NC_005831.2;YP_003766.2;HCoV-NL63_1ab;4692 | 0 | 0 |
| NC_005831.2;YP_003766.2;HCoV-NL63_1ab;4693 | 0 | 0.025 |
| NC_005831.2;YP_003766.2;HCoV-NL63_1ab;4694 | 0 | 0.025 |
| NC_005831.2;YP_003766.2;HCoV-NL63_1ab;4695 | 0.05 | 0.025 |
| NC_005831.2;YP_003766.2;HCoV-NL63_1ab;4696 | 0 | 0.025 |

| | | |
|---|---|---|
| NC_005831.2;YP_003766.2;HCoV-NL63_1ab;4697 | 0 | 0 |
| NC_005831.2;YP_003766.2;HCoV-NL63_1ab;4698 | 0 | 0.025 |
| NC_005831.2;YP_003766.2;HCoV-NL63_1ab;4699 | 0 | 0 |
| NC_005831.2;YP_003766.2;HCoV-NL63_1ab;4700 | 0 | 0 |
| NC_005831.2;YP_003766.2;HCoV-NL63_1ab;4701 | 0 | 0 |
| NC_005831.2;YP_003766.2;HCoV-NL63_1ab;4702 | 0 | 0.025 |
| NC_005831.2;YP_003766.2;HCoV-NL63_1ab;4703 | 0 | 0.025 |
| NC_005831.2;YP_003766.2;HCoV-NL63_1ab;4704 | 0 | 0.025 |
| NC_005831.2;YP_003766.2;HCoV-NL63_1ab;4705 | 0 | 0.05 |
| NC_005831.2;YP_003766.2;HCoV-NL63_1ab;4706 | 0.05 | 0 |
| NC_005831.2;YP_003766.2;HCoV-NL63_1ab;4707 | 0.05 | 0 |
| NC_005831.2;YP_003766.2;HCoV-NL63_1ab;4708 | 0 | 0 |
| NC_005831.2;YP_003766.2;HCoV-NL63_1ab;4709 | 0 | 0 |
| NC_005831.2;YP_003766.2;HCoV-NL63_1ab;4710 | 0 | 0.025 |
| NC_005831.2;YP_003766.2;HCoV-NL63_1ab;4711 | 0 | 0 |
| NC_005831.2;YP_003766.2;HCoV-NL63_1ab;4712 | 0.05 | 0.025 |
| NC_005831.2;YP_003766.2;HCoV-NL63_1ab;4713 | 0.05 | 0.025 |
| NC_005831.2;YP_003766.2;HCoV-NL63_1ab;4714 | 0 | 0 |
| NC_005831.2;YP_003766.2;HCoV-NL63_1ab;4715 | 0 | 0 |
| NC_005831.2;YP_003766.2;HCoV-NL63_1ab;4716 | 0 | 0.025 |
| NC_005831.2;YP_003766.2;HCoV-NL63_1ab;4717 | 0 | 0 |
| NC_005831.2;YP_003766.2;HCoV-NL63_1ab;4718 | 0.05 | 0 |
| NC_005831.2;YP_003766.2;HCoV-NL63_1ab;4719 | 0 | 0 |
| NC_005831.2;YP_003766.2;HCoV-NL63_1ab;4720 | 0 | 0 |
| NC_005831.2;YP_003766.2;HCoV-NL63_1ab;4721 | 0 | 0 |
| NC_005831.2;YP_003766.2;HCoV-NL63_1ab;4722 | 0 | 0 |
| NC_005831.2;YP_003766.2;HCoV-NL63_1ab;4723 | 0 | 0 |
| NC_005831.2;YP_003766.2;HCoV-NL63_1ab;4724 | 0 | 0 |
| NC_005831.2;YP_003766.2;HCoV-NL63_1ab;4725 | 0 | 0 |
| NC_005831.2;YP_003766.2;HCoV-NL63_1ab;4726 | 0 | 0 |
| NC_005831.2;YP_003766.2;HCoV-NL63_1ab;4727 | 0 | 0 |
| NC_005831.2;YP_003766.2;HCoV-NL63_1ab;4728 | 0 | 0 |
| NC_005831.2;YP_003766.2;HCoV-NL63_1ab;4729 | 0 | 0 |
| NC_005831.2;YP_003766.2;HCoV-NL63_1ab;4730 | 0 | 0 |
| NC_005831.2;YP_003766.2;HCoV-NL63_1ab;4731 | 0 | 0 |
| NC_005831.2;YP_003766.2;HCoV-NL63_1ab;4732 | 0 | 0 |
| NC_005831.2;YP_003766.2;HCoV-NL63_1ab;4733 | 0 | 0 |
| NC_005831.2;YP_003766.2;HCoV-NL63_1ab;4734 | 0.05 | 0 |
| NC_005831.2;YP_003766.2;HCoV-NL63_1ab;4735 | 0 | 0 |
| NC_005831.2;YP_003766.2;HCoV-NL63_1ab;4736 | 0.05 | 0.05 |
| NC_005831.2;YP_003766.2;HCoV-NL63_1ab;4737 | 0.2 | 0.05 |
| NC_005831.2;YP_003766.2;HCoV-NL63_1ab;4738 | 0.05 | 0.025 |
| NC_005831.2;YP_003766.2;HCoV-NL63_1ab;4739 | 0 | 0.025 |

| | | |
|---|---|---|
| NC_005831.2;YP_003766.2;HCoV-NL63_1ab;4740 | 0 | 0 |
| NC_005831.2;YP_003766.2;HCoV-NL63_1ab;4741 | 0 | 0 |
| NC_005831.2;YP_003766.2;HCoV-NL63_1ab;4742 | 0 | 0 |
| NC_005831.2;YP_003766.2;HCoV-NL63_1ab;4743 | 0.05 | 0.025 |
| NC_005831.2;YP_003766.2;HCoV-NL63_1ab;4744 | 0 | 0 |
| NC_005831.2;YP_003766.2;HCoV-NL63_1ab;4745 | 0 | 0 |
| NC_005831.2;YP_003766.2;HCoV-NL63_1ab;4746 | 0 | 0 |
| NC_005831.2;YP_003766.2;HCoV-NL63_1ab;4747 | 0 | 0 |
| NC_005831.2;YP_003766.2;HCoV-NL63_1ab;4748 | 0 | 0 |
| NC_005831.2;YP_003766.2;HCoV-NL63_1ab;4749 | 0 | 0 |
| NC_005831.2;YP_003766.2;HCoV-NL63_1ab;4750 | 0 | 0.025 |
| NC_005831.2;YP_003766.2;HCoV-NL63_1ab;4751 | 0.05 | 0.05 |
| NC_005831.2;YP_003766.2;HCoV-NL63_1ab;4752 | 0 | 0.025 |
| NC_005831.2;YP_003766.2;HCoV-NL63_1ab;4753 | 0 | 0 |
| NC_005831.2;YP_003766.2;HCoV-NL63_1ab;4754 | 0 | 0 |
| NC_005831.2;YP_003766.2;HCoV-NL63_1ab;4755 | 0 | 0 |
| NC_005831.2;YP_003766.2;HCoV-NL63_1ab;4756 | 0 | 0 |
| NC_005831.2;YP_003766.2;HCoV-NL63_1ab;4757 | 0 | 0 |
| NC_005831.2;YP_003766.2;HCoV-NL63_1ab;4758 | 0 | 0 |
| NC_005831.2;YP_003766.2;HCoV-NL63_1ab;4759 | 0 | 0 |
| NC_005831.2;YP_003766.2;HCoV-NL63_1ab;4760 | 0 | 0 |
| NC_005831.2;YP_003766.2;HCoV-NL63_1ab;4761 | 0 | 0 |
| NC_005831.2;YP_003766.2;HCoV-NL63_1ab;4762 | 0 | 0 |
| NC_005831.2;YP_003766.2;HCoV-NL63_1ab;4763 | 0 | 0.025 |
| NC_005831.2;YP_003766.2;HCoV-NL63_1ab;4764 | 0 | 0.05 |
| NC_005831.2;YP_003766.2;HCoV-NL63_1ab;4765 | 0 | 0.025 |
| NC_005831.2;YP_003766.2;HCoV-NL63_1ab;4766 | 0 | 0 |
| NC_005831.2;YP_003766.2;HCoV-NL63_1ab;4767 | 0 | 0 |
| NC_005831.2;YP_003766.2;HCoV-NL63_1ab;4768 | 0 | 0 |
| NC_005831.2;YP_003766.2;HCoV-NL63_1ab;4769 | 0 | 0 |
| NC_005831.2;YP_003766.2;HCoV-NL63_1ab;4770 | 0 | 0 |
| NC_005831.2;YP_003766.2;HCoV-NL63_1ab;4771 | 0 | 0 |
| NC_005831.2;YP_003766.2;HCoV-NL63_1ab;4772 | 0.05 | 0 |
| NC_005831.2;YP_003766.2;HCoV-NL63_1ab;4773 | 0.05 | 0 |
| NC_005831.2;YP_003766.2;HCoV-NL63_1ab;4774 | 0.1 | 0 |
| NC_005831.2;YP_003766.2;HCoV-NL63_1ab;4775 | 0.05 | 0 |
| NC_005831.2;YP_003766.2;HCoV-NL63_1ab;4776 | 0 | 0 |
| NC_005831.2;YP_003766.2;HCoV-NL63_1ab;4777 | 0.1 | 0.025 |
| NC_005831.2;YP_003766.2;HCoV-NL63_1ab;4778 | 0.05 | 0 |
| NC_005831.2;YP_003766.2;HCoV-NL63_1ab;4779 | 0 | 0 |
| NC_005831.2;YP_003766.2;HCoV-NL63_1ab;4780 | 0 | 0 |
| NC_005831.2;YP_003766.2;HCoV-NL63_1ab;4781 | 0 | 0 |
| NC_005831.2;YP_003766.2;HCoV-NL63_1ab;4782 | 0 | 0 |

| | | |
|---|---|---|
| NC_005831.2;YP_003766.2;HCoV-NL63_1ab;4783 | 0 | 0 |
| NC_005831.2;YP_003766.2;HCoV-NL63_1ab;4784 | 0 | 0 |
| NC_005831.2;YP_003766.2;HCoV-NL63_1ab;4785 | 0 | 0 |
| NC_005831.2;YP_003766.2;HCoV-NL63_1ab;4786 | 0 | 0 |
| NC_005831.2;YP_003766.2;HCoV-NL63_1ab;4787 | 0 | 0 |
| NC_005831.2;YP_003766.2;HCoV-NL63_1ab;4788 | 0 | 0 |
| NC_005831.2;YP_003766.2;HCoV-NL63_1ab;4789 | 0 | 0 |
| NC_005831.2;YP_003766.2;HCoV-NL63_1ab;4790 | 0 | 0 |
| NC_005831.2;YP_003766.2;HCoV-NL63_1ab;4791 | 0 | 0 |
| NC_005831.2;YP_003766.2;HCoV-NL63_1ab;4792 | 0 | 0 |
| NC_005831.2;YP_003766.2;HCoV-NL63_1ab;4793 | 0 | 0.025 |
| NC_005831.2;YP_003766.2;HCoV-NL63_1ab;4794 | 0 | 0.025 |
| NC_005831.2;YP_003766.2;HCoV-NL63_1ab;4795 | 0 | 0.025 |
| NC_005831.2;YP_003766.2;HCoV-NL63_1ab;4796 | 0.05 | 0.025 |
| NC_005831.2;YP_003766.2;HCoV-NL63_1ab;4797 | 0 | 0.025 |
| NC_005831.2;YP_003766.2;HCoV-NL63_1ab;4798 | 0 | 0 |
| NC_005831.2;YP_003766.2;HCoV-NL63_1ab;4799 | 0 | 0.025 |
| NC_005831.2;YP_003766.2;HCoV-NL63_1ab;4800 | 0 | 0.05 |
| NC_005831.2;YP_003766.2;HCoV-NL63_1ab;4801 | 0 | 0 |
| NC_005831.2;YP_003766.2;HCoV-NL63_1ab;4802 | 0 | 0 |
| NC_005831.2;YP_003766.2;HCoV-NL63_1ab;4803 | 0 | 0 |
| NC_005831.2;YP_003766.2;HCoV-NL63_1ab;4804 | 0 | 0.025 |
| NC_005831.2;YP_003766.2;HCoV-NL63_1ab;4805 | 0 | 0 |
| NC_005831.2;YP_003766.2;HCoV-NL63_1ab;4806 | 0.1 | 0 |
| NC_005831.2;YP_003766.2;HCoV-NL63_1ab;4807 | 0.1 | 0.025 |
| NC_005831.2;YP_003766.2;HCoV-NL63_1ab;4808 | 0 | 0 |
| NC_005831.2;YP_003766.2;HCoV-NL63_1ab;4809 | 0 | 0 |
| NC_005831.2;YP_003766.2;HCoV-NL63_1ab;4810 | 0 | 0 |
| NC_005831.2;YP_003766.2;HCoV-NL63_1ab;4811 | 0 | 0 |
| NC_005831.2;YP_003766.2;HCoV-NL63_1ab;4812 | 0 | 0 |
| NC_005831.2;YP_003766.2;HCoV-NL63_1ab;4813 | 0 | 0 |
| NC_005831.2;YP_003766.2;HCoV-NL63_1ab;4814 | 0 | 0 |
| NC_005831.2;YP_003766.2;HCoV-NL63_1ab;4815 | 0 | 0 |
| NC_005831.2;YP_003766.2;HCoV-NL63_1ab;4816 | 0 | 0 |
| NC_005831.2;YP_003766.2;HCoV-NL63_1ab;4817 | 0 | 0 |
| NC_005831.2;YP_003766.2;HCoV-NL63_1ab;4818 | 0 | 0 |
| NC_005831.2;YP_003766.2;HCoV-NL63_1ab;4819 | 0 | 0 |
| NC_005831.2;YP_003766.2;HCoV-NL63_1ab;4820 | 0 | 0 |
| NC_005831.2;YP_003766.2;HCoV-NL63_1ab;4821 | 0 | 0 |
| NC_005831.2;YP_003766.2;HCoV-NL63_1ab;4822 | 0 | 0 |
| NC_005831.2;YP_003766.2;HCoV-NL63_1ab;4823 | 0 | 0 |
| NC_005831.2;YP_003766.2;HCoV-NL63_1ab;4824 | 0 | 0 |
| NC_005831.2;YP_003766.2;HCoV-NL63_1ab;4825 | 0 | 0 |

| | | |
|---|---|---|
| NC_005831.2;YP_003766.2;HCoV-NL63_1ab;4826 | 0 | 0 |
| NC_005831.2;YP_003766.2;HCoV-NL63_1ab;4827 | 0 | 0 |
| NC_005831.2;YP_003766.2;HCoV-NL63_1ab;4828 | 0 | 0 |
| NC_005831.2;YP_003766.2;HCoV-NL63_1ab;4829 | 0 | 0 |
| NC_005831.2;YP_003766.2;HCoV-NL63_1ab;4830 | 0 | 0.025 |
| NC_005831.2;YP_003766.2;HCoV-NL63_1ab;4831 | 0 | 0.025 |
| NC_005831.2;YP_003766.2;HCoV-NL63_1ab;4832 | 0.05 | 0.025 |
| NC_005831.2;YP_003766.2;HCoV-NL63_1ab;4833 | 0 | 0.05 |
| NC_005831.2;YP_003766.2;HCoV-NL63_1ab;4834 | 0 | 0.025 |
| NC_005831.2;YP_003766.2;HCoV-NL63_1ab;4835 | 0 | 0.025 |
| NC_005831.2;YP_003766.2;HCoV-NL63_1ab;4836 | 0 | 0.025 |
| NC_005831.2;YP_003766.2;HCoV-NL63_1ab;4837 | 0 | 0.025 |
| NC_005831.2;YP_003766.2;HCoV-NL63_1ab;4838 | 0.05 | 0.05 |
| NC_005831.2;YP_003766.2;HCoV-NL63_1ab;4839 | 0 | 0.025 |
| NC_005831.2;YP_003766.2;HCoV-NL63_1ab;4840 | 0 | 0 |
| NC_005831.2;YP_003766.2;HCoV-NL63_1ab;4841 | 0.05 | 0 |
| NC_005831.2;YP_003766.2;HCoV-NL63_1ab;4842 | 0 | 0 |
| NC_005831.2;YP_003766.2;HCoV-NL63_1ab;4843 | 0 | 0 |
| NC_005831.2;YP_003766.2;HCoV-NL63_1ab;4844 | 0 | 0 |
| NC_005831.2;YP_003766.2;HCoV-NL63_1ab;4845 | 0 | 0 |
| NC_005831.2;YP_003766.2;HCoV-NL63_1ab;4846 | 0.2 | 0.15 |
| NC_005831.2;YP_003766.2;HCoV-NL63_1ab;4847 | 0.05 | 0.025 |
| NC_005831.2;YP_003766.2;HCoV-NL63_1ab;4848 | 0 | 0.025 |
| NC_005831.2;YP_003766.2;HCoV-NL63_1ab;4849 | 0 | 0.025 |
| NC_005831.2;YP_003766.2;HCoV-NL63_1ab;4850 | 0 | 0.05 |
| NC_005831.2;YP_003766.2;HCoV-NL63_1ab;4851 | 0 | 0.05 |
| NC_005831.2;YP_003766.2;HCoV-NL63_1ab;4852 | 0 | 0.025 |
| NC_005831.2;YP_003766.2;HCoV-NL63_1ab;4853 | 0 | 0.025 |
| NC_005831.2;YP_003766.2;HCoV-NL63_1ab;4854 | 0 | 0.025 |
| NC_005831.2;YP_003766.2;HCoV-NL63_1ab;4855 | 0 | 0.05 |
| NC_005831.2;YP_003766.2;HCoV-NL63_1ab;4856 | 0 | 0.025 |
| NC_005831.2;YP_003766.2;HCoV-NL63_1ab;4857 | 0.05 | 0.025 |
| NC_005831.2;YP_003766.2;HCoV-NL63_1ab;4858 | 0 | 0.05 |
| NC_005831.2;YP_003766.2;HCoV-NL63_1ab;4859 | 0 | 0 |
| NC_005831.2;YP_003766.2;HCoV-NL63_1ab;4860 | 0 | 0 |
| NC_005831.2;YP_003766.2;HCoV-NL63_1ab;4861 | 0 | 0 |
| NC_005831.2;YP_003766.2;HCoV-NL63_1ab;4862 | 0 | 0 |
| NC_005831.2;YP_003766.2;HCoV-NL63_1ab;4863 | 0 | 0 |
| NC_005831.2;YP_003766.2;HCoV-NL63_1ab;4864 | 0 | 0 |
| NC_005831.2;YP_003766.2;HCoV-NL63_1ab;4865 | 0 | 0 |
| NC_005831.2;YP_003766.2;HCoV-NL63_1ab;4866 | 0 | 0 |
| NC_005831.2;YP_003766.2;HCoV-NL63_1ab;4867 | 0 | 0 |
| NC_005831.2;YP_003766.2;HCoV-NL63_1ab;4868 | 0 | 0 |

| | | |
|---|---|---|
| NC_005831.2;YP_003766.2;HCoV-NL63_1ab;4869 | 0 | 0 |
| NC_005831.2;YP_003766.2;HCoV-NL63_1ab;4870 | 0 | 0 |
| NC_005831.2;YP_003766.2;HCoV-NL63_1ab;4871 | 0 | 0 |
| NC_005831.2;YP_003766.2;HCoV-NL63_1ab;4872 | 0 | 0 |
| NC_005831.2;YP_003766.2;HCoV-NL63_1ab;4873 | 0 | 0 |
| NC_005831.2;YP_003766.2;HCoV-NL63_1ab;4874 | 0 | 0 |
| NC_005831.2;YP_003766.2;HCoV-NL63_1ab;4875 | 0 | 0 |
| NC_005831.2;YP_003766.2;HCoV-NL63_1ab;4876 | 0 | 0 |
| NC_005831.2;YP_003766.2;HCoV-NL63_1ab;4877 | 0 | 0 |
| NC_005831.2;YP_003766.2;HCoV-NL63_1ab;4878 | 0 | 0 |
| NC_005831.2;YP_003766.2;HCoV-NL63_1ab;4879 | 0 | 0 |
| NC_005831.2;YP_003766.2;HCoV-NL63_1ab;4880 | 0 | 0.025 |
| NC_005831.2;YP_003766.2;HCoV-NL63_1ab;4881 | 0 | 0 |
| NC_005831.2;YP_003766.2;HCoV-NL63_1ab;4882 | 0 | 0.025 |
| NC_005831.2;YP_003766.2;HCoV-NL63_1ab;4883 | 0 | 0.025 |
| NC_005831.2;YP_003766.2;HCoV-NL63_1ab;4884 | 0 | 0.025 |
| NC_005831.2;YP_003766.2;HCoV-NL63_1ab;4885 | 0 | 0.025 |
| NC_005831.2;YP_003766.2;HCoV-NL63_1ab;4886 | 0 | 0.025 |
| NC_005831.2;YP_003766.2;HCoV-NL63_1ab;4887 | 0 | 0.025 |
| NC_005831.2;YP_003766.2;HCoV-NL63_1ab;4888 | 0 | 0.025 |
| NC_005831.2;YP_003766.2;HCoV-NL63_1ab;4889 | 0 | 0.025 |
| NC_005831.2;YP_003766.2;HCoV-NL63_1ab;4890 | 0 | 0.025 |
| NC_005831.2;YP_003766.2;HCoV-NL63_1ab;4891 | 0 | 0.025 |
| NC_005831.2;YP_003766.2;HCoV-NL63_1ab;4892 | 0 | 0.025 |
| NC_005831.2;YP_003766.2;HCoV-NL63_1ab;4893 | 0 | 0 |
| NC_005831.2;YP_003766.2;HCoV-NL63_1ab;4894 | 0 | 0 |
| NC_005831.2;YP_003766.2;HCoV-NL63_1ab;4895 | 0 | 0 |
| NC_005831.2;YP_003766.2;HCoV-NL63_1ab;4896 | 0 | 0 |
| NC_005831.2;YP_003766.2;HCoV-NL63_1ab;4897 | 0 | 0 |
| NC_005831.2;YP_003766.2;HCoV-NL63_1ab;4898 | 0 | 0.025 |
| NC_005831.2;YP_003766.2;HCoV-NL63_1ab;4899 | 0 | 0 |
| NC_005831.2;YP_003766.2;HCoV-NL63_1ab;4900 | 0 | 0 |
| NC_005831.2;YP_003766.2;HCoV-NL63_1ab;4901 | 0 | 0 |
| NC_005831.2;YP_003766.2;HCoV-NL63_1ab;4902 | 0.05 | 0.025 |
| NC_005831.2;YP_003766.2;HCoV-NL63_1ab;4903 | 0.05 | 0.025 |
| NC_005831.2;YP_003766.2;HCoV-NL63_1ab;4904 | 0 | 0 |
| NC_005831.2;YP_003766.2;HCoV-NL63_1ab;4905 | 0 | 0 |
| NC_005831.2;YP_003766.2;HCoV-NL63_1ab;4906 | 0 | 0 |
| NC_005831.2;YP_003766.2;HCoV-NL63_1ab;4907 | 0 | 0 |
| NC_005831.2;YP_003766.2;HCoV-NL63_1ab;4908 | 0 | 0 |
| NC_005831.2;YP_003766.2;HCoV-NL63_1ab;4909 | 0 | 0 |
| NC_005831.2;YP_003766.2;HCoV-NL63_1ab;4910 | 0 | 0 |
| NC_005831.2;YP_003766.2;HCoV-NL63_1ab;4911 | 0 | 0 |

| | | |
|---|---|---|
| NC_005831.2;YP_003766.2;HCoV-NL63_1ab;4912 | 0 | 0 |
| NC_005831.2;YP_003766.2;HCoV-NL63_1ab;4913 | 0 | 0 |
| NC_005831.2;YP_003766.2;HCoV-NL63_1ab;4914 | 0 | 0 |
| NC_005831.2;YP_003766.2;HCoV-NL63_1ab;4915 | 0 | 0 |
| NC_005831.2;YP_003766.2;HCoV-NL63_1ab;4916 | 0 | 0 |
| NC_005831.2;YP_003766.2;HCoV-NL63_1ab;4917 | 0 | 0 |
| NC_005831.2;YP_003766.2;HCoV-NL63_1ab;4918 | 0 | 0 |
| NC_005831.2;YP_003766.2;HCoV-NL63_1ab;4919 | 0.05 | 0 |
| NC_005831.2;YP_003766.2;HCoV-NL63_1ab;4920 | 0.05 | 0.025 |
| NC_005831.2;YP_003766.2;HCoV-NL63_1ab;4921 | 0.2 | 0.1 |
| NC_005831.2;YP_003766.2;HCoV-NL63_1ab;4922 | 0.1 | 0.125 |
| NC_005831.2;YP_003766.2;HCoV-NL63_1ab;4923 | 0.2 | 0.125 |
| NC_005831.2;YP_003766.2;HCoV-NL63_1ab;4924 | 0.2 | 0.125 |
| NC_005831.2;YP_003766.2;HCoV-NL63_1ab;4925 | 0.15 | 0.15 |
| NC_005831.2;YP_003766.2;HCoV-NL63_1ab;4926 | 0.05 | 0.05 |
| NC_005831.2;YP_003766.2;HCoV-NL63_1ab;4927 | 0 | 0 |
| NC_005831.2;YP_003766.2;HCoV-NL63_1ab;4928 | 0 | 0 |
| NC_005831.2;YP_003766.2;HCoV-NL63_1ab;4929 | 0 | 0.025 |
| NC_005831.2;YP_003766.2;HCoV-NL63_1ab;4930 | 0 | 0 |
| NC_005831.2;YP_003766.2;HCoV-NL63_1ab;4931 | 0 | 0 |
| NC_005831.2;YP_003766.2;HCoV-NL63_1ab;4932 | 0 | 0 |
| NC_005831.2;YP_003766.2;HCoV-NL63_1ab;4933 | 0 | 0 |
| NC_005831.2;YP_003766.2;HCoV-NL63_1ab;4934 | 0 | 0 |
| NC_005831.2;YP_003766.2;HCoV-NL63_1ab;4935 | 0 | 0 |
| NC_005831.2;YP_003766.2;HCoV-NL63_1ab;4936 | 0 | 0.025 |
| NC_005831.2;YP_003766.2;HCoV-NL63_1ab;4937 | 0 | 0.05 |
| NC_005831.2;YP_003766.2;HCoV-NL63_1ab;4938 | 0 | 0.05 |
| NC_005831.2;YP_003766.2;HCoV-NL63_1ab;4939 | 0 | 0.025 |
| NC_005831.2;YP_003766.2;HCoV-NL63_1ab;4940 | 0.05 | 0 |
| NC_005831.2;YP_003766.2;HCoV-NL63_1ab;4941 | 0 | 0 |
| NC_005831.2;YP_003766.2;HCoV-NL63_1ab;4942 | 0 | 0 |
| NC_005831.2;YP_003766.2;HCoV-NL63_1ab;4943 | 0 | 0 |
| NC_005831.2;YP_003766.2;HCoV-NL63_1ab;4944 | 0 | 0 |
| NC_005831.2;YP_003766.2;HCoV-NL63_1ab;4945 | 0 | 0.025 |
| NC_005831.2;YP_003766.2;HCoV-NL63_1ab;4946 | 0 | 0 |
| NC_005831.2;YP_003766.2;HCoV-NL63_1ab;4947 | 0 | 0 |
| NC_005831.2;YP_003766.2;HCoV-NL63_1ab;4948 | 0 | 0 |
| NC_005831.2;YP_003766.2;HCoV-NL63_1ab;4949 | 0 | 0 |
| NC_005831.2;YP_003766.2;HCoV-NL63_1ab;4950 | 0 | 0 |
| NC_005831.2;YP_003766.2;HCoV-NL63_1ab;4951 | 0 | 0 |
| NC_005831.2;YP_003766.2;HCoV-NL63_1ab;4952 | 0 | 0 |
| NC_005831.2;YP_003766.2;HCoV-NL63_1ab;4953 | 0 | 0 |
| NC_005831.2;YP_003766.2;HCoV-NL63_1ab;4954 | 0 | 0 |

| | | |
|---|---|---|
| NC_005831.2;YP_003766.2;HCoV-NL63_1ab;4955 | 0 | 0 |
| NC_005831.2;YP_003766.2;HCoV-NL63_1ab;4956 | 0 | 0.025 |
| NC_005831.2;YP_003766.2;HCoV-NL63_1ab;4957 | 0.05 | 0.025 |
| NC_005831.2;YP_003766.2;HCoV-NL63_1ab;4958 | 0 | 0 |
| NC_005831.2;YP_003766.2;HCoV-NL63_1ab;4959 | 0.05 | 0 |
| NC_005831.2;YP_003766.2;HCoV-NL63_1ab;4960 | 0 | 0 |
| NC_005831.2;YP_003766.2;HCoV-NL63_1ab;4961 | 0 | 0 |
| NC_005831.2;YP_003766.2;HCoV-NL63_1ab;4962 | 0 | 0 |
| NC_005831.2;YP_003766.2;HCoV-NL63_1ab;4963 | 0 | 0 |
| NC_005831.2;YP_003766.2;HCoV-NL63_1ab;4964 | 0 | 0 |
| NC_005831.2;YP_003766.2;HCoV-NL63_1ab;4965 | 0 | 0 |
| NC_005831.2;YP_003766.2;HCoV-NL63_1ab;4966 | 0 | 0 |
| NC_005831.2;YP_003766.2;HCoV-NL63_1ab;4967 | 0 | 0 |
| NC_005831.2;YP_003766.2;HCoV-NL63_1ab;4968 | 0 | 0 |
| NC_005831.2;YP_003766.2;HCoV-NL63_1ab;4969 | 0 | 0 |
| NC_005831.2;YP_003766.2;HCoV-NL63_1ab;4970 | 0 | 0 |
| NC_005831.2;YP_003766.2;HCoV-NL63_1ab;4971 | 0 | 0 |
| NC_005831.2;YP_003766.2;HCoV-NL63_1ab;4972 | 0 | 0 |
| NC_005831.2;YP_003766.2;HCoV-NL63_1ab;4973 | 0 | 0 |
| NC_005831.2;YP_003766.2;HCoV-NL63_1ab;4974 | 0 | 0 |
| NC_005831.2;YP_003766.2;HCoV-NL63_1ab;4975 | 0 | 0 |
| NC_005831.2;YP_003766.2;HCoV-NL63_1ab;4976 | 0 | 0 |
| NC_005831.2;YP_003766.2;HCoV-NL63_1ab;4977 | 0 | 0 |
| NC_005831.2;YP_003766.2;HCoV-NL63_1ab;4978 | 0 | 0 |
| NC_005831.2;YP_003766.2;HCoV-NL63_1ab;4979 | 0 | 0 |
| NC_005831.2;YP_003766.2;HCoV-NL63_1ab;4980 | 0 | 0 |
| NC_005831.2;YP_003766.2;HCoV-NL63_1ab;4981 | 0 | 0 |
| NC_005831.2;YP_003766.2;HCoV-NL63_1ab;4982 | 0 | 0 |
| NC_005831.2;YP_003766.2;HCoV-NL63_1ab;4983 | 0 | 0 |
| NC_005831.2;YP_003766.2;HCoV-NL63_1ab;4984 | 0 | 0 |
| NC_005831.2;YP_003766.2;HCoV-NL63_1ab;4985 | 0 | 0 |
| NC_005831.2;YP_003766.2;HCoV-NL63_1ab;4986 | 0 | 0 |
| NC_005831.2;YP_003766.2;HCoV-NL63_1ab;4987 | 0 | 0 |
| NC_005831.2;YP_003766.2;HCoV-NL63_1ab;4988 | 0 | 0 |
| NC_005831.2;YP_003766.2;HCoV-NL63_1ab;4989 | 0 | 0 |
| NC_005831.2;YP_003766.2;HCoV-NL63_1ab;4990 | 0 | 0 |
| NC_005831.2;YP_003766.2;HCoV-NL63_1ab;4991 | 0 | 0 |
| NC_005831.2;YP_003766.2;HCoV-NL63_1ab;4992 | 0 | 0 |
| NC_005831.2;YP_003766.2;HCoV-NL63_1ab;4993 | 0 | 0 |
| NC_005831.2;YP_003766.2;HCoV-NL63_1ab;4994 | 0 | 0 |
| NC_005831.2;YP_003766.2;HCoV-NL63_1ab;4995 | 0 | 0 |
| NC_005831.2;YP_003766.2;HCoV-NL63_1ab;4996 | 0 | 0 |
| NC_005831.2;YP_003766.2;HCoV-NL63_1ab;4997 | 0 | 0 |

| | | |
|---|---|---|
| NC_005831.2;YP_003766.2;HCoV-NL63_1ab;4998 | 0 | 0 |
| NC_005831.2;YP_003766.2;HCoV-NL63_1ab;4999 | 0 | 0 |
| NC_005831.2;YP_003766.2;HCoV-NL63_1ab;5000 | 0 | 0 |
| NC_005831.2;YP_003766.2;HCoV-NL63_1ab;5001 | 0 | 0 |
| NC_005831.2;YP_003766.2;HCoV-NL63_1ab;5002 | 0 | 0 |
| NC_005831.2;YP_003766.2;HCoV-NL63_1ab;5003 | 0 | 0 |
| NC_005831.2;YP_003766.2;HCoV-NL63_1ab;5004 | 0 | 0 |
| NC_005831.2;YP_003766.2;HCoV-NL63_1ab;5005 | 0 | 0 |
| NC_005831.2;YP_003766.2;HCoV-NL63_1ab;5006 | 0 | 0 |
| NC_005831.2;YP_003766.2;HCoV-NL63_1ab;5007 | 0 | 0 |
| NC_005831.2;YP_003766.2;HCoV-NL63_1ab;5008 | 0 | 0 |
| NC_005831.2;YP_003766.2;HCoV-NL63_1ab;5009 | 0 | 0 |
| NC_005831.2;YP_003766.2;HCoV-NL63_1ab;5010 | 0 | 0 |
| NC_005831.2;YP_003766.2;HCoV-NL63_1ab;5011 | 0 | 0 |
| NC_005831.2;YP_003766.2;HCoV-NL63_1ab;5012 | 0 | 0 |
| NC_005831.2;YP_003766.2;HCoV-NL63_1ab;5013 | 0 | 0 |
| NC_005831.2;YP_003766.2;HCoV-NL63_1ab;5014 | 0 | 0 |
| NC_005831.2;YP_003766.2;HCoV-NL63_1ab;5015 | 0 | 0 |
| NC_005831.2;YP_003766.2;HCoV-NL63_1ab;5016 | 0 | 0.05 |
| NC_005831.2;YP_003766.2;HCoV-NL63_1ab;5017 | 0 | 0.025 |
| NC_005831.2;YP_003766.2;HCoV-NL63_1ab;5018 | 0 | 0.025 |
| NC_005831.2;YP_003766.2;HCoV-NL63_1ab;5019 | 0.05 | 0.025 |
| NC_005831.2;YP_003766.2;HCoV-NL63_1ab;5020 | 0 | 0.025 |
| NC_005831.2;YP_003766.2;HCoV-NL63_1ab;5021 | 0 | 0.025 |
| NC_005831.2;YP_003766.2;HCoV-NL63_1ab;5022 | 0 | 0 |
| NC_005831.2;YP_003766.2;HCoV-NL63_1ab;5023 | 0 | 0 |
| NC_005831.2;YP_003766.2;HCoV-NL63_1ab;5024 | 0 | 0 |
| NC_005831.2;YP_003766.2;HCoV-NL63_1ab;5025 | 0 | 0 |
| NC_005831.2;YP_003766.2;HCoV-NL63_1ab;5026 | 0.1 | 0.025 |
| NC_005831.2;YP_003766.2;HCoV-NL63_1ab;5027 | 0 | 0 |
| NC_005831.2;YP_003766.2;HCoV-NL63_1ab;5028 | 0 | 0 |
| NC_005831.2;YP_003766.2;HCoV-NL63_1ab;5029 | 0 | 0 |
| NC_005831.2;YP_003766.2;HCoV-NL63_1ab;5030 | 0 | 0 |
| NC_005831.2;YP_003766.2;HCoV-NL63_1ab;5031 | 0 | 0 |
| NC_005831.2;YP_003766.2;HCoV-NL63_1ab;5032 | 0 | 0 |
| NC_005831.2;YP_003766.2;HCoV-NL63_1ab;5033 | 0 | 0 |
| NC_005831.2;YP_003766.2;HCoV-NL63_1ab;5034 | 0 | 0 |
| NC_005831.2;YP_003766.2;HCoV-NL63_1ab;5035 | 0 | 0 |
| NC_005831.2;YP_003766.2;HCoV-NL63_1ab;5036 | 0 | 0 |
| NC_005831.2;YP_003766.2;HCoV-NL63_1ab;5037 | 0 | 0 |
| NC_005831.2;YP_003766.2;HCoV-NL63_1ab;5038 | 0 | 0 |
| NC_005831.2;YP_003766.2;HCoV-NL63_1ab;5039 | 0 | 0 |
| NC_005831.2;YP_003766.2;HCoV-NL63_1ab;5040 | 0 | 0 |

| | | |
|---|---|---|
| NC_005831.2;YP_003766.2;HCoV-NL63_1ab;5041 | 0 | 0 |
| NC_005831.2;YP_003766.2;HCoV-NL63_1ab;5042 | 0 | 0 |
| NC_005831.2;YP_003766.2;HCoV-NL63_1ab;5043 | 0 | 0 |
| NC_005831.2;YP_003766.2;HCoV-NL63_1ab;5044 | 0 | 0 |
| NC_005831.2;YP_003766.2;HCoV-NL63_1ab;5045 | 0 | 0 |
| NC_005831.2;YP_003766.2;HCoV-NL63_1ab;5046 | 0 | 0 |
| NC_005831.2;YP_003766.2;HCoV-NL63_1ab;5047 | 0 | 0 |
| NC_005831.2;YP_003766.2;HCoV-NL63_1ab;5048 | 0 | 0 |
| NC_005831.2;YP_003766.2;HCoV-NL63_1ab;5049 | 0 | 0 |
| NC_005831.2;YP_003766.2;HCoV-NL63_1ab;5050 | 0 | 0 |
| NC_005831.2;YP_003766.2;HCoV-NL63_1ab;5051 | 0 | 0 |
| NC_005831.2;YP_003766.2;HCoV-NL63_1ab;5052 | 0 | 0.025 |
| NC_005831.2;YP_003766.2;HCoV-NL63_1ab;5053 | 0.05 | 0 |
| NC_005831.2;YP_003766.2;HCoV-NL63_1ab;5054 | 0.05 | 0 |
| NC_005831.2;YP_003766.2;HCoV-NL63_1ab;5055 | 0.05 | 0.05 |
| NC_005831.2;YP_003766.2;HCoV-NL63_1ab;5056 | 0.05 | 0.025 |
| NC_005831.2;YP_003766.2;HCoV-NL63_1ab;5057 | 0.1 | 0.1 |
| NC_005831.2;YP_003766.2;HCoV-NL63_1ab;5058 | 0.1 | 0.05 |
| NC_005831.2;YP_003766.2;HCoV-NL63_1ab;5059 | 0.05 | 0.05 |
| NC_005831.2;YP_003766.2;HCoV-NL63_1ab;5060 | 0.05 | 0.05 |
| NC_005831.2;YP_003766.2;HCoV-NL63_1ab;5061 | 0.05 | 0.05 |
| NC_005831.2;YP_003766.2;HCoV-NL63_1ab;5062 | 0.05 | 0.05 |
| NC_005831.2;YP_003766.2;HCoV-NL63_1ab;5063 | 0.1 | 0.05 |
| NC_005831.2;YP_003766.2;HCoV-NL63_1ab;5064 | 0.1 | 0.05 |
| NC_005831.2;YP_003766.2;HCoV-NL63_1ab;5065 | 0.05 | 0.075 |
| NC_005831.2;YP_003766.2;HCoV-NL63_1ab;5066 | 0.05 | 0.05 |
| NC_005831.2;YP_003766.2;HCoV-NL63_1ab;5067 | 0.05 | 0.025 |
| NC_005831.2;YP_003766.2;HCoV-NL63_1ab;5068 | 0 | 0 |
| NC_005831.2;YP_003766.2;HCoV-NL63_1ab;5069 | 0 | 0.025 |
| NC_005831.2;YP_003766.2;HCoV-NL63_1ab;5070 | 0 | 0.025 |
| NC_005831.2;YP_003766.2;HCoV-NL63_1ab;5071 | 0 | 0 |
| NC_005831.2;YP_003766.2;HCoV-NL63_1ab;5072 | 0 | 0 |
| NC_005831.2;YP_003766.2;HCoV-NL63_1ab;5073 | 0 | 0.025 |
| NC_005831.2;YP_003766.2;HCoV-NL63_1ab;5074 | 0 | 0.025 |
| NC_005831.2;YP_003766.2;HCoV-NL63_1ab;5075 | 0 | 0 |
| NC_005831.2;YP_003766.2;HCoV-NL63_1ab;5076 | 0 | 0 |
| NC_005831.2;YP_003766.2;HCoV-NL63_1ab;5077 | 0 | 0 |
| NC_005831.2;YP_003766.2;HCoV-NL63_1ab;5078 | 0 | 0 |
| NC_005831.2;YP_003766.2;HCoV-NL63_1ab;5079 | 0 | 0 |
| NC_005831.2;YP_003766.2;HCoV-NL63_1ab;5080 | 0 | 0.075 |
| NC_005831.2;YP_003766.2;HCoV-NL63_1ab;5081 | 0.05 | 0 |
| NC_005831.2;YP_003766.2;HCoV-NL63_1ab;5082 | 0.25 | 0.125 |
| NC_005831.2;YP_003766.2;HCoV-NL63_1ab;5083 | 0.15 | 0.1 |

| | | |
|---|---|---|
| NC_005831.2;YP_003766.2;HCoV-NL63_1ab;5084 | 0.25 | 0.15 |
| NC_005831.2;YP_003766.2;HCoV-NL63_1ab;5085 | 0.15 | 0.15 |
| NC_005831.2;YP_003766.2;HCoV-NL63_1ab;5086 | 0.15 | 0.125 |
| NC_005831.2;YP_003766.2;HCoV-NL63_1ab;5087 | 0.15 | 0.125 |
| NC_005831.2;YP_003766.2;HCoV-NL63_1ab;5088 | 0.15 | 0.025 |
| NC_005831.2;YP_003766.2;HCoV-NL63_1ab;5089 | 0.1 | 0.05 |
| NC_005831.2;YP_003766.2;HCoV-NL63_1ab;5090 | 0.15 | 0.05 |
| NC_005831.2;YP_003766.2;HCoV-NL63_1ab;5091 | 0 | 0.025 |
| NC_005831.2;YP_003766.2;HCoV-NL63_1ab;5092 | 0 | 0 |
| NC_005831.2;YP_003766.2;HCoV-NL63_1ab;5093 | 0 | 0 |
| NC_005831.2;YP_003766.2;HCoV-NL63_1ab;5094 | 0 | 0 |
| NC_005831.2;YP_003766.2;HCoV-NL63_1ab;5095 | 0 | 0 |
| NC_005831.2;YP_003766.2;HCoV-NL63_1ab;5096 | 0 | 0 |
| NC_005831.2;YP_003766.2;HCoV-NL63_1ab;5097 | 0.1 | 0 |
| NC_005831.2;YP_003766.2;HCoV-NL63_1ab;5098 | 0 | 0 |
| NC_005831.2;YP_003766.2;HCoV-NL63_1ab;5099 | 0.05 | 0 |
| NC_005831.2;YP_003766.2;HCoV-NL63_1ab;5100 | 0 | 0 |
| NC_005831.2;YP_003766.2;HCoV-NL63_1ab;5101 | 0 | 0 |
| NC_005831.2;YP_003766.2;HCoV-NL63_1ab;5102 | 0 | 0.05 |
| NC_005831.2;YP_003766.2;HCoV-NL63_1ab;5103 | 0 | 0.025 |
| NC_005831.2;YP_003766.2;HCoV-NL63_1ab;5104 | 0 | 0 |
| NC_005831.2;YP_003766.2;HCoV-NL63_1ab;5105 | 0 | 0.1 |
| NC_005831.2;YP_003766.2;HCoV-NL63_1ab;5106 | 0 | 0.025 |
| NC_005831.2;YP_003766.2;HCoV-NL63_1ab;5107 | 0 | 0.025 |
| NC_005831.2;YP_003766.2;HCoV-NL63_1ab;5108 | 0.05 | 0 |
| NC_005831.2;YP_003766.2;HCoV-NL63_1ab;5109 | 0 | 0.025 |
| NC_005831.2;YP_003766.2;HCoV-NL63_1ab;5110 | 0 | 0 |
| NC_005831.2;YP_003766.2;HCoV-NL63_1ab;5111 | 0.05 | 0.025 |
| NC_005831.2;YP_003766.2;HCoV-NL63_1ab;5112 | 0.05 | 0.025 |
| NC_005831.2;YP_003766.2;HCoV-NL63_1ab;5113 | 0 | 0 |
| NC_005831.2;YP_003766.2;HCoV-NL63_1ab;5114 | 0 | 0 |
| NC_005831.2;YP_003766.2;HCoV-NL63_1ab;5115 | 0 | 0 |
| NC_005831.2;YP_003766.2;HCoV-NL63_1ab;5116 | 0.05 | 0 |
| NC_005831.2;YP_003766.2;HCoV-NL63_1ab;5117 | 0.05 | 0 |
| NC_005831.2;YP_003766.2;HCoV-NL63_1ab;5118 | 0 | 0 |
| NC_005831.2;YP_003766.2;HCoV-NL63_1ab;5119 | 0.1 | 0.075 |
| NC_005831.2;YP_003766.2;HCoV-NL63_1ab;5120 | 0.1 | 0.05 |
| NC_005831.2;YP_003766.2;HCoV-NL63_1ab;5121 | 0 | 0 |
| NC_005831.2;YP_003766.2;HCoV-NL63_1ab;5122 | 0 | 0 |
| NC_005831.2;YP_003766.2;HCoV-NL63_1ab;5123 | 0 | 0.025 |
| NC_005831.2;YP_003766.2;HCoV-NL63_1ab;5124 | 0.05 | 0.025 |
| NC_005831.2;YP_003766.2;HCoV-NL63_1ab;5125 | 0 | 0 |
| NC_005831.2;YP_003766.2;HCoV-NL63_1ab;5126 | 0 | 0.025 |

| | | |
|---|---|---|
| NC_005831.2;YP_003766.2;HCoV-NL63_1ab;5127 | 0 | 0.025 |
| NC_005831.2;YP_003766.2;HCoV-NL63_1ab;5128 | 0 | 0 |
| NC_005831.2;YP_003766.2;HCoV-NL63_1ab;5129 | 0 | 0.025 |
| NC_005831.2;YP_003766.2;HCoV-NL63_1ab;5130 | 0 | 0.025 |
| NC_005831.2;YP_003766.2;HCoV-NL63_1ab;5131 | 0 | 0.025 |
| NC_005831.2;YP_003766.2;HCoV-NL63_1ab;5132 | 0 | 0.025 |
| NC_005831.2;YP_003766.2;HCoV-NL63_1ab;5133 | 0.05 | 0.025 |
| NC_005831.2;YP_003766.2;HCoV-NL63_1ab;5134 | 0 | 0.025 |
| NC_005831.2;YP_003766.2;HCoV-NL63_1ab;5135 | 0 | 0.05 |
| NC_005831.2;YP_003766.2;HCoV-NL63_1ab;5136 | 0 | 0.05 |
| NC_005831.2;YP_003766.2;HCoV-NL63_1ab;5137 | 0 | 0.025 |
| NC_005831.2;YP_003766.2;HCoV-NL63_1ab;5138 | 0.05 | 0.025 |
| NC_005831.2;YP_003766.2;HCoV-NL63_1ab;5139 | 0 | 0 |
| NC_005831.2;YP_003766.2;HCoV-NL63_1ab;5140 | 0 | 0 |
| NC_005831.2;YP_003766.2;HCoV-NL63_1ab;5141 | 0 | 0 |
| NC_005831.2;YP_003766.2;HCoV-NL63_1ab;5142 | 0 | 0 |
| NC_005831.2;YP_003766.2;HCoV-NL63_1ab;5143 | 0 | 0 |
| NC_005831.2;YP_003766.2;HCoV-NL63_1ab;5144 | 0 | 0 |
| NC_005831.2;YP_003766.2;HCoV-NL63_1ab;5145 | 0 | 0 |
| NC_005831.2;YP_003766.2;HCoV-NL63_1ab;5146 | 0 | 0 |
| NC_005831.2;YP_003766.2;HCoV-NL63_1ab;5147 | 0 | 0 |
| NC_005831.2;YP_003766.2;HCoV-NL63_1ab;5148 | 0 | 0 |
| NC_005831.2;YP_003766.2;HCoV-NL63_1ab;5149 | 0 | 0 |
| NC_005831.2;YP_003766.2;HCoV-NL63_1ab;5150 | 0 | 0 |
| NC_005831.2;YP_003766.2;HCoV-NL63_1ab;5151 | 0 | 0 |
| NC_005831.2;YP_003766.2;HCoV-NL63_1ab;5152 | 0 | 0 |
| NC_005831.2;YP_003766.2;HCoV-NL63_1ab;5153 | 0 | 0 |
| NC_005831.2;YP_003766.2;HCoV-NL63_1ab;5154 | 0 | 0 |
| NC_005831.2;YP_003766.2;HCoV-NL63_1ab;5155 | 0 | 0 |
| NC_005831.2;YP_003766.2;HCoV-NL63_1ab;5156 | 0 | 0 |
| NC_005831.2;YP_003766.2;HCoV-NL63_1ab;5157 | 0.05 | 0.025 |
| NC_005831.2;YP_003766.2;HCoV-NL63_1ab;5158 | 0 | 0 |
| NC_005831.2;YP_003766.2;HCoV-NL63_1ab;5159 | 0 | 0 |
| NC_005831.2;YP_003766.2;HCoV-NL63_1ab;5160 | 0 | 0 |
| NC_005831.2;YP_003766.2;HCoV-NL63_1ab;5161 | 0 | 0 |
| NC_005831.2;YP_003766.2;HCoV-NL63_1ab;5162 | 0 | 0 |
| NC_005831.2;YP_003766.2;HCoV-NL63_1ab;5163 | 0 | 0 |
| NC_005831.2;YP_003766.2;HCoV-NL63_1ab;5164 | 0.05 | 0 |
| NC_005831.2;YP_003766.2;HCoV-NL63_1ab;5165 | 0 | 0 |
| NC_005831.2;YP_003766.2;HCoV-NL63_1ab;5166 | 0 | 0 |
| NC_005831.2;YP_003766.2;HCoV-NL63_1ab;5167 | 0 | 0 |
| NC_005831.2;YP_003766.2;HCoV-NL63_1ab;5168 | 0 | 0.075 |
| NC_005831.2;YP_003766.2;HCoV-NL63_1ab;5169 | 0 | 0 |

| | | |
|---|---|---|
| NC_005831.2;YP_003766.2;HCoV-NL63_1ab;5170 | 0 | 0.075 |
| NC_005831.2;YP_003766.2;HCoV-NL63_1ab;5171 | 0 | 0 |
| NC_005831.2;YP_003766.2;HCoV-NL63_1ab;5172 | 0 | 0 |
| NC_005831.2;YP_003766.2;HCoV-NL63_1ab;5173 | 0 | 0 |
| NC_005831.2;YP_003766.2;HCoV-NL63_1ab;5174 | 0 | 0.025 |
| NC_005831.2;YP_003766.2;HCoV-NL63_1ab;5175 | 0 | 0 |
| NC_005831.2;YP_003766.2;HCoV-NL63_1ab;5176 | 0 | 0 |
| NC_005831.2;YP_003766.2;HCoV-NL63_1ab;5177 | 0 | 0 |
| NC_005831.2;YP_003766.2;HCoV-NL63_1ab;5178 | 0 | 0 |
| NC_005831.2;YP_003766.2;HCoV-NL63_1ab;5179 | 0 | 0 |
| NC_005831.2;YP_003766.2;HCoV-NL63_1ab;5180 | 0 | 0 |
| NC_005831.2;YP_003766.2;HCoV-NL63_1ab;5181 | 0.05 | 0.025 |
| NC_005831.2;YP_003766.2;HCoV-NL63_1ab;5182 | 0 | 0 |
| NC_005831.2;YP_003766.2;HCoV-NL63_1ab;5183 | 0 | 0 |
| NC_005831.2;YP_003766.2;HCoV-NL63_1ab;5184 | 0 | 0 |
| NC_005831.2;YP_003766.2;HCoV-NL63_1ab;5185 | 0 | 0 |
| NC_005831.2;YP_003766.2;HCoV-NL63_1ab;5186 | 0 | 0 |
| NC_005831.2;YP_003766.2;HCoV-NL63_1ab;5187 | 0 | 0 |
| NC_005831.2;YP_003766.2;HCoV-NL63_1ab;5188 | 0 | 0 |
| NC_005831.2;YP_003766.2;HCoV-NL63_1ab;5189 | 0 | 0 |
| NC_005831.2;YP_003766.2;HCoV-NL63_1ab;5190 | 0 | 0 |
| NC_005831.2;YP_003766.2;HCoV-NL63_1ab;5191 | 0 | 0 |
| NC_005831.2;YP_003766.2;HCoV-NL63_1ab;5192 | 0 | 0 |
| NC_005831.2;YP_003766.2;HCoV-NL63_1ab;5193 | 0 | 0 |
| NC_005831.2;YP_003766.2;HCoV-NL63_1ab;5194 | 0 | 0 |
| NC_005831.2;YP_003766.2;HCoV-NL63_1ab;5195 | 0 | 0 |
| NC_005831.2;YP_003766.2;HCoV-NL63_1ab;5196 | 0 | 0 |
| NC_005831.2;YP_003766.2;HCoV-NL63_1ab;5197 | 0 | 0 |
| NC_005831.2;YP_003766.2;HCoV-NL63_1ab;5198 | 0 | 0 |
| NC_005831.2;YP_003766.2;HCoV-NL63_1ab;5199 | 0 | 0 |
| NC_005831.2;YP_003766.2;HCoV-NL63_1ab;5200 | 0 | 0 |
| NC_005831.2;YP_003766.2;HCoV-NL63_1ab;5201 | 0 | 0 |
| NC_005831.2;YP_003766.2;HCoV-NL63_1ab;5202 | 0 | 0 |
| NC_005831.2;YP_003766.2;HCoV-NL63_1ab;5203 | 0.05 | 0 |
| NC_005831.2;YP_003766.2;HCoV-NL63_1ab;5204 | 0.05 | 0 |
| NC_005831.2;YP_003766.2;HCoV-NL63_1ab;5205 | 0.05 | 0.075 |
| NC_005831.2;YP_003766.2;HCoV-NL63_1ab;5206 | 0 | 0 |
| NC_005831.2;YP_003766.2;HCoV-NL63_1ab;5207 | 0 | 0 |
| NC_005831.2;YP_003766.2;HCoV-NL63_1ab;5208 | 0 | 0 |
| NC_005831.2;YP_003766.2;HCoV-NL63_1ab;5209 | 0.05 | 0 |
| NC_005831.2;YP_003766.2;HCoV-NL63_1ab;5210 | 0 | 0 |
| NC_005831.2;YP_003766.2;HCoV-NL63_1ab;5211 | 0 | 0 |
| NC_005831.2;YP_003766.2;HCoV-NL63_1ab;5212 | 0 | 0 |

| | | |
|---|---|---|
| NC_005831.2;YP_003766.2;HCoV-NL63_1ab;5213 | 0 | 0 |
| NC_005831.2;YP_003766.2;HCoV-NL63_1ab;5214 | 0 | 0 |
| NC_005831.2;YP_003766.2;HCoV-NL63_1ab;5215 | 0 | 0 |
| NC_005831.2;YP_003766.2;HCoV-NL63_1ab;5216 | 0 | 0 |
| NC_005831.2;YP_003766.2;HCoV-NL63_1ab;5217 | 0 | 0 |
| NC_005831.2;YP_003766.2;HCoV-NL63_1ab;5218 | 0 | 0.025 |
| NC_005831.2;YP_003766.2;HCoV-NL63_1ab;5219 | 0 | 0 |
| NC_005831.2;YP_003766.2;HCoV-NL63_1ab;5220 | 0 | 0 |
| NC_005831.2;YP_003766.2;HCoV-NL63_1ab;5221 | 0 | 0.025 |
| NC_005831.2;YP_003766.2;HCoV-NL63_1ab;5222 | 0 | 0.025 |
| NC_005831.2;YP_003766.2;HCoV-NL63_1ab;5223 | 0 | 0.05 |
| NC_005831.2;YP_003766.2;HCoV-NL63_1ab;5224 | 0 | 0.025 |
| NC_005831.2;YP_003766.2;HCoV-NL63_1ab;5225 | 0 | 0.025 |
| NC_005831.2;YP_003766.2;HCoV-NL63_1ab;5226 | 0 | 0.025 |
| NC_005831.2;YP_003766.2;HCoV-NL63_1ab;5227 | 0 | 0.025 |
| NC_005831.2;YP_003766.2;HCoV-NL63_1ab;5228 | 0 | 0.025 |
| NC_005831.2;YP_003766.2;HCoV-NL63_1ab;5229 | 0 | 0.025 |
| NC_005831.2;YP_003766.2;HCoV-NL63_1ab;5230 | 0 | 0 |
| NC_005831.2;YP_003766.2;HCoV-NL63_1ab;5231 | 0 | 0 |
| NC_005831.2;YP_003766.2;HCoV-NL63_1ab;5232 | 0 | 0 |
| NC_005831.2;YP_003766.2;HCoV-NL63_1ab;5233 | 0 | 0 |
| NC_005831.2;YP_003766.2;HCoV-NL63_1ab;5234 | 0 | 0 |
| NC_005831.2;YP_003766.2;HCoV-NL63_1ab;5235 | 0 | 0 |
| NC_005831.2;YP_003766.2;HCoV-NL63_1ab;5236 | 0 | 0 |
| NC_005831.2;YP_003766.2;HCoV-NL63_1ab;5237 | 0 | 0 |
| NC_005831.2;YP_003766.2;HCoV-NL63_1ab;5238 | 0 | 0 |
| NC_005831.2;YP_003766.2;HCoV-NL63_1ab;5239 | 0 | 0 |
| NC_005831.2;YP_003766.2;HCoV-NL63_1ab;5240 | 0 | 0.025 |
| NC_005831.2;YP_003766.2;HCoV-NL63_1ab;5241 | 0 | 0.025 |
| NC_005831.2;YP_003766.2;HCoV-NL63_1ab;5242 | 0 | 0 |
| NC_005831.2;YP_003766.2;HCoV-NL63_1ab;5243 | 0 | 0 |
| NC_005831.2;YP_003766.2;HCoV-NL63_1ab;5244 | 0 | 0 |
| NC_005831.2;YP_003766.2;HCoV-NL63_1ab;5245 | 0 | 0 |
| NC_005831.2;YP_003766.2;HCoV-NL63_1ab;5246 | 0 | 0 |
| NC_005831.2;YP_003766.2;HCoV-NL63_1ab;5247 | 0 | 0 |
| NC_005831.2;YP_003766.2;HCoV-NL63_1ab;5248 | 0 | 0 |
| NC_005831.2;YP_003766.2;HCoV-NL63_1ab;5249 | 0 | 0 |
| NC_005831.2;YP_003766.2;HCoV-NL63_1ab;5250 | 0 | 0 |
| NC_005831.2;YP_003766.2;HCoV-NL63_1ab;5251 | 0 | 0 |
| NC_005831.2;YP_003766.2;HCoV-NL63_1ab;5252 | 0 | 0 |
| NC_005831.2;YP_003766.2;HCoV-NL63_1ab;5253 | 0 | 0 |
| NC_005831.2;YP_003766.2;HCoV-NL63_1ab;5254 | 0 | 0 |
| NC_005831.2;YP_003766.2;HCoV-NL63_1ab;5255 | 0 | 0 |

| | | |
|---|---|---|
| NC_005831.2;YP_003766.2;HCoV-NL63_1ab;5256 | 0 | 0 |
| NC_005831.2;YP_003766.2;HCoV-NL63_1ab;5257 | 0 | 0 |
| NC_005831.2;YP_003766.2;HCoV-NL63_1ab;5258 | 0 | 0 |
| NC_005831.2;YP_003766.2;HCoV-NL63_1ab;5259 | 0 | 0 |
| NC_005831.2;YP_003766.2;HCoV-NL63_1ab;5260 | 0 | 0 |
| NC_005831.2;YP_003766.2;HCoV-NL63_1ab;5261 | 0 | 0 |
| NC_005831.2;YP_003766.2;HCoV-NL63_1ab;5262 | 0 | 0 |
| NC_005831.2;YP_003766.2;HCoV-NL63_1ab;5263 | 0 | 0 |
| NC_005831.2;YP_003766.2;HCoV-NL63_1ab;5264 | 0 | 0 |
| NC_005831.2;YP_003766.2;HCoV-NL63_1ab;5265 | 0 | 0 |
| NC_005831.2;YP_003766.2;HCoV-NL63_1ab;5266 | 0 | 0 |
| NC_005831.2;YP_003766.2;HCoV-NL63_1ab;5267 | 0 | 0 |
| NC_005831.2;YP_003766.2;HCoV-NL63_1ab;5268 | 0 | 0 |
| NC_005831.2;YP_003766.2;HCoV-NL63_1ab;5269 | 0 | 0 |
| NC_005831.2;YP_003766.2;HCoV-NL63_1ab;5270 | 0 | 0 |
| NC_005831.2;YP_003766.2;HCoV-NL63_1ab;5271 | 0 | 0 |
| NC_005831.2;YP_003766.2;HCoV-NL63_1ab;5272 | 0 | 0 |
| NC_005831.2;YP_003766.2;HCoV-NL63_1ab;5273 | 0 | 0.025 |
| NC_005831.2;YP_003766.2;HCoV-NL63_1ab;5274 | 0 | 0 |
| NC_005831.2;YP_003766.2;HCoV-NL63_1ab;5275 | 0 | 0 |
| NC_005831.2;YP_003766.2;HCoV-NL63_1ab;5276 | 0 | 0 |
| NC_005831.2;YP_003766.2;HCoV-NL63_1ab;5277 | 0 | 0 |
| NC_005831.2;YP_003766.2;HCoV-NL63_1ab;5278 | 0 | 0 |
| NC_005831.2;YP_003766.2;HCoV-NL63_1ab;5279 | 0 | 0 |
| NC_005831.2;YP_003766.2;HCoV-NL63_1ab;5280 | 0 | 0 |
| NC_005831.2;YP_003766.2;HCoV-NL63_1ab;5281 | 0 | 0 |
| NC_005831.2;YP_003766.2;HCoV-NL63_1ab;5282 | 0 | 0 |
| NC_005831.2;YP_003766.2;HCoV-NL63_1ab;5283 | 0 | 0 |
| NC_005831.2;YP_003766.2;HCoV-NL63_1ab;5284 | 0 | 0 |
| NC_005831.2;YP_003766.2;HCoV-NL63_1ab;5285 | 0 | 0 |
| NC_005831.2;YP_003766.2;HCoV-NL63_1ab;5286 | 0 | 0 |
| NC_005831.2;YP_003766.2;HCoV-NL63_1ab;5287 | 0 | 0 |
| NC_005831.2;YP_003766.2;HCoV-NL63_1ab;5288 | 0 | 0 |
| NC_005831.2;YP_003766.2;HCoV-NL63_1ab;5289 | 0 | 0 |
| NC_005831.2;YP_003766.2;HCoV-NL63_1ab;5290 | 0 | 0 |
| NC_005831.2;YP_003766.2;HCoV-NL63_1ab;5291 | 0 | 0 |
| NC_005831.2;YP_003766.2;HCoV-NL63_1ab;5292 | 0 | 0 |
| NC_005831.2;YP_003766.2;HCoV-NL63_1ab;5293 | 0 | 0 |
| NC_005831.2;YP_003766.2;HCoV-NL63_1ab;5294 | 0 | 0 |
| NC_005831.2;YP_003766.2;HCoV-NL63_1ab;5295 | 0 | 0 |
| NC_005831.2;YP_003766.2;HCoV-NL63_1ab;5296 | 0 | 0 |
| NC_005831.2;YP_003766.2;HCoV-NL63_1ab;5297 | 0 | 0 |
| NC_005831.2;YP_003766.2;HCoV-NL63_1ab;5298 | 0 | 0 |

| | | |
|---|---|---|
| NC_005831.2;YP_003766.2;HCoV-NL63_1ab;5299 | 0 | 0 |
| NC_005831.2;YP_003766.2;HCoV-NL63_1ab;5300 | 0 | 0 |
| NC_005831.2;YP_003766.2;HCoV-NL63_1ab;5301 | 0 | 0 |
| NC_005831.2;YP_003766.2;HCoV-NL63_1ab;5302 | 0 | 0 |
| NC_005831.2;YP_003766.2;HCoV-NL63_1ab;5303 | 0 | 0 |
| NC_005831.2;YP_003766.2;HCoV-NL63_1ab;5304 | 0 | 0 |
| NC_005831.2;YP_003766.2;HCoV-NL63_1ab;5305 | 0 | 0 |
| NC_005831.2;YP_003766.2;HCoV-NL63_1ab;5306 | 0.05 | 0 |
| NC_005831.2;YP_003766.2;HCoV-NL63_1ab;5307 | 0 | 0 |
| NC_005831.2;YP_003766.2;HCoV-NL63_1ab;5308 | 0 | 0 |
| NC_005831.2;YP_003766.2;HCoV-NL63_1ab;5309 | 0.1 | 0.1 |
| NC_005831.2;YP_003766.2;HCoV-NL63_1ab;5310 | 0.1 | 0.125 |
| NC_005831.2;YP_003766.2;HCoV-NL63_1ab;5311 | 0.2 | 0.125 |
| NC_005831.2;YP_003766.2;HCoV-NL63_1ab;5312 | 0.25 | 0.175 |
| NC_005831.2;YP_003766.2;HCoV-NL63_1ab;5313 | 0.1 | 0 |
| NC_005831.2;YP_003766.2;HCoV-NL63_1ab;5314 | 0.05 | 0.025 |
| NC_005831.2;YP_003766.2;HCoV-NL63_1ab;5315 | 0.1 | 0.025 |
| NC_005831.2;YP_003766.2;HCoV-NL63_1ab;5316 | 0 | 0 |
| NC_005831.2;YP_003766.2;HCoV-NL63_1ab;5317 | 0.1 | 0 |
| NC_005831.2;YP_003766.2;HCoV-NL63_1ab;5318 | 0 | 0 |
| NC_005831.2;YP_003766.2;HCoV-NL63_1ab;5319 | 0 | 0 |
| NC_005831.2;YP_003766.2;HCoV-NL63_1ab;5320 | 0 | 0.025 |
| NC_005831.2;YP_003766.2;HCoV-NL63_1ab;5321 | 0.05 | 0.025 |
| NC_005831.2;YP_003766.2;HCoV-NL63_1ab;5322 | 0.05 | 0 |
| NC_005831.2;YP_003766.2;HCoV-NL63_1ab;5323 | 0.05 | 0.025 |
| NC_005831.2;YP_003766.2;HCoV-NL63_1ab;5324 | 0.05 | 0.025 |
| NC_005831.2;YP_003766.2;HCoV-NL63_1ab;5325 | 0 | 0.025 |
| NC_005831.2;YP_003766.2;HCoV-NL63_1ab;5326 | 0 | 0 |
| NC_005831.2;YP_003766.2;HCoV-NL63_1ab;5327 | 0 | 0.1 |
| NC_005831.2;YP_003766.2;HCoV-NL63_1ab;5328 | 0.05 | 0 |
| NC_005831.2;YP_003766.2;HCoV-NL63_1ab;5329 | 0 | 0 |
| NC_005831.2;YP_003766.2;HCoV-NL63_1ab;5330 | 0.05 | 0 |
| NC_005831.2;YP_003766.2;HCoV-NL63_1ab;5331 | 0.05 | 0 |
| NC_005831.2;YP_003766.2;HCoV-NL63_1ab;5332 | 0.05 | 0 |
| NC_005831.2;YP_003766.2;HCoV-NL63_1ab;5333 | 0.1 | 0 |
| NC_005831.2;YP_003766.2;HCoV-NL63_1ab;5334 | 0.05 | 0 |
| NC_005831.2;YP_003766.2;HCoV-NL63_1ab;5335 | 0.1 | 0 |
| NC_005831.2;YP_003766.2;HCoV-NL63_1ab;5336 | 0.1 | 0.025 |
| NC_005831.2;YP_003766.2;HCoV-NL63_1ab;5337 | 0.1 | 0.025 |
| NC_005831.2;YP_003766.2;HCoV-NL63_1ab;5338 | 0.05 | 0.025 |
| NC_005831.2;YP_003766.2;HCoV-NL63_1ab;5339 | 0 | 0.05 |
| NC_005831.2;YP_003766.2;HCoV-NL63_1ab;5340 | 0 | 0 |
| NC_005831.2;YP_003766.2;HCoV-NL63_1ab;5341 | 0.05 | 0 |

| | | |
|---|---|---|
| NC_005831.2;YP_003766.2;HCoV-NL63_1ab;5342 | 0.05 | 0.075 |
| NC_005831.2;YP_003766.2;HCoV-NL63_1ab;5343 | 0.05 | 0.025 |
| NC_005831.2;YP_003766.2;HCoV-NL63_1ab;5344 | 0.05 | 0.025 |
| NC_005831.2;YP_003766.2;HCoV-NL63_1ab;5345 | 0 | 0 |
| NC_005831.2;YP_003766.2;HCoV-NL63_1ab;5346 | 0 | 0 |
| NC_005831.2;YP_003766.2;HCoV-NL63_1ab;5347 | 0 | 0 |
| NC_005831.2;YP_003766.2;HCoV-NL63_1ab;5348 | 0 | 0 |
| NC_005831.2;YP_003766.2;HCoV-NL63_1ab;5349 | 0 | 0 |
| NC_005831.2;YP_003766.2;HCoV-NL63_1ab;5350 | 0 | 0 |
| NC_005831.2;YP_003766.2;HCoV-NL63_1ab;5351 | 0 | 0 |
| NC_005831.2;YP_003766.2;HCoV-NL63_1ab;5352 | 0 | 0 |
| NC_005831.2;YP_003766.2;HCoV-NL63_1ab;5353 | 0 | 0 |
| NC_005831.2;YP_003766.2;HCoV-NL63_1ab;5354 | 0 | 0 |
| NC_005831.2;YP_003766.2;HCoV-NL63_1ab;5355 | 0 | 0 |
| NC_005831.2;YP_003766.2;HCoV-NL63_1ab;5356 | 0 | 0 |
| NC_005831.2;YP_003766.2;HCoV-NL63_1ab;5357 | 0 | 0 |
| NC_005831.2;YP_003766.2;HCoV-NL63_1ab;5358 | 0 | 0 |
| NC_005831.2;YP_003766.2;HCoV-NL63_1ab;5359 | 0 | 0 |
| NC_005831.2;YP_003766.2;HCoV-NL63_1ab;5360 | 0 | 0 |
| NC_005831.2;YP_003766.2;HCoV-NL63_1ab;5361 | 0 | 0 |
| NC_005831.2;YP_003766.2;HCoV-NL63_1ab;5362 | 0 | 0 |
| NC_005831.2;YP_003766.2;HCoV-NL63_1ab;5363 | 0 | 0 |
| NC_005831.2;YP_003766.2;HCoV-NL63_1ab;5364 | 0 | 0 |
| NC_005831.2;YP_003766.2;HCoV-NL63_1ab;5365 | 0 | 0 |
| NC_005831.2;YP_003766.2;HCoV-NL63_1ab;5366 | 0 | 0 |
| NC_005831.2;YP_003766.2;HCoV-NL63_1ab;5367 | 0 | 0 |
| NC_005831.2;YP_003766.2;HCoV-NL63_1ab;5368 | 0 | 0 |
| NC_005831.2;YP_003766.2;HCoV-NL63_1ab;5369 | 0 | 0 |
| NC_005831.2;YP_003766.2;HCoV-NL63_1ab;5370 | 0 | 0 |
| NC_005831.2;YP_003766.2;HCoV-NL63_1ab;5371 | 0 | 0 |
| NC_005831.2;YP_003766.2;HCoV-NL63_1ab;5372 | 0 | 0 |
| NC_005831.2;YP_003766.2;HCoV-NL63_1ab;5373 | 0 | 0 |
| NC_005831.2;YP_003766.2;HCoV-NL63_1ab;5374 | 0 | 0 |
| NC_005831.2;YP_003766.2;HCoV-NL63_1ab;5375 | 0 | 0 |
| NC_005831.2;YP_003766.2;HCoV-NL63_1ab;5376 | 0 | 0 |
| NC_005831.2;YP_003766.2;HCoV-NL63_1ab;5377 | 0 | 0 |
| NC_005831.2;YP_003766.2;HCoV-NL63_1ab;5378 | 0 | 0.025 |
| NC_005831.2;YP_003766.2;HCoV-NL63_1ab;5379 | 0 | 0 |
| NC_005831.2;YP_003766.2;HCoV-NL63_1ab;5380 | 0 | 0 |
| NC_005831.2;YP_003766.2;HCoV-NL63_1ab;5381 | 0 | 0 |
| NC_005831.2;YP_003766.2;HCoV-NL63_1ab;5382 | 0 | 0 |
| NC_005831.2;YP_003766.2;HCoV-NL63_1ab;5383 | 0.05 | 0 |
| NC_005831.2;YP_003766.2;HCoV-NL63_1ab;5384 | 0 | 0 |

| | | |
|---|---|---|
| NC_005831.2;YP_003766.2;HCoV-NL63_1ab;5385 | 0 | 0.025 |
| NC_005831.2;YP_003766.2;HCoV-NL63_1ab;5386 | 0 | 0 |
| NC_005831.2;YP_003766.2;HCoV-NL63_1ab;5387 | 0 | 0 |
| NC_005831.2;YP_003766.2;HCoV-NL63_1ab;5388 | 0.05 | 0 |
| NC_005831.2;YP_003766.2;HCoV-NL63_1ab;5389 | 0 | 0 |
| NC_005831.2;YP_003766.2;HCoV-NL63_1ab;5390 | 0 | 0 |
| NC_005831.2;YP_003766.2;HCoV-NL63_1ab;5391 | 0 | 0 |
| NC_005831.2;YP_003766.2;HCoV-NL63_1ab;5392 | 0 | 0 |
| NC_005831.2;YP_003766.2;HCoV-NL63_1ab;5393 | 0 | 0 |
| NC_005831.2;YP_003766.2;HCoV-NL63_1ab;5394 | 0 | 0 |
| NC_005831.2;YP_003766.2;HCoV-NL63_1ab;5395 | 0 | 0.05 |
| NC_005831.2;YP_003766.2;HCoV-NL63_1ab;5396 | 0 | 0.05 |
| NC_005831.2;YP_003766.2;HCoV-NL63_1ab;5397 | 0 | 0.05 |
| NC_005831.2;YP_003766.2;HCoV-NL63_1ab;5398 | 0 | 0 |
| NC_005831.2;YP_003766.2;HCoV-NL63_1ab;5399 | 0 | 0 |
| NC_005831.2;YP_003766.2;HCoV-NL63_1ab;5400 | 0 | 0 |
| NC_005831.2;YP_003766.2;HCoV-NL63_1ab;5401 | 0 | 0 |
| NC_005831.2;YP_003766.2;HCoV-NL63_1ab;5402 | 0 | 0 |
| NC_005831.2;YP_003766.2;HCoV-NL63_1ab;5403 | 0 | 0.05 |
| NC_005831.2;YP_003766.2;HCoV-NL63_1ab;5404 | 0 | 0.025 |
| NC_005831.2;YP_003766.2;HCoV-NL63_1ab;5405 | 0 | 0 |
| NC_005831.2;YP_003766.2;HCoV-NL63_1ab;5406 | 0 | 0 |
| NC_005831.2;YP_003766.2;HCoV-NL63_1ab;5407 | 0 | 0 |
| NC_005831.2;YP_003766.2;HCoV-NL63_1ab;5408 | 0 | 0 |
| NC_005831.2;YP_003766.2;HCoV-NL63_1ab;5409 | 0 | 0 |
| NC_005831.2;YP_003766.2;HCoV-NL63_1ab;5410 | 0 | 0 |
| NC_005831.2;YP_003766.2;HCoV-NL63_1ab;5411 | 0 | 0 |
| NC_005831.2;YP_003766.2;HCoV-NL63_1ab;5412 | 0 | 0 |
| NC_005831.2;YP_003766.2;HCoV-NL63_1ab;5413 | 0 | 0 |
| NC_005831.2;YP_003766.2;HCoV-NL63_1ab;5414 | 0 | 0 |
| NC_005831.2;YP_003766.2;HCoV-NL63_1ab;5415 | 0 | 0 |
| NC_005831.2;YP_003766.2;HCoV-NL63_1ab;5416 | 0 | 0 |
| NC_005831.2;YP_003766.2;HCoV-NL63_1ab;5417 | 0.1 | 0.15 |
| NC_005831.2;YP_003766.2;HCoV-NL63_1ab;5418 | 0 | 0 |
| NC_005831.2;YP_003766.2;HCoV-NL63_1ab;5419 | 0.05 | 0.025 |
| NC_005831.2;YP_003766.2;HCoV-NL63_1ab;5420 | 0.05 | 0.1 |
| NC_005831.2;YP_003766.2;HCoV-NL63_1ab;5421 | 0 | 0.025 |
| NC_005831.2;YP_003766.2;HCoV-NL63_1ab;5422 | 0 | 0 |
| NC_005831.2;YP_003766.2;HCoV-NL63_1ab;5423 | 0 | 0.05 |
| NC_005831.2;YP_003766.2;HCoV-NL63_1ab;5424 | 0.05 | 0.225 |
| NC_005831.2;YP_003766.2;HCoV-NL63_1ab;5425 | 0 | 0.1 |
| NC_005831.2;YP_003766.2;HCoV-NL63_1ab;5426 | 0 | 0.025 |
| NC_005831.2;YP_003766.2;HCoV-NL63_1ab;5427 | 0 | 0.025 |

| | | |
|---|---|---|
| NC_005831.2;YP_003766.2;HCoV-NL63_1ab;5428 | 0 | 0.05 |
| NC_005831.2;YP_003766.2;HCoV-NL63_1ab;5429 | 0.05 | 0 |
| NC_005831.2;YP_003766.2;HCoV-NL63_1ab;5430 | 0 | 0.025 |
| NC_005831.2;YP_003766.2;HCoV-NL63_1ab;5431 | 0 | 0 |
| NC_005831.2;YP_003766.2;HCoV-NL63_1ab;5432 | 0 | 0 |
| NC_005831.2;YP_003766.2;HCoV-NL63_1ab;5433 | 0 | 0 |
| NC_005831.2;YP_003766.2;HCoV-NL63_1ab;5434 | 0 | 0 |
| NC_005831.2;YP_003766.2;HCoV-NL63_1ab;5435 | 0 | 0 |
| NC_005831.2;YP_003766.2;HCoV-NL63_1ab;5436 | 0 | 0 |
| NC_005831.2;YP_003766.2;HCoV-NL63_1ab;5437 | 0 | 0 |
| NC_005831.2;YP_003766.2;HCoV-NL63_1ab;5438 | 0 | 0 |
| NC_005831.2;YP_003766.2;HCoV-NL63_1ab;5439 | 0.05 | 0 |
| NC_005831.2;YP_003766.2;HCoV-NL63_1ab;5440 | 0.05 | 0 |
| NC_005831.2;YP_003766.2;HCoV-NL63_1ab;5441 | 0.05 | 0 |
| NC_005831.2;YP_003766.2;HCoV-NL63_1ab;5442 | 0.05 | 0.025 |
| NC_005831.2;YP_003766.2;HCoV-NL63_1ab;5443 | 0.1 | 0.025 |
| NC_005831.2;YP_003766.2;HCoV-NL63_1ab;5444 | 0.1 | 0.075 |
| NC_005831.2;YP_003766.2;HCoV-NL63_1ab;5445 | 0.1 | 0.05 |
| NC_005831.2;YP_003766.2;HCoV-NL63_1ab;5446 | 0.1 | 0.1 |
| NC_005831.2;YP_003766.2;HCoV-NL63_1ab;5447 | 0.05 | 0.05 |
| NC_005831.2;YP_003766.2;HCoV-NL63_1ab;5448 | 0 | 0 |
| NC_005831.2;YP_003766.2;HCoV-NL63_1ab;5449 | 0 | 0.025 |
| NC_005831.2;YP_003766.2;HCoV-NL63_1ab;5450 | 0.05 | 0.025 |
| NC_005831.2;YP_003766.2;HCoV-NL63_1ab;5451 | 0.1 | 0.025 |
| NC_005831.2;YP_003766.2;HCoV-NL63_1ab;5452 | 0.05 | 0.025 |
| NC_005831.2;YP_003766.2;HCoV-NL63_1ab;5453 | 0 | 0 |
| NC_005831.2;YP_003766.2;HCoV-NL63_1ab;5454 | 0 | 0.025 |
| NC_005831.2;YP_003766.2;HCoV-NL63_1ab;5455 | 0 | 0 |
| NC_005831.2;YP_003766.2;HCoV-NL63_1ab;5456 | 0 | 0 |
| NC_005831.2;YP_003766.2;HCoV-NL63_1ab;5457 | 0 | 0 |
| NC_005831.2;YP_003766.2;HCoV-NL63_1ab;5458 | 0 | 0 |
| NC_005831.2;YP_003766.2;HCoV-NL63_1ab;5459 | 0 | 0 |
| NC_005831.2;YP_003766.2;HCoV-NL63_1ab;5460 | 0 | 0 |
| NC_005831.2;YP_003766.2;HCoV-NL63_1ab;5461 | 0 | 0 |
| NC_005831.2;YP_003766.2;HCoV-NL63_1ab;5462 | 0 | 0 |
| NC_005831.2;YP_003766.2;HCoV-NL63_1ab;5463 | 0 | 0 |
| NC_005831.2;YP_003766.2;HCoV-NL63_1ab;5464 | 0 | 0 |
| NC_005831.2;YP_003766.2;HCoV-NL63_1ab;5465 | 0 | 0 |
| NC_005831.2;YP_003766.2;HCoV-NL63_1ab;5466 | 0 | 0 |
| NC_005831.2;YP_003766.2;HCoV-NL63_1ab;5467 | 0 | 0 |
| NC_005831.2;YP_003766.2;HCoV-NL63_1ab;5468 | 0 | 0 |
| NC_005831.2;YP_003766.2;HCoV-NL63_1ab;5469 | 0 | 0 |
| NC_005831.2;YP_003766.2;HCoV-NL63_1ab;5470 | 0 | 0 |

| | | |
|---|---|---|
| NC_005831.2;YP_003766.2;HCoV-NL63_1ab;5471 | 0 | 0 |
| NC_005831.2;YP_003766.2;HCoV-NL63_1ab;5472 | 0 | 0 |
| NC_005831.2;YP_003766.2;HCoV-NL63_1ab;5473 | 0 | 0 |
| NC_005831.2;YP_003766.2;HCoV-NL63_1ab;5474 | 0 | 0 |
| NC_005831.2;YP_003766.2;HCoV-NL63_1ab;5475 | 0.15 | 0 |
| NC_005831.2;YP_003766.2;HCoV-NL63_1ab;5476 | 0 | 0 |
| NC_005831.2;YP_003766.2;HCoV-NL63_1ab;5477 | 0 | 0 |
| NC_005831.2;YP_003766.2;HCoV-NL63_1ab;5478 | 0.05 | 0 |
| NC_005831.2;YP_003766.2;HCoV-NL63_1ab;5479 | 0.05 | 0 |
| NC_005831.2;YP_003766.2;HCoV-NL63_1ab;5480 | 0 | 0 |
| NC_005831.2;YP_003766.2;HCoV-NL63_1ab;5481 | 0 | 0 |
| NC_005831.2;YP_003766.2;HCoV-NL63_1ab;5482 | 0 | 0 |
| NC_005831.2;YP_003766.2;HCoV-NL63_1ab;5483 | 0 | 0 |
| NC_005831.2;YP_003766.2;HCoV-NL63_1ab;5484 | 0 | 0 |
| NC_005831.2;YP_003766.2;HCoV-NL63_1ab;5485 | 0 | 0 |
| NC_005831.2;YP_003766.2;HCoV-NL63_1ab;5486 | 0 | 0 |
| NC_005831.2;YP_003766.2;HCoV-NL63_1ab;5487 | 0 | 0 |
| NC_005831.2;YP_003766.2;HCoV-NL63_1ab;5488 | 0 | 0 |
| NC_005831.2;YP_003766.2;HCoV-NL63_1ab;5489 | 0 | 0 |
| NC_005831.2;YP_003766.2;HCoV-NL63_1ab;5490 | 0 | 0 |
| NC_005831.2;YP_003766.2;HCoV-NL63_1ab;5491 | 0.05 | 0.025 |
| NC_005831.2;YP_003766.2;HCoV-NL63_1ab;5492 | 0.1 | 0.05 |
| NC_005831.2;YP_003766.2;HCoV-NL63_1ab;5493 | 0.1 | 0.05 |
| NC_005831.2;YP_003766.2;HCoV-NL63_1ab;5494 | 0.1 | 0.075 |
| NC_005831.2;YP_003766.2;HCoV-NL63_1ab;5495 | 0.15 | 0.1 |
| NC_005831.2;YP_003766.2;HCoV-NL63_1ab;5496 | 0.1 | 0.125 |
| NC_005831.2;YP_003766.2;HCoV-NL63_1ab;5497 | 0.1 | 0.1 |
| NC_005831.2;YP_003766.2;HCoV-NL63_1ab;5498 | 0.15 | 0.1 |
| NC_005831.2;YP_003766.2;HCoV-NL63_1ab;5499 | 0.1 | 0.1 |
| NC_005831.2;YP_003766.2;HCoV-NL63_1ab;5500 | 0.15 | 0.075 |
| NC_005831.2;YP_003766.2;HCoV-NL63_1ab;5501 | 0.05 | 0 |
| NC_005831.2;YP_003766.2;HCoV-NL63_1ab;5502 | 0 | 0 |
| NC_005831.2;YP_003766.2;HCoV-NL63_1ab;5503 | 0 | 0.025 |
| NC_005831.2;YP_003766.2;HCoV-NL63_1ab;5504 | 0 | 0.025 |
| NC_005831.2;YP_003766.2;HCoV-NL63_1ab;5505 | 0 | 0 |
| NC_005831.2;YP_003766.2;HCoV-NL63_1ab;5506 | 0 | 0 |
| NC_005831.2;YP_003766.2;HCoV-NL63_1ab;5507 | 0 | 0 |
| NC_005831.2;YP_003766.2;HCoV-NL63_1ab;5508 | 0 | 0 |
| NC_005831.2;YP_003766.2;HCoV-NL63_1ab;5509 | 0 | 0 |
| NC_005831.2;YP_003766.2;HCoV-NL63_1ab;5510 | 0 | 0 |
| NC_005831.2;YP_003766.2;HCoV-NL63_1ab;5511 | 0 | 0 |
| NC_005831.2;YP_003766.2;HCoV-NL63_1ab;5512 | 0 | 0 |
| NC_005831.2;YP_003766.2;HCoV-NL63_1ab;5513 | 0 | 0 |

| | | |
|---|---|---|
| NC_005831.2;YP_003766.2;HCoV-NL63_1ab;5514 | 0 | 0 |
| NC_005831.2;YP_003766.2;HCoV-NL63_1ab;5515 | 0 | 0 |
| NC_005831.2;YP_003766.2;HCoV-NL63_1ab;5516 | 0 | 0.025 |
| NC_005831.2;YP_003766.2;HCoV-NL63_1ab;5517 | 0 | 0 |
| NC_005831.2;YP_003766.2;HCoV-NL63_1ab;5518 | 0.05 | 0 |
| NC_005831.2;YP_003766.2;HCoV-NL63_1ab;5519 | 0.05 | 0 |
| NC_005831.2;YP_003766.2;HCoV-NL63_1ab;5520 | 0.05 | 0.05 |
| NC_005831.2;YP_003766.2;HCoV-NL63_1ab;5521 | 0 | 0 |
| NC_005831.2;YP_003766.2;HCoV-NL63_1ab;5522 | 0 | 0 |
| NC_005831.2;YP_003766.2;HCoV-NL63_1ab;5523 | 0 | 0 |
| NC_005831.2;YP_003766.2;HCoV-NL63_1ab;5524 | 0 | 0 |
| NC_005831.2;YP_003766.2;HCoV-NL63_1ab;5525 | 0 | 0 |
| NC_005831.2;YP_003766.2;HCoV-NL63_1ab;5526 | 0 | 0 |
| NC_005831.2;YP_003766.2;HCoV-NL63_1ab;5527 | 0 | 0 |
| NC_005831.2;YP_003766.2;HCoV-NL63_1ab;5528 | 0 | 0 |
| NC_005831.2;YP_003766.2;HCoV-NL63_1ab;5529 | 0.05 | 0 |
| NC_005831.2;YP_003766.2;HCoV-NL63_1ab;5530 | 0.05 | 0 |
| NC_005831.2;YP_003766.2;HCoV-NL63_1ab;5531 | 0 | 0 |
| NC_005831.2;YP_003766.2;HCoV-NL63_1ab;5532 | 0 | 0 |
| NC_005831.2;YP_003766.2;HCoV-NL63_1ab;5533 | 0 | 0 |
| NC_005831.2;YP_003766.2;HCoV-NL63_1ab;5534 | 0 | 0 |
| NC_005831.2;YP_003766.2;HCoV-NL63_1ab;5535 | 0 | 0 |
| NC_005831.2;YP_003766.2;HCoV-NL63_1ab;5536 | 0 | 0 |
| NC_005831.2;YP_003766.2;HCoV-NL63_1ab;5537 | 0 | 0 |
| NC_005831.2;YP_003766.2;HCoV-NL63_1ab;5538 | 0 | 0 |
| NC_005831.2;YP_003766.2;HCoV-NL63_1ab;5539 | 0 | 0 |
| NC_005831.2;YP_003766.2;HCoV-NL63_1ab;5540 | 0 | 0 |
| NC_005831.2;YP_003766.2;HCoV-NL63_1ab;5541 | 0 | 0.025 |
| NC_005831.2;YP_003766.2;HCoV-NL63_1ab;5542 | 0 | 0 |
| NC_005831.2;YP_003766.2;HCoV-NL63_1ab;5543 | 0 | 0 |
| NC_005831.2;YP_003766.2;HCoV-NL63_1ab;5544 | 0 | 0 |
| NC_005831.2;YP_003766.2;HCoV-NL63_1ab;5545 | 0 | 0 |
| NC_005831.2;YP_003766.2;HCoV-NL63_1ab;5546 | 0 | 0 |
| NC_005831.2;YP_003766.2;HCoV-NL63_1ab;5547 | 0 | 0 |
| NC_005831.2;YP_003766.2;HCoV-NL63_1ab;5548 | 0 | 0 |
| NC_005831.2;YP_003766.2;HCoV-NL63_1ab;5549 | 0 | 0 |
| NC_005831.2;YP_003766.2;HCoV-NL63_1ab;5550 | 0 | 0 |
| NC_005831.2;YP_003766.2;HCoV-NL63_1ab;5551 | 0 | 0 |
| NC_005831.2;YP_003766.2;HCoV-NL63_1ab;5552 | 0 | 0 |
| NC_005831.2;YP_003766.2;HCoV-NL63_1ab;5553 | 0 | 0 |
| NC_005831.2;YP_003766.2;HCoV-NL63_1ab;5554 | 0.05 | 0 |
| NC_005831.2;YP_003766.2;HCoV-NL63_1ab;5555 | 0 | 0 |
| NC_005831.2;YP_003766.2;HCoV-NL63_1ab;5556 | 0 | 0 |

| | | |
|---|---|---|
| NC_005831.2;YP_003766.2;HCoV-NL63_1ab;5557 | 0 | 0 |
| NC_005831.2;YP_003766.2;HCoV-NL63_1ab;5558 | 0 | 0.025 |
| NC_005831.2;YP_003766.2;HCoV-NL63_1ab;5559 | 0 | 0.025 |
| NC_005831.2;YP_003766.2;HCoV-NL63_1ab;5560 | 0 | 0 |
| NC_005831.2;YP_003766.2;HCoV-NL63_1ab;5561 | 0 | 0 |
| NC_005831.2;YP_003766.2;HCoV-NL63_1ab;5562 | 0 | 0 |
| NC_005831.2;YP_003766.2;HCoV-NL63_1ab;5563 | 0 | 0 |
| NC_005831.2;YP_003766.2;HCoV-NL63_1ab;5564 | 0 | 0 |
| NC_005831.2;YP_003766.2;HCoV-NL63_1ab;5565 | 0 | 0 |
| NC_005831.2;YP_003766.2;HCoV-NL63_1ab;5566 | 0 | 0 |
| NC_005831.2;YP_003766.2;HCoV-NL63_1ab;5567 | 0 | 0 |
| NC_005831.2;YP_003766.2;HCoV-NL63_1ab;5568 | 0 | 0 |
| NC_005831.2;YP_003766.2;HCoV-NL63_1ab;5569 | 0 | 0 |
| NC_005831.2;YP_003766.2;HCoV-NL63_1ab;5570 | 0 | 0 |
| NC_005831.2;YP_003766.2;HCoV-NL63_1ab;5571 | 0 | 0 |
| NC_005831.2;YP_003766.2;HCoV-NL63_1ab;5572 | 0 | 0 |
| NC_005831.2;YP_003766.2;HCoV-NL63_1ab;5573 | 0 | 0 |
| NC_005831.2;YP_003766.2;HCoV-NL63_1ab;5574 | 0 | 0 |
| NC_005831.2;YP_003766.2;HCoV-NL63_1ab;5575 | 0 | 0 |
| NC_005831.2;YP_003766.2;HCoV-NL63_1ab;5576 | 0 | 0 |
| NC_005831.2;YP_003766.2;HCoV-NL63_1ab;5577 | 0 | 0 |
| NC_005831.2;YP_003766.2;HCoV-NL63_1ab;5578 | 0 | 0 |
| NC_005831.2;YP_003766.2;HCoV-NL63_1ab;5579 | 0 | 0 |
| NC_005831.2;YP_003766.2;HCoV-NL63_1ab;5580 | 0 | 0 |
| NC_005831.2;YP_003766.2;HCoV-NL63_1ab;5581 | 0 | 0 |
| NC_005831.2;YP_003766.2;HCoV-NL63_1ab;5582 | 0 | 0 |
| NC_005831.2;YP_003766.2;HCoV-NL63_1ab;5583 | 0 | 0 |
| NC_005831.2;YP_003766.2;HCoV-NL63_1ab;5584 | 0 | 0 |
| NC_005831.2;YP_003766.2;HCoV-NL63_1ab;5585 | 0 | 0 |
| NC_005831.2;YP_003766.2;HCoV-NL63_1ab;5586 | 0 | 0 |
| NC_005831.2;YP_003766.2;HCoV-NL63_1ab;5587 | 0 | 0 |
| NC_005831.2;YP_003766.2;HCoV-NL63_1ab;5588 | 0 | 0 |
| NC_005831.2;YP_003766.2;HCoV-NL63_1ab;5589 | 0 | 0 |
| NC_005831.2;YP_003766.2;HCoV-NL63_1ab;5590 | 0 | 0 |
| NC_005831.2;YP_003766.2;HCoV-NL63_1ab;5591 | 0 | 0 |
| NC_005831.2;YP_003766.2;HCoV-NL63_1ab;5592 | 0.05 | 0.025 |
| NC_005831.2;YP_003766.2;HCoV-NL63_1ab;5593 | 0.1 | 0.05 |
| NC_005831.2;YP_003766.2;HCoV-NL63_1ab;5594 | 0.05 | 0 |
| NC_005831.2;YP_003766.2;HCoV-NL63_1ab;5595 | 0.05 | 0 |
| NC_005831.2;YP_003766.2;HCoV-NL63_1ab;5596 | 0.05 | 0.05 |
| NC_005831.2;YP_003766.2;HCoV-NL63_1ab;5597 | 0.05 | 0.05 |
| NC_005831.2;YP_003766.2;HCoV-NL63_1ab;5598 | 0.05 | 0.075 |
| NC_005831.2;YP_003766.2;HCoV-NL63_1ab;5599 | 0.1 | 0.15 |

| | | |
|---|---|---|
| NC_005831.2;YP_003766.2;HCoV-NL63_1ab;5600 | 0.1 | 0.15 |
| NC_005831.2;YP_003766.2;HCoV-NL63_1ab;5601 | 0.15 | 0.15 |
| NC_005831.2;YP_003766.2;HCoV-NL63_1ab;5602 | 0 | 0.075 |
| NC_005831.2;YP_003766.2;HCoV-NL63_1ab;5603 | 0 | 0.05 |
| NC_005831.2;YP_003766.2;HCoV-NL63_1ab;5604 | 0 | 0.05 |
| NC_005831.2;YP_003766.2;HCoV-NL63_1ab;5605 | 0 | 0.05 |
| NC_005831.2;YP_003766.2;HCoV-NL63_1ab;5606 | 0 | 0.025 |
| NC_005831.2;YP_003766.2;HCoV-NL63_1ab;5607 | 0 | 0.025 |
| NC_005831.2;YP_003766.2;HCoV-NL63_1ab;5608 | 0 | 0.025 |
| NC_005831.2;YP_003766.2;HCoV-NL63_1ab;5609 | 0.05 | 0.025 |
| NC_005831.2;YP_003766.2;HCoV-NL63_1ab;5610 | 0 | 0.05 |
| NC_005831.2;YP_003766.2;HCoV-NL63_1ab;5611 | 0 | 0 |
| NC_005831.2;YP_003766.2;HCoV-NL63_1ab;5612 | 0 | 0 |
| NC_005831.2;YP_003766.2;HCoV-NL63_1ab;5613 | 0 | 0 |
| NC_005831.2;YP_003766.2;HCoV-NL63_1ab;5614 | 0 | 0 |
| NC_005831.2;YP_003766.2;HCoV-NL63_1ab;5615 | 0 | 0 |
| NC_005831.2;YP_003766.2;HCoV-NL63_1ab;5616 | 0 | 0 |
| NC_005831.2;YP_003766.2;HCoV-NL63_1ab;5617 | 0 | 0 |
| NC_005831.2;YP_003766.2;HCoV-NL63_1ab;5618 | 0 | 0 |
| NC_005831.2;YP_003766.2;HCoV-NL63_1ab;5619 | 0 | 0 |
| NC_005831.2;YP_003766.2;HCoV-NL63_1ab;5620 | 0 | 0 |
| NC_005831.2;YP_003766.2;HCoV-NL63_1ab;5621 | 0 | 0 |
| NC_005831.2;YP_003766.2;HCoV-NL63_1ab;5622 | 0 | 0 |
| NC_005831.2;YP_003766.2;HCoV-NL63_1ab;5623 | 0 | 0 |
| NC_005831.2;YP_003766.2;HCoV-NL63_1ab;5624 | 0 | 0 |
| NC_005831.2;YP_003766.2;HCoV-NL63_1ab;5625 | 0 | 0 |
| NC_005831.2;YP_003766.2;HCoV-NL63_1ab;5626 | 0 | 0 |
| NC_005831.2;YP_003766.2;HCoV-NL63_1ab;5627 | 0 | 0 |
| NC_005831.2;YP_003766.2;HCoV-NL63_1ab;5628 | 0 | 0 |
| NC_005831.2;YP_003766.2;HCoV-NL63_1ab;5629 | 0 | 0 |
| NC_005831.2;YP_003766.2;HCoV-NL63_1ab;5630 | 0 | 0 |
| NC_005831.2;YP_003766.2;HCoV-NL63_1ab;5631 | 0 | 0 |
| NC_005831.2;YP_003766.2;HCoV-NL63_1ab;5632 | 0 | 0 |
| NC_005831.2;YP_003766.2;HCoV-NL63_1ab;5633 | 0 | 0 |
| NC_005831.2;YP_003766.2;HCoV-NL63_1ab;5634 | 0 | 0 |
| NC_005831.2;YP_003766.2;HCoV-NL63_1ab;5635 | 0 | 0 |
| NC_005831.2;YP_003766.2;HCoV-NL63_1ab;5636 | 0 | 0 |
| NC_005831.2;YP_003766.2;HCoV-NL63_1ab;5637 | 0 | 0 |
| NC_005831.2;YP_003766.2;HCoV-NL63_1ab;5638 | 0 | 0 |
| NC_005831.2;YP_003766.2;HCoV-NL63_1ab;5639 | 0 | 0 |
| NC_005831.2;YP_003766.2;HCoV-NL63_1ab;5640 | 0 | 0 |
| NC_005831.2;YP_003766.2;HCoV-NL63_1ab;5641 | 0 | 0 |
| NC_005831.2;YP_003766.2;HCoV-NL63_1ab;5642 | 0 | 0.05 |

| | | |
|---|---|---|
| NC_005831.2;YP_003766.2;HCoV-NL63_1ab;5643 | 0 | 0 |
| NC_005831.2;YP_003766.2;HCoV-NL63_1ab;5644 | 0 | 0 |
| NC_005831.2;YP_003766.2;HCoV-NL63_1ab;5645 | 0 | 0 |
| NC_005831.2;YP_003766.2;HCoV-NL63_1ab;5646 | 0 | 0.025 |
| NC_005831.2;YP_003766.2;HCoV-NL63_1ab;5647 | 0 | 0 |
| NC_005831.2;YP_003766.2;HCoV-NL63_1ab;5648 | 0 | 0 |
| NC_005831.2;YP_003766.2;HCoV-NL63_1ab;5649 | 0 | 0 |
| NC_005831.2;YP_003766.2;HCoV-NL63_1ab;5650 | 0 | 0.025 |
| NC_005831.2;YP_003766.2;HCoV-NL63_1ab;5651 | 0 | 0.025 |
| NC_005831.2;YP_003766.2;HCoV-NL63_1ab;5652 | 0 | 0.05 |
| NC_005831.2;YP_003766.2;HCoV-NL63_1ab;5653 | 0 | 0 |
| NC_005831.2;YP_003766.2;HCoV-NL63_1ab;5654 | 0 | 0 |
| NC_005831.2;YP_003766.2;HCoV-NL63_1ab;5655 | 0 | 0 |
| NC_005831.2;YP_003766.2;HCoV-NL63_1ab;5656 | 0 | 0 |
| NC_005831.2;YP_003766.2;HCoV-NL63_1ab;5657 | 0 | 0.025 |
| NC_005831.2;YP_003766.2;HCoV-NL63_1ab;5658 | 0.05 | 0.025 |
| NC_005831.2;YP_003766.2;HCoV-NL63_1ab;5659 | 0.05 | 0.025 |
| NC_005831.2;YP_003766.2;HCoV-NL63_1ab;5660 | 0.05 | 0.025 |
| NC_005831.2;YP_003766.2;HCoV-NL63_1ab;5661 | 0.05 | 0.025 |
| NC_005831.2;YP_003766.2;HCoV-NL63_1ab;5662 | 0.05 | 0.025 |
| NC_005831.2;YP_003766.2;HCoV-NL63_1ab;5663 | 0.05 | 0.025 |
| NC_005831.2;YP_003766.2;HCoV-NL63_1ab;5664 | 0.05 | 0.075 |
| NC_005831.2;YP_003766.2;HCoV-NL63_1ab;5665 | 0.05 | 0.025 |
| NC_005831.2;YP_003766.2;HCoV-NL63_1ab;5666 | 0.05 | 0.025 |
| NC_005831.2;YP_003766.2;HCoV-NL63_1ab;5667 | 0.05 | 0.025 |
| NC_005831.2;YP_003766.2;HCoV-NL63_1ab;5668 | 0.05 | 0.25 |
| NC_005831.2;YP_003766.2;HCoV-NL63_1ab;5669 | 0 | 0 |
| NC_005831.2;YP_003766.2;HCoV-NL63_1ab;5670 | 0.05 | 0 |
| NC_005831.2;YP_003766.2;HCoV-NL63_1ab;5671 | 0 | 0 |
| NC_005831.2;YP_003766.2;HCoV-NL63_1ab;5672 | 0 | 0 |
| NC_005831.2;YP_003766.2;HCoV-NL63_1ab;5673 | 0 | 0 |
| NC_005831.2;YP_003766.2;HCoV-NL63_1ab;5674 | 0 | 0 |
| NC_005831.2;YP_003766.2;HCoV-NL63_1ab;5675 | 0 | 0 |
| NC_005831.2;YP_003766.2;HCoV-NL63_1ab;5676 | 0 | 0 |
| NC_005831.2;YP_003766.2;HCoV-NL63_1ab;5677 | 0 | 0 |
| NC_005831.2;YP_003766.2;HCoV-NL63_1ab;5678 | 0 | 0 |
| NC_005831.2;YP_003766.2;HCoV-NL63_1ab;5679 | 0 | 0 |
| NC_005831.2;YP_003766.2;HCoV-NL63_1ab;5680 | 0 | 0 |
| NC_005831.2;YP_003766.2;HCoV-NL63_1ab;5681 | 0 | 0 |
| NC_005831.2;YP_003766.2;HCoV-NL63_1ab;5682 | 0 | 0 |
| NC_005831.2;YP_003766.2;HCoV-NL63_1ab;5683 | 0 | 0 |
| NC_005831.2;YP_003766.2;HCoV-NL63_1ab;5684 | 0 | 0 |
| NC_005831.2;YP_003766.2;HCoV-NL63_1ab;5685 | 0 | 0 |

| | | |
|---|---|---|
| NC_005831.2;YP_003766.2;HCoV-NL63_1ab;5686 | 0 | 0 |
| NC_005831.2;YP_003766.2;HCoV-NL63_1ab;5687 | 0 | 0 |
| NC_005831.2;YP_003766.2;HCoV-NL63_1ab;5688 | 0 | 0.025 |
| NC_005831.2;YP_003766.2;HCoV-NL63_1ab;5689 | 0 | 0 |
| NC_005831.2;YP_003766.2;HCoV-NL63_1ab;5690 | 0 | 0 |
| NC_005831.2;YP_003766.2;HCoV-NL63_1ab;5691 | 0 | 0 |
| NC_005831.2;YP_003766.2;HCoV-NL63_1ab;5692 | 0 | 0 |
| NC_005831.2;YP_003766.2;HCoV-NL63_1ab;5693 | 0 | 0 |
| NC_005831.2;YP_003766.2;HCoV-NL63_1ab;5694 | 0 | 0 |
| NC_005831.2;YP_003766.2;HCoV-NL63_1ab;5695 | 0 | 0 |
| NC_005831.2;YP_003766.2;HCoV-NL63_1ab;5696 | 0 | 0 |
| NC_005831.2;YP_003766.2;HCoV-NL63_1ab;5697 | 0 | 0.025 |
| NC_005831.2;YP_003766.2;HCoV-NL63_1ab;5698 | 0 | 0 |
| NC_005831.2;YP_003766.2;HCoV-NL63_1ab;5699 | 0 | 0.025 |
| NC_005831.2;YP_003766.2;HCoV-NL63_1ab;5700 | 0 | 0 |
| NC_005831.2;YP_003766.2;HCoV-NL63_1ab;5701 | 0 | 0 |
| NC_005831.2;YP_003766.2;HCoV-NL63_1ab;5702 | 0 | 0 |
| NC_005831.2;YP_003766.2;HCoV-NL63_1ab;5703 | 0 | 0 |
| NC_005831.2;YP_003766.2;HCoV-NL63_1ab;5704 | 0 | 0 |
| NC_005831.2;YP_003766.2;HCoV-NL63_1ab;5705 | 0 | 0 |
| NC_005831.2;YP_003766.2;HCoV-NL63_1ab;5706 | 0 | 0 |
| NC_005831.2;YP_003766.2;HCoV-NL63_1ab;5707 | 0 | 0 |
| NC_005831.2;YP_003766.2;HCoV-NL63_1ab;5708 | 0 | 0 |
| NC_005831.2;YP_003766.2;HCoV-NL63_1ab;5709 | 0 | 0 |
| NC_005831.2;YP_003766.2;HCoV-NL63_1ab;5710 | 0 | 0 |
| NC_005831.2;YP_003766.2;HCoV-NL63_1ab;5711 | 0 | 0 |
| NC_005831.2;YP_003766.2;HCoV-NL63_1ab;5712 | 0 | 0 |
| NC_005831.2;YP_003766.2;HCoV-NL63_1ab;5713 | 0 | 0 |
| NC_005831.2;YP_003766.2;HCoV-NL63_1ab;5714 | 0 | 0 |
| NC_005831.2;YP_003766.2;HCoV-NL63_1ab;5715 | 0 | 0 |
| NC_005831.2;YP_003766.2;HCoV-NL63_1ab;5716 | 0 | 0 |
| NC_005831.2;YP_003766.2;HCoV-NL63_1ab;5717 | 0 | 0 |
| NC_005831.2;YP_003766.2;HCoV-NL63_1ab;5718 | 0 | 0 |
| NC_005831.2;YP_003766.2;HCoV-NL63_1ab;5719 | 0 | 0 |
| NC_005831.2;YP_003766.2;HCoV-NL63_1ab;5720 | 0 | 0 |
| NC_005831.2;YP_003766.2;HCoV-NL63_1ab;5721 | 0 | 0 |
| NC_005831.2;YP_003766.2;HCoV-NL63_1ab;5722 | 0 | 0 |
| NC_005831.2;YP_003766.2;HCoV-NL63_1ab;5723 | 0 | 0 |
| NC_005831.2;YP_003766.2;HCoV-NL63_1ab;5724 | 0 | 0 |
| NC_005831.2;YP_003766.2;HCoV-NL63_1ab;5725 | 0 | 0 |
| NC_005831.2;YP_003766.2;HCoV-NL63_1ab;5726 | 0 | 0 |
| NC_005831.2;YP_003766.2;HCoV-NL63_1ab;5727 | 0 | 0 |
| NC_005831.2;YP_003766.2;HCoV-NL63_1ab;5728 | 0 | 0 |

| | | |
|---|---|---|
| NC_005831.2;YP_003766.2;HCoV-NL63_1ab;5729 | 0 | 0 |
| NC_005831.2;YP_003766.2;HCoV-NL63_1ab;5730 | 0 | 0 |
| NC_005831.2;YP_003766.2;HCoV-NL63_1ab;5731 | 0 | 0 |
| NC_005831.2;YP_003766.2;HCoV-NL63_1ab;5732 | 0 | 0 |
| NC_005831.2;YP_003766.2;HCoV-NL63_1ab;5733 | 0 | 0.025 |
| NC_005831.2;YP_003766.2;HCoV-NL63_1ab;5734 | 0 | 0 |
| NC_005831.2;YP_003766.2;HCoV-NL63_1ab;5735 | 0 | 0 |
| NC_005831.2;YP_003766.2;HCoV-NL63_1ab;5736 | 0 | 0 |
| NC_005831.2;YP_003766.2;HCoV-NL63_1ab;5737 | 0 | 0 |
| NC_005831.2;YP_003766.2;HCoV-NL63_1ab;5738 | 0 | 0 |
| NC_005831.2;YP_003766.2;HCoV-NL63_1ab;5739 | 0 | 0 |
| NC_005831.2;YP_003766.2;HCoV-NL63_1ab;5740 | 0 | 0 |
| NC_005831.2;YP_003766.2;HCoV-NL63_1ab;5741 | 0 | 0 |
| NC_005831.2;YP_003766.2;HCoV-NL63_1ab;5742 | 0 | 0 |
| NC_005831.2;YP_003766.2;HCoV-NL63_1ab;5743 | 0 | 0 |
| NC_005831.2;YP_003766.2;HCoV-NL63_1ab;5744 | 0 | 0 |
| NC_005831.2;YP_003766.2;HCoV-NL63_1ab;5745 | 0 | 0 |
| NC_005831.2;YP_003766.2;HCoV-NL63_1ab;5746 | 0 | 0 |
| NC_005831.2;YP_003766.2;HCoV-NL63_1ab;5747 | 0 | 0 |
| NC_005831.2;YP_003766.2;HCoV-NL63_1ab;5748 | 0 | 0 |
| NC_005831.2;YP_003766.2;HCoV-NL63_1ab;5749 | 0 | 0 |
| NC_005831.2;YP_003766.2;HCoV-NL63_1ab;5750 | 0 | 0 |
| NC_005831.2;YP_003766.2;HCoV-NL63_1ab;5751 | 0 | 0 |
| NC_005831.2;YP_003766.2;HCoV-NL63_1ab;5752 | 0 | 0 |
| NC_005831.2;YP_003766.2;HCoV-NL63_1ab;5753 | 0 | 0 |
| NC_005831.2;YP_003766.2;HCoV-NL63_1ab;5754 | 0 | 0 |
| NC_005831.2;YP_003766.2;HCoV-NL63_1ab;5755 | 0 | 0 |
| NC_005831.2;YP_003766.2;HCoV-NL63_1ab;5756 | 0 | 0 |
| NC_005831.2;YP_003766.2;HCoV-NL63_1ab;5757 | 0 | 0 |
| NC_005831.2;YP_003766.2;HCoV-NL63_1ab;5758 | 0 | 0 |
| NC_005831.2;YP_003766.2;HCoV-NL63_1ab;5759 | 0 | 0 |
| NC_005831.2;YP_003766.2;HCoV-NL63_1ab;5760 | 0 | 0 |
| NC_005831.2;YP_003766.2;HCoV-NL63_1ab;5761 | 0 | 0 |
| NC_005831.2;YP_003766.2;HCoV-NL63_1ab;5762 | 0 | 0 |
| NC_005831.2;YP_003766.2;HCoV-NL63_1ab;5763 | 0 | 0 |
| NC_005831.2;YP_003766.2;HCoV-NL63_1ab;5764 | 0 | 0 |
| NC_005831.2;YP_003766.2;HCoV-NL63_1ab;5765 | 0 | 0 |
| NC_005831.2;YP_003766.2;HCoV-NL63_1ab;5766 | 0.05 | 0.025 |
| NC_005831.2;YP_003766.2;HCoV-NL63_1ab;5767 | 0 | 0 |
| NC_005831.2;YP_003766.2;HCoV-NL63_1ab;5768 | 0 | 0 |
| NC_005831.2;YP_003766.2;HCoV-NL63_1ab;5769 | 0 | 0 |
| NC_005831.2;YP_003766.2;HCoV-NL63_1ab;5770 | 0 | 0 |
| NC_005831.2;YP_003766.2;HCoV-NL63_1ab;5771 | 0 | 0 |

| | | |
|---|---|---|
| NC_005831.2;YP_003766.2;HCoV-NL63_1ab;5772 | 0 | 0 |
| NC_005831.2;YP_003766.2;HCoV-NL63_1ab;5773 | 0.05 | 0.025 |
| NC_005831.2;YP_003766.2;HCoV-NL63_1ab;5774 | 0.05 | 0.025 |
| NC_005831.2;YP_003766.2;HCoV-NL63_1ab;5775 | 0.05 | 0.025 |
| NC_005831.2;YP_003766.2;HCoV-NL63_1ab;5776 | 0.05 | 0 |
| NC_005831.2;YP_003766.2;HCoV-NL63_1ab;5777 | 0 | 0 |
| NC_005831.2;YP_003766.2;HCoV-NL63_1ab;5778 | 0 | 0 |
| NC_005831.2;YP_003766.2;HCoV-NL63_1ab;5779 | 0 | 0 |
| NC_005831.2;YP_003766.2;HCoV-NL63_1ab;5780 | 0 | 0 |
| NC_005831.2;YP_003766.2;HCoV-NL63_1ab;5781 | 0 | 0 |
| NC_005831.2;YP_003766.2;HCoV-NL63_1ab;5782 | 0 | 0 |
| NC_005831.2;YP_003766.2;HCoV-NL63_1ab;5783 | 0 | 0.025 |
| NC_005831.2;YP_003766.2;HCoV-NL63_1ab;5784 | 0 | 0 |
| NC_005831.2;YP_003766.2;HCoV-NL63_1ab;5785 | 0 | 0 |
| NC_005831.2;YP_003766.2;HCoV-NL63_1ab;5786 | 0 | 0 |
| NC_005831.2;YP_003766.2;HCoV-NL63_1ab;5787 | 0 | 0 |
| NC_005831.2;YP_003766.2;HCoV-NL63_1ab;5788 | 0 | 0 |
| NC_005831.2;YP_003766.2;HCoV-NL63_1ab;5789 | 0 | 0 |
| NC_005831.2;YP_003766.2;HCoV-NL63_1ab;5790 | 0 | 0 |
| NC_005831.2;YP_003766.2;HCoV-NL63_1ab;5791 | 0 | 0 |
| NC_005831.2;YP_003766.2;HCoV-NL63_1ab;5792 | 0 | 0 |
| NC_005831.2;YP_003766.2;HCoV-NL63_1ab;5793 | 0 | 0 |
| NC_005831.2;YP_003766.2;HCoV-NL63_1ab;5794 | 0 | 0 |
| NC_005831.2;YP_003766.2;HCoV-NL63_1ab;5795 | 0 | 0 |
| NC_005831.2;YP_003766.2;HCoV-NL63_1ab;5796 | 0 | 0 |
| NC_005831.2;YP_003766.2;HCoV-NL63_1ab;5797 | 0 | 0 |
| NC_005831.2;YP_003766.2;HCoV-NL63_1ab;5798 | 0 | 0 |
| NC_005831.2;YP_003766.2;HCoV-NL63_1ab;5799 | 0 | 0 |
| NC_005831.2;YP_003766.2;HCoV-NL63_1ab;5800 | 0 | 0 |
| NC_005831.2;YP_003766.2;HCoV-NL63_1ab;5801 | 0 | 0 |
| NC_005831.2;YP_003766.2;HCoV-NL63_1ab;5802 | 0 | 0 |
| NC_005831.2;YP_003766.2;HCoV-NL63_1ab;5803 | 0 | 0 |
| NC_005831.2;YP_003766.2;HCoV-NL63_1ab;5804 | 0 | 0 |
| NC_005831.2;YP_003766.2;HCoV-NL63_1ab;5805 | 0 | 0 |
| NC_005831.2;YP_003766.2;HCoV-NL63_1ab;5806 | 0 | 0 |
| NC_005831.2;YP_003766.2;HCoV-NL63_1ab;5807 | 0 | 0 |
| NC_005831.2;YP_003766.2;HCoV-NL63_1ab;5808 | 0 | 0 |
| NC_005831.2;YP_003766.2;HCoV-NL63_1ab;5809 | 0 | 0 |
| NC_005831.2;YP_003766.2;HCoV-NL63_1ab;5810 | 0 | 0 |
| NC_005831.2;YP_003766.2;HCoV-NL63_1ab;5811 | 0 | 0 |
| NC_005831.2;YP_003766.2;HCoV-NL63_1ab;5812 | 0 | 0 |
| NC_005831.2;YP_003766.2;HCoV-NL63_1ab;5813 | 0 | 0 |
| NC_005831.2;YP_003766.2;HCoV-NL63_1ab;5814 | 0 | 0 |

| | | |
|---|---|---|
| NC_005831.2;YP_003766.2;HCoV-NL63_1ab;5815 | 0 | 0 |
| NC_005831.2;YP_003766.2;HCoV-NL63_1ab;5816 | 0 | 0 |
| NC_005831.2;YP_003766.2;HCoV-NL63_1ab;5817 | 0 | 0 |
| NC_005831.2;YP_003766.2;HCoV-NL63_1ab;5818 | 0 | 0 |
| NC_005831.2;YP_003766.2;HCoV-NL63_1ab;5819 | 0 | 0 |
| NC_005831.2;YP_003766.2;HCoV-NL63_1ab;5820 | 0 | 0 |
| NC_005831.2;YP_003766.2;HCoV-NL63_1ab;5821 | 0 | 0 |
| NC_005831.2;YP_003766.2;HCoV-NL63_1ab;5822 | 0 | 0 |
| NC_005831.2;YP_003766.2;HCoV-NL63_1ab;5823 | 0 | 0 |
| NC_005831.2;YP_003766.2;HCoV-NL63_1ab;5824 | 0 | 0 |
| NC_005831.2;YP_003766.2;HCoV-NL63_1ab;5825 | 0 | 0 |
| NC_005831.2;YP_003766.2;HCoV-NL63_1ab;5826 | 0 | 0 |
| NC_005831.2;YP_003766.2;HCoV-NL63_1ab;5827 | 0 | 0 |
| NC_005831.2;YP_003766.2;HCoV-NL63_1ab;5828 | 0 | 0 |
| NC_005831.2;YP_003766.2;HCoV-NL63_1ab;5829 | 0 | 0 |
| NC_005831.2;YP_003766.2;HCoV-NL63_1ab;5830 | 0 | 0 |
| NC_005831.2;YP_003766.2;HCoV-NL63_1ab;5831 | 0 | 0 |
| NC_005831.2;YP_003766.2;HCoV-NL63_1ab;5832 | 0 | 0 |
| NC_005831.2;YP_003766.2;HCoV-NL63_1ab;5833 | 0 | 0 |
| NC_005831.2;YP_003766.2;HCoV-NL63_1ab;5834 | 0 | 0 |
| NC_005831.2;YP_003766.2;HCoV-NL63_1ab;5835 | 0 | 0 |
| NC_005831.2;YP_003766.2;HCoV-NL63_1ab;5836 | 0 | 0 |
| NC_005831.2;YP_003766.2;HCoV-NL63_1ab;5837 | 0 | 0 |
| NC_005831.2;YP_003766.2;HCoV-NL63_1ab;5838 | 0 | 0 |
| NC_005831.2;YP_003766.2;HCoV-NL63_1ab;5839 | 0 | 0 |
| NC_005831.2;YP_003766.2;HCoV-NL63_1ab;5840 | 0 | 0 |
| NC_005831.2;YP_003766.2;HCoV-NL63_1ab;5841 | 0 | 0 |
| NC_005831.2;YP_003766.2;HCoV-NL63_1ab;5842 | 0 | 0 |
| NC_005831.2;YP_003766.2;HCoV-NL63_1ab;5843 | 0 | 0 |
| NC_005831.2;YP_003766.2;HCoV-NL63_1ab;5844 | 0 | 0 |
| NC_005831.2;YP_003766.2;HCoV-NL63_1ab;5845 | 0 | 0 |
| NC_005831.2;YP_003766.2;HCoV-NL63_1ab;5846 | 0 | 0 |
| NC_005831.2;YP_003766.2;HCoV-NL63_1ab;5847 | 0 | 0 |
| NC_005831.2;YP_003766.2;HCoV-NL63_1ab;5848 | 0 | 0 |
| NC_005831.2;YP_003766.2;HCoV-NL63_1ab;5849 | 0 | 0 |
| NC_005831.2;YP_003766.2;HCoV-NL63_1ab;5850 | 0 | 0 |
| NC_005831.2;YP_003766.2;HCoV-NL63_1ab;5851 | 0 | 0 |
| NC_005831.2;YP_003766.2;HCoV-NL63_1ab;5852 | 0 | 0 |
| NC_005831.2;YP_003766.2;HCoV-NL63_1ab;5853 | 0 | 0 |
| NC_005831.2;YP_003766.2;HCoV-NL63_1ab;5854 | 0 | 0.025 |
| NC_005831.2;YP_003766.2;HCoV-NL63_1ab;5855 | 0 | 0 |
| NC_005831.2;YP_003766.2;HCoV-NL63_1ab;5856 | 0 | 0 |
| NC_005831.2;YP_003766.2;HCoV-NL63_1ab;5857 | 0 | 0.05 |

| | | |
|---|---|---|
| NC_005831.2;YP_003766.2;HCoV-NL63_1ab;5858 | 0 | 0.025 |
| NC_005831.2;YP_003766.2;HCoV-NL63_1ab;5859 | 0 | 0.05 |
| NC_005831.2;YP_003766.2;HCoV-NL63_1ab;5860 | 0 | 0.025 |
| NC_005831.2;YP_003766.2;HCoV-NL63_1ab;5861 | 0 | 0.025 |
| NC_005831.2;YP_003766.2;HCoV-NL63_1ab;5862 | 0 | 0.025 |
| NC_005831.2;YP_003766.2;HCoV-NL63_1ab;5863 | 0.1 | 0.025 |
| NC_005831.2;YP_003766.2;HCoV-NL63_1ab;5864 | 0.05 | 0.025 |
| NC_005831.2;YP_003766.2;HCoV-NL63_1ab;5865 | 0.05 | 0 |
| NC_005831.2;YP_003766.2;HCoV-NL63_1ab;5866 | 0.05 | 0 |
| NC_005831.2;YP_003766.2;HCoV-NL63_1ab;5867 | 0.05 | 0 |
| NC_005831.2;YP_003766.2;HCoV-NL63_1ab;5868 | 0 | 0.025 |
| NC_005831.2;YP_003766.2;HCoV-NL63_1ab;5869 | 0 | 0 |
| NC_005831.2;YP_003766.2;HCoV-NL63_1ab;5870 | 0 | 0 |
| NC_005831.2;YP_003766.2;HCoV-NL63_1ab;5871 | 0.05 | 0 |
| NC_005831.2;YP_003766.2;HCoV-NL63_1ab;5872 | 0 | 0 |
| NC_005831.2;YP_003766.2;HCoV-NL63_1ab;5873 | 0 | 0 |
| NC_005831.2;YP_003766.2;HCoV-NL63_1ab;5874 | 0 | 0 |
| NC_005831.2;YP_003766.2;HCoV-NL63_1ab;5875 | 0 | 0 |
| NC_005831.2;YP_003766.2;HCoV-NL63_1ab;5876 | 0 | 0 |
| NC_005831.2;YP_003766.2;HCoV-NL63_1ab;5877 | 0 | 0 |
| NC_005831.2;YP_003766.2;HCoV-NL63_1ab;5878 | 0 | 0 |
| NC_005831.2;YP_003766.2;HCoV-NL63_1ab;5879 | 0 | 0 |
| NC_005831.2;YP_003766.2;HCoV-NL63_1ab;5880 | 0 | 0 |
| NC_005831.2;YP_003766.2;HCoV-NL63_1ab;5881 | 0 | 0 |
| NC_005831.2;YP_003766.2;HCoV-NL63_1ab;5882 | 0 | 0 |
| NC_005831.2;YP_003766.2;HCoV-NL63_1ab;5883 | 0 | 0 |
| NC_005831.2;YP_003766.2;HCoV-NL63_1ab;5884 | 0 | 0 |
| NC_005831.2;YP_003766.2;HCoV-NL63_1ab;5885 | 0 | 0 |
| NC_005831.2;YP_003766.2;HCoV-NL63_1ab;5886 | 0 | 0 |
| NC_005831.2;YP_003766.2;HCoV-NL63_1ab;5887 | 0 | 0 |
| NC_005831.2;YP_003766.2;HCoV-NL63_1ab;5888 | 0 | 0 |
| NC_005831.2;YP_003766.2;HCoV-NL63_1ab;5889 | 0 | 0 |
| NC_005831.2;YP_003766.2;HCoV-NL63_1ab;5890 | 0 | 0 |
| NC_005831.2;YP_003766.2;HCoV-NL63_1ab;5891 | 0 | 0 |
| NC_005831.2;YP_003766.2;HCoV-NL63_1ab;5892 | 0 | 0 |
| NC_005831.2;YP_003766.2;HCoV-NL63_1ab;5893 | 0 | 0 |
| NC_005831.2;YP_003766.2;HCoV-NL63_1ab;5894 | 0 | 0 |
| NC_005831.2;YP_003766.2;HCoV-NL63_1ab;5895 | 0 | 0 |
| NC_005831.2;YP_003766.2;HCoV-NL63_1ab;5896 | 0 | 0 |
| NC_005831.2;YP_003766.2;HCoV-NL63_1ab;5897 | 0 | 0 |
| NC_005831.2;YP_003766.2;HCoV-NL63_1ab;5898 | 0 | 0 |
| NC_005831.2;YP_003766.2;HCoV-NL63_1ab;5899 | 0 | 0 |
| NC_005831.2;YP_003766.2;HCoV-NL63_1ab;5900 | 0 | 0 |

| | | |
|---|---|---|
| NC_005831.2;YP_003766.2;HCoV-NL63_1ab;5901 | 0 | 0 |
| NC_005831.2;YP_003766.2;HCoV-NL63_1ab;5902 | 0 | 0 |
| NC_005831.2;YP_003766.2;HCoV-NL63_1ab;5903 | 0 | 0 |
| NC_005831.2;YP_003766.2;HCoV-NL63_1ab;5904 | 0 | 0 |
| NC_005831.2;YP_003766.2;HCoV-NL63_1ab;5905 | 0 | 0 |
| NC_005831.2;YP_003766.2;HCoV-NL63_1ab;5906 | 0 | 0 |
| NC_005831.2;YP_003766.2;HCoV-NL63_1ab;5907 | 0 | 0.05 |
| NC_005831.2;YP_003766.2;HCoV-NL63_1ab;5908 | 0.05 | 0.025 |
| NC_005831.2;YP_003766.2;HCoV-NL63_1ab;5909 | 0 | 0.075 |
| NC_005831.2;YP_003766.2;HCoV-NL63_1ab;5910 | 0 | 0.025 |
| NC_005831.2;YP_003766.2;HCoV-NL63_1ab;5911 | 0 | 0 |
| NC_005831.2;YP_003766.2;HCoV-NL63_1ab;5912 | 0 | 0 |
| NC_005831.2;YP_003766.2;HCoV-NL63_1ab;5913 | 0 | 0 |
| NC_005831.2;YP_003766.2;HCoV-NL63_1ab;5914 | 0.05 | 0.05 |
| NC_005831.2;YP_003766.2;HCoV-NL63_1ab;5915 | 0.05 | 0.025 |
| NC_005831.2;YP_003766.2;HCoV-NL63_1ab;5916 | 0.05 | 0.025 |
| NC_005831.2;YP_003766.2;HCoV-NL63_1ab;5917 | 0.05 | 0.05 |
| NC_005831.2;YP_003766.2;HCoV-NL63_1ab;5918 | 0.05 | 0.05 |
| NC_005831.2;YP_003766.2;HCoV-NL63_1ab;5919 | 0 | 0.05 |
| NC_005831.2;YP_003766.2;HCoV-NL63_1ab;5920 | 0.05 | 0.1 |
| NC_005831.2;YP_003766.2;HCoV-NL63_1ab;5921 | 0.05 | 0.05 |
| NC_005831.2;YP_003766.2;HCoV-NL63_1ab;5922 | 0 | 0 |
| NC_005831.2;YP_003766.2;HCoV-NL63_1ab;5923 | 0.1 | 0.075 |
| NC_005831.2;YP_003766.2;HCoV-NL63_1ab;5924 | 0.05 | 0.025 |
| NC_005831.2;YP_003766.2;HCoV-NL63_1ab;5925 | 0.1 | 0 |
| NC_005831.2;YP_003766.2;HCoV-NL63_1ab;5926 | 0.05 | 0 |
| NC_005831.2;YP_003766.2;HCoV-NL63_1ab;5927 | 0.05 | 0 |
| NC_005831.2;YP_003766.2;HCoV-NL63_1ab;5928 | 0.05 | 0 |
| NC_005831.2;YP_003766.2;HCoV-NL63_1ab;5929 | 0 | 0 |
| NC_005831.2;YP_003766.2;HCoV-NL63_1ab;5930 | 0 | 0 |
| NC_005831.2;YP_003766.2;HCoV-NL63_1ab;5931 | 0 | 0 |
| NC_005831.2;YP_003766.2;HCoV-NL63_1ab;5932 | 0 | 0 |
| NC_005831.2;YP_003766.2;HCoV-NL63_1ab;5933 | 0 | 0 |
| NC_005831.2;YP_003766.2;HCoV-NL63_1ab;5934 | 0 | 0 |
| NC_005831.2;YP_003766.2;HCoV-NL63_1ab;5935 | 0 | 0 |
| NC_005831.2;YP_003766.2;HCoV-NL63_1ab;5936 | 0 | 0 |
| NC_005831.2;YP_003766.2;HCoV-NL63_1ab;5937 | 0 | 0 |
| NC_005831.2;YP_003766.2;HCoV-NL63_1ab;5938 | 0 | 0 |
| NC_005831.2;YP_003766.2;HCoV-NL63_1ab;5939 | 0 | 0 |
| NC_005831.2;YP_003766.2;HCoV-NL63_1ab;5940 | 0 | 0 |
| NC_005831.2;YP_003766.2;HCoV-NL63_1ab;5941 | 0 | 0 |
| NC_005831.2;YP_003766.2;HCoV-NL63_1ab;5942 | 0 | 0 |
| NC_005831.2;YP_003766.2;HCoV-NL63_1ab;5943 | 0 | 0 |

| | | |
|---|---|---|
| NC_005831.2;YP_003766.2;HCoV-NL63_1ab;5944 | 0 | 0 |
| NC_005831.2;YP_003766.2;HCoV-NL63_1ab;5945 | 0 | 0 |
| NC_005831.2;YP_003766.2;HCoV-NL63_1ab;5946 | 0 | 0 |
| NC_005831.2;YP_003766.2;HCoV-NL63_1ab;5947 | 0 | 0 |
| NC_005831.2;YP_003766.2;HCoV-NL63_1ab;5948 | 0 | 0 |
| NC_005831.2;YP_003766.2;HCoV-NL63_1ab;5949 | 0 | 0 |
| NC_005831.2;YP_003766.2;HCoV-NL63_1ab;5950 | 0 | 0 |
| NC_005831.2;YP_003766.2;HCoV-NL63_1ab;5951 | 0 | 0 |
| NC_005831.2;YP_003766.2;HCoV-NL63_1ab;5952 | 0 | 0 |
| NC_005831.2;YP_003766.2;HCoV-NL63_1ab;5953 | 0 | 0 |
| NC_005831.2;YP_003766.2;HCoV-NL63_1ab;5954 | 0 | 0 |
| NC_005831.2;YP_003766.2;HCoV-NL63_1ab;5955 | 0 | 0 |
| NC_005831.2;YP_003766.2;HCoV-NL63_1ab;5956 | 0 | 0 |
| NC_005831.2;YP_003766.2;HCoV-NL63_1ab;5957 | 0 | 0 |
| NC_005831.2;YP_003766.2;HCoV-NL63_1ab;5958 | 0 | 0 |
| NC_005831.2;YP_003766.2;HCoV-NL63_1ab;5959 | 0 | 0 |
| NC_005831.2;YP_003766.2;HCoV-NL63_1ab;5960 | 0.05 | 0 |
| NC_005831.2;YP_003766.2;HCoV-NL63_1ab;5961 | 0 | 0 |
| NC_005831.2;YP_003766.2;HCoV-NL63_1ab;5962 | 0.05 | 0.025 |
| NC_005831.2;YP_003766.2;HCoV-NL63_1ab;5963 | 0.05 | 0.025 |
| NC_005831.2;YP_003766.2;HCoV-NL63_1ab;5964 | 0 | 0 |
| NC_005831.2;YP_003766.2;HCoV-NL63_1ab;5965 | 0 | 0 |
| NC_005831.2;YP_003766.2;HCoV-NL63_1ab;5966 | 0.1 | 0.1 |
| NC_005831.2;YP_003766.2;HCoV-NL63_1ab;5967 | 0 | 0.025 |
| NC_005831.2;YP_003766.2;HCoV-NL63_1ab;5968 | 0 | 0 |
| NC_005831.2;YP_003766.2;HCoV-NL63_1ab;5969 | 0 | 0 |
| NC_005831.2;YP_003766.2;HCoV-NL63_1ab;5970 | 0 | 0 |
| NC_005831.2;YP_003766.2;HCoV-NL63_1ab;5971 | 0 | 0 |
| NC_005831.2;YP_003766.2;HCoV-NL63_1ab;5972 | 0 | 0.025 |
| NC_005831.2;YP_003766.2;HCoV-NL63_1ab;5973 | 0 | 0 |
| NC_005831.2;YP_003766.2;HCoV-NL63_1ab;5974 | 0 | 0 |
| NC_005831.2;YP_003766.2;HCoV-NL63_1ab;5975 | 0 | 0 |
| NC_005831.2;YP_003766.2;HCoV-NL63_1ab;5976 | 0 | 0 |
| NC_005831.2;YP_003766.2;HCoV-NL63_1ab;5977 | 0 | 0 |
| NC_005831.2;YP_003766.2;HCoV-NL63_1ab;5978 | 0 | 0 |
| NC_005831.2;YP_003766.2;HCoV-NL63_1ab;5979 | 0 | 0 |
| NC_005831.2;YP_003766.2;HCoV-NL63_1ab;5980 | 0 | 0 |
| NC_005831.2;YP_003766.2;HCoV-NL63_1ab;5981 | 0 | 0 |
| NC_005831.2;YP_003766.2;HCoV-NL63_1ab;5982 | 0 | 0 |
| NC_005831.2;YP_003766.2;HCoV-NL63_1ab;5983 | 0 | 0 |
| NC_005831.2;YP_003766.2;HCoV-NL63_1ab;5984 | 0 | 0 |
| NC_005831.2;YP_003766.2;HCoV-NL63_1ab;5985 | 0 | 0 |
| NC_005831.2;YP_003766.2;HCoV-NL63_1ab;5986 | 0 | 0 |

| | | |
|---|---|---|
| NC_005831.2;YP_003766.2;HCoV-NL63_1ab;5987 | 0 | 0 |
| NC_005831.2;YP_003766.2;HCoV-NL63_1ab;5988 | 0.05 | 0 |
| NC_005831.2;YP_003766.2;HCoV-NL63_1ab;5989 | 0 | 0 |
| NC_005831.2;YP_003766.2;HCoV-NL63_1ab;5990 | 0 | 0 |
| NC_005831.2;YP_003766.2;HCoV-NL63_1ab;5991 | 0 | 0 |
| NC_005831.2;YP_003766.2;HCoV-NL63_1ab;5992 | 0 | 0 |
| NC_005831.2;YP_003766.2;HCoV-NL63_1ab;5993 | 0 | 0 |
| NC_005831.2;YP_003766.2;HCoV-NL63_1ab;5994 | 0 | 0 |
| NC_005831.2;YP_003766.2;HCoV-NL63_1ab;5995 | 0 | 0 |
| NC_005831.2;YP_003766.2;HCoV-NL63_1ab;5996 | 0 | 0 |
| NC_005831.2;YP_003766.2;HCoV-NL63_1ab;5997 | 0 | 0 |
| NC_005831.2;YP_003766.2;HCoV-NL63_1ab;5998 | 0 | 0.025 |
| NC_005831.2;YP_003766.2;HCoV-NL63_1ab;5999 | 0 | 0.025 |
| NC_005831.2;YP_003766.2;HCoV-NL63_1ab;6000 | 0 | 0 |
| NC_005831.2;YP_003766.2;HCoV-NL63_1ab;6001 | 0 | 0 |
| NC_005831.2;YP_003766.2;HCoV-NL63_1ab;6002 | 0 | 0.025 |
| NC_005831.2;YP_003766.2;HCoV-NL63_1ab;6003 | 0.05 | 0 |
| NC_005831.2;YP_003766.2;HCoV-NL63_1ab;6004 | 0.05 | 0 |
| NC_005831.2;YP_003766.2;HCoV-NL63_1ab;6005 | 0.05 | 0.025 |
| NC_005831.2;YP_003766.2;HCoV-NL63_1ab;6006 | 0.05 | 0 |
| NC_005831.2;YP_003766.2;HCoV-NL63_1ab;6007 | 0.1 | 0.025 |
| NC_005831.2;YP_003766.2;HCoV-NL63_1ab;6008 | 0.05 | 0 |
| NC_005831.2;YP_003766.2;HCoV-NL63_1ab;6009 | 0.05 | 0.025 |
| NC_005831.2;YP_003766.2;HCoV-NL63_1ab;6010 | 0.05 | 0.025 |
| NC_005831.2;YP_003766.2;HCoV-NL63_1ab;6011 | 0.05 | 0.025 |
| NC_005831.2;YP_003766.2;HCoV-NL63_1ab;6012 | 0.1 | 0.05 |
| NC_005831.2;YP_003766.2;HCoV-NL63_1ab;6013 | 0.05 | 0 |
| NC_005831.2;YP_003766.2;HCoV-NL63_1ab;6014 | 0.05 | 0 |
| NC_005831.2;YP_003766.2;HCoV-NL63_1ab;6015 | 0.05 | 0 |
| NC_005831.2;YP_003766.2;HCoV-NL63_1ab;6016 | 0.2 | 0 |
| NC_005831.2;YP_003766.2;HCoV-NL63_1ab;6017 | 0.05 | 0 |
| NC_005831.2;YP_003766.2;HCoV-NL63_1ab;6018 | 0 | 0 |
| NC_005831.2;YP_003766.2;HCoV-NL63_1ab;6019 | 0 | 0 |
| NC_005831.2;YP_003766.2;HCoV-NL63_1ab;6020 | 0 | 0 |
| NC_005831.2;YP_003766.2;HCoV-NL63_1ab;6021 | 0 | 0 |
| NC_005831.2;YP_003766.2;HCoV-NL63_1ab;6022 | 0.15 | 0.05 |
| NC_005831.2;YP_003766.2;HCoV-NL63_1ab;6023 | 0.15 | 0.1 |
| NC_005831.2;YP_003766.2;HCoV-NL63_1ab;6024 | 0 | 0 |
| NC_005831.2;YP_003766.2;HCoV-NL63_1ab;6025 | 0 | 0 |
| NC_005831.2;YP_003766.2;HCoV-NL63_1ab;6026 | 0 | 0 |
| NC_005831.2;YP_003766.2;HCoV-NL63_1ab;6027 | 0 | 0 |
| NC_005831.2;YP_003766.2;HCoV-NL63_1ab;6028 | 0 | 0 |
| NC_005831.2;YP_003766.2;HCoV-NL63_1ab;6029 | 0 | 0 |

| | | |
|---|---|---|
| NC_005831.2;YP_003766.2;HCoV-NL63_1ab;6030 | 0 | 0 |
| NC_005831.2;YP_003766.2;HCoV-NL63_1ab;6031 | 0.05 | 0.025 |
| NC_005831.2;YP_003766.2;HCoV-NL63_1ab;6032 | 0.05 | 0.025 |
| NC_005831.2;YP_003766.2;HCoV-NL63_1ab;6033 | 0 | 0.025 |
| NC_005831.2;YP_003766.2;HCoV-NL63_1ab;6034 | 0 | 0.025 |
| NC_005831.2;YP_003766.2;HCoV-NL63_1ab;6035 | 0 | 0.025 |
| NC_005831.2;YP_003766.2;HCoV-NL63_1ab;6036 | 0 | 0.025 |
| NC_005831.2;YP_003766.2;HCoV-NL63_1ab;6037 | 0 | 0.025 |
| NC_005831.2;YP_003766.2;HCoV-NL63_1ab;6038 | 0 | 0.025 |
| NC_005831.2;YP_003766.2;HCoV-NL63_1ab;6039 | 0 | 0.025 |
| NC_005831.2;YP_003766.2;HCoV-NL63_1ab;6040 | 0 | 0.025 |
| NC_005831.2;YP_003766.2;HCoV-NL63_1ab;6041 | 0 | 0.05 |
| NC_005831.2;YP_003766.2;HCoV-NL63_1ab;6042 | 0 | 0 |
| NC_005831.2;YP_003766.2;HCoV-NL63_1ab;6043 | 0 | 0 |
| NC_005831.2;YP_003766.2;HCoV-NL63_1ab;6044 | 0 | 0 |
| NC_005831.2;YP_003766.2;HCoV-NL63_1ab;6045 | 0.05 | 0 |
| NC_005831.2;YP_003766.2;HCoV-NL63_1ab;6046 | 0 | 0 |
| NC_005831.2;YP_003766.2;HCoV-NL63_1ab;6047 | 0 | 0 |
| NC_005831.2;YP_003766.2;HCoV-NL63_1ab;6048 | 0.05 | 0.025 |
| NC_005831.2;YP_003766.2;HCoV-NL63_1ab;6049 | 0 | 0 |
| NC_005831.2;YP_003766.2;HCoV-NL63_1ab;6050 | 0 | 0 |
| NC_005831.2;YP_003766.2;HCoV-NL63_1ab;6051 | 0.05 | 0 |
| NC_005831.2;YP_003766.2;HCoV-NL63_1ab;6052 | 0.05 | 0 |
| NC_005831.2;YP_003766.2;HCoV-NL63_1ab;6053 | 0 | 0 |
| NC_005831.2;YP_003766.2;HCoV-NL63_1ab;6054 | 0 | 0 |
| NC_005831.2;YP_003766.2;HCoV-NL63_1ab;6055 | 0 | 0 |
| NC_005831.2;YP_003766.2;HCoV-NL63_1ab;6056 | 0 | 0 |
| NC_005831.2;YP_003766.2;HCoV-NL63_1ab;6057 | 0 | 0 |
| NC_005831.2;YP_003766.2;HCoV-NL63_1ab;6058 | 0 | 0 |
| NC_005831.2;YP_003766.2;HCoV-NL63_1ab;6059 | 0 | 0 |
| NC_005831.2;YP_003766.2;HCoV-NL63_1ab;6060 | 0 | 0 |
| NC_005831.2;YP_003766.2;HCoV-NL63_1ab;6061 | 0 | 0 |
| NC_005831.2;YP_003766.2;HCoV-NL63_1ab;6062 | 0 | 0 |
| NC_005831.2;YP_003766.2;HCoV-NL63_1ab;6063 | 0 | 0 |
| NC_005831.2;YP_003766.2;HCoV-NL63_1ab;6064 | 0 | 0 |
| NC_005831.2;YP_003766.2;HCoV-NL63_1ab;6065 | 0 | 0 |
| NC_005831.2;YP_003766.2;HCoV-NL63_1ab;6066 | 0 | 0 |
| NC_005831.2;YP_003766.2;HCoV-NL63_1ab;6067 | 0 | 0 |
| NC_005831.2;YP_003766.2;HCoV-NL63_1ab;6068 | 0 | 0 |
| NC_005831.2;YP_003766.2;HCoV-NL63_1ab;6069 | 0 | 0 |
| NC_005831.2;YP_003766.2;HCoV-NL63_1ab;6070 | 0 | 0 |
| NC_005831.2;YP_003766.2;HCoV-NL63_1ab;6071 | 0 | 0 |
| NC_005831.2;YP_003766.2;HCoV-NL63_1ab;6072 | 0.05 | 0 |

| | | |
|---|---|---|
| NC_005831.2;YP_003766.2;HCoV-NL63_1ab;6073 | 0 | 0 |
| NC_005831.2;YP_003766.2;HCoV-NL63_1ab;6074 | 0 | 0 |
| NC_005831.2;YP_003766.2;HCoV-NL63_1ab;6075 | 0 | 0 |
| NC_005831.2;YP_003766.2;HCoV-NL63_1ab;6076 | 0 | 0 |
| NC_005831.2;YP_003766.2;HCoV-NL63_1ab;6077 | 0 | 0 |
| NC_005831.2;YP_003766.2;HCoV-NL63_1ab;6078 | 0.1 | 0.025 |
| NC_005831.2;YP_003766.2;HCoV-NL63_1ab;6079 | 0.05 | 0 |
| NC_005831.2;YP_003766.2;HCoV-NL63_1ab;6080 | 0 | 0 |
| NC_005831.2;YP_003766.2;HCoV-NL63_1ab;6081 | 0.05 | 0 |
| NC_005831.2;YP_003766.2;HCoV-NL63_1ab;6082 | 0 | 0 |
| NC_005831.2;YP_003766.2;HCoV-NL63_1ab;6083 | 0 | 0 |
| NC_005831.2;YP_003766.2;HCoV-NL63_1ab;6084 | 0 | 0 |
| NC_005831.2;YP_003766.2;HCoV-NL63_1ab;6085 | 0.05 | 0 |
| NC_005831.2;YP_003766.2;HCoV-NL63_1ab;6086 | 0.05 | 0 |
| NC_005831.2;YP_003766.2;HCoV-NL63_1ab;6087 | 0.05 | 0 |
| NC_005831.2;YP_003766.2;HCoV-NL63_1ab;6088 | 0.05 | 0 |
| NC_005831.2;YP_003766.2;HCoV-NL63_1ab;6089 | 0.05 | 0 |
| NC_005831.2;YP_003766.2;HCoV-NL63_1ab;6090 | 0.05 | 0 |
| NC_005831.2;YP_003766.2;HCoV-NL63_1ab;6091 | 0.05 | 0 |
| NC_005831.2;YP_003766.2;HCoV-NL63_1ab;6092 | 0.05 | 0 |
| NC_005831.2;YP_003766.2;HCoV-NL63_1ab;6093 | 0 | 0 |
| NC_005831.2;YP_003766.2;HCoV-NL63_1ab;6094 | 0.05 | 0 |
| NC_005831.2;YP_003766.2;HCoV-NL63_1ab;6095 | 0.05 | 0.075 |
| NC_005831.2;YP_003766.2;HCoV-NL63_1ab;6096 | 0 | 0.025 |
| NC_005831.2;YP_003766.2;HCoV-NL63_1ab;6097 | 0 | 0.025 |
| NC_005831.2;YP_003766.2;HCoV-NL63_1ab;6098 | 0 | 0 |
| NC_005831.2;YP_003766.2;HCoV-NL63_1ab;6099 | 0 | 0.025 |
| NC_005831.2;YP_003766.2;HCoV-NL63_1ab;6100 | 0.1 | 0.05 |
| NC_005831.2;YP_003766.2;HCoV-NL63_1ab;6101 | 0.1 | 0.05 |
| NC_005831.2;YP_003766.2;HCoV-NL63_1ab;6102 | 0.1 | 0.075 |
| NC_005831.2;YP_003766.2;HCoV-NL63_1ab;6103 | 0.1 | 0.05 |
| NC_005831.2;YP_003766.2;HCoV-NL63_1ab;6104 | 0.1 | 0.05 |
| NC_005831.2;YP_003766.2;HCoV-NL63_1ab;6105 | 0.1 | 0.05 |
| NC_005831.2;YP_003766.2;HCoV-NL63_1ab;6106 | 0.1 | 0.025 |
| NC_005831.2;YP_003766.2;HCoV-NL63_1ab;6107 | 0.05 | 0 |
| NC_005831.2;YP_003766.2;HCoV-NL63_1ab;6108 | 0 | 0 |
| NC_005831.2;YP_003766.2;HCoV-NL63_1ab;6109 | 0.05 | 0.025 |
| NC_005831.2;YP_003766.2;HCoV-NL63_1ab;6110 | 0 | 0 |
| NC_005831.2;YP_003766.2;HCoV-NL63_1ab;6111 | 0 | 0.025 |
| NC_005831.2;YP_003766.2;HCoV-NL63_1ab;6112 | 0 | 0 |
| NC_005831.2;YP_003766.2;HCoV-NL63_1ab;6113 | 0 | 0 |
| NC_005831.2;YP_003766.2;HCoV-NL63_1ab;6114 | 0 | 0 |
| NC_005831.2;YP_003766.2;HCoV-NL63_1ab;6115 | 0 | 0 |

| | | |
|---|---|---|
| NC_005831.2;YP_003766.2;HCoV-NL63_1ab;6116 | 0.05 | 0 |
| NC_005831.2;YP_003766.2;HCoV-NL63_1ab;6117 | 0.05 | 0.025 |
| NC_005831.2;YP_003766.2;HCoV-NL63_1ab;6118 | 0.05 | 0.05 |
| NC_005831.2;YP_003766.2;HCoV-NL63_1ab;6119 | 0.05 | 0.025 |
| NC_005831.2;YP_003766.2;HCoV-NL63_1ab;6120 | 0.05 | 0.025 |
| NC_005831.2;YP_003766.2;HCoV-NL63_1ab;6121 | 0.05 | 0.075 |
| NC_005831.2;YP_003766.2;HCoV-NL63_1ab;6122 | 0.05 | 0.025 |
| NC_005831.2;YP_003766.2;HCoV-NL63_1ab;6123 | 0.1 | 0.05 |
| NC_005831.2;YP_003766.2;HCoV-NL63_1ab;6124 | 0.05 | 0.05 |
| NC_005831.2;YP_003766.2;HCoV-NL63_1ab;6125 | 0.05 | 0.025 |
| NC_005831.2;YP_003766.2;HCoV-NL63_1ab;6126 | 0.1 | 0.025 |
| NC_005831.2;YP_003766.2;HCoV-NL63_1ab;6127 | 0.05 | 0.05 |
| NC_005831.2;YP_003766.2;HCoV-NL63_1ab;6128 | 0.05 | 0.05 |
| NC_005831.2;YP_003766.2;HCoV-NL63_1ab;6129 | 0 | 0 |
| NC_005831.2;YP_003766.2;HCoV-NL63_1ab;6130 | 0 | 0 |
| NC_005831.2;YP_003766.2;HCoV-NL63_1ab;6131 | 0 | 0 |
| NC_005831.2;YP_003766.2;HCoV-NL63_1ab;6132 | 0 | 0 |
| NC_005831.2;YP_003766.2;HCoV-NL63_1ab;6133 | 0 | 0 |
| NC_005831.2;YP_003766.2;HCoV-NL63_1ab;6134 | 0 | 0 |
| NC_005831.2;YP_003766.2;HCoV-NL63_1ab;6135 | 0 | 0.025 |
| NC_005831.2;YP_003766.2;HCoV-NL63_1ab;6136 | 0.05 | 0.025 |
| NC_005831.2;YP_003766.2;HCoV-NL63_1ab;6137 | 0 | 0.025 |
| NC_005831.2;YP_003766.2;HCoV-NL63_1ab;6138 | 0 | 0.025 |
| NC_005831.2;YP_003766.2;HCoV-NL63_1ab;6139 | 0 | 0.05 |
| NC_005831.2;YP_003766.2;HCoV-NL63_1ab;6140 | 0 | 0.05 |
| NC_005831.2;YP_003766.2;HCoV-NL63_1ab;6141 | 0 | 0.025 |
| NC_005831.2;YP_003766.2;HCoV-NL63_1ab;6142 | 0 | 0.025 |
| NC_005831.2;YP_003766.2;HCoV-NL63_1ab;6143 | 0 | 0.025 |
| NC_005831.2;YP_003766.2;HCoV-NL63_1ab;6144 | 0 | 0.025 |
| NC_005831.2;YP_003766.2;HCoV-NL63_1ab;6145 | 0 | 0.05 |
| NC_005831.2;YP_003766.2;HCoV-NL63_1ab;6146 | 0 | 0 |
| NC_005831.2;YP_003766.2;HCoV-NL63_1ab;6147 | 0 | 0.025 |
| NC_005831.2;YP_003766.2;HCoV-NL63_1ab;6148 | 0 | 0 |
| NC_005831.2;YP_003766.2;HCoV-NL63_1ab;6149 | 0 | 0 |
| NC_005831.2;YP_003766.2;HCoV-NL63_1ab;6150 | 0 | 0 |
| NC_005831.2;YP_003766.2;HCoV-NL63_1ab;6151 | 0 | 0 |
| NC_005831.2;YP_003766.2;HCoV-NL63_1ab;6152 | 0 | 0 |
| NC_005831.2;YP_003766.2;HCoV-NL63_1ab;6153 | 0 | 0 |
| NC_005831.2;YP_003766.2;HCoV-NL63_1ab;6154 | 0 | 0 |
| NC_005831.2;YP_003766.2;HCoV-NL63_1ab;6155 | 0 | 0 |
| NC_005831.2;YP_003766.2;HCoV-NL63_1ab;6156 | 0 | 0 |
| NC_005831.2;YP_003766.2;HCoV-NL63_1ab;6157 | 0 | 0 |
| NC_005831.2;YP_003766.2;HCoV-NL63_1ab;6158 | 0 | 0 |

| | | |
|---|---|---|
| NC_005831.2;YP_003766.2;HCoV-NL63_1ab;6159 | 0 | 0 |
| NC_005831.2;YP_003766.2;HCoV-NL63_1ab;6160 | 0 | 0 |
| NC_005831.2;YP_003766.2;HCoV-NL63_1ab;6161 | 0 | 0 |
| NC_005831.2;YP_003766.2;HCoV-NL63_1ab;6162 | 0 | 0 |
| NC_005831.2;YP_003766.2;HCoV-NL63_1ab;6163 | 0 | 0 |
| NC_005831.2;YP_003766.2;HCoV-NL63_1ab;6164 | 0 | 0 |
| NC_005831.2;YP_003766.2;HCoV-NL63_1ab;6165 | 0 | 0 |
| NC_005831.2;YP_003766.2;HCoV-NL63_1ab;6166 | 0 | 0 |
| NC_005831.2;YP_003766.2;HCoV-NL63_1ab;6167 | 0 | 0 |
| NC_005831.2;YP_003766.2;HCoV-NL63_1ab;6168 | 0.05 | 0 |
| NC_005831.2;YP_003766.2;HCoV-NL63_1ab;6169 | 0.05 | 0 |
| NC_005831.2;YP_003766.2;HCoV-NL63_1ab;6170 | 0.1 | 0 |
| NC_005831.2;YP_003766.2;HCoV-NL63_1ab;6171 | 0.05 | 0 |
| NC_005831.2;YP_003766.2;HCoV-NL63_1ab;6172 | 0.05 | 0 |
| NC_005831.2;YP_003766.2;HCoV-NL63_1ab;6173 | 0 | 0 |
| NC_005831.2;YP_003766.2;HCoV-NL63_1ab;6174 | 0 | 0 |
| NC_005831.2;YP_003766.2;HCoV-NL63_1ab;6175 | 0 | 0 |
| NC_005831.2;YP_003766.2;HCoV-NL63_1ab;6176 | 0 | 0 |
| NC_005831.2;YP_003766.2;HCoV-NL63_1ab;6177 | 0 | 0 |
| NC_005831.2;YP_003766.2;HCoV-NL63_1ab;6178 | 0 | 0 |
| NC_005831.2;YP_003766.2;HCoV-NL63_1ab;6179 | 0 | 0 |
| NC_005831.2;YP_003766.2;HCoV-NL63_1ab;6180 | 0 | 0 |
| NC_005831.2;YP_003766.2;HCoV-NL63_1ab;6181 | 0 | 0.025 |
| NC_005831.2;YP_003766.2;HCoV-NL63_1ab;6182 | 0 | 0 |
| NC_005831.2;YP_003766.2;HCoV-NL63_1ab;6183 | 0 | 0 |
| NC_005831.2;YP_003766.2;HCoV-NL63_1ab;6184 | 0 | 0 |
| NC_005831.2;YP_003766.2;HCoV-NL63_1ab;6185 | 0 | 0 |
| NC_005831.2;YP_003766.2;HCoV-NL63_1ab;6186 | 0 | 0 |
| NC_005831.2;YP_003766.2;HCoV-NL63_1ab;6187 | 0 | 0 |
| NC_005831.2;YP_003766.2;HCoV-NL63_1ab;6188 | 0 | 0 |
| NC_005831.2;YP_003766.2;HCoV-NL63_1ab;6189 | 0 | 0 |
| NC_005831.2;YP_003766.2;HCoV-NL63_1ab;6190 | 0 | 0 |
| NC_005831.2;YP_003766.2;HCoV-NL63_1ab;6191 | 0 | 0 |
| NC_005831.2;YP_003766.2;HCoV-NL63_1ab;6192 | 0 | 0.025 |
| NC_005831.2;YP_003766.2;HCoV-NL63_1ab;6193 | 0 | 0 |
| NC_005831.2;YP_003766.2;HCoV-NL63_1ab;6194 | 0 | 0 |
| NC_005831.2;YP_003766.2;HCoV-NL63_1ab;6195 | 0 | 0 |
| NC_005831.2;YP_003766.2;HCoV-NL63_1ab;6196 | 0 | 0 |
| NC_005831.2;YP_003766.2;HCoV-NL63_1ab;6197 | 0 | 0 |
| NC_005831.2;YP_003766.2;HCoV-NL63_1ab;6198 | 0 | 0 |
| NC_005831.2;YP_003766.2;HCoV-NL63_1ab;6199 | 0 | 0 |
| NC_005831.2;YP_003766.2;HCoV-NL63_1ab;6200 | 0 | 0 |
| NC_005831.2;YP_003766.2;HCoV-NL63_1ab;6201 | 0 | 0 |

| | | |
|---|---|---|
| NC_005831.2;YP_003766.2;HCoV-NL63_1ab;6202 | 0 | 0 |
| NC_005831.2;YP_003766.2;HCoV-NL63_1ab;6203 | 0 | 0 |
| NC_005831.2;YP_003766.2;HCoV-NL63_1ab;6204 | 0 | 0 |
| NC_005831.2;YP_003766.2;HCoV-NL63_1ab;6205 | 0 | 0 |
| NC_005831.2;YP_003766.2;HCoV-NL63_1ab;6206 | 0 | 0 |
| NC_005831.2;YP_003766.2;HCoV-NL63_1ab;6207 | 0 | 0 |
| NC_005831.2;YP_003766.2;HCoV-NL63_1ab;6208 | 0 | 0 |
| NC_005831.2;YP_003766.2;HCoV-NL63_1ab;6209 | 0 | 0 |
| NC_005831.2;YP_003766.2;HCoV-NL63_1ab;6210 | 0 | 0 |
| NC_005831.2;YP_003766.2;HCoV-NL63_1ab;6211 | 0 | 0 |
| NC_005831.2;YP_003766.2;HCoV-NL63_1ab;6212 | 0 | 0 |
| NC_005831.2;YP_003766.2;HCoV-NL63_1ab;6213 | 0 | 0 |
| NC_005831.2;YP_003766.2;HCoV-NL63_1ab;6214 | 0 | 0 |
| NC_005831.2;YP_003766.2;HCoV-NL63_1ab;6215 | 0 | 0 |
| NC_005831.2;YP_003766.2;HCoV-NL63_1ab;6216 | 0 | 0 |
| NC_005831.2;YP_003766.2;HCoV-NL63_1ab;6217 | 0 | 0.025 |
| NC_005831.2;YP_003766.2;HCoV-NL63_1ab;6218 | 0 | 0.025 |
| NC_005831.2;YP_003766.2;HCoV-NL63_1ab;6219 | 0 | 0.025 |
| NC_005831.2;YP_003766.2;HCoV-NL63_1ab;6220 | 0 | 0.025 |
| NC_005831.2;YP_003766.2;HCoV-NL63_1ab;6221 | 0 | 0.075 |
| NC_005831.2;YP_003766.2;HCoV-NL63_1ab;6222 | 0 | 0.05 |
| NC_005831.2;YP_003766.2;HCoV-NL63_1ab;6223 | 0 | 0.025 |
| NC_005831.2;YP_003766.2;HCoV-NL63_1ab;6224 | 0 | 0.075 |
| NC_005831.2;YP_003766.2;HCoV-NL63_1ab;6225 | 0 | 0.075 |
| NC_005831.2;YP_003766.2;HCoV-NL63_1ab;6226 | 0.05 | 0.075 |
| NC_005831.2;YP_003766.2;HCoV-NL63_1ab;6227 | 0 | 0.075 |
| NC_005831.2;YP_003766.2;HCoV-NL63_1ab;6228 | 0.05 | 0.075 |
| NC_005831.2;YP_003766.2;HCoV-NL63_1ab;6229 | 0 | 0.05 |
| NC_005831.2;YP_003766.2;HCoV-NL63_1ab;6230 | 0.05 | 0 |
| NC_005831.2;YP_003766.2;HCoV-NL63_1ab;6231 | 0 | 0.025 |
| NC_005831.2;YP_003766.2;HCoV-NL63_1ab;6232 | 0 | 0 |
| NC_005831.2;YP_003766.2;HCoV-NL63_1ab;6233 | 0 | 0 |
| NC_005831.2;YP_003766.2;HCoV-NL63_1ab;6234 | 0 | 0 |
| NC_005831.2;YP_003766.2;HCoV-NL63_1ab;6235 | 0 | 0 |
| NC_005831.2;YP_003766.2;HCoV-NL63_1ab;6236 | 0 | 0 |
| NC_005831.2;YP_003766.2;HCoV-NL63_1ab;6237 | 0 | 0 |
| NC_005831.2;YP_003766.2;HCoV-NL63_1ab;6238 | 0 | 0 |
| NC_005831.2;YP_003766.2;HCoV-NL63_1ab;6239 | 0 | 0 |
| NC_005831.2;YP_003766.2;HCoV-NL63_1ab;6240 | 0 | 0 |
| NC_005831.2;YP_003766.2;HCoV-NL63_1ab;6241 | 0 | 0 |
| NC_005831.2;YP_003766.2;HCoV-NL63_1ab;6242 | 0 | 0 |
| NC_005831.2;YP_003766.2;HCoV-NL63_1ab;6243 | 0 | 0 |
| NC_005831.2;YP_003766.2;HCoV-NL63_1ab;6244 | 0 | 0.05 |

| | | |
|---|---|---|
| NC_005831.2;YP_003766.2;HCoV-NL63_1ab;6245 | 0 | 0.025 |
| NC_005831.2;YP_003766.2;HCoV-NL63_1ab;6246 | 0 | 0 |
| NC_005831.2;YP_003766.2;HCoV-NL63_1ab;6247 | 0 | 0 |
| NC_005831.2;YP_003766.2;HCoV-NL63_1ab;6248 | 0 | 0 |
| NC_005831.2;YP_003766.2;HCoV-NL63_1ab;6249 | 0 | 0 |
| NC_005831.2;YP_003766.2;HCoV-NL63_1ab;6250 | 0 | 0 |
| NC_005831.2;YP_003766.2;HCoV-NL63_1ab;6251 | 0 | 0 |
| NC_005831.2;YP_003766.2;HCoV-NL63_1ab;6252 | 0 | 0 |
| NC_005831.2;YP_003766.2;HCoV-NL63_1ab;6253 | 0 | 0 |
| NC_005831.2;YP_003766.2;HCoV-NL63_1ab;6254 | 0 | 0 |
| NC_005831.2;YP_003766.2;HCoV-NL63_1ab;6255 | 0 | 0 |
| NC_005831.2;YP_003766.2;HCoV-NL63_1ab;6256 | 0 | 0 |
| NC_005831.2;YP_003766.2;HCoV-NL63_1ab;6257 | 0 | 0 |
| NC_005831.2;YP_003766.2;HCoV-NL63_1ab;6258 | 0.05 | 0 |
| NC_005831.2;YP_003766.2;HCoV-NL63_1ab;6259 | 0.05 | 0.025 |
| NC_005831.2;YP_003766.2;HCoV-NL63_1ab;6260 | 0.1 | 0.025 |
| NC_005831.2;YP_003766.2;HCoV-NL63_1ab;6261 | 0.1 | 0.025 |
| NC_005831.2;YP_003766.2;HCoV-NL63_1ab;6262 | 0.05 | 0.075 |
| NC_005831.2;YP_003766.2;HCoV-NL63_1ab;6263 | 0.05 | 0.05 |
| NC_005831.2;YP_003766.2;HCoV-NL63_1ab;6264 | 0.1 | 0.025 |
| NC_005831.2;YP_003766.2;HCoV-NL63_1ab;6265 | 0 | 0 |
| NC_005831.2;YP_003766.2;HCoV-NL63_1ab;6266 | 0 | 0 |
| NC_005831.2;YP_003766.2;HCoV-NL63_1ab;6267 | 0 | 0 |
| NC_005831.2;YP_003766.2;HCoV-NL63_1ab;6268 | 0 | 0 |
| NC_005831.2;YP_003766.2;HCoV-NL63_1ab;6269 | 0 | 0 |
| NC_005831.2;YP_003766.2;HCoV-NL63_1ab;6270 | 0 | 0 |
| NC_005831.2;YP_003766.2;HCoV-NL63_1ab;6271 | 0 | 0 |
| NC_005831.2;YP_003766.2;HCoV-NL63_1ab;6272 | 0.05 | 0 |
| NC_005831.2;YP_003766.2;HCoV-NL63_1ab;6273 | 0.05 | 0 |
| NC_005831.2;YP_003766.2;HCoV-NL63_1ab;6274 | 0 | 0 |
| NC_005831.2;YP_003766.2;HCoV-NL63_1ab;6275 | 0.05 | 0 |
| NC_005831.2;YP_003766.2;HCoV-NL63_1ab;6276 | 0.05 | 0.05 |
| NC_005831.2;YP_003766.2;HCoV-NL63_1ab;6277 | 0.05 | 0 |
| NC_005831.2;YP_003766.2;HCoV-NL63_1ab;6278 | 0.05 | 0 |
| NC_005831.2;YP_003766.2;HCoV-NL63_1ab;6279 | 0.05 | 0.025 |
| NC_005831.2;YP_003766.2;HCoV-NL63_1ab;6280 | 0.05 | 0 |
| NC_005831.2;YP_003766.2;HCoV-NL63_1ab;6281 | 0 | 0 |
| NC_005831.2;YP_003766.2;HCoV-NL63_1ab;6282 | 0.05 | 0.025 |
| NC_005831.2;YP_003766.2;HCoV-NL63_1ab;6283 | 0.05 | 0.025 |
| NC_005831.2;YP_003766.2;HCoV-NL63_1ab;6284 | 0 | 0.025 |
| NC_005831.2;YP_003766.2;HCoV-NL63_1ab;6285 | 0.05 | 0.025 |
| NC_005831.2;YP_003766.2;HCoV-NL63_1ab;6286 | 0 | 0.05 |
| NC_005831.2;YP_003766.2;HCoV-NL63_1ab;6287 | 0 | 0.05 |

| | | |
|---|---|---|
| NC_005831.2;YP_003766.2;HCoV-NL63_1ab;6288 | 0.05 | 0.05 |
| NC_005831.2;YP_003766.2;HCoV-NL63_1ab;6289 | 0 | 0.05 |
| NC_005831.2;YP_003766.2;HCoV-NL63_1ab;6290 | 0 | 0.05 |
| NC_005831.2;YP_003766.2;HCoV-NL63_1ab;6291 | 0 | 0.05 |
| NC_005831.2;YP_003766.2;HCoV-NL63_1ab;6292 | 0 | 0.05 |
| NC_005831.2;YP_003766.2;HCoV-NL63_1ab;6293 | 0 | 0.025 |
| NC_005831.2;YP_003766.2;HCoV-NL63_1ab;6294 | 0 | 0 |
| NC_005831.2;YP_003766.2;HCoV-NL63_1ab;6295 | 0 | 0 |
| NC_005831.2;YP_003766.2;HCoV-NL63_1ab;6296 | 0 | 0.05 |
| NC_005831.2;YP_003766.2;HCoV-NL63_1ab;6297 | 0 | 0.025 |
| NC_005831.2;YP_003766.2;HCoV-NL63_1ab;6298 | 0 | 0.025 |
| NC_005831.2;YP_003766.2;HCoV-NL63_1ab;6299 | 0 | 0.025 |
| NC_005831.2;YP_003766.2;HCoV-NL63_1ab;6300 | 0.05 | 0.075 |
| NC_005831.2;YP_003766.2;HCoV-NL63_1ab;6301 | 0.05 | 0 |
| NC_005831.2;YP_003766.2;HCoV-NL63_1ab;6302 | 0.05 | 0.05 |
| NC_005831.2;YP_003766.2;HCoV-NL63_1ab;6303 | 0 | 0.05 |
| NC_005831.2;YP_003766.2;HCoV-NL63_1ab;6304 | 0 | 0.05 |
| NC_005831.2;YP_003766.2;HCoV-NL63_1ab;6305 | 0 | 0 |
| NC_005831.2;YP_003766.2;HCoV-NL63_1ab;6306 | 0 | 0 |
| NC_005831.2;YP_003766.2;HCoV-NL63_1ab;6307 | 0 | 0 |
| NC_005831.2;YP_003766.2;HCoV-NL63_1ab;6308 | 0.1 | 0.3 |
| NC_005831.2;YP_003766.2;HCoV-NL63_1ab;6309 | 0.1 | 0.025 |
| NC_005831.2;YP_003766.2;HCoV-NL63_1ab;6310 | 0 | 0 |
| NC_005831.2;YP_003766.2;HCoV-NL63_1ab;6311 | 0 | 0 |
| NC_005831.2;YP_003766.2;HCoV-NL63_1ab;6312 | 0 | 0 |
| NC_005831.2;YP_003766.2;HCoV-NL63_1ab;6313 | 0 | 0 |
| NC_005831.2;YP_003766.2;HCoV-NL63_1ab;6314 | 0.05 | 0 |
| NC_005831.2;YP_003766.2;HCoV-NL63_1ab;6315 | 0 | 0 |
| NC_005831.2;YP_003766.2;HCoV-NL63_1ab;6316 | 0 | 0 |
| NC_005831.2;YP_003766.2;HCoV-NL63_1ab;6317 | 0 | 0 |
| NC_005831.2;YP_003766.2;HCoV-NL63_1ab;6318 | 0 | 0 |
| NC_005831.2;YP_003766.2;HCoV-NL63_1ab;6319 | 0 | 0 |
| NC_005831.2;YP_003766.2;HCoV-NL63_1ab;6320 | 0 | 0 |
| NC_005831.2;YP_003766.2;HCoV-NL63_1ab;6321 | 0 | 0 |
| NC_005831.2;YP_003766.2;HCoV-NL63_1ab;6322 | 0 | 0 |
| NC_005831.2;YP_003766.2;HCoV-NL63_1ab;6323 | 0 | 0 |
| NC_005831.2;YP_003766.2;HCoV-NL63_1ab;6324 | 0 | 0 |
| NC_005831.2;YP_003766.2;HCoV-NL63_1ab;6325 | 0 | 0 |
| NC_005831.2;YP_003766.2;HCoV-NL63_1ab;6326 | 0 | 0 |
| NC_005831.2;YP_003766.2;HCoV-NL63_1ab;6327 | 0 | 0 |
| NC_005831.2;YP_003766.2;HCoV-NL63_1ab;6328 | 0 | 0 |
| NC_005831.2;YP_003766.2;HCoV-NL63_1ab;6329 | 0 | 0 |
| NC_005831.2;YP_003766.2;HCoV-NL63_1ab;6330 | 0 | 0 |

| | | |
|---|---|---|
| NC_005831.2;YP_003766.2;HCoV-NL63_1ab;6331 | 0.05 | 0 |
| NC_005831.2;YP_003766.2;HCoV-NL63_1ab;6332 | 0.05 | 0 |
| NC_005831.2;YP_003766.2;HCoV-NL63_1ab;6333 | 0.05 | 0 |
| NC_005831.2;YP_003766.2;HCoV-NL63_1ab;6334 | 0.05 | 0 |
| NC_005831.2;YP_003766.2;HCoV-NL63_1ab;6335 | 0.1 | 0 |
| NC_005831.2;YP_003766.2;HCoV-NL63_1ab;6336 | 0.1 | 0 |
| NC_005831.2;YP_003766.2;HCoV-NL63_1ab;6337 | 0.1 | 0 |
| NC_005831.2;YP_003766.2;HCoV-NL63_1ab;6338 | 0.1 | 0 |
| NC_005831.2;YP_003766.2;HCoV-NL63_1ab;6339 | 0.1 | 0.025 |
| NC_005831.2;YP_003766.2;HCoV-NL63_1ab;6340 | 0.05 | 0 |
| NC_005831.2;YP_003766.2;HCoV-NL63_1ab;6341 | 0.1 | 0.025 |
| NC_005831.2;YP_003766.2;HCoV-NL63_1ab;6342 | 0 | 0 |
| NC_005831.2;YP_003766.2;HCoV-NL63_1ab;6343 | 0.05 | 0 |
| NC_005831.2;YP_003766.2;HCoV-NL63_1ab;6344 | 0.05 | 0 |
| NC_005831.2;YP_003766.2;HCoV-NL63_1ab;6345 | 0.05 | 0.025 |
| NC_005831.2;YP_003766.2;HCoV-NL63_1ab;6346 | 0 | 0.025 |
| NC_005831.2;YP_003766.2;HCoV-NL63_1ab;6347 | 0 | 0 |
| NC_005831.2;YP_003766.2;HCoV-NL63_1ab;6348 | 0 | 0 |
| NC_005831.2;YP_003766.2;HCoV-NL63_1ab;6349 | 0 | 0.025 |
| NC_005831.2;YP_003766.2;HCoV-NL63_1ab;6350 | 0 | 0 |
| NC_005831.2;YP_003766.2;HCoV-NL63_1ab;6351 | 0 | 0.025 |
| NC_005831.2;YP_003766.2;HCoV-NL63_1ab;6352 | 0 | 0 |
| NC_005831.2;YP_003766.2;HCoV-NL63_1ab;6353 | 0 | 0 |
| NC_005831.2;YP_003766.2;HCoV-NL63_1ab;6354 | 0 | 0 |
| NC_005831.2;YP_003766.2;HCoV-NL63_1ab;6355 | 0 | 0 |
| NC_005831.2;YP_003766.2;HCoV-NL63_1ab;6356 | 0 | 0 |
| NC_005831.2;YP_003766.2;HCoV-NL63_1ab;6357 | 0 | 0.025 |
| NC_005831.2;YP_003766.2;HCoV-NL63_1ab;6358 | 0 | 0 |
| NC_005831.2;YP_003766.2;HCoV-NL63_1ab;6359 | 0 | 0 |
| NC_005831.2;YP_003766.2;HCoV-NL63_1ab;6360 | 0 | 0 |
| NC_005831.2;YP_003766.2;HCoV-NL63_1ab;6361 | 0 | 0 |
| NC_005831.2;YP_003766.2;HCoV-NL63_1ab;6362 | 0 | 0 |
| NC_005831.2;YP_003766.2;HCoV-NL63_1ab;6363 | 0 | 0.025 |
| NC_005831.2;YP_003766.2;HCoV-NL63_1ab;6364 | 0 | 0 |
| NC_005831.2;YP_003766.2;HCoV-NL63_1ab;6365 | 0 | 0 |
| NC_005831.2;YP_003766.2;HCoV-NL63_1ab;6366 | 0 | 0 |
| NC_005831.2;YP_003766.2;HCoV-NL63_1ab;6367 | 0 | 0 |
| NC_005831.2;YP_003766.2;HCoV-NL63_1ab;6368 | 0 | 0 |
| NC_005831.2;YP_003766.2;HCoV-NL63_1ab;6369 | 0 | 0 |
| NC_005831.2;YP_003766.2;HCoV-NL63_1ab;6370 | 0 | 0 |
| NC_005831.2;YP_003766.2;HCoV-NL63_1ab;6371 | 0 | 0 |
| NC_005831.2;YP_003766.2;HCoV-NL63_1ab;6372 | 0 | 0 |
| NC_005831.2;YP_003766.2;HCoV-NL63_1ab;6373 | 0 | 0 |

| | | |
|---|---|---|
| NC_005831.2;YP_003766.2;HCoV-NL63_1ab;6374 | 0 | 0 |
| NC_005831.2;YP_003766.2;HCoV-NL63_1ab;6375 | 0 | 0 |
| NC_005831.2;YP_003766.2;HCoV-NL63_1ab;6376 | 0 | 0 |
| NC_005831.2;YP_003766.2;HCoV-NL63_1ab;6377 | 0 | 0 |
| NC_005831.2;YP_003766.2;HCoV-NL63_1ab;6378 | 0 | 0 |
| NC_005831.2;YP_003766.2;HCoV-NL63_1ab;6379 | 0 | 0 |
| NC_005831.2;YP_003766.2;HCoV-NL63_1ab;6380 | 0 | 0 |
| NC_005831.2;YP_003766.2;HCoV-NL63_1ab;6381 | 0 | 0 |
| NC_005831.2;YP_003766.2;HCoV-NL63_1ab;6382 | 0 | 0 |
| NC_005831.2;YP_003766.2;HCoV-NL63_1ab;6383 | 0 | 0 |
| NC_005831.2;YP_003766.2;HCoV-NL63_1ab;6384 | 0 | 0 |
| NC_005831.2;YP_003766.2;HCoV-NL63_1ab;6385 | 0 | 0 |
| NC_005831.2;YP_003766.2;HCoV-NL63_1ab;6386 | 0 | 0 |
| NC_005831.2;YP_003766.2;HCoV-NL63_1ab;6387 | 0 | 0 |
| NC_005831.2;YP_003766.2;HCoV-NL63_1ab;6388 | 0 | 0 |
| NC_005831.2;YP_003766.2;HCoV-NL63_1ab;6389 | 0 | 0 |
| NC_005831.2;YP_003766.2;HCoV-NL63_1ab;6390 | 0 | 0 |
| NC_005831.2;YP_003766.2;HCoV-NL63_1ab;6391 | 0 | 0 |
| NC_005831.2;YP_003766.2;HCoV-NL63_1ab;6392 | 0 | 0 |
| NC_005831.2;YP_003766.2;HCoV-NL63_1ab;6393 | 0.1 | 0.05 |
| NC_005831.2;YP_003766.2;HCoV-NL63_1ab;6394 | 0 | 0 |
| NC_005831.2;YP_003766.2;HCoV-NL63_1ab;6395 | 0 | 0.05 |
| NC_005831.2;YP_003766.2;HCoV-NL63_1ab;6396 | 0 | 0 |
| NC_005831.2;YP_003766.2;HCoV-NL63_1ab;6397 | 0 | 0 |
| NC_005831.2;YP_003766.2;HCoV-NL63_1ab;6398 | 0 | 0 |
| NC_005831.2;YP_003766.2;HCoV-NL63_1ab;6399 | 0 | 0 |
| NC_005831.2;YP_003766.2;HCoV-NL63_1ab;6400 | 0 | 0 |
| NC_005831.2;YP_003766.2;HCoV-NL63_1ab;6401 | 0 | 0 |
| NC_005831.2;YP_003766.2;HCoV-NL63_1ab;6402 | 0 | 0 |
| NC_005831.2;YP_003766.2;HCoV-NL63_1ab;6403 | 0 | 0 |
| NC_005831.2;YP_003766.2;HCoV-NL63_1ab;6404 | 0 | 0 |
| NC_005831.2;YP_003766.2;HCoV-NL63_1ab;6405 | 0.05 | 0 |
| NC_005831.2;YP_003766.2;HCoV-NL63_1ab;6406 | 0 | 0.025 |
| NC_005831.2;YP_003766.2;HCoV-NL63_1ab;6407 | 0 | 0 |
| NC_005831.2;YP_003766.2;HCoV-NL63_1ab;6408 | 0 | 0 |
| NC_005831.2;YP_003766.2;HCoV-NL63_1ab;6409 | 0 | 0 |
| NC_005831.2;YP_003766.2;HCoV-NL63_1ab;6410 | 0 | 0 |
| NC_005831.2;YP_003766.2;HCoV-NL63_1ab;6411 | 0 | 0 |
| NC_005831.2;YP_003766.2;HCoV-NL63_1ab;6412 | 0 | 0 |
| NC_005831.2;YP_003766.2;HCoV-NL63_1ab;6413 | 0 | 0 |
| NC_005831.2;YP_003766.2;HCoV-NL63_1ab;6414 | 0 | 0 |
| NC_005831.2;YP_003766.2;HCoV-NL63_1ab;6415 | 0 | 0 |
| NC_005831.2;YP_003766.2;HCoV-NL63_1ab;6416 | 0 | 0 |

| | | |
|---|---|---|
| NC_005831.2;YP_003766.2;HCoV-NL63_1ab;6417 | 0.05 | 0.075 |
| NC_005831.2;YP_003766.2;HCoV-NL63_1ab;6418 | 0 | 0.025 |
| NC_005831.2;YP_003766.2;HCoV-NL63_1ab;6419 | 0 | 0.025 |
| NC_005831.2;YP_003766.2;HCoV-NL63_1ab;6420 | 0 | 0.025 |
| NC_005831.2;YP_003766.2;HCoV-NL63_1ab;6421 | 0 | 0 |
| NC_005831.2;YP_003766.2;HCoV-NL63_1ab;6422 | 0 | 0.025 |
| NC_005831.2;YP_003766.2;HCoV-NL63_1ab;6423 | 0 | 0 |
| NC_005831.2;YP_003766.2;HCoV-NL63_1ab;6424 | 0 | 0 |
| NC_005831.2;YP_003766.2;HCoV-NL63_1ab;6425 | 0 | 0 |
| NC_005831.2;YP_003766.2;HCoV-NL63_1ab;6426 | 0.05 | 0 |
| NC_005831.2;YP_003766.2;HCoV-NL63_1ab;6427 | 0 | 0 |
| NC_005831.2;YP_003766.2;HCoV-NL63_1ab;6428 | 0 | 0 |
| NC_005831.2;YP_003766.2;HCoV-NL63_1ab;6429 | 0 | 0 |
| NC_005831.2;YP_003766.2;HCoV-NL63_1ab;6430 | 0 | 0 |
| NC_005831.2;YP_003766.2;HCoV-NL63_1ab;6431 | 0 | 0 |
| NC_005831.2;YP_003766.2;HCoV-NL63_1ab;6432 | 0 | 0 |
| NC_005831.2;YP_003766.2;HCoV-NL63_1ab;6433 | 0 | 0 |
| NC_005831.2;YP_003766.2;HCoV-NL63_1ab;6434 | 0 | 0 |
| NC_005831.2;YP_003766.2;HCoV-NL63_1ab;6435 | 0 | 0 |
| NC_005831.2;YP_003766.2;HCoV-NL63_1ab;6436 | 0 | 0 |
| NC_005831.2;YP_003766.2;HCoV-NL63_1ab;6437 | 0 | 0 |
| NC_005831.2;YP_003766.2;HCoV-NL63_1ab;6438 | 0 | 0 |
| NC_005831.2;YP_003766.2;HCoV-NL63_1ab;6439 | 0 | 0 |
| NC_005831.2;YP_003766.2;HCoV-NL63_1ab;6440 | 0 | 0 |
| NC_005831.2;YP_003766.2;HCoV-NL63_1ab;6441 | 0 | 0 |
| NC_005831.2;YP_003766.2;HCoV-NL63_1ab;6442 | 0 | 0 |
| NC_005831.2;YP_003766.2;HCoV-NL63_1ab;6443 | 0 | 0 |
| NC_005831.2;YP_003766.2;HCoV-NL63_1ab;6444 | 0 | 0 |
| NC_005831.2;YP_003766.2;HCoV-NL63_1ab;6445 | 0 | 0 |
| NC_005831.2;YP_003766.2;HCoV-NL63_1ab;6446 | 0 | 0 |
| NC_005831.2;YP_003766.2;HCoV-NL63_1ab;6447 | 0 | 0 |
| NC_005831.2;YP_003766.2;HCoV-NL63_1ab;6448 | 0 | 0.025 |
| NC_005831.2;YP_003766.2;HCoV-NL63_1ab;6449 | 0 | 0 |
| NC_005831.2;YP_003766.2;HCoV-NL63_1ab;6450 | 0 | 0.05 |
| NC_005831.2;YP_003766.2;HCoV-NL63_1ab;6451 | 0 | 0.05 |
| NC_005831.2;YP_003766.2;HCoV-NL63_1ab;6452 | 0 | 0.05 |
| NC_005831.2;YP_003766.2;HCoV-NL63_1ab;6453 | 0 | 0.05 |
| NC_005831.2;YP_003766.2;HCoV-NL63_1ab;6454 | 0 | 0.05 |
| NC_005831.2;YP_003766.2;HCoV-NL63_1ab;6455 | 0 | 0.025 |
| NC_005831.2;YP_003766.2;HCoV-NL63_1ab;6456 | 0 | 0.05 |
| NC_005831.2;YP_003766.2;HCoV-NL63_1ab;6457 | 0 | 0.025 |
| NC_005831.2;YP_003766.2;HCoV-NL63_1ab;6458 | 0 | 0 |
| NC_005831.2;YP_003766.2;HCoV-NL63_1ab;6459 | 0 | 0 |

| | | |
|---|---|---|
| NC_005831.2;YP_003766.2;HCoV-NL63_1ab;6460 | 0 | 0 |
| NC_005831.2;YP_003766.2;HCoV-NL63_1ab;6461 | 0 | 0 |
| NC_005831.2;YP_003766.2;HCoV-NL63_1ab;6462 | 0 | 0.025 |
| NC_005831.2;YP_003766.2;HCoV-NL63_1ab;6463 | 0 | 0 |
| NC_005831.2;YP_003766.2;HCoV-NL63_1ab;6464 | 0 | 0 |
| NC_005831.2;YP_003766.2;HCoV-NL63_1ab;6465 | 0 | 0 |
| NC_005831.2;YP_003766.2;HCoV-NL63_1ab;6466 | 0 | 0 |
| NC_005831.2;YP_003766.2;HCoV-NL63_1ab;6467 | 0 | 0 |
| NC_005831.2;YP_003766.2;HCoV-NL63_1ab;6468 | 0 | 0 |
| NC_005831.2;YP_003766.2;HCoV-NL63_1ab;6469 | 0 | 0 |
| NC_005831.2;YP_003766.2;HCoV-NL63_1ab;6470 | 0 | 0 |
| NC_005831.2;YP_003766.2;HCoV-NL63_1ab;6471 | 0 | 0 |
| NC_005831.2;YP_003766.2;HCoV-NL63_1ab;6472 | 0 | 0 |
| NC_005831.2;YP_003766.2;HCoV-NL63_1ab;6473 | 0 | 0 |
| NC_005831.2;YP_003766.2;HCoV-NL63_1ab;6474 | 0.05 | 0 |
| NC_005831.2;YP_003766.2;HCoV-NL63_1ab;6475 | 0.05 | 0.025 |
| NC_005831.2;YP_003766.2;HCoV-NL63_1ab;6476 | 0.05 | 0 |
| NC_005831.2;YP_003766.2;HCoV-NL63_1ab;6477 | 0.05 | 0 |
| NC_005831.2;YP_003766.2;HCoV-NL63_1ab;6478 | 0.05 | 0 |
| NC_005831.2;YP_003766.2;HCoV-NL63_1ab;6479 | 0.05 | 0 |
| NC_005831.2;YP_003766.2;HCoV-NL63_1ab;6480 | 0.05 | 0 |
| NC_005831.2;YP_003766.2;HCoV-NL63_1ab;6481 | 0.05 | 0 |
| NC_005831.2;YP_003766.2;HCoV-NL63_1ab;6482 | 0 | 0 |
| NC_005831.2;YP_003766.2;HCoV-NL63_1ab;6483 | 0 | 0.025 |
| NC_005831.2;YP_003766.2;HCoV-NL63_1ab;6484 | 0 | 0 |
| NC_005831.2;YP_003766.2;HCoV-NL63_1ab;6485 | 0 | 0 |
| NC_005831.2;YP_003766.2;HCoV-NL63_1ab;6486 | 0 | 0 |
| NC_005831.2;YP_003766.2;HCoV-NL63_1ab;6487 | 0.05 | 0 |
| NC_005831.2;YP_003766.2;HCoV-NL63_1ab;6488 | 0 | 0.025 |
| NC_005831.2;YP_003766.2;HCoV-NL63_1ab;6489 | 0 | 0.025 |
| NC_005831.2;YP_003766.2;HCoV-NL63_1ab;6490 | 0 | 0 |
| NC_005831.2;YP_003766.2;HCoV-NL63_1ab;6491 | 0 | 0.025 |
| NC_005831.2;YP_003766.2;HCoV-NL63_1ab;6492 | 0.05 | 0.05 |
| NC_005831.2;YP_003766.2;HCoV-NL63_1ab;6493 | 0.05 | 0.05 |
| NC_005831.2;YP_003766.2;HCoV-NL63_1ab;6494 | 0 | 0 |
| NC_005831.2;YP_003766.2;HCoV-NL63_1ab;6495 | 0 | 0.025 |
| NC_005831.2;YP_003766.2;HCoV-NL63_1ab;6496 | 0 | 0.05 |
| NC_005831.2;YP_003766.2;HCoV-NL63_1ab;6497 | 0.05 | 0.025 |
| NC_005831.2;YP_003766.2;HCoV-NL63_1ab;6498 | 0 | 0.025 |
| NC_005831.2;YP_003766.2;HCoV-NL63_1ab;6499 | 0 | 0.025 |
| NC_005831.2;YP_003766.2;HCoV-NL63_1ab;6500 | 0 | 0.025 |
| NC_005831.2;YP_003766.2;HCoV-NL63_1ab;6501 | 0.05 | 0.025 |
| NC_005831.2;YP_003766.2;HCoV-NL63_1ab;6502 | 0 | 0 |

| | | |
|---|---|---|
| NC_005831.2;YP_003766.2;HCoV-NL63_1ab;6503 | 0 | 0.025 |
| NC_005831.2;YP_003766.2;HCoV-NL63_1ab;6504 | 0 | 0.05 |
| NC_005831.2;YP_003766.2;HCoV-NL63_1ab;6505 | 0 | 0 |
| NC_005831.2;YP_003766.2;HCoV-NL63_1ab;6506 | 0 | 0 |
| NC_005831.2;YP_003766.2;HCoV-NL63_1ab;6507 | 0 | 0 |
| NC_005831.2;YP_003766.2;HCoV-NL63_1ab;6508 | 0 | 0 |
| NC_005831.2;YP_003766.2;HCoV-NL63_1ab;6509 | 0 | 0 |
| NC_005831.2;YP_003766.2;HCoV-NL63_1ab;6510 | 0 | 0 |
| NC_005831.2;YP_003766.2;HCoV-NL63_1ab;6511 | 0 | 0 |
| NC_005831.2;YP_003766.2;HCoV-NL63_1ab;6512 | 0 | 0 |
| NC_005831.2;YP_003766.2;HCoV-NL63_1ab;6513 | 0 | 0 |
| NC_005831.2;YP_003766.2;HCoV-NL63_1ab;6514 | 0.05 | 0.1 |
| NC_005831.2;YP_003766.2;HCoV-NL63_1ab;6515 | 0 | 0.025 |
| NC_005831.2;YP_003766.2;HCoV-NL63_1ab;6516 | 0.05 | 0.025 |
| NC_005831.2;YP_003766.2;HCoV-NL63_1ab;6517 | 0 | 0.05 |
| NC_005831.2;YP_003766.2;HCoV-NL63_1ab;6518 | 0 | 0.05 |
| NC_005831.2;YP_003766.2;HCoV-NL63_1ab;6519 | 0 | 0.05 |
| NC_005831.2;YP_003766.2;HCoV-NL63_1ab;6520 | 0 | 0.05 |
| NC_005831.2;YP_003766.2;HCoV-NL63_1ab;6521 | 0 | 0.05 |
| NC_005831.2;YP_003766.2;HCoV-NL63_1ab;6522 | 0 | 0.05 |
| NC_005831.2;YP_003766.2;HCoV-NL63_1ab;6523 | 0 | 0.05 |
| NC_005831.2;YP_003766.2;HCoV-NL63_1ab;6524 | 0 | 0.025 |
| NC_005831.2;YP_003766.2;HCoV-NL63_1ab;6525 | 0 | 0.025 |
| NC_005831.2;YP_003766.2;HCoV-NL63_1ab;6526 | 0 | 0.025 |
| NC_005831.2;YP_003766.2;HCoV-NL63_1ab;6527 | 0 | 0.05 |
| NC_005831.2;YP_003766.2;HCoV-NL63_1ab;6528 | 0 | 0 |
| NC_005831.2;YP_003766.2;HCoV-NL63_1ab;6529 | 0 | 0 |
| NC_005831.2;YP_003766.2;HCoV-NL63_1ab;6530 | 0 | 0 |
| NC_005831.2;YP_003766.2;HCoV-NL63_1ab;6531 | 0 | 0 |
| NC_005831.2;YP_003766.2;HCoV-NL63_1ab;6532 | 0 | 0 |
| NC_005831.2;YP_003766.2;HCoV-NL63_1ab;6533 | 0 | 0 |
| NC_005831.2;YP_003766.2;HCoV-NL63_1ab;6534 | 0 | 0 |
| NC_005831.2;YP_003766.2;HCoV-NL63_1ab;6535 | 0 | 0 |
| NC_005831.2;YP_003766.2;HCoV-NL63_1ab;6536 | 0 | 0 |
| NC_005831.2;YP_003766.2;HCoV-NL63_1ab;6537 | 0 | 0 |
| NC_005831.2;YP_003766.2;HCoV-NL63_1ab;6538 | 0 | 0 |
| NC_005831.2;YP_003766.2;HCoV-NL63_1ab;6539 | 0 | 0 |
| NC_005831.2;YP_003766.2;HCoV-NL63_1ab;6540 | 0 | 0 |
| NC_005831.2;YP_003766.2;HCoV-NL63_1ab;6541 | 0 | 0 |
| NC_005831.2;YP_003766.2;HCoV-NL63_1ab;6542 | 0 | 0 |
| NC_005831.2;YP_003766.2;HCoV-NL63_1ab;6543 | 0 | 0 |
| NC_005831.2;YP_003766.2;HCoV-NL63_1ab;6544 | 0 | 0 |
| NC_005831.2;YP_003766.2;HCoV-NL63_1ab;6545 | 0 | 0 |

| | | |
|---|---|---|
| NC_005831.2;YP_003766.2;HCoV-NL63_1ab;6546 | 0 | 0.025 |
| NC_005831.2;YP_003766.2;HCoV-NL63_1ab;6547 | 0.1 | 0.025 |
| NC_005831.2;YP_003766.2;HCoV-NL63_1ab;6548 | 0.15 | 0.025 |
| NC_005831.2;YP_003766.2;HCoV-NL63_1ab;6549 | 0.3 | 0.025 |
| NC_005831.2;YP_003766.2;HCoV-NL63_1ab;6550 | 0.35 | 0.05 |
| NC_005831.2;YP_003766.2;HCoV-NL63_1ab;6551 | 0 | 0 |
| NC_005831.2;YP_003766.2;HCoV-NL63_1ab;6552 | 0 | 0 |
| NC_005831.2;YP_003766.2;HCoV-NL63_1ab;6553 | 0 | 0 |
| NC_005831.2;YP_003766.2;HCoV-NL63_1ab;6554 | 0 | 0 |
| NC_005831.2;YP_003766.2;HCoV-NL63_1ab;6555 | 0 | 0 |
| NC_005831.2;YP_003766.2;HCoV-NL63_1ab;6556 | 0 | 0 |
| NC_005831.2;YP_003766.2;HCoV-NL63_1ab;6557 | 0 | 0 |
| NC_005831.2;YP_003766.2;HCoV-NL63_1ab;6558 | 0 | 0 |
| NC_005831.2;YP_003766.2;HCoV-NL63_1ab;6559 | 0 | 0 |
| NC_005831.2;YP_003766.2;HCoV-NL63_1ab;6560 | 0 | 0 |
| NC_005831.2;YP_003766.2;HCoV-NL63_1ab;6561 | 0 | 0 |
| NC_005831.2;YP_003766.2;HCoV-NL63_1ab;6562 | 0 | 0 |
| NC_005831.2;YP_003766.2;HCoV-NL63_1ab;6563 | 0 | 0 |
| NC_005831.2;YP_003766.2;HCoV-NL63_1ab;6564 | 0.05 | 0.025 |
| NC_005831.2;YP_003766.2;HCoV-NL63_1ab;6565 | 0 | 0 |
| NC_005831.2;YP_003766.2;HCoV-NL63_1ab;6566 | 0 | 0 |
| NC_005831.2;YP_003766.2;HCoV-NL63_1ab;6567 | 0 | 0 |
| NC_005831.2;YP_003766.2;HCoV-NL63_1ab;6568 | 0.05 | 0 |
| NC_005831.2;YP_003766.2;HCoV-NL63_1ab;6569 | 0.05 | 0 |
| NC_005831.2;YP_003766.2;HCoV-NL63_1ab;6570 | 0.05 | 0 |
| NC_005831.2;YP_003766.2;HCoV-NL63_1ab;6571 | 0 | 0 |
| NC_005831.2;YP_003766.2;HCoV-NL63_1ab;6572 | 0 | 0 |
| NC_005831.2;YP_003766.2;HCoV-NL63_1ab;6573 | 0 | 0 |
| NC_005831.2;YP_003766.2;HCoV-NL63_1ab;6574 | 0 | 0 |
| NC_005831.2;YP_003766.2;HCoV-NL63_1ab;6575 | 0 | 0 |
| NC_005831.2;YP_003766.2;HCoV-NL63_1ab;6576 | 0 | 0 |
| NC_005831.2;YP_003766.2;HCoV-NL63_1ab;6577 | 0 | 0 |
| NC_005831.2;YP_003766.2;HCoV-NL63_1ab;6578 | 0 | 0 |
| NC_005831.2;YP_003766.2;HCoV-NL63_1ab;6579 | 0 | 0 |
| NC_005831.2;YP_003766.2;HCoV-NL63_1ab;6580 | 0 | 0 |
| NC_005831.2;YP_003766.2;HCoV-NL63_1ab;6581 | 0.05 | 0 |
| NC_005831.2;YP_003766.2;HCoV-NL63_1ab;6582 | 0 | 0 |
| NC_005831.2;YP_003766.2;HCoV-NL63_1ab;6583 | 0 | 0 |
| NC_005831.2;YP_003766.2;HCoV-NL63_1ab;6584 | 0 | 0 |
| NC_005831.2;YP_003766.2;HCoV-NL63_1ab;6585 | 0 | 0 |
| NC_005831.2;YP_003766.2;HCoV-NL63_1ab;6586 | 0 | 0 |
| NC_005831.2;YP_003766.2;HCoV-NL63_1ab;6587 | 0 | 0 |
| NC_005831.2;YP_003766.2;HCoV-NL63_1ab;6588 | 0 | 0 |

| | | |
|---|---|---|
| NC_005831.2;YP_003766.2;HCoV-NL63_1ab;6589 | 0 | 0 |
| NC_005831.2;YP_003766.2;HCoV-NL63_1ab;6590 | 0 | 0 |
| NC_005831.2;YP_003766.2;HCoV-NL63_1ab;6591 | 0 | 0 |
| NC_005831.2;YP_003766.2;HCoV-NL63_1ab;6592 | 0 | 0 |
| NC_005831.2;YP_003766.2;HCoV-NL63_1ab;6593 | 0 | 0 |
| NC_005831.2;YP_003766.2;HCoV-NL63_1ab;6594 | 0 | 0 |
| NC_005831.2;YP_003766.2;HCoV-NL63_1ab;6595 | 0 | 0 |
| NC_005831.2;YP_003766.2;HCoV-NL63_1ab;6596 | 0 | 0 |
| NC_005831.2;YP_003766.2;HCoV-NL63_1ab;6597 | 0 | 0 |
| NC_005831.2;YP_003766.2;HCoV-NL63_1ab;6598 | 0 | 0 |
| NC_005831.2;YP_003766.2;HCoV-NL63_1ab;6599 | 0 | 0 |
| NC_005831.2;YP_003766.2;HCoV-NL63_1ab;6600 | 0 | 0 |
| NC_005831.2;YP_003766.2;HCoV-NL63_1ab;6601 | 0 | 0 |
| NC_005831.2;YP_003766.2;HCoV-NL63_1ab;6602 | 0 | 0 |
| NC_005831.2;YP_003766.2;HCoV-NL63_1ab;6603 | 0 | 0 |
| NC_005831.2;YP_003766.2;HCoV-NL63_1ab;6604 | 0 | 0 |
| NC_005831.2;YP_003766.2;HCoV-NL63_1ab;6605 | 0 | 0 |
| NC_005831.2;YP_003766.2;HCoV-NL63_1ab;6606 | 0 | 0 |
| NC_005831.2;YP_003766.2;HCoV-NL63_1ab;6607 | 0 | 0 |
| NC_005831.2;YP_003766.2;HCoV-NL63_1ab;6608 | 0 | 0 |
| NC_005831.2;YP_003766.2;HCoV-NL63_1ab;6609 | 0 | 0 |
| NC_005831.2;YP_003766.2;HCoV-NL63_1ab;6610 | 0 | 0 |
| NC_005831.2;YP_003766.2;HCoV-NL63_1ab;6611 | 0 | 0 |
| NC_005831.2;YP_003766.2;HCoV-NL63_1ab;6612 | 0 | 0 |
| NC_005831.2;YP_003766.2;HCoV-NL63_1ab;6613 | 0 | 0 |
| NC_005831.2;YP_003766.2;HCoV-NL63_1ab;6614 | 0 | 0 |
| NC_005831.2;YP_003766.2;HCoV-NL63_1ab;6615 | 0 | 0 |
| NC_005831.2;YP_003766.2;HCoV-NL63_1ab;6616 | 0 | 0 |
| NC_005831.2;YP_003766.2;HCoV-NL63_1ab;6617 | 0 | 0.025 |
| NC_005831.2;YP_003766.2;HCoV-NL63_1ab;6618 | 0 | 0 |
| NC_005831.2;YP_003766.2;HCoV-NL63_1ab;6619 | 0 | 0 |
| NC_005831.2;YP_003766.2;HCoV-NL63_1ab;6620 | 0 | 0 |
| NC_005831.2;YP_003766.2;HCoV-NL63_1ab;6621 | 0 | 0 |
| NC_005831.2;YP_003766.2;HCoV-NL63_1ab;6622 | 0 | 0 |
| NC_005831.2;YP_003766.2;HCoV-NL63_1ab;6623 | 0.05 | 0 |
| NC_005831.2;YP_003766.2;HCoV-NL63_1ab;6624 | 0 | 0 |
| NC_005831.2;YP_003766.2;HCoV-NL63_1ab;6625 | 0 | 0 |
| NC_005831.2;YP_003766.2;HCoV-NL63_1ab;6626 | 0 | 0 |
| NC_005831.2;YP_003766.2;HCoV-NL63_1ab;6627 | 0 | 0 |
| NC_005831.2;YP_003766.2;HCoV-NL63_1ab;6628 | 0 | 0 |
| NC_005831.2;YP_003766.2;HCoV-NL63_1ab;6629 | 0 | 0 |
| NC_005831.2;YP_003766.2;HCoV-NL63_1ab;6630 | 0 | 0 |
| NC_005831.2;YP_003766.2;HCoV-NL63_1ab;6631 | 0 | 0 |

| | | |
|---|---|---|
| NC_005831.2;YP_003766.2;HCoV-NL63_1ab;6632 | 0 | 0 |
| NC_005831.2;YP_003766.2;HCoV-NL63_1ab;6633 | 0 | 0 |
| NC_005831.2;YP_003766.2;HCoV-NL63_1ab;6634 | 0 | 0 |
| NC_005831.2;YP_003766.2;HCoV-NL63_1ab;6635 | 0 | 0 |
| NC_005831.2;YP_003766.2;HCoV-NL63_1ab;6636 | 0 | 0.025 |
| NC_005831.2;YP_003766.2;HCoV-NL63_1ab;6637 | 0 | 0 |
| NC_005831.2;YP_003766.2;HCoV-NL63_1ab;6638 | 0 | 0 |
| NC_005831.2;YP_003766.2;HCoV-NL63_1ab;6639 | 0 | 0.025 |
| NC_005831.2;YP_003766.2;HCoV-NL63_1ab;6640 | 0 | 0.025 |
| NC_005831.2;YP_003766.2;HCoV-NL63_1ab;6641 | 0 | 0 |
| NC_005831.2;YP_003766.2;HCoV-NL63_1ab;6642 | 0 | 0 |
| NC_005831.2;YP_003766.2;HCoV-NL63_1ab;6643 | 0 | 0 |
| NC_005831.2;YP_003766.2;HCoV-NL63_1ab;6644 | 0 | 0 |
| NC_005831.2;YP_003766.2;HCoV-NL63_1ab;6645 | 0 | 0 |
| NC_005831.2;YP_003766.2;HCoV-NL63_1ab;6646 | 0 | 0 |
| NC_005831.2;YP_003766.2;HCoV-NL63_1ab;6647 | 0 | 0 |
| NC_005831.2;YP_003766.2;HCoV-NL63_1ab;6648 | 0 | 0 |
| NC_005831.2;YP_003766.2;HCoV-NL63_1ab;6649 | 0 | 0 |
| NC_005831.2;YP_003766.2;HCoV-NL63_1ab;6650 | 0 | 0 |
| NC_005831.2;YP_003766.2;HCoV-NL63_1ab;6651 | 0 | 0 |
| NC_005831.2;YP_003766.2;HCoV-NL63_1ab;6652 | 0 | 0 |
| NC_005831.2;YP_003766.2;HCoV-NL63_1ab;6653 | 0 | 0 |
| NC_005831.2;YP_003766.2;HCoV-NL63_1ab;6654 | 0 | 0 |
| NC_005831.2;YP_003766.2;HCoV-NL63_1ab;6655 | 0 | 0 |
| NC_005831.2;YP_003766.2;HCoV-NL63_1ab;6656 | 0 | 0 |
| NC_005831.2;YP_003766.2;HCoV-NL63_1ab;6657 | 0 | 0 |
| NC_005831.2;YP_003766.2;HCoV-NL63_1ab;6658 | 0 | 0 |
| NC_005831.2;YP_003766.2;HCoV-NL63_1ab;6659 | 0 | 0 |
| NC_005831.2;YP_003766.2;HCoV-NL63_1ab;6660 | 0 | 0 |
| NC_005831.2;YP_003766.2;HCoV-NL63_1ab;6661 | 0 | 0 |
| NC_005831.2;YP_003766.2;HCoV-NL63_1ab;6662 | 0 | 0 |
| NC_005831.2;YP_003766.2;HCoV-NL63_1ab;6663 | 0 | 0 |
| NC_005831.2;YP_003766.2;HCoV-NL63_1ab;6664 | 0 | 0 |
| NC_005831.2;YP_003766.2;HCoV-NL63_1ab;6665 | 0 | 0 |
| NC_005831.2;YP_003766.2;HCoV-NL63_1ab;6666 | 0 | 0 |
| NC_005831.2;YP_003766.2;HCoV-NL63_1ab;6667 | 0 | 0 |
| NC_005831.2;YP_003766.2;HCoV-NL63_1ab;6668 | 0 | 0 |
| NC_005831.2;YP_003766.2;HCoV-NL63_1ab;6669 | 0 | 0 |
| NC_005831.2;YP_003766.2;HCoV-NL63_1ab;6670 | 0 | 0 |
| NC_005831.2;YP_003766.2;HCoV-NL63_1ab;6671 | 0 | 0 |
| NC_005831.2;YP_003766.2;HCoV-NL63_1ab;6672 | 0 | 0 |
| NC_005831.2;YP_003766.2;HCoV-NL63_1ab;6673 | 0 | 0 |
| NC_005831.2;YP_003766.2;HCoV-NL63_1ab;6674 | 0 | 0 |

| | | |
|---|---|---|
| NC_005831.2;YP_003766.2;HCoV-NL63_1ab;6675 | 0 | 0 |
| NC_005831.2;YP_003766.2;HCoV-NL63_1ab;6676 | 0 | 0 |
| NC_005831.2;YP_003766.2;HCoV-NL63_1ab;6677 | 0 | 0 |
| NC_005831.2;YP_003766.2;HCoV-NL63_1ab;6678 | 0 | 0 |
| NC_005831.2;YP_003766.2;HCoV-NL63_1ab;6679 | 0 | 0 |
| NC_005831.2;YP_003766.2;HCoV-NL63_1ab;6680 | 0 | 0 |
| NC_005831.2;YP_003766.2;HCoV-NL63_1ab;6681 | 0 | 0 |
| NC_005831.2;YP_003766.2;HCoV-NL63_1ab;6682 | 0 | 0 |
| NC_005831.2;YP_003766.2;HCoV-NL63_1ab;6683 | 0 | 0 |
| NC_005831.2;YP_003766.2;HCoV-NL63_1ab;6684 | 0 | 0 |
| NC_005831.2;YP_003766.2;HCoV-NL63_1ab;6685 | 0 | 0 |
| NC_005831.2;YP_003766.2;HCoV-NL63_1ab;6686 | 0 | 0.025 |
| NC_005831.2;YP_003766.2;HCoV-NL63_1ab;6687 | 0 | 0.025 |
| NC_005831.2;YP_003766.2;HCoV-NL63_1ab;6688 | 0 | 0 |
| NC_005831.2;YP_003766.2;HCoV-NL63_1ab;6689 | 0 | 0 |
| NC_005831.2;YP_003766.2;HCoV-NL63_1ab;6690 | 0 | 0 |
| NC_005831.2;YP_003766.2;HCoV-NL63_1ab;6691 | 0 | 0 |
| NC_005831.2;YP_003766.2;HCoV-NL63_1ab;6692 | 0 | 0.025 |
| NC_005831.2;YP_003766.2;HCoV-NL63_1ab;6693 | 0 | 0 |
| NC_005831.2;YP_003766.2;HCoV-NL63_1ab;6694 | 0 | 0 |
| NC_005831.2;YP_003766.2;HCoV-NL63_1ab;6695 | 0 | 0 |
| NC_005831.2;YP_003766.2;HCoV-NL63_1ab;6696 | 0 | 0 |
| NC_005831.2;YP_003766.2;HCoV-NL63_1ab;6697 | 0 | 0 |
| NC_005831.2;YP_003766.2;HCoV-NL63_1ab;6698 | 0 | 0 |
| NC_005831.2;YP_003766.2;HCoV-NL63_1ab;6699 | 0 | 0 |
| NC_005831.2;YP_003766.2;HCoV-NL63_1ab;6700 | 0 | 0 |
| NC_005831.2;YP_003766.2;HCoV-NL63_1ab;6701 | 0 | 0 |
| NC_005831.2;YP_003766.2;HCoV-NL63_1ab;6702 | 0 | 0 |
| NC_005831.2;YP_003766.2;HCoV-NL63_1ab;6703 | 0 | 0 |
| NC_005831.2;YP_003766.2;HCoV-NL63_1ab;6704 | 0 | 0 |
| NC_005831.2;YP_003766.2;HCoV-NL63_1ab;6705 | 0 | 0 |
| NC_005831.2;YP_003766.2;HCoV-NL63_1ab;6706 | 0 | 0 |
| NC_005831.2;YP_003766.2;HCoV-NL63_1ab;6707 | 0 | 0 |
| NC_005831.2;YP_003766.2;HCoV-NL63_1ab;6708 | 0 | 0 |
| NC_005831.2;YP_003766.2;HCoV-NL63_1ab;6709 | 0 | 0 |
| NC_005831.2;YP_003766.2;HCoV-NL63_1ab;6710 | 0 | 0 |
| NC_005831.2;YP_003766.2;HCoV-NL63_1ab;6711 | 0 | 0 |
| NC_005831.2;YP_003766.2;HCoV-NL63_1ab;6712 | 0 | 0 |
| NC_005831.2;YP_003766.2;HCoV-NL63_1ab;6713 | 0 | 0 |
| NC_005831.2;YP_003766.2;HCoV-NL63_1ab;6714 | 0 | 0 |
| NC_005831.2;YP_003766.2;HCoV-NL63_1ab;6715 | 0 | 0 |
| NC_005831.2;YP_003766.2;HCoV-NL63_1ab;6716 | 0 | 0 |
| NC_005831.2;YP_003766.2;HCoV-NL63_1ab;6717 | 0 | 0 |

| | | |
|---|---|---|
| NC_005831.2;YP_003766.2;HCoV-NL63_1ab;6718 | 0 | 0 |
| NC_005831.2;YP_003767.1;HCoV-NL63_spike;1 | 0 | 0 |
| NC_005831.2;YP_003767.1;HCoV-NL63_spike;2 | 0 | 0 |
| NC_005831.2;YP_003767.1;HCoV-NL63_spike;3 | 0 | 0 |
| NC_005831.2;YP_003767.1;HCoV-NL63_spike;4 | 0 | 0 |
| NC_005831.2;YP_003767.1;HCoV-NL63_spike;5 | 0 | 0 |
| NC_005831.2;YP_003767.1;HCoV-NL63_spike;6 | 0 | 0 |
| NC_005831.2;YP_003767.1;HCoV-NL63_spike;7 | 0 | 0 |
| NC_005831.2;YP_003767.1;HCoV-NL63_spike;8 | 0 | 0 |
| NC_005831.2;YP_003767.1;HCoV-NL63_spike;9 | 0 | 0 |
| NC_005831.2;YP_003767.1;HCoV-NL63_spike;10 | 0 | 0 |
| NC_005831.2;YP_003767.1;HCoV-NL63_spike;11 | 0 | 0 |
| NC_005831.2;YP_003767.1;HCoV-NL63_spike;12 | 0 | 0 |
| NC_005831.2;YP_003767.1;HCoV-NL63_spike;13 | 0 | 0 |
| NC_005831.2;YP_003767.1;HCoV-NL63_spike;14 | 0 | 0 |
| NC_005831.2;YP_003767.1;HCoV-NL63_spike;15 | 0 | 0 |
| NC_005831.2;YP_003767.1;HCoV-NL63_spike;16 | 0 | 0.025 |
| NC_005831.2;YP_003767.1;HCoV-NL63_spike;17 | 0 | 0.025 |
| NC_005831.2;YP_003767.1;HCoV-NL63_spike;18 | 0.1 | 0.05 |
| NC_005831.2;YP_003767.1;HCoV-NL63_spike;19 | 0 | 0.075 |
| NC_005831.2;YP_003767.1;HCoV-NL63_spike;20 | 0 | 0.075 |
| NC_005831.2;YP_003767.1;HCoV-NL63_spike;21 | 0 | 0.025 |
| NC_005831.2;YP_003767.1;HCoV-NL63_spike;22 | 0 | 0 |
| NC_005831.2;YP_003767.1;HCoV-NL63_spike;23 | 0 | 0 |
| NC_005831.2;YP_003767.1;HCoV-NL63_spike;24 | 0 | 0.05 |
| NC_005831.2;YP_003767.1;HCoV-NL63_spike;25 | 0 | 0.025 |
| NC_005831.2;YP_003767.1;HCoV-NL63_spike;26 | 0 | 0 |
| NC_005831.2;YP_003767.1;HCoV-NL63_spike;27 | 0 | 0 |
| NC_005831.2;YP_003767.1;HCoV-NL63_spike;28 | 0 | 0 |
| NC_005831.2;YP_003767.1;HCoV-NL63_spike;29 | 0 | 0 |
| NC_005831.2;YP_003767.1;HCoV-NL63_spike;30 | 0 | 0 |
| NC_005831.2;YP_003767.1;HCoV-NL63_spike;31 | 0 | 0.025 |
| NC_005831.2;YP_003767.1;HCoV-NL63_spike;32 | 0 | 0 |
| NC_005831.2;YP_003767.1;HCoV-NL63_spike;33 | 0 | 0.025 |
| NC_005831.2;YP_003767.1;HCoV-NL63_spike;34 | 0 | 0 |
| NC_005831.2;YP_003767.1;HCoV-NL63_spike;35 | 0 | 0 |
| NC_005831.2;YP_003767.1;HCoV-NL63_spike;36 | 0 | 0 |
| NC_005831.2;YP_003767.1;HCoV-NL63_spike;37 | 0 | 0.025 |
| NC_005831.2;YP_003767.1;HCoV-NL63_spike;38 | 0 | 0 |
| NC_005831.2;YP_003767.1;HCoV-NL63_spike;39 | 0 | 0 |
| NC_005831.2;YP_003767.1;HCoV-NL63_spike;40 | 0 | 0 |
| NC_005831.2;YP_003767.1;HCoV-NL63_spike;41 | 0 | 0 |
| NC_005831.2;YP_003767.1;HCoV-NL63_spike;42 | 0 | 0 |

| | | |
|---|---|---|
| NC_005831.2;YP_003767.1;HCoV-NL63_spike;43 | 0 | 0 |
| NC_005831.2;YP_003767.1;HCoV-NL63_spike;44 | 0 | 0 |
| NC_005831.2;YP_003767.1;HCoV-NL63_spike;45 | 0 | 0 |
| NC_005831.2;YP_003767.1;HCoV-NL63_spike;46 | 0 | 0.025 |
| NC_005831.2;YP_003767.1;HCoV-NL63_spike;47 | 0 | 0 |
| NC_005831.2;YP_003767.1;HCoV-NL63_spike;48 | 0 | 0 |
| NC_005831.2;YP_003767.1;HCoV-NL63_spike;49 | 0 | 0 |
| NC_005831.2;YP_003767.1;HCoV-NL63_spike;50 | 0 | 0 |
| NC_005831.2;YP_003767.1;HCoV-NL63_spike;51 | 0 | 0 |
| NC_005831.2;YP_003767.1;HCoV-NL63_spike;52 | 0 | 0 |
| NC_005831.2;YP_003767.1;HCoV-NL63_spike;53 | 0 | 0 |
| NC_005831.2;YP_003767.1;HCoV-NL63_spike;54 | 0 | 0 |
| NC_005831.2;YP_003767.1;HCoV-NL63_spike;55 | 0 | 0 |
| NC_005831.2;YP_003767.1;HCoV-NL63_spike;56 | 0 | 0 |
| NC_005831.2;YP_003767.1;HCoV-NL63_spike;57 | 0 | 0 |
| NC_005831.2;YP_003767.1;HCoV-NL63_spike;58 | 0 | 0 |
| NC_005831.2;YP_003767.1;HCoV-NL63_spike;59 | 0 | 0 |
| NC_005831.2;YP_003767.1;HCoV-NL63_spike;60 | 0 | 0 |
| NC_005831.2;YP_003767.1;HCoV-NL63_spike;61 | 0 | 0 |
| NC_005831.2;YP_003767.1;HCoV-NL63_spike;62 | 0 | 0 |
| NC_005831.2;YP_003767.1;HCoV-NL63_spike;63 | 0 | 0 |
| NC_005831.2;YP_003767.1;HCoV-NL63_spike;64 | 0 | 0 |
| NC_005831.2;YP_003767.1;HCoV-NL63_spike;65 | 0 | 0 |
| NC_005831.2;YP_003767.1;HCoV-NL63_spike;66 | 0 | 0 |
| NC_005831.2;YP_003767.1;HCoV-NL63_spike;67 | 0 | 0 |
| NC_005831.2;YP_003767.1;HCoV-NL63_spike;68 | 0 | 0 |
| NC_005831.2;YP_003767.1;HCoV-NL63_spike;69 | 0 | 0 |
| NC_005831.2;YP_003767.1;HCoV-NL63_spike;70 | 0 | 0 |
| NC_005831.2;YP_003767.1;HCoV-NL63_spike;71 | 0 | 0 |
| NC_005831.2;YP_003767.1;HCoV-NL63_spike;72 | 0 | 0 |
| NC_005831.2;YP_003767.1;HCoV-NL63_spike;73 | 0 | 0 |
| NC_005831.2;YP_003767.1;HCoV-NL63_spike;74 | 0 | 0 |
| NC_005831.2;YP_003767.1;HCoV-NL63_spike;75 | 0 | 0 |
| NC_005831.2;YP_003767.1;HCoV-NL63_spike;76 | 0 | 0 |
| NC_005831.2;YP_003767.1;HCoV-NL63_spike;77 | 0 | 0 |
| NC_005831.2;YP_003767.1;HCoV-NL63_spike;78 | 0 | 0 |
| NC_005831.2;YP_003767.1;HCoV-NL63_spike;79 | 0 | 0 |
| NC_005831.2;YP_003767.1;HCoV-NL63_spike;80 | 0 | 0 |
| NC_005831.2;YP_003767.1;HCoV-NL63_spike;81 | 0 | 0 |
| NC_005831.2;YP_003767.1;HCoV-NL63_spike;82 | 0 | 0 |
| NC_005831.2;YP_003767.1;HCoV-NL63_spike;83 | 0 | 0 |
| NC_005831.2;YP_003767.1;HCoV-NL63_spike;84 | 0 | 0 |
| NC_005831.2;YP_003767.1;HCoV-NL63_spike;85 | 0 | 0.05 |

| | | |
|---|---|---|
| NC_005831.2;YP_003767.1;HCoV-NL63_spike;86 | 0 | 0.05 |
| NC_005831.2;YP_003767.1;HCoV-NL63_spike;87 | 0.1 | 0.1 |
| NC_005831.2;YP_003767.1;HCoV-NL63_spike;88 | 0 | 0.05 |
| NC_005831.2;YP_003767.1;HCoV-NL63_spike;89 | 0 | 0.05 |
| NC_005831.2;YP_003767.1;HCoV-NL63_spike;90 | 0 | 0.025 |
| NC_005831.2;YP_003767.1;HCoV-NL63_spike;91 | 0 | 0.025 |
| NC_005831.2;YP_003767.1;HCoV-NL63_spike;92 | 0 | 0.025 |
| NC_005831.2;YP_003767.1;HCoV-NL63_spike;93 | 0 | 0 |
| NC_005831.2;YP_003767.1;HCoV-NL63_spike;94 | 0 | 0 |
| NC_005831.2;YP_003767.1;HCoV-NL63_spike;95 | 0 | 0 |
| NC_005831.2;YP_003767.1;HCoV-NL63_spike;96 | 0 | 0 |
| NC_005831.2;YP_003767.1;HCoV-NL63_spike;97 | 0 | 0.025 |
| NC_005831.2;YP_003767.1;HCoV-NL63_spike;98 | 0 | 0 |
| NC_005831.2;YP_003767.1;HCoV-NL63_spike;99 | 0 | 0 |
| NC_005831.2;YP_003767.1;HCoV-NL63_spike;100 | 0 | 0 |
| NC_005831.2;YP_003767.1;HCoV-NL63_spike;101 | 0 | 0 |
| NC_005831.2;YP_003767.1;HCoV-NL63_spike;102 | 0 | 0 |
| NC_005831.2;YP_003767.1;HCoV-NL63_spike;103 | 0 | 0 |
| NC_005831.2;YP_003767.1;HCoV-NL63_spike;104 | 0 | 0 |
| NC_005831.2;YP_003767.1;HCoV-NL63_spike;105 | 0 | 0 |
| NC_005831.2;YP_003767.1;HCoV-NL63_spike;106 | 0 | 0 |
| NC_005831.2;YP_003767.1;HCoV-NL63_spike;107 | 0 | 0 |
| NC_005831.2;YP_003767.1;HCoV-NL63_spike;108 | 0 | 0.075 |
| NC_005831.2;YP_003767.1;HCoV-NL63_spike;109 | 0 | 0.025 |
| NC_005831.2;YP_003767.1;HCoV-NL63_spike;110 | 0 | 0 |
| NC_005831.2;YP_003767.1;HCoV-NL63_spike;111 | 0 | 0 |
| NC_005831.2;YP_003767.1;HCoV-NL63_spike;112 | 0 | 0 |
| NC_005831.2;YP_003767.1;HCoV-NL63_spike;113 | 0 | 0 |
| NC_005831.2;YP_003767.1;HCoV-NL63_spike;114 | 0 | 0 |
| NC_005831.2;YP_003767.1;HCoV-NL63_spike;115 | 0 | 0 |
| NC_005831.2;YP_003767.1;HCoV-NL63_spike;116 | 0 | 0 |
| NC_005831.2;YP_003767.1;HCoV-NL63_spike;117 | 0.05 | 0 |
| NC_005831.2;YP_003767.1;HCoV-NL63_spike;118 | 0 | 0.075 |
| NC_005831.2;YP_003767.1;HCoV-NL63_spike;119 | 0 | 0 |
| NC_005831.2;YP_003767.1;HCoV-NL63_spike;120 | 0 | 0 |
| NC_005831.2;YP_003767.1;HCoV-NL63_spike;121 | 0 | 0 |
| NC_005831.2;YP_003767.1;HCoV-NL63_spike;122 | 0 | 0 |
| NC_005831.2;YP_003767.1;HCoV-NL63_spike;123 | 0 | 0 |
| NC_005831.2;YP_003767.1;HCoV-NL63_spike;124 | 0 | 0 |
| NC_005831.2;YP_003767.1;HCoV-NL63_spike;125 | 0 | 0 |
| NC_005831.2;YP_003767.1;HCoV-NL63_spike;126 | 0 | 0 |
| NC_005831.2;YP_003767.1;HCoV-NL63_spike;127 | 0 | 0 |
| NC_005831.2;YP_003767.1;HCoV-NL63_spike;128 | 0 | 0 |

| | | |
|---|---|---|
| NC_005831.2;YP_003767.1;HCoV-NL63_spike;129 | 0 | 0 |
| NC_005831.2;YP_003767.1;HCoV-NL63_spike;130 | 0 | 0 |
| NC_005831.2;YP_003767.1;HCoV-NL63_spike;131 | 0 | 0 |
| NC_005831.2;YP_003767.1;HCoV-NL63_spike;132 | 0 | 0 |
| NC_005831.2;YP_003767.1;HCoV-NL63_spike;133 | 0 | 0 |
| NC_005831.2;YP_003767.1;HCoV-NL63_spike;134 | 0 | 0 |
| NC_005831.2;YP_003767.1;HCoV-NL63_spike;135 | 0 | 0 |
| NC_005831.2;YP_003767.1;HCoV-NL63_spike;136 | 0 | 0 |
| NC_005831.2;YP_003767.1;HCoV-NL63_spike;137 | 0 | 0 |
| NC_005831.2;YP_003767.1;HCoV-NL63_spike;138 | 0 | 0 |
| NC_005831.2;YP_003767.1;HCoV-NL63_spike;139 | 0 | 0 |
| NC_005831.2;YP_003767.1;HCoV-NL63_spike;140 | 0 | 0 |
| NC_005831.2;YP_003767.1;HCoV-NL63_spike;141 | 0 | 0 |
| NC_005831.2;YP_003767.1;HCoV-NL63_spike;142 | 0 | 0 |
| NC_005831.2;YP_003767.1;HCoV-NL63_spike;143 | 0 | 0 |
| NC_005831.2;YP_003767.1;HCoV-NL63_spike;144 | 0 | 0 |
| NC_005831.2;YP_003767.1;HCoV-NL63_spike;145 | 0 | 0 |
| NC_005831.2;YP_003767.1;HCoV-NL63_spike;146 | 0 | 0 |
| NC_005831.2;YP_003767.1;HCoV-NL63_spike;147 | 0 | 0 |
| NC_005831.2;YP_003767.1;HCoV-NL63_spike;148 | 0 | 0 |
| NC_005831.2;YP_003767.1;HCoV-NL63_spike;149 | 0 | 0 |
| NC_005831.2;YP_003767.1;HCoV-NL63_spike;150 | 0.05 | 0 |
| NC_005831.2;YP_003767.1;HCoV-NL63_spike;151 | 0 | 0 |
| NC_005831.2;YP_003767.1;HCoV-NL63_spike;152 | 0 | 0 |
| NC_005831.2;YP_003767.1;HCoV-NL63_spike;153 | 0 | 0 |
| NC_005831.2;YP_003767.1;HCoV-NL63_spike;154 | 0 | 0 |
| NC_005831.2;YP_003767.1;HCoV-NL63_spike;155 | 0 | 0 |
| NC_005831.2;YP_003767.1;HCoV-NL63_spike;156 | 0 | 0 |
| NC_005831.2;YP_003767.1;HCoV-NL63_spike;157 | 0 | 0 |
| NC_005831.2;YP_003767.1;HCoV-NL63_spike;158 | 0 | 0 |
| NC_005831.2;YP_003767.1;HCoV-NL63_spike;159 | 0 | 0 |
| NC_005831.2;YP_003767.1;HCoV-NL63_spike;160 | 0 | 0 |
| NC_005831.2;YP_003767.1;HCoV-NL63_spike;161 | 0 | 0 |
| NC_005831.2;YP_003767.1;HCoV-NL63_spike;162 | 0 | 0 |
| NC_005831.2;YP_003767.1;HCoV-NL63_spike;163 | 0 | 0 |
| NC_005831.2;YP_003767.1;HCoV-NL63_spike;164 | 0 | 0 |
| NC_005831.2;YP_003767.1;HCoV-NL63_spike;165 | 0 | 0 |
| NC_005831.2;YP_003767.1;HCoV-NL63_spike;166 | 0 | 0 |
| NC_005831.2;YP_003767.1;HCoV-NL63_spike;167 | 0 | 0 |
| NC_005831.2;YP_003767.1;HCoV-NL63_spike;168 | 0 | 0 |
| NC_005831.2;YP_003767.1;HCoV-NL63_spike;169 | 0 | 0 |
| NC_005831.2;YP_003767.1;HCoV-NL63_spike;170 | 0 | 0 |
| NC_005831.2;YP_003767.1;HCoV-NL63_spike;171 | 0 | 0 |

| | | |
|---|---|---|
| NC_005831.2;YP_003767.1;HCoV-NL63_spike;172 | 0 | 0 |
| NC_005831.2;YP_003767.1;HCoV-NL63_spike;173 | 0 | 0 |
| NC_005831.2;YP_003767.1;HCoV-NL63_spike;174 | 0 | 0 |
| NC_005831.2;YP_003767.1;HCoV-NL63_spike;175 | 0 | 0 |
| NC_005831.2;YP_003767.1;HCoV-NL63_spike;176 | 0 | 0 |
| NC_005831.2;YP_003767.1;HCoV-NL63_spike;177 | 0 | 0 |
| NC_005831.2;YP_003767.1;HCoV-NL63_spike;178 | 0 | 0 |
| NC_005831.2;YP_003767.1;HCoV-NL63_spike;179 | 0 | 0 |
| NC_005831.2;YP_003767.1;HCoV-NL63_spike;180 | 0 | 0 |
| NC_005831.2;YP_003767.1;HCoV-NL63_spike;181 | 0 | 0 |
| NC_005831.2;YP_003767.1;HCoV-NL63_spike;182 | 0 | 0 |
| NC_005831.2;YP_003767.1;HCoV-NL63_spike;183 | 0 | 0 |
| NC_005831.2;YP_003767.1;HCoV-NL63_spike;184 | 0.05 | 0 |
| NC_005831.2;YP_003767.1;HCoV-NL63_spike;185 | 0 | 0 |
| NC_005831.2;YP_003767.1;HCoV-NL63_spike;186 | 0.05 | 0 |
| NC_005831.2;YP_003767.1;HCoV-NL63_spike;187 | 0 | 0 |
| NC_005831.2;YP_003767.1;HCoV-NL63_spike;188 | 0 | 0 |
| NC_005831.2;YP_003767.1;HCoV-NL63_spike;189 | 0 | 0 |
| NC_005831.2;YP_003767.1;HCoV-NL63_spike;190 | 0 | 0 |
| NC_005831.2;YP_003767.1;HCoV-NL63_spike;191 | 0 | 0 |
| NC_005831.2;YP_003767.1;HCoV-NL63_spike;192 | 0 | 0 |
| NC_005831.2;YP_003767.1;HCoV-NL63_spike;193 | 0 | 0 |
| NC_005831.2;YP_003767.1;HCoV-NL63_spike;194 | 0 | 0 |
| NC_005831.2;YP_003767.1;HCoV-NL63_spike;195 | 0 | 0 |
| NC_005831.2;YP_003767.1;HCoV-NL63_spike;196 | 0 | 0 |
| NC_005831.2;YP_003767.1;HCoV-NL63_spike;197 | 0 | 0 |
| NC_005831.2;YP_003767.1;HCoV-NL63_spike;198 | 0 | 0 |
| NC_005831.2;YP_003767.1;HCoV-NL63_spike;199 | 0 | 0 |
| NC_005831.2;YP_003767.1;HCoV-NL63_spike;200 | 0 | 0 |
| NC_005831.2;YP_003767.1;HCoV-NL63_spike;201 | 0 | 0.05 |
| NC_005831.2;YP_003767.1;HCoV-NL63_spike;202 | 0 | 0.025 |
| NC_005831.2;YP_003767.1;HCoV-NL63_spike;203 | 0 | 0.025 |
| NC_005831.2;YP_003767.1;HCoV-NL63_spike;204 | 0 | 0.025 |
| NC_005831.2;YP_003767.1;HCoV-NL63_spike;205 | 0 | 0 |
| NC_005831.2;YP_003767.1;HCoV-NL63_spike;206 | 0 | 0.05 |
| NC_005831.2;YP_003767.1;HCoV-NL63_spike;207 | 0 | 0.05 |
| NC_005831.2;YP_003767.1;HCoV-NL63_spike;208 | 0 | 0.05 |
| NC_005831.2;YP_003767.1;HCoV-NL63_spike;209 | 0 | 0.05 |
| NC_005831.2;YP_003767.1;HCoV-NL63_spike;210 | 0 | 0.025 |
| NC_005831.2;YP_003767.1;HCoV-NL63_spike;211 | 0 | 0.025 |
| NC_005831.2;YP_003767.1;HCoV-NL63_spike;212 | 0.3 | 0.1 |
| NC_005831.2;YP_003767.1;HCoV-NL63_spike;213 | 0 | 0.05 |
| NC_005831.2;YP_003767.1;HCoV-NL63_spike;214 | 0 | 0 |

| | | |
|---|---|---|
| NC_005831.2;YP_003767.1;HCoV-NL63_spike;215 | 0 | 0 |
| NC_005831.2;YP_003767.1;HCoV-NL63_spike;216 | 0 | 0.025 |
| NC_005831.2;YP_003767.1;HCoV-NL63_spike;217 | 0.05 | 0.025 |
| NC_005831.2;YP_003767.1;HCoV-NL63_spike;218 | 0.05 | 0 |
| NC_005831.2;YP_003767.1;HCoV-NL63_spike;219 | 0.1 | 0 |
| NC_005831.2;YP_003767.1;HCoV-NL63_spike;220 | 0.05 | 0 |
| NC_005831.2;YP_003767.1;HCoV-NL63_spike;221 | 0 | 0.025 |
| NC_005831.2;YP_003767.1;HCoV-NL63_spike;222 | 0 | 0 |
| NC_005831.2;YP_003767.1;HCoV-NL63_spike;223 | 0 | 0 |
| NC_005831.2;YP_003767.1;HCoV-NL63_spike;224 | 0 | 0 |
| NC_005831.2;YP_003767.1;HCoV-NL63_spike;225 | 0 | 0 |
| NC_005831.2;YP_003767.1;HCoV-NL63_spike;226 | 0 | 0 |
| NC_005831.2;YP_003767.1;HCoV-NL63_spike;227 | 0 | 0 |
| NC_005831.2;YP_003767.1;HCoV-NL63_spike;228 | 0 | 0 |
| NC_005831.2;YP_003767.1;HCoV-NL63_spike;229 | 0 | 0 |
| NC_005831.2;YP_003767.1;HCoV-NL63_spike;230 | 0 | 0 |
| NC_005831.2;YP_003767.1;HCoV-NL63_spike;231 | 0 | 0 |
| NC_005831.2;YP_003767.1;HCoV-NL63_spike;232 | 0 | 0 |
| NC_005831.2;YP_003767.1;HCoV-NL63_spike;233 | 0 | 0 |
| NC_005831.2;YP_003767.1;HCoV-NL63_spike;234 | 0 | 0 |
| NC_005831.2;YP_003767.1;HCoV-NL63_spike;235 | 0 | 0 |
| NC_005831.2;YP_003767.1;HCoV-NL63_spike;236 | 0 | 0 |
| NC_005831.2;YP_003767.1;HCoV-NL63_spike;237 | 0 | 0 |
| NC_005831.2;YP_003767.1;HCoV-NL63_spike;238 | 0 | 0 |
| NC_005831.2;YP_003767.1;HCoV-NL63_spike;239 | 0 | 0 |
| NC_005831.2;YP_003767.1;HCoV-NL63_spike;240 | 0 | 0 |
| NC_005831.2;YP_003767.1;HCoV-NL63_spike;241 | 0 | 0 |
| NC_005831.2;YP_003767.1;HCoV-NL63_spike;242 | 0 | 0 |
| NC_005831.2;YP_003767.1;HCoV-NL63_spike;243 | 0 | 0 |
| NC_005831.2;YP_003767.1;HCoV-NL63_spike;244 | 0.1 | 0.025 |
| NC_005831.2;YP_003767.1;HCoV-NL63_spike;245 | 0 | 0.025 |
| NC_005831.2;YP_003767.1;HCoV-NL63_spike;246 | 0 | 0 |
| NC_005831.2;YP_003767.1;HCoV-NL63_spike;247 | 0 | 0 |
| NC_005831.2;YP_003767.1;HCoV-NL63_spike;248 | 0 | 0 |
| NC_005831.2;YP_003767.1;HCoV-NL63_spike;249 | 0 | 0 |
| NC_005831.2;YP_003767.1;HCoV-NL63_spike;250 | 0 | 0 |
| NC_005831.2;YP_003767.1;HCoV-NL63_spike;251 | 0 | 0 |
| NC_005831.2;YP_003767.1;HCoV-NL63_spike;252 | 0 | 0 |
| NC_005831.2;YP_003767.1;HCoV-NL63_spike;253 | 0 | 0 |
| NC_005831.2;YP_003767.1;HCoV-NL63_spike;254 | 0 | 0 |
| NC_005831.2;YP_003767.1;HCoV-NL63_spike;255 | 0 | 0 |
| NC_005831.2;YP_003767.1;HCoV-NL63_spike;256 | 0 | 0 |
| NC_005831.2;YP_003767.1;HCoV-NL63_spike;257 | 0 | 0 |

| | | |
|---|---|---|
| NC_005831.2;YP_003767.1;HCoV-NL63_spike;258 | 0 | 0 |
| NC_005831.2;YP_003767.1;HCoV-NL63_spike;259 | 0 | 0 |
| NC_005831.2;YP_003767.1;HCoV-NL63_spike;260 | 0 | 0 |
| NC_005831.2;YP_003767.1;HCoV-NL63_spike;261 | 0 | 0 |
| NC_005831.2;YP_003767.1;HCoV-NL63_spike;262 | 0 | 0 |
| NC_005831.2;YP_003767.1;HCoV-NL63_spike;263 | 0 | 0 |
| NC_005831.2;YP_003767.1;HCoV-NL63_spike;264 | 0 | 0 |
| NC_005831.2;YP_003767.1;HCoV-NL63_spike;265 | 0 | 0.025 |
| NC_005831.2;YP_003767.1;HCoV-NL63_spike;266 | 0 | 0.025 |
| NC_005831.2;YP_003767.1;HCoV-NL63_spike;267 | 0.05 | 0.025 |
| NC_005831.2;YP_003767.1;HCoV-NL63_spike;268 | 0.05 | 0.025 |
| NC_005831.2;YP_003767.1;HCoV-NL63_spike;269 | 0.05 | 0.025 |
| NC_005831.2;YP_003767.1;HCoV-NL63_spike;270 | 0.05 | 0.025 |
| NC_005831.2;YP_003767.1;HCoV-NL63_spike;271 | 0.05 | 0.025 |
| NC_005831.2;YP_003767.1;HCoV-NL63_spike;272 | 0.05 | 0.025 |
| NC_005831.2;YP_003767.1;HCoV-NL63_spike;273 | 0.05 | 0.025 |
| NC_005831.2;YP_003767.1;HCoV-NL63_spike;274 | 0.05 | 0.025 |
| NC_005831.2;YP_003767.1;HCoV-NL63_spike;275 | 0.05 | 0.025 |
| NC_005831.2;YP_003767.1;HCoV-NL63_spike;276 | 0.05 | 0.025 |
| NC_005831.2;YP_003767.1;HCoV-NL63_spike;277 | 0.1 | 0.025 |
| NC_005831.2;YP_003767.1;HCoV-NL63_spike;278 | 0 | 0 |
| NC_005831.2;YP_003767.1;HCoV-NL63_spike;279 | 0 | 0 |
| NC_005831.2;YP_003767.1;HCoV-NL63_spike;280 | 0 | 0 |
| NC_005831.2;YP_003767.1;HCoV-NL63_spike;281 | 0 | 0 |
| NC_005831.2;YP_003767.1;HCoV-NL63_spike;282 | 0 | 0 |
| NC_005831.2;YP_003767.1;HCoV-NL63_spike;283 | 0 | 0 |
| NC_005831.2;YP_003767.1;HCoV-NL63_spike;284 | 0 | 0 |
| NC_005831.2;YP_003767.1;HCoV-NL63_spike;285 | 0 | 0 |
| NC_005831.2;YP_003767.1;HCoV-NL63_spike;286 | 0 | 0.025 |
| NC_005831.2;YP_003767.1;HCoV-NL63_spike;287 | 0 | 0 |
| NC_005831.2;YP_003767.1;HCoV-NL63_spike;288 | 0 | 0 |
| NC_005831.2;YP_003767.1;HCoV-NL63_spike;289 | 0 | 0 |
| NC_005831.2;YP_003767.1;HCoV-NL63_spike;290 | 0 | 0 |
| NC_005831.2;YP_003767.1;HCoV-NL63_spike;291 | 0.05 | 0.05 |
| NC_005831.2;YP_003767.1;HCoV-NL63_spike;292 | 0 | 0.025 |
| NC_005831.2;YP_003767.1;HCoV-NL63_spike;293 | 0.2 | 0.2 |
| NC_005831.2;YP_003767.1;HCoV-NL63_spike;294 | 0.15 | 0.25 |
| NC_005831.2;YP_003767.1;HCoV-NL63_spike;295 | 0.1 | 0.225 |
| NC_005831.2;YP_003767.1;HCoV-NL63_spike;296 | 0.05 | 0.1 |
| NC_005831.2;YP_003767.1;HCoV-NL63_spike;297 | 0 | 0.075 |
| NC_005831.2;YP_003767.1;HCoV-NL63_spike;298 | 0.1 | 0.25 |
| NC_005831.2;YP_003767.1;HCoV-NL63_spike;299 | 0.1 | 0.225 |
| NC_005831.2;YP_003767.1;HCoV-NL63_spike;300 | 0.05 | 0 |

| | | |
|---|---|---|
| NC_005831.2;YP_003767.1;HCoV-NL63_spike;301 | 0 | 0 |
| NC_005831.2;YP_003767.1;HCoV-NL63_spike;302 | 0 | 0 |
| NC_005831.2;YP_003767.1;HCoV-NL63_spike;303 | 0 | 0 |
| NC_005831.2;YP_003767.1;HCoV-NL63_spike;304 | 0 | 0 |
| NC_005831.2;YP_003767.1;HCoV-NL63_spike;305 | 0 | 0 |
| NC_005831.2;YP_003767.1;HCoV-NL63_spike;306 | 0 | 0 |
| NC_005831.2;YP_003767.1;HCoV-NL63_spike;307 | 0 | 0 |
| NC_005831.2;YP_003767.1;HCoV-NL63_spike;308 | 0 | 0 |
| NC_005831.2;YP_003767.1;HCoV-NL63_spike;309 | 0 | 0 |
| NC_005831.2;YP_003767.1;HCoV-NL63_spike;310 | 0 | 0 |
| NC_005831.2;YP_003767.1;HCoV-NL63_spike;311 | 0 | 0 |
| NC_005831.2;YP_003767.1;HCoV-NL63_spike;312 | 0 | 0 |
| NC_005831.2;YP_003767.1;HCoV-NL63_spike;313 | 0 | 0 |
| NC_005831.2;YP_003767.1;HCoV-NL63_spike;314 | 0 | 0 |
| NC_005831.2;YP_003767.1;HCoV-NL63_spike;315 | 0 | 0 |
| NC_005831.2;YP_003767.1;HCoV-NL63_spike;316 | 0 | 0 |
| NC_005831.2;YP_003767.1;HCoV-NL63_spike;317 | 0 | 0 |
| NC_005831.2;YP_003767.1;HCoV-NL63_spike;318 | 0 | 0 |
| NC_005831.2;YP_003767.1;HCoV-NL63_spike;319 | 0 | 0 |
| NC_005831.2;YP_003767.1;HCoV-NL63_spike;320 | 0 | 0.025 |
| NC_005831.2;YP_003767.1;HCoV-NL63_spike;321 | 0 | 0.025 |
| NC_005831.2;YP_003767.1;HCoV-NL63_spike;322 | 0 | 0.025 |
| NC_005831.2;YP_003767.1;HCoV-NL63_spike;323 | 0 | 0.025 |
| NC_005831.2;YP_003767.1;HCoV-NL63_spike;324 | 0 | 0.025 |
| NC_005831.2;YP_003767.1;HCoV-NL63_spike;325 | 0 | 0.025 |
| NC_005831.2;YP_003767.1;HCoV-NL63_spike;326 | 0 | 0.05 |
| NC_005831.2;YP_003767.1;HCoV-NL63_spike;327 | 0 | 0.025 |
| NC_005831.2;YP_003767.1;HCoV-NL63_spike;328 | 0 | 0.05 |
| NC_005831.2;YP_003767.1;HCoV-NL63_spike;329 | 0.05 | 0.075 |
| NC_005831.2;YP_003767.1;HCoV-NL63_spike;330 | 0.05 | 0.05 |
| NC_005831.2;YP_003767.1;HCoV-NL63_spike;331 | 0.05 | 0.025 |
| NC_005831.2;YP_003767.1;HCoV-NL63_spike;332 | 0.1 | 0.05 |
| NC_005831.2;YP_003767.1;HCoV-NL63_spike;333 | 0.1 | 0.025 |
| NC_005831.2;YP_003767.1;HCoV-NL63_spike;334 | 0.1 | 0.075 |
| NC_005831.2;YP_003767.1;HCoV-NL63_spike;335 | 0 | 0 |
| NC_005831.2;YP_003767.1;HCoV-NL63_spike;336 | 0 | 0 |
| NC_005831.2;YP_003767.1;HCoV-NL63_spike;337 | 0 | 0 |
| NC_005831.2;YP_003767.1;HCoV-NL63_spike;338 | 0 | 0 |
| NC_005831.2;YP_003767.1;HCoV-NL63_spike;339 | 0 | 0 |
| NC_005831.2;YP_003767.1;HCoV-NL63_spike;340 | 0 | 0 |
| NC_005831.2;YP_003767.1;HCoV-NL63_spike;341 | 0 | 0 |
| NC_005831.2;YP_003767.1;HCoV-NL63_spike;342 | 0 | 0 |
| NC_005831.2;YP_003767.1;HCoV-NL63_spike;343 | 0.05 | 0.05 |

| | | |
|---|---|---|
| NC_005831.2;YP_003767.1;HCoV-NL63_spike;344 | 0.05 | 0 |
| NC_005831.2;YP_003767.1;HCoV-NL63_spike;345 | 0.05 | 0.025 |
| NC_005831.2;YP_003767.1;HCoV-NL63_spike;346 | 0.05 | 0 |
| NC_005831.2;YP_003767.1;HCoV-NL63_spike;347 | 0.05 | 0 |
| NC_005831.2;YP_003767.1;HCoV-NL63_spike;348 | 0.05 | 0 |
| NC_005831.2;YP_003767.1;HCoV-NL63_spike;349 | 0.05 | 0 |
| NC_005831.2;YP_003767.1;HCoV-NL63_spike;350 | 0 | 0 |
| NC_005831.2;YP_003767.1;HCoV-NL63_spike;351 | 0 | 0 |
| NC_005831.2;YP_003767.1;HCoV-NL63_spike;352 | 0.05 | 0 |
| NC_005831.2;YP_003767.1;HCoV-NL63_spike;353 | 0.05 | 0 |
| NC_005831.2;YP_003767.1;HCoV-NL63_spike;354 | 0 | 0 |
| NC_005831.2;YP_003767.1;HCoV-NL63_spike;355 | 0 | 0.025 |
| NC_005831.2;YP_003767.1;HCoV-NL63_spike;356 | 0 | 0 |
| NC_005831.2;YP_003767.1;HCoV-NL63_spike;357 | 0 | 0 |
| NC_005831.2;YP_003767.1;HCoV-NL63_spike;358 | 0 | 0 |
| NC_005831.2;YP_003767.1;HCoV-NL63_spike;359 | 0 | 0 |
| NC_005831.2;YP_003767.1;HCoV-NL63_spike;360 | 0 | 0 |
| NC_005831.2;YP_003767.1;HCoV-NL63_spike;361 | 0 | 0 |
| NC_005831.2;YP_003767.1;HCoV-NL63_spike;362 | 0 | 0 |
| NC_005831.2;YP_003767.1;HCoV-NL63_spike;363 | 0 | 0 |
| NC_005831.2;YP_003767.1;HCoV-NL63_spike;364 | 0 | 0 |
| NC_005831.2;YP_003767.1;HCoV-NL63_spike;365 | 0 | 0 |
| NC_005831.2;YP_003767.1;HCoV-NL63_spike;366 | 0 | 0 |
| NC_005831.2;YP_003767.1;HCoV-NL63_spike;367 | 0 | 0 |
| NC_005831.2;YP_003767.1;HCoV-NL63_spike;368 | 0 | 0 |
| NC_005831.2;YP_003767.1;HCoV-NL63_spike;369 | 0 | 0 |
| NC_005831.2;YP_003767.1;HCoV-NL63_spike;370 | 0 | 0 |
| NC_005831.2;YP_003767.1;HCoV-NL63_spike;371 | 0 | 0 |
| NC_005831.2;YP_003767.1;HCoV-NL63_spike;372 | 0 | 0 |
| NC_005831.2;YP_003767.1;HCoV-NL63_spike;373 | 0 | 0 |
| NC_005831.2;YP_003767.1;HCoV-NL63_spike;374 | 0 | 0 |
| NC_005831.2;YP_003767.1;HCoV-NL63_spike;375 | 0 | 0 |
| NC_005831.2;YP_003767.1;HCoV-NL63_spike;376 | 0 | 0 |
| NC_005831.2;YP_003767.1;HCoV-NL63_spike;377 | 0 | 0 |
| NC_005831.2;YP_003767.1;HCoV-NL63_spike;378 | 0 | 0 |
| NC_005831.2;YP_003767.1;HCoV-NL63_spike;379 | 0 | 0 |
| NC_005831.2;YP_003767.1;HCoV-NL63_spike;380 | 0 | 0 |
| NC_005831.2;YP_003767.1;HCoV-NL63_spike;381 | 0 | 0 |
| NC_005831.2;YP_003767.1;HCoV-NL63_spike;382 | 0 | 0 |
| NC_005831.2;YP_003767.1;HCoV-NL63_spike;383 | 0 | 0 |
| NC_005831.2;YP_003767.1;HCoV-NL63_spike;384 | 0 | 0 |
| NC_005831.2;YP_003767.1;HCoV-NL63_spike;385 | 0 | 0 |
| NC_005831.2;YP_003767.1;HCoV-NL63_spike;386 | 0 | 0 |

| | | |
|---|---|---|
| NC_005831.2;YP_003767.1;HCoV-NL63_spike;387 | 0 | 0 |
| NC_005831.2;YP_003767.1;HCoV-NL63_spike;388 | 0 | 0 |
| NC_005831.2;YP_003767.1;HCoV-NL63_spike;389 | 0 | 0 |
| NC_005831.2;YP_003767.1;HCoV-NL63_spike;390 | 0 | 0 |
| NC_005831.2;YP_003767.1;HCoV-NL63_spike;391 | 0 | 0 |
| NC_005831.2;YP_003767.1;HCoV-NL63_spike;392 | 0 | 0 |
| NC_005831.2;YP_003767.1;HCoV-NL63_spike;393 | 0 | 0 |
| NC_005831.2;YP_003767.1;HCoV-NL63_spike;394 | 0 | 0 |
| NC_005831.2;YP_003767.1;HCoV-NL63_spike;395 | 0.05 | 0 |
| NC_005831.2;YP_003767.1;HCoV-NL63_spike;396 | 0.05 | 0 |
| NC_005831.2;YP_003767.1;HCoV-NL63_spike;397 | 0.15 | 0 |
| NC_005831.2;YP_003767.1;HCoV-NL63_spike;398 | 0.1 | 0 |
| NC_005831.2;YP_003767.1;HCoV-NL63_spike;399 | 0.1 | 0 |
| NC_005831.2;YP_003767.1;HCoV-NL63_spike;400 | 0 | 0 |
| NC_005831.2;YP_003767.1;HCoV-NL63_spike;401 | 0 | 0 |
| NC_005831.2;YP_003767.1;HCoV-NL63_spike;402 | 0 | 0 |
| NC_005831.2;YP_003767.1;HCoV-NL63_spike;403 | 0 | 0 |
| NC_005831.2;YP_003767.1;HCoV-NL63_spike;404 | 0 | 0 |
| NC_005831.2;YP_003767.1;HCoV-NL63_spike;405 | 0 | 0 |
| NC_005831.2;YP_003767.1;HCoV-NL63_spike;406 | 0 | 0 |
| NC_005831.2;YP_003767.1;HCoV-NL63_spike;407 | 0 | 0 |
| NC_005831.2;YP_003767.1;HCoV-NL63_spike;408 | 0 | 0 |
| NC_005831.2;YP_003767.1;HCoV-NL63_spike;409 | 0 | 0 |
| NC_005831.2;YP_003767.1;HCoV-NL63_spike;410 | 0 | 0 |
| NC_005831.2;YP_003767.1;HCoV-NL63_spike;411 | 0 | 0 |
| NC_005831.2;YP_003767.1;HCoV-NL63_spike;412 | 0 | 0 |
| NC_005831.2;YP_003767.1;HCoV-NL63_spike;413 | 0 | 0 |
| NC_005831.2;YP_003767.1;HCoV-NL63_spike;414 | 0 | 0 |
| NC_005831.2;YP_003767.1;HCoV-NL63_spike;415 | 0 | 0 |
| NC_005831.2;YP_003767.1;HCoV-NL63_spike;416 | 0.05 | 0 |
| NC_005831.2;YP_003767.1;HCoV-NL63_spike;417 | 0 | 0 |
| NC_005831.2;YP_003767.1;HCoV-NL63_spike;418 | 0 | 0.025 |
| NC_005831.2;YP_003767.1;HCoV-NL63_spike;419 | 0 | 0 |
| NC_005831.2;YP_003767.1;HCoV-NL63_spike;420 | 0 | 0 |
| NC_005831.2;YP_003767.1;HCoV-NL63_spike;421 | 0 | 0 |
| NC_005831.2;YP_003767.1;HCoV-NL63_spike;422 | 0 | 0 |
| NC_005831.2;YP_003767.1;HCoV-NL63_spike;423 | 0 | 0 |
| NC_005831.2;YP_003767.1;HCoV-NL63_spike;424 | 0 | 0 |
| NC_005831.2;YP_003767.1;HCoV-NL63_spike;425 | 0 | 0.025 |
| NC_005831.2;YP_003767.1;HCoV-NL63_spike;426 | 0 | 0 |
| NC_005831.2;YP_003767.1;HCoV-NL63_spike;427 | 0 | 0 |
| NC_005831.2;YP_003767.1;HCoV-NL63_spike;428 | 0 | 0 |
| NC_005831.2;YP_003767.1;HCoV-NL63_spike;429 | 0 | 0 |

| | | |
|---|---|---|
| NC_005831.2;YP_003767.1;HCoV-NL63_spike;430 | 0 | 0.025 |
| NC_005831.2;YP_003767.1;HCoV-NL63_spike;431 | 0 | 0.025 |
| NC_005831.2;YP_003767.1;HCoV-NL63_spike;432 | 0.05 | 0.05 |
| NC_005831.2;YP_003767.1;HCoV-NL63_spike;433 | 0.05 | 0.1 |
| NC_005831.2;YP_003767.1;HCoV-NL63_spike;434 | 0 | 0.05 |
| NC_005831.2;YP_003767.1;HCoV-NL63_spike;435 | 0.05 | 0.05 |
| NC_005831.2;YP_003767.1;HCoV-NL63_spike;436 | 0.1 | 0.075 |
| NC_005831.2;YP_003767.1;HCoV-NL63_spike;437 | 0.05 | 0.075 |
| NC_005831.2;YP_003767.1;HCoV-NL63_spike;438 | 0.1 | 0.075 |
| NC_005831.2;YP_003767.1;HCoV-NL63_spike;439 | 0.05 | 0.075 |
| NC_005831.2;YP_003767.1;HCoV-NL63_spike;440 | 0.05 | 0.05 |
| NC_005831.2;YP_003767.1;HCoV-NL63_spike;441 | 0 | 0 |
| NC_005831.2;YP_003767.1;HCoV-NL63_spike;442 | 0 | 0 |
| NC_005831.2;YP_003767.1;HCoV-NL63_spike;443 | 0 | 0 |
| NC_005831.2;YP_003767.1;HCoV-NL63_spike;444 | 0 | 0 |
| NC_005831.2;YP_003767.1;HCoV-NL63_spike;445 | 0 | 0 |
| NC_005831.2;YP_003767.1;HCoV-NL63_spike;446 | 0 | 0 |
| NC_005831.2;YP_003767.1;HCoV-NL63_spike;447 | 0 | 0 |
| NC_005831.2;YP_003767.1;HCoV-NL63_spike;448 | 0 | 0 |
| NC_005831.2;YP_003767.1;HCoV-NL63_spike;449 | 0 | 0 |
| NC_005831.2;YP_003767.1;HCoV-NL63_spike;450 | 0.05 | 0.075 |
| NC_005831.2;YP_003767.1;HCoV-NL63_spike;451 | 0.15 | 0.175 |
| NC_005831.2;YP_003767.1;HCoV-NL63_spike;452 | 0.25 | 0.2 |
| NC_005831.2;YP_003767.1;HCoV-NL63_spike;453 | 0.4 | 0.375 |
| NC_005831.2;YP_003767.1;HCoV-NL63_spike;454 | 0.3 | 0.3 |
| NC_005831.2;YP_003767.1;HCoV-NL63_spike;455 | 0.4 | 0.275 |
| NC_005831.2;YP_003767.1;HCoV-NL63_spike;456 | 0.4 | 0.275 |
| NC_005831.2;YP_003767.1;HCoV-NL63_spike;457 | 0.35 | 0.275 |
| NC_005831.2;YP_003767.1;HCoV-NL63_spike;458 | 0.35 | 0.325 |
| NC_005831.2;YP_003767.1;HCoV-NL63_spike;459 | 0.15 | 0.125 |
| NC_005831.2;YP_003767.1;HCoV-NL63_spike;460 | 0.15 | 0.15 |
| NC_005831.2;YP_003767.1;HCoV-NL63_spike;461 | 0 | 0.075 |
| NC_005831.2;YP_003767.1;HCoV-NL63_spike;462 | 0 | 0.075 |
| NC_005831.2;YP_003767.1;HCoV-NL63_spike;463 | 0 | 0.1 |
| NC_005831.2;YP_003767.1;HCoV-NL63_spike;464 | 0 | 0.075 |
| NC_005831.2;YP_003767.1;HCoV-NL63_spike;465 | 0 | 0.075 |
| NC_005831.2;YP_003767.1;HCoV-NL63_spike;466 | 0 | 0.05 |
| NC_005831.2;YP_003767.1;HCoV-NL63_spike;467 | 0 | 0.025 |
| NC_005831.2;YP_003767.1;HCoV-NL63_spike;468 | 0 | 0.05 |
| NC_005831.2;YP_003767.1;HCoV-NL63_spike;469 | 0 | 0.05 |
| NC_005831.2;YP_003767.1;HCoV-NL63_spike;470 | 0 | 0 |
| NC_005831.2;YP_003767.1;HCoV-NL63_spike;471 | 0 | 0.025 |
| NC_005831.2;YP_003767.1;HCoV-NL63_spike;472 | 0 | 0 |

| | | |
|---|---|---|
| NC_005831.2;YP_003767.1;HCoV-NL63_spike;473 | 0 | 0 |
| NC_005831.2;YP_003767.1;HCoV-NL63_spike;474 | 0 | 0 |
| NC_005831.2;YP_003767.1;HCoV-NL63_spike;475 | 0 | 0.025 |
| NC_005831.2;YP_003767.1;HCoV-NL63_spike;476 | 0 | 0.05 |
| NC_005831.2;YP_003767.1;HCoV-NL63_spike;477 | 0 | 0 |
| NC_005831.2;YP_003767.1;HCoV-NL63_spike;478 | 0 | 0 |
| NC_005831.2;YP_003767.1;HCoV-NL63_spike;479 | 0 | 0 |
| NC_005831.2;YP_003767.1;HCoV-NL63_spike;480 | 0.05 | 0 |
| NC_005831.2;YP_003767.1;HCoV-NL63_spike;481 | 0 | 0 |
| NC_005831.2;YP_003767.1;HCoV-NL63_spike;482 | 0 | 0 |
| NC_005831.2;YP_003767.1;HCoV-NL63_spike;483 | 0 | 0 |
| NC_005831.2;YP_003767.1;HCoV-NL63_spike;484 | 0 | 0 |
| NC_005831.2;YP_003767.1;HCoV-NL63_spike;485 | 0 | 0 |
| NC_005831.2;YP_003767.1;HCoV-NL63_spike;486 | 0 | 0 |
| NC_005831.2;YP_003767.1;HCoV-NL63_spike;487 | 0 | 0 |
| NC_005831.2;YP_003767.1;HCoV-NL63_spike;488 | 0 | 0 |
| NC_005831.2;YP_003767.1;HCoV-NL63_spike;489 | 0 | 0 |
| NC_005831.2;YP_003767.1;HCoV-NL63_spike;490 | 0 | 0 |
| NC_005831.2;YP_003767.1;HCoV-NL63_spike;491 | 0 | 0 |
| NC_005831.2;YP_003767.1;HCoV-NL63_spike;492 | 0 | 0 |
| NC_005831.2;YP_003767.1;HCoV-NL63_spike;493 | 0 | 0 |
| NC_005831.2;YP_003767.1;HCoV-NL63_spike;494 | 0 | 0 |
| NC_005831.2;YP_003767.1;HCoV-NL63_spike;495 | 0.05 | 0.025 |
| NC_005831.2;YP_003767.1;HCoV-NL63_spike;496 | 0.05 | 0 |
| NC_005831.2;YP_003767.1;HCoV-NL63_spike;497 | 0.05 | 0.05 |
| NC_005831.2;YP_003767.1;HCoV-NL63_spike;498 | 0.05 | 0.075 |
| NC_005831.2;YP_003767.1;HCoV-NL63_spike;499 | 0.1 | 0.1 |
| NC_005831.2;YP_003767.1;HCoV-NL63_spike;500 | 0.1 | 0.125 |
| NC_005831.2;YP_003767.1;HCoV-NL63_spike;501 | 0.05 | 0.075 |
| NC_005831.2;YP_003767.1;HCoV-NL63_spike;502 | 0.05 | 0.05 |
| NC_005831.2;YP_003767.1;HCoV-NL63_spike;503 | 0.05 | 0.05 |
| NC_005831.2;YP_003767.1;HCoV-NL63_spike;504 | 0.05 | 0.05 |
| NC_005831.2;YP_003767.1;HCoV-NL63_spike;505 | 0.05 | 0.05 |
| NC_005831.2;YP_003767.1;HCoV-NL63_spike;506 | 0 | 0.025 |
| NC_005831.2;YP_003767.1;HCoV-NL63_spike;507 | 0 | 0 |
| NC_005831.2;YP_003767.1;HCoV-NL63_spike;508 | 0.05 | 0 |
| NC_005831.2;YP_003767.1;HCoV-NL63_spike;509 | 0 | 0 |
| NC_005831.2;YP_003767.1;HCoV-NL63_spike;510 | 0 | 0 |
| NC_005831.2;YP_003767.1;HCoV-NL63_spike;511 | 0 | 0 |
| NC_005831.2;YP_003767.1;HCoV-NL63_spike;512 | 0 | 0 |
| NC_005831.2;YP_003767.1;HCoV-NL63_spike;513 | 0 | 0 |
| NC_005831.2;YP_003767.1;HCoV-NL63_spike;514 | 0 | 0 |
| NC_005831.2;YP_003767.1;HCoV-NL63_spike;515 | 0 | 0 |

| | | |
|---|---|---|
| NC_005831.2;YP_003767.1;HCoV-NL63_spike;516 | 0 | 0 |
| NC_005831.2;YP_003767.1;HCoV-NL63_spike;517 | 0 | 0 |
| NC_005831.2;YP_003767.1;HCoV-NL63_spike;518 | 0 | 0 |
| NC_005831.2;YP_003767.1;HCoV-NL63_spike;519 | 0 | 0 |
| NC_005831.2;YP_003767.1;HCoV-NL63_spike;520 | 0 | 0 |
| NC_005831.2;YP_003767.1;HCoV-NL63_spike;521 | 0 | 0 |
| NC_005831.2;YP_003767.1;HCoV-NL63_spike;522 | 0 | 0 |
| NC_005831.2;YP_003767.1;HCoV-NL63_spike;523 | 0 | 0 |
| NC_005831.2;YP_003767.1;HCoV-NL63_spike;524 | 0 | 0 |
| NC_005831.2;YP_003767.1;HCoV-NL63_spike;525 | 0 | 0.05 |
| NC_005831.2;YP_003767.1;HCoV-NL63_spike;526 | 0.15 | 0.05 |
| NC_005831.2;YP_003767.1;HCoV-NL63_spike;527 | 0.1 | 0.05 |
| NC_005831.2;YP_003767.1;HCoV-NL63_spike;528 | 0.1 | 0.05 |
| NC_005831.2;YP_003767.1;HCoV-NL63_spike;529 | 0 | 0.05 |
| NC_005831.2;YP_003767.1;HCoV-NL63_spike;530 | 0 | 0.025 |
| NC_005831.2;YP_003767.1;HCoV-NL63_spike;531 | 0 | 0 |
| NC_005831.2;YP_003767.1;HCoV-NL63_spike;532 | 0 | 0.025 |
| NC_005831.2;YP_003767.1;HCoV-NL63_spike;533 | 0 | 0 |
| NC_005831.2;YP_003767.1;HCoV-NL63_spike;534 | 0 | 0 |
| NC_005831.2;YP_003767.1;HCoV-NL63_spike;535 | 0 | 0 |
| NC_005831.2;YP_003767.1;HCoV-NL63_spike;536 | 0 | 0 |
| NC_005831.2;YP_003767.1;HCoV-NL63_spike;537 | 0 | 0.025 |
| NC_005831.2;YP_003767.1;HCoV-NL63_spike;538 | 0.05 | 0 |
| NC_005831.2;YP_003767.1;HCoV-NL63_spike;539 | 0 | 0 |
| NC_005831.2;YP_003767.1;HCoV-NL63_spike;540 | 0 | 0 |
| NC_005831.2;YP_003767.1;HCoV-NL63_spike;541 | 0 | 0 |
| NC_005831.2;YP_003767.1;HCoV-NL63_spike;542 | 0 | 0 |
| NC_005831.2;YP_003767.1;HCoV-NL63_spike;543 | 0 | 0 |
| NC_005831.2;YP_003767.1;HCoV-NL63_spike;544 | 0.05 | 0 |
| NC_005831.2;YP_003767.1;HCoV-NL63_spike;545 | 0.05 | 0 |
| NC_005831.2;YP_003767.1;HCoV-NL63_spike;546 | 0.05 | 0 |
| NC_005831.2;YP_003767.1;HCoV-NL63_spike;547 | 0 | 0 |
| NC_005831.2;YP_003767.1;HCoV-NL63_spike;548 | 0 | 0.025 |
| NC_005831.2;YP_003767.1;HCoV-NL63_spike;549 | 0 | 0 |
| NC_005831.2;YP_003767.1;HCoV-NL63_spike;550 | 0 | 0 |
| NC_005831.2;YP_003767.1;HCoV-NL63_spike;551 | 0 | 0 |
| NC_005831.2;YP_003767.1;HCoV-NL63_spike;552 | 0 | 0 |
| NC_005831.2;YP_003767.1;HCoV-NL63_spike;553 | 0 | 0 |
| NC_005831.2;YP_003767.1;HCoV-NL63_spike;554 | 0 | 0 |
| NC_005831.2;YP_003767.1;HCoV-NL63_spike;555 | 0 | 0 |
| NC_005831.2;YP_003767.1;HCoV-NL63_spike;556 | 0 | 0 |
| NC_005831.2;YP_003767.1;HCoV-NL63_spike;557 | 0 | 0 |
| NC_005831.2;YP_003767.1;HCoV-NL63_spike;558 | 0 | 0 |

| | | |
|---|---|---|
| NC_005831.2;YP_003767.1;HCoV-NL63_spike;559 | 0 | 0 |
| NC_005831.2;YP_003767.1;HCoV-NL63_spike;560 | 0 | 0 |
| NC_005831.2;YP_003767.1;HCoV-NL63_spike;561 | 0 | 0 |
| NC_005831.2;YP_003767.1;HCoV-NL63_spike;562 | 0 | 0 |
| NC_005831.2;YP_003767.1;HCoV-NL63_spike;563 | 0 | 0 |
| NC_005831.2;YP_003767.1;HCoV-NL63_spike;564 | 0 | 0 |
| NC_005831.2;YP_003767.1;HCoV-NL63_spike;565 | 0 | 0 |
| NC_005831.2;YP_003767.1;HCoV-NL63_spike;566 | 0 | 0 |
| NC_005831.2;YP_003767.1;HCoV-NL63_spike;567 | 0 | 0 |
| NC_005831.2;YP_003767.1;HCoV-NL63_spike;568 | 0 | 0 |
| NC_005831.2;YP_003767.1;HCoV-NL63_spike;569 | 0 | 0 |
| NC_005831.2;YP_003767.1;HCoV-NL63_spike;570 | 0 | 0 |
| NC_005831.2;YP_003767.1;HCoV-NL63_spike;571 | 0 | 0 |
| NC_005831.2;YP_003767.1;HCoV-NL63_spike;572 | 0 | 0 |
| NC_005831.2;YP_003767.1;HCoV-NL63_spike;573 | 0 | 0 |
| NC_005831.2;YP_003767.1;HCoV-NL63_spike;574 | 0 | 0 |
| NC_005831.2;YP_003767.1;HCoV-NL63_spike;575 | 0 | 0 |
| NC_005831.2;YP_003767.1;HCoV-NL63_spike;576 | 0 | 0 |
| NC_005831.2;YP_003767.1;HCoV-NL63_spike;577 | 0 | 0 |
| NC_005831.2;YP_003767.1;HCoV-NL63_spike;578 | 0 | 0 |
| NC_005831.2;YP_003767.1;HCoV-NL63_spike;579 | 0 | 0 |
| NC_005831.2;YP_003767.1;HCoV-NL63_spike;580 | 0 | 0 |
| NC_005831.2;YP_003767.1;HCoV-NL63_spike;581 | 0 | 0 |
| NC_005831.2;YP_003767.1;HCoV-NL63_spike;582 | 0 | 0 |
| NC_005831.2;YP_003767.1;HCoV-NL63_spike;583 | 0 | 0 |
| NC_005831.2;YP_003767.1;HCoV-NL63_spike;584 | 0 | 0 |
| NC_005831.2;YP_003767.1;HCoV-NL63_spike;585 | 0 | 0 |
| NC_005831.2;YP_003767.1;HCoV-NL63_spike;586 | 0 | 0 |
| NC_005831.2;YP_003767.1;HCoV-NL63_spike;587 | 0 | 0 |
| NC_005831.2;YP_003767.1;HCoV-NL63_spike;588 | 0 | 0 |
| NC_005831.2;YP_003767.1;HCoV-NL63_spike;589 | 0 | 0 |
| NC_005831.2;YP_003767.1;HCoV-NL63_spike;590 | 0 | 0 |
| NC_005831.2;YP_003767.1;HCoV-NL63_spike;591 | 0 | 0 |
| NC_005831.2;YP_003767.1;HCoV-NL63_spike;592 | 0 | 0 |
| NC_005831.2;YP_003767.1;HCoV-NL63_spike;593 | 0 | 0 |
| NC_005831.2;YP_003767.1;HCoV-NL63_spike;594 | 0 | 0 |
| NC_005831.2;YP_003767.1;HCoV-NL63_spike;595 | 0 | 0.05 |
| NC_005831.2;YP_003767.1;HCoV-NL63_spike;596 | 0 | 0 |
| NC_005831.2;YP_003767.1;HCoV-NL63_spike;597 | 0 | 0 |
| NC_005831.2;YP_003767.1;HCoV-NL63_spike;598 | 0 | 0 |
| NC_005831.2;YP_003767.1;HCoV-NL63_spike;599 | 0.2 | 0.05 |
| NC_005831.2;YP_003767.1;HCoV-NL63_spike;600 | 0.1 | 0.075 |
| NC_005831.2;YP_003767.1;HCoV-NL63_spike;601 | 0 | 0 |

| | | |
|---|---|---|
| NC_005831.2;YP_003767.1;HCoV-NL63_spike;602 | 0 | 0 |
| NC_005831.2;YP_003767.1;HCoV-NL63_spike;603 | 0 | 0 |
| NC_005831.2;YP_003767.1;HCoV-NL63_spike;604 | 0 | 0 |
| NC_005831.2;YP_003767.1;HCoV-NL63_spike;605 | 0 | 0 |
| NC_005831.2;YP_003767.1;HCoV-NL63_spike;606 | 0 | 0 |
| NC_005831.2;YP_003767.1;HCoV-NL63_spike;607 | 0 | 0 |
| NC_005831.2;YP_003767.1;HCoV-NL63_spike;608 | 0 | 0 |
| NC_005831.2;YP_003767.1;HCoV-NL63_spike;609 | 0 | 0 |
| NC_005831.2;YP_003767.1;HCoV-NL63_spike;610 | 0 | 0 |
| NC_005831.2;YP_003767.1;HCoV-NL63_spike;611 | 0 | 0 |
| NC_005831.2;YP_003767.1;HCoV-NL63_spike;612 | 0 | 0 |
| NC_005831.2;YP_003767.1;HCoV-NL63_spike;613 | 0.05 | 0 |
| NC_005831.2;YP_003767.1;HCoV-NL63_spike;614 | 0.05 | 0 |
| NC_005831.2;YP_003767.1;HCoV-NL63_spike;615 | 0.2 | 0.05 |
| NC_005831.2;YP_003767.1;HCoV-NL63_spike;616 | 0.05 | 0 |
| NC_005831.2;YP_003767.1;HCoV-NL63_spike;617 | 0 | 0 |
| NC_005831.2;YP_003767.1;HCoV-NL63_spike;618 | 0.05 | 0 |
| NC_005831.2;YP_003767.1;HCoV-NL63_spike;619 | 0.1 | 0.075 |
| NC_005831.2;YP_003767.1;HCoV-NL63_spike;620 | 0.1 | 0.05 |
| NC_005831.2;YP_003767.1;HCoV-NL63_spike;621 | 0.1 | 0.025 |
| NC_005831.2;YP_003767.1;HCoV-NL63_spike;622 | 0.1 | 0.075 |
| NC_005831.2;YP_003767.1;HCoV-NL63_spike;623 | 0.15 | 0.1 |
| NC_005831.2;YP_003767.1;HCoV-NL63_spike;624 | 0.2 | 0.125 |
| NC_005831.2;YP_003767.1;HCoV-NL63_spike;625 | 0.1 | 0.1 |
| NC_005831.2;YP_003767.1;HCoV-NL63_spike;626 | 0.05 | 0.05 |
| NC_005831.2;YP_003767.1;HCoV-NL63_spike;627 | 0.05 | 0.05 |
| NC_005831.2;YP_003767.1;HCoV-NL63_spike;628 | 0.1 | 0.075 |
| NC_005831.2;YP_003767.1;HCoV-NL63_spike;629 | 0.1 | 0.075 |
| NC_005831.2;YP_003767.1;HCoV-NL63_spike;630 | 0.1 | 0 |
| NC_005831.2;YP_003767.1;HCoV-NL63_spike;631 | 0.1 | 0.025 |
| NC_005831.2;YP_003767.1;HCoV-NL63_spike;632 | 0.1 | 0 |
| NC_005831.2;YP_003767.1;HCoV-NL63_spike;633 | 0.1 | 0.025 |
| NC_005831.2;YP_003767.1;HCoV-NL63_spike;634 | 0.1 | 0.025 |
| NC_005831.2;YP_003767.1;HCoV-NL63_spike;635 | 0.1 | 0.05 |
| NC_005831.2;YP_003767.1;HCoV-NL63_spike;636 | 0.15 | 0.05 |
| NC_005831.2;YP_003767.1;HCoV-NL63_spike;637 | 0.1 | 0.075 |
| NC_005831.2;YP_003767.1;HCoV-NL63_spike;638 | 0.1 | 0.1 |
| NC_005831.2;YP_003767.1;HCoV-NL63_spike;639 | 0.05 | 0.05 |
| NC_005831.2;YP_003767.1;HCoV-NL63_spike;640 | 0.1 | 0.025 |
| NC_005831.2;YP_003767.1;HCoV-NL63_spike;641 | 0.05 | 0.025 |
| NC_005831.2;YP_003767.1;HCoV-NL63_spike;642 | 0.05 | 0 |
| NC_005831.2;YP_003767.1;HCoV-NL63_spike;643 | 0 | 0 |
| NC_005831.2;YP_003767.1;HCoV-NL63_spike;644 | 0.05 | 0 |

| | | |
|---|---|---|
| NC_005831.2;YP_003767.1;HCoV-NL63_spike;645 | 0 | 0.025 |
| NC_005831.2;YP_003767.1;HCoV-NL63_spike;646 | 0 | 0 |
| NC_005831.2;YP_003767.1;HCoV-NL63_spike;647 | 0 | 0 |
| NC_005831.2;YP_003767.1;HCoV-NL63_spike;648 | 0 | 0.025 |
| NC_005831.2;YP_003767.1;HCoV-NL63_spike;649 | 0 | 0.025 |
| NC_005831.2;YP_003767.1;HCoV-NL63_spike;650 | 0 | 0.05 |
| NC_005831.2;YP_003767.1;HCoV-NL63_spike;651 | 0.2 | 0.225 |
| NC_005831.2;YP_003767.1;HCoV-NL63_spike;652 | 0.05 | 0.05 |
| NC_005831.2;YP_003767.1;HCoV-NL63_spike;653 | 0.05 | 0.05 |
| NC_005831.2;YP_003767.1;HCoV-NL63_spike;654 | 0.05 | 0.05 |
| NC_005831.2;YP_003767.1;HCoV-NL63_spike;655 | 0 | 0.05 |
| NC_005831.2;YP_003767.1;HCoV-NL63_spike;656 | 0.15 | 0.05 |
| NC_005831.2;YP_003767.1;HCoV-NL63_spike;657 | 0 | 0.05 |
| NC_005831.2;YP_003767.1;HCoV-NL63_spike;658 | 0 | 0.075 |
| NC_005831.2;YP_003767.1;HCoV-NL63_spike;659 | 0 | 0 |
| NC_005831.2;YP_003767.1;HCoV-NL63_spike;660 | 0 | 0 |
| NC_005831.2;YP_003767.1;HCoV-NL63_spike;661 | 0 | 0 |
| NC_005831.2;YP_003767.1;HCoV-NL63_spike;662 | 0 | 0 |
| NC_005831.2;YP_003767.1;HCoV-NL63_spike;663 | 0 | 0 |
| NC_005831.2;YP_003767.1;HCoV-NL63_spike;664 | 0 | 0 |
| NC_005831.2;YP_003767.1;HCoV-NL63_spike;665 | 0 | 0 |
| NC_005831.2;YP_003767.1;HCoV-NL63_spike;666 | 0 | 0 |
| NC_005831.2;YP_003767.1;HCoV-NL63_spike;667 | 0 | 0 |
| NC_005831.2;YP_003767.1;HCoV-NL63_spike;668 | 0 | 0 |
| NC_005831.2;YP_003767.1;HCoV-NL63_spike;669 | 0 | 0 |
| NC_005831.2;YP_003767.1;HCoV-NL63_spike;670 | 0 | 0.025 |
| NC_005831.2;YP_003767.1;HCoV-NL63_spike;671 | 0 | 0 |
| NC_005831.2;YP_003767.1;HCoV-NL63_spike;672 | 0 | 0 |
| NC_005831.2;YP_003767.1;HCoV-NL63_spike;673 | 0 | 0 |
| NC_005831.2;YP_003767.1;HCoV-NL63_spike;674 | 0 | 0 |
| NC_005831.2;YP_003767.1;HCoV-NL63_spike;675 | 0 | 0.025 |
| NC_005831.2;YP_003767.1;HCoV-NL63_spike;676 | 0 | 0.025 |
| NC_005831.2;YP_003767.1;HCoV-NL63_spike;677 | 0 | 0 |
| NC_005831.2;YP_003767.1;HCoV-NL63_spike;678 | 0.05 | 0.05 |
| NC_005831.2;YP_003767.1;HCoV-NL63_spike;679 | 0.05 | 0.05 |
| NC_005831.2;YP_003767.1;HCoV-NL63_spike;680 | 0.1 | 0.1 |
| NC_005831.2;YP_003767.1;HCoV-NL63_spike;681 | 0.1 | 0.125 |
| NC_005831.2;YP_003767.1;HCoV-NL63_spike;682 | 0.15 | 0.1 |
| NC_005831.2;YP_003767.1;HCoV-NL63_spike;683 | 0.05 | 0.025 |
| NC_005831.2;YP_003767.1;HCoV-NL63_spike;684 | 0.05 | 0.05 |
| NC_005831.2;YP_003767.1;HCoV-NL63_spike;685 | 0.1 | 0.05 |
| NC_005831.2;YP_003767.1;HCoV-NL63_spike;686 | 0.05 | 0.025 |
| NC_005831.2;YP_003767.1;HCoV-NL63_spike;687 | 0.1 | 0.05 |

| | | |
|---|---|---|
| NC_005831.2;YP_003767.1;HCoV-NL63_spike;688 | 0.1 | 0.05 |
| NC_005831.2;YP_003767.1;HCoV-NL63_spike;689 | 0.05 | 0.05 |
| NC_005831.2;YP_003767.1;HCoV-NL63_spike;690 | 0 | 0.025 |
| NC_005831.2;YP_003767.1;HCoV-NL63_spike;691 | 0 | 0.05 |
| NC_005831.2;YP_003767.1;HCoV-NL63_spike;692 | 0 | 0.05 |
| NC_005831.2;YP_003767.1;HCoV-NL63_spike;693 | 0.05 | 0.1 |
| NC_005831.2;YP_003767.1;HCoV-NL63_spike;694 | 0 | 0.075 |
| NC_005831.2;YP_003767.1;HCoV-NL63_spike;695 | 0 | 0.05 |
| NC_005831.2;YP_003767.1;HCoV-NL63_spike;696 | 0 | 0.025 |
| NC_005831.2;YP_003767.1;HCoV-NL63_spike;697 | 0 | 0.025 |
| NC_005831.2;YP_003767.1;HCoV-NL63_spike;698 | 0.05 | 0.025 |
| NC_005831.2;YP_003767.1;HCoV-NL63_spike;699 | 0 | 0.025 |
| NC_005831.2;YP_003767.1;HCoV-NL63_spike;700 | 0.15 | 0.075 |
| NC_005831.2;YP_003767.1;HCoV-NL63_spike;701 | 0 | 0 |
| NC_005831.2;YP_003767.1;HCoV-NL63_spike;702 | 0 | 0 |
| NC_005831.2;YP_003767.1;HCoV-NL63_spike;703 | 0 | 0 |
| NC_005831.2;YP_003767.1;HCoV-NL63_spike;704 | 0 | 0 |
| NC_005831.2;YP_003767.1;HCoV-NL63_spike;705 | 0 | 0 |
| NC_005831.2;YP_003767.1;HCoV-NL63_spike;706 | 0.05 | 0 |
| NC_005831.2;YP_003767.1;HCoV-NL63_spike;707 | 0 | 0 |
| NC_005831.2;YP_003767.1;HCoV-NL63_spike;708 | 0 | 0.025 |
| NC_005831.2;YP_003767.1;HCoV-NL63_spike;709 | 0 | 0.025 |
| NC_005831.2;YP_003767.1;HCoV-NL63_spike;710 | 0 | 0.025 |
| NC_005831.2;YP_003767.1;HCoV-NL63_spike;711 | 0 | 0.025 |
| NC_005831.2;YP_003767.1;HCoV-NL63_spike;712 | 0 | 0 |
| NC_005831.2;YP_003767.1;HCoV-NL63_spike;713 | 0 | 0 |
| NC_005831.2;YP_003767.1;HCoV-NL63_spike;714 | 0 | 0 |
| NC_005831.2;YP_003767.1;HCoV-NL63_spike;715 | 0 | 0 |
| NC_005831.2;YP_003767.1;HCoV-NL63_spike;716 | 0 | 0 |
| NC_005831.2;YP_003767.1;HCoV-NL63_spike;717 | 0 | 0.025 |
| NC_005831.2;YP_003767.1;HCoV-NL63_spike;718 | 0 | 0 |
| NC_005831.2;YP_003767.1;HCoV-NL63_spike;719 | 0 | 0 |
| NC_005831.2;YP_003767.1;HCoV-NL63_spike;720 | 0 | 0 |
| NC_005831.2;YP_003767.1;HCoV-NL63_spike;721 | 0 | 0 |
| NC_005831.2;YP_003767.1;HCoV-NL63_spike;722 | 0 | 0 |
| NC_005831.2;YP_003767.1;HCoV-NL63_spike;723 | 0 | 0 |
| NC_005831.2;YP_003767.1;HCoV-NL63_spike;724 | 0 | 0 |
| NC_005831.2;YP_003767.1;HCoV-NL63_spike;725 | 0 | 0 |
| NC_005831.2;YP_003767.1;HCoV-NL63_spike;726 | 0 | 0 |
| NC_005831.2;YP_003767.1;HCoV-NL63_spike;727 | 0 | 0 |
| NC_005831.2;YP_003767.1;HCoV-NL63_spike;728 | 0 | 0 |
| NC_005831.2;YP_003767.1;HCoV-NL63_spike;729 | 0 | 0 |
| NC_005831.2;YP_003767.1;HCoV-NL63_spike;730 | 0 | 0 |

| | | |
|---|---|---|
| NC_005831.2;YP_003767.1;HCoV-NL63_spike;731 | 0 | 0 |
| NC_005831.2;YP_003767.1;HCoV-NL63_spike;732 | 0 | 0.025 |
| NC_005831.2;YP_003767.1;HCoV-NL63_spike;733 | 0 | 0 |
| NC_005831.2;YP_003767.1;HCoV-NL63_spike;734 | 0 | 0 |
| NC_005831.2;YP_003767.1;HCoV-NL63_spike;735 | 0 | 0.025 |
| NC_005831.2;YP_003767.1;HCoV-NL63_spike;736 | 0 | 0 |
| NC_005831.2;YP_003767.1;HCoV-NL63_spike;737 | 0 | 0 |
| NC_005831.2;YP_003767.1;HCoV-NL63_spike;738 | 0.25 | 0.15 |
| NC_005831.2;YP_003767.1;HCoV-NL63_spike;739 | 0.25 | 0.175 |
| NC_005831.2;YP_003767.1;HCoV-NL63_spike;740 | 0.25 | 0.2 |
| NC_005831.2;YP_003767.1;HCoV-NL63_spike;741 | 0.3 | 0.3 |
| NC_005831.2;YP_003767.1;HCoV-NL63_spike;742 | 0.35 | 0.2 |
| NC_005831.2;YP_003767.1;HCoV-NL63_spike;743 | 0.4 | 0.25 |
| NC_005831.2;YP_003767.1;HCoV-NL63_spike;744 | 0.25 | 0.15 |
| NC_005831.2;YP_003767.1;HCoV-NL63_spike;745 | 0.2 | 0.125 |
| NC_005831.2;YP_003767.1;HCoV-NL63_spike;746 | 0.15 | 0.125 |
| NC_005831.2;YP_003767.1;HCoV-NL63_spike;747 | 0.05 | 0.125 |
| NC_005831.2;YP_003767.1;HCoV-NL63_spike;748 | 0 | 0.1 |
| NC_005831.2;YP_003767.1;HCoV-NL63_spike;749 | 0 | 0.05 |
| NC_005831.2;YP_003767.1;HCoV-NL63_spike;750 | 0 | 0 |
| NC_005831.2;YP_003767.1;HCoV-NL63_spike;751 | 0 | 0 |
| NC_005831.2;YP_003767.1;HCoV-NL63_spike;752 | 0 | 0 |
| NC_005831.2;YP_003767.1;HCoV-NL63_spike;753 | 0 | 0 |
| NC_005831.2;YP_003767.1;HCoV-NL63_spike;754 | 0 | 0 |
| NC_005831.2;YP_003767.1;HCoV-NL63_spike;755 | 0 | 0 |
| NC_005831.2;YP_003767.1;HCoV-NL63_spike;756 | 0 | 0.025 |
| NC_005831.2;YP_003767.1;HCoV-NL63_spike;757 | 0.05 | 0.05 |
| NC_005831.2;YP_003767.1;HCoV-NL63_spike;758 | 0 | 0 |
| NC_005831.2;YP_003767.1;HCoV-NL63_spike;759 | 0.05 | 0 |
| NC_005831.2;YP_003767.1;HCoV-NL63_spike;760 | 0 | 0 |
| NC_005831.2;YP_003767.1;HCoV-NL63_spike;761 | 0 | 0 |
| NC_005831.2;YP_003767.1;HCoV-NL63_spike;762 | 0 | 0 |
| NC_005831.2;YP_003767.1;HCoV-NL63_spike;763 | 0 | 0 |
| NC_005831.2;YP_003767.1;HCoV-NL63_spike;764 | 0 | 0 |
| NC_005831.2;YP_003767.1;HCoV-NL63_spike;765 | 0 | 0 |
| NC_005831.2;YP_003767.1;HCoV-NL63_spike;766 | 0 | 0 |
| NC_005831.2;YP_003767.1;HCoV-NL63_spike;767 | 0 | 0 |
| NC_005831.2;YP_003767.1;HCoV-NL63_spike;768 | 0 | 0 |
| NC_005831.2;YP_003767.1;HCoV-NL63_spike;769 | 0 | 0 |
| NC_005831.2;YP_003767.1;HCoV-NL63_spike;770 | 0 | 0 |
| NC_005831.2;YP_003767.1;HCoV-NL63_spike;771 | 0 | 0 |
| NC_005831.2;YP_003767.1;HCoV-NL63_spike;772 | 0 | 0 |
| NC_005831.2;YP_003767.1;HCoV-NL63_spike;773 | 0 | 0 |

| | | |
|---|---|---|
| NC_005831.2;YP_003767.1;HCoV-NL63_spike;774 | 0 | 0 |
| NC_005831.2;YP_003767.1;HCoV-NL63_spike;775 | 0 | 0 |
| NC_005831.2;YP_003767.1;HCoV-NL63_spike;776 | 0 | 0 |
| NC_005831.2;YP_003767.1;HCoV-NL63_spike;777 | 0 | 0 |
| NC_005831.2;YP_003767.1;HCoV-NL63_spike;778 | 0 | 0 |
| NC_005831.2;YP_003767.1;HCoV-NL63_spike;779 | 0 | 0 |
| NC_005831.2;YP_003767.1;HCoV-NL63_spike;780 | 0 | 0 |
| NC_005831.2;YP_003767.1;HCoV-NL63_spike;781 | 0 | 0 |
| NC_005831.2;YP_003767.1;HCoV-NL63_spike;782 | 0 | 0 |
| NC_005831.2;YP_003767.1;HCoV-NL63_spike;783 | 0 | 0 |
| NC_005831.2;YP_003767.1;HCoV-NL63_spike;784 | 0 | 0 |
| NC_005831.2;YP_003767.1;HCoV-NL63_spike;785 | 0 | 0 |
| NC_005831.2;YP_003767.1;HCoV-NL63_spike;786 | 0 | 0 |
| NC_005831.2;YP_003767.1;HCoV-NL63_spike;787 | 0 | 0 |
| NC_005831.2;YP_003767.1;HCoV-NL63_spike;788 | 0 | 0 |
| NC_005831.2;YP_003767.1;HCoV-NL63_spike;789 | 0 | 0 |
| NC_005831.2;YP_003767.1;HCoV-NL63_spike;790 | 0 | 0 |
| NC_005831.2;YP_003767.1;HCoV-NL63_spike;791 | 0 | 0 |
| NC_005831.2;YP_003767.1;HCoV-NL63_spike;792 | 0 | 0 |
| NC_005831.2;YP_003767.1;HCoV-NL63_spike;793 | 0 | 0 |
| NC_005831.2;YP_003767.1;HCoV-NL63_spike;794 | 0 | 0.025 |
| NC_005831.2;YP_003767.1;HCoV-NL63_spike;795 | 0 | 0 |
| NC_005831.2;YP_003767.1;HCoV-NL63_spike;796 | 0 | 0 |
| NC_005831.2;YP_003767.1;HCoV-NL63_spike;797 | 0 | 0 |
| NC_005831.2;YP_003767.1;HCoV-NL63_spike;798 | 0 | 0 |
| NC_005831.2;YP_003767.1;HCoV-NL63_spike;799 | 0 | 0 |
| NC_005831.2;YP_003767.1;HCoV-NL63_spike;800 | 0.05 | 0 |
| NC_005831.2;YP_003767.1;HCoV-NL63_spike;801 | 0.05 | 0 |
| NC_005831.2;YP_003767.1;HCoV-NL63_spike;802 | 0 | 0 |
| NC_005831.2;YP_003767.1;HCoV-NL63_spike;803 | 0 | 0 |
| NC_005831.2;YP_003767.1;HCoV-NL63_spike;804 | 0 | 0 |
| NC_005831.2;YP_003767.1;HCoV-NL63_spike;805 | 0 | 0 |
| NC_005831.2;YP_003767.1;HCoV-NL63_spike;806 | 0 | 0 |
| NC_005831.2;YP_003767.1;HCoV-NL63_spike;807 | 0 | 0 |
| NC_005831.2;YP_003767.1;HCoV-NL63_spike;808 | 0 | 0 |
| NC_005831.2;YP_003767.1;HCoV-NL63_spike;809 | 0 | 0 |
| NC_005831.2;YP_003767.1;HCoV-NL63_spike;810 | 0 | 0.025 |
| NC_005831.2;YP_003767.1;HCoV-NL63_spike;811 | 0 | 0.05 |
| NC_005831.2;YP_003767.1;HCoV-NL63_spike;812 | 0 | 0.45 |
| NC_005831.2;YP_003767.1;HCoV-NL63_spike;813 | 0 | 0.4 |
| NC_005831.2;YP_003767.1;HCoV-NL63_spike;814 | 0 | 0.025 |
| NC_005831.2;YP_003767.1;HCoV-NL63_spike;815 | 0 | 0 |
| NC_005831.2;YP_003767.1;HCoV-NL63_spike;816 | 0 | 0.025 |

| | | |
|---|---|---|
| NC_005831.2;YP_003767.1;HCoV-NL63_spike;817 | 0 | 0 |
| NC_005831.2;YP_003767.1;HCoV-NL63_spike;818 | 0 | 0 |
| NC_005831.2;YP_003767.1;HCoV-NL63_spike;819 | 0 | 0.025 |
| NC_005831.2;YP_003767.1;HCoV-NL63_spike;820 | 0 | 0.025 |
| NC_005831.2;YP_003767.1;HCoV-NL63_spike;821 | 0 | 0.025 |
| NC_005831.2;YP_003767.1;HCoV-NL63_spike;822 | 0 | 0 |
| NC_005831.2;YP_003767.1;HCoV-NL63_spike;823 | 0 | 0 |
| NC_005831.2;YP_003767.1;HCoV-NL63_spike;824 | 0 | 0 |
| NC_005831.2;YP_003767.1;HCoV-NL63_spike;825 | 0 | 0 |
| NC_005831.2;YP_003767.1;HCoV-NL63_spike;826 | 0 | 0 |
| NC_005831.2;YP_003767.1;HCoV-NL63_spike;827 | 0 | 0 |
| NC_005831.2;YP_003767.1;HCoV-NL63_spike;828 | 0 | 0 |
| NC_005831.2;YP_003767.1;HCoV-NL63_spike;829 | 0 | 0 |
| NC_005831.2;YP_003767.1;HCoV-NL63_spike;830 | 0 | 0 |
| NC_005831.2;YP_003767.1;HCoV-NL63_spike;831 | 0 | 0 |
| NC_005831.2;YP_003767.1;HCoV-NL63_spike;832 | 0 | 0 |
| NC_005831.2;YP_003767.1;HCoV-NL63_spike;833 | 0 | 0 |
| NC_005831.2;YP_003767.1;HCoV-NL63_spike;834 | 0 | 0 |
| NC_005831.2;YP_003767.1;HCoV-NL63_spike;835 | 0 | 0 |
| NC_005831.2;YP_003767.1;HCoV-NL63_spike;836 | 0.05 | 0 |
| NC_005831.2;YP_003767.1;HCoV-NL63_spike;837 | 0.05 | 0 |
| NC_005831.2;YP_003767.1;HCoV-NL63_spike;838 | 0 | 0 |
| NC_005831.2;YP_003767.1;HCoV-NL63_spike;839 | 0 | 0 |
| NC_005831.2;YP_003767.1;HCoV-NL63_spike;840 | 0 | 0 |
| NC_005831.2;YP_003767.1;HCoV-NL63_spike;841 | 0 | 0 |
| NC_005831.2;YP_003767.1;HCoV-NL63_spike;842 | 0 | 0 |
| NC_005831.2;YP_003767.1;HCoV-NL63_spike;843 | 0 | 0 |
| NC_005831.2;YP_003767.1;HCoV-NL63_spike;844 | 0.05 | 0 |
| NC_005831.2;YP_003767.1;HCoV-NL63_spike;845 | 0.05 | 0 |
| NC_005831.2;YP_003767.1;HCoV-NL63_spike;846 | 0 | 0 |
| NC_005831.2;YP_003767.1;HCoV-NL63_spike;847 | 0 | 0.05 |
| NC_005831.2;YP_003767.1;HCoV-NL63_spike;848 | 0 | 0.025 |
| NC_005831.2;YP_003767.1;HCoV-NL63_spike;849 | 0 | 0.05 |
| NC_005831.2;YP_003767.1;HCoV-NL63_spike;850 | 0 | 0.05 |
| NC_005831.2;YP_003767.1;HCoV-NL63_spike;851 | 0.05 | 0.025 |
| NC_005831.2;YP_003767.1;HCoV-NL63_spike;852 | 0 | 0.025 |
| NC_005831.2;YP_003767.1;HCoV-NL63_spike;853 | 0 | 0.025 |
| NC_005831.2;YP_003767.1;HCoV-NL63_spike;854 | 0.05 | 0.025 |
| NC_005831.2;YP_003767.1;HCoV-NL63_spike;855 | 0 | 0.025 |
| NC_005831.2;YP_003767.1;HCoV-NL63_spike;856 | 0.05 | 0.025 |
| NC_005831.2;YP_003767.1;HCoV-NL63_spike;857 | 0 | 0.025 |
| NC_005831.2;YP_003767.1;HCoV-NL63_spike;858 | 0 | 0 |
| NC_005831.2;YP_003767.1;HCoV-NL63_spike;859 | 0.05 | 0 |

| | | |
|---|---|---|
| NC_005831.2;YP_003767.1;HCoV-NL63_spike;860 | 0 | 0 |
| NC_005831.2;YP_003767.1;HCoV-NL63_spike;861 | 0 | 0 |
| NC_005831.2;YP_003767.1;HCoV-NL63_spike;862 | 0 | 0 |
| NC_005831.2;YP_003767.1;HCoV-NL63_spike;863 | 0.05 | 0.45 |
| NC_005831.2;YP_003767.1;HCoV-NL63_spike;864 | 0.1 | 0.5 |
| NC_005831.2;YP_003767.1;HCoV-NL63_spike;865 | 0.05 | 0.7 |
| NC_005831.2;YP_003767.1;HCoV-NL63_spike;866 | 0 | 0.55 |
| NC_005831.2;YP_003767.1;HCoV-NL63_spike;867 | 0 | 0.55 |
| NC_005831.2;YP_003767.1;HCoV-NL63_spike;868 | 0 | 0.475 |
| NC_005831.2;YP_003767.1;HCoV-NL63_spike;869 | 0 | 0.5 |
| NC_005831.2;YP_003767.1;HCoV-NL63_spike;870 | 0 | 0.45 |
| NC_005831.2;YP_003767.1;HCoV-NL63_spike;871 | 0 | 0.25 |
| NC_005831.2;YP_003767.1;HCoV-NL63_spike;872 | 0.1 | 0.4 |
| NC_005831.2;YP_003767.1;HCoV-NL63_spike;873 | 0 | 0.25 |
| NC_005831.2;YP_003767.1;HCoV-NL63_spike;874 | 0 | 0.05 |
| NC_005831.2;YP_003767.1;HCoV-NL63_spike;875 | 0.05 | 0 |
| NC_005831.2;YP_003767.1;HCoV-NL63_spike;876 | 0.05 | 0 |
| NC_005831.2;YP_003767.1;HCoV-NL63_spike;877 | 0.05 | 0 |
| NC_005831.2;YP_003767.1;HCoV-NL63_spike;878 | 0 | 0 |
| NC_005831.2;YP_003767.1;HCoV-NL63_spike;879 | 0.05 | 0 |
| NC_005831.2;YP_003767.1;HCoV-NL63_spike;880 | 0 | 0 |
| NC_005831.2;YP_003767.1;HCoV-NL63_spike;881 | 0 | 0 |
| NC_005831.2;YP_003767.1;HCoV-NL63_spike;882 | 0 | 0 |
| NC_005831.2;YP_003767.1;HCoV-NL63_spike;883 | 0 | 0.025 |
| NC_005831.2;YP_003767.1;HCoV-NL63_spike;884 | 0 | 0 |
| NC_005831.2;YP_003767.1;HCoV-NL63_spike;885 | 0 | 0.05 |
| NC_005831.2;YP_003767.1;HCoV-NL63_spike;886 | 0 | 0 |
| NC_005831.2;YP_003767.1;HCoV-NL63_spike;887 | 0 | 0 |
| NC_005831.2;YP_003767.1;HCoV-NL63_spike;888 | 0 | 0.025 |
| NC_005831.2;YP_003767.1;HCoV-NL63_spike;889 | 0 | 0.025 |
| NC_005831.2;YP_003767.1;HCoV-NL63_spike;890 | 0.1 | 0 |
| NC_005831.2;YP_003767.1;HCoV-NL63_spike;891 | 0.15 | 0.05 |
| NC_005831.2;YP_003767.1;HCoV-NL63_spike;892 | 0.1 | 0.05 |
| NC_005831.2;YP_003767.1;HCoV-NL63_spike;893 | 0.1 | 0.05 |
| NC_005831.2;YP_003767.1;HCoV-NL63_spike;894 | 0.1 | 0.025 |
| NC_005831.2;YP_003767.1;HCoV-NL63_spike;895 | 0.1 | 0.025 |
| NC_005831.2;YP_003767.1;HCoV-NL63_spike;896 | 0.1 | 0.025 |
| NC_005831.2;YP_003767.1;HCoV-NL63_spike;897 | 0.15 | 0.05 |
| NC_005831.2;YP_003767.1;HCoV-NL63_spike;898 | 0.15 | 0.075 |
| NC_005831.2;YP_003767.1;HCoV-NL63_spike;899 | 0 | 0 |
| NC_005831.2;YP_003767.1;HCoV-NL63_spike;900 | 0.05 | 0.325 |
| NC_005831.2;YP_003767.1;HCoV-NL63_spike;901 | 0 | 0.05 |
| NC_005831.2;YP_003767.1;HCoV-NL63_spike;902 | 0 | 0 |

| | | |
|---|---|---|
| NC_005831.2;YP_003767.1;HCoV-NL63_spike;903 | 0 | 0 |
| NC_005831.2;YP_003767.1;HCoV-NL63_spike;904 | 0 | 0 |
| NC_005831.2;YP_003767.1;HCoV-NL63_spike;905 | 0 | 0 |
| NC_005831.2;YP_003767.1;HCoV-NL63_spike;906 | 0.05 | 0.05 |
| NC_005831.2;YP_003767.1;HCoV-NL63_spike;907 | 0 | 0 |
| NC_005831.2;YP_003767.1;HCoV-NL63_spike;908 | 0.05 | 0.025 |
| NC_005831.2;YP_003767.1;HCoV-NL63_spike;909 | 0 | 0 |
| NC_005831.2;YP_003767.1;HCoV-NL63_spike;910 | 0 | 0 |
| NC_005831.2;YP_003767.1;HCoV-NL63_spike;911 | 0 | 0 |
| NC_005831.2;YP_003767.1;HCoV-NL63_spike;912 | 0 | 0 |
| NC_005831.2;YP_003767.1;HCoV-NL63_spike;913 | 0 | 0 |
| NC_005831.2;YP_003767.1;HCoV-NL63_spike;914 | 0 | 0 |
| NC_005831.2;YP_003767.1;HCoV-NL63_spike;915 | 0 | 0 |
| NC_005831.2;YP_003767.1;HCoV-NL63_spike;916 | 0 | 0 |
| NC_005831.2;YP_003767.1;HCoV-NL63_spike;917 | 0 | 0 |
| NC_005831.2;YP_003767.1;HCoV-NL63_spike;918 | 0 | 0 |
| NC_005831.2;YP_003767.1;HCoV-NL63_spike;919 | 0 | 0 |
| NC_005831.2;YP_003767.1;HCoV-NL63_spike;920 | 0 | 0 |
| NC_005831.2;YP_003767.1;HCoV-NL63_spike;921 | 0 | 0 |
| NC_005831.2;YP_003767.1;HCoV-NL63_spike;922 | 0 | 0.025 |
| NC_005831.2;YP_003767.1;HCoV-NL63_spike;923 | 0 | 0 |
| NC_005831.2;YP_003767.1;HCoV-NL63_spike;924 | 0 | 0 |
| NC_005831.2;YP_003767.1;HCoV-NL63_spike;925 | 0 | 0 |
| NC_005831.2;YP_003767.1;HCoV-NL63_spike;926 | 0.1 | 0 |
| NC_005831.2;YP_003767.1;HCoV-NL63_spike;927 | 0 | 0 |
| NC_005831.2;YP_003767.1;HCoV-NL63_spike;928 | 0 | 0 |
| NC_005831.2;YP_003767.1;HCoV-NL63_spike;929 | 0 | 0 |
| NC_005831.2;YP_003767.1;HCoV-NL63_spike;930 | 0 | 0 |
| NC_005831.2;YP_003767.1;HCoV-NL63_spike;931 | 0.05 | 0.025 |
| NC_005831.2;YP_003767.1;HCoV-NL63_spike;932 | 0.05 | 0.025 |
| NC_005831.2;YP_003767.1;HCoV-NL63_spike;933 | 0 | 0 |
| NC_005831.2;YP_003767.1;HCoV-NL63_spike;934 | 0 | 0 |
| NC_005831.2;YP_003767.1;HCoV-NL63_spike;935 | 0 | 0 |
| NC_005831.2;YP_003767.1;HCoV-NL63_spike;936 | 0 | 0 |
| NC_005831.2;YP_003767.1;HCoV-NL63_spike;937 | 0 | 0 |
| NC_005831.2;YP_003767.1;HCoV-NL63_spike;938 | 0 | 0 |
| NC_005831.2;YP_003767.1;HCoV-NL63_spike;939 | 0 | 0 |
| NC_005831.2;YP_003767.1;HCoV-NL63_spike;940 | 0 | 0 |
| NC_005831.2;YP_003767.1;HCoV-NL63_spike;941 | 0 | 0 |
| NC_005831.2;YP_003767.1;HCoV-NL63_spike;942 | 0 | 0 |
| NC_005831.2;YP_003767.1;HCoV-NL63_spike;943 | 0 | 0 |
| NC_005831.2;YP_003767.1;HCoV-NL63_spike;944 | 0 | 0 |
| NC_005831.2;YP_003767.1;HCoV-NL63_spike;945 | 0 | 0 |

| | | |
|---|---|---|
| NC_005831.2;YP_003767.1;HCoV-NL63_spike;946 | 0 | 0 |
| NC_005831.2;YP_003767.1;HCoV-NL63_spike;947 | 0 | 0 |
| NC_005831.2;YP_003767.1;HCoV-NL63_spike;948 | 0 | 0 |
| NC_005831.2;YP_003767.1;HCoV-NL63_spike;949 | 0 | 0 |
| NC_005831.2;YP_003767.1;HCoV-NL63_spike;950 | 0 | 0 |
| NC_005831.2;YP_003767.1;HCoV-NL63_spike;951 | 0 | 0 |
| NC_005831.2;YP_003767.1;HCoV-NL63_spike;952 | 0 | 0 |
| NC_005831.2;YP_003767.1;HCoV-NL63_spike;953 | 0 | 0 |
| NC_005831.2;YP_003767.1;HCoV-NL63_spike;954 | 0 | 0 |
| NC_005831.2;YP_003767.1;HCoV-NL63_spike;955 | 0 | 0 |
| NC_005831.2;YP_003767.1;HCoV-NL63_spike;956 | 0 | 0 |
| NC_005831.2;YP_003767.1;HCoV-NL63_spike;957 | 0 | 0 |
| NC_005831.2;YP_003767.1;HCoV-NL63_spike;958 | 0 | 0 |
| NC_005831.2;YP_003767.1;HCoV-NL63_spike;959 | 0.05 | 0 |
| NC_005831.2;YP_003767.1;HCoV-NL63_spike;960 | 0 | 0 |
| NC_005831.2;YP_003767.1;HCoV-NL63_spike;961 | 0 | 0 |
| NC_005831.2;YP_003767.1;HCoV-NL63_spike;962 | 0.05 | 0.025 |
| NC_005831.2;YP_003767.1;HCoV-NL63_spike;963 | 0 | 0 |
| NC_005831.2;YP_003767.1;HCoV-NL63_spike;964 | 0 | 0 |
| NC_005831.2;YP_003767.1;HCoV-NL63_spike;965 | 0.05 | 0.05 |
| NC_005831.2;YP_003767.1;HCoV-NL63_spike;966 | 0.3 | 0.1 |
| NC_005831.2;YP_003767.1;HCoV-NL63_spike;967 | 0.2 | 0.125 |
| NC_005831.2;YP_003767.1;HCoV-NL63_spike;968 | 0.2 | 0.1 |
| NC_005831.2;YP_003767.1;HCoV-NL63_spike;969 | 0.2 | 0.125 |
| NC_005831.2;YP_003767.1;HCoV-NL63_spike;970 | 0.2 | 0.15 |
| NC_005831.2;YP_003767.1;HCoV-NL63_spike;971 | 0.15 | 0.1 |
| NC_005831.2;YP_003767.1;HCoV-NL63_spike;972 | 0.2 | 0.1 |
| NC_005831.2;YP_003767.1;HCoV-NL63_spike;973 | 0.2 | 0.075 |
| NC_005831.2;YP_003767.1;HCoV-NL63_spike;974 | 0.25 | 0.1 |
| NC_005831.2;YP_003767.1;HCoV-NL63_spike;975 | 0.25 | 0.075 |
| NC_005831.2;YP_003767.1;HCoV-NL63_spike;976 | 0.25 | 0.1 |
| NC_005831.2;YP_003767.1;HCoV-NL63_spike;977 | 0.1 | 0 |
| NC_005831.2;YP_003767.1;HCoV-NL63_spike;978 | 0.05 | 0 |
| NC_005831.2;YP_003767.1;HCoV-NL63_spike;979 | 0.1 | 0 |
| NC_005831.2;YP_003767.1;HCoV-NL63_spike;980 | 0.1 | 0 |
| NC_005831.2;YP_003767.1;HCoV-NL63_spike;981 | 0.1 | 0 |
| NC_005831.2;YP_003767.1;HCoV-NL63_spike;982 | 0 | 0 |
| NC_005831.2;YP_003767.1;HCoV-NL63_spike;983 | 0 | 0 |
| NC_005831.2;YP_003767.1;HCoV-NL63_spike;984 | 0 | 0 |
| NC_005831.2;YP_003767.1;HCoV-NL63_spike;985 | 0 | 0 |
| NC_005831.2;YP_003767.1;HCoV-NL63_spike;986 | 0 | 0 |
| NC_005831.2;YP_003767.1;HCoV-NL63_spike;987 | 0.05 | 0 |
| NC_005831.2;YP_003767.1;HCoV-NL63_spike;988 | 0.05 | 0 |

| | | |
|---|---|---|
| NC_005831.2;YP_003767.1;HCoV-NL63_spike;989 | 0.05 | 0 |
| NC_005831.2;YP_003767.1;HCoV-NL63_spike;990 | 0.05 | 0.025 |
| NC_005831.2;YP_003767.1;HCoV-NL63_spike;991 | 0.05 | 0.05 |
| NC_005831.2;YP_003767.1;HCoV-NL63_spike;992 | 0.15 | 0.125 |
| NC_005831.2;YP_003767.1;HCoV-NL63_spike;993 | 0.05 | 0 |
| NC_005831.2;YP_003767.1;HCoV-NL63_spike;994 | 0 | 0 |
| NC_005831.2;YP_003767.1;HCoV-NL63_spike;995 | 0 | 0 |
| NC_005831.2;YP_003767.1;HCoV-NL63_spike;996 | 0 | 0 |
| NC_005831.2;YP_003767.1;HCoV-NL63_spike;997 | 0 | 0 |
| NC_005831.2;YP_003767.1;HCoV-NL63_spike;998 | 0 | 0 |
| NC_005831.2;YP_003767.1;HCoV-NL63_spike;999 | 0 | 0 |
| NC_005831.2;YP_003767.1;HCoV-NL63_spike;1000 | 0 | 0 |
| NC_005831.2;YP_003767.1;HCoV-NL63_spike;1001 | 0 | 0 |
| NC_005831.2;YP_003767.1;HCoV-NL63_spike;1002 | 0.05 | 0 |
| NC_005831.2;YP_003767.1;HCoV-NL63_spike;1003 | 0.05 | 0 |
| NC_005831.2;YP_003767.1;HCoV-NL63_spike;1004 | 0.05 | 0 |
| NC_005831.2;YP_003767.1;HCoV-NL63_spike;1005 | 0 | 0 |
| NC_005831.2;YP_003767.1;HCoV-NL63_spike;1006 | 0 | 0.025 |
| NC_005831.2;YP_003767.1;HCoV-NL63_spike;1007 | 0 | 0.05 |
| NC_005831.2;YP_003767.1;HCoV-NL63_spike;1008 | 0 | 0.025 |
| NC_005831.2;YP_003767.1;HCoV-NL63_spike;1009 | 0 | 0.025 |
| NC_005831.2;YP_003767.1;HCoV-NL63_spike;1010 | 0 | 0.025 |
| NC_005831.2;YP_003767.1;HCoV-NL63_spike;1011 | 0.05 | 0.075 |
| NC_005831.2;YP_003767.1;HCoV-NL63_spike;1012 | 0 | 0.025 |
| NC_005831.2;YP_003767.1;HCoV-NL63_spike;1013 | 0 | 0.05 |
| NC_005831.2;YP_003767.1;HCoV-NL63_spike;1014 | 0 | 0.075 |
| NC_005831.2;YP_003767.1;HCoV-NL63_spike;1015 | 0 | 0.05 |
| NC_005831.2;YP_003767.1;HCoV-NL63_spike;1016 | 0 | 0.025 |
| NC_005831.2;YP_003767.1;HCoV-NL63_spike;1017 | 0 | 0.025 |
| NC_005831.2;YP_003767.1;HCoV-NL63_spike;1018 | 0 | 0.025 |
| NC_005831.2;YP_003767.1;HCoV-NL63_spike;1019 | 0 | 0.025 |
| NC_005831.2;YP_003767.1;HCoV-NL63_spike;1020 | 0 | 0 |
| NC_005831.2;YP_003767.1;HCoV-NL63_spike;1021 | 0 | 0 |
| NC_005831.2;YP_003767.1;HCoV-NL63_spike;1022 | 0 | 0 |
| NC_005831.2;YP_003767.1;HCoV-NL63_spike;1023 | 0 | 0 |
| NC_005831.2;YP_003767.1;HCoV-NL63_spike;1024 | 0 | 0 |
| NC_005831.2;YP_003767.1;HCoV-NL63_spike;1025 | 0 | 0 |
| NC_005831.2;YP_003767.1;HCoV-NL63_spike;1026 | 0 | 0 |
| NC_005831.2;YP_003767.1;HCoV-NL63_spike;1027 | 0 | 0 |
| NC_005831.2;YP_003767.1;HCoV-NL63_spike;1028 | 0 | 0 |
| NC_005831.2;YP_003767.1;HCoV-NL63_spike;1029 | 0 | 0.025 |
| NC_005831.2;YP_003767.1;HCoV-NL63_spike;1030 | 0 | 0 |
| NC_005831.2;YP_003767.1;HCoV-NL63_spike;1031 | 0 | 0 |

| | | |
|---|---|---|
| NC_005831.2;YP_003767.1;HCoV-NL63_spike;1032 | 0 | 0 |
| NC_005831.2;YP_003767.1;HCoV-NL63_spike;1033 | 0 | 0 |
| NC_005831.2;YP_003767.1;HCoV-NL63_spike;1034 | 0 | 0 |
| NC_005831.2;YP_003767.1;HCoV-NL63_spike;1035 | 0 | 0 |
| NC_005831.2;YP_003767.1;HCoV-NL63_spike;1036 | 0 | 0 |
| NC_005831.2;YP_003767.1;HCoV-NL63_spike;1037 | 0 | 0 |
| NC_005831.2;YP_003767.1;HCoV-NL63_spike;1038 | 0.1 | 0.025 |
| NC_005831.2;YP_003767.1;HCoV-NL63_spike;1039 | 0.1 | 0 |
| NC_005831.2;YP_003767.1;HCoV-NL63_spike;1040 | 0.05 | 0 |
| NC_005831.2;YP_003767.1;HCoV-NL63_spike;1041 | 0.05 | 0.05 |
| NC_005831.2;YP_003767.1;HCoV-NL63_spike;1042 | 0.05 | 0.05 |
| NC_005831.2;YP_003767.1;HCoV-NL63_spike;1043 | 0.05 | 0.05 |
| NC_005831.2;YP_003767.1;HCoV-NL63_spike;1044 | 0.05 | 0.025 |
| NC_005831.2;YP_003767.1;HCoV-NL63_spike;1045 | 0.2 | 0.075 |
| NC_005831.2;YP_003767.1;HCoV-NL63_spike;1046 | 0.05 | 0.025 |
| NC_005831.2;YP_003767.1;HCoV-NL63_spike;1047 | 0 | 0.025 |
| NC_005831.2;YP_003767.1;HCoV-NL63_spike;1048 | 0 | 0 |
| NC_005831.2;YP_003767.1;HCoV-NL63_spike;1049 | 0 | 0 |
| NC_005831.2;YP_003767.1;HCoV-NL63_spike;1050 | 0 | 0.025 |
| NC_005831.2;YP_003767.1;HCoV-NL63_spike;1051 | 0 | 0 |
| NC_005831.2;YP_003767.1;HCoV-NL63_spike;1052 | 0 | 0 |
| NC_005831.2;YP_003767.1;HCoV-NL63_spike;1053 | 0 | 0 |
| NC_005831.2;YP_003767.1;HCoV-NL63_spike;1054 | 0.05 | 0.025 |
| NC_005831.2;YP_003767.1;HCoV-NL63_spike;1055 | 0.05 | 0 |
| NC_005831.2;YP_003767.1;HCoV-NL63_spike;1056 | 0 | 0 |
| NC_005831.2;YP_003767.1;HCoV-NL63_spike;1057 | 0 | 0 |
| NC_005831.2;YP_003767.1;HCoV-NL63_spike;1058 | 0 | 0 |
| NC_005831.2;YP_003767.1;HCoV-NL63_spike;1059 | 0 | 0 |
| NC_005831.2;YP_003767.1;HCoV-NL63_spike;1060 | 0 | 0 |
| NC_005831.2;YP_003767.1;HCoV-NL63_spike;1061 | 0 | 0 |
| NC_005831.2;YP_003767.1;HCoV-NL63_spike;1062 | 0 | 0 |
| NC_005831.2;YP_003767.1;HCoV-NL63_spike;1063 | 0 | 0 |
| NC_005831.2;YP_003767.1;HCoV-NL63_spike;1064 | 0 | 0 |
| NC_005831.2;YP_003767.1;HCoV-NL63_spike;1065 | 0 | 0 |
| NC_005831.2;YP_003767.1;HCoV-NL63_spike;1066 | 0 | 0 |
| NC_005831.2;YP_003767.1;HCoV-NL63_spike;1067 | 0 | 0 |
| NC_005831.2;YP_003767.1;HCoV-NL63_spike;1068 | 0 | 0.025 |
| NC_005831.2;YP_003767.1;HCoV-NL63_spike;1069 | 0 | 0.025 |
| NC_005831.2;YP_003767.1;HCoV-NL63_spike;1070 | 0 | 0 |
| NC_005831.2;YP_003767.1;HCoV-NL63_spike;1071 | 0 | 0.025 |
| NC_005831.2;YP_003767.1;HCoV-NL63_spike;1072 | 0 | 0.025 |
| NC_005831.2;YP_003767.1;HCoV-NL63_spike;1073 | 0 | 0.025 |
| NC_005831.2;YP_003767.1;HCoV-NL63_spike;1074 | 0 | 0 |

| | | |
|---|---|---|
| NC_005831.2;YP_003767.1;HCoV-NL63_spike;1075 | 0 | 0 |
| NC_005831.2;YP_003767.1;HCoV-NL63_spike;1076 | 0 | 0 |
| NC_005831.2;YP_003767.1;HCoV-NL63_spike;1077 | 0 | 0 |
| NC_005831.2;YP_003767.1;HCoV-NL63_spike;1078 | 0 | 0 |
| NC_005831.2;YP_003767.1;HCoV-NL63_spike;1079 | 0 | 0 |
| NC_005831.2;YP_003767.1;HCoV-NL63_spike;1080 | 0 | 0 |
| NC_005831.2;YP_003767.1;HCoV-NL63_spike;1081 | 0 | 0 |
| NC_005831.2;YP_003767.1;HCoV-NL63_spike;1082 | 0 | 0.025 |
| NC_005831.2;YP_003767.1;HCoV-NL63_spike;1083 | 0.05 | 0.025 |
| NC_005831.2;YP_003767.1;HCoV-NL63_spike;1084 | 0 | 0.025 |
| NC_005831.2;YP_003767.1;HCoV-NL63_spike;1085 | 0 | 0 |
| NC_005831.2;YP_003767.1;HCoV-NL63_spike;1086 | 0.05 | 0 |
| NC_005831.2;YP_003767.1;HCoV-NL63_spike;1087 | 0 | 0 |
| NC_005831.2;YP_003767.1;HCoV-NL63_spike;1088 | 0 | 0 |
| NC_005831.2;YP_003767.1;HCoV-NL63_spike;1089 | 0 | 0 |
| NC_005831.2;YP_003767.1;HCoV-NL63_spike;1090 | 0 | 0 |
| NC_005831.2;YP_003767.1;HCoV-NL63_spike;1091 | 0 | 0 |
| NC_005831.2;YP_003767.1;HCoV-NL63_spike;1092 | 0 | 0 |
| NC_005831.2;YP_003767.1;HCoV-NL63_spike;1093 | 0 | 0 |
| NC_005831.2;YP_003767.1;HCoV-NL63_spike;1094 | 0 | 0 |
| NC_005831.2;YP_003767.1;HCoV-NL63_spike;1095 | 0 | 0 |
| NC_005831.2;YP_003767.1;HCoV-NL63_spike;1096 | 0 | 0 |
| NC_005831.2;YP_003767.1;HCoV-NL63_spike;1097 | 0.05 | 0 |
| NC_005831.2;YP_003767.1;HCoV-NL63_spike;1098 | 0 | 0 |
| NC_005831.2;YP_003767.1;HCoV-NL63_spike;1099 | 0 | 0 |
| NC_005831.2;YP_003767.1;HCoV-NL63_spike;1100 | 0.05 | 0 |
| NC_005831.2;YP_003767.1;HCoV-NL63_spike;1101 | 0.05 | 0 |
| NC_005831.2;YP_003767.1;HCoV-NL63_spike;1102 | 0.05 | 0 |
| NC_005831.2;YP_003767.1;HCoV-NL63_spike;1103 | 0.05 | 0 |
| NC_005831.2;YP_003767.1;HCoV-NL63_spike;1104 | 0 | 0 |
| NC_005831.2;YP_003767.1;HCoV-NL63_spike;1105 | 0 | 0 |
| NC_005831.2;YP_003767.1;HCoV-NL63_spike;1106 | 0 | 0 |
| NC_005831.2;YP_003767.1;HCoV-NL63_spike;1107 | 0 | 0 |
| NC_005831.2;YP_003767.1;HCoV-NL63_spike;1108 | 0 | 0 |
| NC_005831.2;YP_003767.1;HCoV-NL63_spike;1109 | 0 | 0 |
| NC_005831.2;YP_003767.1;HCoV-NL63_spike;1110 | 0 | 0 |
| NC_005831.2;YP_003767.1;HCoV-NL63_spike;1111 | 0 | 0 |
| NC_005831.2;YP_003767.1;HCoV-NL63_spike;1112 | 0 | 0 |
| NC_005831.2;YP_003767.1;HCoV-NL63_spike;1113 | 0 | 0 |
| NC_005831.2;YP_003767.1;HCoV-NL63_spike;1114 | 0 | 0 |
| NC_005831.2;YP_003767.1;HCoV-NL63_spike;1115 | 0 | 0 |
| NC_005831.2;YP_003767.1;HCoV-NL63_spike;1116 | 0 | 0.025 |
| NC_005831.2;YP_003767.1;HCoV-NL63_spike;1117 | 0 | 0 |

| | | |
|---|---|---|
| NC_005831.2;YP_003767.1;HCoV-NL63_spike;1118 | 0 | 0 |
| NC_005831.2;YP_003767.1;HCoV-NL63_spike;1119 | 0 | 0 |
| NC_005831.2;YP_003767.1;HCoV-NL63_spike;1120 | 0 | 0 |
| NC_005831.2;YP_003767.1;HCoV-NL63_spike;1121 | 0 | 0 |
| NC_005831.2;YP_003767.1;HCoV-NL63_spike;1122 | 0 | 0 |
| NC_005831.2;YP_003767.1;HCoV-NL63_spike;1123 | 0 | 0 |
| NC_005831.2;YP_003767.1;HCoV-NL63_spike;1124 | 0 | 0 |
| NC_005831.2;YP_003767.1;HCoV-NL63_spike;1125 | 0 | 0 |
| NC_005831.2;YP_003767.1;HCoV-NL63_spike;1126 | 0 | 0 |
| NC_005831.2;YP_003767.1;HCoV-NL63_spike;1127 | 0 | 0 |
| NC_005831.2;YP_003767.1;HCoV-NL63_spike;1128 | 0 | 0 |
| NC_005831.2;YP_003767.1;HCoV-NL63_spike;1129 | 0 | 0 |
| NC_005831.2;YP_003767.1;HCoV-NL63_spike;1130 | 0 | 0 |
| NC_005831.2;YP_003767.1;HCoV-NL63_spike;1131 | 0 | 0 |
| NC_005831.2;YP_003767.1;HCoV-NL63_spike;1132 | 0 | 0 |
| NC_005831.2;YP_003767.1;HCoV-NL63_spike;1133 | 0 | 0 |
| NC_005831.2;YP_003767.1;HCoV-NL63_spike;1134 | 0 | 0 |
| NC_005831.2;YP_003767.1;HCoV-NL63_spike;1135 | 0.05 | 0 |
| NC_005831.2;YP_003767.1;HCoV-NL63_spike;1136 | 0.05 | 0 |
| NC_005831.2;YP_003767.1;HCoV-NL63_spike;1137 | 0 | 0 |
| NC_005831.2;YP_003767.1;HCoV-NL63_spike;1138 | 0 | 0 |
| NC_005831.2;YP_003767.1;HCoV-NL63_spike;1139 | 0 | 0 |
| NC_005831.2;YP_003767.1;HCoV-NL63_spike;1140 | 0 | 0 |
| NC_005831.2;YP_003767.1;HCoV-NL63_spike;1141 | 0 | 0 |
| NC_005831.2;YP_003767.1;HCoV-NL63_spike;1142 | 0 | 0 |
| NC_005831.2;YP_003767.1;HCoV-NL63_spike;1143 | 0 | 0 |
| NC_005831.2;YP_003767.1;HCoV-NL63_spike;1144 | 0 | 0 |
| NC_005831.2;YP_003767.1;HCoV-NL63_spike;1145 | 0 | 0 |
| NC_005831.2;YP_003767.1;HCoV-NL63_spike;1146 | 0 | 0 |
| NC_005831.2;YP_003767.1;HCoV-NL63_spike;1147 | 0 | 0 |
| NC_005831.2;YP_003767.1;HCoV-NL63_spike;1148 | 0 | 0 |
| NC_005831.2;YP_003767.1;HCoV-NL63_spike;1149 | 0 | 0 |
| NC_005831.2;YP_003767.1;HCoV-NL63_spike;1150 | 0 | 0 |
| NC_005831.2;YP_003767.1;HCoV-NL63_spike;1151 | 0 | 0 |
| NC_005831.2;YP_003767.1;HCoV-NL63_spike;1152 | 0 | 0 |
| NC_005831.2;YP_003767.1;HCoV-NL63_spike;1153 | 0 | 0 |
| NC_005831.2;YP_003767.1;HCoV-NL63_spike;1154 | 0 | 0 |
| NC_005831.2;YP_003767.1;HCoV-NL63_spike;1155 | 0 | 0 |
| NC_005831.2;YP_003767.1;HCoV-NL63_spike;1156 | 0 | 0 |
| NC_005831.2;YP_003767.1;HCoV-NL63_spike;1157 | 0 | 0 |
| NC_005831.2;YP_003767.1;HCoV-NL63_spike;1158 | 0 | 0 |
| NC_005831.2;YP_003767.1;HCoV-NL63_spike;1159 | 0 | 0 |
| NC_005831.2;YP_003767.1;HCoV-NL63_spike;1160 | 0 | 0 |

| | | |
|---|---|---|
| NC_005831.2;YP_003767.1;HCoV-NL63_spike;1161 | 0 | 0 |
| NC_005831.2;YP_003767.1;HCoV-NL63_spike;1162 | 0 | 0 |
| NC_005831.2;YP_003767.1;HCoV-NL63_spike;1163 | 0 | 0 |
| NC_005831.2;YP_003767.1;HCoV-NL63_spike;1164 | 0 | 0 |
| NC_005831.2;YP_003767.1;HCoV-NL63_spike;1165 | 0 | 0 |
| NC_005831.2;YP_003767.1;HCoV-NL63_spike;1166 | 0 | 0 |
| NC_005831.2;YP_003767.1;HCoV-NL63_spike;1167 | 0 | 0 |
| NC_005831.2;YP_003767.1;HCoV-NL63_spike;1168 | 0 | 0 |
| NC_005831.2;YP_003767.1;HCoV-NL63_spike;1169 | 0 | 0 |
| NC_005831.2;YP_003767.1;HCoV-NL63_spike;1170 | 0 | 0 |
| NC_005831.2;YP_003767.1;HCoV-NL63_spike;1171 | 0 | 0 |
| NC_005831.2;YP_003767.1;HCoV-NL63_spike;1172 | 0.05 | 0 |
| NC_005831.2;YP_003767.1;HCoV-NL63_spike;1173 | 0 | 0 |
| NC_005831.2;YP_003767.1;HCoV-NL63_spike;1174 | 0 | 0 |
| NC_005831.2;YP_003767.1;HCoV-NL63_spike;1175 | 0 | 0 |
| NC_005831.2;YP_003767.1;HCoV-NL63_spike;1176 | 0 | 0 |
| NC_005831.2;YP_003767.1;HCoV-NL63_spike;1177 | 0 | 0 |
| NC_005831.2;YP_003767.1;HCoV-NL63_spike;1178 | 0 | 0 |
| NC_005831.2;YP_003767.1;HCoV-NL63_spike;1179 | 0 | 0 |
| NC_005831.2;YP_003767.1;HCoV-NL63_spike;1180 | 0 | 0 |
| NC_005831.2;YP_003767.1;HCoV-NL63_spike;1181 | 0 | 0 |
| NC_005831.2;YP_003767.1;HCoV-NL63_spike;1182 | 0 | 0 |
| NC_005831.2;YP_003767.1;HCoV-NL63_spike;1183 | 0 | 0 |
| NC_005831.2;YP_003767.1;HCoV-NL63_spike;1184 | 0 | 0 |
| NC_005831.2;YP_003767.1;HCoV-NL63_spike;1185 | 0 | 0 |
| NC_005831.2;YP_003767.1;HCoV-NL63_spike;1186 | 0 | 0 |
| NC_005831.2;YP_003767.1;HCoV-NL63_spike;1187 | 0 | 0 |
| NC_005831.2;YP_003767.1;HCoV-NL63_spike;1188 | 0 | 0 |
| NC_005831.2;YP_003767.1;HCoV-NL63_spike;1189 | 0 | 0 |
| NC_005831.2;YP_003767.1;HCoV-NL63_spike;1190 | 0 | 0 |
| NC_005831.2;YP_003767.1;HCoV-NL63_spike;1191 | 0 | 0 |
| NC_005831.2;YP_003767.1;HCoV-NL63_spike;1192 | 0 | 0 |
| NC_005831.2;YP_003767.1;HCoV-NL63_spike;1193 | 0 | 0 |
| NC_005831.2;YP_003767.1;HCoV-NL63_spike;1194 | 0 | 0 |
| NC_005831.2;YP_003767.1;HCoV-NL63_spike;1195 | 0 | 0 |
| NC_005831.2;YP_003767.1;HCoV-NL63_spike;1196 | 0 | 0 |
| NC_005831.2;YP_003767.1;HCoV-NL63_spike;1197 | 0 | 0 |
| NC_005831.2;YP_003767.1;HCoV-NL63_spike;1198 | 0 | 0 |
| NC_005831.2;YP_003767.1;HCoV-NL63_spike;1199 | 0.05 | 0.025 |
| NC_005831.2;YP_003767.1;HCoV-NL63_spike;1200 | 0 | 0 |
| NC_005831.2;YP_003767.1;HCoV-NL63_spike;1201 | 0 | 0 |
| NC_005831.2;YP_003767.1;HCoV-NL63_spike;1202 | 0 | 0 |
| NC_005831.2;YP_003767.1;HCoV-NL63_spike;1203 | 0 | 0 |

| | | |
|---|---|---|
| NC_005831.2;YP_003767.1;HCoV-NL63_spike;1204 | 0 | 0 |
| NC_005831.2;YP_003767.1;HCoV-NL63_spike;1205 | 0 | 0 |
| NC_005831.2;YP_003767.1;HCoV-NL63_spike;1206 | 0 | 0.025 |
| NC_005831.2;YP_003767.1;HCoV-NL63_spike;1207 | 0 | 0 |
| NC_005831.2;YP_003767.1;HCoV-NL63_spike;1208 | 0 | 0 |
| NC_005831.2;YP_003767.1;HCoV-NL63_spike;1209 | 0.05 | 0.025 |
| NC_005831.2;YP_003767.1;HCoV-NL63_spike;1210 | 0 | 0.05 |
| NC_005831.2;YP_003767.1;HCoV-NL63_spike;1211 | 0.05 | 0.1 |
| NC_005831.2;YP_003767.1;HCoV-NL63_spike;1212 | 0.1 | 0.1 |
| NC_005831.2;YP_003767.1;HCoV-NL63_spike;1213 | 0.15 | 0.175 |
| NC_005831.2;YP_003767.1;HCoV-NL63_spike;1214 | 0.2 | 0.25 |
| NC_005831.2;YP_003767.1;HCoV-NL63_spike;1215 | 0.2 | 0.15 |
| NC_005831.2;YP_003767.1;HCoV-NL63_spike;1216 | 0.15 | 0.125 |
| NC_005831.2;YP_003767.1;HCoV-NL63_spike;1217 | 0.25 | 0.125 |
| NC_005831.2;YP_003767.1;HCoV-NL63_spike;1218 | 0 | 0.05 |
| NC_005831.2;YP_003767.1;HCoV-NL63_spike;1219 | 0 | 0.05 |
| NC_005831.2;YP_003767.1;HCoV-NL63_spike;1220 | 0.05 | 0.025 |
| NC_005831.2;YP_003767.1;HCoV-NL63_spike;1221 | 0.05 | 0 |
| NC_005831.2;YP_003767.1;HCoV-NL63_spike;1222 | 0.1 | 0.025 |
| NC_005831.2;YP_003767.1;HCoV-NL63_spike;1223 | 0.05 | 0 |
| NC_005831.2;YP_003767.1;HCoV-NL63_spike;1224 | 0.05 | 0 |
| NC_005831.2;YP_003767.1;HCoV-NL63_spike;1225 | 0.1 | 0 |
| NC_005831.2;YP_003767.1;HCoV-NL63_spike;1226 | 0.05 | 0 |
| NC_005831.2;YP_003767.1;HCoV-NL63_spike;1227 | 0.05 | 0 |
| NC_005831.2;YP_003767.1;HCoV-NL63_spike;1228 | 0.05 | 0 |
| NC_005831.2;YP_003767.1;HCoV-NL63_spike;1229 | 0.05 | 0 |
| NC_005831.2;YP_003767.1;HCoV-NL63_spike;1230 | 0 | 0 |
| NC_005831.2;YP_003767.1;HCoV-NL63_spike;1231 | 0 | 0 |
| NC_005831.2;YP_003767.1;HCoV-NL63_spike;1232 | 0.05 | 0 |
| NC_005831.2;YP_003767.1;HCoV-NL63_spike;1233 | 0 | 0 |
| NC_005831.2;YP_003767.1;HCoV-NL63_spike;1234 | 0 | 0 |
| NC_005831.2;YP_003767.1;HCoV-NL63_spike;1235 | 0 | 0 |
| NC_005831.2;YP_003767.1;HCoV-NL63_spike;1236 | 0 | 0 |
| NC_005831.2;YP_003767.1;HCoV-NL63_spike;1237 | 0 | 0 |
| NC_005831.2;YP_003767.1;HCoV-NL63_spike;1238 | 0 | 0 |
| NC_005831.2;YP_003767.1;HCoV-NL63_spike;1239 | 0 | 0 |
| NC_005831.2;YP_003767.1;HCoV-NL63_spike;1240 | 0 | 0 |
| NC_005831.2;YP_003767.1;HCoV-NL63_spike;1241 | 0 | 0 |
| NC_005831.2;YP_003767.1;HCoV-NL63_spike;1242 | 0 | 0 |
| NC_005831.2;YP_003767.1;HCoV-NL63_spike;1243 | 0 | 0 |
| NC_005831.2;YP_003767.1;HCoV-NL63_spike;1244 | 0 | 0.05 |
| NC_005831.2;YP_003767.1;HCoV-NL63_spike;1245 | 0 | 0.025 |
| NC_005831.2;YP_003767.1;HCoV-NL63_spike;1246 | 0.15 | 0.075 |

| | | |
|---|---|---|
| NC_005831.2;YP_003767.1;HCoV-NL63_spike;1247 | 0.05 | 0 |
| NC_005831.2;YP_003767.1;HCoV-NL63_spike;1248 | 0.05 | 0 |
| NC_005831.2;YP_003767.1;HCoV-NL63_spike;1249 | 0.1 | 0.025 |
| NC_005831.2;YP_003767.1;HCoV-NL63_spike;1250 | 0.1 | 0.025 |
| NC_005831.2;YP_003767.1;HCoV-NL63_spike;1251 | 0.1 | 0.025 |
| NC_005831.2;YP_003767.1;HCoV-NL63_spike;1252 | 0.1 | 0 |
| NC_005831.2;YP_003767.1;HCoV-NL63_spike;1253 | 0.05 | 0 |
| NC_005831.2;YP_003767.1;HCoV-NL63_spike;1254 | 0 | 0 |
| NC_005831.2;YP_003767.1;HCoV-NL63_spike;1255 | 0 | 0 |
| NC_005831.2;YP_003767.1;HCoV-NL63_spike;1256 | 0 | 0 |
| NC_005831.2;YP_003767.1;HCoV-NL63_spike;1257 | 0 | 0 |
| NC_005831.2;YP_003767.1;HCoV-NL63_spike;1258 | 0 | 0 |
| NC_005831.2;YP_003767.1;HCoV-NL63_spike;1259 | 0 | 0 |
| NC_005831.2;YP_003767.1;HCoV-NL63_spike;1260 | 0 | 0 |
| NC_005831.2;YP_003767.1;HCoV-NL63_spike;1261 | 0 | 0 |
| NC_005831.2;YP_003767.1;HCoV-NL63_spike;1262 | 0 | 0.05 |
| NC_005831.2;YP_003767.1;HCoV-NL63_spike;1263 | 0 | 0.05 |
| NC_005831.2;YP_003767.1;HCoV-NL63_spike;1264 | 0 | 0.05 |
| NC_005831.2;YP_003767.1;HCoV-NL63_spike;1265 | 0 | 0.225 |
| NC_005831.2;YP_003767.1;HCoV-NL63_spike;1266 | 0 | 0 |
| NC_005831.2;YP_003767.1;HCoV-NL63_spike;1267 | 0 | 0 |
| NC_005831.2;YP_003767.1;HCoV-NL63_spike;1268 | 0 | 0 |
| NC_005831.2;YP_003767.1;HCoV-NL63_spike;1269 | 0 | 0.05 |
| NC_005831.2;YP_003767.1;HCoV-NL63_spike;1270 | 0 | 0.05 |
| NC_005831.2;YP_003767.1;HCoV-NL63_spike;1271 | 0.2 | 0.175 |
| NC_005831.2;YP_003767.1;HCoV-NL63_spike;1272 | 0.3 | 0.225 |
| NC_005831.2;YP_003767.1;HCoV-NL63_spike;1273 | 0.05 | 0.025 |
| NC_005831.2;YP_003767.1;HCoV-NL63_spike;1274 | 0 | 0 |
| NC_005831.2;YP_003767.1;HCoV-NL63_spike;1275 | 0 | 0.025 |
| NC_005831.2;YP_003767.1;HCoV-NL63_spike;1276 | 0 | 0 |
| NC_005831.2;YP_003767.1;HCoV-NL63_spike;1277 | 0.05 | 0 |
| NC_005831.2;YP_003767.1;HCoV-NL63_spike;1278 | 0 | 0.025 |
| NC_005831.2;YP_003767.1;HCoV-NL63_spike;1279 | 0 | 0 |
| NC_005831.2;YP_003767.1;HCoV-NL63_spike;1280 | 0 | 0 |
| NC_005831.2;YP_003767.1;HCoV-NL63_spike;1281 | 0.05 | 0 |
| NC_005831.2;YP_003767.1;HCoV-NL63_spike;1282 | 0 | 0 |
| NC_005831.2;YP_003767.1;HCoV-NL63_spike;1283 | 0 | 0 |
| NC_005831.2;YP_003767.1;HCoV-NL63_spike;1284 | 0 | 0 |
| NC_005831.2;YP_003767.1;HCoV-NL63_spike;1285 | 0 | 0 |
| NC_005831.2;YP_003767.1;HCoV-NL63_spike;1286 | 0 | 0 |
| NC_005831.2;YP_003767.1;HCoV-NL63_spike;1287 | 0 | 0 |
| NC_005831.2;YP_003767.1;HCoV-NL63_spike;1288 | 0 | 0 |
| NC_005831.2;YP_003767.1;HCoV-NL63_spike;1289 | 0 | 0.025 |

| | | |
|---|---|---|
| NC_005831.2;YP_003767.1;HCoV-NL63_spike;1290 | 0 | 0 |
| NC_005831.2;YP_003767.1;HCoV-NL63_spike;1291 | 0 | 0 |
| NC_005831.2;YP_003767.1;HCoV-NL63_spike;1292 | 0 | 0 |
| NC_005831.2;YP_003767.1;HCoV-NL63_spike;1293 | 0 | 0 |
| NC_005831.2;YP_003767.1;HCoV-NL63_spike;1294 | 0 | 0 |
| NC_005831.2;YP_003767.1;HCoV-NL63_spike;1295 | 0 | 0 |
| NC_005831.2;YP_003767.1;HCoV-NL63_spike;1296 | 0 | 0 |
| NC_005831.2;YP_003767.1;HCoV-NL63_spike;1297 | 0 | 0 |
| NC_005831.2;YP_003767.1;HCoV-NL63_spike;1298 | 0 | 0 |
| NC_005831.2;YP_003767.1;HCoV-NL63_spike;1299 | 0 | 0 |
| NC_005831.2;YP_003767.1;HCoV-NL63_spike;1300 | 0 | 0 |
| NC_005831.2;YP_003767.1;HCoV-NL63_spike;1301 | 0 | 0 |
| NC_005831.2;YP_003767.1;HCoV-NL63_spike;1302 | 0 | 0 |
| NC_005831.2;YP_003767.1;HCoV-NL63_spike;1303 | 0 | 0 |
| NC_005831.2;YP_003767.1;HCoV-NL63_spike;1304 | 0 | 0 |
| NC_005831.2;YP_003767.1;HCoV-NL63_spike;1305 | 0 | 0 |
| NC_005831.2;YP_003767.1;HCoV-NL63_spike;1306 | 0 | 0 |
| NC_005831.2;YP_003767.1;HCoV-NL63_spike;1307 | 0 | 0 |
| NC_005831.2;YP_003767.1;HCoV-NL63_spike;1308 | 0 | 0 |
| NC_005831.2;YP_003767.1;HCoV-NL63_spike;1309 | 0 | 0 |
| NC_005831.2;YP_003767.1;HCoV-NL63_spike;1310 | 0 | 0 |
| NC_005831.2;YP_003767.1;HCoV-NL63_spike;1311 | 0 | 0 |
| NC_005831.2;YP_003767.1;HCoV-NL63_spike;1312 | 0 | 0 |
| NC_005831.2;YP_003767.1;HCoV-NL63_spike;1313 | 0 | 0 |
| NC_005831.2;YP_003767.1;HCoV-NL63_spike;1314 | 0 | 0 |
| NC_005831.2;YP_003767.1;HCoV-NL63_spike;1315 | 0 | 0 |
| NC_005831.2;YP_003767.1;HCoV-NL63_spike;1316 | 0 | 0 |
| NC_005831.2;YP_003767.1;HCoV-NL63_spike;1317 | 0 | 0 |
| NC_005831.2;YP_003767.1;HCoV-NL63_spike;1318 | 0 | 0 |
| NC_005831.2;YP_003767.1;HCoV-NL63_spike;1319 | 0 | 0 |
| NC_005831.2;YP_003767.1;HCoV-NL63_spike;1320 | 0 | 0 |
| NC_005831.2;YP_003767.1;HCoV-NL63_spike;1321 | 0 | 0 |
| NC_005831.2;YP_003767.1;HCoV-NL63_spike;1322 | 0 | 0 |
| NC_005831.2;YP_003767.1;HCoV-NL63_spike;1323 | 0 | 0 |
| NC_005831.2;YP_003767.1;HCoV-NL63_spike;1324 | 0 | 0 |
| NC_005831.2;YP_003767.1;HCoV-NL63_spike;1325 | 0 | 0 |
| NC_005831.2;YP_003767.1;HCoV-NL63_spike;1326 | 0 | 0 |
| NC_005831.2;YP_003767.1;HCoV-NL63_spike;1327 | 0 | 0 |
| NC_005831.2;YP_003767.1;HCoV-NL63_spike;1328 | 0 | 0 |
| NC_005831.2;YP_003767.1;HCoV-NL63_spike;1329 | 0 | 0 |
| NC_005831.2;YP_003767.1;HCoV-NL63_spike;1330 | 0 | 0 |
| NC_005831.2;YP_003767.1;HCoV-NL63_spike;1331 | 0 | 0 |
| NC_005831.2;YP_003767.1;HCoV-NL63_spike;1332 | 0 | 0 |

| | | |
|---|---|---|
| NC_005831.2;YP_003767.1;HCoV-NL63_spike;1333 | 0 | 0 |
| NC_005831.2;YP_003767.1;HCoV-NL63_spike;1334 | 0 | 0.025 |
| NC_005831.2;YP_003767.1;HCoV-NL63_spike;1335 | 0 | 0.025 |
| NC_005831.2;YP_003767.1;HCoV-NL63_spike;1336 | 0.05 | 0.1 |
| NC_005831.2;YP_003767.1;HCoV-NL63_spike;1337 | 0.35 | 0.225 |
| NC_005831.2;YP_003767.1;HCoV-NL63_spike;1338 | 0.35 | 0.25 |
| NC_005831.2;YP_003767.1;HCoV-NL63_spike;1339 | 0.35 | 0.225 |
| NC_005831.2;YP_003767.1;HCoV-NL63_spike;1340 | 0.3 | 0.2 |
| NC_005831.2;YP_003767.1;HCoV-NL63_spike;1341 | 0.45 | 0.25 |
| NC_005831.2;YP_003767.1;HCoV-NL63_spike;1342 | 0.4 | 0.225 |
| NC_005831.2;YP_003767.1;HCoV-NL63_spike;1343 | 0.4 | 0.225 |
| NC_005831.2;YP_003767.1;HCoV-NL63_spike;1344 | 0.45 | 0.225 |
| NC_005831.2;YP_003767.1;HCoV-NL63_spike;1345 | 0.35 | 0.175 |
| NC_005831.2;YP_003768.1;HCoV-NL63_3;1 | 0 | 0 |
| NC_005831.2;YP_003768.1;HCoV-NL63_3;2 | 0.05 | 0 |
| NC_005831.2;YP_003768.1;HCoV-NL63_3;3 | 0 | 0 |
| NC_005831.2;YP_003768.1;HCoV-NL63_3;4 | 0 | 0 |
| NC_005831.2;YP_003768.1;HCoV-NL63_3;5 | 0.05 | 0 |
| NC_005831.2;YP_003768.1;HCoV-NL63_3;6 | 0 | 0 |
| NC_005831.2;YP_003768.1;HCoV-NL63_3;7 | 0 | 0 |
| NC_005831.2;YP_003768.1;HCoV-NL63_3;8 | 0 | 0.05 |
| NC_005831.2;YP_003768.1;HCoV-NL63_3;9 | 0 | 0 |
| NC_005831.2;YP_003768.1;HCoV-NL63_3;10 | 0.05 | 0.025 |
| NC_005831.2;YP_003768.1;HCoV-NL63_3;11 | 0.05 | 0 |
| NC_005831.2;YP_003768.1;HCoV-NL63_3;12 | 0.05 | 0.1 |
| NC_005831.2;YP_003768.1;HCoV-NL63_3;13 | 0.05 | 0 |
| NC_005831.2;YP_003768.1;HCoV-NL63_3;14 | 0.1 | 0 |
| NC_005831.2;YP_003768.1;HCoV-NL63_3;15 | 0.1 | 0.025 |
| NC_005831.2;YP_003768.1;HCoV-NL63_3;16 | 0.15 | 0.05 |
| NC_005831.2;YP_003768.1;HCoV-NL63_3;17 | 0.05 | 0.05 |
| NC_005831.2;YP_003768.1;HCoV-NL63_3;18 | 0.05 | 0 |
| NC_005831.2;YP_003768.1;HCoV-NL63_3;19 | 0.2 | 0.025 |
| NC_005831.2;YP_003768.1;HCoV-NL63_3;20 | 0.2 | 0.025 |
| NC_005831.2;YP_003768.1;HCoV-NL63_3;21 | 0.2 | 0.025 |
| NC_005831.2;YP_003768.1;HCoV-NL63_3;22 | 0.2 | 0.1 |
| NC_005831.2;YP_003768.1;HCoV-NL63_3;23 | 0.3 | 0.125 |
| NC_005831.2;YP_003768.1;HCoV-NL63_3;24 | 0.15 | 0.05 |
| NC_005831.2;YP_003768.1;HCoV-NL63_3;25 | 0.15 | 0.05 |
| NC_005831.2;YP_003768.1;HCoV-NL63_3;26 | 0.05 | 0.05 |
| NC_005831.2;YP_003768.1;HCoV-NL63_3;27 | 0.05 | 0.05 |
| NC_005831.2;YP_003768.1;HCoV-NL63_3;28 | 0.05 | 0.05 |
| NC_005831.2;YP_003768.1;HCoV-NL63_3;29 | 0.05 | 0.075 |
| NC_005831.2;YP_003768.1;HCoV-NL63_3;30 | 0.05 | 0.025 |

| | | |
|---|---|---|
| NC_005831.2;YP_003768.1;HCoV-NL63_3;31 | 0.05 | 0.025 |
| NC_005831.2;YP_003768.1;HCoV-NL63_3;32 | 0 | 0.05 |
| NC_005831.2;YP_003768.1;HCoV-NL63_3;33 | 0 | 0.025 |
| NC_005831.2;YP_003768.1;HCoV-NL63_3;34 | 0 | 0 |
| NC_005831.2;YP_003768.1;HCoV-NL63_3;35 | 0 | 0 |
| NC_005831.2;YP_003768.1;HCoV-NL63_3;36 | 0 | 0 |
| NC_005831.2;YP_003768.1;HCoV-NL63_3;37 | 0 | 0 |
| NC_005831.2;YP_003768.1;HCoV-NL63_3;38 | 0 | 0 |
| NC_005831.2;YP_003768.1;HCoV-NL63_3;39 | 0 | 0 |
| NC_005831.2;YP_003768.1;HCoV-NL63_3;40 | 0 | 0 |
| NC_005831.2;YP_003768.1;HCoV-NL63_3;41 | 0 | 0 |
| NC_005831.2;YP_003768.1;HCoV-NL63_3;42 | 0 | 0 |
| NC_005831.2;YP_003768.1;HCoV-NL63_3;43 | 0 | 0 |
| NC_005831.2;YP_003768.1;HCoV-NL63_3;44 | 0 | 0 |
| NC_005831.2;YP_003768.1;HCoV-NL63_3;45 | 0 | 0 |
| NC_005831.2;YP_003768.1;HCoV-NL63_3;46 | 0 | 0 |
| NC_005831.2;YP_003768.1;HCoV-NL63_3;47 | 0 | 0 |
| NC_005831.2;YP_003768.1;HCoV-NL63_3;48 | 0 | 0 |
| NC_005831.2;YP_003768.1;HCoV-NL63_3;49 | 0 | 0 |
| NC_005831.2;YP_003768.1;HCoV-NL63_3;50 | 0 | 0 |
| NC_005831.2;YP_003768.1;HCoV-NL63_3;51 | 0 | 0 |
| NC_005831.2;YP_003768.1;HCoV-NL63_3;52 | 0 | 0 |
| NC_005831.2;YP_003768.1;HCoV-NL63_3;53 | 0 | 0 |
| NC_005831.2;YP_003768.1;HCoV-NL63_3;54 | 0 | 0 |
| NC_005831.2;YP_003768.1;HCoV-NL63_3;55 | 0 | 0 |
| NC_005831.2;YP_003768.1;HCoV-NL63_3;56 | 0 | 0 |
| NC_005831.2;YP_003768.1;HCoV-NL63_3;57 | 0 | 0 |
| NC_005831.2;YP_003768.1;HCoV-NL63_3;58 | 0 | 0 |
| NC_005831.2;YP_003768.1;HCoV-NL63_3;59 | 0 | 0 |
| NC_005831.2;YP_003768.1;HCoV-NL63_3;60 | 0 | 0 |
| NC_005831.2;YP_003768.1;HCoV-NL63_3;61 | 0 | 0 |
| NC_005831.2;YP_003768.1;HCoV-NL63_3;62 | 0 | 0 |
| NC_005831.2;YP_003768.1;HCoV-NL63_3;63 | 0 | 0 |
| NC_005831.2;YP_003768.1;HCoV-NL63_3;64 | 0 | 0 |
| NC_005831.2;YP_003768.1;HCoV-NL63_3;65 | 0 | 0 |
| NC_005831.2;YP_003768.1;HCoV-NL63_3;66 | 0 | 0 |
| NC_005831.2;YP_003768.1;HCoV-NL63_3;67 | 0 | 0 |
| NC_005831.2;YP_003768.1;HCoV-NL63_3;68 | 0 | 0 |
| NC_005831.2;YP_003768.1;HCoV-NL63_3;69 | 0 | 0 |
| NC_005831.2;YP_003768.1;HCoV-NL63_3;70 | 0 | 0 |
| NC_005831.2;YP_003768.1;HCoV-NL63_3;71 | 0 | 0 |
| NC_005831.2;YP_003768.1;HCoV-NL63_3;72 | 0 | 0 |
| NC_005831.2;YP_003768.1;HCoV-NL63_3;73 | 0 | 0 |

| | | |
|---|---|---|
| NC_005831.2;YP_003768.1;HCoV-NL63_3;74 | 0 | 0 |
| NC_005831.2;YP_003768.1;HCoV-NL63_3;75 | 0 | 0 |
| NC_005831.2;YP_003768.1;HCoV-NL63_3;76 | 0 | 0 |
| NC_005831.2;YP_003768.1;HCoV-NL63_3;77 | 0 | 0 |
| NC_005831.2;YP_003768.1;HCoV-NL63_3;78 | 0 | 0 |
| NC_005831.2;YP_003768.1;HCoV-NL63_3;79 | 0 | 0 |
| NC_005831.2;YP_003768.1;HCoV-NL63_3;80 | 0 | 0 |
| NC_005831.2;YP_003768.1;HCoV-NL63_3;81 | 0 | 0 |
| NC_005831.2;YP_003768.1;HCoV-NL63_3;82 | 0 | 0 |
| NC_005831.2;YP_003768.1;HCoV-NL63_3;83 | 0 | 0 |
| NC_005831.2;YP_003768.1;HCoV-NL63_3;84 | 0 | 0 |
| NC_005831.2;YP_003768.1;HCoV-NL63_3;85 | 0 | 0 |
| NC_005831.2;YP_003768.1;HCoV-NL63_3;86 | 0 | 0 |
| NC_005831.2;YP_003768.1;HCoV-NL63_3;87 | 0 | 0 |
| NC_005831.2;YP_003768.1;HCoV-NL63_3;88 | 0 | 0 |
| NC_005831.2;YP_003768.1;HCoV-NL63_3;89 | 0 | 0 |
| NC_005831.2;YP_003768.1;HCoV-NL63_3;90 | 0 | 0 |
| NC_005831.2;YP_003768.1;HCoV-NL63_3;91 | 0 | 0 |
| NC_005831.2;YP_003768.1;HCoV-NL63_3;92 | 0 | 0 |
| NC_005831.2;YP_003768.1;HCoV-NL63_3;93 | 0 | 0 |
| NC_005831.2;YP_003768.1;HCoV-NL63_3;94 | 0 | 0 |
| NC_005831.2;YP_003768.1;HCoV-NL63_3;95 | 0 | 0 |
| NC_005831.2;YP_003768.1;HCoV-NL63_3;96 | 0 | 0 |
| NC_005831.2;YP_003768.1;HCoV-NL63_3;97 | 0 | 0 |
| NC_005831.2;YP_003768.1;HCoV-NL63_3;98 | 0 | 0 |
| NC_005831.2;YP_003768.1;HCoV-NL63_3;99 | 0 | 0 |
| NC_005831.2;YP_003768.1;HCoV-NL63_3;100 | 0 | 0 |
| NC_005831.2;YP_003768.1;HCoV-NL63_3;101 | 0 | 0 |
| NC_005831.2;YP_003768.1;HCoV-NL63_3;102 | 0 | 0 |
| NC_005831.2;YP_003768.1;HCoV-NL63_3;103 | 0 | 0 |
| NC_005831.2;YP_003768.1;HCoV-NL63_3;104 | 0 | 0 |
| NC_005831.2;YP_003768.1;HCoV-NL63_3;105 | 0 | 0 |
| NC_005831.2;YP_003768.1;HCoV-NL63_3;106 | 0 | 0 |
| NC_005831.2;YP_003768.1;HCoV-NL63_3;107 | 0 | 0 |
| NC_005831.2;YP_003768.1;HCoV-NL63_3;108 | 0 | 0 |
| NC_005831.2;YP_003768.1;HCoV-NL63_3;109 | 0 | 0 |
| NC_005831.2;YP_003768.1;HCoV-NL63_3;110 | 0 | 0 |
| NC_005831.2;YP_003768.1;HCoV-NL63_3;111 | 0 | 0 |
| NC_005831.2;YP_003768.1;HCoV-NL63_3;112 | 0 | 0 |
| NC_005831.2;YP_003768.1;HCoV-NL63_3;113 | 0 | 0 |
| NC_005831.2;YP_003768.1;HCoV-NL63_3;114 | 0 | 0 |
| NC_005831.2;YP_003768.1;HCoV-NL63_3;115 | 0 | 0 |
| NC_005831.2;YP_003768.1;HCoV-NL63_3;116 | 0 | 0 |

| | | |
|---|---|---|
| NC_005831.2;YP_003768.1;HCoV-NL63_3;117 | 0 | 0 |
| NC_005831.2;YP_003768.1;HCoV-NL63_3;118 | 0 | 0 |
| NC_005831.2;YP_003768.1;HCoV-NL63_3;119 | 0 | 0 |
| NC_005831.2;YP_003768.1;HCoV-NL63_3;120 | 0 | 0 |
| NC_005831.2;YP_003768.1;HCoV-NL63_3;121 | 0 | 0 |
| NC_005831.2;YP_003768.1;HCoV-NL63_3;122 | 0 | 0 |
| NC_005831.2;YP_003768.1;HCoV-NL63_3;123 | 0 | 0 |
| NC_005831.2;YP_003768.1;HCoV-NL63_3;124 | 0 | 0 |
| NC_005831.2;YP_003768.1;HCoV-NL63_3;125 | 0 | 0 |
| NC_005831.2;YP_003768.1;HCoV-NL63_3;126 | 0 | 0 |
| NC_005831.2;YP_003768.1;HCoV-NL63_3;127 | 0 | 0 |
| NC_005831.2;YP_003768.1;HCoV-NL63_3;128 | 0 | 0 |
| NC_005831.2;YP_003768.1;HCoV-NL63_3;129 | 0 | 0 |
| NC_005831.2;YP_003768.1;HCoV-NL63_3;130 | 0 | 0 |
| NC_005831.2;YP_003768.1;HCoV-NL63_3;131 | 0 | 0 |
| NC_005831.2;YP_003768.1;HCoV-NL63_3;132 | 0 | 0 |
| NC_005831.2;YP_003768.1;HCoV-NL63_3;133 | 0 | 0 |
| NC_005831.2;YP_003768.1;HCoV-NL63_3;134 | 0 | 0 |
| NC_005831.2;YP_003768.1;HCoV-NL63_3;135 | 0 | 0 |
| NC_005831.2;YP_003768.1;HCoV-NL63_3;136 | 0.05 | 0 |
| NC_005831.2;YP_003768.1;HCoV-NL63_3;137 | 0 | 0 |
| NC_005831.2;YP_003768.1;HCoV-NL63_3;138 | 0 | 0 |
| NC_005831.2;YP_003768.1;HCoV-NL63_3;139 | 0 | 0 |
| NC_005831.2;YP_003768.1;HCoV-NL63_3;140 | 0 | 0 |
| NC_005831.2;YP_003768.1;HCoV-NL63_3;141 | 0 | 0 |
| NC_005831.2;YP_003768.1;HCoV-NL63_3;142 | 0 | 0 |
| NC_005831.2;YP_003768.1;HCoV-NL63_3;143 | 0 | 0 |
| NC_005831.2;YP_003768.1;HCoV-NL63_3;144 | 0 | 0 |
| NC_005831.2;YP_003768.1;HCoV-NL63_3;145 | 0 | 0 |
| NC_005831.2;YP_003768.1;HCoV-NL63_3;146 | 0 | 0 |
| NC_005831.2;YP_003768.1;HCoV-NL63_3;147 | 0 | 0 |
| NC_005831.2;YP_003768.1;HCoV-NL63_3;148 | 0 | 0 |
| NC_005831.2;YP_003768.1;HCoV-NL63_3;149 | 0 | 0 |
| NC_005831.2;YP_003768.1;HCoV-NL63_3;150 | 0 | 0.025 |
| NC_005831.2;YP_003768.1;HCoV-NL63_3;151 | 0 | 0 |
| NC_005831.2;YP_003768.1;HCoV-NL63_3;152 | 0 | 0 |
| NC_005831.2;YP_003768.1;HCoV-NL63_3;153 | 0 | 0 |
| NC_005831.2;YP_003768.1;HCoV-NL63_3;154 | 0 | 0 |
| NC_005831.2;YP_003768.1;HCoV-NL63_3;155 | 0 | 0 |
| NC_005831.2;YP_003768.1;HCoV-NL63_3;156 | 0 | 0 |
| NC_005831.2;YP_003768.1;HCoV-NL63_3;157 | 0 | 0 |
| NC_005831.2;YP_003768.1;HCoV-NL63_3;158 | 0 | 0 |
| NC_005831.2;YP_003768.1;HCoV-NL63_3;159 | 0 | 0 |

| | | |
|---|---|---|
| NC_005831.2;YP_003768.1;HCoV-NL63_3;160 | 0 | 0 |
| NC_005831.2;YP_003768.1;HCoV-NL63_3;161 | 0 | 0 |
| NC_005831.2;YP_003768.1;HCoV-NL63_3;162 | 0 | 0 |
| NC_005831.2;YP_003768.1;HCoV-NL63_3;163 | 0 | 0 |
| NC_005831.2;YP_003768.1;HCoV-NL63_3;164 | 0 | 0 |
| NC_005831.2;YP_003768.1;HCoV-NL63_3;165 | 0 | 0 |
| NC_005831.2;YP_003768.1;HCoV-NL63_3;166 | 0 | 0 |
| NC_005831.2;YP_003768.1;HCoV-NL63_3;167 | 0 | 0 |
| NC_005831.2;YP_003768.1;HCoV-NL63_3;168 | 0 | 0 |
| NC_005831.2;YP_003768.1;HCoV-NL63_3;169 | 0 | 0 |
| NC_005831.2;YP_003768.1;HCoV-NL63_3;170 | 0 | 0.025 |
| NC_005831.2;YP_003768.1;HCoV-NL63_3;171 | 0 | 0 |
| NC_005831.2;YP_003768.1;HCoV-NL63_3;172 | 0.05 | 0 |
| NC_005831.2;YP_003768.1;HCoV-NL63_3;173 | 0.05 | 0 |
| NC_005831.2;YP_003768.1;HCoV-NL63_3;174 | 0 | 0 |
| NC_005831.2;YP_003768.1;HCoV-NL63_3;175 | 0 | 0 |
| NC_005831.2;YP_003768.1;HCoV-NL63_3;176 | 0 | 0 |
| NC_005831.2;YP_003768.1;HCoV-NL63_3;177 | 0 | 0 |
| NC_005831.2;YP_003768.1;HCoV-NL63_3;178 | 0 | 0 |
| NC_005831.2;YP_003768.1;HCoV-NL63_3;179 | 0 | 0 |
| NC_005831.2;YP_003768.1;HCoV-NL63_3;180 | 0 | 0 |
| NC_005831.2;YP_003768.1;HCoV-NL63_3;181 | 0 | 0.025 |
| NC_005831.2;YP_003768.1;HCoV-NL63_3;182 | 0.05 | 0.025 |
| NC_005831.2;YP_003768.1;HCoV-NL63_3;183 | 0 | 0 |
| NC_005831.2;YP_003768.1;HCoV-NL63_3;184 | 0.05 | 0 |
| NC_005831.2;YP_003768.1;HCoV-NL63_3;185 | 0.05 | 0 |
| NC_005831.2;YP_003768.1;HCoV-NL63_3;186 | 0 | 0 |
| NC_005831.2;YP_003768.1;HCoV-NL63_3;187 | 0 | 0 |
| NC_005831.2;YP_003768.1;HCoV-NL63_3;188 | 0 | 0 |
| NC_005831.2;YP_003768.1;HCoV-NL63_3;189 | 0 | 0.025 |
| NC_005831.2;YP_003768.1;HCoV-NL63_3;190 | 0 | 0 |
| NC_005831.2;YP_003768.1;HCoV-NL63_3;191 | 0 | 0 |
| NC_005831.2;YP_003768.1;HCoV-NL63_3;192 | 0 | 0 |
| NC_005831.2;YP_003768.1;HCoV-NL63_3;193 | 0 | 0 |
| NC_005831.2;YP_003768.1;HCoV-NL63_3;194 | 0 | 0 |
| NC_005831.2;YP_003768.1;HCoV-NL63_3;195 | 0 | 0 |
| NC_005831.2;YP_003768.1;HCoV-NL63_3;196 | 0 | 0 |
| NC_005831.2;YP_003768.1;HCoV-NL63_3;197 | 0 | 0 |
| NC_005831.2;YP_003768.1;HCoV-NL63_3;198 | 0 | 0 |
| NC_005831.2;YP_003768.1;HCoV-NL63_3;199 | 0 | 0 |
| NC_005831.2;YP_003768.1;HCoV-NL63_3;200 | 0 | 0 |
| NC_005831.2;YP_003768.1;HCoV-NL63_3;201 | 0 | 0 |
| NC_005831.2;YP_003768.1;HCoV-NL63_3;202 | 0 | 0 |

| | | |
|---|---|---|
| NC_005831.2;YP_003768.1;HCoV-NL63_3;203 | 0 | 0 |
| NC_005831.2;YP_003768.1;HCoV-NL63_3;204 | 0 | 0 |
| NC_005831.2;YP_003768.1;HCoV-NL63_3;205 | 0 | 0 |
| NC_005831.2;YP_003768.1;HCoV-NL63_3;206 | 0 | 0 |
| NC_005831.2;YP_003768.1;HCoV-NL63_3;207 | 0 | 0.05 |
| NC_005831.2;YP_003768.1;HCoV-NL63_3;208 | 0 | 0.15 |
| NC_005831.2;YP_003768.1;HCoV-NL63_3;209 | 0 | 0.075 |
| NC_005831.2;YP_003768.1;HCoV-NL63_3;210 | 0 | 0.125 |
| NC_005831.2;YP_003768.1;HCoV-NL63_3;211 | 0 | 0.125 |
| NC_005831.2;YP_003768.1;HCoV-NL63_3;212 | 0 | 0.2 |
| NC_005831.2;YP_003768.1;HCoV-NL63_3;213 | 0.05 | 0.15 |
| NC_005831.2;YP_003768.1;HCoV-NL63_3;214 | 0.05 | 0.125 |
| NC_005831.2;YP_003769.1;HCoV-NL63_envelope;1 | 0 | 0.025 |
| NC_005831.2;YP_003769.1;HCoV-NL63_envelope;2 | 0 | 0.025 |
| NC_005831.2;YP_003769.1;HCoV-NL63_envelope;3 | 0 | 0.025 |
| NC_005831.2;YP_003769.1;HCoV-NL63_envelope;4 | 0 | 0 |
| NC_005831.2;YP_003769.1;HCoV-NL63_envelope;5 | 0 | 0 |
| NC_005831.2;YP_003769.1;HCoV-NL63_envelope;6 | 0 | 0 |
| NC_005831.2;YP_003769.1;HCoV-NL63_envelope;7 | 0 | 0 |
| NC_005831.2;YP_003769.1;HCoV-NL63_envelope;8 | 0 | 0 |
| NC_005831.2;YP_003769.1;HCoV-NL63_envelope;9 | 0 | 0 |
| NC_005831.2;YP_003769.1;HCoV-NL63_envelope;10 | 0 | 0 |
| NC_005831.2;YP_003769.1;HCoV-NL63_envelope;11 | 0 | 0 |
| NC_005831.2;YP_003769.1;HCoV-NL63_envelope;12 | 0 | 0 |
| NC_005831.2;YP_003769.1;HCoV-NL63_envelope;13 | 0 | 0 |
| NC_005831.2;YP_003769.1;HCoV-NL63_envelope;14 | 0 | 0 |
| NC_005831.2;YP_003769.1;HCoV-NL63_envelope;15 | 0 | 0 |
| NC_005831.2;YP_003769.1;HCoV-NL63_envelope;16 | 0 | 0 |
| NC_005831.2;YP_003769.1;HCoV-NL63_envelope;17 | 0 | 0 |
| NC_005831.2;YP_003769.1;HCoV-NL63_envelope;18 | 0 | 0 |
| NC_005831.2;YP_003769.1;HCoV-NL63_envelope;19 | 0 | 0 |
| NC_005831.2;YP_003769.1;HCoV-NL63_envelope;20 | 0 | 0 |
| NC_005831.2;YP_003769.1;HCoV-NL63_envelope;21 | 0 | 0 |
| NC_005831.2;YP_003769.1;HCoV-NL63_envelope;22 | 0 | 0 |
| NC_005831.2;YP_003769.1;HCoV-NL63_envelope;23 | 0 | 0 |
| NC_005831.2;YP_003769.1;HCoV-NL63_envelope;24 | 0 | 0 |
| NC_005831.2;YP_003769.1;HCoV-NL63_envelope;25 | 0 | 0 |
| NC_005831.2;YP_003769.1;HCoV-NL63_envelope;26 | 0 | 0 |
| NC_005831.2;YP_003769.1;HCoV-NL63_envelope;27 | 0 | 0 |
| NC_005831.2;YP_003769.1;HCoV-NL63_envelope;28 | 0 | 0 |
| NC_005831.2;YP_003769.1;HCoV-NL63_envelope;29 | 0 | 0 |
| NC_005831.2;YP_003769.1;HCoV-NL63_envelope;30 | 0 | 0 |
| NC_005831.2;YP_003769.1;HCoV-NL63_envelope;31 | 0 | 0 |

| | | |
|---|---|---|
| NC_005831.2;YP_003769.1;HCoV-NL63_envelope;32 | 0 | 0 |
| NC_005831.2;YP_003769.1;HCoV-NL63_envelope;33 | 0 | 0 |
| NC_005831.2;YP_003769.1;HCoV-NL63_envelope;34 | 0 | 0 |
| NC_005831.2;YP_003769.1;HCoV-NL63_envelope;35 | 0 | 0 |
| NC_005831.2;YP_003769.1;HCoV-NL63_envelope;36 | 0 | 0 |
| NC_005831.2;YP_003769.1;HCoV-NL63_envelope;37 | 0 | 0 |
| NC_005831.2;YP_003769.1;HCoV-NL63_envelope;38 | 0 | 0 |
| NC_005831.2;YP_003769.1;HCoV-NL63_envelope;39 | 0 | 0 |
| NC_005831.2;YP_003769.1;HCoV-NL63_envelope;40 | 0 | 0 |
| NC_005831.2;YP_003769.1;HCoV-NL63_envelope;41 | 0 | 0 |
| NC_005831.2;YP_003769.1;HCoV-NL63_envelope;42 | 0 | 0 |
| NC_005831.2;YP_003769.1;HCoV-NL63_envelope;43 | 0 | 0 |
| NC_005831.2;YP_003769.1;HCoV-NL63_envelope;44 | 0 | 0 |
| NC_005831.2;YP_003769.1;HCoV-NL63_envelope;45 | 0 | 0 |
| NC_005831.2;YP_003769.1;HCoV-NL63_envelope;46 | 0 | 0 |
| NC_005831.2;YP_003769.1;HCoV-NL63_envelope;47 | 0 | 0 |
| NC_005831.2;YP_003769.1;HCoV-NL63_envelope;48 | 0 | 0 |
| NC_005831.2;YP_003769.1;HCoV-NL63_envelope;49 | 0 | 0 |
| NC_005831.2;YP_003769.1;HCoV-NL63_envelope;50 | 0 | 0 |
| NC_005831.2;YP_003769.1;HCoV-NL63_envelope;51 | 0 | 0 |
| NC_005831.2;YP_003769.1;HCoV-NL63_envelope;52 | 0 | 0 |
| NC_005831.2;YP_003769.1;HCoV-NL63_envelope;53 | 0 | 0 |
| NC_005831.2;YP_003769.1;HCoV-NL63_envelope;54 | 0 | 0 |
| NC_005831.2;YP_003769.1;HCoV-NL63_envelope;55 | 0 | 0 |
| NC_005831.2;YP_003769.1;HCoV-NL63_envelope;56 | 0 | 0 |
| NC_005831.2;YP_003769.1;HCoV-NL63_envelope;57 | 0 | 0 |
| NC_005831.2;YP_003769.1;HCoV-NL63_envelope;58 | 0.05 | 0 |
| NC_005831.2;YP_003769.1;HCoV-NL63_envelope;59 | 0 | 0.075 |
| NC_005831.2;YP_003769.1;HCoV-NL63_envelope;60 | 0.05 | 0.05 |
| NC_005831.2;YP_003769.1;HCoV-NL63_envelope;61 | 0.15 | 0.075 |
| NC_005831.2;YP_003769.1;HCoV-NL63_envelope;62 | 0 | 0.05 |
| NC_005831.2;YP_003769.1;HCoV-NL63_envelope;63 | 0 | 0.05 |
| NC_005831.2;YP_003769.1;HCoV-NL63_envelope;64 | 0 | 0.05 |
| NC_005831.2;YP_003769.1;HCoV-NL63_envelope;65 | 0.05 | 0.025 |
| NC_005831.2;YP_003769.1;HCoV-NL63_envelope;66 | 0.05 | 0.05 |
| NC_005831.2;YP_003770.1;HCoV-NL63_membrane;1 | 0.1 | 0.05 |
| NC_005831.2;YP_003770.1;HCoV-NL63_membrane;2 | 0.15 | 0.175 |
| NC_005831.2;YP_003770.1;HCoV-NL63_membrane;3 | 0 | 0.025 |
| NC_005831.2;YP_003770.1;HCoV-NL63_membrane;4 | 0 | 0.025 |
| NC_005831.2;YP_003770.1;HCoV-NL63_membrane;5 | 0 | 0.025 |
| NC_005831.2;YP_003770.1;HCoV-NL63_membrane;6 | 0 | 0.025 |
| NC_005831.2;YP_003770.1;HCoV-NL63_membrane;7 | 0 | 0.025 |
| NC_005831.2;YP_003770.1;HCoV-NL63_membrane;8 | 0 | 0 |

| | | |
|---|---|---|
| NC_005831.2;YP_003770.1;HCoV-NL63_membrane;9 | 0 | 0.025 |
| NC_005831.2;YP_003770.1;HCoV-NL63_membrane;10 | 0 | 0.025 |
| NC_005831.2;YP_003770.1;HCoV-NL63_membrane;11 | 0 | 0 |
| NC_005831.2;YP_003770.1;HCoV-NL63_membrane;12 | 0 | 0 |
| NC_005831.2;YP_003770.1;HCoV-NL63_membrane;13 | 0 | 0 |
| NC_005831.2;YP_003770.1;HCoV-NL63_membrane;14 | 0 | 0 |
| NC_005831.2;YP_003770.1;HCoV-NL63_membrane;15 | 0 | 0 |
| NC_005831.2;YP_003770.1;HCoV-NL63_membrane;16 | 0 | 0 |
| NC_005831.2;YP_003770.1;HCoV-NL63_membrane;17 | 0 | 0 |
| NC_005831.2;YP_003770.1;HCoV-NL63_membrane;18 | 0 | 0 |
| NC_005831.2;YP_003770.1;HCoV-NL63_membrane;19 | 0 | 0 |
| NC_005831.2;YP_003770.1;HCoV-NL63_membrane;20 | 0 | 0 |
| NC_005831.2;YP_003770.1;HCoV-NL63_membrane;21 | 0 | 0 |
| NC_005831.2;YP_003770.1;HCoV-NL63_membrane;22 | 0 | 0 |
| NC_005831.2;YP_003770.1;HCoV-NL63_membrane;23 | 0 | 0 |
| NC_005831.2;YP_003770.1;HCoV-NL63_membrane;24 | 0 | 0 |
| NC_005831.2;YP_003770.1;HCoV-NL63_membrane;25 | 0 | 0 |
| NC_005831.2;YP_003770.1;HCoV-NL63_membrane;26 | 0 | 0 |
| NC_005831.2;YP_003770.1;HCoV-NL63_membrane;27 | 0 | 0 |
| NC_005831.2;YP_003770.1;HCoV-NL63_membrane;28 | 0 | 0 |
| NC_005831.2;YP_003770.1;HCoV-NL63_membrane;29 | 0 | 0 |
| NC_005831.2;YP_003770.1;HCoV-NL63_membrane;30 | 0 | 0 |
| NC_005831.2;YP_003770.1;HCoV-NL63_membrane;31 | 0 | 0 |
| NC_005831.2;YP_003770.1;HCoV-NL63_membrane;32 | 0 | 0 |
| NC_005831.2;YP_003770.1;HCoV-NL63_membrane;33 | 0 | 0 |
| NC_005831.2;YP_003770.1;HCoV-NL63_membrane;34 | 0 | 0 |
| NC_005831.2;YP_003770.1;HCoV-NL63_membrane;35 | 0 | 0 |
| NC_005831.2;YP_003770.1;HCoV-NL63_membrane;36 | 0 | 0 |
| NC_005831.2;YP_003770.1;HCoV-NL63_membrane;37 | 0 | 0 |
| NC_005831.2;YP_003770.1;HCoV-NL63_membrane;38 | 0 | 0 |
| NC_005831.2;YP_003770.1;HCoV-NL63_membrane;39 | 0 | 0 |
| NC_005831.2;YP_003770.1;HCoV-NL63_membrane;40 | 0 | 0 |
| NC_005831.2;YP_003770.1;HCoV-NL63_membrane;41 | 0 | 0 |
| NC_005831.2;YP_003770.1;HCoV-NL63_membrane;42 | 0 | 0 |
| NC_005831.2;YP_003770.1;HCoV-NL63_membrane;43 | 0 | 0 |
| NC_005831.2;YP_003770.1;HCoV-NL63_membrane;44 | 0 | 0 |
| NC_005831.2;YP_003770.1;HCoV-NL63_membrane;45 | 0 | 0 |
| NC_005831.2;YP_003770.1;HCoV-NL63_membrane;46 | 0 | 0 |
| NC_005831.2;YP_003770.1;HCoV-NL63_membrane;47 | 0 | 0 |
| NC_005831.2;YP_003770.1;HCoV-NL63_membrane;48 | 0 | 0 |
| NC_005831.2;YP_003770.1;HCoV-NL63_membrane;49 | 0 | 0 |
| NC_005831.2;YP_003770.1;HCoV-NL63_membrane;50 | 0 | 0 |
| NC_005831.2;YP_003770.1;HCoV-NL63_membrane;51 | 0 | 0 |

| | | |
|---|---|---|
| NC_005831.2;YP_003770.1;HCoV-NL63_membrane;52 | 0 | 0 |
| NC_005831.2;YP_003770.1;HCoV-NL63_membrane;53 | 0 | 0 |
| NC_005831.2;YP_003770.1;HCoV-NL63_membrane;54 | 0 | 0 |
| NC_005831.2;YP_003770.1;HCoV-NL63_membrane;55 | 0 | 0 |
| NC_005831.2;YP_003770.1;HCoV-NL63_membrane;56 | 0 | 0 |
| NC_005831.2;YP_003770.1;HCoV-NL63_membrane;57 | 0 | 0 |
| NC_005831.2;YP_003770.1;HCoV-NL63_membrane;58 | 0 | 0 |
| NC_005831.2;YP_003770.1;HCoV-NL63_membrane;59 | 0 | 0 |
| NC_005831.2;YP_003770.1;HCoV-NL63_membrane;60 | 0 | 0 |
| NC_005831.2;YP_003770.1;HCoV-NL63_membrane;61 | 0 | 0 |
| NC_005831.2;YP_003770.1;HCoV-NL63_membrane;62 | 0 | 0 |
| NC_005831.2;YP_003770.1;HCoV-NL63_membrane;63 | 0 | 0 |
| NC_005831.2;YP_003770.1;HCoV-NL63_membrane;64 | 0 | 0 |
| NC_005831.2;YP_003770.1;HCoV-NL63_membrane;65 | 0 | 0 |
| NC_005831.2;YP_003770.1;HCoV-NL63_membrane;66 | 0 | 0 |
| NC_005831.2;YP_003770.1;HCoV-NL63_membrane;67 | 0 | 0 |
| NC_005831.2;YP_003770.1;HCoV-NL63_membrane;68 | 0 | 0 |
| NC_005831.2;YP_003770.1;HCoV-NL63_membrane;69 | 0 | 0 |
| NC_005831.2;YP_003770.1;HCoV-NL63_membrane;70 | 0 | 0 |
| NC_005831.2;YP_003770.1;HCoV-NL63_membrane;71 | 0 | 0 |
| NC_005831.2;YP_003770.1;HCoV-NL63_membrane;72 | 0 | 0 |
| NC_005831.2;YP_003770.1;HCoV-NL63_membrane;73 | 0 | 0 |
| NC_005831.2;YP_003770.1;HCoV-NL63_membrane;74 | 0 | 0 |
| NC_005831.2;YP_003770.1;HCoV-NL63_membrane;75 | 0 | 0 |
| NC_005831.2;YP_003770.1;HCoV-NL63_membrane;76 | 0 | 0 |
| NC_005831.2;YP_003770.1;HCoV-NL63_membrane;77 | 0 | 0 |
| NC_005831.2;YP_003770.1;HCoV-NL63_membrane;78 | 0 | 0 |
| NC_005831.2;YP_003770.1;HCoV-NL63_membrane;79 | 0 | 0 |
| NC_005831.2;YP_003770.1;HCoV-NL63_membrane;80 | 0 | 0 |
| NC_005831.2;YP_003770.1;HCoV-NL63_membrane;81 | 0 | 0 |
| NC_005831.2;YP_003770.1;HCoV-NL63_membrane;82 | 0 | 0 |
| NC_005831.2;YP_003770.1;HCoV-NL63_membrane;83 | 0 | 0 |
| NC_005831.2;YP_003770.1;HCoV-NL63_membrane;84 | 0 | 0 |
| NC_005831.2;YP_003770.1;HCoV-NL63_membrane;85 | 0 | 0 |
| NC_005831.2;YP_003770.1;HCoV-NL63_membrane;86 | 0 | 0 |
| NC_005831.2;YP_003770.1;HCoV-NL63_membrane;87 | 0 | 0 |
| NC_005831.2;YP_003770.1;HCoV-NL63_membrane;88 | 0 | 0 |
| NC_005831.2;YP_003770.1;HCoV-NL63_membrane;89 | 0 | 0 |
| NC_005831.2;YP_003770.1;HCoV-NL63_membrane;90 | 0 | 0 |
| NC_005831.2;YP_003770.1;HCoV-NL63_membrane;91 | 0 | 0 |
| NC_005831.2;YP_003770.1;HCoV-NL63_membrane;92 | 0 | 0 |
| NC_005831.2;YP_003770.1;HCoV-NL63_membrane;93 | 0 | 0 |
| NC_005831.2;YP_003770.1;HCoV-NL63_membrane;94 | 0 | 0 |

| | | |
|---|---|---|
| NC_005831.2;YP_003770.1;HCoV-NL63_membrane;95 | 0 | 0 |
| NC_005831.2;YP_003770.1;HCoV-NL63_membrane;96 | 0 | 0 |
| NC_005831.2;YP_003770.1;HCoV-NL63_membrane;97 | 0 | 0 |
| NC_005831.2;YP_003770.1;HCoV-NL63_membrane;98 | 0 | 0 |
| NC_005831.2;YP_003770.1;HCoV-NL63_membrane;99 | 0 | 0 |
| NC_005831.2;YP_003770.1;HCoV-NL63_membrane;100 | 0 | 0 |
| NC_005831.2;YP_003770.1;HCoV-NL63_membrane;101 | 0 | 0 |
| NC_005831.2;YP_003770.1;HCoV-NL63_membrane;102 | 0 | 0 |
| NC_005831.2;YP_003770.1;HCoV-NL63_membrane;103 | 0 | 0 |
| NC_005831.2;YP_003770.1;HCoV-NL63_membrane;104 | 0 | 0 |
| NC_005831.2;YP_003770.1;HCoV-NL63_membrane;105 | 0 | 0 |
| NC_005831.2;YP_003770.1;HCoV-NL63_membrane;106 | 0 | 0 |
| NC_005831.2;YP_003770.1;HCoV-NL63_membrane;107 | 0 | 0 |
| NC_005831.2;YP_003770.1;HCoV-NL63_membrane;108 | 0 | 0 |
| NC_005831.2;YP_003770.1;HCoV-NL63_membrane;109 | 0 | 0 |
| NC_005831.2;YP_003770.1;HCoV-NL63_membrane;110 | 0 | 0 |
| NC_005831.2;YP_003770.1;HCoV-NL63_membrane;111 | 0 | 0 |
| NC_005831.2;YP_003770.1;HCoV-NL63_membrane;112 | 0 | 0 |
| NC_005831.2;YP_003770.1;HCoV-NL63_membrane;113 | 0 | 0 |
| NC_005831.2;YP_003770.1;HCoV-NL63_membrane;114 | 0 | 0 |
| NC_005831.2;YP_003770.1;HCoV-NL63_membrane;115 | 0 | 0 |
| NC_005831.2;YP_003770.1;HCoV-NL63_membrane;116 | 0 | 0 |
| NC_005831.2;YP_003770.1;HCoV-NL63_membrane;117 | 0 | 0 |
| NC_005831.2;YP_003770.1;HCoV-NL63_membrane;118 | 0 | 0 |
| NC_005831.2;YP_003770.1;HCoV-NL63_membrane;119 | 0 | 0 |
| NC_005831.2;YP_003770.1;HCoV-NL63_membrane;120 | 0 | 0 |
| NC_005831.2;YP_003770.1;HCoV-NL63_membrane;121 | 0.05 | 0.025 |
| NC_005831.2;YP_003770.1;HCoV-NL63_membrane;122 | 0 | 0 |
| NC_005831.2;YP_003770.1;HCoV-NL63_membrane;123 | 0 | 0 |
| NC_005831.2;YP_003770.1;HCoV-NL63_membrane;124 | 0 | 0 |
| NC_005831.2;YP_003770.1;HCoV-NL63_membrane;125 | 0 | 0 |
| NC_005831.2;YP_003770.1;HCoV-NL63_membrane;126 | 0 | 0 |
| NC_005831.2;YP_003770.1;HCoV-NL63_membrane;127 | 0 | 0 |
| NC_005831.2;YP_003770.1;HCoV-NL63_membrane;128 | 0 | 0 |
| NC_005831.2;YP_003770.1;HCoV-NL63_membrane;129 | 0 | 0 |
| NC_005831.2;YP_003770.1;HCoV-NL63_membrane;130 | 0 | 0 |
| NC_005831.2;YP_003770.1;HCoV-NL63_membrane;131 | 0 | 0 |
| NC_005831.2;YP_003770.1;HCoV-NL63_membrane;132 | 0 | 0 |
| NC_005831.2;YP_003770.1;HCoV-NL63_membrane;133 | 0 | 0 |
| NC_005831.2;YP_003770.1;HCoV-NL63_membrane;134 | 0 | 0 |
| NC_005831.2;YP_003770.1;HCoV-NL63_membrane;135 | 0 | 0 |
| NC_005831.2;YP_003770.1;HCoV-NL63_membrane;136 | 0 | 0 |
| NC_005831.2;YP_003770.1;HCoV-NL63_membrane;137 | 0 | 0 |

| | | |
|---|---|---|
| NC_005831.2;YP_003770.1;HCoV-NL63_membrane;138 | 0 | 0 |
| NC_005831.2;YP_003770.1;HCoV-NL63_membrane;139 | 0 | 0 |
| NC_005831.2;YP_003770.1;HCoV-NL63_membrane;140 | 0 | 0 |
| NC_005831.2;YP_003770.1;HCoV-NL63_membrane;141 | 0 | 0 |
| NC_005831.2;YP_003770.1;HCoV-NL63_membrane;142 | 0 | 0 |
| NC_005831.2;YP_003770.1;HCoV-NL63_membrane;143 | 0 | 0 |
| NC_005831.2;YP_003770.1;HCoV-NL63_membrane;144 | 0 | 0 |
| NC_005831.2;YP_003770.1;HCoV-NL63_membrane;145 | 0 | 0 |
| NC_005831.2;YP_003770.1;HCoV-NL63_membrane;146 | 0 | 0 |
| NC_005831.2;YP_003770.1;HCoV-NL63_membrane;147 | 0 | 0 |
| NC_005831.2;YP_003770.1;HCoV-NL63_membrane;148 | 0 | 0 |
| NC_005831.2;YP_003770.1;HCoV-NL63_membrane;149 | 0 | 0 |
| NC_005831.2;YP_003770.1;HCoV-NL63_membrane;150 | 0 | 0 |
| NC_005831.2;YP_003770.1;HCoV-NL63_membrane;151 | 0 | 0 |
| NC_005831.2;YP_003770.1;HCoV-NL63_membrane;152 | 0 | 0 |
| NC_005831.2;YP_003770.1;HCoV-NL63_membrane;153 | 0 | 0 |
| NC_005831.2;YP_003770.1;HCoV-NL63_membrane;154 | 0 | 0 |
| NC_005831.2;YP_003770.1;HCoV-NL63_membrane;155 | 0 | 0 |
| NC_005831.2;YP_003770.1;HCoV-NL63_membrane;156 | 0 | 0 |
| NC_005831.2;YP_003770.1;HCoV-NL63_membrane;157 | 0 | 0 |
| NC_005831.2;YP_003770.1;HCoV-NL63_membrane;158 | 0 | 0 |
| NC_005831.2;YP_003770.1;HCoV-NL63_membrane;159 | 0 | 0 |
| NC_005831.2;YP_003770.1;HCoV-NL63_membrane;160 | 0 | 0 |
| NC_005831.2;YP_003770.1;HCoV-NL63_membrane;161 | 0 | 0 |
| NC_005831.2;YP_003770.1;HCoV-NL63_membrane;162 | 0 | 0 |
| NC_005831.2;YP_003770.1;HCoV-NL63_membrane;163 | 0 | 0 |
| NC_005831.2;YP_003770.1;HCoV-NL63_membrane;164 | 0 | 0 |
| NC_005831.2;YP_003770.1;HCoV-NL63_membrane;165 | 0 | 0 |
| NC_005831.2;YP_003770.1;HCoV-NL63_membrane;166 | 0 | 0 |
| NC_005831.2;YP_003770.1;HCoV-NL63_membrane;167 | 0 | 0 |
| NC_005831.2;YP_003770.1;HCoV-NL63_membrane;168 | 0 | 0 |
| NC_005831.2;YP_003770.1;HCoV-NL63_membrane;169 | 0 | 0 |
| NC_005831.2;YP_003770.1;HCoV-NL63_membrane;170 | 0 | 0 |
| NC_005831.2;YP_003770.1;HCoV-NL63_membrane;171 | 0 | 0 |
| NC_005831.2;YP_003770.1;HCoV-NL63_membrane;172 | 0 | 0 |
| NC_005831.2;YP_003770.1;HCoV-NL63_membrane;173 | 0 | 0 |
| NC_005831.2;YP_003770.1;HCoV-NL63_membrane;174 | 0 | 0 |
| NC_005831.2;YP_003770.1;HCoV-NL63_membrane;175 | 0 | 0 |
| NC_005831.2;YP_003770.1;HCoV-NL63_membrane;176 | 0 | 0 |
| NC_005831.2;YP_003770.1;HCoV-NL63_membrane;177 | 0.05 | 0 |
| NC_005831.2;YP_003770.1;HCoV-NL63_membrane;178 | 0.05 | 0 |
| NC_005831.2;YP_003770.1;HCoV-NL63_membrane;179 | 0.05 | 0 |
| NC_005831.2;YP_003770.1;HCoV-NL63_membrane;180 | 0.05 | 0 |

| | | |
|---|---|---|
| NC_005831.2;YP_003770.1;HCoV-NL63_membrane;181 | 0.05 | 0 |
| NC_005831.2;YP_003770.1;HCoV-NL63_membrane;182 | 0.05 | 0.05 |
| NC_005831.2;YP_003770.1;HCoV-NL63_membrane;183 | 0.05 | 0.025 |
| NC_005831.2;YP_003770.1;HCoV-NL63_membrane;184 | 0 | 0 |
| NC_005831.2;YP_003770.1;HCoV-NL63_membrane;185 | 0 | 0 |
| NC_005831.2;YP_003770.1;HCoV-NL63_membrane;186 | 0 | 0 |
| NC_005831.2;YP_003770.1;HCoV-NL63_membrane;187 | 0 | 0 |
| NC_005831.2;YP_003770.1;HCoV-NL63_membrane;188 | 0 | 0 |
| NC_005831.2;YP_003770.1;HCoV-NL63_membrane;189 | 0 | 0 |
| NC_005831.2;YP_003770.1;HCoV-NL63_membrane;190 | 0 | 0 |
| NC_005831.2;YP_003770.1;HCoV-NL63_membrane;191 | 0 | 0 |
| NC_005831.2;YP_003770.1;HCoV-NL63_membrane;192 | 0 | 0 |
| NC_005831.2;YP_003770.1;HCoV-NL63_membrane;193 | 0 | 0 |
| NC_005831.2;YP_003770.1;HCoV-NL63_membrane;194 | 0 | 0 |
| NC_005831.2;YP_003770.1;HCoV-NL63_membrane;195 | 0.05 | 0.05 |
| NC_005831.2;YP_003770.1;HCoV-NL63_membrane;196 | 0 | 0 |
| NC_005831.2;YP_003770.1;HCoV-NL63_membrane;197 | 0 | 0 |
| NC_005831.2;YP_003770.1;HCoV-NL63_membrane;198 | 0 | 0 |
| NC_005831.2;YP_003770.1;HCoV-NL63_membrane;199 | 0 | 0 |
| NC_005831.2;YP_003770.1;HCoV-NL63_membrane;200 | 0 | 0 |
| NC_005831.2;YP_003770.1;HCoV-NL63_membrane;201 | 0 | 0 |
| NC_005831.2;YP_003770.1;HCoV-NL63_membrane;202 | 0 | 0 |
| NC_005831.2;YP_003770.1;HCoV-NL63_membrane;203 | 0 | 0 |
| NC_005831.2;YP_003770.1;HCoV-NL63_membrane;204 | 0.05 | 0 |
| NC_005831.2;YP_003770.1;HCoV-NL63_membrane;205 | 0.05 | 0 |
| NC_005831.2;YP_003770.1;HCoV-NL63_membrane;206 | 0.05 | 0.025 |
| NC_005831.2;YP_003770.1;HCoV-NL63_membrane;207 | 0.05 | 0 |
| NC_005831.2;YP_003770.1;HCoV-NL63_membrane;208 | 0.05 | 0 |
| NC_005831.2;YP_003770.1;HCoV-NL63_membrane;209 | 0.05 | 0 |
| NC_005831.2;YP_003770.1;HCoV-NL63_membrane;210 | 0.05 | 0 |
| NC_005831.2;YP_003770.1;HCoV-NL63_membrane;211 | 0 | 0 |
| NC_005831.2;YP_003770.1;HCoV-NL63_membrane;212 | 0 | 0 |
| NC_005831.2;YP_003770.1;HCoV-NL63_membrane;213 | 0 | 0 |
| NC_005831.2;YP_003770.1;HCoV-NL63_membrane;214 | 0.05 | 0.1 |
| NC_005831.2;YP_003770.1;HCoV-NL63_membrane;215 | 0.05 | 0 |
| NC_005831.2;YP_003771.1;HCoV-NL63_nucleocap;1 | 0.05 | 0.025 |
| NC_005831.2;YP_003771.1;HCoV-NL63_nucleocap;2 | 0.1 | 0 |
| NC_005831.2;YP_003771.1;HCoV-NL63_nucleocap;3 | 0.05 | 0 |
| NC_005831.2;YP_003771.1;HCoV-NL63_nucleocap;4 | 0.05 | 0.025 |
| NC_005831.2;YP_003771.1;HCoV-NL63_nucleocap;5 | 0.05 | 0 |
| NC_005831.2;YP_003771.1;HCoV-NL63_nucleocap;6 | 0.05 | 0 |
| NC_005831.2;YP_003771.1;HCoV-NL63_nucleocap;7 | 0 | 0 |
| NC_005831.2;YP_003771.1;HCoV-NL63_nucleocap;8 | 0 | 0.025 |

| | | |
|---|---|---|
| NC_005831.2;YP_003771.1;HCoV-NL63_nucleocap;9 | 0 | 0 |
| NC_005831.2;YP_003771.1;HCoV-NL63_nucleocap;10 | 0 | 0 |
| NC_005831.2;YP_003771.1;HCoV-NL63_nucleocap;11 | 0 | 0 |
| NC_005831.2;YP_003771.1;HCoV-NL63_nucleocap;12 | 0 | 0 |
| NC_005831.2;YP_003771.1;HCoV-NL63_nucleocap;13 | 0 | 0 |
| NC_005831.2;YP_003771.1;HCoV-NL63_nucleocap;14 | 0 | 0 |
| NC_005831.2;YP_003771.1;HCoV-NL63_nucleocap;15 | 0 | 0 |
| NC_005831.2;YP_003771.1;HCoV-NL63_nucleocap;16 | 0.05 | 0 |
| NC_005831.2;YP_003771.1;HCoV-NL63_nucleocap;17 | 0.05 | 0 |
| NC_005831.2;YP_003771.1;HCoV-NL63_nucleocap;18 | 0.05 | 0 |
| NC_005831.2;YP_003771.1;HCoV-NL63_nucleocap;19 | 0.05 | 0 |
| NC_005831.2;YP_003771.1;HCoV-NL63_nucleocap;20 | 0.05 | 0.025 |
| NC_005831.2;YP_003771.1;HCoV-NL63_nucleocap;21 | 0.05 | 0 |
| NC_005831.2;YP_003771.1;HCoV-NL63_nucleocap;22 | 0.05 | 0 |
| NC_005831.2;YP_003771.1;HCoV-NL63_nucleocap;23 | 0.05 | 0 |
| NC_005831.2;YP_003771.1;HCoV-NL63_nucleocap;24 | 0.1 | 0 |
| NC_005831.2;YP_003771.1;HCoV-NL63_nucleocap;25 | 0.1 | 0 |
| NC_005831.2;YP_003771.1;HCoV-NL63_nucleocap;26 | 0.05 | 0 |
| NC_005831.2;YP_003771.1;HCoV-NL63_nucleocap;27 | 0 | 0 |
| NC_005831.2;YP_003771.1;HCoV-NL63_nucleocap;28 | 0 | 0 |
| NC_005831.2;YP_003771.1;HCoV-NL63_nucleocap;29 | 0 | 0 |
| NC_005831.2;YP_003771.1;HCoV-NL63_nucleocap;30 | 0 | 0 |
| NC_005831.2;YP_003771.1;HCoV-NL63_nucleocap;31 | 0 | 0 |
| NC_005831.2;YP_003771.1;HCoV-NL63_nucleocap;32 | 0 | 0 |
| NC_005831.2;YP_003771.1;HCoV-NL63_nucleocap;33 | 0 | 0 |
| NC_005831.2;YP_003771.1;HCoV-NL63_nucleocap;34 | 0 | 0 |
| NC_005831.2;YP_003771.1;HCoV-NL63_nucleocap;35 | 0.1 | 0.05 |
| NC_005831.2;YP_003771.1;HCoV-NL63_nucleocap;36 | 0.05 | 0 |
| NC_005831.2;YP_003771.1;HCoV-NL63_nucleocap;37 | 0 | 0 |
| NC_005831.2;YP_003771.1;HCoV-NL63_nucleocap;38 | 0.05 | 0 |
| NC_005831.2;YP_003771.1;HCoV-NL63_nucleocap;39 | 0 | 0 |
| NC_005831.2;YP_003771.1;HCoV-NL63_nucleocap;40 | 0.05 | 0.025 |
| NC_005831.2;YP_003771.1;HCoV-NL63_nucleocap;41 | 0.05 | 0.05 |
| NC_005831.2;YP_003771.1;HCoV-NL63_nucleocap;42 | 0.05 | 0.025 |
| NC_005831.2;YP_003771.1;HCoV-NL63_nucleocap;43 | 0 | 0.05 |
| NC_005831.2;YP_003771.1;HCoV-NL63_nucleocap;44 | 0 | 0.075 |
| NC_005831.2;YP_003771.1;HCoV-NL63_nucleocap;45 | 0 | 0.025 |
| NC_005831.2;YP_003771.1;HCoV-NL63_nucleocap;46 | 0.05 | 0.025 |
| NC_005831.2;YP_003771.1;HCoV-NL63_nucleocap;47 | 0 | 0.025 |
| NC_005831.2;YP_003771.1;HCoV-NL63_nucleocap;48 | 0 | 0 |
| NC_005831.2;YP_003771.1;HCoV-NL63_nucleocap;49 | 0.05 | 0 |
| NC_005831.2;YP_003771.1;HCoV-NL63_nucleocap;50 | 0 | 0 |
| NC_005831.2;YP_003771.1;HCoV-NL63_nucleocap;51 | 0 | 0 |

| | | |
|---|---|---|
| NC_005831.2;YP_003771.1;HCoV-NL63_nucleocap;52 | 0 | 0 |
| NC_005831.2;YP_003771.1;HCoV-NL63_nucleocap;53 | 0 | 0 |
| NC_005831.2;YP_003771.1;HCoV-NL63_nucleocap;54 | 0 | 0 |
| NC_005831.2;YP_003771.1;HCoV-NL63_nucleocap;55 | 0 | 0 |
| NC_005831.2;YP_003771.1;HCoV-NL63_nucleocap;56 | 0 | 0 |
| NC_005831.2;YP_003771.1;HCoV-NL63_nucleocap;57 | 0 | 0 |
| NC_005831.2;YP_003771.1;HCoV-NL63_nucleocap;58 | 0.05 | 0 |
| NC_005831.2;YP_003771.1;HCoV-NL63_nucleocap;59 | 0.15 | 0.05 |
| NC_005831.2;YP_003771.1;HCoV-NL63_nucleocap;60 | 0.1 | 0 |
| NC_005831.2;YP_003771.1;HCoV-NL63_nucleocap;61 | 0.1 | 0.025 |
| NC_005831.2;YP_003771.1;HCoV-NL63_nucleocap;62 | 0.05 | 0.025 |
| NC_005831.2;YP_003771.1;HCoV-NL63_nucleocap;63 | 0 | 0.025 |
| NC_005831.2;YP_003771.1;HCoV-NL63_nucleocap;64 | 0 | 0.025 |
| NC_005831.2;YP_003771.1;HCoV-NL63_nucleocap;65 | 0 | 0.025 |
| NC_005831.2;YP_003771.1;HCoV-NL63_nucleocap;66 | 0 | 0 |
| NC_005831.2;YP_003771.1;HCoV-NL63_nucleocap;67 | 0 | 0 |
| NC_005831.2;YP_003771.1;HCoV-NL63_nucleocap;68 | 0 | 0 |
| NC_005831.2;YP_003771.1;HCoV-NL63_nucleocap;69 | 0 | 0 |
| NC_005831.2;YP_003771.1;HCoV-NL63_nucleocap;70 | 0 | 0 |
| NC_005831.2;YP_003771.1;HCoV-NL63_nucleocap;71 | 0 | 0 |
| NC_005831.2;YP_003771.1;HCoV-NL63_nucleocap;72 | 0 | 0 |
| NC_005831.2;YP_003771.1;HCoV-NL63_nucleocap;73 | 0 | 0 |
| NC_005831.2;YP_003771.1;HCoV-NL63_nucleocap;74 | 0 | 0 |
| NC_005831.2;YP_003771.1;HCoV-NL63_nucleocap;75 | 0 | 0 |
| NC_005831.2;YP_003771.1;HCoV-NL63_nucleocap;76 | 0 | 0 |
| NC_005831.2;YP_003771.1;HCoV-NL63_nucleocap;77 | 0 | 0 |
| NC_005831.2;YP_003771.1;HCoV-NL63_nucleocap;78 | 0 | 0 |
| NC_005831.2;YP_003771.1;HCoV-NL63_nucleocap;79 | 0 | 0 |
| NC_005831.2;YP_003771.1;HCoV-NL63_nucleocap;80 | 0 | 0 |
| NC_005831.2;YP_003771.1;HCoV-NL63_nucleocap;81 | 0 | 0 |
| NC_005831.2;YP_003771.1;HCoV-NL63_nucleocap;82 | 0 | 0 |
| NC_005831.2;YP_003771.1;HCoV-NL63_nucleocap;83 | 0 | 0 |
| NC_005831.2;YP_003771.1;HCoV-NL63_nucleocap;84 | 0 | 0 |
| NC_005831.2;YP_003771.1;HCoV-NL63_nucleocap;85 | 0 | 0 |
| NC_005831.2;YP_003771.1;HCoV-NL63_nucleocap;86 | 0 | 0 |
| NC_005831.2;YP_003771.1;HCoV-NL63_nucleocap;87 | 0 | 0 |
| NC_005831.2;YP_003771.1;HCoV-NL63_nucleocap;88 | 0 | 0 |
| NC_005831.2;YP_003771.1;HCoV-NL63_nucleocap;89 | 0 | 0 |
| NC_005831.2;YP_003771.1;HCoV-NL63_nucleocap;90 | 0 | 0 |
| NC_005831.2;YP_003771.1;HCoV-NL63_nucleocap;91 | 0 | 0 |
| NC_005831.2;YP_003771.1;HCoV-NL63_nucleocap;92 | 0 | 0 |
| NC_005831.2;YP_003771.1;HCoV-NL63_nucleocap;93 | 0.05 | 0 |
| NC_005831.2;YP_003771.1;HCoV-NL63_nucleocap;94 | 0.05 | 0.025 |

| | | |
|---|---|---|
| NC_005831.2;YP_003771.1;HCoV-NL63_nucleocap;95 | 0.15 | 0.025 |
| NC_005831.2;YP_003771.1;HCoV-NL63_nucleocap;96 | 0.15 | 0.075 |
| NC_005831.2;YP_003771.1;HCoV-NL63_nucleocap;97 | 0.2 | 0.075 |
| NC_005831.2;YP_003771.1;HCoV-NL63_nucleocap;98 | 0.15 | 0.1 |
| NC_005831.2;YP_003771.1;HCoV-NL63_nucleocap;99 | 0.15 | 0.1 |
| NC_005831.2;YP_003771.1;HCoV-NL63_nucleocap;100 | 0.3 | 0.15 |
| NC_005831.2;YP_003771.1;HCoV-NL63_nucleocap;101 | 0.2 | 0.1 |
| NC_005831.2;YP_003771.1;HCoV-NL63_nucleocap;102 | 0.1 | 0.05 |
| NC_005831.2;YP_003771.1;HCoV-NL63_nucleocap;103 | 0.1 | 0.05 |
| NC_005831.2;YP_003771.1;HCoV-NL63_nucleocap;104 | 0.1 | 0.05 |
| NC_005831.2;YP_003771.1;HCoV-NL63_nucleocap;105 | 0.05 | 0.025 |
| NC_005831.2;YP_003771.1;HCoV-NL63_nucleocap;106 | 0.1 | 0.025 |
| NC_005831.2;YP_003771.1;HCoV-NL63_nucleocap;107 | 0.1 | 0.025 |
| NC_005831.2;YP_003771.1;HCoV-NL63_nucleocap;108 | 0.05 | 0.025 |
| NC_005831.2;YP_003771.1;HCoV-NL63_nucleocap;109 | 0.1 | 0.05 |
| NC_005831.2;YP_003771.1;HCoV-NL63_nucleocap;110 | 0.05 | 0.05 |
| NC_005831.2;YP_003771.1;HCoV-NL63_nucleocap;111 | 0 | 0.025 |
| NC_005831.2;YP_003771.1;HCoV-NL63_nucleocap;112 | 0 | 0 |
| NC_005831.2;YP_003771.1;HCoV-NL63_nucleocap;113 | 0 | 0.025 |
| NC_005831.2;YP_003771.1;HCoV-NL63_nucleocap;114 | 0 | 0.025 |
| NC_005831.2;YP_003771.1;HCoV-NL63_nucleocap;115 | 0 | 0 |
| NC_005831.2;YP_003771.1;HCoV-NL63_nucleocap;116 | 0.05 | 0.025 |
| NC_005831.2;YP_003771.1;HCoV-NL63_nucleocap;117 | 0.05 | 0.025 |
| NC_005831.2;YP_003771.1;HCoV-NL63_nucleocap;118 | 0 | 0 |
| NC_005831.2;YP_003771.1;HCoV-NL63_nucleocap;119 | 0.1 | 0.025 |
| NC_005831.2;YP_003771.1;HCoV-NL63_nucleocap;120 | 0.1 | 0.075 |
| NC_005831.2;YP_003771.1;HCoV-NL63_nucleocap;121 | 0 | 0.1 |
| NC_005831.2;YP_003771.1;HCoV-NL63_nucleocap;122 | 0 | 0.025 |
| NC_005831.2;YP_003771.1;HCoV-NL63_nucleocap;123 | 0 | 0.025 |
| NC_005831.2;YP_003771.1;HCoV-NL63_nucleocap;124 | 0 | 0 |
| NC_005831.2;YP_003771.1;HCoV-NL63_nucleocap;125 | 0 | 0 |
| NC_005831.2;YP_003771.1;HCoV-NL63_nucleocap;126 | 0 | 0 |
| NC_005831.2;YP_003771.1;HCoV-NL63_nucleocap;127 | 0 | 0 |
| NC_005831.2;YP_003771.1;HCoV-NL63_nucleocap;128 | 0 | 0 |
| NC_005831.2;YP_003771.1;HCoV-NL63_nucleocap;129 | 0 | 0 |
| NC_005831.2;YP_003771.1;HCoV-NL63_nucleocap;130 | 0 | 0 |
| NC_005831.2;YP_003771.1;HCoV-NL63_nucleocap;131 | 0 | 0.025 |
| NC_005831.2;YP_003771.1;HCoV-NL63_nucleocap;132 | 0 | 0 |
| NC_005831.2;YP_003771.1;HCoV-NL63_nucleocap;133 | 0 | 0 |
| NC_005831.2;YP_003771.1;HCoV-NL63_nucleocap;134 | 0.05 | 0 |
| NC_005831.2;YP_003771.1;HCoV-NL63_nucleocap;135 | 0.1 | 0.125 |
| NC_005831.2;YP_003771.1;HCoV-NL63_nucleocap;136 | 0.1 | 0.1 |
| NC_005831.2;YP_003771.1;HCoV-NL63_nucleocap;137 | 0.1 | 0.15 |

| | | |
|---|---|---|
| NC_005831.2;YP_003771.1;HCoV-NL63_nucleocap;138 | 0.1 | 0.125 |
| NC_005831.2;YP_003771.1;HCoV-NL63_nucleocap;139 | 0.05 | 0.1 |
| NC_005831.2;YP_003771.1;HCoV-NL63_nucleocap;140 | 0.05 | 0.075 |
| NC_005831.2;YP_003771.1;HCoV-NL63_nucleocap;141 | 0.1 | 0.05 |
| NC_005831.2;YP_003771.1;HCoV-NL63_nucleocap;142 | 0 | 0.025 |
| NC_005831.2;YP_003771.1;HCoV-NL63_nucleocap;143 | 0 | 0 |
| NC_005831.2;YP_003771.1;HCoV-NL63_nucleocap;144 | 0 | 0.025 |
| NC_005831.2;YP_003771.1;HCoV-NL63_nucleocap;145 | 0 | 0.025 |
| NC_005831.2;YP_003771.1;HCoV-NL63_nucleocap;146 | 0 | 0.025 |
| NC_005831.2;YP_003771.1;HCoV-NL63_nucleocap;147 | 0 | 0.05 |
| NC_005831.2;YP_003771.1;HCoV-NL63_nucleocap;148 | 0 | 0.125 |
| NC_005831.2;YP_003771.1;HCoV-NL63_nucleocap;149 | 0 | 0.05 |
| NC_005831.2;YP_003771.1;HCoV-NL63_nucleocap;150 | 0 | 0.05 |
| NC_005831.2;YP_003771.1;HCoV-NL63_nucleocap;151 | 0 | 0.075 |
| NC_005831.2;YP_003771.1;HCoV-NL63_nucleocap;152 | 0.05 | 0.05 |
| NC_005831.2;YP_003771.1;HCoV-NL63_nucleocap;153 | 0.05 | 0.05 |
| NC_005831.2;YP_003771.1;HCoV-NL63_nucleocap;154 | 0 | 0.05 |
| NC_005831.2;YP_003771.1;HCoV-NL63_nucleocap;155 | 0.05 | 0 |
| NC_005831.2;YP_003771.1;HCoV-NL63_nucleocap;156 | 0.05 | 0 |
| NC_005831.2;YP_003771.1;HCoV-NL63_nucleocap;157 | 0.05 | 0 |
| NC_005831.2;YP_003771.1;HCoV-NL63_nucleocap;158 | 0.05 | 0.025 |
| NC_005831.2;YP_003771.1;HCoV-NL63_nucleocap;159 | 0.05 | 0 |
| NC_005831.2;YP_003771.1;HCoV-NL63_nucleocap;160 | 0.05 | 0.025 |
| NC_005831.2;YP_003771.1;HCoV-NL63_nucleocap;161 | 0.05 | 0.025 |
| NC_005831.2;YP_003771.1;HCoV-NL63_nucleocap;162 | 0.05 | 0.05 |
| NC_005831.2;YP_003771.1;HCoV-NL63_nucleocap;163 | 0 | 0 |
| NC_005831.2;YP_003771.1;HCoV-NL63_nucleocap;164 | 0 | 0 |
| NC_005831.2;YP_003771.1;HCoV-NL63_nucleocap;165 | 0 | 0.025 |
| NC_005831.2;YP_003771.1;HCoV-NL63_nucleocap;166 | 0 | 0.025 |
| NC_005831.2;YP_003771.1;HCoV-NL63_nucleocap;167 | 0 | 0 |
| NC_005831.2;YP_003771.1;HCoV-NL63_nucleocap;168 | 0 | 0 |
| NC_005831.2;YP_003771.1;HCoV-NL63_nucleocap;169 | 0 | 0.025 |
| NC_005831.2;YP_003771.1;HCoV-NL63_nucleocap;170 | 0 | 0.025 |
| NC_005831.2;YP_003771.1;HCoV-NL63_nucleocap;171 | 0.05 | 0.025 |
| NC_005831.2;YP_003771.1;HCoV-NL63_nucleocap;172 | 0.05 | 0.05 |
| NC_005831.2;YP_003771.1;HCoV-NL63_nucleocap;173 | 0.05 | 0.05 |
| NC_005831.2;YP_003771.1;HCoV-NL63_nucleocap;174 | 0 | 0.025 |
| NC_005831.2;YP_003771.1;HCoV-NL63_nucleocap;175 | 0 | 0.05 |
| NC_005831.2;YP_003771.1;HCoV-NL63_nucleocap;176 | 0 | 0.025 |
| NC_005831.2;YP_003771.1;HCoV-NL63_nucleocap;177 | 0.05 | 0 |
| NC_005831.2;YP_003771.1;HCoV-NL63_nucleocap;178 | 0 | 0 |
| NC_005831.2;YP_003771.1;HCoV-NL63_nucleocap;179 | 0 | 0 |
| NC_005831.2;YP_003771.1;HCoV-NL63_nucleocap;180 | 0 | 0 |

| | | |
|---|---|---|
| NC_005831.2;YP_003771.1;HCoV-NL63_nucleocap;181 | 0 | 0 |
| NC_005831.2;YP_003771.1;HCoV-NL63_nucleocap;182 | 0 | 0 |
| NC_005831.2;YP_003771.1;HCoV-NL63_nucleocap;183 | 0 | 0 |
| NC_005831.2;YP_003771.1;HCoV-NL63_nucleocap;184 | 0 | 0 |
| NC_005831.2;YP_003771.1;HCoV-NL63_nucleocap;185 | 0 | 0 |
| NC_005831.2;YP_003771.1;HCoV-NL63_nucleocap;186 | 0.05 | 0 |
| NC_005831.2;YP_003771.1;HCoV-NL63_nucleocap;187 | 0.05 | 0.025 |
| NC_005831.2;YP_003771.1;HCoV-NL63_nucleocap;188 | 0.1 | 0 |
| NC_005831.2;YP_003771.1;HCoV-NL63_nucleocap;189 | 0.1 | 0 |
| NC_005831.2;YP_003771.1;HCoV-NL63_nucleocap;190 | 0.1 | 0 |
| NC_005831.2;YP_003771.1;HCoV-NL63_nucleocap;191 | 0.1 | 0 |
| NC_005831.2;YP_003771.1;HCoV-NL63_nucleocap;192 | 0.1 | 0 |
| NC_005831.2;YP_003771.1;HCoV-NL63_nucleocap;193 | 0.2 | 0.05 |
| NC_005831.2;YP_003771.1;HCoV-NL63_nucleocap;194 | 0.15 | 0.05 |
| NC_005831.2;YP_003771.1;HCoV-NL63_nucleocap;195 | 0.2 | 0.075 |
| NC_005831.2;YP_003771.1;HCoV-NL63_nucleocap;196 | 0.25 | 0.075 |
| NC_005831.2;YP_003771.1;HCoV-NL63_nucleocap;197 | 0.05 | 0.05 |
| NC_005831.2;YP_003771.1;HCoV-NL63_nucleocap;198 | 0.05 | 0.05 |
| NC_005831.2;YP_003771.1;HCoV-NL63_nucleocap;199 | 0.1 | 0.05 |
| NC_005831.2;YP_003771.1;HCoV-NL63_nucleocap;200 | 0.05 | 0.025 |
| NC_005831.2;YP_003771.1;HCoV-NL63_nucleocap;201 | 0.05 | 0.025 |
| NC_005831.2;YP_003771.1;HCoV-NL63_nucleocap;202 | 0.05 | 0.025 |
| NC_005831.2;YP_003771.1;HCoV-NL63_nucleocap;203 | 0.05 | 0.025 |
| NC_005831.2;YP_003771.1;HCoV-NL63_nucleocap;204 | 0.05 | 0.075 |
| NC_005831.2;YP_003771.1;HCoV-NL63_nucleocap;205 | 0.05 | 0.025 |
| NC_005831.2;YP_003771.1;HCoV-NL63_nucleocap;206 | 0.05 | 0.075 |
| NC_005831.2;YP_003771.1;HCoV-NL63_nucleocap;207 | 0.05 | 0.075 |
| NC_005831.2;YP_003771.1;HCoV-NL63_nucleocap;208 | 0.1 | 0.1 |
| NC_005831.2;YP_003771.1;HCoV-NL63_nucleocap;209 | 0.05 | 0.1 |
| NC_005831.2;YP_003771.1;HCoV-NL63_nucleocap;210 | 0.1 | 0.125 |
| NC_005831.2;YP_003771.1;HCoV-NL63_nucleocap;211 | 0.1 | 0.125 |
| NC_005831.2;YP_003771.1;HCoV-NL63_nucleocap;212 | 0.05 | 0.1 |
| NC_005831.2;YP_003771.1;HCoV-NL63_nucleocap;213 | 0.1 | 0.15 |
| NC_005831.2;YP_003771.1;HCoV-NL63_nucleocap;214 | 0.1 | 0.1 |
| NC_005831.2;YP_003771.1;HCoV-NL63_nucleocap;215 | 0.05 | 0 |
| NC_005831.2;YP_003771.1;HCoV-NL63_nucleocap;216 | 0 | 0.1 |
| NC_005831.2;YP_003771.1;HCoV-NL63_nucleocap;217 | 0 | 0.025 |
| NC_005831.2;YP_003771.1;HCoV-NL63_nucleocap;218 | 0 | 0 |
| NC_005831.2;YP_003771.1;HCoV-NL63_nucleocap;219 | 0 | 0.025 |
| NC_005831.2;YP_003771.1;HCoV-NL63_nucleocap;220 | 0 | 0.05 |
| NC_005831.2;YP_003771.1;HCoV-NL63_nucleocap;221 | 0 | 0.025 |
| NC_005831.2;YP_003771.1;HCoV-NL63_nucleocap;222 | 0 | 0.025 |
| NC_005831.2;YP_003771.1;HCoV-NL63_nucleocap;223 | 0 | 0 |

| | | |
|---|---|---|
| NC_005831.2;YP_003771.1;HCoV-NL63_nucleocap;224 | 0 | 0 |
| NC_005831.2;YP_003771.1;HCoV-NL63_nucleocap;225 | 0 | 0 |
| NC_005831.2;YP_003771.1;HCoV-NL63_nucleocap;226 | 0 | 0 |
| NC_005831.2;YP_003771.1;HCoV-NL63_nucleocap;227 | 0 | 0 |
| NC_005831.2;YP_003771.1;HCoV-NL63_nucleocap;228 | 0 | 0 |
| NC_005831.2;YP_003771.1;HCoV-NL63_nucleocap;229 | 0.1 | 0 |
| NC_005831.2;YP_003771.1;HCoV-NL63_nucleocap;230 | 0.1 | 0 |
| NC_005831.2;YP_003771.1;HCoV-NL63_nucleocap;231 | 0.1 | 0 |
| NC_005831.2;YP_003771.1;HCoV-NL63_nucleocap;232 | 0.05 | 0 |
| NC_005831.2;YP_003771.1;HCoV-NL63_nucleocap;233 | 0.05 | 0 |
| NC_005831.2;YP_003771.1;HCoV-NL63_nucleocap;234 | 0.1 | 0 |
| NC_005831.2;YP_003771.1;HCoV-NL63_nucleocap;235 | 0.05 | 0 |
| NC_005831.2;YP_003771.1;HCoV-NL63_nucleocap;236 | 0 | 0 |
| NC_005831.2;YP_003771.1;HCoV-NL63_nucleocap;237 | 0 | 0.025 |
| NC_005831.2;YP_003771.1;HCoV-NL63_nucleocap;238 | 0 | 0 |
| NC_005831.2;YP_003771.1;HCoV-NL63_nucleocap;239 | 0.05 | 0 |
| NC_005831.2;YP_003771.1;HCoV-NL63_nucleocap;240 | 0 | 0 |
| NC_005831.2;YP_003771.1;HCoV-NL63_nucleocap;241 | 0 | 0.025 |
| NC_005831.2;YP_003771.1;HCoV-NL63_nucleocap;242 | 0 | 0 |
| NC_005831.2;YP_003771.1;HCoV-NL63_nucleocap;243 | 0 | 0 |
| NC_005831.2;YP_003771.1;HCoV-NL63_nucleocap;244 | 0 | 0 |
| NC_005831.2;YP_003771.1;HCoV-NL63_nucleocap;245 | 0 | 0 |
| NC_005831.2;YP_003771.1;HCoV-NL63_nucleocap;246 | 0 | 0 |
| NC_005831.2;YP_003771.1;HCoV-NL63_nucleocap;247 | 0 | 0 |
| NC_005831.2;YP_003771.1;HCoV-NL63_nucleocap;248 | 0 | 0 |
| NC_005831.2;YP_003771.1;HCoV-NL63_nucleocap;249 | 0 | 0 |
| NC_005831.2;YP_003771.1;HCoV-NL63_nucleocap;250 | 0 | 0 |
| NC_005831.2;YP_003771.1;HCoV-NL63_nucleocap;251 | 0 | 0 |
| NC_005831.2;YP_003771.1;HCoV-NL63_nucleocap;252 | 0 | 0 |
| NC_005831.2;YP_003771.1;HCoV-NL63_nucleocap;253 | 0 | 0 |
| NC_005831.2;YP_003771.1;HCoV-NL63_nucleocap;254 | 0 | 0 |
| NC_005831.2;YP_003771.1;HCoV-NL63_nucleocap;255 | 0 | 0 |
| NC_005831.2;YP_003771.1;HCoV-NL63_nucleocap;256 | 0 | 0 |
| NC_005831.2;YP_003771.1;HCoV-NL63_nucleocap;257 | 0 | 0 |
| NC_005831.2;YP_003771.1;HCoV-NL63_nucleocap;258 | 0 | 0 |
| NC_005831.2;YP_003771.1;HCoV-NL63_nucleocap;259 | 0 | 0 |
| NC_005831.2;YP_003771.1;HCoV-NL63_nucleocap;260 | 0 | 0 |
| NC_005831.2;YP_003771.1;HCoV-NL63_nucleocap;261 | 0.05 | 0 |
| NC_005831.2;YP_003771.1;HCoV-NL63_nucleocap;262 | 0.1 | 0.025 |
| NC_005831.2;YP_003771.1;HCoV-NL63_nucleocap;263 | 0.2 | 0.025 |
| NC_005831.2;YP_003771.1;HCoV-NL63_nucleocap;264 | 0.4 | 0.1 |
| NC_005831.2;YP_003771.1;HCoV-NL63_nucleocap;265 | 0.25 | 0.075 |
| NC_005831.2;YP_003771.1;HCoV-NL63_nucleocap;266 | 0.2 | 0.05 |

| | | |
|---|---|---|
| NC_005831.2;YP_003771.1;HCoV-NL63_nucleocap;267 | 0.2 | 0.05 |
| NC_005831.2;YP_003771.1;HCoV-NL63_nucleocap;268 | 0.4 | 0.125 |
| NC_005831.2;YP_003771.1;HCoV-NL63_nucleocap;269 | 0.4 | 0.05 |
| NC_005831.2;YP_003771.1;HCoV-NL63_nucleocap;270 | 0.35 | 0.075 |
| NC_005831.2;YP_003771.1;HCoV-NL63_nucleocap;271 | 0.35 | 0.1 |
| NC_005831.2;YP_003771.1;HCoV-NL63_nucleocap;272 | 0.3 | 0.075 |
| NC_005831.2;YP_003771.1;HCoV-NL63_nucleocap;273 | 0 | 0.025 |
| NC_005831.2;YP_003771.1;HCoV-NL63_nucleocap;274 | 0 | 0 |
| NC_005831.2;YP_003771.1;HCoV-NL63_nucleocap;275 | 0 | 0 |
| NC_005831.2;YP_003771.1;HCoV-NL63_nucleocap;276 | 0 | 0 |
| NC_005831.2;YP_003771.1;HCoV-NL63_nucleocap;277 | 0 | 0 |
| NC_005831.2;YP_003771.1;HCoV-NL63_nucleocap;278 | 0.1 | 0.025 |
| NC_005831.2;YP_003771.1;HCoV-NL63_nucleocap;279 | 0.1 | 0 |
| NC_005831.2;YP_003771.1;HCoV-NL63_nucleocap;280 | 0.05 | 0 |
| NC_005831.2;YP_003771.1;HCoV-NL63_nucleocap;281 | 0.05 | 0 |
| NC_005831.2;YP_003771.1;HCoV-NL63_nucleocap;282 | 0.05 | 0 |
| NC_005831.2;YP_003771.1;HCoV-NL63_nucleocap;283 | 0.05 | 0 |
| NC_005831.2;YP_003771.1;HCoV-NL63_nucleocap;284 | 0.05 | 0 |
| NC_005831.2;YP_003771.1;HCoV-NL63_nucleocap;285 | 0.05 | 0 |
| NC_005831.2;YP_003771.1;HCoV-NL63_nucleocap;286 | 0.05 | 0.025 |
| NC_005831.2;YP_003771.1;HCoV-NL63_nucleocap;287 | 0.05 | 0 |
| NC_005831.2;YP_003771.1;HCoV-NL63_nucleocap;288 | 0.05 | 0 |
| NC_005831.2;YP_003771.1;HCoV-NL63_nucleocap;289 | 0 | 0 |
| NC_005831.2;YP_003771.1;HCoV-NL63_nucleocap;290 | 0 | 0 |
| NC_005831.2;YP_003771.1;HCoV-NL63_nucleocap;291 | 0 | 0 |
| NC_005831.2;YP_003771.1;HCoV-NL63_nucleocap;292 | 0 | 0 |
| NC_005831.2;YP_003771.1;HCoV-NL63_nucleocap;293 | 0 | 0 |
| NC_005831.2;YP_003771.1;HCoV-NL63_nucleocap;294 | 0.05 | 0 |
| NC_005831.2;YP_003771.1;HCoV-NL63_nucleocap;295 | 0 | 0 |
| NC_005831.2;YP_003771.1;HCoV-NL63_nucleocap;296 | 0.05 | 0 |
| NC_005831.2;YP_003771.1;HCoV-NL63_nucleocap;297 | 0 | 0 |
| NC_005831.2;YP_003771.1;HCoV-NL63_nucleocap;298 | 0 | 0 |
| NC_005831.2;YP_003771.1;HCoV-NL63_nucleocap;299 | 0 | 0 |
| NC_005831.2;YP_003771.1;HCoV-NL63_nucleocap;300 | 0 | 0 |
| NC_005831.2;YP_003771.1;HCoV-NL63_nucleocap;301 | 0 | 0 |
| NC_005831.2;YP_003771.1;HCoV-NL63_nucleocap;302 | 0 | 0.025 |
| NC_005831.2;YP_003771.1;HCoV-NL63_nucleocap;303 | 0 | 0.025 |
| NC_005831.2;YP_003771.1;HCoV-NL63_nucleocap;304 | 0 | 0.025 |
| NC_005831.2;YP_003771.1;HCoV-NL63_nucleocap;305 | 0 | 0.025 |
| NC_005831.2;YP_003771.1;HCoV-NL63_nucleocap;306 | 0 | 0.025 |
| NC_005831.2;YP_003771.1;HCoV-NL63_nucleocap;307 | 0 | 0.025 |
| NC_005831.2;YP_003771.1;HCoV-NL63_nucleocap;308 | 0 | 0.075 |
| NC_005831.2;YP_003771.1;HCoV-NL63_nucleocap;309 | 0 | 0.05 |

| | | |
|---|---|---|
| NC_005831.2;YP_003771.1;HCoV-NL63_nucleocap;310 | 0 | 0 |
| NC_005831.2;YP_003771.1;HCoV-NL63_nucleocap;311 | 0 | 0 |
| NC_005831.2;YP_003771.1;HCoV-NL63_nucleocap;312 | 0 | 0 |
| NC_005831.2;YP_003771.1;HCoV-NL63_nucleocap;313 | 0 | 0 |
| NC_005831.2;YP_003771.1;HCoV-NL63_nucleocap;314 | 0 | 0.025 |
| NC_005831.2;YP_003771.1;HCoV-NL63_nucleocap;315 | 0 | 0 |
| NC_005831.2;YP_003771.1;HCoV-NL63_nucleocap;316 | 0 | 0 |
| NC_005831.2;YP_003771.1;HCoV-NL63_nucleocap;317 | 0 | 0 |
| NC_005831.2;YP_003771.1;HCoV-NL63_nucleocap;318 | 0 | 0.025 |
| NC_005831.2;YP_003771.1;HCoV-NL63_nucleocap;319 | 0 | 0.025 |
| NC_005831.2;YP_003771.1;HCoV-NL63_nucleocap;320 | 0 | 0.025 |
| NC_005831.2;YP_003771.1;HCoV-NL63_nucleocap;321 | 0 | 0 |
| NC_005831.2;YP_003771.1;HCoV-NL63_nucleocap;322 | 0 | 0 |
| NC_005831.2;YP_003771.1;HCoV-NL63_nucleocap;323 | 0 | 0 |
| NC_005831.2;YP_003771.1;HCoV-NL63_nucleocap;324 | 0.15 | 0.025 |
| NC_005831.2;YP_003771.1;HCoV-NL63_nucleocap;325 | 0 | 0 |
| NC_005831.2;YP_003771.1;HCoV-NL63_nucleocap;326 | 0 | 0 |
| NC_005831.2;YP_003771.1;HCoV-NL63_nucleocap;327 | 0.05 | 0 |
| NC_005831.2;YP_003771.1;HCoV-NL63_nucleocap;328 | 0.05 | 0 |
| NC_005831.2;YP_003771.1;HCoV-NL63_nucleocap;329 | 0.05 | 0 |
| NC_005831.2;YP_003771.1;HCoV-NL63_nucleocap;330 | 0.05 | 0 |
| NC_005831.2;YP_003771.1;HCoV-NL63_nucleocap;331 | 0.05 | 0.025 |
| NC_005831.2;YP_003771.1;HCoV-NL63_nucleocap;332 | 0.05 | 0.025 |
| NC_005831.2;YP_003771.1;HCoV-NL63_nucleocap;333 | 0.05 | 0.025 |
| NC_005831.2;YP_003771.1;HCoV-NL63_nucleocap;334 | 0.05 | 0 |
| NC_005831.2;YP_003771.1;HCoV-NL63_nucleocap;335 | 0.05 | 0 |
| NC_005831.2;YP_003771.1;HCoV-NL63_nucleocap;336 | 0.05 | 0 |
| NC_005831.2;YP_003771.1;HCoV-NL63_nucleocap;337 | 0.05 | 0 |
| NC_005831.2;YP_003771.1;HCoV-NL63_nucleocap;338 | 0 | 0 |
| NC_005831.2;YP_003771.1;HCoV-NL63_nucleocap;339 | 0.05 | 0.025 |
| NC_005831.2;YP_003771.1;HCoV-NL63_nucleocap;340 | 0 | 0.05 |
| NC_005831.2;YP_003771.1;HCoV-NL63_nucleocap;341 | 0.1 | 0.025 |
| NC_005831.2;YP_003771.1;HCoV-NL63_nucleocap;342 | 0 | 0 |
| NC_005831.2;YP_003771.1;HCoV-NL63_nucleocap;343 | 0.05 | 0 |
| NC_005831.2;YP_003771.1;HCoV-NL63_nucleocap;344 | 0.1 | 0.05 |
| NC_005831.2;YP_003771.1;HCoV-NL63_nucleocap;345 | 0.2 | 0.05 |
| NC_005831.2;YP_003771.1;HCoV-NL63_nucleocap;346 | 0.4 | 0.125 |
| NC_005831.2;YP_003771.1;HCoV-NL63_nucleocap;347 | 0.6 | 0.3 |
| NC_005831.2;YP_003771.1;HCoV-NL63_nucleocap;348 | 0.6 | 0.5 |
| NC_005831.2;YP_003771.1;HCoV-NL63_nucleocap;349 | 0.65 | 0.475 |
| NC_005831.2;YP_003771.1;HCoV-NL63_nucleocap;350 | 0.65 | 0.425 |
| NC_005831.2;YP_003771.1;HCoV-NL63_nucleocap;351 | 0.65 | 0.5 |
| NC_005831.2;YP_003771.1;HCoV-NL63_nucleocap;352 | 0.55 | 0.45 |

| | | |
|---|---|---|
| NC_005831.2;YP_003771.1;HCoV-NL63_nucleocap;353 | 0.5 | 0.425 |
| NC_005831.2;YP_003771.1;HCoV-NL63_nucleocap;354 | 0.5 | 0.4 |
| NC_005831.2;YP_003771.1;HCoV-NL63_nucleocap;355 | 0.45 | 0.3 |
| NC_005831.2;YP_003771.1;HCoV-NL63_nucleocap;356 | 0.1 | 0.075 |
| NC_005831.2;YP_003771.1;HCoV-NL63_nucleocap;357 | 0.05 | 0 |
| NC_005831.2;YP_003771.1;HCoV-NL63_nucleocap;358 | 0.05 | 0 |
| NC_005831.2;YP_003771.1;HCoV-NL63_nucleocap;359 | 0.05 | 0 |
| NC_005831.2;YP_003771.1;HCoV-NL63_nucleocap;360 | 0.05 | 0 |
| NC_005831.2;YP_003771.1;HCoV-NL63_nucleocap;361 | 0 | 0.025 |
| NC_005831.2;YP_003771.1;HCoV-NL63_nucleocap;362 | 0 | 0 |
| NC_005831.2;YP_003771.1;HCoV-NL63_nucleocap;363 | 0 | 0.025 |
| NC_005831.2;YP_003771.1;HCoV-NL63_nucleocap;364 | 0 | 0 |
| NC_005831.2;YP_003771.1;HCoV-NL63_nucleocap;365 | 0 | 0.05 |
| NC_005831.2;YP_003771.1;HCoV-NL63_nucleocap;366 | 0 | 0.05 |
| NC_002645.1;NP_073549.1;HCoV-229E_1ab;1 | 0.05 | 0 |
| NC_002645.1;NP_073549.1;HCoV-229E_1ab;2 | 0 | 0 |
| NC_002645.1;NP_073549.1;HCoV-229E_1ab;3 | 0 | 0 |
| NC_002645.1;NP_073549.1;HCoV-229E_1ab;4 | 0 | 0 |
| NC_002645.1;NP_073549.1;HCoV-229E_1ab;5 | 0 | 0 |
| NC_002645.1;NP_073549.1;HCoV-229E_1ab;6 | 0 | 0 |
| NC_002645.1;NP_073549.1;HCoV-229E_1ab;7 | 0 | 0 |
| NC_002645.1;NP_073549.1;HCoV-229E_1ab;8 | 0.05 | 0 |
| NC_002645.1;NP_073549.1;HCoV-229E_1ab;9 | 0 | 0 |
| NC_002645.1;NP_073549.1;HCoV-229E_1ab;10 | 0 | 0.025 |
| NC_002645.1;NP_073549.1;HCoV-229E_1ab;11 | 0 | 0 |
| NC_002645.1;NP_073549.1;HCoV-229E_1ab;12 | 0 | 0 |
| NC_002645.1;NP_073549.1;HCoV-229E_1ab;13 | 0 | 0.025 |
| NC_002645.1;NP_073549.1;HCoV-229E_1ab;14 | 0 | 0.025 |
| NC_002645.1;NP_073549.1;HCoV-229E_1ab;15 | 0 | 0 |
| NC_002645.1;NP_073549.1;HCoV-229E_1ab;16 | 0 | 0 |
| NC_002645.1;NP_073549.1;HCoV-229E_1ab;17 | 0 | 0 |
| NC_002645.1;NP_073549.1;HCoV-229E_1ab;18 | 0 | 0 |
| NC_002645.1;NP_073549.1;HCoV-229E_1ab;19 | 0 | 0 |
| NC_002645.1;NP_073549.1;HCoV-229E_1ab;20 | 0.05 | 0.025 |
| NC_002645.1;NP_073549.1;HCoV-229E_1ab;21 | 0.05 | 0 |
| NC_002645.1;NP_073549.1;HCoV-229E_1ab;22 | 0 | 0 |
| NC_002645.1;NP_073549.1;HCoV-229E_1ab;23 | 0 | 0 |
| NC_002645.1;NP_073549.1;HCoV-229E_1ab;24 | 0 | 0 |
| NC_002645.1;NP_073549.1;HCoV-229E_1ab;25 | 0 | 0 |
| NC_002645.1;NP_073549.1;HCoV-229E_1ab;26 | 0 | 0 |
| NC_002645.1;NP_073549.1;HCoV-229E_1ab;27 | 0 | 0 |
| NC_002645.1;NP_073549.1;HCoV-229E_1ab;28 | 0 | 0 |
| NC_002645.1;NP_073549.1;HCoV-229E_1ab;29 | 0 | 0 |

| | | |
|---|---|---|
| NC_002645.1;NP_073549.1;HCoV-229E_1ab;30 | 0 | 0 |
| NC_002645.1;NP_073549.1;HCoV-229E_1ab;31 | 0 | 0 |
| NC_002645.1;NP_073549.1;HCoV-229E_1ab;32 | 0 | 0 |
| NC_002645.1;NP_073549.1;HCoV-229E_1ab;33 | 0 | 0 |
| NC_002645.1;NP_073549.1;HCoV-229E_1ab;34 | 0 | 0 |
| NC_002645.1;NP_073549.1;HCoV-229E_1ab;35 | 0 | 0 |
| NC_002645.1;NP_073549.1;HCoV-229E_1ab;36 | 0 | 0 |
| NC_002645.1;NP_073549.1;HCoV-229E_1ab;37 | 0 | 0 |
| NC_002645.1;NP_073549.1;HCoV-229E_1ab;38 | 0 | 0 |
| NC_002645.1;NP_073549.1;HCoV-229E_1ab;39 | 0 | 0 |
| NC_002645.1;NP_073549.1;HCoV-229E_1ab;40 | 0 | 0 |
| NC_002645.1;NP_073549.1;HCoV-229E_1ab;41 | 0 | 0 |
| NC_002645.1;NP_073549.1;HCoV-229E_1ab;42 | 0 | 0 |
| NC_002645.1;NP_073549.1;HCoV-229E_1ab;43 | 0 | 0 |
| NC_002645.1;NP_073549.1;HCoV-229E_1ab;44 | 0 | 0 |
| NC_002645.1;NP_073549.1;HCoV-229E_1ab;45 | 0 | 0 |
| NC_002645.1;NP_073549.1;HCoV-229E_1ab;46 | 0.05 | 0.025 |
| NC_002645.1;NP_073549.1;HCoV-229E_1ab;47 | 0 | 0 |
| NC_002645.1;NP_073549.1;HCoV-229E_1ab;48 | 0 | 0 |
| NC_002645.1;NP_073549.1;HCoV-229E_1ab;49 | 0 | 0.025 |
| NC_002645.1;NP_073549.1;HCoV-229E_1ab;50 | 0 | 0 |
| NC_002645.1;NP_073549.1;HCoV-229E_1ab;51 | 0 | 0 |
| NC_002645.1;NP_073549.1;HCoV-229E_1ab;52 | 0 | 0 |
| NC_002645.1;NP_073549.1;HCoV-229E_1ab;53 | 0 | 0 |
| NC_002645.1;NP_073549.1;HCoV-229E_1ab;54 | 0 | 0 |
| NC_002645.1;NP_073549.1;HCoV-229E_1ab;55 | 0.2 | 0.025 |
| NC_002645.1;NP_073549.1;HCoV-229E_1ab;56 | 0 | 0 |
| NC_002645.1;NP_073549.1;HCoV-229E_1ab;57 | 0 | 0 |
| NC_002645.1;NP_073549.1;HCoV-229E_1ab;58 | 0.05 | 0 |
| NC_002645.1;NP_073549.1;HCoV-229E_1ab;59 | 0 | 0 |
| NC_002645.1;NP_073549.1;HCoV-229E_1ab;60 | 0 | 0 |
| NC_002645.1;NP_073549.1;HCoV-229E_1ab;61 | 0 | 0 |
| NC_002645.1;NP_073549.1;HCoV-229E_1ab;62 | 0 | 0.025 |
| NC_002645.1;NP_073549.1;HCoV-229E_1ab;63 | 0 | 0.025 |
| NC_002645.1;NP_073549.1;HCoV-229E_1ab;64 | 0 | 0 |
| NC_002645.1;NP_073549.1;HCoV-229E_1ab;65 | 0 | 0 |
| NC_002645.1;NP_073549.1;HCoV-229E_1ab;66 | 0 | 0 |
| NC_002645.1;NP_073549.1;HCoV-229E_1ab;67 | 0 | 0 |
| NC_002645.1;NP_073549.1;HCoV-229E_1ab;68 | 0 | 0 |
| NC_002645.1;NP_073549.1;HCoV-229E_1ab;69 | 0 | 0 |
| NC_002645.1;NP_073549.1;HCoV-229E_1ab;70 | 0 | 0 |
| NC_002645.1;NP_073549.1;HCoV-229E_1ab;71 | 0 | 0 |
| NC_002645.1;NP_073549.1;HCoV-229E_1ab;72 | 0 | 0 |

| | | |
|---|---|---|
| NC_002645.1;NP_073549.1;HCoV-229E_1ab;73 | 0 | 0 |
| NC_002645.1;NP_073549.1;HCoV-229E_1ab;74 | 0 | 0 |
| NC_002645.1;NP_073549.1;HCoV-229E_1ab;75 | 0 | 0 |
| NC_002645.1;NP_073549.1;HCoV-229E_1ab;76 | 0 | 0 |
| NC_002645.1;NP_073549.1;HCoV-229E_1ab;77 | 0 | 0 |
| NC_002645.1;NP_073549.1;HCoV-229E_1ab;78 | 0 | 0 |
| NC_002645.1;NP_073549.1;HCoV-229E_1ab;79 | 0 | 0 |
| NC_002645.1;NP_073549.1;HCoV-229E_1ab;80 | 0 | 0 |
| NC_002645.1;NP_073549.1;HCoV-229E_1ab;81 | 0 | 0 |
| NC_002645.1;NP_073549.1;HCoV-229E_1ab;82 | 0 | 0 |
| NC_002645.1;NP_073549.1;HCoV-229E_1ab;83 | 0 | 0 |
| NC_002645.1;NP_073549.1;HCoV-229E_1ab;84 | 0 | 0 |
| NC_002645.1;NP_073549.1;HCoV-229E_1ab;85 | 0 | 0 |
| NC_002645.1;NP_073549.1;HCoV-229E_1ab;86 | 0 | 0 |
| NC_002645.1;NP_073549.1;HCoV-229E_1ab;87 | 0 | 0 |
| NC_002645.1;NP_073549.1;HCoV-229E_1ab;88 | 0 | 0 |
| NC_002645.1;NP_073549.1;HCoV-229E_1ab;89 | 0.2 | 0.05 |
| NC_002645.1;NP_073549.1;HCoV-229E_1ab;90 | 0.05 | 0 |
| NC_002645.1;NP_073549.1;HCoV-229E_1ab;91 | 0 | 0 |
| NC_002645.1;NP_073549.1;HCoV-229E_1ab;92 | 0 | 0 |
| NC_002645.1;NP_073549.1;HCoV-229E_1ab;93 | 0 | 0 |
| NC_002645.1;NP_073549.1;HCoV-229E_1ab;94 | 0 | 0.025 |
| NC_002645.1;NP_073549.1;HCoV-229E_1ab;95 | 0.05 | 0 |
| NC_002645.1;NP_073549.1;HCoV-229E_1ab;96 | 0.1 | 0.05 |
| NC_002645.1;NP_073549.1;HCoV-229E_1ab;97 | 0.05 | 0 |
| NC_002645.1;NP_073549.1;HCoV-229E_1ab;98 | 0 | 0.025 |
| NC_002645.1;NP_073549.1;HCoV-229E_1ab;99 | 0.05 | 0 |
| NC_002645.1;NP_073549.1;HCoV-229E_1ab;100 | 0 | 0 |
| NC_002645.1;NP_073549.1;HCoV-229E_1ab;101 | 0 | 0 |
| NC_002645.1;NP_073549.1;HCoV-229E_1ab;102 | 0 | 0 |
| NC_002645.1;NP_073549.1;HCoV-229E_1ab;103 | 0.05 | 0 |
| NC_002645.1;NP_073549.1;HCoV-229E_1ab;104 | 0 | 0 |
| NC_002645.1;NP_073549.1;HCoV-229E_1ab;105 | 0 | 0 |
| NC_002645.1;NP_073549.1;HCoV-229E_1ab;106 | 0 | 0 |
| NC_002645.1;NP_073549.1;HCoV-229E_1ab;107 | 0.05 | 0.05 |
| NC_002645.1;NP_073549.1;HCoV-229E_1ab;108 | 0 | 0 |
| NC_002645.1;NP_073549.1;HCoV-229E_1ab;109 | 0 | 0 |
| NC_002645.1;NP_073549.1;HCoV-229E_1ab;110 | 0 | 0 |
| NC_002645.1;NP_073549.1;HCoV-229E_1ab;111 | 0 | 0 |
| NC_002645.1;NP_073549.1;HCoV-229E_1ab;112 | 0 | 0 |
| NC_002645.1;NP_073549.1;HCoV-229E_1ab;113 | 0.05 | 0.025 |
| NC_002645.1;NP_073549.1;HCoV-229E_1ab;114 | 0.05 | 0.025 |
| NC_002645.1;NP_073549.1;HCoV-229E_1ab;115 | 0.05 | 0.025 |

| | | |
|---|---|---|
| NC_002645.1;NP_073549.1;HCoV-229E_1ab;116 | 0.05 | 0.025 |
| NC_002645.1;NP_073549.1;HCoV-229E_1ab;117 | 0.05 | 0.025 |
| NC_002645.1;NP_073549.1;HCoV-229E_1ab;118 | 0.05 | 0.05 |
| NC_002645.1;NP_073549.1;HCoV-229E_1ab;119 | 0.05 | 0.075 |
| NC_002645.1;NP_073549.1;HCoV-229E_1ab;120 | 0.05 | 0.025 |
| NC_002645.1;NP_073549.1;HCoV-229E_1ab;121 | 0.05 | 0.025 |
| NC_002645.1;NP_073549.1;HCoV-229E_1ab;122 | 0 | 0 |
| NC_002645.1;NP_073549.1;HCoV-229E_1ab;123 | 0 | 0 |
| NC_002645.1;NP_073549.1;HCoV-229E_1ab;124 | 0 | 0 |
| NC_002645.1;NP_073549.1;HCoV-229E_1ab;125 | 0 | 0 |
| NC_002645.1;NP_073549.1;HCoV-229E_1ab;126 | 0 | 0 |
| NC_002645.1;NP_073549.1;HCoV-229E_1ab;127 | 0 | 0 |
| NC_002645.1;NP_073549.1;HCoV-229E_1ab;128 | 0 | 0 |
| NC_002645.1;NP_073549.1;HCoV-229E_1ab;129 | 0 | 0.025 |
| NC_002645.1;NP_073549.1;HCoV-229E_1ab;130 | 0 | 0.025 |
| NC_002645.1;NP_073549.1;HCoV-229E_1ab;131 | 0 | 0.025 |
| NC_002645.1;NP_073549.1;HCoV-229E_1ab;132 | 0 | 0.025 |
| NC_002645.1;NP_073549.1;HCoV-229E_1ab;133 | 0 | 0.025 |
| NC_002645.1;NP_073549.1;HCoV-229E_1ab;134 | 0 | 0.025 |
| NC_002645.1;NP_073549.1;HCoV-229E_1ab;135 | 0 | 0.025 |
| NC_002645.1;NP_073549.1;HCoV-229E_1ab;136 | 0 | 0.025 |
| NC_002645.1;NP_073549.1;HCoV-229E_1ab;137 | 0 | 0.025 |
| NC_002645.1;NP_073549.1;HCoV-229E_1ab;138 | 0 | 0 |
| NC_002645.1;NP_073549.1;HCoV-229E_1ab;139 | 0 | 0 |
| NC_002645.1;NP_073549.1;HCoV-229E_1ab;140 | 0 | 0 |
| NC_002645.1;NP_073549.1;HCoV-229E_1ab;141 | 0 | 0 |
| NC_002645.1;NP_073549.1;HCoV-229E_1ab;142 | 0 | 0.025 |
| NC_002645.1;NP_073549.1;HCoV-229E_1ab;143 | 0 | 0 |
| NC_002645.1;NP_073549.1;HCoV-229E_1ab;144 | 0 | 0 |
| NC_002645.1;NP_073549.1;HCoV-229E_1ab;145 | 0 | 0 |
| NC_002645.1;NP_073549.1;HCoV-229E_1ab;146 | 0 | 0 |
| NC_002645.1;NP_073549.1;HCoV-229E_1ab;147 | 0 | 0 |
| NC_002645.1;NP_073549.1;HCoV-229E_1ab;148 | 0 | 0 |
| NC_002645.1;NP_073549.1;HCoV-229E_1ab;149 | 0 | 0 |
| NC_002645.1;NP_073549.1;HCoV-229E_1ab;150 | 0 | 0 |
| NC_002645.1;NP_073549.1;HCoV-229E_1ab;151 | 0 | 0 |
| NC_002645.1;NP_073549.1;HCoV-229E_1ab;152 | 0 | 0 |
| NC_002645.1;NP_073549.1;HCoV-229E_1ab;153 | 0 | 0 |
| NC_002645.1;NP_073549.1;HCoV-229E_1ab;154 | 0 | 0 |
| NC_002645.1;NP_073549.1;HCoV-229E_1ab;155 | 0 | 0 |
| NC_002645.1;NP_073549.1;HCoV-229E_1ab;156 | 0.05 | 0 |
| NC_002645.1;NP_073549.1;HCoV-229E_1ab;157 | 0 | 0 |
| NC_002645.1;NP_073549.1;HCoV-229E_1ab;158 | 0 | 0 |

| | | |
|---|---|---|
| NC_002645.1;NP_073549.1;HCoV-229E_1ab;159 | 0 | 0 |
| NC_002645.1;NP_073549.1;HCoV-229E_1ab;160 | 0 | 0.05 |
| NC_002645.1;NP_073549.1;HCoV-229E_1ab;161 | 0 | 0.025 |
| NC_002645.1;NP_073549.1;HCoV-229E_1ab;162 | 0 | 0 |
| NC_002645.1;NP_073549.1;HCoV-229E_1ab;163 | 0.05 | 0.025 |
| NC_002645.1;NP_073549.1;HCoV-229E_1ab;164 | 0.05 | 0.025 |
| NC_002645.1;NP_073549.1;HCoV-229E_1ab;165 | 0 | 0.025 |
| NC_002645.1;NP_073549.1;HCoV-229E_1ab;166 | 0.05 | 0.025 |
| NC_002645.1;NP_073549.1;HCoV-229E_1ab;167 | 0.05 | 0.05 |
| NC_002645.1;NP_073549.1;HCoV-229E_1ab;168 | 0.05 | 0.025 |
| NC_002645.1;NP_073549.1;HCoV-229E_1ab;169 | 0 | 0.025 |
| NC_002645.1;NP_073549.1;HCoV-229E_1ab;170 | 0.05 | 0.05 |
| NC_002645.1;NP_073549.1;HCoV-229E_1ab;171 | 0.05 | 0.05 |
| NC_002645.1;NP_073549.1;HCoV-229E_1ab;172 | 0 | 0.025 |
| NC_002645.1;NP_073549.1;HCoV-229E_1ab;173 | 0 | 0.025 |
| NC_002645.1;NP_073549.1;HCoV-229E_1ab;174 | 0 | 0 |
| NC_002645.1;NP_073549.1;HCoV-229E_1ab;175 | 0 | 0.025 |
| NC_002645.1;NP_073549.1;HCoV-229E_1ab;176 | 0 | 0.025 |
| NC_002645.1;NP_073549.1;HCoV-229E_1ab;177 | 0 | 0 |
| NC_002645.1;NP_073549.1;HCoV-229E_1ab;178 | 0 | 0 |
| NC_002645.1;NP_073549.1;HCoV-229E_1ab;179 | 0 | 0 |
| NC_002645.1;NP_073549.1;HCoV-229E_1ab;180 | 0 | 0 |
| NC_002645.1;NP_073549.1;HCoV-229E_1ab;181 | 0 | 0 |
| NC_002645.1;NP_073549.1;HCoV-229E_1ab;182 | 0 | 0 |
| NC_002645.1;NP_073549.1;HCoV-229E_1ab;183 | 0 | 0 |
| NC_002645.1;NP_073549.1;HCoV-229E_1ab;184 | 0 | 0 |
| NC_002645.1;NP_073549.1;HCoV-229E_1ab;185 | 0 | 0 |
| NC_002645.1;NP_073549.1;HCoV-229E_1ab;186 | 0 | 0.025 |
| NC_002645.1;NP_073549.1;HCoV-229E_1ab;187 | 0 | 0.025 |
| NC_002645.1;NP_073549.1;HCoV-229E_1ab;188 | 0 | 0.025 |
| NC_002645.1;NP_073549.1;HCoV-229E_1ab;189 | 0.15 | 0.05 |
| NC_002645.1;NP_073549.1;HCoV-229E_1ab;190 | 0 | 0.025 |
| NC_002645.1;NP_073549.1;HCoV-229E_1ab;191 | 0 | 0.025 |
| NC_002645.1;NP_073549.1;HCoV-229E_1ab;192 | 0 | 0.025 |
| NC_002645.1;NP_073549.1;HCoV-229E_1ab;193 | 0 | 0.025 |
| NC_002645.1;NP_073549.1;HCoV-229E_1ab;194 | 0 | 0.025 |
| NC_002645.1;NP_073549.1;HCoV-229E_1ab;195 | 0.05 | 0.025 |
| NC_002645.1;NP_073549.1;HCoV-229E_1ab;196 | 0 | 0.025 |
| NC_002645.1;NP_073549.1;HCoV-229E_1ab;197 | 0 | 0 |
| NC_002645.1;NP_073549.1;HCoV-229E_1ab;198 | 0 | 0 |
| NC_002645.1;NP_073549.1;HCoV-229E_1ab;199 | 0 | 0 |
| NC_002645.1;NP_073549.1;HCoV-229E_1ab;200 | 0 | 0 |
| NC_002645.1;NP_073549.1;HCoV-229E_1ab;201 | 0 | 0 |

| | | |
|---|---|---|
| NC_002645.1;NP_073549.1;HCoV-229E_1ab;202 | 0 | 0.025 |
| NC_002645.1;NP_073549.1;HCoV-229E_1ab;203 | 0.05 | 0.025 |
| NC_002645.1;NP_073549.1;HCoV-229E_1ab;204 | 0 | 0.025 |
| NC_002645.1;NP_073549.1;HCoV-229E_1ab;205 | 0 | 0 |
| NC_002645.1;NP_073549.1;HCoV-229E_1ab;206 | 0 | 0.025 |
| NC_002645.1;NP_073549.1;HCoV-229E_1ab;207 | 0.05 | 0 |
| NC_002645.1;NP_073549.1;HCoV-229E_1ab;208 | 0 | 0 |
| NC_002645.1;NP_073549.1;HCoV-229E_1ab;209 | 0 | 0 |
| NC_002645.1;NP_073549.1;HCoV-229E_1ab;210 | 0.05 | 0 |
| NC_002645.1;NP_073549.1;HCoV-229E_1ab;211 | 0 | 0 |
| NC_002645.1;NP_073549.1;HCoV-229E_1ab;212 | 0 | 0 |
| NC_002645.1;NP_073549.1;HCoV-229E_1ab;213 | 0 | 0 |
| NC_002645.1;NP_073549.1;HCoV-229E_1ab;214 | 0 | 0 |
| NC_002645.1;NP_073549.1;HCoV-229E_1ab;215 | 0 | 0 |
| NC_002645.1;NP_073549.1;HCoV-229E_1ab;216 | 0 | 0 |
| NC_002645.1;NP_073549.1;HCoV-229E_1ab;217 | 0 | 0 |
| NC_002645.1;NP_073549.1;HCoV-229E_1ab;218 | 0 | 0 |
| NC_002645.1;NP_073549.1;HCoV-229E_1ab;219 | 0 | 0 |
| NC_002645.1;NP_073549.1;HCoV-229E_1ab;220 | 0 | 0 |
| NC_002645.1;NP_073549.1;HCoV-229E_1ab;221 | 0 | 0.025 |
| NC_002645.1;NP_073549.1;HCoV-229E_1ab;222 | 0.05 | 0.025 |
| NC_002645.1;NP_073549.1;HCoV-229E_1ab;223 | 0 | 0 |
| NC_002645.1;NP_073549.1;HCoV-229E_1ab;224 | 0 | 0 |
| NC_002645.1;NP_073549.1;HCoV-229E_1ab;225 | 0 | 0 |
| NC_002645.1;NP_073549.1;HCoV-229E_1ab;226 | 0 | 0 |
| NC_002645.1;NP_073549.1;HCoV-229E_1ab;227 | 0 | 0 |
| NC_002645.1;NP_073549.1;HCoV-229E_1ab;228 | 0 | 0 |
| NC_002645.1;NP_073549.1;HCoV-229E_1ab;229 | 0 | 0 |
| NC_002645.1;NP_073549.1;HCoV-229E_1ab;230 | 0 | 0 |
| NC_002645.1;NP_073549.1;HCoV-229E_1ab;231 | 0 | 0 |
| NC_002645.1;NP_073549.1;HCoV-229E_1ab;232 | 0 | 0 |
| NC_002645.1;NP_073549.1;HCoV-229E_1ab;233 | 0 | 0 |
| NC_002645.1;NP_073549.1;HCoV-229E_1ab;234 | 0 | 0 |
| NC_002645.1;NP_073549.1;HCoV-229E_1ab;235 | 0 | 0 |
| NC_002645.1;NP_073549.1;HCoV-229E_1ab;236 | 0 | 0 |
| NC_002645.1;NP_073549.1;HCoV-229E_1ab;237 | 0 | 0 |
| NC_002645.1;NP_073549.1;HCoV-229E_1ab;238 | 0 | 0 |
| NC_002645.1;NP_073549.1;HCoV-229E_1ab;239 | 0 | 0 |
| NC_002645.1;NP_073549.1;HCoV-229E_1ab;240 | 0 | 0 |
| NC_002645.1;NP_073549.1;HCoV-229E_1ab;241 | 0 | 0 |
| NC_002645.1;NP_073549.1;HCoV-229E_1ab;242 | 0 | 0 |
| NC_002645.1;NP_073549.1;HCoV-229E_1ab;243 | 0.05 | 0 |
| NC_002645.1;NP_073549.1;HCoV-229E_1ab;244 | 0.05 | 0 |

| | | |
|---|---|---|
| NC_002645.1;NP_073549.1;HCoV-229E_1ab;245 | 0.05 | 0 |
| NC_002645.1;NP_073549.1;HCoV-229E_1ab;246 | 0 | 0 |
| NC_002645.1;NP_073549.1;HCoV-229E_1ab;247 | 0.05 | 0 |
| NC_002645.1;NP_073549.1;HCoV-229E_1ab;248 | 0.05 | 0 |
| NC_002645.1;NP_073549.1;HCoV-229E_1ab;249 | 0.05 | 0 |
| NC_002645.1;NP_073549.1;HCoV-229E_1ab;250 | 0.05 | 0 |
| NC_002645.1;NP_073549.1;HCoV-229E_1ab;251 | 0.05 | 0 |
| NC_002645.1;NP_073549.1;HCoV-229E_1ab;252 | 0 | 0 |
| NC_002645.1;NP_073549.1;HCoV-229E_1ab;253 | 0 | 0 |
| NC_002645.1;NP_073549.1;HCoV-229E_1ab;254 | 0 | 0 |
| NC_002645.1;NP_073549.1;HCoV-229E_1ab;255 | 0 | 0 |
| NC_002645.1;NP_073549.1;HCoV-229E_1ab;256 | 0 | 0 |
| NC_002645.1;NP_073549.1;HCoV-229E_1ab;257 | 0 | 0 |
| NC_002645.1;NP_073549.1;HCoV-229E_1ab;258 | 0 | 0 |
| NC_002645.1;NP_073549.1;HCoV-229E_1ab;259 | 0 | 0 |
| NC_002645.1;NP_073549.1;HCoV-229E_1ab;260 | 0 | 0 |
| NC_002645.1;NP_073549.1;HCoV-229E_1ab;261 | 0 | 0 |
| NC_002645.1;NP_073549.1;HCoV-229E_1ab;262 | 0 | 0 |
| NC_002645.1;NP_073549.1;HCoV-229E_1ab;263 | 0 | 0 |
| NC_002645.1;NP_073549.1;HCoV-229E_1ab;264 | 0 | 0 |
| NC_002645.1;NP_073549.1;HCoV-229E_1ab;265 | 0 | 0 |
| NC_002645.1;NP_073549.1;HCoV-229E_1ab;266 | 0 | 0 |
| NC_002645.1;NP_073549.1;HCoV-229E_1ab;267 | 0 | 0 |
| NC_002645.1;NP_073549.1;HCoV-229E_1ab;268 | 0 | 0 |
| NC_002645.1;NP_073549.1;HCoV-229E_1ab;269 | 0.15 | 0.05 |
| NC_002645.1;NP_073549.1;HCoV-229E_1ab;270 | 0.15 | 0.05 |
| NC_002645.1;NP_073549.1;HCoV-229E_1ab;271 | 0.1 | 0.05 |
| NC_002645.1;NP_073549.1;HCoV-229E_1ab;272 | 0.1 | 0.05 |
| NC_002645.1;NP_073549.1;HCoV-229E_1ab;273 | 0.05 | 0.05 |
| NC_002645.1;NP_073549.1;HCoV-229E_1ab;274 | 0.1 | 0.05 |
| NC_002645.1;NP_073549.1;HCoV-229E_1ab;275 | 0.05 | 0.05 |
| NC_002645.1;NP_073549.1;HCoV-229E_1ab;276 | 0.05 | 0.05 |
| NC_002645.1;NP_073549.1;HCoV-229E_1ab;277 | 0 | 0.05 |
| NC_002645.1;NP_073549.1;HCoV-229E_1ab;278 | 0.05 | 0.025 |
| NC_002645.1;NP_073549.1;HCoV-229E_1ab;279 | 0 | 0.025 |
| NC_002645.1;NP_073549.1;HCoV-229E_1ab;280 | 0 | 0.05 |
| NC_002645.1;NP_073549.1;HCoV-229E_1ab;281 | 0 | 0.025 |
| NC_002645.1;NP_073549.1;HCoV-229E_1ab;282 | 0 | 0 |
| NC_002645.1;NP_073549.1;HCoV-229E_1ab;283 | 0 | 0 |
| NC_002645.1;NP_073549.1;HCoV-229E_1ab;284 | 0 | 0 |
| NC_002645.1;NP_073549.1;HCoV-229E_1ab;285 | 0 | 0.025 |
| NC_002645.1;NP_073549.1;HCoV-229E_1ab;286 | 0 | 0 |
| NC_002645.1;NP_073549.1;HCoV-229E_1ab;287 | 0 | 0 |

| | | |
|---|---|---|
| NC_002645.1;NP_073549.1;HCoV-229E_1ab;288 | 0 | 0 |
| NC_002645.1;NP_073549.1;HCoV-229E_1ab;289 | 0 | 0 |
| NC_002645.1;NP_073549.1;HCoV-229E_1ab;290 | 0 | 0 |
| NC_002645.1;NP_073549.1;HCoV-229E_1ab;291 | 0 | 0 |
| NC_002645.1;NP_073549.1;HCoV-229E_1ab;292 | 0 | 0 |
| NC_002645.1;NP_073549.1;HCoV-229E_1ab;293 | 0 | 0 |
| NC_002645.1;NP_073549.1;HCoV-229E_1ab;294 | 0 | 0 |
| NC_002645.1;NP_073549.1;HCoV-229E_1ab;295 | 0 | 0 |
| NC_002645.1;NP_073549.1;HCoV-229E_1ab;296 | 0 | 0 |
| NC_002645.1;NP_073549.1;HCoV-229E_1ab;297 | 0 | 0 |
| NC_002645.1;NP_073549.1;HCoV-229E_1ab;298 | 0 | 0 |
| NC_002645.1;NP_073549.1;HCoV-229E_1ab;299 | 0 | 0 |
| NC_002645.1;NP_073549.1;HCoV-229E_1ab;300 | 0 | 0 |
| NC_002645.1;NP_073549.1;HCoV-229E_1ab;301 | 0 | 0 |
| NC_002645.1;NP_073549.1;HCoV-229E_1ab;302 | 0 | 0 |
| NC_002645.1;NP_073549.1;HCoV-229E_1ab;303 | 0 | 0 |
| NC_002645.1;NP_073549.1;HCoV-229E_1ab;304 | 0 | 0 |
| NC_002645.1;NP_073549.1;HCoV-229E_1ab;305 | 0 | 0 |
| NC_002645.1;NP_073549.1;HCoV-229E_1ab;306 | 0 | 0 |
| NC_002645.1;NP_073549.1;HCoV-229E_1ab;307 | 0 | 0 |
| NC_002645.1;NP_073549.1;HCoV-229E_1ab;308 | 0 | 0 |
| NC_002645.1;NP_073549.1;HCoV-229E_1ab;309 | 0 | 0 |
| NC_002645.1;NP_073549.1;HCoV-229E_1ab;310 | 0 | 0 |
| NC_002645.1;NP_073549.1;HCoV-229E_1ab;311 | 0 | 0 |
| NC_002645.1;NP_073549.1;HCoV-229E_1ab;312 | 0 | 0 |
| NC_002645.1;NP_073549.1;HCoV-229E_1ab;313 | 0 | 0 |
| NC_002645.1;NP_073549.1;HCoV-229E_1ab;314 | 0 | 0 |
| NC_002645.1;NP_073549.1;HCoV-229E_1ab;315 | 0 | 0 |
| NC_002645.1;NP_073549.1;HCoV-229E_1ab;316 | 0 | 0 |
| NC_002645.1;NP_073549.1;HCoV-229E_1ab;317 | 0 | 0 |
| NC_002645.1;NP_073549.1;HCoV-229E_1ab;318 | 0 | 0 |
| NC_002645.1;NP_073549.1;HCoV-229E_1ab;319 | 0 | 0 |
| NC_002645.1;NP_073549.1;HCoV-229E_1ab;320 | 0 | 0 |
| NC_002645.1;NP_073549.1;HCoV-229E_1ab;321 | 0 | 0 |
| NC_002645.1;NP_073549.1;HCoV-229E_1ab;322 | 0 | 0 |
| NC_002645.1;NP_073549.1;HCoV-229E_1ab;323 | 0 | 0 |
| NC_002645.1;NP_073549.1;HCoV-229E_1ab;324 | 0 | 0 |
| NC_002645.1;NP_073549.1;HCoV-229E_1ab;325 | 0 | 0 |
| NC_002645.1;NP_073549.1;HCoV-229E_1ab;326 | 0 | 0 |
| NC_002645.1;NP_073549.1;HCoV-229E_1ab;327 | 0 | 0 |
| NC_002645.1;NP_073549.1;HCoV-229E_1ab;328 | 0 | 0 |
| NC_002645.1;NP_073549.1;HCoV-229E_1ab;329 | 0 | 0 |
| NC_002645.1;NP_073549.1;HCoV-229E_1ab;330 | 0 | 0.025 |

| | | |
|---|---|---|
| NC_002645.1;NP_073549.1;HCoV-229E_1ab;331 | 0 | 0 |
| NC_002645.1;NP_073549.1;HCoV-229E_1ab;332 | 0 | 0 |
| NC_002645.1;NP_073549.1;HCoV-229E_1ab;333 | 0 | 0 |
| NC_002645.1;NP_073549.1;HCoV-229E_1ab;334 | 0 | 0.025 |
| NC_002645.1;NP_073549.1;HCoV-229E_1ab;335 | 0 | 0 |
| NC_002645.1;NP_073549.1;HCoV-229E_1ab;336 | 0 | 0 |
| NC_002645.1;NP_073549.1;HCoV-229E_1ab;337 | 0 | 0 |
| NC_002645.1;NP_073549.1;HCoV-229E_1ab;338 | 0 | 0 |
| NC_002645.1;NP_073549.1;HCoV-229E_1ab;339 | 0 | 0 |
| NC_002645.1;NP_073549.1;HCoV-229E_1ab;340 | 0 | 0 |
| NC_002645.1;NP_073549.1;HCoV-229E_1ab;341 | 0 | 0 |
| NC_002645.1;NP_073549.1;HCoV-229E_1ab;342 | 0 | 0 |
| NC_002645.1;NP_073549.1;HCoV-229E_1ab;343 | 0 | 0 |
| NC_002645.1;NP_073549.1;HCoV-229E_1ab;344 | 0 | 0 |
| NC_002645.1;NP_073549.1;HCoV-229E_1ab;345 | 0 | 0 |
| NC_002645.1;NP_073549.1;HCoV-229E_1ab;346 | 0 | 0 |
| NC_002645.1;NP_073549.1;HCoV-229E_1ab;347 | 0 | 0 |
| NC_002645.1;NP_073549.1;HCoV-229E_1ab;348 | 0 | 0 |
| NC_002645.1;NP_073549.1;HCoV-229E_1ab;349 | 0 | 0 |
| NC_002645.1;NP_073549.1;HCoV-229E_1ab;350 | 0 | 0 |
| NC_002645.1;NP_073549.1;HCoV-229E_1ab;351 | 0 | 0 |
| NC_002645.1;NP_073549.1;HCoV-229E_1ab;352 | 0 | 0 |
| NC_002645.1;NP_073549.1;HCoV-229E_1ab;353 | 0 | 0.025 |
| NC_002645.1;NP_073549.1;HCoV-229E_1ab;354 | 0 | 0.025 |
| NC_002645.1;NP_073549.1;HCoV-229E_1ab;355 | 0 | 0.05 |
| NC_002645.1;NP_073549.1;HCoV-229E_1ab;356 | 0 | 0.05 |
| NC_002645.1;NP_073549.1;HCoV-229E_1ab;357 | 0 | 0.05 |
| NC_002645.1;NP_073549.1;HCoV-229E_1ab;358 | 0 | 0.025 |
| NC_002645.1;NP_073549.1;HCoV-229E_1ab;359 | 0 | 0.05 |
| NC_002645.1;NP_073549.1;HCoV-229E_1ab;360 | 0 | 0.1 |
| NC_002645.1;NP_073549.1;HCoV-229E_1ab;361 | 0.05 | 0.05 |
| NC_002645.1;NP_073549.1;HCoV-229E_1ab;362 | 0 | 0.075 |
| NC_002645.1;NP_073549.1;HCoV-229E_1ab;363 | 0 | 0.05 |
| NC_002645.1;NP_073549.1;HCoV-229E_1ab;364 | 0.05 | 0.075 |
| NC_002645.1;NP_073549.1;HCoV-229E_1ab;365 | 0.05 | 0.05 |
| NC_002645.1;NP_073549.1;HCoV-229E_1ab;366 | 0 | 0 |
| NC_002645.1;NP_073549.1;HCoV-229E_1ab;367 | 0 | 0 |
| NC_002645.1;NP_073549.1;HCoV-229E_1ab;368 | 0 | 0 |
| NC_002645.1;NP_073549.1;HCoV-229E_1ab;369 | 0 | 0 |
| NC_002645.1;NP_073549.1;HCoV-229E_1ab;370 | 0 | 0 |
| NC_002645.1;NP_073549.1;HCoV-229E_1ab;371 | 0 | 0 |
| NC_002645.1;NP_073549.1;HCoV-229E_1ab;372 | 0 | 0 |
| NC_002645.1;NP_073549.1;HCoV-229E_1ab;373 | 0 | 0 |

| | | |
|---|---|---|
| NC_002645.1;NP_073549.1;HCoV-229E_1ab;374 | 0 | 0 |
| NC_002645.1;NP_073549.1;HCoV-229E_1ab;375 | 0 | 0 |
| NC_002645.1;NP_073549.1;HCoV-229E_1ab;376 | 0 | 0 |
| NC_002645.1;NP_073549.1;HCoV-229E_1ab;377 | 0 | 0 |
| NC_002645.1;NP_073549.1;HCoV-229E_1ab;378 | 0 | 0 |
| NC_002645.1;NP_073549.1;HCoV-229E_1ab;379 | 0 | 0 |
| NC_002645.1;NP_073549.1;HCoV-229E_1ab;380 | 0 | 0 |
| NC_002645.1;NP_073549.1;HCoV-229E_1ab;381 | 0 | 0 |
| NC_002645.1;NP_073549.1;HCoV-229E_1ab;382 | 0 | 0 |
| NC_002645.1;NP_073549.1;HCoV-229E_1ab;383 | 0 | 0 |
| NC_002645.1;NP_073549.1;HCoV-229E_1ab;384 | 0.05 | 0.025 |
| NC_002645.1;NP_073549.1;HCoV-229E_1ab;385 | 0 | 0 |
| NC_002645.1;NP_073549.1;HCoV-229E_1ab;386 | 0 | 0 |
| NC_002645.1;NP_073549.1;HCoV-229E_1ab;387 | 0.05 | 0.025 |
| NC_002645.1;NP_073549.1;HCoV-229E_1ab;388 | 0.05 | 0 |
| NC_002645.1;NP_073549.1;HCoV-229E_1ab;389 | 0.05 | 0 |
| NC_002645.1;NP_073549.1;HCoV-229E_1ab;390 | 0.05 | 0.025 |
| NC_002645.1;NP_073549.1;HCoV-229E_1ab;391 | 0.05 | 0.025 |
| NC_002645.1;NP_073549.1;HCoV-229E_1ab;392 | 0.05 | 0 |
| NC_002645.1;NP_073549.1;HCoV-229E_1ab;393 | 0.1 | 0 |
| NC_002645.1;NP_073549.1;HCoV-229E_1ab;394 | 0.05 | 0.075 |
| NC_002645.1;NP_073549.1;HCoV-229E_1ab;395 | 0.1 | 0.05 |
| NC_002645.1;NP_073549.1;HCoV-229E_1ab;396 | 0 | 0 |
| NC_002645.1;NP_073549.1;HCoV-229E_1ab;397 | 0 | 0 |
| NC_002645.1;NP_073549.1;HCoV-229E_1ab;398 | 0 | 0 |
| NC_002645.1;NP_073549.1;HCoV-229E_1ab;399 | 0 | 0 |
| NC_002645.1;NP_073549.1;HCoV-229E_1ab;400 | 0 | 0 |
| NC_002645.1;NP_073549.1;HCoV-229E_1ab;401 | 0 | 0 |
| NC_002645.1;NP_073549.1;HCoV-229E_1ab;402 | 0 | 0 |
| NC_002645.1;NP_073549.1;HCoV-229E_1ab;403 | 0 | 0 |
| NC_002645.1;NP_073549.1;HCoV-229E_1ab;404 | 0 | 0.025 |
| NC_002645.1;NP_073549.1;HCoV-229E_1ab;405 | 0.05 | 0.05 |
| NC_002645.1;NP_073549.1;HCoV-229E_1ab;406 | 0.05 | 0.05 |
| NC_002645.1;NP_073549.1;HCoV-229E_1ab;407 | 0 | 0 |
| NC_002645.1;NP_073549.1;HCoV-229E_1ab;408 | 0 | 0 |
| NC_002645.1;NP_073549.1;HCoV-229E_1ab;409 | 0 | 0 |
| NC_002645.1;NP_073549.1;HCoV-229E_1ab;410 | 0 | 0 |
| NC_002645.1;NP_073549.1;HCoV-229E_1ab;411 | 0 | 0 |
| NC_002645.1;NP_073549.1;HCoV-229E_1ab;412 | 0 | 0 |
| NC_002645.1;NP_073549.1;HCoV-229E_1ab;413 | 0.1 | 0.025 |
| NC_002645.1;NP_073549.1;HCoV-229E_1ab;414 | 0.1 | 0.075 |
| NC_002645.1;NP_073549.1;HCoV-229E_1ab;415 | 0.1 | 0.075 |
| NC_002645.1;NP_073549.1;HCoV-229E_1ab;416 | 0.1 | 0.025 |

| | | |
|---|---|---|
| NC_002645.1;NP_073549.1;HCoV-229E_1ab;417 | 0 | 0.05 |
| NC_002645.1;NP_073549.1;HCoV-229E_1ab;418 | 0 | 0 |
| NC_002645.1;NP_073549.1;HCoV-229E_1ab;419 | 0 | 0 |
| NC_002645.1;NP_073549.1;HCoV-229E_1ab;420 | 0 | 0 |
| NC_002645.1;NP_073549.1;HCoV-229E_1ab;421 | 0 | 0 |
| NC_002645.1;NP_073549.1;HCoV-229E_1ab;422 | 0 | 0 |
| NC_002645.1;NP_073549.1;HCoV-229E_1ab;423 | 0 | 0 |
| NC_002645.1;NP_073549.1;HCoV-229E_1ab;424 | 0 | 0 |
| NC_002645.1;NP_073549.1;HCoV-229E_1ab;425 | 0 | 0 |
| NC_002645.1;NP_073549.1;HCoV-229E_1ab;426 | 0 | 0 |
| NC_002645.1;NP_073549.1;HCoV-229E_1ab;427 | 0 | 0 |
| NC_002645.1;NP_073549.1;HCoV-229E_1ab;428 | 0 | 0 |
| NC_002645.1;NP_073549.1;HCoV-229E_1ab;429 | 0 | 0 |
| NC_002645.1;NP_073549.1;HCoV-229E_1ab;430 | 0 | 0 |
| NC_002645.1;NP_073549.1;HCoV-229E_1ab;431 | 0 | 0 |
| NC_002645.1;NP_073549.1;HCoV-229E_1ab;432 | 0 | 0 |
| NC_002645.1;NP_073549.1;HCoV-229E_1ab;433 | 0 | 0 |
| NC_002645.1;NP_073549.1;HCoV-229E_1ab;434 | 0 | 0 |
| NC_002645.1;NP_073549.1;HCoV-229E_1ab;435 | 0 | 0 |
| NC_002645.1;NP_073549.1;HCoV-229E_1ab;436 | 0 | 0 |
| NC_002645.1;NP_073549.1;HCoV-229E_1ab;437 | 0 | 0 |
| NC_002645.1;NP_073549.1;HCoV-229E_1ab;438 | 0 | 0 |
| NC_002645.1;NP_073549.1;HCoV-229E_1ab;439 | 0 | 0.025 |
| NC_002645.1;NP_073549.1;HCoV-229E_1ab;440 | 0 | 0 |
| NC_002645.1;NP_073549.1;HCoV-229E_1ab;441 | 0 | 0 |
| NC_002645.1;NP_073549.1;HCoV-229E_1ab;442 | 0 | 0 |
| NC_002645.1;NP_073549.1;HCoV-229E_1ab;443 | 0.05 | 0 |
| NC_002645.1;NP_073549.1;HCoV-229E_1ab;444 | 0 | 0 |
| NC_002645.1;NP_073549.1;HCoV-229E_1ab;445 | 0 | 0 |
| NC_002645.1;NP_073549.1;HCoV-229E_1ab;446 | 0 | 0 |
| NC_002645.1;NP_073549.1;HCoV-229E_1ab;447 | 0 | 0 |
| NC_002645.1;NP_073549.1;HCoV-229E_1ab;448 | 0.05 | 0.05 |
| NC_002645.1;NP_073549.1;HCoV-229E_1ab;449 | 0.05 | 0.075 |
| NC_002645.1;NP_073549.1;HCoV-229E_1ab;450 | 0.05 | 0.05 |
| NC_002645.1;NP_073549.1;HCoV-229E_1ab;451 | 0.05 | 0.025 |
| NC_002645.1;NP_073549.1;HCoV-229E_1ab;452 | 0.05 | 0.025 |
| NC_002645.1;NP_073549.1;HCoV-229E_1ab;453 | 0 | 0.025 |
| NC_002645.1;NP_073549.1;HCoV-229E_1ab;454 | 0 | 0.025 |
| NC_002645.1;NP_073549.1;HCoV-229E_1ab;455 | 0 | 0 |
| NC_002645.1;NP_073549.1;HCoV-229E_1ab;456 | 0 | 0 |
| NC_002645.1;NP_073549.1;HCoV-229E_1ab;457 | 0 | 0 |
| NC_002645.1;NP_073549.1;HCoV-229E_1ab;458 | 0 | 0 |
| NC_002645.1;NP_073549.1;HCoV-229E_1ab;459 | 0 | 0 |

| | | |
|---|---|---|
| NC_002645.1;NP_073549.1;HCoV-229E_1ab;460 | 0 | 0 |
| NC_002645.1;NP_073549.1;HCoV-229E_1ab;461 | 0 | 0 |
| NC_002645.1;NP_073549.1;HCoV-229E_1ab;462 | 0 | 0 |
| NC_002645.1;NP_073549.1;HCoV-229E_1ab;463 | 0 | 0 |
| NC_002645.1;NP_073549.1;HCoV-229E_1ab;464 | 0 | 0 |
| NC_002645.1;NP_073549.1;HCoV-229E_1ab;465 | 0 | 0 |
| NC_002645.1;NP_073549.1;HCoV-229E_1ab;466 | 0 | 0 |
| NC_002645.1;NP_073549.1;HCoV-229E_1ab;467 | 0 | 0 |
| NC_002645.1;NP_073549.1;HCoV-229E_1ab;468 | 0 | 0 |
| NC_002645.1;NP_073549.1;HCoV-229E_1ab;469 | 0 | 0 |
| NC_002645.1;NP_073549.1;HCoV-229E_1ab;470 | 0 | 0 |
| NC_002645.1;NP_073549.1;HCoV-229E_1ab;471 | 0 | 0 |
| NC_002645.1;NP_073549.1;HCoV-229E_1ab;472 | 0 | 0 |
| NC_002645.1;NP_073549.1;HCoV-229E_1ab;473 | 0 | 0 |
| NC_002645.1;NP_073549.1;HCoV-229E_1ab;474 | 0.05 | 0 |
| NC_002645.1;NP_073549.1;HCoV-229E_1ab;475 | 0.05 | 0 |
| NC_002645.1;NP_073549.1;HCoV-229E_1ab;476 | 0 | 0 |
| NC_002645.1;NP_073549.1;HCoV-229E_1ab;477 | 0.2 | 0.025 |
| NC_002645.1;NP_073549.1;HCoV-229E_1ab;478 | 0.15 | 0.025 |
| NC_002645.1;NP_073549.1;HCoV-229E_1ab;479 | 0 | 0.025 |
| NC_002645.1;NP_073549.1;HCoV-229E_1ab;480 | 0 | 0 |
| NC_002645.1;NP_073549.1;HCoV-229E_1ab;481 | 0 | 0 |
| NC_002645.1;NP_073549.1;HCoV-229E_1ab;482 | 0 | 0 |
| NC_002645.1;NP_073549.1;HCoV-229E_1ab;483 | 0 | 0 |
| NC_002645.1;NP_073549.1;HCoV-229E_1ab;484 | 0 | 0 |
| NC_002645.1;NP_073549.1;HCoV-229E_1ab;485 | 0 | 0 |
| NC_002645.1;NP_073549.1;HCoV-229E_1ab;486 | 0 | 0 |
| NC_002645.1;NP_073549.1;HCoV-229E_1ab;487 | 0 | 0 |
| NC_002645.1;NP_073549.1;HCoV-229E_1ab;488 | 0 | 0 |
| NC_002645.1;NP_073549.1;HCoV-229E_1ab;489 | 0 | 0 |
| NC_002645.1;NP_073549.1;HCoV-229E_1ab;490 | 0 | 0 |
| NC_002645.1;NP_073549.1;HCoV-229E_1ab;491 | 0 | 0 |
| NC_002645.1;NP_073549.1;HCoV-229E_1ab;492 | 0 | 0 |
| NC_002645.1;NP_073549.1;HCoV-229E_1ab;493 | 0 | 0 |
| NC_002645.1;NP_073549.1;HCoV-229E_1ab;494 | 0 | 0 |
| NC_002645.1;NP_073549.1;HCoV-229E_1ab;495 | 0 | 0 |
| NC_002645.1;NP_073549.1;HCoV-229E_1ab;496 | 0 | 0 |
| NC_002645.1;NP_073549.1;HCoV-229E_1ab;497 | 0 | 0 |
| NC_002645.1;NP_073549.1;HCoV-229E_1ab;498 | 0 | 0 |
| NC_002645.1;NP_073549.1;HCoV-229E_1ab;499 | 0 | 0 |
| NC_002645.1;NP_073549.1;HCoV-229E_1ab;500 | 0 | 0 |
| NC_002645.1;NP_073549.1;HCoV-229E_1ab;501 | 0 | 0 |
| NC_002645.1;NP_073549.1;HCoV-229E_1ab;502 | 0 | 0 |

| | | |
|---|---|---|
| NC_002645.1;NP_073549.1;HCoV-229E_1ab;503 | 0 | 0 |
| NC_002645.1;NP_073549.1;HCoV-229E_1ab;504 | 0 | 0 |
| NC_002645.1;NP_073549.1;HCoV-229E_1ab;505 | 0 | 0 |
| NC_002645.1;NP_073549.1;HCoV-229E_1ab;506 | 0 | 0 |
| NC_002645.1;NP_073549.1;HCoV-229E_1ab;507 | 0 | 0 |
| NC_002645.1;NP_073549.1;HCoV-229E_1ab;508 | 0 | 0 |
| NC_002645.1;NP_073549.1;HCoV-229E_1ab;509 | 0 | 0 |
| NC_002645.1;NP_073549.1;HCoV-229E_1ab;510 | 0 | 0 |
| NC_002645.1;NP_073549.1;HCoV-229E_1ab;511 | 0 | 0 |
| NC_002645.1;NP_073549.1;HCoV-229E_1ab;512 | 0 | 0 |
| NC_002645.1;NP_073549.1;HCoV-229E_1ab;513 | 0 | 0 |
| NC_002645.1;NP_073549.1;HCoV-229E_1ab;514 | 0 | 0 |
| NC_002645.1;NP_073549.1;HCoV-229E_1ab;515 | 0 | 0 |
| NC_002645.1;NP_073549.1;HCoV-229E_1ab;516 | 0 | 0 |
| NC_002645.1;NP_073549.1;HCoV-229E_1ab;517 | 0 | 0 |
| NC_002645.1;NP_073549.1;HCoV-229E_1ab;518 | 0 | 0 |
| NC_002645.1;NP_073549.1;HCoV-229E_1ab;519 | 0 | 0 |
| NC_002645.1;NP_073549.1;HCoV-229E_1ab;520 | 0 | 0.025 |
| NC_002645.1;NP_073549.1;HCoV-229E_1ab;521 | 0 | 0 |
| NC_002645.1;NP_073549.1;HCoV-229E_1ab;522 | 0 | 0 |
| NC_002645.1;NP_073549.1;HCoV-229E_1ab;523 | 0 | 0 |
| NC_002645.1;NP_073549.1;HCoV-229E_1ab;524 | 0 | 0 |
| NC_002645.1;NP_073549.1;HCoV-229E_1ab;525 | 0 | 0 |
| NC_002645.1;NP_073549.1;HCoV-229E_1ab;526 | 0 | 0 |
| NC_002645.1;NP_073549.1;HCoV-229E_1ab;527 | 0 | 0 |
| NC_002645.1;NP_073549.1;HCoV-229E_1ab;528 | 0 | 0 |
| NC_002645.1;NP_073549.1;HCoV-229E_1ab;529 | 0 | 0.025 |
| NC_002645.1;NP_073549.1;HCoV-229E_1ab;530 | 0 | 0 |
| NC_002645.1;NP_073549.1;HCoV-229E_1ab;531 | 0 | 0 |
| NC_002645.1;NP_073549.1;HCoV-229E_1ab;532 | 0 | 0 |
| NC_002645.1;NP_073549.1;HCoV-229E_1ab;533 | 0 | 0 |
| NC_002645.1;NP_073549.1;HCoV-229E_1ab;534 | 0 | 0 |
| NC_002645.1;NP_073549.1;HCoV-229E_1ab;535 | 0 | 0 |
| NC_002645.1;NP_073549.1;HCoV-229E_1ab;536 | 0 | 0 |
| NC_002645.1;NP_073549.1;HCoV-229E_1ab;537 | 0 | 0 |
| NC_002645.1;NP_073549.1;HCoV-229E_1ab;538 | 0 | 0 |
| NC_002645.1;NP_073549.1;HCoV-229E_1ab;539 | 0 | 0 |
| NC_002645.1;NP_073549.1;HCoV-229E_1ab;540 | 0 | 0.025 |
| NC_002645.1;NP_073549.1;HCoV-229E_1ab;541 | 0 | 0.025 |
| NC_002645.1;NP_073549.1;HCoV-229E_1ab;542 | 0 | 0.025 |
| NC_002645.1;NP_073549.1;HCoV-229E_1ab;543 | 0 | 0.05 |
| NC_002645.1;NP_073549.1;HCoV-229E_1ab;544 | 0.1 | 0.05 |
| NC_002645.1;NP_073549.1;HCoV-229E_1ab;545 | 0 | 0.025 |

| | | |
|---|---|---|
| NC_002645.1;NP_073549.1;HCoV-229E_1ab;546 | 0 | 0.025 |
| NC_002645.1;NP_073549.1;HCoV-229E_1ab;547 | 0 | 0.025 |
| NC_002645.1;NP_073549.1;HCoV-229E_1ab;548 | 0 | 0 |
| NC_002645.1;NP_073549.1;HCoV-229E_1ab;549 | 0 | 0 |
| NC_002645.1;NP_073549.1;HCoV-229E_1ab;550 | 0 | 0 |
| NC_002645.1;NP_073549.1;HCoV-229E_1ab;551 | 0 | 0 |
| NC_002645.1;NP_073549.1;HCoV-229E_1ab;552 | 0 | 0 |
| NC_002645.1;NP_073549.1;HCoV-229E_1ab;553 | 0 | 0 |
| NC_002645.1;NP_073549.1;HCoV-229E_1ab;554 | 0 | 0 |
| NC_002645.1;NP_073549.1;HCoV-229E_1ab;555 | 0 | 0 |
| NC_002645.1;NP_073549.1;HCoV-229E_1ab;556 | 0 | 0 |
| NC_002645.1;NP_073549.1;HCoV-229E_1ab;557 | 0 | 0 |
| NC_002645.1;NP_073549.1;HCoV-229E_1ab;558 | 0 | 0 |
| NC_002645.1;NP_073549.1;HCoV-229E_1ab;559 | 0 | 0 |
| NC_002645.1;NP_073549.1;HCoV-229E_1ab;560 | 0 | 0 |
| NC_002645.1;NP_073549.1;HCoV-229E_1ab;561 | 0 | 0 |
| NC_002645.1;NP_073549.1;HCoV-229E_1ab;562 | 0 | 0 |
| NC_002645.1;NP_073549.1;HCoV-229E_1ab;563 | 0 | 0 |
| NC_002645.1;NP_073549.1;HCoV-229E_1ab;564 | 0 | 0 |
| NC_002645.1;NP_073549.1;HCoV-229E_1ab;565 | 0 | 0 |
| NC_002645.1;NP_073549.1;HCoV-229E_1ab;566 | 0 | 0 |
| NC_002645.1;NP_073549.1;HCoV-229E_1ab;567 | 0 | 0 |
| NC_002645.1;NP_073549.1;HCoV-229E_1ab;568 | 0 | 0 |
| NC_002645.1;NP_073549.1;HCoV-229E_1ab;569 | 0 | 0 |
| NC_002645.1;NP_073549.1;HCoV-229E_1ab;570 | 0 | 0 |
| NC_002645.1;NP_073549.1;HCoV-229E_1ab;571 | 0 | 0 |
| NC_002645.1;NP_073549.1;HCoV-229E_1ab;572 | 0 | 0 |
| NC_002645.1;NP_073549.1;HCoV-229E_1ab;573 | 0 | 0 |
| NC_002645.1;NP_073549.1;HCoV-229E_1ab;574 | 0 | 0 |
| NC_002645.1;NP_073549.1;HCoV-229E_1ab;575 | 0 | 0 |
| NC_002645.1;NP_073549.1;HCoV-229E_1ab;576 | 0 | 0 |
| NC_002645.1;NP_073549.1;HCoV-229E_1ab;577 | 0 | 0 |
| NC_002645.1;NP_073549.1;HCoV-229E_1ab;578 | 0 | 0 |
| NC_002645.1;NP_073549.1;HCoV-229E_1ab;579 | 0 | 0 |
| NC_002645.1;NP_073549.1;HCoV-229E_1ab;580 | 0 | 0 |
| NC_002645.1;NP_073549.1;HCoV-229E_1ab;581 | 0 | 0 |
| NC_002645.1;NP_073549.1;HCoV-229E_1ab;582 | 0 | 0 |
| NC_002645.1;NP_073549.1;HCoV-229E_1ab;583 | 0 | 0 |
| NC_002645.1;NP_073549.1;HCoV-229E_1ab;584 | 0 | 0 |
| NC_002645.1;NP_073549.1;HCoV-229E_1ab;585 | 0 | 0 |
| NC_002645.1;NP_073549.1;HCoV-229E_1ab;586 | 0 | 0 |
| NC_002645.1;NP_073549.1;HCoV-229E_1ab;587 | 0 | 0 |
| NC_002645.1;NP_073549.1;HCoV-229E_1ab;588 | 0 | 0 |

| | | |
|---|---|---|
| NC_002645.1;NP_073549.1;HCoV-229E_1ab;589 | 0 | 0 |
| NC_002645.1;NP_073549.1;HCoV-229E_1ab;590 | 0 | 0.025 |
| NC_002645.1;NP_073549.1;HCoV-229E_1ab;591 | 0 | 0 |
| NC_002645.1;NP_073549.1;HCoV-229E_1ab;592 | 0.1 | 0 |
| NC_002645.1;NP_073549.1;HCoV-229E_1ab;593 | 0 | 0 |
| NC_002645.1;NP_073549.1;HCoV-229E_1ab;594 | 0.05 | 0.05 |
| NC_002645.1;NP_073549.1;HCoV-229E_1ab;595 | 0 | 0 |
| NC_002645.1;NP_073549.1;HCoV-229E_1ab;596 | 0 | 0 |
| NC_002645.1;NP_073549.1;HCoV-229E_1ab;597 | 0 | 0 |
| NC_002645.1;NP_073549.1;HCoV-229E_1ab;598 | 0 | 0 |
| NC_002645.1;NP_073549.1;HCoV-229E_1ab;599 | 0 | 0 |
| NC_002645.1;NP_073549.1;HCoV-229E_1ab;600 | 0 | 0 |
| NC_002645.1;NP_073549.1;HCoV-229E_1ab;601 | 0 | 0 |
| NC_002645.1;NP_073549.1;HCoV-229E_1ab;602 | 0 | 0 |
| NC_002645.1;NP_073549.1;HCoV-229E_1ab;603 | 0 | 0 |
| NC_002645.1;NP_073549.1;HCoV-229E_1ab;604 | 0 | 0 |
| NC_002645.1;NP_073549.1;HCoV-229E_1ab;605 | 0 | 0 |
| NC_002645.1;NP_073549.1;HCoV-229E_1ab;606 | 0 | 0 |
| NC_002645.1;NP_073549.1;HCoV-229E_1ab;607 | 0 | 0 |
| NC_002645.1;NP_073549.1;HCoV-229E_1ab;608 | 0 | 0 |
| NC_002645.1;NP_073549.1;HCoV-229E_1ab;609 | 0 | 0 |
| NC_002645.1;NP_073549.1;HCoV-229E_1ab;610 | 0.05 | 0 |
| NC_002645.1;NP_073549.1;HCoV-229E_1ab;611 | 0.05 | 0 |
| NC_002645.1;NP_073549.1;HCoV-229E_1ab;612 | 0.05 | 0 |
| NC_002645.1;NP_073549.1;HCoV-229E_1ab;613 | 0.05 | 0.05 |
| NC_002645.1;NP_073549.1;HCoV-229E_1ab;614 | 0.05 | 0 |
| NC_002645.1;NP_073549.1;HCoV-229E_1ab;615 | 0.05 | 0 |
| NC_002645.1;NP_073549.1;HCoV-229E_1ab;616 | 0.05 | 0 |
| NC_002645.1;NP_073549.1;HCoV-229E_1ab;617 | 0.05 | 0 |
| NC_002645.1;NP_073549.1;HCoV-229E_1ab;618 | 0.05 | 0 |
| NC_002645.1;NP_073549.1;HCoV-229E_1ab;619 | 0 | 0 |
| NC_002645.1;NP_073549.1;HCoV-229E_1ab;620 | 0 | 0 |
| NC_002645.1;NP_073549.1;HCoV-229E_1ab;621 | 0 | 0 |
| NC_002645.1;NP_073549.1;HCoV-229E_1ab;622 | 0 | 0.025 |
| NC_002645.1;NP_073549.1;HCoV-229E_1ab;623 | 0 | 0.025 |
| NC_002645.1;NP_073549.1;HCoV-229E_1ab;624 | 0 | 0 |
| NC_002645.1;NP_073549.1;HCoV-229E_1ab;625 | 0 | 0.05 |
| NC_002645.1;NP_073549.1;HCoV-229E_1ab;626 | 0 | 0.025 |
| NC_002645.1;NP_073549.1;HCoV-229E_1ab;627 | 0.05 | 0.025 |
| NC_002645.1;NP_073549.1;HCoV-229E_1ab;628 | 0.05 | 0.025 |
| NC_002645.1;NP_073549.1;HCoV-229E_1ab;629 | 0.05 | 0.05 |
| NC_002645.1;NP_073549.1;HCoV-229E_1ab;630 | 0 | 0 |
| NC_002645.1;NP_073549.1;HCoV-229E_1ab;631 | 0.25 | 0.15 |

| | | |
|---|---|---|
| NC_002645.1;NP_073549.1;HCoV-229E_1ab;632 | 0.05 | 0.075 |
| NC_002645.1;NP_073549.1;HCoV-229E_1ab;633 | 0.2 | 0.175 |
| NC_002645.1;NP_073549.1;HCoV-229E_1ab;634 | 0.45 | 0.2 |
| NC_002645.1;NP_073549.1;HCoV-229E_1ab;635 | 0.5 | 0.175 |
| NC_002645.1;NP_073549.1;HCoV-229E_1ab;636 | 0.4 | 0.175 |
| NC_002645.1;NP_073549.1;HCoV-229E_1ab;637 | 0.4 | 0.2 |
| NC_002645.1;NP_073549.1;HCoV-229E_1ab;638 | 0.3 | 0.15 |
| NC_002645.1;NP_073549.1;HCoV-229E_1ab;639 | 0.4 | 0.175 |
| NC_002645.1;NP_073549.1;HCoV-229E_1ab;640 | 0.35 | 0.15 |
| NC_002645.1;NP_073549.1;HCoV-229E_1ab;641 | 0.2 | 0.125 |
| NC_002645.1;NP_073549.1;HCoV-229E_1ab;642 | 0.2 | 0.125 |
| NC_002645.1;NP_073549.1;HCoV-229E_1ab;643 | 0.05 | 0.025 |
| NC_002645.1;NP_073549.1;HCoV-229E_1ab;644 | 0 | 0 |
| NC_002645.1;NP_073549.1;HCoV-229E_1ab;645 | 0 | 0 |
| NC_002645.1;NP_073549.1;HCoV-229E_1ab;646 | 0 | 0 |
| NC_002645.1;NP_073549.1;HCoV-229E_1ab;647 | 0 | 0 |
| NC_002645.1;NP_073549.1;HCoV-229E_1ab;648 | 0 | 0 |
| NC_002645.1;NP_073549.1;HCoV-229E_1ab;649 | 0 | 0 |
| NC_002645.1;NP_073549.1;HCoV-229E_1ab;650 | 0 | 0 |
| NC_002645.1;NP_073549.1;HCoV-229E_1ab;651 | 0 | 0 |
| NC_002645.1;NP_073549.1;HCoV-229E_1ab;652 | 0 | 0.05 |
| NC_002645.1;NP_073549.1;HCoV-229E_1ab;653 | 0.05 | 0.05 |
| NC_002645.1;NP_073549.1;HCoV-229E_1ab;654 | 0 | 0 |
| NC_002645.1;NP_073549.1;HCoV-229E_1ab;655 | 0 | 0 |
| NC_002645.1;NP_073549.1;HCoV-229E_1ab;656 | 0.05 | 0.025 |
| NC_002645.1;NP_073549.1;HCoV-229E_1ab;657 | 0 | 0.025 |
| NC_002645.1;NP_073549.1;HCoV-229E_1ab;658 | 0.05 | 0 |
| NC_002645.1;NP_073549.1;HCoV-229E_1ab;659 | 0.05 | 0.05 |
| NC_002645.1;NP_073549.1;HCoV-229E_1ab;660 | 0 | 0.1 |
| NC_002645.1;NP_073549.1;HCoV-229E_1ab;661 | 0 | 0 |
| NC_002645.1;NP_073549.1;HCoV-229E_1ab;662 | 0.05 | 0 |
| NC_002645.1;NP_073549.1;HCoV-229E_1ab;663 | 0 | 0.025 |
| NC_002645.1;NP_073549.1;HCoV-229E_1ab;664 | 0 | 0.025 |
| NC_002645.1;NP_073549.1;HCoV-229E_1ab;665 | 0.05 | 0.025 |
| NC_002645.1;NP_073549.1;HCoV-229E_1ab;666 | 0.1 | 0.025 |
| NC_002645.1;NP_073549.1;HCoV-229E_1ab;667 | 0.05 | 0.025 |
| NC_002645.1;NP_073549.1;HCoV-229E_1ab;668 | 0.1 | 0.05 |
| NC_002645.1;NP_073549.1;HCoV-229E_1ab;669 | 0.05 | 0.075 |
| NC_002645.1;NP_073549.1;HCoV-229E_1ab;670 | 0.05 | 0 |
| NC_002645.1;NP_073549.1;HCoV-229E_1ab;671 | 0.05 | 0 |
| NC_002645.1;NP_073549.1;HCoV-229E_1ab;672 | 0.05 | 0.025 |
| NC_002645.1;NP_073549.1;HCoV-229E_1ab;673 | 0 | 0 |
| NC_002645.1;NP_073549.1;HCoV-229E_1ab;674 | 0 | 0 |

| | | |
|---|---|---|
| NC_002645.1;NP_073549.1;HCoV-229E_1ab;675 | 0 | 0 |
| NC_002645.1;NP_073549.1;HCoV-229E_1ab;676 | 0 | 0 |
| NC_002645.1;NP_073549.1;HCoV-229E_1ab;677 | 0 | 0 |
| NC_002645.1;NP_073549.1;HCoV-229E_1ab;678 | 0 | 0 |
| NC_002645.1;NP_073549.1;HCoV-229E_1ab;679 | 0 | 0 |
| NC_002645.1;NP_073549.1;HCoV-229E_1ab;680 | 0 | 0 |
| NC_002645.1;NP_073549.1;HCoV-229E_1ab;681 | 0.05 | 0.025 |
| NC_002645.1;NP_073549.1;HCoV-229E_1ab;682 | 0 | 0.025 |
| NC_002645.1;NP_073549.1;HCoV-229E_1ab;683 | 0.1 | 0.05 |
| NC_002645.1;NP_073549.1;HCoV-229E_1ab;684 | 0.05 | 0.025 |
| NC_002645.1;NP_073549.1;HCoV-229E_1ab;685 | 0 | 0.025 |
| NC_002645.1;NP_073549.1;HCoV-229E_1ab;686 | 0.05 | 0.1 |
| NC_002645.1;NP_073549.1;HCoV-229E_1ab;687 | 0.05 | 0.075 |
| NC_002645.1;NP_073549.1;HCoV-229E_1ab;688 | 0.05 | 0.1 |
| NC_002645.1;NP_073549.1;HCoV-229E_1ab;689 | 0.05 | 0.1 |
| NC_002645.1;NP_073549.1;HCoV-229E_1ab;690 | 0.05 | 0.1 |
| NC_002645.1;NP_073549.1;HCoV-229E_1ab;691 | 0.1 | 0.075 |
| NC_002645.1;NP_073549.1;HCoV-229E_1ab;692 | 0.05 | 0.075 |
| NC_002645.1;NP_073549.1;HCoV-229E_1ab;693 | 0.1 | 0.05 |
| NC_002645.1;NP_073549.1;HCoV-229E_1ab;694 | 0.1 | 0.075 |
| NC_002645.1;NP_073549.1;HCoV-229E_1ab;695 | 0.05 | 0.1 |
| NC_002645.1;NP_073549.1;HCoV-229E_1ab;696 | 0 | 0.075 |
| NC_002645.1;NP_073549.1;HCoV-229E_1ab;697 | 0 | 0.05 |
| NC_002645.1;NP_073549.1;HCoV-229E_1ab;698 | 0.05 | 0.025 |
| NC_002645.1;NP_073549.1;HCoV-229E_1ab;699 | 0.15 | 0.1 |
| NC_002645.1;NP_073549.1;HCoV-229E_1ab;700 | 0.15 | 0.025 |
| NC_002645.1;NP_073549.1;HCoV-229E_1ab;701 | 0 | 0 |
| NC_002645.1;NP_073549.1;HCoV-229E_1ab;702 | 0 | 0.025 |
| NC_002645.1;NP_073549.1;HCoV-229E_1ab;703 | 0 | 0 |
| NC_002645.1;NP_073549.1;HCoV-229E_1ab;704 | 0.05 | 0 |
| NC_002645.1;NP_073549.1;HCoV-229E_1ab;705 | 0 | 0 |
| NC_002645.1;NP_073549.1;HCoV-229E_1ab;706 | 0 | 0 |
| NC_002645.1;NP_073549.1;HCoV-229E_1ab;707 | 0 | 0 |
| NC_002645.1;NP_073549.1;HCoV-229E_1ab;708 | 0 | 0 |
| NC_002645.1;NP_073549.1;HCoV-229E_1ab;709 | 0 | 0 |
| NC_002645.1;NP_073549.1;HCoV-229E_1ab;710 | 0 | 0 |
| NC_002645.1;NP_073549.1;HCoV-229E_1ab;711 | 0 | 0 |
| NC_002645.1;NP_073549.1;HCoV-229E_1ab;712 | 0 | 0 |
| NC_002645.1;NP_073549.1;HCoV-229E_1ab;713 | 0 | 0 |
| NC_002645.1;NP_073549.1;HCoV-229E_1ab;714 | 0 | 0 |
| NC_002645.1;NP_073549.1;HCoV-229E_1ab;715 | 0 | 0 |
| NC_002645.1;NP_073549.1;HCoV-229E_1ab;716 | 0 | 0 |
| NC_002645.1;NP_073549.1;HCoV-229E_1ab;717 | 0 | 0 |

| | | |
|---|---|---|
| NC_002645.1;NP_073549.1;HCoV-229E_1ab;718 | 0 | 0.025 |
| NC_002645.1;NP_073549.1;HCoV-229E_1ab;719 | 0 | 0.025 |
| NC_002645.1;NP_073549.1;HCoV-229E_1ab;720 | 0 | 0 |
| NC_002645.1;NP_073549.1;HCoV-229E_1ab;721 | 0 | 0.025 |
| NC_002645.1;NP_073549.1;HCoV-229E_1ab;722 | 0 | 0.025 |
| NC_002645.1;NP_073549.1;HCoV-229E_1ab;723 | 0.05 | 0.025 |
| NC_002645.1;NP_073549.1;HCoV-229E_1ab;724 | 0 | 0 |
| NC_002645.1;NP_073549.1;HCoV-229E_1ab;725 | 0 | 0.025 |
| NC_002645.1;NP_073549.1;HCoV-229E_1ab;726 | 0 | 0.05 |
| NC_002645.1;NP_073549.1;HCoV-229E_1ab;727 | 0 | 0.05 |
| NC_002645.1;NP_073549.1;HCoV-229E_1ab;728 | 0 | 0.025 |
| NC_002645.1;NP_073549.1;HCoV-229E_1ab;729 | 0 | 0 |
| NC_002645.1;NP_073549.1;HCoV-229E_1ab;730 | 0 | 0 |
| NC_002645.1;NP_073549.1;HCoV-229E_1ab;731 | 0 | 0 |
| NC_002645.1;NP_073549.1;HCoV-229E_1ab;732 | 0 | 0 |
| NC_002645.1;NP_073549.1;HCoV-229E_1ab;733 | 0 | 0 |
| NC_002645.1;NP_073549.1;HCoV-229E_1ab;734 | 0 | 0 |
| NC_002645.1;NP_073549.1;HCoV-229E_1ab;735 | 0 | 0 |
| NC_002645.1;NP_073549.1;HCoV-229E_1ab;736 | 0 | 0 |
| NC_002645.1;NP_073549.1;HCoV-229E_1ab;737 | 0 | 0 |
| NC_002645.1;NP_073549.1;HCoV-229E_1ab;738 | 0.05 | 0 |
| NC_002645.1;NP_073549.1;HCoV-229E_1ab;739 | 0 | 0.025 |
| NC_002645.1;NP_073549.1;HCoV-229E_1ab;740 | 0 | 0 |
| NC_002645.1;NP_073549.1;HCoV-229E_1ab;741 | 0 | 0 |
| NC_002645.1;NP_073549.1;HCoV-229E_1ab;742 | 0 | 0 |
| NC_002645.1;NP_073549.1;HCoV-229E_1ab;743 | 0 | 0 |
| NC_002645.1;NP_073549.1;HCoV-229E_1ab;744 | 0 | 0 |
| NC_002645.1;NP_073549.1;HCoV-229E_1ab;745 | 0 | 0 |
| NC_002645.1;NP_073549.1;HCoV-229E_1ab;746 | 0 | 0 |
| NC_002645.1;NP_073549.1;HCoV-229E_1ab;747 | 0 | 0 |
| NC_002645.1;NP_073549.1;HCoV-229E_1ab;748 | 0 | 0 |
| NC_002645.1;NP_073549.1;HCoV-229E_1ab;749 | 0 | 0 |
| NC_002645.1;NP_073549.1;HCoV-229E_1ab;750 | 0 | 0 |
| NC_002645.1;NP_073549.1;HCoV-229E_1ab;751 | 0 | 0 |
| NC_002645.1;NP_073549.1;HCoV-229E_1ab;752 | 0 | 0 |
| NC_002645.1;NP_073549.1;HCoV-229E_1ab;753 | 0 | 0 |
| NC_002645.1;NP_073549.1;HCoV-229E_1ab;754 | 0 | 0 |
| NC_002645.1;NP_073549.1;HCoV-229E_1ab;755 | 0 | 0 |
| NC_002645.1;NP_073549.1;HCoV-229E_1ab;756 | 0 | 0 |
| NC_002645.1;NP_073549.1;HCoV-229E_1ab;757 | 0 | 0 |
| NC_002645.1;NP_073549.1;HCoV-229E_1ab;758 | 0 | 0 |
| NC_002645.1;NP_073549.1;HCoV-229E_1ab;759 | 0 | 0.025 |
| NC_002645.1;NP_073549.1;HCoV-229E_1ab;760 | 0 | 0 |

| | | |
|---|---|---|
| NC_002645.1;NP_073549.1;HCoV-229E_1ab;761 | 0 | 0 |
| NC_002645.1;NP_073549.1;HCoV-229E_1ab;762 | 0 | 0 |
| NC_002645.1;NP_073549.1;HCoV-229E_1ab;763 | 0 | 0.025 |
| NC_002645.1;NP_073549.1;HCoV-229E_1ab;764 | 0 | 0 |
| NC_002645.1;NP_073549.1;HCoV-229E_1ab;765 | 0 | 0 |
| NC_002645.1;NP_073549.1;HCoV-229E_1ab;766 | 0 | 0 |
| NC_002645.1;NP_073549.1;HCoV-229E_1ab;767 | 0 | 0 |
| NC_002645.1;NP_073549.1;HCoV-229E_1ab;768 | 0 | 0 |
| NC_002645.1;NP_073549.1;HCoV-229E_1ab;769 | 0 | 0.05 |
| NC_002645.1;NP_073549.1;HCoV-229E_1ab;770 | 0 | 0 |
| NC_002645.1;NP_073549.1;HCoV-229E_1ab;771 | 0 | 0.025 |
| NC_002645.1;NP_073549.1;HCoV-229E_1ab;772 | 0 | 0.025 |
| NC_002645.1;NP_073549.1;HCoV-229E_1ab;773 | 0 | 0.025 |
| NC_002645.1;NP_073549.1;HCoV-229E_1ab;774 | 0 | 0.025 |
| NC_002645.1;NP_073549.1;HCoV-229E_1ab;775 | 0 | 0.025 |
| NC_002645.1;NP_073549.1;HCoV-229E_1ab;776 | 0 | 0.025 |
| NC_002645.1;NP_073549.1;HCoV-229E_1ab;777 | 0 | 0 |
| NC_002645.1;NP_073549.1;HCoV-229E_1ab;778 | 0 | 0 |
| NC_002645.1;NP_073549.1;HCoV-229E_1ab;779 | 0 | 0.025 |
| NC_002645.1;NP_073549.1;HCoV-229E_1ab;780 | 0 | 0.025 |
| NC_002645.1;NP_073549.1;HCoV-229E_1ab;781 | 0 | 0.025 |
| NC_002645.1;NP_073549.1;HCoV-229E_1ab;782 | 0 | 0 |
| NC_002645.1;NP_073549.1;HCoV-229E_1ab;783 | 0 | 0 |
| NC_002645.1;NP_073549.1;HCoV-229E_1ab;784 | 0 | 0 |
| NC_002645.1;NP_073549.1;HCoV-229E_1ab;785 | 0 | 0 |
| NC_002645.1;NP_073549.1;HCoV-229E_1ab;786 | 0 | 0 |
| NC_002645.1;NP_073549.1;HCoV-229E_1ab;787 | 0 | 0 |
| NC_002645.1;NP_073549.1;HCoV-229E_1ab;788 | 0 | 0 |
| NC_002645.1;NP_073549.1;HCoV-229E_1ab;789 | 0 | 0 |
| NC_002645.1;NP_073549.1;HCoV-229E_1ab;790 | 0 | 0 |
| NC_002645.1;NP_073549.1;HCoV-229E_1ab;791 | 0 | 0 |
| NC_002645.1;NP_073549.1;HCoV-229E_1ab;792 | 0 | 0 |
| NC_002645.1;NP_073549.1;HCoV-229E_1ab;793 | 0 | 0 |
| NC_002645.1;NP_073549.1;HCoV-229E_1ab;794 | 0 | 0 |
| NC_002645.1;NP_073549.1;HCoV-229E_1ab;795 | 0 | 0 |
| NC_002645.1;NP_073549.1;HCoV-229E_1ab;796 | 0 | 0 |
| NC_002645.1;NP_073549.1;HCoV-229E_1ab;797 | 0.05 | 0 |
| NC_002645.1;NP_073549.1;HCoV-229E_1ab;798 | 0 | 0 |
| NC_002645.1;NP_073549.1;HCoV-229E_1ab;799 | 0 | 0.05 |
| NC_002645.1;NP_073549.1;HCoV-229E_1ab;800 | 0 | 0 |
| NC_002645.1;NP_073549.1;HCoV-229E_1ab;801 | 0 | 0.025 |
| NC_002645.1;NP_073549.1;HCoV-229E_1ab;802 | 0 | 0 |
| NC_002645.1;NP_073549.1;HCoV-229E_1ab;803 | 0 | 0 |

| | | |
|---|---|---|
| NC_002645.1;NP_073549.1;HCoV-229E_1ab;804 | 0 | 0 |
| NC_002645.1;NP_073549.1;HCoV-229E_1ab;805 | 0 | 0 |
| NC_002645.1;NP_073549.1;HCoV-229E_1ab;806 | 0 | 0 |
| NC_002645.1;NP_073549.1;HCoV-229E_1ab;807 | 0 | 0 |
| NC_002645.1;NP_073549.1;HCoV-229E_1ab;808 | 0 | 0 |
| NC_002645.1;NP_073549.1;HCoV-229E_1ab;809 | 0 | 0.025 |
| NC_002645.1;NP_073549.1;HCoV-229E_1ab;810 | 0 | 0.025 |
| NC_002645.1;NP_073549.1;HCoV-229E_1ab;811 | 0 | 0 |
| NC_002645.1;NP_073549.1;HCoV-229E_1ab;812 | 0 | 0.075 |
| NC_002645.1;NP_073549.1;HCoV-229E_1ab;813 | 0 | 0.025 |
| NC_002645.1;NP_073549.1;HCoV-229E_1ab;814 | 0 | 0.025 |
| NC_002645.1;NP_073549.1;HCoV-229E_1ab;815 | 0 | 0 |
| NC_002645.1;NP_073549.1;HCoV-229E_1ab;816 | 0 | 0 |
| NC_002645.1;NP_073549.1;HCoV-229E_1ab;817 | 0 | 0.025 |
| NC_002645.1;NP_073549.1;HCoV-229E_1ab;818 | 0 | 0.025 |
| NC_002645.1;NP_073549.1;HCoV-229E_1ab;819 | 0 | 0 |
| NC_002645.1;NP_073549.1;HCoV-229E_1ab;820 | 0 | 0 |
| NC_002645.1;NP_073549.1;HCoV-229E_1ab;821 | 0 | 0 |
| NC_002645.1;NP_073549.1;HCoV-229E_1ab;822 | 0 | 0 |
| NC_002645.1;NP_073549.1;HCoV-229E_1ab;823 | 0 | 0 |
| NC_002645.1;NP_073549.1;HCoV-229E_1ab;824 | 0.05 | 0.025 |
| NC_002645.1;NP_073549.1;HCoV-229E_1ab;825 | 0.1 | 0.025 |
| NC_002645.1;NP_073549.1;HCoV-229E_1ab;826 | 0.1 | 0.025 |
| NC_002645.1;NP_073549.1;HCoV-229E_1ab;827 | 0.05 | 0.075 |
| NC_002645.1;NP_073549.1;HCoV-229E_1ab;828 | 0 | 0 |
| NC_002645.1;NP_073549.1;HCoV-229E_1ab;829 | 0 | 0 |
| NC_002645.1;NP_073549.1;HCoV-229E_1ab;830 | 0 | 0 |
| NC_002645.1;NP_073549.1;HCoV-229E_1ab;831 | 0.1 | 0.025 |
| NC_002645.1;NP_073549.1;HCoV-229E_1ab;832 | 0 | 0 |
| NC_002645.1;NP_073549.1;HCoV-229E_1ab;833 | 0 | 0 |
| NC_002645.1;NP_073549.1;HCoV-229E_1ab;834 | 0 | 0 |
| NC_002645.1;NP_073549.1;HCoV-229E_1ab;835 | 0 | 0.025 |
| NC_002645.1;NP_073549.1;HCoV-229E_1ab;836 | 0 | 0.025 |
| NC_002645.1;NP_073549.1;HCoV-229E_1ab;837 | 0 | 0 |
| NC_002645.1;NP_073549.1;HCoV-229E_1ab;838 | 0 | 0 |
| NC_002645.1;NP_073549.1;HCoV-229E_1ab;839 | 0 | 0 |
| NC_002645.1;NP_073549.1;HCoV-229E_1ab;840 | 0 | 0 |
| NC_002645.1;NP_073549.1;HCoV-229E_1ab;841 | 0 | 0 |
| NC_002645.1;NP_073549.1;HCoV-229E_1ab;842 | 0 | 0 |
| NC_002645.1;NP_073549.1;HCoV-229E_1ab;843 | 0 | 0 |
| NC_002645.1;NP_073549.1;HCoV-229E_1ab;844 | 0 | 0.025 |
| NC_002645.1;NP_073549.1;HCoV-229E_1ab;845 | 0 | 0.025 |
| NC_002645.1;NP_073549.1;HCoV-229E_1ab;846 | 0 | 0 |

| | | |
|---|---|---|
| NC_002645.1;NP_073549.1;HCoV-229E_1ab;847 | 0 | 0 |
| NC_002645.1;NP_073549.1;HCoV-229E_1ab;848 | 0 | 0 |
| NC_002645.1;NP_073549.1;HCoV-229E_1ab;849 | 0 | 0.025 |
| NC_002645.1;NP_073549.1;HCoV-229E_1ab;850 | 0 | 0 |
| NC_002645.1;NP_073549.1;HCoV-229E_1ab;851 | 0 | 0 |
| NC_002645.1;NP_073549.1;HCoV-229E_1ab;852 | 0.05 | 0 |
| NC_002645.1;NP_073549.1;HCoV-229E_1ab;853 | 0 | 0.05 |
| NC_002645.1;NP_073549.1;HCoV-229E_1ab;854 | 0 | 0 |
| NC_002645.1;NP_073549.1;HCoV-229E_1ab;855 | 0 | 0 |
| NC_002645.1;NP_073549.1;HCoV-229E_1ab;856 | 0 | 0 |
| NC_002645.1;NP_073549.1;HCoV-229E_1ab;857 | 0 | 0 |
| NC_002645.1;NP_073549.1;HCoV-229E_1ab;858 | 0 | 0 |
| NC_002645.1;NP_073549.1;HCoV-229E_1ab;859 | 0 | 0 |
| NC_002645.1;NP_073549.1;HCoV-229E_1ab;860 | 0 | 0 |
| NC_002645.1;NP_073549.1;HCoV-229E_1ab;861 | 0 | 0 |
| NC_002645.1;NP_073549.1;HCoV-229E_1ab;862 | 0 | 0 |
| NC_002645.1;NP_073549.1;HCoV-229E_1ab;863 | 0 | 0 |
| NC_002645.1;NP_073549.1;HCoV-229E_1ab;864 | 0 | 0 |
| NC_002645.1;NP_073549.1;HCoV-229E_1ab;865 | 0.05 | 0 |
| NC_002645.1;NP_073549.1;HCoV-229E_1ab;866 | 0.05 | 0 |
| NC_002645.1;NP_073549.1;HCoV-229E_1ab;867 | 0.05 | 0.025 |
| NC_002645.1;NP_073549.1;HCoV-229E_1ab;868 | 0.05 | 0 |
| NC_002645.1;NP_073549.1;HCoV-229E_1ab;869 | 0.05 | 0 |
| NC_002645.1;NP_073549.1;HCoV-229E_1ab;870 | 0.05 | 0 |
| NC_002645.1;NP_073549.1;HCoV-229E_1ab;871 | 0.05 | 0 |
| NC_002645.1;NP_073549.1;HCoV-229E_1ab;872 | 0.15 | 0.025 |
| NC_002645.1;NP_073549.1;HCoV-229E_1ab;873 | 0.05 | 0.025 |
| NC_002645.1;NP_073549.1;HCoV-229E_1ab;874 | 0.2 | 0.1 |
| NC_002645.1;NP_073549.1;HCoV-229E_1ab;875 | 0.05 | 0.025 |
| NC_002645.1;NP_073549.1;HCoV-229E_1ab;876 | 0.1 | 0.05 |
| NC_002645.1;NP_073549.1;HCoV-229E_1ab;877 | 0 | 0 |
| NC_002645.1;NP_073549.1;HCoV-229E_1ab;878 | 0 | 0 |
| NC_002645.1;NP_073549.1;HCoV-229E_1ab;879 | 0.05 | 0 |
| NC_002645.1;NP_073549.1;HCoV-229E_1ab;880 | 0.05 | 0 |
| NC_002645.1;NP_073549.1;HCoV-229E_1ab;881 | 0.05 | 0 |
| NC_002645.1;NP_073549.1;HCoV-229E_1ab;882 | 0 | 0 |
| NC_002645.1;NP_073549.1;HCoV-229E_1ab;883 | 0.05 | 0 |
| NC_002645.1;NP_073549.1;HCoV-229E_1ab;884 | 0 | 0 |
| NC_002645.1;NP_073549.1;HCoV-229E_1ab;885 | 0 | 0 |
| NC_002645.1;NP_073549.1;HCoV-229E_1ab;886 | 0 | 0 |
| NC_002645.1;NP_073549.1;HCoV-229E_1ab;887 | 0 | 0.025 |
| NC_002645.1;NP_073549.1;HCoV-229E_1ab;888 | 0 | 0.025 |
| NC_002645.1;NP_073549.1;HCoV-229E_1ab;889 | 0 | 0.025 |

| | | |
|---|---|---|
| NC_002645.1;NP_073549.1;HCoV-229E_1ab;890 | 0 | 0.025 |
| NC_002645.1;NP_073549.1;HCoV-229E_1ab;891 | 0 | 0.025 |
| NC_002645.1;NP_073549.1;HCoV-229E_1ab;892 | 0 | 0.025 |
| NC_002645.1;NP_073549.1;HCoV-229E_1ab;893 | 0 | 0.025 |
| NC_002645.1;NP_073549.1;HCoV-229E_1ab;894 | 0 | 0.025 |
| NC_002645.1;NP_073549.1;HCoV-229E_1ab;895 | 0 | 0.025 |
| NC_002645.1;NP_073549.1;HCoV-229E_1ab;896 | 0 | 0 |
| NC_002645.1;NP_073549.1;HCoV-229E_1ab;897 | 0 | 0 |
| NC_002645.1;NP_073549.1;HCoV-229E_1ab;898 | 0 | 0 |
| NC_002645.1;NP_073549.1;HCoV-229E_1ab;899 | 0 | 0 |
| NC_002645.1;NP_073549.1;HCoV-229E_1ab;900 | 0.1 | 0 |
| NC_002645.1;NP_073549.1;HCoV-229E_1ab;901 | 0.05 | 0 |
| NC_002645.1;NP_073549.1;HCoV-229E_1ab;902 | 0 | 0 |
| NC_002645.1;NP_073549.1;HCoV-229E_1ab;903 | 0.1 | 0 |
| NC_002645.1;NP_073549.1;HCoV-229E_1ab;904 | 0 | 0 |
| NC_002645.1;NP_073549.1;HCoV-229E_1ab;905 | 0 | 0 |
| NC_002645.1;NP_073549.1;HCoV-229E_1ab;906 | 0.05 | 0 |
| NC_002645.1;NP_073549.1;HCoV-229E_1ab;907 | 0 | 0 |
| NC_002645.1;NP_073549.1;HCoV-229E_1ab;908 | 0 | 0 |
| NC_002645.1;NP_073549.1;HCoV-229E_1ab;909 | 0 | 0 |
| NC_002645.1;NP_073549.1;HCoV-229E_1ab;910 | 0 | 0 |
| NC_002645.1;NP_073549.1;HCoV-229E_1ab;911 | 0 | 0.025 |
| NC_002645.1;NP_073549.1;HCoV-229E_1ab;912 | 0 | 0 |
| NC_002645.1;NP_073549.1;HCoV-229E_1ab;913 | 0 | 0 |
| NC_002645.1;NP_073549.1;HCoV-229E_1ab;914 | 0 | 0 |
| NC_002645.1;NP_073549.1;HCoV-229E_1ab;915 | 0 | 0 |
| NC_002645.1;NP_073549.1;HCoV-229E_1ab;916 | 0 | 0 |
| NC_002645.1;NP_073549.1;HCoV-229E_1ab;917 | 0 | 0 |
| NC_002645.1;NP_073549.1;HCoV-229E_1ab;918 | 0 | 0 |
| NC_002645.1;NP_073549.1;HCoV-229E_1ab;919 | 0 | 0 |
| NC_002645.1;NP_073549.1;HCoV-229E_1ab;920 | 0 | 0 |
| NC_002645.1;NP_073549.1;HCoV-229E_1ab;921 | 0 | 0 |
| NC_002645.1;NP_073549.1;HCoV-229E_1ab;922 | 0 | 0 |
| NC_002645.1;NP_073549.1;HCoV-229E_1ab;923 | 0 | 0 |
| NC_002645.1;NP_073549.1;HCoV-229E_1ab;924 | 0.05 | 0.05 |
| NC_002645.1;NP_073549.1;HCoV-229E_1ab;925 | 0 | 0 |
| NC_002645.1;NP_073549.1;HCoV-229E_1ab;926 | 0 | 0 |
| NC_002645.1;NP_073549.1;HCoV-229E_1ab;927 | 0 | 0 |
| NC_002645.1;NP_073549.1;HCoV-229E_1ab;928 | 0 | 0 |
| NC_002645.1;NP_073549.1;HCoV-229E_1ab;929 | 0 | 0 |
| NC_002645.1;NP_073549.1;HCoV-229E_1ab;930 | 0 | 0 |
| NC_002645.1;NP_073549.1;HCoV-229E_1ab;931 | 0 | 0 |
| NC_002645.1;NP_073549.1;HCoV-229E_1ab;932 | 0 | 0.025 |

| | | |
|---|---|---|
| NC_002645.1;NP_073549.1;HCoV-229E_1ab;933 | 0 | 0.025 |
| NC_002645.1;NP_073549.1;HCoV-229E_1ab;934 | 0 | 0.075 |
| NC_002645.1;NP_073549.1;HCoV-229E_1ab;935 | 0 | 0.025 |
| NC_002645.1;NP_073549.1;HCoV-229E_1ab;936 | 0 | 0 |
| NC_002645.1;NP_073549.1;HCoV-229E_1ab;937 | 0 | 0.025 |
| NC_002645.1;NP_073549.1;HCoV-229E_1ab;938 | 0 | 0.05 |
| NC_002645.1;NP_073549.1;HCoV-229E_1ab;939 | 0 | 0 |
| NC_002645.1;NP_073549.1;HCoV-229E_1ab;940 | 0 | 0 |
| NC_002645.1;NP_073549.1;HCoV-229E_1ab;941 | 0 | 0 |
| NC_002645.1;NP_073549.1;HCoV-229E_1ab;942 | 0 | 0 |
| NC_002645.1;NP_073549.1;HCoV-229E_1ab;943 | 0 | 0 |
| NC_002645.1;NP_073549.1;HCoV-229E_1ab;944 | 0 | 0 |
| NC_002645.1;NP_073549.1;HCoV-229E_1ab;945 | 0 | 0 |
| NC_002645.1;NP_073549.1;HCoV-229E_1ab;946 | 0 | 0 |
| NC_002645.1;NP_073549.1;HCoV-229E_1ab;947 | 0 | 0 |
| NC_002645.1;NP_073549.1;HCoV-229E_1ab;948 | 0 | 0 |
| NC_002645.1;NP_073549.1;HCoV-229E_1ab;949 | 0 | 0 |
| NC_002645.1;NP_073549.1;HCoV-229E_1ab;950 | 0 | 0 |
| NC_002645.1;NP_073549.1;HCoV-229E_1ab;951 | 0 | 0 |
| NC_002645.1;NP_073549.1;HCoV-229E_1ab;952 | 0 | 0 |
| NC_002645.1;NP_073549.1;HCoV-229E_1ab;953 | 0 | 0 |
| NC_002645.1;NP_073549.1;HCoV-229E_1ab;954 | 0 | 0 |
| NC_002645.1;NP_073549.1;HCoV-229E_1ab;955 | 0 | 0 |
| NC_002645.1;NP_073549.1;HCoV-229E_1ab;956 | 0 | 0 |
| NC_002645.1;NP_073549.1;HCoV-229E_1ab;957 | 0 | 0 |
| NC_002645.1;NP_073549.1;HCoV-229E_1ab;958 | 0 | 0 |
| NC_002645.1;NP_073549.1;HCoV-229E_1ab;959 | 0 | 0 |
| NC_002645.1;NP_073549.1;HCoV-229E_1ab;960 | 0 | 0 |
| NC_002645.1;NP_073549.1;HCoV-229E_1ab;961 | 0 | 0 |
| NC_002645.1;NP_073549.1;HCoV-229E_1ab;962 | 0 | 0 |
| NC_002645.1;NP_073549.1;HCoV-229E_1ab;963 | 0 | 0 |
| NC_002645.1;NP_073549.1;HCoV-229E_1ab;964 | 0 | 0 |
| NC_002645.1;NP_073549.1;HCoV-229E_1ab;965 | 0 | 0.025 |
| NC_002645.1;NP_073549.1;HCoV-229E_1ab;966 | 0 | 0 |
| NC_002645.1;NP_073549.1;HCoV-229E_1ab;967 | 0 | 0 |
| NC_002645.1;NP_073549.1;HCoV-229E_1ab;968 | 0 | 0.025 |
| NC_002645.1;NP_073549.1;HCoV-229E_1ab;969 | 0 | 0 |
| NC_002645.1;NP_073549.1;HCoV-229E_1ab;970 | 0 | 0.025 |
| NC_002645.1;NP_073549.1;HCoV-229E_1ab;971 | 0 | 0 |
| NC_002645.1;NP_073549.1;HCoV-229E_1ab;972 | 0.05 | 0 |
| NC_002645.1;NP_073549.1;HCoV-229E_1ab;973 | 0 | 0 |
| NC_002645.1;NP_073549.1;HCoV-229E_1ab;974 | 0.05 | 0 |
| NC_002645.1;NP_073549.1;HCoV-229E_1ab;975 | 0.1 | 0 |

| | | |
|---|---|---|
| NC_002645.1;NP_073549.1;HCoV-229E_1ab;976 | 0 | 0 |
| NC_002645.1;NP_073549.1;HCoV-229E_1ab;977 | 0 | 0 |
| NC_002645.1;NP_073549.1;HCoV-229E_1ab;978 | 0 | 0 |
| NC_002645.1;NP_073549.1;HCoV-229E_1ab;979 | 0 | 0 |
| NC_002645.1;NP_073549.1;HCoV-229E_1ab;980 | 0 | 0 |
| NC_002645.1;NP_073549.1;HCoV-229E_1ab;981 | 0 | 0 |
| NC_002645.1;NP_073549.1;HCoV-229E_1ab;982 | 0 | 0 |
| NC_002645.1;NP_073549.1;HCoV-229E_1ab;983 | 0.05 | 0.05 |
| NC_002645.1;NP_073549.1;HCoV-229E_1ab;984 | 0 | 0.025 |
| NC_002645.1;NP_073549.1;HCoV-229E_1ab;985 | 0 | 0.025 |
| NC_002645.1;NP_073549.1;HCoV-229E_1ab;986 | 0 | 0.05 |
| NC_002645.1;NP_073549.1;HCoV-229E_1ab;987 | 0 | 0 |
| NC_002645.1;NP_073549.1;HCoV-229E_1ab;988 | 0 | 0.15 |
| NC_002645.1;NP_073549.1;HCoV-229E_1ab;989 | 0.05 | 0.125 |
| NC_002645.1;NP_073549.1;HCoV-229E_1ab;990 | 0 | 0.05 |
| NC_002645.1;NP_073549.1;HCoV-229E_1ab;991 | 0 | 0.1 |
| NC_002645.1;NP_073549.1;HCoV-229E_1ab;992 | 0 | 0.075 |
| NC_002645.1;NP_073549.1;HCoV-229E_1ab;993 | 0 | 0.025 |
| NC_002645.1;NP_073549.1;HCoV-229E_1ab;994 | 0 | 0 |
| NC_002645.1;NP_073549.1;HCoV-229E_1ab;995 | 0 | 0 |
| NC_002645.1;NP_073549.1;HCoV-229E_1ab;996 | 0 | 0 |
| NC_002645.1;NP_073549.1;HCoV-229E_1ab;997 | 0.05 | 0 |
| NC_002645.1;NP_073549.1;HCoV-229E_1ab;998 | 0.05 | 0 |
| NC_002645.1;NP_073549.1;HCoV-229E_1ab;999 | 0.05 | 0 |
| NC_002645.1;NP_073549.1;HCoV-229E_1ab;1000 | 0.05 | 0 |
| NC_002645.1;NP_073549.1;HCoV-229E_1ab;1001 | 0.05 | 0 |
| NC_002645.1;NP_073549.1;HCoV-229E_1ab;1002 | 0.05 | 0 |
| NC_002645.1;NP_073549.1;HCoV-229E_1ab;1003 | 0 | 0 |
| NC_002645.1;NP_073549.1;HCoV-229E_1ab;1004 | 0 | 0.025 |
| NC_002645.1;NP_073549.1;HCoV-229E_1ab;1005 | 0.1 | 0.05 |
| NC_002645.1;NP_073549.1;HCoV-229E_1ab;1006 | 0.15 | 0.05 |
| NC_002645.1;NP_073549.1;HCoV-229E_1ab;1007 | 0.2 | 0.05 |
| NC_002645.1;NP_073549.1;HCoV-229E_1ab;1008 | 0.1 | 0.05 |
| NC_002645.1;NP_073549.1;HCoV-229E_1ab;1009 | 0.05 | 0.05 |
| NC_002645.1;NP_073549.1;HCoV-229E_1ab;1010 | 0 | 0.05 |
| NC_002645.1;NP_073549.1;HCoV-229E_1ab;1011 | 0.05 | 0.075 |
| NC_002645.1;NP_073549.1;HCoV-229E_1ab;1012 | 0.1 | 0.075 |
| NC_002645.1;NP_073549.1;HCoV-229E_1ab;1013 | 0.15 | 0.125 |
| NC_002645.1;NP_073549.1;HCoV-229E_1ab;1014 | 0.05 | 0.075 |
| NC_002645.1;NP_073549.1;HCoV-229E_1ab;1015 | 0.1 | 0.1 |
| NC_002645.1;NP_073549.1;HCoV-229E_1ab;1016 | 0.1 | 0.125 |
| NC_002645.1;NP_073549.1;HCoV-229E_1ab;1017 | 0.15 | 0.075 |
| NC_002645.1;NP_073549.1;HCoV-229E_1ab;1018 | 0.15 | 0.075 |

| | | |
|---|---|---|
| NC_002645.1;NP_073549.1;HCoV-229E_1ab;1019 | 0.1 | 0.05 |
| NC_002645.1;NP_073549.1;HCoV-229E_1ab;1020 | 0 | 0.025 |
| NC_002645.1;NP_073549.1;HCoV-229E_1ab;1021 | 0 | 0 |
| NC_002645.1;NP_073549.1;HCoV-229E_1ab;1022 | 0.05 | 0 |
| NC_002645.1;NP_073549.1;HCoV-229E_1ab;1023 | 0 | 0 |
| NC_002645.1;NP_073549.1;HCoV-229E_1ab;1024 | 0.05 | 0 |
| NC_002645.1;NP_073549.1;HCoV-229E_1ab;1025 | 0.05 | 0 |
| NC_002645.1;NP_073549.1;HCoV-229E_1ab;1026 | 0.05 | 0 |
| NC_002645.1;NP_073549.1;HCoV-229E_1ab;1027 | 0.05 | 0.025 |
| NC_002645.1;NP_073549.1;HCoV-229E_1ab;1028 | 0.1 | 0.05 |
| NC_002645.1;NP_073549.1;HCoV-229E_1ab;1029 | 0.05 | 0.025 |
| NC_002645.1;NP_073549.1;HCoV-229E_1ab;1030 | 0.05 | 0.025 |
| NC_002645.1;NP_073549.1;HCoV-229E_1ab;1031 | 0.05 | 0.05 |
| NC_002645.1;NP_073549.1;HCoV-229E_1ab;1032 | 0 | 0.025 |
| NC_002645.1;NP_073549.1;HCoV-229E_1ab;1033 | 0 | 0 |
| NC_002645.1;NP_073549.1;HCoV-229E_1ab;1034 | 0 | 0.025 |
| NC_002645.1;NP_073549.1;HCoV-229E_1ab;1035 | 0 | 0 |
| NC_002645.1;NP_073549.1;HCoV-229E_1ab;1036 | 0 | 0.025 |
| NC_002645.1;NP_073549.1;HCoV-229E_1ab;1037 | 0 | 0 |
| NC_002645.1;NP_073549.1;HCoV-229E_1ab;1038 | 0.05 | 0 |
| NC_002645.1;NP_073549.1;HCoV-229E_1ab;1039 | 0 | 0 |
| NC_002645.1;NP_073549.1;HCoV-229E_1ab;1040 | 0 | 0.025 |
| NC_002645.1;NP_073549.1;HCoV-229E_1ab;1041 | 0 | 0 |
| NC_002645.1;NP_073549.1;HCoV-229E_1ab;1042 | 0 | 0 |
| NC_002645.1;NP_073549.1;HCoV-229E_1ab;1043 | 0 | 0 |
| NC_002645.1;NP_073549.1;HCoV-229E_1ab;1044 | 0 | 0 |
| NC_002645.1;NP_073549.1;HCoV-229E_1ab;1045 | 0.05 | 0 |
| NC_002645.1;NP_073549.1;HCoV-229E_1ab;1046 | 0 | 0 |
| NC_002645.1;NP_073549.1;HCoV-229E_1ab;1047 | 0 | 0 |
| NC_002645.1;NP_073549.1;HCoV-229E_1ab;1048 | 0 | 0 |
| NC_002645.1;NP_073549.1;HCoV-229E_1ab;1049 | 0 | 0 |
| NC_002645.1;NP_073549.1;HCoV-229E_1ab;1050 | 0 | 0 |
| NC_002645.1;NP_073549.1;HCoV-229E_1ab;1051 | 0 | 0 |
| NC_002645.1;NP_073549.1;HCoV-229E_1ab;1052 | 0 | 0 |
| NC_002645.1;NP_073549.1;HCoV-229E_1ab;1053 | 0 | 0 |
| NC_002645.1;NP_073549.1;HCoV-229E_1ab;1054 | 0 | 0 |
| NC_002645.1;NP_073549.1;HCoV-229E_1ab;1055 | 0 | 0 |
| NC_002645.1;NP_073549.1;HCoV-229E_1ab;1056 | 0 | 0 |
| NC_002645.1;NP_073549.1;HCoV-229E_1ab;1057 | 0 | 0 |
| NC_002645.1;NP_073549.1;HCoV-229E_1ab;1058 | 0 | 0 |
| NC_002645.1;NP_073549.1;HCoV-229E_1ab;1059 | 0 | 0 |
| NC_002645.1;NP_073549.1;HCoV-229E_1ab;1060 | 0 | 0 |
| NC_002645.1;NP_073549.1;HCoV-229E_1ab;1061 | 0 | 0 |

| | | |
|---|---|---|
| NC_002645.1;NP_073549.1;HCoV-229E_1ab;1062 | 0 | 0 |
| NC_002645.1;NP_073549.1;HCoV-229E_1ab;1063 | 0 | 0 |
| NC_002645.1;NP_073549.1;HCoV-229E_1ab;1064 | 0.05 | 0.025 |
| NC_002645.1;NP_073549.1;HCoV-229E_1ab;1065 | 0.05 | 0.025 |
| NC_002645.1;NP_073549.1;HCoV-229E_1ab;1066 | 0.05 | 0.025 |
| NC_002645.1;NP_073549.1;HCoV-229E_1ab;1067 | 0.1 | 0.075 |
| NC_002645.1;NP_073549.1;HCoV-229E_1ab;1068 | 0 | 0 |
| NC_002645.1;NP_073549.1;HCoV-229E_1ab;1069 | 0 | 0.025 |
| NC_002645.1;NP_073549.1;HCoV-229E_1ab;1070 | 0 | 0 |
| NC_002645.1;NP_073549.1;HCoV-229E_1ab;1071 | 0 | 0 |
| NC_002645.1;NP_073549.1;HCoV-229E_1ab;1072 | 0 | 0 |
| NC_002645.1;NP_073549.1;HCoV-229E_1ab;1073 | 0 | 0 |
| NC_002645.1;NP_073549.1;HCoV-229E_1ab;1074 | 0 | 0 |
| NC_002645.1;NP_073549.1;HCoV-229E_1ab;1075 | 0 | 0 |
| NC_002645.1;NP_073549.1;HCoV-229E_1ab;1076 | 0 | 0 |
| NC_002645.1;NP_073549.1;HCoV-229E_1ab;1077 | 0 | 0 |
| NC_002645.1;NP_073549.1;HCoV-229E_1ab;1078 | 0 | 0 |
| NC_002645.1;NP_073549.1;HCoV-229E_1ab;1079 | 0 | 0 |
| NC_002645.1;NP_073549.1;HCoV-229E_1ab;1080 | 0 | 0 |
| NC_002645.1;NP_073549.1;HCoV-229E_1ab;1081 | 0 | 0 |
| NC_002645.1;NP_073549.1;HCoV-229E_1ab;1082 | 0 | 0 |
| NC_002645.1;NP_073549.1;HCoV-229E_1ab;1083 | 0 | 0 |
| NC_002645.1;NP_073549.1;HCoV-229E_1ab;1084 | 0 | 0 |
| NC_002645.1;NP_073549.1;HCoV-229E_1ab;1085 | 0 | 0 |
| NC_002645.1;NP_073549.1;HCoV-229E_1ab;1086 | 0 | 0 |
| NC_002645.1;NP_073549.1;HCoV-229E_1ab;1087 | 0 | 0 |
| NC_002645.1;NP_073549.1;HCoV-229E_1ab;1088 | 0.05 | 0 |
| NC_002645.1;NP_073549.1;HCoV-229E_1ab;1089 | 0 | 0 |
| NC_002645.1;NP_073549.1;HCoV-229E_1ab;1090 | 0 | 0 |
| NC_002645.1;NP_073549.1;HCoV-229E_1ab;1091 | 0 | 0.025 |
| NC_002645.1;NP_073549.1;HCoV-229E_1ab;1092 | 0 | 0 |
| NC_002645.1;NP_073549.1;HCoV-229E_1ab;1093 | 0 | 0 |
| NC_002645.1;NP_073549.1;HCoV-229E_1ab;1094 | 0 | 0 |
| NC_002645.1;NP_073549.1;HCoV-229E_1ab;1095 | 0 | 0 |
| NC_002645.1;NP_073549.1;HCoV-229E_1ab;1096 | 0 | 0 |
| NC_002645.1;NP_073549.1;HCoV-229E_1ab;1097 | 0 | 0 |
| NC_002645.1;NP_073549.1;HCoV-229E_1ab;1098 | 0 | 0 |
| NC_002645.1;NP_073549.1;HCoV-229E_1ab;1099 | 0 | 0 |
| NC_002645.1;NP_073549.1;HCoV-229E_1ab;1100 | 0 | 0 |
| NC_002645.1;NP_073549.1;HCoV-229E_1ab;1101 | 0 | 0 |
| NC_002645.1;NP_073549.1;HCoV-229E_1ab;1102 | 0 | 0 |
| NC_002645.1;NP_073549.1;HCoV-229E_1ab;1103 | 0 | 0 |
| NC_002645.1;NP_073549.1;HCoV-229E_1ab;1104 | 0 | 0 |

| | | |
|---|---|---|
| NC_002645.1;NP_073549.1;HCoV-229E_1ab;1105 | 0 | 0 |
| NC_002645.1;NP_073549.1;HCoV-229E_1ab;1106 | 0 | 0 |
| NC_002645.1;NP_073549.1;HCoV-229E_1ab;1107 | 0 | 0 |
| NC_002645.1;NP_073549.1;HCoV-229E_1ab;1108 | 0 | 0 |
| NC_002645.1;NP_073549.1;HCoV-229E_1ab;1109 | 0 | 0 |
| NC_002645.1;NP_073549.1;HCoV-229E_1ab;1110 | 0.05 | 0 |
| NC_002645.1;NP_073549.1;HCoV-229E_1ab;1111 | 0 | 0 |
| NC_002645.1;NP_073549.1;HCoV-229E_1ab;1112 | 0.05 | 0 |
| NC_002645.1;NP_073549.1;HCoV-229E_1ab;1113 | 0 | 0 |
| NC_002645.1;NP_073549.1;HCoV-229E_1ab;1114 | 0 | 0 |
| NC_002645.1;NP_073549.1;HCoV-229E_1ab;1115 | 0 | 0 |
| NC_002645.1;NP_073549.1;HCoV-229E_1ab;1116 | 0 | 0 |
| NC_002645.1;NP_073549.1;HCoV-229E_1ab;1117 | 0 | 0 |
| NC_002645.1;NP_073549.1;HCoV-229E_1ab;1118 | 0 | 0 |
| NC_002645.1;NP_073549.1;HCoV-229E_1ab;1119 | 0 | 0 |
| NC_002645.1;NP_073549.1;HCoV-229E_1ab;1120 | 0 | 0 |
| NC_002645.1;NP_073549.1;HCoV-229E_1ab;1121 | 0 | 0 |
| NC_002645.1;NP_073549.1;HCoV-229E_1ab;1122 | 0 | 0 |
| NC_002645.1;NP_073549.1;HCoV-229E_1ab;1123 | 0 | 0 |
| NC_002645.1;NP_073549.1;HCoV-229E_1ab;1124 | 0 | 0 |
| NC_002645.1;NP_073549.1;HCoV-229E_1ab;1125 | 0 | 0 |
| NC_002645.1;NP_073549.1;HCoV-229E_1ab;1126 | 0 | 0 |
| NC_002645.1;NP_073549.1;HCoV-229E_1ab;1127 | 0 | 0 |
| NC_002645.1;NP_073549.1;HCoV-229E_1ab;1128 | 0 | 0 |
| NC_002645.1;NP_073549.1;HCoV-229E_1ab;1129 | 0 | 0 |
| NC_002645.1;NP_073549.1;HCoV-229E_1ab;1130 | 0 | 0 |
| NC_002645.1;NP_073549.1;HCoV-229E_1ab;1131 | 0 | 0 |
| NC_002645.1;NP_073549.1;HCoV-229E_1ab;1132 | 0 | 0 |
| NC_002645.1;NP_073549.1;HCoV-229E_1ab;1133 | 0 | 0 |
| NC_002645.1;NP_073549.1;HCoV-229E_1ab;1134 | 0 | 0.025 |
| NC_002645.1;NP_073549.1;HCoV-229E_1ab;1135 | 0 | 0 |
| NC_002645.1;NP_073549.1;HCoV-229E_1ab;1136 | 0 | 0.025 |
| NC_002645.1;NP_073549.1;HCoV-229E_1ab;1137 | 0 | 0.025 |
| NC_002645.1;NP_073549.1;HCoV-229E_1ab;1138 | 0 | 0 |
| NC_002645.1;NP_073549.1;HCoV-229E_1ab;1139 | 0 | 0 |
| NC_002645.1;NP_073549.1;HCoV-229E_1ab;1140 | 0 | 0 |
| NC_002645.1;NP_073549.1;HCoV-229E_1ab;1141 | 0 | 0.025 |
| NC_002645.1;NP_073549.1;HCoV-229E_1ab;1142 | 0 | 0.025 |
| NC_002645.1;NP_073549.1;HCoV-229E_1ab;1143 | 0 | 0.05 |
| NC_002645.1;NP_073549.1;HCoV-229E_1ab;1144 | 0 | 0 |
| NC_002645.1;NP_073549.1;HCoV-229E_1ab;1145 | 0 | 0.025 |
| NC_002645.1;NP_073549.1;HCoV-229E_1ab;1146 | 0 | 0 |
| NC_002645.1;NP_073549.1;HCoV-229E_1ab;1147 | 0 | 0 |

| | | |
|---|---|---|
| NC_002645.1;NP_073549.1;HCoV-229E_1ab;1148 | 0 | 0 |
| NC_002645.1;NP_073549.1;HCoV-229E_1ab;1149 | 0.05 | 0 |
| NC_002645.1;NP_073549.1;HCoV-229E_1ab;1150 | 0 | 0 |
| NC_002645.1;NP_073549.1;HCoV-229E_1ab;1151 | 0 | 0 |
| NC_002645.1;NP_073549.1;HCoV-229E_1ab;1152 | 0 | 0 |
| NC_002645.1;NP_073549.1;HCoV-229E_1ab;1153 | 0 | 0 |
| NC_002645.1;NP_073549.1;HCoV-229E_1ab;1154 | 0 | 0 |
| NC_002645.1;NP_073549.1;HCoV-229E_1ab;1155 | 0 | 0 |
| NC_002645.1;NP_073549.1;HCoV-229E_1ab;1156 | 0 | 0 |
| NC_002645.1;NP_073549.1;HCoV-229E_1ab;1157 | 0 | 0 |
| NC_002645.1;NP_073549.1;HCoV-229E_1ab;1158 | 0 | 0 |
| NC_002645.1;NP_073549.1;HCoV-229E_1ab;1159 | 0 | 0 |
| NC_002645.1;NP_073549.1;HCoV-229E_1ab;1160 | 0 | 0 |
| NC_002645.1;NP_073549.1;HCoV-229E_1ab;1161 | 0 | 0 |
| NC_002645.1;NP_073549.1;HCoV-229E_1ab;1162 | 0 | 0 |
| NC_002645.1;NP_073549.1;HCoV-229E_1ab;1163 | 0 | 0 |
| NC_002645.1;NP_073549.1;HCoV-229E_1ab;1164 | 0 | 0 |
| NC_002645.1;NP_073549.1;HCoV-229E_1ab;1165 | 0 | 0 |
| NC_002645.1;NP_073549.1;HCoV-229E_1ab;1166 | 0 | 0 |
| NC_002645.1;NP_073549.1;HCoV-229E_1ab;1167 | 0 | 0 |
| NC_002645.1;NP_073549.1;HCoV-229E_1ab;1168 | 0 | 0 |
| NC_002645.1;NP_073549.1;HCoV-229E_1ab;1169 | 0 | 0 |
| NC_002645.1;NP_073549.1;HCoV-229E_1ab;1170 | 0 | 0 |
| NC_002645.1;NP_073549.1;HCoV-229E_1ab;1171 | 0 | 0.025 |
| NC_002645.1;NP_073549.1;HCoV-229E_1ab;1172 | 0 | 0.025 |
| NC_002645.1;NP_073549.1;HCoV-229E_1ab;1173 | 0.1 | 0.025 |
| NC_002645.1;NP_073549.1;HCoV-229E_1ab;1174 | 0.05 | 0.025 |
| NC_002645.1;NP_073549.1;HCoV-229E_1ab;1175 | 0 | 0 |
| NC_002645.1;NP_073549.1;HCoV-229E_1ab;1176 | 0 | 0 |
| NC_002645.1;NP_073549.1;HCoV-229E_1ab;1177 | 0 | 0 |
| NC_002645.1;NP_073549.1;HCoV-229E_1ab;1178 | 0 | 0 |
| NC_002645.1;NP_073549.1;HCoV-229E_1ab;1179 | 0 | 0 |
| NC_002645.1;NP_073549.1;HCoV-229E_1ab;1180 | 0 | 0 |
| NC_002645.1;NP_073549.1;HCoV-229E_1ab;1181 | 0 | 0 |
| NC_002645.1;NP_073549.1;HCoV-229E_1ab;1182 | 0 | 0 |
| NC_002645.1;NP_073549.1;HCoV-229E_1ab;1183 | 0.05 | 0.025 |
| NC_002645.1;NP_073549.1;HCoV-229E_1ab;1184 | 0 | 0 |
| NC_002645.1;NP_073549.1;HCoV-229E_1ab;1185 | 0 | 0 |
| NC_002645.1;NP_073549.1;HCoV-229E_1ab;1186 | 0 | 0 |
| NC_002645.1;NP_073549.1;HCoV-229E_1ab;1187 | 0 | 0 |
| NC_002645.1;NP_073549.1;HCoV-229E_1ab;1188 | 0 | 0 |
| NC_002645.1;NP_073549.1;HCoV-229E_1ab;1189 | 0 | 0 |
| NC_002645.1;NP_073549.1;HCoV-229E_1ab;1190 | 0 | 0 |

| | | |
|---|---|---|
| NC_002645.1;NP_073549.1;HCoV-229E_1ab;1191 | 0 | 0 |
| NC_002645.1;NP_073549.1;HCoV-229E_1ab;1192 | 0 | 0 |
| NC_002645.1;NP_073549.1;HCoV-229E_1ab;1193 | 0 | 0 |
| NC_002645.1;NP_073549.1;HCoV-229E_1ab;1194 | 0 | 0 |
| NC_002645.1;NP_073549.1;HCoV-229E_1ab;1195 | 0 | 0 |
| NC_002645.1;NP_073549.1;HCoV-229E_1ab;1196 | 0 | 0 |
| NC_002645.1;NP_073549.1;HCoV-229E_1ab;1197 | 0 | 0 |
| NC_002645.1;NP_073549.1;HCoV-229E_1ab;1198 | 0 | 0 |
| NC_002645.1;NP_073549.1;HCoV-229E_1ab;1199 | 0 | 0.025 |
| NC_002645.1;NP_073549.1;HCoV-229E_1ab;1200 | 0 | 0.025 |
| NC_002645.1;NP_073549.1;HCoV-229E_1ab;1201 | 0 | 0.025 |
| NC_002645.1;NP_073549.1;HCoV-229E_1ab;1202 | 0 | 0.025 |
| NC_002645.1;NP_073549.1;HCoV-229E_1ab;1203 | 0 | 0.025 |
| NC_002645.1;NP_073549.1;HCoV-229E_1ab;1204 | 0 | 0.025 |
| NC_002645.1;NP_073549.1;HCoV-229E_1ab;1205 | 0 | 0.025 |
| NC_002645.1;NP_073549.1;HCoV-229E_1ab;1206 | 0 | 0.025 |
| NC_002645.1;NP_073549.1;HCoV-229E_1ab;1207 | 0 | 0.025 |
| NC_002645.1;NP_073549.1;HCoV-229E_1ab;1208 | 0.05 | 0.025 |
| NC_002645.1;NP_073549.1;HCoV-229E_1ab;1209 | 0 | 0.05 |
| NC_002645.1;NP_073549.1;HCoV-229E_1ab;1210 | 0 | 0.025 |
| NC_002645.1;NP_073549.1;HCoV-229E_1ab;1211 | 0 | 0.025 |
| NC_002645.1;NP_073549.1;HCoV-229E_1ab;1212 | 0 | 0 |
| NC_002645.1;NP_073549.1;HCoV-229E_1ab;1213 | 0 | 0.025 |
| NC_002645.1;NP_073549.1;HCoV-229E_1ab;1214 | 0 | 0.05 |
| NC_002645.1;NP_073549.1;HCoV-229E_1ab;1215 | 0.05 | 0.05 |
| NC_002645.1;NP_073549.1;HCoV-229E_1ab;1216 | 0.05 | 0.1 |
| NC_002645.1;NP_073549.1;HCoV-229E_1ab;1217 | 0.05 | 0.15 |
| NC_002645.1;NP_073549.1;HCoV-229E_1ab;1218 | 0.15 | 0.075 |
| NC_002645.1;NP_073549.1;HCoV-229E_1ab;1219 | 0.05 | 0.1 |
| NC_002645.1;NP_073549.1;HCoV-229E_1ab;1220 | 0.1 | 0.075 |
| NC_002645.1;NP_073549.1;HCoV-229E_1ab;1221 | 0.05 | 0.075 |
| NC_002645.1;NP_073549.1;HCoV-229E_1ab;1222 | 0.1 | 0.075 |
| NC_002645.1;NP_073549.1;HCoV-229E_1ab;1223 | 0.05 | 0.025 |
| NC_002645.1;NP_073549.1;HCoV-229E_1ab;1224 | 0.05 | 0.05 |
| NC_002645.1;NP_073549.1;HCoV-229E_1ab;1225 | 0.05 | 0.025 |
| NC_002645.1;NP_073549.1;HCoV-229E_1ab;1226 | 0.05 | 0 |
| NC_002645.1;NP_073549.1;HCoV-229E_1ab;1227 | 0.05 | 0.025 |
| NC_002645.1;NP_073549.1;HCoV-229E_1ab;1228 | 0 | 0.025 |
| NC_002645.1;NP_073549.1;HCoV-229E_1ab;1229 | 0 | 0 |
| NC_002645.1;NP_073549.1;HCoV-229E_1ab;1230 | 0 | 0 |
| NC_002645.1;NP_073549.1;HCoV-229E_1ab;1231 | 0.05 | 0.1 |
| NC_002645.1;NP_073549.1;HCoV-229E_1ab;1232 | 0.1 | 0.15 |
| NC_002645.1;NP_073549.1;HCoV-229E_1ab;1233 | 0.05 | 0.1 |

| | | |
|---|---|---|
| NC_002645.1;NP_073549.1;HCoV-229E_1ab;1234 | 0.1 | 0.1 |
| NC_002645.1;NP_073549.1;HCoV-229E_1ab;1235 | 0.05 | 0.1 |
| NC_002645.1;NP_073549.1;HCoV-229E_1ab;1236 | 0.05 | 0.05 |
| NC_002645.1;NP_073549.1;HCoV-229E_1ab;1237 | 0.05 | 0.05 |
| NC_002645.1;NP_073549.1;HCoV-229E_1ab;1238 | 0.05 | 0.075 |
| NC_002645.1;NP_073549.1;HCoV-229E_1ab;1239 | 0.1 | 0.1 |
| NC_002645.1;NP_073549.1;HCoV-229E_1ab;1240 | 0.05 | 0.1 |
| NC_002645.1;NP_073549.1;HCoV-229E_1ab;1241 | 0.1 | 0 |
| NC_002645.1;NP_073549.1;HCoV-229E_1ab;1242 | 0.1 | 0.025 |
| NC_002645.1;NP_073549.1;HCoV-229E_1ab;1243 | 0 | 0.025 |
| NC_002645.1;NP_073549.1;HCoV-229E_1ab;1244 | 0 | 0.025 |
| NC_002645.1;NP_073549.1;HCoV-229E_1ab;1245 | 0 | 0.05 |
| NC_002645.1;NP_073549.1;HCoV-229E_1ab;1246 | 0 | 0.05 |
| NC_002645.1;NP_073549.1;HCoV-229E_1ab;1247 | 0 | 0.05 |
| NC_002645.1;NP_073549.1;HCoV-229E_1ab;1248 | 0 | 0.05 |
| NC_002645.1;NP_073549.1;HCoV-229E_1ab;1249 | 0 | 0.05 |
| NC_002645.1;NP_073549.1;HCoV-229E_1ab;1250 | 0.05 | 0.05 |
| NC_002645.1;NP_073549.1;HCoV-229E_1ab;1251 | 0 | 0.05 |
| NC_002645.1;NP_073549.1;HCoV-229E_1ab;1252 | 0 | 0.05 |
| NC_002645.1;NP_073549.1;HCoV-229E_1ab;1253 | 0 | 0 |
| NC_002645.1;NP_073549.1;HCoV-229E_1ab;1254 | 0 | 0 |
| NC_002645.1;NP_073549.1;HCoV-229E_1ab;1255 | 0 | 0 |
| NC_002645.1;NP_073549.1;HCoV-229E_1ab;1256 | 0 | 0.025 |
| NC_002645.1;NP_073549.1;HCoV-229E_1ab;1257 | 0.05 | 0.025 |
| NC_002645.1;NP_073549.1;HCoV-229E_1ab;1258 | 0.1 | 0.125 |
| NC_002645.1;NP_073549.1;HCoV-229E_1ab;1259 | 0.1 | 0.175 |
| NC_002645.1;NP_073549.1;HCoV-229E_1ab;1260 | 0.2 | 0.275 |
| NC_002645.1;NP_073549.1;HCoV-229E_1ab;1261 | 0.15 | 0.175 |
| NC_002645.1;NP_073549.1;HCoV-229E_1ab;1262 | 0.05 | 0.1 |
| NC_002645.1;NP_073549.1;HCoV-229E_1ab;1263 | 0.05 | 0.125 |
| NC_002645.1;NP_073549.1;HCoV-229E_1ab;1264 | 0.1 | 0.05 |
| NC_002645.1;NP_073549.1;HCoV-229E_1ab;1265 | 0.05 | 0.075 |
| NC_002645.1;NP_073549.1;HCoV-229E_1ab;1266 | 0.05 | 0.025 |
| NC_002645.1;NP_073549.1;HCoV-229E_1ab;1267 | 0 | 0 |
| NC_002645.1;NP_073549.1;HCoV-229E_1ab;1268 | 0 | 0 |
| NC_002645.1;NP_073549.1;HCoV-229E_1ab;1269 | 0 | 0 |
| NC_002645.1;NP_073549.1;HCoV-229E_1ab;1270 | 0 | 0 |
| NC_002645.1;NP_073549.1;HCoV-229E_1ab;1271 | 0 | 0 |
| NC_002645.1;NP_073549.1;HCoV-229E_1ab;1272 | 0.05 | 0 |
| NC_002645.1;NP_073549.1;HCoV-229E_1ab;1273 | 0.05 | 0.025 |
| NC_002645.1;NP_073549.1;HCoV-229E_1ab;1274 | 0.05 | 0.05 |
| NC_002645.1;NP_073549.1;HCoV-229E_1ab;1275 | 0.05 | 0.075 |
| NC_002645.1;NP_073549.1;HCoV-229E_1ab;1276 | 0.05 | 0.025 |

| | | |
|---|---|---|
| NC_002645.1;NP_073549.1;HCoV-229E_1ab;1277 | 0.05 | 0.05 |
| NC_002645.1;NP_073549.1;HCoV-229E_1ab;1278 | 0.1 | 0.075 |
| NC_002645.1;NP_073549.1;HCoV-229E_1ab;1279 | 0.05 | 0.025 |
| NC_002645.1;NP_073549.1;HCoV-229E_1ab;1280 | 0.05 | 0 |
| NC_002645.1;NP_073549.1;HCoV-229E_1ab;1281 | 0.05 | 0.025 |
| NC_002645.1;NP_073549.1;HCoV-229E_1ab;1282 | 0.05 | 0.05 |
| NC_002645.1;NP_073549.1;HCoV-229E_1ab;1283 | 0.15 | 0.075 |
| NC_002645.1;NP_073549.1;HCoV-229E_1ab;1284 | 0.05 | 0.05 |
| NC_002645.1;NP_073549.1;HCoV-229E_1ab;1285 | 0.05 | 0 |
| NC_002645.1;NP_073549.1;HCoV-229E_1ab;1286 | 0.05 | 0 |
| NC_002645.1;NP_073549.1;HCoV-229E_1ab;1287 | 0.05 | 0.025 |
| NC_002645.1;NP_073549.1;HCoV-229E_1ab;1288 | 0.1 | 0.125 |
| NC_002645.1;NP_073549.1;HCoV-229E_1ab;1289 | 0 | 0 |
| NC_002645.1;NP_073549.1;HCoV-229E_1ab;1290 | 0 | 0 |
| NC_002645.1;NP_073549.1;HCoV-229E_1ab;1291 | 0 | 0.025 |
| NC_002645.1;NP_073549.1;HCoV-229E_1ab;1292 | 0.05 | 0.025 |
| NC_002645.1;NP_073549.1;HCoV-229E_1ab;1293 | 0 | 0.025 |
| NC_002645.1;NP_073549.1;HCoV-229E_1ab;1294 | 0.3 | 0.225 |
| NC_002645.1;NP_073549.1;HCoV-229E_1ab;1295 | 0.05 | 0 |
| NC_002645.1;NP_073549.1;HCoV-229E_1ab;1296 | 0.05 | 0 |
| NC_002645.1;NP_073549.1;HCoV-229E_1ab;1297 | 0.05 | 0 |
| NC_002645.1;NP_073549.1;HCoV-229E_1ab;1298 | 0.1 | 0.05 |
| NC_002645.1;NP_073549.1;HCoV-229E_1ab;1299 | 0.05 | 0.05 |
| NC_002645.1;NP_073549.1;HCoV-229E_1ab;1300 | 0.05 | 0.025 |
| NC_002645.1;NP_073549.1;HCoV-229E_1ab;1301 | 0.05 | 0 |
| NC_002645.1;NP_073549.1;HCoV-229E_1ab;1302 | 0.05 | 0 |
| NC_002645.1;NP_073549.1;HCoV-229E_1ab;1303 | 0 | 0 |
| NC_002645.1;NP_073549.1;HCoV-229E_1ab;1304 | 0 | 0.025 |
| NC_002645.1;NP_073549.1;HCoV-229E_1ab;1305 | 0 | 0 |
| NC_002645.1;NP_073549.1;HCoV-229E_1ab;1306 | 0 | 0 |
| NC_002645.1;NP_073549.1;HCoV-229E_1ab;1307 | 0 | 0 |
| NC_002645.1;NP_073549.1;HCoV-229E_1ab;1308 | 0 | 0.05 |
| NC_002645.1;NP_073549.1;HCoV-229E_1ab;1309 | 0 | 0.05 |
| NC_002645.1;NP_073549.1;HCoV-229E_1ab;1310 | 0 | 0.05 |
| NC_002645.1;NP_073549.1;HCoV-229E_1ab;1311 | 0 | 0.05 |
| NC_002645.1;NP_073549.1;HCoV-229E_1ab;1312 | 0 | 0.05 |
| NC_002645.1;NP_073549.1;HCoV-229E_1ab;1313 | 0 | 0 |
| NC_002645.1;NP_073549.1;HCoV-229E_1ab;1314 | 0 | 0 |
| NC_002645.1;NP_073549.1;HCoV-229E_1ab;1315 | 0 | 0 |
| NC_002645.1;NP_073549.1;HCoV-229E_1ab;1316 | 0 | 0 |
| NC_002645.1;NP_073549.1;HCoV-229E_1ab;1317 | 0 | 0 |
| NC_002645.1;NP_073549.1;HCoV-229E_1ab;1318 | 0 | 0 |
| NC_002645.1;NP_073549.1;HCoV-229E_1ab;1319 | 0 | 0 |

| | | |
|---|---|---|
| NC_002645.1;NP_073549.1;HCoV-229E_1ab;1320 | 0 | 0 |
| NC_002645.1;NP_073549.1;HCoV-229E_1ab;1321 | 0 | 0 |
| NC_002645.1;NP_073549.1;HCoV-229E_1ab;1322 | 0 | 0 |
| NC_002645.1;NP_073549.1;HCoV-229E_1ab;1323 | 0 | 0.025 |
| NC_002645.1;NP_073549.1;HCoV-229E_1ab;1324 | 0 | 0 |
| NC_002645.1;NP_073549.1;HCoV-229E_1ab;1325 | 0.05 | 0 |
| NC_002645.1;NP_073549.1;HCoV-229E_1ab;1326 | 0.05 | 0 |
| NC_002645.1;NP_073549.1;HCoV-229E_1ab;1327 | 0 | 0 |
| NC_002645.1;NP_073549.1;HCoV-229E_1ab;1328 | 0 | 0 |
| NC_002645.1;NP_073549.1;HCoV-229E_1ab;1329 | 0 | 0 |
| NC_002645.1;NP_073549.1;HCoV-229E_1ab;1330 | 0 | 0 |
| NC_002645.1;NP_073549.1;HCoV-229E_1ab;1331 | 0 | 0 |
| NC_002645.1;NP_073549.1;HCoV-229E_1ab;1332 | 0 | 0 |
| NC_002645.1;NP_073549.1;HCoV-229E_1ab;1333 | 0 | 0.025 |
| NC_002645.1;NP_073549.1;HCoV-229E_1ab;1334 | 0 | 0.025 |
| NC_002645.1;NP_073549.1;HCoV-229E_1ab;1335 | 0 | 0.025 |
| NC_002645.1;NP_073549.1;HCoV-229E_1ab;1336 | 0 | 0.025 |
| NC_002645.1;NP_073549.1;HCoV-229E_1ab;1337 | 0 | 0.025 |
| NC_002645.1;NP_073549.1;HCoV-229E_1ab;1338 | 0 | 0.025 |
| NC_002645.1;NP_073549.1;HCoV-229E_1ab;1339 | 0 | 0.025 |
| NC_002645.1;NP_073549.1;HCoV-229E_1ab;1340 | 0 | 0.025 |
| NC_002645.1;NP_073549.1;HCoV-229E_1ab;1341 | 0 | 0.025 |
| NC_002645.1;NP_073549.1;HCoV-229E_1ab;1342 | 0 | 0 |
| NC_002645.1;NP_073549.1;HCoV-229E_1ab;1343 | 0 | 0 |
| NC_002645.1;NP_073549.1;HCoV-229E_1ab;1344 | 0 | 0 |
| NC_002645.1;NP_073549.1;HCoV-229E_1ab;1345 | 0 | 0 |
| NC_002645.1;NP_073549.1;HCoV-229E_1ab;1346 | 0 | 0 |
| NC_002645.1;NP_073549.1;HCoV-229E_1ab;1347 | 0 | 0 |
| NC_002645.1;NP_073549.1;HCoV-229E_1ab;1348 | 0 | 0 |
| NC_002645.1;NP_073549.1;HCoV-229E_1ab;1349 | 0 | 0 |
| NC_002645.1;NP_073549.1;HCoV-229E_1ab;1350 | 0 | 0 |
| NC_002645.1;NP_073549.1;HCoV-229E_1ab;1351 | 0 | 0 |
| NC_002645.1;NP_073549.1;HCoV-229E_1ab;1352 | 0 | 0 |
| NC_002645.1;NP_073549.1;HCoV-229E_1ab;1353 | 0 | 0 |
| NC_002645.1;NP_073549.1;HCoV-229E_1ab;1354 | 0 | 0 |
| NC_002645.1;NP_073549.1;HCoV-229E_1ab;1355 | 0 | 0 |
| NC_002645.1;NP_073549.1;HCoV-229E_1ab;1356 | 0 | 0 |
| NC_002645.1;NP_073549.1;HCoV-229E_1ab;1357 | 0 | 0 |
| NC_002645.1;NP_073549.1;HCoV-229E_1ab;1358 | 0 | 0 |
| NC_002645.1;NP_073549.1;HCoV-229E_1ab;1359 | 0 | 0 |
| NC_002645.1;NP_073549.1;HCoV-229E_1ab;1360 | 0 | 0 |
| NC_002645.1;NP_073549.1;HCoV-229E_1ab;1361 | 0 | 0 |
| NC_002645.1;NP_073549.1;HCoV-229E_1ab;1362 | 0 | 0.025 |

| | | |
|---|---|---|
| NC_002645.1;NP_073549.1;HCoV-229E_1ab;1363 | 0 | 0.05 |
| NC_002645.1;NP_073549.1;HCoV-229E_1ab;1364 | 0 | 0.05 |
| NC_002645.1;NP_073549.1;HCoV-229E_1ab;1365 | 0.1 | 0.05 |
| NC_002645.1;NP_073549.1;HCoV-229E_1ab;1366 | 0.25 | 0.1 |
| NC_002645.1;NP_073549.1;HCoV-229E_1ab;1367 | 0.45 | 0.125 |
| NC_002645.1;NP_073549.1;HCoV-229E_1ab;1368 | 0.3 | 0.125 |
| NC_002645.1;NP_073549.1;HCoV-229E_1ab;1369 | 0.25 | 0.15 |
| NC_002645.1;NP_073549.1;HCoV-229E_1ab;1370 | 0.25 | 0.175 |
| NC_002645.1;NP_073549.1;HCoV-229E_1ab;1371 | 0.45 | 0.25 |
| NC_002645.1;NP_073549.1;HCoV-229E_1ab;1372 | 0.55 | 0.275 |
| NC_002645.1;NP_073549.1;HCoV-229E_1ab;1373 | 0.5 | 0.25 |
| NC_002645.1;NP_073549.1;HCoV-229E_1ab;1374 | 0.2 | 0.1 |
| NC_002645.1;NP_073549.1;HCoV-229E_1ab;1375 | 0.15 | 0 |
| NC_002645.1;NP_073549.1;HCoV-229E_1ab;1376 | 0 | 0 |
| NC_002645.1;NP_073549.1;HCoV-229E_1ab;1377 | 0 | 0 |
| NC_002645.1;NP_073549.1;HCoV-229E_1ab;1378 | 0 | 0 |
| NC_002645.1;NP_073549.1;HCoV-229E_1ab;1379 | 0 | 0 |
| NC_002645.1;NP_073549.1;HCoV-229E_1ab;1380 | 0 | 0 |
| NC_002645.1;NP_073549.1;HCoV-229E_1ab;1381 | 0 | 0 |
| NC_002645.1;NP_073549.1;HCoV-229E_1ab;1382 | 0 | 0 |
| NC_002645.1;NP_073549.1;HCoV-229E_1ab;1383 | 0 | 0 |
| NC_002645.1;NP_073549.1;HCoV-229E_1ab;1384 | 0 | 0 |
| NC_002645.1;NP_073549.1;HCoV-229E_1ab;1385 | 0 | 0.025 |
| NC_002645.1;NP_073549.1;HCoV-229E_1ab;1386 | 0 | 0.025 |
| NC_002645.1;NP_073549.1;HCoV-229E_1ab;1387 | 0 | 0.025 |
| NC_002645.1;NP_073549.1;HCoV-229E_1ab;1388 | 0 | 0.025 |
| NC_002645.1;NP_073549.1;HCoV-229E_1ab;1389 | 0 | 0.025 |
| NC_002645.1;NP_073549.1;HCoV-229E_1ab;1390 | 0 | 0.025 |
| NC_002645.1;NP_073549.1;HCoV-229E_1ab;1391 | 0 | 0 |
| NC_002645.1;NP_073549.1;HCoV-229E_1ab;1392 | 0 | 0 |
| NC_002645.1;NP_073549.1;HCoV-229E_1ab;1393 | 0 | 0 |
| NC_002645.1;NP_073549.1;HCoV-229E_1ab;1394 | 0 | 0 |
| NC_002645.1;NP_073549.1;HCoV-229E_1ab;1395 | 0 | 0 |
| NC_002645.1;NP_073549.1;HCoV-229E_1ab;1396 | 0 | 0 |
| NC_002645.1;NP_073549.1;HCoV-229E_1ab;1397 | 0 | 0 |
| NC_002645.1;NP_073549.1;HCoV-229E_1ab;1398 | 0 | 0 |
| NC_002645.1;NP_073549.1;HCoV-229E_1ab;1399 | 0 | 0 |
| NC_002645.1;NP_073549.1;HCoV-229E_1ab;1400 | 0 | 0 |
| NC_002645.1;NP_073549.1;HCoV-229E_1ab;1401 | 0 | 0 |
| NC_002645.1;NP_073549.1;HCoV-229E_1ab;1402 | 0 | 0.05 |
| NC_002645.1;NP_073549.1;HCoV-229E_1ab;1403 | 0 | 0.025 |
| NC_002645.1;NP_073549.1;HCoV-229E_1ab;1404 | 0 | 0.025 |
| NC_002645.1;NP_073549.1;HCoV-229E_1ab;1405 | 0 | 0 |

| | | |
|---|---|---|
| NC_002645.1;NP_073549.1;HCoV-229E_1ab;1406 | 0 | 0 |
| NC_002645.1;NP_073549.1;HCoV-229E_1ab;1407 | 0 | 0 |
| NC_002645.1;NP_073549.1;HCoV-229E_1ab;1408 | 0 | 0 |
| NC_002645.1;NP_073549.1;HCoV-229E_1ab;1409 | 0 | 0 |
| NC_002645.1;NP_073549.1;HCoV-229E_1ab;1410 | 0 | 0 |
| NC_002645.1;NP_073549.1;HCoV-229E_1ab;1411 | 0 | 0 |
| NC_002645.1;NP_073549.1;HCoV-229E_1ab;1412 | 0 | 0 |
| NC_002645.1;NP_073549.1;HCoV-229E_1ab;1413 | 0 | 0 |
| NC_002645.1;NP_073549.1;HCoV-229E_1ab;1414 | 0 | 0 |
| NC_002645.1;NP_073549.1;HCoV-229E_1ab;1415 | 0 | 0 |
| NC_002645.1;NP_073549.1;HCoV-229E_1ab;1416 | 0 | 0 |
| NC_002645.1;NP_073549.1;HCoV-229E_1ab;1417 | 0 | 0 |
| NC_002645.1;NP_073549.1;HCoV-229E_1ab;1418 | 0 | 0 |
| NC_002645.1;NP_073549.1;HCoV-229E_1ab;1419 | 0 | 0.025 |
| NC_002645.1;NP_073549.1;HCoV-229E_1ab;1420 | 0 | 0 |
| NC_002645.1;NP_073549.1;HCoV-229E_1ab;1421 | 0 | 0 |
| NC_002645.1;NP_073549.1;HCoV-229E_1ab;1422 | 0 | 0 |
| NC_002645.1;NP_073549.1;HCoV-229E_1ab;1423 | 0 | 0 |
| NC_002645.1;NP_073549.1;HCoV-229E_1ab;1424 | 0 | 0 |
| NC_002645.1;NP_073549.1;HCoV-229E_1ab;1425 | 0 | 0 |
| NC_002645.1;NP_073549.1;HCoV-229E_1ab;1426 | 0 | 0 |
| NC_002645.1;NP_073549.1;HCoV-229E_1ab;1427 | 0 | 0 |
| NC_002645.1;NP_073549.1;HCoV-229E_1ab;1428 | 0 | 0.05 |
| NC_002645.1;NP_073549.1;HCoV-229E_1ab;1429 | 0.05 | 0 |
| NC_002645.1;NP_073549.1;HCoV-229E_1ab;1430 | 0 | 0 |
| NC_002645.1;NP_073549.1;HCoV-229E_1ab;1431 | 0 | 0 |
| NC_002645.1;NP_073549.1;HCoV-229E_1ab;1432 | 0.2 | 0.075 |
| NC_002645.1;NP_073549.1;HCoV-229E_1ab;1433 | 0.2 | 0.075 |
| NC_002645.1;NP_073549.1;HCoV-229E_1ab;1434 | 0.05 | 0.05 |
| NC_002645.1;NP_073549.1;HCoV-229E_1ab;1435 | 0.1 | 0.05 |
| NC_002645.1;NP_073549.1;HCoV-229E_1ab;1436 | 0.15 | 0.125 |
| NC_002645.1;NP_073549.1;HCoV-229E_1ab;1437 | 0.2 | 0.125 |
| NC_002645.1;NP_073549.1;HCoV-229E_1ab;1438 | 0.2 | 0.075 |
| NC_002645.1;NP_073549.1;HCoV-229E_1ab;1439 | 0.2 | 0.125 |
| NC_002645.1;NP_073549.1;HCoV-229E_1ab;1440 | 0.15 | 0.075 |
| NC_002645.1;NP_073549.1;HCoV-229E_1ab;1441 | 0.15 | 0.1 |
| NC_002645.1;NP_073549.1;HCoV-229E_1ab;1442 | 0.05 | 0.025 |
| NC_002645.1;NP_073549.1;HCoV-229E_1ab;1443 | 0.05 | 0.025 |
| NC_002645.1;NP_073549.1;HCoV-229E_1ab;1444 | 0.05 | 0 |
| NC_002645.1;NP_073549.1;HCoV-229E_1ab;1445 | 0.05 | 0 |
| NC_002645.1;NP_073549.1;HCoV-229E_1ab;1446 | 0 | 0 |
| NC_002645.1;NP_073549.1;HCoV-229E_1ab;1447 | 0 | 0 |
| NC_002645.1;NP_073549.1;HCoV-229E_1ab;1448 | 0 | 0 |

| | | |
|---|---|---|
| NC_002645.1;NP_073549.1;HCoV-229E_1ab;1449 | 0 | 0 |
| NC_002645.1;NP_073549.1;HCoV-229E_1ab;1450 | 0 | 0 |
| NC_002645.1;NP_073549.1;HCoV-229E_1ab;1451 | 0 | 0.025 |
| NC_002645.1;NP_073549.1;HCoV-229E_1ab;1452 | 0 | 0.025 |
| NC_002645.1;NP_073549.1;HCoV-229E_1ab;1453 | 0 | 0.025 |
| NC_002645.1;NP_073549.1;HCoV-229E_1ab;1454 | 0 | 0 |
| NC_002645.1;NP_073549.1;HCoV-229E_1ab;1455 | 0 | 0 |
| NC_002645.1;NP_073549.1;HCoV-229E_1ab;1456 | 0 | 0 |
| NC_002645.1;NP_073549.1;HCoV-229E_1ab;1457 | 0 | 0 |
| NC_002645.1;NP_073549.1;HCoV-229E_1ab;1458 | 0 | 0 |
| NC_002645.1;NP_073549.1;HCoV-229E_1ab;1459 | 0 | 0 |
| NC_002645.1;NP_073549.1;HCoV-229E_1ab;1460 | 0 | 0 |
| NC_002645.1;NP_073549.1;HCoV-229E_1ab;1461 | 0 | 0 |
| NC_002645.1;NP_073549.1;HCoV-229E_1ab;1462 | 0 | 0 |
| NC_002645.1;NP_073549.1;HCoV-229E_1ab;1463 | 0 | 0 |
| NC_002645.1;NP_073549.1;HCoV-229E_1ab;1464 | 0 | 0 |
| NC_002645.1;NP_073549.1;HCoV-229E_1ab;1465 | 0 | 0 |
| NC_002645.1;NP_073549.1;HCoV-229E_1ab;1466 | 0 | 0 |
| NC_002645.1;NP_073549.1;HCoV-229E_1ab;1467 | 0.05 | 0 |
| NC_002645.1;NP_073549.1;HCoV-229E_1ab;1468 | 0.05 | 0.025 |
| NC_002645.1;NP_073549.1;HCoV-229E_1ab;1469 | 0.05 | 0 |
| NC_002645.1;NP_073549.1;HCoV-229E_1ab;1470 | 0 | 0 |
| NC_002645.1;NP_073549.1;HCoV-229E_1ab;1471 | 0 | 0 |
| NC_002645.1;NP_073549.1;HCoV-229E_1ab;1472 | 0 | 0 |
| NC_002645.1;NP_073549.1;HCoV-229E_1ab;1473 | 0 | 0 |
| NC_002645.1;NP_073549.1;HCoV-229E_1ab;1474 | 0 | 0 |
| NC_002645.1;NP_073549.1;HCoV-229E_1ab;1475 | 0 | 0 |
| NC_002645.1;NP_073549.1;HCoV-229E_1ab;1476 | 0 | 0 |
| NC_002645.1;NP_073549.1;HCoV-229E_1ab;1477 | 0 | 0 |
| NC_002645.1;NP_073549.1;HCoV-229E_1ab;1478 | 0 | 0 |
| NC_002645.1;NP_073549.1;HCoV-229E_1ab;1479 | 0 | 0 |
| NC_002645.1;NP_073549.1;HCoV-229E_1ab;1480 | 0 | 0 |
| NC_002645.1;NP_073549.1;HCoV-229E_1ab;1481 | 0 | 0 |
| NC_002645.1;NP_073549.1;HCoV-229E_1ab;1482 | 0 | 0 |
| NC_002645.1;NP_073549.1;HCoV-229E_1ab;1483 | 0 | 0 |
| NC_002645.1;NP_073549.1;HCoV-229E_1ab;1484 | 0 | 0 |
| NC_002645.1;NP_073549.1;HCoV-229E_1ab;1485 | 0 | 0 |
| NC_002645.1;NP_073549.1;HCoV-229E_1ab;1486 | 0 | 0 |
| NC_002645.1;NP_073549.1;HCoV-229E_1ab;1487 | 0 | 0.025 |
| NC_002645.1;NP_073549.1;HCoV-229E_1ab;1488 | 0 | 0 |
| NC_002645.1;NP_073549.1;HCoV-229E_1ab;1489 | 0 | 0 |
| NC_002645.1;NP_073549.1;HCoV-229E_1ab;1490 | 0.05 | 0.05 |
| NC_002645.1;NP_073549.1;HCoV-229E_1ab;1491 | 0 | 0.025 |

| | | |
|---|---|---|
| NC_002645.1;NP_073549.1;HCoV-229E_1ab;1492 | 0.05 | 0.025 |
| NC_002645.1;NP_073549.1;HCoV-229E_1ab;1493 | 0.05 | 0 |
| NC_002645.1;NP_073549.1;HCoV-229E_1ab;1494 | 0.05 | 0 |
| NC_002645.1;NP_073549.1;HCoV-229E_1ab;1495 | 0.05 | 0 |
| NC_002645.1;NP_073549.1;HCoV-229E_1ab;1496 | 0.05 | 0.025 |
| NC_002645.1;NP_073549.1;HCoV-229E_1ab;1497 | 0.05 | 0.025 |
| NC_002645.1;NP_073549.1;HCoV-229E_1ab;1498 | 0 | 0 |
| NC_002645.1;NP_073549.1;HCoV-229E_1ab;1499 | 0 | 0 |
| NC_002645.1;NP_073549.1;HCoV-229E_1ab;1500 | 0 | 0 |
| NC_002645.1;NP_073549.1;HCoV-229E_1ab;1501 | 0 | 0.025 |
| NC_002645.1;NP_073549.1;HCoV-229E_1ab;1502 | 0 | 0 |
| NC_002645.1;NP_073549.1;HCoV-229E_1ab;1503 | 0 | 0 |
| NC_002645.1;NP_073549.1;HCoV-229E_1ab;1504 | 0 | 0 |
| NC_002645.1;NP_073549.1;HCoV-229E_1ab;1505 | 0 | 0 |
| NC_002645.1;NP_073549.1;HCoV-229E_1ab;1506 | 0.1 | 0.15 |
| NC_002645.1;NP_073549.1;HCoV-229E_1ab;1507 | 0 | 0.025 |
| NC_002645.1;NP_073549.1;HCoV-229E_1ab;1508 | 0 | 0 |
| NC_002645.1;NP_073549.1;HCoV-229E_1ab;1509 | 0 | 0.025 |
| NC_002645.1;NP_073549.1;HCoV-229E_1ab;1510 | 0.05 | 0 |
| NC_002645.1;NP_073549.1;HCoV-229E_1ab;1511 | 0.05 | 0.05 |
| NC_002645.1;NP_073549.1;HCoV-229E_1ab;1512 | 0 | 0.025 |
| NC_002645.1;NP_073549.1;HCoV-229E_1ab;1513 | 0 | 0.025 |
| NC_002645.1;NP_073549.1;HCoV-229E_1ab;1514 | 0.1 | 0.05 |
| NC_002645.1;NP_073549.1;HCoV-229E_1ab;1515 | 0 | 0.025 |
| NC_002645.1;NP_073549.1;HCoV-229E_1ab;1516 | 0.05 | 0.05 |
| NC_002645.1;NP_073549.1;HCoV-229E_1ab;1517 | 0 | 0.05 |
| NC_002645.1;NP_073549.1;HCoV-229E_1ab;1518 | 0 | 0.05 |
| NC_002645.1;NP_073549.1;HCoV-229E_1ab;1519 | 0.05 | 0.075 |
| NC_002645.1;NP_073549.1;HCoV-229E_1ab;1520 | 0 | 0.025 |
| NC_002645.1;NP_073549.1;HCoV-229E_1ab;1521 | 0.1 | 0.075 |
| NC_002645.1;NP_073549.1;HCoV-229E_1ab;1522 | 0 | 0.025 |
| NC_002645.1;NP_073549.1;HCoV-229E_1ab;1523 | 0 | 0 |
| NC_002645.1;NP_073549.1;HCoV-229E_1ab;1524 | 0 | 0 |
| NC_002645.1;NP_073549.1;HCoV-229E_1ab;1525 | 0 | 0 |
| NC_002645.1;NP_073549.1;HCoV-229E_1ab;1526 | 0 | 0 |
| NC_002645.1;NP_073549.1;HCoV-229E_1ab;1527 | 0 | 0 |
| NC_002645.1;NP_073549.1;HCoV-229E_1ab;1528 | 0 | 0 |
| NC_002645.1;NP_073549.1;HCoV-229E_1ab;1529 | 0 | 0 |
| NC_002645.1;NP_073549.1;HCoV-229E_1ab;1530 | 0 | 0 |
| NC_002645.1;NP_073549.1;HCoV-229E_1ab;1531 | 0 | 0 |
| NC_002645.1;NP_073549.1;HCoV-229E_1ab;1532 | 0 | 0 |
| NC_002645.1;NP_073549.1;HCoV-229E_1ab;1533 | 0 | 0 |
| NC_002645.1;NP_073549.1;HCoV-229E_1ab;1534 | 0 | 0.025 |

| | | |
|---|---|---|
| NC_002645.1;NP_073549.1;HCoV-229E_1ab;1535 | 0 | 0 |
| NC_002645.1;NP_073549.1;HCoV-229E_1ab;1536 | 0 | 0 |
| NC_002645.1;NP_073549.1;HCoV-229E_1ab;1537 | 0 | 0 |
| NC_002645.1;NP_073549.1;HCoV-229E_1ab;1538 | 0 | 0 |
| NC_002645.1;NP_073549.1;HCoV-229E_1ab;1539 | 0 | 0 |
| NC_002645.1;NP_073549.1;HCoV-229E_1ab;1540 | 0 | 0 |
| NC_002645.1;NP_073549.1;HCoV-229E_1ab;1541 | 0 | 0 |
| NC_002645.1;NP_073549.1;HCoV-229E_1ab;1542 | 0 | 0 |
| NC_002645.1;NP_073549.1;HCoV-229E_1ab;1543 | 0 | 0.025 |
| NC_002645.1;NP_073549.1;HCoV-229E_1ab;1544 | 0 | 0 |
| NC_002645.1;NP_073549.1;HCoV-229E_1ab;1545 | 0 | 0 |
| NC_002645.1;NP_073549.1;HCoV-229E_1ab;1546 | 0 | 0 |
| NC_002645.1;NP_073549.1;HCoV-229E_1ab;1547 | 0 | 0 |
| NC_002645.1;NP_073549.1;HCoV-229E_1ab;1548 | 0 | 0 |
| NC_002645.1;NP_073549.1;HCoV-229E_1ab;1549 | 0 | 0 |
| NC_002645.1;NP_073549.1;HCoV-229E_1ab;1550 | 0 | 0 |
| NC_002645.1;NP_073549.1;HCoV-229E_1ab;1551 | 0 | 0 |
| NC_002645.1;NP_073549.1;HCoV-229E_1ab;1552 | 0 | 0 |
| NC_002645.1;NP_073549.1;HCoV-229E_1ab;1553 | 0 | 0 |
| NC_002645.1;NP_073549.1;HCoV-229E_1ab;1554 | 0 | 0 |
| NC_002645.1;NP_073549.1;HCoV-229E_1ab;1555 | 0 | 0 |
| NC_002645.1;NP_073549.1;HCoV-229E_1ab;1556 | 0 | 0 |
| NC_002645.1;NP_073549.1;HCoV-229E_1ab;1557 | 0 | 0 |
| NC_002645.1;NP_073549.1;HCoV-229E_1ab;1558 | 0 | 0 |
| NC_002645.1;NP_073549.1;HCoV-229E_1ab;1559 | 0 | 0 |
| NC_002645.1;NP_073549.1;HCoV-229E_1ab;1560 | 0 | 0 |
| NC_002645.1;NP_073549.1;HCoV-229E_1ab;1561 | 0 | 0 |
| NC_002645.1;NP_073549.1;HCoV-229E_1ab;1562 | 0 | 0 |
| NC_002645.1;NP_073549.1;HCoV-229E_1ab;1563 | 0 | 0 |
| NC_002645.1;NP_073549.1;HCoV-229E_1ab;1564 | 0 | 0 |
| NC_002645.1;NP_073549.1;HCoV-229E_1ab;1565 | 0 | 0 |
| NC_002645.1;NP_073549.1;HCoV-229E_1ab;1566 | 0 | 0 |
| NC_002645.1;NP_073549.1;HCoV-229E_1ab;1567 | 0 | 0 |
| NC_002645.1;NP_073549.1;HCoV-229E_1ab;1568 | 0 | 0 |
| NC_002645.1;NP_073549.1;HCoV-229E_1ab;1569 | 0 | 0 |
| NC_002645.1;NP_073549.1;HCoV-229E_1ab;1570 | 0 | 0 |
| NC_002645.1;NP_073549.1;HCoV-229E_1ab;1571 | 0 | 0 |
| NC_002645.1;NP_073549.1;HCoV-229E_1ab;1572 | 0 | 0 |
| NC_002645.1;NP_073549.1;HCoV-229E_1ab;1573 | 0 | 0 |
| NC_002645.1;NP_073549.1;HCoV-229E_1ab;1574 | 0 | 0 |
| NC_002645.1;NP_073549.1;HCoV-229E_1ab;1575 | 0 | 0 |
| NC_002645.1;NP_073549.1;HCoV-229E_1ab;1576 | 0 | 0 |
| NC_002645.1;NP_073549.1;HCoV-229E_1ab;1577 | 0 | 0 |

| | | |
|---|---|---|
| NC_002645.1;NP_073549.1;HCoV-229E_1ab;1578 | 0 | 0 |
| NC_002645.1;NP_073549.1;HCoV-229E_1ab;1579 | 0 | 0 |
| NC_002645.1;NP_073549.1;HCoV-229E_1ab;1580 | 0 | 0 |
| NC_002645.1;NP_073549.1;HCoV-229E_1ab;1581 | 0 | 0 |
| NC_002645.1;NP_073549.1;HCoV-229E_1ab;1582 | 0 | 0 |
| NC_002645.1;NP_073549.1;HCoV-229E_1ab;1583 | 0.05 | 0.025 |
| NC_002645.1;NP_073549.1;HCoV-229E_1ab;1584 | 0 | 0 |
| NC_002645.1;NP_073549.1;HCoV-229E_1ab;1585 | 0 | 0 |
| NC_002645.1;NP_073549.1;HCoV-229E_1ab;1586 | 0.05 | 0.05 |
| NC_002645.1;NP_073549.1;HCoV-229E_1ab;1587 | 0.05 | 0.075 |
| NC_002645.1;NP_073549.1;HCoV-229E_1ab;1588 | 0.05 | 0.075 |
| NC_002645.1;NP_073549.1;HCoV-229E_1ab;1589 | 0.05 | 0.075 |
| NC_002645.1;NP_073549.1;HCoV-229E_1ab;1590 | 0 | 0.05 |
| NC_002645.1;NP_073549.1;HCoV-229E_1ab;1591 | 0 | 0 |
| NC_002645.1;NP_073549.1;HCoV-229E_1ab;1592 | 0 | 0 |
| NC_002645.1;NP_073549.1;HCoV-229E_1ab;1593 | 0 | 0 |
| NC_002645.1;NP_073549.1;HCoV-229E_1ab;1594 | 0 | 0.025 |
| NC_002645.1;NP_073549.1;HCoV-229E_1ab;1595 | 0 | 0 |
| NC_002645.1;NP_073549.1;HCoV-229E_1ab;1596 | 0 | 0 |
| NC_002645.1;NP_073549.1;HCoV-229E_1ab;1597 | 0 | 0 |
| NC_002645.1;NP_073549.1;HCoV-229E_1ab;1598 | 0 | 0 |
| NC_002645.1;NP_073549.1;HCoV-229E_1ab;1599 | 0 | 0.025 |
| NC_002645.1;NP_073549.1;HCoV-229E_1ab;1600 | 0 | 0 |
| NC_002645.1;NP_073549.1;HCoV-229E_1ab;1601 | 0 | 0 |
| NC_002645.1;NP_073549.1;HCoV-229E_1ab;1602 | 0 | 0 |
| NC_002645.1;NP_073549.1;HCoV-229E_1ab;1603 | 0 | 0.025 |
| NC_002645.1;NP_073549.1;HCoV-229E_1ab;1604 | 0 | 0.025 |
| NC_002645.1;NP_073549.1;HCoV-229E_1ab;1605 | 0 | 0.025 |
| NC_002645.1;NP_073549.1;HCoV-229E_1ab;1606 | 0 | 0.025 |
| NC_002645.1;NP_073549.1;HCoV-229E_1ab;1607 | 0.05 | 0 |
| NC_002645.1;NP_073549.1;HCoV-229E_1ab;1608 | 0 | 0 |
| NC_002645.1;NP_073549.1;HCoV-229E_1ab;1609 | 0 | 0 |
| NC_002645.1;NP_073549.1;HCoV-229E_1ab;1610 | 0 | 0 |
| NC_002645.1;NP_073549.1;HCoV-229E_1ab;1611 | 0.05 | 0 |
| NC_002645.1;NP_073549.1;HCoV-229E_1ab;1612 | 0 | 0.025 |
| NC_002645.1;NP_073549.1;HCoV-229E_1ab;1613 | 0 | 0.025 |
| NC_002645.1;NP_073549.1;HCoV-229E_1ab;1614 | 0 | 0 |
| NC_002645.1;NP_073549.1;HCoV-229E_1ab;1615 | 0 | 0.025 |
| NC_002645.1;NP_073549.1;HCoV-229E_1ab;1616 | 0 | 0.025 |
| NC_002645.1;NP_073549.1;HCoV-229E_1ab;1617 | 0 | 0 |
| NC_002645.1;NP_073549.1;HCoV-229E_1ab;1618 | 0 | 0.05 |
| NC_002645.1;NP_073549.1;HCoV-229E_1ab;1619 | 0 | 0 |
| NC_002645.1;NP_073549.1;HCoV-229E_1ab;1620 | 0 | 0.075 |

| | | |
|---|---|---|
| NC_002645.1;NP_073549.1;HCoV-229E_1ab;1621 | 0 | 0.025 |
| NC_002645.1;NP_073549.1;HCoV-229E_1ab;1622 | 0 | 0.075 |
| NC_002645.1;NP_073549.1;HCoV-229E_1ab;1623 | 0 | 0.075 |
| NC_002645.1;NP_073549.1;HCoV-229E_1ab;1624 | 0 | 0.075 |
| NC_002645.1;NP_073549.1;HCoV-229E_1ab;1625 | 0 | 0.075 |
| NC_002645.1;NP_073549.1;HCoV-229E_1ab;1626 | 0 | 0.075 |
| NC_002645.1;NP_073549.1;HCoV-229E_1ab;1627 | 0.05 | 0.1 |
| NC_002645.1;NP_073549.1;HCoV-229E_1ab;1628 | 0 | 0.075 |
| NC_002645.1;NP_073549.1;HCoV-229E_1ab;1629 | 0 | 0.075 |
| NC_002645.1;NP_073549.1;HCoV-229E_1ab;1630 | 0 | 0.075 |
| NC_002645.1;NP_073549.1;HCoV-229E_1ab;1631 | 0.05 | 0.025 |
| NC_002645.1;NP_073549.1;HCoV-229E_1ab;1632 | 0.05 | 0.05 |
| NC_002645.1;NP_073549.1;HCoV-229E_1ab;1633 | 0.1 | 0.025 |
| NC_002645.1;NP_073549.1;HCoV-229E_1ab;1634 | 0.05 | 0.025 |
| NC_002645.1;NP_073549.1;HCoV-229E_1ab;1635 | 0.05 | 0.025 |
| NC_002645.1;NP_073549.1;HCoV-229E_1ab;1636 | 0 | 0 |
| NC_002645.1;NP_073549.1;HCoV-229E_1ab;1637 | 0 | 0 |
| NC_002645.1;NP_073549.1;HCoV-229E_1ab;1638 | 0 | 0 |
| NC_002645.1;NP_073549.1;HCoV-229E_1ab;1639 | 0 | 0.025 |
| NC_002645.1;NP_073549.1;HCoV-229E_1ab;1640 | 0 | 0 |
| NC_002645.1;NP_073549.1;HCoV-229E_1ab;1641 | 0 | 0 |
| NC_002645.1;NP_073549.1;HCoV-229E_1ab;1642 | 0 | 0 |
| NC_002645.1;NP_073549.1;HCoV-229E_1ab;1643 | 0 | 0 |
| NC_002645.1;NP_073549.1;HCoV-229E_1ab;1644 | 0 | 0 |
| NC_002645.1;NP_073549.1;HCoV-229E_1ab;1645 | 0 | 0.025 |
| NC_002645.1;NP_073549.1;HCoV-229E_1ab;1646 | 0 | 0 |
| NC_002645.1;NP_073549.1;HCoV-229E_1ab;1647 | 0 | 0 |
| NC_002645.1;NP_073549.1;HCoV-229E_1ab;1648 | 0 | 0 |
| NC_002645.1;NP_073549.1;HCoV-229E_1ab;1649 | 0 | 0 |
| NC_002645.1;NP_073549.1;HCoV-229E_1ab;1650 | 0 | 0 |
| NC_002645.1;NP_073549.1;HCoV-229E_1ab;1651 | 0 | 0 |
| NC_002645.1;NP_073549.1;HCoV-229E_1ab;1652 | 0 | 0 |
| NC_002645.1;NP_073549.1;HCoV-229E_1ab;1653 | 0 | 0 |
| NC_002645.1;NP_073549.1;HCoV-229E_1ab;1654 | 0 | 0 |
| NC_002645.1;NP_073549.1;HCoV-229E_1ab;1655 | 0 | 0.025 |
| NC_002645.1;NP_073549.1;HCoV-229E_1ab;1656 | 0 | 0 |
| NC_002645.1;NP_073549.1;HCoV-229E_1ab;1657 | 0 | 0 |
| NC_002645.1;NP_073549.1;HCoV-229E_1ab;1658 | 0 | 0 |
| NC_002645.1;NP_073549.1;HCoV-229E_1ab;1659 | 0 | 0 |
| NC_002645.1;NP_073549.1;HCoV-229E_1ab;1660 | 0 | 0 |
| NC_002645.1;NP_073549.1;HCoV-229E_1ab;1661 | 0 | 0 |
| NC_002645.1;NP_073549.1;HCoV-229E_1ab;1662 | 0 | 0 |
| NC_002645.1;NP_073549.1;HCoV-229E_1ab;1663 | 0 | 0 |

| | | |
|---|---|---|
| NC_002645.1;NP_073549.1;HCoV-229E_1ab;1664 | 0 | 0 |
| NC_002645.1;NP_073549.1;HCoV-229E_1ab;1665 | 0 | 0 |
| NC_002645.1;NP_073549.1;HCoV-229E_1ab;1666 | 0 | 0.025 |
| NC_002645.1;NP_073549.1;HCoV-229E_1ab;1667 | 0 | 0 |
| NC_002645.1;NP_073549.1;HCoV-229E_1ab;1668 | 0 | 0 |
| NC_002645.1;NP_073549.1;HCoV-229E_1ab;1669 | 0 | 0 |
| NC_002645.1;NP_073549.1;HCoV-229E_1ab;1670 | 0 | 0 |
| NC_002645.1;NP_073549.1;HCoV-229E_1ab;1671 | 0 | 0 |
| NC_002645.1;NP_073549.1;HCoV-229E_1ab;1672 | 0 | 0 |
| NC_002645.1;NP_073549.1;HCoV-229E_1ab;1673 | 0 | 0 |
| NC_002645.1;NP_073549.1;HCoV-229E_1ab;1674 | 0 | 0 |
| NC_002645.1;NP_073549.1;HCoV-229E_1ab;1675 | 0 | 0 |
| NC_002645.1;NP_073549.1;HCoV-229E_1ab;1676 | 0 | 0 |
| NC_002645.1;NP_073549.1;HCoV-229E_1ab;1677 | 0 | 0 |
| NC_002645.1;NP_073549.1;HCoV-229E_1ab;1678 | 0 | 0 |
| NC_002645.1;NP_073549.1;HCoV-229E_1ab;1679 | 0 | 0 |
| NC_002645.1;NP_073549.1;HCoV-229E_1ab;1680 | 0 | 0 |
| NC_002645.1;NP_073549.1;HCoV-229E_1ab;1681 | 0 | 0.025 |
| NC_002645.1;NP_073549.1;HCoV-229E_1ab;1682 | 0 | 0 |
| NC_002645.1;NP_073549.1;HCoV-229E_1ab;1683 | 0 | 0 |
| NC_002645.1;NP_073549.1;HCoV-229E_1ab;1684 | 0.05 | 0 |
| NC_002645.1;NP_073549.1;HCoV-229E_1ab;1685 | 0 | 0.025 |
| NC_002645.1;NP_073549.1;HCoV-229E_1ab;1686 | 0 | 0 |
| NC_002645.1;NP_073549.1;HCoV-229E_1ab;1687 | 0 | 0 |
| NC_002645.1;NP_073549.1;HCoV-229E_1ab;1688 | 0 | 0 |
| NC_002645.1;NP_073549.1;HCoV-229E_1ab;1689 | 0 | 0 |
| NC_002645.1;NP_073549.1;HCoV-229E_1ab;1690 | 0.05 | 0 |
| NC_002645.1;NP_073549.1;HCoV-229E_1ab;1691 | 0 | 0 |
| NC_002645.1;NP_073549.1;HCoV-229E_1ab;1692 | 0 | 0 |
| NC_002645.1;NP_073549.1;HCoV-229E_1ab;1693 | 0 | 0 |
| NC_002645.1;NP_073549.1;HCoV-229E_1ab;1694 | 0 | 0 |
| NC_002645.1;NP_073549.1;HCoV-229E_1ab;1695 | 0 | 0 |
| NC_002645.1;NP_073549.1;HCoV-229E_1ab;1696 | 0 | 0 |
| NC_002645.1;NP_073549.1;HCoV-229E_1ab;1697 | 0 | 0 |
| NC_002645.1;NP_073549.1;HCoV-229E_1ab;1698 | 0 | 0 |
| NC_002645.1;NP_073549.1;HCoV-229E_1ab;1699 | 0 | 0 |
| NC_002645.1;NP_073549.1;HCoV-229E_1ab;1700 | 0 | 0 |
| NC_002645.1;NP_073549.1;HCoV-229E_1ab;1701 | 0 | 0 |
| NC_002645.1;NP_073549.1;HCoV-229E_1ab;1702 | 0 | 0 |
| NC_002645.1;NP_073549.1;HCoV-229E_1ab;1703 | 0 | 0 |
| NC_002645.1;NP_073549.1;HCoV-229E_1ab;1704 | 0 | 0 |
| NC_002645.1;NP_073549.1;HCoV-229E_1ab;1705 | 0 | 0 |
| NC_002645.1;NP_073549.1;HCoV-229E_1ab;1706 | 0 | 0 |

| | | |
|---|---|---|
| NC_002645.1;NP_073549.1;HCoV-229E_1ab;1707 | 0 | 0 |
| NC_002645.1;NP_073549.1;HCoV-229E_1ab;1708 | 0 | 0 |
| NC_002645.1;NP_073549.1;HCoV-229E_1ab;1709 | 0 | 0 |
| NC_002645.1;NP_073549.1;HCoV-229E_1ab;1710 | 0.05 | 0.05 |
| NC_002645.1;NP_073549.1;HCoV-229E_1ab;1711 | 0 | 0 |
| NC_002645.1;NP_073549.1;HCoV-229E_1ab;1712 | 0 | 0 |
| NC_002645.1;NP_073549.1;HCoV-229E_1ab;1713 | 0 | 0 |
| NC_002645.1;NP_073549.1;HCoV-229E_1ab;1714 | 0 | 0 |
| NC_002645.1;NP_073549.1;HCoV-229E_1ab;1715 | 0 | 0 |
| NC_002645.1;NP_073549.1;HCoV-229E_1ab;1716 | 0 | 0 |
| NC_002645.1;NP_073549.1;HCoV-229E_1ab;1717 | 0 | 0 |
| NC_002645.1;NP_073549.1;HCoV-229E_1ab;1718 | 0.05 | 0 |
| NC_002645.1;NP_073549.1;HCoV-229E_1ab;1719 | 0.05 | 0 |
| NC_002645.1;NP_073549.1;HCoV-229E_1ab;1720 | 0.05 | 0 |
| NC_002645.1;NP_073549.1;HCoV-229E_1ab;1721 | 0.05 | 0.025 |
| NC_002645.1;NP_073549.1;HCoV-229E_1ab;1722 | 0 | 0 |
| NC_002645.1;NP_073549.1;HCoV-229E_1ab;1723 | 0 | 0 |
| NC_002645.1;NP_073549.1;HCoV-229E_1ab;1724 | 0 | 0 |
| NC_002645.1;NP_073549.1;HCoV-229E_1ab;1725 | 0 | 0 |
| NC_002645.1;NP_073549.1;HCoV-229E_1ab;1726 | 0 | 0 |
| NC_002645.1;NP_073549.1;HCoV-229E_1ab;1727 | 0 | 0 |
| NC_002645.1;NP_073549.1;HCoV-229E_1ab;1728 | 0 | 0 |
| NC_002645.1;NP_073549.1;HCoV-229E_1ab;1729 | 0 | 0 |
| NC_002645.1;NP_073549.1;HCoV-229E_1ab;1730 | 0 | 0 |
| NC_002645.1;NP_073549.1;HCoV-229E_1ab;1731 | 0 | 0 |
| NC_002645.1;NP_073549.1;HCoV-229E_1ab;1732 | 0 | 0 |
| NC_002645.1;NP_073549.1;HCoV-229E_1ab;1733 | 0 | 0 |
| NC_002645.1;NP_073549.1;HCoV-229E_1ab;1734 | 0 | 0 |
| NC_002645.1;NP_073549.1;HCoV-229E_1ab;1735 | 0 | 0 |
| NC_002645.1;NP_073549.1;HCoV-229E_1ab;1736 | 0 | 0 |
| NC_002645.1;NP_073549.1;HCoV-229E_1ab;1737 | 0 | 0 |
| NC_002645.1;NP_073549.1;HCoV-229E_1ab;1738 | 0 | 0 |
| NC_002645.1;NP_073549.1;HCoV-229E_1ab;1739 | 0 | 0 |
| NC_002645.1;NP_073549.1;HCoV-229E_1ab;1740 | 0 | 0 |
| NC_002645.1;NP_073549.1;HCoV-229E_1ab;1741 | 0.1 | 0.025 |
| NC_002645.1;NP_073549.1;HCoV-229E_1ab;1742 | 0 | 0 |
| NC_002645.1;NP_073549.1;HCoV-229E_1ab;1743 | 0 | 0.025 |
| NC_002645.1;NP_073549.1;HCoV-229E_1ab;1744 | 0 | 0.025 |
| NC_002645.1;NP_073549.1;HCoV-229E_1ab;1745 | 0 | 0 |
| NC_002645.1;NP_073549.1;HCoV-229E_1ab;1746 | 0 | 0 |
| NC_002645.1;NP_073549.1;HCoV-229E_1ab;1747 | 0 | 0 |
| NC_002645.1;NP_073549.1;HCoV-229E_1ab;1748 | 0 | 0 |
| NC_002645.1;NP_073549.1;HCoV-229E_1ab;1749 | 0 | 0 |

| | | |
|---|---|---|
| NC_002645.1;NP_073549.1;HCoV-229E_1ab;1750 | 0 | 0.025 |
| NC_002645.1;NP_073549.1;HCoV-229E_1ab;1751 | 0 | 0.025 |
| NC_002645.1;NP_073549.1;HCoV-229E_1ab;1752 | 0 | 0.025 |
| NC_002645.1;NP_073549.1;HCoV-229E_1ab;1753 | 0.05 | 0.05 |
| NC_002645.1;NP_073549.1;HCoV-229E_1ab;1754 | 0.1 | 0.025 |
| NC_002645.1;NP_073549.1;HCoV-229E_1ab;1755 | 0.05 | 0.025 |
| NC_002645.1;NP_073549.1;HCoV-229E_1ab;1756 | 0.05 | 0.05 |
| NC_002645.1;NP_073549.1;HCoV-229E_1ab;1757 | 0.1 | 0.025 |
| NC_002645.1;NP_073549.1;HCoV-229E_1ab;1758 | 0.1 | 0.025 |
| NC_002645.1;NP_073549.1;HCoV-229E_1ab;1759 | 0.05 | 0.025 |
| NC_002645.1;NP_073549.1;HCoV-229E_1ab;1760 | 0.1 | 0.025 |
| NC_002645.1;NP_073549.1;HCoV-229E_1ab;1761 | 0 | 0 |
| NC_002645.1;NP_073549.1;HCoV-229E_1ab;1762 | 0.05 | 0 |
| NC_002645.1;NP_073549.1;HCoV-229E_1ab;1763 | 0 | 0.025 |
| NC_002645.1;NP_073549.1;HCoV-229E_1ab;1764 | 0.05 | 0.025 |
| NC_002645.1;NP_073549.1;HCoV-229E_1ab;1765 | 0 | 0 |
| NC_002645.1;NP_073549.1;HCoV-229E_1ab;1766 | 0 | 0 |
| NC_002645.1;NP_073549.1;HCoV-229E_1ab;1767 | 0 | 0 |
| NC_002645.1;NP_073549.1;HCoV-229E_1ab;1768 | 0 | 0 |
| NC_002645.1;NP_073549.1;HCoV-229E_1ab;1769 | 0 | 0 |
| NC_002645.1;NP_073549.1;HCoV-229E_1ab;1770 | 0 | 0 |
| NC_002645.1;NP_073549.1;HCoV-229E_1ab;1771 | 0 | 0 |
| NC_002645.1;NP_073549.1;HCoV-229E_1ab;1772 | 0 | 0 |
| NC_002645.1;NP_073549.1;HCoV-229E_1ab;1773 | 0 | 0 |
| NC_002645.1;NP_073549.1;HCoV-229E_1ab;1774 | 0 | 0 |
| NC_002645.1;NP_073549.1;HCoV-229E_1ab;1775 | 0 | 0 |
| NC_002645.1;NP_073549.1;HCoV-229E_1ab;1776 | 0 | 0 |
| NC_002645.1;NP_073549.1;HCoV-229E_1ab;1777 | 0.05 | 0.125 |
| NC_002645.1;NP_073549.1;HCoV-229E_1ab;1778 | 0.05 | 0.025 |
| NC_002645.1;NP_073549.1;HCoV-229E_1ab;1779 | 0 | 0 |
| NC_002645.1;NP_073549.1;HCoV-229E_1ab;1780 | 0 | 0.025 |
| NC_002645.1;NP_073549.1;HCoV-229E_1ab;1781 | 0 | 0 |
| NC_002645.1;NP_073549.1;HCoV-229E_1ab;1782 | 0 | 0 |
| NC_002645.1;NP_073549.1;HCoV-229E_1ab;1783 | 0.15 | 0.125 |
| NC_002645.1;NP_073549.1;HCoV-229E_1ab;1784 | 0.15 | 0.125 |
| NC_002645.1;NP_073549.1;HCoV-229E_1ab;1785 | 0.15 | 0.075 |
| NC_002645.1;NP_073549.1;HCoV-229E_1ab;1786 | 0.1 | 0.05 |
| NC_002645.1;NP_073549.1;HCoV-229E_1ab;1787 | 0 | 0 |
| NC_002645.1;NP_073549.1;HCoV-229E_1ab;1788 | 0 | 0 |
| NC_002645.1;NP_073549.1;HCoV-229E_1ab;1789 | 0 | 0 |
| NC_002645.1;NP_073549.1;HCoV-229E_1ab;1790 | 0 | 0 |
| NC_002645.1;NP_073549.1;HCoV-229E_1ab;1791 | 0 | 0.025 |
| NC_002645.1;NP_073549.1;HCoV-229E_1ab;1792 | 0 | 0 |

| | | |
|---|---|---|
| NC_002645.1;NP_073549.1;HCoV-229E_1ab;1793 | 0 | 0.05 |
| NC_002645.1;NP_073549.1;HCoV-229E_1ab;1794 | 0 | 0.1 |
| NC_002645.1;NP_073549.1;HCoV-229E_1ab;1795 | 0 | 0.15 |
| NC_002645.1;NP_073549.1;HCoV-229E_1ab;1796 | 0 | 0.1 |
| NC_002645.1;NP_073549.1;HCoV-229E_1ab;1797 | 0.05 | 0.125 |
| NC_002645.1;NP_073549.1;HCoV-229E_1ab;1798 | 0.05 | 0.15 |
| NC_002645.1;NP_073549.1;HCoV-229E_1ab;1799 | 0 | 0.125 |
| NC_002645.1;NP_073549.1;HCoV-229E_1ab;1800 | 0 | 0.125 |
| NC_002645.1;NP_073549.1;HCoV-229E_1ab;1801 | 0 | 0.125 |
| NC_002645.1;NP_073549.1;HCoV-229E_1ab;1802 | 0.05 | 0.125 |
| NC_002645.1;NP_073549.1;HCoV-229E_1ab;1803 | 0 | 0.075 |
| NC_002645.1;NP_073549.1;HCoV-229E_1ab;1804 | 0.2 | 0.275 |
| NC_002645.1;NP_073549.1;HCoV-229E_1ab;1805 | 0 | 0 |
| NC_002645.1;NP_073549.1;HCoV-229E_1ab;1806 | 0 | 0 |
| NC_002645.1;NP_073549.1;HCoV-229E_1ab;1807 | 0 | 0 |
| NC_002645.1;NP_073549.1;HCoV-229E_1ab;1808 | 0 | 0 |
| NC_002645.1;NP_073549.1;HCoV-229E_1ab;1809 | 0 | 0 |
| NC_002645.1;NP_073549.1;HCoV-229E_1ab;1810 | 0 | 0 |
| NC_002645.1;NP_073549.1;HCoV-229E_1ab;1811 | 0 | 0 |
| NC_002645.1;NP_073549.1;HCoV-229E_1ab;1812 | 0 | 0 |
| NC_002645.1;NP_073549.1;HCoV-229E_1ab;1813 | 0 | 0 |
| NC_002645.1;NP_073549.1;HCoV-229E_1ab;1814 | 0 | 0 |
| NC_002645.1;NP_073549.1;HCoV-229E_1ab;1815 | 0 | 0 |
| NC_002645.1;NP_073549.1;HCoV-229E_1ab;1816 | 0 | 0 |
| NC_002645.1;NP_073549.1;HCoV-229E_1ab;1817 | 0 | 0 |
| NC_002645.1;NP_073549.1;HCoV-229E_1ab;1818 | 0 | 0 |
| NC_002645.1;NP_073549.1;HCoV-229E_1ab;1819 | 0 | 0 |
| NC_002645.1;NP_073549.1;HCoV-229E_1ab;1820 | 0 | 0 |
| NC_002645.1;NP_073549.1;HCoV-229E_1ab;1821 | 0 | 0 |
| NC_002645.1;NP_073549.1;HCoV-229E_1ab;1822 | 0.05 | 0.075 |
| NC_002645.1;NP_073549.1;HCoV-229E_1ab;1823 | 0.05 | 0.025 |
| NC_002645.1;NP_073549.1;HCoV-229E_1ab;1824 | 0.1 | 0.175 |
| NC_002645.1;NP_073549.1;HCoV-229E_1ab;1825 | 0.1 | 0.1 |
| NC_002645.1;NP_073549.1;HCoV-229E_1ab;1826 | 0.05 | 0.075 |
| NC_002645.1;NP_073549.1;HCoV-229E_1ab;1827 | 0 | 0.05 |
| NC_002645.1;NP_073549.1;HCoV-229E_1ab;1828 | 0 | 0.025 |
| NC_002645.1;NP_073549.1;HCoV-229E_1ab;1829 | 0 | 0.025 |
| NC_002645.1;NP_073549.1;HCoV-229E_1ab;1830 | 0 | 0 |
| NC_002645.1;NP_073549.1;HCoV-229E_1ab;1831 | 0 | 0 |
| NC_002645.1;NP_073549.1;HCoV-229E_1ab;1832 | 0.05 | 0 |
| NC_002645.1;NP_073549.1;HCoV-229E_1ab;1833 | 0 | 0.075 |
| NC_002645.1;NP_073549.1;HCoV-229E_1ab;1834 | 0 | 0 |
| NC_002645.1;NP_073549.1;HCoV-229E_1ab;1835 | 0 | 0 |

| | | |
|---|---|---|
| NC_002645.1;NP_073549.1;HCoV-229E_1ab;1836 | 0.05 | 0.05 |
| NC_002645.1;NP_073549.1;HCoV-229E_1ab;1837 | 0 | 0 |
| NC_002645.1;NP_073549.1;HCoV-229E_1ab;1838 | 0 | 0 |
| NC_002645.1;NP_073549.1;HCoV-229E_1ab;1839 | 0 | 0 |
| NC_002645.1;NP_073549.1;HCoV-229E_1ab;1840 | 0 | 0 |
| NC_002645.1;NP_073549.1;HCoV-229E_1ab;1841 | 0 | 0 |
| NC_002645.1;NP_073549.1;HCoV-229E_1ab;1842 | 0 | 0 |
| NC_002645.1;NP_073549.1;HCoV-229E_1ab;1843 | 0 | 0 |
| NC_002645.1;NP_073549.1;HCoV-229E_1ab;1844 | 0 | 0 |
| NC_002645.1;NP_073549.1;HCoV-229E_1ab;1845 | 0 | 0 |
| NC_002645.1;NP_073549.1;HCoV-229E_1ab;1846 | 0 | 0 |
| NC_002645.1;NP_073549.1;HCoV-229E_1ab;1847 | 0 | 0 |
| NC_002645.1;NP_073549.1;HCoV-229E_1ab;1848 | 0 | 0 |
| NC_002645.1;NP_073549.1;HCoV-229E_1ab;1849 | 0.05 | 0.025 |
| NC_002645.1;NP_073549.1;HCoV-229E_1ab;1850 | 0.05 | 0 |
| NC_002645.1;NP_073549.1;HCoV-229E_1ab;1851 | 0.05 | 0.025 |
| NC_002645.1;NP_073549.1;HCoV-229E_1ab;1852 | 0 | 0 |
| NC_002645.1;NP_073549.1;HCoV-229E_1ab;1853 | 0 | 0 |
| NC_002645.1;NP_073549.1;HCoV-229E_1ab;1854 | 0 | 0 |
| NC_002645.1;NP_073549.1;HCoV-229E_1ab;1855 | 0.05 | 0 |
| NC_002645.1;NP_073549.1;HCoV-229E_1ab;1856 | 0 | 0.025 |
| NC_002645.1;NP_073549.1;HCoV-229E_1ab;1857 | 0 | 0.075 |
| NC_002645.1;NP_073549.1;HCoV-229E_1ab;1858 | 0 | 0 |
| NC_002645.1;NP_073549.1;HCoV-229E_1ab;1859 | 0 | 0 |
| NC_002645.1;NP_073549.1;HCoV-229E_1ab;1860 | 0 | 0 |
| NC_002645.1;NP_073549.1;HCoV-229E_1ab;1861 | 0 | 0.025 |
| NC_002645.1;NP_073549.1;HCoV-229E_1ab;1862 | 0 | 0.025 |
| NC_002645.1;NP_073549.1;HCoV-229E_1ab;1863 | 0 | 0.025 |
| NC_002645.1;NP_073549.1;HCoV-229E_1ab;1864 | 0 | 0.05 |
| NC_002645.1;NP_073549.1;HCoV-229E_1ab;1865 | 0 | 0.025 |
| NC_002645.1;NP_073549.1;HCoV-229E_1ab;1866 | 0.05 | 0.025 |
| NC_002645.1;NP_073549.1;HCoV-229E_1ab;1867 | 0 | 0.05 |
| NC_002645.1;NP_073549.1;HCoV-229E_1ab;1868 | 0.15 | 0.075 |
| NC_002645.1;NP_073549.1;HCoV-229E_1ab;1869 | 0 | 0.025 |
| NC_002645.1;NP_073549.1;HCoV-229E_1ab;1870 | 0.05 | 0.025 |
| NC_002645.1;NP_073549.1;HCoV-229E_1ab;1871 | 0 | 0.05 |
| NC_002645.1;NP_073549.1;HCoV-229E_1ab;1872 | 0.05 | 0.025 |
| NC_002645.1;NP_073549.1;HCoV-229E_1ab;1873 | 0 | 0.025 |
| NC_002645.1;NP_073549.1;HCoV-229E_1ab;1874 | 0 | 0.025 |
| NC_002645.1;NP_073549.1;HCoV-229E_1ab;1875 | 0 | 0.025 |
| NC_002645.1;NP_073549.1;HCoV-229E_1ab;1876 | 0 | 0.05 |
| NC_002645.1;NP_073549.1;HCoV-229E_1ab;1877 | 0 | 0 |
| NC_002645.1;NP_073549.1;HCoV-229E_1ab;1878 | 0 | 0 |

| | | |
|---|---|---|
| NC_002645.1;NP_073549.1;HCoV-229E_1ab;1879 | 0 | 0 |
| NC_002645.1;NP_073549.1;HCoV-229E_1ab;1880 | 0 | 0 |
| NC_002645.1;NP_073549.1;HCoV-229E_1ab;1881 | 0 | 0 |
| NC_002645.1;NP_073549.1;HCoV-229E_1ab;1882 | 0 | 0 |
| NC_002645.1;NP_073549.1;HCoV-229E_1ab;1883 | 0 | 0 |
| NC_002645.1;NP_073549.1;HCoV-229E_1ab;1884 | 0 | 0 |
| NC_002645.1;NP_073549.1;HCoV-229E_1ab;1885 | 0 | 0 |
| NC_002645.1;NP_073549.1;HCoV-229E_1ab;1886 | 0 | 0 |
| NC_002645.1;NP_073549.1;HCoV-229E_1ab;1887 | 0 | 0 |
| NC_002645.1;NP_073549.1;HCoV-229E_1ab;1888 | 0 | 0 |
| NC_002645.1;NP_073549.1;HCoV-229E_1ab;1889 | 0 | 0 |
| NC_002645.1;NP_073549.1;HCoV-229E_1ab;1890 | 0 | 0 |
| NC_002645.1;NP_073549.1;HCoV-229E_1ab;1891 | 0 | 0 |
| NC_002645.1;NP_073549.1;HCoV-229E_1ab;1892 | 0 | 0 |
| NC_002645.1;NP_073549.1;HCoV-229E_1ab;1893 | 0 | 0 |
| NC_002645.1;NP_073549.1;HCoV-229E_1ab;1894 | 0 | 0 |
| NC_002645.1;NP_073549.1;HCoV-229E_1ab;1895 | 0 | 0 |
| NC_002645.1;NP_073549.1;HCoV-229E_1ab;1896 | 0 | 0 |
| NC_002645.1;NP_073549.1;HCoV-229E_1ab;1897 | 0 | 0 |
| NC_002645.1;NP_073549.1;HCoV-229E_1ab;1898 | 0 | 0 |
| NC_002645.1;NP_073549.1;HCoV-229E_1ab;1899 | 0.05 | 0.025 |
| NC_002645.1;NP_073549.1;HCoV-229E_1ab;1900 | 0 | 0.05 |
| NC_002645.1;NP_073549.1;HCoV-229E_1ab;1901 | 0.05 | 0.075 |
| NC_002645.1;NP_073549.1;HCoV-229E_1ab;1902 | 0 | 0.05 |
| NC_002645.1;NP_073549.1;HCoV-229E_1ab;1903 | 0.05 | 0.075 |
| NC_002645.1;NP_073549.1;HCoV-229E_1ab;1904 | 0 | 0.025 |
| NC_002645.1;NP_073549.1;HCoV-229E_1ab;1905 | 0 | 0.025 |
| NC_002645.1;NP_073549.1;HCoV-229E_1ab;1906 | 0 | 0.075 |
| NC_002645.1;NP_073549.1;HCoV-229E_1ab;1907 | 0.05 | 0.075 |
| NC_002645.1;NP_073549.1;HCoV-229E_1ab;1908 | 0 | 0 |
| NC_002645.1;NP_073549.1;HCoV-229E_1ab;1909 | 0.2 | 0 |
| NC_002645.1;NP_073549.1;HCoV-229E_1ab;1910 | 0.1 | 0 |
| NC_002645.1;NP_073549.1;HCoV-229E_1ab;1911 | 0.2 | 0.075 |
| NC_002645.1;NP_073549.1;HCoV-229E_1ab;1912 | 0.15 | 0.05 |
| NC_002645.1;NP_073549.1;HCoV-229E_1ab;1913 | 0.1 | 0 |
| NC_002645.1;NP_073549.1;HCoV-229E_1ab;1914 | 0.05 | 0.025 |
| NC_002645.1;NP_073549.1;HCoV-229E_1ab;1915 | 0.05 | 0 |
| NC_002645.1;NP_073549.1;HCoV-229E_1ab;1916 | 0.15 | 0 |
| NC_002645.1;NP_073549.1;HCoV-229E_1ab;1917 | 0.05 | 0 |
| NC_002645.1;NP_073549.1;HCoV-229E_1ab;1918 | 0 | 0 |
| NC_002645.1;NP_073549.1;HCoV-229E_1ab;1919 | 0.05 | 0 |
| NC_002645.1;NP_073549.1;HCoV-229E_1ab;1920 | 0 | 0 |
| NC_002645.1;NP_073549.1;HCoV-229E_1ab;1921 | 0 | 0 |

| | | |
|---|---|---|
| NC_002645.1;NP_073549.1;HCoV-229E_1ab;1922 | 0 | 0 |
| NC_002645.1;NP_073549.1;HCoV-229E_1ab;1923 | 0 | 0 |
| NC_002645.1;NP_073549.1;HCoV-229E_1ab;1924 | 0 | 0 |
| NC_002645.1;NP_073549.1;HCoV-229E_1ab;1925 | 0 | 0 |
| NC_002645.1;NP_073549.1;HCoV-229E_1ab;1926 | 0 | 0 |
| NC_002645.1;NP_073549.1;HCoV-229E_1ab;1927 | 0 | 0 |
| NC_002645.1;NP_073549.1;HCoV-229E_1ab;1928 | 0 | 0 |
| NC_002645.1;NP_073549.1;HCoV-229E_1ab;1929 | 0 | 0 |
| NC_002645.1;NP_073549.1;HCoV-229E_1ab;1930 | 0 | 0 |
| NC_002645.1;NP_073549.1;HCoV-229E_1ab;1931 | 0 | 0 |
| NC_002645.1;NP_073549.1;HCoV-229E_1ab;1932 | 0 | 0 |
| NC_002645.1;NP_073549.1;HCoV-229E_1ab;1933 | 0 | 0 |
| NC_002645.1;NP_073549.1;HCoV-229E_1ab;1934 | 0 | 0 |
| NC_002645.1;NP_073549.1;HCoV-229E_1ab;1935 | 0 | 0 |
| NC_002645.1;NP_073549.1;HCoV-229E_1ab;1936 | 0 | 0 |
| NC_002645.1;NP_073549.1;HCoV-229E_1ab;1937 | 0 | 0 |
| NC_002645.1;NP_073549.1;HCoV-229E_1ab;1938 | 0 | 0 |
| NC_002645.1;NP_073549.1;HCoV-229E_1ab;1939 | 0 | 0 |
| NC_002645.1;NP_073549.1;HCoV-229E_1ab;1940 | 0 | 0 |
| NC_002645.1;NP_073549.1;HCoV-229E_1ab;1941 | 0 | 0 |
| NC_002645.1;NP_073549.1;HCoV-229E_1ab;1942 | 0.05 | 0 |
| NC_002645.1;NP_073549.1;HCoV-229E_1ab;1943 | 0.05 | 0 |
| NC_002645.1;NP_073549.1;HCoV-229E_1ab;1944 | 0 | 0 |
| NC_002645.1;NP_073549.1;HCoV-229E_1ab;1945 | 0 | 0 |
| NC_002645.1;NP_073549.1;HCoV-229E_1ab;1946 | 0 | 0 |
| NC_002645.1;NP_073549.1;HCoV-229E_1ab;1947 | 0 | 0 |
| NC_002645.1;NP_073549.1;HCoV-229E_1ab;1948 | 0 | 0 |
| NC_002645.1;NP_073549.1;HCoV-229E_1ab;1949 | 0 | 0 |
| NC_002645.1;NP_073549.1;HCoV-229E_1ab;1950 | 0.05 | 0 |
| NC_002645.1;NP_073549.1;HCoV-229E_1ab;1951 | 0 | 0 |
| NC_002645.1;NP_073549.1;HCoV-229E_1ab;1952 | 0 | 0 |
| NC_002645.1;NP_073549.1;HCoV-229E_1ab;1953 | 0 | 0 |
| NC_002645.1;NP_073549.1;HCoV-229E_1ab;1954 | 0 | 0 |
| NC_002645.1;NP_073549.1;HCoV-229E_1ab;1955 | 0 | 0 |
| NC_002645.1;NP_073549.1;HCoV-229E_1ab;1956 | 0 | 0 |
| NC_002645.1;NP_073549.1;HCoV-229E_1ab;1957 | 0 | 0 |
| NC_002645.1;NP_073549.1;HCoV-229E_1ab;1958 | 0 | 0 |
| NC_002645.1;NP_073549.1;HCoV-229E_1ab;1959 | 0 | 0 |
| NC_002645.1;NP_073549.1;HCoV-229E_1ab;1960 | 0 | 0 |
| NC_002645.1;NP_073549.1;HCoV-229E_1ab;1961 | 0 | 0 |
| NC_002645.1;NP_073549.1;HCoV-229E_1ab;1962 | 0 | 0 |
| NC_002645.1;NP_073549.1;HCoV-229E_1ab;1963 | 0 | 0 |
| NC_002645.1;NP_073549.1;HCoV-229E_1ab;1964 | 0 | 0 |

| | | |
|---|---|---|
| NC_002645.1;NP_073549.1;HCoV-229E_1ab;1965 | 0 | 0 |
| NC_002645.1;NP_073549.1;HCoV-229E_1ab;1966 | 0 | 0.025 |
| NC_002645.1;NP_073549.1;HCoV-229E_1ab;1967 | 0 | 0.025 |
| NC_002645.1;NP_073549.1;HCoV-229E_1ab;1968 | 0.05 | 0 |
| NC_002645.1;NP_073549.1;HCoV-229E_1ab;1969 | 0 | 0 |
| NC_002645.1;NP_073549.1;HCoV-229E_1ab;1970 | 0 | 0 |
| NC_002645.1;NP_073549.1;HCoV-229E_1ab;1971 | 0 | 0 |
| NC_002645.1;NP_073549.1;HCoV-229E_1ab;1972 | 0 | 0 |
| NC_002645.1;NP_073549.1;HCoV-229E_1ab;1973 | 0 | 0 |
| NC_002645.1;NP_073549.1;HCoV-229E_1ab;1974 | 0 | 0 |
| NC_002645.1;NP_073549.1;HCoV-229E_1ab;1975 | 0 | 0 |
| NC_002645.1;NP_073549.1;HCoV-229E_1ab;1976 | 0 | 0 |
| NC_002645.1;NP_073549.1;HCoV-229E_1ab;1977 | 0 | 0 |
| NC_002645.1;NP_073549.1;HCoV-229E_1ab;1978 | 0 | 0 |
| NC_002645.1;NP_073549.1;HCoV-229E_1ab;1979 | 0 | 0 |
| NC_002645.1;NP_073549.1;HCoV-229E_1ab;1980 | 0 | 0 |
| NC_002645.1;NP_073549.1;HCoV-229E_1ab;1981 | 0 | 0.025 |
| NC_002645.1;NP_073549.1;HCoV-229E_1ab;1982 | 0 | 0 |
| NC_002645.1;NP_073549.1;HCoV-229E_1ab;1983 | 0 | 0 |
| NC_002645.1;NP_073549.1;HCoV-229E_1ab;1984 | 0 | 0 |
| NC_002645.1;NP_073549.1;HCoV-229E_1ab;1985 | 0 | 0 |
| NC_002645.1;NP_073549.1;HCoV-229E_1ab;1986 | 0 | 0 |
| NC_002645.1;NP_073549.1;HCoV-229E_1ab;1987 | 0 | 0 |
| NC_002645.1;NP_073549.1;HCoV-229E_1ab;1988 | 0 | 0 |
| NC_002645.1;NP_073549.1;HCoV-229E_1ab;1989 | 0 | 0 |
| NC_002645.1;NP_073549.1;HCoV-229E_1ab;1990 | 0 | 0 |
| NC_002645.1;NP_073549.1;HCoV-229E_1ab;1991 | 0 | 0 |
| NC_002645.1;NP_073549.1;HCoV-229E_1ab;1992 | 0 | 0 |
| NC_002645.1;NP_073549.1;HCoV-229E_1ab;1993 | 0 | 0 |
| NC_002645.1;NP_073549.1;HCoV-229E_1ab;1994 | 0 | 0 |
| NC_002645.1;NP_073549.1;HCoV-229E_1ab;1995 | 0 | 0 |
| NC_002645.1;NP_073549.1;HCoV-229E_1ab;1996 | 0 | 0 |
| NC_002645.1;NP_073549.1;HCoV-229E_1ab;1997 | 0 | 0 |
| NC_002645.1;NP_073549.1;HCoV-229E_1ab;1998 | 0 | 0 |
| NC_002645.1;NP_073549.1;HCoV-229E_1ab;1999 | 0 | 0 |
| NC_002645.1;NP_073549.1;HCoV-229E_1ab;2000 | 0 | 0 |
| NC_002645.1;NP_073549.1;HCoV-229E_1ab;2001 | 0 | 0 |
| NC_002645.1;NP_073549.1;HCoV-229E_1ab;2002 | 0 | 0 |
| NC_002645.1;NP_073549.1;HCoV-229E_1ab;2003 | 0 | 0 |
| NC_002645.1;NP_073549.1;HCoV-229E_1ab;2004 | 0 | 0 |
| NC_002645.1;NP_073549.1;HCoV-229E_1ab;2005 | 0 | 0 |
| NC_002645.1;NP_073549.1;HCoV-229E_1ab;2006 | 0 | 0 |
| NC_002645.1;NP_073549.1;HCoV-229E_1ab;2007 | 0 | 0 |

| | | |
|---|---|---|
| NC_002645.1;NP_073549.1;HCoV-229E_1ab;2008 | 0 | 0 |
| NC_002645.1;NP_073549.1;HCoV-229E_1ab;2009 | 0 | 0 |
| NC_002645.1;NP_073549.1;HCoV-229E_1ab;2010 | 0 | 0 |
| NC_002645.1;NP_073549.1;HCoV-229E_1ab;2011 | 0 | 0 |
| NC_002645.1;NP_073549.1;HCoV-229E_1ab;2012 | 0 | 0 |
| NC_002645.1;NP_073549.1;HCoV-229E_1ab;2013 | 0 | 0 |
| NC_002645.1;NP_073549.1;HCoV-229E_1ab;2014 | 0 | 0 |
| NC_002645.1;NP_073549.1;HCoV-229E_1ab;2015 | 0 | 0.025 |
| NC_002645.1;NP_073549.1;HCoV-229E_1ab;2016 | 0 | 0.025 |
| NC_002645.1;NP_073549.1;HCoV-229E_1ab;2017 | 0 | 0 |
| NC_002645.1;NP_073549.1;HCoV-229E_1ab;2018 | 0.05 | 0 |
| NC_002645.1;NP_073549.1;HCoV-229E_1ab;2019 | 0.05 | 0 |
| NC_002645.1;NP_073549.1;HCoV-229E_1ab;2020 | 0 | 0.05 |
| NC_002645.1;NP_073549.1;HCoV-229E_1ab;2021 | 0 | 0.025 |
| NC_002645.1;NP_073549.1;HCoV-229E_1ab;2022 | 0.05 | 0.1 |
| NC_002645.1;NP_073549.1;HCoV-229E_1ab;2023 | 0.1 | 0.125 |
| NC_002645.1;NP_073549.1;HCoV-229E_1ab;2024 | 0.05 | 0.05 |
| NC_002645.1;NP_073549.1;HCoV-229E_1ab;2025 | 0.05 | 0.05 |
| NC_002645.1;NP_073549.1;HCoV-229E_1ab;2026 | 0 | 0 |
| NC_002645.1;NP_073549.1;HCoV-229E_1ab;2027 | 0 | 0 |
| NC_002645.1;NP_073549.1;HCoV-229E_1ab;2028 | 0 | 0.025 |
| NC_002645.1;NP_073549.1;HCoV-229E_1ab;2029 | 0 | 0 |
| NC_002645.1;NP_073549.1;HCoV-229E_1ab;2030 | 0 | 0 |
| NC_002645.1;NP_073549.1;HCoV-229E_1ab;2031 | 0 | 0.025 |
| NC_002645.1;NP_073549.1;HCoV-229E_1ab;2032 | 0 | 0.025 |
| NC_002645.1;NP_073549.1;HCoV-229E_1ab;2033 | 0 | 0.025 |
| NC_002645.1;NP_073549.1;HCoV-229E_1ab;2034 | 0 | 0.025 |
| NC_002645.1;NP_073549.1;HCoV-229E_1ab;2035 | 0 | 0.025 |
| NC_002645.1;NP_073549.1;HCoV-229E_1ab;2036 | 0.25 | 0.15 |
| NC_002645.1;NP_073549.1;HCoV-229E_1ab;2037 | 0 | 0.025 |
| NC_002645.1;NP_073549.1;HCoV-229E_1ab;2038 | 0 | 0.025 |
| NC_002645.1;NP_073549.1;HCoV-229E_1ab;2039 | 0.1 | 0.025 |
| NC_002645.1;NP_073549.1;HCoV-229E_1ab;2040 | 0.1 | 0.025 |
| NC_002645.1;NP_073549.1;HCoV-229E_1ab;2041 | 0 | 0 |
| NC_002645.1;NP_073549.1;HCoV-229E_1ab;2042 | 0 | 0 |
| NC_002645.1;NP_073549.1;HCoV-229E_1ab;2043 | 0 | 0 |
| NC_002645.1;NP_073549.1;HCoV-229E_1ab;2044 | 0 | 0 |
| NC_002645.1;NP_073549.1;HCoV-229E_1ab;2045 | 0 | 0 |
| NC_002645.1;NP_073549.1;HCoV-229E_1ab;2046 | 0 | 0 |
| NC_002645.1;NP_073549.1;HCoV-229E_1ab;2047 | 0 | 0 |
| NC_002645.1;NP_073549.1;HCoV-229E_1ab;2048 | 0 | 0 |
| NC_002645.1;NP_073549.1;HCoV-229E_1ab;2049 | 0 | 0 |
| NC_002645.1;NP_073549.1;HCoV-229E_1ab;2050 | 0 | 0 |

| | | |
|---|---|---|
| NC_002645.1;NP_073549.1;HCoV-229E_1ab;2051 | 0 | 0 |
| NC_002645.1;NP_073549.1;HCoV-229E_1ab;2052 | 0 | 0 |
| NC_002645.1;NP_073549.1;HCoV-229E_1ab;2053 | 0 | 0 |
| NC_002645.1;NP_073549.1;HCoV-229E_1ab;2054 | 0 | 0 |
| NC_002645.1;NP_073549.1;HCoV-229E_1ab;2055 | 0 | 0 |
| NC_002645.1;NP_073549.1;HCoV-229E_1ab;2056 | 0 | 0 |
| NC_002645.1;NP_073549.1;HCoV-229E_1ab;2057 | 0 | 0 |
| NC_002645.1;NP_073549.1;HCoV-229E_1ab;2058 | 0 | 0 |
| NC_002645.1;NP_073549.1;HCoV-229E_1ab;2059 | 0 | 0 |
| NC_002645.1;NP_073549.1;HCoV-229E_1ab;2060 | 0 | 0 |
| NC_002645.1;NP_073549.1;HCoV-229E_1ab;2061 | 0 | 0 |
| NC_002645.1;NP_073549.1;HCoV-229E_1ab;2062 | 0 | 0.025 |
| NC_002645.1;NP_073549.1;HCoV-229E_1ab;2063 | 0 | 0 |
| NC_002645.1;NP_073549.1;HCoV-229E_1ab;2064 | 0 | 0 |
| NC_002645.1;NP_073549.1;HCoV-229E_1ab;2065 | 0 | 0 |
| NC_002645.1;NP_073549.1;HCoV-229E_1ab;2066 | 0 | 0 |
| NC_002645.1;NP_073549.1;HCoV-229E_1ab;2067 | 0 | 0 |
| NC_002645.1;NP_073549.1;HCoV-229E_1ab;2068 | 0 | 0 |
| NC_002645.1;NP_073549.1;HCoV-229E_1ab;2069 | 0 | 0 |
| NC_002645.1;NP_073549.1;HCoV-229E_1ab;2070 | 0 | 0 |
| NC_002645.1;NP_073549.1;HCoV-229E_1ab;2071 | 0 | 0 |
| NC_002645.1;NP_073549.1;HCoV-229E_1ab;2072 | 0 | 0 |
| NC_002645.1;NP_073549.1;HCoV-229E_1ab;2073 | 0 | 0 |
| NC_002645.1;NP_073549.1;HCoV-229E_1ab;2074 | 0 | 0 |
| NC_002645.1;NP_073549.1;HCoV-229E_1ab;2075 | 0 | 0 |
| NC_002645.1;NP_073549.1;HCoV-229E_1ab;2076 | 0 | 0 |
| NC_002645.1;NP_073549.1;HCoV-229E_1ab;2077 | 0 | 0 |
| NC_002645.1;NP_073549.1;HCoV-229E_1ab;2078 | 0 | 0 |
| NC_002645.1;NP_073549.1;HCoV-229E_1ab;2079 | 0 | 0 |
| NC_002645.1;NP_073549.1;HCoV-229E_1ab;2080 | 0 | 0 |
| NC_002645.1;NP_073549.1;HCoV-229E_1ab;2081 | 0 | 0 |
| NC_002645.1;NP_073549.1;HCoV-229E_1ab;2082 | 0 | 0 |
| NC_002645.1;NP_073549.1;HCoV-229E_1ab;2083 | 0 | 0 |
| NC_002645.1;NP_073549.1;HCoV-229E_1ab;2084 | 0 | 0 |
| NC_002645.1;NP_073549.1;HCoV-229E_1ab;2085 | 0 | 0 |
| NC_002645.1;NP_073549.1;HCoV-229E_1ab;2086 | 0 | 0 |
| NC_002645.1;NP_073549.1;HCoV-229E_1ab;2087 | 0 | 0 |
| NC_002645.1;NP_073549.1;HCoV-229E_1ab;2088 | 0 | 0 |
| NC_002645.1;NP_073549.1;HCoV-229E_1ab;2089 | 0 | 0 |
| NC_002645.1;NP_073549.1;HCoV-229E_1ab;2090 | 0 | 0 |
| NC_002645.1;NP_073549.1;HCoV-229E_1ab;2091 | 0 | 0 |
| NC_002645.1;NP_073549.1;HCoV-229E_1ab;2092 | 0 | 0 |
| NC_002645.1;NP_073549.1;HCoV-229E_1ab;2093 | 0 | 0 |

| | | |
|---|---|---|
| NC_002645.1;NP_073549.1;HCoV-229E_1ab;2094 | 0 | 0 |
| NC_002645.1;NP_073549.1;HCoV-229E_1ab;2095 | 0 | 0 |
| NC_002645.1;NP_073549.1;HCoV-229E_1ab;2096 | 0 | 0 |
| NC_002645.1;NP_073549.1;HCoV-229E_1ab;2097 | 0 | 0 |
| NC_002645.1;NP_073549.1;HCoV-229E_1ab;2098 | 0 | 0 |
| NC_002645.1;NP_073549.1;HCoV-229E_1ab;2099 | 0 | 0 |
| NC_002645.1;NP_073549.1;HCoV-229E_1ab;2100 | 0 | 0 |
| NC_002645.1;NP_073549.1;HCoV-229E_1ab;2101 | 0 | 0 |
| NC_002645.1;NP_073549.1;HCoV-229E_1ab;2102 | 0 | 0 |
| NC_002645.1;NP_073549.1;HCoV-229E_1ab;2103 | 0.05 | 0 |
| NC_002645.1;NP_073549.1;HCoV-229E_1ab;2104 | 0.05 | 0 |
| NC_002645.1;NP_073549.1;HCoV-229E_1ab;2105 | 0 | 0 |
| NC_002645.1;NP_073549.1;HCoV-229E_1ab;2106 | 0 | 0 |
| NC_002645.1;NP_073549.1;HCoV-229E_1ab;2107 | 0 | 0 |
| NC_002645.1;NP_073549.1;HCoV-229E_1ab;2108 | 0 | 0 |
| NC_002645.1;NP_073549.1;HCoV-229E_1ab;2109 | 0 | 0 |
| NC_002645.1;NP_073549.1;HCoV-229E_1ab;2110 | 0 | 0 |
| NC_002645.1;NP_073549.1;HCoV-229E_1ab;2111 | 0 | 0 |
| NC_002645.1;NP_073549.1;HCoV-229E_1ab;2112 | 0 | 0 |
| NC_002645.1;NP_073549.1;HCoV-229E_1ab;2113 | 0 | 0 |
| NC_002645.1;NP_073549.1;HCoV-229E_1ab;2114 | 0 | 0 |
| NC_002645.1;NP_073549.1;HCoV-229E_1ab;2115 | 0 | 0 |
| NC_002645.1;NP_073549.1;HCoV-229E_1ab;2116 | 0 | 0 |
| NC_002645.1;NP_073549.1;HCoV-229E_1ab;2117 | 0 | 0 |
| NC_002645.1;NP_073549.1;HCoV-229E_1ab;2118 | 0 | 0 |
| NC_002645.1;NP_073549.1;HCoV-229E_1ab;2119 | 0 | 0 |
| NC_002645.1;NP_073549.1;HCoV-229E_1ab;2120 | 0 | 0 |
| NC_002645.1;NP_073549.1;HCoV-229E_1ab;2121 | 0 | 0 |
| NC_002645.1;NP_073549.1;HCoV-229E_1ab;2122 | 0 | 0 |
| NC_002645.1;NP_073549.1;HCoV-229E_1ab;2123 | 0 | 0 |
| NC_002645.1;NP_073549.1;HCoV-229E_1ab;2124 | 0 | 0 |
| NC_002645.1;NP_073549.1;HCoV-229E_1ab;2125 | 0 | 0 |
| NC_002645.1;NP_073549.1;HCoV-229E_1ab;2126 | 0 | 0 |
| NC_002645.1;NP_073549.1;HCoV-229E_1ab;2127 | 0 | 0 |
| NC_002645.1;NP_073549.1;HCoV-229E_1ab;2128 | 0 | 0 |
| NC_002645.1;NP_073549.1;HCoV-229E_1ab;2129 | 0 | 0 |
| NC_002645.1;NP_073549.1;HCoV-229E_1ab;2130 | 0 | 0 |
| NC_002645.1;NP_073549.1;HCoV-229E_1ab;2131 | 0 | 0 |
| NC_002645.1;NP_073549.1;HCoV-229E_1ab;2132 | 0 | 0 |
| NC_002645.1;NP_073549.1;HCoV-229E_1ab;2133 | 0 | 0 |
| NC_002645.1;NP_073549.1;HCoV-229E_1ab;2134 | 0 | 0 |
| NC_002645.1;NP_073549.1;HCoV-229E_1ab;2135 | 0 | 0 |
| NC_002645.1;NP_073549.1;HCoV-229E_1ab;2136 | 0 | 0 |

| | | |
|---|---|---|
| NC_002645.1;NP_073549.1;HCoV-229E_1ab;2137 | 0 | 0 |
| NC_002645.1;NP_073549.1;HCoV-229E_1ab;2138 | 0 | 0 |
| NC_002645.1;NP_073549.1;HCoV-229E_1ab;2139 | 0 | 0 |
| NC_002645.1;NP_073549.1;HCoV-229E_1ab;2140 | 0 | 0 |
| NC_002645.1;NP_073549.1;HCoV-229E_1ab;2141 | 0 | 0 |
| NC_002645.1;NP_073549.1;HCoV-229E_1ab;2142 | 0 | 0 |
| NC_002645.1;NP_073549.1;HCoV-229E_1ab;2143 | 0 | 0 |
| NC_002645.1;NP_073549.1;HCoV-229E_1ab;2144 | 0 | 0 |
| NC_002645.1;NP_073549.1;HCoV-229E_1ab;2145 | 0 | 0 |
| NC_002645.1;NP_073549.1;HCoV-229E_1ab;2146 | 0 | 0 |
| NC_002645.1;NP_073549.1;HCoV-229E_1ab;2147 | 0 | 0 |
| NC_002645.1;NP_073549.1;HCoV-229E_1ab;2148 | 0 | 0 |
| NC_002645.1;NP_073549.1;HCoV-229E_1ab;2149 | 0 | 0 |
| NC_002645.1;NP_073549.1;HCoV-229E_1ab;2150 | 0 | 0 |
| NC_002645.1;NP_073549.1;HCoV-229E_1ab;2151 | 0.05 | 0 |
| NC_002645.1;NP_073549.1;HCoV-229E_1ab;2152 | 0 | 0 |
| NC_002645.1;NP_073549.1;HCoV-229E_1ab;2153 | 0 | 0 |
| NC_002645.1;NP_073549.1;HCoV-229E_1ab;2154 | 0 | 0 |
| NC_002645.1;NP_073549.1;HCoV-229E_1ab;2155 | 0 | 0 |
| NC_002645.1;NP_073549.1;HCoV-229E_1ab;2156 | 0 | 0 |
| NC_002645.1;NP_073549.1;HCoV-229E_1ab;2157 | 0 | 0 |
| NC_002645.1;NP_073549.1;HCoV-229E_1ab;2158 | 0 | 0 |
| NC_002645.1;NP_073549.1;HCoV-229E_1ab;2159 | 0 | 0 |
| NC_002645.1;NP_073549.1;HCoV-229E_1ab;2160 | 0 | 0 |
| NC_002645.1;NP_073549.1;HCoV-229E_1ab;2161 | 0 | 0 |
| NC_002645.1;NP_073549.1;HCoV-229E_1ab;2162 | 0 | 0 |
| NC_002645.1;NP_073549.1;HCoV-229E_1ab;2163 | 0 | 0.025 |
| NC_002645.1;NP_073549.1;HCoV-229E_1ab;2164 | 0 | 0 |
| NC_002645.1;NP_073549.1;HCoV-229E_1ab;2165 | 0 | 0.025 |
| NC_002645.1;NP_073549.1;HCoV-229E_1ab;2166 | 0 | 0 |
| NC_002645.1;NP_073549.1;HCoV-229E_1ab;2167 | 0 | 0 |
| NC_002645.1;NP_073549.1;HCoV-229E_1ab;2168 | 0 | 0 |
| NC_002645.1;NP_073549.1;HCoV-229E_1ab;2169 | 0 | 0 |
| NC_002645.1;NP_073549.1;HCoV-229E_1ab;2170 | 0 | 0 |
| NC_002645.1;NP_073549.1;HCoV-229E_1ab;2171 | 0 | 0 |
| NC_002645.1;NP_073549.1;HCoV-229E_1ab;2172 | 0 | 0 |
| NC_002645.1;NP_073549.1;HCoV-229E_1ab;2173 | 0.05 | 0 |
| NC_002645.1;NP_073549.1;HCoV-229E_1ab;2174 | 0 | 0 |
| NC_002645.1;NP_073549.1;HCoV-229E_1ab;2175 | 0.05 | 0 |
| NC_002645.1;NP_073549.1;HCoV-229E_1ab;2176 | 0 | 0 |
| NC_002645.1;NP_073549.1;HCoV-229E_1ab;2177 | 0 | 0 |
| NC_002645.1;NP_073549.1;HCoV-229E_1ab;2178 | 0 | 0 |
| NC_002645.1;NP_073549.1;HCoV-229E_1ab;2179 | 0 | 0 |

| | | |
|---|---|---|
| NC_002645.1;NP_073549.1;HCoV-229E_1ab;2180 | 0 | 0 |
| NC_002645.1;NP_073549.1;HCoV-229E_1ab;2181 | 0 | 0 |
| NC_002645.1;NP_073549.1;HCoV-229E_1ab;2182 | 0 | 0 |
| NC_002645.1;NP_073549.1;HCoV-229E_1ab;2183 | 0 | 0 |
| NC_002645.1;NP_073549.1;HCoV-229E_1ab;2184 | 0 | 0 |
| NC_002645.1;NP_073549.1;HCoV-229E_1ab;2185 | 0 | 0.025 |
| NC_002645.1;NP_073549.1;HCoV-229E_1ab;2186 | 0 | 0 |
| NC_002645.1;NP_073549.1;HCoV-229E_1ab;2187 | 0 | 0 |
| NC_002645.1;NP_073549.1;HCoV-229E_1ab;2188 | 0 | 0 |
| NC_002645.1;NP_073549.1;HCoV-229E_1ab;2189 | 0 | 0 |
| NC_002645.1;NP_073549.1;HCoV-229E_1ab;2190 | 0 | 0 |
| NC_002645.1;NP_073549.1;HCoV-229E_1ab;2191 | 0 | 0 |
| NC_002645.1;NP_073549.1;HCoV-229E_1ab;2192 | 0 | 0 |
| NC_002645.1;NP_073549.1;HCoV-229E_1ab;2193 | 0 | 0 |
| NC_002645.1;NP_073549.1;HCoV-229E_1ab;2194 | 0 | 0 |
| NC_002645.1;NP_073549.1;HCoV-229E_1ab;2195 | 0 | 0 |
| NC_002645.1;NP_073549.1;HCoV-229E_1ab;2196 | 0 | 0 |
| NC_002645.1;NP_073549.1;HCoV-229E_1ab;2197 | 0 | 0 |
| NC_002645.1;NP_073549.1;HCoV-229E_1ab;2198 | 0 | 0 |
| NC_002645.1;NP_073549.1;HCoV-229E_1ab;2199 | 0 | 0 |
| NC_002645.1;NP_073549.1;HCoV-229E_1ab;2200 | 0 | 0 |
| NC_002645.1;NP_073549.1;HCoV-229E_1ab;2201 | 0 | 0 |
| NC_002645.1;NP_073549.1;HCoV-229E_1ab;2202 | 0 | 0 |
| NC_002645.1;NP_073549.1;HCoV-229E_1ab;2203 | 0 | 0 |
| NC_002645.1;NP_073549.1;HCoV-229E_1ab;2204 | 0 | 0 |
| NC_002645.1;NP_073549.1;HCoV-229E_1ab;2205 | 0 | 0 |
| NC_002645.1;NP_073549.1;HCoV-229E_1ab;2206 | 0 | 0 |
| NC_002645.1;NP_073549.1;HCoV-229E_1ab;2207 | 0 | 0 |
| NC_002645.1;NP_073549.1;HCoV-229E_1ab;2208 | 0.05 | 0.025 |
| NC_002645.1;NP_073549.1;HCoV-229E_1ab;2209 | 0.05 | 0 |
| NC_002645.1;NP_073549.1;HCoV-229E_1ab;2210 | 0.05 | 0 |
| NC_002645.1;NP_073549.1;HCoV-229E_1ab;2211 | 0 | 0.025 |
| NC_002645.1;NP_073549.1;HCoV-229E_1ab;2212 | 0 | 0.05 |
| NC_002645.1;NP_073549.1;HCoV-229E_1ab;2213 | 0 | 0.025 |
| NC_002645.1;NP_073549.1;HCoV-229E_1ab;2214 | 0 | 0.025 |
| NC_002645.1;NP_073549.1;HCoV-229E_1ab;2215 | 0 | 0.025 |
| NC_002645.1;NP_073549.1;HCoV-229E_1ab;2216 | 0.05 | 0.05 |
| NC_002645.1;NP_073549.1;HCoV-229E_1ab;2217 | 0.05 | 0.075 |
| NC_002645.1;NP_073549.1;HCoV-229E_1ab;2218 | 0.05 | 0 |
| NC_002645.1;NP_073549.1;HCoV-229E_1ab;2219 | 0.05 | 0 |
| NC_002645.1;NP_073549.1;HCoV-229E_1ab;2220 | 0.05 | 0 |
| NC_002645.1;NP_073549.1;HCoV-229E_1ab;2221 | 0.05 | 0 |
| NC_002645.1;NP_073549.1;HCoV-229E_1ab;2222 | 0.05 | 0 |

| | | |
|---|---|---|
| NC_002645.1;NP_073549.1;HCoV-229E_1ab;2223 | 0.15 | 0 |
| NC_002645.1;NP_073549.1;HCoV-229E_1ab;2224 | 0.05 | 0 |
| NC_002645.1;NP_073549.1;HCoV-229E_1ab;2225 | 0 | 0 |
| NC_002645.1;NP_073549.1;HCoV-229E_1ab;2226 | 0 | 0 |
| NC_002645.1;NP_073549.1;HCoV-229E_1ab;2227 | 0 | 0 |
| NC_002645.1;NP_073549.1;HCoV-229E_1ab;2228 | 0.1 | 0 |
| NC_002645.1;NP_073549.1;HCoV-229E_1ab;2229 | 0 | 0 |
| NC_002645.1;NP_073549.1;HCoV-229E_1ab;2230 | 0 | 0 |
| NC_002645.1;NP_073549.1;HCoV-229E_1ab;2231 | 0 | 0 |
| NC_002645.1;NP_073549.1;HCoV-229E_1ab;2232 | 0.05 | 0 |
| NC_002645.1;NP_073549.1;HCoV-229E_1ab;2233 | 0.1 | 0 |
| NC_002645.1;NP_073549.1;HCoV-229E_1ab;2234 | 0.3 | 0.25 |
| NC_002645.1;NP_073549.1;HCoV-229E_1ab;2235 | 0 | 0 |
| NC_002645.1;NP_073549.1;HCoV-229E_1ab;2236 | 0 | 0.025 |
| NC_002645.1;NP_073549.1;HCoV-229E_1ab;2237 | 0 | 0 |
| NC_002645.1;NP_073549.1;HCoV-229E_1ab;2238 | 0 | 0 |
| NC_002645.1;NP_073549.1;HCoV-229E_1ab;2239 | 0 | 0 |
| NC_002645.1;NP_073549.1;HCoV-229E_1ab;2240 | 0 | 0 |
| NC_002645.1;NP_073549.1;HCoV-229E_1ab;2241 | 0 | 0 |
| NC_002645.1;NP_073549.1;HCoV-229E_1ab;2242 | 0 | 0 |
| NC_002645.1;NP_073549.1;HCoV-229E_1ab;2243 | 0 | 0 |
| NC_002645.1;NP_073549.1;HCoV-229E_1ab;2244 | 0 | 0 |
| NC_002645.1;NP_073549.1;HCoV-229E_1ab;2245 | 0 | 0 |
| NC_002645.1;NP_073549.1;HCoV-229E_1ab;2246 | 0 | 0 |
| NC_002645.1;NP_073549.1;HCoV-229E_1ab;2247 | 0 | 0 |
| NC_002645.1;NP_073549.1;HCoV-229E_1ab;2248 | 0 | 0 |
| NC_002645.1;NP_073549.1;HCoV-229E_1ab;2249 | 0 | 0 |
| NC_002645.1;NP_073549.1;HCoV-229E_1ab;2250 | 0 | 0 |
| NC_002645.1;NP_073549.1;HCoV-229E_1ab;2251 | 0 | 0 |
| NC_002645.1;NP_073549.1;HCoV-229E_1ab;2252 | 0 | 0 |
| NC_002645.1;NP_073549.1;HCoV-229E_1ab;2253 | 0 | 0 |
| NC_002645.1;NP_073549.1;HCoV-229E_1ab;2254 | 0 | 0 |
| NC_002645.1;NP_073549.1;HCoV-229E_1ab;2255 | 0.05 | 0.025 |
| NC_002645.1;NP_073549.1;HCoV-229E_1ab;2256 | 0 | 0 |
| NC_002645.1;NP_073549.1;HCoV-229E_1ab;2257 | 0 | 0 |
| NC_002645.1;NP_073549.1;HCoV-229E_1ab;2258 | 0 | 0 |
| NC_002645.1;NP_073549.1;HCoV-229E_1ab;2259 | 0.05 | 0.025 |
| NC_002645.1;NP_073549.1;HCoV-229E_1ab;2260 | 0.1 | 0.025 |
| NC_002645.1;NP_073549.1;HCoV-229E_1ab;2261 | 0.05 | 0 |
| NC_002645.1;NP_073549.1;HCoV-229E_1ab;2262 | 0 | 0 |
| NC_002645.1;NP_073549.1;HCoV-229E_1ab;2263 | 0 | 0 |
| NC_002645.1;NP_073549.1;HCoV-229E_1ab;2264 | 0 | 0.025 |
| NC_002645.1;NP_073549.1;HCoV-229E_1ab;2265 | 0 | 0 |

| | | |
|---|---|---|
| NC_002645.1;NP_073549.1;HCoV-229E_1ab;2266 | 0.05 | 0 |
| NC_002645.1;NP_073549.1;HCoV-229E_1ab;2267 | 0.05 | 0.05 |
| NC_002645.1;NP_073549.1;HCoV-229E_1ab;2268 | 0 | 0 |
| NC_002645.1;NP_073549.1;HCoV-229E_1ab;2269 | 0 | 0 |
| NC_002645.1;NP_073549.1;HCoV-229E_1ab;2270 | 0 | 0 |
| NC_002645.1;NP_073549.1;HCoV-229E_1ab;2271 | 0 | 0 |
| NC_002645.1;NP_073549.1;HCoV-229E_1ab;2272 | 0 | 0 |
| NC_002645.1;NP_073549.1;HCoV-229E_1ab;2273 | 0 | 0 |
| NC_002645.1;NP_073549.1;HCoV-229E_1ab;2274 | 0 | 0.025 |
| NC_002645.1;NP_073549.1;HCoV-229E_1ab;2275 | 0 | 0 |
| NC_002645.1;NP_073549.1;HCoV-229E_1ab;2276 | 0 | 0 |
| NC_002645.1;NP_073549.1;HCoV-229E_1ab;2277 | 0 | 0 |
| NC_002645.1;NP_073549.1;HCoV-229E_1ab;2278 | 0 | 0 |
| NC_002645.1;NP_073549.1;HCoV-229E_1ab;2279 | 0 | 0.025 |
| NC_002645.1;NP_073549.1;HCoV-229E_1ab;2280 | 0 | 0.025 |
| NC_002645.1;NP_073549.1;HCoV-229E_1ab;2281 | 0.1 | 0.025 |
| NC_002645.1;NP_073549.1;HCoV-229E_1ab;2282 | 0.1 | 0.025 |
| NC_002645.1;NP_073549.1;HCoV-229E_1ab;2283 | 0.1 | 0.075 |
| NC_002645.1;NP_073549.1;HCoV-229E_1ab;2284 | 0.1 | 0 |
| NC_002645.1;NP_073549.1;HCoV-229E_1ab;2285 | 0.1 | 0 |
| NC_002645.1;NP_073549.1;HCoV-229E_1ab;2286 | 0.25 | 0.05 |
| NC_002645.1;NP_073549.1;HCoV-229E_1ab;2287 | 0.25 | 0.025 |
| NC_002645.1;NP_073549.1;HCoV-229E_1ab;2288 | 0.25 | 0.125 |
| NC_002645.1;NP_073549.1;HCoV-229E_1ab;2289 | 0.2 | 0.075 |
| NC_002645.1;NP_073549.1;HCoV-229E_1ab;2290 | 0.1 | 0 |
| NC_002645.1;NP_073549.1;HCoV-229E_1ab;2291 | 0.1 | 0 |
| NC_002645.1;NP_073549.1;HCoV-229E_1ab;2292 | 0.05 | 0 |
| NC_002645.1;NP_073549.1;HCoV-229E_1ab;2293 | 0.05 | 0 |
| NC_002645.1;NP_073549.1;HCoV-229E_1ab;2294 | 0 | 0 |
| NC_002645.1;NP_073549.1;HCoV-229E_1ab;2295 | 0 | 0 |
| NC_002645.1;NP_073549.1;HCoV-229E_1ab;2296 | 0 | 0 |
| NC_002645.1;NP_073549.1;HCoV-229E_1ab;2297 | 0 | 0 |
| NC_002645.1;NP_073549.1;HCoV-229E_1ab;2298 | 0 | 0 |
| NC_002645.1;NP_073549.1;HCoV-229E_1ab;2299 | 0 | 0 |
| NC_002645.1;NP_073549.1;HCoV-229E_1ab;2300 | 0 | 0 |
| NC_002645.1;NP_073549.1;HCoV-229E_1ab;2301 | 0 | 0 |
| NC_002645.1;NP_073549.1;HCoV-229E_1ab;2302 | 0 | 0 |
| NC_002645.1;NP_073549.1;HCoV-229E_1ab;2303 | 0 | 0 |
| NC_002645.1;NP_073549.1;HCoV-229E_1ab;2304 | 0 | 0.025 |
| NC_002645.1;NP_073549.1;HCoV-229E_1ab;2305 | 0 | 0 |
| NC_002645.1;NP_073549.1;HCoV-229E_1ab;2306 | 0 | 0 |
| NC_002645.1;NP_073549.1;HCoV-229E_1ab;2307 | 0 | 0 |
| NC_002645.1;NP_073549.1;HCoV-229E_1ab;2308 | 0 | 0 |

| | | |
|---|---|---|
| NC_002645.1;NP_073549.1;HCoV-229E_1ab;2309 | 0 | 0 |
| NC_002645.1;NP_073549.1;HCoV-229E_1ab;2310 | 0 | 0.025 |
| NC_002645.1;NP_073549.1;HCoV-229E_1ab;2311 | 0 | 0 |
| NC_002645.1;NP_073549.1;HCoV-229E_1ab;2312 | 0 | 0 |
| NC_002645.1;NP_073549.1;HCoV-229E_1ab;2313 | 0 | 0 |
| NC_002645.1;NP_073549.1;HCoV-229E_1ab;2314 | 0.05 | 0 |
| NC_002645.1;NP_073549.1;HCoV-229E_1ab;2315 | 0 | 0 |
| NC_002645.1;NP_073549.1;HCoV-229E_1ab;2316 | 0 | 0 |
| NC_002645.1;NP_073549.1;HCoV-229E_1ab;2317 | 0 | 0.025 |
| NC_002645.1;NP_073549.1;HCoV-229E_1ab;2318 | 0 | 0 |
| NC_002645.1;NP_073549.1;HCoV-229E_1ab;2319 | 0 | 0 |
| NC_002645.1;NP_073549.1;HCoV-229E_1ab;2320 | 0 | 0 |
| NC_002645.1;NP_073549.1;HCoV-229E_1ab;2321 | 0 | 0 |
| NC_002645.1;NP_073549.1;HCoV-229E_1ab;2322 | 0 | 0 |
| NC_002645.1;NP_073549.1;HCoV-229E_1ab;2323 | 0 | 0 |
| NC_002645.1;NP_073549.1;HCoV-229E_1ab;2324 | 0 | 0 |
| NC_002645.1;NP_073549.1;HCoV-229E_1ab;2325 | 0 | 0 |
| NC_002645.1;NP_073549.1;HCoV-229E_1ab;2326 | 0 | 0 |
| NC_002645.1;NP_073549.1;HCoV-229E_1ab;2327 | 0 | 0 |
| NC_002645.1;NP_073549.1;HCoV-229E_1ab;2328 | 0 | 0 |
| NC_002645.1;NP_073549.1;HCoV-229E_1ab;2329 | 0 | 0 |
| NC_002645.1;NP_073549.1;HCoV-229E_1ab;2330 | 0 | 0 |
| NC_002645.1;NP_073549.1;HCoV-229E_1ab;2331 | 0 | 0 |
| NC_002645.1;NP_073549.1;HCoV-229E_1ab;2332 | 0 | 0.025 |
| NC_002645.1;NP_073549.1;HCoV-229E_1ab;2333 | 0 | 0.025 |
| NC_002645.1;NP_073549.1;HCoV-229E_1ab;2334 | 0 | 0.05 |
| NC_002645.1;NP_073549.1;HCoV-229E_1ab;2335 | 0 | 0.1 |
| NC_002645.1;NP_073549.1;HCoV-229E_1ab;2336 | 0.05 | 0.1 |
| NC_002645.1;NP_073549.1;HCoV-229E_1ab;2337 | 0.1 | 0.05 |
| NC_002645.1;NP_073549.1;HCoV-229E_1ab;2338 | 0.1 | 0.075 |
| NC_002645.1;NP_073549.1;HCoV-229E_1ab;2339 | 0.05 | 0.025 |
| NC_002645.1;NP_073549.1;HCoV-229E_1ab;2340 | 0.05 | 0.075 |
| NC_002645.1;NP_073549.1;HCoV-229E_1ab;2341 | 0.1 | 0.075 |
| NC_002645.1;NP_073549.1;HCoV-229E_1ab;2342 | 0.1 | 0.075 |
| NC_002645.1;NP_073549.1;HCoV-229E_1ab;2343 | 0.05 | 0.05 |
| NC_002645.1;NP_073549.1;HCoV-229E_1ab;2344 | 0.05 | 0.05 |
| NC_002645.1;NP_073549.1;HCoV-229E_1ab;2345 | 0.05 | 0.025 |
| NC_002645.1;NP_073549.1;HCoV-229E_1ab;2346 | 0.05 | 0.1 |
| NC_002645.1;NP_073549.1;HCoV-229E_1ab;2347 | 0.05 | 0.075 |
| NC_002645.1;NP_073549.1;HCoV-229E_1ab;2348 | 0 | 0.1 |
| NC_002645.1;NP_073549.1;HCoV-229E_1ab;2349 | 0 | 0.1 |
| NC_002645.1;NP_073549.1;HCoV-229E_1ab;2350 | 0 | 0.075 |
| NC_002645.1;NP_073549.1;HCoV-229E_1ab;2351 | 0 | 0.075 |

| | | |
|---|---|---|
| NC_002645.1;NP_073549.1;HCoV-229E_1ab;2352 | 0 | 0.075 |
| NC_002645.1;NP_073549.1;HCoV-229E_1ab;2353 | 0 | 0.025 |
| NC_002645.1;NP_073549.1;HCoV-229E_1ab;2354 | 0 | 0.05 |
| NC_002645.1;NP_073549.1;HCoV-229E_1ab;2355 | 0.05 | 0.05 |
| NC_002645.1;NP_073549.1;HCoV-229E_1ab;2356 | 0.05 | 0.05 |
| NC_002645.1;NP_073549.1;HCoV-229E_1ab;2357 | 0.05 | 0.075 |
| NC_002645.1;NP_073549.1;HCoV-229E_1ab;2358 | 0.2 | 0.225 |
| NC_002645.1;NP_073549.1;HCoV-229E_1ab;2359 | 0 | 0 |
| NC_002645.1;NP_073549.1;HCoV-229E_1ab;2360 | 0 | 0 |
| NC_002645.1;NP_073549.1;HCoV-229E_1ab;2361 | 0 | 0 |
| NC_002645.1;NP_073549.1;HCoV-229E_1ab;2362 | 0 | 0 |
| NC_002645.1;NP_073549.1;HCoV-229E_1ab;2363 | 0.05 | 0 |
| NC_002645.1;NP_073549.1;HCoV-229E_1ab;2364 | 0 | 0 |
| NC_002645.1;NP_073549.1;HCoV-229E_1ab;2365 | 0 | 0 |
| NC_002645.1;NP_073549.1;HCoV-229E_1ab;2366 | 0 | 0 |
| NC_002645.1;NP_073549.1;HCoV-229E_1ab;2367 | 0 | 0 |
| NC_002645.1;NP_073549.1;HCoV-229E_1ab;2368 | 0 | 0 |
| NC_002645.1;NP_073549.1;HCoV-229E_1ab;2369 | 0 | 0 |
| NC_002645.1;NP_073549.1;HCoV-229E_1ab;2370 | 0 | 0 |
| NC_002645.1;NP_073549.1;HCoV-229E_1ab;2371 | 0 | 0 |
| NC_002645.1;NP_073549.1;HCoV-229E_1ab;2372 | 0 | 0 |
| NC_002645.1;NP_073549.1;HCoV-229E_1ab;2373 | 0 | 0 |
| NC_002645.1;NP_073549.1;HCoV-229E_1ab;2374 | 0.05 | 0 |
| NC_002645.1;NP_073549.1;HCoV-229E_1ab;2375 | 0 | 0 |
| NC_002645.1;NP_073549.1;HCoV-229E_1ab;2376 | 0 | 0 |
| NC_002645.1;NP_073549.1;HCoV-229E_1ab;2377 | 0 | 0 |
| NC_002645.1;NP_073549.1;HCoV-229E_1ab;2378 | 0 | 0 |
| NC_002645.1;NP_073549.1;HCoV-229E_1ab;2379 | 0 | 0 |
| NC_002645.1;NP_073549.1;HCoV-229E_1ab;2380 | 0 | 0 |
| NC_002645.1;NP_073549.1;HCoV-229E_1ab;2381 | 0 | 0 |
| NC_002645.1;NP_073549.1;HCoV-229E_1ab;2382 | 0 | 0 |
| NC_002645.1;NP_073549.1;HCoV-229E_1ab;2383 | 0.05 | 0 |
| NC_002645.1;NP_073549.1;HCoV-229E_1ab;2384 | 0 | 0 |
| NC_002645.1;NP_073549.1;HCoV-229E_1ab;2385 | 0 | 0 |
| NC_002645.1;NP_073549.1;HCoV-229E_1ab;2386 | 0.1 | 0 |
| NC_002645.1;NP_073549.1;HCoV-229E_1ab;2387 | 0 | 0.025 |
| NC_002645.1;NP_073549.1;HCoV-229E_1ab;2388 | 0.05 | 0.1 |
| NC_002645.1;NP_073549.1;HCoV-229E_1ab;2389 | 0.05 | 0.075 |
| NC_002645.1;NP_073549.1;HCoV-229E_1ab;2390 | 0.1 | 0.075 |
| NC_002645.1;NP_073549.1;HCoV-229E_1ab;2391 | 0 | 0.025 |
| NC_002645.1;NP_073549.1;HCoV-229E_1ab;2392 | 0.05 | 0.025 |
| NC_002645.1;NP_073549.1;HCoV-229E_1ab;2393 | 0 | 0.025 |
| NC_002645.1;NP_073549.1;HCoV-229E_1ab;2394 | 0 | 0.05 |

| | | |
|---|---|---|
| NC_002645.1;NP_073549.1;HCoV-229E_1ab;2395 | 0 | 0.025 |
| NC_002645.1;NP_073549.1;HCoV-229E_1ab;2396 | 0 | 0.025 |
| NC_002645.1;NP_073549.1;HCoV-229E_1ab;2397 | 0 | 0.025 |
| NC_002645.1;NP_073549.1;HCoV-229E_1ab;2398 | 0 | 0.025 |
| NC_002645.1;NP_073549.1;HCoV-229E_1ab;2399 | 0 | 0 |
| NC_002645.1;NP_073549.1;HCoV-229E_1ab;2400 | 0 | 0 |
| NC_002645.1;NP_073549.1;HCoV-229E_1ab;2401 | 0 | 0.025 |
| NC_002645.1;NP_073549.1;HCoV-229E_1ab;2402 | 0 | 0 |
| NC_002645.1;NP_073549.1;HCoV-229E_1ab;2403 | 0 | 0 |
| NC_002645.1;NP_073549.1;HCoV-229E_1ab;2404 | 0 | 0 |
| NC_002645.1;NP_073549.1;HCoV-229E_1ab;2405 | 0 | 0 |
| NC_002645.1;NP_073549.1;HCoV-229E_1ab;2406 | 0 | 0 |
| NC_002645.1;NP_073549.1;HCoV-229E_1ab;2407 | 0 | 0 |
| NC_002645.1;NP_073549.1;HCoV-229E_1ab;2408 | 0 | 0 |
| NC_002645.1;NP_073549.1;HCoV-229E_1ab;2409 | 0 | 0 |
| NC_002645.1;NP_073549.1;HCoV-229E_1ab;2410 | 0 | 0 |
| NC_002645.1;NP_073549.1;HCoV-229E_1ab;2411 | 0 | 0 |
| NC_002645.1;NP_073549.1;HCoV-229E_1ab;2412 | 0 | 0 |
| NC_002645.1;NP_073549.1;HCoV-229E_1ab;2413 | 0 | 0 |
| NC_002645.1;NP_073549.1;HCoV-229E_1ab;2414 | 0 | 0 |
| NC_002645.1;NP_073549.1;HCoV-229E_1ab;2415 | 0.05 | 0 |
| NC_002645.1;NP_073549.1;HCoV-229E_1ab;2416 | 0.05 | 0 |
| NC_002645.1;NP_073549.1;HCoV-229E_1ab;2417 | 0 | 0.025 |
| NC_002645.1;NP_073549.1;HCoV-229E_1ab;2418 | 0.05 | 0 |
| NC_002645.1;NP_073549.1;HCoV-229E_1ab;2419 | 0 | 0 |
| NC_002645.1;NP_073549.1;HCoV-229E_1ab;2420 | 0 | 0 |
| NC_002645.1;NP_073549.1;HCoV-229E_1ab;2421 | 0 | 0 |
| NC_002645.1;NP_073549.1;HCoV-229E_1ab;2422 | 0 | 0 |
| NC_002645.1;NP_073549.1;HCoV-229E_1ab;2423 | 0 | 0 |
| NC_002645.1;NP_073549.1;HCoV-229E_1ab;2424 | 0 | 0 |
| NC_002645.1;NP_073549.1;HCoV-229E_1ab;2425 | 0 | 0 |
| NC_002645.1;NP_073549.1;HCoV-229E_1ab;2426 | 0 | 0 |
| NC_002645.1;NP_073549.1;HCoV-229E_1ab;2427 | 0.05 | 0.05 |
| NC_002645.1;NP_073549.1;HCoV-229E_1ab;2428 | 0.05 | 0.05 |
| NC_002645.1;NP_073549.1;HCoV-229E_1ab;2429 | 0.05 | 0.05 |
| NC_002645.1;NP_073549.1;HCoV-229E_1ab;2430 | 0.1 | 0.05 |
| NC_002645.1;NP_073549.1;HCoV-229E_1ab;2431 | 0 | 0.025 |
| NC_002645.1;NP_073549.1;HCoV-229E_1ab;2432 | 0.1 | 0 |
| NC_002645.1;NP_073549.1;HCoV-229E_1ab;2433 | 0.1 | 0 |
| NC_002645.1;NP_073549.1;HCoV-229E_1ab;2434 | 0.15 | 0 |
| NC_002645.1;NP_073549.1;HCoV-229E_1ab;2435 | 0.05 | 0 |
| NC_002645.1;NP_073549.1;HCoV-229E_1ab;2436 | 0.05 | 0 |
| NC_002645.1;NP_073549.1;HCoV-229E_1ab;2437 | 0 | 0 |

| | | |
|---|---|---|
| NC_002645.1;NP_073549.1;HCoV-229E_1ab;2438 | 0 | 0 |
| NC_002645.1;NP_073549.1;HCoV-229E_1ab;2439 | 0 | 0 |
| NC_002645.1;NP_073549.1;HCoV-229E_1ab;2440 | 0 | 0 |
| NC_002645.1;NP_073549.1;HCoV-229E_1ab;2441 | 0 | 0 |
| NC_002645.1;NP_073549.1;HCoV-229E_1ab;2442 | 0 | 0 |
| NC_002645.1;NP_073549.1;HCoV-229E_1ab;2443 | 0 | 0 |
| NC_002645.1;NP_073549.1;HCoV-229E_1ab;2444 | 0 | 0 |
| NC_002645.1;NP_073549.1;HCoV-229E_1ab;2445 | 0 | 0 |
| NC_002645.1;NP_073549.1;HCoV-229E_1ab;2446 | 0 | 0 |
| NC_002645.1;NP_073549.1;HCoV-229E_1ab;2447 | 0 | 0 |
| NC_002645.1;NP_073549.1;HCoV-229E_1ab;2448 | 0 | 0 |
| NC_002645.1;NP_073549.1;HCoV-229E_1ab;2449 | 0 | 0 |
| NC_002645.1;NP_073549.1;HCoV-229E_1ab;2450 | 0 | 0 |
| NC_002645.1;NP_073549.1;HCoV-229E_1ab;2451 | 0 | 0 |
| NC_002645.1;NP_073549.1;HCoV-229E_1ab;2452 | 0 | 0 |
| NC_002645.1;NP_073549.1;HCoV-229E_1ab;2453 | 0 | 0 |
| NC_002645.1;NP_073549.1;HCoV-229E_1ab;2454 | 0 | 0 |
| NC_002645.1;NP_073549.1;HCoV-229E_1ab;2455 | 0 | 0 |
| NC_002645.1;NP_073549.1;HCoV-229E_1ab;2456 | 0 | 0 |
| NC_002645.1;NP_073549.1;HCoV-229E_1ab;2457 | 0 | 0 |
| NC_002645.1;NP_073549.1;HCoV-229E_1ab;2458 | 0 | 0 |
| NC_002645.1;NP_073549.1;HCoV-229E_1ab;2459 | 0 | 0 |
| NC_002645.1;NP_073549.1;HCoV-229E_1ab;2460 | 0 | 0 |
| NC_002645.1;NP_073549.1;HCoV-229E_1ab;2461 | 0 | 0 |
| NC_002645.1;NP_073549.1;HCoV-229E_1ab;2462 | 0 | 0 |
| NC_002645.1;NP_073549.1;HCoV-229E_1ab;2463 | 0 | 0 |
| NC_002645.1;NP_073549.1;HCoV-229E_1ab;2464 | 0 | 0 |
| NC_002645.1;NP_073549.1;HCoV-229E_1ab;2465 | 0 | 0 |
| NC_002645.1;NP_073549.1;HCoV-229E_1ab;2466 | 0 | 0 |
| NC_002645.1;NP_073549.1;HCoV-229E_1ab;2467 | 0 | 0 |
| NC_002645.1;NP_073549.1;HCoV-229E_1ab;2468 | 0 | 0 |
| NC_002645.1;NP_073549.1;HCoV-229E_1ab;2469 | 0 | 0 |
| NC_002645.1;NP_073549.1;HCoV-229E_1ab;2470 | 0 | 0 |
| NC_002645.1;NP_073549.1;HCoV-229E_1ab;2471 | 0 | 0.025 |
| NC_002645.1;NP_073549.1;HCoV-229E_1ab;2472 | 0 | 0.025 |
| NC_002645.1;NP_073549.1;HCoV-229E_1ab;2473 | 0 | 0.025 |
| NC_002645.1;NP_073549.1;HCoV-229E_1ab;2474 | 0 | 0.025 |
| NC_002645.1;NP_073549.1;HCoV-229E_1ab;2475 | 0 | 0.025 |
| NC_002645.1;NP_073549.1;HCoV-229E_1ab;2476 | 0 | 0.025 |
| NC_002645.1;NP_073549.1;HCoV-229E_1ab;2477 | 0 | 0.025 |
| NC_002645.1;NP_073549.1;HCoV-229E_1ab;2478 | 0 | 0.025 |
| NC_002645.1;NP_073549.1;HCoV-229E_1ab;2479 | 0 | 0.025 |
| NC_002645.1;NP_073549.1;HCoV-229E_1ab;2480 | 0 | 0.025 |

| | | |
|---|---|---|
| NC_002645.1;NP_073549.1;HCoV-229E_1ab;2481 | 0 | 0.025 |
| NC_002645.1;NP_073549.1;HCoV-229E_1ab;2482 | 0 | 0 |
| NC_002645.1;NP_073549.1;HCoV-229E_1ab;2483 | 0 | 0 |
| NC_002645.1;NP_073549.1;HCoV-229E_1ab;2484 | 0 | 0 |
| NC_002645.1;NP_073549.1;HCoV-229E_1ab;2485 | 0 | 0 |
| NC_002645.1;NP_073549.1;HCoV-229E_1ab;2486 | 0 | 0 |
| NC_002645.1;NP_073549.1;HCoV-229E_1ab;2487 | 0 | 0 |
| NC_002645.1;NP_073549.1;HCoV-229E_1ab;2488 | 0 | 0 |
| NC_002645.1;NP_073549.1;HCoV-229E_1ab;2489 | 0 | 0 |
| NC_002645.1;NP_073549.1;HCoV-229E_1ab;2490 | 0 | 0 |
| NC_002645.1;NP_073549.1;HCoV-229E_1ab;2491 | 0 | 0 |
| NC_002645.1;NP_073549.1;HCoV-229E_1ab;2492 | 0 | 0 |
| NC_002645.1;NP_073549.1;HCoV-229E_1ab;2493 | 0 | 0 |
| NC_002645.1;NP_073549.1;HCoV-229E_1ab;2494 | 0 | 0 |
| NC_002645.1;NP_073549.1;HCoV-229E_1ab;2495 | 0 | 0 |
| NC_002645.1;NP_073549.1;HCoV-229E_1ab;2496 | 0 | 0 |
| NC_002645.1;NP_073549.1;HCoV-229E_1ab;2497 | 0 | 0 |
| NC_002645.1;NP_073549.1;HCoV-229E_1ab;2498 | 0 | 0 |
| NC_002645.1;NP_073549.1;HCoV-229E_1ab;2499 | 0 | 0 |
| NC_002645.1;NP_073549.1;HCoV-229E_1ab;2500 | 0 | 0 |
| NC_002645.1;NP_073549.1;HCoV-229E_1ab;2501 | 0 | 0 |
| NC_002645.1;NP_073549.1;HCoV-229E_1ab;2502 | 0 | 0 |
| NC_002645.1;NP_073549.1;HCoV-229E_1ab;2503 | 0 | 0 |
| NC_002645.1;NP_073549.1;HCoV-229E_1ab;2504 | 0 | 0 |
| NC_002645.1;NP_073549.1;HCoV-229E_1ab;2505 | 0 | 0 |
| NC_002645.1;NP_073549.1;HCoV-229E_1ab;2506 | 0 | 0 |
| NC_002645.1;NP_073549.1;HCoV-229E_1ab;2507 | 0 | 0 |
| NC_002645.1;NP_073549.1;HCoV-229E_1ab;2508 | 0 | 0 |
| NC_002645.1;NP_073549.1;HCoV-229E_1ab;2509 | 0 | 0 |
| NC_002645.1;NP_073549.1;HCoV-229E_1ab;2510 | 0 | 0 |
| NC_002645.1;NP_073549.1;HCoV-229E_1ab;2511 | 0 | 0 |
| NC_002645.1;NP_073549.1;HCoV-229E_1ab;2512 | 0 | 0 |
| NC_002645.1;NP_073549.1;HCoV-229E_1ab;2513 | 0 | 0 |
| NC_002645.1;NP_073549.1;HCoV-229E_1ab;2514 | 0 | 0.05 |
| NC_002645.1;NP_073549.1;HCoV-229E_1ab;2515 | 0 | 0 |
| NC_002645.1;NP_073549.1;HCoV-229E_1ab;2516 | 0 | 0 |
| NC_002645.1;NP_073549.1;HCoV-229E_1ab;2517 | 0 | 0 |
| NC_002645.1;NP_073549.1;HCoV-229E_1ab;2518 | 0 | 0.025 |
| NC_002645.1;NP_073549.1;HCoV-229E_1ab;2519 | 0 | 0.025 |
| NC_002645.1;NP_073549.1;HCoV-229E_1ab;2520 | 0 | 0.025 |
| NC_002645.1;NP_073549.1;HCoV-229E_1ab;2521 | 0 | 0.075 |
| NC_002645.1;NP_073549.1;HCoV-229E_1ab;2522 | 0.05 | 0.05 |
| NC_002645.1;NP_073549.1;HCoV-229E_1ab;2523 | 0.05 | 0.075 |

| | | |
|---|---|---|
| NC_002645.1;NP_073549.1;HCoV-229E_1ab;2524 | 0.1 | 0.025 |
| NC_002645.1;NP_073549.1;HCoV-229E_1ab;2525 | 0.15 | 0.1 |
| NC_002645.1;NP_073549.1;HCoV-229E_1ab;2526 | 0.1 | 0 |
| NC_002645.1;NP_073549.1;HCoV-229E_1ab;2527 | 0.15 | 0 |
| NC_002645.1;NP_073549.1;HCoV-229E_1ab;2528 | 0.15 | 0 |
| NC_002645.1;NP_073549.1;HCoV-229E_1ab;2529 | 0.1 | 0.025 |
| NC_002645.1;NP_073549.1;HCoV-229E_1ab;2530 | 0.2 | 0 |
| NC_002645.1;NP_073549.1;HCoV-229E_1ab;2531 | 0.05 | 0 |
| NC_002645.1;NP_073549.1;HCoV-229E_1ab;2532 | 0.05 | 0 |
| NC_002645.1;NP_073549.1;HCoV-229E_1ab;2533 | 0.1 | 0 |
| NC_002645.1;NP_073549.1;HCoV-229E_1ab;2534 | 0.05 | 0 |
| NC_002645.1;NP_073549.1;HCoV-229E_1ab;2535 | 0.05 | 0 |
| NC_002645.1;NP_073549.1;HCoV-229E_1ab;2536 | 0.05 | 0 |
| NC_002645.1;NP_073549.1;HCoV-229E_1ab;2537 | 0.05 | 0 |
| NC_002645.1;NP_073549.1;HCoV-229E_1ab;2538 | 0 | 0 |
| NC_002645.1;NP_073549.1;HCoV-229E_1ab;2539 | 0.05 | 0 |
| NC_002645.1;NP_073549.1;HCoV-229E_1ab;2540 | 0.1 | 0.025 |
| NC_002645.1;NP_073549.1;HCoV-229E_1ab;2541 | 0 | 0 |
| NC_002645.1;NP_073549.1;HCoV-229E_1ab;2542 | 0 | 0 |
| NC_002645.1;NP_073549.1;HCoV-229E_1ab;2543 | 0 | 0 |
| NC_002645.1;NP_073549.1;HCoV-229E_1ab;2544 | 0 | 0 |
| NC_002645.1;NP_073549.1;HCoV-229E_1ab;2545 | 0.1 | 0 |
| NC_002645.1;NP_073549.1;HCoV-229E_1ab;2546 | 0.05 | 0 |
| NC_002645.1;NP_073549.1;HCoV-229E_1ab;2547 | 0.05 | 0.05 |
| NC_002645.1;NP_073549.1;HCoV-229E_1ab;2548 | 0 | 0 |
| NC_002645.1;NP_073549.1;HCoV-229E_1ab;2549 | 0.1 | 0.025 |
| NC_002645.1;NP_073549.1;HCoV-229E_1ab;2550 | 0 | 0 |
| NC_002645.1;NP_073549.1;HCoV-229E_1ab;2551 | 0 | 0 |
| NC_002645.1;NP_073549.1;HCoV-229E_1ab;2552 | 0 | 0 |
| NC_002645.1;NP_073549.1;HCoV-229E_1ab;2553 | 0 | 0 |
| NC_002645.1;NP_073549.1;HCoV-229E_1ab;2554 | 0 | 0 |
| NC_002645.1;NP_073549.1;HCoV-229E_1ab;2555 | 0 | 0 |
| NC_002645.1;NP_073549.1;HCoV-229E_1ab;2556 | 0 | 0 |
| NC_002645.1;NP_073549.1;HCoV-229E_1ab;2557 | 0 | 0 |
| NC_002645.1;NP_073549.1;HCoV-229E_1ab;2558 | 0 | 0 |
| NC_002645.1;NP_073549.1;HCoV-229E_1ab;2559 | 0 | 0 |
| NC_002645.1;NP_073549.1;HCoV-229E_1ab;2560 | 0 | 0 |
| NC_002645.1;NP_073549.1;HCoV-229E_1ab;2561 | 0 | 0 |
| NC_002645.1;NP_073549.1;HCoV-229E_1ab;2562 | 0 | 0 |
| NC_002645.1;NP_073549.1;HCoV-229E_1ab;2563 | 0 | 0 |
| NC_002645.1;NP_073549.1;HCoV-229E_1ab;2564 | 0 | 0 |
| NC_002645.1;NP_073549.1;HCoV-229E_1ab;2565 | 0 | 0 |
| NC_002645.1;NP_073549.1;HCoV-229E_1ab;2566 | 0 | 0 |

| | | |
|---|---|---|
| NC_002645.1;NP_073549.1;HCoV-229E_1ab;2567 | 0 | 0 |
| NC_002645.1;NP_073549.1;HCoV-229E_1ab;2568 | 0 | 0 |
| NC_002645.1;NP_073549.1;HCoV-229E_1ab;2569 | 0 | 0 |
| NC_002645.1;NP_073549.1;HCoV-229E_1ab;2570 | 0 | 0 |
| NC_002645.1;NP_073549.1;HCoV-229E_1ab;2571 | 0 | 0 |
| NC_002645.1;NP_073549.1;HCoV-229E_1ab;2572 | 0 | 0 |
| NC_002645.1;NP_073549.1;HCoV-229E_1ab;2573 | 0 | 0 |
| NC_002645.1;NP_073549.1;HCoV-229E_1ab;2574 | 0 | 0 |
| NC_002645.1;NP_073549.1;HCoV-229E_1ab;2575 | 0 | 0 |
| NC_002645.1;NP_073549.1;HCoV-229E_1ab;2576 | 0.1 | 0.025 |
| NC_002645.1;NP_073549.1;HCoV-229E_1ab;2577 | 0.15 | 0 |
| NC_002645.1;NP_073549.1;HCoV-229E_1ab;2578 | 0 | 0.025 |
| NC_002645.1;NP_073549.1;HCoV-229E_1ab;2579 | 0 | 0 |
| NC_002645.1;NP_073549.1;HCoV-229E_1ab;2580 | 0 | 0 |
| NC_002645.1;NP_073549.1;HCoV-229E_1ab;2581 | 0.05 | 0.075 |
| NC_002645.1;NP_073549.1;HCoV-229E_1ab;2582 | 0 | 0 |
| NC_002645.1;NP_073549.1;HCoV-229E_1ab;2583 | 0 | 0 |
| NC_002645.1;NP_073549.1;HCoV-229E_1ab;2584 | 0 | 0 |
| NC_002645.1;NP_073549.1;HCoV-229E_1ab;2585 | 0 | 0 |
| NC_002645.1;NP_073549.1;HCoV-229E_1ab;2586 | 0 | 0 |
| NC_002645.1;NP_073549.1;HCoV-229E_1ab;2587 | 0 | 0 |
| NC_002645.1;NP_073549.1;HCoV-229E_1ab;2588 | 0 | 0 |
| NC_002645.1;NP_073549.1;HCoV-229E_1ab;2589 | 0 | 0 |
| NC_002645.1;NP_073549.1;HCoV-229E_1ab;2590 | 0 | 0 |
| NC_002645.1;NP_073549.1;HCoV-229E_1ab;2591 | 0 | 0 |
| NC_002645.1;NP_073549.1;HCoV-229E_1ab;2592 | 0 | 0 |
| NC_002645.1;NP_073549.1;HCoV-229E_1ab;2593 | 0 | 0 |
| NC_002645.1;NP_073549.1;HCoV-229E_1ab;2594 | 0 | 0 |
| NC_002645.1;NP_073549.1;HCoV-229E_1ab;2595 | 0 | 0 |
| NC_002645.1;NP_073549.1;HCoV-229E_1ab;2596 | 0 | 0 |
| NC_002645.1;NP_073549.1;HCoV-229E_1ab;2597 | 0 | 0 |
| NC_002645.1;NP_073549.1;HCoV-229E_1ab;2598 | 0 | 0 |
| NC_002645.1;NP_073549.1;HCoV-229E_1ab;2599 | 0 | 0 |
| NC_002645.1;NP_073549.1;HCoV-229E_1ab;2600 | 0 | 0 |
| NC_002645.1;NP_073549.1;HCoV-229E_1ab;2601 | 0 | 0 |
| NC_002645.1;NP_073549.1;HCoV-229E_1ab;2602 | 0 | 0 |
| NC_002645.1;NP_073549.1;HCoV-229E_1ab;2603 | 0 | 0 |
| NC_002645.1;NP_073549.1;HCoV-229E_1ab;2604 | 0 | 0 |
| NC_002645.1;NP_073549.1;HCoV-229E_1ab;2605 | 0 | 0 |
| NC_002645.1;NP_073549.1;HCoV-229E_1ab;2606 | 0 | 0 |
| NC_002645.1;NP_073549.1;HCoV-229E_1ab;2607 | 0 | 0 |
| NC_002645.1;NP_073549.1;HCoV-229E_1ab;2608 | 0 | 0 |
| NC_002645.1;NP_073549.1;HCoV-229E_1ab;2609 | 0 | 0 |

| | | |
|---|---|---|
| NC_002645.1;NP_073549.1;HCoV-229E_1ab;2610 | 0 | 0 |
| NC_002645.1;NP_073549.1;HCoV-229E_1ab;2611 | 0 | 0 |
| NC_002645.1;NP_073549.1;HCoV-229E_1ab;2612 | 0 | 0 |
| NC_002645.1;NP_073549.1;HCoV-229E_1ab;2613 | 0 | 0 |
| NC_002645.1;NP_073549.1;HCoV-229E_1ab;2614 | 0 | 0 |
| NC_002645.1;NP_073549.1;HCoV-229E_1ab;2615 | 0 | 0 |
| NC_002645.1;NP_073549.1;HCoV-229E_1ab;2616 | 0 | 0 |
| NC_002645.1;NP_073549.1;HCoV-229E_1ab;2617 | 0 | 0 |
| NC_002645.1;NP_073549.1;HCoV-229E_1ab;2618 | 0 | 0 |
| NC_002645.1;NP_073549.1;HCoV-229E_1ab;2619 | 0 | 0 |
| NC_002645.1;NP_073549.1;HCoV-229E_1ab;2620 | 0 | 0 |
| NC_002645.1;NP_073549.1;HCoV-229E_1ab;2621 | 0 | 0 |
| NC_002645.1;NP_073549.1;HCoV-229E_1ab;2622 | 0 | 0 |
| NC_002645.1;NP_073549.1;HCoV-229E_1ab;2623 | 0.05 | 0 |
| NC_002645.1;NP_073549.1;HCoV-229E_1ab;2624 | 0 | 0 |
| NC_002645.1;NP_073549.1;HCoV-229E_1ab;2625 | 0 | 0 |
| NC_002645.1;NP_073549.1;HCoV-229E_1ab;2626 | 0 | 0 |
| NC_002645.1;NP_073549.1;HCoV-229E_1ab;2627 | 0 | 0 |
| NC_002645.1;NP_073549.1;HCoV-229E_1ab;2628 | 0 | 0 |
| NC_002645.1;NP_073549.1;HCoV-229E_1ab;2629 | 0 | 0 |
| NC_002645.1;NP_073549.1;HCoV-229E_1ab;2630 | 0 | 0.025 |
| NC_002645.1;NP_073549.1;HCoV-229E_1ab;2631 | 0 | 0.025 |
| NC_002645.1;NP_073549.1;HCoV-229E_1ab;2632 | 0 | 0.025 |
| NC_002645.1;NP_073549.1;HCoV-229E_1ab;2633 | 0 | 0.025 |
| NC_002645.1;NP_073549.1;HCoV-229E_1ab;2634 | 0 | 0 |
| NC_002645.1;NP_073549.1;HCoV-229E_1ab;2635 | 0 | 0 |
| NC_002645.1;NP_073549.1;HCoV-229E_1ab;2636 | 0 | 0 |
| NC_002645.1;NP_073549.1;HCoV-229E_1ab;2637 | 0 | 0 |
| NC_002645.1;NP_073549.1;HCoV-229E_1ab;2638 | 0 | 0 |
| NC_002645.1;NP_073549.1;HCoV-229E_1ab;2639 | 0 | 0.025 |
| NC_002645.1;NP_073549.1;HCoV-229E_1ab;2640 | 0 | 0 |
| NC_002645.1;NP_073549.1;HCoV-229E_1ab;2641 | 0 | 0 |
| NC_002645.1;NP_073549.1;HCoV-229E_1ab;2642 | 0 | 0 |
| NC_002645.1;NP_073549.1;HCoV-229E_1ab;2643 | 0 | 0.025 |
| NC_002645.1;NP_073549.1;HCoV-229E_1ab;2644 | 0 | 0 |
| NC_002645.1;NP_073549.1;HCoV-229E_1ab;2645 | 0 | 0 |
| NC_002645.1;NP_073549.1;HCoV-229E_1ab;2646 | 0 | 0 |
| NC_002645.1;NP_073549.1;HCoV-229E_1ab;2647 | 0 | 0 |
| NC_002645.1;NP_073549.1;HCoV-229E_1ab;2648 | 0 | 0.025 |
| NC_002645.1;NP_073549.1;HCoV-229E_1ab;2649 | 0 | 0 |
| NC_002645.1;NP_073549.1;HCoV-229E_1ab;2650 | 0 | 0.025 |
| NC_002645.1;NP_073549.1;HCoV-229E_1ab;2651 | 0 | 0 |
| NC_002645.1;NP_073549.1;HCoV-229E_1ab;2652 | 0 | 0 |

| | | |
|---|---|---|
| NC_002645.1;NP_073549.1;HCoV-229E_1ab;2653 | 0 | 0 |
| NC_002645.1;NP_073549.1;HCoV-229E_1ab;2654 | 0 | 0 |
| NC_002645.1;NP_073549.1;HCoV-229E_1ab;2655 | 0.1 | 0 |
| NC_002645.1;NP_073549.1;HCoV-229E_1ab;2656 | 0.2 | 0.125 |
| NC_002645.1;NP_073549.1;HCoV-229E_1ab;2657 | 0.2 | 0.05 |
| NC_002645.1;NP_073549.1;HCoV-229E_1ab;2658 | 0.1 | 0.05 |
| NC_002645.1;NP_073549.1;HCoV-229E_1ab;2659 | 0 | 0.025 |
| NC_002645.1;NP_073549.1;HCoV-229E_1ab;2660 | 0 | 0.025 |
| NC_002645.1;NP_073549.1;HCoV-229E_1ab;2661 | 0.05 | 0.075 |
| NC_002645.1;NP_073549.1;HCoV-229E_1ab;2662 | 0.1 | 0.125 |
| NC_002645.1;NP_073549.1;HCoV-229E_1ab;2663 | 0.05 | 0.025 |
| NC_002645.1;NP_073549.1;HCoV-229E_1ab;2664 | 0.1 | 0.15 |
| NC_002645.1;NP_073549.1;HCoV-229E_1ab;2665 | 0.15 | 0.175 |
| NC_002645.1;NP_073549.1;HCoV-229E_1ab;2666 | 0.05 | 0.075 |
| NC_002645.1;NP_073549.1;HCoV-229E_1ab;2667 | 0.05 | 0 |
| NC_002645.1;NP_073549.1;HCoV-229E_1ab;2668 | 0 | 0 |
| NC_002645.1;NP_073549.1;HCoV-229E_1ab;2669 | 0 | 0 |
| NC_002645.1;NP_073549.1;HCoV-229E_1ab;2670 | 0 | 0 |
| NC_002645.1;NP_073549.1;HCoV-229E_1ab;2671 | 0 | 0 |
| NC_002645.1;NP_073549.1;HCoV-229E_1ab;2672 | 0 | 0 |
| NC_002645.1;NP_073549.1;HCoV-229E_1ab;2673 | 0 | 0 |
| NC_002645.1;NP_073549.1;HCoV-229E_1ab;2674 | 0 | 0 |
| NC_002645.1;NP_073549.1;HCoV-229E_1ab;2675 | 0 | 0 |
| NC_002645.1;NP_073549.1;HCoV-229E_1ab;2676 | 0 | 0 |
| NC_002645.1;NP_073549.1;HCoV-229E_1ab;2677 | 0 | 0 |
| NC_002645.1;NP_073549.1;HCoV-229E_1ab;2678 | 0 | 0 |
| NC_002645.1;NP_073549.1;HCoV-229E_1ab;2679 | 0 | 0 |
| NC_002645.1;NP_073549.1;HCoV-229E_1ab;2680 | 0 | 0 |
| NC_002645.1;NP_073549.1;HCoV-229E_1ab;2681 | 0 | 0 |
| NC_002645.1;NP_073549.1;HCoV-229E_1ab;2682 | 0 | 0 |
| NC_002645.1;NP_073549.1;HCoV-229E_1ab;2683 | 0 | 0 |
| NC_002645.1;NP_073549.1;HCoV-229E_1ab;2684 | 0 | 0 |
| NC_002645.1;NP_073549.1;HCoV-229E_1ab;2685 | 0 | 0 |
| NC_002645.1;NP_073549.1;HCoV-229E_1ab;2686 | 0 | 0 |
| NC_002645.1;NP_073549.1;HCoV-229E_1ab;2687 | 0 | 0 |
| NC_002645.1;NP_073549.1;HCoV-229E_1ab;2688 | 0 | 0 |
| NC_002645.1;NP_073549.1;HCoV-229E_1ab;2689 | 0 | 0 |
| NC_002645.1;NP_073549.1;HCoV-229E_1ab;2690 | 0 | 0 |
| NC_002645.1;NP_073549.1;HCoV-229E_1ab;2691 | 0 | 0 |
| NC_002645.1;NP_073549.1;HCoV-229E_1ab;2692 | 0.05 | 0 |
| NC_002645.1;NP_073549.1;HCoV-229E_1ab;2693 | 0.05 | 0 |
| NC_002645.1;NP_073549.1;HCoV-229E_1ab;2694 | 0.05 | 0.025 |
| NC_002645.1;NP_073549.1;HCoV-229E_1ab;2695 | 0.05 | 0.025 |

| | | |
|---|---|---|
| NC_002645.1;NP_073549.1;HCoV-229E_1ab;2696 | 0.05 | 0 |
| NC_002645.1;NP_073549.1;HCoV-229E_1ab;2697 | 0 | 0 |
| NC_002645.1;NP_073549.1;HCoV-229E_1ab;2698 | 0 | 0 |
| NC_002645.1;NP_073549.1;HCoV-229E_1ab;2699 | 0 | 0 |
| NC_002645.1;NP_073549.1;HCoV-229E_1ab;2700 | 0 | 0 |
| NC_002645.1;NP_073549.1;HCoV-229E_1ab;2701 | 0 | 0 |
| NC_002645.1;NP_073549.1;HCoV-229E_1ab;2702 | 0.05 | 0 |
| NC_002645.1;NP_073549.1;HCoV-229E_1ab;2703 | 0 | 0 |
| NC_002645.1;NP_073549.1;HCoV-229E_1ab;2704 | 0 | 0 |
| NC_002645.1;NP_073549.1;HCoV-229E_1ab;2705 | 0 | 0 |
| NC_002645.1;NP_073549.1;HCoV-229E_1ab;2706 | 0 | 0 |
| NC_002645.1;NP_073549.1;HCoV-229E_1ab;2707 | 0.05 | 0.05 |
| NC_002645.1;NP_073549.1;HCoV-229E_1ab;2708 | 0 | 0 |
| NC_002645.1;NP_073549.1;HCoV-229E_1ab;2709 | 0.05 | 0 |
| NC_002645.1;NP_073549.1;HCoV-229E_1ab;2710 | 0 | 0 |
| NC_002645.1;NP_073549.1;HCoV-229E_1ab;2711 | 0 | 0 |
| NC_002645.1;NP_073549.1;HCoV-229E_1ab;2712 | 0.05 | 0.025 |
| NC_002645.1;NP_073549.1;HCoV-229E_1ab;2713 | 0 | 0 |
| NC_002645.1;NP_073549.1;HCoV-229E_1ab;2714 | 0 | 0 |
| NC_002645.1;NP_073549.1;HCoV-229E_1ab;2715 | 0.05 | 0.025 |
| NC_002645.1;NP_073549.1;HCoV-229E_1ab;2716 | 0 | 0 |
| NC_002645.1;NP_073549.1;HCoV-229E_1ab;2717 | 0 | 0 |
| NC_002645.1;NP_073549.1;HCoV-229E_1ab;2718 | 0 | 0 |
| NC_002645.1;NP_073549.1;HCoV-229E_1ab;2719 | 0 | 0 |
| NC_002645.1;NP_073549.1;HCoV-229E_1ab;2720 | 0 | 0 |
| NC_002645.1;NP_073549.1;HCoV-229E_1ab;2721 | 0 | 0 |
| NC_002645.1;NP_073549.1;HCoV-229E_1ab;2722 | 0 | 0 |
| NC_002645.1;NP_073549.1;HCoV-229E_1ab;2723 | 0 | 0 |
| NC_002645.1;NP_073549.1;HCoV-229E_1ab;2724 | 0 | 0 |
| NC_002645.1;NP_073549.1;HCoV-229E_1ab;2725 | 0 | 0 |
| NC_002645.1;NP_073549.1;HCoV-229E_1ab;2726 | 0 | 0 |
| NC_002645.1;NP_073549.1;HCoV-229E_1ab;2727 | 0 | 0 |
| NC_002645.1;NP_073549.1;HCoV-229E_1ab;2728 | 0 | 0 |
| NC_002645.1;NP_073549.1;HCoV-229E_1ab;2729 | 0 | 0 |
| NC_002645.1;NP_073549.1;HCoV-229E_1ab;2730 | 0 | 0 |
| NC_002645.1;NP_073549.1;HCoV-229E_1ab;2731 | 0 | 0 |
| NC_002645.1;NP_073549.1;HCoV-229E_1ab;2732 | 0 | 0 |
| NC_002645.1;NP_073549.1;HCoV-229E_1ab;2733 | 0 | 0 |
| NC_002645.1;NP_073549.1;HCoV-229E_1ab;2734 | 0 | 0 |
| NC_002645.1;NP_073549.1;HCoV-229E_1ab;2735 | 0.05 | 0 |
| NC_002645.1;NP_073549.1;HCoV-229E_1ab;2736 | 0 | 0 |
| NC_002645.1;NP_073549.1;HCoV-229E_1ab;2737 | 0 | 0 |
| NC_002645.1;NP_073549.1;HCoV-229E_1ab;2738 | 0 | 0 |

| | | |
|---|---|---|
| NC_002645.1;NP_073549.1;HCoV-229E_1ab;2739 | 0 | 0 |
| NC_002645.1;NP_073549.1;HCoV-229E_1ab;2740 | 0 | 0 |
| NC_002645.1;NP_073549.1;HCoV-229E_1ab;2741 | 0 | 0 |
| NC_002645.1;NP_073549.1;HCoV-229E_1ab;2742 | 0 | 0 |
| NC_002645.1;NP_073549.1;HCoV-229E_1ab;2743 | 0 | 0 |
| NC_002645.1;NP_073549.1;HCoV-229E_1ab;2744 | 0 | 0 |
| NC_002645.1;NP_073549.1;HCoV-229E_1ab;2745 | 0 | 0 |
| NC_002645.1;NP_073549.1;HCoV-229E_1ab;2746 | 0 | 0 |
| NC_002645.1;NP_073549.1;HCoV-229E_1ab;2747 | 0 | 0 |
| NC_002645.1;NP_073549.1;HCoV-229E_1ab;2748 | 0 | 0 |
| NC_002645.1;NP_073549.1;HCoV-229E_1ab;2749 | 0 | 0 |
| NC_002645.1;NP_073549.1;HCoV-229E_1ab;2750 | 0 | 0 |
| NC_002645.1;NP_073549.1;HCoV-229E_1ab;2751 | 0 | 0 |
| NC_002645.1;NP_073549.1;HCoV-229E_1ab;2752 | 0 | 0 |
| NC_002645.1;NP_073549.1;HCoV-229E_1ab;2753 | 0 | 0 |
| NC_002645.1;NP_073549.1;HCoV-229E_1ab;2754 | 0 | 0 |
| NC_002645.1;NP_073549.1;HCoV-229E_1ab;2755 | 0 | 0 |
| NC_002645.1;NP_073549.1;HCoV-229E_1ab;2756 | 0 | 0 |
| NC_002645.1;NP_073549.1;HCoV-229E_1ab;2757 | 0 | 0 |
| NC_002645.1;NP_073549.1;HCoV-229E_1ab;2758 | 0 | 0 |
| NC_002645.1;NP_073549.1;HCoV-229E_1ab;2759 | 0 | 0 |
| NC_002645.1;NP_073549.1;HCoV-229E_1ab;2760 | 0 | 0 |
| NC_002645.1;NP_073549.1;HCoV-229E_1ab;2761 | 0 | 0 |
| NC_002645.1;NP_073549.1;HCoV-229E_1ab;2762 | 0 | 0 |
| NC_002645.1;NP_073549.1;HCoV-229E_1ab;2763 | 0 | 0 |
| NC_002645.1;NP_073549.1;HCoV-229E_1ab;2764 | 0 | 0 |
| NC_002645.1;NP_073549.1;HCoV-229E_1ab;2765 | 0 | 0 |
| NC_002645.1;NP_073549.1;HCoV-229E_1ab;2766 | 0 | 0 |
| NC_002645.1;NP_073549.1;HCoV-229E_1ab;2767 | 0 | 0 |
| NC_002645.1;NP_073549.1;HCoV-229E_1ab;2768 | 0 | 0 |
| NC_002645.1;NP_073549.1;HCoV-229E_1ab;2769 | 0 | 0 |
| NC_002645.1;NP_073549.1;HCoV-229E_1ab;2770 | 0 | 0 |
| NC_002645.1;NP_073549.1;HCoV-229E_1ab;2771 | 0 | 0 |
| NC_002645.1;NP_073549.1;HCoV-229E_1ab;2772 | 0 | 0 |
| NC_002645.1;NP_073549.1;HCoV-229E_1ab;2773 | 0 | 0 |
| NC_002645.1;NP_073549.1;HCoV-229E_1ab;2774 | 0 | 0 |
| NC_002645.1;NP_073549.1;HCoV-229E_1ab;2775 | 0 | 0 |
| NC_002645.1;NP_073549.1;HCoV-229E_1ab;2776 | 0 | 0 |
| NC_002645.1;NP_073549.1;HCoV-229E_1ab;2777 | 0 | 0 |
| NC_002645.1;NP_073549.1;HCoV-229E_1ab;2778 | 0 | 0 |
| NC_002645.1;NP_073549.1;HCoV-229E_1ab;2779 | 0 | 0 |
| NC_002645.1;NP_073549.1;HCoV-229E_1ab;2780 | 0 | 0 |
| NC_002645.1;NP_073549.1;HCoV-229E_1ab;2781 | 0 | 0 |

| | | |
|---|---|---|
| NC_002645.1;NP_073549.1;HCoV-229E_1ab;2782 | 0 | 0 |
| NC_002645.1;NP_073549.1;HCoV-229E_1ab;2783 | 0 | 0 |
| NC_002645.1;NP_073549.1;HCoV-229E_1ab;2784 | 0 | 0 |
| NC_002645.1;NP_073549.1;HCoV-229E_1ab;2785 | 0 | 0 |
| NC_002645.1;NP_073549.1;HCoV-229E_1ab;2786 | 0 | 0 |
| NC_002645.1;NP_073549.1;HCoV-229E_1ab;2787 | 0 | 0 |
| NC_002645.1;NP_073549.1;HCoV-229E_1ab;2788 | 0 | 0 |
| NC_002645.1;NP_073549.1;HCoV-229E_1ab;2789 | 0 | 0 |
| NC_002645.1;NP_073549.1;HCoV-229E_1ab;2790 | 0 | 0 |
| NC_002645.1;NP_073549.1;HCoV-229E_1ab;2791 | 0 | 0 |
| NC_002645.1;NP_073549.1;HCoV-229E_1ab;2792 | 0 | 0 |
| NC_002645.1;NP_073549.1;HCoV-229E_1ab;2793 | 0 | 0 |
| NC_002645.1;NP_073549.1;HCoV-229E_1ab;2794 | 0 | 0 |
| NC_002645.1;NP_073549.1;HCoV-229E_1ab;2795 | 0 | 0 |
| NC_002645.1;NP_073549.1;HCoV-229E_1ab;2796 | 0 | 0 |
| NC_002645.1;NP_073549.1;HCoV-229E_1ab;2797 | 0 | 0 |
| NC_002645.1;NP_073549.1;HCoV-229E_1ab;2798 | 0 | 0 |
| NC_002645.1;NP_073549.1;HCoV-229E_1ab;2799 | 0 | 0 |
| NC_002645.1;NP_073549.1;HCoV-229E_1ab;2800 | 0 | 0 |
| NC_002645.1;NP_073549.1;HCoV-229E_1ab;2801 | 0 | 0 |
| NC_002645.1;NP_073549.1;HCoV-229E_1ab;2802 | 0 | 0 |
| NC_002645.1;NP_073549.1;HCoV-229E_1ab;2803 | 0 | 0 |
| NC_002645.1;NP_073549.1;HCoV-229E_1ab;2804 | 0 | 0 |
| NC_002645.1;NP_073549.1;HCoV-229E_1ab;2805 | 0 | 0 |
| NC_002645.1;NP_073549.1;HCoV-229E_1ab;2806 | 0 | 0 |
| NC_002645.1;NP_073549.1;HCoV-229E_1ab;2807 | 0 | 0 |
| NC_002645.1;NP_073549.1;HCoV-229E_1ab;2808 | 0 | 0 |
| NC_002645.1;NP_073549.1;HCoV-229E_1ab;2809 | 0 | 0 |
| NC_002645.1;NP_073549.1;HCoV-229E_1ab;2810 | 0 | 0 |
| NC_002645.1;NP_073549.1;HCoV-229E_1ab;2811 | 0 | 0 |
| NC_002645.1;NP_073549.1;HCoV-229E_1ab;2812 | 0 | 0 |
| NC_002645.1;NP_073549.1;HCoV-229E_1ab;2813 | 0 | 0 |
| NC_002645.1;NP_073549.1;HCoV-229E_1ab;2814 | 0 | 0 |
| NC_002645.1;NP_073549.1;HCoV-229E_1ab;2815 | 0 | 0 |
| NC_002645.1;NP_073549.1;HCoV-229E_1ab;2816 | 0 | 0 |
| NC_002645.1;NP_073549.1;HCoV-229E_1ab;2817 | 0 | 0 |
| NC_002645.1;NP_073549.1;HCoV-229E_1ab;2818 | 0 | 0 |
| NC_002645.1;NP_073549.1;HCoV-229E_1ab;2819 | 0 | 0 |
| NC_002645.1;NP_073549.1;HCoV-229E_1ab;2820 | 0 | 0 |
| NC_002645.1;NP_073549.1;HCoV-229E_1ab;2821 | 0 | 0 |
| NC_002645.1;NP_073549.1;HCoV-229E_1ab;2822 | 0 | 0 |
| NC_002645.1;NP_073549.1;HCoV-229E_1ab;2823 | 0 | 0 |
| NC_002645.1;NP_073549.1;HCoV-229E_1ab;2824 | 0 | 0 |

| | | |
|---|---|---|
| NC_002645.1;NP_073549.1;HCoV-229E_1ab;2825 | 0 | 0 |
| NC_002645.1;NP_073549.1;HCoV-229E_1ab;2826 | 0 | 0 |
| NC_002645.1;NP_073549.1;HCoV-229E_1ab;2827 | 0 | 0 |
| NC_002645.1;NP_073549.1;HCoV-229E_1ab;2828 | 0 | 0 |
| NC_002645.1;NP_073549.1;HCoV-229E_1ab;2829 | 0 | 0 |
| NC_002645.1;NP_073549.1;HCoV-229E_1ab;2830 | 0 | 0 |
| NC_002645.1;NP_073549.1;HCoV-229E_1ab;2831 | 0 | 0 |
| NC_002645.1;NP_073549.1;HCoV-229E_1ab;2832 | 0 | 0 |
| NC_002645.1;NP_073549.1;HCoV-229E_1ab;2833 | 0 | 0 |
| NC_002645.1;NP_073549.1;HCoV-229E_1ab;2834 | 0 | 0 |
| NC_002645.1;NP_073549.1;HCoV-229E_1ab;2835 | 0 | 0 |
| NC_002645.1;NP_073549.1;HCoV-229E_1ab;2836 | 0 | 0 |
| NC_002645.1;NP_073549.1;HCoV-229E_1ab;2837 | 0 | 0 |
| NC_002645.1;NP_073549.1;HCoV-229E_1ab;2838 | 0 | 0 |
| NC_002645.1;NP_073549.1;HCoV-229E_1ab;2839 | 0 | 0 |
| NC_002645.1;NP_073549.1;HCoV-229E_1ab;2840 | 0 | 0 |
| NC_002645.1;NP_073549.1;HCoV-229E_1ab;2841 | 0 | 0 |
| NC_002645.1;NP_073549.1;HCoV-229E_1ab;2842 | 0 | 0 |
| NC_002645.1;NP_073549.1;HCoV-229E_1ab;2843 | 0 | 0 |
| NC_002645.1;NP_073549.1;HCoV-229E_1ab;2844 | 0 | 0 |
| NC_002645.1;NP_073549.1;HCoV-229E_1ab;2845 | 0 | 0 |
| NC_002645.1;NP_073549.1;HCoV-229E_1ab;2846 | 0 | 0 |
| NC_002645.1;NP_073549.1;HCoV-229E_1ab;2847 | 0 | 0 |
| NC_002645.1;NP_073549.1;HCoV-229E_1ab;2848 | 0 | 0 |
| NC_002645.1;NP_073549.1;HCoV-229E_1ab;2849 | 0 | 0 |
| NC_002645.1;NP_073549.1;HCoV-229E_1ab;2850 | 0 | 0 |
| NC_002645.1;NP_073549.1;HCoV-229E_1ab;2851 | 0 | 0 |
| NC_002645.1;NP_073549.1;HCoV-229E_1ab;2852 | 0 | 0 |
| NC_002645.1;NP_073549.1;HCoV-229E_1ab;2853 | 0 | 0 |
| NC_002645.1;NP_073549.1;HCoV-229E_1ab;2854 | 0 | 0 |
| NC_002645.1;NP_073549.1;HCoV-229E_1ab;2855 | 0 | 0 |
| NC_002645.1;NP_073549.1;HCoV-229E_1ab;2856 | 0 | 0 |
| NC_002645.1;NP_073549.1;HCoV-229E_1ab;2857 | 0 | 0 |
| NC_002645.1;NP_073549.1;HCoV-229E_1ab;2858 | 0 | 0 |
| NC_002645.1;NP_073549.1;HCoV-229E_1ab;2859 | 0.05 | 0 |
| NC_002645.1;NP_073549.1;HCoV-229E_1ab;2860 | 0 | 0.025 |
| NC_002645.1;NP_073549.1;HCoV-229E_1ab;2861 | 0.05 | 0.025 |
| NC_002645.1;NP_073549.1;HCoV-229E_1ab;2862 | 0.05 | 0.025 |
| NC_002645.1;NP_073549.1;HCoV-229E_1ab;2863 | 0.1 | 0.025 |
| NC_002645.1;NP_073549.1;HCoV-229E_1ab;2864 | 0.1 | 0.075 |
| NC_002645.1;NP_073549.1;HCoV-229E_1ab;2865 | 0.1 | 0.025 |
| NC_002645.1;NP_073549.1;HCoV-229E_1ab;2866 | 0.05 | 0.025 |
| NC_002645.1;NP_073549.1;HCoV-229E_1ab;2867 | 0.15 | 0.025 |

| | | |
|---|---|---|
| NC_002645.1;NP_073549.1;HCoV-229E_1ab;2868 | 0 | 0 |
| NC_002645.1;NP_073549.1;HCoV-229E_1ab;2869 | 0.05 | 0.025 |
| NC_002645.1;NP_073549.1;HCoV-229E_1ab;2870 | 0.05 | 0.025 |
| NC_002645.1;NP_073549.1;HCoV-229E_1ab;2871 | 0 | 0 |
| NC_002645.1;NP_073549.1;HCoV-229E_1ab;2872 | 0 | 0 |
| NC_002645.1;NP_073549.1;HCoV-229E_1ab;2873 | 0 | 0 |
| NC_002645.1;NP_073549.1;HCoV-229E_1ab;2874 | 0 | 0 |
| NC_002645.1;NP_073549.1;HCoV-229E_1ab;2875 | 0 | 0 |
| NC_002645.1;NP_073549.1;HCoV-229E_1ab;2876 | 0 | 0 |
| NC_002645.1;NP_073549.1;HCoV-229E_1ab;2877 | 0 | 0 |
| NC_002645.1;NP_073549.1;HCoV-229E_1ab;2878 | 0 | 0 |
| NC_002645.1;NP_073549.1;HCoV-229E_1ab;2879 | 0 | 0 |
| NC_002645.1;NP_073549.1;HCoV-229E_1ab;2880 | 0 | 0 |
| NC_002645.1;NP_073549.1;HCoV-229E_1ab;2881 | 0 | 0 |
| NC_002645.1;NP_073549.1;HCoV-229E_1ab;2882 | 0 | 0 |
| NC_002645.1;NP_073549.1;HCoV-229E_1ab;2883 | 0 | 0 |
| NC_002645.1;NP_073549.1;HCoV-229E_1ab;2884 | 0 | 0 |
| NC_002645.1;NP_073549.1;HCoV-229E_1ab;2885 | 0 | 0 |
| NC_002645.1;NP_073549.1;HCoV-229E_1ab;2886 | 0 | 0 |
| NC_002645.1;NP_073549.1;HCoV-229E_1ab;2887 | 0 | 0.025 |
| NC_002645.1;NP_073549.1;HCoV-229E_1ab;2888 | 0 | 0 |
| NC_002645.1;NP_073549.1;HCoV-229E_1ab;2889 | 0.05 | 0.025 |
| NC_002645.1;NP_073549.1;HCoV-229E_1ab;2890 | 0.05 | 0 |
| NC_002645.1;NP_073549.1;HCoV-229E_1ab;2891 | 0.05 | 0 |
| NC_002645.1;NP_073549.1;HCoV-229E_1ab;2892 | 0 | 0 |
| NC_002645.1;NP_073549.1;HCoV-229E_1ab;2893 | 0 | 0 |
| NC_002645.1;NP_073549.1;HCoV-229E_1ab;2894 | 0 | 0 |
| NC_002645.1;NP_073549.1;HCoV-229E_1ab;2895 | 0.05 | 0 |
| NC_002645.1;NP_073549.1;HCoV-229E_1ab;2896 | 0.05 | 0 |
| NC_002645.1;NP_073549.1;HCoV-229E_1ab;2897 | 0.05 | 0 |
| NC_002645.1;NP_073549.1;HCoV-229E_1ab;2898 | 0 | 0 |
| NC_002645.1;NP_073549.1;HCoV-229E_1ab;2899 | 0 | 0 |
| NC_002645.1;NP_073549.1;HCoV-229E_1ab;2900 | 0 | 0.025 |
| NC_002645.1;NP_073549.1;HCoV-229E_1ab;2901 | 0 | 0 |
| NC_002645.1;NP_073549.1;HCoV-229E_1ab;2902 | 0 | 0 |
| NC_002645.1;NP_073549.1;HCoV-229E_1ab;2903 | 0.05 | 0 |
| NC_002645.1;NP_073549.1;HCoV-229E_1ab;2904 | 0 | 0 |
| NC_002645.1;NP_073549.1;HCoV-229E_1ab;2905 | 0 | 0 |
| NC_002645.1;NP_073549.1;HCoV-229E_1ab;2906 | 0 | 0 |
| NC_002645.1;NP_073549.1;HCoV-229E_1ab;2907 | 0 | 0 |
| NC_002645.1;NP_073549.1;HCoV-229E_1ab;2908 | 0 | 0.025 |
| NC_002645.1;NP_073549.1;HCoV-229E_1ab;2909 | 0 | 0.025 |
| NC_002645.1;NP_073549.1;HCoV-229E_1ab;2910 | 0 | 0.025 |

| | | |
|---|---|---|
| NC_002645.1;NP_073549.1;HCoV-229E_1ab;2911 | 0 | 0.1 |
| NC_002645.1;NP_073549.1;HCoV-229E_1ab;2912 | 0 | 0.1 |
| NC_002645.1;NP_073549.1;HCoV-229E_1ab;2913 | 0.25 | 0.15 |
| NC_002645.1;NP_073549.1;HCoV-229E_1ab;2914 | 0.05 | 0.125 |
| NC_002645.1;NP_073549.1;HCoV-229E_1ab;2915 | 0 | 0.1 |
| NC_002645.1;NP_073549.1;HCoV-229E_1ab;2916 | 0 | 0.1 |
| NC_002645.1;NP_073549.1;HCoV-229E_1ab;2917 | 0.05 | 0.125 |
| NC_002645.1;NP_073549.1;HCoV-229E_1ab;2918 | 0.05 | 0.15 |
| NC_002645.1;NP_073549.1;HCoV-229E_1ab;2919 | 0 | 0.075 |
| NC_002645.1;NP_073549.1;HCoV-229E_1ab;2920 | 0.05 | 0.075 |
| NC_002645.1;NP_073549.1;HCoV-229E_1ab;2921 | 0 | 0 |
| NC_002645.1;NP_073549.1;HCoV-229E_1ab;2922 | 0 | 0 |
| NC_002645.1;NP_073549.1;HCoV-229E_1ab;2923 | 0 | 0 |
| NC_002645.1;NP_073549.1;HCoV-229E_1ab;2924 | 0 | 0 |
| NC_002645.1;NP_073549.1;HCoV-229E_1ab;2925 | 0.05 | 0 |
| NC_002645.1;NP_073549.1;HCoV-229E_1ab;2926 | 0 | 0 |
| NC_002645.1;NP_073549.1;HCoV-229E_1ab;2927 | 0 | 0 |
| NC_002645.1;NP_073549.1;HCoV-229E_1ab;2928 | 0 | 0 |
| NC_002645.1;NP_073549.1;HCoV-229E_1ab;2929 | 0 | 0 |
| NC_002645.1;NP_073549.1;HCoV-229E_1ab;2930 | 0 | 0 |
| NC_002645.1;NP_073549.1;HCoV-229E_1ab;2931 | 0 | 0 |
| NC_002645.1;NP_073549.1;HCoV-229E_1ab;2932 | 0 | 0.025 |
| NC_002645.1;NP_073549.1;HCoV-229E_1ab;2933 | 0 | 0 |
| NC_002645.1;NP_073549.1;HCoV-229E_1ab;2934 | 0 | 0 |
| NC_002645.1;NP_073549.1;HCoV-229E_1ab;2935 | 0.05 | 0.05 |
| NC_002645.1;NP_073549.1;HCoV-229E_1ab;2936 | 0.05 | 0.075 |
| NC_002645.1;NP_073549.1;HCoV-229E_1ab;2937 | 0 | 0.025 |
| NC_002645.1;NP_073549.1;HCoV-229E_1ab;2938 | 0 | 0.025 |
| NC_002645.1;NP_073549.1;HCoV-229E_1ab;2939 | 0 | 0.025 |
| NC_002645.1;NP_073549.1;HCoV-229E_1ab;2940 | 0.05 | 0.025 |
| NC_002645.1;NP_073549.1;HCoV-229E_1ab;2941 | 0 | 0.075 |
| NC_002645.1;NP_073549.1;HCoV-229E_1ab;2942 | 0 | 0.075 |
| NC_002645.1;NP_073549.1;HCoV-229E_1ab;2943 | 0 | 0.075 |
| NC_002645.1;NP_073549.1;HCoV-229E_1ab;2944 | 0 | 0.1 |
| NC_002645.1;NP_073549.1;HCoV-229E_1ab;2945 | 0 | 0.075 |
| NC_002645.1;NP_073549.1;HCoV-229E_1ab;2946 | 0 | 0.05 |
| NC_002645.1;NP_073549.1;HCoV-229E_1ab;2947 | 0 | 0.025 |
| NC_002645.1;NP_073549.1;HCoV-229E_1ab;2948 | 0 | 0 |
| NC_002645.1;NP_073549.1;HCoV-229E_1ab;2949 | 0 | 0 |
| NC_002645.1;NP_073549.1;HCoV-229E_1ab;2950 | 0 | 0 |
| NC_002645.1;NP_073549.1;HCoV-229E_1ab;2951 | 0 | 0 |
| NC_002645.1;NP_073549.1;HCoV-229E_1ab;2952 | 0 | 0 |
| NC_002645.1;NP_073549.1;HCoV-229E_1ab;2953 | 0 | 0 |

| | | |
|---|---|---|
| NC_002645.1;NP_073549.1;HCoV-229E_1ab;2954 | 0 | 0 |
| NC_002645.1;NP_073549.1;HCoV-229E_1ab;2955 | 0 | 0 |
| NC_002645.1;NP_073549.1;HCoV-229E_1ab;2956 | 0 | 0 |
| NC_002645.1;NP_073549.1;HCoV-229E_1ab;2957 | 0 | 0 |
| NC_002645.1;NP_073549.1;HCoV-229E_1ab;2958 | 0 | 0 |
| NC_002645.1;NP_073549.1;HCoV-229E_1ab;2959 | 0 | 0 |
| NC_002645.1;NP_073549.1;HCoV-229E_1ab;2960 | 0 | 0 |
| NC_002645.1;NP_073549.1;HCoV-229E_1ab;2961 | 0 | 0 |
| NC_002645.1;NP_073549.1;HCoV-229E_1ab;2962 | 0 | 0 |
| NC_002645.1;NP_073549.1;HCoV-229E_1ab;2963 | 0 | 0 |
| NC_002645.1;NP_073549.1;HCoV-229E_1ab;2964 | 0 | 0.025 |
| NC_002645.1;NP_073549.1;HCoV-229E_1ab;2965 | 0 | 0 |
| NC_002645.1;NP_073549.1;HCoV-229E_1ab;2966 | 0 | 0 |
| NC_002645.1;NP_073549.1;HCoV-229E_1ab;2967 | 0 | 0 |
| NC_002645.1;NP_073549.1;HCoV-229E_1ab;2968 | 0 | 0 |
| NC_002645.1;NP_073549.1;HCoV-229E_1ab;2969 | 0 | 0 |
| NC_002645.1;NP_073549.1;HCoV-229E_1ab;2970 | 0 | 0 |
| NC_002645.1;NP_073549.1;HCoV-229E_1ab;2971 | 0 | 0 |
| NC_002645.1;NP_073549.1;HCoV-229E_1ab;2972 | 0 | 0 |
| NC_002645.1;NP_073549.1;HCoV-229E_1ab;2973 | 0 | 0 |
| NC_002645.1;NP_073549.1;HCoV-229E_1ab;2974 | 0 | 0 |
| NC_002645.1;NP_073549.1;HCoV-229E_1ab;2975 | 0 | 0 |
| NC_002645.1;NP_073549.1;HCoV-229E_1ab;2976 | 0 | 0 |
| NC_002645.1;NP_073549.1;HCoV-229E_1ab;2977 | 0 | 0 |
| NC_002645.1;NP_073549.1;HCoV-229E_1ab;2978 | 0 | 0 |
| NC_002645.1;NP_073549.1;HCoV-229E_1ab;2979 | 0 | 0 |
| NC_002645.1;NP_073549.1;HCoV-229E_1ab;2980 | 0 | 0 |
| NC_002645.1;NP_073549.1;HCoV-229E_1ab;2981 | 0 | 0 |
| NC_002645.1;NP_073549.1;HCoV-229E_1ab;2982 | 0 | 0 |
| NC_002645.1;NP_073549.1;HCoV-229E_1ab;2983 | 0 | 0 |
| NC_002645.1;NP_073549.1;HCoV-229E_1ab;2984 | 0 | 0 |
| NC_002645.1;NP_073549.1;HCoV-229E_1ab;2985 | 0 | 0 |
| NC_002645.1;NP_073549.1;HCoV-229E_1ab;2986 | 0 | 0 |
| NC_002645.1;NP_073549.1;HCoV-229E_1ab;2987 | 0 | 0 |
| NC_002645.1;NP_073549.1;HCoV-229E_1ab;2988 | 0 | 0 |
| NC_002645.1;NP_073549.1;HCoV-229E_1ab;2989 | 0 | 0 |
| NC_002645.1;NP_073549.1;HCoV-229E_1ab;2990 | 0 | 0 |
| NC_002645.1;NP_073549.1;HCoV-229E_1ab;2991 | 0 | 0 |
| NC_002645.1;NP_073549.1;HCoV-229E_1ab;2992 | 0 | 0 |
| NC_002645.1;NP_073549.1;HCoV-229E_1ab;2993 | 0 | 0 |
| NC_002645.1;NP_073549.1;HCoV-229E_1ab;2994 | 0 | 0 |
| NC_002645.1;NP_073549.1;HCoV-229E_1ab;2995 | 0 | 0 |
| NC_002645.1;NP_073549.1;HCoV-229E_1ab;2996 | 0 | 0 |

| | | |
|---|---|---|
| NC_002645.1;NP_073549.1;HCoV-229E_1ab;2997 | 0 | 0 |
| NC_002645.1;NP_073549.1;HCoV-229E_1ab;2998 | 0 | 0 |
| NC_002645.1;NP_073549.1;HCoV-229E_1ab;2999 | 0 | 0 |
| NC_002645.1;NP_073549.1;HCoV-229E_1ab;3000 | 0 | 0 |
| NC_002645.1;NP_073549.1;HCoV-229E_1ab;3001 | 0 | 0.025 |
| NC_002645.1;NP_073549.1;HCoV-229E_1ab;3002 | 0 | 0.025 |
| NC_002645.1;NP_073549.1;HCoV-229E_1ab;3003 | 0 | 0 |
| NC_002645.1;NP_073549.1;HCoV-229E_1ab;3004 | 0 | 0 |
| NC_002645.1;NP_073549.1;HCoV-229E_1ab;3005 | 0 | 0 |
| NC_002645.1;NP_073549.1;HCoV-229E_1ab;3006 | 0 | 0.025 |
| NC_002645.1;NP_073549.1;HCoV-229E_1ab;3007 | 0 | 0.075 |
| NC_002645.1;NP_073549.1;HCoV-229E_1ab;3008 | 0 | 0.025 |
| NC_002645.1;NP_073549.1;HCoV-229E_1ab;3009 | 0 | 0 |
| NC_002645.1;NP_073549.1;HCoV-229E_1ab;3010 | 0 | 0 |
| NC_002645.1;NP_073549.1;HCoV-229E_1ab;3011 | 0 | 0 |
| NC_002645.1;NP_073549.1;HCoV-229E_1ab;3012 | 0 | 0 |
| NC_002645.1;NP_073549.1;HCoV-229E_1ab;3013 | 0 | 0 |
| NC_002645.1;NP_073549.1;HCoV-229E_1ab;3014 | 0 | 0.025 |
| NC_002645.1;NP_073549.1;HCoV-229E_1ab;3015 | 0 | 0.025 |
| NC_002645.1;NP_073549.1;HCoV-229E_1ab;3016 | 0 | 0 |
| NC_002645.1;NP_073549.1;HCoV-229E_1ab;3017 | 0 | 0 |
| NC_002645.1;NP_073549.1;HCoV-229E_1ab;3018 | 0 | 0 |
| NC_002645.1;NP_073549.1;HCoV-229E_1ab;3019 | 0 | 0 |
| NC_002645.1;NP_073549.1;HCoV-229E_1ab;3020 | 0 | 0 |
| NC_002645.1;NP_073549.1;HCoV-229E_1ab;3021 | 0 | 0 |
| NC_002645.1;NP_073549.1;HCoV-229E_1ab;3022 | 0 | 0 |
| NC_002645.1;NP_073549.1;HCoV-229E_1ab;3023 | 0 | 0 |
| NC_002645.1;NP_073549.1;HCoV-229E_1ab;3024 | 0 | 0 |
| NC_002645.1;NP_073549.1;HCoV-229E_1ab;3025 | 0 | 0 |
| NC_002645.1;NP_073549.1;HCoV-229E_1ab;3026 | 0 | 0 |
| NC_002645.1;NP_073549.1;HCoV-229E_1ab;3027 | 0 | 0 |
| NC_002645.1;NP_073549.1;HCoV-229E_1ab;3028 | 0 | 0 |
| NC_002645.1;NP_073549.1;HCoV-229E_1ab;3029 | 0 | 0 |
| NC_002645.1;NP_073549.1;HCoV-229E_1ab;3030 | 0 | 0 |
| NC_002645.1;NP_073549.1;HCoV-229E_1ab;3031 | 0 | 0 |
| NC_002645.1;NP_073549.1;HCoV-229E_1ab;3032 | 0 | 0 |
| NC_002645.1;NP_073549.1;HCoV-229E_1ab;3033 | 0 | 0 |
| NC_002645.1;NP_073549.1;HCoV-229E_1ab;3034 | 0 | 0 |
| NC_002645.1;NP_073549.1;HCoV-229E_1ab;3035 | 0 | 0 |
| NC_002645.1;NP_073549.1;HCoV-229E_1ab;3036 | 0 | 0 |
| NC_002645.1;NP_073549.1;HCoV-229E_1ab;3037 | 0 | 0 |
| NC_002645.1;NP_073549.1;HCoV-229E_1ab;3038 | 0 | 0 |
| NC_002645.1;NP_073549.1;HCoV-229E_1ab;3039 | 0 | 0 |

| | | |
|---|---|---|
| NC_002645.1;NP_073549.1;HCoV-229E_1ab;3040 | 0 | 0 |
| NC_002645.1;NP_073549.1;HCoV-229E_1ab;3041 | 0 | 0 |
| NC_002645.1;NP_073549.1;HCoV-229E_1ab;3042 | 0 | 0 |
| NC_002645.1;NP_073549.1;HCoV-229E_1ab;3043 | 0 | 0 |
| NC_002645.1;NP_073549.1;HCoV-229E_1ab;3044 | 0 | 0 |
| NC_002645.1;NP_073549.1;HCoV-229E_1ab;3045 | 0 | 0 |
| NC_002645.1;NP_073549.1;HCoV-229E_1ab;3046 | 0 | 0 |
| NC_002645.1;NP_073549.1;HCoV-229E_1ab;3047 | 0 | 0 |
| NC_002645.1;NP_073549.1;HCoV-229E_1ab;3048 | 0 | 0 |
| NC_002645.1;NP_073549.1;HCoV-229E_1ab;3049 | 0 | 0 |
| NC_002645.1;NP_073549.1;HCoV-229E_1ab;3050 | 0 | 0 |
| NC_002645.1;NP_073549.1;HCoV-229E_1ab;3051 | 0 | 0 |
| NC_002645.1;NP_073549.1;HCoV-229E_1ab;3052 | 0 | 0 |
| NC_002645.1;NP_073549.1;HCoV-229E_1ab;3053 | 0 | 0 |
| NC_002645.1;NP_073549.1;HCoV-229E_1ab;3054 | 0 | 0 |
| NC_002645.1;NP_073549.1;HCoV-229E_1ab;3055 | 0 | 0 |
| NC_002645.1;NP_073549.1;HCoV-229E_1ab;3056 | 0 | 0 |
| NC_002645.1;NP_073549.1;HCoV-229E_1ab;3057 | 0 | 0 |
| NC_002645.1;NP_073549.1;HCoV-229E_1ab;3058 | 0 | 0 |
| NC_002645.1;NP_073549.1;HCoV-229E_1ab;3059 | 0 | 0 |
| NC_002645.1;NP_073549.1;HCoV-229E_1ab;3060 | 0 | 0 |
| NC_002645.1;NP_073549.1;HCoV-229E_1ab;3061 | 0 | 0 |
| NC_002645.1;NP_073549.1;HCoV-229E_1ab;3062 | 0.05 | 0 |
| NC_002645.1;NP_073549.1;HCoV-229E_1ab;3063 | 0 | 0 |
| NC_002645.1;NP_073549.1;HCoV-229E_1ab;3064 | 0 | 0 |
| NC_002645.1;NP_073549.1;HCoV-229E_1ab;3065 | 0 | 0 |
| NC_002645.1;NP_073549.1;HCoV-229E_1ab;3066 | 0 | 0 |
| NC_002645.1;NP_073549.1;HCoV-229E_1ab;3067 | 0 | 0 |
| NC_002645.1;NP_073549.1;HCoV-229E_1ab;3068 | 0 | 0 |
| NC_002645.1;NP_073549.1;HCoV-229E_1ab;3069 | 0 | 0 |
| NC_002645.1;NP_073549.1;HCoV-229E_1ab;3070 | 0 | 0 |
| NC_002645.1;NP_073549.1;HCoV-229E_1ab;3071 | 0 | 0 |
| NC_002645.1;NP_073549.1;HCoV-229E_1ab;3072 | 0 | 0 |
| NC_002645.1;NP_073549.1;HCoV-229E_1ab;3073 | 0 | 0 |
| NC_002645.1;NP_073549.1;HCoV-229E_1ab;3074 | 0 | 0 |
| NC_002645.1;NP_073549.1;HCoV-229E_1ab;3075 | 0 | 0 |
| NC_002645.1;NP_073549.1;HCoV-229E_1ab;3076 | 0 | 0 |
| NC_002645.1;NP_073549.1;HCoV-229E_1ab;3077 | 0 | 0 |
| NC_002645.1;NP_073549.1;HCoV-229E_1ab;3078 | 0 | 0 |
| NC_002645.1;NP_073549.1;HCoV-229E_1ab;3079 | 0 | 0 |
| NC_002645.1;NP_073549.1;HCoV-229E_1ab;3080 | 0 | 0.025 |
| NC_002645.1;NP_073549.1;HCoV-229E_1ab;3081 | 0 | 0.025 |
| NC_002645.1;NP_073549.1;HCoV-229E_1ab;3082 | 0 | 0.025 |

| | | |
|---|---|---|
| NC_002645.1;NP_073549.1;HCoV-229E_1ab;3083 | 0 | 0.025 |
| NC_002645.1;NP_073549.1;HCoV-229E_1ab;3084 | 0 | 0.025 |
| NC_002645.1;NP_073549.1;HCoV-229E_1ab;3085 | 0 | 0 |
| NC_002645.1;NP_073549.1;HCoV-229E_1ab;3086 | 0 | 0 |
| NC_002645.1;NP_073549.1;HCoV-229E_1ab;3087 | 0 | 0 |
| NC_002645.1;NP_073549.1;HCoV-229E_1ab;3088 | 0 | 0 |
| NC_002645.1;NP_073549.1;HCoV-229E_1ab;3089 | 0 | 0 |
| NC_002645.1;NP_073549.1;HCoV-229E_1ab;3090 | 0 | 0 |
| NC_002645.1;NP_073549.1;HCoV-229E_1ab;3091 | 0 | 0 |
| NC_002645.1;NP_073549.1;HCoV-229E_1ab;3092 | 0 | 0 |
| NC_002645.1;NP_073549.1;HCoV-229E_1ab;3093 | 0 | 0 |
| NC_002645.1;NP_073549.1;HCoV-229E_1ab;3094 | 0 | 0 |
| NC_002645.1;NP_073549.1;HCoV-229E_1ab;3095 | 0 | 0 |
| NC_002645.1;NP_073549.1;HCoV-229E_1ab;3096 | 0 | 0 |
| NC_002645.1;NP_073549.1;HCoV-229E_1ab;3097 | 0 | 0 |
| NC_002645.1;NP_073549.1;HCoV-229E_1ab;3098 | 0 | 0 |
| NC_002645.1;NP_073549.1;HCoV-229E_1ab;3099 | 0 | 0 |
| NC_002645.1;NP_073549.1;HCoV-229E_1ab;3100 | 0 | 0 |
| NC_002645.1;NP_073549.1;HCoV-229E_1ab;3101 | 0 | 0 |
| NC_002645.1;NP_073549.1;HCoV-229E_1ab;3102 | 0.05 | 0.025 |
| NC_002645.1;NP_073549.1;HCoV-229E_1ab;3103 | 0.05 | 0.05 |
| NC_002645.1;NP_073549.1;HCoV-229E_1ab;3104 | 0.05 | 0.05 |
| NC_002645.1;NP_073549.1;HCoV-229E_1ab;3105 | 0.05 | 0.1 |
| NC_002645.1;NP_073549.1;HCoV-229E_1ab;3106 | 0.1 | 0.1 |
| NC_002645.1;NP_073549.1;HCoV-229E_1ab;3107 | 0.05 | 0.075 |
| NC_002645.1;NP_073549.1;HCoV-229E_1ab;3108 | 0.05 | 0.05 |
| NC_002645.1;NP_073549.1;HCoV-229E_1ab;3109 | 0.05 | 0.05 |
| NC_002645.1;NP_073549.1;HCoV-229E_1ab;3110 | 0.05 | 0.025 |
| NC_002645.1;NP_073549.1;HCoV-229E_1ab;3111 | 0.05 | 0.025 |
| NC_002645.1;NP_073549.1;HCoV-229E_1ab;3112 | 0 | 0.05 |
| NC_002645.1;NP_073549.1;HCoV-229E_1ab;3113 | 0 | 0.05 |
| NC_002645.1;NP_073549.1;HCoV-229E_1ab;3114 | 0 | 0.025 |
| NC_002645.1;NP_073549.1;HCoV-229E_1ab;3115 | 0 | 0 |
| NC_002645.1;NP_073549.1;HCoV-229E_1ab;3116 | 0 | 0 |
| NC_002645.1;NP_073549.1;HCoV-229E_1ab;3117 | 0 | 0 |
| NC_002645.1;NP_073549.1;HCoV-229E_1ab;3118 | 0 | 0 |
| NC_002645.1;NP_073549.1;HCoV-229E_1ab;3119 | 0 | 0 |
| NC_002645.1;NP_073549.1;HCoV-229E_1ab;3120 | 0 | 0 |
| NC_002645.1;NP_073549.1;HCoV-229E_1ab;3121 | 0 | 0 |
| NC_002645.1;NP_073549.1;HCoV-229E_1ab;3122 | 0 | 0 |
| NC_002645.1;NP_073549.1;HCoV-229E_1ab;3123 | 0 | 0 |
| NC_002645.1;NP_073549.1;HCoV-229E_1ab;3124 | 0 | 0 |
| NC_002645.1;NP_073549.1;HCoV-229E_1ab;3125 | 0.05 | 0 |

| | | |
|---|---|---|
| NC_002645.1;NP_073549.1;HCoV-229E_1ab;3126 | 0 | 0 |
| NC_002645.1;NP_073549.1;HCoV-229E_1ab;3127 | 0 | 0 |
| NC_002645.1;NP_073549.1;HCoV-229E_1ab;3128 | 0 | 0 |
| NC_002645.1;NP_073549.1;HCoV-229E_1ab;3129 | 0 | 0.025 |
| NC_002645.1;NP_073549.1;HCoV-229E_1ab;3130 | 0 | 0.025 |
| NC_002645.1;NP_073549.1;HCoV-229E_1ab;3131 | 0 | 0 |
| NC_002645.1;NP_073549.1;HCoV-229E_1ab;3132 | 0 | 0 |
| NC_002645.1;NP_073549.1;HCoV-229E_1ab;3133 | 0.1 | 0.025 |
| NC_002645.1;NP_073549.1;HCoV-229E_1ab;3134 | 0 | 0 |
| NC_002645.1;NP_073549.1;HCoV-229E_1ab;3135 | 0 | 0 |
| NC_002645.1;NP_073549.1;HCoV-229E_1ab;3136 | 0.05 | 0 |
| NC_002645.1;NP_073549.1;HCoV-229E_1ab;3137 | 0.05 | 0.025 |
| NC_002645.1;NP_073549.1;HCoV-229E_1ab;3138 | 0.05 | 0.025 |
| NC_002645.1;NP_073549.1;HCoV-229E_1ab;3139 | 0.05 | 0.025 |
| NC_002645.1;NP_073549.1;HCoV-229E_1ab;3140 | 0.05 | 0.025 |
| NC_002645.1;NP_073549.1;HCoV-229E_1ab;3141 | 0.05 | 0.025 |
| NC_002645.1;NP_073549.1;HCoV-229E_1ab;3142 | 0.05 | 0.025 |
| NC_002645.1;NP_073549.1;HCoV-229E_1ab;3143 | 0.05 | 0.05 |
| NC_002645.1;NP_073549.1;HCoV-229E_1ab;3144 | 0.05 | 0.025 |
| NC_002645.1;NP_073549.1;HCoV-229E_1ab;3145 | 0.05 | 0.025 |
| NC_002645.1;NP_073549.1;HCoV-229E_1ab;3146 | 0.05 | 0.025 |
| NC_002645.1;NP_073549.1;HCoV-229E_1ab;3147 | 0.05 | 0 |
| NC_002645.1;NP_073549.1;HCoV-229E_1ab;3148 | 0.05 | 0.1 |
| NC_002645.1;NP_073549.1;HCoV-229E_1ab;3149 | 0 | 0 |
| NC_002645.1;NP_073549.1;HCoV-229E_1ab;3150 | 0 | 0 |
| NC_002645.1;NP_073549.1;HCoV-229E_1ab;3151 | 0 | 0 |
| NC_002645.1;NP_073549.1;HCoV-229E_1ab;3152 | 0 | 0 |
| NC_002645.1;NP_073549.1;HCoV-229E_1ab;3153 | 0 | 0 |
| NC_002645.1;NP_073549.1;HCoV-229E_1ab;3154 | 0 | 0.025 |
| NC_002645.1;NP_073549.1;HCoV-229E_1ab;3155 | 0.05 | 0.025 |
| NC_002645.1;NP_073549.1;HCoV-229E_1ab;3156 | 0.05 | 0.025 |
| NC_002645.1;NP_073549.1;HCoV-229E_1ab;3157 | 0 | 0.025 |
| NC_002645.1;NP_073549.1;HCoV-229E_1ab;3158 | 0 | 0 |
| NC_002645.1;NP_073549.1;HCoV-229E_1ab;3159 | 0 | 0 |
| NC_002645.1;NP_073549.1;HCoV-229E_1ab;3160 | 0 | 0 |
| NC_002645.1;NP_073549.1;HCoV-229E_1ab;3161 | 0 | 0 |
| NC_002645.1;NP_073549.1;HCoV-229E_1ab;3162 | 0 | 0 |
| NC_002645.1;NP_073549.1;HCoV-229E_1ab;3163 | 0 | 0 |
| NC_002645.1;NP_073549.1;HCoV-229E_1ab;3164 | 0 | 0 |
| NC_002645.1;NP_073549.1;HCoV-229E_1ab;3165 | 0 | 0 |
| NC_002645.1;NP_073549.1;HCoV-229E_1ab;3166 | 0 | 0 |
| NC_002645.1;NP_073549.1;HCoV-229E_1ab;3167 | 0 | 0 |
| NC_002645.1;NP_073549.1;HCoV-229E_1ab;3168 | 0 | 0 |

| | | |
|---|---|---|
| NC_002645.1;NP_073549.1;HCoV-229E_1ab;3169 | 0 | 0 |
| NC_002645.1;NP_073549.1;HCoV-229E_1ab;3170 | 0 | 0 |
| NC_002645.1;NP_073549.1;HCoV-229E_1ab;3171 | 0 | 0 |
| NC_002645.1;NP_073549.1;HCoV-229E_1ab;3172 | 0 | 0 |
| NC_002645.1;NP_073549.1;HCoV-229E_1ab;3173 | 0.05 | 0 |
| NC_002645.1;NP_073549.1;HCoV-229E_1ab;3174 | 0 | 0 |
| NC_002645.1;NP_073549.1;HCoV-229E_1ab;3175 | 0 | 0 |
| NC_002645.1;NP_073549.1;HCoV-229E_1ab;3176 | 0 | 0 |
| NC_002645.1;NP_073549.1;HCoV-229E_1ab;3177 | 0 | 0 |
| NC_002645.1;NP_073549.1;HCoV-229E_1ab;3178 | 0 | 0 |
| NC_002645.1;NP_073549.1;HCoV-229E_1ab;3179 | 0 | 0 |
| NC_002645.1;NP_073549.1;HCoV-229E_1ab;3180 | 0 | 0 |
| NC_002645.1;NP_073549.1;HCoV-229E_1ab;3181 | 0 | 0 |
| NC_002645.1;NP_073549.1;HCoV-229E_1ab;3182 | 0 | 0 |
| NC_002645.1;NP_073549.1;HCoV-229E_1ab;3183 | 0 | 0 |
| NC_002645.1;NP_073549.1;HCoV-229E_1ab;3184 | 0 | 0 |
| NC_002645.1;NP_073549.1;HCoV-229E_1ab;3185 | 0 | 0 |
| NC_002645.1;NP_073549.1;HCoV-229E_1ab;3186 | 0 | 0.025 |
| NC_002645.1;NP_073549.1;HCoV-229E_1ab;3187 | 0 | 0.025 |
| NC_002645.1;NP_073549.1;HCoV-229E_1ab;3188 | 0 | 0 |
| NC_002645.1;NP_073549.1;HCoV-229E_1ab;3189 | 0 | 0.025 |
| NC_002645.1;NP_073549.1;HCoV-229E_1ab;3190 | 0 | 0.05 |
| NC_002645.1;NP_073549.1;HCoV-229E_1ab;3191 | 0 | 0 |
| NC_002645.1;NP_073549.1;HCoV-229E_1ab;3192 | 0 | 0.025 |
| NC_002645.1;NP_073549.1;HCoV-229E_1ab;3193 | 0.05 | 0.025 |
| NC_002645.1;NP_073549.1;HCoV-229E_1ab;3194 | 0 | 0.025 |
| NC_002645.1;NP_073549.1;HCoV-229E_1ab;3195 | 0 | 0 |
| NC_002645.1;NP_073549.1;HCoV-229E_1ab;3196 | 0 | 0 |
| NC_002645.1;NP_073549.1;HCoV-229E_1ab;3197 | 0 | 0 |
| NC_002645.1;NP_073549.1;HCoV-229E_1ab;3198 | 0 | 0 |
| NC_002645.1;NP_073549.1;HCoV-229E_1ab;3199 | 0 | 0 |
| NC_002645.1;NP_073549.1;HCoV-229E_1ab;3200 | 0 | 0 |
| NC_002645.1;NP_073549.1;HCoV-229E_1ab;3201 | 0 | 0 |
| NC_002645.1;NP_073549.1;HCoV-229E_1ab;3202 | 0.05 | 0.025 |
| NC_002645.1;NP_073549.1;HCoV-229E_1ab;3203 | 0 | 0 |
| NC_002645.1;NP_073549.1;HCoV-229E_1ab;3204 | 0 | 0 |
| NC_002645.1;NP_073549.1;HCoV-229E_1ab;3205 | 0 | 0.025 |
| NC_002645.1;NP_073549.1;HCoV-229E_1ab;3206 | 0 | 0 |
| NC_002645.1;NP_073549.1;HCoV-229E_1ab;3207 | 0 | 0 |
| NC_002645.1;NP_073549.1;HCoV-229E_1ab;3208 | 0 | 0 |
| NC_002645.1;NP_073549.1;HCoV-229E_1ab;3209 | 0 | 0 |
| NC_002645.1;NP_073549.1;HCoV-229E_1ab;3210 | 0 | 0 |
| NC_002645.1;NP_073549.1;HCoV-229E_1ab;3211 | 0 | 0 |

| | | |
|---|---|---|
| NC_002645.1;NP_073549.1;HCoV-229E_1ab;3212 | 0 | 0 |
| NC_002645.1;NP_073549.1;HCoV-229E_1ab;3213 | 0 | 0 |
| NC_002645.1;NP_073549.1;HCoV-229E_1ab;3214 | 0 | 0 |
| NC_002645.1;NP_073549.1;HCoV-229E_1ab;3215 | 0 | 0 |
| NC_002645.1;NP_073549.1;HCoV-229E_1ab;3216 | 0 | 0 |
| NC_002645.1;NP_073549.1;HCoV-229E_1ab;3217 | 0 | 0 |
| NC_002645.1;NP_073549.1;HCoV-229E_1ab;3218 | 0 | 0 |
| NC_002645.1;NP_073549.1;HCoV-229E_1ab;3219 | 0 | 0 |
| NC_002645.1;NP_073549.1;HCoV-229E_1ab;3220 | 0 | 0 |
| NC_002645.1;NP_073549.1;HCoV-229E_1ab;3221 | 0 | 0 |
| NC_002645.1;NP_073549.1;HCoV-229E_1ab;3222 | 0 | 0 |
| NC_002645.1;NP_073549.1;HCoV-229E_1ab;3223 | 0 | 0 |
| NC_002645.1;NP_073549.1;HCoV-229E_1ab;3224 | 0.1 | 0 |
| NC_002645.1;NP_073549.1;HCoV-229E_1ab;3225 | 0 | 0 |
| NC_002645.1;NP_073549.1;HCoV-229E_1ab;3226 | 0 | 0 |
| NC_002645.1;NP_073549.1;HCoV-229E_1ab;3227 | 0 | 0 |
| NC_002645.1;NP_073549.1;HCoV-229E_1ab;3228 | 0 | 0 |
| NC_002645.1;NP_073549.1;HCoV-229E_1ab;3229 | 0.05 | 0 |
| NC_002645.1;NP_073549.1;HCoV-229E_1ab;3230 | 0 | 0 |
| NC_002645.1;NP_073549.1;HCoV-229E_1ab;3231 | 0 | 0 |
| NC_002645.1;NP_073549.1;HCoV-229E_1ab;3232 | 0 | 0 |
| NC_002645.1;NP_073549.1;HCoV-229E_1ab;3233 | 0 | 0 |
| NC_002645.1;NP_073549.1;HCoV-229E_1ab;3234 | 0 | 0 |
| NC_002645.1;NP_073549.1;HCoV-229E_1ab;3235 | 0 | 0 |
| NC_002645.1;NP_073549.1;HCoV-229E_1ab;3236 | 0.05 | 0.025 |
| NC_002645.1;NP_073549.1;HCoV-229E_1ab;3237 | 0.2 | 0.05 |
| NC_002645.1;NP_073549.1;HCoV-229E_1ab;3238 | 0.15 | 0.05 |
| NC_002645.1;NP_073549.1;HCoV-229E_1ab;3239 | 0 | 0 |
| NC_002645.1;NP_073549.1;HCoV-229E_1ab;3240 | 0.1 | 0 |
| NC_002645.1;NP_073549.1;HCoV-229E_1ab;3241 | 0.35 | 0.125 |
| NC_002645.1;NP_073549.1;HCoV-229E_1ab;3242 | 0.4 | 0.125 |
| NC_002645.1;NP_073549.1;HCoV-229E_1ab;3243 | 0.05 | 0 |
| NC_002645.1;NP_073549.1;HCoV-229E_1ab;3244 | 0 | 0 |
| NC_002645.1;NP_073549.1;HCoV-229E_1ab;3245 | 0.05 | 0 |
| NC_002645.1;NP_073549.1;HCoV-229E_1ab;3246 | 0.25 | 0.15 |
| NC_002645.1;NP_073549.1;HCoV-229E_1ab;3247 | 0.55 | 0.275 |
| NC_002645.1;NP_073549.1;HCoV-229E_1ab;3248 | 0.05 | 0 |
| NC_002645.1;NP_073549.1;HCoV-229E_1ab;3249 | 0 | 0 |
| NC_002645.1;NP_073549.1;HCoV-229E_1ab;3250 | 0 | 0.025 |
| NC_002645.1;NP_073549.1;HCoV-229E_1ab;3251 | 0 | 0 |
| NC_002645.1;NP_073549.1;HCoV-229E_1ab;3252 | 0.05 | 0 |
| NC_002645.1;NP_073549.1;HCoV-229E_1ab;3253 | 0 | 0 |
| NC_002645.1;NP_073549.1;HCoV-229E_1ab;3254 | 0.05 | 0.025 |

| | | |
|---|---|---|
| NC_002645.1;NP_073549.1;HCoV-229E_1ab;3255 | 0 | 0.025 |
| NC_002645.1;NP_073549.1;HCoV-229E_1ab;3256 | 0 | 0 |
| NC_002645.1;NP_073549.1;HCoV-229E_1ab;3257 | 0 | 0 |
| NC_002645.1;NP_073549.1;HCoV-229E_1ab;3258 | 0 | 0 |
| NC_002645.1;NP_073549.1;HCoV-229E_1ab;3259 | 0 | 0 |
| NC_002645.1;NP_073549.1;HCoV-229E_1ab;3260 | 0 | 0 |
| NC_002645.1;NP_073549.1;HCoV-229E_1ab;3261 | 0 | 0 |
| NC_002645.1;NP_073549.1;HCoV-229E_1ab;3262 | 0 | 0 |
| NC_002645.1;NP_073549.1;HCoV-229E_1ab;3263 | 0.05 | 0 |
| NC_002645.1;NP_073549.1;HCoV-229E_1ab;3264 | 0 | 0 |
| NC_002645.1;NP_073549.1;HCoV-229E_1ab;3265 | 0 | 0 |
| NC_002645.1;NP_073549.1;HCoV-229E_1ab;3266 | 0 | 0 |
| NC_002645.1;NP_073549.1;HCoV-229E_1ab;3267 | 0 | 0 |
| NC_002645.1;NP_073549.1;HCoV-229E_1ab;3268 | 0 | 0 |
| NC_002645.1;NP_073549.1;HCoV-229E_1ab;3269 | 0 | 0 |
| NC_002645.1;NP_073549.1;HCoV-229E_1ab;3270 | 0 | 0 |
| NC_002645.1;NP_073549.1;HCoV-229E_1ab;3271 | 0 | 0 |
| NC_002645.1;NP_073549.1;HCoV-229E_1ab;3272 | 0 | 0 |
| NC_002645.1;NP_073549.1;HCoV-229E_1ab;3273 | 0 | 0 |
| NC_002645.1;NP_073549.1;HCoV-229E_1ab;3274 | 0 | 0 |
| NC_002645.1;NP_073549.1;HCoV-229E_1ab;3275 | 0 | 0 |
| NC_002645.1;NP_073549.1;HCoV-229E_1ab;3276 | 0 | 0 |
| NC_002645.1;NP_073549.1;HCoV-229E_1ab;3277 | 0 | 0 |
| NC_002645.1;NP_073549.1;HCoV-229E_1ab;3278 | 0 | 0 |
| NC_002645.1;NP_073549.1;HCoV-229E_1ab;3279 | 0 | 0 |
| NC_002645.1;NP_073549.1;HCoV-229E_1ab;3280 | 0 | 0 |
| NC_002645.1;NP_073549.1;HCoV-229E_1ab;3281 | 0 | 0 |
| NC_002645.1;NP_073549.1;HCoV-229E_1ab;3282 | 0 | 0 |
| NC_002645.1;NP_073549.1;HCoV-229E_1ab;3283 | 0 | 0 |
| NC_002645.1;NP_073549.1;HCoV-229E_1ab;3284 | 0 | 0 |
| NC_002645.1;NP_073549.1;HCoV-229E_1ab;3285 | 0 | 0 |
| NC_002645.1;NP_073549.1;HCoV-229E_1ab;3286 | 0 | 0 |
| NC_002645.1;NP_073549.1;HCoV-229E_1ab;3287 | 0 | 0 |
| NC_002645.1;NP_073549.1;HCoV-229E_1ab;3288 | 0 | 0 |
| NC_002645.1;NP_073549.1;HCoV-229E_1ab;3289 | 0 | 0 |
| NC_002645.1;NP_073549.1;HCoV-229E_1ab;3290 | 0 | 0 |
| NC_002645.1;NP_073549.1;HCoV-229E_1ab;3291 | 0 | 0 |
| NC_002645.1;NP_073549.1;HCoV-229E_1ab;3292 | 0 | 0 |
| NC_002645.1;NP_073549.1;HCoV-229E_1ab;3293 | 0 | 0 |
| NC_002645.1;NP_073549.1;HCoV-229E_1ab;3294 | 0 | 0 |
| NC_002645.1;NP_073549.1;HCoV-229E_1ab;3295 | 0 | 0 |
| NC_002645.1;NP_073549.1;HCoV-229E_1ab;3296 | 0 | 0 |
| NC_002645.1;NP_073549.1;HCoV-229E_1ab;3297 | 0 | 0 |

| | | |
|---|---|---|
| NC_002645.1;NP_073549.1;HCoV-229E_1ab;3298 | 0 | 0 |
| NC_002645.1;NP_073549.1;HCoV-229E_1ab;3299 | 0 | 0 |
| NC_002645.1;NP_073549.1;HCoV-229E_1ab;3300 | 0 | 0 |
| NC_002645.1;NP_073549.1;HCoV-229E_1ab;3301 | 0 | 0 |
| NC_002645.1;NP_073549.1;HCoV-229E_1ab;3302 | 0 | 0 |
| NC_002645.1;NP_073549.1;HCoV-229E_1ab;3303 | 0 | 0 |
| NC_002645.1;NP_073549.1;HCoV-229E_1ab;3304 | 0 | 0 |
| NC_002645.1;NP_073549.1;HCoV-229E_1ab;3305 | 0 | 0 |
| NC_002645.1;NP_073549.1;HCoV-229E_1ab;3306 | 0 | 0 |
| NC_002645.1;NP_073549.1;HCoV-229E_1ab;3307 | 0 | 0 |
| NC_002645.1;NP_073549.1;HCoV-229E_1ab;3308 | 0 | 0 |
| NC_002645.1;NP_073549.1;HCoV-229E_1ab;3309 | 0 | 0 |
| NC_002645.1;NP_073549.1;HCoV-229E_1ab;3310 | 0 | 0 |
| NC_002645.1;NP_073549.1;HCoV-229E_1ab;3311 | 0 | 0 |
| NC_002645.1;NP_073549.1;HCoV-229E_1ab;3312 | 0 | 0 |
| NC_002645.1;NP_073549.1;HCoV-229E_1ab;3313 | 0 | 0 |
| NC_002645.1;NP_073549.1;HCoV-229E_1ab;3314 | 0 | 0 |
| NC_002645.1;NP_073549.1;HCoV-229E_1ab;3315 | 0 | 0 |
| NC_002645.1;NP_073549.1;HCoV-229E_1ab;3316 | 0 | 0 |
| NC_002645.1;NP_073549.1;HCoV-229E_1ab;3317 | 0 | 0 |
| NC_002645.1;NP_073549.1;HCoV-229E_1ab;3318 | 0 | 0 |
| NC_002645.1;NP_073549.1;HCoV-229E_1ab;3319 | 0 | 0 |
| NC_002645.1;NP_073549.1;HCoV-229E_1ab;3320 | 0 | 0 |
| NC_002645.1;NP_073549.1;HCoV-229E_1ab;3321 | 0 | 0 |
| NC_002645.1;NP_073549.1;HCoV-229E_1ab;3322 | 0 | 0 |
| NC_002645.1;NP_073549.1;HCoV-229E_1ab;3323 | 0 | 0 |
| NC_002645.1;NP_073549.1;HCoV-229E_1ab;3324 | 0 | 0 |
| NC_002645.1;NP_073549.1;HCoV-229E_1ab;3325 | 0 | 0 |
| NC_002645.1;NP_073549.1;HCoV-229E_1ab;3326 | 0 | 0 |
| NC_002645.1;NP_073549.1;HCoV-229E_1ab;3327 | 0 | 0 |
| NC_002645.1;NP_073549.1;HCoV-229E_1ab;3328 | 0 | 0 |
| NC_002645.1;NP_073549.1;HCoV-229E_1ab;3329 | 0 | 0 |
| NC_002645.1;NP_073549.1;HCoV-229E_1ab;3330 | 0 | 0 |
| NC_002645.1;NP_073549.1;HCoV-229E_1ab;3331 | 0 | 0 |
| NC_002645.1;NP_073549.1;HCoV-229E_1ab;3332 | 0 | 0 |
| NC_002645.1;NP_073549.1;HCoV-229E_1ab;3333 | 0 | 0 |
| NC_002645.1;NP_073549.1;HCoV-229E_1ab;3334 | 0 | 0 |
| NC_002645.1;NP_073549.1;HCoV-229E_1ab;3335 | 0 | 0 |
| NC_002645.1;NP_073549.1;HCoV-229E_1ab;3336 | 0 | 0 |
| NC_002645.1;NP_073549.1;HCoV-229E_1ab;3337 | 0 | 0 |
| NC_002645.1;NP_073549.1;HCoV-229E_1ab;3338 | 0 | 0 |
| NC_002645.1;NP_073549.1;HCoV-229E_1ab;3339 | 0 | 0 |
| NC_002645.1;NP_073549.1;HCoV-229E_1ab;3340 | 0 | 0 |

| | | |
|---|---|---|
| NC_002645.1;NP_073549.1;HCoV-229E_1ab;3341 | 0 | 0 |
| NC_002645.1;NP_073549.1;HCoV-229E_1ab;3342 | 0 | 0 |
| NC_002645.1;NP_073549.1;HCoV-229E_1ab;3343 | 0 | 0 |
| NC_002645.1;NP_073549.1;HCoV-229E_1ab;3344 | 0 | 0 |
| NC_002645.1;NP_073549.1;HCoV-229E_1ab;3345 | 0 | 0 |
| NC_002645.1;NP_073549.1;HCoV-229E_1ab;3346 | 0 | 0 |
| NC_002645.1;NP_073549.1;HCoV-229E_1ab;3347 | 0 | 0.025 |
| NC_002645.1;NP_073549.1;HCoV-229E_1ab;3348 | 0 | 0 |
| NC_002645.1;NP_073549.1;HCoV-229E_1ab;3349 | 0 | 0.025 |
| NC_002645.1;NP_073549.1;HCoV-229E_1ab;3350 | 0.05 | 0.025 |
| NC_002645.1;NP_073549.1;HCoV-229E_1ab;3351 | 0.15 | 0.025 |
| NC_002645.1;NP_073549.1;HCoV-229E_1ab;3352 | 0.15 | 0.025 |
| NC_002645.1;NP_073549.1;HCoV-229E_1ab;3353 | 0.1 | 0.025 |
| NC_002645.1;NP_073549.1;HCoV-229E_1ab;3354 | 0.05 | 0.05 |
| NC_002645.1;NP_073549.1;HCoV-229E_1ab;3355 | 0.1 | 0.05 |
| NC_002645.1;NP_073549.1;HCoV-229E_1ab;3356 | 0.05 | 0 |
| NC_002645.1;NP_073549.1;HCoV-229E_1ab;3357 | 0.1 | 0 |
| NC_002645.1;NP_073549.1;HCoV-229E_1ab;3358 | 0.1 | 0 |
| NC_002645.1;NP_073549.1;HCoV-229E_1ab;3359 | 0.1 | 0 |
| NC_002645.1;NP_073549.1;HCoV-229E_1ab;3360 | 0.05 | 0 |
| NC_002645.1;NP_073549.1;HCoV-229E_1ab;3361 | 0.05 | 0 |
| NC_002645.1;NP_073549.1;HCoV-229E_1ab;3362 | 0.05 | 0 |
| NC_002645.1;NP_073549.1;HCoV-229E_1ab;3363 | 0 | 0 |
| NC_002645.1;NP_073549.1;HCoV-229E_1ab;3364 | 0 | 0 |
| NC_002645.1;NP_073549.1;HCoV-229E_1ab;3365 | 0 | 0 |
| NC_002645.1;NP_073549.1;HCoV-229E_1ab;3366 | 0 | 0 |
| NC_002645.1;NP_073549.1;HCoV-229E_1ab;3367 | 0 | 0 |
| NC_002645.1;NP_073549.1;HCoV-229E_1ab;3368 | 0 | 0 |
| NC_002645.1;NP_073549.1;HCoV-229E_1ab;3369 | 0 | 0 |
| NC_002645.1;NP_073549.1;HCoV-229E_1ab;3370 | 0 | 0 |
| NC_002645.1;NP_073549.1;HCoV-229E_1ab;3371 | 0 | 0 |
| NC_002645.1;NP_073549.1;HCoV-229E_1ab;3372 | 0 | 0 |
| NC_002645.1;NP_073549.1;HCoV-229E_1ab;3373 | 0 | 0 |
| NC_002645.1;NP_073549.1;HCoV-229E_1ab;3374 | 0 | 0 |
| NC_002645.1;NP_073549.1;HCoV-229E_1ab;3375 | 0 | 0 |
| NC_002645.1;NP_073549.1;HCoV-229E_1ab;3376 | 0 | 0 |
| NC_002645.1;NP_073549.1;HCoV-229E_1ab;3377 | 0 | 0 |
| NC_002645.1;NP_073549.1;HCoV-229E_1ab;3378 | 0 | 0 |
| NC_002645.1;NP_073549.1;HCoV-229E_1ab;3379 | 0 | 0 |
| NC_002645.1;NP_073549.1;HCoV-229E_1ab;3380 | 0 | 0 |
| NC_002645.1;NP_073549.1;HCoV-229E_1ab;3381 | 0 | 0 |
| NC_002645.1;NP_073549.1;HCoV-229E_1ab;3382 | 0 | 0 |
| NC_002645.1;NP_073549.1;HCoV-229E_1ab;3383 | 0 | 0 |

| | | |
|---|---|---|
| NC_002645.1;NP_073549.1;HCoV-229E_1ab;3384 | 0 | 0 |
| NC_002645.1;NP_073549.1;HCoV-229E_1ab;3385 | 0 | 0 |
| NC_002645.1;NP_073549.1;HCoV-229E_1ab;3386 | 0 | 0 |
| NC_002645.1;NP_073549.1;HCoV-229E_1ab;3387 | 0 | 0 |
| NC_002645.1;NP_073549.1;HCoV-229E_1ab;3388 | 0 | 0 |
| NC_002645.1;NP_073549.1;HCoV-229E_1ab;3389 | 0 | 0 |
| NC_002645.1;NP_073549.1;HCoV-229E_1ab;3390 | 0 | 0 |
| NC_002645.1;NP_073549.1;HCoV-229E_1ab;3391 | 0 | 0 |
| NC_002645.1;NP_073549.1;HCoV-229E_1ab;3392 | 0 | 0 |
| NC_002645.1;NP_073549.1;HCoV-229E_1ab;3393 | 0 | 0 |
| NC_002645.1;NP_073549.1;HCoV-229E_1ab;3394 | 0 | 0 |
| NC_002645.1;NP_073549.1;HCoV-229E_1ab;3395 | 0 | 0 |
| NC_002645.1;NP_073549.1;HCoV-229E_1ab;3396 | 0 | 0 |
| NC_002645.1;NP_073549.1;HCoV-229E_1ab;3397 | 0 | 0 |
| NC_002645.1;NP_073549.1;HCoV-229E_1ab;3398 | 0 | 0 |
| NC_002645.1;NP_073549.1;HCoV-229E_1ab;3399 | 0 | 0 |
| NC_002645.1;NP_073549.1;HCoV-229E_1ab;3400 | 0 | 0 |
| NC_002645.1;NP_073549.1;HCoV-229E_1ab;3401 | 0 | 0 |
| NC_002645.1;NP_073549.1;HCoV-229E_1ab;3402 | 0 | 0 |
| NC_002645.1;NP_073549.1;HCoV-229E_1ab;3403 | 0 | 0 |
| NC_002645.1;NP_073549.1;HCoV-229E_1ab;3404 | 0 | 0 |
| NC_002645.1;NP_073549.1;HCoV-229E_1ab;3405 | 0 | 0 |
| NC_002645.1;NP_073549.1;HCoV-229E_1ab;3406 | 0 | 0 |
| NC_002645.1;NP_073549.1;HCoV-229E_1ab;3407 | 0 | 0 |
| NC_002645.1;NP_073549.1;HCoV-229E_1ab;3408 | 0 | 0 |
| NC_002645.1;NP_073549.1;HCoV-229E_1ab;3409 | 0 | 0 |
| NC_002645.1;NP_073549.1;HCoV-229E_1ab;3410 | 0 | 0 |
| NC_002645.1;NP_073549.1;HCoV-229E_1ab;3411 | 0 | 0 |
| NC_002645.1;NP_073549.1;HCoV-229E_1ab;3412 | 0 | 0 |
| NC_002645.1;NP_073549.1;HCoV-229E_1ab;3413 | 0 | 0 |
| NC_002645.1;NP_073549.1;HCoV-229E_1ab;3414 | 0 | 0 |
| NC_002645.1;NP_073549.1;HCoV-229E_1ab;3415 | 0 | 0 |
| NC_002645.1;NP_073549.1;HCoV-229E_1ab;3416 | 0 | 0 |
| NC_002645.1;NP_073549.1;HCoV-229E_1ab;3417 | 0 | 0 |
| NC_002645.1;NP_073549.1;HCoV-229E_1ab;3418 | 0 | 0 |
| NC_002645.1;NP_073549.1;HCoV-229E_1ab;3419 | 0 | 0 |
| NC_002645.1;NP_073549.1;HCoV-229E_1ab;3420 | 0 | 0 |
| NC_002645.1;NP_073549.1;HCoV-229E_1ab;3421 | 0 | 0 |
| NC_002645.1;NP_073549.1;HCoV-229E_1ab;3422 | 0 | 0 |
| NC_002645.1;NP_073549.1;HCoV-229E_1ab;3423 | 0 | 0 |
| NC_002645.1;NP_073549.1;HCoV-229E_1ab;3424 | 0 | 0 |
| NC_002645.1;NP_073549.1;HCoV-229E_1ab;3425 | 0 | 0 |
| NC_002645.1;NP_073549.1;HCoV-229E_1ab;3426 | 0 | 0 |

| | | |
|---|---|---|
| NC_002645.1;NP_073549.1;HCoV-229E_1ab;3427 | 0 | 0 |
| NC_002645.1;NP_073549.1;HCoV-229E_1ab;3428 | 0 | 0 |
| NC_002645.1;NP_073549.1;HCoV-229E_1ab;3429 | 0 | 0 |
| NC_002645.1;NP_073549.1;HCoV-229E_1ab;3430 | 0 | 0 |
| NC_002645.1;NP_073549.1;HCoV-229E_1ab;3431 | 0 | 0 |
| NC_002645.1;NP_073549.1;HCoV-229E_1ab;3432 | 0 | 0 |
| NC_002645.1;NP_073549.1;HCoV-229E_1ab;3433 | 0 | 0 |
| NC_002645.1;NP_073549.1;HCoV-229E_1ab;3434 | 0 | 0 |
| NC_002645.1;NP_073549.1;HCoV-229E_1ab;3435 | 0 | 0 |
| NC_002645.1;NP_073549.1;HCoV-229E_1ab;3436 | 0 | 0 |
| NC_002645.1;NP_073549.1;HCoV-229E_1ab;3437 | 0 | 0 |
| NC_002645.1;NP_073549.1;HCoV-229E_1ab;3438 | 0 | 0 |
| NC_002645.1;NP_073549.1;HCoV-229E_1ab;3439 | 0 | 0 |
| NC_002645.1;NP_073549.1;HCoV-229E_1ab;3440 | 0 | 0 |
| NC_002645.1;NP_073549.1;HCoV-229E_1ab;3441 | 0 | 0 |
| NC_002645.1;NP_073549.1;HCoV-229E_1ab;3442 | 0 | 0 |
| NC_002645.1;NP_073549.1;HCoV-229E_1ab;3443 | 0 | 0 |
| NC_002645.1;NP_073549.1;HCoV-229E_1ab;3444 | 0 | 0 |
| NC_002645.1;NP_073549.1;HCoV-229E_1ab;3445 | 0 | 0 |
| NC_002645.1;NP_073549.1;HCoV-229E_1ab;3446 | 0 | 0 |
| NC_002645.1;NP_073549.1;HCoV-229E_1ab;3447 | 0 | 0 |
| NC_002645.1;NP_073549.1;HCoV-229E_1ab;3448 | 0 | 0 |
| NC_002645.1;NP_073549.1;HCoV-229E_1ab;3449 | 0 | 0 |
| NC_002645.1;NP_073549.1;HCoV-229E_1ab;3450 | 0 | 0 |
| NC_002645.1;NP_073549.1;HCoV-229E_1ab;3451 | 0 | 0 |
| NC_002645.1;NP_073549.1;HCoV-229E_1ab;3452 | 0 | 0 |
| NC_002645.1;NP_073549.1;HCoV-229E_1ab;3453 | 0 | 0 |
| NC_002645.1;NP_073549.1;HCoV-229E_1ab;3454 | 0 | 0 |
| NC_002645.1;NP_073549.1;HCoV-229E_1ab;3455 | 0.05 | 0 |
| NC_002645.1;NP_073549.1;HCoV-229E_1ab;3456 | 0 | 0 |
| NC_002645.1;NP_073549.1;HCoV-229E_1ab;3457 | 0 | 0 |
| NC_002645.1;NP_073549.1;HCoV-229E_1ab;3458 | 0 | 0 |
| NC_002645.1;NP_073549.1;HCoV-229E_1ab;3459 | 0 | 0 |
| NC_002645.1;NP_073549.1;HCoV-229E_1ab;3460 | 0 | 0 |
| NC_002645.1;NP_073549.1;HCoV-229E_1ab;3461 | 0 | 0 |
| NC_002645.1;NP_073549.1;HCoV-229E_1ab;3462 | 0 | 0 |
| NC_002645.1;NP_073549.1;HCoV-229E_1ab;3463 | 0 | 0 |
| NC_002645.1;NP_073549.1;HCoV-229E_1ab;3464 | 0 | 0 |
| NC_002645.1;NP_073549.1;HCoV-229E_1ab;3465 | 0 | 0 |
| NC_002645.1;NP_073549.1;HCoV-229E_1ab;3466 | 0 | 0 |
| NC_002645.1;NP_073549.1;HCoV-229E_1ab;3467 | 0 | 0 |
| NC_002645.1;NP_073549.1;HCoV-229E_1ab;3468 | 0 | 0 |
| NC_002645.1;NP_073549.1;HCoV-229E_1ab;3469 | 0 | 0 |

| | | |
|---|---|---|
| NC_002645.1;NP_073549.1;HCoV-229E_1ab;3470 | 0 | 0 |
| NC_002645.1;NP_073549.1;HCoV-229E_1ab;3471 | 0 | 0 |
| NC_002645.1;NP_073549.1;HCoV-229E_1ab;3472 | 0 | 0 |
| NC_002645.1;NP_073549.1;HCoV-229E_1ab;3473 | 0 | 0 |
| NC_002645.1;NP_073549.1;HCoV-229E_1ab;3474 | 0 | 0 |
| NC_002645.1;NP_073549.1;HCoV-229E_1ab;3475 | 0 | 0 |
| NC_002645.1;NP_073549.1;HCoV-229E_1ab;3476 | 0 | 0 |
| NC_002645.1;NP_073549.1;HCoV-229E_1ab;3477 | 0 | 0 |
| NC_002645.1;NP_073549.1;HCoV-229E_1ab;3478 | 0 | 0 |
| NC_002645.1;NP_073549.1;HCoV-229E_1ab;3479 | 0 | 0 |
| NC_002645.1;NP_073549.1;HCoV-229E_1ab;3480 | 0 | 0 |
| NC_002645.1;NP_073549.1;HCoV-229E_1ab;3481 | 0 | 0 |
| NC_002645.1;NP_073549.1;HCoV-229E_1ab;3482 | 0 | 0 |
| NC_002645.1;NP_073549.1;HCoV-229E_1ab;3483 | 0 | 0 |
| NC_002645.1;NP_073549.1;HCoV-229E_1ab;3484 | 0 | 0 |
| NC_002645.1;NP_073549.1;HCoV-229E_1ab;3485 | 0 | 0 |
| NC_002645.1;NP_073549.1;HCoV-229E_1ab;3486 | 0 | 0 |
| NC_002645.1;NP_073549.1;HCoV-229E_1ab;3487 | 0 | 0 |
| NC_002645.1;NP_073549.1;HCoV-229E_1ab;3488 | 0 | 0 |
| NC_002645.1;NP_073549.1;HCoV-229E_1ab;3489 | 0 | 0 |
| NC_002645.1;NP_073549.1;HCoV-229E_1ab;3490 | 0 | 0 |
| NC_002645.1;NP_073549.1;HCoV-229E_1ab;3491 | 0 | 0 |
| NC_002645.1;NP_073549.1;HCoV-229E_1ab;3492 | 0 | 0 |
| NC_002645.1;NP_073549.1;HCoV-229E_1ab;3493 | 0 | 0 |
| NC_002645.1;NP_073549.1;HCoV-229E_1ab;3494 | 0 | 0 |
| NC_002645.1;NP_073549.1;HCoV-229E_1ab;3495 | 0 | 0 |
| NC_002645.1;NP_073549.1;HCoV-229E_1ab;3496 | 0.05 | 0.125 |
| NC_002645.1;NP_073549.1;HCoV-229E_1ab;3497 | 0.05 | 0.05 |
| NC_002645.1;NP_073549.1;HCoV-229E_1ab;3498 | 0 | 0 |
| NC_002645.1;NP_073549.1;HCoV-229E_1ab;3499 | 0 | 0 |
| NC_002645.1;NP_073549.1;HCoV-229E_1ab;3500 | 0 | 0 |
| NC_002645.1;NP_073549.1;HCoV-229E_1ab;3501 | 0 | 0 |
| NC_002645.1;NP_073549.1;HCoV-229E_1ab;3502 | 0.05 | 0 |
| NC_002645.1;NP_073549.1;HCoV-229E_1ab;3503 | 0.25 | 0.175 |
| NC_002645.1;NP_073549.1;HCoV-229E_1ab;3504 | 0.1 | 0.125 |
| NC_002645.1;NP_073549.1;HCoV-229E_1ab;3505 | 0.15 | 0.2 |
| NC_002645.1;NP_073549.1;HCoV-229E_1ab;3506 | 0.15 | 0.225 |
| NC_002645.1;NP_073549.1;HCoV-229E_1ab;3507 | 0.1 | 0.1 |
| NC_002645.1;NP_073549.1;HCoV-229E_1ab;3508 | 0.35 | 0.3 |
| NC_002645.1;NP_073549.1;HCoV-229E_1ab;3509 | 0.1 | 0.075 |
| NC_002645.1;NP_073549.1;HCoV-229E_1ab;3510 | 0.05 | 0.025 |
| NC_002645.1;NP_073549.1;HCoV-229E_1ab;3511 | 0.05 | 0.075 |
| NC_002645.1;NP_073549.1;HCoV-229E_1ab;3512 | 0 | 0.025 |

| | | |
|---|---|---|
| NC_002645.1;NP_073549.1;HCoV-229E_1ab;3513 | 0 | 0 |
| NC_002645.1;NP_073549.1;HCoV-229E_1ab;3514 | 0 | 0 |
| NC_002645.1;NP_073549.1;HCoV-229E_1ab;3515 | 0 | 0 |
| NC_002645.1;NP_073549.1;HCoV-229E_1ab;3516 | 0 | 0 |
| NC_002645.1;NP_073549.1;HCoV-229E_1ab;3517 | 0 | 0 |
| NC_002645.1;NP_073549.1;HCoV-229E_1ab;3518 | 0 | 0 |
| NC_002645.1;NP_073549.1;HCoV-229E_1ab;3519 | 0 | 0 |
| NC_002645.1;NP_073549.1;HCoV-229E_1ab;3520 | 0 | 0 |
| NC_002645.1;NP_073549.1;HCoV-229E_1ab;3521 | 0 | 0 |
| NC_002645.1;NP_073549.1;HCoV-229E_1ab;3522 | 0 | 0 |
| NC_002645.1;NP_073549.1;HCoV-229E_1ab;3523 | 0 | 0 |
| NC_002645.1;NP_073549.1;HCoV-229E_1ab;3524 | 0 | 0 |
| NC_002645.1;NP_073549.1;HCoV-229E_1ab;3525 | 0 | 0 |
| NC_002645.1;NP_073549.1;HCoV-229E_1ab;3526 | 0 | 0 |
| NC_002645.1;NP_073549.1;HCoV-229E_1ab;3527 | 0 | 0 |
| NC_002645.1;NP_073549.1;HCoV-229E_1ab;3528 | 0 | 0 |
| NC_002645.1;NP_073549.1;HCoV-229E_1ab;3529 | 0 | 0 |
| NC_002645.1;NP_073549.1;HCoV-229E_1ab;3530 | 0 | 0 |
| NC_002645.1;NP_073549.1;HCoV-229E_1ab;3531 | 0 | 0 |
| NC_002645.1;NP_073549.1;HCoV-229E_1ab;3532 | 0 | 0 |
| NC_002645.1;NP_073549.1;HCoV-229E_1ab;3533 | 0.1 | 0 |
| NC_002645.1;NP_073549.1;HCoV-229E_1ab;3534 | 0.05 | 0 |
| NC_002645.1;NP_073549.1;HCoV-229E_1ab;3535 | 0.05 | 0.025 |
| NC_002645.1;NP_073549.1;HCoV-229E_1ab;3536 | 0.05 | 0.075 |
| NC_002645.1;NP_073549.1;HCoV-229E_1ab;3537 | 0.05 | 0.05 |
| NC_002645.1;NP_073549.1;HCoV-229E_1ab;3538 | 0.05 | 0 |
| NC_002645.1;NP_073549.1;HCoV-229E_1ab;3539 | 0.05 | 0.025 |
| NC_002645.1;NP_073549.1;HCoV-229E_1ab;3540 | 0.05 | 0 |
| NC_002645.1;NP_073549.1;HCoV-229E_1ab;3541 | 0.05 | 0.025 |
| NC_002645.1;NP_073549.1;HCoV-229E_1ab;3542 | 0 | 0 |
| NC_002645.1;NP_073549.1;HCoV-229E_1ab;3543 | 0 | 0 |
| NC_002645.1;NP_073549.1;HCoV-229E_1ab;3544 | 0 | 0 |
| NC_002645.1;NP_073549.1;HCoV-229E_1ab;3545 | 0 | 0 |
| NC_002645.1;NP_073549.1;HCoV-229E_1ab;3546 | 0 | 0 |
| NC_002645.1;NP_073549.1;HCoV-229E_1ab;3547 | 0 | 0 |
| NC_002645.1;NP_073549.1;HCoV-229E_1ab;3548 | 0.1 | 0.025 |
| NC_002645.1;NP_073549.1;HCoV-229E_1ab;3549 | 0.15 | 0.025 |
| NC_002645.1;NP_073549.1;HCoV-229E_1ab;3550 | 0 | 0 |
| NC_002645.1;NP_073549.1;HCoV-229E_1ab;3551 | 0 | 0 |
| NC_002645.1;NP_073549.1;HCoV-229E_1ab;3552 | 0 | 0 |
| NC_002645.1;NP_073549.1;HCoV-229E_1ab;3553 | 0 | 0 |
| NC_002645.1;NP_073549.1;HCoV-229E_1ab;3554 | 0 | 0 |
| NC_002645.1;NP_073549.1;HCoV-229E_1ab;3555 | 0.05 | 0 |

| | | |
|---|---|---|
| NC_002645.1;NP_073549.1;HCoV-229E_1ab;3556 | 0 | 0 |
| NC_002645.1;NP_073549.1;HCoV-229E_1ab;3557 | 0 | 0 |
| NC_002645.1;NP_073549.1;HCoV-229E_1ab;3558 | 0 | 0.025 |
| NC_002645.1;NP_073549.1;HCoV-229E_1ab;3559 | 0.1 | 0.075 |
| NC_002645.1;NP_073549.1;HCoV-229E_1ab;3560 | 0.1 | 0.175 |
| NC_002645.1;NP_073549.1;HCoV-229E_1ab;3561 | 0.1 | 0.275 |
| NC_002645.1;NP_073549.1;HCoV-229E_1ab;3562 | 0.05 | 0.2 |
| NC_002645.1;NP_073549.1;HCoV-229E_1ab;3563 | 0.05 | 0.125 |
| NC_002645.1;NP_073549.1;HCoV-229E_1ab;3564 | 0.05 | 0.075 |
| NC_002645.1;NP_073549.1;HCoV-229E_1ab;3565 | 0 | 0.025 |
| NC_002645.1;NP_073549.1;HCoV-229E_1ab;3566 | 0 | 0 |
| NC_002645.1;NP_073549.1;HCoV-229E_1ab;3567 | 0.05 | 0 |
| NC_002645.1;NP_073549.1;HCoV-229E_1ab;3568 | 0.1 | 0.025 |
| NC_002645.1;NP_073549.1;HCoV-229E_1ab;3569 | 0.1 | 0 |
| NC_002645.1;NP_073549.1;HCoV-229E_1ab;3570 | 0.1 | 0 |
| NC_002645.1;NP_073549.1;HCoV-229E_1ab;3571 | 0 | 0 |
| NC_002645.1;NP_073549.1;HCoV-229E_1ab;3572 | 0 | 0 |
| NC_002645.1;NP_073549.1;HCoV-229E_1ab;3573 | 0 | 0 |
| NC_002645.1;NP_073549.1;HCoV-229E_1ab;3574 | 0 | 0 |
| NC_002645.1;NP_073549.1;HCoV-229E_1ab;3575 | 0 | 0 |
| NC_002645.1;NP_073549.1;HCoV-229E_1ab;3576 | 0 | 0 |
| NC_002645.1;NP_073549.1;HCoV-229E_1ab;3577 | 0 | 0.025 |
| NC_002645.1;NP_073549.1;HCoV-229E_1ab;3578 | 0 | 0.025 |
| NC_002645.1;NP_073549.1;HCoV-229E_1ab;3579 | 0.05 | 0.025 |
| NC_002645.1;NP_073549.1;HCoV-229E_1ab;3580 | 0 | 0.025 |
| NC_002645.1;NP_073549.1;HCoV-229E_1ab;3581 | 0 | 0 |
| NC_002645.1;NP_073549.1;HCoV-229E_1ab;3582 | 0 | 0 |
| NC_002645.1;NP_073549.1;HCoV-229E_1ab;3583 | 0 | 0.025 |
| NC_002645.1;NP_073549.1;HCoV-229E_1ab;3584 | 0 | 0 |
| NC_002645.1;NP_073549.1;HCoV-229E_1ab;3585 | 0 | 0 |
| NC_002645.1;NP_073549.1;HCoV-229E_1ab;3586 | 0 | 0 |
| NC_002645.1;NP_073549.1;HCoV-229E_1ab;3587 | 0 | 0 |
| NC_002645.1;NP_073549.1;HCoV-229E_1ab;3588 | 0 | 0 |
| NC_002645.1;NP_073549.1;HCoV-229E_1ab;3589 | 0 | 0 |
| NC_002645.1;NP_073549.1;HCoV-229E_1ab;3590 | 0 | 0 |
| NC_002645.1;NP_073549.1;HCoV-229E_1ab;3591 | 0 | 0 |
| NC_002645.1;NP_073549.1;HCoV-229E_1ab;3592 | 0 | 0 |
| NC_002645.1;NP_073549.1;HCoV-229E_1ab;3593 | 0 | 0 |
| NC_002645.1;NP_073549.1;HCoV-229E_1ab;3594 | 0 | 0 |
| NC_002645.1;NP_073549.1;HCoV-229E_1ab;3595 | 0 | 0 |
| NC_002645.1;NP_073549.1;HCoV-229E_1ab;3596 | 0 | 0 |
| NC_002645.1;NP_073549.1;HCoV-229E_1ab;3597 | 0 | 0 |
| NC_002645.1;NP_073549.1;HCoV-229E_1ab;3598 | 0 | 0 |

| | | |
|---|---|---|
| NC_002645.1;NP_073549.1;HCoV-229E_1ab;3599 | 0 | 0 |
| NC_002645.1;NP_073549.1;HCoV-229E_1ab;3600 | 0 | 0 |
| NC_002645.1;NP_073549.1;HCoV-229E_1ab;3601 | 0 | 0 |
| NC_002645.1;NP_073549.1;HCoV-229E_1ab;3602 | 0.05 | 0 |
| NC_002645.1;NP_073549.1;HCoV-229E_1ab;3603 | 0.05 | 0 |
| NC_002645.1;NP_073549.1;HCoV-229E_1ab;3604 | 0.15 | 0.2 |
| NC_002645.1;NP_073549.1;HCoV-229E_1ab;3605 | 0.35 | 0.35 |
| NC_002645.1;NP_073549.1;HCoV-229E_1ab;3606 | 0.3 | 0.3 |
| NC_002645.1;NP_073549.1;HCoV-229E_1ab;3607 | 0.1 | 0.1 |
| NC_002645.1;NP_073549.1;HCoV-229E_1ab;3608 | 0.35 | 0.325 |
| NC_002645.1;NP_073549.1;HCoV-229E_1ab;3609 | 0.35 | 0.3 |
| NC_002645.1;NP_073549.1;HCoV-229E_1ab;3610 | 0.25 | 0.275 |
| NC_002645.1;NP_073549.1;HCoV-229E_1ab;3611 | 0.05 | 0.225 |
| NC_002645.1;NP_073549.1;HCoV-229E_1ab;3612 | 0.1 | 0.275 |
| NC_002645.1;NP_073549.1;HCoV-229E_1ab;3613 | 0.55 | 0.625 |
| NC_002645.1;NP_073549.1;HCoV-229E_1ab;3614 | 0.05 | 0 |
| NC_002645.1;NP_073549.1;HCoV-229E_1ab;3615 | 0.05 | 0.025 |
| NC_002645.1;NP_073549.1;HCoV-229E_1ab;3616 | 0.1 | 0.025 |
| NC_002645.1;NP_073549.1;HCoV-229E_1ab;3617 | 0.15 | 0.025 |
| NC_002645.1;NP_073549.1;HCoV-229E_1ab;3618 | 0 | 0.025 |
| NC_002645.1;NP_073549.1;HCoV-229E_1ab;3619 | 0 | 0 |
| NC_002645.1;NP_073549.1;HCoV-229E_1ab;3620 | 0 | 0 |
| NC_002645.1;NP_073549.1;HCoV-229E_1ab;3621 | 0 | 0 |
| NC_002645.1;NP_073549.1;HCoV-229E_1ab;3622 | 0 | 0 |
| NC_002645.1;NP_073549.1;HCoV-229E_1ab;3623 | 0 | 0 |
| NC_002645.1;NP_073549.1;HCoV-229E_1ab;3624 | 0 | 0 |
| NC_002645.1;NP_073549.1;HCoV-229E_1ab;3625 | 0 | 0 |
| NC_002645.1;NP_073549.1;HCoV-229E_1ab;3626 | 0 | 0 |
| NC_002645.1;NP_073549.1;HCoV-229E_1ab;3627 | 0 | 0 |
| NC_002645.1;NP_073549.1;HCoV-229E_1ab;3628 | 0 | 0 |
| NC_002645.1;NP_073549.1;HCoV-229E_1ab;3629 | 0 | 0 |
| NC_002645.1;NP_073549.1;HCoV-229E_1ab;3630 | 0 | 0.025 |
| NC_002645.1;NP_073549.1;HCoV-229E_1ab;3631 | 0.05 | 0.1 |
| NC_002645.1;NP_073549.1;HCoV-229E_1ab;3632 | 0.05 | 0.025 |
| NC_002645.1;NP_073549.1;HCoV-229E_1ab;3633 | 0.05 | 0.025 |
| NC_002645.1;NP_073549.1;HCoV-229E_1ab;3634 | 0.05 | 0.025 |
| NC_002645.1;NP_073549.1;HCoV-229E_1ab;3635 | 0.05 | 0.075 |
| NC_002645.1;NP_073549.1;HCoV-229E_1ab;3636 | 0.1 | 0.075 |
| NC_002645.1;NP_073549.1;HCoV-229E_1ab;3637 | 0.1 | 0.1 |
| NC_002645.1;NP_073549.1;HCoV-229E_1ab;3638 | 0.1 | 0.125 |
| NC_002645.1;NP_073549.1;HCoV-229E_1ab;3639 | 0.1 | 0.1 |
| NC_002645.1;NP_073549.1;HCoV-229E_1ab;3640 | 0.1 | 0.05 |
| NC_002645.1;NP_073549.1;HCoV-229E_1ab;3641 | 0.1 | 0.075 |

| | | |
|---|---|---|
| NC_002645.1;NP_073549.1;HCoV-229E_1ab;3642 | 0 | 0 |
| NC_002645.1;NP_073549.1;HCoV-229E_1ab;3643 | 0 | 0.075 |
| NC_002645.1;NP_073549.1;HCoV-229E_1ab;3644 | 0 | 0.025 |
| NC_002645.1;NP_073549.1;HCoV-229E_1ab;3645 | 0.05 | 0.025 |
| NC_002645.1;NP_073549.1;HCoV-229E_1ab;3646 | 0 | 0 |
| NC_002645.1;NP_073549.1;HCoV-229E_1ab;3647 | 0 | 0 |
| NC_002645.1;NP_073549.1;HCoV-229E_1ab;3648 | 0 | 0 |
| NC_002645.1;NP_073549.1;HCoV-229E_1ab;3649 | 0 | 0 |
| NC_002645.1;NP_073549.1;HCoV-229E_1ab;3650 | 0 | 0.025 |
| NC_002645.1;NP_073549.1;HCoV-229E_1ab;3651 | 0 | 0.025 |
| NC_002645.1;NP_073549.1;HCoV-229E_1ab;3652 | 0 | 0.05 |
| NC_002645.1;NP_073549.1;HCoV-229E_1ab;3653 | 0 | 0 |
| NC_002645.1;NP_073549.1;HCoV-229E_1ab;3654 | 0 | 0 |
| NC_002645.1;NP_073549.1;HCoV-229E_1ab;3655 | 0 | 0 |
| NC_002645.1;NP_073549.1;HCoV-229E_1ab;3656 | 0 | 0 |
| NC_002645.1;NP_073549.1;HCoV-229E_1ab;3657 | 0 | 0 |
| NC_002645.1;NP_073549.1;HCoV-229E_1ab;3658 | 0 | 0 |
| NC_002645.1;NP_073549.1;HCoV-229E_1ab;3659 | 0 | 0 |
| NC_002645.1;NP_073549.1;HCoV-229E_1ab;3660 | 0 | 0 |
| NC_002645.1;NP_073549.1;HCoV-229E_1ab;3661 | 0 | 0 |
| NC_002645.1;NP_073549.1;HCoV-229E_1ab;3662 | 0 | 0 |
| NC_002645.1;NP_073549.1;HCoV-229E_1ab;3663 | 0 | 0 |
| NC_002645.1;NP_073549.1;HCoV-229E_1ab;3664 | 0 | 0.025 |
| NC_002645.1;NP_073549.1;HCoV-229E_1ab;3665 | 0 | 0.025 |
| NC_002645.1;NP_073549.1;HCoV-229E_1ab;3666 | 0 | 0.025 |
| NC_002645.1;NP_073549.1;HCoV-229E_1ab;3667 | 0 | 0.025 |
| NC_002645.1;NP_073549.1;HCoV-229E_1ab;3668 | 0 | 0.025 |
| NC_002645.1;NP_073549.1;HCoV-229E_1ab;3669 | 0.05 | 0.05 |
| NC_002645.1;NP_073549.1;HCoV-229E_1ab;3670 | 0.05 | 0.025 |
| NC_002645.1;NP_073549.1;HCoV-229E_1ab;3671 | 0 | 0 |
| NC_002645.1;NP_073549.1;HCoV-229E_1ab;3672 | 0 | 0 |
| NC_002645.1;NP_073549.1;HCoV-229E_1ab;3673 | 0 | 0 |
| NC_002645.1;NP_073549.1;HCoV-229E_1ab;3674 | 0 | 0 |
| NC_002645.1;NP_073549.1;HCoV-229E_1ab;3675 | 0 | 0 |
| NC_002645.1;NP_073549.1;HCoV-229E_1ab;3676 | 0 | 0.025 |
| NC_002645.1;NP_073549.1;HCoV-229E_1ab;3677 | 0 | 0 |
| NC_002645.1;NP_073549.1;HCoV-229E_1ab;3678 | 0 | 0 |
| NC_002645.1;NP_073549.1;HCoV-229E_1ab;3679 | 0 | 0 |
| NC_002645.1;NP_073549.1;HCoV-229E_1ab;3680 | 0 | 0 |
| NC_002645.1;NP_073549.1;HCoV-229E_1ab;3681 | 0 | 0.05 |
| NC_002645.1;NP_073549.1;HCoV-229E_1ab;3682 | 0 | 0 |
| NC_002645.1;NP_073549.1;HCoV-229E_1ab;3683 | 0 | 0 |
| NC_002645.1;NP_073549.1;HCoV-229E_1ab;3684 | 0 | 0 |

| | | |
|---|---|---|
| NC_002645.1;NP_073549.1;HCoV-229E_1ab;3685 | 0 | 0 |
| NC_002645.1;NP_073549.1;HCoV-229E_1ab;3686 | 0 | 0 |
| NC_002645.1;NP_073549.1;HCoV-229E_1ab;3687 | 0 | 0 |
| NC_002645.1;NP_073549.1;HCoV-229E_1ab;3688 | 0.05 | 0 |
| NC_002645.1;NP_073549.1;HCoV-229E_1ab;3689 | 0.05 | 0 |
| NC_002645.1;NP_073549.1;HCoV-229E_1ab;3690 | 0.05 | 0 |
| NC_002645.1;NP_073549.1;HCoV-229E_1ab;3691 | 0.1 | 0.025 |
| NC_002645.1;NP_073549.1;HCoV-229E_1ab;3692 | 0.15 | 0.025 |
| NC_002645.1;NP_073549.1;HCoV-229E_1ab;3693 | 0.05 | 0 |
| NC_002645.1;NP_073549.1;HCoV-229E_1ab;3694 | 0 | 0 |
| NC_002645.1;NP_073549.1;HCoV-229E_1ab;3695 | 0 | 0 |
| NC_002645.1;NP_073549.1;HCoV-229E_1ab;3696 | 0 | 0 |
| NC_002645.1;NP_073549.1;HCoV-229E_1ab;3697 | 0 | 0 |
| NC_002645.1;NP_073549.1;HCoV-229E_1ab;3698 | 0 | 0 |
| NC_002645.1;NP_073549.1;HCoV-229E_1ab;3699 | 0 | 0 |
| NC_002645.1;NP_073549.1;HCoV-229E_1ab;3700 | 0 | 0 |
| NC_002645.1;NP_073549.1;HCoV-229E_1ab;3701 | 0 | 0 |
| NC_002645.1;NP_073549.1;HCoV-229E_1ab;3702 | 0 | 0 |
| NC_002645.1;NP_073549.1;HCoV-229E_1ab;3703 | 0 | 0 |
| NC_002645.1;NP_073549.1;HCoV-229E_1ab;3704 | 0 | 0 |
| NC_002645.1;NP_073549.1;HCoV-229E_1ab;3705 | 0 | 0 |
| NC_002645.1;NP_073549.1;HCoV-229E_1ab;3706 | 0 | 0 |
| NC_002645.1;NP_073549.1;HCoV-229E_1ab;3707 | 0 | 0 |
| NC_002645.1;NP_073549.1;HCoV-229E_1ab;3708 | 0 | 0 |
| NC_002645.1;NP_073549.1;HCoV-229E_1ab;3709 | 0 | 0 |
| NC_002645.1;NP_073549.1;HCoV-229E_1ab;3710 | 0 | 0 |
| NC_002645.1;NP_073549.1;HCoV-229E_1ab;3711 | 0 | 0 |
| NC_002645.1;NP_073549.1;HCoV-229E_1ab;3712 | 0 | 0 |
| NC_002645.1;NP_073549.1;HCoV-229E_1ab;3713 | 0 | 0 |
| NC_002645.1;NP_073549.1;HCoV-229E_1ab;3714 | 0 | 0 |
| NC_002645.1;NP_073549.1;HCoV-229E_1ab;3715 | 0 | 0 |
| NC_002645.1;NP_073549.1;HCoV-229E_1ab;3716 | 0 | 0 |
| NC_002645.1;NP_073549.1;HCoV-229E_1ab;3717 | 0 | 0 |
| NC_002645.1;NP_073549.1;HCoV-229E_1ab;3718 | 0 | 0 |
| NC_002645.1;NP_073549.1;HCoV-229E_1ab;3719 | 0 | 0 |
| NC_002645.1;NP_073549.1;HCoV-229E_1ab;3720 | 0 | 0 |
| NC_002645.1;NP_073549.1;HCoV-229E_1ab;3721 | 0 | 0 |
| NC_002645.1;NP_073549.1;HCoV-229E_1ab;3722 | 0 | 0 |
| NC_002645.1;NP_073549.1;HCoV-229E_1ab;3723 | 0 | 0 |
| NC_002645.1;NP_073549.1;HCoV-229E_1ab;3724 | 0 | 0.025 |
| NC_002645.1;NP_073549.1;HCoV-229E_1ab;3725 | 0 | 0 |
| NC_002645.1;NP_073549.1;HCoV-229E_1ab;3726 | 0 | 0.025 |
| NC_002645.1;NP_073549.1;HCoV-229E_1ab;3727 | 0 | 0 |

| | | |
|---|---|---|
| NC_002645.1;NP_073549.1;HCoV-229E_1ab;3728 | 0 | 0.025 |
| NC_002645.1;NP_073549.1;HCoV-229E_1ab;3729 | 0 | 0.025 |
| NC_002645.1;NP_073549.1;HCoV-229E_1ab;3730 | 0 | 0.025 |
| NC_002645.1;NP_073549.1;HCoV-229E_1ab;3731 | 0.05 | 0 |
| NC_002645.1;NP_073549.1;HCoV-229E_1ab;3732 | 0.15 | 0.15 |
| NC_002645.1;NP_073549.1;HCoV-229E_1ab;3733 | 0.05 | 0 |
| NC_002645.1;NP_073549.1;HCoV-229E_1ab;3734 | 0 | 0 |
| NC_002645.1;NP_073549.1;HCoV-229E_1ab;3735 | 0 | 0 |
| NC_002645.1;NP_073549.1;HCoV-229E_1ab;3736 | 0 | 0 |
| NC_002645.1;NP_073549.1;HCoV-229E_1ab;3737 | 0 | 0.025 |
| NC_002645.1;NP_073549.1;HCoV-229E_1ab;3738 | 0 | 0.025 |
| NC_002645.1;NP_073549.1;HCoV-229E_1ab;3739 | 0 | 0.025 |
| NC_002645.1;NP_073549.1;HCoV-229E_1ab;3740 | 0.05 | 0.05 |
| NC_002645.1;NP_073549.1;HCoV-229E_1ab;3741 | 0 | 0.05 |
| NC_002645.1;NP_073549.1;HCoV-229E_1ab;3742 | 0 | 0.025 |
| NC_002645.1;NP_073549.1;HCoV-229E_1ab;3743 | 0 | 0 |
| NC_002645.1;NP_073549.1;HCoV-229E_1ab;3744 | 0 | 0 |
| NC_002645.1;NP_073549.1;HCoV-229E_1ab;3745 | 0 | 0 |
| NC_002645.1;NP_073549.1;HCoV-229E_1ab;3746 | 0 | 0 |
| NC_002645.1;NP_073549.1;HCoV-229E_1ab;3747 | 0 | 0 |
| NC_002645.1;NP_073549.1;HCoV-229E_1ab;3748 | 0 | 0 |
| NC_002645.1;NP_073549.1;HCoV-229E_1ab;3749 | 0 | 0 |
| NC_002645.1;NP_073549.1;HCoV-229E_1ab;3750 | 0 | 0 |
| NC_002645.1;NP_073549.1;HCoV-229E_1ab;3751 | 0 | 0 |
| NC_002645.1;NP_073549.1;HCoV-229E_1ab;3752 | 0 | 0 |
| NC_002645.1;NP_073549.1;HCoV-229E_1ab;3753 | 0 | 0.025 |
| NC_002645.1;NP_073549.1;HCoV-229E_1ab;3754 | 0 | 0 |
| NC_002645.1;NP_073549.1;HCoV-229E_1ab;3755 | 0 | 0 |
| NC_002645.1;NP_073549.1;HCoV-229E_1ab;3756 | 0 | 0 |
| NC_002645.1;NP_073549.1;HCoV-229E_1ab;3757 | 0 | 0.075 |
| NC_002645.1;NP_073549.1;HCoV-229E_1ab;3758 | 0.05 | 0.075 |
| NC_002645.1;NP_073549.1;HCoV-229E_1ab;3759 | 0 | 0.075 |
| NC_002645.1;NP_073549.1;HCoV-229E_1ab;3760 | 0 | 0.075 |
| NC_002645.1;NP_073549.1;HCoV-229E_1ab;3761 | 0 | 0.025 |
| NC_002645.1;NP_073549.1;HCoV-229E_1ab;3762 | 0 | 0 |
| NC_002645.1;NP_073549.1;HCoV-229E_1ab;3763 | 0 | 0 |
| NC_002645.1;NP_073549.1;HCoV-229E_1ab;3764 | 0 | 0 |
| NC_002645.1;NP_073549.1;HCoV-229E_1ab;3765 | 0.1 | 0.025 |
| NC_002645.1;NP_073549.1;HCoV-229E_1ab;3766 | 0 | 0 |
| NC_002645.1;NP_073549.1;HCoV-229E_1ab;3767 | 0 | 0 |
| NC_002645.1;NP_073549.1;HCoV-229E_1ab;3768 | 0 | 0 |
| NC_002645.1;NP_073549.1;HCoV-229E_1ab;3769 | 0 | 0 |
| NC_002645.1;NP_073549.1;HCoV-229E_1ab;3770 | 0 | 0 |

| | | |
|---|---|---|
| NC_002645.1;NP_073549.1;HCoV-229E_1ab;3771 | 0 | 0 |
| NC_002645.1;NP_073549.1;HCoV-229E_1ab;3772 | 0 | 0 |
| NC_002645.1;NP_073549.1;HCoV-229E_1ab;3773 | 0 | 0 |
| NC_002645.1;NP_073549.1;HCoV-229E_1ab;3774 | 0 | 0 |
| NC_002645.1;NP_073549.1;HCoV-229E_1ab;3775 | 0 | 0.025 |
| NC_002645.1;NP_073549.1;HCoV-229E_1ab;3776 | 0 | 0 |
| NC_002645.1;NP_073549.1;HCoV-229E_1ab;3777 | 0 | 0 |
| NC_002645.1;NP_073549.1;HCoV-229E_1ab;3778 | 0 | 0 |
| NC_002645.1;NP_073549.1;HCoV-229E_1ab;3779 | 0 | 0.025 |
| NC_002645.1;NP_073549.1;HCoV-229E_1ab;3780 | 0 | 0 |
| NC_002645.1;NP_073549.1;HCoV-229E_1ab;3781 | 0 | 0.05 |
| NC_002645.1;NP_073549.1;HCoV-229E_1ab;3782 | 0 | 0.075 |
| NC_002645.1;NP_073549.1;HCoV-229E_1ab;3783 | 0 | 0.075 |
| NC_002645.1;NP_073549.1;HCoV-229E_1ab;3784 | 0 | 0 |
| NC_002645.1;NP_073549.1;HCoV-229E_1ab;3785 | 0 | 0.025 |
| NC_002645.1;NP_073549.1;HCoV-229E_1ab;3786 | 0 | 0.025 |
| NC_002645.1;NP_073549.1;HCoV-229E_1ab;3787 | 0 | 0.025 |
| NC_002645.1;NP_073549.1;HCoV-229E_1ab;3788 | 0.05 | 0.025 |
| NC_002645.1;NP_073549.1;HCoV-229E_1ab;3789 | 0.05 | 0.025 |
| NC_002645.1;NP_073549.1;HCoV-229E_1ab;3790 | 0 | 0 |
| NC_002645.1;NP_073549.1;HCoV-229E_1ab;3791 | 0 | 0 |
| NC_002645.1;NP_073549.1;HCoV-229E_1ab;3792 | 0 | 0 |
| NC_002645.1;NP_073549.1;HCoV-229E_1ab;3793 | 0 | 0 |
| NC_002645.1;NP_073549.1;HCoV-229E_1ab;3794 | 0 | 0.025 |
| NC_002645.1;NP_073549.1;HCoV-229E_1ab;3795 | 0 | 0 |
| NC_002645.1;NP_073549.1;HCoV-229E_1ab;3796 | 0 | 0 |
| NC_002645.1;NP_073549.1;HCoV-229E_1ab;3797 | 0 | 0 |
| NC_002645.1;NP_073549.1;HCoV-229E_1ab;3798 | 0.05 | 0 |
| NC_002645.1;NP_073549.1;HCoV-229E_1ab;3799 | 0.05 | 0 |
| NC_002645.1;NP_073549.1;HCoV-229E_1ab;3800 | 0.05 | 0 |
| NC_002645.1;NP_073549.1;HCoV-229E_1ab;3801 | 0 | 0 |
| NC_002645.1;NP_073549.1;HCoV-229E_1ab;3802 | 0 | 0 |
| NC_002645.1;NP_073549.1;HCoV-229E_1ab;3803 | 0 | 0 |
| NC_002645.1;NP_073549.1;HCoV-229E_1ab;3804 | 0 | 0 |
| NC_002645.1;NP_073549.1;HCoV-229E_1ab;3805 | 0 | 0 |
| NC_002645.1;NP_073549.1;HCoV-229E_1ab;3806 | 0 | 0 |
| NC_002645.1;NP_073549.1;HCoV-229E_1ab;3807 | 0 | 0 |
| NC_002645.1;NP_073549.1;HCoV-229E_1ab;3808 | 0 | 0 |
| NC_002645.1;NP_073549.1;HCoV-229E_1ab;3809 | 0 | 0 |
| NC_002645.1;NP_073549.1;HCoV-229E_1ab;3810 | 0 | 0 |
| NC_002645.1;NP_073549.1;HCoV-229E_1ab;3811 | 0 | 0 |
| NC_002645.1;NP_073549.1;HCoV-229E_1ab;3812 | 0 | 0 |
| NC_002645.1;NP_073549.1;HCoV-229E_1ab;3813 | 0 | 0 |

| | | |
|---|---|---|
| NC_002645.1;NP_073549.1;HCoV-229E_1ab;3814 | 0 | 0 |
| NC_002645.1;NP_073549.1;HCoV-229E_1ab;3815 | 0 | 0 |
| NC_002645.1;NP_073549.1;HCoV-229E_1ab;3816 | 0 | 0 |
| NC_002645.1;NP_073549.1;HCoV-229E_1ab;3817 | 0 | 0 |
| NC_002645.1;NP_073549.1;HCoV-229E_1ab;3818 | 0 | 0 |
| NC_002645.1;NP_073549.1;HCoV-229E_1ab;3819 | 0 | 0 |
| NC_002645.1;NP_073549.1;HCoV-229E_1ab;3820 | 0.05 | 0 |
| NC_002645.1;NP_073549.1;HCoV-229E_1ab;3821 | 0 | 0 |
| NC_002645.1;NP_073549.1;HCoV-229E_1ab;3822 | 0 | 0 |
| NC_002645.1;NP_073549.1;HCoV-229E_1ab;3823 | 0 | 0 |
| NC_002645.1;NP_073549.1;HCoV-229E_1ab;3824 | 0 | 0 |
| NC_002645.1;NP_073549.1;HCoV-229E_1ab;3825 | 0 | 0 |
| NC_002645.1;NP_073549.1;HCoV-229E_1ab;3826 | 0 | 0 |
| NC_002645.1;NP_073549.1;HCoV-229E_1ab;3827 | 0 | 0 |
| NC_002645.1;NP_073549.1;HCoV-229E_1ab;3828 | 0 | 0 |
| NC_002645.1;NP_073549.1;HCoV-229E_1ab;3829 | 0 | 0 |
| NC_002645.1;NP_073549.1;HCoV-229E_1ab;3830 | 0 | 0 |
| NC_002645.1;NP_073549.1;HCoV-229E_1ab;3831 | 0 | 0 |
| NC_002645.1;NP_073549.1;HCoV-229E_1ab;3832 | 0 | 0 |
| NC_002645.1;NP_073549.1;HCoV-229E_1ab;3833 | 0 | 0.025 |
| NC_002645.1;NP_073549.1;HCoV-229E_1ab;3834 | 0 | 0.025 |
| NC_002645.1;NP_073549.1;HCoV-229E_1ab;3835 | 0 | 0.025 |
| NC_002645.1;NP_073549.1;HCoV-229E_1ab;3836 | 0 | 0 |
| NC_002645.1;NP_073549.1;HCoV-229E_1ab;3837 | 0 | 0.025 |
| NC_002645.1;NP_073549.1;HCoV-229E_1ab;3838 | 0 | 0.05 |
| NC_002645.1;NP_073549.1;HCoV-229E_1ab;3839 | 0 | 0.025 |
| NC_002645.1;NP_073549.1;HCoV-229E_1ab;3840 | 0 | 0.025 |
| NC_002645.1;NP_073549.1;HCoV-229E_1ab;3841 | 0.05 | 0.025 |
| NC_002645.1;NP_073549.1;HCoV-229E_1ab;3842 | 0.05 | 0.025 |
| NC_002645.1;NP_073549.1;HCoV-229E_1ab;3843 | 0.05 | 0.05 |
| NC_002645.1;NP_073549.1;HCoV-229E_1ab;3844 | 0.15 | 0.1 |
| NC_002645.1;NP_073549.1;HCoV-229E_1ab;3845 | 0.1 | 0.05 |
| NC_002645.1;NP_073549.1;HCoV-229E_1ab;3846 | 0 | 0.025 |
| NC_002645.1;NP_073549.1;HCoV-229E_1ab;3847 | 0 | 0.025 |
| NC_002645.1;NP_073549.1;HCoV-229E_1ab;3848 | 0 | 0.025 |
| NC_002645.1;NP_073549.1;HCoV-229E_1ab;3849 | 0 | 0.025 |
| NC_002645.1;NP_073549.1;HCoV-229E_1ab;3850 | 0 | 0 |
| NC_002645.1;NP_073549.1;HCoV-229E_1ab;3851 | 0.05 | 0 |
| NC_002645.1;NP_073549.1;HCoV-229E_1ab;3852 | 0 | 0 |
| NC_002645.1;NP_073549.1;HCoV-229E_1ab;3853 | 0 | 0 |
| NC_002645.1;NP_073549.1;HCoV-229E_1ab;3854 | 0 | 0 |
| NC_002645.1;NP_073549.1;HCoV-229E_1ab;3855 | 0 | 0 |
| NC_002645.1;NP_073549.1;HCoV-229E_1ab;3856 | 0 | 0 |

| | | |
|---|---|---|
| NC_002645.1;NP_073549.1;HCoV-229E_1ab;3857 | 0.05 | 0.025 |
| NC_002645.1;NP_073549.1;HCoV-229E_1ab;3858 | 0 | 0 |
| NC_002645.1;NP_073549.1;HCoV-229E_1ab;3859 | 0 | 0 |
| NC_002645.1;NP_073549.1;HCoV-229E_1ab;3860 | 0 | 0 |
| NC_002645.1;NP_073549.1;HCoV-229E_1ab;3861 | 0 | 0 |
| NC_002645.1;NP_073549.1;HCoV-229E_1ab;3862 | 0 | 0 |
| NC_002645.1;NP_073549.1;HCoV-229E_1ab;3863 | 0 | 0 |
| NC_002645.1;NP_073549.1;HCoV-229E_1ab;3864 | 0 | 0 |
| NC_002645.1;NP_073549.1;HCoV-229E_1ab;3865 | 0 | 0 |
| NC_002645.1;NP_073549.1;HCoV-229E_1ab;3866 | 0.05 | 0.025 |
| NC_002645.1;NP_073549.1;HCoV-229E_1ab;3867 | 0.05 | 0 |
| NC_002645.1;NP_073549.1;HCoV-229E_1ab;3868 | 0.05 | 0 |
| NC_002645.1;NP_073549.1;HCoV-229E_1ab;3869 | 0.05 | 0 |
| NC_002645.1;NP_073549.1;HCoV-229E_1ab;3870 | 0 | 0 |
| NC_002645.1;NP_073549.1;HCoV-229E_1ab;3871 | 0 | 0 |
| NC_002645.1;NP_073549.1;HCoV-229E_1ab;3872 | 0 | 0 |
| NC_002645.1;NP_073549.1;HCoV-229E_1ab;3873 | 0 | 0 |
| NC_002645.1;NP_073549.1;HCoV-229E_1ab;3874 | 0 | 0 |
| NC_002645.1;NP_073549.1;HCoV-229E_1ab;3875 | 0 | 0 |
| NC_002645.1;NP_073549.1;HCoV-229E_1ab;3876 | 0 | 0 |
| NC_002645.1;NP_073549.1;HCoV-229E_1ab;3877 | 0 | 0 |
| NC_002645.1;NP_073549.1;HCoV-229E_1ab;3878 | 0 | 0 |
| NC_002645.1;NP_073549.1;HCoV-229E_1ab;3879 | 0 | 0 |
| NC_002645.1;NP_073549.1;HCoV-229E_1ab;3880 | 0 | 0 |
| NC_002645.1;NP_073549.1;HCoV-229E_1ab;3881 | 0 | 0 |
| NC_002645.1;NP_073549.1;HCoV-229E_1ab;3882 | 0 | 0 |
| NC_002645.1;NP_073549.1;HCoV-229E_1ab;3883 | 0 | 0 |
| NC_002645.1;NP_073549.1;HCoV-229E_1ab;3884 | 0 | 0 |
| NC_002645.1;NP_073549.1;HCoV-229E_1ab;3885 | 0 | 0.025 |
| NC_002645.1;NP_073549.1;HCoV-229E_1ab;3886 | 0 | 0 |
| NC_002645.1;NP_073549.1;HCoV-229E_1ab;3887 | 0.05 | 0 |
| NC_002645.1;NP_073549.1;HCoV-229E_1ab;3888 | 0 | 0 |
| NC_002645.1;NP_073549.1;HCoV-229E_1ab;3889 | 0 | 0.025 |
| NC_002645.1;NP_073549.1;HCoV-229E_1ab;3890 | 0 | 0 |
| NC_002645.1;NP_073549.1;HCoV-229E_1ab;3891 | 0 | 0 |
| NC_002645.1;NP_073549.1;HCoV-229E_1ab;3892 | 0 | 0 |
| NC_002645.1;NP_073549.1;HCoV-229E_1ab;3893 | 0 | 0 |
| NC_002645.1;NP_073549.1;HCoV-229E_1ab;3894 | 0 | 0 |
| NC_002645.1;NP_073549.1;HCoV-229E_1ab;3895 | 0.05 | 0 |
| NC_002645.1;NP_073549.1;HCoV-229E_1ab;3896 | 0 | 0 |
| NC_002645.1;NP_073549.1;HCoV-229E_1ab;3897 | 0 | 0 |
| NC_002645.1;NP_073549.1;HCoV-229E_1ab;3898 | 0 | 0 |
| NC_002645.1;NP_073549.1;HCoV-229E_1ab;3899 | 0 | 0 |

| | | |
|---|---|---|
| NC_002645.1;NP_073549.1;HCoV-229E_1ab;3900 | 0 | 0 |
| NC_002645.1;NP_073549.1;HCoV-229E_1ab;3901 | 0 | 0 |
| NC_002645.1;NP_073549.1;HCoV-229E_1ab;3902 | 0 | 0 |
| NC_002645.1;NP_073549.1;HCoV-229E_1ab;3903 | 0 | 0 |
| NC_002645.1;NP_073549.1;HCoV-229E_1ab;3904 | 0 | 0 |
| NC_002645.1;NP_073549.1;HCoV-229E_1ab;3905 | 0 | 0 |
| NC_002645.1;NP_073549.1;HCoV-229E_1ab;3906 | 0 | 0 |
| NC_002645.1;NP_073549.1;HCoV-229E_1ab;3907 | 0 | 0 |
| NC_002645.1;NP_073549.1;HCoV-229E_1ab;3908 | 0 | 0 |
| NC_002645.1;NP_073549.1;HCoV-229E_1ab;3909 | 0 | 0 |
| NC_002645.1;NP_073549.1;HCoV-229E_1ab;3910 | 0 | 0 |
| NC_002645.1;NP_073549.1;HCoV-229E_1ab;3911 | 0 | 0 |
| NC_002645.1;NP_073549.1;HCoV-229E_1ab;3912 | 0 | 0 |
| NC_002645.1;NP_073549.1;HCoV-229E_1ab;3913 | 0 | 0 |
| NC_002645.1;NP_073549.1;HCoV-229E_1ab;3914 | 0 | 0 |
| NC_002645.1;NP_073549.1;HCoV-229E_1ab;3915 | 0 | 0 |
| NC_002645.1;NP_073549.1;HCoV-229E_1ab;3916 | 0 | 0 |
| NC_002645.1;NP_073549.1;HCoV-229E_1ab;3917 | 0 | 0 |
| NC_002645.1;NP_073549.1;HCoV-229E_1ab;3918 | 0 | 0 |
| NC_002645.1;NP_073549.1;HCoV-229E_1ab;3919 | 0 | 0 |
| NC_002645.1;NP_073549.1;HCoV-229E_1ab;3920 | 0 | 0 |
| NC_002645.1;NP_073549.1;HCoV-229E_1ab;3921 | 0 | 0 |
| NC_002645.1;NP_073549.1;HCoV-229E_1ab;3922 | 0 | 0 |
| NC_002645.1;NP_073549.1;HCoV-229E_1ab;3923 | 0.05 | 0 |
| NC_002645.1;NP_073549.1;HCoV-229E_1ab;3924 | 0.05 | 0.025 |
| NC_002645.1;NP_073549.1;HCoV-229E_1ab;3925 | 0.05 | 0.025 |
| NC_002645.1;NP_073549.1;HCoV-229E_1ab;3926 | 0.05 | 0.025 |
| NC_002645.1;NP_073549.1;HCoV-229E_1ab;3927 | 0.05 | 0.05 |
| NC_002645.1;NP_073549.1;HCoV-229E_1ab;3928 | 0.05 | 0.025 |
| NC_002645.1;NP_073549.1;HCoV-229E_1ab;3929 | 0 | 0.025 |
| NC_002645.1;NP_073549.1;HCoV-229E_1ab;3930 | 0 | 0.025 |
| NC_002645.1;NP_073549.1;HCoV-229E_1ab;3931 | 0 | 0 |
| NC_002645.1;NP_073549.1;HCoV-229E_1ab;3932 | 0 | 0 |
| NC_002645.1;NP_073549.1;HCoV-229E_1ab;3933 | 0 | 0 |
| NC_002645.1;NP_073549.1;HCoV-229E_1ab;3934 | 0 | 0 |
| NC_002645.1;NP_073549.1;HCoV-229E_1ab;3935 | 0 | 0 |
| NC_002645.1;NP_073549.1;HCoV-229E_1ab;3936 | 0 | 0 |
| NC_002645.1;NP_073549.1;HCoV-229E_1ab;3937 | 0 | 0 |
| NC_002645.1;NP_073549.1;HCoV-229E_1ab;3938 | 0 | 0 |
| NC_002645.1;NP_073549.1;HCoV-229E_1ab;3939 | 0 | 0 |
| NC_002645.1;NP_073549.1;HCoV-229E_1ab;3940 | 0 | 0 |
| NC_002645.1;NP_073549.1;HCoV-229E_1ab;3941 | 0 | 0 |
| NC_002645.1;NP_073549.1;HCoV-229E_1ab;3942 | 0 | 0 |

| | | |
|---|---|---|
| NC_002645.1;NP_073549.1;HCoV-229E_1ab;3943 | 0.05 | 0 |
| NC_002645.1;NP_073549.1;HCoV-229E_1ab;3944 | 0.05 | 0 |
| NC_002645.1;NP_073549.1;HCoV-229E_1ab;3945 | 0 | 0 |
| NC_002645.1;NP_073549.1;HCoV-229E_1ab;3946 | 0 | 0 |
| NC_002645.1;NP_073549.1;HCoV-229E_1ab;3947 | 0.05 | 0 |
| NC_002645.1;NP_073549.1;HCoV-229E_1ab;3948 | 0.05 | 0 |
| NC_002645.1;NP_073549.1;HCoV-229E_1ab;3949 | 0.1 | 0 |
| NC_002645.1;NP_073549.1;HCoV-229E_1ab;3950 | 0.15 | 0 |
| NC_002645.1;NP_073549.1;HCoV-229E_1ab;3951 | 0.2 | 0.025 |
| NC_002645.1;NP_073549.1;HCoV-229E_1ab;3952 | 0.2 | 0.025 |
| NC_002645.1;NP_073549.1;HCoV-229E_1ab;3953 | 0.1 | 0.05 |
| NC_002645.1;NP_073549.1;HCoV-229E_1ab;3954 | 0.05 | 0.025 |
| NC_002645.1;NP_073549.1;HCoV-229E_1ab;3955 | 0.15 | 0.025 |
| NC_002645.1;NP_073549.1;HCoV-229E_1ab;3956 | 0.05 | 0.025 |
| NC_002645.1;NP_073549.1;HCoV-229E_1ab;3957 | 0.05 | 0 |
| NC_002645.1;NP_073549.1;HCoV-229E_1ab;3958 | 0.05 | 0.025 |
| NC_002645.1;NP_073549.1;HCoV-229E_1ab;3959 | 0 | 0.025 |
| NC_002645.1;NP_073549.1;HCoV-229E_1ab;3960 | 0 | 0.075 |
| NC_002645.1;NP_073549.1;HCoV-229E_1ab;3961 | 0 | 0.025 |
| NC_002645.1;NP_073549.1;HCoV-229E_1ab;3962 | 0.05 | 0.025 |
| NC_002645.1;NP_073549.1;HCoV-229E_1ab;3963 | 0.05 | 0.025 |
| NC_002645.1;NP_073549.1;HCoV-229E_1ab;3964 | 0 | 0.025 |
| NC_002645.1;NP_073549.1;HCoV-229E_1ab;3965 | 0 | 0.025 |
| NC_002645.1;NP_073549.1;HCoV-229E_1ab;3966 | 0 | 0.025 |
| NC_002645.1;NP_073549.1;HCoV-229E_1ab;3967 | 0 | 0 |
| NC_002645.1;NP_073549.1;HCoV-229E_1ab;3968 | 0 | 0 |
| NC_002645.1;NP_073549.1;HCoV-229E_1ab;3969 | 0 | 0 |
| NC_002645.1;NP_073549.1;HCoV-229E_1ab;3970 | 0 | 0.025 |
| NC_002645.1;NP_073549.1;HCoV-229E_1ab;3971 | 0.05 | 0.025 |
| NC_002645.1;NP_073549.1;HCoV-229E_1ab;3972 | 0 | 0 |
| NC_002645.1;NP_073549.1;HCoV-229E_1ab;3973 | 0 | 0.025 |
| NC_002645.1;NP_073549.1;HCoV-229E_1ab;3974 | 0 | 0 |
| NC_002645.1;NP_073549.1;HCoV-229E_1ab;3975 | 0 | 0 |
| NC_002645.1;NP_073549.1;HCoV-229E_1ab;3976 | 0 | 0 |
| NC_002645.1;NP_073549.1;HCoV-229E_1ab;3977 | 0 | 0 |
| NC_002645.1;NP_073549.1;HCoV-229E_1ab;3978 | 0 | 0 |
| NC_002645.1;NP_073549.1;HCoV-229E_1ab;3979 | 0 | 0.025 |
| NC_002645.1;NP_073549.1;HCoV-229E_1ab;3980 | 0 | 0.025 |
| NC_002645.1;NP_073549.1;HCoV-229E_1ab;3981 | 0 | 0 |
| NC_002645.1;NP_073549.1;HCoV-229E_1ab;3982 | 0 | 0 |
| NC_002645.1;NP_073549.1;HCoV-229E_1ab;3983 | 0 | 0 |
| NC_002645.1;NP_073549.1;HCoV-229E_1ab;3984 | 0 | 0 |
| NC_002645.1;NP_073549.1;HCoV-229E_1ab;3985 | 0 | 0 |

| | | |
|---|---|---|
| NC_002645.1;NP_073549.1;HCoV-229E_1ab;3986 | 0.05 | 0 |
| NC_002645.1;NP_073549.1;HCoV-229E_1ab;3987 | 0 | 0 |
| NC_002645.1;NP_073549.1;HCoV-229E_1ab;3988 | 0.1 | 0 |
| NC_002645.1;NP_073549.1;HCoV-229E_1ab;3989 | 0 | 0 |
| NC_002645.1;NP_073549.1;HCoV-229E_1ab;3990 | 0.05 | 0.025 |
| NC_002645.1;NP_073549.1;HCoV-229E_1ab;3991 | 0 | 0.075 |
| NC_002645.1;NP_073549.1;HCoV-229E_1ab;3992 | 0 | 0.025 |
| NC_002645.1;NP_073549.1;HCoV-229E_1ab;3993 | 0.05 | 0.025 |
| NC_002645.1;NP_073549.1;HCoV-229E_1ab;3994 | 0 | 0 |
| NC_002645.1;NP_073549.1;HCoV-229E_1ab;3995 | 0 | 0 |
| NC_002645.1;NP_073549.1;HCoV-229E_1ab;3996 | 0 | 0 |
| NC_002645.1;NP_073549.1;HCoV-229E_1ab;3997 | 0 | 0 |
| NC_002645.1;NP_073549.1;HCoV-229E_1ab;3998 | 0 | 0 |
| NC_002645.1;NP_073549.1;HCoV-229E_1ab;3999 | 0 | 0 |
| NC_002645.1;NP_073549.1;HCoV-229E_1ab;4000 | 0 | 0 |
| NC_002645.1;NP_073549.1;HCoV-229E_1ab;4001 | 0 | 0 |
| NC_002645.1;NP_073549.1;HCoV-229E_1ab;4002 | 0 | 0 |
| NC_002645.1;NP_073549.1;HCoV-229E_1ab;4003 | 0 | 0 |
| NC_002645.1;NP_073549.1;HCoV-229E_1ab;4004 | 0 | 0 |
| NC_002645.1;NP_073549.1;HCoV-229E_1ab;4005 | 0 | 0 |
| NC_002645.1;NP_073549.1;HCoV-229E_1ab;4006 | 0 | 0 |
| NC_002645.1;NP_073549.1;HCoV-229E_1ab;4007 | 0 | 0 |
| NC_002645.1;NP_073549.1;HCoV-229E_1ab;4008 | 0 | 0 |
| NC_002645.1;NP_073549.1;HCoV-229E_1ab;4009 | 0 | 0 |
| NC_002645.1;NP_073549.1;HCoV-229E_1ab;4010 | 0 | 0 |
| NC_002645.1;NP_073549.1;HCoV-229E_1ab;4011 | 0 | 0 |
| NC_002645.1;NP_073549.1;HCoV-229E_1ab;4012 | 0 | 0 |
| NC_002645.1;NP_073549.1;HCoV-229E_1ab;4013 | 0 | 0 |
| NC_002645.1;NP_073549.1;HCoV-229E_1ab;4014 | 0 | 0 |
| NC_002645.1;NP_073549.1;HCoV-229E_1ab;4015 | 0 | 0 |
| NC_002645.1;NP_073549.1;HCoV-229E_1ab;4016 | 0 | 0 |
| NC_002645.1;NP_073549.1;HCoV-229E_1ab;4017 | 0 | 0 |
| NC_002645.1;NP_073549.1;HCoV-229E_1ab;4018 | 0 | 0 |
| NC_002645.1;NP_073549.1;HCoV-229E_1ab;4019 | 0 | 0 |
| NC_002645.1;NP_073549.1;HCoV-229E_1ab;4020 | 0 | 0 |
| NC_002645.1;NP_073549.1;HCoV-229E_1ab;4021 | 0 | 0 |
| NC_002645.1;NP_073549.1;HCoV-229E_1ab;4022 | 0 | 0.075 |
| NC_002645.1;NP_073549.1;HCoV-229E_1ab;4023 | 0.05 | 0.075 |
| NC_002645.1;NP_073549.1;HCoV-229E_1ab;4024 | 0.05 | 0.075 |
| NC_002645.1;NP_073549.1;HCoV-229E_1ab;4025 | 0 | 0.1 |
| NC_002645.1;NP_073549.1;HCoV-229E_1ab;4026 | 0 | 0.075 |
| NC_002645.1;NP_073549.1;HCoV-229E_1ab;4027 | 0 | 0.025 |
| NC_002645.1;NP_073549.1;HCoV-229E_1ab;4028 | 0 | 0.025 |

| | | |
|---|---|---|
| NC_002645.1;NP_073549.1;HCoV-229E_1ab;4029 | 0 | 0 |
| NC_002645.1;NP_073549.1;HCoV-229E_1ab;4030 | 0 | 0 |
| NC_002645.1;NP_073549.1;HCoV-229E_1ab;4031 | 0.05 | 0.025 |
| NC_002645.1;NP_073549.1;HCoV-229E_1ab;4032 | 0.05 | 0.025 |
| NC_002645.1;NP_073549.1;HCoV-229E_1ab;4033 | 0 | 0.025 |
| NC_002645.1;NP_073549.1;HCoV-229E_1ab;4034 | 0.05 | 0.125 |
| NC_002645.1;NP_073549.1;HCoV-229E_1ab;4035 | 0 | 0 |
| NC_002645.1;NP_073549.1;HCoV-229E_1ab;4036 | 0 | 0 |
| NC_002645.1;NP_073549.1;HCoV-229E_1ab;4037 | 0 | 0 |
| NC_002645.1;NP_073549.1;HCoV-229E_1ab;4038 | 0 | 0.025 |
| NC_002645.1;NP_073549.1;HCoV-229E_1ab;4039 | 0 | 0 |
| NC_002645.1;NP_073549.1;HCoV-229E_1ab;4040 | 0 | 0 |
| NC_002645.1;NP_073549.1;HCoV-229E_1ab;4041 | 0 | 0 |
| NC_002645.1;NP_073549.1;HCoV-229E_1ab;4042 | 0 | 0 |
| NC_002645.1;NP_073549.1;HCoV-229E_1ab;4043 | 0 | 0 |
| NC_002645.1;NP_073549.1;HCoV-229E_1ab;4044 | 0 | 0 |
| NC_002645.1;NP_073549.1;HCoV-229E_1ab;4045 | 0 | 0 |
| NC_002645.1;NP_073549.1;HCoV-229E_1ab;4046 | 0 | 0 |
| NC_002645.1;NP_073549.1;HCoV-229E_1ab;4047 | 0 | 0 |
| NC_002645.1;NP_073549.1;HCoV-229E_1ab;4048 | 0 | 0 |
| NC_002645.1;NP_073549.1;HCoV-229E_1ab;4049 | 0 | 0 |
| NC_002645.1;NP_073549.1;HCoV-229E_1ab;4050 | 0 | 0 |
| NC_002645.1;NP_073549.1;HCoV-229E_1ab;4051 | 0 | 0 |
| NC_002645.1;NP_073549.1;HCoV-229E_1ab;4052 | 0 | 0 |
| NC_002645.1;NP_073549.1;HCoV-229E_1ab;4053 | 0 | 0 |
| NC_002645.1;NP_073549.1;HCoV-229E_1ab;4054 | 0 | 0 |
| NC_002645.1;NP_073549.1;HCoV-229E_1ab;4055 | 0 | 0 |
| NC_002645.1;NP_073549.1;HCoV-229E_1ab;4056 | 0 | 0.025 |
| NC_002645.1;NP_073549.1;HCoV-229E_1ab;4057 | 0 | 0 |
| NC_002645.1;NP_073549.1;HCoV-229E_1ab;4058 | 0 | 0.025 |
| NC_002645.1;NP_073549.1;HCoV-229E_1ab;4059 | 0 | 0.025 |
| NC_002645.1;NP_073549.1;HCoV-229E_1ab;4060 | 0 | 0.025 |
| NC_002645.1;NP_073549.1;HCoV-229E_1ab;4061 | 0.05 | 0.025 |
| NC_002645.1;NP_073549.1;HCoV-229E_1ab;4062 | 0.05 | 0.025 |
| NC_002645.1;NP_073549.1;HCoV-229E_1ab;4063 | 0.05 | 0 |
| NC_002645.1;NP_073549.1;HCoV-229E_1ab;4064 | 0 | 0 |
| NC_002645.1;NP_073549.1;HCoV-229E_1ab;4065 | 0 | 0 |
| NC_002645.1;NP_073549.1;HCoV-229E_1ab;4066 | 0 | 0 |
| NC_002645.1;NP_073549.1;HCoV-229E_1ab;4067 | 0 | 0 |
| NC_002645.1;NP_073549.1;HCoV-229E_1ab;4068 | 0.05 | 0 |
| NC_002645.1;NP_073549.1;HCoV-229E_1ab;4069 | 0.05 | 0.025 |
| NC_002645.1;NP_073549.1;HCoV-229E_1ab;4070 | 0 | 0 |
| NC_002645.1;NP_073549.1;HCoV-229E_1ab;4071 | 0 | 0 |

| | | |
|---|---|---|
| NC_002645.1;NP_073549.1;HCoV-229E_1ab;4072 | 0 | 0 |
| NC_002645.1;NP_073549.1;HCoV-229E_1ab;4073 | 0.05 | 0 |
| NC_002645.1;NP_073549.1;HCoV-229E_1ab;4074 | 0.05 | 0 |
| NC_002645.1;NP_073549.1;HCoV-229E_1ab;4075 | 0 | 0 |
| NC_002645.1;NP_073549.1;HCoV-229E_1ab;4076 | 0 | 0 |
| NC_002645.1;NP_073549.1;HCoV-229E_1ab;4077 | 0 | 0.05 |
| NC_002645.1;NP_073549.1;HCoV-229E_1ab;4078 | 0 | 0.025 |
| NC_002645.1;NP_073549.1;HCoV-229E_1ab;4079 | 0 | 0 |
| NC_002645.1;NP_073549.1;HCoV-229E_1ab;4080 | 0 | 0 |
| NC_002645.1;NP_073549.1;HCoV-229E_1ab;4081 | 0 | 0 |
| NC_002645.1;NP_073549.1;HCoV-229E_1ab;4082 | 0 | 0 |
| NC_002645.1;NP_073549.1;HCoV-229E_1ab;4083 | 0 | 0 |
| NC_002645.1;NP_073549.1;HCoV-229E_1ab;4084 | 0 | 0 |
| NC_002645.1;NP_073549.1;HCoV-229E_1ab;4085 | 0 | 0 |
| NC_002645.1;NP_073549.1;HCoV-229E_1ab;4086 | 0 | 0 |
| NC_002645.1;NP_073549.1;HCoV-229E_1ab;4087 | 0 | 0 |
| NC_002645.1;NP_073549.1;HCoV-229E_1ab;4088 | 0 | 0 |
| NC_002645.1;NP_073549.1;HCoV-229E_1ab;4089 | 0 | 0 |
| NC_002645.1;NP_073549.1;HCoV-229E_1ab;4090 | 0 | 0 |
| NC_002645.1;NP_073549.1;HCoV-229E_1ab;4091 | 0 | 0 |
| NC_002645.1;NP_073549.1;HCoV-229E_1ab;4092 | 0.05 | 0.025 |
| NC_002645.1;NP_073549.1;HCoV-229E_1ab;4093 | 0 | 0.025 |
| NC_002645.1;NP_073549.1;HCoV-229E_1ab;4094 | 0 | 0.025 |
| NC_002645.1;NP_073549.1;HCoV-229E_1ab;4095 | 0 | 0.025 |
| NC_002645.1;NP_073549.1;HCoV-229E_1ab;4096 | 0 | 0.025 |
| NC_002645.1;NP_073549.1;HCoV-229E_1ab;4097 | 0 | 0 |
| NC_002645.1;NP_073549.1;HCoV-229E_1ab;4098 | 0.05 | 0.025 |
| NC_002645.1;NP_073549.1;HCoV-229E_1ab;4099 | 0.05 | 0.05 |
| NC_002645.1;NP_073549.1;HCoV-229E_1ab;4100 | 0.05 | 0.05 |
| NC_002645.1;NP_073549.1;HCoV-229E_1ab;4101 | 0.1 | 0.075 |
| NC_002645.1;NP_073549.1;HCoV-229E_1ab;4102 | 0.15 | 0.075 |
| NC_002645.1;NP_073549.1;HCoV-229E_1ab;4103 | 0.1 | 0.05 |
| NC_002645.1;NP_073549.1;HCoV-229E_1ab;4104 | 0.1 | 0.05 |
| NC_002645.1;NP_073549.1;HCoV-229E_1ab;4105 | 0.05 | 0.05 |
| NC_002645.1;NP_073549.1;HCoV-229E_1ab;4106 | 0.05 | 0.05 |
| NC_002645.1;NP_073549.1;HCoV-229E_1ab;4107 | 0.05 | 0.025 |
| NC_002645.1;NP_073549.1;HCoV-229E_1ab;4108 | 0 | 0 |
| NC_002645.1;NP_073549.1;HCoV-229E_1ab;4109 | 0 | 0 |
| NC_002645.1;NP_073549.1;HCoV-229E_1ab;4110 | 0 | 0 |
| NC_002645.1;NP_073549.1;HCoV-229E_1ab;4111 | 0 | 0 |
| NC_002645.1;NP_073549.1;HCoV-229E_1ab;4112 | 0 | 0 |
| NC_002645.1;NP_073549.1;HCoV-229E_1ab;4113 | 0 | 0.025 |
| NC_002645.1;NP_073549.1;HCoV-229E_1ab;4114 | 0.05 | 0.05 |

| | | |
|---|---|---|
| NC_002645.1;NP_073549.1;HCoV-229E_1ab;4115 | 0.05 | 0 |
| NC_002645.1;NP_073549.1;HCoV-229E_1ab;4116 | 0.05 | 0.025 |
| NC_002645.1;NP_073549.1;HCoV-229E_1ab;4117 | 0.1 | 0.05 |
| NC_002645.1;NP_073549.1;HCoV-229E_1ab;4118 | 0 | 0 |
| NC_002645.1;NP_073549.1;HCoV-229E_1ab;4119 | 0 | 0 |
| NC_002645.1;NP_073549.1;HCoV-229E_1ab;4120 | 0.05 | 0 |
| NC_002645.1;NP_073549.1;HCoV-229E_1ab;4121 | 0.05 | 0 |
| NC_002645.1;NP_073549.1;HCoV-229E_1ab;4122 | 0 | 0.025 |
| NC_002645.1;NP_073549.1;HCoV-229E_1ab;4123 | 0 | 0 |
| NC_002645.1;NP_073549.1;HCoV-229E_1ab;4124 | 0 | 0 |
| NC_002645.1;NP_073549.1;HCoV-229E_1ab;4125 | 0 | 0 |
| NC_002645.1;NP_073549.1;HCoV-229E_1ab;4126 | 0 | 0 |
| NC_002645.1;NP_073549.1;HCoV-229E_1ab;4127 | 0 | 0 |
| NC_002645.1;NP_073549.1;HCoV-229E_1ab;4128 | 0 | 0 |
| NC_002645.1;NP_073549.1;HCoV-229E_1ab;4129 | 0 | 0 |
| NC_002645.1;NP_073549.1;HCoV-229E_1ab;4130 | 0 | 0 |
| NC_002645.1;NP_073549.1;HCoV-229E_1ab;4131 | 0 | 0 |
| NC_002645.1;NP_073549.1;HCoV-229E_1ab;4132 | 0.05 | 0 |
| NC_002645.1;NP_073549.1;HCoV-229E_1ab;4133 | 0 | 0 |
| NC_002645.1;NP_073549.1;HCoV-229E_1ab;4134 | 0 | 0 |
| NC_002645.1;NP_073549.1;HCoV-229E_1ab;4135 | 0.1 | 0.1 |
| NC_002645.1;NP_073549.1;HCoV-229E_1ab;4136 | 0 | 0 |
| NC_002645.1;NP_073549.1;HCoV-229E_1ab;4137 | 0 | 0 |
| NC_002645.1;NP_073549.1;HCoV-229E_1ab;4138 | 0.05 | 0.025 |
| NC_002645.1;NP_073549.1;HCoV-229E_1ab;4139 | 0 | 0.025 |
| NC_002645.1;NP_073549.1;HCoV-229E_1ab;4140 | 0.05 | 0 |
| NC_002645.1;NP_073549.1;HCoV-229E_1ab;4141 | 0 | 0 |
| NC_002645.1;NP_073549.1;HCoV-229E_1ab;4142 | 0 | 0 |
| NC_002645.1;NP_073549.1;HCoV-229E_1ab;4143 | 0 | 0 |
| NC_002645.1;NP_073549.1;HCoV-229E_1ab;4144 | 0 | 0.025 |
| NC_002645.1;NP_073549.1;HCoV-229E_1ab;4145 | 0 | 0.025 |
| NC_002645.1;NP_073549.1;HCoV-229E_1ab;4146 | 0 | 0 |
| NC_002645.1;NP_073549.1;HCoV-229E_1ab;4147 | 0 | 0 |
| NC_002645.1;NP_073549.1;HCoV-229E_1ab;4148 | 0 | 0.025 |
| NC_002645.1;NP_073549.1;HCoV-229E_1ab;4149 | 0 | 0.025 |
| NC_002645.1;NP_073549.1;HCoV-229E_1ab;4150 | 0 | 0 |
| NC_002645.1;NP_073549.1;HCoV-229E_1ab;4151 | 0 | 0 |
| NC_002645.1;NP_073549.1;HCoV-229E_1ab;4152 | 0 | 0 |
| NC_002645.1;NP_073549.1;HCoV-229E_1ab;4153 | 0 | 0.025 |
| NC_002645.1;NP_073549.1;HCoV-229E_1ab;4154 | 0 | 0 |
| NC_002645.1;NP_073549.1;HCoV-229E_1ab;4155 | 0 | 0.025 |
| NC_002645.1;NP_073549.1;HCoV-229E_1ab;4156 | 0 | 0.025 |
| NC_002645.1;NP_073549.1;HCoV-229E_1ab;4157 | 0 | 0.025 |

| | | |
|---|---|---|
| NC_002645.1;NP_073549.1;HCoV-229E_1ab;4158 | 0 | 0 |
| NC_002645.1;NP_073549.1;HCoV-229E_1ab;4159 | 0 | 0 |
| NC_002645.1;NP_073549.1;HCoV-229E_1ab;4160 | 0 | 0 |
| NC_002645.1;NP_073549.1;HCoV-229E_1ab;4161 | 0 | 0 |
| NC_002645.1;NP_073549.1;HCoV-229E_1ab;4162 | 0 | 0.025 |
| NC_002645.1;NP_073549.1;HCoV-229E_1ab;4163 | 0 | 0 |
| NC_002645.1;NP_073549.1;HCoV-229E_1ab;4164 | 0 | 0 |
| NC_002645.1;NP_073549.1;HCoV-229E_1ab;4165 | 0 | 0.05 |
| NC_002645.1;NP_073549.1;HCoV-229E_1ab;4166 | 0 | 0.025 |
| NC_002645.1;NP_073549.1;HCoV-229E_1ab;4167 | 0 | 0.025 |
| NC_002645.1;NP_073549.1;HCoV-229E_1ab;4168 | 0 | 0.075 |
| NC_002645.1;NP_073549.1;HCoV-229E_1ab;4169 | 0 | 0 |
| NC_002645.1;NP_073549.1;HCoV-229E_1ab;4170 | 0 | 0 |
| NC_002645.1;NP_073549.1;HCoV-229E_1ab;4171 | 0 | 0.025 |
| NC_002645.1;NP_073549.1;HCoV-229E_1ab;4172 | 0 | 0 |
| NC_002645.1;NP_073549.1;HCoV-229E_1ab;4173 | 0 | 0 |
| NC_002645.1;NP_073549.1;HCoV-229E_1ab;4174 | 0 | 0.025 |
| NC_002645.1;NP_073549.1;HCoV-229E_1ab;4175 | 0 | 0 |
| NC_002645.1;NP_073549.1;HCoV-229E_1ab;4176 | 0 | 0 |
| NC_002645.1;NP_073549.1;HCoV-229E_1ab;4177 | 0 | 0 |
| NC_002645.1;NP_073549.1;HCoV-229E_1ab;4178 | 0 | 0 |
| NC_002645.1;NP_073549.1;HCoV-229E_1ab;4179 | 0 | 0 |
| NC_002645.1;NP_073549.1;HCoV-229E_1ab;4180 | 0 | 0 |
| NC_002645.1;NP_073549.1;HCoV-229E_1ab;4181 | 0 | 0 |
| NC_002645.1;NP_073549.1;HCoV-229E_1ab;4182 | 0 | 0 |
| NC_002645.1;NP_073549.1;HCoV-229E_1ab;4183 | 0 | 0 |
| NC_002645.1;NP_073549.1;HCoV-229E_1ab;4184 | 0 | 0.375 |
| NC_002645.1;NP_073549.1;HCoV-229E_1ab;4185 | 0.1 | 0.025 |
| NC_002645.1;NP_073549.1;HCoV-229E_1ab;4186 | 0 | 0 |
| NC_002645.1;NP_073549.1;HCoV-229E_1ab;4187 | 0 | 0 |
| NC_002645.1;NP_073549.1;HCoV-229E_1ab;4188 | 0 | 0 |
| NC_002645.1;NP_073549.1;HCoV-229E_1ab;4189 | 0 | 0 |
| NC_002645.1;NP_073549.1;HCoV-229E_1ab;4190 | 0.1 | 0.075 |
| NC_002645.1;NP_073549.1;HCoV-229E_1ab;4191 | 0.05 | 0.05 |
| NC_002645.1;NP_073549.1;HCoV-229E_1ab;4192 | 0.05 | 0.025 |
| NC_002645.1;NP_073549.1;HCoV-229E_1ab;4193 | 0.05 | 0.1 |
| NC_002645.1;NP_073549.1;HCoV-229E_1ab;4194 | 0 | 0 |
| NC_002645.1;NP_073549.1;HCoV-229E_1ab;4195 | 0 | 0 |
| NC_002645.1;NP_073549.1;HCoV-229E_1ab;4196 | 0 | 0 |
| NC_002645.1;NP_073549.1;HCoV-229E_1ab;4197 | 0.05 | 0.05 |
| NC_002645.1;NP_073549.1;HCoV-229E_1ab;4198 | 0 | 0 |
| NC_002645.1;NP_073549.1;HCoV-229E_1ab;4199 | 0 | 0 |
| NC_002645.1;NP_073549.1;HCoV-229E_1ab;4200 | 0 | 0 |

| | | |
|---|---|---|
| NC_002645.1;NP_073549.1;HCoV-229E_1ab;4201 | 0 | 0 |
| NC_002645.1;NP_073549.1;HCoV-229E_1ab;4202 | 0 | 0 |
| NC_002645.1;NP_073549.1;HCoV-229E_1ab;4203 | 0 | 0 |
| NC_002645.1;NP_073549.1;HCoV-229E_1ab;4204 | 0 | 0 |
| NC_002645.1;NP_073549.1;HCoV-229E_1ab;4205 | 0 | 0 |
| NC_002645.1;NP_073549.1;HCoV-229E_1ab;4206 | 0 | 0 |
| NC_002645.1;NP_073549.1;HCoV-229E_1ab;4207 | 0.2 | 0 |
| NC_002645.1;NP_073549.1;HCoV-229E_1ab;4208 | 0.2 | 0 |
| NC_002645.1;NP_073549.1;HCoV-229E_1ab;4209 | 0.15 | 0 |
| NC_002645.1;NP_073549.1;HCoV-229E_1ab;4210 | 0.15 | 0 |
| NC_002645.1;NP_073549.1;HCoV-229E_1ab;4211 | 0.15 | 0 |
| NC_002645.1;NP_073549.1;HCoV-229E_1ab;4212 | 0.2 | 0 |
| NC_002645.1;NP_073549.1;HCoV-229E_1ab;4213 | 0.15 | 0 |
| NC_002645.1;NP_073549.1;HCoV-229E_1ab;4214 | 0.15 | 0 |
| NC_002645.1;NP_073549.1;HCoV-229E_1ab;4215 | 0.2 | 0 |
| NC_002645.1;NP_073549.1;HCoV-229E_1ab;4216 | 0.15 | 0 |
| NC_002645.1;NP_073549.1;HCoV-229E_1ab;4217 | 0 | 0 |
| NC_002645.1;NP_073549.1;HCoV-229E_1ab;4218 | 0.05 | 0 |
| NC_002645.1;NP_073549.1;HCoV-229E_1ab;4219 | 0 | 0 |
| NC_002645.1;NP_073549.1;HCoV-229E_1ab;4220 | 0 | 0 |
| NC_002645.1;NP_073549.1;HCoV-229E_1ab;4221 | 0 | 0 |
| NC_002645.1;NP_073549.1;HCoV-229E_1ab;4222 | 0 | 0.025 |
| NC_002645.1;NP_073549.1;HCoV-229E_1ab;4223 | 0 | 0.025 |
| NC_002645.1;NP_073549.1;HCoV-229E_1ab;4224 | 0.05 | 0.025 |
| NC_002645.1;NP_073549.1;HCoV-229E_1ab;4225 | 0 | 0.025 |
| NC_002645.1;NP_073549.1;HCoV-229E_1ab;4226 | 0 | 0 |
| NC_002645.1;NP_073549.1;HCoV-229E_1ab;4227 | 0 | 0 |
| NC_002645.1;NP_073549.1;HCoV-229E_1ab;4228 | 0 | 0 |
| NC_002645.1;NP_073549.1;HCoV-229E_1ab;4229 | 0 | 0 |
| NC_002645.1;NP_073549.1;HCoV-229E_1ab;4230 | 0 | 0 |
| NC_002645.1;NP_073549.1;HCoV-229E_1ab;4231 | 0 | 0.025 |
| NC_002645.1;NP_073549.1;HCoV-229E_1ab;4232 | 0 | 0 |
| NC_002645.1;NP_073549.1;HCoV-229E_1ab;4233 | 0.05 | 0 |
| NC_002645.1;NP_073549.1;HCoV-229E_1ab;4234 | 0 | 0 |
| NC_002645.1;NP_073549.1;HCoV-229E_1ab;4235 | 0.1 | 0 |
| NC_002645.1;NP_073549.1;HCoV-229E_1ab;4236 | 0 | 0 |
| NC_002645.1;NP_073549.1;HCoV-229E_1ab;4237 | 0 | 0 |
| NC_002645.1;NP_073549.1;HCoV-229E_1ab;4238 | 0 | 0 |
| NC_002645.1;NP_073549.1;HCoV-229E_1ab;4239 | 0 | 0.025 |
| NC_002645.1;NP_073549.1;HCoV-229E_1ab;4240 | 0 | 0 |
| NC_002645.1;NP_073549.1;HCoV-229E_1ab;4241 | 0 | 0 |
| NC_002645.1;NP_073549.1;HCoV-229E_1ab;4242 | 0 | 0 |
| NC_002645.1;NP_073549.1;HCoV-229E_1ab;4243 | 0 | 0 |

| | | |
|---|---|---|
| NC_002645.1;NP_073549.1;HCoV-229E_1ab;4244 | 0 | 0 |
| NC_002645.1;NP_073549.1;HCoV-229E_1ab;4245 | 0 | 0 |
| NC_002645.1;NP_073549.1;HCoV-229E_1ab;4246 | 0 | 0 |
| NC_002645.1;NP_073549.1;HCoV-229E_1ab;4247 | 0 | 0 |
| NC_002645.1;NP_073549.1;HCoV-229E_1ab;4248 | 0 | 0 |
| NC_002645.1;NP_073549.1;HCoV-229E_1ab;4249 | 0.05 | 0 |
| NC_002645.1;NP_073549.1;HCoV-229E_1ab;4250 | 0.05 | 0 |
| NC_002645.1;NP_073549.1;HCoV-229E_1ab;4251 | 0.05 | 0 |
| NC_002645.1;NP_073549.1;HCoV-229E_1ab;4252 | 0 | 0 |
| NC_002645.1;NP_073549.1;HCoV-229E_1ab;4253 | 0 | 0 |
| NC_002645.1;NP_073549.1;HCoV-229E_1ab;4254 | 0 | 0 |
| NC_002645.1;NP_073549.1;HCoV-229E_1ab;4255 | 0 | 0 |
| NC_002645.1;NP_073549.1;HCoV-229E_1ab;4256 | 0 | 0 |
| NC_002645.1;NP_073549.1;HCoV-229E_1ab;4257 | 0 | 0 |
| NC_002645.1;NP_073549.1;HCoV-229E_1ab;4258 | 0 | 0 |
| NC_002645.1;NP_073549.1;HCoV-229E_1ab;4259 | 0 | 0 |
| NC_002645.1;NP_073549.1;HCoV-229E_1ab;4260 | 0 | 0 |
| NC_002645.1;NP_073549.1;HCoV-229E_1ab;4261 | 0.05 | 0 |
| NC_002645.1;NP_073549.1;HCoV-229E_1ab;4262 | 0 | 0 |
| NC_002645.1;NP_073549.1;HCoV-229E_1ab;4263 | 0 | 0 |
| NC_002645.1;NP_073549.1;HCoV-229E_1ab;4264 | 0 | 0 |
| NC_002645.1;NP_073549.1;HCoV-229E_1ab;4265 | 0.1 | 0 |
| NC_002645.1;NP_073549.1;HCoV-229E_1ab;4266 | 0 | 0 |
| NC_002645.1;NP_073549.1;HCoV-229E_1ab;4267 | 0 | 0 |
| NC_002645.1;NP_073549.1;HCoV-229E_1ab;4268 | 0 | 0 |
| NC_002645.1;NP_073549.1;HCoV-229E_1ab;4269 | 0 | 0 |
| NC_002645.1;NP_073549.1;HCoV-229E_1ab;4270 | 0 | 0 |
| NC_002645.1;NP_073549.1;HCoV-229E_1ab;4271 | 0 | 0 |
| NC_002645.1;NP_073549.1;HCoV-229E_1ab;4272 | 0 | 0 |
| NC_002645.1;NP_073549.1;HCoV-229E_1ab;4273 | 0 | 0 |
| NC_002645.1;NP_073549.1;HCoV-229E_1ab;4274 | 0 | 0 |
| NC_002645.1;NP_073549.1;HCoV-229E_1ab;4275 | 0 | 0 |
| NC_002645.1;NP_073549.1;HCoV-229E_1ab;4276 | 0 | 0 |
| NC_002645.1;NP_073549.1;HCoV-229E_1ab;4277 | 0 | 0 |
| NC_002645.1;NP_073549.1;HCoV-229E_1ab;4278 | 0 | 0 |
| NC_002645.1;NP_073549.1;HCoV-229E_1ab;4279 | 0 | 0 |
| NC_002645.1;NP_073549.1;HCoV-229E_1ab;4280 | 0 | 0 |
| NC_002645.1;NP_073549.1;HCoV-229E_1ab;4281 | 0 | 0 |
| NC_002645.1;NP_073549.1;HCoV-229E_1ab;4282 | 0 | 0 |
| NC_002645.1;NP_073549.1;HCoV-229E_1ab;4283 | 0 | 0 |
| NC_002645.1;NP_073549.1;HCoV-229E_1ab;4284 | 0 | 0 |
| NC_002645.1;NP_073549.1;HCoV-229E_1ab;4285 | 0 | 0 |
| NC_002645.1;NP_073549.1;HCoV-229E_1ab;4286 | 0 | 0 |

| | | |
|---|---|---|
| NC_002645.1;NP_073549.1;HCoV-229E_1ab;4287 | 0 | 0 |
| NC_002645.1;NP_073549.1;HCoV-229E_1ab;4288 | 0 | 0 |
| NC_002645.1;NP_073549.1;HCoV-229E_1ab;4289 | 0 | 0 |
| NC_002645.1;NP_073549.1;HCoV-229E_1ab;4290 | 0 | 0 |
| NC_002645.1;NP_073549.1;HCoV-229E_1ab;4291 | 0 | 0 |
| NC_002645.1;NP_073549.1;HCoV-229E_1ab;4292 | 0 | 0 |
| NC_002645.1;NP_073549.1;HCoV-229E_1ab;4293 | 0 | 0 |
| NC_002645.1;NP_073549.1;HCoV-229E_1ab;4294 | 0 | 0 |
| NC_002645.1;NP_073549.1;HCoV-229E_1ab;4295 | 0 | 0 |
| NC_002645.1;NP_073549.1;HCoV-229E_1ab;4296 | 0 | 0 |
| NC_002645.1;NP_073549.1;HCoV-229E_1ab;4297 | 0 | 0 |
| NC_002645.1;NP_073549.1;HCoV-229E_1ab;4298 | 0 | 0 |
| NC_002645.1;NP_073549.1;HCoV-229E_1ab;4299 | 0 | 0 |
| NC_002645.1;NP_073549.1;HCoV-229E_1ab;4300 | 0 | 0 |
| NC_002645.1;NP_073549.1;HCoV-229E_1ab;4301 | 0 | 0 |
| NC_002645.1;NP_073549.1;HCoV-229E_1ab;4302 | 0 | 0 |
| NC_002645.1;NP_073549.1;HCoV-229E_1ab;4303 | 0 | 0 |
| NC_002645.1;NP_073549.1;HCoV-229E_1ab;4304 | 0 | 0 |
| NC_002645.1;NP_073549.1;HCoV-229E_1ab;4305 | 0 | 0 |
| NC_002645.1;NP_073549.1;HCoV-229E_1ab;4306 | 0 | 0 |
| NC_002645.1;NP_073549.1;HCoV-229E_1ab;4307 | 0 | 0 |
| NC_002645.1;NP_073549.1;HCoV-229E_1ab;4308 | 0 | 0 |
| NC_002645.1;NP_073549.1;HCoV-229E_1ab;4309 | 0 | 0 |
| NC_002645.1;NP_073549.1;HCoV-229E_1ab;4310 | 0 | 0 |
| NC_002645.1;NP_073549.1;HCoV-229E_1ab;4311 | 0 | 0 |
| NC_002645.1;NP_073549.1;HCoV-229E_1ab;4312 | 0 | 0.025 |
| NC_002645.1;NP_073549.1;HCoV-229E_1ab;4313 | 0.1 | 0.05 |
| NC_002645.1;NP_073549.1;HCoV-229E_1ab;4314 | 0.2 | 0.175 |
| NC_002645.1;NP_073549.1;HCoV-229E_1ab;4315 | 0.15 | 0.175 |
| NC_002645.1;NP_073549.1;HCoV-229E_1ab;4316 | 0.1 | 0.125 |
| NC_002645.1;NP_073549.1;HCoV-229E_1ab;4317 | 0.1 | 0.15 |
| NC_002645.1;NP_073549.1;HCoV-229E_1ab;4318 | 0.1 | 0.125 |
| NC_002645.1;NP_073549.1;HCoV-229E_1ab;4319 | 0.1 | 0.15 |
| NC_002645.1;NP_073549.1;HCoV-229E_1ab;4320 | 0.1 | 0.125 |
| NC_002645.1;NP_073549.1;HCoV-229E_1ab;4321 | 0 | 0.075 |
| NC_002645.1;NP_073549.1;HCoV-229E_1ab;4322 | 0 | 0.05 |
| NC_002645.1;NP_073549.1;HCoV-229E_1ab;4323 | 0 | 0.025 |
| NC_002645.1;NP_073549.1;HCoV-229E_1ab;4324 | 0 | 0 |
| NC_002645.1;NP_073549.1;HCoV-229E_1ab;4325 | 0.05 | 0.15 |
| NC_002645.1;NP_073549.1;HCoV-229E_1ab;4326 | 0 | 0 |
| NC_002645.1;NP_073549.1;HCoV-229E_1ab;4327 | 0 | 0 |
| NC_002645.1;NP_073549.1;HCoV-229E_1ab;4328 | 0 | 0.025 |
| NC_002645.1;NP_073549.1;HCoV-229E_1ab;4329 | 0 | 0.025 |

| | | |
|---|---|---|
| NC_002645.1;NP_073549.1;HCoV-229E_1ab;4330 | 0 | 0 |
| NC_002645.1;NP_073549.1;HCoV-229E_1ab;4331 | 0 | 0 |
| NC_002645.1;NP_073549.1;HCoV-229E_1ab;4332 | 0 | 0 |
| NC_002645.1;NP_073549.1;HCoV-229E_1ab;4333 | 0.05 | 0.025 |
| NC_002645.1;NP_073549.1;HCoV-229E_1ab;4334 | 0.1 | 0 |
| NC_002645.1;NP_073549.1;HCoV-229E_1ab;4335 | 0.05 | 0 |
| NC_002645.1;NP_073549.1;HCoV-229E_1ab;4336 | 0 | 0 |
| NC_002645.1;NP_073549.1;HCoV-229E_1ab;4337 | 0 | 0 |
| NC_002645.1;NP_073549.1;HCoV-229E_1ab;4338 | 0 | 0 |
| NC_002645.1;NP_073549.1;HCoV-229E_1ab;4339 | 0 | 0 |
| NC_002645.1;NP_073549.1;HCoV-229E_1ab;4340 | 0 | 0 |
| NC_002645.1;NP_073549.1;HCoV-229E_1ab;4341 | 0 | 0 |
| NC_002645.1;NP_073549.1;HCoV-229E_1ab;4342 | 0 | 0 |
| NC_002645.1;NP_073549.1;HCoV-229E_1ab;4343 | 0 | 0 |
| NC_002645.1;NP_073549.1;HCoV-229E_1ab;4344 | 0 | 0 |
| NC_002645.1;NP_073549.1;HCoV-229E_1ab;4345 | 0 | 0 |
| NC_002645.1;NP_073549.1;HCoV-229E_1ab;4346 | 0 | 0 |
| NC_002645.1;NP_073549.1;HCoV-229E_1ab;4347 | 0 | 0 |
| NC_002645.1;NP_073549.1;HCoV-229E_1ab;4348 | 0 | 0 |
| NC_002645.1;NP_073549.1;HCoV-229E_1ab;4349 | 0 | 0.025 |
| NC_002645.1;NP_073549.1;HCoV-229E_1ab;4350 | 0 | 0.025 |
| NC_002645.1;NP_073549.1;HCoV-229E_1ab;4351 | 0 | 0.025 |
| NC_002645.1;NP_073549.1;HCoV-229E_1ab;4352 | 0 | 0.025 |
| NC_002645.1;NP_073549.1;HCoV-229E_1ab;4353 | 0 | 0 |
| NC_002645.1;NP_073549.1;HCoV-229E_1ab;4354 | 0 | 0 |
| NC_002645.1;NP_073549.1;HCoV-229E_1ab;4355 | 0 | 0 |
| NC_002645.1;NP_073549.1;HCoV-229E_1ab;4356 | 0 | 0 |
| NC_002645.1;NP_073549.1;HCoV-229E_1ab;4357 | 0 | 0 |
| NC_002645.1;NP_073549.1;HCoV-229E_1ab;4358 | 0 | 0 |
| NC_002645.1;NP_073549.1;HCoV-229E_1ab;4359 | 0 | 0 |
| NC_002645.1;NP_073549.1;HCoV-229E_1ab;4360 | 0 | 0 |
| NC_002645.1;NP_073549.1;HCoV-229E_1ab;4361 | 0 | 0 |
| NC_002645.1;NP_073549.1;HCoV-229E_1ab;4362 | 0 | 0 |
| NC_002645.1;NP_073549.1;HCoV-229E_1ab;4363 | 0 | 0 |
| NC_002645.1;NP_073549.1;HCoV-229E_1ab;4364 | 0 | 0 |
| NC_002645.1;NP_073549.1;HCoV-229E_1ab;4365 | 0 | 0 |
| NC_002645.1;NP_073549.1;HCoV-229E_1ab;4366 | 0 | 0 |
| NC_002645.1;NP_073549.1;HCoV-229E_1ab;4367 | 0 | 0 |
| NC_002645.1;NP_073549.1;HCoV-229E_1ab;4368 | 0 | 0 |
| NC_002645.1;NP_073549.1;HCoV-229E_1ab;4369 | 0 | 0 |
| NC_002645.1;NP_073549.1;HCoV-229E_1ab;4370 | 0 | 0 |
| NC_002645.1;NP_073549.1;HCoV-229E_1ab;4371 | 0 | 0 |
| NC_002645.1;NP_073549.1;HCoV-229E_1ab;4372 | 0 | 0 |

| | | |
|---|---|---|
| NC_002645.1;NP_073549.1;HCoV-229E_1ab;4373 | 0 | 0 |
| NC_002645.1;NP_073549.1;HCoV-229E_1ab;4374 | 0 | 0 |
| NC_002645.1;NP_073549.1;HCoV-229E_1ab;4375 | 0 | 0.025 |
| NC_002645.1;NP_073549.1;HCoV-229E_1ab;4376 | 0 | 0 |
| NC_002645.1;NP_073549.1;HCoV-229E_1ab;4377 | 0 | 0 |
| NC_002645.1;NP_073549.1;HCoV-229E_1ab;4378 | 0 | 0 |
| NC_002645.1;NP_073549.1;HCoV-229E_1ab;4379 | 0 | 0 |
| NC_002645.1;NP_073549.1;HCoV-229E_1ab;4380 | 0 | 0 |
| NC_002645.1;NP_073549.1;HCoV-229E_1ab;4381 | 0 | 0 |
| NC_002645.1;NP_073549.1;HCoV-229E_1ab;4382 | 0 | 0 |
| NC_002645.1;NP_073549.1;HCoV-229E_1ab;4383 | 0 | 0 |
| NC_002645.1;NP_073549.1;HCoV-229E_1ab;4384 | 0 | 0 |
| NC_002645.1;NP_073549.1;HCoV-229E_1ab;4385 | 0 | 0 |
| NC_002645.1;NP_073549.1;HCoV-229E_1ab;4386 | 0 | 0 |
| NC_002645.1;NP_073549.1;HCoV-229E_1ab;4387 | 0 | 0 |
| NC_002645.1;NP_073549.1;HCoV-229E_1ab;4388 | 0 | 0 |
| NC_002645.1;NP_073549.1;HCoV-229E_1ab;4389 | 0 | 0 |
| NC_002645.1;NP_073549.1;HCoV-229E_1ab;4390 | 0 | 0 |
| NC_002645.1;NP_073549.1;HCoV-229E_1ab;4391 | 0 | 0 |
| NC_002645.1;NP_073549.1;HCoV-229E_1ab;4392 | 0 | 0 |
| NC_002645.1;NP_073549.1;HCoV-229E_1ab;4393 | 0 | 0 |
| NC_002645.1;NP_073549.1;HCoV-229E_1ab;4394 | 0 | 0 |
| NC_002645.1;NP_073549.1;HCoV-229E_1ab;4395 | 0 | 0 |
| NC_002645.1;NP_073549.1;HCoV-229E_1ab;4396 | 0 | 0 |
| NC_002645.1;NP_073549.1;HCoV-229E_1ab;4397 | 0 | 0 |
| NC_002645.1;NP_073549.1;HCoV-229E_1ab;4398 | 0 | 0 |
| NC_002645.1;NP_073549.1;HCoV-229E_1ab;4399 | 0 | 0 |
| NC_002645.1;NP_073549.1;HCoV-229E_1ab;4400 | 0 | 0 |
| NC_002645.1;NP_073549.1;HCoV-229E_1ab;4401 | 0 | 0 |
| NC_002645.1;NP_073549.1;HCoV-229E_1ab;4402 | 0 | 0 |
| NC_002645.1;NP_073549.1;HCoV-229E_1ab;4403 | 0 | 0 |
| NC_002645.1;NP_073549.1;HCoV-229E_1ab;4404 | 0 | 0 |
| NC_002645.1;NP_073549.1;HCoV-229E_1ab;4405 | 0 | 0 |
| NC_002645.1;NP_073549.1;HCoV-229E_1ab;4406 | 0 | 0 |
| NC_002645.1;NP_073549.1;HCoV-229E_1ab;4407 | 0 | 0 |
| NC_002645.1;NP_073549.1;HCoV-229E_1ab;4408 | 0 | 0 |
| NC_002645.1;NP_073549.1;HCoV-229E_1ab;4409 | 0 | 0 |
| NC_002645.1;NP_073549.1;HCoV-229E_1ab;4410 | 0 | 0 |
| NC_002645.1;NP_073549.1;HCoV-229E_1ab;4411 | 0 | 0 |
| NC_002645.1;NP_073549.1;HCoV-229E_1ab;4412 | 0.05 | 0 |
| NC_002645.1;NP_073549.1;HCoV-229E_1ab;4413 | 0 | 0.025 |
| NC_002645.1;NP_073549.1;HCoV-229E_1ab;4414 | 0 | 0 |
| NC_002645.1;NP_073549.1;HCoV-229E_1ab;4415 | 0 | 0 |

| | | |
|---|---|---|
| NC_002645.1;NP_073549.1;HCoV-229E_1ab;4416 | 0 | 0 |
| NC_002645.1;NP_073549.1;HCoV-229E_1ab;4417 | 0 | 0 |
| NC_002645.1;NP_073549.1;HCoV-229E_1ab;4418 | 0 | 0.025 |
| NC_002645.1;NP_073549.1;HCoV-229E_1ab;4419 | 0 | 0 |
| NC_002645.1;NP_073549.1;HCoV-229E_1ab;4420 | 0 | 0 |
| NC_002645.1;NP_073549.1;HCoV-229E_1ab;4421 | 0 | 0 |
| NC_002645.1;NP_073549.1;HCoV-229E_1ab;4422 | 0 | 0 |
| NC_002645.1;NP_073549.1;HCoV-229E_1ab;4423 | 0 | 0 |
| NC_002645.1;NP_073549.1;HCoV-229E_1ab;4424 | 0 | 0 |
| NC_002645.1;NP_073549.1;HCoV-229E_1ab;4425 | 0 | 0 |
| NC_002645.1;NP_073549.1;HCoV-229E_1ab;4426 | 0 | 0 |
| NC_002645.1;NP_073549.1;HCoV-229E_1ab;4427 | 0 | 0 |
| NC_002645.1;NP_073549.1;HCoV-229E_1ab;4428 | 0 | 0 |
| NC_002645.1;NP_073549.1;HCoV-229E_1ab;4429 | 0 | 0.125 |
| NC_002645.1;NP_073549.1;HCoV-229E_1ab;4430 | 0 | 0 |
| NC_002645.1;NP_073549.1;HCoV-229E_1ab;4431 | 0 | 0 |
| NC_002645.1;NP_073549.1;HCoV-229E_1ab;4432 | 0 | 0 |
| NC_002645.1;NP_073549.1;HCoV-229E_1ab;4433 | 0 | 0 |
| NC_002645.1;NP_073549.1;HCoV-229E_1ab;4434 | 0 | 0 |
| NC_002645.1;NP_073549.1;HCoV-229E_1ab;4435 | 0 | 0 |
| NC_002645.1;NP_073549.1;HCoV-229E_1ab;4436 | 0 | 0 |
| NC_002645.1;NP_073549.1;HCoV-229E_1ab;4437 | 0 | 0 |
| NC_002645.1;NP_073549.1;HCoV-229E_1ab;4438 | 0 | 0 |
| NC_002645.1;NP_073549.1;HCoV-229E_1ab;4439 | 0 | 0 |
| NC_002645.1;NP_073549.1;HCoV-229E_1ab;4440 | 0 | 0 |
| NC_002645.1;NP_073549.1;HCoV-229E_1ab;4441 | 0 | 0 |
| NC_002645.1;NP_073549.1;HCoV-229E_1ab;4442 | 0 | 0 |
| NC_002645.1;NP_073549.1;HCoV-229E_1ab;4443 | 0 | 0 |
| NC_002645.1;NP_073549.1;HCoV-229E_1ab;4444 | 0 | 0 |
| NC_002645.1;NP_073549.1;HCoV-229E_1ab;4445 | 0 | 0 |
| NC_002645.1;NP_073549.1;HCoV-229E_1ab;4446 | 0 | 0.025 |
| NC_002645.1;NP_073549.1;HCoV-229E_1ab;4447 | 0 | 0 |
| NC_002645.1;NP_073549.1;HCoV-229E_1ab;4448 | 0.05 | 0 |
| NC_002645.1;NP_073549.1;HCoV-229E_1ab;4449 | 0 | 0 |
| NC_002645.1;NP_073549.1;HCoV-229E_1ab;4450 | 0 | 0 |
| NC_002645.1;NP_073549.1;HCoV-229E_1ab;4451 | 0 | 0 |
| NC_002645.1;NP_073549.1;HCoV-229E_1ab;4452 | 0 | 0 |
| NC_002645.1;NP_073549.1;HCoV-229E_1ab;4453 | 0 | 0 |
| NC_002645.1;NP_073549.1;HCoV-229E_1ab;4454 | 0 | 0.075 |
| NC_002645.1;NP_073549.1;HCoV-229E_1ab;4455 | 0 | 0.05 |
| NC_002645.1;NP_073549.1;HCoV-229E_1ab;4456 | 0 | 0.05 |
| NC_002645.1;NP_073549.1;HCoV-229E_1ab;4457 | 0 | 0.05 |
| NC_002645.1;NP_073549.1;HCoV-229E_1ab;4458 | 0 | 0.05 |

| | | |
|---|---|---|
| NC_002645.1;NP_073549.1;HCoV-229E_1ab;4459 | 0 | 0.05 |
| NC_002645.1;NP_073549.1;HCoV-229E_1ab;4460 | 0 | 0.05 |
| NC_002645.1;NP_073549.1;HCoV-229E_1ab;4461 | 0 | 0.05 |
| NC_002645.1;NP_073549.1;HCoV-229E_1ab;4462 | 0 | 0.05 |
| NC_002645.1;NP_073549.1;HCoV-229E_1ab;4463 | 0.1 | 0.05 |
| NC_002645.1;NP_073549.1;HCoV-229E_1ab;4464 | 0 | 0.075 |
| NC_002645.1;NP_073549.1;HCoV-229E_1ab;4465 | 0 | 0.05 |
| NC_002645.1;NP_073549.1;HCoV-229E_1ab;4466 | 0 | 0 |
| NC_002645.1;NP_073549.1;HCoV-229E_1ab;4467 | 0.15 | 0.05 |
| NC_002645.1;NP_073549.1;HCoV-229E_1ab;4468 | 0.2 | 0.075 |
| NC_002645.1;NP_073549.1;HCoV-229E_1ab;4469 | 0.25 | 0.125 |
| NC_002645.1;NP_073549.1;HCoV-229E_1ab;4470 | 0.25 | 0.125 |
| NC_002645.1;NP_073549.1;HCoV-229E_1ab;4471 | 0.2 | 0.1 |
| NC_002645.1;NP_073549.1;HCoV-229E_1ab;4472 | 0.15 | 0.05 |
| NC_002645.1;NP_073549.1;HCoV-229E_1ab;4473 | 0.15 | 0.05 |
| NC_002645.1;NP_073549.1;HCoV-229E_1ab;4474 | 0.15 | 0.05 |
| NC_002645.1;NP_073549.1;HCoV-229E_1ab;4475 | 0.15 | 0.05 |
| NC_002645.1;NP_073549.1;HCoV-229E_1ab;4476 | 0.15 | 0.05 |
| NC_002645.1;NP_073549.1;HCoV-229E_1ab;4477 | 0 | 0.025 |
| NC_002645.1;NP_073549.1;HCoV-229E_1ab;4478 | 0.05 | 0.05 |
| NC_002645.1;NP_073549.1;HCoV-229E_1ab;4479 | 0 | 0 |
| NC_002645.1;NP_073549.1;HCoV-229E_1ab;4480 | 0 | 0 |
| NC_002645.1;NP_073549.1;HCoV-229E_1ab;4481 | 0 | 0 |
| NC_002645.1;NP_073549.1;HCoV-229E_1ab;4482 | 0 | 0 |
| NC_002645.1;NP_073549.1;HCoV-229E_1ab;4483 | 0 | 0 |
| NC_002645.1;NP_073549.1;HCoV-229E_1ab;4484 | 0 | 0 |
| NC_002645.1;NP_073549.1;HCoV-229E_1ab;4485 | 0 | 0 |
| NC_002645.1;NP_073549.1;HCoV-229E_1ab;4486 | 0.05 | 0.025 |
| NC_002645.1;NP_073549.1;HCoV-229E_1ab;4487 | 0 | 0 |
| NC_002645.1;NP_073549.1;HCoV-229E_1ab;4488 | 0.05 | 0 |
| NC_002645.1;NP_073549.1;HCoV-229E_1ab;4489 | 0.05 | 0 |
| NC_002645.1;NP_073549.1;HCoV-229E_1ab;4490 | 0.1 | 0 |
| NC_002645.1;NP_073549.1;HCoV-229E_1ab;4491 | 0 | 0 |
| NC_002645.1;NP_073549.1;HCoV-229E_1ab;4492 | 0 | 0 |
| NC_002645.1;NP_073549.1;HCoV-229E_1ab;4493 | 0 | 0 |
| NC_002645.1;NP_073549.1;HCoV-229E_1ab;4494 | 0 | 0 |
| NC_002645.1;NP_073549.1;HCoV-229E_1ab;4495 | 0 | 0 |
| NC_002645.1;NP_073549.1;HCoV-229E_1ab;4496 | 0 | 0 |
| NC_002645.1;NP_073549.1;HCoV-229E_1ab;4497 | 0.05 | 0.025 |
| NC_002645.1;NP_073549.1;HCoV-229E_1ab;4498 | 0 | 0 |
| NC_002645.1;NP_073549.1;HCoV-229E_1ab;4499 | 0 | 0 |
| NC_002645.1;NP_073549.1;HCoV-229E_1ab;4500 | 0 | 0.025 |
| NC_002645.1;NP_073549.1;HCoV-229E_1ab;4501 | 0 | 0.025 |

| | | |
|---|---|---|
| NC_002645.1;NP_073549.1;HCoV-229E_1ab;4502 | 0 | 0.025 |
| NC_002645.1;NP_073549.1;HCoV-229E_1ab;4503 | 0 | 0.05 |
| NC_002645.1;NP_073549.1;HCoV-229E_1ab;4504 | 0 | 0.05 |
| NC_002645.1;NP_073549.1;HCoV-229E_1ab;4505 | 0 | 0.05 |
| NC_002645.1;NP_073549.1;HCoV-229E_1ab;4506 | 0 | 0.05 |
| NC_002645.1;NP_073549.1;HCoV-229E_1ab;4507 | 0.05 | 0.05 |
| NC_002645.1;NP_073549.1;HCoV-229E_1ab;4508 | 0 | 0.025 |
| NC_002645.1;NP_073549.1;HCoV-229E_1ab;4509 | 0 | 0.025 |
| NC_002645.1;NP_073549.1;HCoV-229E_1ab;4510 | 0.15 | 0.025 |
| NC_002645.1;NP_073549.1;HCoV-229E_1ab;4511 | 0.05 | 0 |
| NC_002645.1;NP_073549.1;HCoV-229E_1ab;4512 | 0 | 0 |
| NC_002645.1;NP_073549.1;HCoV-229E_1ab;4513 | 0.1 | 0.025 |
| NC_002645.1;NP_073549.1;HCoV-229E_1ab;4514 | 0.1 | 0 |
| NC_002645.1;NP_073549.1;HCoV-229E_1ab;4515 | 0.05 | 0 |
| NC_002645.1;NP_073549.1;HCoV-229E_1ab;4516 | 0 | 0 |
| NC_002645.1;NP_073549.1;HCoV-229E_1ab;4517 | 0 | 0 |
| NC_002645.1;NP_073549.1;HCoV-229E_1ab;4518 | 0 | 0 |
| NC_002645.1;NP_073549.1;HCoV-229E_1ab;4519 | 0 | 0 |
| NC_002645.1;NP_073549.1;HCoV-229E_1ab;4520 | 0.05 | 0.025 |
| NC_002645.1;NP_073549.1;HCoV-229E_1ab;4521 | 0 | 0 |
| NC_002645.1;NP_073549.1;HCoV-229E_1ab;4522 | 0 | 0 |
| NC_002645.1;NP_073549.1;HCoV-229E_1ab;4523 | 0 | 0 |
| NC_002645.1;NP_073549.1;HCoV-229E_1ab;4524 | 0 | 0.025 |
| NC_002645.1;NP_073549.1;HCoV-229E_1ab;4525 | 0.05 | 0.025 |
| NC_002645.1;NP_073549.1;HCoV-229E_1ab;4526 | 0 | 0 |
| NC_002645.1;NP_073549.1;HCoV-229E_1ab;4527 | 0 | 0 |
| NC_002645.1;NP_073549.1;HCoV-229E_1ab;4528 | 0 | 0 |
| NC_002645.1;NP_073549.1;HCoV-229E_1ab;4529 | 0 | 0 |
| NC_002645.1;NP_073549.1;HCoV-229E_1ab;4530 | 0 | 0 |
| NC_002645.1;NP_073549.1;HCoV-229E_1ab;4531 | 0 | 0 |
| NC_002645.1;NP_073549.1;HCoV-229E_1ab;4532 | 0 | 0 |
| NC_002645.1;NP_073549.1;HCoV-229E_1ab;4533 | 0 | 0 |
| NC_002645.1;NP_073549.1;HCoV-229E_1ab;4534 | 0 | 0 |
| NC_002645.1;NP_073549.1;HCoV-229E_1ab;4535 | 0 | 0.025 |
| NC_002645.1;NP_073549.1;HCoV-229E_1ab;4536 | 0 | 0 |
| NC_002645.1;NP_073549.1;HCoV-229E_1ab;4537 | 0 | 0 |
| NC_002645.1;NP_073549.1;HCoV-229E_1ab;4538 | 0 | 0 |
| NC_002645.1;NP_073549.1;HCoV-229E_1ab;4539 | 0 | 0 |
| NC_002645.1;NP_073549.1;HCoV-229E_1ab;4540 | 0 | 0 |
| NC_002645.1;NP_073549.1;HCoV-229E_1ab;4541 | 0 | 0 |
| NC_002645.1;NP_073549.1;HCoV-229E_1ab;4542 | 0 | 0 |
| NC_002645.1;NP_073549.1;HCoV-229E_1ab;4543 | 0 | 0 |
| NC_002645.1;NP_073549.1;HCoV-229E_1ab;4544 | 0 | 0 |

| | | |
|---|---|---|
| NC_002645.1;NP_073549.1;HCoV-229E_1ab;4545 | 0 | 0 |
| NC_002645.1;NP_073549.1;HCoV-229E_1ab;4546 | 0 | 0.05 |
| NC_002645.1;NP_073549.1;HCoV-229E_1ab;4547 | 0.05 | 0 |
| NC_002645.1;NP_073549.1;HCoV-229E_1ab;4548 | 0.15 | 0.05 |
| NC_002645.1;NP_073549.1;HCoV-229E_1ab;4549 | 0.15 | 0.05 |
| NC_002645.1;NP_073549.1;HCoV-229E_1ab;4550 | 0.1 | 0.1 |
| NC_002645.1;NP_073549.1;HCoV-229E_1ab;4551 | 0.1 | 0.1 |
| NC_002645.1;NP_073549.1;HCoV-229E_1ab;4552 | 0.1 | 0.075 |
| NC_002645.1;NP_073549.1;HCoV-229E_1ab;4553 | 0.1 | 0.1 |
| NC_002645.1;NP_073549.1;HCoV-229E_1ab;4554 | 0.2 | 0.05 |
| NC_002645.1;NP_073549.1;HCoV-229E_1ab;4555 | 0.25 | 0.15 |
| NC_002645.1;NP_073549.1;HCoV-229E_1ab;4556 | 0.1 | 0 |
| NC_002645.1;NP_073549.1;HCoV-229E_1ab;4557 | 0.1 | 0 |
| NC_002645.1;NP_073549.1;HCoV-229E_1ab;4558 | 0.1 | 0.025 |
| NC_002645.1;NP_073549.1;HCoV-229E_1ab;4559 | 0.1 | 0 |
| NC_002645.1;NP_073549.1;HCoV-229E_1ab;4560 | 0.1 | 0 |
| NC_002645.1;NP_073549.1;HCoV-229E_1ab;4561 | 0.1 | 0.05 |
| NC_002645.1;NP_073549.1;HCoV-229E_1ab;4562 | 0.1 | 0.025 |
| NC_002645.1;NP_073549.1;HCoV-229E_1ab;4563 | 0.1 | 0.05 |
| NC_002645.1;NP_073549.1;HCoV-229E_1ab;4564 | 0 | 0.025 |
| NC_002645.1;NP_073549.1;HCoV-229E_1ab;4565 | 0.05 | 0.025 |
| NC_002645.1;NP_073549.1;HCoV-229E_1ab;4566 | 0 | 0 |
| NC_002645.1;NP_073549.1;HCoV-229E_1ab;4567 | 0 | 0 |
| NC_002645.1;NP_073549.1;HCoV-229E_1ab;4568 | 0 | 0 |
| NC_002645.1;NP_073549.1;HCoV-229E_1ab;4569 | 0 | 0 |
| NC_002645.1;NP_073549.1;HCoV-229E_1ab;4570 | 0 | 0 |
| NC_002645.1;NP_073549.1;HCoV-229E_1ab;4571 | 0 | 0 |
| NC_002645.1;NP_073549.1;HCoV-229E_1ab;4572 | 0 | 0 |
| NC_002645.1;NP_073549.1;HCoV-229E_1ab;4573 | 0 | 0.025 |
| NC_002645.1;NP_073549.1;HCoV-229E_1ab;4574 | 0 | 0 |
| NC_002645.1;NP_073549.1;HCoV-229E_1ab;4575 | 0 | 0 |
| NC_002645.1;NP_073549.1;HCoV-229E_1ab;4576 | 0 | 0 |
| NC_002645.1;NP_073549.1;HCoV-229E_1ab;4577 | 0 | 0 |
| NC_002645.1;NP_073549.1;HCoV-229E_1ab;4578 | 0 | 0 |
| NC_002645.1;NP_073549.1;HCoV-229E_1ab;4579 | 0 | 0 |
| NC_002645.1;NP_073549.1;HCoV-229E_1ab;4580 | 0.05 | 0 |
| NC_002645.1;NP_073549.1;HCoV-229E_1ab;4581 | 0 | 0 |
| NC_002645.1;NP_073549.1;HCoV-229E_1ab;4582 | 0.05 | 0 |
| NC_002645.1;NP_073549.1;HCoV-229E_1ab;4583 | 0.05 | 0.025 |
| NC_002645.1;NP_073549.1;HCoV-229E_1ab;4584 | 0.05 | 0.025 |
| NC_002645.1;NP_073549.1;HCoV-229E_1ab;4585 | 0 | 0.025 |
| NC_002645.1;NP_073549.1;HCoV-229E_1ab;4586 | 0.05 | 0.025 |
| NC_002645.1;NP_073549.1;HCoV-229E_1ab;4587 | 0 | 0 |

| | | |
|---|---|---|
| NC_002645.1;NP_073549.1;HCoV-229E_1ab;4588 | 0 | 0.025 |
| NC_002645.1;NP_073549.1;HCoV-229E_1ab;4589 | 0 | 0 |
| NC_002645.1;NP_073549.1;HCoV-229E_1ab;4590 | 0 | 0 |
| NC_002645.1;NP_073549.1;HCoV-229E_1ab;4591 | 0 | 0 |
| NC_002645.1;NP_073549.1;HCoV-229E_1ab;4592 | 0 | 0 |
| NC_002645.1;NP_073549.1;HCoV-229E_1ab;4593 | 0 | 0 |
| NC_002645.1;NP_073549.1;HCoV-229E_1ab;4594 | 0.05 | 0.025 |
| NC_002645.1;NP_073549.1;HCoV-229E_1ab;4595 | 0.15 | 0.15 |
| NC_002645.1;NP_073549.1;HCoV-229E_1ab;4596 | 0 | 0 |
| NC_002645.1;NP_073549.1;HCoV-229E_1ab;4597 | 0 | 0 |
| NC_002645.1;NP_073549.1;HCoV-229E_1ab;4598 | 0 | 0.025 |
| NC_002645.1;NP_073549.1;HCoV-229E_1ab;4599 | 0 | 0 |
| NC_002645.1;NP_073549.1;HCoV-229E_1ab;4600 | 0.05 | 0.05 |
| NC_002645.1;NP_073549.1;HCoV-229E_1ab;4601 | 0.05 | 0.025 |
| NC_002645.1;NP_073549.1;HCoV-229E_1ab;4602 | 0.05 | 0.025 |
| NC_002645.1;NP_073549.1;HCoV-229E_1ab;4603 | 0 | 0 |
| NC_002645.1;NP_073549.1;HCoV-229E_1ab;4604 | 0 | 0 |
| NC_002645.1;NP_073549.1;HCoV-229E_1ab;4605 | 0 | 0 |
| NC_002645.1;NP_073549.1;HCoV-229E_1ab;4606 | 0 | 0 |
| NC_002645.1;NP_073549.1;HCoV-229E_1ab;4607 | 0 | 0 |
| NC_002645.1;NP_073549.1;HCoV-229E_1ab;4608 | 0 | 0 |
| NC_002645.1;NP_073549.1;HCoV-229E_1ab;4609 | 0 | 0.025 |
| NC_002645.1;NP_073549.1;HCoV-229E_1ab;4610 | 0 | 0 |
| NC_002645.1;NP_073549.1;HCoV-229E_1ab;4611 | 0.1 | 0 |
| NC_002645.1;NP_073549.1;HCoV-229E_1ab;4612 | 0 | 0 |
| NC_002645.1;NP_073549.1;HCoV-229E_1ab;4613 | 0 | 0 |
| NC_002645.1;NP_073549.1;HCoV-229E_1ab;4614 | 0 | 0 |
| NC_002645.1;NP_073549.1;HCoV-229E_1ab;4615 | 0 | 0 |
| NC_002645.1;NP_073549.1;HCoV-229E_1ab;4616 | 0 | 0 |
| NC_002645.1;NP_073549.1;HCoV-229E_1ab;4617 | 0 | 0 |
| NC_002645.1;NP_073549.1;HCoV-229E_1ab;4618 | 0 | 0 |
| NC_002645.1;NP_073549.1;HCoV-229E_1ab;4619 | 0 | 0 |
| NC_002645.1;NP_073549.1;HCoV-229E_1ab;4620 | 0 | 0 |
| NC_002645.1;NP_073549.1;HCoV-229E_1ab;4621 | 0 | 0 |
| NC_002645.1;NP_073549.1;HCoV-229E_1ab;4622 | 0 | 0 |
| NC_002645.1;NP_073549.1;HCoV-229E_1ab;4623 | 0 | 0 |
| NC_002645.1;NP_073549.1;HCoV-229E_1ab;4624 | 0 | 0 |
| NC_002645.1;NP_073549.1;HCoV-229E_1ab;4625 | 0 | 0 |
| NC_002645.1;NP_073549.1;HCoV-229E_1ab;4626 | 0 | 0 |
| NC_002645.1;NP_073549.1;HCoV-229E_1ab;4627 | 0 | 0 |
| NC_002645.1;NP_073549.1;HCoV-229E_1ab;4628 | 0 | 0 |
| NC_002645.1;NP_073549.1;HCoV-229E_1ab;4629 | 0 | 0 |
| NC_002645.1;NP_073549.1;HCoV-229E_1ab;4630 | 0 | 0 |

| | | |
|---|---|---|
| NC_002645.1;NP_073549.1;HCoV-229E_1ab;4631 | 0 | 0 |
| NC_002645.1;NP_073549.1;HCoV-229E_1ab;4632 | 0 | 0 |
| NC_002645.1;NP_073549.1;HCoV-229E_1ab;4633 | 0 | 0 |
| NC_002645.1;NP_073549.1;HCoV-229E_1ab;4634 | 0 | 0 |
| NC_002645.1;NP_073549.1;HCoV-229E_1ab;4635 | 0 | 0 |
| NC_002645.1;NP_073549.1;HCoV-229E_1ab;4636 | 0 | 0 |
| NC_002645.1;NP_073549.1;HCoV-229E_1ab;4637 | 0 | 0 |
| NC_002645.1;NP_073549.1;HCoV-229E_1ab;4638 | 0 | 0 |
| NC_002645.1;NP_073549.1;HCoV-229E_1ab;4639 | 0 | 0 |
| NC_002645.1;NP_073549.1;HCoV-229E_1ab;4640 | 0 | 0 |
| NC_002645.1;NP_073549.1;HCoV-229E_1ab;4641 | 0 | 0 |
| NC_002645.1;NP_073549.1;HCoV-229E_1ab;4642 | 0 | 0 |
| NC_002645.1;NP_073549.1;HCoV-229E_1ab;4643 | 0 | 0.025 |
| NC_002645.1;NP_073549.1;HCoV-229E_1ab;4644 | 0 | 0 |
| NC_002645.1;NP_073549.1;HCoV-229E_1ab;4645 | 0 | 0 |
| NC_002645.1;NP_073549.1;HCoV-229E_1ab;4646 | 0 | 0 |
| NC_002645.1;NP_073549.1;HCoV-229E_1ab;4647 | 0 | 0 |
| NC_002645.1;NP_073549.1;HCoV-229E_1ab;4648 | 0 | 0 |
| NC_002645.1;NP_073549.1;HCoV-229E_1ab;4649 | 0 | 0 |
| NC_002645.1;NP_073549.1;HCoV-229E_1ab;4650 | 0 | 0 |
| NC_002645.1;NP_073549.1;HCoV-229E_1ab;4651 | 0 | 0 |
| NC_002645.1;NP_073549.1;HCoV-229E_1ab;4652 | 0.05 | 0 |
| NC_002645.1;NP_073549.1;HCoV-229E_1ab;4653 | 0 | 0 |
| NC_002645.1;NP_073549.1;HCoV-229E_1ab;4654 | 0 | 0.025 |
| NC_002645.1;NP_073549.1;HCoV-229E_1ab;4655 | 0 | 0.025 |
| NC_002645.1;NP_073549.1;HCoV-229E_1ab;4656 | 0.05 | 0.05 |
| NC_002645.1;NP_073549.1;HCoV-229E_1ab;4657 | 0.15 | 0.05 |
| NC_002645.1;NP_073549.1;HCoV-229E_1ab;4658 | 0.05 | 0.025 |
| NC_002645.1;NP_073549.1;HCoV-229E_1ab;4659 | 0 | 0.025 |
| NC_002645.1;NP_073549.1;HCoV-229E_1ab;4660 | 0 | 0.025 |
| NC_002645.1;NP_073549.1;HCoV-229E_1ab;4661 | 0 | 0 |
| NC_002645.1;NP_073549.1;HCoV-229E_1ab;4662 | 0 | 0 |
| NC_002645.1;NP_073549.1;HCoV-229E_1ab;4663 | 0.05 | 0 |
| NC_002645.1;NP_073549.1;HCoV-229E_1ab;4664 | 0.05 | 0 |
| NC_002645.1;NP_073549.1;HCoV-229E_1ab;4665 | 0.05 | 0 |
| NC_002645.1;NP_073549.1;HCoV-229E_1ab;4666 | 0.05 | 0 |
| NC_002645.1;NP_073549.1;HCoV-229E_1ab;4667 | 0.05 | 0 |
| NC_002645.1;NP_073549.1;HCoV-229E_1ab;4668 | 0.1 | 0.025 |
| NC_002645.1;NP_073549.1;HCoV-229E_1ab;4669 | 0.1 | 0.025 |
| NC_002645.1;NP_073549.1;HCoV-229E_1ab;4670 | 0.15 | 0.05 |
| NC_002645.1;NP_073549.1;HCoV-229E_1ab;4671 | 0.05 | 0.025 |
| NC_002645.1;NP_073549.1;HCoV-229E_1ab;4672 | 0.05 | 0.05 |
| NC_002645.1;NP_073549.1;HCoV-229E_1ab;4673 | 0.05 | 0.025 |

| | | |
|---|---|---|
| NC_002645.1;NP_073549.1;HCoV-229E_1ab;4674 | 0.05 | 0.025 |
| NC_002645.1;NP_073549.1;HCoV-229E_1ab;4675 | 0 | 0.05 |
| NC_002645.1;NP_073549.1;HCoV-229E_1ab;4676 | 0 | 0.075 |
| NC_002645.1;NP_073549.1;HCoV-229E_1ab;4677 | 0 | 0.025 |
| NC_002645.1;NP_073549.1;HCoV-229E_1ab;4678 | 0 | 0.025 |
| NC_002645.1;NP_073549.1;HCoV-229E_1ab;4679 | 0 | 0.025 |
| NC_002645.1;NP_073549.1;HCoV-229E_1ab;4680 | 0 | 0 |
| NC_002645.1;NP_073549.1;HCoV-229E_1ab;4681 | 0 | 0 |
| NC_002645.1;NP_073549.1;HCoV-229E_1ab;4682 | 0 | 0 |
| NC_002645.1;NP_073549.1;HCoV-229E_1ab;4683 | 0 | 0 |
| NC_002645.1;NP_073549.1;HCoV-229E_1ab;4684 | 0 | 0 |
| NC_002645.1;NP_073549.1;HCoV-229E_1ab;4685 | 0 | 0 |
| NC_002645.1;NP_073549.1;HCoV-229E_1ab;4686 | 0 | 0 |
| NC_002645.1;NP_073549.1;HCoV-229E_1ab;4687 | 0 | 0 |
| NC_002645.1;NP_073549.1;HCoV-229E_1ab;4688 | 0 | 0 |
| NC_002645.1;NP_073549.1;HCoV-229E_1ab;4689 | 0 | 0 |
| NC_002645.1;NP_073549.1;HCoV-229E_1ab;4690 | 0 | 0 |
| NC_002645.1;NP_073549.1;HCoV-229E_1ab;4691 | 0 | 0 |
| NC_002645.1;NP_073549.1;HCoV-229E_1ab;4692 | 0 | 0 |
| NC_002645.1;NP_073549.1;HCoV-229E_1ab;4693 | 0 | 0 |
| NC_002645.1;NP_073549.1;HCoV-229E_1ab;4694 | 0 | 0 |
| NC_002645.1;NP_073549.1;HCoV-229E_1ab;4695 | 0 | 0 |
| NC_002645.1;NP_073549.1;HCoV-229E_1ab;4696 | 0 | 0 |
| NC_002645.1;NP_073549.1;HCoV-229E_1ab;4697 | 0 | 0 |
| NC_002645.1;NP_073549.1;HCoV-229E_1ab;4698 | 0 | 0 |
| NC_002645.1;NP_073549.1;HCoV-229E_1ab;4699 | 0 | 0 |
| NC_002645.1;NP_073549.1;HCoV-229E_1ab;4700 | 0 | 0 |
| NC_002645.1;NP_073549.1;HCoV-229E_1ab;4701 | 0.1 | 0 |
| NC_002645.1;NP_073549.1;HCoV-229E_1ab;4702 | 0 | 0 |
| NC_002645.1;NP_073549.1;HCoV-229E_1ab;4703 | 0 | 0 |
| NC_002645.1;NP_073549.1;HCoV-229E_1ab;4704 | 0 | 0 |
| NC_002645.1;NP_073549.1;HCoV-229E_1ab;4705 | 0 | 0 |
| NC_002645.1;NP_073549.1;HCoV-229E_1ab;4706 | 0 | 0.025 |
| NC_002645.1;NP_073549.1;HCoV-229E_1ab;4707 | 0 | 0 |
| NC_002645.1;NP_073549.1;HCoV-229E_1ab;4708 | 0 | 0.025 |
| NC_002645.1;NP_073549.1;HCoV-229E_1ab;4709 | 0 | 0.025 |
| NC_002645.1;NP_073549.1;HCoV-229E_1ab;4710 | 0 | 0.025 |
| NC_002645.1;NP_073549.1;HCoV-229E_1ab;4711 | 0.05 | 0.025 |
| NC_002645.1;NP_073549.1;HCoV-229E_1ab;4712 | 0 | 0.025 |
| NC_002645.1;NP_073549.1;HCoV-229E_1ab;4713 | 0 | 0.025 |
| NC_002645.1;NP_073549.1;HCoV-229E_1ab;4714 | 0 | 0 |
| NC_002645.1;NP_073549.1;HCoV-229E_1ab;4715 | 0 | 0.025 |
| NC_002645.1;NP_073549.1;HCoV-229E_1ab;4716 | 0 | 0 |

| | | |
|---|---|---|
| NC_002645.1;NP_073549.1;HCoV-229E_1ab;4717 | 0 | 0 |
| NC_002645.1;NP_073549.1;HCoV-229E_1ab;4718 | 0 | 0.025 |
| NC_002645.1;NP_073549.1;HCoV-229E_1ab;4719 | 0 | 0.025 |
| NC_002645.1;NP_073549.1;HCoV-229E_1ab;4720 | 0.05 | 0.025 |
| NC_002645.1;NP_073549.1;HCoV-229E_1ab;4721 | 0 | 0.025 |
| NC_002645.1;NP_073549.1;HCoV-229E_1ab;4722 | 0 | 0 |
| NC_002645.1;NP_073549.1;HCoV-229E_1ab;4723 | 0 | 0.025 |
| NC_002645.1;NP_073549.1;HCoV-229E_1ab;4724 | 0 | 0 |
| NC_002645.1;NP_073549.1;HCoV-229E_1ab;4725 | 0 | 0 |
| NC_002645.1;NP_073549.1;HCoV-229E_1ab;4726 | 0 | 0 |
| NC_002645.1;NP_073549.1;HCoV-229E_1ab;4727 | 0 | 0.025 |
| NC_002645.1;NP_073549.1;HCoV-229E_1ab;4728 | 0 | 0.025 |
| NC_002645.1;NP_073549.1;HCoV-229E_1ab;4729 | 0 | 0.025 |
| NC_002645.1;NP_073549.1;HCoV-229E_1ab;4730 | 0 | 0.05 |
| NC_002645.1;NP_073549.1;HCoV-229E_1ab;4731 | 0.05 | 0 |
| NC_002645.1;NP_073549.1;HCoV-229E_1ab;4732 | 0.05 | 0 |
| NC_002645.1;NP_073549.1;HCoV-229E_1ab;4733 | 0 | 0 |
| NC_002645.1;NP_073549.1;HCoV-229E_1ab;4734 | 0 | 0 |
| NC_002645.1;NP_073549.1;HCoV-229E_1ab;4735 | 0 | 0 |
| NC_002645.1;NP_073549.1;HCoV-229E_1ab;4736 | 0 | 0 |
| NC_002645.1;NP_073549.1;HCoV-229E_1ab;4737 | 0.05 | 0 |
| NC_002645.1;NP_073549.1;HCoV-229E_1ab;4738 | 0.05 | 0 |
| NC_002645.1;NP_073549.1;HCoV-229E_1ab;4739 | 0.05 | 0 |
| NC_002645.1;NP_073549.1;HCoV-229E_1ab;4740 | 0.05 | 0 |
| NC_002645.1;NP_073549.1;HCoV-229E_1ab;4741 | 0.05 | 0.025 |
| NC_002645.1;NP_073549.1;HCoV-229E_1ab;4742 | 0.05 | 0 |
| NC_002645.1;NP_073549.1;HCoV-229E_1ab;4743 | 0.05 | 0 |
| NC_002645.1;NP_073549.1;HCoV-229E_1ab;4744 | 0 | 0 |
| NC_002645.1;NP_073549.1;HCoV-229E_1ab;4745 | 0 | 0 |
| NC_002645.1;NP_073549.1;HCoV-229E_1ab;4746 | 0 | 0 |
| NC_002645.1;NP_073549.1;HCoV-229E_1ab;4747 | 0 | 0 |
| NC_002645.1;NP_073549.1;HCoV-229E_1ab;4748 | 0 | 0 |
| NC_002645.1;NP_073549.1;HCoV-229E_1ab;4749 | 0 | 0 |
| NC_002645.1;NP_073549.1;HCoV-229E_1ab;4750 | 0 | 0 |
| NC_002645.1;NP_073549.1;HCoV-229E_1ab;4751 | 0 | 0 |
| NC_002645.1;NP_073549.1;HCoV-229E_1ab;4752 | 0 | 0 |
| NC_002645.1;NP_073549.1;HCoV-229E_1ab;4753 | 0 | 0 |
| NC_002645.1;NP_073549.1;HCoV-229E_1ab;4754 | 0 | 0 |
| NC_002645.1;NP_073549.1;HCoV-229E_1ab;4755 | 0 | 0 |
| NC_002645.1;NP_073549.1;HCoV-229E_1ab;4756 | 0 | 0 |
| NC_002645.1;NP_073549.1;HCoV-229E_1ab;4757 | 0 | 0 |
| NC_002645.1;NP_073549.1;HCoV-229E_1ab;4758 | 0 | 0 |
| NC_002645.1;NP_073549.1;HCoV-229E_1ab;4759 | 0 | 0 |

| | | |
|---|---|---|
| NC_002645.1;NP_073549.1;HCoV-229E_1ab;4760 | 0 | 0 |
| NC_002645.1;NP_073549.1;HCoV-229E_1ab;4761 | 0 | 0 |
| NC_002645.1;NP_073549.1;HCoV-229E_1ab;4762 | 0.15 | 0.2 |
| NC_002645.1;NP_073549.1;HCoV-229E_1ab;4763 | 0 | 0.025 |
| NC_002645.1;NP_073549.1;HCoV-229E_1ab;4764 | 0 | 0.025 |
| NC_002645.1;NP_073549.1;HCoV-229E_1ab;4765 | 0 | 0.05 |
| NC_002645.1;NP_073549.1;HCoV-229E_1ab;4766 | 0 | 0 |
| NC_002645.1;NP_073549.1;HCoV-229E_1ab;4767 | 0.2 | 0.025 |
| NC_002645.1;NP_073549.1;HCoV-229E_1ab;4768 | 0.05 | 0.05 |
| NC_002645.1;NP_073549.1;HCoV-229E_1ab;4769 | 0.15 | 0.05 |
| NC_002645.1;NP_073549.1;HCoV-229E_1ab;4770 | 0.2 | 0.1 |
| NC_002645.1;NP_073549.1;HCoV-229E_1ab;4771 | 0.05 | 0.025 |
| NC_002645.1;NP_073549.1;HCoV-229E_1ab;4772 | 0.15 | 0 |
| NC_002645.1;NP_073549.1;HCoV-229E_1ab;4773 | 0.05 | 0 |
| NC_002645.1;NP_073549.1;HCoV-229E_1ab;4774 | 0.05 | 0 |
| NC_002645.1;NP_073549.1;HCoV-229E_1ab;4775 | 0 | 0 |
| NC_002645.1;NP_073549.1;HCoV-229E_1ab;4776 | 0 | 0 |
| NC_002645.1;NP_073549.1;HCoV-229E_1ab;4777 | 0 | 0 |
| NC_002645.1;NP_073549.1;HCoV-229E_1ab;4778 | 0.05 | 0 |
| NC_002645.1;NP_073549.1;HCoV-229E_1ab;4779 | 0.05 | 0 |
| NC_002645.1;NP_073549.1;HCoV-229E_1ab;4780 | 0 | 0 |
| NC_002645.1;NP_073549.1;HCoV-229E_1ab;4781 | 0 | 0 |
| NC_002645.1;NP_073549.1;HCoV-229E_1ab;4782 | 0 | 0 |
| NC_002645.1;NP_073549.1;HCoV-229E_1ab;4783 | 0 | 0 |
| NC_002645.1;NP_073549.1;HCoV-229E_1ab;4784 | 0 | 0.05 |
| NC_002645.1;NP_073549.1;HCoV-229E_1ab;4785 | 0 | 0 |
| NC_002645.1;NP_073549.1;HCoV-229E_1ab;4786 | 0 | 0 |
| NC_002645.1;NP_073549.1;HCoV-229E_1ab;4787 | 0 | 0 |
| NC_002645.1;NP_073549.1;HCoV-229E_1ab;4788 | 0 | 0.025 |
| NC_002645.1;NP_073549.1;HCoV-229E_1ab;4789 | 0 | 0 |
| NC_002645.1;NP_073549.1;HCoV-229E_1ab;4790 | 0 | 0 |
| NC_002645.1;NP_073549.1;HCoV-229E_1ab;4791 | 0 | 0 |
| NC_002645.1;NP_073549.1;HCoV-229E_1ab;4792 | 0 | 0 |
| NC_002645.1;NP_073549.1;HCoV-229E_1ab;4793 | 0 | 0 |
| NC_002645.1;NP_073549.1;HCoV-229E_1ab;4794 | 0 | 0 |
| NC_002645.1;NP_073549.1;HCoV-229E_1ab;4795 | 0.05 | 0 |
| NC_002645.1;NP_073549.1;HCoV-229E_1ab;4796 | 0.05 | 0 |
| NC_002645.1;NP_073549.1;HCoV-229E_1ab;4797 | 0 | 0 |
| NC_002645.1;NP_073549.1;HCoV-229E_1ab;4798 | 0.05 | 0.025 |
| NC_002645.1;NP_073549.1;HCoV-229E_1ab;4799 | 0.05 | 0.025 |
| NC_002645.1;NP_073549.1;HCoV-229E_1ab;4800 | 0.05 | 0 |
| NC_002645.1;NP_073549.1;HCoV-229E_1ab;4801 | 0.05 | 0 |
| NC_002645.1;NP_073549.1;HCoV-229E_1ab;4802 | 0.1 | 0.025 |

| | | |
|---|---|---|
| NC_002645.1;NP_073549.1;HCoV-229E_1ab;4803 | 0.1 | 0.025 |
| NC_002645.1;NP_073549.1;HCoV-229E_1ab;4804 | 0 | 0 |
| NC_002645.1;NP_073549.1;HCoV-229E_1ab;4805 | 0 | 0 |
| NC_002645.1;NP_073549.1;HCoV-229E_1ab;4806 | 0 | 0 |
| NC_002645.1;NP_073549.1;HCoV-229E_1ab;4807 | 0 | 0 |
| NC_002645.1;NP_073549.1;HCoV-229E_1ab;4808 | 0 | 0 |
| NC_002645.1;NP_073549.1;HCoV-229E_1ab;4809 | 0 | 0 |
| NC_002645.1;NP_073549.1;HCoV-229E_1ab;4810 | 0 | 0 |
| NC_002645.1;NP_073549.1;HCoV-229E_1ab;4811 | 0 | 0 |
| NC_002645.1;NP_073549.1;HCoV-229E_1ab;4812 | 0 | 0 |
| NC_002645.1;NP_073549.1;HCoV-229E_1ab;4813 | 0 | 0 |
| NC_002645.1;NP_073549.1;HCoV-229E_1ab;4814 | 0 | 0 |
| NC_002645.1;NP_073549.1;HCoV-229E_1ab;4815 | 0 | 0 |
| NC_002645.1;NP_073549.1;HCoV-229E_1ab;4816 | 0 | 0 |
| NC_002645.1;NP_073549.1;HCoV-229E_1ab;4817 | 0 | 0 |
| NC_002645.1;NP_073549.1;HCoV-229E_1ab;4818 | 0.05 | 0 |
| NC_002645.1;NP_073549.1;HCoV-229E_1ab;4819 | 0.05 | 0 |
| NC_002645.1;NP_073549.1;HCoV-229E_1ab;4820 | 0.05 | 0.025 |
| NC_002645.1;NP_073549.1;HCoV-229E_1ab;4821 | 0.05 | 0 |
| NC_002645.1;NP_073549.1;HCoV-229E_1ab;4822 | 0.05 | 0 |
| NC_002645.1;NP_073549.1;HCoV-229E_1ab;4823 | 0.05 | 0 |
| NC_002645.1;NP_073549.1;HCoV-229E_1ab;4824 | 0 | 0 |
| NC_002645.1;NP_073549.1;HCoV-229E_1ab;4825 | 0 | 0 |
| NC_002645.1;NP_073549.1;HCoV-229E_1ab;4826 | 0 | 0 |
| NC_002645.1;NP_073549.1;HCoV-229E_1ab;4827 | 0 | 0 |
| NC_002645.1;NP_073549.1;HCoV-229E_1ab;4828 | 0 | 0 |
| NC_002645.1;NP_073549.1;HCoV-229E_1ab;4829 | 0.05 | 0 |
| NC_002645.1;NP_073549.1;HCoV-229E_1ab;4830 | 0 | 0 |
| NC_002645.1;NP_073549.1;HCoV-229E_1ab;4831 | 0 | 0 |
| NC_002645.1;NP_073549.1;HCoV-229E_1ab;4832 | 0.05 | 0 |
| NC_002645.1;NP_073549.1;HCoV-229E_1ab;4833 | 0 | 0 |
| NC_002645.1;NP_073549.1;HCoV-229E_1ab;4834 | 0 | 0 |
| NC_002645.1;NP_073549.1;HCoV-229E_1ab;4835 | 0 | 0.025 |
| NC_002645.1;NP_073549.1;HCoV-229E_1ab;4836 | 0 | 0 |
| NC_002645.1;NP_073549.1;HCoV-229E_1ab;4837 | 0 | 0 |
| NC_002645.1;NP_073549.1;HCoV-229E_1ab;4838 | 0 | 0 |
| NC_002645.1;NP_073549.1;HCoV-229E_1ab;4839 | 0 | 0 |
| NC_002645.1;NP_073549.1;HCoV-229E_1ab;4840 | 0 | 0 |
| NC_002645.1;NP_073549.1;HCoV-229E_1ab;4841 | 0 | 0 |
| NC_002645.1;NP_073549.1;HCoV-229E_1ab;4842 | 0 | 0 |
| NC_002645.1;NP_073549.1;HCoV-229E_1ab;4843 | 0 | 0 |
| NC_002645.1;NP_073549.1;HCoV-229E_1ab;4844 | 0 | 0 |
| NC_002645.1;NP_073549.1;HCoV-229E_1ab;4845 | 0 | 0 |

| | | |
|---|---|---|
| NC_002645.1;NP_073549.1;HCoV-229E_1ab;4846 | 0 | 0 |
| NC_002645.1;NP_073549.1;HCoV-229E_1ab;4847 | 0 | 0.025 |
| NC_002645.1;NP_073549.1;HCoV-229E_1ab;4848 | 0 | 0 |
| NC_002645.1;NP_073549.1;HCoV-229E_1ab;4849 | 0 | 0 |
| NC_002645.1;NP_073549.1;HCoV-229E_1ab;4850 | 0 | 0.025 |
| NC_002645.1;NP_073549.1;HCoV-229E_1ab;4851 | 0 | 0.025 |
| NC_002645.1;NP_073549.1;HCoV-229E_1ab;4852 | 0 | 0.025 |
| NC_002645.1;NP_073549.1;HCoV-229E_1ab;4853 | 0 | 0.025 |
| NC_002645.1;NP_073549.1;HCoV-229E_1ab;4854 | 0 | 0.025 |
| NC_002645.1;NP_073549.1;HCoV-229E_1ab;4855 | 0 | 0 |
| NC_002645.1;NP_073549.1;HCoV-229E_1ab;4856 | 0 | 0 |
| NC_002645.1;NP_073549.1;HCoV-229E_1ab;4857 | 0 | 0 |
| NC_002645.1;NP_073549.1;HCoV-229E_1ab;4858 | 0 | 0 |
| NC_002645.1;NP_073549.1;HCoV-229E_1ab;4859 | 0 | 0 |
| NC_002645.1;NP_073549.1;HCoV-229E_1ab;4860 | 0 | 0.025 |
| NC_002645.1;NP_073549.1;HCoV-229E_1ab;4861 | 0 | 0.025 |
| NC_002645.1;NP_073549.1;HCoV-229E_1ab;4862 | 0 | 0 |
| NC_002645.1;NP_073549.1;HCoV-229E_1ab;4863 | 0 | 0 |
| NC_002645.1;NP_073549.1;HCoV-229E_1ab;4864 | 0 | 0 |
| NC_002645.1;NP_073549.1;HCoV-229E_1ab;4865 | 0 | 0 |
| NC_002645.1;NP_073549.1;HCoV-229E_1ab;4866 | 0.05 | 0 |
| NC_002645.1;NP_073549.1;HCoV-229E_1ab;4867 | 0 | 0 |
| NC_002645.1;NP_073549.1;HCoV-229E_1ab;4868 | 0 | 0 |
| NC_002645.1;NP_073549.1;HCoV-229E_1ab;4869 | 0 | 0 |
| NC_002645.1;NP_073549.1;HCoV-229E_1ab;4870 | 0 | 0 |
| NC_002645.1;NP_073549.1;HCoV-229E_1ab;4871 | 0 | 0 |
| NC_002645.1;NP_073549.1;HCoV-229E_1ab;4872 | 0 | 0 |
| NC_002645.1;NP_073549.1;HCoV-229E_1ab;4873 | 0 | 0 |
| NC_002645.1;NP_073549.1;HCoV-229E_1ab;4874 | 0 | 0.05 |
| NC_002645.1;NP_073549.1;HCoV-229E_1ab;4875 | 0 | 0.175 |
| NC_002645.1;NP_073549.1;HCoV-229E_1ab;4876 | 0.05 | 0.175 |
| NC_002645.1;NP_073549.1;HCoV-229E_1ab;4877 | 0 | 0.125 |
| NC_002645.1;NP_073549.1;HCoV-229E_1ab;4878 | 0.05 | 0.125 |
| NC_002645.1;NP_073549.1;HCoV-229E_1ab;4879 | 0 | 0.125 |
| NC_002645.1;NP_073549.1;HCoV-229E_1ab;4880 | 0 | 0.175 |
| NC_002645.1;NP_073549.1;HCoV-229E_1ab;4881 | 0.05 | 0.175 |
| NC_002645.1;NP_073549.1;HCoV-229E_1ab;4882 | 0 | 0.125 |
| NC_002645.1;NP_073549.1;HCoV-229E_1ab;4883 | 0.05 | 0.225 |
| NC_002645.1;NP_073549.1;HCoV-229E_1ab;4884 | 0 | 0.025 |
| NC_002645.1;NP_073549.1;HCoV-229E_1ab;4885 | 0 | 0 |
| NC_002645.1;NP_073549.1;HCoV-229E_1ab;4886 | 0 | 0 |
| NC_002645.1;NP_073549.1;HCoV-229E_1ab;4887 | 0 | 0 |
| NC_002645.1;NP_073549.1;HCoV-229E_1ab;4888 | 0 | 0 |

| | | |
|---|---|---|
| NC_002645.1;NP_073549.1;HCoV-229E_1ab;4889 | 0 | 0 |
| NC_002645.1;NP_073549.1;HCoV-229E_1ab;4890 | 0 | 0 |
| NC_002645.1;NP_073549.1;HCoV-229E_1ab;4891 | 0 | 0 |
| NC_002645.1;NP_073549.1;HCoV-229E_1ab;4892 | 0 | 0 |
| NC_002645.1;NP_073549.1;HCoV-229E_1ab;4893 | 0 | 0 |
| NC_002645.1;NP_073549.1;HCoV-229E_1ab;4894 | 0 | 0 |
| NC_002645.1;NP_073549.1;HCoV-229E_1ab;4895 | 0 | 0 |
| NC_002645.1;NP_073549.1;HCoV-229E_1ab;4896 | 0 | 0 |
| NC_002645.1;NP_073549.1;HCoV-229E_1ab;4897 | 0.05 | 0 |
| NC_002645.1;NP_073549.1;HCoV-229E_1ab;4898 | 0.1 | 0 |
| NC_002645.1;NP_073549.1;HCoV-229E_1ab;4899 | 0.1 | 0 |
| NC_002645.1;NP_073549.1;HCoV-229E_1ab;4900 | 0.15 | 0 |
| NC_002645.1;NP_073549.1;HCoV-229E_1ab;4901 | 0.1 | 0 |
| NC_002645.1;NP_073549.1;HCoV-229E_1ab;4902 | 0.05 | 0 |
| NC_002645.1;NP_073549.1;HCoV-229E_1ab;4903 | 0.05 | 0 |
| NC_002645.1;NP_073549.1;HCoV-229E_1ab;4904 | 0 | 0 |
| NC_002645.1;NP_073549.1;HCoV-229E_1ab;4905 | 0 | 0.025 |
| NC_002645.1;NP_073549.1;HCoV-229E_1ab;4906 | 0 | 0 |
| NC_002645.1;NP_073549.1;HCoV-229E_1ab;4907 | 0 | 0 |
| NC_002645.1;NP_073549.1;HCoV-229E_1ab;4908 | 0.05 | 0.025 |
| NC_002645.1;NP_073549.1;HCoV-229E_1ab;4909 | 0 | 0.025 |
| NC_002645.1;NP_073549.1;HCoV-229E_1ab;4910 | 0 | 0.025 |
| NC_002645.1;NP_073549.1;HCoV-229E_1ab;4911 | 0 | 0.025 |
| NC_002645.1;NP_073549.1;HCoV-229E_1ab;4912 | 0 | 0.025 |
| NC_002645.1;NP_073549.1;HCoV-229E_1ab;4913 | 0 | 0.025 |
| NC_002645.1;NP_073549.1;HCoV-229E_1ab;4914 | 0 | 0.025 |
| NC_002645.1;NP_073549.1;HCoV-229E_1ab;4915 | 0 | 0 |
| NC_002645.1;NP_073549.1;HCoV-229E_1ab;4916 | 0 | 0.025 |
| NC_002645.1;NP_073549.1;HCoV-229E_1ab;4917 | 0 | 0.025 |
| NC_002645.1;NP_073549.1;HCoV-229E_1ab;4918 | 0 | 0 |
| NC_002645.1;NP_073549.1;HCoV-229E_1ab;4919 | 0 | 0 |
| NC_002645.1;NP_073549.1;HCoV-229E_1ab;4920 | 0 | 0 |
| NC_002645.1;NP_073549.1;HCoV-229E_1ab;4921 | 0 | 0 |
| NC_002645.1;NP_073549.1;HCoV-229E_1ab;4922 | 0 | 0 |
| NC_002645.1;NP_073549.1;HCoV-229E_1ab;4923 | 0 | 0 |
| NC_002645.1;NP_073549.1;HCoV-229E_1ab;4924 | 0 | 0 |
| NC_002645.1;NP_073549.1;HCoV-229E_1ab;4925 | 0 | 0.025 |
| NC_002645.1;NP_073549.1;HCoV-229E_1ab;4926 | 0 | 0.05 |
| NC_002645.1;NP_073549.1;HCoV-229E_1ab;4927 | 0 | 0.025 |
| NC_002645.1;NP_073549.1;HCoV-229E_1ab;4928 | 0 | 0.025 |
| NC_002645.1;NP_073549.1;HCoV-229E_1ab;4929 | 0 | 0 |
| NC_002645.1;NP_073549.1;HCoV-229E_1ab;4930 | 0 | 0 |
| NC_002645.1;NP_073549.1;HCoV-229E_1ab;4931 | 0 | 0 |

| | | |
|---|---|---|
| NC_002645.1;NP_073549.1;HCoV-229E_1ab;4932 | 0 | 0 |
| NC_002645.1;NP_073549.1;HCoV-229E_1ab;4933 | 0 | 0 |
| NC_002645.1;NP_073549.1;HCoV-229E_1ab;4934 | 0 | 0 |
| NC_002645.1;NP_073549.1;HCoV-229E_1ab;4935 | 0 | 0 |
| NC_002645.1;NP_073549.1;HCoV-229E_1ab;4936 | 0 | 0 |
| NC_002645.1;NP_073549.1;HCoV-229E_1ab;4937 | 0 | 0 |
| NC_002645.1;NP_073549.1;HCoV-229E_1ab;4938 | 0 | 0 |
| NC_002645.1;NP_073549.1;HCoV-229E_1ab;4939 | 0 | 0 |
| NC_002645.1;NP_073549.1;HCoV-229E_1ab;4940 | 0 | 0 |
| NC_002645.1;NP_073549.1;HCoV-229E_1ab;4941 | 0 | 0 |
| NC_002645.1;NP_073549.1;HCoV-229E_1ab;4942 | 0 | 0 |
| NC_002645.1;NP_073549.1;HCoV-229E_1ab;4943 | 0 | 0 |
| NC_002645.1;NP_073549.1;HCoV-229E_1ab;4944 | 0.1 | 0 |
| NC_002645.1;NP_073549.1;HCoV-229E_1ab;4945 | 0.15 | 0.125 |
| NC_002645.1;NP_073549.1;HCoV-229E_1ab;4946 | 0.2 | 0.175 |
| NC_002645.1;NP_073549.1;HCoV-229E_1ab;4947 | 0.15 | 0.15 |
| NC_002645.1;NP_073549.1;HCoV-229E_1ab;4948 | 0.25 | 0.225 |
| NC_002645.1;NP_073549.1;HCoV-229E_1ab;4949 | 0.2 | 0.225 |
| NC_002645.1;NP_073549.1;HCoV-229E_1ab;4950 | 0.2 | 0.2 |
| NC_002645.1;NP_073549.1;HCoV-229E_1ab;4951 | 0.15 | 0.1 |
| NC_002645.1;NP_073549.1;HCoV-229E_1ab;4952 | 0.1 | 0.025 |
| NC_002645.1;NP_073549.1;HCoV-229E_1ab;4953 | 0.1 | 0.075 |
| NC_002645.1;NP_073549.1;HCoV-229E_1ab;4954 | 0.1 | 0.075 |
| NC_002645.1;NP_073549.1;HCoV-229E_1ab;4955 | 0 | 0 |
| NC_002645.1;NP_073549.1;HCoV-229E_1ab;4956 | 0 | 0 |
| NC_002645.1;NP_073549.1;HCoV-229E_1ab;4957 | 0 | 0 |
| NC_002645.1;NP_073549.1;HCoV-229E_1ab;4958 | 0 | 0 |
| NC_002645.1;NP_073549.1;HCoV-229E_1ab;4959 | 0 | 0 |
| NC_002645.1;NP_073549.1;HCoV-229E_1ab;4960 | 0 | 0 |
| NC_002645.1;NP_073549.1;HCoV-229E_1ab;4961 | 0 | 0 |
| NC_002645.1;NP_073549.1;HCoV-229E_1ab;4962 | 0 | 0 |
| NC_002645.1;NP_073549.1;HCoV-229E_1ab;4963 | 0 | 0 |
| NC_002645.1;NP_073549.1;HCoV-229E_1ab;4964 | 0 | 0 |
| NC_002645.1;NP_073549.1;HCoV-229E_1ab;4965 | 0 | 0 |
| NC_002645.1;NP_073549.1;HCoV-229E_1ab;4966 | 0 | 0 |
| NC_002645.1;NP_073549.1;HCoV-229E_1ab;4967 | 0.05 | 0 |
| NC_002645.1;NP_073549.1;HCoV-229E_1ab;4968 | 0 | 0.025 |
| NC_002645.1;NP_073549.1;HCoV-229E_1ab;4969 | 0.05 | 0 |
| NC_002645.1;NP_073549.1;HCoV-229E_1ab;4970 | 0 | 0 |
| NC_002645.1;NP_073549.1;HCoV-229E_1ab;4971 | 0 | 0 |
| NC_002645.1;NP_073549.1;HCoV-229E_1ab;4972 | 0 | 0 |
| NC_002645.1;NP_073549.1;HCoV-229E_1ab;4973 | 0 | 0.025 |
| NC_002645.1;NP_073549.1;HCoV-229E_1ab;4974 | 0 | 0 |

| | | |
|---|---|---|
| NC_002645.1;NP_073549.1;HCoV-229E_1ab;4975 | 0.05 | 0.025 |
| NC_002645.1;NP_073549.1;HCoV-229E_1ab;4976 | 0.05 | 0 |
| NC_002645.1;NP_073549.1;HCoV-229E_1ab;4977 | 0 | 0 |
| NC_002645.1;NP_073549.1;HCoV-229E_1ab;4978 | 0 | 0 |
| NC_002645.1;NP_073549.1;HCoV-229E_1ab;4979 | 0 | 0 |
| NC_002645.1;NP_073549.1;HCoV-229E_1ab;4980 | 0.05 | 0 |
| NC_002645.1;NP_073549.1;HCoV-229E_1ab;4981 | 0 | 0 |
| NC_002645.1;NP_073549.1;HCoV-229E_1ab;4982 | 0.05 | 0.025 |
| NC_002645.1;NP_073549.1;HCoV-229E_1ab;4983 | 0 | 0 |
| NC_002645.1;NP_073549.1;HCoV-229E_1ab;4984 | 0 | 0 |
| NC_002645.1;NP_073549.1;HCoV-229E_1ab;4985 | 0 | 0 |
| NC_002645.1;NP_073549.1;HCoV-229E_1ab;4986 | 0 | 0 |
| NC_002645.1;NP_073549.1;HCoV-229E_1ab;4987 | 0 | 0 |
| NC_002645.1;NP_073549.1;HCoV-229E_1ab;4988 | 0 | 0 |
| NC_002645.1;NP_073549.1;HCoV-229E_1ab;4989 | 0 | 0 |
| NC_002645.1;NP_073549.1;HCoV-229E_1ab;4990 | 0 | 0 |
| NC_002645.1;NP_073549.1;HCoV-229E_1ab;4991 | 0 | 0 |
| NC_002645.1;NP_073549.1;HCoV-229E_1ab;4992 | 0 | 0 |
| NC_002645.1;NP_073549.1;HCoV-229E_1ab;4993 | 0 | 0 |
| NC_002645.1;NP_073549.1;HCoV-229E_1ab;4994 | 0 | 0 |
| NC_002645.1;NP_073549.1;HCoV-229E_1ab;4995 | 0 | 0 |
| NC_002645.1;NP_073549.1;HCoV-229E_1ab;4996 | 0 | 0 |
| NC_002645.1;NP_073549.1;HCoV-229E_1ab;4997 | 0 | 0 |
| NC_002645.1;NP_073549.1;HCoV-229E_1ab;4998 | 0 | 0 |
| NC_002645.1;NP_073549.1;HCoV-229E_1ab;4999 | 0 | 0 |
| NC_002645.1;NP_073549.1;HCoV-229E_1ab;5000 | 0 | 0 |
| NC_002645.1;NP_073549.1;HCoV-229E_1ab;5001 | 0 | 0 |
| NC_002645.1;NP_073549.1;HCoV-229E_1ab;5002 | 0 | 0 |
| NC_002645.1;NP_073549.1;HCoV-229E_1ab;5003 | 0 | 0 |
| NC_002645.1;NP_073549.1;HCoV-229E_1ab;5004 | 0 | 0 |
| NC_002645.1;NP_073549.1;HCoV-229E_1ab;5005 | 0 | 0 |
| NC_002645.1;NP_073549.1;HCoV-229E_1ab;5006 | 0 | 0 |
| NC_002645.1;NP_073549.1;HCoV-229E_1ab;5007 | 0 | 0 |
| NC_002645.1;NP_073549.1;HCoV-229E_1ab;5008 | 0 | 0 |
| NC_002645.1;NP_073549.1;HCoV-229E_1ab;5009 | 0 | 0 |
| NC_002645.1;NP_073549.1;HCoV-229E_1ab;5010 | 0 | 0 |
| NC_002645.1;NP_073549.1;HCoV-229E_1ab;5011 | 0 | 0 |
| NC_002645.1;NP_073549.1;HCoV-229E_1ab;5012 | 0 | 0 |
| NC_002645.1;NP_073549.1;HCoV-229E_1ab;5013 | 0 | 0 |
| NC_002645.1;NP_073549.1;HCoV-229E_1ab;5014 | 0 | 0 |
| NC_002645.1;NP_073549.1;HCoV-229E_1ab;5015 | 0 | 0 |
| NC_002645.1;NP_073549.1;HCoV-229E_1ab;5016 | 0 | 0 |
| NC_002645.1;NP_073549.1;HCoV-229E_1ab;5017 | 0 | 0 |

| | | |
|---|---|---|
| NC_002645.1;NP_073549.1;HCoV-229E_1ab;5018 | 0 | 0 |
| NC_002645.1;NP_073549.1;HCoV-229E_1ab;5019 | 0 | 0 |
| NC_002645.1;NP_073549.1;HCoV-229E_1ab;5020 | 0 | 0 |
| NC_002645.1;NP_073549.1;HCoV-229E_1ab;5021 | 0 | 0 |
| NC_002645.1;NP_073549.1;HCoV-229E_1ab;5022 | 0 | 0 |
| NC_002645.1;NP_073549.1;HCoV-229E_1ab;5023 | 0 | 0 |
| NC_002645.1;NP_073549.1;HCoV-229E_1ab;5024 | 0 | 0 |
| NC_002645.1;NP_073549.1;HCoV-229E_1ab;5025 | 0 | 0 |
| NC_002645.1;NP_073549.1;HCoV-229E_1ab;5026 | 0 | 0 |
| NC_002645.1;NP_073549.1;HCoV-229E_1ab;5027 | 0 | 0 |
| NC_002645.1;NP_073549.1;HCoV-229E_1ab;5028 | 0 | 0 |
| NC_002645.1;NP_073549.1;HCoV-229E_1ab;5029 | 0 | 0 |
| NC_002645.1;NP_073549.1;HCoV-229E_1ab;5030 | 0 | 0 |
| NC_002645.1;NP_073549.1;HCoV-229E_1ab;5031 | 0 | 0 |
| NC_002645.1;NP_073549.1;HCoV-229E_1ab;5032 | 0 | 0 |
| NC_002645.1;NP_073549.1;HCoV-229E_1ab;5033 | 0 | 0 |
| NC_002645.1;NP_073549.1;HCoV-229E_1ab;5034 | 0 | 0 |
| NC_002645.1;NP_073549.1;HCoV-229E_1ab;5035 | 0 | 0 |
| NC_002645.1;NP_073549.1;HCoV-229E_1ab;5036 | 0 | 0 |
| NC_002645.1;NP_073549.1;HCoV-229E_1ab;5037 | 0 | 0 |
| NC_002645.1;NP_073549.1;HCoV-229E_1ab;5038 | 0 | 0 |
| NC_002645.1;NP_073549.1;HCoV-229E_1ab;5039 | 0 | 0.025 |
| NC_002645.1;NP_073549.1;HCoV-229E_1ab;5040 | 0 | 0 |
| NC_002645.1;NP_073549.1;HCoV-229E_1ab;5041 | 0 | 0.025 |
| NC_002645.1;NP_073549.1;HCoV-229E_1ab;5042 | 0.05 | 0 |
| NC_002645.1;NP_073549.1;HCoV-229E_1ab;5043 | 0.05 | 0.05 |
| NC_002645.1;NP_073549.1;HCoV-229E_1ab;5044 | 0.05 | 0.025 |
| NC_002645.1;NP_073549.1;HCoV-229E_1ab;5045 | 0.05 | 0.025 |
| NC_002645.1;NP_073549.1;HCoV-229E_1ab;5046 | 0.05 | 0.025 |
| NC_002645.1;NP_073549.1;HCoV-229E_1ab;5047 | 0.1 | 0.025 |
| NC_002645.1;NP_073549.1;HCoV-229E_1ab;5048 | 0 | 0.025 |
| NC_002645.1;NP_073549.1;HCoV-229E_1ab;5049 | 0 | 0.025 |
| NC_002645.1;NP_073549.1;HCoV-229E_1ab;5050 | 0 | 0 |
| NC_002645.1;NP_073549.1;HCoV-229E_1ab;5051 | 0 | 0 |
| NC_002645.1;NP_073549.1;HCoV-229E_1ab;5052 | 0 | 0 |
| NC_002645.1;NP_073549.1;HCoV-229E_1ab;5053 | 0 | 0 |
| NC_002645.1;NP_073549.1;HCoV-229E_1ab;5054 | 0 | 0 |
| NC_002645.1;NP_073549.1;HCoV-229E_1ab;5055 | 0 | 0 |
| NC_002645.1;NP_073549.1;HCoV-229E_1ab;5056 | 0 | 0 |
| NC_002645.1;NP_073549.1;HCoV-229E_1ab;5057 | 0 | 0 |
| NC_002645.1;NP_073549.1;HCoV-229E_1ab;5058 | 0 | 0 |
| NC_002645.1;NP_073549.1;HCoV-229E_1ab;5059 | 0 | 0 |
| NC_002645.1;NP_073549.1;HCoV-229E_1ab;5060 | 0 | 0 |

| | | |
|---|---|---|
| NC_002645.1;NP_073549.1;HCoV-229E_1ab;5061 | 0 | 0 |
| NC_002645.1;NP_073549.1;HCoV-229E_1ab;5062 | 0 | 0 |
| NC_002645.1;NP_073549.1;HCoV-229E_1ab;5063 | 0 | 0 |
| NC_002645.1;NP_073549.1;HCoV-229E_1ab;5064 | 0 | 0 |
| NC_002645.1;NP_073549.1;HCoV-229E_1ab;5065 | 0 | 0 |
| NC_002645.1;NP_073549.1;HCoV-229E_1ab;5066 | 0 | 0 |
| NC_002645.1;NP_073549.1;HCoV-229E_1ab;5067 | 0 | 0 |
| NC_002645.1;NP_073549.1;HCoV-229E_1ab;5068 | 0 | 0 |
| NC_002645.1;NP_073549.1;HCoV-229E_1ab;5069 | 0 | 0 |
| NC_002645.1;NP_073549.1;HCoV-229E_1ab;5070 | 0 | 0 |
| NC_002645.1;NP_073549.1;HCoV-229E_1ab;5071 | 0 | 0 |
| NC_002645.1;NP_073549.1;HCoV-229E_1ab;5072 | 0 | 0 |
| NC_002645.1;NP_073549.1;HCoV-229E_1ab;5073 | 0 | 0 |
| NC_002645.1;NP_073549.1;HCoV-229E_1ab;5074 | 0 | 0 |
| NC_002645.1;NP_073549.1;HCoV-229E_1ab;5075 | 0 | 0 |
| NC_002645.1;NP_073549.1;HCoV-229E_1ab;5076 | 0 | 0 |
| NC_002645.1;NP_073549.1;HCoV-229E_1ab;5077 | 0 | 0 |
| NC_002645.1;NP_073549.1;HCoV-229E_1ab;5078 | 0 | 0 |
| NC_002645.1;NP_073549.1;HCoV-229E_1ab;5079 | 0 | 0 |
| NC_002645.1;NP_073549.1;HCoV-229E_1ab;5080 | 0 | 0 |
| NC_002645.1;NP_073549.1;HCoV-229E_1ab;5081 | 0 | 0 |
| NC_002645.1;NP_073549.1;HCoV-229E_1ab;5082 | 0.1 | 0 |
| NC_002645.1;NP_073549.1;HCoV-229E_1ab;5083 | 0.1 | 0 |
| NC_002645.1;NP_073549.1;HCoV-229E_1ab;5084 | 0.05 | 0 |
| NC_002645.1;NP_073549.1;HCoV-229E_1ab;5085 | 0.05 | 0 |
| NC_002645.1;NP_073549.1;HCoV-229E_1ab;5086 | 0.05 | 0 |
| NC_002645.1;NP_073549.1;HCoV-229E_1ab;5087 | 0.05 | 0.05 |
| NC_002645.1;NP_073549.1;HCoV-229E_1ab;5088 | 0.15 | 0.15 |
| NC_002645.1;NP_073549.1;HCoV-229E_1ab;5089 | 0.15 | 0.15 |
| NC_002645.1;NP_073549.1;HCoV-229E_1ab;5090 | 0.1 | 0.15 |
| NC_002645.1;NP_073549.1;HCoV-229E_1ab;5091 | 0.05 | 0.175 |
| NC_002645.1;NP_073549.1;HCoV-229E_1ab;5092 | 0.05 | 0.15 |
| NC_002645.1;NP_073549.1;HCoV-229E_1ab;5093 | 0.05 | 0.2 |
| NC_002645.1;NP_073549.1;HCoV-229E_1ab;5094 | 0.1 | 0.15 |
| NC_002645.1;NP_073549.1;HCoV-229E_1ab;5095 | 0.05 | 0.125 |
| NC_002645.1;NP_073549.1;HCoV-229E_1ab;5096 | 0.05 | 0.125 |
| NC_002645.1;NP_073549.1;HCoV-229E_1ab;5097 | 0.05 | 0.025 |
| NC_002645.1;NP_073549.1;HCoV-229E_1ab;5098 | 0 | 0.025 |
| NC_002645.1;NP_073549.1;HCoV-229E_1ab;5099 | 0 | 0 |
| NC_002645.1;NP_073549.1;HCoV-229E_1ab;5100 | 0 | 0 |
| NC_002645.1;NP_073549.1;HCoV-229E_1ab;5101 | 0 | 0 |
| NC_002645.1;NP_073549.1;HCoV-229E_1ab;5102 | 0 | 0.05 |
| NC_002645.1;NP_073549.1;HCoV-229E_1ab;5103 | 0 | 0 |

| | | |
|---|---|---|
| NC_002645.1;NP_073549.1;HCoV-229E_1ab;5104 | 0 | 0 |
| NC_002645.1;NP_073549.1;HCoV-229E_1ab;5105 | 0 | 0.025 |
| NC_002645.1;NP_073549.1;HCoV-229E_1ab;5106 | 0.05 | 0 |
| NC_002645.1;NP_073549.1;HCoV-229E_1ab;5107 | 0.2 | 0.1 |
| NC_002645.1;NP_073549.1;HCoV-229E_1ab;5108 | 0.15 | 0.075 |
| NC_002645.1;NP_073549.1;HCoV-229E_1ab;5109 | 0.15 | 0.1 |
| NC_002645.1;NP_073549.1;HCoV-229E_1ab;5110 | 0.1 | 0.1 |
| NC_002645.1;NP_073549.1;HCoV-229E_1ab;5111 | 0.15 | 0.15 |
| NC_002645.1;NP_073549.1;HCoV-229E_1ab;5112 | 0.15 | 0.075 |
| NC_002645.1;NP_073549.1;HCoV-229E_1ab;5113 | 0.1 | 0.075 |
| NC_002645.1;NP_073549.1;HCoV-229E_1ab;5114 | 0.1 | 0.1 |
| NC_002645.1;NP_073549.1;HCoV-229E_1ab;5115 | 0.1 | 0.075 |
| NC_002645.1;NP_073549.1;HCoV-229E_1ab;5116 | 0 | 0.025 |
| NC_002645.1;NP_073549.1;HCoV-229E_1ab;5117 | 0 | 0 |
| NC_002645.1;NP_073549.1;HCoV-229E_1ab;5118 | 0 | 0 |
| NC_002645.1;NP_073549.1;HCoV-229E_1ab;5119 | 0 | 0 |
| NC_002645.1;NP_073549.1;HCoV-229E_1ab;5120 | 0 | 0 |
| NC_002645.1;NP_073549.1;HCoV-229E_1ab;5121 | 0 | 0 |
| NC_002645.1;NP_073549.1;HCoV-229E_1ab;5122 | 0.05 | 0 |
| NC_002645.1;NP_073549.1;HCoV-229E_1ab;5123 | 0.05 | 0.025 |
| NC_002645.1;NP_073549.1;HCoV-229E_1ab;5124 | 0 | 0 |
| NC_002645.1;NP_073549.1;HCoV-229E_1ab;5125 | 0 | 0 |
| NC_002645.1;NP_073549.1;HCoV-229E_1ab;5126 | 0 | 0 |
| NC_002645.1;NP_073549.1;HCoV-229E_1ab;5127 | 0 | 0.05 |
| NC_002645.1;NP_073549.1;HCoV-229E_1ab;5128 | 0 | 0.025 |
| NC_002645.1;NP_073549.1;HCoV-229E_1ab;5129 | 0 | 0.025 |
| NC_002645.1;NP_073549.1;HCoV-229E_1ab;5130 | 0.1 | 0.1 |
| NC_002645.1;NP_073549.1;HCoV-229E_1ab;5131 | 0 | 0.025 |
| NC_002645.1;NP_073549.1;HCoV-229E_1ab;5132 | 0.05 | 0.025 |
| NC_002645.1;NP_073549.1;HCoV-229E_1ab;5133 | 0.05 | 0 |
| NC_002645.1;NP_073549.1;HCoV-229E_1ab;5134 | 0.05 | 0 |
| NC_002645.1;NP_073549.1;HCoV-229E_1ab;5135 | 0 | 0.025 |
| NC_002645.1;NP_073549.1;HCoV-229E_1ab;5136 | 0.05 | 0.025 |
| NC_002645.1;NP_073549.1;HCoV-229E_1ab;5137 | 0.1 | 0.05 |
| NC_002645.1;NP_073549.1;HCoV-229E_1ab;5138 | 0 | 0 |
| NC_002645.1;NP_073549.1;HCoV-229E_1ab;5139 | 0 | 0 |
| NC_002645.1;NP_073549.1;HCoV-229E_1ab;5140 | 0 | 0 |
| NC_002645.1;NP_073549.1;HCoV-229E_1ab;5141 | 0.05 | 0 |
| NC_002645.1;NP_073549.1;HCoV-229E_1ab;5142 | 0 | 0 |
| NC_002645.1;NP_073549.1;HCoV-229E_1ab;5143 | 0 | 0 |
| NC_002645.1;NP_073549.1;HCoV-229E_1ab;5144 | 0 | 0.025 |
| NC_002645.1;NP_073549.1;HCoV-229E_1ab;5145 | 0.05 | 0 |
| NC_002645.1;NP_073549.1;HCoV-229E_1ab;5146 | 0 | 0 |

| | | |
|---|---|---|
| NC_002645.1;NP_073549.1;HCoV-229E_1ab;5147 | 0 | 0 |
| NC_002645.1;NP_073549.1;HCoV-229E_1ab;5148 | 0 | 0 |
| NC_002645.1;NP_073549.1;HCoV-229E_1ab;5149 | 0 | 0.025 |
| NC_002645.1;NP_073549.1;HCoV-229E_1ab;5150 | 0 | 0 |
| NC_002645.1;NP_073549.1;HCoV-229E_1ab;5151 | 0 | 0.025 |
| NC_002645.1;NP_073549.1;HCoV-229E_1ab;5152 | 0 | 0.025 |
| NC_002645.1;NP_073549.1;HCoV-229E_1ab;5153 | 0 | 0.025 |
| NC_002645.1;NP_073549.1;HCoV-229E_1ab;5154 | 0 | 0.025 |
| NC_002645.1;NP_073549.1;HCoV-229E_1ab;5155 | 0 | 0.025 |
| NC_002645.1;NP_073549.1;HCoV-229E_1ab;5156 | 0 | 0.025 |
| NC_002645.1;NP_073549.1;HCoV-229E_1ab;5157 | 0 | 0.025 |
| NC_002645.1;NP_073549.1;HCoV-229E_1ab;5158 | 0 | 0.025 |
| NC_002645.1;NP_073549.1;HCoV-229E_1ab;5159 | 0 | 0.025 |
| NC_002645.1;NP_073549.1;HCoV-229E_1ab;5160 | 0 | 0.05 |
| NC_002645.1;NP_073549.1;HCoV-229E_1ab;5161 | 0 | 0.075 |
| NC_002645.1;NP_073549.1;HCoV-229E_1ab;5162 | 0 | 0.025 |
| NC_002645.1;NP_073549.1;HCoV-229E_1ab;5163 | 0.05 | 0.025 |
| NC_002645.1;NP_073549.1;HCoV-229E_1ab;5164 | 0 | 0 |
| NC_002645.1;NP_073549.1;HCoV-229E_1ab;5165 | 0 | 0 |
| NC_002645.1;NP_073549.1;HCoV-229E_1ab;5166 | 0 | 0 |
| NC_002645.1;NP_073549.1;HCoV-229E_1ab;5167 | 0 | 0 |
| NC_002645.1;NP_073549.1;HCoV-229E_1ab;5168 | 0 | 0 |
| NC_002645.1;NP_073549.1;HCoV-229E_1ab;5169 | 0 | 0 |
| NC_002645.1;NP_073549.1;HCoV-229E_1ab;5170 | 0 | 0 |
| NC_002645.1;NP_073549.1;HCoV-229E_1ab;5171 | 0 | 0 |
| NC_002645.1;NP_073549.1;HCoV-229E_1ab;5172 | 0 | 0 |
| NC_002645.1;NP_073549.1;HCoV-229E_1ab;5173 | 0 | 0 |
| NC_002645.1;NP_073549.1;HCoV-229E_1ab;5174 | 0 | 0 |
| NC_002645.1;NP_073549.1;HCoV-229E_1ab;5175 | 0 | 0 |
| NC_002645.1;NP_073549.1;HCoV-229E_1ab;5176 | 0 | 0 |
| NC_002645.1;NP_073549.1;HCoV-229E_1ab;5177 | 0 | 0 |
| NC_002645.1;NP_073549.1;HCoV-229E_1ab;5178 | 0 | 0 |
| NC_002645.1;NP_073549.1;HCoV-229E_1ab;5179 | 0 | 0 |
| NC_002645.1;NP_073549.1;HCoV-229E_1ab;5180 | 0 | 0 |
| NC_002645.1;NP_073549.1;HCoV-229E_1ab;5181 | 0 | 0.025 |
| NC_002645.1;NP_073549.1;HCoV-229E_1ab;5182 | 0 | 0.025 |
| NC_002645.1;NP_073549.1;HCoV-229E_1ab;5183 | 0 | 0.05 |
| NC_002645.1;NP_073549.1;HCoV-229E_1ab;5184 | 0 | 0 |
| NC_002645.1;NP_073549.1;HCoV-229E_1ab;5185 | 0 | 0.025 |
| NC_002645.1;NP_073549.1;HCoV-229E_1ab;5186 | 0 | 0 |
| NC_002645.1;NP_073549.1;HCoV-229E_1ab;5187 | 0 | 0 |
| NC_002645.1;NP_073549.1;HCoV-229E_1ab;5188 | 0 | 0 |
| NC_002645.1;NP_073549.1;HCoV-229E_1ab;5189 | 0 | 0 |

| | | |
|---|---|---|
| NC_002645.1;NP_073549.1;HCoV-229E_1ab;5190 | 0.05 | 0 |
| NC_002645.1;NP_073549.1;HCoV-229E_1ab;5191 | 0.05 | 0 |
| NC_002645.1;NP_073549.1;HCoV-229E_1ab;5192 | 0.05 | 0 |
| NC_002645.1;NP_073549.1;HCoV-229E_1ab;5193 | 0.05 | 0.075 |
| NC_002645.1;NP_073549.1;HCoV-229E_1ab;5194 | 0 | 0 |
| NC_002645.1;NP_073549.1;HCoV-229E_1ab;5195 | 0 | 0.075 |
| NC_002645.1;NP_073549.1;HCoV-229E_1ab;5196 | 0 | 0.025 |
| NC_002645.1;NP_073549.1;HCoV-229E_1ab;5197 | 0.05 | 0.025 |
| NC_002645.1;NP_073549.1;HCoV-229E_1ab;5198 | 0.05 | 0 |
| NC_002645.1;NP_073549.1;HCoV-229E_1ab;5199 | 0 | 0.025 |
| NC_002645.1;NP_073549.1;HCoV-229E_1ab;5200 | 0 | 0 |
| NC_002645.1;NP_073549.1;HCoV-229E_1ab;5201 | 0 | 0 |
| NC_002645.1;NP_073549.1;HCoV-229E_1ab;5202 | 0 | 0.025 |
| NC_002645.1;NP_073549.1;HCoV-229E_1ab;5203 | 0 | 0 |
| NC_002645.1;NP_073549.1;HCoV-229E_1ab;5204 | 0 | 0 |
| NC_002645.1;NP_073549.1;HCoV-229E_1ab;5205 | 0.05 | 0.025 |
| NC_002645.1;NP_073549.1;HCoV-229E_1ab;5206 | 0.05 | 0.025 |
| NC_002645.1;NP_073549.1;HCoV-229E_1ab;5207 | 0 | 0 |
| NC_002645.1;NP_073549.1;HCoV-229E_1ab;5208 | 0 | 0 |
| NC_002645.1;NP_073549.1;HCoV-229E_1ab;5209 | 0 | 0 |
| NC_002645.1;NP_073549.1;HCoV-229E_1ab;5210 | 0 | 0 |
| NC_002645.1;NP_073549.1;HCoV-229E_1ab;5211 | 0 | 0 |
| NC_002645.1;NP_073549.1;HCoV-229E_1ab;5212 | 0 | 0 |
| NC_002645.1;NP_073549.1;HCoV-229E_1ab;5213 | 0 | 0 |
| NC_002645.1;NP_073549.1;HCoV-229E_1ab;5214 | 0 | 0 |
| NC_002645.1;NP_073549.1;HCoV-229E_1ab;5215 | 0 | 0 |
| NC_002645.1;NP_073549.1;HCoV-229E_1ab;5216 | 0 | 0 |
| NC_002645.1;NP_073549.1;HCoV-229E_1ab;5217 | 0 | 0 |
| NC_002645.1;NP_073549.1;HCoV-229E_1ab;5218 | 0 | 0 |
| NC_002645.1;NP_073549.1;HCoV-229E_1ab;5219 | 0 | 0 |
| NC_002645.1;NP_073549.1;HCoV-229E_1ab;5220 | 0 | 0 |
| NC_002645.1;NP_073549.1;HCoV-229E_1ab;5221 | 0 | 0 |
| NC_002645.1;NP_073549.1;HCoV-229E_1ab;5222 | 0 | 0 |
| NC_002645.1;NP_073549.1;HCoV-229E_1ab;5223 | 0 | 0 |
| NC_002645.1;NP_073549.1;HCoV-229E_1ab;5224 | 0 | 0 |
| NC_002645.1;NP_073549.1;HCoV-229E_1ab;5225 | 0 | 0 |
| NC_002645.1;NP_073549.1;HCoV-229E_1ab;5226 | 0 | 0 |
| NC_002645.1;NP_073549.1;HCoV-229E_1ab;5227 | 0 | 0 |
| NC_002645.1;NP_073549.1;HCoV-229E_1ab;5228 | 0 | 0 |
| NC_002645.1;NP_073549.1;HCoV-229E_1ab;5229 | 0.05 | 0.025 |
| NC_002645.1;NP_073549.1;HCoV-229E_1ab;5230 | 0.1 | 0.125 |
| NC_002645.1;NP_073549.1;HCoV-229E_1ab;5231 | 0 | 0.05 |
| NC_002645.1;NP_073549.1;HCoV-229E_1ab;5232 | 0 | 0.025 |

| | | |
|---|---|---|
| NC_002645.1;NP_073549.1;HCoV-229E_1ab;5233 | 0 | 0 |
| NC_002645.1;NP_073549.1;HCoV-229E_1ab;5234 | 0 | 0 |
| NC_002645.1;NP_073549.1;HCoV-229E_1ab;5235 | 0 | 0 |
| NC_002645.1;NP_073549.1;HCoV-229E_1ab;5236 | 0 | 0.025 |
| NC_002645.1;NP_073549.1;HCoV-229E_1ab;5237 | 0 | 0 |
| NC_002645.1;NP_073549.1;HCoV-229E_1ab;5238 | 0 | 0.025 |
| NC_002645.1;NP_073549.1;HCoV-229E_1ab;5239 | 0 | 0 |
| NC_002645.1;NP_073549.1;HCoV-229E_1ab;5240 | 0 | 0.025 |
| NC_002645.1;NP_073549.1;HCoV-229E_1ab;5241 | 0 | 0 |
| NC_002645.1;NP_073549.1;HCoV-229E_1ab;5242 | 0 | 0 |
| NC_002645.1;NP_073549.1;HCoV-229E_1ab;5243 | 0 | 0.025 |
| NC_002645.1;NP_073549.1;HCoV-229E_1ab;5244 | 0 | 0 |
| NC_002645.1;NP_073549.1;HCoV-229E_1ab;5245 | 0 | 0 |
| NC_002645.1;NP_073549.1;HCoV-229E_1ab;5246 | 0 | 0.025 |
| NC_002645.1;NP_073549.1;HCoV-229E_1ab;5247 | 0 | 0.025 |
| NC_002645.1;NP_073549.1;HCoV-229E_1ab;5248 | 0 | 0.05 |
| NC_002645.1;NP_073549.1;HCoV-229E_1ab;5249 | 0 | 0.025 |
| NC_002645.1;NP_073549.1;HCoV-229E_1ab;5250 | 0 | 0.025 |
| NC_002645.1;NP_073549.1;HCoV-229E_1ab;5251 | 0 | 0.025 |
| NC_002645.1;NP_073549.1;HCoV-229E_1ab;5252 | 0 | 0.025 |
| NC_002645.1;NP_073549.1;HCoV-229E_1ab;5253 | 0 | 0.025 |
| NC_002645.1;NP_073549.1;HCoV-229E_1ab;5254 | 0 | 0.025 |
| NC_002645.1;NP_073549.1;HCoV-229E_1ab;5255 | 0 | 0 |
| NC_002645.1;NP_073549.1;HCoV-229E_1ab;5256 | 0 | 0 |
| NC_002645.1;NP_073549.1;HCoV-229E_1ab;5257 | 0 | 0 |
| NC_002645.1;NP_073549.1;HCoV-229E_1ab;5258 | 0 | 0 |
| NC_002645.1;NP_073549.1;HCoV-229E_1ab;5259 | 0 | 0 |
| NC_002645.1;NP_073549.1;HCoV-229E_1ab;5260 | 0 | 0 |
| NC_002645.1;NP_073549.1;HCoV-229E_1ab;5261 | 0 | 0 |
| NC_002645.1;NP_073549.1;HCoV-229E_1ab;5262 | 0 | 0 |
| NC_002645.1;NP_073549.1;HCoV-229E_1ab;5263 | 0 | 0 |
| NC_002645.1;NP_073549.1;HCoV-229E_1ab;5264 | 0 | 0 |
| NC_002645.1;NP_073549.1;HCoV-229E_1ab;5265 | 0 | 0 |
| NC_002645.1;NP_073549.1;HCoV-229E_1ab;5266 | 0 | 0.025 |
| NC_002645.1;NP_073549.1;HCoV-229E_1ab;5267 | 0.05 | 0 |
| NC_002645.1;NP_073549.1;HCoV-229E_1ab;5268 | 0 | 0 |
| NC_002645.1;NP_073549.1;HCoV-229E_1ab;5269 | 0 | 0 |
| NC_002645.1;NP_073549.1;HCoV-229E_1ab;5270 | 0 | 0 |
| NC_002645.1;NP_073549.1;HCoV-229E_1ab;5271 | 0.05 | 0 |
| NC_002645.1;NP_073549.1;HCoV-229E_1ab;5272 | 0 | 0 |
| NC_002645.1;NP_073549.1;HCoV-229E_1ab;5273 | 0 | 0 |
| NC_002645.1;NP_073549.1;HCoV-229E_1ab;5274 | 0 | 0 |
| NC_002645.1;NP_073549.1;HCoV-229E_1ab;5275 | 0 | 0 |

| | | |
|---|---|---|
| NC_002645.1;NP_073549.1;HCoV-229E_1ab;5276 | 0 | 0 |
| NC_002645.1;NP_073549.1;HCoV-229E_1ab;5277 | 0 | 0 |
| NC_002645.1;NP_073549.1;HCoV-229E_1ab;5278 | 0 | 0 |
| NC_002645.1;NP_073549.1;HCoV-229E_1ab;5279 | 0 | 0 |
| NC_002645.1;NP_073549.1;HCoV-229E_1ab;5280 | 0 | 0 |
| NC_002645.1;NP_073549.1;HCoV-229E_1ab;5281 | 0 | 0 |
| NC_002645.1;NP_073549.1;HCoV-229E_1ab;5282 | 0 | 0 |
| NC_002645.1;NP_073549.1;HCoV-229E_1ab;5283 | 0 | 0 |
| NC_002645.1;NP_073549.1;HCoV-229E_1ab;5284 | 0 | 0 |
| NC_002645.1;NP_073549.1;HCoV-229E_1ab;5285 | 0 | 0 |
| NC_002645.1;NP_073549.1;HCoV-229E_1ab;5286 | 0 | 0 |
| NC_002645.1;NP_073549.1;HCoV-229E_1ab;5287 | 0 | 0 |
| NC_002645.1;NP_073549.1;HCoV-229E_1ab;5288 | 0 | 0 |
| NC_002645.1;NP_073549.1;HCoV-229E_1ab;5289 | 0 | 0 |
| NC_002645.1;NP_073549.1;HCoV-229E_1ab;5290 | 0 | 0 |
| NC_002645.1;NP_073549.1;HCoV-229E_1ab;5291 | 0 | 0 |
| NC_002645.1;NP_073549.1;HCoV-229E_1ab;5292 | 0 | 0 |
| NC_002645.1;NP_073549.1;HCoV-229E_1ab;5293 | 0 | 0 |
| NC_002645.1;NP_073549.1;HCoV-229E_1ab;5294 | 0.05 | 0 |
| NC_002645.1;NP_073549.1;HCoV-229E_1ab;5295 | 0 | 0 |
| NC_002645.1;NP_073549.1;HCoV-229E_1ab;5296 | 0 | 0 |
| NC_002645.1;NP_073549.1;HCoV-229E_1ab;5297 | 0 | 0 |
| NC_002645.1;NP_073549.1;HCoV-229E_1ab;5298 | 0 | 0.025 |
| NC_002645.1;NP_073549.1;HCoV-229E_1ab;5299 | 0 | 0.025 |
| NC_002645.1;NP_073549.1;HCoV-229E_1ab;5300 | 0 | 0.025 |
| NC_002645.1;NP_073549.1;HCoV-229E_1ab;5301 | 0 | 0.025 |
| NC_002645.1;NP_073549.1;HCoV-229E_1ab;5302 | 0 | 0.025 |
| NC_002645.1;NP_073549.1;HCoV-229E_1ab;5303 | 0 | 0 |
| NC_002645.1;NP_073549.1;HCoV-229E_1ab;5304 | 0 | 0 |
| NC_002645.1;NP_073549.1;HCoV-229E_1ab;5305 | 0 | 0 |
| NC_002645.1;NP_073549.1;HCoV-229E_1ab;5306 | 0 | 0 |
| NC_002645.1;NP_073549.1;HCoV-229E_1ab;5307 | 0 | 0 |
| NC_002645.1;NP_073549.1;HCoV-229E_1ab;5308 | 0 | 0 |
| NC_002645.1;NP_073549.1;HCoV-229E_1ab;5309 | 0 | 0 |
| NC_002645.1;NP_073549.1;HCoV-229E_1ab;5310 | 0 | 0.025 |
| NC_002645.1;NP_073549.1;HCoV-229E_1ab;5311 | 0 | 0 |
| NC_002645.1;NP_073549.1;HCoV-229E_1ab;5312 | 0 | 0 |
| NC_002645.1;NP_073549.1;HCoV-229E_1ab;5313 | 0 | 0 |
| NC_002645.1;NP_073549.1;HCoV-229E_1ab;5314 | 0 | 0 |
| NC_002645.1;NP_073549.1;HCoV-229E_1ab;5315 | 0 | 0 |
| NC_002645.1;NP_073549.1;HCoV-229E_1ab;5316 | 0 | 0 |
| NC_002645.1;NP_073549.1;HCoV-229E_1ab;5317 | 0 | 0 |
| NC_002645.1;NP_073549.1;HCoV-229E_1ab;5318 | 0 | 0 |

| | | |
|---|---|---|
| NC_002645.1;NP_073549.1;HCoV-229E_1ab;5319 | 0 | 0 |
| NC_002645.1;NP_073549.1;HCoV-229E_1ab;5320 | 0 | 0.025 |
| NC_002645.1;NP_073549.1;HCoV-229E_1ab;5321 | 0 | 0 |
| NC_002645.1;NP_073549.1;HCoV-229E_1ab;5322 | 0 | 0 |
| NC_002645.1;NP_073549.1;HCoV-229E_1ab;5323 | 0 | 0 |
| NC_002645.1;NP_073549.1;HCoV-229E_1ab;5324 | 0 | 0 |
| NC_002645.1;NP_073549.1;HCoV-229E_1ab;5325 | 0 | 0 |
| NC_002645.1;NP_073549.1;HCoV-229E_1ab;5326 | 0 | 0 |
| NC_002645.1;NP_073549.1;HCoV-229E_1ab;5327 | 0 | 0 |
| NC_002645.1;NP_073549.1;HCoV-229E_1ab;5328 | 0 | 0 |
| NC_002645.1;NP_073549.1;HCoV-229E_1ab;5329 | 0 | 0 |
| NC_002645.1;NP_073549.1;HCoV-229E_1ab;5330 | 0 | 0 |
| NC_002645.1;NP_073549.1;HCoV-229E_1ab;5331 | 0.05 | 0 |
| NC_002645.1;NP_073549.1;HCoV-229E_1ab;5332 | 0 | 0 |
| NC_002645.1;NP_073549.1;HCoV-229E_1ab;5333 | 0 | 0 |
| NC_002645.1;NP_073549.1;HCoV-229E_1ab;5334 | 0 | 0.05 |
| NC_002645.1;NP_073549.1;HCoV-229E_1ab;5335 | 0 | 0.025 |
| NC_002645.1;NP_073549.1;HCoV-229E_1ab;5336 | 0.1 | 0.075 |
| NC_002645.1;NP_073549.1;HCoV-229E_1ab;5337 | 0.1 | 0.075 |
| NC_002645.1;NP_073549.1;HCoV-229E_1ab;5338 | 0.05 | 0 |
| NC_002645.1;NP_073549.1;HCoV-229E_1ab;5339 | 0.1 | 0 |
| NC_002645.1;NP_073549.1;HCoV-229E_1ab;5340 | 0.05 | 0 |
| NC_002645.1;NP_073549.1;HCoV-229E_1ab;5341 | 0.05 | 0 |
| NC_002645.1;NP_073549.1;HCoV-229E_1ab;5342 | 0.05 | 0 |
| NC_002645.1;NP_073549.1;HCoV-229E_1ab;5343 | 0.05 | 0 |
| NC_002645.1;NP_073549.1;HCoV-229E_1ab;5344 | 0.05 | 0.025 |
| NC_002645.1;NP_073549.1;HCoV-229E_1ab;5345 | 0.05 | 0 |
| NC_002645.1;NP_073549.1;HCoV-229E_1ab;5346 | 0 | 0.025 |
| NC_002645.1;NP_073549.1;HCoV-229E_1ab;5347 | 0 | 0.025 |
| NC_002645.1;NP_073549.1;HCoV-229E_1ab;5348 | 0.1 | 0.025 |
| NC_002645.1;NP_073549.1;HCoV-229E_1ab;5349 | 0.05 | 0.05 |
| NC_002645.1;NP_073549.1;HCoV-229E_1ab;5350 | 0 | 0.025 |
| NC_002645.1;NP_073549.1;HCoV-229E_1ab;5351 | 0 | 0.05 |
| NC_002645.1;NP_073549.1;HCoV-229E_1ab;5352 | 0.05 | 0.1 |
| NC_002645.1;NP_073549.1;HCoV-229E_1ab;5353 | 0 | 0.025 |
| NC_002645.1;NP_073549.1;HCoV-229E_1ab;5354 | 0 | 0 |
| NC_002645.1;NP_073549.1;HCoV-229E_1ab;5355 | 0.05 | 0 |
| NC_002645.1;NP_073549.1;HCoV-229E_1ab;5356 | 0.05 | 0 |
| NC_002645.1;NP_073549.1;HCoV-229E_1ab;5357 | 0.05 | 0 |
| NC_002645.1;NP_073549.1;HCoV-229E_1ab;5358 | 0.1 | 0 |
| NC_002645.1;NP_073549.1;HCoV-229E_1ab;5359 | 0.05 | 0 |
| NC_002645.1;NP_073549.1;HCoV-229E_1ab;5360 | 0.1 | 0 |
| NC_002645.1;NP_073549.1;HCoV-229E_1ab;5361 | 0.1 | 0 |

| | | |
|---|---|---|
| NC_002645.1;NP_073549.1;HCoV-229E_1ab;5362 | 0.1 | 0.025 |
| NC_002645.1;NP_073549.1;HCoV-229E_1ab;5363 | 0.05 | 0.025 |
| NC_002645.1;NP_073549.1;HCoV-229E_1ab;5364 | 0 | 0.05 |
| NC_002645.1;NP_073549.1;HCoV-229E_1ab;5365 | 0 | 0 |
| NC_002645.1;NP_073549.1;HCoV-229E_1ab;5366 | 0.05 | 0 |
| NC_002645.1;NP_073549.1;HCoV-229E_1ab;5367 | 0.05 | 0.075 |
| NC_002645.1;NP_073549.1;HCoV-229E_1ab;5368 | 0.05 | 0.025 |
| NC_002645.1;NP_073549.1;HCoV-229E_1ab;5369 | 0.05 | 0.025 |
| NC_002645.1;NP_073549.1;HCoV-229E_1ab;5370 | 0 | 0 |
| NC_002645.1;NP_073549.1;HCoV-229E_1ab;5371 | 0 | 0 |
| NC_002645.1;NP_073549.1;HCoV-229E_1ab;5372 | 0 | 0 |
| NC_002645.1;NP_073549.1;HCoV-229E_1ab;5373 | 0 | 0 |
| NC_002645.1;NP_073549.1;HCoV-229E_1ab;5374 | 0 | 0 |
| NC_002645.1;NP_073549.1;HCoV-229E_1ab;5375 | 0 | 0 |
| NC_002645.1;NP_073549.1;HCoV-229E_1ab;5376 | 0 | 0 |
| NC_002645.1;NP_073549.1;HCoV-229E_1ab;5377 | 0 | 0 |
| NC_002645.1;NP_073549.1;HCoV-229E_1ab;5378 | 0 | 0 |
| NC_002645.1;NP_073549.1;HCoV-229E_1ab;5379 | 0 | 0 |
| NC_002645.1;NP_073549.1;HCoV-229E_1ab;5380 | 0 | 0 |
| NC_002645.1;NP_073549.1;HCoV-229E_1ab;5381 | 0 | 0 |
| NC_002645.1;NP_073549.1;HCoV-229E_1ab;5382 | 0 | 0 |
| NC_002645.1;NP_073549.1;HCoV-229E_1ab;5383 | 0 | 0 |
| NC_002645.1;NP_073549.1;HCoV-229E_1ab;5384 | 0 | 0 |
| NC_002645.1;NP_073549.1;HCoV-229E_1ab;5385 | 0 | 0 |
| NC_002645.1;NP_073549.1;HCoV-229E_1ab;5386 | 0 | 0 |
| NC_002645.1;NP_073549.1;HCoV-229E_1ab;5387 | 0 | 0 |
| NC_002645.1;NP_073549.1;HCoV-229E_1ab;5388 | 0 | 0 |
| NC_002645.1;NP_073549.1;HCoV-229E_1ab;5389 | 0 | 0 |
| NC_002645.1;NP_073549.1;HCoV-229E_1ab;5390 | 0 | 0 |
| NC_002645.1;NP_073549.1;HCoV-229E_1ab;5391 | 0 | 0 |
| NC_002645.1;NP_073549.1;HCoV-229E_1ab;5392 | 0 | 0 |
| NC_002645.1;NP_073549.1;HCoV-229E_1ab;5393 | 0 | 0 |
| NC_002645.1;NP_073549.1;HCoV-229E_1ab;5394 | 0 | 0 |
| NC_002645.1;NP_073549.1;HCoV-229E_1ab;5395 | 0 | 0 |
| NC_002645.1;NP_073549.1;HCoV-229E_1ab;5396 | 0 | 0 |
| NC_002645.1;NP_073549.1;HCoV-229E_1ab;5397 | 0 | 0 |
| NC_002645.1;NP_073549.1;HCoV-229E_1ab;5398 | 0 | 0 |
| NC_002645.1;NP_073549.1;HCoV-229E_1ab;5399 | 0 | 0 |
| NC_002645.1;NP_073549.1;HCoV-229E_1ab;5400 | 0 | 0 |
| NC_002645.1;NP_073549.1;HCoV-229E_1ab;5401 | 0 | 0 |
| NC_002645.1;NP_073549.1;HCoV-229E_1ab;5402 | 0 | 0.025 |
| NC_002645.1;NP_073549.1;HCoV-229E_1ab;5403 | 0 | 0.025 |
| NC_002645.1;NP_073549.1;HCoV-229E_1ab;5404 | 0 | 0 |

| | | |
|---|---|---|
| NC_002645.1;NP_073549.1;HCoV-229E_1ab;5405 | 0 | 0 |
| NC_002645.1;NP_073549.1;HCoV-229E_1ab;5406 | 0 | 0 |
| NC_002645.1;NP_073549.1;HCoV-229E_1ab;5407 | 0 | 0 |
| NC_002645.1;NP_073549.1;HCoV-229E_1ab;5408 | 0.05 | 0 |
| NC_002645.1;NP_073549.1;HCoV-229E_1ab;5409 | 0 | 0 |
| NC_002645.1;NP_073549.1;HCoV-229E_1ab;5410 | 0 | 0 |
| NC_002645.1;NP_073549.1;HCoV-229E_1ab;5411 | 0 | 0 |
| NC_002645.1;NP_073549.1;HCoV-229E_1ab;5412 | 0 | 0 |
| NC_002645.1;NP_073549.1;HCoV-229E_1ab;5413 | 0.05 | 0 |
| NC_002645.1;NP_073549.1;HCoV-229E_1ab;5414 | 0 | 0 |
| NC_002645.1;NP_073549.1;HCoV-229E_1ab;5415 | 0 | 0.025 |
| NC_002645.1;NP_073549.1;HCoV-229E_1ab;5416 | 0 | 0 |
| NC_002645.1;NP_073549.1;HCoV-229E_1ab;5417 | 0 | 0 |
| NC_002645.1;NP_073549.1;HCoV-229E_1ab;5418 | 0 | 0 |
| NC_002645.1;NP_073549.1;HCoV-229E_1ab;5419 | 0 | 0 |
| NC_002645.1;NP_073549.1;HCoV-229E_1ab;5420 | 0 | 0.05 |
| NC_002645.1;NP_073549.1;HCoV-229E_1ab;5421 | 0 | 0.05 |
| NC_002645.1;NP_073549.1;HCoV-229E_1ab;5422 | 0 | 0.05 |
| NC_002645.1;NP_073549.1;HCoV-229E_1ab;5423 | 0 | 0 |
| NC_002645.1;NP_073549.1;HCoV-229E_1ab;5424 | 0 | 0 |
| NC_002645.1;NP_073549.1;HCoV-229E_1ab;5425 | 0 | 0 |
| NC_002645.1;NP_073549.1;HCoV-229E_1ab;5426 | 0 | 0 |
| NC_002645.1;NP_073549.1;HCoV-229E_1ab;5427 | 0 | 0 |
| NC_002645.1;NP_073549.1;HCoV-229E_1ab;5428 | 0 | 0.05 |
| NC_002645.1;NP_073549.1;HCoV-229E_1ab;5429 | 0 | 0.025 |
| NC_002645.1;NP_073549.1;HCoV-229E_1ab;5430 | 0 | 0 |
| NC_002645.1;NP_073549.1;HCoV-229E_1ab;5431 | 0 | 0 |
| NC_002645.1;NP_073549.1;HCoV-229E_1ab;5432 | 0 | 0 |
| NC_002645.1;NP_073549.1;HCoV-229E_1ab;5433 | 0 | 0 |
| NC_002645.1;NP_073549.1;HCoV-229E_1ab;5434 | 0 | 0 |
| NC_002645.1;NP_073549.1;HCoV-229E_1ab;5435 | 0 | 0 |
| NC_002645.1;NP_073549.1;HCoV-229E_1ab;5436 | 0 | 0 |
| NC_002645.1;NP_073549.1;HCoV-229E_1ab;5437 | 0 | 0 |
| NC_002645.1;NP_073549.1;HCoV-229E_1ab;5438 | 0 | 0 |
| NC_002645.1;NP_073549.1;HCoV-229E_1ab;5439 | 0 | 0 |
| NC_002645.1;NP_073549.1;HCoV-229E_1ab;5440 | 0 | 0 |
| NC_002645.1;NP_073549.1;HCoV-229E_1ab;5441 | 0 | 0 |
| NC_002645.1;NP_073549.1;HCoV-229E_1ab;5442 | 0.1 | 0.15 |
| NC_002645.1;NP_073549.1;HCoV-229E_1ab;5443 | 0 | 0 |
| NC_002645.1;NP_073549.1;HCoV-229E_1ab;5444 | 0.05 | 0.025 |
| NC_002645.1;NP_073549.1;HCoV-229E_1ab;5445 | 0.05 | 0.1 |
| NC_002645.1;NP_073549.1;HCoV-229E_1ab;5446 | 0 | 0.025 |
| NC_002645.1;NP_073549.1;HCoV-229E_1ab;5447 | 0 | 0.15 |

| | | |
|---|---|---|
| NC_002645.1;NP_073549.1;HCoV-229E_1ab;5448 | 0.05 | 0.075 |
| NC_002645.1;NP_073549.1;HCoV-229E_1ab;5449 | 0 | 0.1 |
| NC_002645.1;NP_073549.1;HCoV-229E_1ab;5450 | 0 | 0.125 |
| NC_002645.1;NP_073549.1;HCoV-229E_1ab;5451 | 0 | 0 |
| NC_002645.1;NP_073549.1;HCoV-229E_1ab;5452 | 0 | 0.025 |
| NC_002645.1;NP_073549.1;HCoV-229E_1ab;5453 | 0 | 0.075 |
| NC_002645.1;NP_073549.1;HCoV-229E_1ab;5454 | 0 | 0 |
| NC_002645.1;NP_073549.1;HCoV-229E_1ab;5455 | 0 | 0 |
| NC_002645.1;NP_073549.1;HCoV-229E_1ab;5456 | 0 | 0 |
| NC_002645.1;NP_073549.1;HCoV-229E_1ab;5457 | 0 | 0 |
| NC_002645.1;NP_073549.1;HCoV-229E_1ab;5458 | 0 | 0 |
| NC_002645.1;NP_073549.1;HCoV-229E_1ab;5459 | 0 | 0.025 |
| NC_002645.1;NP_073549.1;HCoV-229E_1ab;5460 | 0 | 0 |
| NC_002645.1;NP_073549.1;HCoV-229E_1ab;5461 | 0 | 0 |
| NC_002645.1;NP_073549.1;HCoV-229E_1ab;5462 | 0 | 0 |
| NC_002645.1;NP_073549.1;HCoV-229E_1ab;5463 | 0 | 0 |
| NC_002645.1;NP_073549.1;HCoV-229E_1ab;5464 | 0 | 0 |
| NC_002645.1;NP_073549.1;HCoV-229E_1ab;5465 | 0 | 0 |
| NC_002645.1;NP_073549.1;HCoV-229E_1ab;5466 | 0 | 0 |
| NC_002645.1;NP_073549.1;HCoV-229E_1ab;5467 | 0 | 0 |
| NC_002645.1;NP_073549.1;HCoV-229E_1ab;5468 | 0 | 0 |
| NC_002645.1;NP_073549.1;HCoV-229E_1ab;5469 | 0 | 0.025 |
| NC_002645.1;NP_073549.1;HCoV-229E_1ab;5470 | 0 | 0.025 |
| NC_002645.1;NP_073549.1;HCoV-229E_1ab;5471 | 0 | 0.05 |
| NC_002645.1;NP_073549.1;HCoV-229E_1ab;5472 | 0 | 0.025 |
| NC_002645.1;NP_073549.1;HCoV-229E_1ab;5473 | 0 | 0.025 |
| NC_002645.1;NP_073549.1;HCoV-229E_1ab;5474 | 0 | 0.075 |
| NC_002645.1;NP_073549.1;HCoV-229E_1ab;5475 | 0 | 0.025 |
| NC_002645.1;NP_073549.1;HCoV-229E_1ab;5476 | 0.1 | 0.025 |
| NC_002645.1;NP_073549.1;HCoV-229E_1ab;5477 | 0 | 0.025 |
| NC_002645.1;NP_073549.1;HCoV-229E_1ab;5478 | 0 | 0.025 |
| NC_002645.1;NP_073549.1;HCoV-229E_1ab;5479 | 0.05 | 0.075 |
| NC_002645.1;NP_073549.1;HCoV-229E_1ab;5480 | 0 | 0 |
| NC_002645.1;NP_073549.1;HCoV-229E_1ab;5481 | 0 | 0 |
| NC_002645.1;NP_073549.1;HCoV-229E_1ab;5482 | 0 | 0 |
| NC_002645.1;NP_073549.1;HCoV-229E_1ab;5483 | 0 | 0 |
| NC_002645.1;NP_073549.1;HCoV-229E_1ab;5484 | 0 | 0 |
| NC_002645.1;NP_073549.1;HCoV-229E_1ab;5485 | 0 | 0.025 |
| NC_002645.1;NP_073549.1;HCoV-229E_1ab;5486 | 0 | 0 |
| NC_002645.1;NP_073549.1;HCoV-229E_1ab;5487 | 0 | 0 |
| NC_002645.1;NP_073549.1;HCoV-229E_1ab;5488 | 0 | 0 |
| NC_002645.1;NP_073549.1;HCoV-229E_1ab;5489 | 0 | 0 |
| NC_002645.1;NP_073549.1;HCoV-229E_1ab;5490 | 0 | 0 |

| | | |
|---|---|---|
| NC_002645.1;NP_073549.1;HCoV-229E_1ab;5491 | 0 | 0 |
| NC_002645.1;NP_073549.1;HCoV-229E_1ab;5492 | 0 | 0 |
| NC_002645.1;NP_073549.1;HCoV-229E_1ab;5493 | 0 | 0 |
| NC_002645.1;NP_073549.1;HCoV-229E_1ab;5494 | 0 | 0 |
| NC_002645.1;NP_073549.1;HCoV-229E_1ab;5495 | 0 | 0 |
| NC_002645.1;NP_073549.1;HCoV-229E_1ab;5496 | 0 | 0 |
| NC_002645.1;NP_073549.1;HCoV-229E_1ab;5497 | 0 | 0 |
| NC_002645.1;NP_073549.1;HCoV-229E_1ab;5498 | 0 | 0 |
| NC_002645.1;NP_073549.1;HCoV-229E_1ab;5499 | 0 | 0 |
| NC_002645.1;NP_073549.1;HCoV-229E_1ab;5500 | 0.05 | 0 |
| NC_002645.1;NP_073549.1;HCoV-229E_1ab;5501 | 0 | 0 |
| NC_002645.1;NP_073549.1;HCoV-229E_1ab;5502 | 0 | 0 |
| NC_002645.1;NP_073549.1;HCoV-229E_1ab;5503 | 0.05 | 0 |
| NC_002645.1;NP_073549.1;HCoV-229E_1ab;5504 | 0.1 | 0 |
| NC_002645.1;NP_073549.1;HCoV-229E_1ab;5505 | 0.1 | 0 |
| NC_002645.1;NP_073549.1;HCoV-229E_1ab;5506 | 0.05 | 0 |
| NC_002645.1;NP_073549.1;HCoV-229E_1ab;5507 | 0 | 0 |
| NC_002645.1;NP_073549.1;HCoV-229E_1ab;5508 | 0 | 0 |
| NC_002645.1;NP_073549.1;HCoV-229E_1ab;5509 | 0 | 0 |
| NC_002645.1;NP_073549.1;HCoV-229E_1ab;5510 | 0 | 0.025 |
| NC_002645.1;NP_073549.1;HCoV-229E_1ab;5511 | 0 | 0 |
| NC_002645.1;NP_073549.1;HCoV-229E_1ab;5512 | 0 | 0 |
| NC_002645.1;NP_073549.1;HCoV-229E_1ab;5513 | 0 | 0 |
| NC_002645.1;NP_073549.1;HCoV-229E_1ab;5514 | 0 | 0 |
| NC_002645.1;NP_073549.1;HCoV-229E_1ab;5515 | 0 | 0 |
| NC_002645.1;NP_073549.1;HCoV-229E_1ab;5516 | 0 | 0 |
| NC_002645.1;NP_073549.1;HCoV-229E_1ab;5517 | 0 | 0 |
| NC_002645.1;NP_073549.1;HCoV-229E_1ab;5518 | 0 | 0.025 |
| NC_002645.1;NP_073549.1;HCoV-229E_1ab;5519 | 0 | 0.05 |
| NC_002645.1;NP_073549.1;HCoV-229E_1ab;5520 | 0 | 0.05 |
| NC_002645.1;NP_073549.1;HCoV-229E_1ab;5521 | 0 | 0 |
| NC_002645.1;NP_073549.1;HCoV-229E_1ab;5522 | 0 | 0 |
| NC_002645.1;NP_073549.1;HCoV-229E_1ab;5523 | 0.05 | 0.025 |
| NC_002645.1;NP_073549.1;HCoV-229E_1ab;5524 | 0.05 | 0 |
| NC_002645.1;NP_073549.1;HCoV-229E_1ab;5525 | 0.05 | 0 |
| NC_002645.1;NP_073549.1;HCoV-229E_1ab;5526 | 0 | 0 |
| NC_002645.1;NP_073549.1;HCoV-229E_1ab;5527 | 0 | 0 |
| NC_002645.1;NP_073549.1;HCoV-229E_1ab;5528 | 0 | 0.025 |
| NC_002645.1;NP_073549.1;HCoV-229E_1ab;5529 | 0 | 0.025 |
| NC_002645.1;NP_073549.1;HCoV-229E_1ab;5530 | 0 | 0 |
| NC_002645.1;NP_073549.1;HCoV-229E_1ab;5531 | 0 | 0 |
| NC_002645.1;NP_073549.1;HCoV-229E_1ab;5532 | 0 | 0 |
| NC_002645.1;NP_073549.1;HCoV-229E_1ab;5533 | 0 | 0 |

| | | |
|---|---|---|
| NC_002645.1;NP_073549.1;HCoV-229E_1ab;5534 | 0 | 0 |
| NC_002645.1;NP_073549.1;HCoV-229E_1ab;5535 | 0 | 0 |
| NC_002645.1;NP_073549.1;HCoV-229E_1ab;5536 | 0 | 0 |
| NC_002645.1;NP_073549.1;HCoV-229E_1ab;5537 | 0 | 0 |
| NC_002645.1;NP_073549.1;HCoV-229E_1ab;5538 | 0 | 0 |
| NC_002645.1;NP_073549.1;HCoV-229E_1ab;5539 | 0 | 0 |
| NC_002645.1;NP_073549.1;HCoV-229E_1ab;5540 | 0 | 0.025 |
| NC_002645.1;NP_073549.1;HCoV-229E_1ab;5541 | 0 | 0.025 |
| NC_002645.1;NP_073549.1;HCoV-229E_1ab;5542 | 0 | 0.025 |
| NC_002645.1;NP_073549.1;HCoV-229E_1ab;5543 | 0.05 | 0.05 |
| NC_002645.1;NP_073549.1;HCoV-229E_1ab;5544 | 0 | 0.025 |
| NC_002645.1;NP_073549.1;HCoV-229E_1ab;5545 | 0.05 | 0.075 |
| NC_002645.1;NP_073549.1;HCoV-229E_1ab;5546 | 0 | 0.025 |
| NC_002645.1;NP_073549.1;HCoV-229E_1ab;5547 | 0 | 0 |
| NC_002645.1;NP_073549.1;HCoV-229E_1ab;5548 | 0 | 0 |
| NC_002645.1;NP_073549.1;HCoV-229E_1ab;5549 | 0 | 0 |
| NC_002645.1;NP_073549.1;HCoV-229E_1ab;5550 | 0 | 0 |
| NC_002645.1;NP_073549.1;HCoV-229E_1ab;5551 | 0 | 0 |
| NC_002645.1;NP_073549.1;HCoV-229E_1ab;5552 | 0 | 0 |
| NC_002645.1;NP_073549.1;HCoV-229E_1ab;5553 | 0 | 0 |
| NC_002645.1;NP_073549.1;HCoV-229E_1ab;5554 | 0.05 | 0 |
| NC_002645.1;NP_073549.1;HCoV-229E_1ab;5555 | 0.05 | 0 |
| NC_002645.1;NP_073549.1;HCoV-229E_1ab;5556 | 0 | 0 |
| NC_002645.1;NP_073549.1;HCoV-229E_1ab;5557 | 0 | 0 |
| NC_002645.1;NP_073549.1;HCoV-229E_1ab;5558 | 0 | 0 |
| NC_002645.1;NP_073549.1;HCoV-229E_1ab;5559 | 0 | 0 |
| NC_002645.1;NP_073549.1;HCoV-229E_1ab;5560 | 0 | 0 |
| NC_002645.1;NP_073549.1;HCoV-229E_1ab;5561 | 0 | 0 |
| NC_002645.1;NP_073549.1;HCoV-229E_1ab;5562 | 0 | 0 |
| NC_002645.1;NP_073549.1;HCoV-229E_1ab;5563 | 0 | 0 |
| NC_002645.1;NP_073549.1;HCoV-229E_1ab;5564 | 0 | 0 |
| NC_002645.1;NP_073549.1;HCoV-229E_1ab;5565 | 0 | 0 |
| NC_002645.1;NP_073549.1;HCoV-229E_1ab;5566 | 0 | 0 |
| NC_002645.1;NP_073549.1;HCoV-229E_1ab;5567 | 0 | 0 |
| NC_002645.1;NP_073549.1;HCoV-229E_1ab;5568 | 0 | 0 |
| NC_002645.1;NP_073549.1;HCoV-229E_1ab;5569 | 0 | 0 |
| NC_002645.1;NP_073549.1;HCoV-229E_1ab;5570 | 0 | 0 |
| NC_002645.1;NP_073549.1;HCoV-229E_1ab;5571 | 0 | 0 |
| NC_002645.1;NP_073549.1;HCoV-229E_1ab;5572 | 0 | 0 |
| NC_002645.1;NP_073549.1;HCoV-229E_1ab;5573 | 0 | 0 |
| NC_002645.1;NP_073549.1;HCoV-229E_1ab;5574 | 0 | 0 |
| NC_002645.1;NP_073549.1;HCoV-229E_1ab;5575 | 0 | 0 |
| NC_002645.1;NP_073549.1;HCoV-229E_1ab;5576 | 0 | 0 |

| | | |
|---|---|---|
| NC_002645.1;NP_073549.1;HCoV-229E_1ab;5577 | 0 | 0 |
| NC_002645.1;NP_073549.1;HCoV-229E_1ab;5578 | 0 | 0 |
| NC_002645.1;NP_073549.1;HCoV-229E_1ab;5579 | 0 | 0 |
| NC_002645.1;NP_073549.1;HCoV-229E_1ab;5580 | 0 | 0 |
| NC_002645.1;NP_073549.1;HCoV-229E_1ab;5581 | 0 | 0 |
| NC_002645.1;NP_073549.1;HCoV-229E_1ab;5582 | 0 | 0 |
| NC_002645.1;NP_073549.1;HCoV-229E_1ab;5583 | 0 | 0.025 |
| NC_002645.1;NP_073549.1;HCoV-229E_1ab;5584 | 0 | 0 |
| NC_002645.1;NP_073549.1;HCoV-229E_1ab;5585 | 0 | 0 |
| NC_002645.1;NP_073549.1;HCoV-229E_1ab;5586 | 0 | 0.025 |
| NC_002645.1;NP_073549.1;HCoV-229E_1ab;5587 | 0 | 0 |
| NC_002645.1;NP_073549.1;HCoV-229E_1ab;5588 | 0 | 0 |
| NC_002645.1;NP_073549.1;HCoV-229E_1ab;5589 | 0.05 | 0.025 |
| NC_002645.1;NP_073549.1;HCoV-229E_1ab;5590 | 0 | 0 |
| NC_002645.1;NP_073549.1;HCoV-229E_1ab;5591 | 0 | 0 |
| NC_002645.1;NP_073549.1;HCoV-229E_1ab;5592 | 0 | 0.05 |
| NC_002645.1;NP_073549.1;HCoV-229E_1ab;5593 | 0 | 0.05 |
| NC_002645.1;NP_073549.1;HCoV-229E_1ab;5594 | 0 | 0.025 |
| NC_002645.1;NP_073549.1;HCoV-229E_1ab;5595 | 0 | 0.05 |
| NC_002645.1;NP_073549.1;HCoV-229E_1ab;5596 | 0.05 | 0.05 |
| NC_002645.1;NP_073549.1;HCoV-229E_1ab;5597 | 0 | 0.05 |
| NC_002645.1;NP_073549.1;HCoV-229E_1ab;5598 | 0.05 | 0.075 |
| NC_002645.1;NP_073549.1;HCoV-229E_1ab;5599 | 0 | 0.05 |
| NC_002645.1;NP_073549.1;HCoV-229E_1ab;5600 | 0 | 0.05 |
| NC_002645.1;NP_073549.1;HCoV-229E_1ab;5601 | 0 | 0.025 |
| NC_002645.1;NP_073549.1;HCoV-229E_1ab;5602 | 0 | 0.025 |
| NC_002645.1;NP_073549.1;HCoV-229E_1ab;5603 | 0.05 | 0.05 |
| NC_002645.1;NP_073549.1;HCoV-229E_1ab;5604 | 0 | 0.025 |
| NC_002645.1;NP_073549.1;HCoV-229E_1ab;5605 | 0 | 0 |
| NC_002645.1;NP_073549.1;HCoV-229E_1ab;5606 | 0.05 | 0.025 |
| NC_002645.1;NP_073549.1;HCoV-229E_1ab;5607 | 0 | 0 |
| NC_002645.1;NP_073549.1;HCoV-229E_1ab;5608 | 0 | 0 |
| NC_002645.1;NP_073549.1;HCoV-229E_1ab;5609 | 0 | 0 |
| NC_002645.1;NP_073549.1;HCoV-229E_1ab;5610 | 0 | 0 |
| NC_002645.1;NP_073549.1;HCoV-229E_1ab;5611 | 0 | 0 |
| NC_002645.1;NP_073549.1;HCoV-229E_1ab;5612 | 0 | 0 |
| NC_002645.1;NP_073549.1;HCoV-229E_1ab;5613 | 0 | 0 |
| NC_002645.1;NP_073549.1;HCoV-229E_1ab;5614 | 0 | 0 |
| NC_002645.1;NP_073549.1;HCoV-229E_1ab;5615 | 0 | 0 |
| NC_002645.1;NP_073549.1;HCoV-229E_1ab;5616 | 0 | 0 |
| NC_002645.1;NP_073549.1;HCoV-229E_1ab;5617 | 0 | 0 |
| NC_002645.1;NP_073549.1;HCoV-229E_1ab;5618 | 0.15 | 0.05 |
| NC_002645.1;NP_073549.1;HCoV-229E_1ab;5619 | 0.15 | 0.025 |

| | | |
|---|---|---|
| NC_002645.1;NP_073549.1;HCoV-229E_1ab;5620 | 0.05 | 0.025 |
| NC_002645.1;NP_073549.1;HCoV-229E_1ab;5621 | 0.1 | 0.05 |
| NC_002645.1;NP_073549.1;HCoV-229E_1ab;5622 | 0.15 | 0.05 |
| NC_002645.1;NP_073549.1;HCoV-229E_1ab;5623 | 0.1 | 0.1 |
| NC_002645.1;NP_073549.1;HCoV-229E_1ab;5624 | 0.2 | 0.125 |
| NC_002645.1;NP_073549.1;HCoV-229E_1ab;5625 | 0.2 | 0.2 |
| NC_002645.1;NP_073549.1;HCoV-229E_1ab;5626 | 0.2 | 0.125 |
| NC_002645.1;NP_073549.1;HCoV-229E_1ab;5627 | 0.05 | 0.075 |
| NC_002645.1;NP_073549.1;HCoV-229E_1ab;5628 | 0.05 | 0.075 |
| NC_002645.1;NP_073549.1;HCoV-229E_1ab;5629 | 0.05 | 0.05 |
| NC_002645.1;NP_073549.1;HCoV-229E_1ab;5630 | 0.05 | 0.05 |
| NC_002645.1;NP_073549.1;HCoV-229E_1ab;5631 | 0 | 0.025 |
| NC_002645.1;NP_073549.1;HCoV-229E_1ab;5632 | 0 | 0.025 |
| NC_002645.1;NP_073549.1;HCoV-229E_1ab;5633 | 0 | 0.025 |
| NC_002645.1;NP_073549.1;HCoV-229E_1ab;5634 | 0 | 0 |
| NC_002645.1;NP_073549.1;HCoV-229E_1ab;5635 | 0 | 0.025 |
| NC_002645.1;NP_073549.1;HCoV-229E_1ab;5636 | 0 | 0 |
| NC_002645.1;NP_073549.1;HCoV-229E_1ab;5637 | 0 | 0 |
| NC_002645.1;NP_073549.1;HCoV-229E_1ab;5638 | 0 | 0 |
| NC_002645.1;NP_073549.1;HCoV-229E_1ab;5639 | 0 | 0 |
| NC_002645.1;NP_073549.1;HCoV-229E_1ab;5640 | 0 | 0 |
| NC_002645.1;NP_073549.1;HCoV-229E_1ab;5641 | 0 | 0 |
| NC_002645.1;NP_073549.1;HCoV-229E_1ab;5642 | 0 | 0 |
| NC_002645.1;NP_073549.1;HCoV-229E_1ab;5643 | 0 | 0 |
| NC_002645.1;NP_073549.1;HCoV-229E_1ab;5644 | 0 | 0 |
| NC_002645.1;NP_073549.1;HCoV-229E_1ab;5645 | 0 | 0 |
| NC_002645.1;NP_073549.1;HCoV-229E_1ab;5646 | 0 | 0 |
| NC_002645.1;NP_073549.1;HCoV-229E_1ab;5647 | 0 | 0 |
| NC_002645.1;NP_073549.1;HCoV-229E_1ab;5648 | 0 | 0 |
| NC_002645.1;NP_073549.1;HCoV-229E_1ab;5649 | 0 | 0 |
| NC_002645.1;NP_073549.1;HCoV-229E_1ab;5650 | 0 | 0 |
| NC_002645.1;NP_073549.1;HCoV-229E_1ab;5651 | 0 | 0 |
| NC_002645.1;NP_073549.1;HCoV-229E_1ab;5652 | 0 | 0 |
| NC_002645.1;NP_073549.1;HCoV-229E_1ab;5653 | 0 | 0 |
| NC_002645.1;NP_073549.1;HCoV-229E_1ab;5654 | 0 | 0 |
| NC_002645.1;NP_073549.1;HCoV-229E_1ab;5655 | 0 | 0 |
| NC_002645.1;NP_073549.1;HCoV-229E_1ab;5656 | 0 | 0 |
| NC_002645.1;NP_073549.1;HCoV-229E_1ab;5657 | 0 | 0 |
| NC_002645.1;NP_073549.1;HCoV-229E_1ab;5658 | 0 | 0 |
| NC_002645.1;NP_073549.1;HCoV-229E_1ab;5659 | 0 | 0 |
| NC_002645.1;NP_073549.1;HCoV-229E_1ab;5660 | 0 | 0 |
| NC_002645.1;NP_073549.1;HCoV-229E_1ab;5661 | 0 | 0 |
| NC_002645.1;NP_073549.1;HCoV-229E_1ab;5662 | 0 | 0 |

| | | |
|---|---|---|
| NC_002645.1;NP_073549.1;HCoV-229E_1ab;5663 | 0 | 0 |
| NC_002645.1;NP_073549.1;HCoV-229E_1ab;5664 | 0 | 0 |
| NC_002645.1;NP_073549.1;HCoV-229E_1ab;5665 | 0 | 0 |
| NC_002645.1;NP_073549.1;HCoV-229E_1ab;5666 | 0 | 0 |
| NC_002645.1;NP_073549.1;HCoV-229E_1ab;5667 | 0 | 0.025 |
| NC_002645.1;NP_073549.1;HCoV-229E_1ab;5668 | 0 | 0 |
| NC_002645.1;NP_073549.1;HCoV-229E_1ab;5669 | 0 | 0 |
| NC_002645.1;NP_073549.1;HCoV-229E_1ab;5670 | 0 | 0.025 |
| NC_002645.1;NP_073549.1;HCoV-229E_1ab;5671 | 0 | 0.025 |
| NC_002645.1;NP_073549.1;HCoV-229E_1ab;5672 | 0 | 0.025 |
| NC_002645.1;NP_073549.1;HCoV-229E_1ab;5673 | 0 | 0 |
| NC_002645.1;NP_073549.1;HCoV-229E_1ab;5674 | 0 | 0 |
| NC_002645.1;NP_073549.1;HCoV-229E_1ab;5675 | 0 | 0 |
| NC_002645.1;NP_073549.1;HCoV-229E_1ab;5676 | 0 | 0.025 |
| NC_002645.1;NP_073549.1;HCoV-229E_1ab;5677 | 0 | 0.025 |
| NC_002645.1;NP_073549.1;HCoV-229E_1ab;5678 | 0 | 0 |
| NC_002645.1;NP_073549.1;HCoV-229E_1ab;5679 | 0 | 0 |
| NC_002645.1;NP_073549.1;HCoV-229E_1ab;5680 | 0 | 0 |
| NC_002645.1;NP_073549.1;HCoV-229E_1ab;5681 | 0 | 0 |
| NC_002645.1;NP_073549.1;HCoV-229E_1ab;5682 | 0 | 0 |
| NC_002645.1;NP_073549.1;HCoV-229E_1ab;5683 | 0.1 | 0.025 |
| NC_002645.1;NP_073549.1;HCoV-229E_1ab;5684 | 0.05 | 0 |
| NC_002645.1;NP_073549.1;HCoV-229E_1ab;5685 | 0.05 | 0 |
| NC_002645.1;NP_073549.1;HCoV-229E_1ab;5686 | 0.05 | 0 |
| NC_002645.1;NP_073549.1;HCoV-229E_1ab;5687 | 0.05 | 0 |
| NC_002645.1;NP_073549.1;HCoV-229E_1ab;5688 | 0.1 | 0 |
| NC_002645.1;NP_073549.1;HCoV-229E_1ab;5689 | 0.05 | 0 |
| NC_002645.1;NP_073549.1;HCoV-229E_1ab;5690 | 0.05 | 0.025 |
| NC_002645.1;NP_073549.1;HCoV-229E_1ab;5691 | 0.05 | 0 |
| NC_002645.1;NP_073549.1;HCoV-229E_1ab;5692 | 0 | 0 |
| NC_002645.1;NP_073549.1;HCoV-229E_1ab;5693 | 0 | 0.075 |
| NC_002645.1;NP_073549.1;HCoV-229E_1ab;5694 | 0 | 0 |
| NC_002645.1;NP_073549.1;HCoV-229E_1ab;5695 | 0.05 | 0 |
| NC_002645.1;NP_073549.1;HCoV-229E_1ab;5696 | 0 | 0.05 |
| NC_002645.1;NP_073549.1;HCoV-229E_1ab;5697 | 0 | 0 |
| NC_002645.1;NP_073549.1;HCoV-229E_1ab;5698 | 0 | 0 |
| NC_002645.1;NP_073549.1;HCoV-229E_1ab;5699 | 0.05 | 0.025 |
| NC_002645.1;NP_073549.1;HCoV-229E_1ab;5700 | 0.1 | 0.025 |
| NC_002645.1;NP_073549.1;HCoV-229E_1ab;5701 | 0.1 | 0.05 |
| NC_002645.1;NP_073549.1;HCoV-229E_1ab;5702 | 0.05 | 0.05 |
| NC_002645.1;NP_073549.1;HCoV-229E_1ab;5703 | 0 | 0.025 |
| NC_002645.1;NP_073549.1;HCoV-229E_1ab;5704 | 0 | 0 |
| NC_002645.1;NP_073549.1;HCoV-229E_1ab;5705 | 0 | 0 |

| | | |
|---|---|---|
| NC_002645.1;NP_073549.1;HCoV-229E_1ab;5706 | 0.15 | 0.1 |
| NC_002645.1;NP_073549.1;HCoV-229E_1ab;5707 | 0 | 0 |
| NC_002645.1;NP_073549.1;HCoV-229E_1ab;5708 | 0 | 0 |
| NC_002645.1;NP_073549.1;HCoV-229E_1ab;5709 | 0 | 0 |
| NC_002645.1;NP_073549.1;HCoV-229E_1ab;5710 | 0 | 0 |
| NC_002645.1;NP_073549.1;HCoV-229E_1ab;5711 | 0 | 0 |
| NC_002645.1;NP_073549.1;HCoV-229E_1ab;5712 | 0 | 0 |
| NC_002645.1;NP_073549.1;HCoV-229E_1ab;5713 | 0 | 0 |
| NC_002645.1;NP_073549.1;HCoV-229E_1ab;5714 | 0 | 0 |
| NC_002645.1;NP_073549.1;HCoV-229E_1ab;5715 | 0 | 0 |
| NC_002645.1;NP_073549.1;HCoV-229E_1ab;5716 | 0 | 0 |
| NC_002645.1;NP_073549.1;HCoV-229E_1ab;5717 | 0 | 0 |
| NC_002645.1;NP_073549.1;HCoV-229E_1ab;5718 | 0 | 0.025 |
| NC_002645.1;NP_073549.1;HCoV-229E_1ab;5719 | 0 | 0.05 |
| NC_002645.1;NP_073549.1;HCoV-229E_1ab;5720 | 0.05 | 0.025 |
| NC_002645.1;NP_073549.1;HCoV-229E_1ab;5721 | 0 | 0.025 |
| NC_002645.1;NP_073549.1;HCoV-229E_1ab;5722 | 0 | 0.05 |
| NC_002645.1;NP_073549.1;HCoV-229E_1ab;5723 | 0 | 0.025 |
| NC_002645.1;NP_073549.1;HCoV-229E_1ab;5724 | 0 | 0.025 |
| NC_002645.1;NP_073549.1;HCoV-229E_1ab;5725 | 0 | 0.05 |
| NC_002645.1;NP_073549.1;HCoV-229E_1ab;5726 | 0 | 0.025 |
| NC_002645.1;NP_073549.1;HCoV-229E_1ab;5727 | 0.05 | 0 |
| NC_002645.1;NP_073549.1;HCoV-229E_1ab;5728 | 0 | 0 |
| NC_002645.1;NP_073549.1;HCoV-229E_1ab;5729 | 0 | 0 |
| NC_002645.1;NP_073549.1;HCoV-229E_1ab;5730 | 0 | 0 |
| NC_002645.1;NP_073549.1;HCoV-229E_1ab;5731 | 0 | 0 |
| NC_002645.1;NP_073549.1;HCoV-229E_1ab;5732 | 0 | 0 |
| NC_002645.1;NP_073549.1;HCoV-229E_1ab;5733 | 0 | 0 |
| NC_002645.1;NP_073549.1;HCoV-229E_1ab;5734 | 0 | 0 |
| NC_002645.1;NP_073549.1;HCoV-229E_1ab;5735 | 0 | 0 |
| NC_002645.1;NP_073549.1;HCoV-229E_1ab;5736 | 0 | 0 |
| NC_002645.1;NP_073549.1;HCoV-229E_1ab;5737 | 0 | 0 |
| NC_002645.1;NP_073549.1;HCoV-229E_1ab;5738 | 0 | 0 |
| NC_002645.1;NP_073549.1;HCoV-229E_1ab;5739 | 0 | 0 |
| NC_002645.1;NP_073549.1;HCoV-229E_1ab;5740 | 0 | 0 |
| NC_002645.1;NP_073549.1;HCoV-229E_1ab;5741 | 0 | 0 |
| NC_002645.1;NP_073549.1;HCoV-229E_1ab;5742 | 0 | 0 |
| NC_002645.1;NP_073549.1;HCoV-229E_1ab;5743 | 0 | 0 |
| NC_002645.1;NP_073549.1;HCoV-229E_1ab;5744 | 0 | 0 |
| NC_002645.1;NP_073549.1;HCoV-229E_1ab;5745 | 0 | 0 |
| NC_002645.1;NP_073549.1;HCoV-229E_1ab;5746 | 0 | 0 |
| NC_002645.1;NP_073549.1;HCoV-229E_1ab;5747 | 0 | 0 |
| NC_002645.1;NP_073549.1;HCoV-229E_1ab;5748 | 0 | 0 |

| | | |
|---|---|---|
| NC_002645.1;NP_073549.1;HCoV-229E_1ab;5749 | 0 | 0 |
| NC_002645.1;NP_073549.1;HCoV-229E_1ab;5750 | 0 | 0 |
| NC_002645.1;NP_073549.1;HCoV-229E_1ab;5751 | 0 | 0 |
| NC_002645.1;NP_073549.1;HCoV-229E_1ab;5752 | 0 | 0 |
| NC_002645.1;NP_073549.1;HCoV-229E_1ab;5753 | 0 | 0 |
| NC_002645.1;NP_073549.1;HCoV-229E_1ab;5754 | 0 | 0 |
| NC_002645.1;NP_073549.1;HCoV-229E_1ab;5755 | 0 | 0 |
| NC_002645.1;NP_073549.1;HCoV-229E_1ab;5756 | 0.05 | 0.075 |
| NC_002645.1;NP_073549.1;HCoV-229E_1ab;5757 | 0 | 0 |
| NC_002645.1;NP_073549.1;HCoV-229E_1ab;5758 | 0 | 0 |
| NC_002645.1;NP_073549.1;HCoV-229E_1ab;5759 | 0 | 0 |
| NC_002645.1;NP_073549.1;HCoV-229E_1ab;5760 | 0 | 0 |
| NC_002645.1;NP_073549.1;HCoV-229E_1ab;5761 | 0 | 0 |
| NC_002645.1;NP_073549.1;HCoV-229E_1ab;5762 | 0 | 0 |
| NC_002645.1;NP_073549.1;HCoV-229E_1ab;5763 | 0 | 0 |
| NC_002645.1;NP_073549.1;HCoV-229E_1ab;5764 | 0 | 0 |
| NC_002645.1;NP_073549.1;HCoV-229E_1ab;5765 | 0 | 0 |
| NC_002645.1;NP_073549.1;HCoV-229E_1ab;5766 | 0 | 0 |
| NC_002645.1;NP_073549.1;HCoV-229E_1ab;5767 | 0 | 0 |
| NC_002645.1;NP_073549.1;HCoV-229E_1ab;5768 | 0 | 0 |
| NC_002645.1;NP_073549.1;HCoV-229E_1ab;5769 | 0 | 0 |
| NC_002645.1;NP_073549.1;HCoV-229E_1ab;5770 | 0 | 0 |
| NC_002645.1;NP_073549.1;HCoV-229E_1ab;5771 | 0 | 0 |
| NC_002645.1;NP_073549.1;HCoV-229E_1ab;5772 | 0 | 0 |
| NC_002645.1;NP_073549.1;HCoV-229E_1ab;5773 | 0 | 0 |
| NC_002645.1;NP_073549.1;HCoV-229E_1ab;5774 | 0 | 0 |
| NC_002645.1;NP_073549.1;HCoV-229E_1ab;5775 | 0 | 0 |
| NC_002645.1;NP_073549.1;HCoV-229E_1ab;5776 | 0 | 0 |
| NC_002645.1;NP_073549.1;HCoV-229E_1ab;5777 | 0 | 0 |
| NC_002645.1;NP_073549.1;HCoV-229E_1ab;5778 | 0 | 0 |
| NC_002645.1;NP_073549.1;HCoV-229E_1ab;5779 | 0 | 0 |
| NC_002645.1;NP_073549.1;HCoV-229E_1ab;5780 | 0 | 0 |
| NC_002645.1;NP_073549.1;HCoV-229E_1ab;5781 | 0 | 0 |
| NC_002645.1;NP_073549.1;HCoV-229E_1ab;5782 | 0 | 0 |
| NC_002645.1;NP_073549.1;HCoV-229E_1ab;5783 | 0 | 0 |
| NC_002645.1;NP_073549.1;HCoV-229E_1ab;5784 | 0 | 0 |
| NC_002645.1;NP_073549.1;HCoV-229E_1ab;5785 | 0 | 0 |
| NC_002645.1;NP_073549.1;HCoV-229E_1ab;5786 | 0 | 0 |
| NC_002645.1;NP_073549.1;HCoV-229E_1ab;5787 | 0 | 0 |
| NC_002645.1;NP_073549.1;HCoV-229E_1ab;5788 | 0 | 0 |
| NC_002645.1;NP_073549.1;HCoV-229E_1ab;5789 | 0 | 0 |
| NC_002645.1;NP_073549.1;HCoV-229E_1ab;5790 | 0 | 0 |
| NC_002645.1;NP_073549.1;HCoV-229E_1ab;5791 | 0 | 0.075 |

| | | |
|---|---|---|
| NC_002645.1;NP_073549.1;HCoV-229E_1ab;5792 | 0 | 0 |
| NC_002645.1;NP_073549.1;HCoV-229E_1ab;5793 | 0 | 0 |
| NC_002645.1;NP_073549.1;HCoV-229E_1ab;5794 | 0 | 0 |
| NC_002645.1;NP_073549.1;HCoV-229E_1ab;5795 | 0 | 0 |
| NC_002645.1;NP_073549.1;HCoV-229E_1ab;5796 | 0 | 0 |
| NC_002645.1;NP_073549.1;HCoV-229E_1ab;5797 | 0 | 0 |
| NC_002645.1;NP_073549.1;HCoV-229E_1ab;5798 | 0 | 0 |
| NC_002645.1;NP_073549.1;HCoV-229E_1ab;5799 | 0 | 0 |
| NC_002645.1;NP_073549.1;HCoV-229E_1ab;5800 | 0 | 0 |
| NC_002645.1;NP_073549.1;HCoV-229E_1ab;5801 | 0 | 0 |
| NC_002645.1;NP_073549.1;HCoV-229E_1ab;5802 | 0 | 0 |
| NC_002645.1;NP_073549.1;HCoV-229E_1ab;5803 | 0 | 0 |
| NC_002645.1;NP_073549.1;HCoV-229E_1ab;5804 | 0 | 0 |
| NC_002645.1;NP_073549.1;HCoV-229E_1ab;5805 | 0 | 0 |
| NC_002645.1;NP_073549.1;HCoV-229E_1ab;5806 | 0 | 0 |
| NC_002645.1;NP_073549.1;HCoV-229E_1ab;5807 | 0.05 | 0 |
| NC_002645.1;NP_073549.1;HCoV-229E_1ab;5808 | 0.05 | 0.025 |
| NC_002645.1;NP_073549.1;HCoV-229E_1ab;5809 | 0 | 0 |
| NC_002645.1;NP_073549.1;HCoV-229E_1ab;5810 | 0 | 0.025 |
| NC_002645.1;NP_073549.1;HCoV-229E_1ab;5811 | 0 | 0 |
| NC_002645.1;NP_073549.1;HCoV-229E_1ab;5812 | 0.05 | 0.075 |
| NC_002645.1;NP_073549.1;HCoV-229E_1ab;5813 | 0 | 0 |
| NC_002645.1;NP_073549.1;HCoV-229E_1ab;5814 | 0 | 0 |
| NC_002645.1;NP_073549.1;HCoV-229E_1ab;5815 | 0 | 0 |
| NC_002645.1;NP_073549.1;HCoV-229E_1ab;5816 | 0 | 0 |
| NC_002645.1;NP_073549.1;HCoV-229E_1ab;5817 | 0 | 0 |
| NC_002645.1;NP_073549.1;HCoV-229E_1ab;5818 | 0 | 0 |
| NC_002645.1;NP_073549.1;HCoV-229E_1ab;5819 | 0 | 0 |
| NC_002645.1;NP_073549.1;HCoV-229E_1ab;5820 | 0 | 0 |
| NC_002645.1;NP_073549.1;HCoV-229E_1ab;5821 | 0 | 0 |
| NC_002645.1;NP_073549.1;HCoV-229E_1ab;5822 | 0 | 0 |
| NC_002645.1;NP_073549.1;HCoV-229E_1ab;5823 | 0 | 0 |
| NC_002645.1;NP_073549.1;HCoV-229E_1ab;5824 | 0 | 0 |
| NC_002645.1;NP_073549.1;HCoV-229E_1ab;5825 | 0 | 0 |
| NC_002645.1;NP_073549.1;HCoV-229E_1ab;5826 | 0 | 0 |
| NC_002645.1;NP_073549.1;HCoV-229E_1ab;5827 | 0 | 0 |
| NC_002645.1;NP_073549.1;HCoV-229E_1ab;5828 | 0 | 0 |
| NC_002645.1;NP_073549.1;HCoV-229E_1ab;5829 | 0 | 0 |
| NC_002645.1;NP_073549.1;HCoV-229E_1ab;5830 | 0 | 0 |
| NC_002645.1;NP_073549.1;HCoV-229E_1ab;5831 | 0 | 0 |
| NC_002645.1;NP_073549.1;HCoV-229E_1ab;5832 | 0 | 0 |
| NC_002645.1;NP_073549.1;HCoV-229E_1ab;5833 | 0 | 0 |
| NC_002645.1;NP_073549.1;HCoV-229E_1ab;5834 | 0 | 0 |

| | | |
|---|---|---|
| NC_002645.1;NP_073549.1;HCoV-229E_1ab;5835 | 0 | 0 |
| NC_002645.1;NP_073549.1;HCoV-229E_1ab;5836 | 0 | 0 |
| NC_002645.1;NP_073549.1;HCoV-229E_1ab;5837 | 0 | 0 |
| NC_002645.1;NP_073549.1;HCoV-229E_1ab;5838 | 0 | 0 |
| NC_002645.1;NP_073549.1;HCoV-229E_1ab;5839 | 0 | 0 |
| NC_002645.1;NP_073549.1;HCoV-229E_1ab;5840 | 0 | 0 |
| NC_002645.1;NP_073549.1;HCoV-229E_1ab;5841 | 0 | 0 |
| NC_002645.1;NP_073549.1;HCoV-229E_1ab;5842 | 0 | 0 |
| NC_002645.1;NP_073549.1;HCoV-229E_1ab;5843 | 0 | 0 |
| NC_002645.1;NP_073549.1;HCoV-229E_1ab;5844 | 0 | 0 |
| NC_002645.1;NP_073549.1;HCoV-229E_1ab;5845 | 0 | 0 |
| NC_002645.1;NP_073549.1;HCoV-229E_1ab;5846 | 0.05 | 0.05 |
| NC_002645.1;NP_073549.1;HCoV-229E_1ab;5847 | 0 | 0 |
| NC_002645.1;NP_073549.1;HCoV-229E_1ab;5848 | 0 | 0 |
| NC_002645.1;NP_073549.1;HCoV-229E_1ab;5849 | 0 | 0 |
| NC_002645.1;NP_073549.1;HCoV-229E_1ab;5850 | 0 | 0 |
| NC_002645.1;NP_073549.1;HCoV-229E_1ab;5851 | 0 | 0 |
| NC_002645.1;NP_073549.1;HCoV-229E_1ab;5852 | 0 | 0.025 |
| NC_002645.1;NP_073549.1;HCoV-229E_1ab;5853 | 0 | 0.025 |
| NC_002645.1;NP_073549.1;HCoV-229E_1ab;5854 | 0 | 0.025 |
| NC_002645.1;NP_073549.1;HCoV-229E_1ab;5855 | 0 | 0 |
| NC_002645.1;NP_073549.1;HCoV-229E_1ab;5856 | 0 | 0 |
| NC_002645.1;NP_073549.1;HCoV-229E_1ab;5857 | 0 | 0 |
| NC_002645.1;NP_073549.1;HCoV-229E_1ab;5858 | 0 | 0 |
| NC_002645.1;NP_073549.1;HCoV-229E_1ab;5859 | 0 | 0 |
| NC_002645.1;NP_073549.1;HCoV-229E_1ab;5860 | 0 | 0 |
| NC_002645.1;NP_073549.1;HCoV-229E_1ab;5861 | 0 | 0 |
| NC_002645.1;NP_073549.1;HCoV-229E_1ab;5862 | 0 | 0 |
| NC_002645.1;NP_073549.1;HCoV-229E_1ab;5863 | 0 | 0 |
| NC_002645.1;NP_073549.1;HCoV-229E_1ab;5864 | 0 | 0 |
| NC_002645.1;NP_073549.1;HCoV-229E_1ab;5865 | 0 | 0 |
| NC_002645.1;NP_073549.1;HCoV-229E_1ab;5866 | 0 | 0 |
| NC_002645.1;NP_073549.1;HCoV-229E_1ab;5867 | 0 | 0 |
| NC_002645.1;NP_073549.1;HCoV-229E_1ab;5868 | 0 | 0 |
| NC_002645.1;NP_073549.1;HCoV-229E_1ab;5869 | 0 | 0 |
| NC_002645.1;NP_073549.1;HCoV-229E_1ab;5870 | 0 | 0 |
| NC_002645.1;NP_073549.1;HCoV-229E_1ab;5871 | 0 | 0 |
| NC_002645.1;NP_073549.1;HCoV-229E_1ab;5872 | 0 | 0 |
| NC_002645.1;NP_073549.1;HCoV-229E_1ab;5873 | 0 | 0 |
| NC_002645.1;NP_073549.1;HCoV-229E_1ab;5874 | 0 | 0 |
| NC_002645.1;NP_073549.1;HCoV-229E_1ab;5875 | 0 | 0 |
| NC_002645.1;NP_073549.1;HCoV-229E_1ab;5876 | 0 | 0.025 |
| NC_002645.1;NP_073549.1;HCoV-229E_1ab;5877 | 0.05 | 0 |

| | | |
|---|---|---|
| NC_002645.1;NP_073549.1;HCoV-229E_1ab;5878 | 0.1 | 0 |
| NC_002645.1;NP_073549.1;HCoV-229E_1ab;5879 | 0.05 | 0.025 |
| NC_002645.1;NP_073549.1;HCoV-229E_1ab;5880 | 0.05 | 0.025 |
| NC_002645.1;NP_073549.1;HCoV-229E_1ab;5881 | 0.05 | 0.025 |
| NC_002645.1;NP_073549.1;HCoV-229E_1ab;5882 | 0.05 | 0.05 |
| NC_002645.1;NP_073549.1;HCoV-229E_1ab;5883 | 0.1 | 0.05 |
| NC_002645.1;NP_073549.1;HCoV-229E_1ab;5884 | 0.1 | 0.05 |
| NC_002645.1;NP_073549.1;HCoV-229E_1ab;5885 | 0.15 | 0.05 |
| NC_002645.1;NP_073549.1;HCoV-229E_1ab;5886 | 0.1 | 0.05 |
| NC_002645.1;NP_073549.1;HCoV-229E_1ab;5887 | 0.1 | 0.05 |
| NC_002645.1;NP_073549.1;HCoV-229E_1ab;5888 | 0.1 | 0.05 |
| NC_002645.1;NP_073549.1;HCoV-229E_1ab;5889 | 0.1 | 0.05 |
| NC_002645.1;NP_073549.1;HCoV-229E_1ab;5890 | 0.1 | 0.05 |
| NC_002645.1;NP_073549.1;HCoV-229E_1ab;5891 | 0.05 | 0.025 |
| NC_002645.1;NP_073549.1;HCoV-229E_1ab;5892 | 0 | 0 |
| NC_002645.1;NP_073549.1;HCoV-229E_1ab;5893 | 0 | 0 |
| NC_002645.1;NP_073549.1;HCoV-229E_1ab;5894 | 0 | 0 |
| NC_002645.1;NP_073549.1;HCoV-229E_1ab;5895 | 0 | 0 |
| NC_002645.1;NP_073549.1;HCoV-229E_1ab;5896 | 0 | 0 |
| NC_002645.1;NP_073549.1;HCoV-229E_1ab;5897 | 0 | 0 |
| NC_002645.1;NP_073549.1;HCoV-229E_1ab;5898 | 0 | 0 |
| NC_002645.1;NP_073549.1;HCoV-229E_1ab;5899 | 0 | 0 |
| NC_002645.1;NP_073549.1;HCoV-229E_1ab;5900 | 0 | 0 |
| NC_002645.1;NP_073549.1;HCoV-229E_1ab;5901 | 0 | 0 |
| NC_002645.1;NP_073549.1;HCoV-229E_1ab;5902 | 0 | 0 |
| NC_002645.1;NP_073549.1;HCoV-229E_1ab;5903 | 0 | 0 |
| NC_002645.1;NP_073549.1;HCoV-229E_1ab;5904 | 0 | 0 |
| NC_002645.1;NP_073549.1;HCoV-229E_1ab;5905 | 0 | 0 |
| NC_002645.1;NP_073549.1;HCoV-229E_1ab;5906 | 0 | 0 |
| NC_002645.1;NP_073549.1;HCoV-229E_1ab;5907 | 0.05 | 0 |
| NC_002645.1;NP_073549.1;HCoV-229E_1ab;5908 | 0 | 0 |
| NC_002645.1;NP_073549.1;HCoV-229E_1ab;5909 | 0 | 0 |
| NC_002645.1;NP_073549.1;HCoV-229E_1ab;5910 | 0 | 0.025 |
| NC_002645.1;NP_073549.1;HCoV-229E_1ab;5911 | 0 | 0 |
| NC_002645.1;NP_073549.1;HCoV-229E_1ab;5912 | 0 | 0 |
| NC_002645.1;NP_073549.1;HCoV-229E_1ab;5913 | 0 | 0 |
| NC_002645.1;NP_073549.1;HCoV-229E_1ab;5914 | 0 | 0 |
| NC_002645.1;NP_073549.1;HCoV-229E_1ab;5915 | 0 | 0 |
| NC_002645.1;NP_073549.1;HCoV-229E_1ab;5916 | 0 | 0 |
| NC_002645.1;NP_073549.1;HCoV-229E_1ab;5917 | 0 | 0 |
| NC_002645.1;NP_073549.1;HCoV-229E_1ab;5918 | 0 | 0 |
| NC_002645.1;NP_073549.1;HCoV-229E_1ab;5919 | 0.05 | 0 |
| NC_002645.1;NP_073549.1;HCoV-229E_1ab;5920 | 0.05 | 0 |

| | | |
|---|---|---|
| NC_002645.1;NP_073549.1;HCoV-229E_1ab;5921 | 0 | 0 |
| NC_002645.1;NP_073549.1;HCoV-229E_1ab;5922 | 0 | 0 |
| NC_002645.1;NP_073549.1;HCoV-229E_1ab;5923 | 0.05 | 0.025 |
| NC_002645.1;NP_073549.1;HCoV-229E_1ab;5924 | 0 | 0 |
| NC_002645.1;NP_073549.1;HCoV-229E_1ab;5925 | 0 | 0 |
| NC_002645.1;NP_073549.1;HCoV-229E_1ab;5926 | 0 | 0 |
| NC_002645.1;NP_073549.1;HCoV-229E_1ab;5927 | 0 | 0 |
| NC_002645.1;NP_073549.1;HCoV-229E_1ab;5928 | 0 | 0 |
| NC_002645.1;NP_073549.1;HCoV-229E_1ab;5929 | 0 | 0 |
| NC_002645.1;NP_073549.1;HCoV-229E_1ab;5930 | 0 | 0 |
| NC_002645.1;NP_073549.1;HCoV-229E_1ab;5931 | 0 | 0 |
| NC_002645.1;NP_073549.1;HCoV-229E_1ab;5932 | 0 | 0.025 |
| NC_002645.1;NP_073549.1;HCoV-229E_1ab;5933 | 0.05 | 0.025 |
| NC_002645.1;NP_073549.1;HCoV-229E_1ab;5934 | 0 | 0.05 |
| NC_002645.1;NP_073549.1;HCoV-229E_1ab;5935 | 0.05 | 0.075 |
| NC_002645.1;NP_073549.1;HCoV-229E_1ab;5936 | 0 | 0.05 |
| NC_002645.1;NP_073549.1;HCoV-229E_1ab;5937 | 0 | 0.05 |
| NC_002645.1;NP_073549.1;HCoV-229E_1ab;5938 | 0.05 | 0.05 |
| NC_002645.1;NP_073549.1;HCoV-229E_1ab;5939 | 0.05 | 0.075 |
| NC_002645.1;NP_073549.1;HCoV-229E_1ab;5940 | 0.05 | 0.075 |
| NC_002645.1;NP_073549.1;HCoV-229E_1ab;5941 | 0.05 | 0.05 |
| NC_002645.1;NP_073549.1;HCoV-229E_1ab;5942 | 0.1 | 0.05 |
| NC_002645.1;NP_073549.1;HCoV-229E_1ab;5943 | 0.15 | 0.075 |
| NC_002645.1;NP_073549.1;HCoV-229E_1ab;5944 | 0.05 | 0.05 |
| NC_002645.1;NP_073549.1;HCoV-229E_1ab;5945 | 0.05 | 0.025 |
| NC_002645.1;NP_073549.1;HCoV-229E_1ab;5946 | 0.05 | 0.025 |
| NC_002645.1;NP_073549.1;HCoV-229E_1ab;5947 | 0.05 | 0 |
| NC_002645.1;NP_073549.1;HCoV-229E_1ab;5948 | 0.1 | 0.075 |
| NC_002645.1;NP_073549.1;HCoV-229E_1ab;5949 | 0.05 | 0 |
| NC_002645.1;NP_073549.1;HCoV-229E_1ab;5950 | 0.05 | 0 |
| NC_002645.1;NP_073549.1;HCoV-229E_1ab;5951 | 0.05 | 0 |
| NC_002645.1;NP_073549.1;HCoV-229E_1ab;5952 | 0.05 | 0 |
| NC_002645.1;NP_073549.1;HCoV-229E_1ab;5953 | 0.05 | 0 |
| NC_002645.1;NP_073549.1;HCoV-229E_1ab;5954 | 0 | 0 |
| NC_002645.1;NP_073549.1;HCoV-229E_1ab;5955 | 0 | 0 |
| NC_002645.1;NP_073549.1;HCoV-229E_1ab;5956 | 0 | 0 |
| NC_002645.1;NP_073549.1;HCoV-229E_1ab;5957 | 0 | 0 |
| NC_002645.1;NP_073549.1;HCoV-229E_1ab;5958 | 0 | 0 |
| NC_002645.1;NP_073549.1;HCoV-229E_1ab;5959 | 0 | 0 |
| NC_002645.1;NP_073549.1;HCoV-229E_1ab;5960 | 0 | 0 |
| NC_002645.1;NP_073549.1;HCoV-229E_1ab;5961 | 0 | 0 |
| NC_002645.1;NP_073549.1;HCoV-229E_1ab;5962 | 0 | 0 |
| NC_002645.1;NP_073549.1;HCoV-229E_1ab;5963 | 0 | 0 |

| | | |
|---|---|---|
| NC_002645.1;NP_073549.1;HCoV-229E_1ab;5964 | 0 | 0 |
| NC_002645.1;NP_073549.1;HCoV-229E_1ab;5965 | 0 | 0 |
| NC_002645.1;NP_073549.1;HCoV-229E_1ab;5966 | 0 | 0 |
| NC_002645.1;NP_073549.1;HCoV-229E_1ab;5967 | 0 | 0 |
| NC_002645.1;NP_073549.1;HCoV-229E_1ab;5968 | 0 | 0 |
| NC_002645.1;NP_073549.1;HCoV-229E_1ab;5969 | 0 | 0 |
| NC_002645.1;NP_073549.1;HCoV-229E_1ab;5970 | 0 | 0 |
| NC_002645.1;NP_073549.1;HCoV-229E_1ab;5971 | 0 | 0 |
| NC_002645.1;NP_073549.1;HCoV-229E_1ab;5972 | 0 | 0 |
| NC_002645.1;NP_073549.1;HCoV-229E_1ab;5973 | 0 | 0 |
| NC_002645.1;NP_073549.1;HCoV-229E_1ab;5974 | 0 | 0 |
| NC_002645.1;NP_073549.1;HCoV-229E_1ab;5975 | 0 | 0 |
| NC_002645.1;NP_073549.1;HCoV-229E_1ab;5976 | 0 | 0 |
| NC_002645.1;NP_073549.1;HCoV-229E_1ab;5977 | 0 | 0 |
| NC_002645.1;NP_073549.1;HCoV-229E_1ab;5978 | 0 | 0 |
| NC_002645.1;NP_073549.1;HCoV-229E_1ab;5979 | 0 | 0 |
| NC_002645.1;NP_073549.1;HCoV-229E_1ab;5980 | 0 | 0 |
| NC_002645.1;NP_073549.1;HCoV-229E_1ab;5981 | 0 | 0 |
| NC_002645.1;NP_073549.1;HCoV-229E_1ab;5982 | 0 | 0 |
| NC_002645.1;NP_073549.1;HCoV-229E_1ab;5983 | 0 | 0 |
| NC_002645.1;NP_073549.1;HCoV-229E_1ab;5984 | 0 | 0 |
| NC_002645.1;NP_073549.1;HCoV-229E_1ab;5985 | 0 | 0 |
| NC_002645.1;NP_073549.1;HCoV-229E_1ab;5986 | 0 | 0 |
| NC_002645.1;NP_073549.1;HCoV-229E_1ab;5987 | 0.1 | 0.025 |
| NC_002645.1;NP_073549.1;HCoV-229E_1ab;5988 | 0 | 0 |
| NC_002645.1;NP_073549.1;HCoV-229E_1ab;5989 | 0 | 0 |
| NC_002645.1;NP_073549.1;HCoV-229E_1ab;5990 | 0 | 0 |
| NC_002645.1;NP_073549.1;HCoV-229E_1ab;5991 | 0 | 0 |
| NC_002645.1;NP_073549.1;HCoV-229E_1ab;5992 | 0.05 | 0 |
| NC_002645.1;NP_073549.1;HCoV-229E_1ab;5993 | 0.1 | 0.025 |
| NC_002645.1;NP_073549.1;HCoV-229E_1ab;5994 | 0 | 0 |
| NC_002645.1;NP_073549.1;HCoV-229E_1ab;5995 | 0 | 0 |
| NC_002645.1;NP_073549.1;HCoV-229E_1ab;5996 | 0 | 0 |
| NC_002645.1;NP_073549.1;HCoV-229E_1ab;5997 | 0 | 0.025 |
| NC_002645.1;NP_073549.1;HCoV-229E_1ab;5998 | 0 | 0 |
| NC_002645.1;NP_073549.1;HCoV-229E_1ab;5999 | 0 | 0 |
| NC_002645.1;NP_073549.1;HCoV-229E_1ab;6000 | 0 | 0 |
| NC_002645.1;NP_073549.1;HCoV-229E_1ab;6001 | 0 | 0 |
| NC_002645.1;NP_073549.1;HCoV-229E_1ab;6002 | 0 | 0 |
| NC_002645.1;NP_073549.1;HCoV-229E_1ab;6003 | 0 | 0.025 |
| NC_002645.1;NP_073549.1;HCoV-229E_1ab;6004 | 0 | 0 |
| NC_002645.1;NP_073549.1;HCoV-229E_1ab;6005 | 0.05 | 0 |
| NC_002645.1;NP_073549.1;HCoV-229E_1ab;6006 | 0.05 | 0.025 |

| | | |
|---|---|---|
| NC_002645.1;NP_073549.1;HCoV-229E_1ab;6007 | 0.05 | 0.025 |
| NC_002645.1;NP_073549.1;HCoV-229E_1ab;6008 | 0.05 | 0 |
| NC_002645.1;NP_073549.1;HCoV-229E_1ab;6009 | 0.05 | 0 |
| NC_002645.1;NP_073549.1;HCoV-229E_1ab;6010 | 0.05 | 0 |
| NC_002645.1;NP_073549.1;HCoV-229E_1ab;6011 | 0 | 0 |
| NC_002645.1;NP_073549.1;HCoV-229E_1ab;6012 | 0.05 | 0 |
| NC_002645.1;NP_073549.1;HCoV-229E_1ab;6013 | 0.05 | 0 |
| NC_002645.1;NP_073549.1;HCoV-229E_1ab;6014 | 0 | 0 |
| NC_002645.1;NP_073549.1;HCoV-229E_1ab;6015 | 0.05 | 0.025 |
| NC_002645.1;NP_073549.1;HCoV-229E_1ab;6016 | 0 | 0 |
| NC_002645.1;NP_073549.1;HCoV-229E_1ab;6017 | 0 | 0 |
| NC_002645.1;NP_073549.1;HCoV-229E_1ab;6018 | 0 | 0.025 |
| NC_002645.1;NP_073549.1;HCoV-229E_1ab;6019 | 0 | 0 |
| NC_002645.1;NP_073549.1;HCoV-229E_1ab;6020 | 0 | 0 |
| NC_002645.1;NP_073549.1;HCoV-229E_1ab;6021 | 0 | 0 |
| NC_002645.1;NP_073549.1;HCoV-229E_1ab;6022 | 0 | 0 |
| NC_002645.1;NP_073549.1;HCoV-229E_1ab;6023 | 0 | 0 |
| NC_002645.1;NP_073549.1;HCoV-229E_1ab;6024 | 0 | 0 |
| NC_002645.1;NP_073549.1;HCoV-229E_1ab;6025 | 0 | 0 |
| NC_002645.1;NP_073549.1;HCoV-229E_1ab;6026 | 0 | 0 |
| NC_002645.1;NP_073549.1;HCoV-229E_1ab;6027 | 0 | 0 |
| NC_002645.1;NP_073549.1;HCoV-229E_1ab;6028 | 0 | 0 |
| NC_002645.1;NP_073549.1;HCoV-229E_1ab;6029 | 0.05 | 0 |
| NC_002645.1;NP_073549.1;HCoV-229E_1ab;6030 | 0.05 | 0 |
| NC_002645.1;NP_073549.1;HCoV-229E_1ab;6031 | 0 | 0.025 |
| NC_002645.1;NP_073549.1;HCoV-229E_1ab;6032 | 0.05 | 0.025 |
| NC_002645.1;NP_073549.1;HCoV-229E_1ab;6033 | 0 | 0 |
| NC_002645.1;NP_073549.1;HCoV-229E_1ab;6034 | 0 | 0.025 |
| NC_002645.1;NP_073549.1;HCoV-229E_1ab;6035 | 0 | 0 |
| NC_002645.1;NP_073549.1;HCoV-229E_1ab;6036 | 0.1 | 0.025 |
| NC_002645.1;NP_073549.1;HCoV-229E_1ab;6037 | 0 | 0.025 |
| NC_002645.1;NP_073549.1;HCoV-229E_1ab;6038 | 0 | 0.025 |
| NC_002645.1;NP_073549.1;HCoV-229E_1ab;6039 | 0 | 0 |
| NC_002645.1;NP_073549.1;HCoV-229E_1ab;6040 | 0 | 0 |
| NC_002645.1;NP_073549.1;HCoV-229E_1ab;6041 | 0 | 0 |
| NC_002645.1;NP_073549.1;HCoV-229E_1ab;6042 | 0 | 0 |
| NC_002645.1;NP_073549.1;HCoV-229E_1ab;6043 | 0 | 0 |
| NC_002645.1;NP_073549.1;HCoV-229E_1ab;6044 | 0 | 0 |
| NC_002645.1;NP_073549.1;HCoV-229E_1ab;6045 | 0 | 0 |
| NC_002645.1;NP_073549.1;HCoV-229E_1ab;6046 | 0 | 0 |
| NC_002645.1;NP_073549.1;HCoV-229E_1ab;6047 | 0 | 0 |
| NC_002645.1;NP_073549.1;HCoV-229E_1ab;6048 | 0 | 0.025 |
| NC_002645.1;NP_073549.1;HCoV-229E_1ab;6049 | 0 | 0 |

| | | |
|---|---|---|
| NC_002645.1;NP_073549.1;HCoV-229E_1ab;6050 | 0 | 0 |
| NC_002645.1;NP_073549.1;HCoV-229E_1ab;6051 | 0 | 0 |
| NC_002645.1;NP_073549.1;HCoV-229E_1ab;6052 | 0 | 0 |
| NC_002645.1;NP_073549.1;HCoV-229E_1ab;6053 | 0 | 0 |
| NC_002645.1;NP_073549.1;HCoV-229E_1ab;6054 | 0 | 0 |
| NC_002645.1;NP_073549.1;HCoV-229E_1ab;6055 | 0 | 0 |
| NC_002645.1;NP_073549.1;HCoV-229E_1ab;6056 | 0 | 0.025 |
| NC_002645.1;NP_073549.1;HCoV-229E_1ab;6057 | 0.05 | 0.025 |
| NC_002645.1;NP_073549.1;HCoV-229E_1ab;6058 | 0 | 0.025 |
| NC_002645.1;NP_073549.1;HCoV-229E_1ab;6059 | 0 | 0.025 |
| NC_002645.1;NP_073549.1;HCoV-229E_1ab;6060 | 0 | 0 |
| NC_002645.1;NP_073549.1;HCoV-229E_1ab;6061 | 0 | 0 |
| NC_002645.1;NP_073549.1;HCoV-229E_1ab;6062 | 0 | 0 |
| NC_002645.1;NP_073549.1;HCoV-229E_1ab;6063 | 0 | 0 |
| NC_002645.1;NP_073549.1;HCoV-229E_1ab;6064 | 0 | 0 |
| NC_002645.1;NP_073549.1;HCoV-229E_1ab;6065 | 0 | 0 |
| NC_002645.1;NP_073549.1;HCoV-229E_1ab;6066 | 0 | 0 |
| NC_002645.1;NP_073549.1;HCoV-229E_1ab;6067 | 0 | 0 |
| NC_002645.1;NP_073549.1;HCoV-229E_1ab;6068 | 0 | 0 |
| NC_002645.1;NP_073549.1;HCoV-229E_1ab;6069 | 0 | 0 |
| NC_002645.1;NP_073549.1;HCoV-229E_1ab;6070 | 0 | 0 |
| NC_002645.1;NP_073549.1;HCoV-229E_1ab;6071 | 0 | 0 |
| NC_002645.1;NP_073549.1;HCoV-229E_1ab;6072 | 0 | 0 |
| NC_002645.1;NP_073549.1;HCoV-229E_1ab;6073 | 0 | 0 |
| NC_002645.1;NP_073549.1;HCoV-229E_1ab;6074 | 0 | 0 |
| NC_002645.1;NP_073549.1;HCoV-229E_1ab;6075 | 0 | 0 |
| NC_002645.1;NP_073549.1;HCoV-229E_1ab;6076 | 0 | 0 |
| NC_002645.1;NP_073549.1;HCoV-229E_1ab;6077 | 0 | 0 |
| NC_002645.1;NP_073549.1;HCoV-229E_1ab;6078 | 0 | 0 |
| NC_002645.1;NP_073549.1;HCoV-229E_1ab;6079 | 0 | 0 |
| NC_002645.1;NP_073549.1;HCoV-229E_1ab;6080 | 0 | 0 |
| NC_002645.1;NP_073549.1;HCoV-229E_1ab;6081 | 0 | 0 |
| NC_002645.1;NP_073549.1;HCoV-229E_1ab;6082 | 0 | 0 |
| NC_002645.1;NP_073549.1;HCoV-229E_1ab;6083 | 0 | 0 |
| NC_002645.1;NP_073549.1;HCoV-229E_1ab;6084 | 0 | 0 |
| NC_002645.1;NP_073549.1;HCoV-229E_1ab;6085 | 0 | 0 |
| NC_002645.1;NP_073549.1;HCoV-229E_1ab;6086 | 0 | 0 |
| NC_002645.1;NP_073549.1;HCoV-229E_1ab;6087 | 0 | 0 |
| NC_002645.1;NP_073549.1;HCoV-229E_1ab;6088 | 0 | 0 |
| NC_002645.1;NP_073549.1;HCoV-229E_1ab;6089 | 0 | 0 |
| NC_002645.1;NP_073549.1;HCoV-229E_1ab;6090 | 0 | 0 |
| NC_002645.1;NP_073549.1;HCoV-229E_1ab;6091 | 0 | 0 |
| NC_002645.1;NP_073549.1;HCoV-229E_1ab;6092 | 0 | 0 |

| | | |
|---|---|---|
| NC_002645.1;NP_073549.1;HCoV-229E_1ab;6093 | 0 | 0 |
| NC_002645.1;NP_073549.1;HCoV-229E_1ab;6094 | 0 | 0 |
| NC_002645.1;NP_073549.1;HCoV-229E_1ab;6095 | 0 | 0 |
| NC_002645.1;NP_073549.1;HCoV-229E_1ab;6096 | 0 | 0 |
| NC_002645.1;NP_073549.1;HCoV-229E_1ab;6097 | 0 | 0 |
| NC_002645.1;NP_073549.1;HCoV-229E_1ab;6098 | 0 | 0 |
| NC_002645.1;NP_073549.1;HCoV-229E_1ab;6099 | 0 | 0 |
| NC_002645.1;NP_073549.1;HCoV-229E_1ab;6100 | 0 | 0 |
| NC_002645.1;NP_073549.1;HCoV-229E_1ab;6101 | 0 | 0 |
| NC_002645.1;NP_073549.1;HCoV-229E_1ab;6102 | 0 | 0 |
| NC_002645.1;NP_073549.1;HCoV-229E_1ab;6103 | 0.05 | 0 |
| NC_002645.1;NP_073549.1;HCoV-229E_1ab;6104 | 0 | 0 |
| NC_002645.1;NP_073549.1;HCoV-229E_1ab;6105 | 0 | 0 |
| NC_002645.1;NP_073549.1;HCoV-229E_1ab;6106 | 0 | 0 |
| NC_002645.1;NP_073549.1;HCoV-229E_1ab;6107 | 0 | 0 |
| NC_002645.1;NP_073549.1;HCoV-229E_1ab;6108 | 0 | 0 |
| NC_002645.1;NP_073549.1;HCoV-229E_1ab;6109 | 0 | 0.025 |
| NC_002645.1;NP_073549.1;HCoV-229E_1ab;6110 | 0 | 0 |
| NC_002645.1;NP_073549.1;HCoV-229E_1ab;6111 | 0.2 | 0.2 |
| NC_002645.1;NP_073549.1;HCoV-229E_1ab;6112 | 0 | 0 |
| NC_002645.1;NP_073549.1;HCoV-229E_1ab;6113 | 0 | 0 |
| NC_002645.1;NP_073549.1;HCoV-229E_1ab;6114 | 0 | 0 |
| NC_002645.1;NP_073549.1;HCoV-229E_1ab;6115 | 0 | 0 |
| NC_002645.1;NP_073549.1;HCoV-229E_1ab;6116 | 0.05 | 0 |
| NC_002645.1;NP_073549.1;HCoV-229E_1ab;6117 | 0.05 | 0 |
| NC_002645.1;NP_073549.1;HCoV-229E_1ab;6118 | 0.05 | 0 |
| NC_002645.1;NP_073549.1;HCoV-229E_1ab;6119 | 0.05 | 0 |
| NC_002645.1;NP_073549.1;HCoV-229E_1ab;6120 | 0.1 | 0.025 |
| NC_002645.1;NP_073549.1;HCoV-229E_1ab;6121 | 0.05 | 0 |
| NC_002645.1;NP_073549.1;HCoV-229E_1ab;6122 | 0.05 | 0.025 |
| NC_002645.1;NP_073549.1;HCoV-229E_1ab;6123 | 0.05 | 0 |
| NC_002645.1;NP_073549.1;HCoV-229E_1ab;6124 | 0.05 | 0 |
| NC_002645.1;NP_073549.1;HCoV-229E_1ab;6125 | 0 | 0.025 |
| NC_002645.1;NP_073549.1;HCoV-229E_1ab;6126 | 0 | 0.025 |
| NC_002645.1;NP_073549.1;HCoV-229E_1ab;6127 | 0.05 | 0.025 |
| NC_002645.1;NP_073549.1;HCoV-229E_1ab;6128 | 0 | 0.025 |
| NC_002645.1;NP_073549.1;HCoV-229E_1ab;6129 | 0.05 | 0.025 |
| NC_002645.1;NP_073549.1;HCoV-229E_1ab;6130 | 0 | 0.025 |
| NC_002645.1;NP_073549.1;HCoV-229E_1ab;6131 | 0 | 0.025 |
| NC_002645.1;NP_073549.1;HCoV-229E_1ab;6132 | 0 | 0 |
| NC_002645.1;NP_073549.1;HCoV-229E_1ab;6133 | 0.05 | 0 |
| NC_002645.1;NP_073549.1;HCoV-229E_1ab;6134 | 0 | 0 |
| NC_002645.1;NP_073549.1;HCoV-229E_1ab;6135 | 0.05 | 0.025 |

| | | |
|---|---|---|
| NC_002645.1;NP_073549.1;HCoV-229E_1ab;6136 | 0 | 0.025 |
| NC_002645.1;NP_073549.1;HCoV-229E_1ab;6137 | 0 | 0.025 |
| NC_002645.1;NP_073549.1;HCoV-229E_1ab;6138 | 0 | 0.025 |
| NC_002645.1;NP_073549.1;HCoV-229E_1ab;6139 | 0 | 0.025 |
| NC_002645.1;NP_073549.1;HCoV-229E_1ab;6140 | 0 | 0 |
| NC_002645.1;NP_073549.1;HCoV-229E_1ab;6141 | 0.05 | 0 |
| NC_002645.1;NP_073549.1;HCoV-229E_1ab;6142 | 0.05 | 0 |
| NC_002645.1;NP_073549.1;HCoV-229E_1ab;6143 | 0.05 | 0.025 |
| NC_002645.1;NP_073549.1;HCoV-229E_1ab;6144 | 0.05 | 0 |
| NC_002645.1;NP_073549.1;HCoV-229E_1ab;6145 | 0.05 | 0 |
| NC_002645.1;NP_073549.1;HCoV-229E_1ab;6146 | 0.05 | 0 |
| NC_002645.1;NP_073549.1;HCoV-229E_1ab;6147 | 0.05 | 0 |
| NC_002645.1;NP_073549.1;HCoV-229E_1ab;6148 | 0.05 | 0.025 |
| NC_002645.1;NP_073549.1;HCoV-229E_1ab;6149 | 0.05 | 0 |
| NC_002645.1;NP_073549.1;HCoV-229E_1ab;6150 | 0 | 0.025 |
| NC_002645.1;NP_073549.1;HCoV-229E_1ab;6151 | 0.1 | 0.025 |
| NC_002645.1;NP_073549.1;HCoV-229E_1ab;6152 | 0 | 0.025 |
| NC_002645.1;NP_073549.1;HCoV-229E_1ab;6153 | 0.05 | 0.025 |
| NC_002645.1;NP_073549.1;HCoV-229E_1ab;6154 | 0 | 0 |
| NC_002645.1;NP_073549.1;HCoV-229E_1ab;6155 | 0 | 0 |
| NC_002645.1;NP_073549.1;HCoV-229E_1ab;6156 | 0 | 0 |
| NC_002645.1;NP_073549.1;HCoV-229E_1ab;6157 | 0 | 0 |
| NC_002645.1;NP_073549.1;HCoV-229E_1ab;6158 | 0 | 0 |
| NC_002645.1;NP_073549.1;HCoV-229E_1ab;6159 | 0 | 0 |
| NC_002645.1;NP_073549.1;HCoV-229E_1ab;6160 | 0 | 0.025 |
| NC_002645.1;NP_073549.1;HCoV-229E_1ab;6161 | 0 | 0.025 |
| NC_002645.1;NP_073549.1;HCoV-229E_1ab;6162 | 0 | 0.025 |
| NC_002645.1;NP_073549.1;HCoV-229E_1ab;6163 | 0 | 0.05 |
| NC_002645.1;NP_073549.1;HCoV-229E_1ab;6164 | 0 | 0.025 |
| NC_002645.1;NP_073549.1;HCoV-229E_1ab;6165 | 0 | 0.1 |
| NC_002645.1;NP_073549.1;HCoV-229E_1ab;6166 | 0 | 0.075 |
| NC_002645.1;NP_073549.1;HCoV-229E_1ab;6167 | 0 | 0.025 |
| NC_002645.1;NP_073549.1;HCoV-229E_1ab;6168 | 0 | 0.05 |
| NC_002645.1;NP_073549.1;HCoV-229E_1ab;6169 | 0 | 0.05 |
| NC_002645.1;NP_073549.1;HCoV-229E_1ab;6170 | 0.15 | 0.2 |
| NC_002645.1;NP_073549.1;HCoV-229E_1ab;6171 | 0 | 0 |
| NC_002645.1;NP_073549.1;HCoV-229E_1ab;6172 | 0 | 0 |
| NC_002645.1;NP_073549.1;HCoV-229E_1ab;6173 | 0 | 0 |
| NC_002645.1;NP_073549.1;HCoV-229E_1ab;6174 | 0 | 0 |
| NC_002645.1;NP_073549.1;HCoV-229E_1ab;6175 | 0 | 0 |
| NC_002645.1;NP_073549.1;HCoV-229E_1ab;6176 | 0 | 0 |
| NC_002645.1;NP_073549.1;HCoV-229E_1ab;6177 | 0 | 0 |
| NC_002645.1;NP_073549.1;HCoV-229E_1ab;6178 | 0 | 0 |

| | | |
|---|---|---|
| NC_002645.1;NP_073549.1;HCoV-229E_1ab;6179 | 0 | 0 |
| NC_002645.1;NP_073549.1;HCoV-229E_1ab;6180 | 0 | 0 |
| NC_002645.1;NP_073549.1;HCoV-229E_1ab;6181 | 0 | 0 |
| NC_002645.1;NP_073549.1;HCoV-229E_1ab;6182 | 0 | 0 |
| NC_002645.1;NP_073549.1;HCoV-229E_1ab;6183 | 0 | 0 |
| NC_002645.1;NP_073549.1;HCoV-229E_1ab;6184 | 0 | 0 |
| NC_002645.1;NP_073549.1;HCoV-229E_1ab;6185 | 0 | 0 |
| NC_002645.1;NP_073549.1;HCoV-229E_1ab;6186 | 0 | 0 |
| NC_002645.1;NP_073549.1;HCoV-229E_1ab;6187 | 0 | 0 |
| NC_002645.1;NP_073549.1;HCoV-229E_1ab;6188 | 0 | 0 |
| NC_002645.1;NP_073549.1;HCoV-229E_1ab;6189 | 0 | 0 |
| NC_002645.1;NP_073549.1;HCoV-229E_1ab;6190 | 0 | 0 |
| NC_002645.1;NP_073549.1;HCoV-229E_1ab;6191 | 0 | 0 |
| NC_002645.1;NP_073549.1;HCoV-229E_1ab;6192 | 0 | 0 |
| NC_002645.1;NP_073549.1;HCoV-229E_1ab;6193 | 0.05 | 0 |
| NC_002645.1;NP_073549.1;HCoV-229E_1ab;6194 | 0 | 0.025 |
| NC_002645.1;NP_073549.1;HCoV-229E_1ab;6195 | 0 | 0 |
| NC_002645.1;NP_073549.1;HCoV-229E_1ab;6196 | 0 | 0 |
| NC_002645.1;NP_073549.1;HCoV-229E_1ab;6197 | 0 | 0 |
| NC_002645.1;NP_073549.1;HCoV-229E_1ab;6198 | 0 | 0 |
| NC_002645.1;NP_073549.1;HCoV-229E_1ab;6199 | 0 | 0 |
| NC_002645.1;NP_073549.1;HCoV-229E_1ab;6200 | 0 | 0 |
| NC_002645.1;NP_073549.1;HCoV-229E_1ab;6201 | 0 | 0 |
| NC_002645.1;NP_073549.1;HCoV-229E_1ab;6202 | 0 | 0 |
| NC_002645.1;NP_073549.1;HCoV-229E_1ab;6203 | 0 | 0 |
| NC_002645.1;NP_073549.1;HCoV-229E_1ab;6204 | 0 | 0 |
| NC_002645.1;NP_073549.1;HCoV-229E_1ab;6205 | 0 | 0 |
| NC_002645.1;NP_073549.1;HCoV-229E_1ab;6206 | 0 | 0 |
| NC_002645.1;NP_073549.1;HCoV-229E_1ab;6207 | 0 | 0.025 |
| NC_002645.1;NP_073549.1;HCoV-229E_1ab;6208 | 0 | 0 |
| NC_002645.1;NP_073549.1;HCoV-229E_1ab;6209 | 0 | 0 |
| NC_002645.1;NP_073549.1;HCoV-229E_1ab;6210 | 0 | 0 |
| NC_002645.1;NP_073549.1;HCoV-229E_1ab;6211 | 0 | 0.025 |
| NC_002645.1;NP_073549.1;HCoV-229E_1ab;6212 | 0 | 0.025 |
| NC_002645.1;NP_073549.1;HCoV-229E_1ab;6213 | 0.1 | 0.075 |
| NC_002645.1;NP_073549.1;HCoV-229E_1ab;6214 | 0 | 0.025 |
| NC_002645.1;NP_073549.1;HCoV-229E_1ab;6215 | 0 | 0 |
| NC_002645.1;NP_073549.1;HCoV-229E_1ab;6216 | 0 | 0 |
| NC_002645.1;NP_073549.1;HCoV-229E_1ab;6217 | 0 | 0.025 |
| NC_002645.1;NP_073549.1;HCoV-229E_1ab;6218 | 0 | 0 |
| NC_002645.1;NP_073549.1;HCoV-229E_1ab;6219 | 0 | 0 |
| NC_002645.1;NP_073549.1;HCoV-229E_1ab;6220 | 0 | 0 |
| NC_002645.1;NP_073549.1;HCoV-229E_1ab;6221 | 0 | 0 |

| | | |
|---|---|---|
| NC_002645.1;NP_073549.1;HCoV-229E_1ab;6222 | 0 | 0 |
| NC_002645.1;NP_073549.1;HCoV-229E_1ab;6223 | 0 | 0 |
| NC_002645.1;NP_073549.1;HCoV-229E_1ab;6224 | 0.05 | 0 |
| NC_002645.1;NP_073549.1;HCoV-229E_1ab;6225 | 0 | 0 |
| NC_002645.1;NP_073549.1;HCoV-229E_1ab;6226 | 0 | 0 |
| NC_002645.1;NP_073549.1;HCoV-229E_1ab;6227 | 0 | 0 |
| NC_002645.1;NP_073549.1;HCoV-229E_1ab;6228 | 0 | 0 |
| NC_002645.1;NP_073549.1;HCoV-229E_1ab;6229 | 0 | 0 |
| NC_002645.1;NP_073549.1;HCoV-229E_1ab;6230 | 0 | 0 |
| NC_002645.1;NP_073549.1;HCoV-229E_1ab;6231 | 0 | 0 |
| NC_002645.1;NP_073549.1;HCoV-229E_1ab;6232 | 0 | 0 |
| NC_002645.1;NP_073549.1;HCoV-229E_1ab;6233 | 0 | 0 |
| NC_002645.1;NP_073549.1;HCoV-229E_1ab;6234 | 0 | 0 |
| NC_002645.1;NP_073549.1;HCoV-229E_1ab;6235 | 0 | 0 |
| NC_002645.1;NP_073549.1;HCoV-229E_1ab;6236 | 0 | 0 |
| NC_002645.1;NP_073549.1;HCoV-229E_1ab;6237 | 0 | 0 |
| NC_002645.1;NP_073549.1;HCoV-229E_1ab;6238 | 0 | 0 |
| NC_002645.1;NP_073549.1;HCoV-229E_1ab;6239 | 0 | 0 |
| NC_002645.1;NP_073549.1;HCoV-229E_1ab;6240 | 0 | 0 |
| NC_002645.1;NP_073549.1;HCoV-229E_1ab;6241 | 0 | 0 |
| NC_002645.1;NP_073549.1;HCoV-229E_1ab;6242 | 0 | 0.025 |
| NC_002645.1;NP_073549.1;HCoV-229E_1ab;6243 | 0 | 0 |
| NC_002645.1;NP_073549.1;HCoV-229E_1ab;6244 | 0 | 0 |
| NC_002645.1;NP_073549.1;HCoV-229E_1ab;6245 | 0 | 0 |
| NC_002645.1;NP_073549.1;HCoV-229E_1ab;6246 | 0 | 0 |
| NC_002645.1;NP_073549.1;HCoV-229E_1ab;6247 | 0 | 0.025 |
| NC_002645.1;NP_073549.1;HCoV-229E_1ab;6248 | 0 | 0 |
| NC_002645.1;NP_073549.1;HCoV-229E_1ab;6249 | 0 | 0 |
| NC_002645.1;NP_073549.1;HCoV-229E_1ab;6250 | 0 | 0 |
| NC_002645.1;NP_073549.1;HCoV-229E_1ab;6251 | 0 | 0 |
| NC_002645.1;NP_073549.1;HCoV-229E_1ab;6252 | 0 | 0 |
| NC_002645.1;NP_073549.1;HCoV-229E_1ab;6253 | 0 | 0 |
| NC_002645.1;NP_073549.1;HCoV-229E_1ab;6254 | 0 | 0 |
| NC_002645.1;NP_073549.1;HCoV-229E_1ab;6255 | 0 | 0 |
| NC_002645.1;NP_073549.1;HCoV-229E_1ab;6256 | 0 | 0.025 |
| NC_002645.1;NP_073549.1;HCoV-229E_1ab;6257 | 0 | 0.05 |
| NC_002645.1;NP_073549.1;HCoV-229E_1ab;6258 | 0 | 0.05 |
| NC_002645.1;NP_073549.1;HCoV-229E_1ab;6259 | 0 | 0.05 |
| NC_002645.1;NP_073549.1;HCoV-229E_1ab;6260 | 0 | 0.075 |
| NC_002645.1;NP_073549.1;HCoV-229E_1ab;6261 | 0 | 0.05 |
| NC_002645.1;NP_073549.1;HCoV-229E_1ab;6262 | 0 | 0.05 |
| NC_002645.1;NP_073549.1;HCoV-229E_1ab;6263 | 0 | 0.075 |
| NC_002645.1;NP_073549.1;HCoV-229E_1ab;6264 | 0 | 0.025 |

| | | |
|---|---|---|
| NC_002645.1;NP_073549.1;HCoV-229E_1ab;6265 | 0 | 0.025 |
| NC_002645.1;NP_073549.1;HCoV-229E_1ab;6266 | 0 | 0.05 |
| NC_002645.1;NP_073549.1;HCoV-229E_1ab;6267 | 0 | 0.075 |
| NC_002645.1;NP_073549.1;HCoV-229E_1ab;6268 | 0 | 0.1 |
| NC_002645.1;NP_073549.1;HCoV-229E_1ab;6269 | 0 | 0.05 |
| NC_002645.1;NP_073549.1;HCoV-229E_1ab;6270 | 0.05 | 0.1 |
| NC_002645.1;NP_073549.1;HCoV-229E_1ab;6271 | 0.1 | 0.1 |
| NC_002645.1;NP_073549.1;HCoV-229E_1ab;6272 | 0.25 | 0.075 |
| NC_002645.1;NP_073549.1;HCoV-229E_1ab;6273 | 0.1 | 0.05 |
| NC_002645.1;NP_073549.1;HCoV-229E_1ab;6274 | 0.05 | 0.025 |
| NC_002645.1;NP_073549.1;HCoV-229E_1ab;6275 | 0 | 0 |
| NC_002645.1;NP_073549.1;HCoV-229E_1ab;6276 | 0 | 0 |
| NC_002645.1;NP_073549.1;HCoV-229E_1ab;6277 | 0 | 0 |
| NC_002645.1;NP_073549.1;HCoV-229E_1ab;6278 | 0 | 0 |
| NC_002645.1;NP_073549.1;HCoV-229E_1ab;6279 | 0 | 0 |
| NC_002645.1;NP_073549.1;HCoV-229E_1ab;6280 | 0 | 0 |
| NC_002645.1;NP_073549.1;HCoV-229E_1ab;6281 | 0 | 0 |
| NC_002645.1;NP_073549.1;HCoV-229E_1ab;6282 | 0 | 0 |
| NC_002645.1;NP_073549.1;HCoV-229E_1ab;6283 | 0 | 0 |
| NC_002645.1;NP_073549.1;HCoV-229E_1ab;6284 | 0 | 0 |
| NC_002645.1;NP_073549.1;HCoV-229E_1ab;6285 | 0 | 0 |
| NC_002645.1;NP_073549.1;HCoV-229E_1ab;6286 | 0 | 0 |
| NC_002645.1;NP_073549.1;HCoV-229E_1ab;6287 | 0 | 0 |
| NC_002645.1;NP_073549.1;HCoV-229E_1ab;6288 | 0.05 | 0 |
| NC_002645.1;NP_073549.1;HCoV-229E_1ab;6289 | 0.05 | 0.15 |
| NC_002645.1;NP_073549.1;HCoV-229E_1ab;6290 | 0.05 | 0 |
| NC_002645.1;NP_073549.1;HCoV-229E_1ab;6291 | 0 | 0.075 |
| NC_002645.1;NP_073549.1;HCoV-229E_1ab;6292 | 0 | 0 |
| NC_002645.1;NP_073549.1;HCoV-229E_1ab;6293 | 0 | 0 |
| NC_002645.1;NP_073549.1;HCoV-229E_1ab;6294 | 0 | 0 |
| NC_002645.1;NP_073549.1;HCoV-229E_1ab;6295 | 0 | 0 |
| NC_002645.1;NP_073549.1;HCoV-229E_1ab;6296 | 0 | 0 |
| NC_002645.1;NP_073549.1;HCoV-229E_1ab;6297 | 0.05 | 0 |
| NC_002645.1;NP_073549.1;HCoV-229E_1ab;6298 | 0 | 0 |
| NC_002645.1;NP_073549.1;HCoV-229E_1ab;6299 | 0 | 0 |
| NC_002645.1;NP_073549.1;HCoV-229E_1ab;6300 | 0 | 0 |
| NC_002645.1;NP_073549.1;HCoV-229E_1ab;6301 | 0 | 0 |
| NC_002645.1;NP_073549.1;HCoV-229E_1ab;6302 | 0 | 0 |
| NC_002645.1;NP_073549.1;HCoV-229E_1ab;6303 | 0 | 0 |
| NC_002645.1;NP_073549.1;HCoV-229E_1ab;6304 | 0 | 0 |
| NC_002645.1;NP_073549.1;HCoV-229E_1ab;6305 | 0.05 | 0.025 |
| NC_002645.1;NP_073549.1;HCoV-229E_1ab;6306 | 0.05 | 0.025 |
| NC_002645.1;NP_073549.1;HCoV-229E_1ab;6307 | 0.05 | 0 |

| | | |
|---|---|---|
| NC_002645.1;NP_073549.1;HCoV-229E_1ab;6308 | 0 | 0 |
| NC_002645.1;NP_073549.1;HCoV-229E_1ab;6309 | 0 | 0 |
| NC_002645.1;NP_073549.1;HCoV-229E_1ab;6310 | 0 | 0 |
| NC_002645.1;NP_073549.1;HCoV-229E_1ab;6311 | 0.05 | 0.025 |
| NC_002645.1;NP_073549.1;HCoV-229E_1ab;6312 | 0 | 0 |
| NC_002645.1;NP_073549.1;HCoV-229E_1ab;6313 | 0 | 0 |
| NC_002645.1;NP_073549.1;HCoV-229E_1ab;6314 | 0 | 0 |
| NC_002645.1;NP_073549.1;HCoV-229E_1ab;6315 | 0 | 0 |
| NC_002645.1;NP_073549.1;HCoV-229E_1ab;6316 | 0 | 0 |
| NC_002645.1;NP_073549.1;HCoV-229E_1ab;6317 | 0 | 0 |
| NC_002645.1;NP_073549.1;HCoV-229E_1ab;6318 | 0 | 0 |
| NC_002645.1;NP_073549.1;HCoV-229E_1ab;6319 | 0 | 0 |
| NC_002645.1;NP_073549.1;HCoV-229E_1ab;6320 | 0 | 0 |
| NC_002645.1;NP_073549.1;HCoV-229E_1ab;6321 | 0 | 0 |
| NC_002645.1;NP_073549.1;HCoV-229E_1ab;6322 | 0 | 0 |
| NC_002645.1;NP_073549.1;HCoV-229E_1ab;6323 | 0 | 0 |
| NC_002645.1;NP_073549.1;HCoV-229E_1ab;6324 | 0 | 0 |
| NC_002645.1;NP_073549.1;HCoV-229E_1ab;6325 | 0 | 0 |
| NC_002645.1;NP_073549.1;HCoV-229E_1ab;6326 | 0 | 0 |
| NC_002645.1;NP_073549.1;HCoV-229E_1ab;6327 | 0 | 0 |
| NC_002645.1;NP_073549.1;HCoV-229E_1ab;6328 | 0 | 0 |
| NC_002645.1;NP_073549.1;HCoV-229E_1ab;6329 | 0 | 0 |
| NC_002645.1;NP_073549.1;HCoV-229E_1ab;6330 | 0 | 0 |
| NC_002645.1;NP_073549.1;HCoV-229E_1ab;6331 | 0 | 0.025 |
| NC_002645.1;NP_073549.1;HCoV-229E_1ab;6332 | 0 | 0 |
| NC_002645.1;NP_073549.1;HCoV-229E_1ab;6333 | 0.05 | 0.025 |
| NC_002645.1;NP_073549.1;HCoV-229E_1ab;6334 | 0 | 0 |
| NC_002645.1;NP_073549.1;HCoV-229E_1ab;6335 | 0 | 0 |
| NC_002645.1;NP_073549.1;HCoV-229E_1ab;6336 | 0 | 0 |
| NC_002645.1;NP_073549.1;HCoV-229E_1ab;6337 | 0.1 | 0.3 |
| NC_002645.1;NP_073549.1;HCoV-229E_1ab;6338 | 0.1 | 0.025 |
| NC_002645.1;NP_073549.1;HCoV-229E_1ab;6339 | 0 | 0 |
| NC_002645.1;NP_073549.1;HCoV-229E_1ab;6340 | 0 | 0 |
| NC_002645.1;NP_073549.1;HCoV-229E_1ab;6341 | 0 | 0 |
| NC_002645.1;NP_073549.1;HCoV-229E_1ab;6342 | 0 | 0 |
| NC_002645.1;NP_073549.1;HCoV-229E_1ab;6343 | 0.05 | 0 |
| NC_002645.1;NP_073549.1;HCoV-229E_1ab;6344 | 0 | 0 |
| NC_002645.1;NP_073549.1;HCoV-229E_1ab;6345 | 0 | 0 |
| NC_002645.1;NP_073549.1;HCoV-229E_1ab;6346 | 0 | 0 |
| NC_002645.1;NP_073549.1;HCoV-229E_1ab;6347 | 0 | 0 |
| NC_002645.1;NP_073549.1;HCoV-229E_1ab;6348 | 0 | 0 |
| NC_002645.1;NP_073549.1;HCoV-229E_1ab;6349 | 0 | 0 |
| NC_002645.1;NP_073549.1;HCoV-229E_1ab;6350 | 0 | 0 |

| | | |
|---|---|---|
| NC_002645.1;NP_073549.1;HCoV-229E_1ab;6351 | 0 | 0 |
| NC_002645.1;NP_073549.1;HCoV-229E_1ab;6352 | 0 | 0 |
| NC_002645.1;NP_073549.1;HCoV-229E_1ab;6353 | 0 | 0 |
| NC_002645.1;NP_073549.1;HCoV-229E_1ab;6354 | 0 | 0 |
| NC_002645.1;NP_073549.1;HCoV-229E_1ab;6355 | 0 | 0 |
| NC_002645.1;NP_073549.1;HCoV-229E_1ab;6356 | 0 | 0 |
| NC_002645.1;NP_073549.1;HCoV-229E_1ab;6357 | 0 | 0 |
| NC_002645.1;NP_073549.1;HCoV-229E_1ab;6358 | 0 | 0 |
| NC_002645.1;NP_073549.1;HCoV-229E_1ab;6359 | 0 | 0 |
| NC_002645.1;NP_073549.1;HCoV-229E_1ab;6360 | 0.05 | 0 |
| NC_002645.1;NP_073549.1;HCoV-229E_1ab;6361 | 0.05 | 0 |
| NC_002645.1;NP_073549.1;HCoV-229E_1ab;6362 | 0.05 | 0 |
| NC_002645.1;NP_073549.1;HCoV-229E_1ab;6363 | 0.05 | 0 |
| NC_002645.1;NP_073549.1;HCoV-229E_1ab;6364 | 0.05 | 0 |
| NC_002645.1;NP_073549.1;HCoV-229E_1ab;6365 | 0.05 | 0 |
| NC_002645.1;NP_073549.1;HCoV-229E_1ab;6366 | 0.1 | 0.025 |
| NC_002645.1;NP_073549.1;HCoV-229E_1ab;6367 | 0.1 | 0.025 |
| NC_002645.1;NP_073549.1;HCoV-229E_1ab;6368 | 0.05 | 0 |
| NC_002645.1;NP_073549.1;HCoV-229E_1ab;6369 | 0 | 0 |
| NC_002645.1;NP_073549.1;HCoV-229E_1ab;6370 | 0.1 | 0 |
| NC_002645.1;NP_073549.1;HCoV-229E_1ab;6371 | 0 | 0 |
| NC_002645.1;NP_073549.1;HCoV-229E_1ab;6372 | 0 | 0 |
| NC_002645.1;NP_073549.1;HCoV-229E_1ab;6373 | 0 | 0 |
| NC_002645.1;NP_073549.1;HCoV-229E_1ab;6374 | 0 | 0 |
| NC_002645.1;NP_073549.1;HCoV-229E_1ab;6375 | 0 | 0 |
| NC_002645.1;NP_073549.1;HCoV-229E_1ab;6376 | 0 | 0 |
| NC_002645.1;NP_073549.1;HCoV-229E_1ab;6377 | 0 | 0 |
| NC_002645.1;NP_073549.1;HCoV-229E_1ab;6378 | 0.1 | 0 |
| NC_002645.1;NP_073549.1;HCoV-229E_1ab;6379 | 0 | 0 |
| NC_002645.1;NP_073549.1;HCoV-229E_1ab;6380 | 0 | 0 |
| NC_002645.1;NP_073549.1;HCoV-229E_1ab;6381 | 0 | 0 |
| NC_002645.1;NP_073549.1;HCoV-229E_1ab;6382 | 0 | 0 |
| NC_002645.1;NP_073549.1;HCoV-229E_1ab;6383 | 0 | 0.025 |
| NC_002645.1;NP_073549.1;HCoV-229E_1ab;6384 | 0 | 0 |
| NC_002645.1;NP_073549.1;HCoV-229E_1ab;6385 | 0 | 0.025 |
| NC_002645.1;NP_073549.1;HCoV-229E_1ab;6386 | 0 | 0 |
| NC_002645.1;NP_073549.1;HCoV-229E_1ab;6387 | 0 | 0 |
| NC_002645.1;NP_073549.1;HCoV-229E_1ab;6388 | 0 | 0 |
| NC_002645.1;NP_073549.1;HCoV-229E_1ab;6389 | 0 | 0 |
| NC_002645.1;NP_073549.1;HCoV-229E_1ab;6390 | 0 | 0 |
| NC_002645.1;NP_073549.1;HCoV-229E_1ab;6391 | 0 | 0 |
| NC_002645.1;NP_073549.1;HCoV-229E_1ab;6392 | 0 | 0 |
| NC_002645.1;NP_073549.1;HCoV-229E_1ab;6393 | 0 | 0.05 |

| | | |
|---|---|---|
| NC_002645.1;NP_073549.1;HCoV-229E_1ab;6394 | 0 | 0.025 |
| NC_002645.1;NP_073549.1;HCoV-229E_1ab;6395 | 0 | 0.025 |
| NC_002645.1;NP_073549.1;HCoV-229E_1ab;6396 | 0 | 0 |
| NC_002645.1;NP_073549.1;HCoV-229E_1ab;6397 | 0 | 0 |
| NC_002645.1;NP_073549.1;HCoV-229E_1ab;6398 | 0 | 0 |
| NC_002645.1;NP_073549.1;HCoV-229E_1ab;6399 | 0 | 0 |
| NC_002645.1;NP_073549.1;HCoV-229E_1ab;6400 | 0 | 0 |
| NC_002645.1;NP_073549.1;HCoV-229E_1ab;6401 | 0 | 0 |
| NC_002645.1;NP_073549.1;HCoV-229E_1ab;6402 | 0 | 0 |
| NC_002645.1;NP_073549.1;HCoV-229E_1ab;6403 | 0 | 0 |
| NC_002645.1;NP_073549.1;HCoV-229E_1ab;6404 | 0 | 0.025 |
| NC_002645.1;NP_073549.1;HCoV-229E_1ab;6405 | 0 | 0 |
| NC_002645.1;NP_073549.1;HCoV-229E_1ab;6406 | 0 | 0 |
| NC_002645.1;NP_073549.1;HCoV-229E_1ab;6407 | 0.05 | 0 |
| NC_002645.1;NP_073549.1;HCoV-229E_1ab;6408 | 0 | 0 |
| NC_002645.1;NP_073549.1;HCoV-229E_1ab;6409 | 0 | 0.05 |
| NC_002645.1;NP_073549.1;HCoV-229E_1ab;6410 | 0 | 0 |
| NC_002645.1;NP_073549.1;HCoV-229E_1ab;6411 | 0 | 0 |
| NC_002645.1;NP_073549.1;HCoV-229E_1ab;6412 | 0 | 0 |
| NC_002645.1;NP_073549.1;HCoV-229E_1ab;6413 | 0 | 0 |
| NC_002645.1;NP_073549.1;HCoV-229E_1ab;6414 | 0 | 0 |
| NC_002645.1;NP_073549.1;HCoV-229E_1ab;6415 | 0 | 0 |
| NC_002645.1;NP_073549.1;HCoV-229E_1ab;6416 | 0 | 0 |
| NC_002645.1;NP_073549.1;HCoV-229E_1ab;6417 | 0 | 0 |
| NC_002645.1;NP_073549.1;HCoV-229E_1ab;6418 | 0 | 0 |
| NC_002645.1;NP_073549.1;HCoV-229E_1ab;6419 | 0 | 0 |
| NC_002645.1;NP_073549.1;HCoV-229E_1ab;6420 | 0 | 0 |
| NC_002645.1;NP_073549.1;HCoV-229E_1ab;6421 | 0 | 0 |
| NC_002645.1;NP_073549.1;HCoV-229E_1ab;6422 | 0.05 | 0.05 |
| NC_002645.1;NP_073549.1;HCoV-229E_1ab;6423 | 0 | 0.025 |
| NC_002645.1;NP_073549.1;HCoV-229E_1ab;6424 | 0 | 0.025 |
| NC_002645.1;NP_073549.1;HCoV-229E_1ab;6425 | 0 | 0 |
| NC_002645.1;NP_073549.1;HCoV-229E_1ab;6426 | 0 | 0 |
| NC_002645.1;NP_073549.1;HCoV-229E_1ab;6427 | 0 | 0 |
| NC_002645.1;NP_073549.1;HCoV-229E_1ab;6428 | 0 | 0 |
| NC_002645.1;NP_073549.1;HCoV-229E_1ab;6429 | 0 | 0 |
| NC_002645.1;NP_073549.1;HCoV-229E_1ab;6430 | 0 | 0 |
| NC_002645.1;NP_073549.1;HCoV-229E_1ab;6431 | 0 | 0 |
| NC_002645.1;NP_073549.1;HCoV-229E_1ab;6432 | 0 | 0 |
| NC_002645.1;NP_073549.1;HCoV-229E_1ab;6433 | 0 | 0 |
| NC_002645.1;NP_073549.1;HCoV-229E_1ab;6434 | 0.05 | 0 |
| NC_002645.1;NP_073549.1;HCoV-229E_1ab;6435 | 0 | 0 |
| NC_002645.1;NP_073549.1;HCoV-229E_1ab;6436 | 0 | 0.025 |

| | | |
|---|---|---|
| NC_002645.1;NP_073549.1;HCoV-229E_1ab;6437 | 0 | 0 |
| NC_002645.1;NP_073549.1;HCoV-229E_1ab;6438 | 0 | 0 |
| NC_002645.1;NP_073549.1;HCoV-229E_1ab;6439 | 0 | 0 |
| NC_002645.1;NP_073549.1;HCoV-229E_1ab;6440 | 0 | 0 |
| NC_002645.1;NP_073549.1;HCoV-229E_1ab;6441 | 0 | 0 |
| NC_002645.1;NP_073549.1;HCoV-229E_1ab;6442 | 0 | 0 |
| NC_002645.1;NP_073549.1;HCoV-229E_1ab;6443 | 0 | 0 |
| NC_002645.1;NP_073549.1;HCoV-229E_1ab;6444 | 0.05 | 0.05 |
| NC_002645.1;NP_073549.1;HCoV-229E_1ab;6445 | 0 | 0.05 |
| NC_002645.1;NP_073549.1;HCoV-229E_1ab;6446 | 0.05 | 0.075 |
| NC_002645.1;NP_073549.1;HCoV-229E_1ab;6447 | 0 | 0.025 |
| NC_002645.1;NP_073549.1;HCoV-229E_1ab;6448 | 0 | 0.025 |
| NC_002645.1;NP_073549.1;HCoV-229E_1ab;6449 | 0 | 0.025 |
| NC_002645.1;NP_073549.1;HCoV-229E_1ab;6450 | 0 | 0.025 |
| NC_002645.1;NP_073549.1;HCoV-229E_1ab;6451 | 0 | 0.025 |
| NC_002645.1;NP_073549.1;HCoV-229E_1ab;6452 | 0 | 0 |
| NC_002645.1;NP_073549.1;HCoV-229E_1ab;6453 | 0.05 | 0 |
| NC_002645.1;NP_073549.1;HCoV-229E_1ab;6454 | 0.05 | 0 |
| NC_002645.1;NP_073549.1;HCoV-229E_1ab;6455 | 0 | 0 |
| NC_002645.1;NP_073549.1;HCoV-229E_1ab;6456 | 0 | 0 |
| NC_002645.1;NP_073549.1;HCoV-229E_1ab;6457 | 0 | 0 |
| NC_002645.1;NP_073549.1;HCoV-229E_1ab;6458 | 0 | 0 |
| NC_002645.1;NP_073549.1;HCoV-229E_1ab;6459 | 0 | 0 |
| NC_002645.1;NP_073549.1;HCoV-229E_1ab;6460 | 0 | 0 |
| NC_002645.1;NP_073549.1;HCoV-229E_1ab;6461 | 0 | 0 |
| NC_002645.1;NP_073549.1;HCoV-229E_1ab;6462 | 0 | 0 |
| NC_002645.1;NP_073549.1;HCoV-229E_1ab;6463 | 0 | 0 |
| NC_002645.1;NP_073549.1;HCoV-229E_1ab;6464 | 0 | 0 |
| NC_002645.1;NP_073549.1;HCoV-229E_1ab;6465 | 0 | 0 |
| NC_002645.1;NP_073549.1;HCoV-229E_1ab;6466 | 0 | 0 |
| NC_002645.1;NP_073549.1;HCoV-229E_1ab;6467 | 0 | 0.025 |
| NC_002645.1;NP_073549.1;HCoV-229E_1ab;6468 | 0 | 0 |
| NC_002645.1;NP_073549.1;HCoV-229E_1ab;6469 | 0 | 0 |
| NC_002645.1;NP_073549.1;HCoV-229E_1ab;6470 | 0 | 0 |
| NC_002645.1;NP_073549.1;HCoV-229E_1ab;6471 | 0 | 0 |
| NC_002645.1;NP_073549.1;HCoV-229E_1ab;6472 | 0 | 0 |
| NC_002645.1;NP_073549.1;HCoV-229E_1ab;6473 | 0 | 0 |
| NC_002645.1;NP_073549.1;HCoV-229E_1ab;6474 | 0 | 0 |
| NC_002645.1;NP_073549.1;HCoV-229E_1ab;6475 | 0 | 0 |
| NC_002645.1;NP_073549.1;HCoV-229E_1ab;6476 | 0 | 0 |
| NC_002645.1;NP_073549.1;HCoV-229E_1ab;6477 | 0 | 0.025 |
| NC_002645.1;NP_073549.1;HCoV-229E_1ab;6478 | 0 | 0 |
| NC_002645.1;NP_073549.1;HCoV-229E_1ab;6479 | 0 | 0 |

| | | |
|---|---|---|
| NC_002645.1;NP_073549.1;HCoV-229E_1ab;6480 | 0 | 0 |
| NC_002645.1;NP_073549.1;HCoV-229E_1ab;6481 | 0 | 0 |
| NC_002645.1;NP_073549.1;HCoV-229E_1ab;6482 | 0 | 0 |
| NC_002645.1;NP_073549.1;HCoV-229E_1ab;6483 | 0 | 0 |
| NC_002645.1;NP_073549.1;HCoV-229E_1ab;6484 | 0 | 0 |
| NC_002645.1;NP_073549.1;HCoV-229E_1ab;6485 | 0 | 0 |
| NC_002645.1;NP_073549.1;HCoV-229E_1ab;6486 | 0 | 0 |
| NC_002645.1;NP_073549.1;HCoV-229E_1ab;6487 | 0 | 0 |
| NC_002645.1;NP_073549.1;HCoV-229E_1ab;6488 | 0 | 0 |
| NC_002645.1;NP_073549.1;HCoV-229E_1ab;6489 | 0 | 0 |
| NC_002645.1;NP_073549.1;HCoV-229E_1ab;6490 | 0 | 0 |
| NC_002645.1;NP_073549.1;HCoV-229E_1ab;6491 | 0 | 0 |
| NC_002645.1;NP_073549.1;HCoV-229E_1ab;6492 | 0 | 0 |
| NC_002645.1;NP_073549.1;HCoV-229E_1ab;6493 | 0 | 0 |
| NC_002645.1;NP_073549.1;HCoV-229E_1ab;6494 | 0 | 0 |
| NC_002645.1;NP_073549.1;HCoV-229E_1ab;6495 | 0 | 0 |
| NC_002645.1;NP_073549.1;HCoV-229E_1ab;6496 | 0 | 0 |
| NC_002645.1;NP_073549.1;HCoV-229E_1ab;6497 | 0 | 0 |
| NC_002645.1;NP_073549.1;HCoV-229E_1ab;6498 | 0 | 0 |
| NC_002645.1;NP_073549.1;HCoV-229E_1ab;6499 | 0 | 0 |
| NC_002645.1;NP_073549.1;HCoV-229E_1ab;6500 | 0 | 0 |
| NC_002645.1;NP_073549.1;HCoV-229E_1ab;6501 | 0 | 0 |
| NC_002645.1;NP_073549.1;HCoV-229E_1ab;6502 | 0 | 0 |
| NC_002645.1;NP_073549.1;HCoV-229E_1ab;6503 | 0 | 0 |
| NC_002645.1;NP_073549.1;HCoV-229E_1ab;6504 | 0 | 0 |
| NC_002645.1;NP_073549.1;HCoV-229E_1ab;6505 | 0 | 0 |
| NC_002645.1;NP_073549.1;HCoV-229E_1ab;6506 | 0.05 | 0 |
| NC_002645.1;NP_073549.1;HCoV-229E_1ab;6507 | 0.05 | 0 |
| NC_002645.1;NP_073549.1;HCoV-229E_1ab;6508 | 0.05 | 0 |
| NC_002645.1;NP_073549.1;HCoV-229E_1ab;6509 | 0.05 | 0 |
| NC_002645.1;NP_073549.1;HCoV-229E_1ab;6510 | 0 | 0 |
| NC_002645.1;NP_073549.1;HCoV-229E_1ab;6511 | 0 | 0 |
| NC_002645.1;NP_073549.1;HCoV-229E_1ab;6512 | 0 | 0 |
| NC_002645.1;NP_073549.1;HCoV-229E_1ab;6513 | 0 | 0 |
| NC_002645.1;NP_073549.1;HCoV-229E_1ab;6514 | 0 | 0 |
| NC_002645.1;NP_073549.1;HCoV-229E_1ab;6515 | 0 | 0 |
| NC_002645.1;NP_073549.1;HCoV-229E_1ab;6516 | 0 | 0 |
| NC_002645.1;NP_073549.1;HCoV-229E_1ab;6517 | 0 | 0 |
| NC_002645.1;NP_073549.1;HCoV-229E_1ab;6518 | 0 | 0 |
| NC_002645.1;NP_073549.1;HCoV-229E_1ab;6519 | 0 | 0 |
| NC_002645.1;NP_073549.1;HCoV-229E_1ab;6520 | 0 | 0 |
| NC_002645.1;NP_073549.1;HCoV-229E_1ab;6521 | 0.1 | 0.05 |
| NC_002645.1;NP_073549.1;HCoV-229E_1ab;6522 | 0.1 | 0 |

| | | |
|---|---|---|
| NC_002645.1;NP_073549.1;HCoV-229E_1ab;6523 | 0.05 | 0 |
| NC_002645.1;NP_073549.1;HCoV-229E_1ab;6524 | 0.05 | 0 |
| NC_002645.1;NP_073549.1;HCoV-229E_1ab;6525 | 0.05 | 0 |
| NC_002645.1;NP_073549.1;HCoV-229E_1ab;6526 | 0.05 | 0 |
| NC_002645.1;NP_073549.1;HCoV-229E_1ab;6527 | 0.05 | 0 |
| NC_002645.1;NP_073549.1;HCoV-229E_1ab;6528 | 0.05 | 0 |
| NC_002645.1;NP_073549.1;HCoV-229E_1ab;6529 | 0 | 0 |
| NC_002645.1;NP_073549.1;HCoV-229E_1ab;6530 | 0 | 0 |
| NC_002645.1;NP_073549.1;HCoV-229E_1ab;6531 | 0 | 0 |
| NC_002645.1;NP_073549.1;HCoV-229E_1ab;6532 | 0 | 0 |
| NC_002645.1;NP_073549.1;HCoV-229E_1ab;6533 | 0 | 0 |
| NC_002645.1;NP_073549.1;HCoV-229E_1ab;6534 | 0 | 0 |
| NC_002645.1;NP_073549.1;HCoV-229E_1ab;6535 | 0 | 0.025 |
| NC_002645.1;NP_073549.1;HCoV-229E_1ab;6536 | 0 | 0 |
| NC_002645.1;NP_073549.1;HCoV-229E_1ab;6537 | 0 | 0 |
| NC_002645.1;NP_073549.1;HCoV-229E_1ab;6538 | 0 | 0 |
| NC_002645.1;NP_073549.1;HCoV-229E_1ab;6539 | 0 | 0 |
| NC_002645.1;NP_073549.1;HCoV-229E_1ab;6540 | 0 | 0 |
| NC_002645.1;NP_073549.1;HCoV-229E_1ab;6541 | 0 | 0 |
| NC_002645.1;NP_073549.1;HCoV-229E_1ab;6542 | 0 | 0 |
| NC_002645.1;NP_073549.1;HCoV-229E_1ab;6543 | 0 | 0.075 |
| NC_002645.1;NP_073549.1;HCoV-229E_1ab;6544 | 0 | 0.025 |
| NC_002645.1;NP_073549.1;HCoV-229E_1ab;6545 | 0 | 0.025 |
| NC_002645.1;NP_073549.1;HCoV-229E_1ab;6546 | 0 | 0.025 |
| NC_002645.1;NP_073549.1;HCoV-229E_1ab;6547 | 0 | 0.025 |
| NC_002645.1;NP_073549.1;HCoV-229E_1ab;6548 | 0.05 | 0.025 |
| NC_002645.1;NP_073549.1;HCoV-229E_1ab;6549 | 0.05 | 0.025 |
| NC_002645.1;NP_073549.1;HCoV-229E_1ab;6550 | 0 | 0 |
| NC_002645.1;NP_073549.1;HCoV-229E_1ab;6551 | 0 | 0 |
| NC_002645.1;NP_073549.1;HCoV-229E_1ab;6552 | 0 | 0.05 |
| NC_002645.1;NP_073549.1;HCoV-229E_1ab;6553 | 0 | 0 |
| NC_002645.1;NP_073549.1;HCoV-229E_1ab;6554 | 0 | 0 |
| NC_002645.1;NP_073549.1;HCoV-229E_1ab;6555 | 0 | 0 |
| NC_002645.1;NP_073549.1;HCoV-229E_1ab;6556 | 0 | 0 |
| NC_002645.1;NP_073549.1;HCoV-229E_1ab;6557 | 0 | 0.025 |
| NC_002645.1;NP_073549.1;HCoV-229E_1ab;6558 | 0 | 0 |
| NC_002645.1;NP_073549.1;HCoV-229E_1ab;6559 | 0 | 0 |
| NC_002645.1;NP_073549.1;HCoV-229E_1ab;6560 | 0 | 0 |
| NC_002645.1;NP_073549.1;HCoV-229E_1ab;6561 | 0 | 0 |
| NC_002645.1;NP_073549.1;HCoV-229E_1ab;6562 | 0 | 0 |
| NC_002645.1;NP_073549.1;HCoV-229E_1ab;6563 | 0 | 0 |
| NC_002645.1;NP_073549.1;HCoV-229E_1ab;6564 | 0 | 0 |
| NC_002645.1;NP_073549.1;HCoV-229E_1ab;6565 | 0 | 0 |

| | | |
|---|---|---|
| NC_002645.1;NP_073549.1;HCoV-229E_1ab;6566 | 0 | 0 |
| NC_002645.1;NP_073549.1;HCoV-229E_1ab;6567 | 0 | 0 |
| NC_002645.1;NP_073549.1;HCoV-229E_1ab;6568 | 0 | 0 |
| NC_002645.1;NP_073549.1;HCoV-229E_1ab;6569 | 0 | 0 |
| NC_002645.1;NP_073549.1;HCoV-229E_1ab;6570 | 0 | 0 |
| NC_002645.1;NP_073549.1;HCoV-229E_1ab;6571 | 0 | 0 |
| NC_002645.1;NP_073549.1;HCoV-229E_1ab;6572 | 0 | 0 |
| NC_002645.1;NP_073549.1;HCoV-229E_1ab;6573 | 0 | 0 |
| NC_002645.1;NP_073549.1;HCoV-229E_1ab;6574 | 0 | 0 |
| NC_002645.1;NP_073549.1;HCoV-229E_1ab;6575 | 0 | 0.025 |
| NC_002645.1;NP_073549.1;HCoV-229E_1ab;6576 | 0.1 | 0.025 |
| NC_002645.1;NP_073549.1;HCoV-229E_1ab;6577 | 0.15 | 0.025 |
| NC_002645.1;NP_073549.1;HCoV-229E_1ab;6578 | 0.35 | 0.025 |
| NC_002645.1;NP_073549.1;HCoV-229E_1ab;6579 | 0.35 | 0.05 |
| NC_002645.1;NP_073549.1;HCoV-229E_1ab;6580 | 0.05 | 0 |
| NC_002645.1;NP_073549.1;HCoV-229E_1ab;6581 | 0 | 0 |
| NC_002645.1;NP_073549.1;HCoV-229E_1ab;6582 | 0 | 0.025 |
| NC_002645.1;NP_073549.1;HCoV-229E_1ab;6583 | 0 | 0.025 |
| NC_002645.1;NP_073549.1;HCoV-229E_1ab;6584 | 0.05 | 0.025 |
| NC_002645.1;NP_073549.1;HCoV-229E_1ab;6585 | 0 | 0.025 |
| NC_002645.1;NP_073549.1;HCoV-229E_1ab;6586 | 0 | 0.025 |
| NC_002645.1;NP_073549.1;HCoV-229E_1ab;6587 | 0 | 0.025 |
| NC_002645.1;NP_073549.1;HCoV-229E_1ab;6588 | 0 | 0.025 |
| NC_002645.1;NP_073549.1;HCoV-229E_1ab;6589 | 0 | 0 |
| NC_002645.1;NP_073549.1;HCoV-229E_1ab;6590 | 0 | 0 |
| NC_002645.1;NP_073549.1;HCoV-229E_1ab;6591 | 0 | 0 |
| NC_002645.1;NP_073549.1;HCoV-229E_1ab;6592 | 0.05 | 0 |
| NC_002645.1;NP_073549.1;HCoV-229E_1ab;6593 | 0.05 | 0.025 |
| NC_002645.1;NP_073549.1;HCoV-229E_1ab;6594 | 0.05 | 0 |
| NC_002645.1;NP_073549.1;HCoV-229E_1ab;6595 | 0 | 0 |
| NC_002645.1;NP_073549.1;HCoV-229E_1ab;6596 | 0.05 | 0 |
| NC_002645.1;NP_073549.1;HCoV-229E_1ab;6597 | 0.05 | 0 |
| NC_002645.1;NP_073549.1;HCoV-229E_1ab;6598 | 0 | 0 |
| NC_002645.1;NP_073549.1;HCoV-229E_1ab;6599 | 0 | 0 |
| NC_002645.1;NP_073549.1;HCoV-229E_1ab;6600 | 0 | 0 |
| NC_002645.1;NP_073549.1;HCoV-229E_1ab;6601 | 0 | 0 |
| NC_002645.1;NP_073549.1;HCoV-229E_1ab;6602 | 0 | 0 |
| NC_002645.1;NP_073549.1;HCoV-229E_1ab;6603 | 0 | 0 |
| NC_002645.1;NP_073549.1;HCoV-229E_1ab;6604 | 0 | 0 |
| NC_002645.1;NP_073549.1;HCoV-229E_1ab;6605 | 0 | 0 |
| NC_002645.1;NP_073549.1;HCoV-229E_1ab;6606 | 0 | 0 |
| NC_002645.1;NP_073549.1;HCoV-229E_1ab;6607 | 0 | 0 |
| NC_002645.1;NP_073549.1;HCoV-229E_1ab;6608 | 0 | 0 |

| | | |
|---|---|---|
| NC_002645.1;NP_073549.1;HCoV-229E_1ab;6609 | 0 | 0 |
| NC_002645.1;NP_073549.1;HCoV-229E_1ab;6610 | 0 | 0 |
| NC_002645.1;NP_073549.1;HCoV-229E_1ab;6611 | 0 | 0 |
| NC_002645.1;NP_073549.1;HCoV-229E_1ab;6612 | 0 | 0 |
| NC_002645.1;NP_073549.1;HCoV-229E_1ab;6613 | 0 | 0 |
| NC_002645.1;NP_073549.1;HCoV-229E_1ab;6614 | 0 | 0 |
| NC_002645.1;NP_073549.1;HCoV-229E_1ab;6615 | 0 | 0 |
| NC_002645.1;NP_073549.1;HCoV-229E_1ab;6616 | 0 | 0 |
| NC_002645.1;NP_073549.1;HCoV-229E_1ab;6617 | 0 | 0 |
| NC_002645.1;NP_073549.1;HCoV-229E_1ab;6618 | 0 | 0 |
| NC_002645.1;NP_073549.1;HCoV-229E_1ab;6619 | 0 | 0 |
| NC_002645.1;NP_073549.1;HCoV-229E_1ab;6620 | 0 | 0 |
| NC_002645.1;NP_073549.1;HCoV-229E_1ab;6621 | 0 | 0.025 |
| NC_002645.1;NP_073549.1;HCoV-229E_1ab;6622 | 0 | 0.025 |
| NC_002645.1;NP_073549.1;HCoV-229E_1ab;6623 | 0 | 0.025 |
| NC_002645.1;NP_073549.1;HCoV-229E_1ab;6624 | 0 | 0.025 |
| NC_002645.1;NP_073549.1;HCoV-229E_1ab;6625 | 0 | 0.025 |
| NC_002645.1;NP_073549.1;HCoV-229E_1ab;6626 | 0.05 | 0.025 |
| NC_002645.1;NP_073549.1;HCoV-229E_1ab;6627 | 0.05 | 0.025 |
| NC_002645.1;NP_073549.1;HCoV-229E_1ab;6628 | 0 | 0.025 |
| NC_002645.1;NP_073549.1;HCoV-229E_1ab;6629 | 0 | 0.025 |
| NC_002645.1;NP_073549.1;HCoV-229E_1ab;6630 | 0 | 0.025 |
| NC_002645.1;NP_073549.1;HCoV-229E_1ab;6631 | 0 | 0 |
| NC_002645.1;NP_073549.1;HCoV-229E_1ab;6632 | 0 | 0 |
| NC_002645.1;NP_073549.1;HCoV-229E_1ab;6633 | 0 | 0 |
| NC_002645.1;NP_073549.1;HCoV-229E_1ab;6634 | 0 | 0 |
| NC_002645.1;NP_073549.1;HCoV-229E_1ab;6635 | 0 | 0 |
| NC_002645.1;NP_073549.1;HCoV-229E_1ab;6636 | 0 | 0 |
| NC_002645.1;NP_073549.1;HCoV-229E_1ab;6637 | 0 | 0 |
| NC_002645.1;NP_073549.1;HCoV-229E_1ab;6638 | 0 | 0 |
| NC_002645.1;NP_073549.1;HCoV-229E_1ab;6639 | 0 | 0 |
| NC_002645.1;NP_073549.1;HCoV-229E_1ab;6640 | 0 | 0 |
| NC_002645.1;NP_073549.1;HCoV-229E_1ab;6641 | 0 | 0 |
| NC_002645.1;NP_073549.1;HCoV-229E_1ab;6642 | 0 | 0 |
| NC_002645.1;NP_073549.1;HCoV-229E_1ab;6643 | 0 | 0 |
| NC_002645.1;NP_073549.1;HCoV-229E_1ab;6644 | 0 | 0 |
| NC_002645.1;NP_073549.1;HCoV-229E_1ab;6645 | 0 | 0 |
| NC_002645.1;NP_073549.1;HCoV-229E_1ab;6646 | 0 | 0.025 |
| NC_002645.1;NP_073549.1;HCoV-229E_1ab;6647 | 0 | 0.025 |
| NC_002645.1;NP_073549.1;HCoV-229E_1ab;6648 | 0 | 0 |
| NC_002645.1;NP_073549.1;HCoV-229E_1ab;6649 | 0 | 0 |
| NC_002645.1;NP_073549.1;HCoV-229E_1ab;6650 | 0 | 0 |
| NC_002645.1;NP_073549.1;HCoV-229E_1ab;6651 | 0 | 0 |

| | | |
|---|---|---|
| NC_002645.1;NP_073549.1;HCoV-229E_1ab;6652 | 0 | 0 |
| NC_002645.1;NP_073549.1;HCoV-229E_1ab;6653 | 0 | 0 |
| NC_002645.1;NP_073549.1;HCoV-229E_1ab;6654 | 0 | 0 |
| NC_002645.1;NP_073549.1;HCoV-229E_1ab;6655 | 0 | 0 |
| NC_002645.1;NP_073549.1;HCoV-229E_1ab;6656 | 0 | 0 |
| NC_002645.1;NP_073549.1;HCoV-229E_1ab;6657 | 0 | 0 |
| NC_002645.1;NP_073549.1;HCoV-229E_1ab;6658 | 0 | 0 |
| NC_002645.1;NP_073549.1;HCoV-229E_1ab;6659 | 0 | 0 |
| NC_002645.1;NP_073549.1;HCoV-229E_1ab;6660 | 0 | 0 |
| NC_002645.1;NP_073549.1;HCoV-229E_1ab;6661 | 0 | 0 |
| NC_002645.1;NP_073549.1;HCoV-229E_1ab;6662 | 0 | 0 |
| NC_002645.1;NP_073549.1;HCoV-229E_1ab;6663 | 0 | 0 |
| NC_002645.1;NP_073549.1;HCoV-229E_1ab;6664 | 0 | 0.025 |
| NC_002645.1;NP_073549.1;HCoV-229E_1ab;6665 | 0.05 | 0.075 |
| NC_002645.1;NP_073549.1;HCoV-229E_1ab;6666 | 0 | 0 |
| NC_002645.1;NP_073549.1;HCoV-229E_1ab;6667 | 0 | 0 |
| NC_002645.1;NP_073549.1;HCoV-229E_1ab;6668 | 0 | 0 |
| NC_002645.1;NP_073549.1;HCoV-229E_1ab;6669 | 0 | 0 |
| NC_002645.1;NP_073549.1;HCoV-229E_1ab;6670 | 0 | 0 |
| NC_002645.1;NP_073549.1;HCoV-229E_1ab;6671 | 0.1 | 0.05 |
| NC_002645.1;NP_073549.1;HCoV-229E_1ab;6672 | 0.05 | 0 |
| NC_002645.1;NP_073549.1;HCoV-229E_1ab;6673 | 0.05 | 0 |
| NC_002645.1;NP_073549.1;HCoV-229E_1ab;6674 | 0.05 | 0 |
| NC_002645.1;NP_073549.1;HCoV-229E_1ab;6675 | 0 | 0 |
| NC_002645.1;NP_073549.1;HCoV-229E_1ab;6676 | 0 | 0 |
| NC_002645.1;NP_073549.1;HCoV-229E_1ab;6677 | 0 | 0 |
| NC_002645.1;NP_073549.1;HCoV-229E_1ab;6678 | 0 | 0 |
| NC_002645.1;NP_073549.1;HCoV-229E_1ab;6679 | 0 | 0 |
| NC_002645.1;NP_073549.1;HCoV-229E_1ab;6680 | 0 | 0 |
| NC_002645.1;NP_073549.1;HCoV-229E_1ab;6681 | 0 | 0 |
| NC_002645.1;NP_073549.1;HCoV-229E_1ab;6682 | 0 | 0 |
| NC_002645.1;NP_073549.1;HCoV-229E_1ab;6683 | 0 | 0 |
| NC_002645.1;NP_073549.1;HCoV-229E_1ab;6684 | 0 | 0 |
| NC_002645.1;NP_073549.1;HCoV-229E_1ab;6685 | 0 | 0 |
| NC_002645.1;NP_073549.1;HCoV-229E_1ab;6686 | 0 | 0 |
| NC_002645.1;NP_073549.1;HCoV-229E_1ab;6687 | 0 | 0 |
| NC_002645.1;NP_073549.1;HCoV-229E_1ab;6688 | 0 | 0 |
| NC_002645.1;NP_073549.1;HCoV-229E_1ab;6689 | 0 | 0 |
| NC_002645.1;NP_073549.1;HCoV-229E_1ab;6690 | 0 | 0 |
| NC_002645.1;NP_073549.1;HCoV-229E_1ab;6691 | 0 | 0 |
| NC_002645.1;NP_073549.1;HCoV-229E_1ab;6692 | 0 | 0 |
| NC_002645.1;NP_073549.1;HCoV-229E_1ab;6693 | 0 | 0 |
| NC_002645.1;NP_073549.1;HCoV-229E_1ab;6694 | 0 | 0 |

| | | |
|---|---|---|
| NC_002645.1;NP_073549.1;HCoV-229E_1ab;6695 | 0 | 0.025 |
| NC_002645.1;NP_073549.1;HCoV-229E_1ab;6696 | 0 | 0 |
| NC_002645.1;NP_073549.1;HCoV-229E_1ab;6697 | 0 | 0 |
| NC_002645.1;NP_073549.1;HCoV-229E_1ab;6698 | 0 | 0 |
| NC_002645.1;NP_073549.1;HCoV-229E_1ab;6699 | 0 | 0 |
| NC_002645.1;NP_073549.1;HCoV-229E_1ab;6700 | 0 | 0 |
| NC_002645.1;NP_073549.1;HCoV-229E_1ab;6701 | 0 | 0 |
| NC_002645.1;NP_073549.1;HCoV-229E_1ab;6702 | 0 | 0 |
| NC_002645.1;NP_073549.1;HCoV-229E_1ab;6703 | 0 | 0 |
| NC_002645.1;NP_073549.1;HCoV-229E_1ab;6704 | 0 | 0 |
| NC_002645.1;NP_073549.1;HCoV-229E_1ab;6705 | 0 | 0 |
| NC_002645.1;NP_073549.1;HCoV-229E_1ab;6706 | 0 | 0 |
| NC_002645.1;NP_073549.1;HCoV-229E_1ab;6707 | 0 | 0 |
| NC_002645.1;NP_073549.1;HCoV-229E_1ab;6708 | 0 | 0 |
| NC_002645.1;NP_073549.1;HCoV-229E_1ab;6709 | 0 | 0 |
| NC_002645.1;NP_073549.1;HCoV-229E_1ab;6710 | 0 | 0 |
| NC_002645.1;NP_073549.1;HCoV-229E_1ab;6711 | 0 | 0 |
| NC_002645.1;NP_073549.1;HCoV-229E_1ab;6712 | 0 | 0.025 |
| NC_002645.1;NP_073549.1;HCoV-229E_1ab;6713 | 0 | 0 |
| NC_002645.1;NP_073549.1;HCoV-229E_1ab;6714 | 0 | 0 |
| NC_002645.1;NP_073549.1;HCoV-229E_1ab;6715 | 0 | 0 |
| NC_002645.1;NP_073549.1;HCoV-229E_1ab;6716 | 0 | 0 |
| NC_002645.1;NP_073549.1;HCoV-229E_1ab;6717 | 0 | 0 |
| NC_002645.1;NP_073549.1;HCoV-229E_1ab;6718 | 0 | 0 |
| NC_002645.1;NP_073549.1;HCoV-229E_1ab;6719 | 0 | 0 |
| NC_002645.1;NP_073549.1;HCoV-229E_1ab;6720 | 0 | 0 |
| NC_002645.1;NP_073549.1;HCoV-229E_1ab;6721 | 0 | 0.025 |
| NC_002645.1;NP_073549.1;HCoV-229E_1ab;6722 | 0 | 0 |
| NC_002645.1;NP_073549.1;HCoV-229E_1ab;6723 | 0.05 | 0.05 |
| NC_002645.1;NP_073549.1;HCoV-229E_1ab;6724 | 0 | 0 |
| NC_002645.1;NP_073549.1;HCoV-229E_1ab;6725 | 0 | 0 |
| NC_002645.1;NP_073549.1;HCoV-229E_1ab;6726 | 0 | 0 |
| NC_002645.1;NP_073549.1;HCoV-229E_1ab;6727 | 0 | 0 |
| NC_002645.1;NP_073549.1;HCoV-229E_1ab;6728 | 0 | 0 |
| NC_002645.1;NP_073549.1;HCoV-229E_1ab;6729 | 0 | 0 |
| NC_002645.1;NP_073549.1;HCoV-229E_1ab;6730 | 0 | 0 |
| NC_002645.1;NP_073549.1;HCoV-229E_1ab;6731 | 0 | 0 |
| NC_002645.1;NP_073549.1;HCoV-229E_1ab;6732 | 0 | 0 |
| NC_002645.1;NP_073549.1;HCoV-229E_1ab;6733 | 0 | 0 |
| NC_002645.1;NP_073549.1;HCoV-229E_1ab;6734 | 0 | 0 |
| NC_002645.1;NP_073549.1;HCoV-229E_1ab;6735 | 0 | 0 |
| NC_002645.1;NP_073549.1;HCoV-229E_1ab;6736 | 0 | 0 |
| NC_002645.1;NP_073549.1;HCoV-229E_1ab;6737 | 0 | 0 |

| | | |
|---|---|---|
| NC_002645.1;NP_073549.1;HCoV-229E_1ab;6738 | 0 | 0 |
| NC_002645.1;NP_073549.1;HCoV-229E_1ab;6739 | 0 | 0 |
| NC_002645.1;NP_073549.1;HCoV-229E_1ab;6740 | 0 | 0 |
| NC_002645.1;NP_073549.1;HCoV-229E_1ab;6741 | 0 | 0 |
| NC_002645.1;NP_073549.1;HCoV-229E_1ab;6742 | 0 | 0.025 |
| NC_002645.1;NP_073549.1;HCoV-229E_1ab;6743 | 0 | 0 |
| NC_002645.1;NP_073549.1;HCoV-229E_1ab;6744 | 0 | 0 |
| NC_002645.1;NP_073549.1;HCoV-229E_1ab;6745 | 0 | 0 |
| NC_002645.1;NP_073549.1;HCoV-229E_1ab;6746 | 0 | 0 |
| NC_002645.1;NP_073549.1;HCoV-229E_1ab;6747 | 0 | 0 |
| NC_002645.1;NP_073550.1;HCoV-229E_1a;1 | 0.05 | 0 |
| NC_002645.1;NP_073550.1;HCoV-229E_1a;2 | 0 | 0 |
| NC_002645.1;NP_073550.1;HCoV-229E_1a;3 | 0 | 0 |
| NC_002645.1;NP_073550.1;HCoV-229E_1a;4 | 0 | 0 |
| NC_002645.1;NP_073550.1;HCoV-229E_1a;5 | 0 | 0 |
| NC_002645.1;NP_073550.1;HCoV-229E_1a;6 | 0 | 0 |
| NC_002645.1;NP_073550.1;HCoV-229E_1a;7 | 0 | 0 |
| NC_002645.1;NP_073550.1;HCoV-229E_1a;8 | 0.05 | 0 |
| NC_002645.1;NP_073550.1;HCoV-229E_1a;9 | 0 | 0 |
| NC_002645.1;NP_073550.1;HCoV-229E_1a;10 | 0 | 0.025 |
| NC_002645.1;NP_073550.1;HCoV-229E_1a;11 | 0 | 0 |
| NC_002645.1;NP_073550.1;HCoV-229E_1a;12 | 0 | 0 |
| NC_002645.1;NP_073550.1;HCoV-229E_1a;13 | 0 | 0.025 |
| NC_002645.1;NP_073550.1;HCoV-229E_1a;14 | 0 | 0.025 |
| NC_002645.1;NP_073550.1;HCoV-229E_1a;15 | 0 | 0 |
| NC_002645.1;NP_073550.1;HCoV-229E_1a;16 | 0 | 0 |
| NC_002645.1;NP_073550.1;HCoV-229E_1a;17 | 0 | 0 |
| NC_002645.1;NP_073550.1;HCoV-229E_1a;18 | 0 | 0 |
| NC_002645.1;NP_073550.1;HCoV-229E_1a;19 | 0 | 0 |
| NC_002645.1;NP_073550.1;HCoV-229E_1a;20 | 0.05 | 0.025 |
| NC_002645.1;NP_073550.1;HCoV-229E_1a;21 | 0.05 | 0 |
| NC_002645.1;NP_073550.1;HCoV-229E_1a;22 | 0 | 0 |
| NC_002645.1;NP_073550.1;HCoV-229E_1a;23 | 0 | 0 |
| NC_002645.1;NP_073550.1;HCoV-229E_1a;24 | 0 | 0 |
| NC_002645.1;NP_073550.1;HCoV-229E_1a;25 | 0 | 0 |
| NC_002645.1;NP_073550.1;HCoV-229E_1a;26 | 0 | 0 |
| NC_002645.1;NP_073550.1;HCoV-229E_1a;27 | 0 | 0 |
| NC_002645.1;NP_073550.1;HCoV-229E_1a;28 | 0 | 0 |
| NC_002645.1;NP_073550.1;HCoV-229E_1a;29 | 0 | 0 |
| NC_002645.1;NP_073550.1;HCoV-229E_1a;30 | 0 | 0 |
| NC_002645.1;NP_073550.1;HCoV-229E_1a;31 | 0 | 0 |
| NC_002645.1;NP_073550.1;HCoV-229E_1a;32 | 0 | 0 |
| NC_002645.1;NP_073550.1;HCoV-229E_1a;33 | 0 | 0 |

| | | |
|---|---|---|
| NC_002645.1;NP_073550.1;HCoV-229E_1a;34 | 0 | 0 |
| NC_002645.1;NP_073550.1;HCoV-229E_1a;35 | 0 | 0 |
| NC_002645.1;NP_073550.1;HCoV-229E_1a;36 | 0 | 0 |
| NC_002645.1;NP_073550.1;HCoV-229E_1a;37 | 0 | 0 |
| NC_002645.1;NP_073550.1;HCoV-229E_1a;38 | 0 | 0 |
| NC_002645.1;NP_073550.1;HCoV-229E_1a;39 | 0 | 0 |
| NC_002645.1;NP_073550.1;HCoV-229E_1a;40 | 0 | 0 |
| NC_002645.1;NP_073550.1;HCoV-229E_1a;41 | 0 | 0 |
| NC_002645.1;NP_073550.1;HCoV-229E_1a;42 | 0 | 0 |
| NC_002645.1;NP_073550.1;HCoV-229E_1a;43 | 0 | 0 |
| NC_002645.1;NP_073550.1;HCoV-229E_1a;44 | 0 | 0 |
| NC_002645.1;NP_073550.1;HCoV-229E_1a;45 | 0 | 0 |
| NC_002645.1;NP_073550.1;HCoV-229E_1a;46 | 0.05 | 0.025 |
| NC_002645.1;NP_073550.1;HCoV-229E_1a;47 | 0 | 0 |
| NC_002645.1;NP_073550.1;HCoV-229E_1a;48 | 0 | 0 |
| NC_002645.1;NP_073550.1;HCoV-229E_1a;49 | 0 | 0.025 |
| NC_002645.1;NP_073550.1;HCoV-229E_1a;50 | 0 | 0 |
| NC_002645.1;NP_073550.1;HCoV-229E_1a;51 | 0 | 0 |
| NC_002645.1;NP_073550.1;HCoV-229E_1a;52 | 0 | 0 |
| NC_002645.1;NP_073550.1;HCoV-229E_1a;53 | 0 | 0 |
| NC_002645.1;NP_073550.1;HCoV-229E_1a;54 | 0 | 0 |
| NC_002645.1;NP_073550.1;HCoV-229E_1a;55 | 0.2 | 0.025 |
| NC_002645.1;NP_073550.1;HCoV-229E_1a;56 | 0 | 0 |
| NC_002645.1;NP_073550.1;HCoV-229E_1a;57 | 0 | 0 |
| NC_002645.1;NP_073550.1;HCoV-229E_1a;58 | 0.05 | 0 |
| NC_002645.1;NP_073550.1;HCoV-229E_1a;59 | 0 | 0 |
| NC_002645.1;NP_073550.1;HCoV-229E_1a;60 | 0 | 0 |
| NC_002645.1;NP_073550.1;HCoV-229E_1a;61 | 0 | 0 |
| NC_002645.1;NP_073550.1;HCoV-229E_1a;62 | 0 | 0.025 |
| NC_002645.1;NP_073550.1;HCoV-229E_1a;63 | 0 | 0.025 |
| NC_002645.1;NP_073550.1;HCoV-229E_1a;64 | 0 | 0 |
| NC_002645.1;NP_073550.1;HCoV-229E_1a;65 | 0 | 0 |
| NC_002645.1;NP_073550.1;HCoV-229E_1a;66 | 0 | 0 |
| NC_002645.1;NP_073550.1;HCoV-229E_1a;67 | 0 | 0 |
| NC_002645.1;NP_073550.1;HCoV-229E_1a;68 | 0 | 0 |
| NC_002645.1;NP_073550.1;HCoV-229E_1a;69 | 0 | 0 |
| NC_002645.1;NP_073550.1;HCoV-229E_1a;70 | 0 | 0 |
| NC_002645.1;NP_073550.1;HCoV-229E_1a;71 | 0 | 0 |
| NC_002645.1;NP_073550.1;HCoV-229E_1a;72 | 0 | 0 |
| NC_002645.1;NP_073550.1;HCoV-229E_1a;73 | 0 | 0 |
| NC_002645.1;NP_073550.1;HCoV-229E_1a;74 | 0 | 0 |
| NC_002645.1;NP_073550.1;HCoV-229E_1a;75 | 0 | 0 |
| NC_002645.1;NP_073550.1;HCoV-229E_1a;76 | 0 | 0 |

| | | |
|---|---|---|
| NC_002645.1;NP_073550.1;HCoV-229E_1a;77 | 0 | 0 |
| NC_002645.1;NP_073550.1;HCoV-229E_1a;78 | 0 | 0 |
| NC_002645.1;NP_073550.1;HCoV-229E_1a;79 | 0 | 0 |
| NC_002645.1;NP_073550.1;HCoV-229E_1a;80 | 0 | 0 |
| NC_002645.1;NP_073550.1;HCoV-229E_1a;81 | 0 | 0 |
| NC_002645.1;NP_073550.1;HCoV-229E_1a;82 | 0 | 0 |
| NC_002645.1;NP_073550.1;HCoV-229E_1a;83 | 0 | 0 |
| NC_002645.1;NP_073550.1;HCoV-229E_1a;84 | 0 | 0 |
| NC_002645.1;NP_073550.1;HCoV-229E_1a;85 | 0 | 0 |
| NC_002645.1;NP_073550.1;HCoV-229E_1a;86 | 0 | 0 |
| NC_002645.1;NP_073550.1;HCoV-229E_1a;87 | 0 | 0 |
| NC_002645.1;NP_073550.1;HCoV-229E_1a;88 | 0 | 0 |
| NC_002645.1;NP_073550.1;HCoV-229E_1a;89 | 0.2 | 0.05 |
| NC_002645.1;NP_073550.1;HCoV-229E_1a;90 | 0.05 | 0 |
| NC_002645.1;NP_073550.1;HCoV-229E_1a;91 | 0 | 0 |
| NC_002645.1;NP_073550.1;HCoV-229E_1a;92 | 0 | 0 |
| NC_002645.1;NP_073550.1;HCoV-229E_1a;93 | 0 | 0 |
| NC_002645.1;NP_073550.1;HCoV-229E_1a;94 | 0 | 0.025 |
| NC_002645.1;NP_073550.1;HCoV-229E_1a;95 | 0.05 | 0 |
| NC_002645.1;NP_073550.1;HCoV-229E_1a;96 | 0.1 | 0.05 |
| NC_002645.1;NP_073550.1;HCoV-229E_1a;97 | 0.05 | 0 |
| NC_002645.1;NP_073550.1;HCoV-229E_1a;98 | 0 | 0.025 |
| NC_002645.1;NP_073550.1;HCoV-229E_1a;99 | 0.05 | 0 |
| NC_002645.1;NP_073550.1;HCoV-229E_1a;100 | 0 | 0 |
| NC_002645.1;NP_073550.1;HCoV-229E_1a;101 | 0 | 0 |
| NC_002645.1;NP_073550.1;HCoV-229E_1a;102 | 0 | 0 |
| NC_002645.1;NP_073550.1;HCoV-229E_1a;103 | 0.05 | 0 |
| NC_002645.1;NP_073550.1;HCoV-229E_1a;104 | 0 | 0 |
| NC_002645.1;NP_073550.1;HCoV-229E_1a;105 | 0 | 0 |
| NC_002645.1;NP_073550.1;HCoV-229E_1a;106 | 0 | 0 |
| NC_002645.1;NP_073550.1;HCoV-229E_1a;107 | 0.05 | 0.05 |
| NC_002645.1;NP_073550.1;HCoV-229E_1a;108 | 0 | 0 |
| NC_002645.1;NP_073550.1;HCoV-229E_1a;109 | 0 | 0 |
| NC_002645.1;NP_073550.1;HCoV-229E_1a;110 | 0 | 0 |
| NC_002645.1;NP_073550.1;HCoV-229E_1a;111 | 0 | 0 |
| NC_002645.1;NP_073550.1;HCoV-229E_1a;112 | 0 | 0 |
| NC_002645.1;NP_073550.1;HCoV-229E_1a;113 | 0.05 | 0.025 |
| NC_002645.1;NP_073550.1;HCoV-229E_1a;114 | 0.05 | 0.025 |
| NC_002645.1;NP_073550.1;HCoV-229E_1a;115 | 0.05 | 0.025 |
| NC_002645.1;NP_073550.1;HCoV-229E_1a;116 | 0.05 | 0.025 |
| NC_002645.1;NP_073550.1;HCoV-229E_1a;117 | 0.05 | 0.025 |
| NC_002645.1;NP_073550.1;HCoV-229E_1a;118 | 0.05 | 0.05 |
| NC_002645.1;NP_073550.1;HCoV-229E_1a;119 | 0.05 | 0.075 |

| | | |
|---|---|---|
| NC_002645.1;NP_073550.1;HCoV-229E_1a;120 | 0.05 | 0.025 |
| NC_002645.1;NP_073550.1;HCoV-229E_1a;121 | 0.05 | 0.025 |
| NC_002645.1;NP_073550.1;HCoV-229E_1a;122 | 0 | 0 |
| NC_002645.1;NP_073550.1;HCoV-229E_1a;123 | 0 | 0 |
| NC_002645.1;NP_073550.1;HCoV-229E_1a;124 | 0 | 0 |
| NC_002645.1;NP_073550.1;HCoV-229E_1a;125 | 0 | 0 |
| NC_002645.1;NP_073550.1;HCoV-229E_1a;126 | 0 | 0 |
| NC_002645.1;NP_073550.1;HCoV-229E_1a;127 | 0 | 0 |
| NC_002645.1;NP_073550.1;HCoV-229E_1a;128 | 0 | 0 |
| NC_002645.1;NP_073550.1;HCoV-229E_1a;129 | 0 | 0.025 |
| NC_002645.1;NP_073550.1;HCoV-229E_1a;130 | 0 | 0.025 |
| NC_002645.1;NP_073550.1;HCoV-229E_1a;131 | 0 | 0.025 |
| NC_002645.1;NP_073550.1;HCoV-229E_1a;132 | 0 | 0.025 |
| NC_002645.1;NP_073550.1;HCoV-229E_1a;133 | 0 | 0.025 |
| NC_002645.1;NP_073550.1;HCoV-229E_1a;134 | 0 | 0.025 |
| NC_002645.1;NP_073550.1;HCoV-229E_1a;135 | 0 | 0.025 |
| NC_002645.1;NP_073550.1;HCoV-229E_1a;136 | 0 | 0.025 |
| NC_002645.1;NP_073550.1;HCoV-229E_1a;137 | 0 | 0.025 |
| NC_002645.1;NP_073550.1;HCoV-229E_1a;138 | 0 | 0 |
| NC_002645.1;NP_073550.1;HCoV-229E_1a;139 | 0 | 0 |
| NC_002645.1;NP_073550.1;HCoV-229E_1a;140 | 0 | 0 |
| NC_002645.1;NP_073550.1;HCoV-229E_1a;141 | 0 | 0 |
| NC_002645.1;NP_073550.1;HCoV-229E_1a;142 | 0 | 0.025 |
| NC_002645.1;NP_073550.1;HCoV-229E_1a;143 | 0 | 0 |
| NC_002645.1;NP_073550.1;HCoV-229E_1a;144 | 0 | 0 |
| NC_002645.1;NP_073550.1;HCoV-229E_1a;145 | 0 | 0 |
| NC_002645.1;NP_073550.1;HCoV-229E_1a;146 | 0 | 0 |
| NC_002645.1;NP_073550.1;HCoV-229E_1a;147 | 0 | 0 |
| NC_002645.1;NP_073550.1;HCoV-229E_1a;148 | 0 | 0 |
| NC_002645.1;NP_073550.1;HCoV-229E_1a;149 | 0 | 0 |
| NC_002645.1;NP_073550.1;HCoV-229E_1a;150 | 0 | 0 |
| NC_002645.1;NP_073550.1;HCoV-229E_1a;151 | 0 | 0 |
| NC_002645.1;NP_073550.1;HCoV-229E_1a;152 | 0 | 0 |
| NC_002645.1;NP_073550.1;HCoV-229E_1a;153 | 0 | 0 |
| NC_002645.1;NP_073550.1;HCoV-229E_1a;154 | 0 | 0 |
| NC_002645.1;NP_073550.1;HCoV-229E_1a;155 | 0 | 0 |
| NC_002645.1;NP_073550.1;HCoV-229E_1a;156 | 0.05 | 0 |
| NC_002645.1;NP_073550.1;HCoV-229E_1a;157 | 0 | 0 |
| NC_002645.1;NP_073550.1;HCoV-229E_1a;158 | 0 | 0 |
| NC_002645.1;NP_073550.1;HCoV-229E_1a;159 | 0 | 0 |
| NC_002645.1;NP_073550.1;HCoV-229E_1a;160 | 0 | 0.05 |
| NC_002645.1;NP_073550.1;HCoV-229E_1a;161 | 0 | 0.025 |
| NC_002645.1;NP_073550.1;HCoV-229E_1a;162 | 0 | 0 |

| | | |
|---|---|---|
| NC_002645.1;NP_073550.1;HCoV-229E_1a;163 | 0.05 | 0.025 |
| NC_002645.1;NP_073550.1;HCoV-229E_1a;164 | 0.05 | 0.025 |
| NC_002645.1;NP_073550.1;HCoV-229E_1a;165 | 0 | 0.025 |
| NC_002645.1;NP_073550.1;HCoV-229E_1a;166 | 0.05 | 0.025 |
| NC_002645.1;NP_073550.1;HCoV-229E_1a;167 | 0.05 | 0.05 |
| NC_002645.1;NP_073550.1;HCoV-229E_1a;168 | 0.05 | 0.025 |
| NC_002645.1;NP_073550.1;HCoV-229E_1a;169 | 0 | 0.025 |
| NC_002645.1;NP_073550.1;HCoV-229E_1a;170 | 0.05 | 0.05 |
| NC_002645.1;NP_073550.1;HCoV-229E_1a;171 | 0.05 | 0.05 |
| NC_002645.1;NP_073550.1;HCoV-229E_1a;172 | 0 | 0.025 |
| NC_002645.1;NP_073550.1;HCoV-229E_1a;173 | 0 | 0.025 |
| NC_002645.1;NP_073550.1;HCoV-229E_1a;174 | 0 | 0 |
| NC_002645.1;NP_073550.1;HCoV-229E_1a;175 | 0 | 0.025 |
| NC_002645.1;NP_073550.1;HCoV-229E_1a;176 | 0 | 0.025 |
| NC_002645.1;NP_073550.1;HCoV-229E_1a;177 | 0 | 0 |
| NC_002645.1;NP_073550.1;HCoV-229E_1a;178 | 0 | 0 |
| NC_002645.1;NP_073550.1;HCoV-229E_1a;179 | 0 | 0 |
| NC_002645.1;NP_073550.1;HCoV-229E_1a;180 | 0 | 0 |
| NC_002645.1;NP_073550.1;HCoV-229E_1a;181 | 0 | 0 |
| NC_002645.1;NP_073550.1;HCoV-229E_1a;182 | 0 | 0 |
| NC_002645.1;NP_073550.1;HCoV-229E_1a;183 | 0 | 0 |
| NC_002645.1;NP_073550.1;HCoV-229E_1a;184 | 0 | 0 |
| NC_002645.1;NP_073550.1;HCoV-229E_1a;185 | 0 | 0 |
| NC_002645.1;NP_073550.1;HCoV-229E_1a;186 | 0 | 0.025 |
| NC_002645.1;NP_073550.1;HCoV-229E_1a;187 | 0 | 0.025 |
| NC_002645.1;NP_073550.1;HCoV-229E_1a;188 | 0 | 0.025 |
| NC_002645.1;NP_073550.1;HCoV-229E_1a;189 | 0.15 | 0.05 |
| NC_002645.1;NP_073550.1;HCoV-229E_1a;190 | 0 | 0.025 |
| NC_002645.1;NP_073550.1;HCoV-229E_1a;191 | 0 | 0.025 |
| NC_002645.1;NP_073550.1;HCoV-229E_1a;192 | 0 | 0.025 |
| NC_002645.1;NP_073550.1;HCoV-229E_1a;193 | 0 | 0.025 |
| NC_002645.1;NP_073550.1;HCoV-229E_1a;194 | 0 | 0.025 |
| NC_002645.1;NP_073550.1;HCoV-229E_1a;195 | 0.05 | 0.025 |
| NC_002645.1;NP_073550.1;HCoV-229E_1a;196 | 0 | 0.025 |
| NC_002645.1;NP_073550.1;HCoV-229E_1a;197 | 0 | 0 |
| NC_002645.1;NP_073550.1;HCoV-229E_1a;198 | 0 | 0 |
| NC_002645.1;NP_073550.1;HCoV-229E_1a;199 | 0 | 0 |
| NC_002645.1;NP_073550.1;HCoV-229E_1a;200 | 0 | 0 |
| NC_002645.1;NP_073550.1;HCoV-229E_1a;201 | 0 | 0 |
| NC_002645.1;NP_073550.1;HCoV-229E_1a;202 | 0 | 0.025 |
| NC_002645.1;NP_073550.1;HCoV-229E_1a;203 | 0.05 | 0.025 |
| NC_002645.1;NP_073550.1;HCoV-229E_1a;204 | 0 | 0.025 |
| NC_002645.1;NP_073550.1;HCoV-229E_1a;205 | 0 | 0 |

| | | |
|---|---|---|
| NC_002645.1;NP_073550.1;HCoV-229E_1a;206 | 0 | 0.025 |
| NC_002645.1;NP_073550.1;HCoV-229E_1a;207 | 0.05 | 0 |
| NC_002645.1;NP_073550.1;HCoV-229E_1a;208 | 0 | 0 |
| NC_002645.1;NP_073550.1;HCoV-229E_1a;209 | 0 | 0 |
| NC_002645.1;NP_073550.1;HCoV-229E_1a;210 | 0.05 | 0 |
| NC_002645.1;NP_073550.1;HCoV-229E_1a;211 | 0 | 0 |
| NC_002645.1;NP_073550.1;HCoV-229E_1a;212 | 0 | 0 |
| NC_002645.1;NP_073550.1;HCoV-229E_1a;213 | 0 | 0 |
| NC_002645.1;NP_073550.1;HCoV-229E_1a;214 | 0 | 0 |
| NC_002645.1;NP_073550.1;HCoV-229E_1a;215 | 0 | 0 |
| NC_002645.1;NP_073550.1;HCoV-229E_1a;216 | 0 | 0 |
| NC_002645.1;NP_073550.1;HCoV-229E_1a;217 | 0 | 0 |
| NC_002645.1;NP_073550.1;HCoV-229E_1a;218 | 0 | 0 |
| NC_002645.1;NP_073550.1;HCoV-229E_1a;219 | 0 | 0 |
| NC_002645.1;NP_073550.1;HCoV-229E_1a;220 | 0 | 0 |
| NC_002645.1;NP_073550.1;HCoV-229E_1a;221 | 0 | 0.025 |
| NC_002645.1;NP_073550.1;HCoV-229E_1a;222 | 0.05 | 0.025 |
| NC_002645.1;NP_073550.1;HCoV-229E_1a;223 | 0 | 0 |
| NC_002645.1;NP_073550.1;HCoV-229E_1a;224 | 0 | 0 |
| NC_002645.1;NP_073550.1;HCoV-229E_1a;225 | 0 | 0 |
| NC_002645.1;NP_073550.1;HCoV-229E_1a;226 | 0 | 0 |
| NC_002645.1;NP_073550.1;HCoV-229E_1a;227 | 0 | 0 |
| NC_002645.1;NP_073550.1;HCoV-229E_1a;228 | 0 | 0 |
| NC_002645.1;NP_073550.1;HCoV-229E_1a;229 | 0 | 0 |
| NC_002645.1;NP_073550.1;HCoV-229E_1a;230 | 0 | 0 |
| NC_002645.1;NP_073550.1;HCoV-229E_1a;231 | 0 | 0 |
| NC_002645.1;NP_073550.1;HCoV-229E_1a;232 | 0 | 0 |
| NC_002645.1;NP_073550.1;HCoV-229E_1a;233 | 0 | 0 |
| NC_002645.1;NP_073550.1;HCoV-229E_1a;234 | 0 | 0 |
| NC_002645.1;NP_073550.1;HCoV-229E_1a;235 | 0 | 0 |
| NC_002645.1;NP_073550.1;HCoV-229E_1a;236 | 0 | 0 |
| NC_002645.1;NP_073550.1;HCoV-229E_1a;237 | 0 | 0 |
| NC_002645.1;NP_073550.1;HCoV-229E_1a;238 | 0 | 0 |
| NC_002645.1;NP_073550.1;HCoV-229E_1a;239 | 0 | 0 |
| NC_002645.1;NP_073550.1;HCoV-229E_1a;240 | 0 | 0 |
| NC_002645.1;NP_073550.1;HCoV-229E_1a;241 | 0 | 0 |
| NC_002645.1;NP_073550.1;HCoV-229E_1a;242 | 0 | 0 |
| NC_002645.1;NP_073550.1;HCoV-229E_1a;243 | 0.05 | 0 |
| NC_002645.1;NP_073550.1;HCoV-229E_1a;244 | 0.05 | 0 |
| NC_002645.1;NP_073550.1;HCoV-229E_1a;245 | 0.05 | 0 |
| NC_002645.1;NP_073550.1;HCoV-229E_1a;246 | 0 | 0 |
| NC_002645.1;NP_073550.1;HCoV-229E_1a;247 | 0.05 | 0 |
| NC_002645.1;NP_073550.1;HCoV-229E_1a;248 | 0.05 | 0 |

| | | |
|---|---|---|
| NC_002645.1;NP_073550.1;HCoV-229E_1a;249 | 0.05 | 0 |
| NC_002645.1;NP_073550.1;HCoV-229E_1a;250 | 0.05 | 0 |
| NC_002645.1;NP_073550.1;HCoV-229E_1a;251 | 0.05 | 0 |
| NC_002645.1;NP_073550.1;HCoV-229E_1a;252 | 0 | 0 |
| NC_002645.1;NP_073550.1;HCoV-229E_1a;253 | 0 | 0 |
| NC_002645.1;NP_073550.1;HCoV-229E_1a;254 | 0 | 0 |
| NC_002645.1;NP_073550.1;HCoV-229E_1a;255 | 0 | 0 |
| NC_002645.1;NP_073550.1;HCoV-229E_1a;256 | 0 | 0 |
| NC_002645.1;NP_073550.1;HCoV-229E_1a;257 | 0 | 0 |
| NC_002645.1;NP_073550.1;HCoV-229E_1a;258 | 0 | 0 |
| NC_002645.1;NP_073550.1;HCoV-229E_1a;259 | 0 | 0 |
| NC_002645.1;NP_073550.1;HCoV-229E_1a;260 | 0 | 0 |
| NC_002645.1;NP_073550.1;HCoV-229E_1a;261 | 0 | 0 |
| NC_002645.1;NP_073550.1;HCoV-229E_1a;262 | 0 | 0 |
| NC_002645.1;NP_073550.1;HCoV-229E_1a;263 | 0 | 0 |
| NC_002645.1;NP_073550.1;HCoV-229E_1a;264 | 0 | 0 |
| NC_002645.1;NP_073550.1;HCoV-229E_1a;265 | 0 | 0 |
| NC_002645.1;NP_073550.1;HCoV-229E_1a;266 | 0 | 0 |
| NC_002645.1;NP_073550.1;HCoV-229E_1a;267 | 0 | 0 |
| NC_002645.1;NP_073550.1;HCoV-229E_1a;268 | 0 | 0 |
| NC_002645.1;NP_073550.1;HCoV-229E_1a;269 | 0.15 | 0.05 |
| NC_002645.1;NP_073550.1;HCoV-229E_1a;270 | 0.15 | 0.05 |
| NC_002645.1;NP_073550.1;HCoV-229E_1a;271 | 0.1 | 0.05 |
| NC_002645.1;NP_073550.1;HCoV-229E_1a;272 | 0.1 | 0.05 |
| NC_002645.1;NP_073550.1;HCoV-229E_1a;273 | 0.05 | 0.05 |
| NC_002645.1;NP_073550.1;HCoV-229E_1a;274 | 0.1 | 0.05 |
| NC_002645.1;NP_073550.1;HCoV-229E_1a;275 | 0.05 | 0.05 |
| NC_002645.1;NP_073550.1;HCoV-229E_1a;276 | 0.05 | 0.05 |
| NC_002645.1;NP_073550.1;HCoV-229E_1a;277 | 0 | 0.05 |
| NC_002645.1;NP_073550.1;HCoV-229E_1a;278 | 0.05 | 0.025 |
| NC_002645.1;NP_073550.1;HCoV-229E_1a;279 | 0 | 0.025 |
| NC_002645.1;NP_073550.1;HCoV-229E_1a;280 | 0 | 0.05 |
| NC_002645.1;NP_073550.1;HCoV-229E_1a;281 | 0 | 0.025 |
| NC_002645.1;NP_073550.1;HCoV-229E_1a;282 | 0 | 0 |
| NC_002645.1;NP_073550.1;HCoV-229E_1a;283 | 0 | 0 |
| NC_002645.1;NP_073550.1;HCoV-229E_1a;284 | 0 | 0 |
| NC_002645.1;NP_073550.1;HCoV-229E_1a;285 | 0 | 0.025 |
| NC_002645.1;NP_073550.1;HCoV-229E_1a;286 | 0 | 0 |
| NC_002645.1;NP_073550.1;HCoV-229E_1a;287 | 0 | 0 |
| NC_002645.1;NP_073550.1;HCoV-229E_1a;288 | 0 | 0 |
| NC_002645.1;NP_073550.1;HCoV-229E_1a;289 | 0 | 0 |
| NC_002645.1;NP_073550.1;HCoV-229E_1a;290 | 0 | 0 |
| NC_002645.1;NP_073550.1;HCoV-229E_1a;291 | 0 | 0 |

| | | |
|---|---|---|
| NC_002645.1;NP_073550.1;HCoV-229E_1a;292 | 0 | 0 |
| NC_002645.1;NP_073550.1;HCoV-229E_1a;293 | 0 | 0 |
| NC_002645.1;NP_073550.1;HCoV-229E_1a;294 | 0 | 0 |
| NC_002645.1;NP_073550.1;HCoV-229E_1a;295 | 0 | 0 |
| NC_002645.1;NP_073550.1;HCoV-229E_1a;296 | 0 | 0 |
| NC_002645.1;NP_073550.1;HCoV-229E_1a;297 | 0 | 0 |
| NC_002645.1;NP_073550.1;HCoV-229E_1a;298 | 0 | 0 |
| NC_002645.1;NP_073550.1;HCoV-229E_1a;299 | 0 | 0 |
| NC_002645.1;NP_073550.1;HCoV-229E_1a;300 | 0 | 0 |
| NC_002645.1;NP_073550.1;HCoV-229E_1a;301 | 0 | 0 |
| NC_002645.1;NP_073550.1;HCoV-229E_1a;302 | 0 | 0 |
| NC_002645.1;NP_073550.1;HCoV-229E_1a;303 | 0 | 0 |
| NC_002645.1;NP_073550.1;HCoV-229E_1a;304 | 0 | 0 |
| NC_002645.1;NP_073550.1;HCoV-229E_1a;305 | 0 | 0 |
| NC_002645.1;NP_073550.1;HCoV-229E_1a;306 | 0 | 0 |
| NC_002645.1;NP_073550.1;HCoV-229E_1a;307 | 0 | 0 |
| NC_002645.1;NP_073550.1;HCoV-229E_1a;308 | 0 | 0 |
| NC_002645.1;NP_073550.1;HCoV-229E_1a;309 | 0 | 0 |
| NC_002645.1;NP_073550.1;HCoV-229E_1a;310 | 0 | 0 |
| NC_002645.1;NP_073550.1;HCoV-229E_1a;311 | 0 | 0 |
| NC_002645.1;NP_073550.1;HCoV-229E_1a;312 | 0 | 0 |
| NC_002645.1;NP_073550.1;HCoV-229E_1a;313 | 0 | 0 |
| NC_002645.1;NP_073550.1;HCoV-229E_1a;314 | 0 | 0 |
| NC_002645.1;NP_073550.1;HCoV-229E_1a;315 | 0 | 0 |
| NC_002645.1;NP_073550.1;HCoV-229E_1a;316 | 0 | 0 |
| NC_002645.1;NP_073550.1;HCoV-229E_1a;317 | 0 | 0 |
| NC_002645.1;NP_073550.1;HCoV-229E_1a;318 | 0 | 0 |
| NC_002645.1;NP_073550.1;HCoV-229E_1a;319 | 0 | 0 |
| NC_002645.1;NP_073550.1;HCoV-229E_1a;320 | 0 | 0 |
| NC_002645.1;NP_073550.1;HCoV-229E_1a;321 | 0 | 0 |
| NC_002645.1;NP_073550.1;HCoV-229E_1a;322 | 0 | 0 |
| NC_002645.1;NP_073550.1;HCoV-229E_1a;323 | 0 | 0 |
| NC_002645.1;NP_073550.1;HCoV-229E_1a;324 | 0 | 0 |
| NC_002645.1;NP_073550.1;HCoV-229E_1a;325 | 0 | 0 |
| NC_002645.1;NP_073550.1;HCoV-229E_1a;326 | 0 | 0 |
| NC_002645.1;NP_073550.1;HCoV-229E_1a;327 | 0 | 0 |
| NC_002645.1;NP_073550.1;HCoV-229E_1a;328 | 0 | 0 |
| NC_002645.1;NP_073550.1;HCoV-229E_1a;329 | 0 | 0 |
| NC_002645.1;NP_073550.1;HCoV-229E_1a;330 | 0 | 0.025 |
| NC_002645.1;NP_073550.1;HCoV-229E_1a;331 | 0 | 0 |
| NC_002645.1;NP_073550.1;HCoV-229E_1a;332 | 0 | 0 |
| NC_002645.1;NP_073550.1;HCoV-229E_1a;333 | 0 | 0 |
| NC_002645.1;NP_073550.1;HCoV-229E_1a;334 | 0 | 0.025 |

| | | |
|---|---|---|
| NC_002645.1;NP_073550.1;HCoV-229E_1a;335 | 0 | 0 |
| NC_002645.1;NP_073550.1;HCoV-229E_1a;336 | 0 | 0 |
| NC_002645.1;NP_073550.1;HCoV-229E_1a;337 | 0 | 0 |
| NC_002645.1;NP_073550.1;HCoV-229E_1a;338 | 0 | 0 |
| NC_002645.1;NP_073550.1;HCoV-229E_1a;339 | 0 | 0 |
| NC_002645.1;NP_073550.1;HCoV-229E_1a;340 | 0 | 0 |
| NC_002645.1;NP_073550.1;HCoV-229E_1a;341 | 0 | 0 |
| NC_002645.1;NP_073550.1;HCoV-229E_1a;342 | 0 | 0 |
| NC_002645.1;NP_073550.1;HCoV-229E_1a;343 | 0 | 0 |
| NC_002645.1;NP_073550.1;HCoV-229E_1a;344 | 0 | 0 |
| NC_002645.1;NP_073550.1;HCoV-229E_1a;345 | 0 | 0 |
| NC_002645.1;NP_073550.1;HCoV-229E_1a;346 | 0 | 0 |
| NC_002645.1;NP_073550.1;HCoV-229E_1a;347 | 0 | 0 |
| NC_002645.1;NP_073550.1;HCoV-229E_1a;348 | 0 | 0 |
| NC_002645.1;NP_073550.1;HCoV-229E_1a;349 | 0 | 0 |
| NC_002645.1;NP_073550.1;HCoV-229E_1a;350 | 0 | 0 |
| NC_002645.1;NP_073550.1;HCoV-229E_1a;351 | 0 | 0 |
| NC_002645.1;NP_073550.1;HCoV-229E_1a;352 | 0 | 0 |
| NC_002645.1;NP_073550.1;HCoV-229E_1a;353 | 0 | 0.025 |
| NC_002645.1;NP_073550.1;HCoV-229E_1a;354 | 0 | 0.025 |
| NC_002645.1;NP_073550.1;HCoV-229E_1a;355 | 0 | 0.05 |
| NC_002645.1;NP_073550.1;HCoV-229E_1a;356 | 0 | 0.05 |
| NC_002645.1;NP_073550.1;HCoV-229E_1a;357 | 0 | 0.05 |
| NC_002645.1;NP_073550.1;HCoV-229E_1a;358 | 0 | 0.025 |
| NC_002645.1;NP_073550.1;HCoV-229E_1a;359 | 0 | 0.05 |
| NC_002645.1;NP_073550.1;HCoV-229E_1a;360 | 0 | 0.1 |
| NC_002645.1;NP_073550.1;HCoV-229E_1a;361 | 0.05 | 0.05 |
| NC_002645.1;NP_073550.1;HCoV-229E_1a;362 | 0 | 0.075 |
| NC_002645.1;NP_073550.1;HCoV-229E_1a;363 | 0 | 0.05 |
| NC_002645.1;NP_073550.1;HCoV-229E_1a;364 | 0.05 | 0.075 |
| NC_002645.1;NP_073550.1;HCoV-229E_1a;365 | 0.05 | 0.05 |
| NC_002645.1;NP_073550.1;HCoV-229E_1a;366 | 0 | 0 |
| NC_002645.1;NP_073550.1;HCoV-229E_1a;367 | 0 | 0 |
| NC_002645.1;NP_073550.1;HCoV-229E_1a;368 | 0 | 0 |
| NC_002645.1;NP_073550.1;HCoV-229E_1a;369 | 0 | 0 |
| NC_002645.1;NP_073550.1;HCoV-229E_1a;370 | 0 | 0 |
| NC_002645.1;NP_073550.1;HCoV-229E_1a;371 | 0 | 0 |
| NC_002645.1;NP_073550.1;HCoV-229E_1a;372 | 0 | 0 |
| NC_002645.1;NP_073550.1;HCoV-229E_1a;373 | 0 | 0 |
| NC_002645.1;NP_073550.1;HCoV-229E_1a;374 | 0 | 0 |
| NC_002645.1;NP_073550.1;HCoV-229E_1a;375 | 0 | 0 |
| NC_002645.1;NP_073550.1;HCoV-229E_1a;376 | 0 | 0 |
| NC_002645.1;NP_073550.1;HCoV-229E_1a;377 | 0 | 0 |

| | | |
|---|---|---|
| NC_002645.1;NP_073550.1;HCoV-229E_1a;378 | 0 | 0 |
| NC_002645.1;NP_073550.1;HCoV-229E_1a;379 | 0 | 0 |
| NC_002645.1;NP_073550.1;HCoV-229E_1a;380 | 0 | 0 |
| NC_002645.1;NP_073550.1;HCoV-229E_1a;381 | 0 | 0 |
| NC_002645.1;NP_073550.1;HCoV-229E_1a;382 | 0 | 0 |
| NC_002645.1;NP_073550.1;HCoV-229E_1a;383 | 0 | 0 |
| NC_002645.1;NP_073550.1;HCoV-229E_1a;384 | 0.05 | 0.025 |
| NC_002645.1;NP_073550.1;HCoV-229E_1a;385 | 0 | 0 |
| NC_002645.1;NP_073550.1;HCoV-229E_1a;386 | 0 | 0 |
| NC_002645.1;NP_073550.1;HCoV-229E_1a;387 | 0.05 | 0.025 |
| NC_002645.1;NP_073550.1;HCoV-229E_1a;388 | 0.05 | 0 |
| NC_002645.1;NP_073550.1;HCoV-229E_1a;389 | 0.05 | 0 |
| NC_002645.1;NP_073550.1;HCoV-229E_1a;390 | 0.05 | 0.025 |
| NC_002645.1;NP_073550.1;HCoV-229E_1a;391 | 0.05 | 0.025 |
| NC_002645.1;NP_073550.1;HCoV-229E_1a;392 | 0.05 | 0 |
| NC_002645.1;NP_073550.1;HCoV-229E_1a;393 | 0.1 | 0 |
| NC_002645.1;NP_073550.1;HCoV-229E_1a;394 | 0.05 | 0.075 |
| NC_002645.1;NP_073550.1;HCoV-229E_1a;395 | 0.1 | 0.05 |
| NC_002645.1;NP_073550.1;HCoV-229E_1a;396 | 0 | 0 |
| NC_002645.1;NP_073550.1;HCoV-229E_1a;397 | 0 | 0 |
| NC_002645.1;NP_073550.1;HCoV-229E_1a;398 | 0 | 0 |
| NC_002645.1;NP_073550.1;HCoV-229E_1a;399 | 0 | 0 |
| NC_002645.1;NP_073550.1;HCoV-229E_1a;400 | 0 | 0 |
| NC_002645.1;NP_073550.1;HCoV-229E_1a;401 | 0 | 0 |
| NC_002645.1;NP_073550.1;HCoV-229E_1a;402 | 0 | 0 |
| NC_002645.1;NP_073550.1;HCoV-229E_1a;403 | 0 | 0 |
| NC_002645.1;NP_073550.1;HCoV-229E_1a;404 | 0 | 0.025 |
| NC_002645.1;NP_073550.1;HCoV-229E_1a;405 | 0.05 | 0.05 |
| NC_002645.1;NP_073550.1;HCoV-229E_1a;406 | 0.05 | 0.05 |
| NC_002645.1;NP_073550.1;HCoV-229E_1a;407 | 0 | 0 |
| NC_002645.1;NP_073550.1;HCoV-229E_1a;408 | 0 | 0 |
| NC_002645.1;NP_073550.1;HCoV-229E_1a;409 | 0 | 0 |
| NC_002645.1;NP_073550.1;HCoV-229E_1a;410 | 0 | 0 |
| NC_002645.1;NP_073550.1;HCoV-229E_1a;411 | 0 | 0 |
| NC_002645.1;NP_073550.1;HCoV-229E_1a;412 | 0 | 0 |
| NC_002645.1;NP_073550.1;HCoV-229E_1a;413 | 0.1 | 0.025 |
| NC_002645.1;NP_073550.1;HCoV-229E_1a;414 | 0.1 | 0.075 |
| NC_002645.1;NP_073550.1;HCoV-229E_1a;415 | 0.1 | 0.075 |
| NC_002645.1;NP_073550.1;HCoV-229E_1a;416 | 0.1 | 0.025 |
| NC_002645.1;NP_073550.1;HCoV-229E_1a;417 | 0 | 0.05 |
| NC_002645.1;NP_073550.1;HCoV-229E_1a;418 | 0 | 0 |
| NC_002645.1;NP_073550.1;HCoV-229E_1a;419 | 0 | 0 |
| NC_002645.1;NP_073550.1;HCoV-229E_1a;420 | 0 | 0 |

| | | |
|---|---|---|
| NC_002645.1;NP_073550.1;HCoV-229E_1a;421 | 0 | 0 |
| NC_002645.1;NP_073550.1;HCoV-229E_1a;422 | 0 | 0 |
| NC_002645.1;NP_073550.1;HCoV-229E_1a;423 | 0 | 0 |
| NC_002645.1;NP_073550.1;HCoV-229E_1a;424 | 0 | 0 |
| NC_002645.1;NP_073550.1;HCoV-229E_1a;425 | 0 | 0 |
| NC_002645.1;NP_073550.1;HCoV-229E_1a;426 | 0 | 0 |
| NC_002645.1;NP_073550.1;HCoV-229E_1a;427 | 0 | 0 |
| NC_002645.1;NP_073550.1;HCoV-229E_1a;428 | 0 | 0 |
| NC_002645.1;NP_073550.1;HCoV-229E_1a;429 | 0 | 0 |
| NC_002645.1;NP_073550.1;HCoV-229E_1a;430 | 0 | 0 |
| NC_002645.1;NP_073550.1;HCoV-229E_1a;431 | 0 | 0 |
| NC_002645.1;NP_073550.1;HCoV-229E_1a;432 | 0 | 0 |
| NC_002645.1;NP_073550.1;HCoV-229E_1a;433 | 0 | 0 |
| NC_002645.1;NP_073550.1;HCoV-229E_1a;434 | 0 | 0 |
| NC_002645.1;NP_073550.1;HCoV-229E_1a;435 | 0 | 0 |
| NC_002645.1;NP_073550.1;HCoV-229E_1a;436 | 0 | 0 |
| NC_002645.1;NP_073550.1;HCoV-229E_1a;437 | 0 | 0 |
| NC_002645.1;NP_073550.1;HCoV-229E_1a;438 | 0 | 0 |
| NC_002645.1;NP_073550.1;HCoV-229E_1a;439 | 0 | 0.025 |
| NC_002645.1;NP_073550.1;HCoV-229E_1a;440 | 0 | 0 |
| NC_002645.1;NP_073550.1;HCoV-229E_1a;441 | 0 | 0 |
| NC_002645.1;NP_073550.1;HCoV-229E_1a;442 | 0 | 0 |
| NC_002645.1;NP_073550.1;HCoV-229E_1a;443 | 0.05 | 0 |
| NC_002645.1;NP_073550.1;HCoV-229E_1a;444 | 0 | 0 |
| NC_002645.1;NP_073550.1;HCoV-229E_1a;445 | 0 | 0 |
| NC_002645.1;NP_073550.1;HCoV-229E_1a;446 | 0 | 0 |
| NC_002645.1;NP_073550.1;HCoV-229E_1a;447 | 0 | 0 |
| NC_002645.1;NP_073550.1;HCoV-229E_1a;448 | 0.05 | 0.05 |
| NC_002645.1;NP_073550.1;HCoV-229E_1a;449 | 0.05 | 0.075 |
| NC_002645.1;NP_073550.1;HCoV-229E_1a;450 | 0.05 | 0.05 |
| NC_002645.1;NP_073550.1;HCoV-229E_1a;451 | 0.05 | 0.025 |
| NC_002645.1;NP_073550.1;HCoV-229E_1a;452 | 0.05 | 0.025 |
| NC_002645.1;NP_073550.1;HCoV-229E_1a;453 | 0 | 0.025 |
| NC_002645.1;NP_073550.1;HCoV-229E_1a;454 | 0 | 0.025 |
| NC_002645.1;NP_073550.1;HCoV-229E_1a;455 | 0 | 0 |
| NC_002645.1;NP_073550.1;HCoV-229E_1a;456 | 0 | 0 |
| NC_002645.1;NP_073550.1;HCoV-229E_1a;457 | 0 | 0 |
| NC_002645.1;NP_073550.1;HCoV-229E_1a;458 | 0 | 0 |
| NC_002645.1;NP_073550.1;HCoV-229E_1a;459 | 0 | 0 |
| NC_002645.1;NP_073550.1;HCoV-229E_1a;460 | 0 | 0 |
| NC_002645.1;NP_073550.1;HCoV-229E_1a;461 | 0 | 0 |
| NC_002645.1;NP_073550.1;HCoV-229E_1a;462 | 0 | 0 |
| NC_002645.1;NP_073550.1;HCoV-229E_1a;463 | 0 | 0 |

| | | |
|---|---|---|
| NC_002645.1;NP_073550.1;HCoV-229E_1a;464 | 0 | 0 |
| NC_002645.1;NP_073550.1;HCoV-229E_1a;465 | 0 | 0 |
| NC_002645.1;NP_073550.1;HCoV-229E_1a;466 | 0 | 0 |
| NC_002645.1;NP_073550.1;HCoV-229E_1a;467 | 0 | 0 |
| NC_002645.1;NP_073550.1;HCoV-229E_1a;468 | 0 | 0 |
| NC_002645.1;NP_073550.1;HCoV-229E_1a;469 | 0 | 0 |
| NC_002645.1;NP_073550.1;HCoV-229E_1a;470 | 0 | 0 |
| NC_002645.1;NP_073550.1;HCoV-229E_1a;471 | 0 | 0 |
| NC_002645.1;NP_073550.1;HCoV-229E_1a;472 | 0 | 0 |
| NC_002645.1;NP_073550.1;HCoV-229E_1a;473 | 0 | 0 |
| NC_002645.1;NP_073550.1;HCoV-229E_1a;474 | 0.05 | 0 |
| NC_002645.1;NP_073550.1;HCoV-229E_1a;475 | 0.05 | 0 |
| NC_002645.1;NP_073550.1;HCoV-229E_1a;476 | 0 | 0 |
| NC_002645.1;NP_073550.1;HCoV-229E_1a;477 | 0.2 | 0.025 |
| NC_002645.1;NP_073550.1;HCoV-229E_1a;478 | 0.15 | 0.025 |
| NC_002645.1;NP_073550.1;HCoV-229E_1a;479 | 0 | 0.025 |
| NC_002645.1;NP_073550.1;HCoV-229E_1a;480 | 0 | 0 |
| NC_002645.1;NP_073550.1;HCoV-229E_1a;481 | 0 | 0 |
| NC_002645.1;NP_073550.1;HCoV-229E_1a;482 | 0 | 0 |
| NC_002645.1;NP_073550.1;HCoV-229E_1a;483 | 0 | 0 |
| NC_002645.1;NP_073550.1;HCoV-229E_1a;484 | 0 | 0 |
| NC_002645.1;NP_073550.1;HCoV-229E_1a;485 | 0 | 0 |
| NC_002645.1;NP_073550.1;HCoV-229E_1a;486 | 0 | 0 |
| NC_002645.1;NP_073550.1;HCoV-229E_1a;487 | 0 | 0 |
| NC_002645.1;NP_073550.1;HCoV-229E_1a;488 | 0 | 0 |
| NC_002645.1;NP_073550.1;HCoV-229E_1a;489 | 0 | 0 |
| NC_002645.1;NP_073550.1;HCoV-229E_1a;490 | 0 | 0 |
| NC_002645.1;NP_073550.1;HCoV-229E_1a;491 | 0 | 0 |
| NC_002645.1;NP_073550.1;HCoV-229E_1a;492 | 0 | 0 |
| NC_002645.1;NP_073550.1;HCoV-229E_1a;493 | 0 | 0 |
| NC_002645.1;NP_073550.1;HCoV-229E_1a;494 | 0 | 0 |
| NC_002645.1;NP_073550.1;HCoV-229E_1a;495 | 0 | 0 |
| NC_002645.1;NP_073550.1;HCoV-229E_1a;496 | 0 | 0 |
| NC_002645.1;NP_073550.1;HCoV-229E_1a;497 | 0 | 0 |
| NC_002645.1;NP_073550.1;HCoV-229E_1a;498 | 0 | 0 |
| NC_002645.1;NP_073550.1;HCoV-229E_1a;499 | 0 | 0 |
| NC_002645.1;NP_073550.1;HCoV-229E_1a;500 | 0 | 0 |
| NC_002645.1;NP_073550.1;HCoV-229E_1a;501 | 0 | 0 |
| NC_002645.1;NP_073550.1;HCoV-229E_1a;502 | 0 | 0 |
| NC_002645.1;NP_073550.1;HCoV-229E_1a;503 | 0 | 0 |
| NC_002645.1;NP_073550.1;HCoV-229E_1a;504 | 0 | 0 |
| NC_002645.1;NP_073550.1;HCoV-229E_1a;505 | 0 | 0 |
| NC_002645.1;NP_073550.1;HCoV-229E_1a;506 | 0 | 0 |

| | | |
|---|---|---|
| NC_002645.1;NP_073550.1;HCoV-229E_1a;507 | 0 | 0 |
| NC_002645.1;NP_073550.1;HCoV-229E_1a;508 | 0 | 0 |
| NC_002645.1;NP_073550.1;HCoV-229E_1a;509 | 0 | 0 |
| NC_002645.1;NP_073550.1;HCoV-229E_1a;510 | 0 | 0 |
| NC_002645.1;NP_073550.1;HCoV-229E_1a;511 | 0 | 0 |
| NC_002645.1;NP_073550.1;HCoV-229E_1a;512 | 0 | 0 |
| NC_002645.1;NP_073550.1;HCoV-229E_1a;513 | 0 | 0 |
| NC_002645.1;NP_073550.1;HCoV-229E_1a;514 | 0 | 0 |
| NC_002645.1;NP_073550.1;HCoV-229E_1a;515 | 0 | 0 |
| NC_002645.1;NP_073550.1;HCoV-229E_1a;516 | 0 | 0 |
| NC_002645.1;NP_073550.1;HCoV-229E_1a;517 | 0 | 0 |
| NC_002645.1;NP_073550.1;HCoV-229E_1a;518 | 0 | 0 |
| NC_002645.1;NP_073550.1;HCoV-229E_1a;519 | 0 | 0 |
| NC_002645.1;NP_073550.1;HCoV-229E_1a;520 | 0 | 0.025 |
| NC_002645.1;NP_073550.1;HCoV-229E_1a;521 | 0 | 0 |
| NC_002645.1;NP_073550.1;HCoV-229E_1a;522 | 0 | 0 |
| NC_002645.1;NP_073550.1;HCoV-229E_1a;523 | 0 | 0 |
| NC_002645.1;NP_073550.1;HCoV-229E_1a;524 | 0 | 0 |
| NC_002645.1;NP_073550.1;HCoV-229E_1a;525 | 0 | 0 |
| NC_002645.1;NP_073550.1;HCoV-229E_1a;526 | 0 | 0 |
| NC_002645.1;NP_073550.1;HCoV-229E_1a;527 | 0 | 0 |
| NC_002645.1;NP_073550.1;HCoV-229E_1a;528 | 0 | 0 |
| NC_002645.1;NP_073550.1;HCoV-229E_1a;529 | 0 | 0.025 |
| NC_002645.1;NP_073550.1;HCoV-229E_1a;530 | 0 | 0 |
| NC_002645.1;NP_073550.1;HCoV-229E_1a;531 | 0 | 0 |
| NC_002645.1;NP_073550.1;HCoV-229E_1a;532 | 0 | 0 |
| NC_002645.1;NP_073550.1;HCoV-229E_1a;533 | 0 | 0 |
| NC_002645.1;NP_073550.1;HCoV-229E_1a;534 | 0 | 0 |
| NC_002645.1;NP_073550.1;HCoV-229E_1a;535 | 0 | 0 |
| NC_002645.1;NP_073550.1;HCoV-229E_1a;536 | 0 | 0 |
| NC_002645.1;NP_073550.1;HCoV-229E_1a;537 | 0 | 0 |
| NC_002645.1;NP_073550.1;HCoV-229E_1a;538 | 0 | 0 |
| NC_002645.1;NP_073550.1;HCoV-229E_1a;539 | 0 | 0 |
| NC_002645.1;NP_073550.1;HCoV-229E_1a;540 | 0 | 0.025 |
| NC_002645.1;NP_073550.1;HCoV-229E_1a;541 | 0 | 0.025 |
| NC_002645.1;NP_073550.1;HCoV-229E_1a;542 | 0 | 0.025 |
| NC_002645.1;NP_073550.1;HCoV-229E_1a;543 | 0 | 0.05 |
| NC_002645.1;NP_073550.1;HCoV-229E_1a;544 | 0.1 | 0.05 |
| NC_002645.1;NP_073550.1;HCoV-229E_1a;545 | 0 | 0.025 |
| NC_002645.1;NP_073550.1;HCoV-229E_1a;546 | 0 | 0.025 |
| NC_002645.1;NP_073550.1;HCoV-229E_1a;547 | 0 | 0.025 |
| NC_002645.1;NP_073550.1;HCoV-229E_1a;548 | 0 | 0 |
| NC_002645.1;NP_073550.1;HCoV-229E_1a;549 | 0 | 0 |

| | | |
|---|---|---|
| NC_002645.1;NP_073550.1;HCoV-229E_1a;550 | 0 | 0 |
| NC_002645.1;NP_073550.1;HCoV-229E_1a;551 | 0 | 0 |
| NC_002645.1;NP_073550.1;HCoV-229E_1a;552 | 0 | 0 |
| NC_002645.1;NP_073550.1;HCoV-229E_1a;553 | 0 | 0 |
| NC_002645.1;NP_073550.1;HCoV-229E_1a;554 | 0 | 0 |
| NC_002645.1;NP_073550.1;HCoV-229E_1a;555 | 0 | 0 |
| NC_002645.1;NP_073550.1;HCoV-229E_1a;556 | 0 | 0 |
| NC_002645.1;NP_073550.1;HCoV-229E_1a;557 | 0 | 0 |
| NC_002645.1;NP_073550.1;HCoV-229E_1a;558 | 0 | 0 |
| NC_002645.1;NP_073550.1;HCoV-229E_1a;559 | 0 | 0 |
| NC_002645.1;NP_073550.1;HCoV-229E_1a;560 | 0 | 0 |
| NC_002645.1;NP_073550.1;HCoV-229E_1a;561 | 0 | 0 |
| NC_002645.1;NP_073550.1;HCoV-229E_1a;562 | 0 | 0 |
| NC_002645.1;NP_073550.1;HCoV-229E_1a;563 | 0 | 0 |
| NC_002645.1;NP_073550.1;HCoV-229E_1a;564 | 0 | 0 |
| NC_002645.1;NP_073550.1;HCoV-229E_1a;565 | 0 | 0 |
| NC_002645.1;NP_073550.1;HCoV-229E_1a;566 | 0 | 0 |
| NC_002645.1;NP_073550.1;HCoV-229E_1a;567 | 0 | 0 |
| NC_002645.1;NP_073550.1;HCoV-229E_1a;568 | 0 | 0 |
| NC_002645.1;NP_073550.1;HCoV-229E_1a;569 | 0 | 0 |
| NC_002645.1;NP_073550.1;HCoV-229E_1a;570 | 0 | 0 |
| NC_002645.1;NP_073550.1;HCoV-229E_1a;571 | 0 | 0 |
| NC_002645.1;NP_073550.1;HCoV-229E_1a;572 | 0 | 0 |
| NC_002645.1;NP_073550.1;HCoV-229E_1a;573 | 0 | 0 |
| NC_002645.1;NP_073550.1;HCoV-229E_1a;574 | 0 | 0 |
| NC_002645.1;NP_073550.1;HCoV-229E_1a;575 | 0 | 0 |
| NC_002645.1;NP_073550.1;HCoV-229E_1a;576 | 0 | 0 |
| NC_002645.1;NP_073550.1;HCoV-229E_1a;577 | 0 | 0 |
| NC_002645.1;NP_073550.1;HCoV-229E_1a;578 | 0 | 0 |
| NC_002645.1;NP_073550.1;HCoV-229E_1a;579 | 0 | 0 |
| NC_002645.1;NP_073550.1;HCoV-229E_1a;580 | 0 | 0 |
| NC_002645.1;NP_073550.1;HCoV-229E_1a;581 | 0 | 0 |
| NC_002645.1;NP_073550.1;HCoV-229E_1a;582 | 0 | 0 |
| NC_002645.1;NP_073550.1;HCoV-229E_1a;583 | 0 | 0 |
| NC_002645.1;NP_073550.1;HCoV-229E_1a;584 | 0 | 0 |
| NC_002645.1;NP_073550.1;HCoV-229E_1a;585 | 0 | 0 |
| NC_002645.1;NP_073550.1;HCoV-229E_1a;586 | 0 | 0 |
| NC_002645.1;NP_073550.1;HCoV-229E_1a;587 | 0 | 0 |
| NC_002645.1;NP_073550.1;HCoV-229E_1a;588 | 0 | 0 |
| NC_002645.1;NP_073550.1;HCoV-229E_1a;589 | 0 | 0 |
| NC_002645.1;NP_073550.1;HCoV-229E_1a;590 | 0 | 0.025 |
| NC_002645.1;NP_073550.1;HCoV-229E_1a;591 | 0 | 0 |
| NC_002645.1;NP_073550.1;HCoV-229E_1a;592 | 0.1 | 0 |

| | | |
|---|---|---|
| NC_002645.1;NP_073550.1;HCoV-229E_1a;593 | 0 | 0 |
| NC_002645.1;NP_073550.1;HCoV-229E_1a;594 | 0.05 | 0.05 |
| NC_002645.1;NP_073550.1;HCoV-229E_1a;595 | 0 | 0 |
| NC_002645.1;NP_073550.1;HCoV-229E_1a;596 | 0 | 0 |
| NC_002645.1;NP_073550.1;HCoV-229E_1a;597 | 0 | 0 |
| NC_002645.1;NP_073550.1;HCoV-229E_1a;598 | 0 | 0 |
| NC_002645.1;NP_073550.1;HCoV-229E_1a;599 | 0 | 0 |
| NC_002645.1;NP_073550.1;HCoV-229E_1a;600 | 0 | 0 |
| NC_002645.1;NP_073550.1;HCoV-229E_1a;601 | 0 | 0 |
| NC_002645.1;NP_073550.1;HCoV-229E_1a;602 | 0 | 0 |
| NC_002645.1;NP_073550.1;HCoV-229E_1a;603 | 0 | 0 |
| NC_002645.1;NP_073550.1;HCoV-229E_1a;604 | 0 | 0 |
| NC_002645.1;NP_073550.1;HCoV-229E_1a;605 | 0 | 0 |
| NC_002645.1;NP_073550.1;HCoV-229E_1a;606 | 0 | 0 |
| NC_002645.1;NP_073550.1;HCoV-229E_1a;607 | 0 | 0 |
| NC_002645.1;NP_073550.1;HCoV-229E_1a;608 | 0 | 0 |
| NC_002645.1;NP_073550.1;HCoV-229E_1a;609 | 0 | 0 |
| NC_002645.1;NP_073550.1;HCoV-229E_1a;610 | 0.05 | 0 |
| NC_002645.1;NP_073550.1;HCoV-229E_1a;611 | 0.05 | 0 |
| NC_002645.1;NP_073550.1;HCoV-229E_1a;612 | 0.05 | 0 |
| NC_002645.1;NP_073550.1;HCoV-229E_1a;613 | 0.05 | 0.05 |
| NC_002645.1;NP_073550.1;HCoV-229E_1a;614 | 0.05 | 0 |
| NC_002645.1;NP_073550.1;HCoV-229E_1a;615 | 0.05 | 0 |
| NC_002645.1;NP_073550.1;HCoV-229E_1a;616 | 0.05 | 0 |
| NC_002645.1;NP_073550.1;HCoV-229E_1a;617 | 0.05 | 0 |
| NC_002645.1;NP_073550.1;HCoV-229E_1a;618 | 0.05 | 0 |
| NC_002645.1;NP_073550.1;HCoV-229E_1a;619 | 0 | 0 |
| NC_002645.1;NP_073550.1;HCoV-229E_1a;620 | 0 | 0 |
| NC_002645.1;NP_073550.1;HCoV-229E_1a;621 | 0 | 0 |
| NC_002645.1;NP_073550.1;HCoV-229E_1a;622 | 0 | 0.025 |
| NC_002645.1;NP_073550.1;HCoV-229E_1a;623 | 0 | 0.025 |
| NC_002645.1;NP_073550.1;HCoV-229E_1a;624 | 0 | 0 |
| NC_002645.1;NP_073550.1;HCoV-229E_1a;625 | 0 | 0.05 |
| NC_002645.1;NP_073550.1;HCoV-229E_1a;626 | 0 | 0.025 |
| NC_002645.1;NP_073550.1;HCoV-229E_1a;627 | 0.05 | 0.025 |
| NC_002645.1;NP_073550.1;HCoV-229E_1a;628 | 0.05 | 0.025 |
| NC_002645.1;NP_073550.1;HCoV-229E_1a;629 | 0.05 | 0.05 |
| NC_002645.1;NP_073550.1;HCoV-229E_1a;630 | 0 | 0 |
| NC_002645.1;NP_073550.1;HCoV-229E_1a;631 | 0.25 | 0.15 |
| NC_002645.1;NP_073550.1;HCoV-229E_1a;632 | 0.05 | 0.075 |
| NC_002645.1;NP_073550.1;HCoV-229E_1a;633 | 0.2 | 0.175 |
| NC_002645.1;NP_073550.1;HCoV-229E_1a;634 | 0.45 | 0.2 |
| NC_002645.1;NP_073550.1;HCoV-229E_1a;635 | 0.5 | 0.175 |

| | | |
|---|---|---|
| NC_002645.1;NP_073550.1;HCoV-229E_1a;636 | 0.4 | 0.175 |
| NC_002645.1;NP_073550.1;HCoV-229E_1a;637 | 0.4 | 0.2 |
| NC_002645.1;NP_073550.1;HCoV-229E_1a;638 | 0.3 | 0.15 |
| NC_002645.1;NP_073550.1;HCoV-229E_1a;639 | 0.4 | 0.175 |
| NC_002645.1;NP_073550.1;HCoV-229E_1a;640 | 0.35 | 0.15 |
| NC_002645.1;NP_073550.1;HCoV-229E_1a;641 | 0.2 | 0.125 |
| NC_002645.1;NP_073550.1;HCoV-229E_1a;642 | 0.2 | 0.125 |
| NC_002645.1;NP_073550.1;HCoV-229E_1a;643 | 0.05 | 0.025 |
| NC_002645.1;NP_073550.1;HCoV-229E_1a;644 | 0 | 0 |
| NC_002645.1;NP_073550.1;HCoV-229E_1a;645 | 0 | 0 |
| NC_002645.1;NP_073550.1;HCoV-229E_1a;646 | 0 | 0 |
| NC_002645.1;NP_073550.1;HCoV-229E_1a;647 | 0 | 0 |
| NC_002645.1;NP_073550.1;HCoV-229E_1a;648 | 0 | 0 |
| NC_002645.1;NP_073550.1;HCoV-229E_1a;649 | 0 | 0 |
| NC_002645.1;NP_073550.1;HCoV-229E_1a;650 | 0 | 0 |
| NC_002645.1;NP_073550.1;HCoV-229E_1a;651 | 0 | 0 |
| NC_002645.1;NP_073550.1;HCoV-229E_1a;652 | 0 | 0.05 |
| NC_002645.1;NP_073550.1;HCoV-229E_1a;653 | 0.05 | 0.05 |
| NC_002645.1;NP_073550.1;HCoV-229E_1a;654 | 0 | 0 |
| NC_002645.1;NP_073550.1;HCoV-229E_1a;655 | 0 | 0 |
| NC_002645.1;NP_073550.1;HCoV-229E_1a;656 | 0.05 | 0.025 |
| NC_002645.1;NP_073550.1;HCoV-229E_1a;657 | 0 | 0.025 |
| NC_002645.1;NP_073550.1;HCoV-229E_1a;658 | 0.05 | 0 |
| NC_002645.1;NP_073550.1;HCoV-229E_1a;659 | 0.05 | 0.05 |
| NC_002645.1;NP_073550.1;HCoV-229E_1a;660 | 0 | 0.1 |
| NC_002645.1;NP_073550.1;HCoV-229E_1a;661 | 0 | 0 |
| NC_002645.1;NP_073550.1;HCoV-229E_1a;662 | 0.05 | 0 |
| NC_002645.1;NP_073550.1;HCoV-229E_1a;663 | 0 | 0.025 |
| NC_002645.1;NP_073550.1;HCoV-229E_1a;664 | 0 | 0.025 |
| NC_002645.1;NP_073550.1;HCoV-229E_1a;665 | 0.05 | 0.025 |
| NC_002645.1;NP_073550.1;HCoV-229E_1a;666 | 0.1 | 0.025 |
| NC_002645.1;NP_073550.1;HCoV-229E_1a;667 | 0.05 | 0.025 |
| NC_002645.1;NP_073550.1;HCoV-229E_1a;668 | 0.1 | 0.05 |
| NC_002645.1;NP_073550.1;HCoV-229E_1a;669 | 0.05 | 0.075 |
| NC_002645.1;NP_073550.1;HCoV-229E_1a;670 | 0.05 | 0 |
| NC_002645.1;NP_073550.1;HCoV-229E_1a;671 | 0.05 | 0 |
| NC_002645.1;NP_073550.1;HCoV-229E_1a;672 | 0.05 | 0.025 |
| NC_002645.1;NP_073550.1;HCoV-229E_1a;673 | 0 | 0 |
| NC_002645.1;NP_073550.1;HCoV-229E_1a;674 | 0 | 0 |
| NC_002645.1;NP_073550.1;HCoV-229E_1a;675 | 0 | 0 |
| NC_002645.1;NP_073550.1;HCoV-229E_1a;676 | 0 | 0 |
| NC_002645.1;NP_073550.1;HCoV-229E_1a;677 | 0 | 0 |
| NC_002645.1;NP_073550.1;HCoV-229E_1a;678 | 0 | 0 |

| | | |
|---|---|---|
| NC_002645.1;NP_073550.1;HCoV-229E_1a;679 | 0 | 0 |
| NC_002645.1;NP_073550.1;HCoV-229E_1a;680 | 0 | 0 |
| NC_002645.1;NP_073550.1;HCoV-229E_1a;681 | 0.05 | 0.025 |
| NC_002645.1;NP_073550.1;HCoV-229E_1a;682 | 0 | 0.025 |
| NC_002645.1;NP_073550.1;HCoV-229E_1a;683 | 0.1 | 0.05 |
| NC_002645.1;NP_073550.1;HCoV-229E_1a;684 | 0.05 | 0.025 |
| NC_002645.1;NP_073550.1;HCoV-229E_1a;685 | 0 | 0.025 |
| NC_002645.1;NP_073550.1;HCoV-229E_1a;686 | 0.05 | 0.1 |
| NC_002645.1;NP_073550.1;HCoV-229E_1a;687 | 0.05 | 0.075 |
| NC_002645.1;NP_073550.1;HCoV-229E_1a;688 | 0.05 | 0.1 |
| NC_002645.1;NP_073550.1;HCoV-229E_1a;689 | 0.05 | 0.1 |
| NC_002645.1;NP_073550.1;HCoV-229E_1a;690 | 0.05 | 0.1 |
| NC_002645.1;NP_073550.1;HCoV-229E_1a;691 | 0.1 | 0.075 |
| NC_002645.1;NP_073550.1;HCoV-229E_1a;692 | 0.05 | 0.075 |
| NC_002645.1;NP_073550.1;HCoV-229E_1a;693 | 0.1 | 0.05 |
| NC_002645.1;NP_073550.1;HCoV-229E_1a;694 | 0.1 | 0.075 |
| NC_002645.1;NP_073550.1;HCoV-229E_1a;695 | 0.05 | 0.1 |
| NC_002645.1;NP_073550.1;HCoV-229E_1a;696 | 0 | 0.075 |
| NC_002645.1;NP_073550.1;HCoV-229E_1a;697 | 0 | 0.05 |
| NC_002645.1;NP_073550.1;HCoV-229E_1a;698 | 0.05 | 0.025 |
| NC_002645.1;NP_073550.1;HCoV-229E_1a;699 | 0.15 | 0.1 |
| NC_002645.1;NP_073550.1;HCoV-229E_1a;700 | 0.15 | 0.025 |
| NC_002645.1;NP_073550.1;HCoV-229E_1a;701 | 0 | 0 |
| NC_002645.1;NP_073550.1;HCoV-229E_1a;702 | 0 | 0.025 |
| NC_002645.1;NP_073550.1;HCoV-229E_1a;703 | 0 | 0 |
| NC_002645.1;NP_073550.1;HCoV-229E_1a;704 | 0.05 | 0 |
| NC_002645.1;NP_073550.1;HCoV-229E_1a;705 | 0 | 0 |
| NC_002645.1;NP_073550.1;HCoV-229E_1a;706 | 0 | 0 |
| NC_002645.1;NP_073550.1;HCoV-229E_1a;707 | 0 | 0 |
| NC_002645.1;NP_073550.1;HCoV-229E_1a;708 | 0 | 0 |
| NC_002645.1;NP_073550.1;HCoV-229E_1a;709 | 0 | 0 |
| NC_002645.1;NP_073550.1;HCoV-229E_1a;710 | 0 | 0 |
| NC_002645.1;NP_073550.1;HCoV-229E_1a;711 | 0 | 0 |
| NC_002645.1;NP_073550.1;HCoV-229E_1a;712 | 0 | 0 |
| NC_002645.1;NP_073550.1;HCoV-229E_1a;713 | 0 | 0 |
| NC_002645.1;NP_073550.1;HCoV-229E_1a;714 | 0 | 0 |
| NC_002645.1;NP_073550.1;HCoV-229E_1a;715 | 0 | 0 |
| NC_002645.1;NP_073550.1;HCoV-229E_1a;716 | 0 | 0 |
| NC_002645.1;NP_073550.1;HCoV-229E_1a;717 | 0 | 0 |
| NC_002645.1;NP_073550.1;HCoV-229E_1a;718 | 0 | 0.025 |
| NC_002645.1;NP_073550.1;HCoV-229E_1a;719 | 0 | 0.025 |
| NC_002645.1;NP_073550.1;HCoV-229E_1a;720 | 0 | 0 |
| NC_002645.1;NP_073550.1;HCoV-229E_1a;721 | 0 | 0.025 |

| | | |
|---|---|---|
| NC_002645.1;NP_073550.1;HCoV-229E_1a;722 | 0 | 0.025 |
| NC_002645.1;NP_073550.1;HCoV-229E_1a;723 | 0.05 | 0.025 |
| NC_002645.1;NP_073550.1;HCoV-229E_1a;724 | 0 | 0 |
| NC_002645.1;NP_073550.1;HCoV-229E_1a;725 | 0 | 0.025 |
| NC_002645.1;NP_073550.1;HCoV-229E_1a;726 | 0 | 0.05 |
| NC_002645.1;NP_073550.1;HCoV-229E_1a;727 | 0 | 0.05 |
| NC_002645.1;NP_073550.1;HCoV-229E_1a;728 | 0 | 0.025 |
| NC_002645.1;NP_073550.1;HCoV-229E_1a;729 | 0 | 0 |
| NC_002645.1;NP_073550.1;HCoV-229E_1a;730 | 0 | 0 |
| NC_002645.1;NP_073550.1;HCoV-229E_1a;731 | 0 | 0 |
| NC_002645.1;NP_073550.1;HCoV-229E_1a;732 | 0 | 0 |
| NC_002645.1;NP_073550.1;HCoV-229E_1a;733 | 0 | 0 |
| NC_002645.1;NP_073550.1;HCoV-229E_1a;734 | 0 | 0 |
| NC_002645.1;NP_073550.1;HCoV-229E_1a;735 | 0 | 0 |
| NC_002645.1;NP_073550.1;HCoV-229E_1a;736 | 0 | 0 |
| NC_002645.1;NP_073550.1;HCoV-229E_1a;737 | 0 | 0 |
| NC_002645.1;NP_073550.1;HCoV-229E_1a;738 | 0.05 | 0 |
| NC_002645.1;NP_073550.1;HCoV-229E_1a;739 | 0 | 0.025 |
| NC_002645.1;NP_073550.1;HCoV-229E_1a;740 | 0 | 0 |
| NC_002645.1;NP_073550.1;HCoV-229E_1a;741 | 0 | 0 |
| NC_002645.1;NP_073550.1;HCoV-229E_1a;742 | 0 | 0 |
| NC_002645.1;NP_073550.1;HCoV-229E_1a;743 | 0 | 0 |
| NC_002645.1;NP_073550.1;HCoV-229E_1a;744 | 0 | 0 |
| NC_002645.1;NP_073550.1;HCoV-229E_1a;745 | 0 | 0 |
| NC_002645.1;NP_073550.1;HCoV-229E_1a;746 | 0 | 0 |
| NC_002645.1;NP_073550.1;HCoV-229E_1a;747 | 0 | 0 |
| NC_002645.1;NP_073550.1;HCoV-229E_1a;748 | 0 | 0 |
| NC_002645.1;NP_073550.1;HCoV-229E_1a;749 | 0 | 0 |
| NC_002645.1;NP_073550.1;HCoV-229E_1a;750 | 0 | 0 |
| NC_002645.1;NP_073550.1;HCoV-229E_1a;751 | 0 | 0 |
| NC_002645.1;NP_073550.1;HCoV-229E_1a;752 | 0 | 0 |
| NC_002645.1;NP_073550.1;HCoV-229E_1a;753 | 0 | 0 |
| NC_002645.1;NP_073550.1;HCoV-229E_1a;754 | 0 | 0 |
| NC_002645.1;NP_073550.1;HCoV-229E_1a;755 | 0 | 0 |
| NC_002645.1;NP_073550.1;HCoV-229E_1a;756 | 0 | 0 |
| NC_002645.1;NP_073550.1;HCoV-229E_1a;757 | 0 | 0 |
| NC_002645.1;NP_073550.1;HCoV-229E_1a;758 | 0 | 0 |
| NC_002645.1;NP_073550.1;HCoV-229E_1a;759 | 0 | 0.025 |
| NC_002645.1;NP_073550.1;HCoV-229E_1a;760 | 0 | 0 |
| NC_002645.1;NP_073550.1;HCoV-229E_1a;761 | 0 | 0 |
| NC_002645.1;NP_073550.1;HCoV-229E_1a;762 | 0 | 0 |
| NC_002645.1;NP_073550.1;HCoV-229E_1a;763 | 0 | 0.025 |
| NC_002645.1;NP_073550.1;HCoV-229E_1a;764 | 0 | 0 |

| | | |
|---|---|---|
| NC_002645.1;NP_073550.1;HCoV-229E_1a;765 | 0 | 0 |
| NC_002645.1;NP_073550.1;HCoV-229E_1a;766 | 0 | 0 |
| NC_002645.1;NP_073550.1;HCoV-229E_1a;767 | 0 | 0 |
| NC_002645.1;NP_073550.1;HCoV-229E_1a;768 | 0 | 0 |
| NC_002645.1;NP_073550.1;HCoV-229E_1a;769 | 0 | 0.05 |
| NC_002645.1;NP_073550.1;HCoV-229E_1a;770 | 0 | 0 |
| NC_002645.1;NP_073550.1;HCoV-229E_1a;771 | 0 | 0.025 |
| NC_002645.1;NP_073550.1;HCoV-229E_1a;772 | 0 | 0.025 |
| NC_002645.1;NP_073550.1;HCoV-229E_1a;773 | 0 | 0.025 |
| NC_002645.1;NP_073550.1;HCoV-229E_1a;774 | 0 | 0.025 |
| NC_002645.1;NP_073550.1;HCoV-229E_1a;775 | 0 | 0.025 |
| NC_002645.1;NP_073550.1;HCoV-229E_1a;776 | 0 | 0.025 |
| NC_002645.1;NP_073550.1;HCoV-229E_1a;777 | 0 | 0 |
| NC_002645.1;NP_073550.1;HCoV-229E_1a;778 | 0 | 0 |
| NC_002645.1;NP_073550.1;HCoV-229E_1a;779 | 0 | 0.025 |
| NC_002645.1;NP_073550.1;HCoV-229E_1a;780 | 0 | 0.025 |
| NC_002645.1;NP_073550.1;HCoV-229E_1a;781 | 0 | 0.025 |
| NC_002645.1;NP_073550.1;HCoV-229E_1a;782 | 0 | 0 |
| NC_002645.1;NP_073550.1;HCoV-229E_1a;783 | 0 | 0 |
| NC_002645.1;NP_073550.1;HCoV-229E_1a;784 | 0 | 0 |
| NC_002645.1;NP_073550.1;HCoV-229E_1a;785 | 0 | 0 |
| NC_002645.1;NP_073550.1;HCoV-229E_1a;786 | 0 | 0 |
| NC_002645.1;NP_073550.1;HCoV-229E_1a;787 | 0 | 0 |
| NC_002645.1;NP_073550.1;HCoV-229E_1a;788 | 0 | 0 |
| NC_002645.1;NP_073550.1;HCoV-229E_1a;789 | 0 | 0 |
| NC_002645.1;NP_073550.1;HCoV-229E_1a;790 | 0 | 0 |
| NC_002645.1;NP_073550.1;HCoV-229E_1a;791 | 0 | 0 |
| NC_002645.1;NP_073550.1;HCoV-229E_1a;792 | 0 | 0 |
| NC_002645.1;NP_073550.1;HCoV-229E_1a;793 | 0 | 0 |
| NC_002645.1;NP_073550.1;HCoV-229E_1a;794 | 0 | 0 |
| NC_002645.1;NP_073550.1;HCoV-229E_1a;795 | 0 | 0 |
| NC_002645.1;NP_073550.1;HCoV-229E_1a;796 | 0 | 0 |
| NC_002645.1;NP_073550.1;HCoV-229E_1a;797 | 0.05 | 0 |
| NC_002645.1;NP_073550.1;HCoV-229E_1a;798 | 0 | 0 |
| NC_002645.1;NP_073550.1;HCoV-229E_1a;799 | 0 | 0.05 |
| NC_002645.1;NP_073550.1;HCoV-229E_1a;800 | 0 | 0 |
| NC_002645.1;NP_073550.1;HCoV-229E_1a;801 | 0 | 0.025 |
| NC_002645.1;NP_073550.1;HCoV-229E_1a;802 | 0 | 0 |
| NC_002645.1;NP_073550.1;HCoV-229E_1a;803 | 0 | 0 |
| NC_002645.1;NP_073550.1;HCoV-229E_1a;804 | 0 | 0 |
| NC_002645.1;NP_073550.1;HCoV-229E_1a;805 | 0 | 0 |
| NC_002645.1;NP_073550.1;HCoV-229E_1a;806 | 0 | 0 |
| NC_002645.1;NP_073550.1;HCoV-229E_1a;807 | 0 | 0 |

| | | |
|---|---|---|
| NC_002645.1;NP_073550.1;HCoV-229E_1a;808 | 0 | 0 |
| NC_002645.1;NP_073550.1;HCoV-229E_1a;809 | 0 | 0.025 |
| NC_002645.1;NP_073550.1;HCoV-229E_1a;810 | 0 | 0.025 |
| NC_002645.1;NP_073550.1;HCoV-229E_1a;811 | 0 | 0 |
| NC_002645.1;NP_073550.1;HCoV-229E_1a;812 | 0 | 0.075 |
| NC_002645.1;NP_073550.1;HCoV-229E_1a;813 | 0 | 0.025 |
| NC_002645.1;NP_073550.1;HCoV-229E_1a;814 | 0 | 0.025 |
| NC_002645.1;NP_073550.1;HCoV-229E_1a;815 | 0 | 0 |
| NC_002645.1;NP_073550.1;HCoV-229E_1a;816 | 0 | 0 |
| NC_002645.1;NP_073550.1;HCoV-229E_1a;817 | 0 | 0.025 |
| NC_002645.1;NP_073550.1;HCoV-229E_1a;818 | 0 | 0.025 |
| NC_002645.1;NP_073550.1;HCoV-229E_1a;819 | 0 | 0 |
| NC_002645.1;NP_073550.1;HCoV-229E_1a;820 | 0 | 0 |
| NC_002645.1;NP_073550.1;HCoV-229E_1a;821 | 0 | 0 |
| NC_002645.1;NP_073550.1;HCoV-229E_1a;822 | 0 | 0 |
| NC_002645.1;NP_073550.1;HCoV-229E_1a;823 | 0 | 0 |
| NC_002645.1;NP_073550.1;HCoV-229E_1a;824 | 0.05 | 0.025 |
| NC_002645.1;NP_073550.1;HCoV-229E_1a;825 | 0.1 | 0.025 |
| NC_002645.1;NP_073550.1;HCoV-229E_1a;826 | 0.1 | 0.025 |
| NC_002645.1;NP_073550.1;HCoV-229E_1a;827 | 0.05 | 0.075 |
| NC_002645.1;NP_073550.1;HCoV-229E_1a;828 | 0 | 0 |
| NC_002645.1;NP_073550.1;HCoV-229E_1a;829 | 0 | 0 |
| NC_002645.1;NP_073550.1;HCoV-229E_1a;830 | 0 | 0 |
| NC_002645.1;NP_073550.1;HCoV-229E_1a;831 | 0.1 | 0.025 |
| NC_002645.1;NP_073550.1;HCoV-229E_1a;832 | 0 | 0 |
| NC_002645.1;NP_073550.1;HCoV-229E_1a;833 | 0 | 0 |
| NC_002645.1;NP_073550.1;HCoV-229E_1a;834 | 0 | 0 |
| NC_002645.1;NP_073550.1;HCoV-229E_1a;835 | 0 | 0.025 |
| NC_002645.1;NP_073550.1;HCoV-229E_1a;836 | 0 | 0.025 |
| NC_002645.1;NP_073550.1;HCoV-229E_1a;837 | 0 | 0 |
| NC_002645.1;NP_073550.1;HCoV-229E_1a;838 | 0 | 0 |
| NC_002645.1;NP_073550.1;HCoV-229E_1a;839 | 0 | 0 |
| NC_002645.1;NP_073550.1;HCoV-229E_1a;840 | 0 | 0 |
| NC_002645.1;NP_073550.1;HCoV-229E_1a;841 | 0 | 0 |
| NC_002645.1;NP_073550.1;HCoV-229E_1a;842 | 0 | 0 |
| NC_002645.1;NP_073550.1;HCoV-229E_1a;843 | 0 | 0 |
| NC_002645.1;NP_073550.1;HCoV-229E_1a;844 | 0 | 0.025 |
| NC_002645.1;NP_073550.1;HCoV-229E_1a;845 | 0 | 0.025 |
| NC_002645.1;NP_073550.1;HCoV-229E_1a;846 | 0 | 0 |
| NC_002645.1;NP_073550.1;HCoV-229E_1a;847 | 0 | 0 |
| NC_002645.1;NP_073550.1;HCoV-229E_1a;848 | 0 | 0 |
| NC_002645.1;NP_073550.1;HCoV-229E_1a;849 | 0 | 0.025 |
| NC_002645.1;NP_073550.1;HCoV-229E_1a;850 | 0 | 0 |

| | | |
|---|---|---|
| NC_002645.1;NP_073550.1;HCoV-229E_1a;851 | 0 | 0 |
| NC_002645.1;NP_073550.1;HCoV-229E_1a;852 | 0.05 | 0 |
| NC_002645.1;NP_073550.1;HCoV-229E_1a;853 | 0 | 0.05 |
| NC_002645.1;NP_073550.1;HCoV-229E_1a;854 | 0 | 0 |
| NC_002645.1;NP_073550.1;HCoV-229E_1a;855 | 0 | 0 |
| NC_002645.1;NP_073550.1;HCoV-229E_1a;856 | 0 | 0 |
| NC_002645.1;NP_073550.1;HCoV-229E_1a;857 | 0 | 0 |
| NC_002645.1;NP_073550.1;HCoV-229E_1a;858 | 0 | 0 |
| NC_002645.1;NP_073550.1;HCoV-229E_1a;859 | 0 | 0 |
| NC_002645.1;NP_073550.1;HCoV-229E_1a;860 | 0 | 0 |
| NC_002645.1;NP_073550.1;HCoV-229E_1a;861 | 0 | 0 |
| NC_002645.1;NP_073550.1;HCoV-229E_1a;862 | 0 | 0 |
| NC_002645.1;NP_073550.1;HCoV-229E_1a;863 | 0 | 0 |
| NC_002645.1;NP_073550.1;HCoV-229E_1a;864 | 0 | 0 |
| NC_002645.1;NP_073550.1;HCoV-229E_1a;865 | 0.05 | 0 |
| NC_002645.1;NP_073550.1;HCoV-229E_1a;866 | 0.05 | 0 |
| NC_002645.1;NP_073550.1;HCoV-229E_1a;867 | 0.05 | 0.025 |
| NC_002645.1;NP_073550.1;HCoV-229E_1a;868 | 0.05 | 0 |
| NC_002645.1;NP_073550.1;HCoV-229E_1a;869 | 0.05 | 0 |
| NC_002645.1;NP_073550.1;HCoV-229E_1a;870 | 0.05 | 0 |
| NC_002645.1;NP_073550.1;HCoV-229E_1a;871 | 0.05 | 0 |
| NC_002645.1;NP_073550.1;HCoV-229E_1a;872 | 0.15 | 0.025 |
| NC_002645.1;NP_073550.1;HCoV-229E_1a;873 | 0.05 | 0.025 |
| NC_002645.1;NP_073550.1;HCoV-229E_1a;874 | 0.2 | 0.1 |
| NC_002645.1;NP_073550.1;HCoV-229E_1a;875 | 0.05 | 0.025 |
| NC_002645.1;NP_073550.1;HCoV-229E_1a;876 | 0.1 | 0.05 |
| NC_002645.1;NP_073550.1;HCoV-229E_1a;877 | 0 | 0 |
| NC_002645.1;NP_073550.1;HCoV-229E_1a;878 | 0 | 0 |
| NC_002645.1;NP_073550.1;HCoV-229E_1a;879 | 0.05 | 0 |
| NC_002645.1;NP_073550.1;HCoV-229E_1a;880 | 0.05 | 0 |
| NC_002645.1;NP_073550.1;HCoV-229E_1a;881 | 0.05 | 0 |
| NC_002645.1;NP_073550.1;HCoV-229E_1a;882 | 0 | 0 |
| NC_002645.1;NP_073550.1;HCoV-229E_1a;883 | 0.05 | 0 |
| NC_002645.1;NP_073550.1;HCoV-229E_1a;884 | 0 | 0 |
| NC_002645.1;NP_073550.1;HCoV-229E_1a;885 | 0 | 0 |
| NC_002645.1;NP_073550.1;HCoV-229E_1a;886 | 0 | 0 |
| NC_002645.1;NP_073550.1;HCoV-229E_1a;887 | 0 | 0.025 |
| NC_002645.1;NP_073550.1;HCoV-229E_1a;888 | 0 | 0.025 |
| NC_002645.1;NP_073550.1;HCoV-229E_1a;889 | 0 | 0.025 |
| NC_002645.1;NP_073550.1;HCoV-229E_1a;890 | 0 | 0.025 |
| NC_002645.1;NP_073550.1;HCoV-229E_1a;891 | 0 | 0.025 |
| NC_002645.1;NP_073550.1;HCoV-229E_1a;892 | 0 | 0.025 |
| NC_002645.1;NP_073550.1;HCoV-229E_1a;893 | 0 | 0.025 |

| | | |
|---|---|---|
| NC_002645.1;NP_073550.1;HCoV-229E_1a;894 | 0 | 0.025 |
| NC_002645.1;NP_073550.1;HCoV-229E_1a;895 | 0 | 0.025 |
| NC_002645.1;NP_073550.1;HCoV-229E_1a;896 | 0 | 0 |
| NC_002645.1;NP_073550.1;HCoV-229E_1a;897 | 0 | 0 |
| NC_002645.1;NP_073550.1;HCoV-229E_1a;898 | 0 | 0 |
| NC_002645.1;NP_073550.1;HCoV-229E_1a;899 | 0 | 0 |
| NC_002645.1;NP_073550.1;HCoV-229E_1a;900 | 0.1 | 0 |
| NC_002645.1;NP_073550.1;HCoV-229E_1a;901 | 0.05 | 0 |
| NC_002645.1;NP_073550.1;HCoV-229E_1a;902 | 0 | 0 |
| NC_002645.1;NP_073550.1;HCoV-229E_1a;903 | 0.1 | 0 |
| NC_002645.1;NP_073550.1;HCoV-229E_1a;904 | 0 | 0 |
| NC_002645.1;NP_073550.1;HCoV-229E_1a;905 | 0 | 0 |
| NC_002645.1;NP_073550.1;HCoV-229E_1a;906 | 0.05 | 0 |
| NC_002645.1;NP_073550.1;HCoV-229E_1a;907 | 0 | 0 |
| NC_002645.1;NP_073550.1;HCoV-229E_1a;908 | 0 | 0 |
| NC_002645.1;NP_073550.1;HCoV-229E_1a;909 | 0 | 0 |
| NC_002645.1;NP_073550.1;HCoV-229E_1a;910 | 0 | 0 |
| NC_002645.1;NP_073550.1;HCoV-229E_1a;911 | 0 | 0.025 |
| NC_002645.1;NP_073550.1;HCoV-229E_1a;912 | 0 | 0 |
| NC_002645.1;NP_073550.1;HCoV-229E_1a;913 | 0 | 0 |
| NC_002645.1;NP_073550.1;HCoV-229E_1a;914 | 0 | 0 |
| NC_002645.1;NP_073550.1;HCoV-229E_1a;915 | 0 | 0 |
| NC_002645.1;NP_073550.1;HCoV-229E_1a;916 | 0 | 0 |
| NC_002645.1;NP_073550.1;HCoV-229E_1a;917 | 0 | 0 |
| NC_002645.1;NP_073550.1;HCoV-229E_1a;918 | 0 | 0 |
| NC_002645.1;NP_073550.1;HCoV-229E_1a;919 | 0 | 0 |
| NC_002645.1;NP_073550.1;HCoV-229E_1a;920 | 0 | 0 |
| NC_002645.1;NP_073550.1;HCoV-229E_1a;921 | 0 | 0 |
| NC_002645.1;NP_073550.1;HCoV-229E_1a;922 | 0 | 0 |
| NC_002645.1;NP_073550.1;HCoV-229E_1a;923 | 0 | 0 |
| NC_002645.1;NP_073550.1;HCoV-229E_1a;924 | 0.05 | 0.05 |
| NC_002645.1;NP_073550.1;HCoV-229E_1a;925 | 0 | 0 |
| NC_002645.1;NP_073550.1;HCoV-229E_1a;926 | 0 | 0 |
| NC_002645.1;NP_073550.1;HCoV-229E_1a;927 | 0 | 0 |
| NC_002645.1;NP_073550.1;HCoV-229E_1a;928 | 0 | 0 |
| NC_002645.1;NP_073550.1;HCoV-229E_1a;929 | 0 | 0 |
| NC_002645.1;NP_073550.1;HCoV-229E_1a;930 | 0 | 0 |
| NC_002645.1;NP_073550.1;HCoV-229E_1a;931 | 0 | 0 |
| NC_002645.1;NP_073550.1;HCoV-229E_1a;932 | 0 | 0.025 |
| NC_002645.1;NP_073550.1;HCoV-229E_1a;933 | 0 | 0.025 |
| NC_002645.1;NP_073550.1;HCoV-229E_1a;934 | 0 | 0.075 |
| NC_002645.1;NP_073550.1;HCoV-229E_1a;935 | 0 | 0.025 |
| NC_002645.1;NP_073550.1;HCoV-229E_1a;936 | 0 | 0 |

| | | |
|---|---|---|
| NC_002645.1;NP_073550.1;HCoV-229E_1a;937 | 0 | 0.025 |
| NC_002645.1;NP_073550.1;HCoV-229E_1a;938 | 0 | 0.05 |
| NC_002645.1;NP_073550.1;HCoV-229E_1a;939 | 0 | 0 |
| NC_002645.1;NP_073550.1;HCoV-229E_1a;940 | 0 | 0 |
| NC_002645.1;NP_073550.1;HCoV-229E_1a;941 | 0 | 0 |
| NC_002645.1;NP_073550.1;HCoV-229E_1a;942 | 0 | 0 |
| NC_002645.1;NP_073550.1;HCoV-229E_1a;943 | 0 | 0 |
| NC_002645.1;NP_073550.1;HCoV-229E_1a;944 | 0 | 0 |
| NC_002645.1;NP_073550.1;HCoV-229E_1a;945 | 0 | 0 |
| NC_002645.1;NP_073550.1;HCoV-229E_1a;946 | 0 | 0 |
| NC_002645.1;NP_073550.1;HCoV-229E_1a;947 | 0 | 0 |
| NC_002645.1;NP_073550.1;HCoV-229E_1a;948 | 0 | 0 |
| NC_002645.1;NP_073550.1;HCoV-229E_1a;949 | 0 | 0 |
| NC_002645.1;NP_073550.1;HCoV-229E_1a;950 | 0 | 0 |
| NC_002645.1;NP_073550.1;HCoV-229E_1a;951 | 0 | 0 |
| NC_002645.1;NP_073550.1;HCoV-229E_1a;952 | 0 | 0 |
| NC_002645.1;NP_073550.1;HCoV-229E_1a;953 | 0 | 0 |
| NC_002645.1;NP_073550.1;HCoV-229E_1a;954 | 0 | 0 |
| NC_002645.1;NP_073550.1;HCoV-229E_1a;955 | 0 | 0 |
| NC_002645.1;NP_073550.1;HCoV-229E_1a;956 | 0 | 0 |
| NC_002645.1;NP_073550.1;HCoV-229E_1a;957 | 0 | 0 |
| NC_002645.1;NP_073550.1;HCoV-229E_1a;958 | 0 | 0 |
| NC_002645.1;NP_073550.1;HCoV-229E_1a;959 | 0 | 0 |
| NC_002645.1;NP_073550.1;HCoV-229E_1a;960 | 0 | 0 |
| NC_002645.1;NP_073550.1;HCoV-229E_1a;961 | 0 | 0 |
| NC_002645.1;NP_073550.1;HCoV-229E_1a;962 | 0 | 0 |
| NC_002645.1;NP_073550.1;HCoV-229E_1a;963 | 0 | 0 |
| NC_002645.1;NP_073550.1;HCoV-229E_1a;964 | 0 | 0 |
| NC_002645.1;NP_073550.1;HCoV-229E_1a;965 | 0 | 0.025 |
| NC_002645.1;NP_073550.1;HCoV-229E_1a;966 | 0 | 0 |
| NC_002645.1;NP_073550.1;HCoV-229E_1a;967 | 0 | 0 |
| NC_002645.1;NP_073550.1;HCoV-229E_1a;968 | 0 | 0.025 |
| NC_002645.1;NP_073550.1;HCoV-229E_1a;969 | 0 | 0 |
| NC_002645.1;NP_073550.1;HCoV-229E_1a;970 | 0 | 0.025 |
| NC_002645.1;NP_073550.1;HCoV-229E_1a;971 | 0 | 0 |
| NC_002645.1;NP_073550.1;HCoV-229E_1a;972 | 0.05 | 0 |
| NC_002645.1;NP_073550.1;HCoV-229E_1a;973 | 0 | 0 |
| NC_002645.1;NP_073550.1;HCoV-229E_1a;974 | 0.05 | 0 |
| NC_002645.1;NP_073550.1;HCoV-229E_1a;975 | 0.1 | 0 |
| NC_002645.1;NP_073550.1;HCoV-229E_1a;976 | 0 | 0 |
| NC_002645.1;NP_073550.1;HCoV-229E_1a;977 | 0 | 0 |
| NC_002645.1;NP_073550.1;HCoV-229E_1a;978 | 0 | 0 |
| NC_002645.1;NP_073550.1;HCoV-229E_1a;979 | 0 | 0 |

| | | |
|---|---|---|
| NC_002645.1;NP_073550.1;HCoV-229E_1a;980 | 0 | 0 |
| NC_002645.1;NP_073550.1;HCoV-229E_1a;981 | 0 | 0 |
| NC_002645.1;NP_073550.1;HCoV-229E_1a;982 | 0 | 0 |
| NC_002645.1;NP_073550.1;HCoV-229E_1a;983 | 0.05 | 0.05 |
| NC_002645.1;NP_073550.1;HCoV-229E_1a;984 | 0 | 0.025 |
| NC_002645.1;NP_073550.1;HCoV-229E_1a;985 | 0 | 0.025 |
| NC_002645.1;NP_073550.1;HCoV-229E_1a;986 | 0 | 0.05 |
| NC_002645.1;NP_073550.1;HCoV-229E_1a;987 | 0 | 0 |
| NC_002645.1;NP_073550.1;HCoV-229E_1a;988 | 0 | 0.15 |
| NC_002645.1;NP_073550.1;HCoV-229E_1a;989 | 0.05 | 0.125 |
| NC_002645.1;NP_073550.1;HCoV-229E_1a;990 | 0 | 0.05 |
| NC_002645.1;NP_073550.1;HCoV-229E_1a;991 | 0 | 0.1 |
| NC_002645.1;NP_073550.1;HCoV-229E_1a;992 | 0 | 0.075 |
| NC_002645.1;NP_073550.1;HCoV-229E_1a;993 | 0 | 0.025 |
| NC_002645.1;NP_073550.1;HCoV-229E_1a;994 | 0 | 0 |
| NC_002645.1;NP_073550.1;HCoV-229E_1a;995 | 0 | 0 |
| NC_002645.1;NP_073550.1;HCoV-229E_1a;996 | 0 | 0 |
| NC_002645.1;NP_073550.1;HCoV-229E_1a;997 | 0.05 | 0 |
| NC_002645.1;NP_073550.1;HCoV-229E_1a;998 | 0.05 | 0 |
| NC_002645.1;NP_073550.1;HCoV-229E_1a;999 | 0.05 | 0 |
| NC_002645.1;NP_073550.1;HCoV-229E_1a;1000 | 0.05 | 0 |
| NC_002645.1;NP_073550.1;HCoV-229E_1a;1001 | 0.05 | 0 |
| NC_002645.1;NP_073550.1;HCoV-229E_1a;1002 | 0.05 | 0 |
| NC_002645.1;NP_073550.1;HCoV-229E_1a;1003 | 0 | 0 |
| NC_002645.1;NP_073550.1;HCoV-229E_1a;1004 | 0 | 0.025 |
| NC_002645.1;NP_073550.1;HCoV-229E_1a;1005 | 0.1 | 0.05 |
| NC_002645.1;NP_073550.1;HCoV-229E_1a;1006 | 0.15 | 0.05 |
| NC_002645.1;NP_073550.1;HCoV-229E_1a;1007 | 0.2 | 0.05 |
| NC_002645.1;NP_073550.1;HCoV-229E_1a;1008 | 0.1 | 0.05 |
| NC_002645.1;NP_073550.1;HCoV-229E_1a;1009 | 0.05 | 0.05 |
| NC_002645.1;NP_073550.1;HCoV-229E_1a;1010 | 0 | 0.05 |
| NC_002645.1;NP_073550.1;HCoV-229E_1a;1011 | 0.05 | 0.075 |
| NC_002645.1;NP_073550.1;HCoV-229E_1a;1012 | 0.1 | 0.075 |
| NC_002645.1;NP_073550.1;HCoV-229E_1a;1013 | 0.15 | 0.125 |
| NC_002645.1;NP_073550.1;HCoV-229E_1a;1014 | 0.05 | 0.075 |
| NC_002645.1;NP_073550.1;HCoV-229E_1a;1015 | 0.1 | 0.1 |
| NC_002645.1;NP_073550.1;HCoV-229E_1a;1016 | 0.1 | 0.125 |
| NC_002645.1;NP_073550.1;HCoV-229E_1a;1017 | 0.15 | 0.075 |
| NC_002645.1;NP_073550.1;HCoV-229E_1a;1018 | 0.15 | 0.075 |
| NC_002645.1;NP_073550.1;HCoV-229E_1a;1019 | 0.1 | 0.05 |
| NC_002645.1;NP_073550.1;HCoV-229E_1a;1020 | 0 | 0.025 |
| NC_002645.1;NP_073550.1;HCoV-229E_1a;1021 | 0 | 0 |
| NC_002645.1;NP_073550.1;HCoV-229E_1a;1022 | 0.05 | 0 |

| | | |
|---|---|---|
| NC_002645.1;NP_073550.1;HCoV-229E_1a;1023 | 0 | 0 |
| NC_002645.1;NP_073550.1;HCoV-229E_1a;1024 | 0.05 | 0 |
| NC_002645.1;NP_073550.1;HCoV-229E_1a;1025 | 0.05 | 0 |
| NC_002645.1;NP_073550.1;HCoV-229E_1a;1026 | 0.05 | 0 |
| NC_002645.1;NP_073550.1;HCoV-229E_1a;1027 | 0.05 | 0.025 |
| NC_002645.1;NP_073550.1;HCoV-229E_1a;1028 | 0.1 | 0.05 |
| NC_002645.1;NP_073550.1;HCoV-229E_1a;1029 | 0.05 | 0.025 |
| NC_002645.1;NP_073550.1;HCoV-229E_1a;1030 | 0.05 | 0.025 |
| NC_002645.1;NP_073550.1;HCoV-229E_1a;1031 | 0.05 | 0.05 |
| NC_002645.1;NP_073550.1;HCoV-229E_1a;1032 | 0 | 0.025 |
| NC_002645.1;NP_073550.1;HCoV-229E_1a;1033 | 0 | 0 |
| NC_002645.1;NP_073550.1;HCoV-229E_1a;1034 | 0 | 0.025 |
| NC_002645.1;NP_073550.1;HCoV-229E_1a;1035 | 0 | 0 |
| NC_002645.1;NP_073550.1;HCoV-229E_1a;1036 | 0 | 0.025 |
| NC_002645.1;NP_073550.1;HCoV-229E_1a;1037 | 0 | 0 |
| NC_002645.1;NP_073550.1;HCoV-229E_1a;1038 | 0.05 | 0 |
| NC_002645.1;NP_073550.1;HCoV-229E_1a;1039 | 0 | 0 |
| NC_002645.1;NP_073550.1;HCoV-229E_1a;1040 | 0 | 0.025 |
| NC_002645.1;NP_073550.1;HCoV-229E_1a;1041 | 0 | 0 |
| NC_002645.1;NP_073550.1;HCoV-229E_1a;1042 | 0 | 0 |
| NC_002645.1;NP_073550.1;HCoV-229E_1a;1043 | 0 | 0 |
| NC_002645.1;NP_073550.1;HCoV-229E_1a;1044 | 0 | 0 |
| NC_002645.1;NP_073550.1;HCoV-229E_1a;1045 | 0.05 | 0 |
| NC_002645.1;NP_073550.1;HCoV-229E_1a;1046 | 0 | 0 |
| NC_002645.1;NP_073550.1;HCoV-229E_1a;1047 | 0 | 0 |
| NC_002645.1;NP_073550.1;HCoV-229E_1a;1048 | 0 | 0 |
| NC_002645.1;NP_073550.1;HCoV-229E_1a;1049 | 0 | 0 |
| NC_002645.1;NP_073550.1;HCoV-229E_1a;1050 | 0 | 0 |
| NC_002645.1;NP_073550.1;HCoV-229E_1a;1051 | 0 | 0 |
| NC_002645.1;NP_073550.1;HCoV-229E_1a;1052 | 0 | 0 |
| NC_002645.1;NP_073550.1;HCoV-229E_1a;1053 | 0 | 0 |
| NC_002645.1;NP_073550.1;HCoV-229E_1a;1054 | 0 | 0 |
| NC_002645.1;NP_073550.1;HCoV-229E_1a;1055 | 0 | 0 |
| NC_002645.1;NP_073550.1;HCoV-229E_1a;1056 | 0 | 0 |
| NC_002645.1;NP_073550.1;HCoV-229E_1a;1057 | 0 | 0 |
| NC_002645.1;NP_073550.1;HCoV-229E_1a;1058 | 0 | 0 |
| NC_002645.1;NP_073550.1;HCoV-229E_1a;1059 | 0 | 0 |
| NC_002645.1;NP_073550.1;HCoV-229E_1a;1060 | 0 | 0 |
| NC_002645.1;NP_073550.1;HCoV-229E_1a;1061 | 0 | 0 |
| NC_002645.1;NP_073550.1;HCoV-229E_1a;1062 | 0 | 0 |
| NC_002645.1;NP_073550.1;HCoV-229E_1a;1063 | 0 | 0 |
| NC_002645.1;NP_073550.1;HCoV-229E_1a;1064 | 0.05 | 0.025 |
| NC_002645.1;NP_073550.1;HCoV-229E_1a;1065 | 0.05 | 0.025 |

| | | |
|---|---|---|
| NC_002645.1;NP_073550.1;HCoV-229E_1a;1066 | 0.05 | 0.025 |
| NC_002645.1;NP_073550.1;HCoV-229E_1a;1067 | 0.1 | 0.075 |
| NC_002645.1;NP_073550.1;HCoV-229E_1a;1068 | 0 | 0 |
| NC_002645.1;NP_073550.1;HCoV-229E_1a;1069 | 0 | 0.025 |
| NC_002645.1;NP_073550.1;HCoV-229E_1a;1070 | 0 | 0 |
| NC_002645.1;NP_073550.1;HCoV-229E_1a;1071 | 0 | 0 |
| NC_002645.1;NP_073550.1;HCoV-229E_1a;1072 | 0 | 0 |
| NC_002645.1;NP_073550.1;HCoV-229E_1a;1073 | 0 | 0 |
| NC_002645.1;NP_073550.1;HCoV-229E_1a;1074 | 0 | 0 |
| NC_002645.1;NP_073550.1;HCoV-229E_1a;1075 | 0 | 0 |
| NC_002645.1;NP_073550.1;HCoV-229E_1a;1076 | 0 | 0 |
| NC_002645.1;NP_073550.1;HCoV-229E_1a;1077 | 0 | 0 |
| NC_002645.1;NP_073550.1;HCoV-229E_1a;1078 | 0 | 0 |
| NC_002645.1;NP_073550.1;HCoV-229E_1a;1079 | 0 | 0 |
| NC_002645.1;NP_073550.1;HCoV-229E_1a;1080 | 0 | 0 |
| NC_002645.1;NP_073550.1;HCoV-229E_1a;1081 | 0 | 0 |
| NC_002645.1;NP_073550.1;HCoV-229E_1a;1082 | 0 | 0 |
| NC_002645.1;NP_073550.1;HCoV-229E_1a;1083 | 0 | 0 |
| NC_002645.1;NP_073550.1;HCoV-229E_1a;1084 | 0 | 0 |
| NC_002645.1;NP_073550.1;HCoV-229E_1a;1085 | 0 | 0 |
| NC_002645.1;NP_073550.1;HCoV-229E_1a;1086 | 0 | 0 |
| NC_002645.1;NP_073550.1;HCoV-229E_1a;1087 | 0 | 0 |
| NC_002645.1;NP_073550.1;HCoV-229E_1a;1088 | 0.05 | 0 |
| NC_002645.1;NP_073550.1;HCoV-229E_1a;1089 | 0 | 0 |
| NC_002645.1;NP_073550.1;HCoV-229E_1a;1090 | 0 | 0 |
| NC_002645.1;NP_073550.1;HCoV-229E_1a;1091 | 0 | 0.025 |
| NC_002645.1;NP_073550.1;HCoV-229E_1a;1092 | 0 | 0 |
| NC_002645.1;NP_073550.1;HCoV-229E_1a;1093 | 0 | 0 |
| NC_002645.1;NP_073550.1;HCoV-229E_1a;1094 | 0 | 0 |
| NC_002645.1;NP_073550.1;HCoV-229E_1a;1095 | 0 | 0 |
| NC_002645.1;NP_073550.1;HCoV-229E_1a;1096 | 0 | 0 |
| NC_002645.1;NP_073550.1;HCoV-229E_1a;1097 | 0 | 0 |
| NC_002645.1;NP_073550.1;HCoV-229E_1a;1098 | 0 | 0 |
| NC_002645.1;NP_073550.1;HCoV-229E_1a;1099 | 0 | 0 |
| NC_002645.1;NP_073550.1;HCoV-229E_1a;1100 | 0 | 0 |
| NC_002645.1;NP_073550.1;HCoV-229E_1a;1101 | 0 | 0 |
| NC_002645.1;NP_073550.1;HCoV-229E_1a;1102 | 0 | 0 |
| NC_002645.1;NP_073550.1;HCoV-229E_1a;1103 | 0 | 0 |
| NC_002645.1;NP_073550.1;HCoV-229E_1a;1104 | 0 | 0 |
| NC_002645.1;NP_073550.1;HCoV-229E_1a;1105 | 0 | 0 |
| NC_002645.1;NP_073550.1;HCoV-229E_1a;1106 | 0 | 0 |
| NC_002645.1;NP_073550.1;HCoV-229E_1a;1107 | 0 | 0 |
| NC_002645.1;NP_073550.1;HCoV-229E_1a;1108 | 0 | 0 |

| | | |
|---|---|---|
| NC_002645.1;NP_073550.1;HCoV-229E_1a;1109 | 0 | 0 |
| NC_002645.1;NP_073550.1;HCoV-229E_1a;1110 | 0.05 | 0 |
| NC_002645.1;NP_073550.1;HCoV-229E_1a;1111 | 0 | 0 |
| NC_002645.1;NP_073550.1;HCoV-229E_1a;1112 | 0.05 | 0 |
| NC_002645.1;NP_073550.1;HCoV-229E_1a;1113 | 0 | 0 |
| NC_002645.1;NP_073550.1;HCoV-229E_1a;1114 | 0 | 0 |
| NC_002645.1;NP_073550.1;HCoV-229E_1a;1115 | 0 | 0 |
| NC_002645.1;NP_073550.1;HCoV-229E_1a;1116 | 0 | 0 |
| NC_002645.1;NP_073550.1;HCoV-229E_1a;1117 | 0 | 0 |
| NC_002645.1;NP_073550.1;HCoV-229E_1a;1118 | 0 | 0 |
| NC_002645.1;NP_073550.1;HCoV-229E_1a;1119 | 0 | 0 |
| NC_002645.1;NP_073550.1;HCoV-229E_1a;1120 | 0 | 0 |
| NC_002645.1;NP_073550.1;HCoV-229E_1a;1121 | 0 | 0 |
| NC_002645.1;NP_073550.1;HCoV-229E_1a;1122 | 0 | 0 |
| NC_002645.1;NP_073550.1;HCoV-229E_1a;1123 | 0 | 0 |
| NC_002645.1;NP_073550.1;HCoV-229E_1a;1124 | 0 | 0 |
| NC_002645.1;NP_073550.1;HCoV-229E_1a;1125 | 0 | 0 |
| NC_002645.1;NP_073550.1;HCoV-229E_1a;1126 | 0 | 0 |
| NC_002645.1;NP_073550.1;HCoV-229E_1a;1127 | 0 | 0 |
| NC_002645.1;NP_073550.1;HCoV-229E_1a;1128 | 0 | 0 |
| NC_002645.1;NP_073550.1;HCoV-229E_1a;1129 | 0 | 0 |
| NC_002645.1;NP_073550.1;HCoV-229E_1a;1130 | 0 | 0 |
| NC_002645.1;NP_073550.1;HCoV-229E_1a;1131 | 0 | 0 |
| NC_002645.1;NP_073550.1;HCoV-229E_1a;1132 | 0 | 0 |
| NC_002645.1;NP_073550.1;HCoV-229E_1a;1133 | 0 | 0 |
| NC_002645.1;NP_073550.1;HCoV-229E_1a;1134 | 0 | 0.025 |
| NC_002645.1;NP_073550.1;HCoV-229E_1a;1135 | 0 | 0 |
| NC_002645.1;NP_073550.1;HCoV-229E_1a;1136 | 0 | 0.025 |
| NC_002645.1;NP_073550.1;HCoV-229E_1a;1137 | 0 | 0.025 |
| NC_002645.1;NP_073550.1;HCoV-229E_1a;1138 | 0 | 0 |
| NC_002645.1;NP_073550.1;HCoV-229E_1a;1139 | 0 | 0 |
| NC_002645.1;NP_073550.1;HCoV-229E_1a;1140 | 0 | 0 |
| NC_002645.1;NP_073550.1;HCoV-229E_1a;1141 | 0 | 0.025 |
| NC_002645.1;NP_073550.1;HCoV-229E_1a;1142 | 0 | 0.025 |
| NC_002645.1;NP_073550.1;HCoV-229E_1a;1143 | 0 | 0.05 |
| NC_002645.1;NP_073550.1;HCoV-229E_1a;1144 | 0 | 0 |
| NC_002645.1;NP_073550.1;HCoV-229E_1a;1145 | 0 | 0.025 |
| NC_002645.1;NP_073550.1;HCoV-229E_1a;1146 | 0 | 0 |
| NC_002645.1;NP_073550.1;HCoV-229E_1a;1147 | 0 | 0 |
| NC_002645.1;NP_073550.1;HCoV-229E_1a;1148 | 0 | 0 |
| NC_002645.1;NP_073550.1;HCoV-229E_1a;1149 | 0.05 | 0 |
| NC_002645.1;NP_073550.1;HCoV-229E_1a;1150 | 0 | 0 |
| NC_002645.1;NP_073550.1;HCoV-229E_1a;1151 | 0 | 0 |

| | | |
|---|---|---|
| NC_002645.1;NP_073550.1;HCoV-229E_1a;1152 | 0 | 0 |
| NC_002645.1;NP_073550.1;HCoV-229E_1a;1153 | 0 | 0 |
| NC_002645.1;NP_073550.1;HCoV-229E_1a;1154 | 0 | 0 |
| NC_002645.1;NP_073550.1;HCoV-229E_1a;1155 | 0 | 0 |
| NC_002645.1;NP_073550.1;HCoV-229E_1a;1156 | 0 | 0 |
| NC_002645.1;NP_073550.1;HCoV-229E_1a;1157 | 0 | 0 |
| NC_002645.1;NP_073550.1;HCoV-229E_1a;1158 | 0 | 0 |
| NC_002645.1;NP_073550.1;HCoV-229E_1a;1159 | 0 | 0 |
| NC_002645.1;NP_073550.1;HCoV-229E_1a;1160 | 0 | 0 |
| NC_002645.1;NP_073550.1;HCoV-229E_1a;1161 | 0 | 0 |
| NC_002645.1;NP_073550.1;HCoV-229E_1a;1162 | 0 | 0 |
| NC_002645.1;NP_073550.1;HCoV-229E_1a;1163 | 0 | 0 |
| NC_002645.1;NP_073550.1;HCoV-229E_1a;1164 | 0 | 0 |
| NC_002645.1;NP_073550.1;HCoV-229E_1a;1165 | 0 | 0 |
| NC_002645.1;NP_073550.1;HCoV-229E_1a;1166 | 0 | 0 |
| NC_002645.1;NP_073550.1;HCoV-229E_1a;1167 | 0 | 0 |
| NC_002645.1;NP_073550.1;HCoV-229E_1a;1168 | 0 | 0 |
| NC_002645.1;NP_073550.1;HCoV-229E_1a;1169 | 0 | 0 |
| NC_002645.1;NP_073550.1;HCoV-229E_1a;1170 | 0 | 0 |
| NC_002645.1;NP_073550.1;HCoV-229E_1a;1171 | 0 | 0.025 |
| NC_002645.1;NP_073550.1;HCoV-229E_1a;1172 | 0 | 0.025 |
| NC_002645.1;NP_073550.1;HCoV-229E_1a;1173 | 0.1 | 0.025 |
| NC_002645.1;NP_073550.1;HCoV-229E_1a;1174 | 0.05 | 0.025 |
| NC_002645.1;NP_073550.1;HCoV-229E_1a;1175 | 0 | 0 |
| NC_002645.1;NP_073550.1;HCoV-229E_1a;1176 | 0 | 0 |
| NC_002645.1;NP_073550.1;HCoV-229E_1a;1177 | 0 | 0 |
| NC_002645.1;NP_073550.1;HCoV-229E_1a;1178 | 0 | 0 |
| NC_002645.1;NP_073550.1;HCoV-229E_1a;1179 | 0 | 0 |
| NC_002645.1;NP_073550.1;HCoV-229E_1a;1180 | 0 | 0 |
| NC_002645.1;NP_073550.1;HCoV-229E_1a;1181 | 0 | 0 |
| NC_002645.1;NP_073550.1;HCoV-229E_1a;1182 | 0 | 0 |
| NC_002645.1;NP_073550.1;HCoV-229E_1a;1183 | 0.05 | 0.025 |
| NC_002645.1;NP_073550.1;HCoV-229E_1a;1184 | 0 | 0 |
| NC_002645.1;NP_073550.1;HCoV-229E_1a;1185 | 0 | 0 |
| NC_002645.1;NP_073550.1;HCoV-229E_1a;1186 | 0 | 0 |
| NC_002645.1;NP_073550.1;HCoV-229E_1a;1187 | 0 | 0 |
| NC_002645.1;NP_073550.1;HCoV-229E_1a;1188 | 0 | 0 |
| NC_002645.1;NP_073550.1;HCoV-229E_1a;1189 | 0 | 0 |
| NC_002645.1;NP_073550.1;HCoV-229E_1a;1190 | 0 | 0 |
| NC_002645.1;NP_073550.1;HCoV-229E_1a;1191 | 0 | 0 |
| NC_002645.1;NP_073550.1;HCoV-229E_1a;1192 | 0 | 0 |
| NC_002645.1;NP_073550.1;HCoV-229E_1a;1193 | 0 | 0 |
| NC_002645.1;NP_073550.1;HCoV-229E_1a;1194 | 0 | 0 |

| | | |
|---|---|---|
| NC_002645.1;NP_073550.1;HCoV-229E_1a;1195 | 0 | 0 |
| NC_002645.1;NP_073550.1;HCoV-229E_1a;1196 | 0 | 0 |
| NC_002645.1;NP_073550.1;HCoV-229E_1a;1197 | 0 | 0 |
| NC_002645.1;NP_073550.1;HCoV-229E_1a;1198 | 0 | 0 |
| NC_002645.1;NP_073550.1;HCoV-229E_1a;1199 | 0 | 0.025 |
| NC_002645.1;NP_073550.1;HCoV-229E_1a;1200 | 0 | 0.025 |
| NC_002645.1;NP_073550.1;HCoV-229E_1a;1201 | 0 | 0.025 |
| NC_002645.1;NP_073550.1;HCoV-229E_1a;1202 | 0 | 0.025 |
| NC_002645.1;NP_073550.1;HCoV-229E_1a;1203 | 0 | 0.025 |
| NC_002645.1;NP_073550.1;HCoV-229E_1a;1204 | 0 | 0.025 |
| NC_002645.1;NP_073550.1;HCoV-229E_1a;1205 | 0 | 0.025 |
| NC_002645.1;NP_073550.1;HCoV-229E_1a;1206 | 0 | 0.025 |
| NC_002645.1;NP_073550.1;HCoV-229E_1a;1207 | 0 | 0.025 |
| NC_002645.1;NP_073550.1;HCoV-229E_1a;1208 | 0.05 | 0.025 |
| NC_002645.1;NP_073550.1;HCoV-229E_1a;1209 | 0 | 0.05 |
| NC_002645.1;NP_073550.1;HCoV-229E_1a;1210 | 0 | 0.025 |
| NC_002645.1;NP_073550.1;HCoV-229E_1a;1211 | 0 | 0.025 |
| NC_002645.1;NP_073550.1;HCoV-229E_1a;1212 | 0 | 0 |
| NC_002645.1;NP_073550.1;HCoV-229E_1a;1213 | 0 | 0.025 |
| NC_002645.1;NP_073550.1;HCoV-229E_1a;1214 | 0 | 0.05 |
| NC_002645.1;NP_073550.1;HCoV-229E_1a;1215 | 0.05 | 0.05 |
| NC_002645.1;NP_073550.1;HCoV-229E_1a;1216 | 0.05 | 0.1 |
| NC_002645.1;NP_073550.1;HCoV-229E_1a;1217 | 0.05 | 0.15 |
| NC_002645.1;NP_073550.1;HCoV-229E_1a;1218 | 0.15 | 0.075 |
| NC_002645.1;NP_073550.1;HCoV-229E_1a;1219 | 0.05 | 0.1 |
| NC_002645.1;NP_073550.1;HCoV-229E_1a;1220 | 0.1 | 0.075 |
| NC_002645.1;NP_073550.1;HCoV-229E_1a;1221 | 0.05 | 0.075 |
| NC_002645.1;NP_073550.1;HCoV-229E_1a;1222 | 0.1 | 0.075 |
| NC_002645.1;NP_073550.1;HCoV-229E_1a;1223 | 0.05 | 0.025 |
| NC_002645.1;NP_073550.1;HCoV-229E_1a;1224 | 0.05 | 0.05 |
| NC_002645.1;NP_073550.1;HCoV-229E_1a;1225 | 0.05 | 0.025 |
| NC_002645.1;NP_073550.1;HCoV-229E_1a;1226 | 0.05 | 0 |
| NC_002645.1;NP_073550.1;HCoV-229E_1a;1227 | 0.05 | 0.025 |
| NC_002645.1;NP_073550.1;HCoV-229E_1a;1228 | 0 | 0.025 |
| NC_002645.1;NP_073550.1;HCoV-229E_1a;1229 | 0 | 0 |
| NC_002645.1;NP_073550.1;HCoV-229E_1a;1230 | 0 | 0 |
| NC_002645.1;NP_073550.1;HCoV-229E_1a;1231 | 0.05 | 0.1 |
| NC_002645.1;NP_073550.1;HCoV-229E_1a;1232 | 0.1 | 0.15 |
| NC_002645.1;NP_073550.1;HCoV-229E_1a;1233 | 0.05 | 0.1 |
| NC_002645.1;NP_073550.1;HCoV-229E_1a;1234 | 0.1 | 0.1 |
| NC_002645.1;NP_073550.1;HCoV-229E_1a;1235 | 0.05 | 0.1 |
| NC_002645.1;NP_073550.1;HCoV-229E_1a;1236 | 0.05 | 0.05 |
| NC_002645.1;NP_073550.1;HCoV-229E_1a;1237 | 0.05 | 0.05 |

| | | |
|---|---|---|
| NC_002645.1;NP_073550.1;HCoV-229E_1a;1238 | 0.05 | 0.075 |
| NC_002645.1;NP_073550.1;HCoV-229E_1a;1239 | 0.1 | 0.1 |
| NC_002645.1;NP_073550.1;HCoV-229E_1a;1240 | 0.05 | 0.1 |
| NC_002645.1;NP_073550.1;HCoV-229E_1a;1241 | 0.1 | 0 |
| NC_002645.1;NP_073550.1;HCoV-229E_1a;1242 | 0.1 | 0.025 |
| NC_002645.1;NP_073550.1;HCoV-229E_1a;1243 | 0 | 0.025 |
| NC_002645.1;NP_073550.1;HCoV-229E_1a;1244 | 0 | 0.025 |
| NC_002645.1;NP_073550.1;HCoV-229E_1a;1245 | 0 | 0.05 |
| NC_002645.1;NP_073550.1;HCoV-229E_1a;1246 | 0 | 0.05 |
| NC_002645.1;NP_073550.1;HCoV-229E_1a;1247 | 0 | 0.05 |
| NC_002645.1;NP_073550.1;HCoV-229E_1a;1248 | 0 | 0.05 |
| NC_002645.1;NP_073550.1;HCoV-229E_1a;1249 | 0 | 0.05 |
| NC_002645.1;NP_073550.1;HCoV-229E_1a;1250 | 0.05 | 0.05 |
| NC_002645.1;NP_073550.1;HCoV-229E_1a;1251 | 0 | 0.05 |
| NC_002645.1;NP_073550.1;HCoV-229E_1a;1252 | 0 | 0.05 |
| NC_002645.1;NP_073550.1;HCoV-229E_1a;1253 | 0 | 0 |
| NC_002645.1;NP_073550.1;HCoV-229E_1a;1254 | 0 | 0 |
| NC_002645.1;NP_073550.1;HCoV-229E_1a;1255 | 0 | 0 |
| NC_002645.1;NP_073550.1;HCoV-229E_1a;1256 | 0 | 0.025 |
| NC_002645.1;NP_073550.1;HCoV-229E_1a;1257 | 0.05 | 0.025 |
| NC_002645.1;NP_073550.1;HCoV-229E_1a;1258 | 0.1 | 0.125 |
| NC_002645.1;NP_073550.1;HCoV-229E_1a;1259 | 0.1 | 0.175 |
| NC_002645.1;NP_073550.1;HCoV-229E_1a;1260 | 0.2 | 0.275 |
| NC_002645.1;NP_073550.1;HCoV-229E_1a;1261 | 0.15 | 0.175 |
| NC_002645.1;NP_073550.1;HCoV-229E_1a;1262 | 0.05 | 0.1 |
| NC_002645.1;NP_073550.1;HCoV-229E_1a;1263 | 0.05 | 0.125 |
| NC_002645.1;NP_073550.1;HCoV-229E_1a;1264 | 0.1 | 0.05 |
| NC_002645.1;NP_073550.1;HCoV-229E_1a;1265 | 0.05 | 0.075 |
| NC_002645.1;NP_073550.1;HCoV-229E_1a;1266 | 0.05 | 0.025 |
| NC_002645.1;NP_073550.1;HCoV-229E_1a;1267 | 0 | 0 |
| NC_002645.1;NP_073550.1;HCoV-229E_1a;1268 | 0 | 0 |
| NC_002645.1;NP_073550.1;HCoV-229E_1a;1269 | 0 | 0 |
| NC_002645.1;NP_073550.1;HCoV-229E_1a;1270 | 0 | 0 |
| NC_002645.1;NP_073550.1;HCoV-229E_1a;1271 | 0 | 0 |
| NC_002645.1;NP_073550.1;HCoV-229E_1a;1272 | 0.05 | 0 |
| NC_002645.1;NP_073550.1;HCoV-229E_1a;1273 | 0.05 | 0.025 |
| NC_002645.1;NP_073550.1;HCoV-229E_1a;1274 | 0.05 | 0.05 |
| NC_002645.1;NP_073550.1;HCoV-229E_1a;1275 | 0.05 | 0.075 |
| NC_002645.1;NP_073550.1;HCoV-229E_1a;1276 | 0.05 | 0.025 |
| NC_002645.1;NP_073550.1;HCoV-229E_1a;1277 | 0.05 | 0.05 |
| NC_002645.1;NP_073550.1;HCoV-229E_1a;1278 | 0.1 | 0.075 |
| NC_002645.1;NP_073550.1;HCoV-229E_1a;1279 | 0.05 | 0.025 |
| NC_002645.1;NP_073550.1;HCoV-229E_1a;1280 | 0.05 | 0 |

| | | |
|---|---|---|
| NC_002645.1;NP_073550.1;HCoV-229E_1a;1281 | 0.05 | 0.025 |
| NC_002645.1;NP_073550.1;HCoV-229E_1a;1282 | 0.05 | 0.05 |
| NC_002645.1;NP_073550.1;HCoV-229E_1a;1283 | 0.15 | 0.075 |
| NC_002645.1;NP_073550.1;HCoV-229E_1a;1284 | 0.05 | 0.05 |
| NC_002645.1;NP_073550.1;HCoV-229E_1a;1285 | 0.05 | 0 |
| NC_002645.1;NP_073550.1;HCoV-229E_1a;1286 | 0.05 | 0 |
| NC_002645.1;NP_073550.1;HCoV-229E_1a;1287 | 0.05 | 0.025 |
| NC_002645.1;NP_073550.1;HCoV-229E_1a;1288 | 0.1 | 0.125 |
| NC_002645.1;NP_073550.1;HCoV-229E_1a;1289 | 0 | 0 |
| NC_002645.1;NP_073550.1;HCoV-229E_1a;1290 | 0 | 0 |
| NC_002645.1;NP_073550.1;HCoV-229E_1a;1291 | 0 | 0.025 |
| NC_002645.1;NP_073550.1;HCoV-229E_1a;1292 | 0.05 | 0.025 |
| NC_002645.1;NP_073550.1;HCoV-229E_1a;1293 | 0 | 0.025 |
| NC_002645.1;NP_073550.1;HCoV-229E_1a;1294 | 0.3 | 0.225 |
| NC_002645.1;NP_073550.1;HCoV-229E_1a;1295 | 0.05 | 0 |
| NC_002645.1;NP_073550.1;HCoV-229E_1a;1296 | 0.05 | 0 |
| NC_002645.1;NP_073550.1;HCoV-229E_1a;1297 | 0.05 | 0 |
| NC_002645.1;NP_073550.1;HCoV-229E_1a;1298 | 0.1 | 0.05 |
| NC_002645.1;NP_073550.1;HCoV-229E_1a;1299 | 0.05 | 0.05 |
| NC_002645.1;NP_073550.1;HCoV-229E_1a;1300 | 0.05 | 0.025 |
| NC_002645.1;NP_073550.1;HCoV-229E_1a;1301 | 0.05 | 0 |
| NC_002645.1;NP_073550.1;HCoV-229E_1a;1302 | 0.05 | 0 |
| NC_002645.1;NP_073550.1;HCoV-229E_1a;1303 | 0 | 0 |
| NC_002645.1;NP_073550.1;HCoV-229E_1a;1304 | 0 | 0.025 |
| NC_002645.1;NP_073550.1;HCoV-229E_1a;1305 | 0 | 0 |
| NC_002645.1;NP_073550.1;HCoV-229E_1a;1306 | 0 | 0 |
| NC_002645.1;NP_073550.1;HCoV-229E_1a;1307 | 0 | 0 |
| NC_002645.1;NP_073550.1;HCoV-229E_1a;1308 | 0 | 0.05 |
| NC_002645.1;NP_073550.1;HCoV-229E_1a;1309 | 0 | 0.05 |
| NC_002645.1;NP_073550.1;HCoV-229E_1a;1310 | 0 | 0.05 |
| NC_002645.1;NP_073550.1;HCoV-229E_1a;1311 | 0 | 0.05 |
| NC_002645.1;NP_073550.1;HCoV-229E_1a;1312 | 0 | 0.05 |
| NC_002645.1;NP_073550.1;HCoV-229E_1a;1313 | 0 | 0 |
| NC_002645.1;NP_073550.1;HCoV-229E_1a;1314 | 0 | 0 |
| NC_002645.1;NP_073550.1;HCoV-229E_1a;1315 | 0 | 0 |
| NC_002645.1;NP_073550.1;HCoV-229E_1a;1316 | 0 | 0 |
| NC_002645.1;NP_073550.1;HCoV-229E_1a;1317 | 0 | 0 |
| NC_002645.1;NP_073550.1;HCoV-229E_1a;1318 | 0 | 0 |
| NC_002645.1;NP_073550.1;HCoV-229E_1a;1319 | 0 | 0 |
| NC_002645.1;NP_073550.1;HCoV-229E_1a;1320 | 0 | 0 |
| NC_002645.1;NP_073550.1;HCoV-229E_1a;1321 | 0 | 0 |
| NC_002645.1;NP_073550.1;HCoV-229E_1a;1322 | 0 | 0 |
| NC_002645.1;NP_073550.1;HCoV-229E_1a;1323 | 0 | 0.025 |

| | | |
|---|---|---|
| NC_002645.1;NP_073550.1;HCoV-229E_1a;1324 | 0 | 0 |
| NC_002645.1;NP_073550.1;HCoV-229E_1a;1325 | 0.05 | 0 |
| NC_002645.1;NP_073550.1;HCoV-229E_1a;1326 | 0.05 | 0 |
| NC_002645.1;NP_073550.1;HCoV-229E_1a;1327 | 0 | 0 |
| NC_002645.1;NP_073550.1;HCoV-229E_1a;1328 | 0 | 0 |
| NC_002645.1;NP_073550.1;HCoV-229E_1a;1329 | 0 | 0 |
| NC_002645.1;NP_073550.1;HCoV-229E_1a;1330 | 0 | 0 |
| NC_002645.1;NP_073550.1;HCoV-229E_1a;1331 | 0 | 0 |
| NC_002645.1;NP_073550.1;HCoV-229E_1a;1332 | 0 | 0 |
| NC_002645.1;NP_073550.1;HCoV-229E_1a;1333 | 0 | 0.025 |
| NC_002645.1;NP_073550.1;HCoV-229E_1a;1334 | 0 | 0.025 |
| NC_002645.1;NP_073550.1;HCoV-229E_1a;1335 | 0 | 0.025 |
| NC_002645.1;NP_073550.1;HCoV-229E_1a;1336 | 0 | 0.025 |
| NC_002645.1;NP_073550.1;HCoV-229E_1a;1337 | 0 | 0.025 |
| NC_002645.1;NP_073550.1;HCoV-229E_1a;1338 | 0 | 0.025 |
| NC_002645.1;NP_073550.1;HCoV-229E_1a;1339 | 0 | 0.025 |
| NC_002645.1;NP_073550.1;HCoV-229E_1a;1340 | 0 | 0.025 |
| NC_002645.1;NP_073550.1;HCoV-229E_1a;1341 | 0 | 0.025 |
| NC_002645.1;NP_073550.1;HCoV-229E_1a;1342 | 0 | 0 |
| NC_002645.1;NP_073550.1;HCoV-229E_1a;1343 | 0 | 0 |
| NC_002645.1;NP_073550.1;HCoV-229E_1a;1344 | 0 | 0 |
| NC_002645.1;NP_073550.1;HCoV-229E_1a;1345 | 0 | 0 |
| NC_002645.1;NP_073550.1;HCoV-229E_1a;1346 | 0 | 0 |
| NC_002645.1;NP_073550.1;HCoV-229E_1a;1347 | 0 | 0 |
| NC_002645.1;NP_073550.1;HCoV-229E_1a;1348 | 0 | 0 |
| NC_002645.1;NP_073550.1;HCoV-229E_1a;1349 | 0 | 0 |
| NC_002645.1;NP_073550.1;HCoV-229E_1a;1350 | 0 | 0 |
| NC_002645.1;NP_073550.1;HCoV-229E_1a;1351 | 0 | 0 |
| NC_002645.1;NP_073550.1;HCoV-229E_1a;1352 | 0 | 0 |
| NC_002645.1;NP_073550.1;HCoV-229E_1a;1353 | 0 | 0 |
| NC_002645.1;NP_073550.1;HCoV-229E_1a;1354 | 0 | 0 |
| NC_002645.1;NP_073550.1;HCoV-229E_1a;1355 | 0 | 0 |
| NC_002645.1;NP_073550.1;HCoV-229E_1a;1356 | 0 | 0 |
| NC_002645.1;NP_073550.1;HCoV-229E_1a;1357 | 0 | 0 |
| NC_002645.1;NP_073550.1;HCoV-229E_1a;1358 | 0 | 0 |
| NC_002645.1;NP_073550.1;HCoV-229E_1a;1359 | 0 | 0 |
| NC_002645.1;NP_073550.1;HCoV-229E_1a;1360 | 0 | 0 |
| NC_002645.1;NP_073550.1;HCoV-229E_1a;1361 | 0 | 0 |
| NC_002645.1;NP_073550.1;HCoV-229E_1a;1362 | 0 | 0.025 |
| NC_002645.1;NP_073550.1;HCoV-229E_1a;1363 | 0 | 0.05 |
| NC_002645.1;NP_073550.1;HCoV-229E_1a;1364 | 0 | 0.05 |
| NC_002645.1;NP_073550.1;HCoV-229E_1a;1365 | 0.1 | 0.05 |
| NC_002645.1;NP_073550.1;HCoV-229E_1a;1366 | 0.25 | 0.1 |

| | | |
|---|---|---|
| NC_002645.1;NP_073550.1;HCoV-229E_1a;1367 | 0.45 | 0.125 |
| NC_002645.1;NP_073550.1;HCoV-229E_1a;1368 | 0.3 | 0.125 |
| NC_002645.1;NP_073550.1;HCoV-229E_1a;1369 | 0.25 | 0.15 |
| NC_002645.1;NP_073550.1;HCoV-229E_1a;1370 | 0.25 | 0.175 |
| NC_002645.1;NP_073550.1;HCoV-229E_1a;1371 | 0.45 | 0.25 |
| NC_002645.1;NP_073550.1;HCoV-229E_1a;1372 | 0.55 | 0.275 |
| NC_002645.1;NP_073550.1;HCoV-229E_1a;1373 | 0.5 | 0.25 |
| NC_002645.1;NP_073550.1;HCoV-229E_1a;1374 | 0.2 | 0.1 |
| NC_002645.1;NP_073550.1;HCoV-229E_1a;1375 | 0.15 | 0 |
| NC_002645.1;NP_073550.1;HCoV-229E_1a;1376 | 0 | 0 |
| NC_002645.1;NP_073550.1;HCoV-229E_1a;1377 | 0 | 0 |
| NC_002645.1;NP_073550.1;HCoV-229E_1a;1378 | 0 | 0 |
| NC_002645.1;NP_073550.1;HCoV-229E_1a;1379 | 0 | 0 |
| NC_002645.1;NP_073550.1;HCoV-229E_1a;1380 | 0 | 0 |
| NC_002645.1;NP_073550.1;HCoV-229E_1a;1381 | 0 | 0 |
| NC_002645.1;NP_073550.1;HCoV-229E_1a;1382 | 0 | 0 |
| NC_002645.1;NP_073550.1;HCoV-229E_1a;1383 | 0 | 0 |
| NC_002645.1;NP_073550.1;HCoV-229E_1a;1384 | 0 | 0 |
| NC_002645.1;NP_073550.1;HCoV-229E_1a;1385 | 0 | 0.025 |
| NC_002645.1;NP_073550.1;HCoV-229E_1a;1386 | 0 | 0.025 |
| NC_002645.1;NP_073550.1;HCoV-229E_1a;1387 | 0 | 0.025 |
| NC_002645.1;NP_073550.1;HCoV-229E_1a;1388 | 0 | 0.025 |
| NC_002645.1;NP_073550.1;HCoV-229E_1a;1389 | 0 | 0.025 |
| NC_002645.1;NP_073550.1;HCoV-229E_1a;1390 | 0 | 0.025 |
| NC_002645.1;NP_073550.1;HCoV-229E_1a;1391 | 0 | 0 |
| NC_002645.1;NP_073550.1;HCoV-229E_1a;1392 | 0 | 0 |
| NC_002645.1;NP_073550.1;HCoV-229E_1a;1393 | 0 | 0 |
| NC_002645.1;NP_073550.1;HCoV-229E_1a;1394 | 0 | 0 |
| NC_002645.1;NP_073550.1;HCoV-229E_1a;1395 | 0 | 0 |
| NC_002645.1;NP_073550.1;HCoV-229E_1a;1396 | 0 | 0 |
| NC_002645.1;NP_073550.1;HCoV-229E_1a;1397 | 0 | 0 |
| NC_002645.1;NP_073550.1;HCoV-229E_1a;1398 | 0 | 0 |
| NC_002645.1;NP_073550.1;HCoV-229E_1a;1399 | 0 | 0 |
| NC_002645.1;NP_073550.1;HCoV-229E_1a;1400 | 0 | 0 |
| NC_002645.1;NP_073550.1;HCoV-229E_1a;1401 | 0 | 0 |
| NC_002645.1;NP_073550.1;HCoV-229E_1a;1402 | 0 | 0.05 |
| NC_002645.1;NP_073550.1;HCoV-229E_1a;1403 | 0 | 0.025 |
| NC_002645.1;NP_073550.1;HCoV-229E_1a;1404 | 0 | 0.025 |
| NC_002645.1;NP_073550.1;HCoV-229E_1a;1405 | 0 | 0 |
| NC_002645.1;NP_073550.1;HCoV-229E_1a;1406 | 0 | 0 |
| NC_002645.1;NP_073550.1;HCoV-229E_1a;1407 | 0 | 0 |
| NC_002645.1;NP_073550.1;HCoV-229E_1a;1408 | 0 | 0 |
| NC_002645.1;NP_073550.1;HCoV-229E_1a;1409 | 0 | 0 |

| | | |
|---|---|---|
| NC_002645.1;NP_073550.1;HCoV-229E_1a;1410 | 0 | 0 |
| NC_002645.1;NP_073550.1;HCoV-229E_1a;1411 | 0 | 0 |
| NC_002645.1;NP_073550.1;HCoV-229E_1a;1412 | 0 | 0 |
| NC_002645.1;NP_073550.1;HCoV-229E_1a;1413 | 0 | 0 |
| NC_002645.1;NP_073550.1;HCoV-229E_1a;1414 | 0 | 0 |
| NC_002645.1;NP_073550.1;HCoV-229E_1a;1415 | 0 | 0 |
| NC_002645.1;NP_073550.1;HCoV-229E_1a;1416 | 0 | 0 |
| NC_002645.1;NP_073550.1;HCoV-229E_1a;1417 | 0 | 0 |
| NC_002645.1;NP_073550.1;HCoV-229E_1a;1418 | 0 | 0 |
| NC_002645.1;NP_073550.1;HCoV-229E_1a;1419 | 0 | 0.025 |
| NC_002645.1;NP_073550.1;HCoV-229E_1a;1420 | 0 | 0 |
| NC_002645.1;NP_073550.1;HCoV-229E_1a;1421 | 0 | 0 |
| NC_002645.1;NP_073550.1;HCoV-229E_1a;1422 | 0 | 0 |
| NC_002645.1;NP_073550.1;HCoV-229E_1a;1423 | 0 | 0 |
| NC_002645.1;NP_073550.1;HCoV-229E_1a;1424 | 0 | 0 |
| NC_002645.1;NP_073550.1;HCoV-229E_1a;1425 | 0 | 0 |
| NC_002645.1;NP_073550.1;HCoV-229E_1a;1426 | 0 | 0 |
| NC_002645.1;NP_073550.1;HCoV-229E_1a;1427 | 0 | 0 |
| NC_002645.1;NP_073550.1;HCoV-229E_1a;1428 | 0 | 0.05 |
| NC_002645.1;NP_073550.1;HCoV-229E_1a;1429 | 0.05 | 0 |
| NC_002645.1;NP_073550.1;HCoV-229E_1a;1430 | 0 | 0 |
| NC_002645.1;NP_073550.1;HCoV-229E_1a;1431 | 0 | 0 |
| NC_002645.1;NP_073550.1;HCoV-229E_1a;1432 | 0.2 | 0.075 |
| NC_002645.1;NP_073550.1;HCoV-229E_1a;1433 | 0.2 | 0.075 |
| NC_002645.1;NP_073550.1;HCoV-229E_1a;1434 | 0.05 | 0.05 |
| NC_002645.1;NP_073550.1;HCoV-229E_1a;1435 | 0.1 | 0.05 |
| NC_002645.1;NP_073550.1;HCoV-229E_1a;1436 | 0.15 | 0.125 |
| NC_002645.1;NP_073550.1;HCoV-229E_1a;1437 | 0.2 | 0.125 |
| NC_002645.1;NP_073550.1;HCoV-229E_1a;1438 | 0.2 | 0.075 |
| NC_002645.1;NP_073550.1;HCoV-229E_1a;1439 | 0.2 | 0.125 |
| NC_002645.1;NP_073550.1;HCoV-229E_1a;1440 | 0.15 | 0.075 |
| NC_002645.1;NP_073550.1;HCoV-229E_1a;1441 | 0.15 | 0.1 |
| NC_002645.1;NP_073550.1;HCoV-229E_1a;1442 | 0.05 | 0.025 |
| NC_002645.1;NP_073550.1;HCoV-229E_1a;1443 | 0.05 | 0.025 |
| NC_002645.1;NP_073550.1;HCoV-229E_1a;1444 | 0.05 | 0 |
| NC_002645.1;NP_073550.1;HCoV-229E_1a;1445 | 0.05 | 0 |
| NC_002645.1;NP_073550.1;HCoV-229E_1a;1446 | 0 | 0 |
| NC_002645.1;NP_073550.1;HCoV-229E_1a;1447 | 0 | 0 |
| NC_002645.1;NP_073550.1;HCoV-229E_1a;1448 | 0 | 0 |
| NC_002645.1;NP_073550.1;HCoV-229E_1a;1449 | 0 | 0 |
| NC_002645.1;NP_073550.1;HCoV-229E_1a;1450 | 0 | 0 |
| NC_002645.1;NP_073550.1;HCoV-229E_1a;1451 | 0 | 0.025 |
| NC_002645.1;NP_073550.1;HCoV-229E_1a;1452 | 0 | 0.025 |

| | | |
|---|---|---|
| NC_002645.1;NP_073550.1;HCoV-229E_1a;1453 | 0 | 0.025 |
| NC_002645.1;NP_073550.1;HCoV-229E_1a;1454 | 0 | 0 |
| NC_002645.1;NP_073550.1;HCoV-229E_1a;1455 | 0 | 0 |
| NC_002645.1;NP_073550.1;HCoV-229E_1a;1456 | 0 | 0 |
| NC_002645.1;NP_073550.1;HCoV-229E_1a;1457 | 0 | 0 |
| NC_002645.1;NP_073550.1;HCoV-229E_1a;1458 | 0 | 0 |
| NC_002645.1;NP_073550.1;HCoV-229E_1a;1459 | 0 | 0 |
| NC_002645.1;NP_073550.1;HCoV-229E_1a;1460 | 0 | 0 |
| NC_002645.1;NP_073550.1;HCoV-229E_1a;1461 | 0 | 0 |
| NC_002645.1;NP_073550.1;HCoV-229E_1a;1462 | 0 | 0 |
| NC_002645.1;NP_073550.1;HCoV-229E_1a;1463 | 0 | 0 |
| NC_002645.1;NP_073550.1;HCoV-229E_1a;1464 | 0 | 0 |
| NC_002645.1;NP_073550.1;HCoV-229E_1a;1465 | 0 | 0 |
| NC_002645.1;NP_073550.1;HCoV-229E_1a;1466 | 0 | 0 |
| NC_002645.1;NP_073550.1;HCoV-229E_1a;1467 | 0.05 | 0 |
| NC_002645.1;NP_073550.1;HCoV-229E_1a;1468 | 0.05 | 0.025 |
| NC_002645.1;NP_073550.1;HCoV-229E_1a;1469 | 0.05 | 0 |
| NC_002645.1;NP_073550.1;HCoV-229E_1a;1470 | 0 | 0 |
| NC_002645.1;NP_073550.1;HCoV-229E_1a;1471 | 0 | 0 |
| NC_002645.1;NP_073550.1;HCoV-229E_1a;1472 | 0 | 0 |
| NC_002645.1;NP_073550.1;HCoV-229E_1a;1473 | 0 | 0 |
| NC_002645.1;NP_073550.1;HCoV-229E_1a;1474 | 0 | 0 |
| NC_002645.1;NP_073550.1;HCoV-229E_1a;1475 | 0 | 0 |
| NC_002645.1;NP_073550.1;HCoV-229E_1a;1476 | 0 | 0 |
| NC_002645.1;NP_073550.1;HCoV-229E_1a;1477 | 0 | 0 |
| NC_002645.1;NP_073550.1;HCoV-229E_1a;1478 | 0 | 0 |
| NC_002645.1;NP_073550.1;HCoV-229E_1a;1479 | 0 | 0 |
| NC_002645.1;NP_073550.1;HCoV-229E_1a;1480 | 0 | 0 |
| NC_002645.1;NP_073550.1;HCoV-229E_1a;1481 | 0 | 0 |
| NC_002645.1;NP_073550.1;HCoV-229E_1a;1482 | 0 | 0 |
| NC_002645.1;NP_073550.1;HCoV-229E_1a;1483 | 0 | 0 |
| NC_002645.1;NP_073550.1;HCoV-229E_1a;1484 | 0 | 0 |
| NC_002645.1;NP_073550.1;HCoV-229E_1a;1485 | 0 | 0 |
| NC_002645.1;NP_073550.1;HCoV-229E_1a;1486 | 0 | 0 |
| NC_002645.1;NP_073550.1;HCoV-229E_1a;1487 | 0 | 0.025 |
| NC_002645.1;NP_073550.1;HCoV-229E_1a;1488 | 0 | 0 |
| NC_002645.1;NP_073550.1;HCoV-229E_1a;1489 | 0 | 0 |
| NC_002645.1;NP_073550.1;HCoV-229E_1a;1490 | 0.05 | 0.05 |
| NC_002645.1;NP_073550.1;HCoV-229E_1a;1491 | 0 | 0.025 |
| NC_002645.1;NP_073550.1;HCoV-229E_1a;1492 | 0.05 | 0.025 |
| NC_002645.1;NP_073550.1;HCoV-229E_1a;1493 | 0.05 | 0 |
| NC_002645.1;NP_073550.1;HCoV-229E_1a;1494 | 0.05 | 0 |
| NC_002645.1;NP_073550.1;HCoV-229E_1a;1495 | 0.05 | 0 |

| | | |
|---|---|---|
| NC_002645.1;NP_073550.1;HCoV-229E_1a;1496 | 0.05 | 0.025 |
| NC_002645.1;NP_073550.1;HCoV-229E_1a;1497 | 0.05 | 0.025 |
| NC_002645.1;NP_073550.1;HCoV-229E_1a;1498 | 0 | 0 |
| NC_002645.1;NP_073550.1;HCoV-229E_1a;1499 | 0 | 0 |
| NC_002645.1;NP_073550.1;HCoV-229E_1a;1500 | 0 | 0 |
| NC_002645.1;NP_073550.1;HCoV-229E_1a;1501 | 0 | 0.025 |
| NC_002645.1;NP_073550.1;HCoV-229E_1a;1502 | 0 | 0 |
| NC_002645.1;NP_073550.1;HCoV-229E_1a;1503 | 0 | 0 |
| NC_002645.1;NP_073550.1;HCoV-229E_1a;1504 | 0 | 0 |
| NC_002645.1;NP_073550.1;HCoV-229E_1a;1505 | 0 | 0 |
| NC_002645.1;NP_073550.1;HCoV-229E_1a;1506 | 0.1 | 0.15 |
| NC_002645.1;NP_073550.1;HCoV-229E_1a;1507 | 0 | 0.025 |
| NC_002645.1;NP_073550.1;HCoV-229E_1a;1508 | 0 | 0 |
| NC_002645.1;NP_073550.1;HCoV-229E_1a;1509 | 0 | 0.025 |
| NC_002645.1;NP_073550.1;HCoV-229E_1a;1510 | 0.05 | 0 |
| NC_002645.1;NP_073550.1;HCoV-229E_1a;1511 | 0.05 | 0.05 |
| NC_002645.1;NP_073550.1;HCoV-229E_1a;1512 | 0 | 0.025 |
| NC_002645.1;NP_073550.1;HCoV-229E_1a;1513 | 0 | 0.025 |
| NC_002645.1;NP_073550.1;HCoV-229E_1a;1514 | 0.1 | 0.05 |
| NC_002645.1;NP_073550.1;HCoV-229E_1a;1515 | 0 | 0.025 |
| NC_002645.1;NP_073550.1;HCoV-229E_1a;1516 | 0.05 | 0.05 |
| NC_002645.1;NP_073550.1;HCoV-229E_1a;1517 | 0 | 0.05 |
| NC_002645.1;NP_073550.1;HCoV-229E_1a;1518 | 0 | 0.05 |
| NC_002645.1;NP_073550.1;HCoV-229E_1a;1519 | 0.05 | 0.075 |
| NC_002645.1;NP_073550.1;HCoV-229E_1a;1520 | 0 | 0.025 |
| NC_002645.1;NP_073550.1;HCoV-229E_1a;1521 | 0.1 | 0.075 |
| NC_002645.1;NP_073550.1;HCoV-229E_1a;1522 | 0 | 0.025 |
| NC_002645.1;NP_073550.1;HCoV-229E_1a;1523 | 0 | 0 |
| NC_002645.1;NP_073550.1;HCoV-229E_1a;1524 | 0 | 0 |
| NC_002645.1;NP_073550.1;HCoV-229E_1a;1525 | 0 | 0 |
| NC_002645.1;NP_073550.1;HCoV-229E_1a;1526 | 0 | 0 |
| NC_002645.1;NP_073550.1;HCoV-229E_1a;1527 | 0 | 0 |
| NC_002645.1;NP_073550.1;HCoV-229E_1a;1528 | 0 | 0 |
| NC_002645.1;NP_073550.1;HCoV-229E_1a;1529 | 0 | 0 |
| NC_002645.1;NP_073550.1;HCoV-229E_1a;1530 | 0 | 0 |
| NC_002645.1;NP_073550.1;HCoV-229E_1a;1531 | 0 | 0 |
| NC_002645.1;NP_073550.1;HCoV-229E_1a;1532 | 0 | 0 |
| NC_002645.1;NP_073550.1;HCoV-229E_1a;1533 | 0 | 0 |
| NC_002645.1;NP_073550.1;HCoV-229E_1a;1534 | 0 | 0.025 |
| NC_002645.1;NP_073550.1;HCoV-229E_1a;1535 | 0 | 0 |
| NC_002645.1;NP_073550.1;HCoV-229E_1a;1536 | 0 | 0 |
| NC_002645.1;NP_073550.1;HCoV-229E_1a;1537 | 0 | 0 |
| NC_002645.1;NP_073550.1;HCoV-229E_1a;1538 | 0 | 0 |

| | | |
|---|---|---|
| NC_002645.1;NP_073550.1;HCoV-229E_1a;1539 | 0 | 0 |
| NC_002645.1;NP_073550.1;HCoV-229E_1a;1540 | 0 | 0 |
| NC_002645.1;NP_073550.1;HCoV-229E_1a;1541 | 0 | 0 |
| NC_002645.1;NP_073550.1;HCoV-229E_1a;1542 | 0 | 0 |
| NC_002645.1;NP_073550.1;HCoV-229E_1a;1543 | 0 | 0.025 |
| NC_002645.1;NP_073550.1;HCoV-229E_1a;1544 | 0 | 0 |
| NC_002645.1;NP_073550.1;HCoV-229E_1a;1545 | 0 | 0 |
| NC_002645.1;NP_073550.1;HCoV-229E_1a;1546 | 0 | 0 |
| NC_002645.1;NP_073550.1;HCoV-229E_1a;1547 | 0 | 0 |
| NC_002645.1;NP_073550.1;HCoV-229E_1a;1548 | 0 | 0 |
| NC_002645.1;NP_073550.1;HCoV-229E_1a;1549 | 0 | 0 |
| NC_002645.1;NP_073550.1;HCoV-229E_1a;1550 | 0 | 0 |
| NC_002645.1;NP_073550.1;HCoV-229E_1a;1551 | 0 | 0 |
| NC_002645.1;NP_073550.1;HCoV-229E_1a;1552 | 0 | 0 |
| NC_002645.1;NP_073550.1;HCoV-229E_1a;1553 | 0 | 0 |
| NC_002645.1;NP_073550.1;HCoV-229E_1a;1554 | 0 | 0 |
| NC_002645.1;NP_073550.1;HCoV-229E_1a;1555 | 0 | 0 |
| NC_002645.1;NP_073550.1;HCoV-229E_1a;1556 | 0 | 0 |
| NC_002645.1;NP_073550.1;HCoV-229E_1a;1557 | 0 | 0 |
| NC_002645.1;NP_073550.1;HCoV-229E_1a;1558 | 0 | 0 |
| NC_002645.1;NP_073550.1;HCoV-229E_1a;1559 | 0 | 0 |
| NC_002645.1;NP_073550.1;HCoV-229E_1a;1560 | 0 | 0 |
| NC_002645.1;NP_073550.1;HCoV-229E_1a;1561 | 0 | 0 |
| NC_002645.1;NP_073550.1;HCoV-229E_1a;1562 | 0 | 0 |
| NC_002645.1;NP_073550.1;HCoV-229E_1a;1563 | 0 | 0 |
| NC_002645.1;NP_073550.1;HCoV-229E_1a;1564 | 0 | 0 |
| NC_002645.1;NP_073550.1;HCoV-229E_1a;1565 | 0 | 0 |
| NC_002645.1;NP_073550.1;HCoV-229E_1a;1566 | 0 | 0 |
| NC_002645.1;NP_073550.1;HCoV-229E_1a;1567 | 0 | 0 |
| NC_002645.1;NP_073550.1;HCoV-229E_1a;1568 | 0 | 0 |
| NC_002645.1;NP_073550.1;HCoV-229E_1a;1569 | 0 | 0 |
| NC_002645.1;NP_073550.1;HCoV-229E_1a;1570 | 0 | 0 |
| NC_002645.1;NP_073550.1;HCoV-229E_1a;1571 | 0 | 0 |
| NC_002645.1;NP_073550.1;HCoV-229E_1a;1572 | 0 | 0 |
| NC_002645.1;NP_073550.1;HCoV-229E_1a;1573 | 0 | 0 |
| NC_002645.1;NP_073550.1;HCoV-229E_1a;1574 | 0 | 0 |
| NC_002645.1;NP_073550.1;HCoV-229E_1a;1575 | 0 | 0 |
| NC_002645.1;NP_073550.1;HCoV-229E_1a;1576 | 0 | 0 |
| NC_002645.1;NP_073550.1;HCoV-229E_1a;1577 | 0 | 0 |
| NC_002645.1;NP_073550.1;HCoV-229E_1a;1578 | 0 | 0 |
| NC_002645.1;NP_073550.1;HCoV-229E_1a;1579 | 0 | 0 |
| NC_002645.1;NP_073550.1;HCoV-229E_1a;1580 | 0 | 0 |
| NC_002645.1;NP_073550.1;HCoV-229E_1a;1581 | 0 | 0 |

| | | |
|---|---|---|
| NC_002645.1;NP_073550.1;HCoV-229E_1a;1582 | 0 | 0 |
| NC_002645.1;NP_073550.1;HCoV-229E_1a;1583 | 0.05 | 0.025 |
| NC_002645.1;NP_073550.1;HCoV-229E_1a;1584 | 0 | 0 |
| NC_002645.1;NP_073550.1;HCoV-229E_1a;1585 | 0 | 0 |
| NC_002645.1;NP_073550.1;HCoV-229E_1a;1586 | 0.05 | 0.05 |
| NC_002645.1;NP_073550.1;HCoV-229E_1a;1587 | 0.05 | 0.075 |
| NC_002645.1;NP_073550.1;HCoV-229E_1a;1588 | 0.05 | 0.075 |
| NC_002645.1;NP_073550.1;HCoV-229E_1a;1589 | 0.05 | 0.075 |
| NC_002645.1;NP_073550.1;HCoV-229E_1a;1590 | 0 | 0.05 |
| NC_002645.1;NP_073550.1;HCoV-229E_1a;1591 | 0 | 0 |
| NC_002645.1;NP_073550.1;HCoV-229E_1a;1592 | 0 | 0 |
| NC_002645.1;NP_073550.1;HCoV-229E_1a;1593 | 0 | 0 |
| NC_002645.1;NP_073550.1;HCoV-229E_1a;1594 | 0 | 0.025 |
| NC_002645.1;NP_073550.1;HCoV-229E_1a;1595 | 0 | 0 |
| NC_002645.1;NP_073550.1;HCoV-229E_1a;1596 | 0 | 0 |
| NC_002645.1;NP_073550.1;HCoV-229E_1a;1597 | 0 | 0 |
| NC_002645.1;NP_073550.1;HCoV-229E_1a;1598 | 0 | 0 |
| NC_002645.1;NP_073550.1;HCoV-229E_1a;1599 | 0 | 0.025 |
| NC_002645.1;NP_073550.1;HCoV-229E_1a;1600 | 0 | 0 |
| NC_002645.1;NP_073550.1;HCoV-229E_1a;1601 | 0 | 0 |
| NC_002645.1;NP_073550.1;HCoV-229E_1a;1602 | 0 | 0 |
| NC_002645.1;NP_073550.1;HCoV-229E_1a;1603 | 0 | 0.025 |
| NC_002645.1;NP_073550.1;HCoV-229E_1a;1604 | 0 | 0.025 |
| NC_002645.1;NP_073550.1;HCoV-229E_1a;1605 | 0 | 0.025 |
| NC_002645.1;NP_073550.1;HCoV-229E_1a;1606 | 0 | 0.025 |
| NC_002645.1;NP_073550.1;HCoV-229E_1a;1607 | 0.05 | 0 |
| NC_002645.1;NP_073550.1;HCoV-229E_1a;1608 | 0 | 0 |
| NC_002645.1;NP_073550.1;HCoV-229E_1a;1609 | 0 | 0 |
| NC_002645.1;NP_073550.1;HCoV-229E_1a;1610 | 0 | 0 |
| NC_002645.1;NP_073550.1;HCoV-229E_1a;1611 | 0.05 | 0 |
| NC_002645.1;NP_073550.1;HCoV-229E_1a;1612 | 0 | 0.025 |
| NC_002645.1;NP_073550.1;HCoV-229E_1a;1613 | 0 | 0.025 |
| NC_002645.1;NP_073550.1;HCoV-229E_1a;1614 | 0 | 0 |
| NC_002645.1;NP_073550.1;HCoV-229E_1a;1615 | 0 | 0.025 |
| NC_002645.1;NP_073550.1;HCoV-229E_1a;1616 | 0 | 0.025 |
| NC_002645.1;NP_073550.1;HCoV-229E_1a;1617 | 0 | 0 |
| NC_002645.1;NP_073550.1;HCoV-229E_1a;1618 | 0 | 0.05 |
| NC_002645.1;NP_073550.1;HCoV-229E_1a;1619 | 0 | 0 |
| NC_002645.1;NP_073550.1;HCoV-229E_1a;1620 | 0 | 0.075 |
| NC_002645.1;NP_073550.1;HCoV-229E_1a;1621 | 0 | 0.025 |
| NC_002645.1;NP_073550.1;HCoV-229E_1a;1622 | 0 | 0.075 |
| NC_002645.1;NP_073550.1;HCoV-229E_1a;1623 | 0 | 0.075 |
| NC_002645.1;NP_073550.1;HCoV-229E_1a;1624 | 0 | 0.075 |

| | | |
|---|---|---|
| NC_002645.1;NP_073550.1;HCoV-229E_1a;1625 | 0 | 0.075 |
| NC_002645.1;NP_073550.1;HCoV-229E_1a;1626 | 0 | 0.075 |
| NC_002645.1;NP_073550.1;HCoV-229E_1a;1627 | 0.05 | 0.1 |
| NC_002645.1;NP_073550.1;HCoV-229E_1a;1628 | 0 | 0.075 |
| NC_002645.1;NP_073550.1;HCoV-229E_1a;1629 | 0 | 0.075 |
| NC_002645.1;NP_073550.1;HCoV-229E_1a;1630 | 0 | 0.075 |
| NC_002645.1;NP_073550.1;HCoV-229E_1a;1631 | 0.05 | 0.025 |
| NC_002645.1;NP_073550.1;HCoV-229E_1a;1632 | 0.05 | 0.05 |
| NC_002645.1;NP_073550.1;HCoV-229E_1a;1633 | 0.1 | 0.025 |
| NC_002645.1;NP_073550.1;HCoV-229E_1a;1634 | 0.05 | 0.025 |
| NC_002645.1;NP_073550.1;HCoV-229E_1a;1635 | 0.05 | 0.025 |
| NC_002645.1;NP_073550.1;HCoV-229E_1a;1636 | 0 | 0 |
| NC_002645.1;NP_073550.1;HCoV-229E_1a;1637 | 0 | 0 |
| NC_002645.1;NP_073550.1;HCoV-229E_1a;1638 | 0 | 0 |
| NC_002645.1;NP_073550.1;HCoV-229E_1a;1639 | 0 | 0.025 |
| NC_002645.1;NP_073550.1;HCoV-229E_1a;1640 | 0 | 0 |
| NC_002645.1;NP_073550.1;HCoV-229E_1a;1641 | 0 | 0 |
| NC_002645.1;NP_073550.1;HCoV-229E_1a;1642 | 0 | 0 |
| NC_002645.1;NP_073550.1;HCoV-229E_1a;1643 | 0 | 0 |
| NC_002645.1;NP_073550.1;HCoV-229E_1a;1644 | 0 | 0 |
| NC_002645.1;NP_073550.1;HCoV-229E_1a;1645 | 0 | 0.025 |
| NC_002645.1;NP_073550.1;HCoV-229E_1a;1646 | 0 | 0 |
| NC_002645.1;NP_073550.1;HCoV-229E_1a;1647 | 0 | 0 |
| NC_002645.1;NP_073550.1;HCoV-229E_1a;1648 | 0 | 0 |
| NC_002645.1;NP_073550.1;HCoV-229E_1a;1649 | 0 | 0 |
| NC_002645.1;NP_073550.1;HCoV-229E_1a;1650 | 0 | 0 |
| NC_002645.1;NP_073550.1;HCoV-229E_1a;1651 | 0 | 0 |
| NC_002645.1;NP_073550.1;HCoV-229E_1a;1652 | 0 | 0 |
| NC_002645.1;NP_073550.1;HCoV-229E_1a;1653 | 0 | 0 |
| NC_002645.1;NP_073550.1;HCoV-229E_1a;1654 | 0 | 0 |
| NC_002645.1;NP_073550.1;HCoV-229E_1a;1655 | 0 | 0.025 |
| NC_002645.1;NP_073550.1;HCoV-229E_1a;1656 | 0 | 0 |
| NC_002645.1;NP_073550.1;HCoV-229E_1a;1657 | 0 | 0 |
| NC_002645.1;NP_073550.1;HCoV-229E_1a;1658 | 0 | 0 |
| NC_002645.1;NP_073550.1;HCoV-229E_1a;1659 | 0 | 0 |
| NC_002645.1;NP_073550.1;HCoV-229E_1a;1660 | 0 | 0 |
| NC_002645.1;NP_073550.1;HCoV-229E_1a;1661 | 0 | 0 |
| NC_002645.1;NP_073550.1;HCoV-229E_1a;1662 | 0 | 0 |
| NC_002645.1;NP_073550.1;HCoV-229E_1a;1663 | 0 | 0 |
| NC_002645.1;NP_073550.1;HCoV-229E_1a;1664 | 0 | 0 |
| NC_002645.1;NP_073550.1;HCoV-229E_1a;1665 | 0 | 0 |
| NC_002645.1;NP_073550.1;HCoV-229E_1a;1666 | 0 | 0.025 |
| NC_002645.1;NP_073550.1;HCoV-229E_1a;1667 | 0 | 0 |

| | | |
|---|---|---|
| NC_002645.1;NP_073550.1;HCoV-229E_1a;1668 | 0 | 0 |
| NC_002645.1;NP_073550.1;HCoV-229E_1a;1669 | 0 | 0 |
| NC_002645.1;NP_073550.1;HCoV-229E_1a;1670 | 0 | 0 |
| NC_002645.1;NP_073550.1;HCoV-229E_1a;1671 | 0 | 0 |
| NC_002645.1;NP_073550.1;HCoV-229E_1a;1672 | 0 | 0 |
| NC_002645.1;NP_073550.1;HCoV-229E_1a;1673 | 0 | 0 |
| NC_002645.1;NP_073550.1;HCoV-229E_1a;1674 | 0 | 0 |
| NC_002645.1;NP_073550.1;HCoV-229E_1a;1675 | 0 | 0 |
| NC_002645.1;NP_073550.1;HCoV-229E_1a;1676 | 0 | 0 |
| NC_002645.1;NP_073550.1;HCoV-229E_1a;1677 | 0 | 0 |
| NC_002645.1;NP_073550.1;HCoV-229E_1a;1678 | 0 | 0 |
| NC_002645.1;NP_073550.1;HCoV-229E_1a;1679 | 0 | 0 |
| NC_002645.1;NP_073550.1;HCoV-229E_1a;1680 | 0 | 0 |
| NC_002645.1;NP_073550.1;HCoV-229E_1a;1681 | 0 | 0.025 |
| NC_002645.1;NP_073550.1;HCoV-229E_1a;1682 | 0 | 0 |
| NC_002645.1;NP_073550.1;HCoV-229E_1a;1683 | 0 | 0 |
| NC_002645.1;NP_073550.1;HCoV-229E_1a;1684 | 0.05 | 0 |
| NC_002645.1;NP_073550.1;HCoV-229E_1a;1685 | 0 | 0.025 |
| NC_002645.1;NP_073550.1;HCoV-229E_1a;1686 | 0 | 0 |
| NC_002645.1;NP_073550.1;HCoV-229E_1a;1687 | 0 | 0 |
| NC_002645.1;NP_073550.1;HCoV-229E_1a;1688 | 0 | 0 |
| NC_002645.1;NP_073550.1;HCoV-229E_1a;1689 | 0 | 0 |
| NC_002645.1;NP_073550.1;HCoV-229E_1a;1690 | 0.05 | 0 |
| NC_002645.1;NP_073550.1;HCoV-229E_1a;1691 | 0 | 0 |
| NC_002645.1;NP_073550.1;HCoV-229E_1a;1692 | 0 | 0 |
| NC_002645.1;NP_073550.1;HCoV-229E_1a;1693 | 0 | 0 |
| NC_002645.1;NP_073550.1;HCoV-229E_1a;1694 | 0 | 0 |
| NC_002645.1;NP_073550.1;HCoV-229E_1a;1695 | 0 | 0 |
| NC_002645.1;NP_073550.1;HCoV-229E_1a;1696 | 0 | 0 |
| NC_002645.1;NP_073550.1;HCoV-229E_1a;1697 | 0 | 0 |
| NC_002645.1;NP_073550.1;HCoV-229E_1a;1698 | 0 | 0 |
| NC_002645.1;NP_073550.1;HCoV-229E_1a;1699 | 0 | 0 |
| NC_002645.1;NP_073550.1;HCoV-229E_1a;1700 | 0 | 0 |
| NC_002645.1;NP_073550.1;HCoV-229E_1a;1701 | 0 | 0 |
| NC_002645.1;NP_073550.1;HCoV-229E_1a;1702 | 0 | 0 |
| NC_002645.1;NP_073550.1;HCoV-229E_1a;1703 | 0 | 0 |
| NC_002645.1;NP_073550.1;HCoV-229E_1a;1704 | 0 | 0 |
| NC_002645.1;NP_073550.1;HCoV-229E_1a;1705 | 0 | 0 |
| NC_002645.1;NP_073550.1;HCoV-229E_1a;1706 | 0 | 0 |
| NC_002645.1;NP_073550.1;HCoV-229E_1a;1707 | 0 | 0 |
| NC_002645.1;NP_073550.1;HCoV-229E_1a;1708 | 0 | 0 |
| NC_002645.1;NP_073550.1;HCoV-229E_1a;1709 | 0 | 0 |
| NC_002645.1;NP_073550.1;HCoV-229E_1a;1710 | 0.05 | 0.05 |

| | | |
|---|---|---|
| NC_002645.1;NP_073550.1;HCoV-229E_1a;1711 | 0 | 0 |
| NC_002645.1;NP_073550.1;HCoV-229E_1a;1712 | 0 | 0 |
| NC_002645.1;NP_073550.1;HCoV-229E_1a;1713 | 0 | 0 |
| NC_002645.1;NP_073550.1;HCoV-229E_1a;1714 | 0 | 0 |
| NC_002645.1;NP_073550.1;HCoV-229E_1a;1715 | 0 | 0 |
| NC_002645.1;NP_073550.1;HCoV-229E_1a;1716 | 0 | 0 |
| NC_002645.1;NP_073550.1;HCoV-229E_1a;1717 | 0 | 0 |
| NC_002645.1;NP_073550.1;HCoV-229E_1a;1718 | 0.05 | 0 |
| NC_002645.1;NP_073550.1;HCoV-229E_1a;1719 | 0.05 | 0 |
| NC_002645.1;NP_073550.1;HCoV-229E_1a;1720 | 0.05 | 0 |
| NC_002645.1;NP_073550.1;HCoV-229E_1a;1721 | 0.05 | 0.025 |
| NC_002645.1;NP_073550.1;HCoV-229E_1a;1722 | 0 | 0 |
| NC_002645.1;NP_073550.1;HCoV-229E_1a;1723 | 0 | 0 |
| NC_002645.1;NP_073550.1;HCoV-229E_1a;1724 | 0 | 0 |
| NC_002645.1;NP_073550.1;HCoV-229E_1a;1725 | 0 | 0 |
| NC_002645.1;NP_073550.1;HCoV-229E_1a;1726 | 0 | 0 |
| NC_002645.1;NP_073550.1;HCoV-229E_1a;1727 | 0 | 0 |
| NC_002645.1;NP_073550.1;HCoV-229E_1a;1728 | 0 | 0 |
| NC_002645.1;NP_073550.1;HCoV-229E_1a;1729 | 0 | 0 |
| NC_002645.1;NP_073550.1;HCoV-229E_1a;1730 | 0 | 0 |
| NC_002645.1;NP_073550.1;HCoV-229E_1a;1731 | 0 | 0 |
| NC_002645.1;NP_073550.1;HCoV-229E_1a;1732 | 0 | 0 |
| NC_002645.1;NP_073550.1;HCoV-229E_1a;1733 | 0 | 0 |
| NC_002645.1;NP_073550.1;HCoV-229E_1a;1734 | 0 | 0 |
| NC_002645.1;NP_073550.1;HCoV-229E_1a;1735 | 0 | 0 |
| NC_002645.1;NP_073550.1;HCoV-229E_1a;1736 | 0 | 0 |
| NC_002645.1;NP_073550.1;HCoV-229E_1a;1737 | 0 | 0 |
| NC_002645.1;NP_073550.1;HCoV-229E_1a;1738 | 0 | 0 |
| NC_002645.1;NP_073550.1;HCoV-229E_1a;1739 | 0 | 0 |
| NC_002645.1;NP_073550.1;HCoV-229E_1a;1740 | 0 | 0 |
| NC_002645.1;NP_073550.1;HCoV-229E_1a;1741 | 0.1 | 0.025 |
| NC_002645.1;NP_073550.1;HCoV-229E_1a;1742 | 0 | 0 |
| NC_002645.1;NP_073550.1;HCoV-229E_1a;1743 | 0 | 0.025 |
| NC_002645.1;NP_073550.1;HCoV-229E_1a;1744 | 0 | 0.025 |
| NC_002645.1;NP_073550.1;HCoV-229E_1a;1745 | 0 | 0 |
| NC_002645.1;NP_073550.1;HCoV-229E_1a;1746 | 0 | 0 |
| NC_002645.1;NP_073550.1;HCoV-229E_1a;1747 | 0 | 0 |
| NC_002645.1;NP_073550.1;HCoV-229E_1a;1748 | 0 | 0 |
| NC_002645.1;NP_073550.1;HCoV-229E_1a;1749 | 0 | 0 |
| NC_002645.1;NP_073550.1;HCoV-229E_1a;1750 | 0 | 0.025 |
| NC_002645.1;NP_073550.1;HCoV-229E_1a;1751 | 0 | 0.025 |
| NC_002645.1;NP_073550.1;HCoV-229E_1a;1752 | 0 | 0.025 |
| NC_002645.1;NP_073550.1;HCoV-229E_1a;1753 | 0.05 | 0.05 |

| | | |
|---|---|---|
| NC_002645.1;NP_073550.1;HCoV-229E_1a;1754 | 0.1 | 0.025 |
| NC_002645.1;NP_073550.1;HCoV-229E_1a;1755 | 0.05 | 0.025 |
| NC_002645.1;NP_073550.1;HCoV-229E_1a;1756 | 0.05 | 0.05 |
| NC_002645.1;NP_073550.1;HCoV-229E_1a;1757 | 0.1 | 0.025 |
| NC_002645.1;NP_073550.1;HCoV-229E_1a;1758 | 0.1 | 0.025 |
| NC_002645.1;NP_073550.1;HCoV-229E_1a;1759 | 0.05 | 0.025 |
| NC_002645.1;NP_073550.1;HCoV-229E_1a;1760 | 0.1 | 0.025 |
| NC_002645.1;NP_073550.1;HCoV-229E_1a;1761 | 0 | 0 |
| NC_002645.1;NP_073550.1;HCoV-229E_1a;1762 | 0.05 | 0 |
| NC_002645.1;NP_073550.1;HCoV-229E_1a;1763 | 0 | 0.025 |
| NC_002645.1;NP_073550.1;HCoV-229E_1a;1764 | 0.05 | 0.025 |
| NC_002645.1;NP_073550.1;HCoV-229E_1a;1765 | 0 | 0 |
| NC_002645.1;NP_073550.1;HCoV-229E_1a;1766 | 0 | 0 |
| NC_002645.1;NP_073550.1;HCoV-229E_1a;1767 | 0 | 0 |
| NC_002645.1;NP_073550.1;HCoV-229E_1a;1768 | 0 | 0 |
| NC_002645.1;NP_073550.1;HCoV-229E_1a;1769 | 0 | 0 |
| NC_002645.1;NP_073550.1;HCoV-229E_1a;1770 | 0 | 0 |
| NC_002645.1;NP_073550.1;HCoV-229E_1a;1771 | 0 | 0 |
| NC_002645.1;NP_073550.1;HCoV-229E_1a;1772 | 0 | 0 |
| NC_002645.1;NP_073550.1;HCoV-229E_1a;1773 | 0 | 0 |
| NC_002645.1;NP_073550.1;HCoV-229E_1a;1774 | 0 | 0 |
| NC_002645.1;NP_073550.1;HCoV-229E_1a;1775 | 0 | 0 |
| NC_002645.1;NP_073550.1;HCoV-229E_1a;1776 | 0 | 0 |
| NC_002645.1;NP_073550.1;HCoV-229E_1a;1777 | 0.05 | 0.125 |
| NC_002645.1;NP_073550.1;HCoV-229E_1a;1778 | 0.05 | 0.025 |
| NC_002645.1;NP_073550.1;HCoV-229E_1a;1779 | 0 | 0 |
| NC_002645.1;NP_073550.1;HCoV-229E_1a;1780 | 0 | 0.025 |
| NC_002645.1;NP_073550.1;HCoV-229E_1a;1781 | 0 | 0 |
| NC_002645.1;NP_073550.1;HCoV-229E_1a;1782 | 0 | 0 |
| NC_002645.1;NP_073550.1;HCoV-229E_1a;1783 | 0.15 | 0.125 |
| NC_002645.1;NP_073550.1;HCoV-229E_1a;1784 | 0.15 | 0.125 |
| NC_002645.1;NP_073550.1;HCoV-229E_1a;1785 | 0.15 | 0.075 |
| NC_002645.1;NP_073550.1;HCoV-229E_1a;1786 | 0.1 | 0.05 |
| NC_002645.1;NP_073550.1;HCoV-229E_1a;1787 | 0 | 0 |
| NC_002645.1;NP_073550.1;HCoV-229E_1a;1788 | 0 | 0 |
| NC_002645.1;NP_073550.1;HCoV-229E_1a;1789 | 0 | 0 |
| NC_002645.1;NP_073550.1;HCoV-229E_1a;1790 | 0 | 0 |
| NC_002645.1;NP_073550.1;HCoV-229E_1a;1791 | 0 | 0.025 |
| NC_002645.1;NP_073550.1;HCoV-229E_1a;1792 | 0 | 0 |
| NC_002645.1;NP_073550.1;HCoV-229E_1a;1793 | 0 | 0.05 |
| NC_002645.1;NP_073550.1;HCoV-229E_1a;1794 | 0 | 0.1 |
| NC_002645.1;NP_073550.1;HCoV-229E_1a;1795 | 0 | 0.15 |
| NC_002645.1;NP_073550.1;HCoV-229E_1a;1796 | 0 | 0.1 |

| | | |
|---|---|---|
| NC_002645.1;NP_073550.1;HCoV-229E_1a;1797 | 0.05 | 0.125 |
| NC_002645.1;NP_073550.1;HCoV-229E_1a;1798 | 0.05 | 0.15 |
| NC_002645.1;NP_073550.1;HCoV-229E_1a;1799 | 0 | 0.125 |
| NC_002645.1;NP_073550.1;HCoV-229E_1a;1800 | 0 | 0.125 |
| NC_002645.1;NP_073550.1;HCoV-229E_1a;1801 | 0 | 0.125 |
| NC_002645.1;NP_073550.1;HCoV-229E_1a;1802 | 0.05 | 0.125 |
| NC_002645.1;NP_073550.1;HCoV-229E_1a;1803 | 0 | 0.075 |
| NC_002645.1;NP_073550.1;HCoV-229E_1a;1804 | 0.2 | 0.275 |
| NC_002645.1;NP_073550.1;HCoV-229E_1a;1805 | 0 | 0 |
| NC_002645.1;NP_073550.1;HCoV-229E_1a;1806 | 0 | 0 |
| NC_002645.1;NP_073550.1;HCoV-229E_1a;1807 | 0 | 0 |
| NC_002645.1;NP_073550.1;HCoV-229E_1a;1808 | 0 | 0 |
| NC_002645.1;NP_073550.1;HCoV-229E_1a;1809 | 0 | 0 |
| NC_002645.1;NP_073550.1;HCoV-229E_1a;1810 | 0 | 0 |
| NC_002645.1;NP_073550.1;HCoV-229E_1a;1811 | 0 | 0 |
| NC_002645.1;NP_073550.1;HCoV-229E_1a;1812 | 0 | 0 |
| NC_002645.1;NP_073550.1;HCoV-229E_1a;1813 | 0 | 0 |
| NC_002645.1;NP_073550.1;HCoV-229E_1a;1814 | 0 | 0 |
| NC_002645.1;NP_073550.1;HCoV-229E_1a;1815 | 0 | 0 |
| NC_002645.1;NP_073550.1;HCoV-229E_1a;1816 | 0 | 0 |
| NC_002645.1;NP_073550.1;HCoV-229E_1a;1817 | 0 | 0 |
| NC_002645.1;NP_073550.1;HCoV-229E_1a;1818 | 0 | 0 |
| NC_002645.1;NP_073550.1;HCoV-229E_1a;1819 | 0 | 0 |
| NC_002645.1;NP_073550.1;HCoV-229E_1a;1820 | 0 | 0 |
| NC_002645.1;NP_073550.1;HCoV-229E_1a;1821 | 0 | 0 |
| NC_002645.1;NP_073550.1;HCoV-229E_1a;1822 | 0.05 | 0.075 |
| NC_002645.1;NP_073550.1;HCoV-229E_1a;1823 | 0.05 | 0.025 |
| NC_002645.1;NP_073550.1;HCoV-229E_1a;1824 | 0.1 | 0.175 |
| NC_002645.1;NP_073550.1;HCoV-229E_1a;1825 | 0.1 | 0.1 |
| NC_002645.1;NP_073550.1;HCoV-229E_1a;1826 | 0.05 | 0.075 |
| NC_002645.1;NP_073550.1;HCoV-229E_1a;1827 | 0 | 0.05 |
| NC_002645.1;NP_073550.1;HCoV-229E_1a;1828 | 0 | 0.025 |
| NC_002645.1;NP_073550.1;HCoV-229E_1a;1829 | 0 | 0.025 |
| NC_002645.1;NP_073550.1;HCoV-229E_1a;1830 | 0 | 0 |
| NC_002645.1;NP_073550.1;HCoV-229E_1a;1831 | 0 | 0 |
| NC_002645.1;NP_073550.1;HCoV-229E_1a;1832 | 0.05 | 0 |
| NC_002645.1;NP_073550.1;HCoV-229E_1a;1833 | 0 | 0.075 |
| NC_002645.1;NP_073550.1;HCoV-229E_1a;1834 | 0 | 0 |
| NC_002645.1;NP_073550.1;HCoV-229E_1a;1835 | 0 | 0 |
| NC_002645.1;NP_073550.1;HCoV-229E_1a;1836 | 0.05 | 0.05 |
| NC_002645.1;NP_073550.1;HCoV-229E_1a;1837 | 0 | 0 |
| NC_002645.1;NP_073550.1;HCoV-229E_1a;1838 | 0 | 0 |
| NC_002645.1;NP_073550.1;HCoV-229E_1a;1839 | 0 | 0 |

| | | |
|---|---|---|
| NC_002645.1;NP_073550.1;HCoV-229E_1a;1840 | 0 | 0 |
| NC_002645.1;NP_073550.1;HCoV-229E_1a;1841 | 0 | 0 |
| NC_002645.1;NP_073550.1;HCoV-229E_1a;1842 | 0 | 0 |
| NC_002645.1;NP_073550.1;HCoV-229E_1a;1843 | 0 | 0 |
| NC_002645.1;NP_073550.1;HCoV-229E_1a;1844 | 0 | 0 |
| NC_002645.1;NP_073550.1;HCoV-229E_1a;1845 | 0 | 0 |
| NC_002645.1;NP_073550.1;HCoV-229E_1a;1846 | 0 | 0 |
| NC_002645.1;NP_073550.1;HCoV-229E_1a;1847 | 0 | 0 |
| NC_002645.1;NP_073550.1;HCoV-229E_1a;1848 | 0 | 0 |
| NC_002645.1;NP_073550.1;HCoV-229E_1a;1849 | 0.05 | 0.025 |
| NC_002645.1;NP_073550.1;HCoV-229E_1a;1850 | 0.05 | 0 |
| NC_002645.1;NP_073550.1;HCoV-229E_1a;1851 | 0.05 | 0.025 |
| NC_002645.1;NP_073550.1;HCoV-229E_1a;1852 | 0 | 0 |
| NC_002645.1;NP_073550.1;HCoV-229E_1a;1853 | 0 | 0 |
| NC_002645.1;NP_073550.1;HCoV-229E_1a;1854 | 0 | 0 |
| NC_002645.1;NP_073550.1;HCoV-229E_1a;1855 | 0.05 | 0 |
| NC_002645.1;NP_073550.1;HCoV-229E_1a;1856 | 0 | 0.025 |
| NC_002645.1;NP_073550.1;HCoV-229E_1a;1857 | 0 | 0.075 |
| NC_002645.1;NP_073550.1;HCoV-229E_1a;1858 | 0 | 0 |
| NC_002645.1;NP_073550.1;HCoV-229E_1a;1859 | 0 | 0 |
| NC_002645.1;NP_073550.1;HCoV-229E_1a;1860 | 0 | 0 |
| NC_002645.1;NP_073550.1;HCoV-229E_1a;1861 | 0 | 0.025 |
| NC_002645.1;NP_073550.1;HCoV-229E_1a;1862 | 0 | 0.025 |
| NC_002645.1;NP_073550.1;HCoV-229E_1a;1863 | 0 | 0.025 |
| NC_002645.1;NP_073550.1;HCoV-229E_1a;1864 | 0 | 0.05 |
| NC_002645.1;NP_073550.1;HCoV-229E_1a;1865 | 0 | 0.025 |
| NC_002645.1;NP_073550.1;HCoV-229E_1a;1866 | 0.05 | 0.025 |
| NC_002645.1;NP_073550.1;HCoV-229E_1a;1867 | 0 | 0.05 |
| NC_002645.1;NP_073550.1;HCoV-229E_1a;1868 | 0.15 | 0.075 |
| NC_002645.1;NP_073550.1;HCoV-229E_1a;1869 | 0 | 0.025 |
| NC_002645.1;NP_073550.1;HCoV-229E_1a;1870 | 0.05 | 0.025 |
| NC_002645.1;NP_073550.1;HCoV-229E_1a;1871 | 0 | 0.05 |
| NC_002645.1;NP_073550.1;HCoV-229E_1a;1872 | 0.05 | 0.025 |
| NC_002645.1;NP_073550.1;HCoV-229E_1a;1873 | 0 | 0.025 |
| NC_002645.1;NP_073550.1;HCoV-229E_1a;1874 | 0 | 0.025 |
| NC_002645.1;NP_073550.1;HCoV-229E_1a;1875 | 0 | 0.025 |
| NC_002645.1;NP_073550.1;HCoV-229E_1a;1876 | 0 | 0.05 |
| NC_002645.1;NP_073550.1;HCoV-229E_1a;1877 | 0 | 0 |
| NC_002645.1;NP_073550.1;HCoV-229E_1a;1878 | 0 | 0 |
| NC_002645.1;NP_073550.1;HCoV-229E_1a;1879 | 0 | 0 |
| NC_002645.1;NP_073550.1;HCoV-229E_1a;1880 | 0 | 0 |
| NC_002645.1;NP_073550.1;HCoV-229E_1a;1881 | 0 | 0 |
| NC_002645.1;NP_073550.1;HCoV-229E_1a;1882 | 0 | 0 |

| | | |
|---|---|---|
| NC_002645.1;NP_073550.1;HCoV-229E_1a;1883 | 0 | 0 |
| NC_002645.1;NP_073550.1;HCoV-229E_1a;1884 | 0 | 0 |
| NC_002645.1;NP_073550.1;HCoV-229E_1a;1885 | 0 | 0 |
| NC_002645.1;NP_073550.1;HCoV-229E_1a;1886 | 0 | 0 |
| NC_002645.1;NP_073550.1;HCoV-229E_1a;1887 | 0 | 0 |
| NC_002645.1;NP_073550.1;HCoV-229E_1a;1888 | 0 | 0 |
| NC_002645.1;NP_073550.1;HCoV-229E_1a;1889 | 0 | 0 |
| NC_002645.1;NP_073550.1;HCoV-229E_1a;1890 | 0 | 0 |
| NC_002645.1;NP_073550.1;HCoV-229E_1a;1891 | 0 | 0 |
| NC_002645.1;NP_073550.1;HCoV-229E_1a;1892 | 0 | 0 |
| NC_002645.1;NP_073550.1;HCoV-229E_1a;1893 | 0 | 0 |
| NC_002645.1;NP_073550.1;HCoV-229E_1a;1894 | 0 | 0 |
| NC_002645.1;NP_073550.1;HCoV-229E_1a;1895 | 0 | 0 |
| NC_002645.1;NP_073550.1;HCoV-229E_1a;1896 | 0 | 0 |
| NC_002645.1;NP_073550.1;HCoV-229E_1a;1897 | 0 | 0 |
| NC_002645.1;NP_073550.1;HCoV-229E_1a;1898 | 0 | 0 |
| NC_002645.1;NP_073550.1;HCoV-229E_1a;1899 | 0.05 | 0.025 |
| NC_002645.1;NP_073550.1;HCoV-229E_1a;1900 | 0 | 0.05 |
| NC_002645.1;NP_073550.1;HCoV-229E_1a;1901 | 0.05 | 0.075 |
| NC_002645.1;NP_073550.1;HCoV-229E_1a;1902 | 0 | 0.05 |
| NC_002645.1;NP_073550.1;HCoV-229E_1a;1903 | 0.05 | 0.075 |
| NC_002645.1;NP_073550.1;HCoV-229E_1a;1904 | 0 | 0.025 |
| NC_002645.1;NP_073550.1;HCoV-229E_1a;1905 | 0 | 0.025 |
| NC_002645.1;NP_073550.1;HCoV-229E_1a;1906 | 0 | 0.075 |
| NC_002645.1;NP_073550.1;HCoV-229E_1a;1907 | 0.05 | 0.075 |
| NC_002645.1;NP_073550.1;HCoV-229E_1a;1908 | 0 | 0 |
| NC_002645.1;NP_073550.1;HCoV-229E_1a;1909 | 0.2 | 0 |
| NC_002645.1;NP_073550.1;HCoV-229E_1a;1910 | 0.1 | 0 |
| NC_002645.1;NP_073550.1;HCoV-229E_1a;1911 | 0.2 | 0.075 |
| NC_002645.1;NP_073550.1;HCoV-229E_1a;1912 | 0.15 | 0.05 |
| NC_002645.1;NP_073550.1;HCoV-229E_1a;1913 | 0.1 | 0 |
| NC_002645.1;NP_073550.1;HCoV-229E_1a;1914 | 0.05 | 0.025 |
| NC_002645.1;NP_073550.1;HCoV-229E_1a;1915 | 0.05 | 0 |
| NC_002645.1;NP_073550.1;HCoV-229E_1a;1916 | 0.15 | 0 |
| NC_002645.1;NP_073550.1;HCoV-229E_1a;1917 | 0.05 | 0 |
| NC_002645.1;NP_073550.1;HCoV-229E_1a;1918 | 0 | 0 |
| NC_002645.1;NP_073550.1;HCoV-229E_1a;1919 | 0.05 | 0 |
| NC_002645.1;NP_073550.1;HCoV-229E_1a;1920 | 0 | 0 |
| NC_002645.1;NP_073550.1;HCoV-229E_1a;1921 | 0 | 0 |
| NC_002645.1;NP_073550.1;HCoV-229E_1a;1922 | 0 | 0 |
| NC_002645.1;NP_073550.1;HCoV-229E_1a;1923 | 0 | 0 |
| NC_002645.1;NP_073550.1;HCoV-229E_1a;1924 | 0 | 0 |
| NC_002645.1;NP_073550.1;HCoV-229E_1a;1925 | 0 | 0 |

| | | |
|---|---|---|
| NC_002645.1;NP_073550.1;HCoV-229E_1a;1926 | 0 | 0 |
| NC_002645.1;NP_073550.1;HCoV-229E_1a;1927 | 0 | 0 |
| NC_002645.1;NP_073550.1;HCoV-229E_1a;1928 | 0 | 0 |
| NC_002645.1;NP_073550.1;HCoV-229E_1a;1929 | 0 | 0 |
| NC_002645.1;NP_073550.1;HCoV-229E_1a;1930 | 0 | 0 |
| NC_002645.1;NP_073550.1;HCoV-229E_1a;1931 | 0 | 0 |
| NC_002645.1;NP_073550.1;HCoV-229E_1a;1932 | 0 | 0 |
| NC_002645.1;NP_073550.1;HCoV-229E_1a;1933 | 0 | 0 |
| NC_002645.1;NP_073550.1;HCoV-229E_1a;1934 | 0 | 0 |
| NC_002645.1;NP_073550.1;HCoV-229E_1a;1935 | 0 | 0 |
| NC_002645.1;NP_073550.1;HCoV-229E_1a;1936 | 0 | 0 |
| NC_002645.1;NP_073550.1;HCoV-229E_1a;1937 | 0 | 0 |
| NC_002645.1;NP_073550.1;HCoV-229E_1a;1938 | 0 | 0 |
| NC_002645.1;NP_073550.1;HCoV-229E_1a;1939 | 0 | 0 |
| NC_002645.1;NP_073550.1;HCoV-229E_1a;1940 | 0 | 0 |
| NC_002645.1;NP_073550.1;HCoV-229E_1a;1941 | 0 | 0 |
| NC_002645.1;NP_073550.1;HCoV-229E_1a;1942 | 0.05 | 0 |
| NC_002645.1;NP_073550.1;HCoV-229E_1a;1943 | 0.05 | 0 |
| NC_002645.1;NP_073550.1;HCoV-229E_1a;1944 | 0 | 0 |
| NC_002645.1;NP_073550.1;HCoV-229E_1a;1945 | 0 | 0 |
| NC_002645.1;NP_073550.1;HCoV-229E_1a;1946 | 0 | 0 |
| NC_002645.1;NP_073550.1;HCoV-229E_1a;1947 | 0 | 0 |
| NC_002645.1;NP_073550.1;HCoV-229E_1a;1948 | 0 | 0 |
| NC_002645.1;NP_073550.1;HCoV-229E_1a;1949 | 0 | 0 |
| NC_002645.1;NP_073550.1;HCoV-229E_1a;1950 | 0.05 | 0 |
| NC_002645.1;NP_073550.1;HCoV-229E_1a;1951 | 0 | 0 |
| NC_002645.1;NP_073550.1;HCoV-229E_1a;1952 | 0 | 0 |
| NC_002645.1;NP_073550.1;HCoV-229E_1a;1953 | 0 | 0 |
| NC_002645.1;NP_073550.1;HCoV-229E_1a;1954 | 0 | 0 |
| NC_002645.1;NP_073550.1;HCoV-229E_1a;1955 | 0 | 0 |
| NC_002645.1;NP_073550.1;HCoV-229E_1a;1956 | 0 | 0 |
| NC_002645.1;NP_073550.1;HCoV-229E_1a;1957 | 0 | 0 |
| NC_002645.1;NP_073550.1;HCoV-229E_1a;1958 | 0 | 0 |
| NC_002645.1;NP_073550.1;HCoV-229E_1a;1959 | 0 | 0 |
| NC_002645.1;NP_073550.1;HCoV-229E_1a;1960 | 0 | 0 |
| NC_002645.1;NP_073550.1;HCoV-229E_1a;1961 | 0 | 0 |
| NC_002645.1;NP_073550.1;HCoV-229E_1a;1962 | 0 | 0 |
| NC_002645.1;NP_073550.1;HCoV-229E_1a;1963 | 0 | 0 |
| NC_002645.1;NP_073550.1;HCoV-229E_1a;1964 | 0 | 0 |
| NC_002645.1;NP_073550.1;HCoV-229E_1a;1965 | 0 | 0 |
| NC_002645.1;NP_073550.1;HCoV-229E_1a;1966 | 0 | 0.025 |
| NC_002645.1;NP_073550.1;HCoV-229E_1a;1967 | 0 | 0.025 |
| NC_002645.1;NP_073550.1;HCoV-229E_1a;1968 | 0.05 | 0 |

| | | |
|---|---|---|
| NC_002645.1;NP_073550.1;HCoV-229E_1a;1969 | 0 | 0 |
| NC_002645.1;NP_073550.1;HCoV-229E_1a;1970 | 0 | 0 |
| NC_002645.1;NP_073550.1;HCoV-229E_1a;1971 | 0 | 0 |
| NC_002645.1;NP_073550.1;HCoV-229E_1a;1972 | 0 | 0 |
| NC_002645.1;NP_073550.1;HCoV-229E_1a;1973 | 0 | 0 |
| NC_002645.1;NP_073550.1;HCoV-229E_1a;1974 | 0 | 0 |
| NC_002645.1;NP_073550.1;HCoV-229E_1a;1975 | 0 | 0 |
| NC_002645.1;NP_073550.1;HCoV-229E_1a;1976 | 0 | 0 |
| NC_002645.1;NP_073550.1;HCoV-229E_1a;1977 | 0 | 0 |
| NC_002645.1;NP_073550.1;HCoV-229E_1a;1978 | 0 | 0 |
| NC_002645.1;NP_073550.1;HCoV-229E_1a;1979 | 0 | 0 |
| NC_002645.1;NP_073550.1;HCoV-229E_1a;1980 | 0 | 0 |
| NC_002645.1;NP_073550.1;HCoV-229E_1a;1981 | 0 | 0.025 |
| NC_002645.1;NP_073550.1;HCoV-229E_1a;1982 | 0 | 0 |
| NC_002645.1;NP_073550.1;HCoV-229E_1a;1983 | 0 | 0 |
| NC_002645.1;NP_073550.1;HCoV-229E_1a;1984 | 0 | 0 |
| NC_002645.1;NP_073550.1;HCoV-229E_1a;1985 | 0 | 0 |
| NC_002645.1;NP_073550.1;HCoV-229E_1a;1986 | 0 | 0 |
| NC_002645.1;NP_073550.1;HCoV-229E_1a;1987 | 0 | 0 |
| NC_002645.1;NP_073550.1;HCoV-229E_1a;1988 | 0 | 0 |
| NC_002645.1;NP_073550.1;HCoV-229E_1a;1989 | 0 | 0 |
| NC_002645.1;NP_073550.1;HCoV-229E_1a;1990 | 0 | 0 |
| NC_002645.1;NP_073550.1;HCoV-229E_1a;1991 | 0 | 0 |
| NC_002645.1;NP_073550.1;HCoV-229E_1a;1992 | 0 | 0 |
| NC_002645.1;NP_073550.1;HCoV-229E_1a;1993 | 0 | 0 |
| NC_002645.1;NP_073550.1;HCoV-229E_1a;1994 | 0 | 0 |
| NC_002645.1;NP_073550.1;HCoV-229E_1a;1995 | 0 | 0 |
| NC_002645.1;NP_073550.1;HCoV-229E_1a;1996 | 0 | 0 |
| NC_002645.1;NP_073550.1;HCoV-229E_1a;1997 | 0 | 0 |
| NC_002645.1;NP_073550.1;HCoV-229E_1a;1998 | 0 | 0 |
| NC_002645.1;NP_073550.1;HCoV-229E_1a;1999 | 0 | 0 |
| NC_002645.1;NP_073550.1;HCoV-229E_1a;2000 | 0 | 0 |
| NC_002645.1;NP_073550.1;HCoV-229E_1a;2001 | 0 | 0 |
| NC_002645.1;NP_073550.1;HCoV-229E_1a;2002 | 0 | 0 |
| NC_002645.1;NP_073550.1;HCoV-229E_1a;2003 | 0 | 0 |
| NC_002645.1;NP_073550.1;HCoV-229E_1a;2004 | 0 | 0 |
| NC_002645.1;NP_073550.1;HCoV-229E_1a;2005 | 0 | 0 |
| NC_002645.1;NP_073550.1;HCoV-229E_1a;2006 | 0 | 0 |
| NC_002645.1;NP_073550.1;HCoV-229E_1a;2007 | 0 | 0 |
| NC_002645.1;NP_073550.1;HCoV-229E_1a;2008 | 0 | 0 |
| NC_002645.1;NP_073550.1;HCoV-229E_1a;2009 | 0 | 0 |
| NC_002645.1;NP_073550.1;HCoV-229E_1a;2010 | 0 | 0 |
| NC_002645.1;NP_073550.1;HCoV-229E_1a;2011 | 0 | 0 |

| | | |
|---|---|---|
| NC_002645.1;NP_073550.1;HCoV-229E_1a;2012 | 0 | 0 |
| NC_002645.1;NP_073550.1;HCoV-229E_1a;2013 | 0 | 0 |
| NC_002645.1;NP_073550.1;HCoV-229E_1a;2014 | 0 | 0 |
| NC_002645.1;NP_073550.1;HCoV-229E_1a;2015 | 0 | 0.025 |
| NC_002645.1;NP_073550.1;HCoV-229E_1a;2016 | 0 | 0.025 |
| NC_002645.1;NP_073550.1;HCoV-229E_1a;2017 | 0 | 0 |
| NC_002645.1;NP_073550.1;HCoV-229E_1a;2018 | 0.05 | 0 |
| NC_002645.1;NP_073550.1;HCoV-229E_1a;2019 | 0.05 | 0 |
| NC_002645.1;NP_073550.1;HCoV-229E_1a;2020 | 0 | 0.05 |
| NC_002645.1;NP_073550.1;HCoV-229E_1a;2021 | 0 | 0.025 |
| NC_002645.1;NP_073550.1;HCoV-229E_1a;2022 | 0.05 | 0.1 |
| NC_002645.1;NP_073550.1;HCoV-229E_1a;2023 | 0.1 | 0.125 |
| NC_002645.1;NP_073550.1;HCoV-229E_1a;2024 | 0.05 | 0.05 |
| NC_002645.1;NP_073550.1;HCoV-229E_1a;2025 | 0.05 | 0.05 |
| NC_002645.1;NP_073550.1;HCoV-229E_1a;2026 | 0 | 0 |
| NC_002645.1;NP_073550.1;HCoV-229E_1a;2027 | 0 | 0 |
| NC_002645.1;NP_073550.1;HCoV-229E_1a;2028 | 0 | 0.025 |
| NC_002645.1;NP_073550.1;HCoV-229E_1a;2029 | 0 | 0 |
| NC_002645.1;NP_073550.1;HCoV-229E_1a;2030 | 0 | 0 |
| NC_002645.1;NP_073550.1;HCoV-229E_1a;2031 | 0 | 0.025 |
| NC_002645.1;NP_073550.1;HCoV-229E_1a;2032 | 0 | 0.025 |
| NC_002645.1;NP_073550.1;HCoV-229E_1a;2033 | 0 | 0.025 |
| NC_002645.1;NP_073550.1;HCoV-229E_1a;2034 | 0 | 0.025 |
| NC_002645.1;NP_073550.1;HCoV-229E_1a;2035 | 0 | 0.025 |
| NC_002645.1;NP_073550.1;HCoV-229E_1a;2036 | 0.25 | 0.15 |
| NC_002645.1;NP_073550.1;HCoV-229E_1a;2037 | 0 | 0.025 |
| NC_002645.1;NP_073550.1;HCoV-229E_1a;2038 | 0 | 0.025 |
| NC_002645.1;NP_073550.1;HCoV-229E_1a;2039 | 0.1 | 0.025 |
| NC_002645.1;NP_073550.1;HCoV-229E_1a;2040 | 0.1 | 0.025 |
| NC_002645.1;NP_073550.1;HCoV-229E_1a;2041 | 0 | 0 |
| NC_002645.1;NP_073550.1;HCoV-229E_1a;2042 | 0 | 0 |
| NC_002645.1;NP_073550.1;HCoV-229E_1a;2043 | 0 | 0 |
| NC_002645.1;NP_073550.1;HCoV-229E_1a;2044 | 0 | 0 |
| NC_002645.1;NP_073550.1;HCoV-229E_1a;2045 | 0 | 0 |
| NC_002645.1;NP_073550.1;HCoV-229E_1a;2046 | 0 | 0 |
| NC_002645.1;NP_073550.1;HCoV-229E_1a;2047 | 0 | 0 |
| NC_002645.1;NP_073550.1;HCoV-229E_1a;2048 | 0 | 0 |
| NC_002645.1;NP_073550.1;HCoV-229E_1a;2049 | 0 | 0 |
| NC_002645.1;NP_073550.1;HCoV-229E_1a;2050 | 0 | 0 |
| NC_002645.1;NP_073550.1;HCoV-229E_1a;2051 | 0 | 0 |
| NC_002645.1;NP_073550.1;HCoV-229E_1a;2052 | 0 | 0 |
| NC_002645.1;NP_073550.1;HCoV-229E_1a;2053 | 0 | 0 |
| NC_002645.1;NP_073550.1;HCoV-229E_1a;2054 | 0 | 0 |

| | | |
|---|---|---|
| NC_002645.1;NP_073550.1;HCoV-229E_1a;2055 | 0 | 0 |
| NC_002645.1;NP_073550.1;HCoV-229E_1a;2056 | 0 | 0 |
| NC_002645.1;NP_073550.1;HCoV-229E_1a;2057 | 0 | 0 |
| NC_002645.1;NP_073550.1;HCoV-229E_1a;2058 | 0 | 0 |
| NC_002645.1;NP_073550.1;HCoV-229E_1a;2059 | 0 | 0 |
| NC_002645.1;NP_073550.1;HCoV-229E_1a;2060 | 0 | 0 |
| NC_002645.1;NP_073550.1;HCoV-229E_1a;2061 | 0 | 0 |
| NC_002645.1;NP_073550.1;HCoV-229E_1a;2062 | 0 | 0.025 |
| NC_002645.1;NP_073550.1;HCoV-229E_1a;2063 | 0 | 0 |
| NC_002645.1;NP_073550.1;HCoV-229E_1a;2064 | 0 | 0 |
| NC_002645.1;NP_073550.1;HCoV-229E_1a;2065 | 0 | 0 |
| NC_002645.1;NP_073550.1;HCoV-229E_1a;2066 | 0 | 0 |
| NC_002645.1;NP_073550.1;HCoV-229E_1a;2067 | 0 | 0 |
| NC_002645.1;NP_073550.1;HCoV-229E_1a;2068 | 0 | 0 |
| NC_002645.1;NP_073550.1;HCoV-229E_1a;2069 | 0 | 0 |
| NC_002645.1;NP_073550.1;HCoV-229E_1a;2070 | 0 | 0 |
| NC_002645.1;NP_073550.1;HCoV-229E_1a;2071 | 0 | 0 |
| NC_002645.1;NP_073550.1;HCoV-229E_1a;2072 | 0 | 0 |
| NC_002645.1;NP_073550.1;HCoV-229E_1a;2073 | 0 | 0 |
| NC_002645.1;NP_073550.1;HCoV-229E_1a;2074 | 0 | 0 |
| NC_002645.1;NP_073550.1;HCoV-229E_1a;2075 | 0 | 0 |
| NC_002645.1;NP_073550.1;HCoV-229E_1a;2076 | 0 | 0 |
| NC_002645.1;NP_073550.1;HCoV-229E_1a;2077 | 0 | 0 |
| NC_002645.1;NP_073550.1;HCoV-229E_1a;2078 | 0 | 0 |
| NC_002645.1;NP_073550.1;HCoV-229E_1a;2079 | 0 | 0 |
| NC_002645.1;NP_073550.1;HCoV-229E_1a;2080 | 0 | 0 |
| NC_002645.1;NP_073550.1;HCoV-229E_1a;2081 | 0 | 0 |
| NC_002645.1;NP_073550.1;HCoV-229E_1a;2082 | 0 | 0 |
| NC_002645.1;NP_073550.1;HCoV-229E_1a;2083 | 0 | 0 |
| NC_002645.1;NP_073550.1;HCoV-229E_1a;2084 | 0 | 0 |
| NC_002645.1;NP_073550.1;HCoV-229E_1a;2085 | 0 | 0 |
| NC_002645.1;NP_073550.1;HCoV-229E_1a;2086 | 0 | 0 |
| NC_002645.1;NP_073550.1;HCoV-229E_1a;2087 | 0 | 0 |
| NC_002645.1;NP_073550.1;HCoV-229E_1a;2088 | 0 | 0 |
| NC_002645.1;NP_073550.1;HCoV-229E_1a;2089 | 0 | 0 |
| NC_002645.1;NP_073550.1;HCoV-229E_1a;2090 | 0 | 0 |
| NC_002645.1;NP_073550.1;HCoV-229E_1a;2091 | 0 | 0 |
| NC_002645.1;NP_073550.1;HCoV-229E_1a;2092 | 0 | 0 |
| NC_002645.1;NP_073550.1;HCoV-229E_1a;2093 | 0 | 0 |
| NC_002645.1;NP_073550.1;HCoV-229E_1a;2094 | 0 | 0 |
| NC_002645.1;NP_073550.1;HCoV-229E_1a;2095 | 0 | 0 |
| NC_002645.1;NP_073550.1;HCoV-229E_1a;2096 | 0 | 0 |
| NC_002645.1;NP_073550.1;HCoV-229E_1a;2097 | 0 | 0 |

| | | |
|---|---|---|
| NC_002645.1;NP_073550.1;HCoV-229E_1a;2098 | 0 | 0 |
| NC_002645.1;NP_073550.1;HCoV-229E_1a;2099 | 0 | 0 |
| NC_002645.1;NP_073550.1;HCoV-229E_1a;2100 | 0 | 0 |
| NC_002645.1;NP_073550.1;HCoV-229E_1a;2101 | 0 | 0 |
| NC_002645.1;NP_073550.1;HCoV-229E_1a;2102 | 0 | 0 |
| NC_002645.1;NP_073550.1;HCoV-229E_1a;2103 | 0.05 | 0 |
| NC_002645.1;NP_073550.1;HCoV-229E_1a;2104 | 0.05 | 0 |
| NC_002645.1;NP_073550.1;HCoV-229E_1a;2105 | 0 | 0 |
| NC_002645.1;NP_073550.1;HCoV-229E_1a;2106 | 0 | 0 |
| NC_002645.1;NP_073550.1;HCoV-229E_1a;2107 | 0 | 0 |
| NC_002645.1;NP_073550.1;HCoV-229E_1a;2108 | 0 | 0 |
| NC_002645.1;NP_073550.1;HCoV-229E_1a;2109 | 0 | 0 |
| NC_002645.1;NP_073550.1;HCoV-229E_1a;2110 | 0 | 0 |
| NC_002645.1;NP_073550.1;HCoV-229E_1a;2111 | 0 | 0 |
| NC_002645.1;NP_073550.1;HCoV-229E_1a;2112 | 0 | 0 |
| NC_002645.1;NP_073550.1;HCoV-229E_1a;2113 | 0 | 0 |
| NC_002645.1;NP_073550.1;HCoV-229E_1a;2114 | 0 | 0 |
| NC_002645.1;NP_073550.1;HCoV-229E_1a;2115 | 0 | 0 |
| NC_002645.1;NP_073550.1;HCoV-229E_1a;2116 | 0 | 0 |
| NC_002645.1;NP_073550.1;HCoV-229E_1a;2117 | 0 | 0 |
| NC_002645.1;NP_073550.1;HCoV-229E_1a;2118 | 0 | 0 |
| NC_002645.1;NP_073550.1;HCoV-229E_1a;2119 | 0 | 0 |
| NC_002645.1;NP_073550.1;HCoV-229E_1a;2120 | 0 | 0 |
| NC_002645.1;NP_073550.1;HCoV-229E_1a;2121 | 0 | 0 |
| NC_002645.1;NP_073550.1;HCoV-229E_1a;2122 | 0 | 0 |
| NC_002645.1;NP_073550.1;HCoV-229E_1a;2123 | 0 | 0 |
| NC_002645.1;NP_073550.1;HCoV-229E_1a;2124 | 0 | 0 |
| NC_002645.1;NP_073550.1;HCoV-229E_1a;2125 | 0 | 0 |
| NC_002645.1;NP_073550.1;HCoV-229E_1a;2126 | 0 | 0 |
| NC_002645.1;NP_073550.1;HCoV-229E_1a;2127 | 0 | 0 |
| NC_002645.1;NP_073550.1;HCoV-229E_1a;2128 | 0 | 0 |
| NC_002645.1;NP_073550.1;HCoV-229E_1a;2129 | 0 | 0 |
| NC_002645.1;NP_073550.1;HCoV-229E_1a;2130 | 0 | 0 |
| NC_002645.1;NP_073550.1;HCoV-229E_1a;2131 | 0 | 0 |
| NC_002645.1;NP_073550.1;HCoV-229E_1a;2132 | 0 | 0 |
| NC_002645.1;NP_073550.1;HCoV-229E_1a;2133 | 0 | 0 |
| NC_002645.1;NP_073550.1;HCoV-229E_1a;2134 | 0 | 0 |
| NC_002645.1;NP_073550.1;HCoV-229E_1a;2135 | 0 | 0 |
| NC_002645.1;NP_073550.1;HCoV-229E_1a;2136 | 0 | 0 |
| NC_002645.1;NP_073550.1;HCoV-229E_1a;2137 | 0 | 0 |
| NC_002645.1;NP_073550.1;HCoV-229E_1a;2138 | 0 | 0 |
| NC_002645.1;NP_073550.1;HCoV-229E_1a;2139 | 0 | 0 |
| NC_002645.1;NP_073550.1;HCoV-229E_1a;2140 | 0 | 0 |

| | | |
|---|---|---|
| NC_002645.1;NP_073550.1;HCoV-229E_1a;2141 | 0 | 0 |
| NC_002645.1;NP_073550.1;HCoV-229E_1a;2142 | 0 | 0 |
| NC_002645.1;NP_073550.1;HCoV-229E_1a;2143 | 0 | 0 |
| NC_002645.1;NP_073550.1;HCoV-229E_1a;2144 | 0 | 0 |
| NC_002645.1;NP_073550.1;HCoV-229E_1a;2145 | 0 | 0 |
| NC_002645.1;NP_073550.1;HCoV-229E_1a;2146 | 0 | 0 |
| NC_002645.1;NP_073550.1;HCoV-229E_1a;2147 | 0 | 0 |
| NC_002645.1;NP_073550.1;HCoV-229E_1a;2148 | 0 | 0 |
| NC_002645.1;NP_073550.1;HCoV-229E_1a;2149 | 0 | 0 |
| NC_002645.1;NP_073550.1;HCoV-229E_1a;2150 | 0 | 0 |
| NC_002645.1;NP_073550.1;HCoV-229E_1a;2151 | 0.05 | 0 |
| NC_002645.1;NP_073550.1;HCoV-229E_1a;2152 | 0 | 0 |
| NC_002645.1;NP_073550.1;HCoV-229E_1a;2153 | 0 | 0 |
| NC_002645.1;NP_073550.1;HCoV-229E_1a;2154 | 0 | 0 |
| NC_002645.1;NP_073550.1;HCoV-229E_1a;2155 | 0 | 0 |
| NC_002645.1;NP_073550.1;HCoV-229E_1a;2156 | 0 | 0 |
| NC_002645.1;NP_073550.1;HCoV-229E_1a;2157 | 0 | 0 |
| NC_002645.1;NP_073550.1;HCoV-229E_1a;2158 | 0 | 0 |
| NC_002645.1;NP_073550.1;HCoV-229E_1a;2159 | 0 | 0 |
| NC_002645.1;NP_073550.1;HCoV-229E_1a;2160 | 0 | 0 |
| NC_002645.1;NP_073550.1;HCoV-229E_1a;2161 | 0 | 0 |
| NC_002645.1;NP_073550.1;HCoV-229E_1a;2162 | 0 | 0 |
| NC_002645.1;NP_073550.1;HCoV-229E_1a;2163 | 0 | 0.025 |
| NC_002645.1;NP_073550.1;HCoV-229E_1a;2164 | 0 | 0 |
| NC_002645.1;NP_073550.1;HCoV-229E_1a;2165 | 0 | 0.025 |
| NC_002645.1;NP_073550.1;HCoV-229E_1a;2166 | 0 | 0 |
| NC_002645.1;NP_073550.1;HCoV-229E_1a;2167 | 0 | 0 |
| NC_002645.1;NP_073550.1;HCoV-229E_1a;2168 | 0 | 0 |
| NC_002645.1;NP_073550.1;HCoV-229E_1a;2169 | 0 | 0 |
| NC_002645.1;NP_073550.1;HCoV-229E_1a;2170 | 0 | 0 |
| NC_002645.1;NP_073550.1;HCoV-229E_1a;2171 | 0 | 0 |
| NC_002645.1;NP_073550.1;HCoV-229E_1a;2172 | 0 | 0 |
| NC_002645.1;NP_073550.1;HCoV-229E_1a;2173 | 0.05 | 0 |
| NC_002645.1;NP_073550.1;HCoV-229E_1a;2174 | 0 | 0 |
| NC_002645.1;NP_073550.1;HCoV-229E_1a;2175 | 0.05 | 0 |
| NC_002645.1;NP_073550.1;HCoV-229E_1a;2176 | 0 | 0 |
| NC_002645.1;NP_073550.1;HCoV-229E_1a;2177 | 0 | 0 |
| NC_002645.1;NP_073550.1;HCoV-229E_1a;2178 | 0 | 0 |
| NC_002645.1;NP_073550.1;HCoV-229E_1a;2179 | 0 | 0 |
| NC_002645.1;NP_073550.1;HCoV-229E_1a;2180 | 0 | 0 |
| NC_002645.1;NP_073550.1;HCoV-229E_1a;2181 | 0 | 0 |
| NC_002645.1;NP_073550.1;HCoV-229E_1a;2182 | 0 | 0 |
| NC_002645.1;NP_073550.1;HCoV-229E_1a;2183 | 0 | 0 |

| | | |
|---|---|---|
| NC_002645.1;NP_073550.1;HCoV-229E_1a;2184 | 0 | 0 |
| NC_002645.1;NP_073550.1;HCoV-229E_1a;2185 | 0 | 0.025 |
| NC_002645.1;NP_073550.1;HCoV-229E_1a;2186 | 0 | 0 |
| NC_002645.1;NP_073550.1;HCoV-229E_1a;2187 | 0 | 0 |
| NC_002645.1;NP_073550.1;HCoV-229E_1a;2188 | 0 | 0 |
| NC_002645.1;NP_073550.1;HCoV-229E_1a;2189 | 0 | 0 |
| NC_002645.1;NP_073550.1;HCoV-229E_1a;2190 | 0 | 0 |
| NC_002645.1;NP_073550.1;HCoV-229E_1a;2191 | 0 | 0 |
| NC_002645.1;NP_073550.1;HCoV-229E_1a;2192 | 0 | 0 |
| NC_002645.1;NP_073550.1;HCoV-229E_1a;2193 | 0 | 0 |
| NC_002645.1;NP_073550.1;HCoV-229E_1a;2194 | 0 | 0 |
| NC_002645.1;NP_073550.1;HCoV-229E_1a;2195 | 0 | 0 |
| NC_002645.1;NP_073550.1;HCoV-229E_1a;2196 | 0 | 0 |
| NC_002645.1;NP_073550.1;HCoV-229E_1a;2197 | 0 | 0 |
| NC_002645.1;NP_073550.1;HCoV-229E_1a;2198 | 0 | 0 |
| NC_002645.1;NP_073550.1;HCoV-229E_1a;2199 | 0 | 0 |
| NC_002645.1;NP_073550.1;HCoV-229E_1a;2200 | 0 | 0 |
| NC_002645.1;NP_073550.1;HCoV-229E_1a;2201 | 0 | 0 |
| NC_002645.1;NP_073550.1;HCoV-229E_1a;2202 | 0 | 0 |
| NC_002645.1;NP_073550.1;HCoV-229E_1a;2203 | 0 | 0 |
| NC_002645.1;NP_073550.1;HCoV-229E_1a;2204 | 0 | 0 |
| NC_002645.1;NP_073550.1;HCoV-229E_1a;2205 | 0 | 0 |
| NC_002645.1;NP_073550.1;HCoV-229E_1a;2206 | 0 | 0 |
| NC_002645.1;NP_073550.1;HCoV-229E_1a;2207 | 0 | 0 |
| NC_002645.1;NP_073550.1;HCoV-229E_1a;2208 | 0.05 | 0.025 |
| NC_002645.1;NP_073550.1;HCoV-229E_1a;2209 | 0.05 | 0 |
| NC_002645.1;NP_073550.1;HCoV-229E_1a;2210 | 0.05 | 0 |
| NC_002645.1;NP_073550.1;HCoV-229E_1a;2211 | 0 | 0.025 |
| NC_002645.1;NP_073550.1;HCoV-229E_1a;2212 | 0 | 0.05 |
| NC_002645.1;NP_073550.1;HCoV-229E_1a;2213 | 0 | 0.025 |
| NC_002645.1;NP_073550.1;HCoV-229E_1a;2214 | 0 | 0.025 |
| NC_002645.1;NP_073550.1;HCoV-229E_1a;2215 | 0 | 0.025 |
| NC_002645.1;NP_073550.1;HCoV-229E_1a;2216 | 0.05 | 0.05 |
| NC_002645.1;NP_073550.1;HCoV-229E_1a;2217 | 0.05 | 0.075 |
| NC_002645.1;NP_073550.1;HCoV-229E_1a;2218 | 0.05 | 0 |
| NC_002645.1;NP_073550.1;HCoV-229E_1a;2219 | 0.05 | 0 |
| NC_002645.1;NP_073550.1;HCoV-229E_1a;2220 | 0.05 | 0 |
| NC_002645.1;NP_073550.1;HCoV-229E_1a;2221 | 0.05 | 0 |
| NC_002645.1;NP_073550.1;HCoV-229E_1a;2222 | 0.05 | 0 |
| NC_002645.1;NP_073550.1;HCoV-229E_1a;2223 | 0.15 | 0 |
| NC_002645.1;NP_073550.1;HCoV-229E_1a;2224 | 0.05 | 0 |
| NC_002645.1;NP_073550.1;HCoV-229E_1a;2225 | 0 | 0 |
| NC_002645.1;NP_073550.1;HCoV-229E_1a;2226 | 0 | 0 |

| | | |
|---|---|---|
| NC_002645.1;NP_073550.1;HCoV-229E_1a;2227 | 0 | 0 |
| NC_002645.1;NP_073550.1;HCoV-229E_1a;2228 | 0.1 | 0 |
| NC_002645.1;NP_073550.1;HCoV-229E_1a;2229 | 0 | 0 |
| NC_002645.1;NP_073550.1;HCoV-229E_1a;2230 | 0 | 0 |
| NC_002645.1;NP_073550.1;HCoV-229E_1a;2231 | 0 | 0 |
| NC_002645.1;NP_073550.1;HCoV-229E_1a;2232 | 0.05 | 0 |
| NC_002645.1;NP_073550.1;HCoV-229E_1a;2233 | 0.1 | 0 |
| NC_002645.1;NP_073550.1;HCoV-229E_1a;2234 | 0.3 | 0.25 |
| NC_002645.1;NP_073550.1;HCoV-229E_1a;2235 | 0 | 0 |
| NC_002645.1;NP_073550.1;HCoV-229E_1a;2236 | 0 | 0.025 |
| NC_002645.1;NP_073550.1;HCoV-229E_1a;2237 | 0 | 0 |
| NC_002645.1;NP_073550.1;HCoV-229E_1a;2238 | 0 | 0 |
| NC_002645.1;NP_073550.1;HCoV-229E_1a;2239 | 0 | 0 |
| NC_002645.1;NP_073550.1;HCoV-229E_1a;2240 | 0 | 0 |
| NC_002645.1;NP_073550.1;HCoV-229E_1a;2241 | 0 | 0 |
| NC_002645.1;NP_073550.1;HCoV-229E_1a;2242 | 0 | 0 |
| NC_002645.1;NP_073550.1;HCoV-229E_1a;2243 | 0 | 0 |
| NC_002645.1;NP_073550.1;HCoV-229E_1a;2244 | 0 | 0 |
| NC_002645.1;NP_073550.1;HCoV-229E_1a;2245 | 0 | 0 |
| NC_002645.1;NP_073550.1;HCoV-229E_1a;2246 | 0 | 0 |
| NC_002645.1;NP_073550.1;HCoV-229E_1a;2247 | 0 | 0 |
| NC_002645.1;NP_073550.1;HCoV-229E_1a;2248 | 0 | 0 |
| NC_002645.1;NP_073550.1;HCoV-229E_1a;2249 | 0 | 0 |
| NC_002645.1;NP_073550.1;HCoV-229E_1a;2250 | 0 | 0 |
| NC_002645.1;NP_073550.1;HCoV-229E_1a;2251 | 0 | 0 |
| NC_002645.1;NP_073550.1;HCoV-229E_1a;2252 | 0 | 0 |
| NC_002645.1;NP_073550.1;HCoV-229E_1a;2253 | 0 | 0 |
| NC_002645.1;NP_073550.1;HCoV-229E_1a;2254 | 0 | 0 |
| NC_002645.1;NP_073550.1;HCoV-229E_1a;2255 | 0.05 | 0.025 |
| NC_002645.1;NP_073550.1;HCoV-229E_1a;2256 | 0 | 0 |
| NC_002645.1;NP_073550.1;HCoV-229E_1a;2257 | 0 | 0 |
| NC_002645.1;NP_073550.1;HCoV-229E_1a;2258 | 0 | 0 |
| NC_002645.1;NP_073550.1;HCoV-229E_1a;2259 | 0.05 | 0.025 |
| NC_002645.1;NP_073550.1;HCoV-229E_1a;2260 | 0.1 | 0.025 |
| NC_002645.1;NP_073550.1;HCoV-229E_1a;2261 | 0.05 | 0 |
| NC_002645.1;NP_073550.1;HCoV-229E_1a;2262 | 0 | 0 |
| NC_002645.1;NP_073550.1;HCoV-229E_1a;2263 | 0 | 0 |
| NC_002645.1;NP_073550.1;HCoV-229E_1a;2264 | 0 | 0.025 |
| NC_002645.1;NP_073550.1;HCoV-229E_1a;2265 | 0 | 0 |
| NC_002645.1;NP_073550.1;HCoV-229E_1a;2266 | 0.05 | 0 |
| NC_002645.1;NP_073550.1;HCoV-229E_1a;2267 | 0.05 | 0.05 |
| NC_002645.1;NP_073550.1;HCoV-229E_1a;2268 | 0 | 0 |
| NC_002645.1;NP_073550.1;HCoV-229E_1a;2269 | 0 | 0 |

| | | |
|---|---|---|
| NC_002645.1;NP_073550.1;HCoV-229E_1a;2270 | 0 | 0 |
| NC_002645.1;NP_073550.1;HCoV-229E_1a;2271 | 0 | 0 |
| NC_002645.1;NP_073550.1;HCoV-229E_1a;2272 | 0 | 0 |
| NC_002645.1;NP_073550.1;HCoV-229E_1a;2273 | 0 | 0 |
| NC_002645.1;NP_073550.1;HCoV-229E_1a;2274 | 0 | 0.025 |
| NC_002645.1;NP_073550.1;HCoV-229E_1a;2275 | 0 | 0 |
| NC_002645.1;NP_073550.1;HCoV-229E_1a;2276 | 0 | 0 |
| NC_002645.1;NP_073550.1;HCoV-229E_1a;2277 | 0 | 0 |
| NC_002645.1;NP_073550.1;HCoV-229E_1a;2278 | 0 | 0 |
| NC_002645.1;NP_073550.1;HCoV-229E_1a;2279 | 0 | 0.025 |
| NC_002645.1;NP_073550.1;HCoV-229E_1a;2280 | 0 | 0.025 |
| NC_002645.1;NP_073550.1;HCoV-229E_1a;2281 | 0.1 | 0.025 |
| NC_002645.1;NP_073550.1;HCoV-229E_1a;2282 | 0.1 | 0.025 |
| NC_002645.1;NP_073550.1;HCoV-229E_1a;2283 | 0.1 | 0.075 |
| NC_002645.1;NP_073550.1;HCoV-229E_1a;2284 | 0.1 | 0 |
| NC_002645.1;NP_073550.1;HCoV-229E_1a;2285 | 0.1 | 0 |
| NC_002645.1;NP_073550.1;HCoV-229E_1a;2286 | 0.25 | 0.05 |
| NC_002645.1;NP_073550.1;HCoV-229E_1a;2287 | 0.25 | 0.025 |
| NC_002645.1;NP_073550.1;HCoV-229E_1a;2288 | 0.25 | 0.125 |
| NC_002645.1;NP_073550.1;HCoV-229E_1a;2289 | 0.2 | 0.075 |
| NC_002645.1;NP_073550.1;HCoV-229E_1a;2290 | 0.1 | 0 |
| NC_002645.1;NP_073550.1;HCoV-229E_1a;2291 | 0.1 | 0 |
| NC_002645.1;NP_073550.1;HCoV-229E_1a;2292 | 0.05 | 0 |
| NC_002645.1;NP_073550.1;HCoV-229E_1a;2293 | 0.05 | 0 |
| NC_002645.1;NP_073550.1;HCoV-229E_1a;2294 | 0 | 0 |
| NC_002645.1;NP_073550.1;HCoV-229E_1a;2295 | 0 | 0 |
| NC_002645.1;NP_073550.1;HCoV-229E_1a;2296 | 0 | 0 |
| NC_002645.1;NP_073550.1;HCoV-229E_1a;2297 | 0 | 0 |
| NC_002645.1;NP_073550.1;HCoV-229E_1a;2298 | 0 | 0 |
| NC_002645.1;NP_073550.1;HCoV-229E_1a;2299 | 0 | 0 |
| NC_002645.1;NP_073550.1;HCoV-229E_1a;2300 | 0 | 0 |
| NC_002645.1;NP_073550.1;HCoV-229E_1a;2301 | 0 | 0 |
| NC_002645.1;NP_073550.1;HCoV-229E_1a;2302 | 0 | 0 |
| NC_002645.1;NP_073550.1;HCoV-229E_1a;2303 | 0 | 0 |
| NC_002645.1;NP_073550.1;HCoV-229E_1a;2304 | 0 | 0.025 |
| NC_002645.1;NP_073550.1;HCoV-229E_1a;2305 | 0 | 0 |
| NC_002645.1;NP_073550.1;HCoV-229E_1a;2306 | 0 | 0 |
| NC_002645.1;NP_073550.1;HCoV-229E_1a;2307 | 0 | 0 |
| NC_002645.1;NP_073550.1;HCoV-229E_1a;2308 | 0 | 0 |
| NC_002645.1;NP_073550.1;HCoV-229E_1a;2309 | 0 | 0 |
| NC_002645.1;NP_073550.1;HCoV-229E_1a;2310 | 0 | 0.025 |
| NC_002645.1;NP_073550.1;HCoV-229E_1a;2311 | 0 | 0 |
| NC_002645.1;NP_073550.1;HCoV-229E_1a;2312 | 0 | 0 |

| | | |
|---|---|---|
| NC_002645.1;NP_073550.1;HCoV-229E_1a;2313 | 0 | 0 |
| NC_002645.1;NP_073550.1;HCoV-229E_1a;2314 | 0.05 | 0 |
| NC_002645.1;NP_073550.1;HCoV-229E_1a;2315 | 0 | 0 |
| NC_002645.1;NP_073550.1;HCoV-229E_1a;2316 | 0 | 0 |
| NC_002645.1;NP_073550.1;HCoV-229E_1a;2317 | 0 | 0.025 |
| NC_002645.1;NP_073550.1;HCoV-229E_1a;2318 | 0 | 0 |
| NC_002645.1;NP_073550.1;HCoV-229E_1a;2319 | 0 | 0 |
| NC_002645.1;NP_073550.1;HCoV-229E_1a;2320 | 0 | 0 |
| NC_002645.1;NP_073550.1;HCoV-229E_1a;2321 | 0 | 0 |
| NC_002645.1;NP_073550.1;HCoV-229E_1a;2322 | 0 | 0 |
| NC_002645.1;NP_073550.1;HCoV-229E_1a;2323 | 0 | 0 |
| NC_002645.1;NP_073550.1;HCoV-229E_1a;2324 | 0 | 0 |
| NC_002645.1;NP_073550.1;HCoV-229E_1a;2325 | 0 | 0 |
| NC_002645.1;NP_073550.1;HCoV-229E_1a;2326 | 0 | 0 |
| NC_002645.1;NP_073550.1;HCoV-229E_1a;2327 | 0 | 0 |
| NC_002645.1;NP_073550.1;HCoV-229E_1a;2328 | 0 | 0 |
| NC_002645.1;NP_073550.1;HCoV-229E_1a;2329 | 0 | 0 |
| NC_002645.1;NP_073550.1;HCoV-229E_1a;2330 | 0 | 0 |
| NC_002645.1;NP_073550.1;HCoV-229E_1a;2331 | 0 | 0 |
| NC_002645.1;NP_073550.1;HCoV-229E_1a;2332 | 0 | 0.025 |
| NC_002645.1;NP_073550.1;HCoV-229E_1a;2333 | 0 | 0.025 |
| NC_002645.1;NP_073550.1;HCoV-229E_1a;2334 | 0 | 0.05 |
| NC_002645.1;NP_073550.1;HCoV-229E_1a;2335 | 0 | 0.1 |
| NC_002645.1;NP_073550.1;HCoV-229E_1a;2336 | 0.05 | 0.1 |
| NC_002645.1;NP_073550.1;HCoV-229E_1a;2337 | 0.1 | 0.05 |
| NC_002645.1;NP_073550.1;HCoV-229E_1a;2338 | 0.1 | 0.075 |
| NC_002645.1;NP_073550.1;HCoV-229E_1a;2339 | 0.05 | 0.025 |
| NC_002645.1;NP_073550.1;HCoV-229E_1a;2340 | 0.05 | 0.075 |
| NC_002645.1;NP_073550.1;HCoV-229E_1a;2341 | 0.1 | 0.075 |
| NC_002645.1;NP_073550.1;HCoV-229E_1a;2342 | 0.1 | 0.075 |
| NC_002645.1;NP_073550.1;HCoV-229E_1a;2343 | 0.05 | 0.05 |
| NC_002645.1;NP_073550.1;HCoV-229E_1a;2344 | 0.05 | 0.05 |
| NC_002645.1;NP_073550.1;HCoV-229E_1a;2345 | 0.05 | 0.025 |
| NC_002645.1;NP_073550.1;HCoV-229E_1a;2346 | 0.05 | 0.1 |
| NC_002645.1;NP_073550.1;HCoV-229E_1a;2347 | 0.05 | 0.075 |
| NC_002645.1;NP_073550.1;HCoV-229E_1a;2348 | 0 | 0.1 |
| NC_002645.1;NP_073550.1;HCoV-229E_1a;2349 | 0 | 0.1 |
| NC_002645.1;NP_073550.1;HCoV-229E_1a;2350 | 0 | 0.075 |
| NC_002645.1;NP_073550.1;HCoV-229E_1a;2351 | 0 | 0.075 |
| NC_002645.1;NP_073550.1;HCoV-229E_1a;2352 | 0 | 0.075 |
| NC_002645.1;NP_073550.1;HCoV-229E_1a;2353 | 0 | 0.025 |
| NC_002645.1;NP_073550.1;HCoV-229E_1a;2354 | 0 | 0.05 |
| NC_002645.1;NP_073550.1;HCoV-229E_1a;2355 | 0.05 | 0.05 |

| | | |
|---|---|---|
| NC_002645.1;NP_073550.1;HCoV-229E_1a;2356 | 0.05 | 0.05 |
| NC_002645.1;NP_073550.1;HCoV-229E_1a;2357 | 0.05 | 0.075 |
| NC_002645.1;NP_073550.1;HCoV-229E_1a;2358 | 0.2 | 0.225 |
| NC_002645.1;NP_073550.1;HCoV-229E_1a;2359 | 0 | 0 |
| NC_002645.1;NP_073550.1;HCoV-229E_1a;2360 | 0 | 0 |
| NC_002645.1;NP_073550.1;HCoV-229E_1a;2361 | 0 | 0 |
| NC_002645.1;NP_073550.1;HCoV-229E_1a;2362 | 0 | 0 |
| NC_002645.1;NP_073550.1;HCoV-229E_1a;2363 | 0.05 | 0 |
| NC_002645.1;NP_073550.1;HCoV-229E_1a;2364 | 0 | 0 |
| NC_002645.1;NP_073550.1;HCoV-229E_1a;2365 | 0 | 0 |
| NC_002645.1;NP_073550.1;HCoV-229E_1a;2366 | 0 | 0 |
| NC_002645.1;NP_073550.1;HCoV-229E_1a;2367 | 0 | 0 |
| NC_002645.1;NP_073550.1;HCoV-229E_1a;2368 | 0 | 0 |
| NC_002645.1;NP_073550.1;HCoV-229E_1a;2369 | 0 | 0 |
| NC_002645.1;NP_073550.1;HCoV-229E_1a;2370 | 0 | 0 |
| NC_002645.1;NP_073550.1;HCoV-229E_1a;2371 | 0 | 0 |
| NC_002645.1;NP_073550.1;HCoV-229E_1a;2372 | 0 | 0 |
| NC_002645.1;NP_073550.1;HCoV-229E_1a;2373 | 0 | 0 |
| NC_002645.1;NP_073550.1;HCoV-229E_1a;2374 | 0.05 | 0 |
| NC_002645.1;NP_073550.1;HCoV-229E_1a;2375 | 0 | 0 |
| NC_002645.1;NP_073550.1;HCoV-229E_1a;2376 | 0 | 0 |
| NC_002645.1;NP_073550.1;HCoV-229E_1a;2377 | 0 | 0 |
| NC_002645.1;NP_073550.1;HCoV-229E_1a;2378 | 0 | 0 |
| NC_002645.1;NP_073550.1;HCoV-229E_1a;2379 | 0 | 0 |
| NC_002645.1;NP_073550.1;HCoV-229E_1a;2380 | 0 | 0 |
| NC_002645.1;NP_073550.1;HCoV-229E_1a;2381 | 0 | 0 |
| NC_002645.1;NP_073550.1;HCoV-229E_1a;2382 | 0 | 0 |
| NC_002645.1;NP_073550.1;HCoV-229E_1a;2383 | 0.05 | 0 |
| NC_002645.1;NP_073550.1;HCoV-229E_1a;2384 | 0 | 0 |
| NC_002645.1;NP_073550.1;HCoV-229E_1a;2385 | 0 | 0 |
| NC_002645.1;NP_073550.1;HCoV-229E_1a;2386 | 0.1 | 0 |
| NC_002645.1;NP_073550.1;HCoV-229E_1a;2387 | 0 | 0.025 |
| NC_002645.1;NP_073550.1;HCoV-229E_1a;2388 | 0.05 | 0.1 |
| NC_002645.1;NP_073550.1;HCoV-229E_1a;2389 | 0.05 | 0.075 |
| NC_002645.1;NP_073550.1;HCoV-229E_1a;2390 | 0.1 | 0.075 |
| NC_002645.1;NP_073550.1;HCoV-229E_1a;2391 | 0 | 0.025 |
| NC_002645.1;NP_073550.1;HCoV-229E_1a;2392 | 0.05 | 0.025 |
| NC_002645.1;NP_073550.1;HCoV-229E_1a;2393 | 0 | 0.025 |
| NC_002645.1;NP_073550.1;HCoV-229E_1a;2394 | 0 | 0.05 |
| NC_002645.1;NP_073550.1;HCoV-229E_1a;2395 | 0 | 0.025 |
| NC_002645.1;NP_073550.1;HCoV-229E_1a;2396 | 0 | 0.025 |
| NC_002645.1;NP_073550.1;HCoV-229E_1a;2397 | 0 | 0.025 |
| NC_002645.1;NP_073550.1;HCoV-229E_1a;2398 | 0 | 0.025 |

| | | |
|---|---|---|
| NC_002645.1;NP_073550.1;HCoV-229E_1a;2399 | 0 | 0 |
| NC_002645.1;NP_073550.1;HCoV-229E_1a;2400 | 0 | 0 |
| NC_002645.1;NP_073550.1;HCoV-229E_1a;2401 | 0 | 0.025 |
| NC_002645.1;NP_073550.1;HCoV-229E_1a;2402 | 0 | 0 |
| NC_002645.1;NP_073550.1;HCoV-229E_1a;2403 | 0 | 0 |
| NC_002645.1;NP_073550.1;HCoV-229E_1a;2404 | 0 | 0 |
| NC_002645.1;NP_073550.1;HCoV-229E_1a;2405 | 0 | 0 |
| NC_002645.1;NP_073550.1;HCoV-229E_1a;2406 | 0 | 0 |
| NC_002645.1;NP_073550.1;HCoV-229E_1a;2407 | 0 | 0 |
| NC_002645.1;NP_073550.1;HCoV-229E_1a;2408 | 0 | 0 |
| NC_002645.1;NP_073550.1;HCoV-229E_1a;2409 | 0 | 0 |
| NC_002645.1;NP_073550.1;HCoV-229E_1a;2410 | 0 | 0 |
| NC_002645.1;NP_073550.1;HCoV-229E_1a;2411 | 0 | 0 |
| NC_002645.1;NP_073550.1;HCoV-229E_1a;2412 | 0 | 0 |
| NC_002645.1;NP_073550.1;HCoV-229E_1a;2413 | 0 | 0 |
| NC_002645.1;NP_073550.1;HCoV-229E_1a;2414 | 0 | 0 |
| NC_002645.1;NP_073550.1;HCoV-229E_1a;2415 | 0.05 | 0 |
| NC_002645.1;NP_073550.1;HCoV-229E_1a;2416 | 0.05 | 0 |
| NC_002645.1;NP_073550.1;HCoV-229E_1a;2417 | 0 | 0.025 |
| NC_002645.1;NP_073550.1;HCoV-229E_1a;2418 | 0.05 | 0 |
| NC_002645.1;NP_073550.1;HCoV-229E_1a;2419 | 0 | 0 |
| NC_002645.1;NP_073550.1;HCoV-229E_1a;2420 | 0 | 0 |
| NC_002645.1;NP_073550.1;HCoV-229E_1a;2421 | 0 | 0 |
| NC_002645.1;NP_073550.1;HCoV-229E_1a;2422 | 0 | 0 |
| NC_002645.1;NP_073550.1;HCoV-229E_1a;2423 | 0 | 0 |
| NC_002645.1;NP_073550.1;HCoV-229E_1a;2424 | 0 | 0 |
| NC_002645.1;NP_073550.1;HCoV-229E_1a;2425 | 0 | 0 |
| NC_002645.1;NP_073550.1;HCoV-229E_1a;2426 | 0 | 0 |
| NC_002645.1;NP_073550.1;HCoV-229E_1a;2427 | 0.05 | 0.05 |
| NC_002645.1;NP_073550.1;HCoV-229E_1a;2428 | 0.05 | 0.05 |
| NC_002645.1;NP_073550.1;HCoV-229E_1a;2429 | 0.05 | 0.05 |
| NC_002645.1;NP_073550.1;HCoV-229E_1a;2430 | 0.1 | 0.05 |
| NC_002645.1;NP_073550.1;HCoV-229E_1a;2431 | 0 | 0.025 |
| NC_002645.1;NP_073550.1;HCoV-229E_1a;2432 | 0.1 | 0 |
| NC_002645.1;NP_073550.1;HCoV-229E_1a;2433 | 0.1 | 0 |
| NC_002645.1;NP_073550.1;HCoV-229E_1a;2434 | 0.15 | 0 |
| NC_002645.1;NP_073550.1;HCoV-229E_1a;2435 | 0.05 | 0 |
| NC_002645.1;NP_073550.1;HCoV-229E_1a;2436 | 0.05 | 0 |
| NC_002645.1;NP_073550.1;HCoV-229E_1a;2437 | 0 | 0 |
| NC_002645.1;NP_073550.1;HCoV-229E_1a;2438 | 0 | 0 |
| NC_002645.1;NP_073550.1;HCoV-229E_1a;2439 | 0 | 0 |
| NC_002645.1;NP_073550.1;HCoV-229E_1a;2440 | 0 | 0 |
| NC_002645.1;NP_073550.1;HCoV-229E_1a;2441 | 0 | 0 |

| | | |
|---|---|---|
| NC_002645.1;NP_073550.1;HCoV-229E_1a;2442 | 0 | 0 |
| NC_002645.1;NP_073550.1;HCoV-229E_1a;2443 | 0 | 0 |
| NC_002645.1;NP_073550.1;HCoV-229E_1a;2444 | 0 | 0 |
| NC_002645.1;NP_073550.1;HCoV-229E_1a;2445 | 0 | 0 |
| NC_002645.1;NP_073550.1;HCoV-229E_1a;2446 | 0 | 0 |
| NC_002645.1;NP_073550.1;HCoV-229E_1a;2447 | 0 | 0 |
| NC_002645.1;NP_073550.1;HCoV-229E_1a;2448 | 0 | 0 |
| NC_002645.1;NP_073550.1;HCoV-229E_1a;2449 | 0 | 0 |
| NC_002645.1;NP_073550.1;HCoV-229E_1a;2450 | 0 | 0 |
| NC_002645.1;NP_073550.1;HCoV-229E_1a;2451 | 0 | 0 |
| NC_002645.1;NP_073550.1;HCoV-229E_1a;2452 | 0 | 0 |
| NC_002645.1;NP_073550.1;HCoV-229E_1a;2453 | 0 | 0 |
| NC_002645.1;NP_073550.1;HCoV-229E_1a;2454 | 0 | 0 |
| NC_002645.1;NP_073550.1;HCoV-229E_1a;2455 | 0 | 0 |
| NC_002645.1;NP_073550.1;HCoV-229E_1a;2456 | 0 | 0 |
| NC_002645.1;NP_073550.1;HCoV-229E_1a;2457 | 0 | 0 |
| NC_002645.1;NP_073550.1;HCoV-229E_1a;2458 | 0 | 0 |
| NC_002645.1;NP_073550.1;HCoV-229E_1a;2459 | 0 | 0 |
| NC_002645.1;NP_073550.1;HCoV-229E_1a;2460 | 0 | 0 |
| NC_002645.1;NP_073550.1;HCoV-229E_1a;2461 | 0 | 0 |
| NC_002645.1;NP_073550.1;HCoV-229E_1a;2462 | 0 | 0 |
| NC_002645.1;NP_073550.1;HCoV-229E_1a;2463 | 0 | 0 |
| NC_002645.1;NP_073550.1;HCoV-229E_1a;2464 | 0 | 0 |
| NC_002645.1;NP_073550.1;HCoV-229E_1a;2465 | 0 | 0 |
| NC_002645.1;NP_073550.1;HCoV-229E_1a;2466 | 0 | 0 |
| NC_002645.1;NP_073550.1;HCoV-229E_1a;2467 | 0 | 0 |
| NC_002645.1;NP_073550.1;HCoV-229E_1a;2468 | 0 | 0 |
| NC_002645.1;NP_073550.1;HCoV-229E_1a;2469 | 0 | 0 |
| NC_002645.1;NP_073550.1;HCoV-229E_1a;2470 | 0 | 0 |
| NC_002645.1;NP_073550.1;HCoV-229E_1a;2471 | 0 | 0.025 |
| NC_002645.1;NP_073550.1;HCoV-229E_1a;2472 | 0 | 0.025 |
| NC_002645.1;NP_073550.1;HCoV-229E_1a;2473 | 0 | 0.025 |
| NC_002645.1;NP_073550.1;HCoV-229E_1a;2474 | 0 | 0.025 |
| NC_002645.1;NP_073550.1;HCoV-229E_1a;2475 | 0 | 0.025 |
| NC_002645.1;NP_073550.1;HCoV-229E_1a;2476 | 0 | 0.025 |
| NC_002645.1;NP_073550.1;HCoV-229E_1a;2477 | 0 | 0.025 |
| NC_002645.1;NP_073550.1;HCoV-229E_1a;2478 | 0 | 0.025 |
| NC_002645.1;NP_073550.1;HCoV-229E_1a;2479 | 0 | 0.025 |
| NC_002645.1;NP_073550.1;HCoV-229E_1a;2480 | 0 | 0.025 |
| NC_002645.1;NP_073550.1;HCoV-229E_1a;2481 | 0 | 0.025 |
| NC_002645.1;NP_073550.1;HCoV-229E_1a;2482 | 0 | 0 |
| NC_002645.1;NP_073550.1;HCoV-229E_1a;2483 | 0 | 0 |
| NC_002645.1;NP_073550.1;HCoV-229E_1a;2484 | 0 | 0 |

| | | |
|---|---|---|
| NC_002645.1;NP_073550.1;HCoV-229E_1a;2485 | 0 | 0 |
| NC_002645.1;NP_073550.1;HCoV-229E_1a;2486 | 0 | 0 |
| NC_002645.1;NP_073550.1;HCoV-229E_1a;2487 | 0 | 0 |
| NC_002645.1;NP_073550.1;HCoV-229E_1a;2488 | 0 | 0 |
| NC_002645.1;NP_073550.1;HCoV-229E_1a;2489 | 0 | 0 |
| NC_002645.1;NP_073550.1;HCoV-229E_1a;2490 | 0 | 0 |
| NC_002645.1;NP_073550.1;HCoV-229E_1a;2491 | 0 | 0 |
| NC_002645.1;NP_073550.1;HCoV-229E_1a;2492 | 0 | 0 |
| NC_002645.1;NP_073550.1;HCoV-229E_1a;2493 | 0 | 0 |
| NC_002645.1;NP_073550.1;HCoV-229E_1a;2494 | 0 | 0 |
| NC_002645.1;NP_073550.1;HCoV-229E_1a;2495 | 0 | 0 |
| NC_002645.1;NP_073550.1;HCoV-229E_1a;2496 | 0 | 0 |
| NC_002645.1;NP_073550.1;HCoV-229E_1a;2497 | 0 | 0 |
| NC_002645.1;NP_073550.1;HCoV-229E_1a;2498 | 0 | 0 |
| NC_002645.1;NP_073550.1;HCoV-229E_1a;2499 | 0 | 0 |
| NC_002645.1;NP_073550.1;HCoV-229E_1a;2500 | 0 | 0 |
| NC_002645.1;NP_073550.1;HCoV-229E_1a;2501 | 0 | 0 |
| NC_002645.1;NP_073550.1;HCoV-229E_1a;2502 | 0 | 0 |
| NC_002645.1;NP_073550.1;HCoV-229E_1a;2503 | 0 | 0 |
| NC_002645.1;NP_073550.1;HCoV-229E_1a;2504 | 0 | 0 |
| NC_002645.1;NP_073550.1;HCoV-229E_1a;2505 | 0 | 0 |
| NC_002645.1;NP_073550.1;HCoV-229E_1a;2506 | 0 | 0 |
| NC_002645.1;NP_073550.1;HCoV-229E_1a;2507 | 0 | 0 |
| NC_002645.1;NP_073550.1;HCoV-229E_1a;2508 | 0 | 0 |
| NC_002645.1;NP_073550.1;HCoV-229E_1a;2509 | 0 | 0 |
| NC_002645.1;NP_073550.1;HCoV-229E_1a;2510 | 0 | 0 |
| NC_002645.1;NP_073550.1;HCoV-229E_1a;2511 | 0 | 0 |
| NC_002645.1;NP_073550.1;HCoV-229E_1a;2512 | 0 | 0 |
| NC_002645.1;NP_073550.1;HCoV-229E_1a;2513 | 0 | 0 |
| NC_002645.1;NP_073550.1;HCoV-229E_1a;2514 | 0 | 0.05 |
| NC_002645.1;NP_073550.1;HCoV-229E_1a;2515 | 0 | 0 |
| NC_002645.1;NP_073550.1;HCoV-229E_1a;2516 | 0 | 0 |
| NC_002645.1;NP_073550.1;HCoV-229E_1a;2517 | 0 | 0 |
| NC_002645.1;NP_073550.1;HCoV-229E_1a;2518 | 0 | 0.025 |
| NC_002645.1;NP_073550.1;HCoV-229E_1a;2519 | 0 | 0.025 |
| NC_002645.1;NP_073550.1;HCoV-229E_1a;2520 | 0 | 0.025 |
| NC_002645.1;NP_073550.1;HCoV-229E_1a;2521 | 0 | 0.075 |
| NC_002645.1;NP_073550.1;HCoV-229E_1a;2522 | 0.05 | 0.05 |
| NC_002645.1;NP_073550.1;HCoV-229E_1a;2523 | 0.05 | 0.075 |
| NC_002645.1;NP_073550.1;HCoV-229E_1a;2524 | 0.1 | 0.025 |
| NC_002645.1;NP_073550.1;HCoV-229E_1a;2525 | 0.15 | 0.1 |
| NC_002645.1;NP_073550.1;HCoV-229E_1a;2526 | 0.1 | 0 |
| NC_002645.1;NP_073550.1;HCoV-229E_1a;2527 | 0.15 | 0 |

| | | |
|---|---|---|
| NC_002645.1;NP_073550.1;HCoV-229E_1a;2528 | 0.15 | 0 |
| NC_002645.1;NP_073550.1;HCoV-229E_1a;2529 | 0.1 | 0.025 |
| NC_002645.1;NP_073550.1;HCoV-229E_1a;2530 | 0.2 | 0 |
| NC_002645.1;NP_073550.1;HCoV-229E_1a;2531 | 0.05 | 0 |
| NC_002645.1;NP_073550.1;HCoV-229E_1a;2532 | 0.05 | 0 |
| NC_002645.1;NP_073550.1;HCoV-229E_1a;2533 | 0.1 | 0 |
| NC_002645.1;NP_073550.1;HCoV-229E_1a;2534 | 0.05 | 0 |
| NC_002645.1;NP_073550.1;HCoV-229E_1a;2535 | 0.05 | 0 |
| NC_002645.1;NP_073550.1;HCoV-229E_1a;2536 | 0.05 | 0 |
| NC_002645.1;NP_073550.1;HCoV-229E_1a;2537 | 0.05 | 0 |
| NC_002645.1;NP_073550.1;HCoV-229E_1a;2538 | 0 | 0 |
| NC_002645.1;NP_073550.1;HCoV-229E_1a;2539 | 0.05 | 0 |
| NC_002645.1;NP_073550.1;HCoV-229E_1a;2540 | 0.1 | 0.025 |
| NC_002645.1;NP_073550.1;HCoV-229E_1a;2541 | 0 | 0 |
| NC_002645.1;NP_073550.1;HCoV-229E_1a;2542 | 0 | 0 |
| NC_002645.1;NP_073550.1;HCoV-229E_1a;2543 | 0 | 0 |
| NC_002645.1;NP_073550.1;HCoV-229E_1a;2544 | 0 | 0 |
| NC_002645.1;NP_073550.1;HCoV-229E_1a;2545 | 0.1 | 0 |
| NC_002645.1;NP_073550.1;HCoV-229E_1a;2546 | 0.05 | 0 |
| NC_002645.1;NP_073550.1;HCoV-229E_1a;2547 | 0.05 | 0.05 |
| NC_002645.1;NP_073550.1;HCoV-229E_1a;2548 | 0 | 0 |
| NC_002645.1;NP_073550.1;HCoV-229E_1a;2549 | 0.1 | 0.025 |
| NC_002645.1;NP_073550.1;HCoV-229E_1a;2550 | 0 | 0 |
| NC_002645.1;NP_073550.1;HCoV-229E_1a;2551 | 0 | 0 |
| NC_002645.1;NP_073550.1;HCoV-229E_1a;2552 | 0 | 0 |
| NC_002645.1;NP_073550.1;HCoV-229E_1a;2553 | 0 | 0 |
| NC_002645.1;NP_073550.1;HCoV-229E_1a;2554 | 0 | 0 |
| NC_002645.1;NP_073550.1;HCoV-229E_1a;2555 | 0 | 0 |
| NC_002645.1;NP_073550.1;HCoV-229E_1a;2556 | 0 | 0 |
| NC_002645.1;NP_073550.1;HCoV-229E_1a;2557 | 0 | 0 |
| NC_002645.1;NP_073550.1;HCoV-229E_1a;2558 | 0 | 0 |
| NC_002645.1;NP_073550.1;HCoV-229E_1a;2559 | 0 | 0 |
| NC_002645.1;NP_073550.1;HCoV-229E_1a;2560 | 0 | 0 |
| NC_002645.1;NP_073550.1;HCoV-229E_1a;2561 | 0 | 0 |
| NC_002645.1;NP_073550.1;HCoV-229E_1a;2562 | 0 | 0 |
| NC_002645.1;NP_073550.1;HCoV-229E_1a;2563 | 0 | 0 |
| NC_002645.1;NP_073550.1;HCoV-229E_1a;2564 | 0 | 0 |
| NC_002645.1;NP_073550.1;HCoV-229E_1a;2565 | 0 | 0 |
| NC_002645.1;NP_073550.1;HCoV-229E_1a;2566 | 0 | 0 |
| NC_002645.1;NP_073550.1;HCoV-229E_1a;2567 | 0 | 0 |
| NC_002645.1;NP_073550.1;HCoV-229E_1a;2568 | 0 | 0 |
| NC_002645.1;NP_073550.1;HCoV-229E_1a;2569 | 0 | 0 |
| NC_002645.1;NP_073550.1;HCoV-229E_1a;2570 | 0 | 0 |

| | | |
|---|---|---|
| NC_002645.1;NP_073550.1;HCoV-229E_1a;2571 | 0 | 0 |
| NC_002645.1;NP_073550.1;HCoV-229E_1a;2572 | 0 | 0 |
| NC_002645.1;NP_073550.1;HCoV-229E_1a;2573 | 0 | 0 |
| NC_002645.1;NP_073550.1;HCoV-229E_1a;2574 | 0 | 0 |
| NC_002645.1;NP_073550.1;HCoV-229E_1a;2575 | 0 | 0 |
| NC_002645.1;NP_073550.1;HCoV-229E_1a;2576 | 0.1 | 0.025 |
| NC_002645.1;NP_073550.1;HCoV-229E_1a;2577 | 0.15 | 0 |
| NC_002645.1;NP_073550.1;HCoV-229E_1a;2578 | 0 | 0.025 |
| NC_002645.1;NP_073550.1;HCoV-229E_1a;2579 | 0 | 0 |
| NC_002645.1;NP_073550.1;HCoV-229E_1a;2580 | 0 | 0 |
| NC_002645.1;NP_073550.1;HCoV-229E_1a;2581 | 0.05 | 0.075 |
| NC_002645.1;NP_073550.1;HCoV-229E_1a;2582 | 0 | 0 |
| NC_002645.1;NP_073550.1;HCoV-229E_1a;2583 | 0 | 0 |
| NC_002645.1;NP_073550.1;HCoV-229E_1a;2584 | 0 | 0 |
| NC_002645.1;NP_073550.1;HCoV-229E_1a;2585 | 0 | 0 |
| NC_002645.1;NP_073550.1;HCoV-229E_1a;2586 | 0 | 0 |
| NC_002645.1;NP_073550.1;HCoV-229E_1a;2587 | 0 | 0 |
| NC_002645.1;NP_073550.1;HCoV-229E_1a;2588 | 0 | 0 |
| NC_002645.1;NP_073550.1;HCoV-229E_1a;2589 | 0 | 0 |
| NC_002645.1;NP_073550.1;HCoV-229E_1a;2590 | 0 | 0 |
| NC_002645.1;NP_073550.1;HCoV-229E_1a;2591 | 0 | 0 |
| NC_002645.1;NP_073550.1;HCoV-229E_1a;2592 | 0 | 0 |
| NC_002645.1;NP_073550.1;HCoV-229E_1a;2593 | 0 | 0 |
| NC_002645.1;NP_073550.1;HCoV-229E_1a;2594 | 0 | 0 |
| NC_002645.1;NP_073550.1;HCoV-229E_1a;2595 | 0 | 0 |
| NC_002645.1;NP_073550.1;HCoV-229E_1a;2596 | 0 | 0 |
| NC_002645.1;NP_073550.1;HCoV-229E_1a;2597 | 0 | 0 |
| NC_002645.1;NP_073550.1;HCoV-229E_1a;2598 | 0 | 0 |
| NC_002645.1;NP_073550.1;HCoV-229E_1a;2599 | 0 | 0 |
| NC_002645.1;NP_073550.1;HCoV-229E_1a;2600 | 0 | 0 |
| NC_002645.1;NP_073550.1;HCoV-229E_1a;2601 | 0 | 0 |
| NC_002645.1;NP_073550.1;HCoV-229E_1a;2602 | 0 | 0 |
| NC_002645.1;NP_073550.1;HCoV-229E_1a;2603 | 0 | 0 |
| NC_002645.1;NP_073550.1;HCoV-229E_1a;2604 | 0 | 0 |
| NC_002645.1;NP_073550.1;HCoV-229E_1a;2605 | 0 | 0 |
| NC_002645.1;NP_073550.1;HCoV-229E_1a;2606 | 0 | 0 |
| NC_002645.1;NP_073550.1;HCoV-229E_1a;2607 | 0 | 0 |
| NC_002645.1;NP_073550.1;HCoV-229E_1a;2608 | 0 | 0 |
| NC_002645.1;NP_073550.1;HCoV-229E_1a;2609 | 0 | 0 |
| NC_002645.1;NP_073550.1;HCoV-229E_1a;2610 | 0 | 0 |
| NC_002645.1;NP_073550.1;HCoV-229E_1a;2611 | 0 | 0 |
| NC_002645.1;NP_073550.1;HCoV-229E_1a;2612 | 0 | 0 |
| NC_002645.1;NP_073550.1;HCoV-229E_1a;2613 | 0 | 0 |

| | | |
|---|---|---|
| NC_002645.1;NP_073550.1;HCoV-229E_1a;2614 | 0 | 0 |
| NC_002645.1;NP_073550.1;HCoV-229E_1a;2615 | 0 | 0 |
| NC_002645.1;NP_073550.1;HCoV-229E_1a;2616 | 0 | 0 |
| NC_002645.1;NP_073550.1;HCoV-229E_1a;2617 | 0 | 0 |
| NC_002645.1;NP_073550.1;HCoV-229E_1a;2618 | 0 | 0 |
| NC_002645.1;NP_073550.1;HCoV-229E_1a;2619 | 0 | 0 |
| NC_002645.1;NP_073550.1;HCoV-229E_1a;2620 | 0 | 0 |
| NC_002645.1;NP_073550.1;HCoV-229E_1a;2621 | 0 | 0 |
| NC_002645.1;NP_073550.1;HCoV-229E_1a;2622 | 0 | 0 |
| NC_002645.1;NP_073550.1;HCoV-229E_1a;2623 | 0.05 | 0 |
| NC_002645.1;NP_073550.1;HCoV-229E_1a;2624 | 0 | 0 |
| NC_002645.1;NP_073550.1;HCoV-229E_1a;2625 | 0 | 0 |
| NC_002645.1;NP_073550.1;HCoV-229E_1a;2626 | 0 | 0 |
| NC_002645.1;NP_073550.1;HCoV-229E_1a;2627 | 0 | 0 |
| NC_002645.1;NP_073550.1;HCoV-229E_1a;2628 | 0 | 0 |
| NC_002645.1;NP_073550.1;HCoV-229E_1a;2629 | 0 | 0 |
| NC_002645.1;NP_073550.1;HCoV-229E_1a;2630 | 0 | 0.025 |
| NC_002645.1;NP_073550.1;HCoV-229E_1a;2631 | 0 | 0.025 |
| NC_002645.1;NP_073550.1;HCoV-229E_1a;2632 | 0 | 0.025 |
| NC_002645.1;NP_073550.1;HCoV-229E_1a;2633 | 0 | 0.025 |
| NC_002645.1;NP_073550.1;HCoV-229E_1a;2634 | 0 | 0 |
| NC_002645.1;NP_073550.1;HCoV-229E_1a;2635 | 0 | 0 |
| NC_002645.1;NP_073550.1;HCoV-229E_1a;2636 | 0 | 0 |
| NC_002645.1;NP_073550.1;HCoV-229E_1a;2637 | 0 | 0 |
| NC_002645.1;NP_073550.1;HCoV-229E_1a;2638 | 0 | 0 |
| NC_002645.1;NP_073550.1;HCoV-229E_1a;2639 | 0 | 0.025 |
| NC_002645.1;NP_073550.1;HCoV-229E_1a;2640 | 0 | 0 |
| NC_002645.1;NP_073550.1;HCoV-229E_1a;2641 | 0 | 0 |
| NC_002645.1;NP_073550.1;HCoV-229E_1a;2642 | 0 | 0 |
| NC_002645.1;NP_073550.1;HCoV-229E_1a;2643 | 0 | 0.025 |
| NC_002645.1;NP_073550.1;HCoV-229E_1a;2644 | 0 | 0 |
| NC_002645.1;NP_073550.1;HCoV-229E_1a;2645 | 0 | 0 |
| NC_002645.1;NP_073550.1;HCoV-229E_1a;2646 | 0 | 0 |
| NC_002645.1;NP_073550.1;HCoV-229E_1a;2647 | 0 | 0 |
| NC_002645.1;NP_073550.1;HCoV-229E_1a;2648 | 0 | 0.025 |
| NC_002645.1;NP_073550.1;HCoV-229E_1a;2649 | 0 | 0 |
| NC_002645.1;NP_073550.1;HCoV-229E_1a;2650 | 0 | 0.025 |
| NC_002645.1;NP_073550.1;HCoV-229E_1a;2651 | 0 | 0 |
| NC_002645.1;NP_073550.1;HCoV-229E_1a;2652 | 0 | 0 |
| NC_002645.1;NP_073550.1;HCoV-229E_1a;2653 | 0 | 0 |
| NC_002645.1;NP_073550.1;HCoV-229E_1a;2654 | 0 | 0 |
| NC_002645.1;NP_073550.1;HCoV-229E_1a;2655 | 0.1 | 0 |
| NC_002645.1;NP_073550.1;HCoV-229E_1a;2656 | 0.2 | 0.125 |

| | | |
|---|---|---|
| NC_002645.1;NP_073550.1;HCoV-229E_1a;2657 | 0.2 | 0.05 |
| NC_002645.1;NP_073550.1;HCoV-229E_1a;2658 | 0.1 | 0.05 |
| NC_002645.1;NP_073550.1;HCoV-229E_1a;2659 | 0 | 0.025 |
| NC_002645.1;NP_073550.1;HCoV-229E_1a;2660 | 0 | 0.025 |
| NC_002645.1;NP_073550.1;HCoV-229E_1a;2661 | 0.05 | 0.075 |
| NC_002645.1;NP_073550.1;HCoV-229E_1a;2662 | 0.1 | 0.125 |
| NC_002645.1;NP_073550.1;HCoV-229E_1a;2663 | 0.05 | 0.025 |
| NC_002645.1;NP_073550.1;HCoV-229E_1a;2664 | 0.1 | 0.15 |
| NC_002645.1;NP_073550.1;HCoV-229E_1a;2665 | 0.15 | 0.175 |
| NC_002645.1;NP_073550.1;HCoV-229E_1a;2666 | 0.05 | 0.075 |
| NC_002645.1;NP_073550.1;HCoV-229E_1a;2667 | 0.05 | 0 |
| NC_002645.1;NP_073550.1;HCoV-229E_1a;2668 | 0 | 0 |
| NC_002645.1;NP_073550.1;HCoV-229E_1a;2669 | 0 | 0 |
| NC_002645.1;NP_073550.1;HCoV-229E_1a;2670 | 0 | 0 |
| NC_002645.1;NP_073550.1;HCoV-229E_1a;2671 | 0 | 0 |
| NC_002645.1;NP_073550.1;HCoV-229E_1a;2672 | 0 | 0 |
| NC_002645.1;NP_073550.1;HCoV-229E_1a;2673 | 0 | 0 |
| NC_002645.1;NP_073550.1;HCoV-229E_1a;2674 | 0 | 0 |
| NC_002645.1;NP_073550.1;HCoV-229E_1a;2675 | 0 | 0 |
| NC_002645.1;NP_073550.1;HCoV-229E_1a;2676 | 0 | 0 |
| NC_002645.1;NP_073550.1;HCoV-229E_1a;2677 | 0 | 0 |
| NC_002645.1;NP_073550.1;HCoV-229E_1a;2678 | 0 | 0 |
| NC_002645.1;NP_073550.1;HCoV-229E_1a;2679 | 0 | 0 |
| NC_002645.1;NP_073550.1;HCoV-229E_1a;2680 | 0 | 0 |
| NC_002645.1;NP_073550.1;HCoV-229E_1a;2681 | 0 | 0 |
| NC_002645.1;NP_073550.1;HCoV-229E_1a;2682 | 0 | 0 |
| NC_002645.1;NP_073550.1;HCoV-229E_1a;2683 | 0 | 0 |
| NC_002645.1;NP_073550.1;HCoV-229E_1a;2684 | 0 | 0 |
| NC_002645.1;NP_073550.1;HCoV-229E_1a;2685 | 0 | 0 |
| NC_002645.1;NP_073550.1;HCoV-229E_1a;2686 | 0 | 0 |
| NC_002645.1;NP_073550.1;HCoV-229E_1a;2687 | 0 | 0 |
| NC_002645.1;NP_073550.1;HCoV-229E_1a;2688 | 0 | 0 |
| NC_002645.1;NP_073550.1;HCoV-229E_1a;2689 | 0 | 0 |
| NC_002645.1;NP_073550.1;HCoV-229E_1a;2690 | 0 | 0 |
| NC_002645.1;NP_073550.1;HCoV-229E_1a;2691 | 0 | 0 |
| NC_002645.1;NP_073550.1;HCoV-229E_1a;2692 | 0.05 | 0 |
| NC_002645.1;NP_073550.1;HCoV-229E_1a;2693 | 0.05 | 0 |
| NC_002645.1;NP_073550.1;HCoV-229E_1a;2694 | 0.05 | 0.025 |
| NC_002645.1;NP_073550.1;HCoV-229E_1a;2695 | 0.05 | 0.025 |
| NC_002645.1;NP_073550.1;HCoV-229E_1a;2696 | 0.05 | 0 |
| NC_002645.1;NP_073550.1;HCoV-229E_1a;2697 | 0 | 0 |
| NC_002645.1;NP_073550.1;HCoV-229E_1a;2698 | 0 | 0 |
| NC_002645.1;NP_073550.1;HCoV-229E_1a;2699 | 0 | 0 |

| | | |
|---|---|---|
| NC_002645.1;NP_073550.1;HCoV-229E_1a;2700 | 0 | 0 |
| NC_002645.1;NP_073550.1;HCoV-229E_1a;2701 | 0 | 0 |
| NC_002645.1;NP_073550.1;HCoV-229E_1a;2702 | 0.05 | 0 |
| NC_002645.1;NP_073550.1;HCoV-229E_1a;2703 | 0 | 0 |
| NC_002645.1;NP_073550.1;HCoV-229E_1a;2704 | 0 | 0 |
| NC_002645.1;NP_073550.1;HCoV-229E_1a;2705 | 0 | 0 |
| NC_002645.1;NP_073550.1;HCoV-229E_1a;2706 | 0 | 0 |
| NC_002645.1;NP_073550.1;HCoV-229E_1a;2707 | 0.05 | 0.05 |
| NC_002645.1;NP_073550.1;HCoV-229E_1a;2708 | 0 | 0 |
| NC_002645.1;NP_073550.1;HCoV-229E_1a;2709 | 0.05 | 0 |
| NC_002645.1;NP_073550.1;HCoV-229E_1a;2710 | 0 | 0 |
| NC_002645.1;NP_073550.1;HCoV-229E_1a;2711 | 0 | 0 |
| NC_002645.1;NP_073550.1;HCoV-229E_1a;2712 | 0.05 | 0.025 |
| NC_002645.1;NP_073550.1;HCoV-229E_1a;2713 | 0 | 0 |
| NC_002645.1;NP_073550.1;HCoV-229E_1a;2714 | 0 | 0 |
| NC_002645.1;NP_073550.1;HCoV-229E_1a;2715 | 0.05 | 0.025 |
| NC_002645.1;NP_073550.1;HCoV-229E_1a;2716 | 0 | 0 |
| NC_002645.1;NP_073550.1;HCoV-229E_1a;2717 | 0 | 0 |
| NC_002645.1;NP_073550.1;HCoV-229E_1a;2718 | 0 | 0 |
| NC_002645.1;NP_073550.1;HCoV-229E_1a;2719 | 0 | 0 |
| NC_002645.1;NP_073550.1;HCoV-229E_1a;2720 | 0 | 0 |
| NC_002645.1;NP_073550.1;HCoV-229E_1a;2721 | 0 | 0 |
| NC_002645.1;NP_073550.1;HCoV-229E_1a;2722 | 0 | 0 |
| NC_002645.1;NP_073550.1;HCoV-229E_1a;2723 | 0 | 0 |
| NC_002645.1;NP_073550.1;HCoV-229E_1a;2724 | 0 | 0 |
| NC_002645.1;NP_073550.1;HCoV-229E_1a;2725 | 0 | 0 |
| NC_002645.1;NP_073550.1;HCoV-229E_1a;2726 | 0 | 0 |
| NC_002645.1;NP_073550.1;HCoV-229E_1a;2727 | 0 | 0 |
| NC_002645.1;NP_073550.1;HCoV-229E_1a;2728 | 0 | 0 |
| NC_002645.1;NP_073550.1;HCoV-229E_1a;2729 | 0 | 0 |
| NC_002645.1;NP_073550.1;HCoV-229E_1a;2730 | 0 | 0 |
| NC_002645.1;NP_073550.1;HCoV-229E_1a;2731 | 0 | 0 |
| NC_002645.1;NP_073550.1;HCoV-229E_1a;2732 | 0 | 0 |
| NC_002645.1;NP_073550.1;HCoV-229E_1a;2733 | 0 | 0 |
| NC_002645.1;NP_073550.1;HCoV-229E_1a;2734 | 0 | 0 |
| NC_002645.1;NP_073550.1;HCoV-229E_1a;2735 | 0.05 | 0 |
| NC_002645.1;NP_073550.1;HCoV-229E_1a;2736 | 0 | 0 |
| NC_002645.1;NP_073550.1;HCoV-229E_1a;2737 | 0 | 0 |
| NC_002645.1;NP_073550.1;HCoV-229E_1a;2738 | 0 | 0 |
| NC_002645.1;NP_073550.1;HCoV-229E_1a;2739 | 0 | 0 |
| NC_002645.1;NP_073550.1;HCoV-229E_1a;2740 | 0 | 0 |
| NC_002645.1;NP_073550.1;HCoV-229E_1a;2741 | 0 | 0 |
| NC_002645.1;NP_073550.1;HCoV-229E_1a;2742 | 0 | 0 |

| | | |
|---|---|---|
| NC_002645.1;NP_073550.1;HCoV-229E_1a;2743 | 0 | 0 |
| NC_002645.1;NP_073550.1;HCoV-229E_1a;2744 | 0 | 0 |
| NC_002645.1;NP_073550.1;HCoV-229E_1a;2745 | 0 | 0 |
| NC_002645.1;NP_073550.1;HCoV-229E_1a;2746 | 0 | 0 |
| NC_002645.1;NP_073550.1;HCoV-229E_1a;2747 | 0 | 0 |
| NC_002645.1;NP_073550.1;HCoV-229E_1a;2748 | 0 | 0 |
| NC_002645.1;NP_073550.1;HCoV-229E_1a;2749 | 0 | 0 |
| NC_002645.1;NP_073550.1;HCoV-229E_1a;2750 | 0 | 0 |
| NC_002645.1;NP_073550.1;HCoV-229E_1a;2751 | 0 | 0 |
| NC_002645.1;NP_073550.1;HCoV-229E_1a;2752 | 0 | 0 |
| NC_002645.1;NP_073550.1;HCoV-229E_1a;2753 | 0 | 0 |
| NC_002645.1;NP_073550.1;HCoV-229E_1a;2754 | 0 | 0 |
| NC_002645.1;NP_073550.1;HCoV-229E_1a;2755 | 0 | 0 |
| NC_002645.1;NP_073550.1;HCoV-229E_1a;2756 | 0 | 0 |
| NC_002645.1;NP_073550.1;HCoV-229E_1a;2757 | 0 | 0 |
| NC_002645.1;NP_073550.1;HCoV-229E_1a;2758 | 0 | 0 |
| NC_002645.1;NP_073550.1;HCoV-229E_1a;2759 | 0 | 0 |
| NC_002645.1;NP_073550.1;HCoV-229E_1a;2760 | 0 | 0 |
| NC_002645.1;NP_073550.1;HCoV-229E_1a;2761 | 0 | 0 |
| NC_002645.1;NP_073550.1;HCoV-229E_1a;2762 | 0 | 0 |
| NC_002645.1;NP_073550.1;HCoV-229E_1a;2763 | 0 | 0 |
| NC_002645.1;NP_073550.1;HCoV-229E_1a;2764 | 0 | 0 |
| NC_002645.1;NP_073550.1;HCoV-229E_1a;2765 | 0 | 0 |
| NC_002645.1;NP_073550.1;HCoV-229E_1a;2766 | 0 | 0 |
| NC_002645.1;NP_073550.1;HCoV-229E_1a;2767 | 0 | 0 |
| NC_002645.1;NP_073550.1;HCoV-229E_1a;2768 | 0 | 0 |
| NC_002645.1;NP_073550.1;HCoV-229E_1a;2769 | 0 | 0 |
| NC_002645.1;NP_073550.1;HCoV-229E_1a;2770 | 0 | 0 |
| NC_002645.1;NP_073550.1;HCoV-229E_1a;2771 | 0 | 0 |
| NC_002645.1;NP_073550.1;HCoV-229E_1a;2772 | 0 | 0 |
| NC_002645.1;NP_073550.1;HCoV-229E_1a;2773 | 0 | 0 |
| NC_002645.1;NP_073550.1;HCoV-229E_1a;2774 | 0 | 0 |
| NC_002645.1;NP_073550.1;HCoV-229E_1a;2775 | 0 | 0 |
| NC_002645.1;NP_073550.1;HCoV-229E_1a;2776 | 0 | 0 |
| NC_002645.1;NP_073550.1;HCoV-229E_1a;2777 | 0 | 0 |
| NC_002645.1;NP_073550.1;HCoV-229E_1a;2778 | 0 | 0 |
| NC_002645.1;NP_073550.1;HCoV-229E_1a;2779 | 0 | 0 |
| NC_002645.1;NP_073550.1;HCoV-229E_1a;2780 | 0 | 0 |
| NC_002645.1;NP_073550.1;HCoV-229E_1a;2781 | 0 | 0 |
| NC_002645.1;NP_073550.1;HCoV-229E_1a;2782 | 0 | 0 |
| NC_002645.1;NP_073550.1;HCoV-229E_1a;2783 | 0 | 0 |
| NC_002645.1;NP_073550.1;HCoV-229E_1a;2784 | 0 | 0 |
| NC_002645.1;NP_073550.1;HCoV-229E_1a;2785 | 0 | 0 |

| | | |
|---|---|---|
| NC_002645.1;NP_073550.1;HCoV-229E_1a;2786 | 0 | 0 |
| NC_002645.1;NP_073550.1;HCoV-229E_1a;2787 | 0 | 0 |
| NC_002645.1;NP_073550.1;HCoV-229E_1a;2788 | 0 | 0 |
| NC_002645.1;NP_073550.1;HCoV-229E_1a;2789 | 0 | 0 |
| NC_002645.1;NP_073550.1;HCoV-229E_1a;2790 | 0 | 0 |
| NC_002645.1;NP_073550.1;HCoV-229E_1a;2791 | 0 | 0 |
| NC_002645.1;NP_073550.1;HCoV-229E_1a;2792 | 0 | 0 |
| NC_002645.1;NP_073550.1;HCoV-229E_1a;2793 | 0 | 0 |
| NC_002645.1;NP_073550.1;HCoV-229E_1a;2794 | 0 | 0 |
| NC_002645.1;NP_073550.1;HCoV-229E_1a;2795 | 0 | 0 |
| NC_002645.1;NP_073550.1;HCoV-229E_1a;2796 | 0 | 0 |
| NC_002645.1;NP_073550.1;HCoV-229E_1a;2797 | 0 | 0 |
| NC_002645.1;NP_073550.1;HCoV-229E_1a;2798 | 0 | 0 |
| NC_002645.1;NP_073550.1;HCoV-229E_1a;2799 | 0 | 0 |
| NC_002645.1;NP_073550.1;HCoV-229E_1a;2800 | 0 | 0 |
| NC_002645.1;NP_073550.1;HCoV-229E_1a;2801 | 0 | 0 |
| NC_002645.1;NP_073550.1;HCoV-229E_1a;2802 | 0 | 0 |
| NC_002645.1;NP_073550.1;HCoV-229E_1a;2803 | 0 | 0 |
| NC_002645.1;NP_073550.1;HCoV-229E_1a;2804 | 0 | 0 |
| NC_002645.1;NP_073550.1;HCoV-229E_1a;2805 | 0 | 0 |
| NC_002645.1;NP_073550.1;HCoV-229E_1a;2806 | 0 | 0 |
| NC_002645.1;NP_073550.1;HCoV-229E_1a;2807 | 0 | 0 |
| NC_002645.1;NP_073550.1;HCoV-229E_1a;2808 | 0 | 0 |
| NC_002645.1;NP_073550.1;HCoV-229E_1a;2809 | 0 | 0 |
| NC_002645.1;NP_073550.1;HCoV-229E_1a;2810 | 0 | 0 |
| NC_002645.1;NP_073550.1;HCoV-229E_1a;2811 | 0 | 0 |
| NC_002645.1;NP_073550.1;HCoV-229E_1a;2812 | 0 | 0 |
| NC_002645.1;NP_073550.1;HCoV-229E_1a;2813 | 0 | 0 |
| NC_002645.1;NP_073550.1;HCoV-229E_1a;2814 | 0 | 0 |
| NC_002645.1;NP_073550.1;HCoV-229E_1a;2815 | 0 | 0 |
| NC_002645.1;NP_073550.1;HCoV-229E_1a;2816 | 0 | 0 |
| NC_002645.1;NP_073550.1;HCoV-229E_1a;2817 | 0 | 0 |
| NC_002645.1;NP_073550.1;HCoV-229E_1a;2818 | 0 | 0 |
| NC_002645.1;NP_073550.1;HCoV-229E_1a;2819 | 0 | 0 |
| NC_002645.1;NP_073550.1;HCoV-229E_1a;2820 | 0 | 0 |
| NC_002645.1;NP_073550.1;HCoV-229E_1a;2821 | 0 | 0 |
| NC_002645.1;NP_073550.1;HCoV-229E_1a;2822 | 0 | 0 |
| NC_002645.1;NP_073550.1;HCoV-229E_1a;2823 | 0 | 0 |
| NC_002645.1;NP_073550.1;HCoV-229E_1a;2824 | 0 | 0 |
| NC_002645.1;NP_073550.1;HCoV-229E_1a;2825 | 0 | 0 |
| NC_002645.1;NP_073550.1;HCoV-229E_1a;2826 | 0 | 0 |
| NC_002645.1;NP_073550.1;HCoV-229E_1a;2827 | 0 | 0 |
| NC_002645.1;NP_073550.1;HCoV-229E_1a;2828 | 0 | 0 |

| | | |
|---|---|---|
| NC_002645.1;NP_073550.1;HCoV-229E_1a;2829 | 0 | 0 |
| NC_002645.1;NP_073550.1;HCoV-229E_1a;2830 | 0 | 0 |
| NC_002645.1;NP_073550.1;HCoV-229E_1a;2831 | 0 | 0 |
| NC_002645.1;NP_073550.1;HCoV-229E_1a;2832 | 0 | 0 |
| NC_002645.1;NP_073550.1;HCoV-229E_1a;2833 | 0 | 0 |
| NC_002645.1;NP_073550.1;HCoV-229E_1a;2834 | 0 | 0 |
| NC_002645.1;NP_073550.1;HCoV-229E_1a;2835 | 0 | 0 |
| NC_002645.1;NP_073550.1;HCoV-229E_1a;2836 | 0 | 0 |
| NC_002645.1;NP_073550.1;HCoV-229E_1a;2837 | 0 | 0 |
| NC_002645.1;NP_073550.1;HCoV-229E_1a;2838 | 0 | 0 |
| NC_002645.1;NP_073550.1;HCoV-229E_1a;2839 | 0 | 0 |
| NC_002645.1;NP_073550.1;HCoV-229E_1a;2840 | 0 | 0 |
| NC_002645.1;NP_073550.1;HCoV-229E_1a;2841 | 0 | 0 |
| NC_002645.1;NP_073550.1;HCoV-229E_1a;2842 | 0 | 0 |
| NC_002645.1;NP_073550.1;HCoV-229E_1a;2843 | 0 | 0 |
| NC_002645.1;NP_073550.1;HCoV-229E_1a;2844 | 0 | 0 |
| NC_002645.1;NP_073550.1;HCoV-229E_1a;2845 | 0 | 0 |
| NC_002645.1;NP_073550.1;HCoV-229E_1a;2846 | 0 | 0 |
| NC_002645.1;NP_073550.1;HCoV-229E_1a;2847 | 0 | 0 |
| NC_002645.1;NP_073550.1;HCoV-229E_1a;2848 | 0 | 0 |
| NC_002645.1;NP_073550.1;HCoV-229E_1a;2849 | 0 | 0 |
| NC_002645.1;NP_073550.1;HCoV-229E_1a;2850 | 0 | 0 |
| NC_002645.1;NP_073550.1;HCoV-229E_1a;2851 | 0 | 0 |
| NC_002645.1;NP_073550.1;HCoV-229E_1a;2852 | 0 | 0 |
| NC_002645.1;NP_073550.1;HCoV-229E_1a;2853 | 0 | 0 |
| NC_002645.1;NP_073550.1;HCoV-229E_1a;2854 | 0 | 0 |
| NC_002645.1;NP_073550.1;HCoV-229E_1a;2855 | 0 | 0 |
| NC_002645.1;NP_073550.1;HCoV-229E_1a;2856 | 0 | 0 |
| NC_002645.1;NP_073550.1;HCoV-229E_1a;2857 | 0 | 0 |
| NC_002645.1;NP_073550.1;HCoV-229E_1a;2858 | 0 | 0 |
| NC_002645.1;NP_073550.1;HCoV-229E_1a;2859 | 0.05 | 0 |
| NC_002645.1;NP_073550.1;HCoV-229E_1a;2860 | 0 | 0.025 |
| NC_002645.1;NP_073550.1;HCoV-229E_1a;2861 | 0.05 | 0.025 |
| NC_002645.1;NP_073550.1;HCoV-229E_1a;2862 | 0.05 | 0.025 |
| NC_002645.1;NP_073550.1;HCoV-229E_1a;2863 | 0.1 | 0.025 |
| NC_002645.1;NP_073550.1;HCoV-229E_1a;2864 | 0.1 | 0.075 |
| NC_002645.1;NP_073550.1;HCoV-229E_1a;2865 | 0.1 | 0.025 |
| NC_002645.1;NP_073550.1;HCoV-229E_1a;2866 | 0.05 | 0.025 |
| NC_002645.1;NP_073550.1;HCoV-229E_1a;2867 | 0.15 | 0.025 |
| NC_002645.1;NP_073550.1;HCoV-229E_1a;2868 | 0 | 0 |
| NC_002645.1;NP_073550.1;HCoV-229E_1a;2869 | 0.05 | 0.025 |
| NC_002645.1;NP_073550.1;HCoV-229E_1a;2870 | 0.05 | 0.025 |
| NC_002645.1;NP_073550.1;HCoV-229E_1a;2871 | 0 | 0 |

| | | |
|---|---|---|
| NC_002645.1;NP_073550.1;HCoV-229E_1a;2872 | 0 | 0 |
| NC_002645.1;NP_073550.1;HCoV-229E_1a;2873 | 0 | 0 |
| NC_002645.1;NP_073550.1;HCoV-229E_1a;2874 | 0 | 0 |
| NC_002645.1;NP_073550.1;HCoV-229E_1a;2875 | 0 | 0 |
| NC_002645.1;NP_073550.1;HCoV-229E_1a;2876 | 0 | 0 |
| NC_002645.1;NP_073550.1;HCoV-229E_1a;2877 | 0 | 0 |
| NC_002645.1;NP_073550.1;HCoV-229E_1a;2878 | 0 | 0 |
| NC_002645.1;NP_073550.1;HCoV-229E_1a;2879 | 0 | 0 |
| NC_002645.1;NP_073550.1;HCoV-229E_1a;2880 | 0 | 0 |
| NC_002645.1;NP_073550.1;HCoV-229E_1a;2881 | 0 | 0 |
| NC_002645.1;NP_073550.1;HCoV-229E_1a;2882 | 0 | 0 |
| NC_002645.1;NP_073550.1;HCoV-229E_1a;2883 | 0 | 0 |
| NC_002645.1;NP_073550.1;HCoV-229E_1a;2884 | 0 | 0 |
| NC_002645.1;NP_073550.1;HCoV-229E_1a;2885 | 0 | 0 |
| NC_002645.1;NP_073550.1;HCoV-229E_1a;2886 | 0 | 0 |
| NC_002645.1;NP_073550.1;HCoV-229E_1a;2887 | 0 | 0.025 |
| NC_002645.1;NP_073550.1;HCoV-229E_1a;2888 | 0 | 0 |
| NC_002645.1;NP_073550.1;HCoV-229E_1a;2889 | 0.05 | 0.025 |
| NC_002645.1;NP_073550.1;HCoV-229E_1a;2890 | 0.05 | 0 |
| NC_002645.1;NP_073550.1;HCoV-229E_1a;2891 | 0.05 | 0 |
| NC_002645.1;NP_073550.1;HCoV-229E_1a;2892 | 0 | 0 |
| NC_002645.1;NP_073550.1;HCoV-229E_1a;2893 | 0 | 0 |
| NC_002645.1;NP_073550.1;HCoV-229E_1a;2894 | 0 | 0 |
| NC_002645.1;NP_073550.1;HCoV-229E_1a;2895 | 0.05 | 0 |
| NC_002645.1;NP_073550.1;HCoV-229E_1a;2896 | 0.05 | 0 |
| NC_002645.1;NP_073550.1;HCoV-229E_1a;2897 | 0.05 | 0 |
| NC_002645.1;NP_073550.1;HCoV-229E_1a;2898 | 0 | 0 |
| NC_002645.1;NP_073550.1;HCoV-229E_1a;2899 | 0 | 0 |
| NC_002645.1;NP_073550.1;HCoV-229E_1a;2900 | 0 | 0.025 |
| NC_002645.1;NP_073550.1;HCoV-229E_1a;2901 | 0 | 0 |
| NC_002645.1;NP_073550.1;HCoV-229E_1a;2902 | 0 | 0 |
| NC_002645.1;NP_073550.1;HCoV-229E_1a;2903 | 0.05 | 0 |
| NC_002645.1;NP_073550.1;HCoV-229E_1a;2904 | 0 | 0 |
| NC_002645.1;NP_073550.1;HCoV-229E_1a;2905 | 0 | 0 |
| NC_002645.1;NP_073550.1;HCoV-229E_1a;2906 | 0 | 0 |
| NC_002645.1;NP_073550.1;HCoV-229E_1a;2907 | 0 | 0 |
| NC_002645.1;NP_073550.1;HCoV-229E_1a;2908 | 0 | 0.025 |
| NC_002645.1;NP_073550.1;HCoV-229E_1a;2909 | 0 | 0.025 |
| NC_002645.1;NP_073550.1;HCoV-229E_1a;2910 | 0 | 0.025 |
| NC_002645.1;NP_073550.1;HCoV-229E_1a;2911 | 0 | 0.1 |
| NC_002645.1;NP_073550.1;HCoV-229E_1a;2912 | 0 | 0.1 |
| NC_002645.1;NP_073550.1;HCoV-229E_1a;2913 | 0.25 | 0.15 |
| NC_002645.1;NP_073550.1;HCoV-229E_1a;2914 | 0.05 | 0.125 |

| | | |
|---|---|---|
| NC_002645.1;NP_073550.1;HCoV-229E_1a;2915 | 0 | 0.1 |
| NC_002645.1;NP_073550.1;HCoV-229E_1a;2916 | 0 | 0.1 |
| NC_002645.1;NP_073550.1;HCoV-229E_1a;2917 | 0.05 | 0.125 |
| NC_002645.1;NP_073550.1;HCoV-229E_1a;2918 | 0.05 | 0.15 |
| NC_002645.1;NP_073550.1;HCoV-229E_1a;2919 | 0 | 0.075 |
| NC_002645.1;NP_073550.1;HCoV-229E_1a;2920 | 0.05 | 0.075 |
| NC_002645.1;NP_073550.1;HCoV-229E_1a;2921 | 0 | 0 |
| NC_002645.1;NP_073550.1;HCoV-229E_1a;2922 | 0 | 0 |
| NC_002645.1;NP_073550.1;HCoV-229E_1a;2923 | 0 | 0 |
| NC_002645.1;NP_073550.1;HCoV-229E_1a;2924 | 0 | 0 |
| NC_002645.1;NP_073550.1;HCoV-229E_1a;2925 | 0.05 | 0 |
| NC_002645.1;NP_073550.1;HCoV-229E_1a;2926 | 0 | 0 |
| NC_002645.1;NP_073550.1;HCoV-229E_1a;2927 | 0 | 0 |
| NC_002645.1;NP_073550.1;HCoV-229E_1a;2928 | 0 | 0 |
| NC_002645.1;NP_073550.1;HCoV-229E_1a;2929 | 0 | 0 |
| NC_002645.1;NP_073550.1;HCoV-229E_1a;2930 | 0 | 0 |
| NC_002645.1;NP_073550.1;HCoV-229E_1a;2931 | 0 | 0 |
| NC_002645.1;NP_073550.1;HCoV-229E_1a;2932 | 0 | 0.025 |
| NC_002645.1;NP_073550.1;HCoV-229E_1a;2933 | 0 | 0 |
| NC_002645.1;NP_073550.1;HCoV-229E_1a;2934 | 0 | 0 |
| NC_002645.1;NP_073550.1;HCoV-229E_1a;2935 | 0.05 | 0.05 |
| NC_002645.1;NP_073550.1;HCoV-229E_1a;2936 | 0.05 | 0.075 |
| NC_002645.1;NP_073550.1;HCoV-229E_1a;2937 | 0 | 0.025 |
| NC_002645.1;NP_073550.1;HCoV-229E_1a;2938 | 0 | 0.025 |
| NC_002645.1;NP_073550.1;HCoV-229E_1a;2939 | 0 | 0.025 |
| NC_002645.1;NP_073550.1;HCoV-229E_1a;2940 | 0.05 | 0.025 |
| NC_002645.1;NP_073550.1;HCoV-229E_1a;2941 | 0 | 0.075 |
| NC_002645.1;NP_073550.1;HCoV-229E_1a;2942 | 0 | 0.075 |
| NC_002645.1;NP_073550.1;HCoV-229E_1a;2943 | 0 | 0.075 |
| NC_002645.1;NP_073550.1;HCoV-229E_1a;2944 | 0 | 0.1 |
| NC_002645.1;NP_073550.1;HCoV-229E_1a;2945 | 0 | 0.075 |
| NC_002645.1;NP_073550.1;HCoV-229E_1a;2946 | 0 | 0.05 |
| NC_002645.1;NP_073550.1;HCoV-229E_1a;2947 | 0 | 0.025 |
| NC_002645.1;NP_073550.1;HCoV-229E_1a;2948 | 0 | 0 |
| NC_002645.1;NP_073550.1;HCoV-229E_1a;2949 | 0 | 0 |
| NC_002645.1;NP_073550.1;HCoV-229E_1a;2950 | 0 | 0 |
| NC_002645.1;NP_073550.1;HCoV-229E_1a;2951 | 0 | 0 |
| NC_002645.1;NP_073550.1;HCoV-229E_1a;2952 | 0 | 0 |
| NC_002645.1;NP_073550.1;HCoV-229E_1a;2953 | 0 | 0 |
| NC_002645.1;NP_073550.1;HCoV-229E_1a;2954 | 0 | 0 |
| NC_002645.1;NP_073550.1;HCoV-229E_1a;2955 | 0 | 0 |
| NC_002645.1;NP_073550.1;HCoV-229E_1a;2956 | 0 | 0 |
| NC_002645.1;NP_073550.1;HCoV-229E_1a;2957 | 0 | 0 |

| | | |
|---|---|---|
| NC_002645.1;NP_073550.1;HCoV-229E_1a;2958 | 0 | 0 |
| NC_002645.1;NP_073550.1;HCoV-229E_1a;2959 | 0 | 0 |
| NC_002645.1;NP_073550.1;HCoV-229E_1a;2960 | 0 | 0 |
| NC_002645.1;NP_073550.1;HCoV-229E_1a;2961 | 0 | 0 |
| NC_002645.1;NP_073550.1;HCoV-229E_1a;2962 | 0 | 0 |
| NC_002645.1;NP_073550.1;HCoV-229E_1a;2963 | 0 | 0 |
| NC_002645.1;NP_073550.1;HCoV-229E_1a;2964 | 0 | 0.025 |
| NC_002645.1;NP_073550.1;HCoV-229E_1a;2965 | 0 | 0 |
| NC_002645.1;NP_073550.1;HCoV-229E_1a;2966 | 0 | 0 |
| NC_002645.1;NP_073550.1;HCoV-229E_1a;2967 | 0 | 0 |
| NC_002645.1;NP_073550.1;HCoV-229E_1a;2968 | 0 | 0 |
| NC_002645.1;NP_073550.1;HCoV-229E_1a;2969 | 0 | 0 |
| NC_002645.1;NP_073550.1;HCoV-229E_1a;2970 | 0 | 0 |
| NC_002645.1;NP_073550.1;HCoV-229E_1a;2971 | 0 | 0 |
| NC_002645.1;NP_073550.1;HCoV-229E_1a;2972 | 0 | 0 |
| NC_002645.1;NP_073550.1;HCoV-229E_1a;2973 | 0 | 0 |
| NC_002645.1;NP_073550.1;HCoV-229E_1a;2974 | 0 | 0 |
| NC_002645.1;NP_073550.1;HCoV-229E_1a;2975 | 0 | 0 |
| NC_002645.1;NP_073550.1;HCoV-229E_1a;2976 | 0 | 0 |
| NC_002645.1;NP_073550.1;HCoV-229E_1a;2977 | 0 | 0 |
| NC_002645.1;NP_073550.1;HCoV-229E_1a;2978 | 0 | 0 |
| NC_002645.1;NP_073550.1;HCoV-229E_1a;2979 | 0 | 0 |
| NC_002645.1;NP_073550.1;HCoV-229E_1a;2980 | 0 | 0 |
| NC_002645.1;NP_073550.1;HCoV-229E_1a;2981 | 0 | 0 |
| NC_002645.1;NP_073550.1;HCoV-229E_1a;2982 | 0 | 0 |
| NC_002645.1;NP_073550.1;HCoV-229E_1a;2983 | 0 | 0 |
| NC_002645.1;NP_073550.1;HCoV-229E_1a;2984 | 0 | 0 |
| NC_002645.1;NP_073550.1;HCoV-229E_1a;2985 | 0 | 0 |
| NC_002645.1;NP_073550.1;HCoV-229E_1a;2986 | 0 | 0 |
| NC_002645.1;NP_073550.1;HCoV-229E_1a;2987 | 0 | 0 |
| NC_002645.1;NP_073550.1;HCoV-229E_1a;2988 | 0 | 0 |
| NC_002645.1;NP_073550.1;HCoV-229E_1a;2989 | 0 | 0 |
| NC_002645.1;NP_073550.1;HCoV-229E_1a;2990 | 0 | 0 |
| NC_002645.1;NP_073550.1;HCoV-229E_1a;2991 | 0 | 0 |
| NC_002645.1;NP_073550.1;HCoV-229E_1a;2992 | 0 | 0 |
| NC_002645.1;NP_073550.1;HCoV-229E_1a;2993 | 0 | 0 |
| NC_002645.1;NP_073550.1;HCoV-229E_1a;2994 | 0 | 0 |
| NC_002645.1;NP_073550.1;HCoV-229E_1a;2995 | 0 | 0 |
| NC_002645.1;NP_073550.1;HCoV-229E_1a;2996 | 0 | 0 |
| NC_002645.1;NP_073550.1;HCoV-229E_1a;2997 | 0 | 0 |
| NC_002645.1;NP_073550.1;HCoV-229E_1a;2998 | 0 | 0 |
| NC_002645.1;NP_073550.1;HCoV-229E_1a;2999 | 0 | 0 |
| NC_002645.1;NP_073550.1;HCoV-229E_1a;3000 | 0 | 0 |

| | | |
|---|---|---|
| NC_002645.1;NP_073550.1;HCoV-229E_1a;3001 | 0 | 0.025 |
| NC_002645.1;NP_073550.1;HCoV-229E_1a;3002 | 0 | 0.025 |
| NC_002645.1;NP_073550.1;HCoV-229E_1a;3003 | 0 | 0 |
| NC_002645.1;NP_073550.1;HCoV-229E_1a;3004 | 0 | 0 |
| NC_002645.1;NP_073550.1;HCoV-229E_1a;3005 | 0 | 0 |
| NC_002645.1;NP_073550.1;HCoV-229E_1a;3006 | 0 | 0.025 |
| NC_002645.1;NP_073550.1;HCoV-229E_1a;3007 | 0 | 0.075 |
| NC_002645.1;NP_073550.1;HCoV-229E_1a;3008 | 0 | 0.025 |
| NC_002645.1;NP_073550.1;HCoV-229E_1a;3009 | 0 | 0 |
| NC_002645.1;NP_073550.1;HCoV-229E_1a;3010 | 0 | 0 |
| NC_002645.1;NP_073550.1;HCoV-229E_1a;3011 | 0 | 0 |
| NC_002645.1;NP_073550.1;HCoV-229E_1a;3012 | 0 | 0 |
| NC_002645.1;NP_073550.1;HCoV-229E_1a;3013 | 0 | 0 |
| NC_002645.1;NP_073550.1;HCoV-229E_1a;3014 | 0 | 0.025 |
| NC_002645.1;NP_073550.1;HCoV-229E_1a;3015 | 0 | 0.025 |
| NC_002645.1;NP_073550.1;HCoV-229E_1a;3016 | 0 | 0 |
| NC_002645.1;NP_073550.1;HCoV-229E_1a;3017 | 0 | 0 |
| NC_002645.1;NP_073550.1;HCoV-229E_1a;3018 | 0 | 0 |
| NC_002645.1;NP_073550.1;HCoV-229E_1a;3019 | 0 | 0 |
| NC_002645.1;NP_073550.1;HCoV-229E_1a;3020 | 0 | 0 |
| NC_002645.1;NP_073550.1;HCoV-229E_1a;3021 | 0 | 0 |
| NC_002645.1;NP_073550.1;HCoV-229E_1a;3022 | 0 | 0 |
| NC_002645.1;NP_073550.1;HCoV-229E_1a;3023 | 0 | 0 |
| NC_002645.1;NP_073550.1;HCoV-229E_1a;3024 | 0 | 0 |
| NC_002645.1;NP_073550.1;HCoV-229E_1a;3025 | 0 | 0 |
| NC_002645.1;NP_073550.1;HCoV-229E_1a;3026 | 0 | 0 |
| NC_002645.1;NP_073550.1;HCoV-229E_1a;3027 | 0 | 0 |
| NC_002645.1;NP_073550.1;HCoV-229E_1a;3028 | 0 | 0 |
| NC_002645.1;NP_073550.1;HCoV-229E_1a;3029 | 0 | 0 |
| NC_002645.1;NP_073550.1;HCoV-229E_1a;3030 | 0 | 0 |
| NC_002645.1;NP_073550.1;HCoV-229E_1a;3031 | 0 | 0 |
| NC_002645.1;NP_073550.1;HCoV-229E_1a;3032 | 0 | 0 |
| NC_002645.1;NP_073550.1;HCoV-229E_1a;3033 | 0 | 0 |
| NC_002645.1;NP_073550.1;HCoV-229E_1a;3034 | 0 | 0 |
| NC_002645.1;NP_073550.1;HCoV-229E_1a;3035 | 0 | 0 |
| NC_002645.1;NP_073550.1;HCoV-229E_1a;3036 | 0 | 0 |
| NC_002645.1;NP_073550.1;HCoV-229E_1a;3037 | 0 | 0 |
| NC_002645.1;NP_073550.1;HCoV-229E_1a;3038 | 0 | 0 |
| NC_002645.1;NP_073550.1;HCoV-229E_1a;3039 | 0 | 0 |
| NC_002645.1;NP_073550.1;HCoV-229E_1a;3040 | 0 | 0 |
| NC_002645.1;NP_073550.1;HCoV-229E_1a;3041 | 0 | 0 |
| NC_002645.1;NP_073550.1;HCoV-229E_1a;3042 | 0 | 0 |
| NC_002645.1;NP_073550.1;HCoV-229E_1a;3043 | 0 | 0 |

| | | |
|---|---|---|
| NC_002645.1;NP_073550.1;HCoV-229E_1a;3044 | 0 | 0 |
| NC_002645.1;NP_073550.1;HCoV-229E_1a;3045 | 0 | 0 |
| NC_002645.1;NP_073550.1;HCoV-229E_1a;3046 | 0 | 0 |
| NC_002645.1;NP_073550.1;HCoV-229E_1a;3047 | 0 | 0 |
| NC_002645.1;NP_073550.1;HCoV-229E_1a;3048 | 0 | 0 |
| NC_002645.1;NP_073550.1;HCoV-229E_1a;3049 | 0 | 0 |
| NC_002645.1;NP_073550.1;HCoV-229E_1a;3050 | 0 | 0 |
| NC_002645.1;NP_073550.1;HCoV-229E_1a;3051 | 0 | 0 |
| NC_002645.1;NP_073550.1;HCoV-229E_1a;3052 | 0 | 0 |
| NC_002645.1;NP_073550.1;HCoV-229E_1a;3053 | 0 | 0 |
| NC_002645.1;NP_073550.1;HCoV-229E_1a;3054 | 0 | 0 |
| NC_002645.1;NP_073550.1;HCoV-229E_1a;3055 | 0 | 0 |
| NC_002645.1;NP_073550.1;HCoV-229E_1a;3056 | 0 | 0 |
| NC_002645.1;NP_073550.1;HCoV-229E_1a;3057 | 0 | 0 |
| NC_002645.1;NP_073550.1;HCoV-229E_1a;3058 | 0 | 0 |
| NC_002645.1;NP_073550.1;HCoV-229E_1a;3059 | 0 | 0 |
| NC_002645.1;NP_073550.1;HCoV-229E_1a;3060 | 0 | 0 |
| NC_002645.1;NP_073550.1;HCoV-229E_1a;3061 | 0 | 0 |
| NC_002645.1;NP_073550.1;HCoV-229E_1a;3062 | 0.05 | 0 |
| NC_002645.1;NP_073550.1;HCoV-229E_1a;3063 | 0 | 0 |
| NC_002645.1;NP_073550.1;HCoV-229E_1a;3064 | 0 | 0 |
| NC_002645.1;NP_073550.1;HCoV-229E_1a;3065 | 0 | 0 |
| NC_002645.1;NP_073550.1;HCoV-229E_1a;3066 | 0 | 0 |
| NC_002645.1;NP_073550.1;HCoV-229E_1a;3067 | 0 | 0 |
| NC_002645.1;NP_073550.1;HCoV-229E_1a;3068 | 0 | 0 |
| NC_002645.1;NP_073550.1;HCoV-229E_1a;3069 | 0 | 0 |
| NC_002645.1;NP_073550.1;HCoV-229E_1a;3070 | 0 | 0 |
| NC_002645.1;NP_073550.1;HCoV-229E_1a;3071 | 0 | 0 |
| NC_002645.1;NP_073550.1;HCoV-229E_1a;3072 | 0 | 0 |
| NC_002645.1;NP_073550.1;HCoV-229E_1a;3073 | 0 | 0 |
| NC_002645.1;NP_073550.1;HCoV-229E_1a;3074 | 0 | 0 |
| NC_002645.1;NP_073550.1;HCoV-229E_1a;3075 | 0 | 0 |
| NC_002645.1;NP_073550.1;HCoV-229E_1a;3076 | 0 | 0 |
| NC_002645.1;NP_073550.1;HCoV-229E_1a;3077 | 0 | 0 |
| NC_002645.1;NP_073550.1;HCoV-229E_1a;3078 | 0 | 0 |
| NC_002645.1;NP_073550.1;HCoV-229E_1a;3079 | 0 | 0 |
| NC_002645.1;NP_073550.1;HCoV-229E_1a;3080 | 0 | 0.025 |
| NC_002645.1;NP_073550.1;HCoV-229E_1a;3081 | 0 | 0.025 |
| NC_002645.1;NP_073550.1;HCoV-229E_1a;3082 | 0 | 0.025 |
| NC_002645.1;NP_073550.1;HCoV-229E_1a;3083 | 0 | 0.025 |
| NC_002645.1;NP_073550.1;HCoV-229E_1a;3084 | 0 | 0.025 |
| NC_002645.1;NP_073550.1;HCoV-229E_1a;3085 | 0 | 0 |
| NC_002645.1;NP_073550.1;HCoV-229E_1a;3086 | 0 | 0 |

| | | |
|---|---|---|
| NC_002645.1;NP_073550.1;HCoV-229E_1a;3087 | 0 | 0 |
| NC_002645.1;NP_073550.1;HCoV-229E_1a;3088 | 0 | 0 |
| NC_002645.1;NP_073550.1;HCoV-229E_1a;3089 | 0 | 0 |
| NC_002645.1;NP_073550.1;HCoV-229E_1a;3090 | 0 | 0 |
| NC_002645.1;NP_073550.1;HCoV-229E_1a;3091 | 0 | 0 |
| NC_002645.1;NP_073550.1;HCoV-229E_1a;3092 | 0 | 0 |
| NC_002645.1;NP_073550.1;HCoV-229E_1a;3093 | 0 | 0 |
| NC_002645.1;NP_073550.1;HCoV-229E_1a;3094 | 0 | 0 |
| NC_002645.1;NP_073550.1;HCoV-229E_1a;3095 | 0 | 0 |
| NC_002645.1;NP_073550.1;HCoV-229E_1a;3096 | 0 | 0 |
| NC_002645.1;NP_073550.1;HCoV-229E_1a;3097 | 0 | 0 |
| NC_002645.1;NP_073550.1;HCoV-229E_1a;3098 | 0 | 0 |
| NC_002645.1;NP_073550.1;HCoV-229E_1a;3099 | 0 | 0 |
| NC_002645.1;NP_073550.1;HCoV-229E_1a;3100 | 0 | 0 |
| NC_002645.1;NP_073550.1;HCoV-229E_1a;3101 | 0 | 0 |
| NC_002645.1;NP_073550.1;HCoV-229E_1a;3102 | 0.05 | 0.025 |
| NC_002645.1;NP_073550.1;HCoV-229E_1a;3103 | 0.05 | 0.05 |
| NC_002645.1;NP_073550.1;HCoV-229E_1a;3104 | 0.05 | 0.05 |
| NC_002645.1;NP_073550.1;HCoV-229E_1a;3105 | 0.05 | 0.1 |
| NC_002645.1;NP_073550.1;HCoV-229E_1a;3106 | 0.1 | 0.1 |
| NC_002645.1;NP_073550.1;HCoV-229E_1a;3107 | 0.05 | 0.075 |
| NC_002645.1;NP_073550.1;HCoV-229E_1a;3108 | 0.05 | 0.05 |
| NC_002645.1;NP_073550.1;HCoV-229E_1a;3109 | 0.05 | 0.05 |
| NC_002645.1;NP_073550.1;HCoV-229E_1a;3110 | 0.05 | 0.025 |
| NC_002645.1;NP_073550.1;HCoV-229E_1a;3111 | 0.05 | 0.025 |
| NC_002645.1;NP_073550.1;HCoV-229E_1a;3112 | 0 | 0.05 |
| NC_002645.1;NP_073550.1;HCoV-229E_1a;3113 | 0 | 0.05 |
| NC_002645.1;NP_073550.1;HCoV-229E_1a;3114 | 0 | 0.025 |
| NC_002645.1;NP_073550.1;HCoV-229E_1a;3115 | 0 | 0 |
| NC_002645.1;NP_073550.1;HCoV-229E_1a;3116 | 0 | 0 |
| NC_002645.1;NP_073550.1;HCoV-229E_1a;3117 | 0 | 0 |
| NC_002645.1;NP_073550.1;HCoV-229E_1a;3118 | 0 | 0 |
| NC_002645.1;NP_073550.1;HCoV-229E_1a;3119 | 0 | 0 |
| NC_002645.1;NP_073550.1;HCoV-229E_1a;3120 | 0 | 0 |
| NC_002645.1;NP_073550.1;HCoV-229E_1a;3121 | 0 | 0 |
| NC_002645.1;NP_073550.1;HCoV-229E_1a;3122 | 0 | 0 |
| NC_002645.1;NP_073550.1;HCoV-229E_1a;3123 | 0 | 0 |
| NC_002645.1;NP_073550.1;HCoV-229E_1a;3124 | 0 | 0 |
| NC_002645.1;NP_073550.1;HCoV-229E_1a;3125 | 0.05 | 0 |
| NC_002645.1;NP_073550.1;HCoV-229E_1a;3126 | 0 | 0 |
| NC_002645.1;NP_073550.1;HCoV-229E_1a;3127 | 0 | 0 |
| NC_002645.1;NP_073550.1;HCoV-229E_1a;3128 | 0 | 0 |
| NC_002645.1;NP_073550.1;HCoV-229E_1a;3129 | 0 | 0.025 |

| | | |
|---|---|---|
| NC_002645.1;NP_073550.1;HCoV-229E_1a;3130 | 0 | 0.025 |
| NC_002645.1;NP_073550.1;HCoV-229E_1a;3131 | 0 | 0 |
| NC_002645.1;NP_073550.1;HCoV-229E_1a;3132 | 0 | 0 |
| NC_002645.1;NP_073550.1;HCoV-229E_1a;3133 | 0.1 | 0.025 |
| NC_002645.1;NP_073550.1;HCoV-229E_1a;3134 | 0 | 0 |
| NC_002645.1;NP_073550.1;HCoV-229E_1a;3135 | 0 | 0 |
| NC_002645.1;NP_073550.1;HCoV-229E_1a;3136 | 0.05 | 0 |
| NC_002645.1;NP_073550.1;HCoV-229E_1a;3137 | 0.05 | 0.025 |
| NC_002645.1;NP_073550.1;HCoV-229E_1a;3138 | 0.05 | 0.025 |
| NC_002645.1;NP_073550.1;HCoV-229E_1a;3139 | 0.05 | 0.025 |
| NC_002645.1;NP_073550.1;HCoV-229E_1a;3140 | 0.05 | 0.025 |
| NC_002645.1;NP_073550.1;HCoV-229E_1a;3141 | 0.05 | 0.025 |
| NC_002645.1;NP_073550.1;HCoV-229E_1a;3142 | 0.05 | 0.025 |
| NC_002645.1;NP_073550.1;HCoV-229E_1a;3143 | 0.05 | 0.05 |
| NC_002645.1;NP_073550.1;HCoV-229E_1a;3144 | 0.05 | 0.025 |
| NC_002645.1;NP_073550.1;HCoV-229E_1a;3145 | 0.05 | 0.025 |
| NC_002645.1;NP_073550.1;HCoV-229E_1a;3146 | 0.05 | 0.025 |
| NC_002645.1;NP_073550.1;HCoV-229E_1a;3147 | 0.05 | 0 |
| NC_002645.1;NP_073550.1;HCoV-229E_1a;3148 | 0.05 | 0.1 |
| NC_002645.1;NP_073550.1;HCoV-229E_1a;3149 | 0 | 0 |
| NC_002645.1;NP_073550.1;HCoV-229E_1a;3150 | 0 | 0 |
| NC_002645.1;NP_073550.1;HCoV-229E_1a;3151 | 0 | 0 |
| NC_002645.1;NP_073550.1;HCoV-229E_1a;3152 | 0 | 0 |
| NC_002645.1;NP_073550.1;HCoV-229E_1a;3153 | 0 | 0 |
| NC_002645.1;NP_073550.1;HCoV-229E_1a;3154 | 0 | 0.025 |
| NC_002645.1;NP_073550.1;HCoV-229E_1a;3155 | 0.05 | 0.025 |
| NC_002645.1;NP_073550.1;HCoV-229E_1a;3156 | 0.05 | 0.025 |
| NC_002645.1;NP_073550.1;HCoV-229E_1a;3157 | 0 | 0.025 |
| NC_002645.1;NP_073550.1;HCoV-229E_1a;3158 | 0 | 0 |
| NC_002645.1;NP_073550.1;HCoV-229E_1a;3159 | 0 | 0 |
| NC_002645.1;NP_073550.1;HCoV-229E_1a;3160 | 0 | 0 |
| NC_002645.1;NP_073550.1;HCoV-229E_1a;3161 | 0 | 0 |
| NC_002645.1;NP_073550.1;HCoV-229E_1a;3162 | 0 | 0 |
| NC_002645.1;NP_073550.1;HCoV-229E_1a;3163 | 0 | 0 |
| NC_002645.1;NP_073550.1;HCoV-229E_1a;3164 | 0 | 0 |
| NC_002645.1;NP_073550.1;HCoV-229E_1a;3165 | 0 | 0 |
| NC_002645.1;NP_073550.1;HCoV-229E_1a;3166 | 0 | 0 |
| NC_002645.1;NP_073550.1;HCoV-229E_1a;3167 | 0 | 0 |
| NC_002645.1;NP_073550.1;HCoV-229E_1a;3168 | 0 | 0 |
| NC_002645.1;NP_073550.1;HCoV-229E_1a;3169 | 0 | 0 |
| NC_002645.1;NP_073550.1;HCoV-229E_1a;3170 | 0 | 0 |
| NC_002645.1;NP_073550.1;HCoV-229E_1a;3171 | 0 | 0 |
| NC_002645.1;NP_073550.1;HCoV-229E_1a;3172 | 0 | 0 |

| | | |
|---|---|---|
| NC_002645.1;NP_073550.1;HCoV-229E_1a;3173 | 0.05 | 0 |
| NC_002645.1;NP_073550.1;HCoV-229E_1a;3174 | 0 | 0 |
| NC_002645.1;NP_073550.1;HCoV-229E_1a;3175 | 0 | 0 |
| NC_002645.1;NP_073550.1;HCoV-229E_1a;3176 | 0 | 0 |
| NC_002645.1;NP_073550.1;HCoV-229E_1a;3177 | 0 | 0 |
| NC_002645.1;NP_073550.1;HCoV-229E_1a;3178 | 0 | 0 |
| NC_002645.1;NP_073550.1;HCoV-229E_1a;3179 | 0 | 0 |
| NC_002645.1;NP_073550.1;HCoV-229E_1a;3180 | 0 | 0 |
| NC_002645.1;NP_073550.1;HCoV-229E_1a;3181 | 0 | 0 |
| NC_002645.1;NP_073550.1;HCoV-229E_1a;3182 | 0 | 0 |
| NC_002645.1;NP_073550.1;HCoV-229E_1a;3183 | 0 | 0 |
| NC_002645.1;NP_073550.1;HCoV-229E_1a;3184 | 0 | 0 |
| NC_002645.1;NP_073550.1;HCoV-229E_1a;3185 | 0 | 0 |
| NC_002645.1;NP_073550.1;HCoV-229E_1a;3186 | 0 | 0.025 |
| NC_002645.1;NP_073550.1;HCoV-229E_1a;3187 | 0 | 0.025 |
| NC_002645.1;NP_073550.1;HCoV-229E_1a;3188 | 0 | 0 |
| NC_002645.1;NP_073550.1;HCoV-229E_1a;3189 | 0 | 0.025 |
| NC_002645.1;NP_073550.1;HCoV-229E_1a;3190 | 0 | 0.05 |
| NC_002645.1;NP_073550.1;HCoV-229E_1a;3191 | 0 | 0 |
| NC_002645.1;NP_073550.1;HCoV-229E_1a;3192 | 0 | 0.025 |
| NC_002645.1;NP_073550.1;HCoV-229E_1a;3193 | 0.05 | 0.025 |
| NC_002645.1;NP_073550.1;HCoV-229E_1a;3194 | 0 | 0.025 |
| NC_002645.1;NP_073550.1;HCoV-229E_1a;3195 | 0 | 0 |
| NC_002645.1;NP_073550.1;HCoV-229E_1a;3196 | 0 | 0 |
| NC_002645.1;NP_073550.1;HCoV-229E_1a;3197 | 0 | 0 |
| NC_002645.1;NP_073550.1;HCoV-229E_1a;3198 | 0 | 0 |
| NC_002645.1;NP_073550.1;HCoV-229E_1a;3199 | 0 | 0 |
| NC_002645.1;NP_073550.1;HCoV-229E_1a;3200 | 0 | 0 |
| NC_002645.1;NP_073550.1;HCoV-229E_1a;3201 | 0 | 0 |
| NC_002645.1;NP_073550.1;HCoV-229E_1a;3202 | 0.05 | 0.025 |
| NC_002645.1;NP_073550.1;HCoV-229E_1a;3203 | 0 | 0 |
| NC_002645.1;NP_073550.1;HCoV-229E_1a;3204 | 0 | 0 |
| NC_002645.1;NP_073550.1;HCoV-229E_1a;3205 | 0 | 0.025 |
| NC_002645.1;NP_073550.1;HCoV-229E_1a;3206 | 0 | 0 |
| NC_002645.1;NP_073550.1;HCoV-229E_1a;3207 | 0 | 0 |
| NC_002645.1;NP_073550.1;HCoV-229E_1a;3208 | 0 | 0 |
| NC_002645.1;NP_073550.1;HCoV-229E_1a;3209 | 0 | 0 |
| NC_002645.1;NP_073550.1;HCoV-229E_1a;3210 | 0 | 0 |
| NC_002645.1;NP_073550.1;HCoV-229E_1a;3211 | 0 | 0 |
| NC_002645.1;NP_073550.1;HCoV-229E_1a;3212 | 0 | 0 |
| NC_002645.1;NP_073550.1;HCoV-229E_1a;3213 | 0 | 0 |
| NC_002645.1;NP_073550.1;HCoV-229E_1a;3214 | 0 | 0 |
| NC_002645.1;NP_073550.1;HCoV-229E_1a;3215 | 0 | 0 |

| | | |
|---|---|---|
| NC_002645.1;NP_073550.1;HCoV-229E_1a;3216 | 0 | 0 |
| NC_002645.1;NP_073550.1;HCoV-229E_1a;3217 | 0 | 0 |
| NC_002645.1;NP_073550.1;HCoV-229E_1a;3218 | 0 | 0 |
| NC_002645.1;NP_073550.1;HCoV-229E_1a;3219 | 0 | 0 |
| NC_002645.1;NP_073550.1;HCoV-229E_1a;3220 | 0 | 0 |
| NC_002645.1;NP_073550.1;HCoV-229E_1a;3221 | 0 | 0 |
| NC_002645.1;NP_073550.1;HCoV-229E_1a;3222 | 0 | 0 |
| NC_002645.1;NP_073550.1;HCoV-229E_1a;3223 | 0 | 0 |
| NC_002645.1;NP_073550.1;HCoV-229E_1a;3224 | 0.1 | 0 |
| NC_002645.1;NP_073550.1;HCoV-229E_1a;3225 | 0 | 0 |
| NC_002645.1;NP_073550.1;HCoV-229E_1a;3226 | 0 | 0 |
| NC_002645.1;NP_073550.1;HCoV-229E_1a;3227 | 0 | 0 |
| NC_002645.1;NP_073550.1;HCoV-229E_1a;3228 | 0 | 0 |
| NC_002645.1;NP_073550.1;HCoV-229E_1a;3229 | 0.05 | 0 |
| NC_002645.1;NP_073550.1;HCoV-229E_1a;3230 | 0 | 0 |
| NC_002645.1;NP_073550.1;HCoV-229E_1a;3231 | 0 | 0 |
| NC_002645.1;NP_073550.1;HCoV-229E_1a;3232 | 0 | 0 |
| NC_002645.1;NP_073550.1;HCoV-229E_1a;3233 | 0 | 0 |
| NC_002645.1;NP_073550.1;HCoV-229E_1a;3234 | 0 | 0 |
| NC_002645.1;NP_073550.1;HCoV-229E_1a;3235 | 0 | 0 |
| NC_002645.1;NP_073550.1;HCoV-229E_1a;3236 | 0.05 | 0.025 |
| NC_002645.1;NP_073550.1;HCoV-229E_1a;3237 | 0.2 | 0.05 |
| NC_002645.1;NP_073550.1;HCoV-229E_1a;3238 | 0.15 | 0.05 |
| NC_002645.1;NP_073550.1;HCoV-229E_1a;3239 | 0 | 0 |
| NC_002645.1;NP_073550.1;HCoV-229E_1a;3240 | 0.1 | 0 |
| NC_002645.1;NP_073550.1;HCoV-229E_1a;3241 | 0.35 | 0.125 |
| NC_002645.1;NP_073550.1;HCoV-229E_1a;3242 | 0.4 | 0.125 |
| NC_002645.1;NP_073550.1;HCoV-229E_1a;3243 | 0.05 | 0 |
| NC_002645.1;NP_073550.1;HCoV-229E_1a;3244 | 0 | 0 |
| NC_002645.1;NP_073550.1;HCoV-229E_1a;3245 | 0.05 | 0 |
| NC_002645.1;NP_073550.1;HCoV-229E_1a;3246 | 0.25 | 0.15 |
| NC_002645.1;NP_073550.1;HCoV-229E_1a;3247 | 0.55 | 0.275 |
| NC_002645.1;NP_073550.1;HCoV-229E_1a;3248 | 0.05 | 0 |
| NC_002645.1;NP_073550.1;HCoV-229E_1a;3249 | 0 | 0 |
| NC_002645.1;NP_073550.1;HCoV-229E_1a;3250 | 0 | 0.025 |
| NC_002645.1;NP_073550.1;HCoV-229E_1a;3251 | 0 | 0 |
| NC_002645.1;NP_073550.1;HCoV-229E_1a;3252 | 0.05 | 0 |
| NC_002645.1;NP_073550.1;HCoV-229E_1a;3253 | 0 | 0 |
| NC_002645.1;NP_073550.1;HCoV-229E_1a;3254 | 0.05 | 0.025 |
| NC_002645.1;NP_073550.1;HCoV-229E_1a;3255 | 0 | 0.025 |
| NC_002645.1;NP_073550.1;HCoV-229E_1a;3256 | 0 | 0 |
| NC_002645.1;NP_073550.1;HCoV-229E_1a;3257 | 0 | 0 |
| NC_002645.1;NP_073550.1;HCoV-229E_1a;3258 | 0 | 0 |

| | | |
|---|---|---|
| NC_002645.1;NP_073550.1;HCoV-229E_1a;3259 | 0 | 0 |
| NC_002645.1;NP_073550.1;HCoV-229E_1a;3260 | 0 | 0 |
| NC_002645.1;NP_073550.1;HCoV-229E_1a;3261 | 0 | 0 |
| NC_002645.1;NP_073550.1;HCoV-229E_1a;3262 | 0 | 0 |
| NC_002645.1;NP_073550.1;HCoV-229E_1a;3263 | 0.05 | 0 |
| NC_002645.1;NP_073550.1;HCoV-229E_1a;3264 | 0 | 0 |
| NC_002645.1;NP_073550.1;HCoV-229E_1a;3265 | 0 | 0 |
| NC_002645.1;NP_073550.1;HCoV-229E_1a;3266 | 0 | 0 |
| NC_002645.1;NP_073550.1;HCoV-229E_1a;3267 | 0 | 0 |
| NC_002645.1;NP_073550.1;HCoV-229E_1a;3268 | 0 | 0 |
| NC_002645.1;NP_073550.1;HCoV-229E_1a;3269 | 0 | 0 |
| NC_002645.1;NP_073550.1;HCoV-229E_1a;3270 | 0 | 0 |
| NC_002645.1;NP_073550.1;HCoV-229E_1a;3271 | 0 | 0 |
| NC_002645.1;NP_073550.1;HCoV-229E_1a;3272 | 0 | 0 |
| NC_002645.1;NP_073550.1;HCoV-229E_1a;3273 | 0 | 0 |
| NC_002645.1;NP_073550.1;HCoV-229E_1a;3274 | 0 | 0 |
| NC_002645.1;NP_073550.1;HCoV-229E_1a;3275 | 0 | 0 |
| NC_002645.1;NP_073550.1;HCoV-229E_1a;3276 | 0 | 0 |
| NC_002645.1;NP_073550.1;HCoV-229E_1a;3277 | 0 | 0 |
| NC_002645.1;NP_073550.1;HCoV-229E_1a;3278 | 0 | 0 |
| NC_002645.1;NP_073550.1;HCoV-229E_1a;3279 | 0 | 0 |
| NC_002645.1;NP_073550.1;HCoV-229E_1a;3280 | 0 | 0 |
| NC_002645.1;NP_073550.1;HCoV-229E_1a;3281 | 0 | 0 |
| NC_002645.1;NP_073550.1;HCoV-229E_1a;3282 | 0 | 0 |
| NC_002645.1;NP_073550.1;HCoV-229E_1a;3283 | 0 | 0 |
| NC_002645.1;NP_073550.1;HCoV-229E_1a;3284 | 0 | 0 |
| NC_002645.1;NP_073550.1;HCoV-229E_1a;3285 | 0 | 0 |
| NC_002645.1;NP_073550.1;HCoV-229E_1a;3286 | 0 | 0 |
| NC_002645.1;NP_073550.1;HCoV-229E_1a;3287 | 0 | 0 |
| NC_002645.1;NP_073550.1;HCoV-229E_1a;3288 | 0 | 0 |
| NC_002645.1;NP_073550.1;HCoV-229E_1a;3289 | 0 | 0 |
| NC_002645.1;NP_073550.1;HCoV-229E_1a;3290 | 0 | 0 |
| NC_002645.1;NP_073550.1;HCoV-229E_1a;3291 | 0 | 0 |
| NC_002645.1;NP_073550.1;HCoV-229E_1a;3292 | 0 | 0 |
| NC_002645.1;NP_073550.1;HCoV-229E_1a;3293 | 0 | 0 |
| NC_002645.1;NP_073550.1;HCoV-229E_1a;3294 | 0 | 0 |
| NC_002645.1;NP_073550.1;HCoV-229E_1a;3295 | 0 | 0 |
| NC_002645.1;NP_073550.1;HCoV-229E_1a;3296 | 0 | 0 |
| NC_002645.1;NP_073550.1;HCoV-229E_1a;3297 | 0 | 0 |
| NC_002645.1;NP_073550.1;HCoV-229E_1a;3298 | 0 | 0 |
| NC_002645.1;NP_073550.1;HCoV-229E_1a;3299 | 0 | 0 |
| NC_002645.1;NP_073550.1;HCoV-229E_1a;3300 | 0 | 0 |
| NC_002645.1;NP_073550.1;HCoV-229E_1a;3301 | 0 | 0 |

| | | |
|---|---|---|
| NC_002645.1;NP_073550.1;HCoV-229E_1a;3302 | 0 | 0 |
| NC_002645.1;NP_073550.1;HCoV-229E_1a;3303 | 0 | 0 |
| NC_002645.1;NP_073550.1;HCoV-229E_1a;3304 | 0 | 0 |
| NC_002645.1;NP_073550.1;HCoV-229E_1a;3305 | 0 | 0 |
| NC_002645.1;NP_073550.1;HCoV-229E_1a;3306 | 0 | 0 |
| NC_002645.1;NP_073550.1;HCoV-229E_1a;3307 | 0 | 0 |
| NC_002645.1;NP_073550.1;HCoV-229E_1a;3308 | 0 | 0 |
| NC_002645.1;NP_073550.1;HCoV-229E_1a;3309 | 0 | 0 |
| NC_002645.1;NP_073550.1;HCoV-229E_1a;3310 | 0 | 0 |
| NC_002645.1;NP_073550.1;HCoV-229E_1a;3311 | 0 | 0 |
| NC_002645.1;NP_073550.1;HCoV-229E_1a;3312 | 0 | 0 |
| NC_002645.1;NP_073550.1;HCoV-229E_1a;3313 | 0 | 0 |
| NC_002645.1;NP_073550.1;HCoV-229E_1a;3314 | 0 | 0 |
| NC_002645.1;NP_073550.1;HCoV-229E_1a;3315 | 0 | 0 |
| NC_002645.1;NP_073550.1;HCoV-229E_1a;3316 | 0 | 0 |
| NC_002645.1;NP_073550.1;HCoV-229E_1a;3317 | 0 | 0 |
| NC_002645.1;NP_073550.1;HCoV-229E_1a;3318 | 0 | 0 |
| NC_002645.1;NP_073550.1;HCoV-229E_1a;3319 | 0 | 0 |
| NC_002645.1;NP_073550.1;HCoV-229E_1a;3320 | 0 | 0 |
| NC_002645.1;NP_073550.1;HCoV-229E_1a;3321 | 0 | 0 |
| NC_002645.1;NP_073550.1;HCoV-229E_1a;3322 | 0 | 0 |
| NC_002645.1;NP_073550.1;HCoV-229E_1a;3323 | 0 | 0 |
| NC_002645.1;NP_073550.1;HCoV-229E_1a;3324 | 0 | 0 |
| NC_002645.1;NP_073550.1;HCoV-229E_1a;3325 | 0 | 0 |
| NC_002645.1;NP_073550.1;HCoV-229E_1a;3326 | 0 | 0 |
| NC_002645.1;NP_073550.1;HCoV-229E_1a;3327 | 0 | 0 |
| NC_002645.1;NP_073550.1;HCoV-229E_1a;3328 | 0 | 0 |
| NC_002645.1;NP_073550.1;HCoV-229E_1a;3329 | 0 | 0 |
| NC_002645.1;NP_073550.1;HCoV-229E_1a;3330 | 0 | 0 |
| NC_002645.1;NP_073550.1;HCoV-229E_1a;3331 | 0 | 0 |
| NC_002645.1;NP_073550.1;HCoV-229E_1a;3332 | 0 | 0 |
| NC_002645.1;NP_073550.1;HCoV-229E_1a;3333 | 0 | 0 |
| NC_002645.1;NP_073550.1;HCoV-229E_1a;3334 | 0 | 0 |
| NC_002645.1;NP_073550.1;HCoV-229E_1a;3335 | 0 | 0 |
| NC_002645.1;NP_073550.1;HCoV-229E_1a;3336 | 0 | 0 |
| NC_002645.1;NP_073550.1;HCoV-229E_1a;3337 | 0 | 0 |
| NC_002645.1;NP_073550.1;HCoV-229E_1a;3338 | 0 | 0 |
| NC_002645.1;NP_073550.1;HCoV-229E_1a;3339 | 0 | 0 |
| NC_002645.1;NP_073550.1;HCoV-229E_1a;3340 | 0 | 0 |
| NC_002645.1;NP_073550.1;HCoV-229E_1a;3341 | 0 | 0 |
| NC_002645.1;NP_073550.1;HCoV-229E_1a;3342 | 0 | 0 |
| NC_002645.1;NP_073550.1;HCoV-229E_1a;3343 | 0 | 0 |
| NC_002645.1;NP_073550.1;HCoV-229E_1a;3344 | 0 | 0 |

| | | |
|---|---|---|
| NC_002645.1;NP_073550.1;HCoV-229E_1a;3345 | 0 | 0 |
| NC_002645.1;NP_073550.1;HCoV-229E_1a;3346 | 0 | 0 |
| NC_002645.1;NP_073550.1;HCoV-229E_1a;3347 | 0 | 0.025 |
| NC_002645.1;NP_073550.1;HCoV-229E_1a;3348 | 0 | 0 |
| NC_002645.1;NP_073550.1;HCoV-229E_1a;3349 | 0 | 0.025 |
| NC_002645.1;NP_073550.1;HCoV-229E_1a;3350 | 0.05 | 0.025 |
| NC_002645.1;NP_073550.1;HCoV-229E_1a;3351 | 0.15 | 0.025 |
| NC_002645.1;NP_073550.1;HCoV-229E_1a;3352 | 0.15 | 0.025 |
| NC_002645.1;NP_073550.1;HCoV-229E_1a;3353 | 0.1 | 0.025 |
| NC_002645.1;NP_073550.1;HCoV-229E_1a;3354 | 0.05 | 0.05 |
| NC_002645.1;NP_073550.1;HCoV-229E_1a;3355 | 0.1 | 0.05 |
| NC_002645.1;NP_073550.1;HCoV-229E_1a;3356 | 0.05 | 0 |
| NC_002645.1;NP_073550.1;HCoV-229E_1a;3357 | 0.1 | 0 |
| NC_002645.1;NP_073550.1;HCoV-229E_1a;3358 | 0.1 | 0 |
| NC_002645.1;NP_073550.1;HCoV-229E_1a;3359 | 0.1 | 0 |
| NC_002645.1;NP_073550.1;HCoV-229E_1a;3360 | 0.05 | 0 |
| NC_002645.1;NP_073550.1;HCoV-229E_1a;3361 | 0.05 | 0 |
| NC_002645.1;NP_073550.1;HCoV-229E_1a;3362 | 0.05 | 0 |
| NC_002645.1;NP_073550.1;HCoV-229E_1a;3363 | 0 | 0 |
| NC_002645.1;NP_073550.1;HCoV-229E_1a;3364 | 0 | 0 |
| NC_002645.1;NP_073550.1;HCoV-229E_1a;3365 | 0 | 0 |
| NC_002645.1;NP_073550.1;HCoV-229E_1a;3366 | 0 | 0 |
| NC_002645.1;NP_073550.1;HCoV-229E_1a;3367 | 0 | 0 |
| NC_002645.1;NP_073550.1;HCoV-229E_1a;3368 | 0 | 0 |
| NC_002645.1;NP_073550.1;HCoV-229E_1a;3369 | 0 | 0 |
| NC_002645.1;NP_073550.1;HCoV-229E_1a;3370 | 0 | 0 |
| NC_002645.1;NP_073550.1;HCoV-229E_1a;3371 | 0 | 0 |
| NC_002645.1;NP_073550.1;HCoV-229E_1a;3372 | 0 | 0 |
| NC_002645.1;NP_073550.1;HCoV-229E_1a;3373 | 0 | 0 |
| NC_002645.1;NP_073550.1;HCoV-229E_1a;3374 | 0 | 0 |
| NC_002645.1;NP_073550.1;HCoV-229E_1a;3375 | 0 | 0 |
| NC_002645.1;NP_073550.1;HCoV-229E_1a;3376 | 0 | 0 |
| NC_002645.1;NP_073550.1;HCoV-229E_1a;3377 | 0 | 0 |
| NC_002645.1;NP_073550.1;HCoV-229E_1a;3378 | 0 | 0 |
| NC_002645.1;NP_073550.1;HCoV-229E_1a;3379 | 0 | 0 |
| NC_002645.1;NP_073550.1;HCoV-229E_1a;3380 | 0 | 0 |
| NC_002645.1;NP_073550.1;HCoV-229E_1a;3381 | 0 | 0 |
| NC_002645.1;NP_073550.1;HCoV-229E_1a;3382 | 0 | 0 |
| NC_002645.1;NP_073550.1;HCoV-229E_1a;3383 | 0 | 0 |
| NC_002645.1;NP_073550.1;HCoV-229E_1a;3384 | 0 | 0 |
| NC_002645.1;NP_073550.1;HCoV-229E_1a;3385 | 0 | 0 |
| NC_002645.1;NP_073550.1;HCoV-229E_1a;3386 | 0 | 0 |
| NC_002645.1;NP_073550.1;HCoV-229E_1a;3387 | 0 | 0 |

| | | |
|---|---|---|
| NC_002645.1;NP_073550.1;HCoV-229E_1a;3388 | 0 | 0 |
| NC_002645.1;NP_073550.1;HCoV-229E_1a;3389 | 0 | 0 |
| NC_002645.1;NP_073550.1;HCoV-229E_1a;3390 | 0 | 0 |
| NC_002645.1;NP_073550.1;HCoV-229E_1a;3391 | 0 | 0 |
| NC_002645.1;NP_073550.1;HCoV-229E_1a;3392 | 0 | 0 |
| NC_002645.1;NP_073550.1;HCoV-229E_1a;3393 | 0 | 0 |
| NC_002645.1;NP_073550.1;HCoV-229E_1a;3394 | 0 | 0 |
| NC_002645.1;NP_073550.1;HCoV-229E_1a;3395 | 0 | 0 |
| NC_002645.1;NP_073550.1;HCoV-229E_1a;3396 | 0 | 0 |
| NC_002645.1;NP_073550.1;HCoV-229E_1a;3397 | 0 | 0 |
| NC_002645.1;NP_073550.1;HCoV-229E_1a;3398 | 0 | 0 |
| NC_002645.1;NP_073550.1;HCoV-229E_1a;3399 | 0 | 0 |
| NC_002645.1;NP_073550.1;HCoV-229E_1a;3400 | 0 | 0 |
| NC_002645.1;NP_073550.1;HCoV-229E_1a;3401 | 0 | 0 |
| NC_002645.1;NP_073550.1;HCoV-229E_1a;3402 | 0 | 0 |
| NC_002645.1;NP_073550.1;HCoV-229E_1a;3403 | 0 | 0 |
| NC_002645.1;NP_073550.1;HCoV-229E_1a;3404 | 0 | 0 |
| NC_002645.1;NP_073550.1;HCoV-229E_1a;3405 | 0 | 0 |
| NC_002645.1;NP_073550.1;HCoV-229E_1a;3406 | 0 | 0 |
| NC_002645.1;NP_073550.1;HCoV-229E_1a;3407 | 0 | 0 |
| NC_002645.1;NP_073550.1;HCoV-229E_1a;3408 | 0 | 0 |
| NC_002645.1;NP_073550.1;HCoV-229E_1a;3409 | 0 | 0 |
| NC_002645.1;NP_073550.1;HCoV-229E_1a;3410 | 0 | 0 |
| NC_002645.1;NP_073550.1;HCoV-229E_1a;3411 | 0 | 0 |
| NC_002645.1;NP_073550.1;HCoV-229E_1a;3412 | 0 | 0 |
| NC_002645.1;NP_073550.1;HCoV-229E_1a;3413 | 0 | 0 |
| NC_002645.1;NP_073550.1;HCoV-229E_1a;3414 | 0 | 0 |
| NC_002645.1;NP_073550.1;HCoV-229E_1a;3415 | 0 | 0 |
| NC_002645.1;NP_073550.1;HCoV-229E_1a;3416 | 0 | 0 |
| NC_002645.1;NP_073550.1;HCoV-229E_1a;3417 | 0 | 0 |
| NC_002645.1;NP_073550.1;HCoV-229E_1a;3418 | 0 | 0 |
| NC_002645.1;NP_073550.1;HCoV-229E_1a;3419 | 0 | 0 |
| NC_002645.1;NP_073550.1;HCoV-229E_1a;3420 | 0 | 0 |
| NC_002645.1;NP_073550.1;HCoV-229E_1a;3421 | 0 | 0 |
| NC_002645.1;NP_073550.1;HCoV-229E_1a;3422 | 0 | 0 |
| NC_002645.1;NP_073550.1;HCoV-229E_1a;3423 | 0 | 0 |
| NC_002645.1;NP_073550.1;HCoV-229E_1a;3424 | 0 | 0 |
| NC_002645.1;NP_073550.1;HCoV-229E_1a;3425 | 0 | 0 |
| NC_002645.1;NP_073550.1;HCoV-229E_1a;3426 | 0 | 0 |
| NC_002645.1;NP_073550.1;HCoV-229E_1a;3427 | 0 | 0 |
| NC_002645.1;NP_073550.1;HCoV-229E_1a;3428 | 0 | 0 |
| NC_002645.1;NP_073550.1;HCoV-229E_1a;3429 | 0 | 0 |
| NC_002645.1;NP_073550.1;HCoV-229E_1a;3430 | 0 | 0 |

| | | |
|---|---|---|
| NC_002645.1;NP_073550.1;HCoV-229E_1a;3431 | 0 | 0 |
| NC_002645.1;NP_073550.1;HCoV-229E_1a;3432 | 0 | 0 |
| NC_002645.1;NP_073550.1;HCoV-229E_1a;3433 | 0 | 0 |
| NC_002645.1;NP_073550.1;HCoV-229E_1a;3434 | 0 | 0 |
| NC_002645.1;NP_073550.1;HCoV-229E_1a;3435 | 0 | 0 |
| NC_002645.1;NP_073550.1;HCoV-229E_1a;3436 | 0 | 0 |
| NC_002645.1;NP_073550.1;HCoV-229E_1a;3437 | 0 | 0 |
| NC_002645.1;NP_073550.1;HCoV-229E_1a;3438 | 0 | 0 |
| NC_002645.1;NP_073550.1;HCoV-229E_1a;3439 | 0 | 0 |
| NC_002645.1;NP_073550.1;HCoV-229E_1a;3440 | 0 | 0 |
| NC_002645.1;NP_073550.1;HCoV-229E_1a;3441 | 0 | 0 |
| NC_002645.1;NP_073550.1;HCoV-229E_1a;3442 | 0 | 0 |
| NC_002645.1;NP_073550.1;HCoV-229E_1a;3443 | 0 | 0 |
| NC_002645.1;NP_073550.1;HCoV-229E_1a;3444 | 0 | 0 |
| NC_002645.1;NP_073550.1;HCoV-229E_1a;3445 | 0 | 0 |
| NC_002645.1;NP_073550.1;HCoV-229E_1a;3446 | 0 | 0 |
| NC_002645.1;NP_073550.1;HCoV-229E_1a;3447 | 0 | 0 |
| NC_002645.1;NP_073550.1;HCoV-229E_1a;3448 | 0 | 0 |
| NC_002645.1;NP_073550.1;HCoV-229E_1a;3449 | 0 | 0 |
| NC_002645.1;NP_073550.1;HCoV-229E_1a;3450 | 0 | 0 |
| NC_002645.1;NP_073550.1;HCoV-229E_1a;3451 | 0 | 0 |
| NC_002645.1;NP_073550.1;HCoV-229E_1a;3452 | 0 | 0 |
| NC_002645.1;NP_073550.1;HCoV-229E_1a;3453 | 0 | 0 |
| NC_002645.1;NP_073550.1;HCoV-229E_1a;3454 | 0 | 0 |
| NC_002645.1;NP_073550.1;HCoV-229E_1a;3455 | 0.05 | 0 |
| NC_002645.1;NP_073550.1;HCoV-229E_1a;3456 | 0 | 0 |
| NC_002645.1;NP_073550.1;HCoV-229E_1a;3457 | 0 | 0 |
| NC_002645.1;NP_073550.1;HCoV-229E_1a;3458 | 0 | 0 |
| NC_002645.1;NP_073550.1;HCoV-229E_1a;3459 | 0 | 0 |
| NC_002645.1;NP_073550.1;HCoV-229E_1a;3460 | 0 | 0 |
| NC_002645.1;NP_073550.1;HCoV-229E_1a;3461 | 0 | 0 |
| NC_002645.1;NP_073550.1;HCoV-229E_1a;3462 | 0 | 0 |
| NC_002645.1;NP_073550.1;HCoV-229E_1a;3463 | 0 | 0 |
| NC_002645.1;NP_073550.1;HCoV-229E_1a;3464 | 0 | 0 |
| NC_002645.1;NP_073550.1;HCoV-229E_1a;3465 | 0 | 0 |
| NC_002645.1;NP_073550.1;HCoV-229E_1a;3466 | 0 | 0 |
| NC_002645.1;NP_073550.1;HCoV-229E_1a;3467 | 0 | 0 |
| NC_002645.1;NP_073550.1;HCoV-229E_1a;3468 | 0 | 0 |
| NC_002645.1;NP_073550.1;HCoV-229E_1a;3469 | 0 | 0 |
| NC_002645.1;NP_073550.1;HCoV-229E_1a;3470 | 0 | 0 |
| NC_002645.1;NP_073550.1;HCoV-229E_1a;3471 | 0 | 0 |
| NC_002645.1;NP_073550.1;HCoV-229E_1a;3472 | 0 | 0 |
| NC_002645.1;NP_073550.1;HCoV-229E_1a;3473 | 0 | 0 |

| | | |
|---|---|---|
| NC_002645.1;NP_073550.1;HCoV-229E_1a;3474 | 0 | 0 |
| NC_002645.1;NP_073550.1;HCoV-229E_1a;3475 | 0 | 0 |
| NC_002645.1;NP_073550.1;HCoV-229E_1a;3476 | 0 | 0 |
| NC_002645.1;NP_073550.1;HCoV-229E_1a;3477 | 0 | 0 |
| NC_002645.1;NP_073550.1;HCoV-229E_1a;3478 | 0 | 0 |
| NC_002645.1;NP_073550.1;HCoV-229E_1a;3479 | 0 | 0 |
| NC_002645.1;NP_073550.1;HCoV-229E_1a;3480 | 0 | 0 |
| NC_002645.1;NP_073550.1;HCoV-229E_1a;3481 | 0 | 0 |
| NC_002645.1;NP_073550.1;HCoV-229E_1a;3482 | 0 | 0 |
| NC_002645.1;NP_073550.1;HCoV-229E_1a;3483 | 0 | 0 |
| NC_002645.1;NP_073550.1;HCoV-229E_1a;3484 | 0 | 0 |
| NC_002645.1;NP_073550.1;HCoV-229E_1a;3485 | 0 | 0 |
| NC_002645.1;NP_073550.1;HCoV-229E_1a;3486 | 0 | 0 |
| NC_002645.1;NP_073550.1;HCoV-229E_1a;3487 | 0 | 0 |
| NC_002645.1;NP_073550.1;HCoV-229E_1a;3488 | 0 | 0 |
| NC_002645.1;NP_073550.1;HCoV-229E_1a;3489 | 0 | 0 |
| NC_002645.1;NP_073550.1;HCoV-229E_1a;3490 | 0 | 0 |
| NC_002645.1;NP_073550.1;HCoV-229E_1a;3491 | 0 | 0 |
| NC_002645.1;NP_073550.1;HCoV-229E_1a;3492 | 0 | 0 |
| NC_002645.1;NP_073550.1;HCoV-229E_1a;3493 | 0 | 0 |
| NC_002645.1;NP_073550.1;HCoV-229E_1a;3494 | 0 | 0 |
| NC_002645.1;NP_073550.1;HCoV-229E_1a;3495 | 0 | 0 |
| NC_002645.1;NP_073550.1;HCoV-229E_1a;3496 | 0.05 | 0.125 |
| NC_002645.1;NP_073550.1;HCoV-229E_1a;3497 | 0.05 | 0.05 |
| NC_002645.1;NP_073550.1;HCoV-229E_1a;3498 | 0 | 0 |
| NC_002645.1;NP_073550.1;HCoV-229E_1a;3499 | 0 | 0 |
| NC_002645.1;NP_073550.1;HCoV-229E_1a;3500 | 0 | 0 |
| NC_002645.1;NP_073550.1;HCoV-229E_1a;3501 | 0 | 0 |
| NC_002645.1;NP_073550.1;HCoV-229E_1a;3502 | 0.05 | 0 |
| NC_002645.1;NP_073550.1;HCoV-229E_1a;3503 | 0.25 | 0.175 |
| NC_002645.1;NP_073550.1;HCoV-229E_1a;3504 | 0.1 | 0.125 |
| NC_002645.1;NP_073550.1;HCoV-229E_1a;3505 | 0.15 | 0.2 |
| NC_002645.1;NP_073550.1;HCoV-229E_1a;3506 | 0.15 | 0.225 |
| NC_002645.1;NP_073550.1;HCoV-229E_1a;3507 | 0.1 | 0.1 |
| NC_002645.1;NP_073550.1;HCoV-229E_1a;3508 | 0.35 | 0.3 |
| NC_002645.1;NP_073550.1;HCoV-229E_1a;3509 | 0.1 | 0.075 |
| NC_002645.1;NP_073550.1;HCoV-229E_1a;3510 | 0.05 | 0.025 |
| NC_002645.1;NP_073550.1;HCoV-229E_1a;3511 | 0.05 | 0.075 |
| NC_002645.1;NP_073550.1;HCoV-229E_1a;3512 | 0 | 0.025 |
| NC_002645.1;NP_073550.1;HCoV-229E_1a;3513 | 0 | 0 |
| NC_002645.1;NP_073550.1;HCoV-229E_1a;3514 | 0 | 0 |
| NC_002645.1;NP_073550.1;HCoV-229E_1a;3515 | 0 | 0 |
| NC_002645.1;NP_073550.1;HCoV-229E_1a;3516 | 0 | 0 |

| | | |
|---|---|---|
| NC_002645.1;NP_073550.1;HCoV-229E_1a;3517 | 0 | 0 |
| NC_002645.1;NP_073550.1;HCoV-229E_1a;3518 | 0 | 0 |
| NC_002645.1;NP_073550.1;HCoV-229E_1a;3519 | 0 | 0 |
| NC_002645.1;NP_073550.1;HCoV-229E_1a;3520 | 0 | 0 |
| NC_002645.1;NP_073550.1;HCoV-229E_1a;3521 | 0 | 0 |
| NC_002645.1;NP_073550.1;HCoV-229E_1a;3522 | 0 | 0 |
| NC_002645.1;NP_073550.1;HCoV-229E_1a;3523 | 0 | 0 |
| NC_002645.1;NP_073550.1;HCoV-229E_1a;3524 | 0 | 0 |
| NC_002645.1;NP_073550.1;HCoV-229E_1a;3525 | 0 | 0 |
| NC_002645.1;NP_073550.1;HCoV-229E_1a;3526 | 0 | 0 |
| NC_002645.1;NP_073550.1;HCoV-229E_1a;3527 | 0 | 0 |
| NC_002645.1;NP_073550.1;HCoV-229E_1a;3528 | 0 | 0 |
| NC_002645.1;NP_073550.1;HCoV-229E_1a;3529 | 0 | 0 |
| NC_002645.1;NP_073550.1;HCoV-229E_1a;3530 | 0 | 0 |
| NC_002645.1;NP_073550.1;HCoV-229E_1a;3531 | 0 | 0 |
| NC_002645.1;NP_073550.1;HCoV-229E_1a;3532 | 0 | 0 |
| NC_002645.1;NP_073550.1;HCoV-229E_1a;3533 | 0.1 | 0 |
| NC_002645.1;NP_073550.1;HCoV-229E_1a;3534 | 0.05 | 0 |
| NC_002645.1;NP_073550.1;HCoV-229E_1a;3535 | 0.05 | 0.025 |
| NC_002645.1;NP_073550.1;HCoV-229E_1a;3536 | 0.05 | 0.075 |
| NC_002645.1;NP_073550.1;HCoV-229E_1a;3537 | 0.05 | 0.05 |
| NC_002645.1;NP_073550.1;HCoV-229E_1a;3538 | 0.05 | 0 |
| NC_002645.1;NP_073550.1;HCoV-229E_1a;3539 | 0.05 | 0.025 |
| NC_002645.1;NP_073550.1;HCoV-229E_1a;3540 | 0.05 | 0 |
| NC_002645.1;NP_073550.1;HCoV-229E_1a;3541 | 0.05 | 0.025 |
| NC_002645.1;NP_073550.1;HCoV-229E_1a;3542 | 0 | 0 |
| NC_002645.1;NP_073550.1;HCoV-229E_1a;3543 | 0 | 0 |
| NC_002645.1;NP_073550.1;HCoV-229E_1a;3544 | 0 | 0 |
| NC_002645.1;NP_073550.1;HCoV-229E_1a;3545 | 0 | 0 |
| NC_002645.1;NP_073550.1;HCoV-229E_1a;3546 | 0 | 0 |
| NC_002645.1;NP_073550.1;HCoV-229E_1a;3547 | 0 | 0 |
| NC_002645.1;NP_073550.1;HCoV-229E_1a;3548 | 0.1 | 0.025 |
| NC_002645.1;NP_073550.1;HCoV-229E_1a;3549 | 0.15 | 0.025 |
| NC_002645.1;NP_073550.1;HCoV-229E_1a;3550 | 0 | 0 |
| NC_002645.1;NP_073550.1;HCoV-229E_1a;3551 | 0 | 0 |
| NC_002645.1;NP_073550.1;HCoV-229E_1a;3552 | 0 | 0 |
| NC_002645.1;NP_073550.1;HCoV-229E_1a;3553 | 0 | 0 |
| NC_002645.1;NP_073550.1;HCoV-229E_1a;3554 | 0 | 0 |
| NC_002645.1;NP_073550.1;HCoV-229E_1a;3555 | 0.05 | 0 |
| NC_002645.1;NP_073550.1;HCoV-229E_1a;3556 | 0 | 0 |
| NC_002645.1;NP_073550.1;HCoV-229E_1a;3557 | 0 | 0 |
| NC_002645.1;NP_073550.1;HCoV-229E_1a;3558 | 0 | 0.025 |
| NC_002645.1;NP_073550.1;HCoV-229E_1a;3559 | 0.1 | 0.075 |

| | | |
|---|---|---|
| NC_002645.1;NP_073550.1;HCoV-229E_1a;3560 | 0.1 | 0.175 |
| NC_002645.1;NP_073550.1;HCoV-229E_1a;3561 | 0.1 | 0.275 |
| NC_002645.1;NP_073550.1;HCoV-229E_1a;3562 | 0.05 | 0.2 |
| NC_002645.1;NP_073550.1;HCoV-229E_1a;3563 | 0.05 | 0.125 |
| NC_002645.1;NP_073550.1;HCoV-229E_1a;3564 | 0.05 | 0.075 |
| NC_002645.1;NP_073550.1;HCoV-229E_1a;3565 | 0 | 0.025 |
| NC_002645.1;NP_073550.1;HCoV-229E_1a;3566 | 0 | 0 |
| NC_002645.1;NP_073550.1;HCoV-229E_1a;3567 | 0.05 | 0 |
| NC_002645.1;NP_073550.1;HCoV-229E_1a;3568 | 0.1 | 0.025 |
| NC_002645.1;NP_073550.1;HCoV-229E_1a;3569 | 0.1 | 0 |
| NC_002645.1;NP_073550.1;HCoV-229E_1a;3570 | 0.1 | 0 |
| NC_002645.1;NP_073550.1;HCoV-229E_1a;3571 | 0 | 0 |
| NC_002645.1;NP_073550.1;HCoV-229E_1a;3572 | 0 | 0 |
| NC_002645.1;NP_073550.1;HCoV-229E_1a;3573 | 0 | 0 |
| NC_002645.1;NP_073550.1;HCoV-229E_1a;3574 | 0 | 0 |
| NC_002645.1;NP_073550.1;HCoV-229E_1a;3575 | 0 | 0 |
| NC_002645.1;NP_073550.1;HCoV-229E_1a;3576 | 0 | 0 |
| NC_002645.1;NP_073550.1;HCoV-229E_1a;3577 | 0 | 0.025 |
| NC_002645.1;NP_073550.1;HCoV-229E_1a;3578 | 0 | 0.025 |
| NC_002645.1;NP_073550.1;HCoV-229E_1a;3579 | 0.05 | 0.025 |
| NC_002645.1;NP_073550.1;HCoV-229E_1a;3580 | 0 | 0.025 |
| NC_002645.1;NP_073550.1;HCoV-229E_1a;3581 | 0 | 0 |
| NC_002645.1;NP_073550.1;HCoV-229E_1a;3582 | 0 | 0 |
| NC_002645.1;NP_073550.1;HCoV-229E_1a;3583 | 0 | 0.025 |
| NC_002645.1;NP_073550.1;HCoV-229E_1a;3584 | 0 | 0 |
| NC_002645.1;NP_073550.1;HCoV-229E_1a;3585 | 0 | 0 |
| NC_002645.1;NP_073550.1;HCoV-229E_1a;3586 | 0 | 0 |
| NC_002645.1;NP_073550.1;HCoV-229E_1a;3587 | 0 | 0 |
| NC_002645.1;NP_073550.1;HCoV-229E_1a;3588 | 0 | 0 |
| NC_002645.1;NP_073550.1;HCoV-229E_1a;3589 | 0 | 0 |
| NC_002645.1;NP_073550.1;HCoV-229E_1a;3590 | 0 | 0 |
| NC_002645.1;NP_073550.1;HCoV-229E_1a;3591 | 0 | 0 |
| NC_002645.1;NP_073550.1;HCoV-229E_1a;3592 | 0 | 0 |
| NC_002645.1;NP_073550.1;HCoV-229E_1a;3593 | 0 | 0 |
| NC_002645.1;NP_073550.1;HCoV-229E_1a;3594 | 0 | 0 |
| NC_002645.1;NP_073550.1;HCoV-229E_1a;3595 | 0 | 0 |
| NC_002645.1;NP_073550.1;HCoV-229E_1a;3596 | 0 | 0 |
| NC_002645.1;NP_073550.1;HCoV-229E_1a;3597 | 0 | 0 |
| NC_002645.1;NP_073550.1;HCoV-229E_1a;3598 | 0 | 0 |
| NC_002645.1;NP_073550.1;HCoV-229E_1a;3599 | 0 | 0 |
| NC_002645.1;NP_073550.1;HCoV-229E_1a;3600 | 0 | 0 |
| NC_002645.1;NP_073550.1;HCoV-229E_1a;3601 | 0 | 0 |
| NC_002645.1;NP_073550.1;HCoV-229E_1a;3602 | 0.05 | 0 |

| | | |
|---|---|---|
| NC_002645.1;NP_073550.1;HCoV-229E_1a;3603 | 0.05 | 0 |
| NC_002645.1;NP_073550.1;HCoV-229E_1a;3604 | 0.15 | 0.2 |
| NC_002645.1;NP_073550.1;HCoV-229E_1a;3605 | 0.35 | 0.35 |
| NC_002645.1;NP_073550.1;HCoV-229E_1a;3606 | 0.3 | 0.3 |
| NC_002645.1;NP_073550.1;HCoV-229E_1a;3607 | 0.1 | 0.1 |
| NC_002645.1;NP_073550.1;HCoV-229E_1a;3608 | 0.35 | 0.325 |
| NC_002645.1;NP_073550.1;HCoV-229E_1a;3609 | 0.35 | 0.3 |
| NC_002645.1;NP_073550.1;HCoV-229E_1a;3610 | 0.25 | 0.275 |
| NC_002645.1;NP_073550.1;HCoV-229E_1a;3611 | 0.05 | 0.225 |
| NC_002645.1;NP_073550.1;HCoV-229E_1a;3612 | 0.1 | 0.275 |
| NC_002645.1;NP_073550.1;HCoV-229E_1a;3613 | 0.55 | 0.625 |
| NC_002645.1;NP_073550.1;HCoV-229E_1a;3614 | 0.05 | 0 |
| NC_002645.1;NP_073550.1;HCoV-229E_1a;3615 | 0.05 | 0.025 |
| NC_002645.1;NP_073550.1;HCoV-229E_1a;3616 | 0.1 | 0.025 |
| NC_002645.1;NP_073550.1;HCoV-229E_1a;3617 | 0.15 | 0.025 |
| NC_002645.1;NP_073550.1;HCoV-229E_1a;3618 | 0 | 0.025 |
| NC_002645.1;NP_073550.1;HCoV-229E_1a;3619 | 0 | 0 |
| NC_002645.1;NP_073550.1;HCoV-229E_1a;3620 | 0 | 0 |
| NC_002645.1;NP_073550.1;HCoV-229E_1a;3621 | 0 | 0 |
| NC_002645.1;NP_073550.1;HCoV-229E_1a;3622 | 0 | 0 |
| NC_002645.1;NP_073550.1;HCoV-229E_1a;3623 | 0 | 0 |
| NC_002645.1;NP_073550.1;HCoV-229E_1a;3624 | 0 | 0 |
| NC_002645.1;NP_073550.1;HCoV-229E_1a;3625 | 0 | 0 |
| NC_002645.1;NP_073550.1;HCoV-229E_1a;3626 | 0 | 0 |
| NC_002645.1;NP_073550.1;HCoV-229E_1a;3627 | 0 | 0 |
| NC_002645.1;NP_073550.1;HCoV-229E_1a;3628 | 0 | 0 |
| NC_002645.1;NP_073550.1;HCoV-229E_1a;3629 | 0 | 0 |
| NC_002645.1;NP_073550.1;HCoV-229E_1a;3630 | 0 | 0.025 |
| NC_002645.1;NP_073550.1;HCoV-229E_1a;3631 | 0.05 | 0.1 |
| NC_002645.1;NP_073550.1;HCoV-229E_1a;3632 | 0.05 | 0.025 |
| NC_002645.1;NP_073550.1;HCoV-229E_1a;3633 | 0.05 | 0.025 |
| NC_002645.1;NP_073550.1;HCoV-229E_1a;3634 | 0.05 | 0.025 |
| NC_002645.1;NP_073550.1;HCoV-229E_1a;3635 | 0.05 | 0.075 |
| NC_002645.1;NP_073550.1;HCoV-229E_1a;3636 | 0.1 | 0.075 |
| NC_002645.1;NP_073550.1;HCoV-229E_1a;3637 | 0.1 | 0.1 |
| NC_002645.1;NP_073550.1;HCoV-229E_1a;3638 | 0.1 | 0.125 |
| NC_002645.1;NP_073550.1;HCoV-229E_1a;3639 | 0.1 | 0.1 |
| NC_002645.1;NP_073550.1;HCoV-229E_1a;3640 | 0.1 | 0.05 |
| NC_002645.1;NP_073550.1;HCoV-229E_1a;3641 | 0.1 | 0.075 |
| NC_002645.1;NP_073550.1;HCoV-229E_1a;3642 | 0 | 0 |
| NC_002645.1;NP_073550.1;HCoV-229E_1a;3643 | 0 | 0.075 |
| NC_002645.1;NP_073550.1;HCoV-229E_1a;3644 | 0 | 0.025 |
| NC_002645.1;NP_073550.1;HCoV-229E_1a;3645 | 0.05 | 0.025 |

| | | |
|---|---|---|
| NC_002645.1;NP_073550.1;HCoV-229E_1a;3646 | 0 | 0 |
| NC_002645.1;NP_073550.1;HCoV-229E_1a;3647 | 0 | 0 |
| NC_002645.1;NP_073550.1;HCoV-229E_1a;3648 | 0 | 0 |
| NC_002645.1;NP_073550.1;HCoV-229E_1a;3649 | 0 | 0 |
| NC_002645.1;NP_073550.1;HCoV-229E_1a;3650 | 0 | 0.025 |
| NC_002645.1;NP_073550.1;HCoV-229E_1a;3651 | 0 | 0.025 |
| NC_002645.1;NP_073550.1;HCoV-229E_1a;3652 | 0 | 0.05 |
| NC_002645.1;NP_073550.1;HCoV-229E_1a;3653 | 0 | 0 |
| NC_002645.1;NP_073550.1;HCoV-229E_1a;3654 | 0 | 0 |
| NC_002645.1;NP_073550.1;HCoV-229E_1a;3655 | 0 | 0 |
| NC_002645.1;NP_073550.1;HCoV-229E_1a;3656 | 0 | 0 |
| NC_002645.1;NP_073550.1;HCoV-229E_1a;3657 | 0 | 0 |
| NC_002645.1;NP_073550.1;HCoV-229E_1a;3658 | 0 | 0 |
| NC_002645.1;NP_073550.1;HCoV-229E_1a;3659 | 0 | 0 |
| NC_002645.1;NP_073550.1;HCoV-229E_1a;3660 | 0 | 0 |
| NC_002645.1;NP_073550.1;HCoV-229E_1a;3661 | 0 | 0 |
| NC_002645.1;NP_073550.1;HCoV-229E_1a;3662 | 0 | 0 |
| NC_002645.1;NP_073550.1;HCoV-229E_1a;3663 | 0 | 0 |
| NC_002645.1;NP_073550.1;HCoV-229E_1a;3664 | 0 | 0.025 |
| NC_002645.1;NP_073550.1;HCoV-229E_1a;3665 | 0 | 0.025 |
| NC_002645.1;NP_073550.1;HCoV-229E_1a;3666 | 0 | 0.025 |
| NC_002645.1;NP_073550.1;HCoV-229E_1a;3667 | 0 | 0.025 |
| NC_002645.1;NP_073550.1;HCoV-229E_1a;3668 | 0 | 0.025 |
| NC_002645.1;NP_073550.1;HCoV-229E_1a;3669 | 0.05 | 0.05 |
| NC_002645.1;NP_073550.1;HCoV-229E_1a;3670 | 0.05 | 0.025 |
| NC_002645.1;NP_073550.1;HCoV-229E_1a;3671 | 0 | 0 |
| NC_002645.1;NP_073550.1;HCoV-229E_1a;3672 | 0 | 0 |
| NC_002645.1;NP_073550.1;HCoV-229E_1a;3673 | 0 | 0 |
| NC_002645.1;NP_073550.1;HCoV-229E_1a;3674 | 0 | 0 |
| NC_002645.1;NP_073550.1;HCoV-229E_1a;3675 | 0 | 0 |
| NC_002645.1;NP_073550.1;HCoV-229E_1a;3676 | 0 | 0.025 |
| NC_002645.1;NP_073550.1;HCoV-229E_1a;3677 | 0 | 0 |
| NC_002645.1;NP_073550.1;HCoV-229E_1a;3678 | 0 | 0 |
| NC_002645.1;NP_073550.1;HCoV-229E_1a;3679 | 0 | 0 |
| NC_002645.1;NP_073550.1;HCoV-229E_1a;3680 | 0 | 0 |
| NC_002645.1;NP_073550.1;HCoV-229E_1a;3681 | 0 | 0.05 |
| NC_002645.1;NP_073550.1;HCoV-229E_1a;3682 | 0 | 0 |
| NC_002645.1;NP_073550.1;HCoV-229E_1a;3683 | 0 | 0 |
| NC_002645.1;NP_073550.1;HCoV-229E_1a;3684 | 0 | 0 |
| NC_002645.1;NP_073550.1;HCoV-229E_1a;3685 | 0 | 0 |
| NC_002645.1;NP_073550.1;HCoV-229E_1a;3686 | 0 | 0 |
| NC_002645.1;NP_073550.1;HCoV-229E_1a;3687 | 0 | 0 |
| NC_002645.1;NP_073550.1;HCoV-229E_1a;3688 | 0.05 | 0 |

| | | |
|---|---|---|
| NC_002645.1;NP_073550.1;HCoV-229E_1a;3689 | 0.05 | 0 |
| NC_002645.1;NP_073550.1;HCoV-229E_1a;3690 | 0.05 | 0 |
| NC_002645.1;NP_073550.1;HCoV-229E_1a;3691 | 0.1 | 0.025 |
| NC_002645.1;NP_073550.1;HCoV-229E_1a;3692 | 0.15 | 0.025 |
| NC_002645.1;NP_073550.1;HCoV-229E_1a;3693 | 0.05 | 0 |
| NC_002645.1;NP_073550.1;HCoV-229E_1a;3694 | 0 | 0 |
| NC_002645.1;NP_073550.1;HCoV-229E_1a;3695 | 0 | 0 |
| NC_002645.1;NP_073550.1;HCoV-229E_1a;3696 | 0 | 0 |
| NC_002645.1;NP_073550.1;HCoV-229E_1a;3697 | 0 | 0 |
| NC_002645.1;NP_073550.1;HCoV-229E_1a;3698 | 0 | 0 |
| NC_002645.1;NP_073550.1;HCoV-229E_1a;3699 | 0 | 0 |
| NC_002645.1;NP_073550.1;HCoV-229E_1a;3700 | 0 | 0 |
| NC_002645.1;NP_073550.1;HCoV-229E_1a;3701 | 0 | 0 |
| NC_002645.1;NP_073550.1;HCoV-229E_1a;3702 | 0 | 0 |
| NC_002645.1;NP_073550.1;HCoV-229E_1a;3703 | 0 | 0 |
| NC_002645.1;NP_073550.1;HCoV-229E_1a;3704 | 0 | 0 |
| NC_002645.1;NP_073550.1;HCoV-229E_1a;3705 | 0 | 0 |
| NC_002645.1;NP_073550.1;HCoV-229E_1a;3706 | 0 | 0 |
| NC_002645.1;NP_073550.1;HCoV-229E_1a;3707 | 0 | 0 |
| NC_002645.1;NP_073550.1;HCoV-229E_1a;3708 | 0 | 0 |
| NC_002645.1;NP_073550.1;HCoV-229E_1a;3709 | 0 | 0 |
| NC_002645.1;NP_073550.1;HCoV-229E_1a;3710 | 0 | 0 |
| NC_002645.1;NP_073550.1;HCoV-229E_1a;3711 | 0 | 0 |
| NC_002645.1;NP_073550.1;HCoV-229E_1a;3712 | 0 | 0 |
| NC_002645.1;NP_073550.1;HCoV-229E_1a;3713 | 0 | 0 |
| NC_002645.1;NP_073550.1;HCoV-229E_1a;3714 | 0 | 0 |
| NC_002645.1;NP_073550.1;HCoV-229E_1a;3715 | 0 | 0 |
| NC_002645.1;NP_073550.1;HCoV-229E_1a;3716 | 0 | 0 |
| NC_002645.1;NP_073550.1;HCoV-229E_1a;3717 | 0 | 0 |
| NC_002645.1;NP_073550.1;HCoV-229E_1a;3718 | 0 | 0 |
| NC_002645.1;NP_073550.1;HCoV-229E_1a;3719 | 0 | 0 |
| NC_002645.1;NP_073550.1;HCoV-229E_1a;3720 | 0 | 0 |
| NC_002645.1;NP_073550.1;HCoV-229E_1a;3721 | 0 | 0 |
| NC_002645.1;NP_073550.1;HCoV-229E_1a;3722 | 0 | 0 |
| NC_002645.1;NP_073550.1;HCoV-229E_1a;3723 | 0 | 0 |
| NC_002645.1;NP_073550.1;HCoV-229E_1a;3724 | 0 | 0.025 |
| NC_002645.1;NP_073550.1;HCoV-229E_1a;3725 | 0 | 0 |
| NC_002645.1;NP_073550.1;HCoV-229E_1a;3726 | 0 | 0.025 |
| NC_002645.1;NP_073550.1;HCoV-229E_1a;3727 | 0 | 0 |
| NC_002645.1;NP_073550.1;HCoV-229E_1a;3728 | 0 | 0.025 |
| NC_002645.1;NP_073550.1;HCoV-229E_1a;3729 | 0 | 0.025 |
| NC_002645.1;NP_073550.1;HCoV-229E_1a;3730 | 0 | 0.025 |
| NC_002645.1;NP_073550.1;HCoV-229E_1a;3731 | 0.05 | 0 |

| | | |
|---|---|---|
| NC_002645.1;NP_073550.1;HCoV-229E_1a;3732 | 0.15 | 0.15 |
| NC_002645.1;NP_073550.1;HCoV-229E_1a;3733 | 0.05 | 0 |
| NC_002645.1;NP_073550.1;HCoV-229E_1a;3734 | 0 | 0 |
| NC_002645.1;NP_073550.1;HCoV-229E_1a;3735 | 0 | 0 |
| NC_002645.1;NP_073550.1;HCoV-229E_1a;3736 | 0 | 0 |
| NC_002645.1;NP_073550.1;HCoV-229E_1a;3737 | 0 | 0.025 |
| NC_002645.1;NP_073550.1;HCoV-229E_1a;3738 | 0 | 0.025 |
| NC_002645.1;NP_073550.1;HCoV-229E_1a;3739 | 0 | 0.025 |
| NC_002645.1;NP_073550.1;HCoV-229E_1a;3740 | 0.05 | 0.05 |
| NC_002645.1;NP_073550.1;HCoV-229E_1a;3741 | 0 | 0.05 |
| NC_002645.1;NP_073550.1;HCoV-229E_1a;3742 | 0 | 0.025 |
| NC_002645.1;NP_073550.1;HCoV-229E_1a;3743 | 0 | 0 |
| NC_002645.1;NP_073550.1;HCoV-229E_1a;3744 | 0 | 0 |
| NC_002645.1;NP_073550.1;HCoV-229E_1a;3745 | 0 | 0 |
| NC_002645.1;NP_073550.1;HCoV-229E_1a;3746 | 0 | 0 |
| NC_002645.1;NP_073550.1;HCoV-229E_1a;3747 | 0 | 0 |
| NC_002645.1;NP_073550.1;HCoV-229E_1a;3748 | 0 | 0 |
| NC_002645.1;NP_073550.1;HCoV-229E_1a;3749 | 0 | 0 |
| NC_002645.1;NP_073550.1;HCoV-229E_1a;3750 | 0 | 0 |
| NC_002645.1;NP_073550.1;HCoV-229E_1a;3751 | 0 | 0 |
| NC_002645.1;NP_073550.1;HCoV-229E_1a;3752 | 0 | 0 |
| NC_002645.1;NP_073550.1;HCoV-229E_1a;3753 | 0 | 0.025 |
| NC_002645.1;NP_073550.1;HCoV-229E_1a;3754 | 0 | 0 |
| NC_002645.1;NP_073550.1;HCoV-229E_1a;3755 | 0 | 0 |
| NC_002645.1;NP_073550.1;HCoV-229E_1a;3756 | 0 | 0 |
| NC_002645.1;NP_073550.1;HCoV-229E_1a;3757 | 0 | 0.075 |
| NC_002645.1;NP_073550.1;HCoV-229E_1a;3758 | 0.05 | 0.075 |
| NC_002645.1;NP_073550.1;HCoV-229E_1a;3759 | 0 | 0.075 |
| NC_002645.1;NP_073550.1;HCoV-229E_1a;3760 | 0 | 0.075 |
| NC_002645.1;NP_073550.1;HCoV-229E_1a;3761 | 0 | 0.025 |
| NC_002645.1;NP_073550.1;HCoV-229E_1a;3762 | 0 | 0 |
| NC_002645.1;NP_073550.1;HCoV-229E_1a;3763 | 0 | 0 |
| NC_002645.1;NP_073550.1;HCoV-229E_1a;3764 | 0 | 0 |
| NC_002645.1;NP_073550.1;HCoV-229E_1a;3765 | 0.1 | 0.025 |
| NC_002645.1;NP_073550.1;HCoV-229E_1a;3766 | 0 | 0 |
| NC_002645.1;NP_073550.1;HCoV-229E_1a;3767 | 0 | 0 |
| NC_002645.1;NP_073550.1;HCoV-229E_1a;3768 | 0 | 0 |
| NC_002645.1;NP_073550.1;HCoV-229E_1a;3769 | 0 | 0 |
| NC_002645.1;NP_073550.1;HCoV-229E_1a;3770 | 0 | 0 |
| NC_002645.1;NP_073550.1;HCoV-229E_1a;3771 | 0 | 0 |
| NC_002645.1;NP_073550.1;HCoV-229E_1a;3772 | 0 | 0 |
| NC_002645.1;NP_073550.1;HCoV-229E_1a;3773 | 0 | 0 |
| NC_002645.1;NP_073550.1;HCoV-229E_1a;3774 | 0 | 0 |

| | | |
|---|---|---|
| NC_002645.1;NP_073550.1;HCoV-229E_1a;3775 | 0 | 0.025 |
| NC_002645.1;NP_073550.1;HCoV-229E_1a;3776 | 0 | 0 |
| NC_002645.1;NP_073550.1;HCoV-229E_1a;3777 | 0 | 0 |
| NC_002645.1;NP_073550.1;HCoV-229E_1a;3778 | 0 | 0 |
| NC_002645.1;NP_073550.1;HCoV-229E_1a;3779 | 0 | 0.025 |
| NC_002645.1;NP_073550.1;HCoV-229E_1a;3780 | 0 | 0 |
| NC_002645.1;NP_073550.1;HCoV-229E_1a;3781 | 0 | 0.05 |
| NC_002645.1;NP_073550.1;HCoV-229E_1a;3782 | 0 | 0.075 |
| NC_002645.1;NP_073550.1;HCoV-229E_1a;3783 | 0 | 0.075 |
| NC_002645.1;NP_073550.1;HCoV-229E_1a;3784 | 0 | 0 |
| NC_002645.1;NP_073550.1;HCoV-229E_1a;3785 | 0 | 0.025 |
| NC_002645.1;NP_073550.1;HCoV-229E_1a;3786 | 0 | 0.025 |
| NC_002645.1;NP_073550.1;HCoV-229E_1a;3787 | 0 | 0.025 |
| NC_002645.1;NP_073550.1;HCoV-229E_1a;3788 | 0.05 | 0.025 |
| NC_002645.1;NP_073550.1;HCoV-229E_1a;3789 | 0.05 | 0.025 |
| NC_002645.1;NP_073550.1;HCoV-229E_1a;3790 | 0 | 0 |
| NC_002645.1;NP_073550.1;HCoV-229E_1a;3791 | 0 | 0 |
| NC_002645.1;NP_073550.1;HCoV-229E_1a;3792 | 0 | 0 |
| NC_002645.1;NP_073550.1;HCoV-229E_1a;3793 | 0 | 0 |
| NC_002645.1;NP_073550.1;HCoV-229E_1a;3794 | 0 | 0.025 |
| NC_002645.1;NP_073550.1;HCoV-229E_1a;3795 | 0 | 0 |
| NC_002645.1;NP_073550.1;HCoV-229E_1a;3796 | 0 | 0 |
| NC_002645.1;NP_073550.1;HCoV-229E_1a;3797 | 0 | 0 |
| NC_002645.1;NP_073550.1;HCoV-229E_1a;3798 | 0.05 | 0 |
| NC_002645.1;NP_073550.1;HCoV-229E_1a;3799 | 0.05 | 0 |
| NC_002645.1;NP_073550.1;HCoV-229E_1a;3800 | 0.05 | 0 |
| NC_002645.1;NP_073550.1;HCoV-229E_1a;3801 | 0 | 0 |
| NC_002645.1;NP_073550.1;HCoV-229E_1a;3802 | 0 | 0 |
| NC_002645.1;NP_073550.1;HCoV-229E_1a;3803 | 0 | 0 |
| NC_002645.1;NP_073550.1;HCoV-229E_1a;3804 | 0 | 0 |
| NC_002645.1;NP_073550.1;HCoV-229E_1a;3805 | 0 | 0 |
| NC_002645.1;NP_073550.1;HCoV-229E_1a;3806 | 0 | 0 |
| NC_002645.1;NP_073550.1;HCoV-229E_1a;3807 | 0 | 0 |
| NC_002645.1;NP_073550.1;HCoV-229E_1a;3808 | 0 | 0 |
| NC_002645.1;NP_073550.1;HCoV-229E_1a;3809 | 0 | 0 |
| NC_002645.1;NP_073550.1;HCoV-229E_1a;3810 | 0 | 0 |
| NC_002645.1;NP_073550.1;HCoV-229E_1a;3811 | 0 | 0 |
| NC_002645.1;NP_073550.1;HCoV-229E_1a;3812 | 0 | 0 |
| NC_002645.1;NP_073550.1;HCoV-229E_1a;3813 | 0 | 0 |
| NC_002645.1;NP_073550.1;HCoV-229E_1a;3814 | 0 | 0 |
| NC_002645.1;NP_073550.1;HCoV-229E_1a;3815 | 0 | 0 |
| NC_002645.1;NP_073550.1;HCoV-229E_1a;3816 | 0 | 0 |
| NC_002645.1;NP_073550.1;HCoV-229E_1a;3817 | 0 | 0 |

| | | |
|---|---|---|
| NC_002645.1;NP_073550.1;HCoV-229E_1a;3818 | 0 | 0 |
| NC_002645.1;NP_073550.1;HCoV-229E_1a;3819 | 0 | 0 |
| NC_002645.1;NP_073550.1;HCoV-229E_1a;3820 | 0.05 | 0 |
| NC_002645.1;NP_073550.1;HCoV-229E_1a;3821 | 0 | 0 |
| NC_002645.1;NP_073550.1;HCoV-229E_1a;3822 | 0 | 0 |
| NC_002645.1;NP_073550.1;HCoV-229E_1a;3823 | 0 | 0 |
| NC_002645.1;NP_073550.1;HCoV-229E_1a;3824 | 0 | 0 |
| NC_002645.1;NP_073550.1;HCoV-229E_1a;3825 | 0 | 0 |
| NC_002645.1;NP_073550.1;HCoV-229E_1a;3826 | 0 | 0 |
| NC_002645.1;NP_073550.1;HCoV-229E_1a;3827 | 0 | 0 |
| NC_002645.1;NP_073550.1;HCoV-229E_1a;3828 | 0 | 0 |
| NC_002645.1;NP_073550.1;HCoV-229E_1a;3829 | 0 | 0 |
| NC_002645.1;NP_073550.1;HCoV-229E_1a;3830 | 0 | 0 |
| NC_002645.1;NP_073550.1;HCoV-229E_1a;3831 | 0 | 0 |
| NC_002645.1;NP_073550.1;HCoV-229E_1a;3832 | 0 | 0 |
| NC_002645.1;NP_073550.1;HCoV-229E_1a;3833 | 0 | 0.025 |
| NC_002645.1;NP_073550.1;HCoV-229E_1a;3834 | 0 | 0.025 |
| NC_002645.1;NP_073550.1;HCoV-229E_1a;3835 | 0 | 0.025 |
| NC_002645.1;NP_073550.1;HCoV-229E_1a;3836 | 0 | 0 |
| NC_002645.1;NP_073550.1;HCoV-229E_1a;3837 | 0 | 0.025 |
| NC_002645.1;NP_073550.1;HCoV-229E_1a;3838 | 0 | 0.05 |
| NC_002645.1;NP_073550.1;HCoV-229E_1a;3839 | 0 | 0.025 |
| NC_002645.1;NP_073550.1;HCoV-229E_1a;3840 | 0 | 0.025 |
| NC_002645.1;NP_073550.1;HCoV-229E_1a;3841 | 0.05 | 0.025 |
| NC_002645.1;NP_073550.1;HCoV-229E_1a;3842 | 0.05 | 0.025 |
| NC_002645.1;NP_073550.1;HCoV-229E_1a;3843 | 0.05 | 0.05 |
| NC_002645.1;NP_073550.1;HCoV-229E_1a;3844 | 0.15 | 0.1 |
| NC_002645.1;NP_073550.1;HCoV-229E_1a;3845 | 0.1 | 0.05 |
| NC_002645.1;NP_073550.1;HCoV-229E_1a;3846 | 0 | 0.025 |
| NC_002645.1;NP_073550.1;HCoV-229E_1a;3847 | 0 | 0.025 |
| NC_002645.1;NP_073550.1;HCoV-229E_1a;3848 | 0 | 0.025 |
| NC_002645.1;NP_073550.1;HCoV-229E_1a;3849 | 0 | 0.025 |
| NC_002645.1;NP_073550.1;HCoV-229E_1a;3850 | 0 | 0 |
| NC_002645.1;NP_073550.1;HCoV-229E_1a;3851 | 0.05 | 0 |
| NC_002645.1;NP_073550.1;HCoV-229E_1a;3852 | 0 | 0 |
| NC_002645.1;NP_073550.1;HCoV-229E_1a;3853 | 0 | 0 |
| NC_002645.1;NP_073550.1;HCoV-229E_1a;3854 | 0 | 0 |
| NC_002645.1;NP_073550.1;HCoV-229E_1a;3855 | 0 | 0 |
| NC_002645.1;NP_073550.1;HCoV-229E_1a;3856 | 0 | 0 |
| NC_002645.1;NP_073550.1;HCoV-229E_1a;3857 | 0.05 | 0.025 |
| NC_002645.1;NP_073550.1;HCoV-229E_1a;3858 | 0 | 0 |
| NC_002645.1;NP_073550.1;HCoV-229E_1a;3859 | 0 | 0 |
| NC_002645.1;NP_073550.1;HCoV-229E_1a;3860 | 0 | 0 |

| | | |
|---|---|---|
| NC_002645.1;NP_073550.1;HCoV-229E_1a;3861 | 0 | 0 |
| NC_002645.1;NP_073550.1;HCoV-229E_1a;3862 | 0 | 0 |
| NC_002645.1;NP_073550.1;HCoV-229E_1a;3863 | 0 | 0 |
| NC_002645.1;NP_073550.1;HCoV-229E_1a;3864 | 0 | 0 |
| NC_002645.1;NP_073550.1;HCoV-229E_1a;3865 | 0 | 0 |
| NC_002645.1;NP_073550.1;HCoV-229E_1a;3866 | 0.05 | 0.025 |
| NC_002645.1;NP_073550.1;HCoV-229E_1a;3867 | 0.05 | 0 |
| NC_002645.1;NP_073550.1;HCoV-229E_1a;3868 | 0.05 | 0 |
| NC_002645.1;NP_073550.1;HCoV-229E_1a;3869 | 0.05 | 0 |
| NC_002645.1;NP_073550.1;HCoV-229E_1a;3870 | 0 | 0 |
| NC_002645.1;NP_073550.1;HCoV-229E_1a;3871 | 0 | 0 |
| NC_002645.1;NP_073550.1;HCoV-229E_1a;3872 | 0 | 0 |
| NC_002645.1;NP_073550.1;HCoV-229E_1a;3873 | 0 | 0 |
| NC_002645.1;NP_073550.1;HCoV-229E_1a;3874 | 0 | 0 |
| NC_002645.1;NP_073550.1;HCoV-229E_1a;3875 | 0 | 0 |
| NC_002645.1;NP_073550.1;HCoV-229E_1a;3876 | 0 | 0 |
| NC_002645.1;NP_073550.1;HCoV-229E_1a;3877 | 0 | 0 |
| NC_002645.1;NP_073550.1;HCoV-229E_1a;3878 | 0 | 0 |
| NC_002645.1;NP_073550.1;HCoV-229E_1a;3879 | 0 | 0 |
| NC_002645.1;NP_073550.1;HCoV-229E_1a;3880 | 0 | 0 |
| NC_002645.1;NP_073550.1;HCoV-229E_1a;3881 | 0 | 0 |
| NC_002645.1;NP_073550.1;HCoV-229E_1a;3882 | 0 | 0 |
| NC_002645.1;NP_073550.1;HCoV-229E_1a;3883 | 0 | 0 |
| NC_002645.1;NP_073550.1;HCoV-229E_1a;3884 | 0 | 0 |
| NC_002645.1;NP_073550.1;HCoV-229E_1a;3885 | 0 | 0.025 |
| NC_002645.1;NP_073550.1;HCoV-229E_1a;3886 | 0 | 0 |
| NC_002645.1;NP_073550.1;HCoV-229E_1a;3887 | 0.05 | 0 |
| NC_002645.1;NP_073550.1;HCoV-229E_1a;3888 | 0 | 0 |
| NC_002645.1;NP_073550.1;HCoV-229E_1a;3889 | 0 | 0.025 |
| NC_002645.1;NP_073550.1;HCoV-229E_1a;3890 | 0 | 0 |
| NC_002645.1;NP_073550.1;HCoV-229E_1a;3891 | 0 | 0 |
| NC_002645.1;NP_073550.1;HCoV-229E_1a;3892 | 0 | 0 |
| NC_002645.1;NP_073550.1;HCoV-229E_1a;3893 | 0 | 0 |
| NC_002645.1;NP_073550.1;HCoV-229E_1a;3894 | 0 | 0 |
| NC_002645.1;NP_073550.1;HCoV-229E_1a;3895 | 0.05 | 0 |
| NC_002645.1;NP_073550.1;HCoV-229E_1a;3896 | 0 | 0 |
| NC_002645.1;NP_073550.1;HCoV-229E_1a;3897 | 0 | 0 |
| NC_002645.1;NP_073550.1;HCoV-229E_1a;3898 | 0 | 0 |
| NC_002645.1;NP_073550.1;HCoV-229E_1a;3899 | 0 | 0 |
| NC_002645.1;NP_073550.1;HCoV-229E_1a;3900 | 0 | 0 |
| NC_002645.1;NP_073550.1;HCoV-229E_1a;3901 | 0 | 0 |
| NC_002645.1;NP_073550.1;HCoV-229E_1a;3902 | 0 | 0 |
| NC_002645.1;NP_073550.1;HCoV-229E_1a;3903 | 0 | 0 |

| | | |
|---|---|---|
| NC_002645.1;NP_073550.1;HCoV-229E_1a;3904 | 0 | 0 |
| NC_002645.1;NP_073550.1;HCoV-229E_1a;3905 | 0 | 0 |
| NC_002645.1;NP_073550.1;HCoV-229E_1a;3906 | 0 | 0 |
| NC_002645.1;NP_073550.1;HCoV-229E_1a;3907 | 0 | 0 |
| NC_002645.1;NP_073550.1;HCoV-229E_1a;3908 | 0 | 0 |
| NC_002645.1;NP_073550.1;HCoV-229E_1a;3909 | 0 | 0 |
| NC_002645.1;NP_073550.1;HCoV-229E_1a;3910 | 0 | 0 |
| NC_002645.1;NP_073550.1;HCoV-229E_1a;3911 | 0 | 0 |
| NC_002645.1;NP_073550.1;HCoV-229E_1a;3912 | 0 | 0 |
| NC_002645.1;NP_073550.1;HCoV-229E_1a;3913 | 0 | 0 |
| NC_002645.1;NP_073550.1;HCoV-229E_1a;3914 | 0 | 0 |
| NC_002645.1;NP_073550.1;HCoV-229E_1a;3915 | 0 | 0 |
| NC_002645.1;NP_073550.1;HCoV-229E_1a;3916 | 0 | 0 |
| NC_002645.1;NP_073550.1;HCoV-229E_1a;3917 | 0 | 0 |
| NC_002645.1;NP_073550.1;HCoV-229E_1a;3918 | 0 | 0 |
| NC_002645.1;NP_073550.1;HCoV-229E_1a;3919 | 0 | 0 |
| NC_002645.1;NP_073550.1;HCoV-229E_1a;3920 | 0 | 0 |
| NC_002645.1;NP_073550.1;HCoV-229E_1a;3921 | 0 | 0 |
| NC_002645.1;NP_073550.1;HCoV-229E_1a;3922 | 0 | 0 |
| NC_002645.1;NP_073550.1;HCoV-229E_1a;3923 | 0.05 | 0 |
| NC_002645.1;NP_073550.1;HCoV-229E_1a;3924 | 0.05 | 0.025 |
| NC_002645.1;NP_073550.1;HCoV-229E_1a;3925 | 0.05 | 0.025 |
| NC_002645.1;NP_073550.1;HCoV-229E_1a;3926 | 0.05 | 0.025 |
| NC_002645.1;NP_073550.1;HCoV-229E_1a;3927 | 0.05 | 0.05 |
| NC_002645.1;NP_073550.1;HCoV-229E_1a;3928 | 0.05 | 0.025 |
| NC_002645.1;NP_073550.1;HCoV-229E_1a;3929 | 0 | 0.025 |
| NC_002645.1;NP_073550.1;HCoV-229E_1a;3930 | 0 | 0.025 |
| NC_002645.1;NP_073550.1;HCoV-229E_1a;3931 | 0 | 0 |
| NC_002645.1;NP_073550.1;HCoV-229E_1a;3932 | 0 | 0 |
| NC_002645.1;NP_073550.1;HCoV-229E_1a;3933 | 0 | 0 |
| NC_002645.1;NP_073550.1;HCoV-229E_1a;3934 | 0 | 0 |
| NC_002645.1;NP_073550.1;HCoV-229E_1a;3935 | 0 | 0 |
| NC_002645.1;NP_073550.1;HCoV-229E_1a;3936 | 0 | 0 |
| NC_002645.1;NP_073550.1;HCoV-229E_1a;3937 | 0 | 0 |
| NC_002645.1;NP_073550.1;HCoV-229E_1a;3938 | 0 | 0 |
| NC_002645.1;NP_073550.1;HCoV-229E_1a;3939 | 0 | 0 |
| NC_002645.1;NP_073550.1;HCoV-229E_1a;3940 | 0 | 0 |
| NC_002645.1;NP_073550.1;HCoV-229E_1a;3941 | 0 | 0 |
| NC_002645.1;NP_073550.1;HCoV-229E_1a;3942 | 0 | 0 |
| NC_002645.1;NP_073550.1;HCoV-229E_1a;3943 | 0.05 | 0 |
| NC_002645.1;NP_073550.1;HCoV-229E_1a;3944 | 0.05 | 0 |
| NC_002645.1;NP_073550.1;HCoV-229E_1a;3945 | 0 | 0 |
| NC_002645.1;NP_073550.1;HCoV-229E_1a;3946 | 0 | 0 |

| | | |
|---|---|---|
| NC_002645.1;NP_073550.1;HCoV-229E_1a;3947 | 0.05 | 0 |
| NC_002645.1;NP_073550.1;HCoV-229E_1a;3948 | 0.05 | 0 |
| NC_002645.1;NP_073550.1;HCoV-229E_1a;3949 | 0.1 | 0 |
| NC_002645.1;NP_073550.1;HCoV-229E_1a;3950 | 0.15 | 0 |
| NC_002645.1;NP_073550.1;HCoV-229E_1a;3951 | 0.2 | 0.025 |
| NC_002645.1;NP_073550.1;HCoV-229E_1a;3952 | 0.2 | 0.025 |
| NC_002645.1;NP_073550.1;HCoV-229E_1a;3953 | 0.1 | 0.05 |
| NC_002645.1;NP_073550.1;HCoV-229E_1a;3954 | 0.05 | 0.025 |
| NC_002645.1;NP_073550.1;HCoV-229E_1a;3955 | 0.15 | 0.025 |
| NC_002645.1;NP_073550.1;HCoV-229E_1a;3956 | 0.05 | 0.025 |
| NC_002645.1;NP_073550.1;HCoV-229E_1a;3957 | 0.05 | 0 |
| NC_002645.1;NP_073550.1;HCoV-229E_1a;3958 | 0.05 | 0.025 |
| NC_002645.1;NP_073550.1;HCoV-229E_1a;3959 | 0 | 0.025 |
| NC_002645.1;NP_073550.1;HCoV-229E_1a;3960 | 0 | 0.075 |
| NC_002645.1;NP_073550.1;HCoV-229E_1a;3961 | 0 | 0.025 |
| NC_002645.1;NP_073550.1;HCoV-229E_1a;3962 | 0.05 | 0.025 |
| NC_002645.1;NP_073550.1;HCoV-229E_1a;3963 | 0.05 | 0.025 |
| NC_002645.1;NP_073550.1;HCoV-229E_1a;3964 | 0 | 0.025 |
| NC_002645.1;NP_073550.1;HCoV-229E_1a;3965 | 0 | 0.025 |
| NC_002645.1;NP_073550.1;HCoV-229E_1a;3966 | 0 | 0.025 |
| NC_002645.1;NP_073550.1;HCoV-229E_1a;3967 | 0 | 0 |
| NC_002645.1;NP_073550.1;HCoV-229E_1a;3968 | 0 | 0 |
| NC_002645.1;NP_073550.1;HCoV-229E_1a;3969 | 0 | 0 |
| NC_002645.1;NP_073550.1;HCoV-229E_1a;3970 | 0 | 0.025 |
| NC_002645.1;NP_073550.1;HCoV-229E_1a;3971 | 0.05 | 0.025 |
| NC_002645.1;NP_073550.1;HCoV-229E_1a;3972 | 0 | 0 |
| NC_002645.1;NP_073550.1;HCoV-229E_1a;3973 | 0 | 0.025 |
| NC_002645.1;NP_073550.1;HCoV-229E_1a;3974 | 0 | 0 |
| NC_002645.1;NP_073550.1;HCoV-229E_1a;3975 | 0 | 0 |
| NC_002645.1;NP_073550.1;HCoV-229E_1a;3976 | 0 | 0 |
| NC_002645.1;NP_073550.1;HCoV-229E_1a;3977 | 0 | 0 |
| NC_002645.1;NP_073550.1;HCoV-229E_1a;3978 | 0 | 0 |
| NC_002645.1;NP_073550.1;HCoV-229E_1a;3979 | 0 | 0.025 |
| NC_002645.1;NP_073550.1;HCoV-229E_1a;3980 | 0 | 0.025 |
| NC_002645.1;NP_073550.1;HCoV-229E_1a;3981 | 0 | 0 |
| NC_002645.1;NP_073550.1;HCoV-229E_1a;3982 | 0 | 0 |
| NC_002645.1;NP_073550.1;HCoV-229E_1a;3983 | 0 | 0 |
| NC_002645.1;NP_073550.1;HCoV-229E_1a;3984 | 0 | 0 |
| NC_002645.1;NP_073550.1;HCoV-229E_1a;3985 | 0 | 0 |
| NC_002645.1;NP_073550.1;HCoV-229E_1a;3986 | 0.05 | 0 |
| NC_002645.1;NP_073550.1;HCoV-229E_1a;3987 | 0 | 0 |
| NC_002645.1;NP_073550.1;HCoV-229E_1a;3988 | 0.1 | 0 |
| NC_002645.1;NP_073550.1;HCoV-229E_1a;3989 | 0 | 0 |

| | | |
|---|---|---|
| NC_002645.1;NP_073550.1;HCoV-229E_1a;3990 | 0.05 | 0.025 |
| NC_002645.1;NP_073550.1;HCoV-229E_1a;3991 | 0 | 0.075 |
| NC_002645.1;NP_073550.1;HCoV-229E_1a;3992 | 0 | 0.025 |
| NC_002645.1;NP_073550.1;HCoV-229E_1a;3993 | 0.05 | 0.025 |
| NC_002645.1;NP_073550.1;HCoV-229E_1a;3994 | 0 | 0 |
| NC_002645.1;NP_073550.1;HCoV-229E_1a;3995 | 0 | 0 |
| NC_002645.1;NP_073550.1;HCoV-229E_1a;3996 | 0 | 0 |
| NC_002645.1;NP_073550.1;HCoV-229E_1a;3997 | 0 | 0 |
| NC_002645.1;NP_073550.1;HCoV-229E_1a;3998 | 0 | 0 |
| NC_002645.1;NP_073550.1;HCoV-229E_1a;3999 | 0 | 0 |
| NC_002645.1;NP_073550.1;HCoV-229E_1a;4000 | 0 | 0 |
| NC_002645.1;NP_073550.1;HCoV-229E_1a;4001 | 0 | 0 |
| NC_002645.1;NP_073550.1;HCoV-229E_1a;4002 | 0 | 0 |
| NC_002645.1;NP_073550.1;HCoV-229E_1a;4003 | 0 | 0 |
| NC_002645.1;NP_073550.1;HCoV-229E_1a;4004 | 0 | 0 |
| NC_002645.1;NP_073550.1;HCoV-229E_1a;4005 | 0 | 0 |
| NC_002645.1;NP_073550.1;HCoV-229E_1a;4006 | 0 | 0 |
| NC_002645.1;NP_073550.1;HCoV-229E_1a;4007 | 0 | 0 |
| NC_002645.1;NP_073550.1;HCoV-229E_1a;4008 | 0 | 0 |
| NC_002645.1;NP_073550.1;HCoV-229E_1a;4009 | 0 | 0 |
| NC_002645.1;NP_073550.1;HCoV-229E_1a;4010 | 0 | 0 |
| NC_002645.1;NP_073550.1;HCoV-229E_1a;4011 | 0 | 0 |
| NC_002645.1;NP_073550.1;HCoV-229E_1a;4012 | 0 | 0 |
| NC_002645.1;NP_073550.1;HCoV-229E_1a;4013 | 0 | 0 |
| NC_002645.1;NP_073550.1;HCoV-229E_1a;4014 | 0 | 0 |
| NC_002645.1;NP_073550.1;HCoV-229E_1a;4015 | 0 | 0 |
| NC_002645.1;NP_073550.1;HCoV-229E_1a;4016 | 0 | 0 |
| NC_002645.1;NP_073550.1;HCoV-229E_1a;4017 | 0 | 0 |
| NC_002645.1;NP_073550.1;HCoV-229E_1a;4018 | 0 | 0 |
| NC_002645.1;NP_073550.1;HCoV-229E_1a;4019 | 0 | 0 |
| NC_002645.1;NP_073550.1;HCoV-229E_1a;4020 | 0 | 0 |
| NC_002645.1;NP_073550.1;HCoV-229E_1a;4021 | 0 | 0 |
| NC_002645.1;NP_073550.1;HCoV-229E_1a;4022 | 0 | 0.075 |
| NC_002645.1;NP_073550.1;HCoV-229E_1a;4023 | 0.05 | 0.075 |
| NC_002645.1;NP_073550.1;HCoV-229E_1a;4024 | 0.05 | 0.075 |
| NC_002645.1;NP_073550.1;HCoV-229E_1a;4025 | 0 | 0.1 |
| NC_002645.1;NP_073550.1;HCoV-229E_1a;4026 | 0 | 0.075 |
| NC_002645.1;NP_073550.1;HCoV-229E_1a;4027 | 0 | 0.025 |
| NC_002645.1;NP_073550.1;HCoV-229E_1a;4028 | 0 | 0.025 |
| NC_002645.1;NP_073550.1;HCoV-229E_1a;4029 | 0 | 0 |
| NC_002645.1;NP_073550.1;HCoV-229E_1a;4030 | 0 | 0 |
| NC_002645.1;NP_073550.1;HCoV-229E_1a;4031 | 0.05 | 0.025 |
| NC_002645.1;NP_073550.1;HCoV-229E_1a;4032 | 0.05 | 0.025 |

| | | |
|---|---|---|
| NC_002645.1;NP_073550.1;HCoV-229E_1a;4033 | 0 | 0.025 |
| NC_002645.1;NP_073550.1;HCoV-229E_1a;4034 | 0.05 | 0.125 |
| NC_002645.1;NP_073550.1;HCoV-229E_1a;4035 | 0 | 0 |
| NC_002645.1;NP_073550.1;HCoV-229E_1a;4036 | 0 | 0 |
| NC_002645.1;NP_073550.1;HCoV-229E_1a;4037 | 0 | 0 |
| NC_002645.1;NP_073550.1;HCoV-229E_1a;4038 | 0 | 0.025 |
| NC_002645.1;NP_073550.1;HCoV-229E_1a;4039 | 0 | 0 |
| NC_002645.1;NP_073550.1;HCoV-229E_1a;4040 | 0 | 0 |
| NC_002645.1;NP_073550.1;HCoV-229E_1a;4041 | 0 | 0 |
| NC_002645.1;NP_073550.1;HCoV-229E_1a;4042 | 0 | 0 |
| NC_002645.1;NP_073550.1;HCoV-229E_1a;4043 | 0 | 0 |
| NC_002645.1;NP_073550.1;HCoV-229E_1a;4044 | 0 | 0 |
| NC_002645.1;NP_073550.1;HCoV-229E_1a;4045 | 0 | 0 |
| NC_002645.1;NP_073550.1;HCoV-229E_1a;4046 | 0 | 0 |
| NC_002645.1;NP_073550.1;HCoV-229E_1a;4047 | 0 | 0 |
| NC_002645.1;NP_073550.1;HCoV-229E_1a;4048 | 0 | 0 |
| NC_002645.1;NP_073550.1;HCoV-229E_1a;4049 | 0 | 0 |
| NC_002645.1;NP_073550.1;HCoV-229E_1a;4050 | 0 | 0 |
| NC_002645.1;NP_073550.1;HCoV-229E_1a;4051 | 0 | 0 |
| NC_002645.1;NP_073550.1;HCoV-229E_1a;4052 | 0 | 0 |
| NC_002645.1;NP_073550.1;HCoV-229E_1a;4053 | 0 | 0 |
| NC_002645.1;NP_073550.1;HCoV-229E_1a;4054 | 0 | 0 |
| NC_002645.1;NP_073550.1;HCoV-229E_1a;4055 | 0 | 0 |
| NC_002645.1;NP_073550.1;HCoV-229E_1a;4056 | 0 | 0.025 |
| NC_002645.1;NP_073550.1;HCoV-229E_1a;4057 | 0 | 0 |
| NC_002645.1;NP_073550.1;HCoV-229E_1a;4058 | 0 | 0.025 |
| NC_002645.1;NP_073550.1;HCoV-229E_1a;4059 | 0 | 0.025 |
| NC_002645.1;NP_073550.1;HCoV-229E_1a;4060 | 0 | 0.025 |
| NC_002645.1;NP_073550.1;HCoV-229E_1a;4061 | 0.05 | 0.025 |
| NC_002645.1;NP_073550.1;HCoV-229E_1a;4062 | 0 | 0.025 |
| NC_002645.1;NP_073550.1;HCoV-229E_1a;4063 | 0 | 0.075 |
| NC_002645.1;NP_073550.1;HCoV-229E_1a;4064 | 0.05 | 0.05 |
| NC_002645.1;NP_073550.1;HCoV-229E_1a;4065 | 0.05 | 0.025 |
| NC_002645.1;NP_073550.1;HCoV-229E_1a;4066 | 0 | 0.05 |
| NC_002645.1;NP_073550.1;HCoV-229E_1a;4067 | 0 | 0.1 |
| NC_002645.1;NP_073550.1;HCoV-229E_1a;4068 | 0 | 0.075 |
| NC_002645.1;NP_073550.1;HCoV-229E_1a;4069 | 0 | 0.05 |
| NC_002645.1;NP_073550.1;HCoV-229E_1a;4070 | 0 | 0.075 |
| NC_002645.1;NP_073550.1;HCoV-229E_1a;4071 | 0 | 0.075 |
| NC_002645.1;NP_073550.1;HCoV-229E_1a;4072 | 0 | 0.1 |
| NC_002645.1;NP_073550.1;HCoV-229E_1a;4073 | 0 | 0.05 |
| NC_002645.1;NP_073550.1;HCoV-229E_1a;4074 | 0 | 0.1 |
| NC_002645.1;NP_073551.1;HCoV-229E_surface;1 | 0 | 0 |

| | | |
|---|---|---|
| NC_002645.1;NP_073551.1;HCoV-229E_surface;2 | 0 | 0 |
| NC_002645.1;NP_073551.1;HCoV-229E_surface;3 | 0 | 0 |
| NC_002645.1;NP_073551.1;HCoV-229E_surface;4 | 0 | 0 |
| NC_002645.1;NP_073551.1;HCoV-229E_surface;5 | 0 | 0 |
| NC_002645.1;NP_073551.1;HCoV-229E_surface;6 | 0 | 0 |
| NC_002645.1;NP_073551.1;HCoV-229E_surface;7 | 0 | 0 |
| NC_002645.1;NP_073551.1;HCoV-229E_surface;8 | 0 | 0.025 |
| NC_002645.1;NP_073551.1;HCoV-229E_surface;9 | 0.05 | 0.025 |
| NC_002645.1;NP_073551.1;HCoV-229E_surface;10 | 0 | 0.025 |
| NC_002645.1;NP_073551.1;HCoV-229E_surface;11 | 0 | 0.025 |
| NC_002645.1;NP_073551.1;HCoV-229E_surface;12 | 0 | 0.025 |
| NC_002645.1;NP_073551.1;HCoV-229E_surface;13 | 0 | 0.025 |
| NC_002645.1;NP_073551.1;HCoV-229E_surface;14 | 0.05 | 0 |
| NC_002645.1;NP_073551.1;HCoV-229E_surface;15 | 0.05 | 0.025 |
| NC_002645.1;NP_073551.1;HCoV-229E_surface;16 | 0.05 | 0.025 |
| NC_002645.1;NP_073551.1;HCoV-229E_surface;17 | 0.05 | 0 |
| NC_002645.1;NP_073551.1;HCoV-229E_surface;18 | 0 | 0 |
| NC_002645.1;NP_073551.1;HCoV-229E_surface;19 | 0 | 0 |
| NC_002645.1;NP_073551.1;HCoV-229E_surface;20 | 0 | 0 |
| NC_002645.1;NP_073551.1;HCoV-229E_surface;21 | 0 | 0 |
| NC_002645.1;NP_073551.1;HCoV-229E_surface;22 | 0 | 0 |
| NC_002645.1;NP_073551.1;HCoV-229E_surface;23 | 0 | 0 |
| NC_002645.1;NP_073551.1;HCoV-229E_surface;24 | 0 | 0.025 |
| NC_002645.1;NP_073551.1;HCoV-229E_surface;25 | 0 | 0 |
| NC_002645.1;NP_073551.1;HCoV-229E_surface;26 | 0 | 0 |
| NC_002645.1;NP_073551.1;HCoV-229E_surface;27 | 0 | 0 |
| NC_002645.1;NP_073551.1;HCoV-229E_surface;28 | 0 | 0 |
| NC_002645.1;NP_073551.1;HCoV-229E_surface;29 | 0 | 0 |
| NC_002645.1;NP_073551.1;HCoV-229E_surface;30 | 0 | 0 |
| NC_002645.1;NP_073551.1;HCoV-229E_surface;31 | 0.05 | 0 |
| NC_002645.1;NP_073551.1;HCoV-229E_surface;32 | 0 | 0 |
| NC_002645.1;NP_073551.1;HCoV-229E_surface;33 | 0 | 0 |
| NC_002645.1;NP_073551.1;HCoV-229E_surface;34 | 0 | 0 |
| NC_002645.1;NP_073551.1;HCoV-229E_surface;35 | 0 | 0.025 |
| NC_002645.1;NP_073551.1;HCoV-229E_surface;36 | 0 | 0 |
| NC_002645.1;NP_073551.1;HCoV-229E_surface;37 | 0.05 | 0.025 |
| NC_002645.1;NP_073551.1;HCoV-229E_surface;38 | 0.05 | 0.05 |
| NC_002645.1;NP_073551.1;HCoV-229E_surface;39 | 0 | 0.025 |
| NC_002645.1;NP_073551.1;HCoV-229E_surface;40 | 0.05 | 0.075 |
| NC_002645.1;NP_073551.1;HCoV-229E_surface;41 | 0.05 | 0.075 |
| NC_002645.1;NP_073551.1;HCoV-229E_surface;42 | 0.05 | 0.075 |
| NC_002645.1;NP_073551.1;HCoV-229E_surface;43 | 0.05 | 0.05 |
| NC_002645.1;NP_073551.1;HCoV-229E_surface;44 | 0 | 0.025 |

| | | |
|---|---|---|
| NC_002645.1;NP_073551.1;HCoV-229E_surface;45 | 0 | 0 |
| NC_002645.1;NP_073551.1;HCoV-229E_surface;46 | 0 | 0 |
| NC_002645.1;NP_073551.1;HCoV-229E_surface;47 | 0 | 0 |
| NC_002645.1;NP_073551.1;HCoV-229E_surface;48 | 0 | 0 |
| NC_002645.1;NP_073551.1;HCoV-229E_surface;49 | 0 | 0.025 |
| NC_002645.1;NP_073551.1;HCoV-229E_surface;50 | 0 | 0.025 |
| NC_002645.1;NP_073551.1;HCoV-229E_surface;51 | 0 | 0.025 |
| NC_002645.1;NP_073551.1;HCoV-229E_surface;52 | 0 | 0 |
| NC_002645.1;NP_073551.1;HCoV-229E_surface;53 | 0 | 0.025 |
| NC_002645.1;NP_073551.1;HCoV-229E_surface;54 | 0.05 | 0.05 |
| NC_002645.1;NP_073551.1;HCoV-229E_surface;55 | 0 | 0.025 |
| NC_002645.1;NP_073551.1;HCoV-229E_surface;56 | 0 | 0.025 |
| NC_002645.1;NP_073551.1;HCoV-229E_surface;57 | 0 | 0.025 |
| NC_002645.1;NP_073551.1;HCoV-229E_surface;58 | 0.05 | 0.025 |
| NC_002645.1;NP_073551.1;HCoV-229E_surface;59 | 0.05 | 0.05 |
| NC_002645.1;NP_073551.1;HCoV-229E_surface;60 | 0.1 | 0.05 |
| NC_002645.1;NP_073551.1;HCoV-229E_surface;61 | 0.1 | 0.05 |
| NC_002645.1;NP_073551.1;HCoV-229E_surface;62 | 0 | 0 |
| NC_002645.1;NP_073551.1;HCoV-229E_surface;63 | 0 | 0 |
| NC_002645.1;NP_073551.1;HCoV-229E_surface;64 | 0 | 0 |
| NC_002645.1;NP_073551.1;HCoV-229E_surface;65 | 0 | 0 |
| NC_002645.1;NP_073551.1;HCoV-229E_surface;66 | 0 | 0 |
| NC_002645.1;NP_073551.1;HCoV-229E_surface;67 | 0 | 0 |
| NC_002645.1;NP_073551.1;HCoV-229E_surface;68 | 0 | 0 |
| NC_002645.1;NP_073551.1;HCoV-229E_surface;69 | 0 | 0 |
| NC_002645.1;NP_073551.1;HCoV-229E_surface;70 | 0 | 0 |
| NC_002645.1;NP_073551.1;HCoV-229E_surface;71 | 0 | 0 |
| NC_002645.1;NP_073551.1;HCoV-229E_surface;72 | 0 | 0 |
| NC_002645.1;NP_073551.1;HCoV-229E_surface;73 | 0 | 0 |
| NC_002645.1;NP_073551.1;HCoV-229E_surface;74 | 0 | 0 |
| NC_002645.1;NP_073551.1;HCoV-229E_surface;75 | 0 | 0 |
| NC_002645.1;NP_073551.1;HCoV-229E_surface;76 | 0 | 0 |
| NC_002645.1;NP_073551.1;HCoV-229E_surface;77 | 0 | 0 |
| NC_002645.1;NP_073551.1;HCoV-229E_surface;78 | 0 | 0 |
| NC_002645.1;NP_073551.1;HCoV-229E_surface;79 | 0 | 0 |
| NC_002645.1;NP_073551.1;HCoV-229E_surface;80 | 0 | 0 |
| NC_002645.1;NP_073551.1;HCoV-229E_surface;81 | 0 | 0 |
| NC_002645.1;NP_073551.1;HCoV-229E_surface;82 | 0 | 0 |
| NC_002645.1;NP_073551.1;HCoV-229E_surface;83 | 0 | 0 |
| NC_002645.1;NP_073551.1;HCoV-229E_surface;84 | 0 | 0 |
| NC_002645.1;NP_073551.1;HCoV-229E_surface;85 | 0 | 0 |
| NC_002645.1;NP_073551.1;HCoV-229E_surface;86 | 0 | 0 |
| NC_002645.1;NP_073551.1;HCoV-229E_surface;87 | 0 | 0 |

| | | |
|---|---|---|
| NC_002645.1;NP_073551.1;HCoV-229E_surface;88 | 0 | 0 |
| NC_002645.1;NP_073551.1;HCoV-229E_surface;89 | 0 | 0 |
| NC_002645.1;NP_073551.1;HCoV-229E_surface;90 | 0 | 0 |
| NC_002645.1;NP_073551.1;HCoV-229E_surface;91 | 0 | 0 |
| NC_002645.1;NP_073551.1;HCoV-229E_surface;92 | 0 | 0 |
| NC_002645.1;NP_073551.1;HCoV-229E_surface;93 | 0 | 0 |
| NC_002645.1;NP_073551.1;HCoV-229E_surface;94 | 0 | 0 |
| NC_002645.1;NP_073551.1;HCoV-229E_surface;95 | 0 | 0 |
| NC_002645.1;NP_073551.1;HCoV-229E_surface;96 | 0 | 0.025 |
| NC_002645.1;NP_073551.1;HCoV-229E_surface;97 | 0 | 0.025 |
| NC_002645.1;NP_073551.1;HCoV-229E_surface;98 | 0 | 0.025 |
| NC_002645.1;NP_073551.1;HCoV-229E_surface;99 | 0 | 0.025 |
| NC_002645.1;NP_073551.1;HCoV-229E_surface;100 | 0 | 0.025 |
| NC_002645.1;NP_073551.1;HCoV-229E_surface;101 | 0 | 0.025 |
| NC_002645.1;NP_073551.1;HCoV-229E_surface;102 | 0 | 0 |
| NC_002645.1;NP_073551.1;HCoV-229E_surface;103 | 0 | 0 |
| NC_002645.1;NP_073551.1;HCoV-229E_surface;104 | 0 | 0 |
| NC_002645.1;NP_073551.1;HCoV-229E_surface;105 | 0 | 0 |
| NC_002645.1;NP_073551.1;HCoV-229E_surface;106 | 0 | 0 |
| NC_002645.1;NP_073551.1;HCoV-229E_surface;107 | 0 | 0 |
| NC_002645.1;NP_073551.1;HCoV-229E_surface;108 | 0 | 0 |
| NC_002645.1;NP_073551.1;HCoV-229E_surface;109 | 0 | 0 |
| NC_002645.1;NP_073551.1;HCoV-229E_surface;110 | 0 | 0.025 |
| NC_002645.1;NP_073551.1;HCoV-229E_surface;111 | 0 | 0.025 |
| NC_002645.1;NP_073551.1;HCoV-229E_surface;112 | 0.05 | 0.05 |
| NC_002645.1;NP_073551.1;HCoV-229E_surface;113 | 0 | 0.075 |
| NC_002645.1;NP_073551.1;HCoV-229E_surface;114 | 0 | 0.025 |
| NC_002645.1;NP_073551.1;HCoV-229E_surface;115 | 0 | 0.025 |
| NC_002645.1;NP_073551.1;HCoV-229E_surface;116 | 0 | 0.025 |
| NC_002645.1;NP_073551.1;HCoV-229E_surface;117 | 0 | 0 |
| NC_002645.1;NP_073551.1;HCoV-229E_surface;118 | 0 | 0 |
| NC_002645.1;NP_073551.1;HCoV-229E_surface;119 | 0 | 0 |
| NC_002645.1;NP_073551.1;HCoV-229E_surface;120 | 0 | 0 |
| NC_002645.1;NP_073551.1;HCoV-229E_surface;121 | 0 | 0 |
| NC_002645.1;NP_073551.1;HCoV-229E_surface;122 | 0 | 0 |
| NC_002645.1;NP_073551.1;HCoV-229E_surface;123 | 0 | 0 |
| NC_002645.1;NP_073551.1;HCoV-229E_surface;124 | 0 | 0 |
| NC_002645.1;NP_073551.1;HCoV-229E_surface;125 | 0 | 0 |
| NC_002645.1;NP_073551.1;HCoV-229E_surface;126 | 0 | 0 |
| NC_002645.1;NP_073551.1;HCoV-229E_surface;127 | 0 | 0 |
| NC_002645.1;NP_073551.1;HCoV-229E_surface;128 | 0 | 0 |
| NC_002645.1;NP_073551.1;HCoV-229E_surface;129 | 0 | 0 |
| NC_002645.1;NP_073551.1;HCoV-229E_surface;130 | 0 | 0 |

| | | |
|---|---|---|
| NC_002645.1;NP_073551.1;HCoV-229E_surface;131 | 0 | 0 |
| NC_002645.1;NP_073551.1;HCoV-229E_surface;132 | 0 | 0 |
| NC_002645.1;NP_073551.1;HCoV-229E_surface;133 | 0 | 0 |
| NC_002645.1;NP_073551.1;HCoV-229E_surface;134 | 0 | 0 |
| NC_002645.1;NP_073551.1;HCoV-229E_surface;135 | 0 | 0 |
| NC_002645.1;NP_073551.1;HCoV-229E_surface;136 | 0 | 0 |
| NC_002645.1;NP_073551.1;HCoV-229E_surface;137 | 0 | 0 |
| NC_002645.1;NP_073551.1;HCoV-229E_surface;138 | 0 | 0 |
| NC_002645.1;NP_073551.1;HCoV-229E_surface;139 | 0 | 0 |
| NC_002645.1;NP_073551.1;HCoV-229E_surface;140 | 0 | 0 |
| NC_002645.1;NP_073551.1;HCoV-229E_surface;141 | 0 | 0 |
| NC_002645.1;NP_073551.1;HCoV-229E_surface;142 | 0 | 0.025 |
| NC_002645.1;NP_073551.1;HCoV-229E_surface;143 | 0.05 | 0.025 |
| NC_002645.1;NP_073551.1;HCoV-229E_surface;144 | 0.1 | 0.05 |
| NC_002645.1;NP_073551.1;HCoV-229E_surface;145 | 0.1 | 0.05 |
| NC_002645.1;NP_073551.1;HCoV-229E_surface;146 | 0.15 | 0.1 |
| NC_002645.1;NP_073551.1;HCoV-229E_surface;147 | 0.15 | 0.125 |
| NC_002645.1;NP_073551.1;HCoV-229E_surface;148 | 0.1 | 0.1 |
| NC_002645.1;NP_073551.1;HCoV-229E_surface;149 | 0.1 | 0.1 |
| NC_002645.1;NP_073551.1;HCoV-229E_surface;150 | 0.15 | 0.1 |
| NC_002645.1;NP_073551.1;HCoV-229E_surface;151 | 0.1 | 0.1 |
| NC_002645.1;NP_073551.1;HCoV-229E_surface;152 | 0.05 | 0.075 |
| NC_002645.1;NP_073551.1;HCoV-229E_surface;153 | 0 | 0 |
| NC_002645.1;NP_073551.1;HCoV-229E_surface;154 | 0 | 0 |
| NC_002645.1;NP_073551.1;HCoV-229E_surface;155 | 0 | 0 |
| NC_002645.1;NP_073551.1;HCoV-229E_surface;156 | 0 | 0 |
| NC_002645.1;NP_073551.1;HCoV-229E_surface;157 | 0 | 0 |
| NC_002645.1;NP_073551.1;HCoV-229E_surface;158 | 0 | 0 |
| NC_002645.1;NP_073551.1;HCoV-229E_surface;159 | 0 | 0 |
| NC_002645.1;NP_073551.1;HCoV-229E_surface;160 | 0 | 0 |
| NC_002645.1;NP_073551.1;HCoV-229E_surface;161 | 0 | 0 |
| NC_002645.1;NP_073551.1;HCoV-229E_surface;162 | 0 | 0 |
| NC_002645.1;NP_073551.1;HCoV-229E_surface;163 | 0 | 0 |
| NC_002645.1;NP_073551.1;HCoV-229E_surface;164 | 0 | 0.025 |
| NC_002645.1;NP_073551.1;HCoV-229E_surface;165 | 0 | 0.025 |
| NC_002645.1;NP_073551.1;HCoV-229E_surface;166 | 0 | 0.025 |
| NC_002645.1;NP_073551.1;HCoV-229E_surface;167 | 0 | 0.025 |
| NC_002645.1;NP_073551.1;HCoV-229E_surface;168 | 0 | 0.025 |
| NC_002645.1;NP_073551.1;HCoV-229E_surface;169 | 0.05 | 0.05 |
| NC_002645.1;NP_073551.1;HCoV-229E_surface;170 | 0 | 0.05 |
| NC_002645.1;NP_073551.1;HCoV-229E_surface;171 | 0 | 0 |
| NC_002645.1;NP_073551.1;HCoV-229E_surface;172 | 0 | 0.025 |
| NC_002645.1;NP_073551.1;HCoV-229E_surface;173 | 0 | 0.025 |

| | | |
|---|---|---|
| NC_002645.1;NP_073551.1;HCoV-229E_surface;174 | 0.05 | 0 |
| NC_002645.1;NP_073551.1;HCoV-229E_surface;175 | 0.05 | 0 |
| NC_002645.1;NP_073551.1;HCoV-229E_surface;176 | 0 | 0 |
| NC_002645.1;NP_073551.1;HCoV-229E_surface;177 | 0 | 0.075 |
| NC_002645.1;NP_073551.1;HCoV-229E_surface;178 | 0 | 0 |
| NC_002645.1;NP_073551.1;HCoV-229E_surface;179 | 0 | 0 |
| NC_002645.1;NP_073551.1;HCoV-229E_surface;180 | 0 | 0 |
| NC_002645.1;NP_073551.1;HCoV-229E_surface;181 | 0 | 0 |
| NC_002645.1;NP_073551.1;HCoV-229E_surface;182 | 0 | 0 |
| NC_002645.1;NP_073551.1;HCoV-229E_surface;183 | 0 | 0 |
| NC_002645.1;NP_073551.1;HCoV-229E_surface;184 | 0 | 0 |
| NC_002645.1;NP_073551.1;HCoV-229E_surface;185 | 0 | 0 |
| NC_002645.1;NP_073551.1;HCoV-229E_surface;186 | 0 | 0 |
| NC_002645.1;NP_073551.1;HCoV-229E_surface;187 | 0 | 0 |
| NC_002645.1;NP_073551.1;HCoV-229E_surface;188 | 0 | 0 |
| NC_002645.1;NP_073551.1;HCoV-229E_surface;189 | 0 | 0 |
| NC_002645.1;NP_073551.1;HCoV-229E_surface;190 | 0 | 0 |
| NC_002645.1;NP_073551.1;HCoV-229E_surface;191 | 0 | 0 |
| NC_002645.1;NP_073551.1;HCoV-229E_surface;192 | 0 | 0 |
| NC_002645.1;NP_073551.1;HCoV-229E_surface;193 | 0 | 0 |
| NC_002645.1;NP_073551.1;HCoV-229E_surface;194 | 0 | 0 |
| NC_002645.1;NP_073551.1;HCoV-229E_surface;195 | 0 | 0 |
| NC_002645.1;NP_073551.1;HCoV-229E_surface;196 | 0 | 0 |
| NC_002645.1;NP_073551.1;HCoV-229E_surface;197 | 0 | 0 |
| NC_002645.1;NP_073551.1;HCoV-229E_surface;198 | 0 | 0 |
| NC_002645.1;NP_073551.1;HCoV-229E_surface;199 | 0 | 0 |
| NC_002645.1;NP_073551.1;HCoV-229E_surface;200 | 0 | 0 |
| NC_002645.1;NP_073551.1;HCoV-229E_surface;201 | 0 | 0 |
| NC_002645.1;NP_073551.1;HCoV-229E_surface;202 | 0 | 0 |
| NC_002645.1;NP_073551.1;HCoV-229E_surface;203 | 0 | 0 |
| NC_002645.1;NP_073551.1;HCoV-229E_surface;204 | 0 | 0 |
| NC_002645.1;NP_073551.1;HCoV-229E_surface;205 | 0 | 0 |
| NC_002645.1;NP_073551.1;HCoV-229E_surface;206 | 0 | 0 |
| NC_002645.1;NP_073551.1;HCoV-229E_surface;207 | 0.05 | 0 |
| NC_002645.1;NP_073551.1;HCoV-229E_surface;208 | 0 | 0 |
| NC_002645.1;NP_073551.1;HCoV-229E_surface;209 | 0 | 0 |
| NC_002645.1;NP_073551.1;HCoV-229E_surface;210 | 0 | 0.05 |
| NC_002645.1;NP_073551.1;HCoV-229E_surface;211 | 0.05 | 0.025 |
| NC_002645.1;NP_073551.1;HCoV-229E_surface;212 | 0 | 0.05 |
| NC_002645.1;NP_073551.1;HCoV-229E_surface;213 | 0 | 0.125 |
| NC_002645.1;NP_073551.1;HCoV-229E_surface;214 | 0 | 0.15 |
| NC_002645.1;NP_073551.1;HCoV-229E_surface;215 | 0 | 0 |
| NC_002645.1;NP_073551.1;HCoV-229E_surface;216 | 0 | 0.025 |

| | | |
|---|---|---|
| NC_002645.1;NP_073551.1;HCoV-229E_surface;217 | 0 | 0 |
| NC_002645.1;NP_073551.1;HCoV-229E_surface;218 | 0 | 0 |
| NC_002645.1;NP_073551.1;HCoV-229E_surface;219 | 0 | 0 |
| NC_002645.1;NP_073551.1;HCoV-229E_surface;220 | 0 | 0 |
| NC_002645.1;NP_073551.1;HCoV-229E_surface;221 | 0 | 0 |
| NC_002645.1;NP_073551.1;HCoV-229E_surface;222 | 0 | 0 |
| NC_002645.1;NP_073551.1;HCoV-229E_surface;223 | 0 | 0 |
| NC_002645.1;NP_073551.1;HCoV-229E_surface;224 | 0 | 0 |
| NC_002645.1;NP_073551.1;HCoV-229E_surface;225 | 0 | 0.025 |
| NC_002645.1;NP_073551.1;HCoV-229E_surface;226 | 0 | 0 |
| NC_002645.1;NP_073551.1;HCoV-229E_surface;227 | 0 | 0 |
| NC_002645.1;NP_073551.1;HCoV-229E_surface;228 | 0 | 0 |
| NC_002645.1;NP_073551.1;HCoV-229E_surface;229 | 0 | 0 |
| NC_002645.1;NP_073551.1;HCoV-229E_surface;230 | 0 | 0 |
| NC_002645.1;NP_073551.1;HCoV-229E_surface;231 | 0 | 0 |
| NC_002645.1;NP_073551.1;HCoV-229E_surface;232 | 0 | 0 |
| NC_002645.1;NP_073551.1;HCoV-229E_surface;233 | 0 | 0 |
| NC_002645.1;NP_073551.1;HCoV-229E_surface;234 | 0 | 0 |
| NC_002645.1;NP_073551.1;HCoV-229E_surface;235 | 0 | 0 |
| NC_002645.1;NP_073551.1;HCoV-229E_surface;236 | 0 | 0 |
| NC_002645.1;NP_073551.1;HCoV-229E_surface;237 | 0 | 0 |
| NC_002645.1;NP_073551.1;HCoV-229E_surface;238 | 0 | 0 |
| NC_002645.1;NP_073551.1;HCoV-229E_surface;239 | 0 | 0 |
| NC_002645.1;NP_073551.1;HCoV-229E_surface;240 | 0 | 0 |
| NC_002645.1;NP_073551.1;HCoV-229E_surface;241 | 0 | 0 |
| NC_002645.1;NP_073551.1;HCoV-229E_surface;242 | 0 | 0 |
| NC_002645.1;NP_073551.1;HCoV-229E_surface;243 | 0 | 0 |
| NC_002645.1;NP_073551.1;HCoV-229E_surface;244 | 0 | 0 |
| NC_002645.1;NP_073551.1;HCoV-229E_surface;245 | 0 | 0 |
| NC_002645.1;NP_073551.1;HCoV-229E_surface;246 | 0 | 0 |
| NC_002645.1;NP_073551.1;HCoV-229E_surface;247 | 0 | 0 |
| NC_002645.1;NP_073551.1;HCoV-229E_surface;248 | 0 | 0 |
| NC_002645.1;NP_073551.1;HCoV-229E_surface;249 | 0 | 0 |
| NC_002645.1;NP_073551.1;HCoV-229E_surface;250 | 0 | 0.025 |
| NC_002645.1;NP_073551.1;HCoV-229E_surface;251 | 0 | 0 |
| NC_002645.1;NP_073551.1;HCoV-229E_surface;252 | 0 | 0 |
| NC_002645.1;NP_073551.1;HCoV-229E_surface;253 | 0 | 0.025 |
| NC_002645.1;NP_073551.1;HCoV-229E_surface;254 | 0 | 0 |
| NC_002645.1;NP_073551.1;HCoV-229E_surface;255 | 0 | 0.025 |
| NC_002645.1;NP_073551.1;HCoV-229E_surface;256 | 0 | 0.025 |
| NC_002645.1;NP_073551.1;HCoV-229E_surface;257 | 0 | 0.025 |
| NC_002645.1;NP_073551.1;HCoV-229E_surface;258 | 0.05 | 0 |
| NC_002645.1;NP_073551.1;HCoV-229E_surface;259 | 0.05 | 0 |

| | | |
|---|---|---|
| NC_002645.1;NP_073551.1;HCoV-229E_surface;260 | 0.05 | 0 |
| NC_002645.1;NP_073551.1;HCoV-229E_surface;261 | 0.05 | 0 |
| NC_002645.1;NP_073551.1;HCoV-229E_surface;262 | 0.05 | 0 |
| NC_002645.1;NP_073551.1;HCoV-229E_surface;263 | 0.05 | 0 |
| NC_002645.1;NP_073551.1;HCoV-229E_surface;264 | 0.05 | 0.025 |
| NC_002645.1;NP_073551.1;HCoV-229E_surface;265 | 0 | 0.025 |
| NC_002645.1;NP_073551.1;HCoV-229E_surface;266 | 0 | 0.05 |
| NC_002645.1;NP_073551.1;HCoV-229E_surface;267 | 0 | 0.05 |
| NC_002645.1;NP_073551.1;HCoV-229E_surface;268 | 0 | 0.05 |
| NC_002645.1;NP_073551.1;HCoV-229E_surface;269 | 0 | 0.075 |
| NC_002645.1;NP_073551.1;HCoV-229E_surface;270 | 0 | 0.05 |
| NC_002645.1;NP_073551.1;HCoV-229E_surface;271 | 0 | 0.025 |
| NC_002645.1;NP_073551.1;HCoV-229E_surface;272 | 0 | 0 |
| NC_002645.1;NP_073551.1;HCoV-229E_surface;273 | 0 | 0 |
| NC_002645.1;NP_073551.1;HCoV-229E_surface;274 | 0 | 0 |
| NC_002645.1;NP_073551.1;HCoV-229E_surface;275 | 0 | 0 |
| NC_002645.1;NP_073551.1;HCoV-229E_surface;276 | 0 | 0 |
| NC_002645.1;NP_073551.1;HCoV-229E_surface;277 | 0 | 0 |
| NC_002645.1;NP_073551.1;HCoV-229E_surface;278 | 0 | 0 |
| NC_002645.1;NP_073551.1;HCoV-229E_surface;279 | 0 | 0.025 |
| NC_002645.1;NP_073551.1;HCoV-229E_surface;280 | 0 | 0 |
| NC_002645.1;NP_073551.1;HCoV-229E_surface;281 | 0 | 0 |
| NC_002645.1;NP_073551.1;HCoV-229E_surface;282 | 0 | 0 |
| NC_002645.1;NP_073551.1;HCoV-229E_surface;283 | 0 | 0 |
| NC_002645.1;NP_073551.1;HCoV-229E_surface;284 | 0 | 0 |
| NC_002645.1;NP_073551.1;HCoV-229E_surface;285 | 0 | 0 |
| NC_002645.1;NP_073551.1;HCoV-229E_surface;286 | 0 | 0 |
| NC_002645.1;NP_073551.1;HCoV-229E_surface;287 | 0 | 0 |
| NC_002645.1;NP_073551.1;HCoV-229E_surface;288 | 0 | 0 |
| NC_002645.1;NP_073551.1;HCoV-229E_surface;289 | 0 | 0 |
| NC_002645.1;NP_073551.1;HCoV-229E_surface;290 | 0 | 0 |
| NC_002645.1;NP_073551.1;HCoV-229E_surface;291 | 0 | 0 |
| NC_002645.1;NP_073551.1;HCoV-229E_surface;292 | 0 | 0 |
| NC_002645.1;NP_073551.1;HCoV-229E_surface;293 | 0 | 0 |
| NC_002645.1;NP_073551.1;HCoV-229E_surface;294 | 0 | 0 |
| NC_002645.1;NP_073551.1;HCoV-229E_surface;295 | 0 | 0 |
| NC_002645.1;NP_073551.1;HCoV-229E_surface;296 | 0 | 0 |
| NC_002645.1;NP_073551.1;HCoV-229E_surface;297 | 0 | 0 |
| NC_002645.1;NP_073551.1;HCoV-229E_surface;298 | 0 | 0 |
| NC_002645.1;NP_073551.1;HCoV-229E_surface;299 | 0 | 0 |
| NC_002645.1;NP_073551.1;HCoV-229E_surface;300 | 0.05 | 0 |
| NC_002645.1;NP_073551.1;HCoV-229E_surface;301 | 0 | 0 |
| NC_002645.1;NP_073551.1;HCoV-229E_surface;302 | 0 | 0 |

| | | |
|---|---|---|
| NC_002645.1;NP_073551.1;HCoV-229E_surface;303 | 0 | 0 |
| NC_002645.1;NP_073551.1;HCoV-229E_surface;304 | 0.05 | 0.05 |
| NC_002645.1;NP_073551.1;HCoV-229E_surface;305 | 0.05 | 0.025 |
| NC_002645.1;NP_073551.1;HCoV-229E_surface;306 | 0 | 0.025 |
| NC_002645.1;NP_073551.1;HCoV-229E_surface;307 | 0 | 0.025 |
| NC_002645.1;NP_073551.1;HCoV-229E_surface;308 | 0 | 0.025 |
| NC_002645.1;NP_073551.1;HCoV-229E_surface;309 | 0 | 0.025 |
| NC_002645.1;NP_073551.1;HCoV-229E_surface;310 | 0 | 0.025 |
| NC_002645.1;NP_073551.1;HCoV-229E_surface;311 | 0 | 0.05 |
| NC_002645.1;NP_073551.1;HCoV-229E_surface;312 | 0 | 0.05 |
| NC_002645.1;NP_073551.1;HCoV-229E_surface;313 | 0.1 | 0.05 |
| NC_002645.1;NP_073551.1;HCoV-229E_surface;314 | 0 | 0.025 |
| NC_002645.1;NP_073551.1;HCoV-229E_surface;315 | 0 | 0 |
| NC_002645.1;NP_073551.1;HCoV-229E_surface;316 | 0 | 0.025 |
| NC_002645.1;NP_073551.1;HCoV-229E_surface;317 | 0 | 0 |
| NC_002645.1;NP_073551.1;HCoV-229E_surface;318 | 0 | 0 |
| NC_002645.1;NP_073551.1;HCoV-229E_surface;319 | 0 | 0 |
| NC_002645.1;NP_073551.1;HCoV-229E_surface;320 | 0 | 0.05 |
| NC_002645.1;NP_073551.1;HCoV-229E_surface;321 | 0.1 | 0.025 |
| NC_002645.1;NP_073551.1;HCoV-229E_surface;322 | 0.1 | 0.05 |
| NC_002645.1;NP_073551.1;HCoV-229E_surface;323 | 0.15 | 0.075 |
| NC_002645.1;NP_073551.1;HCoV-229E_surface;324 | 0.1 | 0.05 |
| NC_002645.1;NP_073551.1;HCoV-229E_surface;325 | 0.1 | 0.05 |
| NC_002645.1;NP_073551.1;HCoV-229E_surface;326 | 0.05 | 0 |
| NC_002645.1;NP_073551.1;HCoV-229E_surface;327 | 0.05 | 0.025 |
| NC_002645.1;NP_073551.1;HCoV-229E_surface;328 | 0.05 | 0.025 |
| NC_002645.1;NP_073551.1;HCoV-229E_surface;329 | 0.1 | 0.025 |
| NC_002645.1;NP_073551.1;HCoV-229E_surface;330 | 0.05 | 0.05 |
| NC_002645.1;NP_073551.1;HCoV-229E_surface;331 | 0.05 | 0.025 |
| NC_002645.1;NP_073551.1;HCoV-229E_surface;332 | 0.05 | 0.025 |
| NC_002645.1;NP_073551.1;HCoV-229E_surface;333 | 0.05 | 0 |
| NC_002645.1;NP_073551.1;HCoV-229E_surface;334 | 0 | 0.025 |
| NC_002645.1;NP_073551.1;HCoV-229E_surface;335 | 0 | 0 |
| NC_002645.1;NP_073551.1;HCoV-229E_surface;336 | 0 | 0.025 |
| NC_002645.1;NP_073551.1;HCoV-229E_surface;337 | 0 | 0 |
| NC_002645.1;NP_073551.1;HCoV-229E_surface;338 | 0 | 0 |
| NC_002645.1;NP_073551.1;HCoV-229E_surface;339 | 0 | 0 |
| NC_002645.1;NP_073551.1;HCoV-229E_surface;340 | 0 | 0 |
| NC_002645.1;NP_073551.1;HCoV-229E_surface;341 | 0 | 0 |
| NC_002645.1;NP_073551.1;HCoV-229E_surface;342 | 0 | 0 |
| NC_002645.1;NP_073551.1;HCoV-229E_surface;343 | 0 | 0 |
| NC_002645.1;NP_073551.1;HCoV-229E_surface;344 | 0 | 0 |
| NC_002645.1;NP_073551.1;HCoV-229E_surface;345 | 0 | 0 |

| | | |
|---|---|---|
| NC_002645.1;NP_073551.1;HCoV-229E_surface;346 | 0 | 0 |
| NC_002645.1;NP_073551.1;HCoV-229E_surface;347 | 0 | 0 |
| NC_002645.1;NP_073551.1;HCoV-229E_surface;348 | 0 | 0 |
| NC_002645.1;NP_073551.1;HCoV-229E_surface;349 | 0 | 0 |
| NC_002645.1;NP_073551.1;HCoV-229E_surface;350 | 0 | 0 |
| NC_002645.1;NP_073551.1;HCoV-229E_surface;351 | 0 | 0.025 |
| NC_002645.1;NP_073551.1;HCoV-229E_surface;352 | 0 | 0.025 |
| NC_002645.1;NP_073551.1;HCoV-229E_surface;353 | 0 | 0.025 |
| NC_002645.1;NP_073551.1;HCoV-229E_surface;354 | 0 | 0.025 |
| NC_002645.1;NP_073551.1;HCoV-229E_surface;355 | 0 | 0.075 |
| NC_002645.1;NP_073551.1;HCoV-229E_surface;356 | 0 | 0.05 |
| NC_002645.1;NP_073551.1;HCoV-229E_surface;357 | 0 | 0.025 |
| NC_002645.1;NP_073551.1;HCoV-229E_surface;358 | 0.05 | 0 |
| NC_002645.1;NP_073551.1;HCoV-229E_surface;359 | 0 | 0 |
| NC_002645.1;NP_073551.1;HCoV-229E_surface;360 | 0 | 0.025 |
| NC_002645.1;NP_073551.1;HCoV-229E_surface;361 | 0 | 0.05 |
| NC_002645.1;NP_073551.1;HCoV-229E_surface;362 | 0.05 | 0 |
| NC_002645.1;NP_073551.1;HCoV-229E_surface;363 | 0.1 | 0.025 |
| NC_002645.1;NP_073551.1;HCoV-229E_surface;364 | 0.05 | 0 |
| NC_002645.1;NP_073551.1;HCoV-229E_surface;365 | 0 | 0 |
| NC_002645.1;NP_073551.1;HCoV-229E_surface;366 | 0 | 0 |
| NC_002645.1;NP_073551.1;HCoV-229E_surface;367 | 0 | 0 |
| NC_002645.1;NP_073551.1;HCoV-229E_surface;368 | 0 | 0 |
| NC_002645.1;NP_073551.1;HCoV-229E_surface;369 | 0 | 0 |
| NC_002645.1;NP_073551.1;HCoV-229E_surface;370 | 0 | 0 |
| NC_002645.1;NP_073551.1;HCoV-229E_surface;371 | 0 | 0 |
| NC_002645.1;NP_073551.1;HCoV-229E_surface;372 | 0 | 0 |
| NC_002645.1;NP_073551.1;HCoV-229E_surface;373 | 0 | 0 |
| NC_002645.1;NP_073551.1;HCoV-229E_surface;374 | 0 | 0 |
| NC_002645.1;NP_073551.1;HCoV-229E_surface;375 | 0 | 0 |
| NC_002645.1;NP_073551.1;HCoV-229E_surface;376 | 0 | 0 |
| NC_002645.1;NP_073551.1;HCoV-229E_surface;377 | 0 | 0 |
| NC_002645.1;NP_073551.1;HCoV-229E_surface;378 | 0 | 0 |
| NC_002645.1;NP_073551.1;HCoV-229E_surface;379 | 0 | 0 |
| NC_002645.1;NP_073551.1;HCoV-229E_surface;380 | 0 | 0 |
| NC_002645.1;NP_073551.1;HCoV-229E_surface;381 | 0 | 0 |
| NC_002645.1;NP_073551.1;HCoV-229E_surface;382 | 0 | 0 |
| NC_002645.1;NP_073551.1;HCoV-229E_surface;383 | 0 | 0 |
| NC_002645.1;NP_073551.1;HCoV-229E_surface;384 | 0 | 0 |
| NC_002645.1;NP_073551.1;HCoV-229E_surface;385 | 0 | 0 |
| NC_002645.1;NP_073551.1;HCoV-229E_surface;386 | 0 | 0 |
| NC_002645.1;NP_073551.1;HCoV-229E_surface;387 | 0 | 0.025 |
| NC_002645.1;NP_073551.1;HCoV-229E_surface;388 | 0 | 0 |

| | | |
|---|---|---|
| NC_002645.1;NP_073551.1;HCoV-229E_surface;389 | 0 | 0 |
| NC_002645.1;NP_073551.1;HCoV-229E_surface;390 | 0.05 | 0 |
| NC_002645.1;NP_073551.1;HCoV-229E_surface;391 | 0 | 0.025 |
| NC_002645.1;NP_073551.1;HCoV-229E_surface;392 | 0 | 0 |
| NC_002645.1;NP_073551.1;HCoV-229E_surface;393 | 0 | 0 |
| NC_002645.1;NP_073551.1;HCoV-229E_surface;394 | 0 | 0 |
| NC_002645.1;NP_073551.1;HCoV-229E_surface;395 | 0 | 0 |
| NC_002645.1;NP_073551.1;HCoV-229E_surface;396 | 0 | 0 |
| NC_002645.1;NP_073551.1;HCoV-229E_surface;397 | 0 | 0 |
| NC_002645.1;NP_073551.1;HCoV-229E_surface;398 | 0 | 0 |
| NC_002645.1;NP_073551.1;HCoV-229E_surface;399 | 0 | 0 |
| NC_002645.1;NP_073551.1;HCoV-229E_surface;400 | 0 | 0 |
| NC_002645.1;NP_073551.1;HCoV-229E_surface;401 | 0 | 0 |
| NC_002645.1;NP_073551.1;HCoV-229E_surface;402 | 0 | 0 |
| NC_002645.1;NP_073551.1;HCoV-229E_surface;403 | 0 | 0 |
| NC_002645.1;NP_073551.1;HCoV-229E_surface;404 | 0 | 0 |
| NC_002645.1;NP_073551.1;HCoV-229E_surface;405 | 0 | 0 |
| NC_002645.1;NP_073551.1;HCoV-229E_surface;406 | 0 | 0 |
| NC_002645.1;NP_073551.1;HCoV-229E_surface;407 | 0 | 0 |
| NC_002645.1;NP_073551.1;HCoV-229E_surface;408 | 0.05 | 0 |
| NC_002645.1;NP_073551.1;HCoV-229E_surface;409 | 0 | 0 |
| NC_002645.1;NP_073551.1;HCoV-229E_surface;410 | 0.05 | 0 |
| NC_002645.1;NP_073551.1;HCoV-229E_surface;411 | 0.05 | 0 |
| NC_002645.1;NP_073551.1;HCoV-229E_surface;412 | 0.05 | 0 |
| NC_002645.1;NP_073551.1;HCoV-229E_surface;413 | 0.05 | 0 |
| NC_002645.1;NP_073551.1;HCoV-229E_surface;414 | 0.05 | 0.05 |
| NC_002645.1;NP_073551.1;HCoV-229E_surface;415 | 0.05 | 0.025 |
| NC_002645.1;NP_073551.1;HCoV-229E_surface;416 | 0.05 | 0 |
| NC_002645.1;NP_073551.1;HCoV-229E_surface;417 | 0.05 | 0 |
| NC_002645.1;NP_073551.1;HCoV-229E_surface;418 | 0.05 | 0 |
| NC_002645.1;NP_073551.1;HCoV-229E_surface;419 | 0 | 0 |
| NC_002645.1;NP_073551.1;HCoV-229E_surface;420 | 0 | 0 |
| NC_002645.1;NP_073551.1;HCoV-229E_surface;421 | 0 | 0 |
| NC_002645.1;NP_073551.1;HCoV-229E_surface;422 | 0 | 0 |
| NC_002645.1;NP_073551.1;HCoV-229E_surface;423 | 0 | 0 |
| NC_002645.1;NP_073551.1;HCoV-229E_surface;424 | 0 | 0 |
| NC_002645.1;NP_073551.1;HCoV-229E_surface;425 | 0 | 0 |
| NC_002645.1;NP_073551.1;HCoV-229E_surface;426 | 0 | 0 |
| NC_002645.1;NP_073551.1;HCoV-229E_surface;427 | 0 | 0 |
| NC_002645.1;NP_073551.1;HCoV-229E_surface;428 | 0 | 0 |
| NC_002645.1;NP_073551.1;HCoV-229E_surface;429 | 0 | 0 |
| NC_002645.1;NP_073551.1;HCoV-229E_surface;430 | 0.05 | 0 |
| NC_002645.1;NP_073551.1;HCoV-229E_surface;431 | 0 | 0 |

| | | |
|---|---|---|
| NC_002645.1;NP_073551.1;HCoV-229E_surface;432 | 0 | 0 |
| NC_002645.1;NP_073551.1;HCoV-229E_surface;433 | 0 | 0 |
| NC_002645.1;NP_073551.1;HCoV-229E_surface;434 | 0 | 0 |
| NC_002645.1;NP_073551.1;HCoV-229E_surface;435 | 0 | 0 |
| NC_002645.1;NP_073551.1;HCoV-229E_surface;436 | 0 | 0 |
| NC_002645.1;NP_073551.1;HCoV-229E_surface;437 | 0.05 | 0 |
| NC_002645.1;NP_073551.1;HCoV-229E_surface;438 | 0.35 | 0.075 |
| NC_002645.1;NP_073551.1;HCoV-229E_surface;439 | 0.35 | 0.15 |
| NC_002645.1;NP_073551.1;HCoV-229E_surface;440 | 0.3 | 0.125 |
| NC_002645.1;NP_073551.1;HCoV-229E_surface;441 | 0.3 | 0.2 |
| NC_002645.1;NP_073551.1;HCoV-229E_surface;442 | 0.3 | 0.2 |
| NC_002645.1;NP_073551.1;HCoV-229E_surface;443 | 0.3 | 0.25 |
| NC_002645.1;NP_073551.1;HCoV-229E_surface;444 | 0.15 | 0.2 |
| NC_002645.1;NP_073551.1;HCoV-229E_surface;445 | 0.1 | 0.075 |
| NC_002645.1;NP_073551.1;HCoV-229E_surface;446 | 0.05 | 0.025 |
| NC_002645.1;NP_073551.1;HCoV-229E_surface;447 | 0.1 | 0.05 |
| NC_002645.1;NP_073551.1;HCoV-229E_surface;448 | 0.1 | 0.025 |
| NC_002645.1;NP_073551.1;HCoV-229E_surface;449 | 0 | 0 |
| NC_002645.1;NP_073551.1;HCoV-229E_surface;450 | 0 | 0 |
| NC_002645.1;NP_073551.1;HCoV-229E_surface;451 | 0 | 0 |
| NC_002645.1;NP_073551.1;HCoV-229E_surface;452 | 0 | 0 |
| NC_002645.1;NP_073551.1;HCoV-229E_surface;453 | 0 | 0 |
| NC_002645.1;NP_073551.1;HCoV-229E_surface;454 | 0 | 0 |
| NC_002645.1;NP_073551.1;HCoV-229E_surface;455 | 0 | 0 |
| NC_002645.1;NP_073551.1;HCoV-229E_surface;456 | 0 | 0 |
| NC_002645.1;NP_073551.1;HCoV-229E_surface;457 | 0 | 0 |
| NC_002645.1;NP_073551.1;HCoV-229E_surface;458 | 0 | 0.025 |
| NC_002645.1;NP_073551.1;HCoV-229E_surface;459 | 0 | 0 |
| NC_002645.1;NP_073551.1;HCoV-229E_surface;460 | 0 | 0 |
| NC_002645.1;NP_073551.1;HCoV-229E_surface;461 | 0 | 0 |
| NC_002645.1;NP_073551.1;HCoV-229E_surface;462 | 0 | 0 |
| NC_002645.1;NP_073551.1;HCoV-229E_surface;463 | 0 | 0 |
| NC_002645.1;NP_073551.1;HCoV-229E_surface;464 | 0 | 0.025 |
| NC_002645.1;NP_073551.1;HCoV-229E_surface;465 | 0 | 0 |
| NC_002645.1;NP_073551.1;HCoV-229E_surface;466 | 0 | 0.025 |
| NC_002645.1;NP_073551.1;HCoV-229E_surface;467 | 0 | 0.025 |
| NC_002645.1;NP_073551.1;HCoV-229E_surface;468 | 0 | 0.025 |
| NC_002645.1;NP_073551.1;HCoV-229E_surface;469 | 0 | 0.05 |
| NC_002645.1;NP_073551.1;HCoV-229E_surface;470 | 0.1 | 0.1 |
| NC_002645.1;NP_073551.1;HCoV-229E_surface;471 | 0.1 | 0.05 |
| NC_002645.1;NP_073551.1;HCoV-229E_surface;472 | 0.15 | 0.075 |
| NC_002645.1;NP_073551.1;HCoV-229E_surface;473 | 0.05 | 0.075 |
| NC_002645.1;NP_073551.1;HCoV-229E_surface;474 | 0 | 0.025 |

| | | |
|---|---|---|
| NC_002645.1;NP_073551.1;HCoV-229E_surface;475 | 0 | 0.025 |
| NC_002645.1;NP_073551.1;HCoV-229E_surface;476 | 0 | 0.025 |
| NC_002645.1;NP_073551.1;HCoV-229E_surface;477 | 0 | 0.025 |
| NC_002645.1;NP_073551.1;HCoV-229E_surface;478 | 0 | 0 |
| NC_002645.1;NP_073551.1;HCoV-229E_surface;479 | 0 | 0 |
| NC_002645.1;NP_073551.1;HCoV-229E_surface;480 | 0 | 0 |
| NC_002645.1;NP_073551.1;HCoV-229E_surface;481 | 0 | 0 |
| NC_002645.1;NP_073551.1;HCoV-229E_surface;482 | 0 | 0 |
| NC_002645.1;NP_073551.1;HCoV-229E_surface;483 | 0 | 0 |
| NC_002645.1;NP_073551.1;HCoV-229E_surface;484 | 0 | 0 |
| NC_002645.1;NP_073551.1;HCoV-229E_surface;485 | 0 | 0 |
| NC_002645.1;NP_073551.1;HCoV-229E_surface;486 | 0 | 0 |
| NC_002645.1;NP_073551.1;HCoV-229E_surface;487 | 0 | 0 |
| NC_002645.1;NP_073551.1;HCoV-229E_surface;488 | 0 | 0.025 |
| NC_002645.1;NP_073551.1;HCoV-229E_surface;489 | 0 | 0 |
| NC_002645.1;NP_073551.1;HCoV-229E_surface;490 | 0 | 0 |
| NC_002645.1;NP_073551.1;HCoV-229E_surface;491 | 0 | 0 |
| NC_002645.1;NP_073551.1;HCoV-229E_surface;492 | 0 | 0 |
| NC_002645.1;NP_073551.1;HCoV-229E_surface;493 | 0 | 0 |
| NC_002645.1;NP_073551.1;HCoV-229E_surface;494 | 0 | 0 |
| NC_002645.1;NP_073551.1;HCoV-229E_surface;495 | 0 | 0 |
| NC_002645.1;NP_073551.1;HCoV-229E_surface;496 | 0 | 0 |
| NC_002645.1;NP_073551.1;HCoV-229E_surface;497 | 0 | 0 |
| NC_002645.1;NP_073551.1;HCoV-229E_surface;498 | 0 | 0 |
| NC_002645.1;NP_073551.1;HCoV-229E_surface;499 | 0 | 0 |
| NC_002645.1;NP_073551.1;HCoV-229E_surface;500 | 0.05 | 0 |
| NC_002645.1;NP_073551.1;HCoV-229E_surface;501 | 0 | 0 |
| NC_002645.1;NP_073551.1;HCoV-229E_surface;502 | 0 | 0 |
| NC_002645.1;NP_073551.1;HCoV-229E_surface;503 | 0 | 0.025 |
| NC_002645.1;NP_073551.1;HCoV-229E_surface;504 | 0 | 0.05 |
| NC_002645.1;NP_073551.1;HCoV-229E_surface;505 | 0 | 0.075 |
| NC_002645.1;NP_073551.1;HCoV-229E_surface;506 | 0 | 0.075 |
| NC_002645.1;NP_073551.1;HCoV-229E_surface;507 | 0 | 0.075 |
| NC_002645.1;NP_073551.1;HCoV-229E_surface;508 | 0 | 0.075 |
| NC_002645.1;NP_073551.1;HCoV-229E_surface;509 | 0.05 | 0.025 |
| NC_002645.1;NP_073551.1;HCoV-229E_surface;510 | 0.05 | 0.05 |
| NC_002645.1;NP_073551.1;HCoV-229E_surface;511 | 0.05 | 0.025 |
| NC_002645.1;NP_073551.1;HCoV-229E_surface;512 | 0.05 | 0.05 |
| NC_002645.1;NP_073551.1;HCoV-229E_surface;513 | 0.05 | 0.025 |
| NC_002645.1;NP_073551.1;HCoV-229E_surface;514 | 0.1 | 0.025 |
| NC_002645.1;NP_073551.1;HCoV-229E_surface;515 | 0.05 | 0 |
| NC_002645.1;NP_073551.1;HCoV-229E_surface;516 | 0.05 | 0 |
| NC_002645.1;NP_073551.1;HCoV-229E_surface;517 | 0.15 | 0 |

| | | |
|---|---|---|
| NC_002645.1;NP_073551.1;HCoV-229E_surface;518 | 0.15 | 0 |
| NC_002645.1;NP_073551.1;HCoV-229E_surface;519 | 0.05 | 0 |
| NC_002645.1;NP_073551.1;HCoV-229E_surface;520 | 0 | 0 |
| NC_002645.1;NP_073551.1;HCoV-229E_surface;521 | 0 | 0 |
| NC_002645.1;NP_073551.1;HCoV-229E_surface;522 | 0.1 | 0 |
| NC_002645.1;NP_073551.1;HCoV-229E_surface;523 | 0.15 | 0.05 |
| NC_002645.1;NP_073551.1;HCoV-229E_surface;524 | 0.1 | 0.05 |
| NC_002645.1;NP_073551.1;HCoV-229E_surface;525 | 0.15 | 0.05 |
| NC_002645.1;NP_073551.1;HCoV-229E_surface;526 | 0.05 | 0.05 |
| NC_002645.1;NP_073551.1;HCoV-229E_surface;527 | 0.05 | 0.05 |
| NC_002645.1;NP_073551.1;HCoV-229E_surface;528 | 0.05 | 0.05 |
| NC_002645.1;NP_073551.1;HCoV-229E_surface;529 | 0.05 | 0.025 |
| NC_002645.1;NP_073551.1;HCoV-229E_surface;530 | 0 | 0.05 |
| NC_002645.1;NP_073551.1;HCoV-229E_surface;531 | 0 | 0.025 |
| NC_002645.1;NP_073551.1;HCoV-229E_surface;532 | 0 | 0.025 |
| NC_002645.1;NP_073551.1;HCoV-229E_surface;533 | 0 | 0 |
| NC_002645.1;NP_073551.1;HCoV-229E_surface;534 | 0 | 0.025 |
| NC_002645.1;NP_073551.1;HCoV-229E_surface;535 | 0 | 0 |
| NC_002645.1;NP_073551.1;HCoV-229E_surface;536 | 0 | 0 |
| NC_002645.1;NP_073551.1;HCoV-229E_surface;537 | 0 | 0 |
| NC_002645.1;NP_073551.1;HCoV-229E_surface;538 | 0 | 0 |
| NC_002645.1;NP_073551.1;HCoV-229E_surface;539 | 0 | 0 |
| NC_002645.1;NP_073551.1;HCoV-229E_surface;540 | 0 | 0 |
| NC_002645.1;NP_073551.1;HCoV-229E_surface;541 | 0 | 0 |
| NC_002645.1;NP_073551.1;HCoV-229E_surface;542 | 0 | 0 |
| NC_002645.1;NP_073551.1;HCoV-229E_surface;543 | 0.05 | 0 |
| NC_002645.1;NP_073551.1;HCoV-229E_surface;544 | 0.05 | 0 |
| NC_002645.1;NP_073551.1;HCoV-229E_surface;545 | 0.05 | 0 |
| NC_002645.1;NP_073551.1;HCoV-229E_surface;546 | 0 | 0 |
| NC_002645.1;NP_073551.1;HCoV-229E_surface;547 | 0 | 0 |
| NC_002645.1;NP_073551.1;HCoV-229E_surface;548 | 0 | 0 |
| NC_002645.1;NP_073551.1;HCoV-229E_surface;549 | 0 | 0 |
| NC_002645.1;NP_073551.1;HCoV-229E_surface;550 | 0 | 0 |
| NC_002645.1;NP_073551.1;HCoV-229E_surface;551 | 0 | 0 |
| NC_002645.1;NP_073551.1;HCoV-229E_surface;552 | 0 | 0 |
| NC_002645.1;NP_073551.1;HCoV-229E_surface;553 | 0 | 0 |
| NC_002645.1;NP_073551.1;HCoV-229E_surface;554 | 0 | 0 |
| NC_002645.1;NP_073551.1;HCoV-229E_surface;555 | 0 | 0 |
| NC_002645.1;NP_073551.1;HCoV-229E_surface;556 | 0 | 0 |
| NC_002645.1;NP_073551.1;HCoV-229E_surface;557 | 0 | 0 |
| NC_002645.1;NP_073551.1;HCoV-229E_surface;558 | 0 | 0 |
| NC_002645.1;NP_073551.1;HCoV-229E_surface;559 | 0 | 0 |
| NC_002645.1;NP_073551.1;HCoV-229E_surface;560 | 0 | 0.025 |

| | | |
|---|---|---|
| NC_002645.1;NP_073551.1;HCoV-229E_surface;561 | 0 | 0 |
| NC_002645.1;NP_073551.1;HCoV-229E_surface;562 | 0.1 | 0 |
| NC_002645.1;NP_073551.1;HCoV-229E_surface;563 | 0 | 0 |
| NC_002645.1;NP_073551.1;HCoV-229E_surface;564 | 0 | 0 |
| NC_002645.1;NP_073551.1;HCoV-229E_surface;565 | 0.05 | 0 |
| NC_002645.1;NP_073551.1;HCoV-229E_surface;566 | 0 | 0 |
| NC_002645.1;NP_073551.1;HCoV-229E_surface;567 | 0 | 0 |
| NC_002645.1;NP_073551.1;HCoV-229E_surface;568 | 0 | 0 |
| NC_002645.1;NP_073551.1;HCoV-229E_surface;569 | 0 | 0 |
| NC_002645.1;NP_073551.1;HCoV-229E_surface;570 | 0 | 0 |
| NC_002645.1;NP_073551.1;HCoV-229E_surface;571 | 0 | 0 |
| NC_002645.1;NP_073551.1;HCoV-229E_surface;572 | 0 | 0 |
| NC_002645.1;NP_073551.1;HCoV-229E_surface;573 | 0 | 0.025 |
| NC_002645.1;NP_073551.1;HCoV-229E_surface;574 | 0 | 0 |
| NC_002645.1;NP_073551.1;HCoV-229E_surface;575 | 0 | 0.025 |
| NC_002645.1;NP_073551.1;HCoV-229E_surface;576 | 0.05 | 0.05 |
| NC_002645.1;NP_073551.1;HCoV-229E_surface;577 | 0.05 | 0.05 |
| NC_002645.1;NP_073551.1;HCoV-229E_surface;578 | 0.05 | 0 |
| NC_002645.1;NP_073551.1;HCoV-229E_surface;579 | 0 | 0 |
| NC_002645.1;NP_073551.1;HCoV-229E_surface;580 | 0 | 0 |
| NC_002645.1;NP_073551.1;HCoV-229E_surface;581 | 0 | 0 |
| NC_002645.1;NP_073551.1;HCoV-229E_surface;582 | 0 | 0 |
| NC_002645.1;NP_073551.1;HCoV-229E_surface;583 | 0 | 0 |
| NC_002645.1;NP_073551.1;HCoV-229E_surface;584 | 0 | 0 |
| NC_002645.1;NP_073551.1;HCoV-229E_surface;585 | 0 | 0 |
| NC_002645.1;NP_073551.1;HCoV-229E_surface;586 | 0 | 0 |
| NC_002645.1;NP_073551.1;HCoV-229E_surface;587 | 0 | 0 |
| NC_002645.1;NP_073551.1;HCoV-229E_surface;588 | 0 | 0 |
| NC_002645.1;NP_073551.1;HCoV-229E_surface;589 | 0 | 0 |
| NC_002645.1;NP_073551.1;HCoV-229E_surface;590 | 0 | 0 |
| NC_002645.1;NP_073551.1;HCoV-229E_surface;591 | 0 | 0 |
| NC_002645.1;NP_073551.1;HCoV-229E_surface;592 | 0 | 0 |
| NC_002645.1;NP_073551.1;HCoV-229E_surface;593 | 0 | 0 |
| NC_002645.1;NP_073551.1;HCoV-229E_surface;594 | 0 | 0 |
| NC_002645.1;NP_073551.1;HCoV-229E_surface;595 | 0 | 0 |
| NC_002645.1;NP_073551.1;HCoV-229E_surface;596 | 0 | 0 |
| NC_002645.1;NP_073551.1;HCoV-229E_surface;597 | 0 | 0 |
| NC_002645.1;NP_073551.1;HCoV-229E_surface;598 | 0 | 0 |
| NC_002645.1;NP_073551.1;HCoV-229E_surface;599 | 0 | 0 |
| NC_002645.1;NP_073551.1;HCoV-229E_surface;600 | 0 | 0 |
| NC_002645.1;NP_073551.1;HCoV-229E_surface;601 | 0 | 0 |
| NC_002645.1;NP_073551.1;HCoV-229E_surface;602 | 0 | 0.025 |
| NC_002645.1;NP_073551.1;HCoV-229E_surface;603 | 0 | 0 |

| | | |
|---|---|---|
| NC_002645.1;NP_073551.1;HCoV-229E_surface;604 | 0 | 0 |
| NC_002645.1;NP_073551.1;HCoV-229E_surface;605 | 0 | 0 |
| NC_002645.1;NP_073551.1;HCoV-229E_surface;606 | 0 | 0.025 |
| NC_002645.1;NP_073551.1;HCoV-229E_surface;607 | 0 | 0 |
| NC_002645.1;NP_073551.1;HCoV-229E_surface;608 | 0 | 0 |
| NC_002645.1;NP_073551.1;HCoV-229E_surface;609 | 0 | 0 |
| NC_002645.1;NP_073551.1;HCoV-229E_surface;610 | 0.05 | 0.025 |
| NC_002645.1;NP_073551.1;HCoV-229E_surface;611 | 0 | 0.025 |
| NC_002645.1;NP_073551.1;HCoV-229E_surface;612 | 0.05 | 0.05 |
| NC_002645.1;NP_073551.1;HCoV-229E_surface;613 | 0 | 0.075 |
| NC_002645.1;NP_073551.1;HCoV-229E_surface;614 | 0 | 0.025 |
| NC_002645.1;NP_073551.1;HCoV-229E_surface;615 | 0 | 0 |
| NC_002645.1;NP_073551.1;HCoV-229E_surface;616 | 0 | 0 |
| NC_002645.1;NP_073551.1;HCoV-229E_surface;617 | 0 | 0 |
| NC_002645.1;NP_073551.1;HCoV-229E_surface;618 | 0 | 0 |
| NC_002645.1;NP_073551.1;HCoV-229E_surface;619 | 0 | 0 |
| NC_002645.1;NP_073551.1;HCoV-229E_surface;620 | 0 | 0 |
| NC_002645.1;NP_073551.1;HCoV-229E_surface;621 | 0 | 0 |
| NC_002645.1;NP_073551.1;HCoV-229E_surface;622 | 0 | 0 |
| NC_002645.1;NP_073551.1;HCoV-229E_surface;623 | 0 | 0 |
| NC_002645.1;NP_073551.1;HCoV-229E_surface;624 | 0 | 0 |
| NC_002645.1;NP_073551.1;HCoV-229E_surface;625 | 0 | 0.025 |
| NC_002645.1;NP_073551.1;HCoV-229E_surface;626 | 0 | 0.025 |
| NC_002645.1;NP_073551.1;HCoV-229E_surface;627 | 0.05 | 0.05 |
| NC_002645.1;NP_073551.1;HCoV-229E_surface;628 | 0 | 0.05 |
| NC_002645.1;NP_073551.1;HCoV-229E_surface;629 | 0.05 | 0.05 |
| NC_002645.1;NP_073551.1;HCoV-229E_surface;630 | 0.05 | 0.1 |
| NC_002645.1;NP_073551.1;HCoV-229E_surface;631 | 0.05 | 0.275 |
| NC_002645.1;NP_073551.1;HCoV-229E_surface;632 | 0.05 | 0.125 |
| NC_002645.1;NP_073551.1;HCoV-229E_surface;633 | 0 | 0 |
| NC_002645.1;NP_073551.1;HCoV-229E_surface;634 | 0 | 0 |
| NC_002645.1;NP_073551.1;HCoV-229E_surface;635 | 0 | 0 |
| NC_002645.1;NP_073551.1;HCoV-229E_surface;636 | 0 | 0 |
| NC_002645.1;NP_073551.1;HCoV-229E_surface;637 | 0 | 0 |
| NC_002645.1;NP_073551.1;HCoV-229E_surface;638 | 0 | 0 |
| NC_002645.1;NP_073551.1;HCoV-229E_surface;639 | 0 | 0 |
| NC_002645.1;NP_073551.1;HCoV-229E_surface;640 | 0 | 0 |
| NC_002645.1;NP_073551.1;HCoV-229E_surface;641 | 0.1 | 0 |
| NC_002645.1;NP_073551.1;HCoV-229E_surface;642 | 0.05 | 0.05 |
| NC_002645.1;NP_073551.1;HCoV-229E_surface;643 | 0.1 | 0.025 |
| NC_002645.1;NP_073551.1;HCoV-229E_surface;644 | 0.05 | 0.025 |
| NC_002645.1;NP_073551.1;HCoV-229E_surface;645 | 0.1 | 0.075 |
| NC_002645.1;NP_073551.1;HCoV-229E_surface;646 | 0.05 | 0.025 |

| | | |
|---|---|---|
| NC_002645.1;NP_073551.1;HCoV-229E_surface;647 | 0.05 | 0.075 |
| NC_002645.1;NP_073551.1;HCoV-229E_surface;648 | 0.05 | 0.025 |
| NC_002645.1;NP_073551.1;HCoV-229E_surface;649 | 0.05 | 0.025 |
| NC_002645.1;NP_073551.1;HCoV-229E_surface;650 | 0.05 | 0.025 |
| NC_002645.1;NP_073551.1;HCoV-229E_surface;651 | 0.2 | 0.05 |
| NC_002645.1;NP_073551.1;HCoV-229E_surface;652 | 0 | 0 |
| NC_002645.1;NP_073551.1;HCoV-229E_surface;653 | 0 | 0 |
| NC_002645.1;NP_073551.1;HCoV-229E_surface;654 | 0 | 0 |
| NC_002645.1;NP_073551.1;HCoV-229E_surface;655 | 0 | 0.025 |
| NC_002645.1;NP_073551.1;HCoV-229E_surface;656 | 0.05 | 0 |
| NC_002645.1;NP_073551.1;HCoV-229E_surface;657 | 0 | 0 |
| NC_002645.1;NP_073551.1;HCoV-229E_surface;658 | 0 | 0 |
| NC_002645.1;NP_073551.1;HCoV-229E_surface;659 | 0 | 0.025 |
| NC_002645.1;NP_073551.1;HCoV-229E_surface;660 | 0 | 0 |
| NC_002645.1;NP_073551.1;HCoV-229E_surface;661 | 0 | 0 |
| NC_002645.1;NP_073551.1;HCoV-229E_surface;662 | 0 | 0 |
| NC_002645.1;NP_073551.1;HCoV-229E_surface;663 | 0.05 | 0.025 |
| NC_002645.1;NP_073551.1;HCoV-229E_surface;664 | 0 | 0 |
| NC_002645.1;NP_073551.1;HCoV-229E_surface;665 | 0.05 | 0.05 |
| NC_002645.1;NP_073551.1;HCoV-229E_surface;666 | 0.05 | 0.05 |
| NC_002645.1;NP_073551.1;HCoV-229E_surface;667 | 0.05 | 0 |
| NC_002645.1;NP_073551.1;HCoV-229E_surface;668 | 0 | 0 |
| NC_002645.1;NP_073551.1;HCoV-229E_surface;669 | 0.15 | 0.075 |
| NC_002645.1;NP_073551.1;HCoV-229E_surface;670 | 0 | 0 |
| NC_002645.1;NP_073551.1;HCoV-229E_surface;671 | 0 | 0 |
| NC_002645.1;NP_073551.1;HCoV-229E_surface;672 | 0 | 0.025 |
| NC_002645.1;NP_073551.1;HCoV-229E_surface;673 | 0 | 0 |
| NC_002645.1;NP_073551.1;HCoV-229E_surface;674 | 0 | 0 |
| NC_002645.1;NP_073551.1;HCoV-229E_surface;675 | 0 | 0 |
| NC_002645.1;NP_073551.1;HCoV-229E_surface;676 | 0 | 0 |
| NC_002645.1;NP_073551.1;HCoV-229E_surface;677 | 0 | 0 |
| NC_002645.1;NP_073551.1;HCoV-229E_surface;678 | 0 | 0 |
| NC_002645.1;NP_073551.1;HCoV-229E_surface;679 | 0 | 0.025 |
| NC_002645.1;NP_073551.1;HCoV-229E_surface;680 | 0 | 0.025 |
| NC_002645.1;NP_073551.1;HCoV-229E_surface;681 | 0 | 0.025 |
| NC_002645.1;NP_073551.1;HCoV-229E_surface;682 | 0 | 0.475 |
| NC_002645.1;NP_073551.1;HCoV-229E_surface;683 | 0 | 0.5 |
| NC_002645.1;NP_073551.1;HCoV-229E_surface;684 | 0.05 | 0.7 |
| NC_002645.1;NP_073551.1;HCoV-229E_surface;685 | 0 | 0.6 |
| NC_002645.1;NP_073551.1;HCoV-229E_surface;686 | 0 | 0.5 |
| NC_002645.1;NP_073551.1;HCoV-229E_surface;687 | 0 | 0.4 |
| NC_002645.1;NP_073551.1;HCoV-229E_surface;688 | 0 | 0.4 |
| NC_002645.1;NP_073551.1;HCoV-229E_surface;689 | 0 | 0.375 |

| | | |
|---|---|---|
| NC_002645.1;NP_073551.1;HCoV-229E_surface;690 | 0 | 0.225 |
| NC_002645.1;NP_073551.1;HCoV-229E_surface;691 | 0.15 | 0.25 |
| NC_002645.1;NP_073551.1;HCoV-229E_surface;692 | 0 | 0.2 |
| NC_002645.1;NP_073551.1;HCoV-229E_surface;693 | 0 | 0.025 |
| NC_002645.1;NP_073551.1;HCoV-229E_surface;694 | 0.05 | 0 |
| NC_002645.1;NP_073551.1;HCoV-229E_surface;695 | 0.05 | 0 |
| NC_002645.1;NP_073551.1;HCoV-229E_surface;696 | 0.05 | 0 |
| NC_002645.1;NP_073551.1;HCoV-229E_surface;697 | 0 | 0 |
| NC_002645.1;NP_073551.1;HCoV-229E_surface;698 | 0.05 | 0 |
| NC_002645.1;NP_073551.1;HCoV-229E_surface;699 | 0 | 0 |
| NC_002645.1;NP_073551.1;HCoV-229E_surface;700 | 0.1 | 0 |
| NC_002645.1;NP_073551.1;HCoV-229E_surface;701 | 0.1 | 0 |
| NC_002645.1;NP_073551.1;HCoV-229E_surface;702 | 0 | 0 |
| NC_002645.1;NP_073551.1;HCoV-229E_surface;703 | 0 | 0 |
| NC_002645.1;NP_073551.1;HCoV-229E_surface;704 | 0.05 | 0 |
| NC_002645.1;NP_073551.1;HCoV-229E_surface;705 | 0 | 0 |
| NC_002645.1;NP_073551.1;HCoV-229E_surface;706 | 0 | 0 |
| NC_002645.1;NP_073551.1;HCoV-229E_surface;707 | 0 | 0 |
| NC_002645.1;NP_073551.1;HCoV-229E_surface;708 | 0.05 | 0.025 |
| NC_002645.1;NP_073551.1;HCoV-229E_surface;709 | 0.1 | 0 |
| NC_002645.1;NP_073551.1;HCoV-229E_surface;710 | 0.15 | 0.05 |
| NC_002645.1;NP_073551.1;HCoV-229E_surface;711 | 0.15 | 0.05 |
| NC_002645.1;NP_073551.1;HCoV-229E_surface;712 | 0.1 | 0.025 |
| NC_002645.1;NP_073551.1;HCoV-229E_surface;713 | 0.15 | 0.05 |
| NC_002645.1;NP_073551.1;HCoV-229E_surface;714 | 0.1 | 0.05 |
| NC_002645.1;NP_073551.1;HCoV-229E_surface;715 | 0.1 | 0.025 |
| NC_002645.1;NP_073551.1;HCoV-229E_surface;716 | 0.15 | 0.05 |
| NC_002645.1;NP_073551.1;HCoV-229E_surface;717 | 0.15 | 0.075 |
| NC_002645.1;NP_073551.1;HCoV-229E_surface;718 | 0 | 0 |
| NC_002645.1;NP_073551.1;HCoV-229E_surface;719 | 0.05 | 0.325 |
| NC_002645.1;NP_073551.1;HCoV-229E_surface;720 | 0 | 0.05 |
| NC_002645.1;NP_073551.1;HCoV-229E_surface;721 | 0 | 0 |
| NC_002645.1;NP_073551.1;HCoV-229E_surface;722 | 0 | 0 |
| NC_002645.1;NP_073551.1;HCoV-229E_surface;723 | 0 | 0 |
| NC_002645.1;NP_073551.1;HCoV-229E_surface;724 | 0 | 0 |
| NC_002645.1;NP_073551.1;HCoV-229E_surface;725 | 0.05 | 0.05 |
| NC_002645.1;NP_073551.1;HCoV-229E_surface;726 | 0 | 0 |
| NC_002645.1;NP_073551.1;HCoV-229E_surface;727 | 0.05 | 0.025 |
| NC_002645.1;NP_073551.1;HCoV-229E_surface;728 | 0 | 0 |
| NC_002645.1;NP_073551.1;HCoV-229E_surface;729 | 0 | 0 |
| NC_002645.1;NP_073551.1;HCoV-229E_surface;730 | 0 | 0 |
| NC_002645.1;NP_073551.1;HCoV-229E_surface;731 | 0 | 0 |
| NC_002645.1;NP_073551.1;HCoV-229E_surface;732 | 0 | 0 |

| | | |
|---|---|---|
| NC_002645.1;NP_073551.1;HCoV-229E_surface;733 | 0 | 0 |
| NC_002645.1;NP_073551.1;HCoV-229E_surface;734 | 0 | 0 |
| NC_002645.1;NP_073551.1;HCoV-229E_surface;735 | 0 | 0 |
| NC_002645.1;NP_073551.1;HCoV-229E_surface;736 | 0 | 0 |
| NC_002645.1;NP_073551.1;HCoV-229E_surface;737 | 0 | 0 |
| NC_002645.1;NP_073551.1;HCoV-229E_surface;738 | 0 | 0 |
| NC_002645.1;NP_073551.1;HCoV-229E_surface;739 | 0 | 0 |
| NC_002645.1;NP_073551.1;HCoV-229E_surface;740 | 0 | 0 |
| NC_002645.1;NP_073551.1;HCoV-229E_surface;741 | 0 | 0.025 |
| NC_002645.1;NP_073551.1;HCoV-229E_surface;742 | 0 | 0.025 |
| NC_002645.1;NP_073551.1;HCoV-229E_surface;743 | 0 | 0 |
| NC_002645.1;NP_073551.1;HCoV-229E_surface;744 | 0 | 0 |
| NC_002645.1;NP_073551.1;HCoV-229E_surface;745 | 0 | 0 |
| NC_002645.1;NP_073551.1;HCoV-229E_surface;746 | 0 | 0 |
| NC_002645.1;NP_073551.1;HCoV-229E_surface;747 | 0 | 0 |
| NC_002645.1;NP_073551.1;HCoV-229E_surface;748 | 0 | 0 |
| NC_002645.1;NP_073551.1;HCoV-229E_surface;749 | 0 | 0 |
| NC_002645.1;NP_073551.1;HCoV-229E_surface;750 | 0 | 0 |
| NC_002645.1;NP_073551.1;HCoV-229E_surface;751 | 0 | 0 |
| NC_002645.1;NP_073551.1;HCoV-229E_surface;752 | 0 | 0 |
| NC_002645.1;NP_073551.1;HCoV-229E_surface;753 | 0 | 0 |
| NC_002645.1;NP_073551.1;HCoV-229E_surface;754 | 0 | 0 |
| NC_002645.1;NP_073551.1;HCoV-229E_surface;755 | 0 | 0 |
| NC_002645.1;NP_073551.1;HCoV-229E_surface;756 | 0 | 0 |
| NC_002645.1;NP_073551.1;HCoV-229E_surface;757 | 0 | 0 |
| NC_002645.1;NP_073551.1;HCoV-229E_surface;758 | 0 | 0 |
| NC_002645.1;NP_073551.1;HCoV-229E_surface;759 | 0 | 0 |
| NC_002645.1;NP_073551.1;HCoV-229E_surface;760 | 0 | 0 |
| NC_002645.1;NP_073551.1;HCoV-229E_surface;761 | 0 | 0 |
| NC_002645.1;NP_073551.1;HCoV-229E_surface;762 | 0 | 0 |
| NC_002645.1;NP_073551.1;HCoV-229E_surface;763 | 0 | 0 |
| NC_002645.1;NP_073551.1;HCoV-229E_surface;764 | 0 | 0 |
| NC_002645.1;NP_073551.1;HCoV-229E_surface;765 | 0 | 0 |
| NC_002645.1;NP_073551.1;HCoV-229E_surface;766 | 0 | 0 |
| NC_002645.1;NP_073551.1;HCoV-229E_surface;767 | 0 | 0 |
| NC_002645.1;NP_073551.1;HCoV-229E_surface;768 | 0 | 0 |
| NC_002645.1;NP_073551.1;HCoV-229E_surface;769 | 0 | 0 |
| NC_002645.1;NP_073551.1;HCoV-229E_surface;770 | 0 | 0 |
| NC_002645.1;NP_073551.1;HCoV-229E_surface;771 | 0 | 0 |
| NC_002645.1;NP_073551.1;HCoV-229E_surface;772 | 0 | 0 |
| NC_002645.1;NP_073551.1;HCoV-229E_surface;773 | 0 | 0 |
| NC_002645.1;NP_073551.1;HCoV-229E_surface;774 | 0 | 0 |
| NC_002645.1;NP_073551.1;HCoV-229E_surface;775 | 0 | 0 |

| | | |
|---|---|---|
| NC_002645.1;NP_073551.1;HCoV-229E_surface;776 | 0 | 0 |
| NC_002645.1;NP_073551.1;HCoV-229E_surface;777 | 0 | 0 |
| NC_002645.1;NP_073551.1;HCoV-229E_surface;778 | 0.05 | 0 |
| NC_002645.1;NP_073551.1;HCoV-229E_surface;779 | 0 | 0 |
| NC_002645.1;NP_073551.1;HCoV-229E_surface;780 | 0 | 0 |
| NC_002645.1;NP_073551.1;HCoV-229E_surface;781 | 0.05 | 0.025 |
| NC_002645.1;NP_073551.1;HCoV-229E_surface;782 | 0 | 0 |
| NC_002645.1;NP_073551.1;HCoV-229E_surface;783 | 0 | 0 |
| NC_002645.1;NP_073551.1;HCoV-229E_surface;784 | 0.05 | 0.05 |
| NC_002645.1;NP_073551.1;HCoV-229E_surface;785 | 0.3 | 0.1 |
| NC_002645.1;NP_073551.1;HCoV-229E_surface;786 | 0.2 | 0.125 |
| NC_002645.1;NP_073551.1;HCoV-229E_surface;787 | 0.2 | 0.1 |
| NC_002645.1;NP_073551.1;HCoV-229E_surface;788 | 0.2 | 0.125 |
| NC_002645.1;NP_073551.1;HCoV-229E_surface;789 | 0.2 | 0.1 |
| NC_002645.1;NP_073551.1;HCoV-229E_surface;790 | 0.15 | 0.1 |
| NC_002645.1;NP_073551.1;HCoV-229E_surface;791 | 0.2 | 0.1 |
| NC_002645.1;NP_073551.1;HCoV-229E_surface;792 | 0.15 | 0.1 |
| NC_002645.1;NP_073551.1;HCoV-229E_surface;793 | 0.15 | 0.1 |
| NC_002645.1;NP_073551.1;HCoV-229E_surface;794 | 0.15 | 0.1 |
| NC_002645.1;NP_073551.1;HCoV-229E_surface;795 | 0.15 | 0.075 |
| NC_002645.1;NP_073551.1;HCoV-229E_surface;796 | 0.05 | 0 |
| NC_002645.1;NP_073551.1;HCoV-229E_surface;797 | 0 | 0 |
| NC_002645.1;NP_073551.1;HCoV-229E_surface;798 | 0.05 | 0 |
| NC_002645.1;NP_073551.1;HCoV-229E_surface;799 | 0.05 | 0 |
| NC_002645.1;NP_073551.1;HCoV-229E_surface;800 | 0.05 | 0 |
| NC_002645.1;NP_073551.1;HCoV-229E_surface;801 | 0.05 | 0 |
| NC_002645.1;NP_073551.1;HCoV-229E_surface;802 | 0 | 0 |
| NC_002645.1;NP_073551.1;HCoV-229E_surface;803 | 0.05 | 0 |
| NC_002645.1;NP_073551.1;HCoV-229E_surface;804 | 0.1 | 0 |
| NC_002645.1;NP_073551.1;HCoV-229E_surface;805 | 0 | 0 |
| NC_002645.1;NP_073551.1;HCoV-229E_surface;806 | 0 | 0 |
| NC_002645.1;NP_073551.1;HCoV-229E_surface;807 | 0.05 | 0 |
| NC_002645.1;NP_073551.1;HCoV-229E_surface;808 | 0 | 0 |
| NC_002645.1;NP_073551.1;HCoV-229E_surface;809 | 0 | 0 |
| NC_002645.1;NP_073551.1;HCoV-229E_surface;810 | 0 | 0 |
| NC_002645.1;NP_073551.1;HCoV-229E_surface;811 | 0.05 | 0.05 |
| NC_002645.1;NP_073551.1;HCoV-229E_surface;812 | 0 | 0.025 |
| NC_002645.1;NP_073551.1;HCoV-229E_surface;813 | 0 | 0.025 |
| NC_002645.1;NP_073551.1;HCoV-229E_surface;814 | 0 | 0.025 |
| NC_002645.1;NP_073551.1;HCoV-229E_surface;815 | 0 | 0.025 |
| NC_002645.1;NP_073551.1;HCoV-229E_surface;816 | 0 | 0.025 |
| NC_002645.1;NP_073551.1;HCoV-229E_surface;817 | 0.05 | 0.025 |
| NC_002645.1;NP_073551.1;HCoV-229E_surface;818 | 0.05 | 0 |

| | | |
|---|---|---|
| NC_002645.1;NP_073551.1;HCoV-229E_surface;819 | 0.05 | 0.025 |
| NC_002645.1;NP_073551.1;HCoV-229E_surface;820 | 0.15 | 0 |
| NC_002645.1;NP_073551.1;HCoV-229E_surface;821 | 0.15 | 0.075 |
| NC_002645.1;NP_073551.1;HCoV-229E_surface;822 | 0.15 | 0.05 |
| NC_002645.1;NP_073551.1;HCoV-229E_surface;823 | 0.15 | 0.05 |
| NC_002645.1;NP_073551.1;HCoV-229E_surface;824 | 0.1 | 0.075 |
| NC_002645.1;NP_073551.1;HCoV-229E_surface;825 | 0.15 | 0.05 |
| NC_002645.1;NP_073551.1;HCoV-229E_surface;826 | 0.1 | 0.025 |
| NC_002645.1;NP_073551.1;HCoV-229E_surface;827 | 0.1 | 0.025 |
| NC_002645.1;NP_073551.1;HCoV-229E_surface;828 | 0 | 0.025 |
| NC_002645.1;NP_073551.1;HCoV-229E_surface;829 | 0.1 | 0.075 |
| NC_002645.1;NP_073551.1;HCoV-229E_surface;830 | 0.1 | 0.1 |
| NC_002645.1;NP_073551.1;HCoV-229E_surface;831 | 0.05 | 0.05 |
| NC_002645.1;NP_073551.1;HCoV-229E_surface;832 | 0.05 | 0.075 |
| NC_002645.1;NP_073551.1;HCoV-229E_surface;833 | 0.05 | 0.1 |
| NC_002645.1;NP_073551.1;HCoV-229E_surface;834 | 0.05 | 0.025 |
| NC_002645.1;NP_073551.1;HCoV-229E_surface;835 | 0.05 | 0.05 |
| NC_002645.1;NP_073551.1;HCoV-229E_surface;836 | 0.05 | 0.05 |
| NC_002645.1;NP_073551.1;HCoV-229E_surface;837 | 0.05 | 0.025 |
| NC_002645.1;NP_073551.1;HCoV-229E_surface;838 | 0.05 | 0.075 |
| NC_002645.1;NP_073551.1;HCoV-229E_surface;839 | 0.05 | 0.05 |
| NC_002645.1;NP_073551.1;HCoV-229E_surface;840 | 0 | 0 |
| NC_002645.1;NP_073551.1;HCoV-229E_surface;841 | 0 | 0 |
| NC_002645.1;NP_073551.1;HCoV-229E_surface;842 | 0 | 0 |
| NC_002645.1;NP_073551.1;HCoV-229E_surface;843 | 0 | 0 |
| NC_002645.1;NP_073551.1;HCoV-229E_surface;844 | 0 | 0 |
| NC_002645.1;NP_073551.1;HCoV-229E_surface;845 | 0 | 0 |
| NC_002645.1;NP_073551.1;HCoV-229E_surface;846 | 0 | 0 |
| NC_002645.1;NP_073551.1;HCoV-229E_surface;847 | 0 | 0 |
| NC_002645.1;NP_073551.1;HCoV-229E_surface;848 | 0 | 0 |
| NC_002645.1;NP_073551.1;HCoV-229E_surface;849 | 0 | 0 |
| NC_002645.1;NP_073551.1;HCoV-229E_surface;850 | 0 | 0 |
| NC_002645.1;NP_073551.1;HCoV-229E_surface;851 | 0 | 0 |
| NC_002645.1;NP_073551.1;HCoV-229E_surface;852 | 0 | 0 |
| NC_002645.1;NP_073551.1;HCoV-229E_surface;853 | 0 | 0 |
| NC_002645.1;NP_073551.1;HCoV-229E_surface;854 | 0 | 0 |
| NC_002645.1;NP_073551.1;HCoV-229E_surface;855 | 0 | 0 |
| NC_002645.1;NP_073551.1;HCoV-229E_surface;856 | 0 | 0 |
| NC_002645.1;NP_073551.1;HCoV-229E_surface;857 | 0.05 | 0 |
| NC_002645.1;NP_073551.1;HCoV-229E_surface;858 | 0 | 0 |
| NC_002645.1;NP_073551.1;HCoV-229E_surface;859 | 0 | 0 |
| NC_002645.1;NP_073551.1;HCoV-229E_surface;860 | 0.05 | 0.025 |
| NC_002645.1;NP_073551.1;HCoV-229E_surface;861 | 0.05 | 0.05 |

| | | |
|---|---|---|
| NC_002645.1;NP_073551.1;HCoV-229E_surface;862 | 0.05 | 0.05 |
| NC_002645.1;NP_073551.1;HCoV-229E_surface;863 | 0 | 0.025 |
| NC_002645.1;NP_073551.1;HCoV-229E_surface;864 | 0.15 | 0.05 |
| NC_002645.1;NP_073551.1;HCoV-229E_surface;865 | 0.05 | 0.025 |
| NC_002645.1;NP_073551.1;HCoV-229E_surface;866 | 0 | 0.025 |
| NC_002645.1;NP_073551.1;HCoV-229E_surface;867 | 0 | 0 |
| NC_002645.1;NP_073551.1;HCoV-229E_surface;868 | 0 | 0 |
| NC_002645.1;NP_073551.1;HCoV-229E_surface;869 | 0 | 0 |
| NC_002645.1;NP_073551.1;HCoV-229E_surface;870 | 0 | 0 |
| NC_002645.1;NP_073551.1;HCoV-229E_surface;871 | 0 | 0 |
| NC_002645.1;NP_073551.1;HCoV-229E_surface;872 | 0 | 0 |
| NC_002645.1;NP_073551.1;HCoV-229E_surface;873 | 0.05 | 0.025 |
| NC_002645.1;NP_073551.1;HCoV-229E_surface;874 | 0.05 | 0 |
| NC_002645.1;NP_073551.1;HCoV-229E_surface;875 | 0 | 0 |
| NC_002645.1;NP_073551.1;HCoV-229E_surface;876 | 0 | 0 |
| NC_002645.1;NP_073551.1;HCoV-229E_surface;877 | 0 | 0 |
| NC_002645.1;NP_073551.1;HCoV-229E_surface;878 | 0 | 0 |
| NC_002645.1;NP_073551.1;HCoV-229E_surface;879 | 0 | 0 |
| NC_002645.1;NP_073551.1;HCoV-229E_surface;880 | 0 | 0 |
| NC_002645.1;NP_073551.1;HCoV-229E_surface;881 | 0 | 0 |
| NC_002645.1;NP_073551.1;HCoV-229E_surface;882 | 0 | 0 |
| NC_002645.1;NP_073551.1;HCoV-229E_surface;883 | 0 | 0 |
| NC_002645.1;NP_073551.1;HCoV-229E_surface;884 | 0 | 0 |
| NC_002645.1;NP_073551.1;HCoV-229E_surface;885 | 0 | 0 |
| NC_002645.1;NP_073551.1;HCoV-229E_surface;886 | 0 | 0 |
| NC_002645.1;NP_073551.1;HCoV-229E_surface;887 | 0.05 | 0 |
| NC_002645.1;NP_073551.1;HCoV-229E_surface;888 | 0.05 | 0 |
| NC_002645.1;NP_073551.1;HCoV-229E_surface;889 | 0 | 0 |
| NC_002645.1;NP_073551.1;HCoV-229E_surface;890 | 0 | 0 |
| NC_002645.1;NP_073551.1;HCoV-229E_surface;891 | 0 | 0 |
| NC_002645.1;NP_073551.1;HCoV-229E_surface;892 | 0 | 0 |
| NC_002645.1;NP_073551.1;HCoV-229E_surface;893 | 0 | 0 |
| NC_002645.1;NP_073551.1;HCoV-229E_surface;894 | 0 | 0 |
| NC_002645.1;NP_073551.1;HCoV-229E_surface;895 | 0 | 0 |
| NC_002645.1;NP_073551.1;HCoV-229E_surface;896 | 0 | 0 |
| NC_002645.1;NP_073551.1;HCoV-229E_surface;897 | 0 | 0 |
| NC_002645.1;NP_073551.1;HCoV-229E_surface;898 | 0.1 | 0.025 |
| NC_002645.1;NP_073551.1;HCoV-229E_surface;899 | 0.1 | 0.05 |
| NC_002645.1;NP_073551.1;HCoV-229E_surface;900 | 0 | 0 |
| NC_002645.1;NP_073551.1;HCoV-229E_surface;901 | 0 | 0 |
| NC_002645.1;NP_073551.1;HCoV-229E_surface;902 | 0 | 0.025 |
| NC_002645.1;NP_073551.1;HCoV-229E_surface;903 | 0 | 0.025 |
| NC_002645.1;NP_073551.1;HCoV-229E_surface;904 | 0 | 0 |

| | | |
|---|---|---|
| NC_002645.1;NP_073551.1;HCoV-229E_surface;905 | 0 | 0 |
| NC_002645.1;NP_073551.1;HCoV-229E_surface;906 | 0 | 0 |
| NC_002645.1;NP_073551.1;HCoV-229E_surface;907 | 0.1 | 0 |
| NC_002645.1;NP_073551.1;HCoV-229E_surface;908 | 0 | 0 |
| NC_002645.1;NP_073551.1;HCoV-229E_surface;909 | 0 | 0 |
| NC_002645.1;NP_073551.1;HCoV-229E_surface;910 | 0 | 0 |
| NC_002645.1;NP_073551.1;HCoV-229E_surface;911 | 0 | 0 |
| NC_002645.1;NP_073551.1;HCoV-229E_surface;912 | 0 | 0 |
| NC_002645.1;NP_073551.1;HCoV-229E_surface;913 | 0 | 0 |
| NC_002645.1;NP_073551.1;HCoV-229E_surface;914 | 0 | 0 |
| NC_002645.1;NP_073551.1;HCoV-229E_surface;915 | 0 | 0 |
| NC_002645.1;NP_073551.1;HCoV-229E_surface;916 | 0 | 0 |
| NC_002645.1;NP_073551.1;HCoV-229E_surface;917 | 0 | 0 |
| NC_002645.1;NP_073551.1;HCoV-229E_surface;918 | 0 | 0 |
| NC_002645.1;NP_073551.1;HCoV-229E_surface;919 | 0 | 0 |
| NC_002645.1;NP_073551.1;HCoV-229E_surface;920 | 0 | 0 |
| NC_002645.1;NP_073551.1;HCoV-229E_surface;921 | 0 | 0 |
| NC_002645.1;NP_073551.1;HCoV-229E_surface;922 | 0 | 0 |
| NC_002645.1;NP_073551.1;HCoV-229E_surface;923 | 0 | 0 |
| NC_002645.1;NP_073551.1;HCoV-229E_surface;924 | 0 | 0 |
| NC_002645.1;NP_073551.1;HCoV-229E_surface;925 | 0 | 0 |
| NC_002645.1;NP_073551.1;HCoV-229E_surface;926 | 0 | 0 |
| NC_002645.1;NP_073551.1;HCoV-229E_surface;927 | 0 | 0 |
| NC_002645.1;NP_073551.1;HCoV-229E_surface;928 | 0 | 0 |
| NC_002645.1;NP_073551.1;HCoV-229E_surface;929 | 0 | 0 |
| NC_002645.1;NP_073551.1;HCoV-229E_surface;930 | 0 | 0 |
| NC_002645.1;NP_073551.1;HCoV-229E_surface;931 | 0 | 0 |
| NC_002645.1;NP_073551.1;HCoV-229E_surface;932 | 0 | 0 |
| NC_002645.1;NP_073551.1;HCoV-229E_surface;933 | 0 | 0 |
| NC_002645.1;NP_073551.1;HCoV-229E_surface;934 | 0 | 0 |
| NC_002645.1;NP_073551.1;HCoV-229E_surface;935 | 0 | 0 |
| NC_002645.1;NP_073551.1;HCoV-229E_surface;936 | 0 | 0 |
| NC_002645.1;NP_073551.1;HCoV-229E_surface;937 | 0 | 0 |
| NC_002645.1;NP_073551.1;HCoV-229E_surface;938 | 0 | 0 |
| NC_002645.1;NP_073551.1;HCoV-229E_surface;939 | 0 | 0 |
| NC_002645.1;NP_073551.1;HCoV-229E_surface;940 | 0 | 0 |
| NC_002645.1;NP_073551.1;HCoV-229E_surface;941 | 0 | 0 |
| NC_002645.1;NP_073551.1;HCoV-229E_surface;942 | 0 | 0 |
| NC_002645.1;NP_073551.1;HCoV-229E_surface;943 | 0 | 0 |
| NC_002645.1;NP_073551.1;HCoV-229E_surface;944 | 0 | 0 |
| NC_002645.1;NP_073551.1;HCoV-229E_surface;945 | 0 | 0 |
| NC_002645.1;NP_073551.1;HCoV-229E_surface;946 | 0 | 0 |
| NC_002645.1;NP_073551.1;HCoV-229E_surface;947 | 0 | 0.025 |

| | | |
|---|---|---|
| NC_002645.1;NP_073551.1;HCoV-229E_surface;948 | 0 | 0 |
| NC_002645.1;NP_073551.1;HCoV-229E_surface;949 | 0 | 0 |
| NC_002645.1;NP_073551.1;HCoV-229E_surface;950 | 0 | 0.1 |
| NC_002645.1;NP_073551.1;HCoV-229E_surface;951 | 0 | 0 |
| NC_002645.1;NP_073551.1;HCoV-229E_surface;952 | 0 | 0 |
| NC_002645.1;NP_073551.1;HCoV-229E_surface;953 | 0 | 0 |
| NC_002645.1;NP_073551.1;HCoV-229E_surface;954 | 0 | 0.05 |
| NC_002645.1;NP_073551.1;HCoV-229E_surface;955 | 0 | 0 |
| NC_002645.1;NP_073551.1;HCoV-229E_surface;956 | 0 | 0 |
| NC_002645.1;NP_073551.1;HCoV-229E_surface;957 | 0 | 0 |
| NC_002645.1;NP_073551.1;HCoV-229E_surface;958 | 0 | 0 |
| NC_002645.1;NP_073551.1;HCoV-229E_surface;959 | 0 | 0 |
| NC_002645.1;NP_073551.1;HCoV-229E_surface;960 | 0 | 0 |
| NC_002645.1;NP_073551.1;HCoV-229E_surface;961 | 0 | 0 |
| NC_002645.1;NP_073551.1;HCoV-229E_surface;962 | 0 | 0.025 |
| NC_002645.1;NP_073551.1;HCoV-229E_surface;963 | 0 | 0 |
| NC_002645.1;NP_073551.1;HCoV-229E_surface;964 | 0 | 0 |
| NC_002645.1;NP_073551.1;HCoV-229E_surface;965 | 0 | 0 |
| NC_002645.1;NP_073551.1;HCoV-229E_surface;966 | 0 | 0.025 |
| NC_002645.1;NP_073551.1;HCoV-229E_surface;967 | 0 | 0 |
| NC_002645.1;NP_073551.1;HCoV-229E_surface;968 | 0 | 0 |
| NC_002645.1;NP_073551.1;HCoV-229E_surface;969 | 0 | 0 |
| NC_002645.1;NP_073551.1;HCoV-229E_surface;970 | 0 | 0 |
| NC_002645.1;NP_073551.1;HCoV-229E_surface;971 | 0 | 0 |
| NC_002645.1;NP_073551.1;HCoV-229E_surface;972 | 0 | 0 |
| NC_002645.1;NP_073551.1;HCoV-229E_surface;973 | 0 | 0 |
| NC_002645.1;NP_073551.1;HCoV-229E_surface;974 | 0 | 0 |
| NC_002645.1;NP_073551.1;HCoV-229E_surface;975 | 0.05 | 0 |
| NC_002645.1;NP_073551.1;HCoV-229E_surface;976 | 0 | 0 |
| NC_002645.1;NP_073551.1;HCoV-229E_surface;977 | 0 | 0 |
| NC_002645.1;NP_073551.1;HCoV-229E_surface;978 | 0 | 0 |
| NC_002645.1;NP_073551.1;HCoV-229E_surface;979 | 0 | 0 |
| NC_002645.1;NP_073551.1;HCoV-229E_surface;980 | 0 | 0 |
| NC_002645.1;NP_073551.1;HCoV-229E_surface;981 | 0 | 0 |
| NC_002645.1;NP_073551.1;HCoV-229E_surface;982 | 0 | 0 |
| NC_002645.1;NP_073551.1;HCoV-229E_surface;983 | 0 | 0 |
| NC_002645.1;NP_073551.1;HCoV-229E_surface;984 | 0 | 0 |
| NC_002645.1;NP_073551.1;HCoV-229E_surface;985 | 0 | 0 |
| NC_002645.1;NP_073551.1;HCoV-229E_surface;986 | 0 | 0 |
| NC_002645.1;NP_073551.1;HCoV-229E_surface;987 | 0 | 0 |
| NC_002645.1;NP_073551.1;HCoV-229E_surface;988 | 0 | 0 |
| NC_002645.1;NP_073551.1;HCoV-229E_surface;989 | 0 | 0 |
| NC_002645.1;NP_073551.1;HCoV-229E_surface;990 | 0 | 0 |

| | | |
|---|---|---|
| NC_002645.1;NP_073551.1;HCoV-229E_surface;991 | 0.05 | 0 |
| NC_002645.1;NP_073551.1;HCoV-229E_surface;992 | 0 | 0 |
| NC_002645.1;NP_073551.1;HCoV-229E_surface;993 | 0 | 0 |
| NC_002645.1;NP_073551.1;HCoV-229E_surface;994 | 0 | 0.025 |
| NC_002645.1;NP_073551.1;HCoV-229E_surface;995 | 0 | 0 |
| NC_002645.1;NP_073551.1;HCoV-229E_surface;996 | 0 | 0 |
| NC_002645.1;NP_073551.1;HCoV-229E_surface;997 | 0 | 0.025 |
| NC_002645.1;NP_073551.1;HCoV-229E_surface;998 | 0 | 0 |
| NC_002645.1;NP_073551.1;HCoV-229E_surface;999 | 0 | 0 |
| NC_002645.1;NP_073551.1;HCoV-229E_surface;1000 | 0 | 0 |
| NC_002645.1;NP_073551.1;HCoV-229E_surface;1001 | 0 | 0 |
| NC_002645.1;NP_073551.1;HCoV-229E_surface;1002 | 0 | 0 |
| NC_002645.1;NP_073551.1;HCoV-229E_surface;1003 | 0 | 0 |
| NC_002645.1;NP_073551.1;HCoV-229E_surface;1004 | 0 | 0 |
| NC_002645.1;NP_073551.1;HCoV-229E_surface;1005 | 0 | 0 |
| NC_002645.1;NP_073551.1;HCoV-229E_surface;1006 | 0 | 0 |
| NC_002645.1;NP_073551.1;HCoV-229E_surface;1007 | 0 | 0 |
| NC_002645.1;NP_073551.1;HCoV-229E_surface;1008 | 0 | 0 |
| NC_002645.1;NP_073551.1;HCoV-229E_surface;1009 | 0 | 0 |
| NC_002645.1;NP_073551.1;HCoV-229E_surface;1010 | 0.05 | 0 |
| NC_002645.1;NP_073551.1;HCoV-229E_surface;1011 | 0.05 | 0 |
| NC_002645.1;NP_073551.1;HCoV-229E_surface;1012 | 0.05 | 0 |
| NC_002645.1;NP_073551.1;HCoV-229E_surface;1013 | 0.05 | 0 |
| NC_002645.1;NP_073551.1;HCoV-229E_surface;1014 | 0 | 0 |
| NC_002645.1;NP_073551.1;HCoV-229E_surface;1015 | 0 | 0 |
| NC_002645.1;NP_073551.1;HCoV-229E_surface;1016 | 0 | 0 |
| NC_002645.1;NP_073551.1;HCoV-229E_surface;1017 | 0.05 | 0 |
| NC_002645.1;NP_073551.1;HCoV-229E_surface;1018 | 0.05 | 0.025 |
| NC_002645.1;NP_073551.1;HCoV-229E_surface;1019 | 0.05 | 0 |
| NC_002645.1;NP_073551.1;HCoV-229E_surface;1020 | 0 | 0 |
| NC_002645.1;NP_073551.1;HCoV-229E_surface;1021 | 0.05 | 0.025 |
| NC_002645.1;NP_073551.1;HCoV-229E_surface;1022 | 0.05 | 0 |
| NC_002645.1;NP_073551.1;HCoV-229E_surface;1023 | 0.05 | 0.025 |
| NC_002645.1;NP_073551.1;HCoV-229E_surface;1024 | 0.05 | 0.05 |
| NC_002645.1;NP_073551.1;HCoV-229E_surface;1025 | 0.05 | 0.025 |
| NC_002645.1;NP_073551.1;HCoV-229E_surface;1026 | 0.1 | 0.025 |
| NC_002645.1;NP_073551.1;HCoV-229E_surface;1027 | 0 | 0.025 |
| NC_002645.1;NP_073551.1;HCoV-229E_surface;1028 | 0 | 0.05 |
| NC_002645.1;NP_073551.1;HCoV-229E_surface;1029 | 0 | 0.05 |
| NC_002645.1;NP_073551.1;HCoV-229E_surface;1030 | 0 | 0.1 |
| NC_002645.1;NP_073551.1;HCoV-229E_surface;1031 | 0 | 0.05 |
| NC_002645.1;NP_073551.1;HCoV-229E_surface;1032 | 0.05 | 0.075 |
| NC_002645.1;NP_073551.1;HCoV-229E_surface;1033 | 0 | 0.05 |

| | | |
|---|---|---|
| NC_002645.1;NP_073551.1;HCoV-229E_surface;1034 | 0 | 0.05 |
| NC_002645.1;NP_073551.1;HCoV-229E_surface;1035 | 0 | 0.05 |
| NC_002645.1;NP_073551.1;HCoV-229E_surface;1036 | 0.05 | 0.075 |
| NC_002645.1;NP_073551.1;HCoV-229E_surface;1037 | 0 | 0 |
| NC_002645.1;NP_073551.1;HCoV-229E_surface;1038 | 0 | 0.025 |
| NC_002645.1;NP_073551.1;HCoV-229E_surface;1039 | 0.05 | 0.075 |
| NC_002645.1;NP_073551.1;HCoV-229E_surface;1040 | 0.1 | 0.05 |
| NC_002645.1;NP_073551.1;HCoV-229E_surface;1041 | 0.1 | 0.05 |
| NC_002645.1;NP_073551.1;HCoV-229E_surface;1042 | 0.1 | 0.05 |
| NC_002645.1;NP_073551.1;HCoV-229E_surface;1043 | 0.1 | 0.025 |
| NC_002645.1;NP_073551.1;HCoV-229E_surface;1044 | 0.25 | 0.05 |
| NC_002645.1;NP_073551.1;HCoV-229E_surface;1045 | 0.2 | 0.025 |
| NC_002645.1;NP_073551.1;HCoV-229E_surface;1046 | 0.35 | 0.075 |
| NC_002645.1;NP_073551.1;HCoV-229E_surface;1047 | 0.3 | 0.125 |
| NC_002645.1;NP_073551.1;HCoV-229E_surface;1048 | 0.3 | 0.15 |
| NC_002645.1;NP_073551.1;HCoV-229E_surface;1049 | 0.2 | 0.1 |
| NC_002645.1;NP_073551.1;HCoV-229E_surface;1050 | 0.15 | 0.025 |
| NC_002645.1;NP_073551.1;HCoV-229E_surface;1051 | 0.15 | 0.075 |
| NC_002645.1;NP_073551.1;HCoV-229E_surface;1052 | 0.1 | 0.025 |
| NC_002645.1;NP_073551.1;HCoV-229E_surface;1053 | 0.05 | 0 |
| NC_002645.1;NP_073551.1;HCoV-229E_surface;1054 | 0 | 0 |
| NC_002645.1;NP_073551.1;HCoV-229E_surface;1055 | 0 | 0.025 |
| NC_002645.1;NP_073551.1;HCoV-229E_surface;1056 | 0.05 | 0.025 |
| NC_002645.1;NP_073551.1;HCoV-229E_surface;1057 | 0.05 | 0.075 |
| NC_002645.1;NP_073551.1;HCoV-229E_surface;1058 | 0 | 0.025 |
| NC_002645.1;NP_073551.1;HCoV-229E_surface;1059 | 0 | 0 |
| NC_002645.1;NP_073551.1;HCoV-229E_surface;1060 | 0 | 0 |
| NC_002645.1;NP_073551.1;HCoV-229E_surface;1061 | 0 | 0 |
| NC_002645.1;NP_073551.1;HCoV-229E_surface;1062 | 0 | 0 |
| NC_002645.1;NP_073551.1;HCoV-229E_surface;1063 | 0 | 0 |
| NC_002645.1;NP_073551.1;HCoV-229E_surface;1064 | 0 | 0.025 |
| NC_002645.1;NP_073551.1;HCoV-229E_surface;1065 | 0 | 0 |
| NC_002645.1;NP_073551.1;HCoV-229E_surface;1066 | 0 | 0 |
| NC_002645.1;NP_073551.1;HCoV-229E_surface;1067 | 0 | 0.025 |
| NC_002645.1;NP_073551.1;HCoV-229E_surface;1068 | 0 | 0 |
| NC_002645.1;NP_073551.1;HCoV-229E_surface;1069 | 0 | 0 |
| NC_002645.1;NP_073551.1;HCoV-229E_surface;1070 | 0 | 0 |
| NC_002645.1;NP_073551.1;HCoV-229E_surface;1071 | 0 | 0.025 |
| NC_002645.1;NP_073551.1;HCoV-229E_surface;1072 | 0.05 | 0.05 |
| NC_002645.1;NP_073551.1;HCoV-229E_surface;1073 | 0.05 | 0.025 |
| NC_002645.1;NP_073551.1;HCoV-229E_surface;1074 | 0.05 | 0.05 |
| NC_002645.1;NP_073551.1;HCoV-229E_surface;1075 | 0.05 | 0.075 |
| NC_002645.1;NP_073551.1;HCoV-229E_surface;1076 | 0 | 0 |

| | | |
|---|---|---|
| NC_002645.1;NP_073551.1;HCoV-229E_surface;1077 | 0 | 0.025 |
| NC_002645.1;NP_073551.1;HCoV-229E_surface;1078 | 0 | 0 |
| NC_002645.1;NP_073551.1;HCoV-229E_surface;1079 | 0 | 0 |
| NC_002645.1;NP_073551.1;HCoV-229E_surface;1080 | 0 | 0 |
| NC_002645.1;NP_073551.1;HCoV-229E_surface;1081 | 0 | 0.025 |
| NC_002645.1;NP_073551.1;HCoV-229E_surface;1082 | 0 | 0.025 |
| NC_002645.1;NP_073551.1;HCoV-229E_surface;1083 | 0.05 | 0.05 |
| NC_002645.1;NP_073551.1;HCoV-229E_surface;1084 | 0.05 | 0.2 |
| NC_002645.1;NP_073551.1;HCoV-229E_surface;1085 | 0.05 | 0.125 |
| NC_002645.1;NP_073551.1;HCoV-229E_surface;1086 | 0.05 | 0.175 |
| NC_002645.1;NP_073551.1;HCoV-229E_surface;1087 | 0.05 | 0.075 |
| NC_002645.1;NP_073551.1;HCoV-229E_surface;1088 | 0 | 0.075 |
| NC_002645.1;NP_073551.1;HCoV-229E_surface;1089 | 0 | 0.075 |
| NC_002645.1;NP_073551.1;HCoV-229E_surface;1090 | 0 | 0.025 |
| NC_002645.1;NP_073551.1;HCoV-229E_surface;1091 | 0.15 | 0.025 |
| NC_002645.1;NP_073551.1;HCoV-229E_surface;1092 | 0.05 | 0.075 |
| NC_002645.1;NP_073551.1;HCoV-229E_surface;1093 | 0.05 | 0.05 |
| NC_002645.1;NP_073551.1;HCoV-229E_surface;1094 | 0.05 | 0.025 |
| NC_002645.1;NP_073551.1;HCoV-229E_surface;1095 | 0.05 | 0.075 |
| NC_002645.1;NP_073551.1;HCoV-229E_surface;1096 | 0 | 0 |
| NC_002645.1;NP_073551.1;HCoV-229E_surface;1097 | 0 | 0 |
| NC_002645.1;NP_073551.1;HCoV-229E_surface;1098 | 0.05 | 0.05 |
| NC_002645.1;NP_073551.1;HCoV-229E_surface;1099 | 0.1 | 0.025 |
| NC_002645.1;NP_073551.1;HCoV-229E_surface;1100 | 0 | 0 |
| NC_002645.1;NP_073551.1;HCoV-229E_surface;1101 | 0 | 0 |
| NC_002645.1;NP_073551.1;HCoV-229E_surface;1102 | 0 | 0 |
| NC_002645.1;NP_073551.1;HCoV-229E_surface;1103 | 0 | 0 |
| NC_002645.1;NP_073551.1;HCoV-229E_surface;1104 | 0 | 0 |
| NC_002645.1;NP_073551.1;HCoV-229E_surface;1105 | 0 | 0 |
| NC_002645.1;NP_073551.1;HCoV-229E_surface;1106 | 0.05 | 0 |
| NC_002645.1;NP_073551.1;HCoV-229E_surface;1107 | 0 | 0 |
| NC_002645.1;NP_073551.1;HCoV-229E_surface;1108 | 0 | 0.025 |
| NC_002645.1;NP_073551.1;HCoV-229E_surface;1109 | 0 | 0.025 |
| NC_002645.1;NP_073551.1;HCoV-229E_surface;1110 | 0 | 0 |
| NC_002645.1;NP_073551.1;HCoV-229E_surface;1111 | 0 | 0 |
| NC_002645.1;NP_073551.1;HCoV-229E_surface;1112 | 0 | 0 |
| NC_002645.1;NP_073551.1;HCoV-229E_surface;1113 | 0 | 0 |
| NC_002645.1;NP_073551.1;HCoV-229E_surface;1114 | 0 | 0 |
| NC_002645.1;NP_073551.1;HCoV-229E_surface;1115 | 0 | 0 |
| NC_002645.1;NP_073551.1;HCoV-229E_surface;1116 | 0 | 0 |
| NC_002645.1;NP_073551.1;HCoV-229E_surface;1117 | 0 | 0 |
| NC_002645.1;NP_073551.1;HCoV-229E_surface;1118 | 0 | 0 |
| NC_002645.1;NP_073551.1;HCoV-229E_surface;1119 | 0 | 0 |

| | | |
|---|---|---|
| NC_002645.1;NP_073551.1;HCoV-229E_surface;1120 | 0 | 0 |
| NC_002645.1;NP_073551.1;HCoV-229E_surface;1121 | 0 | 0 |
| NC_002645.1;NP_073551.1;HCoV-229E_surface;1122 | 0 | 0 |
| NC_002645.1;NP_073551.1;HCoV-229E_surface;1123 | 0 | 0 |
| NC_002645.1;NP_073551.1;HCoV-229E_surface;1124 | 0 | 0 |
| NC_002645.1;NP_073551.1;HCoV-229E_surface;1125 | 0 | 0 |
| NC_002645.1;NP_073551.1;HCoV-229E_surface;1126 | 0 | 0 |
| NC_002645.1;NP_073551.1;HCoV-229E_surface;1127 | 0 | 0 |
| NC_002645.1;NP_073551.1;HCoV-229E_surface;1128 | 0 | 0 |
| NC_002645.1;NP_073551.1;HCoV-229E_surface;1129 | 0 | 0 |
| NC_002645.1;NP_073551.1;HCoV-229E_surface;1130 | 0 | 0 |
| NC_002645.1;NP_073551.1;HCoV-229E_surface;1131 | 0 | 0 |
| NC_002645.1;NP_073551.1;HCoV-229E_surface;1132 | 0 | 0 |
| NC_002645.1;NP_073551.1;HCoV-229E_surface;1133 | 0 | 0 |
| NC_002645.1;NP_073551.1;HCoV-229E_surface;1134 | 0 | 0 |
| NC_002645.1;NP_073551.1;HCoV-229E_surface;1135 | 0 | 0 |
| NC_002645.1;NP_073551.1;HCoV-229E_surface;1136 | 0 | 0 |
| NC_002645.1;NP_073551.1;HCoV-229E_surface;1137 | 0 | 0 |
| NC_002645.1;NP_073551.1;HCoV-229E_surface;1138 | 0 | 0 |
| NC_002645.1;NP_073551.1;HCoV-229E_surface;1139 | 0 | 0 |
| NC_002645.1;NP_073551.1;HCoV-229E_surface;1140 | 0 | 0 |
| NC_002645.1;NP_073551.1;HCoV-229E_surface;1141 | 0 | 0 |
| NC_002645.1;NP_073551.1;HCoV-229E_surface;1142 | 0 | 0 |
| NC_002645.1;NP_073551.1;HCoV-229E_surface;1143 | 0 | 0 |
| NC_002645.1;NP_073551.1;HCoV-229E_surface;1144 | 0 | 0 |
| NC_002645.1;NP_073551.1;HCoV-229E_surface;1145 | 0 | 0 |
| NC_002645.1;NP_073551.1;HCoV-229E_surface;1146 | 0 | 0 |
| NC_002645.1;NP_073551.1;HCoV-229E_surface;1147 | 0 | 0 |
| NC_002645.1;NP_073551.1;HCoV-229E_surface;1148 | 0 | 0 |
| NC_002645.1;NP_073551.1;HCoV-229E_surface;1149 | 0 | 0 |
| NC_002645.1;NP_073551.1;HCoV-229E_surface;1150 | 0 | 0.05 |
| NC_002645.1;NP_073551.1;HCoV-229E_surface;1151 | 0 | 0.15 |
| NC_002645.1;NP_073551.1;HCoV-229E_surface;1152 | 0 | 0.025 |
| NC_002645.1;NP_073551.1;HCoV-229E_surface;1153 | 0 | 0.05 |
| NC_002645.1;NP_073551.1;HCoV-229E_surface;1154 | 0.05 | 0.075 |
| NC_002645.1;NP_073551.1;HCoV-229E_surface;1155 | 0.05 | 0.075 |
| NC_002645.1;NP_073551.1;HCoV-229E_surface;1156 | 0.05 | 0.05 |
| NC_002645.1;NP_073551.1;HCoV-229E_surface;1157 | 0.05 | 0.05 |
| NC_002645.1;NP_073551.1;HCoV-229E_surface;1158 | 0.05 | 0.2 |
| NC_002645.1;NP_073551.1;HCoV-229E_surface;1159 | 0.05 | 0.025 |
| NC_002645.1;NP_073551.1;HCoV-229E_surface;1160 | 0.05 | 0.05 |
| NC_002645.1;NP_073551.1;HCoV-229E_surface;1161 | 0.1 | 0.05 |
| NC_002645.1;NP_073551.1;HCoV-229E_surface;1162 | 0 | 0.025 |

| | | |
|---|---|---|
| NC_002645.1;NP_073552.1;HCoV-229E_4a;1 | 0 | 0 |
| NC_002645.1;NP_073552.1;HCoV-229E_4a;2 | 0 | 0 |
| NC_002645.1;NP_073552.1;HCoV-229E_4a;3 | 0 | 0 |
| NC_002645.1;NP_073552.1;HCoV-229E_4a;4 | 0.05 | 0.025 |
| NC_002645.1;NP_073552.1;HCoV-229E_4a;5 | 0.1 | 0.075 |
| NC_002645.1;NP_073552.1;HCoV-229E_4a;6 | 0.1 | 0.1 |
| NC_002645.1;NP_073552.1;HCoV-229E_4a;7 | 0 | 0.025 |
| NC_002645.1;NP_073552.1;HCoV-229E_4a;8 | 0.05 | 0.025 |
| NC_002645.1;NP_073552.1;HCoV-229E_4a;9 | 0 | 0.075 |
| NC_002645.1;NP_073552.1;HCoV-229E_4a;10 | 0.1 | 0.075 |
| NC_002645.1;NP_073552.1;HCoV-229E_4a;11 | 0.05 | 0.125 |
| NC_002645.1;NP_073552.1;HCoV-229E_4a;12 | 0.05 | 0.075 |
| NC_002645.1;NP_073552.1;HCoV-229E_4a;13 | 0.05 | 0.1 |
| NC_002645.1;NP_073552.1;HCoV-229E_4a;14 | 0 | 0.075 |
| NC_002645.1;NP_073552.1;HCoV-229E_4a;15 | 0 | 0.075 |
| NC_002645.1;NP_073552.1;HCoV-229E_4a;16 | 0 | 0.025 |
| NC_002645.1;NP_073552.1;HCoV-229E_4a;17 | 0 | 0.05 |
| NC_002645.1;NP_073552.1;HCoV-229E_4a;18 | 0.05 | 0.05 |
| NC_002645.1;NP_073552.1;HCoV-229E_4a;19 | 0.1 | 0.075 |
| NC_002645.1;NP_073552.1;HCoV-229E_4a;20 | 0.15 | 0.1 |
| NC_002645.1;NP_073552.1;HCoV-229E_4a;21 | 0.15 | 0.125 |
| NC_002645.1;NP_073552.1;HCoV-229E_4a;22 | 0.2 | 0.075 |
| NC_002645.1;NP_073552.1;HCoV-229E_4a;23 | 0.15 | 0.075 |
| NC_002645.1;NP_073552.1;HCoV-229E_4a;24 | 0.15 | 0.1 |
| NC_002645.1;NP_073552.1;HCoV-229E_4a;25 | 0.15 | 0.05 |
| NC_002645.1;NP_073552.1;HCoV-229E_4a;26 | 0.15 | 0.075 |
| NC_002645.1;NP_073552.1;HCoV-229E_4a;27 | 0.15 | 0.05 |
| NC_002645.1;NP_073552.1;HCoV-229E_4a;28 | 0.2 | 0.05 |
| NC_002645.1;NP_073552.1;HCoV-229E_4a;29 | 0.2 | 0.05 |
| NC_002645.1;NP_073552.1;HCoV-229E_4a;30 | 0.05 | 0 |
| NC_002645.1;NP_073552.1;HCoV-229E_4a;31 | 0.1 | 0.025 |
| NC_002645.1;NP_073552.1;HCoV-229E_4a;32 | 0.1 | 0.025 |
| NC_002645.1;NP_073552.1;HCoV-229E_4a;33 | 0 | 0 |
| NC_002645.1;NP_073552.1;HCoV-229E_4a;34 | 0 | 0 |
| NC_002645.1;NP_073552.1;HCoV-229E_4a;35 | 0 | 0 |
| NC_002645.1;NP_073552.1;HCoV-229E_4a;36 | 0 | 0 |
| NC_002645.1;NP_073552.1;HCoV-229E_4a;37 | 0 | 0 |
| NC_002645.1;NP_073552.1;HCoV-229E_4a;38 | 0 | 0 |
| NC_002645.1;NP_073552.1;HCoV-229E_4a;39 | 0 | 0 |
| NC_002645.1;NP_073552.1;HCoV-229E_4a;40 | 0 | 0 |
| NC_002645.1;NP_073552.1;HCoV-229E_4a;41 | 0 | 0 |
| NC_002645.1;NP_073552.1;HCoV-229E_4a;42 | 0 | 0 |
| NC_002645.1;NP_073552.1;HCoV-229E_4a;43 | 0 | 0 |

| | | |
|---|---|---|
| NC_002645.1;NP_073552.1;HCoV-229E_4a;44 | 0 | 0 |
| NC_002645.1;NP_073552.1;HCoV-229E_4a;45 | 0 | 0 |
| NC_002645.1;NP_073552.1;HCoV-229E_4a;46 | 0 | 0 |
| NC_002645.1;NP_073552.1;HCoV-229E_4a;47 | 0 | 0 |
| NC_002645.1;NP_073552.1;HCoV-229E_4a;48 | 0 | 0 |
| NC_002645.1;NP_073552.1;HCoV-229E_4a;49 | 0 | 0 |
| NC_002645.1;NP_073552.1;HCoV-229E_4a;50 | 0 | 0 |
| NC_002645.1;NP_073552.1;HCoV-229E_4a;51 | 0 | 0 |
| NC_002645.1;NP_073552.1;HCoV-229E_4a;52 | 0 | 0 |
| NC_002645.1;NP_073552.1;HCoV-229E_4a;53 | 0 | 0 |
| NC_002645.1;NP_073552.1;HCoV-229E_4a;54 | 0 | 0 |
| NC_002645.1;NP_073552.1;HCoV-229E_4a;55 | 0 | 0 |
| NC_002645.1;NP_073552.1;HCoV-229E_4a;56 | 0 | 0 |
| NC_002645.1;NP_073552.1;HCoV-229E_4a;57 | 0 | 0 |
| NC_002645.1;NP_073552.1;HCoV-229E_4a;58 | 0 | 0 |
| NC_002645.1;NP_073552.1;HCoV-229E_4a;59 | 0 | 0 |
| NC_002645.1;NP_073552.1;HCoV-229E_4a;60 | 0 | 0 |
| NC_002645.1;NP_073552.1;HCoV-229E_4a;61 | 0 | 0 |
| NC_002645.1;NP_073552.1;HCoV-229E_4a;62 | 0 | 0 |
| NC_002645.1;NP_073552.1;HCoV-229E_4a;63 | 0 | 0 |
| NC_002645.1;NP_073552.1;HCoV-229E_4a;64 | 0 | 0 |
| NC_002645.1;NP_073552.1;HCoV-229E_4a;65 | 0 | 0 |
| NC_002645.1;NP_073552.1;HCoV-229E_4a;66 | 0 | 0 |
| NC_002645.1;NP_073552.1;HCoV-229E_4a;67 | 0 | 0 |
| NC_002645.1;NP_073552.1;HCoV-229E_4a;68 | 0 | 0 |
| NC_002645.1;NP_073552.1;HCoV-229E_4a;69 | 0 | 0 |
| NC_002645.1;NP_073552.1;HCoV-229E_4a;70 | 0 | 0 |
| NC_002645.1;NP_073552.1;HCoV-229E_4a;71 | 0 | 0 |
| NC_002645.1;NP_073552.1;HCoV-229E_4a;72 | 0 | 0 |
| NC_002645.1;NP_073552.1;HCoV-229E_4a;73 | 0 | 0 |
| NC_002645.1;NP_073552.1;HCoV-229E_4a;74 | 0 | 0 |
| NC_002645.1;NP_073552.1;HCoV-229E_4a;75 | 0 | 0 |
| NC_002645.1;NP_073552.1;HCoV-229E_4a;76 | 0 | 0 |
| NC_002645.1;NP_073552.1;HCoV-229E_4a;77 | 0 | 0 |
| NC_002645.1;NP_073552.1;HCoV-229E_4a;78 | 0 | 0 |
| NC_002645.1;NP_073552.1;HCoV-229E_4a;79 | 0 | 0 |
| NC_002645.1;NP_073552.1;HCoV-229E_4a;80 | 0 | 0 |
| NC_002645.1;NP_073552.1;HCoV-229E_4a;81 | 0 | 0 |
| NC_002645.1;NP_073552.1;HCoV-229E_4a;82 | 0 | 0 |
| NC_002645.1;NP_073552.1;HCoV-229E_4a;83 | 0 | 0 |
| NC_002645.1;NP_073552.1;HCoV-229E_4a;84 | 0 | 0 |
| NC_002645.1;NP_073552.1;HCoV-229E_4a;85 | 0 | 0 |
| NC_002645.1;NP_073552.1;HCoV-229E_4a;86 | 0 | 0 |

| | | |
|---|---|---|
| NC_002645.1;NP_073552.1;HCoV-229E_4a;87 | 0 | 0 |
| NC_002645.1;NP_073552.1;HCoV-229E_4a;88 | 0 | 0 |
| NC_002645.1;NP_073552.1;HCoV-229E_4a;89 | 0 | 0 |
| NC_002645.1;NP_073552.1;HCoV-229E_4a;90 | 0 | 0 |
| NC_002645.1;NP_073552.1;HCoV-229E_4a;91 | 0 | 0 |
| NC_002645.1;NP_073552.1;HCoV-229E_4a;92 | 0 | 0 |
| NC_002645.1;NP_073552.1;HCoV-229E_4a;93 | 0 | 0 |
| NC_002645.1;NP_073552.1;HCoV-229E_4a;94 | 0 | 0 |
| NC_002645.1;NP_073552.1;HCoV-229E_4a;95 | 0 | 0 |
| NC_002645.1;NP_073552.1;HCoV-229E_4a;96 | 0 | 0 |
| NC_002645.1;NP_073552.1;HCoV-229E_4a;97 | 0 | 0 |
| NC_002645.1;NP_073552.1;HCoV-229E_4a;98 | 0 | 0 |
| NC_002645.1;NP_073552.1;HCoV-229E_4a;99 | 0 | 0 |
| NC_002645.1;NP_073552.1;HCoV-229E_4a;100 | 0 | 0 |
| NC_002645.1;NP_073552.1;HCoV-229E_4a;101 | 0 | 0 |
| NC_002645.1;NP_073552.1;HCoV-229E_4a;102 | 0 | 0 |
| NC_002645.1;NP_073552.1;HCoV-229E_4a;103 | 0 | 0 |
| NC_002645.1;NP_073552.1;HCoV-229E_4a;104 | 0 | 0 |
| NC_002645.1;NP_073552.1;HCoV-229E_4a;105 | 0 | 0 |
| NC_002645.1;NP_073552.1;HCoV-229E_4a;106 | 0 | 0 |
| NC_002645.1;NP_073552.1;HCoV-229E_4a;107 | 0 | 0 |
| NC_002645.1;NP_073552.1;HCoV-229E_4a;108 | 0 | 0 |
| NC_002645.1;NP_073552.1;HCoV-229E_4a;109 | 0 | 0 |
| NC_002645.1;NP_073552.1;HCoV-229E_4a;110 | 0 | 0 |
| NC_002645.1;NP_073552.1;HCoV-229E_4a;111 | 0 | 0 |
| NC_002645.1;NP_073552.1;HCoV-229E_4a;112 | 0 | 0 |
| NC_002645.1;NP_073552.1;HCoV-229E_4a;113 | 0 | 0 |
| NC_002645.1;NP_073552.1;HCoV-229E_4a;114 | 0 | 0 |
| NC_002645.1;NP_073552.1;HCoV-229E_4a;115 | 0 | 0 |
| NC_002645.1;NP_073552.1;HCoV-229E_4a;116 | 0 | 0 |
| NC_002645.1;NP_073552.1;HCoV-229E_4a;117 | 0 | 0 |
| NC_002645.1;NP_073552.1;HCoV-229E_4a;118 | 0 | 0 |
| NC_002645.1;NP_073552.1;HCoV-229E_4a;119 | 0 | 0 |
| NC_002645.1;NP_073552.1;HCoV-229E_4a;120 | 0 | 0 |
| NC_002645.1;NP_073552.1;HCoV-229E_4a;121 | 0 | 0 |
| NC_002645.1;NP_073552.1;HCoV-229E_4a;122 | 0 | 0 |
| NC_002645.1;NP_073553.1;HCoV-229E_4b;1 | 0 | 0 |
| NC_002645.1;NP_073553.1;HCoV-229E_4b;2 | 0 | 0 |
| NC_002645.1;NP_073553.1;HCoV-229E_4b;3 | 0 | 0 |
| NC_002645.1;NP_073553.1;HCoV-229E_4b;4 | 0 | 0 |
| NC_002645.1;NP_073553.1;HCoV-229E_4b;5 | 0 | 0 |
| NC_002645.1;NP_073553.1;HCoV-229E_4b;6 | 0 | 0 |
| NC_002645.1;NP_073553.1;HCoV-229E_4b;7 | 0 | 0 |

| | | |
|---|---|---|
| NC_002645.1;NP_073553.1;HCoV-229E_4b;8 | 0 | 0 |
| NC_002645.1;NP_073553.1;HCoV-229E_4b;9 | 0 | 0 |
| NC_002645.1;NP_073553.1;HCoV-229E_4b;10 | 0 | 0 |
| NC_002645.1;NP_073553.1;HCoV-229E_4b;11 | 0 | 0 |
| NC_002645.1;NP_073553.1;HCoV-229E_4b;12 | 0 | 0 |
| NC_002645.1;NP_073553.1;HCoV-229E_4b;13 | 0 | 0 |
| NC_002645.1;NP_073553.1;HCoV-229E_4b;14 | 0 | 0 |
| NC_002645.1;NP_073553.1;HCoV-229E_4b;15 | 0 | 0 |
| NC_002645.1;NP_073553.1;HCoV-229E_4b;16 | 0 | 0 |
| NC_002645.1;NP_073553.1;HCoV-229E_4b;17 | 0 | 0 |
| NC_002645.1;NP_073553.1;HCoV-229E_4b;18 | 0 | 0 |
| NC_002645.1;NP_073553.1;HCoV-229E_4b;19 | 0 | 0 |
| NC_002645.1;NP_073553.1;HCoV-229E_4b;20 | 0 | 0 |
| NC_002645.1;NP_073553.1;HCoV-229E_4b;21 | 0 | 0 |
| NC_002645.1;NP_073553.1;HCoV-229E_4b;22 | 0 | 0 |
| NC_002645.1;NP_073553.1;HCoV-229E_4b;23 | 0 | 0 |
| NC_002645.1;NP_073553.1;HCoV-229E_4b;24 | 0 | 0 |
| NC_002645.1;NP_073553.1;HCoV-229E_4b;25 | 0 | 0 |
| NC_002645.1;NP_073553.1;HCoV-229E_4b;26 | 0 | 0 |
| NC_002645.1;NP_073553.1;HCoV-229E_4b;27 | 0 | 0 |
| NC_002645.1;NP_073553.1;HCoV-229E_4b;28 | 0 | 0 |
| NC_002645.1;NP_073553.1;HCoV-229E_4b;29 | 0 | 0 |
| NC_002645.1;NP_073553.1;HCoV-229E_4b;30 | 0 | 0 |
| NC_002645.1;NP_073553.1;HCoV-229E_4b;31 | 0 | 0 |
| NC_002645.1;NP_073553.1;HCoV-229E_4b;32 | 0 | 0 |
| NC_002645.1;NP_073553.1;HCoV-229E_4b;33 | 0 | 0 |
| NC_002645.1;NP_073553.1;HCoV-229E_4b;34 | 0 | 0 |
| NC_002645.1;NP_073553.1;HCoV-229E_4b;35 | 0 | 0 |
| NC_002645.1;NP_073553.1;HCoV-229E_4b;36 | 0 | 0.025 |
| NC_002645.1;NP_073553.1;HCoV-229E_4b;37 | 0.05 | 0.1 |
| NC_002645.1;NP_073553.1;HCoV-229E_4b;38 | 0 | 0.025 |
| NC_002645.1;NP_073553.1;HCoV-229E_4b;39 | 0 | 0.025 |
| NC_002645.1;NP_073553.1;HCoV-229E_4b;40 | 0.05 | 0.025 |
| NC_002645.1;NP_073553.1;HCoV-229E_4b;41 | 0 | 0.075 |
| NC_002645.1;NP_073553.1;HCoV-229E_4b;42 | 0 | 0.025 |
| NC_002645.1;NP_073553.1;HCoV-229E_4b;43 | 0 | 0.025 |
| NC_002645.1;NP_073553.1;HCoV-229E_4b;44 | 0 | 0 |
| NC_002645.1;NP_073553.1;HCoV-229E_4b;45 | 0 | 0.025 |
| NC_002645.1;NP_073553.1;HCoV-229E_4b;46 | 0 | 0.025 |
| NC_002645.1;NP_073553.1;HCoV-229E_4b;47 | 0 | 0 |
| NC_002645.1;NP_073553.1;HCoV-229E_4b;48 | 0 | 0 |
| NC_002645.1;NP_073553.1;HCoV-229E_4b;49 | 0 | 0 |
| NC_002645.1;NP_073553.1;HCoV-229E_4b;50 | 0 | 0 |

| | | |
|---|---|---|
| NC_002645.1;NP_073553.1;HCoV-229E_4b;51 | 0 | 0 |
| NC_002645.1;NP_073553.1;HCoV-229E_4b;52 | 0 | 0 |
| NC_002645.1;NP_073553.1;HCoV-229E_4b;53 | 0 | 0 |
| NC_002645.1;NP_073553.1;HCoV-229E_4b;54 | 0 | 0 |
| NC_002645.1;NP_073553.1;HCoV-229E_4b;55 | 0 | 0 |
| NC_002645.1;NP_073553.1;HCoV-229E_4b;56 | 0 | 0 |
| NC_002645.1;NP_073553.1;HCoV-229E_4b;57 | 0 | 0 |
| NC_002645.1;NP_073553.1;HCoV-229E_4b;58 | 0 | 0 |
| NC_002645.1;NP_073553.1;HCoV-229E_4b;59 | 0 | 0 |
| NC_002645.1;NP_073553.1;HCoV-229E_4b;60 | 0 | 0 |
| NC_002645.1;NP_073553.1;HCoV-229E_4b;61 | 0 | 0 |
| NC_002645.1;NP_073553.1;HCoV-229E_4b;62 | 0 | 0 |
| NC_002645.1;NP_073553.1;HCoV-229E_4b;63 | 0 | 0 |
| NC_002645.1;NP_073553.1;HCoV-229E_4b;64 | 0 | 0 |
| NC_002645.1;NP_073553.1;HCoV-229E_4b;65 | 0 | 0 |
| NC_002645.1;NP_073553.1;HCoV-229E_4b;66 | 0 | 0 |
| NC_002645.1;NP_073553.1;HCoV-229E_4b;67 | 0 | 0 |
| NC_002645.1;NP_073553.1;HCoV-229E_4b;68 | 0 | 0 |
| NC_002645.1;NP_073553.1;HCoV-229E_4b;69 | 0.05 | 0.025 |
| NC_002645.1;NP_073553.1;HCoV-229E_4b;70 | 0.05 | 0.025 |
| NC_002645.1;NP_073553.1;HCoV-229E_4b;71 | 0.05 | 0.025 |
| NC_002645.1;NP_073553.1;HCoV-229E_4b;72 | 0.05 | 0.025 |
| NC_002645.1;NP_073553.1;HCoV-229E_4b;73 | 0.05 | 0 |
| NC_002645.1;NP_073553.1;HCoV-229E_4b;74 | 0.05 | 0 |
| NC_002645.1;NP_073553.1;HCoV-229E_4b;75 | 0.05 | 0.025 |
| NC_002645.1;NP_073553.1;HCoV-229E_4b;76 | 0.05 | 0.025 |
| NC_002645.1;NP_073553.1;HCoV-229E_4b;77 | 0.05 | 0.025 |
| NC_002645.1;NP_073554.1;HCoV-229E_envelope;1 | 0 | 0 |
| NC_002645.1;NP_073554.1;HCoV-229E_envelope;2 | 0 | 0 |
| NC_002645.1;NP_073554.1;HCoV-229E_envelope;3 | 0 | 0 |
| NC_002645.1;NP_073554.1;HCoV-229E_envelope;4 | 0 | 0 |
| NC_002645.1;NP_073554.1;HCoV-229E_envelope;5 | 0 | 0 |
| NC_002645.1;NP_073554.1;HCoV-229E_envelope;6 | 0 | 0 |
| NC_002645.1;NP_073554.1;HCoV-229E_envelope;7 | 0 | 0 |
| NC_002645.1;NP_073554.1;HCoV-229E_envelope;8 | 0 | 0 |
| NC_002645.1;NP_073554.1;HCoV-229E_envelope;9 | 0 | 0 |
| NC_002645.1;NP_073554.1;HCoV-229E_envelope;10 | 0 | 0 |
| NC_002645.1;NP_073554.1;HCoV-229E_envelope;11 | 0 | 0 |
| NC_002645.1;NP_073554.1;HCoV-229E_envelope;12 | 0 | 0 |
| NC_002645.1;NP_073554.1;HCoV-229E_envelope;13 | 0 | 0 |
| NC_002645.1;NP_073554.1;HCoV-229E_envelope;14 | 0 | 0 |
| NC_002645.1;NP_073554.1;HCoV-229E_envelope;15 | 0 | 0 |
| NC_002645.1;NP_073554.1;HCoV-229E_envelope;16 | 0 | 0 |

| | | |
|---|---|---|
| NC_002645.1;NP_073554.1;HCoV-229E_envelope;17 | 0 | 0 |
| NC_002645.1;NP_073554.1;HCoV-229E_envelope;18 | 0 | 0 |
| NC_002645.1;NP_073554.1;HCoV-229E_envelope;19 | 0 | 0 |
| NC_002645.1;NP_073554.1;HCoV-229E_envelope;20 | 0 | 0 |
| NC_002645.1;NP_073554.1;HCoV-229E_envelope;21 | 0 | 0 |
| NC_002645.1;NP_073554.1;HCoV-229E_envelope;22 | 0 | 0 |
| NC_002645.1;NP_073554.1;HCoV-229E_envelope;23 | 0 | 0 |
| NC_002645.1;NP_073554.1;HCoV-229E_envelope;24 | 0 | 0 |
| NC_002645.1;NP_073554.1;HCoV-229E_envelope;25 | 0 | 0 |
| NC_002645.1;NP_073554.1;HCoV-229E_envelope;26 | 0 | 0 |
| NC_002645.1;NP_073554.1;HCoV-229E_envelope;27 | 0 | 0 |
| NC_002645.1;NP_073554.1;HCoV-229E_envelope;28 | 0 | 0 |
| NC_002645.1;NP_073554.1;HCoV-229E_envelope;29 | 0 | 0 |
| NC_002645.1;NP_073554.1;HCoV-229E_envelope;30 | 0 | 0 |
| NC_002645.1;NP_073554.1;HCoV-229E_envelope;31 | 0 | 0 |
| NC_002645.1;NP_073554.1;HCoV-229E_envelope;32 | 0 | 0 |
| NC_002645.1;NP_073554.1;HCoV-229E_envelope;33 | 0 | 0 |
| NC_002645.1;NP_073554.1;HCoV-229E_envelope;34 | 0 | 0 |
| NC_002645.1;NP_073554.1;HCoV-229E_envelope;35 | 0 | 0 |
| NC_002645.1;NP_073554.1;HCoV-229E_envelope;36 | 0 | 0 |
| NC_002645.1;NP_073554.1;HCoV-229E_envelope;37 | 0 | 0 |
| NC_002645.1;NP_073554.1;HCoV-229E_envelope;38 | 0 | 0 |
| NC_002645.1;NP_073554.1;HCoV-229E_envelope;39 | 0 | 0 |
| NC_002645.1;NP_073554.1;HCoV-229E_envelope;40 | 0 | 0 |
| NC_002645.1;NP_073554.1;HCoV-229E_envelope;41 | 0 | 0 |
| NC_002645.1;NP_073554.1;HCoV-229E_envelope;42 | 0 | 0.025 |
| NC_002645.1;NP_073554.1;HCoV-229E_envelope;43 | 0 | 0 |
| NC_002645.1;NP_073554.1;HCoV-229E_envelope;44 | 0 | 0 |
| NC_002645.1;NP_073554.1;HCoV-229E_envelope;45 | 0 | 0 |
| NC_002645.1;NP_073554.1;HCoV-229E_envelope;46 | 0 | 0 |
| NC_002645.1;NP_073554.1;HCoV-229E_envelope;47 | 0 | 0 |
| NC_002645.1;NP_073554.1;HCoV-229E_envelope;48 | 0 | 0 |
| NC_002645.1;NP_073554.1;HCoV-229E_envelope;49 | 0 | 0 |
| NC_002645.1;NP_073554.1;HCoV-229E_envelope;50 | 0 | 0 |
| NC_002645.1;NP_073554.1;HCoV-229E_envelope;51 | 0 | 0 |
| NC_002645.1;NP_073554.1;HCoV-229E_envelope;52 | 0 | 0 |
| NC_002645.1;NP_073554.1;HCoV-229E_envelope;53 | 0 | 0 |
| NC_002645.1;NP_073554.1;HCoV-229E_envelope;54 | 0 | 0 |
| NC_002645.1;NP_073554.1;HCoV-229E_envelope;55 | 0 | 0 |
| NC_002645.1;NP_073554.1;HCoV-229E_envelope;56 | 0 | 0 |
| NC_002645.1;NP_073554.1;HCoV-229E_envelope;57 | 0 | 0 |
| NC_002645.1;NP_073554.1;HCoV-229E_envelope;58 | 0 | 0 |
| NC_002645.1;NP_073554.1;HCoV-229E_envelope;59 | 0 | 0 |

| | | |
|---|---|---|
| NC_002645.1;NP_073554.1;HCoV-229E_envelope;60 | 0 | 0 |
| NC_002645.1;NP_073554.1;HCoV-229E_envelope;61 | 0 | 0 |
| NC_002645.1;NP_073554.1;HCoV-229E_envelope;62 | 0 | 0 |
| NC_002645.1;NP_073554.1;HCoV-229E_envelope;63 | 0 | 0 |
| NC_002645.1;NP_073554.1;HCoV-229E_envelope;64 | 0 | 0 |
| NC_002645.1;NP_073554.1;HCoV-229E_envelope;65 | 0 | 0.025 |
| NC_002645.1;NP_073554.1;HCoV-229E_envelope;66 | 0 | 0 |
| NC_002645.1;NP_073555.1;HCoV-229E_membrane;1 | 0 | 0 |
| NC_002645.1;NP_073555.1;HCoV-229E_membrane;2 | 0 | 0.05 |
| NC_002645.1;NP_073555.1;HCoV-229E_membrane;3 | 0 | 0 |
| NC_002645.1;NP_073555.1;HCoV-229E_membrane;4 | 0 | 0 |
| NC_002645.1;NP_073555.1;HCoV-229E_membrane;5 | 0 | 0 |
| NC_002645.1;NP_073555.1;HCoV-229E_membrane;6 | 0 | 0 |
| NC_002645.1;NP_073555.1;HCoV-229E_membrane;7 | 0 | 0 |
| NC_002645.1;NP_073555.1;HCoV-229E_membrane;8 | 0 | 0 |
| NC_002645.1;NP_073555.1;HCoV-229E_membrane;9 | 0 | 0 |
| NC_002645.1;NP_073555.1;HCoV-229E_membrane;10 | 0 | 0 |
| NC_002645.1;NP_073555.1;HCoV-229E_membrane;11 | 0 | 0 |
| NC_002645.1;NP_073555.1;HCoV-229E_membrane;12 | 0 | 0 |
| NC_002645.1;NP_073555.1;HCoV-229E_membrane;13 | 0 | 0 |
| NC_002645.1;NP_073555.1;HCoV-229E_membrane;14 | 0 | 0 |
| NC_002645.1;NP_073555.1;HCoV-229E_membrane;15 | 0 | 0 |
| NC_002645.1;NP_073555.1;HCoV-229E_membrane;16 | 0 | 0 |
| NC_002645.1;NP_073555.1;HCoV-229E_membrane;17 | 0 | 0 |
| NC_002645.1;NP_073555.1;HCoV-229E_membrane;18 | 0 | 0 |
| NC_002645.1;NP_073555.1;HCoV-229E_membrane;19 | 0 | 0 |
| NC_002645.1;NP_073555.1;HCoV-229E_membrane;20 | 0 | 0 |
| NC_002645.1;NP_073555.1;HCoV-229E_membrane;21 | 0 | 0 |
| NC_002645.1;NP_073555.1;HCoV-229E_membrane;22 | 0 | 0 |
| NC_002645.1;NP_073555.1;HCoV-229E_membrane;23 | 0 | 0 |
| NC_002645.1;NP_073555.1;HCoV-229E_membrane;24 | 0 | 0 |
| NC_002645.1;NP_073555.1;HCoV-229E_membrane;25 | 0 | 0 |
| NC_002645.1;NP_073555.1;HCoV-229E_membrane;26 | 0 | 0 |
| NC_002645.1;NP_073555.1;HCoV-229E_membrane;27 | 0 | 0 |
| NC_002645.1;NP_073555.1;HCoV-229E_membrane;28 | 0 | 0 |
| NC_002645.1;NP_073555.1;HCoV-229E_membrane;29 | 0 | 0 |
| NC_002645.1;NP_073555.1;HCoV-229E_membrane;30 | 0 | 0 |
| NC_002645.1;NP_073555.1;HCoV-229E_membrane;31 | 0 | 0 |
| NC_002645.1;NP_073555.1;HCoV-229E_membrane;32 | 0 | 0 |
| NC_002645.1;NP_073555.1;HCoV-229E_membrane;33 | 0 | 0 |
| NC_002645.1;NP_073555.1;HCoV-229E_membrane;34 | 0 | 0 |
| NC_002645.1;NP_073555.1;HCoV-229E_membrane;35 | 0 | 0 |
| NC_002645.1;NP_073555.1;HCoV-229E_membrane;36 | 0 | 0 |

| | | |
|---|---|---|
| NC_002645.1;NP_073555.1;HCoV-229E_membrane;37 | 0 | 0 |
| NC_002645.1;NP_073555.1;HCoV-229E_membrane;38 | 0 | 0 |
| NC_002645.1;NP_073555.1;HCoV-229E_membrane;39 | 0 | 0 |
| NC_002645.1;NP_073555.1;HCoV-229E_membrane;40 | 0 | 0 |
| NC_002645.1;NP_073555.1;HCoV-229E_membrane;41 | 0 | 0 |
| NC_002645.1;NP_073555.1;HCoV-229E_membrane;42 | 0 | 0 |
| NC_002645.1;NP_073555.1;HCoV-229E_membrane;43 | 0 | 0 |
| NC_002645.1;NP_073555.1;HCoV-229E_membrane;44 | 0 | 0 |
| NC_002645.1;NP_073555.1;HCoV-229E_membrane;45 | 0 | 0 |
| NC_002645.1;NP_073555.1;HCoV-229E_membrane;46 | 0 | 0 |
| NC_002645.1;NP_073555.1;HCoV-229E_membrane;47 | 0 | 0 |
| NC_002645.1;NP_073555.1;HCoV-229E_membrane;48 | 0 | 0 |
| NC_002645.1;NP_073555.1;HCoV-229E_membrane;49 | 0 | 0 |
| NC_002645.1;NP_073555.1;HCoV-229E_membrane;50 | 0 | 0 |
| NC_002645.1;NP_073555.1;HCoV-229E_membrane;51 | 0 | 0 |
| NC_002645.1;NP_073555.1;HCoV-229E_membrane;52 | 0 | 0 |
| NC_002645.1;NP_073555.1;HCoV-229E_membrane;53 | 0 | 0 |
| NC_002645.1;NP_073555.1;HCoV-229E_membrane;54 | 0 | 0 |
| NC_002645.1;NP_073555.1;HCoV-229E_membrane;55 | 0 | 0 |
| NC_002645.1;NP_073555.1;HCoV-229E_membrane;56 | 0 | 0 |
| NC_002645.1;NP_073555.1;HCoV-229E_membrane;57 | 0 | 0 |
| NC_002645.1;NP_073555.1;HCoV-229E_membrane;58 | 0 | 0 |
| NC_002645.1;NP_073555.1;HCoV-229E_membrane;59 | 0 | 0 |
| NC_002645.1;NP_073555.1;HCoV-229E_membrane;60 | 0 | 0.025 |
| NC_002645.1;NP_073555.1;HCoV-229E_membrane;61 | 0 | 0 |
| NC_002645.1;NP_073555.1;HCoV-229E_membrane;62 | 0 | 0 |
| NC_002645.1;NP_073555.1;HCoV-229E_membrane;63 | 0 | 0 |
| NC_002645.1;NP_073555.1;HCoV-229E_membrane;64 | 0 | 0 |
| NC_002645.1;NP_073555.1;HCoV-229E_membrane;65 | 0 | 0 |
| NC_002645.1;NP_073555.1;HCoV-229E_membrane;66 | 0 | 0 |
| NC_002645.1;NP_073555.1;HCoV-229E_membrane;67 | 0 | 0 |
| NC_002645.1;NP_073555.1;HCoV-229E_membrane;68 | 0 | 0 |
| NC_002645.1;NP_073555.1;HCoV-229E_membrane;69 | 0 | 0 |
| NC_002645.1;NP_073555.1;HCoV-229E_membrane;70 | 0 | 0 |
| NC_002645.1;NP_073555.1;HCoV-229E_membrane;71 | 0 | 0 |
| NC_002645.1;NP_073555.1;HCoV-229E_membrane;72 | 0 | 0 |
| NC_002645.1;NP_073555.1;HCoV-229E_membrane;73 | 0 | 0 |
| NC_002645.1;NP_073555.1;HCoV-229E_membrane;74 | 0 | 0 |
| NC_002645.1;NP_073555.1;HCoV-229E_membrane;75 | 0 | 0 |
| NC_002645.1;NP_073555.1;HCoV-229E_membrane;76 | 0 | 0 |
| NC_002645.1;NP_073555.1;HCoV-229E_membrane;77 | 0 | 0 |
| NC_002645.1;NP_073555.1;HCoV-229E_membrane;78 | 0 | 0 |
| NC_002645.1;NP_073555.1;HCoV-229E_membrane;79 | 0 | 0 |

| | | |
|---|---|---|
| NC_002645.1;NP_073555.1;HCoV-229E_membrane;80 | 0 | 0 |
| NC_002645.1;NP_073555.1;HCoV-229E_membrane;81 | 0 | 0 |
| NC_002645.1;NP_073555.1;HCoV-229E_membrane;82 | 0 | 0 |
| NC_002645.1;NP_073555.1;HCoV-229E_membrane;83 | 0 | 0 |
| NC_002645.1;NP_073555.1;HCoV-229E_membrane;84 | 0 | 0 |
| NC_002645.1;NP_073555.1;HCoV-229E_membrane;85 | 0 | 0 |
| NC_002645.1;NP_073555.1;HCoV-229E_membrane;86 | 0 | 0 |
| NC_002645.1;NP_073555.1;HCoV-229E_membrane;87 | 0 | 0 |
| NC_002645.1;NP_073555.1;HCoV-229E_membrane;88 | 0 | 0 |
| NC_002645.1;NP_073555.1;HCoV-229E_membrane;89 | 0 | 0 |
| NC_002645.1;NP_073555.1;HCoV-229E_membrane;90 | 0 | 0 |
| NC_002645.1;NP_073555.1;HCoV-229E_membrane;91 | 0 | 0 |
| NC_002645.1;NP_073555.1;HCoV-229E_membrane;92 | 0 | 0 |
| NC_002645.1;NP_073555.1;HCoV-229E_membrane;93 | 0 | 0 |
| NC_002645.1;NP_073555.1;HCoV-229E_membrane;94 | 0 | 0 |
| NC_002645.1;NP_073555.1;HCoV-229E_membrane;95 | 0 | 0 |
| NC_002645.1;NP_073555.1;HCoV-229E_membrane;96 | 0 | 0 |
| NC_002645.1;NP_073555.1;HCoV-229E_membrane;97 | 0 | 0 |
| NC_002645.1;NP_073555.1;HCoV-229E_membrane;98 | 0 | 0 |
| NC_002645.1;NP_073555.1;HCoV-229E_membrane;99 | 0 | 0 |
| NC_002645.1;NP_073555.1;HCoV-229E_membrane;100 | 0 | 0 |
| NC_002645.1;NP_073555.1;HCoV-229E_membrane;101 | 0 | 0 |
| NC_002645.1;NP_073555.1;HCoV-229E_membrane;102 | 0 | 0 |
| NC_002645.1;NP_073555.1;HCoV-229E_membrane;103 | 0 | 0 |
| NC_002645.1;NP_073555.1;HCoV-229E_membrane;104 | 0 | 0 |
| NC_002645.1;NP_073555.1;HCoV-229E_membrane;105 | 0 | 0 |
| NC_002645.1;NP_073555.1;HCoV-229E_membrane;106 | 0 | 0 |
| NC_002645.1;NP_073555.1;HCoV-229E_membrane;107 | 0 | 0 |
| NC_002645.1;NP_073555.1;HCoV-229E_membrane;108 | 0 | 0 |
| NC_002645.1;NP_073555.1;HCoV-229E_membrane;109 | 0 | 0 |
| NC_002645.1;NP_073555.1;HCoV-229E_membrane;110 | 0 | 0 |
| NC_002645.1;NP_073555.1;HCoV-229E_membrane;111 | 0 | 0 |
| NC_002645.1;NP_073555.1;HCoV-229E_membrane;112 | 0 | 0.025 |
| NC_002645.1;NP_073555.1;HCoV-229E_membrane;113 | 0 | 0 |
| NC_002645.1;NP_073555.1;HCoV-229E_membrane;114 | 0 | 0 |
| NC_002645.1;NP_073555.1;HCoV-229E_membrane;115 | 0 | 0 |
| NC_002645.1;NP_073555.1;HCoV-229E_membrane;116 | 0 | 0 |
| NC_002645.1;NP_073555.1;HCoV-229E_membrane;117 | 0 | 0 |
| NC_002645.1;NP_073555.1;HCoV-229E_membrane;118 | 0 | 0 |
| NC_002645.1;NP_073555.1;HCoV-229E_membrane;119 | 0 | 0 |
| NC_002645.1;NP_073555.1;HCoV-229E_membrane;120 | 0 | 0 |
| NC_002645.1;NP_073555.1;HCoV-229E_membrane;121 | 0 | 0.025 |
| NC_002645.1;NP_073555.1;HCoV-229E_membrane;122 | 0 | 0 |

| | | |
|---|---|---|
| NC_002645.1;NP_073555.1;HCoV-229E_membrane;123 | 0 | 0 |
| NC_002645.1;NP_073555.1;HCoV-229E_membrane;124 | 0 | 0 |
| NC_002645.1;NP_073555.1;HCoV-229E_membrane;125 | 0 | 0 |
| NC_002645.1;NP_073555.1;HCoV-229E_membrane;126 | 0 | 0 |
| NC_002645.1;NP_073555.1;HCoV-229E_membrane;127 | 0 | 0 |
| NC_002645.1;NP_073555.1;HCoV-229E_membrane;128 | 0 | 0 |
| NC_002645.1;NP_073555.1;HCoV-229E_membrane;129 | 0 | 0 |
| NC_002645.1;NP_073555.1;HCoV-229E_membrane;130 | 0 | 0 |
| NC_002645.1;NP_073555.1;HCoV-229E_membrane;131 | 0 | 0 |
| NC_002645.1;NP_073555.1;HCoV-229E_membrane;132 | 0 | 0 |
| NC_002645.1;NP_073555.1;HCoV-229E_membrane;133 | 0 | 0 |
| NC_002645.1;NP_073555.1;HCoV-229E_membrane;134 | 0 | 0 |
| NC_002645.1;NP_073555.1;HCoV-229E_membrane;135 | 0 | 0 |
| NC_002645.1;NP_073555.1;HCoV-229E_membrane;136 | 0 | 0 |
| NC_002645.1;NP_073555.1;HCoV-229E_membrane;137 | 0 | 0 |
| NC_002645.1;NP_073555.1;HCoV-229E_membrane;138 | 0 | 0 |
| NC_002645.1;NP_073555.1;HCoV-229E_membrane;139 | 0 | 0 |
| NC_002645.1;NP_073555.1;HCoV-229E_membrane;140 | 0 | 0 |
| NC_002645.1;NP_073555.1;HCoV-229E_membrane;141 | 0 | 0 |
| NC_002645.1;NP_073555.1;HCoV-229E_membrane;142 | 0 | 0 |
| NC_002645.1;NP_073555.1;HCoV-229E_membrane;143 | 0 | 0 |
| NC_002645.1;NP_073555.1;HCoV-229E_membrane;144 | 0 | 0 |
| NC_002645.1;NP_073555.1;HCoV-229E_membrane;145 | 0 | 0 |
| NC_002645.1;NP_073555.1;HCoV-229E_membrane;146 | 0.05 | 0 |
| NC_002645.1;NP_073555.1;HCoV-229E_membrane;147 | 0 | 0 |
| NC_002645.1;NP_073555.1;HCoV-229E_membrane;148 | 0 | 0 |
| NC_002645.1;NP_073555.1;HCoV-229E_membrane;149 | 0 | 0.025 |
| NC_002645.1;NP_073555.1;HCoV-229E_membrane;150 | 0 | 0.025 |
| NC_002645.1;NP_073555.1;HCoV-229E_membrane;151 | 0 | 0.05 |
| NC_002645.1;NP_073555.1;HCoV-229E_membrane;152 | 0 | 0.05 |
| NC_002645.1;NP_073555.1;HCoV-229E_membrane;153 | 0 | 0.025 |
| NC_002645.1;NP_073555.1;HCoV-229E_membrane;154 | 0 | 0.025 |
| NC_002645.1;NP_073555.1;HCoV-229E_membrane;155 | 0 | 0 |
| NC_002645.1;NP_073555.1;HCoV-229E_membrane;156 | 0.05 | 0 |
| NC_002645.1;NP_073555.1;HCoV-229E_membrane;157 | 0 | 0 |
| NC_002645.1;NP_073555.1;HCoV-229E_membrane;158 | 0 | 0 |
| NC_002645.1;NP_073555.1;HCoV-229E_membrane;159 | 0 | 0.025 |
| NC_002645.1;NP_073555.1;HCoV-229E_membrane;160 | 0 | 0 |
| NC_002645.1;NP_073555.1;HCoV-229E_membrane;161 | 0 | 0 |
| NC_002645.1;NP_073555.1;HCoV-229E_membrane;162 | 0 | 0 |
| NC_002645.1;NP_073555.1;HCoV-229E_membrane;163 | 0 | 0 |
| NC_002645.1;NP_073555.1;HCoV-229E_membrane;164 | 0 | 0 |
| NC_002645.1;NP_073555.1;HCoV-229E_membrane;165 | 0 | 0 |

| | | |
|---|---|---|
| NC_002645.1;NP_073555.1;HCoV-229E_membrane;166 | 0 | 0 |
| NC_002645.1;NP_073555.1;HCoV-229E_membrane;167 | 0 | 0 |
| NC_002645.1;NP_073555.1;HCoV-229E_membrane;168 | 0 | 0 |
| NC_002645.1;NP_073555.1;HCoV-229E_membrane;169 | 0 | 0 |
| NC_002645.1;NP_073555.1;HCoV-229E_membrane;170 | 0 | 0 |
| NC_002645.1;NP_073555.1;HCoV-229E_membrane;171 | 0 | 0 |
| NC_002645.1;NP_073555.1;HCoV-229E_membrane;172 | 0 | 0 |
| NC_002645.1;NP_073555.1;HCoV-229E_membrane;173 | 0 | 0 |
| NC_002645.1;NP_073555.1;HCoV-229E_membrane;174 | 0 | 0 |
| NC_002645.1;NP_073555.1;HCoV-229E_membrane;175 | 0 | 0 |
| NC_002645.1;NP_073555.1;HCoV-229E_membrane;176 | 0 | 0 |
| NC_002645.1;NP_073555.1;HCoV-229E_membrane;177 | 0 | 0 |
| NC_002645.1;NP_073555.1;HCoV-229E_membrane;178 | 0 | 0 |
| NC_002645.1;NP_073555.1;HCoV-229E_membrane;179 | 0 | 0 |
| NC_002645.1;NP_073555.1;HCoV-229E_membrane;180 | 0.05 | 0.025 |
| NC_002645.1;NP_073555.1;HCoV-229E_membrane;181 | 0 | 0.025 |
| NC_002645.1;NP_073555.1;HCoV-229E_membrane;182 | 0 | 0.025 |
| NC_002645.1;NP_073555.1;HCoV-229E_membrane;183 | 0 | 0 |
| NC_002645.1;NP_073555.1;HCoV-229E_membrane;184 | 0 | 0 |
| NC_002645.1;NP_073555.1;HCoV-229E_membrane;185 | 0 | 0 |
| NC_002645.1;NP_073555.1;HCoV-229E_membrane;186 | 0 | 0 |
| NC_002645.1;NP_073555.1;HCoV-229E_membrane;187 | 0 | 0 |
| NC_002645.1;NP_073555.1;HCoV-229E_membrane;188 | 0 | 0 |
| NC_002645.1;NP_073555.1;HCoV-229E_membrane;189 | 0 | 0 |
| NC_002645.1;NP_073555.1;HCoV-229E_membrane;190 | 0 | 0 |
| NC_002645.1;NP_073555.1;HCoV-229E_membrane;191 | 0 | 0 |
| NC_002645.1;NP_073555.1;HCoV-229E_membrane;192 | 0 | 0 |
| NC_002645.1;NP_073555.1;HCoV-229E_membrane;193 | 0 | 0.025 |
| NC_002645.1;NP_073555.1;HCoV-229E_membrane;194 | 0 | 0.025 |
| NC_002645.1;NP_073555.1;HCoV-229E_membrane;195 | 0 | 0.025 |
| NC_002645.1;NP_073555.1;HCoV-229E_membrane;196 | 0 | 0.025 |
| NC_002645.1;NP_073555.1;HCoV-229E_membrane;197 | 0 | 0.025 |
| NC_002645.1;NP_073555.1;HCoV-229E_membrane;198 | 0 | 0.025 |
| NC_002645.1;NP_073555.1;HCoV-229E_membrane;199 | 0 | 0.025 |
| NC_002645.1;NP_073555.1;HCoV-229E_membrane;200 | 0 | 0.025 |
| NC_002645.1;NP_073555.1;HCoV-229E_membrane;201 | 0 | 0.025 |
| NC_002645.1;NP_073555.1;HCoV-229E_membrane;202 | 0 | 0 |
| NC_002645.1;NP_073555.1;HCoV-229E_membrane;203 | 0.3 | 0.325 |
| NC_002645.1;NP_073555.1;HCoV-229E_membrane;204 | 0.35 | 0.3 |
| NC_002645.1;NP_073555.1;HCoV-229E_membrane;205 | 0.35 | 0.325 |
| NC_002645.1;NP_073555.1;HCoV-229E_membrane;206 | 0.3 | 0.275 |
| NC_002645.1;NP_073555.1;HCoV-229E_membrane;207 | 0.3 | 0.225 |
| NC_002645.1;NP_073555.1;HCoV-229E_membrane;208 | 0.15 | 0.175 |

| | | |
|---|---|---|
| NC_002645.1;NP_073555.1;HCoV-229E_membrane;209 | 0.2 | 0.2 |
| NC_002645.1;NP_073555.1;HCoV-229E_membrane;210 | 0.2 | 0.15 |
| NC_002645.1;NP_073555.1;HCoV-229E_membrane;211 | 0.1 | 0.05 |
| NC_002645.1;NP_073555.1;HCoV-229E_membrane;212 | 0.2 | 0.225 |
| NC_002645.1;NP_073555.1;HCoV-229E_membrane;213 | 0.2 | 0.175 |
| NC_002645.1;NP_073555.1;HCoV-229E_membrane;214 | 0.1 | 0.025 |
| NC_002645.1;NP_073556.1;HCoV-229E_nucleocapsi;1 | 0.05 | 0.05 |
| NC_002645.1;NP_073556.1;HCoV-229E_nucleocapsi;2 | 0 | 0 |
| NC_002645.1;NP_073556.1;HCoV-229E_nucleocapsi;3 | 0 | 0.025 |
| NC_002645.1;NP_073556.1;HCoV-229E_nucleocapsi;4 | 0.05 | 0.025 |
| NC_002645.1;NP_073556.1;HCoV-229E_nucleocapsi;5 | 0.05 | 0.025 |
| NC_002645.1;NP_073556.1;HCoV-229E_nucleocapsi;6 | 0.05 | 0.025 |
| NC_002645.1;NP_073556.1;HCoV-229E_nucleocapsi;7 | 0.1 | 0.025 |
| NC_002645.1;NP_073556.1;HCoV-229E_nucleocapsi;8 | 0.1 | 0.025 |
| NC_002645.1;NP_073556.1;HCoV-229E_nucleocapsi;9 | 0.05 | 0 |
| NC_002645.1;NP_073556.1;HCoV-229E_nucleocapsi;10 | 0.05 | 0 |
| NC_002645.1;NP_073556.1;HCoV-229E_nucleocapsi;11 | 0.05 | 0 |
| NC_002645.1;NP_073556.1;HCoV-229E_nucleocapsi;12 | 0.05 | 0 |
| NC_002645.1;NP_073556.1;HCoV-229E_nucleocapsi;13 | 0.05 | 0 |
| NC_002645.1;NP_073556.1;HCoV-229E_nucleocapsi;14 | 0.05 | 0 |
| NC_002645.1;NP_073556.1;HCoV-229E_nucleocapsi;15 | 0 | 0 |
| NC_002645.1;NP_073556.1;HCoV-229E_nucleocapsi;16 | 0 | 0 |
| NC_002645.1;NP_073556.1;HCoV-229E_nucleocapsi;17 | 0 | 0 |
| NC_002645.1;NP_073556.1;HCoV-229E_nucleocapsi;18 | 0 | 0 |
| NC_002645.1;NP_073556.1;HCoV-229E_nucleocapsi;19 | 0 | 0 |
| NC_002645.1;NP_073556.1;HCoV-229E_nucleocapsi;20 | 0 | 0 |
| NC_002645.1;NP_073556.1;HCoV-229E_nucleocapsi;21 | 0 | 0 |
| NC_002645.1;NP_073556.1;HCoV-229E_nucleocapsi;22 | 0 | 0 |
| NC_002645.1;NP_073556.1;HCoV-229E_nucleocapsi;23 | 0.05 | 0 |
| NC_002645.1;NP_073556.1;HCoV-229E_nucleocapsi;24 | 0 | 0 |
| NC_002645.1;NP_073556.1;HCoV-229E_nucleocapsi;25 | 0 | 0 |
| NC_002645.1;NP_073556.1;HCoV-229E_nucleocapsi;26 | 0 | 0 |
| NC_002645.1;NP_073556.1;HCoV-229E_nucleocapsi;27 | 0 | 0 |
| NC_002645.1;NP_073556.1;HCoV-229E_nucleocapsi;28 | 0.05 | 0 |
| NC_002645.1;NP_073556.1;HCoV-229E_nucleocapsi;29 | 0 | 0 |
| NC_002645.1;NP_073556.1;HCoV-229E_nucleocapsi;30 | 0 | 0.025 |
| NC_002645.1;NP_073556.1;HCoV-229E_nucleocapsi;31 | 0.05 | 0 |
| NC_002645.1;NP_073556.1;HCoV-229E_nucleocapsi;32 | 0.05 | 0 |
| NC_002645.1;NP_073556.1;HCoV-229E_nucleocapsi;33 | 0 | 0.025 |
| NC_002645.1;NP_073556.1;HCoV-229E_nucleocapsi;34 | 0 | 0 |
| NC_002645.1;NP_073556.1;HCoV-229E_nucleocapsi;35 | 0 | 0 |
| NC_002645.1;NP_073556.1;HCoV-229E_nucleocapsi;36 | 0 | 0 |
| NC_002645.1;NP_073556.1;HCoV-229E_nucleocapsi;37 | 0 | 0 |

| | | |
|---|---|---|
| NC_002645.1;NP_073556.1;HCoV-229E_nucleocapsi;38 | 0 | 0 |
| NC_002645.1;NP_073556.1;HCoV-229E_nucleocapsi;39 | 0 | 0 |
| NC_002645.1;NP_073556.1;HCoV-229E_nucleocapsi;40 | 0 | 0 |
| NC_002645.1;NP_073556.1;HCoV-229E_nucleocapsi;41 | 0 | 0 |
| NC_002645.1;NP_073556.1;HCoV-229E_nucleocapsi;42 | 0 | 0 |
| NC_002645.1;NP_073556.1;HCoV-229E_nucleocapsi;43 | 0 | 0 |
| NC_002645.1;NP_073556.1;HCoV-229E_nucleocapsi;44 | 0 | 0 |
| NC_002645.1;NP_073556.1;HCoV-229E_nucleocapsi;45 | 0 | 0 |
| NC_002645.1;NP_073556.1;HCoV-229E_nucleocapsi;46 | 0 | 0 |
| NC_002645.1;NP_073556.1;HCoV-229E_nucleocapsi;47 | 0 | 0 |
| NC_002645.1;NP_073556.1;HCoV-229E_nucleocapsi;48 | 0 | 0 |
| NC_002645.1;NP_073556.1;HCoV-229E_nucleocapsi;49 | 0 | 0 |
| NC_002645.1;NP_073556.1;HCoV-229E_nucleocapsi;50 | 0 | 0 |
| NC_002645.1;NP_073556.1;HCoV-229E_nucleocapsi;51 | 0 | 0 |
| NC_002645.1;NP_073556.1;HCoV-229E_nucleocapsi;52 | 0 | 0 |
| NC_002645.1;NP_073556.1;HCoV-229E_nucleocapsi;53 | 0 | 0 |
| NC_002645.1;NP_073556.1;HCoV-229E_nucleocapsi;54 | 0 | 0 |
| NC_002645.1;NP_073556.1;HCoV-229E_nucleocapsi;55 | 0.05 | 0 |
| NC_002645.1;NP_073556.1;HCoV-229E_nucleocapsi;56 | 0.05 | 0 |
| NC_002645.1;NP_073556.1;HCoV-229E_nucleocapsi;57 | 0.05 | 0 |
| NC_002645.1;NP_073556.1;HCoV-229E_nucleocapsi;58 | 0.05 | 0 |
| NC_002645.1;NP_073556.1;HCoV-229E_nucleocapsi;59 | 0.05 | 0 |
| NC_002645.1;NP_073556.1;HCoV-229E_nucleocapsi;60 | 0.05 | 0 |
| NC_002645.1;NP_073556.1;HCoV-229E_nucleocapsi;61 | 0 | 0 |
| NC_002645.1;NP_073556.1;HCoV-229E_nucleocapsi;62 | 0 | 0 |
| NC_002645.1;NP_073556.1;HCoV-229E_nucleocapsi;63 | 0.05 | 0 |
| NC_002645.1;NP_073556.1;HCoV-229E_nucleocapsi;64 | 0 | 0 |
| NC_002645.1;NP_073556.1;HCoV-229E_nucleocapsi;65 | 0 | 0 |
| NC_002645.1;NP_073556.1;HCoV-229E_nucleocapsi;66 | 0 | 0 |
| NC_002645.1;NP_073556.1;HCoV-229E_nucleocapsi;67 | 0 | 0 |
| NC_002645.1;NP_073556.1;HCoV-229E_nucleocapsi;68 | 0 | 0 |
| NC_002645.1;NP_073556.1;HCoV-229E_nucleocapsi;69 | 0 | 0 |
| NC_002645.1;NP_073556.1;HCoV-229E_nucleocapsi;70 | 0 | 0 |
| NC_002645.1;NP_073556.1;HCoV-229E_nucleocapsi;71 | 0 | 0 |
| NC_002645.1;NP_073556.1;HCoV-229E_nucleocapsi;72 | 0 | 0 |
| NC_002645.1;NP_073556.1;HCoV-229E_nucleocapsi;73 | 0 | 0 |
| NC_002645.1;NP_073556.1;HCoV-229E_nucleocapsi;74 | 0 | 0 |
| NC_002645.1;NP_073556.1;HCoV-229E_nucleocapsi;75 | 0 | 0 |
| NC_002645.1;NP_073556.1;HCoV-229E_nucleocapsi;76 | 0 | 0 |
| NC_002645.1;NP_073556.1;HCoV-229E_nucleocapsi;77 | 0 | 0 |
| NC_002645.1;NP_073556.1;HCoV-229E_nucleocapsi;78 | 0 | 0 |
| NC_002645.1;NP_073556.1;HCoV-229E_nucleocapsi;79 | 0 | 0 |
| NC_002645.1;NP_073556.1;HCoV-229E_nucleocapsi;80 | 0 | 0 |

| | | |
|---|---|---|
| NC_002645.1;NP_073556.1;HCoV-229E_nucleocapsi;81 | 0 | 0 |
| NC_002645.1;NP_073556.1;HCoV-229E_nucleocapsi;82 | 0 | 0 |
| NC_002645.1;NP_073556.1;HCoV-229E_nucleocapsi;83 | 0 | 0 |
| NC_002645.1;NP_073556.1;HCoV-229E_nucleocapsi;84 | 0 | 0 |
| NC_002645.1;NP_073556.1;HCoV-229E_nucleocapsi;85 | 0 | 0 |
| NC_002645.1;NP_073556.1;HCoV-229E_nucleocapsi;86 | 0 | 0 |
| NC_002645.1;NP_073556.1;HCoV-229E_nucleocapsi;87 | 0 | 0 |
| NC_002645.1;NP_073556.1;HCoV-229E_nucleocapsi;88 | 0 | 0 |
| NC_002645.1;NP_073556.1;HCoV-229E_nucleocapsi;89 | 0 | 0 |
| NC_002645.1;NP_073556.1;HCoV-229E_nucleocapsi;90 | 0 | 0 |
| NC_002645.1;NP_073556.1;HCoV-229E_nucleocapsi;91 | 0 | 0 |
| NC_002645.1;NP_073556.1;HCoV-229E_nucleocapsi;92 | 0 | 0 |
| NC_002645.1;NP_073556.1;HCoV-229E_nucleocapsi;93 | 0 | 0 |
| NC_002645.1;NP_073556.1;HCoV-229E_nucleocapsi;94 | 0 | 0 |
| NC_002645.1;NP_073556.1;HCoV-229E_nucleocapsi;95 | 0.05 | 0 |
| NC_002645.1;NP_073556.1;HCoV-229E_nucleocapsi;96 | 0 | 0 |
| NC_002645.1;NP_073556.1;HCoV-229E_nucleocapsi;97 | 0 | 0 |
| NC_002645.1;NP_073556.1;HCoV-229E_nucleocapsi;98 | 0 | 0 |
| NC_002645.1;NP_073556.1;HCoV-229E_nucleocapsi;99 | 0 | 0 |
| NC_002645.1;NP_073556.1;HCoV-229E_nucleocapsi;100 | 0 | 0.025 |
| NC_002645.1;NP_073556.1;HCoV-229E_nucleocapsi;101 | 0 | 0.05 |
| NC_002645.1;NP_073556.1;HCoV-229E_nucleocapsi;102 | 0 | 0.025 |
| NC_002645.1;NP_073556.1;HCoV-229E_nucleocapsi;103 | 0.05 | 0.025 |
| NC_002645.1;NP_073556.1;HCoV-229E_nucleocapsi;104 | 0 | 0 |
| NC_002645.1;NP_073556.1;HCoV-229E_nucleocapsi;105 | 0 | 0 |
| NC_002645.1;NP_073556.1;HCoV-229E_nucleocapsi;106 | 0 | 0 |
| NC_002645.1;NP_073556.1;HCoV-229E_nucleocapsi;107 | 0 | 0 |
| NC_002645.1;NP_073556.1;HCoV-229E_nucleocapsi;108 | 0 | 0 |
| NC_002645.1;NP_073556.1;HCoV-229E_nucleocapsi;109 | 0 | 0 |
| NC_002645.1;NP_073556.1;HCoV-229E_nucleocapsi;110 | 0 | 0 |
| NC_002645.1;NP_073556.1;HCoV-229E_nucleocapsi;111 | 0 | 0.025 |
| NC_002645.1;NP_073556.1;HCoV-229E_nucleocapsi;112 | 0 | 0.025 |
| NC_002645.1;NP_073556.1;HCoV-229E_nucleocapsi;113 | 0 | 0.025 |
| NC_002645.1;NP_073556.1;HCoV-229E_nucleocapsi;114 | 0 | 0.025 |
| NC_002645.1;NP_073556.1;HCoV-229E_nucleocapsi;115 | 0 | 0.025 |
| NC_002645.1;NP_073556.1;HCoV-229E_nucleocapsi;116 | 0 | 0.025 |
| NC_002645.1;NP_073556.1;HCoV-229E_nucleocapsi;117 | 0.05 | 0 |
| NC_002645.1;NP_073556.1;HCoV-229E_nucleocapsi;118 | 0 | 0.025 |
| NC_002645.1;NP_073556.1;HCoV-229E_nucleocapsi;119 | 0 | 0 |
| NC_002645.1;NP_073556.1;HCoV-229E_nucleocapsi;120 | 0 | 0 |
| NC_002645.1;NP_073556.1;HCoV-229E_nucleocapsi;121 | 0.05 | 0 |
| NC_002645.1;NP_073556.1;HCoV-229E_nucleocapsi;122 | 0 | 0 |
| NC_002645.1;NP_073556.1;HCoV-229E_nucleocapsi;123 | 0 | 0.05 |

| | | |
|---|---|---|
| NC_002645.1;NP_073556.1;HCoV-229E_nucleocapsi;124 | 0 | 0.05 |
| NC_002645.1;NP_073556.1;HCoV-229E_nucleocapsi;125 | 0 | 0.025 |
| NC_002645.1;NP_073556.1;HCoV-229E_nucleocapsi;126 | 0 | 0.025 |
| NC_002645.1;NP_073556.1;HCoV-229E_nucleocapsi;127 | 0 | 0 |
| NC_002645.1;NP_073556.1;HCoV-229E_nucleocapsi;128 | 0 | 0.025 |
| NC_002645.1;NP_073556.1;HCoV-229E_nucleocapsi;129 | 0 | 0 |
| NC_002645.1;NP_073556.1;HCoV-229E_nucleocapsi;130 | 0 | 0 |
| NC_002645.1;NP_073556.1;HCoV-229E_nucleocapsi;131 | 0 | 0.025 |
| NC_002645.1;NP_073556.1;HCoV-229E_nucleocapsi;132 | 0.05 | 0 |
| NC_002645.1;NP_073556.1;HCoV-229E_nucleocapsi;133 | 0.05 | 0 |
| NC_002645.1;NP_073556.1;HCoV-229E_nucleocapsi;134 | 0.05 | 0 |
| NC_002645.1;NP_073556.1;HCoV-229E_nucleocapsi;135 | 0.05 | 0 |
| NC_002645.1;NP_073556.1;HCoV-229E_nucleocapsi;136 | 0.05 | 0 |
| NC_002645.1;NP_073556.1;HCoV-229E_nucleocapsi;137 | 0.05 | 0 |
| NC_002645.1;NP_073556.1;HCoV-229E_nucleocapsi;138 | 0.05 | 0 |
| NC_002645.1;NP_073556.1;HCoV-229E_nucleocapsi;139 | 0.05 | 0 |
| NC_002645.1;NP_073556.1;HCoV-229E_nucleocapsi;140 | 0.05 | 0 |
| NC_002645.1;NP_073556.1;HCoV-229E_nucleocapsi;141 | 0.05 | 0 |
| NC_002645.1;NP_073556.1;HCoV-229E_nucleocapsi;142 | 0 | 0 |
| NC_002645.1;NP_073556.1;HCoV-229E_nucleocapsi;143 | 0 | 0 |
| NC_002645.1;NP_073556.1;HCoV-229E_nucleocapsi;144 | 0.05 | 0 |
| NC_002645.1;NP_073556.1;HCoV-229E_nucleocapsi;145 | 0 | 0 |
| NC_002645.1;NP_073556.1;HCoV-229E_nucleocapsi;146 | 0 | 0 |
| NC_002645.1;NP_073556.1;HCoV-229E_nucleocapsi;147 | 0 | 0.025 |
| NC_002645.1;NP_073556.1;HCoV-229E_nucleocapsi;148 | 0.05 | 0.025 |
| NC_002645.1;NP_073556.1;HCoV-229E_nucleocapsi;149 | 0 | 0 |
| NC_002645.1;NP_073556.1;HCoV-229E_nucleocapsi;150 | 0 | 0 |
| NC_002645.1;NP_073556.1;HCoV-229E_nucleocapsi;151 | 0 | 0.025 |
| NC_002645.1;NP_073556.1;HCoV-229E_nucleocapsi;152 | 0 | 0 |
| NC_002645.1;NP_073556.1;HCoV-229E_nucleocapsi;153 | 0 | 0 |
| NC_002645.1;NP_073556.1;HCoV-229E_nucleocapsi;154 | 0 | 0 |
| NC_002645.1;NP_073556.1;HCoV-229E_nucleocapsi;155 | 0 | 0.05 |
| NC_002645.1;NP_073556.1;HCoV-229E_nucleocapsi;156 | 0 | 0.025 |
| NC_002645.1;NP_073556.1;HCoV-229E_nucleocapsi;157 | 0 | 0 |
| NC_002645.1;NP_073556.1;HCoV-229E_nucleocapsi;158 | 0 | 0 |
| NC_002645.1;NP_073556.1;HCoV-229E_nucleocapsi;159 | 0 | 0.025 |
| NC_002645.1;NP_073556.1;HCoV-229E_nucleocapsi;160 | 0 | 0 |
| NC_002645.1;NP_073556.1;HCoV-229E_nucleocapsi;161 | 0 | 0 |
| NC_002645.1;NP_073556.1;HCoV-229E_nucleocapsi;162 | 0.1 | 0.15 |
| NC_002645.1;NP_073556.1;HCoV-229E_nucleocapsi;163 | 0 | 0.025 |
| NC_002645.1;NP_073556.1;HCoV-229E_nucleocapsi;164 | 0 | 0 |
| NC_002645.1;NP_073556.1;HCoV-229E_nucleocapsi;165 | 0 | 0 |
| NC_002645.1;NP_073556.1;HCoV-229E_nucleocapsi;166 | 0 | 0 |

| | | |
|---|---|---|
| NC_002645.1;NP_073556.1;HCoV-229E_nucleocapsi;167 | 0 | 0.025 |
| NC_002645.1;NP_073556.1;HCoV-229E_nucleocapsi;168 | 0 | 0.05 |
| NC_002645.1;NP_073556.1;HCoV-229E_nucleocapsi;169 | 0 | 0 |
| NC_002645.1;NP_073556.1;HCoV-229E_nucleocapsi;170 | 0 | 0 |
| NC_002645.1;NP_073556.1;HCoV-229E_nucleocapsi;171 | 0 | 0 |
| NC_002645.1;NP_073556.1;HCoV-229E_nucleocapsi;172 | 0 | 0 |
| NC_002645.1;NP_073556.1;HCoV-229E_nucleocapsi;173 | 0.05 | 0.025 |
| NC_002645.1;NP_073556.1;HCoV-229E_nucleocapsi;174 | 0.05 | 0.05 |
| NC_002645.1;NP_073556.1;HCoV-229E_nucleocapsi;175 | 0.05 | 0 |
| NC_002645.1;NP_073556.1;HCoV-229E_nucleocapsi;176 | 0.05 | 0 |
| NC_002645.1;NP_073556.1;HCoV-229E_nucleocapsi;177 | 0.05 | 0.025 |
| NC_002645.1;NP_073556.1;HCoV-229E_nucleocapsi;178 | 0.05 | 0.05 |
| NC_002645.1;NP_073556.1;HCoV-229E_nucleocapsi;179 | 0 | 0.025 |
| NC_002645.1;NP_073556.1;HCoV-229E_nucleocapsi;180 | 0 | 0 |
| NC_002645.1;NP_073556.1;HCoV-229E_nucleocapsi;181 | 0.05 | 0 |
| NC_002645.1;NP_073556.1;HCoV-229E_nucleocapsi;182 | 0.05 | 0 |
| NC_002645.1;NP_073556.1;HCoV-229E_nucleocapsi;183 | 0.05 | 0.025 |
| NC_002645.1;NP_073556.1;HCoV-229E_nucleocapsi;184 | 0.05 | 0.025 |
| NC_002645.1;NP_073556.1;HCoV-229E_nucleocapsi;185 | 0.05 | 0.025 |
| NC_002645.1;NP_073556.1;HCoV-229E_nucleocapsi;186 | 0.05 | 0.025 |
| NC_002645.1;NP_073556.1;HCoV-229E_nucleocapsi;187 | 0.05 | 0.025 |
| NC_002645.1;NP_073556.1;HCoV-229E_nucleocapsi;188 | 0.1 | 0.025 |
| NC_002645.1;NP_073556.1;HCoV-229E_nucleocapsi;189 | 0.05 | 0.05 |
| NC_002645.1;NP_073556.1;HCoV-229E_nucleocapsi;190 | 0.05 | 0.025 |
| NC_002645.1;NP_073556.1;HCoV-229E_nucleocapsi;191 | 0.05 | 0.025 |
| NC_002645.1;NP_073556.1;HCoV-229E_nucleocapsi;192 | 0.05 | 0.025 |
| NC_002645.1;NP_073556.1;HCoV-229E_nucleocapsi;193 | 0.05 | 0.025 |
| NC_002645.1;NP_073556.1;HCoV-229E_nucleocapsi;194 | 0.05 | 0.025 |
| NC_002645.1;NP_073556.1;HCoV-229E_nucleocapsi;195 | 0.05 | 0.025 |
| NC_002645.1;NP_073556.1;HCoV-229E_nucleocapsi;196 | 0.1 | 0.025 |
| NC_002645.1;NP_073556.1;HCoV-229E_nucleocapsi;197 | 0.05 | 0.025 |
| NC_002645.1;NP_073556.1;HCoV-229E_nucleocapsi;198 | 0.05 | 0.025 |
| NC_002645.1;NP_073556.1;HCoV-229E_nucleocapsi;199 | 0.05 | 0.05 |
| NC_002645.1;NP_073556.1;HCoV-229E_nucleocapsi;200 | 0.05 | 0 |
| NC_002645.1;NP_073556.1;HCoV-229E_nucleocapsi;201 | 0.05 | 0 |
| NC_002645.1;NP_073556.1;HCoV-229E_nucleocapsi;202 | 0.05 | 0 |
| NC_002645.1;NP_073556.1;HCoV-229E_nucleocapsi;203 | 0.05 | 0.025 |
| NC_002645.1;NP_073556.1;HCoV-229E_nucleocapsi;204 | 0.05 | 0.025 |
| NC_002645.1;NP_073556.1;HCoV-229E_nucleocapsi;205 | 0.05 | 0.025 |
| NC_002645.1;NP_073556.1;HCoV-229E_nucleocapsi;206 | 0.05 | 0.025 |
| NC_002645.1;NP_073556.1;HCoV-229E_nucleocapsi;207 | 0.05 | 0.025 |
| NC_002645.1;NP_073556.1;HCoV-229E_nucleocapsi;208 | 0 | 0.025 |
| NC_002645.1;NP_073556.1;HCoV-229E_nucleocapsi;209 | 0 | 0.025 |

| | | |
|---|---|---|
| NC_002645.1;NP_073556.1;HCoV-229E_nucleocapsi;210 | 0 | 0 |
| NC_002645.1;NP_073556.1;HCoV-229E_nucleocapsi;211 | 0 | 0 |
| NC_002645.1;NP_073556.1;HCoV-229E_nucleocapsi;212 | 0 | 0 |
| NC_002645.1;NP_073556.1;HCoV-229E_nucleocapsi;213 | 0.05 | 0 |
| NC_002645.1;NP_073556.1;HCoV-229E_nucleocapsi;214 | 0 | 0 |
| NC_002645.1;NP_073556.1;HCoV-229E_nucleocapsi;215 | 0 | 0 |
| NC_002645.1;NP_073556.1;HCoV-229E_nucleocapsi;216 | 0 | 0.025 |
| NC_002645.1;NP_073556.1;HCoV-229E_nucleocapsi;217 | 0 | 0.025 |
| NC_002645.1;NP_073556.1;HCoV-229E_nucleocapsi;218 | 0 | 0.175 |
| NC_002645.1;NP_073556.1;HCoV-229E_nucleocapsi;219 | 0.05 | 0.025 |
| NC_002645.1;NP_073556.1;HCoV-229E_nucleocapsi;220 | 0 | 0.025 |
| NC_002645.1;NP_073556.1;HCoV-229E_nucleocapsi;221 | 0 | 0.05 |
| NC_002645.1;NP_073556.1;HCoV-229E_nucleocapsi;222 | 0 | 0.075 |
| NC_002645.1;NP_073556.1;HCoV-229E_nucleocapsi;223 | 0 | 0 |
| NC_002645.1;NP_073556.1;HCoV-229E_nucleocapsi;224 | 0 | 0.025 |
| NC_002645.1;NP_073556.1;HCoV-229E_nucleocapsi;225 | 0 | 0.025 |
| NC_002645.1;NP_073556.1;HCoV-229E_nucleocapsi;226 | 0 | 0 |
| NC_002645.1;NP_073556.1;HCoV-229E_nucleocapsi;227 | 0 | 0 |
| NC_002645.1;NP_073556.1;HCoV-229E_nucleocapsi;228 | 0 | 0 |
| NC_002645.1;NP_073556.1;HCoV-229E_nucleocapsi;229 | 0 | 0 |
| NC_002645.1;NP_073556.1;HCoV-229E_nucleocapsi;230 | 0 | 0 |
| NC_002645.1;NP_073556.1;HCoV-229E_nucleocapsi;231 | 0 | 0 |
| NC_002645.1;NP_073556.1;HCoV-229E_nucleocapsi;232 | 0.05 | 0.1 |
| NC_002645.1;NP_073556.1;HCoV-229E_nucleocapsi;233 | 0.1 | 0 |
| NC_002645.1;NP_073556.1;HCoV-229E_nucleocapsi;234 | 0.1 | 0.025 |
| NC_002645.1;NP_073556.1;HCoV-229E_nucleocapsi;235 | 0 | 0 |
| NC_002645.1;NP_073556.1;HCoV-229E_nucleocapsi;236 | 0 | 0.025 |
| NC_002645.1;NP_073556.1;HCoV-229E_nucleocapsi;237 | 0 | 0 |
| NC_002645.1;NP_073556.1;HCoV-229E_nucleocapsi;238 | 0 | 0 |
| NC_002645.1;NP_073556.1;HCoV-229E_nucleocapsi;239 | 0 | 0 |
| NC_002645.1;NP_073556.1;HCoV-229E_nucleocapsi;240 | 0 | 0 |
| NC_002645.1;NP_073556.1;HCoV-229E_nucleocapsi;241 | 0.05 | 0 |
| NC_002645.1;NP_073556.1;HCoV-229E_nucleocapsi;242 | 0 | 0.025 |
| NC_002645.1;NP_073556.1;HCoV-229E_nucleocapsi;243 | 0.1 | 0.025 |
| NC_002645.1;NP_073556.1;HCoV-229E_nucleocapsi;244 | 0.1 | 0.05 |
| NC_002645.1;NP_073556.1;HCoV-229E_nucleocapsi;245 | 0.05 | 0.025 |
| NC_002645.1;NP_073556.1;HCoV-229E_nucleocapsi;246 | 0.1 | 0.025 |
| NC_002645.1;NP_073556.1;HCoV-229E_nucleocapsi;247 | 0.2 | 0.05 |
| NC_002645.1;NP_073556.1;HCoV-229E_nucleocapsi;248 | 0.2 | 0.15 |
| NC_002645.1;NP_073556.1;HCoV-229E_nucleocapsi;249 | 0.15 | 0.125 |
| NC_002645.1;NP_073556.1;HCoV-229E_nucleocapsi;250 | 0.15 | 0.05 |
| NC_002645.1;NP_073556.1;HCoV-229E_nucleocapsi;251 | 0.05 | 0.075 |
| NC_002645.1;NP_073556.1;HCoV-229E_nucleocapsi;252 | 0.05 | 0.05 |

| | | |
|---|---|---|
| NC_002645.1;NP_073556.1;HCoV-229E_nucleocapsi;253 | 0.05 | 0.05 |
| NC_002645.1;NP_073556.1;HCoV-229E_nucleocapsi;254 | 0.05 | 0.025 |
| NC_002645.1;NP_073556.1;HCoV-229E_nucleocapsi;255 | 0.05 | 0 |
| NC_002645.1;NP_073556.1;HCoV-229E_nucleocapsi;256 | 0.1 | 0 |
| NC_002645.1;NP_073556.1;HCoV-229E_nucleocapsi;257 | 0 | 0 |
| NC_002645.1;NP_073556.1;HCoV-229E_nucleocapsi;258 | 0.05 | 0 |
| NC_002645.1;NP_073556.1;HCoV-229E_nucleocapsi;259 | 0.1 | 0 |
| NC_002645.1;NP_073556.1;HCoV-229E_nucleocapsi;260 | 0.05 | 0 |
| NC_002645.1;NP_073556.1;HCoV-229E_nucleocapsi;261 | 0 | 0 |
| NC_002645.1;NP_073556.1;HCoV-229E_nucleocapsi;262 | 0 | 0 |
| NC_002645.1;NP_073556.1;HCoV-229E_nucleocapsi;263 | 0 | 0 |
| NC_002645.1;NP_073556.1;HCoV-229E_nucleocapsi;264 | 0 | 0 |
| NC_002645.1;NP_073556.1;HCoV-229E_nucleocapsi;265 | 0 | 0.025 |
| NC_002645.1;NP_073556.1;HCoV-229E_nucleocapsi;266 | 0.05 | 0.1 |
| NC_002645.1;NP_073556.1;HCoV-229E_nucleocapsi;267 | 0 | 0 |
| NC_002645.1;NP_073556.1;HCoV-229E_nucleocapsi;268 | 0.05 | 0 |
| NC_002645.1;NP_073556.1;HCoV-229E_nucleocapsi;269 | 0 | 0.025 |
| NC_002645.1;NP_073556.1;HCoV-229E_nucleocapsi;270 | 0 | 0 |
| NC_002645.1;NP_073556.1;HCoV-229E_nucleocapsi;271 | 0 | 0 |
| NC_002645.1;NP_073556.1;HCoV-229E_nucleocapsi;272 | 0 | 0 |
| NC_002645.1;NP_073556.1;HCoV-229E_nucleocapsi;273 | 0 | 0 |
| NC_002645.1;NP_073556.1;HCoV-229E_nucleocapsi;274 | 0 | 0.025 |
| NC_002645.1;NP_073556.1;HCoV-229E_nucleocapsi;275 | 0 | 0.025 |
| NC_002645.1;NP_073556.1;HCoV-229E_nucleocapsi;276 | 0 | 0.025 |
| NC_002645.1;NP_073556.1;HCoV-229E_nucleocapsi;277 | 0 | 0.025 |
| NC_002645.1;NP_073556.1;HCoV-229E_nucleocapsi;278 | 0 | 0.025 |
| NC_002645.1;NP_073556.1;HCoV-229E_nucleocapsi;279 | 0.1 | 0.05 |
| NC_002645.1;NP_073556.1;HCoV-229E_nucleocapsi;280 | 0.05 | 0.05 |
| NC_002645.1;NP_073556.1;HCoV-229E_nucleocapsi;281 | 0.05 | 0.025 |
| NC_002645.1;NP_073556.1;HCoV-229E_nucleocapsi;282 | 0.15 | 0.075 |
| NC_002645.1;NP_073556.1;HCoV-229E_nucleocapsi;283 | 0.1 | 0.075 |
| NC_002645.1;NP_073556.1;HCoV-229E_nucleocapsi;284 | 0.1 | 0.025 |
| NC_002645.1;NP_073556.1;HCoV-229E_nucleocapsi;285 | 0.1 | 0.025 |
| NC_002645.1;NP_073556.1;HCoV-229E_nucleocapsi;286 | 0.2 | 0.025 |
| NC_002645.1;NP_073556.1;HCoV-229E_nucleocapsi;287 | 0.05 | 0.05 |
| NC_002645.1;NP_073556.1;HCoV-229E_nucleocapsi;288 | 0.05 | 0.025 |
| NC_002645.1;NP_073556.1;HCoV-229E_nucleocapsi;289 | 0.05 | 0 |
| NC_002645.1;NP_073556.1;HCoV-229E_nucleocapsi;290 | 0 | 0 |
| NC_002645.1;NP_073556.1;HCoV-229E_nucleocapsi;291 | 0 | 0 |
| NC_002645.1;NP_073556.1;HCoV-229E_nucleocapsi;292 | 0 | 0 |
| NC_002645.1;NP_073556.1;HCoV-229E_nucleocapsi;293 | 0 | 0.05 |
| NC_002645.1;NP_073556.1;HCoV-229E_nucleocapsi;294 | 0 | 0 |
| NC_002645.1;NP_073556.1;HCoV-229E_nucleocapsi;295 | 0 | 0 |

| | | |
|---|---|---|
| NC_002645.1;NP_073556.1;HCoV-229E_nucleocapsi;296 | 0 | 0 |
| NC_002645.1;NP_073556.1;HCoV-229E_nucleocapsi;297 | 0 | 0 |
| NC_002645.1;NP_073556.1;HCoV-229E_nucleocapsi;298 | 0 | 0 |
| NC_002645.1;NP_073556.1;HCoV-229E_nucleocapsi;299 | 0 | 0.025 |
| NC_002645.1;NP_073556.1;HCoV-229E_nucleocapsi;300 | 0.1 | 0.075 |
| NC_002645.1;NP_073556.1;HCoV-229E_nucleocapsi;301 | 0.1 | 0.1 |
| NC_002645.1;NP_073556.1;HCoV-229E_nucleocapsi;302 | 0.15 | 0.1 |
| NC_002645.1;NP_073556.1;HCoV-229E_nucleocapsi;303 | 0.1 | 0.1 |
| NC_002645.1;NP_073556.1;HCoV-229E_nucleocapsi;304 | 0.05 | 0.05 |
| NC_002645.1;NP_073556.1;HCoV-229E_nucleocapsi;305 | 0.05 | 0.05 |
| NC_002645.1;NP_073556.1;HCoV-229E_nucleocapsi;306 | 0.05 | 0.025 |
| NC_002645.1;NP_073556.1;HCoV-229E_nucleocapsi;307 | 0.05 | 0.025 |
| NC_002645.1;NP_073556.1;HCoV-229E_nucleocapsi;308 | 0.05 | 0.025 |
| NC_002645.1;NP_073556.1;HCoV-229E_nucleocapsi;309 | 0 | 0.025 |
| NC_002645.1;NP_073556.1;HCoV-229E_nucleocapsi;310 | 0 | 0 |
| NC_002645.1;NP_073556.1;HCoV-229E_nucleocapsi;311 | 0 | 0 |
| NC_002645.1;NP_073556.1;HCoV-229E_nucleocapsi;312 | 0 | 0 |
| NC_002645.1;NP_073556.1;HCoV-229E_nucleocapsi;313 | 0 | 0 |
| NC_002645.1;NP_073556.1;HCoV-229E_nucleocapsi;314 | 0 | 0 |
| NC_002645.1;NP_073556.1;HCoV-229E_nucleocapsi;315 | 0 | 0.025 |
| NC_002645.1;NP_073556.1;HCoV-229E_nucleocapsi;316 | 0 | 0 |
| NC_002645.1;NP_073556.1;HCoV-229E_nucleocapsi;317 | 0 | 0 |
| NC_002645.1;NP_073556.1;HCoV-229E_nucleocapsi;318 | 0 | 0 |
| NC_002645.1;NP_073556.1;HCoV-229E_nucleocapsi;319 | 0 | 0 |
| NC_002645.1;NP_073556.1;HCoV-229E_nucleocapsi;320 | 0 | 0 |
| NC_002645.1;NP_073556.1;HCoV-229E_nucleocapsi;321 | 0 | 0 |
| NC_002645.1;NP_073556.1;HCoV-229E_nucleocapsi;322 | 0 | 0.025 |
| NC_002645.1;NP_073556.1;HCoV-229E_nucleocapsi;323 | 0 | 0.025 |
| NC_002645.1;NP_073556.1;HCoV-229E_nucleocapsi;324 | 0 | 0.025 |
| NC_002645.1;NP_073556.1;HCoV-229E_nucleocapsi;325 | 0 | 0 |
| NC_002645.1;NP_073556.1;HCoV-229E_nucleocapsi;326 | 0.05 | 0 |
| NC_002645.1;NP_073556.1;HCoV-229E_nucleocapsi;327 | 0.05 | 0 |
| NC_002645.1;NP_073556.1;HCoV-229E_nucleocapsi;328 | 0 | 0 |
| NC_002645.1;NP_073556.1;HCoV-229E_nucleocapsi;329 | 0 | 0 |
| NC_002645.1;NP_073556.1;HCoV-229E_nucleocapsi;330 | 0 | 0 |
| NC_002645.1;NP_073556.1;HCoV-229E_nucleocapsi;331 | 0.05 | 0 |
| NC_002645.1;NP_073556.1;HCoV-229E_nucleocapsi;332 | 0.05 | 0 |
| NC_002645.1;NP_073556.1;HCoV-229E_nucleocapsi;333 | 0.05 | 0 |
| NC_002645.1;NP_073556.1;HCoV-229E_nucleocapsi;334 | 0.1 | 0.025 |
| NC_002645.1;NP_073556.1;HCoV-229E_nucleocapsi;335 | 0.15 | 0.175 |
| NC_002645.1;NP_073556.1;HCoV-229E_nucleocapsi;336 | 0.1 | 0.1 |
| NC_002645.1;NP_073556.1;HCoV-229E_nucleocapsi;337 | 0.05 | 0.075 |
| NC_002645.1;NP_073556.1;HCoV-229E_nucleocapsi;338 | 0.05 | 0 |

| | | |
|---|---|---|
| NC_002645.1;NP_073556.1;HCoV-229E_nucleocapsi;339 | 0.05 | 0 |
| NC_002645.1;NP_073556.1;HCoV-229E_nucleocapsi;340 | 0.05 | 0 |
| NC_002645.1;NP_073556.1;HCoV-229E_nucleocapsi;341 | 0.1 | 0 |
| NC_002645.1;NP_073556.1;HCoV-229E_nucleocapsi;342 | 0.05 | 0.025 |
| NC_002645.1;NP_073556.1;HCoV-229E_nucleocapsi;343 | 0.05 | 0.025 |
| NC_002645.1;NP_073556.1;HCoV-229E_nucleocapsi;344 | 0.05 | 0.025 |
| NC_002645.1;NP_073556.1;HCoV-229E_nucleocapsi;345 | 0.05 | 0.025 |
| NC_002645.1;NP_073556.1;HCoV-229E_nucleocapsi;346 | 0.05 | 0.05 |
| NC_002645.1;NP_073556.1;HCoV-229E_nucleocapsi;347 | 0.1 | 0.025 |
| NC_002645.1;NP_073556.1;HCoV-229E_nucleocapsi;348 | 0.05 | 0.075 |
| NC_002645.1;NP_073556.1;HCoV-229E_nucleocapsi;349 | 0.05 | 0.025 |
| NC_002645.1;NP_073556.1;HCoV-229E_nucleocapsi;350 | 0.05 | 0.05 |
| NC_002645.1;NP_073556.1;HCoV-229E_nucleocapsi;351 | 0.05 | 0.075 |
| NC_002645.1;NP_073556.1;HCoV-229E_nucleocapsi;352 | 0.05 | 0.075 |
| NC_002645.1;NP_073556.1;HCoV-229E_nucleocapsi;353 | 0.05 | 0.075 |
| NC_002645.1;NP_073556.1;HCoV-229E_nucleocapsi;354 | 0.05 | 0.1 |
| NC_002645.1;NP_073556.1;HCoV-229E_nucleocapsi;355 | 0.05 | 0.075 |
| NC_002645.1;NP_073556.1;HCoV-229E_nucleocapsi;356 | 0.05 | 0.025 |
| NC_002645.1;NP_073556.1;HCoV-229E_nucleocapsi;357 | 0.05 | 0.05 |
| NC_002645.1;NP_073556.1;HCoV-229E_nucleocapsi;358 | 0.05 | 0.05 |
| NC_002645.1;NP_073556.1;HCoV-229E_nucleocapsi;359 | 0.05 | 0.05 |
| NC_002645.1;NP_073556.1;HCoV-229E_nucleocapsi;360 | 0.15 | 0.05 |
| NC_002645.1;NP_073556.1;HCoV-229E_nucleocapsi;361 | 0.05 | 0 |
| NC_002645.1;NP_073556.1;HCoV-229E_nucleocapsi;362 | 0.05 | 0 |
| NC_002645.1;NP_073556.1;HCoV-229E_nucleocapsi;363 | 0.05 | 0 |
| NC_002645.1;NP_073556.1;HCoV-229E_nucleocapsi;364 | 0.1 | 0 |
| NC_002645.1;NP_073556.1;HCoV-229E_nucleocapsi;365 | 0.05 | 0 |
| NC_002645.1;NP_073556.1;HCoV-229E_nucleocapsi;366 | 0.05 | 0 |
| NC_002645.1;NP_073556.1;HCoV-229E_nucleocapsi;367 | 0.05 | 0 |
| NC_002645.1;NP_073556.1;HCoV-229E_nucleocapsi;368 | 0.05 | 0 |
| NC_002645.1;NP_073556.1;HCoV-229E_nucleocapsi;369 | 0.05 | 0.025 |
| NC_002645.1;NP_073556.1;HCoV-229E_nucleocapsi;370 | 0.05 | 0.025 |
| NC_002645.1;NP_073556.1;HCoV-229E_nucleocapsi;371 | 0.05 | 0.025 |
| NC_002645.1;NP_073556.1;HCoV-229E_nucleocapsi;372 | 0.1 | 0.025 |
| NC_002645.1;NP_073556.1;HCoV-229E_nucleocapsi;373 | 0.05 | 0.025 |
| NC_002645.1;NP_073556.1;HCoV-229E_nucleocapsi;374 | 0.05 | 0.025 |
| NC_002645.1;NP_073556.1;HCoV-229E_nucleocapsi;375 | 0.05 | 0 |
| NC_002645.1;NP_073556.1;HCoV-229E_nucleocapsi;376 | 0.05 | 0.025 |
| NC_002645.1;NP_073556.1;HCoV-229E_nucleocapsi;377 | 0.05 | 0 |
| NC_002645.1;NP_073556.1;HCoV-229E_nucleocapsi;378 | 0.05 | 0 |
