## Extended data 3 for "The landscape of antibody binding in SARS-CoV-2 infection"

| Extended data 3 | | | | | |
|---|---|---|---|---|---|
| **Protein** | **First aa position** | **Sequence** | **Specificity** | **Sensitivity** | **F1** |
| M | 8 | ITVEELKKLLEQWNLV | 1 | 0.975 | 0.98734177 |
| M | 7 | TITVEELKKLLEQWNL | 1 | 0.95 | 0.97435897 |
| N | 390 | QTVTLLPAADLDDFSK | 1 | 0.95 | 0.97435897 |
| N | 388 | KQQTVTLLPAADLDDF | 1 | 0.9 | 0.94736842 |
| N | 391 | TVTLLPAADLDDFSKQ | 1 | 0.9 | 0.94736842 |
| S | 570 | ADTTDAVRDPQTLEIL | 1 | 0.875 | 0.93333333 |
| S | 571 | DTTDAVRDPQTLEILD | 1 | 0.875 | 0.93333333 |
| S | 574 | DAVRDPQTLEILDITP | 1 | 0.85 | 0.91891892 |
| S | 576 | VRDPQTLEILDITPCS | 1 | 0.85 | 0.91891892 |
| S | 1253 | CCKFDEDDSEPVLKGV | 1 | 0.85 | 0.91891892 |
| S | 572 | TTDAVRDPQTLEILDI | 1 | 0.825 | 0.90410959 |
| S | 573 | TDAVRDPQTLEILDIT | 1 | 0.825 | 0.90410959 |
| S | 577 | RDPQTLEILDITPCSF | 1 | 0.825 | 0.90410959 |
| S | 1252 | SCCKFDEDDSEPVLKG | 1 | 0.825 | 0.90410959 |
| M | 162 | KDLPKEITVATSRTLS | 1 | 0.825 | 0.90410959 |
| S | 1250 | CGSCCKFDEDDSEPVL | 1 | 0.8 | 0.88888889 |
| S | 814 | KRSFIEDLLFNKVTLA | 0.95 | 0.9 | 0.93506494 |
| M | 5 | NGTITVEELKKLLEQW | 0.95 | 0.9 | 0.93506494 |
| N | 392 | VTLLPAADLDDFSKQL | 0.95 | 0.9 | 0.93506494 |
| S | 626 | ADQLTPTWRVYSTGSN | 0.95 | 0.875 | 0.92105263 |
| S | 811 | KPSKRSFIEDLLFNKV | 0.95 | 0.875 | 0.92105263 |
| M | 4 | SNGTITVEELKKLLEQ | 0.95 | 0.875 | 0.92105263 |
| N | 389 | QQTVTLLPAADLDDFS | 0.95 | 0.875 | 0.92105263 |
| S | 575 | AVRDPQTLEILDITPC | 0.95 | 0.85 | 0.90666667 |
| N | 386 | QKKQQTVTLLPAADLD | 0.95 | 0.85 | 0.90666667 |
| S | 1251 | GSCCKFDEDDSEPVLK | 0.95 | 0.825 | 0.89189189 |
| S | 1255 | KFDEDDSEPVLKGVKL | 0.95 | 0.825 | 0.89189189 |
| M | 9 | TVEELKKLLEQWNLVI | 0.95 | 0.825 | 0.89189189 |
| M | 160 | DIKDLPKEITVATSRT | 0.95 | 0.825 | 0.89189189 |
| M | 161 | IKDLPKEITVATSRTL | 0.95 | 0.825 | 0.89189189 |
| M | 184 | SQRVAGDSGFAAYSRY | 0.95 | 0.8 | 0.87671233 |
| orf1ab | 4514 | YTMADLVYALRHFDEG | 0.95 | 0.775 | 0.86111111 |
| S | 1257 | DEDDSEPVLKGVKLHY | 0.95 | 0.75 | 0.84507042 |
| N | 213 | NGGDAALALLLLDRLN | 0.9 | 0.925 | 0.93670886 |
| N | 215 | GDAALALLLLDRLNQL | 0.9 | 0.9 | 0.92307692 |
| S | 554 | ESNKKFLPFQQFGRDI | 0.9 | 0.875 | 0.90909091 |
| S | 795 | KDFGGFNFSQILPDPS | 0.9 | 0.875 | 0.90909091 |
| N | 214 | GGDAALALLLLDRLNQ | 0.9 | 0.875 | 0.90909091 |
| N | 220 | ALLLLDRLNQLESKMS | 0.9 | 0.875 | 0.90909091 |
| S | 557 | KKFLPFQQFGRDIADT | 0.9 | 0.85 | 0.89473684 |
| S | 787 | QIYKTPPIKDFGGFNF | 0.9 | 0.85 | 0.89473684 |

| | | | | | |
|---|---|---|---|---|---|
| M | 3 | DSNGTITVEELKKLLE | 0.9 | 0.85 | 0.89473684 |
| M | 181 | LGASQRVAGDSGFAAY | 0.9 | 0.85 | 0.89473684 |
| N | 212 | GNGGDAALALLLLDRL | 0.9 | 0.85 | 0.89473684 |
| N | 216 | DAALALLLLDRLNQLE | 0.9 | 0.85 | 0.89473684 |
| S | 556 | NKKFLPFQQFGRDIAD | 0.9 | 0.825 | 0.88 |
| S | 558 | KFLPFQQFGRDIADTT | 0.9 | 0.825 | 0.88 |
| S | 559 | FLPFQQFGRDIADTTD | 0.9 | 0.825 | 0.88 |
| S | 568 | DIADTTDAVRDPQTLE | 0.9 | 0.825 | 0.88 |
| M | 159 | CDIKDLPKEITVATSR | 0.9 | 0.825 | 0.88 |
| M | 182 | GASQRVAGDSGFAAYS | 0.9 | 0.825 | 0.88 |
| N | 11 | NAPRITFGGPSDSTGS | 0.9 | 0.825 | 0.88 |
| N | 210 | MAGNGGDAALALLLLD | 0.9 | 0.825 | 0.88 |
| S | 792 | PPIKDFGGFNFSQILP | 0.9 | 0.8 | 0.86486486 |
| S | 793 | PIKDFGGFNFSQILPD | 0.9 | 0.8 | 0.86486486 |
| S | 1254 | CKFDEDDSEPVLKGVK | 0.9 | 0.8 | 0.86486486 |
| M | 183 | ASQRVAGDSGFAAYSR | 0.9 | 0.8 | 0.86486486 |
| M | 185 | QRVAGDSGFAAYSRYR | 0.9 | 0.775 | 0.84931507 |
| S | 625 | HADQLTPTWRVYSTGS | 0.9 | 0.75 | 0.83333333 |
| S | 1256 | FDEDDSEPVLKGVKLH | 0.9 | 0.75 | 0.83333333 |
| S | 1249 | SCGSCCKFDEDDSEPV | 0.9 | 0.725 | 0.81690141 |
| S | 810 | SKPSKRSFIEDLLFNK | 0.85 | 0.9 | 0.91139241 |
| S | 812 | PSKRSFIEDLLFNKVT | 0.85 | 0.9 | 0.91139241 |
| N | 393 | TLLPAADLDDFSKQLQ | 0.85 | 0.9 | 0.91139241 |
| N | 394 | LLPAADLDDFSKQLQQ | 0.85 | 0.9 | 0.91139241 |
| S | 553 | TESNKKFLPFQQFGRD | 0.85 | 0.875 | 0.8974359 |
| S | 788 | IYKTPPIKDFGGFNFS | 0.85 | 0.875 | 0.8974359 |
| S | 794 | IKDFGGFNFSQILPDP | 0.85 | 0.875 | 0.8974359 |
| S | 813 | SKRSFIEDLLFNKVTL | 0.85 | 0.875 | 0.8974359 |
| N | 9 | QRNAPRITFGGPSDST | 0.85 | 0.875 | 0.8974359 |
| N | 10 | RNAPRITFGGPSDSTG | 0.85 | 0.875 | 0.8974359 |
| N | 387 | KKQQTVTLLPAADLDD | 0.85 | 0.875 | 0.8974359 |
| S | 560 | LPFQQFGRDIADTTDA | 0.85 | 0.85 | 0.88311688 |
| S | 569 | IADTTDAVRDPQTLEI | 0.85 | 0.85 | 0.88311688 |
| S | 1141 | LQPELDSFKEELDKYF | 0.85 | 0.85 | 0.88311688 |
| S | 1143 | PELDSFKEELDKYFKN | 0.85 | 0.85 | 0.88311688 |
| N | 396 | PAADLDDFSKQLQQSM | 0.85 | 0.85 | 0.88311688 |
| S | 561 | PFQQFGRDIADTTDAV | 0.85 | 0.825 | 0.86842105 |
| S | 692 | IIAYTMSLGAENSVAY | 0.85 | 0.825 | 0.86842105 |
| S | 789 | YKTPPIKDFGGFNFSQ | 0.85 | 0.825 | 0.86842105 |
| N | 211 | AGNGGDAALALLLLDR | 0.85 | 0.825 | 0.86842105 |
| M | 188 | AGDSGFAAYSRYRIGN | 0.85 | 0.775 | 0.83783784 |
| M | 6 | GTITVEELKKLLEQWN | 0.8 | 0.9 | 0.9 |
| N | 395 | LPAADLDDFSKQLQQS | 0.8 | 0.9 | 0.9 |

| | | | | | |
|---|---|---|---|---|---|
| orf1ab | 5999 | ITREEAIRHVRAWIGF | 0.8 | 0.85 | 0.87179487 |
| S | 549 | TGVLTESNKKFLPFQQ | 0.8 | 0.85 | 0.87179487 |
| S | 686 | SVASQSIIAYTMSLGA | 0.8 | 0.85 | 0.87179487 |
| S | 688 | ASQSIIAYTMSLGAEN | 0.8 | 0.85 | 0.87179487 |
| S | 791 | TPPIKDFGGFNFSQIL | 0.8 | 0.85 | 0.87179487 |
| S | 1142 | QPELDSFKEELDKYFK | 0.8 | 0.85 | 0.87179487 |
| S | 1144 | ELDSFKEELDKYFKNH | 0.8 | 0.85 | 0.87179487 |
| N | 221 | LLLLDRLNQLESKMSG | 0.8 | 0.85 | 0.87179487 |
| S | 622 | VAIHADQLTPTWRVYS | 0.8 | 0.825 | 0.85714286 |
| S | 790 | KTPPIKDFGGFNFSQI | 0.8 | 0.825 | 0.85714286 |
| M | 155 | HLGRCDIKDLPKEITV | 0.8 | 0.825 | 0.85714286 |
| M | 157 | GRCDIKDLPKEITVAT | 0.8 | 0.825 | 0.85714286 |
| N | 37 | SKQRRPQGLPNNTASW | 0.8 | 0.8 | 0.84210526 |
| N | 250 | SAAEASKKPRQKRTAT | 0.8 | 0.8 | 0.84210526 |
| N | 399 | DLDDFSKQLQQSMSSA | 0.8 | 0.8 | 0.84210526 |
| M | 211 | SSSSDNIALLVQ | 0.8 | 0.775 | 0.82666667 |
| M | 187 | VAGDSGFAAYSRYRIG | 0.8 | 0.75 | 0.81081081 |
| M | 189 | GDSGFAAYSRYRIGNY | 0.8 | 0.75 | 0.81081081 |
| N | 7 | QNQRNAPRITFGGPSD | 0.75 | 0.9 | 0.88888889 |
| M | 2 | ADSNGTITVEELKKLL | 0.75 | 0.875 | 0.875 |
| N | 397 | AADLDDFSKQLQQSMS | 0.75 | 0.875 | 0.875 |
| S | 555 | SNKKFLPFQQFGRDIA | 0.75 | 0.85 | 0.86075949 |
| S | 693 | IAYTMSLGAENSVAYS | 0.75 | 0.85 | 0.86075949 |
| M | 158 | RCDIKDLPKEITVATS | 0.75 | 0.85 | 0.86075949 |
| N | 155 | AAIVLQLPQGTTLPKG | 0.75 | 0.85 | 0.86075949 |
| N | 161 | LPQGTTLPKGFYAEGS | 0.75 | 0.85 | 0.86075949 |
| N | 219 | LALLLLDRLNQLESKM | 0.75 | 0.85 | 0.86075949 |
| S | 689 | SQSIIAYTMSLGAENS | 0.75 | 0.825 | 0.84615385 |
| M | 154 | HHLGRCDIKDLPKEIT | 0.75 | 0.825 | 0.84615385 |
| M | 205 | KLNTDHSSSSDNIALL | 0.75 | 0.825 | 0.84615385 |
| N | 14 | RITFGGPSDSTGSNQN | 0.75 | 0.825 | 0.84615385 |
| N | 208 | ARMAGNGGDAALALLL | 0.75 | 0.825 | 0.84615385 |
| M | 208 | TDHSSSSDNIALLVQ | 0.75 | 0.8 | 0.83116883 |
| N | 12 | APRITFGGPSDSTGSN | 0.75 | 0.8 | 0.83116883 |
| N | 34 | GARSKQRRPQGLPNNT | 0.75 | 0.8 | 0.83116883 |
| N | 154 | NAAIVLQLPQGTTLPK | 0.75 | 0.8 | 0.83116883 |
| S | 562 | FQQFGRDIADTTDAVR | 0.75 | 0.775 | 0.81578947 |
| S | 627 | DQLTPTWRVYSTGSNV | 0.75 | 0.775 | 0.81578947 |
| S | 615 | VNCTEVPVAIHADQLT | 0.75 | 0.75 | 0.8 |
| S | 816 | SFIEDLLFNKVTLADA | 0.7 | 0.875 | 0.86419753 |
| S | 940 | STASALGKLQDVVNQN | 0.7 | 0.875 | 0.86419753 |
| N | 160 | QLPQGTTLPKGFYAEG | 0.7 | 0.875 | 0.86419753 |
| N | 218 | ALALLLLDRLNQLESK | 0.7 | 0.875 | 0.86419753 |

| | | | | | |
|---|---|---|---|---|---|
| N | 400 | LDDFSKQLQQSMSSAD | 0.7 | 0.875 | 0.86419753 |
| S | 624 | IHADQLTPTWRVYSTG | 0.7 | 0.85 | 0.85 |
| S | 815 | RSFIEDLLFNKVTLAD | 0.7 | 0.85 | 0.85 |
| S | 1260 | DSEPVLKGVKLHYT | 0.7 | 0.85 | 0.85 |
| N | 223 | LLDRLNQLESKMSGKG | 0.7 | 0.85 | 0.85 |
| S | 690 | QSIIAYTMSLGAENSV | 0.7 | 0.825 | 0.83544304 |
| S | 1140 | PLQPELDSFKEELDKY | 0.7 | 0.825 | 0.83544304 |
| N | 156 | AIVLQLPQGTTLPKGF | 0.7 | 0.825 | 0.83544304 |
| N | 158 | VLQLPQGTTLPKGFYA | 0.7 | 0.825 | 0.83544304 |
| N | 222 | LLLDRLNQLESKMSGK | 0.7 | 0.825 | 0.83544304 |
| M | 207 | NTDHSSSSDNIALLVQ | 0.7 | 0.8 | 0.82051282 |
| orf3a | 258 | PVMEPIYDEPTTTTSV | 0.7 | 0.775 | 0.80519481 |
| M | 206 | LNTDHSSSSDNIALLV | 0.7 | 0.775 | 0.80519481 |
| M | 210 | HSSSSDNIALLVQ | 0.7 | 0.775 | 0.80519481 |
| N | 33 | SGARSKQRRPQGLPNN | 0.7 | 0.775 | 0.80519481 |
| S | 621 | PVAIHADQLTPTWRVY | 0.7 | 0.75 | 0.78947368 |
| S | 1248 | CSCGSCCKFDEDDSEP | 0.7 | 0.75 | 0.78947368 |
| S | 290 | DCALDPLSETKCTLKS | 0.65 | 0.9 | 0.86746988 |
| orf1ab | 1720 | KTVGELGDVRETMSYL | 0.65 | 0.875 | 0.85365854 |
| S | 550 | GVLTESNKKFLPFQQF | 0.65 | 0.875 | 0.85365854 |
| N | 8 | NQRNAPRITFGGPSDS | 0.65 | 0.875 | 0.85365854 |
| orf1ab | 2309 | ITISSFKWDLTAFGLV | 0.65 | 0.85 | 0.83950617 |
| S | 687 | VASQSIIAYTMSLGAE | 0.65 | 0.85 | 0.83950617 |
| N | 159 | LQLPQGTTLPKGFYAE | 0.65 | 0.85 | 0.83950617 |
| N | 209 | RMAGNGGDAALALLLL | 0.65 | 0.85 | 0.83950617 |
| S | 563 | QQFGRDIADTTDAVRD | 0.65 | 0.825 | 0.825 |
| S | 619 | EVPVAIHADQLTPTWR | 0.65 | 0.825 | 0.825 |
| S | 1155 | YFKNHTSPDVDLGDIS | 0.65 | 0.825 | 0.825 |
| S | 1258 | EDDSEPVLKGVKLHYT | 0.65 | 0.825 | 0.825 |
| S | 1259 | DDSEPVLKGVKLHYT | 0.65 | 0.825 | 0.825 |
| N | 157 | IVLQLPQGTTLPKGFY | 0.65 | 0.825 | 0.825 |
| orf1ab | 6058 | FSRVSAKPPPGDQFKH | 0.65 | 0.8 | 0.81012658 |
| S | 808 | DPSKPSKRSFIEDLLF | 0.65 | 0.8 | 0.81012658 |
| S | 1157 | KNHTSPDVDLGDISGI | 0.65 | 0.8 | 0.81012658 |
| S | 616 | NCTEVPVAIHADQLTP | 0.65 | 0.725 | 0.76315789 |
| orf1ab | 1241 | TTLEETKFLTENLLLY | 0.6 | 0.9 | 0.85714286 |
| S | 1156 | FKNHTSPDVDLGDISG | 0.6 | 0.9 | 0.85714286 |
| S | 1147 | SFKEELDKYFKNHTSP | 0.6 | 0.875 | 0.84337349 |
| orf1ab | 1551 | ITFDNLKTLLSLREVR | 0.6 | 0.85 | 0.82926829 |
| S | 289 | VDCALDPLSETKCTLK | 0.6 | 0.85 | 0.82926829 |
| S | 630 | TPTWRVYSTGSNVFQT | 0.6 | 0.85 | 0.82926829 |
| M | 156 | LGRCDIKDLPKEITVA | 0.6 | 0.85 | 0.82926829 |
| N | 15 | ITFGGPSDSTGSNQNG | 0.6 | 0.85 | 0.82926829 |

| | | | | | |
|---|---|---|---|---|---|
| N | 401 | DDFSKQLQQSMSSADS | 0.6 | 0.85 | 0.82926829 |
| S | 1148 | FKEELDKYFKNHTSPD | 0.6 | 0.825 | 0.81481481 |
| N | 32 | RSGARSKQRRPQGLPN | 0.6 | 0.825 | 0.81481481 |
| N | 377 | DETQALPQRQKKQQTV | 0.6 | 0.825 | 0.81481481 |
| M | 209 | DHSSSSDNIALLVQ | 0.6 | 0.775 | 0.78481013 |
| N | 30 | GERSGARSKQRRPQGL | 0.6 | 0.775 | 0.78481013 |
| S | 404 | GDEVRQIAPGQTGKIA | 0.55 | 0.95 | 0.87356322 |
| S | 807 | PDPSKPSKRSFIEDLL | 0.55 | 0.9 | 0.84705882 |
| M | 1 | MADSNGTITVEELKKL | 0.55 | 0.9 | 0.84705882 |
| S | 656 | VNNSYECDIPIGAGIC | 0.55 | 0.875 | 0.83333333 |
| S | 657 | NNSYECDIPIGAGICA | 0.55 | 0.875 | 0.83333333 |
| S | 1158 | NHTSPDVDLGDISGIN | 0.55 | 0.85 | 0.81927711 |
| S | 551 | VLTESNKKFLPFQQFG | 0.55 | 0.825 | 0.80487805 |
| orf3a | 259 | VMEPIYDEPTTTTSVP | 0.55 | 0.825 | 0.80487805 |
| N | 231 | ESKMSGKGQQQQGQTV | 0.55 | 0.825 | 0.80487805 |
| S | 639 | GSNVFQTRAGCLIGAE | 0.55 | 0.8 | 0.79012346 |
| S | 809 | PSKPSKRSFIEDLLFN | 0.55 | 0.8 | 0.79012346 |
| M | 186 | RVAGDSGFAAYSRYRI | 0.55 | 0.8 | 0.79012346 |
| N | 29 | NGERSGARSKQRRPQG | 0.55 | 0.8 | 0.79012346 |
| N | 35 | ARSKQRRPQGLPNNTA | 0.55 | 0.8 | 0.79012346 |
| N | 233 | KMSGKGQQQQGQTVTK | 0.55 | 0.8 | 0.79012346 |
| N | 398 | ADLDDFSKQLQQSMSS | 0.55 | 0.8 | 0.79012346 |
| N | 31 | ERSGARSKQRRPQGLP | 0.55 | 0.775 | 0.775 |
| N | 38 | KQRRPQGLPNNTASWF | 0.55 | 0.775 | 0.775 |
| N | 162 | PQGTTLPKGFYAEGSR | 0.55 | 0.775 | 0.775 |
| N | 217 | AALALLLLDRLNQLES | 0.55 | 0.775 | 0.775 |
| orf3a | 255 | VVNPVMEPIYDEPTTT | 0.55 | 0.75 | 0.75949367 |
| S | 1262 | EPVLKGVKLHYT | 0.5 | 0.925 | 0.85057471 |
| S | 691 | SIIAYTMSLGAENSVA | 0.5 | 0.9 | 0.8372093 |
| orf8 | 12 | TVAAFHQECSLQSCTQ | 0.5 | 0.9 | 0.8372093 |
| orf8 | 67 | SKSPIQYIDIGNYTVS | 0.5 | 0.9 | 0.8372093 |
| orf1ab | 6057 | DFSRVSAKPPPGDQFK | 0.5 | 0.875 | 0.82352941 |
| S | 636 | YSTGSNVFQTRAGCLI | 0.5 | 0.875 | 0.82352941 |
| S | 1149 | KEELDKYFKNHTSPDV | 0.5 | 0.875 | 0.82352941 |
| orf8 | 60 | LCVDEAGSKSPIQYID | 0.5 | 0.825 | 0.79518072 |
| N | 36 | RSKQRRPQGLPNNTAS | 0.5 | 0.825 | 0.79518072 |
| N | 242 | QGQTVTKKSAAEASKK | 0.5 | 0.825 | 0.79518072 |
| N | 243 | GQTVTKKSAAEASKKP | 0.5 | 0.825 | 0.79518072 |
| S | 578 | DPQTLEILDITPCSFG | 0.5 | 0.8 | 0.7804878 |
| S | 614 | DVNCTEVPVAIHADQL | 0.5 | 0.8 | 0.7804878 |
| orf3a | 260 | MEPIYDEPTTTTSVPL | 0.5 | 0.8 | 0.7804878 |
| M | 194 | AAYSRYRIGNYKLNTD | 0.5 | 0.8 | 0.7804878 |
| S | 617 | CTEVPVAIHADQLTPT | 0.5 | 0.775 | 0.7654321 |

| | | | | | |
|---|---|---|---|---|---|
| orf3a | 253 | SGVVNPVMEPIYDEPT | 0.5 | 0.775 | 0.7654321 |
| N | 163 | QGTTLPKGFYAEGSRG | 0.5 | 0.775 | 0.7654321 |
| N | 245 | TVTKKSAAEASKKPRQ | 0.5 | 0.775 | 0.7654321 |
| orf3a | 254 | GVVNPVMEPIYDEPTT | 0.5 | 0.75 | 0.75 |
| orf3a | 256 | VNPVMEPIYDEPTTTT | 0.5 | 0.75 | 0.75 |
| S | 410 | IAPGQTGKIADYNYKL | 0.45 | 0.925 | 0.84090909 |
| S | 817 | FIEDLLFNKVTLADAG | 0.45 | 0.925 | 0.84090909 |
| N | 118 | EAGLPYGANKDGIIWV | 0.45 | 0.925 | 0.84090909 |
| N | 119 | AGLPYGANKDGIIWVA | 0.45 | 0.9 | 0.82758621 |
| N | 230 | LESKMSGKGQQQQGQT | 0.45 | 0.9 | 0.82758621 |
| N | 235 | SGKGQQQQGQTVTKKS | 0.45 | 0.9 | 0.82758621 |
| N | 340 | DDKDPNFKDQVILLNK | 0.45 | 0.9 | 0.82758621 |
| S | 1164 | VDLGDISGINASVVNI | 0.45 | 0.875 | 0.81395349 |
| N | 381 | ALPQRQKKQQTVTLLP | 0.45 | 0.875 | 0.81395349 |
| N | 382 | LPQRQKKQQTVTLLPA | 0.45 | 0.875 | 0.81395349 |
| S | 844 | IAARDLICAQKFNGLT | 0.45 | 0.85 | 0.8 |
| S | 1145 | LDSFKEELDKYFKNHT | 0.45 | 0.85 | 0.8 |
| S | 1146 | DSFKEELDKYFKNHTS | 0.45 | 0.85 | 0.8 |
| S | 1159 | HTSPDVDLGDISGINA | 0.45 | 0.85 | 0.8 |
| M | 153 | GHHLGRCDIKDLPKEI | 0.45 | 0.85 | 0.8 |
| N | 339 | LDDKDPNFKDQVILLN | 0.45 | 0.85 | 0.8 |
| M | 190 | DSGFAAYSRYRIGNYK | 0.45 | 0.825 | 0.78571429 |
| S | 620 | VPVAIHADQLTPTWRV | 0.45 | 0.8 | 0.77108434 |
| S | 761 | TQLNRALTGIAVEQDK | 0.45 | 0.8 | 0.77108434 |
| orf8 | 61 | CVDEAGSKSPIQYIDI | 0.45 | 0.775 | 0.75609756 |
| N | 94 | IRGGDGKMKDLSPRWY | 0.45 | 0.775 | 0.75609756 |
| orf1ab | 1240 | TTTLEETKFLTENLLL | 0.4 | 0.95 | 0.84444444 |
| orf1ab | 4451 | KDEDDNLIDSYFVVKR | 0.4 | 0.95 | 0.84444444 |
| orf8 | 62 | VDEAGSKSPIQYIDIG | 0.4 | 0.925 | 0.83146067 |
| S | 406 | EVRQIAPGQTGKIADY | 0.4 | 0.9 | 0.81818182 |
| S | 1161 | SPDVDLGDISGINASV | 0.4 | 0.9 | 0.81818182 |
| N | 126 | NKDGIIWVATEGALNT | 0.4 | 0.9 | 0.81818182 |
| S | 405 | DEVRQIAPGQTGKIAD | 0.4 | 0.875 | 0.8045977 |
| S | 694 | AYTMSLGAENSVAYSN | 0.4 | 0.875 | 0.8045977 |
| S | 798 | GGFNFSQILPDPSKPS | 0.4 | 0.875 | 0.8045977 |
| S | 1162 | PDVDLGDISGINASVV | 0.4 | 0.875 | 0.8045977 |
| orf3a | 257 | NPVMEPIYDEPTTTTS | 0.4 | 0.875 | 0.8045977 |
| orf8 | 66 | GSKSPIQYIDIGNYTV | 0.4 | 0.85 | 0.79069767 |
| orf8 | 68 | KSPIQYIDIGNYTVSC | 0.4 | 0.85 | 0.79069767 |
| N | 234 | MSGKGQQQQGQTVTKK | 0.4 | 0.85 | 0.79069767 |
| N | 244 | QTVTKKSAAEASKKPR | 0.4 | 0.85 | 0.79069767 |
| N | 356 | HIDAYKTFPPTEPKKD | 0.4 | 0.85 | 0.79069767 |
| S | 618 | TEVPVAIHADQLTPTW | 0.4 | 0.825 | 0.77647059 |

| | | | | | |
|---|---|---|---|---|---|
| N | 153 | NNAAIVLQLPQGTTLP | 0.4 | 0.825 | 0.77647059 |
| N | 232 | SKMSGKGQQQQGQTVT | 0.4 | 0.825 | 0.77647059 |
| S | 768 | TGIAVEQDKNTQEVFA | 0.4 | 0.8 | 0.76190476 |
| S | 637 | STGSNVFQTRAGCLIG | 0.4 | 0.775 | 0.74698795 |
| N | 122 | PYGANKDGIIWVATEG | 0.35 | 1 | 0.86021505 |
| S | 631 | PTWRVYSTGSNVFQTR | 0.35 | 0.9 | 0.80898876 |
| S | 644 | QTRAGCLIGAEHVNNS | 0.35 | 0.9 | 0.80898876 |
| S | 796 | DFGGFNFSQILPDPSK | 0.35 | 0.9 | 0.80898876 |
| orf3a | 252 | SSGVVNPVMEPIYDEP | 0.35 | 0.9 | 0.80898876 |
| N | 385 | RQKKQQTVTLLPAADL | 0.35 | 0.9 | 0.80898876 |
| S | 1261 | SEPVLKGVKLHYT | 0.35 | 0.875 | 0.79545455 |
| N | 378 | ETQALPQRQKKQQTVT | 0.35 | 0.875 | 0.79545455 |
| N | 384 | QRQKKQQTVTLLPAAD | 0.35 | 0.875 | 0.79545455 |
| S | 537 | KCVNFNFNGLTGTGVL | 0.35 | 0.85 | 0.7816092 |
| S | 785 | VKQIYKTPPIKDFGGF | 0.35 | 0.85 | 0.7816092 |
| orf3a | 261 | EPIYDEPTTTTSVPL | 0.35 | 0.85 | 0.7816092 |
| M | 152 | AGHHLGRCDIKDLPKE | 0.35 | 0.825 | 0.76744186 |
| orf8 | 64 | EAGSKSPIQYIDIGNY | 0.35 | 0.825 | 0.76744186 |
| N | 28 | QNGERSGARSKQRRPQ | 0.35 | 0.775 | 0.73809524 |
| orf3a | 236 | IVDEPEEHVQIHTIDG | 0.3 | 0.975 | 0.83870968 |
| orf1ab | 1239 | VTTTLEETKFLTENLL | 0.3 | 0.925 | 0.81318681 |
| N | 227 | LNQLESKMSGKGQQQQ | 0.3 | 0.925 | 0.81318681 |
| N | 380 | QALPQRQKKQQTVTLL | 0.3 | 0.925 | 0.81318681 |
| orf1ab | 1681 | LTLQQIELKFNPPALQ | 0.3 | 0.9 | 0.8 |
| orf1ab | 2584 | AEVAVKMFDAYVNTFS | 0.3 | 0.9 | 0.8 |
| N | 124 | GANKDGIIWVATEGAL | 0.3 | 0.9 | 0.8 |
| M | 197 | SRYRIGNYKLNTDHSS | 0.3 | 0.875 | 0.78651685 |
| N | 249 | KSAAEASKKPRQKRTA | 0.3 | 0.875 | 0.78651685 |
| S | 628 | QLTPTWRVYSTGSNVF | 0.3 | 0.85 | 0.77272727 |
| M | 177 | SYYKLGASQRVAGDSG | 0.3 | 0.85 | 0.77272727 |
| M | 179 | YKLGASQRVAGDSGFA | 0.3 | 0.85 | 0.77272727 |
| S | 613 | QDVNCTEVPVAIHADQ | 0.3 | 0.825 | 0.75862069 |
| S | 1178 | NIQKEIDRLNEVAKNL | 0.3 | 0.825 | 0.75862069 |
| N | 336 | AIKLDDKDPNFKDQVI | 0.3 | 0.825 | 0.75862069 |
| M | 178 | YYKLGASQRVAGDSGF | 0.3 | 0.8 | 0.74418605 |
| N | 338 | KLDDKDPNFKDQVILL | 0.3 | 0.8 | 0.74418605 |
| S | 685 | RSVASQSIIAYTMSLG | 0.25 | 0.925 | 0.80434783 |
| S | 629 | LTPTWRVYSTGSNVFQ | 0.25 | 0.9 | 0.79120879 |
| N | 117 | PEAGLPYGANKDGIIW | 0.25 | 0.875 | 0.77777778 |
| N | 251 | AAEASKKPRQKRTATK | 0.25 | 0.875 | 0.77777778 |
| S | 638 | TGSNVFQTRAGCLIGA | 0.25 | 0.85 | 0.76404494 |
| S | 786 | KQIYKTPPIKDFGGFN | 0.25 | 0.85 | 0.76404494 |
| M | 195 | AYSRYRIGNYKLNTDH | 0.25 | 0.85 | 0.76404494 |

| | | | | | |
|---|---|---|---|---|---|
| orf8 | 63 | DEAGSKSPIQYIDIGN | 0.25 | 0.85 | 0.76404494 |
| S | 661 | ECDIPIGAGICASYQT | 0.25 | 0.8 | 0.73563218 |
| S | 769 | GIAVEQDKNTQEVFAQ | 0.25 | 0.775 | 0.72093023 |
| S | 770 | IAVEQDKNTQEVFAQV | 0.25 | 0.775 | 0.72093023 |
| S | 536 | NKCVNFNFNGLTGTGV | 0.2 | 0.95 | 0.80851064 |
| N | 341 | DKDPNFKDQVILLNKH | 0.2 | 0.95 | 0.80851064 |
| N | 127 | KDGIIWVATEGALNTP | 0.2 | 0.9 | 0.7826087 |
| S | 541 | FNFNGLTGTGVLTESN | 0.2 | 0.875 | 0.76923077 |
| M | 175 | TLSYYKLGASQRVAGD | 0.2 | 0.875 | 0.76923077 |
| N | 376 | ADETQALPQRQKKQQT | 0.2 | 0.875 | 0.76923077 |
| S | 804 | QILPDPSKPSKRSFIE | 0.2 | 0.85 | 0.75555556 |
| S | 241 | LLALHRSYLTPGDSSS | 0.15 | 0.975 | 0.8125 |
| orf1ab | 1572 | TTVDNINLHTQVVDMS | 0.15 | 0.95 | 0.8 |
| S | 799 | GFNFSQILPDPSKPSK | 0.15 | 0.925 | 0.78723404 |
| N | 379 | TQALPQRQKKQQTVTL | 0.15 | 0.925 | 0.78723404 |
| S | 635 | VYSTGSNVFQTRAGCL | 0.15 | 0.875 | 0.76086957 |
| M | 176 | LSYYKLGASQRVAGDS | 0.15 | 0.875 | 0.76086957 |
| S | 805 | ILPDPSKPSKRSFIED | 0.15 | 0.85 | 0.74725275 |
| orf1ab | 1546 | LDGEVITFDNLKTLLS | 0.1 | 1 | 0.81632653 |
| S | 306 | FTVEKGIYQTSNFRVQ | 0.1 | 1 | 0.81632653 |
| N | 228 | NQLESKMSGKGQQQQG | 0.1 | 1 | 0.81632653 |
| M | 196 | YSRYRIGNYKLNTDHS | 0.1 | 0.975 | 0.80412371 |
| N | 120 | GLPYGANKDGIIWVAT | 0.1 | 0.95 | 0.79166667 |
| M | 192 | GFAAYSRYRIGNYKLN | 0.1 | 0.925 | 0.77894737 |
| orf6 | 9 | VTIAEILLIIMRTFKV | 0.1 | 0.925 | 0.77894737 |
| M | 193 | FAAYSRYRIGNYKLNT | 0.1 | 0.875 | 0.75268817 |
| M | 191 | SGFAAYSRYRIGNYKL | 0.05 | 1 | 0.80808081 |
| orf3a | 16 | KQGEIKDATPSDFVRA | 0.05 | 0.925 | 0.77083333 |
| N | 96 | GGDGKMKDLSPRWYFY | 0.05 | 0.925 | 0.77083333 |
| S | 1247 | CCSCGSCCKFDEDDSE | 0.05 | 0.875 | 0.74468085 |
| orf8 | 53 | KSAPLIELCVDEAGSK | 0.05 | 0.875 | 0.74468085 |
| orf3a | 21 | KDATPSDFVRATATIP | 0 | 1 | 0.8 |
| orf3a | 18 | GEIKDATPSDFVRATA | 0 | 0.975 | 0.78787879 |
| N | 40 | RRPQGLPNNTASWFTA | 0 | 0.975 | 0.78787879 |
| N | 97 | GDGKMKDLSPRWYFYY | 0 | 0.975 | 0.78787879 |
| S | 172 | SQPFLMDLEGKQGNFK | 0 | 0.95 | 0.7755102 |
| orf3a | 235 | KIVDEPEEHVQIHTID | 0 | 0.95 | 0.7755102 |
| N | 39 | QRRPQGLPNNTASWFT | 0 | 0.95 | 0.7755102 |
