## Extended data 4 for "The landscape of antibody binding in SARS-CoV-2 infection"

| Extended data 4 | | | |
|---|---|---|---|
| **Epitope identifier** | **AUC-ROC** | **Sensitivity** | **Specificity** |
| 384-N-33 | 1 | 0.95 | 1 |
| 568-S-26 | 0.999375 | 0.95 | 1 |
| 1247-S-27 | 0.995 | 0.875 | 1 |
| 208-N-31 | 0.995625 | 0.95 | 1 |
| 807-S-26 | 1 | 0.925 | 1 |
| 553-S-26 | 0.995625 | 0.95 | 1 |
| 785-S-27 | 0.989375 | 0.975 | 1 |
| 1140-S-25 | 0.990625 | 0.925 | 1 |
| 624-S-23 | 0.99125 | 0.925 | 1 |
| 181-M-32 | 0.99125 | 0.925 | 1 |
| 28-N-28 | 0.999375 | 0.925 | 1 |
| 4514-orf1ab-16 | 0.996875 | 0.925 | 1 |
| 152-M-26 | 0.995625 | 0.95 | 1 |
| 549-S-18 | 0.994375 | 0.925 | 1 |
| 685-S-25 | 0.996875 | 0.925 | 1 |
| 249-N-18 | 0.994375 | 0.925 | 1 |
| 205-M-18 | 0.996875 | 0.925 | 1 |
| 5999-orf1ab-16 | 0.996875 | 0.925 | 1 |
| 1239-orf1ab-18 | 0.996875 | 0.925 | 1 |
| 2309-orf1ab-16 | 0.99625 | 0.95 | 1 |
| 613-S-25 | 0.996875 | 0.9 | 1 |
| 1551-orf1ab-16 | 0.99625 | 0.9 | 1 |
| 6057-orf1ab-17 | 1 | 0.9 | 1 |
| 153-N-26 | 0.99875 | 0.925 | 1 |
| 1720-orf1ab-16 | 0.9925 | 0.95 | 1 |
| 635-S-20 | 0.995625 | 0.925 | 1 |
| 14-N-17 | 0.996875 | 0.925 | 1 |
| 7-N-21 | 0.996875 | 0.925 | 0.95 |
| 940-S-16 | 0.994375 | 0.925 | 1 |
| 1155-S-20 | 0.996875 | 0.9 | 1 |
| 338-N-19 | 0.995 | 0.9 | 1 |
| 404-S-18 | 0.998125 | 0.925 | 1 |
| 60-orf8-20 | 0.996875 | 0.925 | 1 |
| 376-N-22 | 0.996875 | 0.95 | 1 |
| 252-orf3a-24 | 0.996875 | 0.925 | 1 |
| 230-N-21 | 0.999375 | 0.875 | 1 |
| 94-N-16 | 0.9875 | 0.9 | 1 |
| 356-N-16 | 0.99125 | 0.95 | 1 |
| 536-S-17 | 0.9925 | 0.925 | 1 |
| 12-orf8-16 | 0.996875 | 0.925 | 1 |
| 798-S-17 | 0.998125 | 0.95 | 1 |

| Extended data 4 | | | |
|---|---|---|---|
| **Epitope identifier** | **AUC-ROC** | **Sensitivity** | **Specificity** |
| 227-N-17 | 1 | 0.9 | 1 |
| 66-orf8-18 | 0.994375 | 0.925 | 1 |
| 4451-orf1ab-16 | 1 | 0.95 | 1 |
| 289-S-17 | 0.995 | 0.925 | 1 |
| 117-N-19 | 0.999375 | 0.925 | 1 |
| 175-M-20 | 0.996875 | 0.875 | 1 |
| 644-S-16 | 0.996875 | 0.925 | 1 |
| 9-orf6-16 | 0.99375 | 0.95 | 1 |
| 242-N-19 | 0.99625 | 0.925 | 1 |
| 656-S-17 | 0.995 | 0.9 | 1 |
| 541-S-16 | 0.999375 | 0.925 | 1 |
| 844-S-16 | 0.996875 | 0.95 | 1 |
| 804-S-17 | 0.994375 | 0.925 | 1 |
| 126-N-17 | 0.994375 | 0.875 | 1 |
| 96-N-17 | 0.999375 | 0.95 | 1 |
| 235-orf3a-17 | 0.996875 | 0.95 | 1 |
| 122-N-16 | 0.995625 | 0.925 | 1 |
| 53-orf8-16 | 0.998125 | 0.925 | 1 |
| 124-N-16 | 0.996875 | 0.925 | 1 |
| 336-N-16 | 0.995 | 0.9 | 1 |
| 1546-orf1ab-16 | 0.996875 | 0.925 | 1 |
| 306-S-16 | 0.99625 | 0.9 | 1 |
| 241-S-16 | 0.993125 | 0.9 | 1 |
| 768-S-18 | 0.9925 | 0.9 | 1 |
| 16-orf3a-16 | 0.996875 | 0.925 | 1 |
| 1164-S-16 | 0.999375 | 0.925 | 1 |
| 172-S-16 | 0.995625 | 0.925 | 1 |
| 21-orf3a-16 | 0.996875 | 0.95 | 1 |
| 2584-orf1ab-16 | 0.998125 | 0.95 | 1 |
| 1178-S-16 | 0.98875 | 0.925 | 1 |
| 661-S-16 | 0.99125 | 0.925 | 1 |
| 18-orf3a-16 | 0.994375 | 0.95 | 1 |
| 410-S-16 | 0.9925 | 0.95 | 1 |
| 1161-S-17 | 0.999375 | 0.9 | 1 |
| 761-S-16 | 0.983125 | 0.95 | 1 |
| 1681-orf1ab-16 | 0.995 | 0.925 | 1 |
| 1572-orf1ab-16 | 0.98875 | 0.925 | 1 |
