## Supplementary figures and images for "The landscape of antibody binding in SARS-CoV-2 infection"

### Extended data 5

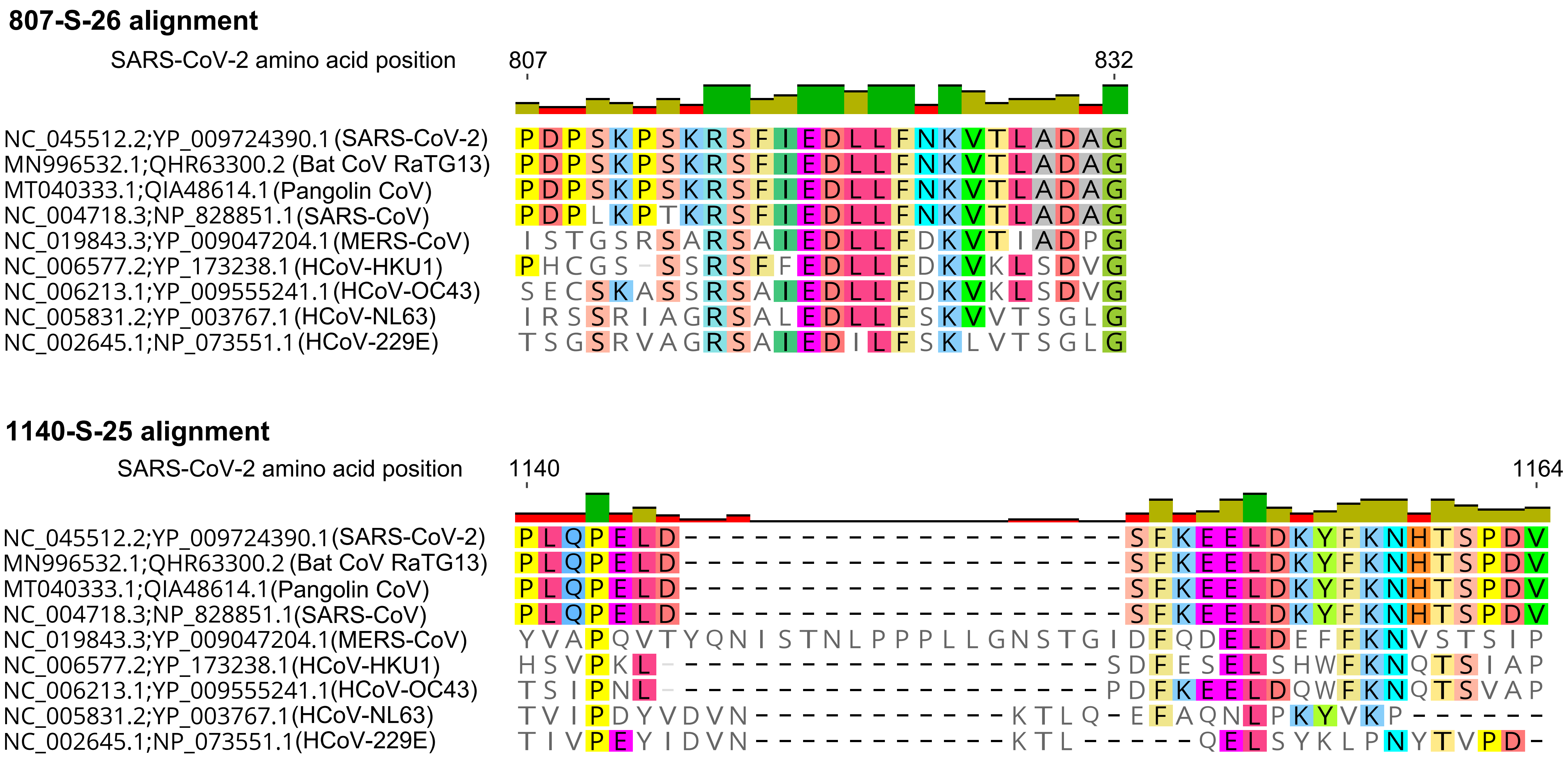
