## Extended data 6 for "The landscape of antibody binding in SARS-CoV-2 infection"

| Virus | Protein | Start position | Sequence | Identical to SARS-CoV-2? | Min.Pvalue | Max.Pvalue | Mean.Pvalue | Min.Signal | Max.Signal | Mean.Signal | Minimum fold change | Maximum fold change | Average fold change |
|---|---|---|---|---|---|---|---|---|---|---|---|---|---|
| RaTG13 bat coronavirus | orf1ab | 4513 | YTMADLVYALRHFDEG | Y | 3.44E-08 | 3.44E-08 | 3.44E-08 | 2.882777378 | 2.882777378 | 2.882777378 | 3.840173961 | 3.840173961 | 3.840173961 |
| | orf1ab | 1238 | VTTTLEETKFLTENLLLY | Y | 6.69E-06 | 0.067158556 | 0.022668343 | 1.375000984 | 1.550166538 | 1.486060242 | 1.162966962 | 2.03537696 | 1.597073998 |
| | orf1ab | 2308 | ITISSFKWDLTAFGLV | Y | 6.79E-06 | 6.79E-06 | 6.79E-06 | 1.054295457 | 1.054295457 | 1.054295457 | 2.075117706 | 2.075117706 | 2.075117706 |
| | orf1ab | 6056 | DFSRVSAKPPPGDQFKH | Y | 6.02E-05 | 0.009276928 | 0.004668546 | 2.850704005 | 3.319341123 | 3.085022564 | 1.237010975 | 1.602201149 | 1.419606062 |
| | orf1ab | 5998 | ITREEAVRHVRAWIGF | N | 0.002251724 | 0.002251724 | 0.002251724 | 1.028856465 | 1.028856465 | 1.028856465 | 1.346000579 | 1.346000579 | 1.346000579 |
| | orf1ab | 1720 | TVGELGDVRETMNYLF | N | 0.002353468 | 0.002353468 | 0.002353468 | 1.739692029 | 1.739692029 | 1.739692029 | 1.546393445 | 1.546393445 | 1.546393445 |
| | orf1ab | 4450 | KDEDDNLIDSYFVVKR | Y | 0.004331827 | 0.004331827 | 0.004331827 | 0.96557138 | 0.96557138 | 0.96557138 | 1.191567751 | 1.191567751 | 1.191567751 |
| | orf1ab | 1330 | GQGLNGYTVEEARTVL | N | 0.026789404 | 0.026789404 | 0.026789404 | 1.514422096 | 1.514422096 | 1.514422096 | 1.134638201 | 1.134638201 | 1.134638201 |
| | orf1ab | 2072 | GDIILKPANDGLKITE | N | 0.036983953 | 0.036983953 | 0.036983953 | 1.045574693 | 1.045574693 | 1.045574693 | 1.021571183 | 1.021571183 | 1.021571183 |
| | orf1ab | 1680 | LTLQQIELKFNPPALQ | Y | 0.082797327 | 0.082797327 | 0.082797327 | 1.636274231 | 1.636274231 | 1.636274231 | 1.189205696 | 1.189205696 | 1.189205696 |
| | orf1ab | 1571 | TTVDNINLHTQVVDMS | Y | 0.085996286 | 0.085996286 | 0.085996286 | 2.28420266 | 2.28420266 | 2.28420266 | 1.403214111 | 1.403214111 | 1.403214111 |
| | S | 568 | DIADTTDAVRDPQTLEILDITPCSFG | Y | 1.61E-14 | 0.000149677 | 1.36E-05 | 0.641875345 | 4.438674675 | 3.547247207 | 1.300346605 | 5.937648135 | 4.828993759 |
| | S | 1243 | CCSCGSCCKFDEDDSEPVLKGVKLHYT | Y | 7.33E-13 | 0.046944992 | 0.003885534 | 2.061703942 | 4.981158638 | 3.767558665 | 1.80188453 | 5.691348595 | 3.96345523 |
| | S | 803 | PDPSKPSKRSFIEDLLFNKVTLADAG | Y | 4.68E-12 | 0.025438608 | 0.00415206 | 3.750858671 | 6.291024386 | 5.329647697 | 1.994346878 | 4.828357464 | 3.618150919 |
| | S | 553 | TESNKKFLPFQQFGRDIADTTDAVRD | Y | 7.22E-12 | 0.000367517 | 4.21E-05 | 2.012189068 | 5.786836915 | 4.40914918 | 1.335175158 | 4.72916528 | 3.660377334 |
| | S | 781 | VKQIYKTPPIKDFGGFNFSQILPDPSK | Y | 3.42E-11 | 0.026129673 | 0.003057208 | 2.218296198 | 4.948993553 | 3.80261023 | 1.113887985 | 3.872141913 | 2.988014448 |
| | S | 1136 | PLQPELDSFKEELDKYFKNHTSPDV | Y | 1.08E-09 | 0.027133519 | 0.005751614 | 3.404069954 | 6.709830772 | 5.839000544 | 1.965130905 | 4.538846053 | 3.163294698 |
| | S | 624 | IHADQLTPTWRVYSTGSNVFQTR | Y | 8.75E-09 | 0.017672625 | 0.005496831 | 0.844750877 | 2.249419745 | 1.502687078 | 1.111159084 | 2.655638641 | 1.97660505 |
| | S | 549 | TGVLTESNKKFLPFQQFG | Y | 1.14E-07 | 0.002595492 | 0.00170456 | 4.350836323 | 4.959213016 | 4.716295354 | 2.525264751 | 3.636617916 | 3.001255932 |
| | S | 681 | RSVASQSIIAYTMSLGAENSVAYSN | Y | 3.26E-07 | 0.044796282 | 0.008264212 | 1.182794065 | 2.946218683 | 2.277405557 | 1.298100687 | 2.686638698 | 2.15574528 |
| | S | 613 | QDVNCTEVPVAIHADQLTPTWRVYS | Y | 8.65E-06 | 0.025201644 | 0.005261101 | 1.52247736 | 2.898127744 | 2.346066294 | 1.577945922 | 2.482387112 | 2.119005817 |
| | S | 635 | VYSTGSNVFQTRAGCLIGAE | Y | 9.25E-05 | 0.046657457 | 0.013871249 | 0.975750784 | 1.913725864 | 1.348463904 | 1.188509857 | 1.533997809 | 1.332618267 |
| | S | 936 | STASALGKLQDVVNQN | Y | 0.000174862 | 0.000174862 | 0.000174862 | 2.53060297 | 2.53060297 | 2.53060297 | 2.051292423 | 2.051292423 | 2.051292423 |
| | S | 1151 | YFKNHTSPDVDLGDISGINA | Y | 0.000216151 | 0.0718081 | 0.022778077 | 2.444300697 | 4.067703063 | 3.268706262 | 1.780824846 | 2.52762732 | 2.159830138 |
| | S | 403 | TGDEVRQIAPGQTGKIADY | N | 0.00037711 | 0.033259779 | 0.011798906 | 2.003095742 | 2.71978386 | 2.315224064 | 1.268238466 | 1.715586822 | 1.50424506 |
| | S | 536 | NKCVNFNFNGLTGTGVL | Y | 0.002097616 | 0.08962105 | 0.045859333 | 1.808445746 | 1.947513063 | 1.877979404 | 1.031084896 | 1.266391603 | 1.148738249 |
| | S | 794 | GGFNFSQILPDPSKPSK | Y | 0.002978263 | 0.0254641 | 0.014221182 | 2.746544032 | 3.16577222 | 2.956158126 | 1.194730019 | 1.561768943 | 1.378249481 |
| | S | 289 | VDCALDPLSETKCTLKS | Y | 0.005494354 | 0.00861176 | 0.007053057 | 2.187937149 | 2.197782173 | 2.192859661 | 1.223863868 | 1.383428156 | 1.303646012 |
| | S | 644 | QTRAGCLIGAEHVNNS | Y | 0.006883358 | 0.006883358 | 0.006883358 | 1.660777167 | 1.660777167 | 1.660777167 | 1.219148559 | 1.219148559 | 1.219148559 |
| | S | 656 | VNNSYECDIPIGAGICA | Y | 0.008913453 | 0.040619754 | 0.024766604 | 3.320680986 | 3.357477767 | 3.339079377 | 1.648243034 | 1.924824879 | 1.786533957 |
| | S | 541 | FNFNGLTGTGVLTESN | Y | 0.009092157 | 0.009092157 | 0.009092157 | 1.682464191 | 1.682464191 | 1.682464191 | 1.355527047 | 1.355527047 | 1.355527047 |
| | S | 840 | IAARDLICAQKFNGLT | Y | 0.009631276 | 0.009631276 | 0.009631276 | 1.934507286 | 1.934507286 | 1.934507286 | 1.187008674 | 1.187008674 | 1.187008674 |
| | S | 800 | QILPDPSKPSKRSFIED | Y | 0.01416558 | 0.082438627 | 0.048302104 | 2.609973692 | 2.967270272 | 2.788621982 | 1.362094837 | 1.543749885 | 1.452922361 |
| | S | 306 | FTVEKGIYQTSNFRVQ | Y | 0.039964939 | 0.039964939 | 0.039964939 | 1.922949885 | 1.922949885 | 1.922949885 | 1.382185206 | 1.382185206 | 1.382185206 |
| | S | 241 | LLALHRSYLTPGDSSS | Y | 0.040647898 | 0.040647898 | 0.040647898 | 1.123709348 | 1.123709348 | 1.123709348 | 1.063450683 | 1.063450683 | 1.063450683 |
| | S | 764 | TGIAVEQDKNTQEVFAQV | Y | 0.043278766 | 0.072374822 | 0.061360863 | 3.231291248 | 4.151098014 | 3.764004324 | 1.902117341 | 2.548039355 | 2.159231495 |
| | S | 1160 | VDLGDISGINASVVNI | Y | 0.050414892 | 0.050414892 | 0.050414892 | 3.288057911 | 3.288057911 | 3.288057911 | 1.548259579 | 1.548259579 | 1.548259579 |
| | S | 172 | SQPFLMDLEGKQGNFK | Y | 0.051046528 | 0.051046528 | 0.051046528 | 2.688603341 | 2.688603341 | 2.688603341 | 1.092667365 | 1.092667365 | 1.092667365 |
| | S | 1174 | NIQKEIDRLNEVAKNL | Y | 0.070452364 | 0.070452364 | 0.070452364 | 4.189173886 | 4.189173886 | 4.189173886 | 1.633966032 | 1.633966032 | 1.633966032 |
| | S | 661 | ECDIPIGAGICASYQT | Y | 0.071744324 | 0.071744324 | 0.071744324 | 2.349375887 | 2.349375887 | 2.349375887 | 1.318253895 | 1.318253895 | 1.318253895 |
| | S | 410 | IAPGQTGKIADYNYKL | Y | 0.077164884 | 0.077164884 | 0.077164884 | 3.229047366 | 3.229047366 | 3.229047366 | 1.351729641 | 1.351729641 | 1.351729641 |
| | S | 1157 | SPDVDLGDISGINASVV | Y | 0.078396556 | 0.091260595 | 0.084828575 | 3.876221731 | 4.493734401 | 4.184978066 | 1.651883749 | 1.838878379 | 1.745381064 |
| | S | 757 | TQLNRALTGIAVEQDK | Y | 0.081244846 | 0.081244846 | 0.081244846 | 3.337645681 | 3.337645681 | 3.337645681 | 2.102133406 | 2.102133406 | 2.102133406 |
| | orf3 | 253 | SGVVNPAMEPIYDEPTTTTSV | N | 1.37E-05 | 0.002429345 | 0.000813474 | 2.973432623 | 3.844470371 | 3.275557943 | 2.065743722 | 2.562768111 | 2.243809591 |
| | orf3 | 235 | KIVDEPEEHVQIHTIDG | Y | 0.015608587 | 0.03747391 | 0.026541248 | 1.195194322 | 1.196190879 | 1.1956926 | 1.419905444 | 1.499188048 | 1.459546746 |
| | orf3 | 260 | MEPIYDEPTTTTSVPL | Y | 0.024292619 | 0.079047032 | 0.051669825 | 3.648568559 | 3.648568559 | 3.648568559 | 2.054172097 | 2.054172097 | 2.054172097 |
| | M | 1 | MADNGTITVEELKKLLEQWNLVI | N | 2.19E-23 | 1.12E-06 | 1.40E-07 | 3.45107702 | 5.969414382 | 4.533319853 | 4.03136054 | 6.804550691 | 5.031049213 |
| | M | 180 | LGASQRVAGDSGFAAYSRYRIGNYKLNTD | Y | 1.35E-08 | 0.008850894 | 0.002361658 | 1.439919597 | 3.860438524 | 2.340234436 | 1.483219803 | 4.160525836 | 2.605255671 |
| | M | 151 | AGHHLGRCDIKDLPKEITVATSRTLS | Y | 6.33E-08 | 0.043789648 | 0.006095071 | 2.964983937 | 4.25674342 | 3.440166979 | 1.961929208 | 3.418942102 | 2.95926923 |
| | M | 204 | KLNTDHSSSSDNIALLVQ | Y | 2.64E-06 | 0.001531138 | 0.000275345 | 2.141560861 | 3.315080245 | 2.411855062 | 2.675262618 | 3.222673693 | 2.926653207 |
| | M | 174 | TLSYYKLGASQRVAGDSGFA | Y | 0.005781236 | 0.020018989 | 0.014866107 | 1.883011253 | 2.652089195 | 2.350120084 | 1.238256256 | 1.354633688 | 1.29578082 |
| | orf6 | 9 | VTIAEILLIIMRTFKV | Y | 0.006984144 | 0.006984144 | 0.006984144 | 1.066558888 | 1.066558888 | 1.066558888 | 1.086903432 | 1.086903432 | 1.086903432 |
| | orf8 | 64 | EVGSKSPIQYIDIGNYTVSC | N | 0.003991613 | 0.073512789 | 0.02204642 | 2.002154366 | 3.120247355 | 2.638651901 | 1.44486258 | 1.587153935 | 1.546539453 |
| | orf8 | 60 | LCVDEVGSKSPIQYIDI | N | 0.016164801 | 0.083331229 | 0.049748015 | 2.20015474 | 2.244468082 | 2.222311411 | 1.549067415 | 1.57301957 | 1.561043493 |
| | N | 384 | QRQKKQQTVTLLPAADLDDFSKQLQQSMS | Y | 9.10E-18 | 0.023356029 | 0.002451109 | 2.629074865 | 5.63270403 | 4.481566394 | 1.814070173 | 5.945368552 | 3.869521069 |
| | N | 210 | MAGNGSDAALALLLLDRLNQLESKMSGKG | N | 2.04E-12 | 0.005353431 | 0.001017316 | 0.871404698 | 3.928054236 | 2.282742936 | 1.317635374 | 3.388815253 | 2.373432392 |
| | N | 249 | KSAAEASKKPRQKRTATK | Y | 2.03E-06 | 0.087810918 | 0.058378101 | 2.098073104 | 3.394016162 | 2.72143097 | 1.010944384 | 2.568675219 | 1.601595882 |

**Extended data 6**

| Virus | Protein | Start position | Sequence | Identical to SARS-CoV-2? | Min.Pvalue | Max.Pvalue | Mean.Pvalue | Min.Signal | Max.Signal | Mean.Signal | Minimum fold change | Maximum fold change | Average fold change |
|---|---|---|---|---|---|---|---|---|---|---|---|---|---|
| | N | 30 | GERSGARPKQRRPQGLPNNTASWFTA | N | 2.28E-06 | 0.06337612 | 0.015966055 | 1.754841917 | 3.955736822 | 3.214076311 | 1.276881818 | 2.510866979 | 2.006162136 |
| | N | 153 | NNAAIVLQLPQGTTLPKGFYAEGSRG | Y | 7.56E-05 | 0.012414363 | 0.003510001 | 1.91066604 | 6.01560407 | 4.699895747 | 1.825709421 | 3.643803384 | 3.173890314 |
| | N | 14 | RITFGGPSDSTGSNQNG | Y | 9.46E-05 | 0.008997088 | 0.004545863 | 3.348608432 | 3.823385006 | 3.585996719 | 1.969875555 | 2.344659254 | 2.157267404 |
| | N | 7 | QNQRNAPRITFGGPSDSTGSN | Y | 0.000169707 | 0.00427168 | 0.001445535 | 3.23703526 | 3.698161125 | 3.385369177 | 2.124786099 | 2.706394076 | 2.335381416 |
| | N | 338 | KLDDKDPNFKDQVILLNKH | Y | 0.000361728 | 0.004381829 | 0.002000941 | 1.888590687 | 2.453447688 | 2.223691867 | 1.433295359 | 2.136694966 | 1.76243273 |
| | N | 376 | ADETQALPQRQKKQQTVTLLPA | Y | 0.000597719 | 0.037200054 | 0.013331027 | 2.309177862 | 2.922942754 | 2.667322022 | 1.473478931 | 2.319955597 | 1.925174373 |
| | N | 94 | IRGGDGKMKDLSPRWY | Y | 0.001332372 | 0.001332372 | 0.001332372 | 2.643480008 | 2.643480008 | 2.643480008 | 1.595240784 | 1.595240784 | 1.595240784 |
| | N | 356 | HIDAYKTFPPTEPKKD | Y | 0.002063175 | 0.002063175 | 0.002063175 | 3.297846371 | 3.297846371 | 3.297846371 | 1.84459208 | 1.84459208 | 1.84459208 |
| | N | 227 | LNQLESKMSGKGQQQQS | N | 0.003531491 | 0.005276791 | 0.004404141 | 3.388720006 | 3.415751095 | 3.402235551 | 2.029064152 | 2.152955309 | 2.09100973 |
| | N | 117 | PEAGLPYGANKDGIIWVAT | Y | 0.005733913 | 0.020529033 | 0.010132879 | 0.945657613 | 2.725917288 | 1.670521645 | 1.257724522 | 1.78848254 | 1.557416012 |
| | N | 242 | QSQTVTKKSAAEASKKPRQ | N | 0.006839298 | 0.027602996 | 0.018689137 | 2.552801031 | 3.188228773 | 2.954942246 | 1.22981961 | 1.54491229 | 1.363099667 |
| | N | 230 | LESKMSGKGQQQQSQTVTKK | N | 0.009133956 | 0.067051405 | 0.038175105 | 2.092369523 | 3.844299729 | 3.118470835 | 1.013431668 | 2.184451085 | 1.758471016 |
| | N | 126 | NKDGIIWVATEGALNTP | Y | 0.01431483 | 0.04668202 | 0.030498425 | 1.161357955 | 1.40100272 | 1.281180337 | 1.066865664 | 1.15001003 | 1.108437847 |
| | N | 96 | GGDGKMKDLSPRWYFYY | Y | 0.014873814 | 0.07645573 | 0.045664772 | 0.847216319 | 1.409055572 | 1.128135946 | 1.008940135 | 1.361803991 | 1.185372063 |
| | N | 122 | PYGANKDGIIWVATEG | Y | 0.019519431 | 0.019519431 | 0.019519431 | 0.846191795 | 0.846191795 | 0.846191795 | 1.121821679 | 1.121821679 | 1.121821679 |
| | N | 124 | GANKDGIIWVATEGAL | Y | 0.025536657 | 0.025536657 | 0.025536657 | 0.907214716 | 0.907214716 | 0.907214716 | 1.026788095 | 1.026788095 | 1.026788095 |
| | N | 336 | AIKLDDKDPNFKDQVI | Y | 0.025765456 | 0.025765456 | 0.025765456 | 3.308735848 | 3.308735848 | 3.308735848 | 1.427493675 | 1.427493675 | 1.427493675 |
| | N | 28 | QNGERSGARPKQRRPQ | N | 0.083538557 | 0.083538557 | 0.083538557 | 3.004810297 | 3.004810297 | 3.004810297 | 1.44629088 | 1.44629088 | 1.44629088 |
| Pangolin coronavirus | orf1ab | 4506 | YTMADLVYALRHFDEG | Y | 3.44E-08 | 3.44E-08 | 3.44E-08 | 2.882777378 | 2.882777378 | 2.882777378 | 3.840173961 | 3.840173961 | 3.840173961 |
| | orf1ab | 5991 | ITREEAIKHVRAWVGF | N | 2.00E-06 | 2.00E-06 | 2.00E-06 | 1.85111002 | 1.85111002 | 1.85111002 | 2.438627591 | 2.438627591 | 2.438627591 |
| | orf1ab | 1230 | VTTTLEETKFLTENLLLYA | N | 6.69E-06 | 0.067158556 | 0.024801846 | 0.754114726 | 1.550166538 | 1.303073863 | 1.043503424 | 2.03537696 | 1.458681354 |
| | orf1ab | 6050 | FSRVSAKPPPGDQFKH | Y | 6.02E-05 | 6.02E-05 | 6.02E-05 | 2.850704005 | 2.850704005 | 2.850704005 | 1.602201149 | 1.602201149 | 1.602201149 |
| | orf1ab | 1877 | DGVVCTEIDPKLDGYYK | N | 0.002492102 | 0.071656415 | 0.037074258 | 0.999146616 | 2.318157079 | 1.658651847 | 1.043767157 | 1.362098425 | 1.202932791 |
| | orf1ab | 1712 | KTVGELGDVRETMSHL | N | 0.047692722 | 0.047692722 | 0.047692722 | 1.568448722 | 1.568448722 | 1.568448722 | 1.181151311 | 1.181151311 | 1.181151311 |
| | orf1ab | 6135 | CTASDTYACWHHSVGF | N | 0.05720529 | 0.05720529 | 0.05720529 | 0.984745672 | 0.984745672 | 0.984745672 | 1.087279806 | 1.087279806 | 1.087279806 |
| | orf1ab | 2576 | AEVAVKMFDAYVNTFS | Y | 0.065722225 | 0.065722225 | 0.065722225 | 1.543624493 | 1.543624493 | 1.543624493 | 1.043177157 | 1.043177157 | 1.043177157 |
| | orf1ab | 516 | FKKNAWNIGEPKSILS | N | 0.071720273 | 0.071720273 | 0.071720273 | 2.264332391 | 2.264332391 | 2.264332391 | 1.021135301 | 1.021135301 | 1.021135301 |
| | orf1ab | 1673 | LTLQQIELKFNPPALQ | Y | 0.082797327 | 0.082797327 | 0.082797327 | 1.636274231 | 1.636274231 | 1.636274231 | 1.189205696 | 1.189205696 | 1.189205696 |
| | orf1ab | 1564 | TTVDNINLHTQVVDMS | Y | 0.085996286 | 0.085996286 | 0.085996286 | 2.28420266 | 2.28420266 | 2.28420266 | 1.403214111 | 1.403214111 | 1.403214111 |
| | orf1ab | 518 | KNAWNIGEPKSILSPLY | N | 0.086712551 | 0.095776005 | 0.091244278 | 1.642529758 | 3.145086888 | 2.393808323 | 1.20654872 | 1.67904735 | 1.442798035 |
| | S | 568 | DISDTTDAVRDPQTLEILDITPCSFG | N | 1.61E-14 | 0.000149677 | 1.36E-05 | 0.641875345 | 4.604402693 | 3.578544963 | 1.300346605 | 5.937648135 | 4.88129758 |
| | S | 1243 | CCSCGSCCKFDEDDSEPVLKGVKLHYT | Y | 7.33E-13 | 0.046944992 | 0.003885534 | 2.061703942 | 4.981158638 | 3.767558665 | 1.80188453 | 5.691348595 | 3.96345523 |
| | S | 803 | PDPSKPSKRSFIEDLLFNKVTLADAG | Y | 4.68E-12 | 0.025438608 | 0.00415206 | 3.750858671 | 6.291024386 | 5.329647697 | 1.994346878 | 4.828357464 | 3.618150919 |
| | S | 549 | TGVLTTSKKQFLPFQQFGRDISDTTDAVR | N | 1.59E-10 | 0.085049641 | 0.012684086 | 1.614025715 | 4.881499682 | 3.415203079 | 1.401556563 | 4.18964913 | 2.880673785 |
| | S | 781 | VKQIYKTPPIKDFGGFNFLQILPDPSK | N | 1.77E-10 | 0.026129673 | 0.003621874 | 1.46883516 | 3.784235386 | 2.7232966 | 1.113887985 | 3.72108484 | 2.683518451 |
| | S | 1136 | PLQPELDSFKEELDKYFKNHTSPDV | Y | 1.08E-09 | 0.027133519 | 0.005751614 | 3.404069954 | 6.709830772 | 5.839000544 | 1.965130905 | 4.538846053 | 3.163294698 |
| | S | 624 | IHAEQLTPAWRVYSAGANVFQT | N | 5.50E-07 | 0.001504662 | 0.00052437 | 0.927998749 | 2.585654306 | 1.486756811 | 1.462095003 | 2.847472396 | 1.932215589 |
| | S | 682 | SVNQRSIIAYTMSLGAENSVAYSN | N | 5.77E-06 | 0.044796282 | 0.007656846 | 1.26969743 | 2.946218683 | 2.319120921 | 1.448297239 | 2.514807294 | 2.125486741 |
| | S | 619 | EVPMAIHAEQLTPAWRVYS | N | 9.46E-06 | 0.000585429 | 0.000154718 | 1.406369855 | 1.735403991 | 1.540472762 | 1.557598251 | 2.248367868 | 1.89045439 |
| | S | 936 | STASALGKLQDVVNQN | Y | 0.000174862 | 0.000174862 | 0.000174862 | 2.53060297 | 2.53060297 | 2.53060297 | 2.051292423 | 2.051292423 | 2.051292423 |
| | S | 1151 | YFKNHTSPDVDLGDISGINA | Y | 0.000216151 | 0.0718081 | 0.022778077 | 2.444300697 | 4.067703063 | 3.268706262 | 1.780824846 | 2.52762732 | 2.159830138 |
| | S | 536 | DKCVNFNFNGLTGTGVL | N | 0.002097616 | 0.04668202 | 0.024389818 | 1.808445746 | 2.395161868 | 2.101803807 | 1.266391603 | 1.317185656 | 1.29178863 |
| | S | 794 | GGFNFLQILPDPSKPS | N | 0.004023965 | 0.004023965 | 0.004023965 | 1.583549061 | 1.583549061 | 1.583549061 | 1.510997662 | 1.510997662 | 1.510997662 |
| | S | 657 | NNSYECDIPVGAGICA | N | 0.004133539 | 0.004133539 | 0.004133539 | 2.457648421 | 2.457648421 | 2.457648421 | 1.450592552 | 1.450592552 | 1.450592552 |
| | S | 840 | IAARDLICAQKFNGLT | Y | 0.009631276 | 0.009631276 | 0.009631276 | 1.934507286 | 1.934507286 | 1.934507286 | 1.187008674 | 1.187008674 | 1.187008674 |
| | S | 796 | FNFLQILPDPSKPSKR | N | 0.012728812 | 0.012728812 | 0.012728812 | 1.269427962 | 1.269427962 | 1.269427962 | 1.091269469 | 1.091269469 | 1.091269469 |
| | S | 798 | FLQILPDPSKPSKRSFIED | N | 0.01416558 | 0.091444743 | 0.053658361 | 1.355609609 | 2.967270272 | 2.097310799 | 1.011767431 | 1.543749885 | 1.299710367 |
| | S | 764 | TGIAVEQDKNTQEVFAQV | Y | 0.043278766 | 0.072374822 | 0.061360863 | 3.231291248 | 4.151098014 | 3.764004324 | 1.902117341 | 2.548039355 | 2.159231495 |
| | S | 1160 | VDLGDISGINASVVNI | Y | 0.050414892 | 0.050414892 | 0.050414892 | 3.288057911 | 3.288057911 | 3.288057911 | 1.548259579 | 1.548259579 | 1.548259579 |
| | S | 172 | SQPFLMDLEGKQGNFK | Y | 0.051046528 | 0.051046528 | 0.051046528 | 2.688603341 | 2.688603341 | 2.688603341 | 1.092667365 | 1.092667365 | 1.092667365 |
| | S | 1174 | NIQKEIDRLNEVAKNL | Y | 0.070452364 | 0.070452364 | 0.070452364 | 4.189173886 | 4.189173886 | 4.189173886 | 1.633966032 | 1.633966032 | 1.633966032 |
| | S | 303 | LKSLTVEKGIYQTSNF | N | 0.070999816 | 0.070999816 | 0.070999816 | 2.089181643 | 2.089181643 | 2.089181643 | 1.329396106 | 1.329396106 | 1.329396106 |
| | S | 310 | KGIYQTSNFRVQPTIS | N | 0.074362926 | 0.074362926 | 0.074362926 | 2.307595033 | 2.307595033 | 2.307595033 | 1.620990306 | 1.620990306 | 1.620990306 |
| | S | 1157 | SPDVDLGDISGINASVV | Y | 0.078396556 | 0.091260595 | 0.084828575 | 3.876221731 | 4.493734401 | 4.184978066 | 1.651883749 | 1.838878379 | 1.745381064 |
| | S | 757 | TQLNRALTGIAVEQDK | Y | 0.081244846 | 0.081244846 | 0.081244846 | 3.337645681 | 3.337645681 | 3.337645681 | 2.102133406 | 2.102133406 | 2.102133406 |
| | orf3 | 234 | SRIVDEPEDHVQIHTI | N | 0.003757461 | 0.003757461 | 0.003757461 | 1.911963242 | 1.911963242 | 1.911963242 | 1.737246792 | 1.737246792 | 1.737246792 |
| | orf3 | 254 | GVVNPAMDPIYDEPTTT | N | 0.019795468 | 0.032801977 | 0.026298723 | 2.674564255 | 2.674564255 | 2.674564255 | 1.482053411 | 1.482053411 | 1.482053411 |

**Extended data 6**

| Virus | Protein | Start position | Sequence | Identical to SARS-CoV-2? | Min.Pvalue | Max.Pvalue | Mean.Pvalue | Min.Signal | Max.Signal | Mean.Signal | Minimum fold change | Maximum fold change | Average fold change |
|---|---|---|---|---|---|---|---|---|---|---|---|---|---|
| | M | 1 | MSANNGTITVEELKKLLEQWNLVI | N | 2.19E-23 | 0.000159604 | 1.77E-05 | 3.45107702 | 5.969414382 | 4.568237141 | 2.408249901 | 6.804550691 | 4.78156831 |
| | M | 181 | LGASQRVAGDSGFAAYSRYRIGNYKLNTD | N | 1.35E-08 | 0.056636448 | 0.005289566 | 1.439919597 | 3.860438524 | 2.359151021 | 1.483219803 | 4.160525836 | 2.656888104 |
| | M | 152 | AGHHLGRCDIKDLPKEITVATSRTLS | Y | 6.33E-08 | 0.043789648 | 0.006095071 | 2.964983937 | 4.25674342 | 3.440166979 | 1.961929208 | 3.418942102 | 2.95926923 |
| | M | 175 | TLSYYKLGASQRVAGDSGFA | Y | 0.005781236 | 0.020018989 | 0.014866107 | 1.883011253 | 2.652089195 | 2.350120084 | 1.238256256 | 1.354633688 | 1.29578082 |
| | N | 382 | QRQKKQQTVTLLPAADLDDFSKQLQQSMS | Y | 9.10E-18 | 0.023356029 | 0.002451109 | 2.629074865 | 5.63270403 | 4.481566394 | 1.814070173 | 5.945368552 | 3.869521069 |
| | N | 206 | ARIAGNGGDAALALLLLDRLNALESKMSGK | N | 8.99E-12 | 0.053424974 | 0.003713924 | 0.88369067 | 4.083870949 | 2.455515672 | 1.120017381 | 3.680488275 | 2.643671721 |
| | N | 247 | KSAAEASKKPRQKRTATK | Y | 2.03E-06 | 0.087810918 | 0.058378101 | 2.098073104 | 3.394016162 | 2.72143097 | 1.010944384 | 2.568675219 | 1.601595882 |
| | N | 28 | GDRSGARPKQRRPQGLPNNTASWFTA | N | 2.28E-06 | 0.074305203 | 0.022866567 | 1.754841917 | 3.955736822 | 3.223017531 | 1.276881818 | 2.510866979 | 1.924348005 |
| | N | 5 | GPQARAPRITFGGPSDSTDNNQN | N | 5.68E-06 | 0.054793174 | 0.012774627 | 2.734425416 | 3.952740813 | 3.481896508 | 1.826708572 | 2.905707727 | 2.281232232 |
| | N | 152 | NAAIVLQLPQGTALPKGFYAEGS | N | 4.89E-05 | 0.001560483 | 0.000376963 | 2.251809591 | 4.408021472 | 3.113813883 | 1.960517285 | 2.745158834 | 2.340670634 |
| | N | 92 | VRGGDGKMKDLSPRWY | N | 0.000366276 | 0.000366276 | 0.000366276 | 2.601646662 | 2.601646662 | 2.601646662 | 1.745702387 | 1.745702387 | 1.745702387 |
| | N | 378 | QPLPQRQKKQQTVTLLPA | N | 0.001680856 | 0.008928167 | 0.004384668 | 2.435346934 | 2.900726155 | 2.698403867 | 1.937900408 | 2.319955597 | 2.182652985 |
| | N | 354 | HIDAYKTFPPTEPKKD | Y | 0.002063175 | 0.002063175 | 0.002063175 | 3.297846371 | 3.297846371 | 3.297846371 | 1.84459208 | 1.84459208 | 1.84459208 |
| | N | 240 | QSQTVTKKSAAEASKKPRQ | N | 0.006839298 | 0.027602996 | 0.018689137 | 2.552801031 | 3.188228773 | 2.954942246 | 1.22981961 | 1.54491229 | 1.363099667 |
| | N | 225 | LNALESKMSGKGSQQQSQTVTK | N | 0.008086483 | 0.041782704 | 0.02387664 | 2.599250044 | 3.599339966 | 3.043328667 | 1.472562874 | 2.123581859 | 1.850758848 |
| | N | 94 | GGDGKMKDLSPRWYFYY | Y | 0.014873814 | 0.07645573 | 0.045664772 | 0.847216319 | 1.409055572 | 1.128135946 | 1.008940135 | 1.361803991 | 1.185372063 |
| | N | 336 | KLDDKDPSFKDNVILL | N | 0.024731598 | 0.024731598 | 0.024731598 | 1.981331186 | 1.981331186 | 1.981331186 | 1.142448531 | 1.142448531 | 1.142448531 |
| | N | 161 | QGTALPKGFYAEGSRG | N | 0.026972572 | 0.026972572 | 0.026972572 | 2.908066532 | 2.908066532 | 2.908066532 | 1.628781763 | 1.628781763 | 1.628781763 |
| | N | 126 | EGIIWVATEGALNTPK | N | 0.041397952 | 0.041397952 | 0.041397952 | 1.724069616 | 1.724069616 | 1.724069616 | 1.11835473 | 1.11835473 | 1.11835473 |
| | N | 375 | DESQPLPQRQKKQQTVT | N | 0.05720529 | 0.076207317 | 0.066706304 | 2.665903986 | 2.666254953 | 2.666079469 | 1.358949593 | 1.422030324 | 1.390489959 |
| | N | 338 | DDKDPSFKDNVILLNKH | N | 0.063904587 | 0.079684888 | 0.071794738 | 1.842263373 | 2.335531316 | 2.088897345 | 1.127526241 | 1.166872567 | 1.147199404 |
| SARS-CoV | orf1ab | 4491 | YTMADLVYALRHFDEG | Y | 3.44E-08 | 3.44E-08 | 3.44E-08 | 2.882777378 | 2.882777378 | 2.882777378 | 3.840173961 | 3.840173961 | 3.840173961 |
| | orf1ab | 5976 | ITREEAIRHVRAWIGF | Y | 4.02E-06 | 4.02E-06 | 4.02E-06 | 1.563287127 | 1.563287127 | 1.563287127 | 2.170975661 | 2.170975661 | 2.170975661 |
| | orf1ab | 1217 | TTLEETKFLTNKLLLF | N | 1.11E-05 | 1.11E-05 | 1.11E-05 | 1.667904307 | 1.667904307 | 1.667904307 | 1.902326585 | 1.902326585 | 1.902326585 |
| | orf1ab | 1697 | KTVGELGDVRETMTHL | N | 0.004183233 | 0.004183233 | 0.004183233 | 1.629246856 | 1.629246856 | 1.629246856 | 1.288394748 | 1.288394748 | 1.288394748 |
| | orf1ab | 2228 | GVLLSNFGAPSYCNGV | N | 0.01539261 | 0.01539261 | 0.01539261 | 1.011483051 | 1.011483051 | 1.011483051 | 1.119869818 | 1.119869818 | 1.119869818 |
| | S | 1229 | ACSCGSCCKFDEDDSEPVLKGVKLHYT | N | 7.33E-13 | 0.02312273 | 0.002396642 | 2.061703942 | 4.981158638 | 3.768434487 | 1.893803548 | 5.691348595 | 3.969583164 |
| | S | 789 | PDPLKPTKRSFIEDLLFNKVTLADAG | N | 4.68E-12 | 0.084289032 | 0.009498743 | 3.644442968 | 6.327094018 | 5.189641573 | 1.805575202 | 4.828357464 | 3.684242105 |
| | S | 1122 | PLQPELDSFKEELDKYFKNHTSPDV | Y | 1.08E-09 | 0.027133519 | 0.005751614 | 3.404069954 | 6.709830772 | 5.839000544 | 1.965130905 | 4.538846053 | 3.163294698 |
| | S | 539 | TPSSKRFQPFQQFGRDVSDFTDSVR | N | 1.94E-07 | 0.069661716 | 0.007072101 | 3.005860353 | 4.360992023 | 3.429030872 | 1.413704496 | 2.62264757 | 2.204667749 |
| | S | 610 | IHADQLTPAWRIYSTGNNV | N | 6.12E-07 | 0.088953012 | 0.022446631 | 0.567958538 | 2.399548393 | 1.664724168 | 1.448451375 | 2.199568387 | 1.895549552 |
| | S | 668 | STSQKSIVAYTMSLGADSSIAY | N | 4.63E-06 | 0.075329431 | 0.016879573 | 1.494269551 | 2.366567556 | 2.065506646 | 1.242004329 | 2.354702223 | 1.92637735 |
| | S | 599 | QDVNCTDVSTAIHADQLTPAWRIYS | N | 5.16E-05 | 0.023356029 | 0.007670506 | 1.228575664 | 2.312155218 | 1.609695034 | 1.181830283 | 2.186249413 | 1.635982796 |
| | S | 1137 | YFKNHTSPDVDLGDISGINA | Y | 0.000216151 | 0.0718081 | 0.022778077 | 2.444300697 | 4.067703063 | 3.268706262 | 1.780824846 | 2.52762732 | 2.159830138 |
| | S | 776 | LKYFGGFNFSQILPDPL | N | 0.000375336 | 0.002282253 | 0.001328794 | 1.34794881 | 1.876951485 | 1.612450147 | 1.454771614 | 1.71156843 | 1.583170022 |
| | S | 647 | ECDIPIGAGICASYHT | N | 0.021737563 | 0.021737563 | 0.021737563 | 1.724174955 | 1.724174955 | 1.724174955 | 1.421369277 | 1.421369277 | 1.421369277 |
| | S | 643 | DTSYECDIPIGAGICA | N | 0.027016454 | 0.027016454 | 0.027016454 | 2.698315952 | 2.698315952 | 2.698315952 | 1.801140583 | 1.801140583 | 1.801140583 |
| | S | 1146 | VDLGDISGINASVVNI | Y | 0.050414892 | 0.050414892 | 0.050414892 | 3.288057911 | 3.288057911 | 3.288057911 | 1.548259579 | 1.548259579 | 1.548259579 |
| | S | 1160 | NIQKEIDRLNEVAKNL | Y | 0.070452364 | 0.070452364 | 0.070452364 | 4.189173886 | 4.189173886 | 4.189173886 | 1.633966032 | 1.633966032 | 1.633966032 |
| | S | 1143 | SPDVDLGDISGINASVV | Y | 0.078396556 | 0.091260595 | 0.084828575 | 3.876221731 | 4.493734401 | 4.184978066 | 1.651883749 | 1.838878379 | 1.745381064 |
| | orf3a | 24 | SPASTVHATATIPLQA | N | 0.0001358 | 0.0001358 | 0.0001358 | 2.315778326 | 2.315778326 | 2.315778326 | 2.042244392 | 2.042244392 | 2.042244392 |
| | orf3a | 253 | GVANPAMDPIYDEPTT | N | 0.069894753 | 0.069894753 | 0.069894753 | 2.820683684 | 2.820683684 | 2.820683684 | 1.272920565 | 1.272920565 | 1.272920565 |
| | orf3b | 20 | ITVSQIQLSLLKVTAF | N | 0.000312565 | 0.000312565 | 0.000312565 | 1.526480555 | 1.526480555 | 1.526480555 | 1.716912228 | 1.716912228 | 1.716912228 |
| | M | 1 | MADNGTITVEELKQLLEQWNLVI | N | 4.79E-15 | 4.34E-07 | 8.67E-08 | 2.945925501 | 5.96080996 | 4.442772502 | 3.523598895 | 6.355094961 | 4.671632585 |
| | M | 152 | GHSLGRCDIKDLPKEITVATSRTLS | N | 6.33E-08 | 0.010639875 | 0.001451217 | 2.964983937 | 4.25674342 | 3.576013162 | 2.645584668 | 3.418942102 | 3.12448893 |
| | M | 180 | LGASQRVGTDSGFAAYNRYRIGNYKLNTD | N | 1.84E-07 | 0.04576054 | 0.010655557 | 1.092137728 | 3.560404557 | 2.284943155 | 1.27472962 | 3.35962305 | 2.404502463 |
| | M | 204 | KLNTDHAGSNDNIALLVQ | N | 7.25E-05 | 0.038392365 | 0.005652127 | 1.466519438 | 2.390937799 | 1.749636622 | 1.611174866 | 2.626175904 | 2.382864794 |
| | M | 174 | TLSYYKLGASQRVGTDSGFA | N | 0.000479187 | 0.082902925 | 0.020309445 | 1.735936531 | 2.633022465 | 2.173237457 | 1.08199326 | 1.62272666 | 1.375747616 |
| | N | 30 | NGGRNGARPKQRRPQGLPNNTASWFTA | N | 7.52E-07 | 0.06337612 | 0.020775039 | 1.754841917 | 3.955736822 | 3.194443014 | 1.276881818 | 2.510866979 | 1.914023371 |
| | N | 250 | KSAAEASKKPRQKRTATK | Y | 2.03E-06 | 0.087810918 | 0.058378101 | 2.098073104 | 3.394016162 | 2.72143097 | 1.010944384 | 2.568675219 | 1.601595882 |
| | N | 216 | GETALALLLLDRLNQLESKVSGKG | N | 7.52E-06 | 0.025806808 | 0.00553581 | 0.820810198 | 3.66023139 | 2.35854049 | 1.060927226 | 3.443479411 | 2.041439819 |
| | N | 388 | KKQPTVTLLPAADMDDFSRQLQNSM | N | 2.08E-05 | 0.067051405 | 0.008235775 | 1.702544206 | 2.98773267 | 2.448847101 | 1.257739965 | 2.657874655 | 1.887686634 |
| | N | 154 | NNAATVLQLPQGTTLPKGFYAEGSRG | N | 7.56E-05 | 0.012414363 | 0.003420922 | 3.175478362 | 6.01560407 | 5.002658237 | 2.171073798 | 3.619921991 | 3.207257265 |
| | N | 357 | HIDAYKTFPPTEPKKD | Y | 0.002063175 | 0.002063175 | 0.002063175 | 3.297846371 | 3.297846371 | 3.297846371 | 1.84459208 | 1.84459208 | 1.84459208 |
| | N | 9 | NQRSAPRITFGGPTDSTDNN | N | 0.004435256 | 0.050890331 | 0.019795702 | 3.210946324 | 3.523280343 | 3.371314232 | 1.671132852 | 2.16789219 | 1.971640117 |
| | N | 400 | DMDDFSRQLQNSMSGAS | N | 0.005610197 | 0.061438887 | 0.033524542 | 2.633690644 | 2.633690644 | 2.633690644 | 1.686017222 | 1.686017222 | 1.686017222 |
| | N | 95 | VRGGDGKMKELSPRWY | N | 0.005805698 | 0.005805698 | 0.005805698 | 2.55854887 | 2.55854887 | 2.55854887 | 1.475753722 | 1.475753722 | 1.475753722 |

**Extended data 6**

| Virus | Protein | Start position | Sequence | Identical to SARS-CoV-2? | Min.Pvalue | Max.Pvalue | Mean.Pvalue | Min.Signal | Max.Signal | Mean.Signal | Minimum fold change | Maximum fold change | Average fold change |
|---|---|---|---|---|---|---|---|---|---|---|---|---|---|
| | N | 129 | EGIVWVATEGALNTPKD | N | 0.007393829 | 0.07095814 | 0.039175984 | 1.650179139 | 1.887108192 | 1.768643666 | 1.178172544 | 1.344705982 | 1.261439263 |
| | N | 243 | QGQTVTKKSAAEASKKPRQ | Y | 0.007498694 | 0.027602996 | 0.017633245 | 2.976489998 | 3.145298981 | 3.0882613 | 1.316046976 | 1.601036545 | 1.420829855 |
| | N | 233 | SKVSGKGQQQQGQTVTKKS | N | 0.013480103 | 0.060892559 | 0.026337643 | 2.776572428 | 2.884411284 | 2.81386683 | 1.527737272 | 1.688543957 | 1.610072208 |
| | N | 341 | DDKDPQFKDNVILLNK | N | 0.024852038 | 0.024852038 | 0.024852038 | 2.343279295 | 2.343279295 | 2.343279295 | 1.095406844 | 1.095406844 | 1.095406844 |
| | N | 339 | KLDDKDPQFKDNVILL | N | 0.033435812 | 0.033435812 | 0.033435812 | 1.923890803 | 1.923890803 | 1.923890803 | 1.059720958 | 1.059720958 | 1.059720958 |
| | N | 377 | TDEAQPLPQRQKKQPT | N | 0.042460351 | 0.042460351 | 0.042460351 | 2.764284899 | 2.764284899 | 2.764284899 | 1.57751202 | 1.57751202 | 1.57751202 |
| | N | 181 | SQASSRSSSRSRGNSR | N | 0.064354665 | 0.064354665 | 0.064354665 | 1.963862083 | 1.963862083 | 1.963862083 | 1.114712519 | 1.114712519 | 1.114712519 |
| | N | 97 | GGDGKMKELSPRWYFY | N | 0.068609528 | 0.068609528 | 0.068609528 | 1.555904506 | 1.555904506 | 1.555904506 | 1.344957647 | 1.344957647 | 1.344957647 |
| | N | 228 | LNQLESKVSGKGQQQQ | N | 0.082902925 | 0.082902925 | 0.082902925 | 2.686565052 | 2.686565052 | 2.686565052 | 1.408442792 | 1.408442792 | 1.408442792 |
| MERS-CoV | orf1ab | 3507 | VAIEQLLYAIQQLYTG | N | 1.86E-09 | 1.86E-09 | 1.86E-09 | 2.436042867 | 2.436042867 | 2.436042867 | 3.804147793 | 3.804147793 | 3.804147793 |
| | orf1ab | 1693 | VQTVEDLRARMTYVCQC | N | 2.49E-09 | 0.002900434 | 0.001450218 | 2.534585074 | 2.909546035 | 2.722065555 | 2.231882346 | 3.356975218 | 2.794428782 |
| | orf1ab | 4501 | YTMMDLVYALRHFDQN | N | 3.52E-07 | 3.52E-07 | 3.52E-07 | 2.574647351 | 2.574647351 | 2.574647351 | 3.057732924 | 3.057732924 | 3.057732924 |
| | orf1ab | 1599 | MTLDLLKDIKFVIPAL | N | 7.52E-07 | 7.52E-07 | 7.52E-07 | 2.174700995 | 2.174700995 | 2.174700995 | 2.718538646 | 2.718538646 | 2.718538646 |
| | orf1ab | 5529 | LTVGDIFVLTSHSVAT | N | 1.11E-06 | 1.11E-06 | 1.11E-06 | 1.222339316 | 1.222339316 | 1.222339316 | 2.353659398 | 2.353659398 | 2.353659398 |
| | orf1ab | 1125 | IVLGDAIQVAKCYGES | N | 8.37E-05 | 8.37E-05 | 8.37E-05 | 1.077083722 | 1.077083722 | 1.077083722 | 1.122406865 | 1.122406865 | 1.122406865 |
| | orf1ab | 795 | TVVGQLEQTNMHSPDVI | N | 0.000414334 | 0.000835839 | 0.000625086 | 0.923651528 | 1.06736544 | 0.995508484 | 1.108171927 | 1.269254536 | 1.188713231 |
| | orf1ab | 517 | VTLDKLRDYLADYDVA | N | 0.007393829 | 0.007393829 | 0.007393829 | 3.735508612 | 3.735508612 | 3.735508612 | 2.226705591 | 2.226705591 | 2.226705591 |
| | orf1ab | 903 | LSIEEFADVVKEQVSD | N | 0.011184166 | 0.011184166 | 0.011184166 | 1.761038341 | 1.761038341 | 1.761038341 | 1.380695156 | 1.380695156 | 1.380695156 |
| | orf1ab | 5566 | ITVPEEFASHVANFQK | N | 0.02397666 | 0.02397666 | 0.02397666 | 2.930193702 | 2.930193702 | 2.930193702 | 1.519834193 | 1.519834193 | 1.519834193 |
| | orf1ab | 5843 | VDSSQGSEYQYVIFCQ | N | 0.031833524 | 0.031833524 | 0.031833524 | 0.41601362 | 0.41601362 | 0.41601362 | 1.071160643 | 1.071160643 | 1.071160643 |
| | orf1ab | 1271 | LTIVDIPQSLTFSYDG | N | 0.043733671 | 0.043733671 | 0.043733671 | 0.586360158 | 0.586360158 | 0.586360158 | 1.040066111 | 1.040066111 | 1.040066111 |
| | orf1ab | 2670 | AASVNQIVLRNSNGAC | N | 0.0509719 | 0.0509719 | 0.0509719 | 1.684727975 | 1.684727975 | 1.684727975 | 1.169922318 | 1.169922318 | 1.169922318 |
| | orf1ab | 793 | VETVVGQLEQTNMHSP | N | 0.07589902 | 0.07589902 | 0.07589902 | 1.635629191 | 1.635629191 | 1.635629191 | 1.047098561 | 1.047098561 | 1.047098561 |
| | S | 1223 | LGNSTGIDFQDELDEFFKNVSTSIP | N | 6.23E-09 | 0.09400429 | 0.018597717 | 2.030301784 | 5.872708544 | 4.482786092 | 1.425554379 | 3.794438087 | 3.09004074 |
| | S | 882 | GSRSARSAIEDLLFDKVTIAD | N | 2.51E-07 | 0.000275584 | 7.53E-05 | 4.847278331 | 6.624386201 | 5.864766775 | 2.762090638 | 3.759808897 | 3.312708281 |
| | S | 1046 | AISASIGDIIQRLDVLEQD | N | 4.27E-05 | 0.028772774 | 0.008403883 | 0.658005868 | 1.706736357 | 1.202282631 | 1.11063994 | 1.73319283 | 1.376546456 |
| | S | 880 | STGSRSARSAIEDLLF | N | 0.024322029 | 0.024322029 | 0.024322029 | 5.28963489 | 5.28963489 | 5.28963489 | 2.420043697 | 2.420043697 | 2.420043697 |
| | M | 208 | ITADIELALLRA | N | 6.36E-06 | 6.36E-06 | 6.36E-06 | 2.270071573 | 2.270071573 | 2.270071573 | 2.629392843 | 2.629392843 | 2.629392843 |
| | N | 114 | RAVKDGIVWVHEDGATD | N | 0.003737679 | 0.017498085 | 0.010617882 | 1.461216931 | 2.037673541 | 1.749445236 | 1.213431078 | 1.224392222 | 1.21891165 |
| | N | 173 | SRASSLSRNSSRSSSQ | N | 0.06260626 | 0.06260626 | 0.06260626 | 2.772902299 | 2.772902299 | 2.772902299 | 1.349208405 | 1.349208405 | 1.349208405 |
| HCoV-HKU1 | orf1ab | 5616 | SVASLSAPTLVPQENY | N | 4.55E-05 | 4.55E-05 | 4.55E-05 | 3.635204612 | 3.635204612 | 3.635204612 | 2.697825758 | 2.697825758 | 2.697825758 |
| | orf1ab | 6061 | ITKDEAIKRVRGWVGF | N | 6.37E-05 | 6.37E-05 | 6.37E-05 | 1.436441711 | 1.436441711 | 1.436441711 | 1.926382393 | 1.926382393 | 1.926382393 |
| | orf1ab | 1951 | LTQTFTSMLTNYFLDDVEM | N | 7.48E-05 | 0.007308104 | 0.003534995 | 0.471761596 | 1.740243787 | 1.134484582 | 1.033213515 | 1.772781422 | 1.392907074 |
| | orf1ab | 3395 | VTLGDFTIMSGRMSLT | N | 0.000329744 | 0.000329744 | 0.000329744 | 1.490420574 | 1.490420574 | 1.490420574 | 1.369034421 | 1.369034421 | 1.369034421 |
| | orf1ab | 2541 | ITVEAAIELSKELKRP | N | 0.000471939 | 0.000471939 | 0.000471939 | 3.493402634 | 3.493402634 | 3.493402634 | 2.666354458 | 2.666354458 | 2.666354458 |
| | orf1ab | 857 | VTVDDFVAVVCDAIEN | N | 0.002003674 | 0.002003674 | 0.002003674 | 0.892832695 | 0.892832695 | 0.892832695 | 1.285410019 | 1.285410019 | 1.285410019 |
| | orf1ab | 686 | VTVDVLKDMPVLKTIN | N | 0.002014909 | 0.002014909 | 0.002014909 | 2.492398472 | 2.492398472 | 2.492398472 | 1.764882002 | 1.764882002 | 1.764882002 |
| | orf1ab | 3591 | VSIETLLAAIKRLYMG | N | 0.007270328 | 0.007270328 | 0.007270328 | 1.192052443 | 1.192052443 | 1.192052443 | 1.1673651 | 1.1673651 | 1.1673651 |
| | orf1ab | 1668 | LTVGEVFGKILGNVFC | N | 0.007654986 | 0.007654986 | 0.007654986 | 0.588705795 | 0.588705795 | 0.588705795 | 1.054108285 | 1.054108285 | 1.054108285 |
| | orf1ab | 5577 | KTVLGEYVFDKSELTN | N | 0.078355906 | 0.078355906 | 0.078355906 | 2.698004145 | 2.698004145 | 2.698004145 | 1.114442844 | 1.114442844 | 1.114442844 |
| | orf1ab | 3880 | ISVQELRYMNANGLRP | N | 0.080077475 | 0.080077475 | 0.080077475 | 2.557296182 | 2.557296182 | 2.557296182 | 1.609468357 | 1.609468357 | 1.609468357 |
| | orf1ab | 2042 | VATGDVVLASDDLYVK | N | 0.099898533 | 0.099898533 | 0.099898533 | 1.023111555 | 1.023111555 | 1.023111555 | 1.137047978 | 1.137047978 | 1.137047978 |
| | S | 897 | PHCGSSSRSFFEDLLFDKVKLSDVGF | N | 4.29E-07 | 0.03807346 | 0.004824521 | 2.293577435 | 5.582720832 | 4.348873298 | 1.997603214 | 3.922955993 | 3.065556771 |
| | S | 814 | VTIDCSLFVCSNYAAC | N | 0.000237433 | 0.000237433 | 0.000237433 | 0.832894223 | 0.832894223 | 0.832894223 | 1.700157872 | 1.700157872 | 1.700157872 |
| | S | 12 | LAVIGDFNCTNFAIND | N | 0.000788163 | 0.000788163 | 0.000788163 | 2.375695738 | 2.375695738 | 2.375695738 | 2.046973398 | 2.046973398 | 2.046973398 |
| | S | 1227 | SVPKLSDFESELSHWF | N | 0.015240727 | 0.015240727 | 0.015240727 | 1.994119904 | 1.994119904 | 1.994119904 | 2.03017788 | 2.03017788 | 2.03017788 |
| | S | 1230 | KLSDFESELSHWFKNQTSI | N | 0.030540593 | 0.05401528 | 0.040623089 | 3.636800359 | 3.831769625 | 3.765698594 | 1.963950097 | 2.178166131 | 2.104441735 |
| | S | 1105 | FGAALAMEKVNECVKS | N | 0.061438672 | 0.061438672 | 0.061438672 | 2.152803794 | 2.152803794 | 2.152803794 | 1.116140456 | 1.116140456 | 1.116140456 |
| | M | 161 | YTLSDLPVYVTVAKVQ | N | 0.001132425 | 0.001132425 | 0.001132425 | 1.233532077 | 1.233532077 | 1.233532077 | 1.470249492 | 1.470249492 | 1.470249492 |
| HCoV-OC43 | orf1ab | 3471 | CSVEDFNVWALSNGFS | N | 6.02E-05 | 6.02E-05 | 6.02E-05 | 2.539705355 | 2.539705355 | 2.539705355 | 2.285867892 | 2.285867892 | 2.285867892 |
| | orf1ab | 4603 | ADSYYSYIMPMLTMCH | N | 9.75E-05 | 9.75E-05 | 9.75E-05 | 1.211609753 | 1.211609753 | 1.211609753 | 1.439801037 | 1.439801037 | 1.439801037 |
| | orf1ab | 2453 | ITVEAALDLSKELKRP | N | 0.000100286 | 0.000100286 | 0.000100286 | 3.342319555 | 3.342319555 | 3.342319555 | 2.607384034 | 2.607384034 | 2.607384034 |
| | orf1ab | 5973 | ITKEEAVKRVRAWVGF | N | 0.001172565 | 0.001172565 | 0.001172565 | 1.22986641 | 1.22986641 | 1.22986641 | 1.573387499 | 1.573387499 | 1.573387499 |
| | orf1ab | 6747 | ILLDDFVALVKSLNLNC | N | 0.001255817 | 0.078506035 | 0.039880926 | 1.45215926 | 1.45215926 | 1.45215926 | 1.822741882 | 1.822741882 | 1.822741882 |
| | orf1ab | 789 | ADSVVEVVTTSLTPCG | N | 0.004381829 | 0.004381829 | 0.004381829 | 0.717148911 | 0.717148911 | 0.717148911 | 1.055892691 | 1.055892691 | 1.055892691 |
| | orf1ab | 833 | VVVDDHVGLLDQAWRV | N | 0.011306393 | 0.011306393 | 0.011306393 | 1.24988491 | 1.24988491 | 1.24988491 | 1.085367085 | 1.085367085 | 1.085367085 |
| | orf1ab | 1956 | TATGDVVLATDDLYVK | N | 0.013362631 | 0.013362631 | 0.013362631 | 1.030272488 | 1.030272488 | 1.030272488 | 1.177914102 | 1.177914102 | 1.177914102 |

**Extended data 6**

| Virus | Protein | Start position | Sequence | Identical to SARS-CoV-2? | Min.Pvalue | Max.Pvalue | Mean.Pvalue | Min.Signal | Max.Signal | Mean.Signal | Minimum fold change | Maximum fold change | Average fold change |
|---|---|---|---|---|---|---|---|---|---|---|---|---|---|
| | orf1ab | 5449 | ATIQEIVSERELILSW | N | 0.035607819 | 0.035607819 | 0.035607819 | 0.625904524 | 0.625904524 | 0.625904524 | 1.093508279 | 1.093508279 | 1.093508279 |
| | orf1ab | 6594 | FAVRKEGQDVIFSQFD | N | 0.068668937 | 0.068668937 | 0.068668937 | 1.059032259 | 1.059032259 | 1.059032259 | 1.004118111 | 1.004118111 | 1.004118111 |
| | orf1ab | 5489 | KTVLGEYVFDKSELTN | N | 0.078355906 | 0.078355906 | 0.078355906 | 2.698004145 | 2.698004145 | 2.698004145 | 1.114442844 | 1.114442844 | 1.114442844 |
| | orf1ab | 3792 | ISVQELRYMNANGLRP | N | 0.080077475 | 0.080077475 | 0.080077475 | 2.557296182 | 2.557296182 | 2.557296182 | 1.609468357 | 1.609468357 | 1.609468357 |
| | orf1ab | 5487 | NGKTVLGEYVFDKSEL | N | 0.084099276 | 0.084099276 | 0.084099276 | 2.730902834 | 2.730902834 | 2.730902834 | 1.240457187 | 1.240457187 | 1.240457187 |
| | ns2 | 130 | CTIAQLTDAALSIKEN | N | 6.59E-06 | 6.59E-06 | 6.59E-06 | 2.406950734 | 2.406950734 | 2.406950734 | 2.466762841 | 2.466762841 | 2.466762841 |
| | S | 1225 | TSIPNLPDFKEELDQWFKNQTSVA | N | 6.33E-09 | 0.000199248 | 2.34E-05 | 2.82059038 | 5.791212963 | 4.945558021 | 2.119506153 | 4.135173602 | 3.656200989 |
| | S | 896 | ECSKASSRSAIEDLLFDKVKLSDV | N | 8.19E-08 | 0.068966836 | 0.013024802 | 4.806391274 | 6.649110128 | 5.719781997 | 2.153728539 | 3.962354064 | 3.186988228 |
| | S | 394 | ITIDKFAIPNGRKVDL | N | 3.66E-06 | 3.66E-06 | 3.66E-06 | 3.959581434 | 3.959581434 | 3.959581434 | 3.010625893 | 3.010625893 | 3.010625893 |
| | S | 812 | VTIDCAAFVCGDYAAC | N | 0.000147968 | 0.000147968 | 0.000147968 | 1.280910425 | 1.280910425 | 1.280910425 | 1.502435733 | 1.502435733 | 1.502435733 |
| | S | 579 | ADSCLQGDKCNIFANF | N | 0.000584567 | 0.000584567 | 0.000584567 | 1.856269591 | 1.856269591 | 1.856269591 | 1.640513321 | 1.640513321 | 1.640513321 |
| | M | 13 | WTADEAIKFLKEWNFS | N | 0.000191165 | 0.000191165 | 0.000191165 | 5.157704719 | 5.157704719 | 5.157704719 | 3.674105025 | 3.674105025 | 3.674105025 |
| | N | 11 | SRASSGNRSGNGILKW | N | 6.16E-05 | 6.16E-05 | 6.16E-05 | 2.5148809 | 2.5148809 | 2.5148809 | 1.95270531 | 1.95270531 | 1.95270531 |
| HCoV-NL63 | orf1ab | 4404 | LTITELLQFVTDPSLI | N | 1.81E-08 | 1.81E-08 | 1.81E-08 | 1.959617 | 1.959617 | 1.959617 | 3.637672073 | 3.637672073 | 3.637672073 |
| | orf1ab | 3605 | SVASSFVSMPSYIAYE | N | 2.93E-08 | 2.93E-08 | 2.93E-08 | 1.249413987 | 1.249413987 | 1.249413987 | 2.473242207 | 2.473242207 | 2.473242207 |
| | orf1ab | 3196 | VSVEQLLASIQHLHEG | N | 1.02E-06 | 1.02E-06 | 1.02E-06 | 2.428786859 | 2.428786859 | 2.428786859 | 2.872757212 | 2.872757212 | 2.872757212 |
| | orf1ab | 1749 | VTLEQYSTCDICKSTV | N | 6.47E-06 | 6.47E-06 | 6.47E-06 | 1.902575332 | 1.902575332 | 1.902575332 | 1.940556126 | 1.940556126 | 1.940556126 |
| | orf1ab | 2341 | LTVSDDDFVSAVANAH | N | 2.16E-05 | 2.16E-05 | 2.16E-05 | 1.643952367 | 1.643952367 | 1.643952367 | 1.62160613 | 1.62160613 | 1.62160613 |
| | orf1ab | 874 | YVVDDIYYPASCNGVL | N | 0.000292071 | 0.000292071 | 0.000292071 | 1.279895149 | 1.279895149 | 1.279895149 | 1.504282678 | 1.504282678 | 1.504282678 |
| | orf1ab | 1091 | VDSIVQKCYELSHLIS | N | 0.000814638 | 0.000814638 | 0.000814638 | 0.74952685 | 0.74952685 | 0.74952685 | 1.043742174 | 1.043742174 | 1.043742174 |
| | orf1ab | 43 | RFVAFGLQDCVTGIND | N | 0.004122514 | 0.004122514 | 0.004122514 | 0.580910527 | 0.580910527 | 0.580910527 | 1.069171499 | 1.069171499 | 1.069171499 |
| | orf1ab | 4159 | YTMMDLVYAMRNFDEQ | N | 0.005794406 | 0.005794406 | 0.005794406 | 3.380957649 | 3.380957649 | 3.380957649 | 2.606478894 | 2.606478894 | 2.606478894 |
| | orf1ab | 434 | SIASTVLTVSNGVIIM | N | 0.017254733 | 0.017254733 | 0.017254733 | 0.754879284 | 0.754879284 | 0.754879284 | 1.229462672 | 1.229462672 | 1.229462672 |
| | orf1ab | 4557 | ISYEEQDALFALTKRN | N | 0.052297053 | 0.052297053 | 0.052297053 | 1.74935607 | 1.74935607 | 1.74935607 | 1.021957876 | 1.021957876 | 1.021957876 |
| | orf1ab | 6138 | VAFELFAKRKMGLTPP | N | 0.068668937 | 0.068668937 | 0.068668937 | 1.197852002 | 1.197852002 | 1.197852002 | 1.155970963 | 1.155970963 | 1.155970963 |
| | orf1ab | 5421 | NTVSELVYENKFVPVK | N | 0.081022685 | 0.081022685 | 0.081022685 | 2.556390404 | 2.556390404 | 2.556390404 | 1.146981684 | 1.146981684 | 1.146981684 |
| | S | 809 | SACKTIEDALRLSAHLETND | N | 2.39E-09 | 0.018864546 | 0.004327241 | 1.414280646 | 3.84619817 | 2.613186507 | 1.019627561 | 3.764639897 | 2.128362786 |
| | S | 863 | RSSRIAGRSALEDLLFSKVVTSGL | N | 4.60E-07 | 0.003988237 | 0.000996819 | 3.422598676 | 5.659010583 | 4.510428264 | 2.566430665 | 4.188699911 | 3.503490264 |
| | S | 1264 | QTTVELQGLIDQINSTY | N | 7.30E-07 | 0.001869843 | 0.000935286 | 2.004480571 | 2.004480571 | 2.004480571 | 1.612191441 | 1.612191441 | 1.612191441 |
| | S | 118 | SNASSSFDCIVNLLFT | N | 0.000270868 | 0.000270868 | 0.000270868 | 1.437130102 | 1.437130102 | 1.437130102 | 2.209168332 | 2.209168332 | 2.209168332 |
| | S | 900 | LSIADLACAQYYNGIM | N | 0.004348974 | 0.004348974 | 0.004348974 | 2.901260115 | 2.901260115 | 2.901260115 | 2.642385725 | 2.642385725 | 2.642385725 |
| | S | 1262 | LFQTTVELQGLIDQIN | N | 0.009319009 | 0.009319009 | 0.009319009 | 1.205482012 | 1.205482012 | 1.205482012 | 1.233921898 | 1.233921898 | 1.233921898 |
| | S | 873 | LEDLLFSKVVTSGLGT | N | 0.064874364 | 0.064874364 | 0.064874364 | 2.680977439 | 2.680977439 | 2.680977439 | 1.964081178 | 1.964081178 | 1.964081178 |
| HCoV-229E | orf1ab | 3630 | SVASSFVGMPSFVAYE | N | 1.35E-08 | 1.35E-08 | 1.35E-08 | 1.628400992 | 1.628400992 | 1.628400992 | 2.759706208 | 2.759706208 | 2.759706208 |
| | orf1ab | 4429 | LTITELLQFVTDPTLI | N | 7.42E-08 | 7.42E-08 | 7.42E-08 | 1.805868746 | 1.805868746 | 1.805868746 | 3.132645462 | 3.132645462 | 3.132645462 |
| | orf1ab | 1833 | VSVEQLEPCAQSRLLS | N | 4.19E-06 | 4.19E-06 | 4.19E-06 | 1.939530724 | 1.939530724 | 1.939530724 | 2.171435179 | 2.171435179 | 2.171435179 |
| | orf1ab | 4184 | YTMMDLCFALRNFDEK | N | 4.92E-06 | 4.92E-06 | 4.92E-06 | 3.598754868 | 3.598754868 | 3.598754868 | 3.050448403 | 3.050448403 | 3.050448403 |
| | orf1ab | 1790 | SVASINSAIVCASVKR | N | 0.000424455 | 0.000424455 | 0.000424455 | 1.456886264 | 1.456886264 | 1.456886264 | 1.313831781 | 1.313831781 | 1.313831781 |
| | orf1ab | 3885 | VTVELEPPCRFVIDTP | N | 0.002831917 | 0.002831917 | 0.002831917 | 0.717553713 | 0.717553713 | 0.717553713 | 1.218222078 | 1.218222078 | 1.218222078 |
| | orf1ab | 5446 | NTVSELVYENKFVPVKE | N | 0.007100249 | 0.081022685 | 0.044061467 | 2.556390404 | 3.284096396 | 2.9202434 | 1.146981684 | 1.690871416 | 1.41892655 |
| | orf1ab | 6165 | FELFAKRKVGLTPPLS | N | 0.052198974 | 0.052198974 | 0.052198974 | 1.289224027 | 1.289224027 | 1.289224027 | 1.354317458 | 1.354317458 | 1.354317458 |
| | S | 682 | SGSRVAGRSAIEDILFSKLVTSGL | N | 8.53E-08 | 0.013645547 | 0.001525746 | 2.677477572 | 5.631105658 | 4.018550847 | 2.011734598 | 4.434920589 | 3.694222772 |
| | S | 1080 | ELNYTVQKLQTLIDNI | N | 0.000942039 | 0.000942039 | 0.000942039 | 1.377127359 | 1.377127359 | 1.377127359 | 1.186917297 | 1.186917297 | 1.186917297 |
| | S | 631 | KTIEDALRNSARLESA | N | 0.002845292 | 0.002845292 | 0.002845292 | 3.735051844 | 3.735051844 | 3.735051844 | 2.165313409 | 2.165313409 | 2.165313409 |
| | S | 719 | LSIADLACAQYYNGIM | N | 0.004348974 | 0.004348974 | 0.004348974 | 2.901260115 | 2.901260115 | 2.901260115 | 2.642385725 | 2.642385725 | 2.642385725 |
| | N | 218 | SPASSQTSAKSLARSQ | N | 2.75E-06 | 2.75E-06 | 2.75E-06 | 2.729637946 | 2.729637946 | 2.729637946 | 2.308508164 | 2.308508164 | 2.308508164 |
