## Extended data 7 for "The landscape of antibody binding in SARS-CoV-2 infection"

| S | SARS-CoV-2 epitope | 172-S-16 | 241-S-16 | 289-S-17 | 306-S-16 | 404-S-18 | 410-S-16 | 536-S-17 | 541-S-16 | 549-S-18 | 553-S-26 | 568-S-26 | 613-S-25 | 624-S-23 | 635-S-20 | 644-S-16 |
|---|---|---|---|---|---|---|---|---|---|---|---|---|---|---|---|---|
| | SARS-CoV-2 | 172-172 | 241-241 | 289-290 | 306-306 | 404-406 | 410-410 | 536-537 | 541-541 | 549-551 | 553-563 | 568-578 | 613-622 | 624-631 | 635-639 | 644-644 |
| | Bat CoV (RaTG13) | 172-172 | 241-241 | 289-290 | 306-306 | 404-406 | 410-410 | 536-537 | 541-541 | 549-551 | 553-563 | 568-578 | 613-622 | 624-631 | 635-639 | 644-644 |
| | Pangolin | 172-172 | | | | | | 536-537 | | 549-551 | 553-562 | 568-578 | 619-622 | 624-630 | | |
| | SARS-CoV | | | | | | | | | | 539-548 | | 599-608 | 610-613 | | |
| | MERS-CoV | | | | | | | | | | | | | | | |
| | HCoV-HKU1 | | | | | | | | | | | | | | | |
| | HCoV-OC43 | | | | | | | | | | | | | | | |
| | HCoV-NL63 | | | | | | | | | | | | | | | |
| | HCoV-229E | | | | | | | | | | | | | | | |

| S | SARS-CoV-2 epitope | 656-S-17 | 661-S-16 | 685-S-25 | 761-S-16 | 768-S-18 | 785-S-27 | 798-S-17 | 804-S-17 | 807-S-26 | 844-S-16 | 940-S-16 | 1140-S-25 | 1155-S-20 | 1161-S-17 | 1164-S-16 | 1178-S-16 | 1247-S-27 |
|---|---|---|---|---|---|---|---|---|---|---|---|---|---|---|---|---|---|---|
| | SARS-CoV-2 | 656-657 | 661-661 | 685-694 | 761-761 | 768-770 | 785-796 | 798-799 | 804-805 | 807-817 | 844-844 | 940-940 | 1140-1149 | 1155-1159 | 1161-1162 | 1164-1164 | 1178-1178 | 1247-1262 |
| | Bat CoV (RaTG13) | 656-657 | 661-661 | 681-690 | 757-757 | 764-766 | 781-792 | 794-795 | 800-801 | 803-813 | 840-840 | 936-936 | 1136-1145 | 1151-1155 | 1157-1158 | 1160-1160 | 1174-1174 | 1243-1258 |
| | Pangolin | 657-657 | | 682-690 | 757-757 | 764-766 | 781-792, 794, 796 | 798-799 | | 803-813 | 840-840 | 936-936 | 1136-1145 | 1151-1155 | 1157-1158 | 1160-1160 | 1174-1174 | 1243-1258 |
| | SARS-CoV | 643-643 | 647-647 | 668-674 | | | 776-777 | | | 789-799 | | | 1122-1131 | 1137-1141 | 1143-1144 | 1146-1146 | 1160-1160 | 1230-1244 |
| | MERS-CoV | | | | | | | | | 880, 882-887 | | | 1223-1232 | | | | | |
| | HCoV-HKU1 | | | | | | | | | 897-907 | | | 1230-1233 | | | | | |
| | HCoV-OC43 | | | | | | | | | 896-904 | | | 1225-1233 | | | | | |
| | HCoV-NL63 | | | | | | | | | 863-871 | | | | | | | | |
| | HCoV-229E | | | | | | | | | 682-690 | | | | | | | | |

| M | SARS-CoV-2 epitope | 1-M-24 | 152-M-26 | 175-M-20 | 181-M-32 | 205-M-18 |
|---|---|---|---|---|---|---|
| | SARS-CoV-2 | 1-9 | 152-162 | 175-179 | 181-197 | 205-211 |
| | Bat CoV (RaTG13) | 1-9 | 152-162 | 175-179 | 181-197 | |
| | Pangolin | 1-8 | 151-161 | 174-178 | 180-196 | 204-210 |
| | SARS-CoV | 1-8 | 152-161 | 174-178 | 180-196 | 204-210 |
| | MERS-CoV | | | | | 208 |
| | HCoV-HKU1 | | 161 | | | |
| | HCoV-OC43 | | | | | |
| | HCoV-NL63 | | | | | |
| | HCoV-229E | | | | | |

| N | SARS-CoV-2 epitope | 7-N-21 | 14-N-17 | 28-N-28 | 94-N-16 | 96-N-17 | 117-N-19 | 122-N-16 | 124-N-16 | 126-N-17 | 153-N-26 | 208-N-31 | 227-N-17 | 230-N-21 | 242-N-19 | 249-N-18 | 336-N-16 | 338-N-19 | 356-N-16 | 376-N-22 | 384-N-33 |
|---|---|---|---|---|---|---|---|---|---|---|---|---|---|---|---|---|---|---|---|---|---|
| | SARS-CoV-2 | 7-12 | 14-15 | 28-40 | 94-94 | 96-97 | 117-120 | 122-122 | 124-124 | 126-127 | 153-163 | 208-223 | 227-228 | 230-235 | 242-245 | 249-251 | 336-336 | 338-341 | 356-356 | 376-382 | 384-401 |
| | Bat CoV (RaTG13) | 7-12 | 14-15 | 28, 30-40 | 94-94 | 96-97 | 117-120 | 122-122 | 124-124 | 126-127 | 153-163 | 210-223 | 227-228 | 230-234 | 242-245 | 249-251 | 336-336 | 338-341 | 356-356 | 376-382 | 384-401 |
| | Pangolin | 5-10 | | 28-38 | 92-92 | 94-95 | | | | 126-126 | 152-159, 161 | 206-221 | 225-226 | 228-231 | 240-243 | 247-249 | | 336,338-339 | 354-354 | 375-376,378-380 | 382-399 |
| | SARS-CoV | 9-13 | | 30-41 | 95-95 | 97-97 | | | | | 154-164 | 216-224 | 228-228 | 233-236 | 243-246 | 250-252 | 337-337 | 339, 341 | 357-357 | 377-377 | 388-397, 400-401 |
| | MERS-CoV | | | | | | | | 115-115 | | 173-173 | | | | | | | | | | |
| | HCoV-HKU1 | | | | | | | | | | | | | | | | | | | | |
| | HCoV-OC43 | 11-11 | | | | | | | | | | | | | | | | | | | |
| | HCoV-NL63 | | | | | | | | | | | | | | | | | | | | |
| | HCoV-229E | | | | | | | | | | | | | | | | | | | | |

| ORF1ab | SARS-CoV-2 epitope | 1239-orf1ab-18 | 1546-orf1ab-16 | 1551-orf1ab-16 | 1572-orf1ab-16 | 1681-orf1ab-16 | 1720-orf1ab-16 | 2309-orf1ab-16 | 2584-orf1ab-16 | 4451-orf1ab-16 | 4514-orf1ab-16 | 5999-orf1ab-16 | 6057-orf1ab-17 |
|---|---|---|---|---|---|---|---|---|---|---|---|---|---|
| | SARS-CoV-2 | 1239-1241 | 1546-1546 | 1551-1551 | 1572-1572 | 1681-1681 | 1720-1720 | 2309-2309 | 2584-2584 | 4451-4451 | 4514-4514 | 5999-5999 | 6057-6058 |
| | Bat CoV (RaTG13) | 1238-1240 | | | 1571-1571 | 1680-1680 | 1720-1720 | 2308-2308 | | 4450-4450 | 4513-4513 | 5998-5998 | 6056-6057 |
| | Pangolin | | | | 1564-1564 | 1673-1673 | 1712-1712 | | 2576-2576 | | 4506-4506 | 5991-5991 | 6050-6050 |
| | SARS-CoV | 1217-1217 | | | | | 1697-1697 | | | | 4491-4491 | 5976-5976 | |
| | MERS-CoV | | | | | | | | | | 4501-4501 | | |
| | HCoV-HKU1 | | | | | | | | | | | 6061-6061 | |
| | HCoV-OC43 | | | | | | | | | | | | |
| | HCoV-NL63 | | | | | | | | | | 4159-4159 | | |
| | HCoV-229E | | | | | | | | | | 4184-4184 | | |

| ORF3a | SARS-CoV-2 epitope | 16-orf3a-16 | 18-orf3a-16 | 21-orf3a-16 | 235-orf3a-17 | 252-orf3a-24 |
|---|---|---|---|---|---|---|
| | SARS-CoV-2 | 16-16 | 18-18 | 21-21 | 235-236 | 252-264 |
| | Bat CoV (RaTG13) | | | | 235-236 | 253-258, 260-261 |
| | Pangolin | | | | 234 | 254-255 |
| | SARS-CoV | | | 24 | | 253 |
| | MERS-CoV | | | | | |
| | HCoV-HKU1 | | | | | |
| | HCoV-OC43 | | | | | |

| | | | |
|---|---|---|---|
| | HCoV-NL63 | | |
| | HCoV-229E | | |
| **ORF6** | **SARS-CoV-2 epitope** | **9-orf6-16** | |
| | SARS-CoV-2 | 9-9 | |
| | Bat CoV (RaTG13) | 9-9 | |
| | Pangolin | | |
| | SARS-CoV | | |
| | MERS-CoV | | |
| | HCoV-HKU1 | | |
| | HCoV-OC43 | | |
| | HCoV-NL63 | | |
| | HCoV-229E | | |
| **ORF8** | **SARS-CoV-2 epitope** | **60-orf8-20** | **66-orf8-18** |
| | SARS-CoV-2 | 60-64 | 66-68 |
| | Bat CoV (RaTG13) | 60-61 | 64-68 |
| | Pangolin | | |
| | SARS-CoV | | |
| | MERS-CoV | | |
| | HCoV-HKU1 | | |
| | HCoV-OC43 | | |
| | HCoV-NL63 | | |
| | HCoV-229E | | |
