## Extended data 8 for "The landscape of antibody binding in SARS-CoV-2 infection"

| Epitope identifier | Adjusted p-value, intubated vs never-hospitalized | Log2 odds ratio, intubated vs never-hospitalized |
|---|---|---|
| 21-orf3a-16 | <0.01 | 0.73 |
| 18-orf3a-16 | <0.01 | 0.48 |
| 613-S-25 | <0.01 | 1.2 |
| 16-orf3a-16 | <0.01 | 0.45 |
| 336-N-16 | <0.01 | 0.72 |
| 376-N-22 | 0.01 | 1 |
| 289-S-17 | 0.02 | 1.22 |
| 252-orf3a-24 | 0.02 | 1.83 |
| 1-M-24 | 0.02 | 1.32 |
| 1140-S-25 | 0.08 | 3.36 |
| 553-S-26 | 0.08 | 1.09 |
| 28-N-28 | 0.09 | 0.75 |
| 5999-orf1ab-16 | 0.09 | 0.5 |
| 1247-S-27 | 0.09 | 0.53 |
| 798-S-17 | 0.11 | 0.82 |
| 568-S-26 | 0.12 | 0.5 |
| 761-S-16 | 0.12 | 0.47 |
| 338-N-19 | 0.12 | 0.39 |
| 181-M-32 | 0.24 | 0.31 |
| 785-S-27 | 0.26 | 1.43 |
| 241-S-16 | 0.26 | 0.35 |
| 1155-S-20 | 0.26 | 0.67 |
| 306-S-16 | 0.26 | 0.48 |
| 768-S-18 | 0.26 | 0.46 |
| 9-orf6-16 | 0.26 | 0.3 |
| 1239-orf1ab-18 | 0.26 | 0.64 |
| 1720-orf1ab-16 | 0.36 | 0.5 |
| 126-N-17 | 0.36 | 0.34 |
| 661-S-16 | 0.42 | 0.44 |
| 410-S-16 | 0.42 | -0.4 |
| 624-S-23 | 0.42 | 0.36 |
| 635-S-20 | 0.42 | 0.55 |
| 644-S-16 | 0.42 | 0.35 |
| 235-orf3a-17 | 0.42 | 0.27 |
| 124-N-16 | 0.42 | 0.29 |
| 356-N-16 | 0.42 | 0.21 |
| 1681-orf1ab-16 | 0.43 | 0.27 |
| 2309-orf1ab-16 | 0.43 | 0.18 |
| 1178-S-16 | 0.43 | 0.39 |
| 807-S-26 | 0.43 | 0.24 |

| | | |
|---|---|---|
| 12-orf8-16 | 0.43 | 0.53 |
| 66-orf8-18 | 0.43 | -0.39 |
| 122-N-16 | 0.43 | 0.33 |
| 153-N-26 | 0.43 | 0.14 |
| 208-N-31 | 0.43 | 0.36 |
| 7-N-21 | 0.45 | 0.36 |
| 656-S-17 | 0.45 | 0.39 |
| 6057-orf1ab-17 | 0.46 | -0.39 |
| 549-S-18 | 0.48 | 0.25 |
| 1551-orf1ab-16 | 0.48 | 0.22 |
| 2584-orf1ab-16 | 0.5 | 0.84 |
| 249-N-18 | 0.52 | 0.25 |
| 1164-S-16 | 0.54 | 0.37 |
| 242-N-19 | 0.54 | 0.23 |
| 4514-orf1ab-16 | 0.55 | 0.14 |
| 1161-S-17 | 0.55 | 0.3 |
| 4451-orf1ab-16 | 0.58 | 0.17 |
| 60-orf8-20 | 0.59 | -0.22 |
| 227-N-17 | 0.59 | -0.29 |
| 53-orf8-16 | 0.61 | -0.33 |
| 1572-orf1ab-16 | 0.65 | -0.09 |
| 230-N-21 | 0.67 | -0.39 |
| 404-S-18 | 0.68 | -0.1 |
| 94-N-16 | 0.68 | 0.2 |
| 804-S-17 | 0.69 | 0.36 |
| 940-S-16 | 0.69 | 0.04 |
| 384-N-33 | 0.69 | 0.07 |
| 172-S-16 | 0.7 | -0.14 |
| 844-S-16 | 0.7 | -0.37 |
| 96-N-17 | 0.7 | -0.24 |
| 175-M-20 | 0.72 | -0.29 |
| 685-S-25 | 0.73 | 0.17 |
| 205-M-22 | 0.75 | -0.02 |
| 152-M-26 | 0.77 | -0.04 |
| 541-S-16 | 0.81 | -0.02 |
| 536-S-17 | 0.85 | 0.02 |
| 117-N-19 | 0.87 | -0.02 |
| 1546-orf1ab-16 | 0.92 | -0.03 |
| 14-N-17 | 1 | -0.02 |
